# Prediction of *Burkholderia pseudomallei* DsbA substrates identifies potential virulence factors and vaccine targets

**DOI:** 10.1101/2020.07.03.186213

**Authors:** Ben Vezina, Guillaume A. Petit, Jennifer L. Martin, Maria A. Halili

## Abstract

Identification of bacterial virulence factors is critical for understanding disease pathogenesis, drug discovery and vaccine development. In this study we used two approaches to predict virulence factors of *Burkholderia pseudomallei*, the Gram-negative bacterium that causes melioidosis. *B. pseudomallei* is naturally antibiotic resistant and there are no melioidosis vaccines. To identify *B. pseudomallei* protein targets for drug discovery and vaccine development, we chose to search for substrates of the *B. pseudomallei* periplasmic disulfide bond forming protein A (DsbA). DsbA introduces disulfide bonds into extra-cytoplasmic proteins and is essential for virulence in many Gram-negative organism, including *B. pseudomallei*. The first approach to identify *B. pseudomallei* DsbA virulence factor substrates was a large-scale genomic analysis of 511 unique *B. pseudomallei* disease-associated strains. This yielded 4,496 core gene products, of which we hypothesise 263 are DsbA substrates. Manual curation of the 263 mature proteins yielded 73 associated with disease pathogenesis or virulence. These were screened for structural homologues to predict potential B-cell epitopes. In the second approach, we searched the *B. pseudomallei* genome for homologues of the more than 90 known DsbA substrates in other bacteria. Using this approach, we identified 15 potential *B. pseudomallei* DsbA virulence factor substrates. Two putative *B. pseudomallei* virulence factors were identified by both methods: homologues of PenI family β-lactamase and of succinate dehydrogenase flavoprotein subunit. These two proteins could serve as high priority targets for future *B. pseudomallei* virulence factor characterization.

## Introduction

*Burkholderia pseudomallei* is a Gram-negative soil dwelling saprophyte, and an opportunistic pathogen responsible for the severe tropical disease melioidosis [1]. *B. pseudomallei* infections are difficult to treat [2-4] and are intrinsically resistant to almost all available antibiotics [5-8]. Predominant resistance factors utilised by *B. pseudomallei* include a thick, impermeable cell wall combined with efficient efflux pumps that interfere with drug activity [9]. Furthermore, *B. pseudomallei* infections are difficult to diagnose as melioidosis symptoms vary significantly, ranging from fever, pneumonia, urinary tract infections, and on rare occasions encephalomyelitis [4]. Standard treatment consists of a combination of intravenous antibiotic for two weeks to stop septicaemia, followed by a second eradication phase that can last for up to six months, with no guarantee of success [10].

More generally, antibiotic resistance is increasing at an accelerating rate among pathogenic bacteria [11]. New approaches and treatment strategies are needed including vaccination [12], novel antimicrobial compounds [13] and antivirulence strategies [14]. There is currently no successful, persistent vaccine against *B. pseudomallei* [15]. However Outer Membrane Protein A (OmpA) has been used as a subunit vaccination against melioidosis in mice [16].

Identification of *B. pseudomallei* virulence factors would contribute towards understanding pathogenesis and could aid in drug discovery and vaccine development [17]. Targeting virulence rather than viability is an approach that is hypothesized to have a number of benefits including an increased range of possible anti-virulence mechanisms compared to antimicrobial compounds, as well as the possibility of reducing selection pressure [18, 19]. Both vaccine development and anti-virulence approaches could reduce selection pressure and potentially reduce resistance development [14, 18, 19].

The formation of correct disulfide bonds is critical for the proper folding and function of proteins [20]. In bacteria, the introduction of disulfide bonds is mediated by the DiSufide Bond-forming proteins (DSB). The DSB proteins are of particular interest as an antivirulence strategy, because many virulence factors contain disulfide bonds [19, 21-23]. The Disulfide bond forming protein A (DsbA) is a periplasmic protein found in most Gram-negative bacteria and incorporates a thioredoxin fold with two cysteines which introduce disulfide bonds into substrate proteins via a redox transfer reaction [24].

Mice infected with *B. pseudomallei* DsbA knockouts (or of its redox partner DsbB) have an increased rate of survival compared with mice infected with wild type *B. pseudomallei* [25, 26]. These findings suggest that many *B. pseudomallei* virulence factors are substrates of DsbA, as is also observed in *Escherichia coli* [27, 28], *Klebsiella pneumoniae* [29], *Salmonella enterica* [30], *Francisella tularensis* [31] and many more [22, 23, 32]. However, the full extent of *B. pseudomallei* DsbA substrates has not been investigated. Identification of *B. pseudomallei* DsbA substrates would help identification of infection mechanisms, and could lead to the discovery of key virulence factors and potential drug and vaccine targets. Finding potential DsbA substrates is assisted by the observation that: (i) DsbA is located in the periplasm, and thus its substrates are likely to have a secretion signal sequence; and (ii) proteins containing disulfide bonds may have an even rather than an odd number of cysteines in their sequence. This last point is thought to have evolved to limit formation of mis-matched disulfide bonds and therefore misfolded proteins [33, 34].

In the present study, we used two approaches to identify potential *B. pseudomallei* DsbA substrates for further study as virulence factors. In one approach, we used computational methods to generate a curated list of 263 putatively extra-cytoplasmic proteins from the core genome of 511 disease-associated isolates of *B. pseudomallei*, 73 of which were predicted to be virulence-associated. In the second approach, 15 candidate DsbA virulence factor substrates were identified by sequence homology to known DsbA virulence factor substrates in other bacteria.

## Results

### Genomic analysis to predict *B. pseudomallei* DsbA virulence factor substrates

In this approach, our strategy was to cast a wide net initially, by determining the pangenome of disease-associated isolates of *B. pseudomallei*, and then filtering from that the core genome (i.e. the highly conserved genes). The disease-associated *B. pseudomallei* core genome should then be enriched in conserved virulence factors. At the time of this analysis the NCBI database [35] contained 1577 *B. pseudomallei* isolates. Metadata notation allowed selection of 512 isolates associated with disease (i.e. isolates from swabs/clinical isolates: accession numbers of these are given in S1 Fig); other genomes were discarded. We note that only 355 of the 512 isolates were tagged ‘pathogen’ in the NCBI database indicating a discrepancy between NCBI assignment and user-uploaded metadata. Analysis of the pangenome, that is the core, accessory and unique genes of these 512 *B. pseudomallei* isolates (see Table 1), revealed two identical strains. Therefore for the remainder of this analysis, only the 511 unique strains were used.

**Table 1:**
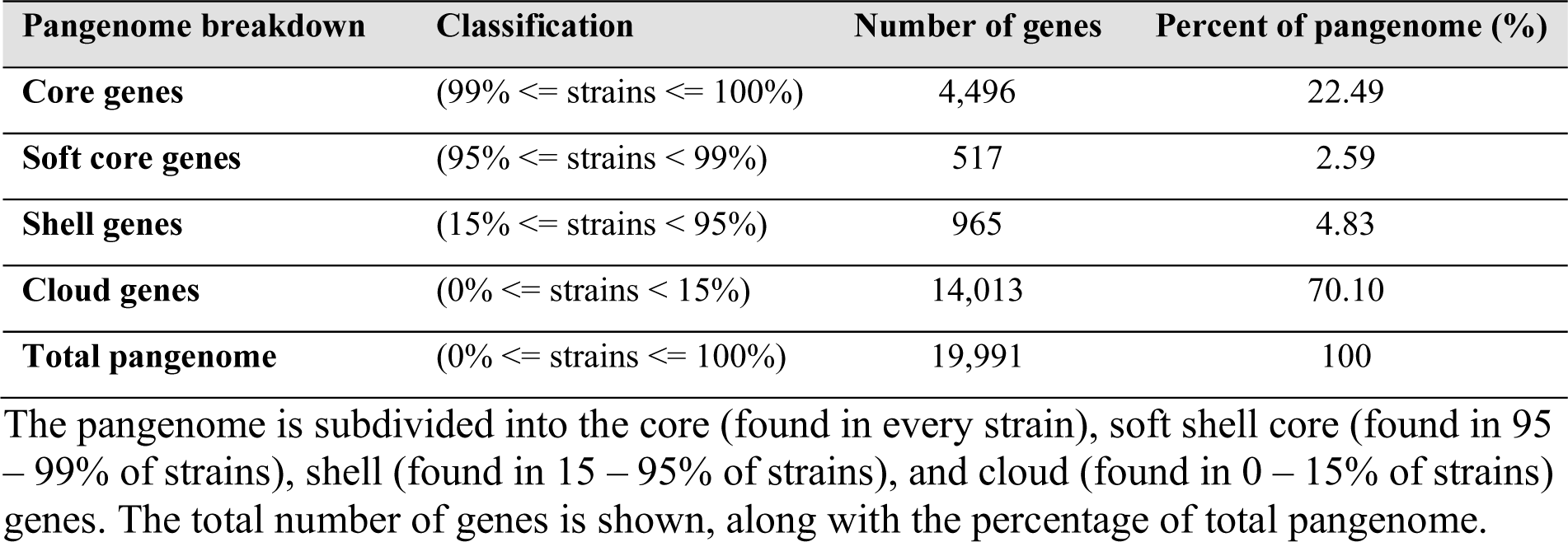
Pangenome results of 511 disease-associated *B. pseudomallei* strains.

We found that the core genome consisted of 4,496 genes (see S2 Fig) or 22.49% of the total 19,991 pangenome. This analysis largely agrees with a previous pangenomic analysis which extrapolated a modelled core genome of 4,568±16 from a much smaller set of 37 isolate genomes [36]. In that approach, modelling was used to predict the core genome if the number of isolates was expanded. Our approach gives an exact number because all 4,496 genes were found in all 511 genomes. Notably, the dithiol oxidase redox enzyme pair DsbA and DsbB and the disulfide isomerase redox relay enzymes DsbC and DsbD were all identified as core genes.

We then used the *B. pseudomallei* core genome for further analysis, because it encodes highly conserved proteins - a key criteria for selecting vaccine or anti-virulence targets.

From these 4,496 core genes, 726 were predicted to encode proteins with a signal sequence and which are therefore likely to be exported out of the cytoplasm and into the periplasm where DsbA is localised. Of these 726 proteins, 263 have an even number of cysteines, indicating the likelihood that the proteins form intramolecular disulfide bonds (see S3 Fig). We predict that these 263 proteins are substrates of *B. pseudomallei* DsbA. The workflow for this analysis is shown in Fig 1.

**Fig 1:**
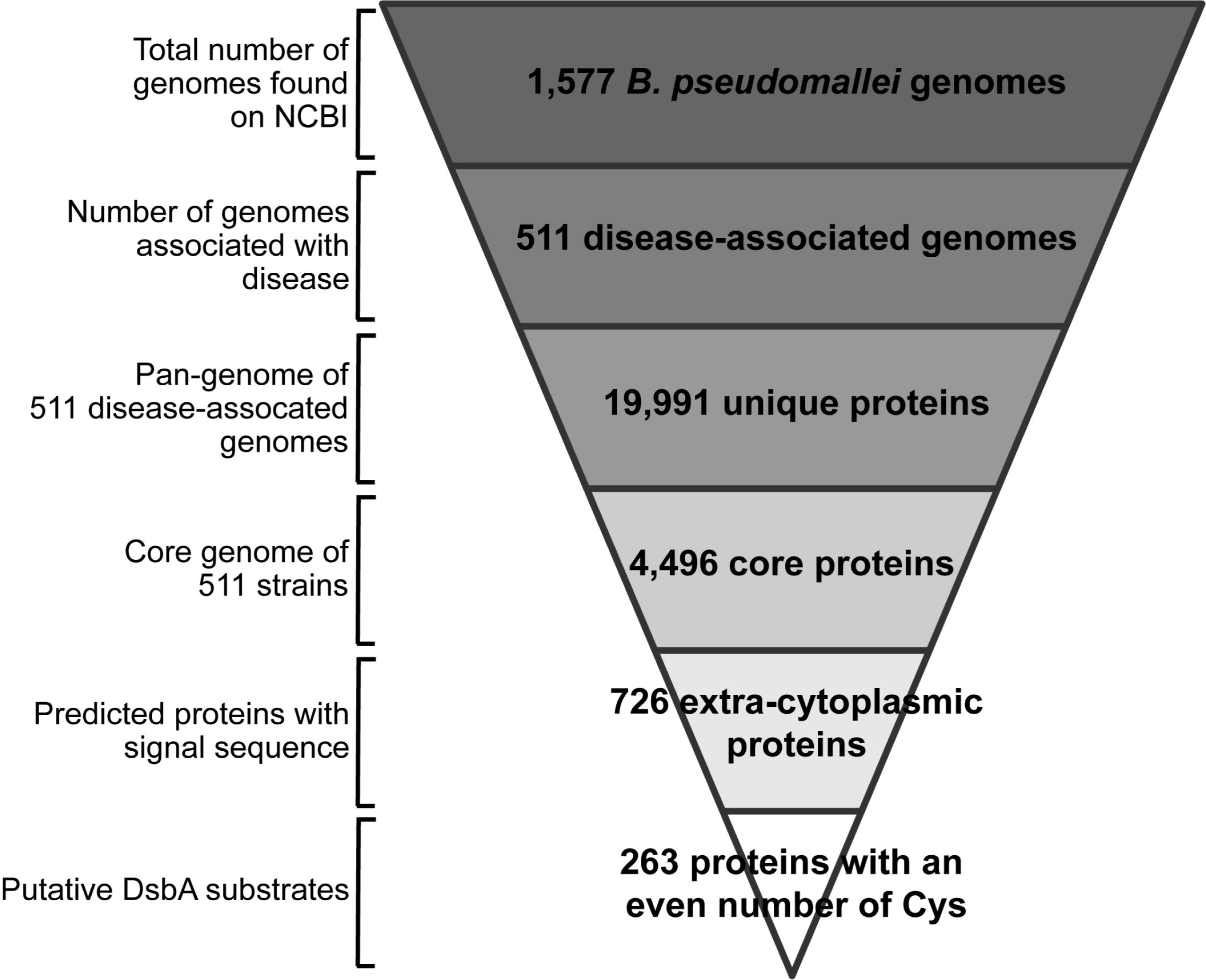
Bioinformatic workflow. From the 1,577 *B. pseudomallei* genomes found on NCBI, 511 were unique and associated with disease and these were used for further analysis. The pangenome of these 511 genomes comprised 19,991 unique genes. 4,496 of these were classified as core genes. Predicted translation of these genes gave 726 predicted extra-cytoplasmic proteins. Of these extra-cytoplasmic proteins, 263 were predicted to contain an even number of cysteines. We predict that these 263 proteins are substrates of *B. pseudomallei* DsbA.

### Distribution of cysteines in the core genome of disease-related *B. pseudomallei*

Many bacterial extra-cytoplasmic (periplasmic and extracellular) proteins have a strong preference for an even number of cysteines, which is thought to reduce the chances of non-native disulfide bond formation [33]. We examined the cysteine distribution of encoded proteins in the *B. pseudomallei* pangenome to investigate whether the previously demonstrated enrichment of an even number of cysteines in extra-cytoplasmic proteins in other Gram-negative bacteria [33] was also true for *B. pseudomallei*.

The distribution of cysteines in *B. pseudomallei* cytoplasmic and extra-cytoplasmic proteins was calculated for the pangenome (total of 19,991 genes) and the core genome (4,496 genes) (refer to Table 1). In cytoplasmic *B. pseudomallei* proteins, cysteine distribution followed a Poisson law peaking at zero for the pangenome and at one for the core genome (denoted by the orange lines in the histograms on Figs 2A and 2B). This distribution changed for extra-cytoplasmic *B. pseudomallei* proteins. For the core genome (blue bars Fig 2B), *B. pseudomallei* proteins with an even number of cysteines were over-represented compared to a typical Poisson distribution. As extra-cytoplasmic proteins represent a small fraction of the total number of the translated core genome and pangenome (16% and 11.5% of all proteins, respectively), we also analysed the normalised frequency (Figs 2C and 2D). The core genome normalised cysteine distribution reveals a sawtooth pattern with a preference for even number of cysteines with peaks for two, four, six and eight cysteines (Fig 2D). In contrast, the pangenomic normalised cysteine distribution for extra-cytoplasmic *B. pseudomallei* proteins does not indicate a strong preference for even number of cysteines (Fig 2C). Overall, the saw-tooth pattern observed in Figs 2B and 2D is similar to that described for *E. coli* exported proteins [33] although not as pronounced.

**Fig 2:**
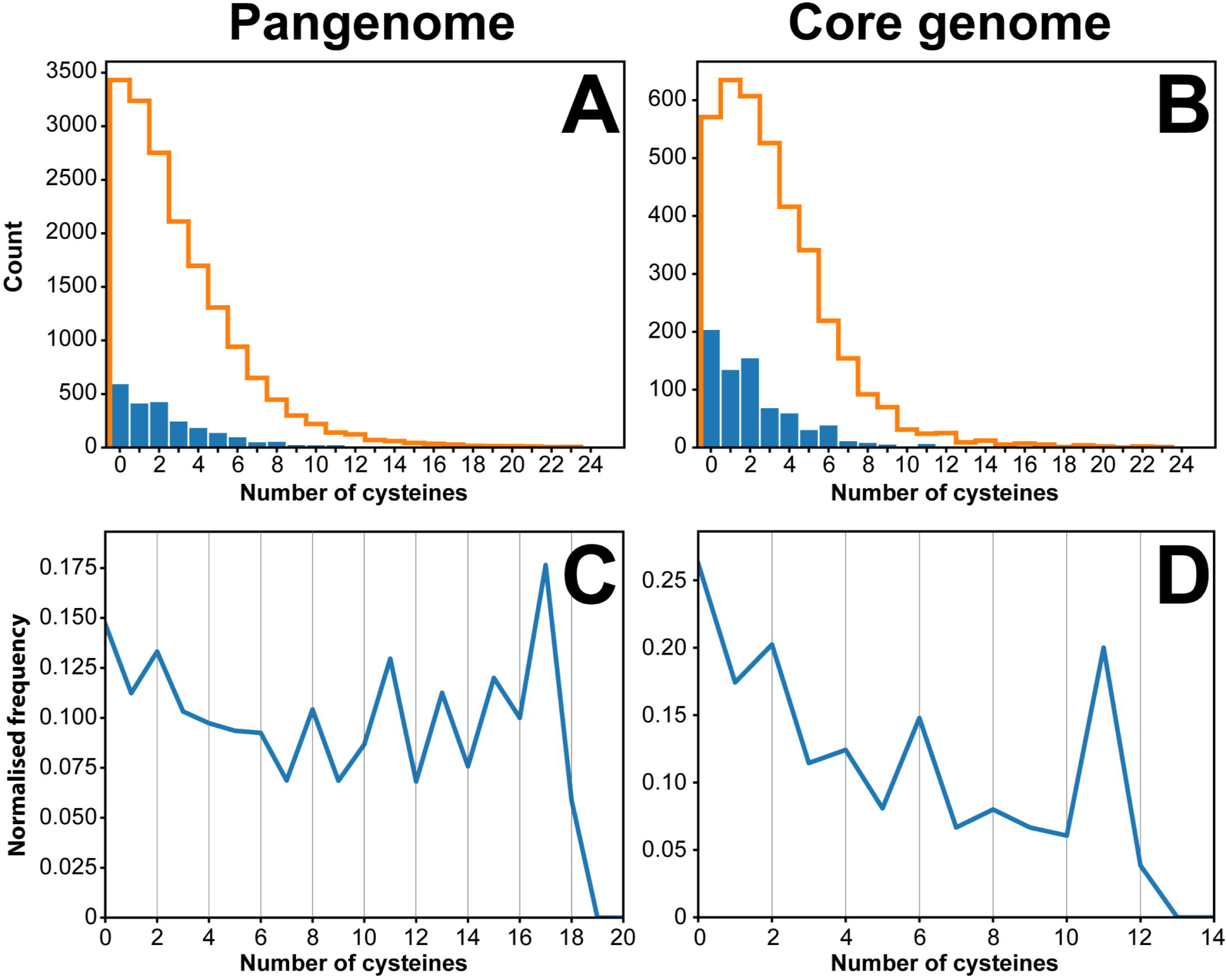
Cysteine distribution in the translated genome of *B. pseudomallei*. Panel **A** shows the distribution of cysteines in the pangenome (19,991 proteins). Panel **B** represents the same analysis for the core genome, comprising 4,496 translated genes. Predicted number of extra-cytoplasmic proteins for each number of cysteines are represented as blue bars. Similarly, predicted cytoplasmic proteins are represented as orange lines. Panels **C** and **D** represent the normalised frequency of cysteine-containing extra-cytoplasmic proteins. The blue line in panel **D** peaks for proteins with 2, 4, 6 and 8 cysteines suggesting a preference for an even number of cysteines. This trend is not observed as strongly in panel **C**, where a clear peak can only be seen for two and eight cysteines. The normalised frequency was calculated by dividing the number of extra-cytoplasmic proteins (having *N* number of cysteines) by the total number of proteins with *N* cysteines (*N* being a number between 0 - 20 as per the data points in **C** and **D** above).

### Functional assignment of core, extra-cytoplasmic, putative DsbA substrates

The next step in the genomic analysis was to predict which of the 263 putative DsbA substrates are associated with virulence. Of the 263 selected proteins, 44 were annotated as hypothetical/uncharacterised. The remaining 219 proteins include ABC transporter-related proteins, housekeeping proteins like cytochrome C, proteins required for motility such as flagellar and fimbrial proteins, enzymes such as collagenase, peptidases and proteases, as well as antibiotic resistance enzymes, β-lactamases. Many oxidoreductases were also present including DsbA, DsbD and others such as Gfo/Idh/MocA family, glycerol-3-phosphate dehydrogenase GpsA and thioredoxin-like TlpA oxidoreductases. Redox enzymes such as DsbB and DsbC are core genes with signal sequences, and they have catalytic rather than structural disulfides. These two enzymes are not identified as DsbA substrates in our filter as they have an odd number of cysteines.

Gene Ontology (GO) classification of the gene and gene-product function of the 263 proteins reveals a variety of functions, totalling 223 GO descriptions (Fig 3) (see S4 File for a full list). The highest frequency are integral components of the membrane (66 proteins), followed by proteins involved in redox processes (25 proteins). Of particular interest due to their putative involvement in virulence, are proteins associated with: proteolysis (20), heme binding (15), hydrolase activity (9), carbohydrate metabolism (8), serine-type endopeptidase activity (7), cell adhesion (6), metallo-endopeptidase activity (6), pilus formation and organisation (6), copper binding (5), lipid catabolism (4), choline binding (3), triglyceride lipase activity (3), aminopeptidase activity (2), porin activity (OmpA family proteins) (2), chitin catabolism (1), *N*-carbamoylputrescine amidase activity (1) and toxin activity (Tat pathway signal protein) (1).

**Fig 3:**
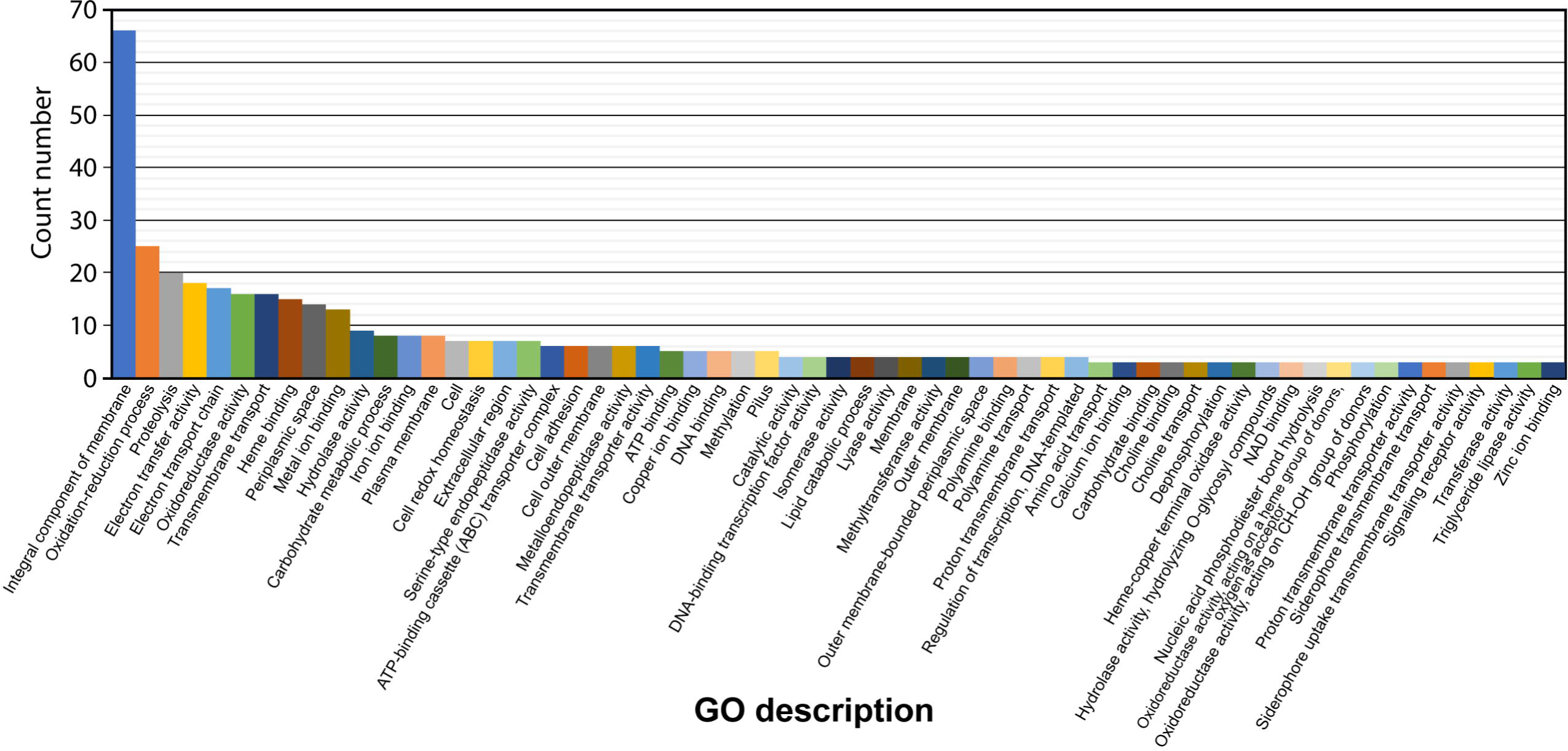
Gene Ontology (GO) descriptions of predicted extra-cytoplasmic proteins with an even number of cysteines. The highest frequency of proteins with an even number of cysteines are integral components of membranes (66 proteins), followed by proteins involved in redox (oxidation-reduction) processes (25 proteins) and proteolysis (20 proteins). For ease of representation and clarity, GO descriptors with less than three counts were excluded from this graph. A complete graph, along with raw values can be found in S4 File.

By further inspection of the 263 core, putatively extra-cytoplasmic DsbA substrates, and by using the GO descriptions to aid in predicting protein functions,73 sequences were identified which were virulence-associated (Table 2). These include serine-type endopeptidases [37] associated with adherence, choline binding proteins N-carbamoylputrescine amidase, essential for production of putrescine, a component of Gram-negative cell walls of pathogens and key virulence [39-42], many proteases and peptidases.

**Table 2:**
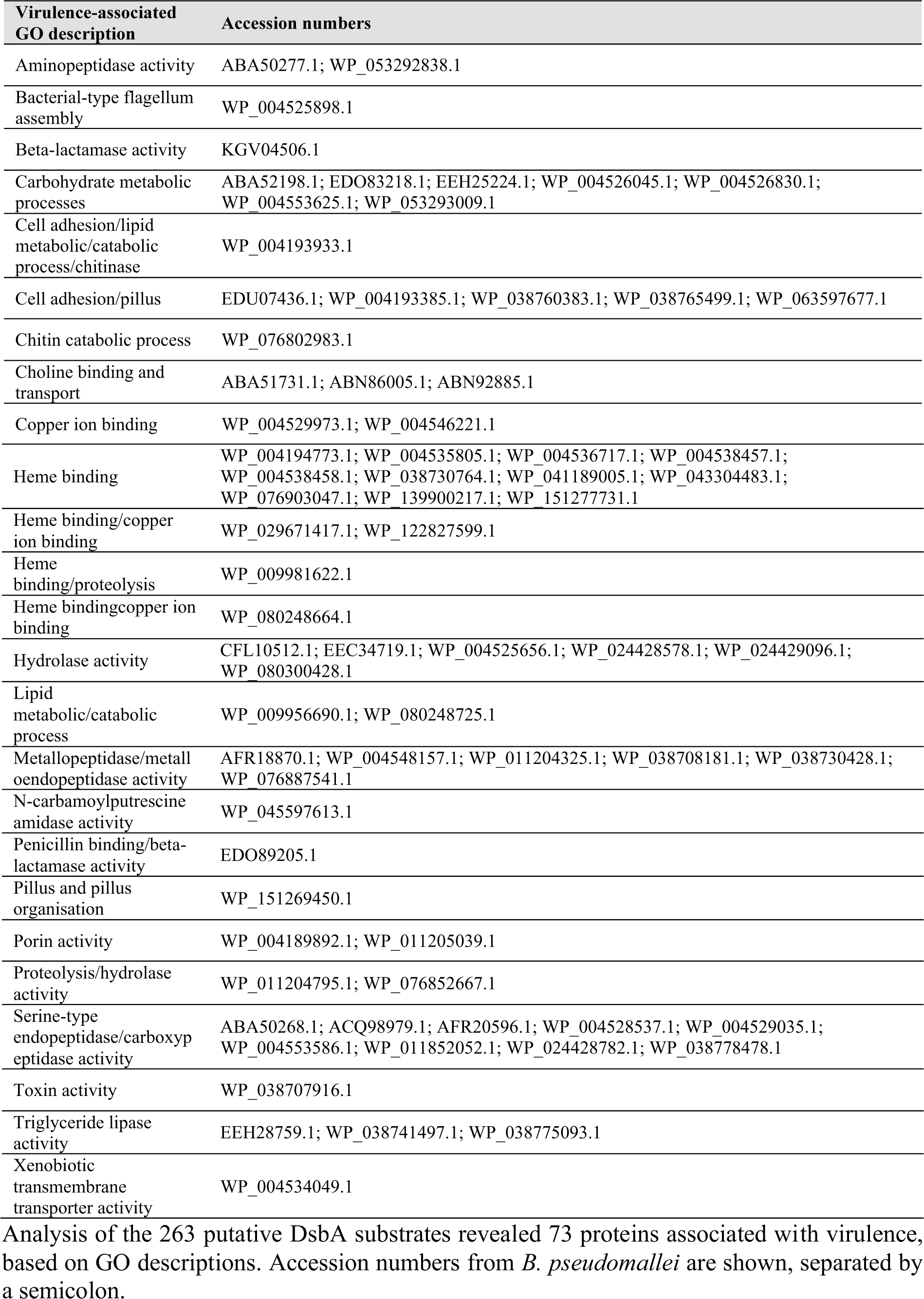
Predicted virulence-associated core, extra-cytoplasmic proteins.

### Sequence homology prediction of *B. pseudomallei* DsbA virulence factor substrates

To complement the genomic analysis described above we used a second approach to identify DsbA substrates, by screening all *B. pseudomallei* genomes uploaded on NCBI [43] (taxid 28450) for homologues of known DsbA substrates. We implemented this approach because some DsbA substrates might be filtered out using the genomic approach described above if the substrates are not encoded by core genes, or if the gene product has an odd number of cysteines.

Over 90 DsbA substrates have been reported in the literature. We searched for *B. pseudomallei* homologues of these DsbA substrates using the following criteria: (i) presence of secretion signal, (ii) at least two cysteines in the mature sequence, (iii) at least 20% identity and (iv) 50% coverage to a known DsbA substrate sequence. After removing duplicates, our analysis found that *B. pseudomallei* encodes homologues of 15 DsbA substrates (Table 3). Two of these 15 are DsbA substrates in other *Burkholderia* species *B. cepacia* and *B. cenocepacia* [44-47]: a metalloproteases, ZmpA and a sulfatase-like hydrolase transferase. In *B. cenocepacia*, ZmpA is a wide spectrum metalloprotease, thought to cause tissue damage during infection [48].

**Table 3:** List of *B. pseudomallei* proteins homologous to previously reported DsbA substrates.

| Accession Number (DsbA substrate) | Organism | Reference | *B. pseudomallei* homologue | Identity / coverage (%) | Protein function | Cys # |
| --- | --- | --- | --- | --- | --- | --- |
| WP_059237834 | *B. cepacia* | [44] | WP_076835606.1 | 89 /100 | Sulfatase like hydrolase /transferase | 3 |
| WP_006481898 | *B. cenocepacia* | [45, 46] | WP_139900467 | 87/100 | M4 family metallopeptidase | 4 |
| gi\|89255876 | *F. tularensis* | [34] | WP_050859308 | 24/92 | lytic transglycosylase | 3 |
| gi\|89255615 | *F. tularensis* | [34] | WP_080367462 | 40/51 | Pilin | 2 |
| gi\|89255615 | *F. tularensis* | [34] | WP_076953316 | 27/92 | Pilin | 2 |
| gi\|89256194 | *F. tularensis* | [34] | WP_041862011 | 30/83 | Molybdopterin synthase adenyl transferase (MoeB) | 13 |
| gi\|89256236 | *F. tularensis* | [34] | WP_064459078 | 34/53 | DNA/RNA endonuclease | 2 |
| gi\|89256237 | *F. tularensis* | [34] | WP_050772403 | 31/90 | PenI family Beta-lactamase | 4 |
| gi\|89256856 | *F. tularensis* | [34] | WP_044360358 | 21/80 | hypothetical protein | 4 |
| gi\|89256859 | *F. tularensis* | [34] | WP_058035453 | 39/80 | Polyamine ABC transporter substrate binding protein | 3 |
| gi\|89257049 | *F. tularensis* | [34] | WP_009915682 | 54/99 | Succinate dehydrogenase | 6 |
| WP_001363619 | *E. coli* | [22] | WP_102811167 | 38/88 | Molecular chaperone | 3 |
| AAC38377 | *E. coli* | [22] | WP_082252625 | 44/93 | T3SS outer membrane ring protein | 4 |
| AAA24962 | *Haemophilus Influenza* | [22] | WP_053293022 | 47/92 | ABC transporter substrate binding protein | 4 |
| CAA43967 | *Yersinia pestis* | [22] | WP_085538626 | 32/83 | Pilus assembly protein PapD | 2 |

The accession number of the known DsbA substrate (in an organism other than *B. pseudomallei)*, the organism and the publication reference are given in the first three columns. The corresponding *B. pseudomallei* homologue is given in the fourth column. The identity and coverage (number of residues in the result sequence that overlap with the search sequence) is given in percent in the column “identity/coverage”. The final two columns provide the protein function and the number of cysteines in the predicted mature sequence. All proteins in this table are known or predicted to be secreted or periplasmic.

Over 50 DsbA substrates in *Francisella tularensis* were identified by trapping and co-purifying substrates bound to a DsbA variant [34]. Of these 50, we found nine homologues encoded in *B. pseudomallei* (see Table 3). These include homologues of the lytic transglycosylase domain containing protein (implicated in peptidoglycan rearrangement) and homologues of two pilin proteins involved in the formation of pilus and flagella. Also present is an MoeB homologue; MoeB is a molybdopterin synthase adenyl transferase (cytoplasmic in *E. coli* but likely periplasmic in *B. pseudomallei* due to the twin-arginine translocation (TAT) signal sequence). A PenI family β-lactamase homologue is also found in *B. pseudomallei*; this is a class A β-lactamase that confers resistance to β-lactams including, in rare cases, ceftazidime (commonly used to treat melioidosis) [49]. A succinate dehydrogenase flavoprotein subunit homologue, found in the bacterial inner membrane and part of the electron transport chain, is also encoded in *B. pseudomallei*. This protein is cytoplasmically oriented in *E. coli*, though again the *B. pseudomallei* version has a TAT signal sequence suggesting a possible periplasmic localisation.

A number of DsbA substrates identified in *E. coli* (reviewed in [22]) have *B. pseudomallei* homologues including a molecular chaperone homologous to PapD and EscC, involved in the formation of the Type III secretion system (T3SS). The T3SS assembly requires DsbA activity in many Gram-negative bacteria, including *E. coli* and *S. typhimurium*. [50, 51]. Finally, a *B. pseudomallei* protein homologous to the *Y. pestis* pilus assembly protein Caf1M (a molecular chaperone involved with assembly of the surface capsule of the bacterium) was also identified.

Of the 15 putative *B. pseudomallei* DsbA substrates identified using this substrate homology method, two were also identified in the genomic pipeline method. These are the PenI and succinate dehydrogenase flavoprotein subunit homologues.

We then aligned the sequences of the Table 3 *B. pseudomallei* proteins to identify any possible sequence conservation around the cysteine residues, but no pattern was identified. This lack of peptide sequence motif in DsbA substrates has also been observed in *E*.*coli*, demonstrating the difficulty of DsbA substrate prediction [52].

### Epitope prediction of virulence-associated proteins

To determine whether the DsbA substrates identified in the two methods above could contribute to vaccination efforts against *B. pseudomallei*, we also predicted B-cell epitopes, using a structure-informed approach. The sequences of the 73 putative, extra-cytoplasmic DsbA substrates (predicted virulence factors, Table 2) along with the 15 homologous DsbA substrates (Table 3) were screened against the Protein Data Bank (PDB) [53], to identify structurally characterised homologues (see S6 File). Six of the 73 proteins were found to have at least 80% similarity to a structurally characterised protein. Three of these six protein structures were from *Pseudomonas* species, while the other three were from *Burkholderia* species. Similarity was used rather than identity to account for mutations of functionally similar residues. The six protein structures were then used as models to predict structurally-informed B-cell epitopes of length 10-20 residues (Table 4 and Fig 4) using the SEPPA3 server.

**Table 4:**
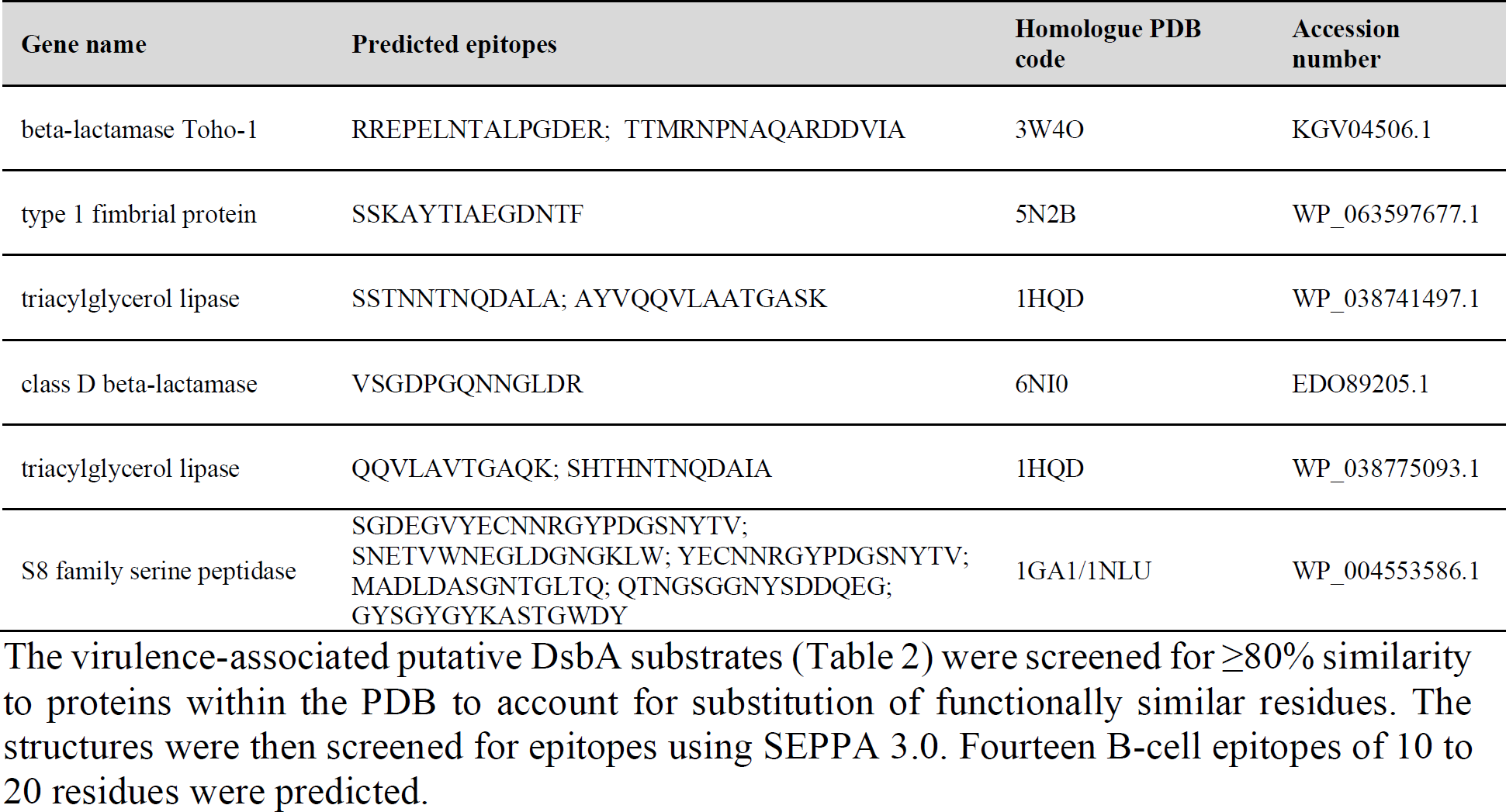
B-cell epitope prediction.

**Fig 4:**
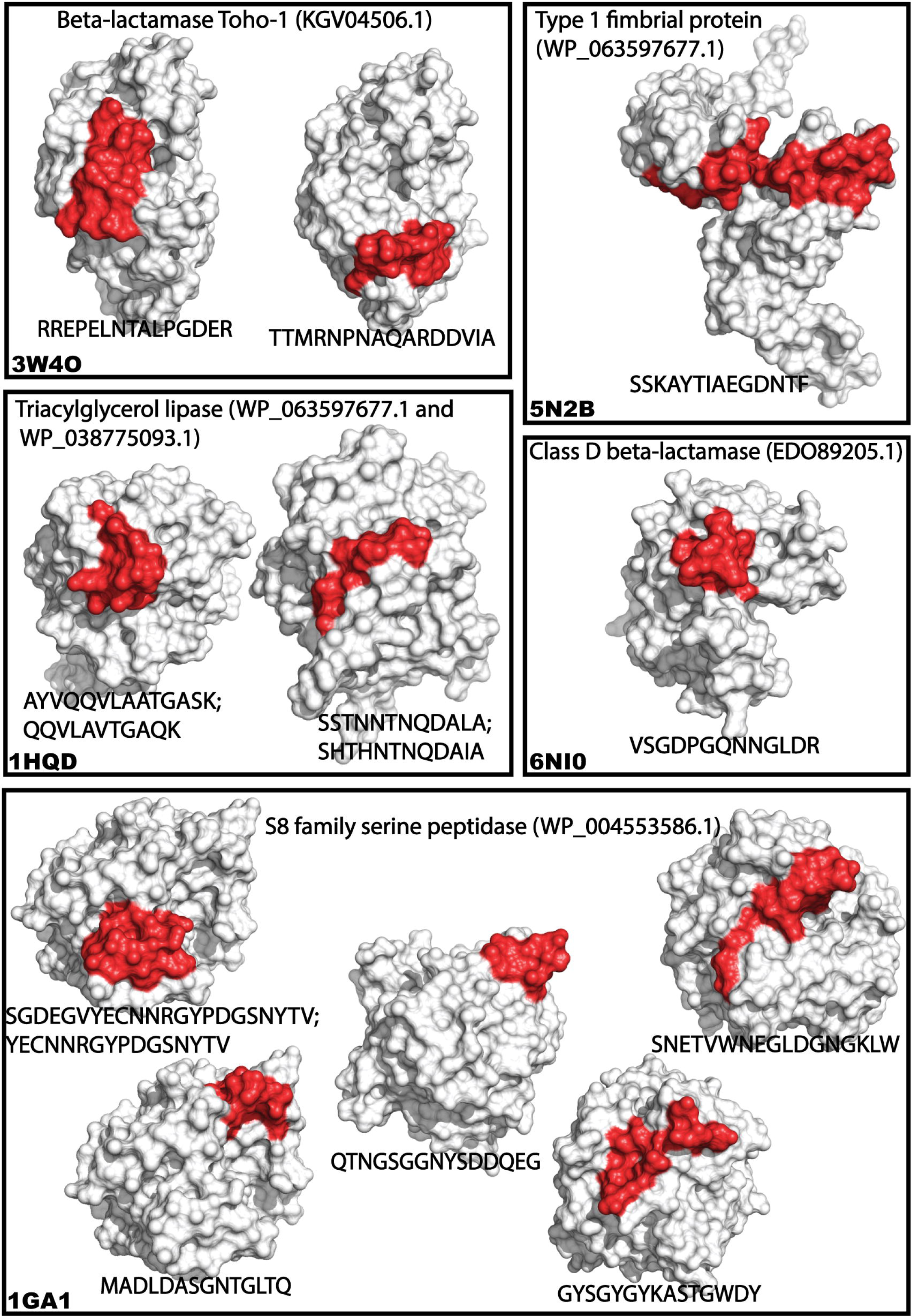
Predicted B-cell epitopes. Graphical representation of B-cell epitopes found in Table 4. Proteins are shown as white surfaces and their respective PDB ID is given in the bottom left corner of each box. The epitope region is highlighted in red and the corresponding homologous sequences found in *B. pseudomallei* are given in one letter code under each respective structure and separated by semicolon when more than one sequence pointed to the same epitope.

These epitopes provide an interesting list for further evaluation. For example, epitopes from beta-lactamase Toho-1 and class D beta-lactamase could provide a useful vaccination approach for *B. pseudomallei* because these directly target antibiotic resistance proteins. Similar approaches have conferred protection against other bacteria in animal models [54-57].

Vaccination targeting adhesion proteins and essential virulence factors such as FimA [58, 59] and type 1 fimbrial protein is a commonly used approach due to the external localisation of these proteins and their exposure to host immune systems. Anti-fimbrial antibodies have been shown to interfere with function and reduce disease [60, 61] and a FimA vaccine provided protection against *Streptococcus parasanguis, Streptococcus mitis, Streptococcus mutans and Streptococcus salivarius* in rats [62-64].

Vaccination against conserved, secreted enzymes such as the triacylglycerol lipase (EstA) and S8 family serine peptidase enzymes may also be a useful strategy. Secreted peptidases are known virulence factors in many pathogenic bacteria [37, 65] and vaccines targeting them have attenuated disease in animal models [66, 67]. Two triacylglycerol lipases (WP_038741497.1 and WP_038775093.1) were identified as having a structural homologue in the PDB. These two lipases are both core genes and share 78% similarity (72% identity, 87% query cover). and their sequences were both aligned to the same PDB code, resulting in epitope variants of similar sequences.

## Discussion

In the present study, we analysed genomes from 512 *B. pseudomallei* isolates specifically associated with disease to identify core putative DsbA substrates and virulence factors. Pangenomic analysis of *B. pseudomallei* has previously been performed utilising 37 isolates from a variety of isolation sources [36] and concluded the pangenome to be ‘open’, indicating that new isolates will continually increase the number of total genes, which we found to be the case, based on a pangenome of 19,991 genes from 512 isolates. Previous studies comparing the *B. pseudomallei* genome with the obligate pathogen *Burkholderia mallei* (responsible for glanders) and the generally non-pathogenic *Burkholderia thailandensis* [68-71], identified several loci likely to be involved in *B. pseudomallei* virulence. These include the capsular polysaccharide gene cluster and Type III secretion needle complex [71], which were not considered core genes, demonstrating the importance of large-scale analysis.

In the present study, we used two orthogonal approaches to identify a total of 278 putative DsbA substrates, with 86 predicted to be virulence factors (S5 File). Of these, 73 were identified by the genome analysis approach and 15 were identified by the DsbA substrate homology approach. Two of the putative 86 DsbA virulence factor substrates were identified in both approaches. These two are the experimentally validated bacterial virulence factors and DsbA substrates succinate dehydrogenase flavoprotein subunit, and a PenI family β-lactamase (both reported to be *F. tularensis* DsbA substrates) [34].

Delving deeper into the results presents some curious outcomes. For example, the well-characterised *E*.*coli* DsbA substrate and virulence factor FlgI [27, 72] was not picked up as a potential *B. pseudomallei* DsbA substrate by either method, though *B. pseudomallei* encodes FlgI. The *B. pseudomallei* FlgI sequence has 4 cysteines in the translated gene product but the predicted mature sequence after cleavage of the signal sequence has just one cysteine. Generally, DsbA does not interact with proteins having just one cysteine. If *B. pseudomallei* FlgI is a DsbA substrate (that is yet to be tested), then the most likely reasons that it was not identified as a substrate by either of the two methods we used are that (i) the predicted signal peptide is incorrect and/or (ii) the single cysteine of *B. pseudomallei* FlgI forms an inter-molecular disulfide bond.

The finding that the two orthogonal approaches identified the same two target proteins suggests that there is merit in using different theoretical approaches to select high priority targets for further evaluation (in this case, the PenI family Beta-lactamase and succinate dehydrogenase flavoprotein subunit). On the other hand, the fact that there were so few overlaps in the predicted substrates from the two methods raises questions about the filters we applied. Specifically, we found that of the 15 potential substrates identified by the substrate homology method, 5 had an odd numbers of cysteines, whereas the genomic analysis filtered these proteins out of consideration. We applied the even cysteine filter because previous reports showed that *E. coli* exported proteins have a strong preference for an even number of cysteines. This even number of cysteine preference is present in *B. pseudomallei* exported proteins (Fig 2) though is not as pronounced as in *E. coli*. By restricting our genomic analysis to core, extra-cytoplasmic *B. pseudomallei* proteins with an even number of cysteines, some DsbA substrates may therefore have been missed. There is considerable evidence that many virulence factors such as adhesion and motility proteins, toxins and enzymes are extra-cytoplasmic proteins in both Gram-positive and Gram-negative bacteria [21, 22, 73]. Given that extra-cytoplasmic proteins in the translated core genome *of B. pseudomallei* have a slight preference for even number of cysteines (Fig 2) and the identification of many virulence-associated proteins within the 263 proteins in the list, the approach taken in this analysis (Fig 1) to identify DsbA substrates was justified. Further, the genomic analysis focused on highly conserved proteins from the core genome; accessory proteins associated with virulence would not be identified using this approach. Nevertheless, the genomic analysis identified homologues of known DsbA substrates in other bacteria, such as the OmpA porin, supporting the use of this approach. However, attempting to identify epitopes from proteins which are not found in every disease-causing isolate may present challenges for anti-virulence and vaccination attempts.

In addition, the genomic analysis identified several proteins of unknown function which could represent novel virulence factors for future studies. Importantly, our theoretical approach was extended to predict structurally-informed surface epitopes for several core gene DsbA substrates for potential vaccine or antibody development (Table 4).

In summary, our *in silico* analysis combined a substrate homology approach and a genomic analysis approach to identify more than 80 potential *B. pseudomallei* DsbA virulence factor substrates, two of which we mark as high priority for experimental validation. Future characterization of these proteins will aid our understanding of *B. pseudomallei* virulence and could provide new targets for antivirulence drug discovery and vaccine development. The approaches we report here could also be applied to identify potential DsbA virulence factor substrates in other pathogenic bacteria.

## Methods

### Data acquisition and filtering of core, extra-cytoplasmic, putative DsbA substrates

1577 *B. pseudomallei* genomes were obtained from the genome information table from NCBI (https://www.ncbi.nlm.nih.gov/genome/genomes/476) (date accessed: 1/2/20).The biosample accession numbers were batch downloaded using Entrez. A list of assembly accession numbers can be found in S1 Fig. Metadata was then scraped for disease association using grep with the following command:

grep -A 1 “disease”

The assemblies were then downloaded using Entrez and annotated using a prokka (version 1.14.5) [74] for loop with the following command:

for file in *.fna; do tag=${file%.fna}; prokka --prefix “$tag” --locustag “$tag” --genus Burkholderia -- species pseudomallei --strain “$tag” --outdir “$tag”_prokka --force --addgenes “$file”; done

The .gff files were used as input for roary (version 3.11.2) [75] without splitting paralogues via the following command:

roary -e --mafft -i 90 -v -p 72 -z -s -o output -f *.gff

The roary output file was altered from interleaved fasta to one line per sequence

awk ‘{if(NR==1) {print $0} else {if($0 ∼ /^>/) {print “\n”$0} else {printf $0}}}’ input.fa > output.fa

The core genome was then used in the remaining analysis and core DNA sequences were translated into protein sequences using transeq [76] with the following command:

transeq -sequence input.fasta -outseq output.fasta -table 11 -frame 1

The core genome was then filtered based on signal sequence and then the sequence of the mature exported protein, as predicted utilising SignalP 5.0 [77, 78]

signalp -fasta prot_core_genome_complete.fasta -format short -mature -org gram--verbose

These sequences were then filtered for genes containing even numbers of cysteines

awk -F \C ‘NF % 2’ < input.fasta | awk “/C.*C/” | sed ‘/>/{$!N;/\n.*>/!P;D}’ > output.fasta

This list was then annotated via screening sequences against NCBI and Gene Ontology [79] using the PANNZER2 server [80].

### Identification of DsbA substrate homologues in *B. pseudomallei*

DsbA substrates were also predicted using a substrate homology search. This approach may identify proteins not encoded in the core genome. The *B. pseudomallei* genome was screened for homologues of known DsbA substrates using BLASTP. A starting list of confirmed DsbA substrates was extracted from the literature [22, 34, 45-48, 81], and their amino acid sequences used in BLAST searches [82] against the NCBI protein database [43] for homologues in *B. pseudomallei* using default search parameters. In some cases two search proteins identified the same homologue in *B. pseudomallei*. In these cases only the search protein most similar to the *B. pseudomallei* homologue is given in Table 3. The results were filtered to select proteins with at least 20% sequence identity and a sequence coverage of at least 50%. Protein sequences with fewer than two cysteines were removed. Exported proteins were selected on the basis of predicted signal sequence (SignalP 5.0 [77]) or experimental evidence of extra-cytoplasmic localisation for the reported DsbA substrate in another *Burkholderia* species.

### Cysteine distribution analysis

Fasta files containing either the 19,991 pan genes or the 4,496 core gene of *B. pseudomallei* with their corresponding amino acid sequences and descriptors were utilised to calculate the distribution of cysteines with a custom Python 3.0 script (available on Github : (https://github.com/gpetit99/cysteineCount_bPseudomallei/blob/master/CysCountFrequency.py“). Briefly, lists of the extra-cytoplasmic protein sequences with signal peptides removed were compared to lists of the protein sequences from the whole genome to create dataframes with either cytoplasmic or extra-cytoplasmic proteins. Proteins were grouped based on the presence or absence of SP, and based on the number of cysteines in the mature protein. To calculate the normalised frequency of cysteines for extra-cytoplasmic proteins, we divided the number of extra-cytoplasmic proteins having N cysteines by the total number of proteins having N cysteines (N being an integer from 0 to 73 – No protein has more than 73 cysteines in the *B. pseudomallei* translated genome). This analysis was run for the core genome and pangenome independently. Other statistics (e.g. number of proteins in each group) were extracted from the dataframes.

### Epitope prediction

The metadata for each of the 263 proteins in the annotated list was manually inspected to select for further analysis a total of 73 proteins likely related to virulence. The sequences of these 73 selected proteins were combined with the 15 selected proteins from the homology analysis (to give 86 unique protein sequences). These were screened against the protein data bank using BLAST (criteria: ≥80% positive substitutions/similarity used as a threshold) to find structurally characterised homologues. These structural homologues were then used to predict B-cell epitopes using SEPPA 3.0 (http://www.badd-cao.net/seppa3/index.html) with a threshold of 0.1 [83]. Similarity was used rather than identity to account for mutations of functionally similar residues. Predicted B-cell epitopes were accepted if they were 10 – 20 residues in length, as described in [84].

## Acknowledgments

We gratefully acknowledge the support of the Griffith University eResearch Services Team and the use of the High Performance Computing Cluster “Gowonda” to complete this research.

## Supporting Information

**S1 Fig**. Accession numbers for disease related genomes of *B. pseudomallei* used in this analysis

**S2 Fig**. Core genome (4,496 gene products) of disease related B. pseudomallei (fasta format).

**S3 Fig** *B. pseudomallei* proteins from the core genome with a signal peptide (removed before counting cysteines) and even number of cysteines (263 proteins, fasta format).

**S4 File Gene Ontology (GO) classification of the gene and gene-product descriptions**.

**S5 File Predicted virulence-associated substrates of DsbA**

**S6 File Predicted B-cell epitopes**

