## Supplemental Figures 01-03 for "Prediction of *Burkholderia pseudomallei* DsbA substrates identifies potential virulence factors and vaccine targets"

Supplementary data

S1:

Accession numbers for disease related genomes of *B. pseudomallei* used in this analysis

| | | | | |
|---|---|---|---|---|
| GCA_000410895.1 | GCA_001207785.1 | GCA_001321105.1 | GCA_001976275.1 | GCA_001977605.1 |
| GCA_000452945.1 | GCA_001208265.1 | GCA_001321125.1 | GCA_001976325.1 | GCA_001977615.1 |
| GCA_000452965.1 | GCA_001208545.1 | GCA_001321145.1 | GCA_001976335.1 | GCA_001977645.1 |
| GCA_000452985.1 | GCA_001209045.1 | GCA_001321165.1 | GCA_001976345.1 | GCA_001977655.1 |
| GCA_000453005.1 | GCA_001209405.1 | GCA_001321185.1 | GCA_001976385.1 | GCA_001977675.1 |
| GCA_000511895.1 | GCA_001209505.1 | GCA_001321205.1 | GCA_001976395.1 | GCA_001977695.1 |
| GCA_000511915.1 | GCA_001209525.1 | GCA_001321225.1 | GCA_001976405.1 | GCA_001977725.1 |
| GCA_000520895.1 | GCA_001210025.1 | GCA_001321245.1 | GCA_001976415.1 | GCA_001977735.1 |
| GCA_000521645.1 | GCA_001210645.1 | GCA_001321265.1 | GCA_001976465.1 | GCA_001977745.1 |
| GCA_000714875.1 | GCA_001210705.1 | GCA_001321285.1 | GCA_001976475.1 | GCA_001977785.1 |
| GCA_000932075.1 | GCA_001210965.1 | GCA_001321305.1 | GCA_001976485.1 | GCA_001977805.1 |
| GCA_000932085.1 | GCA_001211245.1 | GCA_001321325.1 | GCA_001976495.1 | GCA_001977815.1 |
| GCA_000932095.1 | GCA_001211305.1 | GCA_001321345.1 | GCA_001976545.1 | GCA_001977825.1 |
| GCA_000932105.1 | GCA_001211345.1 | GCA_001321365.1 | GCA_001976565.1 | GCA_001977865.1 |
| GCA_000932145.1 | GCA_001211385.1 | GCA_001321385.1 | GCA_001976575.1 | GCA_001977875.1 |
| GCA_000953095.1 | GCA_001211405.1 | GCA_001321405.1 | GCA_001976585.1 | GCA_001977885.1 |
| GCA_000954175.1 | GCA_001211505.1 | GCA_001321425.1 | GCA_001976625.1 | GCA_001977915.1 |
| GCA_000959305.1 | GCA_001211565.1 | GCA_001321445.1 | GCA_001976645.1 | GCA_001977945.1 |
| GCA_000961535.1 | GCA_001211705.1 | GCA_001321465.1 | GCA_001976655.1 | GCA_001977955.1 |
| GCA_000981285.2 | GCA_001211865.1 | GCA_001321485.1 | GCA_001976675.1 | GCA_001977975.1 |
| GCA_001182265.1 | GCA_001212025.1 | GCA_001321505.1 | GCA_001976685.1 | GCA_001978005.1 |
| GCA_001182285.1 | GCA_001212065.1 | GCA_001321525.1 | GCA_001976725.1 | GCA_001978015.1 |
| GCA_001182305.1 | GCA_001212185.1 | GCA_001321545.1 | GCA_001976735.1 | GCA_001978045.1 |
| GCA_001182325.1 | GCA_001212265.1 | GCA_001321565.1 | GCA_001976755.1 | GCA_001978055.1 |
| GCA_001191815.1 | GCA_001212325.1 | GCA_001321585.1 | GCA_001976785.1 | GCA_001978085.1 |
| GCA_001192415.1 | GCA_001212405.1 | GCA_001321605.1 | GCA_001976805.1 | GCA_001978105.1 |
| GCA_001192815.1 | GCA_001212465.1 | GCA_001321625.1 | GCA_001976815.1 | GCA_001978115.1 |
| GCA_001193035.1 | GCA_001212705.1 | GCA_001326895.1 | GCA_001976825.1 | GCA_001978125.1 |
| GCA_001193375.1 | GCA_001233045.1 | GCA_001326915.1 | GCA_001976865.1 | GCA_001978165.1 |
| GCA_001193615.1 | GCA_001262275.1 | GCA_001326935.1 | GCA_001976885.1 | GCA_001978175.1 |
| GCA_001193655.1 | GCA_001262315.1 | GCA_001326955.1 | GCA_001976895.1 | GCA_001978185.1 |
| GCA_001194765.1 | GCA_001262355.1 | GCA_001326975.1 | GCA_001976905.1 | GCA_001978205.1 |
| GCA_001194905.1 | GCA_001270725.1 | GCA_001326995.1 | GCA_001976925.1 | GCA_001978245.1 |
| GCA_001195465.1 | GCA_001275585.1 | GCA_001327015.1 | GCA_001976965.1 | GCA_001978265.1 |
| GCA_001196125.1 | GCA_001277975.1 | GCA_001327035.1 | GCA_001976975.1 | GCA_001978285.1 |
| GCA_001196775.1 | GCA_001320065.1 | GCA_001327075.1 | GCA_001977005.1 | GCA_001978295.1 |
| GCA_001197015.1 | GCA_001320105.1 | GCA_001327095.1 | GCA_001977015.1 | GCA_001978325.1 |
| GCA_001197235.1 | GCA_001320145.1 | GCA_001327115.1 | GCA_001977045.1 | GCA_001978345.1 |
| GCA_001197775.1 | GCA_001320185.1 | GCA_001327135.1 | GCA_001977055.1 | GCA_001978365.1 |
| GCA_001197795.1 | GCA_001320215.1 | GCA_001327155.1 | GCA_001977085.1 | GCA_001978385.1 |
| GCA_001197955.1 | GCA_001320255.1 | GCA_001327175.1 | GCA_001977095.1 | GCA_001978405.1 |
| GCA_001199035.1 | GCA_001320285.1 | GCA_001327195.1 | GCA_001977125.1 | GCA_001978415.1 |
| GCA_001199195.1 | GCA_001320325.1 | GCA_001327275.1 | GCA_001977135.1 | GCA_001978445.1 |
| GCA_001199695.1 | GCA_001320365.1 | GCA_001327315.1 | GCA_001977165.1 | GCA_001978455.1 |
| GCA_001199735.1 | GCA_001320395.1 | GCA_001327395.1 | GCA_001977185.1 | GCA_001978485.1 |
| GCA_001199815.1 | GCA_001320425.1 | GCA_001327415.1 | GCA_001977195.1 | GCA_001978505.1 |
| GCA_001199855.1 | GCA_001320465.1 | GCA_001327455.1 | GCA_001977225.1 | GCA_001978515.1 |
| GCA_001200615.1 | GCA_001320505.1 | GCA_001327515.1 | GCA_001977245.1 | GCA_001978525.1 |
| GCA_001200935.1 | GCA_001320545.1 | GCA_001327535.1 | GCA_001977265.1 | GCA_001978565.1 |
| GCA_001202135.1 | GCA_001320585.1 | GCA_001327575.1 | GCA_001977275.1 | GCA_001978585.1 |
| GCA_001202415.1 | GCA_001320615.1 | GCA_001885195.1 | GCA_001977285.1 | GCA_001978605.1 |
| GCA_001202895.1 | GCA_001320645.1 | GCA_001887555.1 | GCA_001977325.1 | GCA_001978615.1 |
| GCA_001203375.1 | GCA_001320685.1 | GCA_001887575.1 | GCA_001977345.1 | GCA_001978635.1 |
| GCA_001203735.1 | GCA_001320725.1 | GCA_001905265.1 | GCA_001977365.1 | GCA_001978665.1 |
| GCA_001204155.1 | GCA_001320755.1 | GCA_001974745.1 | GCA_001977375.1 | GCA_001978675.1 |
| GCA_001204275.1 | GCA_001320795.1 | GCA_001975065.1 | GCA_001977385.1 | GCA_001978705.1 |
| GCA_001204455.1 | GCA_001320825.1 | GCA_001975085.1 | GCA_001977425.1 | GCA_001978725.1 |
| GCA_001204515.1 | GCA_001320865.1 | GCA_001975105.1 | GCA_001977435.1 | GCA_001978745.1 |
| GCA_001205635.1 | GCA_001320905.1 | GCA_001976165.1 | GCA_001977465.1 | GCA_001978765.1 |
| GCA_001205935.1 | GCA_001320945.1 | GCA_001976175.1 | GCA_001977475.1 | GCA_001978785.1 |
| GCA_001206455.1 | GCA_001320985.1 | GCA_001976185.1 | GCA_001977495.1 | GCA_001978795.1 |
| GCA_001206495.1 | GCA_001321015.1 | GCA_001976195.1 | GCA_001977525.1 | GCA_001978825.1 |
| GCA_001207035.1 | GCA_001321045.1 | GCA_001976245.1 | GCA_001977545.1 | GCA_001978835.1 |
| GCA_001207055.1 | GCA_001321065.1 | GCA_001976255.1 | GCA_001977565.1 | GCA_001978865.1 |
| GCA_001207675.1 | GCA_001321085.1 | GCA_001976265.1 | GCA_001977575.1 | GCA_001978875.1 |

GCA_001978905.1
GCA_001978925.1
GCA_001978935.1
GCA_001978965.1
GCA_001978985.1
GCA_001978995.1
GCA_001979015.1
GCA_001979045.1
GCA_001979065.1
GCA_001979085.1
GCA_001979105.1
GCA_001979115.1
GCA_001979135.1
GCA_001979165.1
GCA_001979175.1
GCA_001979195.1
GCA_001979215.1
GCA_001979245.1
GCA_001979255.1
GCA_001979275.1
GCA_001979285.1
GCA_001979325.1
GCA_001979335.1
GCA_001979345.1
GCA_001979385.1
GCA_001979405.1
GCA_001979415.1
GCA_001979435.1
GCA_001979455.1
GCA_001979485.1
GCA_001979495.1
GCA_001979505.1
GCA_001979545.1
GCA_001979565.1
GCA_001979585.1
GCA_001979595.1
GCA_001979615.1
GCA_001979645.1
GCA_001979665.1
GCA_001979675.1
GCA_001979695.1
GCA_001979725.1
GCA_001979745.1
GCA_001979755.1
GCA_001979765.1
GCA_001979805.1
GCA_001979815.1
GCA_001979835.1
GCA_001979865.1
GCA_001979885.1
GCA_001979895.1
GCA_001979905.1
GCA_001979945.1
GCA_001979965.1
GCA_001979975.1
GCA_001979995.1
GCA_001980025.1
GCA_001980045.1
GCA_001980065.1
GCA_001980075.1
GCA_001980105.1
GCA_001980125.1
GCA_001980145.1
GCA_001980155.1
GCA_001980175.1
GCA_001980205.1
GCA_001980215.1
GCA_001980245.1
GCA_001980265.1
GCA_001980275.1
GCA_001980305.1
GCA_001980315.1
GCA_001980335.1
GCA_001980365.1
GCA_001980385.1
GCA_001980395.1
GCA_001980425.1
GCA_001980435.1
GCA_001980465.1
GCA_001980485.1
GCA_001980495.1
GCA_001980515.1
GCA_001980545.1
GCA_001980565.1
GCA_001980585.1
GCA_001980605.1
GCA_001980625.1
GCA_001980645.1
GCA_001980655.1
GCA_001980675.1
GCA_001980695.1
GCA_001980725.1
GCA_001980735.1
GCA_001980755.1
GCA_001980775.1
GCA_001980805.1
GCA_001980815.1
GCA_001980835.1
GCA_001980845.1
GCA_001980885.1
GCA_001980895.1
GCA_001980905.1
GCA_001980915.1
GCA_001980965.1
GCA_001980985.1
GCA_001980995.1
GCA_001981025.1
GCA_001981045.1
GCA_001981055.1
GCA_001981085.1
GCA_001981105.1
GCA_001981125.1
GCA_001981135.1
GCA_001981165.1
GCA_001981185.1
GCA_001981195.1
GCA_002110925.1
GCA_002110945.1
GCA_002110965.1
GCA_002110985.1
GCA_002111005.1
GCA_002111025.1
GCA_002111045.1
GCA_002111065.1
GCA_002111085.1
GCA_002111105.1
GCA_002111125.1
GCA_002111165.1
GCA_002111185.1
GCA_002111205.1
GCA_002111225.1
GCA_002111245.1
GCA_002111265.1
GCA_002111285.1
GCA_002111305.1
GCA_002111325.1
GCA_002111345.1
GCA_002111365.1
GCA_002111385.1
GCA_002113945.1
GCA_002115385.1
GCA_002245325.2
GCA_002843645.1
GCA_002860065.1
GCA_002900605.1
GCA_002900625.1
GCA_002900645.1
GCA_002900665.1
GCA_002920945.1
GCA_002920995.1
GCA_002921005.1
GCA_002921015.1
GCA_002921055.1
GCA_002921075.1
GCA_002921105.1
GCA_003268455.1
GCA_003268465.1
GCA_003546995.3
GCA_003547015.1
GCA_003547035.1
GCA_003547055.1
GCA_003583425.1
GCA_003583435.1
GCA_003584055.1
GCA_003584065.1
GCA_004323015.1
GCA_004323035.1
GCA_004348055.1
GCA_004348075.1
GCA_004360045.1
GCA_004360055.1
GCA_004360065.1
GCA_004360075.1
GCA_004360125.1
GCA_004360205.1
GCA_004367665.1
GCA_004367685.1
GCA_004367705.1
GCA_004367725.1
GCA_004526325.1
GCA_005853645.1
GCA_005862325.1
GCA_006542565.1
GCA_006542585.1
GCA_007995115.1
GCA_900006245.1
GCA_900446265.1

### S2 Core genome (4496 gene products) of disease related *B. pseudomallei* (fasta format):

>GCA_000452945.1_BPMSHR5858_genomic_00014_1 yhcG
VEWPQGDSRTLSLYLKPVSQIMYCEQCGARCQQIHETTVRRVRDLPLFEYRVVLHVPRRR
VWCERCGAARLEKLDWLGRYQRVTERFAKACEKLLQAASVQAVAAFYELGWHTVKSIDKM
RLRARVAEPDWSTIRYLAMDEFALHKGHRYATVVVDPIGRQVLWVGPGRSRETARAFFEQ
LPEGVAERIEAVAIDMTTAYELEIKEQCPQAEIVFDLYHVVAKYGREVIDRVRVDQANQL
RHDKPARKVLKSSRWLLLRNRHNLKPEQAVHLKELLAANQSLLCVYVLRDELKRLWFYRK
PAWAEKAWGQWFEQAQQSGIAALQKFAQRLQGYWHGIVARCRHPLNTSVVEGINNTIKVI
KRRAYGYRDEQYFFLKIRAAFPGIPR*
>GCA_000452945.1_BPMSHR5858_genomic_00015_1 syrM1_3
MSQQREAIDTYLLRVLHTLLMERSVTRAAVKLNQSQPAISAALRRLRDITGDPLLVRGKS
GMVPTEYGLRLLEPVQNALREIERIKFQQHNFDPATSIRCYRIGCPDYLNVLFVPTVVER
FRQAAPNATLEFHSLGPAFDYELALEDGKLDIVVGNWPEPPEQLHLSNLFVDEIVCLMSN
SHPFAKRGGLTLDQYLNAPHLAPTPYSVGQRGAIDVHLARERLKRHVVVTLPYFNLAPYV
LVKSDLIFTTTRLFADHYAKFLPLSVVPAPLDFPPMQYYQLWHERCHYSDEVRWLRGLVA
EATRTLIDA*
>GCA_000452945.1_BPMSHR5858_genomic_00016_1 dadD_1
MERHPSARAGAHSLSQPPSLSPNRSKTLVVKHADVLVTMDGARRELRDAGLYVEDNRIVA
VGPSAELPEQADEVLDLRGHLVIPGLVNTHHHMYQSLTRAIPAAQNAELFGWLTNLYRIW
AHLTPEMIEVSALTAMAELLLSGCTTSSDHLYIYPNGSRLDDSIAAARRIGMRFHASRGS
MSVGQRDGGLPPDAVVEREADILRDTQRVIETYHDEGRYAMLRVAVAPCSPFSVSRGLMR
DAAALAREHRVSLHTHLAENVNDVAYSREKFGMTPAEYAEDLGWVGRDVWHAHCVRLDEP
GIALFARTGTGVAHCPCSNMRLASGIAPIARMRRAGVPVGLGVDGCASNDGAQMVAEARQ
ALLLQRVGFGPDALSARDALEIATLGGARVLNRDDIGALAPGMAADFVAFDLRTPQFAGA
LHDPVAALVFCAPPQAAYSVVNGRVVVREGRLTTLEIEPLVERHNALARALCDAAR*
>GCA_000452945.1_BPMSHR5858_genomic_00019_1 allA_2
MKTLSIEPLTRAAFAPFGDVIETQGAKQIPINLGTTMRFHDLAKIDVADEGGRPLVNLFR
GQPRTLPFEVTMLERHPLGSQAFVPLTDRPYIVVVAPAGDLDASKIRAFVTSGWQGVNYA
KGVWHHPLIALGEVSDFIVVDRGGDGRNLNEQNLPESLWLTDDALLAVGA*
>GCA_000452945.1_BPMSHR5858_genomic_00021_1 pucL
MMKAMHYTLEQLNAMSTDAFVAALSGIFEHSPWVAEAAAAARPFGSIDALHGTMKDAVEA
AGDARQLALINAHPELAGKAAVRGELTAESTREQSGAGLDRCTQAEFDRLQQLNRAYREK
FGFPFILAVRGFDRHGIIANFEARVGHTRDEELRASLAQIYRIARFRLDDLIGA*
>GCA_000452945.1_BPMSHR5858_genomic_00027_1 tadA
VRLAGAPASGGELDDSTQSPPHAALLSAAAERDRRYMRLALAAAEEARAAGEVPVGAVIV
RGDEVIARGFNHPIGGHDPSAHAEMAALRAAARALRNYRMPGCELYVTLEPCLMCSGAIM
HARIARVVFGAPDPKTGACGSVVDAFADARLNHHTTVEGGVLADECGAVLKSFFAQRRQA
VREARRAQQAPGAPPRDE*
>GCA_000452945.1_BPMSHR5858_genomic_00043_1 guaA
MHDKILILDFGSQVTQLIARRVREAHVYCEIHPNDVSDDFVREFAPKGVILSGSHASTYE
DHQLRAPQAVWDLGVPVLGICYGMQTMAVQLGGKVEWSDHREFGYAEVRVHGRTRLLDGI
QDFATPEGHGMLKVWMSHGDKVGEMPPGFALMASTPSCPIAGMADEARGYYAVQFHPEVT
HTVQGRKLLERFVLDIADAKPDWIMRDHIEEAVARIREQVGDEEVILGLSGGVDSSVAAA
LIHRAIGDQLTCVFVDHGLLRLNEGKMVLDMFEGRLHAKVVHVDASEQFLGHLAGVADPE
HKRKIIGREFVEVFQAEAKKLTNAKWLAQGTIYPDVIESGGAKTKKATTIKSHHNVGGLP
ETLGLKLLEPLRDLFKDEVRELGVALGLPAEMVYRHPFPGPGLGVRILGEVKRDYAELLR
RADAIFIEELRGTLATEQDAAAGLCEPSQVGKSWYDLTSQAFAVFLPVKSVGVMGDGRTY
DYVAALRAVQTTDFMTAHWAHLPYALLGRASNRIINEVRGINRVVYDVSGKPPATIEWE*
>GCA_000452945.1_BPMSHR5858_genomic_00053_1 qmcA
MDSLIVWVVLLVIAFVIVSQTVKIVPQQHAWVLERFGRYHATLSPGLNIVLPFVDRIAYR
HVLKEIPLDVPSQICITRDNTQLQVDGVLYFQVTDPMKASYGSSNFVLAITQLAQTTLRS
VIGKLELDKTFEERDFINHSIVSALDEAASNWGVKVLRYEIKDLTPPKEILHAMQAQITA
EREKRALIAASEGRKQEQINLASGAREAAIQKSEGEKQAAINQAQGEAAAILAVAEANSQ
AIQKIAQAIQSQGGMDAVNLKVAEQYVGAFGNLAKAGNTLIVPSNLSDLSTAIASALTIV
NRSAPGALAPGASKS*
>GCA_000452945.1_BPMSHR5858_genomic_00059_1 ppsR
MLPTVFIVSDGTGITAETFAHSILSQFDQKFRLVRVPFIDSIEKAYDTVRKINDAAQHDG
RRPIVFTTLVDGESNEIVKRSNALVLDMFQRFVEPLEQELQLKSSHAMGRVHQNADTEEY
KTRIEAINFSLAHDDGQSNRNLADADVILIGVSRSGKTPTSLYLAMQYGVKAANYPLIPE
DFERGKLPTPLHPHRDKLFGLSIDPMRLSEIRNERRPGSKYAAPENCRYEINEAEAMMRR
EGVKWLSSTHKSIEEIATTILQEIKLERQSY*
>GCA_000452945.1_BPMSHR5858_genomic_00060_1 aviRb
MKAITSRDNPLYKRLKALAGSTHQQRRGEQALLEGFHLASAYLDAAGQPQLCVVTDGALA
HEEARGIVARIEPARVVTLPDVLFGQLSNVVNGVGLLLLVARPAPALPARVAGTSVVLDG
VQDAGNVGSILRSAAAAGVERVFCAPGTAYAWSSKVLRSAMGAHFLLDIHEDVDAGTLIE
RLAIPVALTDSHGARAIYDCDLAGPLAWVFGNEGAGVSARWREAATHRVTIPQPGGMESL
NVAAAAAVCLFEQCRQQRALRA*
>GCA_000452945.1_BPMSHR5858_genomic_00063_1 lpxA
MSRIHPTAIIEPGAQLHETVEVGPYAIVGSHVTIGARTTIGSHSVIEGHTTIGEDNRIGH
YASVGGRPQDMKYKDEPTRLVIGDRNTIREFTTIHTGTVQDTGVTTLGDDNWIMAYVHIG
HDCRVGSHVILSSNAQMAGHVEIGDWAIVGGMSGVHQFVRIGAHSMLGGASALVQDIPPF
VIAAGNKAEPHGINVEGLRRRGFSPDAISALRSAYRILYKNSLSLEEAKVQLSELAQAGG
DGDAAVKSLVDFVESSQRGIIR*
>GCA_000452945.1_BPMSHR5858_genomic_00066_1 group_19257
LLTGKFSKRVMCALALAAALGATAARAQDVARIAAVNSDRILRESAPAKAAQTKLEAEFA
KRDKDLQDMAARLKSMSDALDRNGPTMSANDRAQKQRDLAQLDTDFQRKQREFREDLNQR
RNEELAAVLDKANKVIKQIAEQQNYDLIVQEAVYVSPRIDITDKVLKALASPSSLSN*
>GCA_000452945.1_BPMSHR5858_genomic_00067_1 bamA_2
MLFKPHRFVPKTVAAAALAAHGLAAHATAPFVVQDIKIEGLQRVEAGSVFAYLPIKQGDT
FTDDKASEAIRALYATGFFNDVRIATQGGVVIVQVQERPAIASIDFTGIKEFDKDNLNKA
LKAVGLSQGRYYDKALVDKAEQELKRQYLTRGFYAAEVSTTVTPVDANRVSILFAVAEGP
SAKIRQINFIGNKAFKTSTLRDEMQLSTPNWFSWYTKNDLYSKEKLTGDLENVRSYYLNR
GYLEFNIESTQVSISPDKKDMYLTVALHEGEPYTVSSVKLAGNLLDRQAELEKLVKIKPG
DRFSAEKLQQTTKAIVDKLGQYGYAFATVNAQPEIDQATHKVGLTLVVDPSRRVYVRRIN
IVGNTRTRDEVVRREMRQLESSWFDSSRLALSKDRVNRLGYFTDVDVTTVPVEGTNDQVD
VNVKVAEKPTGAITLGAGFSSTDKVVLSAGISQDNVFGSGTSLAVNVNTAKSYRTLTVTQ
VDPYFTVDGIKRITDVFYRTYQPLYYSTNSSFRIITAGGNLKFGIPFSETDTVYFGAGFE
QNRLDVDSNTPQSYQDYVNEFGRVSNTVPLTIGWSRDARDSALIPSRGYFTQANAEYGVP
VGKIQYYKMDVQGQYYYSFARGFILGLNFQAGYGNGIGNPYPIFKNYYAGGIGSVRGYEP
SSLGPRDTKTNDPIGGSKMVVGNIELTFPLPGTGYDRTLRVFTFLDGGNVWGNAPGGTST
GANGLRYGYGIGLAWISPIGPLKLSLGFPLQKHEGDQYQKFQFQIGTAF*
>GCA_000452945.1_BPMSHR5858_genomic_00068_1 dxr
MNVLVELIAFAVAIGVLVVVHEYGHYRVARWCGVKVLRFSIGFGAPVARWVSKKTGTEWT
LSALPLGGYVKMLDERDPGDGIRADELPHAFNRQPVGKRIAIVAAGPVANFLLAIALFSA
VFATGVTEPAAIVAPPAAGTPAAVAGFDGGETIVSIRIPGAGGAQGGQAEPVRSWSELRW
KLLGAAFDHRDVVLGARNRADGATYDFRVDLHGIADRDVDDDFMSRLGFEPGGGSLTVTS
VLPGGAAQRAGLQAGDKLVALDGARIGGSTRFIDDVKAHAGRALALRIERAGAERRVSIV
PQAQRDDETGKQVGRIGAALALQTPSVDVRYGVLESVELGARRTWDISVYSLKMFGRMVT
GEASLKNLSGPVTIADYAGKSARLGLSAFLSFLALVSISLGVLNLLPIPVLDGGHLLYYL
VEAATGKAVSERWQLILQRAGLICIVALSAIALFNDLARLIHF*
>GCA_000452945.1_BPMSHR5858_genomic_00070_1 cdsA
MLKTRVITAVVLLAVLLPVTLFAPLAAFGALIALALVFAAWEWGRLLKLGGAGPLGYAAV
AALALGASTRLGIGAAAARPLFAAAGVFWLLAGPYALWRKPALAERAWKPFLLVAGLVVF
SACWHALVAARAQGVPFVLSLLVVVWLADIGAYFAGKAFGKHKLAPSVSPGKTWEGAAGG
WLAVMIVAGAAVAAHAFEPTLYSTFVARYGAAGAFAALTVLVAFSVVGDLFESLLKRQAG
VKDSSGLLPGHGGVLDRVDALLPVLPLAMLLLG*
>GCA_000452945.1_BPMSHR5858_genomic_00071_1 ispU
MTYTSSTVRVPDVAAVPRHIAIIMDGNGRWATERRLPRVAGHTRGVDAVRSVVEGCARAG
VEYLTLFAFSSENWRRPNDEVSFLMRLFITALEREVGKLHANGIRLRVVGDLTKFEPRIQ
ALIRRGETKTARNTRLTLTIAANYGGRWDILQATRKLVEEAAREGRAPEIDEDAFARHLA
MAYAPEPDLFIRTGGEQRISNFLLWQLAYTEFYFTDTFWPDFDAGALAEALASYTHRERR
FGRTSAQLEPQAQDADSLSC*
>GCA_000452945.1_BPMSHR5858_genomic_00072_1 frr
MSVADIKKSVEQKMQRSIEAFKNDLAKIRTGRAHTGLLDHVQVDYYGSMVPISQVANLTL
VDARTIGVQPWEKTMVAKVEKAIREADLGLNPATSGDLIRVPMPPLTEERRRELTKVVKS
EGETAKVAVRNLRRDANEQLKKLVKDKEISEDDERRASDDVQKLTDKHVAEIDKLVQAKD
AEIMTV*
>GCA_000452945.1_BPMSHR5858_genomic_00073_1 pyrH
MPNAYKRVLLKLSGEALMGDDAFGINRATIERMVADIAEVVRLGTQLAVVIGGGNIFRGV
AGGAAGMDRATADYMGMLATMMNALALQDAMRHAGIEARVQSALRMDQVVEPYIRPRAIR
QLEEGKVVIFAAGTGNPFFTTDTAAALRGSEVGAEVVLKATKVDGVYSADPKKDPSATRY
SSISFDEAIGRNLQVMDATAFALCRDQKLPIRVFSINKPGALKRIVQGEDEGTLVHV*
>GCA_000452945.1_BPMSHR5858_genomic_00074_1 tsf
MAAITASMVAELRAKTDAPMMECKKALTEADGDMAKAEELLRVKLGNKASKAASRVTAEG
VVASFVGANAGALVELNCETDFVAKNDDFNAFAKTVAELVATQNPADVAALSALPLDGKT
VDEVRLALVGKIGENISIRRFVRFETSNKLATYLHGSRIGVIVEYTGAQEQVGKDVAMHV
AAMKPVSLSADEVPADLIEKERRVAEQKAAESGKPAEIVAKMVDGSVQKFLKEVSLLNQP
FVKNDKQTIEQMLKAADAAVQKFALFVVGEGIEKRQDDFAAEVAAQVAAAKQQ*
>GCA_000452945.1_BPMSHR5858_genomic_00079_1 def
MIREILKMGDPRLLEVARPVEAFNTPELHALVADMFETMHHANGAGLAAPQIGVGLQVII
FGFGSSERYPEAPPVPETVLVNPSIEYLPPDLEEGWEGCLSVPGLRGVVSRYRRVRYSGF
DQYGAKLERIAEGFHARVVQHEYDHLIGKLYPMRITDFSKFGFADVLFPGLDAQADD*
>GCA_000452945.1_BPMSHR5858_genomic_00080_1 ligA
MARSPVEPPASQPAKRAAWLRAELERANYAYYVLDQPDLPDAEYDRLFVELQRIEAEHPD
LVTPDSPTQRVGGEAASGFTPVVHDKPMLSLNNGFADEDVIAFDKRVADGLDKATDLAGT
VTEPVEYACELKFDGLAISLRYENGRFVQASTRGDGTTGEDVTENIRTIRAIPLTLKGKR
IPRMLDVRGEVLMFKRDFARLNERQRAAGQREFANPRNAAAGSLRQLDSKITASRPLSFF
AYGIGVLDGADMPDTHSGLLDWYETLGLPVNRERAVVRGAAGLLAFFHSVGERRESLPYD
IDGVVYKVNRRDEQDRLGFVSRAPRFALAHKFPAQEALTKLIAIDVQVGRTGAITPVARL
EPVFVGGATVTNATLHNEDEVRRKDIRIGDTVIVRRAGDVIPEVVSAVLDRRPADAQEFV
MPTECPECGSRIERLPDEAIARCTGGLFCPAQRKQALWHFAQRRALDIDGLGEKIIDQLV
EQNLVRTPADLFNLGFSTLVALDRFAEKSARNLIDSLEKAKHTTLARFIYALGIRHVGES
TAKDLAKHFGSLDPIMDAPIDALLEVNDVGPIVAESIHQFFAEEHNRTVIEQLRARGKVT
WPEGPPAPRAPQGVLAGKTVVLTGTLPTLTREAAKEMLEAAGAKVAGSVSKKTDYVVAGA
DAGSKLAKAEELGIPVLDEAGMHTLLEGHAR*
>GCA_000452945.1_BPMSHR5858_genomic_00148_1 group_19547
MDPIFYGFVLLLFVAVVLAFEGVYQWWNARHGPAARRIEARIRAMSAGGQVQKERLSILK
ERMLNDAKPFDRLLMRVPRVQTLDLMIQQSGLTWSVVKLAWISLAIPFAAFVALGFTPLP
LWATATASVAAVCLPTLFVLRCRTRRLRKLERQLPDACDMIARALRAGHAFSSAISMVGG
EFAEPMGGEFRITFDEINYGVSLQEALLNLATRVPVSDLRYFVIAVLIQRETGGNLAELL
DGIAQLIRERFKLFDKIRVLSAEGKLSALILTLLPFGTAALLSIMNPDFISVLWEDPAGV
LLVGVTIASMAFGILWMRHIIRIRV*
>GCA_000452945.1_BPMSHR5858_genomic_00152_1 group_1747
MNRFSLPPRPSRMRGAVAVEFAIVMIPLVLLATGVAEFGRAIYQYEALTKATRDAARYLS
TYLPTDPAYPLAQAQCLAVYGSTTCGSTGSELAPGLATSMVVVCDAAHAPDCSDSSDPAQ
FANVPTYDTNNGSPDPASLAGSMNLVEVKIKGYQYRPIPAFPGLPNLSFGNIVTVMRQVS
*
>GCA_000452945.1_BPMSHR5858_genomic_00155_1 sctC_4
MDTKPQLSRPTPARSRAVRGAALAAACCAGLFTVFAAAAQDAVENAAIAKGTAAAPMALG
RAPMQMTISMAPAPAAGAAAAAAAVPLRGPSCVGEMRESTSVSVPLGKSLLVPLDEPVRN
RTIGNPAVAQATMVSPRTLYLVGMSVGTTNMIVQGKSGACRIIDVAVGADAGGLQASLRQ
LMPRERDIRVSTAADTIVLAGNVSSAQAAQQAVAIAKAYSNRTNAGAAGSGGGARADVLN
MLTVTSPQQVMLEVKVAEVSKTLINQMGSAVNIQGGFGSWTGALVTSLLTGVGSGIAASK
SNKLPLNGSIDAQNTDQPVKILAEPRLVTISGQEATFLAGGKIFIPVPQSSGIGTNTITL
QEEEFGVGLKFTPTVLSNGRISLKVAPEVSELSPTGVTLSAANVGNVSILPLITTRRAST
TVQMGDGETFAIGGLIKDNAQGTLKALPGIGELPVLGALFRSTSFQQDRTELVFIITPHL
VQPLETADVPLPTDSFTKPNEADVYATGNMEGRGGLRQHGAAPAAAPANAAPGAAAPTGA
APAAAVPAPAAPAPAAPAAPAAAAAPAKDAAASPAPAKQAPIAAQAPAPQPAKPAASAPA
LANARPPATVQAPPPALPSAAALAQRQGAHAAAPDTAHAGPAESAAATASPAPAAALAAP
AAAGATAKRQGAQPPQQLANAGAIAPAADR*
>GCA_000452945.1_BPMSHR5858_genomic_00173_1 group_19439
MSAAPTDPRAARPARRKMRPLPAVIFMSRWLQVPLYLGLIVAQAIYVFLFLKEVWHLLSH
ATGLDETNIMLAVLGLIDVVMISNLLIMVIVGGYETFVSRLGVEGHPDEPEWLDHVNAGV
LKVKLSMALISISSIHLLKTFINPDQHTTHAIMWQVLIHVAFLVSALVMAWVDRLTTHTH
PEHFHEASTPPSAPREPAQQSA*
>GCA_000452945.1_BPMSHR5858_genomic_00178_1 group_1749
MQVSNSLPGASAPAFSRMNPRIVTAAVTVLAGHALLLTGALLMRNDVPHRPLESKTITAQ
LLSAPVAQPAGIQSAPAPTPPQPVPKAKPKPAPTPVPRPVAKPSPTPLPVTHEPAPNAIT
APEPAPAAPVAPAETSAKAAPPAGAPTNRPTMEIVAPKEGAHLSCQIAKAAYPSMSKRRG
ETGVVKVRFVVGLTGKIESAQVVQSSGFARLDDAALDAIRSSPCQPYLQNGQPMRAAYTQ
PYDFTLTD*
>GCA_000452945.1_BPMSHR5858_genomic_00183_1 group_19409
LNFEHLIQINAADNPAVPTLTRAQLWEGLVLRAEQPQLFVLGLDSCIVHERTETTLEREL
HYGNATVRDHVTFTPNEQVRYDIHAADGEIGGSLTMTIEERDDQQLFLRFEYRTTLPVSN
DSEDARQTQEIVKEAYRTSDIDTVRLIREYAQGRKDPDPLH*
>GCA_000452945.1_BPMSHR5858_genomic_00184_1 group_1750
MKPAAASAGQRVVSSDALLQGQSHVSIAHNGETYQLRATRLGKLILTK*
>GCA_000452945.1_BPMSHR5858_genomic_00185_1 group_19054
MRTKPVLTSDDVNKMAAAAEAFAAANQWAVTVAIFDDGAHLLHLHRMDGAAPSTVEMAIG
KGRTAALGRRESKVYEDIVLQGRISFLSVPLVGLVEGGVPIVVDGETVGAIGVSGVKSEQ
DAAVARAGIAALTDAG*
>GCA_000452945.1_BPMSHR5858_genomic_00188_1 group_19389
MTFPRAYAFAAAILVAGAAHAADLPTFRLEMADGKLNPARIEVPAGQRIKIQIRNTGKGA
VEFESVQLRKEKVLAPGGESFVVIAPLSPGEYKFFDDFHQQAQGVIVAK*
>GCA_000452945.1_BPMSHR5858_genomic_00189_1 tpd
MLRSSFVRSGVALAAACAALSATAAEYPIGKHQIQGGMEIGAVYLQPITMDPEGMMRKAS
DSDIHLEADIHAVKNNPTGFAEGDWMPYLQVTYKLTKQGDAKWAAQGDLMGMVASDGPHY
GDNVKLNGPGKYHLTLVVKPPMQTGHMAFGRHVDKETGVGAWFKPITLEYDFPFAGIGKK
GGY*
>GCA_000452945.1_BPMSHR5858_genomic_00190_1 uvrB
MSEHHSDTRDDLDESKFVTFEGSPFQLYQPYPPSGDQPTAIATLVEGVEDGLSFQTLLGV
TGSGKTYTMANTIARLGRPAIVFAPNKTLAAQLYAEFREFFPRNAVEYFVSYYDYYQPEA
YVPQRDLFIEKDSSINEHIEQMRLSATKSLMERRDVVIVATVSAIYGIGNPSEYHQMILT
LRTGDKIGQREVIARLIAMQYTRNEQDFQRGTFRVRGDTIDIFPAEHAEMAVRVELFDDE
VDTLHLFDPLTGRVRQKIPRFTVYPSSHYVTPRETVMRAVETIKDELRERLEFFHRDGKL
VEAQRLEQRTRFDLEMLQELGFCKGIENYSRHFSGAAPGEPPPTLVDYLPPDALMLLDES
HVLIGQLNGMYNGDRARKENLVDYGFRLPSALDNRPLKFPEFERKMRQVVFVSATPADYE
QRVSGQTAEQVVRPTGLVDPQIEVRPASTQVDDVLSEITERVKANERVLITVLTKRMAEQ
LTEFLADHGVKVRYLHSDIDTVERVEIIRDLRLGTFDVLVGINLLREGLDIPEVSLVAIL
DADKEGFLRAERSLIQTIGRAARNVNGKALLYADRITDSMRRAIDETERRRAKQIAYNEK
MGITPRGVVKRIKDIIDGVYNADEARAELKEAQQRAKFEDMSEKQIAKEIKRLEKQMADY
AKNLEFEKAAQTRDQLALLRERVFGANVGDHVSGGE*
>GCA_000452945.1_BPMSHR5858_genomic_00203_1 group_19347
MDAESIAGLIGLGIGLLVLVSLSVYESREYKRAHNGEGMIHHWMAEHHLLDWRRKH*
>GCA_000452945.1_BPMSHR5858_genomic_00204_1 group_19609
MRELPTPTLPFASLPLDLPRRRLPRAAETVTAEFRLRAAWARTEDELREAQRLRHSVFAE
EMGAHVSGPAGLDVDPFDPYCDHLLVRDLDTLKVVGTYRVLPPHQAARVGRLYAEGEFDL
SRLTHLRSKMVEVGRSCVHRDYRSGAVIMALWGGLGTYMLQNGYETMLGCASVSMADGGH
YAANLYQSLGDALTAPEYRAFPHTPLPVDELQTGAKVAPPPLIKGYLRLGAKICGAPAWD

PDFNCADFLTLFRLSDINARYARHFLSDPLPR*
>GCA_000452945.1_BPMSHR5858_genomic_00212_1 group_19342
MNNPLRLIARLLIPFATLAAAPAASADSQALDAALDCSSTGHAFVAPLVASGAVRSQPMH
VEANSMNAFRTNRSLSAYGFSVYVVLGYQANDPLFAHGDGEPIGDWAYGVVVRGSKQAVE
AKVRAAGSQAVVKDAFPFLTAIVCTSP*
>GCA_000452945.1_BPMSHR5858_genomic_00218_1 pstB
MNMAESHLDPSKLATGPAGFGAAAGQRPLAPLNAKIEVKNLNFFYNQFHALKNINLSIPE
GKVTAFIGPSGCGKSTLLRTFNKMYALYPEQRAEGEIVMDGENLLQSKLDISLLRARIGM
VFQKPTPFPMSIYDNIAFGVKMFERLTRSEMDDRVEWALTKAALWNEVKDKLSQSGYGLS
GGQQQRLCIARGIAIRPEVLLLDEPCSALDPISTGRIEELIAELKSDYTVVIVTHNMQQA
ARCSDYTAYMYLGELIEFGETEKIFIKPARKETEDYITGRFG*
>GCA_000452945.1_BPMSHR5858_genomic_00219_1 pstA
MSAPIVNMPGPDGAALDAMRARLQRRRKATNVVALTASLAAMAFGLLWLVWILYTTLHLG
VGGLSLALFTESTPPPNTAGGGLANAIVGSLLLVALATFVGTPIGILAGVYLAEYGQKSW
LASVMRFINDILLSAPSIVVGLFVYALVVAKSGHFSGWAGVIALALLQIPIVIRTTENML
KLVPNALREAAFALGTPKWRMILKITLRASVGGIVTGVLLAVARIAGETAPLLFTALSNQ
FFSADMSQPMANLPVTIFKFAMSPFAEWQSLAWAGVFLITLGVLGLNILARSIFSKK*
>GCA_000452945.1_BPMSHR5858_genomic_00225_1 rlmE
MAKNRFNQSWLHDHINDPYVKMAQREGYRARAAYKLKEIDEQDKLIRPGQVIVDLGAAPG
SWSQYARNKLAQGKRRDAVREGGIDGTIIALDMLPMEPVADVHFIQGDFREESVLHQLEE
VLAGRAVDLVISDMAPNLSGVAVADAARIEHVCDLALEFAQNHLKPDGALLVKCFHGSGY
SQIVEKFKHQFKTVAPRKPKASRDKSSETFILGRHLKQPR*
>GCA_000452945.1_BPMSHR5858_genomic_00227_1 group_1392
MAGAAAPHRLFRLLATVWVGSLLTIGYAVAPVLFATLDRATAGAVAARLFSIEATIGAVC
GVLLIGLSNLLVRRGHDAYRPLRWLLAAMLGCVLVGYFALQPFMNALRVAALEAGTDVGH
SAYAGRFALLHGVSSVFYLVESLVGIGLIWRLPGQGGASPLGGERRA*
>GCA_000452945.1_BPMSHR5858_genomic_00234_1 sauU
MQFRVFALFAIGYFVSYVFRGVNLGFAPLITRELGLSAADLGLLTSLYFLGFACAQIPAG
VLLDRYGPRRVSAALLLVAAAGAWVFGAAHDIGAMMLGRLLIGVGVSVCLGGAFKALAQH
FPVARLPLLNGFVMAVGGLGGVAVGSPLDWLLGFAGWRAVCAGLGAFTVAVAAALWAFAP
DARQSRHDASVAAQFAGTWRIVTNRTFWKIASFSVVTQGVFYAMQSLWVGAYLRDVSGFA
PRDAAVLVSVLGFAMMAGCVGFGAAARGLERRGMSLYAFCGFGMALFVATQLAIMLRAPL
PAAVLWAAYGVFGGVGILSYAVLAEYFPVHVIGRANTTLTLVIFVLIFGFQVGVGAVLSH
WPMRAGHYPAEAHLSAWGALVALQIASAVWYVLPDRALDKSARHAG*
>GCA_000452945.1_BPMSHR5858_genomic_00235_1 mltD
MRFLLSALLVLTLAACAGTGPTAQNPAASDPQAASDYLRKSASAKETVDVDKQSVGDLTT
ADSDLWARIRRGFQMPDLQSDLVDMQASWYAQRPDYVQRMTERSQKYLYHIVEELESRHM
PTELALLPFIESAYNPQALSVAKAAGMWQFMPATGRTFNLKRNMWQDERRDVLASTSAAL
DYLSRLHDMFGDWYLALAAYNWGEGNLQRAIARNQAAGLPTDYQSLRMPNETRNYVPKLQ
AVKNIVTNPQQYGLTLPEIPNHPYFVTVTTSRDIDVTVAAKLANLPLDEFKSLNPSFSKP
VILGATQPQILLPFDNAAAFEKGLKSYDGQLSSWTAYTVTERARPAAIAEKIGVDADTLM
QVNKIPAGMRLKPGSTIVVPRTDDDDEDISADVAENGVLAMEPDVPDTRKMLIRVRRKQS
MEALASRYGVSVAQLRGWNRTRRAVVMPGQTMVLHVPVGRAVPAEPGPERIATSVGGGRI
ERASLHTGGKASSRGRKAGPAAKASAKAATKSGAKAAPAKTAGRKTTKKK*
>GCA_000452945.1_BPMSHR5858_genomic_00238_1 rnhA
MTLQTIDIYTDGACKGNPGPGGWGALLRYGAQEKELFGGEPGTTNNRMELTAVIAALEAL
KRPCKVVVHTDSQYVQKGISEWIHGWKKKGWVTAAKTPVKNADLWQRLDALVAQHDVEWR
WVKGHAGHPENERADALANRGVESLAQA*
>GCA_000452945.1_BPMSHR5858_genomic_00254_1 group_1751
MIIYGTALLAFCHLAGLFLGDLLGAAIGAKTNVGGVGIAMLLLIGLRLALHRRGWLPKET
EAGIAFWGAMYIPVVVAMAANQNVVAALKGGPVALLSAVGAVAVCVLSIAALARVGHDAT
SLAALPRPEQN*
>GCA_000452945.1_BPMSHR5858_genomic_00260_1 group_19326
MSDLYFGADLFSEFDRLHRQMANLFGGFPSSIRASRLGGFPALNVGTTEDSIEIVVFAPG
MRAADFDVSIDKGLLAISGERKAAQRAEGEVRAYAQERFSGAFRRVVELPQNADPDKIRA
RYENGCLLISVGKREASKPRAITVQ*
>GCA_000452945.1_BPMSHR5858_genomic_00272_1 group_1135
MCGIVGAVARRDVVPNLVDGLKRLEYRGYDSCGVVVYQDRRLVRARSVDRVAALQREISA
RALSGYTGIAHTRWATHGAPVTANAHPHFSSGVATSGVATSGVATSGVATPGVATSGAPA
ASAAASGGTAQADASPEARARIALSHNGIIENYEALRADLERHGYAFASQTDSEAIAHLV
DHLYDGDLFDAVRRALARLRGSYAIAVMCRDEPHRIVGARDGMPLVVGIGDGEHFLASDA
IALSNLTDRIAYLENGDVVDIQLHRHWIVDASGRRVERTVHTIAAHVAAADLGGYRYYMQ
KEIFEQPRAVADTLLDVTSIMPELFGDRAWRALSAADSVLLLACGGSYHAALTAKYWIES
IAGLPANVEIASEFRYRDGVPNPKALVVAVSQSGETADVLGAVHAAKRLGMPNTLAICNV
PTSALVRECASSFITRAGIEVGVASTKAFTTQLVALFLLTLALAQVRARLSDEDEKRHLH
ALRHLPEAMSKVLALEPRIIAWSERLARRENMLFLGRGMHYPIALEGALKMKEISYIHAE
AYAAGELKHGPLALVSDEMPVIAVAPNDRLLEKLRSNMHEVSARNGRLFVFADVDCAIAS
APGLDVIRLTEYYGPLSPILHAVPMQLLAFHAALARGTDIDKPRNLAKSVTVE*
>GCA_000452945.1_BPMSHR5858_genomic_00280_1 group_1368
MTSPAQPTWRDTLSAFAYRHLRLPPPKLETGRFGWATRWFLDWLVPTGASAWAAYHLAAA
FVAHYRSVLPANVGPDVPLGVMSELMSLANSKPFAYTGISLVAVLVALMFHRYLTWPAPG
RWLNRRKRPVIHALPFMLGFLAGKLLAFAVPALAVAVAAYASEGAIGAIAPLAAPIALLS
AAYYFLLYDEVRPPAIVSLSFHRSRQRSAGAGATHQAREPDGAAAVGIGAWRDAHDELRD
DVLAIADAAGGMCGQNVFVSHDIDDIAIAIALSIDEAHRSDRQKRSLDAQIDRVLRKWRD
RGYASGSLHDAKATRASPGAPEARRADVAFWLPESERIVRHGSVLLKARDGRDVRIVEES
IGSVQADASAVRPRPRRMRTFEFV*
>GCA_000452945.1_BPMSHR5858_genomic_00307_1 cysA_2
MKLDSVPITLTRCAKTFHGTRVLKPLDLAIGAGETLVLLGPSGCGKTTTLRLIAGLDTPD
AGGAIAFGADDVTALPIEKRGVGMVFQNYALFPNLSVRGNVGYGLRIRRTEPRVLRERVD
ALLAMMRLTAHADKPIDALSGGQRQRVALARALAARPRVLLLDEPLTALDAQLRDALRRE
MNALLRELGVTTVYVTHDQAEAMALGDRIVVMSAGAIEQIGTPRDVYYRPASRAVARFIG
TLNRLEGVWRDGALVTEGGALAVASPANELFFRPEDARLVDPSDAPLRGAVAHCAFLGER
TRVTIEHAAPDPLVIDVPGRVDLARGTAVGVAIAAEGLIALA*
>GCA_000452945.1_BPMSHR5858_genomic_00314_1 yedZ1
MSATPTTSASTTAMAGARASGAADAPARAPTGADVDAGATAWADATANATANATTDPTID
ATADATTDAMANVTTDAATRVTANPPIHETANVPASTAAAATTEAVAGARAGAALDAAHP
GATQTMSQTMSQTMAQTMTQAGTEAASPAGTDAAASGACITRATRARRPIHPLWVRASHW
LNALAAVTMALSGWRIYNASPIYAGFVFPRGATLGGWLGGALQWHFAAMWLLFFNGVFYL
TMGVATRRFTRKLLPLSASGVARDLRDALRGRLSHDDLSVYNALQRAAYVAAIADLALLV
LSGLVIWKSVQFPLLREWLGGYDRARVVHFWAMAFLVAFFVVHVAMAALVPRSLLAMLRG
H*
>GCA_000452945.1_BPMSHR5858_genomic_00317_1 thiC
MNANPKFLSADARVDAAAVAPLPNSRKVYVTGSQPDIRVPMREITQADTPTSFGGEKNPP
IYVYDTSGPYTDPDAKIDIRAGLPALRQRWIDARGDTETLAGLTSDYGRERAADPATAEL
RFPGLHRHPRRAKAGKNVTQMHYARQGIITPEMEYIAIRENQRRAEYLESLKASGPNGAK
LAAMMGRQHAGQAFGAAAFGANAPAEITPEFVRDEVARGRAIIPANINHPETEPMIIGRN
FLVKINANIGNSAVTSSIGEEVDKMTWAIRWGGDTVMDLSTGKHIHETREWIIRNSPVPI
GTVPIYQALEKVNGKAEDLTWEIFRDTLIEQAEQGVDYFTIHAGVRLQYVPLTANRMTGI
VSRGGSIMAKWCLAHHKESFLYEHFEEICEIMKAYDVSFSLGDGLRPGSIYDANDEAQLG
ELKTLGELTQIAWKHDVQVMIEGPGHVPMQLIKENMDLQLDWCKEAPFYTLGPLTTDIAP
GYDHITSGIGAAMIGWFGTAMLCYVTPKEHLGLPNKDDVKEGIITYKLAAHAADLAKGHP
GAQVRDNALSKARFEFRWQDQFNLGLDPDKAREFHDETLPKDSAKVAHFCSMCGPHFCSM
KITQDVREFAAQQGVSENDALKKGMEVKAVEFVKSGSEIYHRQ*
>GCA_000452945.1_BPMSHR5858_genomic_00318_1 group_18925
MKSIRRIGVCAMLIATMASLSACDTMTTRQRNAAIGAGIGGVAGAAIGGNALSTIGGAAA
GGLVGSQVGK*
>GCA_000452945.1_BPMSHR5858_genomic_00369_1 group_1292
MPFLKQAAGALRRNLSDTARHTLSGPHRGWKIALYAALLIVPGGSLAALGFAWFDQRRHR
AARDAGMRAASAQEPSAPMLCLAATHGEPVRCS*
>GCA_000452945.1_BPMSHR5858_genomic_00370_1 slyA_1
MSGLYDPEHIELESSLGYYLTKARQALVERLDRALGPLELTAQQISVILLLARGYARTPF
ELSRKLSYDSGSMTRMLDRLEKKGFVVRARSESDRRVIELALTERGAHAARALPALIATE
LNAQLEGFSADELALLTDLLRRFIANAPGAADAACAEPPPDQR*
>GCA_000452945.1_BPMSHR5858_genomic_00376_1 fliM
MGHEEFMSQEEVDALLKGVTGEDDSADEPAEASGIRPYNIATQERIVRGRMPGLEIINDR
FARLLRIGIFNFMRRTAEISVSQVKVQKYSEFTRNLPIPTNLNLVHVKPLRGTSLFVFDP
NLVFFVVDNLFGGDGRFHTRVEGRDFTATEQRIIGKLLNLVFEHYASAWKSVRPLQFEFV
RSEMHTQFANVATPNEIVIVTQFSIEFGPTGGTLHICMPYSMIEPIRDVLSSPIQGEALE
VDRRWVRVLSQQVQSAEVELIADLAEVPTTFEKILNLRTGDVLPLDITDSITAKVDGVPV
MECGYGIFNGQYALRVQRMISASDTMKEGGYD*
>GCA_000452945.1_BPMSHR5858_genomic_00378_1 group_1143
MKPRFAPLAARAVPVCVPAFLSPRRAAERAPRGRRRAPAALAVLASGTAALPVSAADMNA
VNHAASLASGVVVGSAAPSLGVGAVLQTLVGLAVVIGLVFGCAWLARRFGLQPQRRGGAL
KIVASVAVGGKESATVVEIGDTWLVLGVAPGNVRLLHTLPAGSAGVIGAPAGGGLTRSPG
APGGTAMSGPLPEGASFGARFRDAMLGEAAKRFKRDGGKDR*
>GCA_000452945.1_BPMSHR5858_genomic_00379_1 fliP
MPRLRFVAPRADAAACAPSDSRSMKTACLRGAARWLPAILIGLAPALACAQAAGLPAFNS
APGPNGGTTYSLSVQTMLLLTMLSFLPAMLLMMTSFTRIIIVLSLLRQAIGTASTPPNQV
LVGLALFLTLFVMSPVIDRAYNDAYKPFSEGTLQMDQAVQRGTAPFKAFMLKQTRETDLA
LFAKISKAAPMQGPEDVPLSLLVPAFVTSELKTGFQIGFTIFIPFLIIDMVVASVLMSMG
MMMVSPATVSLPFKLMLFVLVDGWQLLIGSLAQSFT*
>GCA_000452945.1_BPMSHR5858_genomic_00383_1 COQ5_2
MTATDRNDNGQAAHWNGRAGRAWADAQPMLDRMFEPFAQRLVEAARTGRRVLDVGCGAGA
TTLAVAQRLGARGRCVGVDVSRPLIAAARARAERGGVPASFVHADAQTHAFVPASFDTII
SRFGVMFFENAVDAFANLLRAATSDASLAFVAWRTAAENPFMTTAERAAAPLVPNLPARQ
PDAPGQFFFGDARRIETVLAQSGWCGIDVRPIDVECTLPERELIGHFSRLGPLGQLFGDL
DDATRARVVDTVRAAFAPYVHGAEVRFTAACWLVGARAPAKWSKRKEAAGV*
>GCA_000452945.1_BPMSHR5858_genomic_00386_1 nrtD
MNPHVSAHAPAIEFRNVSCRFISPDGRATVALHDFTMSVARGEFVAVVGPTGCGKSTTLN
LITGLLKPVSGEVRVMGKPVDGIDPRIGFVFQADAVFPWRTVIDNVAAGPLFRGRSKESA
YAQAEEWIRRVGLAKFAKHYPHQLSGGMRKRVALAQTFINQPEILLMDEPFSALDMQTRT
LMQDELLQLWSANKGSVVFVTHDLEEAIALADRVFVLTSRPATLKRVYEIDLPRPRVTSE
VRYEPRFIEISKDIWHDLREEVQIG*
>GCA_000452945.1_BPMSHR5858_genomic_00388_1 group_19107
MSHSLRGRLLWWLLLPLAVFVAIAGAMSYDTARKTADLVQDGALVASARVIAEDVDWEGG
ALVANVPPAALELFASPAQDHVYYKVRTGGGRLLAGNPDLDGPAAPAASGAQPVLFDTAL
GGLAIRAVAYTRELYNAGNTETVTVVVGKTQTSRQMMIAAIWHPQLWRLALMLALAMALV
YLGLTFELRPLMKLKEDVADRGPMELEPIRTERLHFELRPIVDAINQCIAQLNLHAATQR
RFIADAAHQLRTPIAVIDTQIQCARQRENGDAALAALLASMQRSSRRMADVTDKLLLLAH
AEAASPARLAARVDIAAVVSGVLEEAIVLAERRRIDLGAELDDDLQVAGSESLLSALLMN
LVDNAVRYAHEGGRVTVSARRDGDAVVLEVVDDGPGIPAEARPHVFKRFYRVARDEEGTG
LGLAIVEEIAQSHGGAVSLATGPGNRGVRMTVRLPAYRN*
>GCA_000452945.1_BPMSHR5858_genomic_00389_1 tctD_2
MKLLLVEDNAELAHWIVDLLRGEGFGVDSAPDGESADTVLKAQRYDALLLDMRLPGMSGK
ELLARLRRRGDNVPVLMLTAHGSVDDKVDCFSAGADDYVVKPFESRELVARIRALIRRQC
GVGATQLACGDLVYLFATREFQCGGAPLALRRREHAILETLMLQQGKTVSKARLMDSVYG
LDDEPSADAIDIYIHRLRKHLSGSLAQIITLRGLGYILRTQDAAE*
>GCA_000452945.1_BPMSHR5858_genomic_00391_1 group_19400
MQKLDAASPEAQSADLVAANVERLKALFPDVVTEGPDGASVNLDALAALVGASAAAAADA
DEKYGLNWHGKRRARRLALTPSTGTLRPCPRESAGWASTRNLMIEGENLEVLKLLQKSYA
GRVKLVYIDPPYNTGKDFVYPDNFTDSLRHYLELTGQTTGGKRVTSHTDASGRFHTDWLN
MIYPRLKLARDLLTEDGVIAVHIDEHEQHALVLVMREIFGEDNELGVAVWDKRNPKGDAR
GIAYQHESIVLFARDAERLFERAPLKRPKRNAQRMLGAAREAVAGAATIADANAAYRGWV
KSQTTLSGGEAMYDRISADGRVYRLVSMAWPNKKKAPDDYFVPLVHPVTGKPCPVPERGW
RNPPATMRALIDKGLVEFGADETTQPQRIYFLDENMYENVPSVLPFGGSDDALMKSLGIP
FDQPKPVEFAASIIGWCTDGDDLIVDFFGGSGTTAHAVMALNAADGGHRRYVLVQLPEPL
DADSKDQKAAADFCAAQRVPLNLAELTKERLRRAAARIAAEHPGTRADLGFRVFRLDSTN
VSEWDPRGDDIQQSLFAAVEHIKPNRSEEDLLYELMLKLGLDLCAPIDARMIAGKAVYMI
DGAIVACFDAHIDRASTDALGEGIVGLIAEAADAREVTCVFRDSGFADDVAKVNLSAILE
QHGVKRIRSL*
>GCA_000452945.1_BPMSHR5858_genomic_00399_1 group_19366
MKPLSLNMKRALAGACIGAAGFLAQQAALAQSCGLANGKPATGAPIPIGAVVGKTGPDDF
SSSARAAAAYFKCVNANGGINGRPVQYLVEDDQWNPETASQVASKLVRDRKVLALAGNAS
FVECGANAKFYEQENVIAIAGVGVPRECYFARNYAPLNMGPRLSMTEAALYAKQQYKATR
MVCIAPNIPSLGAWSCEGPALWGKRNGVSVDTIVMDPDSADPTSVVLQAASKNPQAILLG
LPKGLMVPILSAAEQQNLGRRIHFVSAASGYDLGVPKAIGPYWKGNFDVNLEFQPLDAQT
PDNRNWLAVMDKYGDRKDPRDTFSQAGYLAARLVTDTLLKLPANQLDRAHVTAALREVKD
FRSDILCGPFYVGAGERHNANNAGRMAQSTGTGWKTVSTCQAVDDPQLADIRAAEKKMH*
>GCA_000452945.1_BPMSHR5858_genomic_00411_1 fadA_1
MEAYIFDAVRTPRGKGKKDGSLHGVTPLALAATALRAIRDRNGLDTRAVDDVVLGCVEPV
GEQGACIGRIAVLAAGYAETTAGVQINRFCASGLEACNMAAAQVMSGQSEMAIGGGVESM
SRVPMGSSGGAWPVDPAIAIPSYFVPQGVSADTIATKWGYSRADVDAYAAESHRRAHAAA
QAGWFARSIVPVRDANGLTILDHDETIRPQTTPETLATLKPSFAELGEMYGFDAVIRQRY
PELERIEHVHHAGNSSGIVDGASAVLIGSLEAGKRAGLAPRARIRSFASIGSEPSIMLTG
PAYAAQKALERAGMRAADIDLYELNEAFASVVLRFMDAMAIAHDRINVNGGAIAMGHPLG
ATGAMILGTLLDELERRGAATGLATLCVGAGMGTATIIERI*
>GCA_000452945.1_BPMSHR5858_genomic_00412_1 fadJ_2
MIDYTLDDDGIATLSWNLPARSQNVLNGESCEAFFAAAARALQDAAVKGMLVTSAKPDFI
AGGDLEWLQASDSAETLFARTCELHRALRALETGGKPVAIAMPGSALGGGLEIALAGHYR
VAADNPKARFGLPEVTLGLLPGGGGTQRLPRLVGVQASLPLLLEGKRLKAADALAAGILH
AVVPAGEEIAAARAWLLDASRRTATQPWDAKGFRIPGGALTHPSVQQVFMAANALARQKT
YGNYPAVASILSCVYEGLLTDLDTGLKTEARYFVKAVLSPEAKAMIRTLFFGMNEANKLA
ARPAGVPTQRYRKVGVLGAGMMGAGIAYVSAKAGLDVVLIDTGEEAAARGKDYSRKLVDK
QVQRGRLAREKADALLAKIVPTTDFARLDGAQLVIEAVFEDRAIKADVTRRSEAVLAPDA
LFASNTSTLPITGLAQASARPANFIGLHFFSPVDKMPLVEVIVGRDTSEATLARALDYVK
TIGMTPIVVNDSRGFYTSRVFSTYVLEGLAMLAEGVAPALIENAGLLAGMPVGPLALTDE
VSSELIHKITRQTRADLGDAYVARPGEDVAARMVELGRLGRKAGLGYYDYPASGGKKALW
PGLAQEFPLAAEQPDVAALIERLVTVQAVETARCLEERVLTTARDADVGAILGWGFPAFR
GGPASYIHGVGVDAFVATCDRLAARHGARFAAPALLREMAAQGRSFY*
>GCA_000452945.1_BPMSHR5858_genomic_00413_1 uctC_2
MGPLHGIRIVEIAGIGPAPFCGMLLADMGADVILVERTAGRDGDPLDLGRRAIFNRGKRS
LALDLKQPEAVGAVLRLVEHADALIEGMRPGVMERLGLGPDTCFARNPKLVYGRMTGWGQ
DGPLAHAAGHDINYIGLSGALWYAGQPGDAPFAPPTLVGDLGGGALYLAMGVLAGIVDAR
LRGKGQIVDAAIVDGSANLMNLLLSIHAAGFMPMARGQGLLDGPHWYGSYRCACGGYVSV
GALEPQFYALLVDKLGLADDDDFRNPYDASRWPRLRERLAEIFASRPRAHWVELMEGTDA
CFAPVLSPAEAQSHPHLAARGVYAQRDGILQAAPAPRFSGSPNAQPGAVPKRGADSAAIL
REAGFAEDEIAALAAKE*
>GCA_000452945.1_BPMSHR5858_genomic_00414_1 group_1761
MRTIRAWLALTFVASIACLIATTPAARAAPAGKALTFDGVAFFQRDAAGNRLEYTPKGED
DLSAWRTMITLVRYPEATTADGLASIADSVLRLYKQNGAMILKVDAVPAARGKPAEYFIS
VAFPTRDFIEAVFTRIALVDGSGAAAIYSHREYGRAVGNEVSAWIEKNGAAKEHALKQWN
PLPTVGGK*
>GCA_000452945.1_BPMSHR5858_genomic_00419_1 group_1062
MNIPPDEHDLHAYVDGRLDADARTAVKRWLALHPARAEQVRRWRQDAQQLRAALDSVPTP
APTRTLDPAVLRARRAGRARARLAMAASLVLCVALGAIGGWQAHGWSGAPAAPMSDAVAA
YKLMVVDRSARVDVAQKQPGELQAWVARALGAAARLPDLSSAGFAPVGGRLFATERGAAA
MVLYEDAAGRTLSFYIRPPESANRLLAAGERADGALLARYGSARGFNYAVVGRADSVGEA
AVARALEVEGRS*
>GCA_000452945.1_BPMSHR5858_genomic_00583_1 pcpR_2
MPDLDLNLIPYLVALNEARNVSRAGELLGVSQPRVSAALGRLREHFGDPLFVRTSRGMEP
TPRALALIPAARDALASIERGLGAPHEFDPAKSARTFSIALSDVGEIVFLPRLLQAFAQH
APHANLRSASLSHAEVAHALEDGGVDLAVGYFPDLGGTNFFQQRLFTHRFICLLRRGHPL
AERPLTLERFLACGHAVVRAEGRSQEVLEQYLTKRRLTRRAVLETPHFMSLPFILSRTDL
IATVPHAIGYAYAAEHASITLVEPPLPLPRFDLKQHWHRKFHNDPSITWLRGVVAQLFND
ALDEWPK*
>GCA_000452945.1_BPMSHR5858_genomic_00587_1 group_18955
MHSTTTAFTHRGYLLNCAPARASDGSFQPYVVISRSSDGELVANRFFPSDLHFNDEDSAI
AHARDWAVRWIDASSLTI*

>GCA_000452945.1_BPMSHR5858_genomic_00589_1 gnl
MSEIGSRRYPDPAVRVLDPRFGALRLPSASVECLFQGARWSEGPVWFGDGRYLLWSDIPN
NRMLRWDEQSGTVGVFRQPSNNANGNTRDRAGRLVTCEHLTRRVTRTEYDGSITVLAERF
AGKRFNSPNDVIVKSDGSIWFSDPDFGIQSFYEGEKQASEVPERVYRIDPQTGAVDIVVD
GVLGPNGLAFSPDEAMLYVVESKGRPRTIRAYDVAADGRTLRNERVLIDAAQGTPDGFRI
DVNGNLWCGWGMGTDELDGVRVFTPEGDAIGHIALPERCANVCFGGRHRNRLFMAASHGL
YSLYVNTQGVPGG*
>GCA_000452945.1_BPMSHR5858_genomic_00591_1 group_19657
MSYVALEVLTEDANRYSLPELIGVGGVSPDVPHICEMLLADAQWPTIQAYLDRQELPYKF
ARPSTGRRVGRNNPCW*
>GCA_000452945.1_BPMSHR5858_genomic_00605_1 group_1767
MTRHHTHAGIPRTLKIVGYCGLAAIALALFVAMCAILKDSALDREAARQLQNGNVELHGG
ADVEPGSTNDAAQARTPG*
>GCA_000452945.1_BPMSHR5858_genomic_00613_1 group_1769
MNAPQVFDPHGAAAAVAADPAPRLREIPYNYTSFSDREIVIRLLGDEAWAALDELRGERR
TGRSARMLYEVLGDVWVVRRNPYLQDDLLDNPKRRALLIEALNHRLSEIEKRRRADLSEH
GDAAGRERAARVEMLAAAAQRAVDEFAQEFEKMAELRRRATKVLGRCTQKDNVRFDGLAR
VSHVTDATDWRVEYPFVVLTPDTEAEIAGLVKACFELGLTVIPRGGGTGYTGGAVPLTPF
SAVINTEKLEQLGAVELTELPGVARKVPTIFSGAGVVTRRVTEAAEAAGYVFAVDPTSLD
ASCIGGNVAMNAGGKKAVLWGTALDNLAWWRMVDPDGNWLEVTRIEHNLGKIHDIPVARF
ELKWFDGARAPGEKLLRTETLDIEGRRFRKEGLGKDVTDKFLAGLPGVQKEGCDGLITSA
RWVLHKMPAHTRTVCLEFFGQAREAIPSIVEIKDYLFETSKQGGAILAGLEHLDERYLRA
VGYATKSKRQAFPKMVLIGDIVGDDADAVAHATSEVVRMANGKSGEGFVAVSAEARKRFW
LDRSRTAAIAKHTNAFKINEDVVIPLNRMGEYTDGIERINIELSIKNKLQLVDALEAFFA
AGNLPLGKSDDASEIPSAELLEDRVQQALELLKRVRARWSFLREKLDMSLREAQHYLVTL
GYAALAEKFADRVDAQPDATVFHVTQDRTVRVSWKQEIRAELRQIFNGGAFKPILDEAQA
IHRQVLRGRVFVALHMHAGDGNVHTNIPVNSDNYAMLQDAHAAVARIMTLARSLDGVISG
EHGIGITKLEFLTDDEIAEFRAYKQRVDPQGRFNKGKLLEGADLRNAYTPSFGLMGYESL
IMQQSDIGAIAESVKDCLRCGKCKPVCATHVPRANLLYSPRNKILATSLLVEAFLYEEQT
RRGVSIKHWDEFNDVADHCTVCHKCATPCPVKIDFGDVTMNMRNLLRKMGKKKFNPGSAA
GMFFLNATNPQTINATRAVMMGLGYKAQRFANDVLKKVAQKQTAHPPSTTGKRPAVEQVI
HFVNKKMPGNLPKKTARALLDIEDNKIVPIIRDPKTTTVDSEAVFYFPGCGSERLFSQVG
LATQAMLWEAGVQTVLPPGYLCCGYPQRGSGQYDKAEKIVTDNRVLFHRVANTLNYLDIK
TVVVSCGTCYDQLAGYEFDKIFPGCRIIDIHEFLLEKGLKLDGVTGTRYMYHDPCHTPIK
TMDPVKLVNDLMGAHNDGYRIEKNDRCCGESGTLAVARPDVSTQVRFRKEEEIKKGAAKL
RNIPVVSGNGANGPEGAAAGASDVKILTSCPACLQGLSRYNEDADIEADYIVVEIARHML
GENWMADYVARANNGGIERVLV*
>GCA_000452945.1_BPMSHR5858_genomic_00616_1 ubiE
MSKTHFGFETVEENEKAKKVAGVFHSVASNYDLMNDLMSAGLHRAWKAFTIAQANVRPGG
KVLDIAAGTGDLTKAFAKAAGPTGEVWHTDINESMLRVGRDRLLDKGVVTPSLLCDAEKL
PFPDNYFDVVTVAFGLRNMTHKDSALAEMRRVAKPGGRVMVLEFSKVWEPLKKAYDVYSF
KVLPWLGDKFAKDADSYRYLAESIRMHPDQETLKTMMEQAGLDAVKYYNLSGGVVALHVG
TKY*
>GCA_000452945.1_BPMSHR5858_genomic_00620_1 tpm
MSQGDGVTNEANQPEAAGQATGDAQPASPAGPAHIANPANPANPPALPSFSPPAAASSSA
APFSSRDPGDASFWDERFEQGVTPWDSARVPDAFAAFVARHARVPVLIPGCGSAYEARWL
ARAGWPVRAIDFSAQAVAAARRELGEDAGLVEQADFFTYAPPFVPQWIYERAFLCAIPRS
RRADYARRMAELLPPGGFLAGFFFIGATPKGPPFGIERAELDALLCPHFALVEDEPVADS
LPVFAGRERWLAWRRS*
>GCA_000452945.1_BPMSHR5858_genomic_00622_1 group_19364
MKKTTLKSVFLTGLLVLVPLAITLWVLGLIIGTMDQTLLLLPASWQPERLFGFRLPGIGA
VLTLAFIFVVGLATQNFIGQKLVTWWNAVVRHIPVVGPIYTSVKQVSDTLLSSSGNAFRK
ALLIEYPRRGSYTIAFLTGTPGGDVVNHLKEEHVSVYVPTTPNPTSGFFLMVPKSEVVEL
DMSVDAALKYIVSMGVVAPPAPVAAPARRPVEPPM*
>GCA_000452945.1_BPMSHR5858_genomic_00623_1 aspS
MSMRTEYCGLVTEHLLGQTVSLCGWVHRRRDHGGVIFIDLRDREGLVQVVCDPDRAEMFA
TAEGVRNEFCIQVKGLVRGRPEGTINAGLKSGRIEVLCHELNVLNASVTPPFQLDDDNLS
ETTRLTHRVLDLRRPQMQHNLRLRYRVAIEARKYLDEQGFIDIETPMLTKSTPEGARDYL
VPSRVNAGQFFALPQSPQLFKQLLMVANFDRYYQITKCFRDEDLRADRQPEFTQIDCETS
FLGEQEIRDLFEDMIRHIFKTTIGVELDATFPVMPYSEAMARFGSDKPDLRVKLEFTELT
DAMKDVDFKVFSTPANTKDGRVAALRVPKGGELTRGDIDGYTEFVRIYGAKGLAWIKVNE
RAKGRDGLQSPIVKNLHDASIAAILERTGAQDGDIIFFAADRAKVVNDSLGALRLKIGHS
EFGKANGLVEAGWKPLWVVDFPMFEYDDEEARYVAAHHPFTSPKDEHLEYLETDPGRCLA
KAYDMVLNGWEIGGGSVRIHREEVQSKVFRALKIGPEEAQAKFGFLLDALQYGAPPHGGI
AFGLDRIVTMMAGADSIRDVIAFPKTQRAQCLLTQAPSPVDERQLRELHIRLRQPEQPKA
*
>GCA_000452945.1_BPMSHR5858_genomic_00624_1 nudB
MTKPPKIPESVLVVIHTPALDVLVIKRADQPDFWQSVTGSKDALDEPIAQVAAREVAEET
GIVVGGPGLAPAALVDWHHAIEYTIYPQYLHRYAPGVTRNVEHWFSLEVPARVDVTLSPR
EHTDYLWLPYREAAARCYSPSNAEAILQLPERLASRAA*
>GCA_000452945.1_BPMSHR5858_genomic_00630_1 group_19376
MDIQIENAGGVLTITLARPAKKNAITAAMYQTMADALAAAQDDKSVRAILIRGSDGNFSA
GNDLDDFMKAPPKDEHAPVFQFLRQIAGAHKPIVAAVAGVAVGIGVTMLLHCDLVYAADT
AQLSLPFAQLGLCPEAASSLLLPRVAGHQVAAEKLLLGEPFDALEAHRIGLVNRVLPAAD
LDAFAAQQAAKLVALPASSLRATKAFLKKGSAADIAARMSEEGDQFGKMLRSPEVREAIT
AFFEKRKPDFRQFD*
>GCA_000452945.1_BPMSHR5858_genomic_00633_1 group_19218
MTDSPMQLPQKPPALRVVPQPSDANVHGDVFGGWIMAQVDIAGSIPASRRANGRVATVAV
NSFLFKQPVFVGDLLSFYASIVKTGNTSVTVDVEVYAQRMSLMSEIVKVTEATLTYVATD
SDRRPRALPPLE*
>GCA_000452945.1_BPMSHR5858_genomic_00639_1 pdxA
MSDRPQPLRIAITTGEPAGVGPELTARALADAATRWPDARFTVLGDASLIAARAAAVGLD
WTRMTAAGGGAHVADAHVANAHVANAHVAVAHRALAAPAEAGKLNPANGRYVLDLLDAAI
DGALAGEYDAIVTAPLQKSTINDAGVPFTGHTEYLAERTRTPHVVMMLAGTGERPLRVAL
ATTHLPLRDVSAALTIDGLADTLSIIDRDLRGSFGLAAPRILVTGLNPHAGEHGYLGREE
IDVIEPALERARAAGIDARGPYPADTLFQPRYLEHADCVLAMFHDQGLPVLKYATFGEGI
NVTLGLPIIRTSVDHGTALDLAGTGRADPGSLVAAIDTAVTMARHRRAG*
>GCA_000452945.1_BPMSHR5858_genomic_00645_1 group_19338
MPTSLSKKLVVLDRDGVINVDSDAFIKSPDEWIPLPGSLEAIARLNHAGYRVVVATNQSG
IGRGLFDMATLNAMHLKMHRAAAAVGGRIDAVFFCPHTADDHCDCRKPKPGMMKLIAERF
EIDPADTPVVGDSLRDLQAGEALGFRPHLVLTGKGKKALAAGGLPEGTRVHDDLRAFALD
FLSEEHE*
>GCA_000452945.1_BPMSHR5858_genomic_00647_1 glyQ
MLTFQQIILTLQSYWDRQGCALLQPIDMEVGAGTSHVHTFLRAIGPEPWRAAYVQPSRRP
KDGRYGENPNRLQHYYQYQVVLKPAPENILDLYLGSLEALGFDLKQNDVRFVEDDWENPT
LGAWGLGWEVWLNGMEVTQFTYFQQVGGLDCKPVLGEITYGLERLAMYLQKVENVYDLVW
TEWEEPGPNGPELRRLTYGDVYHQNEVEQSTYNFEQANVELLFTFFNSYEAEAKRMIEAQ
LALPAYELVLKAGHTFNLLDARGAISVTERAAYIGRIRALSRLVAQAYYDSREKLGFPMI
GNPVPGVPGLTTDAQEAAQPAWAPPLKAERKIDQD*
>GCA_000452945.1_BPMSHR5858_genomic_00658_1 gph_2
MLDHLICDCDGVLVDSEVIADRVLFDTLSATFPAIDFTADAKAAFGQQTSRFLADLEARH
GIAMPANFLDTIEHNIELGLAQWLAPIAGVRDALTRIALPAAVVSNSRLARVRNSLKRAA
LTDVFGERVFSAEQVARPKPYPDVYLHAARTLGVEPARCVVVEDSVSGLNAARAAGMKTI
AFVGASHIPGGYEDALRKMGITRIMRSMGELPALVEAGMRGEFGSQQS*
>GCA_000452945.1_BPMSHR5858_genomic_00661_1 asnB_2
MCGIGAIFSTQRASVDGIERSLSALMSKIAHRGDASRFNENLIRERFGLATNRLAIVERD
HARQPVSDHERDLHVVLNGQIYNYKELRDELAARGHRFSHAGDAEVLLRAYLEYGPSFAT
KLDGMFSFVLFDNLNGKFLIGRDHVGIKPLYYAQRDGVWYVASELKALTHVNARIETLAP
GTLFDGKETIRYVSYDASRPVSRDSFDEAKARVRTLLEESVRRRVDTDLPICVMFSGGID
SAIVLQLAHRYHPDVTAISFGLPGSNDIEIAHRYCAEHGIRHVIYTFTKQDLLENIDDAV
YYGEFFEKIDAIDSAIAHFGYYIANKLGLKVALCGEGSDEIFGGYDLFKTHAQPGALSLY
RLNNLHRTDLQRVDRASMRNTVEARVPFMDAELIDYVLSLDFSYKLHGGVEKHILREAFR
DVLPGYIIDRPKIRMPDGSGVKNVIIDHIDSLGDAPPDDAARRLGEIGLSDRSALFFARK
YLECNFPFPTQRFKQAGKDFSESGYFDFIS*
>GCA_000452945.1_BPMSHR5858_genomic_00664_1 rutC_2
MSTIDTSEFRKTVFSPSFWEDKMGYARGKRVGNHVYIAGCVASDGNAAVIGNDAYEQTAF
IIAKIEKYLNELGAELSDVVSTVTHLTGFEHFDDYCRAFSERFRDIRPVNTTVAVKSLVE
PRHYVEITATAIVGD*
>GCA_000452945.1_BPMSHR5858_genomic_00666_1 glpK
MQDQYILALDQGTTSSRAMLFDRQGNIVSIAQKEFEQIYPQPGWVEHDPQEIWSTQAGVA
AEAVTRVGLNGTAIAAIGITNQRETTIVWDRETGHPIYNAIVWQDRRTADFCDKLKAQGL
SEKVRAKTGLPIDSYFSATKIRWLLDNVEGAREKARQGKLAFGTVDSWLVWNFTKHELHV
TDVTNASRTMLFNIHTLDWDDELLDALEIPRSMLPQVRASSEIYGPTKTTVFASKIPLAG
IAGDQQAALFGQMCTSSGMVKNTYGTGCFLMMNTGETPIESRNNLVTTIAWQVDGKVSYA
LEGSIFIAGAVVQWLRDGLGIIKSASEIEALAAGVPHTDGVYLVPAFAGLGAPHWNARAR
GSLFGVTRGTTSAHLARAALDSIAYQSLDVLKAMEADSGIRIGELRVDGGASANNLLMQF
QADLLGVDAVRPRITETTALGAAYLAGLAIGYWQNVDELHSQWQLERRFAPSMQAEQVGS
CLAGWQRAVRAAKAWADDTQ*
>GCA_000452945.1_BPMSHR5858_genomic_00668_1 group_1404
MRTMIAAGALALLAGCAGGPAGFGSWGGPSFAQLQRGCGDVRDYGDDARSVYSAVFDAWV
AKCHGKLTDARFCAFENELAQRHAALGTNADAAARGRWVSYLNDARARALSWRAAVDPSL
RGG*
>GCA_000452945.1_BPMSHR5858_genomic_00669_1 group_18960
MLSPHEFATLLLVKDAPDQADMDRDELDALLERQLVKLEALGSGRKYCVTEIGDAALRSI
KLRYS*
>GCA_000452945.1_BPMSHR5858_genomic_00670_1 glpR_3
MTRDPRLTLNARQQELLEWVQRDGFVTVDDLAAHFNVTPQTIRRDVNWLADLNLLRRYHG
GASLPTSSENVSYTARQRMFHDEKRRIAALAASHIPDQASLFINLGTTTEEVARALNRHH
GLRVITNNLNVASMMSGYPECEVLITGGIVRPWDKGIVGELAIDFIRQFKVDYAIIGTSS
IESDGTLRDFDTREVRVAEAIIQHARTVYLVTDHSKFGRPALVRQGHLSQIHALFTDKPL
PDDMAETLVQAGTQVYVAD*
>GCA_000452945.1_BPMSHR5858_genomic_00671_1 ywrD
MNLHSNLSGAPFSWRNPYPTTRVPVFARNVVSTSHPLAAQAGLRMLWKGGNAVDAAIAAA
AAITVVEPVSCGLGGDAFALVWDGAKLHGLNASGVAPAAWSVDYFRRRHGEAGNGLARQP
VRGWDTVTVPGVIAGWEALHAKFGTLPFADLLEPAIELAERGHAVAAVVAHKWAAAVPEL
SGQPGFAQTFMPHGRAPDVSERVRLPGHARTLRTIAAEGARAFYEGSLAESIAAFFRDGG
GALTLDDLRAYRPEWVEPIGKDFRGYTVHEIPPNGQGIAALIALGIVERFGLDDLPLDSA
QAQHVQIEAMKLAFADVYRYVADPRAMELTPAQMLDDAYLAARAKLIDPRRATHFSFGMP
RAGGTIYLSAADERGMMVSFIQSNYMGFGSGLVVPGTGIALQNRGCGFSMDPASPNVVAG
GKRPFHTIIPAFVTQQAGSARHAVMSFGVMGGDMQPQGHLQTIVRMLGYGQQPQAACDAP
RWKVSRSFTLDVEATFDASVVEALAARGHTIQAIDDPYMDFGSGQFIWRLDRDEPDRGYV
AASDSRRDGLAAGF*
>GCA_000452945.1_BPMSHR5858_genomic_00676_1 group_19010
MGAAGLLSTAHAQTKPTEPAAAKAPLKAPDTMAERVRGCTACHGTHGQGTDNDYFPRLAG
KPAEYLYNQLVNFRDGRRKYPPMNYLLTYLSDDYLREIAQHFSDERPPYPAPTKPTVPAA
VVERGKQLALHGDPARKLPACVACHGNGLTGMQPAIPGLVGLHSDYLSAQIGAWRSGTRH
AKAPDCMHDVASKLSDEDVTAVTAWLAAQPAPANPVPAPARSMKTPLACGSEPQ*
>GCA_000452945.1_BPMSHR5858_genomic_00678_1 dgoD
MKITRLETFVVPPRWLFLKIETDAGIVGWGEPIVEGRAHTVEAAVHELADYLVGQDPLRI
EDHWQVMYRAGFYRGGPITMSAIAGIDQALWDIKGKHHGAPVHALLGGPVRERIKVYSWI
GGDRPSDVANNARAVVERGFQAVKMNGSEELQIVDTFDKVDKVIANVAAVRDAVGPYVGI
GVDFHGRVHKPMAKVLARELDPYKLMFIEEPVLSENAEALRDIANQTSTPIALGERLYSR
WDFKRILEGGYVDIVQPDASHAGGITECRKIATLAESYDVALALHCPLGPIALAACLQLD
AVSYNAFIQEQSLGIHYNQGSDLLDYLRNPDVFRYADGFVAIPQGPGLGIDVDEDKVCEM
AKTGHRWRNPVWRHADGSVAEW*
>GCA_000452945.1_BPMSHR5858_genomic_00733_1 phnE
MNAREPAGAALGEPLATAAAARRGWRAWLAWALAFAILGGTWRSADMRPLDLLTESANMG
EFARGFFPPDFSEWRYYASEMGVTLSIALWGTALAIVSAVPFGLMSARNIAPAWLLHPVR
RAMDSCRAINEMVFAMLFIVAVGLGPFSGVLALWVHTTGVLAKLFAEAVEAVDPRPAEGV
RAAGATPLDEIVYAVLPQVLPLWISYALYRFESNVRSAMVVGMVGAGGIGVVLYEAIRSF
NYAQTAALLVIVIAVVTLIDMSSAWLRARAI*
>GCA_000452945.1_BPMSHR5858_genomic_00736_1 phnM
MLISNARVVTRDDVFIGTVQIDGESIRAVSRGTTAARDADDWGGDYLLPGLVELHTDNLE
KHLAPRPGVYWNVDAAFVTHDAQVAAAGITTVFDALAIGTRLNVGVRGRDVQTQCADALA
RFASHGLLRAEHFLHLRCEIATEDVVDVFDSLARHPRLRLASVMDHTPGQRQWKDPQQWR
HYQERHRKWSDEEAATTIAELADEQQRYADAHRRAIVARCRALGVPVASHDDTVIEHVEQ
AAREGIALAEFPTTHAAAEAAHARGIATVMGAPNLVRGGSHSGNVSALALARAGLLDILS
SDYVPSSLLCAAFDLVHQAHWSLPRAIDTVSGSPARHTGLRDRGEIAPGLRADFVRVALH
GTLPVPRATYVRGRRVV*
>GCA_000452945.1_BPMSHR5858_genomic_00737_1 phnL
MQTIESSRRACDHDHARAFAGNAALMLRAVGVGKTFTLHAHGGMTIDALENVSLDVEAGE
CVALTGPSGAGKSTLLRCLYGNYLANRGTIAVRAGARAAEHVVLTASEPHEVIALRRDVI
GYVSQFLRAIPRVPALALVAAPLVARGMPEGDAADRAAHLLARLNVPRRLWTLAPATFSG
GEQQRVNIARGLIAEHPVLLLDEPTASLDAENRDVVAELIGDVRRRGAAIVGIFHDEATR
ERVATRCFPLRPPSRARLPFDEVNA*
>GCA_000452945.1_BPMSHR5858_genomic_00738_1 phnK
MTPLLSARGLTKRFGGRNGFANASFDLYPGEVLCIVGESGSGKTTLLNVLSLQTQADAGA
LRYTGGGGERIDLFALDEPQRRALMRTEWGFVTQNPRDGLRGGVSAGANIGEPLMAVGAR
HYGRLRETAVDWMARVELDPARIDDLPSTFSGGMQQRLQIARNLVIGPRLVFMDEPTAGL
DVSVQARLLDLLRALTSTLALSMLIVTHDIGVARLLAHRLMVMQRGEIVEAGLTDQVLDD
PQHPYTQMLVSSVLPV*
>GCA_000452945.1_BPMSHR5858_genomic_00740_1 phnI
MYVAVKGGERAIEASWRLLDQARRGDPAVPELSVAQIREQQRLAVARVMTEGSLYDEALA
ALAIKQAAGDLVEAIFLLRAYRTTLPRFGYTAPIDTGAMRLERRISATFKDIPGGQLLGP
TYDYTQRLLDFALLAEGGAARHEPHEAESARAAADAGSPPPHATHAPPPAAARVIALLND
EGLIEEERPTAVGAEPGDLSREPLAFPADRATRLQNLARGDEGFLLAMGYATQRGYGHSH
PFAGELRFGAVAVEMALDELDGETIEIGELDVTECQMINHFSGGDGEPPRFTQGYGLAFG
HSERKAIAMALVDRALRASELGEAAHSPPQDQEFVLSHSDNVEASGFVQHLKLPHYVDFQ
SELELVRRLRAGHAAQAGAGANAHENAPARAERADTHTEESR*
>GCA_000452945.1_BPMSHR5858_genomic_00748_1 dps2
MAKKSNATQINIGISDKDRKKIAAGLSRLLADTYTLYLKTHNFHWNVTGPMFNTLHLMFE
EQYNELWLAVDLVAERIRTLGVVAPGTYREFAKLSSIPEADGVPAAEEMVRQLVEGHEAV
VRTARAIFPDADAASDEPTADLLTQRLQTHEKTAWMLRSLLA*
>GCA_000452945.1_BPMSHR5858_genomic_00765_1 group_1105
MKRRDFLALASLAGAAGVSLPVPYAFAAAQGETSATGAMGAVAAARAARYSNLLILVELK
GGNDGLNTVIPYANPLYRTLRPAIGVKREQVVQLDERAALHPALEPLMPIWRDGRLAIVE
GVGYPQPNLSHFRSIEIWDTASRANEYLREGWLTRAFAQASVPPGFAADGIVLGSAEMGP
LANGARAIALVNPAQFARAARLAQPVSLRERNPALAHVIDIENDIVKAADRLRPHAGTPA
LATAFPGGPFGASVKTAMQVLAACDTPQRTPAPGQGVAVLRLTLNGFDTHQNQPGQQAGL
LGQLAQGLVAMRSALIELGRWNDTLVMTYAEFGRRARENQSNGTDHGTAAPHFVMGGRVR
GGLYGAPPALDALDGNGNLPVAVDFRQLYATVLGPWWGLDAASVLRQRFEPLPLLRA*
>GCA_000452945.1_BPMSHR5858_genomic_00769_1 envZ_1
MRIDRRLLTLVFGGLFWRTFLLIALLIAVSLAAWFQSFRVIEREPRAQRVALQLVAIVKL
TRTALLYSDPDLRRALLQDLESNEGVRVYPREKTDKFKLQPDESVNRLIEHDIRSRLGDD
TVIAQSVNDIPGVWISFKIDDDDYWVALDRDQLDTVTGLQWAGWGLFALALSLIGSAFIT
SLVNQPFSRLALAARKVGSGQTPEPLPERGMGVAAETNRSFNQMVRDLEQLEADRALMLA
GISHDLRTPLARLRLETEMSPSDQATKDAMVDDIEQMDRIIAAFIDYARPAQRTPEPVDL
SQVAREVAARIVSEDGVQLDIRLAPAAVIEADETDMRRVIGNLVENARKYGQSKQDGIAR
ILLETRVTHARVELVVADEGPGIPEDQLPLVMRPFYRVDTARSKADGTGLGMAIVLRLVN
RYRGALRLRNRTPDAGLEVTLEFPSAKARAAA*
>GCA_000452945.1_BPMSHR5858_genomic_00770_1 ahpC_1
MKTVGDKLEAFTVVAAKPGFNNHEENGQSAFETVTEASFPGKWKIIYFYPKDFTFVCPTE
IVEFAKLAKQFEERDAVLLGGSSDNEFVKLAWRREHKDLDKLNHYSFGDVKGELIDQLGV
RDKEAGVALRATFIVDPDNTIQHVSVNNLNVGRSPEEILRILDGLQTDELCPCNRAIGGA
TL*
>GCA_000452945.1_BPMSHR5858_genomic_00776_1 argE
MSVLDATAPSASSSSSLAEPISLSWVKQLVSIDTTSRVPNLGLIEMVRDALAAAGVESTL
THDARDGWANLFATIPAHDGTTNGGIVLSGHTDVVPVDGQQWDSDPFKPQVRDGKLYGRG

TCDMKGFIGAALALLPEMQAAKLAQPLHFALSFDEEIGCVGAPLLLADLKARGVAPAGCI
VGEPTGMRPVIAHKGINVYRCCVRGHAAHSSLTPKGLNAIEYAARLICHIRDLADEFRAQ
GPFDALYDVPFTTAQTSLIQGGNAVNTVPAECQFSFEFRNLPTLDPDAIFARIDAYARDT
LLPRMRREHPNAAIEFSKVAAAPGLDAAEQAAITELVRSLTADQEKRKVAYGTEAGLFQR
AGIPSVVCGPGYIEQAHKPNEFVELEQLASCERFVRKLVQGMTLA*
>GCA_000452945.1_BPMSHR5858_genomic_00781_1 group_1641
MKKLLAAVGLSLILLSAAANAAVPSLQQIQQSIAQGNWQRADAQLSQVIDAYPDNARARY
LYGQVLDREGRPAEALAQIERAKSLDPQLRFTDPSRFAQTEARVRADARRATAAQDSRSA
TSGGMLAAPQAPAQARAPFSAAPVAPAAPVHRGPSVGMWIGFAVLIGVIVIVLRKTLRRA
RSTDDQRADDERRAQLKRATDILNEVRPLKLDARLSTAPGAAALNGEIEGLEAQARELVE
TLSNGKNPAPPYRLDELEKQFASLKARVEGRPDPNAAAPGGPGQTGSVFAQEADRLTGAQ
GQPPYSPYPPQPQQPPPVVIQQGGGGFGGGMGGLLTGVLLGQAMSHSRDRVIERDVIVDD
EARRRAGADPGIDFGQGDSWDSGGSDGGGSIDLGSSGDDWSNNG*
>GCA_000452945.1_BPMSHR5858_genomic_00785_1 group_18923
MNDHVYKLIELTGSSRQSSDDAIRNAINKAAKTVRNLHWFEVTDMRGHIEGDQVIHWQVT
LKVGLRIED*
>GCA_000452945.1_BPMSHR5858_genomic_00801_1 sfnC_3
MSHAPADPTRSTGADYASLAARFRPIFARIAEGAIERDRTRALPHEPIRWLKEAGFGALR
VPVHAGGAGASVPQLVQLLIELAAADSNLPQALRGHFAFVEDWLNAPPDAARRAWFDRFA
SGQLVGNAWTEVGDVALGEVRTKVAKRDGGWVVNGEKYYSTGAIFADWIDVYAQRTDDGG
PVIAAVAARQDGVILGDDWDGFGQATTGSGTTRFVDAHVDEANVIDFARRFKYQTAFYQL
FHLATLAGIGRAVERDASALVRGRRRVYSHGNAPRVSDDAQILQVVGEISAWAYAAEAIA
LRAAQPSQRAYEARVGGDAAAEHDANVAAEIESAQGQLVVSELVLRAATHLFDALGASAT
RATNALDRHWRNARTVASHNPLVYKARIVGDRAVNGTEPPYVWQIGAGPGGPREPEQAA*
>GCA_000452945.1_BPMSHR5858_genomic_00820_1 hscA_2
MSDARYSIGIDLGTTHCALSYVDLAASDGEKTSLRVLPIAQLTAPGAIDAPNLLPSFLYL
PHPSELAPGELALPWAAERDFAVGELARSRGAGTPIRLVSSAKSWLCHPGVDRRAGILPS
DAPPEVARVSPLDASVRYLSHLREAWNHAHPDAPFDRQDVTVTIPASFDPAARELTAEAA
QAAGYAHMTLLEEPQAALYSWIEKSGGAWRKDVRVGDIILVVDVGGGTTDLSLIAVVERD
GNLELHRIAVGEHILLGGDNMDLALAHVVARKLAAQGTQADPWQLRALTYACRGAKETLL
GDPSAQAVPLVVPSRGAKLIGGSIRTELTRAELTQTILEGFFPAVDASARPVSRTRAGLM
QLGLPYAQDAGITRHLAAFLGRQAGALAELDGLRHAHDASASFLHPSAVLFNGGVFKSPL
LVERMLDTLNGWLAAERAAPARLLGGADLDLAVARGAAYYGYVKRGRGVRIRGGTARAYY
IAVESAMPAVPGLEPPIQALCVAPFGMEEGTDAALPPQEFGLVVGEPVRFRFFGSSVRRQ
DQVGTLLDFWSPDELQELDAIEATLPTAGRTPGEVVPVRLHARVTEAGTLELEAVQGGTH
ERWKVEFDVRGGANA*
>GCA_000452945.1_BPMSHR5858_genomic_00874_1 tycC_1
MTAFGNFVALCTHRAAANGDALAYHYLSPHGEDRALSFGALDRAARRVAARLAASGAPGD
RVLIVCPQSLDYVSAFFGCLYGGFIAVPAYAPRNNHHFARLSKIIEDARPRVVMLCRKQY
AAVHAFIERNPPLREVELVVVDELDDVEPGGCRPHAAARDDVAFLQYTSGSTGQAKGIMV
SHGNLLANEEMIRTTCGNTPDSRAVFWLPLFHDMGLMTLLQGVYVGYPTYLMAPMDFLAN
PLRWLQAVSRVRATLTVAPNFAWQLCVEKIPPEQLDGLDLSSVTAAVNGSEPISVRTLDG
FVARFGACGFRREAFRPSYGLAEATLLVTGSSGHAPDGVIEETMVSSAGRRTVFQRVGCG
RPAAGCEVAIVERDTRAVRRDGELGEIWVKGPHVAQGYWNNPEQTAQTFGNRTADGAGPY
LATGDLGFLHDGALVVTGRCKDVIILRGDNYYPSDLEASATAAHPALVPDGAAAFTLAGD
EAGPDAAAAQALAVVAEVRRNTPPAQFAQIAAAIVERISAGYGLALERLVLIKERSIPKT
SSGKVQRSAVRAELDAGQLKTLHDVRPGETGARAAAQPFAELHAMLAAARGDGLARIVDG
FVLSQFAELMKTPFYALDLTRSVAALGLDSLALVNVQHRLGAALKIDVPAQLLFGDGTLG
ELAAQIGALARRAWTDAPCAAASTDAAESADSADSAAAPQVPAVSDEAARAARNSPEAAA
PALPAQADTDTGSDSTSTSTSPSAAAGWDATPGQAALWLIQQAARESHDYNEGFVAAVDG
AFDPELFMLALARLAARRDALRVRFAEVDGRPVARDAHALEQSTLRCADDAAALQAARDA
LAAPFDLSRATWRAVLAKGPNATYVALALHHAIVDMWSFDILIAELKQLYDALARGADAS
LGAARGYREFAAWQHGWLNSPQATGHAAYWREQLRGMPARSAFPSTPIATPASGARAHRF
AIDARTMGAVRALARRHGLTAYHVLFAAGALLLGRYAGQRDVVVGMPAHGRPRAADADTI
GYFANVLPIRVALDDRRALADWLHDVKGAIVGGLLHQALPLPVLTEALAAERGAPDGALV
QTVISLLAPSAARAVEAGADAGDRLSPFVLGEAGGRLALGDATLTSCESRHDRAPFDVAF
TLIDDGGELAASIGYRADRHSAAMIERLAANYRCVLEALAHGACARLGDVPHVSAAEHAL
LRRSLTHRARAHAPAAGADGDSVVDRFRRTAARQPDALALAAAGRALTYGELARATDALG
HMLVEHGVTPGDRVALAVGERAMQTRVALAILKVGAAYVPVDLANPPERLAYLLDDCGAK
LVLTTRDDRPRLPATGANVVCADALDDAAAARHAGRPLPRVAIAAGQPAYCIYTSGSTGQ
PKGVLVTHGGLANLVDWHVDAFALDAGARASMLAGPGFDAAVWEIWPALCAGASLAEPAP
DARHDVAELARWLDAHAISHCFMPTPLAEAFIAAAARPRALRFLLTGGDQLKARGRAGDG
FQLINAYGPTENTVVTTSGAVEPASESAQGPAGPARLAPLPDIGAPIRGQALHILDAQLR
PTPLGVSGELYVSGAGLALGYLNRPALTAERFVPAPDADAPDVPTPGARLYATGDLVRLD
ENGRLHFVGRADDQVQIRGFRVEPGEVEAVLATHPGVAQCKVIAFEREPSGKLLAAYVAG
DASLTEAALRAFVDSRLPSYMRPAGFVIADALPLDANGKISRRALPPPTFAAADEAAFAD
PLEQAVANVYQEALGCGPVRADDDFFALGGHSLLVARVAAGIRARLGITVSIADLFAHSK
VAALAAHLRGAAAAPIPPAVRLTRAGRPARVPLSPAQQRMWLLQALEPDSTDYHVCGALR
VHGALDLERLAAALTQTVARHEALRTCFPRQDGQACQVVLPPAAVAVERLSVDIDADRAS
PAAHDAAVQARIDAWEAQRFDLETGPLLKAAYLPLAADDGYLLLNLHHIIADGRSIAILL
DDIAARYADLREAGAGADAPGAPEAPHYADYCVWSRSEAAREREAASLPFWRTQLAGAWP
AGLAALRPARTAAAQHAPGAARGDEIGDTLPADLARNVRAVAARWGVTESTVYLAAYGLL
VAKLGGDADVLIGIAYAGRDLPDTADMIGMFVNVLPIRVALGDADAFEQQVRGWRDACVA
AFRHAHVSYERMVELAGGDRRDGPGELVKTMFDFEEPGPPPRALGDARLSVERRADRSAK
FDLTLRCRPDALGRMRIALNFRRAALSVERARGLLDAYRCVLEQVLQAPELTPARTRLVD
AAQADALLRLGGGPAAPWAGAPVHRQIERIAHADPQALALAGEDGARIGYAVLNATANRL
ARRLRTLGVAPRDAVALCMRPGPSFALAALAVLKLGAAYVPIDPRYPDPRKLRIVADSGA
RLVVAEPDAAPAQAPDAVAPVWWDALAAQAATLPDSDLDIETAPDDLAYIVYTSGTTGAP
KGVEIPHRGLANLCAWHARAYGLHDAPQSIRASQTAGIGFDAAVWEIWPYLCTGASVWFA
PDAARQSSRHLEEWLSTRRITHCFAATPLAHAVLADGWLGSPSLAYLLTGGERLTRRPPA
GARYRLFNHYGPSENSVVATAGEVAQGAGEPPSIGAALDNVRVYVLDRHGQLAPRGVPGE
LYIGGASLMRGYRSNDALTHARLVADPFAGVPDARMYRTGDLVCWNDAGELDYVGRADDQ
VKIRGHRIEPSEILHAVKSHAGVYDAAVTTLDHPQAGPQLVAYVVFDPAHGAGAPDRAAR
LKRAIAAQLPAFMVPTHVVELDALPLTSNGKIDYAALPAPRAANADARRTPLRTETERAL
AAIWSALLGEPVDDAHANFFALGGHSLLAARAVALIETRMAREIALKDFFAADDLAALAR
ALDAASAHAGIPVAPAGAAIPLSPSQQRLWLVQAFNRGSVDYNVVGALRVSGALDAARLR
AAVAATVERHEILRTRIVDTEDGPRQRAPGADAREADLLQMNLTGNPPSMQDDFVRDYLA
RLAVEPFDLAAGPLFKLVLIQTAVDSGVLAASFHHIVVDAWSANVFLRDLLECYAAQCAQ
RAPRLAPLPVQYRDYSVWAQRELARSEDALRAFWRDYLHALPTHAPLPVAHHRAGAAPRD
GRRLSIAWPADTLARLKALARREQASLYEVLIAAYFAWLHRLSGQSDLVVGMPYNDRGRA
ELENLVGFFVNVLPVRARVRPHLSYAALLRQVRDDLRRVYRHHALPFDRIAEQCANASGS
LFQTMFDLDAQPLAHDARTADGDGLRAELLDGAPQAPIADLSVTLRETAQGLTLSCVYAA
ERFDAPAVERWLENFRALLDSVAADADQAVGRLAFLTANEQAIVERLRNPASVDHRGVLA
QAGGANGPALLHRLVDHWATAAPANPALVSPARTLSFAALARSTDALAAVLREEGVRPGM
VVGVEATHSVDAVLGIIATIKAGAICLPVDTRLPPERLDAMIADSGCRHVLASAGAPLGR
FDGKRLALDGAARHAGDAPAPAAEATPDHGVFLTYTSGTTGAPKASVLHHRGIVNYIGTV
IERFGYTCGDRAMLFAPLTFDASLEEIFAPLCAGASLYIGDENVKRSVPALVDACRAQRI
SVLTLPTAYWRVLGEHLAANGGAAGLGAVRLVSIGGEKVTLEAIRQWHRATAGRIALYNI
YGPSECSIGSIVDRIDVARALEDGEVYLHRPVANAHLHVLDACLNPVPADMPGELYIGGV
GVAHGYHGRPALTAQRFVADPFAAAPGARLYRSGDRVRYDLEGRLHYLGRTDFQVKVDGI
RVEPEEIQAVLEAHPDVAQAVVLAGEARHARNPLIGYVILEKAPARARAATGTDGAAFVD
YLRARLPAHMVPAQVVVMDAFPLTTNQKVDRRALLLAANDAVVAAPALSPTETALLALWR
ELLGNDAFGVDDNFFSLGGSSLLAIRINSRVEAMYAVRLPVAALFDCPTVRALAVLVERL
RASRDAAAPDQAGPPIERIDARATPLSGAQYRLWYLHQRDPDSTAYHLPDLLKLAGPLDA
AALRTALAHTLDEHESLRTTFVVEGETPLQVIAARAQPRLDGHDLSALAPDARDAALDAL
VRRALDTPFDLARGPLATFALVTLAPDEHLLLSVFHHIVVDGWSVGLFQRSLARHYNAAR
GGRAAEPAADASARALQYRDFAAWHRRVLDGGERERQLDYWRAQLDGTLPVLQLPTDFKR
PPQASGGGEAFSFAFDDVLAARLQAVAQRHHVSMFMLLLAAYAALLARYARQDDLLIGVP
ALGRTRPEFESVVGFFVNTLVIRVRANPSTTFASLLEHVRATCLDAFDHQDVALEEIAAD
VNARRERDGTGLFQTMFSYQEADALEPAAFDGLVATSVEPEHRSAKFDLYLATWTHDGRL
HGGFEFSTDLFEAATVRRMADHYRNLLAGVAARPDGALHALPLLGDDERAYQVRALNQAS
RPLPDGVYVRDLFARQAALHPARVAASCGEAALTYGELDRASDRVARNLLAAGARGEDLV
GLLIGRNLDYLIAMLGVLKAGVAFTPMNPDDPAHKLDRIAELGDVRYVVHDAASAGRAQA
LTARATRLALDALSREPAAAPEFLPLTPASLAYVIYTSGSTGLPKGAMIEQRGVLNHLLA
KIDDLAIGEDDVVAEMAVTTFDVSIWQYLVALLVGGRTAVMPGDAAWDPQQLFAQLDADG
VTVFESVPSHMKILIDELEARPGRHRLGRVRVYVSNAEALTPALCARWFACAPHVPVVNT
YGATECSDDTSHLWIREPLSGALPYVPIQGTLPNLTTYLLDERLEPVPIGVTGEVHIGGV
GVGRGYLGDPARTARAFVPDPFSPRPGRRLYKTGDLARYRPDGTLEFLGREDFQVKIRGQ
RVEIGEVERAIGDHDNVRQAVVVAARDGKDRLYLLGYVIPHRHPAPTVPELRAFVAGRVA
SYMVPASFVLMDQFPLNANGKVDRKRLPKPADQDIFRRNEHVAPATQTEARLAELWRELL
DVDEIGALDSFFELGGHSLLAAELTLRMRTLFGVALPLRTLFQSPQLRDVAAALDTLRTS
ADAAGPARVERLPARAHYELAPCQVPEWYAYQMDPSSPVYNISVADLFFTGKLNRDAFIA
AWRAILERHDVLRVKFDYRDGAPIQIVDPAIDIRAEDVFLDRTTLAGPDAIDEANRLGAR
FGTAPFDFANGPLFRLHVASYAGDFHQLIFVVHHIIWDETSLINLMLELSELYNAHAAGR
AANVPALEVGYFDYVQWTHQQLRCGAFDEHKRYWLDMYRTLPPPLDLPTDRARPNLMSYR
GDALRTWLPRGIVRKIEAYLKQHDVTLFMLQLAILDCYLSRISGQRDFVIGCPIAGRADE
RLKPLLGLFATPMPIRCTIGENMTFGELLAQVSQRTLEAFEHYHYPSNQVIEQLQHEKDL
SRPKLFSIMYGVQNNKTDLMGRLKFDGLTLSLENVVDTENKSSRFDLNFVVDQFGSDIMF
SCIYNVDLFDASTIEQMLENMTALMDQVLDDPDQPLSRYTMVGANGNPPAALHGPRVDYD
EAATMHALMAEQAARTPAATALVVEGVAYDYETLNRQANRLAHYLLSLGVTSGENVAVML
RPSFEMVVALYAILKVGGAFVPIGPQYPQKRVDAILRHAHARWALTHSGLRRAFAAFPYD
VVCVDDVMGALGGYSDRNPPSVDPRQLAYILHTSGSTGTPKGIEIEHRGVVSMLADLQRT
YCLDAHDRVLFHTPFTFDVFIQDVFWPLAYGARVVVMGDDALKSAHGLADVIERERVTLA
QFVPAMLETLVDARERGEIAGLASLRQVICGAAALYRGLAERFARAFGCRLANHYGPTEV
TVDASRFDCAEPYAGDTVPIGRPVGNASLHVLDAHLQPVPRGVIGEICVASPGLARRYLN
DDEGTARAFVEIVVDGAPLRLYRTGDLGHCDRTGVVYFHGRADKQLKIRGNRVELDEIGS
VLRSHPAIAAAALAYREDAAHGGRLVAYIEQTARDHQVPAEPGDAPLYGFTLEQRPELAA
AAFALPPGADEAARELARRFPACQLVVTDRASTVTAFCQAVPLRIDATAQAGAPALERDA
ALRLAFAQQDDGVEPNALYVLDDAATGAPATSPAVRAALLAPWRRLAAAFGLARLLRASA
AAEPAEPADAGGQGAPAHAYLTRDAVRAWLRQSLPDYMIPDQVHFLPAIALTDSGKVDAR
NLPVIEVDEQPGRQAAGTALQRELADLWARLLQVGQIGVTDDFFVLGGQSLKAIEMVAEV
GRRYGVKIQLRQFYENPTIRYLETLLTERA*
>GCA_000452945.1_BPMSHR5858_genomic_00875_1 novN_1
VYILGINAAYHESAACLVKDGQVVMAIEEERLNRIKHGKLSTPETTGQLPWKAIRKCLDA
AGIGLDACAHIGYSFSPDILQKGVASWRSANGGAHAWPLDKESYQTLEGGLNFAEGVLSS
RRQLIEDAGFAGEFHFLPHHDCHAASAFHVSPFEKAAVLVVDGIGEWASTSLYHGDGTTL
RKVHDFLFPNSLGFVWEKLSKYLGFSQYDASKVMGLAAYGDGAKTIEPFSRIVTDKENMS
VDLAVLRHESPDFTTLEALFGLPRRHTPVDFDTDNWQAYVDVAAGLQLLTEQVLLHILRK
FDRTTYRHLCMAGGVALNCAANGVIVREKLFDDVFIQPASNDAGTAMGAAFLIWNQVLGH
PREYVFNHAYLGPEYSDGEIEAVLRRTNLAYRKVDIEKEVAQLIANGSIVGWFNGRMEWG
PRALGGRSLLADPRNKAVRELMNVKVKHRELYRPFCPSILGERAEEWFEGSASVAAGKYM
LTTSRVRAEKAERIPAVVHFDGTVRAQTVYAQDNPRYHRLISEFDALTGVPLLLNTSFND
SEPIVCTPQDAINTFMKTDIDYLALGSYLISKH*
>GCA_000452945.1_BPMSHR5858_genomic_00913_1 group_19219
MPMTWPMRVRRVAPWILIGGVAILVTLVALLPASWVTPQFGRATGGHVNLVDPDGSLWRG
SATLLLAPGADRSASTLLPGRVEWRTQFWPLFAGRVRMRLRQTDAMPDGVTLDATLRGAV
LSAGSMAVPASLLAGLGTPFNTLDLQGDVRLDWTDWRLFGKDAFGQLTVTITDMSSRVSR
VKPLGSYRAVLEARGANSTLDLSTSKGPLMMSGQGNFGATGASFRGTASAEPEQRDNLAG
LLNLLGHPIGNGAVSLIYGDAAR*
>GCA_000452945.1_BPMSHR5858_genomic_00920_1 xcpT
MQTWITRRSQAARRQRGFTLIEIMVVVAILGILAALIVPKIMSRPDEARRIAAKQDIGTI
MQALKLYRLDNGRYPTQDQGLNALIQKPTTDPIPNNWKDGGYLERLPNDPWGNSYKYLNP
GVHGEIDVFSYGADGKEGGESNDSDIGSWQ*
>GCA_000452945.1_BPMSHR5858_genomic_00927_1 hupB
MNKQELIDAVAAQTGASKAQTGETLDTLLEVIKKAVSKGDSVQLIGFGSFGSGKRAARTG
RNPKTGETIKIPAAKTVKFTAGKAFKDAVNKR*
>GCA_000452945.1_BPMSHR5858_genomic_00930_1 ybdK
MALETFVNSEPFTFGVELEIQIVNTHNYDLTKAASDLMRLIKDAKFPGNITPEITESMIE
LSTGICRTHDQALGELHAIRDTLVSAADQLNVGLCGGGTHAFQQWSERQIFDAPRFQYIS
ELYGYLAKQFTVFGQHVHIGCPDADSALFLLHSMSRFIPHFIALSASSPYVQNVDTGFHS
ARLNSVFAFPLSGRAPFVLTWHGFEEYFTKMVNTGVVNSMKDFYWDIRPKPGYGTIEVRV
MDTPLSVDRAAAIACYIQTLARYLLIDRPLKLSEDDYLVYTFNRFEACRFGLEGTCVNPQ
TGERRTIAEDILDTLDRIAPHAAALGSRAALDEIGALAKARVNDASWLRTIFKQEKSLNE
TVRQQCLRWRE*
>GCA_000452945.1_BPMSHR5858_genomic_00936_1 tctD_3
MRLLLIEDDRPIARGIQSSLEQAGFTVDMVHDGIFAEQALAQNRHELVILDLGLPGIDGM
TLLSRFRQTNRHTPVIVLTARDELNDRVQGLNSGADDYMLKPFEPAELEARIRAVMRRSG
PHSDMPRPEVSLGGVRLSGVDRRIFNDDKPLELSPREFAVLEMLLLRHGRVVSKAQLQDH
LTHFGGDLGDTAIEVYVHRVRKKLEQCRVEIVTVRGFGYLLQEIRQTASV*
>GCA_000452945.1_BPMSHR5858_genomic_00938_1 phhB
MIHKLTSEERKTQLESLHHWTAVPGRDAIQRSLRFADFNEAFGFMTRVAIKAQEMNHHPE
WFNVYNRVDVTLSTHDANGLTERDIKLAHFIDEVGKHAKAA*
>GCA_000452945.1_BPMSHR5858_genomic_00945_1 proV
MILGDTILETRGLTKEFKGFTAVNGVNLRVRRGSIHALIGPNGAGKTTCFNLLTKFLKPT
SGQIVYNGIDITDERPSQIARRGVIRSFQISAVFPHLSALQNVRIGLQRSLGTAFHFWRS
ERLLKRLDDRAMDLLTQVGLTDFAHVPTVELAYGRKRALEIATTLAMEPELMLLDEPTQG
MGHEDVDRVTALIKKVANGRTILMVEHNMNVIAGISDTITVLQRGEVLAQGTYAEVSKNP
LVVEAYMGSADAALAGAHA*
>GCA_000452945.1_BPMSHR5858_genomic_00946_1 livF_3
MNHSEREERELNSVESGTPALALAGLEAWYGESHILHGVDLTVHRGEVVTLLGRNGAGRT
TTLRAIMGLTGRREGSIRIAGHETIGLPTHKIAHYGVGYCPEERGIFSSLSCEENLLLPP
VLGDRKRAMSLEDIYEMFPNLASRRASQGTRLSGGEQQMLAVARILRTGANLLLLDEISE
GLAPVIVQALAKMIVTLKARGYTIVMVEQNFRFAAPLADRFYVMEHGRIVEHFGAAELEG
KMPVLHDLLGV*
>GCA_000452945.1_BPMSHR5858_genomic_00948_1 group_19537
MDIFGIPVSAMLSQLLLGLVNGSFYAILSLGLAVIFGLLNVINFAHGALFMLGAMLAWMG
LNYLGLPYWAMLVLAPVVVGLFGILIERSMLRWLYKLDHLYGLLLTFGLTLVVEGVFRAI
YGSSGQPYDVPALLEGATNVGFMFLPNYRAWVVVASLAVCLATWFVIEKTRLGAYLRAGT
ENPKLVEAFGVNVPMMVTLTYGFGVALAAFAGVLAAPVIQVSPLMGQPMIITVFAVVVIG
GMGSILGSIVTGLMLGVVEGLTRVFYPEASATVVFVIMAIVLLIRPAGLFGKER*
>GCA_000452945.1_BPMSHR5858_genomic_00951_1 group_19346
MKMNRWIEVAFAAGCLCTAGLASAQVKIGVTVSATGAAASLGIPEKNTVALLPKEIGGKR
VEYIVLDDASDTSRAVQNTRKLIDEDHVDAIVGSSITPNALAMIDVVAQGKTPAISLAAS
AHIIAPMDAKRAWVFKTPQNDRLMADALAGYMAKHGVKTVGFIGFADAYGDGWYGVFSAA
AAANGLRIVANERYNRTDASVTGQVLKTLGARPDAVLIAGAGTPAALPAKTLKARGYTGK
VYQTHGVANNDFLRVCGKDCDGELLPAGPVLVADQLPDSNPAKQPALAYKAAYEKAYGAA
AVSTFGGHVWDAGLMLQRAIPDALKKAQPGTPAFREALRGALENVKDLPVSHGVINTTPA
DHNGFDTRARVIVQIVGDKWKLQAD*
>GCA_000452945.1_BPMSHR5858_genomic_00952_1 livH_4
MDFSIAAILAQDGITTGAIYALLSLALVLVFSVTRVIFIPQGEFVTYGALTLAALQAQKF
PATCWLLFVAGIACFMLEVGGLIRHRARRRRLARTLTTLASRYVLLPLAIYAVTRSAAAQ
PLPMIAQIALTLAIVVPLGPFIYRLAYQPIAEGTTLLLLIVSVAVHFALVGLGLVMFGAE
GSRTTAFSDTSLALGSLSISMQSAWVVGTALVLIAALYLYFERTLPGKALRATSVNRLGA
RLVGIGTTEAGRLAFTLAAGLGALSGVLVGPLTTIYYDSGFLIGLKGFVGAIIGGLVSYP
LAAAGSLLVGVLESYSSFWASAYKEVIVFTLIVPVLLWRSLAAPQTEDEEA*
>GCA_000452945.1_BPMSHR5858_genomic_00957_1 soj_1
MAKIFCVANQKGGVGKTTTTVNLAASLAAQGQRVLLIDLDPQGNATMGSGIDKAECESTV
YEVLVDGVTVADARVRPEAVKYDVLPANRELAGAEIELVSVENRERQLKAALAKVADDYD
FVLIDCPPALSLLTLNGLCAAHGVVIPMQCEYFALEGLSDLVNTIKQVHANLNRDLKVIG
LLRVMFDPRITLQQQVSDQLKAHFGDKVFDAVIPRNVRLAEAPSYGLPGVVFDRGSRGAQ
AYIQFGAEMIERVRAL*
>GCA_000452945.1_BPMSHR5858_genomic_00958_1 parB
MNAVAKKKGLGRGLEALLGGSADITEAVKIEGAPNVLALGKLQAGKYQPRTRMDEGSLQE
LAASIRAQGVMQPILVRPISSDKYEIIAGERRFRAARLAGLEEVPVLVKDVSDQAAAAMA
LIENIQREDLNPLEEAHGIQRLLDEFGFTHEQAAESVGRSRSAVSNLLRLLNLAMPVQTM
LLAGDLDMGHARALLAVDAATQITLAHQVVNKRMSVRETEKLVAHTTKAEPAVKARAKDD
GGRDTRRLEEELSDLLASNVKIKVGRRGRGQLTIDFGDLDALEGILARLRGNVAA*
>GCA_000452945.1_BPMSHR5858_genomic_00960_1 group_19323

MVLSLVATLAWWLFSRSPGAAAQSAFLGGAIGWVPSALFVARLRLSGAGTVMGWVAGEAL
KLGVTIAMFAAVAFGYPGVHWVPLLVTYLVVLKTYWIALAWR*
>GCA_000452945.1_BPMSHR5858_genomic_00961_1 atpB_1
MAASEGTRAPDPSEYIAHHLQNFSTSHQTSIFDIHVWNLDTLFWSIVCGLITILLLRLAA
RKATSGVPGRFQCAVEMLVEMVEDQSKAIVHGNRAFIAPLALTVFVWVALMNSLDFLPVD
LPGRVIGWLGLSGIISHHRIVPTADLNGTLGIALGVFVLMLYYSIKIKGAGGFAHELVSA
PFGSHPLLWIPNLALNIVEFVAKTVSLGMRLFGNMYAGELVFLLIALLGSMWSFGGDATF
LGFVGHVIAGSVWAIFHILIVLLQAFIFMMLTLVYLGQAHDAH*
>GCA_000452945.1_BPMSHR5858_genomic_00963_1 atpF_1
VNLNATLFAQMVVFLVLAWFTMKFVWPPLINALDERSKKIADGLAAAEKGKAELEAAHKR
VDQELAQARNDGQQRIADAEKRALAVADEIKTNAQAEAARIIAQAKAEAEQQIVKARETL
RGEVAALAVKGAEQILKREVDQTAHAELLNQLKAEL*
>GCA_000452945.1_BPMSHR5858_genomic_00967_1 atpD1
MSTAALVEGKIVQCIGAVIDVEFPRESMPKIYDALILEGSELTLEVQQQLGDGVVRTICL
GASDGLRRGVVVKNTGNPISVPVGKPTLGRIMDVLGRPIDEAGPIESENKRSIHQKAPAF
DELSPSTELLETGIKVIDLICPFAKGGKVGLFGGAGVGKTVNMMELINNIAKEHGGYSVF
AGVGERTREGNDFYHEMKDSNVLDKVALVYGQMNEPPGNRLRVALTGLTMAEHFRDEGLD
VLFFVDNIYRFTLAGTEVSALLGRMPSAVGYQPTLAEEMGKLQERITSTKKGSITSVQAV
YVPADDLTDPSPATTFGHLDATVVLSRDIASLGIYPAVDPLDSTSRQIDPNVIGEEHYSI
TRRVQQTLQRYKELRDIIAILGMDELSPEDKLSVARARKIQRFLSQPFHVAEVFTGSPGK
YVPLKETIRGFKMIVDGECDHLPEQAFYMVGTIDEAFEKAKKIQ*
>GCA_000452945.1_BPMSHR5858_genomic_00971_1 pheC
MKMNQTLLALACCAGLAHAQTAPQPGAPASRLDDVLARGTLRVCTTGDYKPYTYRREDGA
FEGIDIDMAASLAKSLGVKTAFVKTTWPTLTDDFVAKCDIAVGGISTTLERQKRVFFTQP
YVSDGKTPIVRCADAERYQTIAQIDQPQTRVIVNPGGTNERFAKQHLTRAKLTVFPDNVT
IFKQILAGNADVMVTDASETLLQQKLNPGLCSVHPDKPFQFGEKAYMVPRGDVVFQQYVD
QWLHLARETGELRAISDKWLK*
>GCA_000452945.1_BPMSHR5858_genomic_00980_1 ybdL_1
MQSAAAPRSKLPDVGTTIFTVIGQLAAEHEALNLSQGAPNFAPDPALVERVARAMRDGHN
QYAPMAGIAALREALAVKTERLYGERYDPDSEVTIVASASEGLYAAISALVHPGDEVIYF
EPSFDSYAPIVRLQGATPVAIRLSPERFRVNWDEVAAKITPRTRMLIVNTPHNPSATILG
EADVARLAQLVAGTDIVVLSDEVYEHVVFDGARHHSMARDRALAERSVIVSSFGKSYHVT
GWRVGYCLAPAELTRELRKVHQFMTFAADTPMQHAFAAALAEPASYLELGAFYERKRDLL
VRELAGSRFELLPSEGSFFMLARFGHFSDESDADFVLRAIRDARVATIPLSAFYADGANT
GCIRLSFSKDDATLVEGARRLCSL*
>GCA_000452945.1_BPMSHR5858_genomic_00985_1 zntR
MKIGELAKAARCTPETIRFYEKEGLMPDAERTDSNYRHYTDAHVERLRFIRNCRALDMTH
DEIRALLRFTDDPADRCDSVNALLDEHIGHVNTRLAELQHLRMQLIELRERCQGEHAVED
CGIVHGLATMETPDMPGKRSHVG*
>GCA_000452945.1_BPMSHR5858_genomic_00988_1 group_19506
MDSSSFLTSMQSTLGVYLPKIAGALGILVIGWLIAVIVRAGVRRLLSALKVDARIAESTG
QGTQVERIIAGGLFWLVLLVTAVGIFNVLNLYAISNPFSLLVTKIINYLPNLLGGAALTL
VAWLIASLLRSLADKTLKASRIDSKLSETAGMRPMSSYLGDVLFWLVILMFIPAILAAFD
LNGLLSPVQGMIDKILAIVPNVFAAAVIGFVGWLVARILRGLVTNLLVAAGADRLTQSVE
SPTPVKLSNLVGTVVFIFVFVPTLISALDALKIEAISRPATDLLGQFLNAVPDIVAAVVI
VLVTFYVARFVGALVQKLLVAAGVDGLPAMLGVERVFTGMLQPSVLAGRLIVFFAMLFAT
VEAANRLGFTQVRDVVTLFIEFGGHVLMGGVILVIGFWLAGLARRVIEQADNEHSRLLSR
IAQFAILGLVFAMGLRAMGIANEIVQLAFGLVLGAIAVAVALSFGLGGREAAGKLLDRWF
NQRGKQE*
>GCA_000452945.1_BPMSHR5858_genomic_00989_1 group_1785
MRDSYLDTARGAGIILVVYGHVLRGLFSAGLVPAGWPSALLAATDYTIYTFHMPLFFLLS
GLHVPKSLRRAGDVFLLTKLRTIVYPYFVWSLLQGAVQIALSARGTNHAFTPNDLLAIGW
RPFGQFWFLYALFICMLIAWAVSAITLRARTHGGASGANGENGGTEVSGENGEPSGAPCA
VPAMPIVLVLLAIGGLAAVAFVAGSATRWGIVSMTLAYFPFFVLGMLIGERLPAFLERVS
SGPALVAVAATFAASVAFAHRFGESDSIWALPAALSGSALVLLVAHRAARRGDAARHAPR
ASWLEYLGFASMPIYLAHILATAATRIALVTLGIVDVGAQLALGTFAGVLGPTLLYALAL
RAGTARLAGFPPLPAGYAMTPDKGRIGRADAA*
>GCA_000452945.1_BPMSHR5858_genomic_00991_1 eutC
MRYTETIGSRTYRFADLKTLLAKASPLRSGDQLAGVAASTEEERVAAKMALASAPLTAFL
NEAVIPYEDDEVTRLVVDTHSRDAFAEISHLTVGDFRNWLLSPAADAAALERIAPGLTPE
MVAAVSKLMRNQDLIAAARKRRVVTRFRNTVGLPGRMSVRLQPNHPTDDVKGIAASMLDG
LMYGCGDAMIGINPATDSLAAITKLLAMIDAFRERYRVPTQSCVLTHVTNTIAAIDKGAP
VDLVFQSIAGTEKANASFGISLALLAEAREAALALKRGTVGNNLMYFETGQGSALSANAH
HGVDQQTCEVRAYAVARAFEPFLVNTVVGFIGPEYLYDGKQIIRAGLEDHFCGKLLGVPM
GCDICYTNHAQADQDDMDNLLTLLGAAGINFIMGIPGADDVMLNYQSTSFHDQLYVREVL
GLRRAPEFEEWLETMEITDAHGALRAASARVPLLAGANDWMGRCA*
>GCA_000452945.1_BPMSHR5858_genomic_00994_1 acoD
MNHADMQHLNIEFPYRKQYGNFIGGEWVAPVGGEYFDNVSPVTGRPFTAIPRSREADIEL
ALDAAHAAKAGWAAKGAAERANVLLRIADRMEANLTRLAVAETIDNGKPLRETTAADVPL
AIDHFRYFAGCIRAQEGSIADIGGDMVAYHFHEPLGVVGQIIPWNFPLLMAAWKLAPALA
AGNCVVLKPAEQTPASILVFAELIQDLLPPGVLNIVNGFGLEAGKPLASSKRIAKIAFTG
ETSTGRLIMQYASENLIPVTLELGGKSPNIFFADVMDRDDSYFDKALEGFAMFALNQGEV
CTCPSRALVEESIYDRFIERALKRVEAIKQGHPLDSQTMIGAQASAEQLEKILSYIDIGR
GEGAQCLTGGERNVLGGELAEGYYVKPTVFRGHNKMRIFQEEIFGPVLAVTTFKTEEEAL
EIANDTLYGLGAGVWTRDGNRAYRFGRGIQAGRVWTNCYHAYPAHAAFGGYKQSGIGRET
HKMMLDHYQQTKNLLVSYSEKPLGFF*
>GCA_000452945.1_BPMSHR5858_genomic_00995_1 rhaS_4
MDHEHTGAETTAEDTGRSADGDGGAAGRALVSVAHDADEQARNLIGWRQTYDQLAAGRFV
GTLTELPLDTMKVFRETTSHTLRQACEVRGDAYWFGIPLARDGAARIDARPIAADALAFR
PGNVEFELLTPAQFSIYGVVVRGAVLRRYAQEVERCGLDERLPLVPVVRVGEARLTRLCA
LLAQRLDDADAMSAAGEPLSDCARNDLQAEVLAALFDLCASPAADASVEHSSRRRKIVAA
ARDYVLAHRSRPVGVPELCEQLHVSRRTLQYCFQDVLGMAPATYLRALRLNGVRRDLRGR
AAASVQDAAAAWGFWHLSQFATDYRRMFGARPSETLRDALAC*
>GCA_000452945.1_BPMSHR5858_genomic_00997_1 ilvG
MSISPDREQRATTGARLLVDALLANHVERVFCVPGESFLAVLDSLADETERIRTIVCRHE
AAAANMAEAVGKLTGRPGIAFATRGPGATHASIGVHTAFQDSTPMILFIGQCAREHLDRE
AFQEIDYRRMFGQLAKWVAQIDDPRRIPEYLSHAFHVAMSGRPGPVVLALPEDVLSEPCA
AQPVAPAARRVAAAPSAAQMSELRERLARAQRPLVLAGGSGWTPGACADLRTFVERWQLP
IACAFRYQDTFDNAHPQYAGDVGLGVNPELARRIREADLLFALGPRLGEATTGGYTLIDI
PRPKQVLIHVHQGADELGRVYAADLPIASGMPEIAAQLAALEPPASLAWAGSAADAHRAY
LDWHAPRPMPGEVQLGDVMRQLRERLPHDAIVTNGAGNYAAWLHRHFAYRHFRSQLAPTS
GAMGYGVPAALAAKSLYPQRTVVALAGDGCFMMAGNELATAIQYGLAIIVLVVNNGHFGT
IRMHQERHYPGRVHGTGLTNPDFAAYARAFGAHGETVERTADFLPALERALASGAPALIE
IRVPQDACTPAATLEQVREQGLRARRA*
>GCA_000452945.1_BPMSHR5858_genomic_01001_1 gcvP
MKLEHPDRLMNRTPLSLAALETHDAFAERHIGPDAASQQAMLDTLGFATRAALIDAVIPA
SIRRAETLPLGPFAQPKSEAEALAALRALADKNQVFRSYIGQGYYDTHTPAVILRNVLEN
PAWYTAYTPYQPEISQGRLEALLNFQQMVADLTGLEISNASLLDEATAAAEAMTLLQRVG
KPQSNVFYVADDVLPQTLEVIKTRAKPIGIEVKSGPAADAAAANAFGVLLQYPGANGDVR
DYRALADAIHAAGGHVVVAADILALTVLMPPGEWGADVAVGNTQRFGVPMGFGGPHAAYM
AVRDEFKRQMPGRLVGVTVDAQGKPALRLALQTREQHIRREKATSNVCTAQALLAIMASM
YAVYHGPRGLKTIALRVNRIAALLAAGIRHLGYATVNDTFFDTLTIDTGARTAQLHAFAQ
AKRINLRRAGDTRVGVSVDETTTRADLADLLTIFAQAAGATAPDIDALDAGLLPAPALPP
SLERTSAYLTHHVFNRHHSETEMLRYLRSLSDKDLALDRSMIPLGSCTMKLNATSEMLPV
TWPEFGRIHPFAPAEQTVGYREMIDQLEQMLVAATGYAAVSLQPNAGSQGEYAGLLIIHA
YHESRGESHRDVCLIPASAHGTNPASAHMAGMKVVVVACDAQGNVDIADLKAKADAHSHD
LAAIMITYPSTHGVFEQNVREICEIVHAHGGQVYVDGANMNAMVGLTAPGQFGGDVSHLN
LHKTFCIPHGGGGPGVGPVAVGPHLAKFLPNQRSTGYARGEDGIGAVSAAPYGSASILPI
SWMYIAMMGAKNLTAATETAILNANYIAKRLAPHYPVLYSGPGGLVAHECILDLRPIKDS
SGITVDDVAKRLMDYGFHAPTMSFPVPGTLMVEPTESESQEELDRFIAAMIAIRDEIRAV
EEGRADREDNPLRHAPHTAAVVTANEWPHAYSREQAAFPVASLVANKYWPPVGRADNAYG
DRNLFCSCVPVSDYA*
>GCA_000452945.1_BPMSHR5858_genomic_01018_1 group_19310
MSTTDSRSATKSSSISPRHGRVITPESRAVYLYEAGRLDFGQVNELEGGKFFPATQSGLR
DPDAPDDVASGMPPRDGEIASGGRTADARAQLNEPDSVAHWQKHAVRSGQSLQISWSYSM
PHKTRRWTYWITKPGWDTQARLARAHFEPDPLKVYLNTYQPYWGPDADKELIPQGETIHE
FNLPTRTGYHVLLAVWDVADTANAFYQVIDLNFA*
>GCA_000452945.1_BPMSHR5858_genomic_01019_1 aknOx
MSYVYVDGADPRHSTLQKGFNLRWPPTAEAGAAAIYVCRTADDVIAASRDALAKGCRITV
RSGGHCYEGFVANRLPDDHGRPLAIIDLGLMSGFDIDDRGQIASPFDAAGAPRYRVRVAA
GRQNWDGYVGLYKTMGKTLPGGSCYSVGAGGHIVGGGYGLLSRLHGLTVDWLTGVDILAP
AAGGALGAKHVHSGSTDPADRALFIACRGGGGGNFGVILNYYFNALPNAPRTAYWLTLAY
DWSHFTQASFGRFLQTYWDWFSRNDANWNAADPALANGGLFTLLKLQHRSAGPIVLAIQY
TGKDGTVGGPEDAPFIDFVDTMNAAANRLPVVWDRFVAPNLPALGHPLAGLRIAHAARSA
LKMDWLYLTQTINTSGANRRGKYKSAYQIGNFGETEVDALWRYLNGADDPRLSQTLVQID
SYGGCVNANDESANPTSVYQRRSLLKSQYQTYWTNPADDAFHVEWLHDLFSAVHADQGGK
PYRDASGRYEGCYINYPDIDMKYLGSDPAQIDPRWLDLYYGEKAASLIATKRSVDPSNLF
MHEMSIPLTPPLG*
>GCA_000452945.1_BPMSHR5858_genomic_01025_1 group_19466
MFKFITARRLAAASVLALLAACAAPAKHVDYTAFKKSQPRAILLLPPQNDTSDVKATYGL
LSQMTRPVAEAGYYVVPVAVMEETFKQNGLTNAADIQTVSPAKLRDIFGADAALYTTVTQ
YGSVYTVLDSTTVVSASAKLVDLRTGDLLWQGSGRATGKELGSNVDFSGFGLIATLAQAA
VKQIAHTLSDDAVDVAGLTSGRMLSAGQPNGLLYGPRSPKYGTD*
>GCA_000452945.1_BPMSHR5858_genomic_01050_1 motB
MSKGKDQAIIVKRVNQQKKGHHGGAWKLAYADFMTAMMAFFLLMWLLSAVTPVQLKGIAE
YFNTPLKAAIFGSGDRSADDASIIKGGGRDISSLDAGTTRRTDGSTHLAERVAKKSEDDA
LAQAQGSLERREQVRLHDLQVKLMAAIEANPTLRQFKQQIRIDSTLMGLRIEIVDTQKRP
MFAMSSDAVEPYMRDILREIGKTLNDVPNRIIVQGHTDAVPYAGGEKGYSNWELSADRAN
ASRRELIAGGMDEAKVLRVLGLASTQNLNKADPLDPENRRISIIVLNRKSELALMRDDAT
TTTLSSDAAGSARLAQQITPPGAAAMPALPGASGLAVQKP*
>GCA_000452945.1_BPMSHR5858_genomic_01057_1 cheB_2
VQKKIKVLCVDDSALIRSLMTEIINSQPDMEVCATAPDPLVARELIKQHNPDVLTLDVEM
PRMDGLDFLEKLMRLRPMPVVMVSSLTERGSEITLRALELGAVDFVTKPRVGIRDGMLDY
SEKLADKVRAASRARVRQNPQPHAAAAAAAHGHAGAAAPLINNPLVSTEKLIIVGASTGG
TEAIREVLTPLPPDAPAVLIAQHMPPGFTRSFAQRLNGLCRISVKEAEHGERVLPGHAYI
APGHAHLLLARSGANYIAHLSDEPPVNRHRPSVDVLFRSAAQHAGKNALGVILTGMGRDG
AAGLLEMKKAGAYTFAQDEASCVVFGMPREAIAMGGVDDVAPLSDMSRRIMARLASMGDR
VQRV*
>GCA_000452945.1_BPMSHR5858_genomic_01058_1 cheY_2
MDKSMKILVVDDFPTMRRIVRNLLKELGYSNVDEAEDGLAGLARLRGGGYDFVISDWNMP
NLDGLAMLKEIRADASLTHLPVLMVTAESKKENIIAAAQAGASGYVVKPFTAATLDEKLN
KILEKMAKAGS*
>GCA_000452945.1_BPMSHR5858_genomic_01063_1 flhB_2
VAEDSDLDRTEAATPKRREKAREEGQVARSRELASFALLSAGFYGAWMLSGPIGEHLRTM
LHAAFSFDRAAAFDTNRMLSHAGTLSLEGLYALAPVLALTGVAALAAPMAMGGWLVSTKT
FELKFERLNPITGLGRIFSIQGPIQLGMSIAKTLVVGGIGGIAIWRSKDELLGLATQPLH
AALADALHLVAVCCGMTVAGMLVVAGLDVPYQLWQYNKKLRMTKEEVKREHRENEGDPHV
KGRIRQQQRAMARRRMMANVPTADVVVTNPTHFAVALKYTDGEMRAPKVVAKGVNLVAAR
IRELAAEHHVPLLEAPPLARALYHNVELEREIPGTLYSAVAEVLAWVYQLKRFRAEGGIA
PETPVDLDVPAELDKGAHVAPQDEREEAEDALGGRAAGGAA*
>GCA_000452945.1_BPMSHR5858_genomic_01065_1 ftsY_2
LNIRKFIGPTSRDALRLVREAMGADAAVLSNRTLEDGRVEIVALPAAELAEISTRRAADA
AHGAAQPHAPRQGAQHAPSGPSFASPAASVVVPSAASGASAPASRAAANPYAAGGIPDVF
SSVFGASVDASADDDARPAVSPPVPPALPLAAPSEPAPWLVEHAKRLTKQHEELIARPRA
TAALPAQNAPQAPRERASADEAPEWARDIVRNAARRLPADEPAAAKPALRLPEDTAAAVA
DAVKARIERIVNDTVMQELGSLRELMEEQFAGLMWNERQRRNPVHGALTKYLFAAGFSAQ
LVRMIVDNLPEGEGYDTLDAAADWAQSVLAANLPVLDSEDALMERGGVFALMGPTGVGKT
TTTAKLAARCVMRFGASKVALLTTDSYRIGGHEQLRIFGKILGVPVHAVKDGGDLQLALA
ELRNKHMVLIDTIGMSQRDRTVSDQIAMLHGADTPVQRLLLLNATSHGDTLNEVVQAYRS
AAGQPKAALPDLAGCILTKLDEASNLGGVLDTVIRYKLPVHYVSTGQKVPENLYVATKKF
LLKSAFCAPREDSPFVPHDDDLPAMLSALSTRTGNELHEVRFG*
>GCA_000452945.1_BPMSHR5858_genomic_01068_1 ahcY
MNAAVIDSHSAQDYVVADIALAGWGRKELNIAETEMPGLVQIRDEYKAQQPLKGARIAGS
LHMTIQTGVLIETLKALGADVRWASCNIFSTQDHAAAAIVEAGTPVFAFKGESLDEYWEF
SHRIFEWPNGEFANMILDDGGDATLLLILGSKAEKDRSVIARPTNEEEVALFKSIERHLE
IDGSWYSKRLAHIKGVTEETTTGVHRLYQMEKDGRLPFPAFNVNDSVTKSKFDNLYGCRE
SLVDGIKRATDVMIAGKIAVVAGYGDVGKGCAQSLRGLGATVWVTEIDPICALQAAMEGY
RVVTMEYAADKADIFVTATGNYHVINHDHMKAMRHNAIVCNIGHFDSEIDVASTRQYQWE
NIKPQVDHIIFPDGKRVILLAEGRLVNLGCATGHPSFVMSNSFTNQTLAQIELFTRGGEY
ANKVYVLPKHLDEKVARLHLARIGAQLSELSDDQAAYIGVSKAGPFKPDHYRY*
>GCA_000452945.1_BPMSHR5858_genomic_01074_1 braC_2
MNIKMQKLLPISAAAMLLAAAATNAAADQVVKIGHVAPLTGGIAHLGKDNENGARLAVEE
INAKGLTIGGQKVTLQLDAQDDAADPRTATQVAQKLVDDKVVAVVGHLNSGTSIPASKIY
SDAGIVQISPSATNPAYTQQGFKTTYRVVATDAQQGPALANYARSKGVKSVAVVDDSTAY
GQGLANEFEKKAKALGLKVVSHDATNDKAVDFRAILTKIKGENPDAVMYGGMDATGGPFA
KQAKQLGLRAKIFAGDGVCTEKLADLAGDATDNVVCSEAGASLEKMPGGAAFKAKYEKRF
GQPIQIYAPFTYDAVYIIVDAMKRANSTDPAKIVAAMPKTNYTGVIGTTTFDSKGDLQHG
VISLYDYKGGKKTFLDEVKM*
>GCA_000452945.1_BPMSHR5858_genomic_01076_1 group_19452
VNISAALPLHLLHRNVLGTLATHSRAPEGYPYPTVVPYAVDARHRPIVLVSGLAEHTRNL
AADPRAGFLVVDGLGDAAGASDSVLEAERATLVGRFEPAGADPHVVARYLRYHPDGERYL
ALGDFTFWALACERLRYIGGFGRMGWLDATELDALAPVGYDDERALWDEYGASRERRAGL
ELLGVDRYGADWRLNGLKRRTPFDAPKMDVESLLAALREAAAVSLM*
>GCA_000452945.1_BPMSHR5858_genomic_01077_1 group_18959
MSSYKDLLAQREKLEKQIEEAKSREYAEVLSDVKQKIADYGFTLAELGLGRGKAGKAGRP
RAGVAAKYRDPESGATWSGRGKPPRWIAGKNREQFAI*
>GCA_000452945.1_BPMSHR5858_genomic_01079_1 lrp_8
VTLDTFSQKILRLLQLDARRSVQEISDQVGLSSTPCWRRIKDMEQSGVIQRYTALLDREK
LGLHICALAHVHLTRHNEGGVEQFEREIATCPEVTECYSTTGEADYILKIVAPDIKAYDI
FLHERIFKIPAVSQVRTSVVLREIKFDTQLPL*
>GCA_000452945.1_BPMSHR5858_genomic_01084_1 group_1792
LCHAREASPGTGEPEAAPRDNFPREAGSKRGIGAAFETRAQRFLERAGLALVARNVTVRG
GEIDLVMRERDGTLVFVEVRARANSRYGGAAASIGARKRMRLLLAAHAFWARTGGANACR
FDVVAFEGGRLVWLRDAFRADDAG*
>GCA_000452945.1_BPMSHR5858_genomic_01085_1 gmhA_1
MSVERIQQHIRDSAALVHEALEALSLPIAAAVDAMFMALANGNKIVACGDGPSAAAAERL
AAALIGGFERERPGLPGLALDTDASRSGTGADGDQIFAQQVRTLGQTGDVLLVLDPLGES
PRVLAAVEEAREREMIVIALTGNGGGRLGASLADTDIQICVPSERAVRIQEVHLLTIHCL
CDGIDAMLLGED*
>GCA_000452945.1_BPMSHR5858_genomic_01359_1 group_19620
MPTFDSRLDTASAAFRDNAEHMRALIAQWHALNARAADESAKAGPRFDKRGQLLPRERLA
LLVDAGAPFLELASLAGYLVGASDPDQSVPGAGLIGGIGYVSGTRCMIVVADSGIDAGAL
QPMGLEKFQRLQRIALEHRLPFVHLVESAGANLMQYRVEEFIHGGRHFFQLAKLSAAGIP
VFTLVHGSSTAGGAYMPGLSDYVVMVRDRAKAFLAGPPLLKAATGEIATDEELGGAAMHA
SVSGLAEYVADDDADGIRILRELVDKLGWREREAALPTGAEPALDPEGLLGILPRDGKKP
VDMREVIARLVDASDLLEFQPAYGPATVCVHARIHGMPVGIVTNNGPLDPAGATKATHFI
QACCQSDVPLVYLQNTTGFIVGKASERAGMIKHGAKMIQAVSNATVPQITILCGASFGAG
NYGMCGRGFDPAFVFSWPNARTAVMGAEQAALTMAIVMEGAARAKGIEPERARIDALREK
IVANFERQTHAFYTSGRMLDDGVIDPRDTRRVLAITLAVCRDGRRKSLQPLTFGVARP*
>GCA_000452945.1_BPMSHR5858_genomic_01375_1 group_19185
MKAIPFSSLKTMCEFVLRTSKRTVLPIVLASALAVPSFGAQARSSVSKRHAAAAASIRTA
SFHPQSGHAPLARHALASRHLKAGRYRLAGGMRRVAFSPRAPLGGRAFEPRDASQSFGLR
SNIVYMVDQHTGETLFDRNSRSVVPIASVTKLMTAMVVLDSNAPLDERIEVTDDDRDQDK
FTGSRLAIGSALSRDDMLHIALMASENRAAAALSRYYPGGREAFVDAMNRKARSLGMVDT
YFENPTGLSKHNVSSARDLAKMVEAAYQYPLIRQYSTDLKYDVDTGLRTLYYHSTNVLLR
DNAWNIGVQKTGYIKESGICLVMQTTIDGRDVVMVLLDSAGRHSDFFDAARLRTMLTSGA
AHVTEANSATSGN*
>GCA_000452945.1_BPMSHR5858_genomic_01383_1 group_1804

MIARRAVALGGWALMSVLAAARAHAADAMDAPAAAAAAAAPGSRFARVSATSESAPSSAF
EFASPAASASTSASALAVVPMSALTLASAPAPAPAPSAAPSPSPTSSPVQPGVRPGASCA
KTRPAILFNRWQEDWSALADPCVPRRPLDALKYVPLFGRVDSYLSLGAGLRERLELNDAP
LFGLGRARGDTYVLQRVQMHADLRIAGHVQAFVQLEDARPFGKDNVGPVDRNRVDLRQAF
VTYVDAIGSGAFKARVGRQEMAFDLQRFVSVRDGPNVRQAFDGIWADWEQGPWRLIGYAT
QPVQYRDDGAFDDVSNRNLTFSGVRIERQRVGPGDLSAYYSRYNRTQAQFPDGAGGEHRD
VFDVRYAGKRRNVDWDIEGMYQTGRVGAQRIEAWAVGSLAGYTFAGVGWMPRIGLQVDAA
SGDRRPRDGRIETFNPLFPNGYYFALAGYTGYTNLIHVKPSLTLKPSSALTLLAAVGLQW
RATTADAVYAQGATPVPGTAGRGGNWTGFYTQLRADWAVTANLAAALEVVHFQIGDALRA
AGGRNADYVGAELKFGW*
>GCA_000452945.1_BPMSHR5858_genomic_01384_1 nfdA
MFGLAQAPAAFADAAAEAAAPARAAASAPAVSVGPQYDTTHVYVAASDLDAFVDSFVATF
GGKASPRVTLTVTPTPSETLSQYVQTPVGMLSVFGFKTPVPYPFGAERTGYLVTDLDAAV
KAARAAGADLVVDSFDDPIGRDAVVRWPGGVMMQLYWHRKAPSYAPLATVPDNRVYLSPY
EADRFVTSWVRFSHGKVVADDRRGDGGAIGRPGTDVRRIVIESGFGRVVAFVTDGKLPYP
YGRETTGYAVADLDAVLDKAKAAGVEVLAQPYRTRAGRTAMLAFPGGYLAEVHDAK*
>GCA_000452945.1_BPMSHR5858_genomic_01426_1 group_1336
MALTRTLGKTAAWIVGIFAVLTAAVGVFIFTFDWNRAKPWINEQASASIGRPFEIRGDIK
VGWRRPDGETGWRAWVPWPVFSASRIVIGNPPWAKAPNFATLDEARFELAVLPLLAHEVV
IPTIELVNPALDIERLADGRNNWTFEFKQSAGPSPWKVQLRDFGFAKGTVVYRDAITKAD
LSVAVDTLGQPIPLGDVLRQQEETSRAASAQRVGKRGAAQLAAKAQAQIASDASAASAAG
APNASGAAAASGASGVNAAASAAKVSGNDRSRAAWATAGASAASKPAAAASGGSAAAVAA
GKAPAAGAGGAVASASASAASSASAVSTSAAVSGASGARGASGASAASAAEQRSAGPTYA
FGLTVKGRYKGVPIDGTGKVGGVLSLRDASRPFPLQADVKAGDTRLAIVGTLTDPVHLAA
IDLRLWLQGSSMSHLYPLTGVTLPDTPPYATEGRLVGRFKRAASTFRYENFNGRVGGSDL
RGTLVYQQRAPRPLLSGELVSNLLQFSDLAPIIGADTAASKARRGDTTKQPPGRVLPVET
FRTDRWRALDADVKFTGRKLIKSAALPISDLYTHIVMTDGVLSLEPLKFGVAGGSLATTA
HLDGSAAPLKGRFSVAARHLKLKQLFPTQKVMQSALGEINGDAALSATGNSPAALAATSN
GEVKMLMTDGAISRLLMEAAGLNVANVLYEKMFGSHDVKINCAAVDFVATNGLLDTKLFA
FDTDDALINIDGPIDLRNESMNLKVHPHTKGFRVFSLRSPLYVKGTFKNPDVGVDAAALA
LRAGAMVGLGLVNPFAALIPLIAPSNNRDVPCSELFAQLKAPQKAANARK*
>GCA_000452945.1_BPMSHR5858_genomic_01427_1 ybiT
VLSTANITMQFGPKPLFENISVKFGEGNRYGLIGANGCGKSTFMKILGGDLEPSAGNVAL
EPNVRLGKLRQDQFAYEDVRVLDVVMMGHTEMWAAMTERDAIYANPEATDDDYMHAAELE
AKFAEYGGYDAEARAGALLLGIGIEEKFHSGPMRDVAPGWKLRVLLAQALFSNPDVLLLD
EPTNNLDINSIRWLEDTLNNYDSTMIIISHDRHFLNSVCTHMADMDYGTLKVWPGNYDDY
MLASAQARERQAAANARAKERVAELQDFVRRFSANKSKARQATSRAKQIDKIKIEEFKPS
SRQNPFIRFEFEKKLHNIAVVADSITKKYERTIFENFNLSVQPGERIAIIGENGAGKTTL
LRALFGNLTLDHGTVKWAENANVGYMPQDTYEEFPNDVTLMDWIDQYRKDGDDETMVRGT
LGRLLFSADDIKKSVKVLSGGEKGRMIWGKLMLGRHNVLLMDEPTNHMDMESIESLQIAL
EKFEGTLIFVSHDREFVSGLANRIIEVTTEGKLVDFGGNYEDFLASKGLQ*
>GCA_000452945.1_BPMSHR5858_genomic_01431_1 ompW_1
MIPSLSHAASPGEGINQGDIIARVRGISIMPDERTSNTLSALNVGVNNAIVPELDFTYMI
RDYLGVELILGTSRHQITSSLGDLGGVGVLPPTLLLQYHFNHAGKVRPYVGAGINYTLFY
NNGLHAGGEGIGINNHSFGPALQFGVDVQVTKKVFVNVDVKKIWMHTDATLGGQPLGRLN
IDPLVVGVGVGMKF*
>GCA_000452945.1_BPMSHR5858_genomic_01433_1 mntH_2
MNSRPLAARSPSSRRPSDSGGPGGGRRWAAFVGAGVLVAVGYMDPGNWATALAGGAGYGY
ALLGVVIAASLMAMLLQWLAARIGVVSGRDLAQLCRERFDPRATVLLWIAAEIAIVACDV
AEVVGAAVALQMLLGVSLSTGVLVSAVGTLAMLALERHGRRTLEAAIASLIFFVGLCFVI
EVALARPDWHAALAGAAPSPRLLREPGMLWLAAGILGATVMPHNLYLHSMLVKHHASARD
EPSIRRALFGVNLDTFVSLGFAFVINAALLIVAAAVFHASGHDDVTDLADAHRLIAPLVG
NRWAGPLFAAALLACGLSATVTGTLAGQAIMEGFLKLTLPRWQRALLTRALAIGPALAAV
GLFGQHGSNQLLVASQVVLSLQLPLAMVPLVVFASSRALMGRWRVRGLPLAAAWMCAAAI
VALNGALVWQVVAAG*
>GCA_000452945.1_BPMSHR5858_genomic_01434_1 potF
MKAKVAGRLTALALCAGATVAAAKDTQLNVYNWSDYIAKDTIPNFEKQTGVKVRYDNYDS
DDTLQAKLLTGNSGYDIVVPTSNYAGKQIQAGIFTPLDKSKLPNLKYLDAQLMALVAGAD
PGNKYVVPWAYGTTGLGYNVDKAQKVLGKVPLDNWDILFKPENLSKLKTCGVSVLDAPDQ
MFAATLHYIGKDPMSTNPADYQAAMQVLKKIRPYITQFNSSGYINDMVGGDICFAFGWSG
DVVIAKHRALEAKKPYKLEYYVPKGGAPVWFDVMAIPKDAKNKDAALQWINYIEDPKVHA
SITNAVYYPSANAQARKYVRPDVANDPAVYPPPDVVKTLFLLKPLPPEIQRLQTRLWTEL
KSGR*
>GCA_000452945.1_BPMSHR5858_genomic_01440_1 group_19004
VTTASTMTVEGFFDPATCTISYLLFDSGSGECALIDSVLDYDPKSGRTRTASADQLIARV
AALGARVRWLLETHVHADHLSAAPYLKTRVGGEIAIGRHVTRVQDVFGKLFNAGPAFAHD
GSQFDRLLDDGDTLALGALSIRAMHTPGHTPACMTYVVTEAHAAHDARDAAAFVGDTLFM
PDYGTARCDFPGGDARSLYRSIRKVLSLPPATRLYMCHDYQPNGRAIQYASTVADELREN
VHIREGVTEDDFVAMRTARDATLDMPVLMLPSVQVNMRAGRLPEPEDNGVRYLKIPLDAI
*
>GCA_000452945.1_BPMSHR5858_genomic_01450_1 clcB_1
VLSFLLKLRTRAQTLFRLSDAHTMLVWSVIVGIGGALATVAFREGIELIQHAISGTSGSF
TEMAKRLPWQVRFWLPAAGGFLAGCVLLVAQRGARKESKTDYMEAVALGDGVVPVRQSLW
RSASSLLTIGSGGSIGREGPMVQLAALAASLVGRFAHFDPPRLRLLVACGAAAGITSAYN
APIAGAFFVCEIVLGSIAMESFGPVVVASVVANIVMREFAGYKPPYQMPVFPAVAGWEVL
FFVALGLLCGAAAPQFLRLLELSKAQFKRLPVPLPVRLAAGGLIVGVISVWVPDVWGNGY
SVVNAILHAPWTWQALVAVLVLKLIATAATAGSGAIGGIFTPTLFVGAVVGSLFGIALHA
LLPAHVSAWYAYAMVGMGAFLAGATQAPLMAILMIFEMTLSYQVVLPLMVSCVVAYFAAR
ALGKTSMYEITLRRHRDDEARTRLRTAQMRDLIQPAATVVPLTASVADMTRVFLEYPVKY
LYVTDDAGRFRGAVALKDITSDLLEKRDTTHKSAANYLHTPFPLLTPDMPIGAALEHFMR
FQGERLPVVERADDPTLAGVVYKTSLLDAYRRMNMDLER*
>GCA_000452945.1_BPMSHR5858_genomic_01532_1 davD_1
MYLEHALKDPSLLRHQAYIGGEWQAADSDATFEVFDPATGESLGTVPKMGAAETARAIDA
AQAAWAGWRMKTAKERAAILRRWFDLVIANSDDLALILTTEQGKPLAEAKGEIAYAASFI
EWFAEEGKRVAGDTLPTPDANKRIVVVKEPIGVCAAITPWNFPAAMIARKVGPALAAGCP
IVVKPAESTPFSALAMAFLAERAGVPKGVLSVVIGDPKAIGTEITSNPIVRKLSFTGSTA
VGRLLMAQSAPTVKKLTLELGGNAPFIVFDDADLDAAVEGAIASKYRNNGQTCVCTNRFF
VHERVYDAFADKLAAAVSKLKVGRGTESGATLGPLINEAAVKKVESHIADALAKGASLMT
GGKRHALGHGFFEPTVLTGVKPDMDVAKEETFGPLAPLFRFASEEELVRLANDTEFGLAA
YLYSRDIGRVWRVAEALEYGMVGINTGLISNEVAPFGGVKQSGLGREGSHYGIDDYVVIK
YLCVAV*
>GCA_000452945.1_BPMSHR5858_genomic_01567_1 group_19091
MRNFLSLGRRLALAAAVLAPVAAFANNVIVLNSAEATLSLIDQQSRQVVGQYPTGKEPHH
LMATPDNASLIVANSVSNSLMFLDPKTGKPQRTLEGIDDPYQLGFSPDRKWFVAAGLRLD
RVDIYSYDGRDLKLAKRLPLAVMPSHLAFTKDSKTVLVSLQVSGELAAIDLATQTLKWKM
KVGKVPAGLWLTPGDKYVLIGMTGADYVAVVDWRNQKVVKTIPTGKGAHNFRSLADGTHV
AVTNRVANTISILDENALTNVGDITGLLPGPDDMELSADRKTLWVTFRFAKKVGVIDLPS
RKLIQTISVGRSPHGIYFYDRAPWTAPNGA*
>GCA_000452945.1_BPMSHR5858_genomic_01570_1 cinA
MTIGIIIIGDEILSGRRQDKHLAKAIELLAARGLALGWAEYVGDDPARITATLARTFASG
DIVFSTGGIGATPDDHTRQCAAAALGVPLELHPQARALIEERIRETHADPAKPVDPDSPE
NRHRLNMGVFPRGATIIPNGYNRIPGFSCGDHHFVPGFPVMAWPMIEWVLDTKYAHLHHA
SPHAERSLYVFELPESTLTPLMEKIERDFPGVRVFSLPSVGDAEKGGIYARRHIDLGVKG
EPEAVAAAFVKLREGVHLLGGDIVEPAEPQHG*
>GCA_000452945.1_BPMSHR5858_genomic_01585_1 dinB
MPPSDPPPPPSGDAAHLARKIIHCDCDCFYASVEMRDDPSLRGRPLAVGGRPDQRGVIAT
CNYDARRYGVHSAMSSALAMRKCPDLLILPPAMDKYRAASRLIMAIYRDYTPDVEPLSLD
EAYLDVSRATRCQGSGTLIAREIRERVRDTVGVTVSAGVAPNKFIAKIASDWNKPDGLFV
VRPHEVDAFVAALPVRKLHGVGKVTAARLDKLGIRTCAQLRDWSLFDLHRTFGAFGRRLH
ELARGIDERPVRADQARKSVSVETTYVQDLAALDACAAEIRRLTGLLDARIERAGAARSI
RKLYVKIRFADFQRTTVECVADATDADTAVALLAKGLARRRQAVRLLGVGVRIDEDTAER
HGQIALFDDEDMSDEDAEGDEVGEDGEDGEGDEGDASPGA*
>GCA_000452945.1_BPMSHR5858_genomic_01590_1 aceE_2
MSAVPNEVMKYVAAERDDDAQETVEWLEALDGVISSVGPGRAHYLIEKQIEFARMHGEHL
PFSANTPYINTIPVEAQAKIPGDQDIEHRIRSYTRWNAIAMVLRAGKHTNVGGHIASFAS
AATLYDVGYNHFWHAPSAEHGGDLVFVQGHSSPGIYSRAFLLGRLTEDQLDNFRQEVGGN
GISSYPHPWLMPDFWQFPTVSMGLGPIMAIYQARFMKYLQARGIVKTEGRKVWAFLGDGE
TDEPESLGAIGMASREKLDNLVFVINCNLQRLDGPVRGNGKIIQELESEFRGAGWNVIKV
IWGSRWDALFARDKTGALMRRMMEAVDGEYQTYKSESGAYVREHFFNTPELKALVADWSD
DDIWNLNRGGHDPHKIYAAFHEASNSKGAPTVILAKTIKGYGMGEAGQAMNITHQQKKLP
VEQLKKFRDQFRLPIADDVIADVPYLKFDEGSKELEYMRAHRQALGGYLPQRRQKAQSLP
VPALDAFEPLLKGTGEGREISTTMAFVRILNILLKDKALGKRVVPIVPDESRTFGMEGLF
RQIGIWNQEGQKYVPEDSDQLMFYKESETGQILQEGINEAGGMCDWIAAATSYSTHGEIM
VPFYIFYSMFGFQRIGDLAWAAGDMRSRGFLLGGTAGRTTLNGEGLQHEDGHSLMWAASV
PNCVSYDPTFGYELAVIVQDGLRRMVQEQEDVYYYVTVMNENYEHPAIPQGEHVAADIIK
GMYAFRKADADKKAPRVQLLGAGTIFNEVIAAADLLKNDWGVAADLWSVPSFTELAREGH
DVERWNLLHPAEARRLSHVQTCLKDTQGPVIASTDYVRALADQIRGQIDRRYVVLGTDGF
GRSDTRGALRHFFEVDRYWVTVAALNALADEGTIERKVVADAIAKYNLDPAKPNPMTV*
>GCA_000452945.1_BPMSHR5858_genomic_01601_1 iscS_2
MNNDIPHLPIYMDYSATTPVDPRVVDKMVPYLREQFGNPASRSHAYGWDAERAVEEAREQ
VAALVNADPREIIWTSGATESDNLAIKGAAHFYQGKGKHIVTVKTEHKAVLDTCRELERE
GFEVTYLDVKDDGLVDLDVFKAALRPDTILVSVMHVNNEIGVIQDIATIGEICREKGIIF
HVDAAQATGKVEIDLAKLKVDLMSFSAHKTYGPKGIGALYVRRKPRVRIEAQMHGGGHER
GMRSGTLPTHQIVGMGEAFRIAREEMATENERIRMLRDKLLRGLSEIDETYVNGDLEHRI
PHNLNISFNFVEGESLIMAIKDVAVSSGSACTSASLEPSYVLRALGRNDELAHSSIRFTV
GRFTTEQEVDYVIDLLKSKIAKLRDLSPLWEMHQEGIDLSTIEWAAH*
>GCA_000452945.1_BPMSHR5858_genomic_01602_1 iscU
MSYSNKVLDHYENPRNVGSFAKDDDTVGTGMVGAPACGDVMKLQIRVGADGVIEDAKFKT
YGCGSAIASSSLVTEWVKGKTLDEALAIKNTQIAEELALPPVKIHCSILAEDAIKAAVAD
YKKRHETAEDGKAAA*
>GCA_000452945.1_BPMSHR5858_genomic_01604_1 hscB
MVSLKDSHFDLFHLPARFALDEPTLDAAYRAVQSQVHPDRFAAAGDAQKRIAMQWATRAN
EAYQTLRDPLKRATYLLHLRGVDVGAENNTAMEPAFLMQQMEWRERIEDAAGAKNVDALD
ALLAELRDERRARLAKLGALLDSGSDQGAAEAVRQLMFVERVSAEIGAQIERLEH*
>GCA_000452945.1_BPMSHR5858_genomic_01606_1 fdx_2
MPQLVVLPHVELCPDGAVIDAMPGKSICDTLLENGVEIEHACEKSCACTTCHVIIREGFN
ELDPSEEDEDDLLDKAWGLEPTSRLSCQAIVKEGVDLVVEIPKYSINHAKENH*
>GCA_000452945.1_BPMSHR5858_genomic_01607_1 iscX
MKWTDSREIAIALADQHPDVDPKTIRFTELRQWVLDLDGFDDDPDHSGEKILEAIQAHWI
DESDFDDED*
>GCA_000452945.1_BPMSHR5858_genomic_01609_1 lysS
MTEPTQPQAAVAADENQIVAERRDKLRALRDQGIAYPNDFQPTHHAAGLQTEYADADKEA
LDAKALDVAVAGRMMLKRVMGKASFATVQDGSGQIQFFVTPADVGAETYDAFKKWDLGDI
VAARGVLFRTNKGELSVKCTELRLLAKALRPLPDKFHGLADQETRYRQRYVDLIVTPETR
ATFRARTKAIASIRKFMSDADFMEVETPMLHPIPGGAAAKPFVTHHNALDMQMFLRIAPE
LYLKRLIVGGFERVFEINRNFRNEGVSPRHNPEFTMMEFYAAYTDYRWLMDFTERLIRQA
AVDALGTATIRYQGRELDLAKPFRRLTITQAIQKYAPNYTDGQLSDDAFLRGELKRLGVD
VTQPAFLNAGIGALQLALFEETAEAQLWEPTFIIDYPIEVSPLARESDTVAGITERFELF
VTGREIANGFSELNDPEDQAARFRKQVEQKDAGDEEAMFFDADYIRALEYGMPPTGGCGI
GIDRLVMLLTDSPTIRDVLLFPHLRRED*
>GCA_000452945.1_BPMSHR5858_genomic_01611_1 recJ
MTRIVTRSAVPAALDALARHGLHPVLARLYAARGIASPADVETALSRLVPPAQLKGCAQA
AALLADALEAGRKMLVVADYDCDGATACAVAVRGLRMFGAKIDYLVPNRFEYGYGLTPEI
VELAAARKPDLLITVDNGIASVDGVAAANARGIDVLVTDHHLPGDTLPAARAIVNPNQPG
CAFPSKHIAGVGVMFYVLLALRAELRARGAFGDARPEPRLDGLLDLVALGTVADVVRLDG
NNRVLVAQGLQRIRNGRMQPGIAALFRAAARDARSASGFDLGFGLGPRLNAAGRLSDMSL
GIECLTTDDVGRAWELAQQLDAMNRERREIEAGMQQQALADLASVDPAEAATITLFNPEW
HQGVIGIVAGRLKERFHRPAFTFAHADDGGARVKGSGRSIPGFHLRDAIDLISKREPDLI
VTFGGHAMAAGVTLETANVPRFAAAFETVAREWLSDDALARVIETDGDLEDAYFTPQVVE
LLDAAVWGQGFPAPLFSGEFDVASQALVKDKHLKLQLTRGRQRFNAIWFNHAEPLPARAT
LAFRLASDTWNGTTRVQLIVEHAA*
>GCA_000452945.1_BPMSHR5858_genomic_01613_1 lolE
LKLPYEWQIGWRYTRAGKRTTGNGFISFIALVSMLGIALGVAALIVVLSVMNGFQKEVRD
RMLSVLAHVEIFSPTGSMPDWQLTAKEARLNRSVIGAAPYVDAQALLTRQDAVSGVMLRG
VEPSLEPQVSDIGKDMKAGALTALAPGQFGIVLGNALAGNLGVGVGDKVTLVAPEGTITP
AGMMPRLKQFTVVGIFESGHYEYDSTLAMIDIQDAQALFRLPAPTGVRLRLTDMQKAPQV
ARELAHTLSGDLYIRDWTQQNKTWFSAVQIEKRMMFIILTLIIAVAAFNLVSSLVMTVTN
KQADIAILRTLGAQPGSIMKIFVVQGVTIGFVGTATGVALGCLIAWSIPWLIPMIEHAFG
VQFLPPSVYFISELPSELVAGDVIKIGVIAFALSALATLYPSWRGAKVRPAEALRYE*
>GCA_000452945.1_BPMSHR5858_genomic_01618_1 pyrG
MTKYVFVTGGVVSSLGKGIAAASLAAILESRGLKVTLLKLDPYINVDPGTMSPFQHGEVF
VTEDGAETDLDLGHYERFISTKMRKANNFTTGQIYESVIRKERRGDYLGKTVQVIPHITN
EIQAFIERGAASATCGEPDVAIVEIGGTVGDIESLPFLEAARQMSLRLGRNSACFVHLTL
VPFIATAGELKTKPTQHSVQKLREIGISPHVLLCRADRPIPDDESKKISLFSNVPEDAVI
SVWDVDSIYKIPQMLHDQGLDRLICEELRLDPQPADLRMWAALVEKLQNPKHEVTIGMVG
KYVDLTESYKSLIEALRHASIHTSTKVNIEYIDSEELETNGTASLAHLDAVLVPGGFGRR
GTEGKIAAVRYAREAKVPYLGICLGMQLAVIEFARDVVGLKQANSTEFDPNTPERVVALI
TEWYDREGKVEKRTEDSDLGGTMRLGSQRCPIKPGTLAEAIYGKDVNERHRHRYEVNNRF
VPQLEAGGLVISARTPSEDLPEMMELPSTMHPWFVGVQFHPEFTSTPRDGHPLFKSFVQA
ALACQQTRAGAKA*
>GCA_000452945.1_BPMSHR5858_genomic_01620_1 eno
MSAIVDIIGREILDSRGNPTVECDVLLESGTMGRAAVPSGASTGSREAIELRDGEAGRYG
GKGVLKAVEHINTEISEAIMGLDASEQAFLDKTLLELDGTDNKSRLGANAMLAVSMAVAK
AAAEEAGLPLYRYFGGSGAMQLPVPMMNIVNGGAHANNSLDIQEFMIVPVSQPTFREALR
CGAEVFHALKKILGDRGMSTAVGDEGGFAPNFGSNDECLSTILQAIEKAGYRAGEDVLLA
LDCAASEFYHDGKYQLAGEGLQLSSAEFTDYLATLADKFPIVSIEDGMHEGDWDGWKLLT
ERLGKKVQLVGDDLFVTNTRILKEGIEKGIANSILIKINQIGTLTETFAAIEMAKRARYT
AVISHRSGETEDSTIADIAVGLNAGQIKTGSLSRSDRISKYNQLLRIEEDLGDIASYPGK
SAFYNLR*
>GCA_000452945.1_BPMSHR5858_genomic_01623_1 hslO
VSDQLQKFMFNAAPVRGEIVSLRSTWQEVLTRRDYPTPVRNVLGEMMAACALLSANLKFD
GTLIMQIFGDGPVKMLVVQCSSDLAMRATAKFSGDAARTVGDGTSFAELINASGHGRCVI
TLDPADKRPGQQPYQGIVPLNGEDGPLASIADVLEHYMRHSEQLDTRLWLAADHDRAVGV
LLQKLPGDGGIVPRVEQTDTDTWERVCTLGGTLSSKELLEVEPETVFRRLFWQENVQHFE
PTSTRFQCTCSREKVGGMLRMLGRVEIDGVIEERGHVEIHCEFCNQRYEFDPVDVAQLFS
TPELGTGVAPAAAQRH*
>GCA_000452945.1_BPMSHR5858_genomic_01628_1 yciO
MSQFFRIHPDDPQPRLVRQAAEIVRGGGVIALPTDSSYALACHLDDKDAVERVRRIRGLD
DKQHLSLLVRDLSELATFAMVDNRQYRLIKSVTPGPYVFILQATKEVPRRLSHPSRKTIG
LRVPAHAITLALLEALGQPLLGTTLILPPDDEPLNDPEEIRARLEKQVDLVIDGGACPRE
PSTVIDLTGDEPQLVRAGRGPLEPFGLAA*
>GCA_000452945.1_BPMSHR5858_genomic_01629_1 group_19079
MDASSLIQTIAVYALPVVFAITLHEAAHGYVARLLGDNTAYMMGRVSFNPMRHIDPFGTI
VIPLVLYFLTSGAFLFGYARPVPVTFRNLRNPRWGSLWVALAGPGCNFVQALIWGFVSIG
LAALAVDEPFFTRMAGAGVGVNLVLAVLNLFPLPPLDGGRVLAALLPPKQSIALSRLEPY
GFFIVLALVATGLLTKLWLRPLVGAGYAVVTAILTPFASLF*
>GCA_000452945.1_BPMSHR5858_genomic_01632_1 dapA_3
MANGTQDGIQIRGSVPAIVTPMLEDGGLDLAAFRKLIDWHIEEGTDALVVVGTSGESATL
SVDEHVLMIETAVKHAAKRIPIVAGAGGNSTTEAIELSKHAKAVGADATLQVVPYYNKPT
QEGIYRHFKAIAEAVDLPVILYNVPGRTVADMSNETTLRLAQVPGIIGVKDATGNIDRAA
QLIKAAPAHFSIYSGDDPTAIALMLLGGHGNISVTANVAPRAMSELCRAALAADVKTARE
IHMKLLSLHKHLFIEANPIPVKWALQQMGKIAGGIRLPLTPLDERCHETVRGALREAGLL
*
>GCA_000452945.1_BPMSHR5858_genomic_01646_1 group_19331
MKPSRGRAPSAATLVATAAVSAALALLPGAAALAQKAPAPTDGTPEIDASITNRNWAQAL
AQLDARIASNPRDVQAQFKRGTVLARLNRDDDAIKQFVELTQAYPELPEPYNNLAALYAK
HGRYDDARSALVTATHANPNYGLAYENLGDLYLRLAAESYKRAQSLGRTTGASAQRLSDL
QKIIAPPKTRAAAAPAPAARDYSARAAANVTTTTLPLSPTFQFGGPSGMLAMPPYAAPSQ

*
>GCA_000452945.1_BPMSHR5858_genomic_01648_1 alkA_1
MATARKAPAKRAASQPPMAAKRAGARAPAVHKAGAKRAAPNGAANGAAHQPGLRAAAGKP
ARAKVDGERAVDALPREAAAVAAEPAARKARVSEAAVPVQLSDAETVARPPYWDKACADL
VKRDRILKKLIPKFGPAHLVKRGDSFVTLARSVVGQQISVAAAQSVWVKIETACPKLAPP
QIIKLGQEKLIACGLSKRKSEYILDLAQHFVSGALHVDKWASMDDEDVIAELTQIRGIGR
WTAEMFLIFNLSRPDVLPLDDLGLIRAISVNYFSGEPVTRSEAREVAANWEPWRTVATWY
MWRSLDPLTANS*
>GCA_000452945.1_BPMSHR5858_genomic_01649_1 accA
MKTTFLDFEQPIAELEAKIEELRFVQDDSAVDISEEIERLSKKSQQLTKDLYANLTPWQV
SQIARHPQRPYTLDYVSELFTDFHELHGDRAFADDQSIVGGLARFNGHACMVIGHQKGRD
TKERAARNFGMPRPEGYRKAERLMRVAEKFGLPIFTFVDTPGAYPGVGAEERGQSEAIGH
NLYVMAELKTPIIATVIGEGGSGGALAIAVADTVMMLQFSTYSVISPEGCASILWKSAAK
APEAAEALGLTAHRLKALGLIDKIVNEPLGGAHRDPKGMAALLRRALGDSLRQFQGMSVD
ALRERRFERLMAYGKFKETTPRA*
>GCA_000452945.1_BPMSHR5858_genomic_01650_1 tilS
VFDALGVALSALPDDTPIAIAYSGGLDSTVLLHAAARIAGAGRCIALHVHHGLSANADAW
LAHCAETAQALGARFDAARVDVPRASGQGIEASARDARYRALETMCARYGARTLWLAQHA
DDQAETVLLQLLRGAGIAGLAAMAPQYRPALADVVRMRPLLHLLRAQLERYAQQHALRWI
DDESNTDTRYARNALRVDVLPALAPHFPGFRDALARTAQHAAAAQRLLDDLAAIDLRAVA
RADVRVLSRDALVALDDERGANLLRYWMRSLGLPGASAARLAEMVKQLRAARDAHALRVD
HAGWRLRLYRDDVQWEAGDGAASEAARAARADVADDDATDARDDRADASAAARLPACALA
WRGHEVWRLPGWRGSFVFSPVAAHEHDAVPEALLSSAALRACARAGGERMRTRQGGPGRT
LKNLFQERGVPAWQRDVPLLYVGERLLFVPRIGVNRATHDGADAPGGWRRIEWRPDMLIA
*
>GCA_000452945.1_BPMSHR5858_genomic_01651_1 group_19244
MGSVERIKNVAKRVAKWHQAGHQMVVVPSAMSGETNRLLGLAKEISSQPSPRELDMIAST
GEQVSVGLLSIALQDIGVEAVSYAGWQVPIKTDSAYTKARIHSIDDERVRRDLDAGKVVI
ITGFQGVDPDGHITTLGRGGSDTSAVAVAAALEADECLIYTDVDGVYTTDPRVVEEARRL
DSVTFEEMLEMASLGSKVLQIRSVEFAGKYRVKTRVLSSLTDPQIPLDEEMRSGTLITFE
EDETMEKAVISGIAFQRDEARIAVMGVPDKPGIAYQILGPVADANIDVDMIIQNQSVEGK
TDFTFTVGRGDYQKAMDILTNQVKGHVNAEQVLGDPKVSKVSVVGVGMRSHVGVASTMFR
TLSEEGINIQMISTSEIKISVLIDEKYMELAVRALHKAFELDQA*
>GCA_000452945.1_BPMSHR5858_genomic_01683_1 group_1326
MVETRRIHVVLLAAAITAIAAIAPAGHAQNATDWLANTYGTLAAHVGNTARSMWVAPEGV
IYTASMWDEYEGGVAIYQNGKSVGSIGSHAEFQGGAITGNAASVFVALQYDKSHGSGAVG
RYNRVTKARELSIQVSASTDQPRVDVVTGLATVGSLLYASDFYGNRVRLFTTDGVWQRDI
GVASPGALAVDRAGNVWVARKRAGEIVEFSAAGALLNTLRMPGGSQPSALYFDAPSGQLM
VGDEGPDMNIKRYAVTGAPALVGTFGIQGGYLDTTTGIKGQVGARRFTRVAGIGKDAAGN
LYVLNNPWGGSWDLGRNGATDIHAYDSAGNLQWTLQSLNFEGIAAADPATDGALFYGGTH
IYAGSAGGTFVANTVDPFSYPSDPRIDMNDTQRDEHFGQLVAVGANRILVASGQNPPIFY
FFHFNKANGYVAIPDASLPGAAFNTAQRVTSGFCIDSEGGVWAGLEKTGAIYHYPLAGFD
ASGKPTWGAGVPTRIPASIQPLTRIVYLAESDTMMLAQGIVGSADWTSIGTRIEVYHGWR
AGNTAAPDPVITLANAGAKSIDAAGNHLFVGYWFGGGGPARPNVDAFNLATGKLDATLVN
TSRATVDASSAVDSMYGVRAYQRSTGEYVVTKNNVKGNSITVYRWKP*
>GCA_000452945.1_BPMSHR5858_genomic_01687_1 dctB_2
MRLRGIRRLHRTAAVLAVIACCVATAWLGYSVALRYYFRAHMNEVAQRSSLYALTLESQL
ARYESIPRIAALEPVLSRLLNDPGERALRDEANAYLNAVQKNAELSAAYVMNARGETLAA
SNSNEPGSFIGSNYAFRPYFADAMKRGFGRFYGVGNTTGQAGYFLAVPIRNPGQPARAPI
GVVAIKASLDRYEAALTKSGDVVLLVDRDGVAFLSSVASWRYRTLTPIPAATLRKLAATR
QYGDAALEPLSDTPPAPAGAGRSSTQADSAPTPESLHLSLSGGRPAWYHVVYRAAGPLGW
RVAVLVDSTDDERRALFASGGVAAATALVFALIMVAWLRHSREKERQRARTALLAAQRDL
ERRIAERTAELTAANAALEEKVDALDAARRILRDTRDTAIQAGKLAALGQMAAGITHELN
QPLAAIMTLSSNAVRMAALGRSADLTRNLELVNDLAGRMGKIVAHMKGFARHESVSREAV
SISEALQQALTLIEAHRKAAKVSVRVEPVPGQGIVLASSIRIEQLLVNLLTNAIDASAAG
MDNRQVVVTTDVAGDIVRISVADSGKGIPADVMPRLFEPFFTTKQTGKGLGLGLAISQAI
VDEFGGRLHARNGSGVEPELGGAVFIVELHRVSRKFSSHDERR*
>GCA_000452945.1_BPMSHR5858_genomic_01688_1 kgtP_3
MEPTFTETGARASEPHATTLAGRAASSGDKLQRLKSIFSASAGNLIEWYDWYVYSAFALY
FAHSFFPAGNQTAQLLNTAAVFAVGFLARPVGGWLMGIYADRYGRRPALLVSVVLMCVGS
LIIALCPGYDVIGTAAPVLLVVARLLQGLSVGGEYGTSATYLSEVATARDRGFYSSFQYV
TLVAGQLVALALLIVLQQFVLTTQQLESWGWRIPFFIGAACALAAMRLRSSMEETGEFKR
TLNARDKRGTLAELSKHPRAVLTVVGLTMGGTIAFYTYSIYMQKFLVNTVGMSKHDATLV
SAASLALFAILQPIVGSISDRIGRRPVLIAFGVLGTLFTVPIMTAISRTHDVWTAFFLNM
AALVIVSGYTSINAVVKAELFPAKIRALGVGFPYALTVSIFGGTAEYLALWLKQAGHESL
FYWYVTAAIFCSLLVYVCMRDTGKHSLIKD*
>GCA_000452945.1_BPMSHR5858_genomic_01715_1 group_19515
MKPAAQRSTKPTDAARDAAKTRAPQRLTYVIGNLDRLLRRHMTDALAPLGITLAQYTALS
VLEARGPSSNAKVAERSFITPQSANEVTSAMAGRGWIMREADPNHGRVILLRLTPEGAAM
LRECEAALRPLESRMLDGVPRADADAAQHWLEIFVRNLRG*
>GCA_000452945.1_BPMSHR5858_genomic_01721_1 plcN_1
MFRRAWIATACAGSAIALYACGGTDSNAPVTSQNALRTATPIKHLVVIYGENVSFDHYFG
TYPNAANPPGEPAFSAASGTPSINGLSGLLLTANPNATNPANGAGASNPFRLDRTQAATA
DQNHAYTAEEQAYDNGLADLFPKYTGNGSSGGAGAFGTTGQVMGYFDGNTVTALWNYAQR
FAMSDNAYTSTYGPSTPGALNVIAGQTDGMQIVKTSKAVSTLAKTSYYINDGQGGLTMIN
DVDPGYDVCSSTTDQAMMSGKNIGDLLNARNVTWGGFMGGFNLSTTNANGTTGCKRSTIA
TAVNAATTDYIPHHNWFQYYASTANPQHTRPSSIAAIGSSVQTDGKTPEPANHQYDSDDF
FAAVKAGNFPSVSFLKAPAAQDAHAGYSDPLDEQQFVTKVVNFLQQQPDWASTAVIVTYD
DSDGWYDHAYTAPTRASFDAVDQLNGNGACGSGAATTGINGAAVNGRCGPGTRIPFVVVS
PWAKQNYVSHTLIDQASVVRFIEDNWLGGQRIGGGSFDATAGDMRDLFDFSGAVNKTPLY
LDPTLGTKLATAPSI*
>GCA_000452945.1_BPMSHR5858_genomic_01723_1 group_19500
MTQKFVGTHVVGPREKLPSGKPWINAPLTVKVPFPAAFNAIPIVVASALQDPKHTSTYPD
TFAVTVISVTKTDFTVNICRADYVRDNYTTSGWGQNLHLSYIAETPA*
>GCA_000452945.1_BPMSHR5858_genomic_01725_1 mcbR
LAPRIVPPALEAIEQETMTDKVYQQLREALMSGRFAPGQALSLRSVAEAVGSSTMPVRAA
LTRLRAERALADGPNRALVVPPMTIEMLDELRDVRIALEGCVAERAATRMSVSQIAEVRR
ICEAMNAHVEAGRSRAYLQSNFAFHSAIYAHGASENTLAIIRNLWMRIGPFLNMIALDVP
HMRRSMDAHRAIVDALERRDGAGARAGIALDIGGAAHDLAETLAAERARLGPRRLANE*
>GCA_000452945.1_BPMSHR5858_genomic_01728_1 ttuB_1
MSTPHAEAFCAPAGSARASLNRLLFRKLMPILITAYVISFLDRTNIAFAKHSMGIDLGLS
SAAYGLGAGLFFVTYALFEIPSNLIMHRVGARFWIARIMITWGALSVAMALVRGETSFYA
MRLLLGAAEAGLFPGVMLYLTYWFGHEERARATGYFLLGVCVANIVGGPLAGALLALDGA
LGLHGWQWLFIVEGVPAIALAFVVWARLPERPSDARWLAPELGRALERALASEQAQSAAA
HGAQAFGVALRDPQIWLAIFVYFCHQLTIYTMIFFLPGIIGESGRFSPIVVGLLAAMPWL
AAALGAATLPRFARDSSRSRRILCAGLGVMAAGLVCAAYAPPAAALASVCVAASMFFVVQ
SIVFTFPASRLAGSALAGGLGLVNTCGLLGGFVGPTVVGTIEQTTGHAKNGLALLAIALG
VAALASLRLRHGHERAHPARPSRAR*
>GCA_000452945.1_BPMSHR5858_genomic_01733_1 group_1440
VKNIRFACATAGVLAATAAHAQSSVTLYGLIDAGIMYTNNVASGNSHGGLVQATTGAVNG
SRFGVRGTEDLGGGLKALFVLENGFNAENGKLGQDGRLFGRFAYVGLSDNRFGTLTIGRQ
YDSLVDFVAPLSATAGTFGDASFAHPFDNDNLNHSLRINNAIKYTSNTYAGLKFGGMYAM
SNSTDFATNRAYSFGASYTRGPLNVAAGYLQINGSKGTTAGSPGAVDIVESAANGKGGFS
LGADRMRSYGGGLNYAFGPATLGFVYTRAEYENTASFGSTGGTVRFNNYELNGKYQLTPA
FSVGGAYTYTDGHVENTVKYGSDPKWHQVDLMAVYRLSVRTDVYLEGMYQHASGRNYVAM
INTAGGASSTGNQVVAAVGLRTRF*
>GCA_000452945.1_BPMSHR5858_genomic_01739_1 group_1171
MRTRDLGIRIGRGKPGRLNAITDVAGVRVGHHTVHVEAGDASAHTGVTVIEPRAARARDE
PCFAGVHVLNGNGDATGLEWIREAGLLTTPIAYTNTHSVGIVRDALVAAERAQGGARERE
HVYWCMPVVMETFDGLLNDIWGQHVCVGHVAQALAAARSGPVAEGCVGGGTGMICHEFKG
GIGTASRVVAEAAGGWTVGALVQANYGQRAALRVAGYPVGEVLRDAHSPFDEAGGAGEPG
MGSIVVTLATDAPLLPHQCTRLAQRASVGLARVGGGTDNSSGDIFVAFATGNTGLPIASY
GRPGPTTVGVRMVADAHISALFDAAAEAVEEAIVNALVAATDLAARGVRVEALGAARLVD
ALRETGWRPRAGDAQL*
>GCA_000452945.1_BPMSHR5858_genomic_01748_1 bauA_1
MRNIFIQSRIAAPTDMNAPTEPTAPDTTVRTDAEWLDAHWMPFTANRQFKRDPRIIVEGK
GAYYTAADGRQIFDGLSGLWCCGLGHGRAEISDAVARQVARLDYSPAFQFGHPQSFELAN
RIKAMTPAGLDYVFFTGSGSEAADTSLKLARAYWRAKGKGTKTRLIGREKGYHGVNFGGI
SVGGIGPNRKLFGQGIDADFLPHTQLPENRFSRGLPERGAELAERLLDLIALHDASNIAA
VIVEPFSGSAGVVVPPKGYLQRLREICTAHDILLIFDEVITGFGRTGAMTGAEAFGVVPD
IMNVAKQVTNGAQPLGAVIASKDVYDAFMAAGGPDYMLEFPHGYTYSAHPVACAAGVAAL
DLLVKEDALGRVRELAPYFEAAVHGLKGVRHVADIRNCGLAAGLTIEHAPGEPARRPYEI
AMHCWKNGFYVRYGGDTIQLAPPFIAEKREIDSLIGALNDALLATD*
>GCA_000452945.1_BPMSHR5858_genomic_01752_1 group_19194
MDTLPNQSDAQFQAFLAKLLEQPQPAVWGEKQQIELDMARELAVEMLQLAERMNAAHTPD
LENCLVLLRHAKVLDFILSSLAARRDIHPRTLRTIFKLAKLKVHDSYPE*
>GCA_000452945.1_BPMSHR5858_genomic_01754_1 group_1172
MPMRFHLVRRLSRAASFAIGPMASLLFVAHGAAGAEPPRAALLAAGCKTLAERVHATPGS
GPVFLGSYEPAPGGAALAGALTQAAFVYDNALASIALIACHRPGDARRIADAILQASRQD
RHYRDARVRNAYRAGALPPGPAPLPGWWDTPSKRWFEDAYQAGTATGNVAWAALSLLAVY
EATRERRYLDGAAALMGWVDPGRLDATAPAGYVGGEFGHEPQPLRQGWKSTEHNVDAYAV
FRWLAACTGDARWHAAAGRARRFVSAMWEPGDGRFLIGTRDDGHTPNTGPSALDASLWPL
LAMPDAPADWRRSLAWVERAHRIDGGYGFNAHPDGVWTEGTAQAALALQAAGRSDDARPL
WALLMSQRAPSGLLFATPEPSIRTGLSIGPTSKTDDFRYFHLPHLGATAWAVLAAAGWNP
FRPGGCLAAGYPGDAAPACGA*
>GCA_000452945.1_BPMSHR5858_genomic_01759_1 group_1595
MQATDSPDTTARIPYPRKPCVMTDDVQAPHRPPLAADEAARADEAAAEPASAPAEAPVEA
PVETSAAAPVRAPAAAAGAGELRALFSRCGWFNTLAPEHQALVVAQSHAEYRDAGDWVAR
RQAPSEYWIGVHRGLVKLAIYNASGRGCTFSGVPSGGWFGEGSVIKRELRKYDVIAIQRS
LVLFVPSATFHALLDSSLPFTGFVIHQLNNRMGEFIASIQNSRLLDVNARVAQSLAQLFN
PDLYPDTGATLAISQEELGMLVGVSRQRINQALQHLERLGAVRLAYNQIDVLDLPKLAAF
GMEQI*
>GCA_000452945.1_BPMSHR5858_genomic_01760_1 sasA_1
MMPTGRGHARPALSPPPSAPAHRAAAHTEPVSILIGRTQVALARERSAKRLRETFESNLE
AFERLRIIVNDMLFLARRDRGERATDLADVSLAAELTRMLAFLEMSLDDAQLRAELTGDA
RASVDPSLFGRAMTNLLINAIQHTRAENTIRATSTPRGETVEIAVSHPGEPIDAAARASV
RARLPDRAGAREQRRESRARPVDREGSRRDARRRRVRVVRGRLERDRLIGRRATGRHGPA
RARPAAGGGRAAAGVEDDLSGGASLHRCAGASAHQCIGGS*
>GCA_000452945.1_BPMSHR5858_genomic_01762_1 group_1657
MDYGVLLSNFERDTARDALARASQQHRLYACLRAAILNGTLEAGTYLMSSRALAETLRIA
RNTVLYAYERLAAEGFVVARRQGTMVARVGLPAASATASPAHARPALARRVTGLPDIDAD
DEREPLPFLPGMPALDQFPLAPWRRAVERAWRRIGPAQLGHAPLGGNLRLRQAIAEYLRV
SRGIGCDARQVFITDGTQHGLDLCARTLADAGDTVWIEHPGYAGARAAFEAADLRLVPIP
VDADGLAPSAEHWRAHPPRLVYITPSHQYPLGAVMSVERRVALVANARAAGAWIVEDDYD
SEFRHFGAPLAALQSLGDDAPVVYLGTFSKTMFPTLRIGFVVASAALAPQLRHTIGALAP
RGRLAEQLALADFIEAGHFTRHLRRMRRLYEERRDALQDALARHLGGALTVSGGAGGMHL
SARLDAPVADVDVARAALARAITVRPLSRFCLPGTDRAAYNGLVLGYGAVPIEQIDACVR
RLGAAIDDALREVTRAPRDAAR*
>GCA_000452945.1_BPMSHR5858_genomic_01763_1 fabG_1
MHIDLTGKTALVTASTAGIGLAIAEGLARAGAQLFINGRSDASVKAALAHLRVTAPGAHA
QGVVADLSDADGARRVIDAAPSADILVNNAGIYGPKPFFDIDDAEWEHFFQMNVMSGVRL
ARHYLKGMLERDWGRVVFISSESALNVPADMIHYGFTKTAQLSISRGLAKLAAGSRVTVN
AVLPGPTMSDGVKAMLKATADAEHTSIEQAGVDFVRTHRSSSIIQRPATCDEVANLVVYV
CSPQASATTGAALRVDGGVVDTIA*
>GCA_000452945.1_BPMSHR5858_genomic_01765_1 ydiK_1
MVQNASVPPASAGRPKAQSIALGALYAALVALALWVIRDFMPAIAWACVVAIALWPALRR
IDALPALRGRATLVAALLTSAVALLVVVPIAAGLVEAVDQSHELLGWLHRVEQTGIPEPD
VLSHLPFGSQQASAWWQANLAKPLHPATAVKGVDGASVVTFGRHFGSRLAHAAVLFGFML
VTLFVIFQAGPRLSGSLLTGVRRAFGDSGAALLQRMAAAVYGTVTGLVVVGFGEGVLLGL
AYAFAGLPHAALLGLVTAIAAMLPFCAPLVFGGAALWLFVQGALGWAIGLAAFGSVVVFV
AEHFVRPVLIGNSARLPFLLVLFGILGGAETFGLLGLFVGPALMTVLTVLWAEWVD*
>GCA_000452945.1_BPMSHR5858_genomic_01770_1 hchA
MKILVVLTSHDELGTTGKKTGFWLEELAAPYYALKDAGAELTLASPRGGLPPVDPKSDDP
ASQTDATRRFAKDAEAQAALASTRKLADVSIDDYDAVFYPGGHGPLWDLAEDRHSIALIE
RAIATNKPVAAVCHAPGVLRNVKDPSGASFVKGLNVTGFSNSEEAAVELTEVVPFLVEDM
LKTNGGNYSRAADWAPHVVVDGLLVTGQNPASSEPVAEALLNLLET*
>GCA_000452945.1_BPMSHR5858_genomic_01773_1 group_19350
MNTKRSRPELVAALLWAALYLASGYLSHALNGPVRLTGYIWLPAGVTVGAFMLRPVREWP
MLAAAFVAAQLALTGIEHGNPFNAALFAIDELGAAALAVGFVRRIRFSLEGLYFLRSVIL
AGVIASVLGALGGAAWYTVVNGASFVDVGLVWAASDFIGVLLVTPVLASWSRFRAHRSGD
HERFDLMLGIAAFALVAIGAFAIFDGDSASKFGIGAGFAMTYIPLFLTVAVTLLLGGRAG
SSSVLVLALIVIMQTAQGEGPFASLDANHGRSLLEAQLYLAVASLLVLTVSTLKTTRERV
HEHAQVLRNNMELALASAGQIAYVLDPSSGRIDWSGDVERVFGVGVDAAQIASVPLVLER
VHADDRDALRDYWRAEIAGEDRASLSLRVVQRDGGTRTITDHGAPLLDSNVDVAVVAGVW
QIERVWPADE*
>GCA_000452945.1_BPMSHR5858_genomic_01777_1 egtB_1
MTKNEESARGLASELARGFTDVRRHSVELARPLSAEDQALQSMPDASPTKWHLAHTTWFF
ETVILARHARGYRLFDSRYPYLFNSYYEALGPRHARPQRGMLSRPSQDDVRRYRRHVDDA
LLELLRTADLPTLIAIEPEITLGLHHEQQHQELILTDILHAFSLNPLLPAYRSDDAVPTA
DARRANGAMRWLPGPIGVVEIGHDGRGFSFDNERPRHRTMLHPYEIAERLVTNGEYAAFI
DDGGYARPEFWLSDGWAIVQREGWKAPLYWIASDGGEGLGWREFGLGGLQPLTRDAPVSH
VSFYEAAAYAEWARARLPTEAEWEAAFDAPGIAQMTGCVWQWTRSSYEPYPGFRPMAGVA
AEYNGKFMVGQQVLRGGSVATPPGHARATYRNFFPPAARWQFTGVRLARDL*
>GCA_000452945.1_BPMSHR5858_genomic_01782_1 group_19544
MKKRTAVAMTAAGLAAVATAHAQSSVTLYGIVDNGIAYQSSSTSLGSTTGGRSAVKMSTG
VWAGSRFGLKGSEDLGGGSKAIFQLESGFSTANGTSQFAGGIFTRQAWVGLTNPTYGTLT
AGRQYTAYYTLLSPYSPTTWLTGYFGAHPGDIDSLDTSYRTNNSLVYLSPKFYGFTFGGS
YAFGGQPGSVNAGSTWSAGIQYMNGPLGIAAAFQRVNNSTSGGGDWGANSTTSNGGAQTA
VSAINNGYKTAQAQQRVAVTAGYQFSSAWDISVSYSNVQYIPGVNSAFRNTAIFNTAGAV
LHFKPSAQWDFAGGYAYTRATQSNGITSAAQYHQFTLSQYYSLSKRTGLYAVEAYQRANG
KTLAGGKIIDATASIGDGFNTSPSSSRSQVGVGVGLIHRF*
>GCA_000452945.1_BPMSHR5858_genomic_01785_1 tctD_1
MRVLLVEDNPILAQSLADALSSARFAVDHMADGEAADHVLHTQDYALVILDLGLPKLDGL
DVLRRLRARRNPVPVLILTAHGSVEDRVRGLDLGADDYLAKPFELTELEARARALIRRSL
GHEHGRVECGPLAFDSVDRSFRLNNDALALTPRERSVLEVLILRNGRAINKETLSEKIFG
LDESVNADAIEIYVHRLRKKLEGSGVAIVTLRGLGYLLEAQPS*
>GCA_000452945.1_BPMSHR5858_genomic_01789_1 bdhA_2
VTGAASGIGKEIALELARAGAAVAIADLNQDGANAVAELIEQAGGRAIGVAMDVTNEDAV
NAGIDKVAQTFGSVDILVSNAGIQIVNPIENYAFADWKKMQAIHVDGAFLTTKAALKHMY
KDDRGGVVIYMGSVHSHEASPLKSAYVTAKHGLLGLARVLAKEGAKHNVRSHVVCPGFVR
TPLVDKQIPEQAKELGISEEEVVKKVMLGQTVDGVFTTVEDVAKTVLFLSAFPSAALTGQ
SFVVSHGWFMQ*
>GCA_000452945.1_BPMSHR5858_genomic_01796_1 group_18958
MSDLLDQYPMLIFALESLLALALLVFIVVWTASGKKKQPSRHDKPQR*
>GCA_000452945.1_BPMSHR5858_genomic_01801_1 trpG
MLLMIDNYDSFTYNLVQYFGELGEAVRTYRNDEITLDEIARLNPDAICLSPGPSNPQHAG
ITLSVLREFAGKKPILGVCLGHQAIGEAFGGRVVRAKTIMHGKVSRIETDGRGVFADLPK
HFDVTRYHSLAIERESLPDCLEISAWTDDGEIMGVRHKTLPIEGVQFHPESILSEHGHAL
LENFLKEARQAAAHSA*
>GCA_000452945.1_BPMSHR5858_genomic_01802_1 trpD
MTITPQEALQRTIEHREIFHDEMLHLMRLIMRGDMSPVMAAAIITGLRVKKETIGEIAAA
ATVMREFARRVEVEDNANFVDIVGTGGDGSHTFNISTATMFVAAAAGAKVAKHGNRGVSS
KSGSADVLEALGVNIDLQPEQVAASIAETGMGFMFAPNHHPAMRNIAPVRRELGVRTIFN
ILGPLTNPADAPNQLMGVFHPDLVGIQVRVMQRLGAQHVLVVYGKDGMDEVSLGAATLVG
ELRDGEVREYEIHPENFGMQMVSNRTLKVESADESRVMLLEALGNKPGVAREIVTLNAGT
ALYSADVAGSIADGIQLARDAIASGRAREKVDELVRFTQQFKR*
>GCA_000452945.1_BPMSHR5858_genomic_01803_1 trpC
MSDILDKIIAVKREEIAAALESAPLEELKVQASARDSRDFVGALRDKHAAGHAAVIAEVK

KASPSKGVLREHFVPADIARSYAQHGAACLSVLTDERFFQGSARYLEQARAACALPVLRK
DFIVDAYQVLEARAMGADAILLIAAALDTPLMIDLEAYAHSLGLAVLVEVHNRGELDEAL
KLKTPLVGINNRNLRTFETTIDTTLGMLDAIPDDRIVVTESGILSRADVERMEAAGVHTF
LVGEAFMRAENPGAELARMFF*
>GCA_000452945.1_BPMSHR5858_genomic_01809_1 ubiI_2
MTAYHQTFDVAVVGGGLVGKTAALALTQAGHKTALLAQPAVPRPADLAFDSRIYALSASS
QALLERLRVWQALDHGRLAPVYDMRVYGDAHAELHFSAFQSAVSQLAWIGESSLIECALD
AALRFQPNLTWFDARAQGLDVTPGAATLALANGDVIECDLVVGADGAHSWVRSQIGSKVE
RRDYRQTGVVANFKAAQPHRETAYQWFRDGEIVALLPLPDGHVSLVWSAHTAHADELLAL
DPARLAAEVERVTLNRLGALECVTPAQGFPLALQTVDRLVAPRVALVGDAAHLIHPLAGQ
GMNLGLRDVAALADVLANREAFRDLGDTVLLRRYERARREDIRALMVATDGLQRLFSLPG
ALARIVRNAGMALVGAQPLVKRWLVASALG*
>GCA_000452945.1_BPMSHR5858_genomic_01813_1 group_1000
VRGGLPASEVSAPLAACDSNRSSMYFCAWRDRLVVERERQRGVDRQASAHPAISTARYRP
FHSRSLRARLHALALRYGMSAGGTAALPEYRSS*
>GCA_000452945.1_BPMSHR5858_genomic_01824_1 group_19679
MNWFGIVVLGFAVGLLGWALNPLRRLSRASLWLAAAAGVLGAAAAKMAGNVAGLFYDGET
LEWPVCTAAALLAVAVTVGLSARR*
>GCA_000452945.1_BPMSHR5858_genomic_01827_1 prfA
MKTSMQSKLDQLTTRLAELNDLLSRENVTADLDQYRKLTREHAEIGPVVEHYAQWRQARA
DELAAQELLADASMRDFAEDELRGARDRMGRLAAELQTMLLPKDPNDERNIFVEIRAGTG
GDESALFAGDLLRMYLRYAERQRWQVEMMSESPSDLGGYKEVIVRIAGYGAYSRLKFESG
GHRVQRVPATETQGRIHTSACTVAVMPEADEIGEVEINPADLRIDTFRASGAGGQHINKT
DSAVRVTHIPTGIVVECQDDRSQHKNKDRALKVLAARIKDKQYHEQHAKEAATRKSLIGS
GDRSERIRTYNFPQGRMTDHRINLTLYKLEQIMDGDLDELIAALVSEHQAELLASLGDAE
*
>GCA_000452945.1_BPMSHR5858_genomic_01829_1 grxD
MDTQQRIKQIVDENPVVLFMKGTAQFPMCGFSGRAVQVLKACGVDQFKTVNVLEDEEIRQ
GIKEFSNWPTIPQLYVKGEFVGGSDIMMEMYQSGELQQLFTA*
>GCA_000452945.1_BPMSHR5858_genomic_01831_1 ygiD
MNRLPSLYLSHGAPTLPIDPTLPSGAFTELGAELPQPRAVLMLSAHWLTQQPVVSTAERP
DTIHDFYGFPRALYEIRYPAPGAPDVAAHAAALLRSAGIETAATPHGLDHGAWVPMLLMF
PHADVPVAQLSIQPRADAAHHFRVGRALRPLRDEGVMVIGSGQITHNLRAADFSAPPDDV
DPRVAEFTGWFEEALAARDVDALVDYRRRAPHAALMHPTDEHLLPVFAALGAADDDYRLR
IQSLGTYQRMLAMTNYVFDSAA*
>GCA_000452945.1_BPMSHR5858_genomic_01832_1 yveA_1
VKSSIQRNIGPFALMLTGLGSIIGSGWLFGAWKAAKIAGPAAICAWIIGAVVILAIALTY
AELGAMFPESGGMVRYARYSHGSLVGFISAWANWIAIVSVIPIEAEASIQYMSTWPYAWA
HALFVNGELTTPGLLLSAILVVIYFLLNYWGVKLFARANTAITIFKFLIPGLTIAGLLLS
SFHSENLGTAGGSSGFAPYGWSAVLTAVATSGIVFAFNGFQSPVNLAGEARNPSRSVPFA
VISSILIALVIYVLLQVAYIGAVNPADVAKGWAHFNFASPFAELAIALNLNWLAILLYVD
AFVSPSGTGTTYMATTTRMIYAMERNNTMPKIFGSVHPLYGVPRPAMWFNLMVSFVFLFF
FRGWSSLAAVISVATVISYLTGPISLMALRRAATDLERPLAIPCMKLIAPFAFVCASLIL
YWAKWPLTGEIILLMIVALPVYFYYQGKAGWSGWGRDLKAAWWLVAYLPTMAVLSLIGSK
EFGGHGVLPYGWDMLVVAAIALVFYYWGVHTGYRTEYLDERESRDEILEGIGA*
>GCA_000452945.1_BPMSHR5858_genomic_01833_1 cspD
MATGTVKWFNDAKGFGFITPEGGGEDLFAHFSEIRVDGFKTLQENQKVEFEVKTGPKGLQ
AANIKPL*
>GCA_000452945.1_BPMSHR5858_genomic_01834_1 hscA_3
MTYCAIDFGTSNSAVALPRGGGTTGMRLAPVEGEHLTLPTAIFFNTDESTREYGRAALAS
YIDGFDGRLMRSMKSILGSPLAETTTDLGDGSAIAYTDVIALFLMHLKQKAEACAGGAIG
CAVLGRPVFFVDDDPRADRLAQQQLEAAAHAVGLADVQFQYEPIAAAFDYESRQDAERLV
LVADIGGGTSDFSLVRVGPERMRRLERKDDVLAHHGVHVAGTDYDRRVELSAVLSAFGYR
ALDPEGRELPNRIYFDLATWHLINTVYTPKRVGELKLMKHLYADARHFDRLLRVVEQRLG
HALAARAEEAKIGVSAGGETMIDLNDVEEDLLIAFDADQLIDASRDETARIVDAARETVR
LAGIAPRDVGALYFTGGSTGLAFLSGALAAAFPDAQPVYGDRLASVATGLGIHARRVYGG
*
>GCA_000452945.1_BPMSHR5858_genomic_01875_1 group_19330
METYLDLLAEREALMIKLDAARMQARSAALSEVKRIVEEFELTTQEVFGRAGVLRRRGKL
KPKYRDPKTGATWNGRGRPPSWIAGKDRAAFVA*
>GCA_000452945.1_BPMSHR5858_genomic_01878_1 benM_2
MESKHLRYFLAIADTGSLTHAAIRLGVAQPALSHALTKMERELGVKLFDRSRRGATLTTA
GEAIVDDIRLSLARMDAAARRASEIGAQRAGRLSIGFVSAALFDTLPSAVCALRAASPRV
DLVMREMSNAEQVVALERGEIDIGILHTPVAIGGPVHEKLIRRDPLIAVIPKSFPRQPDG
SITLRELSRAGLIWFPHEQLPSIRAAILSAFRLAGHPIEVVQDANRTLTVIACVAAGCGV
SLLPSSIRTLTFEGVDFSPVSDGHALPEFPLTAIWPKRSRRTLADRFAELLPRFQPN*
>GCA_000452945.1_BPMSHR5858_genomic_01885_1 group_1666
MKSDGFCGGKFCRRILVCLCAAALAFLTLPQLSNRMQESMLIATEPCKWTHVYPVLWGSH
VLVALLSYGWAHRDLKRLQAALRASRGSLATASHELRAPVNGMLALVELVRQGLLAPEQQ
RLLGLVQDSGQSLARVLNDILDHASIEAGQFAIEKTRVDVRELVDSVSAMLAVHAHRKGL
RLRLHVDANVPAAVLTDGNRLRQILVNLIGNGIKYTERGGVSVRVRAADDVDGMVELVAT
IVDTGIGIAADELRRLFTPFARATRGARRMESGTGLGLAIARKLARLLGGDVTLTSDEGR
GTTATARIRCAVAARGYANDALAGRVVSIDVGDAALAATLAAYARAAGVRVAARGAPDVE
AVLCDDARVDASRTQLGWVGVTDAPALAGWTDSGTHVRLSSNPLGWRAFIDGLRAALDAR
DARDAAGAHTAARAEPGAYRVLVVDDQEMNRIVLRYQLDALGHHARLCASGDEALRALGT
AAYDVVLTDCRMPGMDGIALTAAIRAHPDARVRATPIVGVTALVSDAEHARCVDAGMTLC
IGKPTTLDALERALVEAIEPVPALRFDPSRIDAARLFAGFGASSAPGAARDAILRACGRA
LERDRDTLTRHLRDGLSERELRVWCHTARGTYSLFAQAHVDALLDEFHHLLTAGDARAIR
GATRDILAMVNHLLRCVAHSAQSRDRGDPKGGR*
>GCA_000452945.1_BPMSHR5858_genomic_01892_1 mauG_1
MRRLPRYARQHRSFFVAPRAFAAAAALAAGVAACDANGPGASAAAAVAPAALAVPAASAA
SAASAARPAPLAQPAAPAVVDSQPQTRAQVYEAVKQMTALGRQLFFDPSLSGSGKLACAS
CHSPQHAFGPPNALPAQFGGDDLRQQGFRAVPTLKYLQKVPAFSEHYHESDDEGDESVDA
GPTGGLTWDGRVDSGAEQARAPLTSPFEMNGTPEKVARAVRAAPYAPAFRAAFGARVLDD
DRATFEAVLQALGTFEQAPDVFYPYTSKYDAYLAGRARLTRAELHGLQVFNDEKKGNCAS
CHVSRRGLDGSPPQFSDFGLIALGVPRNRALAVNRNPNFYDLGACGPERRDLKGRDEFCG
LFRTPTLRNVALKKTFFHNGVYHSLDDVLRFYAERDTHPEKFYPVKRGVVQKFDDLPKRY
WKNLNDEPPFERKRGDPPAMTDAEIRDVIAFLGTLTDGYDPRAKPAGGAR*
>GCA_000452945.1_BPMSHR5858_genomic_01896_1 rppH_1
MSAMKPPGRTRARTVSCGVVLLDSDGRVLLAHATDTTHWDIPKGQGEPGETAQQAALREL
AEETGIVLDPARLVDLGLFAYRRDKDLHLFAARAAAGETDLSRCTCTSMFPSRRDGTMIP
EMDAFRWTAPADVDAYASRSLARLFGTTLSLAALHRTLAG*
>GCA_000452945.1_BPMSHR5858_genomic_01919_1 murR_1
MDDTTPPETPAVDDLLQRIAGAYDTLPRQLKRIASYIDEHRSSVMMDRTSDIAQRCGVHP
SAVVRFAQRFGFSGFSDLQAVFKDAYTGQNPTVQSYQQRIRSLIDGEAGPMTGGAVARQF
IDASRAGLDELAAGLDDDQFDAAVTMLERAENIYVVGVRRSFPVASYIVYALQHTAKRVH
LVSGLGGMYREQIRSVKKGDVVIAISFAPYGKETQYCLRAARHNHAQTLVVTDSRLSPLA
RDASAHLLVKEGSAFAFRSLTSTICLCQALFIALAYRLELNVEEVKDIGGYDD*
>GCA_000452945.1_BPMSHR5858_genomic_01921_1 iolX
MTVSHSAAAVRIGIAGLGRLGRRHAENLARRVAGARLAAACSPLADECAWARETLAVPRV
YEDFDALAADPELDALWLVTPSALHADQIVAALRAGKHVFCEKPLSLDLGECERVLAEAR
ARPHLVATIGFMRRFDPSYRDAYARVAAGEIGRPFLVRSQTCDRNDPEGFFVRFAPSSGG
IFLDCTVHDIDVARWLLGAPRATRVYAAGAVALHEGLRACGDVDNGVAICEFEGGRLAMF
YASRTMAHGNDTHSEVIGTAGALAIGRNPRANRVEIYDAAGIRNTCTPTFFDRFEDAFLI
EAQAFVAAVRGEGAAGGATLADALEATRIGHALRESLQTGRAIEL*
>GCA_000452945.1_BPMSHR5858_genomic_01925_1 group_1827
VNARGFVTKYVLTACCAAATALAAQHAAAQSAKNRSVATVDYTTRSGDTLYDVSARYLQG
TDDWPLVAQLNDVPVPKHLQPGVVLKLPAARLRKERLSARVIAAHGTVESAGRGSAQFAP
VAVDATLTEGDRLRTGSNAFVTLELSDGTHLSLPPDSQIDLATLRRTVLTGTLERVIDLR
RGSVDSEVTHLKKKDDRFQIRSPSVVAGVRGTRFRVNYDKDGRASTTVEVLDGTVGVAPS
AKRSADTLVHANFGNVTSASGVVGSPIALLDAPQLANPAKIQDDPQVAFDLVPLGGAQSY
HVQIARDAGLYDLFKEVQVPAPRATFADVPDGTYFVRIAAIDSHGLEGQPRIYAFERRRF
GVDASAAPADGGYAFRWSTTQDGAAAGATRFRFVLSRSKDLSNPIVDQVDAQGGRIAVSN
LTPGDYYWSVIAERYEGGRFHEKASAVNAFTIAR*
>GCA_000452945.1_BPMSHR5858_genomic_01933_1 tap_1
MKGVSLHGKHFLPDDATGNRGRTPKPGKPRRARRAWSVKTTLRAAFALLLVGTLAVGLFS
LAQISRLNSSIRSVYEQGHVASRAAEEVRASVLRASRAQKMLITATTAKERDELGADIDK
GLAAIGAELATLQRHADGAADDAARLKAFGAAVGAWSAHLRDFVRLVREQPLDLSQMSWQ
VGSQDVSLLVETGKLEKLVDGLVDARGDASKATLDASGAIFRESFAMLAAMTAALVVLAF
VIAAWVVRRLGAQLGGEPAYAKEIAVSISRGDLSNVIRLDTRDHDSMLHALRDMQEGLAG
TVREISASAEAIASAAGEIAMGNLDLSQRTEQQAVALERTATSMGQLTSTVHQNAENARQ
ASTLAANASSVAEAGGTVVGRVVSTMNEIDESAKSIRDIIGVIESIAFQTNILALNAAVE
AARAGEEGRGFSVVASEVRSLAQRSASAAKEIKALIGASVERVANGAVLAQDAGRTMDEV
VRAVKRVTDIMGEISAASSEQSAGIDAIERAVTQMDAGTQQNAALVEEAAAAARSLDEQA
QMLKEMVGRFHLPAHAAG*
>GCA_000452945.1_BPMSHR5858_genomic_01935_1 aaeA
MFLRKLFGFVATAVILIVAILIGRTLWVHYMDEPWTRDGRVRAEIVNVAPDVSGAVVELP
VRDNQFVKQGDLVMQIDPSHFRIAVEQAQAAVAARRAELQMRRADAARRADLDALVVSKE
SRENSAQTASSAEAQYQQALAALDAAKLNLERTRVVAPVDGYVTNLQVFKGDYASAGQAK
LAIVDSHSFWVYGYFEETKLPRVKIGAKAEMRLMSGGVLKGHVESISRGIYDRDNPQSRD
LVADVNPTFNWVRLAQRVPVRIHIDEVPKDLVLSAGTTCTVIVEPDGGGKRKS*
>GCA_000452945.1_BPMSHR5858_genomic_01944_1 group_19273
MKDILVLYYSRHGATRELALAIAAGIDSVPGMQARIRTVPPVSTVCEATQPDIPDDGPPY
AEPRDLDECAGLALGSPTRFGNMAAPLKYFLDGTTPQWLSGALAGKPACVFTSTGSLHGG
QESTLLSMMLPLLHHGMLIVGIPYTETALSVTQTGGTPYGASHFARVADPARERISADEK
TLAHALGARLARTASQLGAAAA*
>GCA_000452945.1_BPMSHR5858_genomic_01945_1 group_19674
LPKLSVDLDTIKRLARFAAKRSAEDRIPQVAGSLTFTTMLALVPLVTVAFALFTAFPMFA
SFQNSLQGFLADHLMPAQFNSQIFKYLNQFASKAKGLTTAGLIVLVFTAVMMMMTIESAF
NLIWRVRKPRPFAQRVLAYWALITLGPLLFGVSLSMSSYLFTKSLAFAGSTSTPPVFEWL
LTAASLPLTVLAFTLLYVYLPNCTVAWRDAVVGGVCAALAFELAKRGFGYYVRRIPTYTA
VYGAFATVPMFLLWLYLSWFVTLAGAMIASALPAIRVGQFHRIHYPGSDLLDSLELLARL
ADAQQAGKPGHTAARLALMARCDMETAQRLLSTLEEREWVARLQSGNDAAPRYVMLANPE
RLTVANLFDVLVIDRSELVYQAQRGRTEVDCATLVDAFTSEKLDLSLAELIACRRQGASR
DAPGAAHSTSALPPRTA*
>GCA_000452945.1_BPMSHR5858_genomic_01955_1 group_19578
MKQRNLILAMACAAPFISACSGGGGGSESQPLVEAALCPSSVDYNTVYTGGGGDGELVKL
QIDTTKMTWQITFVESPVPKTTGTAAPTRAGTTQSGTLTQETLLPTQKLNNCAFRLNGAS
LDPNRPARIFLGMGIVGGTIPGAEIQYGGLLGQGAIPDTTFPYYPFIAFSSLETDVTKLA
GTYSQLGYGQVGSQNFAPTTIDAKVTINPDGTWVRCDTTGIYAGTCRQPGTNIVQSADGT
GAFETDHYQGQVKPTLATIPQAKGYIIVGKLRNQLVPIMIRTGVANPNPTPDANGVPGLT
ADDESGISILAPQTAIAVGSQNGEYIGVDSQFDYRTTALVNAQATLLDPFQASQASLATP
LNLDYTQAVPGTVTSTHVGASSTTPTGKFIFTGGVFGFLDMSNSSSPYFTVGAFVQ*
>GCA_000452945.1_BPMSHR5858_genomic_01960_1 pyrE
MTGFDRQTISDTTAKMLLEVQAVHFNADKPFIFTSGWASPVYIDCRKLISFPRVRRGLME
MAEATILRDVGFEQIDAVAGGETAGIPFAAWIADRMMLPMQYVRKKPKGFGRNAQIEGYL
EEGSRVLLVEDLTTDSRSKINFVNALRTAGAKVNHCFVLFHYDIFKESVSVLKDIDVDLH
ALATWWDVLRVAKASGYFETKTLDEVEKFLHAPAEWSAAHGGATGAQE*
>GCA_000452945.1_BPMSHR5858_genomic_01961_1 maeB_2
MDEQLKQSALAYHQNPKPGKISVTPTKPLSNQLDLSLAYSPGVAAACEAIHADPLDAQKY
TSRGNLVGVVTNGTAVLGLGNIGPLAAKPVMEGKGCLFKKFAGIDVFDIELAESDPDKLV
EAIAMLEPTLGGINLEDIKAPECFYIEQKLRERMKIPVFHDDQHGTAIIASAAILNGLKV
VGKDLSSVKLVCSGAGAAAIACLDLLVKLGLSKSNVLVADSKGVIYEGRGNLDPSKQRYA
ATTGARTLADAIAGADVFLGCSSAGVLKQDMVKTMADRPLILALANPEPEIRPEDAKAVR
PDAIVATGRSDYPNQVNNVLCFPFIFRGALDVGATTITEEMKLACVRAIAELAQETDQSE
EVAKAYEGHSLEFGPEYLIPKPFDPRLIIKIAPAVAQAAMDSGVATRPIADMDAYREQLG
TTVYRTGMVMRPVFATAKTKAARIVFAEGEDERVLRAAQFVLQERIAKPIIVGRPSVVEM
RLKKIGSKLVGGVDFEIVNPEDDPRYQQCWQAYHEIGARDGVTPEVAKAALRKFNTLIGA
MLVHLGDAHGMICGMIDTYHSHLKFIEQVLGRAKGAEHFAAMNLLMLPGRNLFMCDTYVN
EQPSAEQLADMTIQAAAEIERFGITPKAALLSNSNFGSAPSASSRRMAEARKLIVERAPE
LEVDGEMHGDAALSEVVRKAAFPGTTLTGEANLLIMPNVEAANIAYNLLKMVGGEGVTVG
PFLLGAAKPVHILTPAATVRRIINMTAVAAANVQVK*
>GCA_000452945.1_BPMSHR5858_genomic_01964_1 hpd
MQIPTWDNPVGTDGFEFIEYTAPDPKALGQLFERMGFTAVARHRHKDVTLYRQGDINFII
NAEPDSFAQRFARLHGPSICAIAFRVQDAAKAYRHALELGAWGFDNKTGPMELNIPAIKG
IGDSLIYFVDRWRGKNGAKPGAIGDISIYDVDFEPIPGADPNPAGHGLTYIDHLTHNVHR
GRMQEWAEFYERLFNFREVRYFDIEGKVTGVKSKAMTSPCGKIRIPINEEGSDTAGQIQE
YLDAYRGEGIQHIALGAADIYRAVDGLRAKGVTLLDTIDTYYELVDRRVPNHGEPLDELR
KRKILIDGAHDDLLLQIFTENQIGPIFFEIIQRKGNQGFGEGNFKALFESIELDQIRRGV
VQDKA*
>GCA_000452945.1_BPMSHR5858_genomic_01966_1 group_19278
MNKPDLRPAESVGIAGPAPVLVRELASGDRQQLLTHFLALGEEDRLLRFGQAVPDHVIEN
YVRTMDFGRDTVFGVFDHALELIGVGHLAYLPAEGDKRTAEFGVSVLERARGQGVGTKLF
ERAAIRSRNTHVTTLYMHCLSRNATMMHIAKKSGMRIEYAYGEADAYLSLPPADHSTIIA
EMMQEQAAVFDYALKRQARRATQIFESLLPAGLTA*
>GCA_000452945.1_BPMSHR5858_genomic_01967_1 group_19241
MARTRAPDHESQREQILDLAAAKFAQTSYPSTSMTDLANASGTSKARLYHYYEGKEAILF
DLLDRYTKRLMLIIAEVEGASQRRGLTEREAFAELVRAFLAEYETSHSRHVALLNDVKYL
EDTQRQIVLDRQRDIVAAFARQLARAYPERISKDNQTPVTMMVFGMINWTFTWLKPGGRL
GYRDFAEQVIGMIEHGLGG*
>GCA_000452945.1_BPMSHR5858_genomic_01979_1 fadH_2
VLDPKGAVGAAEKSLIATSTWAMLIVVVPVILLTLLFAWRYRASNRNATYAPKWAHSTAI
EIVIWTVPSLIILFLGILTWKTTHALDPYRPLESSVKPIDVEVVALDWKWLFVYPDLGIA
SVNQLAVPVGTPINFRITSDSVMNSFFIPQLGTQVYAMAGMQTRLHLIADEPGDFAGISA
NYSGRGFSDMKFRTLATSRDAFDAWVAKVRASADRLDMTAYGQLAQPSEKQPVRYYSTVD
PRLFNNIIAKYNDGRVLDLTDAACRTKR*
>GCA_000452945.1_BPMSHR5858_genomic_01984_1 catA2
MNKEAIDALLKSFDDAATRPGNPRVRAIVNRIVKDLFYTIEDFDVQPSEFWTALNYLNDA
GKEFGLIAAGLGFERFLDVRMDEAEAKAGLQGGTPRTIEGPLYVAGAPESVGHARLDDGT
DPGETLVMRGRVLGGNGAPLANALVEVWHANHLGNYSYFDPSQPAFNLRRSIRTDAEGRY
SFRSVLPVGYSVPPGSKTEQLLDQLGRHGHRPAHIHFFVSAPGHRKLTTQINIEGDPHIW
DDFAFATREGLIPKIAQAEGAQGKPYGIDGRFALIDFDFTLTRERGDVPASEVERVRAQA
*
>GCA_000452945.1_BPMSHR5858_genomic_01985_1 catB
MIATGITIDRIDTLLVDVPTVRPHKLSVATMNCQTLVLVRVRCSDGIEGVGEGTTIGGLA
YGEESPESIKTNIDAYFAPMLRGADASRPGAAMARVRKLLQGNRFAKCALETALFDAHAR
RLGVPLSELLGGRTTDALDVAWTLASGDTARDIAEAEAMLEARRHRAFKLKIGARAVADD
VAHVVAIKRALGERGDVRVDVNQAWTESEAVWAGARLADAGVSLVEQPIAAANRAGLKRL
TALAHIPIMADEALHGPVDAFALARERAADVFAVKIAQSGGLQGAAAVAAIAAAAGIELY
GGTMLEGAAGTIASAQLFSTFGTLEWGTELFGPLLLTEEILVEPLRYEDFKLHLPSAPGL
GIAFDWARIERMQRRAR*
>GCA_000452945.1_BPMSHR5858_genomic_01986_1 benM_5
MELRQLRYFIAVAEEMNITRAAQRLHMTQPPLSRQLQLIEDEIGLPLFERGARPLKLTDA
GRVLYAQARRVLEQADELAPLTRRLAQAAERIVIGFVPSTLYGALPEVIRAFREAAPAVE
LSLIEMFTLEQLGALKGGRIDIGFGRLRFDDDRLVREVLVEERLIAALPDGHPLAAPDAS
ISLADIAGQTLIVYPSTPRPSFADQQLSALRDGGLAPAAVHEVRELQTALGLVAAQVGVS
LVPESVEGVRVKGVAYRRLPEPVATSPIILSRRLHDESRATALFCSLARELMAGH*
>GCA_000452945.1_BPMSHR5858_genomic_01987_1 rhaR_3
MPPTPFDPLALREHRLFESRDLDETRERISRVMQPHALLPDGSRHGPSHMDYVRLGGLGI
GTIAFGDAMRVHLDAVDGYHLLMFCLTGSAQVRTMGRAFDVDAHTGVLCAPGEPFDAHLS
RDCEQFVLRIDAATLAAHAGDAAAALDPVIGIDDSALSAWMQQLQLVARSPELLASASAN
PRVATRLEQLLLDLLIDGHPPAAPPARRADPAPGFVRRAQEFIGAQLAQPLQLADIAQAA
GVPERTLRDGFLQFRGTSPMQYLRQRRLERARELLRTAAPERRIAEIALDCGFAHFGRFA
IAYRERFGELPSATLADRRDA*
>GCA_000452945.1_BPMSHR5858_genomic_01990_1 andAb
MSNETAAAWHTLGALDEFSEDEPAARVVGNKPIAVFRVGDDVFALHDLCTHGHARLSEGF
VENGCVECPLHQGLFDLRTGAPKCAPVTQPVRAYPIRIVDGQVEIHVDG*
>GCA_000452945.1_BPMSHR5858_genomic_01991_1 andAa

MSTADPYLIAGGGHAARRAAETLRERDPAARIVMIGAEPELPYDRPVLSKEALVGGDAGE
RRAFVRDAAWYRERDIELRLGVRVEAIERGARRVRLSDGARLGYARLLIATGSRVRRFAG
PVDAGVHVHYVRTLADTRALRAALAPGKRVAVLGGGFIGLEVAASAVRLGCRATVVDPAP
RLLQRGMPEAVGAFALALHARHGVDVRLGALPERIRRAANGAAVVETSAGGIVADVVVAG
IGVVPNVELAQAAGLDVDDGVCVDEMCRTADPAIFAAGEVTRHFNPLVGRALRIESWQIA
ENQPAVAAANMLGAAETYAQWPWLWSDQYDSNLQTLGLFGGERTLVLRGSPERDRAFCVF
ALNARRELAAVAAVNAGREIAVCRRLMTAGVALDPARLADPSDALRAMLPRGV*
>GCA_000452945.1_BPMSHR5858_genomic_01992_1 group_19139
MAYDQSNIFAKILRGEVPCIRLCETDTTLAFMDIMPQSRGHALVVPKEAAETFYELSEAA
AAEAMKMTKRVALALRRTLEPEGLFIGQFNGAAAGQTVPHVHFHVIPRWADEPLKMHARE
MADAAELEALAARIRTAL*
>GCA_000452945.1_BPMSHR5858_genomic_01993_1 clsC
VPDRPALRGLAPRHPPRIVKAAARALVLCAALALSGCATHPPATTLERTVSHALPPDAST
PLADALAVQARAHPGESGFVVLPRGDEALQMRIAVARAATKTLDIQYYIAAEDTTGKLLL
GAALYAADRGVRVRMLVDALNFKDIDKLMAALDAHANLEVRVFNPFGAPRLGMFARTANV
FTRIDNFTRRMHNKAMISDNQIAIVGGRNLGDEYFNASPTLQFRDLDVLAAGPVTRAVSA
SFDAYWSSALTYPLPALNRRRYDAKDLDAARDALRAHWRANATPYNAKPLNATPLAAQIA
RNELGLVWASAEFTADSPEKIAAPDDSYKSPPMQRLFALTRDAQREFLVLSPYFVPHDAG
VNALGRLTARGVRVAILTNSLAATDAIAVQAGYAPYRVPMLERGVELYEYKPDPGRSRIG
MLGSRSRASLHAKAYVIDRKILVIGSMNLDPRSAHLNTELALVIHSPRLANEIANLFDEV
TKPTISYRVTLAPDTPGAAQTTGAGAPAWPLVWTEIADGQVRTYSVDPNAGFYRNLLTGL
CLLLPIDDQL*
>GCA_000452945.1_BPMSHR5858_genomic_02003_1 group_19247
MLFLLFHLDGERYALDAAEIAEVLPLAATKTIPGAPAWAAGILMYRGEPVPVIDVPRLAL
GRRAQPLRSTRLVLVRYRLPAAAEPAFVPPRVRLLGLIVERATQTARIAREAFRDSGFAT
PHARWLGPIASDADGVVQWVALRHIVDGEARALLFDYAASLDAHGAPAGGGR*
>GCA_000452945.1_BPMSHR5858_genomic_02006_1 rcsC_10
MSTDDDFGRASLLELFREETHTQTQALSERLLALERGAQDAATLEACMRAAHSLKGAARI
VGVPQGVDIAGRMEDCFVAAQHGRQPLTPCHVDALLTGVDLLVRVGDPQTAASVAPHEID
AFAAALAAADAGVDPGADARGANAYESDGNAHERGGAARASVAAAPGEAADASRESGARE
AAGPNRAGAAASAADVRASADAGDARGASDTRDVRHTAGARGANRSTEVFGAGEATGIGA
STHANDANGMNATNLATSAAHPPRGAFAAAPDAPPSSAPSDAPRAPSQMRRVRADTLNRL
LSLSGESLVESRWLKPFAESMLRVKRAQRDAARSLDSAVEALADDADPRVRGALNEARQM
FTDLQRTFAARLDELDRFERRSSHIAEQLYDEALQCRMRPFGDATRAYPRVVRDLARSLG
KRVRFSIIGEATQVDRDILDMLDAPLGHLLRNALDHGVEPPEVRLARGKPAEATITLEAR
HSAGSLLVSVSDDGPGADLAAVRAAIVRHHLTDEDTAARLSDQELLEFLLLPGFSMRERV
TDVSGRGVGLDAVQEMVKSVRGAVRIFNEPGLGMRFVLQLPLTLSVIRSLIVDVGGEPYA
FPLVQVRRTLELERADIDVLEGQQHFPLDDRRVGLVTAHQLLDAGELDESRPATAVVVVG
GEPETYGVAVDRFLGERMLVVQPLDGRLNKIQNIAAGALLENGDPVLIVDVEDLIRSIDK
LIRGGQLAKVRRGDRDALARRAKRVLVVDDSLTVRELERKLLEKRGYDVTIAIDGMDGWN
AIRGDAFDLVVTDIDMPRMDGIELVTLIKGDPLLKSVPVMIVSYKDRDEDRRRGLEAGAD
YYLAKGGFHDEALLDAVHDLIGDA*
>GCA_000452945.1_BPMSHR5858_genomic_02012_1 group_1003
MNTKPIQTTTTKRAAFTKTTLLLALASAAALSLSACVAPNAYGPYGSQPQYGTPAYAQPS
YAQPAYTQPSYAQPDYSQPGYPQQGYTQPGYQQGYQPGYQPGYSGYSTQYGTIAGIRPIG
GATSPSGVAGTVVGALVGGVLGNQIGRGHGRDAATVIGALGGAVAGNQIGQQMGAQPSAY
RVDVQLSDGSMRSFDLQTPGDLRPGDRVRVDGNQISRY*
>GCA_000452945.1_BPMSHR5858_genomic_02013_1 trpS2
MHDDTSNPSRAPIALTGDRPTGALHLGHYIGSLKTRIALQDTHRQFILVADTQALTDNAH
DPDKVRRNVIEVALDYLAVGIDPRRSTIAVQSGLPALAELTLFYLNFVTVARLERNPTIK
DEIHARGFGRDIPAGFLCYPVAQAADITAFKADTVPVGDDQLPLIEQTNEIVRRVNRQAG
ADVLPEASALISPTGRLPGVDGKAKMSKSQGNAIALSSSPDAIADAVRRMYTDPNHLRVS
DPGTVEGNVVFTYLDAFDGEPDELERLKAAYRAGGLGDMVLKRRLEDRLQALIAPIRERR
AQLARDPGYVVDVLKDGTAHARDVTQRTLDEVRDALGAFSFRRFVG*
>GCA_000452945.1_BPMSHR5858_genomic_02015_1 inhA_1
MTIEVGLLLFPQVQQLDLTGPHDVFASLPGVTVRLVWKTREPVASSSGLMLAPNATFDDC
PPLDVICVPGGIGVADLMEDADTLAFLRRHAAHARYVSSVCTGALVLGAAGLLRGRRATT
HWAFHSLLAPFGATPVRERVVRDGNLVTGGGVTAGIDFALTLAAELVGEDDAQAVQLQLE
YAPAPPFAAGDPDTAPARIVERMRERSKPVLARREQVVARAVAALERRRADG*
>GCA_000452945.1_BPMSHR5858_genomic_02017_1 yfeX
MSDFQPGILAPIPAASRYLSFRISHPEHAASVLAALRDALDGVDTVIGLGPSLVQALDRK
IDGLKEFPELSVPGAPVPSTPAALWVWLRADDLGAATVRSRDIERLVAPAFELDAAANAF
RYADDRDLSGYEDGTENPEGDEALEVAFVSGKGAGLDGASFVAVQQWVHDFGKMREIPGD
EMDRVIGRRKDDNEELEDAPAYAHVKRTEQESFKPEAKVLRRSSPWADERRAGLYFVAFG
HSFRAFEVQMRRMIGATDGVHDGLFRFTQPITGSFFWCPPVKGGKLDLSALGL*
>GCA_000452945.1_BPMSHR5858_genomic_02018_1 group_1831
LKFNCIAATILAAVAADATAAGACLNGSTIASTTRAPLVARQGSVFSSTLYDPAITSNNR
THNPVMLTVKVTNNGRPVAGCDVAWQPRGAGGASGWLFPASASTDANGIASAWWVAGSGA
AQTAVASIRRFDGTTQGVAIGGSAQPHATRANSIHLNYEPASDWTAFRVDVTPEALAPTT
YWEAIGWPGAYTGIQSIDGKQDGLVLFSVWDVNGKSPQIIAKGPGVDCTQFGGEGTGYKC
AKRHAPVAGRTYRFMASIAPVAGQNQTDYSVWFTDTSTNARELIATLRYQKAVQSANYAN
SFVEDWATQGASCLGATQRAGQYGNVWALDRASAQWRTVKRASTSAVYTPDHNEVCSNYQ
FSVVNGRFRMSTGGHAVGQPLNLPNGPKSFPLTLP*
>GCA_000452945.1_BPMSHR5858_genomic_02023_1 group_1832
MTELLLIRHAQASFDAADYDCLSPLGDEQSARLGAWMARGARRPALIATGTLRRHAQTAD
GCARAAGVDAPRLALAGLDELDADELIARHRPALASRDALLRAMAAEADPRRAFQTLFAA
AVARWTGGAHDGEYGCAWPEFRARTLAAWDALASQPAREIWAFTSGGPIGVIVAALFGVP
AERSFELAWPLVNTSVTRIRIGRGGARVTTYNGWPHLDGERDERLVTHR*
>GCA_000452945.1_BPMSHR5858_genomic_02038_1 group_1004
MMDAAVHRAIDAVWRIEAARIIAHVARLVRDVGVAEELAQDALVAALEHWPSGGVPDNPG
AWLMTAAKRRALDHLRQHALHARKREQIGLDLDALGAHVAPDVADVFEAARDDDIGDDLL
RLVFTACHPVLSTDARVALTLRLLGGLTTGEIARAFLTPEPTIAQRIVRAKRTLSAAKVP
FEVPRAPERAARLASVLEVIYLVFNEGYSATAGDDWMRPALTDEALRLGRVLAGLAPDES
EVHGLVALMEIQASRMHARVDAQGRPVLLLDQDRSRWDPLLIRRGLAALARSEALGGASG
PYALQAALAACHARARHADDTDWEQIVALYDALAQVAPSPVVELNRAVAVGMAFGPAAGL
EIVDALAADPALARYHWLPSVRGDLLAKLGRRAEAQAEFQRAADMTLNAREREMLLARAT
QR*
>GCA_000452945.1_BPMSHR5858_genomic_02041_1 rbr
MAQLKGSKTEENLKYAFAGESQANRRYLYFASKADVEGQNDIAALFRSTAEGETGHAHGH
LEYLEAVGDPATGLPFGTSRQNLQSAIAGETHEYTDMYPGMAKTARDEGFEEIANWFETL
AKAERSHANRYTKALDGLVD*
>GCA_000452945.1_BPMSHR5858_genomic_02045_1 algA_1
MNATLAADARSACAADTRVAVQPVVLAGGSGTRLWPMSRERHPKQLINLLGDESLLQSTA
RRLDGLSGVHPVSERMIVVCGEDHRFTTAEQLRLAGKDATLVLEPLGRDTAPALTVAALR
AVGGGDDAVLTVMPADHAVADVARFQQAIAAAVRYAEQGMIATLGIVPTHAETGYGYIRV
GAAANAADAGAEAGVAVRKLDRFVEKPHLELARQYVASGEYWWNSGIFIVRASVWLKAIR
HFQPAIHDACARAVAQGSEDGPFFRVDRDSFAASPSNSIDYAVMEPLSNLPAVCEGVVVP
LDAGWSDVGSWDTLWQISEKDDEGNVARGRVCFEQASSTFAHSEGRLIACVGTQDLVVVE
TPDAILVADRAHVQDVKKIVGRLKAESGTEASDHRKVHRPWGHYDSVDSGERFQVKRIVV
KPGARLSLQMHHHRAEHWIVVRGTARITRGDETFLLSENESTYIPLGVSHRLENPGKMPL
ELIEVQSGAYLGEDDIVRFEDNYGRH*
>GCA_000452945.1_BPMSHR5858_genomic_02058_1 group_19385
MNQPAKSTDPVSVMPVRRDLRFDLPVERATDWHGQGSQVSHFFNALSLLFPAGERFFMDS
VRNYRDRIDDPVLKKQVLGFIGQEAMHTREHVEYNELMHANRLPARTVDRRVWKVTGWMK
KVLPHSVQLAHTVAAEHYTAMLANWLLSEPAALAGSHEGYKQMWLWHALEETEHKAVSFD
VWNTVMKPGPKRYLIRIGVYLLTTMTFWPNVFLIHSQLIRWDRKSDKSVRGLWRVVSLLY
GPRHGMFPAIAREWLSFFRPDFHPWDYDNHHHLARVDAIVEATQAREARKLAAVELDGAK
PA*
>GCA_000452945.1_BPMSHR5858_genomic_02069_1 group_19683
MNACEARWFLFGADERKPDARIRLFCFHYAGSGGSIFRHWGEALPDDVELVAVQMPGREN
RLNEPLLYSMDEAVPPLVDALAPLLDRPFAFFGHSTGALIGFEVARVLRARGYPQPRLLI
ASAQNAPDVKPEVIRHRLSDAEFVEVLRGCNGTPDAILQDPALLELLLPRIRADGAVFET
YRYERQAPLDCRIVVFHGAADALVHDAGLARWAGETQHSFARYRFAGDHFFIHDEEASVL
DHINRELEPLLSDASSTLIG*
>GCA_000452945.1_BPMSHR5858_genomic_02070_1 ligK
MAQWNSDDELFALMRAELSTCPVSDVMEVLGFAYPMLPPEIRPLRRDMVMIGRAMPVQDE
PPVPHGGLKRYDAKPFGLLFESVEALRPGEVYIASGGPTAVARLGDLLVTRARNLGAAGV
VLHAHVRDANAILDLNLPAFAHGTYAYGLQGRHNVVDYRCSITVGHVRIRPGDLIFGDGD
GVCVIPREAEREVITRAIAKNRLERKVRGAIAEGRSVVEAFNQYNVM*
>GCA_000452945.1_BPMSHR5858_genomic_02073_1 bphJ
MKSKSSRTRVAILGSGSIGLDLMFKVKASEQFDLKFVVGRNANSDGLRLARSCNVETSSD
GLDFLKAHEDAYDLVFDATSAAAHKVNNRFFSDAGKFVIDLTPAKVGRLCVPCINLDDMG
AEQNVNLITCGGQASLPLAYALKQAVDEIDYLEVVSAIASRSAGIATRENIDEYMTTTEY
ALAKFSGAKKTKAILNINPAEPGVRMQTTLYAYARYRDFDRVRASVADMVEKVREYVPGY
RLVVEPLESQGRITIGLTVRGRGDYLPEYAGNLDIINCAALAVASHRHATARLGATQ*
>GCA_000452945.1_BPMSHR5858_genomic_02075_1 group_1005
VTGLNERGFGAWNRIERETLTERIERQVVSGDALTMARLYLKKGAFVGTHSHPNEQLTYI
LEGRLLFRYGEHLEHEIEVGPGEILHLPANVPHNALCLEDAVDLDVFTPVRTDWLAPGGN
RYFAGASAATSPTPVPSASR*
>GCA_000452945.1_BPMSHR5858_genomic_02088_1 group_19412
VIPSAEPAAALLRERAARYAAEVALFVRDHALSVASHDLRSPLNAMHSWAYVLERRLTAS
DENLARALDGIRIGIEQQTKLLETTIDAPRAETRSLPIARAGVALDALADECAALARIAL
GDARGTAVTIASPMRPASLECDRERVAQALWSMLTFAIEASAPGGEVALGCDGPADAARL
NVTFLARLPALTDATLPHVFETFARREALAERQGERPAAVFALAQRVARAHGGAFAQEPS
PLADGARATLALTIPAARA*
>GCA_000452945.1_BPMSHR5858_genomic_02114_1 ynfE
MHFEFFEESEVILIWGANPIASNLHFWTRAQEAKRRGARLIAIDPYRSLTAEKCHQHIAL
KPGTDGALALGIMHVLISEDLLDRDYIRDHTLGFDALKARALTYTPERVAEICGVTADEV
AELARLYGRTRKASIRLNYGMQRVRGGGNAVRAIACLPALTGAWRDRAGGLLLSSSEWAP
VDAAAIGRPDLLPGWPNKLPRSINMNAIGDALLHSGDAEFGPRIEAIIVYNSNPVAVAPD
SEKVAAGFAREDLFTVVLEHFKTDTADYADILLPATTQLEHLDVHKSYGHTYVMANLPSI
APIGEVRPNTEIFRGIARSMGLDEPALYEDDETLARTSLRWDDASLQSDWDTLKRAGWLK
LKLADAPFANGGFRTPSGKCEFYSERLVEMGLDPLPDYLPPYESADGSPELAVRYPLAMI
SPPARHFLNSTFVNVASLRATEGEPHLDIHPADAARRGVADGDAVRIFNDRGSLRAKARV
TDRAREGLVVGLSIWWKKLAPDGRNANQVTSQALTDLGRGATFYDCLVEVERG*
>GCA_000452945.1_BPMSHR5858_genomic_02115_1 fadD_2
MDKIWLKSYPPGVPAEIDASPYSSVPDLLEESFGLYRNQQAFICMGKAITYGELDTLSRK
LGAWFQSRGLPRGARVAIMMPNVLQYPVTIAAVLRAGYTVVNVNPLYTPRELEHQLKDSG
AEAIVILENFATTLQAVIARTAVKHVVVAAMGDLLGLKGHLVNYVVRRVKKMVPAWSLPS
YTRFNAALATGGRQTFKPAKPAPDDVAFLQYTGGTTGVAKGATLLHRNIVSNVLQAQAWH
EPAHSKRPEVKQFITVIALPLYHVFALTVCGLLTLRTGGTGILIPNPRDIAGMIKELKGY
PIITIPAVNTLYNALLNHPDFDQLDFSKLAVANGGGMAIQENVAKRWYEKTKTPIVEGYG
LSETSPVATCNPVTATEYSGTIGLPLPSTEIAIRDDAGDDVPLGQPGEICIRGPQVMAGY
WNRPDETAKVMMPDGFFKTGDVGVMDARGYVKIVDRKKDMILVSGFNVYPNEVEDVVASH
PGVFEVAAVGVPDEHSGEAVKLFVVKKDPALTDKDLLAYCKDRLTGYKRPKYVEFRADLP
KTNVGKILRRELRDGKA*
>GCA_000452945.1_BPMSHR5858_genomic_02118_1 mraZ
VFQGASALTLDAKGRMSVPSRYREALQGQAEGRVTVTKHPDGCLLLFPRPEWEVFRAKIA
ALPMDAHWWRRIFLGNAMDVDLDSAGRILVSPELRMAAGLEKEVMLLGMGSHFELWDAQT
YTAKEQAAMAQGMPEALKNFTF*
>GCA_000452945.1_BPMSHR5858_genomic_02121_1 penA_1
MKPASKRTQGVKFASSPVLSVHLPMWRSKLVVFMLFIAFVALAMRAFWIQGPGNAFYQKQ
GESRYQRTLELPATRGKILDRNGLVLATSLPVRAIWAIPADVPDDLGADKLDALGKLLGM
TTKELKAKLSEDKTFVYVKRQVSLDVAEQVAKLDIPGVYQRGEYKRFYPEGEITAHLIGF
TNVEDEGQEGVELADQKLLVGTPGSRHVIKDRVGHIVEDVDAQVPPHNGADVGLSIDSKI
QYITYTNLKAAVEKFHAKAGAAMVIDVQTGEVLSLVNYPTYNPNDRSHLTGEQLRNRVLT
DVFEPGSIMKPFTVSLALDLHRVTPNTLVETGNGHFVLDGAPITDDSGFGTLTVGGVIQK
SSNIGATKIAMTMKPEEMWNMYTGIGLGQAPKVGFPGAAAGRLRPWKNWRRIEQATMSYG
YGLSVSLFQLARAYTAIAHDGELMPVTIFKTNDAQPPTGPRVFSPTTAREVRTMLESVVS
PQGTSPNAAVPGYRVGGKSGTAYKQVGRGYDHKKYRASFVGMAPMPNPRIVVAVSVDEPT
TGGHFGGQVSGPVFSAIVGDTLRALNVPPNMPVKQLVVSDDATNAAAAQKLAANAPAKRL
IVSSTTHNRPGVVR*
>GCA_000452945.1_BPMSHR5858_genomic_02122_1 murE
MSAARSSHPAHQQIAAALAWLRAHVAPGAQLHADTRSLRAGDVFVAYAVDGADNRPFIAD
ALARGAAAVLYQPEGFPAEGLDSATSLAVPALDALAGDIASGWYGDPSDAMFSVGVTGTN
GKTSCTQWIAAALTALREPCAVIGTLGSGMPGQLVHTGFTTPDAPQLQRSLAQLRDAGAR
AVAMEVSSHALHQGRVNGTAFDVAVFTNLTQDHLDYHRTFEAYEAAKAKLFAWRGLRCAV
VNRDDPAGRRLLANLAGRVRTIAYGIDSPTQAADADRELVATNVRATATGTAFRVTSAWG
EADVEVGTLGAFNVSNLLAVLGTLLAADVPFDAALAEIGKLESVNGRMQRLGGRLQNDEP
LVVVDYAHTPDALEKTLDALRPIAQARGGELICMFGCGGDRDATKRPLMGAIAERLADKV
VVTSDNPRSEDPQKIIDQIVAGMRDASGARRVEDRASAILQAVRGAAREDVVLLAGKGHE
ATQEIMGKKRAFSDQDHARLALAARTTQTRGGGE*
>GCA_000452945.1_BPMSHR5858_genomic_02126_1 ftsW
MSWSDRLVSRFNGVRDAGGAAAPRTAARAASGARATAGGLASVVNGARPTRSRMLDFDYS
LLWVSIALLGLGVVMVYSASIAMPDSPKYASYHDYAFLLRHCVSLVVAFVAAVIAFRVPV
STWDKYAPHLFLIALVGLVIVLIPHVGKGVNGARRWIPLGITNMQPSEIMKLAVTIYAAN
YTVRKQEYMQSFAKGFLPMACAVGLVGALLLLEPDMGAFMVIAAIAMGVLFLGGVNGKLF
GGLVATAVGTFTMLVWLSPWRRERIFAYLDPWDERYAQGKAYQLTHSLIAFGRGEWFGVG
LGGSVEKLNYLPEAHTDFILAVIGEELGFVGVLVVILLFYWIVRRAFEIGRQALALDRTF
AGLMAKGVGIWFGAQAFINMGVNLGLLPTKGLTLPLVSYGGSGILLNCVALAVLLRVDYE
NRVLMRGGKV*
>GCA_000452945.1_BPMSHR5858_genomic_02130_1 ftsQ
MWNNVRQLNLAASALYALLLLVLAAAGCYWLIQRPAFALREIRIDGDTEHINAPTVRAGV
VGRLKGNFFTVDLDLARVAFEQMPWVRHASVRRVWPNALAVTLEEYKPLGTWGNDQLVSV
DGELFTANQGELDAELPSFDGPEGSAKEVVARYRDFAKWFAPIHATPEEVTLSPRYAWTV
KLSNGMQVELGRERNSDTLPDRIQRLVAAWPSVTQRWGGDIEYADLRYPNGFAIRAAGMR
FLTDTDKGKK*
>GCA_000452945.1_BPMSHR5858_genomic_02132_1 ftsZ
MEFEMLETETNGTIIKVIGVGGAGGNAVQHMINKGVQGVDFVVMNTDAQALSRSRAPSVI
QLGNTGLGAGAKPEMGRAAAEEARERIADALRGAHMVFITAGMGGGTGTGAAPVVAQIAK
EMGILTVGVVSKPFEFEGGKRMRVAEAGSQQLEDHVDSLIVVLNDKLFEVMGDDAEMDKC
FQCADDVLNNAVAGIAEIINVDGLVNVDFEDVKTVMGEQGKAMMGTATVAGVDRARLAAE
QAVASPLLEGVDLSGARGVLVNITSSRSLRLSETREVMNTIKSYAADDATVIFGAVYDDA
MGDALRVTVVATGLGRAAKKQQSAPMTLLRTGTDNQPISAAPQGYAASHHVSTGDYGAFD
TPAVWRNSRETAASHVQALQEKGVDTYDIPAFLRKQAD*
>GCA_000452945.1_BPMSHR5858_genomic_02133_1 group_19525
MIQVGDTLPDAQLFEYLDDARAGCTPGPNAFGVREQTAGKRVVIFGLPGAFTPTCSAQHV
PGYVAHAEPLRSAGIDEIWCVAVNDAFVMGAWGRDLHTAGKVRMMADGSAAFTHALGLTQ
DLSARGMGIRSRRYAMVVDDGVVKTLFVEAPGKFEVSDAASVLAGLTR*
>GCA_000452945.1_BPMSHR5858_genomic_02134_1 lpxC_1
MLKQRTIKSIVKTVGIGVHSGRKVELTLRPAAPDTGIVFSRVDLPTPVDIPASALSIGDT
RLASVLQKDGVRVSTVEHLMSACAGLGIDNLYVDVTAEEIPIMDGSAATFVFLIQSAGIE
EQNAAKKFIKVTKPVEIRDGDKFARLDPYFGFKLKFTIDFRHPAVDKTGQELEVDFANTS
YVREIARARTFGFAHEVEMMRELGLARGGSMDNAIVLDEYRILNNDGLRYDDEFVKHKML
DAIGDLYVIGHPLLASYTAYKSGHGLNNALLRELLAHEQAYEIVTFDDPKTAPTGFGFDA
QTAFA*
>GCA_000452945.1_BPMSHR5858_genomic_02137_1 adh_2
MKAVVFHGIGDIRIDTVPDPGLAQTTDAVVRLTSSAICGTDLHMVRGTLGGMKPGTILGH
EGVGLVEEVGRDVRNLRRGDRVLIPSTISCGHCAYCRAGYTAQCDTANPGGPLAGTAFFG
GPQASGAFDGLQAEYARTPFAHASLIRLPDSLDDERAILMSDIFPTGYFGAQLAEVRMGD
TVAVFGAGPVGQFAIASAKLMGAGRVIAIDRVVSRLEMACAQGAEVINFDDDDPVDTILQ
LTGGIGVDRVIDAVGVDAVRPTRGPAAPHDAQSRRAFDDEVRELAPRTRADGGHWVPGDA
PSQVLQWAVRAVAKAGTVSIIGVYPMNDRFFPIGAAMNRNLTVKMGNCNHRAITPKLIEI
VRSGAFDPITVLTQAEPLTSAIDAYRAFDTREPGWIKVKLEP*
>GCA_000452945.1_BPMSHR5858_genomic_02138_1 group_1835
MATKKFPPDENAPRRPQTDKDTGVGTFVPDPGLDDDADDVGYASGDEAQTAPLEDEESGT
PGRGERSRNGGGDSGDGGGHDNGRRRR*
>GCA_000452945.1_BPMSHR5858_genomic_02139_1 group_19463

MKKYALALWFMLVSSMSAMAGAIIPSGATLVAGQYAISNNGNFAFGVDAASGKLYFHYRF
PTQAWGASAMSQMYGTQPEGALYVGKGDRLVMQTDGRLVFYSGNDVVWSNWYYGPAPIPG
SYAVVEDFGVVAIYPPQGGYPTFTWPFASVTPNSGALATVVGYPFGVNFPVLNGRQWLQI
FNNVTYYLWNKSFAADSVAMQTDGNLVVYNKGQPVWSTGTSGNPGAYMLVTPVGFVLGKQ
YGGILANVPYPVSVDQSSRASGPNDPKPTPAAPPPPSLNLPINASWKCYYVKDWLACYAP
G*
>GCA_000452945.1_BPMSHR5858_genomic_02164_1 ybiB
VADTRRQVQVDWLHDGHCETIVEPARASSDAPPVELPASRDAATTAAWTDSVMRGEIPVP
DAIAMQVDTIVRLARLA*
>GCA_000452945.1_BPMSHR5858_genomic_02168_1 pagL
MNDKNGGRVGRAIARTALALALVGASGSAFADRWGLQLGGGVADHDMKKGDIAVVWDPNW
TWWEIGGWHFAFVAEGHLSYWRYTGDRAINSSIWEVGATPIIRFIKSAGYVRPFVELGAG
VRFLSHPTISQNYSMSTSFQFADMVGVGAQFGNRQQYQAGFRFQHVSNAGIKDPNPGINF
SQLYVQYNF*
>GCA_000452945.1_BPMSHR5858_genomic_02169_1 rpoH
VSNALTLPNTLRPASAKAVSAGSLTLASHSMLPGHLGNIDAYIQAVNRIPLLSAEEERQY
ATEYREQNNLDAARRLVLSHLRLVVSVARNYLGYGLPHGDLIQEGNIGLMKAVKRFDPAQ
NVRLVSYAIHWIKAEIHEYILRNWRMVKVATTKAQRKLFFNLRSHKKGTQAFTPEEIDGL
AQELNVKREEVAEMETRLSGGDIALEGQIDDGEESYAPIAYLADSHNEPTAVLAARQRDM
LQTDGIARALESLDARSRRIIEARWLNVDDDGSGGSTLHDLAAEFGVSAERIRQIEASAM
KKMRTALAAYA*
>GCA_000452945.1_BPMSHR5858_genomic_02171_1 cydA_1
MISSEVVDLSRLQFGITALYHFLFVPLTLGMSWLLVIMEAVYVMTGKQVYKDMTQFWGKL
FGINFAMGVTTGITLEFQFGTNWSYYSHYVGDIFGVPLAVEGLMAFFLESTFVGLFFFGW
NRLSKVQHLVVTFLVALGSNLSALWILVANGWMNNPVGAEFNYQTMRMELANLFDVLFNP
VAQVKFVHTVSAGYVTAAMFVLGVSAWYLLKKRDVDFALRSFAIAAGFGLASTLCVIVLG
DESGYTTGEVQKMKLAAIESEWETQPAPASFTLIGLPNQEAERTDYAIKIPYALGLIATR
SVSEPVIGLRELAKRSEEHIQSGMIAYGALQKLKAGDTSAETKATFDQHKQYLGYGLMLK
QFTQNVTDATPDQIKAAAKQTIPPVAPVFFSFRIMVFLGFLFLATFIAAFWFCARRQLLQ
DNRRWFLRYAVWAIPLPWLAAEFGWVVAEVGRQPWTIAGILPTHLSASSLTATDLYLSLA
GFIVFYTALFIIEITLMFKYARLGPSSLHTGRYHHEQAGRPLEKAVAGTAA*
>GCA_000452945.1_BPMSHR5858_genomic_02173_1 group_18949
MWYFSWILGIGVALAFGIINAMWLEARQKRDA*
>GCA_000452945.1_BPMSHR5858_genomic_02176_1 nagE
MDGNPFLKIQSLGRALMLPIAVLPVAGILLRLGQQDVLNIKMIADAGGAIFENLPLLFAI
GVAVGFAKDNNGVAALAGAIGYLIEVAIMKDIDPKLNMGVLSGIIAGVVAGLLYNRYKDI
KLPDYLAFFGGKRFVPIITGLACVVLGIVFGYVWQPVQHAIDAVGQWLLTAGAIGTFVYG
FLNRLLLVTGLHHIINSLVWFVFGTFTPAGGAAVTGDLHRFFAGDPSAGGFMAGFFPIMM
FGLPAACLAMFHEAPKARRAIVGGLLFSMALTSFLTGVTEPIEFSFMFLAPVLYVIHAVL
TGLSLAICQLLGVKLGFTFSAGAIDYVLNYGLSTKGWIAIPLGLAYGLAYYGLFRFFIRK
FNMATPGREPAGADAQAQSFASGGFVAPTAGASVPRAQRYIAALGGAANLSVVDACTTRL
RLSVVDPEKVSEADLRTIGARGVLKRGGSSVQVIIGPEADLIADEIRATLGSGAAAPAAA
AAAAPAAAATAAGAQSGPLDPEPTRWLAVFGGATNVASLDAVAATRLRVVVRDPSAVDRQ
RLATLDVAWVASDTFHIVCGQSAPRYAQQLAARLPSSDGGTAAQPA*
>GCA_000452945.1_BPMSHR5858_genomic_02177_1 ptsI_2
MRRAEESQLKQQASHDQIVLVAPLTGPVVPLADVPDPVFSGGMFGDGIGIDPLEGRLLAP
CAGVVSHVARTGHAVTIAADGGAEILLHIGIDTVELNGLGFTAKIAEGARVAAGDLLIEF
DQDAIARAAHSLVSVIAIANSDAFEVVERAGAGVVKAGETPLLALRARGADASADASADA
SASASAGAAADASCAQPAAEARKSITLTQPGGLHARPAARAREAARGLDAHVDVHFEGRK
AALQSVVGLLGLGAGEHATIELVATGRDAAKALERVAHELLREAHGEAEEKPARIVSPAP
AAAGIARAPLEPNTLAGVCAAPGIAVGTLVRWDDAQIVPTELASGTPAAESRLLDRALAE
VDAQLETTVREASRRGAIGEAGIFAVHRVLLEDPALVDAARDLISLGKSAGYAWRETIRA
QTAVLADVDDTLLAERAADLRDIDKRVLRALGYASASARELPAEAVLAAEEFTPSDLASL
DRERVAALVMARGGATSHAAIIARQLGIPALVAVGDALYAIAQRTQVVVDASAGRLEYAP
SALDVERARHERQRLAGVREANRRMSGEAALTRDGHRIEVAANIATLDDARVALDNGADA
VGLLRTELMFIHRQAAPTASEHQQSYQSIVDALQGRTAIIRTLDVGADKEVDYLTLPPEP
NPALGLRGIRLAQVRPDLLDDQLRGLLAVKPYGSVRILLPMVTDVGELVRIRKRIDDFAR
AMGRAQAVEVGVMIEVPSAALLADQLAQHADFLSIGTNDLTQYTLAMDRCQADLAAQADG
LHPAVLRLVDATVRGAEKHGKWVGVCGALGGDPVAVPVLAGLGVTELSVDPVSVPGIKAQ
VRRLDYQLCRQRAQDLLALESAQAVRAASREIWPAE*
>GCA_000452945.1_BPMSHR5858_genomic_02179_1 nagA
MLTGNILTAEGWLHGTLEFENGRITTLSGDAADPSNNDAPYILPGFIDLHVHGAGGADVM
EGGDAIETIARTHARYGTTSLLATTMTAPRDELMRVVAELGDVARTRTPGGSRVLGVHLE
GPYINPGKLGAQPDAAVSAALDEVLKYLSIAPIRVVTLAPEIAGHIEIISEMAARGVRVQ
LGHSLGTYDDAVAALKHGACGFTHLFNAMSPLHHRNPGIVGAALAHAEYAEIIPDLLHVH
PGAIRAALRAIPRLYVVTDSTSATGMPDGEYRLGSQRVTKCLGGVRLADGTLAGSTLTMD
QALRNLVSLGLPIADVSSRMSRYAADYLGLADRGRIARGAWADLAVFDRELNLTATFVEG
ESIVEYA*
>GCA_000452945.1_BPMSHR5858_genomic_02180_1 nagR_1
MDIGWSTLAPDARSDTPLYLQLARNLAGAIHGGAWRAGEALPSERGLSAAAGVSRITARR
ALALLVEQGLIRRVRGAGSFITPRVADPLSRLVGFTAKMRQRGFAPDSVWLARSLRAASR
DELARFGLSPGATVARLERLRRADGVVMAYERSTLPASCVPEPQALEGSLYGYLQSRGLD
VVRASQRFRAVNASADVAQWLGLAPGAALLVITRIGYGADRRAIEATETYCRDDYYDFVA
ELKR*
>GCA_000452945.1_BPMSHR5858_genomic_02181_1 benM_1
MIDLRRLRYFIVVAEELHFGKAAQRLHLAQPPLTRHISALEGELGLRLFDRSTRSVKLTA
EGELFLPHARDVLDAVHRAEIASQRAALGKEGKVALGYTSSVPLCDAFGTLIRNFARSFP
DIELSLVEASSAQQGRSIKEGLLDIGIGWKNSFEDYEGCRVKTIAAEPLVVAVSQERALA
SEASLAVEQLADETFVLFPPGYGSTLDRKAFDLCATAGFTPRRGPSASQMTTMIALVAAG
RGVAIVPEAVSTLRKPGVAYVPLTDESALIELVLMWREAGLSLAARSFVEWHKAHVAQRV
DGMVGVG*
>GCA_000452945.1_BPMSHR5858_genomic_02184_1 group_19662
MSTPAAETLPLLAREVAKILHVETVETNVGIGELGIDSLNIVELIVFCEQLYGSIDPETL
NITQYTTLAQLDAELRAPQTAA*
>GCA_000452945.1_BPMSHR5858_genomic_02190_1 lcfB_1
MNPHIRLADLLDSAAAQRPDGIAIAEPRRRVRYAQLAADVRRAAAALAAAGIAPGERVAT
YAPKAYETIVTMLAANLAGAIVVPINPQLRDHQVRHILADSGARLLLTTAPRLARLAERP
AGLVTWLVEDVAALAGAGDDVRDAGRAHAVDSDPAALLYTSGSTGQPKGVVLSQRNLTAG
ADSVAAYQRLSHDDVILGALPLSFDAGLSQLTSALAAQACYAPLDFLRAEEVPAWCAQVG
VTSITGVPPLWMQLAAVAWPDAARLPVRRIANTGGTMPQPLLHRLRQIFPNAAPYLMYGL
TEAFRSTYLPPEEAAARPGSIGKAVPNAQILVLRADGSECDAGEPGELVHRGAFVTLGYW
NAPELTAQRFRALPHARHPVPLADVAVWSGDIVTRDDEGFLYFVSRADEMIKTSGYRVSP
TEIEDILFECPQTLEAVAFGVPHPALGQAVVACVYGGGDPASCRQALLDACRARLPSYMV
PQHIEIAGAPLPRNPNGKIDRPLLKRAHLSRFEAEPRAAALG*
>GCA_000452945.1_BPMSHR5858_genomic_02211_1 group_1008
MSQVCYFAYGSNMSAARLSERLSRFGEALIERRPGIVEGYRLTFNKVSSRQDWVGFANIE
AAADCRVEGTLNAMSARALDALDSIELVPLHYRRAGVLVRDGATGRLTFAVTYVANPAMV
RPNLRPTRDYLDHLLAAADVLPRAYLDHVRAVECWT*
>GCA_000452945.1_BPMSHR5858_genomic_02216_1 tauD_1
MSDGPALGIGPLSAALGARIDGIDARAELDADTVRAIRAAWLRFGLLVFRGQALDPPRLV
AFTRRFGEPVVYTRACNACDGQPEVLVLSNVVKDGKPIGAALSGRYWHTDGHYLACPPAG
TLLFGAEIPAVGGDTHFVNMTSAYRALPAWVRTQIDGRTFVMDRVQTLPFHYPQRPAPPP
DQKQAWPDMPQPVVRTHPETGENALYIGGVVPWRIVGMEQRLSDALMAHLHAIAFDEARF
GYRHRWRAGDLLMWDNRCLAHRATDYDMVRHRRTMYRTTIAGDRPRYVRGAAQPGVS*
>GCA_000452945.1_BPMSHR5858_genomic_02223_1 ubiI_1
MKNNPVDVLINGSGIAGVALAHLLGARGHSVTVIERAARNRAQNGADLLKPSGIGVVRAM
GLLDDVFAAGGLRRDAMRLYHDKAPIASLDYRSASALGYFILIPCEQLRRLLLAKLDGLP
NVRLRFETSIERIERDADGTVTSLTLSDGERVAPTVLVGADGARSMIRDDVLRMPAERVP
YATPMAFGTIALTDSVRECNRLYVDSNQGLAYFYPIGDQATRLVVSFPADEMQGYLADTT
RTKLLARLNEFVGDESADAMAAIGAGTAFPLIPLGRMNLDRYHKRNAVLLGDAAHSIHPI
TGQGMNLAIEDAGELGACLDLHLSGQITLADALERFERIRRPVNEAVISYGHTLATTYHD
RAALVANFRSQLQTSGRDAAWLGAAVEV*
>GCA_000452945.1_BPMSHR5858_genomic_02226_1 group_19108
MTLKANLEGVRDTYRLCFVRNPWAYFTCLPLDRQCGDRWSEAPYELYAGAPYGDSPEQIL
RVAFDGPLLPPEAGRGGLTCSVVDINEGRAPWLRTESYFGGEPLAIAAGATLGAFVETVE
KAGGTVFVPLTWGALPDAGKRAHAMAATATN*
>GCA_000452945.1_BPMSHR5858_genomic_02229_1 group_19328
MIKVLIADDHALVRDGLRHILQNASGYEVAGEAFDSATTIALVRSTPAQVLVLDLSMPGR
NGVELIKQIKEEKPALRILVLTMHAEQQYAVRAFRAGASGYLTKESASSELVGAVTKVAS
GGVYVSLAMAERFAQSLNEPAETLPHHRLSDREFDVFRRIASGQTLTQIADELCVSAKTV
STYKTRILEKMQMPHEAALVRYALRHKLIDDPDED*
>GCA_000452945.1_BPMSHR5858_genomic_02234_1 adh_1
MKALVYHGPGRKSLDERPMPELAAPTDVIVRVTRTTICGTDLHILKGDVPSCEPGRVLGH
EGVGVVHRAGGAVSTFKEGDRVLISCISSCGQCEYCRRGMYSHCATGGWILGNRIDGTQA
EYVRIPHAQTSLYRIAPGADEEALVMLSDILPTGFECGVLNGRVQPGGTVAIIGAGPIGL
AALLTAQFYSPAQIVMIDPDPNRLALARRFGATACIDNGQVDAVAGVKALTGDGVDCAIE
AVGIPASFELCQRIIAPGGVIANVGVHGMKANLHLETLWDRNITITTRLVDTSSTPMLLK
TVGAGRLDPKQLITHRFSLADIGQAYETFGHAAHTRALKVIVEVSE*
>GCA_000452945.1_BPMSHR5858_genomic_02244_1 group_19675
MPAIVRRDVRFALPPDRIGDWHVNGVANTHYFNALSLMFPAGERFFIDAVRHYRDRIQDP
ELLRQVQGFIGQEAMHSREHVEFNDVAEAAGYPAHRLDRGFWRFTGLMQKILPPPLRLAQ
TIAFEHYTAIMTDMLLGNFERFHDSADAYANMWLWHSMEETEHKAVAFDVWNAVMKPGPA
RYLMRTGSMLLTSVVFWGTNFYFHLAFMNAHRRMHGKVTGKWAFLRFFLKGMVRIAPKTL
AYFKPGFHPWQHDNRRHFAQLDQLLANIDASNARYAAQAAPRRIPLHPMTIQPS*
>GCA_000452945.1_BPMSHR5858_genomic_02245_1 group_19182
MSRDLLLLLEPGFTDPRHPGERFVCPGSVPIEGLLASDPSKQPRLDIRRLPFARPREAAI
AALDADHQGLPVLILGDDAPPPDDALALGQTRFVTDSRRILELLAERHGFPKLH*
>GCA_000452945.1_BPMSHR5858_genomic_02246_1 ntaA_1
MTSLSKRPQRRQLHLNVNILHSGFAPSAWRIDDADPRAFVDVSHYVRIARIAEAAKFDAV
FLADNASIVDQIDFRPITALESTIVLASIAAATARIGLIGTASTSYNEPFNLARRFASLD
HVSGGRAGWNVVTTADAGSARNFGLDAAPDHAARYARAAEFVDVVKALWDSWDDDALVGD
KASGRFVDTAKVRPIAHRGTHFRVHGPLNLPRPPQGHPVLVQAGGSADGRAFAARHAQAV
FSASQSFDEALAFARALKAAAAGCGRGDIMVLPGLTTIVGATEADALRRRDELVDLIPLR
YGLNRLAGTLGVPVERLAPDAPLPDDLALPDGGNGNHTFFHATLARARRRGYTARELIRA
MAGGGGHRVIAGTPEQIADDIAHWFGAGAADGFNLMPDVLPGGLRDFVDGVVPILQRRGL
FRTEYEGTTLRDHFGLPRPAKPGACERA*
>GCA_000452945.1_BPMSHR5858_genomic_02247_1 rihB
MSLHKIIYDTDPGVDDSMALVFQALHPDIELLGVTSVFGNATIDTTTRNALYLAGRFAPG
VPVARGAAAPLRRPAPEPLGGIHGDDGLGNTGLSVSVDVAAAPNLDARPAHRFIIDTVRA
HPHEITLLAVGPLTNLAHALAEDPQVAMLVKQVVIMGGAFGTAGVLGNVSPAAEANIAGD
PDAADIVMSAPWPLAVVGLDVTQATIMTTEYLAALRDDAGEAGRFVWDVSRHYEAFHRAS
AGLAGIYVHDSSAVAYVVAPQLYRTRTGPVRVLTSGIAVGETIQKPAAMTVPAPDWDGRP
PRDVCVGVDANALLALYRKTLVG*
>GCA_000452945.1_BPMSHR5858_genomic_02252_1 group_181
MGCARAAGARRAYLAQVPRAACVSYVLMRAMHAIRGAAERADSTDAARLPAVLHTLKRQG
VTGAIAFGGEGNLRDPAFTIYRAGGRTWAVVGVLGGVGGVGGVGASGNAAACVAAGVSGL
SGGKAARLARFRESRERIERRIEMATSPGGRPMTTIPIHPAAGGAAHDEPLGILARRRIE
AEIIKPIYEIMKREFGLERAQAVIAEAVRGAALDAGRAFAAKEPGGTSIASFVALQVLWE
KDDALDVDVHRADDAHYDYDVRRCAYAQMYREMGLAEIGHLLSCARDSVFIEGYDARIAL
TRTRTLMQGGTHCDFRYRLAQPPRDGAADATHAAPGGLAGGMREARVRENEPAGQPEPAP
GDRAGPSPESPPRSRAESRTETGTEPGTASQASGDAPGIRRASSPTFDRRAVAASPAGEH
TRPEAGDAAR*
>GCA_000452945.1_BPMSHR5858_genomic_02262_1 feaB_1
MTQTDLVAVADTVRAFTEREFGVFIDGAMRAAHSPRRLDVFDPATGARLSRVPDADAHDV
DAAVASAQRAFDARAWSGLRPAERERILLKLADVLEAHAEELAQLETLNQGKSILVSRGV
EVGATIEYVRYMAGWATKITGQTLDVSIPFPPGARYTAYTRKEPVGVVAAIVPWNFPLMI
AVWKLVPALAAGCTVVLKPSPETPLTALRLAELALEAGVPAGVFNVVTGARACGAALASH
PAVRKISFTGSTATGKLVGAAAVQNMTRFSLELGGKNPIVMLEDVDVDAALGGVAAGAFF
NQGQVCAAASRIYVHRSRFRRLAEGLAGVASAMRLGPGLDPAAQINPLVSAHHRDTVARH
IDAARREGLTFLAGGTRADDLPGYFVRPAVIADAAHDSAIVRDEVFGPVVVVLPFDDPAE
AVRLANASPYGLAASLWSNDLKAVMDLVPQIEAGTVWVNCHIPLDPSMPFGGYKQSGIGR
EFGQYAIEGFTETKSVCIAH*
>GCA_000452945.1_BPMSHR5858_genomic_02288_1 nagAb
MSEQWVCAGHAGALSEDSPIEFKRTDGVEIGIYRVGDDVYALENVCPHAYALLTQGFVDE
GTVECPLHEAVFDIKTGQCLKGPGGRALKTYAVRLAGEEIQIKVE*
>GCA_000452945.1_BPMSHR5858_genomic_02293_1 todF
MSTISTTTTEAGAAPFAALLARFPERRAAVGGAGVIGYREAGEAHARRAGCEPVVLLHGI
GSGAASWVRQLDALGATRRALAWDAPGYGASSRVAAYSPVAADYAASLAAWLDALRIERC
VLVGHSLGAIMAGAFARIAGERLAGLLLISPAGGYGSAPADVRAGRRDSRLALLASLGPQ
GLAAERSANMLSAQASGAAREWVRWNMARIAPAGYAQATHLLANADLASDLAHCRGRIAV
AVGGDDAITPPAACERIARAAGVGLQVIARAGHAGYVEAPDAYSALIDAFCRQCDGSRGT
P*
>GCA_000452945.1_BPMSHR5858_genomic_02296_1 nadX
MSNPTTTTVGELIAAFLEQCGVKTAFGVISIHNMPILDAIHTRGKIRYVGARGEAGAVNM
ADGLARVSGGLGVAFTSTGTAAGNAAGAMVEALTAGTALVHITGQIETPYLDQDLAYIHE
APDQLSMLNAISKAAYRVRTVETVLPTIREAVRVAMTAPSGPVSVEIPIDIQAAEIEWPD
DLAPPHVAVREHDSARVAQLAERLAGARRPLLWLGGGARHARAEVERLVALGFGVVTSVQ
GRGVLPEDHPATLGAFNVHAAVEHFYKTCDAVVVVGSRLRGNETLKYKLALPRPLFRVDA
DALADNRGYRNDLFVHGDAKRVLAELADRLEGRLAIDPRFADDLAAARAAAVADVGKGLG
PYKRLVDTLQQALGRDYNWVRDVTISNSTWGNRLLRIFEPRAGVHALGGGIGQGMQMAIG
AALAGNAAKTVCLVGDGGLMVNVGELATAVQENANVMIVLMNDQCYGVIRNIQDAQYGGR
RCYVELHQPDFAQFCASLKLAHHRITSLDDAERIVREGLAHEGPVLVEVDMLSVGSFAAA
FAGPPVKKDTPAERQYA*
>GCA_000452945.1_BPMSHR5858_genomic_02298_1 aldA
VTAFDSSLVPSGDILIGGEWRRGRGATTPSFYPADGSLNTEIHMADAADAREAVQAADAA
WRRADWAGLKPHQRADVLYRIADLIHAHREALAQLQRRDNGKPINETRALVASAASTFRY
FAACAQTLDEALTPSRGDYLSMSVHEPLGVVAAITPWNSPIASDAQKLGPALAAGNAVVL
KPAEVTPLASLALARLCEQAGVPRGVISVLPGKGSVIGDALVRDPLVKKVSFTGGTEVGR
GIARLAAEKLMPLSLELGGKSPTIVFDDAELDHAVNGVLYGIFSSSGESCIAGSRLFVQR
SIYGAFVARLVEAARKLRVGDPASERTQMGPLITARHRDTVERYVALGRDEGARVLCGGE
RPTGEGRDAGFFYLPTILDGLSNHARICREEIFGPVLVALPFDDEAALVADANDSVFGLA
AGIWTRDYKRAWRVARALDAGTVWINTYKQFSISTPFSGRKESGMGREKGSLGIREYMQQ
KSLYWGLNDSPLPWAN*
>GCA_000452945.1_BPMSHR5858_genomic_02299_1 group_19365
MSILGIEQITYGVTDLATCRRFFADWGLREIAHDDAHAVFETMNGCTVRVVDARDPSLPP
AFEEGPTLREVTWGVGTQAELDALRGRLAGQPGHYETDDAIGCIDPNGMAIRIEVTRRRA
LDVKGSLPNVWGATLRVDQPAAIYARAEPVEVGHVVFFTNRLAEQEAFYHTLLGFETSDR
YPGRGAFMRCAPHGGHHDLFLLALPTAKRGLNHVAFTVRDIHEVFGGGMHVSRCGWRTQL
GPGRHPVSSAYFWYFRNPAGGLIEYYADEDMLTPAWQPREFEPGPTVFAEWAIDGGLDGD
TRRQKNAQAPQGKFMTERDDD*
>GCA_000452945.1_BPMSHR5858_genomic_02301_1 naiP_1
MSTLETAIAGAPAADAAAGAATPGAIIARIERLPANAMQIRARVLIGTATFFDGFDVITI
AATLPLLIHKWGLSPNQIGMLIASGAIGQLLGAFLFPALAQKHGRVRAIAWSSAVIGVTS
IACGFASTFETFVLLRIVQGLGLGGELPVAATYINEITRAHGRGRFVLLYEIVFPIGLLA
SMAFGAWLVPRFGWEVMYFIGGMPLLLALVLTRLVPESPRWLASRGRLDEAGRALHAFEA
SVKEPLPPVAHAGEFDTLVLSHPKRRMVDLFGPAYLKRTLAVATLWATCGFIQYGLSTWL
PTIYKNFYHAPLQLALNLAVIGSVMGVAGSLVSALAVDVVGRKPVIVWSFVLCALSLVFA
GVYHASSVYVVAAFCSLALGLMASGFITSYVYTPEQYPTSIRAAGCGLGSAWLKIASFVA
PMVVPRAIVGGDLSPAFYLLAAVPLIAALTVHSVGIETKGQVLEKLEA*
>GCA_000452945.1_BPMSHR5858_genomic_02302_1 sadA
MSLAVSAAMGCMATDAAAQVSYAAGENAYAGPGGNTGPWAFYNPAFSAGTLLYGTAVGNY
AYANGEGSSAYGDHATVKGRIGSAFGAYSEAAGDGSTAIGASARALPDFSIAIGTNAQAL
KDTGQSIPGREDIGTIAIGAGALAQGDNSDPLHVSAPNAFGGYSSATASGAVALGEGAAS
SGYYANALGSYSKASGAGAVAVGGGAQASAQGAVAIGGATSVDNATALSGYASASGVNAI
AIGSGAQATGARSISIGTGNVVSGASSGAFGDPSTVTGTGSYSFGNNNTINSNNAFVLGN
NVTIGPGFDGSVALGSGTTVAAANPTGSATITTSSGGQLTLSGFAGANPTSVVSVGAPGA
ERQITNVAAGRITPTSTDAVNGSQLYAVASTIDNAVNGGGIKYFHANSTLADSTAAGTDS
VAVGPAALAYGNDSIAEGTNATAGVSGNPAVAGDVALGSGAQATGGRSLALGANASVNTA

GGVALGAGSVANRAAGTYTDPITGSSFTTAFGAVSVGLEGSLRQITNVAAGTQATDAVNV
GQLQGAIAQLNQTIQNITNGSNSGNTGNNGNNTGQTVSGQWITGNPSTYTPPVASGIGST
AAGSGSVASGANSVAIGDGASASGNNSVALGAHSVASAPNTVSVGSVGNERTISNVAPGV
NGTDAVNVNQLNSGIGNAVGQANQYTDQKVDHLRREMNGGVAAAMAVAGLPQPTAPGKSM
VAIAGSTWQGQQGFALGVSTISENGKWLYKGSLTTSTRGGTGAVLGAGYQW*
>GCA_000452945.1_BPMSHR5858_genomic_02303_1 ompA_1
MKMNRMSKYAAAAGICAALLTGCGMAPTSAPDGSVGFPARDSAWLKEGTFVNVENLRQIG
PGLDKNQVYALIGEPHFSEGIIGVHVWNYVFNFRTGPGASDVVTCQYQIQYDKHYRVKST
YWKEPACKALVDGPKPAPAVQAPPGDDVKQFTLDYQLLFPFDKSALRDLLPAGRAELDRI
AATLEQQYQHIRSVRVTGYTDRLGTDAYNRQLSLARAETIRAYLTTRGLPGDLIVAQGLG
KADPVSNGCPAGRSAEAIACLAPDRRVRIDVAGERR*
>GCA_000452945.1_BPMSHR5858_genomic_02311_1 group_1842
MREVCWASLEHDGIEHLAFERHARGSVAESVVVGWSGGRAYGLAYRVVCDARWRAKHVIV
TMMGGGTLELRGDGEGRWRNAADERLGALDGCIDVDIAATPYTNTLPIRRLGLARGERRP
VDVVYVSIPDLAVSAMQQAYQCIVPDRVYRYESVVSGFTAHLEVDRDGLVVDYEALFRRA
SDDAR*
>GCA_000452945.1_BPMSHR5858_genomic_02313_1 ahpF_1
VNFDVVIIGGSFAGQAAALQLARARRGVLLIDAKRPRNRFARTSHGFLGQDGHAPSAINA
TAGRQLAAYPTVQFDAGDAIDAAAVSDGFRVSLADGRQAQGRRLILATGVRDILPVLPGL
QERWGVSVLHCPYCHGYELERRPMGVLASSEQAVHQAMLVSDWGPTTLFTQGAFVPTREQ
SAMLSARGVTVEQTPVAALVGAAPALEAVCLADGRTAALCGLFIAPRTVPACDLPERLGC
VFDDGPTGPYLSVDERQQTSVPGVFAAGDVARPMANATLAAAAGVIAAAGAHHSLIYGAG
ERG*
>GCA_000452945.1_BPMSHR5858_genomic_02315_1 dapE_1
VEEPQNTREAAIAAAYQAFDSGAFFADLQRRVAFRTSSQDGVNPALSAYLTDEISATLAR
LSFTARVVENPVAGGPPFLIAERHESDDLPTVLTYGHGDVVRGYDEQWRAALKPWEIVVD
ADRWYGRGTADNKGQHSVNFAALAAVLQARDGKLGYNVKVIFETGEEIGSPGLNEVCAAH
KLALAADVFIASDGPRVSAERPTLFLGSRGSVPFELTVNLREGGHHSGNWGGVLRNPAVV
LSHAIASLIDRDGRIVVEGLRPPPISGAVREALTDIELGRDDSSPDIDPTWGEPGLTPTE
RVIGWNALEVLAMKSGNADAPVNAIPPFAKAVCQLRFVVGTDWENVQAHLERHFAENGFA
QVAVEVARASPATRLDLDDPWVRWALDSMRATTGKKPALLPNLGGTVPNDVFAQTLGLPT
LWVPHSYPACGQHAPNEHVLAPVMRESLGMMAGLWWDLGEMGAAIVAARHAH*
>GCA_000452945.1_BPMSHR5858_genomic_02319_1 sadH_1
MKSFSNKVAAITGAGSGMGRSLAVQLAQQHCHVALADRNAVGLAETERIIRALAPSVRVT
SRVLDVGDRAAMHAWADETAAAHGRVNLVFNNAGVALSSTIDGMEYSDLEWIVNINFWGV
VHGTKAFLPYLKASGDGHVINTSSIFGIFAQPGMSGYNATKYAVRGFTESLRQELDMMKC
GVSATCVHPGGIRTNIAQSARLAQNMVGFMVESEQQGKADFEKFFITTADEAARVILHGV
RKNKRRVLIGRDARGADWLARTLPAAYQPLVVLATKLQSARARRAGRATPAPAHATYNN
AGNQGEQS*
>GCA_000452945.1_BPMSHR5858_genomic_02320_1 group_19030
MNARNMSFGPLDSTNREETLDIAIIGTGFSGLGMAIRLRQTGNQDFAVFEKASSVGGTWR
DNHYPGCACDVQSHVYSFSFAPNPRWTRMFAPQPEIRAYMEDCVQRFNVGSHLRFDHELV
NATYDETAHRWHLTFANGKRVCARVLVSGMGGLSRAAYPNIPGIETFKGEAFHSQHWKHD
YALEGKRVAVIGTGASAIQFVPQIAPRVAKLALFQRTPPWIMPKPDRAVSGVEQWLFRNL
PFTQKLMRAGLYWMLESRVLGFAVHPSTMKLVQKLALRHIDKQISEPELRAAVTPDYTIG
CKRILMSNDYYPALTRKNVDVITTGIERIEEDAVVTTDGKRHEVDCLIYGTGFQINNPFP
RGTIVGRGGVDVVDAWRDGLHAYLGATVPGYPNWFILMGPNTGLGHNSMVYMIESQIEYV
LGALKAMRTERAEAIEVRPLVELEYNNRIQNKLKGTVWSTGGCSSWYLDPRTGKNTTLWP
TFTWRFRQETAFFSTADYHLYYANPRQSPQPVPALASAAAATTTAETTA*
>GCA_000452945.1_BPMSHR5858_genomic_02323_1 group_19141
MSNEYTVDELARVSDTTVRNVRAYQDRGLLAPPRKRGRVGIYDDTHVARLKVINHLLARG
YTLSNIQDLIKAIDDGHDLRSILGLENAIGGRWSNERPQTFSLAQLIQMFGPQTPASLGR
VAELGLLERRGVSFVAKSPALIEAAAAMVREGIPARELLDAISITRPYFDAIARVLVELV
VRRLDRYDEGSLPPPTDVPALVDAIWRLRPLSSVFVEGEMNRALEAASSAYLGGRVATIL
DKKLSSQAAREGEPELERVTAKGAPKKR*
>GCA_000452945.1_BPMSHR5858_genomic_02352_1 vgb
MIHIRLKRASFLLLLGAQAVSLGAAAGTLSGTVSSAGTPLAGAMVTVFDAAQARRDTVYT
DHNGRYRITVDFAGELRVRARTPYFKDAAQDLALAPDASKTLDFSLARQTVADELSASLP
ASAHLATLPWSSQDSRTAFISQCNYCHQVGNALTRTPRDEAAWGATVRRMEGYAALLTDR
QARDITHTLYQGMNAHAVAAVEKYLYDDRLAPAKIREWAAGDGLTFIHDADVGFDDHLYG
ADEGHDKIWELDRKTGRLTEWKEPDVDLPVGGIFSGVQLPIGVFSGKHGPHSLAQAPDGR
FWITNALSSTLASFDPATKRFKLYELGHSHLYPHTLRIDRAGIVWFTIVASNEVARFDPK
TGRFTIIHLPDGGIWRAMSHYLFPLVVKMAAWFPGQNLHLALTHHKWAFQGRDAFPFPYG
IDVNPVDGSIWYAKLYANKIGRIDPKTLAVTEFDTPLGGPRRLRFDPQGNLWIPAFDDGG
LMRFDTRTHRFETFKLPLLAHNEYEVPYALNVQPKTGDIWITSNMSDRIFRFVPSTQTFI
TYPLPTRVTWLRDMTFTQDGAVCSSSSNLPAYGIEGGRASFICLYPDGERGGTAANAAGN
ANAAANLARVRAAAARRTPERTAPARAALARAGSNRDGAIQTAAIQADGTRTDRSAH*
>GCA_000452945.1_BPMSHR5858_genomic_02371_1 nirD
MNRQPQASAWTPVCALDEIVPNTGVCALVNGEQIAVFHVDDGGGGRAFAIGNVDPHSQAA
VLSRGLIGSLGERIVVASPLYKHHFDLRTGECVEAPEKSVSAYPSRVEDGYVWVAA*
>GCA_000452945.1_BPMSHR5858_genomic_02441_1 group_19470
MSENRPLLTALRRAALAFGMCATLVANGASAEPLYAGEDALYAKAADEGLVVSFDTGPEW
ANWKALFAAFRKRYPKVELTYNDIGSAATVVALDKSRRRPQADTAYYFAASALDAAGKDV
VAPFKPVNFDKLPPVFREADGRWFTVHSLNVAFLVNRKLVKNVPRRWSDLLKPEYRNAVV
YLDPRSTGQGQVVVFAAASALGGGVDDPKPGAEFFGKLKHAGNVLRIEGTTPYAKFVKGE
IPILIGYENDGLKAKYADGLGDAVDVVIPQDGSVCAPYAMSLVKNGPNPAAAQLWLNFVM
SDAGQALFAHGYVRPAVPGVALAPDVAAKMPNAPQVRALDVAKAAARKAEVDRLWSQAAL
GQ*
>GCA_000452945.1_BPMSHR5858_genomic_02446_1 rpiB
MKVAIGCDEAAYALKESIKRHLRATHPELELVDFGTHSADEAVLYPDIAIEVAQRVAAGE
FPRAILLCGTGIGVAISANKVPGIRAAQCHDTYSAERARRSNDAQIVTMGARVIGTELAN
SIVDAWLRAEFDGGRSQPKVQRIADYERRAGLTQAGPNDAHRGS*
>GCA_000452945.1_BPMSHR5858_genomic_02450_1 golD_2
MTNLWDRAFDLTGRVALVTGGAAGIGHACARLLAQRGASVALVDRHPETAGIAATLEGGA
ARHSGMSLDLRDCSAAQAGVALAASRFGGVDMLVNSAGVALLDKALDVGEAAWDATMAIN
VKASFFVAQAAARQMIAGARGGRIVNLASQASVVGLERHAAYCASKAAIVGMTKALALEW
APHGITVNAVSPTIVETALGKQAWAGEVGERAKREIPAGRFAQPDEIAALVLYLLSDAAA
MMTGENVVIDGGYTVR*
>GCA_000452945.1_BPMSHR5858_genomic_02453_1 dhaL-2
MNTAAIQCLPLADAGFVVRELVDVIRRNRDHLSEIDAAIGDGDHGINMSKGFGQCGARLD
ARGATSLPDALDVLSTALMDGIGGSMGPLYGSFFMDFAAPLKGRDTLDAALFGEALAAGF
AGVQAISDAKIGDKTLIDTLAPAAAAFHEALRAGDDFRHALAAMSAAAERGKESTRWLQA
RVGRASRLGERSVGTLDAGAASCCLILCSLASSIGARLN*
>GCA_000452945.1_BPMSHR5858_genomic_02457_1 group_19622
LNKKLLTTAILAATASAAHAQSSVTLYGLIDAGVSYVNHSKSASGSDKLFKYDDGIAQGS
RWGLRGAEDLGGGLKAIFVLENGFNSGNGTIGQGGAIFGRQAYVGLSHKDAGTLTFGRQY
SFTTDILGANYSTGGNTVAGNYAYHVNDIDQLTSSRINNAVKFQSANYAGFTFGALYGFS
NSTSFAGAPATTTGTTTTPGSSRAYSFGLNYANGPIGLGAAYTDIRYPSQATPAFTTNIA
NLTISNVRDLRSFGVGGRYIWGPATLWALWTNTRLEPVTGGSTTYNAYEAGGKYAFTTAL
SAGVGYTYSRVSGANSGHWNQVNGALDYALSKRTDVYLLAIYQEASGKNLQAQIGSSTSY
FNTSGTGSKNQFAARVGIRHKF*
>GCA_000452945.1_BPMSHR5858_genomic_02470_1 fusA_2
VPRKTPIERYRNIGISAHIDAGKTTTTERILFYTGVSHKIGEVHDGAATMDWMEQEQERG
ITITSAATTAFWKGMAGNYPEHRINIIDTPGHVDFTIEVERSMRVLDGACMVYDSVGGVQ
PQSETVWRQANKYKVPRIAFVNKMDRVGADFFRVQRQIGERLKGVAVPIQIPVGAEEHFQ
GVVDLVKMKAIVWDDESQGVKFTYEDIPANLVELAHEWREKMVEAAAEASEELLEKYLTD
HNSLTEDEIKAALRKRTIANEIVPMLCGSAFKNKGVQAMLDAVIDYLPSPADVPAILGHD
LDDKEAERHPSDDEPFSALAFKIMTDPFVGQLIFFRVYSGVVESGDTLLNATKDKKERLG
RILQMHANERKEIKEVRAGDIAAAVGLKEATTGDTLCDPGKPIILEKMEFPEPVISQAVE
PKTKADQEKMGLALNRLAQEDPSFRVQTDEESGQTIISGMGELHLEIIVDRMKREFGVEA
TVGKPQVAYRETVRTVAEDVEGKFVKQSGGRGQYGHAVIKLEPNPGKGYEFLDEIKGGVI
PREFIPAVNKGIEETLKSGVLAGYPVVDVKVHLTFGSYHDVDSNENAFRMAGSMAFKEAM
RRAKPVLLEPMMAVEVETPEDFMGNVMGDLSSRRGIVQGMEDIAGGGGKLVRAEVPLAEM
FGYSTSLRSATQGRATYTMEFKHYAETPSNVSEAVINAKQIGRG*
>GCA_000452945.1_BPMSHR5858_genomic_02475_1 cspA
MATGIVKWFNDAKGFGFITPDEGGEDLFAHFSAINMQGFKTLKEGQKVSFEVVQGPKGKQ
ASNIQAA*
>GCA_000452945.1_BPMSHR5858_genomic_02477_1 clpA
MIAQELEVSLHMAFMEARQARHEFITVEHLLLALLDNPTAAEVLRACAANIEDLRQNLRN
FIHDNTPTVPGTDDVDTQPTLGFQRVIQRAIMHVQSTSNGKKEVTGANVLVAIFGEKDSH
AVYYLQQQGVTRLDVVNFISHGIAKTSSGEAAKPADANAEGEDAGAQKETPLAQFTQNLN
QMAKDGRIDPLIGRESEVERVVQVLCRRRKNNPLLVGEAGVGKTAIAEGLAYRITRGEVP
DILANAQVYSLDMGALLAGTKYRGDFEQRLKTVLKELKERPHAILFIDEIHTLIGAGAAS
GGTLDASNLLKPALSSGTLKCIGATTFTEYRGIFEKDAALSRRFQKIDVTEPSVEQTVAI
LRGLKSRFEEHHGVKYSSGALSAAAELSARFITDRHLPDKAIDVIDEAGAAQRVLPKSKQ
KKTIGKSEIEEIISKIARVPPQSVSQDDRSKLQTLDRDLKSVVFGQDPAIDALAAAIKMA
RAGLGKLDKPIGAFLFSGPTGVGKTEVARQLAFTLGIELIRFDMSEYMERHAVSRLIGAP
PGYVGFDQGGLLTEAVTKKPHCVLLLDEIEKAHPDIFNVLLQVMDHGTLTDNNGRKADFR
NVIIIMTTNAGAESMQKATIGFTTRRETGDEMADIKRLFTPEFRNRLDATISFRSLDEEI
IMRVVDKFLIQLEEQLHEKKVDALFTDALRKHLAKHGFDPLMGARPMQRLIQDTIRRALA
DELLFGKLVNGGHVTVDVDENDKVLLSFDKTATPPSKPNEEAVEVE*
>GCA_000452945.1_BPMSHR5858_genomic_02478_1 pheP
LTQQQRTFDSIVEREKGLRRGLSSAQMAMIAIGGAIGTGLFLGSGFAIGLAGPGVLVSYA
IGALIALLLMGALAEMTVAHPTSGSFGAYAEHYVGPLAGFLVRYAYWSAVVFAVGTEVSA
IAVFMKYWYPGVPGWYWVVGFSAVLIAVNMTSVTLYGVVEYVFSMLKIAAIVAFIALGAY
FVAAAPVSSGIGFANYSAHGGFFPKGFSGVWVAVIVAIFSYMSIETVAIAAGEARDPQRA
VSRAFRSTVFRLVLFYLLTLALMLAIVPWTQAGTDESPFVKVMAATHVPYAAGVINFVVL
VAALSAMNSQLYVTTRMMFSLSRARLAPAMFGRVGANGVPLAALAVSSSGVAVAAVLVAL
YPQTAFTLLMAIAMFGALFTWLMIFVTHLRFRARYRGPTPVFRMWLHPAGSLTGAALVGA
VLVTTAFTREFRMTMMVGVAFIVLLTLAYRWHYRARHA*
>GCA_000452945.1_BPMSHR5858_genomic_02481_1 dut
MKLDLKILDARMRDYLPKYATTGSAGLDLRACLDAPVTLKPGDTALVPTGLAIHLADPGY
AALILPRSGLGHKHGIVLGNLVGLIDSDYQGELMISTWNRGQTEFALNPFERLAQLVIVP
VVQARFNLVDDFAQSERGAGGFGSTGRG*
>GCA_000452945.1_BPMSHR5858_genomic_02482_1 coaBC
LAHAELAGKHLVLGLTGGIACYKIAELTRLLTKAGATVQVAMTEAATQFITPVTMQALSG
RPVYTSQWDARMPNNMAHIDLSREADAIVIAPASTDFLAKLAHGFADDLLSTLCIARDCP
LLVVPAMNRQMWQNPATQRNVAQLRHDGVAVLGPDSGAQACGEVGDGRMLEPEAIYEAIV
THFQPKALAHRRVLITAGPTFEPLDPVRGLTNRSSGKMGFALARAAQQAGADVHLVAGPV
HLPTPWGVYREDVQTAQQMYDAVMNAVPDADVFIAVAAVADWRAAQPAEHKIKKTADHKM
PALTFVENPDILASVAALPDAPYCVGFAAESGDLDVHGEEKRVRKRVPLLIGNLGPLTFG
LDDNEVVLFEASGTTRLPRADKMTLAHALVAEIAKRLPDTRLI*
>GCA_000452945.1_BPMSHR5858_genomic_02483_1 lspA
MAKTLSKSSGGALAPWLGISLIVILFDQLTKIAVLKTFAYGAMHALTPFFNLTLIYNRGA
AFGFLATAGGWQRWAFTALGIGATLVICYLLKRHGHQRLFSLSLALILGGALGNVIDRLI
YGHVIDFLDFHVGAWHWPAFNLADSAITVGAVLLIYDELRRVRGTR*
>GCA_000452945.1_BPMSHR5858_genomic_02484_1 ileS_1
MSNKKADSKPQAKYPVNLLDTPFPMRGDLPKREPQWVEDWEARGVYEKIRAASQGRPKFI
LHDGPPYANGDIHLGHAVNKILKDMVVKSRNMAGFDAPYVPGWDCHGMPIEIQIEKRFGK
SLPAAEVMAKARAYATEQIEKQKVGFKRLGVLGEWGNPYKTMNFQNEAEEIRALGKIIEK
GYVYRGLKPVNWCFDCGSALAEAEVEYKDRTDPTIDVLFAFAEPEKTAHAFGLAELPRAE
GGIVIWTTTPWTIPANQALNLHPEIVYALVDTERGLLVMAEERVEACMKDFGLTGRVIAS
TPGEKLANLRFHHPLAAAHPGYKRTSPVYLGDYVTTDTGTGVVHSSPAYGVEDFTSCKAH
GMTDSDIINPVMGDGRYIESLPLFGGLTIWDANPKIVDALKAAGSLLRNERYAHSYMHCW
RHKTPIIYRATSQWFAGMDTQPADGGKTLRETALDAVDATAFYPSWGKQRLHAMIANRPD
WTLSRQRQWGVPMAFFVHKETGELHPRTLELLEEVAKRIERQGIEAWQTLDARELIGDDA
NLYEKNRDTLDVWFDSGTTHWHVLRGSHKDQLQFPADLYLEGSDQHRGWFHSSLLTASML
DGRAPYKGLLTHGFTVDGEGRKMSKSLGNGIDPHEVANRLGAEIIRLWIASTDYSGELAI
SEEILKRVTEGYRRIRNTLRFLLANLSDFDYAKDALPAGQWLEIDRYAVAFAAQLQAELL
AHYEKYEFHPVVAKLQTFCSEDLGGFYLDVLKDRLYTSAPASPARRSAQTALYHVTQGLL
RVLAPFLSFTAEEAWRVFQPQSDTIFTETYYAYPEIAGAEALIAKWTLLRDVRGDVTKAL
EEARTANRIGSSLQAQVEVRASGARYDALASLGDDLKFVLITSAATVVKVDAQGDESVDV
AASTYPKCERCWHYREDVGAHADHPTLCGRCFSNLFENGEIRSAA*
>GCA_000452945.1_BPMSHR5858_genomic_02485_1 ribF
VKVFRGLPNAESRAPCALTIGNFDGVHRGHQALLARVRAAADARGLPVCVMTFEPHPREF
FNPASAPPRIAMLRDKLEALREHGVDRVVVEHFNHTFASQSPEAFVERTLVHGLHTRWIM
VGDDFCYGARRAGDFASLKAAGERFGFEVEQMETLADADGARISSSGVRAALAAGDLDAA
ARALGHGYAISGHVAHGLKLGRDLGFPTLNLPIAHKRPALSGIFVVRVHGLAPEPLPGVA
SLGVRPTVDDSGRVLLEVHLLDWHGDAYGKLVRVEFLKKLRDEAKFVDLETLSRAIAQDV
VDTRAYFAEHGRAPGSRATGFATSATDRIS*
>GCA_000452945.1_BPMSHR5858_genomic_02495_1 group_1268
MQYEIVSAGEDVDDERARGRRAAAPAAPSSAVPSSAALSSAALSSAAQPTGAPPATAAAR
RGRDLPRERLLARGPAALSDAELVALLLGSGLPGHDVFALAHTLLARFGSLRALLDAAPD
DFKGLRGIGPARTAILVAVVELARRALAEKARERPLVDSPGAVDDYLRLLIGTRPREVFV
CLFLDARHRLVQTEETAHGSLTRMAVYPREIVRRALALNAAALIVAHNHPSGAVRPSAAD
RRLTRVLRDALALVDIKLIDHFVVGASDTFSFAQAGWI*
>GCA_000452945.1_BPMSHR5858_genomic_02499_1 livH_6
MDIFVQQILNGLVLGSVYAIIALGYTMVYGILGIINFAHGDVLMVGAMVALSAITVLQNH
FPELGHVATLTIGLLIAAVVCACVGFTIERVAYRPLRRAPRLAPLITAIGVSILLQTAAM
MIWSRNPLPFPQLLPTDPINVIKAGENNPGAVISMTEITIIIVAFLVMAGLLLLVHKTKL
GRAMRAIAENPNTASLMGVNPNFVISATFMIGSALAALAGVMIASEYGNVHFYMGFIPGM
KAFTAAVLGGIGNLGGAMVGGVLLGLIEQLGAGYIGNLTGGVFGSNYQDVFAFVVLIIVL
VFRPSGLLGERVADRA*
>GCA_000452945.1_BPMSHR5858_genomic_02500_1 group_19587
MTSIQPIETSASLVEERHPAKTATVGILIAAFVIAAPLVIGAAGGNYWVRVLDFAMLYVM
LALGLNVVVGFAGLLDLGYIAFYAVGAYTAALLSSPHLTSHFDWIAQMVPGGLHVPFLLI
VPIAMALAAIFGILLGAPTLRLRGDYLAIVTLGFGEIVRIFMNNLDRPVNVTNGPQGITG
IDPVQVAGFNLSQTHEIFGFSLPSVYMYYYLFVLCALLVIWVCTRLQHSRIGRAWAAIRE
DEIAAKAMGINTRNVKLLAFAMGASFGGLSGAMFGAFQGFVSPESFTFWESVVVLACVVL
GGMGHIPGVILGAVLLAVFPEFLRSTMSPLQHMLFGHDIVDTEVIRQALYGLAMVVIMLY
RSEGLWPAPKHEDRIAKLAKRGGKKPVRA*
>GCA_000452945.1_BPMSHR5858_genomic_02501_1 lptB_4
MSEQIRLSVKGVNKRFGGLQALSDVGLEIREGQIYGLIGPNGAGKTTFFNVITGLYTPDS
GEFKLDGTEYTPTAVHQVAKAGIARTFQNIRLFGGMTALENVMVGRHVRTKHGLLGAVFQ
TPAERREEREIKERAIELLDYVGVLQYADYTARNLSYGHQRRLEIARALATDPKLLALDE
PAAGMNATEKVELTRLLDKIRSDGRTILLIEHDVKLVMGLCNRMTVLDYGKVIAEGLPQD
VQKNPKVIEAYLGAGVH*
>GCA_000452945.1_BPMSHR5858_genomic_02502_1 livF_7
MAAAMLKIKGLQVNYGGIQAVKGVDMEVRQGELVTLIGANGAGKTTTMKAITGLKPYSAG
DIEYDGQSIKGVPPHELLKRGLAMVPEGRGIFARMSIIENMQMGAYLRNDNEQIKKDVER
MFGFFPRLKERATQLAGTLSGGEQQMLAMSRAILSKPKLLLLDEPSMGLSPIMVEKIFEV
VREISKEGITVLLVEQNARLALQAADRGYVMDSGTVTMEGDAKQMLDDPKVRAAYLGE*
>GCA_000452945.1_BPMSHR5858_genomic_02506_1 ascD
MAFNVTLKQSGRQFQVEADETVLAAALRQNVHLPYGCKNGACGSCKGTIVQGQFEQGPHS
ASALSNDERTRGLALLCCSKPQSDLEVDVREIAGVDGVQVKKLPCRVAALERRADDVIVL
KLQLPANERLQYLAGQYIEFILKDGARRSYSMATAPHEEGPIELHIRHMPGGKFTDHVFG
PMKERDILRFEGPLGTFFLREDSDKPIVLLASGTGFAPIKAIIEHAHHVKLERPMTLYWG
ARRKKDLYLLDAAEQWAKEIPNFKFVPVLSEPDASDAWTGRTGFVHRAVIEDLPDLSGHQ
VYACGAPVMVESAQRDFTQHHRLPADEFYADSFTSAADLAHTV*
>GCA_000452945.1_BPMSHR5858_genomic_02509_1 rsuA
MDLESILFTQGFGSRRQCRALVEAGHVAVGGATCADAHASFDTTQLVFEVDGVAWPYRAR
AYVALNKPAGYECSREPQHHSSVFALLPPQFAERGVQCVGRLDQDTTGLLLLSDDGQFVH
AYTSPKRKVPKTYVATVRHPLDDAQLDALRAGVLLHGEPKPVAALAAVARGERLLEMTIA
EGKYHQVKRMVAAASNRVEALHRSRIGGFALPEDLAEGAWRWLDERDLAALRDTAETLSG
*
>GCA_000452945.1_BPMSHR5858_genomic_02511_1 pgrR_5
MREVNRFSEMAVFVQVIESCGFSAAGRRLDMTPSAISKFVQRLETRLGVRLLNRTTRKLQ
LTPEGAAFYERCVRILEDIADAEREATHGATPQGRVRINSHVPFGKHHLMPLLPEFQRRY

PDITLDIVLSDSLVDPLDERSDIAIRSGALDDSRLVMRKLGGSRMVVVGAPAYLERNGVP
ATPDDLDRHNCIGLSGNAKAWPFVAEEGQRLLPSAGNLLLGDNESIREAAVSGLGLARLA
RFHVDADLDAGRLVPVLEAFNPGDEEEINAVFVGPSEHLPMRVHALLDFLKERIRIGPRA
AGERGEASALGARAFVTRLR*
>GCA_000452945.1_BPMSHR5858_genomic_02512_1 hdfR_3
MNAAPDSAAMSLDIVLLRTFLEVADRRGFAPAAERLALTPSAVSGHIKRLEQATGAVLLA
RTTRRVELTQAGETLYAYARNIVEMEREARARLRGSPLHGRLRIGASEDFASAWLPQVLH
TFGRLHPRATIELKVGITAELLRQQASGRLDVVFGKHCSRLDDEAGELLWEEPLVWAFAS
DAVLDANDALPLALFPQPCVYREAALAALNGALRPFRIAFESGSMAGCISAALAGFALTV
VARSQLRHGLRECGPDDALPALPAARFYAFPHRPTPAGTALIDAVRETGRRRGFAPKP*
>GCA_000452945.1_BPMSHR5858_genomic_02514_1 yiiM
MHPTTAKLDALLTGAIRPLAGTARTSAIDKQPVRERSWLARTGFAGDEQADRKHHGGPDK
AVHHYAFDHYPQWIAEIGPRDVLARPGAFGENLSTRGVTERDVCVGDVFTLGGATLQVSQ
SRQPCWKLNARFGLPKMSARVQQSGRTGWYYRVLDEGWVEPDAELALRERPYPQWPLSNV
LDVLYRRTLDVGALEALSRIDALPPSWRQLFERRLKAGAVEDWTKRLDE*
>GCA_000452945.1_BPMSHR5858_genomic_02519_1 cysB
MNLHQFRFVREAVRQNFNLTEAAKALYTSQPGVSKAIIELEDELGVEIFTRHGKRVRSLT
EPGRIILASVERILQEVESLKRVGKDYAAQDQGNLTIAATHTQARYSLPAAIAEFKKRFP
KVHLSILQGSPTQVAEMVIHDQADLAIATEAIADYKELVSLPCFQWHHAAVVPADHPLLE
RKPLSLDDLAQFPLITYDDAFAGRKKINQAFALHRLTPDIVLEAIDADVIKTYVELGLGV
GIMADIAFNPERDRNLRAIAVGHLFGSNVTRVALKQGAYLRGYIYTLVELLSPTLNRKLV
ETALKGEAETYEL*
>GCA_000452945.1_BPMSHR5858_genomic_02521_1 group_19355
MALIIKNREVVDDAWQLVRAAEDGTLPAPDALPAGQVLVPLALWQTARDALAASRTKDAL
GVWLSPDSEPADLAADFDRIAVIGVEFPRFADGRGYSIARLLRERHGWKGELRAIGDVLR
DQLLYMSRCGFDAFAVRADKDVRDALNAFGEFSQRYQGAFDEPAPLFRRRAAAAEAKVSA
*
>GCA_000452945.1_BPMSHR5858_genomic_02528_1 moeA_1
MSHPTPQTPRVPMLSTADALTALLAAARPLAEIETVPTLEALNRVLAADVASGLDVPPMN
TSAMDGYAVRIADLTRGERRLPVSQRIAAGHPAAPLAPGTAARIFTGASVPAGADAVVMQ
EQTEAAGDQVDILHTPTVGEWITAQGADIRRGSVILPAGTRMTPQAMGLAASVGCASLAV
TRRVKVAVFFTGDELTMPGEPLKPGAIYNSNRFTLRGLLERLGCDVTDYGIVPDRLDATR
AALREAAREHDVIVTSGGVSVGDEDHVKPAVEAEGRLALWQIAMKPGKPLAFGAIRRDGA
ASGAAAVEANAHGASDASDTRAMHATSGGEAYFIGLPGNPVSSFVTFLLFVRPFLLRVAG
AAQVAPRAVSLRADFTQNKADRRNEFLRARVNAAGGLDLFPNQSSAVLTSTVWGDGLIDN
PPNHAISAGETVRFIPFSELLG*
>GCA_000452945.1_BPMSHR5858_genomic_02610_1 relA
MTADSASTAAAPSLDDVLAFVREHAGDARLSSGERLADHAAGTSTIMQRLNVDPPATQAA
ALFALAPYLSDPDKQIAGHFGDEVAHLVADVRKLLRLGTVSLRAAQSAPVDAGRDAQAER
RAQIEALRKMLLAFAQDIRVVLIRLASRLQSLRYYAAAKIEPPPDVARETLEIYAPLANR
LGIWQLKWELEDLAFRFEDPVTYKRIAKLLDEKRVEREAYVAGAIERLQHELAAAHIQAE
VSGRPKHIYSIWRKMRGKELDFSELYDVRAFRVIVPDIKDCYTVLGIVHHLWQPVPKEFD
DYISRPKPNGYKSLHTVVIGDDGRAFEVQIRTQEMHRFAEYGIAAHWRYKEAGARGYGGQ
FSASEKYDEKIAWLRQLLAWKDEIADGGGAEVSGQQAWAQLREATLDDDHIYVLTPQARV
IALPQGATPVDFAYHLHSELGHRCRGARVDGAMVPLNTQLANGQTVEIVAVKEGGPSRDW
LNPQLGYLHSSRARQKVRAWFNAVDQQENVAHGRALVEKTLQREGKTSVNLEHLAAKLGF
KSPDELFSVVGKEEFSLRNVEQALSDAPAPEPEPEAPANFEKRASGANVAHGASAGVLVV
GVDALLTQLARCCRPAPPDPISGFVTRGKGMSIHRSDCATFRRMAERAPERVLQTTWSAD
VLGGRGASVYPVDLMIEATDRQGLLRDISEVFAREKMNVIGVKTQSRRNAAFMQFTVEVS
SASQVQRACTLLGEIQGVVRAARKA*
>GCA_000452945.1_BPMSHR5858_genomic_02614_1 rpmI
MPKMKTKKSAAKRFVVRPGGTVKRGQAFKRHILTKKTTKNKRHLRGATAVHDSDLNSVRA
MLPFA*
>GCA_000452945.1_BPMSHR5858_genomic_02616_1 pheS
MGSMDLDQIVADAQQSFEGAADITTLENEKARFLGKSGALTELLKGLGKLDPETRKTEGA
RINVAKQQVEAALNARRQALADALLNQRLAAEAIDVTLPGRGAGAGSLHPVMRTWERVEQ
IFRSIGFDVADGPEIETDWYNFTALNSPENHPARSMQDTFYVDGKDADGRPLLLRTHTSP
MQVRYARMNRPPIKVIAPGRTYRVDSDATHSPMFNQVEGLWIDENVSFADLKGVYTDFLK
KFFERDDILVRFRPSYFPFTEPSAEIDMMFEHGKNAGKWLEISGSGQVHPTVIRNMGLDP
ERYIGFAFGSGLERLTMLRYGVQDLRLFFENDLRFLRQFA*
>GCA_000452945.1_BPMSHR5858_genomic_02617_1 pheT
MQFPESWLRTFVDPQLTTDELSHALTMAGLEVESLRPAAPPTEKIVVGRVLEVVKHPDAD
KLNVCQVDAGTGATLQIVCGAPNVAPGIKVPVALVGAKLPPAEEGGAPFAIKLSKLRGVE
SQGMLCSARELKLSEDHSGLMILPEGTPVGQDIREALNLDDTVFEIKLTPNKADCLSVFG
IARETAAITGAPLAAPDIRPVLAELTETLPVKISAPDLCGRFSGRVIRGVNARAKTPHWM
VERLERAGQRSVSALVDISNYVMFELGRPSHVFDLDKIHGGIDVRWGKRGESLKLLNGNT
IELDETVGVISDGAQVESLAGIMGGDSTAVTLDTTNIYLEAAFWWPDSIRGRARKYNFST
DAAHRFERGVDYSTTVEHVERITQLILDICGGQAGPVDDQIVNLPQRAPVSMRASRANRI
IGVEIGEDEIAQIFTRLGLAFERDGDVFRVTPPPHRFDIEIEEDLIEEVARIYGFEKIPA
RPPVAKSEMRATDETRRSVHAIRHALAARDYAETVNFSFVDAEWERDFAGNDNPVRLLNP
IASQLSVMRTTLFGSLVGVLRHNLNRRAERVRVFEAGRVFVADPSVKAGELAVEGYAQPK
RIGALAYGPVVEEQWGTATRQVDYFDVKGDLEALLAPVPARFVKAEHPALHPGRSARIEV
DGHAVGWIGELHPRLMQKYELPHAPVMFEIDTDALVARALPAPSEVSKFPPVRRDIAVVV
DQKVEVQALFDEMKKALADDACKFVQRVALFDEFRAKSNTSGGLSAHEKSLAFRVTLQDA
AGTLQDETVDQAIQTLVDRMARVYGARLRG*
>GCA_000452945.1_BPMSHR5858_genomic_02619_1 group_19063
MTSTVEKVVLPPIPAKRYFTIGEVSELCGVKPHVLRYWEQEFTQLRPVKRRGNRRYYQHH
EVLLIRRIRELLYEQGFTINGARNRLDAVGSERGAPPEPDELEQHAAEHAARETVDVAQL
RRALLDVIGALKRD*
>GCA_000452945.1_BPMSHR5858_genomic_02623_1 group_1106
MPLKIAWYTAALAATLAACGGGGGGGGSTTSNPNGTGVTPSSAANVQPVTVSAGITGAAN
MLMTSVTVCVPNTNQCQTIDNVQVDTGSDGLRILAAALPSSVALPVVPAANAPITGACAI
FGTGFTWGAVRRADVRIAGEIAANIPIQVIADSSTPNAPIDCTQSAGPMQTQGALRANGI
LGIGLFVTDCGTACANSALPGWYYGCTTSGTCASTSQPIAQQIANPIGAFASDNNGSIIV
LPAISPTGAATVNGSLIFGIGTQANNGLGTATPLVTTASDAYVTSVLNGNTNARTFFDSG
SNGIFFANASLPQCGSWFCPASTQTFDITLSSGNGASKATTFQVASSQVLFSTFNFAFND
LAGPIGNVVDLGLSFFYGRNVYTALEGSATPAGPGPYYAY*
>GCA_000452945.1_BPMSHR5858_genomic_02627_1 group_19032
MSNIQLDIEWTEAATRKIKQLMPRGSDDAFLALPPIECLPMEGDVLFLGPQGKQQPFIVA
ERQYHHDGDADWTIILILDVPNASH*
>GCA_000452945.1_BPMSHR5858_genomic_02632_1 group_19110
MMLFDDLRDNEWALLEGLFCSEPARSERRGRPRVEARSVVNAVLWVLSTGEGWSKLPGRY
PSPPTCRRRFDEWQADGTLAEVVKRLGTSGRQISLRGRIGASAVKPPAPPSRDRLRGAFW
TNPESWRAPVKMA*
>GCA_000452945.1_BPMSHR5858_genomic_02633_1 gltC_3
MNITLRQLRVFIEVARLKSFSRAGDEIGLTQSAVSRSVRELEGEIGLKLIDRTTRDVQLT
DVGTNLIASVSRLLGDLDDALREIREIGEQRRGRVIVAASPTIACRLMPRVVAACERRFP
FVTLGLRDDVQSDVLRKVKSGEVDFGVVIGPLATSELISEPLMTDSFCLVARADHPLAAR
DRVPWTALAGERLVLLDYASGSRPLIDAALAAHRVAAGVVQELGHSATVFGLVEAGVGVS
VLPWLSLPLPAGSSLVARPLAPRAERTVELVRRRDRSLSPAADAVWGLVKEMPARVEDLN
*
>GCA_000452945.1_BPMSHR5858_genomic_02639_1 rbfA
MSKKRSSPNRNVQIADQIQRDLSELIMREVKDPRIGIVTIQSVELTPDYAHAKVYFTALT
GTPADTQEALNHAAGHLHNLLFKRLHIHTVPTLHFHYDQTIEKAVAMSRLIDEANATRAK
DD*
>GCA_000452945.1_BPMSHR5858_genomic_02644_1 slyA_2
MSDPSSPQPDQDLALYQINDSVGYLMSRVKSLMTNMVTQRTQTELGITGTQATMLFMLAV
GKCSTAAELAREYGIDASAITRLLDRVEKRGLLQRVRSSEDRRVVRLELTDEGRALTKRM
PAIFRSVLDEVLDGFTPEEVGFLKSMLRRILVNSGECPGATGSTS*
>GCA_000452945.1_BPMSHR5858_genomic_02645_1 typA
MTRALRNIAIIAHVDHGKTTLVDQLLRQSGTFRENQQVAERVMDSNDIEKERGITILAKN
CAVEYEGTHINIVDTPGHADFGGEVERVLSMVDSVLLLVDAVEGPMPQTRFVTKKALALG
LKPIVVINKIDRPGARIDWVINQTFDLFDKLGATEEQLDFPIVYASGLNGYASLDPAARD
GDMRPLFEAILQHVPVRPADPDAPLQLQITSLDYSTYVGRIGVGRITRGRIKPGQPVVMR
FGLEGDVLNRKINQVLSFQGLERVQVDSAEAGDIVLINGIEDVGIGATICAVEAPEALPM
ITVDEPTLTMNFLVNSSPLAGREGKFVTSRQIRDRLMKELNHNVALRVKDTGDETVFEVS
GRGELHLTILVENMRREGYELAVSRPRVVMQEIDGVKHEPYELLTVDLEDEHQGGVMEEL
GRRKGEMLDMVSDGRGRTRLEYRIPARGLIGFQSEFLTLTRGTGLMSHIFDSYAPVKEGS
VGERRNGVLISQDDGAAVAYALWKLQDRGRMFVKPGDALYEGMIIGIHSRDNDLVVNPIK
GKQLTNVRASGTDEAVRLVPPIQMSLEYAVEFIDDDELVEVTPQSIRLRKRHLKEHERRR
ASREAEAG*
>GCA_000452945.1_BPMSHR5858_genomic_02646_1 sucA
MSDVMKQFQLNSYLFGGNASYVEELYEAYLDNPASVPDNWREYFDALQNVPATDGSNASD
VAHNPIVESFAQRAKANAFIPRESGGNLATARKQVHVQSLISAYRFLGSQWANLDPLKRR
ERPAIPELEPAFYDFSEADLDQTFSASNLYFGFEQASLRDIVKGLRDTYCGTIGAEFMYI
SDPEQKRWWQERLESTRATPNFSADKKKHILNRLTAGEGLERYLHTKYVGQKRFSLEGGE
SFIAAMDEVVQHAGSKGVQEIVIGMAHRGRLNVLVNTLGKMPADLFAEFEGKHVDDLPAG
DVKYHKGFSSDIATEGGPVHLSLAFNPSHLEIVNPVVEGSAKARMDRRGDADGLQVLPVQ
IHGDAAFAGQGVVMETLNLAQTRGYGTHGTLHIVINNQIGFTTSDPRDARSTLYCTDVVK
MIEAPVLHVNGDDPEAVVLATQIAIDYRMQFHKDVVIDIVCFRKLGHNEQDTPAVTQPLM
YKKIAQHPGTRALYAEKLVQQGVISADDADGFVKAYRKAMDDGHHTVDPVLSNYKSKYAV
DWMPFLNRKWTDAADTAVPLAELKRLGERITTVPENFKVHPLVERVINDRRNMARGDQPL
DWGMGEHLAYASLVASGYAVRLTGQDSGRGTFTHRHAVLHDQNRERWNDGTYVPLQNVSE
GQAKFNVIDSVLSEEAVLGFEYGYSTAEPNTLVLWEAQFGDFANGAQVVIDQFISSGEVK
WGRVSGLTMLLPHGYEGQGPEHSSARIERYLQLCAEHNMQVVQPTTPAQIFHLLRRQMIR
LFRKPLIVATPKSLLRHKEAVSDLSELAKGSFQPVLGEVDGGIDAKKVKRVIACSGRVYY
DLVAHRREAKANDVAIVRIEQLYPFAHKQFEAEMKKYENATEVVWVQDEPQNQGPWFYVE
HHLKEGMKEGQKLAYSGRPASASPAVGYYAKHYEQQKALVEGAFGRLKSASIAK*
>GCA_000452945.1_BPMSHR5858_genomic_02648_1 lpdG
MSKEFDVVVIGAGPGGYIAAIRAAQLGKTVACIEKWKNPAGALKLGGTCLNVGCIPSKAL
LASSEEFENAQHHLADHGISVDGVKMDVAKMLARKDGIVEKMTGGIEFLFKKNKITWLKG
HGKFTGKSDAGVQIEVSGEGETQVVTAKNVIIATGSKARHLPNVPVDNKIVADNEGALAF
DSVPKKLAVIGAGVIGLELGSVWRRLGAEVTVLEALPAFLGAADEALAKEAAKLFKKQGL
DIHLGVKIDEVKTGANGVSIAYTDKDGNAKTLDADRLIVSVGRVPNTDNLGLEAIGLKAN
ERGFIDVDDHCRTAVPNVYAIGDVVRGPMLAHKAEDEGVLVAEVIDGQKPHIDYNCIPWV
IYTYPEIAWVGKTEQQLKAEGREIKSGKFPFSINGRALGMNAPDGFVKMIADAKTDELLG
VHVIGANASDLIAEAVVAMEFKAASEDIARICHPHPSMSEVMREAALAVDKRSLNS*
>GCA_000452945.1_BPMSHR5858_genomic_02649_1 zapE
MNVTEYYEHELNTRGYQSDAAQRAAVERLQRCYGEWVAYKARRSNAFKKLVVRPDLPRGV
YMWGGVGRGKSFLMDSFYAVVPVQRKTRLHFHEFMREVHRELEELKGQADPLDELARRIA
KRYRLICFDEFHVSDIADAMILYRLLDRLFSNGVQFVMTSNYDPDDLYPDGLHRDRMLPA
IALLKERLDVLNVDAGVDYRQRTLTQVRMYHTPLGADADRELRHAFAQLAAVPDESPILH
IEKREIKALRRADGVVWFDFATLCGGPRSQNDYLELASRFHAIVLSDVPQMSPRMASEAR
RFTWLIDVLYDHKVKLLMSAAVPADDLYVEGPMANEFSRTVSRIVEMQSKEYIEAPRRLI
DTSLT*
>GCA_000452945.1_BPMSHR5858_genomic_02652_1 group_1190
MKSRHDAWMLLLALVAAAGAAHAQSRSGGNPLEALPQINTPQKPSVTVQVAPQEVQVQAL
LARHLTPSSFQVEGVKSIPFEEISQRFTPLVGKDITIGQLIETANGVTKLYQERGYALSF
AFVPAQTFEGGVVRVTVVEGYVANLKITGRPGAMEPKVRAIAAHIMADRPLRRATFERYV
NTFGLLPGVTVKANVPPPQNTDGATTLELNVDRKPFNVSAGLNTNNPGLQGLFTVTENGL
TSLGEQMSISALFPKGPNNQTYVSFNGAVPIGSNGLVTRLDASHYRGNPSVDQTVLPNVQ
RTVINDKLGLSASYPLMLSNQRSLLGTVSGYASHSEDRYQNQSTGATIGMRSQVRVLQMQ
FDYTSVQPKQVQKLSFNVAKAFDILGASKSGFTNLPGVIATNPASTTFVRTGATFVQTNE
WPFKIGSTVQLTGQYSPDSLPSTEQISFGAQRFALGYQPGETSGDSGWGASLELNRAFAP
GFTYLKNITPYIVYDMARVYLHSGTPVPRRLSSAGFGVRLTDSRFYNLDVSIAKPVGDAP
IESASRSPRVNASFSYQLY*
>GCA_000452945.1_BPMSHR5858_genomic_02655_1 group_19548
MKQLMHRFFKEEAGVTAIEYGLIAGLIAVAIATTVGTVGTDLSALFSTIASKLPAA*
>GCA_000452945.1_BPMSHR5858_genomic_02656_1 group_1191
MIHLFSIGFFFAWAAAVAIADCRDRRIPNELVLVGLAAVIIFTVCRQNPFGTTLSGALIG
GAVGLVSLFPFFALRVMGAADVKVFAVLGAWCGLSALPRLWVVASVAAGVHALALMLLTR
TPPGSLGRGGAPAFALGARRAAPYAAFLVAPAAAWLAYLIHTGGTR*
>GCA_000452945.1_BPMSHR5858_genomic_02659_1 group_19017
MKNKLIAWAVAFGVLLSFAARAAETGTIELAAGTQRQIAVGHGVQRVAIADPNVADVLVI
KGGRGGVLLVAKAAGTTSLMVWERGRDEPAAYTVNVVSGAARALLDGGSPSVKAYNGTAV
VAGSAATLDAHARALAVGKGMAGKDGGVLDVSTVGGKNVVQVDVRVVEFSRSVLKQAGLN
FFKQNNGFTFGSFAPAGLASVTGGGTSSMSVSANIPIASAFNLVVGSATRGLFADLSILE
ANNLARVLAQPTLVALSGQSASFLAGGEIPVPVPQSLGTISIDWKPYGVGLTLTPTVLSP
RRIALKVAPESSQLDFVHSITINGVTVPALTTRRADTTVELGDGESFAIGGLIDRETTSN
VDKVPFLGDLPIIGTFFKHLSYQQNDKELVIIVTPHLVAPIAKNASLPATPGELSEQRDG
PVWRSYLGGVLSPDAGPGFSK*
>GCA_000452945.1_BPMSHR5858_genomic_02667_1 group_19520
MRNTNAIEGLDLYVWEGKADIVDRVARCMSSFDVEVIRADNEELSAERTAMRPSLAIISV
SMIESGAAFLRTWQAEIGMPVVWVGAARDHDPSLYPPEYSHILPLDFTCAELRGMISKLA
VQLRAHAAKALEPSTLVAHSDCMQALLQEVDTFADCDTNVLLHGETGVGKERIAQLLHEK
HSRYGMGEFVPVNCGAIPDGLFESLFFGHAKGSFTGAVGTHKGYFEQAAGGTLFLDEVGD
LPLYQQVKLLRVLEDGAVLRIGATAPVKVDFRLVAASNKKLPQLVKDGLFRADLYYRLAV
IELSIPSLEERGPVDKIALFKSFVASIVGEDRLAALPELPYWLAEAVADSYFPGNVRELR
NLAERVGVTVRQTGGWDTARLQRLIAHARSAAQPAPAESAPDVFVDRSKWDMTERNRVIA
ALDANGWRRQDTAQHLGISRKVLWEKMRKYQIFDEEPEARESE*
>GCA_000452945.1_BPMSHR5858_genomic_02672_1 fadD_1
MTAPVAPAHAHTHAPNTDGIWYASYPKGVPHDIDVTRYESLAQYFDECTTRYAQRVAFIS
AGARMTYAALARKAAAFASYLQSLGVKPGDRVAIMLPNTFQYPVTLFGTLKAGAIVVNVN
PLYTARELAHQLKDSGAQTIIVFENFAKTLQDALPETQIKHVVVTALGDLLADGLNPKGR
LINFVLKHVKKLVPAYRLPQAVRLRAALAAGAKKAPAPVALKRDDLAFLQYTGGTTGVAK
GAMLTHGNLIANLLQAKAWIEDQLTGDVETVLTPLPLYHIYSLTVNAFIFLGLGGRNILI
ANPRDTKTMMKILRHETFTGITGINTLYNAFLDNEEFRKRDFSKLKLAMAGGMAMQRAVA
ERFEQVTGCPIVEGYGLTECSPIVTMNPYDAHEKRAFSGTIGLPAPSTIVRFRTEDGAWA
NVGEPGELCVHGPQVMRGYWQRPDETAKVIDADGWLATGDIGVMDEQGFIRLIDRKKDMI
LVSGFNVYPNEIEDVLVSHPGIREAAAIGIPDPVHGERIKVFVVPRDASLTVEDVLTHCR
KNLTGYKMPKAVEFRDALPQTNVGKILRRALRDEELAKLANAPADPHAKH*
>GCA_000452945.1_BPMSHR5858_genomic_02676_1 group_18997
MTIGQDVHLIPAKLDALRPTQITVGYREVKAKRKHWKALGKKARKAAIESHWFPAVLGPG
GDYFITDHHHLGLALLEEGVTEVKAMLLKDLSWLDETIFWRMMEHNQWVHPFGANGARHD
YDHLPTSLVGLKDDPYRSLAGELRAAGGYAKDATPFSEFLWADFLRPRIALAQIRKEFAK
ALEAALGYAHTQEARYLPGWSGVIAAPR*
>GCA_000452945.1_BPMSHR5858_genomic_02680_1 group_19288
VGFLNTFSLRRYASLLLGLLAVAAVAIGLCQWRLDAANHRVAQAYQQRYVSTQLANELRR
SSDDLTRLARTYVATGDAKWEQQYNEILAIRSGKAPRPLDYDRIYWDFRAADEPLRAEQG
ETISLQEMMKRAGFTDEEFAKLHDAEQNSNDLVKTETVAMNLVKGLTPDDAGHFTKQGPP
DLEKARALMFDANYHRFKAKIMHPIDDFLKLLDARTEGAIARAQASAQTWKIISAAVAIG
ILAFFALMLHMMFKRVLAGLEAAASTASRVAAGDLTSHFDAHRVDPQAKDEISRVMRALQ
TMNDGLVRIVTDVRSGTDTIATASHEIMAGNNDLSARTEQQAASLQETAASMSELTATVK
LNLENARQANMIGSNAVSTVEKGSVSVEQLVTTVNAISTNSGKIADIISLIEGIAFQTNI
LALNAAVEAARAGEQGRGFAVVASEVRSLAQRSSSAAKEIKDLIETSIDTVRDGVSKADE
VGQHIVEVKQAIRRVADLVGEITAASEEQTRGIEQVDAAVSQMDRVTQQNAALVEQAAAA
SKAMDDQAGNLRAAASIFKLPGRASLFAHA*
>GCA_000452945.1_BPMSHR5858_genomic_02683_1 rlmB
MSGAPVRLRLPRSRKPNATRKIIDKPLSKYPYPIPSREEILGVLRTSDAPLAANDIAEAL
SIKRQEREGFFRRVAAMERDGQIRLDKRGHYQLTHPSNFVAGRVQGHRDGYGFVIRDDGQ
DDLFLPNAEMQKVMHNDRVLARIVGYDRRGRPEGHVVEVTERANKRVIGRLLNENGALIV
APEDKRISHDILVTQNAKKAKVGQVVVVELTDFPSRHSQPLGRVCEVLGDIDDPGMEIEI
AVRKYGVPHEFGAQALGEAAALPDKVRPADLRYRVDLRDVPLVTIDGEDARDFDDAVYCE
PAAVGRGEGFRLIVAIADVSHYVQPGSPLDADAVERSTSVYFPRRVIPMLPEKLSNGLCS
LNPQVDRCVLACDMIINARGDIKAYQFYPAVIHSAARLTYTEVAAVLSNTKGPEAARRAE
LMPHLQHLYGVYKALFVARQKRGAIDFDTTETYIVCNAQGKIEQILPRQRNDAHRLIEEC
MLAANVCAADFLKRNKHPGLYRVHAGPTPEKLENLRAFLRDMGLTLGGGDTPHASDYAAL
MAHIRDRPDAQMLQPMLLRSMQQAVYSPDNIGHFGLAYDAYAHFTSPIRRYPDLLTHRAI

YAILSGKKYTPKSPEGVELNTALSPRARAMQREDDEARGRARSNAAIWEELGLHCSSNER
RADEASRDVEAWLKCYFMRDKLGEEYGGMVNGVTSFGIFVQLDALFIEGLVHVTELGADY
FQYDEVKNELRGERTGIRYRLSDRVRVQVSRVDLDARKIDFRLVRETPVKAPRPASAVAA
EAGAGGPRVRALPPAEGAARRKKAASAPSAAVKEARAARKKGAASKPAAKKARSRKKY*
>GCA_000452945.1_BPMSHR5858_genomic_02687_1 pdhR_1
MEQGKDRSLVSKVMDGLVTGIVEDKYGGVLPPQDVLSKEFDVSRTVMREALSMLLARDML
DVRPKVGTRVRPMSHWRMIDEDVVNWRFRAKPDRQFIRDVIEFRTLIEPRATAQAASRAT
TAEIAGIREAYELFKLAEPGTPAYDEADELLHTRIVHASGNQFFQQMAAIIRGALRVVNP
RMSQRDDIRALALKSHGRVVDAIERRDARDAEAAALELIDYTAEEMRDFAVDMPERN*
>GCA_000452945.1_BPMSHR5858_genomic_02688_1 iolU
MTDHQPTPIRFGIVGAGSIARRFAHSIAHVRVAELSRIWARRADAAAAFCAAHGGAPCDS
VDALLASDIDAVYIATLNDSHAHYARTALAAGKAVLCEKPATVNARELETVLDAAHAAKR
LFMEAMKPPFYPLYRKLRAHLERDPIGEIALVRAGCASSRVPADHPVYRFEQAGGALLDI
GVYEMFLAVDWLGAPLDVQTLGRVGATGVDVFASVNARHARGVSQLFCGLDVMGRGDALL
AAPGGYVTIHENWWNPARATIRYADGRVVELDEPFDGGGLNYEIAHFCELLRAGELESPV
MTHDHSRRMIAMTDAARAALGVRYPMD*
>GCA_000452945.1_BPMSHR5858_genomic_02691_1 pyrD
VFSSLYPLARASLFKMDAEDAHHLTLRMLGAAGRTGLACALSPRVPDAPRTVMGLSFRNP
VGLAAGLDKDGAAIDGFAALGFGFIEVGTVTPRAQPGNPRPRMFRLPEADAIINRMGFNN
SGVDQFVKNVQAARYRGVLGLNIGKNADTPIERAADDYLYCLERVYPFASYVTINISSPN
TKNLRQLQGAGELDALLAALKDKQRRLADLHGKLVPLALKIAPDLDDEQVKEIAATLLRH
DIEGVIATNTTLSREAVKGLPHADEAGGLSGRPVFDASNAVIRKLRAELGDAVPIIGVGG
IFSGEDARAKLAAGAALVQLYTGFIYRGPALVAECVKAIARG*
>GCA_000452945.1_BPMSHR5858_genomic_02692_1 bpt
MTHPTELPLSPLSALQFYATAPYPCSYLDGRVARSQVATPSHLINSDIYTELVKAGFRRS
GVFTYRPYCDGCRACVPVRVPVDAFAPNRTQRRTWKRHRALVATVAALHYDEEHYALYMR
YQSARHAGGGMDRDSRDQYEQFLLQSRINSRLVEFRDLDPAENGASTLRMVSMIDILGDG
LSSVYTFFDPDESHASYGTYNILWQIEQAKSLRLPYVYLGYWIRESPKMAYKANFHPLEG
LVDGRWKVLDPTLADLPPVDAALARAPLPGGHSGTR*
>GCA_000452945.1_BPMSHR5858_genomic_02711_1 rpsG
MPRRREVPKREVLPDPKYGNVDVAKFMNMLMLSGKKSVAERIVYGAFEQIQTKGGKDPLE
VFTVALNNVKPVVEVKSRRVGGANYQVPVEVRPSRRMALAMRWLREAAKKRSEKSMALRL
AGELSEAAEGRGGAMKKRDEVHRMAEANRAFSHFRF*
>GCA_000452945.1_BPMSHR5858_genomic_02712_1 rpsL
MPTINQLVRKGRASETTKSKSPALQDCPQRRGVCTRVYTTTPKKPNSALRKVAKVRLTNG
FEVISYIGGEGHNLQEHSVVLIRGGRVKDLPGVRYHMVRGSLDTQGVKDRKQARSKYGAK
RAKAAK*
>GCA_000452945.1_BPMSHR5858_genomic_02713_1 recQ
MSRSLEILNEVFGYPAFRGQQGEIVEHVAAGGDCLVLMPTGGGKSLCYQIPALVRSEAGR
GAGIVVSPLIALMQDQVAALSEVGVRAAYLNSTLSGAEAAATERALRDGDIDLLYVAPER
LMTPRFLDLLERARIGLFAIDEAHCVSQWGHDFRPEYIQLSVLHERFPDVPRIALTATAD
AITRDEIVQRLALDDARIFVSSFDRPNIRYRIVEKDNARSQLLDFIRAEHTNADGTTDAG
VVYCLSRRKVEETAEWLKAQGVRALPYHAGMEFEIRQKHQEMFQREEGVVMCATIAFGMG
IDKPDVRFVAHLDLPKSVEGYYQETGRAGRDGLPANAWMAYGLGDVVQQRKMIDESEADD
THKRVQTSKLDALLGLCETASCRRVRLLAYFGETSGPCGNCDTCLEPPATWDATREAQMA
LSCVFRAQRASGFNFGASHLIEVLRGARTEKVQQRGHDSLSTFGIGAELSEAEWRAIFRQ
LVAFGYLAVDHGGFGALVLTEASKPVLKGEEKVTLRRYVKPARTRQSSGRSGARVDPTAG
MSARERACWERLRAWRAETAKSDGVPAYVIFHDATLAEIARNAPESIDDLRHIPGIGARK
LDRFGDEILEVVESA*
>GCA_000452945.1_BPMSHR5858_genomic_02716_1 rplL
MAIAKEDILAAVEGMTVLELNELVKAFEEKFGVSAAAVAVAGPAAGGAAAAAEEKTEFTV
VLAEAGSNKVAVIKAVREITGLGLKEAKDLVDGAPKPVKEGVDKASADEAKKKLEDAGAK
VELK*
>GCA_000452945.1_BPMSHR5858_genomic_02717_1 rplJ
VPLNREDKQAVVAEVAAQVAKAQTVVLAEYRGIAVGDLTTLRAKAREQKVYLRVLKNTLA
RRAVEGTPFAPLAEQMTGPLIYGISEDAIAAAKVVHDFSKSNDKLVIKAGSYDGKVMDKA
GVQALASIPSREELLSKLLFVMQAPVSGFARALAALAEKKQAEAA*
>GCA_000452945.1_BPMSHR5858_genomic_02718_1 rplA
MAKISKRRQAFAAKVDRQKLYPIDDALALVKECASAKFDESIDVAVQLGIDAKKSDQVVR
GSVVLPAGTGKSVRVAVFAQGEKAEQARAAGAEVVGMEDLAEQIKAGQMDFDIVIASPDT
MRIVGTLGQILGPRGLMPNPKVGTVTPDVATAVKNAKAGQVQFRVDKAGIIHATIGRASF
EPTALRTNLSALIEALQKAKPATSKGVYLRKIALSSTMGVGVRVDQGSLAAQ*
>GCA_000452945.1_BPMSHR5858_genomic_02730_1 group_1193
MKTYSELLAQWDDLKNEIDRAREREAQRLAQRVLELLAESGVDVRDMMNPRPRGQGRGRR
ITPKYWNPQTGATWSGRGRMPKWLIGQNPRQFLILHPDDERDGCAQAHPGGSN*
>GCA_000452945.1_BPMSHR5858_genomic_02740_1 lrp_2
MQEPLLDRIDRHLLDILQENGRISNLELAKAVGLSPAQTLRRHRRLEEIGAIKRYETRLD
ADTLGFGVTAFVQVTMERGHIRDLSTFQKLVSELAQIQECFSVTGDIDYVLKVVAADLKG
LSTFLLDTLMRIPGVSGVHSSVCLDEIKCTSAMPL*
>GCA_000452945.1_BPMSHR5858_genomic_02743_1 group_19146
MSISIVRLGTPRADDEGVRIGTVRRPPRGVPKDEFASRDFYDVWLPTLSPSPELVAEAQA
AQSDADWKAFARKFRAEMKHGDASRMLDVLAALSATSNFAIGCYCENEAHCHRSVLRELL
EERGASIRS*
>GCA_000452945.1_BPMSHR5858_genomic_02745_1 group_19101
METAELHFRCNEGGMADYAAQLREVGTVMLPAYVAFDAHELARIDALQARLPEEPVTAGD
AGDTHDIYVRRIMVDRAGERPQLVNLPHSETILNLLGDARRTRFFGDMFGTRAEYFIRRC
QINRMLKDSFIGMHLDAASNPDYEFSVVIQLGRAFDGGEFVVHPQGRPPNVFAPAYGTVI
VTSCAHRHEVRTVRANERTSLVYFYSRHNGANRRAA*
>GCA_000452945.1_BPMSHR5858_genomic_02746_1 group_19685
MFESFNKDVIDAFARNKRRLTKNAASRQVRALLDEVGASAFKLLPHPFLFPADEFEMLAR
RARALLRAQNKIVRHLCRHRPRADLLAQCSLNPALARYVDWDELEAGRRRVARVDILPLP
DGYAVCELNFSAAVGGAELFDCHRRFMQASGWPACERGGSPYANLRALYEDTLRSMEIDR
IVLLDWSSHAARGYATPELARRYLAGIGNGMPVDVHTELTYPSKWLGNGRGKGILIHRNF
VYDDVTVNADRFDAIHRSGATFSNGLEAELLMSKAWLALLCDARFHPLLDEDEIAAIRNH
VPHTFRLDEGTFAEALAHKDEWVFKLNVSYGGVDVLIGKDHPADALADRLRRQGIANWIC
QAFRAVQTIRHPFDDTFVAAEHKPVLGMYLHRDDASGMALRSNRFSSVVNAGSGAGVHWA
AVVTEAEKAALLAAMDG*
>GCA_000452945.1_BPMSHR5858_genomic_02834_1 nth
MNATKRRAIFETLQSLNPHPTTELEYTTPFELLIAVMLSAQATDVSVNKAMRKMFPVANT
PKKIVALGEEGVADYIKTIGLYRTKAKNVVAASRILLEQYGGEVPAEREALESLPGVGRK
TANVVLNTAFGQPTIAVDTHIFRVANRTGLAPGKDVRAVEAALEKLTPKEFLHDAHHWLI
LHGRYVCKARKPECWHCAIEPLCEYRPKTPAPMQ*
>GCA_000452945.1_BPMSHR5858_genomic_02835_1 group_1854
MPFIADSSRAALRGVLYVALSAVAFGAMAIFGRYAYAGGADVLGLLIVRFSIAGALLVAV
ARRRRVRWPRGRALAAIVGMGALGYVGQSLCYFSALQHAQASLVALLLYLYPAFVALLAA
WWLGERLTRAKAVALALCVAGSALMVGGGRGEPLGIALALGAAVVYSLYIVVGAKAARGV
DPLATVAVICCAAAAMLAMLALARAAAFDAPPHWPRAAAGWAALVAIALVSTVAAMLAFF
AGLARLGAARTSMLSTLEPVVTVALAAELFGETLTPLQWAGGVAILAAVLWLVRAGDAAD
SRGAGDDRERRRLGRRDDEPSAPGGSGAGGGLAGFVDPNECGIRRVRSADENARPM*
>GCA_000452945.1_BPMSHR5858_genomic_02837_1 cc4
MKKPQTALKTAAALALAAGFAIGTAHAANVAKGKELVESHNCAACHGAKLDNPINAEYPR
LAGQHADYLVWAMRQYQMGLTNPLLGRNNAIMQAQVQSLSIADMKDIAAYLESLQGSLVF
KK*
>GCA_000452945.1_BPMSHR5858_genomic_02844_1 mctP
MNLTATFVFVLFFVGVTIMGFLAANWRRGNLAHLDEWGLGGRRFGTVVTWFLLGGDLYTA
YTFVAVPALVFGAGAMGFFALPYTILIYPFAFVVFPKLWSIAKRHGYVTAADFVSARYGS
RSLALAVAVTGIVATMPYIALQLVGIEVVIGGLGFDTKGFIGDLPLIIAFAILAAYTYTS
GLRAPAMIAIVKDILIYITIAAAVIVIPAKLGGFGHIFGAVPPAKLLLKAPDAASLNGFS
AYTTLAIGSALALFLYPHSVTAILSSSSGNTIRRNMAMLPAYSFVLGLLALLGYMALASG
VKDMPEYAPYFKAFGPNFAVPALFLHFFPSWFVGVAFAAIGIGALVPAAIMSIAAANLYT
RNIHREFVNRNMTHDQETHVAKLVSLIVKVGAVAFILGLPLTYAIQLQLLGGIWIIQTLP
AIVLGLYTRVLDYRGLLAGWAAGLVCGTWMAISLKLASSIFTIHLFGHAIPGYAAVWALA
VNLVVSIVVSVLVRAFGIAHAEDRTRPEDYLDVVES*
>GCA_000452945.1_BPMSHR5858_genomic_02848_1 ubiC_1
MARMRFDAADAHWRETPRPGASSAQKDWLTRGGSLTAHLARLGRVTVRVTRETVAAPWAD
EHRALSCASRAPVWVREVVLAVDGAPFVAAHSIAPLAASKGVWQAMRRLRTRPLAELLYS
DPEVTRSALVSRRVLAGHPLFSLASLALVRAHATEHAFAARRSVFERRGTPLMVTECMLP
ALWRHLDAHGERRARGLEQT*
>GCA_000452945.1_BPMSHR5858_genomic_02855_1 yddE
MAAARRAAHTVRFKQVDVFTSVPFKGNPLAVVFDADALSDADMLAIARWTNLSETAFLVA
PSDPAADYRVRIFTTGGELPFAGHPTLGAAHAFLDAGAQPKAPGRLVQQCGAGLVELAKR
DGGWAFAAPSARVTPLAEREWPALAAALRTGAIDFGVPPRAVDNGAPWLVVRLASAADCL
ALAPDHAALAPIVAALGAAGLAAYGPHPADGPATFEVRCLMTGDAFGAAGEDPVTGSANA
AIAGVLAQAGRRPGRRYTARQGTVLGRDGRISVDYDDDAGKIWIGGAAVTLVDGTIRLR*
>GCA_000452945.1_BPMSHR5858_genomic_02856_1 group_19406
MSSSRSHAARAARPVRATRKSRRRRALFAIPLLGTLALALLWVAIIARLSVEKDSAFRDA
GASAAILSTALEQHTVKAIHQVDQITRFVKYEFEKSPTRFNLASTVEKGVVPSDTLIQVS
ILDARGRLVANTAEAHPKPIDLSDREHFKVHMQRNDDDLYISKPVFGRVSRHWTLQMTRR
LNHPDGSFAGVVVVSEDPAYFTNDFYNNATLGKDGVIAVISDTGAVLARSTGSTQKTPGT
FSASGVYPVSERASGTYVDPIDGIVRIVSYRHLDGYPLGVMVGLSQAEEFADYHHTRNVY
LLMTGFITLAMLAFFGVATGLIGKLLGREREMTQLAEFDLLTGLANRYATLRSLRNDVAL
PSNLSRLAILFIDLDNFKTVNDTLGHNAGDIVLQMTAARLSDAVANEGALARIGGDEFVV
VVKGENVERRAERLAEAIVRMFAEPFDVRGSSFVLHASIGIALHSVENESEIDLLKKADL
AMYSAKDAGKNCYQFYAPHLSHRADHLMRWEQQLRVALSDGQLFLAYQPKIDLTYRCITG
FEALVRWDHPEHGVISANEFISIAESTGLIVPIGDFVIRTACEQLARWRNEGHDKLTLAV
NISPVQFWRGDLIDTISRALAETGIEARRLELEITETAMMEYPELVSEKIVALKRLGIRV
ALDDFGTGYSSLSYLHRFAVDTLKVDRSFIQAIPADRSVCVMVSSIVHLARSLGLTVVVE
GTETEEQIAWLAALGRIEAQGFLFSRPVPSDGIAALLARFGVCGGDSARAAGRDIANAAK
QSFGSD*
>GCA_000452945.1_BPMSHR5858_genomic_02857_1 srpC_1
MTIASVEAACCGERESLWALFKTVTGVSAVSWGGLAMMAQLERHYVEHERRIDPLSFADL
VALAWMMPGPVGCNVAVQVGHALRGRAGAWIAGVASVLPFSAAMTVFAIFYQTPLVRSLA
SPVLLHHFAMVLAALIGLTWFRQVRALVHAPLERVIAALATALLALAHNPAAFVAILAAA
FAVGWLASGRKQGEALRLALPAREWRLLASLALLIALFALPLPNEYESSLLWPRLAGAGM
TLFGGGFSALPVLKSLFVTRSTGITEQDFMLAFTLSPVSPGPLLNVVPFLGYLEDGWRGA
LLSTVALFVPSGCLVIFARRHVERLKRHPRFASGMRVLRAATTAFLAIAAVRLVAKTPAE
PMYWATGVIAWLCLARFKVPVYALYGAVAAACGGWLILAAHG*
>GCA_000452945.1_BPMSHR5858_genomic_02859_1 hyuC_1
MMASALGGPTLAIDGERLWRSLMEMARIGGTAHGGVRRLAATDDDRRGRDRFAQWCREAG
LALSVDEIGNLFARRAGAEAGAPPVLIGSHLDTQPEGGRFDGTYGVLAALEVVRSLNDAG
IVTRKPIEIVSWTNEEGARFAPAMLGSAVAAGVMPLAQALAVRDAAGVTLGDALERIGYR
GARRAVRPAIDAYFEAHIEQGPVLEANGVEIGVVTGGQAIRWLDVTIAGQPAHAGTTPVA
HRRDALFGFAQIADAIERMLAEFAPHGLATIGCVEIPNASRNTIAGAVAFTLDLRHPDDA
TLDAMARAATDACARIAARRGLRADVSTHWASPSTRFDAACIASVEAAAARFGYRHERIV
SGAGHDAIHLAKHGPGAMVFIPCVGGLSHNEAEAVLPADATRGANVLLGAVLDRAGRAAR
*
>GCA_000452945.1_BPMSHR5858_genomic_02860_1 abaF_2
MTAPQPCARQPARAAAAAFVGTMIEWYDFYIYATAAALVFGELYFPSGDPFVSTMASFGT
FAVGFFARPLGGVVFGHLGDRIGRKKALMTTLAMMGAATVCIGLLPSHASIGALAPVLLV
LLRIVQGVAVGGEWGGAVLMAGEHAPDGRRTFFASFAQLGSPAGLILSLVAFRAATSLEK
ADFLAWGWRLPFLASAVLLIVGLAIRLGVNESPEFEAVKRTRATAKLPIADVLRSASGLV
LLCIGANTIGIAGVYFTNTFMIAYTTQQLGIPRATILDCLFAVAIIQFVAQPAAAWLAEK
LGGARFLKLAAVAAMASPYPMFMLVTSGRPGAIVAGIALAVVCMAGFYSVIAGFVSGVFA
ARIRYSAISLSYQVCGALAGGLTPLVGTWLAHRYAGAWLPLALFYTCLAGLSLVSVVALD
ARRAARGASGREALEAR*
>GCA_000452945.1_BPMSHR5858_genomic_02865_1 parC
MDDNTPDLFAEPAAPEGDFLTLGRYAERQYLDYAVSVVKGRALPDVSDGQKPVQRRILYA
MNEMGLGDNAKPVKSARVVGDVLGKYHPHGDQSAYDALVRLAQDFSMRYPLIDGQGNFGS
RDGDGAAAMRYTEARLTPIAKLLLDEIDEGTVEFMPNYDGSFDEPKLLPARLPFVLLNGA
SGIAVGLATEIPSHNLREVAAAAVALIRHPALPHAELMRLVPGPDFPGGGQIISSDAEIS
AAYETGRGSLKVRARWKIEDLARGQWQLVVTELPPNTSCQKVLEEIEELTNPKLKAGKKT
LTQEQINTKKTLLDLLDAVRDESGKDAPVRLVFEPKTRAIDQTEFVNSLLAHTSLESNAA
LNLVMIGEDGRPAQKGLASILGEWVRFRQATVTRRCRHRLGKVNDRIHILEGRMIVFLNI
DEVIRIIREADEPKAALISAFGLTERQADDILEIRLRQLARLEKIKIEKELEELRAEKAK
LEELLANESAMKRLMIKEIEADAKQYGDERRTLIQQEKRATFEAKVVDEPVTVVVSQKGW
VRALKGHGLDPASFSFKAGDGLYAAFQCRTPDTLIAWGSTGRVYSVAVQVLPGGRGDGVP
VTSLIELESGSHLLHYYAAPAEQPLLLASSNGFGFVAKVGDMVSRVKAGKSFMTIDDGAA
PLAPMPVLPDASQVACLSSGGRLLVFGMDEMKTLSGGGRGVILMALDAKETLVQALAIDA
AGVVLTGTGRGGKAQEEALAGRALEPHVGKRARKGRAPDTRLKVNGLRPVLG*
>GCA_000452945.1_BPMSHR5858_genomic_02866_1 parE
MSTKKPAAAYSEASIKVLKGLEPVKQRPGMYTRTENPLHIIQEVIDNASDEALGGFGKQI
TVTLHPDQSVSVEDDGRGIPFGLHPEEKVPVVEIVFTRLHAGGKFDKAKGGAYTFSGGLH
GVGVSVTNALATRLDVTVWRDGRIAQLGFADGDVVKPLATQGAGRGEKKSGTRVTVWPNP
KYFDSPNLPLGELQRLLRSKAVLLPGVEVVLVNEKTGERQSWKYDDGLRGYLLDEMNGSE
LLIPLFEGERFADSRSGDDTFAEGEGASWVVAWSEEGSLVRESYVNLIPTPAGGTHESGL
RDGLYQAVKSFVELHNLQPKGVKLLAEDVFARVSFVLSAKVLDPQFQGQIKERLNSRDAV
KLVSSFTRPALELWLNQHVEHGKKLAELVIKQAQARTRAGQKVEKKKSSGVAVLPGKLTD
CETEDIARNELFLVEGDSAGGSAKMGRDKEYQAILPLRGKVLNTWETERDRLFANNEVHD
ISVAIGVDPHGPDDSVDVSNLRYGKICILSDADVDGAHIQVLLLTLFFKHFPQLIERGHV
FVARPPLFRVDAPARGKKPAQKLYALDEGELEAILDKLRKDGVRESQWTISRFKGLGEMS
AEQLWDTTMNPDTRRLMPVALGELDFEATVARMTMLMGKGEAAARRNWLEEKGNEVEADI
*
>GCA_000452945.1_BPMSHR5858_genomic_02868_1 group_19290
MLYNKWLAGAFTAAALFASASAHAEARVYFVEPADGASVSSPVHVKFGLEGDIVLRPAGD
MTPHTGHHHLLIDGKPVARGDVIPANDHSLHFGKPQTETEVRLSPGRHTLTLQFGDGAHR
SYGPEMSQTITVNVK*
>GCA_000452945.1_BPMSHR5858_genomic_02869_1 rubA2
MSEVVEYKSWVCLICGWIYNEAEGLPDEGIAAGTRFADIPHGWRCPLCDVGKEDFVVVEF
*
>GCA_000452945.1_BPMSHR5858_genomic_02872_1 infA_1
MAKEELIELDGIVDEVLPDSRYRVTLDNGVVVGAYASGQMRRHRIRILAGDRVTLELSVY
DLTKGRINFRHKDERRSDAAPRASARRR*
>GCA_000452945.1_BPMSHR5858_genomic_02892_1 group_1475
MQNPNAVNAPVQTPPAPPRVGEEILRVENVSRGFNKTQGELLVLDGANLSLREGEIVGLL
GRSGSGKSTLLRIIAGLIEPTGGEVTYLGKPLSGPAEGVAMVFQTFALFPWLTVLQNVEA
GLEALGVGARERRERALAAIDLIGLDGFENAYPRELSGGMRQRVGFARALVVDPTLLLMD
EPFSALDVLTAETLRTDLLDLWTQGRMPIKSVLIVTHNIEEAVFMCDRILVLSSNPGRVI
AEIKVPFKHPRNRLDPAFRRLVDDIYAKMTARQVGEATKKGLELGSWLPQVSTNLMAGLI
ETLAAPPYHGRADMPEIARTLHLEVDDLFPIAEVLQYLGFADVREGDVFLTPPGRVFAEF
GTQERKMMFAEHLLRHVPLAARIKKVLNERPGHRAPRVRFEQELEDFLSDGAAEETLDAV
IDWGRYGEIFSYNDQTEIFSLEDVES*
>GCA_000452945.1_BPMSHR5858_genomic_02896_1 envZ_2
MRKPIDSLFGRLALLVVCVLLLSHFAWYFAIRLERNQFQTRYAVEEAAFLVDAVRQHAER
SPDQPLPSRVRLVAPGSGDVPEAKQDLPPPLKRFADELRERLPAGTQVRIGRPGHPPVLW
VKQPSDRDWIVVPVQPLRPPRSLDRMVLWLVMIFSAAVMAALFAAWQLQQPLSSLARAVA
RFGRGMAVPPLPERGPRELRQLTHGFNQMVQQVSRAENDRAVMLAGVAHDLRTPLARMRL
RAEMMDDARLRDGVVRDVDSMTHIVDQFLVFAHGDADRSEAVPVDQACERITRTYRAVSP
NAPTVRTELAAGPGFRLPAATLDRVLSNLLDNAHAYGAPPVVIATARTATGYTLTVSDHG
KGIAPRDLADATRPFVRLDPARGGNGHSGLGLAIVERLVQRTGGACEIGNREEGGLRVAM
TFAFDVVPKAEPQTEAP*
>GCA_000452945.1_BPMSHR5858_genomic_02902_1 hisM_2
MIELIQEYWRNYLYTDGYRFTGLAITMWLLVVSIGIGFCLSVPLAVARVSKKKWLAGAVW
LYTYVFRGTPLYVQLLLCYTGLYSLQAVRGTPMLDAFFRDGMNCTLLAFTLNTCAYTTEI
FAGAIKATSYGEIEAARAYGMSTFTLYRRIVLPSALRRALPLYSNEVILMLHATTVAFTA
TVPDILKIARDVNSATYMSFHAFGIAALLYLAISFTLVWLFRRAERRWLAYLRPQGK*
>GCA_000452945.1_BPMSHR5858_genomic_02937_1 btuF_2
MADAPITHGGNLHEAALRYGIPRDAWLDLSTGINPHGFPVPPVPADAWRRLPEDDGVLAA
HAARYYRAPGAAHVLPVAGSQAAIRALPALFARGTVGVAPLAYSEYAPAFARHGHSGAPL

DCGADTLPAALTYAIVANPNNPTAERIDRTRLLRWHAQLVARGGALIVDEAFADAESAAH
ASLAADTHRDGLVVLRSVGKFFGLAGVRAGFALAAPALLARLRDALGAWTVSGPARHAVL
AAFADAAWQHAMRERLAHDGARLAALLRAHGFVTHATPLFSWSADPRAHALHDALAARGI
WTRYFAHAPSVRIGLPAGDDDWRRLERTLAECVPTLAAAAAHPSESTTRD*
>GCA_000452945.1_BPMSHR5858_genomic_02940_1 cobC
MDVVLIRHPAVALGTGVCYGASDVPLAAGAAAGAAGIRRRLRSLGVARASRVWSSPLARC
ADVARALAGGVRAPLEIDPRLREIDFGAWEMQSWDAIGRDALDRWSADLMHAREHGGESA
AQFVARVRPAFATLAVAARTSWVLTHAGVIRALASHALDAPLDAMLARPIAMGGIVWLRC
DEGGDSDCARGWRIAHWDE*
>GCA_000452945.1_BPMSHR5858_genomic_02947_1 group_19444
MSTKQIEVSILGQTYRLACSAETEAALLEAVARVDAEMSKIRANSSVRGTDRIAVMAALS
LASELLRLQQSVRHGEAFPAEEIRRTMRQMNEQLGAVLAQHEVQ*
>GCA_000452945.1_BPMSHR5858_genomic_02950_1 lgt
MIIHPNFDPVAIHLGPLAVRWYGLMYLVGFILAIVVGRLRLKLPHVAAQGWSAKDIDDMM
FYGVLGVVLGGRLGYVLFYKAGYYFSHPLDIFRVWEGGMSFHGGFLGVTLAMALFAWQRK
RHWLEVTDFVAPMVPTGLAAGRLGNFINGELWGRVTSPDAPWAMLFPGASRDDAAWLAAH
QDIAAKWNLNEVFLSHQMLPRHPSQLYEIALEGIALFFVLWFFSRKPRPMGAISALFLIG
YGAARFTVEFAREPDDFLGLLTFGLSMGQWLSLPMIVAGVLMMIWAYRRGGVAKQA*
>GCA_000452945.1_BPMSHR5858_genomic_02952_1 ilvD
MSYNRRSKNITQGVARSPNRSMYYALGYQKEDFDKPMIGIANGHSTITPCNAGLQRLSDA
AVAAVKDAGANPQIFGTPTISDGMSMGTEGMKYSLVSREVIADCIETCVQGQWMDGVVVV
GGCDKNMPGGMIALARINVPGIYVYGGTIRPGHWKGHDLTIVSSFEAVGEFTAGRMSQED
FEGVEKNACPTTGSCGGMYTANTMSSSFEALGMSLLYSSTMANPDQEKVDSAAESARVLV
EAVKKDLKPRDIITKQSIENAVSVIMATGGSTNAVLHYLAIAHAAEIDWSIEDFERIRKR
VPVICDLKPSGQYVATDLHAAGGIPQVMKLLLDAGLLHGDCMTITGRTLAEELKDVPSVP
RADQKVIHPIDQALYKEGHLAILKGNLAEDGAVAKITGLKNPVITGPARVFDDEQSALAA
ILDDRIRAGDVVVLRYLGPQGGPGMPEMLAPTSAIIGKGLGESVGLITDGRFSGGTWGMV
VGHVAPEAFVGGTIALVQEGDSITIDAHKLLLQLNVDDAELARRRAAWKQPAPRYTRGVL
AKYAALARPANQGAVTG*
>GCA_000452945.1_BPMSHR5858_genomic_02953_1 group_1859
MALTRGDKKRQDEEGGATYSRPLSFYNARTAKPDTVPAETSMKRTTLRLTAFALVAAAAN
TQAQQADGLALAQRKNCMACHAIDKTLMGPSFHAIADKYAARGDAAGYLAQTIVKGSVGV
WGNVPMPANTQLTSAEAGTLAHWILSSK*
>GCA_000452945.1_BPMSHR5858_genomic_02956_1 pepA
MDFSIKGCDWSKGTANGFLTGKSDCIVLGVFEAQTLSGAALDIDEATKGLVSRVIKAGDI
DGKLGKTLFLHEVSGIGASRVLLVGLGRQDAFSQKAYGDAAKAAWRALLGTKVVQVTFTL
AQLPVPERASDWGVRAAILALRNETYKFTQMKSKPDAGAPALKRVVFSVDPADDKAAKVA
AKQAVALANGMDLTRDLGNLPGNVCTPTYLANTAKKIAKDWGLKVDVLGLKQIQALKMGS
FLSVAKGSVEPPQFIVLQYRGAAAKAAPVVLVGKGITFDSGGISLKPGEGMDEMKYDMCG
AGSVLGTMRAVAEMGLKVNVVAIVPTCENMPAGNANKPGDIVTSMKGLTIEVLNTDAEGR
LILCDALTYAERFKPAAVIDVATLTGACIIALGHHNTGLFSKDDALAGELLDASREAGDP
AWRLPLDDEYQDQLKSNFADLANIGGRPAGSVTAACFLSRFAENYPWAHLDIAGTAWKSG
AAKGATGRPVPLLAQFLIDRAGA*
>GCA_000452945.1_BPMSHR5858_genomic_02957_1 lptF
MIFERSLQRELAYTAGAVFMVLLTIMLTSMMIRIVGYAASGEVNPKDVLVLIGLTVIGYL
AMILVGTLFVSILFVLTRWYKDSEMVVWLASGVSLTRLIKPIGVFATPIVILIAFFAFVG
WPWSNQQSKLIKARFQQRDEVSLLAPGQFRESAANHRVFFIEKMSPDQAKVQNVFVTTTE
HGKVNVVVSQTGHTETMKNGDRFIVLENGRRYDGVPGQPNFKIMEFERYGVKIQSKQIVS
QPTTTGTSTPDLLRNPTNQNLAEFAWRAGLPLIAINLMILAIPLAYQNPRRGRTVNLVMA
VLIYLTYSNLLNVVQSQMERGKIAFGLGLIGLHVIVAGIVALLFWLRVRNRPLFSRAAAG
RSQGA*
>GCA_000452945.1_BPMSHR5858_genomic_02958_1 lptG
MRLYEKYFARQIYVTFIFVLFAFSGLFFFFDLISELNSVGHGNYKFGYAVLRVALQAPSR
FYEIIPVAALISAIYVFAQMAANSEFTIFRVSGLATNQALRSLVKIGVPIVIATYLIGEF
IGPYSDQLSERVRLEALGSSVSTNFASGVWVKDTLTARDNGEPVTRFVNVGTLSPDSTIS
DVRIYEFDSKFNLQNVRIAKRGHYQPPGHWLLTDVTDTQLTSLAGNGTASPVDTLNPVYR
AQQVTLPQYSLRSDLTPQILSVLLVSPERMSLFNLFRYIQHLKENQQDTQRYDIALWRKL
LYPFAVFVMLVLSLPFAYLHTRAGVVGVKVFGGIMLGMSFQLFNTLFSHIGTLNTWPAPL
TAALPGCIYLALGLFALKWVDRH*
>GCA_000452945.1_BPMSHR5858_genomic_02980_1 group_19616
MRFDLTDLRLFLHVCEAGSITGGAERAHMTLQSASERIRGMEEELGVPLLQRAKRGTRAT
EAGRALEHHARVVLQQIDHMRGELQQFGAGLRGHIRLLSNTAALSEYLPDALAEYLPRHP
KLSVSVEERSSQEIVHAIRGKTADFGIVADSVGLDGLEQMPFREDWLIAVAAVDHSLAAR
ERVAFAELVDADFIGMTDGSALQVHLADQAKALGKRIDYRVQLKSFDAICRLIERGVGIG
IVSRHAALRAQQTMQIRLIELTDPWAHRRLTICARSFDELPKYTREFIAFLAHDPGKDES
FAA*
>GCA_000452945.1_BPMSHR5858_genomic_02985_1 group_19158
MQRPSLIASSLLTASMFALSFAAHAQQTGIVRFTGMIVEPPCSFSVAGGAGTHAQLQPAC
PRPAKGSVTFVDPQSGAALRTVTFTELSRAIALPAGQTGDARPIVAVVSYL*
>GCA_000452945.1_BPMSHR5858_genomic_03072_1 bfpB
MKQPAPAECDAPPPVAPFHDGELAAQRRAGVAAAAAAFGGRGIRTFMPDQHRAFFAQLPL
LVVGGVDGGGQPWATLRAGAPGFVSAPQPRTLDIAGGTLRGDPLEGSWRIGSLLGGLGIE
LHTRRRNRVNGVVASIAGDAMSVTVEQSFGNCAKYIQGRTPTPVPPQAGGAPPQRRAARL
DAADRALLARADTFFIATANTSDAAGGARGVDVSHRGGPPGFVRVDDDSTLTTPDYSGNN
FFNTIGNLLHDPRAGLLFVDFERGDLLYIAADAQIVWDGPALAAFDGARRLVRLRIREVR
RSAAVLPFRWSAPRFAPQFAAWGDAGARSLASRAESAAVGGSATALVSVTQAASAGRGES
AEWPESSVSLKVLKASEASEAPEVSETPETPGVAGACVR*
>GCA_000452945.1_BPMSHR5858_genomic_03101_1 group_1867
MSILIRTASFKATVLCAALAGLVSAAQAETAAAPQVPGPADAVNQLIVKLRAVKTPPGAT
AAKAERADVQAVIDRVLAARNARAAGRAFGAAAASAPGNPDDPAAGIRIKRDMSGGATVL
SLQRHVSLAQAEALARDFAADGAIEYAEPDARMHPFVVPNDTRYSEQWGYFNPTAGANLP
KAWDRTTGSARVVVAVIDTGYRPHADLAANLLPGYDFISDIPSANDGNGRDSDASDPGDW
VSAQEDGDPSGPFYGCGASDSSWHGTHVAGTIGAVTNNGVGVAGISWVGKVLPVRVLGKC
GGMLSDIADGMRWAAGLPVPGAPSNPNPAKVLNLSLGGYGRTCSSTYQNAINEITSRGAN
VVVAAGNNGGSVSTTQPANCRGVIAVGAIDSRGVRASFSNTGAAVKISAPGVGILSTLNA
GKTSPGADSYASYSGTSMATPHVAGTVALMLAVNSTLSPSQILQRLQSSARPFSSGSSCS
TSTCGAGLLDAGNAVDAAAQ*
>GCA_000452945.1_BPMSHR5858_genomic_03110_1 group_19264
MKPTEFKITVSGAALDGRQIAHASAQLNIALLTALVGENATQIKGPAWNVVLPNGGRIIV
AEEAERLLRVTEQERIGE*
>GCA_000452945.1_BPMSHR5858_genomic_03111_1 gltI_3
VAIAAIASSLIAALHARTLKKIADTNRMTVSYRESAVPFCYLLTPHKAVGFSADLTDAVV
DDVRKKLKMPGLQIAYIPVTGQNRNPLLVNDTYDLEGGSTTNTAARGQDVSFSISLFYAG
TRVLVKKASRIRGYADLAGKTVATAAGSTNENVIRKYAADHHLDIDVVPAKDYAAGPRMV
ESDRTAVLALDDVLLFGMRANAADPELLEVAGDALQVEPRGCMVRKGDPASGLSAARSRG
*
>GCA_000452945.1_BPMSHR5858_genomic_03119_1 group_1481
MEGGEAARQPGPISSFAALDALLAHFANRALYPSLSAVVMMGHSAGAQLLQRYAVAGREG
EKLARDGISVRYIVANPSSYLYFDDERPNGEVTAGGACPAATEWKYGLKSAPPYVVSQDL
QDLESRYVRRSVVYLLGEADTDPYTHFIDRSCAAMAQGPYRLARGLAYFDYLKKRHPSDL
AHRVVEVPGVGHDNLRMFTSDCGLAVLFDKATPQSCPIFTASAPSVRAEPR*
>GCA_000452945.1_BPMSHR5858_genomic_03120_1 group_1199
VCTGKLLDIGSPAGDRWRVRYPNAPKSEAAPTPLAQRGPRWFAIQPAIGDQIHDEGLSHR
RVECRRALAAALRSIYAAASEEAARQALQDFADGPWGAKYPTIVQPWQRAWEHAVPFYMF
PPEILRVAYTTNAIESLNMPLRKFIKTRGHCPNDEAAITLLWLALRNVLAKTVRAAFDWK
SAMNQFAILFGKRFMQARGSSVVQPSRP*
>GCA_000452945.1_BPMSHR5858_genomic_03124_1 group_19686
MHIDRVPPPDDRSLRTHAHPRRWLVSAALLTPALYALLWLALPYAWRYWRAVAGWGARQL
DPSLTAVVLGYPAHAPHVSLLSINLAARMPSGTLLFATAAACAAGFAATFVRRTTWLPVA
YLLRIAAFAQLMICMYFWAMYEPADTVASGVAAPGVAAPAAFPYDALLHLRDVFVMNVAA
IVLIPLVMAALYYPLDFSFAQKAFASLLVLGYFVLALPFVLLLHAIVIYHGSLLFVPFCY
FVLGAPLLIGLLVTLYSYCASWRGALTREPR*
>GCA_000452945.1_BPMSHR5858_genomic_03131_1 nhaR
MLNFRHLYYFWVVVKEGGFARAAGRLDMAVQTISAQVRELERSIGHQLLKPAGRGVTMTD
AGQAAFARAEEIFRIGQLIEDEVREAARGEGVRLAVGLTDGISKLTAHAVLEPALAAPSL
RLLCHDGEYEELLGELALHRLDLVLAGEPAPRRANLRLASERIVASPVDWYGPTAIAAAA
RGRFPQCLAELPVLLPTGHAALRARLDRWFDAHGIRPRIVGEFEDSALMAVFAARGLGVF
ALSEPGAGDVALLRGLRRLGRAQGVTEEIYAIRSHRGQHHPLVAQVLAAASR*
>GCA_000452945.1_BPMSHR5858_genomic_03134_1 group_1482
MKCPVCVTPDLLMTERQSIEIDYCPTCRGVWLDRGELDKLIARADDDASERRRDPARDAG
RDAPPARDEHDEHGRRRHDARERDASYRSQQGGYRKKKSLFDMFDFD*
>GCA_000452945.1_BPMSHR5858_genomic_03140_1 lipA
MTDLTATPAPAEPAASAYDPTAKQKAQAKTARIPIKIVPIEKLKKPEWIRVKAATGSSRF
NEIKTILREHNLHTVCEEASCPNIGECFGKGTATFMIMGDKCTRRCPFCDVGHGRPDPLD
ADEPKNLARTIAALKLKYVVITSVDRDDLRDGGAGHFVECIREVREQSPATRIEILTPDF
RGRLDRALAILNAAPPDVMNHNLETVPRLYKEARPGSDYAHSLKLLKDFKALHPDVATKS
GLMVGLGETTDEILQVMRDLRAHDVDMLTIGQYLQPSEHHLPVREYVHPDTFKMYEEEAY
KMGFTHAAVGAMVRSSYHADLQAHGAGVV*
>GCA_000452945.1_BPMSHR5858_genomic_03142_1 group_19368
MKEISSSITFEIPPGETVPMKVARSTRLAVRGAPVWATRSNDVHDYFLSPGATLKLRRGE
RLWLSADGATSACVSFSAIAPPQQAALRGVARFASWLSAHWRDGWRTV*
>GCA_000452945.1_BPMSHR5858_genomic_03143_1 gcvA_2
MNIHRLPMLNALRVFEAAARHESFSRAADELSVTHGAVSHQMRALEAELGVPLFVRHGKR
LALTDAGGRYAQQVRAALALLADATREVRASERDKRLVVSTLPSFAARWITPRIGPFIER
HPEIDLELRASDSLVDFARDDVDVAIRFGHGVYPGLHVEPLLDETFFPVCAPTLNGGMLP
ETPADLVRYPLLRSDDELWRPWFDAAGLDTLTEPKRGVLYQDSSNLLQAAIDGQGIALVR
RSLAVPEVAAGRIVRLFDIAGPSPWHYFFVCPPSLAQTPRVQALRSWLLDEIARFRALCA
AQEAQHTAAYAAARARGKEEN*
>GCA_000452945.1_BPMSHR5858_genomic_03144_1 group_19259
MTEPNKTVELTGEIATPKETLLEFPCDFPIKVMGRAHPEFKDTIFKVVSVHDNEIDLEKI
EERASSGGNYTGLTITVRATSQAQLDNIYRALTGHPMVKVVL*
>GCA_000452945.1_BPMSHR5858_genomic_03145_1 dat_1
MNQADIEPIVYLSVASHEELVPLSEARVPVLDRGFIFGDGVYEVVPVYADGARRAPFRIA
QHLARLARSLKRIGIADPHDEAGWRALVAQVVDANAAALGDGRHAIVYIQVTRGVAKRGH
AFPANAVPTVFAMASPLALPSDAQRAQGVRCVTAEDRRWLHCDIKSVSLLGNVLMAQHAA
EHDAIETIQLRDGNVTEGSSSNVWIVKNGELIAPPRSNRILEGIRYALVEELAEECGIRF
VAREINEAELRAADEILLTSATKEVLPVTRLDDLPVQGGRPGPVFDALYAAYQRAKAREM
EASERAVPPTRGETT*
>GCA_000452945.1_BPMSHR5858_genomic_03152_1 mlaF_1
MSARDDDFVIEVRDLTKRYGRNVVHEHLDFDVRAGEIVSIVGGSGSGKTTLVRQILGLER
PSSGTIRVFGEDTSKIDADTARVMRSRSGMLFQHGALFSSLTVFDNVAQPLRELGRVPDD
LLHDIVMLKLEMVGLPCKHASKMPAALSGGMVKRVGIARAIALEPELLFLDEPTAGLDPG
ASDEFVELIATLHRTLGLTVVMVTHDLDTMVALSTRVAVIAERKVLVAASVEEAAGVDHP
FIREYFLGRRGRRALQALPPERRARLPKAALEPAPSDVEL*
>GCA_000452945.1_BPMSHR5858_genomic_03154_1 cpdB
MPFPLRFRRRRLAAALVACAALVAGCNDDVDSPGAQSGASAPAGTKATLAVLETTDLHTN
VLSYDYFKLAADKSLGFERVATLIAQARAQYPNTLLLDNGDTIQGTALADYQALVKPVSC
GETLAIYKVMNAAKFDGGGIGNHEFNYGLPYLSQVTGNAFAVDGLPDPARQKKCAGPDFP
QVLANVISAKTNAPLFTPYTILTKNVTATTPDGRTVTAPVKVGIIGFTPPAIMNWDKRWL
DGKVYTTGLKEAAEKYIPEMRAKGADLVVAISHGGLDNSPYSPTMENGSWWLSTVPGIDA
MLIGHSHQVFPDATSTVAQFNLPGVDKVKGTVNGVPTVMANYWGKHLGVIKLGLAFDGKT
WRVDKSQTTVEARSIQNPDKSYVDADPSVAAAIAAEHQATIDYVKTPIGSTDYRMTSYFA
DVGDPGAIQLVNEAQADYVANYVQTNLPQYASLPVLSVSAPFKSGFGGGTDFTDVAAGAL
AINNAADLYLYPNTVYAVKVSGADIKNWLETAAKRFNTIDPTKATVQKLVSTFPGYNFDM
FTSADLRYEIDVTQPAGSRIRNLAYKGAPIDPNAQFIVATNNYRASGGGNFPGLDGSKTI
FASPDANRDVLIAYIKKRGRITRAADGMQRSWRFTKLAGSVAHVQFASAPNLAALASTAG
LAGITQVAADDGSGKGLATYEIDLTQ*
>GCA_000452945.1_BPMSHR5858_genomic_03163_1 group_19664
MTDLALYAGYEALRIQRRPHGIVEIVMSGEGANRSGLAVASEAMHRELADIWRDVDRDPD
ARVAVIRGEGKGFSAGGDLALVERMADDHAVRERVWREARDLVYNVINCSKPIVSAMHGP
AVGAGLVAGLLADVSIAAKHARIIDGHTRLGVAAGDHAAIVWPLLCGMAKAKYHLLLCEP
VSGEEAERIGLVSLALDDHELLPKAYEIAERLAQGSQSAIRWTKYALNNWLRLAGPTFDA
SLALEFMGFSGPDVREGIRSLRERRAPDFTGGAPS*
>GCA_000452945.1_BPMSHR5858_genomic_03168_1 metH_1
MSEPTPIAPFASSAAPAAPYTRGAALPQLLRQRILILDGAMGTMIQRYKLDEAAYRGERF
KDFPRDVKGNNELLSITQPRIIREIHDQYFAAGADIVETNTFGATAVAQADYGMEALVVE
MNVASAALARESAAKYATPEKPRFVAGAIGPTPKTASISPDVNDPGARNVTFDELRDAYY
QQAKALLDGGVDLFLVETIFDTLNAKAALFALDQLFDDTGERLPIMISGTVTDASGRILS
GQTVEAFWNSLRHAKPLTFGLNCALGAALMRPYIAELAKLCDTYVSCYPNAGLPNPMSDT
GFDETPDVTSGLLKEFAQAGLVNLAGGCCGTTPEHIAAIAKALAEVKPRRWPSQYSEAA*
>GCA_000452945.1_BPMSHR5858_genomic_03172_1 ftsN
MAKPRRTSKQSKQAGGTFLGIVLGLIVGLAIAVVVALYITRAPSPFVSKVAPPADNNASQ
PQQFDPNRALQGKTPGQPVTPQAAQPAPPNTAPGQAANPSQPPLLPEPQIVEVPSSNNNG
NGSPSASNNAADNGVAVAPKPAEPAPPPAKKPQTAANGSSAPHVANNNAQASAAATPPKA
AQAPKGASSATTTAAKPTSGADANTGYFLQVGAYKTEADAEQQRARLGFQGFESKVSKRD
VSGVTYFRVRIGPFSKFEDMNSARQRLSDAGVDTAVIRFTKQ*
>GCA_000452945.1_BPMSHR5858_genomic_03173_1 dsbA
MKKLLSSLFLSLSLVAGFAQASPSAPVAGKDFEVMKSPQPVSAPAGKVEVIEFFWYGCPH
CYEFEPTIEAWVKKQGDKIAFKRVPVAFRDDFVPHSKLFYALAALGVSEKVTPAVFNAIH
KEKNYLLTPQAQADFLATQGVDKKKFLDAYNSFSVQGQVKQSAELLKNYNIDGVPTIVVQ
GKYKTGPAYTNSLEGTAQVLDFLVKQVQDKKL*
>GCA_000452945.1_BPMSHR5858_genomic_03178_1 dppA_1
MRFTHIAAAALVAAASVAAAKPLTVCTESSPDGFDVVQYNSLVTTNASADVVFNTLVSYD
EATKKVVPALADKWDVSADGLAYTFHLRPNVAFQTTDSFKPTRALDADDVVFTFGRMLDD
ANPWHKVAGASGFPHAQSMGLAKLVKAVTKIDESTVKFELNEPNATFVPILTMGFASIYS
AEYADQLLKAGKQADLNAKPVGTGPFVLKSYTKDAVIRYEANPTYWGPKPKVERLIYAIT
PDPSVRAQKVKAGECQIALSPKPQDVAAAKADRALRVVETPAFMTAFVALNTQKKPLDND
KVREALNLAFDRATYLKVVFDNTATPATNPYPPNTWSYAKSIAPYPHDPAKAKQLLAQAG
FPNGFSTTIWVRPTGSVLNPNPKAGAELLQADLAKIGVKADVRVIEWGELIKQAKLGQHD
LLFMGFAGDNGDPDNILTPQFSCNSVKSGLNFARYCDPKLDKLIADAKETADQAKRAKLY
EAAQKIIHDEALWIPLGYPTAAAITRPNVAGYRVSPFGRQHFDAVSVQ*
>GCA_000452945.1_BPMSHR5858_genomic_03179_1 expZ
MARTAPAGALVALHHVSFRFDDGVTLFDSLDLTFNRAPSAIVGRNGAGKSVLARLIAGRL
APTAGAIERHASIAFVAQHDADAMHDSRCTAAAVARLDAPLAALERVARGDARADDFDLL
DGRWDLAERWQAALDEAGLPPLDARTPAHALSGGQLARVAMIGALLSGADLLVADEPTNH
LDAPGREWLRAALAGWRGGLVVVSHDRALLADVERIVELTPRGARVYGGNYAAYRAQRDA
EAQATQAALDHAHAERERERRRLAREHDTIQRHAAATRRYAETANLPSGKRVSLKNSARE
IMGRVRRDHRDTKTALGDAVQAAAARIEPDAPVLVSLPGTEIAARRRLFTLDAARLPWLP
AHARAATVTWSAHGPARIALTGPNGCGKSTLLRMLAGECAPRAGRCDTHVPLAYLDQRLA
LLDPRRSVVEQLAALRTPLGQGELRSRLALLQLDATRATQPSARLSGGERLKAALACALW
RETPAQLLLLDEPTNHLDLESVRAFEAALADFPGAIVAVSHDAAFIDALAPTHAMRWTSE
GWRFEPVA*
>GCA_000452945.1_BPMSHR5858_genomic_03181_1 aceK
MNHFPKLLSSQIGFDVAQTILENFDRHYRIFREAAVEAKDLFERADWHGLQRLARERITS
YDDRVRECVELLEDEYDAENIDNEVWPQIKLHYIGLLTSHRQPECAETFFNSVCCKILHR
AYFNNDFIFVRPAISTEYIENDEPAAKPTYRAYYPGSEGLAATLERIVTNFQLNPPFEDL
ERDIACIMQAIHDEFGAFDEAVNFQIHVLSSLFYRNKTAYVVGRIINGDRVLPFAVPIRH
ARAGILALDTVLLRRDQLKIIFSFSHSYFLVDMNVPSAYVQFLRSIMPGKPKAEIYTSVG
LQKQGKNLFYRDLLHHLSHSSDRFIVAPGIKGLVMLVFTLPSFPYVFKMIKDHFPPPKDT
TREQIMAKYLLVKRHDRLGRMADTLEYSSVALPLARLDDALVRELEKEVPSLIEYEGENL
VIKHLYIERRMVPLNLYLQNGSDAEIEHGVREYGNAVKELMQANIFPGDMLYKNFGVTRH
GRVVFYDYDEIEYLTDCNVRRVPPPRNDEDEMSGEPWYTVGPHDIFPETYAPFLLGDPRV
REHFLAHHADFFDPQLWQDSKDRLLRGELPDFFAYEPALRFCIRYPERFAPGDAADGGKL
AAA*
>GCA_000452945.1_BPMSHR5858_genomic_03182_1 can_1
MNKNDHPLSHLFDNNDAWVKRKLADDPQYFSRLADQQAPEYLWIGCSDSRVPANQIIGLP
PGEVFVHRNIANVVVHTDLNCLSVIQFAVDLLKVKHVMVVGHYGCSGVNAALHNRRVGLA
DNWLHHVQDVREKHAALLEDWPLGEARYRRLIELNAIEQVVNVCRTTIVNDAWARGQPLT

VHALVYGVHDGRMRNLGMAVSHAEQLDATYRRAVAALSASGAHSADNDVVAADAAQLAGA
VDLIAQTIKETKHDGC*
>GCA_000452945.1_BPMSHR5858_genomic_03193_1 phoA_1
MSTIKRIAAAAFAAAALSGGFGHAAHAAGQAKNVIFFLGDGMGPATVTASRLYKVGEAGQ
LTMEKLPRTARIKTFSNDAQTTDSAPSMAAYMTGVKMNNEVLSMSPDTRAIAPGSDANGN
KTVNRCGVGNGTPAATLLELAKARGKAVGAITTTELTHATPAATYSHICHRDAQYDIAAQ
AVPGGAGYNAALGDGVDVLMGGGRNHWTPYDPLANRRGRADGRNLLAELQAKGYAVVATK
DQLAQAGAGKLIGLFSTTSHLEYELDRVAGKGEGATQPSLAEMTAKAIDVLRKNPNGYFL
MVEGGRIDHALHGTNAKRALEDTVAFDEAIRTALAKVDLSDTLIVVTADHDHTMTINGYS
KRGNPVLDISRNYRDGQPNKDADGNPYTTLVFGNGANRPNARVPVDSSTATNDAYLQEVG
VRMGSAGSETHGGGDVMLFADGAGAKAFKGTLDNTKVFGLVKAAFGF*
>GCA_000452945.1_BPMSHR5858_genomic_03195_1 group_19481
MKRPLSLTVLAWFVIVSNVIGLISMPFSLRDPNAVALLRYSPLPLEATVAIAVVSEGLHL
VAGTAILKRRIWGRTLYAIVALAGFAISVFTTPFKGMLIIAAALIALFLFILYRPAATAY
FRQRAS*
>GCA_000452945.1_BPMSHR5858_genomic_03197_1 nhaK_1
MAIVFTVLILLLAVALSGVATRVLPLRLPLPLMQIAIGAVLAWPRFNLHVTFDPEIFMLL
FIPPLLFADGWRIPKRELYLQRRAILMLAFGLVFMTVLAVGYFAHGLIPGLPLPIAFALA
AVLSPTDAVALAGIAGKGRIPPQLMHILEGEALMNDASGLVALKFAIAAALTGMFSLRDA
SISFVIVASGGLAAGAALAWLFSALSTRFLNAAQEGDPAPGIVMTLLVPFASYLIAEHLG
LSGILSAVASGMMMNYTSFSKSSTVAARVRAESTWAMIEFVFNGMVFIMLGLQLPHIIGR
ALVDAHHTSDALVGAMIGYVAAMLAALYAIRFTWVWLLRWIASRRAAKQGLAGTMAGLRT
IAVMTVGGVRGAITLAGVLSIPVALGDGTPLAGRDTAIFVASGVILGSLVVAVIALPLLL
RGVRPMRNPHAEEERAARAAAAQAAIRAVDAAHDALAVDLDESAAARCADVTARVMDLYR
RRLAGLAADGATPREEARQTEKMELALKIAAIRAERAVLYRLRGEQKINDETLAKLMREV
DFSETALSTRKKGIV*
>GCA_000452945.1_BPMSHR5858_genomic_03201_1 group_1872
MKKSMHGGYGGLARPCGRMPATAAVAAALLLALGGCGDDLQSTTTPAQLNQPYTDTTAYS
PKAGDGLPASQVSERAAVMSHQWTANGASVDYLTTTGHLTATDPNGNAEATMSYVAYTAP
SRDGSPRPVTFFYNGGPGSSSVWLRLGSFAPTRVATPDPLMTNWPNFPLVDNPESLIATT
DMVFIDPPGTGLSEAIQPNTNQTFWGADADVKVMRDFIRRYLSVNGRGGSPIYLYGESYG
TPRTDMLALALESAGVPLTGIVLQSSILNYMAAAGDQAVGTFPSYAQVAAYFNQVSPSPT
NLGAYAQRIENFVTAQYAPIVHYATASSPISPDAGTLAAWSSQTGMATASIGAYFQYFYD
TEPSPGQTTLVPGYTIGRYDGRVSLPNGDARLASDDDPSDILISKPFTSALASQMPNYLG
YTAPNATYQTLNPDIIGVWNFSHAGQPYPDTIPDLLAALQLNPKLKVLASNGYHDLATPY
FETEKELARLQTVSGLAPNLQVTFYQGGHMIYLDDVARPQMQADLVAFYQNRPVANALTL
AALPSPWPDESPANTPTAKIARAAAAR*
>GCA_000452945.1_BPMSHR5858_genomic_03210_1 menH_1
MTNVDACGHAAADRWVDTGRGRLFARCWPAPHRAHMPDDAPPIVLLHDSLGCVRMWRTFP
RTLAACAGRRVIAYDRLGFGQSDPRTDALGFDFVRDEARRFFPMLRDAFGFDRFVAFGHS
VGGGMAVHCAAEFASACDALITESAQAFVEDRTRAGIVDAREQFKQRDAFERLRAYHGEK
ARWVLDAWIDTWLSPEFAGWSLAAVLPRVTCPTLAIHGGDDEYGSALHPETIARLASGPA
QIEILPGVRHVPHREREQWVAQRVAAFIGAGSHGIAE*
>GCA_000452945.1_BPMSHR5858_genomic_03211_1 group_19546
MIDVPTLLTIVLMASSTYLSRILGYVALRNRTLSPRMMSVMENVPGCVLVSVIAPAFVSD
KPANLLALGITLLAATRLSILPTVIIGIVSTGVLRHLLGQ*
>GCA_000452945.1_BPMSHR5858_genomic_03215_1 plcN_2
MTSESRRRFLHTVAQSAGAAAALTVFPESIRRALAIPAASRTGTLRDVEHIVVFMQENRS
FDHYFGHLRGVRGYNDRFPIPLPNGKPVWYQPSKANPGKPVLPFRLNTLTTSAQCIGDLD
HSWYKTHAAIDGGRYDQWPANKTDMTMGYHVREDIPFHYALADAFTVCDNYFCSLPGPTH
PNRSYLMTGTVDPTGRYGGPLLDNSDYVDGDVPPKYDLLTWTTYPERLEANGISWQIYQQ
GTTGADPLNGNYGTNILQNFANFINAKPGSSLYRRAQTARTLDDLKADVLANKLPQVSWL
LPPAAFSEHPRYTPAYGANYTSQILDALTSNPDVWRKTVLFIMYDENDGFFDHVVPPQPA
TTRAQGLSTVTVDGEIHDVVNPGRGGSYTADGLPYGLGPRVPMTVVSPWTKGGFVCSQVF
DHTSVIRFIAARFGIDEPNITPWRRAVCGDLTSAFDFRTPDATLPPLPDTSGYRSIADQL
CSTKPAPAVPAAPSPVDPQEPGVRPARAALRAARERRGARRRAPAHRVRESRRARRALLC
VRIESQRRPVALHGRRAPLVARNVRSRRDERRVRVRRVWSERLRARVRRPRRGRRPRPAS
GRAPRGEGGLRRRERQPVSEARESWRTERATDARRQRVRRAGAPDHAARRR*
>GCA_000452945.1_BPMSHR5858_genomic_03220_1 liuE
MTLPQSVKIVEVGPRDGLQNEQTFVPTGVKIELVNRLSQAGFRNVEAASFVSPKWVPQMA
DGADVMAGIARRAGTIYSALTPNLKGVENALAARADEVVIFGAASEAFSRRNINCSIAES
IARFEPVARAAKDAGVRLRASVSCALGCPYQGDVPLEAVVDVVERLAALGCDEIDIADTI
GVGTPGRTRAVLDAVAKAFPRERLSGHFHDTYGQALANIYAALLEGIAIFHASVAGLGGC
PYAKGATGNVATEDVLYMMRGLNIETGVDLDQVVAAGDFISNAIGRANISRAGRALLAKQ
RAAAEPPSACA*
>GCA_000452945.1_BPMSHR5858_genomic_03225_1 group_19287
MSFEEFAPFRVTAQDVDIFGVKGGAGPPILLLHGHPQTHMIWHRVAATLARHFTVIATDL
RGYGASGKPPSDARHAPYSKRAMAADQVAVMRHFGFPRFYVCAHDRGARVAHRMALDHPD
AVERMLLLDIAPTLAMYERTDRAFATAYFHWFFLIQPEPLPETLIGGQSDAYVEAVMGNR
SAGLAPFSPEALHAYREALRQPGAVHAMCEDYRASATIDLDHDRADLERGHKLACPLRVL
WGEQGVVARCFEPLDEWRRVARDVSGRALACGHYIPEEAPVALIDEIVSFFEAREAA*
>GCA_000452945.1_BPMSHR5858_genomic_03227_1 dmlR_3
MDTLVSMNVFRYVVEVGSFVGAAERMEMSAAMASKHVMHLEHQLGARLLHRTTRRVAPTE
AGREYYERLVQALTELDEAGQAVGAASVVPQGRLRVTSLSAFGLRHVMDAVTSYAERYAD
VTVEITLSDRVVELIDEGYDVAVRAAPAGMKSSSLVARQIATAHILLVASPDYLAKHGTP
ATIADLQHHSFIRREPNETLLDAAISEVAAASRVPMSGNVIVNHLEGLRVAVLGGAGIAL
LGTEVVGEDIESGRLVPLLLDALPPRELPIYAVYASRRHVSAKVRSFVDFLADRFAGQSL
CPSIDERLRQLAMPRMKRLG*
>GCA_000452945.1_BPMSHR5858_genomic_03242_1 glmU
MNIVILAAGTGKRMRSALPKVLHPLAGRPLLSHVIDTARALAPSRLVVVIGHGAEQVRAA
VAAPDVQFAVQEQQLGTGHAVRQALPLLDPSQPTLVLYGDVPLTRTATLKRLADAATDAR
YGVLTVTLDDPTGYGRIVRDQAGCVTRIVEQKDASPDELRIDEINTGIVVAPTAQLSMWL
GALGNDNAQGEYYLTDVVEQAIEAGFEIVTTQPDDEWETLGVNSKAQLAELERIHQRNLA
DALLAAGVTLADPARIDVRGTLACGRDVSIDVNCVFEGDVTLADGVTIGANCVIRHAAIA
AGARVDAFSHLDGATVGANAVVGPYARLRPGAVLAADAHVGNFVEVKNATLGQGSKANHL
TYLGDADIGARVNVGAGTITCNYDGANKFRTVIEDDVFVGSDTQFVAPVRVGRGVTVAAG
TTVWKDVAADMLVLNDKTQTAKSGYVRPVKKKS*
>GCA_000452945.1_BPMSHR5858_genomic_03262_1 group_1875
MTGAKHRENNFRPAHAESFGAMQAARKLTGELARKPNRKESPYFMGFSGLIGRARSCASA
QRLCHFYLPIFMRPAYNPPQFIKESHVDTSPGSSRPMPVGIGAAFAN*
>GCA_000452945.1_BPMSHR5858_genomic_03266_1 group_19656
MQNEPQRTRSDAHTPRLIKHAGRIAALAGLAISLWLVARDDPHAVLALLRAAGIGLVVAA
ASHLLPMLANAKDWQTLIAPRCRPSLPAMLRLVWIRESVNGLLPVARIGGEFASFRLLRA
HGLTAPAAVASLVSDMQLTLISQALFALAGVGYLLVHATSDTARVAGMFAWGLAVLAPLL
LLFALVQHAKPFERAMHALNRVAGGQLASLVERSAQIDDALKAVWRQRGTVVRYLFFWQP
LQHLATSLEIWLALHFLGASVSLVDALVIEALIQALSSAAFFVPGGLGVQEGGFVLIGGA
LGLDPATSLGIAGARRIRDLLIFVPGLFAWQHAEASVAASTP*
>GCA_000452945.1_BPMSHR5858_genomic_03267_1 group_1206
MRWQYPVAAALAIAAFVWACARIVSTDAAPATAPAGQTASAAPAALASAFTAASSAAPRS
APPPLPPRVTEHIAREYADSAVTREALTQIAYGWNQAVNDVHDPGDAKIAGDAIAKGIAC
ALSARVLAKAGIDQQTMLDRIKGTRAVMLDTETDTLAYIRFQSLAGGQFFDDPGPASCGF
NPTAMPN*
>GCA_000452945.1_BPMSHR5858_genomic_03268_1 group_19694
MKKLVTLMIAGALVALPCTTNLNAAQAATRAPAVAGSPGSAIVFVNGINNTFDDAIASLQ
VLKTQLNARDANNAYVYGNAYNATQGTFSDLWQVFKQKSVEGASPADFWRAIDGGTVPAG
GMDAALQQKYIDILTKSETPELPDHLNQYRAYLQQNRKVVLVGHSQGTLYANFEMNLLIT
GPERAQGRISSVNVGNAARYLLPGSSYLTSSSDTVIGSLGLVQTVMPSNFSLGLHLFTDP
LGHSFTKIYLNPSFSAARQILTQVSQQANSAAKAAVR*
>GCA_000452945.1_BPMSHR5858_genomic_03270_1 group_1487
MNLDAASRNLTVASLLTLSGGYLDAYTYVGHGHVFANTMTGNVALLGINVSAGDWAQALH
HVPPLAGFVIAVFVAHTLGLAAQKGWVRHTAFISLIVEIGFLAFASAGWPAMSSAWLIPG
ISFVATLQTLSFTHLENLSYTSVMTTGNLRRAAQLLFAGLIPRYDALALHDSGLLAVISL
CFLMGAVAGGFVTRLVGDGALWGAVLLLAAAFAEIVRRARHDSGAGAARAERAARDAPAE
RAAHER*
>GCA_000452945.1_BPMSHR5858_genomic_03272_1 group_19163
MTAAKFFLRYTLVPFVAIVLVVVWAFNRESDEIDARQYAALVAAYPGFTPDLRDEVNRAM
RDGKLAKSDYASIVRDSLNHGYVLDWSSTNVDLNAERGKLLGLVKSDAHAPD*
>GCA_000452945.1_BPMSHR5858_genomic_03273_1 group_19446
MSDTLAALAVIFGQLSLLAFGGGNTILPEMQRQVVDVHHWMSAQAFTALFALAQAAPGPN
MMIVPLVGWHVASWPGLLVSSIAKFGPSSIVTMAALHAWERFKDKPWRRYVQQGMMPVTA
GLVVASAVLISEASNRSALQWGITAAVAALAYRTRVHPLWLLAGGALAGLVGGWFA*
>GCA_000452945.1_BPMSHR5858_genomic_03275_1 gcvA_1
VTHGAISRQIAAFEAWLGKPVFHRHGKRVKLTDEGRRYLATVEAAFDSIAGATVQLRETG
AARVLRINALPTFAMKWLLPRLSRFQREAPNVELKLSTSNAPLGSLDGFDVAIRRGPGRW
PNCTSGRFLDESVIPVCSAALLKRAPIASADDLARHVLLHSDTRPDAWREWFAAAGAPLK
GRKKQSFDHFYLALQAAVDGLGVALGPLPLIADELASGRLVTPLPGPRIATRSYWWIAPR
ETANDVLVGRFCAWLDAQAQAS*
>GCA_000452945.1_BPMSHR5858_genomic_03278_1 group_19627
MSNTLMNLGVSGLNAALWGLTTTGQNISNAATPGYSVERPVYAEASGQYTSSGYLPQGVS
TVTVERQYNQYLSNQLNAAQTQGSSLSTYYTLVAQLNNYVGSPTAGIATAITNYFTGLQT
VANNAADPSARQTAMSNAQTLASQLVAAGQQYSQLRQSVNSQLTDTVTQINSYTSQIAQL
NEQIASASSQGQPPNQLLDQRDLAVSKLSQLAGVQVVQSNGNYSVFLSGGQPLVVGNASY
QLATVASPSDPSELTIVSKGVAGSAQPGPTQYLPDVSLTGGALGGLLAFRSQTLDPAQAQ
LGALAVSFASQVNAQNALGVDMSGNPGGSLFAVGAPAVYANQNNTGSATLSVSFVDGTQP
TTSDYALSYDGAKYTLTDRATGSVVGTATPSSTPPTMTIGGLKLSLSSTPNAGDSFTVLP
TRGALDGFSLATANGSAIAAASPVLAAGVATNSGTGVISQGSVSAGYQLPSGTTTLAYNA
ASKTLSGFPVGTTVTIAGTPPTSINITSATTPVPYDPSKGASMTISSTTQPAPSGVMNGV
SVSLSGTPADGDQFTIGANKGTNDGRNALALSQLVNSKTMNNGTTTLTGAYAGYVNAIGN
AASQLKASSAAQTALVGQITQAQQSVSGVNQNEEAANLMQYQQLYQANAKVIQTANSVFQ
TVLGLFN*
>GCA_000452945.1_BPMSHR5858_genomic_03279_1 ycgR
MNTEQSTSPAASAAAHSGHDYGRRNPLEIGVQLRNLVNRGDFLTVQYQGGQLVTRILDVD
VGARTFVFDWGALADQNAGILAAPRCVFDASPDGVRVEFSTATPRETRYENLPAFEADFP
DVLYCMQRREYFRVDAPILDPYVCRGRLPDGETFLFEVHNLSLGGLGLRTSDDRVASLEP
GMTLPDVELNLNGHGMLSLDLQLVSHRASETPSGARRYQLGFRFVSLPGSAENTLQRIIT
QLEMKRRQLARA*
>GCA_000452945.1_BPMSHR5858_genomic_03280_1 flgJ
VTTFTKPNDLSQRFALDVQGFDALRSKATAAAPREGVKMVAGQFDAMFTQMMLKSMRDAT
PSDGLLDSSSSKMYTSMLDQQLAQQMSSKGIGVADALTKQLLRNANVAPDAQGEGGLAAM
NALAKAYANSNGAPGNGALAGTRGYSAASALTPPLKGNGNSAQADAFVEKMALAAQAASA
TTGIPARFIVGQAALESGWGKREIRGANGESSYNVFGIKATKGWTGRTVSAVTTEYVNGK
PHRVVAQFRAYDSYEHAMTDYANLLKNNPRYASVLNAGHNAEGFAHGMQKAGYATDPHYA
KKLISIMQQIG*
>GCA_000452945.1_BPMSHR5858_genomic_03285_1 flgE
MGYQQGLSGLAGASSDLDVIGNNIANANTVGFKGSTAQFSDMYANSVASAVNNPIGIGTM
LASVQQQFSQGTITSSTSSLNVAINGNGFFQMSNNGVTTYSRDGTFQRDKNGYIVNSQGL
NLMGYAANANGVINTAATVPLQAPTTNIAPTATTKITGQFNLNSQDAVPATTPFNYTDPT
SYNYTTSVQVFDTLGGSQNVNLYFVKSATSGQWEAYAGPAGKTPTDLGSVKFSTAGTITG
TSTPAGVPTTNVGQFSFSIPTTTGAANPQNLTLDLTGTTQYGGKNGINNLAQDGFASGVL
TTFSIGADGKVTGNYSNGQTSTLGQIVLANFNNPNGLVNVGNNQYVESAASGVPQISAPG
STNHGTLQGSALENSNVDLTSQLVKLITAQRNYQANAQTIKTQQTVDQTLINL*
>GCA_000452945.1_BPMSHR5858_genomic_03287_1 flgC
MPSLMNIFDVAGSAMSAQSQRLNVTASNLANADSVTGPDGKPYKAKQVVFATAPMGRART
ASGQGVGGVQVSKVIDDPSPMKTAYDPSNPAADQNGYVTMPNVDPVQEMVNMISASRSYQ
ANVETLNTAKQLMLKTLTIGS*
>GCA_000452945.1_BPMSHR5858_genomic_03292_1 group_1488
MAFPTSVNDQITDSVTQANTKVLGDAPAIAMGNLYQATAQALANTAHNATNAQQQSYVTA
LAERRDGGAATTI*
>GCA_000452945.1_BPMSHR5858_genomic_03293_1 cca
MKIYAVGGAIRDALLGLPVRDRDYVVVGATPEQMAAQRFRPVGKDFPVFLHPDTHEEYAL
ARTERKTAAGYHGFQFYYAPDVTLEQDLVRRDLTINAMAREVSPDGALVGPVVDPFGGQA
DLRAKLFRHVGDAFVEDPVRILRVARFAARFAEFAVAPDTAALMRAMVDAGEVDALVPER
VWQELARGLMEAKPSRMFAVLRECGALARILPEIDALFGVPQRADYHPEVDTGVHVMMVI
DHAAKQGYSLAVRFAALTHDLGKATTPADVLPRHIGHEGRSVDLLKPLCERLRVPNECRD
LALVVAREHGNLHRVMEMGAAALVRLLERADALRKPARFAEALQASEADARGRLGLETKP
YPQAERLRQALVAARAVDAGAIAQGLAGEPAKIKDAVHRARVRAVAQAVGVAD*
>GCA_000452945.1_BPMSHR5858_genomic_03299_1 pxpB_1
MTTPRIYPLGDTALVCEAPPPATLECQRRVWAVASAAAGWPHVVDVVPGMNNLTIVFDPL
ETTADTLAPLMKAAWRDADGVAESGREIEIPVEYGGAFGPDLDAVARHTGLARDEVVRRH
AAGTYVVFFLGFQPGFTYMGGLDGSLHTPRRAAPRLEVPAGSVGIGGEQTGIYPAAAPGG
WQLIGRTPLALFDPARTPPTLLQPGDRVRFIVAGVHAG*
>GCA_000452945.1_BPMSHR5858_genomic_03302_1 group_19065
MQGTVSLWPLIGVAVIVVGFALRFNPMLIVATAAIVTAASAHFPPERILAAIGAGFLKTR
NIPLIILLPLAVIGLLERHGLRERAQAWIASIRAATAGRLLILYLLVRELTAAVGLTGLG
GHPQMVRPLIAPMAEGATENRFGPLPDAIRHRLRAYAAATDNVGLFFGEDIFVAFGAIVL
MVTFLKEAGIAVEPMRVALWGIPTAVSAFVIHGARLWLLDRRLERELRASAAPHAASNGA
NAAADNGGRA*
>GCA_000452945.1_BPMSHR5858_genomic_03303_1 group_19109
MTLTIGYLFWLVGLVLLAVGATIVTDRAHPRRFTAGGFWLLYALVFLVGDKLPVELVGAL
VIVMALIAGFGGVTAARPKLPSDDARRASAARIGNRLFVPALTIPFVTVAITLAASHLHV
GGAPLVDPKNVTLIGFGIGCVIALGFACWITRDSVGQSLQEARRLIDALSWAAVLPQMLG
MLGLVFSDAGVGKAVAHVTTAYVNLDYRLIAVAVYCIGMALFTMVMGNGFAAFPVMTGGV
GVPILVDVFHGDPAVMVAIGMFSGYCGTLMTPMAANFNMVPAALLELPDKNGVIKVQIPT
ALALLATNIVLLNVLMFR*
>GCA_000452945.1_BPMSHR5858_genomic_03306_1 dppD
MSDLLTIRNLAVNFSGLPAVDRINLSIARGEVVGVVGESGSGKSVTMMALMGLIDAPGIV
TADEVTFDGVDLLKASAKARRRIIGKDVAMVFQDALTSLNPSYTVGYQIKEVLKLHEGLR
GDALNKRALELLDQVGIPDARNRIGTFPHQMSGGMNQRVMIAMAVACNPKLLIADEPTTA
LDVTIQAQIMDLLVTLQKERGMALVLISHDLAVVSEVAQRVAVMYAGEIIETNRVPDIFA
HPHHPYTEALLAAIPEHNAGAKRLAALPGMVPGRDDRPSGCLFAPRCKYVVDDCMKARPT
LAELAPGTGMRARCIKPLNLEAPHTQGGAR*
>GCA_000452945.1_BPMSHR5858_genomic_03308_1 dppB_2
MFRFVLRRVGMVIPTFIGITVLAFALIHLIPGDPIEVMMGERGVDPTMHAQALHRLGLDE
PLPMQYLHYVGRALHGDLGTSIITNTSVMGEFLARFPATVELSLCALAFALALGLPAGVF
AALRRGTIVDHGVMGTALTGYSMPIFWWGLILIMVFSATLGWTPVSGRIAVEYEIPHTTG
FMLIDTLLSGEEGAFASAVSHLILPAIVLGTIPLAVIARMTRSSMLEVLREDYIRTARAK
GLSPARIVVVHALRNALIPVVTVIGLQVGTLLAGAVLTETLFSWPGVGKWLIDAIGRRDY
PVVQGGILMIATLVIVVNLVVDLLYGVLNPRIRHTR*
>GCA_000452945.1_BPMSHR5858_genomic_03316_1 rhtC
MNLLPILLQIAAVYLVALVTPGPNIFMISQLSLSGRRGLGAVSALGVGTASLTWATLAML
GVAAVLHQLAWLYEAIRIGGAVYLAYFGIKLLRASAQRDPAPAAGLADDAGALPSPPATR
DYLRAYRTGLFTCLTNPKSCVFWTSVFAAMMPAHVPLWFSGATLLTIGAMSGGWYCGVAY
LFANPRAQRGYRRVRRPLDALCGAALVGLGAKLAADR*
>GCA_000452945.1_BPMSHR5858_genomic_03320_1 yhdG_2
MSLFRKKNVDRMIAGAQAAGLKKALGAVDLTFLGIGAIIGTGIFVLTGTGAVQAGPALML
SFVIAAIACGLAALSYAEFASTIPVAGSIYTYSYATLGELVAWIIGWDLMLEYGLAASAV
SVGWSGYLQSLLQGFGVSLPTALTAAPGALPGVVTYFNLPAFVVMLVITTLLSIGIREST
RVNNIMVFIKVAVVLLVIAVGVFHVTPANWKPFMPHGWNGVFGAAAVMFFAFIGFDAVSS
AAEEVKNPKRDLPVGIIASLGVCAFLYVAVAAIATGIVPWAQYANVSHPISYALQVAGEK
WVAGFIDLGAVIGMLTVILVMSYGQTRIIFAMSRDGLLPAALSRVHPRFATPFLTTWLVG
LFFGLIAALIPLNVLAELINIGTLAAFSMVSIAVLVLRRTHPDLPRAFRCPGVPLVPILA
VAACLFLMLNLQPVTWIAFLTWLVIGLFVYFAYSRSRSKLAHGAQQH*
>GCA_000452945.1_BPMSHR5858_genomic_03323_1 flhB_1
MSGPPRKRAAVLAYDAKGGDTAPRVVAKGYGLVAERIIERARDAGLYVHTAPEMVSLLMQ
VDLDARIPPQLYQAVAELLAWLYALERDAGDGGGAQPAFPPLPKR*
>GCA_000452945.1_BPMSHR5858_genomic_03325_1 group_18986
MNRKDEYFARYEAIAAVSGRMLVAARDAAWHAMGGLQEEYLRLVDGLKEAEQGVLLDDAE
RTRKFELIRRILADDAAIRDLASPEVARLSAFFHAGRSTLKDLYRARG*

>GCA_000452945.1_BPMSHR5858_genomic_03327_1 fliE
MVAPVNGIASALQQMQAMAAQAAGGASPATSLAGSGAASAGSFASAMKASLDKISGDQQK
ALGEAHAFEIGAQNVSLNDVMVDMQKANIGFQFGLQVRNKLVSAYNEIMQMSV*
>GCA_000452945.1_BPMSHR5858_genomic_03329_1 fliG
MNAEGLNKSALLLMSIGEEEAAQVFKFLAPREVQKIGAAMAALKNVTREQVEDVLNDFVQ
EAEKHTALSLDSSEYIRTVLTKALGEDKAGVLIDRILQGSDTSGIEGLKWMDSAAVAELI
KNEHPQIIATILVHLDRDQASEIASCFTERLRNDVLLRIATLDGIQPTALRELDDVLTGL
LSGSDNLKRAPMGGIRTAAEILNFMTSVHEEAVIENVKQYDPDLAQKIIDQMFVFENLLD
LEDRAIQLLLKEVESEALIIALKGAPPALRQKFLSNMSQRAAELLAEDLDARGPVRVSEV
ETQQRKILQVVRNLAESGQIVIGGKAEDAYV*
>GCA_000452945.1_BPMSHR5858_genomic_03330_1 fliH
MSDRASASDKPVTAYQRWEMASFDPPPPPPPDDGGAAAAALAAELQRVRDAAHAEGLAAG
HVEGQALGYQAGYEQGRAKGFDEGRAEAHTHAAQLAALAASFRDALAGVERDLADDIATL
ALEIAQQVVRQHVQHDPAALIAAAREVLAAEPALAGAPHLIVNPADLPVVEAYLKDELDT
LGWSVRTDTSIERGGCRAHASTGEIDATLTTRWERVAAALGKVSAW*
>GCA_000452945.1_BPMSHR5858_genomic_03332_1 group_19579
MAQSFPLQLLLERAQDDLDTAAKQLGRAQRERTDAQAQLDALMRYRDEYRVRFAESAQSG
MPAGNWRNFQAFLDTLDAAIEQQRRVLAAAQTRIDAARPEWQAKKRTLGSYEILQARGAR
QDAQRAAKREQRDADEHAAKVLRMRADAAK*
>GCA_000452945.1_BPMSHR5858_genomic_03335_1 aidB_1
MQQDLMTATHRVTNQAPPLSDYNAFETDAALVEAVRRYGASWRQTALSHDGAALTTPDVL
ALADLANRHAPELATHSPRGERIDALEFHPSWHQLLARLRAEGLHALPFSDPRPGAMAAR
CAGYFLHAQLESGSLCPLTMTFASIPVLQREPALFATLRDKLYAREHDARDAPLPQKRSM
MVGMGMTEKQGGSDVRSNETHAYPLGAGGRGQAYRLVGHKWFFSAPQCDAHLVLARTAEQ
AGISCFYVPRFSPDGSKNAVHVQRLKDKLGNRSNASSEVEFLNAYGVMIGDEGRGVPTII
EMANYTRLDCVIGSAALMRAALVQAIHHARHRSAFGRLLADQPLMRNVLADLALESEAAT
VLFMRLAHAFEEDAAARSPQARGWRRIVTPAAKFWVCKRTLEFTGEAMEVWGGNGYVETG
PMARFYREAPVNSIWEGSGNVMCIDVLRAIEREPDAAAALFDEWRDAARHSAPLAGALAE
LTRLLSLEPEPREACARRIAQRIALVAQASLLLRDGPYAVAQAFVATRFGHDSSETGRVF
GTLPATIDHAALVERAFPA*
>GCA_000452945.1_BPMSHR5858_genomic_03338_1 srkA
MRSMNDATSESNAAGACAGLPFAGLTPERVLDALDSVLIPAGSRTDGRLLALNSYENRVY
QAGIEDGAPIVAKFYRPQRWSNDAILEEHTFVAELAAREIPAVPPLAFDGRTLHEFDGFR
FAIFERRGGRAPELDRRDTLEWLGRFIGRIHAVGATKPYAARPTLDLRTFGYEPRDFLMS
HDFVPDDVRPAYEAAVALALEGVERAYERAGDVRMLRAHGDCHPSNVLWTDAGPHFVDFD
DSRMAPAVQDLWLLLPGDRPGASRALTDLLAGYEDFCEFDPRELHLIEALRTLRLIHYAA
WLARRWDDPAFPAAFPWFNTHRYWEARVLELREQIGAMQEGPLWPV*
>GCA_000452945.1_BPMSHR5858_genomic_03340_1 ynfM_1
VTVPPGAPRPAAVGRANSAHAAQGASPVGAAPYLERGSRGYWHASVALLFAGYATFSLLY
YVQPLLPEFSKAFGVSPAQSSLALSFATAALAAAVFVAGFVSEALSRHRLMTASLTASSL
LTLAAAFAPHWHQLLILRALTGLALGGVPAVAMAYLAEEVHPDGLGLAMGLYVGGTAIGG
MAGRVITGILTDLFSWRIAVGAIGVLGLASMLAFRMLLPPSRHFVPRRGLNLAHHRTSLA
HHLRGQRELPVLFAMAFVLMGSFVTLYNYIGYRLLAPPYSMGQATIGAIFVVYLVGVVAS
PLSGRLADTLGRGRVLIASLAVMLGGVALTLLHPVAAIVAGIACVTFGFFAGHAVASGWV
GRLAQHGKGQAAALYLLAYYLGSSVVGSLGGRFWSTLGWPGVAALVGALLALGVVAAAWL
RARERGGAA*
>GCA_000452945.1_BPMSHR5858_genomic_03341_1 group_1207
MASVRPALAAHGATSLFVLLWSSGAIFAELGLRHASAFAFLIARFALASLVLLVLSLARR
RWLPPRGARRATLATGVLLTGGYSILYLLALERGLAPGILATILGVQPILTLAVLERRAS
PARLAGLALALTGLSLVVYRSVAGGAAPAAGTACALGALLAITVGALLQKRVRAAPIDVL
PLQNALGLALCALIAPFEPISFEASWAFALPLAWLGIVISVFAQLLFYRLMQRGDLVNVT
SLFYLVPVVTALMDAAWLGNRLAPLEIAGMIAILAGLALAFRRAARD*
>GCA_000452945.1_BPMSHR5858_genomic_03344_1 group_19671
MTRAISVALIVGGVVLLYFGGQSFHSINDGVARFFTGSPSTKTIMLIAGGVVATLVGLIG
LSMPSGKR*
>GCA_000452945.1_BPMSHR5858_genomic_03355_1 hslV
MEQFHGTTILSVRRGDKVALGGDGQVTLGNIVMKGGARKVRRIYNNQVLVGFAGGTADAF
SLLDRFEAKLEKHQGNLTRAAVELAKDWRTDRLLRRLEAMLIAADATTTLVITGNGDVLD
PEGGICAIGSGGSYAQAAARALAENTDLSPREIVEKALGIAGDMCIYTNHNRIIETIE*
>GCA_000452945.1_BPMSHR5858_genomic_03358_1 regB
MQRITTTGRVNLSHLFWLRNLAIIGQLVTIAVVQTYFGVHLPLPAMLMVIALEIVFNALT
WVRVLRARPETNFELLGQLWVDLGALSALLFLSGGTTNPFVSLYLPSLAIAAAVLPWHLM
IWLAAFAVACYLALGFDSVPLNLDNPANLFDYFRAGMWVNFMVSVGLIAWFVARMSSALR
QRDAALGEAQQALLRDERAVALGVQAATVAHEMGTPLSTIAMLTEELREAARTDAGLARY
DADLKVLEEQMSQCTSALARLRSRASERPSREPVGEWLDTFVEHWRLRHPHVLFEQRGPQ
PVGVALDDTVAVGQILTILLDNAARASRDRVTLAATIAHDGARDEIEFEVCDNGPGIPAA
LRDTLGAAPVDSTQGGHGVGLYLAFSAAARLGGSIELADAQPRGTRAILRLPVARQVAAE
MANENG*
>GCA_000452945.1_BPMSHR5858_genomic_03362_1 slmA
MQPTHPHDPAVTAAAENERSAATANRTRPKPGERRVHILQTLASMLEAPKSEKITTAALA
ARLDVSEAALYRHFSSKAQMFEGLIEFIEETFFGLVNQIAANEPNGVLQARSIALMLLNF
SAKNPGMTRVLTGEALVGEHERLAERVNQMLERVEASIKQCLRVALLEAQAHAAGGAPPP
VPLPDDYDPALRASLVISYVLGRWHRYAKSGFTKAPGEHADAQLRLILQ*
>GCA_000452945.1_BPMSHR5858_genomic_03363_1 metXS
MESIGVVAPHTMHFAEPLRLQSGSVLGNYQLVVETYGELNAARSNAVLVCHALNASHHVA
GVYADDPRSTGWWDNMVGPGKPLDTNRFFVIGVNNLGSCFGSTGPMSIDPATGTPYGARF
PVVTVEDWVHAQARVADAFGIERFAAVMGGSLGGMQALAWSLLYPERVAHCIDIASTPKL
SAQNIAFNEVARSAILSDPDFHGGDYYAHGVKPRRGLRVARMIGHITYLSDDDMAEKFGR
ALRRADGALDAYNFNFDVEFEVESYLRYQGDKFADYFDANTYLLITRALDYFDPAKAFNG
DLSAALAHTKAKYLIASFMTDWRFAPARSREIVKALLDNRRSVSYAEIDAPHGHDAFLLD
DARYHNLVRAYYERIAEEVGA*
>GCA_000452945.1_BPMSHR5858_genomic_03364_1 metXA
MNQQALNTLSARADFRAIARWVEPRSTVLDLGCGDGSLLSLLTEELEVSGYGVEINDAGV
LASVKNGINVIQQNLEDGLRLFEDHSFDFAILSQTLQTIHQTAAILRETARVGRECIVSF
PNFGYWAHRLSVLRGRMPVSKSLPYQWHNTPNVRVLTIKDFEALAPEVGVEILDRVVLHE
GQPIRWGVNWRGSLAVYRVKRRQP*
>GCA_000452945.1_BPMSHR5858_genomic_03370_1 group_19641
MAKQPSLDDYRVPYHTREKEAAAFTFDAFDPAAKPFSSGSKETDRERLSAVALQLDTLQE
RLHTQRHKRMLLVLQGMDSSGKDGTVRAVFHEVDPLGLRIVSFKAPTPVERSHDFLWRVH
AQAPAAGELTIFNRSHYEDVLVPRVTAEIDKAECERRYRQIRQFEEMLAESGTRLVKCFL
HISKDEQRARLQARIDDPNKHWKFDISDLEARKHWDAYQAAYRDALAATSAEHAPWYVIP
ADSKTHRNVMIAELLLRELTAMKLEYPPAKPELAGIQIQ*
>GCA_000452945.1_BPMSHR5858_genomic_03371_1 gatB
MTQWEVVIGLETHAQLSTVSKIFSGASTQFGAQPNTQACPVDLALPGVLPVLNRGAVERA
IRFGLAIGATVAPRSVFARKNYFYPDLPKGYQISQYEIPVVQGGQITIQVPANEKAGKQA
YSKTVNLTRAHLEEDAGKSLHEDFAGMTGIDLNRAGTPLLEIVTEPEMRSAAEAVAYAKA
LHGLVMWLGICDGNMQEGSFRCDANVSVRPVGQEKFGTRAEIKNLNSFRFLEDAINYEVR
RQIELIEDGGEVVQETRLYDPDKRETRSMRSKEDAHDYRYFPDPDLMPLVIGADWIARVK
GEMPELPAAMQQRFVEQYGVSAYDAGVLTSTKAMAEYFEALVAKAGAANAKLAANWLMGD
VSSQLNRDGIDIDACPVSAAQLALVLQRIADGTISNKIAKEIFVTIWDEKAADEGAADRI
IEAKGLKQISDTGALEAIIDEVLAANAKSVEEFRAGKDKAFNALVGQAMKATKGKANPQQ
VNELLKKKLG*
>GCA_000452945.1_BPMSHR5858_genomic_03377_1 mrdB
MTEFKDTQQQLSKFRLRVAAAGVFVFVCFGLLATRFFYLQVWQHGKYALQAEENRISVAP
IVPNRGIITDRNGVVLAKNYSAYTLEITPSKLADTLDNTINALAEVVQIDARDRRRFKKL
QEDSKNFESLPIRTRLTDDEVARFTAQRWRFPGVDVRARLFRQYPLGPTAAHVIGYIGRI
SKRDQDRIDAMSDENDSDPETYDPRRDANNYKGTDYIGKVGVEQSYETELHGITGFEEVE
VTAGGRPVRTLSRTQATPGSNLVLSLDIGLQQVAEQAFAGKRGALVAIEPKTGDVLAFVS
SPSFDPNSFVDGIDQQTWDELNNSPDKPLLNRPLHGTYPPGSTYKPFMALAGLALGKRTP
GWGFQDPGYFTFGGHTFRNDVRSGQGWVDMNKAIMVSNDTYFYMLARDLGVTAIANFMKP
FGFGQLTGIDIQGEARGILPSPEWKKKTFKKAAQQKWFDGETISLGIGQGYNSFTILQLA
HATATLANDGVIMKPHLVKEVENPISRARHLTVPKESGVIPLKQNDLDVVKRGMENVVES
PSGTAFKVFRGAPYLAAGKTGTAQVFSLQGGNYRGHLLAEHLRDHALFIAYAPVDHPQIA
LALIVENGGWGAQSAGPIARRVLDFYLLERKNPSTEAAAVAAAASATEPVNAPVIGDATK
PVGIAAGFKALPQPVPPASASGADGASGASGAGGEPTEHANAGGNPAGGGIAGGAAGTAN
NGSGAAAPGGMPGANGAAMGTPPASRRLPPRKPRRTPASDAAISAAASRDDPRKPSSGAP
KTPAQNDNHAVSAARPAAGGIDE*
>GCA_000452945.1_BPMSHR5858_genomic_03381_1 group_19335
VLITRKLEFDAGHRIPDHRSQCRNLHGHRYVLEITLRGDLVETEGAPDRGMVMDFADVKA
LAMEHLVSKWDHAFLVYARDEIVRSFLEKMADHKTVVLDRIPTVENLAAIAFDILANVYD
AHYGVNLRLERVRLYETPNCWADVERDPRV*
>GCA_000452945.1_BPMSHR5858_genomic_03383_1 queC
VIRTDAKDGALVLFSGGQDSATCVAWALERYQTVETLGFDYGQRHRVELECREGVRDALK
RRFPQWSHKLGDDHLIDLSVLGSISDTAMTRAIEIETASNGLPNTFVPGRNLLFMTIAAA
IAYRRGLRALVGGMCETDFSGYPDCRDDTMKALQVALNLGMDTRFVLETPLMWLDKADTW
RLAEQLGGAPLVELIRVETHTCYVGERSELHDWGFGCGECPACKLRKRGYDAYLRGESVT
EAPA*
>GCA_000452945.1_BPMSHR5858_genomic_03390_1 leuE_3
MFGITHLGLFMAAVFLLNITPGPDTAYIVGRSVAQGRGAGLMSALGISAGCCVHVLACAF
GLTAVLAASATAFTVIKVVGAAYLVYLGVRLIFAKQEPAAERPARRGADKSLRQLFMQGF
LTNVLNPKVVLFFVSFFPQFVSVDSNHKVLAFLTLGAVFLAMSTVWSCTVAWVAGTVTRR
FAGKPGVKKWLDRGVGSAFVGLGVKLATSTR*
>GCA_000452945.1_BPMSHR5858_genomic_03391_1 fmt
MTHSLRVIFAGTPEFAAAALAAIHEAGFPVPLVLTQPDRPAGRGMKLQASAVKRYAFERG
MAVAQPPSLRRAGKYPAEAVAALDLLHATPHDVMVVAAYGLLLPQEVLELPRHGCINIHA
SLLPRWRGAAPIHRAIEAGDAETGVTLMQMDAGLDTGAMLHEARVAIAPDDTTATLHDKL
AAAGARLVVDALVELERTGALAATPQPADGVTYAEKIGKHEAALDWRKPAAALARQVRAF
DPFPGGAGTLDGATLKLWAADAVPGRDDAAPGTIVDIGPDGVVIACGEGALRVTQLQKPG
GKRLPAREFLAGAPLAVGQRFAPADAA*
>GCA_000452945.1_BPMSHR5858_genomic_03394_1 group_18995
MPALNLDTDEGRIAERLRNPDTLLVACLCAEWCGTCREYRATFDRLADTHPGACFAWIDV
ETHADRLDELDVENFPTLLIEDANAVRFFGTVLPHAAIVERMLADLTALPGAPHAPKLRN
LFD*
>GCA_000452945.1_BPMSHR5858_genomic_03396_1 dmlR_18
MDHLQSMRVFVKVADLGSFARAASAMDISNAVATRHVADLEGRLGTRLLNRTTRSLSLTE
SGQVYLERARQILDELEDVEQMVVARNHEPVGTLRVVAPVVFGLHNLAPVLQTYTQRFPK
VIPDLTLVDRQVDLVEEGFDVGIVITRQMRSASIVTRRLTTGCMVVCATPSYLEKHGMPT
HPEQLVEHPCLSLPSEYWGDERVFTGPDGEVRVRPSNVVVANNTEMLRQFAMLGMGVAIL
PSYLIGSDIARGKLVRLLSDYRLPQVEINIAYPSRRHLPAKVRTFIDHLVEHFSQSPDTM
LGEQWATQGVAGQHVDALPADALAERSSADPALSPRLPRAPRARTVVPSPL*
>GCA_000452945.1_BPMSHR5858_genomic_03397_1 ghrB_2
MKIAILDDYQDAVRKLNCFEMLADHEVKIFNNTVRGLGQLASRLAEVEALVLIRERTHIT
SQLLGKLPHLRMISQTGRVSTHIDLEACTERGIAVLEGTGSPTAPAELTWALIMAAQRRI
PQYVANLKQGAWQQSGLKTSAMPPNFGLGQVLRGQTLGIWGYGKVGRLVAGYGKAFGMNV
LIWGREHSLEAARADGFEAAESREAFFEQADVLSLHLRLHDDTRGVVKLDDLLRMKPTSL
LVNTSRAELLEENALVSALARNRPGMVAIDVYESEPILQGYSLLRMENVICTPHIGYVER
ESYELYFSAAFRNILAFDAGDLSSVANPDALTQGRVRR*
>GCA_000452945.1_BPMSHR5858_genomic_03407_1 hdaH
MPVMNRAEILDYVECWSNGEWFEPPVSFAGLAKSFRASTHHSSALFFKANVLASTFRPHR
WLSRHAFERWALDFLTFGNGYLERRRNMVGGTLRLEPALAKYVRRKADFSGFVYVNGWQE
RHEFEPDSVFQLVRPDINQEVYGLPEYLSSLHSAWLNESSTLFRRKYYENGSHAGFILYM
TDAAQKQDDVDNMRDALKNAKGPGNFRNVFMYAPGGKKDGIQLIPVSEVAAKDEFFNIKN
VTRDDLLAAHRVPPQLLGIVPSNSGGFGTPDTAARVFGRNEIRPLQARFAELNDWLGDEV
VRFDDYEIPPAPVAA*
>GCA_000452945.1_BPMSHR5858_genomic_03415_1 rpoD_1
MKVSARELRYEQGIDRLLELAGARDDTEYADVADEFASAPSSAFAHDLGEPDSLDHQASD
ESAGAGSFGDKPTRPAHGDTDVDTDWRDDDAPPSGSNDAAPVDTAASTDPLALFVRRMHA
VPLLTHAQEIALAAEIEAGREQVLQALAEALDRLDLASLPADPTLGRARAALQRARARGD
EGARAFARARTAAYAALAAHGWPSAEVDAVRRAMRRLPTSPKKSDAGAQRAFADGEARIS
AATARMAEANLRLVLSIARKYASRGLDLADLLQEGNLGLLRAIEKFEHRRGFKFSTYATW
WIRQAISRAIADRARTIRVPVHVGDEARRVRKAADRIERRSGVSASLAELSEATGLPAER
LRAVLTLPREPRSLNAPLGDDDQAEFGDIVEDTSTPSLFDALAHGQMCDVVTSLLKTMSA
QEADILRRRFGIGGDEPWSYEQIAARAGVSREQVRRIEKRALQTLRVTAEACNAHDFLDV
QP*
>GCA_000452945.1_BPMSHR5858_genomic_03416_1 gshA
MPNNMTSRQTDLLQQRLAVLSTSPTREQLPDGLRGIEKESLRVTRDGMIAFTPHPRALGS
ALTHPSLTTDYSEALVELITPAERDAATTLERLDELHRYVYASLGDEMLWTDSMPGLLPA
DDEIPIADYGTSNIGRLKTVYRRGLAYRYGRTMQCIAGIHYNYSLSEEVWRRLHAEEGST
AGIVDYQSERYLALIRNFRRTSWLLMYLFGASPALDAGFLRGRAHKLDTFDAATLYRPHA
TSLRMSDLGYSNTTAQAALQADYNTLSGYLDALSKAVSEPYPPYEAIGTHRDGEWIQINT
NVLQIENEFYSTIRPKRVTYSGERPLHALASRGVQYIEVRCLDIDPFEPTGIALDTARFL
DAYLLVCALDESAPLTCDAYRESNANFGRVTMEGRKPGLELTRDGAAVPMRAWADEIFAQ
IEAAGRVLDDIRGDDAHARAIAAQRAKLDDPELTSSARVLRAMRENGQSFLEFARAQSAA
HAAYFHGRPLDEAATRDARALAERSLAEQAELEGKNAGSFDAFVAAYRAYTLNRFSV*
>GCA_000452945.1_BPMSHR5858_genomic_03443_1 group_18966
MSSLVQYVKQFVRDEGGVSAIEYGLIAALIAIVIIGAVKTVGTNLNSVFSTIGNDL*
>GCA_000452945.1_BPMSHR5858_genomic_03450_1 group_1493
MSDNLCRTDLSSDAAAAVERVYRQWDRAWAADDLDAMIALYAPDAVLESPLVTHLLGTES
GVVEGRERIRALLEQAAPRKPEARMFHRIGYFTDGKRLIWEYPRATPTGEQMDFVEMMEI
EHGLIRRHRVYWGWRGVSVIKADAYRKGRR*
>GCA_000452945.1_BPMSHR5858_genomic_03452_1 cynR_2
MREINHQRLRYFHEVLTHGTIRGAADHINTSPSVITRQIRLLEEELGAALFERQARGVRP
TEAAAHLLEFWRGYRSQQEKLEDQLHALKGLQQGHIRIVVSEGFVDTLVDDVLAPFCAKY
PKLNVDVDMLAVDTILDEVALSRAHIGLAYNPPPHPLIAHRASSRQPVVLLLRRDHPLAR
RRRPATVDDLHAYPLALMPPTFGIGHVVKMLELAENIEIRPTLTTNSLTALKRIVTAENF
ITLIGAFAAYREIAAGELTTVPIAHPIFEGTHARLIVKSGRPLAPAPVELIRWIETRLAV
FAAATGGGGKTRGRRGTR*
>GCA_000452945.1_BPMSHR5858_genomic_03454_1 dhmA
MSHPINARRRRLLGTTLAGISLVDLGLGGFAHAQSAPRATSAAGVARAAGGASFDTIHQI
DAGVLNVGYADVGPKGGPAVILLHGWPYDIHSFAEVAPLLAAAGYRVIVPYLRGYGTTRF
LSADTPRNGQQAVTAVDIIALMDALKIDRAVLGGFDWGARTAGIIAAAWPQRCHALVSVS
GYLIGSQEANRKPLAPKAELAWWYQFYFATERGAAGYAANRDDFNRLIWQLASPRWRFDD
ETFARSAASFRNPDHVAIVVHNYRWRLGLAQGEPQYDALEKRLAAAPAIAAPTITLEGDA
NGAPHPEPAAYAKKFTGKYRHRDIAGGIGHNLPQEAPKAFADAILQAAHL*
>GCA_000452945.1_BPMSHR5858_genomic_03469_1 group_1675
MNRRHFLAAAVAAGLPLAATLAPRGARAQPNATQAAAPTLARQLAEYAAGLRYEDLDSAT
IDIVKSHLIDALGCALAALDEPPVRIARDAARSAGGGGPSTIIGTAERTSPDLATFATGT
ALRYFDFNDAYAGREIGHPSDNIAACVAVAEAQHASGRELILSIALAYEIACRLMDAAAI
SPRGWDHTCYSLPAAALAAGKLMRMPVEALTQAVNLSLNSHLALNQTRVQQLSNWKALAD
ADAARNAVFSTQLARAGLTGPSPIFEGEAGFFRQVSGPFELETSRFGGRGEPFRIARCFV
KYYPAQGFTQTAIPAALDVASQAGDLSRIRRIDVHTTRVGYVTAGSEPEKWKPSTRETAD
HSLPYVVARAMLDGDIRTTSFSDAALRDPVLHALIAKIRVEEDPALTAGYPARAANRVTA
HCRDGAVYAKQVDDLPGSPTRPMRREDFEAKFVKNGGARLSEQRMRAALDRLWRLDELQD
VAALPPLFVAG*
>GCA_000452945.1_BPMSHR5858_genomic_03470_1 group_1494
MDKRLSALCGLGLASAGACAQTSVTLYGVADAYVEYATNQADAKGEPAALARMGSGGKSG
SRWGIKGTEVLGGGWRAAFRLESGVNLNNGAGTGAVGFDRSAWVGLEHPRWGALRFGRQY
TTMFDIMEHYSPTGAYSTLYEPDGAIVGISFRENNVVKYLATAGPLTFEAHYAFSNEPGA
FQASAAHGAGFEYTGGALSFAFAYDDVHTPQAGGFAHLRRYAAAAMLTVEATQLIAGAAH
GQGNVATPSVVTRYTFWWIGVRQAITPVVQLIGALYAERVRAQHPASPPAARHASGTPQQ
ATLQLNYFVSKTTTLYAATGYARHAALDFDNYNYGFLHYSLAGARAGSAGVAVGVRKLF*
>GCA_000452945.1_BPMSHR5858_genomic_03473_1 smtB_2
MDVRQTQVEERASPRRRADTGPIHAKRFDADARGPLAGVRVVDLSRLMAGNMLSLQLADF
GADVVKVENERGDTLRAVGAGGVSTNWKVYGRNKRSVCVDLRSADGIDIVKRLVRDADVF
VESFKPGVLEQMGLGPDVLLELRPSLVIARISGWGQTGPYRHKPGFGTLAEGYAGFASIN
GFADREPVLPPMFLGDMTAGLSGAIAVLVALHACRDGARGQVIDVSLFEPLFSILGPAAA
NYVLTGRVKERTGSRSSNTAPRNAYRTRDDKWLCLSSSTQGMAERLFRAIGRADLIDDPR
YATNVQRVQHAGELDAIVGAFVAARDLADNLAFFEKTGVTVGPILDIAQIVQDHYVIERE

ALVELPDDDVGSLPMHGVVPRLSGTPGGFRRAAPKLGEHNREVLLPLLGAHEYARLARNG
VIRPE*
>GCA_000452945.1_BPMSHR5858_genomic_03502_1 rplS
MNLIAKLEQEEIERALAGKTIPEFAPGDTVIVNVNVVEGNRKRVQAYEGVVIAKRNRGLN
SSFIVRKISSGEGVERTFQTYSPLLASIVVKRRGDVRRAKLYYLRERSGKSARIKEKLVS
KDRAAAAQQ*
>GCA_000452945.1_BPMSHR5858_genomic_03505_1 rpsP
MVIIRLARGGSKKRPFYNIVATDSRNRRDGRFIERVGFYNPVATKGEALRIAQDRLTYWQ
GVGAQLSPTVERLVKQAQKAQPAA*
>GCA_000452945.1_BPMSHR5858_genomic_03506_1 group_1496
LPRAAPAACLVLATIVTVATAPAASAALPIDELRVPPGFRVQVLADDVPTAREMAWSPRG
ILYVGSMNGRVHALVVRDGHVREHHVIASGLEMPVGVAYRSGALFVSAVSRILRLDRIDE
RLAAPPKPVVVTNALPTDRHHGWKFIAFGPDGKLYVPTGAPCNICVADRDRYAMIGRMNA
DGSGYEVYARGVRNTVGFAWHPATRELWFTDNGRDLMGDDRPDDKLNRAPRAGLDFGYPF
CHGGDVLDPQFGRGHTCSSYAPPVLKLGAHVAALGMRFYTGGMFPPEYRDNIFIAEHGSW
NRSRKVGYRVVRVIASPDGRAAREETFVHGWLRPDESVWGRPADVLPLPDGSLLVSDDYA
GAIYRITYDATH*
>GCA_000452945.1_BPMSHR5858_genomic_03507_1 group_19668
MSSSTSTPPRFRSKTLTALLAFLFGSLGAHRFYLYGWKDLAGWAHLLATIVGVPGFMLLA
ASGRAAALGWWLAVPGAISLLAAFLAALVYGLRPDDKWDAQFNASTGRHSRSGWGVVFVV
IFSLLIGAFLLMTGLALAFQTYFESQVEAAKQISQ*
>GCA_000452945.1_BPMSHR5858_genomic_03509_1 lrp_9
MRTQRQPVRALDKLDRRILKLLQEDGRMAMKDLAERVGLTVTPCIERVRRMERDGVITGY
HARVDPSQLGASLLVFVEITLDHKNGNMFEQFRREVMKIDEVLECHLVSGDFDYLIKARI
GEMADYRKLLGDILLQLPGAVQSKSYVVMEEIKETLMIAVDE*
>GCA_000452945.1_BPMSHR5858_genomic_03511_1 dmdC_1
MSYTAPVKDMLFVMKELAGIDAVAKLPGYEDAGFDTAQAVLDESAKFCGEVLAPLNVEGD
KNPSSWKDGAVTATPGFKDAFRQFVEGGWQGLQHPTEYDGQGLPKLIATPCIEMLNASNL
SFALCPLLTDGAIEALLTAGTDEQKSRYVPKLISGEWTGTMNLTEPQAGSDLALVRSRAE
PQPDGSYKVFGTKIFITWGEHDMADNIVHLVLARTPDAPEGVKGISLFIVPKFLVNDDGS
LGARNDVHCVSIEHKLGIKASPTAVLQYGDHGGAIGYLVGEENRGLEYMFIMMNAARFGV
GMQGIGVAERAYQKAAEFAKERVQSRPVDGSAKQSVTIIHHPDVRRMLATMRALTEGARA
LSYVAAAHSDHAHRAADEGVRAKHQAIYEYLVPVVKGWSTEMVNEVASLGIQVHGGMGFI
EETGAAQYYRDARILAIYEGTTAIQANDLVGRKTARDGGAVAKALIGEIRQTAEALGAHD
GPAFTAMKTQLENGARALSAVVDFVVANVKGDPNAVFAGSVPYLKLAGVVLCGWQMARAL
LVAQQKRADDPRFYDAKIALAQFYAEHILAQAGGLEASIVGAKGGQGVFALTEDQF*
>GCA_000452945.1_BPMSHR5858_genomic_03512_1 etfA_2
MTILVIAEHDNASIKAATLNTVAAATKIGGDIHVLVAGHNAQGAADAAAKIAGVSKVLLA
DAPQLADGLAENVEATALNVAKDYSHILAPATAYGKNIAPRIAAKLDVAQISDITAVDSA
DTFERPIYAGNAIATVQSSDPIKVITVRATGFDPVAAQGGSASVEKIDAAADAGVSQFVS
REVTKLDRPELTSASIIVSGGRGLGSGENYTKVLEPLADKLQAALGASRAAVDAGYVPND
YQVGQTGKIVAPQLYIAVGISGAIQHLAGMKDSKVIVAINKDPEAPIFSVADYGLVGDLF
TLVPELVNELG*
>GCA_000452945.1_BPMSHR5858_genomic_03515_1 metI
MLSEMFDMFVQSFWETLIMVGISGAVGALVGLPLGVLLYLTDRQGVLQNLAVNRVLGGVV
NAVRSTPFIILLVAVIPFTRLAVGSSIGTAAAVVPLMIAAAPFIARLVETALREVDRGLI
EAAQAMGATTGQIVFKVLLPESLPGIVAGLTITFVSLVGYSAMAGAIGGGGLGDLGIRYG
YQRYLPEVMWTVVAILIVFVQIVQSFGDWLVRRLSHK*
>GCA_000452945.1_BPMSHR5858_genomic_03516_1 metN_1
MIEIRNLSQRFEGPRGWIEALHNVNLTIPQGEVFGIIGRSGAGKSTLVRTINLLTRPSEG
SVFVGGRDLTQLSAAELRGARRDIGMIFQHFNLLSSRTVFDNVALPLELAGVKRAQIEAT
VLPLLDLVGLAAQKDRYPAQISGGQKQRVGIARALASKPKVLLSDEATSALDPETTRAIL
DLLRRINRELGLTIVLITHQMEVIKDVCDRVAVLDAGRVVEEGNVIDVFMRPHHEVTRAL
IGDVIAQELPPAMKARVAERLKTGSGHLLRLAFTGSGVDQPILSETIRRYELDFNILHGQ
IDEIQGRAFGSLAVLATGEPGKVGQAFAYLREQGVVVEELSYVE*
>GCA_000452945.1_BPMSHR5858_genomic_03518_1 group_19295
MPTAFYTHPDCLLHEMGEWHPESPARLSAIQDQLIASRIDDLIVHETAPFASETALARVH
TQAHIDYIRSRIPKDGYVEIDPDTSLNRYTWRAALRAAGAAVEATDAVLEGRYDNAFCSV
RPPGHHAEPARAMGFCFFNNVAIAARHALEIHGLERVAIIDFDVHHGNGTEAAFANDERV
LMCSFFQHPLYPFSGVDHQAPNMVNLPMAARTNGMAVREAVDLMWLPRLDAFRPQMLFVS
AGFDAHREDDIGGMGLVEADYEWLTQQICEVARRHAQGRIVSCLEGGYNLSALGRSVVAH
LRVLAGV*
>GCA_000452945.1_BPMSHR5858_genomic_03519_1 mltB
MIFHQPAAPASLAFRLRFPLAALSLAATLFASAAVAQTQPAAAVVAQEPAQSQSQSQSQS
QLQLQPQPQPQPQPQPSVQQGQTFEEEIIPQRYANNAKIDAFIADMVARHDFDANALHAL
FARVSYSATAAKLVMPAPSPAVKNWRVYQSRFLDAVRVNAGVKFWRANQGTLQRASTEFG
VPPEVIVGIIGVETIYGRYMGNFRTLDALTTLAFDYPNTPNRDARQATFRKNLEDFLVWT
RDSQLDPTGVLGSYTGAVGIPQFLPSSIRDYAVDYDGNGHIDLRASQADAIGSVANYLKQ
HGWETGRPVVWNIAPDTGSQGVAQAAADGRPEPHWVLSQLLRAGLVLDEPSVNIASEAST
PVTVVDLPTPGRATEYKLGLQNFYVLTRYNRSFFYALAVYQLGERVKAQMEASGALTPSP
ADAATGSPAAQPPSE*
>GCA_000452945.1_BPMSHR5858_genomic_03581_1 group_19393
MIYSMTGYASATRDLATASGNGGTSVSVELRTVNSRFLDLNFRMPDDVRACEPALREMLM
NKLSRGKVDVRINLQRGDQSVNAGALNHAALGQLAELERAVLDAFPGVGRLRAGEILRWP
GVLAESGVSADALREAVLACGKEAVAELVVVRSREGAQLATMLLGNVAEMEAIVARIAPL
VPELIAKHQQKIVERLQEALGVAAPEGGAPLVSREEAAERIRQEVTMYGIRIDIAEELSR
LTAHLNETRHVIEKGGRVGKRLDFMMQELNREANTLGSKAAAKELADASMALKLLIEQMR
EQVQNLE*
>GCA_000452945.1_BPMSHR5858_genomic_03582_1 rph
MTNSSLRPSGRRADQLRDVRITRHYTKHAEGAVLVEFGDTKVICTASVAERVPEFLRERG
QGWLTAEYGMLPRATHTRSDREAARGKQTGRTQEIQRLIGRALRAVFDLNALGPRTLHLD
CDVIQADGGTRTASITGAFVAAHDAVTKLVAAGRIARSPITDYVAAISVGVFGGTPVLDL
DYDEDSACDTDMNVVMTGAGGFVEVQGTAEGAPFSRTEMNALLDLAQAGIGELVRLQRAA
LEA*
>GCA_000452945.1_BPMSHR5858_genomic_03584_1 hemW
MSDAATNGARVVATFAAPGRIRLASLPPLALYVHFPWCVRKCPYCDFNSHEWKDGGALPE
RDYLDALRADLEHALPLVWGRQVHTVFIGGGTPSLLSAAGLDRLLSDVRALLPLDADAEI
TLEANPGTFEAAKFAQFRASGVNRLSIGIQSFNEAHLKALGRIHDAAQARAAVEIAARTF
ENFNLDLMFALPNQTLDECRADLETALAFAPPHLSLYHLTLEPNTYFAKFPPTVPDDDAS
ADMQDWLHERTAQAGYARYEVSAYTKPHRQSKHNLNYWRFGDYLGIGAGAHTKLSFPNRI
LRQARYKHPATFIEQAKAGTPVQEEREVGPRDLPFEFMLNALRLVDGVPAYSFEARTGMS
LAAIGPALAQAEQRGLLVHDHTRIAPTPLGQRFLNDLQALFLRDA*
>GCA_000452945.1_BPMSHR5858_genomic_03594_1 group_19037
MSFSLHRVVLPLVAAGAGVCALAAPLAASADEIVVEQAVPGVVVAPAMREVVVVAPSAPP
PVRYEVVPEPRVGYVWDRGHWRWDHDRYVWIAGHWEVERIGMHWAPGHWSQRGPGWVWVR
GHWA*
>GCA_000452945.1_BPMSHR5858_genomic_03598_1 group_19349
MSSTLTVRRIAADQGGVYRELRAASLREPYASGETPEAELAVDAARAAAIAQQRAASDES
TTFVLYTEGHPAGMIGAYFDGTPARRAFVSELWVAHAVRHLRGGVLLVNTARAWLATRGA
GEIYAWIADQNRSAIRFYEHVGFADTGERAPITRVPGAMKSLFVWREHA*
>GCA_000452945.1_BPMSHR5858_genomic_03602_1 group_19681
MAIELDKDVRDRAIASLQRYFTENLEEPIGNLQAGALLHFFVEEVGPAIYNLAIADAQSS
LHARVAELDIDCHQAPFGYWKKTPVRKR*
>GCA_000452945.1_BPMSHR5858_genomic_03603_1 rarA
MSDLFAVEPRRPLAEALRPKTLADVIGQTHLLGEGKPLRLAFESGKPHSMILWGPPGVGK
TTLARLTALAFDCEFIALSAVLGGVKDIREAMEQARDTLNRTGRHTILFVDEIHRFNKAQ
QDALLPFVESGLVTFIGATTENPSFEVNSALLSRAQVYVLKSLAEDELRQLLKRAQDVAL
GGLAFDDKAVDTLVGYADGDARRFLNLLEQAQTAASSAGVTTIDAAFVESAMTMNARRFD
KGGDNFYDQISALHKSVRGSSPDGALYWLCRMLDGGADPKYLARRVVRMAWEDIGLADPR
AMQIANDAAQTYERLGSPEGELALAQAVLYLACAAKSNAGYNAYNEAMAFVRQDKSREVP
VHLRNAPTKLMKELGYGHAYRYAHDEPNAYAAGETYLPDGMREPRWYKPVPRGLETKIAD
KLAWLRELDRDAGKND*
>GCA_000452945.1_BPMSHR5858_genomic_03607_1 mutS2
MAKNQPHPSDPAKRQAPARATEPAPASPPAAPAALRGQGLAGLGSLRAALTGAAERRARE
AAQAQQSARQAAADADLFRREIGAIKPLAAPPRAASGRLAPAPVPKHTRQDEEAVLSETL
SDEFDPETLLDTDETLYYHRPGVSRDVVRKLRSGAWIVQAQLDLHGMRRDEAREALAEFI
RESGKKGLRCLRVIHGKGLGSIGKEPVLKGKVRAWLVQKDEVIAFCQARGHDGGAGAVLV
LLQPS*
>GCA_000452945.1_BPMSHR5858_genomic_03610_1 ngcG
MQPKMTLSRAVIYAALILFALYFLFPIYVMLSTSFKDLDQLRTGNLLTPPSSWTVAPWAK
AWGEACTGVRCEGMKPFFFNSIKMVIPAVLISSLIGAFNGYVLTHWRFRGADALFTMLLV
GCFIPFQVILLPMARLQGYFGMANTIPGLVLVHVVYGIAFTTMFFRNFYVSVPAELVKAA
RIDGAGFFTIFTKILLPVSLPIFMVCLIWQFTQIWNDFLFGIVFSGVDSMPITVALNNLV
NTSTGVKEYNVDMAGAIIAALPTLLVYVIAGRYFVRGLTAGAVKG*
>GCA_000452945.1_BPMSHR5858_genomic_03614_1 pgl_1
VIEVHAFETPRAQTEALAQAVGEALAATLAQRARATLAVSGGTSPRPFLQTLSGAALDWP
RIDVTLVDDRWVPDTDDASNAKLVRETLLRNAAAHARFAPLVDLRESLDAQIAALNQSPA
HPLPDVAVLGMGEDGHTASIFADAPEWDHAIATTERFVGVHPGAAPHARVSLSLGALKRI
GRVFLLIAGSRKRAVLEAAAAAPQKNAISQLANDKGTKLDVYWCAN*
>GCA_000452945.1_BPMSHR5858_genomic_03615_1 glk
MSTGAQTKAAAASQHADGPRLLADVGGTNARFALETGPGEITQIRVYPGAEYPTLTDAIR
KYLKDAKIGRVNHAAIAIANPVDGDQVRMTNHNWSFSIEATRRALGFDTLLVVNDFTALA
MALPGLTDAQRVQIGGGTRRQNSVIGLMGPGTGLGVSGLIPADDRWIALGSEGGHATFAP
MDEREDLVLQYARRKYPHVSFERVCAGPGMEIIYRALAARDKKRIAANVDTADIVERAHA
GDALALEAVECFCAILGTFAGNLAVTLGALGGIYIGGGVVPKLGELFMRSPFRARFEAKG
RFEAYLANIPTYLITAEYPAFLGVSAILAEQLSNRTGGASSAVFERIRQMRDALTPAERR
VADLALNHPRSIINDPIVDIARKADVSQPTVIRFCRSLGCQGLSDFKLKLATGLTGTIPM
SHSQVHLGDTATDFGAKVLDNTVSAILQLREHLNFEHVEQAIDILNNARRIEFYGLGNSN
IVAQDAHYKFFRFGIPTIAYGDLYMQAASAALLGKGDVIVAVSKSGRAPELLRVLDVAMQ
AGAKVIAITSSNTPLAKRATVALETDHIEMRESQLSMISRILHLVMIDILAVGVAIRRAA
PNAELAEAMARAKARAGASAGDEAADVLDWLSHGAAPAAKD*
>GCA_000452945.1_BPMSHR5858_genomic_03617_1 yecS_3
MQAFDLVVHTLPVLAKGAVLTMKFAVASMVLGLIVGLLVALVRIGSNRLLSAIAQGYVSL
MRGTPLLVQMFVVYYGLPDIGISLDPTAAGIFTLTLNAGAYLSESMRGAILGIGRGQWAA
SHSLGLTHVQTLRHVICPQALRLAVPSLGNTLISLIKDTSLVSVITVTELLRSTQEVIAS
TFQPLPLYLAAAAIYWVLSTLLARLQGRIETRCALPSAH*
>GCA_000452945.1_BPMSHR5858_genomic_03625_1 ribD_1
MFSESDFTHMQRALALAARGMYTTTPNPRVGCVIVKDDVVIGEGFTQPAGQDHAEVQALK
DARARGNDLRDATVYVTLEPCSHFGRTPPCAHTLIDARVGKVVAAMEDPNPQVSGRGLAM
LRDAGIDVRCGLLAHEAHEMNIGFVSRMTRGRPWVRMKTAASLDGRTALASGESKWITGD
AARDDGHKWRARACAILTGIGTVREDDPQLNVRGVDTPRQPRRVLIDSRLDIPLAARLLE
GGSLLVFCGALDAQSEPRADALRARGAEIVPLANARGKVDLPAMLAALGERGINELHVEA
GHKLNGSLLRERCVDELLVYLAPSLLGADAAAMFDLAAPASLVDRTRLAFHSVERVGDDL
RILARFANAHASASPTHS*
>GCA_000452945.1_BPMSHR5858_genomic_03626_1 ribE
MFTGIVAAVGRIEAITPIAASPDAGVRLTLRAGGLDLADVALGDSIAIQGACMTVVAKTA
DTFDVEVSRESLNRTVGLAEPGDVNLEKALRAHDRLGGHIVSGHVDGLGTVTRFAPVGES
HELRVLAPRELGRYLAYKGSITVNGVSLTVNSVDDRDDGCEFSINLIPHTVEVTTLRHLT
AGAKVNLEIDMIARYVERMTNAPK*
>GCA_000452945.1_BPMSHR5858_genomic_03627_1 ribBA_2
MTLASTPDIIAELKAGRMVILVDEEDRENEGDLVLAAEFVTPEAINFMARYGRGLICLTL
TQERCKQLNLPLMTYRNGTQYGTAFTVSIEAAEGVTTGISAADRARTIATAVAHDVRPEH
IVQPGHVFPIMAQPGGVLVRAGHTEAGCDLTALAGLTPAAVICEIIKDDGTMARLPDLLE
FGKEHGLKIGTIADLIHYRSRTESIVERVAERTMQTAHGPFKTVLYRDHPTGSPHIALVR
GAPTPNDETPVRVHEPLSVLDLLETGVSTHSWTLDAAMKEIAARELGVIVLLNCGDTKEH
LVDVFLAFDEKEKAAELKRRPVDFKTFGIGAQILRDVGVGRMQVLSNPRKLGSMSGYGLE
VTGFAPMPGSAAPASGAA*
>GCA_000452945.1_BPMSHR5858_genomic_03629_1 nusB
MKKSARRQSRELATQGLYQWLLSNAAPGEIDAQLRGALGYDKADKTLLDTILHGVIREHA
TLAEAISPSLDRPIDQLSPVERAVLLIATYELTHQIETPYRVIINEAVELAKTFGGSDGY
KYVNGVLDKLAVKLRPAETQARRGA*
>GCA_000452945.1_BPMSHR5858_genomic_03651_1 glpE_1
MQILSASALAEWLRDPARPAPIVLDVREPWEIATAKIDGSVSIPMQQIPARSEELDDEAA
IVCVCHHGMRSAQVAMFLESRGFTKLYNLQGGIDAWSRDVDPAVPRY*
>GCA_000452945.1_BPMSHR5858_genomic_03656_1 cbiO
MNENTMIREPAQPDGADRLAVSGAAGMSHVVTPGAIDTSHGTILYLDDIEVSFDGFRALK
KLSLSIDVGELRCVIGPNGAGKTTMMDVITGKTRPDSGHAFLGQTLDLTRMNEPQIARAG
IGRKFQKPTVFEQHPVWENLELAMAADKRWFASLRARLDRAAQARIEQTLALIRLEDSAY
RLAGELSHGQKQRLEIGMLLMQRPALLLLDEPAAGMTDHETMELAELLNTLRGTCSMMVV
EHDMEFVAALAGDAGRVTVMAEGAVLAQGTLDQVKRDEAVIESYLGR*
>GCA_000452945.1_BPMSHR5858_genomic_03657_1 livF_5
MLTIESLNQYYGGSHILRDVSLTAETGRLTVLLGRNGVGKTTLLRCLMGVVPTKSGGIAW
CGKPLGALPPHARVASGLAYVPQGRDIFARLTVEENLLVGAASRKASAKVPDRIYALFPV
LKDMRARRGGDLSGGQQQQLAIGRALMSEPQLLILDEPTEGIQPSIIQDIGRTLRQLVDE
TKMTVLLVEQYYDFARSIADRYWVMSRGEIVAGGAGGEMDAHGVRELIAV*
>GCA_000452945.1_BPMSHR5858_genomic_03658_1 ureD
MSAHEPHTSLVRPAAKAWHARLELGFERQPGGRTALAHRRHVGPLRVQRALYPEGDAICH
AVIVHPPGGVAGGDRLEIDVRLDAGTHAVLTTPGATKWYKSNGLDARQRIDIDVGAHAKL
DWLPQNNLFFDAAHASLEFVLALGDGASVLGWDATQLGRQAAGEAWSAGSIASFSKIVGP
SGRPLWVERARLDAGDPLRAAPQGLGGFAVYGTLWALGAACTEALAESIAPALPFDDALR
AGVTCVAPGTLLIRALAHSMEALQRLLAEQWLALRPIVHGVDPKPLRLWQT*
>GCA_000452945.1_BPMSHR5858_genomic_03662_1 ureE
MRTIDKRIAPNVRLAATLVARAPALTLAYDARCKSRLAATLDTGEDVALVLPRGTVLRDG
DVLVADDGALVRVAAAHEAVLLVRAPDALTLTRAAYHLGNRHTPVEVGAGCLKLEYDPVL
ADMLTRLGATVERASAPFQPEAGAYGGGHRHGHDATFAEDYALAQQVFDEHHGHSHSHDH
DHDHDHQHGPCCSHGHHHGHR*
>GCA_000452945.1_BPMSHR5858_genomic_03669_1 group_19011
MLKSRLSNPDVAKAVANLAWLGLERLTQIGVAIVVSGLLARYFGPDTFGKWQYANTLLLV
LAPLTWVCGAEILVPTIVQRTGAQLGAVLGSAFALRFTVSVAALALTWAAIALGAFDPLV
GAMLAGLAVTLALREPFVGIVNAWLQSMTYSKPQLLASIAAALAKMALVWLLVRAAAAPA
RFGWLWALEAAAIAAALIWYYRARNGGTLGWRVERPLVREFATTGTVFWLGLVCMYLFLK
LDRLMLERYVSFAELGRYAAAQQLNENWITLALMLAQTIAPAFVYRVQNAPQLRRNVWRL
IGMTAALMIAGALVLDLLAGFIIRKVFGPGFETSVDVFRWAVWLSVPAGIEAIGNLVVLK
YQAKFVLLAKWSAALALAALVNLIAIPALGLYGALAGLAAGYLAAAAVNFHYIRLKLRP*
>GCA_000452945.1_BPMSHR5858_genomic_03670_1 group_19082
MTPTFAPPGAARALDDVAVLIPAYNAHDDLLRTLMSLREDAPVRVLVVDDGSTPPIAAPE
LPGLSVEVLRMPQNGGIERALAAGIDALAARGVRYAARIDAGDLAAPGRLAKQRAYFDTH
PSVAGLGTWTQVVSRDGRPLFMLTPSADPATLRRTRFLRSPFVHPSMMLDIAAVKEVGNY
RIKYRAAEDLDLFLRLMERYDCANLPELGLYYELNEGGISATKRRRQIASTLALALRYFD
PLNPCDWLGVAKNLLHFVTPYSTLQRAKRLLFARRGAA*
>GCA_000452945.1_BPMSHR5858_genomic_03688_1 rfbB
MILVTGGAGFIGANFVLDWLAQSDEAVLNVDKLTYAGNLGTLKSLQGNPKHVFARVDICD
RAAIDALLAQHKPRAIVHFAAESHVDRSIHGPADFVQTNVVGTFTLLEAARQYWSALGPD
AKAAFRFLHVSTDEVFGSLSPADPQFSETTPYAPNSPYSATKAGSDHLVRAYHHTYGLPV
LTTNCSNNYGPYQFPEKLIPLMIANALGGKPLPVYGDGQNVRDWLYVGDHCSAIREVLAR
GVPGETYNVGGWNEKKNLDVVHTLCDLLDEARPKAGGSYRDQITYVTDRPGHDRRYAIDA
RKLERELGWKPAETFETGLAKTVRWYLDHQEWVDEVVSGDYRKWVETNYAQRA*
>GCA_000452945.1_BPMSHR5858_genomic_03689_1 apaH
MTNFSSSPPIAFGDLQGCHAAYRQLFDTLAPAADTPLWFAGDLVNRGPASLATLREIVAL
GERAIAVLGNHDLHLLAVAAGIRTLKPGDTIGEILDAPDADDLIEWVRHRPFAHFERGML
MVHAGLLPQWDAALALELADELQRALRAPNWRDTLRSLYGNDPNCWSPDLKHADRLRVAF
NAFTRIRFCTPEGAMEFRANGGPAAAPAGYLPWFDAPGRKTADVTVVFGHWAALGLMLRE
NLVALDSGCVWGNRLSAVRLADDPAARVVTQVACERCGAADE*
>GCA_000452945.1_BPMSHR5858_genomic_03692_1 pyrB_1
MTTDTSGRTGNPAAAAPAERFRYGFLKGNPQLTKNGELKHLLTIEGLPRAILNQILDTAE
QFVSVTDREVKKVPLLRGKSVFNLFFENSTRTRTTFEIAAKRLSADVINLNINASSTSKG
ESLLDTINNLSAMHADLFVVRHASSGAPYLIAEHCAPHVHVINAGDGRHAHPTQGLLDMY
TIRHYKRDFTKLRVAIVGDILHSRVARSDIHALTTLGVPEVRAIGPRTLLPGGLEQMGVR

VFHNLDEGLRDVDVIIMLRLQNERMSGALLPSAQEYFKSWGLTPERLALAAPDAIVMHPG
PMNRGVEIDSQVADGPQSVILNQVTFGIAVRMAVMGIVAGTSD*
>GCA_000452945.1_BPMSHR5858_genomic_03693_1 pyrR
MSSIDAEALYRVLLDQIRAAYGDAAFAQAGGPVLAGIHSGGAWLAERLARDLGAPGFGVV
NVALHRDDYAKKGLHSQASPTSLPFEVEGRRIVLVDDVLYTGRTVRAALNELYDYGRPAA
VELAVLADRGGRELPVAARFVGGALAVDADATLVLERADAGRFTLRTEARAG*
>GCA_000452945.1_BPMSHR5858_genomic_03695_1 group_1500
MSKSSDRINLTNQFLIAMPNMADPTFSGTVVYLCDHSERGALGLVINRPTDIDLESLFNR
IDLKLEIEPLLHIPVYFGGPVQTERGFVLHEPVEGSAYNSSMTVEGGLEMTTSKDVLEAV
ATGTGPKRFLLTLGHAGWGAGQLEEEISKNGWLTVAADPRIVFDTPAEERFEAALGLLGV
SSSMLSGEAGHA*
>GCA_000452945.1_BPMSHR5858_genomic_03707_1 group_1886
MYWRKSLAAALLAPVLTACGGGDDPPPAPVVRLCPKTIDYNTVFTGGSGSGELVRVQLDT
TKMAFQITYLASPVPATTGTVQPTRDAAPNNIVTGTLTDETGLPTEKLNQCTFRLNNASL
DPNRPARVFLGEGVLGGAIPGATIQFGGVIGVGQIPKTTFPYYPFISFSDQETDLSKIAG
NYNQLGYHQVPSQNFAQAAVDAKVTINADGTYVETDNFGRKNGGQPLASSATVNQKLTLR
ADAPVFQSLNYQPQVPPTLASLDPSKAGKGILIVGKLRNQLVPIFIRTGAANADLTQGAP
VADDESGISILSPQTAIASGSQDGEYTGVDSVLDYRATALVGAQATLLDPFHASQAALTR
SLNLDYTQTVPGVVTTVQTNAASGPPTGKFVFTGGVFGLLDMSDVNNPYFTVGAFVQ*
>GCA_000452945.1_BPMSHR5858_genomic_03709_1 group_1887
MRKNAPEGEQITDRKPSLLDAPRKGNVYASECPTRLVLDRIADKWTVLILALLSHEPLRF
NTLLRHIEGLSQKVLSQTLKRLERDGLVARTVYATVPVSVEYALTPLGHTLAHSITPIIV
WSETHIDAVLDARTAYDARQGGRAPE*
>GCA_000452945.1_BPMSHR5858_genomic_03710_1 group_1501
MFLRSLVALAAAAASMFGAGPAHATGAGAPLQIDVYNPGEKSLFPVSSEIVTGKTGTILI
DAQFQRNDAEALVNKIKATGKPLKLVYVSHSDPDYYFGLDTIKAAFPDAKIVATPQTVAA
IKANKDDKLAYWGPILKDNAPRTLVVPQPLHGDRLTLDGHELRIVGLDGPSPDRTFVSIP
SARAVVGGIPVAANIHVWIADTQTPESRRNWIKTLDRIAALHPSRVVPGHYLANPDGSEP
YTLASVKFTRDYLVAFDAEAAKARNSGELITAMKARYPNLADASSLEMSAKVIKGEMQWP
VVGTASAFPAAGKKALVQFGDIGFRLDFKDDHTMTFVGTAGQYQGATDTVEYKATQIRPQ
VFMVYWHEPKSGDNVVHVEDFENGIVYTNIAHRNGEFQHLKGTIRIEDAK*
>GCA_000452945.1_BPMSHR5858_genomic_03711_1 group_1888
MDINQRIARRVRALRDLRGYSLDALAERSKVSRSNISLIERAQSNPTAVVLERLANALGV
SLASLFEDDRAARAASPLSRATEQPVWKDPASGYVRRSLSPAVASPLQLVEVKFPPGGRV
AYDGGERNADVHQQIWLLEGEMDIASGDDTWRIAAGDCLAMRLDRPAVFFNPGRKAARYV
VALAAANAVRPGWIE*
>GCA_000452945.1_BPMSHR5858_genomic_03715_1 stp_1
MTHGLHGPKRWYALVVLCLGVLMIVLDSTIVNVALPSIGADLHFTETALVWIVNAYMLTF
GGCLLLGGRLGDLYGQRRMFLAGLTLFTLASLACGLAPTQFVLIAARAVQGFGGAVVSAV
ALSLIMNLFTEPGERAKAMGVYSFVCAGGGSLGVLLGGVLTSVLSWHWIFLVNLPIGVAA
YALSAALLPKVRPQAADARLDVAGAIAVTASLMLAVYGIVNGNEAGWLSTQTVATLAGAA
ALLAAFIAIETRAAHPLMPLALATQRNVAVANVIGVLWAAAMFAWFFLSALYMQRVLGYG
PLQVGLAFLPANLIMAAFSLGLSARIVMRFGIRRAIGAGLVIAAAGLALFARAPADGGFV
AHVLPGMILVGVGAGVAFNPVLLAAMSDVAPSDSGLASGIVNTSFMMGGALGLAVLASVA
SARTDALAAARAALPVALNGGYHAAFACGAAFAALAAGLAYALRIRAQSNAAAELHGAAH
*
>GCA_000452945.1_BPMSHR5858_genomic_03718_1 accC_1
MTAIRGGSRRAPGLALALLGGVLLGACHGDENAQVNALPGFVSGSVRKTAYDGASDDLLT
AGLGKTGLGSDTRPGFANPAQPSAAELRRLAIYSNYRALVDITPNGGYGRFWGPNVDLAG
NDTLGEGKIAGTEYLAYSDDGSGRKNVTLLVQVPASFDPANPCIVTATASGSRGVYGAIA
AAGEWGLKRGCAVAYNDKGGGNGAHEIGTGVVTLIDGTLATASSAGSSSLFTASESSSTL
AAFNSAFPNRYAYKHAHSQQNPEQDWGRVTLQAVEFAYWALNEQFGPVVDGTRHGIRYRP
GDITTIAASVSNGGGSALAAAEQDTRGWITAVVVGEPQINVRMTPGVTVEQGGAPVPSFG
RPLADYATLANLLQPCAAAAVAATGAPYLSALPMGVTQSIRTQRCATLAAAGLVSGADTA
SQASDALAQLHAAGYLADSDLLQAPMWDSQAMPAIAVTYANAYTRSRVTDNLCNFSFATT
NPVTGAVAAPAVSPMTNLFGAGNGVPPTNGINLVFNGASGGVDHRLATPDASFAGAFCLR
QLWAANQLGIGTNVDAVRVAANLQHKPAIIVHGRSDALVPVNHASRAYVAQNSATEGRAS
QLSFYEVTNGQHFDAFLSVPGFDTRFVPVHYYDEQALNLMWNHLKSGAPLPPSQVIRTVP
RGGVPGAAPALSTANLPPIVQSPGANAIAVNAGVIDVPL*
>GCA_000452945.1_BPMSHR5858_genomic_03720_1 group_1503
MKPEPTRRRRPHRIVARFVAVSWRDLAMSIGPTVVLSIAAVWLAIALIQPAPPTSLTISA
GPPGSTNWRSAQRYKQILAKNGVTLRVLESEGSAENLARLSDPAQKVDVGFVQSGIEQKG
KHEALVSLGSVGYVPLAILYRGPVIERLSQFKGKRLALGAEGAGAHELGLALLKMNGIVP
GGPTPLLPLSGEDAARALTEGRIDAAFLSGDSTQIPVMAKLFRTPGVHFYSFTQAEAYTR
RVAYLTDITLPMGVYDPGTNLPPSDIHTLSPTVELIARDTLHPALSDLLIEAARDVHGRA
TILQRAGEFPSPVTHSSFLLSDDAARYYKSGKTFLYRKLPFWVASLVDRLLFIVVPLVVV
LIPGLRLVPTLYGWRVRSRIYRWYGALIALERSALGEHTAQERVVLLDKLDDVEESVNRM
KMPLAYAGQFYVLREHIGFVRARLLARDYETPQPAAAKPAGAPPAGSRQPGDA*
>GCA_000452945.1_BPMSHR5858_genomic_03722_1 syrM1_1
MQPDQDRAASLDIWLVRVLRTLLVERSVTQAALRLNQTQPAISAALRKLRDALGDPLLVR
GKTGMVPTEYGASLLDAAERALRDVDFVATPHGPFDPDTARRTFRIAAPDYLNDFFMPTL
VARFRAAAPHARIEIDSLNPSLDHAGALDSGALDLVIGNWPKPDPRFVRTDLFSDTVVCL
MRRDHPLARAPLTRDAYANALHLAPAPYIGARNNAIELGLARAGIARRVAATLPYFGLVP
QVLLQSDLIFTTTRRFAIHYARLLPLVVVAPPVPFPRIRCYQLRHPAPDRPTDADWLSAL
MADVSAELTMRGKRR*
>GCA_000452945.1_BPMSHR5858_genomic_03725_1 fecE
MSATLDIERLNWSPAGAGALLRELSLTVRAGELAGLIGPNGSGKTSALRCAFRFARPHAG
TVTLDGRDVWRAPPRWSAQRIAVLQQEAPDDFGLTVEELARMGRTPHKRLFDADTADDAR
VVEAALRDVDLWALRERGFASLSGGEKQRALLARALVQQPALLLLDEPTNHLDVRHQLEL
LARVRALRVTTLATIHDLNLAAAYCDRLHVLAGGRIVASGAPADVLTRALIADVFGVDAI
VDAHPVSGRPRVTLIYPEDAERC*
>GCA_000452945.1_BPMSHR5858_genomic_03745_1 mhpA
MTIDYQTLKFDYRPRAALADAGEPHPAVVVGAGPVGLAAAIDLAQQGVPVVLLDDDDTLS
SGSRAICFAKRTLEIFDRLGCGERVAQKGVGWHVGKVFLQDELIYAFDLLPEVGHARPAF
VNLQQYYVEGYLTERALELPNLDLRWKNRVTGIRQTPEHAELDVDTPEGPHTLRARYVIA
ADGSKSPLRAMMGLDSRGRTFKDRFLIADVKMKAPFPAERWFWFDPPFHRNQSVLLHRQP
DDMWRIDFQLGWDADPVAEKAPERVIPRVRALLGPDVEFELEWVSVYTFRCQRMDSFRHG
RVLFAGDAAHGVSPFGARGANSGVQDADNLAWKLKLVLDGRAPDRLLDTYAREREFAADE
NIRNSTRSTDFITPKSPISRVFRDATLKLARDCEFARRLVNSGRLSVPAVLADSPLNTPD
RAGEAFERAPVPGAVAPDAPVRANGEPGWLLQHLQGGFVGVLFGSIGDAGALVDATRGHA
LPVAPVLIVPKGEARAVAGVDVIEDSEGLVARRYDARPGTFYLLRPDQHVCARFRTAQAS
AVRAALARAICDA*
>GCA_000452945.1_BPMSHR5858_genomic_03746_1 group_19160
MAGDKKFASQADLAEKKVTFERLSEHAYAYTAEGDPNTGIIVGDDAVLVADTQATPVMAE
DVIRRIREVTDKPIKYVLLTHYHAVRVLGASAYGADHVIASQDTYDLIVERGAADMKSEI
ERFPRLFRSVESVPGLTWPTLTFKGEMTLWLGKLEVKIMQLGRGHTKGDTVVWLPQEKVL
LSGDLVEYGATPYAGDAYFQDWPATLDAIAALEPEKLVPGRGAALKTPREVADGLAGTRA
FVSELYAKVKAGAAAGRDLNAIYKETYAALKPTFGDWVIFDHCMPFDVTRAFDEATQYPD
PRIWTAERDKDMWHTLEG*
>GCA_000452945.1_BPMSHR5858_genomic_03751_1 group_19407
MSLRLGDIAPDFEQDSSLGRIKFHEWLGNSWGVLFSHPADYTPVCTTELGLTAKLKGEFE
KRNVKVIALSVDSVESHKGWIDDINETQATSVGFPIIADGDRKVSELYDMIHPNANETLT
VRSLFVIDPNKKVRLIITYPASTGRNFDEVLRVIDSLQLTDHYKVATPGNWKDGDDVVIV
PSLQDPDELKQRFPKGFKAVRPYLRLTPQPNK*
>GCA_000452945.1_BPMSHR5858_genomic_03752_1 yqjD
MSEVNKEKLMSDIKTVLADAEDLLKQAASSTGDRATELREKALARLKQAKEKATDVQVVV
VEKGKKAARATDDYVHEHPWTSIGIAAGVGVLIGLLINRK*
>GCA_000452945.1_BPMSHR5858_genomic_03753_1 group_19198
MTTETSSHQSGHGPLRRLLGSVLALLQTRLELVGIELAEEKERLMGVLFVGLAAMMLATM
ALISLTVLIAIAFWDTYRWQSLAVVTALYALGALACWLKARSGLRDAPSVFEATLNELEK
DRELFRGKP*
>GCA_000452945.1_BPMSHR5858_genomic_03754_1 group_19099
MSAAQYRALRKELLILRSDVERLELAQAGAELRQSITHFKWLKLLVPGVSGGSLGKSARN
VNATLGMLVSQYPLLSSLASVVLAKPVRALLRASARPALKWGAVGFALWEGYQIWKQAKH
DDASDAGARQRTRP*
>GCA_000452945.1_BPMSHR5858_genomic_03755_1 group_19696
MPCTGRNSPSRGFTLIEVVVAIAIVAVLAAFAVPSYRSYVERVNRLTAVAALYRAAQYVD
AFGDAPPTALPEGVNRAPESGKLVYVLRIMFDDARGGYALEARPAADGAMRDDRCGVYVL
HADGTRENRVAGGVALDGGAAEGDACWRTG*
>GCA_000452945.1_BPMSHR5858_genomic_03760_1 nrdR
MRCPFCRHDDTQVVDSRVSEDGAAIRRRRRCSACDKRFTTYERVELALPAVVKKDGSRTE
FDRRKIVASMQLALRKRPVAADAIDAAVARIEYQLLASGEREVRSEKLGELVMNELRQLD
TIAYVRFASVYRRFEDVSEFEDVIEEFRRAAPAKTPRKR*
>GCA_000452945.1_BPMSHR5858_genomic_03764_1 tolQ_1
MNTSQDLSIISLVLNASVLAQAVMGLLLVLSLMSWTFIFRKWFAIRRARAQTERFERDFW
SGGDLQALYQSAANNRHTIGALERIFESGMREFLKAKEKRLSDPALLLDGARRAMRASFQ
REMDVLESNLSFLASVGSVSPYIGLFGTVWGIMNSFRGLANVQQATLANVAPGIAEALVA
TAIGLFAAIPAVVAYNRYAHDIDRLAIRFETFIEEFSNILQRQAQ*
>GCA_000452945.1_BPMSHR5858_genomic_03765_1 exbD_1
MAGTPIRSSMRGGRSRRSMADINVVPYIDVMLVLLVIFMVTAPLVAPSIVNLPTVGNAAP
QEQTPPVVVNIQADGKISVRYKGDSGASQEDTMTKAELDGFIAARQADHPDQPVVIAADK
TVQYDAVMTVMSDLKARGVKRVGLLVKSQ*
>GCA_000452945.1_BPMSHR5858_genomic_03768_1 pal_2
MMSKKLRLAFAMLMIGALAACKSGVKLDEHANQGDAVSTQPNPENVAQVTVDPLNDPNSP
LAKRSVYFDFDSYSVQDQYQALLQQHAQYLKSHPQRHILIQGNTDERGTSEYNLALGQKR
AEAVRRALSLLGVGDAQMEAVSLGKEKPVALGHDEASWAQNRRADLVYQQ*
>GCA_000452945.1_BPMSHR5858_genomic_03769_1 cpoB_2
MTHRFPWLRTAAALCVAGAALLAAPAHAGLFDDDEARRAVLDLRSKSDSQASQLAAAQRT
ILDQTNRLDQLNQQVATLRGENEDLTNRLTTLERQQKEYYTDLDTRLKKFEPQKTTVDGV
EGTVQPGETDAFNAASQQFRNGNFKGAAASFRAFIAKYPQSPYQPTAQYWLGNAQYALRD
YKGSTATWQGLVKNYPQHPRAADALIAIGTNQLEQGQKAAAKKTFEQVVSQFPGSNAAET
AQSKLDAIK*
>GCA_000452945.1_BPMSHR5858_genomic_03772_1 gtaB_3
MLKVTKAVFPVAGLGTRFLPATKASPKEMLPIVDKPLIQYAVEEAIAAGITEMIFVTGRS
KRAIEDHFDKSYEIEAELEARGKEKLLTLVRGIKPRHVDCFYVRQAEALGLGHAVLCAQK
LVGDSPFAVVLADDLLDGRPPVLRQMIDVFNHYHTSVIGVEQIAPQDSKSYGVIDGKRWE
DNLFKLSGIVEKPEPAAAPSNFGVVGRYVLTPRIFDHLRALKPGAGGELQLTDALQSLLK
DEQVLAHRYDGLRFDCGSKIGYLKATVEFALRHPEVAPEFERYLSERFPELVAA*
>GCA_000452945.1_BPMSHR5858_genomic_03785_1 rhaS_7
MQAGRMAEIFNLANRFRHAGEDSGGYEMLLLSATGGAVYSSSGIPVWTQPLHERAIERVH
AIFAIGEPDPADRDDEKVSEWLRSARAHVRASGGARRLMGLAALERGDAAADDLSIDAML
SGAASAEKVRPTAAERAAFSAALAIIRHDCGDSAAHEVAERLCPALSARPAESAFAAREA
RASKLIRASVQQLRDNSANRISIADTAHAAAMSERNFLRRFKQEIGVTPSEFVQKVRLEH
ACHMLVHTDLPVDKIARRTGLGSGDRLAKLFRQHLSMSPTEYRAIERSRGADADLACGDF
VSRLSGSVS*
>GCA_000452945.1_BPMSHR5858_genomic_03793_1 tdh_2
MVNIKIAIIGASCRFPGGVAGLEEYWQLLKSERDAVTEIPPERFGTDFYRHPSKREPGKS
YTFSAGVLDNVAGFDAAFFGISPREATQMDPQQRLLLELAWEAFEDAGVRPADMRGSNCG
VYVGVAGTDYGNRSMDDLNAIDPYSATGNTLSIASNRVSYLFDLRGPSMSVDTACSSSLV
ALHQAVQALQSGEAETALAGGVNLLLHPFGFVSFSKASMLSPRGRCRAFDATGDGYVRAE
GGAFVLLKPLDRAIADGDTIHAVIAGSGVNSVGHSPGGISVPGAAAQAALLRSVYARAGV
DPQSLAYLEAHGTGTAVGDPIEARALIDVVSGARPADRPLLIGSVKTNIGHLETASGMAG
LLKAVLCLKHRAVPRSLHFSIPNPAIDFDGGRLRVVDRYMPLEAGNAPLTVGVNSFGFGG
TNAHIVLTEAPVSAVVADDVPTSATQAARAPSALVLTARSPNALGALANRYLAALENGGD
WQALAAAAARRRQWLEHRAIVAPADAAEGRAALAALARPAPEGGPSCVTTGRAPADAPRT
ALVFSGNGCQWVGMGNALYAEDAVFRAALDEVDALWCADGSPSLVDVMRGGPGAQWLAGA
GAEWLAATENAQPLLFAIQVGMIRVIDARGMRYDAAIGHSVGEVAAAWVTGALSLTDAVR
VIKIRSRAQAMTRGCGRMAAVGIGDAAARELIARHGLARRVEIAGVNSPDAVTLAGEPQG
LQVLEAALRGSGKFFQMLDLDYAFHSSHMDRIEPVVLAGLAGLRPQPGNGAFVSTVTGGA
LAGSELDARYWWRNIREPVRFGDGIAHLIEQGVRLFVEVSPHSILRTYVKQALAAAGVTG
VALPTLKRDHGSAATLRQAILAAIAHGASVDPDRFAPGASRAALPSYPWQRERFWLTPTV
EGYGLVNRRREHPLLGYRLHEHAFAWENQLDPAKLPMLADHVVDGGVAFPGAGYVEMALA
AACTFFDTPDAALENVEIRMPVVFQPQQAKLFRLVIEPRTATFTIETRDRMSDGAWTLNV
TGRMLESGNALGAAGVVPSDMLERLLALPAADGDMLYANTAAIGLGYGPAFRWVRTVRVA
EGDDAALADVAAPDACGDARTLSAYRLHPALMDSGFHPLFALLAAHARDGEHPAYVPVQI
GQVDYLRGDTVARVLARIDRRSPHSIVAHFEFLDAQGAIVARLGACRFRRVDLVGRRQNL
PARFVYRLEAMPLPGDDDAAGLPAPDALLARAVARLDAGHDGRRTRHLTEILPLLDVLAS
LYVLQAFDALGAFAGTWRPAHGCAALAARLADMLVEDGLAHRDGDRLVRDDAACAALPPL
DELWRGLLAESPAHVAELTLLAHCGSALPGVLSGAKDGTRLLSPTGHSLVEHLLAASPTW
QHVYALLTASVEQAVDAWQPARRLRVLELGATDGDVLQTLGMHLAPARCDHTIAATAGQL
AGFGADAQAAVTTVALQPGERLSLAADEAGPYDVIVANRALSGRRDVADALNAMRSWLAP
GGLLLLAESRRGRFSDIVFGMQAPSTQTDAPAPLAPADLEAALAHAGYVGVVRHVEQSLD
LEGTPMFVIARNPASAVAAPSGGDSSDLDTLARSEQWLVLHAPDVVGGFGARLADALAHA
GYPADIVEITHAADAIASLPAGQPAHVVFCAPETPLAEAATGDDLMAAQRNGVIALAALV
RVLGAQPNATTRLTIVTRGGAPFAGTAHAHPEQATLWGLGRVLANEHPELACRLIDVDCA
ADRIPEALIRELTGAALEEEVVLGAHGRRVPRMLTAAQDAARNDGAIARAAVLAFDAPGS
LRNLEWFALAESALGADEVEIEPVATGLNFRDVMYAMGLLSDEAVETGFAGATIGMELSG
RVVRVGSAVTEFAPGDAILGFAPASFATRVRTRAQAIAPKPERLSFEEAATVPTTFFTAY
YALVELARLRRGERVLVHGGAGGVGIAAIQLARYFGAEVFATAGSDEKREFVRLLGADHV
LDSRSLAFADEIRAMTGGEGIDIVLNSLAGEAMVRSIDTLRPFGRFLELGKRDFYENSHI
GLRPFRNNISYFGIDADQLMGALPELTARLFGDVMGLFAAGVLHPLPYRAFPAERAEDAF
RYMQQARQIGKVLVTYPCGTPAPTHGAARADSLALDPHGAYLVVGGTGGLGFASARWMVE
RGARRLTLASRSGELAVAARDEIECWRATLGVAVDIVSCDVIDAAAVDAMIAAIVRRDIP
LKGVLHSAMTIDDSLVRNLDDARMAAVLAPKVAGAWNLHRATRSLPLDLFVVYSSATTYL
GNPGQSNYVAANSFLEALVEHRRAAGLPGTFMAWGPLEDVGFLARHADTREALQSRIGGA
SITSGEAMIALERALVAGAAGEAVVRLDWHAVARGMPAAKARRYSLLQSHANGGDARDGG
TQLREEVLALPRDEAIALVAGTLQAQIARILHMTPDRIALDKSVLDMGMDSLMGMELGLA
VEEAFEVKLSVMAIAEGASAMTLAGRIVDSIGASADAGAGAAADSAHEAVAALAAKHAID
GDARAVLDIRPAPVAGQVSPTLEVAR*
>GCA_000452945.1_BPMSHR5858_genomic_03794_1 group_1649
VASSMPITFVFAIALSFAADAIAIPRAAVRRPFLAGALHTASVLFVACVILALTRRPHFA
AFLALALVALAGAVSNAKFSSLREPFVFTDLSLFSQLFSHPRLYLPFLSATTVVAMVLGS
VALAAGFFLDPAVSAQTAWQAAMAALICFATGGLCAARLPLTLDPLDDQRRHGFFAVFVA
YLLNGMRPATFRSVAGLTCAGPFASEDELPGHPDVIVIQSESFFDPRPLSAAIDSSILGH
FDRVCRESVEHGQLAVPAWGANTMRSEFAFLTGLPSSHLGYARFYPYAFVRRISASLAGW
FRRAGYRTTAIHPYYADFFGRDRVFPLLQFDRFFDIRAFGDAPRAGPYISDAAVLDQIVA
VLDEKRTQPAFLFAMTMENHGPLHLEPVEAGEAARYHTLGDDATWRDLTAYLRHVANADA
MIGRLVAYLRQCRRDTILCFYGDHVPALSHVFERFGNIPDQSNYFIWRNFGEHAPRKQDR
AVEELGTALLRAMKTTGRQAVSTGASEITT*
>GCA_000452945.1_BPMSHR5858_genomic_03795_1 group_19654
VSDSAGPRHIAITGASAGLGRALARAYARPGVVLSLGGRDAVRLEESAADCRARGATVFV
ASIDVRDADAMRRWLEQFDDAHPIHLLIANAGVASTLAHGGDWEARERTAAIVDTNFYGA
MNAVLPVIDRMRARGSGQVALISSLAALRGMAISPAYCASKAALKAWGDSVRPVLKRDGI
RLSVVLPGFVKTAMSDVFPADKPLLWLPDKAAQYIQRGIAARRAEIAFPALLALGMRLLP
LLPAVMADAILGRLSYLPREER*
>GCA_000452945.1_BPMSHR5858_genomic_03800_1 hddA
MNPTIIRARAPLRLGLAGGGTDVAPYADTFGGYVLNATIDRYAYAVIKTLTIPAVRFVST
DQQVEKHQLISEPLELNGTLNLHKAVYNHMIRNYNHGKPIALELSTFCDAPAGSGLGSSS
TLVVVMIKAFVELLNLPLDDYAIAQLAYRIERVDCGLAGGRQDQYSATFGGFNFMEFYEE
ERTIVNPLRIKNWVLCELEASLVLFYTGVSRESAKIIQDQSDNVVSHKTAAIEAMHGIKR
EALVMKEALLKGDFKAFVASMRLGWDNKKNSARTVSNAHIDEIYDAAIRAGAQAGKVSGA
GGGGFMLFFVPTEKRMDLIRTLGEYDGQVSNCHFTKNGTQAWRIAN*
>GCA_000452945.1_BPMSHR5858_genomic_03804_1 group_1506
MKLAFFTPFHPACGPGFMSKAVVEHLKDYFDVTVFYQWHAQHQPYEVEDVSVRIIEDDMD
MEPVAREFDAVIYNLGNNEENHYSIFQCLKRLPGIVVLHDFVMQHGIVNDLFNRKQKSDL
YYWLLAALYGKDGVRACHASALSYAGGVRGGWDSSSVASFPMFEVFSGLASACVVHSKFF
ENQLKPHFAGPILMTRNPYDLKRVPTEDVIETFYKVRGKAGGKVVFAAFGHMSPSKCIDR
VLRVFGHSERLRKEARLILCGGASPEYDQYLRRLCADGDLGHCVEFRGSVSDTELYALQV

EADVFVNLRQPNTEGQSGSLIEQLAAGKPVICYDTGCYGDLSEDVAYKVHSTTDFNELQR
VFERLLNGVAERREVGLRGRAAALEYDCARYARDMFEFVFENREYLNAFQRRNFDGLRPL
GVSGTTIDTRHPENYSRTQSIWGVLDGFVHHASSEALSLIPGDARHHYLSFLTQPYRPLV
DSLLELSANDVPEVELEADGTSLYSTASAPVLNTSFWTKLKCPVSEHFAYIGYKKLLCRD
PDLAGLRDYAEKLASETISRKEMILAMLRSTEFAERFGDIRKSSDYVSLIRWAGAQVA*
>GCA_000452945.1_BPMSHR5858_genomic_03807_1 group_1507
LLFRTLLGLKKMAVTGLLQQGGRDITAKMVYAAESERPAAQIRAMSKIVVSFTDDGRRGV
AGAISRAISIVMRGISLRASVIAGLSEKLGFFAPVEFEDFVWRRFFAKTLPVDDRPHVLA
AQYRVLPLAMATLHRLKISLSGLLRVKRFKKIETRDFDVFISQTPFPGVLSSNTKLIVRY
HDAIPIFLPHTIKDRAFHHGAHYDALARNVRSGAYFACVSAATRNDLLKIFPEIEDRAVV
IHNTVSPIYSEIDSPKELVKDIVSSRLFRPLAKKYKALSSSHVDVFTRSRLNADQQDFEY
LLMVSTVEPRKNHTSLVAAWELLRSRHNEALKLVLVGDLGWDYAEFVDAIVPWVERGELF
LLSNVAAPDLRDLYKHAKVTICPSYAEGFDYSGVEAMMSGGVVAASDIPVHREVYEDACI
YFNPYSVESIVGGIREAISLRNASETFAEYRRCAKRISARYTAEAISPRWDEFIQSVGSA
P*
>GCA_000452945.1_BPMSHR5858_genomic_03809_1 kpsM
MTNFGRSLAVQRRVIFALLMREVLTRYGRHNIGFLWLFVEPMIFTVGITILWNATGATHG
SNLPISAFALTGYSTVLLWRNMPGRCSMAVTPNLALMYHRNVKVIDIFIARILLEVVGNT
ASFFVLMITFHALGLVDYPEDILEVMFAWVMIIWFGASLGFIIGALSEKTELVEKLWHPV
TYLMFPLSGAIFMVDWLSPAFQKIVLWLPMVHGVEMLREGYFGSLVAAHYDIGYMACACL
AMSLVGLALVRDVGREVTPE*
>GCA_000452945.1_BPMSHR5858_genomic_03811_1 group_19619
MGAVTLAGCSSIPTSGASGAQIARAAQSPSGIQIVDVTEDVARQLFADRNTADFVTALGG
GASFRQQLGVGDTIQVSIWEAPPATLFGAAQSEGSSGPANARVTVLPDQAIDGDGNVNIP
FAGQVKAAGRSPTQLAREIAARLKSMAHDPQVLVKLSRNETSYVTVVGDVAENARMALTA
RGERLLDALASAGGAKHPVDKVTIQITRGKTVASLPLDMVIRDPRQNVPLHAGDVVTVLF
QPYSFTVLGATGKNDEINFEAKGITLAQALARAGGLQDSRADAKGVFIFRLEDANALKWP
TAPVRTTADGKVPVVYRVNLRDPNSFFVAQSFRVDNNDLLYVSNAPIAELQKFLNVVFSV
AYPVITGVQTVRY*
>GCA_000452945.1_BPMSHR5858_genomic_03813_1 group_1889
VSEFRAPPRLPGETRVLLIDERKYSQGIGAVATRNNRGAFERMIQAARAAHPDAEFWLAR
TRDRGSGVWLSASAADILPPDIHRLGEHESLCAALEHVDHVYTVGASEGMQALLAGRRVH
VFGAPYYAGWGLTDDAVQLPGRHARPTLAALFDVVFLRFARYLNPATHAPGRIDDLLDAI
EWQNTVRRRFADLRQVAGIRFQWWKRPFATPYLTAGGGTLRWTRDASRLREGEHAALWGA
RGTNDLPPGTRVIRIEDGFLHSTGLGSDHVAPCSQVIDRSGLYFDPSRPSDLTTILNETD
FDDAELVRANRLRREIARLGLTKYNLGRRKPAWSPPPGKRVVLVPGQVADDASIRLGTRG
ITTAEDLLREVRARRPDAFIVYKPHPDVLSGNRRGAIEVNAWADLIEQDADLISLIEVAD
EIHTLSSLSGFEALIRGKAVHTYGLPFYAGWGLTQDALAQPWRKRTLSLDMLTAGVLLRY
PVYWDWSLRLFASPELVVRQLAIPAARPLTSIRGDRLRPVRKASRWIASCLRHLLWQCGK
*
>GCA_000452945.1_BPMSHR5858_genomic_03816_1 mscS
VDLASVQTFIVTRGLDFGLQVIASIALWIFGRWAIRIVTNLMGKLIRRSGKVDSTLSHYL
ESVVSVLLTVLLILAILQVFGVQTTSFAALLAGLGLAVGTAWGGLLAHFAAGVFMQVLRP
FKVGDVISAGGVTGTVKELGLFGTTIVTADNVVTLVGNNKIFSDNIANYSATPTRRVDLT
AKIANGVDAADAIKRLKAALELIPNVVKHPAPDVGVLSFTPEGPLLFVRPSTQPANYWQV
YCDTNRAILDTFREAGYPTPETPVAHRTTADVAKS*
>GCA_000452945.1_BPMSHR5858_genomic_03822_1 hda
VTVTRQLTLDLGTPPPSTFDNFYSGTNAELVTRLRELDLALAAGPVADRTFYVWGEAGSG
RSHLLQALVHDTTYGHARYVSPQGGLDALAFDPRVSLYAVDDCDALNDAQQIALFNLFNE
VRAHPMTALVVAGPAAPLALDVREDLRTRLGWGLVFHLAPLTDEGKAAVLKHAAKERGIA
LADDVPSYLLTHFRRDMPSLMSLLDALDRFSLEQKRAVTLPLLRAMLAAPEREEPAPGRF
K*
>GCA_000452945.1_BPMSHR5858_genomic_03823_1 serB1
MTNLALFDLDHTLIPTDSDHEWGRFMVRLGIVDADSFSRQNDRFYADYKAGKLDIHAYLS
AMLTPLAKYSRAQLAQWHDRYMHEVIRPAMLPAAIELVRRHQDAGDLCCIVTATNEFITR
PIATAFGVDTLIACEVETVDGHPDSAFTGRPTGTPSYREGKITRTQAWLASLGKRWDDFD
RSYFYSDSHNDIPLLEKVTDPIATNPDDTLRAYAREHGWRILDLFQPS*
>GCA_000452945.1_BPMSHR5858_genomic_03826_1 group_19564
MNSTPLTVTAPDWRPPCRYLAIEGPIGVGKTTLATLLAERWSMRALLERPQDNPFLERFY
RDTARYALPTQLAFALQRERQAREIAAAHEANTPIVADFMPQKNEIFARLTLPDDEWQLY
RSLAAHLDAPAPAPDLVVYLQASPEVLFSRIQKRGEPMELQIGDAYLRSLCDAYNEFFYH
YDRTPVLTVAAEHLNPLDSPDDLALLVDRIASMRGRKEFFVKGGSAR*
>GCA_000452945.1_BPMSHR5858_genomic_03827_1 panB
MTYLQESSRPAVTVPKLQAMREAGEKIAMLTSYDASFAALLDRANVDVQLIGDSLGNVLQ
GQATTLPVTLDDIAYHTACVARAQPRGLVVADLPFGTYGTPADAFASAVKLMRAGAQMVK
LEGGEWLAETVRFLVERAVPVCAHVGLTPQSVHAFGGFKVQGKTEAGAAQLLRDARAVEE
AGAQLIVLEAVPTLVAAEVTRELSIPTIGIGAGAECSGQVLVLHDMLGVFPGKRPRFVKD
FMQGQPSIFAAVEAYVRAVKDGSFPGPEHSF*
>GCA_000452945.1_BPMSHR5858_genomic_03876_1 group_1308
MLDDFSRVLRFKPHLLVRDAGSGALYVVDEFRRSVLPGDVFPAIAACMRDRLTIAQTFAA
LAARFSQWEVLAALDQLVRRGYVRADAPGERDAELAFHERAGVDGDAASGVASRLTVAVE
AFGVDPRAQLDAFAACGIGVAPDAPLTVALTDGYDRAELIVAAERAAARGGALLVVVADR
VEPLIGPLLGAAAGLAASAMSTAPTAPSTSPAPQGAADAPPCIECVRYWTALNHPVETLL
ARLHGGDAARLPPARSRASAAAVAAVVASFVEQIAVNAQRRRHAGSHIVSLRVDTLATAA
HRVVRRPQCPRCAHAGWMREQAERPVTLASADAGARRDGGYRTLAAAELFKRYGHLISPV
SGPIAYLHPMPGRNAGMRHVYVAGYLVCPPSAPRENRFDKLCSGKGASDAQARASALAEA
LERFSGVYQGDEAALRGSLAQLSAHAPPGSGPIDVNALQQYSDRQFERRERHNATTDDPR
KQVPRRFTRDSVIDWTPAWSIATGARRLVPLAYCYAETPASSGADYCVHNPNGCAAGACI
EEAILQGLLELVERDAVAIWWYNMLRRPAVDIESFGDPYFDALAADYASLGWRLWALDIT
HDLRIPVFVALARETATGRFSIGFGCHPDSRIALQRALTEVNQLLDVGAAAPPPWDVDKL
PDDAFLHPDPALPPVRAPARAPHGRCDLKRDIEDCVARLAAAGIDTLVVDKTRPDIGLPV
VQVIAPGLCHFWPRFGAPRLYSVPVAQRWRERPRDEDALNRALLFL*
>GCA_000452945.1_BPMSHR5858_genomic_03877_1 group_19183
MAENNAVPTHQSLLDFPEVYLRAIALSWENEQFKRELLADPLDALERYFDYRCPWILNLK
VAEVPPDESHYGWDAKRQRWNLPVNTLRVGIPTPPKHLAEEGIALAAYNDAGPAYLFTCC
*
>GCA_000452945.1_BPMSHR5858_genomic_03878_1 group_19458
MTDRSPNVPTYDQFLEYRAVIVQAIAAAWHDAEFREELIAHPVDALHKRFDYRFPMKMHL
KVHENSATWTPLTNGGWTTNEVNGLDLVLPPAPPPEQRAAALAAYNARHISLFGPDRKEI
*
>GCA_000452945.1_BPMSHR5858_genomic_03880_1 group_19196
MALLENGLSGEASLLRARHVLGRDPQRCDTVIADPYVSRIHASICWTAGRWELHDHGRNG
TFVSGRFVGEGECVVLRDGDLIQFGSAGSVRWRARELGEPVDMLWPLRAPARPIALDRTH
ALPGAAFTVSRSAQGDWLCNDTTPARVLHDGDAVICGEFAWQLVLAHRGVTAALPRATQV
ATLPQRIDFTVSRDEEHVTATLHTRGGAVDLGARAHHYCLVTLARARFADAQAGYDTASR
GWIELDVLARMLGLDESHINVQIHRARTQFLPLLSPGSAELVERRRGGVRFGALAFRVVR
GDRLECQSADADASAPLPDPLARGAATFVSPAALG*
>GCA_000452945.1_BPMSHR5858_genomic_03881_1 group_19165
MGFYGPEPFERATATYVWLGLRVPGALIVEVEGNAPRYTTGIQLVRDPRFVGGLKIDVMG
WTGPLSSGTQSYKVRHTFQGVFHPTIVVHGSNKTEVVEVKQIPHEEADAFLQALDAA*
>GCA_000452945.1_BPMSHR5858_genomic_03882_1 group_1033
MSSDIVESRIVLVPVTRAYATYTSFSLHLLGTFHVYAEGEKPLNIRNPHLTPQMWYPPGG
LAVVGVVGRPRLGPITSYRTPVYSYQGQYQDTIYVQGANGIVTVKVRLLKDEEAESFLDT
LDAAA*
>GCA_000452945.1_BPMSHR5858_genomic_03884_1 group_19193
MNRELLTHIRADQLDGSIFDASCHAVTVGPVTVEFRTSAEPPSSTAELYISRQLVGGCTV
NAAHPSGKFGGSVGKLKAEVDLTLDVPGGKLDYKLTVCAPIIGGTSKSGTLGL*
>GCA_000452945.1_BPMSHR5858_genomic_03886_1 group_1509
MLSLTQRVLTYSFIDRPPVNPRAIGTSSADAALLAQIDALLASAAASFKARAYDAALDDY
FACESLIYSHLDAQWNPDLGGRLRSRLPRDAALFDSLLSATSQWLNVLPVPAPASPVRPA
TPPPAQALAGVAALRGAGLAPVSPNPAATAQALSDMQLASLYTSQGNSAASSVAVTRAKA
VDAAVVGAFSPPQMPNPSALPAANPNAAPSTTPGFHPAPGVMPPRGIDLAPAALTPLKIQ
PMPKLPIALLAQKQVGLLTGSGAQTAVKAIQWAASGAPDIASIKTILYAPHASAAALPDA
LTNANSLWERSVLLPHDYFYTIPLAIAHCYQALGDYANAETYYLQAAGYAYLNTATEGPY
IWVALAQLYRAWGDSLYLQGDRAGATNAYGKVVTPGSPAAPATALYQLAGLATAAKRATA
LLPQLATLAQTGTGGVTADDVAIATVLLEVYAKLVQIGAGLDYWGNYAAAVPIWSFSYLQ
QVAINFAQLAQQAENQVVNFWNQADQAKLTRTELANQVSQASGQINAAQQQLAVAQAQAQ
AYQAGVALAQTRATNAAKNAQEYGSLNSQVIVIQATGQQVSGGDDGDYNGVSAMANQYLS
GQRISGDSATVAAATNLAANRLSQQFQIDSMNRTTAEMQQALAQAQAQLAAANAQVSAAG
ANLAVAQLNAQAAAQTLGVFDADTFTPQVWKAMGNFVDQIYERYMNMALRAAKLMQQAYN
FENDVSVSFIKASYQGVVDGLLAADALMADIQSFTDDLVNAKRGKKQYLKQSISLASRYG
YLFETQLRKTGTMTFETTLDDFDSAYPGTYQGRIRRVLVSVQGIVPPTGISGTLGNEGIS
FYRLPADVATPAAPSKVRVQSAETQVISDYDPVQDAVLAPPPENQTGIFEGAGVASSWTL
SLPPALNDINYGTLTDVVLTFLYEARFDPRLVQPVLAQLASRPGFYNRERAIPLAWLYPD
LFYGFVSTGTLTLNLSAADFPIDQTAPAVTAVSLLVAMKPGTPASNVTIALAAPGKGALS
GVTDATGAISSQSAGSAWAGAVGGAALGDWTLTLGAAANPSLAPGGKLDLSPLINLVLVI
DYAFKPRG*
>GCA_000452945.1_BPMSHR5858_genomic_03891_1 group_19691
MRLIEPNEHAGYLAALHARGLSGNEFTLRETDTTDPKGDENIGQTGFVTITRHSTHATRE
YPLGDAGDWLQHFKKDLEAGAFD*
>GCA_000452945.1_BPMSHR5858_genomic_03896_1 uvrA_1
MEQIRIRGARTHNLKNVNLDLPRHKLIVITGLSGSGKSSLAFDTLYAEGQRRYVESLSAY
ARQFLQLMEKPDVDLIEGLSPAISIEQKATSHNPRSTVGTVTEIHDYLRLLYARVGTPYC
PDHEIPLEAQSVSQMVDAALALPEETRLMILAPVVANRKGEHAELFDEMQAQGFVRFRVR
SGGGAANEGAAKIYEVDSLPKLKKNDKHTIDVVVDRLKVRPDMKQRLAESFETALRLADG
RAIALEMDTDREHLFSSKFACPICSYSLQELEPRLFSFNNPMGACPECDGLGQITFFDPK
RVVAHPSLSLAAGAVKGWDRRNQFYFQMLQSLAAFYDFDIDTAFEDLPEKIRKILLYGSG
KQTIPFSYINERGRTTIREHVFEGIIPNLERRYRETDSAAVREELAKYQNNQPCPSCDGT
RLRREARFVRVGQGDYARAIFEVSSWPLRDALGYFDGLTLDGAKREIADKVIKEIVARLT
FLNNVGLDYLSLERSAETLSGGEAQRIRLASQIGSGLTGVMYVLDEPSIGLHQRDNDRLI
GTLKHLRDLGNSVIVVEHDEDMIRTADYVVDMGPGAGEHGGMIVAEGTPKQVQANGDSLT
GQYLVGKRTIEYPDERISPDPQRMLRIVDAHGNNLKHVDLDLPVGLLTCITGVSGSGKST
LINDTLYNAVARHLYGSATEPAPHESIEGLEHFDKVINVDQSPIGRTPRSNPATYTGLFT
PIRELFAGVPASKERGYDPGRFSFNVKGGRCESCQGDGVLKVEMHFLPDVYVPCDVCHGK
RYNRETLEVQYKGRNISEVLDMTVEHAYEFFSAVPVVARKLKTLLDVGLGYIRLGQSATT
LSGGEAQRVKLSLELSKRDTGRTLYILDEPTTGLHFHDIALLLEVIHRLRDQGNTVVIIE
HNLDVIKTADWVIDLGPEGGAGGGQIIAQGTPEQVAKTKASFTGKYLAPLLKRKK*
>GCA_000452945.1_BPMSHR5858_genomic_03898_1 purU_2
MPADHSFILKLSCPDRRGIVHAVSGFLLERGSNILDSAQFGDSRTGEFFMRVHFEQAGAC
GAAAALDALRAEFAPLAEAFAMRWELHDAAVKPRVVILVSKIGHCLNDLLFRYRTGQLPI
EISAIVSNHKDFYQLAASYDIPFHHLPLAAGASADAKAAQEARVLEVIDGHAADLVVLAR
YMQILSPALCERLAGRAINIHHSFLPSFKGAKPYYQAFDRGVKLIGATAHYVTTDLDEGP
IIEQEVERVDHSMTPDELTAVGRDVECVTLARAVKWHVEHRIVLNGTKTVVFR*
>GCA_000452945.1_BPMSHR5858_genomic_03900_1 rhtB_2
MPNFMLFLATSIAITVAPGPDNLQVLARGISQGRAAGIVAALGFASGVLFHTTLAAFGVA
ALLRSSPVAFHLLKLAGGAYLIWIGIKALRSKGLATAHARAPQPLSTIFRQSVIGNLLNP
KVTLFFVVFLPQFVDPHGAQPVALQMLELGALFMAQTAAVFSVFGVGAGAIGVWLKRRPR
VGVWLDRLAGATFVALGLRVALKD*
>GCA_000452945.1_BPMSHR5858_genomic_03902_1 kefC_2
VISPLEMTLLLLLASVVGVVVFRSLNLPPMLGYLTVGILVGPHALGFAADLERAEHLAEF
GVVFLMFSIGLEFSLSKLRAMRRLVFGLGLAQVLGTIAIALVLGALFERWMHITWEGSVA
LGGALAMSSTAIVSKMLAERLEIETEHGRNIFGVLLFQDLAVVPLLIVIAAFGGESSKDL
AMSLGFAAVKIVVALALLLIVGQRFMTRWLNVVARRRSQELFVLNLLLVTLGAAFITDRF
GLSLALGAFIAGMLISETPYRHQVEEDIKPFRDVLLGLFFVTTGMLLDPSVIWRHPLLVL
AFFVGQVLLKSVMITGLARAFGATPGVAMRTGLGLAQAGEFGFVLLNLILDRHLVDATLL
QSILASMLLSMLAAPFLIQNADRIVMRLSAAEWMQQSLQMTRIATQSIKQKGHVIICGYG
RAGQNLARMLEQEGIAYVALDLDPDRVAAAAAAGESVVFGDAARRESLLAAGIHRAAAVA
VTYANTPSALRVLHNVHELEPTLPAIVRTVDDADLEKLLAAGATEVIPEIVEGSLMLASH
TLVLMGVPMRRVVRRVEEMRDARYSLLRGYFHGADDADDDDHEQVRLQSVPVDARADAVG
RSLSELGLFALGLEVTAIRRHGIRGVEPDPETKLRASDIVVLRGLPEALAQAEERLSRHR
REGGSAAVA*
>GCA_000452945.1_BPMSHR5858_genomic_03903_1 kdsD_1
MIAKINDDRALALARDVLDIEANAVRALAEQLDGEFVAAVGLLLNCRGRVVVSGIGKSGH
IARKIAATLASTGTPAFFVHPAEASHGDLGMVTKDDVFVAISNSGESEELVAILPLIKRL
GAKLIAMTGRPASSLAMLSDVHLNAGVAKEACPLNLAPTASTTAALALGDALAVAVLDAR
GFGSDDFARSHPGGALGRRLLTYVRDVMRTGDEVPAVPLDATLSDALFQITAKRMGMTAV
IDDANRVAGIFTDGDLRRVLERDGDFRRLPIVDVMTRHPHTIAPDHLAVEAVELMERHRI
NQMLVVDERGALIGALNMHDLFSKKVI*
>GCA_000452945.1_BPMSHR5858_genomic_03905_1 lptC
MDGKFRLTSMIPLAAMAALAGGTYWLLQATLPPPGEAVVRPKSHTPDYFADNFSVTELDQ
TGSTQYRLTAAGLVHYEDDETSDLTRPALRAFQPGKPTVTATAKRGAVNGDVSIVDLYDD
ARILRAAGAGDPPMQADSQHFRVLVNDDVIETEKPVKLQRGLSIANAAAGMKYNNVTRVI
ELYGNVRGTIAASDTSGGAAGQPK*
>GCA_000452945.1_BPMSHR5858_genomic_03907_1 lptB_2
VNALPNRQPAGTTSSLVVRNLKKRYGSRTVVKDVSLDVKSGEVVGLLGPNGAGKTTSFYM
IVGLVPLDAGDISLNGSPISLMPIHKRASLGLSYLPQEASVFRKLTVEQNVRAVLELQHD
ENGKRLSKDAIGARTEALLEELQIAHLRENPALSLSGGERRRVEIARALASNPSFILLDE
PFAGVDPIAVLEIQKIVKFLKQRNIGVLITDHNVRETLGICDHAYIISDGSVLASGAPKE
IIENESVRRVYLGEHFRM*
>GCA_000452945.1_BPMSHR5858_genomic_03910_1 hpf
MNLKISGHHLEVTPAIREYVITKLDRVLRHSDQVIDGAVILTVDNHKEKDKRQKAEITLH
LKGKDIFVESANGDMYAAIDLLIDKLDRQVVKHMERLQTHAHEPLKHQQAAEQIELPPQ*
>GCA_000452945.1_BPMSHR5858_genomic_03913_1 rapZ
MRIVLITGISGSGKSVALNALEDAGYYCVDNLPPHVLPELARYLAHEGQNRLAVAIDARS
SASLDEMPGLIRALSHEHDVRVLFLNASTQALIQRFSETRRRHPLSGSPSHDADVGLLVS
LEEAIERERELVAPLAEFGHQIDTSNLRANVLRTWVKRFIEQKNDDLVLMFESFGFKRGV
PLDADFMFDVRALPNPYYDHELRPLTGLDQPVVAFLDALPVVHQMLDDIETFLVKWLPHF
REDNRSYLTVAIGCTGGQHRSVFLAETLAARLSRQASVIVRHRDAPVAVDASSRLVT*
>GCA_000452945.1_BPMSHR5858_genomic_03914_1 group_19100
MSSLSTTLIDLPLFPLHTVLFPGGLLPLKVFEARYLDMARACLRDDAPFGVCLLKSGPEV
AQEGEVSVPETIGCMARIVECDTGEFGMLLLRTIGTQRFELLSHRVEANGLLVGIAEPMQ
EDIPLEGDSALAQFGACAEALERIVEVLRRSDAELPFAEPFRFDDPTWVSNRLAEVLPLD
LRARQKLMEFPDVGARIDAVHRELNRHGWL*
>GCA_000452945.1_BPMSHR5858_genomic_03915_1 mutM
LKPPRIPPAPPVITPLHSAFAPTLIAWQRKHGRHDLPWQNTRDPYRIWLSEIMLQQTQVS
TAVPYYVRFLERYPDVAALAAAPIDDVMALWAGLGYYSRARNLHRCAQAVVERHGGAFPA
SPEALAELPGIGRSTAAAIASFAFGARATILDGNVKRVLARVFGVEGFPGDKRIENEMWA
LAEALLPDAAEPTDVTAYTQGLMDLGATLCVRGKPECGRCPFAGECVAQRSGRQRELPAA
RPKKAVPTRRTWMLVLRDGDAVLLQRRPPAGIWGGLWSLPEADGDAALARRAREFGGAAL
VPLAPLTHTFTHFRLEIEPRLAELDGAAGGAPPAADDETAWVLLDRLDAYGVPAPVRKLL
DGLSGPLL*
>GCA_000452945.1_BPMSHR5858_genomic_03919_1 ispE
MTDTTRSLRDCLAPAKLNLFLHITGRRPDGYHALQSVFQLLDWGDRLHFTLRDDGKVSRV
TDVPGVPEESDLVVRAASLLKAHAGATLGVDIEIDKRLPMGAGLGGGSSDAATTLLALNR
LWRLDLPRTTLQSLAVKLGADVPFFVFGKNAFAEGIGEALQAVELPARWFLVVTPRVHVP
TAAIFSEKSLTRDSKPITITDFLAQRGIDAGWPDSFGRNDMQPVVTSKYAEVAKVVEWFY
NLTPARMTGSGASVFAAFKSKADAEAAQAKLPAGWNSAVAESMSEHPLFAFAS*
>GCA_000452945.1_BPMSHR5858_genomic_03921_1 prs
MSSHDGLMVFTGNANPALAQEVVKILGIPLGKAMVSRFSDGEIQVEIQENVRGKDVFVLQ
STCAPTNDNLMELMIMVDALKRASAGRITAAIPYFGYARQDRRPRSARVAISAKVVANML
EIAGVERIITMDLHADQIQGFFDIPVDNIYATPILLGDLRKQNYPDLLVVSPDVGGVVRA
RALAKQLNCDLAIIDKRRPKANVAEVMNIIGEVEGRTCVIMDDMVDTAGTLCKAAQVLKE
RGAKQVFAYATHPVLSGGAADRIAASALDELVVTDTIPLSAESLACPKIRALSSAGLLAE
TFSRIRRGDSVMSLFAES*
>GCA_000452945.1_BPMSHR5858_genomic_03923_1 pth
MIKLIVGLGNPGAEYTATRHNAGFWLVDQLAREAGATLRDERRFHGFYAKARLFGEEVHL
LEPQTYMNRSGQAVVALAHFFKILPTEILVAHDELDLPPGAAKLKLGGGSGGHNGLKDIS
AHLSSQQYWRLRIGIGHPRDLIPESARAGAKPDVANFVLKPPRKDEQDLINAAIERALAV

MPTAIKGETERAMMQLHRNGA*
>GCA_000452945.1_BPMSHR5858_genomic_03924_1 hisC_2
VSRYWSDIVRQLEPYVPGEQPALAHPVKLNTNENPYPPSPRALDAIRRELGDTGEALRRY
PDPVARRLRETVAAYHGIAPEQVFAGNGSDEVLAHAFQALLQHDRPLRFPDITYSFYPTY
ARLYRVAYETVPLADDFSIVVDDYLDDAGCVLFPNPNAPTGRALPLADIERIVAANPSSV
VVIDEAYVDFGAESAVSLIARYPNLLVVHTVSKARSLAGMRVGFAFGDAALIDALTRVKD
SFNSYPLDRLAQVATQASYEDEAWFQATRKQVIASRERLVGALAALGFDVVPSAANFVFA
RPRSHDAATLAAQLKQREIFVRHFKLPRIDQHLRITVGSDAECDALVAALRELLAA*
>GCA_000452945.1_BPMSHR5858_genomic_03925_1 fdx_1
MALMITDECINCDVCEPECPNGAISMGPEIYVIDPNKCTECVGHFDEPQCQQVCPVECIP
QDPGRPETKDQLMAKYRALIAAEQKDDA*
>GCA_000452945.1_BPMSHR5858_genomic_03926_1 coaD
MVVAVYPGTFDPLTRGHEDLVRRASSIFDTLVVGVADSRAKKPFFSLEERLKIANEVLGH
YPNVKVMGFTGLLKDFVRANDARVIVRGLRAVSDFEYEFQMAGMNRYLLPDVETMFMTPS
DQYQFISGTIVREIAQLGGDVSKFVFPSVEKWLTEKVAAMAQGPSA*
>GCA_000452945.1_BPMSHR5858_genomic_03927_1 rsmD
MSRSSPARPQAQRAPAAGRGKPHTIRIIGGDWKRTPLPVLDLDGLRPTPDRVRETLFNWL
GQDLEGRRCLDLFAGSGALGFEAASRGAASVVMVERHPRAAQQLRALKDKLAARAIEIAE
ADALRIAAGLAPRSFDVVFVDPPFGDAAAFARALDLAPNLVAPGGFLYVETGAPLDPSAH
ASLAGWQVARRGKAGAVHYHLLQRENDE*
>GCA_000452945.1_BPMSHR5858_genomic_03928_1 ftsY_1
MFSFFKRFKKAPDAAPAEPQQAPDAPRDASRAPASPDAARVDTPAAPQPRDERSENHDRP
EHDETPTAEHSRSGAQPAARATPPASAAASEAPVGQPRAAAARASEPAPSVVMTVTPSSN
GRHGVVETVEIVPPPAPEPAAKKSWIARLKSGLAKTSSSITNVFVNTKIDDALYEELETA
LLMSDAGIDATEHLLDALREKVRAGRLTDPQQVKDALRGLLVELLAPLEKSLVLGRAQPL
VMMIAGVNGAGKTTSIGKLAKHLQSFDQSVLLAAGDTFRAAAREQLAIWGERNNVTVVQQ
ESGDPAAVIFDAVSAARARNIDVMMADTAGRLPTQLHLMEELRKVKRVIGKAHDGAPHEV
LLVIDANTGQNALAQVKAFDDALGLTGLIVTKLDGTAKGGILAAIARQRPIPVYFIGIGE
KVEDLQPFSAEEFADALLG*
>GCA_000452945.1_BPMSHR5858_genomic_03930_1 nasD_1
MRKMKLVMVGNGMAGVRTLEELFKLAPDLYDVTVFGAEPHPNYNRILLSPVLAGEQTLEQ
IVLNDYAWYASHGVTLHVGKKIVKIDRVKRVVIADDGTEAAYDRLLLATGSNPFMLPVPG
AQLDGVLGYRDIADTQAMIDAAARHTHAVVIGGGLLGLEAANGLKLRGMDVTVVHLAPTL
LERQLDAKAGGLLRASLEARGLTFLMPKQTQALVGDEHGRVRAVHFKDGGECKADLVVMA
VGIRPNTALAESAGLYCNRGIVVNDTMQTYDPRIYAVGECVSHRGIAYGLVAPLFEQAKV
AANHLAQFGIGRYTGSVTSTKLKVTGIDLFSAGDFLGGDDTEDITLSDPIAGVYKKLVIK
DDRIVGACLYGDTADGAWYFKLLREGRNIADIRETLMFGETSLGDTGHSGATRAMTMADD
AEVCGCNGVCKGTIVSAIKEKGFFTLDDVRRHTKASASCGSCTGLVEQILMSTLGGDYSA
SPKAKPVCGCTDHTHAEVRAAIREHKLLSVPEAMRFLEWRTPNGCSSCRPALNYYCISTW
PHDAKDDPQSRFINERAHANIQKDGTYSVVPRMWGGVTSADELRRIADVVDKYAIPTVKV
TGGQRIDLLGVKKEDLPGVWRDLGMPSGHAYAKALRTVKTCVGSEWCRFGTQDSTLMGQQ
LERALWRMYAPHKVKLAVSGCPRNCAESGIKDVGVIGVDSGWEIYVGGNGGIKTEVAQFF
CKVKTHEEVLEYAGAFLQLYREEGWYLERTVHYLERVGLDHVKARVLGDAENRRALWDRL
QFALKDEPDPWHDTQDAKVDLRQFIPIAVAPADA*
>GCA_000452945.1_BPMSHR5858_genomic_03932_1 group_1892
MTVDTLAQPIAPSTPGAVRETRSTCCYCGVGCGVVIETRRDANGRDTIVGVRGDPDHPAN
FGRLCSKGSTLHLTATPERYAQTRATHPELRASRDAARTRVSWDDALEHVAQRLARTVER
HGPDSVGFYISGQLLTEDYYVFNKLAKGLVGTNNIDSNSRLCMSSAVVGYKKTLGADAPP
CSYEDLDHALTVLIAGANPAYAHPILYRRLEAARARNPQMKVIVADPRRTDSASDATLHL
ALQPGTDTMLFNAMLHVLIWDGALDSAFIAAHTHGFDALRDAVRDATPAAAAQVCGLRAD
DIVQAARWFGAGPSLSLYCQGLNQSSSGTAKNVALINLHLATGQIGKPGAGPFSLTGQPN
AMGGREVGGMATLASAHRDLANPAERAEIARLWGVPDLPAKPGLTAVEMFERLAEGALKA
IWIVCTNPAHSIPSQALARAGLERAEFVVLQDAYAHTGTAPYADVLLPAATWGEKEGTVT
NSERRISRVRAATQPYGDARADWRIAADVGRRLEARLAPGRPTLFPYEDAEAIWNEHRAT
TLGRDCDIGGLSWPLLARDGPQQWPFPAGATQGLARLYTDGRFPSADGRARFVPTPYQPV
AEAVDARYRFALTTGRLRDQWHGGSRTGSVAKLFAHAPQPCVELNPGDCARLGLGAHDFV
HLTSRRGSALMPARADDAVAPGQAFAPMHWGDEYVSGVAGNRAAMGINALTTPARDPHSQ
QPELKHAAIKLLKADLPWRWLIAARVPPGELLARQRAARAFFARFPYASCVPFGTDAYAG
LAWRAAAYEPAPAELQREIEALFELPAQAADVMSYGDARRGTGRRVRIVDGALVAFSVAG
AAGATEDSAAVLRDYVDTGASVGALGRLLLFAGRVPLQTAPARGRTICNCVGVSEREIGA
VLAADAPSAAPVERLANLQERLKCGTQCGSCIPELRRLIAETATAAAPPPR*
>GCA_000452945.1_BPMSHR5858_genomic_03957_1 puuC
MDKTTLADWQDKAATLAIEGRAFIDGAYRDAHGGKTFDCVSPIDGRVLAKVADCGAADVD
AAVAAARRAFDAQAWAGLNPRERKAILLRWAALMRAHLDELALLETLDAGKPIGDTTSVD
VPGAAYCVEWFAEAIDKVGGEVVPADHHLVGLVTREPLGVVAAVVPWNFPILMASWKFGP
ALAAGNSVVLKPSEKSPLTAIRVARLAHEAGIPAGVFNVVPGGGEPGKLLALHRDVDCLA
FTGSTGVGKLIMQYAGQSNLKRVWLELGGKSPNIVLPDCPDLDRAAKAAAGAIFYNMGEM
CTAGSRLLVHREIKDAFVEKLVAAARAYKPGNPLDPNVSMGAIVDAIQLERVLGYIEAGR
AEARLLLGGARVNEASGGFYIEPTVFDTAPDTRIAREEIFGPVLSMITFDSVDEAVRIAN
DSEYGLGAAVWTANLTTAHELARRLRAGTVWVNCYDEGGDMNFPFGGYKQSGNGRDKSLH
ALEKYTELKSTLVRLR*
>GCA_000452945.1_BPMSHR5858_genomic_03959_1 group_1894
MAYAIKADVSDPRAKTMTFVAQKTMYGGKRIARGDAVYVFASENEGGQGLVARGVVTDAR
AVAKTPGVARQTPRVSVAIACAAFAVARPGRGELERFADWRDGRPQTELNFKRYRQATNK
IVGISNEAAAFLDTFLR*
>GCA_000452945.1_BPMSHR5858_genomic_03967_1 pqsD
MRPNLEIAGLGHALPQRIVDNDEVARVIDTSDAFIRTRTGVRHRRYLAPGQQLADLACPA
AERAMAEAGVAPADIDLLIVNTLSPDHHDPSQACYMQPRLGLREIPCFDIRAQCSGGLYG
IEIARHFVASGLYRNVLLICAEALSRRIDTSNAGRNLSILLSDGAAAMVLRAQPDATRGL
IDLALGADGTQFDLLCTEAPGAKRPNFIDADDIAAGRHYFRMRGADMFADATRRIVDACR
RMLDKHRLALGDIGLVVPHQPNLRIIDAVIEQLGLPRERCMISVEDLGNMASAAFPVALS
LAREQGRMPAGQLNLLVTYGAGATWACALYRS*
>GCA_000452945.1_BPMSHR5858_genomic_03969_1 menE_4
MTTTNLPKTINELLRVRAATRGEKVAYLFLSGQPDDEQRQSITFAELDRTARRVALLLQR
QSVGVGERVLLLCRPGLDYIAGFLGCLYAGAVAVPAYPPRNRQHASRVVGIVESAGAKAI
LSTGDDFARCTKLLEDTAASHVALLDLDAAKPLDATLEPADVAPSHVAFLQYTSGTTGKP
KGVMVTHGNLIHNLALIGQWMGYHEESTMVSWLPPYHDMGLIGGILTSLFGGFRCVLMAP
ERFIQHPFLWLRAISDYRADVTGAPDFAYRMCSRRVPDEQLATLDLSCLKTAYSGAESVR
YGTLAEFAQRFARTGFDSERFKPCYGLAECTLLVAGRSAPRPLRTVCVDQAALQQQKVVI
RRAFEGLAPQLDERDGERVLVSVGVTIGEQRVVVRDLNTNERCADGEIGEICVAGPSVAP
GYWQQYEQTLATFQRGIGGETGQAFAGTGDLGFHHRGDLYVTGRLKDMIIIAGRNYYSED
IEYAVIGSRPELVPNGCAAFTVDAGDEERLVVVAEIERTHRKGDLDALLKGIREAIWLRH
DLSPGAVLLVSPGSVPKTSSGKVRRGECRKRLGDGELTVLARWDADDCIAAATRGDTAAN
APSNVPSNVPSNVPSSAPASAAATAPAAPAEKTPAADAAGAASPNAAKVEQLKDWLRHYA
RTRIDSRTIDERRTIPPHIVLDFGNEGLLGMQIDRAYGGLGLTNHEMLQVVSQLATIDST
LAFFVGLNNTLGIRPIMLHAQPALRDELLPLLATGRTLAAFALTEPDAGSNARAIASVAQ
RADGGHWVVSGRKSWSGSSAWAGVINVFAKQADGAGMVGLALRQGTPGLRIGAEDLTMGV
RGMIQNTLHLERARVSDACRLGAPGQGMAVAQQTMNFARLGIGAVCVGAMKRCAQLMHRY
AARRRIGTGLLLDHPLSRQRLGDLRHRIDALSALIEQLAADFDAGRDAPEDGLLIAKILG
SECLSQTSDELMQMLGGRGYIETNVAPQLFRDARLPRIFEGPTETLLAHLGSRLLNGGDD
LLGYLGERSGALALAAELRGLGEQLLEDGLANADALGGAAHAANWVNYWLGSVAQWALLL
AVVEQAAKRRGVDGATLEWAQSQYELALEAAQRQVGRRRVLASAAQLGEWAQRVEREIGP
IEQTVPGATQRVDPLLRAECDVRSQPSDIGALDDEPPAAGVPPAPVPSPASASAEQADPE
LKREVERWLLTWLGARLRNRRIALTAETTFADIGLDSILAVELTMAFGDAFRTTVDASAV
WDYASIDALATHLAARMDRHAPAQADAASSAPSASSSL*
>GCA_000452945.1_BPMSHR5858_genomic_03970_1 group_19475
MQSLCELATSAGALGAQLENYLRSHADDSCKLLGVPLGMPLRVEIVDAGAPIETRRDERG
ATLRIGADDAQRVMAYTRSCNWANGIVLVPTLTRLVLAGAFDGLKAGEPRAMRAFAATRK
ADPTRNLGLLWGALNLLGLAGWVTLGSGDEHADYALTPAGARVVECVNAQRPLFTRLADA
TAMLQHLHALCHRRRENADESARYAALVRICIDGWPLPAARDDLERHADAQLRTAMDGLL
LGPTWVALDMPVFEKHGAQQGKVAASFFEVFDAHREGVAIGDAWPHADRVALDAAWALLR
HAGVADSEAGGARVRLNETGRIHRPIAAPYAGLAASYLRSYALLDELLFGNPDPLDVDRD
GHIDRVMNVYASSGAGSGPASKEITTKIIRRLFDETPLDQQPAGISDMGCGDGSALRRLA
QYVIESTRRGRHLADHPLIVIGADYNESARSRAADTLAALSDVPGVHVRVIDADISRPDR
YDEAVAASGLTVKGSDGGPRAARLSDLLHTFMFLVHNRRLGVRSGEAADAILERHLRQVD
RAHLRGVVERYYPGQLTVSDDAALPVALDTIKRAFRVAYSDAEGLVPGYVAAADLIDFVA
RWKPHAKHGFLVVEGHSPWAASLLDDAVAEPGRWTRTEQLPSVFNWGMHFLSRQFMAPFD
EFMLAMCIAGLSPRDAIHGRIHPEGFPGPDLLDAYRFFSIADYVAYDAAHA*
>GCA_000452945.1_BPMSHR5858_genomic_03974_1 ahpF_2
MLDAHLKTQLKSYLERITRPVELVASLDAGDQSRELRALLDEIAALSPQIFVTERRDAAE
RAPSFSVGEPGKAAGIRFAGMPMGHEFTSLVLALLQTGGHPIKLDDATIEQIRALDGDYA
FETYISLTCQNCPEVVQALNVMALVNPRIRHVMIDGALFQDEVEARRIMAVPTIFLNGET
FGQGRSSVKEILAKLDSGASRRAAQALAEKPVFDMLIVGGGPAGAAAAIYAARKGISTGV
VAERFGGQVLDTMAIENFVSVQETEGPKFATALEQHVKQYDVDVIDVERADRLVPGRVHE
IRLASGAVLKAKTVVLATGARWREIGVPGEREYRNRGVAYCPHCDGPLFKRKRVAVVGGG
NSGVEAAIDLAGLAAAVTLIEYGEQLRADDVLQRKLRSLANVTVITQARTTELTGDGAKL
NGLVYEDRRTGEAKRIALEGVFVQIGLVPNTEWLKGTVELSRHGEIVVDARGATSVPGVF
AAGDVTTVPYKQIVIAVGEGAKASLGAFDHLIRSEAHAVQAADIEASANAEAMAA*
>GCA_000452945.1_BPMSHR5858_genomic_03982_1 group_1283
MSIHVALHHVTRYRYDRLVRLGPQIVRLRPAPHCRTPVLSYSMKVEPAQHFVNWQQDPFA
NYMARLVFPERTRTLEIAIDLVAEMSVYNPFDFFLEEHAQTFPFDYGDALRRELAPYLAC
DSATGACDAFRSYIESIDRAPAGTVDFLVALNRRLQHDIRYVVRLEPGVQTPEQTLALAS
GSCRDSGWLLVQLCRHLGLAARFVSGYLIQLTPDVKSLDGPSGAEADFTDLHAWCEVYLP
GAGWIGFDPTSGLLAGEGHIPLAATPQPTSAAPVEGLVDECEVQFEHEMRVTRIYESPRV
TKPYTDEAWQRVLRLGAQVDAALNAGDVRLTQGGEPTFVSIDDRDGAEWNTDALGPTKRG
HATSLVQKLRAEYGVGGFLHFGQGKWYPGEQLPRWALSIFWRADGQPVWRDPARFADERE
PSAYTSADAERFIRALAARLGLAGDYVTPGYEDVWYYLWRERRLPVNVDPFDARLDDELE
RARLRKVFSQQLDSVIGYVLPLKPLAPNPALAGPRWETGPWFFRDERMYLVPGDSPMGYR
LPLDSLPWASRGDYPYLVERDPFAPRDALPDADALRARHAGGGLGAPRDPGAAARDADAP
ARPVMQARAGERAFARPAGEAARFPQRFESAGWITRTALCVEARGGVLYVFMPPIAALED
YLDLLAAIELTAESLDAKLVLEGYPPPRDARLKMLQVTPDPGVIEVNIHPAHDFDELVQH
TEFLYDAAYRSRLSSEKFMVDGRHVGTGGGNHFVLGGATPADSPFLRRPDLLASLIAYWH
NHPSLSYLFSGLFIGPTSQAPRVDEARDDQVYELDIAFAELRRNTLRAGEDMPPWLVDRV
LRNLLIDVTGNTHRSEFCIDKLYSPDSATGRLGLLELRAFEMPPHARMSIVQQLLLRALI
ARFWRAPYTAPLARWGTALHDRFLLPSFVRMDFDDVLTELREAGFGFDAAWFAPHFEFRF
PLFGQIAARGVALALRGALEPWHVMGEEGAIGGTVRYVDSSLERLEVRVSGLNDSRYVVT
VNGRALPLQPTGTAGEYVAGVRYKAWAPPSALHPTIGVHAPLTFDIVDTWMRRSLGGCRY
HVAHPGGRHYDTFPVNAYEAESRRLARFVSMGHTPGAMTVEPAAPSREFPFTLDLRHG*
>GCA_000452945.1_BPMSHR5858_genomic_03984_1 group_1350
MGMTTRPQANAKARARKAAARGEPARGALLRVTHDTRYRYAARVESAQHQARLRPLETPR
QRVIEFSLEIEPRAEGLVVDIDSFGNERASFALNQPHEELFVRSRSVVRVTPPALAAGKR
GEPPPAVAAPRDGCASAWEAVRERLTFRAGRPFDPASEFVFASPHVACHSDLAAYAAASF
TPGRPLVQAAWELMRRIHADFAYAPNSTDVGTTALDALALRQGVCQDFAHVMIGALRSLG
LAARYVSGYLLTQPPPGQPRLIGADASHAWVEVYDPAWPEDGGWLPLDPTNDRAPGDDYV
MLSIGRDYADVTPLRGVIRGGGADQVLTVGVTVEPLDSAS*
>GCA_000452945.1_BPMSHR5858_genomic_03985_1 gabR_6
MARGKTAIPLDLPRPPGLSVRRAQSKQDCIGDALRDAILRKLLPGGSPLPSTRTLAARWG
VARGTVEAAFDRLCAEGYIARAHGSGTRVSAVVPDAYLSPARHPAKPAPHARGKPADALA
RERAAEADGSVRAGVPFVARLADPGLLPMRTWKRHIGIGLSEATHAALCALPPQGAEPLR
ARIADYLGQYRGIACDAADIFITTGIRHSLDLIARTIAGSGSRIAIEDPGYRTARRIFEL
AGAESICVAVGPEGIDTHALSAHRRVAAAYVTPAHQSPLGVTMSISRRLDLLDWAHARRT
WVIEDDYDGEFGYQSAPLPALKSLDVRERVIYCGSFNKTLFSGLRVGFMVVPAALRAPIS
ALVQAVGRPVGTTEQLTLASFIGSGDFAKHLRTSRHAYLQRRDAVLEQLRRYADGQYAIS
GEHTGFHFVLWLGRGIDERTFAEEAANIGLALEPLRAFCTQAALPAGVVVGYSALSVAQA
RYAGKKLGQLLAGVRRDAAA*
>GCA_000452945.1_BPMSHR5858_genomic_03986_1 bcr_3
MNTTETNTLSDRGWLSLLMLVVCLPRMTIDAYLPSLPAMADALHGTDAQMQLTLTMYMAG
YALSMLLSGPLSDRYGRRPVLLAGMLVYLGASVACATATSVQGIVVARIGQALGGCCGTV
IGRVIVRERFGTAMQAAMLSRISAGMALSPVVAPLAGHVVAQWLGWRGVFASLAFGGAAA
ALMVHRFLPETRVRAAASERGAGLVKTYLSLLRDRRFVRYSLAIGFVYCTYFPFIAESSA
LFQRTLRASGAVYAAIFGITVLGYLIGSSVFRRASGRFGIDAIIASAAFVNLVGAAALWI
GTSVAPPSVWSIALPMFVVMVSVGVSIPACQFAVLQPFAKLAGTASGLFFFIQMAMTAGC
GGMLSRLSDGSARPMAVVTAASSAAFMAVVMLVRTRGVNGRRG*
>GCA_000452945.1_BPMSHR5858_genomic_03987_1 yeeA
MTSTLQSRLPGLVRNALRPVLDPYRRYRHAKLIHAARVALSVLASIALTTGLHVPHGEWA
TITVLIVIGGLQHHGNIRKKAAERALGTLIGAIAGLSLILLQTTVHLSPLTFLVMSVACG
LCAYHAIGKAGYIALLSAITMVIVAGHGDNEIADGLWRAVNVLVGIVIALAFSFALPLYA
TYSWRYKLADALRGCAAVHARIAGERYVSDNERLKDMAALNALLVQLRSLMPSVSKEISV
SMPQLEAIQRSLRLCMSALEILSSRQPGEDDAAGRRFVQIQMKADNRRIQEMLVGAARAL
KFGTPGRLGPLHGPAAAVGENVPPADLSGYVSLTAKLSHEIEQLRQRLRDTAPQWNI*
>GCA_000452945.1_BPMSHR5858_genomic_03990_1 phnW_2
MLLLNPGPVTLTERVRQSLLQPDLCHRESEFFDLQDEARARLVAAYALDPAEWSAVLMTG
SGTAAVESMIAALVPENGKLLVIENGVYGERITQIATQYRIAHDVLKHDWMQAPDLARVA
EKLDADRAITHVAVIHHETTTGRLNDLDALAAVCRARGVRMLVDGVSSFGAEAIDFAGGD
IDALAATANKCLHGVPGAAFVIVRRRALASAASRTYYLDLGRLAKLQDARNTPFTPSVHA
YYALVEALREFDEAGGWRARHARYAALAEQVRAGLAARGMPLVLPDGESSVVLRAYRLPA
GVAYEQLHDGLKARGFVIYAGQGGLSSALFRVSTMGAIEPADVERLLDGFTALAR*
>GCA_000452945.1_BPMSHR5858_genomic_03993_1 group_19605
MRAIILAAGLGLRLQLPPEAQFPKCLLRFDDTSLLERHLRVLDAAGVDEIVLALGFQSEK
VEAELARLNRRAEIVLNPRYDLGSVLTVHTAADALTRGGDVLLMDADVLYDEAILHALVA
DPSRAIDRLLIDRDFEAGDEPVKLCVKDGVPVELRKQLAAGLEYDTIGESVGFFRFSEGT
ARRLAEIVAGYVDGGRANMPHEEAVRDLLLEGGRTFDIADVTGSPWIEIDFPGDVARARD
EVLPHIQQRTIGAIR*
>GCA_000452945.1_BPMSHR5858_genomic_03995_1 group_19534
MMMDLEAPAAHGSADAAVAARVAVLDRARLRGEFARQDAFLYLDEFLPKDLAGRVADCAR
ALVPEINRNYLPGHKQGGSVSRHTIDEKAPLIAELYRSKALVGFLETLTGDKLLPSPDDD
PHAYALYYYTKPGDHIGWHYDTSYYDGRRYTLLFGVIDDSTSRLDYALHTRNPAVPDEPG
SVQIAHGGIVFFDGDKLRHRVTPLGENEIRVSLTFEYVTDPGMRPWKRFVSNMKDAIAYF
GFRQVFKQTAARRSPRS*
>GCA_000452945.1_BPMSHR5858_genomic_03997_1 group_19125
MPTSRILAPSLRSLVRTADEAAALIGPNMTVAMSGFTGSGYPKAVPAALAARIAASHARG
EDFRINVLTGASTAPELDGALAKADGISMRLPYQSDPTLREKINAGELDYQDIHLSHVAQ
YAWFGLFGELDVAIVEVAGIREDGRLIPSASVGNNKTWLDQAKRVILEVNARQPLGLDGM
HDIYYGTALPPNRKPIPLVKSDDRIGEPYLRCPAEKIVAIVETDAPDRSNAFAAPDETSK
QIAGLLIDFLRHEIKRGRLPEHLLPLQSGVGNITNAVLAGLGEGGFSNLSAYTEVIQDGM
LDLLANGTLSFASATALSLSPDAVKRFADEIDTFRSKIVLRPQEISNHPELVRRLGIIAM
NGMIEADIYGNVNSTHVMGTRIQNGIGGSGDFARNGYLSCFMSASTAKGGAISRIVPMAS
HVDHTEHDVAVVVTEQGLADLRGLSPKQRARKVIANCAHPDYRPMLEDYFERASRESFGK
QTPHLLGEALSWHERYVRTGSMKA*
>GCA_000452945.1_BPMSHR5858_genomic_03999_1 group_19504
MVRQKDGQKFIGESRAPRVQIEYDVEVYGSQKKVELPFVAGVMADLSGDNVEPLGPVEDR
RFQEIDVENFDERMAQIAPSLSYHVKNVLTNDGTLIPIDLTFTSMESFEPADVVKRIPEL
STLLEARNRLKELLTYMDGKAAAEDVIQELLKSPQWANEADAAEQSGGAGEGGDHQPEEG
AK*
>GCA_000452945.1_BPMSHR5858_genomic_04000_1 group_19072
MSMQQLESSAEKVVVDQNNVNEDLKDILRRSFRPRTNEAAEAVQNAVETLLTYARRSRVV
VREDVAQTIEQLVAELDKKISEQLTLVLHNKRFQSLEGAWRGLHYLVSNTDTSENLKIRY
LNISKADLGKTLRRFKGVVWDQSPIFKMIYEQEYGQFGGEPFGCLIGDFYFDHSMQDVSI
LTEMSKISAAAHAPFIAAAAPGLLQMDDWSELSNPRDVSKIFTATEYAFWRRLRESNDSR
YLALTLPRFLARVPYGPKTQPVEEFGFEEKVDPNRAEDFCWANSAYAMGANITRAFKTYG
WCTKIRGVESGGAVEVLPKFVLPSQDREVDLHCPTEIAISDRREHELSESGLMPLVYRKN
SDTAAFIGAKTVHRPAIYEDDDATANSNLSSRLPYIFATCRFAHYLKCIVRDKIGSFKSA
EDTQRWLNDWLMNYVDGDPSISSEVTKSQRPLSAAEVVVDEIPENPGYYRAQFFLRPHFQ
LEGLTVSLRLVSKLPSTKHEVTT*
>GCA_000452945.1_BPMSHR5858_genomic_04001_1 hcp1_4
MANALVDYFLQIDGVEGESTDQQYPGLIQIQSWQWAEENSGRWGFGSGGGAGKVEMKDFE
FRMVSNKASPKLFLMCATGEHIQNAKLICRKSGKGQQEFLTISFASGLVSSFRTLGNMPI
SQLGHASGEVDGVLPTDQIRINFAQIEFEYREQRNDGTMGAVIKAGYDLKQNAPI*

>GCA_000452945.1_BPMSHR5858_genomic_04004_1 group_19038
MTPLERVSRALTPRRTFFELMRRVEALQRRHDGRSARKRRMPKWLRIEQPAQMHFASTEV
ERVQVTLARFVEDDDHPQVTVAQRHFGLFAPYGPLPLHVTEHAMQEKRFERNAAFERFVN
VACGDLAWLHYSAWSSMHPVLGYERARNPFVERVTALADACRAQQDDGEPYGRHALACRR
AFPGIYCAPRRSLADLQRLLRAYFGVALQVVPRHGRWVPVPAAASHARRLGGWRLGARIW
DVQHSIEIVVGPIEADEFYRWQRRAAAVMALSAVVTDFVDGRIYPVIKVQVWTRPELAGR
VGCMRVGVDAWSRPNRALRTLTVFESFRD*
>GCA_000452945.1_BPMSHR5858_genomic_04005_1 clpV1_4
VNRERIFNCLGRTTYAALVDATALGRSRRHAFIDLDHWALCLLQREQSDLARLFELFGSD
AGEAKRRMEKALDGFDVSGDSLRDISSSLERSVGPAVIWSQIAARAGKVRSGHLLLAWLD
EDLTRRWLQQRVPSGITSVALDDVVKRYEALAAGWPEADEAPAALDGAALGAQAGEAGAD
GTGDALAKWATCVTEQAARGELDPVVGRDDELRTVIDILSRRRQNNPILVGEAGVGKTAV
VEALAQKIHAGAVPPGLVGAQVWALDLARMQAGAGVRGEFEQRLKSLIDAVIASPAPIIL
FCDETHTLIGAGGAAGTGDAANLIKPMLARGQLRMVAATTWSEYKQYIEPDAALVRRFQA
VAVDEPSDDAAVDMLRTIAPRFAAHHGVRIVDSALRGAVELSRRYLPARQLPDKAISLLD
TACARVAMSQSCAPAELERLQHQAFAIGQTLDWRASDRRMGVRTPGDEAELEGRQASLAQ
QAATLETVVDAQRDEVRAWLARLNDATPQAADGDGAAFAARIGANRWVRPWVDEHVVSEV
LAEWTGVPVAQLAQDDAQRVVELEAALNAGIHGQTGAMRSIAQALQVSHSGLNDPRRPLG
VMLLAGPTGTGKSQAAAKLAELLFGGERNLLQFNMNEFQEAHTVSTLKGAPPGYVGYGKG
GRLTEAVRKKPYSVLLLDEFDRAHPDIHEVFYQVFDQGWMEDGEGRRISFRNCLILLTSN
LGEAEIEAACKADPRISQAKLDKLVGERLQGRFSPALLARIQLVAFRTLDVDALTGIATQ
ALDELGERLAQNDLQWRADEGVASWIAHAVSQHPANGRAVRDLLRQHVMPAVARGVLVAR
AEGRALQTVRLTANEKLSLVFDEDAWELSGTDAASLGEQAQAVAMAREAEAVAAAVAARE
AHGARGTNEADGGGGAPHAAKADAHVDSDDERPGGAPHPDETASANAGTTGEPSCV*
>GCA_000452945.1_BPMSHR5858_genomic_04011_1 group_19299
MRLKFKIFLLAIVPFLVTIAAIGLGVRQQATVLARTQHATIEAAYLSSKEIELKHYVDLA
TSAVAPLYEAGRANARDDALLRLQALAMLQRMEFGPDGYFFVYDLHGNALMHPREPERVG
HNYWTLRDPRGSPTIQQLIAAASRGGGYVRYVWQRPSTGKLAPKLGYVVSLPRWGWMIGT
GIYLDDVDNALARIDARASANIERTMTWLSAIALAGAAVIALCALVLNVSESRSADAKLK
RLAQQVVESQEQERARLARELHDGISQMMVSAKLMFESALERLAHLPARENAAQAALSKG
IARLGDTLREVRRISHALRPTMLDDLGLAAALDQLVRELGAETGVELGFTQVDHSGAPPL
PAPVKTALFRIAQEALTNILRHAHATRAAVTLELSPREVALTIADNGRGFDAGRAQADAR
GGIGLRNMRERLDALGGTLEISSQIGHTVVAARVPLRGTDSPGTPR*
>GCA_000452945.1_BPMSHR5858_genomic_04012_1 liaR
MTPASSTPRTARLLLVDDHPLVRDGLKMRLEAAPGLAVVGEAGNADEALTLAEALAPDLA
LMDIGMQGMNGIALAGVFHERFPAIRVLMLSMHDNVEYVTQAVRAGASGYLLKDSPATEI
VRAIGAVLAGQTFFSEGIAARMIHASAEASPLDRLTPRERDILDALAQGLSSKQIAQRNG
LSVRTVETHRLNLKRKLDIEGQAELIKFAVEHRRR*
>GCA_000452945.1_BPMSHR5858_genomic_04040_1 group_19232
MSETVLTAEQRDALQEIANLAMGRAAARLALLLGAFIELSVPRVRVVRAADVGRVLRDMT
GIHDNVTAVRQGFRSDIKGEALMLCRSAGVGQLVSLVDGAYADDEFGTVTQAELVFDMAN
VLMGACVSSILDELGRTPVFSPPGLLGEDIALDDVFQPDVLAWSETLLLEVNFGVADDGA
RTHFVMLLAEDSIRHMSDALDLLLSSL*
>GCA_000452945.1_BPMSHR5858_genomic_04041_1 spo0A
MPLPIVIADDSLLARKLLTKALPADWDVDVNYASNGREALALYRDGKASVMFLDLTMPDM
SGYEVLETLRGEDLNTFVIVVSADIQPMAQARVRELGAAAFVAKPVTSEALLPILKEYGL
YV*
>GCA_000452945.1_BPMSHR5858_genomic_04046_1 yfgD
MITIYHNPRCSKSRETLALVETLNPTGEAINVVEYLKQPPTAAQLKVLHRQLGRPVRDMI
RDNEAPYKDLNLADAALSDTQLYDAIAEHPILLQRPIVVHGDKAAIGRPPEAVRSLFE*
>GCA_000452945.1_BPMSHR5858_genomic_04051_1 smc_1
MTLDEIRQSIREELDALRVSGARRQDLSLHACKRLFFDLGIRPSTANVRELTQTGSASDI
PKDIDHFWERIRSASKVRLEGAALPKPLEEKAGALLGALYEEALKQARESLDGERQQASS
EVAAAEQKYRDALIRQEALEAAVARSDARAEQLQSRLTDVEVQLASASTQGLAHQDSLQG
LIRRMENENALLNQRLEAEQSQNAALRERIDALHAELRENTEHYAQQIKDAIAEAERRVK
PMLVELDSLRSMASTYQQGLRDVNRKEFDFLQQLSAAKARADRLDAQLREQGDELTAALN
EVQILRASQGMTPEIAALIRRLANAGNLDAAAFDTIGTSLDQHVTLPARCPKCGDGEPEL
SHDAHGYELLCPECEHASGTQPSRFAAAACFTRTT*
>GCA_000452945.1_BPMSHR5858_genomic_04058_1 group_19560
MDTKPDLAPAAGARERLLDAAETLIYAGGIHATGVDAIVKRSGAARKSFYAHFESKEALV
AAALERRDARWMAWFVEATKKRGKTPRARLVGMFDVLREWFEQADFHGCAFLNAAGEIER
ADDPIRVVVREHKARLLAFVREQLDAHAAQAGADRRFVARLARQWLVLIDGAIGVALVSG
DASAARDARAAAELLLDATLPNRSS*
>GCA_000452945.1_BPMSHR5858_genomic_04059_1 group_19644
MSDSTEIRPPVPPFTRETAIQKVRAAEDGWNTRDPERVSLAYTPQSKWRNRAEFATGRAE
IVALLRRKWTREIDYRLIKELWAFTGNRIAVRFAYEWRDDAGNWFRSYGNENWEFDENGL
MAHRHACINDMPIREADRLFHWPLGRRPDDHPGLSDLGL*
>GCA_000452945.1_BPMSHR5858_genomic_04060_1 leuB_2
MTESAPVFPHYSIEVDFPDLEAHRTGNAGVDYVHRFDSGVPGPRVMINALTHGNEVCGAI
VVDALLKRGLRPRRGVLTVSFANVTAYERFDPARPDAARFVDQDFNRVWAPAVLDDLSQH
SVELDRARAIRPFVDEADWLLDLHSMHERSAPLIVAGPLAKGTALALRIGAPATVIRDEG
HPEGKRMRDYGGFGDPASAKNALLVECGQHWEARAVAVARDSTARFLMASGIVDERDLPA
DWFLPLATTMRIVQVTQPVVATSRDFRFADAYTGLEHFAEAGAVIGWSDGKPVTTPYPDC
VLVMPSLRQLHPGVTVVRLGRVEREVAQRAARGTRGKC*
>GCA_000452945.1_BPMSHR5858_genomic_04061_1 dmlA
MSEKVYRIAVIPGDGIGVEVMPEGLRALDAVSRRFGLRFAYEPIEWASCDYYAKHGQMMP
DDWKTQLSGMDALLFGAVGWPETVPDHISLWGSLIKFRREFDQYVNLRPARLFDGVPCPL
AGRKAGDIDFMIVRENTEGEYSAVGGTMFEGTEREFVVQQAVFTRHGTERVLKFAFELAQ
RRAKRLTVATKSNGIAISMPWWDARAAEMAARYPDVAWDKQHIDILCARFVMQPDRFDVV
VASNLFGDILSDLGPACTGTIGIAPSANLNPERNFPSLFEPVHGSAPDIAGKHIANPIAM
IWSAAMMVDFLGAGTGPAREAHDAIVAAIEHVLKHGPRTRDMGGRASTREVGDAIVAHLL
G*
>GCA_000452945.1_BPMSHR5858_genomic_04062_1 dmlR_19
VNNSPNLDDLRVFSVVVRLASFSAAAEQLAVSPAYVSKRVALLEKQLGTRLLHRSTRRVA
VTEAGERVYAWTEKILDDVDHLVEDVSTTRSVPRGTLRISSSFGFGRHVLAPALLGFNER
YPQLNVRLDLFDRLVDVAGEGFDLDIRIGDEIADHLIAKRLATNYRVLCASPGYLARYGT
PRQLADLGAHQCLAIKERDHPFGVWRLTVRGETSTVKVGGALSTNHGEVAVQWALAGRGI
VLRSIWEAGPLLASGELRRVLPEASQPANVWAVYPARLAASAKVRVCVDFLVDAFAHLNE
RANGG*
>GCA_000452945.1_BPMSHR5858_genomic_04071_1 astD_2
MTHPLFTKHEDTLKHALSTIETRGYWSPFAEMPSPKVYGESANTDGEAAFKAQLDKPFEL
DQPASGGTVGAERSPYGFALGVRYPKSTPDELIAAAAQAECAWRKAGPTAWAGVCLEILA
RLNRASFEIAYSVMHTTGQAFMMAFQAGGPHAQDRALEAVAYAWQELQRIPAEAHWEKPQ
GKNPPLAMRKRYTIVPRGTGLVLGCCTFPTWNGYPGLFADLATGNTVIVKPHPGAILPLA
ITVRIARDVLREAGFDPNIVTLLATEGNDGALVQDLARRPEIKLIDFTGSSQNGTWLERN
AYQAQVYTEKAGVNQIVIDSVDDLKAAVKNIAFSLALYSGQMCTAPQNIYVPRDGIRTAE
GHVGFDDVAQAIADAVQKLTGDPARSVELIGALQNEGVAARIDEARKLGRILADSQALEH
PAFKDARVRTPLVLQLDVADRAKYTQEWFGPISFVIATDSTAQSLDLAGSIAAEHGALTL
SVYSTDDAVVEAAHEAAVRGGVALSINLTGGVFVNQSAAFSDFHGTGANPAANASLADAA
FVANRFRVVQSRHHVAPKAAPAEAGQTA*
>GCA_000452945.1_BPMSHR5858_genomic_04072_1 echA8
MAYENILVETRGRVGLITLNRPKALNALNDALMDELGLALKAFDADEGIGAIVLTGSEKA
FAAGADIGMMATYSYMDVFKGDYITRNWETVRQIRKPIIAAVAGFALGGGCELAMMCDII
FAADTAKFGQPEIKLGVLPGAGGTQRLPRAVSKAKAMDMCLTARFMDAEEAERAGLVSRI
LPADKLLDEAIAAATTIAEFSLPAVMMVKEAVNRAYETTLSEGVHFERRLFHSAFATEDQ
KEGMAAFVEKRKPVFKHR*
>GCA_000452945.1_BPMSHR5858_genomic_04078_1 hpcH
MKAAVVHRAGERPTYAEFEPPRALPGHRLIDVSASALSRLAQARASGAHYSSTGGFPFVA
GVDGVGRLDDGRRVYFFGPPAPFGALAERTLVPAAQCIPLPDSIDDATAAAIAIPGMSSW
AALTERARLAAGETVLVNGATGASGRLAVRIAKHLGAASVIATGRNAHALDALSSAGADA
TISLAQDDEQVARAFEAHFRAGVDVVLDYLWGASARAALLAAAKAPQQARPVRFVQIGTI
GGAELPLPGAVLRASAITLMGSGLGSIALPRLLNAARAVLGAACEARLRIDTRTVPLADV
DAHWGDTGSTLRPVFTMRAPG*
>GCA_000452945.1_BPMSHR5858_genomic_04085_1 hpcE_2
MRTARVVFNGALHAAEPAGDALVRLDTGRVLAETEVSWLPPVVPRTAFALGLNYADHAKE
LAFSAPAEPLIFLKGPNTFIGHRAQTVRPADATHMHYECELAVVIGREARRVTRARALDH
VLGYTIGNDYAIRDYLENFYRPNLRVKNRDTCTPLGPWLVSRDEVGDASDLALRTTVNGV
ETQRGNTRDMIFDVASLIEYISGFMTLSPGDLILTGTPEGLADTQPGDEVVTEIEGIGRL
VNTIVGEADYYRAG*
>GCA_000452945.1_BPMSHR5858_genomic_04087_1 farR
MNRTFDHRNLAMLLLEARETLMGRFRPILKEFALTEQQWRIIRVLDGEPSHELEAGQIAK
RCCILSPSLTGVLERMERDGLIRRTRAQEDQRRLIVSLTPQSCQLVTQISPRIDEQYRLI
EQRFGADALGEVYAALDKLIDVGGS*
>GCA_000452945.1_BPMSHR5858_genomic_04088_1 group_19449
VLIWKTALSLTWKAGLVSPASAPAAAAHVDRVVLTGATGFIGGAVLVSLVNAGLLDRVVC
IVRACDRAHALARLRAAALRSGLAPYWAERLSEANVIAGELDGALADADAAHIALASHVI
HCAGVASLADARIVNETNVGATLRFARRFAGSRRLQRFVHVGAAFACGLRARGTIREDDT
PARGREIDFAPYTRGKRDAEAQLRALGLPLVVVRPSCVVGHTLLGTQPSASTFWMFRIVH
AARRFTARPMARIDVIAVDDCARALMLLALKPSLAHDTYHVSAGDEAPTVTQIVRAMDEA
VGLDDEPRYALCSPAEFPSIARDVLGRRDAPRERVIRRALQSYAAFAELDHVFDNARVRR
EIDFEPLPFVDYVNECMRTSRGIDVLAQMPRTAAR*
>GCA_000452945.1_BPMSHR5858_genomic_04089_1 group_1899
VTDAQAAATRFAPYSTKGGIDSASMSVAWTYTFSKHWSVDTTLGLTRLVGRYGNSPIVQR
KTQYFGLSALTYRF*
>GCA_000452945.1_BPMSHR5858_genomic_04099_1 iorA
MSTSFVLNGKNLTLDADPAMPLLWAIREDAGLHGTKFGCGAAQCGACTVHLEGQAVRSCV
LPLAAVAGKRVTTIEGLASRPAKAVQAAWVKLQVPQCGYCQSGQIMSATALLEQNPAPTD
ADIDAAMNGNICRCATYTRIRAAIHEAAATLKA*
>GCA_000452945.1_BPMSHR5858_genomic_04102_1 asnB_1
MCGITGFVGPVACAPQTARATLARMTDSLAHRGPDDTGLWIDHDAGVALGHRRLSIVDLS
EHGHQPMLSACGRYAMTFNGEIYNFRALRSELEREHRAPEWRGHSDSEVLLAAFCAWGVD
RALQKAVGMFALALWNRESRVLTLARDRIGEKPLYYGRIGGSFVFGSELKALRRFPGFDA
HPIDRDALCLYLRHASVPAPYTIYEGIRKLPPGTYAQLEDLAEAPAPRRYWSLERAIAAG
RANPFDGDANAAVAQLDHVLRDAIAEQSVADVPLGAFLSGGVDSSVIVALMQTQTAQPVQ
TFTIGFHEAGYDEAGYAKAVARHLGTSHTELYVKPADVLATVPRMPCVYDEPFADSSQIP
TLLVSELTRRHVKVSLSGDGGDELFGGYTRYFLTPRLWGRLRRVPAAVRMRIAAALHALR
PDHADQIASLVQGAWSASAQRDASPRVGDRLHKLANVMTADSRASLYRLLMSSVHHPERF
TLAGHEPQTLIDTPGAWPPGLSFAEQAMAVDTLTYLPTDILTKVDRAAMAVSLETRMPFL
DHRVVEFAWRLPEALRIPDGQDKWLLRKLLYRYVPRPLIDRPKQGFCAPVGAWLRGELSD
WAAELLRPARLRDEGYLDAARVERLWRQHQSGRMNWQHPLWTVLMFQAWLERQRAE*
>GCA_000452945.1_BPMSHR5858_genomic_04103_1 group_19425
MTIADSAPRIEWFSHAELAASRGGFAPRDTQPSVSSFAAPPDGPRRATFDVADCDARLRA
AGFTTLCYGAFELIGQQMLAAYLLRDFAPASFLRAYVEGRLFEVDPRFAEWRQGVSPIAW
RLDELDAHARRTGERRMHALVNSLRAHAMHSGVIFCWPVPHLDLRVAVSLASEAHDGSLD
DRVIGCALALSLDVHRAAQPHLDARVGGARTFVLERDEAAVLERLVHGLSDQEIANALST
SLHRVGAQIRTLEKRFNARNRAQLAYLAARRLRG*
>GCA_000452945.1_BPMSHR5858_genomic_04104_1 group_19123
MSDIAVQEDAAVAASARPASRANPGGTARDAQTIDVVWCGHARRAADADPRFAPRLGVAE
MLASARGTAECGRIATSLLRLMGFATFAYFALEFMRDDAECLYLHEAFTPAAYRGDYVRR
RHHDVDPRMLGTRASSMPVVWDLRQLAREHAARGCDALDGFLRVMHDDDMCSGVMYSMPV
PGTRVHAFASFTAPRRSRDWISSATLEQVLSLGLSVHRFAAPQLIASARERAAERLTPFE
RELLVGIAEGASDKEIGRRLDTSAHNVDYHLRKLRKRFGVANRIQLTYFASARGLI*
>GCA_000452945.1_BPMSHR5858_genomic_04109_1 hsrA_1
MPSALYAFIAPLIVACALFMESVDANIIVTALPAMARDFGQNPVTLNIAITSYVVGLGVF
IPICGWLADRFGARTVFRTAIGIFVAGSLLCAASNNLELFTFARFVQGVGGAMMVPVGRI
IIFRAVPRTELVRAMNYLSVPALFGPAAGPLLGGFITTYLHWRLIFFINVPIGILGIYLA
NRHIANTHELDPGPLDGFGFVLSAAGAALLLMGLTLLDGALVTRGAALAMGATGAALLGG
YVLYARRVERPVLDLRFLRIPTYHASVVGGSLFRIGLGAVPFLLPLALQEGLGMSAFHSG
AITCASAVGGALTRMLAPRTLKRFGFRTVLMYNAAFSGLAIAAYGVFHPGMSTLAIWLIV
LVGGIFPALQFTSLNSMIYAEIAARDAGRATSLGSVVQQMSLGLGVTVAALVLHVSHWAQ
GHPAMVWSDFWPAFVVVGLCSFASIPITRRLSPNAGDEVARGKRG*
>GCA_000452945.1_BPMSHR5858_genomic_04113_1 group_1519
MKNSHNVVNRFIVAASIIIGVVLYSSAWANPQPMHTKQARMPRIPQNLPLSPDQAKYDLP
LSKYDRATLMEPLRRKQSAKPDRRTRPGADCRDMSIMTQYHGTALADYIANLPDYECHYG
LFSIDRAMAAQIFNSENVWAVASRLTQEINRYDATNITLVNLLIYLRAAYFQYDAAQLAD
PIPGLVVWLRPYILQSLSGDALYLENSRAPSTANELMILITNMKDEAYYLPTLKDRIAFY
TASATNPQAAAPLLQRSAAGGFTGLLTVFFYAHQRSGAQPMLDSDATLPETLNRFVTANR
AYLSNTSAAYQLADAARETYRFLRYPSQKPRVKKMIQDMLASTTMTGPDNDLWLAAAEAA
DYGDPGNCADYGTCDYQKRLIEAVLTHRYSCNANVRILAQDMTVPQFQSACQSVAQEEDY
FHRMMKTGHVPVANDHNDTIEIVVFGDYDNYRKYASVIYGISTDNGGMYVEGDPSAPGNQ
ARFIAHEASWLRPEFKVWNLEHEFTHYLDGRYDMAGDFAASTAKPTVWWIEGLAEYISRK
NDDQESIDAVRTNAYRLSDVLQTTYSSGDYVTRAYRWGYMATRFMFERHRADVDAIVSRF
RVGDYDGYADYVAYMGNRYDSEFVDWARGATTTGEPPLPPTKAGH*
>GCA_000452945.1_BPMSHR5858_genomic_04122_1 group_19527
MKIVTYLGFTVAATIISMSNLGAAFAYDDLPKPPPPPEGWVQPPKPSEGTVTRDVENQIA
QRFTAATGGNPNGLLSKQQAKAAGWGLVSDRFSDIDRAGTGYVRLDDVLRFMSERTPQRV
MRMRNATKPNP*
>GCA_000452945.1_BPMSHR5858_genomic_04123_1 group_19576
MAKRIPIGTTVAALATVTLVACGGDDPSSGAVSAGNSPAIVPSVPAATPATPTATAAESE
QDVFARTAPADKVLDDSNVYDTTKDGSIALSKVNEDSSVKRHTITINGKVLPYVARAGHL
VAYRQNGAGKKAEAAIFYTAYTRDALPKEHRPVTFLWNGGPGSASIWLHMGSWGPKRLKS
DAPNMADPTKQPDSFPFEDNAISLLDQSDIVFVDPPGTGLSTAIAPLKNGDLWGTDDDAQ
VVADFITSYTNKYNRQSSPKYLYGESYGGIRTPIVANLLEQAGTSGYVPDPSGKPARVLD
GFILNSPLVDYNSNCDMMGGRVTCEGYIPSYAMTADYFKKSVKRGTRTQEQYLGELRTLA
RTTFQTTYGTYFTNGKPNSQWNAYAASTAGQALLNRIADYTGIPASTWNGSFNYTPRPFR
NALVPGYELGRYDARMKVPNGNSFAADSYIDVAFLNQLKTYFPDFVNYKTQSIYEPLNNA
TIRNWKWKRAGSKYDYPQSITDIQAVLTANPDAKLLILHGYEDIATPGFQTELDLEGVNL
SDRIPVKWFEGGHMIYNTEASRVPLKQAIDSYYKSPTRIADGSLL*
>GCA_000452945.1_BPMSHR5858_genomic_04124_1 pepX_1
VSSRASNEGSQPGTPSQSIPSGTVTTAPAPAPFAPPTPVAEVGAQFERSPTGLPYPKLAT
LYPGHDGPIVDNGMILPWLSMRPPLKSNVMVQTPFDTDQDGKLDRIALRIVQPAEVAEGL
KTPVIVRPSVYYADPTYATQTRAPFLGEAEYLRMGYTIVYADSIGTNQSDGCWSVMDRTE
REAMASVVRWLTNDPGAPGFDAQGKQVAASWSTGHVAMEGVSYGGTLPTMVAATGVPGLE
AIVPVEGISSGYDYFRYNGVIADIDNTVSLGSYMKSQQAFARSSICEPARVAAVTASDDA
TYAYNDFWKVRNTVSLVDRIQAATLIAQGQADNNVKTKNATQLYDALYRAKKPVQLWLHS
RNHDDPAWQKEWQKQIAMWYSRYLFGVNNGVETQPTYVRETPTGDIPVGATLPPDPNDTS
DTLIGHCHSGHNPRDCIPTGELFIKEDAWPKTVDTFYHLRGDGRVGGLLTPSPADGTQAA
SVDLSNATAVTYETKSLANVTRYAGAIRVAMRGRFAPAVSNIKATLSVDGHDVTYGWANP
RFYKGLDVAQLIVPNTDYDFTLEMMPRDFTVLPGSKVMLKLEGYQGTSLVTLDLSHTVLA
MPVVPKAHVAAVMVGK*
>GCA_000452945.1_BPMSHR5858_genomic_04125_1 group_1034
MGDRGLRGGHRRTIAHDTAAGVALPSLAALPPGVLPADLAAASSGALLSVGGGPSQQIDD
HAFTSRNYTALVCSSGLRQEFTTPHCSR*
>GCA_000452945.1_BPMSHR5858_genomic_04131_1 group_1520
MSHVTEIPAADSAAAVAHFAASLAFETDCRDVHDALACGAPDFVLLDVRGPEQYATAHVS
GALNLPHRKIIESKLSEFARDTCFVVYCAGPHCNGAARAALRLARLGRPVKLMLGGVTGW
LDEGFALARGNGASRRQGGVRQRTAGNDPLRNRNN*
>GCA_000452945.1_BPMSHR5858_genomic_04132_1 group_19203
MQMEILIQIVGSVVAIALYFLPAIVADRRHRDDKLTIALFNALFGWTGIGWLLTLYWSLQ
PNATDDFVSDVRLKRRALSMKTFSTGLVERVQRRIAAQEQWADKQGTR*
>GCA_000452945.1_BPMSHR5858_genomic_04135_1 basR
MRVLLVEDDPLIGSGLEQGLKQEGFAVDWAQNADAATLALRTTSYGLLLLDLGLPDKDGL
AVLAALRRRDDALPVIVVTARDALPDRIAGLDCGADDYLVKPFALEELLARIRAVNRRQA
GRAQTVLAVGALRLDPARHQVWRGDEEVALSPKEFVLLHELMREPGAVISREQFEERLYS
WGEEIESNAVQVHIHNLRKKLGHDTILTVRGVGYRIGDGA*
>GCA_000452945.1_BPMSHR5858_genomic_04138_1 group_19075
MSLKFYGHPFSLYCQKVLIALYEYDVPFEWCLLAGDTPQASAEFAARWPLKHMPMLVDGE
RTIVESTILIEYLGLRRPASVRLVPADAGVALDARMMDRFFDCYVSTPVQKIVFDALRPQ

AERDARGVADARRMLDTSYAWLERTLDGRDWAAGGAFTLADCAAAPSLFYADWAHPIDPA
LQRVTAYRERLLARPSFARAVDEARPYRHLFPLGAPERD*
>GCA_000452945.1_BPMSHR5858_genomic_04139_1 group_19422
MPTAVKPIPEGMHSLTPHLICDGAAAAIEFYKKAFDAVEISRLPSRDAGRLMHAAVRIGD
STLMLVDEMGQCGALSPKALKGSPVFIHLYVPDVDAVIARAVEAGAKLTMPPADMFWGDR
YGQLEDPFGHRWSVATHRQDLTPEQIREGMASCVPPAQ*
>GCA_000452945.1_BPMSHR5858_genomic_04148_1 rstB
MNSTQAHRLSPFRYLKWRWLHFRRSWTDTRADRIPSWSRLYLRTYLHLVGLVLVTVGGTI
AVLCSALGPRTVWRALDALPGGALPLAAFVLAVPALAGYRWMRPVWSDLVMVRARAIDFT
GGRFNTRARESRSVIIGPLARTLNALAMRMERLIATQRDLTNGISHELRTPLARVRFALE
SLREPGSAAEYQNAIDSIEQDVSELDELIDMSLTYARLEYSSLQSNLEQTALVAWFDKQV
ADAALLYPSKAIEARVALPAELRVKMDKRLMSYAMRNLLRNASKHARGRISVGLRIRHGN
IEIDVEDDGPGVPPDERERIFDAFVRLDRHTAGYGLGLAITRLVLQAHNGRVAVVDPLRL
PGARFEMSWPV*
>GCA_000452945.1_BPMSHR5858_genomic_04151_1 glsA
MDYQPILERIHTELAPWIGAGRVADYIPELAKVPAERFGMAVVTLDGRIHTVGDARERFS
IQSISKLFACTLAFQLLGDALWERVGREPSGNAFNSLVQLESERGIPRNPFINAGALIVT
DVLCRRFVGAETALVQFMRRLTGVAELDYDLRVASSELAHAERNRAMAHFMASFGNLHMP
PETVIDAYCRQCAIEMNCIELATAALFLANAGVAPVSGERILDASSAKRLSALMLTCGTY
DAAGDFVYRVGLPAKSGVGGGIVAVLPGEMAVCVWSPGLDANGNSVAGVQALEQLTTLTG
RSIF*
>GCA_000452945.1_BPMSHR5858_genomic_04154_1 sfnG
MSHEPDPLKFAYWVPNVSGGLVVSKIEQRTSWDIDYNRRLARLAEQSGFDYALSQIRFTA
GYGAEYQHESVAFSHALLAATERLNVIAAILPGPWHPAVVAKQLATIDQLNQGRVAINVV
SGWFKGEFTAIGEPWLEHDERYRRSEEFIRAVKGVWTQDNFTFKGDFYRFNDYTLKPKPL
RQPHPEIFQGGNSAAARRMAAAVSDWYFMNGNTPDGHRAQIDEIRAAAAAHGRRVKFGVN
AFIIARDTEREARDVLDEIVRHADVDAVNAFGHAVQQAGKAAPEGRGMWADSKFADLVQY
NDGFKTNLIGTPEQIAERIVALKAIGVDLVLGGFLHYLEDVEYFGKRVLPLVRELERRRD
AQPA*
>GCA_000452945.1_BPMSHR5858_genomic_04155_1 macA
MKKSQRRWIAVAAGGVVLVSLALLYHRFVSKDKAPHYLTANVARADLENAVLATGTLHAF
KQVDVGAQVSGQLKSLKVKLGDKVTKDQWLAEIDPVISRNELRQAEANVENLVAQKRSSS
AQFKQAELAFRRQQQMLPDDATSRQDYEAAAASLDVQRASLAALDAQIRAARIKIETARA
NLGYTRIVAPMDGEVVAIVTQEGQTVIAQQQAPVILKLAQLDTMTVKAQVSEADVIRIHP
EQAAYFTILGEPDKRYYGKLRAIEPAPQNFLDTQGGMGGAGGGGSAKTNTAVFYNALFEV
PNPGHRLRISMTAQVNILLGTARNALNIPVAALGAKGKDGAYAVRVVGADGKVATRQVRT
GINNNVKVEVLSGLTEGERVIIGDAAEPAGAGAAASDARN*
>GCA_000452945.1_BPMSHR5858_genomic_04156_1 macB
MTGPLLQLTRVTRRFPAGEKDVVVLDDVSLSIDAGEIVAIVGASGSGKSTLMNILGCLDH
PSSGSYTVGGRETSELESDELARLRREHFGFIFQRYHLLPHLCAAENVEMPAVYAGSAQA
QRRERALALLARLGLSDRASHRPSQLSGGQQQRVSIARALMNGGEVILADEPTGALDSKS
GRDVIRVLRELNALGHTVIIVTHDEQVAAHARRIIEISDGRIVGDRLNPHADAADAAPDA
SGGAQPRRARRLSAGVGRFAEAFRMAWIALVSHRLRTLLTMLGIIIGITSVVSIVAIGEG
AKRYMLDEIGSIGTNTINVYPGADWGDSRADAIQTLVAADAAALAEQIYIDSATPETSRS
LLLRYRNVDVNALVSGVGERFFQVRGMKLAQGIAFGADEVRRQAQVAVIDENTRRKLFGA
NPNPLGEVILIDNLPCVVIGVTASKKSAFGDMKNLNVWVPYTTASGRLFGQRHLDSITVR
VRDGQPSDAAERSLTKLMLQRHGRKDFFTYNMDSVVKTVEKTGQSLTLLLSLIAVISLVV
GGIGVMNIMLVSVTERTREIGIRMAVGARQTDIMQQFLVEAVTVCLMGGAIGIVLSFGMS
FVFSLFVDQWKMVFSAASIASAFLCSTLIGVVFGFMPARNASRLDPIDALARD*
>GCA_000452945.1_BPMSHR5858_genomic_04161_1 mmsA
MNATPSSRKGHHVPTVKLLIAGEFVESHATEWRDIVNPATQELLARVPFSTVAEVGAAVE
AAHAAFAKWKSTPISARMRIMLKFQDLVRANLPQIAKTLTAEQGKTLPDAEGDVFRGLEV
VEHACSVGTLQLGEFAENVAGGVDTYTLRQPLGVCVGITPFNFPAMIPLWMFPMAIVCGN
TFVLKPSEQDPLSTMQLVELAIEAGVPKGVLNVVHGGKEVVDALCSHPLVKAISFVGSTA
VGTHVYRLGSEHGKRVQSMMGAKNHAVILPDANREQTVNALVGAAFGAAGQRCMATSVAV
LVGAARDWLPDIVAKAKTLKVNAGAEAGTDVGPLVSRAAKQRVLGLIETGEQEGARLVLD
GRGVSVPGYEHGNFVGPTIFADVRPEMSVYTHEIFGPVLCVMSVDTLDEAIALVNANPFG
NGVGLFTQSGAAARKFQSEIDIGQVGINIPIPVPVPFFSFTGSRGSKLGDLGPYGKQVVQ
FYTQTKTVTARWFDDDATAGAVNTTIRLH*
>GCA_000452945.1_BPMSHR5858_genomic_04162_1 acsA_1
MTAQAFLDARDFLLRHRTDYDTAYRDFRWPALDEFNWALDYFDAIARGNDKPALWIVDAA
SGDGARYSFAQMSERSARIANWLREIGVARGERILLMLPNRVELWDTMLAAMKLGAVVLP
ATTQLSADDVRERVQIGGARYAIVDEHEAEKFEQPGLDVTKIVAGAPRAGWLALADGYRA
PAEFAPDARTRASDPMLLYFTSGTTSKPKLVEHTHRTYPVGSLSTMYWVGLQPGDIHWNI
SSPGWAKHAWSCFYAPWNAQACVFAFNYARFEPKVVLDALVKYQVTTMCAPPTVWRMLVQ
QPLSTFAVKLREIVGAGEPLNPEIIERVRKAWGVTIRDGYGQTETTCLIGNSPGQPVVPG
SMGRPLPGYAIALLDPDGAHASEGEIALPVGPDVERPVGLMKGYASNPEATAHAMRDGHY
RTSDIALRRDDGYFVYVGRADDVFKSSDYRLSPFELESVLIEHEAIAEAAVVPSPDPVRL
SVPKTFVMLRAGYEPSETLAREIFRFSREKLAPYKRIRRLQFAELPKTISGKIRRVELRR
RELERGDDASSRMPGEYWEEDFAADGK*
>GCA_000452945.1_BPMSHR5858_genomic_04163_1 mmgC_1
MDDLYTEDQRMILDAARAFSAEVLAPNAAQWDRESHLPDEIVAQMGELGFLGMIVPADWG
GSYTDYVAYALALEEIAAGCASCATLVSVHNSVGCGPVLNYGTAEQKARWLRDLASGKTV
GAFSLTEPHAGSEAHNLRTRAELRDGKWILNGSKQFVTNGARAGLAIVFAMSDPDEGKRG
LSAFVVPTDTPGFIVGKPEKKMGIRASDTCPITLENCAIAQENLLGKRGEGLKIALSNLE
GGRIGIAAQATGIARAAFDRARRYARERVQFGKPIAEHQAIAEKLANMATRINAARLLTH
HAARLRTAGLPCLSEASQAKLFASEMAEAVCSDAIQIHGGYGFLADYEVERHYRDARITQ
IYEGTSEVQRMVIARQL*
>GCA_000452945.1_BPMSHR5858_genomic_04167_1 group_19148
MAEHPLPEVHIDELDSFRSVQQLAYRCVETVGDMLYPGITEKQAAQLMTEWARDHGVHDW
LHKPFAWFGDRTAFEGFSGLKHLGGFNLAFFPSNRVLEPDMPVILDVAPVVDGIVADVGY
ATCLGHNPILEQLQDDLIAHRDMIVKLVKERRTLAEVAQEVDALCRRQGVEPRHKAYPFK
VLAHRVAKLHKLSKPRFVARFGLNSTRNLLLEQARAGKEQGWSPLWSIDARSDHAPTPGL
WAVEPHLGFAGVGAKFEELLVITDDDAYWLDDDLPHVRRWQARAASKALVHEQAA*
>GCA_000452945.1_BPMSHR5858_genomic_04170_1 pphA_1
VPIRELISAPITNEQRRRSFKEKLATGRALRFIESHSPISAMISEKVYVAHGASQSEFDG
FWSSSLTDSTLRGRPDIEILDIASRLANVQHIFDVTTKPLIIDGDTGGKPEHFALNVQSL
ERSGVSAVIIEDKTGLKKNSLLGNDVIQHQESIEDFCEKIRVGKAAQITDDFQIIARVES
LILDKGMPDALARAAAYCEAGADGVMIHSRKKDADEVIEFAQRFRACRRDAYLVCVPTSF
NAISFSELARHFSVVIYANHLLRAAYPAMLAVAEGILAHGRTLEIEERCLPVNEILKLIP
GTA*
>GCA_000452945.1_BPMSHR5858_genomic_04205_1 group_1216
MMLSRLTSMSLAAMLVAAGAAAHAETVKIAIAGPMSGSVAQYGDMVKAGALTAVEQVNAA
GGAGGNKLEVVMMDDACEPKQAVAVANKIVSQKIKFVIGHVCSGSTIPASDIYENEGVVM
ITPSATAPQLTEGKKRKFIFRTIGRDDQQGPAAAQYIINKVKPKKVAVLHDKQSYGQGIA
SAVKKELDAAKVPVVIFEGINAGDTDYSAIVTKLKSQGIDFVYFGGYHPEMGLLMRQARE
QGVKAVFMGPEGVGNKDVTAIAGPASEGMLVTLPADFTIDPANAGVMKAFAQKKRDPNGT
FQMPAYSGVKIIADSIAGAKSTDPAKVAAYMHQHTFDTPIGKVAYDAQGDLKSFKFVVYK
WHKDATKTAAN*
>GCA_000452945.1_BPMSHR5858_genomic_04206_1 dadA1
MDFDVIVLGAGIVGVSAALHLRDRGRRVALVDRGAPGDGTSFGNAGLIERSSVAPYAFAR
SPLALLRYAANRSTELYWDRAALPSFAPWLARFWWESSPRRHAAAARDMLPLIERSVAEH
DALIARAGAGELVRACGWIEAYRTPARLEHATRDALLDASRHGLRVTALDAAALAAQEPA
LGPGFCGALHWLDPKSVVDPGALVKAYARLFARDGGTLLEGDAATLEAGDGGWRVRTAEG
VAAARDVVVALGPWSDTVFAKLGYRLPLREKHGYHMHYAPPDGAPLSAPVVDLEHGYVVA
PMRRGLRLTTGVEIAARTLPPTGVQLARAERVAQPTFRLGRRLDAAPWLGFRPCTPDMRP
VIGAAPRHRGIWFAFGHNHHGLTLGPITGRLLAEMMCGGPTVANVAPFRAERFL*
>GCA_000452945.1_BPMSHR5858_genomic_04207_1 group_18970
MKSSKSQPVLDPADVHVEILERSDTLLVVRWVEPGRCHYGEQRWRRRFAQRTGTCALSRQ
VIQRGDEVFRPAERPAPANAGAMISAAEVLGHTGGK*
>GCA_000452945.1_BPMSHR5858_genomic_04209_1 group_19249
MKSLVYAVVAASALTASFGAFAQSSQPLTRAQVRAELVQLEKAGYHPGVSSPYYPEDIQA
AQARVQGADTSGYGAQAAPVAHVGAPVAAAPRSPRESIYFGQ*
>GCA_000452945.1_BPMSHR5858_genomic_04214_1 opuAC
MKSTTTFVFGAALAAACALPSQSFAQDSAACRNVRFADIGWTDITSTTALASLLFDGLGY
KPTTTIASVPISFAGLKNRQLDVSLGYWWPVQQHQLQPFLDSKSISVVEPPNLSGAKATL
AVPSYVYQAGLKSFDDIAKHRAELDGKIYGIEPGSSANATIQKMIDTNQYGLGGFKLVES
SEAGMLVTVERAIRDKKWVVFLGWEPHPMNIQIGMNYLSGGDAAFGPNYGEARVYTLTSP
DYMARCPNAGKLVGNLRFTTQMENQLMQAVMNKVKPAEAAKAYIRKNPQVLDAWLAGVKT
YDGKDGLAAVKAYLGL*
>GCA_000452945.1_BPMSHR5858_genomic_04216_1 group_19138
MQTSRKALPLALGLAIGLGAALPAWADSKAPSPQEESRRASLTRGVVAPAEQAGKTGQFR
PGAVAVTLASPAFHAKKADAAAMAREYVTARAAQLGLDKAALANLVVASERADAAFTVVR
FQQRAAGLPVYDSDIAVTVAPDGRVLYVASKAVSGVAAVSSKTQAVDEQQALDRARAYLG
VGGFVNVQSQLVAFVDGAGTHTAWKVSGRPQDSLHGDWELIIDAGSGEVLRAQDKASYAT
DGSGLVFRPDPLSPTKSSYGSPGFKDNNDADSPQLSAARVRVTLKDLTQTSGGYKLSGPY
ASCIDFDAPLDKACPVQASTTFDFTRSNLYFEAVNAYYHIDTFLRYVNLTLGIKALPYQY
AGGVQYDPHGQSGDDNSSYSPSSGRLSFGQGGVDDAEDADVVIHELGHGIHDWITNGGLS
QVEGLSEGTGDYLAAAYSRDFNQWSPSDAQYHWVFNWDGHNEFWAGRVTNYNVGRTYAQI
RNAAIHTAGQYWASCNMVARDAIGGAAMDKAFLKGLSMTNGSTNQKAAAQAVLTAAAALG
YSSAQLNAIGDAYNKSCTYGVTVPQKL*
>GCA_000452945.1_BPMSHR5858_genomic_04223_1 group_19612
MSQAGLRARSTSEIEATISHEERGHTLHRGLSWKDAFWVTSGVPAGVLFTIGGVCATVGQ
PAWAVWIAAITMGLIQSATYAEISGLFPHKSGGASVYGAIGWVRYGKLIAPVSVWCNWLA
WSPMLALGCGLAASYALTSLFPADAAILHWQWTLADLGFVKPGLTLRVNATFVIATILLL
ITFKLQHSGASKAARTQRILGIASLTPLLIVGIVPFVTGDVPMSNLLPLLPLGHDAQGNL
TAAAFGSWNGQGVVMALGAMFMAGWASYGFETAVCYTREFRDPRRDTAKAIFWSGALCLI
VMTLVPMAFQGALGTAAMLDPRIGDGTGVAAAMARIVGGGAWVANAIVVMLMLSILLIVM
TSMMGSSRTLYQASVDGWLPKYLSHVNEHGAPTRAMWTDLGFNLVLLLMSDYMTVLSISN
VCYMLFVFLNLQSGWIHRMDRGQWQRPFRCPTWLLVLGSICGYANLVYVGAGADLQGAGT
LRNGLIAMLLIVPVFVYRHYWQDRGRFPAQMQRDMELAVAARGMWLNVAPYAALAAAALT
IAASYYFAWLR*
>GCA_000452945.1_BPMSHR5858_genomic_04231_1 group_19565
MQPQLPIDVDPNTGVWTTDALPMLYVPRHFFTNNHTAVEEALGRDAYADILYRAGYKSAY
HWCDKEAKQHGIAGMAVFEHYLKRLSQRGWGLFSIVEAEPAAARAKIELRHSSFVLAQPG
KEGKLCYMFAGWFAGALDWVNDTTPEGRGAPRARSAEAQCAAEGHDHCVFEVSPIAR*
>GCA_000452945.1_BPMSHR5858_genomic_04233_1 glyA2
MSNANPFFSQSLAERDASVRGAILKELERQQSQVELIASENIVSRAVLDAQGSVLTNKYA
EGYPGKRYYGGCEFADEVEALAIERVKRLFNAGHANVQPHSGAQANGAVMLALAKPGDTV
LGMSLDAGGHLTHGAKPALSGKWFNALQYGVSRDTMLLDYDQVEALAQQHKPSLIIAGFS
AYPRKLDFARFRAIADSVGAKLMVDMAHIAGVIAAGRHANPVEHAHVVTSTTHKTLRGPR
GGFVLTNDEEIAKKINSAVFPGLQGGPLMHVIAGKAVAFGEALTDDFKTYIDRVLANAQA
LGDVLKAGGVDLVTGGTDNHLLLVDLRPKGLKGAQVEQALERAGITCNKNGIPFDPEKPT
ITSGIRLGTPAGTTRGFGAAEFREVGRLILEVFEALRTNPEGDHATEQRVRREIFALCER
FPIY*
>GCA_000452945.1_BPMSHR5858_genomic_04235_1 fdhA
MSSNRGVVYQGPGKVEVRKIDYPKMVDPSGRAIGHGVILKVVSTNICGSDQHMVRGRTTA
PVGLVLGHEITGEVIEVGRDVETLKIGDLVSVPFNVACGRCAMCKEQHTGVCLNVNPARA
GGAYGYVDMGGWIGGQAEYVLVPYADFNLLKFPDRDRAMAKIRDLTCLSDILPTGYHGAV
TAGVKPGSTVYVAGAGPVGMAAAASARLLGAACTIVGDMNAERLAHAKAMGFEVVDLSKD
ATLGEQIEQILGKPEIDCAVDCVGFEAHGHGSSGHAQEAPATVLNSLMEITRPAGAIGIP
GLYVTDDPGAVDVAAKHGSLSIRFGLGWAKSHSFHTGQTPVLKYSRNLMQAILFDRLPIA
EIVNVAVISLDDAPEGYKKFDGGAPRKFVIDPHGMLKAA*
>GCA_000452945.1_BPMSHR5858_genomic_04237_1 lsdA
MTYSKQAVSRRHKRLALSAAALAAAACMSAHAQSDGAGPAPTPHTQQAYDPESHFTMRWT
RADMRQLVKQSHTAGADKNSLPPALTMPDIAQNFPLVDSNVWVWDTWPLADMRANQLSYK
GWEVIFSLSADPHAGYTFDDRHVHARIGFFYRRAGIPASQRPANGGWTWGGHLFPDGASA
KVFGTAPMTNNAEWSGSARLTHGENVSLYYTALSFNRSAPGGADITPPIAIITRADGHIH
ADDKHVWFSGFDDHKALLQPEGKMYQTGQQNTYYSFRDPFVFTDPAHPGNTYMVFEGNTG
GPRGARTCTEADLGYAPNDPYREDLNAVMNLGAVYQKANVGLAIATNPQLTEWKFLPPLL
SANCVDDQTERPQIYLKDGKYYLFTISHRTTMAAGIDGPDGVYGFVGNGIRSDFLPLNGG
SGLVLGNPTDFSAPAGAPYAQDPNQNPRAFQSYSHYVMPGGRVESFIDAIGARRGGTLAP
TVKIDIHGDSTTVDRAYGAGGLGGYGDIPANLPAVGAGHHD*
>GCA_000452945.1_BPMSHR5858_genomic_04238_1 sacC
MSNPRLRFSSVVYGLIALAACASFAAHAAGARCAASPAKSADGTPQWRPALHYTPRRNWM
NDPNGLVYENGVYHLFYQYNPHGNFWGDMSWGHATSRDLVHWDEQPVAMPANAREDIFSG
SIVADVRNTSGLGTPNAPPLVALYTSVYKAGSGHEPGTQAQSLAYSIDHGKTWRPYAHNP
VLTLAPESRHFRDPNVTWYAPGGYWMMTAVVADAPVVKLYRSSDLIRWDFLSDFTLPDVP
HRGALWEMPELLPMPLDGDAARMKWVMIVNVNPWSIAGGSGAMYFIGEFDGRTFTPDRAA
PANADPAQYSWLDHGADYYAAGTFANAPGEGPVAIAWMSNWDYAERIPTTPWKGAMALPR
ELALKTIDGRPQVTVAPARAFDAFARTRPAVRIGSLAVASATRELGADARGTVQRIAVTI
EPRSARRAGLIVRRSANGRVGTRIVYDSSAHTLSVDRSASGETNFSNAFSQRHIVALPLV
NGKLRLDVIVDRDSVEVFDGDGRTVITDLVFPSPADNRLAVFAEGGDATFGDVVVTNLDE
TAGERRGCPTS*
>GCA_000452945.1_BPMSHR5858_genomic_04271_1 pmbA
MAANLDAQAHYFPHTQDELKAIATDILRHAKALGATDAATEISEGDGLSVSVRRGEVETI
EHNRDKTVGVTVFIGKKRGNASTSDFSPTALKDTVAAAYNIARFTAEDDAAGLAEAELLE
TEPRDLDLYHPWRLSADEAVDLARRSEDAAFAVSPQIRNSEGASVSAQHSQFVLATTRGF
LAGYPYSRHYIACAPIAGSGRHMQRDDWYTSKRRADELASPESVGRYAAQRALARMGARR
LDTRKVPVLFEAPLAAGLLGAFVQAVSGGALYRKTSFLIDSLGKEVFAPHVQIVEDPHVP
RAMGSAPFDEEGVRTRARNVVRDGVVEGYFLSTYSARKLGTQTTGNAGGSHNIALRSALT
APGDDFDAMLKKLGTGLLLTELMGQGVNYVTGDYSRGAAGFWVENGEIQYPVEEITVAST
LQEMFRHIVAIGADSIVRGTKETGSVLIEQMTIAGQ*
>GCA_000452945.1_BPMSHR5858_genomic_04278_1 cbiB
MTFFSVLLALIIEQVRALSPSNPVFALFQSHAESAAHGFDAGKKKHGVLAWLVVVLPWVL
AVGLVYFLLYKVSFVLAFLWNVVVVYFTLGFRQFSHYFTDIHLALNNDDVPRARDILREW
TGIDTVDMPVGEIVRHTLIHAVVASHRHVFGVFFWFVLPVGPAGAVLYRIAEYLSRSWSV
PADDRTAAFSTFAQRAFFVIDWVPARLTALGFAIVGNFEDAIYAWRNHTRQWPDPNDGVL
LAAGSGALGARLAGPLAEPSSVDALASGDSGPLTVGDDCTPRTLQSAVGLVWRAVILWML
LLLMLTIAVWVS*
>GCA_000452945.1_BPMSHR5858_genomic_04285_1 dnaN
MQLVKTERDTLLRPLQTVSGIVERRHTLPILANLLITKNGPDVSFLSTDLELQITTRADF
GVGGDQVATTVAARKLLDILRAMPDGQVTLSLADKRLTVQSGKSRFALQTLAADEFPTVA
QSKDFGASLAVPQKAFRQLLGMVYFAMAQQDIRYYLNGMLLVVDGDRLMAVATDGHRLAF
SSMKIEGSFGRQEVIVPRKTILELQRLLEDIDDTVKIDIAQTQAKFTFGQVELVSKLVEG
KFPDFQRVIPKAHKNTFEIGREELQRSLQRAAILTSDKFKGVRCIIAPGQLKIMSTNADQ
EEAQEELEIAYQGDTVDIGFNVTYLLDVLANLKVDTVQVSLGDASSSALITVPENDEFKY
VVMPMRI*
>GCA_000452945.1_BPMSHR5858_genomic_04290_1 group_19476
MPTHEEKAAFTERLKFALLRSPEKVTGATELALHFNLRHHGEHPVSPQTAHKWLTGRTIP
TPDKLHTLAEWLRVDLHWLHYGPPPSAARTTPQPLPRDEKYPPTPETIELASKIEALSPH
HRYLVQELIEQFYGDTSKR*
>GCA_000452945.1_BPMSHR5858_genomic_04299_1 group_1912
MRRSRESVVLRRPVKVECVPSGAPLELPAGTAAEITQALGTNFTLVAGGQMVRLRGEDAD
AIGKPPPGPLDVPDGPVTPDGARACVMRTLKTCYDPEIPVDIVELGLIYGCEIEPAGHDR
LKVSITMTLTAPGCGMGEALADEVADKVLALPFVGEVNVDMVFDPPWDRSRMSEAAMLTL
GL*
>GCA_000452945.1_BPMSHR5858_genomic_04302_1 group_19655
MTTTATDRAAARAQLAQHGWISRHHESFRHLPPPAADTWLDAEADRADAARPPGAAAHAA
WLLAPVGSGSLEQVDACWLGTADRAERAALFADLPTSPDDDAAPFFWAHRALCETGLRLR
IGSGPAQRAPTMLQLRLPAQGRAEAPLVVIDVMPGVHCVLVESHEHAAGRDAPLTQNLHA
YVRLGAGASLDHLRIATPGARDRIAHHVHVCLQRDAQYRQSLLASGSEYHLQRTVLELRG
ERAVARASGVLFASSQAKLEQQVRAAHAAPRTRSAIDTLALASGHAQIVANAFSSIAAGA
SGADLRQRLSGIPTSGEPRVVLRPHLEIHHDDVQAAHGATWGALPEDALFYACQRGLDRR
DALAMIVAGMARAALAHGIDTPGRLEALGGDALLAREVSRHLAEQTEARHG*
>GCA_000452945.1_BPMSHR5858_genomic_04304_1 sufB
MLNVDQPTPLARALEQTYRHGFVTDIESDSLPPGLDEGVVREISRRKREPAFLLKWRLAA
FERWLAMSPPDWGKLRIAPIDFQAQSYYSAPKSLKDQPKSLDEVDPKLLETYEKLNVPLH

ERARLAGVAVDAVFDSVSVGTTFREQLAAAGVIFCSFSHAVEHHPELIERYLGTVVPPAD
NFYAALNSAVFSDGSFVYVPQGVRCPMELSSYFRINAQNTGQFERTLIIAEPGSHVSYLE
GCTAPQRDEHQLHAAVVELVAHDDAHIKYSTVQNWYPGDADGVGGIYNFVTKRGLCAGAR
SRIAWTQIETGSAITWKYPSVVLRGDESSGEFHSIAVSNHCQQADTGTKMIHLGRNTRSR
IVSKGISAGRAQNSYRGLVRVARTAEGARNHTQCDSLLIGPQCGAHTFPYVESARGDAVV
EHEATTTRISEDRLFYCQQRGIDPQEATALIVGGFCQAVLEALPMEFALEAKALLAVSLE
GAVG*
>GCA_000452945.1_BPMSHR5858_genomic_04311_1 yhbU
MTQSSHFATGAAPIELVCPAGSLPALKAAVDNGADCVYLGFRDATNARNFAGLNFDAQAI
AAGIRYARERGRKVLVALNTYPQPDGWAAWREAVSRAADAGVDAIIVADPGLMRFARERY
PELRLHLSVQGSATNYEAINFYHEHFGVSRAVLPRVLSLAQVEQVAENTPVEIEVFGFGS
LCVMVEGRCALSSYATGESPNTRGVCSPAKAVRWQKTPDGLESRLNGVLIDRYEDGENAG
YPTLCKGRFTVADESYYAIEEPTSLNTLELLPKLMQIGIRAIKIEGRQRSPAYVAQVTRV
WRDAIDQCTANLARYYVKPAWMTELNKVAEGQQHTLGAYHRPWK*
>GCA_000452945.1_BPMSHR5858_genomic_04313_1 group_19239
MNAPVFQLPPTLGKLLSKLPAYPGALLFASGINLVLRRHLPAETLALLEDRPLRIQVKDA
GVAFDFVCRRGAFSALSGGRDVDLTIGATAYDFYLLSQRREDPDTLFFSRRLTMEGDTEL
GLLVKNSLDAIDLSVFSLERWLPTRLFRQRPLERDAMPD*
>GCA_000452945.1_BPMSHR5858_genomic_04315_1 queE_2
MNLALPRRRPLKVGGLVPFTATDYPGQFAAVVFVQGCPWRCGYCHNPHLQPRSQPAEIEW
DALLAFLARRVGLIDAVVFSGGEPSIDPALAASIDDVRRLGFKVGLHSAGTHPRRLAHLL
PSLDWIGLDVKAPFDDYAHTTRVRASGGHARQSLEAVLASGIAYECRTTAHPDLLPEPAL
LRVAHELADLGVEHYVLQVFRAQGCGSGALNAASLAGYPSDAVLGRLDRLFANFAIRRG*
>GCA_000452945.1_BPMSHR5858_genomic_04323_1 phrA
LSSSISIAPAVVWFRDDLRVSDHPALMRAAESGRPVIGVYVFDERAGGRPLGGAARWWLH
GSLRALGAALAALGGTLALLRGDEHDTIVSFAAALGAGAVYWSRRYALAARAVDNAVQTA
LDAQGIAVETFNASLLFEPGDLVNADGRPYQVFSAYWRAAMRRGVPAPPLGAPRRLNGGA
LPAALARQCVALDALGLEPTGPDWAGGLRDAWRFGEDAAHRRLRAFVDSRLGGYAAERDR
PGIASTSRLSPYVRFGNLSARQLWHAAADAYAHGAATQADLDKFLSELGWREFGWTQLHH
FPDLPRRNLRHTLDGMPWRDDPAALAAWRRGATGYPFVDAGMRELWATGWMHNRVRMVCA
SFLVKHLLIDWRAGEAWFWDTLVDADAASNAANWQWVAGCGADAAPYFRIFNPVAQGRKF
DPDGAYVRRWVPELAGLPPATIHEPWAAPGAQLAAAGVRLGETYPLPIVAHDAARRRALD
AFGAIR*
>GCA_000452945.1_BPMSHR5858_genomic_04328_1 hutH
MITLTPGRLTLPQLRRIARENVQIALDPASFAAIDRGAQAVADIAAKGEPAYGINTGFGR
LASTHIPHDQLELLQKNLVLSHAVGVGEPMARPVVRLLMALKLSSLGRGHSGIRRVVMDA
LVTLFNADVLPLIPVKGSVGASGDLAPLAHMSAVLLGIGDVFIRGERASAAEGLRVAGLA
PLTLEAKEGLALLNGTQASTALALDNLFAIEDLYRTALVSGALSVDAAAGSVKPFDARIH
ELRGHRGQIDAAAAYRSLLDGSAINVSHRDCDKVQDPYSLRCQPQVMGACLDQIRHAAGV
LLIEANAVSDNPLIFPDTGEVLSGGNFHAEPVAFAADNLAIAAAEIGALAERRIALLIDA
TLSGLPPFLVKDGGVNSGFMIAHVTAAALASENKTLAHPASVDSLPTSANQEDHVSMATF
AARKLADIAENVANILAIELLAAAQGVDLRAPHATSPALQHAMKTIRADVAHYDLDHYFA
PDIAVVARRVRERAFATLSPLSFESEQ*
>GCA_000452945.1_BPMSHR5858_genomic_04332_1 hutI
MKSILWHNLKLCAHGDPNDTIADAAIAVNGDGTIAWTGRASDVPAGYVHWPREDLRGAWV
TPGLVDCHTHLVYGGQRADEFAQRLAGASYEEIAQRGGGIVSTVRATRDASEAALFEQAC
ARLRPLLAEGVTAIEIKSGYGLELASERRMLRVARQLGERFPVSVYTTFLGAHALPPEYA
GRADEYIDEVCERMLPALADEGLVDAVDVFCERIGFTLAQSERVFEAAARRGLPVKMHAE
QLSNGGGSALAARYRALSADHLEYLDAAGVAAMRASGTTAVLLPGAYYFIRETKLPPIDL
LRRHGVPIALATDHNPGTSPLTSLLLTMNMGCTVFKLTVQEALLGVTRHAAAALGASDRH
GSLAPGRQADFAVWSVSTLAELAYWFGRPLCERVVKGGVTVFTRDAR*
>GCA_000452945.1_BPMSHR5858_genomic_04397_1 group_19581
MRFLVLNSDAERRDGLKTLLRQIDRHAGYNDAKDWRQAQRLIKQQSFDLIVVDWQDIKRL
GDLHALCQACAPTPVAALVDDASPVHAQALFEAGVLGVISRTTRPHLIVRALEMVLLGGH
YVPPSALNLTEPAARARDGRVEAIVSQLPRRKRSPCSLLSPRQAQIMRFVHMGSTNKMIA
RTLGISEGTVKIHLASIFQQLGATNRAAAVAIYNGWLPPHLEVLLSARETSPKPVHGAAG
PVPLRPAKSRHPYPLAAENESSAQPLIAAEPRTPFKANE*
>GCA_000452945.1_BPMSHR5858_genomic_04398_1 group_19459
MGFLYTPLPFWIAVGGWIAAVGLLVLALWKRPFGRLQDATLQHVWLALVTTITVLWASNA
WLDDGLVMHLLGATLLVTLFDWTLALVGMAAVIGVAAIVFDATWQGVGLTYLVYGALPVA
VSTLLQRAALAWLPHNLPALIVGQGFVSPAIAVGIVAAAAAGVQIAIAEGTLVVVPSGYA
LNALLLALGEAWFTGMATALIAVYRPAWVTTSDVRRYRLGGPRA*
>GCA_000452945.1_BPMSHR5858_genomic_04399_1 group_19682
MDSFPASLRLLGALGKLTACAAGLSLALAAPAFADLLDQRAELVNKYVNEMHADPLVADC
AAHGNFIASTSSAFDHVEFAPNAFDSGSASITPWNDSFDEGKQRVKVDNIVTVEGLGVRA
SGGDPEPLKFRCGYVGTQMLAFSWNDPVPPAKPRVERATSSKKKLRAHSGKGKVKAKATG
KSGKRAVSKKSSTTKKAAKKRSTRKS*
>GCA_000452945.1_BPMSHR5858_genomic_04401_1 yaaA
MIIVLSPAKSLDYETPPHVSHHTQPQFADDAAALIDELRRLSPQQIATLMSISDPLARLN
FQRYADWSRASTPANAKQAVLAFNGDVYEGLDARSLSPDDLDYAQRHVRVLSGLYGLLRP
LDLLQPYRLEMGTRFSNARGKDLYAFWGERITHALNAELKTRVGASRVLVNCASAEYFKS
VKPKLLDARVVTPVFEDWKDGRYKIISFHAKRARGLMARYVVEGRIDSPDALKDFASEGY
AFDASASNDDTYVFRRRAGA*
>GCA_000452945.1_BPMSHR5858_genomic_04412_1 group_19168
MGISYEDLEVGSRTEVGKHRFTADEIVTFAQAYDPQPFHVDAQAAKSSMFGGLVASGWHT
CSVFMGMLVRGVLQGSTSMGSPGIEEIRWIKPVRAGDTITMYNKILDKRLSESKPDRGIV
TTEWEGVDEQGETVITVRSKVIFGLRSPQVRA*
>GCA_000452945.1_BPMSHR5858_genomic_04421_1 nuoK
MLTLAHYLVLGAILFAIAIVGIFLNRRNIIIILMAIELMLLAVNTNFVAFSHYLGDVHGQ
IFVFFVLTVAAAEAAIGLAILVTLFRKLDTINVEDLDQLKG*
>GCA_000452945.1_BPMSHR5858_genomic_04422_1 nuoJ
MDFTTVLFYIFSLLLVVSGLKVITSRNPVASALFLVLAFFNAAAIWMLLEAEFLAILLVL
VYVGAVMVLFLFVVMMLDVNIDVLRRDFKRFVPMATAVGAIIVVETALILWRGYGATASP
VRDMAAGALAGMPNTKLIGKVIYTDYIFAFEIAGLVLLVAIIAAVALTAQKGKDSKRQRV
SEQVKVRREDRVRLVKMPADKPQPEAAASDAASGSNNG*
>GCA_000452945.1_BPMSHR5858_genomic_04425_1 nqo3
MVELEIDGKKVEVPEGSMVIQAAHKADTYIPHFCYHKKLSVAANCRMCLVEVEKMPKAVP
ACATPVSAGMIVHTNSEKAVKAQQSVMEFLLINHPLDCPICDQGGECQLQDLAVGYGKST
SRYAEEKRVVFHKEVGPLISMEEMSRCIHCTRCVRFGQEIAGVMEFGMLGRGEHSEITTF
VGKTVDSELSGNMIDLCPVGALTSKPFRYSARTWELSRRKSVSPHDSVGANLVVQVKNNR
VMRVLPFENEAVNECWISDKDRFSYEGLNSEERLTKPMIKQGGEWREVDWQTALEYVARG
LKGIGDEHGPAAIAALGSAHSTVEELFLLKQLAHELKTPNVDFRLRQTDFSAAAQGAPWL
GMPIADLSAVDAAFVVGSFLRRDHPLFAARLRQAAKNGAKLHFLHATGDDALIPTAQRIV
AAPSAWLDTLAGVAAAVAQARGVALPEALAGVEASDAARSVAQSLANGERRAVLLGNSVV
RHPQFAQIHAIAQWIAENTGATLGFLTEAANTVGAHLVGALPGANGLNARDAFAQPRKGY
VLLNVEPEFDTADPAQALAALKQAETVVVLSPFKYGLDYADVLLPISPFTETAGTYVNAE
GRAQSFNGVVRPLGDTRPAWKVLRVLGSLLGLPNFEYETSEEVRLAALGDGEITSRLSNK
TAAVPARAAARAANGALERLADVPIYHADALSRRAGALHLTAASKAAHQVGLPAALFDKL
GLKNGDAVRVRQGDRTVQLPAVRDENLAEAVVRVSAATPAGAALGSLSGELVVEKA*
>GCA_000452945.1_BPMSHR5858_genomic_04426_1 nuoF
MTSLHDRHIKPLILAGLTGENWHLEDYVARGGYKQLRRILEEKIPPEQVIADVKASGLRG
RGGAGFPTGLKWSFMPRQFPGQKYLVCNSDEGEPGTFKDRDILRWNPHALIEGMAIGAYA
MGITVGYNYIHGEIFEVYRRFEAALDEARRAGFLGEKIMGSDFSFQLHAHHGYGAYICGE
ETALLESLEGKKGQPRFKPPFPASFGVYGKPTTINNTETFAAVPFLLAVGPQNYLEMGKP
NNGGTKIFSVSGDVERPGNYEIPLGTPFATLMELAGGMRGGKKIKAVIPGGSSAPVIPGD
IMMQTDMDYDSIAKAGSMLGSGAVIVMDETRCMVRSLLRLSYFYYEESCGQCTPCREGTG
WLYRVVNRIEHGEGRQEDLDLLNSVAENIMGRTICALGDAAAMPVRGMLKHYWDEFAYHV
EHKHCMVGGHAHAAAA*
>GCA_000452945.1_BPMSHR5858_genomic_04427_1 nqo2
MISAEGLKEIDRAIAKYPADQKQSAVMSALAVAQEEHGWLSPELMQFVADYLGMPAIAVQ
EVATFYTMYELAPVGKHKITLCTNLPCQLGPHGGAEATADYLKQKLGIDFGETTPDGKFT
LKEGECFGACGDAPVLLLNNHKMCSFMSREKIDQLLEELSK*
>GCA_000452945.1_BPMSHR5858_genomic_04428_1 nuoD
MAEIKNYTLNFGPQHPAAHGVLRLVLELDGEVIQRADPHIGLLHRATEKLAENKTFIQSV
PYMDRLDYVSMMVNEHGYVLAIEKLLGIEVPERAQYIRVLFDEITRVLNHLMWIGAHALD
VGAMAVFLYAFREREDLMDVYEAVSGARMHAAYYRPGGVYRDLPEAMPQYKASKIRNERA
LAKMNEARSGSVLDFIDDFFTRFPKCVDEYETLLTDNRIWKQRLVGIGVVSPERALQLGL
TGPMIRGSGIAWDLRKKQPYEVYDRLDFDIPVGVNGDCYDRYLVRVEEMRQSTRIAKQCI
EWLRKNPGPVITDNHKVAPPSRVGMKTNMEDLIHHFKLFTEGFHVPEGETYAAVEHPKGE
FGIYLVSDGANKPYRLKIRAPGYAHLSALDEMARGHMIADAVTIIGTQDIVFGEIDR*
>GCA_000452945.1_BPMSHR5858_genomic_04429_1 ndhJ
MASKIETLKANLEAALGARAVSLVEAVGELTLVVKASDYLEVAKQLRDDRSLGFEQLIDL
CGVDYQTYGDGAYDGPRFAAVLHLLSVANNWRLRVRVFASDDDLPIVPSVVDIWNSANWY
EREAFDLYGIVFEGHPDLRRILTDYGFIGHPFRKDFPVSGYVEMRYDPQEKRVVYQPVTI
EPREITPRVIREDRYGGLKH*
>GCA_000452945.1_BPMSHR5858_genomic_04430_1 nuoB_1
MSIEGVLKEGFVTTTADKLINWTRTGSLWPMTFGLACCAVEMMHAGAARYDLDRFGVVFR
PSPRQSDVMIVAGTLCNKMAPALRRVYDQMAEPRWVISMGSCANGGGYYHYSYSVVRGCD
RIVPVDVYVPGCPPTAEALVYGVIQLQAKIRRTSTIARQ*
>GCA_000452945.1_BPMSHR5858_genomic_04431_1 ndhC
LNLAAYYPVLLFLLVGTGLGIALVSIGKILGPNKPDSEKNAPYECGFEAFEDARMKFDVR
YYLVAILFIIFDLETAFLFPWGVALREIGWPGFIAMMIFLLEFLLGFAYIWKKGGLDWE*
>GCA_000452945.1_BPMSHR5858_genomic_04436_1 pnp
MSLFNKIVKEFQWGQHKVRLETGEIARQASGAVIVDIEDTVVLATVVGAKSAKPGQDFFP
LTVDYIEKTYSAGKIPGGFFRREGRPSEHETLTSRLIDRPLRPLFPEGFYNEVQVVIHVL
SVNPEIPADIPALIGASAALAVSGLPFNGPVGAARVAYVNNEYVLNPTREQIKASRLDLV
VAGTERAVLMVESEADQLPEDVMLGAVVFGHEQMQTAIDAIHELVREGGKPEWDWQPAPK
DEALNARVTELAQPELLAAYQIRDKQARSTKLKEVYAATSAKLEEEAVAAGTVAADKATV
GNILFDLEAKIVRGQILNGEPRIDGRDTRTVRPIEIRTGVLPRTHGSALFTRGETQALVV
ATLGTKGDEQIIDALEGEYRERFMLHYNMPPFATGETGRVGSPKRREIGHGRLAKRALVA
CLPSADEFGYSIRVVSEITESNGSSSMASVCGGCLALMDAGVPMKAHVAGIAMGLILEGN
KFAVLTDILGDEDHLGDMDFKVAGTADGVTALQMDIKIQGITKEIMQVALAQAKEGRMHI
LGKMKDAVAGANTQLSEFAPRMITIKINPEKIRDVIGKGGSVIRALTEETGTTIDISDDG
VVTIASTNSEGMAEAKKRIENITAEIEVGHVYEGTVLKLLDFGAIVNLLPGKDGLLHISE
IVNERVKDINDYLKEGQQVKVKVIQTDEKGRVRLSAKALLNEAAAQADTPPQQ*
>GCA_000452945.1_BPMSHR5858_genomic_04439_1 group_19090
MFARLPVLFACAAALLAGCAQPWQQFHAGDDSSAIVARLGPPREIYDLPNGGKRLMWPTQ
PMGEVTTAADIDAHDRIVSVRQVLQPLEFYRAEIGKWTKTDVLVNFGRPVETSYFPLMKR
EVWTYRYLEDNVWYMLYSFYFDDDGILRLTQKTPDPLHDPDRRTLF*
>GCA_000452945.1_BPMSHR5858_genomic_04519_1 group_19697
MTTNLVVQSTAPLSDAHHKPLAALAHGARVVALDAQSIRIERANPAQRADIDAYCGTHAL
DYAFVDAARKLADFGLVAMDMDSTLITIECIDEIADFCGLKAEVSAITEAAMRGEIKDFN
ESLTRRVALLAGLEASALERVYEERLRLSPGAESMLAGVKAAGLKTLLVSGGFTFFTERL
KARLGLDFAHSNTLEIVDGKLTGKVVGEIVNADVKARAVRETCAALGIEPARAIVIGDGS
NDLKMMAAGGFSIAFRAKPIVRSAASAAFDHVGLDGLLRLF*
>GCA_000452945.1_BPMSHR5858_genomic_04526_1 group_19074
MTTTKKTGERLIKKYPNRRLYDTETSTYITLTDVKQLVLEQEDFKVIDAKSSEDLTRSIL
LQIILEEESGGVPMFSSSMLSQIIRFYGHAMQGMMGTYLEKNIQAFIDIQSKLADQSKNL
YENNAMNPEVWSQFMNMQAPMMQGMMTSYIEQSKNMFVQMQEQMQNQAKTMFSSFPFKPA
TPPGSEPEKK*
>GCA_000452945.1_BPMSHR5858_genomic_04528_1 phaA
MTDVVIVSAARTAVGKFGGSLAKIAAPELGASVIRAVLERAGVKPEQVSEVILGQVLTAG
SGQNPARQALIAAGLPNAVPGMTINKVCGSGLKAVMLAANAVVAGDAEIVVAGGQENMSA
APHVLPGSRDGFRMGDAKLVDSMIVDGLWDVYNKYHMGITAENVAKEYGITREAQDQFAA
LSQNKAEAAQKAGRFDDEIVPIEIPQRKGEPLRFATDEFVRHGVTAESLAGLKPAFAKEG
TVTAANASGINDGAAAVLVMSAKKAEALGLEPLARIKAYANAGVDPSVMGMGPVPASRRC
LERAGWSVGDLDLMEINEAFAAQALAVHKQMGWDTSKVNVNGGAIAIGHPIGASGCRILV
TLLHEMLKRDAKRGLASLCIGGGMGVALALERP*
>GCA_000452945.1_BPMSHR5858_genomic_04550_1 ctaB
MDTTLSHTPGSRLSQYLALTKPRVTQLAVFCAVIGMFLATPGMVPWKVLLGGTIGIGLLA
GSAFAINCLVEQKIDAMMRRTAWRPSARGEITTLQILAFSTVLGGLGAWTLYTFTNPLTM
WLTIATFVGYAVIYTLLLKPMTPQNIVIGGASGAMPPALGWAAVTGAVPGDAWILVLIIF
VWTPPHFWVLALYRRKDYENAGLPMLPVTHGEQFTRLHILLYTVILFAVTMMPFISGMSG
AVYLTSAVLLGALFLAYAWKIYRDYSDALARRAFRYSIVYLSLLFAALLVDHYARPVIGM
*
>GCA_000452945.1_BPMSHR5858_genomic_04552_1 group_19529
MQTSRPTRPQDGRQAEPAQRGSWRRGRWVLLALALVCAAPVIASYFTYYVIKPRGGATNY
GTLIEPQRPIPPDLSVVDETGRTIPLASLRGVWLFVMEDGGACNDACAKKLYFMRQVRAT
QAGERHRITMVWLKSDAANVPAAIADAYPDTRKLRADPAAVAAWLPADAGTRVTDHLYLV
DPNGNLMMRFPKNPDPSRIKQDVTKLLKWSSIG*
>GCA_000452945.1_BPMSHR5858_genomic_04553_1 group_19094
MRIRWLPALLILAVIAITVRLGFWQRERAHQKEALDAQITRYEHANPVDVPRERIALKDI
EFHRVRATGRFMPEHAVFLDNRPYNDQPGFYVVMPMKLANGGYVLVNRGWLPRNYADRTA
IEPFATPAGDVTIEGIARGNASRAFELGEGGSAARQKIRQNLDAAEYAQETGLPLQPFVI
QQTSDDGDKLVRDWPAPTTGVERNYGYMFQWWGMAAAAAGFGLYAARRAARKQSGDA*
>GCA_000452945.1_BPMSHR5858_genomic_04557_1 ctaG
MSKSEANVDRAFNRSMLLKLVVVAGLMFGFGFALVPMYRAICQITGINNLLQRDVSAREA
KNTQVDYTRTVSIELDANARGPLGFKPRRNSIDVHPGELATVVYEVTNGQGRTVVAQAIP
SYAPKQATEFFKKIECFCFTQQTLAANETRDMPVVFVIDPKLPKDVKTITLSYTFFELNT
PAPAVKGAAVQTGAAAAKPAA*
>GCA_000452945.1_BPMSHR5858_genomic_04560_1 group_19356
MEAANGLHDAEPVLADWLMKRNCSVSPRQFVAFYVSLAAFSLLIAVLLLWRGAWLVLPFT
GIELLAVGVAFAIYARHAVDYERIRLFPHRLVIERMSAERLTQIELNPRWVRVEPGASPR
DPITLVSRGESVVVGQHLAQYRRAQFARELRASLSRYG*
>GCA_000452945.1_BPMSHR5858_genomic_04565_1 gpsA
MKVAVLGAGAWGTALAAHLAVRHDTLLWARDAALVAELAVRRENARYLGGVALPPGLRYE
ADLATALSHAQADDALCVIAAPVAGLRALCRAMRDARRVPAHFVWVCKGFEADTRRLPHQ
MVAEELPDHASYGVLSGPSFAREVAQGLPVALTVASASAACRERTLAAFHHGAMRIYTGD
DVVGVEVGGAVKNVLAIATGIADGLGLGLNARAALVTRGLAEMSRLGVALGGRAETFTGL
TGLGDLILTATGDLSRNRSVGLQLAAGRSLDDILAALGHVAEGVRCARAVLSIARERGVD
MPITEAVCAVLFDGVAPRDAVSGLLRRDAKAE*
>GCA_000452945.1_BPMSHR5858_genomic_04568_1 yibN
VTFFTDYTNLALIAILVVSGGLLAWPALRRGGGGLSAAEATQLINRRNALVVDLRPAAEY
GAGHLPSARSVEFGELQAKAGQLAKNKATPVLLVCQNGQQSQKARKIVEEAGYQDVHVLQ
GGVAAWQQAGMPVVK*
>GCA_000452945.1_BPMSHR5858_genomic_04569_1 gpmA
MYKLVLIRHGESTWNKENRFTGWVDVDLTEQGNREARQAGQLLKEAGYTFDIAYTSVLKR
AIRTLWHVQDQMDLMYVPVVHSWRLNERHYGALSGLNKAETAAKYGDEQVLVWRRSYDTP
PPALEPGDERAPYADPRYAKVPREQLPLTECLKDTVARVLPLWNESIAPAVKAGKQVLIA
AHGNSLRALIKYLDGISDADIVGLNIPNGVPLVYELDESLTPIRHYYPGDQEAIAKAQAA
VAQQGKSAA*
>GCA_000452945.1_BPMSHR5858_genomic_04573_1 ptsH
MLQQETTIVNKLGLHARASAKLTQLAGNFQAEVWMTRNGRRINAKSIMGVMMLAAGIGST
VTIETEGPDEREAMDALLKLIADKFGEGQ*
>GCA_000452945.1_BPMSHR5858_genomic_04576_1 group_19399
MVPHLVTALNGPLLELERKILDATPAIERWFRLEWQEHTPPFYCSVDLRNAGFKLAPVDA
NLFPGAFNNLPPEVLPLAVQAAMAAIEKICPDAKNLLVIPELPTRNAFYLENVARLATIM
RQAGLNVRFGSLDPSIAEVTPITLADGQKIVLEPLERTPRRLGLKNFDPCSILLNNDLSA
GIPSVLENLHEQYLLPPLHAGWAVRRKSTHFSCYDDVAKKFAKMVGVDPWMLNPYFSPVE
GVDWHASDGGEALAEAIDAVLKKIARKYREYGIGEKPYVVVKADAGTAGRGVMTVHDASE
IGRMTKHERAQMAESKAGVPVSDVIVQEGVYTFERIGDEVAEPVVYMIDRYVVGGFYRTH
GARERDQNLNAPGMHYVPLGFEHTALPDTHAKPGAAPPNRFYMYGVVARLGLLASSVELE
KTDPEAIQV*
>GCA_000452945.1_BPMSHR5858_genomic_04578_1 glnB_1
MKLITAIIKPFKLDETREALSALGVSGITVTEVKGFGRQKGHTELYRGAEYVVDFLPKMK
IEAAVSDDLVDQAVEAIERAARTGKIGDGKIFVTSIEQVIRIRTGETGADAL*
>GCA_000452945.1_BPMSHR5858_genomic_04582_1 lgrE_1
MALLPGTLTIPDRREHARHRLLMFHHACGSATNYFRWAAAFAPHIEVWLVELPGRGRSLR

HGAFDALPPLRDYLRELAPELPADYSIFGHSMGALVAYCFAQDMMQLGRAPRWLGVSGAD
SPFAASRRRAFPVHQRPDAAIVEHLATLGGTPPEVFAHDELRAMLLQLAKADFRIVEAFF
PLASATPLPVPVTVFAGAQDTVLSEGGLQDWQRASTLPIERETFDGGHFYLLDTPCAIHD
AIGAALARHERQASVPAVL*
>GCA_000452945.1_BPMSHR5858_genomic_04592_1 group_18979
MDLNFTAEEEAFRAQVQRFLADELPPRISRKVKGGLRLTRDDMREWHAILNARGWLASHW
PREWGGPGWSVAQKFLFDNECAIAGAPRIVPFGVNMLGPVLIKYGSPAQKRHWLPRILDG
TDWWCQGYSEPGAGSDLAAVSTSAVRGVDARGDHYIVNGQKTWTTLGHYANMIFCLVRTA
TDVRKQEGISFLLVDMNSPGVEVRPIVTLDGEHEVNEVFFTDVRVPAENRVGEENQGWTC
AKYLLTYERTNIAGVGFSVAALDKLRAVAAKVTKNGRPLADDPLFAARLARVAIELDNMK
TTNLRVLAAVAGGGAPGAESSMLKIRGTQIRQEISSLMRRAMGPYAQPFVDAALDADDGE
PPGGLPEAASAATQYFNNRKLSIFGGSNEIQKNIISKMMLGL*
>GCA_000452945.1_BPMSHR5858_genomic_04593_1 fadJ_1
MNQQAFPATGTVTRERRDKVLVVTIDHPPVNALSADVRRGLVDALDAADADAAVGAVLIV
GAGRNFIAGADIREFGKLPVPPLLPDVCNRIETSGKPVVVALHGAALGGGLEVALAAHYR
LAVPGAKLGLPEVQLGLLPGAGGTQRTPRLIGAKAALELMLSGRHASAEEALALGLVDRL
AHSDDTLAEGLAYVHELLAAGAGVRRTRDAQGLADRAAAEAALDAARADVARHSRGLFSP
ARIVDAVAAALAQPFDEGLRTERRLFVECLDSPQRAGLVHAFFAQREAAKSPETRRGAPR
AIERIGVVGGGTMGAGIAVAALDAGFPVTMIERDDASLERGRAHVEKVYAGLVAKGRMTP
AAQAARLARFKGGTSYEALAQADVVIEAVFEDMAVKKAVFAELARACKPGAVLATNTSYL
DIDELAASVERAPDVIGLHFFSPANIMKLLEVVVPAKVSADVVATAFELARKLGKIPVRA
GVCDGFIGNRVLAVYRTAADYMMEDGASPYQIDRAIREFGFPMGPYQVVDLAGGDIGWAT
RKRRAPTRDPRLRYVEIGDRLCERGWFGQKTGRGYYLYPDGARVGTPDPEVEAIIAQARA
AKGVSPRAFTDEEIVRRYLAAMINEGANVVHEKIALRPLDVDAVFLHGYGFPRHRGGPMH
YADTVGLANVLADIRAFAGEDPVFWKPSPLLVDLVARGETFASLNRID*
>GCA_000452945.1_BPMSHR5858_genomic_04597_1 dapL
VNPRLDTLQPYPFEKLRALFAGVTPSTPLPPISFGIGEPKHATPAPIRDAAAAALDGLAS
YPATAGTDALRDTVARWLERRYGLPAIDATTQVLAASGSREALFSLAQAVIDSSPRENGA
TQDGERPIVLCPNPFYQIYEGAALLAGAAPYFVNSDPARNFAPDYSSVPADVWARTQLVY
VCSPGNPTGAVLTLDDWRELFALSDEHGFVIASDECYSEIYFDETKPPLGGLEAAHRLGR
DFTRLVMLSSLSKRSNVPGMRSGFVAGDAALLKRFLLYRTYHGAALSPVWQKASVAAWGD
EAHVRENRALYAQKFATVTPMLAEVLDVRLPDAAFYLWANVARTGLSDTEFARRLYADYN
VTVLPGSYLARDAHGANPGRDFVRIALVAGTAECVEGAQRIVDFCRGLAR*
>GCA_000452945.1_BPMSHR5858_genomic_04598_1 dapD
MSQQLQQIIDNTWENRAELSPKAASAEIREAVAHAIEQLDRGALRVAEKIDGAWTVHQWL
KKAVLLSFRLEDNAPMPAGGYSQFYDKVPSKFANYTAEDFAAGGFRVVPPAIARRGSFIA
KNVVLMPSYTNIGAYVDEGTMVDTWATVGSCAQIGKNVHLSGGVGIGGVLEPLQANPVII
EDNCFIGARSEVVEGVIVEENSVISMGVYLGQSTKIYDRETGEVTYGRIPAGSVVVAGNL
PAKDGTHSLYCAVIVKKVDAKTRAKVGLNELLRGD*
>GCA_000452945.1_BPMSHR5858_genomic_04602_1 ywiE
MTLDWLHLGTLIGLAHVLGVIAACHAILNTRTSQGAIAWAVSLTAMPYLTLIPYLFLGRS
KFSGYVDARRHETAALRTRTHPAPWDAEGSTLGAPKEALGAARVRALTRLVGMPFVAGNA
VRTLVNGDATFSAILAAIDAARSYVIVQFFIVRDDALGAMLRDTLIARAKAGVRCYLLYD
SIGSFDLPSSYVHALREGGVEVHPFATNKQFVNRFQLNFRNHRKIVVVDGERAFVGGHNV
GVEYLGGNPRLSPWRDTHIEVRGPAAANIQYVFAEDWHWATQSLPPQAPPPAPVPDADMH
CLVVPMGPADKQETGSLFFAEAINAARERVWITTPYLVPDEAVISALKLAVMRGVDVRVM
IPSRRDHYVVFEASKLYARDLVDAGVRIFRYRPGFLHQKVVLIDRAAAAVGSANLDNRSF
RLNFEIMVLTVDAGFAAEVEAMLARDFDQAFEVDLSEYRRSPAWRRVAMHVARLFAPIL*
>GCA_000452945.1_BPMSHR5858_genomic_04604_1 yheS
VIRFNQFSLARGTKPLFDATSFTLNPGEKAGLVGANGAGKSTLFAVLRGELHADAGDFSM
PPAWRIAHVSQETPAVERSALDYTLDGDAALRAIEARIAQASAAHDGAAEADAHAAFADA
DGYTAPARAEALLLGLGFTLAQTREPVASFSGGWRMRLNLAQALMCRSDLLLLDEPTNHL
DLDAIVWLEDWLHRYSGTLVIISHDREFLDAVCDVTLHLENRQVKRYGGNYSQFEVLRAQ
QLELQQSAYEKQQKTIAHLQSFVDRFKAKASKAKQAQSRVKALEKMERIAPAHIASPFTF
EFRTPDSAPNPMLVMENVRCGYRADDGGEIPIVERVALSIQNGQRIGLLGANGQGKSTLI
KTLAGTLAPLSGELRTGRGLTIGYFAQHQLETLREDDSALAHLARLAPDTREQELRDFLG
GFNFSGEMATAPIAPFSGGEKARLALALIIWRKPNLLLLDEPTNHLDLETRHALTMALAQ
FEGTLILVSHDRHLLRATTDQFMLVAKHRLQPFDGDLDDYRDWLLQHAAEQRAAAKAACG
PAGDADGGAPAVNRKDQKRQEAEARQRLSQLKKPLQARITKIEKEMERLHAQKAELDAFV
ADPASYAVDQKTRLTEAIRKLGDVNGQLETLEADWLGAQDELEKIG*
>GCA_000452945.1_BPMSHR5858_genomic_04616_1 aceA
MSRQQQAQELQKQWETDPRWKGIKRAFTAEDVVRLRGSIQQEHTLAKRGAEKLWTLINNE
PFVNALGALTGNQAMQQVKAGLKAIYLSGWQVAGDANVAGEMYPDQSLYPANSVPLVVKR
INNTLTRADQIQWSEGKNPGDEGYVDFFAPIVADAEAGFGGVLNAFELMKAMIEAGASGV
HFEDQLASVKKCGHMGGKVLVPTREAVAKLTAARLAADVMGTPTVLVARTDAEAADLITS
DIDDNDKPYLTGERTVEGFFRTKPGLEQAISRGLAYAPYADLIWCETGKPDLEYAKKFAE
AIHKQFPGKLLSYNCSPSFNWKKNLDDATIAKFQKELGAMGYKFQFITLAGFHALNYSMF
NLAHGYARTQMSAFVELQQAEFAAADKGFTAVKHQREVGTGYFDAVTQTVEREASTTALH
GSTEDEQFFDGQKVA*
>GCA_000452945.1_BPMSHR5858_genomic_04618_1 dmlR_9
MDRFKQIETFAAVAAKGSLSAAAHAEGVAPAIIGRRLDALEERLGVKLLVRTTRKLTLTF
EGSAFLEDCQRIINDMQNAEASVSAGGVKASGHLRVSAPAGFGRRHVAPLVPPFTGAHPD
VSVTLDLSDRMVDLVNEGFDCAVRLGELPDSSLVSLKLGENRRVCVASPAYLARAGMPGT
LDDLARHNCLALAANANQQRGWSFVDGGKVVSIRVSGTMECSDGAVLHEWCLEGHGLAWR
SWWEVGADIAAGRLVSVLDAFAAPPVGIHAVFPQRRHLPLRVRLFLDFLKHTYERPGYWG
*
>GCA_000452945.1_BPMSHR5858_genomic_04707_1 fumC
MSEAVRMERDTFGEIAVPAARLWGAQTQRSLQNFKISTEKQSPELIHALALIKRAAAAVN
LELGVLAQDKANAIVAAADEIIAGRHADEFPLAVWQTGSGTQTNMNLNEVIANRASELLG
GEHGESRAVHPNDDVNRGQSSNDVFPTAMHIAAARAIVDHLLPALRTLRATLDAKAAAFA
DIVKIGRTHLQDATPLTLGQEFSGYVAQLDQGIRHVEAALPHLYELAQGGTAVGTGLNAH
PKFAAGVAAEIGRLTGLPFVSAPNKFEVMAAADALVFAHGALKTVAASLMKIANDIRWLA
SGPRCGLGELSIPENEPGSSIMPGKVNPTQSEAVTMLCCQVFGNDVAVNFGGASGNFELN
VFRPMIAHNVLQSVRLLADGAQGFNDHCAVGIEPNRARIDALLNESLMLVTALNPHIGYD
KAAQIAKKAHKEGTTLKAAALALGYVTDAQFDEWVRPEHMVGNR*
>GCA_000452945.1_BPMSHR5858_genomic_04716_1 group_19045
MTHVVTEACIKCKYTDCVDVCPVDCFREGPNFLAIDPDECIDCAVCVAECPTNAIYAEED
VPGDQQHFTALNAELAKDWPSITKTKPAPADADEWKDVQEKLHLLER*
>GCA_000452945.1_BPMSHR5858_genomic_04719_1 group_19122
MKRHAAWAGMALGGALGAAPALASAATLDGAALSAFWAVPFAGILLSIAVFPLIAPVFWH
HHFGKIAAAWAVAFLVPFAATFGFGTAFGTLMHALFEEYIPFIVLLSALYTVAGGICVRG
NLHGTPRLNTGILALGTVLASVMGTTGAAMLLIRPLLRANDNRKHVVHVVVFFIFLVANA
GGSLSPLGDPPLFLGFLNGVDFFWTTIHLALPMLFICAILLTLFFVLDTYFYRKGGEEGK
PFLDPTPDSHGVSIEGKVNFVLLGAVIALVLMSGLWKPGIAFDLFGTHVALQNAVRDVAL
VAVALVSLAVTPRSAREGNAFNWAPIEEVAKLFAGIFVTIAPVIVILRAGGDGVFAPIVH
LVTGADGKPVDAMYFWATGVLSSFLDNAPTYLVFFNLAGGDAQTLMTTGATTLAAISAGA
VFMGANSYIGNAPNFMVKAIAESRGVRMPSFFAYLGWALAILVPVFLLTTWVFFSAQAAQ
*
>GCA_000452945.1_BPMSHR5858_genomic_04720_1 ghrB_3
MTAARAMRGAMRARRPRGATETAMQKILVARPIFPDVIERLKQYFDVDWNDGDALAPDAL
KARLADKDGALTAGDMIDASVLAAAPRLRVVSNMAVGYNNFDIGAFDAAHVLGTNTPDVL
TETTADFGWALMMAAARRITESEHWLRAGQWRKWSYDSFLGADIHGATLGVLGMGRIGQA
LARRARGFGMRVIYHNRSRVAPEIEAELNAEYVPKAALLAQADHVVLVLPYSAQSHHTIG
AAELALMKPSATLTNIARGGIVDDAALADALREKRIAAAGLDVFEGEPSVHPALLDVPNV
VLTPHIASASEGTRRAMANLAADNLIAALGAGPRAGRPPNPINPGVLGKARI*
>GCA_000452945.1_BPMSHR5858_genomic_04723_1 group_18956
MKKQISSIAPGQTAKALILVYLTFSVPIVLLGIVNAFFREQSVGLGTIFSTIFSALLLNA
IVGFVLLWIACHAYNWVASRFGGIEIVLSDAPEEA*
>GCA_000452945.1_BPMSHR5858_genomic_04728_1 rluC
MNELGKKSHNSVASGQVSLIEIDESAAGQRIDNFLLRVCKGVPKSHIYRILRSGEVRVNK
GRIDAQYRLAFGDVVRVPPVRVAAADLARAAGPAPVPAAEFEILFEDDAIIVLNKPAGVA
VHGGSGVAFGVIEQMRHARPHAKFLELAHRLDRETSGILMLAKKRSALVGLHEQIRENRM
DKRYFACVHGDWAADWGRRRVVKAPLFKYATPDGERRVRVQEDGLPSHTVFNLVDRWPDY
ALVEAELKTGRTHQIRVHLAHLGLPIVGDAKYGDFALNKALARANAVPSIKRMFLHAHRL
RLAHPLTGEPLQFDAPLPAECRQFIDQLSDLRDTA*
>GCA_000452945.1_BPMSHR5858_genomic_04733_1 yceF
MQHHACSPPRLILASSSRYRRELLERLRVPFDVVAPEIDETPLPDETPCATALRLAAAKA
RAAAERARAPHGALVIGSDQVATFDGLQIGKPGTHARALAQLQAMRGRDVEFHSALCLYD
SRSGATQSEDIVTRVRFRTLTDVELDAYLRAETPYDVAGSAKSEGLGIALLDAIDSDDPT
ALVGLPLIALTRMLRAAGYPLFGAPAPAADGVNGR*
>GCA_000452945.1_BPMSHR5858_genomic_04735_1 rpmF
MAVQQNKKSPSKRGMHRSHDFLTTSPLAVEPSTGEVHLRHHISPNGYYRGKKVVKTKND*
>GCA_000452945.1_BPMSHR5858_genomic_04739_1 fabG_9
MDKTLDKQVAIVTGASRGIGRAIALELARRGAMVIGTATTEAGAEGIGAAFKQAGLEGRG
AVLNVNDATAVDALVESTLKEFGALNVLVNNAGITQDQLAMRMKDDEWDAVIDTNLKAVF
RLSRAVLRPMMKARGGRIVNITSVVGSAGNPGQVNYAAAKAGVAGMTRALAREIGSRGIT
VNCVAPGFIDTDMTKVLPQEQQTALKTQIPLGRLGSPEDIAHAVAFLASPQAGYITGTTL
HVNGGMFMS*
>GCA_000452945.1_BPMSHR5858_genomic_04740_1 acpP_1
MDNIEQRVKKIVAEQLGVAEAEIKNEASFVNDLGADSLDTVELVMALEDEFGMEIPDEEA
EKITTVQQAIDYARANVKA*
>GCA_000452945.1_BPMSHR5858_genomic_04744_1 group_19306
MGSVSTQSQPCSQRERVSALVDGEAPEGLSLKQILAGLADAERATWAQYHLIGDALRSDE
LTLEPAESSAFMARFSASLAAEPHLLAPAAAPIARRLLAMRRRVVPAFAIAAAAATLTWI
VVPQMQGVGAPGAVQVASVSPVQQGPLQRVAAAQTGAQDVNIIRDASLDQYLEAHQQFAQ
QPVVTGSMPVIRTAVTTQGQ*
>GCA_000452945.1_BPMSHR5858_genomic_04745_1 mucB
MRTLQLNQAILGWRRLPALLLCAAALLSAQPRAFAQPDDPVATRRQAAEWLDRIQQAAQQ
QNYEGTFVYQRGAYVQSSRIVHVATKGDGEYERVESLDGKPRKLLRHNDDLYTFVPERRL
CVVERRQNKDSFPALLGASGEQVLSVYEPKSLGRDRVAGLDAQVVELVPKDAYRFAYKLW
TDAKTGLLLRSQTLDTDDRVLEQIAFSQLQLGAAAPSQKTAIAAGMRNLGGWTVVRPPVA
PVDIEAQGWQIAPTVAGFRKIREVRRPMAARDANDPPIAVDQAVFTDGLATISVFIEPAE
KNSRKEGAGGTGATHVLVKRRGDFWITVLGEAPPSTLQQFASAIEYKASK*
>GCA_000452945.1_BPMSHR5858_genomic_04746_1 group_1536
MMNPSLRTWLVAAAVTALTPLAAQSATAAPNVTTTPAATGAVPAARAGLPDFADLVERVG
PAVVNIRTTANVPADTRGALPPGLDNGDMSEFFRRFFGIPLPQGPGGQKNAPSTPDAPDT
EQNRGVGSGFILSPDGYVMTNAHVVDDADTIYVTLTDKREFKAKLIGVDERTDVAIVKIN
ASSLPTVAIGDSNRVRVGEWVVAIGSPFGLDNTVTAGIVSAKGRNTGDYLPFIQTDVAVN
PGNSGGPLINMQGEVIGINSQIYSRTGGFMGISFAIPIDEAMRVAEQLKASGKVTRGRIA
VAIGEVTKEVADSIGLPKAEGALVSSVEPGGPADKAGLQPGDIILKFNGRPVEAASDLPR
MVGDTKPGAKATVTVWRKGQSRDLPITIAEFPADKAAKADSRQAPQQKPRSSALGLTVSD
LSPEQLKTLKLRNGVQIDAVDGPASRAGLQRGDIVLRVGDVDITSAKQFVDVTSKLDPQR
AVAVLVRRGENTQFIPIRPRQK*
>GCA_000452945.1_BPMSHR5858_genomic_04753_1 pdxJ
MSFFLTTPAAIDLGVNIDHVATLRNARGTAYPDPVRAALAAEDAGADAITLHLREDRRHI
VDADVRTLRPRVKTRMNLECAVTPEMLDIACEIRPHDACLVPEKRSELTTEGGLDVVGHF
DAVRAACKQLADAGVRVSLFIDPDEAQIRAAHETGAPVIELHTGRYADAHDAAEQQREFE
RIATGVDAGIALGLKVNAGHGLHYTNVQAIAALPGIAELNIGHAIVAHAVFVGWDNAVRE
MKAIMVAARVAALHGGR*
>GCA_000452945.1_BPMSHR5858_genomic_04758_1 group_1231
MTSSAPLPPPVPPPADTASPLQAASPVACDIFCAVVDNFGDIGVCWRLARQLALEHGWQV
RIFVDALATFARLQPAALPDAARQTVDGIVVEHWRAPAHAGDTLEIADIVIEAFACELPG
AYVAAMARRARPPVWINLEYLSAEDWVGEFHLRPSPHPRYPLTKTFFFPGLGPGTGGVLK
ERDLDARRAAFETGDDARRTWWQNVAGAPIPAPDTTVVSLFAYENPALDALLEQWRDGRE
PVALLVPEGRISARVARFFGAGAFGAGAHAARGSLVAHGLAFVAQPDYDRLLWASDANFV
RGEDSFVRAQWARRPFVWQIYPQADDAHLPKLDAALAHVTARVDHATRAATERFWHAWNG
AGTPDWTDFWRHRAALAARAASWADELAAVGDLAGNLANFAKTQLK*
>GCA_000452945.1_BPMSHR5858_genomic_04759_1 uvrC
MTSPDAPESRFEPKPILAQLPHLPGVYRYYDAQDAVLYVGKARDLKKRVSSYFTKTQLSP
RIAMMITRIARIETTVTRSEAEALLLENNLIKALAPRYNILFRDDKSYPYLKLTGHRFPR
MAYYRGAVDKKNQYFGPFPSAWAVRESIQILQRVFQLRTCEDSVFNNRTRPCLLHQIGRC
SAPCVGAIGEEDYARDVDNASRFLLGRQGEVMGELERKMHAFAAELKFEQAAAVRNQMSS
LAKVLHQQAIDVGGDSDVDILAVVAQGGRVCVNLAMVRGGRHLGDKAYFPAHVETALALA
GDIEALAGEGAGDGVQAAAQPAQAPLATDADATDAAATEAKTVTAAAAARAGARTAQAAG
ARAAASAEGDVERRAEGETHARADAREAAALPDGAAAAQEADADVDAAPLETEVLEAFIA
QHYLGNRVPPVLVVSHAPANRELIDLLVEQAGHKVAVVRQPQGQKRAWLTMAEQNARLAL
ARLLSEQGSQQARTRSLADVLGYESDDLAQLRIECFDISHTMGEATQASCVVYHHHRMQS
SEYRRYNIAGITPGDDYAAMRQVLTRRYEKMVEEAAAEASADEAAGIDGNAVHAAASAGR
LPNVVLIDGGRGQVEIARQVFSELGLDISMLVGVAKGEGRKVGLETLIFADGRAPLELGK
ESAALMLVAQIRDEAHRFAITGMRAKRAKTRQTSRLEELEGVGAKRRQRLLARFGGLRGV
VAASVDELASVEGISRALAEQIYRQLH*
>GCA_000452945.1_BPMSHR5858_genomic_04760_1 pgsA
MPFNFPIFLTWVRIVLIPLVVGVFYLPDTMLSADHRNLAAAVIFILAALTDWFDGYLARK
WNQTSAFGAFLDPVADKLMVTAALLMLVQLARIDAAIALVIVGREIAISALREWMAQIGA
SKSVAVNSLGKFKTACQMVAIPMLLFYGRLPFGGGVAIDTRGWGEWLIYLAAVLTIWSML
YYMKLAWPQIRERGGAA*
>GCA_000452945.1_BPMSHR5858_genomic_04770_1 group_1922
MAVAHDCFEPNSPMPFPFFTARAQAASSPASARRTRFLRFAAACAAAAALAGCTPRGEPW
RLANVTGHLPDLTFSLTGEDGRPIDASAFRGRAALVYFGYTHCPDVCPETMARLMRVLAL
LSPQARSEVRILFVSVDPARDTPAAMRDYVGAFDAEHAYGLTGSERQIESLAKRYRVAYQ
MEKRDPNGSYEVTHSSAVYVFDAQGRARLLATDQDSPDAIAADLRRIIVNRS*
>GCA_000452945.1_BPMSHR5858_genomic_04779_1 group_1923
MTNQNRRDFLRLAAGTAGAAALQLFPPVIREALAIPANRRTGTIRDVEHIVILMQENRSF
DHYFGKLRGVRGFGDPRPLALQNGKSVFHQPVLLGPAELLPFHPDASNLGMQFLQDLPHG
WQDMHGAWNKGRYDRWIANKGTTTMAYLERDDIPFHYQLADAFTICDAYHCSIPSSTDPN
RYYMWTGYVGNDGAGGGPVLGNEEAGYGWSTYPETLEQAGVSWKIYQDIGTGLDAAGSWG
WTQNPYIGNYGDNSLLYFNQYRNAQPGSPLYDKARTGTNVSAGGTLFDVLQQDVKNGTLP
QVSWICAPEAYSEHPNWPTNYGAWYVEQVLKALTSNPDVWSKTALFITYDENDGFFDHVA
PPFAPQSRENGLSTVSTACEIFAGDATHMAGPYGLGPRVPMLVVSPWTKGGWVCSQTFDH
TSLLQFIEARFNDRYSVRAPNVTPWRRAVCGDLTSAFNFSSPDGSWPQLPDTSGYAPPDR
NRHPSYVPVPPAAQSMPKQEAGLRAARALPYELFVLGRIDQSTGKFKLTFANTGRAGAAF
QVTAGNRLDGPWAYTVEARKRLSDEWSTALTLSIYDLTVYGPNGFLCQFRGSTAAALGLS
ANPEVIYGYDVANGNITLRLSNRGRAAVRLTVTNAYGNAAPRVYELKPGQRINDYWDLRD
SHSWYDLSVSDGAPNGFLRRFAGHVETGRPSTSDPLIATA*
>GCA_000452945.1_BPMSHR5858_genomic_04785_1 chbG
VAACDAAPRALIFTADDFGLHERVNAAVERAHRDGVLSAASLMVGAPAARDAIERAKRLP
SLAVGLHIVLADGPATLPAHAIAALVGPDGRFRDAMAKDGCRFFFLPHVRAQLRREIRAQ
FEAFASTGLVLDHVNTHKHFHLHPTVLSMIVDIGREFGLAAMRLPYEPSTPFWLKPWIAL
VRARLDRAGIAHNDYVVGIEHTGAMDEAALLAALAKLPAGVGEIYSHPAEAGDGPITPSM
RDYRPADELAALLSPRVAAAVQAAGVACGGFADVFGGGARARISGAQPS*
>GCA_000452945.1_BPMSHR5858_genomic_04797_1 group_1108
MNFKGIDLGGTPPKPGSFGDIKLRANSGVGALKLIAAEQNGVELTWDLDTNWAQLKNELD
SSKWNAAVQEWDAKRRKVLLDRRAQLKARIGDLLVKGKPKLQRIRLYVFGFSRGAAEART
FSNWLADALDPDFSLCGVPVSYDFLGIFDTVASVGIAQSAAATLFDGHGGWGRKELMAVP
HYVRRCVHMVAAHEPRGSFPLDLIDCSMDGREEIVYPGVHSDVGGGYGPGEQGRGRDDAD
KLSQVPLLDMYREARKAGVPLDINGPGIGADVIAAFKVSAGLKQAFAAYVKKSENYYYEK
DHGTPGLMRCHYGLYLRWRKMRLKDMHAQPSFKAAQARYPQDAADLDSANAELKEEWNDL
LEVEKAGGPTITRYVEKFVTKWASENPKTAAVVSTSLLPAAVAAEVFVPGVSAATVLFGD
KVVKLVQAQLKEKWEQWLQVRSDWNMGPPEPAISALYDNYMHDSRAWFKPLGDDDDVWNH
RQIQELKTKQASFERAHAAWKKRVEMGMPSLLQVTQAMGVGSTGWGPVGGLPPEPEPRSP
LTAQQADLLKRYDAAMQSAKQARAAKDPNAPTDSAVLTDPKVTGGLALQTSGREFYFLWG
FLRWRTVFVNGVRWDQPRVPTVQEEMEGMRMQMQRQVDMKGIGVLFQ*
>GCA_000452945.1_BPMSHR5858_genomic_04802_1 group_1925
MARPFIVLGDKHSHGGTVTTAAAASAIHGKGIARRNDLVSCPIHGPNHIVEGDDTMIIDN
EGVARDGYRTACGAVLIASQSSTGGDAA*
>GCA_000452945.1_BPMSHR5858_genomic_04803_1 group_1118
MKQRAWLSKAAFVAAVFVVVWIATIAYWKATYHPPSATELLTHGIALPALLAAGLAWARR
AASRLREASAATAQTAAPSAPQASTAQAADGAAMRTWTLALLDSSVRFPTGATTADIDGA

ARDETVVGLHPALRRADGTGVFASGVASVSIDHFDERLLPPGADAALNDEHARALLLAAD
VLDELLERHAVLATDGASRPAEPLRLHLLLPERWQPMAGALAAWLDTHVARERWMPGVER
VQTRMVANPVQAWAVVDELIVSLHREPSNASHVVLACDSSLSDATVHALDHAGKLYGHNR
PDGRIPGEGACALLLAHPAAADAADAPRVHRLVAARAPTPDGAPAKAPADTLAPLLEAAR
TQTAAAEIAPAGCGLVSDADQRGGSRTEIAAVAERNWPDDARERCRHLGVANGESGAALA
LGAIAVAGWRASEEQMPMLAASIADPVARGVMLVAPAPIAPVTASELAAAA*
>GCA_000452945.1_BPMSHR5858_genomic_04804_1 group_19588
MDELLSYYERELAFLRRHVRDFAERYPKIASRMQLASGADGSSEDPQVERLFESFALTGA
RASRHIDDDYSEFTKAFIEVLYPHYLRVFPSCSIASFDIDTSRAAQMSAAVVVPRGTQLY
SRPVRGAKLFFRTAYDVTLSPLQLTAARFHAMPQAPRSFRLPPNASAQLSLSFAIRSPHA
SVADLKLDSVRLYTRGEPLMSAALRDALSIHALQAYVEPEQGGRWVALERVPFAAVGVSR
EDSLIPCPQGVHPAYPLLTEYFAFAEKFGFFDCDLREAGRLGRRHFTLHLLLKGIPADSA
KAGVLESLSAEHVLLGCTPVINLFETTGKLGQQPSAASGAHMQPLVVDKQNAYAYEVYSV
DAVTQVQETPQGERVTTFPSLHSLYHGGQAARASLYWRMRRDALVARSEPGHELSLGFVD
GALDPVAAPAGLDFKLTCSNRDLPEHLPHGAPGGDLMMEGGTLASRIGLLQRPTRPLRLR
EDRGVLWRLVSQLSSNSLLLAGGAGAVRDLLGLHDVQESPATVRQIAGIVDVSQKPVTAW
VSEKPFASVVRGLEIRITVDEECFAGTGVHTFAQLMDCLLSRYVAPNGFTQLVLVSSRTG
DVLCTCARRAGGGFLI*
>GCA_000452945.1_BPMSHR5858_genomic_04809_1 group_1926
MNPPSLIRRRLVLAGAALVSLPACAAESREIAHRPWGAIEPAEAAPDIEVSLADGTRGRL
VDLLRGKTSAIQFVFTGCSATCSLQGAIFQSLQAQLARHPIGGAQLLSISIDPANDTAVA
MTAWLRRFGAQPGWQAAIPASGDLARLARLYRDERNPADSHIDQAFIASRSARFVWKTDH
LPTPESVHDALRYYASH*
>GCA_000452945.1_BPMSHR5858_genomic_04936_1 group_1540
MRFMQIMRRARMLAPLAFAALQACGGDGVDPALPRLAAAAPAAMSGTCGALAAKLRYPRT
AFTSVATAAAGALSVAGKPVGEHCVIAGKMNERVSAIDGETYAIGFEMRLPKAWNGRFFY
QANGGLDGDVVAATGDIGGGGPLDNALRMGFAVISSDAGHSAAQNPLFGIDPDARLDYGY
RAVDALTPMAKQVIRLAYGKAPDRSYFGGCSNGGRHAMVAAVRNAGDYDGIVAGDPGLHL
PKAAIAEMAGAQQFARIATATAAAGRPDIRSGFTDTERRFVASRILAKCDALDGVADGLV
EDIAACQARFDLNADVPGCTNGVRTGQCLTGAQKTALANVFAGARNSAGEALYASFPFDA
GVSGDNWAAWKQGNAMTLDPAAAAFTFTTPPRRPSALADLGAYALGFDMDRDAPAIFATS
GVYAASAWSFMTPPDEADLSALKRRGGKLIVYHGTSDPVFSSTDTTDWYVRVMQANGGDA
SDFARFFRVAGMNHCSGGPAADQFDMLTPLVAWVEAGKAPDAIVATARDASNAAPNPEVP
ASWGAARTRPLCPYPRVARYDGGDVNAAASFSCR*
>GCA_000452945.1_BPMSHR5858_genomic_04941_1 group_19556
MSAPAIDARLLLLSPADNCLIAATRLCAGQAVTIDGERVVLAQTVELGHKIARAPLAAND
KVLRYGAPIGHVTREVARGEHLHTHNLVSDYLPTYTRDAGRAFVAH*
>GCA_000452945.1_BPMSHR5858_genomic_04942_1 suyB
MSMTVSTPASPELSADGVRRVLLAGYARSDGRKGIRNVVAVAYLVECAHHVAREIVAQFR
APPRLGDAAPAAEAPPVHLIGFPGCYPNGYAEKMIERLATHPNVGAVLFVSLGCESMNKH
YLVDAVRASGRPAEVLTIQEAGGTLNTIGAGVDWVRDARATLARQAKVPMGVGELIVGTI
CGGSDGTSGITANPAVGRAFDLLVEQGAACLFEETGELVGCEYHMQSRAARPELGREIVA
CVKKAERYYSILGHGSFAVGNADGGLTTQEEKSLGAYAKSGASPIVGIVKPGDVPPTPGL
YLLDVVPDGEPRFGFPNISDNAEIAELIACGAHVILFTTGRGSVVGSALAPVVKVCANPA
TYRKLEGDMDVDAGRILEGRATLDEVGREIFAKTLAVASGAASKSEALGHQEFILTYKAF
EPLGPACLPVAGPLRAVRPA*
>GCA_000452945.1_BPMSHR5858_genomic_04947_1 group_18988
VRYCLALDLKDDPDAIARYEAHHERIWPEVAAYLRAHGVVAMEIYRLGTRMTMVMDTDDA
CFDAARFDADARADAKIVEWEALMGAFQQPTPWTPAGVKWTPMTRIFDLSKQ*
>GCA_000452945.1_BPMSHR5858_genomic_04958_1 phnU
MSSLSHLEPRAALAADPAAAARIRASAAAHDAAARRRERRASARIVVLAAVVLGPLVLYP
LVRLVLLSAFGAHGPTLAAYRAFFASHDVRDVLGTTLWVLFASAGTASIVGVLLAALLFF
RPFPGASLVTRFLELYVAFPSFLVAFTLIFLYGSQGSVSIALQRVLGLDAPPLDFLFGPG
GVILAQTVFYTPFVVRPALASFATLDARLVEAAASLGANGWTLARRVVLPLAWPGIAAGV
ILCFLLTLNEFGILLVLGSARLVTLPVAIYSSATVDIDLPAAAAGAVVMLAMSLTLYALY
RRMSRRANEGGRDVR*
>GCA_000452945.1_BPMSHR5858_genomic_04961_1 phnW_1
MPERDPILLTPGPLTTSRMTRDAMLRDWGSWDAAFNRLTKSVCADLVRIAGGGDAYVCVP
LQGSGTFAVEATLGTLVPRDARVLVPNNGAYCARIAAILRRLGVAHVELPFAEDEPASAH
AIDAALARDARLTHVALVHLETSAGLLNPLDDIAAVCRARGKALIVDAMSSFGALPIALA
ASGIDALISASGKCLEGVPGMGFAIVRRSALEAAEGRSPSVALDLHDQYAYMQRTSQWRF
TPPTHVLAALRAALDQFFDEGGQPARGARYARNCATLVDGMRALGFEPFLDARAQASVIV
TFYAPADPAYAFPAFYAAVRDAGYVLYPGKLTTADTFRVGCIGALGADEMRGAVAAIGGA
LESLGIAMR*
>GCA_000452945.1_BPMSHR5858_genomic_04967_1 group_1932
MSAVDTSAAVRPAAPSAKPKTMLRRLLSHHEIATLLILLHAPIGANAKPELPALQEAGLV
EMIRSDMDAAPSFRLTEDGSAVLKGLGYR*
>GCA_000452945.1_BPMSHR5858_genomic_05103_1 group_1937
MRSSDAYRLGVDIGIDVRIDARNGANPFDVVARRRHARPDIQRNPFSHLPMAIRQSPVRA
CSRRATATRTRFPAWPTPSPGCMRRDAVTGESRGMPRAHNATR*
>GCA_000452945.1_BPMSHR5858_genomic_05104_1 group_1938
MSNVNAPRVRRSVRDLQKLYDNGEKKPLEDLVRAWAGIQALPPSDPKSFFALGGYHGEPF
QYRKPVDALPQSDIYPYWGGYCNHGNVLFPTWHRMYVYKLEEALQSIVPGVSMPFWDETD
EYTLRHGIPSILTQETFELDGTPIDNPLRSFVLPDALSDRLPGDGSIYEKPKGYLTVRYP
LSGLVGTPEALEQTKLHNAKFPLPEKNTELLNGNVRAWLRGDSPTPDDPDPTRNGVYAKY
VRCLSAPNYTVFSNTTSASVWSSSNPGLVTAVESPHNDIHLAVGGFDYGGGETGQIAGAN
GDMGENNTAGMDPIFFFHHCNVDRMFWVWQKQTGHTDRLDIIRNYPGTNASDSQGPTPGF
APGESLNLKTPLNPFKKASGEAYTSEDCINIERQLGFTYGPGSLDDVTPELKSLLAVPSG
NSTKKLTVTGIDRALIQGSFIMKAYASVTDANGKTREYYLGHKSILSRWNVVHCANCLTH
LDVVAHFPLSAMPADDVPKAEFRVKIIHRGGGVPSASKAAIGVVSGLQPNFEVSD*
>GCA_000452945.1_BPMSHR5858_genomic_05108_1 fhaC
MTVLLAGRSAPASAFGPTVQDRERDLGEKARRQQMIEQHRRGHAAEPAAGLPAPVQAPEL
GGARFPITQIEVRGGARHSREIDRIVERYRGATMGQAEIFALLRDLMNFYYGRGYITTGV
SLAEQNLSSGKLIVVVHWGYVKGWRVNGKAPEGMRERLMLSSAMPGIEGSVLNIHDIDQA
IENLNGLGRAAKINVVPAEEAGYSYLDVVLQRSKPFAVSASVDNSGLGDRAGDGLYRYGV
DIAVAHLLGWNDVLTLSAGRRYYQDPAHHAYDSLAVSYGWRVGRWSADLRYIAQPTKSLL
SGTYGDYQLSGNTQDVNLRVSRVLRRWRTGTDAVYVAIDRKRVRNYIDDTLIEINSNTHT
SVTAGVNRLDSLFGGVVFADGGWTRGVGWLGASEDPVSAGQGPLSRYDKFNLNLNWSRDF
TAGSLRARYTAAMGAQYSKDDLYYDSKFVIGDQYTVRGFKIRSAYGDSGVSVSNTLSFPF
PTSIGTVTPLVGVDWGAVGNNGEADNGGWIGAVAGGLRVSTKWANAAVTIGKPFRALPGM
GYGVVWYLSASATY*
>GCA_000452945.1_BPMSHR5858_genomic_05111_1 prpD
MSAPIPNVRPAPDSVLVEIADYALNHRIDSALALETARHCLIDTLGCGLEALSYPACTKL
LGPIVPGTIVPHGAKVPGTSLQLDPVQAAFNIGAAIRWLDFNDTWLAAEWGHPSDNLGGI
LATADWLSRTAVASGKRPLAMKDVLVAMIHAHEIQGCLALENSFNAVGLDHVLLVKVAST
ATVGCLLGLTRDELINALSLAFVDGQALRTYRHAPNTGSRKSWAAGDATSRAVRLALIAK
TGEMGYPRALTAKTWGFYDVSFGGKPFRFQRPYGTYVMENVLFKISYPAEFHAQTAVEAA
MTLHARLAAAGRTADEIRRVTVRTHAAAMRIIDKQGPLANPADRDHCIQYMIAVPLLFGR
LTAADYEDAVAADPRIDALRAKIACVEDPQFTQDYHDPAKRSIANALTIEFDDGSTLAEV
AVEYPIGHMRRRAEGMALLIDKFRTNLARRFPAKQQQAILDVSLDQAKLEAMPVDEYVDL
YVI*
>GCA_000452945.1_BPMSHR5858_genomic_05116_1 sdhC
MSETVRKPRPEYRNIGFGDITLKYRMPLAARVSILHRVSGALLFLFLPFLLYLFDQSLTS
ELSFEVFKAFLSNIIVKLIVLALSWAFLFHFCAGIRHLMMDMNHDAVSKERGKKTSVVVF
AVSTLLTIAVALKLFGAF*
>GCA_000452945.1_BPMSHR5858_genomic_05117_1 sdhD
MAANNRIGSKRLVVGAHYGLRDWLAQRVTASVMAIYTVILLVWFFIARDFSYEGWASIFA
TQWMKLATFVTLLSLFYHAWVGVRDIWMDYIKPVGVRLLLQSLTIVWLLACAGYAAQILW
RV*
>GCA_000452945.1_BPMSHR5858_genomic_05119_1 sdhB_1
MAKRIFEIYRYDPDKDAAPRMQTYELEIQHERMLLDALVKLKSVDETLSFRRSCREGVCG
SDAMNINGKNGLACLTNLNDLPQKIVLRPLPGLPVVRDLICDFTQFFNQYHSIRPYLIND
EPPPEKERLQSPEERDELDGLYECILCASCSTSCPSFWWNPDKFVGPAGLLQAYRFIADS
RDRATGERLDNLDDPYRLFRCHTIMNCVDVCPKGLNPTKAIGKIKELMVRRAV*
>GCA_000452945.1_BPMSHR5858_genomic_05120_1 group_19677
MSESSHQSDPHRRARLRWRARRGLLENDLIFERFFARHEHDLSDADVSALSRLLDLSDND
LMDLLLARKEPEGDLAGPDMSRLLEMLRSV*
>GCA_000452945.1_BPMSHR5858_genomic_05122_1 group_1632
MKPRYERVAIPDGCSVRVYRRRLAQIPFEWHHHPEYELTLTLNSLGKRFVGDHVADYAGD
DLVLVPPNLPHTWVSDERLDPGEPLVALVLWFDGDWAQRVADCCPEFAGLRTLLRRAAPG
LAFAPAAAADMRARLPALVDRAPRVRLAAALDVLACLADAPATPLATAAAYRAASGAALA
PEAERLDRVLDLLDRRFHEPLRVAELAALAHLSERSLQRRFARHVGESIGSYLQRLRLAH
AARLLASTDWPVSLVATRSGYANLANFNRQFLAARRVTPRAYRRFLAEHGRAPDDMPAHE
ASIDVRPPSLDHGPPRAGKNKIAR*
>GCA_000452945.1_BPMSHR5858_genomic_05124_1 ybiU
MPFPIDDLPAAIRTTKAVLRAAMPNRAPVFRALEGDIARQADAILTDRAQGRDTIPVLRF
ADIAADRVDPASLAALRTRGACVIRGVFDARQASDWNDEIAAYVEANRLDDKLARRAEDR
YFGTLASSRPQIYGIYWSKPQIAARQSPALTQARVFLNRLWRAQSEGRVHFDPARAPAYA
DRIRRRPPGSSSLGLSPHVDGGSVERWLEPNYRRVYRHVLAGDWRAYDPFDAAFRPDVEE
IPSPAVCSMFRTFQGWTALTPQGPGDGTLQLIPVANAMAYVVLRALQDDVPDDDLCGAQP
GRALSVLPAWHAPLLAALVPIPPMEPGDAVFWHGDVVHAVEDAHRGTGYSNVMYIASAPG
CAKNDAYLKRQLPSFLRGESPPDFPADHFETDFTGRARADDLSALGREQMGFGP*
>GCA_000452945.1_BPMSHR5858_genomic_05129_1 ntaA_2
MHEHRQMHLNCHIVGVGQHPAGWRTLRNTRAIVDPDFYRRVARVAEQGKFDALFFSDSLS
LHGHAHGPSQMLDPLVVASALALVTERIGLICTASTTFSDPFSLARRFLSLDQMSGGRAG
WNAVTTYNPAAAANFGPVALPSAAERYARADEFIDVTIKLWESWGERALVADAASGVFAD
PAQVRRIDHHGRHFDIAGPLNVPRSPQGRPLLVQAGASEAGLELAARHADMVFTSQHSLE
GAQRFYADLKSRVHAKGRDPDQFGILPGLYPVIGSTMAEARARKDEMDALRDPGGDIEML
ARQFGIRPEDLALDAPLPYERIARASRDHVSHGFAAQMIGFARGKNLTVRELIGHNLGMH
RIVVGTPESIADDMTQWFEQGAADGFNLNFDVFPDGLQTMVEHVVPLLQKRGLFRSDYSA
RTLKGHLGVRS*
>GCA_000452945.1_BPMSHR5858_genomic_05132_1 leuD1
MEKFNVHTGVVAPLDRENVDTDAIIPKQFLKSIKRTGFGPNAFDEWRYLDHGEPGQDNSK
RPLNPDFVLNQPRYQGASVLLARKNFGCGSSREHAPWALQQYGFRAIVAPSFADIFFNNC
YKNGLLPIVLTEQQVDHLFNDTYAFNGYQLTIDLDAQVVRAPDGREYPFEITAFRKYCLL
NGFDDIGLTLRHADKIRQFEAERLAKQPWLDNRLVG*
>GCA_000452945.1_BPMSHR5858_genomic_05133_1 leuB
MKIAVLPGDGIGPEIVNEAVKVLNALDEKFELEQAPVGGAGYEASGHPLPDATLALAKEA
DAILFGAVGDWKYDSLERALRPEQAILGLRKHLELFANFRPAICYPQLVDASPLKPELVA
GLDILIVRELNGDIYFGQPRGVRAAPDGPFAGAREGFDTMRYSEPEVRRIAHVAFQAARK
RAKKLLSVDKSNVLETSQFWRDVMIDVSKEYADVELSHMYVDNAAMQLAKAPKQFDVIVT
GNMFGDILSDEASMLTGSIGMLPSASLDQRNKGLYEPSHGSAPDIAGKGIANPLATILSA
AMLLRYSLNRAEQADRIERAVKAVLEQGYRTGDIATPGCKQVGTAAMGDAVVAAL*
>GCA_000452945.1_BPMSHR5858_genomic_05141_1 trpA
MSRIQNTFAALAAQGRKGLIPFITAGDPDPAKTVELMHALAEGGADVIELGVPFSDPMAD
GPVIQRSSERALAKGVTLHSVLDDVKRFRARDQKTPVVLMGYANPIERMGADAFAAAARD
AGVDGVLVVDYPPEESHDFAAKMRAAGIDPIFLLAPTSTDDRIAAVGQVASGYVYYVSLK
GVTGAANLDVSSIAGKIPAIKSRVPLPVGVGFGIRDAATARAVAEVADAVVIGSRLVQLL
EQAVPERAAAELAGFVAELRAAIDGAAKPAA*
>GCA_000452945.1_BPMSHR5858_genomic_05142_1 accD
MSWLDKLLPPKIKQTDPKSRKGIPEGLWVKCPSCEAVLYRNDVDANQHVCPKCDHHMRIG
ARERLDALLDPEGRYEIGQEIVPVDTLKFKDSRKYPDRLKEAMDETGETDAMVVMGGAIH
TLPVVAACFEFSFMGGSMGSVVGERFARGAQNALEQQVPFICFASSGGARMQESLLSLMQ
MAKTTAMLTKLSEAKLPFISVLTDPTMGGVSASFAFLGDVVIAEPKALIGFAGPRVIEQT
VREKLPEGFQRAEFLLQKGAIDMIVDRRKMRDEIAQLLALLQRQPADALA*
>GCA_000452945.1_BPMSHR5858_genomic_05145_1 cvpA
MLTAFDYAVLLVIGLSALRGAWRGFVGEIFGLIGWIAAFLIACRFVGVVLPYIPATWPGG
ALTQWVIAFALIIIGVVLVAGVANALLSRIAQATGLGGIDRSLGMMFGLVRGVLLVLLLV
AAAGLTELPKQEFWRDALLRPFAEQGVRELKPLLPDGLAAYIHV*
>GCA_000452945.1_BPMSHR5858_genomic_05146_1 purF
MCGIVGVISQSPVNQLVYDSLLLLQHRGQDAAGIATADGSNFHMYKANGMVRDVFRTRNM
RSLPGTAGIGQVRYPTAGSASSEAEAQPFYVNAPFGIILAHNGNLTNWQQLKDEMFRIDR
RHINTNSDSEVLLNVLAHELQLSTTGLELDPDSLFKAVAGVHRRAHGSYAIVSLIAGYGL
LAFRDPFGIRPLCIGKLETEQGTEWMVASESVAVEGIGFEFVRDVQPGEAVFIDVNGKFH
SRQCADKPSLNPCMFEYVYLARPDSCLDGVPVYNVRLRMGDYLAEKIRRELPNVPIDVVM
PIPDSSRPAAMQVAAKLGVEYREGFFKNRYVGRTFIMPGQAVRKKSVRQKLNAMSIEFKD
KHVLIVDDSIVRGTTSHEIVQMARDAGAKSVIFASAAPPVKFPNVYGIDMPTRGELVAHG
RTDEEVAKIIGADYLIYQDVGDLRRAVRDINPSLEGFEASCFDGDYITGDVTPEYLDAIE
RARLAPASQADRDPSGEGAERSQMNLQLSVE*
>GCA_000452945.1_BPMSHR5858_genomic_05176_1 group_18981
MSKIRSAVPPPPLPEVVEGQRYATPQRGVTLADTMFVDCHFERVEWTGCRLANLRFVNCT
FDANRFDRCELEKLSYDSSRIRAGAWTQSALQRVSFNECELDGGTWTGSLVKDVVCTQSK
GGAWTFDAVRGAHVSLVAGDYAGVTLRGGRWSDTSWIGSRLADLRLESVELENLIAGQSG
FERVVLVECRGVNVRWIDSRIERMTVHGCELKQAAWSHSTWATGEIHASRLPIASFDHAS
VNGLTVTNSELPQAIFDSASVADSALQGVRAPRIALRDAWLTRVNLSGAQMQQLDARGVH
LERVDLRGADCRGGNLIGQLSHTWAAADTRDAVFEEATSADDRLWWQRVQPGARGV*
>GCA_000452945.1_BPMSHR5858_genomic_05178_1 group_19153
MFANCSAGGMAIAPGNDVCKTPPLAIPIPYPNIANKPEAVPNVPNILINGGPAHNLNTII
PVTHSDEGGSMGGVASGTVSGPSRHAKGSSKVMIQGAPETRLTDTNLPNNQNTAGFSAVP
SQTITMTLS*
>GCA_000452945.1_BPMSHR5858_genomic_05182_1 group_19332
MIRTSLRVFAAILIAILIWWVGPLFAFGIYHPLGPVWVREILVALVLIWGFWPTLARLWA
RLAMSPRQVKVAPKAKQLDFVDKHLRTLDQQLKERWRKEPRGRWKRWVGALTREHRTMLP
WYLVLGSEGSGKSSLVAKAVSVSGSLQDRVLGSDATYGRGDDLNFRITREAVWFDVGGRW
SLRAGADEAEFDAWRKLLRGMRRLRRGAPISGVVLCVDGLEMIDAPLDARKRLADSVRAR
LEEMREAFGQQVQVYVALNGLDRLDGAVSTLSLLEASKWVKGVGFSLPDDGAHADAARAD
ASWQHALQGLQQRVQQQVLYSAPAATEVSMNHAQLRFVETLSRLQKALVAWLHVALAPGE
PHTAARLRGVWLGSMAELAEPHPAGAGGVGSAELPVPSRPLSELWTPLIRQVALERDAVR
PSGPKSWRGRLGEALRWGAVPLVALSLLLWFGWGYVTERDYLDGVWAQFTEAKRLAQAEA
SYGNDGGSTLIEIANQMRYAQLQAEDAAQGMATPYFEHGLVAETARETYYRHLQKMLMPE
LYNEVRRTLVSQVDGSPGDIYQTLKVYLMLCRPARRSADDVVRWLDGRWDALSGGQYSDD
DRRSLLAHVRTLMSLKEVPATPEDANLVRSARAKAAQIPLVTRVLQHIHAQGLPQQVNDI
SLSRAAGFEASMSLRMRSNVPSTDTAVSGWFTRAGYTDVFLPRLQKSARAMLEEESWVLR
DETLSGNSFQIDGLVQKLADSARNQFLQDYISAWQNFLNDVTVRGVTGLDDASQLAAAMM
EAQSPLANLLRFAARETTLTGASDEGNIDSWIDRQKYRFEKGRRQVVGELSGQHYRTVLL
PEHVVEEHFQAIRQLAAQLNRNNTIANNPLSRLFEPLYRQLGLVNGALQAGQVLPAQYDA
FSRLKETAARQPEPVRGIMLDLVSSGSTMTTRESGALLNRGAAGATKMVCDQGFTGRYPM
RRGAQADAGVEDFERLFSAQGLMATYFRDHLAAYVDTSAKPWQALRSNGGPNGMVSQSVL
NSYETAERIRGAMLDDSGHLRVSTVLRFIDMDSQLSEAQLSVAGQTVRFAHGVTSPHRVD
WTNQNTQLAIKLQLKSVDGRMTTLQFDGPWALFRFFDAGQAVGGTGTADRRERLYQTSLG
TVRIEWQALTLPSPIWSGILQSFRCPS*
>GCA_000452945.1_BPMSHR5858_genomic_05183_1 group_19382
MIDPKEIFAACNNAHQYGQDASGSMAKPFMDSIPDLSTASSSNASNLIGNVQQVGSKMLE
HMSTIKSAVDKQVMIHQDNQQRQKTYDFNVDMRDLRPTNAVGFPEEMPSIFFAPFRTGK*
>GCA_000452945.1_BPMSHR5858_genomic_05185_1 group_19557
MFQTSIISVVGVLVAIVLIFVVVTTTRTLSTVMSSVQAQTVAKNGYEAMADVLSIGQTGV
TLNNVPMMRIELRVHHNGASWDVTIKQFIDLGNIPRAGERVRVMVDPADNGHVAYVGLAG
AGR*
>GCA_000452945.1_BPMSHR5858_genomic_05205_1 tig
MANVVENLGKLERRVTISLPKDVVQKEIDARIQKLAKNVRMPGFRPGKVPLKMVAQQYAG
QVEAEVLSDKIGQEFFTISRAENLRVAGQPSFAPKEDTQQESAYAFDATFEVYPEVKIGD
LATAEVERSTTTIGDAEIDRTLDILRKQRVHFHARGEGGEHGDGGADTAAQNGDRVTVDF
VGKIDGVAFQGGTAEDFVFVLGEGRMLPEFETAALGLKAGESREFDLKFPDDYHGKDVAG
KTAQFKVTLKKVEWPHLPEIDADFAKSLGVEDGDLTKMRAEIKENLEREAKRRTQSIVKN
QVMDALLKISELDVPKALIEQDQQRLVEMARQDLAQRGVPNAKDAPIPAEMFADQAERRV
KLGLVLAELVKANGLEAKPEQIRAEVDEFAKSYEDPKEVVRWYYSNQQRLAEMEAFVVES
NVVDFVLGKAKVTDKEVSFEALASATAQA*
>GCA_000452945.1_BPMSHR5858_genomic_05206_1 clpP

MINRAQLLDTLASQAPRDFEAQALGLVPIVVETSGRGERSYDIYSRLLKERIVFMVGEVN
DQTANLVVAQLLFLESENPDKDISLYINSPGGSVSAGMAIYDTMQFVKPDVSTLCMGLAA
SMGAFLLASGAKGKRYALPNARVMIHQPLGGARGQASDIEIQAREILYLRDRLNHLLAHH
TGQDVERIARDTDRDNFMSSEDAKAYGLIDHVSTKRP*
>GCA_000452945.1_BPMSHR5858_genomic_05233_1 hcaD
MREFDYLLVGGGIASVTAARTLRGEDASASIAILCGEPVLPYQRPPLSQEWLMGAVQPAS
ITLHDAAFYAAQRIDIALGVRAEQLDRAKRIVRTSSGATFRYGKLLIATGASAKAPALPG
FGLDGVHVLHTFAQAQALKDAAAHARRATVLGGGFLGVEIAATLQALGLQVTLVERAPGV
MPSLRAPALGGHFEALCRARGIDVLTRCEVRRVLGAQCVEAVETSDGGTRPCDLFVAAVG
VTPNCGWLDGSGLALGDGVEVDAFLQTADPDVFAAGDVAHFDDPIFGVRRRIEHWDNAVR
QGRIAARNMLGHRLPYRDVSIFYGSVFGLSYNLLGYPVGATETIERGAFDEASYMQLYLA
DDVLRAAFTIDRPAVEIAALNDAIRLHVNVAAHKTKLSDPNYPLDKLPVQTILILQGGGA
LGAFECGAVKALEQHGVRPDVISAVSIGAFNGAIVASHPKGATKALEAFWRELSIALPLP
YESQWREAWLSSYVLSFGVPNFLKPQWSQLGCDADRLRAKWTSFFDATAIEQLIARYVDF
DSLGMSPTRLLIGAVDVETGEPRIFDSYVDRLSPAHVLASGSLPPGMPWTVVDGRPYWDG
GVVSNSPLDLVIARCGQIGGRVFIVDLFSGSKALPTNLVEVMLRREEIAYMDRVRNDLRF
EEYASDFRDLVDGILAHVDAPSANEIRQQPLYIRLMGNRAAIDIARITLNREGRLSFIED
YDFSSDAIAQLQQQGYDAALVALANGRSRRRRGRRRA*
>GCA_000452945.1_BPMSHR5858_genomic_05240_1 group_1299
LTERERAARGRNDARRGGGDSGGDDEAWRRWLPGIATLRAYRRAWLARDLYAGVALTAVL
VPVGMSYAQAAGLPVIAGLNASIAALVGYAIFGPSRILVLGPDSALAALIAGAIAPLAHG
EPAHAVALAAALALMSGGFCVLAGLLKLGFVTDLLSKPIQYGYLNGLALTLIASQLPSLL
GTVPLGGTFVDQVASLAATVAQGRIDFASLALGGGCLAGIALLRRVAPAWPGMLIAVAGA
SIVAAWLGAAPDAGGAHAHVAYAHVAYAHVANAHVALVGSLAGTLPPLGLPSISLADASR
LIAGALAIAMVSVADISVLSHVFAQHDGSETDRNQELCALGAANLLAGMLRGCAVSSSAS
RTPVALAAGARTQLTSLVAAACIALLLVAPTLLARVPLAALAAVVVYSANALVDVRAIVR
LYRVRRGECAVSVLCFAGVVLLGVVPGILLAVGLSLLSFVWRAWHPYDAVLGRVEGMHGY
HDVSRHPGAALTRGLVAFRWDAPLFHANATIFRDHVRDAIAEADAPVRCVVIAAEPITDV
DVTAADMLATLRDELAARRIALVFAEMKGPVKDRLRTYGLFEKIGADHFFPTVTDAIEHF
TRMRKDVATARRARR*
>GCA_000452945.1_BPMSHR5858_genomic_05254_1 soeA
MDDRARSGGEPREVKTTTCYMCACRCGIRVHLRNGEVRYIDGNPDHPLNQGVICAKGASG
IMKQYSPARLTQPLMRKAGAERGSAQFEPVSWDVAFSVLEQRLAHLRATDPKRFALFTGR
DQMQALTGLFAKQYGTPNYAAHGGFCSANMAAGMIYTVGGSFWEFGGPDLDRAKLFFMIG
TAEDHHSNPLKIAISKFKRAGGRFVAINPVRTGYAAIADEWVPIRPGTDGALFMAMIREL
IETGGYDRDFVTRYTNAAELLDMRAEADTFGLFVRDASRPERNPLFPQNHLWWDLGSGRA
VAHHTRGATPALDGRYALDDGTPVAPSFALLRERVAECTPQWAERITGIPAATIRRLAHE
MADVARDHKITLPIRWTDAWGETHDTVTGNPVAFHAMRGLAAHSNGFQSIRALAVLMSLL
GTIDRPGGFRHKSPYPRAVPPSAKPPNGPDAVRPNTPLAAGPLGWPAAPEDLFVDEQGGP
VRIDKAFSWEYPLAVHGLMHSVITNAWRGDPYPIDTLMIFMANMAWNSSMNTVEVRRMLA
DRHDNGDYKIPFIVVCDAFQSEMTAFADLILPDTTYLERHDAMSMLDRPISEFDGPVDSV
RIPVVPPTGECKPFQEVLIELASRLKLPAFTNADGTRKFRDYPDFVINYQTAPDSGVGFL
IGWRGEDGGDALVGAPNPRQWDEYEKHGCVFHYTLPDTLQYMRGCNGPYLKWAVEKGFRK
YDAPIVIHLYSDVLQKFRLAAQGRTRGRQPPEHLRARIARHFDPLPFWYEPLELGATDLQ
RYPLAAVTQRPMAMYHSWDSQNAWLRQIHGENALFVNPKVARDAGIDDGGWIYVESQWGK
VRCRARYSEVVEPGTVWTWNAIGKAAGAWNLGPDANESQRAFLLNHVITDELPGEGAHAP
RISNSDPITGQAAWYDVRVRIYPAEADADHTLPQFAPMPALPGVTGAVRRIVQTYFAGRG
EFAARLRDAAKRR*
>GCA_000452945.1_BPMSHR5858_genomic_05264_1 group_19648
MKQTKHLAALAVPAGFLLSAGAHAQSSVTLYGIVDAGIAYVHNVQGADGRNEASLFKFSS
GNLSGSRWGLRGTEDLGGGLAALFQLENGFNLGTGQQNGNREFGRKAIVGLASNTWGSVT
LGRQYDPVVDLVQGLTEDNYFGGVFATPGDLDNYDNSLRVSNSVKYTSPVLSGFQFEALY
GFGGVAGATGSGRTYSFAASYANGPLSLGAGYLYADGGANATNGVRTWTGSSDTLFNTVI
NQGFASAKSIQIVRAGGQYVLGSATFGLAYSNTQYRRDSLSAFSDTAKFNSGSAFFNYQF
SPALRAGVGYNYTSLTGPASAHYHQVNLGAGYALSKRTDLYALFGYQKASGHTLGANGAV
IDAQASVGSYGANSGTDSQELAIVGIRHKF*
>GCA_000452945.1_BPMSHR5858_genomic_05276_1 astC
MKSPNVSRQTFDEVMVPVFSPAPFIPDRGEGSRVWDTEGRDYVDFAGGIAVTALGHGHPE
LLKVLDEQSRKLWHIGNGYTNEPVLRLAKRLESLTFADRAFFANSGAEANEAALKLARRV
AFERHGADKYEIVSFLQSFHGRTFFTVSVGGQSKYSEGFGPVPEGIKHLPFNDIAAAQAA
IGPKTCAVIVEPIQGEGGVIPADPAFLKALREACDAHGALLIFDEVQTGVGRTGHFYAYM
DTGVTPDILTTAKALGNGLPIGAMLTTEALAAHFKVGVHGTTYGGNPLASAIAEKVVELV
GDPALLEGVKQRSARLTATLEKINARFKLFKEIRGKGLLIGAELVGALDGRAKDFVSAAA
EHGVIMLIAGPNVLRFAPSLVIPLDVLDEGLARFEKAVEQVLAQAEAAPR*
>GCA_000452945.1_BPMSHR5858_genomic_05277_1 astA_1
MLFVRPGRLADLDALAQMARTAQPVLHSLPHDRAALEARVALSEDSFRAEVDFPGEEFYL
FVLEDGATGKLLGTASIVAAAGYAEPFYAFRNDALIHASRELHVNRKIHALTMSHELTGK
SRLAGFYVDPSLRGDAAAHLISRARMMYIAANRNRFTPEVFSLLLGVTDETGASPFWEAV
GRKFFGRDFAAVEIESGGRSRTFIAEVMPTYPLYVPLLPEAAQRVLGEPNEHALLAYDIH
LEEGFEPDRYVDIFDAGPVLTAQVDRTACVTKHAERTVREAAHAHGGTTYLLSGGRGDAF
RCVLADLPADGADAPLPADARAALGVADGDTVRCVPLHQRDGQPTDQPGDAA*
>GCA_000452945.1_BPMSHR5858_genomic_05278_1 astA_2
MIVVRVVQTGDVDALVALAQETGPGLTTFKPDRDALAARIERTRRTLGNDATPAEAGYFF
VMEDSQTGDIAGVCGIETEVGLEQPFYNYRVSTVVHASQDLGVWTKMSLLNISHDLTGYA
EVCSLFLSPRYRTGGVGGLLSRSRFMFIAQFRERFPERICAELRGHFDDDGTSPFWRAVG
SHFYQIDFNAADYLSSHGRKSFLAELMPRFPVYVDLLPRDAQDVIGVTHRDTLPARKMLE
AEGLRYQNHVDIFDAGPVLECHVADLRTVRESVVVPVAIGVPAARENAPRSLVSNTSLAD
FRVGVAPGLVDAGVFTLSAEDAAALRVSEGELVRVLPLKPKQG*
>GCA_000452945.1_BPMSHR5858_genomic_05279_1 astD_1
MTELFIDGAWRDGAGPVFASRNPGTGEPVWEGAGASADDVERAVASARRAFAAWSALDLD
ARCAIVKRFAALLVERKEALATMIGRETGKPLWEARTEVASMAAKVDVSIAAYHERTGER
RSPTADGVAVLRHRPHGVVAVFGPYNFPGHLPNGHIVPALIAGNTVVFKPSELAPGVARA
TVEIWREAGLPAGVLNLVQGEKDTGVALANHRQIDGLFFTGSSDTGTLLHRQFGGRPEIV
LALEMGGNNPLVVADVEDIDAAVHHAIQSAFLSAGQRCTCARRILVPRGAFGDRFLERFA
DVASRITADVYDADPQPFMGAVISARAASRLVAAQAKLLELGAAPIIEMRQRDPALGFVN
ASILDVTPVRELPDEEHFGPLAQIVRYTDLDDAIARANDTAFGLSAGLLADDETVWNTFR
RAIRAGIVNWNRPTNGASSAAPFGGAGRSGNHRPSAYYAADYCAYPMASVESAQLQMPAN
LSPGLHF*
>GCA_000452945.1_BPMSHR5858_genomic_05280_1 astB
MNAKEANFDGLVGPTHNYAGLSFGNVASLSNEKSDANPKAAAKQGLRKMKQLADLGFAQG
VLPPQERPSLRLLRELGFSGKDADVIAKAARQAPELLAAASSASAMWTANAATVSPSADT
SDARVHFTPANLCSKLHRAIEHESTRRTLAAIFADEARFAVHDALPGTPALGDEGAANHT
RFCAEYGAPGVEFFVYGRAEYRRGPEPTRFPARQTFEASRAVAHRHGLREEATIYAQQRP
DVIDAGVFHNDVIAVGNRDTLFCHEHAFVDRQAVYDALAASLGALGAQLNVIEVPDRAVS
VADAVGSYLFNSQLLAREDGTQMLVVPQECRENANVAAYLDALVAGNGPIRDVRVFDLRE
SMKNGGGPACLRLRVVLNDAERAAVKPNVWIGDALFASLDAWIDKHYRDRLSPVDLADPA
LLDESRTALDELTQILGLGSLYDFQR*
>GCA_000452945.1_BPMSHR5858_genomic_05282_1 gstA
MRRNSRKVAIVPPRARRAVEPALGRRRGGCAARTARNPQEDVAIAQGRSVRRAPAPLRAR
AVVGERVGQRRAQMRVDGTRHTETAALLQRIGDLAGAGTLIPAHGTPARYEVIEWLTFVS
SELHKTYSPWLFHADTADSTRRQCLEKLDRRLQEVDAHLATRDYLTGAFTVADAYLFAVA
NWSNFLRIPLAPYPHLVAFMARVGARAKVGEALAAEGLGR*
>GCA_000452945.1_BPMSHR5858_genomic_05283_1 argT_4
MKLNWRKMAAHAVVTATALAAGSAFAADLKEIRFGVEASYAPFEYKTPDGKLAGFDIDIG
NAVCAKLKVKCVWVENAFDGLIPALQARKFDAINSDMTITEQRRKAIDFTDPIYTIPNQL
IAKKGSGLLPTTASLKGKRIGVLQGTIQEAYAKKRWAPAGVEVVPYQTQDLAYEDLKSGR
LDATFQDSEAGAKGFLSKPQGAGFAFAGDHVSDAEILGTGVGFGMRKNDAQLKSAVNQAL
KELKADGTIDGLAKKYFSVPVTLK*
>GCA_000452945.1_BPMSHR5858_genomic_05309_1 yajC
VSFIPNAYAQGAAGGAESSLMSFLPLILMFAVLYFIMIRPQMKRQKEHRNMLAAMAKGDE
VVTSGGLVGKVTKVSEAYIGVEIAEGTEITVQKAAVTTILPKGTIKSL*
>GCA_000452945.1_BPMSHR5858_genomic_05312_1 group_1050
MHAADAASPATDTVPPRVWRAVVAASIGNALEWFDLVVYGFFAITISRLFFPAGNDTVSL
LLTLGTFGVSFFMRPLGAIVLGAYADRHGRKAALTLSILLMMAGTFVIAVLPTYATIGIA
APVVLVGARLMQGFSAGGEFGSATAFLAEHVPGRRGFFASWQVASQGLTTLLAAIFGTAL
NAGLTSAQMAAWGWRVPFCFGLLLGPVAYYIRSKVDETPEFLASTPAANPLRDTFAAHKG
RLAAAIGAVVLGTVATYLVLFMPTYGVKELKLAPSAAFAAILVVGVIQMGFAPVVGHWSD
RHGRVRTMCVPALAILVLIYPAFAWLAAHPTFGALIAVQIVFAFLMTGYFAALPGLLSEI
FPVATRTTGMSLAYNVAVTVFGGFGPFIIAWLIRATGTKAAPSFYLIFAALASLAALVAL
RRRFGLR*
>GCA_000452945.1_BPMSHR5858_genomic_05313_1 yceI_2
LNKQLMIAAGALAAALSFSAHAAPATYQFDPSHTYPSFEADHFGGLSVWRGKFDRSSGTV
TLDRAAKTGTVDVTTKVASIATGSQKLDEHLQTADFFDSAKYPDATYKGTIKFEGDRPAE
VVGNLTLHGVTKPVTLKIDSFKCMPHPMLKKEVCGVDAVGEFNRDDFGLDYGKQYGFKMK
TKLLITAEAVKQ*
>GCA_000452945.1_BPMSHR5858_genomic_05316_1 yebS
MDRNALIACHECDALLHKPRLSGKDTARCPRCDALLYRSGSAQIDRICALTTGALITFLI
AQGFPILEMDVNGMRVQTTLFGALEAMWNQGMPLVAVMVFCSTLLFPLVEMSALLYVLFP
LRSGAIAPGFNGVLRAIQLVRPWGMIEVFMLGILVTIVKMVSLARVVPEAALFAFAALTL
MLAVVVMFDPRTLWDIADSLRERAAGAAPGSADESSAR*
>GCA_000452945.1_BPMSHR5858_genomic_05317_1 pqiA
MTGIVTAARAGMASCHACGHVQKLARPAQQALGSHDRPERCARCGAALHLRTPDSIARTL
ALLIAAAILYIPANLLPIMRTTSIVGSQEDTIMSGVVYFWTSGEWPLAVVVFVASILVPM
MKLGVLAILVYTAQRRSAWRPMQRTRLYRIVERIGRWSMLDIFVVTLTVALVHFRSLAVI
TAGPGALAFGSVVILTMLASMQFDPRLIWDNVEHSGNAHE*
>GCA_000452945.1_BPMSHR5858_genomic_05318_1 pqiB
MNRPQGPQHDPNAAPPTPPGGEPPVLPEPVITRRSGWLPSLVWLVPLVAALIGIGLVVKS
VLERGPEITISFKSAEGLEPGKTQVKYKDVEIGMVETIKLSKDLSHVLVEVQLKKEAEDF
AVKGTRFWVVRPRIGATGVSGLGTLLSGAYIGVDAGRSDETEKSFTGLETPPAVTGDQKG
TQYVLRGDSLGSIDIGSPVYYRRVQVGQVVGFSLDKDGTGVTFRVFVNAPYDQYVGLNSR
WWQASGVDLRLDSSGFKLNTQSLATVILGGIAFQSPPNQAAGPPAPNNMTFRLGADEGDA
MREPDGEPVQVVMNFNQSLRGLSVGAPVDFRGIVLGQVTNIGIDFDPKTKNFTMPVTMNV
YPDRLGRKFREASQDKGYAARREVLSRLVQHGLRGQLRTGNLLTSQLYVALDFFPKAPPV
KIDLAREPVELPTVPNTLDELQLQVADIAKKLDKVPFDQIGANLNSALANADKLFKRLDT
EIAPEARDTLSAAKQTFTTAEATLQQDSPLQSDVRGALKELTRTLQSLNALADYLERHPE
SLLKGKPGDQK*
>GCA_000452945.1_BPMSHR5858_genomic_05322_1 rlmL
MSSPTLFDFFAPCPRGLEAALAAELAQIAARHAIAPYDAGAQVPGGVHFRGAWAAGMAAN
LHSRIASRVLLKIAHRPYRNEQDVYALALEQPWERWFASTQTLRVDITAIKSPLKSLEFA
TLRVKDAICDRLREKTGARPSIDTAQPDVRVFAFLTAGDCTLYLDTSGEPLFKRGWRLDK
GAAPLRENLAAGILRLAGWTPGTPLYDPMCGSGTFVAEAAQIALGVAPGVERRFGFEKLK
QYDITAWQTLKVAAMDAKRAARGKRDDLLIFGSDISGDMLEKARANLERAGVPSLWLKQI
DARNMTPPAAAPGVIVTNPPYGERIEVRGRGPRGDVRETGKNRGGDEAFRRTHADSPDSE
FFRALGDALKQRFAGWQAFMLSSDRTLPGQLRLRESAKTPLFNGALECRLFRFDLIAGSI
RQRPAAADGE*
>GCA_000452945.1_BPMSHR5858_genomic_05326_1 pntB
MSMNVVTLLYLVASVCFIQALKGLSNPKSARRGNLFGMVGMAIAILTTVALIAKQAAWLG
ANLPLGLALVFGALVIGGAVGAVIAARVEMTKMPELVAAMHSLIGLAAVCIAIAVVAEPE
AFGLVPQDASAPNFIPYGNRIELFIGTFVGAITFSGSVIAFGKLSGKYRFRLFQGAPVVY
PGQHLINLMLALAMLGFGILFFITQSWLPFGIMTAIAFALGVLIIIPIGGADMPVVVSML
NSYSGWAAAGIGFSLNNAMLIIAGSLVGSSGAILSYIMCHAMNRSFFNVILGGFGGDAAA
ATAGGAKEQRPVKSGSAEDASFMLGNAETVVIVPGYGLAVARAQHALKELTDKLVEKGID
VKYAIHPVAGRMPGHMNVLLAEAEVPYEIVHEMEDINGEFGQVDVVLVLGANDVVNPAAK
NDPKSPIAGMPIIEAYKARTVIVNKRSMAAGYAGLDNELFYMDKTMMVFGDAKKVIEDMV
KSVE*
>GCA_000452945.1_BPMSHR5858_genomic_05328_1 pntAA
MHIGVPAETRANEARVAATPETVRKYAAAGHRVTVERGAGAGASFPDDAYAAAGAELADA
AAAFGADIVLKVQAPTDAECGQLKRGSVLIGMLEPFNAEQAARLAAAGVTGFALEAAPRT
TRAQSLDVLSSQANIAGYKAVLVAAALYPRFFPMLMTAAGTVKAARVLVLGAGVAGLQAI
ATAKRLGAVIEASDVRPAVKEQIESLGAKFLDVPYETQDERDAAQGVGGYARPMPASWLG
RQAALVHERAKQADIVITTALIPGRPAPTLISVETVQSMKPGSVLVDLAAGRGPEFDGRK
SGNCPLTVADQVIVHHGVTIAGHTNLAAMVPADASSLYARNLFDFLKLIVTKEGALNIDL
SDDIVAATLLCRDGEVVRK*
>GCA_000452945.1_BPMSHR5858_genomic_05334_1 ubiH
MTTAASTPAFDFDIAIVGAGPVGLALAGWLARRSATRALSVALVDARDPDASENDPRAIA
VSQGSRMLLDTLGWPGDATPIEHIHVSQRGHFGRTLIERDEHDLAALGYVARYGSIVRAL
ARAVRGTPARWLTSTSAHSPAQDADGVTIALDTPQGARTLRARVLVNAEGGLFDERKRKL
TDSATNRDYGQTALVGTVTVSAPRPRVAWERFTAEGPLALLPLGGPQQADYALVWCCAPD
VAQRRGALPDDAFLQELGATFGTRMGRFTRIVGRAAFPLGLAAAHTLVEGRIAIVGNAAQ
TLHPVAGQGLNLGLRDAHTLAETLSEHGATPLALATFNARRALDRRMTIGATDTLARLFT
VDSRPLAMLRGAALTALEFVPPLKTAIARQMMFGQRR*
>GCA_000452945.1_BPMSHR5858_genomic_05336_1 fis
MSKHNIEQCVRDSLDGYFRDLDGSNPHDVYEMVMSCVEKPMLEVVLEQAGGNQSLAAEYL
GINRNTLRKKLQQHGLL*
>GCA_000452945.1_BPMSHR5858_genomic_05337_1 purH
MIKQALISVSDKTGIVDFAKALSALGVKLLSTGGTAKLLADAGLPVTEVADYTGFPEMLD
GRVKTLHPKVHGGILARRDLPEHMQALEAHGIPTIDLLVVNLYPFVQTIAKDDCTLADAI
ENIDIGGPTMLRSAAKNHRDVTVVVDPADYAVVLDEMKANGNTLGYKTNFRLATKVFAHT
AQYDGAITNYLTSLGDDLQHGSRSAYPATLNLAFDKVQDLRYGENPHQSAAFYRDVATPA
GALANYRQLQGKELSYNNIADSDAAWECVKTFDAPACVIIKHANPCGVAVGEDAGEAYAK
AFQTDPTSAFGGIIAFNREVDEAAAQAVAKQFVEVLIAPSFSDAAKQVFAAKQNVRLLEI
ALGEGHNAFDLKRVGGGLLVQSLDSKNVQPRELRVVTKRHPTPKEMDDLLFAWRVAKYVK
SNAIVFCGNGMTLGVGAGQMSRVDSARIASIKAQNAGLTLAGSAVASDAFFPFRDGLDVV
VAAGATCVIQPGGSVRDDEVIAAADEHNIAMVVTGVRHFRH*
>GCA_000452945.1_BPMSHR5858_genomic_05344_1 dtd
MIALIQRVKRADVRVGERVTGEIGPGLLALVCAERGDTEAAADKLLAKVLGYRVFSDAAG
KMNLPVSNLDGAGRAGGLLLVSQFTLAADTNSGLRPSFTPAAPPDEGERLFDYFVRRARE
RHPIVATGEFGADMQVSLVNDGPVTFWLQTRA*
>GCA_000452945.1_BPMSHR5858_genomic_05348_1 yfcG_3
MLRFFYHPSPNPAKVALFLEEAGVPYEFVPVDTRKGEQHSDAYKAINPNAKTPAIADGDA
IVFDSNAILLYLAEKTGQFLPDDTPAARGELLSWLMFVATGIGPYCGQAVHFKHFAPEPK
AYAVNRYDFEAWRHWRIVDERLAGRRYMVGERYTIADMAVWGWARAIPRVLGDDAWAQLP
HVKRLVDEIDARPAAQRADALKARHAFKVEFDDEARRALFPQNARLAATAGTGA*
>GCA_000452945.1_BPMSHR5858_genomic_05349_1 rhaR_8
MIDRLKDLLARFELHARVFHFGALPGASTFEIGADGFHMHLVRTGAVNVTGEALGRHAVP
EPGAVFIGCPGKYRIEARGDAPVEVLSAAIEFGLGDENPLLRGLPDLLAIPLASMSPLGG
VQQALFAEASAPACGHDTVINRLTEVLVVQLLRFVMRNRLVASGSLAGLSDARLAKALNA
MHADPALPWSLERMAAIAGMSRSRFAAHFAGTVGLPPGEYLLQWRVGLAKTLLRRGYAVK
EIAPEVGYGSASALTRAFAQSTGQAPTDWLARAGDAPGAIGAAADSMPDVGVRAA*
>GCA_000452945.1_BPMSHR5858_genomic_05350_1 erpA
MNAVTESAATTEMPAPFVFTDAAADKVKQLIDEEGNPDLKLRVFVQGGGCSGFQYGFTFD
EEVNEDDTVLNKNGVVLLVDAMSYQYLVGAEIDYKDDLNGAQFVIKNPNATTTCGCGSSF
SV*
>GCA_000452945.1_BPMSHR5858_genomic_05351_1 rpsI
MIGNWNYGTGRRKSAVARVFIKAGKGDIVVNGKPISDYFSRETSLMIVRQPLELTNHAQT
FDIKVNVSGGGETGQAGAVRHGITRALIDYDATLKPALSNAGFVTRDAREVERKKVGLHK
ARRAKQFSKR*
>GCA_000452945.1_BPMSHR5858_genomic_05352_1 rplM
MKTFSAKAHEVTREWYVIDATDKVLGRVASEVARRLRGKHKPEFTPHVDTGDFIIVINAS
KLKVTGNKTLDKKYYRHSGYPGGIYETTFGKMQERFPGRALEKAVKGMLPKGPLGYAMIK
KLKVYAEATHPHSAQQPKALEI*
>GCA_000452945.1_BPMSHR5858_genomic_05354_1 group_1947
MLNDEARAERLATGRGLPLRFIAQAALPAGIAYETHIAETGAVPTRHNLHDFFNALVWFA
YPRIKAALNARQAAAIDAAGVGAVRGGVRDALTLLDENGALFATSDPALAAALRGFDWPT
LMRASRDAWGARCDARIVGHALCEKLVDPYKGCTAHAWIVEVPAAYFDWPDARRRAWLDE
RVAAALAATDPASRGFAPLPVLGVPGWWPANASPAFYDDPQVFRRGRRARAE*
>GCA_000452945.1_BPMSHR5858_genomic_05362_1 group_19700
MHLNLTIDSKPLAIEIDDVIVGLLAIRLNLPENADPRDAISRHLSEAGGPWILDEEHMRK

RVLRRLILEVADPALVIRYLMVEQ*
>GCA_000452945.1_BPMSHR5858_genomic_05364_1 gltK
MHQFDWSSIPGALPTLWTGAVVTFKITLLAIVIGIVWGTLLALMRLSGVKPLAWFAQGYV
TVFRSIPLVMVLLWFFLIVPQLLQGVLGLSPTIDIRLASAMVAFSLFEAAYYSEIIRAGI
QAVPRGQVNAAFALGMNYAQAMRLVILPQAFRAMVPLLLTQAIVLFQDTSLVYVISLADF
FRTAANIGDRDGTSVEMILFAGACYFVVCSVASALVKGLQKKVTR*
>GCA_000452945.1_BPMSHR5858_genomic_05368_1 yjiE
MELKWLEDFVSLAETRSFSRSAELRHVSQPAFSRRIQALEAWLATELIDRSVYPTRLTQA
GQVFYEQALAMLSQAHEARTLLRGHVGAPVPTIEFAVPHTLSLTYFPRWLQRIEAKMGPV
HTRLRALNVHDAVLSLVEGGCDLVMGYHHPSHPVALDPARYDMLTLGTEPISAFSAPGRG
GRARYALPGTADAPVPYLSYTPNAYLGRMTEVIIANARAPLFLDKVYETDMAEALKAMAL
AGHGVAFLPHSAVEDALASGRLVRLGRAAKGAAEPFMLTMEIRLYRDKLAAQGDEPRQKL
VRTLWDVVREELGATA*
>GCA_000452945.1_BPMSHR5858_genomic_05373_1 eda
MKTIGEIVKLGPVIPVLAFDSVEQGEQVSRALHAGGVKVLEITLRTAAGLEAIRRASELA
EDIVVGVGTITKPEQCALAKQAGAKFGVSPGLTRQMHAAAVDAGLPLLPGVMTPTDIIVA
LELGYEIVKFFPAQQAGGVPMLQAFHGPFPTLRFCPTGGITAETAPSFLKLPNVVCVGGS
WLTPKAALAAQDWAEITRLAQAASGLAR*
>GCA_000452945.1_BPMSHR5858_genomic_05378_1 holA
MQLRLDALEPHLSKGLAGLYVVYGDEPLLAQEACDKIRAAARAAGFTERSVHTVERGFDW
STLIGASQAMSLFGERQLVELRIPSGKPGKDGADALKTLAGAANPDVLMLVTLPRLDAAT
QKSAWFTALANGGVALKIDPVERAQLPNWIGQRLALQGQRVAPGDDGRRALAFVAERVEG
NLLAAHQEIQKLGLLYPAGALTFEQIHDAVLNVARYDVFKLNEAMLAGDAARLSRMIDGL
KGEGEALVLVLWAVVEELRTLLRIKRGVAAGKPLAVLVRENRVWGPRERLVGPALSRVSE
AALEHALAFAARLDRQVKGLSAVSRGAARGEPPPDPWDGLFQLAMTVARAAGPGGDAPRR
PA*
>GCA_000452945.1_BPMSHR5858_genomic_05380_1 leuS_2
MHERYVPADVEAAAQSDWRAADAYRSKEDANRKKFYCVSMLPYPSGKLHMGHVRNYTIND
VMYRYLRMNGYNTLMPMGWDAFGMPAENAAMANGVPPAQWTYENIAYMKKQMQSMGLAID
WSREVTTCKPDYYKWNQWLFLKMLEKGVAYKKTGTVNWDPVDQTVLANEQVIDGRGWRSG
ALVEKREIPMYYMRITQYADELLNDLDGLGWPERVKVMQHNWIGKSFGVNFGFPYELDGE
KKLLRVFTTRADTIMGVTFCAIAAEHPLAARLARDKPALQAFIDECKRGGVAEADIATME
KKGVATGFSVSHPLTGEPVEVWIGNYVLMSYGEGAVMGVPAHDERDFAFAKKYGLPIRQV
IAVEGETYSTDAWQEWYGDKTRAVCVNSGKYDGLAHDAAVDAIAAELKAGGLGDKQITYR
LRDWGISRQRYWGTPIPIIHCPSCGDVPVPEQDLPVVLPEDLVPDGTGNPLAKSDAFLNC
TCPKCGAAAKRETDTMDTFVDSAWYFSRYAAPDAQTMVDARTDYWMPMDQYIGGIEHAIL
HLLYSRFWAKVMRDLGLVAFGEPAKNLLTQGMVLNETFYREDAAGKKTWYNPADVTVSFD
DKGRPVGAVLKSDGQPVELGGIEKMSKSKNNGVDPQMLIDHYGADTARLFTMFAAPPEQQ
LEWSGAGVDGASRFLRRVWAFGFANREALAVRAPFDAAQLAEADKTLRREIHGVLKQADF
DYQRLQYNTVVSAAMKMLNAIEGAKGATPAVLRETYGVLLRVLYPVVPHVTFELWKALGY
ADEFGPLLDAPWPKVDEAALEQAEIELVLQVNGKVRGALKVAKDASREAIEAAAVADGMF
AKFAEGRPAKKIIVVPGRLVNVVV*
>GCA_000452945.1_BPMSHR5858_genomic_05382_1 tolQ_2
MATPTGILHYLASGDAVTHAVAYVLLAMSIASWCFLLVKAWVLVRAKRQGPAALAAFWRA
ATLDDGIAALRAADKERVFAPLAEAAREADAASRVEPHALAARVERGERVLRALRHALRA
SQRRLEFGQVLLASIGSTAPFVGLLGTVWGIYHALGSIAASGQAQIENVAGPVGEALIMT
AFGLVVAIPAVLAYNILGRLVRQLVEELDGFARDLHVFVCGEPAAAAADDAATAA*
>GCA_000452945.1_BPMSHR5858_genomic_05384_1 bamE
VRSTVIAAVAVVGLAACSSYDSVTQRIAQSITPYRITVVQGNFVSQEKASQLQVGMSREQ
VRALLGTPLLTDMFHADRWDYLFYFKRGSTSVVQQRDLVVNFAGDRLAGWSGAENLPSEL
DLLADIDGDRRGKKARAAAAARAASAPEAAAAAPEAMQAAEQNDANAQAARAANRATAQV
PGQSGAASRFAPSAQMAPNAPTPGGLPPGAAPAIQPQFQFHRPPPPPAQGGANQPVGPQG
ADALPNQPLTAPASGAQGTGG*
>GCA_000452945.1_BPMSHR5858_genomic_05386_1 allA_1
MQTLHTLRLERLTREAFAPFGDVIELAGARRILINGGTTERFHDLASIDVTEAGGRPLVS
LFRAQPRAWPIEIDMMERHPLGSQAFVPLAAVARYALVVAPAGEFDPTRMRAFLAQGWQG
VNYAKGVWHHPLLALDALSDFVVIDRGGAQPNCDELALDARWRLVTDDAHASAA*
>GCA_000452945.1_BPMSHR5858_genomic_05387_1 alc_1
MAAPILDPNAPAFTRRYMNLADPRLGAKALFASDEFFAPKERMLDPEPAVFIPGKYDDHG
KWMDGWETRRKRTTGHDFCVVRLARPGVVYGVDLDTSHFTGNFPPAASIDACVSDADTPP
DDAVWETLVPATTLAGNQHHYVDVSNPRAYTHLRVNLYPDGGLARLRVYGQPQHDWSRAA
RGELVDLAAIENGAYLVAANNEHFGPASRMLMPGRGANMGDGWETRRRREPGNDWAIVAL
ARPGVIRRVEVDTAHFKGNFPDRCSLQAARVAGGTDASLVTQAMFWPMLLGEQPLGMDSV
HTFETQLAALGPVTHVRLNIHPDGGVSRLRLWGELA*
>GCA_000452945.1_BPMSHR5858_genomic_05388_1 dctA_2
VSKFLNSLFGRVVIALLIGVALGAFFPHFAQSLRPLGDGFLKLIKMVIGPIVFCVVVSGM
ANAGDLKKVGRVGLKAVIYFEAMTTLALGIGLVLAYLTKPGVGMNVDLHTLDPASLADYA
KNAQSLKDTAGYLLKIIPETAFDAFAKGDILQILVFSVLFGSALSLLGEKAKRVNTLIDE
FAQVFFRVMGFIIKLAPLGVLGAIAFTTGKYGVASLKQLGLLVIVFYASCIAFVVIVLGA
VMKLAGFSVFKLIRYLREELLIVLGTASSDAVLPQVMRKLEWMGVKDSTVGLVIPTGYSF
NLDGFSIYLTLAVLFIAQATNTPLSVHDLIVVLLVSLVTSKGAHGIPGSAIVILAATLSA
IPALPVLGLVLILPVDWFVGIARALTNLLGNCVATIVVAVWEHDIDRARAQRVLNEELRY
VPAGDESPRGAISDGGAHAL*
>GCA_000452945.1_BPMSHR5858_genomic_05389_1 lldR_1
MKNVPHTVTDAAIATIRERIERGFYPVGSLLPAQRQLSEELEISRASLREALSTLEALGM
LRIRPGKGVYVESAQASAAHAWRFAEQSSLPDTYQMRYALEGFAARMAALAVSEEDLAWF
EENVAALQEALANDELDSASQLDFDFHMRIVSLAGNAAIESILRGSADIMKESQRMPFYR
RELVLSTYREHRAILDALIAKDPAAAGAAIEAHISNAAQRAGVFFPTPKR*
>GCA_000452945.1_BPMSHR5858_genomic_05394_1 epd
MTIRVAINGYGRIGRNTLRAFYENGKKHDLEIVAINDLGDAKTNAHLTQYDTAHGKFPGE
VSVDGDYLVVNGDRIRVLANRNPAELPWGELGVDVVMECTGFFTSKEKASAHLKGGAKKV
IISAPGGKDVDATIVYGVNHDVLKAEHTVISNASCTTNCLAPLVKPLNDKIGLETGLMTT
IHAYTNDQVLTDVYHEDLRRARSATHSQIPTKTGAAAAVGLVLPELNGKLDGYAIRVPTI
NVSIVDLSFIAKRDTTAAEVNAIMKEASEGALKGILGYNEAPLVSIDFNHNPASSTFDAT
LTKVSGRLVKVSSWYDNEWGFSNRMLDTAIALANAK*
>GCA_000452945.1_BPMSHR5858_genomic_05397_1 group_19575
MTDPRPADVPWLTPYLTVRSAQASIQFFSDAFGFEKRDLIDEDGAIMHVEMSYRGQLIVM
FAPEGAFGSTARTPRSADAIAPQSFYLYVDDVDATYRRALDAGAKSLSAPQDQFWGDRFA
QIEDLDGYRWALARRIGA*
>GCA_000452945.1_BPMSHR5858_genomic_05399_1 group_19561
MSDNTYARDTSAAADLFAAGDGNLFQRVMQMRRAAHAGGAPDGAASPGLSSIEEPMSLFA
TVRPNLVQSIRAFRVQDLADEANQLGQHFLYAYCGNAQSKQEVLETIATSFLFPKHFGKN
YDALYDCLTDLVHKAGAQPGFVIVLEALPIAHKFDKEGRETLLDVFREAAEFWAERKVAF
RVFYSFA*
>GCA_000452945.1_BPMSHR5858_genomic_05400_1 group_19404
MARKWLRNGALASVFAIVAMGNIGATPGSLVSAAYAREAATVDGAAAQVDTIPASRLPRE
AATTLSLIAAGGPYPYGKDGAVFGNYERILPKMRRGYYHEYTVPTPRARNRGARRIVCGG
PLRRVDNCYYTGDHYNSFKRIVD*
>GCA_000452945.1_BPMSHR5858_genomic_05401_1 maeB_1
MSTSSSSSSSKEKLREAALDYHEFPTPGKVAIAPTKQMINQRDLALAYSPGVAFACEEIV
ENPLNAARFTARSNLVGVVTNGTAVLGLGNIGPLASKPVMEGKAVLFKKFAGIDVFDIEL
NESDPHKLVDVIAALEPTFGGINLEDIKAPDCFIVERECRKRMKIPVFHDDQHGTAIVVA
AAVTNGLKVVGKSIKEVKLVASGAGAAALACLDLLVDLGLPLEHITVTDLAGVVYKGRAE
LMDPDKERFARETDARTLSEVIDGADIFLGLSAAGVLKADMVKKMAERPLILALANPTPE
ILPEVALEARPDAVLATGRTDYPNQVNNVLCFPFIFRGALDVGATTITREMEIAAVNAIA
ELAQQEQSDIVATAYGIQDLSFGPEYLIPKPFDPRLIVKIAPAVAQAAMDGGVATRPIED
MEAYKVHLQQFVYHSGTTMKPVFQIARGAPAEKKRVVFAEGEEERVLRAVQIVVDEKLAK
PILIGRPAVIEHRIQRYGLRLTPGVDFTIVNTEHDERYRDFWQTYFKMMARKGISEQLAR
VEMRRRTTLIGSMLVKKGEADGMICGTISTTHRHLHFIDQVIGKRAGCSVYGAMNALVLP
GRQIFLVDTHVNVDPTPTQLAEITIMAAEEVRRFGIEPKVALLSHSNFGTSNAPSAQKMR
DTLAILQERAPDLHVDGEMHGDVALDAALRKEILPESTLEGEANLLVLPNIDAANIAYNL
LKTAAGNNIAIGPILLGAAQPVHVLTESATVRRIVNMTALLVADVNAVR*
>GCA_000452945.1_BPMSHR5858_genomic_05406_1 yphB
MTATHSRSTATDAVQTLQTRTARRAAAAHRVSPGPQTSQIARGAGVTETAIATLSNDMLR
LDVAPHLGGGVTRFDWRGDGALTPIFRRCDAPGARTDPNELACYSLLPYSNRIGGGRFEC
DGRLVRVPRNRSAEPLPIHGDGWLAHWQLDDATDTQLGLSLDRSNGAPYAYRATQVYALD
GATLTIALGIENTGATRLPFGLGVHPFIVRDASTELAAAASGLWLSTPDWLPSRHVGAPP
AWRFGIAYPLPDTLVNHAFTGWGGGATIAWPQRRLGLTVTADADCYVLYTPPGEPFFCFE
PVDHPIDAVNLPGGGAAHGMTLLAPGERLMRRFRFTVARTDARAAPVARQSRRRAIA*
>GCA_000452945.1_BPMSHR5858_genomic_05407_1 group_19078
MASHRESNGALDAQHERYARYPSLVGRAVLITGGATGIGASFVEHFARQGARVAFVDLDE
KAGRALVARLADAAHEPVFVVCDLTDIGALRGAIDAIRVRIGPIAVLVNNAANDVRHAVA
DVTPESFDASIAVNLRHQFFAAQAVIDDMKRLGGGAIVNLGSIGWMLKNAGYPVYATAKA
AVQGLTRALARELGPFGIRVNTLVPGWVMTDKQRRLWLDDAGRAAIKAGQCIDAELLPGD
LARMALFLAADDSRLITAQDVVVDGGWA*
>GCA_000452945.1_BPMSHR5858_genomic_05408_1 araH_2
MQARENLPPAAAHAAAVPTEDRQRWRQHAADYSLVAIFAAMFVAMSLTVDHFFSIDNMLG
LALSISQIGMVACTMMFCLASRDFDLSIGSTVAFAGVLCAMVLNATDNTFVAIAAAVAAG
AVIGFVNGAVIAYLRINALITTLATMEIVRGLGFIVSKGQAVGVSSETFIALGGLTFFGV
SLPIWVTLACFVVFGVLLNQTVYGRNTLAIGGNPEASRLAGINVERTRVYIFLIQGAVTA
LAGVILASRITSGQPNAAQGFELNVISACVLGGVSLAGGRASISGVVIGVLIMGTVENVM
NLLNIDAFYQYLVRGAILLAAVLLDQLKNRGARD*
>GCA_000452945.1_BPMSHR5858_genomic_05413_1 dgoK1
MTAQASTPALIGLDWGTTSLRAYLFAADGSVLDTRALAAGVMSLPDAPGASPQQTFDAAF
ERACGAWLDRAPHVPALAAGMVGSAQGWREAPYVTMPAAADTFASSLVRFRTARGSALAI
VPGALEPGELPDVMRGEETQIVGALAAEPALAAERAGALIGLPGTHAKWAWVRGGRIEWF
RTYMTGELFATLRGHTILGRTMQASDAPDWSAFARGVRTARRKRDEGLLATLFSTRTLGL
TGQLAPHAQGDYLSGLLIGHELNALEALLGGQHASLAALAPRLVGEGALCDRYRRALAEF
GCGDARIVERASERGLWRIAVQAGLVGAACDAPLAPQ*
>GCA_000452945.1_BPMSHR5858_genomic_05415_1 group_19016
MKIQSAFAWSALVVAGLAFFTQTAFAQEAASDAQSDRHAQAQLGDPYVPPAARKPTAGTQ
TSGAALHAQVVRKLQRQFAAADAGNAGLTEAQAKAAGLGYVSKNFKQIDVNHTGRVSFSD
VQRYIQSQSATQK*
>GCA_000452945.1_BPMSHR5858_genomic_05416_1 group_19036
MPARKAQLLLAAVFSSLLIAACNGDDAGSPASAAASENSSGNIDASSGADKPYRDPNRYS
SSGSASLSPSAVTENAAITHHQITVNGKTINYTATAGHLSARNPQSGAAEASFFYVAYTA
DNQPLGKRPVTFFYNGGPGSSTVWLHLGSFGPKRLVTGDPNANSPTPFPFVDNQETLLDT
TDLVFVDAIGTGFSEAIAPNTNQTFWGVDQDAGAFRDFVIRYLQVNQRNDSPKYLFGESY
GTPRTDVLANLLETAGVKLDGIVLQSSILNYNSNCDMASNSVGGSNSGSSAVSCAGFVPS
YGAIGAYYQLDNPNPTNLPQYADQMRLLTAGTYSPAVDAYLANHTPPSSSLVTTMSNATG
ATVPLWRANFNLLPTSYDNSYQLALIPGTLIGRYDARVNVPTNSPLASDGDPSSSFITKP
FTDTIGTYLPNVLKYTAKSAYAVQSNAIASWDWSHDGLDLPDTIPDLAAALTLNPALKVL
SLNGYHDIATPFYQTELDLARLGAQPNLTIKDYQGGHMVYLDDTSRPQEKADLATFYAAA
PAAR*
>GCA_000452945.1_BPMSHR5858_genomic_05417_1 aroE_1
MRNSPVSPGPGYAPARGSERARRRGVARWLAYVGGVFAGAWLATQLYYAVQIAMWSVFDP
GSSAFVRADAWRLSNAQPATTIRHRWVPYDQIARTLKRAVIASEDADFANNSGYEVDAIL
QAWEKNRARGRIVSGGSTITQQLARNLFLSGERSYIRKGQELIITWMLETLLDKERIFEI
YLNSVEFGRGVYGAQAAAQYYYRIPASRLSAWQSARLAVMLPNPKYFDAHRGSPYLAQRA
GVIARRMGAAELPASQ*
>GCA_000452945.1_BPMSHR5858_genomic_05421_1 mpl
MHIHILGICGTFMGGLAVLARAAGHTVTGCDAGVYPPMSTQLEAQGIQLIEGYGAEQIDL
KPDLFVIGNVVSRGNPLMEAILDRGLPYVSGPQWLGEHVLAGKWVLAVAGTHGKTTTSSM
LAWVLEDAGLNPGFLIGGVPLNFGVSARLTDSSFFVIEADEYDTAFFDKRSKFVHYRPRT
AILNNLEFDHADIFPDLAAIETQFHHLVRTVPGVGRLVTNGRDDALERVLSRGCWSDVER
FGVDGGWQALPVENGVPVDGRFAVYWRSERVGAVDWQVQGEHNRMNALAAIAAARHVGVP
PAQAAAALAAFRNVKRRMEVRGSVDGVTVYDDFAHHPTAIETTIAGLRTRIGRENTRILA
VLEPRSNTMKLGVMKAQLPASLADADLVFGYGAPSGRDALGWSLPDALAPLGGKARAFDD
LHALVKAVTASARPGDHVLVMSNGGFGGVHQKLLDALSARGDAAPARSGA*
>GCA_000452945.1_BPMSHR5858_genomic_05424_1 aroQ_1
MTRLLVLHGPNLNLLGTREPEVYGRVTLEQIDQALAARAQEAGAELESFQSNHEGALVDR
VQAAGSDGTAFILINPAAYTHTSVAIRDALAGVGIPFVEIHLSNVHRREPFRHHSYFSDQ
AEGVICGLGWKGYLYALEYALDKLQGASRG*
>GCA_000452945.1_BPMSHR5858_genomic_05428_1 group_1674
MLLATRCPHCETVFRLQREQLALHDGLVRCGHCQQVFDAARALVPTEPEAAAAQAAPRAP
EQPAARRLFDATSPDRRPLEAGHRDFTPGAWDMRGPWLDGAIDPKLQMTSASVSTAAGGA
VAAHAAAHETRNAAADTPAQEASKASDALSVSEASQASRASDTSTHDTLPVPETPETPAG
GTPAPGERPAHAWPEANAGAPAIRPSDADLPHAGAREFAPPPRPTLDLDLDRSATAKHKP
DTARASASPPGDAVEPMLASHAAAAAALAGVPHDREPRFGAAPPARADAEPFAAAPEADN
REHFAMTRETRTDAARGGLARALVALVALALAALLAAQLAWWQRETITIYWPSSEPLFKQ
ACAALGCAVTPPRAIDGLRLDASDLRQLDGPRLLELKVPLTNRYRVALAYPSIELTLLDE
ANNITARRVLAPRDYVRPGTRVEAGMPAGATQTMIVRIDTDGIAASNFRVQIFYP*
>GCA_000452945.1_BPMSHR5858_genomic_05429_1 tpx
MSKVTLGGNPIDIAGTFPTVGSPAPDFKLVGKDLADVSLATFAGKRKVLNIVPSLDTPTC
AASTRKFNEAASKLDNTVVVVVSADLPFAATRFCTTEGLANVITASTFRGGRAFADAYGV
NVTSGPLDGLAARAVIVIDENDKVTYTELVGEIKNEPNYDAALAALK*
>GCA_000452945.1_BPMSHR5858_genomic_05442_1 proS
MKASRFFIGTLKEAPADAEIVSHKLMVRAGMIRRVAGGIYNYLPVGLRSIRKVEAIVREE
MNRAGAIELLMPAVQPAELWQESGRWEQYGPELLRFKDRKQNEFVIGPTHEEVVTDIARN
QIKSYRQMPVNFYQIQTKFRDEIRPRFGVMRGREFIMKDAYSFDKDHESLKESYKKMYDA
YVRIFTRIGLEFRPVAADNGSIGGSGSHEFHVIADTGEDAIAYCPTSDFAANVEAAEALP
LLASRAAPAEAMQKVATPGKAKCEAVAELMGIPLERTIKSIVLATDNEGAEPTIWLLMLR
GDHVLNEIKTAKLPGLAGHRFATEAEIVEWFGTPPGYLGPIGTKKPVRVVADRTVANMSD
FVVGANEVDYHIAGVNWGRDLPEPVVADIRNVKAGDPSPDGKGALDICRGIEVGHVFQLG
TKYSDAMGATFIDESGKAQPMVMGCYGIGITRILGAAIEQNFDDKGIVWPEAIAPFEVVL
CPMGYDRSDAVREAADKLYADLAAAGIDVILDDRGERPGVMFADWELIGVPHRLVIGERG
LKDGKIEYQGRRDAEATLLPADSAAAAVAEKVRAALAR*
>GCA_000452945.1_BPMSHR5858_genomic_05443_1 rppH_2
MLDREGFRPNVGIILLNAHNEVFWGKRLREHSWQFPQGGIKYGETPMQAMYRELHEETGL
LPEHVKIIGRTRDWLRYEVPDKFIKREVRGHYRGQKQIWFLLRMVGRDCDICLRATDHPE
FDAWRWNEYWVPLDAVIEFKRDVYQLALTELSRFLRRPAQRTDKSRGPRAPRYPRVANGH
AASEAPAAIDTSAVCSEVEPGANALDETPPRVSLRD*
>GCA_000452945.1_BPMSHR5858_genomic_05444_1 group_19643
LKAIALAAASIAAAAALAGCAHSNTPSNKDDSEFVYLLDRQPQWTENKVDKLPPLPQTSD
LLPFNVSQNTPLKFFVDSKSLDVGTDGVVRYIVVVTSPAGARNVNYEGIRCDTYEWRQYA
GLNADHSGWDRTIETDWQRIENGELNAYHAALYQDFFCANKMPAAKRPTIIENIRYNRTQ
LNQIR*
>GCA_000452945.1_BPMSHR5858_genomic_05446_1 obg
MKFIDEARIEVIAGDGGDGSASMRREKFVPFGGPDGGDGGRGGSVYVIADRNINTLIDYR
YAKKHMARNGENGRGSDCYGKGGDDITLRMPVGTVINDMDTGELIADLTEHDQKVLVAKG
GAGGLGNLHFKSSTNRAPRQKTDGKPGERRMLKLELKVLADVGLLGMPNAGKSTFISSVS
NAKPKIADYPFTTLAPNLGVVRVGPGKSFVIADIPGLIEGAAEGAGLGHQFLRHLQRTGL
LLHLVDLAPFDERVDPVAEARAIVGELRKYDESLYEKPRWLVLNKLDMVPEDERRARVAD
FIERFGWTGPVFEISALTGQGCEGLVYAIHDYLVEHSDAHRAELAEDLASDVRFRDAPGA
GGEPHERDAGAH*
>GCA_000452945.1_BPMSHR5858_genomic_05448_1 rplU
MYAVIKTGGKQYKVAVGEKLKVEQIPADIDAEITLDQVLAVGEGESIQFGTPLVSGASVK
ATVVSHGRHAKVTIFKMRRRKHYQKHGGHRQNYTELRIDAINA*
>GCA_000452945.1_BPMSHR5858_genomic_05451_1 group_19285
MMALNRHRLKHLASHGALGAKTTQRLLARTDQLLSVILIGNNLFNTIIPVLTTSIALRTF
GHDNFVLSIATGIVAFLIIVFAEITPKIVGATFPERIALPASVVIAPLMRVMMPVVWFVN
TLSGAILAALRINTKAGRDQRLSTEELRSLVLESGSFMPTKHRSILLNLFDLENITVDDV
MIPRRQIESLNILAPLDDVLHQLETCYHNRLVVYEGDIDRVLGVLHVRKTLAAFHNQDFD
RGTLRELLAQPYYVPSGTPVFQQLQYFQESRHRTALVVNEYGELEGLVTPEDIIEELIGE
FTTTMPRSENSKSGWDANGECIVAASMPLRELNRWLHLGLPTNGPKTLNGLILEELEEIP
EDDVCLKIGDVMIEVMRTEDQGVRTVKLFKPRAPRGGRAAGA*
>GCA_000452945.1_BPMSHR5858_genomic_05452_1 epsE_2
MSATPSNPSFPAVRSAASALPRPRAPELADDAPAVGIVADMLRAAHERDASDIHVEPGES

GWRVRLRIDGVLHEFARPPAHLRDACVTRIKVLARMDIAERRVPQDGRLRLPLAPGRAGD
YRVSSLPTLHGEKLVLRRLETLPADLSLAALGFDAAQARGVEAAIGAPHGLILVTGPTGS
GKTLSLYCFLQMLNDVSRNVCTVEDPAEIELDGINQVGVREKAGLTFAVALRAFLRQDPD
VIMVGEIRDAQTADVALKAAQTGHLVLSTLHTNDAPAAVARLLDIGVAPYNLAAALRLVT
AQRLVRRLCPACRAPSDAPPATLRAAGFDAASIDAGWRPYAANGCAACHGIGYRGRIGIH
QVMPLSAELRNLTVARASSAELARQARAEGMASLRDAALARVRDGTTSLSEALATTEAA*
>GCA_000452945.1_BPMSHR5858_genomic_05455_1 coaE
MFSVGLTGGIGSGKTTVADLFGKLGATIVDTDLIAHRITAPQGLAMPLIAREFGAEFVAA
DGSLDRAKMRALVFSDESARKRLEAITHPLIREETEREARTAQGAYVVFVVPLLVESGTW
KTRVDRVLVVDCDVETQIARVTARNGFTRAQVEAIVARQASRDARLAAADDVIANDNASV
AELAAEVAALHQRYLECAAAARN*
>GCA_000452945.1_BPMSHR5858_genomic_05456_1 zapD
LILYEYPFNERIRTLLRLEDLFERFTFFVAQEDAREHHVALTTLFEISEVAGRADLKSDL
MKELERQRQTLAPFRGNPGIEQNALEAVLGEIEQTLANLAQMQGKTGQHLIDNEWLASIR
SRAVIPGGTCKFDLPSYYAWQQWPAEQRRHDIAKWAMPLLPLRDAAMIVLRLARESGQAS
KVMAMQGSYQQMLSGRTYQLMQVRVPPELRVIPEASANKYMLWVRFTAQDGDVRPRAVDI
DVPFQLTLCNL*
>GCA_000452945.1_BPMSHR5858_genomic_05459_1 argJ
MAVNFPSIDPAQLHPVAGVTLGWAEANIRKPNRKDVLVVSVEEGATVSGVFTENRFCAAP
VTVCREHLAKVRAGGAGIRALVVNTGNANAGTGEPGLAHARETCAELARLAGIAPGQVLP
FSTGVILEPLPIERLKAGLPAALANRAAANWHDAAQAIMTTDTLPKAASRQVTIDGHTIT
LTGISKGAGMIKPNMATMLGFLAFDAKVAQPVLDALVKDVADRSFNCITIDGDTSTNDSF
ILIASGKASLPQIASTDSPAYAALREAVTAVAQALAQLIVRDGEGATKFITVTVEGGKSA
AECRQIAYAIGHSPLVKTAFYASDPNLGRILAAIGYAGVADLDVGKIDLYLDDVLVAKAG
GRNPAYLEEDGQRVMKQSEIAVRVLLGRGDAQATIWTCDLSHDYVSINADYRS*
>GCA_000452945.1_BPMSHR5858_genomic_05472_1 group_19660
VDARLLEILVCPICKGPLHYDRGAQELVCHADKLAYPIRDGIPVMLVDEARQTVEGTPVD
PAGPARGR*
>GCA_000452945.1_BPMSHR5858_genomic_05473_1 kdsB
MTSPLPFVAVVPARLASTRLPNKPLADLGGKPMVVRVAERAREAGAQQVLVASDAQRVLD
AVREHGFDAVLTRADHPSGTDRLAEVAAKLGFDDDTIVVNVQGDEPLIDPQLVRDVASHL
AAHPSCAIATAAHPIHEAHEVFNPNYVKVVLDAHGVALYFSRAPIPWSRDAYLPHWPNVA
AMPAPTCPVYRHIGLYAYRARFLRTYPTLAQAPIEAAEQLEQLRAMWHGERIAVRVTEHA
PEAGIDTPADLERVQALFRSRAK*
>GCA_000452945.1_BPMSHR5858_genomic_05474_1 adk
MRLILLGAPGAGKGTQANFIKEKFGIPQISTGDMLRAAVKAGTPLGVEAKTYMDEGKLVP
DSLIIGLVKERLKEADCANGYLFDGFPRTIAQADAMKEAGVAIDYVLEIDVPFSEIIERM
SGRRTHPASGRTYHVKFNPPKVEGKDDVTGEPLVQRDDDKEETVKKRLDVYEAQTKPLIT
YYGDWARRGAENGLKAPAYRKISGLGAVEEIRARVFDALK*
>GCA_000452945.1_BPMSHR5858_genomic_05475_1 group_19531
MEIRDNVFLITGGASGLGAGTARLLTEAGGKVVLADLNQQAGEALARELGGVFVKCDVAR
EEDAQAAVAAAAKLGTLRGLVNCAGIAPAAKTVGKDGPHPLELFAKTITVNLIGTFNMIR
VAAAAMAANEPAQTGERGVIVSTASVAAFDGQIGQAAYAASKAGVAGMTLPIARDLSRNA
IRVMTIAPGIFETPMLLGMPQDVQDALGAMVPFPPRLGKPAEYAMLVKQIFENPMLNGEV
IRLDGAIRMQPK*
>GCA_000452945.1_BPMSHR5858_genomic_05477_1 murJ
MNLFRALLTVSGFTLLSRVTGLARETLIARAFGASQFTDAFYVAFRIPNLLRRLSAEGAF
SQAFVPILAEFKNQKGHDATKALVDAMSTVLAWALALLSLAGIAGASWVVFAVASGLRTD
GQAFPLAVAMTRIMFPYIVFISLTTLASGVLNTYKRFSLPAFAPVLLNVAFIVAAVFVAP
HLKVPVYALAWAVIAGGALQFAVQLPGLKKIDMMPAIGVNPLRALAHPGVKRVLAKMVPA
TFAVSVAQLSLIINTNIASRLGQGAVSWINYADRLMEFPTALLGVALGTILLPSLSKAHV
DADSTEYSALLDWGLRVTFLLAAPSALALFFFAEPLTATLFNYGKFDAHTVTMVARALAT
YGIGLVGIILIKILAPGFYAKQDIKTPVKIAIGVLVVTQLSNYVFVPIIGHAGLTLSIGV
GACLNSLLLFIGLRRRGIYQPSSGWLRFFAQLAGASLVLAGVMRWLAINFDWTAMRAAPL
ERIALMAACLVLFAALYFGMLWLMGFKYAYFRRRAK*
>GCA_000452945.1_BPMSHR5858_genomic_05485_1 lcfB_4
MTIEFSPGAAIDADALLDTLPQRIAALPARVAEHAPDHVALIEDGRRLTYAQLARAVEAA
AERLRGLGVRGGDRVMIVAENSIAQIVLLFAATRLDAWAIVSNARLSAAELDAIATHAQP
RLIAFVTGTSPDARAHADRRNAHAAAPFELDIGAWSHTVEAGTRAEPVEAHGARQCAALV
YTTGTTGAPKGVMLSHRNLLFVAAVSSMQRRVAPTDVVYAVLPISHVYGLASVCLGSLYA
GATLRLAPRFSPETLRRALADEGVTIFQGVPAMHAKLLEHLRAHGHAWRAPHLRFVYSGG
SPLDADLKARVERAYGLPLHNGYGMTESSPTIAQTPIDAPRADCSVGAPIPGVAVRFCSA
DGADVAPGEVGELWVRGPNVMLGYYRDPEGTRAAVTAHGWLKTGDLARAGTDGAMTIVGR
SKELIIRSGFNVYPSEVEQALNAHPDVVQSAVIGRAAEAGNEDVIAFVELAPNAAATESD
LKEWCVGRLAPYKRPVQIRALDALPAASTGKVLKHRLRAML*
>GCA_000452945.1_BPMSHR5858_genomic_05487_1 ogt
MFNAVIDAPFGKVGIRTDASVVREIVYLPESMKRVAPGTPLAKQAARQIELYFERASARF
DLPLAEVGTPFQHRVWHAICAIPPGVVLTYGQIAKQIGSAPRAVGQACGANYFPLVIPCH
RVVASGGLGGFANHDDEGYFLKVKRWLLAHEGVRY*
>GCA_000452945.1_BPMSHR5858_genomic_05490_1 amiC_2
MSRKMLIKPFRSIESAATATHNWRRRQILRAGASTLVLGLALPRLAHASSVLGVRVWPAR
DYTRVTIESDQPLQNTQQLLQGPDRLVVDLNGLELDQALRDLVSKIAPNDPQIQSVRVGQ
YQPHVVRMVFDLKGSVKPQVFTLPPVGTYKYRLVFDLYPAVAPDPLSDLIAQTERKEQQL
NDTLRAQQAQPPTAALNGPTAAPPAATDNSDAFFQRFAQNNPPAAHSARPAPANPPVAQA
SPPAAPPRPAVKPPPVIARRNDDDSGDTDTYAFTAPKSGRGTVRLLTVAIDPGHGGEDPG
AIGGGGTYEKHIALDIAKKLRAKIDAAPNMRAMMTRDADFFVPLNVRVQKARRVGADLFV
SIHADAFTTPSARGSSVFALSDHGASSAAARWLANKENSSDLIGGINIKTQDVAVNRALF
DMSTTAQIRDSLRYGNYVLREVGGINKLHKGSVEQAGFAVLKAPDIPSILVETAFISNPD
EERRLNDDAYRDQMADAIFRGIKRYFSANPPLAKSRMT*
>GCA_000452945.1_BPMSHR5858_genomic_05494_1 tcdA
MSRTDAIATPHDVTPQPSGELDADRARRFGGVARLYGADALAAFERARVAVIGIGGVGSW
AAEALARSAVGELTLIDLDNVAESNTNRQIHALDGNYGKPKVDAMAERIALIDPACRVVK
IEDFVEPDNLDALLGGGFDYIVDAIDSVRTKVALIAWCVARAQPLVTVGGAGGQLDPTRI
RIDDLAQTIQDPLLSKVRAQLRKQHGFPRGPKARFKVSAVYSDEPLIYPEAAVCDVDDVA
MHTATDAQAPGPTGLNCAGFGSSVCVTASFGFAAAAHALRALAARAGR*
>GCA_000452945.1_BPMSHR5858_genomic_05495_1 pdxH
MTTLADLRTNYSRASLDAADVNPNPFVQFDVWFKEALDAQLPEPNTMTLATVDESGRPSA
RIVLIKGADERGFVFFTNYESRKGRELAHNPNAALLFYWIELERQVRVEGRIEKTSEEES
DRYFASRPLGSRIGAWASEQSAVIESRALLEAREKEIGARFGENPPRPPHWGGYRLVPSS
IEFWQGRPSRLHDRLLYTRDAASASGWKIARLAP*
>GCA_000452945.1_BPMSHR5858_genomic_05500_1 decR_4
MHAITLDATDCRILAVLQEEGRISNLDLAERISLSPSACLRRMRLLEEQGVIERYRACLS
REKLGFELEAFVQVSMRNDQTQWHERFAQAVREWPEVVGAFVVTGETHYLLRVLAHNLKH
YSDFVLNQLYKTPGVLDIRSNIVLQTLKDEAGPPVALARTGPAIKAV*
>GCA_000452945.1_BPMSHR5858_genomic_05504_1 pcaK_2
MSARHGGAPLALDIGATLDHGPFTAMQRVAVLLAAFAIVLDGFDSQLIGFAIPVLIKEWR
ITRDAFAPAVAAGLFGMGVGSACAGLFADRFGRRWAIIGSVFVFGAATCAIGFAPNVATI
AALRFVAGLGIGGALPTATTMTAEYTPARRRTMMVTATIVCVPAGGMLAGLFAHEVLPAY
GWRSLFWLGGALPLALGLLLVRALPESPRYLARNPARWRELGALLARMGRPVADGTAFTD
LAEARAHEGQRRGVRALFSAAYARDTIALWCAFCMCLLAVYSAFSWLPTMLTSQGLSVSV
AGSGLTAYNLGGVLGALGCALAIGRFGSRWPLAFCCAGGAASAAWLLGVDAGSHAGWLIV
GLAAHGFFVNAVQSTMYALCTFIYPTPVRATGTAGAVAFGRVGAILSAFAGAYVISAGGA
NAYLAMLAAAMAVVLVALLALRRHIPRLRRGGLPAADELARTAP*
>GCA_000452945.1_BPMSHR5858_genomic_05505_1 panE
MATIEMTALRIAMLGAGAMGSLFGGLLAECGHDVTLVDVNAAHVDAIARDGLRLDTDRGE
RRVARLAALRPDGVAAHAGMPPDLVIVFTKTLHTRAALAGARALFGPRTHALTLQNGLGN
VETLAEYVPLERILVGVTTWPADLAGPAHVRSHGAGWVRLMSADGAMRPIVQASAHALDR
AGLNCAIDTGVWATIREKVGFNAALNTLCALTRGTVDALGAREDGPALALAIVAEVAAVA
RAKGVGADERRMRENVLHAIREHRGHRPSMLQDVLAGRRTEIDAINGAVVAAAGELGVAV
PHVRTLLQLVRLIDAQGG*
>GCA_000452945.1_BPMSHR5858_genomic_05506_1 vdh_2
MQDISMLIGGERRPATGGATFERRNPLDGELATRAPAATAADARAAVDAASAAFAPWAAL
GPSARRALLLKAAAALEGKRDAFIAAMAAETGASAIWARFNVELAANGLVEAAALTTQIG
GELIPSDVPGSLAMGVRQPAGVVLGIAPWNAPVILGVRALALPLACGNTVVFKGSELCPA
THGLIADALHEAGLPRGVVNFVTNAPADAGAVVDAMIAHPAVRRVNFTGSTRVGRIIAER
CARHLKPAVLELGGKAPFVVLDDADLDAAVAAAAFGAFANSGQICMSTERIIVDERIADA
FVAKLADKAASLPLGDPRNGPVVLGSVIDAQTVERCNALIDDALAKGAVLRCGGKADSTL
MPATLVDRVTPAMRLYAEESFGPVKGIVRVAGEEAAIACANDNAFGLSSAVFSRDVARAM
RVAARIEAGICHVNGPTVHDEAQMPFGGMKDSGFGHFGGKAGIAEFTDLRWITVQTAPRH
YPF*
>GCA_000452945.1_BPMSHR5858_genomic_05521_1 kbaZ
MRGVSARAEPPRSGDHGRWLREIFEANRAGRASGIYSVCSAHRLVLEAAFEAARADESPL
LVEATCNQVNHHGGYTGMTPADFRRDVDALARAAGFPARALILGGDHLGPNPWRHLAARD
AMREARAMVAAYVAAGFTKIHLDASMACADDAAPLSDATIAERAAQLCAAAEEAAEAAGA
LPVYVIGTEVPTPGGEVSAQAGGSAADTASGAAQAAARKAAVGAAHGMAHEHGGAANLPA
DDARGAAAGHRGAFPQIEVTRADSVSATLAAHRDAFARHGLQHAWSRVIALVAQPGVDFD
DRRVLDYDPARAAALGASILRVPSLVFEAHSTDYQTESALAALVRDHFAILKVGPALTFA
LREALFALTYIEDALFDDASERSQLRDVIDAAMRERPEYWAPYYRGDALAQRIARQFSYS
DRIRYYWLQPAVAAALERLFVNLARRAPPETLVAQWLPDVYAACRRGELAREPLAWVRHR
IREVISRYARACAMQHNA*
>GCA_000452945.1_BPMSHR5858_genomic_05523_1 polS
VRLQEKVAILTGAASGIGEAVARRYLEEGARCVLVDLKPADGALARLIDAHAGRAIAVSA
DVTRRDDIARIVATAVERFGGVDILFNNAALFDMRPLLDESWDVFDRLFAVNVKGLFFLM
QAVAQRMVEQGRGGKIVNMSSQAGRRGEALVSHYCATKAAVLSYTQSAALALARHGINVN
GIAPGVVDTPMWEQVDALFARYENRPLGEKKRLVGEAVPLGRMGVPDDLTGAALFLASAD
ADYITAQTLNVDGGNWMS*
>GCA_000452945.1_BPMSHR5858_genomic_05527_1 mntB_2
MGMRIHMAAGTHTMFEYEFMINAFAAAGIVAVLAGIVGYFLVLRGQTFAGHALSHVGFAG
ATGAALFGLSPIWGMLGFTLAAGIGMGALGERLAGRDVAIGVILSVALGCGLLFLHFYTN
YATQVTALLFGNVLGVSHATLVVLAAIGAVSLAALAAIARPLLFASLQPELAEAKGVSLR
LVSILFLSIGALAVAACTQIVGVLLVFTLLVGPAAAAQNLATRLSTGVWLAAAFALGEAW
LGITLAYYTDWPTSFWITALSALVYGASLVGRRHHA*
>GCA_000452945.1_BPMSHR5858_genomic_05529_1 flhA_2
MKTKAAIAWKAGEPLTIEEVDLEGPRAGEVLIEVKATGICHTDYYTLSGADPEGIFPAIL
GHEGAGVVVDVGPGVGTLKKGDHVIPLYTPECRECKFCLSRKTNLCQKIRATQGRGLMPD
ATSRFSVGGKPIFHYMGTSTFSNYIVVPEIAVAKVREDAPFDKICYIGCGVTTGVGAVVY
SAKVEAGANVVVFGLGGIGLNVIQGARMVGADKIIGVDLNPKRVELARKFGMTHFINPNE
VENVVDHIVQLTDGGADYSFECIGNVKVMRQALECTHKGWGQSFIIGVAAAGEEISTRPF
QLVTGREWKGSAFGGARGRTDVPKIVDWYMEGKINIDDLITHTLPLEKINEGFELMKKGE
SIRSVVLY*
>GCA_000452945.1_BPMSHR5858_genomic_05530_1 cdhR_5
MAAQTPLRHRTTTTVDVVIYPGFKVIEAVGVINVFDYANARLAAAGLAPVYDLQIAAPAK
GAVKSDTLIVLEATKALDTLAVPDTAIVVGARDIERALRDTSMLVGWCRDVSARIGRMVG
LCSGCFFLAEAGMLDGRRATTHWSVAPLLRARYPAVKVEPDAIFVREGNVWTSAGVTAGL
DLALAMVEEDLGREIALAVARDLVIYLKRPGGQSQFSVYLASQMTAHASIRDIQDWILNA
LDARLSIAQLARRAAMSERNFIRVFVRETGYRPAEFIEIARLEKARRLLEQEALPLKTVA
VRSGFRSDDQLRRVFMRRLGVTPGAYRERFSGTGVREARGSGDVD*
>GCA_000452945.1_BPMSHR5858_genomic_05538_1 hdfR_4
MGSEIGWELYRSFLGVLREGSLSGAARALGLTQPTVGRHVAALEAALRVPLFTRSSSGLM
PTDVALALRAHAEAMESTADALARAATSFGEDVRGVVRISASDVVGVEVLPPIVARLRQR
HPALTVELALTNRVQDLLRREADIAVRMTRPGQTQLIARHIGGIELGLHAHRDYLARCGT
PRDAGELVRHALIGHDRPTAFIRQIAKSFPGFDRGAFALRTDSDLAQLALIRCGAGIGAC
QAALAKRDPALVRVLPKAFAGRLDMWVTMHEDLRGSPRCRAAFDALAEGLDAYVDEQRAP
AIARRRRPLPGTRSA*
>GCA_000452945.1_BPMSHR5858_genomic_05596_1 rpsJ
MQQQKIRIRLKAFDYRLIDQSAAEIVDTAKRTGAIVRGPVPLPTRIQRFDILRSPHVNKT
SRDQLEIRTHQRLMDIVDPTDKTVDALMKLDLPAGVDVEIKLQ*
>GCA_000452945.1_BPMSHR5858_genomic_05598_1 rplD
MELKLLNSNGQEGAVVNASDVVFGRDYNEALIHQVVVAYQANARQGNRAQKDREQVKHTT
KKPWRQKGTGRARAGMSSSPLWRGGGRIFPNSPDENFSHKVNKKMHRAGLCSIFSQLARE
GRLSVVEDIVLEAPKTKLLADKFKAMGLDSVLVITDTVDENLYLASRNLPHVAVVEPRYA
DPLSLIYFKKVLVTKAAVAQIEELLS*
>GCA_000452945.1_BPMSHR5858_genomic_05599_1 rplW
MSEIRKNDHRLMQVLLAPVISEKATLVADKNEQVVFEVAPDATKQEVKAAVELLFKVEVD
SVNVLVQKGKQKRFGRSMGRRKDVKKAYVCLKPGQEINFEAEAK*
>GCA_000452945.1_BPMSHR5858_genomic_05600_1 rplB
MAIVKVKPTSPGRRAMVKVVNKDLHKGKPHAALLDTQSSKAGRNNNGRITTRHQGGGHKQ
HYRVIDFRRTKDGIPAKVERLEYDPNRSANIALVLYADGERRYIIAPKGVTVGQQLMSGS
EAPIRAGNTLPIRNIPVGTTIHCIEMLPGKGAQMARSAGTSAMLLAREGLYAQVRLRSGE
IRRVHIECRATIGEVGNEEHSLRQIGKAGANRWRGIRPTVRGVAMNPIDHPHGGGEGRTA
AGRDPVSPWGTPTKGFRTRRNKRTTTMIVQRRHKR*
>GCA_000452945.1_BPMSHR5858_genomic_05601_1 rpsS
MARSVKKGPFCDAHLLKKVEAAAASRDKKPIKTWSRRSTILPDFIGLTIAVHNGRQHVPV
YISENMVGHKLGEFALTRTFKGHAADKKAKK*
>GCA_000452945.1_BPMSHR5858_genomic_05602_1 rplV
MEVKAIHRGARISAQKTRLVADQIRGLPVDKALNVLTFSPKKAAGIVKKVVLSAIANAEH
NEGADIDELKIKSIYVDKAASLKRFTARAKGRGNRIEKQSCHITVTVGN*
>GCA_000452945.1_BPMSHR5858_genomic_05603_1 group_19178
MGQKIHPTGFRLAVSRNWASRWYANNNNFAAMLQEDIGVREYLKKKLKNASVGRVVIERP
AKNARITIFSSRPGVVIGKKGEDIELLKTELQRRMGVPVHVNIEEIRKPETDAQLIADSI
TQQLERRIMFRRAMKRAMQNAMRLGAQGIKIMSAGRLNGIEIARTEWYREGRVPLHTLRA
DIDYATSEAKTTYGIIGVKVWVYKGDTLGRNDAPVVEEVTEDKRPRRNARPGDRRPRRDG
EGGAPGARRGGPRRGAGKPEDGKTGE*
>GCA_000452945.1_BPMSHR5858_genomic_05604_1 rplP
MLQPKRRKYRKEQKGRNTGIATRGNAVSFGEFGLKAVGRGRLTARQIEAARRAMTRHIKR
GGRIWIRIFPDKPISQKPAEVRMGNGKGNPEYYVAEIQPGKMLYEMDGVSEELAREAFRL
AAAKLPLKTTFIVRQLGA*
>GCA_000452945.1_BPMSHR5858_genomic_05605_1 rpmC
MKASELLQKDQAALNKELSDLLKAQFGLRMQLATQQLTNTSQLKKVRRDIARVRTVLTQK
ANQK*
>GCA_000452945.1_BPMSHR5858_genomic_05607_1 rplN
MIQTESRLEVADNTGAREVMCIKVLGGSKRRYASIGDIIKVSVKEATPRGRVKKGEIYNA
VVVRTAKGVRRQDGSLIKFDGNAAVLLNNKLEPIGTRIFGPVTRELRSERFMKIVSLAPE
VL*
>GCA_000452945.1_BPMSHR5858_genomic_05609_1 rplE
MARFQEFYKEKVVPGLIEKFGYKSVMEVPRITKITLNMGLGEAVADKKIIENAVGDLTKI
AGQKPVVTKARKAIAGFKIRQGYPIGAMVTLRGRAMYEFLDRFVTVALPRVRDFRGVSGR
AFDGRGNYNIGVKEQIIFPEIDYDKIDALRGLNISITTTAKTDDEAKALLASFKFPFRN*
>GCA_000452945.1_BPMSHR5858_genomic_05610_1 rpsN
VAKLALIEREKKRARLAQKYAPKRAELKAIIDDASKSDEERYAARLELQQLPRNANPTRK
RNRCAITGRPRGTFRKFGLARNKIREIAFRGEIPGLTKASW*
>GCA_000452945.1_BPMSHR5858_genomic_05611_1 rpsH
MSMSDPIADMLTRIRNAQMVEKVSVSMPSSKVKVAIAQVLKDEGYIDDFAVKADGAKAEL
NIALKYYAGRPVIERLERVSKPGLRVYRGRNEIPQVMNGLGVAIVSTPKGVMTDRKARAT
GVGGEVICYVA*
>GCA_000452945.1_BPMSHR5858_genomic_05612_1 rplF
MSRVGKSPIALQGAEVKLADGAITVKGPLGTITQAVNPLVNVANNDGTLNLSPVDDSREA
NALSGTMRAIIANAVHGVTKGFERKLTLVGVGYRAQAQGDKLNLSLGFSHPVVHQMPEGI
KAETPTQTEIVIKGIDKQKVGQVAAEVRGYRPPEPYKGKGVRYADEVVILKETKKK*
>GCA_000452945.1_BPMSHR5858_genomic_05613_1 rplR
MDKTQSRLRRARQTRIKIAELQVARLAVHRTNTHIYAQVFSPCGTKVLASASTLEAEVRA
QLADKSGKGGNVAAATLIGKRIAEKAKAAGIESVAFDRSGFRYHGRVKALAEAAREAGLK
F*
>GCA_000452945.1_BPMSHR5858_genomic_05614_1 rpsE
MAKMQAKVQADERDDGLREKMISVNRVTKVVKGGRILGFAALTVVGDGDGRVGMGKGKAK
EVPVAVQKAMEQARRNMFKVPLKNGTLQHEVHGKHGASTVLLAPAKDGTGVIAGGPMRAV
FDVMGVQNVVAKSHGSTNPYNLVRATLDGLRKQSTPADIAAKRGKSVEEILG*
>GCA_000452945.1_BPMSHR5858_genomic_05615_1 rpmD

MSEKTVKVQLVKSLIGTRESHRATVRGLGLRRLNSVSELQDTPAVRGMINKVSYLVKVIG
*
>GCA_000452945.1_BPMSHR5858_genomic_05616_1 rplO
MELNNLKPAEGAKHAKRRVGRGIGSGLGKTAGRGHKGQKSRSGGFHKVGFEGGQMPLQRR
LPKRGFTSLTKEFVGEVRLGDLEKLPVDEIDLLALKQAGLVGELIKSAKIIATGELKRKI
VVKGLGATKGARAAIEAAGGSFAE*
>GCA_000452945.1_BPMSHR5858_genomic_05618_1 infA_2
MAKDDVIQMQGEVIENLPNATFRVKLENGHVVLGHISGKMRMHYIRILPGDKVTVELTPY
DLSRARIVFRAK*
>GCA_000452945.1_BPMSHR5858_genomic_05620_1 rpsK
MAKASNTAAQRVRKKVKKNVAEGVVHVHASFNNTIITITDRQGNALAWATSGGQGFKGSR
KSTPFAAQVAAESAGRVAMEYGVKNLEVRIKGPGPGRESAVRALHGLGIKITAISDVTPI
PHNGCRPPKRRRI*
>GCA_000452945.1_BPMSHR5858_genomic_05621_1 rpsD
VARYIGPKAKLSRREGTDLFLKSARRSLADKCKLDSKPGQHGRISGARTSDYGTQLREKQ
KVKRIYGVLERQFRRYFAEADRRKGNTGETLLQLLESRLDNVVYRMGFGSTRAEARQLVS
HKAITVNGIVANIPSQQVKAGDVVAIREKAKKQARIVEALSLAEQGGMPSWVAVDAKKFE
GTFKQVPERADIAGDINESLIVELYSR*
>GCA_000452945.1_BPMSHR5858_genomic_05622_1 rpoA
MQTSLLKPKIIAVESLGENHAKVVMEPFERGYGHTLGNALRRVLLSSMVGYAPTEVTIAG
VVHEYSTLDGVQEDVVNLLLNLKGVVFKLHNRDEVTVTLRKEGEGVVTAGDIELAHDCEV
INPNHVIAHLSKGGKLDVQIKVEKGRGYVPGNVRRYGDETAKIIGRIVLDASFSPVRRVS
YTVESARVEQRTDLDKLVMNIETSGVITPEEAIRQSARILVDQLSVFAALEGTETAAEAP
SRAPQIDPILLRPVDDLELTVRSANCLKAENIYYIGDLIQRTENELLKTPNLGRKSLNEI
KEVLASRGLTLGMKLENWPPAGLDK*
>GCA_000452945.1_BPMSHR5858_genomic_05623_1 rplQ
MRHRHGLRKLNRTSSHRLAMLRNMSNSLIEHEVIKTTLPKAKELRKVVEPLITLGKKPSL
ANRRLAFNRLRDRDSVAKLFDVLGPRFANRPGGYLRILKFGFRVGDNAPMALVELLDRPE
VEETENVQEAE*
>GCA_000452945.1_BPMSHR5858_genomic_05624_1 cutA
MVIVMMLTTVPDAAVARALAEGALSARLAACVSELGAIRSSYHWQGKVETADEIQLLFKT
SAVRALELERYIQSHHPYDVPEIVSWQATASAAYGQWVATETQRLFHV*
>GCA_000452945.1_BPMSHR5858_genomic_05626_1 hemB
MSIHPLHRPRRMRRDDFSRRLMRENILTTNDLIYPVFVVEGTNVRQPVPSMPGVERVSID
LLMGVAEQCVELGVPVLSLFPAIEPSLKTPDGREAANPEGLIPRAVRELKRRFPELGVLT
DVALDPYTSHGQDGVLDEAGYVLNDETLEILVEQARAQAEAGVDIVAPSDMMDGRIGAVR
EMLEREGHIYTRIMAYSAKYASAFYGPFRDAVGSASNLGKGNKMTYQMDPANSDEALREV
RLDIDEGADMVMVKPGMPYLDIVRRVKDEFRYPTYVYQVSGEYAMLKAAAQNGWLDHDKV
VLESLLAFKRAGADGILTYFALDAARLLRAQK*
>GCA_000452945.1_BPMSHR5858_genomic_05628_1 cycA
MNRLSKSLVVLQFAVAGLAGFMAEARAADAAKPDLDRGKAIATQVCAACHGVDGNSATGS
FPKLAGQHPDYLVKQLHDFKTQPGAKGPVRVNSVMVGFASALNDQDARNVAAYYGSQSAK
PGAARNAATVPVGQKIYRGGIAEKGVPACASCHGPTGQGIPVQYPRLSGQWADYTVAQLT
AFQQGTRSNDAMHQIALRLTDSEVKAVADYIAGLH*
>GCA_000452945.1_BPMSHR5858_genomic_05635_1 cyaY
MSDTDYLTRAEAVLAAVERSVDAANDGDADIDLERNGSVLTLTFENGSKIIVNLQPPMKE
VWIAAKAGGFHYRFVDGAWRDTRSGDEFFAALTGYATQQAGMPIAFSA*
>GCA_000452945.1_BPMSHR5858_genomic_05640_1 sctC_9
MRRGLVRFLLGCALGAAGEAVASLPPLPVGAPSGWSASASVGAAGRAPLPEAAAPQWRFD
SARDPVAGAPSPDVDGGAPAAEFAGEAMPERMPAAPTAEPARSTSADAGTSSAVASAGLQ
AQEAALEGPPVPLAPAQRMSDESDEHRSSPPAAGAVSTASVAGTGTETGDPSGDNRPISI
NLQQASLAAVFDAFARFTGLNIVVSERVRGTVSLRLNNVRWRSAFDALLDAHGLAMARRG
SVIWVAPVAELAERERQRFDAHARAAQLEPLASRGFVLRYARAADVQRLLAGSAGQRILS
KRGSVLADPRTNLLFVTDLSGRLAQIADLIGKLDTPSRQVLIEARIVEGDRGFSRNLGAR
LALRAPDAGERATGIVAGRNGTLAELAARPISGFDAATAGLTLFAARASRLLDVELSALE
AEGRGQIVSSPRVVTADRTKAVVEQGAELPYQAKVGNGVSGVQFRRATLKLEVEPQITPD
GRVILDLDVAKDSVGEETASGPAIHTKHVQTRVEVENGGTVSIGGIFESDDRDDVTRVPL
LGKIPVLGALFRHRAQRAQRSELVVYITPTVVIGP*
>GCA_000452945.1_BPMSHR5858_genomic_05641_1 aroK
LQARDPHVNVIFVGLMGAGKTTVGRAVARRLDRPFFDSDHEIEARTGARIPVIFELEGEA
GFRDREAQMIAELTQRENIVLATGGGAILRPENRKLLHERGLVVYLRANPHDLWLRTRKD
KNRPLLQTDDPKAKLEALYEARDPLYRECAHFVIETGRPSVNGLVNMVLMQLEMAGIVAK
PLQA*
>GCA_000452945.1_BPMSHR5858_genomic_05646_1 araQ_2
MIENRKGFDLFCHAVLIAGIAVIVFPVYVGFCAATMNAREVFTVPLSLVPSTHLFENVAA
IWRRGSGGMTTPFGTLLVNSVVMALVIAAGKISASILSAYAIVYFRFPLRNTAFWLIFVT
LMLPVEVRIFPTVQVVSTLHLTNSYAGLTLPLIASATATFLFRQFFMTLPDELVDAARID
GAGPLRFFWDVVLPLSKTSIAALFVITFIYGWNQYLWPILITTDASLTTAVVGIKSMIAS
GDAATEWHLVMAATLVAMLPPLVVVLAMQRWFVRGLVDSEK*
>GCA_000452945.1_BPMSHR5858_genomic_05648_1 ugpQ
MTNLQTWPYPRVVAHRGGGALAPENTLAALDAGAHYGHKMVEFDAKLSADGVAFLLHDDT
VERTSNGAGAARDMRYAALEALDAGAWYDARFSGERMPTLAQAAARCLALGLAANVEIKP
CPERDAETGSVVAALAAELWRDAAVPPLLSSFSAAALDAAREAAPALPRGMLYEEIPAGW
LEQTQRLGCVSLHAHHAQFNAALVRDLKASGLRLLAYTVNDPVRARELADWGVDLICTDR
IDLIGADFADREG*
>GCA_000452945.1_BPMSHR5858_genomic_05649_1 group_19127
MKQKMAITGAVALAFAAAATGAHAQSAGSFYVTTGWFHLAPQDSSDPLKIMNVGGSPVNH
EVPNTGAGIDNADTIGLAAGYFLTDHIATELVAGIPPRFNLNGKGSLEQFGVLGHAYQWS
PALLLKYYFNDAKAKFRPYVGIGASYIWFTGAKITNTAFERGALGGPTSVQTSNQWAPVL
NAGFTYNFTDHWFGGLSVSYIPVSLTATLTTQRPTPIGNVTQTSQAKITLNPIVTYLNIG
YRF*
>GCA_000452945.1_BPMSHR5858_genomic_05650_1 group_19433
MARLARLYVPDQPQHVILRGLDQQPAFVDDQDYELFIDCLKAAARDHHLAVHAYVLLPRQ
VQLLVTPSDEASLPKAMQAVGRRYVAHFNRRYSRRGTLWEGRYRATVIEGERYFLLASRV
VEMSPVRAQLVTSADAYRWSSYRHHVGLTVDSLITDHPLYWALGNTPFERQRAYKELCEQ
PLDERQADQLQQATLKGWVLGGETYREWAARTANRRVSPLPRGRPRKVRENTPPIQQ*
>GCA_000452945.1_BPMSHR5858_genomic_05651_1 gltB
MIDHQQPKSALPAAQGLYDPQNEHDACGVGFVAHIKGKKSHEIIEQGLKILENLDHRGAV
GADPLMGDGAGILIQIPDAFYREEMARQDVTLPPAGEYGVGMIFLPKESASRIACEQELE
RTVKAEGQVVLGWRDVPVDHAMPISPAVKASEPVIRQIFIGRGKDVMVTDALERKLYVIR
KTASHRIQALKLKHGKEYFVPSMSARTVVYKGLLLAGQVGVYYRDLQDARTVSALALVHQ
RFSTNTFPAWELAHPYRMIAHNGEINTVKGNVNWLNARTGAIASHVLGDDLPKLWPLIYP
GQSDTASFDNCLELLVMAGYPLVHAVMMMIPEAWEQHTLMDDNRRAFYEYHAAMMEPWDG
PAAIAFTDGRQIGATLDRNGLRPARYIITDDDLVIMASEAGVLPIPESKIVKKWRLQPGK
MFLIDMEHGRIIDDKELKDNLANAKPYKSWIDAVRIKLDEIEPEGEDVAAERRSAAALLD
RQQAFGYTQEDVKFLMAPMAQQGEEAVGSMGNDSPLAVMSNKNKTLYHYFKQLFAQVTNP
PIDPIRENMVMSLVSFIGPKPNLLDTNNINPPMRLEVSQPVLDFKDIAKIRSIDRYTGGK
FSAYELNICYPVAWGKEGIEARLASLCAEAVDAVKSGYNILIVSDRKTDAENVAIPALLA
TSAIHTHLVQQGLRTSTGLVVETGSARETHHFALLAGYGAEAVHPYLAMETLAKMAHGLP
GDLSPEKAIYNFTKAVGKGLQKVMSKMGISTYMSYTGAQIFEALGLATDLVEKYFKGTAS
KVGGIGLFEVAEEAIRLHRDAFGENPVLADMLDAGGEYAFRVRGEDHMWTPDSIAKLQHA
TRGNSYQTYKEYAHLINDQTKRHMTLRGLFEFRVDPAKAIPIDEVEPAKEIVKRFATGAM
SLGSISTEAHTTLAIAMNRIGGKSNTGEGGEDENRYRNELRGIPIKVGDTLKSVIGDEIV
RDIPLKEGDSLRSKIKQVASGRFGVTAEYLASADQIQIKMAQGAKPGEGGQLPGHKVSEY
IGKLRYSVPGVGLISPPPHHDIYSIEDLAQLIHDLKNVNPVASISVKLVSEVGVGTVAAG
VAKAKADHVVIAGHDGGTGASPLSSVKHAGTPWELGLAETQQTLVLNRLRGRIRVQADGQ
MKTGRDIVIGALLGADEFGFATAPLVVEGCIMMRKCHLNTCPVGVATQDPVLRAKFSGQP
EHVVNYFFFVAEEAREIMAQLGIAKFDDLVGRADLLDMRRGVEHWKAKGLDFSRVFYQPE
GCEGIARRHVESQDHGLERALDHTLIEKAKAAIENGEHVSFIQPVRNVNRTVGAMLSGAI
AKKHGHDGLADDAVHIQLKGTAGQSFGAFLAKGVTLDLVGDGNDYVGKGLSGGRIIIRPT
NDFRGKSEENIICGNTVMYGALEGEAFFRGVAGERFCVRNSGATAVVEGTGDHGCEYMTG
GTVVVLGETGRNFAAGMSGGIAYVYDADGTFAAKCNKSMVALDPVLQQAEQERTVDPALW
HAGTTDEALLKGLVERHFQFTGSPRAKSLLENWDAARRQFVKVFPHEYKRALGEIGAKKK
AGEVLAA*
>GCA_000452945.1_BPMSHR5858_genomic_05656_1 mnmC_2
MNMRASESDFAVLGGGLVGRLIAWRLAGAGYRVSLYERGDAAGSGSAAWVAAAMLAPLAE
AASAERLITDLGVASFDLWPSWLAELPEPVYFQRNGTLVVWHHADRAEAPLFERRVRANA
SAELLDGGLIALAGAQLDAAEPALAGRFARGLLLPREGQLDNRQALRALAAGLAERGVDV
HWNAAVAPDAAPAARITIDCRGLGAKAQMPALRGIRGEVARVHAPGIGLTRPVRLLHPRY
PLYVAPKENDLYVIGATEIEGEDMSPVSVRSALELLSAAFSVHPAFGEARILELNTQCRP
TLPDHRPALVWDGGATLAVNGLYRHGFMIAPEIADAAARFAQALVERTVKDADTFAAWRR
DARWPALLQQRAAHASA*
>GCA_000452945.1_BPMSHR5858_genomic_05658_1 thiG
MTPLSSADTLTLHGQTFASRVLLGTSRYPSLQSLSDSIAAARPGMVTVALRRQMNAGAAE
AGFFELLKRHDVPLLPNTAGCQTIAEAVTTAQMAREVFETDWIKLELIGDDYTLQPDPVG
LIEAATLLVKDGFKVLPYCTEDLVIARRLLDAGCEALMPWGAPIGTGKGIVNPYGLRVLR
ERLPDVPLIVDAGLGVPSHACQVMEWGFDGVLLNTAVSQATHPEIMARAFARGVEAGRAA
YLAGPMDARESAHASTPVVGMPFWHQDGSHA*
>GCA_000452945.1_BPMSHR5858_genomic_05660_1 mlaF_2
VSSSSESLLELRDVDFGYGERLVLSNLNMRFARGQVVAVMGGSGCGKTTVLRLIGGLVRA
RRGQVLFDGADVGAQTRDGLYALRRKMGMLFQFGALFTDMSVYDNVAFALREHTDLPEAL
IRDLVLMKLNAVGLRGARELMPSEVSGGMARRIALARAIALDPQLIMYDEPFAGLDPISL
GITANLIRTLNHALGATSILVTHDVPESFAIADYVYFLANGGVLAEGTPDALRASTDPSV
RQFIDGAPDGPFKFHYMSQPLAADFGLGGGRR*
>GCA_000452945.1_BPMSHR5858_genomic_05664_1 mlaC_2
MKKLFLIPVLAALFSFGAAAAHAEVDQSNPQALIKSATQQVLDEVRTQTIKQGDTARIMT
IVNKDILPYTDFRRTTQLAMGRNWRTATPAQQQQVIEQFKQLLIRTYSGALAQLKPDQQI
QYPPFRADADATDVVVRTVAMNNGQPVQIDYRLYKTAQGWRVYDLNVLGAWLIQTYQQQF
SEKIQQSGVEGLIQFLTERNQQLASGKQAS*
>GCA_000452945.1_BPMSHR5858_genomic_05666_1 btuD_18
MSAIEIRHVKKRYKSLQALKGVSLSVEEGEFFGLLGPNGAGKTTLISILAGLARADEGTI
AVRGHDVVKDFRAARRALGVVPQELVFDPFFTVRETLRIQSGYFGLHRNDDWIDEVMANL
DLTDKADANMRALSGGMKRRVLVAQALVHRPPVIVLDEPTAGVDVELRQTLWKFISRLNR
EGHTIVLTTHYLEEAESLCDRIAMLRRGEVVALDHTRTLLQRFSGLQLFVRLSHGALPAE
LRALEAEPAPAAQPEHLLRITDYDEVERILSLCRAAGCGFDEIEIRKADLEDVFVQVMNG
ADVIEGLA*
>GCA_000452945.1_BPMSHR5858_genomic_05667_1 yadH_2
MSGFRTLFYKEILRFWKVSFQTVLAPVVTALLYLTIFGHALTGRVNVYPGVEYVSFLVPG
LVMMSVLQNAFANSSSSLIQSKITGNLVFMLLPPLSSADIFGAYVLASVVRGLAVGAGVF
VVTVWFIPMSFAAPLYIVAFALFGSAILGTLGLIAGIWAEKFDQLAAFQNFLIMPLTFLS
GVFYSTHSLPPVWREVSRLNPFFYMIDGFRYGFFGIADVNPLASLSVVAGFFVLLALIAM
RLLATGYKLRH*
>GCA_000452945.1_BPMSHR5858_genomic_05668_1 group_19667
MLPTPELVKQYIEAGLACTHLEVEGDGQHFFATIVSSAFEGKRPIQRHQLVYAALGDRMK
QEIHALSMKTLTPAEWQNA*
>GCA_000452945.1_BPMSHR5858_genomic_05669_1 murA_2
VQVTVNEHDAVERVATATPAGNREAHAHGTDKLAIEGGRRLAGEIAVSGAKNAALPILCA
GLLSAEPVRLDNVPDLKDVRTTLALLGQMGMREETDGARVVLDASRVDNPVAPYELVKTM
RASILVLGPLLARFGYAKVSLPGGCAIGARPVDQHIKGLQAMGAEIHIEHGYIEARAKRL
SGARIVTDMITVTGTENLLMAATLADGETVIENAAREPEVTDLAHLLVAMGAKIDGIGTD
RLVIQGVERLHGATHAVIPDRIEAGTFLCAVAAAGGDVTLTGMRAHILDAVIDKLREAGA
TIDEGVDTLRVRMDGRPSAVAIRTSEYPAFPTDMQAQFMALNAVAQGVAQVTETIFENRF
MHVQELNRLGANIAVDGNTALVTGVPKLSGASVMATDLRASASLVIAGLCAQGETLVERI
YHLDRGYDRMETKLTAVGANVRRISGSEA*
>GCA_000452945.1_BPMSHR5858_genomic_05670_1 hisG
MSAPLTLALSKGRIFEETVPLLAAAGVTVAEDPETSRKLILPTTDPNLRVIVVRATDVPT
YVEYGAADFGVAGKDVLLEHGGGGLYQPIDLNIARCRMSVAVPAGFDYANAVRQGARLRV
ATKYVETAREHFAAKGVHVDLIKLYGSMELAPLVGLADAIVDLVSSGGTLKANNLVEVEE
IMPISSRLVVNQAALKLKRAALKPFLDAFERASLGSGA*
>GCA_000452945.1_BPMSHR5858_genomic_05673_1 hisB_2
MRVAQVVRNTSETQISVKIDLDGTGRQKLATGVPFLDHMLDQIARHGLVDLDIEAHGDTH
IDDHHTVEDVGITLGQAVAKAVGDRKGIRRYGHSYVPLDEALSRVVIDFSGRPGLEFHVP
FTRARIGTFDVDLSIEFFRGFVNHAGVTLHIDNLRGVNAHHQLETVFKAFGRALRMAVEL
DERAAGQIPSTKGSL*
>GCA_000452945.1_BPMSHR5858_genomic_05674_1 hisH
MDLLKSFISLLALINPLGAVPFFLSLTAQQTEAERRHTIRIAAVSVFCVIAVTTLLGQQI
IDFFGISVGSLEVGGGIIMLLMAISMLNAQIGNTRSTPEERDEAESKNSIAVVPLAIPLL
TGPGSISTVIVYAANARHWYDRLGLVAIGAALALICFFAMRLAEPIAHWIGRTGINIVTR
LMGLMLSALAVEFIVDGLKALLPALT*
>GCA_000452945.1_BPMSHR5858_genomic_05676_1 hisA
MLLIPAIDLKDGQCVRLKQGDMDQATIFSEDPAAMARKWVDLGARRLHLVDLNGAFAGKP
KNLEAIEAILGEVGDEIPVQLGGGIRSLETIEKYLDAGLSYVIIGTAAVKDPGFLQDACS
AFAGNIIVGLDAKDGKVATDGWSKLTGHEVIDLARKFEDYGVESIVYTDIGRDGMLQGIN
IEATVKLAQAVGIPVIASGGLSNIVDIEKLCEVEDEGIEGVICGRAIYSGDLDFAAAQKR
ADELNGELDDA*
>GCA_000452945.1_BPMSHR5858_genomic_05678_1 hisI
MNAEAKPGDWLGKVRWDANGLVPVIAQDAATNDVLMFAWMNRDALAKTIELKRAVYYSRS
RQRLWFKGEESGHVQHVHEVRLDCDEDVVLLKVEQVEGIACHTGRRSCFFQKFEGTVDDG
EWVAVDPVLKDPEHIYK*
>GCA_000452945.1_BPMSHR5858_genomic_05679_1 hisE
MTQSTTEDTLLRLAAVIDSRKGGDPEQSYVSRLFHKGDDAILKKIGEEATEVVLAAKDVR
QGGAPSALVGEVADLWFHCLVALSHFDLSPADVIAELERREGMSGIEEKALRKRREREEN
GG*
>GCA_000452945.1_BPMSHR5858_genomic_05680_1 group_19170
MPESPNQFPPSYQSATESERQQSLRTLTHILYLLYAIHWLTGGITGIVAIIINYVKRDDA
VGTAYQAHFEWQIRTFWRALIAYLIGFALLLVGIGFIVLGAVWIWTLYRIIKGWLYLNDN
KTLDPQAWF*
>GCA_000452945.1_BPMSHR5858_genomic_05682_1 tatA
MGGLSIWHWLIVLLIVALVFGTKKLRNIGNDLGSAVKGFKDGMKESEAPADAQQLPRSGS
VNVDAKDAARSSDSNKA*
>GCA_000452945.1_BPMSHR5858_genomic_05684_1 tatC
VSDPQHNPDDGPEETFISHLVELRDRIIRAGAAVIVVFLGLVYWAPDIFRLLARPLMENL
PKGGRMIVTDVTGSFFVPMKVTMLVALVIALPVVLYQIWAFVAPGLYQHEKKLVAPLVGS
SYVLFLCGMAFAYFLVFPTIFRVMAHYNAPLGAEMSTDIDNYLSFVLGMFVAFGVTFEVP
IVVVLLVRMGVLTVQKLKEIRPYVIVGAFVVAAVVTPPDVFSQLMLALPLVLLYEAGIIA
ARLFVKPPAKEEDESKAAAN*
>GCA_000452945.1_BPMSHR5858_genomic_05685_1 degQ
MLRRFWLFFAQAVTVLLALMFIVATLKPQWLQRQGQLGKQLAAPIVALREVAPGVGGAPA
QSSYADAAQKAMPAVVNVFSSKDGSLPPDPRAKDPLFRYFFGDRNARRQQEEPAANLGSG
VIVSSEGYILTNQHVVDGADQIEVALADGRTATAKVIGSDPETDLAVLKINMTNLPTITL
GRSDQSRVGDVVLAIGNPFGVGQTVTMGIISALGRNHLGINTFENFIQTDAPINPGNSGG
ALVDVNGNLLGINTAIYSRSGGSLGIGFAIPVSTARNVLESIITTGTVTRGWIGVEPQDV
TPEIAESFSLAQKSGAIVAGVLQGGPADKAGIKPGDILMSIDGEDITDTTKLLNVVAQIK
PGTPAKVHVVRKGKELDVTVVIGKRPPPPKQALDDQNSDEEE*
>GCA_000452945.1_BPMSHR5858_genomic_05687_1 petA
MRDKEEEHVDSGRRTWLIATSVAGGVGGVATVIPFAASLAPSTKAKAAGAPVEVDIGALK
PGEMLTVPWRGKPVWVLNRTDAMLADVVKADKEVADPQSKSPYSMPLPEYCANEYRSRAD
RKNILVVMAVCTHLGCTPSQRFTPGPQPNLPDDWPGGFLCPCHGSTYDLAGRVFKNKPAP
QNLDVPPYMFTSATTLVIGKDEKGEA*
>GCA_000452945.1_BPMSHR5858_genomic_05689_1 petC
MKKLLSTLAMFIVSACLLTSVSVRAEEGEFPLDRAPDNTENLVSLQHGAQLFVNYCLNCH
SANLMRYNRLTDLGISQKEIEKNLLFATDKVGNTMTVAMRPEDAKTWLGVAPPDLSVEAR
ARSRDWLYTYLRTFYRDDTRPTGWNNAVFPNVGMPHVLWQLQGERIAKFEEKTDEETGEK
THKLVGFQQVTPGTLAPADYDAAVGDLVAYLNWMSEPAQQTRKRLGVWVLLFLGVLTFLA
WRLNAAYWKDIK*
>GCA_000452945.1_BPMSHR5858_genomic_05690_1 sspA
MMVLYSGTTCPFSQRCRLVLFEKGMDFEIRDVDLFNKPEDIAVMNPYGQVPILVERDLIL
YESNIINEYIDERFPHPQLMPADPVQRARARLFLLNFEKELFVHVSTLENEKGKAAEKSH

EKARLAIRDRLTQLAPIFLKNKYMLGEEFSMLDVAIAPLLWRLDHYGIELSKNAAPLMKY
AERIFSRPAYIEALTPSEKVMRR*
>GCA_000452945.1_BPMSHR5858_genomic_05732_1 group_18942
MSEIKEMPLIELTAKDLPAYCPNPAMPRWSAHPRVFIDVSHGEARCPYCGTRYKLRDGEV
LRGHH*
>GCA_000452945.1_BPMSHR5858_genomic_05733_1 ilvE_2
MSMADRDGKIWMDGKLIEWRDAKIHVLTHTLHYGMGVFEGVRAYKTADGGTAIFRLKEHT
KRLLNSAKIFQMDVPFDQETLEAAQRDVVRENKLESCYLRPIIWIGSEKLGVSAKGNTIH
VAIAAWPWGAYLGEEGLAKGIRVKTSSFTRHHVNVSMVRAKASGWYVNSILANQEATADG
YDEALLLDVDGYVSEGSGENFFLVNRGKLYTPDLASCLDGITRDTVITLAKEAGIEVIEK
RITRDEVYTADEAFFTGTAAEVTPIRELDNRTIGGGARGPITEKLQSAFFDVVNGKSAKH
ADWLTKI*
>GCA_000452945.1_BPMSHR5858_genomic_05734_1 group_1960
MLARLSATDRFALAQGFRDFSPTLMAILSWGLVTGIAMSKSVLTTGQALGMSILVYAGSS
QLAVLPLFAAKLPVWTILLTAAMVNMRFVIFSAGLAPHFSYLPLWRRLAIGYFNGDVIYL
LFQKQGFAAGHVPGKEAYFWGMALTSWFAWQASSLVGIALASAFPDSWGLALAGTLALIP
IMVSAISNRSTLAAVAVAGIVSLVAFDLPYRLALPLAVVAALAAGSAADWFVERADWRRI
RAAHTEDAE*
>GCA_000452945.1_BPMSHR5858_genomic_05741_1 purK
MTALPTPNSPILPGAWLGMVGGGQLGRMFCFAAQAMGYRVAVLDPDPTSPAGAVADKHLR
AAYDDEAALAELAQLCDAVSTEFENVPAASLELLAQSTFVAPAGRCVAIAQDRIAEKRFI
AASGVPVAPHVVIESHAQLAALADADLAAVLPGILKTARLGYDGKGQVRVATVREARDAY
ASLGGVPCVLEKRLPLEYEVSALIARGANGASAVFPLAQNTHHGGILSLSVVPAPAASDA
LVREAQQAAVRIADSLGYVGVLCVEFFVLEDGSLVANEMAPRPHNSGHYTVDACETSQFE
QQVRAMTRLPLGSTRQHSPAAMLNVLGDVWFANGVSGEPVTPPWDEVAAMPTARLHLYGK
EEARAGRKMGHVNFTAATRDEAVAGATACARLLRIALD*
>GCA_000452945.1_BPMSHR5858_genomic_05744_1 group_19132
MQFNVELLLLALAPVFLLCIGWEAWHLARTRPGEHVYNLRDTLCNAALALMHQGADKIAW
IFVIPVYAYCYAHYRLVTWDATWIAFAVLFVAQDLLYYVFHRCSHRVRWLWAAHVVHHSS
ERLNFSTAMRQSLMYPIAGMWAFWLPLAFLGFPPQQIVGIVLINLAFQFFVHTQTIPKLG
WLEYVLNTPSIHRAHHARNPRYIDRNYAGVLVIWDRLFGSYVDEDPRDPPEYGIVEPLRS
NNPLVATFHEWRSMAADAIGVEGWRNKLRAVFGPPEWASAYHARVAHARAQSDEPPALAA
APRER*
>GCA_000452945.1_BPMSHR5858_genomic_05745_1 dacB
MNHHACRIAASPSPPPPAAVRARRTLARGAMLVAACAALAFAAPADARRKPKPPRYPAAV
SAARNVLPASVLVALQRARVPASSLSVVVERIGDRTPAVAWNASRPMQPASTMKLVTTYA
GLSLLGADYRWRTSAYADGDVDENGTLHGTLYIKGTGDPKLVPEELIDLVNKIRRAGIVN
VDGALVLDKTFFAPQTRDLPPLDDDASAPYNVGPDPLLYAFKALSFTVTPTDSGAIAVDV
VPALANLNVDNRLVEGQGSCDAARPTLVTDANGELTASFAGDYPPSCGPTTTNLAVLNHS
TFFARGFLALWRQTGGSFSGTIAEGKVPGRARPVASHHGPVLSSVVHDINKFSNNVMARN
LFLTIGAVEHRPPATPAQSADTIRAFLAHSGLPTDGLALENGSGLSRDERVSALGLADML
QAANASPVAQAFVDSLPIAGVDGTMRNRLTNAPVGGNAHIKTGTLRDVRAIAGYVASADG
SSYVVVSFINDDRASAARAAHDALLEWVYEGPR*
>GCA_000452945.1_BPMSHR5858_genomic_05746_1 qseB_2
MRILLVEDDRMIAEGVRKALKADGCAVDWVQDGDAALTALGGEAYDLLLLDLGLPKRDGI
DVLRTLRARGLALPVLILTARDAVADRVKGLDAGADDYLVKPFDLDELAARMRALIRRQS
GRSESMIRHGALTLDPASHQVTLDGAPVALSAREFALLEALLARPGAVLSKSQLEEKMYG
WGEEIGSNTVEVYIHALRKKLGSDLIRNVRGLGYMVVKEA*
>GCA_000452945.1_BPMSHR5858_genomic_05759_1 nsrR_1
MRLTDYTDYSLRVMLFLAVRGEGLSTIQEISDAYGISKNHLMKVVQQLGELGWIETVRGR
NGGLRLNAQSVTLTVGEVVRKTENDFALVGCFAGESESHRSCVIQSACRLRGVFAAAREA
FFAELDKHTIGELAQPASQLATLLGLAPVRIVSAADIRAAAPVVG*
>GCA_000452945.1_BPMSHR5858_genomic_05761_1 group_19253
MGLLDIVGGLIGGQGGNQNALVSAALEFINNQPGGLNGLIEKFNAGGAGGIIGSWIGTGE
NQPISPETLQNVLGSDTIGALAQKVGIDPQQASGILAQVLPHVVNGATPNGEVPAGGQLD
TSNVLGTLSQLASLLGNKQA*
>GCA_000452945.1_BPMSHR5858_genomic_05774_1 ybaB
MMKGQLAGLMKQAQQMQENMKKMQEQLALIEVEGQSGAGLVKVTMTCRNEVRRVAIDPSL
LADDKDMLEDLVAAAFNDAVRKAEATSQEKMSGMTSGLPLPPGFKLPF*
>GCA_000452945.1_BPMSHR5858_genomic_05776_1 smtB_3
MSATPGPLAGVKVLELGTLIAGPFAARLLAEFGAQVIKIEDPNGGDPLRKWRKLYPEAGG
TSLWWAVQARNKKSVTINLKSDEGKEIVRRLARDADIVVENFRPGLLEKLGLGYDVLSAD
NPGLVMVRLSGYGQTGPYRDRPGFGAIAESMGGLRHITGYPELPPPRIGISIGDSIAALH
GVIGALMALHHRNVNGGAGQVVDVALYEAVFNMMESVVPEYGVYGLVRERTGASLPGIVP
SNTYPCRDGSVVIGGNSDPIFKRLMKAIGRDDLADDPALAHNDGRVPRTGEIDAAIGAWL
AERTIDDALAVLNAADVPASRIYSVADMFDDPQFVARQMIQRFKLADGAQIPLPNVTPKL
SGTPGETRWLGPALGEHTDEVLGTLGYDAPAIAALRARRAI*
>GCA_000452945.1_BPMSHR5858_genomic_05778_1 pcm_2
MSGERAKRFPLALEDLKREPRKPEGRVAERQAAGDAARQRLTAAAAAPAAASPIVPERRA
PHGGVFAAKPARAKQHAPAAPGAAKRAPQGGAKQGDRSAAPNVALSGALALTSERVRERM
VERLRANGVADPRVLAAMSAVPRHMFVDPGLAAQAYEDAALPIGHQQTISKPSVVARMIE
LAAAGRALERVLEIGTGCGYQAAVLSRVARDVYSIERVKPLYERAKLNLRPLRVPNIRLH
YGDGRVGLPAAAPFDAIVIAAAGLDVPRALLEQLAIGGRLVAPVGEQAGEQVLTLVERVA
PAQWRESRLDRVFFVPLKSGVI*
>GCA_000452945.1_BPMSHR5858_genomic_05781_1 group_19309
MTPILVFDIETIPDVDGIRRLEDLPADLADAEVAEHAFAARREKTGSDFLPHHLQRIAAI
SCVFRDNNGFRVRSLGTPDDPEATLIQSFYRVIEKYTPQLVSWNGGGFDLPVLHYRALVH
GVAAPRYWDMGEDDRDFKWNNYLSRYHTRHLDLMDLLALYQPRASAPLDALAKLCGFPGK
LGMDGSQVWTAFREGRIEEIRNYCETDVVNTYLLYCRFQQMRGGLSPAEYTEEIALVKNT
LVQEPAPHWAEYLAAFDA*
>GCA_000452945.1_BPMSHR5858_genomic_05788_1 ispG
MQSEAQSPRSSQICSTEPVFGGHAPRRVSHAVDVRWGGTLVTIGGAAPVRVQSMTNTDTA
DAIGTAIQVKELANAGSELVRITVNTPEAAAAVPAIREQLDRMGVTVPLVGDFHYNGHLL
LRDYPDCAQALSKYRINPGNVGQGAKRDSQFAQMIEAAIKYDKPVRIGVNWGSLDQDLLA
RMMDENGARAEPWEAQSVMYEALIQSAIGSAERAVELGLGRDKIVLSCKVSGVQDLVAVY
RELSRRCGFALHLGLTEAGMGSKGIVASTAAIGLLLQEGIGDTIRISLTPEPGAPRTGEV
VVGQEILQTMGLRSFAPMVVACPGCGRTTSTLFQELALRIQTYLREQMPVWRSEYPGVEK
MNVAVMGCIVNGPGESKHANIGISLPGSGENPAAPVFVDGEKVKTLRGEHIAEEFQQIVS
DYVARTYGRAAAQN*
>GCA_000452945.1_BPMSHR5858_genomic_05789_1 hisS
MTEQKRKLEKLTGVKGMNDILPQDAGLWEFFEATVKSLLRAYGYQNIRTPIVEHTQLFTR
GIGEVTDIVEKEMYSFVDALNGENLTLRPENTAAVVRAAIEHNMLYDGPKRLWYLGPMFR
HERPQRGRYRQFHQVGVEALGFAGPDADAEIIMMCQRLWDDLGLTGIKLEINSLGLAEER
AAHRVELIKYLEQHVDKLDDDAQRRLYTNPLRVLDTKNPALQEIVRNAPQLIDFLGDVSR
AHFDGLQQLLKANNLPFTINPRLVRGLDYYNLTVFEWVTDKLGAQGTVAAGGRYDPLIEQ
LGGKPTAACGWAMGVERILELLKEEHLVPEQEGVDVYVVHQGDAAREQAFIVAERLRDTG
LDVILHCSADGAGASFKSQMKRADASGAAFAVIFGEDEVANGTVSVKPLRGTGAEGEKNV
QQSVPVESLTEFLINAMVATAEDGDD*
>GCA_000452945.1_BPMSHR5858_genomic_05790_1 group_19430
MSYHDEQESIESLKAWWARWGNLTTWVAIAVLAAAAAWNGWNYWQRRQATQAAVLYEQAQ
QAVRSNDKAKISRVASDMEDKFGGTAYAQMTALEAAKALYGAGDAAGAKAQLQWAADHAK
DDEYKQIAKLRLASLLLDEKAYDAGLALLSGTPSDAFKGAVADRRGDLLAAQGKSDDAKA
AYKLALDALPKDDASARQLVQFKLDALGG*
>GCA_000452945.1_BPMSHR5858_genomic_05791_1 bamB
MNLLKRYAAPVACAAAVLVFAACSSTKDARRVPTPLTEFKPVLDVQQVWTASVGKGGRYS
FSPVAVGDAVYVAGANGSVEKIDAKTGQQVWRTKIGSDLSAGVGSDGNLTAVGALKGGVF
VLGPDGKQLWKATVQGEIFSPPLVGNGLVIVRTIDGQVIAFAAQTGEQKWTYRNRAVPLN
LRVSAGMTFAGDAAVLAGFPGGGLVALNLQTGEPFWQTPVSFPKGVTEVERINDVTGAPT
LVGAETCAVTFQGQLGCFDANSGRPLWEKPFSSRSGVAQDDTVVVGGDDWSVVSAYDVAT
GKALWRNDKLKSRDVGVPYLLGRAVVLGDYKGFVHFLSRDDGTFVARMKTDGSAIAAAPV
LAGNTLVVLTQDGGVYGFRPR*
>GCA_000452945.1_BPMSHR5858_genomic_05792_1 der
MKPVIALVGRPNVGKSTLFNRLTRSRDALVADLPGLTRDRHYGEGRVGARPYLVVDTGGF
EPVAKDGILHEMARQTRQAVEEADVVVFIVDGRNGLAPQDKSIADYLRKTGRPIFLVVNK
AEGMKYTAVASDFYELGLGDPRAISAAHGDGVNDMINEALEVAYAGEPQESEEAAAARGI
KIAIVGRPNVGKSTLVNTLIGEDRVIAFDMPGTTRDSIYVDFERNGKHYTLIDTAGLRRR
GKVFEAIEKFSVVKTLQSISDANVVILLLDARQDISDQDAHIAGFVVEQGRALVVGVNKW
DGLDPHVRERTKADLARKLKFLEFAKFHFISAAEKTGIGALMRSVDDAYAAAMKKLPTPK
LTRALIEAVEFQQPRRRGPVRPKLRYAHQGGQNPPIIVIHGNALDAVTETYKRYLENRFR
ETFSLTGTPLRIEFRSSTNPYADKD*
>GCA_000452945.1_BPMSHR5858_genomic_05793_1 hfq_2
MSNKGQLLQDPFLNALRKEHVPVSIYLVNGIKLQGNIESFDQYVVLLRNTVTQMVYKHAI
STVVPARPVNFHPDAEAAS*
>GCA_000452945.1_BPMSHR5858_genomic_05800_1 gpt
MTQNLTASITMSDPRNDDKNLWVGWDEYHRLIEMLALAVHDSGWKFDKILCLARGGLRVG
DQLSRIYDLPLAILATSSYREAAGTQQGDLDIAQYITMTRGELTGNVLLVDDLVDSGVTL
ARVQQHLKERYPAVTAVRSAVLWYKGCSKVKPDYHVQYLPTNPWIHQPFEEWDTVRPHNL
AAWIKRGNAQRDDASAA*
>GCA_000452945.1_BPMSHR5858_genomic_05810_1 group_1564
MNDDVRAAGGVRAAHDRLQPALLDRLTDAERERRTEPPDAQAIGGERLRAAVLRDLAWLL
NTRNGEDGFVDWSAFAHAQASVLNYGMRPLVGKPMSGVERMSVEASIRDAIVRFEPRIAP
DSVEVRSVLDAPGGAAGERRHNVLMFEIKGTLWSIPHPVEFVLRSDLDLETGAMALHPAA
GG*
>GCA_000452945.1_BPMSHR5858_genomic_05811_1 hcp1_3
MGVAMFMKVDGVTGESADAQHKGWTDIQSFSWGASQPGAMASGSGGNAGKASFNDLVVAA
YMDKGATAIIKNCASGKHLPTVEISACKTGGSQIEFMRVTLQEVLVTSAQIAGVDPGDAA
DRLMMQYGFQAAKVKKQYWQQNDNGGKGAEVSVGWNIKENTEM*
>GCA_000452945.1_BPMSHR5858_genomic_05813_1 group_19403
MTSKYKASASGQKFIARNRAPRVQIEYDVETYGAERKVQLPFVMGVIADLAGKRAEPLPD
LPERKFLEIDVDNFDERMKSIAPRVAFQVPNTLSGDGMLSVDMTFESIDDFSPAAIARNV
DALRRLLDARTELSNLLSYMDGKHGAEQLIENAINDPALLASLVRQPLAADAEGGARQPV
TGTEDDPEIRHE*
>GCA_000452945.1_BPMSHR5858_genomic_05817_1 group_1565
MSHAVARIVRPLPSRDIWTFAGLVVLACFVWLAGPLFAFAEFRPFESGAVRAATIVALFV
AWGARIAWRGWRAGQLNAQLLNQLREAAPRPAATGDPAQAQLDELRSRFDEAATLLKKVR
FGEADGARKGLPRWLEQMSRQYLYQLPWYVFIGAPGSGKTTALVNSGLSFPLAEQFGRAA
IRGVGGTRHCDWWFTNDAVLIDTAGRYTTHESNRALDEAEWKGFVDLLKKYRARQPLNGA
MLTISVADLLGASEAERTQHAMVLRKRLLELRAQLGIRFPVYLLVTKADLLAGFAEYFGG
FGRAECAQVWGFTFPLAESEAPGFELRAAFDREYRLLHQRLNDGLPELLASQTDARQREM
TYLLPQQIADLQDMLGQFVAEVFSVSSFEPMPMLRGVYLTSGTQEGTAFDRVMSGIKRFL
KIEGVPPAAQTGSSGRSFFLKSLLQDHIFREAALAGSNLRWHQRQRALQIVGYAAIALLC
VAVLFAWLRSYSRNRDYLDEVAARVPAVDAQIGRAKFTGAADIVQLLPVLDELSGLPNAG
GVDLRHPPLAYRWGLFQGEKIEEASDAVYRRALDDVLLPIAASRMEQALRDARPDEVEYA
YAALKAYLMLYDSAHYDPAFVQAVVDLEMERALPADFSSAQRSALRAHLGALFGNRVAVS
PFPMNERLVADVRERLRQVPFSQRLYRQLARTLHASTASYDFSVARAVGPDASLVFRRQS
GKSLADGVPGLYTRSGYRNVFAPRLPGAIDSYGREEVWVLNLGASEIPNPADAAAWARDI
RQLYLNDYIKTWDDYLADIRLQRTSTLAQSIQVARTLSSADSPLTRLMVALARDTPLGDA
PGGARNLASRAQDKVDEARNSLAQIFAGQPGGEAGAAAAPPASPEQIVDSHFAGLRAFAP
GGGDQAASFDAVLKAIDALYTYLTATDDALRSGAAPPPSDAPARLRAQAGRLPTPVREVL
DDLSNVANGSIASVEQRNVAQRAGANVGDFCRQAIAGRYPFARGAARDVAPSDFAQLFAA
GGLMDDFFQKNLQTLVDTTAHPWRFNNRNAEADPSAAAMLGSFEKAAVIRDVYFGGGART
AQIKVEIVPLEMDPSISEMLLDVDGQIVRYAHGPQVPTAVQWPGTRGSNQVRLQVTEQSG
ATGGFTTEGPWALHRLFDRAGVSGGRGPEQMVARFAVDGKPIVLQVTASSVRNPFRLPQM
ESFTCPPKQ*
>GCA_000452945.1_BPMSHR5858_genomic_05819_1 group_1964
MNEPVLSATPAAALRQRVIWTEGMFLRPQHFQQLERHWERYVGMRCLPLQGFYWGYDELQ
IDRELLALGKVALLAATGVMRDGTPFDLSHPDDRPEPLDVPADAKDQLVVLALPLWRGGA
QEVSFGAGGNGEGHGGDGNAHAGFARYVVREYEVADANEVALGPALLQTGRLNVRLMLES
ELTGDWQALGVARVVERRTDGRLLVDDGYIPPRLVAQRDPVLLRHTRELHGLLTQRSEAL
GERLSEPGRGGVSEVADFLLLQLVNRYLALTWHAQQDVAAHPETLFCDWLKLACDLSTFT
AAGRRPQSLAVYRHDDLRASFGELMAELRRSLSTVLEQNAIQIELRDVGNGMKVATIADP
ALRDTAGFVLAVRADVPADSLRARFPAQAKLGPVERIRDLVQLQLPGIAMRQLPVAPRQI
PYHAGHTYFEIDKGGEMWKQLERSGGLAFHFAGEFPGLSMEFWAIRG*
>GCA_000452945.1_BPMSHR5858_genomic_05823_1 group_1734
MSILIRLSCAACASLFIAAAGASPLDADSADADAAPAAVVSDVHVFVVQRDGSVDEHDDS
TLRANTASGIDDVAQRYVWFNKDIDRVELLRAETIDRAGVAHPVGPEAIRDVQEPRSAGA
PFFEDGVLRSVIFPGVDAGARTRLVFRKSRTKPRDPGYFGYFAAPSRMPVEAQRLIFDLP
ADMPLYADARGYVARAPVTENGRTRYAFDYRHGPYPRIEEGSVGYTTYGDRLMVSTLRDF
AAFAGRYRAAAADASAADPAVARLAQAIVANANAAGPRAKARAIYDWVRTHVRYVALFLG
ETAAAPHKVTDILRNRYGDCKDHVALFGALLAAAGIRSEPALLNLGAVYTLPDVPGYGGG
AISHAITWLPDLGLFADTTVGGVEFGYLPPVVMDRPVLLVDDGVLSRTPAAQPRTRDARL
RIDVAPDGDARYQYHVEDGGWPAEFERGAFRQAARERVRQLAADRLRQSGLRGTARLGTS
ERDVTGGPFSTSMTGALERFAWPDGTTALPALSSLAGGIATQVQGWLAEPVRTQPWLCIG
ADFDETAQIALPENLRVTDLPADAGVHDRFFDYESHYVFDAPARVVQITRRLRARFAHQV
CAPEDFAAARASLEHIERDALAQIVVRAKPRD*
>GCA_000452945.1_BPMSHR5858_genomic_05824_1 phzS
MKKIVIVGAGIAGLTLALTLHSRGLAVRVFEAAPRIEPLGVGLNIQPYAVRVLHELGLLE
TLARQSITTKEMVFFTRYGQRIYSLPLGKYGGHNFPQLSIHRGILQLALLKAATERLGPD
FLVNGHSFVSCTQNSRSVNAVFSGAPGGADLVSVECDALIGADGVHSKVYRSLYPDGPRT
KGNGVLMWRGVTIAPPFLTGASMVRMGVPSHGKLVVYPIRSDVNAQGDQLINWVAELDEA
VYRKGANAAPGNGKDFIRFFASRTFDWLDVPALLERAASPIIAMPMADRDPLPRWQTGRV
TLIGDAAHPMVPVGSNGAGQAMLDAHSLGTRLAETADIEAALATYEHERRAYTSNIVLGD
RLDLPDRLIQEVDCRSHGKPFSDIGEIIASDEASRLALPDYHGRPLAPTQAGGA*
>GCA_000452945.1_BPMSHR5858_genomic_05841_1 group_1966
MKPRKDAPAGAAQAPHDCEEIERYRAPALDKGLDILELLAEQKEGLTRTEITKELGRNAS
EIYRMLERLVARQYVIRSPGGDRYTLSLKLYSLAHRHPPLKRLIGEALPPMQRFADAAEQ
SCHLCVYDRGNLLVIAQVDGPGAWGISVRLGSRVGLVDTASGPVLLAFQTPEQRARMLVE
HTKVEGEIALDMREFDALLHEIRATGRLRQKSRQAFGVTDIAYPVLGPSGHAIAALTCPY
VRRIDTHAAPCAEDVDAQLRATARHLSMLKEAAA*
>GCA_000452945.1_BPMSHR5858_genomic_05843_1 group_19245
MDVMAKLLTDLEFQRFSELQQKQASFTITPEEADELRDIVARAQKKRDDRASAMRAIEAY
IAQFDITPDELFSPEQIGEAARTYGLIAAGKKERSLPPSLTFNGKPYQWTKTLPDEVRLP
LFDAFKAGESIKRFIAMPKDAARCALTVARLERETGAVYADTLLDELAISRKHVEEAAAK
LAA*
>GCA_000452945.1_BPMSHR5858_genomic_05844_1 pat
VTVRNLDALFHPTSVAVVGASPRPGSVGAMVWACVLDGRFGGAIWPVNPKYGELNGHKVY
PYVDQLPSAPSVALICTPPATWPGIVRKLGGLGVRAAIIVGETRSGADRAALERALAAAK
PYLLRVVGPGSLGVVSPALGAHFGAPACIVKAGGVAWVSQSNALTNAVLGWAHARGLGFS
HAVALGGEADVDAADVLDYLASDAETRAILLELDTVKSARKFMSAARAAARNKPVLALRA
GRGDSGDLLYTAAFQRAGMVRVDALDDLLDEIEALGVGRAAATSGGVTLVTSDRGVAKLA
VDALAAAGETLAQWPRAAVDEVGGALPAGIVAGNPLLLGDDARPEYFGAALKALAQHPPT
GTVFVVHATSHSAPAVDVARVLIESRKFARRGMLACFFGGVDAATRDALHVHGIPVHTTP
QRLARAHARLVDYQLGRELLMQTPEGTPPQPAASISAARHTARAALAQGRDGFEGDAALE
WLAGFGIERATDADVDMGDTIVDITVGMYDDPNFGPVFRYSVPPADGVSAPFVVYGLPPF
NTVLARAVVARSPYAHRAPPEPLLQALTALSQAVCDVREIVEMSLVLRVRPTRVVALGPH
IRLATGRSRLAIVPYPRYLEQQLDWRGERITVRPIRPEDEAAHRELLSAMTPDDLRMRFF
GAIRNFDHSQIARMTQIDYDREMALIATLDDADGRAHTLGAVRAVTDPDNEATEFAIAVR
PDQKGKGLGRMLMTRIIDYARSRGTAWMIGEALRENTAMISLAKDSGFAVSSTEEPGVVA
FRLKLQP*
>GCA_000452945.1_BPMSHR5858_genomic_05846_1 group_18932
MATYRQLTAQLEKLQQKIDKEREKAITDAIADIKAKIEEYDITPEELGFRSVKAAAKPKR
ALPPKYLNPKTGETWSGRGRAPAWLGKNRNRFLIKD*
>GCA_000452945.1_BPMSHR5858_genomic_05850_1 group_19645
MRTTGSSGAMTLLTEFDQADGRELRSLRLVATEDGKSVLLIEVDERKPGIHREVRYEITP
AELIAAIRSHGAELPGETHGTVPLARTTAP*
>GCA_000452945.1_BPMSHR5858_genomic_05856_1 oppB
MLAYALRRTLWAVPTILAVVTVCYLLLHFTPGGPFDSEKQLSAATLANLNAKYHLDEPLW
KQYLLYLGSLLHGDLGPSFRYTDWSVNDLVRKAFPVSLGVGGVSIPLSIVLGVLLGTLAA

VRRDSFVDRFVMLIGNFGNVIPPFVLGPVLVLIFAIGLKTSEGAGWLPAGGWGDGGWRYR
VLPVTLLTLINVSLLARVMRGSMIETLSSNYIRTARAKGLPGRTIVLRHALKPALMPVVS
LFGTVCISSITAAVVTESVFALPGLGQLVVNGAINRDYTLVLGLVVLTTVCAVLFNLLVD
LAYAWLDPRIRY*
>GCA_000452945.1_BPMSHR5858_genomic_05868_1 group_1968
MNSQHARFIRPVLGTLCIVSLAALSGCRGDACFGFDGCFDDGGVQTVTLSGTAATGPALA
SATVNVDCARGSVATLSDGGGHYSVTFDAALPCAITVTSGGTSLHSLAYAGGTFNTTPQT
ELLLVYLAAQLGTSPARLIGDLPNSSRLQQALENQTNVQAAQTAVVANLQQRYAVMFAVP
GFLTTPFFVGQPGVDSDLGALANAGAIDANGMPDALAVSLLQQAGAAHPL*
>GCA_000452945.1_BPMSHR5858_genomic_05881_1 ompA_4
MSRQIRFWLVWGAIPFVVLASLCLPLRMQPEWSHVLTIVLVIAAIVLAIEIFEHSRETAQ
DRTAEAEIDDHELAVIIVAGPHASSLFNSSGDDTAFRREAGALWLRADSVEQLRDTMMRV
KASRGRFPDAAVVPLVSESDDESDMRQAFAKWRTELREALCYPDAAYVLPCYLAVYTCLG
FKGADGARPKWTGDAIEIGTRQMNAASKVGELLASIRQQLLRPSPTGSSAQCALGLGVFD
WLDATALLSSMTMLANTPPFMLKGLTLANVGRRPIRPGAWTRWLTVRTGLHIRQTKSSMA
PLPVPRVAKHAESGIREGWSSARHLVSTRASFVPALVFSTVSTLGIAVAVSGWMNSRMIQ
RVSNDIADYRSGADEWIGRKRFAQARLEHDRTAIVQTLLDGVPTGFGWGLYRGQELLVKL
DTVLDSPRTASIAMQLDTVSLFAAGKATFSPGAAQRELQHALRMILANPEQRVLIVGHAD
SSGSDAANFQLSEARARAIRDWFVDQGVLKTRFIVQGAGDTHPIADNDSPSGRAQNRRVE
VLLIPDVASDRKASGTEL*
>GCA_000452945.1_BPMSHR5858_genomic_05882_1 group_19631
MAISNSSQKFIARNRAPRVQIEYDVEIYGSEKKVELPFVMGVLADLSGKPLEPLPAVGDR
KFFSIDIDNFDERMKAMKPRVAFSVPNTLSGDGQLMVDITFESMDDFSPAAIAKKVDALS
QLLDARTQLANLQTYMDGKSGAENLVSKVLKDPALLSALAKAPKPVAEQAENRESKEKH*
>GCA_000452945.1_BPMSHR5858_genomic_05888_1 group_1254
MSYLKRFLRFVFSWQMLACIAVLLVCAAVWFVGPLLAFDELRPLAGVVVRVAVIVLLVAL
LAFWLLRWPLSPIGVAALCLLIWHAGPLFAFGDHRPFGPAWVRVLIVAAMLFCYAVYGLY
RLWQAVRTNDALLKRILEPSAGKPDAAAQANIRAVSVAVSKAIGQLKRLRGGAFGWRRLL
ESGRYLYELPWYMVVGAPGAGKSAAIARSGLKFPLADQMEASSERARGGTANCEWWFANE
AVLIDTAGRYVRHEVPGDEEATVANGAEWKGFLGLLRKHRPRAPVNGVVLSVSVEELVGR
TPAERTAHAATLRARLGELHQELGIHFPVYVIVTKLDLLPGFPEYFQSLTAEGRTQIWGF
TLPYDAENRRSAVGALREHCADELKRLEMRIDAGLNNRLLEEYENDRRKRLYALPQEFRS
LSAALSDMLALVFLDSRYDDAQLQNTLRGVYFTSAEQTDQVLAADRETILQRLKRQLGRM
LGGDAGAQTRDSGAVSGSRGYFLRDVFQHVIVPEAHLVRPNVRWEVRFRLMRWAGHLLAV
ALLVWLASALTVSFDNNRGYLDAISEKTAALAARVNAYNKAPKPAQIGGVLDGARDLPQH
GNLDLDAPGASFRYGLYVAPGIVDASDATYRSLLRRSLLPQIVRRVENALSAQIDAKHAD
EVYRTLTVYLMLDDAARHDAKAVKDWVMRDWERSDSAAEMGGRNLMARHLDALFVDARPF
EPSGHRNAALVQRARLFLNANPAPRRLYERAMAAIEKEAPENFTLARAVGLQGAGIFRLA
DGSRFQRGMPGLYTYDGYHQVFSVRLPEFLARAQSDDAWVMGNAGPAARWGDAIRNTQAV
AGRSSLADDIRRQYLTDYGNYWQQYLADIRPASSGEDGAGGTLAFDLATLRALAAPDSPL
VRLARAVVRETSLSVVDAREDASLTDAALSALGRRSGAAKEVADGAQKLAARRPEQRLEK
EVVDNRFAALREVVTGQADTGSGPAMTDVPVAAGGRALQLDAILTLINEQYTRLVVADNA
LSSQSMPPALDIGTTLQMEAEKLPAPLRTVLGGIATQAADKVGREVGSLLAMQVDSSVGN
ACRAAVDGKYPFARSAQEVDIEDFNRLFAAGGLFDAFFQKALAPHVDTHSKPWRYKALNP
GMPPIRGPSLEPFERAAAIRDVFFREPGAKRMAWKMDAKVASIDPEITELIVDVDGQSQR
YVHGPVLPFSVNWPGPRGGAIAEITAKPRIRPDTSTIATTGPWALFRLIERGRLTGTTSA
SRLMLDFDFDGRHAALELRTNGQMNPLTSGLLTHFRCPGSVG*
>GCA_000452945.1_BPMSHR5858_genomic_05905_1 rcsB_3
MTVFSIKVVIADDHPAVLTGLLHALAPVSSIEVVSACKNSTELLTALNEYPCDVVISDYN
MPDDNFGDGISLFSYIQRRYPSTRLIALTMISNPAIIHNLISHGISTVLSKSDEIGHIIA
AIHASYAKGSYLSPTMETIVNSGHGKSPDGEVQLSPREIEVVRLFTSGLSVNEIANRLKR
SKQTISSQKANAMRKLGVNSDIDLLKYALKAKLVEDTNSY*
>GCA_000452945.1_BPMSHR5858_genomic_05921_1 gcvA_7
MRATRGASRRNGSSQDSGAQMSEQSKNSQAGHAPLAYPLAALTRALPTLAALQSFVAAAQ
LGSLSKAAAHLCRTQGAVSRQIQQLEAHYRCALFVRQPTGLTLTADGDALFAVAADVLAR
LVRHARERDEAAAAVTVRVPSTFAIRWLLPRLPAIRRALGATQLCLSTSANDTPDFSEPD
IDAIVARGDGRWPGVEAIPLFAETLAPMCGPALAASLKSVDDLAHVTLLHPGRGRAEWRC
WLDAVGAAHIDATRGPVFDTLELTLSAAAEGHGVAIGDPRMAGDRLRAGTLTMPLRETAA
NGLSYYLVYPAQRATQPKIRALADVLVKLARAR*
>GCA_000452945.1_BPMSHR5858_genomic_05925_1 group_1572
MGERRVDTGGGSHGQSQSRRGWRQDMKKILVAMLALGVLIGVVAIVSICVAWTLADRLCY
*
>GCA_000452945.1_BPMSHR5858_genomic_05928_1 thpA_1
MLEIKHKLVQTAAACALAFAACAAHAADKPLKSIGITVGSLGNPYFVTVVKGAQAQAKEI
NPSAKVTAVSADYDLNKQFTQIDNFISAHVDMILLNAADPKAIEPAVRKAQAAGIAVIAV
DVAASGADATVQTNNVKAGELACDYLAKKLNGKGNVIIENGPQVSAVVDRVNGCKAVLAK
NPGLRLLSSDQDGKGSREGGMNAMQGYLTRFPKVDGLFAINDPQAIGSDLAAKQLRRTGI
VITSVDGAPDIETALTSDTQVQASSSQDPFAMAKKAVSVGYGIMNGHKPANAMILLEPTL
ITRENVKRYKGWSAH*
>GCA_000452945.1_BPMSHR5858_genomic_05929_1 rbsC_7
MNQQIKQSETAAAPPGAAQPRAPRRAADHRARMQALMRTAGMLPVLVLLCVGFGFMTDGF
FTLQNLSIVTQQASINIVLAAGMTFVILTGGIDLSVGSVLAAAAVASLLASNLPGWGWLG
IPAALAVGLAFGVVNGGLISFLRLPPFIVTLGALTAVRGIARLMGNDTTIFNPQLPFAFI
GNGSILGVPWLVVIALAVVAISWFILRRTVLGMRIYAVGGNPEAARLSGINVRGIELFVY
ASSGLLAGLGALMSAARLYAANGLQLGQSYELDAIAAVILGGTSFVGGVGSIVGTLIGAL
IIAVLSNGLVLLGVSDIWQYIIKGLVIIGAVALDRYRQHGSART*
>GCA_000452945.1_BPMSHR5858_genomic_05945_1 group_19640
MRLRHIEVFHAIMRTGSLSKAAQLLCVSQPAVSKVLAHAEQNLGIRLFSRAHGRLLPTRE
AELLFGETQKLQSNLERIRALSRNLALQPEGQLRVGCLPSLGLSLIPQAVKAFRDDYPRV
ALKIQTLHTDALLNALLTRDLDVAVAIDPPARPGITSAELGRTAVVSVGPPDDDAPDAPL
SLHAFMEGESIGIGTEDPLGDAIGNALEAFGEDRVTMVEAHTWYVARALAARGVGRVLLD
ELTARAPGEPVTIRPIEPALSVGVFALWRDGGLDSHAGATFVDALRAPFGEPSGKPPSMR
APGAPKRRAR*
>GCA_000452945.1_BPMSHR5858_genomic_05951_1 hpaB
MNDLFLQRAVSLEPLTSDAGAPCSFGVLVKPRRARTHVGELSIEWLRALVRSQQLVVLRG
FDSFADAPGMTRYCAAFGEIMMWPFGAVLELREQANPVDHVFASSYVPLHWDGMYLETVP
EFQVFQCVQAIGDAHGGRTTFSSTTEALRVATPETRALWHRAHGRYRRTVELYSNTVEAP
IVERHPRREFPILRFCEPPIADDPTFINPSSYAFGGIADSERDALLGSLTRALYDPRAHY
AHRWRTGDVVLTDNFTLLHGRERFTSRSGRHLRRVHIHGDPPLRNPHLSKQPHDAAAADA
AASA*
>GCA_000452945.1_BPMSHR5858_genomic_05959_1 group_19545
MSWHNKVVWSEGLFMRPQLFQQQERYLEHYAHKRAAPLSPFFFGFSHFSLDSEAPALGKI
IVKSASGVFADGTPFDAPGSTPPPAPLTIRPEHLDQVIYLAVPIRVPNGEETAFDRAAES
LARYAVFETDLRDTNSIGQGPKTVQLSNLRLRLLPEKELTDAWIGLALTRVKTIRADASI
ELDDMLIPPVVGYGASDTLASWLAKIHDLTRLRANALAERLTGSDGRAGTTAEVSDYLLL
QTLNRYEPLLKHLQRVPTTSPAELYALLIGMAGELSTYVRTDTRRPLDTHPPYQHVAPHL
CLKPVVDDTHRLLNAVLVRSAQRLALADLGHGMLNAVVDPVDMQGFTAVVLAVHAQMPPD
LLQQQFAAQAKAGPSERLPDLVRSHLSGIALQALPVPPRQIPFNAGYVYYELARGGPLWD
EVARHGGLALHIAGEFPSLKLELWGIRG*
>GCA_000452945.1_BPMSHR5858_genomic_05969_1 group_19177
MARDMTQAAVADTALSPQALARLRAEPWRYGFLALLRRIGADARIDPIGKARRPQAEPFR
LGQQPSLAFAPREIASVGDANGRLKVRLFGLGMLGPNGPLPIHVTEIARDREESRRDPTL
GNFLDIFHHRYLTLLYRAWASAQAAAGLDRPDDERFSFYVASLAGQDLDEVGARPLPAHA
RLSASPHLVREARNADGLRMTLERYFGVPVTLEENVFHWIAVDPLEHSRLGRPGDASTMA
AGALLGELVPDRQHKFRLVFGPLDIDAYLRFTPRGEDLPRLVEWVRAFVGYEFEWELELR
IKPNGAPPAVMGGPHQLGWSGWLGRSPSGEPVTGMRFEPEHYAHGFARGTARDTRGMRGE
R*
>GCA_000452945.1_BPMSHR5858_genomic_05970_1 group_1257
MTPSRKPAGRAAAARTPKPDGWMAPVDAAASCGADLEYDPEFVVLAAKVAPRAEAQYGDF
VGSPEPVNWSDVERDCRRLMMRSKDMRLAVLFARSRTRLAGATGLAEGIGLLAAWLAAFP
DAIHPQADVDADRDAALEIRRNALQALTDADGLLADVREIALTRSSATRLQVRDVERAFA
QPRPGDALAPESVVRQLDDLHAQQPETLAGFGDALAGLAAIDAWSGEHLGDYAPDLSALD
ALLRRIAGANARGDRAEAEPIAPAEADAPPASEAAAAHAHASPPRRREPAAQALAAAIAG
EPAAQPVDRYAARELIRQARQWFEQHEPSSPIPILLRRAEHFVGKRYADVVQAIPAELLA
LWSADET*
>GCA_000452945.1_BPMSHR5858_genomic_05972_1 group_19205
MKIVKPLAISPLTRVYRMHGREHLGVAALLIATLGDEPKLLAESALWRLAGDELRGYPLD
MALPKACPEFLVSGYAYGKYASDPHACACEVGVRIAGLEKRLRVCGDRQWAGARITAPRP
FERLPIDWDLAYGGAGCADNPRGRGAHAREGAPRDLPNVEYAHSPMRFAHEQPAPAGFCP
VDAAWPARAGLYGALDRQWQEEDCPGFPRTLDPRYFNIAPADQQLPELRAFPDGARYELT
HMHPDHATLAGNLPALRARSFVVRRGSDAPEEMPMRLTTAWFVPHRERVILIYHGVTPVR
AFDASDVQTVLFGAEASGHARPADWYRQVIEWRTRDDRAALYALRDRDLLPEHALAPEAA
ATPEPTQQSAKQRQLRERLSVFPDAPRAQTPAPDRLAEFVEQQQALADEKRAALEAMRRE
LATSEVFSVGRRRGPPGRIAPADEDPARHAGASAESPDIRALERDADERLRGLYQQCAQH
QDAPARLHGAAARARRECVASAAAAGQSLQVADLTGADLSGMDLRGARLAGAMLENADLS
DADLTGADLSRTVLVRADLTRAKLVDARLTAANLSLAHCERTDFSGSDLSDGIFEQVHLR
DCRFNGSVLASTRFDACRFDAVDFGRATLRELIFIEQSFSGVSFSDATIRKMLLMRCAFA
DVRFSAASIDGFGIVETQASGQLRFDRASVNKACFVGRCDIGRADFSFATLTEVNFRETQ
LVEANFGGARIGNCDFTDACLRAADLRGAKAEGSPFVRADLTRADLRDTDLIAAYLRGAK
LDGADLRRANLFRANLSQILTDADTRWQGAYLNRAVRFPLAEART*
>GCA_000452945.1_BPMSHR5858_genomic_05974_1 group_19692
MSEIEAKPDIRGRSSVPDKAGSVVGALPDGVYVVECGGAKLRCRRAFSCLVEPRIGDRVV
VCGADSRCVYVIAILARPNAHGVQLRVEGDLVLEATRRVCVTSGDALRLASREQLSIEAQ
RLTMSARQAALTSEKTTLTSAAFDGRLGKIRLIGRLLETVMEHVAQSCRSSFRTVETVEH
LRASHIDHAAAESMRMHAKHTLLTAETLTKIDAAQIHLG*
>GCA_000452945.1_BPMSHR5858_genomic_05977_1 syrM1_2
MDNLRRLDLNLLITLDALLSEHNVTRAAARLNLSQPSVSIHLAKLRDIFGDPLLLPGPRG
MRPTARADELREPLRRALETLGRAVSPGAPFDPAASAHTWRVAASDYSESTVVLPALNGL
RAAAPGTRLAVVETVPSRIAKQAEQGDIDLAFHITECAPGNLRHRALFTERYVLVGRAGH
PRLKRRPTLAQFCKLDHVIVSPDGGGFQGGTDHALAKLGLARRVVLSVPHFLFMMSAVAN
SDLVAMMPSRLVRDSRALQVVEPPVDVPGFRMAMLWHERSHRDPAHQWLREHIAASV*
>GCA_000452945.1_BPMSHR5858_genomic_05982_1 group_1706
MSSAPTATSASAPAAPSRGASPKPGSAASALASPASAASAPSPVSNAPAPRTLTVPPLPP
LTEAESADVTRRALALRAAFAQDVARRLNVPDAEQRAYGERLQQTLDANGLGELAREYVV
LVDRAPNVQAVFLYFRTTRSNAWQMIGASPVATGLPGQYDHFVTPLGVFEHTPENMDFRA
EGTTNDNGIRGYGQRDMRIYDFGWTDAERGWGKGGVSQMRFQMHATDPEYLEPLLGIRHS
KGCVRIPASLNVFLDQHGILDAEYEARAAGGDPPWVLRAHRQVTPWAGRYLVVIDSQRKT
RPAWSPAPGKKAQAKLPKGGDTAD*
>GCA_000452945.1_BPMSHR5858_genomic_05988_1 pxpA_2
MRKHSVDLNSDMGEGFGPWKIGDGVDEEIMPLISSANIATGFHAGDPNIIARTVQLAKNA
GVGVGAHPGFRDLVGFGRRDIGETPQALVNDIVYQLGALREFARFHGVSVQHVKPHGALY
MRAARDEALSRLLVETLQQLDPALRLYCMEASVTYRIARELGQPVVREFYADRDYGRNGS
IVFTRRAGRLDPRQVADKVLRACVDGRVATVDGEDIDIDFDSICLHSDTPGALALARATR
DALTAHGIRIAAPATAEANGIRIDEA*
>GCA_000452945.1_BPMSHR5858_genomic_05991_1 quiP_2
MSIRHATRSVALCSMLALASTLGCTSGAHDAPAAHAPRTAQTAHATAVAAGIAGLREPVR
IVRDSVGVSHIYARNTHDLFFAQGYNAARDRLWQLDLWRRQGEGKMAEQFGPRFAAQDRA
ARLFLYRGDLEAEYASYHPEAKQILQAFADGINAYVREAKADPAKMPPEFRLTGAEPGEW
RPESSLIRIYGITRNVTGEVALARQVAAVGLSKALSLNAFEPPVRQTAIPAGVDVGLIDK
RILADYLLARNGLPFRAEDFPRSPLGAAQRDALAARLNAARLTLADDSANPLAPRYESNN
WVVAGARTASGKPLLANDPHRRIGMPSLRYMVHLHAPGWNVIGAGEPALPGVSVGHNDRV
AFGLTIFAFGDEEDLYVYDTRPGHPDEYRYRGRWEKMRIVEETVAVRGQANRVERLAFTR
HGPVLYEDAAHRKAYALRAAYLEYPGTAAYLASLRLDQARDWPSFVAGMRRHYVPGENMV
YADVDGNIGWFGGALAPIRREADWSGALPVPGDGRYEWAGLTEGEALPHVLNPAAGYIAT
ANAYDLPDGYRYVDRSAHVWAEPFRQRRLDEVLRGASGLTVADMQRLQYDDRSLPAQAML
GFASQLKADEPGTRDALARLARWSGDMAIDSVPATIYEFWMREVMRRVRDVEVPAAARDA
FPKLEVRQVLRYLQQPDGAFGADPVAGRDALLLAALRDGLANARAKLGENVDAWRWGRLH
HAQFDHQLAGVLPAFGAFGTERYAVGGNDDTVHRGTYRPSDFRQTSGASYRQVIDVADWD
RSMIQNTPGQSGDPRSPFHANLLKGWASGEYWPMAFSDRAVDALAHDTLILEPAPAR*
>GCA_000452945.1_BPMSHR5858_genomic_05998_1 prpB_1
MGNPQLISAGAKFRAAVAAEQPLQVVGAITAYAAKMAEAVGFKAVYLSGGGVAANSLGIP
DLGISTMDDVLVDANRITNATNLPLLVDIDTGWGGAFNIARTIRSFIKAGVGAVHLEDQV
GQKRCGHRPGKECVPAGEMVDRIKAAVDARTDETFVIMARTDAAAAEGIDAAIERAIAYV
EAGADMIFPEAMKTLDDYRRFKEAVKVPILANLTEFGSTPLFTLDELKGANVDIALYCCG
AYRAMNKAALNFYETVRRDGTQKAAVPTMQTRAQLYDYLGYYAYEEKLDQLFNQGRN*
>GCA_000452945.1_BPMSHR5858_genomic_05999_1 prpC
MSETKEAAATGAAGGFKPKKSVALSGVAAGNTALCTVGKTGNDLHYRGYDILDIAGSSEF
EEIAHLLVHETLPNVTELAAYKAKLRAMRGLPANVKAALEWIPASAHPMDVMRTGVSVLG
TVLPEKDDHNLPGARDIADRLMASLGSMLLYWYHYSHNGRRIETETDDDSIGGHFLHLLH
GKTPSKSWIDAMHTSLILYAEHEFNASTFTARVIAGTGSDMYSSITGAIGALRGPKHGGA
NEVAYEIQNRYRTPDEAQADIRRRVENKEVVIGFGHPVYTISDPRNKVIKEVARKLSKEA
GDLKLFNIAERLESVMWEIKKMFPNLDWFSAVSYHMMGVPTAMFTPLFVIARTAGWSAHI
IEQRVDNKIIRPSANYTGPEDLKFVPIEKR*
>GCA_000452945.1_BPMSHR5858_genomic_06001_1 group_19200
MAHVPQIKIPATYLRGGTSKGVFFRLQDLPEAARTPGAARDALLSRVIGSPDPYGKQIDG
MGGASSSTSKTVIVSKSARPDHDVDYLFGQVSIDKPFVDWSGNCGNLSAAVGPFAIAGGL
IDPARVPENGVATVRIWQANIGKTIVAHVPITNGAVQETGDFELDGVTFPAAEVQLEFMD
PAADEDGAGGAMFPTGNLVDDLDVPGVGTLKATMINAGIPTIFVDAAAIGYTGTELQDAI
NSDAKALAMFETIRAHGALRMGLIASLDEIATRQHTPKVAFVAKPADYVASSGKRVAAAD
VDLLVRALSMGKLHHAMMGTAAVAIGTAAAIPGTLVNLAAGGGARHAVRFGHPSGTLRVG
AEAKQDGGEWVVTKAIMSRSARVLMEGWVRVPGDAF*
>GCA_000452945.1_BPMSHR5858_genomic_06002_1 group_19298
MTDIVDHARGALRAQPFSMLLGTELVHIGGEEASLRLPIRDELRQQYGFVHGGVISYLAD
NALTFAGALALGPRVVTGEYKINYLRPAVVGTLIARAKLIYAGRNQATCQCHVFVIDGDH
ERLVAVAQGTINRVGDGREPGAAEEKA*
>GCA_000452945.1_BPMSHR5858_genomic_06004_1 group_19438
MSEDLRVGLFPVRYLVGTGLPGAPQLVLDLMVDTVDHSVVGRAAVSQAVSPPLNFHADVW
GSYVFRLGPPPRRDGSGAIVQISLQGNQGGPQSNSMITFYGELLLKGDGKTGVASYRYYS
NGSWHEVENVPVKADPELVPIEPGPVIGQSSMSAIGSAAMYGVAIQSAAASGDLAHMRTL
SAYARQQLESRDEIAAALSELKAEIAKLESRQ*
>GCA_000452945.1_BPMSHR5858_genomic_06011_1 group_19595
MLNNVELIMRLMLAAALGSVIGIERERLSWAAGLRTHMLVCVGSALIMIVSAFGFADVLG
QAHVDLDPSRMAAQVVSGIGFLGAGSILLRGEIVRGLTTAASLWSVAAVGLAVGGGLYAA
AIAATIIILVILAGIKPLERRYLSVRQRRHLMLLVERGTLTFDSLHAALGLDSARVKRFI
VQQSEDSADSDEVTIALGRVSDVEYESICARLRQLPGVKGFSERKGGLPDD*
>GCA_000452945.1_BPMSHR5858_genomic_06014_1 group_18947
MSLIFALLQKIEDLFTSPHLPLDADYSYKARHAEVERVRKSHVSQVGLIRR*
>GCA_000452945.1_BPMSHR5858_genomic_06015_1 yycN
MLQMRPMTQTEFREYRERAARGYARDLIESGQSAPDEADARALACIDTLLPDGLLTDDQV
LLTLSESADGSVLGHLWYGVVAEGPHRSLFIYDLEIEPAFRRQGWATRVLQALEDDAREL
RVSEIGLSVFNHNAAALALYRELGFAAVTTTFVKSIESP*
>GCA_000452945.1_BPMSHR5858_genomic_00017_1 pucM
MGKLTTHVLDTAHGRPGAAIKLELHAISGDGSRLLKTLHTNDDGRCDAPLLEGDALAAGT
YELVFHAGDYFAAQGVALPEPRFVDLVVLRFGIADPTAHYHVPLLVSPWSYSTYRGS*
>GCA_000452945.1_BPMSHR5858_genomic_00018_1 group_3046
MEGFITDWLNLAIRWLHVVAAIAWIGESFYFVALDNSLKPPRDAQLKSRGVFGELWHVHG
GGFYNMQKYLVAPPEMPEDLHWSKWPSYTTWMSGFGLFFVLYLLAPNTYLIDRNVLDMGP
VVAVASALGFLLAGWIVYDCACRLLGGRDKLLGVFVGLYVIAAAYLACHVFSGRAAYLIV
GAMLATIMTANVFFVIIPGQRKMVDKLLKGETPDPIYGKRGKQRSVHNTYFTLPVVFAML
SNHYAMTYANRYNWIVLVVIMLAGALIRQFFVMRHRGQQLWYLPLGGVALMLLALVTTMP
RPIAPEARAANAPNVPKLAINDIAPILQQRCVACHSAKPTLMGSAPAGVMFDTPDEIARN
APRIYQQAVQLKAMPIGNVTHMTDDERTKIAAWFEGGATK*
>GCA_000452945.1_BPMSHR5858_genomic_00020_1 alc_2
MAFPLSDPNAPEFTRRYVNLADPRLGAQALEASDDFFAPKERMLNPEPAVFIPGKYDDHG
KWMDGWETRRKRTTGYDWCVVKLARPGVIKGVDIDTSHFTGNFPPAASIEAAHVPDGAPN
EATKWVEIVPATTLQGNSHHYVEARDANAYTHLRVNLYPDGGIARLRVYGQPQLDWAGAS
RSALFDLAAMENGGYVVAANNQHFGLASNVLLPGRGVNMGDGWETRRRREPGNDWAIVAL
AQPGVIRKVEIDTAHFKGNYPDRCSIQAAYVQGGTDSSLVTQAMFWPVLLGEQKLQMDKQ

HAFEAELAALGPVTHVRLNIIPDGGVSRLRVWGTLDK*
>GCA_000452945.1_BPMSHR5858_genomic_00022_1 group_19784
MAFDPNYPRDLIGYGRHPVQANWPGRARVAVQFVLNYEEGGENCVLHGDPASEQFLSEIV
GAAAYPARHMSMESIYEYGSRAGVWRILREFDKRGLPLTVFGVGMAIERHPELARAFVEL
GHEIACHGWRWIHYQDMTPEREAEHMRLGMEAIERVTGVRPLGWYTGRDSPNTHRLVAEY
GGFLYDSDHYGDDLPFWMDVEVSGGASVPQLIVPYTLDANDMRFATPQGFNTADHFFHYL
RDAFDVLYEEGDEAPKMMSIGMHCRLLGRPGRFRALQRFLDHIERHDRVWVARRVEIARH
WREHHPYRPHARREGDA*
>GCA_000452945.1_BPMSHR5858_genomic_00023_1 hyuE
MKIKLINPNTTRRMTDAMGRCAREVASAGTAIVAVSPAMGPPSIEGYYDEALATPGLLAE
VAAGERDGFDGYVIACFGDPGLYAARELARGPVIGIAEAAMHAASVLAPGFSVVTTLART
CAMAWHLAERYGMKRFCRNVRATDVAVLDLDRPGSAARRVIVDECRRALDEDGADAIVLG
CAGMAEFAHEIEAAIGAPVVEGVTAAVKWTEALIALKLATAKRGDFARPLAKRYDGEFAR
FSPDGAFAREVAAAGAAGAPAASAAPAASAAPAVSAVPAVPAVPATSATSAAPATSAAGS
DAAPSGAPATGVFPQAHIHSV*
>GCA_000452945.1_BPMSHR5858_genomic_00024_1 pucI
MAQFSVARQSASYRPNEDRAGGPDGGAQMPAGYSSRLYNEDLAPLASQRWGAYNIFAFWM
SDVHSVGGYVFAGSLFALGLTSWQVLVALIVGISIVNVLCNLIARPSQQLGVPYPVACRA
TFGVLGANVPAVIRGLIAIAWYGIQTYLASSALVIVVLKFFPHWMPYADVHRYGFLGLSA
LGWAGFMLLWVLQAFVFWNGMETIKKFIDFAGPAVYVVMFALAGYMVWRAGWRNIGLNLG
GVRYHGAEVIPVMVTAISLVVSYFSGPMLNFGDFSRYCSSYGGVKRGNFWGLPVNFLAFS
LVTVITTAATLPVFGELITDPVETVGRIDHPSAVILGALTFTIATIGINIVANFVSPAFD
FSNVAPRLISWRAGGMLAAVASVFITPWNLFNNPAVIHYTLDVLGGFIGPLYGVLIVDFY
LVKRGALRRDDLYTTSADGAYWYRDGVNRRAIAALLPAAAIAVACVMAPALSGLANFSWF
IGAALGGAFYRALAKA*
>GCA_000452945.1_BPMSHR5858_genomic_00025_1 rspR_2
MTKKDAAPTGASAESIAERIRTAILEHRLAPGTKLTEAQLCEVFGVKRGAIRQALALLAT
DRLVDLEPNRGAFVASPTLQDVHEVFEMRRIVELAVMERLATGPGAKRLKGVAAMIDKER
HAFERRDFPAWIRLSGEFHTALAALTGNTVLRDCLGGLVARSTLMSALYESHGRSPCSFD
DHAQILAALEAGDAKRAAQLMAHHLQHVELKMLDRPAQGAVDLREVFGDPG*
>GCA_000452945.1_BPMSHR5858_genomic_00026_1 ldcA
MISTPCTPRTIRLLAPSGYPHDPAAIERALERLSDEQHRIQNLDATQRRYQRFAGTDGER
AGDLNRLADPSLPLPDIGLAVRGGYGAARILHGLDYRGLERRLRDQPIALVGHSDFTAIQ
LALYAKARVKTFGGPMLSADFGAPTPSEFTLAHFWQTLTQPTTTIVSDVPQTQVVNASGT
LWGGNLAILASLVGTPYMPQIEGGILFVEDVNEQPFRIERMIYQLHLAGILARQQALVLG
QFTGARPFDYDNGYDMQAMIEQVRGVIGIPIVTGLQFGHVPDLLTLPFGARAELVANEHG
FRLTMSDYPHLAA*
>GCA_000452945.1_BPMSHR5858_genomic_00028_1 group_2221
MKLLDALVEQRIAAAAARGAFDDLPGAGAPMELDDDLLVPEEVRVANRILKNAGFVPPAV
EQLRALRNLQDELRAVSDRATRCRLQAKMLALDMALESLRGGPMVVPREYCRRIAERLSE
RVLGDAQGEAGAM*
>GCA_000452945.1_BPMSHR5858_genomic_00029_1 group_19764
MALSMYDVSIPALIRGLNNLSAVLDKGDAHARAQGIDPAELVRARLRDDMYALDAQVQRA
SDTAKGCGARLAGIEVPSFADTETTFAELRARIGKTIDFLKTIQPAQLDGSESRTIELPL
RDGVMTFDGKSYLVGFALPNFYFHTTTAYDILRHKGVPLGKMDYLGGR*
>GCA_000452945.1_BPMSHR5858_genomic_00045_1 guaB
MRLIQKALTFDDVLLVPAFSDVLPRDTSLKTQLTRNISLNMPLVSAAMDTVTEGRLAIAM
AQQGGVGIVHKNLTPVEQAREVAKVKRFESGVVRDPITVPPSMKVRDVIALSRQHGISGF
PVVEGPKLVGIVTNRDLRFETRLDEPVKSIMTPRERLVTVAEGTPLAEAKALMHSHRLER
VLVVNDAFELRGLMTVKDITKQTEHPEACKDEHGKLRVGAAVGVGADNEERVELLVQAGV
DVIVVDTAHGHSKGVLERVRWVKQNFPKVEVIGGNIATAAAAKALVEYGADAVKVGIGPG
SICTTRIVAGVGVPQISAIANVSDALRGTGVPCVADGGIRFSGDVSKALAAGANAVMMGS
MFAGTEEAPGDVFLYQGRQYKSYRGMGSVGAMKDGAADRYFQDNSANIDKLVPEGIEGRV
AYKGSVNAIIFQLIGGVRASMGYCGCKTIAELHEKAEFVQITAAGMRESHVHDVQITKEA
PNYHVD*
>GCA_000452945.1_BPMSHR5858_genomic_00049_1 ytfF
MQRGVVYGVLAGALWGMVFLVPRLLTDFSPLLLSVGRYAMYGLVSLAAALPAARSLASRL
THDDLVALVKLALVGNVAYYMLLSSAVHLIGIAPSSLIVGVLPVTVTLAGLGDHGAVPLK
RLAGPLALVVAGIACINVDLFTSEAAHATTLAQKLAGIACATGALASWTWYAVANARYLQ
RHHHFDGNEWSVLWGVVTGLIGGLCWIGIAALPAGALQPALPASRWQLFWLLNLGLAIGA
SWLGNGLWNAASKRLPLTLSGQLIVFETVFALLYGFVFDHRLPRVLEIAALVLLLAGVYG
SVRRHGDKRAADDPLNANLGAH*
>GCA_000452945.1_BPMSHR5858_genomic_00050_1 group_19990
MPKVQVCYALPERQTLVAVDVPAGASVRDAIAASGVLALHPDIDASALKTGIFGKLAPLD
APVADGDRVEIYRPLIVDPKLARQRRVDKTRREGSIEGRKWLPKDSR*
>GCA_000452945.1_BPMSHR5858_genomic_00051_1 pasT
MFDLVTDVDDYPNFLPWCGGVEIRRRDETGMEARIDINFKGIKQHFATRNTQERPTRIDM
EFADGPFRKFTGYWRFTPLRADACKIEFALHYEFSSIILEKIIGPVFTHIANTFVESFVK
RADQRYGKG*
>GCA_000452945.1_BPMSHR5858_genomic_00052_1 smpB
MSIIDNRKAFFDYHIEERYEAGLVLEGWEVKALRAGRGQIKEGYVVVKHAEIFLIGTHIS
PLPEASTHIKPDPVRTRKLLLHRDEIKKLIGKVEQRGYTLVPLNFHYKGGRVKCEIGLAK
GKKLHDKRETEKKRDWEREKARIMRSAT*
>GCA_000452945.1_BPMSHR5858_genomic_00054_1 group_3050
MVSGQFFWWIGVGVFVVAELLTGTFYLLMIALGFLAGGLAHLAGVPAAWQLAAAALVALV
AVIALRRSGLGRKQKRDAGANPDVNLDIGATIEVTHWRDRRARAQYRGAPWDVELAAGER
EDAHLYEVRAVRGNCLIVAAKPPA*
>GCA_000452945.1_BPMSHR5858_genomic_00055_1 ppsA
MTNAANVAKDQAYVIPFEQLRMTDVEIVGGKNASLGEMISQLAEAGVRVPTGFATTALAF
RDFLQHNNLTERIAQRLESLDIDDVKALAEAGAEIRKWIVDAPLQPRLEQEIRAQFEILK
NGSPGELSFAVRSSATAEDLPDASFAGQQESYLNVVGIEDVLDRMKHVFASLYNDRAISY
RVHKGFTHAEVALSAGVQRMVRSDVGAAGVMFTIDTESGFKDAVFITASYGLGETVVQGA
VNPDEFYVFKTTLAQDKYPIIRRSIGSKLIKMEFTQPGEPGRVKTVDVPHEQRNRYSITD
DDVIELAKYAVIIEKHYQRPMDIEWGKDGRDGKIFILQARPETVKSQAHGKVEQRFKLKG
QSQVLATGRAIGQKIGAGPVRVIHDPSEMERVQPGDVLVADMTDPNWEPVMKRASAIVTN
RGGRTCHAAIIARELGVPAVVGCGDATDVLKDGALVTVSCAEGDEGKIYDGLLETEVSEV
QRGELPSVPVKIMMNVGNPQLAFDFSQLPNAGVGLARLEFIINNNIGVHPKAILEYPNVD
ADLKKAVESVARGHASPRAFYVDKLTEGIATIAAAFYPKPVIVRLSDFKSNEYKKLIGGS
RYEPDEENPMLGFRGASRYIAEDFAQAFEMECMALKRVRDEMGLTNVEIMVPFVRTVKQA
ERVVGLLGKFGLKRGDNGLRLIMMCEVPSNAILAEEFLQHFDGFSIGSNDLTQLTLGLDR
DSGMELLAVDFDERDPAVKFMLKRAIDTCRKLDKYVGICGQGPSDHPDFAKWLADEGIAS
ISLNPDTVIETWQALAAKQ*
>GCA_000452945.1_BPMSHR5858_genomic_00056_1 group_3919
VNTRLAGAVLAALALIGPAAAQPLPLPAGLIDLGSLAGEQMLAESGARSAYASLGSHFVT
QKTQSYCGVASLVMVLNALRVPAPAAAQYPPFHYFTQDNVLGDATERIRPRALIERHGMT
LDQLGALANALGASGDVRHASDVSVDVFRADAIAHLGRPGRYVLVNYLRSRLGQQTGGHI
SPLGAYDAAADRFLILDVSRYKYPPVWVTTADLYAAMNTPDADSGGRSRGYVLIDGVVAG
ADS*
>GCA_000452945.1_BPMSHR5858_genomic_00058_1 pcp_4
MTSRKWARPRASQAKHAIYAATFFAAAALSAHAAAAWVDTQTGAYPALAQQALAASQASA
AATAAGKAIDTAPGEPVRVVVSLNLNDEARLDRFLRDLHTPGSAAYGRHLTPAEFAAQYA
PTPQQVALVEAHLRRAGFRDIEVAPNRLLISATGTAAAVKTAFNTRLKRFTLEGRRVYAN
QDAAQVPAELGRIVGAVLGLDNATLARTYNRQAAVTGAVGGAKASLAARASDASAAASGA
PVLTGHDPLEFSRIYRAGSTPTASQTTVGVIMAGDAAPVLRDLDTFAAKAGLARVAATVT
RTGPPGSDYSDNSGLSEWDMDSQAIVGAAGGAVKGLVLYAAPSMLLSDITSAYNRAVVDN
VAKVINVSLGVCEADARASGTQAADDRIFKSAVAQGQTFVVAAGDAGAYECSVSRVSGGQ
GVPARSNYSVSEPATSPYVVAVGGTTLSTDRTTLAYAGEVAWNEGLQPIGVYDAYGSYDG
TRRLWATGGGYSRSEAAPAWQRSVLGASAKARALPDVAFDADGRSGAHVYVNGRTEQWGG
TSLAAPIFTGIWARVQSDNGNRLGFPLASLYRYAPANGAFAHDVKSGNNGSGGYGYKAGA
GWDPVTGFGSLDIANFAAFVKQTADFAR*
>GCA_000452945.1_BPMSHR5858_genomic_00061_1 rnhB
MATTRKPRGGAGGATQPALDFDAPGEIVCGVDEAGRGPLAGPVVAAAVVLDPARPIVGLD
DSKALSAKKRERLFDEIVAHALAYSVASASVEEIDSLNILHATMLAMKRAVEGLSVLPTL
AKIDGNRCPMLAIRSEAIVGGDALVPSISAASILAKVTRDRMLVELHQQFPMYGFDAHAG
YGTPQHLAALREHGPCEHHRRSFAPVREAFDLIR*
>GCA_000452945.1_BPMSHR5858_genomic_00062_1 lpxB
MAFQLTPLRVALVAGEPSGDLLGASLLGGLHARLPASSRYYGIGGPRMSAVEFDAHWPME
KLAVRGYVEALKHIPEILRIRGELKRQLLAEPPDAFVGIDAPDFNFGLEQALRGAGIPTI
HFVCPSIWAWRGGRIKKIVKAVDHMLCLFPFEPELLEKAGVAATFVGHPLADEIPLEPDT
HGARIALGLPGGGPVIAVLPGSRRSEIELIGPTFFDAMELMQQREPGVRFVVPAATPALR
ALLQPLVDAHPSLSVTLTEGRAQVAMTAADAILVKSGTVTLEAALLKKPMVISYKVPWLT
GQIMRRQGYLPYVGLPNILAGRFVVPELLQHFATPDALADATLTQLRDDANRRALTDIFT
DMHLALRQNTAQRAAEAVARVIDSRKPR*
>GCA_000452945.1_BPMSHR5858_genomic_00065_1 lpxD
MALTLEALAARFGGEIVGDGRCEVGALAPLDQAGPRQLAFLANPKYLAQVETTGAGAVLI
APGDLEKLGAAAHGRNFIVTPNPYAYFARVAQMFIDLAAPPRAAGVHPSATIDPAAQVAA
SAVIGPHVTVEAGAVIGERAQLDANVFVGRGTRIGDDSHLYPNVAIYHGCTLGPRAIVHS
GAVIGSDGFGFAPDFVGEGDARTGAWVKIPQVGGVKVGPDVEIGANTTIDRGAMADTVID
ECVKIDNLVQIGHNCRIGAYTVIAGCAGIAGSTTIGKHCMIGGAVGIAGHVTLGDYVIVT
AKSGVSKSLPKAGIYTSAFPAVEHGDWNRSAALVRNLDKLRDRIKALETALAAREGDAGG
A*
>GCA_000452945.1_BPMSHR5858_genomic_00075_1 rpsB
MAITMRQMLEAGVHFGHQTRFWNPKMAPFIFGHRNKIHIINLEKTLPMYNDALKYVRQLA
ANRGTILFVGTKRQSRDTIAQEALRAGMPYVNARWLGGMLTNFKTLKVSIKRLKDMEAAV
EAGELEKMSKKEALLFEREIAKLQKSIGGVKDMGGIPDAIFVVDVGYHKIAVTEANKLGV
PVIAVVDTNHSPEGVDYVIPGNDDSSKAVALYAQGVADAILEGRANAVNEVVQAVRGDDE
YVEENA*
>GCA_000452945.1_BPMSHR5858_genomic_00077_1 glnD
MSASVAEPPPALSRKAEFKAAKAELLARFKSANHVTPLMHALSRATDDALRSLWQECGLP
ATLALVAVGGFGRGELSPHSDVDILVLLPDAHASELDERIERFIGMAWDLGLEIGSSVRT
VDQCIEEASHDVTVQTSLLEARRIVGSTALFERFMLRYREALDARAFFQAKVLEMRQRHA
KFQDTPYSLEPNVKESPGGLRDLQTILWIARAAGFGSSWRELDTRGLITDREARELRRNE
GFLKTLRARLHVIAGRRQDILVFDLQTQAAESFGYQPTSAKRASEQLMRRYYWAAKAVTQ
LATILIQNIEAQLFPATSGVTRVLSPGRFVEKQGMLEIAADDVFERHPDAILEAFLLYEA
TRGVKGLSARTLRALYNSRDVMNNAWRRDLRNRHTFMQILQQPEGITHAFRLMNQTSVLG
RYLLNFRRIVGQMQHDLYHVYTVDQHILMVLRNIRRFAVAEHAHEYPFCSQLIVNFERPW
VLYVAALFHDIAKGRGGDHSALGMADARRFCREHGIEGDDAALVVWLVQHHLTMSQVAQK
QDTSDPVVIKRFAELVGSERRLTALYLLTVADIRGTSPKVWNTWKGKLLEDLYRATLAVL
GGAQPDAHSELKTRQEEALALLRLETVPPDAHRALWDQLDVGYFLRHDAADIAWQTRVLY
RHVAADTAIVRARPSPVGDALQVLVYVKDRSDLFAGICAYFDRNGLSVLDARVNTTRHGY
ALDNFIVTQTEHDVQYRDIANLVEQQLAARLAESAPLPEPSKGRLSRLSRTFPITPRVDL
RADERGQYYILSVSANDRPGLLYSIARVLAEHRVGVHAARINTLGERVEDVFMLDGTGLS
DNRLQIQVETELLRAIAV*
>GCA_000452945.1_BPMSHR5858_genomic_00083_1 btuD_12
MTTPADPAARANHAERAGRGLRARIALDLWRAIRRYRLRVLAALALLVLAKAAAVAVPLL
LKQIVDELGRVAAHPFALPVLLLFGYAVVRFASNALNEVRDMTFVHVTQRTVADFTVRTF
AHLHRLGARFHARRETGAVVRDLEKGTAGIGYLLGIAVFTIVPTVLEIGAVLVVVIGKYG
GGFTAIIFVTFAVYAAYTFVFTRRRMRLQRRVNALEARSNGRIVDSLLNYDTVKYFARED
FERERLDTVLDDWREAGIDNQFALSTLHIGQSACIGAGIAAVMLLAGERVVSGVMTVGDL
VLINAYIIQISLPLNALGFVFREANDALTNVERLFALLDARGKPGEDGDAPFAQPLAVRG
GEIRFEHVDFGYEPGRQILWDVSLAIGAGETVAVVGGSGSGKSTLARLLFRLYQPDAGSI
SIDGQDLRLVTERSLRDAIGIVPQDTILFNDTLAYNIGYGRRDATRADIVAAARGAQLDA
LIERLPDSYDTRVGERGVRLSGGERQRVAIARALLKAPPIVVFDEATSALDTRSERAIQD
ELMRIAEHRTSLVIAHRLSTIVDADRIVVMEHGRLVEQGTHDELLDRGGVYAQMWALQAK
QREFERTEARIALHAVRINPLVAHVLDSLREVADTRGVPLFSHLAEDDLVVTADPAALRR
IVWDLCRGGIDASRAGGRVEIVTARAGNDARISVSSVGADAPELSLPNLETMQAALESQG
GYVARGRDDVGIVVHLALPLNAIDTAAPPAAAPGGARAGANGNADGGPSGHPDGAVPGAT
RGAANASPNGAASGGTNGRVNGGANGGQNGVATSADIAPPSVAAAATAGRPLDRVTIACV
DDHDDARDALSALLSDAGARVLAFGSGQALLDRLWQMRRRDWPAALVCDIDLGDEDGCTV
MTQVRQLEHARRDDGARAVEALALSGYASEHDRARAIEAGFRGYLTKPVKAVDLIAALRA
LAFHRGESDEHVEYPATEREGADERARR*
>GCA_000452945.1_BPMSHR5858_genomic_00115_1 group_2227
MTQHSQRQPTPRAPNSPPMKDKDEIRTHRSEDRIDKALEDTFPASDPPATGGVTRVDPSK
PTRGDRPHGPAHDTHRRDDAKH*
>GCA_000452945.1_BPMSHR5858_genomic_00122_1 group_4398
MQNAESNPTPQKQREIAAKSPVKTPAAGRDMPDAATQEQETRAHERKLTAEAIAGKTPTG
LPPIDVDQYASGDGDPVRRAASRIVTLDNANMAATDNTVDVDGKGLEARTGASEWHDNVI
YSNASLDEAVDTPDDSLAGIESRPHGYRPEVEPSEGWRVVYLGTVDIEHTLGARTEHKFS
FERST*
>GCA_000452945.1_BPMSHR5858_genomic_00123_1 yciE
MTTPQEHLEDWLRDAYAMERHAETMLKAQAARLDNYPVLRDRIQQHVEETLEQQRLLESC
LQRLDTSPSAIKDLAARAAAYGQAASGLFVADEVVKGAMAGYVFEHVEIASYKTLIAAAQ
LAGENEIRDCCERILKQEVDMAEWMSQHLPDIAVAFLERSAAERRDAKR*
>GCA_000452945.1_BPMSHR5858_genomic_00130_1 group_3058
MDAVLAFFAPLFGSGAPLGAPQMAARAAAAFFIALALIRIAGRRSFGQRSPFDYVVGILL
GATLSRAIVGVSPFWPTVAASLAIVLLHRALAWACVRSARLERFFVGAQRELYRNGRFDE
AQMSAALITMTDIYESVRQKTGECDLSGVAAAILERNGHVSIIRRKS*
>GCA_000452945.1_BPMSHR5858_genomic_00147_1 group_2231
MENLNVMHILLLLGLFAIVVAGVMATWLAFAPRGLARRLEQIGGAGAVAAGDGRAPPAVW
IEKLAEISQPISKLSLPKSGWENSPLRLRLINAGWRSPGAAHLYFAAKTVLALALPAAAM
PLLIGTAIADSPYSVSAMLVLIGAFGFYLPNAMLARRIAARQRMIVEDFPDALDLLTVCV
EAGLGLDAALTKVADELRLRSPVVADELGLMLLEMRSGFTKETALRNLALRTGAEDVDAF
STMLIQADRFGTSIGASLRVLSDTLRTQRRMRAEERAAKIALKLLFPLIFCIFPALLTVL
LGPAFIHIYRTLLPTMAGG*
>GCA_000452945.1_BPMSHR5858_genomic_00149_1 group_19771
MSLRDQMSVRRAQPLMTRSDAIGAATAMAREAYQKLRRQMHGAVLERVELERLSRLPAEQ
VRNEIATLIARILDEEKLLANDLERRQLTIDIYDEMFGFGPLESLLRDPSVSDILVNTAS
AVYVERYGRLELTDVTFYDDAHLMKVIEKIVSRVGRRIDESSPMVDARLPDGSRVNAIIP
PSAIDGPLMSIRRFAVNPLKMDDLVNFQTLTPPMAQLLEALARAKVNVLVSGGTGSGKTT
LLNILSGFIPDDERIVTIEDAAELQLQQHHVLRLETRPPNIEGKGEITQRTLVRNALRMR
PDRIILGEVRGAEALDMLNAMNTGHEGSLATIHANTPRDALTRLENMIGVAGLALPPKTM
RQQISSALSVVVQTARLTDGKRKIISIQELTGMEGEIINMQEIFTFKRTGLDASGNVLGY
FTATGVRPKFTERLTAFGIQLPDAMYDPARRFEVA*
>GCA_000452945.1_BPMSHR5858_genomic_00169_1 actP_2
MLTNRLVRAYALYTIGFAAFVLLLWRIERATGPGVWIGYVFLFVPIAVYAVIGLLSRTSD
LVEYYVAGRRVPSAFNGMATAADWLSAASFIGLAGSLYATGYDALAYLMGWTGGFCLVAF
LLAPYVRKLARYTIPDFLGTRFSSTLVRALAAIAAILCSFVYLVAQIQGIGLIATRFIGV
DFSIGIFCGLAGILVCSFLGGMRAVTWTQVAQYIILIIAFLLPVSLIAMKNGLGPVPQFN
YGRLMSRVETLEGEMRDAPQERQVRETYRRQAGAIQARLDRLPASYDEARAKLVDQVAEL
RRHNGPLREINQRERALAEFPRDPAAARVVWEQARDELLARAAAPVPMHEPFPAASGDDR
RPRGRNFLALLLCLSLGTASLPHILTRYNTTTSVAAARRSVGWTLFFIALFYLTVPVLAV
LIKYEILTNLVGRPFADLPAWITQWHRFEPGLIGVTDLLRDGIVHWSEIQMQPDIVVLAA
PEIAGLPYVVSGLIAAGALAAALSTADGLLLTIANALSHDVYYHMVAPDASSQRRVTISK
VLLLGVALFASYVASLNTGKILFLVGAAFSLAASSFFPVLVLGVFWKRTTTRGAVAGMMT
GLGVCVYYIVSTYPFFTQITGFAGPGWLGIEPISSGVFGVPAGFATAIVVSLLDRRPDAY
TNALVDYIRHP*
>GCA_000452945.1_BPMSHR5858_genomic_00170_1 group_4404
MAASSGTSSPTPSHAAEPPFVPAPLARAHARYWRFNVALIAVLMTIGFAVSFLVPLFAPA
LAHVRFAGFSLPFYVGAQGAILVYLGLIGAYIVLMQRADRVLRRDYDAYAAEAKRRAAAS
TDADAC*
>GCA_000452945.1_BPMSHR5858_genomic_00171_1 acs
MSAIESVLHERRQFAPPAAVEKAAAISGMAAYRALAEEAERDYEGFWARLARETLEWRKP
FGKVLDETNAPFYKWFEDGELNASYNCLDRHVAAGLGERVAVIFEADDGTVTRVTYADLL
ARVSRFANALKKRGVGRGDRVVIYIPMSVEGIVAMQACARIGATHSVVFGGFSSKSLHER
IVDVGATALVTADEQMRGGKTLPLKSIADEALAMGGCDAVKTVVVYRRTGGNVDWHAGRD
VWMHEMVANESDACEPEWVGAEHPLFILYTSGSTGKPKGVQHSTAGYLLWVAQTMKWTFD
WKPDDVFWCTADIGWVTGHSYITYGPLAVGATQVVFEGVPTYPNAGRFWKMIGDHKVTVF
YTAPTAIRSLIKAAEADDRVHPRSYDLSSLRIIGTVGEPINPEAWIWYHKNVGQARCPIV

DTWWQTETGGHMITPLPGATPTVPGSCTLPLPGIMAAVVDETGQDVPNGQGGILVVKRPW
PAMARTIWGDPERFKKSYFPEELGGRLYLAGDGTVRDKETGYFTIMGRIDDVLNVSGHRL
GTMEIESALVSHELVAEAAVVGRPDDTTGEAVVAFVVLKRSRPEGEEAAALAKTLRDWVG
KEIGPIAKPKDIRFGDNLPKTRSGKIMRRLLRSLAKGEAITQDTSTLENPAILEQLAEVR
*
>GCA_000452945.1_BPMSHR5858_genomic_00172_1 group_3064
LNRYAFFLIASMLLVGSNVGIGKSIVAFVPVVILATLRFVIANAVLWPLFSPVKMRAVKR
GEWLNLFLQAFFGTFMFTLLMLNGVQRTSAVAAGVITSTIPAVVALFAWLFLREKPNGRA
LVSIALAIVGVVTINVANGTASEGGAHASSLAGNLLMLGAVCCESIYVILSRRLTQTLAP
IDICAYTHLFGLLLMLPLGAGALWHFDYASVPAGIWALVVWYGVSASIFSFWLWMKGIRH
VPGSLAGVFSAVLPIAAAIYGILFLGERPTLAHGFALACVVTGIVLASLRVRRAPPVSP*
>GCA_000452945.1_BPMSHR5858_genomic_00175_1 bfr
MQGDKKVIEYLNAQLKNELTAINQYFLHARMYKHWGLEKLGKHEYDESIGEMKHADWLIE
RVFMLDGLPNLQDLHKLLVGEETEEILKCDLKLEQVSQATCKEAIAYCESVRDYVSREIF
EKILDDTEEHIDWLETQIDLIGKVGIQNYQQSMMGSPE*
>GCA_000452945.1_BPMSHR5858_genomic_00179_1 tolQ_3
MQSYGIAHVWAQGDFVTRFIAIALLVMSILSWMVIVIKGWNVIRLKRLSKDAEQSFWHSD
DFDEGVDKLGRASSSSQDNPFLALALSGKEAADHHHQTQPHLHDRMDVSDWVTRCLKDTM
DEGIARMQGGLAILASIGSTAPFVGLFGTVWGIYHALLVIGATGQTSIDQVAGPVGESLI
MTAFGLFVAIPAVLGYNALTRANKSVVSRLRRFAHGLHAYFVTGTRLASSQRDGLRLAAR
AN*
>GCA_000452945.1_BPMSHR5858_genomic_00180_1 exbD_2
MAMNTSFGNDDDDGLMNEINMTPLVDVMLVLLIIFLVTIPAMHHAVKIDLPRASSQPVEV
KPKTIDVAIESDGTVLWDDHPVSAADLQARIAQAAQTNPQPELHLRADRKVAYERVAEVM
SAAQAGGLTKLGFVTEPKSNAK*
>GCA_000452945.1_BPMSHR5858_genomic_00181_1 group_4407
MTIDAHNLNDLMYFSQVVEHGGFSAAERVLGISKSRLSRRLTELEATLGVRLLQRSTRKL
ALTEAGQLFYQHCQAMLSEAQAAMNAVQQLRSAPRGTVRVSVPVTVSQTMLSRLLPEFLR
RYPEVRVQIRVTNRVIDLFEDSIDVALRVRSEPPQSANIVVRPLWRTEQMLVGAPSLLQQ
NAPPLAPDDLAGFDTLDTPSGDGRHVFNLIAPDGTRHTHEHEPRLVTADLMTIREAVLDG
VGIAALPESMYGNALRAGQLSPVMPGWTLPVPQLYAVFVSRQGMPPAVRAFVDYLVEKLD
NERYEEPGCPERGAKDTQTKISI*
>GCA_000452945.1_BPMSHR5858_genomic_00182_1 group_3066
MTIARSIQRTYPAHRTTEGGGFVVHRPFPTRLLTDFDPFLLLDEMGPVDYAPNDAKGAPD
HPHRGFETVTYVLEGWFRHRDSAGHAGALGPGDVQWMTAGAGVVHSEMPDPEFARRGGRA
HAFQLWVNLPRRDKMIAPRYQDIPAARIPSAKSPDGRAKVRVIAGEAFGACATIETRTPM
LYLHFTLSPGAVVEQPVPAGFRVFAYPFEGGGFYGGDREAVDARHVVVYAEDGDTAVFAA
GEAPLDLLLIGGAPLNEPIVRYGPFVMNTEDEIREAVADYQTGRMGRISA*
>GCA_000452945.1_BPMSHR5858_genomic_00186_1 efeU_2
MSAVAGCRRTSLVAQAGHWMQRHGALIRGIQWVVVLVYAFLILVPAFTPLPDDTAHLWSN
LTLAAQFVFWGIWWPFVLLSMVMLGRVWCGVLCPEGALSEFASKFGRGRAIPRWMRWGGW
PFVAFGLTTIYGQMVSVYQYPRAVLLVLGGSTLAAIVIGFLYGREKRVWCRYLCPVNGVF
SLLARLSPLRYKVDEDAWRRSYRKGEDGHRVIPINCAPLVPLRNMKGAAACHMCGRCSGH
RGAIELTARSPSEEVVALGDRQASSWDTALILYGLLGIAIGAFHWTVSPWFVQIKQTLAG
WLIDHDIMWPLDTNAPWFIFTHYPEHNDVFSWLDGSLVIAYIVGTGLAYGTALAVLLAGA
AAMLGRFERVRLHHLAQALIPLAGAGVFLGLSATTLSLLRAEHVPLGWAPDVRIAILVGA
NLWSAWLAWLVTGRYASWPRRLLAMLWFGAALAVVDSAWWLMFWGF*
>GCA_000452945.1_BPMSHR5858_genomic_00200_1 group_3067
MPSNTPPSASHSPLYLNLLGETAKIDWCDLERFFAQGKLLSVARDLDLVSVAEAIASDDK
EQVTRWLSAGHVERMQAGTAQDYATRNPELWAVVVSPWVCVQARS*
>GCA_000452945.1_BPMSHR5858_genomic_00201_1 dinF
MSASSPPTHADEDPEPRAAANGHRRVLGLALPIIAANLTQPILGAVDTAVAGHLNGPQYL
GGVALGGLFFNFVYWGFGFLRMGTTGLVAQAFGANDLAGIRINVLRALLLAFTLGAAVLA
LQTPLLGIAFAALGGSDSVRSTALEYAHARIWAAPFALANYVVLGYLLGMQRVRLALVIQ
VLVNVVNIGAVLLYVYRFDWGVAGIGAATATADIAGFAASALLLRWLNPRGLAKIRRVEL
FDRAALARLVALNRDIFIRTLCLLSAFGWFAHLGARQGDATLAANALLLNFQTFMAYALD
GFAHAAEALVGAAAGARNRDAFRQALRTTFFWAGIGALAFALAYWCAGGWIVAQLTDQPS
VRGVAERYLPWAALSPVISVWGFMLDGVFIGATHTRALMRAMATSFALFVAATLALVGPL
GNHGLWIALLVFMAARGATLARRLPGLERAVAPDAARAPSAAG*
>GCA_000452945.1_BPMSHR5858_genomic_00202_1 group_3068
MTRIVIIALVAAVLAGCASDPRQGRRGTPPQDPPDYQGVPTDMTPPSMLGEPGTGGTAAT
PAK*
>GCA_000452945.1_BPMSHR5858_genomic_00205_1 group_19890
MNLILWRHAEAEDYAGTDLARQLTARGRKDAQTVAKWLRARIERHATVLASPAARTVQTA
EALTDHYRTANELAPGCDVGDVLAAAGWPDGIAPTVVVVGHQPTLGSVAAHLLAGSPESW
SVKKGGLLWLSSRTREGNGQTVLRAAISPDLV*
>GCA_000452945.1_BPMSHR5858_genomic_00210_1 group_19735
MASPVAHKIMIVRHAEKPAGDGAPYGVNAKGEQDVEALTVFGWQRAGALAVLFAPSRGPL
QHAELATPDWLFAAAVAKHAKSERPIDTITPLAHKLGLAIGDAYTKGQEKDLAQAAVASK
GCVLIAWQHEDIPAIADALPLGKQTVPQKWPDDRFDIVWVFDLDTGSGTYRFSQVPQMLL
KGDRNSVIE*
>GCA_000452945.1_BPMSHR5858_genomic_00213_1 ppx
MVTSPHLLAAVDLGSNSFRLIVGRVEETPAGSQIYPVDALREPVRLAAGLSRDKMLDRAS
QERGWEALKRFGERLRDFHPDHVRAVATNTLRVAKNAGEFLAEAQAALGFPIEVIAGREE
ARLIYAGAAHSVPANAGKRLVVDIGGGSTEFIIGQHYTPLVMESLYIGCVSHSRAFFPAG
NVDEYTMRQAELAAKREIQIISGEYKKAGWDQAIGSSGTARALAELVEANGFNDAGVSHG
ISRGGLERLKRALIKAENVNRLKLVALKPDRVPVLAGGLSIMLAVFEELGVDYVDTTDGA
LRLGVLYDLLGRTQHEDMRAVTVDGFTRRYGVDRAQAARIGALAARFYDQLDEPEGEDRD
ESRTFLGWAAALHEIGLSISHSAYHKHSAYIASNADMPGFSRTDQARLAALVLGHAGKLG
KLSQSREVEWPLLFCLRLAALLCRRRNDAGLPDINVAQMKKGGYEVHLPSAWVQQNPLTD
YSLSQEAAEWEKVGIPYRVVYTDA*
>GCA_000452945.1_BPMSHR5858_genomic_00214_1 ppk
MSLRYPLLNRELGILGFNERVLAQAADPHVPLLERLRFICITSSNLDEFFEVRMAGLQEQ
MRDNPGALAPDGMSLQHVYDLVVERAQRLVHRQYTMLHETILPALEQEGIYFHGTETWSD
AQIEWARRYFFDELLPVLTPIGLDPAHPFPRVLNKSLNFVIELEGRDAFGRQAVMGIVQA
PRALPRLVRMPHALSGFQHGFVLLSSLMQRFVGELFPKLVVKSCNQFRITRNSELFVDED
EITNLRVALQGELPARHLGNAVRLEVSADTPAHLVRRLLDESGLTEKDCYRVDGPVNLVR
LMQLPDLVDLPELKFTPFLPAISPAIANAPSMFDAIDHGDILLHHPYESFQPVLELLQQA
AQDPSVVAIKQTIYRTGTDSPLMDALMEAARNGKEVTVVVELLARFDEETNINWAAQLES
VGAHVVYGVVGHKCHAKMMLIVRRVTQGGKSVLRRYVHLGTGNYHPRTARLYTDFGLMTS
EPTICEDVHHVFQQLTGIGGELKLHEMWQSPFTLHPRLIEHIRAEAGNARAGKRARIVAK
MNALLEPTVIAELYEASQAGVKIDLIVRGVCALQPGVPGLSENITVRSIVGRFLEHHRIY
YFHANGEELVYLSSADWMDRNLFRRVEVAFPVRERKLKRRVIAEGLSVFLGDNQSAWLMQ
SDGHYRRRRAGKAPRNAQLGLLAKFC*
>GCA_000452945.1_BPMSHR5858_genomic_00220_1 pstC
MRAPSRYGDIVFGGLTRVAAIVTLLLLGGIIVSLIVASMPTIEKFGLSFLWHAEWDPNSD
TYGALVPIYGTIVTSIIALVIAVPVSFGIALFLTELSPVWLRRPLGVAIELLAAIPSIVY
GMWGLLVFAPIFATYFQKPVGALLGPIPVIGALFQGPPIGIGILSAGVILAIMIIPYIAS
VMRDVFEVTPVLLKESAYGIGCTTWEVMWKIVLPYTKTGVIGGVMLGLGRALGETMAVTF
VIGNTNLLDNVSLYSPGNSITSALANEFAEAAPGLHTSALMELGLILFVITFVVLAVSKL
MLLRLERGEGRK*
>GCA_000452945.1_BPMSHR5858_genomic_00223_1 folP
MGILNVTPDSFSDGGRYVARDAALAQAERMLAEGADILDIGGESTRPGAPPVPLNEELER
VIPLVEALRGRNVPLSIDTYKPAVMRASLAAGADLINDIWGLRQPGAIDAVRDSACGLCV
MHMLGEPRTMQVHAPEYRDVIDDVRAFLAERADALVQAGIDARRISVDPGFGFGKSVIEH
NYALLAHLPRTAPARPDGRAYPILAGMSRKSMLGAVIGRPAPERIAASVAAAVCAAERGA
AIVRVHDVAATVDALKVWSAVRDAARRA*
>GCA_000452945.1_BPMSHR5858_genomic_00228_1 greA
MSTIPLTKRGADQLRDELQRLKSVERPAVINAIAEARAQGDLSENAEYDAAKEKQGFIEG
RIAEIESKLSAAQIIDPATLDTDGRVVFSSTVELEDLETGAKVKYQIVGDDEADLDHGLI
SVSSPIARALIGKSEGDVAAVQAPSGVREYEIISVQFV*
>GCA_000452945.1_BPMSHR5858_genomic_00229_1 carB
MPKRTDINSILIIGAGPIIIGQACEFDYSGAQACKALREEGYKVILVNSNPATIMTDPNT
ADVTYIEPITWEVVERIIAKERPDAILPTMGGQTALNCALDLFHHGVLEKYGVELIGASP
EAIDKAEDRQKFKDAMTKIGLGSAKSGIAHSMEEALKVHADIAAATGGSGYPVVIRPSFT
LGGSGGGIAYNREEFEEICKRGLDLSPTRELLIEESLLGWKEYEMEVVRDRADNCIIVCS
IENLDPMGVHTGDSITVAPAQTLTDKEYQILRNASLAVLREIGVDTGGSNVQFSINPKDG
RMVVIEMNPRVSRSSALASKATGFPIAKVAAKLAVGYTLDELKNEITGGATPASFEPTID
YVVTKIPRFAFEKFREADSRLTTQMKSVGEVMAIGRTFQESFQKALRGLEVGVDGLDEKT
TSRDEIIREIGEPGPDRIWYVGDAFRVGMSREQIFEETAIDPWFLAQIEQIVLKEKALAG
RTLASLTKDELHYLKQSGFSDRRLAKLLGATPADVRARRLELNVRPVYKRVDTCAAEFAT
KTAYMYSTYEEECEAQPTGNRKIMVLGGGPNRIGQGIEFDYCCVHAALAMREDGYETIMV
NCNPETVSTDYDTSDRLYFEPLTLEDVLEIVDKEKPVGVIVQYGGQTPLKLALDLEAHGV
PIIGTSPDMIDAAEDRERFQKLLQDLGLRQPPNRTARAEDEALALADEIGYPLVVRPSYV
LGGRAMEIVHEPRDLERYMREAVKVSNDSPVLLDRFLNDAIECDVDCICDGDAVFIGGVM
EHIEQAGVHSGDSACSLPPYSLSTQTVAELKRQTGAMAKALNVVGLMNVQFAIQQVPRPD
GSKEDIIYVLEVNPRASRTVPYVSKATSLPLAKIAARAMVGQKLAQQGVTKEVEPPYFSV
KEAVFPFIKFPTVDPVLGPEMRSTGEVMGVGATFGEALFKSQLAAGSRLPESGTVLLTVM
DADKPKAVEVAKMLHDLGYPLVATRGTAAAIEAAGVPVKVVNKVKDGRPHIVDMIKNGEI
ALVFTTVDETRAAIADSRSIRMSAQAHKVTYYTTMSGARAAVEGLRYLKDLEVYDLQGLH
ARLN*
>GCA_000452945.1_BPMSHR5858_genomic_00232_1 group_19848
LTAKLAHWDFVRPVATTRVLVDVGVEQGLAVGQCLGGSGVCADLLDKPDATVAAAQELRI
VRNLMRLLGPDFPLGIEIGRRYHATAYGVWGFALMSSATFGDAVSVGLRYLQLTSTFCEI
RPTLRGDDATLVIDDRDLPGDVRDVLVEITVAALITLQFDLDAANLPVKRLALKMKPPSY
AGRFRTLFDATPEFGAAHNALTVDARCLDLKLPQRNALTRRRCEDECRRVLERHRRSEGW
AGRVRRQLAGDPARGPTMDVLAAELGVSPRTLRRRLANEGTDYETVADELRQALAEALLA
TTTLTLAEVSERLGYSEPSAFARAFRRWKGMSPSEYRRAA*
>GCA_000452945.1_BPMSHR5858_genomic_00237_1 group_3072
MSDRSIIDWPAWTDSPPGRYVLGWEQAQLDRTVSDVFGFHALQLGLPQLDALRENRMPCR
GLVLDPASGASAPYHYPWAREARSAEHAPAGRSTVWCDLLDLPFESQSVDLIVMPHTLEF
TSDPHRLLREAERVLMPEGQLVITGFNSLSLWGARHSVGKMAKRPFVPATRDQITFIRLK
DWIKLLGFDLERGRFGCYRPPLATDKWLSRYAFMEAAGDRWWPIFGAVYMVTAIKRVRGM
RLVGPIKMKKPVLAPGLTPAATPTTHQEHS*
>GCA_000452945.1_BPMSHR5858_genomic_00242_1 yibF_2
MKLIGMLDSPFVRRVAISAKLLGLPFEHHSISVFRQFDAFKSFNPVVKAPTLLTDDGTML
VDSSLIVDYLDRLVAPQRRLVPDDLPARVRVLELTGLALAAAEKTVQVVYEHGLRPGEKQ
HQPWLDRVLGQLDAAYGALDARIAGASGWLVGERITQADVTVAVAWRFTQFMQGDYPDLA
RVDPARYPALAALSARAEALPEFEGTPLD*
>GCA_000452945.1_BPMSHR5858_genomic_00251_1 mdcC
MEQLNYRHTARARAKGDAEWALVGVVASGNLEVLVERVLPGNECEVDIRTAAVGFGDVWQ
AVVADFVERRSPGGLRLSVNDGGARPDMVSLRLAQAVRAIEANAR*
>GCA_000452945.1_BPMSHR5858_genomic_00252_1 madA
MTGWNEARAARAARLAAAAPLARGKRVAARDATALLEAVLRPGERVCIEGDNQKQADLLA
AALAAVDSAKVHGLHMVQSGVVLPEHLDVFERGIAKRLDYAYSGPQSQRIAKLLFGGGIE
LGAVHTYLELFARYFIDLTPHVALIAAVSADADGNLYTGPNTEDTPTVVEATAFKDGIVI
AQVEKIVERVPRVDIPGDRVHFVVEAGRPFYVEPLFTRDPAGITETQILTAMLAIKGIYA
RYGVKRLNHGIGFNTAAIELLLPTYGERLGLKGQVCTHWALNPHPTLIPAIESGWVEQIH
CFGSEVGMDDYIRARSDIFFTGADGSLRSNRAFCQTAGLYACDLFIGSTLQIDLAGHSST
VTAGRIAGFGGAPNMGSDARGRRHPSEPWLKAGEEADARTPAALRRGRKLVVQIGETFGE
KNAPMFVEQLDALRLADKLALDLAPVMVYGDDVTHVVTEEGIANLLMCRDADEREHAIRG
VAGYTEIGRGRDRRLVERLRERGVIRRPEDLGIDALDADRRLLAARSIKDLVHWSEGLYA
PPARFRNW*
>GCA_000452945.1_BPMSHR5858_genomic_00253_1 group_2208
MIDMLGKVVAHNGLVASFALVGCVMFASAALSRKLTLGRVHGSAIAIVIGLALAYAGGAY
TGGEKGIADVPLFAGVGLMGGAMLRDFAIVATAFEVQPAQARKAGLVGAVSLLLGTVLPF
IVGASVARAFGYSDAVSMTTIGAGAVTYIVGPVTGAAIGASSDVIALSIATGLVKAVIVM
VGTPFAAGFMGLRTPRAAMIFGGLAGTVSGVSAGLAATDRRLVPYGALIATFHTGVGCLL
GPSLLFFATKAAMGG*
>GCA_000452945.1_BPMSHR5858_genomic_00255_1 hdfR_2
MRPLPPELLRSFVAVAQTGSFTAASERVSLSQSTVSQHIRRLEELLDRPLFERDTRNVRL
APSGEALFRYAVRILALMDEAVASVCGPPLSGKVRLGLSEDFALTHLTAALASFVQRNPE
VELEIATGLSGDLFDALDAGRHDLVFAKRIAGSRRGRVIRTEPLYWCAGPASPLTGREPV
LPLALHPQPSVSRQRVLETLDAIGRPYRIAVASTSIAVLRAAASAGLGISAFAGYVIPHG
LAKLEAGLPPLGELEYVIDRPANASRPALALEATLVAAAHEF*
>GCA_000452945.1_BPMSHR5858_genomic_00256_1 lcdH_3
MKIQRIAVVGAGVIGASWTAFYLSKGFDVAVTDPAPGARERLDAALANFLGERAGTLASR
VAFEPTLDAALEGADFVQENGPERLDFKRELYRRIDARLPAHVLVASSSSGLKMSDIQTA
CDAHPQRCLVAHPFNPPHLIPLVELVGGAATSADAIARAKRFYDSLGKVTIVLNKERAGH
VANRLAAALFREVYYLVGEGVVSVEDADKAVSWGPGLRWGLMGQSLVYHLGGGEGGIAHF
LEHLSGPMASWWNDLGAPSFSPDVDRQLIDALREIQGAHSMRELGAERDRLLVELIDARR
NSFLP*
>GCA_000452945.1_BPMSHR5858_genomic_00257_1 pckG
MTRSNVVAATRTVPIDVPEYVKHRGLIDWVARIAELTEPDRVVWCDGSQQEYDRLCDAMV
EQRTMVRLNPAKRPNSFLALSDPSDVARVEDRTFICSEHRDDAGPTNHWVAPAEMRATLN
GLFRGAMRGRTLYVVPFSMGPLGSPIAHIGVELSDSPYVVVNMRIMTRMGRAVLDALGER
GEYVPCVHSVGRPLAAGEQDVPWPCNPTKYIVHFPESREIWSFGSGYGGNALLGKKCFAL
RIASTMGRDEGWLAEHMLILGVTSPEGRKYHIAAAFPSACGKTNFAMLIPPKGFEGWRVT
TIGDDIAWLKPGRDGRLYAINPEAGYFGVAPGTGEKTNPNALATLRENVIFTNVALTEDG
DVWWEGLTDTPPARLTDWQGNAWTPEIGRETGRKAAHPNSRFTAPASQCPSIDDDWENPG
GVPIDAFIFGGRRSTTVPLVTEARDWIEGVYMAATMGSETTAAAAGQQGIVRRDPFAMLP
FCGYNMSDYFSHWLALGEKLAAAGATLPKIYCVNWFRKDADGRFAWPGFGENMRVLKWML
DRIDGRGEGVEHAFGVTPRYEDLHWAGLAFSPAQYAQVTSMNPDEWRAELALHAELFNKL
SARLPDALAETKARIEKRLGG*
>GCA_000452945.1_BPMSHR5858_genomic_00261_1 group_2238
VNTRNPAAERPSAVPARRDAGDAPRALTVTPAVDVYEDKHGVTLWADLPGVSKERLEVKI
QDGHLSIDAHAVLPMPASLRAQHVEVREPHFARTFQLSPDFDTTKIEANLQDGVLKLTIP
RRDEARPRRIDVQ*
>GCA_000452945.1_BPMSHR5858_genomic_00263_1 norG_2
MKRYETLARAMADDIRSGNITTGTRMPSLRQMIAQHRVSQSTVFRAYYLLEQWGLIRARE
RSGYYVAHGAQASVRANGMNGMNGAHATQDGRLTRDAAPAESRKVDISDLVFSVLEAATH
PRIVPLGSAFASPHLFPLPRLAKSLAQAARLVSPWSAVVDLPPGSEQLRRQIALRYLGIG
VSQPIDEIVVTNGALEALNLCLTAVTRPGDVVAVEAPGFYAALQAIERLDLRAVEIPVDP
RTGLDLGALAAALERHDVRACWFMTNFQNPTGVTLSADKKKALVDMLAAREVPLIEDDVY
GELHHGPDYPLPARAFDKHGLVMHCSSFSKTLAPGYRIGWAAAGRFAEKVQRLKLMTTLS
ASIPAQVGIADYLEHGGYDRHLRKLRGALRGQMERMNDALMRHLPPDVHWTLPDGGYFVW
LEFPDAIDAMGLHREAIRRGVSVAPGPLFSAAHAFERCVRINFGHPWSAAVDDAIRTVGE
LVAQPSVRKRR*
>GCA_000452945.1_BPMSHR5858_genomic_00264_1 group_19943
MSLARMMRMVLWSFFGVRKGAGHQADLASVKLPWVPVMAILLAACFGGLLFGLAKVAITV
IH*
>GCA_000452945.1_BPMSHR5858_genomic_00270_1 group_3783
MTSFALPSHIAFDGYRRLAAGPLPTVALAVKRAIANDAIGGALLIFDNATGRSIDIDTRG
SDDEVAARYAPAASADIAPGAAAGAHRERAADLADAAGTSNAADADPLAPEPPRGRGRPK
LGVVAREVTLLPRHWAWLATQPGGASVALRRLVDEARRAHAHKDRSRRAQERAYHFMSAI
AGDMPGFEEASRALFANDAARLRAQIAGWPADVRDHALALASPDDGAPLPEVR*
>GCA_000452945.1_BPMSHR5858_genomic_00276_1 paiB
MYVPAHFEENRTDALHALIAQHPFGILVTHGDGGLDANHIPFELAPGEGPLGVLRAHVAR
ANPVWQQVSDGDEVLAVFRAGDAYISPNWYPSKHELHRQVPTWNYVVVHAHGRIAVRDEA
SFVRGVVARLTRTHEAAQPVPWKIGDAPRDYIDTMLQAIVGLQIDITRLVGKCKLGQNKD
ARDIRGAGQALKARGDDAIGNRMLAKAAEKPE*
>GCA_000452945.1_BPMSHR5858_genomic_00308_1 cpdA_1
MLLAHISDLHIKRPGALAYRRVDTAAHLARCVARLNALEPRPDAVLVTGDLTDFGDDDEY
AHLKALLAPLAMPYYLLVGNHDDRAALRRAFPARVELQSGEFVQYALDLGPLRVIALDSQ
IPGASGGTLCDARLAWLADQLDAARDRPVIVALHHPPFACGIGHMDAMRLAPDASSRLDA
LLRRHPNVERVLCGHVHRTMFARFGGTIASAVPAPAHQVTFDLRDGGPSSFSMEPPAFAV
HRHAPGAGLASHHVYVDASDGPHPFFEPTGTLID*
>GCA_000452945.1_BPMSHR5858_genomic_00319_1 group_3084

MNKTLVAAALASVALAGCYYPYGYYPASGYYAVPVQTAPVYVAPAYPAYYYPAPSLSLNF
GYWSGGGRGYWRHH*
>GCA_000452945.1_BPMSHR5858_genomic_00320_1 group_2244
MNLKNALLLIVLAALWGASFLFIRVGVAEFGVAPLMALRVGIGALFLVALMLTRYAPREL
IALLRQHAWPLFVVGLLNSAAPFCLFAFAELTLSAGVTSVINATTPLWGALVAYLWLKDS
LSLPRALGLVIGFAGVLTLVWDQVFSPHGANPASPATAALAAAAALGATLLYGIAANFTK
RKLTGVDPLVSATGSMVGATILLLPFALAT*
>GCA_000452945.1_BPMSHR5858_genomic_00324_1 arfB
MTLRYAFSPDEVELTAVRAQGAGGQNVNKVSSAIHLRFDIRASSLPDDVKARLFARADQR
ITREGVIVIKAQEYRTQEKNRAAALARLDTLIRSAGVTPRKRVATRPTRASKERRLAEKG
RRSEVKSGRGRVVE*
>GCA_000452945.1_BPMSHR5858_genomic_00340_1 tsaQ1_2
VTTRSNAQAIPREREATTDEITALARGLAVLRRIATSDAPVSNRELAELTGIPKPTVSRI
TATLVSAGFLFQLPDSERFVLTASVLELSNGFLRNFDIRARSRPFLVELAEKTSLSVHLA
VRDRLDMVAIDVIRPRSAVLVTRLETGSRMDIARTAVGRAYLAALEDDARRELIGALQAA
AGDDWPFVVSRLNAALADIAQHGYAIAIGEWREELNAIAAGFVAPTGQCYAVNCGGSAHQ
CTPDFLRSVAVPALRECIAKITREIGGAAWPNLAR*
>GCA_000452945.1_BPMSHR5858_genomic_00373_1 cidA
MSRISATAATATRPTTAAPARAGRVARIALQTAALGALWAAIDWTVRAFGVPVPSGVVGL
AVLLALLLSGRVAPGWVKDGANWLLSDMLLFFIPATVAAVQYGGLFKADGWRLALVVVAG
TTFVMLSVAIAVDVAAGFERRLAVMRVRAGRRRARA*
>GCA_000452945.1_BPMSHR5858_genomic_00374_1 yohK
MTTPDLSFLTAPSSAAISAGCFALTIALYFASKRLYACRKTLFFSPLVLVPAVLVAFVAL
TGIPYAVYFRDTRWLMWLLGPATIAFAVPIYEYRELMKRHWFSLAVGVVVGIAVAICGSL
LLSKLLHLSPELQRSLVTRSVSTPFALAVSDRIHAPRDLTALFVIATGICGMLVGEFVLA
LVPLRSRLARGALFGAAAHAVGTAKAREIGSEEGVVSSLTMMIAGVAMVLIAPLLTLLPI
*
>GCA_000452945.1_BPMSHR5858_genomic_00375_1 group_19866
MATTANPTVDKPASSGKLKRLVLFLLIGTVAAAAAAGGTYFMLSKEGAHSAAPSAPAPLA
VPAFFPLEPLTVNLLSDDGIQHYLRVGLSLKLTDPKAQEYLTQHMPELRSRILLALSNKH
PEQLATLEGKHALADELKTLIEQPTQPGNQSARVDDVLFTEFVVQ*
>GCA_000452945.1_BPMSHR5858_genomic_00377_1 fliN
MTELNSTPEADGIDEPAFADTAMSAAAAGNAAGAAPQEEMAMDDWAAALAEQNQQPIETG
ATGAGVFRPLSKATASSTHNDIDLILDIPVKMTVELGRTKIAIRNLLQLAQGSVVELDGL
AGEPMDVLVNGCLIAQGEVVVVNDKFGIRLTDIITPSERIRKLNR*
>GCA_000452945.1_BPMSHR5858_genomic_00380_1 fliQ
MTPENVMTLAHQAMYIGLLLAAPLLLVALAVGLVVSLFQAATQINEATLSFIPKLLAVAA
TMVIAGPWMLSTMIDYLRETLLRVATLGAG*
>GCA_000452945.1_BPMSHR5858_genomic_00387_1 ssuC_1
MTLPTPLGTLTSLEDEERAAQRRLRRRRQLIVGLRIAVLVAVLGGWEIAARLKWIDPFFF
SMPSLIAAQIQDWFVNGTSQGPLLLQVWVTLEETIAGFLIGSVAGIFCGIVLGRNKLLAD
VFGLYIQIANSIPRVVLGSVFVIALGLGMASKIALAVVMVFFVVFGNAFQGVREADRYLI
ANAQILGASRRQITTSVVIPSALSWILASLHVSFGFALVGAVVGEFLGSKQGIGLLISTA
QGAFNASGVFAAMIVLAVVALAADFLLTRLEKRLLKWRPAAF*
>GCA_000452945.1_BPMSHR5858_genomic_00390_1 group_2251
LKRQYLALSIATAACAAPQAHAQSSVQLYGLIDLSVPTYRSHANAKGDHVIGMGLGGEPW
FSGSRWGLKGAEDIGGGTKVIFRLESEYTVADGNMEDPGQIFDRDAWVGVENDTFGKLTA
GFQNTIARDAAAIYGDPYGSAKLTTEEGGWTNANNFKQMIFYAAGATGTRYNNGLAWKKL
FGNGIFASAGYAFSNSTSFGQNSTYQVALGYNGGPFNVSGFFSHVNHAGYANKSFSVGGN
YTFDIFRVNAGYFRYLGDQGALGQRQDNAWTVSFKVAPKGALDYELGYQQMRVHNAAYNS
DGNVPNANIGDFSLTSGVGNGFKETLYGSVFYHLSKRTELYLAGDYMRLHGGYTVASTHG
ATNQLELTTGIRTRF*
>GCA_000452945.1_BPMSHR5858_genomic_00393_1 group_3092
MKKFAVAAAGLAVATGAHASDGSVTLFGLIDAGVSYVSNEGGKRNVYFDDGIAVPNLWGL
RGTEDLGGGAKAIFELTSQYALGNGAALPTPGSMFSRTALVGLWSERLGSVTLGQQYDFM
TDSLTFGSFDGAFRYGGLYNFRQGPFSKLGIPDNPTGSFDFDRLAGSSRVPNSVKYTSAN
LNGLVFGLMYGFGNQAGGGLAANSTVSAGLKYETGSFALGAAYVDVKYPQMNNGHDGLRN
WGLGARYALSAFDLNLLYTNTRNTLTGAAIDVIQAGVRYVGAPWTIGANYEYMKGNAQLD
RNYAHQVTAAAQYALSKRTSAYVETVYQYAGGSAGAHAWINGVMGPDAQSSSRSQFLARI
GMLTRF*
>GCA_000452945.1_BPMSHR5858_genomic_00394_1 feaR
VNGRFFTTAGESPAFRGRAWGRVVTQYFGGLDACCDGDDAFDAQLSQYEIGPMRVFTIAA
PAHRIVRPAAALHDHGSDFFKLILQLSGVSEIEQRGKTFRLHSGDWSLYDPRVPYSIANL
THVEQLAIQIPRKQLGGFAVPDLHTSDVREFELKGLFSLLSSFLVSLSEQLPSLPGTTGT
ALSETILGLIVSTLTAQRDAQGEHVALPAVLRMRVKQYIHGHLADADLSIDRIARELRCS
KRYLHRIFEEEGVTIDRYIWSSRLERCKDALDNARAAKPAISEIAFSWGFSSSAHFCRSF
KQRYGMTPREFVRRRAWS*
>GCA_000452945.1_BPMSHR5858_genomic_00396_1 lptB_3
MIEISNLTVQFGGTRVLDALDAKLAAPICGLIGPNGAGKTTLLNVLSGFLRPRAGSVALD
GRALLALSVAERVRAGVRRTFQTEQIVEDLSVHDNVLALAEHVMPALAARDDTRRALELV
GLADVAHAQGAALNLYQRRMLELGKALVGAPRLLLLDEPGAGLNETEAARLADVIVRIPE
CVGAQVLLIDHDVDLIDAVCEATLVLDFGKRLALGPTRAVLDDPLVRSAYLGQAFDEHEA
*
>GCA_000452945.1_BPMSHR5858_genomic_00397_1 group_2253
MKTPLAFAAPPRDAAARAKPAGARLALVATLGTLAAIALVVPAVADAYWIKTLTSALTVS
IAAAGVALLYRQLGLVCLSQHALFGVGGWIALRLAHLGAPFELCVLAGAAGGSAIGMIAG
LPALRLRGLYLALVTLMMAGGFQVVVSAIGFPDGGGGFTGHLSFGARQMMARPLVGQSDA
AYFRYVAAWAALAFALIELHRRSKAGRSWALIRRGEMPALAAGVNVVLYQTWAFGLAGLL
AGLSGGLLAGSVGQLDGRAFAASESVLLFALSVVGGVYHWFGALITGLLLRAVPALLTDF
GVNGYLAMIFFGAALLHALVTAPAGIAGQLAGLASRVGRAFRARDGGAR*
>GCA_000452945.1_BPMSHR5858_genomic_00398_1 livH_3
MNAIDFVPYLISGLGVGAVYALSGVGLVVLYRASGVLNFAFGATGALGAYVAAACLDADY
PQALAWGAAVAASTAASLVYGLVLAPRLAARERVVRSVATLGFALVLLGFCEWYWGDTPR
RLVLPTDSEALDFGDVRFTYTRIVGLALACAMMAAIGVVLARTRLGLHMRALSNDRHLSG
VLGIRVLSVDAAAWAISGVFAGITGLLLANLVRLQAAVLTFLVIPAFAAAIVGRLASLPA
TVAAGVAIGLAEALAITVPGFAPYRSATPFLIALVAMLFIGSPSSGRAANES*
>GCA_000452945.1_BPMSHR5858_genomic_00400_1 feaB_2
MNLAALSTQHQRQSGFLARRQFGNWIDGRAAEPRSGRYLPVVDPATEMTIAEVAASDARD
VDAAVAAARRAFDSGDWPRMRPASREKLLHQLAERLERYADELAALETLETGKLIGVARA
IDVLGGAEYVRYMAGWATKLEGSTLDTSIAAPAGAEYFAYTRREAVGVVGAIVPWNFPLA
IALWKVATALACGCTVVLKPSEETPLTALRLGELAQEAGLPDGVLNIVTGAGAEAGAALA
AHPGIDKITFTGSVGVGRAIGHAAVERMARFTLELGGKSPLIVLDDADPDFAAHGAAQGI
FFNQGQVCTAGSRVYVQKRLFERVVAGIAAAAEAMKIGSGFDPNTQIGPLVSKRHFERVL
GHVGAAKEEGATLVTGGTRALDGGYFVKPTVFVDAAPAMRIVREEVFGPVVTVTPFDTVD
DAVRLANDTDFGLAASVWSQNLSHVHRVVPRLKAGIVWVNTHNMLDPNLPFGGFKQSGYG
RELGRAALEQFTELKSVCIAH*
>GCA_000452945.1_BPMSHR5858_genomic_00401_1 group_19991
MRTPYQIVADHYAASDRRDPAAMMADIAPTIEWTEMAGFPCAGTYRSADEIVRNVFRRLG
EEWDGYTFALDALHDAGDTVIGIGRYSGTYRRTGKSFACRVAHVWRVDAGKIVRFEQFTD
TLLVAQAMQP*
>GCA_000452945.1_BPMSHR5858_genomic_00403_1 ucpA_1
MHTFNDKVALITGGGTGIGAAVARKFVEAGGRVVLLGRRREPLDAVAKPLGAAAIAVAGD
AADAHDVRRALDDARAAFGRVDVLVANAGGHGVGSALDTDDASWAQSTRVNLDTAFVCAR
ELLPELIERRGSIVILSSLAGHFAGPNVIGYVTTKHALIGLTRSLARDYGRAGVRVNAVC
PGWVRTAMADEQMDALRDAHGLATREDAYRLVTRDVPLGRPAQPDEVADTVLYLASPYAS
MVTGTSLLVDGGASAVDLPTIEFAR*
>GCA_000452945.1_BPMSHR5858_genomic_00406_1 group_2854
MKQDAFIQRLRQALGSLPKRDVDEIVADYREYIGDALSAGRTEEDVIAALGEPEKLAREL
KAQANFRQWEERRSFGNLMRVVGSIAGLGLLHLVLLVPFLLYMLVLTLGYVFFGALTVAG
LVTLVAFGSHYVFGTPVPGVLPFGGGAAQLGAAGASGKADGKAATDGKAAQTGKIAQDPK
SASAAAADALNDAKELRVEGERYVLDLHDGSRVSIVTRKGVIDARKQGGKLSIDATGDGA
RELLTTGKGDTLGIARADVIALDLKSTNGDQMTVANTGGSVPGVNVPGIAGGNVQMTPDG
RGGMNLSVTNGENSVSIVGGRITVDNGKQHVSVAAPAGFALGGIAFGYGLAMLPIGIVGL
LVCVWLTRVTWRALGRYVRRRIDAVSAKLERERAN*
>GCA_000452945.1_BPMSHR5858_genomic_00407_1 group_19888
MNDRIRIRDLFTRDEIRALTARSNWRGAWAVGSTWAVIALAFAGLAYARAHFPAWGFALA
LAAGMTIIAGRQLCLGILQHDAAHGTLFRTRWANDVLVDWLCARPIWNALHKYRPYHLMH
HATTSSRADPDLSLVAGLPTTRASLARKCLRDLSGVTGVKFLIGRALMDAGVLQWSLTND
IRRLPQTGRRWWDYPRAFLRAAAGMLITNAILLGALWASGHPWLYGVWVLAYVTPFPLFI
RIRSMAEHACLETSDDPLRNTRTTHAGWLARATVAPIRVNYHVEHHLMASVPYFRLPKMH
RMLRARGCVARPPSYWQVLRTVSAKTAA*
>GCA_000452945.1_BPMSHR5858_genomic_00415_1 group_3097
MNRLILRCTTALIGSGAFVCACAQTYDGTTAPIDTSDCRAITGQADIDGTMQQVVGRACL
QSDGTWRLVQNPDGSVPWYPLAAYPYPDPWYWGAPFFVGAGTTFVFVDHFHHFHRFHRFH
PMTPGHVGPPAGGGVHHGPSRSGGMGGWGGTARH*
>GCA_000452945.1_BPMSHR5858_genomic_00416_1 group_19767
MPYVSTHYSNNASAPVGRWTCAPTSKLAPFDKAPTGSVTSGVDLCGQCVSYVKRVCPTLP
LTGQWRKGAPVKGNATIVAGTVIATFNAAGKYDGHAAIYVSQTKDGGILVYDQFVTPPTP
QPVRQRRLRWGAHGRSNNGDNFYVVE*
>GCA_000452945.1_BPMSHR5858_genomic_00547_1 sasA_2
MARQAGRDEGVGRIGRIGRIGREAGDAREAAGAGEACGRRHARADAWARRASRASRGWRT
FWRGGGRGRVRAVVASVVASFSVMAFAAATLPVSWRVAAAAERTRSGGERAGGLRDTAGL
IEISAAAPASTPIPAAPRRFAQPFAQPARAFAVASACAPSWPRWDRFKRDFVSADGRVID
VGSADERTVSEGQAYGLFFALVANDRAAFDALLRWTEDNLAQGDLSARLPAWLWGRAADG
AWRVLDANAASDADLWLAYALLEAGRLWRERSYTARGALLAKRVLDEETATLPGLGLVLL
PGPTGFRPARDAWRLNPSYSPPQAIRGIGAHVPDDARWARLAAGVGRVLTDSAPRGFAPD
WALYRAGRGFEPDAETHAASAYNAIRVYLWAGMLDAGDPLARPLVAHFAPFAEHVAAHGA
PPEAVDATTGAAAPRDGNAGFSAAAVPFLEARGERASADAQLARVARLERETASGYYANV
LTLFGLGWRDGRYRFAADGTLRVRWSEPCSTPAR*
>GCA_000452945.1_BPMSHR5858_genomic_00576_1 pcaK_3
MTDLARQLDVQQFIDERRFSPYQWLILILCFLIVAADGFDTAAIGFVAPALAQQWHVTKA
ALGPVMSAALVGLALGALAAGPLADRIGRKRVLVGSVVLFGLFSIGCAFTQSVTELAVLR
LLTGLGLGAAMPNATTLMAEYAPHAKRSFLVNTMFCGFTLGSSAGGLVAAALIPEHGWRS
VFIVGGIAPLALGALLIALPESIRFMVLRGAPHERIAAVLRRIAPDAAFDGVRFVLHDDR
GEQRRSGAAVVLSPRYRAGTAMLWLTYFMGLLVYYLMTSWLPTLIRDTGFTVRDAALVTA
LFPLGGGIGAIACGWLMDRIEPHRVIAFTYALTALFVWMIGLQAGHLALLATVVFVAGVC
MNGAQSSLPALAAAFYPTSGRATGVAWMLGVGRFGGILGAFSGGLLLQAQLGFGTIFSML
AVPALIAAGALMVKRSAASRETQPDAPTRAA*
>GCA_000452945.1_BPMSHR5858_genomic_00577_1 group_19985
MNDDQRYEAGMNVRRAVLGDAHVDRSLANRTDLTDEFQNLITRYAWGEIWTRDGLPRHTR
SLLTIAMMVALNRGEELALHLRAAKNNGVTRDEIKEVLLQTAIYCGVPAANSAFHLAQRI
FGEEDSAS*
>GCA_000452945.1_BPMSHR5858_genomic_00580_1 pcaJ
MKRLTRDDMAKRVAQDIPEGAYVNLGIGVPTLVANHLDPSREIFLHSENGLLGMGPAPAP
GAEDDELINAGKQHVTLLTGGAYFHHADSFAMMRGGHLDYCVLGAFQVSAHGDLANWHTG
APDAIPAVGGAMDLAIGAKHVFVMMEHLTKQGESKIVAECSYPVTGVRCVDRIYTDLAVL
DVISDGLAVREICAGLSFDELQQLTGVPLIDATQKAAA*
>GCA_000452945.1_BPMSHR5858_genomic_00582_1 praI
MRTQVAIIGAGPSGLLLSHLLRLQGVDSVVLETRSREYCENRIRAGVLEQGTVDTLNDAG
LGARMRREGLVHRGIELLFDGRRHRIDFGELTPGRAITVYSQHEVVRDLIAAALEHGQPI
HFDVRDVALHDVATERPSVTFMHADGRPDRIDCDYVAGCDGFHGVARQAIPAERRRTFER
VYPYAWLGILADAAPSLDELVYAHHARGFALFSMRSPTVTRLYLQCRPDENLAEWPDARI
WEELGTRFENDGGWTPTPGRITQKSVTPMRSFVSETMQHGRLLLAGDAAHIVPPTGAKGM
NLAVADVRALSRALVARYRHGRGDLLDAYSSTCLERIWRAEHFSYSMTNMLHPCADDSPF
VNRLKLAELKYVTRSRAAAQSLAENYVGLPFDETAGDAPSFGEAAAA*
>GCA_000452945.1_BPMSHR5858_genomic_00585_1 fabG_8
MQSSAKPATLPVVLVTGAARRAGRAFALRFAARGYRVAVHYDRSADAAQAVVREIESSGG
EAVALQADLAHADAITQLVDAVYTRFGRLDVLVNNASVFWQDHLPSFDLAAFDNAWAINC
RAPILLTRAYYERAKAALAQGVVINVVDQKIKENFHRDHFSYTVAKAALGNLTQMLALSA
SPVLRVNAVFPGLMLPSDDQTPADFEHASRASTPLARIAGPNDVADAILLLTGPAYNGAD
FVVDAGQNLIRVDQDVLYKHRSPDGKQ*
>GCA_000452945.1_BPMSHR5858_genomic_00594_1 group_2269
MLDYALLDALAEVIRHGSFERAAKALNVSPSAVSQRVKLLEERVGSVLVKRGQPCVATTS
GALLCRHTERVRMLEAELSGHMPAFAGAQAGAWPTLRVAVNDDSVATWFVDAVGPFCAQS
GTLLDLVIDDQDYTAERIRDGSVQGAVTALAEPIQGCRSTRLGRMRYHAVCSPAFHARYF
GNGVNRDALRRAPCVMFNPKDSLQARFIRRVTRVDLDPPQHWIPHVAGYLRACEAGLGWG
MCPERMVERQIAEGGLIDLSDGRSIDIELYWQSWRLSIGWLDEFSAALKASAAQYLD*
>GCA_000452945.1_BPMSHR5858_genomic_00595_1 argO
MNWIAFSHGAMLCASLIVTIGAQNAFILRQGIMRSHVGKIVLLCTLSDMLLIGAGVGGAS
VLVERYPAFVHAVLYVGLAYLAWFGLSALRRAFKPGHETLDATGAVAAPPAQRAWPIVLM
TLAFTWLNPHVYLDTFLLIGTAGAREPEGARGAFAIGAMAVSAVWFVALGYGARLLAPWF
KRALAWRVLDGAIGGMVLFLAAVQLR*
>GCA_000452945.1_BPMSHR5858_genomic_00600_1 group_3114
MATKPKKTKGVSVSIEGKLPKMTLDMPVDAKKIKAIQKCLENGKLTITMSKVDLAGGRMG
EGYLYD*
>GCA_000452945.1_BPMSHR5858_genomic_00603_1 group_2270
MRHAHRFSRRACGMSDSFGHSPFEMFRQMFGGHRRGGGRHGFGPFGGFGGGGPGGFGGFG
DDAMPRGRQFSADDLQLLLLALLAEQPSHGYELIKALDTRSSGFYAPSPGMVYPALTYLE
ELGYVTMQAEGNRKRYALADAGRAHLDAQRERVEMLFAKLAHLGRKMEFMRRAFAGEAPA
APEDDTRGWLPEFVEARMALKQALFRRSGANADEQRRIAAILRRAVADIEGRAQG*
>GCA_000452945.1_BPMSHR5858_genomic_00615_1 group_19790
MSGQAAAAPVPSGIVVHAVSRVLELQYPSGECFRVPFELMRVYSPSAEVRGHGPGQETLQ
TGKREVTVTALEPVGNYALKPTFSDGHSTGIYSWALLYELATRQDALWREYFDKLKAAGV
ERDAPMPADSLPRGHHH*
>GCA_000452945.1_BPMSHR5858_genomic_00617_1 group_2193
MSVSRSSFNRSREPKSLARRIGTFLMVGLLAAGTFASLDAEAKRMGGGRSVGRQSQSLQQ
RQATPPAQQPMQQAAPTQAPRQQPTPAAQPNRSRWLGPIAGLAAGLGIAALLSHLGLGGA
FAGMMANVIVIALLAMAGIWLVRKFMNRRGAQQPAYAGGQPTLNRGHDGYSQDAGVQRQA
APGGAFPGSGSAYAGEAQRVFGGAAATTAAAAAPAVPAGFDTEAFLRNAKVYFVRLQAAW
DEGNLADIREFTTPEMFAEVKIDLDGRGAGANQTDVVQLDAELLAVEDRGGEHLASVRFH
GLIRETQGAAAEPFDEVWNLSKSGSQGWLLAGIQQASTH*
>GCA_000452945.1_BPMSHR5858_genomic_00619_1 ubiB
MRIFRFVKIVFTVIRFGLDEVMLSRIENPRVKLLLRITTIGRRFADPPAVRLRRALESLG
PIFVKFGQVLSTRRDLLPVDFANELAKLQDQVPPFDSAVAIAIVEKSLGARIDVLFDEFE
RVPVASASIAQVHFAKLKQGEHKGKAVAVKVLRPNMLPVIDSDLALMRDIATWAERLWAD
GRRLKPREVVAEFDKYLHDELDLMREAANGSQLRRNFAGLDLLLVPEMFWDYSTPAVLVM
ERMTGVPISQVDTLRAAGVDIPKLAREGVEIFFTQVFRDGFFHADMHPGNIQVSLDPKHF
GRYIALDFGIVGALSDFDKNYLAQNFLAFFKRDYHRVATLHLESGWVPPDTRVEELESAI
RAVCEPYFDRALKDISLGQVLMRLFSTSRRFNVEIQPQLVLLQKTMLNVEGLGRSLDPEL
DLWKTAKPYLERWMTEQIGLRGWYERFKVEAPQWSKTLPQLPRLVHQALISHHEAPRAIS
DDLIRQILVEQRRTNRLLQALLVFGLAVGAGAVIARVLIVLAYGG*
>GCA_000452945.1_BPMSHR5858_genomic_00627_1 dmdC_2
MGQYAAPLRDMQFVLHELLNVEAEVKQMPKHADLDADTINQVLEEAGKFCSEVLFPLNQV
GDREGCTYVGDGVVKTPTGFKEAYQQYIDAGWPALGCDPDYGGQGLPAFVNNALYEMLNS
ANQAWTMYPGLSHGAYECLHAHGAPELQKLYLPKLVSGEWTGTMCLTEPHCGTDLGILRT
KAEPNGDGTYSISGTKIFISSGEHDFSKNIVHLVLARLPDAPKGTKGISLFIVPKFVPDA
SGEPGERNGVKCGSIEHKMGIHGNATCVMNLDNAKGWLIGEPNKGLNAMFVMMNAARLGV
GMQGLGLTEVAYQNSLTYAKERLQMRSLTGPKAPDKPADPIIVHPDVRRMLLTQKAYAEG
ARAFTYWSALQIDKALSHADEAVRKEAEDLVALLTPIIKAFLTDNAFECTNHAMQIYGGH
GFISEWGMEQYVRDARINMIYEGTNSVQALDLLGRKVLGDMGAKLKKFGKIVAEFAEAEG
VKPEMSEFITPLADIGEKVQKLTMEIGMKAMQNPDEVGAAAVPYLRTVGHLVFSYFWARM
ARIALDNEASGDPFYKSKLATARFYFARLLPETAASIRLARAGAKSMMDVDEALF*
>GCA_000452945.1_BPMSHR5858_genomic_00629_1 fadA_3
MSKQLQDAYIVAASRTPIGKAPRGAFKNTRPDELLVHAIKSAIAQVPEFDTKLIEDAIVG
CAIPEAEQGLNVARMGALLAGLPNTVGGVTVNRFCASGITALAMAADRIRVGESDALLAA
GVESMSMVPMMGNKPSMSPHIFDRSEDFGIAYGMGLTAERVAEQWKVSREDQDAFSVESH
RKAIAAQQAGEFKDEIAPYAIVERFPNLATGEIDTKTREIALDEGPRAETSLEGLAKLKT
VFANKGSVTAGNSSQTSDGSGALLVVSEKVLKQFNLTPLARFVSFAVRGVPPEIMGIGPK

EAIPAALKAAGLKQDDIDWIELNEAFAAQSLAVIRDLGLDPSKVNPLGGAIALGHPLGAT
GAIRAATVVNGLRRRNLKYGMVTMCVGTGMGAAGIIERL*
>GCA_000452945.1_BPMSHR5858_genomic_00631_1 fdhD
VSLSETVEPSGIVELAVRRRRGDAAETAVDRVGQEWPVALVFNGISHAVMMCTPRDLEAF
AVGFAVSEGIVERGSDVKDIEVALHGAGPLPHAEVQLTVVQQAFAALKEKRRALAGRTGC
GVCGIESIDLLDLVPQRLPDTGFLARLAPDAIARAARELPAHQALTRQTGGLHAAAWCDA
SGAIVHAFEDIGRHNALDKLIGTLTLTRADMANGFVFLSSRASYELVRKSARVGIPMVAT
ISAPSSLAIEIARQAGLRLVSFCREAGYVDYGTA*
>GCA_000452945.1_BPMSHR5858_genomic_00635_1 murU
MTHTLDTAMIFAAGRGERMRPLTDTTPKPLLAAGGKPLIVWQIERLAAAGIRTIVVNHAW
LGAQIEAALGDGARWGVRLRYSAEREALETAGGIARALPLIEDGAGAKVFVGVSGDIYTG
FDYAALSARADALAGAREPGMHLVMVPNPPFHPAGDFALGDDGVLSLDGAPRATFGNIGL
YDTRMFRDLPAGTRRALTPYYRETIAARRATGELYTGAWENVGTPAQLGELDAALKAGR*
>GCA_000452945.1_BPMSHR5858_genomic_00637_1 lptD
MPPKTLFPLVPACDAAPRKKRLAVALLAVPGLVPAVSQAQLSGAAAEPQAFGSPWDLRLA
PQLDEHPQKQGGKPATFVLADHTNGTAEQDLAAKGAAEIRRGNAAVKADAIHYDQDTDMA
DAYGKVTVANGGTTFSGPEAHLKVEANQGFMTTPKYRFTATGGTGSAERVQLLDSERSVF
TNGTYTGCQCSTNPAWYIKGSEFDFDTGADEGVARNGVLFFQGVPLFGSPWLTFPLSGDR
RSGFLPPTFSPFSSTNGFELSLPYYFNIAPNRDLTITPHIISKRGIFTQATFRYLSTNYS
GTLTGEYLPDDRVAHRNRYAIFWQHQQNFGNGFGGYVYYNKVSDNLYPEELGSTNQFVNG
VQTVYQQEAGLTYNNGPWSVLGRYQHWQTLPPSAAPYGREPQLNVKYTKYNVGGFDFGAE
ADYSRFRITTADQPEGDRVMFNPYVSYGLYGPGYFFVPKAQLHMASYDLTTTTGGVPGQP
KRFTYSIPTLSLDTGLVFDRSVRLFGQDFIQTLEPRLFYVYTPYRNQSNAPLFDTAVSDF
GLAEIFTPNTFVGNDRIADANRLTAALTTRFINPTTGDERARFVIAQQYYFTDQRVTLLP
TEAPATARHSDLILGASVKLGAGFASETAFQYNVDNNQLVKSSVGFGYSPGERRVINVGY
RYTRQNPTLSNEPINQILMSAQWPLTRRLYAVGRLNYDLASSRVVDGLVGFQYDADCWAF
GVGVQRYANGLNSSGQQNSSTRVLAQLVLKGLTSIDNGLVTAFRAGVQGYTPLPPAPAPL
SRFSNYD*
>GCA_000452945.1_BPMSHR5858_genomic_00640_1 rsmA
MSNSRQHQGHFARKRFGQNFLVDHGVIDAIVAAIRPERGERMVEIGPGLGALTGPVIARL
ATPGSPLHAVELDRDLIGRLEQRFGELLELHAGDALTFDFGSIARPGDEPSLRIIGNLPY
NISSPLLFHLMSFAPVVIDQHFMLQNEVVERMVAEPGTKAFSRLSVMLQYRYVMDKLIDV
PPESFQPPPKVDSAIVRMIPHAPHELPAVDPAVLGEVVTAAFSQRRKMLRNTLGGYRDLV
DFDALGFDLARRAEDIGVDEYVRVAQAVASARASG*
>GCA_000452945.1_BPMSHR5858_genomic_00641_1 eamA_3
MSAVACAPARPSLDLRAIGVMILLCAIWGFQQVAIKSATHAIPPMLQAGLRSAIAAVGVW
AWARARGTPIFRTDGTLGAGIVAGTLFAGEFVCLFFGLTLTSAARMAIFLYTAPCFTALG
LHLFAPGETMRRQQWAGVAIAFAGIAVAFADGFARPAAGGASALAGLAGDALGVLGGVMW
AATTVVVRSTSLAHASASKTLFYQLTVSSAVLLGLAVVTRQTTFANVTPLAVASLAYQGV
IVAFASYLAWYWLLTRYSAARLSVFTFLAPLFGVSFGVLLLGDAIGPRFVAAAALVLAGI
ALVNAPPRGARN*
>GCA_000452945.1_BPMSHR5858_genomic_00643_1 group_2274
MPKRPRPRPAVVALDHRQLDLPLFDGPAASPSPASPPAAPPASSATPPEADRSRRRTLAL
DSRALEYKLKRSARRTIGFAIDGSGLTITAPRWVTLADIEAAIAEKRRWIFNKLAEWQTR
AEQRALPQIDWKEGAQIPYLGKPVSIALASAKGALSFDPLAATLELGLPAHASEQQIKDR
VQGWLQGEAKRIFGERLVVYAEKLGVTYSMYALSSAATRWGSCSSDGKIRLNWRLIHFPM
SIVDYVVAHELSHLREMNHSPAFWQTVESIFPEFREARHTLKHHPPELLPAL*
>GCA_000452945.1_BPMSHR5858_genomic_00644_1 plsC
MRFVRSLLLFVYFIVYTVPYATACFIAFPFMRPNARYWMAVGWCKSTLFVARWLTGIRYR
VEGYENLPDGPAVLLPKHQSAWETIALPALMPKPLCYVFKRELLYVPFFGWALGMLHMVH
INRKDGKNAFDSVIRQGRKRMAEGAWVIMFPEGTRTPVGSQGKYKTGGARFAIGAGAPVV
PIAHNAGRVWPRNSFTKYPGTVTVSIGQPIDTQGLTPDQLNERVENWIEAEMRRIDPHAY
RHERDGARGAARASDAAHP*
>GCA_000452945.1_BPMSHR5858_genomic_00646_1 glyS
MTQNHSASLLVELLTEELPPKALARLGDAFAEGIAQRLAARDLIEGELAFERYATPRRLA
VVVQNVRAVAPERQVREKVLPVSVALDAQGAPSAPLAKKLAALGHPNLSIADLERAHDGK
AEAFFVNYAAPGATLADGLQAALDETLAKLPIPKVMTYQRPDGTDVQFVRPVHRLTVLHG
DRIVPVAAFGIDAGDTTLGHRFLSDGLVAIRHANAYADTLREKGRVLAHFAQRRETIRTQ
LEANANGDRVVMPEALLDEVTSLVEWPVVYPCRFEDEFLQVPQECLILTMQTNQKYFALT
DAAGKLRSRFLIVSNIETKTPGEIVEGNERVVRPRLADAKFFFEQDKKKPLAERVPLLAN
VVYHNKLGSQLARVERLETLAGEIAPAIGADATVAIRAARLAKADLLTDMVGEFPELQGT
MGTYYARHDGEPEDVAIACSEHYQPRFSGDALPTTPVSTAVALADKLETIVGIWGIGLAP
TGEKDPFALRRHALGVLRLLLEKQLPLDLVWLLRTAYERFAAVPGVAESTDAIHAFFIDR
LRGLLRERGYTAGEIDAVLSLNPTRLDDIVARLNAVREFTRLAEAEALAAANKRISNILK
KSESGANGAVQPALFVEAAEKALAEQLADVTPRVQSQLEARAYTGALTALAALRAPVDAF
FDGVMVNADDPALRANRLALLSALHQQMNCVADISKLAA*
>GCA_000452945.1_BPMSHR5858_genomic_00649_1 corC
MNDSYPSRKSSDKPHEKRSLLERLTDIISPEPDSRAELLEILQDAHERNLIDADSLSMIE
GVFQVSDLCARDIMVPRAQMDALNIADKPEDFIPFVLEKAHSRYPVYEDNRDNVIGVLLA
KDLLRFYAEEEFDVRGMLRPAVFIPESKRLNVLLHDFRVNRNHLAIVVDEYGGVAGLITI
EDVLEQIVGDIEDEYDFDEEAGNIIAAPDGRYRVRALTEIEQFNETFGTDFSDDEVDTIG
GLITHHFGRVPHRGEKVRLGELLFEIQRGDARQIHVLLVRRAPLAGRRSGSAGED*
>GCA_000452945.1_BPMSHR5858_genomic_00652_1 ybeZ
LKTAQALEFTAPRDDNARLANLCGPLDENLRQIEQALDVTLSRRGHRIAIRGRGAKIALA
ALEDFYNRAREPLSVDDIQLALVEARHPGNPRRNGNGDGEPEIDVRFRGDPDHPFDEPVV
RLGDEPPADEAAPKLYTRRADLRGRTPAQREYLKQILSHDVTFGVGPAGTGKTYLAVACA
VDALERDQVKRIVLTRPAVEAGERLGFLPGDLAQKVDPYLRPLYDALYDLLGFDKTAKMF
ERQMIEIAPLAYMRGRTLNHAFIILDEAQNTTPEQMKMFLTRIGFGSKAVVTGDTSQVDL
PRGHKSGLVEAQQVLGGVRGIALTRFTSADVVRHPLVARIVEAYDEFHAQHKDE*
>GCA_000452945.1_BPMSHR5858_genomic_00653_1 miaB
MTKKVYVKTFGCQMNEYDSDKMVDVLNAAEGLEKTDTPEDADIILFNTCSVREKAQEKVF
SDLGRVRELKEAKPDLLIGVGGCVASQEGASIVARAPYVDLVFGPQTLHRLPQMIDARRE
SGRAQVDITFPEIEKFDHLPPARVEGPSAFVSIMEGCSKYCSYCVVPYTRGDEVSRPLDD
VLTEVAGLADQGVREVTLLGQNVNAYRGAIAAGSAEIADFATLIEYVADIPGIERIRYTT
SHPKEFTQRLLDVYAKVPKLVDHLHLPVQHGSDRILMAMKRGYTVLEYKSVIRKLRAIRP
NLSLSTDIIVGFPGETDADFDKTMALVHEMSYDTSFSFIYSPRPGTPAANLADDTPRELK
LKRLQHLQATIEENVARISQSMLGKVERILVEGPSRKDPNELAGRTENNRVVNFPAPSAA
HPRLIGQMIDVKINHAYPHSLRGELVLAHGDASAATH*
>GCA_000452945.1_BPMSHR5858_genomic_00657_1 ribB
MSFPLPSKSAPADAFSDFPLLDAEPVPPRIAAALAALREGRAVVLQDDHDRENEADLIVA
AERLTDETMALLIRECSGIVCLCLTDEKVRALELPPMAQLNESKNGTAFTVSIEARAGVT
TGVSAADRVTTIRAAIADNASPHDIVRPGHVFPLRAAPGGVLARRGHTEGTVDLAILAGL
KPAGVLCELMNPDGTMARGADVERFAAQHGLPMLTIAELVAFRERLASLRARDACCADLA
*
>GCA_000452945.1_BPMSHR5858_genomic_00663_1 aroA_2
MSHATVHPSGALKGTVTLPASKPHVQRALLVALLNGATTRIENVSWCAETELQFAALRQF
GLTVVERGDASLTLRGAGAGVEATGTIDAAGSGMLFRMSAALASLSERPVTIRCNDSLFS
RDSVFDDGFFSHLGIDARRRAGNLVTISRKPHPERIALTTRKSTQFISFALFVAPFSADR
TLRVADDGSQAGYIDMTIKAMSLLHATVTRAPGRLLAGEYRADDIVIGIPTDFTSLSYLA
SAALSVGARSAIDIANYRPGDTLNEAALFDVYRALGVRLTRDDARHALRIECERDRAARR
APRELSLRELPSAATNIIAAASNLGDSIRFGGVDGINNHKCQRAFVIHENLRAMGGRSAL
VFNDVGRFDKIDVMSDGPLQGGAELPSYRDHRICAANIVAALGARRKTVVCDTDKLDDGF
PKFIDTLRALGAEIA*
>GCA_000452945.1_BPMSHR5858_genomic_00667_1 eryB
VTQQNRYDLLVVGGGINGAGIARDAAGRGLSVLLCEQDDLASHTSSSSTKLIHGGLRYLE
YKEFGLVRKALQERETLLRAAPHIMWPLRFVMPHMPNLRPAWLIRVGLFLYDHLAKRELL
PGSRGIDMRRHPAGAPLVDSIKRGFVYSDGWVDDARLVVLNALDAQERGARILTRTKLVS
AERRDGQWHARLQRADGSTLDVLARAIANAAGPWVGEVLHGALGRGAQHSVRLVKGSHII
TPRLFDHDHAYIFQNPDKRIIFAIPYERDFTLIGTTDVEYRDDPSRVAIDRDETRYLCES
INRYFKRKISPADVCWTYSGVRPLLEDENADNPSAVTRDYRLELDDGEGAPLLSVFGGKI
TTFRKLAEEATDMLGGALGAARGAWTAGVPLPGGDIANARFAPFAEAFAKRHPWLPAALA
RRYARAYGTRAERVIGRARSLAELDAQLAPGLYEAELRYLRDVEWAGCADDVLWRRSKLG
LHVAPGTLEPVTAALDAWFGAAREAASAAH*
>GCA_000452945.1_BPMSHR5858_genomic_00726_1 cobO_2
MGNRLSKIATRTGDDGTTGLGDGSRVRKDDARIAAIGDVDELNSQLGVLLAEPLPDDVRA
ALSSIQHDLFDLGGELCIPGHAAITDAHLARLDDWLAHYNAQLPPLEEFILPGGARGAAL
AHVCRTVCRRAERAIVALGVHAPLDAAPRRYVNRLSDLLFVLARVLNRAAGGADVLWDRT
RAHGTPHGTR*
>GCA_000452945.1_BPMSHR5858_genomic_00730_1 group_3130
MQTNLADFIRHTPDGDDADAILRRCVHCGFCTATCPTYQLLGDELDGPRGRIYLIKQMLE
GAEVTRSTQQHLDRCLTCRSCETTCPSGVEYGRLVDIGRKVVETKVSRPLSQRIVRRLLA
SFVPNAAVFAPVMRLGQHVRPLLPKRLRDKVPPRARLLAWPSARRERRVLMLAGCVQPSM
MPNINIATARVLDALGIETVVAPEAGCCGAIRLHLGYHDEALDDARRNIDAWWPHIERGV
EAIVMNASGCGATVREYAHLLRGDPAYADKARRIVELTKDASELLGGFEAELIALTRRRG
IHTVAYHPPCTLQHGQQLRGKVERLLEGLGIEVRLPADSHLCCGSAGTYSLTQPSLAYKL
RKRKLAKLQETEPQMIVSANIGCIAHLQSGTAVPVVHWVQLVENLLYG*
>GCA_000452945.1_BPMSHR5858_genomic_00732_1 proC
MKIAFIGGGNMAAALIGGLVKRGVDASGLLAVDVNEDARQRAQAQFGVRTAAAVDATLAG
YDAIVLAVKPQVLKDVALALAPHLTTQLVVSIAAGIRGADLSRWLGDYPRVVRTMPNTPA
LVGLGVTGLSALPGVDAAGRDLASKVLGAVGETVWFDDESQLDAVTAISGSGPAYVFYFI
EALQEAARQLGMNDEQGRALAVATFAGAAQLAVQSGEPAGVLRERVTSKGGTTAAALASF
DAQHVKDAIVRGVLAAQARAREMGDELGGA*
>GCA_000452945.1_BPMSHR5858_genomic_00741_1 phnH
MIDTRFSLADVAPGFADPVQATQRVFRTLLDAMARPGTPGRIEPIGANGSDAAPAMRVAP
AAFASLLTLCDASTPVFLGAPDAALASALRFHTGAPIAAEPRAACFAYLDDANALPPLQC
FASGAPQTPEHAVTLFVRVASLADGEPRTLRGPGIRDARTIAPAGLPDAFWRERAQFAPR
FPCGIDCYLVCGDMLIGLPRTTIVEAG*
>GCA_000452945.1_BPMSHR5858_genomic_00747_1 group_2674
MSDDACPVARAVDAVGDRWSLLIVRDAFDGVRRFSEFQRNLGIARNMLADRLKTLVEAGV
LAAAPASDGSAHHEYVLTDKGRALFPVVVALRQWGESQLFGAGEPHSRLVERRSGLPVAP
MSPRTQSDRALRPEDAVVDKRGADDAAT*
>GCA_000452945.1_BPMSHR5858_genomic_00756_1 gtfC
MAKVIVTAIGSAGDVHPLLGVSRALAARGHDVVFCTHAPFEAAVRASGFAFVPVGTAEAY
AQAMADPALWDPRTSFRTLWRVIAPVLRPHFDTLRALSDADTVLVGTLWAFSARLMQERF
GARYVSVQVSPSTLLSAHAPPTHKRLTIPKGLPLAVKAGLMTLIERQVLDRVCGPELNAA
RRALGLAPARRILGRWLHSTDGVLCLFPSWFAPAQPDWPANHLQSGFALFNDVGPVSADA
ELDAFVASGEAPVVFTAGSTLVDGRAYERAVTQVLRATGVRGILLAPDAPAASDGTTGPM
ERTAETTARANGAALLKRRYVPLAALLPRCRALVHHGGIGTASLAYAAGVPQVVTPFAHD
QFDNAQRVAASGCGVRLDAPVRGEPLARALARVLGDAAMAARCAEVRARMAAQPDGCDEA
ARFIERFAPGVAARQAQPA*
>GCA_000452945.1_BPMSHR5858_genomic_00761_1 emrA_2
VSGRDDARTSASGECRERENVNPNASFTVYTRDSTETEHRSVPRRAAAAPGPVRSAPAGA
TDITHATILNDTATDTAHAKAPTDPAVSDSAPASGHGLPASPETAATLAARRATRRRRFA
WFFGVLAIVALGAALYWFIVGRFSEETDDAYVAGNVVQIAAQIQGTVTDVLVADTQQVKA
GQALVRLDDAEAAAAFAQAQAQLAQAVRQVANTRLSMQTYEQTVKAREADLKLAQQAYRA
RAGASVEVVAPEELARAKSSLANAQAALAGAQAQREAARALGSERPVGQNPAVQQAAAQF
KLAYRNLKRTTIVSPVDGTVGQRSVQIGQQVGPGVPLMSVVQLRQVWVEANFKEGQLRHM
RVGQPVRLESDLYGARVTYHGRVEGFSAGTGSAFSMLPSQNAAGNWIKVVQRLPVVISLE
PSELAAHPLRVGLSMRATVQTKARGGHLLDGDAPLPAQRTRVHEMHAGEAEAAAEAVIRA
NDGGQ*
>GCA_000452945.1_BPMSHR5858_genomic_00764_1 crtB
MPGRPAPALPAPSRSRLTRSVPVNFDEYCQQKAAPPGSSFYYALRQAPLARQPLVAALFA
LRRELEQTVKEVSDPTVGHTKLAWWQKELAALASGHPSHPVSQALAAHHPDIARETDALR
TLVAGYGMDLEQARYLDFANLRRYADQTGGMFAALVASASASAGRPTANGARWAQPLGQA
LTLAQFVQDLGQDARQGRIYVPIDELQRYGVTAADLLNRRYSPAYTELMRFQTSRARDAL
NASLSGIPADERRAQRTLRALGALALAVLDEIERDGFQVLHQRIALTPIRKLWIAWRAAR
RR*
>GCA_000452945.1_BPMSHR5858_genomic_00766_1 group_19724
MSDSRTPGRPPAKPQRMPAAMQTLLDADDARFLLTRTGFSPPPRALARFVGMTRAQALAE
LLDGARTQSVTPPPDWVREPPPSRARRAAFSPDERRAEQQLRNRRYDELRAWWVREMIVT
PSPLTERMTLFWHNHFTSGQDKVPFPQTIAAQHALLRANALGNFGAMLHGVAKDPAMLQY
LDGASNRKGRPNENFAREAMELFTLGEGHYTQRDVSEAARAYTGWGLDPDALTYVFRPNV
HDDGVKTVLGETGRFDGDAVLDILLGRPETARFVVAKLWREFVSDAPDAGEVERIAARLR
QSDYDIRAALTELFSSDAFWAERNRGVLVKSPAEFVVGTVRLFDVDYTDAAPFANTLRAL
GQNLFYPPNVKGWPGGVSWINSATLLARKQFVEQMMRATEAPGMRAAPVSRDMAGQPAPT
RRGAMRFDLDAWLAAYRTKPQAQPDLSTELQLQHAVLPISPVAAIEAGATSGAYLQALLM
DPAYQLK*
>GCA_000452945.1_BPMSHR5858_genomic_00771_1 ahpD
MEFIESIKAQIPDYAKDIRLNLDGTISRSSLEGTDAVGAALAAAFAAKSPVLVKAIREAG
VLSPEETQAALTAAALMGMNNAWYPYVEMADDADLKTQRAELRMGAYATHGGVDKRKFEM
YALAASIVGKCHFCVKSHYALLKNEQGMTVTQLRDVGRIAAVINAAAQVISAEGK*
>GCA_000452945.1_BPMSHR5858_genomic_00772_1 ispF
MDFRIGQGYDVHQLVPGRPLIIGGVTIPYERGLLGHSDADVLLHAITDALFGAAALGDIG
RHFSDTDPRFKGADSRALLRECASRVAQAGFAIRNVDSTIIAQAPKLAPHIDAMRANIAA
DLDLPLDRVNVKAKTNEKLGYLGRGEGIEAQAAALVVREAAA*
>GCA_000452945.1_BPMSHR5858_genomic_00773_1 ispD
VTSRLFALIPCAGTGSRSGSALPKQYRTLAGRALLHYTLAAFDACSEFAQTLVVISPDDA
HFDARRFAGLRFAVRRCGGASRQASVMNGLIQLAEFGATDADWVLVHDAARPGITPALIR
TLIGALKGDPVGGIVALPVADTLKRVPAGGDAIERTESRNGLWQAQTPQMFRIGMLRDAI
RRAQLDGHDLTDEASAIEWAGHTPRVVQGSLRNFKVTYPEDFDLAEAILAQPARAS*
>GCA_000452945.1_BPMSHR5858_genomic_00779_1 mrcA_1
VRRPSDMPIIKRPPSSRATNEHHYSLRRSPAGAYYTDDDDDDERAPRRTGRDGGSRTFGS
RVALWFAGLFATLLVVGALIVGYALVVMGPQLPSLDALTNYQPKVPLRVYSADHVLLGEF
GEERRSLVRFADIPDVMKKAVLAIEDYRFYEHGGVDFVGILRAGVADLLHGGARQGASTI
TMQVARNFFLSSEKTYTRKIYEMLLAYKIEKALTKDQILELYMNQIYLGQRAYGFAAAAR
VYFGKDLKDVTLAEAAMLAGLPKAPSAYNPVVNPKRAKVRQEYILKRMLEIGYITQQQYD
QAVKEEIHVRTPGNQYAVHGEYVAEMVRQMMYAQYKDETYTRGLTVTTTINAADQEAAYQ
AVRRGIMDYERRHGYRGPEGFVALPPAGDERDEAIDDALADHPDNGDLQSAVVLAVSPTA
VDVQFVGGATATINGAGLRFAAGALSARASAALKIKPGSIVRVMKDARSAWQIVQLPQVE
GALVAVAPQDGAIRSLVGGFDFNKSKFNHVTQAWRQPGSSFKPFIYSASLEKGLGPATII
NDAPLYFPPSVPGGQPWEPKDDDQPDGPMPMRTALQRSKNLVSIRILASIGTSYAQDYVT
QRFGFDPAKTPPYLPMALGAGLVTPLQLAAGYAVFANGGFKVDPYLIAEVDDARGQALQK
AQPVVAGASATRTIDARNAYVMNSLLHTVATAGTGAGTNALGRGDLQGKTGTTNEAKDGW
FAGYQQSLVAVAWMGFDQPKSLGSREFGAQLALPIWVNYMRTALNGVPEQQMAMPDGLTT
IDGELYYADRTPGAGFVASVDFNPAASPTVSANDALGSAGAAGLTPPPVTPEEKRQIIDM
FEGNKP*
>GCA_000452945.1_BPMSHR5858_genomic_00799_1 group_3135
MATTRRHATATALAAAARPCAAPHADSFLGILIEVFARVVAALALGVFAYAAYLQWRLDP
SRITLILLAVSTTLTVGLSLFAEPPKKRDWSPLALFFSVGGTFYYLVFQLTASRQLLPEA
AGAAIQLIGLFWQLFAKVSLRKSFGILPANRGIVSRGAYRFVRHPMYLGYLIIDAGFLLT
NFSARNLIAVALQIGLQLGRIRREERLLSEDAAYRAYRRSVRYRVIPGLF*
>GCA_000452945.1_BPMSHR5858_genomic_00804_1 group_3136
MNTKTTIVWATAAALTALALSPAAFAQGKTRDEVRQELIRAQHEGVVPAGKNDYPPSAAL
VARNKELHAISVHGGKTDLALDQHDQTVAR*
>GCA_000452945.1_BPMSHR5858_genomic_00807_1 czcA_2
MFERLIRFAIAHRWLVMLAIVAAASLGVYSYQKLPIDAVPDITNVQVQVNTSAPGYSPLE
AEQRITYPVETAMAGLPNLEQTRSISRYGLSQVTVIFKDGTDIYFARQLVNERIQEAKDK
LPPGVAPAMGPTSTGLGEIYLWTVEADAGARKPDGTRYTSADLRELQDWVVRPQLRNVPG
VTEVNSIGGYVKEYRVAPNPTKLMSYGLTLADVVRALERNNDNVGAGYIEKRGEQYLVRV
PGQARSAEDIANIVLANAGGVPVRMKDIGAVDLGRELRTGAATANGEEVVLGTVFMLMGE
NSRTVAKAVAARMADVNRTLPAGVRAIAVYDRTVLVEKAVRTVKKNLLEGAVLVIAVLFL
FLGNIRAAVITALVIPLSMLMTFTGMVNAKVSANLMSLGALDFGIIVDGAVVIVENCIRR
LAHAQQAAGRPLTRDERFAEVFAASQEARRALIFGQLIIMVVYLPIFALTGVEGKMFHPM
AITVVMALAAAMALTVTFIPAAVALFIGRRVEEKENRLMSAAKRLYAPVLDAFVARPGQV
LAGATAVVALSLVLATRLGSEFIPSLNEGDLAVSAQRIPGTSLTQSVAMQQSIEKTLKAR
FPEIDRVFARTGTAEIAADPMPPNLSDGYIMLKPADQWPDPRKSRDALRGEIEAVLAQLP
GNAYEFSQPIQLRFNELISGVRSDVAVKIFGDDMATLNATGEQIAAALQKVPGASEVKVE
QTTGLPVLTVNLDRDKLARYGVTVGDVQDTVAAAVGGRKAGLLFQGDRRFDIVVRLPDEL

RTDVEAIKRLPIALPLAAPAGASGPLAQAPYVPLAELATVDTSPGPNQISREDGKRRVVV
SANVRGRDVGSFVADARERIASDVRVPAGYWLSWGGQFEQLQSASERLKIVVPLALAMVF
VLLFVMFNNAKDGLLVFTGIPFALTGGVVSLWLRGIPLSITAAVGFIALSGVAVLNGLVM
ISFIRNLRDEGAPLDAAVREGALTRLRPVLMTALVASLGFLPMAFATGTGAEVQRPLATV
VIGGILSSTALTLLVLPVLYRVAHSAPLRATLGARAAGWLARLAPAAVLRRRRESSR*
>GCA_000452945.1_BPMSHR5858_genomic_00808_1 czcB
MQRKPLILAGAAAAGALAIVLALARAPSTSSAPAASGAAAAAPASGATGPHGGAMFAQGA
LSAEIVLSEKPDDARLVVYPFVDGKPVEQGAAVSATLTRHDGERVALQFAQAGAKYTSAQ
PIARPHVFDASIDVRIGARSTSFPFSRADGAIALDDAQLGAARIGIARAGPAQIITSFQL
PGEIKFNEDRTAHVVPRVAGIVERVAVSIGENVKPGQVLAVIASTDLADRRSELMTAERR
LQAARTSYAREKTLWEERISAEQDYLQAQVQLREAEIAAANARQKLAALNASAASGALNR
YELRAPFAGTIVEKHIAPGEAIAADANVFVVSDLSSVWAEMAVPAQRLNDVRVGRHATVS
AAAFESKSSGPIAYVGALLGEQTRTAPARVVLPNPDGAWRPGMFVNVTVDAGAQNVPVAI
ESDALQDVDGAPAVFVRTPKGFVAQAIETGRRDERRVEIVKGLAPGQTYAAANSFVLKAE
LGKGVADEH*
>GCA_000452945.1_BPMSHR5858_genomic_00869_1 group_19979
MMLRRLLYRETPFEPLTDAELRRLEAAFGEMVAGNPLIYYWVHRVDGARWLITDFFHPSM
LRYRGLEFVLVERGTVSYYRLPGARVGGTGHVAAGDYRVSITSPADAAFLIEIRKNALGR
LELLGVSAAPASGAAPSHVELPRHALEPSKFADEMKAAIAGGVEWVYRRYRSADDPARAA
LARELRDARWPRAVRGASVDADTYLWMLEQSIA*
>GCA_000452945.1_BPMSHR5858_genomic_00914_1 group_19839
MKTTEFNEALSQFWEARTAREKLLLGWGGAVLALAIAYSVLWSPAQEGRARIQRELPTMR
HELAEMTAQANEARSLAGAAQGVAPTGVALKDALAASLSDHGLAAAAPQVVGGGVQIQLK
NASFPAWTQWLDDVRRQFKAQVVEAHVSALKEDGQVDLTAVLQPASVK*
>GCA_000452945.1_BPMSHR5858_genomic_00916_1 xcpX
MRAFALLARFASPGRPRAALGPHAPDAAQPAPARSPHAPHGGSAVADVRVCPPARRPGAP
VHRERGVAIINALLVVALSAILVSGMLWRQQVQVRRIENQRLLAQAQWVARGALDWTRMV
LRSEGDTAPGITYLGGIWAVPIAKTKLSDFLGRIGVTDSAEGSDTYLSGSIEDAQAKFNL
RNLVSSPAPGALQLNAAQVQTFQRLLATLGLDQQLAKIVARRVRAGLKQSATRFQAAQLP
GAASGVPVPLPDGGTESGGNFTDEPGIAGGEGRGPAPLMMTSVDSLLDVEGFTPEMVERL
RPFVTVLPTVTPVNLNTAPAEVIAALVPGMSVSSAQALVSRRETVFFRNVGDVQLALRGS
GAPNVNLDSSQIDVTSSYFIVHGRIEHDRAVVDRTSLVYRDPTTHTTRIVRIRDQL*
>GCA_000452945.1_BPMSHR5858_genomic_00917_1 atpF_2
MKPAVRSERRAPGEKRPRRARRAFRRGARHGARGFTLIEMMIAITILAVIAILSWRGLDQ
IIRGREKVAAAMEDERVFAQMFDQMRIDARRAATDDEAGQPAVRVAGDTLQIVREFDAPG
AAPRLQVVRYRISNGRVVRYASPPIGDVNALRDALRGGDTEGWSEVALMRGVGMINARLY
VPKVGWTTSMPDADNALEQNNNALKVPMLGNAPPPRAVTGLEVSIGATSLRVPITRIFLI
GE*
>GCA_000452945.1_BPMSHR5858_genomic_00918_1 group_19884
MMRAPAPPRSPARSRGFTMIEVLVALAIIAVALAASIRAVGSMATGASDLHARLLAGWSA
DNALAQLRLAHAWPDIGMQTFDCSQGNVALTCTQRVSSTPNPVFRRVEIAVSMNGRAGVL
AQMVTVVANETSRPL*
>GCA_000452945.1_BPMSHR5858_genomic_00919_1 group_3150
MYARSGTIFVCDTRRHAGAALGGAAKARAGACAGVSSGRPAGTASARRARGFTLLEMLVV
LVIAGILVSVASLTLRRNPRTDLREEAQRIALLFETAGDEAQVRARPIAWRATEHGFRFD
IRTGDGWRPLRDDVLRARDWDGGVTGAAIDYPGSDTHTDAVVFGTESIDVPVRVTLYSAV
GSATIVGTGNGRYEVR*
>GCA_000452945.1_BPMSHR5858_genomic_00922_1 epsF
MPAFRFEAIDASGRAQKGVIEADSARNARGQLRTQGLTPLVVEPAASAQRGARSQRLALG
RKLSQREQAILTRQLASLLVAGLPLDEALAVLTEQAERDYIRELMAAIRAEVLGGHSLAN
ALTQHPRDFPEIYRALVAAGEHTGKLGIVLSRLADYIEERNALKQKILLAFTYPAIVTVI
AFGIVTFLLSYVVPQVVNVFASTKQQLPVLTIVMMALSDFVRHWWWAILIGIAAVVYLVK
ATLSRDGPRLAFDRWLLTAPLAGKLVRGYNTVRFASTLGILTAAGVPILRALQAAGETLS
NRAMRGNIEDAIVRVREGSALSRALNNVKTFPPVLVHLIRSGEATGDVTTMLDRAAEGES
RELERRTMFLTSLLEPLLILAMGGIVLVIVLAVMLPIIELNNMVQ*
>GCA_000452945.1_BPMSHR5858_genomic_00924_1 sctC_1
MTRTRFALRRVATALVVAGIVAAQAAHAQVTLNFVNADIDQVAKAIGAATGKTIIVDPRV
KGQLNLVAERPVPEDQALKTLQSALRMQGFALVQDHGVLKVVPEADAKLQGVPTYIGNAP
QARGDQVVTQVFELRNESANNLLPVLRPLISPNNTITAYPANNTIVVTDYADNVRRIAQI
IAGVDSAAGSQVAVVPLKNANAIDIAAQLTKLLDPGAIGNTDATLKVTVQADPRTNALLL
RASNAQRLAAAKKIAQQLDAPSGVPGNMHVVPLRNAEAVKLAKTLRGMLGKGGGESGSSA
SSNDANAFNQGGSQSGSNFSTGASGTPPLPSGLSSNSSGGAGGTTGGGGLGNAGLLGGDK
DKGDDNQPGGMIQADAASNSLIITASDPVYRNLRAVIDQLDARRAQVYIEALVVELQATT
SANLGIQWQVANNALYAGTNLVTGQTGLGNSIVNLTAGAVTNPGGTLGSLGSITNGLNIG
WLHNMFGVQGLGALLQFFAGSSDANVLSTPNLVTLDNEEAKIVVGQNVPIPTGSYSNLTS
GTTANAFNTYDRRDVGLTLHVKPQITEGGILKLQLYTEDSAVVPGTNTTSANSPGPTFTK
RSIQSTVLADNGEIIVLGGLMQDNYQVSNTKVPLLGDIPWIGQLFRSEGKTRQKTNLMVF
LRPVIINDRETAQAVTSNRYDYIQGVTGAYKSDNNVIRDKDDPIVPPMPLGPSQGGSPAM
NLFDLDKMRRQQLAPPAPASAPAAASDAGTASPGARP*
>GCA_000452945.1_BPMSHR5858_genomic_00925_1 group_19986
MEKRFVIVAMFAAGAWFASAPARADCYDEAAKYQKVNPLVLRAIAWQESHNRPEALNKNA
NGSTDYGLMQINSIHLPTLSRYGITKDTLMEPCKSVYIAAWHLRRKMDKYGNTWQAIGAY
HSETPSLRDKYARQIADILARWKLLPASAASATQDTQSAQR*
>GCA_000452945.1_BPMSHR5858_genomic_00926_1 yciC
MNQPLPVTVLSGFLGAGKTTLLNHILANRAGLKVAVIVNDLAAANVDATFVRGATELSHV
EAHLVEMSNGCICCTLRDDLLVEIRRLAAENRFDAIVVESTGIAEPMPIAETFTFVDDDG
STLEDVARLDTMVTVIDAFNFLHDYARDDALAEHGLAATDEDDRTLVELLIEQIEFCDVL
VINKADLVDADALARLQRILANLNPRAQQIVSRFGDVPLAEVINTGRFDFDAAANAPGWL
ASLEHRRDADEAECGQGQSQGDGRVHSEADEYGIGHFVYRARRPFHPQRLWALLHEEWKG
VLRSKGFFWLATRNDIAGSLSQAGGVCRHGPAGHWWAAQDRTEWPEAGDELYDEIVADWH
GELADTSIGDRRQELVLIGIGLDAAAWRAKFDACLLTGAEYAQGKQAWAGYADPFPAWDV
DDHDHDHAHDHHDHDHGDDSEIVHRH*
>GCA_000452945.1_BPMSHR5858_genomic_00928_1 mnmC
MTDRIVPATLVFREDGTVVSPLYGDIYHSAAGALAQADHVFIRGNGLPERWRHERAFTII
ETGFGTGCNFLATWAAWRADPSHCERLHFVSVEKHPFAREDLRRAAAHIVAYTTITTITP
IAPLVDELANAWPALTPGVHRLEFDDGRVTLTLVFGDALDVLPNLALRAHAFYLDGFAPS
KNADLWSPAIFKSLAKLADERATFATYTSSGAVKRALDEAGFAYRKVDGFAGKRAMLVGE
FAPRWRVRRHEPPRAFSTDRRDAIVIGAGLAGCAVVERLAARGWHVTLIERRERIASEAS
GNPAGVFHPMIARDDNLAARLSRAGFLHALHRWRALERAGHAFSRSTHGLVQLATSDDEF
ERMRESIDALGVPAELASALSRDDARALLRTDVAHGGWLFAQGGSISPAALAAAQCAAAG
DRLSRIVGVEIARLERGGDGRWRALDASGATIAQASVVVVANAADAARIAGLRHAPTQRV
RGQLTLLPPGSAPAVPLPVIGDGYVVPLANGVTLTGATYEPDDTDATPREAGHRENLERL
ERLLPAFSANALDAGALAGRVGFRCVASDRLPLVGELGDEAAAAREAAALTGARLRDVPR
ATGLYGAFGYGSRGLVWAALGAELIAAQIDGEPWPLERELAEAIDPARFLVRALRHGRVA
*
>GCA_000452945.1_BPMSHR5858_genomic_00931_1 group_19865
MRSQAWDWAEGAGLTPTQGEILVLLLQRKGPMRLGEIARETQLTAATTSDAVSTLEHKGL
VEKRRALDDGRALAVRLSARGRTAAKKALQWPDFLSKAVGTLGGDEQGVLYRALLKTLRE
LQVNGDIPPHRMCVTCKHFQPGKAARKPTYRCSLLDLTMTDSDLRLDCAVHEEADVLTQK
KTWKLFAQA*
>GCA_000452945.1_BPMSHR5858_genomic_00932_1 group_19796
MNAEVVAIRHVHFEDLGSFEQVLGERGRRVRYVDVGGSRVEVLDALEPSLLVVLGGPISA
YDDELYPTTAPLAALVGKRIEAGLPTLGVCLGSQLIARALGARVYPAAAKELGWIPLALT
DAGRASPLRHVDGAITSMMHWHGDTFDLPAGAIHLASTPACRNQAFSWGTHVLALQCHPE
IRADRFEPWLIGHAGEIAATPGTDAKRLREQTAQLGPTLECAARRMFGEWLDSVGL*
>GCA_000452945.1_BPMSHR5858_genomic_00935_1 qseC_1
MSPDPAVTSLRRTLLRRLAAPLSMLALMSGLIAYWLAWQYTQHVVDRSLADLASAISKQI
QIAGPDAPFTVPPLAQAMFSDPTEQLIYRISDGEHELAGDPKLPLQGTSVRRMHVAYVFE
AEYDNQAVRVAQVRVAQTESGNPMIVEVAQPVRYRYRIAAEFLIAIMMPLLLLLLAGWGI
VWRVVNQQLGPLTHLADSLNRQTHTSLEPVDETEVPLEIRPLTSAMNALLGRLKTALDAQ
RKFIADAAHQLRTPLTAIKLHAEQAAVARDPQQALTAVRELRSAADRAVRLSNQLLSLAR
AEPGEQAARFVDVDLAALAFETGAEWVPRALAAHVDLGFQRGDGRGDDEPLLVRGNPVLL
REVIANLLDNALKYVPLARPQGARITMDVSRIALDEGHAAAEIVVEDNGSGVPAAQQADL
FKRFFRGDAQSGNGVETGAGLGLAIVHDIIAMHGGTVSYADAPEGGSCFIVRVPLADTPS
TPAPPPCESPAASA*
>GCA_000452945.1_BPMSHR5858_genomic_00937_1 group_19737
MSDISIHELEAAINFWRARSPASGDELELCEEASALSKPYALLIVQRQSALQLEGLDPKA
RNAWETYVRLNNGLEG*
>GCA_000452945.1_BPMSHR5858_genomic_00939_1 phhA
MSTVVTAKLKEQFDAGLETRADFTIDQPLARYGDVDHAVWTQLYTRQAALLRGRACDAFI
EGLARIGLAPDRVPSFADVNRRLEPATGWRIVAVPGLVPDAVFFEHLANRRFPVTWWMRR
PDQLDYLQEPDCFHDLFGHVPLLIDPVFADYMHAYGRAALRVADDASALALLARLYWYTV
EFGLIRDTRGENGLRIYGAGIVSSKGETLYSLESASPNRIGFDLERVMRTRYRIDTFQKT
YFVIDDFAQLFALADLDARALAARLAGAPEHAAGAVLDGDHVLTRGTGEGWAADADA*
>GCA_000452945.1_BPMSHR5858_genomic_00944_1 alkJ_2
VTTERTLEGEFDYVIVGAGTAGCVLANRLTEDPDVTVLLLEAGGRDDYHWIHIPVGYLYC
IGNPRTDWLYKTEPEAGLNGRALSYPRGRVLGGSSSINGMIYMRGQRGDYDDWARATGDA
GWSWDSVLPVFRRSEDHHAGATDMHGAGGMWRVEKQRLRWEILEAFSQAAQQTGIPATDD
FNRGDNTGVGYFEVNQKRGIRWNASKAFLRPALARPNLTVITGAQAERLVFDGKRCAGVE
YRGGGAPFVARARVEVLVASGAVNSPQLLELSGIGDGSRLQALGIGVVADLRGVGENLQD
HLQLRMAFRVRGVRTLNTLSAHWWGKLWIGAQYALMQRGPMSMAPSQLGAFAKSDPNDPA
LAQPDLEYHVQPLSLERFGEPLHRFNAFTASVCHLRPTSRGSVHAASPDPARAPSIAPNY
LSTDYDRHVAANALRLTRRIASAPALARYAPEEILPGARYVSEAELIAAAGAVGTTIFHP
VGTCRMGRADDPDAVVDSRLRVRGVTGLRVVDASVMPTITSGNTNSPTLMIAERASDMIR
ADRRGASERGASARAEAVLPT*
>GCA_000452945.1_BPMSHR5858_genomic_00947_1 group_3153
MKLKTLAHACLAVAAAWSVGAAQAADSVKIGFITDMSGLYADIDGQGGLEAIKMAVADFG
GKVNGKPIEVVYADHQNKADIAASKAREWMDRGGLDLLVGGTNSATALSMNQVAAEKKKV
YINIGAGADTLTNEQCTPYTVHYAYDTMALAKGTGSAVVKQGGKTWFFLTADYAFGKALE
KNTADVVKANGGKVLGEVRHPLSASDFSSFLLQAQSSKAQILGLANAGGDTVNAIKAAKE
FGITKTMKLAALLMFINDVHALGLETTQGLVLTDSWYWNRDQASRQWAQRYFAKMKKMPS
SLQAADYSSVTTYLKAVQAAGSTDSDKVMAQLKKMKIDDFYAKGSIRTDGSMIHDMYLME
VKKPSESKEPWDYYKVVATIPGEQAFTTKQETRCALWK*
>GCA_000452945.1_BPMSHR5858_genomic_00949_1 group_3154
MQRKALYGLLLLGLLVAPFVGAYPVFVMKVLCFALFAAAFNLLIGFTGLLSFGHAMFLAT
AGYVTGYAMQSLGTAPELGVLAGTAAATLLGLVVGLFAIRRQGIYFAMITLAFAQMVYFI
YLQAPFTHGEDGLQGVPRGRLFGLVDLSSDLALYYVVLVVIACACFFIVRIVHSPFGQVL
VAIKENEPRAISLGYDTDRFKLLAFILSATLAGLAGALKVVVLGFETLSDAYWTMSGLVV
LMTLVGGMGTLFGPLVGAALIVALEDRLGDIGGWLASVTHIDWFRSLGESATIVTGLIFI
VCVLAFRRGIVGEVIARVRPLRAS*
>GCA_000452945.1_BPMSHR5858_genomic_00953_1 btuD_11
MKAILRNKAFWLFLVALFALPALPGAARVPEYWITLLNYIGLYAIVAVGLVLLTGIGGMT
SFGQAAFVGVGAYATAFLTTAYGVSPWLGLIAGVVLTALVALVLGVVTMRLSGHFLPLGT
IAWGLALFFLFGNLDWLGKYDGINGIPALALFGHAFDTGRSLYYLIWIVVLAAIVSVQNL
LNSRPGRAIRALRGGGMMAEAMGVNTGWLRVVIFVYAAVLASISGFLYAHLQRAVNPTPF
GLSHGIEFLFMAVVGGVAHVWGAVLGAAIVTVLQDYLQTLLPKLLGAEGNFEIIVFGVLM
VLLLQYARQGVWPFVAKLMPREPAARAPGQAAPLPRRARPQAGEALLVVDNVRKQFGGLV
AVNDVSFDVLAGQIVGLIGPNGAGKSTTFNLVTGVLRPTSGAITFRGERIDKLVSREIVK
RGIGRTFQHVKLLPGMTVLENVAIGAHLRGSAGIWRSVVRLNAHEEARLMSEAARQLRRV
GLDGHMYDEAGSLALGQQRILEIARALCSDPTLLLLDEPAAGLRYQEKQQLAELLRKLKA
EGMSVLLVEHDMDFVMNLTDRLVVMEFGTRIAEGLPQDVQKDPAVLEAYLGGVE*
>GCA_000452945.1_BPMSHR5858_genomic_00955_1 mnmG
MLYPTEFDVIVVGGGHAGTEAALASARMGAKTLLLTHNIETLGQMSCNPSIGGIGKGHLV
KEVDALGGAMAAATDEGGIQFRILNSSKGPAVRATRAQADRVLYKQAIRRRLENQPNLWL
FQQAVDDLMVEGDRVVGAVTQVGVRFRARAVVLTAGTFLDGKIHVGLNHYTGGRAGDPAA
VSLSSRLKELNLPQGRLKTGTPPRIDGRTIDFSKLDEQPGDLDPIPVFSFLGRAEQHPQQ
LPCWVTHTNERTHDIIRSGLDRSPMYTGVIEGVGPRYCPSIEDKIHRFASKDSHQIFLEP
EGLTTNEFYPNGISTSLPFDVQLALVHSMRGLEQAHILRPGYAIEYDYFDPRALKSSLET
KAIGGLFFAGQINGTTGYEEAAAQGLLAGINAGRYAQEKDAWCPRRDQAYLGVLVDDLVT
RGVSEPYRMFTSRAEYRLSLREDNADMRLTEIGRELGVVDDVRWDAFNRKRDAVSRETER
LRTTWVTPKTLPVDEATALLGKPIDHEYSLAELLRRPGVSYDGVCGLRGGECGPSEPLAE
DELLLAQIKEQIEIGIKYQGYIERQAGEIERNGANENTRLPDGIDYTEVRGLSFEVSQKL
NQFRPETIGQASRISGMTPAAISLLMVHLKKRGLGRRKGTDSVPGADVQADNTAAQQ*
>GCA_000452945.1_BPMSHR5858_genomic_00956_1 rsmG
MTVQQRRRPPIASRETLQALLSEGAQALGVALSDAQRGALLDYVALLAKWNAVYNLTAIR
DPRQMLIQHILDSLSIVPHLGAHGAAAAALDVGSGGGLPGVVLAIALPGWRVTLNDIVHK
KSAFQNQAKAELKLGNLSVVTGRVETLRPGADVPAKFDVIVSRAFADLADFVTLARHLVA
PGGSIWAMKGVRPDEEIGRLPDGARVKQMIRLTVPSLDAERHLIEVELDEAI*
>GCA_000452945.1_BPMSHR5858_genomic_00959_1 ybiR
VSNSVVGRAPARIIAFVRHEPVLSILVAALIALQAVHSLSAIALVRLVDWQTVATLAGLL
MLTKALELSGFLMWLAHRIVHRVRSERALAALLVVFAAALSVWLTNDVALFVVIPLVLSL
RELTPLPFKRLVIYIALAVNAGAIATPLGNPQNLFLWQLSGVSFGRFVIALGPLALVLMA
LLLAMVVFAFDGRPLDLSRDSTERPVDRAQAFATVAMFAGFVLLADAHHALAGVLAVAAV
FLIVRRDAVLKIDWLLLLIFVLMFIVLRSAASLPAVHQAIAGIGLDTPLRAYAAGALLSQ
IISNVPAAILLSEFTHDWRALAFGVSVGGFGIAFGSLANLIAIRLSRARGMWLPFHLVSI
PFGIASAVLGALLLHYF*
>GCA_000452945.1_BPMSHR5858_genomic_00964_1 atpH_1
MAELATIARPYAEALFRVAEGGDISAWSTLVQELAQVAQLPEVLSVASSPKVSRTQVAEL
LLAALKSPLASGAQAKNFVQMLVDNHRIALLPEIAEQFEALKNAREGAADVQIVSAFPLE
GAQLAELVTSLERKFKRKLKPAVEVDSSLIGGVRVTVGDEVLDTSVRARLAGMQAALTA*
>GCA_000452945.1_BPMSHR5858_genomic_00965_1 atpA_1
MQLNPSEISELIKSRIQGLEASADVRNQGTVISVTDGIVRIHGLSDVMQGEMLEFPGNTF
GLALNLERDSVGAVILGEYEHISEGDIVKTTGRILEVPVGPELVGRVVDALGNPIDGKGP
VNAKLTDAIEKIAPGVIWRKSVSQPVQTGLKSIDSMVPIGRGQRELIIGDRQCGKTAVAI
DTIINQKGKDLICIYVAIGQKASSIMNVVRKLEETGALEYTIVVAASASESAAMQYLAPY
AGCTMGEYFRDRGQDALIIYDDLTKQAWAYRQISLLLRRPPGREAYPGDVFYLHSRLLER
AARVSEEYVEKFTNGEVKGKSGSLTALPVIETQAGDVTAFVPTNVISITDGQIFLETDLF
NAGIRPAINAGVSVSRVGGAAQTKVVKKLSGGIRTDLAQYRELAAFAQFASDLDEATRKQ
LERGRRVTELLKQPQYQPLQVWELAVSLFSANNGYLDDLDVKDVLPFEKGLREYLKTSHA
DLIKRIEDTKDLSKDDESALHAALKDFKKSGAY*
>GCA_000452945.1_BPMSHR5858_genomic_00973_1 priA
MADAFVRVALDHPLPTLFDYRYRGDMPAAPGMLVQVPFGRRSVVGLVCEVTAHTDVPAAK
LKAVDAVCTELPPLSRHWLELAAFAADYYQRGLGEVALPALPQALRDVPRWGRLLAPELR
YRLLDAGRAALPDALPARGAALRRLAQALADAPALSLAEVRALHPKAVAVLDEWQAQGWA
ACDEIAITHADAQLVDNPAPASARRPTALPPQLTDQQAEALDAIAAADGFAPFLLHGVTG
SGKTEVYLRALAALLERQPEAQALVLVPEINLTPQFEAAFRARFAATLPADAIVTLHSGL
AEGERARHWLAAHTGRARIVLGTRLAVLASLPALAMIVVDEEHEPAYKQQEGLRYSARDL
AVWRAKQLAIPVVLGSATPSLETWWQAEQGRYRRLTLTKRAVADAALPTVRLVDLEEERR
RGRGSMGGLSGPLVAALKARLERGEQSLVFLNRRGYAPALACDACGWVAGCPRCSAYVVM
HKPEHAVRCHHCGWEARIPRACPECGNVDIAPLGRGTQRVEETLALAVPGARILRIDADS
TRRKGSAQALFSDVHAGEVDILVGTQMIAKGHDFQRVSLVGVLNADTALFSHDFRASERL
FAQLMQVSGRAGRAGLPGEVLIQTRYPRHALYHALARQDYVGFAGATLGERRDAHLPPFV
YQALLRAEGRTLDAALAFLQQAADAFSGLPGAERVTVYDAVPMTIVKVANVHRAQLLIES
ASRAALQHTLRAWQPLLRELKGVLRWSVEVDPLDI*
>GCA_000452945.1_BPMSHR5858_genomic_00974_1 group_19981
MDGWVVIGIVLAAWIAGAIVFVHRASAVFFQALGCAQRVKKDERRARLSGLREAHE*
>GCA_000452945.1_BPMSHR5858_genomic_00976_1 braC_4
MQHTMKKLAGATFVAVMSLAGTAHADDVKIGFAAPMTGAQAHYGKDMQNGIVLAIEDINA
TKPTIGGKPVKFVLDTQDDQADPRTGTTVAQKLVDDGIKGMLGHFNSGTTIPASRIYANA
GIPQIAMATAPEYTTQGYKTTFRMMTSDTQQGSVAGTFAVKDLGMKKIAIVDDRTAYGQG
LADQFEKAAKAAGATIVDREFTNDKAVDFKAILTKLKATKPDLVYYGGADSQAAPMAKQM
KSLGVTAPLMGGEMVHTPTFLKIAGDAAEGSIASLAGLPLAEMPGGKAYADKYKKRFGED
VQTYSPYAYDGAMAMFSAMKKANSTDPAKYLPLLAKTDMAGVTSTHIAYDAKGDLRNGGI
TMYKVEKGEWKPLKSIGGK*
>GCA_000452945.1_BPMSHR5858_genomic_00981_1 argT_2
MKRLQFWFALACALGAATGAGAARADASQTLRFGLEAQYPPFESKAANGALQGFDIDVGN
AVCNAAKLTCKWVETSFDGLIPALQGRKFDAINSAMNATDQRRQAIDFTTVIYRVPTQLI
ARADSGLEPTPASLKGKRVGVLQGSIQEAYANAHWAGAGVQVVAYQDQNQAYADLTAGRL

DGTLVLAPAGQRGFLSRPEAKGFAFVGPPVRDDKILGSGIAFGLRKGDDALKARLNAAID
KLKADGAVKALGRKYFGDIDISAK*
>GCA_000452945.1_BPMSHR5858_genomic_00996_1 group_2292
MTPTLDETALPPADALAYPGTLLSPDGRLVAPYDLERPGADARGHVGAAARAGLLHAASG
PFTLDYATLSTVHVINGMGVALGDSVIGLTALAAIKARHPHVHFVLYRPAHAPAYVDALY
RLAADAIAPIRALPWPAAELPARDTRIDLGNHLYWPAFAARPMIDFFLASLGVPPESISA
DAKCNRWLARLTLPPLPPEWRDRPYALFCPQASTPLRSIPAAQRAALVERLARTYRLPVL
GFGPIDHPDYVDVSGHSADTAQFVSWVGRARVALCSDTSAVHIAAGFDVPTLACFSSIAP
ELRVRDYPLCTAIELDMPEMLRNRHKSDAANDIAQVDAAYRRIDWDAIAWPPARE*
>GCA_000452945.1_BPMSHR5858_genomic_00999_1 sdaA
MAVSVFDLFKIGIGPSSSHTVGPMRAALMFAQGLERDALLAATASVKVELYGSLGATGKG
HGTDRGVMLGLMGDAPDTVDPSTIAARLDAVRTSKTLALLGTHPVPFVPKEHIAFYRQAL
PEHPNAMKLRAADTAGAVLREATYLSVGGGFVVTSGAPNTKVLAAAEQMPHPFRTGAELL
ALTASTGKSIAQLMWENERAWHTEEETRTGLLKIWDVMQSCVARGCGIGNPDADGNLPGP
FQVKRRAPQLYRTLTGHPERALQDPLSMVDWINLYAIAVNEENAAGGRVVTAPTNGAAGI
IPAVLHYYTRFTPGANEQGVIDFLMTAAAIGILYKLNASISGAEVGCQGEVGVACSMAAG
ALAAVLGGTPQQVENAAEIGMEHNLGLTCDPVGGMVQIPCIERNAMASVKAVNAARMALR
GDGSHYVSLDSVIKTMRETGADMKTKYKETSRGGLAVNIVEC*
>GCA_000452945.1_BPMSHR5858_genomic_01000_1 group_3986
MVRQTFPGRAQALRQRLSALAPALVAAAALAAAGPARAAMNFCAAPALQSSEATHAEPGV
QALIKSVDAHLNDEPKALPRVHTEGTLPHEGIYDQSAEALNDMELIRNAALAWRVTNQSR
YLALVDRFLSTWVNTYRPSFNPIDETRFESLILAYDMTASALPVKTRNAAAAFIAALGNG
YVQQIDAQKRPLKGTWRNNWQSHRIKLIALAAFTLGDRRMMNAAQRLFVEHLADNIEPDG
TTYDFLERDALHYAVYDLQPLATAALAARRFNRNWLRERAPNGATLAAALDWLAPYARGE
KTHEEFVHSPVPFDAKRREAGLPGYSGMWEPKNATELFHLAARLDGRYAGIAQQLSPMPP
AWLAACLPLPAR*
>GCA_000452945.1_BPMSHR5858_genomic_01002_1 gcvH
MSNVPAELKYTDEHEWIRTEADGTLTIGITDHAQTTLGDIVFLELPKVGSQVKAGDAVGV
VESVKAASDIYSPVTGEVIAINTDVADAPEEVNGDAYGAWLFKIKLAADASTDSLLDAAA
YEKLVG*
>GCA_000452945.1_BPMSHR5858_genomic_01004_1 group_220
MQRLIRAAACCVLVSSLAACVVTPPRPAPAPAPRPSPQVVGYERMQQIQGRIDNLSHRID
ARVNAGYYPPPQGAALHRRLDVIRQESTDMAAQHGGGLSGDEQRVLNQELDVAARAIGE*
>GCA_000452945.1_BPMSHR5858_genomic_01005_1 ydgJ
MSSTLKIGLMGFGFAGATFHAPVITHSGRTRLAAIATGQPERATAAYPEAAVVPDLDALA
ARDDVECVVIATPNDTHVDLAKRALEAGKHVVVDKPVALSAADALVLARLAAARGRVFAP
FHNRRWDGDFLTVRQLVESGELGRIVCFESHFDRFRPHVRTRWREEAARGGGLLFDLGPH
LIDQALALFGPPETVSATVKTRRDGGDAPDFVHLQLGYPDKDVVLHASALAAIEPARFTV
LGTQGGYQKHGLDTQEDQLKAGLTPDDVEFGGGNPPGVLRGLDGELEVERPVPTLDGQYA
EFYRALAASIHDGAPFPVAPQDAVDVMTIIETAMRSEREGRRLPFVREAV*
>GCA_000452945.1_BPMSHR5858_genomic_01009_1 petJ
MSEAPHGAPIKTPGQLIAVVIASFVIPIAIIVLFATYANHAFRTGAGTDGLSDEAVAKRI
APLAQVDIKDANAPHVYKTGEEVYKAVCVTCHGTGAAGAPKFGDAAAWAPRIAAGYDEVL
HLALTGKGAMPPRGGTNPDDYSDYEIARAVVYMANQGGAKFAEPAQPAANAAPASGAAAA
SAPGASDAAANSQAAAAMAAIAALPKAGEAPAAGANAESSASAGKALYESTCQACHATGV
LNAPKFGNKADWAPRLKDSMDTVYNFALHGKGAMPPKGGSNASDADVKAAVDYMVNAAK*
>GCA_000452945.1_BPMSHR5858_genomic_01023_1 group_2297
LKTSKTVLACLAASAIVALSGCATEASRTLPVPPSASAQKPFAGKPVGIAVGKFDNRSSY
MRGIFTDGVDRLGGQAKTILVTQLQQSRRFNVLDRENLDEIKQEAAFMKKAQAVKGANYV
VTGDVTEFGRKEVGDQQLFGILGRGKSQIAYAKVNLNIVDTTTSEVVLSSQGAGEYSLSN
REIIGFGGTAGYDSTLNGKVLDLAIQDAVNHLVEQVDAGALASGK*
>GCA_000452945.1_BPMSHR5858_genomic_01024_1 group_2298
MKKTFKTGAWLPVAAAGVLLAGCAANQPKPLYQWDGYQPQVYEYFKGQKSPEDQIDALEK
ALQQIRAKGNRPPPGFEAHLGMLYASVGKEQQAEQSFQAEKASFPESSPFMDFLLKKKAA
APQAKPSAPAQPQTQTQTQAQQ*
>GCA_000452945.1_BPMSHR5858_genomic_01026_1 group_3166
VTDKHPPSTDLPEDPDRRRVLGGIAALGASVALAGCETAVPAAPRSAAELRLDAALRREV
RNIVVIYAENRSFANLFGDFPGVQQPLGAVSPERYAQLDRDGKTPLATLPKIWGGLVPQA
QVVDGKRYMIGEHDIVGLPNAPFVIPDAQGKPLPNGVITRDLWHRFYQNQMQINGGRNDQ
FAAWADSGGLVMGHYRNSAETLRLWNLARGYTLCDNFFMAAFGGSWLNHIFLISAQAPLY
PDVHQSPAKHLVSVVEGDDPAGTRLKLAADSPASALDGPPKFENDGLFTPDGYAVNTMAP
PYQPSSVRPPVEGNPAYADPSNPRVLPPQHYATIGDRLSEKGVDWAWYGGAWQYALEHRD
TGSVPDFQYHHQPFNYFASYAPGTEARRRHLRDAGLGDDPSTNHLIADIDAGRLPTVTFY
KPQGNLNMHAGYADVASGDRHIATVIEHIMRGPQWAHTVVILTVDENGGWWDPVSPPKGD
RWGPGSRIPAMVISPFAKKGYVDHTLYDTNSILRFISRVHGLAPLDGVLARERAFAARGA
QPPGDLTEALDLG*
>GCA_000452945.1_BPMSHR5858_genomic_01036_1 bepA_4
MSESTTSDTLTPEQQFEEDIALVLKNAIELHHQGDFENARALYEAIVDAKPDHADAQYDL
GVLKVQIGRAADALPHFEIALGGAPNNGQYWVSYVNALIDAGQIAAAWTALGLCQQRGVR
GPAVDGLILRLAHSEEGKPAYIGTAAAAPEAAAPADAAANAAEPVRADARKPSEQDVRRF
ATLYNKRRMAEAIKFARALAQRYPGSGVAWKSLGFALHRDGQYGPACEALTKGAAMLPDD
AECNTLYADTLRVLNRLADAEAQVRRVLDGTPDYAEAHRVLNMTLSARGRYQEAIEAGRR
SVELAPNSVNAHGSLAVTLSDYGQTDEAEIHFRRAHELNPKDPMAYSNLLFCQSHKIDVS
IRELFDAHRAFGELYEAPLRGATPKHANSRDPERRLRIGFVSGDLFLHAVSSYLLPLVDE
LAKDPSVSLHFYYTFAREDSVTERIRSYAQGWHMVMPLSDEQFAERVRHDRIDILVDLSG
HSGRNRLPMFARKPAPIQVSWIGYPGTTGLEAIDYYLADPYAVPFGPMQAQFTEKIVHLS
SGATFSPDGNAPPVNMLPALHNGHVSFGSFNRLNKLRGDVIAVWARIMRAVPGSRIVLGS
IPKDGGGAAMIEWFEQEGIARERLSFQPRSVTAVYLQQHHHVDVCLDTFPYTGSTTVLNA
LWMGVPTLTMRGETLPSRAGLTWMSHVGLESFIADDIDDFVAKGIALASDIPALARIRGE
LRERCMRSPAFQPQRVAQDVSDAFRIMWRRWCDGQPPAPFAVPPRDATTTVAGDRS*
>GCA_000452945.1_BPMSHR5858_genomic_01039_1 fliC
MLGINSNINSLVAQQNLNGSQGALSQAITRLSSGKRINSAADDAAGLAIATRMQTQINGL
NQGVSNANDGVSILQTASSGLTSLTNSLQRIRQLAVQASNGPLSASDASALQQEVAQQIS
EVNRIASQTNYNGKNILDGSAGTLSFQVGANVGQTVSVDLTQSMSAAKIGGGMVQTGQTL
GTIKVAIDSSGAAWSSGSTGQETTQINVVSDGKGGFTFTDQNNQALSSTAVTAVFGSSTA
GTGTAASPSFQTLALSTSATSALSATDQANATAMVAQINAVNKPQTVSNLDISTQTGAYQ
AMVSIDNALATVNNLQATLGAAQNRFTAIATTQQAGSNNLAQAQSQIQSADFAQETANLS
RAQVLQQAGISVLAQANSLPQQVLKLLQ*
>GCA_000452945.1_BPMSHR5858_genomic_01041_1 tag
MPKQGRCSWVKTEADAHYHDTEWGVPSRDDRHLFEMLVLEGAQAGLSWSTILNKRAGYRA
AFAGFDIDEVARFTPKRIDELMLDASIVRNRAKIEAAVANARAVQQIRAEHGSLAAFLWS
FVDHSPLQNAWASYRDAPASTERSDALSQALKRYGCKFVGSTICYALMQATGMVNDHEST
CPCHARCAALGEKARRRKKPAA*
>GCA_000452945.1_BPMSHR5858_genomic_01044_1 aqpZ2
MKLPQRLAAEVFGTFWLVLGGCGSAVLAAAFPGLGIGFAGVALAFGLTVLTMAFAIGHVS
GCHLNPAVSVGLTVAGRFPARDLVPYVIAQVIGATLGAFVLYLIATGKPGFDVAGSGFAT
NGYDARSPGHYSLAAAFICEVVLTGFFLFVILGSTDKRAPAGFAPIAIGLCLTLIHLISI
PVTNTSVNPARSTGPALFVGSEAIGQLWLFWVAPIIGAAIAGVVYPLVAGRDES*
>GCA_000452945.1_BPMSHR5858_genomic_01046_1 pimA
MRIAQIAPLYEAVPPKLYGGTERVVSYLTEALVEMGHDVTLFASGDSVTSARLEAAWPRA
LRLDPSIRDAMAPHMRLLEQVARAAHTFDILHFHLDYLPFPLLSRLDAPFVTTLHGRLDL
PELQPVFDAFPNAPVVSISDSQRTPLPQAGWAGTVYHGLPDTLLTPQAGRKPEYLAFLGR
ICPEKRVDTAIRIAAQSGLPLKIAAKVDKVDEDYFKAEIEPLLDSAHVEFIGEINEAQKP
AFLSGAKALLFPIDWPEPFGLVMIEAMACGTPVVAFNRGSVPEVIDDGLTGFIVEDVQGA
VGALHRIDELSRDAIRAQFEQRFSSHAMARRYIDIYETLRGATKQPQWQRVAAG*
>GCA_000452945.1_BPMSHR5858_genomic_01049_1 motA
VLIFVGTLVTLLSVFGGYALAGGHLGALVQPVEILMIVGAGVGAFIIGNGIKTIKATLRV
LPTLFKGSKYNKDIYMELMALLYVLLAKARKEGTLTLEADIDDPEKSPIFTQYPKILADH
HIVEFLTDYLRLMVGGNMNAFEIESLMDEEIETHHAEGEGPAHALMRVGDAMPAFGIVAA
VMGVVHTMASADKPPAVLGAMIAQALVGTFLGILLSYGLIGPLASLAEQRVAESTKMFQC
IKVTILASLNGYAPAIAVEFGRKVLFSTERPSFAELEEHVRRVKAK*
>GCA_000452945.1_BPMSHR5858_genomic_01051_1 group_19946
MIRTILAIDDSATMRTLLSATLGEAGYDVTVASDGEVGLDVALATRFDLVLTDHHMPRKN
GLELIVALRRQLGYEATPILVLTTENGDAFKDAARAAGATGWIEKPIDPDALIELVAALS
ASSSLS*
>GCA_000452945.1_BPMSHR5858_genomic_01053_1 cheW_2
VSEVQTNHPAAPNAASRRDAEQGDAAGQELLVFTLGDEEYGIDILKVQEIRGYDSVTRIA
NAPDFIKGVINLRGIIVPIVDMRIKFHLGRVEYDHQTVVIILNVAHRVVGMVVDGVSDVL
TLSTEQIMPAPEFGGVLTTEYLTGLGTVDGRMLILMDIEKLMTSKEMALIETLGA*
>GCA_000452945.1_BPMSHR5858_genomic_01061_1 group_2112
LRFQHTLTRWLGALSLAAATAALVAACGGGGDGGSNASVNTGTGGGDTSAGGGSNGGTGG
TGGSGSTPLASNQAAITVSTGVATIINMPTVSVTICAPGTSACQTIGNVLVDTASYGLRI
VNTAASGVLGSLPVSSVSGGTLTECGKFVTSYTWGTVRTADVRIAGEQASSLPIQIIGDL
GSSGVPSSCSNGNAPSNTAADLGANGILGIGPAPYDCGANCTTSTLYSNYYACPNGNTSC
QVTTVPLAQQVANPVPRFATDNNGVIVAMNPPAGGSATGTLTFGINTQSNNALPAGVTKL
TSTTAGDLTGTLLGRAITTAFFDTGSNAYFFDTGGSTAVSLPVCSNNSSFYCPSSTQTTT
ATVVGLNGAQASQSISIGNASALFSSYKSAVPNLGGPFGNTRVLDFGLPHFFGKTLYFGM
DLTGSGGAAPYVAF*
>GCA_000452945.1_BPMSHR5858_genomic_01062_1 group_3168
MTARVQRITPFLWFDGNAEEAATFYVSVFDNARLTRVARYGKEGAQASGQGEGAVMTVAF
ELDGQSFVALNGGPVFQFNPAVSFVVNCATQDEIDYYWGRLSEGGDASAQQCGWLRDRFG
LSWQIVPVQLPELLAGDDPAKASRTMQALMQMKKIDLGALRRAAAD*
>GCA_000452945.1_BPMSHR5858_genomic_01071_1 nylA
MTVPSDYLCHDAIGLAQLVAQREVSARELLDAAIGRAQALNPAINAIVLNDYAAARERAA
SGTLAGPLAGVPYLVKDLGSAVAGLPLSLGSRHYRHHVPAEDSPLIVRTKAAGLNIFGKT
NTPELGQMPYTESALLGVCRNPWNLDHTPGGSSGGAAAAVAAGIVPLAHASDGGGSIRIP
ASCCGLFGFKPSRDPALTPWPVPGEIVVEHAVSRSVRDSALLLDITTGRTAPGTLEGVGA
PGTYLGALDAPPPALKIGYVTDPMLAPALSADVRDALDRAAALAASLGHNVEPATLNLDF
AQIGEVFLTIWAAIADELVLGAERITGRKPARAEFEPATWAMAQVGRRLVRERLPHALEL
QRQITARVAGLVSRYDVLLCATLAAPPIKIGEMQPTPLETRQMELLGVLPIKPLLKRMLA
EVSHQAFAWAGCTELFNLTGQPAMSVPLHWSARGLPVGVQFVARHGDDARLFALARQLEL
AQPWFDKRPPLARAQA*
>GCA_000452945.1_BPMSHR5858_genomic_01073_1 pnoA_2
MTSRIAPTPFAARFDLRLPLVQAPMVGATTPALVAAASNAGALGSLGGASFAPEKLAAEI
AAVRAATRRAFAVNLFVLPDAQPDDAAVRRALDAIDPLRARFGLPPGAPLPRYAPDFRAQ
LDALVDARVPVASFTFGVLDKKDVVRLQAAGTYVIGTATHVAEGLAWQAAGADAICAQGA
EAGGHRGTFIGSAEDALVGTIALVPQLVDATNLPVLAAGGIMDGRGIAAALALGAQAAQL
GTAFLTCAESAIPACWKARLLASDDTSTSVTRAITGRHARGIRNALMAQLAGRPDSVAPY
PVQNALTQELRQTAARAGDAEYLSLWSGQGAPLGKHRDGAQTTAQLIDALDAEWRAALSR
SVISLV*
>GCA_000452945.1_BPMSHR5858_genomic_01075_1 ybjJ_1
MALFFVAGMMYASWGVHVPTVRDKFALSPALLSFALLAVAIGSIVAMTTNARWIARVGSR
TACLAGGLAMSASGALILVVPTYPLLLAVLALFGFSMATLDVAMNAEASAVESAFGKPIM
SMLHGMFSVGGMAGAAAGGALLSAGMASAVHLGLAALASALVLALACPAVLPHVPHTAAA
DGAPRVNRWRSPALWALGAIALVALIAEGAMYDWATVYMRDVVLASPAFSSAAYAAFSGG
MAAARFAGDAVRARFGAPQLVCASATLACVGMIAALALPSPFVALAGFTLMGLGLANMMP
VLFAAAARIDGIHAAEGLAHVAGLAYFGLLFGPVAIGAVTQAANLSVGLSIVALCAALVA
IVAPKVLSRLKI*
>GCA_000452945.1_BPMSHR5858_genomic_01081_1 gloC
MKVTLIPVTPFQQNCSLLVCEKTGRAAVVDPGGDLELIEQEVARQHVQVEKVLLTHGHVD
HCAGAKRLAAHYGVPIEGPHEDERFWIEKLPMQSERFGFPAAEAFEPDRWLADGDTLAFG
EETLEVYHCPGHTPGHVVFFSRAHRLAIVGDVLFAGSIGRTDFPRGNHADLIRSIREKLW
PLGDDVTFVPGHGPVSTFGDERRTNPFVADEVAG*
>GCA_000452945.1_BPMSHR5858_genomic_01082_1 rlpA
MKFRLPRHLGTLAVFFALTGCAVPPNAPTSEQAKSAASKDALRTAASGAASNNKESAGAP
LNFDTALASMPAAGSQDASRQSLADAKPIDAADISDFRQTGRASWYGKGFHGRRTANGER
FNMNEFTAAHRTLPLASYVRVTNQANGKSVVVKINDRGPFSRGRILDLSYAAAKVIGLVH
AGTARVKIQGLSPEEARVARDEMLASNSTTK*
>GCA_000452945.1_BPMSHR5858_genomic_01083_1 rsmI_1
MTSLLDLAQAQHYPAGALYIVATPIGNAADITLRALHVLTLADRIAAEDTRNTGQLLVRY
GISKPLVAVHEHNERAAAAKLIDHLRAGERIAYVSDAGTPGISDPGAKLVDAVRAAGFGV
IPLPGANAAAAAVSVAGDWAGAFTFAGFLPPKPKQRDAALQPLKTHPYALVFYEAPHRIV
ETVEALAAALGGERRLLIARELTKLHEELFEGTLADGPTWLRADANRQRGEFVLVVEGAP
QGAQDEDERAHDALLELLLDEVPVKSAAKLAAALTGASRNALYARALALKKEEEE*
>GCA_000452945.1_BPMSHR5858_genomic_01086_1 group_2302
MSNSRVRQTLVRTTLVIGLSAGLAATLQGCVLAVAGAAAGGGALVATDRRTLGAQTEDRE
IQVKALTQMNNGLPDGSHVNVTVFNRRVLLTGEVPSDAAKQRAEEIVRAINNVNAIVNEL
AVGPASSLSDRANDSYLEGRVKTAMIAEKGISANNYKVVCERGNVYLMGLVTTDEGSRGA
DVASRVPGVEKVVKVFQYIKPQDAQALTAATPASGASAAAAPATAPEGATVGAVPDSSVT
ATPLSAPAPVSNSSNVHPGNPKAGTP*
>GCA_000452945.1_BPMSHR5858_genomic_01087_1 group_19945
VTIAKRVKRTMSAAAAAMAVVSCAMAAAPAAHADAGDGLKVARSNACMGCHAVDRKLVGP
SFQQIAERYKNDKQAEPKLAKKVKDGGSGVWGAIPMPAHPRMSDADVRSVVQWVLAGAPS
K*
>GCA_000452945.1_BPMSHR5858_genomic_01090_1 group_4009
MPINSDWVASTGLPAAVYLFCLGYALLRAFGPRHARYPKLMRRYAMACAGAVLCGMLATI
GAALRTPAAPVLASFGSTVTFAGIVGVLIGCMGEAAVSFGKRWER*
>GCA_000452945.1_BPMSHR5858_genomic_01092_1 group_2965
MKKIIASVATAAIAFASASVFAQASAPAASAPAAASAAKQDKSKAKHELKKHGSTKAKAK
AEAASAAGTNDKGAQQ*
>GCA_000452945.1_BPMSHR5858_genomic_01094_1 garR
MDIGFIGAGEMGSAIAGNLLKAGHRVRVWNRSAERVAPLVALGAQQVATVADAFAGDAVF
SMLADDAAARDVFDAALLEQAPRGLIHVNMATISVALAEQLAHAHAQRAIHYVAAPVMGR
PHVAAAARLTIIAAGPAEAIDRMQPAFDAIGQRTWRLGSLPQHANAVKIAANFTIASAIE
TMGEAGALLAAHGVAMNDYLDVITNSVFPGPVYQGYGAMIAESRYEPALFKARLGLKDVR
LALEAGDALSVPLPVASVVRDSLIEAVAHGHGDKDFAVLGQVAARRAGR*
>GCA_000452945.1_BPMSHR5858_genomic_01095_1 rhtB_3
MTALVAMLSVSAAGLGVFLEAWPAMFDALRFAGAAYLIYLGVKAWRARVDEPSDAQSNAH
AEAAAPRAGRSGSAGQGSRWALFRNGFLVAGSNPKAILFAAALLPQFIDASAPTLPQFGI
LVATFAVIEVSWYIVYASFGTRIGVTLRSANVAKAFNRLTGGLFVGFGAMMALVRH*
>GCA_000452945.1_BPMSHR5858_genomic_01096_1 group_3021
MNCPADLRTGPAAGRPRTAPAPALTPALTMFFSATVGVIVLNLFAAQPLTGPIAAELRLP
ASLTGLVAMLPQLGYAAGLVLLVPLVDLLENRRLIVTTLAVCAATLALPAVTRSGAVYLA
AVFAAGAASSVIQMLVPMAASMAPDERRGRAVGNVMSGLMLGILLSRPLASLIAGAAGWR
AFYGAAAAADIAIAAVLAAKLPLRAPQLSTRYAALLRSLWTLVATERVLQRRALSAALSM
AAFSAFWTAIGLRLAAAPFDLGLHGIAMFAFAGATGAIVTPFAGLAGDRGRERDALRGAH
VAMLAALAALGVAGAGWGGFDTAAHPALALALLVAGAAALDAGVVADQTLGRRAINLLDP
AMRGRLNGLFVGLFFVGGSLGAVLAGAAWAWAGWGAVCAVGLVFAGAAFALDWIGAHGPA
PR*
>GCA_000452945.1_BPMSHR5858_genomic_01131_1 eryD
MTAAPAIVILGPGALETARRIQARFDGARVHALAGRAEADLAFAELGPHLRELYARGAPI
VALFAAGIAIRCVAPCLADKGAEPPVLAVAEDGSAVVPLLGGLAGANALAREIADALGVA
AAITTSGELRFGACMLNPPEGYALADLDAGKRFVSDLLAGESTRVEGDARWLDDVALPRD
AHAARVIRVTPHASRGDAHALLIHPRSVVVALGDGDGDNDGEGLRDDARSVSAPDATGRA
RANAAPDLPPLAARIAAALAAHDLAPLALAAVVAPARRIAEHALTDAARALRAPLRFVET
NARDAAGVLDAALPGALAPRVDASRAAAGSPRTTPPEPRDVAIAVARAPVDADALGIARG
RLTVLGLGPGRADLMTPAARAALADATDIVGYATYVNMAGPLRADQQLHVSDNREELQRA
RHAFELAARGRRVAVVSSGDPGVFAMAAAVLEALDGADDARWAAVELDVVPGVSAALATA
AEAGAPLGHDFCLISLSDNLKPWAIIEKRIDHAAAADFALAFYNPVSRARPVQFDRALDI
VRRHRAPETVVVLGRDIGRPGATLATTTLAALSAQQVDMRTMVIVGSSTTRRVARDGARD
WVYTPRWYR*
>GCA_000452945.1_BPMSHR5858_genomic_01234_1 ilvA_2
MSEKDTRRDTAPLHERQRRFEEDLVDAYDEELEMELDDRRFDDESLFSPERREARKRYFR
ELFRLQGELVKLQDWVVSTGHRLVVIFEGRDAAGKGGVIKRITQRLNPRVCRVAALPAPS
NRERTQWYFQRYVAHLPAGGEIVLFDRSWYNRAGVERVMNFCTDAEYEEFFRSVPEFEKM
LVRSGIQIVKYWFSITDEEQEVRFQNRIEDPLKQWKLSPMDLESRRRWEAYTAAKEEMLL
RTHIAEAPWWVVQAVDKKRARLNCIHHLLGLVPYYEIERDSVHLPRREHHDDYIRRPVPA
EMIVPEIY*

>GCA_000452945.1_BPMSHR5858_genomic_01310_1 yihX
MTQPIKLVLFDMEGVLSDYDRAARTARLADLTGRPPEAVRYAIWESGLEARADAGRIGVD
QYLRELAQLLGAPVSRDDWLIARKASITPNLEVVALAERVARCSRIAVLTNNCRLVTDYI
DYLNPPVARVFGAHVYPSATFGAAKPDAQAYLGCVGWLGVTAQETLFIDDADANVAGAVS
AGLLGCKFVGAGALAQELARRRLA*
>GCA_000452945.1_BPMSHR5858_genomic_01354_1 xylB_2
MTEPCVLAIDLGTSGPKAAIISLDGRVVAAARDAVATLRMPDGGVEQDPLAVWRAVKRAC
GGALRAAGIASRDVLAVACASQYSSIVPVGADGAPVANMMLWLDRRGAPRARRGAADGPL
RADSPLRQWRWLRVHGLPPVEGGISLTHMRYLKHAKPDVYARTATFLEPMDYLNLCFTGR
ACANQCTAFMSLVVDNRRLNVACYDPRLVRYSRIDASKLPELLPVDAIVGRVLPDVADEL
GLPRGTPVVAGLNDTQAGGIGAHAFAGEHAALSIGSSSVMIAHVRFKRTDIRHAILSMPS
PIPDTYFVMAENGIGGAALKHFIEQQVYADDPFGALPRDDCFARLQKAIDATPPGSGGVM
FMPWLAGSLAPHADASMRGGFVNLGLDATRSHLARAVLEGVAMNLRWLRGPVEAFAKRRF
SHFVFYGGGAESDAWSQIVADVLDAPVHRIEQPQYTTCVGVALLAFQRLGLLGFDDFASR
VRIRGVCEPNRANARVYAEMSAQFVEAFRRNRPIFRALQRNLRRPASAAANGLRPSG*
>GCA_000452945.1_BPMSHR5858_genomic_01355_1 group_2328
MKHDAAHHGERAISMNHSTITWAAVLALCTGFAVTNAHARDSAECRRLSDATGGADDNFR
PPLTATVTGAGRAYFHSAPASACIARRVFVVPGDTVTVYKPYRHWYQVTYVNGRTGEDVE
GWIEAGRLRLGGHLGGGQ*
>GCA_000452945.1_BPMSHR5858_genomic_01356_1 kstR2
MSAIPQAARKRGRPVKGESATLRDELILKSAKLFRTQGYERTTVRDIAAAAGVQAGSWFY
YFKTKQDILVAVMEQGMSNALARIEALDVEHLPARDAFRALVHTHLHTLVSPDHDFIPVL
LYEWKSLDEAMQAKVLKLKDRYEAVWDGVIERLQAAGEWPAPTPIDRLLMFGALNWVAQW
YKPDGALGLDALAEHAVRFLLRTAETPAPAPAGTAKRRRGVKAS*
>GCA_000452945.1_BPMSHR5858_genomic_01358_1 fadH_3
MGYRSQLRAGSFEGKVALVTGGGSGIGRCCAHELASLGAHVALLGRNADKLANVAAEIHA
DGGRADALACDIRDEDGVRHAIAQIVERHGRVDLLVNNAGGQFPAPLEQISAKGWQAVLN
TNLLGGFLVARECYVQSMKPRKSGAIVNIIADMWHGMTGMGHSGAARAGMLNFTETAAAE
WAPVRVNAVAPGWIASSGMDTYPDAIKPMLRGLPKMVPLGRIGNEAEVSAAIAFLLSDAA
SFISGACLRVDGGAPNARRHFAMQPTDTNPPFDGFHRATVPELFKR*
>GCA_000452945.1_BPMSHR5858_genomic_01365_1 group_2330
LTSIAPENLPLERLQHWERARADDVWLVQPGADGGVRRFTWRDAVDEARRVAAYLRAFDL
PRGSNIAILSKNCAHWVLADFAIWLSGHVSVPLYPTLGAESIRQVLTHCEAAALFVGKLD
AWETMRAGVPPHVRSIGLPYLHDASSVGATWDDIVRDTAPLAEHVTRAADALATIVYTSG
TTGEPKGVMLTFGALGWCVEPVFDLVAIGPDDRMISYLPLSHVAERGYVEMLSVRAGFTV
YFSDSPDTFIADLQRARPTFFISVPRLWAKFRHAVASRLPPGPVPDAMKPMILRQLGLDQ
VRLAASAAAAIEPALLRWYRDLGLELLEGYGMSEVCGVSHSCRQHDMRLGYVGTPVRDVE
SRLADTGEIEIRSPGNTIGYYKRPDLSAALFTPDGFVRSGDKGELDEAGRLKITGRVKEI
FKTSKGKYVAPSPIESRLATHPFVDACCVVGSGLAQPCALVSLSDEGRRFDGDGRRAALE
TSLGEHLAHVNASLDDHEKLRFVVVVDSVWNETSGFVTPTFKVRRNRVEARYAPFLDAWD
ARGATIVWESDTA*
>GCA_000452945.1_BPMSHR5858_genomic_01367_1 group_19880
MKSATLKAVEIEARSHTDIVDWMTGYLAARLPADSGPIDVNRQFIEYGLDSADAMKMVGD
LEDYLGFELSPSLPYQYPTIDALARALAELSAGR*
>GCA_000452945.1_BPMSHR5858_genomic_01368_1 cmaA
MTVHLESREAAAPGRDAPYGASADAIQYHYDIGNAFLTLAQERGRNYSCAMYEEGDTHEQ
AQIRKLDYHIAQIRARGAARVLDIGCGWGALLDRLVTTAGVKQAVGLTLSNEQIRYIGEQ
YPHPNVDVLLRNWQDYEPEEPFDGIISLGAFEHFAKIDEDKVQAYRHFFRKCHDFLKPGG
RLSLQTMGYGDVPRERRHSDLFIAREVFPESDLPYLADIVRACEMLFEVELVRNDRHDYV
KTMRAWFENLRAHRREALELAPLNVVERYERLYRTMSYSFDLGAFVLYRITFRRIEPNRI
DAGTQHRAALA*
>GCA_000452945.1_BPMSHR5858_genomic_01369_1 group_3198
MKHASAESPDSAGCAETPARAPAGSAADTAAPPAPDERASGHLSRASSARHLGVAALPAA
GTAAAIALWAGFGLAPRVQDIAMLAVFYVLNILGMELALHRYFAHRTFKAKPAVKIALAI
LGSLAYMGPLMWWVAIHRLHHANADRPGDPHTPQLGGRGFAGRAKGILHGHVGWLFDPSS
ARPKGWNQYANDMYRDPTLLRIHLAYDYWLLLGLLLPAALGWLLDPSWRGALLGLLWGGT
VRIFLATNAIWAVNSIGHALGGRRPFPGRDQSRNAAWLAFVTLGAGWHNNHHAFPQYAST
RLTRWQIDVTGMLIALLERLGLVWDVQHPDRDAVRERLANARRDDA*
>GCA_000452945.1_BPMSHR5858_genomic_01373_1 group_3199
MSTQHLKDAPHAAARGRRDGLDMTWLDPLRLGPRAARIARGWAARSRPDIRFCDVPAATI
RARVAGTRASGRPTIVLVCDPPSVVEHFDPLISLVAPHARIVCFEPPGFGFSVPNRHFTF
SFDDYRAAIEAMLADLNEGPYLLAFSCVWAHIALQIAAAQPALVGRLLLWQSPTWDEQVA
WARHVDARNLLATPFVGQCACAFAARKIGAGWFRQAMAKDRYREFTPMLDTSFDHGAFCC
LASLWQRFYSGTPRAVQVTQPTLLTWGAADRTHRHSDKWSIARQVPHAIRHPGFEHAGHS
PELEESSAFSQLLLSWVDAKDPR*
>GCA_000452945.1_BPMSHR5858_genomic_01376_1 group_19875
MHIAILTFEGFNELDSLIALGVLNCVKKPGWRVSIASPTPRVCSMNGVAIDAQASLREAN
DADAVLVGSGMRTREIVADAALLAQLRLDASRQLLGAQCSGTLVLAKLGLLDGVPACTDL
TTKPWVQEAGVDVLNRPFYANGNVATAGGCLASHYLAAWVIARLEGRQAAERALHYVAPV
GEKEAYVSRAMAHVTPYLAVSATAA*
>GCA_000452945.1_BPMSHR5858_genomic_01377_1 hisC_5
MAHARYKRLVDTLAADIRSGRLPPGARLPTHRKLAAAEGLALVTATRVYAELEAMGLVSG
ETGRGTFVRETALPRGLGVDQHASAAGVVDLAFNYPSLPEQAELLRGALRQLASSGDLDA
LLRYQPHGGRWHERASVARHLARRGLSVDAQRVAIVNGAQHGLAVTAMALLRPGDVVAVD
ALTYPGFKVVADAQHLELAPLPASGQGPDPDALERLCRTRRVRAVYTMPTLHNPLGWVTS
AHRRRRLVAIAHRHGLLIIEDGAYAFLADDPPAPIAALAPEATVYVSGLSKNVATGLRVG
FVAAPEPWAPAIERAIRGTTWNTPGVMTAIACGWLDDGTVERLEADKRRDAAARQAIASE
AFAGLRCIRHPASYFVWLPLADDARADRVAMTLMRERVAVSTAEPFATSAHAPHAIRVAL
GSVDPPTLRDALGKVRRAIDAHSY*
>GCA_000452945.1_BPMSHR5858_genomic_01382_1 cpo
MDTITTKDGTRIFYKDWGAGRPVVFSHGWPLNADAWDAQMLFLAQRGYRAIAHDRRGHGR
SGQPATGNDMDTYADDLAALLDALDLREATLVGHSTGGGEVARYIGRHGSKRVAKAVLIG
AVPPIMLQTEANPGGLPMEAFDGIRRGVAGNRSQFYRDVALPFFGFNRPDAQVSQATIDA
FWAQGMTGGILGQYACIREFSEVDYTADLKKIDVPTLILHGDDDQMVPIDASARLSAKLV
AGATLKVYPGGSHGICVTHADEVNADLLAFLES*
>GCA_000452945.1_BPMSHR5858_genomic_01386_1 group_19937
MKVYVLSLAMGLLVGVIYGVFNVRSPAPPVIALVGLLGILAGEQVPPLVKRLAGRAETST
TWFQHQVKPHMFGQLPSCPKAPDAPAQVAAARPPASRSGDTHHG*
>GCA_000452945.1_BPMSHR5858_genomic_01387_1 ycaC
MSDPKLEVLTPHNSQLIFIDQQPQMAFGVQSIDRQTLKNNVVGLAKAAKIFNVPTTITTV
ESESFSGFTYPELLDVYPDHKLLERSSMNSWDDQKVRDALKANGRKKVVVSGLWTEVCNT
TFALCAMAEGGYEIYMVADASGGTSKDAHDYAMQRMIQAGVVPVTWQQVLLEWQRDWAHR
DTYDAVMTLVKEHSGAYGMGVDYAYTMVHKAAQRTANAHDMLAPVPAR*
>GCA_000452945.1_BPMSHR5858_genomic_01429_1 ald_1
MKTCEQFYIGGAWRPSTGKGMIDVIDSGTEAVIGRVPEGTAEDAKAAVAAARAAFDGWAA
TPPAERARYLHKITENLKARSEELAQSITGEVGMPIKLSRAIQVGGPIYNWGAYAKLAET
FAFEEQVGNSLVVREPVGVVAAITPWNYPLNQITLKVAAALAAGCTVALKPSEVAPLNAF
ILAEAIHDAGLPAGVFNLVCGYGPVVGEALASDADVDMVSFTGSTRAGKRVAELAAAGVK
RVALELGGKSASVILDDADFAAAVKGTVSACYLNAGQTCSAHTRMLVPESRYEEARDLAK
AAAQAYVAGDPRDDATRLGALASSVQQKRVQDYIRRGIDEGAELVTGGLGMPEGITRGFF
VKPTVFGRVKPNATIAQEEIFGPVLSILTYRDEDDAVRIANDSPYGLGGAVWAGSDERAM
RVARRIRTGQVDINGGAWNMAAPFGGFKQSGIGRENGVYGLDEYLEYKSMQLRPHKRA*
>GCA_000452945.1_BPMSHR5858_genomic_01430_1 group_3206
MTGSIPFPPLMIRGRSLLPIVQGGMGVGISAHRLAGSVAREGALGTIASIDLRHHHTDLI
ERCKQHPDRETMEAANLEALAREIQRAKTWGEGRGMIAVNVMKAVRSHADYVRIACEFGA
DAIVMGAGLPLDLPDMTQGHDIALIPILSDSRGIALVLKKWMKKGRLPDAIVIEHPARAG
GHLGVTSLDDMDDPRFEFARVIDETRQTFATLGLERERIALVVAGGINSHEAVRAALAEG
ANGVQVGTPFAVTEEGDAHPNFKRVLANAKPDDIVEFLSVTGLPARAVKTPWLERYLRHE
TRIRAKIGALKQRCPSALECLSVCGLRDGIERFGHFCIDTRLAAALRGDVANGLFFRGRE
ALPFGQAIRSVRDLLELLLTGTAPEAAANRPTFSLS*
>GCA_000452945.1_BPMSHR5858_genomic_01436_1 potH_1
MNALSSLFAWPVRKFGLTGRTAVIAGPFFWLVLFFLVPFVLVVKISFAQLQLGIPPYTEL
TALADGVLSIKLNLQHYAFLFSDSLYFATYVNSVVVAAVTTLLCLLIGYPMAYYIARSNP
GTRNLLMMAVMLPFWTSFLIRVYAWIGILKNNGLLNNFLLWTGLISTPIELYRTNVGVYI
GMVYSYLPFLVMPLYAHLVKMDLRLLEAAYDLGAKPWKAFVQITLPLSKNGIIAGCLLVF
IPAVGEYVIPELLGGANTLMIGRVMWNEFFNNADWPMASAVTCAMVLLLLVPMAMFQHFQ
AKEQEGHRR*
>GCA_000452945.1_BPMSHR5858_genomic_01439_1 group_3207
MEIRKLTDTLSVSPQLVAADLPAIAQAGFRAIICNRPDGEGPDQPTVAEIDAAAQRFGIA
VHYLPVDSGKATDAQGEQFGVLLDTLASPVLAYCRTGMRSATLWGLSQAGLRPLNDIVAT
GRAAGYDLAALGRRIMQRSQSATAQTVEASHDIVIVGAGAAGIAVASSLLAREPSLDIAV
VDPADTHYYQPGWTMVGAGVFQPGTTARRMAAVLPRGVHWIKAAVAAFEPDTHTVVLDGC
RLVRYRQLVVCPGLKLDWHGIEGLPETLGRNGVTSNYRYDLAPYTWQLVRAFAGRRALFT
QPPMPIKCAGAPQKAMYLSADHWRRTNRLDAIDIEFFNAGAALFGVAEYVPALMEYVKRY
GIALNFGHNLIAIDGERKRATFARALPEGGTETLTREFDMIHVVPPQKAPDFVRASPLAD
AAGWIDVDPATLRHRRHADVFALGDVTNTTNAKTAAAARKQAPVVAHNVLAALGKAQGDA
AYDGYGSCPLTVERGKIVLAEFLYGGKLAPTFPAWLIDGKRPSRLAWLLKERLLPPLYWQ
AMLKGREWLAKPEIVG*
>GCA_000452945.1_BPMSHR5858_genomic_01448_1 mdtA_1
VNRSGSRAALLIGVALIAAACHRKEAAPSAPRPVVAVPAQADGAAAAVSLPGEIQPRYAT
PLSFRIAGKLVERKVRLGDIVKKGQVVALLDTSDVARSAASAQAQLDAATHALTFAQQQR
ERDRAQARENLIAPAQLEQTENAYASARAQRDQAAQQLALAKNQLQYATLVADHAGYITA
EQADTGQNVSAGQPVYQLAWSGDIDVVTDAPESELPRLAPGRTARVTLPALPGRAFDARV
REIAPAADPQSRTYRVKLTLAQPDAAVRLGMTANVALSPDAPSAANAQPRFTLPSTALFH
AGNAPALWIVRKQDDTLELRRVDVVRYNERTVTVSGGIQPGERVVLQGVHTVSAGEKVRA
VPPLHPEDVAS*
>GCA_000452945.1_BPMSHR5858_genomic_01451_1 group_2337
MSTLTSPPLSVAAPSMRGRLNGVLFVGLFALAVTSVAQLPAVAHLGLSPLIVGIVAGALY
GNLLREGMPASWAAGVDFSARKLLRIAVAFFGLRVSLQEIAQVGVPGLAVSALVVASTLA
VGTWAGIKLMKLDRDTALLTAAGSAICGAAAVLAFESALRSKPHQSAMAVGSVVLFGTLS
MFAYPLAYRAGWLHLDATGLGLFFGGTIHEVAQVVGAASDVGPEVARVATIVKMTRVMLL
VPVLLVLGAWLARSARRSTAAGRGAPRKLAVPWFALGFLGFVIVNSLQMLPAAAIGTLNA
LDTFALTMAMTALGIETRMSQIRAAGPRALMTGLILYAWLVFGGYAIVWATQRWLG*
>GCA_000452945.1_BPMSHR5858_genomic_01566_1 group_213
MSNVPKSPAQKRAGTPGEPQPAGAAAPRPPRPRQAPPAQARAPRTERRDAGPEAARAPHA
PRTRCAPNPVPPITFAESLPVSGKRDEIARAIAAHPVVIVCGETGSGKTTQLPKICLALG
RGLGAGGAGLIGHTQPRRLAASSTGRRIAEELGTPFGEVVGYKVRFTDNLAPGASVKLMT
DGILLAETQTDPLLKAYDTLIIDEAHERSLNIDFLLGYLRQILPRRPDLKLIVTSATIDA
ERFARHFGSDERPAPVIEVSGRLYPVEVRYRPIADDRPAAVRHAEGASSGRDRAKSAREA
ERDLMDGIVDAVDELCREGPGDVLVFLPGEREIRDAAEALRKHHPPHTEILPLFARLSAA
EQERVFKASNARRIVLATNVAETSLTVPGIRYVVDTGLARVKRYSYRNKVEQLQIEPISQ
AAANQRAGRCGRVADGICIRLYEESDFAGRARFTDPEILRSSLASVILRMKSLHLSAIES
FPFIEPPPGRAIADGYQLLNELGAVDDENALTPLGRELARLPLDPRVGRMILAARDQQAL
REVLVIASALSVQDPRERPVDAQEQADQAHRRFADERSEFLQWLRIWAWFEEAVAHKKSN
RQLVDACRQHFLSHLRLREWRDVHSQLLTVVREHGWRLNEADATFEQIHLSLLTGLLGNI
GFKAEDEPHYLGARGIKFHLWPGSALVKKAGRWVMAAELVETSRLYARCIAKIEPEWIER
IGAHLLKKSLSEPHWEKRPAQVAAFERATLYGLTIYHRRRVAFGRQDPARARELFIRGAL
VDGEFDTKLAFFAHNRKLLADIEQLEHKSRRQDVLVDDELIHAFYDQAIPAGIHTGAAFE
RWYRDEVSKSGQPEDKLRLLYLSRDDLMRHEAAGVTTELFPKRVTMAGVEMALAYHFEPG
SPRDGVTLAVPLFALNQVDARRAEWLVPGMLKEKAHLLLKSLPQKLRRHCVPLPEYAAGF
VERAGRERFGAGGLVDALIADVREQTQVATKTSDFKLETLPAHLFMNFKVIDEHGRQLAM
GRNLAQLRAELGAQAQQHFQKIAAAATLAPAGEPAAAAAGASGARARRVPLGAPPRAAEP
AAQAGAAAGATALYENLTTWNFGKLPELLEIRRRGETLFGYPALVDRGTHCDVEVFDSPD
EAARIHRAGLRRLFALQLKEPIKYLEKNLPGLREMAIQYMSLGTQDELRDQLIATALDRA
CLQEPLPADDASFHARRDEGRSRLNLLAQEIARLVGQILAEYAGLAKKLAQAKPFPAAHA
DMQGQLAALVGKRFVVDTPYAQLAHFPRYLKGIALRIDKLKADPARDARQAAELQPLAQH
YQRSVAQRGGVADARLAEFRWLLEELRISLFAQELRTPMPVSVKRLYKVWESMQR*
>GCA_000452945.1_BPMSHR5858_genomic_01569_1 group_3223
MNDLLRSFGRALASALHPRMLWLTFLPFIVSAVVWGVLLWFSWQPLVDAARAALDGFALT
ALLYKFFNAIGASQLHAVVAPFVVVSLAIPLIVLTVLLLIATLSMPAVVSHLSKRQFASL
EMKRGGSLVGSVFNSLWAAFASIALLIVTVPLWLIPPFFALLPPVIWGWLTYRVMTYDAL
ALHASAGERRALVRKHRWPLFGIGVASGLLGTVPTFIWVSSIWMMVLFPFVAAAMIWVYA
FILVFSALWFGFYCLRALQDERAAAANAISTDSTRQA*
>GCA_000452945.1_BPMSHR5858_genomic_01571_1 group_19955
MSTLEQLYAQADTRRAQGNLNYAGALSPAEAFELLQLDPNARLVDVRTRAELDWVGRPLV
GDGQYAHVEWSRYPGGVPNVEFIDQLRTLAQPDQPILFLCRSAARSKLAAVAAAQAGFTK
AFDLLEGFEGAKDAQGHRKTVEGWCHRGLPWIGA*
>GCA_000452945.1_BPMSHR5858_genomic_01574_1 glnG
MKPIWIVDDDQSIRWVLEKALARESFATRSFANVRDALAALDQETPQVLVSDIRMPGGSG
LELLQAMHDRLPGLPVIIMTAFSDLDSAVAAFQGGAFEYLAKPFDVDKAVELIRRAVEES
LRGGVPDDDKLAEAPEMLGQAPAMQDMFRAIGRLSHSAATVLITGESGTGKELVARALHR
HSPRANGPFIALNTAAIPKDLLESELFGHERGAFTGAQTTRQGRFEQAENGTLFLDEIGD
MPFDLQTRLLRVLSDGQFYRVGGHSPLRANVRVIAATHQNLEARVRQGLFREDLYHRLNV
IRLRLPPLRERSEDIALLTRHFLQKSARDLGVEPKRVSDEALAYLTSLPFPGNVRQLENL
ANWLTVMAPAQTVEIKDLPPDLVATQTASGSMMPVEATVEADAQASGLGVGAAEPHAPSA
VAAVPATAGAPTAVGAPAGHPLWEHGLRTEVARLLRENSADVMDALARRFEAAVIREALD
FTRGRKVEAAERLGIGRNTITRKIQELHLES*
>GCA_000452945.1_BPMSHR5858_genomic_01579_1 narJ_2
MSIYPILSALLTYPEQPLLDALPEIERALDAQPGARAALAPLVESLRTAPLIELQERYVA
TFDRTPSHSLHLFEHVHGESRDRGQAMVDLLDEYRRHGFEPVGNELPDYVPLFVEFLGAI
AGDGDGAHAAHLLGDAIDVLAALGERLARAQSPYAGAFDVLRACSPVEPRPVAEPPPRTM
DEALERFGPGPDGVEPLLSPAADASLAQPIRFHPPRRAATPAPRA*
>GCA_000452945.1_BPMSHR5858_genomic_01580_1 narH
MKIRAQVAMVLNLDKCIGCHTCSVTCKNVWTSRDGVEYAWFNNVETKPGIGYPREWENQD
KWQGGWKRNDAGKLEPRQGGKLKILANLFANPNLPAIDDYYEPFTYDYAHLQNAPLSQTP
PTARPVSAITGQKMEKIEWGPNWEDDLGGEFKSRSRDKLFEAVQKEMYSTFENTFMMYLP
RLCEHCLNPTCVASCPSGSVYKREDDGIVLVDQDKCRGWRMCVSGCPYKKIYFNWQTGKA
EKCVFCFPRIEAGQPTVCSETCVGRIRYLGVMLYDADRIEQAASVADPQKLYEEQLRVFV
DPHDPAIERQALADGVPQSWLDAAKRSPVYRMACEWKVAFPLHPEYRTLPMVWYVPPLSP
IQGAAQSGHMGMNGVIPDVRSLRIPLRYLANLLTAGDEAPVAAALDRMLAMRAYKRAQTV
HGAHDAQVLEQVGLTAEQVEDMYQLMAIANYEDRFVIPSSHKEMAEDSFDEKGSCGFSFG
NGCSGGTSDGTLFGRKKETAVSYAALPTVTRKKASAA*
>GCA_000452945.1_BPMSHR5858_genomic_01586_1 prlC_2
MSVSANANPLLDFSGLPRFGEIRPEHVTPALDTLLANANDAVEQAARPDTPATWADVVET
IEQATEPLGRAWGVVGHLNAVADTPELRAVYGENLPRVTEFWSSVGQNLALYEKYKAIAA
SPEYAALSAERKKILDNALRDFRLSGAELPEDRKPRFAQLQEQQAALSKSFSDHVLDATN
GYAYYARSEDELAGLPDDAIAAAREAARKEDKAGWKFTLHFPSYFPVLQYARNRAMREAM
YRAYVTRASELGPQYGDGKADWDNTTIIADQLALRREEATMLGYHDFAEVSLAPKMAESP
QQVIAFLEDLAKRARPFAEKDWDELRAFASSELGLAALEPWDVAYAAEKLRERRYAFSEN
EVKQYFPEPAVLKGLFTVAETLFGVRITRDDAPVWHEDVRFFRVENRDGSLVAQFYLDLY
AREGKRGGAWMDDARSRAKRGDGRVQTPVAYLTCNFSAPIGGKPACFTHDEVITLFHEFG
HGLHHMLTRVDEIGVSGINGVEWDAVELPSQFMENFCWEWDVLKSMSSHVDTGDALPRVL
FDKMLAAKNFQSGLGTLRQIVFSMFDMLLHVDFDPAGKTSVNEFAREINERFHVIAQAPF
SRWPNTFSHIFAGGYAAGYYSYKWAEVLSADAYAAFEEAAGSAGSVLDATTGARYRREIL
EVGGSRPAMDSFKAFRGREPNIDALLRHSGMADDAQARA*
>GCA_000452945.1_BPMSHR5858_genomic_01589_1 sasA_11
MWNIAAPHRLDPRLPCVVYRQFRFRFNLSCTPFVQDALSYNPAMLTDRLFARSARPSGSP
AESQPSRWHHGPWWSNSYLLTPLLSILVFLVVMSLILWSLNRREEQQQEDTLFRNVAWAQ
QQIRLSMTSAQEQLQAFSRDIAAGRIDEHAFQATVGDVMQAHPEILYLNWYTSPGTKRWP
TMQLPLLGQRLAKPNDAQMDEVVRGAYAQARGTRRQSYSPLVYDDFGNGYLTLQTPVIRE
REYLGSIAAVFSVEGILKHDIPPELSAKYKISITDANNRELASTSSRPRLPRDAHYDLPL
DPPGQGLTVRVYAYPQTTNLTNNTLVWLVAGLSCFVLWSLWSLWKHTRQRFEAQQALYAE
AFFRRAMENSVLIGMRVLDMHGRITHVNPAFCRMTGWDESDLVGKTAPFPYWPRDAYPEM
QRQLDMTLRGKAPSSGFELRVRRKDGSLFHARLYVSPLIDSAGRQTGWMSSMTDITEPKR
AREELAAAHERFTTVLESLDAAVSVLAADEAELLFANRYYRHLFGIRPDGHLELSGGGFD
TAQASSDSIDMVDAYAGLPAAALTESTADAQEVYVESIQKWFEVRRQYIQWVDGHLAQMQ
IATDITTRKKAQELAHQQEEKLQFTSRLMTMGEMASSIAHELNQPLAAINNYCSGTLALV
KSGRASPETLAPALEKTAQQALRAGMIVKRIREFVKRSEPKRQPSRVADIVADAVGLAEI

EARKRRIRIVTEIRARMPIIYVDPVLIEQVLVNLMKNAAEAMQEARPQAENGVIRVVADL
EAGFVDIRVIDQGPGVDEATAERLFEPFYSTKSDGMGMGLNICRSIIESHRGRLWVVNNV
EPDGLVSGATFHCSLPIGEPEDLGRGSETSPSQTVTGEI*
>GCA_000452945.1_BPMSHR5858_genomic_01592_1 lpdA
MSLIEVKVPDIGDFSGVDVIEVNVKPGDVIEKEQTLITLESDKASMEVPSDVAGTVKEIK
IKAGDKVSQGTVIAIVEASAGAAAPAPAKAPEAPKPAAAAPAPAAAPAPAPQAGSYSGSA
DIECDMLVLGAGPGGYSAAFRAADLGMKTVLVERYSTLGGVCLNVGCIPSKALLHTALVV
EEAQALASHGITFGKPQVELDKLRDFKGGVVKKLTTGLAGMAKARKVEVVTGVGAFVDPY
HMEVQGENGKKIVKFKQAIIAAGSQAVKLPFMPEDPRVVDSTGALELRQLPKRMLVIGGG
IIGLEMATVYSTLGAEIDVVEMMDGLMMGADRDLVKVWEKYNAKRFGNVMLKTKTVGAQA
KEDGIYVTFEGEKAPAQAQRYDLVLVAVGRSPNGKKIGADKAGVAVTDRGFIDVDKQMRT
NVPHIFAIGDIVGQPMLAHKAVHEGHVAAEAAHGEKAYFDALQIPSVAYTDPEVAWAGKT
EDQCKAEGIKYGKAVFPWAASGRAIANGRDEGFTKLLFDEETHRVIGGGIVGLNAGDLIS
EVCLAVEMGADAEDIGKTIHPHPTLGESIGMAAELYEGVCTDLPPQRKK*
>GCA_000452945.1_BPMSHR5858_genomic_01595_1 xynR_3
MSNPNPDSKTSIQVIERMMRLLDALAAHSDPVSLKELAQRTELHPSTAHRILNDMVTCRL
VDRSDPGTYRLGMRLLELGNLVKARLSVRDAALMPMRELHRLTGQTVNLSVRQGDEIVYI
ERAYSERSGMQVVRAIGGRAPLHLTSVGKLFLAADETSRVRAYATRTGLAGHTQNSITDL
PKLERELTLVRQQACARDNEELELGVRCIAAGIYDDSGKLVAGLSLSAPADRLQDAWLGQ
LSRTALTISESLGYRPETAKEMDAPAQKQA*
>GCA_000452945.1_BPMSHR5858_genomic_01599_1 yfkJ
MKSVAICFVCLGNICRSPTAEGVMRHQVAAAGLDGAIEIDSAGTGDWHVGEAPDARAQQA
ARARGYDLSALRARQIGDADFERFDLVLAMDGANLAALRKRCPPQYRGKVRLLMEFAGDG
SAGDIADPYFGGARGFEQVLDQCEAACRGLLDSLRETAR*
>GCA_000452945.1_BPMSHR5858_genomic_01600_1 iscR
MRLTTKGRFAVTAMIDLALRQEQGPVTLAGISQRQRISLSYLEQLFGKLRRHEIVESVRG
PGGGYNLARRAQDVTVADIIIAVDEPLDATQCGGKGACEGSKQPDGHCMTHELWSTLNQK
MVEYLDSVSLQDLVDQQRAREGTPAVLRDKRKPEPVAASVEPVRSVPLGPNSVFNIANS*
>GCA_000452945.1_BPMSHR5858_genomic_01603_1 iscA
MAITLTEKAAQHVQKYLTRRGKGLGLRLGVRTTGCSGLAYKLEYVDELTPEDQMFESHGV
KVFVDPKSLAYIDGTELDFAREGLNEGFKFNNPNVKDECGCGESFRV*
>GCA_000452945.1_BPMSHR5858_genomic_01608_1 group_3232
MDNQNTTTQQRQRLHPLVATAAGAVIVASLAATAAITGVFPKASSTNAQNAQTQAALIAS
QPAVDTAAAASAALAAQAAQQAAQQPQQTVAQAAPKPTSHQTTRHHHKTSAAPQPPQYAQ
QPYPQQQQSTYCASCGTVASIVPVRTAGTSSGLGAIGGAAAGGLVGNQFGRGNGRTAMTI
IGALGGGLAGNAVEKQVRSETDYQVQVQMQDGSTRTFTYHNPPPFGQGQRVRVENGTLVG
A*
>GCA_000452945.1_BPMSHR5858_genomic_01610_1 prfB
MFSNPADPNNCFIDIQAGAGGTEACDWASMLLRQYLRYCERKGFKAEVLEESDGDVAGIK
NATIKVSGEYAYGYLRTETGIHRLVRKSPFDSSGGRHTSFSSVFVYPEIDDSIEVEINPA
DLRIDTYRASGAGGQHINKTDSAVRITHMPTGIVVQCQNDRSQHRNRAEAMAMLKSRLFE
AELRKRQAEQDKLESSKTDVGWGHQIRSYVLDQSRVKDLRTNVEMSNTKAVLDGDLDDFI
SASLKQGV*
>GCA_000452945.1_BPMSHR5858_genomic_01625_1 eno_2
MSSVPISSSEPLRCARSAALAALGESDPARKVALVAELHAALADGRAAVFPERELSAPAR
GVPGRPARPELVEPRRLERRSMRSPQGRAVLLHALAHIEFNAINLALDAVWRFARMPTAF
YADWLKVAAEEAHHFSLLAARLAEFGHAYGDFPAHDGLWEMCERTAGDVLARMALVPRTL
EARGLDASPPIRARLQQAGDHASAAILDVILRDEIGHVWIGNRWFRHLCDAAGLDPHPTY
ERLAGQYRAPRLRGPFNFDARRAAGFNDDELNALVAQDADPNG*
>GCA_000452945.1_BPMSHR5858_genomic_01626_1 menH_3
MHSTQSVSDFVTVRGVKLHVRRWGRPDAPTLYMLHGWMDVAASFQFVVDALAGDWQVIAP
DARGFGLSDWPVAAQGGGHYWFHEYLVDLEALIDHYTPDGEVNLVGHSMGANVVCLYAGA
RPQRVRRVVDLEGFGLAPARAEQAPRRLAQWLDELRAPPELKRYASLEDVAARLVKTNPR
LEPRRAAFLAAHWSTRDADGWYRLLADPAHKLRGPLLYRLDEVMAIWSKVRAKVLHVEAV
DSPTLAFLAGEIPIAEFKARFAAFADWREKRVEDAGHMVHHDQPEQIAALIEAFCA*
>GCA_000452945.1_BPMSHR5858_genomic_01631_1 group_2351
LKTPAIEIERVPGAGAHGAATEPSAWVRRWSHLVPAGGAVLDIAAGHGRHAHWFARRGHP
VRALEREPAALASLGALPGVRAQAADLEGAPWPLADDARFAAVVVTNYLHRPLLPRLVAA
IAPGGVLLYETFAQGNQTVGKPSNPAFLLAPGELLDAVRGQLRVVAFEDGFAAAPREAFV
QRICAVREHAAPQEGAGFPRYGLAG*
>GCA_000452945.1_BPMSHR5858_genomic_01633_1 group_19819
MKHSAFSSRAIQVSVLALALAALAGCDTLNDYLAPDRVNYKSTGSAPPLQVPQDLTAMPL
SPSYVAPPTNSGLGSAPTRAVTAAGNATEGQPSAQDPFGMHVERDGDRRWLVVDGRTPEQ
LWPQLKEFWQDNGFALKTDAPSTGIMATDWAENRANIPDDWFRRTIGRVIDFAYSSGTRD
RFRTLVTRTADGNTDISITHSAMEEKLTGAQGGTSSRWEERPRNPVLEAVFLAKLMEKFG
LTDAQAKQLLADARPATAPATVVGSGAASTLDLAESFDRAWLRVGLALDRTNFTVDNRDR
AKGVYYVRYANSMEELKRDGLFGKLFYGGPTAAKPGKEFLVNVRAQGDAKTQVAVIDANG
QIDTSSDAQRIISLLHAQLN*
>GCA_000452945.1_BPMSHR5858_genomic_01634_1 yycJ
VRFASLGSGSEGNALVVEASNGATTTRVLLDCGFSAKELERRLARIDLRVDDLDAILVTH
EHGDHVGCALTLARRASLPLYTSWGTARAVGADEADVDLHVLWSDETAAVGDLCVLPYTV
PHDAREPLQFVFSDGATRLGVLTDVGMSTPHIGAVLSGCDGLVLEANHDVAMLAASRYPA
SLKARIGGSHGHLSNDAAAAILAELDRSRLRHLVAAHLSQQNNAPALAQAALAAALGGQA
DDVLVASQDDGFGWLSLS*
>GCA_000452945.1_BPMSHR5858_genomic_01636_1 slyD
MKIAKNTVVSVAYKLSDAQGNLIEESDEPMVYLHGGYDGTFPKIEEQLDGQEPGYQAQIQ
LEPQDAFGDYDPELVKIEPRDRFPEPLEVGMQFEGTPEDGDEEIDSLIYTVTDIAEDKVV
LDGNHPLAGMALRFALTVKDVREATEDEIEHEHAHGAEGLEIVDEDEDDDNEGNGSARTL
H*
>GCA_000452945.1_BPMSHR5858_genomic_01643_1 lpxH
MLQETPPRSVFAGVPGERGLAHAARPFLFVSDLHLSDAIPHTVAAFERFVRVTADSADSV
FILGDLFEYWIGDDILEHDPFAARMAALLHTFSERGIALYVMHGNRDFLLGKRFMKAAGA
LLIPDPSVIVAFGNRIVLAHGDAQCTADRGYQWFRRFARNGVAQRLFLAWPFAWRRALAQ
RMRASSETGRIRPASPRYDVTRKGVAALLRRSGARTIIHGHTHKPARHVEADGVRWVLPD
WELDHGLRRGGYLRVDAEGIRALPLD*
>GCA_000452945.1_BPMSHR5858_genomic_01644_1 group_3235
MVELHTNHGVIKLELDEAKAPKTVENFLNYVKKGHYDGTIFHRVINGFMIQGGGFEPGLK
QKPTDAPIANEANNGLKNDTYTIAMARTNDPHSATAQFFINVNDNEFLNHSSPTPQGWGY
AVFGKVVEGQDIVDKIKAVKTGSKGFHQDVPNDDVVIEKAVVV*
>GCA_000452945.1_BPMSHR5858_genomic_01653_1 group_19709
LNDTPFTPDSREAQDSTAEAAKTAARAATEEIFVFNGDADGLCALQQLRLAEGAGAMLVT
GVKRDIRLLSRVSAGPGCRVTVLDVSHERNRGDVARLLAAGAALRYFDHHFAGELPNHSR
FHAYIDTAPDVCTSVLVDRYLRGRHAGWAVVAAFGDALPDVGRALASARGIAPGALESLA
QLGRCLNYNAYGEDLDDLHFSPYALADAMLPYADPLDFIAATDVMPVLADGYRDDLQRAR
AIEPALVAPGATMLVLPAEKWARRVSGVLANERMAEDGCARALAILSPRRDGGYVVSVRV
RAGSALGADEFCRRFETGGGRKLAAGIDRLPETDVERFAAHFRATFAA*
>GCA_000452945.1_BPMSHR5858_genomic_01654_1 group_2693
MTIGSKTKLALGIALAMPAAAWAYTFYALGDSLTDNGRVVRLTGILPNATSTIFRGGRSS
NGPVWAEYLPGLIGARFAPDDDYAINGALSGHGGYLNIIPTRPTWRTLPGFVDQVEQLIA
AHPRLRRDDLVGVWIGTNDQDLTKASLNGIEPFLGVPRPANIAEMSAYTLTNLNAQLQRL
IGAGARQFVILNLNDAGGTRPGYIDYNGKLPEDLARFSRQGVNVHLFDVSALLNQMRRNP
SAYGLNDTPDVQCRYVPSCSGGSIELQNTYLTADGTHMMTSVHEYIARYLANQLNAPAAI
STAPGLGLDVARAGALSALNAADGGPLGPARVAWSDRLSLFADVGYTRNFHGAAGGMNAF
DSDVEMFSIGADYKLSEASRAGALLSSGNANGSLAGGQGRIGLHAYRLGVYHAFERASLF
VRAYAGAGWSRYRLDRAAVLPGAVRASTSGFDFGALVKAGYLFALGGVRLGPVADVGYTQ
LVARGYTEDGDPILAQNVGVQRLKGVSAGAGVRFAAPLAAIGLRAGELSAEAQLRHDAFG
DRTLVTAQRYAPGLPIGTAVDGASATYERLSVALTVNPAKRWRGKLVVQTDLGTAQRRSY
TLLALLGGTF*
>GCA_000452945.1_BPMSHR5858_genomic_01681_1 plcN_3
MNARRRFLQGTATTGIAAATLAAFPLSIRRALAIPANNRTGTIRDVEHIVILMQENRSFD
NYFGTLRGVRGFGDRFGIPLPNALTVWQQRNATGALVLPYHLDGSKGNAQRVSGTPHSWD
DGQNAWDGGRMYQWPRYKNTASMGYFRESELPFQFALANAFTICDAYHCSMHAGTNSNRM
FMWTGTNGPTGAGVASVVNEWDDIGPSTFGYEWKTYPERLQEAGVSWKVYQNMPDNFTDN
ALAGFRQYRRANEASGKPVSNSDKVVSPAYDPASDDVRNPLYKGIANTMPDGGFLGAFKE
DIRAGKLPQVSWVVAPAAYSEHPGPSSPVQGAWYIQEVLDALTATPDVWSKTVLLVNFDE
NDGYFDHVPSPSAPSLNPDGSPAGKTTLPEADIAFERFTHPKPPGTKSQPQPDARVYGPG
VRVPMYVISPWSRGGWVNSQVFDHTSTLRFIEARFGVREPNISAFRRAVCGDLTSAFNFV
RPNNEPLPTLAGRKTRAQADALRAAQQQMPQIVPPANGLLPRQETGTRPSRALPYELHVN
AVARTLDGTLRLVFANTGQAAAVFHVYDKLNLGRLPRRYMVEAGKQLDDVWAAMTDNGGK
YDLWVLGPNGLHRHFTGDMNRLRAGGAPVPEVRAEYDRQHGNVQLLLRNEGARECVFTVR
AKAYRDDGPWTFKVKGGAERKHHWKLDGSGNWYDFVVTCDADTGYSRRFAGRVETGEHSV
SDPAMGFGDRF*
>GCA_000452945.1_BPMSHR5858_genomic_01686_1 dctD_2
MMNADNAYGREAVYVIEDDAAVRLGCSQALALEGIGVREFEDAESALAALKRDPPAAIVS
DVRLPGMGGLALLDSMKAHPRHADIPVILMTGHGDVAMAVGAMRCGAYDFIEKPFHSDRL
VDTVRRALEHRKLVIENRSLRAQLSGERPLLGSAPAMQRVHALIDAIGPTSADVLIIGET
GTGKEVLARALHAASRRTGPFVALNCAALPEAVFESEIFGHEPGAFTGAQQRRIGKFEYA
SGGTLFLDELESMPLSLQAKLLRALQERSIERLGSNVSVAVDVRVIAAVKQDLKQLAADG
LFRSDLYFRLNVASIELPPLRRRTDDIPELFSHFLRAAAVRFEKPEPVWTQEDMMRWQLY
DWPGNVRELKNTAERFCLGLEDGLPRSPAGFDSLASRMVSAERAYIEEALRNAGGQVAKA
AELLGLPRKTLYDKITRHGIDMTAFRCT*
>GCA_000452945.1_BPMSHR5858_genomic_01690_1 group_2354
MRTTRTLSISLPYEMADYIRAKVSTGEFATESEVIRDALRTLIARDRVVEAWLRDAVNSG
EFALDPDAQWAADKDEAKPARPRRRP*
>GCA_000452945.1_BPMSHR5858_genomic_01710_1 group_349
MSRSVFEAVRLDDVVLQPSPIDPGWIIEGAPVARSGRWSCSPDSTTSMWVWDCTAGRFNW
YFDCDETIHVIEGEVIVSSDGDAPRTLRAGDAALFYAGSRTEWHVPKYVRKHAILRPHIA
GPALFVHKAFRKLASALR*
>GCA_000452945.1_BPMSHR5858_genomic_01712_1 araB
MESSNRPARTHEASAATLLVDCRNALGEGATWCDAAHALYWVDIEGARLWRWRAAGAHGG
ERCDSWEMPERIACFALTGDPDVLLVGLASRLAFFDTRRRALTPIVDVEPDRPTRLNDGR
CDRAGAFVFGTKDESGGASPRAIGGYYRLNADLSLQRLALPPAAIANGIAFSPDGSAMYF
CDSPTREIQVCDYRPGGDVDRVRSFVRLADAHGEPDGSTVDASGGVWNAQWGGARVVRYD
AQGVETDRIAVPTPQPSCVTLDAAGRLYVTSARVGLDDGALAGNSGAGGVFVAHTRDPGG
ATPRFALARHA*
>GCA_000452945.1_BPMSHR5858_genomic_01714_1 gbpR_1
VRYEPLTGEPVAAVVRAGHPLLERAPLARADAQRAAWVVPPAGSVLRRRVERMFQRASLT
PPANVVETAALLFVTRVLERSDMIAVLAEDIARYYATHGIVAMLPLEMDGRMDDFGIITR
TRCTRPRRA*
>GCA_000452945.1_BPMSHR5858_genomic_01718_1 uvrY_2
MIRLLMVDRHAVLRDGLRHILEKSGEFEIDGEAADAASAFELAGRGCAEIALLDQTTLGR
ECAELIRRLRRAAPSTRILVLAARASQRHAAAAFQAGASGYLTKQSGCEELLAALRKIAA
GGVYVSMHVAEQFAENLHSPSDAPPHERLSRREFEVFRRLAAGDTPADIAQALGISAKTV
RIYKAHVLDKLELPNETALVRYAVAHRIGDPL*
>GCA_000452945.1_BPMSHR5858_genomic_01724_1 clcB_2
MPNPIACASPVSPIRSAPAAARGPLARIAAVTLLTGAGAGLGGMLLALLLHAIQHLAYGY
DVARAIGDEGFLEGVTAASPLRRFIVLTGCGVLAGGGWWALYRFGAPLVGIRRAVRAEDP
RMPVASTTAHALLQIATVALGSPLGREVAPREIGAMWAGWLAHRAGLPVADARMMVACGA
GAGLAAVYNVPLGGALFVLEGLLGTFAWRALVPAVATSAIAALVARIGLGDAQQYHVAPI
AAGAPLFAWSAVCGPLFGVAAHGFVRLTAHARARAPKDGRLPALALVNFAIVGALATVFP
QLLGNGKGPAALGFDGQLAIGFAATLLVLKIAITWSSLRAGAEGGLLTPGLAHGALLAIV
LGGLWRAAWPGEPLGAFAVVGASAFLAASMRMPLTAIVLVVEFTRVGHDVLIPMLIAMVG
AVAVFRLLERRRERKQSRLHEAIGEVAR*
>GCA_000452945.1_BPMSHR5858_genomic_01727_1 kce_2
VASPRKVIITCAPTGAIHTPSMSPHLPVTPREIEEAAIAAAEAGAAILHLHARDPADGRP
TQDPAVFAEFLPRIKARTDAVINITTGGSPHMTVAERLRPAHHFQPEVASLNMGSMNFGL
YPMLERFREFRHPWEREHLEKSRDLVFKNTFADIETILASCGANGTRFEFECYDISHLYN
LAHFVERGLAKPPFFVQSVFGLLGGIGAHPEDLAHMRRTADRLFGADYVWSILGAGKHQI
PLATIGAAQGANVRVGLEDSLWIAPGRLAESSAAQVRKIRQVLEGLSLEIATPAEARAML
ALKGGGAVNF*
>GCA_000452945.1_BPMSHR5858_genomic_01732_1 group_2995
MHAVSRRPSLSSAVFYRDPRAALEWLERAFGFARGIVVSDAQGRIAHAELSFGDGVVMIG
AAGWSEFAFSPESLDGKNTQCVHVQLESDLDAHCERARAAGAVILQEPADQFYGDRTYRA
RDPERHVWTFARTVRRVSREEAERHGGFSIEGGR*
>GCA_000452945.1_BPMSHR5858_genomic_01734_1 group_3246
LKSIVVTAAAALVLAAPVVSFAQSSNGSLTRAQVIQELVDLESVGYQPSRGNDNTYPDDI
QAAQRRLDAKRLATRKTAETAYGPAVAGTAQAGKPAAGAR*
>GCA_000452945.1_BPMSHR5858_genomic_01736_1 aaeB_1
MKHEPALQRFPIDPHRWQERLRSAGRLARDWAGRDGLVWLHLAKTVAAALLAMGIAMLLD
LSQPRIAMTTVFVLMQPMSGMVLAKSFYRVVGTGVGLVAALALGGLFAQQPELYMAGITL
WVAGCIAAAVRNRHFRWYGYVLAGYTAALIGLPAVMAPNTLFLSALTRAAEVAVGIFCSG
AVSALVLPLSSSDALMRTLSARHADFVAFAASTTAGAVERGEFERRFADFVDGIVGFEAT
RAFATFEDPYIRARSRRLARLNSEFMNACARLHAFHQLLKRLRANHAAEVLTAIAPHVDA
LSGVLYALRADLARPRARSAPSPSPSPALVALSAYLAALPKRARASRRGLETLAPAGVLD
FDTAIELLYRFVDEFLGYAETHASLALDSHVLERSVTRYAVKTNRYFVGFTFLRTLVAMG
AMSAFWIASEWSSGSLAVIGTAIACALSSTAPRASRFVAQMAAGAALATLTGYLYVCRVY
PNIDGFPLLCAALAPVLAAGAYLATRPGKSGYGIGFVVFFCLLAGPDNVIVYAPDVLINN
GLALVVSMLAASIAFAVVFPAEMPWLTGRIARDLRRQIALACDGPLQGLDQRFQSSSHDL
MSQLRNLLMKRSRRHRDALRWMLATLEIGHAVIDLRREMQAFTAAQPAQALRWRALIDAV
REALPRLFETPDAHRLARALKSVNLAIRAVRHTQHLWYAVPDERRRMQRIVSCLHFIRSA
LIDQDAPFNRGSRARERVRARRM*
>GCA_000452945.1_BPMSHR5858_genomic_01737_1 group_2356
MKSLQIAVVALSLSAVMAAAHAQPAAADAGRQPANRTEAVQAAGRVPAAHQDRQDPNACV
GPVSFCNIYFGS*
>GCA_000452945.1_BPMSHR5858_genomic_01740_1 yhdG_1
MAIISESSSAAAAAAPGGARAPDTLARSLNVRDVVMITVSGVTPASSIFVIAPFAIQQAG
SGAVMSFVLGGLLAFAFALCYAELSAAHRSAGGEYVMVKRVFGALPGYLTFVAVLAVNVF
IPAVLASGAAPYLNAALGTQLGNQSVALAIVLASYALGILNIRTNAWITGAFLVIEILVL
ALIAYLGFSEPHRSPLALVHPVVANGAALVPATLAAIMPAVGTAIFCYNGFGAAVFLAED
LEGGNRDIAKAVICSLAVILVVELVPLTAIVLGAPSFVELAKSADPIGYVVRALSNPGVS
RIVSGGIFLSVFNAIIAIVIMVGRFLYSSGRHALWSHACNGAFTRIHPRLESPWIATLTL
AVPSTLLVFVSSLDALTAFTVDLLLLVYLAVGVAALASRVMRRDVDHHYRMPWWPLPPIV
AIAGAGYTLYTTIAAATKPTDLYIIAGLAAVSLAAYARWARRSAAFREL*
>GCA_000452945.1_BPMSHR5858_genomic_01741_1 group_3247
MARSLGSQIARRHSPLGLRQTGAHMDRLFSEVAMHQAFGRALDHLGEPRFWRLLILFLNE
MVPFDNALATALGRDGAPVVLDEYDTGGASGPSPVPLYLNGLYLLDPFLQAAHDGLADGC
YRLEEVAPDLFRQSEYFLSYFRDAVGEDELQIIVRTRPDEMLSLSLGAHARFDTDTLGRL
AACSPWLLAAIRQHRRLQGDASRARASDDLAERVERALASFGAPLLTEREMAIARLVLRG
NSSKAIAERLAISPETVKVHRRHLYSKLGISSQPELFSLFLQALGHDAPR*
>GCA_000452945.1_BPMSHR5858_genomic_01747_1 bauC_1
MKHDSNVTSPLGHLIDGKRVDGGERVQPVFDPATGASTKRVRMADRPSVEAAIAAAQAAY
PAWRNTPPLKRARVMSRFKTLLEEHANELCALITAEHGKVLADAMGELQRGIENVEYASY
APELLKGEHSKNVGPAIDSWSEFQALGVVAGITPFNFPIMVPLWMWPMAVACGNTFVLKP
SERTPSSTLRMAELALEAGLPPGVLNVVNGDKEAVDTILTDSRVKAVSFVGSTPIAESIY
TTGCAHGKRVQALGGAKNFAIVMPDADIGNAVNALMGAAYGSCGERCMAIPLVVAIGDDT
AEQVVDGLKAEIAKMKVGPGTGEQVDMGPLVTRQHFEKVTGFVEAGIAAGATLVVDGRGV
KVDGHEGGYYLGPCLFDHVKPGMPIYQHEIFGPVLGVVRVASLAEAMALVDAHEYGNGTC
LFTRDGEAARFFSDNIQVGMVGINVPLPVPVAYHSFGGWKRSLFGDLHAYGPDAVRFYTK
RKTITQRWPSAGVREGTVFSFPSSR*
>GCA_000452945.1_BPMSHR5858_genomic_01750_1 group_3248
MNPKVTGGSGVDPTYAATSTTAEPQAASHASGTSRTIPSELRELQRTKFRLGGSPVFQRP
VQPNRDGSGVGVVVPENSVGRIPPGEAANLCYPNISSCVSVSALSRNGSLRGVHLTTPEG
FEQKDMETMFEAIDAKNCAQVVVAGPIQEAKGRTENSEFNTRANITARIHKLAPDAQVGF
VETEATVRPHNIYVRHDPQTADGKLEVHVAPSDVSSEDGVKTGRIPQDASIPGHAHRVAD
GDIVQSRQGGLRRLFSFSKNG*
>GCA_000452945.1_BPMSHR5858_genomic_01755_1 acsAB
MFDFFLANRSLLDINGGVLLVVLLLTRAGRRERAADRIVFGGVTVVLLLVYLAWRATQTL
PEPRMDFPSAWAHVFFGFECMALAYTLISVAVLTRTTERTPQADAGEAALRRAAAPPRVD

IFIATYNEGLDILEKTIVAALAIDYPDFRVWVLDDTRRDWLKAFCDQVGARYATRADNAH
AKAGNLNNGLRLSARSEGGGAPYIMVLDADFAPHRKILLRTVGLFADRTVGIVQTPQFYY
NADPVQYNLRSAECWVDEQRAFFDVMQPAKDAWGAAFCIGTSFVVRRDLVEHIGGFPTGT
VTEDIHLTYRLMRHGYVTRWLNERLSVGLSAEGLPEYISQRSRWALGTIQVALTPDGPLR
GRGYTFAQRLHYVHGMLHWLSRPFTLMLLLGPLLYWYFDIPTLYGEPLQFLAYGLPALIA
YWGYSMWITGRRALPVFTEVTQIVCALAVSLSLASALVRPFGRPFKVTNKGLDRSKLIVH
GRFAAFYAGLLVLSALGLGRALASAPGAPGLAFNAAWTAISLALYLASLLVCIELPRPRR
EERFPYRARARLRVGDREYAVATRDLSCNGAAVTTAQAASLPLHAAGALWLAPIGWIPCR
IVRRDGALLGVALGADTAARHGLIRLLFGDPPHNIAIRGRPRLAISRLMQRALSG*
>GCA_000452945.1_BPMSHR5858_genomic_01758_1 group_3250
MTQKFEEGLGRREANYVPLTPIDFIARAAEVYGERLAVVHGEIRRTWSETYARARRLASA
LERAGVGRGDTVAALLPNIPAMIEAHFGVPMAGAVLNALNTRLDVASMLFMLRHGEAKVL
IVDTEYAEFAHRAALEVPGLAIVSVADAMPADPARFPAALDYEAFLAGGDPAYAWTPPSD
EWEAIALNYTSGTTGDPKGVVYHHRGAYLAAISNLLEWDMPKHAVYLWTLPMFHCNGWCF
PWAVAARAGVNVCLRKFDAKRVLELIRRERVTHYSGAPIVQSAIANAPAEWRAGIAHEVH
AFVAGAAPAPAVIAKMKEIGFDLTHVYGLTEVYGPATVCAKQAHWASLPDDELAQLNARQ
GVRYHLQAGATVLDPDTMAPVPADGETLGEIMFRGNLCMKGYLKNPHATDEAFAGGWFHT
GDLGVLTPDGYIRIKDRRKDIIISGGENISSIEVEDALYRHPAVAVAAVVALPDPKWGEV
PCAFVELRDGMSATEEEIIAHCRLLLAAYKIPKTVRFGELPKTSTGKIQKFQLRAQVGSS
SAIDLVAAARKPG*
>GCA_000452945.1_BPMSHR5858_genomic_01774_1 group_2702
MTGRTGILLIHGLGGTQYDLGSLHKAMRRAGGDTHMITLPGHGTRPEDLVDVRAEAWLDA
VTAQYRALEREYDTVHVAGMCMGALVARLLCRRVRHARGRLALLAAPMFIDGWSTPWYRS
LRHLLYRVPGVPARLRVDEDEPFGIKNPTIRAIVKKKFERQDNFHYPWVPLACIRQVDRL
RSWARAAAADTHCPTLVLHAREDELTSLRSADFLLKHLPDARGIVLENSYHMICADNDRD
EVARQVLTFFGFDPSHAMSPAMARRMGRADP*
>GCA_000452945.1_BPMSHR5858_genomic_01775_1 group_2359
MRHVDRQDRSRLARRLTAALLTSATMAAHAAHAAPPPPPELVPRDAARDRAFAAAGIQVY
SCEYDGNHRLAWTFQHPEATLYDAAGVAVIKHGAGPAWEAQDGSRIVGEKLADAPSPNAD
SIAQLLLSTRASASGSLASVRYVQRLDTKGGTAPAAACTAEHQTGGSPYYARYVFWK*
>GCA_000452945.1_BPMSHR5858_genomic_01776_1 egtD
LREISERMDEAAAGVANETMRDSAFGRDLLAGLRRSPRSIAPKYFYDAAGSALFDRICEL
PEYYPTRTELAILKRRAHEIAEHIGRDANLIEFGAGSLSKIRVLLDACAASNPPARYLPV
DISADHLAQSAAALRDAYAWLDVRPVVADYLQSEQLRAIERLDGRRVGCFLGSTIGNFSP
DEASAFLRRAASLLKGGGLLIGVDLVKDVSILHRAYNDAAGVTAAFNLNLLRRANAELGA
DFALDAWAHRAFYDAGRQRIEMHLVSRRAQMVRVAGYAFRFDAGETLHTENSHKFTVDGF
RALAQSAGFVPGTVWIDDARLFSVHWLESRA*
>GCA_000452945.1_BPMSHR5858_genomic_01778_1 group_3253
MTPSTHPVKMPGPDHPITIEATGARVVVKAGGHTIADTRDALTLREASYPAVQYVPRKDV
DLARLERTTHESYCPYKGDASYYSIKGVGERGVNAVWSYETPHDAVEQIAGRLAFYPDRV
DSITIG*
>GCA_000452945.1_BPMSHR5858_genomic_01780_1 dpgD_1
MDRPHPAPLLRDDRDGVATLRLNRPSQFNALSEALLDALQRELAVLAGDPHVRCVVLAAE
GRAFCAGHDLREMRGTPDLAHYRGLFGQCSRVMQAIRALPVPVVARVHGIATAAGCQLVA
SCDLAIAADTARFAVSGINVGLFCSTPAVALSRNVSAKRAFDMLVTGRFIDAATAVDWGL
VNEAVPQDALDAAVARTVAAIASKSPRAVRYGKAMFYRQRQMPLDDAYAYACDVMARNMM
EEDACEGIDAFLDKRPPRWRT*
>GCA_000452945.1_BPMSHR5858_genomic_01787_1 group_19923
MLTTGSGRPCAPTLGALGGAQPTPNAGGLWPLAGMLRWIVVAVLTISVALPIAFIVFQSL
LSAPFFDANKTLGVEGFRFIFDDPDFWSAVKNSFVIAGGMLFISIPLGGILAFLMVRTDL
PGRRWLEPLVLTPVFVSPMVLAFGYVVAAGPVGFYSVWFKALFGVASVPWNVYSVLAITI
IVGLTHVPHVYLYASAALRNLGSDVEEAARVAGARPFRVALDVSLPMTLPALLFSGVLVF
FLGFEVFGLPLVLGDPEGHLVLATYLYKLTNKLGVPSYHLMAAVAVCIIAITFPLVLLQR
HLLKSANRYVTVKGKAGRATVLPLRAWRWVALAIVAAWLALTVIVPISGIVLRAFVTHWG
EGVPLADVLTVANFVELFEQDNLVRAIVNTLGIGVIGGALAIAFYSLVAFAGYRRHDWAA
RLLDYIVLLPRAVPGLLAGLAFLWIFLFVPGLRELKGSMWSIWIAYTVVWLAYGMRVIQS
ALLQVGPELEEAGRSVGATRARVSLDVTLPLVRFGLLAAWLLIFMIFEREYSTAVYLLSP
GTEVIGALLVSLWATGAVDQVAALSVINIAMVGAGLAVALRFGVKLHG*
>GCA_000452945.1_BPMSHR5858_genomic_01794_1 paaK
MATSLPLDPIERASRDELLALQLDRLKWSLAHAYENSPVYRRKFDEAGVHPADLKTLADL
SRFPFTTKNDLRDNYPFGMFAVPQERISRIHASSGTTGKPTVVGYTARDIDTWAGLVARS
IRAAGARPGDKVHVSYGYGLFTGGLGAHYGAERAGLTVIPFGGGQTEKQVQLIQDFQPDI
IMVTPSYMLSIADEMERQHLDPAQCSLRIGIFGAEPWTNDMRVAIEQRMGIDAVDIYGLS
EVIGPGVASECVETKDGPTIWEDHFYPEIIDPDTGEVLPDGAFGELVFTSLTKEALPIVR
YRTRDLTRLLPGTARTMRRMEKITGRSDDMMIVRGVNVFPTQLEEQLLKQRALAPHYQVV
LTKDGPLDVLTLNVEPCPETAPDAAAIDAARRALAHDIKSLIGVTAIINVLPVNGIERSV
GKARRIVDKRRP*
>GCA_000452945.1_BPMSHR5858_genomic_01797_1 apaG
MSQYRFSVSVKTSYLPEQSDPEHRQYAFAYTLTIRNTGQVAAQLIARHWIITDSESQVQE
VKGLGVVGHQPLLQPGEQFEYTSWAVIATPVGTMRGAYFCVAEDGERFEAPVDEFALHMP
RTLH*
>GCA_000452945.1_BPMSHR5858_genomic_01798_1 rpe
MTQFRIAPSILSADFARLGEEVRNVVAAGADWIHFDVMDNHYVPNLTIGPLVCEAIRPHV
QVPIDVHLMVRPVDRIVPDFAKAGANLISFHPEASDHIDRTLGLIRDHGCKAGLVFNPAT
PLNYLDHVMDRVDLVLIMSVNPGFGGQSFIPEALNKLREARARIDAYTARTGREIHLEID
GGVKAENIAEIAAAGADTFVAGSAIFGKPDYRQVIGEMRDALATVGRACAR*
>GCA_000452945.1_BPMSHR5858_genomic_01800_1 trpE
MTELEFQSLANEGYNRIPLIAEALADLETPLSLYLKLAQPERGGANSFLLESVVGGERFG
RYSFIGLPAHTLVRTKNGVSEVVTDGQVTETHDGDPFAFIATFQSRFKVAQRPGLPRFCG
GLAGYFGYDAVRYIEKKLAHTAPRDDLGLPDIQLLLTEEVAVIDNLAGKLYLIVYADPTK
PEAYTKAKQRLRELKQRLRASVVPPVTSASVRTEIYREFKKDDYLAAVRTAKEYIAAGEL
MQIQVGQRLTKPYRDNPLSLYRALRSLNPSPYMYYYNFGEFHVVGASPEILVRQEKRGDD
QIVTIRPLAGTRPRGNTPERDAELATELLNDPKEIAEHVMLIDLARNDVGRIAEIGSVHV
TDKMVIEKYSHVQHIVSSVEGKLKPGVTNYDVLRATFPAGTLSGAPKVRAMELIDELEPI
KRGLYGGAVGYLSFSGEMDLAIAIRTGLIHNGNLYVQAAAGIVADSVPESEWQETENKAR
AVLRAAEQVQDGLDSDF*
>GCA_000452945.1_BPMSHR5858_genomic_01806_1 azoR
MTTILQINSAARSQGAQSTLLADELTAKLQQGNPGATVKVRNLLADALPHLDDAVLGAFF
TPADQRSAEQNAIVAKSDELVDELRSADVIVIGAPMYNFGVSSQLKAYFDWIARAGVTFR
YTSEGPEGLIKGKKAYVVSARGGKHVGMPTDSQTPFLKTFLGFIGLTDVTFVYAEGLALG
PDAATEALASAREAIAAV*
>GCA_000452945.1_BPMSHR5858_genomic_01807_1 group_2098
MKPIRYTIVPKDPAAHLFEVTLTLADPDPAGQRFALPVWIPGSYMVREFARNIVTLRAFN
EAGRKLRIGKLDKQTWQAAPAPGPITLRYDVYAWDLSVRAAHLDDTGGFFNGTSVFLAPL
GREDAPCEVMIERPAGDAYRRWRVATALPEARGTKRYGFGAYRAENYDELIDHPVTLGEF
ALASFDAHGVPHDIAIAGRVTGLDLERLAADLKRVCEAQIALFEPKTRRAPMSRYVFMTQ
AVSDGYGGLEHRASTALVCNRTDLPVKGRPEKTDGYRTYLGLCSHEYFHTWNVKRIKPAA
FAPYDLSQENYTSLLWLFEGFTSYYDDLMLARSGLISQDDYFALVGRTIAGVQRGAGRLR
QSVAESSFDAWIKYYRQDENATNAIVSYYTKGSLVALAFDLAIRARSRHRKSLDDVMRLL
WQRFGRDFYHGKPQGVGEDDVKALIAEATGVDLGRLFDEAVSGTRDLPLAELFEPFGVTL
APDGGAGGAGGAADAPAKPTLGARTRGGAECTLAAVYEGGAAHRAGLSAGDALVAIDGLR
VTGSNLDALLARYRVGDKVEIHAFRRDELRVVQLKLDGPDIARYKLAAQPKPAAARARRD
AWLGLPAARGGR*
>GCA_000452945.1_BPMSHR5858_genomic_01810_1 group_3258
MKRIAVLAALAVALGSVLAGCIVVPDGGYGYAHHRDYYYYRY*
>GCA_000452945.1_BPMSHR5858_genomic_01811_1 group_19935
MSDLRHHLNQQDRVELLCWLVVGNLGAFYLNESWPGAAFQVQSAHKWLDRHAREADWLTL
AKLSLIALDIAKKHADFVNAPWARDAVEEIIDTDDLNYEARLAQLVLDDCRAALADRRIA
D*
>GCA_000452945.1_BPMSHR5858_genomic_01812_1 ychF
MSLKCGIVGLPNVGKSTLFNALTKAGIAAENYPFCTIEPNVGVVEVPDARLKALAEIIKP
ERVVPAVVEFVDIAGLVAGASKGEGLGNQFLANIRETDAITHVVRCFEDENVIHVAGKVS
PLDDIEVINTELALADLGTVEKALARYSKAAKSGNDKEAAKLVAVLDKARAHLDQGKAVR
GLDLSDDERALLKPFCLITAKPAMYVANVKEDGFENNPHLDAVRRYAESEKSPVVAVCAA
IEAEIADLDDADKEAFLADMGMDEPGLDRVIRAGFKLLGLQTYFTAGVKEVRAWTIHIGD
TAPQAAGVIHTDFERGFIRAQTIAYDDFIAYKGEQGAKEAGKMRAEGKEYVVHDGDVMNF
LFNV*
>GCA_000452945.1_BPMSHR5858_genomic_01821_1 group_2361
MTVKAVVFDFGGVLIDWSPEYLYRKLIPDEAERRWFLTHVCGMDWVVRQDGGQTIEEGTA
ERIALFPEHEALIRAFYARWHEMIGGELADGAALVDALDARGVPLFGLTNWSAQTFPYAW
ENFPVLRRFRDIVVSGRVKLAKPDPAIYREMHARIEPHLPGIAPHELVFIDDNAKNAAAA
AALGWHGIHHTSAAATRARLRELGALG*
>GCA_000452945.1_BPMSHR5858_genomic_01823_1 group_2362
MNARLPDPSPVPARLALLRGAMTREDLAAYVVPSADPHLSEYLPERWQARQWLSGFTGSV
GTLVVTADFAGLWVDSRYWMQAEAQLAGTGVALMKMVGGQQTQPHVEWLAEHVPEGTTVG
VDGAVLGVAAARALTSALTPRGIVLRTDLDLLDAIWPQRPSLPGDAVFEHAAPQADTARA
GKLAQVRRAMHEQGAQWHFVSTLDDLAWLFNLRGADVNYNPVFVAHALVGLERATLFVAD
GKVSAELATSLARDGVDVKPYDAAAAALAALPEGAGLLIDPRRVTYGLLQAVPQQVRVIE
AVNPSTFAKSRKTPAEIEHVRATMEHDGAALAEFFAWFERALGRETITELTIDEQLTAAR
ARRPGYVSPSFATIAGFNANGAMPHYRATRAAHATIEGDGLLLVDSGGQYLSGTTDITRV
VPVGAIGDAHRRDFTIVLKAMMALSRARFPRGIRSPMLDAIARAPMWAAGLDYGHGTGHG
VGYFLNVHEGPQVISHYAPAEPYTAMEEGMITSIEPGVYRPGNWGVRIENLVVNRAAGQT
EFGDFLEFETLTLCPIDTRCVLPALLDDVERAWLNAYHATVRERVGKHVSGDARAWLDAR
TQPI*
>GCA_000452945.1_BPMSHR5858_genomic_01826_1 hemA
MGLNDSLLDMQLLTIGINHHTAPVALRERVAFPLEQIKPALSTFKSVFLGHPAPNAPEAA
ILSTCNRTELYCATNDRAARDAAIRWMSDYHRIPADELAPHVYALPQSEAVRHAFRVASG
LDSMVLGETQILGQMKNAVRTASEAGSLGTYLNQLFQRTFAVAKEVRGTTEIGAQSVSMA
AAAVRLAQRIFEQIAQQRVLFIGAGEMIELCATHFAAQGPRELVVANRTAERGAKLAERF
GGRAMPLADLPARMHEFDIIVSCTASTLPIIGLGAVERAVKARRHRPIFMVDLAVPRDIE
PEVGKLKDVFLYTVDDLGAIVREGNASRQAAVAQAEAIIETRVQNFMQWLDARSIVPVIR
HMHTQADALRRAEVERARKMLARGDDPDAVLDALSQALTNKLIHGPTSALNRANGADRDS
LIDLMRGFYQHAPRSSDTSDR*
>GCA_000452945.1_BPMSHR5858_genomic_01830_1 ubiX
MEPRPAPPRRLIVAITGATGAIYGVRMLDMLRASGGVETHLLISSAGWLNIRHELQLSKD
DVLARADVVHSVRDVGASIASGSFATEGMIVAPCSMKTLASIAHGLSDNLITRAADVALK
ERRRLVLLVREAPFNLAHLRNMTAVTEMGGVIFPPVPAFYAMPKTIDEIVDHTVARVLDM
FALGAPLTAPWQGLR*
>GCA_000452945.1_BPMSHR5858_genomic_01876_1 pmdD
MDQASNDPAVETTGRPFATRTHEARKAHEAREAHDKRAPLVDAWYGSGAAPIEAVDSHAH
VFLRSLPRIPSARHSPEYDATLESYVAHLSACGITHAVLVQPSFLGTDNHFFVDALARYP
QRFRGIAVVNPCTAEDEFARLEATDVVGIRLNLVGLPIPDFTAPRWRALLARANALGWHV
EVHRRAADLPAIIPALLDQSCRVVVDHFGRPAPHLGTLDPGFRFLLSIAGTGQVWVKLSA
AYRNIGSGDGTAFGTRAARALLGAFAPNRLVWGSDWPHTQHRDRTDYQTTRSALDDWVPD
PSLRRIILCDSARALFRFDRQTPARDT*
>GCA_000452945.1_BPMSHR5858_genomic_01877_1 group_19734
VNWYKEISPVERRTFKGCFSGWALDGVDTQMFSLVIPALLTAWGIGKGQAGLIGGATLAA
GAIGGLLAGMIADRFGRVRALQITVCWFSLFTFLSAFAQNFEQLLVLKTLQGLGFGGEWT
AGAVLLSETIRARHRGKAMGIVQSAWGFGWGGAVLLYTLVFSWLPPEWAWRVLFAIGVLP
ALLVLYIRRAIPEPPRDDARVAVSTSAAVAQTAPARASAKSIFDPSVLRMTIVGGLIGVG
AHGGYHAITTWLPTYLKTERHLSVLGTGAYLAVIIVAFIIGCMTSAYLQDRIGRRRNLML
FSACCVVTVNLYVMLPLDNVAMLLLGFPLGFFAAGIPATLGALFNELYPRNVRGRGVGFC
YNFGRVASAIFPVLVGHMGTSMPLGVALGIDAGIAYGLVFVAVWFLPETRARDIQPSQPE
ARIREAAL*
>GCA_000452945.1_BPMSHR5858_genomic_01879_1 group_3263
MHKREVQVVSGAAFMLFFATVSAGQLTGTMQVNLQVSRGCEVAGVAASGDLGRLDFGAQG
PLWSDYLTADGRATSSGAVRVVCSPDVNGFLVSIDGGRNGDQSTRYLVKRGANGRVAGRI
PYNVYRDAARSVPYVPLMPQSFFVDGGRDDVTLPVYGVVNGMTRAVPSGTYEDLLGITLD
W*
>GCA_000452945.1_BPMSHR5858_genomic_01880_1 group_19838
MKAHWKVWAALACVSACAGVQAQTSPLTGTVNSQLVLTTGCAVDTGGGSVNSANFGTLDF
GTQPSGFTGRLTSAAKGGGSTSTQVTCSPDVTSIQVTIDGGQNASKGATVGTGTRALANG
ASFVPYEVYADAGHSQQYVSGTAQSVAVPTPGAAFELPLYGVVNKTNASALAAGTYTDVL
NVTLGW*
>GCA_000452945.1_BPMSHR5858_genomic_01882_1 group_4019
MKVSRSPGAPARAARALRCALLGAAAAAPFVAQHAHAAASVMIWPIDPVIESGDRAAALW
LENRDRRPVTLQVRVLGWREANGEDVYDEDQPRVAGSPPMATVEPGKRQLIRLTRLADTA
PGTEQAYRVLIDEIPQPDDDAQPAASGTSLGVKFQMHYSVPLFVYGDGLWTKEHPDRRRD
PATAGRPALRWRVEHDGGKRWLVVSNRGPVHARITHAAFEASGARADFARGLLGYVLPGA
RMRWALPETLKLHSNSKLIATVNGVADLAIDVDGAASNR*
>GCA_000452945.1_BPMSHR5858_genomic_01883_1 fimD
MSRRGNALKRYNDDRTSGLPRRRAAAWAVGIAFAAAGHARAGETATLADSFGRALPPVGG
TAAHGTLYLELVVNALSTGRIVPVRYRDGVYYARAGDLAQASVRTGAQPDALVDLSRLDG
VQVEYESAEQRLKLTVPPDWLPRQTLGSPRLYDRTPAAVSFGLLFNYDVYANSPTLGTSY
TSAWTEQRLFDRWGTVTNTGVYRRDYGGGAGGVGSNRYLRYDTFWRYSDQDRLRTYTAGD
VITGALSWSSAVRLGGVSVERDFKVRPDIVTYPLPQFSGQAAVPTAVDLFINGSKTTTGQ
VNPGPFTMNNVPFINGAGEATVVTTDALGRQVATTIPFYVANTLLQKGLSDYSLSAGAMR
RDYGIRSFSYGKFAASGTARHGLTDYLTLEGHVEGGERFALGGLGFDLGIGMFGVLGVAA
TQSRLAGASGRQYAFGYSYASQRFSVSLQRIQRTNGFRDLSVYDLPANVAYRLVRSSTQA
TGALNLGALGGTLGAGYFDVRGADGARTRIANLSYTRPLWRRATLYASVNKTVGEHGVAA
QLQLIVPLGEPGVVTGALARDANNSFSERVQYSRSVPSDGGLGWNLAYAGGGSHYQQADA
TWRNRYFQAQGGVYGYGAGRGYARWGEVQGSVVVMDGAVLPANRVDDAFVLIDTQGRGGV
PVRYENQLVGKTDGGGHLLVPWAPSYYAGKYEIDPLDLPSNVRVPIVERRVAVRDHGGAL
VTFPIRRIVCAQIALVDAAGRPVAIGSRVLHEESGETALVGWQGETYLEGLSALNHLRVR
TPDGRTCRATFAADVDAAQMSRVGPLACGE*
>GCA_000452945.1_BPMSHR5858_genomic_01886_1 rcsB_1
MSVQVLIADDHPLVLLGVRHMLAGMGDVSIVGEAHDPAGLLALLAATPCDIVITDFAMPE
QPAADGLAMLTAIRDGYPSVRVIVLTMLDNPVLMHTMRQAGALAVLSKRGDLDELPRALA
AVYQGRPFVGTHAGAAGGGAMRGTDAPRQLSPREIEVVRLCATGMTMTDIANRYGRSIKT
VSTHKHNAMGKLGLKSDVDLFMYATENGLV*
>GCA_000452945.1_BPMSHR5858_genomic_01888_1 emrB_2
MNGSTPTPPAAPRRDRNGNGDGDGTWRPAANPWLIAVVVTLAAFMEVLDTTIVNVALPHI
AGTMSASYDEATWTLTSYLVANGIVLPISGFLGRLLGRKRYFVLCIVAFTICSFLCGIAT
DLGQLIVFRVLQGLFGGGLQPNQQSIILDTFPPEQRNRAFSISAVAIVVAPVLGPTLGGW
ITDNFSWRWVFLLNVPIGVLTSLAVIQLVEDPPWKRGRARGLSIDYIGITLIAIGLGCLQ
VMLDRGEDEDWFASTFIRTFAVLTVAGLVGATFWLLYAKKPVVDLSCLKDRNFALGCVTI
ATFAVVLYGSAVLVPQLAQQRLGYTAMLAGLVLSPGALLITLEIPIVSKLMPYVQTRFLV
CFGFLLLAASLVYSHTLVPDIDYPNLLKIRSAQSLAIGFLFVPITTLAYLTVPRRLNDDA
SALFTMFRNVAGSIGISLSTALIRERAQARMAHLSTHLSPLSQNFDDTLRQNAQTIASLS
GMPPPAALQTASGQLYETFVSQATILAYLDVFAMLAIFCFVCTPLTLLFSPVKAAGGPGG
H*
>GCA_000452945.1_BPMSHR5858_genomic_01889_1 emrK
MPDPNDAAERGNDERSEHDEPGRERARGRREEADAHARDARPGDAPSGERKRPGKKPLVV
LAIIVVLLLVGAFVWWFATRNQVSTDDAYTDGNAITIAPKVSGYVVALAIDDNVYVHRGD
LLLVIDQRDYQAQVDAARAQLGLAQAQLDAAQVQLDIAHVQFPAQYRQAQAQIEAAQASF
RQALAAYERQHAVDARATSQQAIDVADAQRLTADANVATTRAQARTASLVPQQIRQAQTA
VEQRRQQVLQAQAQLEAAQLALSYCEVRAPSDGWITRRNVQLGSFLQAGAALFAIVTPQL
WVTANFKESQLERMRAGDRVSVSVDAYPNLELHGHVDSIQLGSGSRFSAFPPENATGNFV
KIVQRVPVKIAIDGGLPRDPPLGIGLSVVPKVYLK*
>GCA_000452945.1_BPMSHR5858_genomic_01891_1 group_2367
MNKHWIRVSPIALAAAIGLTACGGDDVTSPGLSAVKNVVVIYAENRSFDNLYGNFPGANG
LQNVTAASAAQVDRDGTPLATLPKIWGGLTAAGVTPAVTEAMTANLPNAPFAIDDPNGFN
TPMSVTTRDLVHRFYENQMQIDGGKNDKYAAWSDAGGLVMGHYTADPSKLPLWKLAQQYT
LADNFFMGTFGGSFLNHQYLICACAPFYPNADKSPAASQISVVNPDGVTLTTASNSPASA
LSGPPKFVKSGGLTPDFYAVNTMQPPYQPSANAAPTGGDPNLADPANPSTLPPQTQQNIG
DLLTSAGVSWAWYAGAWGAALQAAQSGTSGVIYGPNMTSPNFQPHHQPFNYYANQAPGSA
NRAQHLLDAGSNGAALIQAIDAGKLPQVTFYKPQGNLNEHPGYTDVASGDQHIADVIAHL
QKSPQWNNMVVIVTYDENGGFWDHVAPPKGDRWGPGTRIPAFVISPFAKQGFVDHTQYDT

ASILRFITRRFALPKLAGLKQRDDALIANGFKPMGDLTNALTLSTGN*
>GCA_000452945.1_BPMSHR5858_genomic_01895_1 group_19928
MTTRARVSILPRVLVGAACAAAALGAHAQNNLNFLNDTPISYFSKADTASLAKAVQKVRD
EGKDGETVDWVNDGRGTKLAAKLTPSTTEQEGRTCREIKTEIEAKGQSMTLRPLYCKTAA
GKWQLQKR*
>GCA_000452945.1_BPMSHR5858_genomic_01897_1 glnS
MPGHPPKNAPRRPPQGAPLLMTVVRQVELTYSGGAPCAPSPFLPLFRMSTERNDATAPSN
FIRNIIDDDNRTGKWSGRVETRFPPEPNGYLHIGHAKSICLNFSVARDYGGVCHLRFDDT
NPEKESVEYVNSIVDAVRWLGFDWQKAGVDHQYFASDYYDKLYEFAELLIQRGCAYVDSQ
SADEMRTNRGSLTEPGTPSPFRDRSPDENLALFRRMKAGEFAEGEHVLRAKIDMSSPNMN
MRDPVIYRIRYAHHYRTGDKWCVYPMYDYTHCISDALEGITHSLCTLEFEDHRPLYDWVL
NQLADAGVFARPLPQQIEFSRLNLTYAITSKRKLLQLVTEGHVDDWDDPRMPTIVGVRRR
GFTPESIQLFCERIGVTKVDSWIDMSVFEGALRDDLDDKAPRTVAVLDPLKLVIDNYPQG
QSEECTAPVHPHHPERGQRTFPISRELWIEREDFSEHPPKGYFRLFPGNKVRLRYGYVIE
CTGADKDEHGNVTAVHCNYYPDSKSGTEGANNYKVKGNIHWVSAAHACPAEVRIYDRLFK
EPHPDAGGRNFLEALNPDSKKIVQAYLEPGARDALPEARYQFERHGYFVADRVDSKPGQP
VFNRIVGLRDSWGKPA*
>GCA_000452945.1_BPMSHR5858_genomic_01923_1 group_19929
MLFRSGTIRIPFEPNDKARERERPIQRVELDPEHDRHARAALEAYADSSAHEMPWLAEWM
REALGSPQPELTRCAQTVERVFDLAHAWAAGQPDYAGAAWEHVRGRLLEALQHAPRAIAP
DALETDPA*
>GCA_000452945.1_BPMSHR5858_genomic_01929_1 valS
MSDTTLAKSFEPQTIESQWGPEWEKRGYATPALDPSRPDFSIQLPPPNVTGTLHMGHAFN
QTIMDGLVRYHRMLGHNTLWVPGTDHAGIATQIVVERQLDAQGVSRHDLGREKFVERVWE
WKERSGSTITGQVRRIGASPDWSREYFTMNDKMSEAVREVFVRLYEQGLIYRGKRLVNWD
PVLLTAVSDLEVVSEEENGHLWHIRYPLADGSGHLSVATTRPETMLGDVAVMVHPEDERY
RHLVGRHVKLPLCEREIPIIADDYVDREFGTGVVKVTPAHDFNDYQVGLRHALAPIEILT
LDAKINDNAPAAYRGLDRFDARKAIVDELDAQGLLESVKPHKLMVPRGDRTGVVIEPMLT
DQWFVAMTKPAPQGTFHPGKSITEVSLEVVRRGEIKFVPENWTTTYYQWLENIQDWCISR
QLWWGHQIPAWYGENGEIFVARNEEDARAQAAAKGYTGALKRDDDVLDTWFSSALVPFSS
LGWPNETPEMKHFLPSSVLVTGFDIIFFWVARMVMMTTHFTGKVPFGTVYVHGLVRDAEG
QKMSKSKGNTLDPIDIVDGIGLDALVAKRTTGLMNPKQAATIEKKTRKEFPDGIPAFGTD
ALRFTMASMATLGRNVNFDLARCEGYRNFCNKLWNATRFVLMNCEGHDCGFDKPDVCGAG
DCGPGGYLDFSPADRWIVSLMQRVEADIAKGFADYRFDNIANAIYKFVWDEYCDWYLELA
KVQIQNGTPEQQRATRRTLLRVLETVLRLAHPIIPFITEALWQKVAPLAGRYPAGKAEGE
ASLMVQAYPVAEPKKLDEACEQWAAELKAVVDACRNLRGEMNLSPATKVPLLAAGDAAQL
QAFAPYVQALARLSEVRVLPDEAALDADAHGAPIAIVGGNKLVLKVEIDVAAERERLSKE
IARLEGEIAKCNAKLGNEAFVAKAPPAVVAQEQKRLAEFQSTLTKLGAQLARLPA*
>GCA_000452945.1_BPMSHR5858_genomic_01932_1 fsr
METSLDNRPSVARAAASPAPAAVSRTVYPVLGAISFSHLLNDMIQSLILAIYPMLKSEFS
LSFAQIGLITLTYQITASLLQPVIGLYTDKRPQPFSLPVGMGFTLTGLLLMAFAPTFPFL
LVAAALVGCGSSVFHPESSRVARMASGGRHGLAQSLFQVGGNAGSSLGPLLAALIVIPHG
QRSIAWFSAAALVAIFVLVQIGRWYRRHPAAKKKAAHAAHPTLSRRQVGLALGVLVMLVF
SKYFYLASINSYFTFYLIDKFHLSVQAAQIHLFVFLAAVAAGTIIGGPVGDRIGRKYVIW
TSILGVAPFTLMLPYANLFWTTVLTIVIGVVLASAFAAIIVYGQELIPGKVGTVAGLFFG
LSFGLGGVGAAVLGQLADATSIAFVYKVCSFLPLIGVLTVFLPNVEGRTTLDKQRA*
>GCA_000452945.1_BPMSHR5858_genomic_01936_1 aaeX
MMPRDLAILDAYVPTVVLMFVIGAFATWLIDRLLAYTGLYRVVWHPSLFRASLLVCICGG
LSLAVYR*
>GCA_000452945.1_BPMSHR5858_genomic_01937_1 aaeB_2
MPAASPPSAATPQSAGAAFVASMSDWARTDGAAWLYLAKALAAAFLALGVSMVLDLPAPK
TAMTTVFIVMQPQSGAVFAKSFYRLIGTFVGLTATLALVGLFPQQPVLFLLAVALWVAAC
TAGAARNRNFRSYGFLLAGYTAALIGLPASQAPNGAFMTAMTRVSEISVGILSAGIVSAL
IFPQYTGEQMRTTVRRRFGAFVDYVAAALAGALDRSKIESVHTRFVADVVGFEAARSMAV
FENPDTRMRSGRLSRLNSEFMTASSRFHSLHQLMNRLHATSAQAVDALEPYFREIAPLLS
NDGDHVRTSADAARAAEQLLAYRDALPRRIRATRAEHETQPAFPLLDFDTASELLYRFIT
DLHAYAATYASLAASTHARERWIERYEPRTNATAALIAGIRTASVMLLLSVFWIYSAWPS
GVMLVLNAAAVCALASSSPQPTRMAAQMALGTAFAVATGFVLMFGVYPHIDGFVLLCAAL
APFLAPGVLMSLKPKYAGCGAGYLIFFCTLAGPENLTHYDPIGFMNDALALVLSMIVSAI
AFAVLFPPSAPWLKKRLFADLRHQAVAACHARLPGLRTRFESGARDLMFQAHGLSADQPD
AQRDALRWMFAVLETGNAVIDLRHELAALPRDPRYAATMPWRRKIEAARDALAALFDKPA
PARFDAALAAAVDAIAATQQALVAFAPPREERHQMQRILSQLHFVRTALLDPESPLAALN
GARTPTDRPGVSS*
>GCA_000452945.1_BPMSHR5858_genomic_01940_1 group_3270
MNTPLLMGFGALALFGAIGIAASRELLRAIDAQQRRAPVPVRMRTRRPHRDGRRTVL*
>GCA_000452945.1_BPMSHR5858_genomic_01946_1 group_19976
MAHSAHTEHFASFAEFYPFYLDEHRHLVSRRLHFVGSLGVIGCLAMALAMGLWWWLPAAV
VCGYGFAWIGHFFFEKNRPATFRHPIYSLMGDWVMFKDICTGKISI*
>GCA_000452945.1_BPMSHR5858_genomic_01948_1 lplT_1
MSDPSITSTARREKRRAHPSQFDLLRERRFAPFFATQFMGALNDNVFKIGFTSLVTYQTA
KFSGVDPKTAAFLISAIFILPFVLFSATAGQLADKYDKARLTRFVKSFEIALMAVGAAGF
VTHGAPLLYLCTFMMGMHSTLFGPVKYSYLPQHLNDHELVGGNGLVEMGTFVAILIGTIV
GGAAAGISGRGELLLAVCCVGIALVGRVAAAGVPATPAPQPKLAINWNPFSETWRNLKLA
RENRTVFLSLLGISWLWFVGATFLTSFFGFAKDVLSANPDVVTVLLATFSFGIGLGSLLC
ERLSRRRVEIGLVPLGSLGISVFAIELYFASHALPTQTHLLTIGEFLGHAAHWRILADLF
LLAMFGGFYSVPLYALIQSRSAPTHRARIIAANNILNALFMIVSALMAMLLTKAGVSIPG
IFLVTALLNVVVASYIYSLVPEFLLRFIAWALVHTFYRIRLVHAERIPDEGAALLVCNHV
SYVDALVLAAASPRPIRFVMDHRIFNTRFASWAFRHAKAIPIAPRHEDPEMLARAYDACE
AALREGELVCIFPEGKLTKTGDVNTFHHGVTEILTRARAPVVPMALRGLWGSVFSRHEDA
RWPRPVKRGVMSRLTLAVGEPIPAAAATPEALQAVVAELRGARK*
>GCA_000452945.1_BPMSHR5858_genomic_01957_1 argC
MSTRVFVDGQEGTTGLKIFEYLLARSDIDILRIDEAKRKDVDERRRLINASDVTFLCLPD
VASRESATLVENPNTTLIDASTAFRTSADWAYGLPELNRAQRERIRGAKRIAVPGCHASA
FVLAMRPLVDAGVVAPGFAAHSYSITGYSGGGKKMIADYEAGGAKLASPRPYALGLAHKH
LPEMAAHTGLTSAPIFTPIVGPFYKGLAVTTYFTPGQLAKRVTPQDVQRVLAEHYAGEAF
VRVAPYDAAENLDDGFFDVQANNDTNRVDLFVFGTAERFVTVARLDNLGKGASGAAIQCM
NLNVGAAEGTGLAR*
>GCA_000452945.1_BPMSHR5858_genomic_01959_1 group_19851
MRVGRRAVPRGSSRRLLSLALVCACALAPVGGSRALAEDAFMLTSDDLHPGGRVGAPHVF
NQGDCKGGNRSPQLAWRNPPPGTKGFAITMFDPDAPGRGWWHWAVVGIPASVSSVPADAS
ASGFLRKLGAAEARNDFGTDGYGGPCPPPGKPHRYVITVHALKATDLRVATGRPAQMFDH
EIGTESIGSAQLVVTYGR*
>GCA_000452945.1_BPMSHR5858_genomic_01963_1 group_4000
MKTRNIQNFLMVSAAFFAMSGPARSEGAGVLATASTQMLQLAAAAGADAPTFAGVATNGR
GAMQK*
>GCA_000452945.1_BPMSHR5858_genomic_01965_1 lrp_11
LSNQDIAERVNLSPSPCLRRIRRLEEMGVITGYVALLDPQKLGLDLLAYVSVRLEKRGGV
APVRGDETSARAGATHAELFRAAVQAWPEVVACHAMTGDMDYLLRVQVEDMAHFSRFMQE
QLLHHPSVIDVKTSFSLERFKETTALPIR*
>GCA_000452945.1_BPMSHR5858_genomic_01968_1 paaE
MSTIHVTMGGSAAASLRIALADAGRDEGVVELFDDLSVGPLRGADDSPDVRAAFWERAIG
DVQQDWTAGLGGDFAKLETLAADTGQVVVWHAQSVADQLMLRRVAYHLRNAPQRLNEVRL
SSADIDDPQSWVRLRADQATATGIFSSAQLRAKLPEAAPISVLRISRLALEWQEAKQANA
ELRYWICNTFKSGHYADIDALVLEHAPDEWGPAARTIGAVMAFADRGHLFVGDVIAFWRC
RELAAAGRIELQGDARSIDDLRTATLRATPAVALPR*
>GCA_000452945.1_BPMSHR5858_genomic_01973_1 paaA
MYTQSLDIPGNIAPIDAAADSPEQAGFDAAIAADGKIEPQDWMPDAYRKTLVRQISQHAH
SEVVGMLPEGNWITRAPSLKRKAILLAKVQDEAGHGVYLYSAAETLGVSRDALIDALHAG
KAKYSSIFNYPTLTWADVGVIGWLVDGAAIMNQVPLCRCTYGPYARAMIRVCKEESFHQR
QGFDALLAMMKGTDAQRAMVQEAVDRWWWPVLMMFGPPDADSVHSNQSAKWGIKRISNDD
LRQKFVDATVEQAKVLGVTLPDPDLKWNDARGHFDYGALDWDEFWRVVNGDGPCNKERLA
TRVKAHENGAWVRDAALAHEEKRRRRAQQQAA*
>GCA_000452945.1_BPMSHR5858_genomic_01974_1 group_2371
MTSRRWFSALSCCLALAASQPGSAAQADAPARWVASWATALQPIPDLAAPPPLYRAPDVA
GRTVRQIVYPTLAGKAVRIRVSNAYGKTPLAIGEMNIGRSAGGAAVAAGSSTAVTFGGRR
ETEVPPGQERDSDPVAYDVRAGEPYALSLYLGSRQTMTVWHRVSNQVNYVSAPGNHTGDA
SPDAFRTRFTQSAWIAELAVEVQQPGAAAIAAVGDSITDGLRSSLNRNRRWPDALAARLE
RAGAGDIGVANLGISGNRLLSDSRCYGIALERRFERDVLTRAGVKVAVLLIGINDINFAA
MPARSGLDCDAPHTRVDAQALIAGYRRVIAAAHARGVAVFGATLTPASLPPAREAIRREV
NEWIRTSGAFDGVVDFDAALRDPAKPSTLLRRYNSGDDIHPSDAGYAAMAEAVPLERLAA
AAGRR*
>GCA_000452945.1_BPMSHR5858_genomic_01980_1 cyaA_1
MFGKLTLSAIPFDQPIIMGAGAFMALVVLGILGALTYTKRWKWLWCEWLTTVDHKKLGVM
YIIVALIMLLRGFADAVMMRLQLALAYNGPGYLPPHHYDQVFTAHGVIMIFFMAMAFMIG
LMNIIVPLQIGARDVAFPFINSLSFWMTAVSAILINISLVVGEFAQTGWLAYPPLSELQF
SPGVGVDYYLWALQISGVGTLLTGVNFFVTILRMRAPGMTLMKMPVFTWTALCTNVLIMA
SFPILTVTLALLGLDRYLGMHFFTNDAGGNAMLYLNLIWAWGHPEVYILILPAFGIFSEV
VATYAKKPLFGYKTMVYASCAIMVLSFLVWLHHFFTMGSGANVNAFFGMMTMVIAIPTGV
KVFNWLFTIYRGRLEFSTPILWTIGFMVTFTLGGMTGVMMAIPGADFVLHNSLFLIAHFH
NVIIGGVLFGYLAGFNYWFPKVFGFKLNETLGKAAFWFWQVGFYVAFVPLYVLGFMGMTR
RLNHYDNPAWHPWLIVAACGAALIAIGIACQLLQLAVSLRDRRLLENRDPTGDPWGGRTL
EWATASPPPAYNFATIPTVHALDEFAYRKERGLGIGKPAEYRDIHMPSNTSAGLFVGVFS
LVLGFAAVWHIWWLAIAALAGILVTVIAYSAGENDGYYIPADAVRRIEEKRGGARFVARP
AQVELEAN*
>GCA_000452945.1_BPMSHR5858_genomic_01988_1 andAc
MEMTESPVALIDRADSSDVRFPRDDGSRVPYKVFSSQAVYEREQERIFRGPTWNFVALEA
EIPKPGDFKSTFVGDTPVVVTRGEDGALAAWVNRCAHRGAQVCRKARGNASSHTCVYHQW
SFDSRGDLLGVPFRRGQKGMAGMPADFDPKNHGLRKLRVDSYKGLVFATFSDEVAPLPDY
LGAQMRPWIDRIFHKPIEYLGCTRQFSKSNWKLYFENVKDPYHASMLHLFHTTFNIFRVG
MKARSIPDATHGLHSIITMTKTQDDADTASAYKQQNIRSFDEGFSLEDDSILGLVSEYDE
DTTNHIQPIFPQLVIQQIHNTLVARQLLPKGPKNFELIFHFFGYADDTPELRDLRIKQAN
LVGPAGYISMEDTEATELVQRGTVRDANAASVIEMSRGNPDQQDTAITESLIRRFWVGYQ
QLMGY*
>GCA_000452945.1_BPMSHR5858_genomic_01989_1 andAd
MTDITDGLTGDIRTWFELHMLQNRYIGHLDNDRLERWPELFTEDCLYEIVPKENADLGLP
IGIVYCTNRRMLRDRVVSLRHANIYEAHTYRHMTSGLTIVGGEGGEIETESSYVVVQTRS
DGESNVYQAGKYYDTVVRTAEGLRYRKKRVIYDTSRVQTLLATPI*
>GCA_000452945.1_BPMSHR5858_genomic_01994_1 group_206
MFVAPAARRPGVGAALLDAALAAAARTAGQVALAAVDGRGAARAFCERHGFVAYGVEPRA
LDAANGCADALPMVKFPPLAPPHA*
>GCA_000452945.1_BPMSHR5858_genomic_01996_1 nhaK_2
MSPVSAFKLVLLSFLAIVALELLAKRLRLPPAAALLVGGAGIAFLPGLPPVNLDPDLVLI
VFLPPLLMDGAYFSVWEEFKRNVGGIMMLAIGAVAFTTLAVGVAVHLVAPGLPWAACFAL
GAIVSPPDAVAAKAVLERVALPRRLMVLLEGESLLNDAAGLVLFRFAVAAALTGAFSTGD
AVVRFAQLGVGGVAVGFIVGWLIVRFLKLVEDDYLVITVAVLAAWISYIAGELFEVSGVI
ATVTTGMMLGWHQHEVFSASVRNRGTAFWQVIVFLLEALVFVLIGLSLRGVIERLGGLGE
VFATMTPAVAAVVAAVVASRFVWVFAVELLKAPVARLRGRGPRGDWKAATVMGWAGMRGV
VTLAIALSLPDAMPGRDLILVAAFAVILVTVLLQGTTIGPLIRLLKLRDPGPAAHHLTEP
QTWARVEAAQLSAIQPLVHDAEGNVIHPRLLEQYTYRASVTARHQDEPAFPQRERDAHYD
VVLAAIAAGRAELLRLHRGGQIHDEMLYLLERDLDLQEIAAQHAKG*
>GCA_000452945.1_BPMSHR5858_genomic_01997_1 budB
MKASDLFVKALEAEGVQYVFGIPGEENLDLLESLRRSRIKLVLTRHEQAAGFMAATYGRL
TGKTGVCLATLGPGATNFVTAAAYAQLGGMPMLMITGQKPIKSSKQGHFQIVDVVGMMQP
LTKLTRQIVSIGNIPSAVREAFRRAEEERPGATHLELPEDIAHEEGDGKPIPASYSRRPI
AEEKALARAVDAIQAARHPLLMIGAGGNRKTTCRMLKEFVDTTGIPFFTTQMGKGVIDET
HPLWLGNATLSDGDFVHRAIEHADCIINVGHDVIEKPPFFMRADDKTVIHVNFLGAQVDP
VYFPQIEVVGDIANAVWQMKEKLARQPHWDFSRFMLIKAHFDAHLKKGQDDPRFPMYPVR
IVNDLYRSLPEDGIVCLDNGMYKIWFARYWRAHEPNSLLLDNALASMGAGLPSAIATKIV
HPQRKVIAVCGDGGFMMNSQELETAVRLKLDLVVMILRDDAFGMIRWKQENMNFPDYAMT
LANPDFVAYAQSYGAHGHRIASADALEPLVRECFASPGVHLIDVPIDYSDNERVLNREIK
RLSAQL*
>GCA_000452945.1_BPMSHR5858_genomic_01998_1 safD
MLKETYPYYLANEAVYANTGLEVTDKYSGKVATRVALADARAIDAAIAAAVAAQRPLRAL
PAFRRQAILEHCVARFRERFDELAEALCIEAGKPINDSKGEVTRLIDTFRVAAEESVRIE
GGLVNLEISPRAQGYSGYYRRVPIGPCSFISPFNFPLNLAAHKVAPALAAGCPFVLKPAS
RTPIGALIIGEVLAQTDLPKGAFSILPAHRDGADLFTTDERFKLLSFTGSPAVGWELKKR
AGKKKVVLELGGNAAAIVDADQRDRLDYVVDRLAFGAYYQSGQSCIGVQRIIAHADLYDA
LREKLIAKTRSLKMGDPKDPATFVGPMISEAEARRLAGWMDAAVAAGARIIAGGNVDGAM
FEATLLEGVGRDQDLYRKEAFGPVALLERFADFDDALARVNDSDFGLQAGVFTDSLSHAQ
RAWDELEVGGVVINDVPSFRVDNMPYGGVKDSGLGREGIRYAIEDMTELRLMVVRQR*
>GCA_000452945.1_BPMSHR5858_genomic_02002_1 yoaH
MPTVTPRASNETLRKSAAGSRLTLGNRILFSFGVLFVLMLFMAALSYQRLRAINDEAISI
ERDSLPGVYLASSLRASANESYIVLQRAVFVDAEAEAVQRDLAKVPGLLEEFDKLSSAYQ
ASTFRSDDQERFNAFRAAYERYLPLLNDAVQKARGARPDALAAYARVTPAWEEVIRHANV
LVQENRRFADQSATLIRESVHGTEITLAVALGVVLVVALMLGYLLHRAVTVPMARLIDVH
DVMRTGNLTQRLNLGRSDEFGTLESGFNRMADELTALVAQAQQSSLQVTTSVAEIAATSR
EQQATANETAATTTEIGATSREIFATSRDLLRTMNEVSAVAEQSATLAGVGQSGLTRMGE
TMRSVMDAAGSVNAKLAILNEKAININQVVATITKVADQTNLLSLNAAIEAEKAGEYGRG
FAVVATEIRRLADQTAVATYDIEQTVKEIQSAVSAGVMGMDKFSEEVRRGMRDVQQVGGQ
LSQIIAEVQTLAPRFQMVNEGMQTQASGAEQITQALAQLSEAAQQTAESLRQSSQAIDDL
TLVANQLRTGVSRFKVDA*
>GCA_000452945.1_BPMSHR5858_genomic_02005_1 group_19852
MERDVVERPRIAFDIDACWNRIGTRGDRSCERLDTYRRCLNCPVFERHAALLLDRPLTDA
DLADAARVAAERAPGANAPASKASAAARDGDADAVHSALAFRVADEWLALPIRVLREITD
TRAIHPLPHRRNRAVLGIVNVRGMLRVAVSLAELLSLDARADRAAPRMSFTRMLVVAHRG
DPVVFPVDEVEGVLRFASADWMPVPATVARTGAVHSRGVFAWRGKSIGLLDEDRLFDSLT
RSLR*
>GCA_000452945.1_BPMSHR5858_genomic_02021_1 bbsG_2
MDFSPSARCRELSERIAAFMRDEIAPVEARYAEQLTGDADWRRWRQPDVMETLKAKARAA
GLWNLFLPEAEHGGAGLSNAEYAPLAELMGHSFIAPEAFNCNAPDTGNMEVLARYGSPEQ
RRRWLEPLLAGEIRSAFCMTEPEVASSDATNMRATAKLEGDEIVLNGRKWWSTGIGHPLA
RVVIFMGLTDPQAEPHRRHTMVLCPLDAPGVRIERMLPVFNAYDEPSGHGEVSFTNVRLP
ASNVILGPGRGFEIAQGRLGPGRIHHCMRALGAAEKALTLLCARATARTAFGKPLVKLGG
NGDVVANLRMAIEQARLLTLKAAWTIDTQGVKAALSLISQIKVVVPAVAQQAADAAIQIH
GGAGLSNDFPLAALYAYARVLRIADGPDEVHRAVVAKLEVKRQLAAAEAA*
>GCA_000452945.1_BPMSHR5858_genomic_02022_1 group_2809
MSVRLDDARDVRDEDRLDAAKLDAYLKPRIAGLSGEPRIRQFHGGASNLTYLIGYGDREL
VLRRPPAGAKAGAAHDMLREAAVMAALGPDYRYVPAILARCDDPAVLGSEFYVMERIAGV
ILRRELPAELKLDRADVRKLCERFVDRLIELHAIDASRPEIAALGKGEGYVARQLSGWAQ
RWRNALTDGTNPCDDVLAWLERHRPAGERRICVIHNDFRFDNVVLDPADPLSIVGVLDWE
MATLGDPLMDLGGSLAYWAQADDDPAFVAMRRQPTHADGMMTRRELVEYYGARTGLDVGG
FVFYEVFGLFRLMVIAQQIYRRFVLGHTTNEQFAGLGAVVRYLGERCRRVIDAAGGAR*
>GCA_000452945.1_BPMSHR5858_genomic_02025_1 linX
MTKTKHLAPRLAGKRAFITGAAGGLGRAIARRMAEQGAQVFLTDIVDAAVLEAFAAELND
TAGERIAWAAVQDVADEAQWASRLSQAADAMGGLSVLVHNAGVGSFGAIGQIARDEWRRV
MAINVESIVLGTKHALPYLEAGAPASIVNISSVAAFKQEPDYTAYNASKAAVASLTKSIA
VDCARRQTEVRCNSIHPSFIMTGIVAPIVRLVGEKEAARKLARGVPMRRLGEPDDVAHAA
VYLASDESRYVTGAELVIDGGMCAV*
>GCA_000452945.1_BPMSHR5858_genomic_02026_1 yfmJ
MSTPVNRQLRLKARPDGRVGHEHFTLAEAPLPALGPGEMLVRVLYLSMDPTNRVWMSDIP
QYLPPVAIGDVMRALGIGRVVASNAPGFAEGDLVQGLVGWQDYAHVRADEIAQYTKLPAA
LGLPLPRLLGACGMSGLTAYYGLTEIAPVQPGETLVVSAAAGSVGSVAGQIGKIHGARVV
GIAGGADKCRYLTDELGFDAAVDYKSDDWKRALKDATPDGVHVSFENVGGEIMRAVLSRM
AIGGRVALCGVIANYNNGRPADDVSVLIAKRLTMRGFLILDYRKSREAIATLAGWLRDGR
LKAEETVADGLTNAPDVLNRLFDGSHRGKLVLRVDPQA*
>GCA_000452945.1_BPMSHR5858_genomic_02027_1 bcr_2
MHPLRSILPLALFTAVGLLATDLYLPAVPSLPQQLGGSIESAQATLAAFSAALAVSQLVW
GAAADRFGHRRTLAFAVLLQLVAGAACALAPSMGALIGARLAQGFGVGAAMVIVPALVRQ

SFGDGGAVRALAWLGIVESAVPGLAPLVGAALLVVADWRTSFWIIVALSAIAAPLVFRVI
PTARAMRACAPANVGAHAGGYRRLLRSPVYLGYALGHALCFAALLAFVASAPQVVEIWLG
AGPSTFSLMQACGVAAFMLSAARSGKWSDALGLDRIIALGALLQFAASAAFLLLAYADWR
STPLVVASWMLFCGSLGLRGPASMARALAAEPAVAGRAAGLLMFFGLGGAALATQAVAPF
LRLGLAPVAWMCAGFTLASGAVVLWGIAIRGRHRAAATEIA*
>GCA_000452945.1_BPMSHR5858_genomic_02030_1 ydiK_2
MDSGNDHQKFFHLLLLVVTVGLCWILTPFFGAIFWGTILAILFQPVQRWLAARFGKRRNL
AALVTLSLIILIVILPLAFVTATLVQEIAYAYQQIKTMQPNMTQYFQEFMHALPSSVHRV
LHNYGLTDIAGIQKKLTDGAAAISQFVAAQALSIGQNTFQFVVSFGVMLYLVFFLLRDGG
EIGRRVRRALPLDEEHKQHLLTKFTTVVRATVKGNIAVAAVQGALGGLIFWILGIEGVIL
WGALMAFLSLLPAIGAGLVWVPAAGYFAVTGQIWKCVILVAFCVGVIGLVDNLLRPILVG
KDTKMPDWVVLISTLGGMALFGINGFVIGPLVAALFMASWDIFARTEQTDWE*
>GCA_000452945.1_BPMSHR5858_genomic_02032_1 group_19963
MALSDTVEYKLDRGLSEARRAGRRFARDARSAARDLNGDVKDNMRSLVDELDALLKEDVD
VDALRKRLHGRLEAARDTLDDASWHASRRLRQSAERVTQAVHDNPWQTAGIVAGLAFAAG
ILLARR*
>GCA_000452945.1_BPMSHR5858_genomic_02034_1 group_3280
MRTPTPLLAALSLLSTVAAAPAFAAAASPVSTNCGGSATPIAEIQGAGAPSPLAGQNVSI
EAVVTADFGGADGFGGFFVQQADPQRRNQPGVSEGLFVYSPKARARAGELVHVTGKVEEK
YGQTQLTLSGAVAVCASGQSVTPATLALPVDSPNAFAAYEGMLVRLPQTLTVTEVYELGR
YGSVLLSNGRLRTPTSVVPPAQAKAVAEANARNRLILDDGSNKQNPATVPYPAPALSAAN
TLRAGYTAADVEGVLELRYGAWRLQPVPGARTPTFGASANPRTAAPARDPRANLRVASFN
VLNYFNGDGAGGGFDDPSNRGAKNYDEFVRQDAKIVSALKALDADVIGLMEIENDGYGPL
SAVRQLAAKLGENWRVVDPGSARLGGDAIAVALIYDSRKVKPIGNAATLAIDDKNRQPLA
QTFRPLGGSRAVTVAVNHLKSKNCPDATGDDLDQGDGQGCWNATRSRAAAKIADWLARNP
TGAPSEGVLLIGDLNSYTYEDPVRTLESRGYVNLVSSKVGSGAYSYVYNGEAGYLDHALA
TNALAWRVKAVHDWHINADEPIALQYTLAYKSAEQQRTYYAPDAYRSSDHDPVLIDIALA
DEYAAAASRRGAEMAAAHSRRPWH*
>GCA_000452945.1_BPMSHR5858_genomic_02036_1 group_19745
MRFMIMVKANAASESGAMPDASLIAAMAAYHEELAKAGVLLDAAGLQPSSKGWRVRYSGG
KRAVVDGPFAETKELIAGYTLIQVRSRDEALEWTRRFPAPFGEHDGGEIEVRQLFELDDF
ESGDGIDRFRELESKLG*
>GCA_000452945.1_BPMSHR5858_genomic_02037_1 group_3281
MVQIVVIAGAAALAFALPLMLYAASRPGTRRVEHRARIEAGAARIFPAPADLRRFDACNP
CERKAPALRGAYGAVRSGVGASDAWTREKAGVGRFGTTGLGEPSHVRTRLDFVRRFDAHD
VAEFTLRPDGDATVVTWAMRGPSPFLSKLMRACLSIGKMVGKDFYTRLANPKALAEGRA*
>GCA_000452945.1_BPMSHR5858_genomic_02042_1 lutA_2
MSHREGSLEAPTRHPLDWRSDAFYDQAAIDAELARVFDICAGCRRCVSLCGAFPTLFDLV
DETETGEAHAVDPKSFGKVVDQCYLCDLCYMTKCPYVPPHPWNVDFPHLMLRARAARYRR
NEVRLRDKLLSNTDMLGHFAGIPIVTQAVNAANRTPAMRGALEATFGVDRRAWLPPFASR
KFRRAAPRSPATPPRDGERTPGRVAIYATCYVNYNEPGIGHDLLAVLAHNGIPYELVASE
ACCGMPLLEQGNLEGVAAKKAANIPVLARHAREGHALIGAIPSCVLMYKSELPLMFPDDA
DVRAVADAFWDPFEYMIARHRDGLLKTDFKHELGVVSYHVPCHARVQNIGRKAADALSLV
PGTKVKVVERCSGHAGTFGVKREFHAQALKIGAPVFNAIAEQAPDYMSSDCALAGHHIEE
GIGAKGVEPPPFAHPITLLRRAYGI*
>GCA_000452945.1_BPMSHR5858_genomic_02043_1 group_19798
MTLTRDSLLTLEAYAKVRRQEHARVIAHKKRRAVSIGNHLRLLFEDETTIRYQIHEMLHI
EKIFDEDGIQAELDAYLPLVPDGSNLKATLQIEYENETQRRAALARLVGIEDRVFLRVDD
EAPVYAIADEDLERDTAEKTSAVHFLRFELGDAMKAKLKAGAPLSIGCDHPHYPIQAARI
DPDVAASLAGDLD*
>GCA_000452945.1_BPMSHR5858_genomic_02044_1 group_3283
MFFDELNDEEWARLSTLIADEPIRLNRRGRPRAENRVVANAVLWILTTGEPWSKLPGRYP
SGPTCRRRFEEWLASGTLGEMVRVLSEESGRAFAYVPPPPESVAPVRRAEPAADCDRLRG
VFWQNPESWQLPVAQTDVCHPGAGVNVRRFASEDDEDGAHDADAHPHGFNVPGAPPVRDH
AHGRAHAASFSFATHAPQTETYRGYTVYGIAQPVQNLMYRAWAEIVQDDRRVERSGLIGP
RFTDAESAEQYALDWARQWIDRRCAGAPSVRETPAPVAGQAGKSVPTIAGLSALAMAETN
IKHFAAERHRAPAEGRGDASQVQIDRHEYAYRVG*
>GCA_000452945.1_BPMSHR5858_genomic_02046_1 wcaJ_2
MLSVLARMIDIAMAVAGAMIAAMIHRGGFVWLDDLQRTVVVFDCMLVVVFFPAFGIYQSW
RGKPVFGLMWRVSVAWLVVEGAGILMSFSVHHAGELSRLWLGYWAMATITLLVASKACVY
AVLRQLRRGGYNHKSVAIVGRAQASATKRLIAHMRARPDAGFNPVCIYDEHDDASTGETD
GIAIERRFGHLVRRVRSREIGELWLVLPLSEEPQIHRIVTEFRNDFVNIRFVPDVRSLSF
FNQEVIELLGVPAINLAASPITDVRILPKFVFDRLFALVALMVLSPAMLVIATLVKLTSP
GPVFFRQKRKGIDGNEFEIYKFRSMKVHQEMGGKVTQARKNDSRVTPVGRFLRRTSLDEL
PQFINVLKGEMSVVGPRPHALEHDDIYKDLVKGYMFRYRIKPGITGWAQINGFRGETDRI
EKMMGRVKLDLYYMQNWSFWLDIKIVALTLWKGFTGSNAY*
>GCA_000452945.1_BPMSHR5858_genomic_02047_1 ptp_2
MNLTIIGSGYVGLVTGACLADIGHDVFCLDVDSKKIDVLNNGGVPIHEPGLGEIIARNRA
AGRLQFSTDVEAAVAHGDVQFIAVGTPPDEDGSADLQYVLAAARNIGRHMNGFKVIVDKS
TVPVGTAERVRAAVADELKARGEDSMFSVVSNPEFLKEGAAVEDFTRPDRIVIGCDDDVP
GERARELMKKLYAPFNRNHERTLYMDVRSAEFTKYAANAMLATRISFMNELANLAERFGA
DIEAVRRGIGSDPRIGYHFLYAGCGYGGSCFPKDVEALIRTAAEHGQALRVLQAVSAVND
EQKHVLARKVVARFGGDLTGRRFGVWGLAFKPNTDDMREAPSRALIAALLSRGAHVVAYD
PVAQDEARRVLAIDLGEYPDALGRLAFADDEAQVAQDADALVIVTEWKAFKSPDFAALGK
RWKTPVIFDGRNLYEPIAMRELGIEYYPIGRPGAAPRTAAGESDAARAARAGA*
>GCA_000452945.1_BPMSHR5858_genomic_02054_1 mshA_5
VRIAIVTHVVRHNDGQGRVNYEIARAALAENCQVTLVASHVAPELLADARVRWIAVKAGR
FWPSNLVKQQVFAIKSAWWLRRHREAYDVLHVNGFISWVRADVNTAHFVHSGWFASRYYP
FGLSKGLWSAYQYVYTRVNTRLERWAYRRARAITAVSQKVADEIRRIGIDGGRIGVIYNG
VDAQAFANAAPDRRAFGLPAEPFMLLFVGDLRTPRKNLGTVLKALAHLPPNVHLAVAGYL
PGSPYPDEARALKIDSRVHFLGLVKNMPTLMSSVDAYVFPSRYEAMSLSLLEAMAAGLPV
VTARTAGGAEIITPECGIVLDDPDDPAALAAAIERLARSRDVCRAMGEAARRLMEGFGWA
RMGAQYIALYRRLRQSSQPSPLAGTEHVVTQERS*
>GCA_000452945.1_BPMSHR5858_genomic_02057_1 fbp_1
MTVVTTESGLKYEDLTEGSGAEARAGQTVSVHYTGWLTDGQKFDSSKDRNDPFAFVLGGG
MVIKGWDEGVQGMKVGGVRRLTIPPQLGYGARGAGGVIPPNATLVFEVELLDV*
>GCA_000452945.1_BPMSHR5858_genomic_02060_1 group_3286
MSRPALKPFSTPPAATVEQMSAAGLRAFFNIARDWNLSTDEQLVLLGSPGRSTFFKWKAS
PQTARLTHDTLERLSLVLGIYKALQILLPQPAAADSWVKRPNDAAPFNGKRALDRMLAGN
VSDLVAVRQYLDAARGGWA*
>GCA_000452945.1_BPMSHR5858_genomic_02061_1 group_19795
VTSPERQTNWPTASLDWAPAYRVIPTRFPAINLFDRVASAEDFDALYALEALTNDRVRTE
VGMLDLVPPQERRFGPGYGPIMAAFTHLNPNGSRFSDGSYGVFYCARARNTAIAETRYHT
SLFLAATKEPPMRQQMRLYTVAAQGEVADVRAWRERDPALLDLVDYSAGQALGRAVRDAG
GAGIAYPSVRDPGGECLAAFRTTLLHDCRHAAYLEYNWNGAAIDAVFELNQVG*
>GCA_000452945.1_BPMSHR5858_genomic_02063_1 group_19921
MTFYPQVLRNRPRMVIAVAAGVALGLLFPYPLRPFARVLIGWDCTIWLYLVLMWVRMVRA
HHHKVREIAMREDENATMVLTIICFATVASIAAIVLELVSAKSVGFRSGLGHYAVTGATM
FGAWFLIPTIFTLHYARLYYLSPKEARAMAFPDRELEPDYWDFLYFSFTIAVASQTADVS
LRGRSIRRAALAQSILSFYFNMAVLGLSVNVAAGLLS*
>GCA_000452945.1_BPMSHR5858_genomic_02064_1 group_19931
MNTSTDRIEKRIVLNAPRSRVWRALTNADEFGAWFRVDLAGQTFEAGRCVEGRITYPGYE
HLVLRMRIERIEPEHRFSYHWHPAAIDPAVDYAQETPTLVVFELADVAGGTLLTVVESGF
DKLPIERRAEAFRMNDGGWGEQMTNIAAHVDAR*
>GCA_000452945.1_BPMSHR5858_genomic_02065_1 group_19914
MSTRAEGRTSRAPLKRAQLRDCAEVFAALGDETRLRIVAALCAGTALSIAQITAGTDITR
QSVTQHLLVLAQAGLVHDVKVGRERRWSFDPARIDAARAALDAIGRQWDQALLRLRQAVE
G*
>GCA_000452945.1_BPMSHR5858_genomic_02067_1 group_19770
MKTETMLIQILESVIGVKKVTGDTRFLDIGGNSLNLVEVLKQIKAKTGVAPSPRLFFDKT
RSTVAALSAEIDTLREMSSAAAPVAG*
>GCA_000452945.1_BPMSHR5858_genomic_02068_1 group_3287
MQKRFHLSEAEKKFFYENGYAGPFTLYEPEEMTRVWEEVRMDLLDTEKAAFPNSKLNYDR
HLDMLSLNQIISHPKIVDRISSILGPDVLSWRTEWFPKYPGDEGTDWHQAESFVEFEGTE
KLEPTAAEEGRPWELTAWIAMSEATKENGCLKLMPGSHRTWFFDEKRNIPFEPENFNKRV
LDDGQKSGFYGYDYEKLKLDPSWKPDEARAVHMEVKPGQFFIFTSRCMHGSNPNTSKDSV
RFGWATRFVPTHVKVYAGQDSFHHFGEVLPLDRYSTVLVAGEDRYGHNRVTKPLAAVAAL
A*
>GCA_000452945.1_BPMSHR5858_genomic_02071_1 ilvE_1
MKSSYVWKNGELVPWEHAQVHVMSHALHYGTSVFEGVRAYEIGEKAAILCGREHFERLLF
SCKVARIASPMSVGQWMEATVETLRANGQRSAYIRPLVYRGAGESLGLDARQCPAEALLI
TTPWGAYLGDEALQQGVDVQVSSWRRNGAGAASTLAKIGGQYVNGQAIVMEAHENGMSEG
IALDTNGFVSEGSGENVFLVYKNEIFTPSIGSSILSGITRHCVIRIARDLGYTVTETSIP
REMLYLADEIFFTGTAVEICPVRSVDRMPVGSGKRGAVTKAIQDQYFGIVRGTHADKWNW
LTPVAVPSRDGARA*
>GCA_000452945.1_BPMSHR5858_genomic_02074_1 group_19830
MILISDATLRDGNHAIRHQLSAAQIHAYARAADEAGIDVVEVGHGNGLGGSSCLLGQTPI
SDRLMLETARAALRTSRLGVHFIPGLGKAADISLALEIGVDVVRVATHCTEANVSARFIE
QTRAAGRTAFGVLMMSHMAPPDALLAQAKLMERYGAQAVVLMDSAGYSTPSLVRAKVERL
VDGLDIDVGFHAHNNLGLAVANSLVALEAGARIVDACVKGFGAGAGNTQLETLVAAMERE
GHDTRTTFERVMTLARGTETFLNPKTPHIQPANIASGLYGLFSGYVPHIQKAAQEFGVNE
FELYKRLAERKLVAGQEDIIIEEASRLARERDVQRATGGVRVRELSA*
>GCA_000452945.1_BPMSHR5858_genomic_02077_1 group_19749
MARNRTVLPLERAQVDVVQERSGVVGDRRISLRVCGEATQGYLLERYHAAPGEPSFVQRV
NLFDDATIASFVEHDPYSAELESLYRAVIDVPKKALKFAGHETIQPEFASECVSESGLLT
VMRNTILECGATNCFYHYFRIDEKTGNLKNHELLIGGTPVWPHRYVHRHWYLNDPAMAHA
RNDTRPLRASTLAPLPHDHWLNQQAQFLGLTSNVFFPAHRRDDDTIGLLHVSSSLPAAQG
EEIIWRNRRTLRGLATEMLEWKVTAQRNTLAREMSLSHHELVALRLVGRGANARHVAEEL
KLQEHAVYQLFTSINKKMNSSHIRTSANKAKQFGFLAEGYISE*
>GCA_000452945.1_BPMSHR5858_genomic_02078_1 group_19881
LNDCARDIAPDRDQIRIGSGAPGIERVEAHFLDHAFTPHRHDTYAIGITLSGLQTFGYLG
EIHHCLPGQCHILHPDELHDGRAGTDEGFGYRIIYVDPALVQEALGGRMLPFVRSPIFQA
PAVSEALAAGIWNLDEEIDNVSRIDIAVAVANLLTAAAAKGSAPKAGPLAMVELTRIRDI
IASCPREPISMDALEHASGLDRWTIARQFRRLFGTSPSRFRTQRQLNLVRRLLMEGESLS
TASTDAGFSDQSHMSRHFKSTYGITPGAWISAVRTRHAQHPSDH*
>GCA_000452945.1_BPMSHR5858_genomic_02079_1 group_2380
MVNQVDLVSETLRSLVALLFAAAVVMGSPGPSTVSATAMGASYGVRRSLKYAWGLIAGTV
AVLLLVSTGVTAFVMSMPSAARILNIVSALYILYLAYKIATAPPLDKRAENLSSPAFKGG
FLLAVANPKAYIAIGAVFAGTTLVATDYGLDAAAKIVLLSVMIVVIHLGWLLAGVSLSRF
LHDPVASRIINVSLAAILAIVSLIALFE*
>GCA_000452945.1_BPMSHR5858_genomic_02080_1 group_19750
MDTKHDGGSPIAERRTEGFRLLESGMPQADVARRLVVSRQSVCRWAKRLAERDGRIAKAP
GRPRRLNSAQCATLRAMLDDGALAAGFASPQWTLARVRKLIEREFGVAYSKTGGWELLRT
LGVSLQRTERHPHIAR*
>GCA_000452945.1_BPMSHR5858_genomic_02086_1 thiE_1
VSARDGDRAPDHDMHDDLALPPYYLITPEPASGSDADLAAFLDRLSDALATGLTLVQLRV
KTLDAPAYAALAAGALARCRAQRARMIVNGPIAVEAALALGAAGVHLGSAALRAATARPL
GSEGLLSAACHSLDELRHAQRIGADLATLSPVLPTLTHPGAPTLGWTRFAECAAHTRVPV
YALGGMTRTHLETARAHHAHGIASIRGLW*
>GCA_000452945.1_BPMSHR5858_genomic_02087_1 group_2381
LIQIFALIGALLLVALNGFFVAAEFGLVKLRATRVKTLARQHGLRGRILRIVHARLDAYL
SACQLGITLASLGLGWIGEPAFAALLSPVLAFAGVESEKLVHLISLVFAFSVISFLHIVV
GELAPKSMAIRQAEKTGLWVALPLYAFYWAMYPAIWVLNTSANAVLRLAGLSADHGGDAH
YSTDELKLILRSRRNAAGAAKAPKHAYSNDEWNTLAHSLDFSSMTVSDLMRPAHEMVGLR
RDLPLADNMEVVARHRFSRYPLFDDQSRETVNGLIHLKDLLLARHAGAALADLSDYVRPV
QYVKPDTPALDLFRRFRKGAPHFALVGKKGEKPIGFLTLDNLLGALVGQIHDEFRQGDSD
WSRLDDGTLIGKGSLPVVSLEQALGIDIDEGRAESVGGLVIQALNDLPTEGRRVSFDRFD
IVVKKMIGPRIVLVRVYPKSPKEADE*
>GCA_000452945.1_BPMSHR5858_genomic_02089_1 group_2382
MKHDARKHLRANLLMLAAAAIWGSAFVAQRLSLAVIGPFLFTGLRFLLGAAVLVPLLCAN
GAARAHCATLARERTRLLPGLALGGLLAVSISLQQIGLQYTKIANAGFISSLYVVLVPVI
GVFFRHRTGIGTWLGALLAAIGLYFLSVDAHFSMLYGDWFQLAGAIVIAFHVIAVGHLVR
RHDPLVLSFMQFVVCGALCLALGLAIEPLDRATLTRALPTLLYGGLLSVGVGYTLQVVAQ
RDAAPAHAAVIFSMEGVFAAIAGWAALGETLSLRALAGCALMIAGLLVCQLLPGHARRAS
DNALAA*
>GCA_000452945.1_BPMSHR5858_genomic_02116_1 group_19844
MKKSLLALVALSAFAGAAHAQSSVTLYGIIDEGFNINTNAGGKHLYNLSSGVMQGSRWGL
RGTEDLGGGLKALFVLENGFDVNSGKLNQGGLEFGRQAYVGLSSGFGTVTLGRQYDSVVD
FVGPLEAGDQWGGYIAAHPGDLDNFNNAYRVNNAVKFTSANYGGFTFGGLYSFGGVAGDF
SRNQTWSLGAGYTNGPLVLGVGYLNARTPSTAGGLFGNNTTSSTPAAVTTPVYAGYASAH
TYQVIGAGGAYSFGAATVGITYSNIKFMNFASTVFPNQTATFNNAEINFKYQLTPTLLAG
AAYDYTQGSKIAGSSAAKYHQGSVGVDYFLSKRTDVYAIGVYQHASGNVIEADGNTVGPA
TAAINGLTPSSNRNQFAARVGIRHKF*
>GCA_000452945.1_BPMSHR5858_genomic_02117_1 coq7
MVFDELITEFDRGLRSIAGVSRMSRPVPKPAAAAPAELSAAERKHAAGLMRVNHVGEVCA
QALYQAQKLTTSSAGLKEMFEHAAREEEDHLAWTAHRLKDLDSRPSLLNPLWYAGALAIG
VVAGRLGDKMSLGFMAETERQVESHLDGHLSELPAADVESRAIVEQMRADEVKHGKSATD
AGGIELPMPARMLMRAASKVMTSTAYYL*
>GCA_000452945.1_BPMSHR5858_genomic_02120_1 ftsL
MSRLNIFLLIIVMGCALSVVNSTNQQRQIFIQLQRAQSQEHQLQQDYAQLQYQQSALSKT
SRIEQLATSSLKMQPITTGRTQYLTLSPGAAKAVDAPLPASAAPTGGAR*
>GCA_000452945.1_BPMSHR5858_genomic_02123_1 murF
MTMLTLREAAAMIPGATVLGDERASFERVSTDSRSAGPGDLFVALKGERFDAHDFVGDVA
ARGASAAIVSRSPAEWNLPALKVADTRAALGALAHGWRMRFSMPLVAVTGSNGKTTVKEM
IASIFAAAVGEHARLATAGNFNNDVGLPLTLLRLNAAHRLAVVELGMNHPGETEMLAKIA
APTVALVNNAQREHQEFMATVEAVALEHAAVIHALGPDGVAVFPADDAYAGIWRVAATGN
RIVDFALHGAAGGAPAAVTGALDGSLLAIDTPGARIDVRLNALGEHNARNALAATAAALA
AGVPLAAIKVGLEAFAPVKGRLQVARAAAGSLAGATVIDDTYNANPDSMRAAIDVLAAQS
APRVLVIGEMGEVGDEGPAFHREIGAYARERGIDALFALGAATRPACDAYGAGARHFDDA
AALVAALLAANFGAHATLLVKGSRFMKMERVVDALANETASGAAPGAH*
>GCA_000452945.1_BPMSHR5858_genomic_02124_1 mraY
MLLALAQWLQGDASFLRLFTYLTFRAVMATITALVIGLVCGPWVIRKLTQMKVGQAVRKD
GPQTHLVKSGTPTMGGVLILIGIAVATLLWGDLTNRFIWIVMLVTFGFGVIGWVDDYRKV
VYKDPRGMSSREKYFWQSVIGLFAAVYLAFSVSEANNVRVFDLFMAWVRSGLSMGLPARA
DLMLPFLKSISYPLGVWGFIALTYFVIVGASNAVNLTDGLDGLVIMPVVLVGASLGVFAY
VMGSAVYSKYLLFPHIPGAGELLIFCSAMGGAGLAFLWYNTHPAQVFMGDVGALALGGAL
GTVAVIVRQEIVLFIMGGIFVAETLSVMLQVTWFKYTKRRYGEGRRIFKMAPLHHHFELS
GWKETQVVVRFWIITLMLCLFGLSTLKLR*
>GCA_000452945.1_BPMSHR5858_genomic_02128_1 murC
MKHIVKHIHFVGIGGAGMSGIAEVLVNLGYQVSGSDLARNAVTERLEALGARVSIGHDAA
NIEGANAVVVSTAVRSDNPEVLAARRLRVPIVPRAVMLAELMRLKQGIAIAGTHGKTTTT
SLVASVLAAGGLDPTFVIGGRLTSAGANARLGTGDFIVAEADESDASFLNLYPVIEVITN
IDADHMDTYGHDFARLKQAFIEFTQRLPFYGSAVVCIDDANVRQIVPLISKPVVRYGFAA
DAQVRAENVEARDGRMHFTVRREGREPLPVVLNLPGLHNVQNALAAIAIATDLDVADAAI
QQALAEFNGVGRRFQRYGEIAAAGGGAYTLIDDYGHHPVEMAATIAAARGAFPGRRLVLA
FQPHRYTRTRDCFDDFVNVLSTVDALVLTEVYAAGEAPISTANGDALSRALRAAGKVEPV
FVATVDEVPDALAKLARDGDVVITMGAGSIGGVPGKLAQDTQQKG*
>GCA_000452945.1_BPMSHR5858_genomic_02166_1 leuA_1
MKRNPAEKYRPFEPVRINGRRWPSRAIERAPIWMSTDLRDGNQSLIEPMTIEQKLEYFEM
LVAIGFKEIEVGFPSASQTDFDFVRKLIVERRIPDDVTIEVLVQSREDLIARTFDALEGA
PRAIVHLYNAICPSFRRIVFGMSKDDVKALAVDGTRIIKAHAQARPATHWTFQYSPETFS
MTELPFAREVCDAVAQTWRPTRDHKMIVNLPATVEAATPNVFADQIEWMDRNLGYRDSIV
LSVHPHNDRGTAVAAAELALLAGAERIEGCLFGNGERTGNVDLVTLALNLYTQGIDPGLD
FSDIDAVRRVVERCNRIPVHPRHPYAGDLVFTAFSGSHQDAIRKGLAQQRPDAIWEVPYL
PIDPADLGRSYDAVIRVNSQSGKGGATYLLERAMGFAPPRRVQIEFSHAVQTLADTSGAE
VTADAICALFTREYLQTIGPAARAGASASWHDREVALAHAEDAPSAARQAAAAFASAAGV
AIDITSCEVERTVNGETAVFVGARIGDAALRHGAGVHRDPVAAATDAVASAINRSAWSRD
KRRAAA*

>GCA_000452945.1_BPMSHR5858_genomic_02172_1 cydB_1
MDYATLKVIWWLLVGILLIGFALTDGFDMGVAALLPFLGKTDDERRIIVNTVGATWEGNQ
VWLITAGGAMFAAWPLVYAASFSGFYFAMLLVLFALFFRPVGFDYRSKRDDPRWRTSWDW
ALFAGGFVPALVFGVAFGNLLQGVPFAFDTDMRVTYHGSFWALLNPFALLCGLVSLTMLV
AHGAAFVKMKTDGAIARRASIALRVSSLVAVVLFVLAGALIASFIGGFQITQAAPLDTVA
NPLLKQASAAPGLWLVNYGLYPWMIAAPAVGIAGGVLAMLLAGSKREKTAFICTGLMITG
VILTAGFSMFPFIMPSSLDPKSSLTVWDSTSSQLTLEVMLVAVIVFLPIVLLYTGWVYRV
LRGKVTPHVLDENRHSMY*
>GCA_000452945.1_BPMSHR5858_genomic_02182_1 menE_1
MRDSSPLIERLLAPLAEGRNCRAHGVGTLDESGIERVLSYRQLSEAGLRMSAALNALGAR
AGDIALVALPATGDYLALLVGCVLSGVVPCTVPPPLKRASGSGPQVLEVACRLYRPRFVF
TASAHVAELEAQLRDTGARVIDVGILHAASTGAEPRLALRGAGDTHHIQLTSGSTSHPKA
AVISHRNVAANIAGIANACGYSKHAADNTVIWLPLHHDMGLVSLLLHLYYRTSLRLMPSM
SFVRNPLGWLRRIAHARSTIAVAPTFALRYCVRRFNAATMDGADFSHLRTFLVGAERVDR
ATLSDFASTFAPYGFHASALQPCYGMAEATLAVTLHRSFDTPNAPAFTHVIADRIDTATL
ARHRRAAPAPDGHTGATDTLLAVGAPLDDMAYEIRDAQGRLAAEREAGEIFIRGASVMQG
YLPSAGAPAPQPFDAHGWFATGDIGYVSGGQLFILGRKKEIIIIRGTNYFPHEIEETIDS
HPAVYKGACIAVGIHDETQGTENLVVLVEAHANHVSRDSRAELQARMQQRLGYCAQRIAF
VEPGALPRTTSGKPQRLKARAMFVSGELREIALPPSRRADADRVAPYP*
>GCA_000452945.1_BPMSHR5858_genomic_02189_1 btrK
MNTDTIGSAPHAAADAGNAEWLDGLDTPCYVFDPQCVLAGYAALRAALGTRLVVSLKANS
HADLFLRCAHAFEDGIELASQGELDTVVGRGKHVKYLNNPAMGDALMRAGIASRCHFILD
GPDMVRRFIALASGKETEDVMLRINAGALLGAHAPKSLRDHFGMTPAEALGMANLLTQAG
RRVAGLHAFVGSGNFRAGPTKGGDADSADLAWALAQLAEQIEALVRAPVTHLNLGGGFSE
KGHDDATFERYRARIAPLAKRYTLVHEAGRAIFANAGVFVTRVRAVKVWDDRVIAVCDGG
MSHNFLLAKTEAVLKTWAAPTLVPAGAADAHRAPNARPITFVGNTCNRADVIGRLERHPR
LPQPGDFVVFSQCGAYHHSYTVSGFLSHKPAQVYIRQA*
>GCA_000452945.1_BPMSHR5858_genomic_02191_1 group_3298
MLTIHHTAIRQPSRRSVDAAQASGRYRYDGRLAPHAIRFAMPYERLDYDTHTFAHLARPA
PASLPVETRGRTLSDLALEAVRDVVGAAGPGACASISQIVVAQASLNEHAGESVAGRMQH
ALGLKHIVPFALGQCGTLGLHAALPLAHGLLRDGGRMLFVAADKWVYPFLRTYGEFVAYG
DGAAAILLGHAETDAEADAAQARVLGHALAHGEPIADPWARQPAELERALVAPTVDAARA
ALADAGVGAARIDCFAPGGFSASFRTTLARALAIPPSRLQVRDGVEHLSTADTPRALAQA
QASLAPGERRLALFCDTALAGGAGALVAQLRGAHASSSSTRTRYPS*
>GCA_000452945.1_BPMSHR5858_genomic_02192_1 group_4110
MSAYQVSLRELRFFLWELCDAEAAFLSHAPYRTHDRAYYDRLLERARDFALEIGESYRAS
DIESCSLRDDGTVRIPADFHALWPRFRDEWAGLLFDRHSAEAGDAPHADVPMVVKQAVFE
MLMGANPSFMTYGGFTHPACKLLTLHGTPEQKALRGPLLRYDWDACFCATEPQAGSDMTA
VRTVATPLGDGVYAVTGEKVYISAGMHDLTENTLYFVLGRIGSTAPDSFSLSCLIVPRFW
RDDATGKLESNHVECVALPRKMGLNGCANAHLVFGRSGTTRAHLLGNRRNVGLLQLVPLM
NQARMGTGLFGIGVASSAYLQSVSYACRRVQGRPIDAASDTAAPRVRIVEHGDVQRMLVD
MKARVEGCRGLLGKLTAAATRATILEATPGADPADIERARKLQLLLTPICKAFISDQAWR
VCETAIQVHGGVGYTDASPVEQNARDVKILSIWEGTNYIQAQDLVREKLGFGRKPLLLRY
FRDALDASLARIEPTAPATFAPWFAQLRGAADALARALESIARAVRDGHMHASSQVYTRF
LEMFGLVACAWALLEAACIAERRLATREVAAEPAEAAFYRGKVKAARYCFANLLPLAAQH
AAAIDALPDIACAITAEELAEVE*
>GCA_000452945.1_BPMSHR5858_genomic_02197_1 group_2393
MLEVFLILTRNELLGQLRSKARLFWTFVFPILLMSVMLVAFGKHSSLGVVELAFDGNAAA
PQARACRAAIEAAFAGNGTVGARVVAPNAAGAAGGDRVRVSWPARAADPVRVSYDFNGPL
AARAAARTIEIALVRCAASEAGLPAAYQVRFENDGHALAPLDYGEFFATGILIMAFMSIG
VVSTATTIATLRERNTFKMYVCFPVSRFVFLASLIVSRVILMLAASVTLMLAARYLFQVP
LPLWSLRALRAIPVVLLGAAMLLSLGTLLASRARSLAAAEAWCNLIYFPLLFFSDLTIPL
RAAPHWLRVVLLVLPTNQFAVALRGVFVRDLGYAQAAWPLAVVAGWTLVFLAGAALTFRW
HQD*
>GCA_000452945.1_BPMSHR5858_genomic_02201_1 cysD
MNATLEPSSPGGAAGASTSRMGHLDWLEAESIHILRELVAECSKPALLFSGGKDSVVVLH
LALKAFGLGANRRTTLPFPLVHIDTGHNYDEVIDFRDRRAKQIGAELVVGHVEDSIARGT
VVLRRETDSRNAAQAVTLLETIERHGYTAMIGGARRDEEKARAKERIFSFRDEFGQWDPK
AQRPELWSLYNARLHRGEHLRVFPISNWTELDVWQYIAREKLELPSIYYAHRREIVRRNG
LLVPVTPLTPMREGETSEQALVRFRTVGDISCTCPVESDADDVEKIIAETAVTEITERGA
TRMDDQASEAAMEQRKKQGYF*
>GCA_000452945.1_BPMSHR5858_genomic_02213_1 group_2394
MTTVTFSGPIFDLVDGVLRDAPEYRGFRAGAQSRAEFAAFLERVRTQLVGRGVPANDVHF
AKRFPVSIDAAVEHAFAELAPMGLAWPAGFPAVCARAAARIGRGFDHQGLGTYIYPEEGR
LLLAIALAFRPRNAVFLGSYYGYWAAWALPAIVAGGARAVLVDPDPRVAEVARRSLARLY
PGARVEIVCDTGEHYLAGGGGPFDLVVLDAELPRDHVDPTRRGKGVYAHLLRAALPRLAE
RSLLVCHNILFRDHSGCAFFDDVIARNREELAPFLALVAREYDCFVECPTTEGVGVGMRT
APRGRA*
>GCA_000452945.1_BPMSHR5858_genomic_02214_1 group_2061
MDAYRSPFDDWHQAASVRSMPADAWLAGGAGAWIAPQAGGVLAGLPARDARRERAAGGLL
IAHLSFTVALENTLISPVARDIAARALHADYDDGVVADALRVQCDEAYHALLAQELMTRV
RAATGADKPRRTPGFLRHVERLAATLPGVDAPLLRFCAAVVAETLITHTLRDDWRDEGLQ
PDVRTFLRRHYLDEARHSAYFSRLLTLLWPQWPAGVRDALRPLWSGLIDAFLFADARIAA
EALAAAGLTACEIDRAMSACGAPDAMQRRRAQSARQTLHALRAAGALEAGESAAIALGNA
*
>GCA_000452945.1_BPMSHR5858_genomic_02215_1 group_3300
MSAAGIRAFPGFAFGGGDRALAFVGGGLILDSGEAENTFAFRCQDRFLVYTGLLGLPSVL
LVLDDAHSRHYARLLATLDERLQGLAAVHRVAAPRLPPPVDVRCDLRTLSLSASASFDRA
AHALARDGARIGFVANQVAPSTARMLGELADLLRGRGVEVDFDRAACERVAALAHAWHNK
ARFLAFMREAAIRGGPPLAPSAVLAPEAFLRMREWAELAARFAAECDRRAPSSGLFLKSS
QDSSGNVSAILSSANFAERAPAFAAEVRRWLLADGFDDPAHVRELRKECEQPPSMEPVPL
SDATLAELRRRQARRRTGLPLIVQPALQPPADACGSPASVGASFCIEDDGRHRVVAVAAQ
LYRDAQRRRFLGLHVDDGLLDDRRVREWVARCGALVDALAARGYRGPVNFDGCMGADGRY
WFVGDCNPRLTAVYVPLAVRTWLRAAGIDVRSVSSFGYRGEFATPAIRACVDTWSRAGLI
FSRERPRGFVVLPNLACRDGHDVLAINLGVAQATAGLARMRELVAGAVPAHLGAIHV*
>GCA_000452945.1_BPMSHR5858_genomic_02217_1 group_2922
MLENPVGNLAAEARRLASAWRRGGPGLAARVALDRLCDAALEWRLRIRSGGLVPIESLVA
EWHGCHDYYPSSIRAFSRVLRDLHVTPDDVFVDYGSGMGRAIVIAARFPFRALIGVETSD
RLHRAAQANVGRALSDADRRRIRLLHCDARRFRLPGSASVLYFYNPFHGDILRAVFADIE
RSLRQHSRRLRIVFNNPAHFRAIERDYGWLRRSREYAFEYPIAVYETPGCAAPRT*
>GCA_000452945.1_BPMSHR5858_genomic_02220_1 group_2395
MPMTEVFRKTRRWSAVAALSAFVGLAGAASANTQPMQPTQQKQARMPRLPQNLPVSPEQA
EYNLPLSEQDRAALTRPSPLKQPAKRGKRSAPGADCRDMSVMTQYRGAALADYIANLPDY
ECHYGLFSVDKTLAAQIFSAENVHAVASRFVQDIYRYDASNLILVNLLIYLRSAYYQYDV
SGIANPIPNLAVWLRPYIKQSLEGAALYRENARAPSTANELMKLITNMKDEAFYLPTLKA
RIAFYTASATNPQAAAPLLQPSAAGGFTGLLTVFFYAHQRSGAQPMLDSDATLPETLNRF
VTANRASLSNTSAAYQLADAARETFRFLRYPAQKPRVKKMIQDMLASTSMTGADSDLWLA
AAEAVDYGDPGNCADYGTCDYKKRLTDAVLTHRYACNAGVRILAQDMTLPQLQSVCTSVA
QQDDYFHRMMKTGRKPVAGDRNDTIELVIFDDYANYRKYASVIYGISTDNGGMYLEGDPS
APGNQARFIAHEASWLRPEFKVWNLEHEFTHYLDGRYDMAGDFAASTAKPTVWWIEGLAE
YLSRKNDNQESIDAARTGAYRFSDVLGTLYSSSDYVARAYRWGYMATRFMFERHRADVDT
IVSRFRVGDYDGYANYVAYIGNRYDGEFVDWARAATTAGEPPLPTKR*
>GCA_000452945.1_BPMSHR5858_genomic_02225_1 group_3301
MVRKYRVVLATIAVFISLAGMMEAVRGLLFDSNAEFLYGVIALSIGVAAFVVLLNPTAND
ER*
>GCA_000452945.1_BPMSHR5858_genomic_02227_1 group_19877
MSVMKISFPQDPPPEYCARDLVVAFPALVDGEYVQCAITAEALEDHFGAASLKEDDLLTA
FDLHRGKIEHAARRMLDEIGGKPVLLHSGFFRFLYE*
>GCA_000452945.1_BPMSHR5858_genomic_02228_1 group_19807
MTLSGNIKDGDWTVEVTTSPVQGGYVCDIEVMHGAPGGAFRHAFRHGGTYPAERDAMIEG
LRAGMTWIELKMSKAFNL*
>GCA_000452945.1_BPMSHR5858_genomic_02236_1 group_2396
MPARSASSPRPPIPRTVWALGFVSLCMDVSSELIHALLPIYLVTTMGMSVAALGVLEGAA
EATAMIVKIFSGALSDWLGRRKALLLLGYGLAALTKPLFPLAAGPATVAAARLLDRVGKG
IRGAPRDALVADVAPPEIRGACFGLRQSMDTVGAFAGPLLAIALMLAFADHIRAVLWFAV
VPAFAAVALILFGVEEPASAPAAARAFRSPLHWRALRAFSGRYWFVVLIGTAFTLARFSE
AFLVLRAQQVGLDIAWIPAVMVVMSVAYAASAYPVGIVSDKFGARAPLAAGMLLLIAADL
LLGASASRTALFAGVAVWGLHMGFTQGMLAALVAQTAPAALRGTAFGVFNLAGGIAMLAA
SALAGWLWEHHGAPTTFFTGAALAAVALAMCGFVRRRPGPAA*
>GCA_000452945.1_BPMSHR5858_genomic_02238_1 group_3303
MSTVIEARPPVREGGEIEAQVNYLAKGVARPVAYTFEPPPGTPWTTGELAPRRVKIRDAR
PLAAAGALGLDASGFELVGHRSALADFADDAAIRSTYYAEVDALLRAATGAEKIVIFDHT
LRDSAHGSRATAALREPVRRVHNDQTFSSAPRRVRDHLPEDEAAARLRQRFAIINVWRAL
DVVERLPLALCDARTIAPDDLVPSDLVYRDKVGETYSVVANPAHRWFYFPRLRADEVLLL
KIYDSLDDGRARLSAHTAFDDPTTPDTAAPRRSIEVRALVFWPAADAKR*
>GCA_000452945.1_BPMSHR5858_genomic_02239_1 proY_1
VTSRDERLHRGLEERHINLMALGATIGVGLFLGSATAIRAAGPAILLTYILGGVAIFLIM
RALGEMAIDNPVAGTFSRYARDYIGPLAGYLTGWTYWFMWIVTCMAEITAVGVYMHMWFP
DVPNWIWTFAALIVMGAVNFIAVKLYGEFEFWFALIKIVTIVLMIAGGALMIVAGVGNGG
VPTGVSNLWAHGGFLPNGWPGVVDALPIVMFAYLGVEMLGLTAGEARNPEKSLAKAVNSV
FWRVVIFYVGALFVIMSVYPWNQIGTQGSPFVMTFSRLGIPAAAGIINFVVLTAALSSCN
SGIFSTARMLFNLAEQGQAPERLASVNRRGVPVHGVLISVALLFVGVVLNYFAPQKVFVW
LTSVSTFGAIWTWCVILIAQLRFRRRVSAERLARLPLRVPCYPLSSFVALAFLAFVVVLM
AFSPDTRVALVIGPLWIAALTVYYYASQAGRRSPLGAPTDG*
>GCA_000452945.1_BPMSHR5858_genomic_02240_1 group_19823
MLPSIHHPRYVVLRTHLRALRRAAGLTQTQLAERLSIDQSYLSKIERGERYVDILLYLDW
CRHCGVEPNHAVSELIDAGV*
>GCA_000452945.1_BPMSHR5858_genomic_02242_1 sfnC_2
MAISSPDSTLRRATAPAAPPQADADASTHAHDGKIADAALLFDDDPLIRRFAPLFERIAH
GAAARDRDRTLPYEPVEWLRAAGFTKLRVPRAAGGAGIGLAPFFALIARLGEADPNLPQI
LRVHGGFIEMLHESADDALRSRWFARIAQGTIVGGATAERSAVTSNTVRLSRENGKLYLD
GEKYYTTGTLYADWIDVSANDGDTDLRVLVPAATPGVERLDDWDGFGQRLTGSGTTRFTR
VEVEPGDIYRRFDASRPRGNSLLTAYFQTLHLANLAGIARAVLRDAVAFTRDRTRTFGVP
GASSPRHDPLVQRVIGRLASLAYSTQSLVATIARTLDDVCAARAAGRATEDAYVRVDIQA
FQAQQIVLAQTLEAATLLFEVGGASATSETRRFDRHWRNARVLASHNPAIVREAVIGNYY
LNGVGHNERFGIARTGVR*
>GCA_000452945.1_BPMSHR5858_genomic_02243_1 group_2397
MSSISSRPHKRPPASPWLSLVIRLLGVSFVLFFGAAIVTILLGIDHRAAGDPIGLLVLRL
VRWGSVHGGGEHYELMISTVYVVWGAFLWKAADAPFEHALFLDFTVVANVAHFGLMFVQG
LTMPGEHLHLVGDVALGWLALALFAATWLPARGNTRKRHTAGVDR*
>GCA_000452945.1_BPMSHR5858_genomic_02249_1 group_3305
MKTFSHALKRVVALVIAMPAAAGWASGQHRHGAADADELPRAFAAKEMGRSAAGARSYGS
PVTSVFEEAKGALPAFDAKRSGAAGAAGFAHELGAAERRQRLAEEDRAVQRLRDSRHDGR
GRWPGGAMPPRVPGRPDIAIEKPVPGVTCASGSDNSIGCGTGF*
>GCA_000452945.1_BPMSHR5858_genomic_02250_1 fan1
VPPDPPSPPAFYYLSNFERALAWLVERYDDVLDAEEHAFVAAFGALPRASRALLVRMLMR
KGPMFRASKLVYDEIGCPFAAAAPLVALGWIDPQPMLALDALFALATKAELRDAFSDAPA
SGALRKADWLDALRARHDGERPWAQWLPSIDDRVLRVTVDALCNRLRLMFFGNLHQDWSE
FVLADLGLLQYEAVAFAPSSRAFQRRGDVDAYLQLHACREQLDAWPDDAPLAPLVDAAAA
VDCGNAWLAMRRAKLTYAIGRACERRADWGGALDAYASSAWPGSRQRRVRVLERCERFDA
ALALADEAAREPENEAQAQQIARMLPRLRRRAGLPTARAPRAQEIPRGCVELARPGVPYP
VEYVARDHLSRADAPVFYVENALVNSLFGLLCWEPVFAAVPGAFFHPFQRGPADLHAPDF
RARRAAQFDACLAQLDGAQYRDTIRRHYAQKRGVQSPFVFWAALDETLLEHALACLPAEH
LRLWFERLLDDVRGNRSGLPDLVRFWPAERRYELIEVKGPGDRLQDNQIRWLDYCVRHRM
PVRVLDVRWTGDARASSQGEEALA*
>GCA_000452945.1_BPMSHR5858_genomic_02255_1 fliD_1
MAITQTGTTSQSDVWSQIQDAAQSLINGATGKTSMDVSGLVSVLVNAKTAGQAAEIKNQA
AWNSTQISALGALKLALSNLKTGVEPLSDGTFAQKFTSKASGKGLGATMDKGTVAGSYQV
EVKQVARSQTLVSAGFDPKHTLGSGTLTLKLGDRSTSIDIDATNNTPAGIAAAINSAKNN
PGVTATVVTGTDGAHLVLRSTASGSANVISMSVSNVKDDAGLSGLAVQSTADDKGGKSAI
TSAGDAWKQSDFAQDAIVTVGGVITARSADNAVKGVIAGVTINVTEEAIGAPQTLSIARD
IDGQARAVTNFVDLYNSMIGTMAQLTSFDKTAKPGQQGGPMIGDSMLNGIRNSLAHIVGG
GVPHGENKRASLAALGITFARPGDKQPEGSLIVDKAKLNEALQNDPQAVEALFNKTNGIG
TQITKALDVHLRKDGSFDVRSNAIDRDMKSIAQRQARLETYASQLTAQYKAQFTALDALM
ARMQQNTNYLTQLFGGANSSGALANNK*
>GCA_000452945.1_BPMSHR5858_genomic_02256_1 dmlR_1
VSAAAQHLDMSRAMASRYVAFVEQWTGARLLHRTTRRLTLTAAGAQMLPLCRDMLGLAEH
VATVVAEPGDAPRGALRITASAIFAQTHVTDAVLDYLARYPTVSVDLLVTDRTADLVDER
IDLAIRITNAVDPSLIARRLGTCRSALCASPGYLAEHGAPKRPHDLTRHNCLTYAYFGQS
LWHLTRDGEPATVPVQGNLSANDALVLLRAATAGGGIALLPTFAAAEPIRTGALVRVLPD
CAAPELGIYAVYASRKQMPLAMRTMIDFLAERFGDTPAWDA*
>GCA_000452945.1_BPMSHR5858_genomic_02257_1 group_2399
MTAMNPFRLQYFFDPLCGWCYASAPALAGLDAAHPGVLELMPSGLFADEGARELTPEWGE
YAWRNDQRIEQMTGQRFTHAYREQVLRRGGVRFDSGPANRALSALRGVDARLERPLLEAI
QLARYVDGLDTARPDVLARIAAGVAANAGVTTVDADALARRVEGDAALASETAGRIARTQ
RAMRQLGASGVPQLLLTVGERGYVLHGASLYGGAQAAVAAVERVLQEA*
>GCA_000452945.1_BPMSHR5858_genomic_02259_1 moaF
MGADPVFIQVGALAEGFAPDSHILAPVDDLAGRTLALAATHGATREYTFAGRSTLRWRER
QAARDIDGPARDTSDTRDARGEAAYRATQLREGVYFVDYVDPARRATSVSLVLDLRRGVW
TSVEGVLPAEADVRIDSFSRVASGLPLTGVDTRFRHGAIVGAAASGPLHAPTRELIGKRT
MYRYSPTECYEHIYLNDDFYAWHCLAGVERGLADVDRCHSFKLAEALYLFVWREKVVPTL
GVVLIDLDRCKTDGKIFGYREGDFGALSNFAIGAHAQVLNETMHPLAR*
>GCA_000452945.1_BPMSHR5858_genomic_02284_1 rutE_1
MTLSDSALDQIFRTARTHNAWQPKPVDDALLRELIDLVKWGPTSANSSPARFVFVKSPEA
KAKLKPALSEGNLAKTMAAPVTVIVGMDMAFHDHLPKLFPHADARSWFAGNDALIEATAF
RNASLQGAYLIVAARALGLDAGPMSGFDQAAVDAAFFAGTSIKSNFLVNLGYGDAAGLFP
RSPRFSFDEIARIE*
>GCA_000452945.1_BPMSHR5858_genomic_02285_1 group_3310
MDDLFDDLPRPDADWHPDWLAPADADRFHARLVDEVAWRQDTMRTPRGLLPLPRLTAWQG
EPDALYVYSGIRNEPAPWTPAVLELKRRVEAASRARFNSVLLNRYRNGFDSMGWHADDEP
ELGAEPVIASLSLGATRVFDLRHNETGVTHAYRLTNGSLLVMRGRTQHAWRHRVPKEPAV
RGERINLTFRWVSAPPAAERRARPRSRG*
>GCA_000452945.1_BPMSHR5858_genomic_02286_1 group_3311
MAKTTTTAAGRPAADAASAKSRRAAAGARPELDAADDEAAGGASSYLVPGLERGLRILSE
FSAREPVLSAPELSKRIGIPRTTTFRLLQTLEALGFIERANGDRHFRLGIGVLRLGFEYL
NSLELTDLGAPVLERLRDAAGLSTHLLIRDRRDVVFVAKAQSNAPMFGSVKVHVGTRLPA
HATVHGHVLMGDLTRDAMRQLYPEKRLEAFTEHTPATVDELYERVRHYARLGYALSEGAF
ESGISAVTAPVRDHSGSIVAAITATVPRSEIGAVDEKERLVELVCGAAVDLSQRLNYRPL
DSDPTVAHARHKVALL*
>GCA_000452945.1_BPMSHR5858_genomic_02287_1 group_3312
MVKHAVATLLGALAVAGAWAQSSVTLYGSLDAGVAYVNNVGGGAKWSMIQGNTQPDRWGL
KGVEDLGGGLKAIFQLENGFYTNNGQMAAAGTMWNRQAFVGLNSDRLGALTLGHQTPFNF
DWLDPLSSAFLAQSWYAFHPGNLDQLADTSTVPFNNSVKYRSPVFAGFTVGAMLGFGNTT
NFSTGRTMSFGVNYANGPFKAAAVYSNEHDQAFPMATVGGIAGPGGTFQGMPVASYVAKK
AQNMGAGLSYRFGPLLVHGLYTRVKLQANGHSDTFQSYDAGANYQSSPFNVIAGGAATST
LAGRRWSQFELGDTYSLSKRTQLYVNVLYEHASGNAKAAFFTAGASSTANQVIVLTGIHH
SF*
>GCA_000452945.1_BPMSHR5858_genomic_02290_1 group_3313
MTSSDQTNSGADPVRAYLDRGIRNYWYPVAPSWQVSSAPVGITRLSEQIVLWRDREGNVH
ALEDRCPHRGARLSLGWNLGDRVACWYHGVEVNGGGTVTKVPAVSNCPLEGQACVKSYPV
EERAGAIFLWFGDDAHREPAPLVLPDELVAPEYAQFVCVSNWTCNYQYAIDNVMDPMHGA
YLHATSHSMAEGDKQADMRVRRTETGLMFEKVNQRDVNFDWVELGETGCIWLRLAIPYQR
KFGPGGSFGIVGFATPIDDEHCQVYFWRTRKVAGWQRDVWRFLYRNRLEGLHWAVLEQDR
YVLESMAPNARDHEYLYQHDIGVTRVRRMLRQRAQEHLAALDAHRAAQADAAERA*
>GCA_000452945.1_BPMSHR5858_genomic_02294_1 xynR_1
MSEPLLKDADDARNDSNYVVPGLERGLRILAEFSPREPVLGAPELSKRLGIPRTTVFRLL
QTLESLGFLERADKDRNYRLGVAVLRLGFEYLSSLELTDLGLPVIESLRDATGFTAHIVI
RDGRDVVFVAKAQNAGNAFGSVRVNVGTRLPAHATTHGHVLMGDLSLAQLRALYPERALE

RVTKATPETVDALFNVIRDDARRGYGIGNSAFERDISVVTAPVRNEASRIVAVVTVTVPR
PEIDAALVAEGLVEKVVRAAAELSRRLNYRSDDENTYLKALGLR*
>GCA_000452945.1_BPMSHR5858_genomic_02295_1 group_19783
MIQIDLSGQVAVVTGGSSGIGYASAELFLRAGASVAICGRGDERLASAHARLVRQFPRER
VLAVRCDVLDEADVTAFAQAVAERFGGADMLVNNAGQGRVSTFADTTDDAWRDELELKYF
SIIRPTRAFLPLLRASSAPSITCVNSLLALQPEPHMVATSSARAGALSLVKSLATELAPE
RVRVNSILIGIVESGQWRRRYDAQAAPGESWEAWTAALAAKKNIPLARFGRPDEAAWALF
YLATNLSSYTTGSHIDVSGGFARHV*
>GCA_000452945.1_BPMSHR5858_genomic_02300_1 cmtAa
MTEHAGADATAMPRTVVVIGGGQAAGWVVKTLRDEGFAGRIVMIADEAHLPYERPALSKA
VLAGEADIATVRVMQPDAFGALDVEAWQPERAASIDRARRVVRTASGREIEYDRLVIATG
GAPRRLPDAIVGTAHLHYLRTLDEALALGEKLRASRRVLVIGGGWIGLEVAATARKLGVD
ALVVEGAPRLCARSVPGAVSDFLLDLHRANGVDVRLNASLASLDAHPDDARRVRATLAGG
GVVDADFAVAGIGLALNTSLATEAGLHVDDGIVVDEYGATSDPAIFACGDVANHPNAWLK
RRVRLESWANAQNQAIAAAKAVLGVRAPYAEIPWFWSDQYDVNLQILGDLPADARCIVRG
ELAAKRATLFFVADGALRGVIAINAARELKLARKWMNQGRAIDVAALADTAKALA*
>GCA_000452945.1_BPMSHR5858_genomic_02309_1 group_3314
MKTINRLAKRAALVCCAGSVWVSGAALAQVDWQSLGASLLGGGESQQAGASGLSQLLQSY
VGANQQVLAGQASLASAMGLTGAAGQAQHAAGLLTNGGTGALTPGALSQVGGAQQSVAQA
LTQAFASGGANGAASGAGVDKAAFTRGLASLGEGVTQYSRLQSGLGKLGESNPASLLQSG
LDPNNLKAASYIAQSAPNQFQSIGGTLGAAVQYAMSHGISVPSVASSALKLLP*
>GCA_000452945.1_BPMSHR5858_genomic_02314_1 hdfR_1
MLKESVVLHGIGLKYFVEVARTGSLAAASQQLHVAVSAISRQIAKLEEQVGTPLFERMPR
GMILTEAGELLVQHARRALMDGDAVLAEISEIHALGLGIVRIGCTEGFTRCFLPSVMATH
YRQHPRTRFVMRSGTPAQVEHWVASGEVDMGLAFSTSTSTALSVEYSVKAPVCALLPPTH
PLAGRTSVTLDDILQYPTAVLDRGTTVRQLIDLCCSARGVHLEPILTSNNSSAMHHFAAL
AGTITLGSQVALEGNPGPPTLVAMPIDEPLLNERLLQVTVMRERRLPPAVVQFLKTLSNA
LRESQGASAARARRKKR*
>GCA_000452945.1_BPMSHR5858_genomic_02322_1 ssuA_1
MIRPARWFARTVTAALASLCVASAFAQDAAGDKVLRVGYQKAGLLAVVKAQGTLETRVKP
LGYTVKWFEFPAGPQLLEALNADSVDIGYTGAPPAVFAQASGVRFVYVGAEPPSPHNEAI
VVKADSPLRRVADLRGRKVALQKGSSANYLLLEALRQAGVRYDEIRPIYLPPADARAAFE
SGTVDAWVVWDPYYAAAQHALKARTLADYTGLSATYNFYEATRDFATRHPDVVGAVLQQL
RETGLWVNAHSAETAALIAPKIGLDRALVETWVKRVPYGTTPLDDKIVASQQRVADAFLA
AKLIPRKLTVADNVWTERRAADALAAK*
>GCA_000452945.1_BPMSHR5858_genomic_02338_1 group_19942
MTHASLATGVLLAAGLGTRFDPGGVQSKLLAKLPDGTPVAVAAAQRLTAVLPDVVAVVRP
GAERLAHLLNDVGCQVVFAPDAERGMGASLAAGVRARADSPGWIVALADMPWIAPATIES
VACALDAGLASTVAPVYRGQRGHPVGFASRHYDALAALDGDTGARALLAAAPPALVDVTD
PAVLRDIDTRADLHAPPRD*
>GCA_000452945.1_BPMSHR5858_genomic_02342_1 araF_2
MGLRWPQAALVCASLAAGLSAAAPAHAQGAAPVKIGFVVKQPDDPWFQDEWRFAEQAAKD
KHFTLVKIAAPSGEKVSTALDSLAAQKAQGVIICAPDVKLGPGIAAKARRYGMKLMSVDD
QLVDGRGAPLADVPHMGISAYRIGRQVGDAIAAEAKRRGWNPAEVGVLRLAYDQLPTARE
RTTGAVDALKAAGFAAANVVDAPEMTADTEGAFNAANIAFTKHRNFKHWVAFGSNDDTTV
GAVRAGEGRGIGADDMIAVGINGSQVALNEFAKPKPTGFFGSILLNPRLHGYDTSVNMYD
WITQNRAPPPVVLTSGTLITRANEKTARAQLGL*
>GCA_000452945.1_BPMSHR5858_genomic_02343_1 bacC_3
MTTSRTTLITGAAGGIGQALVRGFLAAGDRVLALDRDRAALAAFVDALGDAAVAPIIDDL
TDAERLAAALANEPVDVLVANAGTAASATLRTTTSASWRADLDANLTATYASVEAVLAGM
RARRRGAITIIGSVNGVAALGHPAYSAAKAGLISYAKSLAIEYGRDGVRANVVCPGTVKT
PAWEARVRQNPQVFEQLKKWYPLEDFATPDDVANAALFLSSDAARAITGAMLPVDGGLLA
GNRVMAQELTLESFY*
>GCA_000452945.1_BPMSHR5858_genomic_02349_1 ybaL
MPHDVSLIALLAAGFGLAMIFGYLASLLKMPPLVGYLLAGIVMGPGTPGFVGDLALAQQL
AEIGVMLLMFGVGLHFSLGDLLSVRKIALPGAIVQIAVATALGAGLALWWGWSVGGALVF
GLSLSVASTVVLLRALEGRGLIESVNGRIAVGWLVVEDLVMVLVLVLLPPLAALLGGGAP
THGDAAAHAGEAGGSLWAALGVTLLKVAAFVALMLVVGKRVFPRILWLVARTGSRELFTL
CMIAAAVGVAFGAAKLFDVSFALGAFFAGMMMRESEFSRRAADETLPLRDAFSVLFFISV
GMLFDPRVLIDEPLHVLEVAAIVVLGKTLAAVALVLAFRYPLNTALTVGVSLAQIGEFSF
ILASLGRGLGLLSAEGQSLILAVALLSIALNTLLFAAVDPVLAWIRKRSAFARRLESRDD
PLAALPMSTPQAHLTGQVVIVGYGRVGTRIAHALDARGIAYVVVEQNRETVEKLRADGVA
AVSGDAVEPIVLVQAHIARAGMLVVTLPDVFDVRQIVDISRTLNPAIEIALCTNSDEEAA
LLANEGMGEVFVSETELAHGMTEHVLARMGAGEAHARRSAAH*
>GCA_000452945.1_BPMSHR5858_genomic_02351_1 group_2411
MAVGNHHPIKARHVKFDFGETPLQWIPGDPESTHIIDTLNLLFPEGELWFCRVYNKALPL
IDDPALHADADGFLRQEAVHSRSHHAVAKQCYARHGVDTQPYTRRLTWLFSKVLGEAPLG
LKIGHTRFWLRQQLGIIAALEHFFAYLGRWVLNARELDEARADPAMLDLLRWHGAEEVEH
GTVAFDLHRYMGGGYFGRAFHMLLAIALLVLFVSMGSKFMHTRDPGAGRYPGFVRAWTRG
SRRRHLPSFWKTIGAALRYFNPAYTPHGEGSTELALDYLARSPAAQAAAHGGNWGAKKTT
*
>GCA_000452945.1_BPMSHR5858_genomic_02361_1 lacF
MFNHGKTSLPWLFLGPPLALMAVLGLVPTIAAINLALKNRVLRYADSDYVGLRNFVRLAS
DRRFLNAIEVSALWEAVTVAGAVAVGIVLAVFLFERVHGRWRHAAALILIMPVLLPRVSA
AFIWKFMYAPLTGILGWLLGALGMTNVAWLADPRLALAAVALVDVWQWGLFFCVVVLKLL
ETLPPEPLEAARLDYASTWQVYAYIALPMLKAPIMSLVFIKMVESLRSFDLIYVMTKGGP
GIATETLDMYAYAQGIGLSGKVSYASSMAVLMMVATTLVFTFIWKRVDKWDD*
>GCA_000452945.1_BPMSHR5858_genomic_02364_1 msmX_2
MARIVCTGLTKRYDGGAPVLHPLDLEIGDGEFIVLLGPSGCGKSTMLRMIAGLETITGGT
LAIGGAVVNDLAARERNVAMVFQNYALYPHMSVYENIAFGLRRLKVPAAEIDRRVREVAR
VLGLDALLDRKPRAMSGGQQQRAAIARAMIKTPDVFLFDEPLSNLDAKLRAQLRGDIKRL
HQRLKTTTLYVTHDQLEAMTLADRIVLMRGGRIEQLGTPAELYGCPHTVFAAGFVGTPAM
NFADGVIERTAGHVALAAGGARWTLAARRFAGLPDGLRVKLAIRPNYLRIAPGAHAPAAT
LALEGRVELVELLGAEALVTFGCNGAPFAALVPASAAPALGAVVTFTFDERDLHLFDAAT
GRNVMLPEAGAPADAERPGAAMRPSPGAPHRFTMSRP*
>GCA_000452945.1_BPMSHR5858_genomic_02372_1 nasD_2
MKIVVIGHGMVGHKLLECIAAGAPGALAAPRVAVICEEPRPAYDRVHLSEFFAGKSADDL
SLVEPGFFERHPNFRLLLNTKAVAIDRAARTVALSTGETLAYDKLVLATGSTPFVPNVPG
RERAGCFVYRTIEDLEAMRACGARANSGVVVGGGLLGLECAKALRDMGLAAHVVEFAPRL
MAVQVDEGGGRMLREKIEALGVQVHTGKNTLEIVDGETGAHRMAFADGTHLDADMIVFSA
GIRPRDEIARACGLEIGPRGGVAIDERCRTSDEDIYAIGECAAWNGQTFGLVAPGYDMAR
VVARQLAGGEGAFAGADLSTKLKLMGVDVASIGDAHGRTPGSRAYQFADERRGVYKKLVV
SECGKALLGAVMVGDAADYGTLLQMMLNRIELPRSPEFLILPQADGAAKPALGVDALPDA
AQICSCNNVTKAQLREAVAGGATSLGALKGCTGAGTSCGGCVPLVTQVLKAEMKKQGLAV
NNHLCEHFPYSRQELYHMVRVERFKTFGALLKKHGRGLGCDVCKPAVAGILASCWNEFVL
KKEHAALQDSNDYYLANIQRDGTYSVVPRMPGGEVTPEGLIAVGEVARKYGLYTKITGGQ
RVDLFGARVEQLPLIWEELIAAGFESGHAYGKSVRTVKSCVGSTWCRYGVGDSVGLAIEL
ENRYKGLRAPHKLKFGVSGCTRECAEAQGKDIGVIATEKGWNLYVCGNGGMKPRHAELLA
ADLDRATLVRYVDRFLMFYIRTADRLQRTSVWRDNLEGGLDYLIDVVVHDKLGLAAELEA
EMQHVVDTYECEWKRAVTDPATRRRFRHFVNSDAPDSTIAFVEARGQIRPVTPDERDAEG
QADRQPETETA*
>GCA_000452945.1_BPMSHR5858_genomic_02406_1 group_3323
MSADAILTGRERAHRRVDAAGPHADLEQALGDPFDAANPLGYARILDADEREALSPDAEA
VLERWGMNAEFVPARLGGRWDSVTEMVRRLRPVFRRDAALGIGHGVSSLVAAVGVWAAGS
PAQQRELSELLLRGGKVSAAYHELDHGNDFLRNAFEAYPGADGALVLSGTKEVINNADRA
HAWVVFARTDARAGGRSHSVLLLDRAALQPAIDAGELEILRRHRTAGVRGCRLAGLRFSR
CRVPAERLVAARGAGVEISLRAFQVTRIALPGMAIGMLDTALRTVVAFARERTLYGTRVI
EMPYVRALLSDAMADLLAADCLVRAAARGLHLAPAHACVHAAAVKYLVPQLLRDSMNALS
VVLGARAYLRGGRHAIFGKMMRDMPIVGLAHAGGTACLLTVLSQLPALARRAAHRARDAG
APFDERVALPPLDFDALTIAAHGEDWLLDALPNALDAFEAAGERATPLARAVRHATAALD
ALRASAKRLPPSETGPAAHPESFELARRYAVLAAAAACVGVWRAERMRAGASFVGAPVWL
EAALSRLSERLGMPKQPAPPHVCDAIVSELVARADEHRSFCLYHARLNGGADVPASV*
>GCA_000452945.1_BPMSHR5858_genomic_02417_1 group_352
VLSSEPFLDLTPLIGHAARLRLATGAGESSWNGYVTYAAYADSDGEITRYRLTMESWLAP
LRLRRNCLYFVDVDTKDICERVFGD*
>GCA_000452945.1_BPMSHR5858_genomic_02438_1 gcvA_4
MKHLYPNVAELHAFASSAKHLNFSYAARELGLTPSAVSRQIASLEALLGVKLFVREGRNL
ALTRAGQVYQARVAGPLREIGNASLELLSAREDSNLLTIASVPTFTTKWLVPRLPRFLET
APDITLSFRRHLAPGDLFPLGLDAAIRYGDGRWEGVQCDYLDGRTFVPVCAPGFAERHAV
REPADIAAAPRLVHEQAECAWLAWADRHRATQMNALAGPRFEQYSVLIQAAQAGLGIALI
PAFLIRAPLAAGTLVQPLDAPVDVDEQSHYLCYAPERLQASASLRLLREWMLAECACA*
>GCA_000452945.1_BPMSHR5858_genomic_02439_1 phnV_2
MRSSTTAGFAGGIPFRRGKRRPSFGAIVVNGVVALFALYLLMPIVLLLIGSIGQSWTNTL
LPAGFTWRWFAELAADPSLRRAFGVSFVVALACCALDTLAGVPLAYALHHRALRGRGVAA
RVVMLTPIGVPALTLGFGYIVVFGGDALPWLGTLPLMIAAHAVLTLPYLLQTLLSDLRHL
DLAKLEACAATLGAPPLRQFFTIVLPNLRQSLFSGLVMVAALSIGEFQVSNLIAGFRYRN
FPVVLLQAFYGASGIACAASVVLLLLAVLATLASTHTVQRLK*
>GCA_000452945.1_BPMSHR5858_genomic_02440_1 potA_3
MSLEFDNVSFRYPGGAHGVDAVTLHAQPGELLAVIGRSGSGKSTLLRLSAGLLDGYAGRI
AIGGADMAGVPVWRREVGMVFQQYALFPHLSVADNVAYGLRMRGVATAERRRRALEMLER
VGLAAHAARSTTQLSGGQQQRVALARALVIEPALLLLDEPLAALDAGIRHQLRDEIRALQ
RACRATTLFVTHDQDEALSMADRVAIVDGGRVLQVGTPRSLYERPVSAQVARFIGHSTIL
RGRVQAGGTIDVRFATLCADTGAHRPGAEVDVLVRPEHVEPDPPAQCVNRIEGALGEVRY
FGATHRYDFLPHGAAHPLLCEGRVLAARGVAIEPRHLLVLPASGGAR*
>GCA_000452945.1_BPMSHR5858_genomic_02444_1 group_2417
MKKGAAAWLVAGSMCAGAHAQGTVTLYGIVDAGLGYTSEQRVAQTKGALGSPVGYRNESS
YGFASGIWSGSRWGLKGKEELGGGLAAVFQLENGFNIGTGQAGQGGRMFGRQAWMGLSSE
RYGTLTMGRQYDPIVDFVGTIGAGAFLTGMGAHPGDLDNIDNQARENNSIKYVSPKFGGL
ALGALYGFGNQAGSVKNQNTWSVGGQYVNGPFSLGAAYLYATNAYGANGGAWTGSYDGTF
ASSINEGFASAKSMQIVAAASTYQIGAVTLGLSYSNTRYKSGAFSTFNGTATYNSIGGTV
SWQATPELRVAAGYDFTRGSSIDGQSAPKYHQVNFASYYYLSKRTALYGLVGYQKASGKT
LDAYGNVVAATASVGDVGNGISSAGDTQTLVRIGVRHTF*
>GCA_000452945.1_BPMSHR5858_genomic_02447_1 mglA
VTPVLLEASQVAKRFNGVSALRDGRLSLAAGRVHALCGGNGTGKSTFLNILMGLLRRDEG
TIRLNGRDVDFASPAEALASRMAIITQELSPVPGMTVAENLYLGREPTRAGVVVDFRALK
RRAQALLDRLGFAIDAGAPMHRLSLAQTQLVEIAKAFSHDCQVMIMDEPTSAIGERETET
LFAAVRNVTAHGAGIIYVSHRLSELFDIADDYTVFRDGAYVESGRLADIDRAHLVRAIVG
REMPVVDKARRPAHGEPCLKVEGLTRANEFEDVSLDVRRGEILGIYGLMGSGRSEFLNCV
YGLTRPDAGAATLGGKPLPRGEPARSIRAGIALVTEDRKDSGLVLCGSVQENIAMAAYRR
LSRGGVIRRSLVRRLAQAMVERLRIKAASLRMPVSAMSGGNQQKVVLAKCLSTEPVLLLC
DEPTRGIDEGAKQEIYRLLDAFARDGGAVIVVSSEALELLYLSDRIAVFKGGRVAAVRPG
DGATQESLLHLAS*
>GCA_000452945.1_BPMSHR5858_genomic_02448_1 rbsC_4
MNKTMNTQTLSRLDAAPRRVNVAELASRFGIPIVFVALCVVLAFASPYFLTWRNWSDILR
QTSINGILAIGMTYVILTKGIDLSVGSVLALAGIVSGLAGAAGHGLAVSLAAGVACGAAL
GAINGVAIARLNVPPFVATLGMLSVARGVTYIANDGSPVANLPDDYLSLGIGRLGPLGMP
VLIFAGVALACWWVLRYTTYGRYLYAVGGNEKSARTSGIGVRKVVFSVYVVSGALAGLAG
MILAARTTSALPQAGVSYELDAIAAVVIGGTSLSGGQGGVVGTLFGALLIGVINNGLNLL
GVSSYYQQIAKGLIIVLAVLIDVARKQQR*
>GCA_000452945.1_BPMSHR5858_genomic_02449_1 thpA
MQTRKTWFGRFAAAAALGATLAIAAAGSAAHAASGPYRIGAAVYGMKGQFMQNWVREIKA
HPAVKSGAVQLTVFDGNYDALTQNNQIETMLTQQYSGILFVPIDTKAGIGVAARAGASDT
PLVASNTMLATPKVPYIGNDDVEGGRLQAEALARRIGGKGNVVIIQGPIGQSAQIDREKG
EMEVLAKYPGIKVIEKKTANWSRAEAMNLMEDWLNAHPKQINGVIAQNDDMALGALQAIK
NRGLTPKDIPITSIDGMPDAIQAAKRGEITTFLQDAQAQSQGALDLVLRQLVGAGYKPQS
VIWQRYAKDLKWDGGTAKRYILPWVPVTPANADQLYKQVTGG*
>GCA_000452945.1_BPMSHR5858_genomic_02452_1 dhaK-2
MNRVINHPDYVVEDMLRGIVAAHPALALDADNPRVIGVAHPVPGKVGVVTGGGSGHEPAF
VGYTGPGLVDAVAIGEIFSSPTAKSFLDAFRRADRGAGVACLYGNYAGDNMNVKMAIKMA
AAQGIDVKTVVANDDVASAPREERAKRRGVAGEILMWKAGGARAAAGGDLDAVIASAKKA
IDNTRSVGIGLSACTIPANGKANFHIADGEMEVGIGHHGEHGVRVMRTVSAKDMAAMMLD
IVLPDFPLERGEEVAVLVSGLGATPLMEQYILYAEVSQRLAAAGLKIGFRLVGNLFTSLE
MMGVTLTVTRLDDELKQLFAAPCSSIGLTVGERA*
>GCA_000452945.1_BPMSHR5858_genomic_02458_1 metQ_3
MRRLASIIRRVAGGAVWALAATLAHAGGPALMIGTMSGPDAQIWAEVTKVAARDGLAIKV
IEFNDYVQPNAALDAGDLDANGFQHQPFLDSQIRQRGYRIVNVGLTYTSPMGFYSKRLKS
LKDLPVGAKVGIQNDPSNGNRALLLLQKYGIVKLKPGAGVNGVNATPLDVAENPRKIRLV
ELDAAQLPRALPDLDAASINTDYAVKAGLSPVKDAIAIEDLKGPYANLIAVRAQDKDKPW
VKKLVAAYESNDVRKFIETKFNGAIIPAF*
>GCA_000452945.1_BPMSHR5858_genomic_02460_1 group_242
MSVSELQWAGSQAGGPAAHGKTAYLPYIDGLRAWAVLSVMLYHLDARWLPGGFAGVDIFF
VISGFVVSVSVANRGPMSLGPFLLFFYARRMRRILPALVAALLATALVSTLFIPASYLSD
RNRTTGLLAFFGLSNVILGAPGGDYFSPKAEFNPYTHTWSLGVEEQFYLLFPLLFHAWRA
GGKRRYASAALFAAGLCASLAWSVWARPAMHAQAFYLIFGRFWELAAGVLLYQCVARAGR
PAGRPAAAFGIEAGVWASAVAVLAGFVASKPALYPFPGALVPVAGALGLLGLLHGRARAS
PVRAALEQPALRFVGRISYSLYLWHWPVFVVFRWTLGLDSAGGKASALLVACALATASYR
FIETPLRRGGAAARRAPRVAALAGGAGVLAAGAALSSMLAAAQPQLSLSTVARHRLDWYP
YGVETNAAYPGCVADVGARDVRGGKLWIYARKGCGRPPESRQRVFVIGDSHAMAYAGMFK
QFVVRTGIAVYSYNNAGCPFVSLQPWRERGEPGCRRYGDAALDDMLSKLAPGDIVFLPSL
RLPRFVTQWAFVGDDDARRLSSGREAAQVRERAIADAVAVLREIGRRGARIVLEAPPPVF
RTVPYRCSDWFNRRNAICARGPLVGRGEIDALRAPVLTAYARIAAQVPDVHTWDPLPLLC
DEKACSAYDGARPLFFDGDHLSGYGNLRVLPGFEAFVRELIEAA*
>GCA_000452945.1_BPMSHR5858_genomic_02472_1 rluE
MRLIALNKPFGTICQFSAHETRASLGDWVKTPGVYPAGRLDADSEGLLLLTDDGALQAKI
AEPRHKLVKRYWAQVEGAPDAAALAALERGVDLGDYVSRPCRARFITPPGALWPRNPPIR
HRAAIPTAWIELEIAEGKNRQVRRMTAAVGFPTLRLVRVGIGALDVFALGLAPGETRELP
PRAPWERAQRPG*
>GCA_000452945.1_BPMSHR5858_genomic_02473_1 icd
MPYQHIKVPEGGDKITVNKDFSLNVSDQPIIPYIEGDGTGFDITPVMIKVVDAAVEKAYG
GKKKIHWMEIYAGEKATKVYGPDVWLPEETLQVLKEYVVSIKGPLTTPVGGGIRSLNVAL
RQELDLYVCLRPIQYFKGVPSPVREPEKTNMVIFRENSEDIYAGIEWAAESEQAKKVIKF
LQEEMGVKKIRFPQTSGIGIKPVSKEGTERLVRKAIQYAIDNDRKSVTLVHKGNIMKFTE
GAFRDAGYALAQKEFGAELIDGGPWMKFKNPKTGNEIVVKDSIADAFLQQILLRPAEYDV
IATLNLNGDYISDALAAQVGGIGIAPGANLSDSVAMFEATHGTAPKYAGKDYVNPGSEIL
SAEMMLRHLGWTEAADVIISAMEKSIKQKRVTYDFARLMEGATQVSCSGFGQVLIENME*
>GCA_000452945.1_BPMSHR5858_genomic_02476_1 clpS
MAIIPDKQDSSVLERKEQKLKPPSMYKVVLLNDDFTPMEFVVMVVQEYFKKDRETATQIM
LKVHREGRGVCGVYTRDIASTKVEQVVTHARQAGHPLQCVMEEA*
>GCA_000452945.1_BPMSHR5858_genomic_02479_1 group_3335
MTTIKAGDASGAGCAAAARARVAARIMRRALAVLGGAAFAGWFGAAFAASSGTTVPPLDT
TPWLASVQAPKTPFFTRASDAQGVSFHFANSGHDVERGREVVVSETADGDMWQGGMVHAT
YIDAAGVETSGWLVRSHLRRVKKPVPPPPTWDGRWRSSAGARRLIVHGERISYSFAGGGG
TGGAQPRVEMLLRMRPVSDDEAVLSRMQAPDGGMCDLAVRRLGDYLIVSARDCFIPGVSP
EGILRKQR*
>GCA_000452945.1_BPMSHR5858_genomic_02496_1 fkpB
MGFALRCPPFFGRRLNEYPMSLIDLAEVKPGSHVTLHYRLALADGADIVNTFSDKPATLL
LGAGQLAPSLEEILIGLKAGDHSTFRLAPEHAFGPRNPDMIQRVSLATLRENGMIGDDFA
PGDLIEFNAPDGGRYAGVLKEIGETSALFDFNHPLAGQALTFEVNIIGIL*
>GCA_000452945.1_BPMSHR5858_genomic_02510_1 yobA_1
MTMTITSLARRGLAAALALALSSAAFAHAKPEKSDPPANAAVAAPQAVSIDFTETLEPAF
SSIVVVDAAGAPAQAAKAVVDASDRKRMSVALKPLQAGVYTVKWVALATDGHRTQGRYTF
TVK*
>GCA_000452945.1_BPMSHR5858_genomic_02515_1 dapA_2
MSNLLQGIIAYPVTPFSPDGRLDTAALGALIERLIASGVHGIAPLGSTGESAYLSDAEWE
AAASASIRAVERRVPTVVGISDLTTANAVRRAKFAEQAGADAVMVLPVSYWRLDDEAIVG
HYRAIGDAIGIPIMLYNNPATSGIDMSPELIARIFRTVDNVTMVKESTGDIKRMHRLAQL
GDGAIPFYNGSNPMALAALAAGAAGWCTAAPNLNARLPLALYDAMRASDLDTARAVFHRQ
LPLLQFIVSGGLPVTVKAGLRLAGFDAGEPRKPLRPLDEARTRELAAILDALRDTAHA*

>GCA_000452945.1_BPMSHR5858_genomic_02520_1 sir_3
MAGSARAVQDSGAAWLGRRAVISSKPWGPRMYQYDQYDQTIVDERVAQYRDQVRRRLSGE
LSEDEFRPLRLQNGLYMQRHAYMHRIAIPYGNLRSVQLRMLARIAREHDRGYGHFSTRSN
IQYNWVKLEETPEILAKLASVQMHAIQTSGNCIRNITADQFAGVAQDEELDPRPWAEILR
QWSTFHPEFAWLPRKFKIAVSGSKADRAAVQIHDLGVYLKKNDAGEVVASILAGGGLGRT
PIVGAIIRENLPWQHLLTYCEAVLRVYNRYGRRDNLYKARIKILVKALSPAKFAQQVEAE
WQHLKDGPSTLTQAELERVSQFFQPPAYEKLADTDASFEQHLLENRAFARWVERNVAPHK
VPGYAAVTLSLKNHRVAPGDASAEQMEQVADWADAYSFGELRVSHEQNLILANVKKRDLF
AVWEKAKAAGFATPNVGLLTDIIACPGGDFCSLANAKSIPIALAIQQRFDDLDYVYDLGD
VSLNISGCMNSCGHHHIGNIGILGVDKDGAEWYQVSLGGEQGTAAGGARLGRVIGPSFSA
EEVPDVISKLIDTFVESRIDGERFIDTYERIGIAPFKERVYAARQTAHA*
>GCA_000452945.1_BPMSHR5858_genomic_02522_1 cysH
MSAAHAPALTPALAEKSARLDALLAQIAARHPKVKFASSLAAEDMLLAHAILSKGVAIGI
FSLNTGRLHAETLDMIERVRERYGYEIEQFHPRQDAVDAYVAEHGLNAFYESVELRKRCC
EIRKVEPLNRALAGVDAWVTGQRREQSVTRAELHEQEHDAARGIAKYNPLADWTESDVWA
YLNAFDVPVNPLHARGYPSIGCEPCTRAIRPGEDSRAGRWWWESRDTKECGLHITITPVP
ADGAESACA*
>GCA_000452945.1_BPMSHR5858_genomic_02525_1 group_3339
MEPALRIAQSLQLSRFGWDYRIPAEQQAILDLIAPKRPDDAAHRPRGEPFPTKIIGRTAR
DDT*
>GCA_000452945.1_BPMSHR5858_genomic_02527_1 moaD
MKIELKFFASVREALGVASETVTVPDGVATVGDVRAWLRARGGAWADTLAEGRALRMACN
HTMTDAGTRITEGCEVAFFPPVTGG*
>GCA_000452945.1_BPMSHR5858_genomic_02606_1 dmlR_7
MLIDDLPALATFARIVAAGSLSAAARELDLSLSVVSKRLAQLETRLGVRLLQRTTRQQTL
TDEGALFHAQVVRILAEIEQAESLMRERRGTVSGALRITAPGELGRRRIAPIVAAFQRQH
PQVTVQMQLTDAVVDLIAHDVDVAVRIGMLADSSLIARELAPNYRVLCASPAYLAEHGMP
AQPADLCAHRCIVLGEQPRVEWRFDGSHPAAVEVSAAFLTNDGGAARTLALEGAGIALKS
IWDVGPDLAAGRLRRVLPECASSAAPLHAVYPSGRHLALRVRTFVDYLGVELRRAWRWDE
R*
>GCA_000452945.1_BPMSHR5858_genomic_02608_1 menH_2
MSSNPATSVSPRERRVQCANPLGLHHIAYTEWGDPANPRVLVCVHGLTRSGRDFDRLAAA
LSDVYRVVCPDVAGRGRSDWLADPKRYAVPQYVSDIVTLIARLDVERVDWFGTSMGGLIG
LALGGLPDTPIGRMLVNDIGPHIEPAALERIGEYVGLDKRFATEQEGVDYLTSLSLSFGP
LTPDEWRALNRPLLVRDADGSWRARYDPRIAEPFKTATPEANAQGEAWLWHALGAFAGPL
LVVRGVQSDLLSRETLEAMKARGQAVTSVEIAEAGHAPAFVSAEQIAIARQFFVGDAPHA
S*
>GCA_000452945.1_BPMSHR5858_genomic_02618_1 ihfA
MNDMNSSEFEALLTAQRSAMSRDVPASTPAGDTPTLTKAELAELLFDSVGLNKREAKDMV
EAFFEVIRDALENGESVKLSGFGNFQLRDKPQRPGRNPKTGEAIPIAARRVVTFHASQKL
KALVENGAEPDLAR*
>GCA_000452945.1_BPMSHR5858_genomic_02626_1 group_19707
MSEVAVVALIVAKPGAEAKLLEQLEGIVGPTRSEEGALQYDLHRDLQEPRRFIFIERWAS
EAALAAHARSAHIDAYKRAAADWIESSEIRVASKVA*
>GCA_000452945.1_BPMSHR5858_genomic_02628_1 group_2429
MKKMTMAILVSMSALAGAAQAQESTEIKTITDPAKIAEIEQRAQALQQQQAAAPAAEAMP
MHEQHKPKHHAKKAHKGAKKAATAAPADAASQ*
>GCA_000452945.1_BPMSHR5858_genomic_02629_1 group_3731
LILFNSFGLLFVVFLLCRKRRGAFAILAAGAALLWLIASGWLAAPLVALAQAGMQPVAHP
TMTGDTVIVMLGAGIERRDGVAIPPRDALARIDKTAELYSACRRTAQRCTVVVSGGDPQV
RGIAEADTYAPYLLARGVASADLIREKRSHTTYENARFTAPILRAQHYDDLILVTSSYQM
KRALLDFERFGFAPQPVYANRRDARPGWLPGAANLANAELALHEIVGVAQFHVYRRLGWF
*
>GCA_000452945.1_BPMSHR5858_genomic_02631_1 group_3354
MKPMSLLTACGVVVSLALAPPSRADQFSPRQHAQREQFARSQANAAAPADARPNVPRGNL
RGDIASNARARNNDAQRPPGDSNHH*
>GCA_000452945.1_BPMSHR5858_genomic_02642_1 emrA
MSDPQQNAAPAQPQNNGKRKRMMTLLVVVIAIAAIAYGLYYLLVARFHETTDDAYVNGNV
VQITPQVTGTVIAVKADDTQTVKSGDPLVVLDPADSQVALQQAEANLAQTVRQVRGLYVN
DDQYRAQVALRQSDLSKAQDDLRRRLAVAQTGAVSQEEISHARDAVKAAQASLDAAGQQL
ASNRALTANTTVADHPNVLAAAAKVRDAYLNNARNTLPAPVTGYVAKRSVQVGQRVSPGT
PLMSVVPLNAVWVDANFKEVQLKHMRIGQPVELTADIYGSSVKYHGKVIGFSAGTGAAFS
LLPAQNATGNWIKVVQRLPVRVELDPKELKEHPLRIGLSMQVDVDIKDESGNQLGNVQNT
VYETDVFAKYGDEADAEIARIIAQNAGGGAAGGASAPTPAAKLVGRASNLM*
>GCA_000452945.1_BPMSHR5858_genomic_02647_1 sucB
MAIVEVKVPQLSESVSEATMLQWKKKPGEAVAQDEILIELETDKVVLEVPAPAAGVLAQV
LQNDGDTVVADQVIATIDTEAKAGAAAAAAGAADVQPAAAPVAAPAPAAQPAAAAASSTA
AASPAASKLMAEKGIGAGDVAGSGRDGRITKGDVLSAGSAPAAAPAAAPAKAAAKPALPE
VKVPASATTWLNDRPEQRVPMSRLRARIAERLLESQQTNAILTTFNEVNMQPVMDLRAKY
KDKFEKEHGVKLGFMSFFVKAAVHALKKFPLVNASIDGNDIVYHGYFDIGIAVGSPRGLV
VPILRNADQLSLAEIEKKIAEFGQKAKDGKLSIEEMTGGTFSISNGGVFGSMLSTPIINP
PQSAILGVHATKERPVVENGQIVIRPINYLALSYDHRIIDGREAVLSLVAMKDALEDPAR
LLLDL*
>GCA_000452945.1_BPMSHR5858_genomic_02653_1 group_2047
MTQQRFAPSFALAAWRVPLTALATVCVLAACGGSDVTAPPSAGGGTSGTSGTSGTSGTSG
TSGTSGTAGTSGVVNTLGKTATDLGNTISSTSVPGLGSGVTQGVGATVGSTGGIINAAAD
ALSNGLGQTGSTQNPVGTTVAGLGNVVSATSNTVQGLSQTVQALGTGPLAPLSPVTTPVA
GALATAANGVNAAGTMLGNALSTGPVQQVTQPISSALTPLVITAGQVTQTVGTTTGVGQP
VSGLLQQVGGAISSAGTQIAGTSPSQPLVGDVGQLVSAVGNTVTNAGGLVNPAAPNGAAP
IPGLITSVVGGSTATVATGSSSGSTLGSPLNGLLSSLGGGALPSGPLGGSAVSGGASNPL
APITSLASGGTGASTNPLAPVTGLLNTVTGTLSGATSSTGGSTGLLAPVTGSLGTLGSFG
K*
>GCA_000452945.1_BPMSHR5858_genomic_02660_1 group_2431
MNARTLSLAEPAVTDYFVCASPHGEHVSWLAQTLVSAGTVEPAPLEPTALAQRIAGLNPV
LVFVDFSGGQAQAASAAAAALRVSHPGLPIVALGSLGEPESALAALRAGVRDFIDFSAPA
EDALRITRGLLDHVGDQPSRHGRLIALLGARAGMGASTLAANLSVLVQKRSAALGRQTAL
VDLGLPAGDGALFLNTRCEFDFVEAVRNLRRFDRTFVNTALARHASGVALTSLPPNLAGL
RDVSYASCVGLMNRLRAFFDCQIVDLGGFSNRDFVAQTANAADESWLLCDQGVASVVSAV
ELLESLRDTGVDTENVRLVVNQYDPALGLAPAQIAERLGLALAATLPSRRVPIGHAANQG
KLIVDVAERDPYVRALEPLVERATGGAGAAAPRPAGGLSALRRFIQPTSKRS*
>GCA_000452945.1_BPMSHR5858_genomic_02668_1 group_2432
MSHIRGMGRVFVAIAAAGGLCGAHAQALEGNGGAASPALTASGAAASPGVSAPAVVNLSA
PAGAPAAGASAEAGGKSTIDELQRQIQARELTEMRTTYNGSYGASLLLNIQEGTYFVALF
QQKAFWRVIRTASEARAEAVYRDFAKQSETLAVNELQAAKLESQKALTDRQIAVAQERAS
RLQADLSIAREQRAAVATRQKDKLDETVALREQKSERQDQLRKLQQQVRSLQRQAEAGLP
KVR*
>GCA_000452945.1_BPMSHR5858_genomic_02670_1 group_3355
MKKWLTITMLGVALLGAAGAATAGGVDLSIGVGVPVAPVYVAPQPVYVAPQPAVVGYPAY
GWRGDDEDDDWKHYRKHWRKWHRHHDDDDD*
>GCA_000452945.1_BPMSHR5858_genomic_02674_1 yjhB
MAMSWTREQRNVTIAAYLGWTLDAFDFFLMVFVLKDIAAEFNTKIPAVAFAITLTLAARP
IGALIFGRLADHFGRRPTLMINIACYSLLELASGFAPSLAALLVLRTLFGVAMGGEWGVG
SALTMETVPPRARGAVSGLLQAGYPSGYLLASVVFGLLYPYIGWRGMFMIGVLPALLVLY
VRAKVPESPAWKQMEKRARPGLVATLKQNWKLSIYAVVLMTAFNFFSHGTQDLYPTFLRE
QHHFDPHTVSWITIVLNIGAIVGGLTFGWLSERIGRRRAIFIAAMIALPVLPLWAFSTGA
LALAAGAFLMQISVQGAWGVIPVHLNEISPDEIRATFPGFVYQLGNLLASGNATLQAQFA
VDHGNNYGMALATVAGIVAVVICVLIVFSRERRGIDMTQTAAMSPTSG*
>GCA_000452945.1_BPMSHR5858_genomic_02675_1 group_3357
MNPDAQAHSSLPTLETDRLWLRPRVLADLDACIEMDRDPDVTRHIAGPWADPVEHRRFVE
HRITRAYPPGLGYWSIFEKARPDRFVGWVLLIPDYVEGSRDVEIGWRLVRASWGRGIASE
AARRVVRHAFETVRLPRIVADIAAANSASLNVARKLGMRRVGIVQDGIPYVRHRLERADL
AARG*
>GCA_000452945.1_BPMSHR5858_genomic_02686_1 rpiA
MTQDELKRLVGEAAARYVTENVPQGAVIGVGTGSTANCFIDALAAVKDRYRGAVSSSVAT
TERLKSHGIKVFDLNEIESLQVYVDGADEIDGSGAMIKGGGGALTREKIVASVAETFVCI
ADASKRVAVLGQFPLPVEVVPMARTAIGRRLAALGGVPVLRVKQDGAPYVTDNGNEILDV
KGLRIDDPRALEAAINGWPGVVTVGLFAQRGADLCLLGTERGVETLRYAAH*
>GCA_000452945.1_BPMSHR5858_genomic_02689_1 group_3358
MPITSLLAQSLPVLLQGALLTVKFAVLSMLFGLVGAVALALMGIGRSRVLAGLARAYVSV
MRGTPLLVQIFVVYYGLPSLGVSLDPTPAGVIALSANVAAYMSESMRGAINGIDKGQWLA
AYSLGLSWRQTLRYVIGPQALRIAVPSLSNSLISLIKDTSLVSVITVTELLRSAQEIIAS
TYQPLPLYLAAAAVYWVLCQILEWVQRWYERRLALPSRH*
>GCA_000452945.1_BPMSHR5858_genomic_02715_1 rpoB
MQYSFTEKKRIRKSFAKRSIVHQVPFLLATQLESFSTFLQADVPTAQRKSEGLQAAFTSV
FPIVSHNGFARLEFVSYALSSPAFNIKECQQRGLTYCSALRAKVRLVLLDKESPSKPVVK
EVKEQEVYMGEIPLMTPTGSFVINGTERVIVSQLHRSPGVFFEHDKGKTHSSGKLLFSAR
IIPYRGSWLDFEFDPKDVLYFRVDRRRKMPVTILLKAIGLTPEQILANFFVFDNFTLMDE
GAQMEFVPERLRGEVARFDITDREGKVIVQKDKRINAKHIRDLEAAKTKYISVPEDYLLG
RVLAKNVVDGDTGEVIANANDEITEGVLEKLREAKIKEIQTLYTNDLDQGPYISSTLRVD
ETVDKTAARIAIYRMMRPGEPPTEEAVEALFNRLFYSEDAYDLSKVGRMKFNRRVGRDEI
TGPMTLQDDDILATIKILVELRNGKGEVDDIDHLGNRRVRCVGELAENQFRAGLVRVERA
VKERLGQAESENLMPHDLINSKPISSAIREFFGSSQLSQFMDQTNPLSEITHKRRVSALG
PGGLTRERAGFEVRDVHPTHYGRVCPIETPEGPNIGLINSLALYAHLNEYGFLETPYRKV
VDSKVTDQIDYLSAIEEGRYMIAQANAAIGDDGALVDELVSSREAGETMMVTPDRIQYMD
VAPSQIVSVAASLIPFLEHDDANRALMGSNMQRQAVPCLRPEKPVVGTGIERTVAVDSGT
TVQALRGGVVDYVDAGRIVIRVNDDEAVAGEVGVDIYNLIKYTRSNQNTNINQRPIVKMG
DKVSRGDVLADGASTDLGELALGQNMLIAFMPWNGYNFEDSILISERVVADDRYTSIHIE
ELNVVARDTKLGPEEITRDISNLAEVQLGRLDESGIVYIGAEVEAGDVLVGKVTPKGETQ
LTPEEKLLRAIFGEKASDVKDTSLRVPSGMSGTVIDVQVFTREGIQRDKRAQQIIDDELK
RYRLDLNDQLRIVEGDAFQRLARMLVGKVANGGPKKLAKGTKIDQAYLEDLDHYHWFDIR
LADDEAAVQLEAIKNSIEEKRHQFDLAFEEKRKKLTQGDELPPGVLKMVKVYLAVKRRLQ
PGDKMAGRHGNKGVVSKIVPVEDMPYMADGRPADVVLNPLGVPSRMNVGQVLEVHLGWAA
KGLGWRIGEMLARQTKIEELRVFLTKIYNESGRAEDLESFSDDEILELAKNLREGVPFAT
PVFDGATEEEMSKMLDLAFPDDIAEQLDMNPSKNQVRLYDGRTGEPFERRVTVGYMHYLK
LHHLVDDKMHARSTGPYSLVTQQPLGGKAQFGGQRFGEMEVWALEAYGASYVLQEMLTVK
SDDVTGRTKVYENLVKGDHVIDAGMPESFNVLVKEIRSLGIDIDLDRN*
>GCA_000452945.1_BPMSHR5858_genomic_02720_1 nusG
MSDTPASPSGKRWYVVHAYSGMEKSVQRALQERIERAGMQDKFGQILVPTEEVVEVKGGH
KAVTERRFFPGYVLVEMEMTDETWHLVKNTAKVTGFVGGARNRPSPISPREVEKIMSQMQ
EGVEKPRPKTLFEVGEMVRVKEGPFTDFNGTVEEVNYEKSRVRVSVTIFGRATPVELEFG
QVEKV*
>GCA_000452945.1_BPMSHR5858_genomic_02721_1 secE
MANPSVETVNTSGDKLMLALGVLLVLAGFAGFFLLGGKEWYFRGAALAVGVIAGVGVALM
SLPGKSFIAFAKDSYREVRKVVWPTRKEATQTTLVVFGFVLVMALFLWISDKSIEWAIFS
VILGWK*
>GCA_000452945.1_BPMSHR5858_genomic_02733_1 group_3362
MVELQNHSHLCAAYGKRFGVEAMRKLGWRAYRWGGSGMILTAIGDNRFIVALPARDSSPR
TAERPVAESCLLALSGTPISHGGVHALAVVDVHTIDAESGQGSNRTREPADQVGPVPPAQ
HTAGWRNQYMSDMHAAVAFRQSLRSNLSYLSFQPIVRRNSAPAVLYEQALLSIVGDDDQS
GEHPASIVAALERLGLARLFDYRVVHAVVARLLRTSDARLGCRISPRSAMLDVWWASLVD
WLASHPCVARRFTVEIPCNTPVTRRAAALAFVRKLQTLGCSVAVTDFDTGGCDLEFALLA
RPQVVKISAGALRALPAGNSLAERLARLVRPCSSFDPQIVVEGVNDPHDLNVLEAVDAAW
VQGLLQSAAVNGSRAEPLVVWQDQ*
>GCA_000452945.1_BPMSHR5858_genomic_02735_1 group_3363
MSQVLEMFNAAGSAQFSKMVCQMAPYFGTIEPEVVGLRPGRAEAKVGFRRDITNHIGTIH
AIALCNAAELVAGLMTQVSIPDGMRWIPSGMTVEYLAKAKTDVTAVADGSAVDWQSEGDK
IVPVEITDSEGQTVFTARITMNLKA*
>GCA_000452945.1_BPMSHR5858_genomic_02737_1 group_3364
VHVGERFNSISHLVGAVLSVAGLATLVTMGALDGDAYKVVSFSVYGAMLCVLYAISTLYH
SVRGPRLKAILQKCDHAAIYLLIAGSYTPFTLVTLRGPWGWSLFGVSWGLAAFGIVQELT
LGRRTRIVSMALYVLMGWLALVAIRPLVHALPAAGTAWLVAGGVIYSAGIYFFVNDERIR
HGHGIWHLFVLAGSLCQFVSVAGYVA*
>GCA_000452945.1_BPMSHR5858_genomic_02738_1 group_19859
MGGSISKSSDAVSTASIPATQSHADTDTASKASSHSAQRRPSSGAFEGLMPLAQKGTVGE
LQNFAAAAFAATTGASPALVDAAHHNPAMQTLREQPMTQENVTQLCQMETDAARKGDPAN
WQNRLAASSVADVARHAVAQTVALNVSLGRDGPIKTQEEGGWNVHEHVIGVHGS*
>GCA_000452945.1_BPMSHR5858_genomic_02739_1 group_2774
MHGTYNLWLVALSLAVATLASYTALDLSGRIALLAHGRLRHAWLAGSAVAMGIGIGSMHV
IGVLSVTLPIRIGFDAAITGLSLALAIVVSGFALYTIAGAQLTRLRLGIGGVLMGLGIAG
MHYTGDAAMRMKPAIEYDPLLFTASIVIALVASTVALWIAQTLIGIGRSRLARIGAAGVM
GAAITGMHYTAYAAAIFAPGSTCGAATGIDLRWLAATVALFTVAILIVALILSHFDVRTT
LLAGSVSRLSGQIVRMAAFDTLTDLPNRHTLNEHLGRALAAARARGGKLAILFMDLDGFK
TINDSLGHPAGDEVLKTFSQRLLRCVRDGDIVARLGGDEFVVLLESVSSDDEAARVARDV
LERMRKGRWNDGQALQVTPSIGIALYPKDGDTADTLLRHADAAMYAAKRSGRCTYRFFEA
TMDEAAARALKLQHALHEALDSDYFSLCFQPKFRCDGHRLAGAEALLRLNHPTLGKLAPM
EFIPVAERSGQIIEIGYWVIRETCRHLRRWRSAHRAACRIAINLSPRQLAQPDLVANILE
IVRAEHVEPRQLMFEITESVAMEDPPRTIEMIRLFQAHGFEIAIDDFGTGYSSLAYLQRF
GAKQLKIDRFFTHALDEQGHEGGAIVSAIIALAHSLAMDVVAEGVETRSQLDRLKTLMCD
EIQGFLLGRPVPPEEFEAMVPLHSEAPA*
>GCA_000452945.1_BPMSHR5858_genomic_02741_1 aroP
MATDQDNDGLKRGLKNRHIQLIALGGAIGTGLFLGIAQTIRMAGPSVLLGYALAGVVAFF
IMRQLGEMVVDEPVAGSFSYFASKYCGHFTGFLSGWNYWVLYVLVSMAELSAVGIYVQYW
WPGVPTWISALVFFVVINAVNLASVKSYGETEFWFSIVKVAAIIGMIGFGGYLLLSGHAG
PDAGVANLWRHGGFFPNGIGGLAMAMAVIMFSFGGLELVGITAAEADDPSHSIPRATNQV
IYRILIFYIGALGVLLSLYPWQKVVTGGSPFVLIFHALSSDIAASVLNVVVLTAALSVYN
SCVYCNSRMLYGLAQQGNAPQALARVNRRGIPIAALGVSAFATALCVVINYFMPGKAFEL
LMGLVVSAIIINWAMISVIHLRFRQAKRAAGETTRFRSLGYPLTNYFCLAFMAAILVVMF
RTPELRLSVYLIPVWLVVLALGYRFRQRGAGYALGGGVSAR*
>GCA_000452945.1_BPMSHR5858_genomic_02744_1 group_3365
MQQQVDGGADSLAPVIETARLILSGPRLEDFDGSRALWADPDVTRLIRDEPFTSEECWLR
LLRYVGHWRLLGFGYWMAREKDTGRFIGEVGFADAKRDIVPPLDDAPELGGALAPAAWGQ
GIAREALHAVLEWADANLTARRTVCLIGPKNRFSMRLAAEFGYREVAVAEYKGEPIRVFE
RPAAGPRAGETRPPADAAERGGRA*
>GCA_000452945.1_BPMSHR5858_genomic_02836_1 group_2438
MFNPSRDDVRRFFIDTWRKQRSGEILTPLEAMAADWIVEHPEYHAELEDAGRSAAHDYTP
DEGRTNPFLHLSMHLAISEQLSIDQPPGIRAAHEKLAARCDSAHDAQHAIMECLGETIWE
AQRTHTPPDSDAYLQRILRRASRG*
>GCA_000452945.1_BPMSHR5858_genomic_02840_1 group_3369
MLIDFFYSLRAAKLPVSVKEYLTLLEALKAQVIEPSLDAFYFLARLTLVKDEQYFDKFDK
AFGAYFHGVSALPADAFDIPLDWLEKRLQRELTPEEKAQIQALGGPDKLMERLKQLLDEQ
KARHEGGNKWIGTGGTSPFGHGGYNPEGVRIGGPSNGNRTAVKVWEARAYRDYDDSVEIG
TRNIKVALRRLRRFAREGAAEELDLPGTIRSTAANAGWLDLRMVPERHNNVKVLMLLDVG
GSMDDHIKRTEELFSAAKAEFKHLEFYYFHNCVYDHLWKNNRRRHAERTPTWDVLHKFTP
DYKLIFVGDATMSPYEVLQPGGSVEYNNPEAGAVWLRRLADQFPHYAWLNPEPERLWEYR
QSIAIIRDVLGDRMYPLTLAGLESAMRTLSK*
>GCA_000452945.1_BPMSHR5858_genomic_02842_1 talB
MTTALDQLKQYTTVVADTGDFQQLAQYKPQDATTNPSLILKAVQKDAYRPILEKTVRDHA
GESVGFIIDRLLIAFGTEILKLIPGRVSTEVDARLSFDTQRSIDKGREIIKLYEAAGVGR
ERVLIKLASTWEGIRAAEVLQREGIRCNMTLLFSLVQAAACAEAGAQLISPFVGRIYDWY
KKQKGADWDEAQDGGANDPGVQSVRRIYTYYKHFGYRTEVMGASFRTTSQITELAGCDLL
TISPELLQKLHDSTEAVARKLSPDEARDARLERVAIDESSFRFQLNDDAMATEKLAEGIR
LFSADAVKLEKMIEALR*
>GCA_000452945.1_BPMSHR5858_genomic_02845_1 group_19757
MAHDADANRAAKRWLWLLVLPLVAMVWVPSYNRTEPQWFGFPFFYWYQLLWVFVSAVITA
FVYQKTKHCWKGVPRGDGK*
>GCA_000452945.1_BPMSHR5858_genomic_02847_1 group_19712
MLRGFPPVASPGTHTLILGSFPGEASLAAAQYYAHPRNQFWRLLGAVLGEPQLHALPYDA
RLARVLAHGFGIWDVLAACHREGSLDAAIRHAQPNDFASLREVAPKLTKVCFNGKTAGRF

EPTIRAAGFDTLVLPSSSPANAMLSFEQKLVFWQRIVG*
>GCA_000452945.1_BPMSHR5858_genomic_02851_1 group_19918
MAAHSLTLDHLVVAARTLEEGVAYVADTLGIEPAGGGAHPSMRTHNRLFGLWGRAYLEVI
AADPDAPAPADGRPRPRLFGLDDPATHARLEQGPYLAHWVARVDRPRELGLWQRQYPARI
ARVVPMTRGDFGWRLTVPDDGALPAWQGAGDGVLPSLIQWSDARHPCDALPHGDVALTAL
KAAHPRADTVREQLVWLNAAHLLELDAGETPALAAEFDTPRGARTLR*
>GCA_000452945.1_BPMSHR5858_genomic_02852_1 group_2439
MDSRETRGMLLGLIGVVIFSLTLPMTRVVVTELHPLLNGLGRALAASVPAGLLLWLRRER
LPTRAQLKSLAVVSAGVIVAFPVFSAWAMKTVPAAHGAVVNGLQPLLVALYAAWLARERP
SRAFWASAVLGSALVIAFALRDGGGALQPGDLLMLVAVGIGALGYAQGARLAREIGGWQV
ICWALVVSAPFLVLPVGWLAWAHHAAHPGPVSPRVWLAFGYVTLFSQFIGFFAWYAGLAV
GGIARVGQVQLLQIFFTIAFSALFFGETVAPSTWLFAAAVIATVVLGRRAAVGAAPHPAR
AA*
>GCA_000452945.1_BPMSHR5858_genomic_02853_1 lysN_1
MNPSDLKPPHWALSERARKLTSSAIREILKVTERPEVISFAGGLPSPATFPAERMREAAE
RVLRDSPAAALQYSATEGFLPLREWIAERYRVRTTQVLVTTGSQQALDLLGKVLIDPASR
VLVETPTYLGALQSFSLYEPIYAQVPTDDAGLLPEALTPELTKDARLLYAQPNFQNPTGR
RLSVERRRALAAFARTSPFPVLEDDPYGALNYAGEPLPTMLSMAPDHIVHLGTFSKVLAP
GLRIGYIIAPEELHFKLVQAKQATDLHTPSLTQRIAHEVIQDGFLDAHIPTIRKLYGAQC
EAMLASLARHMPQGVSWNRPEGGMFIWVTLPAQIDSMQLLETAVANNVAFVPGAPFFAND
AQKNTLRLSFVTVPPEKIEEGVARLGKLLRERL*
>GCA_000452945.1_BPMSHR5858_genomic_02864_1 group_19917
MSHAIAVALFLHLLAVAVWVGGMVFANFCLRPALSDLSPQLRLPLVEAVFGRFFNWVAGA
VIVILLTGGFLLVKFGGAHATWPLHAMAGLGVVMMLIYGHIRFALFPRIRRAVQAQNWPD
GARAVNAVRLLVMVNLVLGVVTIGAAVLSRGF*
>GCA_000452945.1_BPMSHR5858_genomic_02871_1 ephD_3
MHTPDLFVQSSNVRLAVYTWGDKPSADKPRDIVVLAHGFPDRALFWEQVAAALQRDFYVV
AYDMRGCANSTHIKGARHYRFALLLADLYAVIDAVSAGRPVHLVGHDWGGVYGWDAIADP
EGARRIASLTTLSPSLDQIGFYLRRRLLRPTPRHLAQLVGQLMRNSLMTFFTAPLLPELL
FASGLAMAMFRRIIAHYEPRITFRKNDGMEGDAIRYLGIYRANLLQRVLRPRKRVSTTPV
HALMAIHDPFLPPALFEGCREFTTRYSESTVDAAHWAPLSRPQEIAETVGAFVRQASRRP
DVALQSAS*
>GCA_000452945.1_BPMSHR5858_genomic_02873_1 group_19811
MRFNSTVFLIDPTPRRADGEYMAHARISANRSDGREYDVCRSGDLAGFDLRADAIAHARR
WAETWLEDHFG*
>GCA_000452945.1_BPMSHR5858_genomic_02875_1 group_19746
MSMHLYRGFEIHPLIYPRLASVDGRAHNYEAGFDAAVRICFPRTDATASASRVFKLDNAQ
AFDSAGDARRASLRHAEYLIDRHGGERWSSAGGA*
>GCA_000452945.1_BPMSHR5858_genomic_02893_1 group_3376
MDLGFFDPNRTANASAWRVLPNRWDFVAFPLIICVIAMAAIGFHETMAPIATLETQRISL
DPANLPEYALRTTLRMLAAMVAALAFTLVYGTLAAKSRRAGMVLVPILDILQSVPVLGYI
SFTVTFFLALIPSRVLGAELAAIFAIFTSQAWNMTFSFYQSLRTVPRDLDEVSRGFHLTA
WQRFWKLEVPFSMPGLIWNMMMSMSGGWFFVVASEAITVGNHTITLPGIGAYLAQAIVEK
NLGAVGWVILAMTVVILAYDQLLFRPLVAWADKFRMETTSSGNAPESWLLDLVRRTRLIH
QLLVPAGWFFARLARIPLRVPSLDAVRFSMPRVEKKASRAADIAWAIAVIVGTVYVVWRV
FAYVSTGVTLAEVGHVFVLGLITLLRVALLIAIASLVWVPIGVWIGLRPAIAEKVQPLAQ
FLAAFPANLLFPVFVIVIARFHLNADIWLSPLIVLGTQWYILFNVIAGATAYPNDYREAA
TNFRIRGWQWWRQAILPGIFPYYVTGAITASGGAWNASIVSEFVQWGDTKIEAHGLGAYI
AQTTAAGDYPKIILGIAVMSLFVTLFNRLLWRPLYAYAEAKLRLD*
>GCA_000452945.1_BPMSHR5858_genomic_02899_1 yhhW_1
MFEIRRAGDRGHADHGWLDTHHSFSFADYRDPEHMHFGALRVLNDDRIAPTRGFGMHPHR
DMEIVTYVLEGALAHRDSMGNGSIVRAGDVQRMSAGTGIVHSEYNASRDAPLHLLQIWLL
PVEPGGRPGYQEARFTDADKRGRLRLVASPDGRDGAVTVRADASIYAGLVDGDERAEFAL
PAGRRAYVHVARGCVVVNGEVLAAGDGARIAEVGRVVFERGERAEVLLFDLA*
>GCA_000452945.1_BPMSHR5858_genomic_02900_1 argT_3
MKKLALCAALALAAGGAFAKEWKTVRIGVDASYPPFESTAPSGEIVGFDVDLAKEVCKRI
NAKCTWTPQDLDGIIPALKAKKFDVIVSSLTVTDKRREQIDFSDKLYDAPARMIAKAGSP
LAPSVESLKGKRVGVEQGSTQETFAKAHWEPKGVTIVPYQNQDQVYADLGSGRLDATLQD
ELQADYGFLRTPRGKGFAWAGTAVKDPRTLGDGTAMGLRKEDADLKDAINRALASMHKDG
TYDRLSRKYFPYSVYSAK*
>GCA_000452945.1_BPMSHR5858_genomic_02959_1 cbiX
MNKHGIVLFGHGARDARWAGPFERLAAKLRAARGAEASVVLAFLELMEPSLAAATAALAA
QGCDAITVIPVFFGEGGHVRRDLPGLIDACRAAHPGVDIRCATAVGEDDAVLDAIVAYCM
RASADPTS*
>GCA_000452945.1_BPMSHR5858_genomic_02961_1 cysN_2
MSIIENNEDLGVLRFITAGSVDDGKSTLIGRLLYDSKAVLSDQLSALSRAKNKRTVGDEL
DLALLTDGLEAEREQGITIDVAYRYFATAKRKFIIADTPGHEQYTRNMVTGASTAHAAII
LIDATRVTFDAGAAQLLPQTKRHSAIVKLLDLQHVIVAINKMDLVDYSETRFNEIRDAYV
KLAQQLGLADVRFVPVSALKGDNIVAASERMPWYAGEPLLNVLETLPVETQAHDALRFPV
QWVARQDGSSADDFRGYMGRIEAGEVKVGDEIVVLPSNRTATIAEIIAPVPGGTAAVERA
FAGQAVTIRLAEDVDVSRGDTFVPRAQGVEPAKKLEADLCWFDETPLSSQRKYLLKQTTN
TVFTKIGAVKQVLDVHTLSHATDRHELKMNDIGRVALTLQKPIVCDTYDAHPGTGAFVLI
DEATHHTVAAGMIRAFSA*
>GCA_000452945.1_BPMSHR5858_genomic_02964_1 group_375
MAFNKETVQSNLQKLGVAARSRRAKRAGIGALIFLVAFGLLGFFAAPPLIRHVAEQQLSQ
QLERPATIARIAFNPYTLRLEADGVRVGERGGQGAFVDIGKLVVRASWTSLARFAPIIDE
VRLDQPRFRIVRYDAQRFNFTDLVDKFSKPSPQPSAKPARFSVSNIQVNDGRIEFDDRLL
NARHVVDQWTVGIPFIATLPSKTDIFVEPKLRARLDGSPIAIDGKTKPFAQSRESSINLK
FDGLDVPRLLSYAPTKLPVDVRSGRLSSNLDVNFAMAGDTPSLRVSGTLDLNDAQVTSRA
NEPLFAAHAVHVAAAQLEPLRNVLHFDDIRIDRPVVSLARDKAGVLNVMKLAGGEKAGAP
PAARAAAPTSAGASAAAAGSSVSAVSPGSPAAARPASGAASQPAAAASSAAAAHAAAEKA
PPLDLTIKRFAINDGALNLNDASLATPAAISLQHVATTLADFTLAGKTPARYTLAADVAS
GGSLKAEGAFSLAARQADTKLAIDALALPAVQPYLAGASSARVLDGVLSTSVNAKADWSK
TPLDAQVSDSELGLKSLKIALPNAKTPAVALPDARVKVTKIDLAARTAEIASVDATGLAL
DVARLQDGRIDLASLAGDGGRAAGAKPAASNASAARGARPAAPAWRYRIDELNVKDAKAN
FTDRSTPRPVKLAIAPLALNVRQISEDLSKPLPVRLTATLNRKGSLELSGNVTAQPLKVG
LKLNGNRLDAAAFEPYFGSMLNATVASALLNAKGELSVEQAKQSTKASYRGDAALVDVRM
LDKATSDPFAGWRSLALANLKASYDDARGTDVDASRVTFSNFYGRVLLDAQGKLNLREVV
AKESGPAQSLTRDASGKEPIPLTPQAASQAKAAIAASAVEPASAPVLASAAAASAAAGAS
AVAASPASGAAVVRAAAPPQHPVRLHFGQLLLQRGRVTYTDNFIKPNYTANLVAINGTIG
AFGTDSTAPAPVDVGANLAGNGPITIKGTVNPLIEKPSLDLTATAHDIELTNLTPYSAKY
AGYPITKGKLNVDLHYKLDNDLLSANNHIFIDQLTFGEHVENDTATKLPVRLAIALLKNS
RGEIDVNVPVSGSLSNPEFSIGGLIWRAVLNLIAKAVTSPFTLLAHAFGGDGEALGYVEF
EPGRSKLSDASQKKLDTISKMLAEKPSIRLDLIGRVDPDKDLPGLRDAYVERLVRQQKLK
DVVGQGESVDPMTVKVDPAEYTKYLTKAYKAADFKKPRNMIGFTKTLPDDDMRKALADHA
SVDDSSLRALAQARAQAVRQYLEGKVDSSRMFIVAPKLDAKGIEDKGATTRVDFGLK*
>GCA_000452945.1_BPMSHR5858_genomic_02967_1 yycE
MNPPVLRVARPTNDIDTLVRFYTKALGFDVVARFENHEGFDGAIVGRAGYPWQIEFTHRH
GVTVARAPTAEHLLVFYLPDRGAWTEAVARMRAYGATPCRSENPYWDREGVTFEDPDGYR
IVLQNAQPA*
>GCA_000452945.1_BPMSHR5858_genomic_02969_1 group_2448
MRLYALPRGADIGRHAPCGAPGRTLRGRPCAKRRRPARICRYNAAAIRMTDLIRSMGNST
RAFLLGPLLKGVSRSFYLTLRVLPQGMRDPIGLAYLLARAADTIADTSLIPPAQRLELLL
SLRAQINGAPDPHALARIAGEVAGQQAQTDEKTLLESLGPALAVLSQLSASDRAAVRGVV
TTLTEGMEFDLRTFPDESSGRIVALREWPALDRYTYLVAGCVGEFWTKMTCAHLPGLLTE
REDTMSERGIRFGKALQMTNVLRDCGRDLRIGRCYLPQAMLDEYGLRAEDLLGPGASLRA
RPLMHALVRTTLAHFHAGIDYTFAIPRRAPRLRLACFWPIAIGLRTLALLASNDAWLDPA
RASKVRRGDVYRILAYSLPLAMSNGALRAWLDRLIADVEARLIA*
>GCA_000452945.1_BPMSHR5858_genomic_02973_1 group_3383
MKTSITTTTEPRAVARGARLPGWMWFALLWMGGVAGAVTLGYAFKLFMYATLFAVK*
>GCA_000452945.1_BPMSHR5858_genomic_02976_1 ocd
MTTPVFDAAQTARVTPFAELVDALRTAALDADAGRIESPARLVVPLNEGGVMLSMPASAA
DLAIHKLVTVCPRNRGGDLPTIQGQVSVVDPATGAPLFALDGPTVTGRRTAAVTLLALRT
FVAAAPRDVLLIGTGTQAAHHVEALAALFPGVRVRVKARAPAQAAAFCARLRAVLPTLEP
LGGDTLPETLDAVITATTSATPVYDDAARAGRLVVGVGAFTPGAAEVGARTIAGSALYVD
DPAGARHEAGDLIVAGVDWARVRSLADALRATVSTPASNASPAPGAPVFFKSVGCAAWDL
AACRVARAVTSRTRACT*
>GCA_000452945.1_BPMSHR5858_genomic_02977_1 xynZ
MRASNHYAVMRRFVSPLAALLAAALTAAAHAAPASSIVTRTFRSPALHRDWSYTVYLPAG
YNPEGARYPVLYLLHGNAGNANDWITQGRLQLTADALIERRDIAPVVIVMPQGGTDWYVD
RKEKMQSAFLDDLIPDVEAHYAVSNQRAGRAIGGVSMGGYGALRFAFLEPERFCGAMLLS
PAIYANEPPASSAARYVGVFGDRQFDPKVWHELNYPALWRGYFAQPLRLRMFIAAGDDDL
SIQAESSALYTSLRRAQNPAALRIVDGAHTWDVWRRLIGPALKYTLECVK*
>GCA_000452945.1_BPMSHR5858_genomic_02979_1 group_3385
MKKHVISAAALLAFAAPVFAQSSVTLYGVIDEGFNYTSNVNVNGVGKSNYQLASGFVQGS
RWGLRGSEDLGGGLKAIFTLENGFDVNNGRLGQGGRMFGRQAFVGLSHAQYGSLTLGRQY
DSLVDYLAPLTANGNWGGTLFSHPFDNDNTDNSFRVNNTVKYASPDWNGLQVGGTYSFSN
ATGFSNNRQYSIGAAYTLGGLQLAAAYLQANNPGKTAGGAIADNDANFTADRLRIFGGGV
NYTFGPATVGFVYTKTDVKNPVSTVYLPTATFAGLGLSATKFQNFEINGKYQLTPALFIG
AQYVYTDGKFDAAAGSVKPKYHTVGLMADYNLSKRTDVYLQGAYQKVAGDKTGTIADGGY
VVGTDGPSASANQFAVRAAIRHKF*
>GCA_000452945.1_BPMSHR5858_genomic_02981_1 group_3386
MRSNSSWLLAFKIAVHIFFLCCGIAFAYSTLSAP*
>GCA_000452945.1_BPMSHR5858_genomic_02983_1 bla_1
MNHSPLRRSLLVAAISTPLIGACAPLRGQAKNVAAAERQLRELESTFDGRLGFVALDTAT
GARIAHRGDERFPFCSTSKMMLCAAVLARSAGEPALLQRRIAYAKGDLIRYSPITEQHVG
AGMSVAELCAATLQYSDNTAANLLIALLGGPQTVTAYARSIGDATFRLDRREPELNTALP
GDERDTTTPAAMAASVHRLLVGDALGAAQRAQLNAWMLGNKTGDARIRAGVPADWRVADK
TGTGDYGTANDIGVAYPPNRAPIVFIVYTTMRNPNAQARDDVIASATRIAARAFA*
>GCA_000452945.1_BPMSHR5858_genomic_02984_1 group_3387
MRRLPPLNALQIFATVARHRSFTRAADALCVTQGAISRQIQSLEAHYGFALFMRHARGLT
LTVEGEQLLPVVVESFARIEDISLKLTRQRTDLALKVPTCVMHWVLPRIMRFQREHPDLH
VQMTTTWRHDVDFQSEPFDAAIVYGMSPGPDVAAVPLFDERLTPVCAPDLLEGRPLARVE
DLACHTLLHPTRDHRDWRRWLDYAGAAGVDPDRGPSFDSLDLATSAATQGFGVALGDLTL
SEEDFAARRLAMPLDIVQRTGARYYFVYPESVAQQQKIQRFSGWLDANRD*
>GCA_000452945.1_BPMSHR5858_genomic_02991_1 ddpX
LKTSAHRLVEIAPHTHGVDVDLVYATERNLTGQPIYRRAHCLLLEPAEAALRRAVAVAAR
SGLRLRIYDAYRPPRAQQILWDFLPDPNFVADLGRGSNHSRGTALDLTLVDANGDALDMG
TGFDEMIAASNHFHDGLPEPVQRNRLLLLGVMHAAGFTHIASEWWHYELPGSRALPLIDE
EAGGPWRLM*
>GCA_000452945.1_BPMSHR5858_genomic_02992_1 group_3974
MPPAFVQTVETRFAELTPTAKRIASYMLANLERLGLETADQIARQTGTSGISVGRFLRSV
GYRNLDDLKRELRGTAERSWMITDRLDEYRRVTGANAPARDEAAPGGAPGALDDALAREL
DAIRHVYQLARTPGFAQVADRIAQADAVFILGIQSTRGISNAFHSYLEYLRPRVFYSDGM
SGSYVDSLNSEFADPYCIVTDTRAYSRIARRYCEAAGARGVRFALVTDVYCPWARELPCD
LLQVKTDVGQFWDSLAPLTCLFNLLISALVDRLGGTIDARVARNRELQRELDQFEP*
>GCA_000452945.1_BPMSHR5858_genomic_03045_1 tnaA_2
MSFDQTLIADLLDQWLGGQSPDWDRLYPAGRPPIEHAPTYPFARNVYWVHAADRTPEPGL
AAAPPRSRDAAPPLEAGTMPAAAIAVVGATAAATATTAVTARAATTVTNAIDATDATNAT
NATNATAATTAAAATTAVTASGATGASAAIVLKSAADEARRHFTPSRERAPVVLRALAPA
AADVADVTDVAASRAAQTQVFDGAAVEAARLAPDAIGAALIDSLARRLLVSPQSIGARDA
FDALGVDSLIGQEWLRELNRTYGTSIDGATLAECGHIGALASRIAGARADAAAPAWVPRA
EPPQRAAQCAPDAGTVSASVLASISTAASTLASASASASAAAAAATTHAGDRSAPRITRD
ALVHALAASLAKALYMDVADIDIEQPFMEMGLDSIVGVEWVHQVNRAYGIGINAIQVYDY
PNIVTFAGLVESLTNAGGAAGGDVASGVGGSDSNADTDRDRDRDRDGDGDVDVDVDVDVN
SDGVLTRPTGAAEPARAAAAATSPATAEIAAATAAAASSAIAAPPTTAETMAAASAACAP
RGELRREVIDSLVAALYMDASEIDVLQPFVEMGLDSIIGVEWIHALNRRYGTSIEAIQVY
DYPNVEKLTELLSKSIDALSPPHARCESDSDVLPGEPMTAATAANDTNDTNDTNDTNNAN
NANNANNANNANNANNANKCGVVAAEAAFAAPPAAPAAACAAAAIAAPIAPIAPAPLAAP
AAKPATLLDELVASLAQALFRPADTIDAERGFEAAGLDPIVADEWLARVHRRYGVRVRAD
EALACLRIADFAALVAARQEAGGAQAAGAAPVAAHDARCAARASADARAGDREPIAIVGM
SGRYPGARDLDAYWENLAAGRSAIGEIPASRWDVARHFDAHPATPGKVYSKWIGLLDDVD
CFDPAFFRISPAEAQEMDPQHRLFLQEGYRAFENAGYSADTLDGRNCGVYLGIMNQEYRQ
LGAGGAVTMLEKSNSFAIGAARLAYHLNLKGPAIPVDTACSSALVAIHLACQALRAGEID
MALAGGVTLYLSPDAYIEMCSSGMLSPDGRCKVFDDSADGFVPGEGVGAVVLKRLGDAQR
DGDPIIATIIGSGINQDGKTNGITAPNMASQFELVSGVHGRYGIDPATIRYVEAHGTGTK
LGDPIELTALGDAFRVRTAQTGFCALGSVKSNIGHTSAAAGVAGLHKVLLCMRHRTLVPT
LHFAVPNRHFDFAASPFYVNTERAPWAPLAASPRRAAVSSFGFSGTNAHLVVEEYVHPAA
AAPEAGGPFLFPLSARTREQLAAYAAQLRDHVRRAAHEDAGLADLAYTLQVARKPMAERV
GLIARTKHELAALLDAFVDGRDGGDGLIAGRRDRAGGTPPAPSPEALRALVDAGESRTIL
QRWALGATIDWACLYRDLDAAARPRRIAAPSYPFARERYWLPDPGTRRTPAARAARAAGP
HPLLREIDSAQSAACFGATLAGDEPFLRDHRIDGRPVMPSSAYLEMVREAAARALGEPAD
AMLVIEGIAWRNPLTVSGGARRLQLRAQAEPGARALRFDVSSQAAGDAAAEPLAHCDGVA
RYVPRAPAGAPDLAALRARLASGPRAADEAACAALHARFARLGIEYGATHRVLLRLRVDG
DEALAELAPAADGIAQHELHPGTLDAALQPMLALLGERVGDGVPVVPYRIERAEIHAPTH
GARWAWLRMRPDAHEWIFDVDLCDARGALCVALRGIAVTAWRRPDEVVRLEPVWRAAPVE
ADLHDERAGLDAQRVVFVCGAHGAPRAWPADGIAPVRYAALAAGAPPGEPDALAGWFEAH
ALALFDEVRRLLAPGMRQATLVQIVVPAAGPGAILHALGALLQTAHLENPLLHGQVIAID
DIDAPDLPRRLARDARRAADTRIRYVNGERQVACFDEAAAPDGARALPWRQGGVYLVTGA
AGGLARALADAIARGMGGDAPRATLVLTGRSPARDDMRALVASLCALGVAADYRVLDVAD
RDAVARMVEAIVGEFGALHGVVHCAGVLRDNYLLRKSADEFAQVLAPKVRGTVNLDFATR
DVRSLDFFVTFSSGAGVVGNPGQADYAVANAFMDAFAAHRASLGAARPGVSVSIAWPIWQ
AGGMRIDRQTEAELERRLAMRPMPTALGLDALHACLLGASPCPTVIHGARARILALARQG
FAAPPAAPPGFGAADRGADAPGADADADAVKARVRAAIDGALSAVLKLPDARLREPEYFE
SYGIDSINAIRLTVELERTFGPLPKTLFFEYRHADELERYLVSVHGAAVGARIRSAARGA
PAGPACARAGESGEPPGAPTRPPASEGGAAGAPRASAAPPLAERDIAVIGMAGRYPQADD
LQQYWDNLRDGRDCIEEIPPHRWDWRKHYDPARGHGAHHSKWGGFINDVDAFDPLFFNIS
PKEAVSMDPKERLFLEQVWTAMEDAGLRPEDLRRDAQRGTGVYVGLMYENYQLLAAEAAA
AGSDVGMAGGSYASIANRVSFFLDLRGPSLAVDTMCSSSMVAVHLACRDLLAGEIGVAIA
GGVNLSLHPNKYRMLSAARFMSGDGRCASFGSGGEGYVPGEGVGVLLLKRRADAERDGDR
ILGLIKASAINHGGRSNGYTVPNAAAQGGVIANAIRAAGIDARTINYVEAHGTGTALGDP
IELAGLARGFADSGANGPCRIGSVKSNIGHCEGAAGIAALTKVLLQLAHRQIVPSLHSRE
LNPDLPLDGSRWIVNQSLCDWERVVVDGVPLPRTAGVSAFGAGGTNAHLILSEYPADACA
APAGVIEPAGRDAHDMHDTQDMQDMQDMQDMQDALVVPLSARNAQRLHAYAQRLRAFVAA
HARGERGAPPRLVDLAFTYQRGRIAMPERLAIVARSLAELERALTAYVAGQRTGDGIYAG
RADRAAAGDARGAASERTAADFADRLAARWVAGEAVDWHALFDGRAPRRIAAPTYPFERG
RYWIGASRAAAGAAEAAMRAGGGASRATTAPRALSESGGEVRQPDQTIRRVRLAPTSSFT
ATARAPSPTQATQATQATQATQATQATQATQATQATQATQATQATQAAQAAQAAQAAQAA
QATRATRAAAAFAPATKASLEAALRDSLASALFVGVDEIDPSRPFSELGLDSIVGVEWIR
DVNRRYGVSIRTTDVYDYPSVGEFAGLLERLLRESSVAGAPPAPATEPTMEPTMEPVTAP
RPEAAGAAPCEPPPDEGRGGAPGNVERIRRELMRSLADALFVDIAEIDVDRPFAQIGMDS
IVGVEWIKGINQRYRVALKATDVYDHPTIASIAALVDARGASAASTASTASTASTASTAS
TASTASTASTASTASTARPEADVRLAADARIASAASGAPESEPARAAHPGAAASRARADV
APDAGPAPRRLDAAAPDARAARAEPVPERIAIVGMSGRYPGAPDLDAFWDNLAAGRDAIA
EIPPSRWPVGAFYDPEPGKPGKVYCTRIGLLDDVDRFDPDFFRISPAEAEEMDPQHRLFL
QEGYRAIEQMGCAPASLSRRKCGVYLGVMNHEYGELAMRHRGAASGIGSSYAIGAARLAY
YLNLKGPAIPVDTACSSALVATHLACQALRNGEIDLALVGGVTVYLTPESYVAMCAAGML
SPEGRCKTFDDAADGFVPGEGVGALVLKRLADAERDRDPILGVIVGSGLNQDGRTNGITA
PSGSSQTELLRDVYRRHRIDPAGIGYVEAHGTGTKLGDPIELTALSAVFGDYTDRRGFCA
LGSVKTNIGHTSAAAGVASIHKVLLCLAHRELVPTLNYANPNRHFDFADSPFYVNTDRRA
WDAAGDAPRRAAVSSFGFSGTNAHVVIEEYRPAAAAAPDASPPRVIVPLSARHPERLRAY
ARNLADWLAQAAARGAPERLAAHLAYTMQVGRDAMAERVAFVADGRDELERQLRRYADTG
ETSDGVYAGRAEPHAQASNALMLDEAFGAAIDGWMRTGKHEPLAKLWAGGFDLDWARLYD
GVPAAAMPRRIAAPTYPFASGRYWIDVEPDGRAAGPDADAASPEADSGSDSEHAHEHEHE
HEPAATLAYLPVWEELPPAQPRAAPDAQAGGVLVVHRGGAWGLVDAIERECVDGRHAGAT
CMTLDLSGHAPSPEGRAWRNAAPGAARLAAWLGEFGPVRAVFFAAGCSEARHDASGAHGW
ASAPDAHGEDERALLQLAQALMRSQAADASIEFVVLSLDHHRTDGTPSNPAGGGVAGIAY
AIAQGDHRFRVTNVDVSLDELRAARHAPAPHPVLAAVLRLAPSDRGALVRLRAGRGYRQA

FVRLDWAAEAGASGLKQGGVYVMLGGAGRVGRALTRRLIERYRANVAWIGRSPADSASVA
HALRALGPAGPAPYYAQADATDAAQMRRAIEAVRQRHGRIDGAVFCGMVFDANHAIASVP
AHRFDEILDVKARGSRIFYEALAHEPLDFLCYCSSAQSFSFSGAARLGAYAAATTAGDAI
VRSIAPVAAFPVGTIHWGFWETSVEDSALGSRHLGALSDDEGFACFERFVGQCMRGNPLR
EVVCMRASPEVEHLMQVLPGETATLAAPGQPAQPAPLRAAPDGAAASEAAPPDGAADVSA
DIDAWLARLTFATLRPMLDGPRPARACHARWWDETLRIFAARGWLRIVDGAPRVIAEPDA
GEHVWRDWARYRFDTPAARGRRAQIDLADVCVRALPDVLAGRLPAADVLFPGGSMERVEG
VYRDNPISDYFNAVQADALIRHVRAWIDAGRREPIRILEVGAGTGGTTALALERLRPYAA
AIGEYCFTDVSQAFLQHAQAAFGARAGYLRTALFDVERPLDAQRMPAGRYDIVIATNVLH
ATRQVRGALRNVKACLRAGGVLLLNEISEKSLFAHLTFGLLEGWWLHEDSSLREPGSPVL
APATWRRLLEDEGFGAIAFAARDAHALGQQVVCATSDGVIRQRAGEPSGHSSRQGHRNRQ
DHQGRQESSTHAGEAGAPAPASAGRAGAAPAASPAGTAREPVVAAIHRALQQSLKLPEAR
IGDHTPFLDYGIDSILGVRFVDSLKQALDVPLNTAVLFDYPTVERLADFIVATYGARLAA
RGVSAAPASVATASATLAASTASTASTAPTAPTAPTAPTASAAPAAPAAPNAPAAPAMPA
EAVSRDAAAPRAEPAGARPADIAVIGMAGQFPDAPDVDAFRALLEHARDGAPGVSGGMLE
NRDRFDHAFFHITPDEADAMHPYQRLVLQESWKALEDAGYNPAALAGARVGVFVGAEPAD
YRSTTFSGSSDALIASRVSYHLNLRGPAYVVNTGCSSGAVAIHLACESLRRNESDVVLAC
GIFAAMGPRMLGALGQAGMLSAGGRCRSFDAGADGTAFAEGIGVVALKRLADAIADGDPI
HGIVKASGVNQDGTSNGIMAPNGVAQEELIVDVYERFGIDPADIRYVEAHGTGTLFGDAV
EANALVRAFRRFTERSAYCALGTVKATIGHTAAAAGVIGLIRILLSMRARRLPGMPGLGR
ANPMIDLDASAFSLGLVSREWPAGRDGRPRLAALNTFGHSGTNVHIVVQEPPQARARPAR
AADGPRVAVPLSAMDREALRRYAARLCERLEAEGAALCVGDVAHTLRVAREPMAQRIVLF
ASTTGELAALLRAFVDGRDSPCLLDGAVTAAARAAGLDAAQLAQAARWLAGERVDWPPAG
GTPMRVHLPAYPFAGRRCGAAGWARAEAGASRDCAAAGPCEPPAGVAAAAMTVAAAGPRV
DATPSAAADRARGAARPAEWLAARVAARLGVPAARVDRRRSLLDLGLTSQDLVSLAGQLR
DATGEALLPSVLFDYPTIERLAAHLADTCPAAFGAAEPAEPAEPAETGRAAAGDAASGPA
PGVIALLERLEGGGLSLEETIYLIENTK*
>GCA_000452945.1_BPMSHR5858_genomic_03123_1 group_2814
MLAKSGKFLRVVFALFALAFAFSPGGARAATNQLLVLVPDTLTLPDPRVSAWLDSAQEEG
LQIKVMTDSQFMQAGAALSQYPGLILPDQVHTVADDTLVTAIQNYALNGGKVMLVYDFGV
LNSAGFYPIPKSRFSSMVGVDYVLYDQLGANMIGLGAVTGMSSWLRTIQVPPGKSMPWTA
ASATTSSALSSTATGTSSTSSTTSSGTVMFLSASPSNPGGLKNYNHNAWFGYTLPNGGKG
TFTSTSPKLTGFVTGKPVKSGTYGASSFTSTSTATANTFTDTLEGISGYVYGFLTYPSFV
TQGTYTGTTILTSPNFGLVSGYNTYGNGGVLFVNLPLAYLDGQTDGMLIHGYLHYFGTNL
LKLPSLSALPKAKAGLVFNWHFCAEDQIQPALYLKSLGVWTNGPFSIVFTAGPDEATFGD
GNGINLDYNATALNLVKYFLTKGHHVGSHGGWIHDYWGAYASETNQSTFQQYLTMNHSAV
QTATGLPDVEYAAPEGNTPTWSVNWLESNHFNGYYFTGHTGSEPTRAYRNGALLNPGLWA
YPVMPFGKYATFEEFQQYGVSTTDVTNWYTNLINFVVANRTSRLIYAHPYGASFYPSVLT
SIFTQANSLKQLGQFTWYTMSDVTNFDRARLNVAWTATDMGASWAFAASHPTSLANMTWL
LPKSAYAQPIVTLGSAAVVGTDATNWLVIANGGTSLKFTSAKVQ*
>GCA_000452945.1_BPMSHR5858_genomic_03127_1 group_2101
MSLCVQCVLLLIVFACSLSARGTALGPASASLATAPFARTAQARAFAQPSLPHAEAVAFA
FAQAEPFDAERPAAAGRPAAADAPPAPGKGIVISLCAAYLWVFALLTLVYASRHYVFGLD
RLFKPQRAPYRAITHADWPEITVFVAAHNEEAVVADCLTALLATTYPRERLTIVPVNDRS
TDNTRALIDEVQARAPELIKPFHRESGKPGKAAALKDALREIRGDIMVVFDADYLPRPGL
LKELVAPFFDPEVGAVMGRVVPQNADRNLLARLLDLERAGGYQVNQQARNNLGLVPQYGG
TVGGVRKSALDAVGGWRDDTLAEDTDMTYRLLLSNWRTVYLNHAECYEEVPERWPVRARQ
LTRWAKGHNQTLFRYLIPLLRSPVTPRRCRLDGALLLGVFVMPALLALAWGIALALYLTN
GIDSLVLGLLVSVFALFAFSTFGNFGVFFEIVVAARLDGRATRLRLVPVNVVGFCVTIAA
VVAALWGLALDALLRRELRWDKTERFRRPLNSGR*
>GCA_000452945.1_BPMSHR5858_genomic_03132_1 group_3991
MDYLLTLAADPAVWAALVTLVAMEIVLGIDNLIFISILSNKLPEAQRERTRRLGIGLALI
LRLALLGTVAWIARLTEPVFTLFDHAFSWRDLILIAGGLFLVWKATKEMHHHVSHDAAPE
GGATSRAGLTTWAAVSQILMLDIVFSIDSIVTAVGMTTHIPIMFVAVIAAVTAMLFAAGP
LARFIDRNPTIVILALSFLMVIGMTLITEGFGSHVPKGYIYAAMAFSAFVEGMNMLARRA
KLRRAKRTAAQ*
>GCA_000452945.1_BPMSHR5858_genomic_03141_1 lipB
MSAPPVSIVPSAPAAPDAAASVAPNPPAALPVTVRWLGETPYDACFDAMRAFTDARTPDT
DDEIWVVEHPPVYTLGQAGNPAHLLVADSGVPLVKVDRGGQITYHGPGQIVAYLLVDLRR
RKLMVRTLVTRIEEAVIETLAAYNLASARKAGAPGIYVESGPHRGAKIAALGLKIRNGCS
YHGLSVNVKMDLRPFLAINPCGYAGLETIDMASLGATADWHEVAQTLVRRLIAHLDGATA
AAALPQQALEQSND*
>GCA_000452945.1_BPMSHR5858_genomic_03147_1 group_19753
MNAHTQKSLIAGPVGHIEIAIDLPDAVRDGSAAPRGIALVAHPHPLFGGTMDNKVAQTLA
RIFVQLNYAVIRSNFRGAGATEGEHDNGAGEVDDLLAVLAHMRALPGHADLPLVLAGFSF
GTFVLSHVGKRLRDAGQAIERMVFVGTAASRWQVAAVPEDTIVIHGENDDTVPIASVYDW
ARPQELPVIVIPGAEHFLHRKLHILKRIVVGAWR*
>GCA_000452945.1_BPMSHR5858_genomic_03148_1 group_3400
MGHIMDSYYRHHVFFCLNQREKGAERPSCANCGSQEMQEYAKKRVKELGLAGAGKVRVNK
AGCLDRCEEGPVVVVYPEGTWYTYVDKNDIDEIVESHLRDGQVVERLRI*
>GCA_000452945.1_BPMSHR5858_genomic_03153_1 mlaE_1
LNFQTPPGLSIDAGSQGRTVRLCGQWTALALARDRGNVARRIARLAKEAVGEWDLSGVER
LDHVGGQALWRVWGRRLPDGIALTDNQRVVFERIERLDAGRESPEPVVRIDPVTRFGQGL
FTFGEHLYGGIALLGGLIVDLLAVLRRPRTMPWTEISANVYAAGTKALPITALVAFLIGI
VLSYLSAQQLQLFGANRYIVNILGLSVIRELGPVLSAILVAGRSGSAITAQIGVMRVTEE
LDAMRVMGIPHGLRITLPRVLALGVAMPLLVMWTNVIALTGGALAAKFVLGIDLNFFVRS
LPSVVPIANLFIGLGKGVVFGMLIALVACHFGFRIKANSQSLGEGTTTSVVTSITVVILA
DAVFAILFQNVGLG*
>GCA_000452945.1_BPMSHR5858_genomic_03161_1 group_3829
MRLALVLMGGGARAAYQAGVLKGLAEIAHDVDPKRRTSPFSVICGSSAGAINATSLASHA
DDFEHGVRRLLEFWEQLRAERVYRTDWLGIAAAGARWLAAMSIGWAARRSPRGLLDNAPL
AYLLRRELDFHRIELMLEARKLHALSVTALSYSSGRHLTFYQASEPIQAWRRAERTARMV
DLSAEHLLASSAIPFVFPAVPLVLDGQIEYFGDGSIRQIAPLSPAIHFGSDRIVVVGAAD
PRPEVPAANGNGRGYPSLAQIGQQVLASVFLDSIGADIERIDHVNRMIEHLPAYVEPESG
WRHVDVLAIAPSERIELIASKHLKRLPLTVRGLLGAVGGNKPAGASFASYLLFEAEFTRE
LVELGYRDAHGQRERLAQWIASAERRGGPAERGPGAGTDARTGAGAHAARRTTT*
>GCA_000452945.1_BPMSHR5858_genomic_03166_1 group_363
MPLSSIVSSNPRGAARRVRRAACVLALVLALHWLAALWLVRFREPFRPVEPDHVPVQVEL
LKPQPIERAPAPEKPAADRPRAAPKRAARASAPPAHAPRASAPVSSAAESSTESSAESPA
AASGTEPASAAGGQAAGATSGAAAGASGASAPPGEAAQGMKFALPPSADLQYDTFYNGMQ
NMPGTIHWRTDGGGYSLYVSMPVPFVGPYTYESRGRVDAFGVAPARYVETRGRRPPDFAI
FNRQTKQIVFTGTPNSLALPDGAQDRFSMLMQLAGLVGGDPDAYRPGVTREFFVVDRDSG
ETWPITTIGDETISTGMGSLDARHFMRLPRRAGDTRRIDIWLAPSLGWLPVRMVQTEPNG
AQIELLLHRRTNANGDADVHSDTSANTDADANADGGAAPAAAPASASGAPLNANEPVNST
EKADGSPRPPPADPGEPQP*
>GCA_000452945.1_BPMSHR5858_genomic_03167_1 group_2461
MQMIYNSPNYCVVEFPPQDGHHAMNSGGYEIVDKNAQREIFIDGELAARFREHVKQLIQA
EPTLDEVDEFLGQFDSLMTQPVVLH*
>GCA_000452945.1_BPMSHR5858_genomic_03174_1 group_3401
MNAPAPLKVFITGASSGLGLALAAEYARHGATLGLVARRADALAEFAPRFPKASISIYPA
DVRDADALALAASRFVAAHGCPDVVIANAGISKGAITGEGDLAAFREIMDVNYYGMIATF
EPFIAPMTAARRGTLVGIASVAGVRGLPGSGAYSASKAAAIKYLEALRVELRPAQVAVVT
IAPGYIRTPLTEHNPYPMPFLMDADRFAKRAALAISQKTAFRVIPWQMGIVAKLLHVMPR
WLYDRLFEKAPRKPRAKAG*
>GCA_000452945.1_BPMSHR5858_genomic_03175_1 group_4360
MNLNDAVVLVTGANRGLGLAFVEGLKAAGAKKIYAAARDPARVTTPGVQPVRLDVTRAQD
IAAAARELRDVNLLVNNAGIFRMGSLLAEADGGGLQAQLDTNFFGPLAMARAFAPVLREN
GGGAIINVLSVLSWLGLPNTGAYGISKAAAWAATNAIRNELREQRTRVLALHSAYIDTDM
VAHAQGVPKNRPEDVVRQTLDALAAGRDEVLVDALTRDVKAGLSAEPAVYTQPPTGA*
>GCA_000452945.1_BPMSHR5858_genomic_03185_1 group_2462
MQPTTNAPERPLDCPCGGAAPGANANARAPRYADCCARLIDGHATAATALELMRSRYSAY
VLGATDYLRATWDPSTCPAELDADTGRAHAPRWLGLVIKRHAQLDATHAEVEFVARYKIG
GRAHRMHETSRFTRDAQGAWRYVDGDVSER*
>GCA_000452945.1_BPMSHR5858_genomic_03186_1 group_3402
LVQFAWVGSGMAFGKVMTGWIAACVLSAAQADPLPSDAPSAPAAAPYAPRVSHAPRAAGA
LANAERFDYGDSGLPPVAANLNETIIRIPVDAAGAITLEATVYKPDGPGPFPLVVFNHGK
NPGDLRAQPRSRPLSFAREFVRRGYAVVAPNREGFAGSGGTYIQEGCDVERNGVAQARDV
AATIGYMSKLSYVDARHVVVAGTSHGGLVSLAYGTEAARGVRGIINFSGGLRQDLCEGWQ
KNLVDAFDTYGSRTHVPSLWLYGENDSVWSPALVAQLRDAYMSHGASTLFVDFGRYKDDA
HRIIVDRDGVPIWWPPVASFLAQLSLPTSVRYAVANPHEPKASGYAAIESVDAVPFIDDA
GRAAYRRFLAQHPSRAFAVSSEGAWSWAEGGDDPMALALEGCRKQGAGACQLYAVDERVV
WRDAGTQTADESTSAAHALASR*
>GCA_000452945.1_BPMSHR5858_genomic_03194_1 phoA
MKRVRLAALLGASVFAAAGCGSDEPKTPGASDNGAQTGAARNVIFFLGDGMGMTTLTAAR
IYALGEDGALTLDTLPETAFVKTYSNDAQVTDSAPSMSAYMTGVKTNNEVISMTPDTKAI
EPSASLTGNCGANNGKPVPTLLEIAKAKGLATGVVTTTRVTHATPAATYAHVCHRDAEND
IAAQLVPGGAGYNAALGGGVDVVLGGGAQFFVPKEAGGKRADGRHLVNELKAKGYAIAQN
RDELLAADATKRGKLAGLFASSHMSYDLDRGATKEPSLADMATRALDVLQKNPNGYFLMV
EGGRIDHALHDTNAKRALQDTVAFDNAIKATLDKVRQTDPELKNTLVVVTADHDHTLVLN
GYAARTGKTESGKPGVLGVLRNYQTGAVAKDADGAPYTIIGFGNGENRVQGSRAGTSLTD
AVTGADDYRQEAVVRMAKGGETHGGTDVFLGAIGRGADGFHGVIENNKVFELVRGAAQL*
>GCA_000452945.1_BPMSHR5858_genomic_03198_1 group_3403
MNARLPIDGPPALADYRLTDNLSATRGRIFLTGTQALVRLLLMQRALDAAQGVNTAGFVS
GYRGSPLGMVDQQLWKAKKQLDASGVRFLPAINEELGGTAVLGTQRVEADPQRTVDGVFA
MWYGKGPGVDRAGDALKHGNAYGSSPHGGVLVVAGDDHGCVSSSMPHQSDFAMIAWHMPV
VNPSNIADMLEFGLYGWALSRYSGAWVGFKAISETVESGSTVDLDALAMRWPAPEGFAPP
EGGLHNRWPDLPSLTIEARLAAKLDAVRHFARANSIDKWIAKSARANVGIVTCGKAHLDL
MEALRRLDLTAADLDAAGVRIYKVGLSFPLEMTRIDTFVEGLAEVLVIEEKGPVIEQQIK
DHLYNRAAGARPVVLGKHDAAGAPLLSELGELRPSRILPVFAEWLARHKPALDRRARVVD
LVAPSILSNEADAVRRTPYFCSGCPHNTSTKVPEGSIAQAGIGCHFMASWMERDTTGLIQ
MGGEGVDWAAHAMFTNTRHVFQNLGDGTYFHSGILAIRQAVAARANITYKILYNDAVAMT
GGQPVDGSISVPQIARQVEAEGVSRFVVVSDEPEKYDGHHDLFPKGTTFHHRRELDAVQR
ELRETSGVTVLIYDQTCAAEKRRRRKKGEFPDPDRRLFINEAVCEGCGDCGVQSNCLSVE
PVETALGRKRRIDQSSCNKDFSCANGFCPSFVTVEGATLKKAAGVAFDEAALAARVGALP
RPATHLADAPYDMLVTGVGGTGVVTVGALVSMAAHLEGKSASVLDFMGFAQKGGAVLSFV
RIASSDAWLNQVRIDTQQADVLLACDMVVGASADALQTVRRERTRIVVNTHAIPNAAFVR
DPDASLHADALVEKMRHAAGDGFLARCDAQALATKFLGDTIGANILMLGFAWQQGLVPVS
LDAMMRAIELNNVAVPMNKLAFSIGRMAAADAAGLDALWDARHPVHAPPLPQTLDAIVAD
REARLADYGGARYVARYRALVGAARATGDERLARAVATTFYRLLAVKDEYEVARLYTDGA
FRTALEAQFEGAAGKDFRVKFNLAPPATAKAGRHGAAPRKQMFGQWLWPVLGVLARVRGL
RGTWLDPFGGTVERKMERALADDYETTMHRAFAAHSPQTAELVARLAELHAKVRGYGHVK
LANVAGVKRSERELAARLSIDAATSDAVRHALDEMKGAGALRGIAVVVAKS*
>GCA_000452945.1_BPMSHR5858_genomic_03199_1 group_3404
MNVDPFSPHSSDSFAQAASPARKPPRGGRRIWSGTREVIAYGMPASVWRDLYYWALKVSW
PVFFASLAALFVVNNTLFALLYQLGDAPIANQSPPGFVGAFFFSVETLATVGYGDMHPQT
VYAHAIATLEIFVGMSGIALSTGLVFARFARPRAKIMFARHAIVRPFNGRMTLMVRAANA
RQNVIAEARAKMRLMRREHSSEGYSLMKIHDLKLVRNEHPIFLLGWNMMHVIDESSPLFG
ETPESLAEGRAMLLVMIEGSDETTAQVMQARHAWEHDDIRWHHRYVDLMSDVDGMTHIDY
TRFNDTEPVEPPGAAPDAQAFAAKPGEGDARPV*
>GCA_000452945.1_BPMSHR5858_genomic_03200_1 group_19862
MFKILQSRRVVTAAAFALAAAYSVGVPAETVQAVSSVAPLHGGVDGPFFPKAASAVPASA
SATTGAQLQQQAQSRLAASLGANGALRNGAALTKAQAQASGLGYVAQHFDEIDTARTGKV
TLKQVQQFIEQRRQ*
>GCA_000452945.1_BPMSHR5858_genomic_03217_1 group_19940
MRSCSKSTWPPRLVTAPGLHGSEGAHWQTWLERQFARALRVEQDDWDAPHVERWAQKVRD
LLARERGPFVIAAHSFGCLATAHALAQAAQTRAADVVGVLFVAPANPRKFAFAGDFDARR
LNVPSIVVGSESDPWMTLADAREFAQRLGSAFVNLGDAGHINTAAGYGPWPRAKYFVDTL
VHCAAPLRLRDDDAPASGAPARQPLVTAA*
>GCA_000452945.1_BPMSHR5858_genomic_03218_1 group_19760
MSATPRGALRPFHLAFPVTSLEHARRFYGGLLGCPEGRSSDHWVDFDFFGHQLVAHLAPG
ETGRSAVNPVDGDDVPVRHFGVVLSMDEWHALADTLKAAGTRFVIEPHIRFKGEAGEQAT
MFFVDPCGNALEFKAFADIGQLFAK*
>GCA_000452945.1_BPMSHR5858_genomic_03219_1 ghrA
MRILLYTPHRDAHDWKRDVERALPGAQLRTWTPGDDAGADYALVWRAPREFFAPRAGLKA
VFNLGAGVDAILALEREHPGLLPRDVPLVRLEDAGMARQMVEYATHAALRYLRRFDEYAL
LQRERRWQVLAPHARETFVVGVLGLGALGAEVARALAALGLPVRGYSRAPKAIDGVTTFA
GAAQLDAFLGGAKLLVNLLPSTAATDGILNARTFSKLAHGAYLVNLARGAHLVEADLLDA
LASGRVAAATLDVFATEPLPPDHPFWREPRITITPHCSADTLRAEAVGQIAAKIGALERG
EPIGGIVDRDRGY*
>GCA_000452945.1_BPMSHR5858_genomic_03223_1 group_3407
MASNPASSDARRGALEMVVAMLMSGTIGWLVVSSRQHLMNVVFFRCLFGGATMLIICAAL
GLLRRDLLTKRTIAFALLGSVAIVANWVLLFAAYSRASISMATAVYNTQPFMLVALGALV
FRERLTASAAAWLVLAFGGLALVVRVEPAVLAVPGEYLEGVALALGAAFLYAVSSIVTKR
LKGTPPHLIALFQTGVGVLALAPFVHYDALPATAGQWLDLVALGVVNTGIMYVLLYGAIQ
KLPTATIGALSFVYPVVAIVVDRFAFGQTLAWSQVGGAALILLAAAGVNLGWRIVPARRV
AARG*
>GCA_000452945.1_BPMSHR5858_genomic_03224_1 lrpC
MQKRLSTPLAAANPLDAIDRELLRTLADDARQPVSELARRVGLSAPSTADRLRRLEAQGV
IARFTVELDPRALGYTLQAIVRVKPLPGQLHLVEELLRRIPEFVECDKVTGDDCFVCRLY
LHSIEQLDGILAKVTERAETSTAIVKSTPVARRLPPLAREED*
>GCA_000452945.1_BPMSHR5858_genomic_03226_1 pnoA_1
MALPAVLQNLALPVIASPMFIVSYPELVLAQCKAGIVGSFPALNARPAELLDEWLTQLQT
QLAEHKAANPDAVIGPIAVNQIVHQSNVRLEQDIRVCVEHKVPIFITSLRAPAREIVDAV
HGYGGIVLHDVINLRHAQKALEAGVDGLILVAAGAGGHAGTTSPFALVGEVRRIFDGPIV
LSGSIANGGSILAAQAMGADLAYMGTRFIATQEAHAVDAYKRAILEAKSADIIYTNLFTG
VHGNYIRESIVNAGLDPDALPESDKTKMNFGGDKAKAWKDIWGAGQGVGLMDDVPSVAEL
VARLKREYDDAKARLGIRA*
>GCA_000452945.1_BPMSHR5858_genomic_03228_1 group_19957
MKSSFSRALQTTFACAAVAALAAGASGACAQSSVQLYGQVDEWIGAQKFPGGQRAWGVQG
GGMSTSYWGLRGTEDLGGGYQAIFTLEDFFRAQNGHYGRFDGDTFFGRNAYVGLATPYGT
VRAGRLTTQLFISTILFNPFIDSYVFSPMVYHVFLGLGTFPTYTTDQGVVGDSGWNNAID
YTSPSFGGLNAAAMYALGNTAGDNRSKKWSGQLNYSSGPFAATAVYQYVNFNGGPGDLGA
LVSGMKSQGVAQLGLSYDFKLAKIYAQYMYTKNELNTGSWHVNTAQGGVSVPLGPGSALA
SYAYSRDSGGHDQTRRTWALGYDYPLSKRTDVYAAYMNDRYSGMSGGDTFGAGIRAKF*
>GCA_000452945.1_BPMSHR5858_genomic_03229_1 group_2466
MMTERAIRKIGQWAAGVTCALAAVLVFAQPPHGGGPGFGGGRGGVPAMAGGWRGAPGYLQ
AAPHGGEARIGVNGYRQRWGLRPTPSYGRYAAQSPYRPISAEARPAPHASGPGNVPMRAG
SIRADVARYNEERGGRPAPAPRQDDSQRLPFFAPFSRN*
>GCA_000452945.1_BPMSHR5858_genomic_03234_1 yihS
MTTSATPAPRPAPAAPPAAAPYVASFRDPQFLLAHVEDTLRFYAPHAFDPTGGFYHFFRD
DGSVYDRTTRHLVSSCRFVFNHAMAYRHFGDARDLAYARHGLRFLRDAHWDAEHEGYDWE
IEWRDGAKRATRDGTRHCYGLAFVLLAAAHAAMAGVEEAKPLIAHTFDLMEHRFWDRAAG
LYADDATPNWTVSSYRGQNANMHATEALLAAYEATGHLTYLDRAEKIATNIVQRQAALSQ
GLVWEHFHADWSVDWEYNKEDSSNIFRPWGFQPGHQTEWAKLLLILERHRPLDWLVPRAI
ELFDAAFAHAWDSDHGGLCYGFGPDYTVCDHDKYFWVQAESFAAAALLAQRTGSERFWDW
YDEIWRYSWAHFVDHRHGAWYRILTCDNRKYSDEKSPAGKTDYHTMGACYEVLHALASTR
DDSGRGANAGGGKGDDDRAPRAAHDSGRGGGVKGGVKGGGKGGGTDDHGHRVDKGYKGGE
GNAADGAGRRRNGGAR*
>GCA_000452945.1_BPMSHR5858_genomic_03239_1 mcl1_1
MTVPADTCDTSAARTARVVLITGAVQTGAAPARAAEAARAGARQAIGAGRPGDAQPGASA
PAGAPAHPPRAGEQQAQAPSAPPAARPAPDAAAVGRALAVGFARRGWNVALAAAAGDEAR
AAAALAAEVEALGRRAAVLAADLAIEADVARLVADCGAALGRPACVIAQAAPVADTARDV
GYASLAGAMARSVAAPLVLARTLADATPDAARDDERERAVVIHLLDETLFHPAPERLSHS
LAQAALHRATAAQALALAPKVRVVGLVGGRAPRADDIADAACYLADAPGVTGATLTVDGG
EHLAPPADERN*
>GCA_000452945.1_BPMSHR5858_genomic_03259_1 group_19739
MGILNTVGEIAGAVAAVEAAEKVDPDAGLLTKAAAAVAGFKGAEALENLVEKKKDDEAAD
GENTAAPDSDASQA*

>GCA_000452945.1_BPMSHR5858_genomic_03261_1 group_2467
MQFGLLLSQLPHVDESRVHYDAMLSLKAAGEPASASFWQRLKALIS*
>GCA_000452945.1_BPMSHR5858_genomic_03264_1 oprB
MKKKTLHQMLVKSGVASGLPLLLSLGIVQPALAQAPSPAAAPAENAAPPAPATRAPDGVP
GGNGATAAQTDAAPTGFWDRSNLFGDMGGLRTKLGDHGITLNLQETSEYLRNLSGGTSRG
GAYDGLTQFGFSVDTEKAIGLPGGTFNVSGLQIHGTSLTARNLQLLQTASGIEAEATTRL
WELWYQQSFANGRADVKVGQQSLDQEFMVSQYASTFINATFGWPVLPAVDMPAGGPAYPL
SSLGVRLRAKPSDAWTVMAGVFDGNPAGGVGDAQQLNRHGTNFNLRNGALFIGELQYALN
APPADPKAPQAGLPGMYKLGVWYNSERFADPRYDTNGVPLADPASNGVAATHRGNYGFYA
VADQMVWRPGADSPRSLNVFARVMGAPGDRNAVDFTLNAGVTLKAPFAGRDNDTAGLAVS
YAKIGSRARGADGDTGVFQTPGYPVRRAETLIEATYQYQVTPWWQLQGDFQYAFRPGGGI
PNPNEPGSRIGNEAIVGVRTTITF*
>GCA_000452945.1_BPMSHR5858_genomic_03265_1 group_2976
MKALAWLTGAVLALLAAELISFGDIGRFVTLLAALLAALCALAAAFGCVYTLVAAALTHR
FFARAPREPHACPPVTIVKPLHGVERTLFANLASFCEQRYDGPIQFLFGVHDRDDPALRA
VDALRTAFPRAHVTIVADARLYGPNRKIANLVNMLPAAAHDVLIFADSDVSVGPDYVRHI
VGELGEPGVGLVTCVYRGRPDPGFWPRVEALVTSHQFLPGVVTGLALKLARPCFGQTIAM
RRAMLDAIGGLAQFAHHLAEDHAIGEAVRARGARVVVPPFAVEHGCVETRVAQLVEHELR
WSRTIRAVDPRGHLGSLLTHPLALALLAGVLSSGAAWAWPLVPAALVARVAAKRIVDRAT
KRPVRDLWLLPLADLIAFGIFVASFSSSRVIWRGFSFDVDRDGRLCPAPEKRPNA*
>GCA_000452945.1_BPMSHR5858_genomic_03269_1 group_3413
MKQTTKITGLIGGAVLAIAGQQAMAQSSVTLWGVADVSLRYLSNSNAQNDGRFFMTNGAI
TNSRIGLHGSEDLGGGLKAIFNLESGVNLQDGAFSDSKRIFNRAAYVGLTSQYGTVTLGR
QKTVLFDLLSDTYDPLTVGNYLENAWLPVALGAGLYADNSVKYRGTFGGLTIGAMYSFGT
DSTATGAGGFSGQLPGHMGAGNMYGFSLSYVAGPVSVAAGVQQNSDNSNRKQTIYHANVV
YAFSKAKIYAGYLHSKDDTGFVDTVLAQQPGLFTSGAPKRTGRIDDGPFAGVSWQATPAL
TLTGAFYYDRMRNATIANGQLDSGNRYTVVGLVEYALSKRTEIYGTVDFNKVNGAATVEM
PGRSNQTGVAIGLRNIF*
>GCA_000452945.1_BPMSHR5858_genomic_03274_1 group_3414
MQSRSEPGGGDARGVSVADLFAGFLGLGLISFGGALPFARRALVEERRWLSADEFTDLLG
LCQFLPGGNVINLSVAVGMRFRGVPGAIAGLLGLVAGPTLVVIALGVLYAKTQQNPAVHH
LFGGLAAAAAGLLIAMAVKVAKPLRHMRAGACVAVLAFVAIAVLRVPLLPTMLVVTPVSI
WLASRAPRGANAGAQGVRR*
>GCA_000452945.1_BPMSHR5858_genomic_03284_1 flgF
MDRLIYTAMTGATQSLEQQSVVANNLANASTTGFRAQLATFRAVPMNFEDGSGNVDPATT
RTYTLSSTPGADFAPGPIERTGNPLDVAVQGPGWLSVMLPDGTEAYTRAGNLHVDQNGQL
VTANNLPVVGNGGPLAVPPNAQLTIGADGTVSALMPGDPPTAVAMIDRLKLVNPDPATMK
RGDDGLFRTADGNPADADPAVKVVPNSLEGSNVNPVAAMVSMIDNARAFELQTKLIQSAD
QNEQAANQLLNFS*
>GCA_000452945.1_BPMSHR5858_genomic_03286_1 group_4187
MTSSFTTIGGSGTTVNSLPYDTMSSNGQSSGTSGANTTNGTAGTNGTSGANSGINVASSL
STTSAADLQTTFLKLLVTQLQNQDPTSPVDSSQMTSQLAQINTVSGIAQLNTALTSLSSQ
LTAGQQTQAAMLIGSNVLAPGNTVPVKSGAASPFGVQLTSAVSNLTITVKNASGVVVNTI
NAGAQSAGTVPFNWTPTDTAGNKLPDGTYTISASYTDTSGKQYAPTTLSSAQVLSVIKQA
DGTPGLVLSNGSTVGFSQVASIFPNTTKSASTGNASSSTN*
>GCA_000452945.1_BPMSHR5858_genomic_03290_1 group_19989
VKVDSTPTSNARTLSNASAGAARTQAGQPAAAQTPAGAAGAPTGGDANVSLSGLSSTLRS
LAASGSADIDTAQVDAIRDAIKSGTLSIDTGKIADGILQTARDLLKQPSQAGNR*
>GCA_000452945.1_BPMSHR5858_genomic_03291_1 group_3897
MAHGRSARRHVERNEMRDELLATVNDEHATVEAFASLLAYEEKALTTAAPLDALPGIVER
KTELLEKLAQLERRRDTLLASLGLPAGKKGMDLAAENDARLANGWQLLQHSAERARHANA
INGMLIRIRMDYNERTLSVLRAAPQRNGFYGPDGRVAAVAR*
>GCA_000452945.1_BPMSHR5858_genomic_03297_1 group_3985
VSESIARAPAPNPKRTLRETLARRRLDAQRAPRAHDALARRVLAILAQCAPRTVGFYWPL
PGEFDARDAIASWLAGDASRQAALPVIGKPRTPLEFHAWTPDAPMREGRHRIAEPAAGAR
LVPELLLIPCVGFDHGCFRLGYGGGYYDRTLAAWPGATPPVTVGVAYDACGVRELPREAH
DLPLDWIVTETRLYRSDD*
>GCA_000452945.1_BPMSHR5858_genomic_03298_1 group_3415
MQRHPTKIVSSEHLVSETSAELSELEYALIMAGNAFNRWMVRCMAAAGAKDMTAVEVSLL
HHVSHRDRKKKLADICFVLNIEDTHVATYALRKLVARGYVKSEKTGKEVFFDATDAGRAL
CLRYRDVRERCLIETLKDSGLTNEQIGDAAQLLRHASGLYDTAARAAASL*
>GCA_000452945.1_BPMSHR5858_genomic_03300_1 pxpC_1
MNSRQATGGIEVLRAGPLSTIQDLGRRGYRHLGVAQSGALDSLALEVGNRLVGNRPDAAA
IEITLGPASFRFPRATRIAITGTEFGATLDGAPIYSWWSVPVAAGQTLSLPVAKRGMRGY
LCIAGGIDVLPMLGSRSTDLASRFGGLGGRVLRDGDRLPTGAPPAGAPACVAPDAPEFGV
KAPAWCAFARVDEAPRRHKHAHAVWAMPVRVLPGPQYASFTAAAKHAFWDEEWIVTPNSN
RMGYRLAGVKLERAESADLLSHAVLPGTIQVPGNGQPIVLMNDAQTTGGYPRIGAVIRAD
LWKLAQARLNLPIRFVRVTDKAARDALAAERAYLRQIDIAIEMHEEALQRALAARAAAA*
>GCA_000452945.1_BPMSHR5858_genomic_03312_1 lhgD
MTIERIDCVVIGAGVVGLAIARELAARGRETLILEAADAIGTGMSSRNSEVIHAGLYYPR
GSLKASLCLHGRDLLYDFCETHGVPHRRCGKLVVATSPAQAKQLKAIAARAEENGVLDLL
TLSRDEVQALEPELECLEALFSPSTGIIDSHQLMLALLGDAERDGASCALRSPVESIDAA
GGRFVVRTGGEAPAAIAAACVINSAGLDAQALAKRIRGLDSRWVPPLYLARGNYFSLSGR
APFAHLVYPVPDRAGLGVHLTLDLAGDARFGPDVEWIDALRYDVDPRRAESFCTSIRAYW
PGLPEGSLRPAYAGIRAKVSGPGEPAADFVIQGTAQHGVRGLVNLFGIESPGLTAALAIA
QRVGDMAARTRHT*
>GCA_000452945.1_BPMSHR5858_genomic_03314_1 group_3419
MIPFKRKMFASPASRPARAGKFVLEACFAGALAGAAHAQGAPQPVVDWEIQVVRDGQTID
AFEQKTTVGQARSDTHSLPIAPAAGCAATSAAASAADAPGSGLSRTITVAPLYVEDGAVA
LAIDAQETLADEGATPPAGAPCTFALPRRIVASHPGLNARAAQWTDWTLAERNPQLVYRV
RARVVED*
>GCA_000452945.1_BPMSHR5858_genomic_03318_1 uvrY_3
MDTPSVAPSIASAPASSSSGLAPEVGIPFVSRCASRAPTAERLGARVAQLSAELVAADES
ARRHLAGELHDGLGADLTAARFALANVETWLPDDAPDAARRALALAQQALDAATAANRRL
IDGLRTPALEAGLVGTLAAWIGAHGARTGLRTSFVCAADARVTQLSADGALAVFRVAQEA
LANVAKHAHASAADVRIDVDDTHLSLVVTDDGVGVGVGVGVGVGAAAARRRGGYGLGSMR
ARCEALGGTLELALPRAGGAGTALRARFAWHALIAARMPAAARKVRKEAR*
>GCA_000452945.1_BPMSHR5858_genomic_03331_1 fliI
MVSDVCGGALGRDELDALERELALASRGPEHEDAAAHGDGFGAHHHAPPRPLAMGAPSSR
ATLANPHLAHWRAHLDGIRERNARALPLRPCGRLTRAAGLVLEAIGLRLSVGAECTIELP
AGSALPYAQAEVVGFAGERLFLMPTTAVAGVLPGARVWPLESAPVADPLAGAKRLPVGWE
MLGRVVDASGRPLDNLGPLASKVDAPLTAPSINPLDREPIHHVLDVGVRAINGLLTVGRG
QRMGLFAGSGVGKSVLLGTMARYTSAEVIVIGLIGERGREVKEFIEQILGEDGLARSVVV
AAPADVSPLLRMQGAAYATTLAEYFRDQGKHVLLLMDSLTRYAMAQREIALAIGEPPATK
GYPPSVFAKLPALVERTGNGPEGGGSITAFYTVLTEGDDQQDPIADSARAILDGHVVLSR
SLAEAGHYPAIDIEASISRAMTALIDDAHLDRVRQFKQMLSRYQRNRDLIAVGAYAPGRD
AQLDRAIALYPRIESFLQQGFRECAPYAPSLAALDALFESEGG*
>GCA_000452945.1_BPMSHR5858_genomic_03336_1 calB
MKNDLPGLATLDALLRDQRAAYLRAPYPSWATRADHLRALRKMLLENRDALAAAINADFG
HRAKEEVLMSEIWLAKEEIDEALKHGKRWIKPKSRTMNKWLRPARAKVMPQPLGVVGIVV
PWNYPVLLAAGPLICALAAGNRAIVKMSELTPRTSQLFEELISKTFARDHVAVVNGDAQI
GAAFSGLPFDHLLFTGSTNVGRHVMRAAAEHLTPVTLELGGKSPVIVGPRARFDAAVDAV
ITGKTLNAGQTCIAPDYVLVPRGKEAEFVARARARMARLYPNLSTNPDYTSIISERHFAR
LQRLASEAQQAGAQLHPLTDAAPDPALRRLPPVLVTQAPDASQLMQEEIFGPLLPIVPYD
TLDDAIAYVNARPRPLALYLFDEDRTTIERVMRDTISGGVTVNDTLMHIACGTLPFGGVG
ASGMGAYHGYDGFVTFSKMKPVLTQPRLNTRAMIAPPYGKRFAAILKLMLKF*
>GCA_000452945.1_BPMSHR5858_genomic_03349_1 dapF
MKLSFTKMHGAGNDFVVLDGYTRALPPLTGAQVRALADRHFGIGADQLLLVEKPTVDGAD
FKYRIFNCDGGEVEHCGNGARCFVKFVRDHGLTGKASVRVEVKHGVITLTMQDNGEVVVD
MGAPVFEPARVPFDASGLDGRREGADTLWPLPVNGVTRWISVVSMGNPHAVQIVDDAEAF
AVRVDGPAIECDPRFPQRVNAGFMQIVSRHEVNLRVYERGAGETLACGTGACAAVAAGIR
RGRLDSPVTVHTHGGTLTISWNGACDERAPLMMAGPATTVFEGVIELPA*
>GCA_000452945.1_BPMSHR5858_genomic_03351_1 xerC_1
MTAKPIADPIVDYLSNLRHVRKLSDHTLRAYAHELDELKRLANGRPLDSLTAVDMRSAVA
RAHAGGLSARSISHRLSAWRAFYRWFSQHVEMNANPVAAVRAPKRAKTLPKALSVDDTAA
LMDAPTAGTAESLRDHAILELFYSSGLRLAELIGLDIEYVKDGAYRSAGWLDLAEAEVTV
RGKGDKERKVPVGRKALDALHAWLAVRGEFVKHDPRPLFLSVRGNRMSPGVVRERVKRAA
LAAGIPANVHPHVLRHSFATHVLQSSGDLRAVQELLGHASISATQVYTSLDFQHLAKIYD
SAHPRAKKRD*
>GCA_000452945.1_BPMSHR5858_genomic_03352_1 rlmI
MHTVTLKPSKDKSLLRRHPWVYANAIDRVDGKPAPGATVIVRAHDGRFLARAAYSPHSQI
RLRVWSFDENEPIDHAFFKRRVQRALAHRRAMISGTDAVRLVFGEADGLPGLIVDYYVAA
RGAAHTGDAAARAAEGGAAAPVAPGDGEGRGQLVCQFMAAGVEHWKGAIVAALVAATGCP
NVYERSDVSIREKEGLEQTTGVLAGDAPPDTLIANENGVLYHVDVRNGHKTGFYVDQREN
RALVAQYARDRDVLNCFCYTGGFSLAALKGGAKRVVSIDSSGDALALAQRNVAANGFDAA
RAQWLDADAFKTLRRLVDEGERFDLIVLDPPKFAPTRDSVDRAARAYKDINLSGLKLLRP
GGLLFTYSCSGAIDMDLFQKIVAGAAADAKVDARILKRLGAGVDHPLLTAFPEGEYLKGL
LLQIA*
>GCA_000452945.1_BPMSHR5858_genomic_03353_1 yjiA
MAIPVTILTGFLGSGKTTLLKRILNEQHGMKIAVIENEFGEENIDNEILVQDSTEQIIQM
SNGCICCTIRGDLARALADLAAKKRDGTFDFDRVVIETTGLANPGPVAQTFFIDSEIADE
FLLDAVITLVDAKHANAQLDEHEVVQRQVGFADRLFITKADLVDDEAVSELKHRLLHMNP
KAAIKVVNFGDADIKEIFDLRGFNLNAKLEIDPDFLAEDDHAHAHHDDHDHDHAHCDHDH
GHCEHDHGHGHHDHHHAHHDDKIKSFVYRNDRPFDPNKLEDFLGGILQIYGERMLRYKGV
LYMKGVDRKVVFQGVHQMMGSDLAAKWLPAEKKTNKMVFIGVDLPRDLITDGLDACLA*
>GCA_000452945.1_BPMSHR5858_genomic_03356_1 hslU
VSTMTPAEIVSELDKHIIGQAKAKKAVAVALRNRWRRQQVAEPLRQEITPKNILMIGPTG
VGKTEIARRLAKLADAPFIKIEATKFTEVGYVGRDVDSIVRDLIEISVKQTRETEMRKVR
SKATDLAEDRILDVLLPQPRAVGFGASAEHANDDNNATRQTFRKRLREGQLDDKEIELDI
EQPAVGMDIMAPPGMEEMTEQIRSMFSNLGSGKKQRRKVKIKEALKLLTDEEAAKMLNDE
EVKTKAVQNVEQNGIVFLDEIDKITSRNHEGGGGEVSRQGVQRDLLPLVEGTTINTKYGM
VKTDHILFIASGAFHLAKPSDLIPELQGRFPIRVELDSLSVKDFEAILVATDASLVKQYQ
ALLATEDVKLEFADDGIRRLAEIAYAVNEKTENIGARRLYTVIEKLLEEVSFAAGNHAGQ
SVTIDSAYVDHALGEVSKDEDLSRYVL*
>GCA_000452945.1_BPMSHR5858_genomic_03360_1 argB
MSEPIDLSQISPALKAEILAEALPYIRRYHGKTVVIKYGGNAMTEERLKQGFARDVILLK
LVGINPVIVHGGGPQIDQALKKIGKQGTFIQGMRVTDEETMEVVEWVLGGEVQQDIVTLI
NHFGGHAVGLTGKDGGLIHARKLMMPDRDNPGEYVDIGQVGEVEAINPAVVKALQDDAFI
PVISPIGFGEDGLSYNINADLVAGKLATVLNAEKLVMMTNIPGVMDKEGNLLTDLSAREI
DALFEDGTISGGMLPKISSALDAAKSGVKSVHIVDGRIEHSVLLEILTEQPFGTMIRSH*
>GCA_000452945.1_BPMSHR5858_genomic_03366_1 bepA_2
VQAKLRVARAALTFVGGVALGVAGPVLGFELCMAEPMLVADNGAPPVPSAPAVATPAAPA
AAPAAGNPQPYALGDQQVRYGNAIVFRSLIPSPLLEQLTDNEYRQTVQDAAQRKRLLPPN
NARVKRLRTIVMRLAPYAVKWSERVKGWNWEIEVLRSRSIRAFCLPGGKVLVDSGLLERL
RLTDDELGVLFAHEIAHALREHARSSLGEQQAASLGTGATPLPPLFGLSEPLPAPLGVVE
RFASVRYDPTDETEADVIGGDIAARAGFDPRAAITLWDKLAVATRADKDNGFIHAHPYDT
RRRNDLRKRLADLMPLYRKALVKNADARANAAGAGAAAGAKQRGAATANR*
>GCA_000452945.1_BPMSHR5858_genomic_03369_1 exoA
MLRVITANLNGIRSAAKKGFFEWLGEQNADCVCVQEIKVSADDLPAEFVEPHGLRSYFHH
AQKKGYSGAGLYSRREPDDVIIGYGSSEFDAEGRYVEARFGKLSVVSVYVPSGSSGDERQ
QAKYRFMDEFMPHLAELKAKREVILCGDVNIVHKEIDIKNWKSNQKNSGCLPEERAWLTK
LFDDVGYVDVFRTLDPRPEQYTWWSNRGQAYAKNVGWRIDYQIATPGVAGTAKNTSIFRD
IKFSDHAPLTVDYDYK*
>GCA_000452945.1_BPMSHR5858_genomic_03373_1 gatC
MALTLTDVTRIAHLARLEMADADAERTLTQLNEFFGLVEQMQAVDTTGIAPLAHPIEQIL
EVAQRLREDAVTEHVNRDDNQRPAPAVQDGLYLVPKVIE*
>GCA_000452945.1_BPMSHR5858_genomic_03388_1 rsmB_1
MTQTRPSRSRASGAPSRAPRLSALHLVPDSLGFALDGAAQAVDAVDRGAALPAALAAVHA
ALPSNAQAASRGAIQDIGYRAMRRRGTADWLIAQLVAKAPPAHVHALLVCAFALLVDDEH
AAYAPFTVVDQAVTAIGARRECSFAKGLVNAVLRRFLRERAAWLERLQADRRALWNYEPW
WIDAVERAWPDAWQRILAAGDAPGPLTLRVNARRMTADAYLDTLRAERIDAEKIGLHAIR
LASARPVERIPGFADGIVSVQDAGAQLSAEWLGARDGMRVLDACAAPGGKTGHILELAHA
EVVALESDPARAPRIGENLARLSLTADVRVGDAADPAQWHDGRPFDRVLADVPCSASGIV
RRHPDIRWLRRAADIPALVAEQRRILSALWPLVKPGGELLYVTCSIFPEEGELQARWFGD
ACQDAVRLDAPGQLLPQGATGGAPAVGGSGESGRHVDHDGFFYARFQKR*
>GCA_000452945.1_BPMSHR5858_genomic_03392_1 def1
MALLNILHYPDKRLHKVAKPVAKVDDRIRKLVADMAETMYAAPGIGLAATQVDVHERVIV
IDVSEDKNELRVFINPEIVWTGDGKQVYEEGCLSVPGVYDEVERPDRVRVRALDGQGEPF
ELDCEGLLAVCIQHEMDHLMGRVFVQYLSPLKQTRIKTKMKKLERAM*
>GCA_000452945.1_BPMSHR5858_genomic_03393_1 group_19755
MSPRALTRAELSGWLRLAGASGVPAAACAALLGAFASLEALFAASHAELAALVGDAAAQS
VLAPPAADFEQRVDAALAWLDEPGNALVTRHDPAYPGPLAELYDPPPLLYIKGRVALLHA
RAVAIVGSRGATPQGLADATRFARELSDAGLAIVSGLARGIDGAAHRGGLDGASGTVAVI
GTGADLVYPACHHALAHEIAERGALVSEWPLGTPARSAHFPQRNRLIAALSGGVLIVEAA
LRSGSLITARLANEMGRDVFAIPGSIHAPLSQGCHALIRDGAKLVETAADVLDEFGLDPA
RPVNSKRGAPAAASADADLDDDTRRLLDAIGYGPVPLELLAQRTSLPSGTLHRLLLQLEL
AGRVAALPGGRYTRIDGAQAEPAQARGTSAVLHSGA*
>GCA_000452945.1_BPMSHR5858_genomic_03411_1 spuE_1
MKRIAWLAAVLASLACAAARAADGNVLNIYNWAEYFAPDTIAGFEKETGIKVRLDVYDSN
EALQTKLTTGNSGYDLVFPSNDFLARQIQAGLYRKLDKSRLPNLTNLDPAIVARAAEVDP
GNQYSVPYMQGTFGLGLNVAKVKQALGGPLPANTLELIFNPAYAAKLERCGIAFNDAGSE
VFPLALRYIGRDPNTTDPRDYEAALDMMKKIRPTIRQFIATPVMNDLATGDVCVVTGYSG
AVLVAARRAAEAKNGQQIVYSLPSAGAPFWFDSMAIPKGAAHADHALRFIDYILRPDVVA
KISNKVMYPNPNRVATPLVDRRLTANPAIYPDAATMRTLWVKRPMPPQAMRMQTRYWTRF
KTGY*
>GCA_000452945.1_BPMSHR5858_genomic_03420_1 group_19874
MKFHIHEIDHVVLRAADLAAMTRFYCDVLGCHVEKEQRELGLVQLRAGRSLIDLLAVGAP
IDRAGSGAPGKGRNMDHLCLRVEPFDAAALRTHLAAHGARPGDEARRYGADGYGPSLYLF
DPEGNMVELKGPPEAVAP*
>GCA_000452945.1_BPMSHR5858_genomic_03421_1 yaeQ
MALKSTIYKAELQIADMDRHYYADHSLTVARHPSETDERMMVRIAAFALFAHERLEFCKG
LSDTDEPDLWQKDLTGAIEVWVDVGQPDERRIAKASGRAGRVNVIAYGGRTSDIWWQGAR
GKVERLRNVQVTTLADGVAAALGGLAERTMRLQCTIQDGAAWLSSATHDPVAIEWTTLKA
REDA*
>GCA_000452945.1_BPMSHR5858_genomic_03425_1 group_2471
MKKIIIALAVSAGILSGCAVYVPDQPGVIVAPQGGPGNGFCPPGQAKKGNC*
>GCA_000452945.1_BPMSHR5858_genomic_03426_1 group_3431
MKTIWKFFGLAAAVSVLLAGCGGGGDNSGQTGTLHVAMTDAPSCGFDHIYVTVAKVRVNM
SAQAGDNDSGWTDVALAAPQKVDLLSLTNGVLADLGRNALPAGQYQQVRLVLAQNQGNTL
ANSIVPTGGTEQPLATPSATQSGYKIITPFTVQPNTLVDLVLDFNACKSIVQRGNGTYAL
KPVVTATPTVVSGAIDGYVSPTEAGATVYAEQNGHVIKGTLADSTGHFVLTPLVQSSTNG
NYDIVITQNNVSTGIVRSVPVVVNTTTSVSTSSAPITLPASTMNMVSGTATASADAILRA
LQMVNSLPYEIASTNANLDTGAYALTLPTAAPIVGTYSGSLPVAMSAAPSAAGQYTIEAD
AANGATQQQPANITAGSVSNVNFGF*
>GCA_000452945.1_BPMSHR5858_genomic_03442_1 group_19869
MRTQSFSRKSPRAHGIGARPARAFMRWLRDESAVSAIEYALIASLIAIVIIGAVQVVGTN
LQSVFSTVASDV*
>GCA_000452945.1_BPMSHR5858_genomic_03451_1 group_19919
MSAQLNHTIVWCRDKRRSTHFLTNILGLPAPVSFGPMLVVSLSNGVSLDFYERDEPISTQ
HYAFLVGDGEFDRAFARIREQGLTYWADPAKRRAGEINTHGGGRGVYFNDPDGHFLELLT
RPYDLGAAS*
>GCA_000452945.1_BPMSHR5858_genomic_03453_1 group_2679
MNILIAGFQHETNTFAPTRASYQSFVRGEGFPPLARDVNVPIGGFIRAAQASGHALLPVV
WAGACPSAHVTSGAFERIGGEIVAAVQAGGFDAIYLDLHGAMVTEQFDDGEGELLARVRR
IVGERMPIVVSLDLHANVTARMAAHASALVAYRTYPHVDMAQTGERAARVLERLAAEARP
LHCAIRRLPFLIPINGMCTHAEPASGAYRLLAQLERDGVVSMSFAPGFPAADFPECGPTV
WAHAFEADAAQRAADALFAKLVGDEARWSVPLLAPDAAAAEAIRMSRTATRPIVIADTQD
NPGAGGDADTMGMVRALLRCGARDAAVGVIWDPDAAAAAHRAGVGARIGLRLGGRSRVRG

DAPLDAEFEVEHLSDGRFRFDGPMFNGAHGELGPVACLRIDGVRIAVSTNKMQTFERNQF
RVAGIEPERTKIVVNKSSVHFRADFEAIADAILVAKSPGPMAADPADLAWARLDPDIRVR
PNGPTLRSLRAMAR*
>GCA_000452945.1_BPMSHR5858_genomic_03456_1 cstA
MNRASGYLVWIAVALLGAFAFGTIALAHGERISALWIVIAAVCVYLIAYRFYSRFIASKV
VQLDGLRMTPAVKYNDGLDYVPTNKYVLFGHHFAAIAGAGPLVGPVLAAQMGYTPGMLWI
LAGVVFAGAVQDFIVLFISTRRDGRSLGDLVKMELGTVPGVIALFGAFLIMVIILAVLAL
IVVKALTNSPWGTFTVAATIPIALFMGVYTRYIRPGRIGEVSIIGFVLLMASIAYGQHVH
DSAALAAWFTFSGTQLTWILIGYGFVASVLPVWLLLAPRDYLSTFLKIGTIVGLAIGILI
VAPELKMPALTKFVDGTGPVWSGNLFPFLFITIACGAVSGFHSLISSGTTPKLLDNETNA
RFIGYGAMLMESFVAIMALVAACVIEPGVYFAMNTPAAVLGTTPEAVAHTVTQWGFMLTP
DMLTQTAKAVGETTIIARAGGAPTLAVGMAQILHQVIGGEAMMAFWYHFAILFEALFILT
AVDAGTRAGRFMLQDLLGTFHPALKRTESLPANLTATALCVAAWGYFLYQGVVDPLGGIN
TLWPLFGISNQMLAAIALVLGTVVLFKMKRERYAWVTIVPTAWLLICTLTAGWQKVFDSN
PKVSFLAHAAKLQAAAADGKVLAPAKSLAQMQRIIFNDYVDAALSALFILVVVSIAVYGV
VAVVRARRAVQPTSRETPYEPMPVAQALGSGR*
>GCA_000452945.1_BPMSHR5858_genomic_03457_1 group_19855
MFSDLGGEWRTAGRYLGQALRLMVGLPDYDGYVAHMRATHPDRPVMTYEEFFRERQNARY
GSGAGKCC*
>GCA_000452945.1_BPMSHR5858_genomic_03459_1 hemL_2
LEIRHGINLARARALLARERDAFERAMPKSRARSAEAAQHLLFGVPLHWMRDWSTPFSLY
VDHARGAHFTDIDGHRYTDFCLGDTGAMFGHAPEPVARALAEQAARGYTTMLPSEDAAWV
AAELARRFRLPYWQFALSASDANRFVLRWARAATGRKHIVVFNGCYHGTVDDVFVDLADG
RPVQRDSLLGQAHDLSAYTRVVEFNDLAALEAALKDGEVACVLAEPALTNIGMVLPDAGF
WREARALTRRHGTLLAIDETHTISSGPGGYALAHGLEPDLLVVGKPIGGGVPCAVYGFSA
SCAERAQHAKASAPPGHSGIGTTLTANMLAMRAIRATLADVMTDTAYAHMFDLAARLAAG
LEQTIARRGLPWCVTRIGARTEFQFAPAPPRNGTIAGLQLDGELEHIVHLYLLNRGVLIT
PFHNMMLVCPQTSAEDVDRLVAAFDACLGELL*
>GCA_000452945.1_BPMSHR5858_genomic_03461_1 moaA_1
LSIPLLQQVRVGAYIMRQHLSGNKRYPLALMLEPLFRCNLACNGCGKIDYPDPILNQRLS
VEECLQAVDECGAPVVSIAGGEPLLHKEMPEIVKGIMKRKKFVYLCTNALLMEKKMDDYA
PSPYFVWSVHLDGDREMHDHSVSQEGVYDKAVAAIREAKRRGFRVNINCTLFNDALPERV
AKFFDTLGPIGVDGITVSPGYAYERAPDQQHFLNRDKTKNLFREVFKRGEGGKRWSFSQS
SLFLDFLAGNQTYKCTPWGNPARTVFGWQKPCYLVGEGYVKTFKELMESTDWDNYGVGNY
EKCADCMVHCGFEATAVMDTIAHPLKALKVSMSGIRTEGAFAPDIPIDNQRPAEYVFSRH
VEIKLEEIQRAGKGKLQKAPKPAATA*
>GCA_000452945.1_BPMSHR5858_genomic_03462_1 ispH_1
MRVILAQPRGFCAGVVRAIEIVERALQQNGAPVYVRHEIVHNRHVVENLRNKGARFVEEL
DEVPHGAVAIFSAHGVAQTVEQDAQARGLDVLDATCPLVTKVHVQGRQYVAAGRRLILIG
HAGHPEVEGTIGQIPAEVILVQSEAEVDTLTLPADTPVAYVTQTTLSVDDTRGIIEALQR
RFTDIVGPDTRDICYATQNRQAAVRELSEQVDVLLVVGATNSSNSNRLREIGTESGVPSY
LVADGSEVRAEWFANARTVGLTAGASAPEEMVEDVIAALRALGPLEVATMSGREEKVEFK
LPAKLTQAVAREV*
>GCA_000452945.1_BPMSHR5858_genomic_03464_1 group_332
MNLNDQIDTLDSRFDQLIRTVCEVASAARHTAATAAADKTAAQAAFEAAAEKHTQAASMS
VEQATLEALAAQAAADKEAADALARQAAAVQAAAEAAAAQAVTDKTAAEALAMQAAAERA
AAERAAAQAAADKSAAEALAARAAGDRDAAERASGERSAADAAAAQAAEDKAAAEALAAQ
AAADRTAAEHAATRAAADKTAAESLVAQAAAERATAEHAVTQAAAERAASEQAAARAAAE
KTAAETLAAQAAAERAAAEHAAEQAAADKAQAQALAQQAAADRAAAEQAAAQAEADRAAA
QHAAAQAATDRAAAEARATEQAADAKAAAESLAAQAAADRAAAEQAAAQATADKDAAESL
FKQAAAERSAAEAAAARAASYKATAEALAAQAERNRAMIEAEPAAPVATAQTANGTLSLH
VARPSQDEISLALGEHSVSLPPHQLSQLIEELAHARASMTIEQPSGLPNGWRFVTTRNPL
MAVQKQANGDRLLVARHSGYGWVPFTFSPDVVIQLYMLLTQK*
>GCA_000452945.1_BPMSHR5858_genomic_03466_1 yccX
MSGDDLDERIETYYVRVRGVVQGVGFRHATVREAHALKLRGWVANLDDGSVEAMLQGSAP
QIDRMLAWLRHGPPAAHVTEVTFEEHRTDKRFERFQQH*
>GCA_000452945.1_BPMSHR5858_genomic_03467_1 rarD
LTGAGDDPQAARGIALSVAASALFALLSAYAKLLAPLTGLDIFAWRIIWTAPGALAVVAL
RGRWPALRELLARAFASRRLALSLAASAGLLGLQLWLFLWAPLHGRMLEVSLGYFLLPLT
MVLVGRFHYHERLSALQWLALACAAAGVAHEIWATRAFAWPTLVVALGYPPYFVLRRRVN
ADSLALFSVEMLALLPIAAAMLAAGGTSVAGRPLLWAMLLPGLGAISTLALASYLKASRM
LPMALFGILGYVEPVLLVAVAVLVLGEALSWQQLATYAPIWAAVALTAWHGFLIARR*
>GCA_000452945.1_BPMSHR5858_genomic_03472_1 xynR_2
MTNQTDGGVAAVNRALAALVAFGEAPGGLTLAQVSEASGVNMSTLLRMFESLERFRFIKR
LNDGRYVLGPAVFQLGMMYRESFQLREHVMPVLDALGAETGETTAFYVREGDQRVCLFRV
HPRRAVRTYLREGDRYPLDVGAAGRVLLAFSGARGGSFDKTAAQGYAVSLGERDPDSAAI
ACPVFGVGRALIGALSLGVPRFRFNKKVQADYLPRVQAAANALTHALGGDLPPAAGVAQV
VDLFGAAHE*
>GCA_000452945.1_BPMSHR5858_genomic_03510_1 dadA_1
MRVVILGSGVVGVASAYYLARAGHEVTVIDREAGPALDTSFANAGQISPGYAAPWAAPGV
PLKAVKWMFEKHAPLAIRLDGTRFQLQWMWQMLRNCTTERYALNKGRMVRLAEYSRDCLQ
ALRAETAIQYEGRTGGTLQVFRTQQQLDGAAKDIAVLREANVPFELLSSDELKKAEPALA
AVSHKLTGGLRLPGDETGDCQLFTTRLAALAEQLGVKFRFNTRIDALAVAGGKIAGVQCG
GEMVRADAYVVALGSYSTNLVASLVKIPVYPLKGYSITAPIVDAAKAPVSTVLDETYKIA
ITRFDDRIRVGGMAEIVGFDKRLRDARRGTLEMCVNDLFPGGGDTEKATFWTGLRPMTPD
GTPIVGRTPVPNLFLNTGHGTLGWTMSCGSGQLLADLMSGKKPVIRADDLSVHRYLSETD
GEHRPAYA*
>GCA_000452945.1_BPMSHR5858_genomic_03513_1 etfB
MKILVPVKRVVDYNVKVRVKSDGTGVDIANVKMSMNPFDEIAVEEAVRLKEAGVATEVIA
VSVGVAQAQETLRTALAIGADRAILVESNDEVQPLSVAKILKALVDKEQPQLVILGKQAI
DDDSNQTGQMLAALANLPQATFASKVTVADGKATVAREVDGGAETLSLTLPAVVTTDLRL
NEPRYVTLPNIMKAKKKPLETVKPEDLGVDVAPRLKTLKVVEPPKRAAGVKVADVKTLVE
KLKTEAKVL*
>GCA_000452945.1_BPMSHR5858_genomic_03514_1 metQ_2
MQRRTMLKLAAALGAAACVAGAQAQTQTQTIKVGVTGGPHAQVMEEVKKVAAKHGLDIRI
VEFSDYVQPNAALAAGDLDANSYQHAPYLDAQVKDRGYRIVKVADTVTFPMGVYSKRVNS
LAALRAGARIAIPNDPTNGGRALLLLQKQGLLKLRDGAGLKATRFDVVDNPKQLKLVELD
AAQIPRSLADVDAAAINTNYAMEAGLKPKQDAIAIEGPNGPYANILAVRAEDKDKPWVAK
LVAAYRSADVKRFIERQFGGAVIAAW*
>GCA_000452945.1_BPMSHR5858_genomic_03578_1 spoT
MSTTPSSASSEAGHAEATASSHARQYIDAVLEQSFRHLFGPTATPEQPRKHGVVSIAKLT
AALADYLGAEEIKEVKAAFHFSDEAHLGQYRQSGEPYITHPVAVAEICAGWKLDAQAIMA
ALLHDVMEDQGVTKSELAERFGPKVAELVDGLSKLDKMEFRSREEAQAENFRKMLLAMAR
DVRVILVKLADRLHNMRTLGAVPMEKRRRVARETLDIYAPIAHRLGLNNTYRELQDMSFA
NFNPHRYATLEKAVKAARGNRREVIGKILESVQRAMADAKIDAEITGREKTIYSIYKKMR
DKQLSFSQVLDVYGFRIVVEHPLDCYTCIGVLHALYKPVPGKFKDYIAIPKVNGYQSLHT
TLVGPFGAPIEFQVRTRKMHEIAEAGVAAHWLYKNGGADLNDVQKRAHQWLKSLLDIQSE
AGDSSEFLEHVKIDLFPDAVYVFTPKSKIMALPRGATALDFAYSIHSDLGNQCVAVKINN
ELLPLRTELKSGDIVEVITAPYSKPNPAWLGFVRTGKARSAIRHYLKTMRLNESVQLGER
LVDQSLKGYGLALADVTPEVWEKLVQWTGNKSRQEIFADIGLGRRVAAVMAKRIEVLMSG
RDADDDLPRAERHAANHAPPVVITGTEGMSVQLSACCRPIPGDDIMGYIGIGLGMAIHTT
ACRVAQRIHRRDPGRWIDVAWAPQPGRLFDVAVKALVKNTKGIFARVAADITSADANIVH
IAMDEDLTHESTVLRFVIQVSDRVHLANVMRRVRTNPDVMRIMRERSNDDVVHARHDGGM
RIDRERQDY*
>GCA_000452945.1_BPMSHR5858_genomic_03579_1 rpoZ
MARITVEDCLKQIPNRFELALAATYRARQLAQGHTPKIESRDKPTVVALREIAAGQVGVE
MLKKVPA*
>GCA_000452945.1_BPMSHR5858_genomic_03583_1 rdgB
MSHASPDAARSRIVLASNNPGKLREFAALFSTAGIDIVPQGELGVSEADEPHATFVENAL
AKARHASRATGLPAVADDSGLCVPALLGAPGVYSARYAQRAGREKSDAANNAYLVEQLRE
VADRRAYYYCVLALVRHANDPEPLIAEGRWAGEIVDAPRGAHGFGYDPHFFVPALGATAA
ELDPAAKNAASHRALALKALVARLGEIR*
>GCA_000452945.1_BPMSHR5858_genomic_03588_1 group_2477
MTTRSELFRFRAAWRGAALCACAAVLAACTITLPPPHGAVVVPGVSSMSATEYRAAIAQR
IADSNSDRILHGTPQAMLRSLVVVAFTVDRNGQLVKSAVYRSNGDDEAEGIALASLRRAA
PLPPPPAKMLDGRGQLELMESWLFNDNGQFQLRTKAAPQAQTLN*
>GCA_000452945.1_BPMSHR5858_genomic_03589_1 yhjE
MSDTRLSASAAAPRAANTTNHRRIVFASFIGTAIEFYDFYVYATAAALVIGPVFFPHGSA
TAQALSAFVTFGIAFIARPIGSFLFGHFGDRIGRKSTLVASLLVMGVSTTAIGFVPGYDA
IGALAPVLLCVLRFGQGIGLGGEWGGAALLATEHAPAGKRGWFGMFPQLGPSIGFLASNG
LFFALALTLSDEQFRSWGWRIPFLVSAALVAVGLYVRLKIAETPAFQAALDRHERVRVPL
AALVAHHWRPTLLGALAMVVCYTLFYISTVFSLSYGVSTLHFSRQSFLGLLCFAVLFMAL
ATPLAAAAADRFGRKPVLIAGAIAALLSGFTMAPLLGSGSMPLVALFLTIELFLMGMTFA
PMGALLPELFPTNVRYTGAGVAYNLGGILGASIAPYIAQLLAARGGLAWVGAYVSAAAAI
SLVGVLLMRETRDAKLT*
>GCA_000452945.1_BPMSHR5858_genomic_03590_1 group_19956
MDPIRLQNADLVLILALSLGGALLLAMRFGPKSWLGIAVEALVANLAAIAAVIAFELLMS
*
>GCA_000452945.1_BPMSHR5858_genomic_03591_1 rfaC_2
MKRILIVKVTSLGDVVQTLPVVADLHRAFPGVQVDWAVDESCADVVRWNTGVSRVLCAPL
RRFKKARSPADLKAIAASISELRAHRYDAVIDLHGVYKSAIISFLARARRRYGYRNQDLG
ELGAMFAYNGRFGPRPACDAWHGMRVSAGQALGYEPQGRADYLLDVPPESREPHTAPLVA
LAEPGGPYALFFHATSNDDKRWPTDRWSAVAREMRARGLRVLLPWGSEREREEAQRIAAR
APGAVVLPRLTLADVARKIDRAALVIGVDTGFVHMAHALEKPTVMIFCATSRQHLGVSGA
PHSLSIGDEGATPGVDDVLGAIEHVYPTGAVEQPRRVAAM*
>GCA_000452945.1_BPMSHR5858_genomic_03601_1 serS
MLDIQLLRKDLDGVAKRLADRGYPLDVAAFSALEAERRAIQTRTEELQARRNSLSKQIGA
MKGRGEDTSAVMAEVGGIGDEMKASAVKLDEIQARLSELMLEMPNVPHESVPVGRDETEN
VEVRRWGAPRQFDFDVKDHVDVGTPLGLDFETGAKLSGARFTVLRGPIARLHRALAQFML
DTHTQQHGYSETYTPYIVNPDVLYGTGQLPKFAEDMFRVEKGGAENTVTQYLISTSEISL
TNTVRDSIVEASALPIKLTAHSPCFRSEAGSYGRDTRGMIRQHQFDKVEMVQIVAPEASY
AALDEMVGHAEAILQKLELPYRVVALCTGDMGFSAAKTFDLEVWLPAQNTYREISSCSNT
ESFQARRMQARFRNAQGKPELVHTLNGSGLAVGRTLVAVLENYQNADGSVTVPVALRPYM
GGVERIDAPSSAA*
>GCA_000452945.1_BPMSHR5858_genomic_03604_1 lolA
MQHQSLAVHTGSGSGFAPKHGFARALRAALAGAACALVASHAFASGTEQLKAFVAKVHAA
KGEFTQQIVKAPPKAASGAVPTVKPTDNSSGSFVFSRPGKFIWTYQKPYQQVLQADGDKL
YVYDKDLNQVTERKLAGALGASPAAILFGSNDLDKNYTLRDAGEKGGIDWLEMVPKAQDT
QFQRIGIGFRNGMLAAMELHDVFGNVTLLTFTNIQTNPPLKADQFKFVVPKGADVVTG*
>GCA_000452945.1_BPMSHR5858_genomic_03618_1 group_3451
MITLDNVSKWYGSHQVLADCSAVVSKGEVVVVCGPSGSGKSTLVKTINGLEPFQKGSITV
DGARLGDPSVKLAQLRARVGMVFQHFELFPHLSVLENLTLAQTKVLRRERDEACDIGLRL
LDRVGLKAHACKFPGQLSGGQQQRVAIARALSMNPVAMLFDEPTSALDPEMINEVLDVMV
ELAREGMTMVCVTHEMGFARKVAHRVLFMDRGAIVEDAPSDAFFAAPRSGRAREFLDKIL
H*
>GCA_000452945.1_BPMSHR5858_genomic_03624_1 hemL_1
MSNNQTLFERAQRTIPGGVNSPVRAFRSVGGTPRFVARAQGAYFWDADGKRYIDYIGSWG
PMIVGHVHPDVLAAVQRVLADGFSFGAPTEAEIEIAEEICKLVPSIEQVRMVSSGTEATM
SALRLARGFTGRSRIVKFEGCYHGHADSLLVKAGSGLLTFGNPTSAGVPADVAKHTTVLE
YNNVAALEEAFAAFGGEIAAVIVEPVAGNMNLVRGTPEFLNALRALTAKHGAVLIFDEVM
CGFRVALGGAQQHYGITPDLTCLGKVIGGGMPAAAFGGRGDIMSHLAPLGGVYQAGTLSG
NPVAVAAGLATLRLIQAPGFHDALADKTRRLADGLAAEARAAGVPFSADAIGGMFGLYFT
EQVPASFADVTKSDIERFNRFFHLMLDAGVYFAPSAYEAGFVSSAHDDATLDATLDAARR
AFAALRA*
>GCA_000452945.1_BPMSHR5858_genomic_03628_1 ribH
MEIGQYQPNLEGDGLRIGIVQSRFNEPVCNGLADACVEELERLGVSGEDVLLVTVPGALE
IPLALQKLAESNQFDALIALGAVIRGETYHFELVSNESGAGITRIALDFNTPIANAVLTT
ETDEQAIARMTEKGRDAARVAVEMANLTMTLDQLSDDEEDEEDEDDEDEEERA*
>GCA_000452945.1_BPMSHR5858_genomic_03632_1 ubiD
MKYKDLRDFIHGLEQRGELRRVTQPVSPVLEMTELCDRVLRAGGPALLFDAPAGHRFPVL
GNLFGTPRRVALGMGVDADDEAALASLRDIGRLLSALKEPDPPKRLKDAGKLLSLAKAVW
DMSPKTVSAPPCQEIVWEGDDVDLHKLPIQTCWPGDAGPLLTWGLTVTRGPNKTRQNLGI
YRQQLIGRNKLIMRWLAHRGGALDFREFALKHPGQPYPVAVVLGADPATMLGAVTPVPDS
LSEYQFAGLLRGARTELAKCVTPGVDALQVPARAEIVLEGFIHPQQGAPAPAPEGAPPRP
AAGAAAGYEHALEGPYGDHTGYYNEQEWFPVFTVERITMRRDAIYHSTYTGKPPDEPAVL
GVALNEVFVPLLQKQFAEITDFYLPPEGCSYRMAIVQMKKSYAGHAKRVMFGVWSFLRQF
MYTKFIVVVDEDVNVRDWKEVIWAITTRVDPARDTVLVENTPIDYLDFASPVAGLGSKMG
LDATNKWPGETQREWGRPIEMDAAVKARVDRLWTEIGLS*
>GCA_000452945.1_BPMSHR5858_genomic_03650_1 pcm_1
MNIEKARFNMIEQQIRPWDVLDQDVLSLLSVVKRENFVPAAYRDLAFADVELPLPSGQKM
LFPRVEARILQELAVKKHENVLEIGAGGGYMAALLAARGQRVTTVEIDPDLAKFAEENLK
KNGVTNAEVVLGDGSRGWPAKAPYDVICVSGGLPVVPQELLEQLKVGGRLAAFVGGRPVM
KAQIITRIDDTQYRVADVFETYVDHLVNAIEPSRFKF*
>GCA_000452945.1_BPMSHR5858_genomic_03652_1 bauA_2
MTEHVSQTPDLSAFWMPFTANRQFKEAPRLLVAAKGMYYTSHDGRRILDGTAGLWCVNAG
HGRDEIVAAVQAQAEEMDFAPTFQMGHPKAFEAATKIARHTPGDLKHIFFTNSGSEAVDT
ALKIALAYHRARGEGQRTRFIGRERGYHGVGFGGISVGGIAPNRNAYSGALLPSVDHLPH
TLNLKEAAFSKGQPAWGAQLADELERLVALHDASTIAAVIVEPVAGSTGVLIPPRGYLER
LRELCDKHGILLIFDEVITGWGRLGAPFAAQFFGVTPDLLTMAKGTNNAAVPMGAVAASG
AIHDAIVGGAPAGIELFHGYTYSGHPLAAAAAVATIDLYEREGLLTRAARIAPIFEREIH
KLKDARHVIDVRNLGIVGGVELTPRDGKPGARAYEVFVKCFEKGAMIRYTGDILAFSPPL
IVEEAQIVELFSIVAQALKETE*
>GCA_000452945.1_BPMSHR5858_genomic_03653_1 amiC
MKRRSLLKFGSMAGVMALAGQSPVARAADSGKGPIKVGILHSLSGTMAISETSLKDTALM
TIADINKNGGVLGRPLQPVVVDPASNWPLFAEKARQLLTQEKVACVFGCWTSVSRKSVLP
VFEELNGLLYYPVQYEGEEMSRNVFYTGAAPNQQAIPAVEYMMSAEGGGAKRFFLLGTDY
VYPRTTNKILRAFLKSKGVKDSDIQEVYTPFGHSDYQTIVANIKTFAQGGKTTVISTING
DSNVPFYKELGNQGLKATDVPVVAFSVGEEELRGIDTKPLVGHLAAWNYFMSVKGPANAK
FKEQFAAWVKSQNLPGGAKRVTNDPMEATFVGIHMWKQAVEKAKSTDVDRVRTAMIGQSV
AAPSGFTLTMDGNHHLHKPVMIGEIRGDGQFNVVWKTKTAIRAQPWSPFIAGNQGKPDVV
GSIPEFLRRRRAALA*
>GCA_000452945.1_BPMSHR5858_genomic_03654_1 group_3457
MPMPLSRAARALAALAACAAFSFAAPRAALAVTAADVAALAGDDFDAKRAAIDRLAAGHD
AAAAALLNALANGDALATDAGRILIQHGDAARDALTNAPAQAGDAQPVMLNNLLRTKIAN
ALSGLDLASPDIDTRRRAIDALLKRPDAALKPMIDAARAKETDPVLKRRLDALWAIAALR
DADPAKRLEAVRLVAARSDLDMIEQLRPLVAKRPDGGDAEPDARVREAAQQGLGALYAIQ
RRGEIAGTLFAGLSLGSVLLLAALGLAITYGLIGVINMAHGEFLMIGAYATYVVQTLVQR
YLPGAFDWYPLAAIPVSFAAAAALGIVLERTVLRHLYGRPLETLLATFGVSLILIQATRM
IFGAQNVQVVNPSWMSGGVTVMQNLILPYNRLAILAFALVVVGIAWAVLTKTRLGLFVRA
VTQNRRMAACVGVKTARVDSYAFAFGAGIAGLGGCALSQIGNVGPDLGQSYIVDSFMAVV
LGGVGQIAGTVLGGFGLGLVSKAIEPFWGAVLAKIAVLVMIVLFIQKRPQGMFALKGRSA
EA*
>GCA_000452945.1_BPMSHR5858_genomic_03659_1 ureA
MKLTLREKDKLLIFTAALLAERRRARGLKLNYPETVAFITAALMEAARDGRTVAEVMHYG
TTLLTRDDVMEGVPEMIPDIQVEATFPDGTKLVTVHHPIP*
>GCA_000452945.1_BPMSHR5858_genomic_03660_1 ureB
MIPGELVIDDGEHALNAGRHTIALVVANTGDRPVQVGSHYHFHEVNDALSFDRAAARGFR
LNIAAGTAVRFEPGQTRTVELVELGGARAVYGFQGKVMGPL*
>GCA_000452945.1_BPMSHR5858_genomic_03663_1 ureF
MDTAELVALLHLASPALPIGAFSYSQGLEAALDAPLIRDADGARDWIASGLADVLAQGEL
PFLAHQLARWHAHDAAALADANDEFVASRESFELRRETEQMGWSLAQLCASLEWGDAARR
ATLASIPSVALPSAFAFAAAAHGATPDAALAAYAFGWVENQTAAAIKAVPLGQLAGQKII
VALREPIRDAVRRALATPPEAINTFAPQLGILSARHESQYSRLFRS*
>GCA_000452945.1_BPMSHR5858_genomic_03666_1 group_3458
MSESQIIEIASADWSGQQLSVPREQLLAGLEDGKVLFFAHLRFAIEGGEEALLDPALADP
KRKNISLAPNGGALAGVAGDAVTQSAVRALIARFQQQAGALVDGLFPEYRGKLRVAPTSL
RLMQVETRETSWRKDDSRLHVDAFPSRPNYGERILRVFTNVNPAGVPRVWRVGEPFEDVA
KRFLPHIKPQVPGVAWLLELLHVTKSRRSAYDHLMLKLHDSMKADLDYQKNSPQQTMPFP

SGSVWVCFSDQASHAVMSGQFMLEQTFFLPVGAMARPQRAPLGILERLQGRALV*
>GCA_000452945.1_BPMSHR5858_genomic_03667_1 rfaC_1
VQKILIVRVSSLGDVVHNMPVIADIRRRHPDAQIDWLVEEGFADLVRLVDGVRDVLPFSL
RRWRKRLSASQTWREIRAFRRRLAEERYDLVIDCQGLIKTAWVASWARGPLVGLGNRTDG
AGYEWPVRFFYDRRVPIAPRTHVVERSRQLVAAALGDPAPAPGEPIDFGLDTHGAARALA
ALDLNLPVPYVVFVHATSRADKQWPDEAWTGLGEALVRRGASLVLPWGSDAERATSERLA
KAFGAAAIVPPKLSLPAVVGLVDGAAATVGVDTGLVHIAAALKRPTVELYNFATAWRTGG
YWSPNVVNLGTAGAPPSLSQAKDALASFGLL*
>GCA_000452945.1_BPMSHR5858_genomic_03668_1 algC
MISQSIFKAYDIRGVIGKTLDADVARSIGRAFGSEVRAQGGDAVVVARDGRLSGPELVGA
LADGLRAAGVDVVDVGMVPTPVGYFAASVPLALSGGERRVDSCIVVTGSHNPPDYNGFKM
VLRGAAIYGDQIQGLYKRIVDARFETGNGSYERYDVADQYVERIVGDIRLARPLKLVVDA
GNGVAGPLATRLFKALGCELVELFTDIDGNFPNHHPDPAHPENLQDVIAKLKTTDAEIGF
AFDGDGDRLGVVTKDGQIIYPDRQLMLFAEEVLSRNPGAQIIYDVKCTRNLAQWVREKGG
EPLMWKTGHSLVKAKLRETGAPLAGEMSGHVFFKDRWYGFDDGLYTGARLLEILARAADP
SALLNGLPNAVSTPELQLKLDEGENVKLIDKLRADAKFDGADEVVTIDGLRVEYPDGFGL
ARSSNTTPVVVLRFEASSDAALARIQDDFRRALKAAKPDANLPF*
>GCA_000452945.1_BPMSHR5858_genomic_03671_1 pglA
MISFQSDMSSASAPRIVLVCNTAWAIYTYRQGLLRMLIARGAQVTVLAPRDRTVEPLVRM
GCRYAELPVASKGTSPREDLRTLIALYRHYRAIRPDLVFHYTIKPNIYGSIAAWLARVPS
IAVTTGLGYVFIQQSHAARVAKQLYRFALRFPREVWFLNRDDLHTFTHEQLLAHPARARL
LHGEGVDLEQFALAPLPARDTFTFVLIGRLLWDKGVREYVDAARMLRARYPHARFALLGP
VGVDNPSAISQADVDAWVREGVIDYLGEAHDVRPHIARADCVVLPSYREGVPRTLMEASA
MGRPIVATDVPGCRDVVADGSTGLLCAARDSASLAAQLARMLDMSAAERRAMGERGRRKI
VAEFDEAKVVERYHQTISALTGITL*
>GCA_000452945.1_BPMSHR5858_genomic_03694_1 yqgF
MSAALSRDATLLAFDYGEKRIGVAVGNLLTRTARALVIVRNLNREHRFKAVGELIAEWKP
DALVVGLPLHPDGAPHEMTQRAMRFGNQLNGRFNLPVSWVDERYSSVEARAGLRARGDAA
DRVDAEAARVILQQYLDGLPDHHEFN*
>GCA_000452945.1_BPMSHR5858_genomic_03703_1 adhT
MKAIMLRQPAGLEHLERVTLADPGAPAAGEIRVRIHASSLNYHDLGVVTGRLNPADKRIP
MSDGAGVVEAIGEGVTEFAPGDAVVSTFFPYWLDGGPTVADFATVPGDGVDGYARESVVR
PAHWFTHAPQGYTHEQAATLTTAGLTAWRALVVDGGLKAGDSVLVLGTGGVSIFALQFAK
RMGATVIATSSSDEKLERARALGADHVVNYRRDANWAAKVLALTNGRGVDHVIEVGGPDT
LPQSIQACRIGGHIALIGVLTGIAGPVPTVALMARQQNLQGLIVGSRRHQIDMVRAIDAT
GIKPVVDRTFALDDIADAFAHQAAGKHFGKICLSI*
>GCA_000452945.1_BPMSHR5858_genomic_03704_1 group_3464
MPDKHNHRADAEGLSLGSLRTAAWLRYLLDHAERFAAAAASSREARVAAAARDTLAKAGA
HARREPAADAGLLLNELLAALEGDGASVDEARGNAYGELARTPPKRLLAAVDRYLPGSKA
VFDGAHGVADVLGYDDAELGWAALDSIELGIGRALASDGKAADDAMRGIAAFEGPAAGRV
IDHAIGRMWGADEKRADALSALAAFSATLAMNAPITLDAYGGEHNGVAAAILRERGDLAS
APGLLAALAVCRLDRLTGAGSFWAYYLLAGVMIAAPQAVASIAARFHRDALSAWLDMLLA
VPARSPIGDAGEAALARLAASMSFTSRHNPIVHTRHRSMG*
>GCA_000452945.1_BPMSHR5858_genomic_03706_1 ompW_3
MKKLIAACVVAGASTGAFAQQAGDLVATLGWLHVMPQDSTNGLTTNVVNSPINGPLRLPG
SFTSPGTSLTVNNADTVGLTFTYFFTDHIAVTSVLGIPPEFALTGHGVIRPPGPSGSLGN
VDLDKPANQPAVKNARQWSPTIILNYYFNAPTAKFRPFVGIGVAYSWFSNIELNGNFAKD
INDNLGSVLAAGAGKPEPTSVEGKASSSFTPVYNIGASYAIAKHWALTATLSYMPLKTYS
SLVIKAADGTTLATTRTRLKADPLITFVGISYKF*
>GCA_000452945.1_BPMSHR5858_genomic_03708_1 group_2484
MLCIPNLRATVLSCMVAVMPFVAGCGGGDDPGPIQVGQCTGGSCPPSGPSTTPPTVTKLC
PDALDYSTTYTGGSGSGEYVKVKFDTAKKTYQMQFIASSVPTSAGQINNTRAGLTISGTF
HNATGLPTAEQNRCAFVLDNGATSDGAYSVTINRADPPMLFVGMGVVGGGIPGATIAFPG
LELFPGFTIGTVPSRSFDFYPFIGFTDTETDFTKVAGVYNEVGVHLQPTGTSFQTAAPQG
WQPDVVNWTQTLNADGSCTITPGSDYSCRTTGTPWTLRKNTDGTSDNVFVSRPPPSQPYP
SAGQGQPLVIVTPSQAQGVMIVGKLNGQLVPVVIRVGYANVDAGNPLASVADAEIGISVL
APAAAVAANSLQGGYIGATSAVACGVVSFGGSSNAPASSGSAFDNTAPHPELPGTYRGGF
FYPSAGNCTDGSAVSTLAANYTSTLFQGATAAFLDPMTSAVSAQFSLDYTQTAPGKIKVT
ATRDFNAQGSAGTVAVFKKGDYGWLIKVGNVYGMVVNNSQYNPFFTVGAFVQ*
>GCA_000452945.1_BPMSHR5858_genomic_03714_1 glpQ
MFLKRSFAFSPVALACAATLALAACGGDDPDYTQPISAKVQVVGHRGASALRPEHTLASY
RKAIEDGADVIEPDLVATRDGVLVARHENEISGTTNVSTLPQFASRKTTKTIDGAQLTGW
FTEDFTLAELKTLRARERIPQIRPANTAYDDQFEIPTFDEIVALAKQMSAQVGRTIHLYP
ETKHPTYFQSIGLPLEDRLVDALRKDPYTARNATVYIQSFEVANLKAIRARIGASQPNWK
LVQLMDEPKQRPYDFVKAGDARTYGDLSTQGGMREIATYANGVGPYKTSIIPVGPDGSLQ
RPTGYVRFAHEAGLTVHPYTFRPENNFLPASLKDGGAPSARHTAGSVREIQAYLRAGIDG
FFTDDPAVGRTAVDTFRR*
>GCA_000452945.1_BPMSHR5858_genomic_03717_1 group_2085
MKIQKSLKDGFTLGWCRAARPVAAAALAALLVAACGGDDGGGGSPSLAAANVANTSTSTN
ATTAADATTNAALPPDQPYIDNDVYGTGPNDSVSDATEGTAVVHRQVKIGDQILTYTATV
GHLVTIDPITSKPNAKMFYVAYTLDNPNPGKPRPVTFFYNGGPGSSSVYLLLGSFGPKRL
QSSFPNFTPPAPYRLRDNPESLLDRSDLVFINPVGTGYSAAIAPAKNKDFWGVDQDAHSI
DRFIQRYLTKYARWNSPKFLFGESYGTARSAVTAWVLHEDGIELNGITLQSSILDYANAV
SAIGIFPTLAADAFYWNKTTISPKPADLDAYMAQARSYADNVLAPLAQAPNPQDGGFVNV
RLNLNVATAQQMGAYIGTDPISLVQTFGNPAALGNVPSSNDNPPYTFFLTLVPGIQIGQY
DGRANYTGKGIAPYILPNSGSNDPSISNVGGAYTVLWNDYINNDLKYVSTSSFVDLNDQV
FNNWDFSHTDPTGANRGGGNTLYTAGDLAATMSLNPDLKVLSANGYFDAVTPFHQTELTL
AQMPLDPSLKSANLTMKYYPSGHMIYLNDHSRIAMKADLATFYDGILADRTAMRRVLLRQ
QKALQLKQQKQQQGQ*
>GCA_000452945.1_BPMSHR5858_genomic_03723_1 btuC
MSGAAPADAPPASRGRRWRVRLVLPALAALVAISIVASTALGATTIAPARVVAIVLSHLA
DLRVTNAHVAHAHVAHAHVADTLARSGASFAEDSIVWQIRLPRALLAALVGATLAMVGVA
LQAATANRLADPHLLGVSSGAMLGAVAATFGLGAAFGPFTLSFAAFAGALAATALVIALA
YRRGRLEPERLLLAGVAVSFMMTALANLLLYLGDPRAASSVLFWMLGGVGLARWDLLAAP
ACCAAFAALLLFARRRELNALMSGDVAAVSLGVPVARMRREVFVIASFATGAMVAVSGAI
GFVGLVTPHLCRRLVGAEHARLLPVAGLSGAALLVWADLAARTLAAPEDLPIGIVTAGLG
GAFFVALMRRR*
>GCA_000452945.1_BPMSHR5858_genomic_03724_1 btuF_1
MLKRTAAERGAPALRAAFTLPFAAAAMAAAAIVATVATAAEANAYPVTVRSCGRDVTFER
APARAVSNDVNLTEMMIALGLQTRMAGYTGIAGWKTGNARLRAALAGVPELATHYPSLEA
LVDARADFYFAGWNYGMHPGGPVTPASLERFGIRTYELTESCSLVMRRPPASLDDVYRDL
ANLGRIFGVDARAAQVAGAMRARIDAVRRTLASAAGGGTPPRVFVYDSGTDKPMTAGALA
MPTALIAAAGGRNVMDDVPRGWTQVGWESVVARDPQAIVIVDYSAVTAEQKKQFLLSQPA
LAQVEAIRARRFIVIPYDAATPGVENAAAVETIARGLHPRAFAKAAR*
>GCA_000452945.1_BPMSHR5858_genomic_03727_1 ais
MQAHQPYFDEIAAGGDAIGNWLSGDVSGPAALDALMARFAPHFTMIGTDGAAYDRAATRA
LFARLAGCKPGLKITFSQMHALVADAAHGVISYCECQTDATGALPTRRSTAVFERDPQTG
AVRWTHLQETFCAA*
>GCA_000452945.1_BPMSHR5858_genomic_03730_1 group_19902
MSEPIRFYHRRAIREVSGANVTRTVLQYLREDVHCTGTKEGCAEGDCGACTVVVGELNDA
GGVEFKAVNACIQFLPTLDGRALFTVEDLRQPDGALHPAQEALVECHGSQCGFCTPGFAM
SMWALYERHGHEAQRADRTTPSRAQIADALTGNLCRCTGYRPIVDAAVRMFDARPPKAPV
DVAALARALQALRRDDTFHYEHAGQAFDAPRTLDALARLKAEKPAARVLAGSTDVGLWVT
KQLRDLGDVLYVGQVPELRRIVEHDDWIEIGAGASVEAAYAALAAHYPELVEMWKRFASL
PIRNAGTIGGNVANGSPIGDSMPGLIALGARVVLRGGRAARELPLEDLYVGYQKKDMAEH
ELVVALKVPKRTGARANLRFRTYKLSKRFDSDISAVCAAFAFEADGEVLRAPRIAFGGMA
ATPKRAAHAEAALAGAVWDEAAALAAMRALDHDYAPLTDMRATSAYRLETAKNLLYRFWL
ETRPHDPLPASAVNVREVAAEPAAS*
>GCA_000452945.1_BPMSHR5858_genomic_03733_1 eamA_1
MNLSLYAVTVLIWGTTWIAIKWQLGSVPPPVSIAWRFWLAAAAMFVLLRALRRPVRPPRD
AWRLLVAQGFALFCVNFLCFYYAEQVVPSGLVAVVFSTAPLLNSINGRLFMGRPLRQSAI
AGALLGLIGIGCLFWQQMAGHVDDRATWIGLAIALAGTMSFSAGNLLSSRMQTMGLHPLA
TNGWAMLIGAAILTAGSIAAGLPLAPDTSPRYLAALVYLAVPGSVIGFTAYLTLIGRIGP
ERAAYCTVLFPIVALAVSTVFEGYRWSPVAVVGLMLVVAGNLVAFDLTRRRFVRTA*
>GCA_000452945.1_BPMSHR5858_genomic_03741_1 group_19806
MSAIPDTLRASLDPSRKSWLDTANAAACDFPIQNLPFGIFSDARDASRRAGVALGDQIID
LAALARAGLLTVDGGAAVFARPALNDFISLGRDAWRSVRAQLSALFERGEARLRDDAALR
AKVLVAQRDAALHLPVDIPGYTDFYSSKEHATNVGSMFRDPKNALLPNWSEMPIGYNGRA
SSVVVSGTPVRRPNGQLKLPDSERPVFGACRKLDIELETGFIVGRGNALGEPIACEDAES
HIFGMVLLNDWSARDIQQWEYVPLGPFNAKTFATSISPWIVTLDALEPFRTAQPRQEPEP
LAYLRHGGAHAFDIELEVRLRPEGAADATTIARTNFRHMYWTMAQQLAHHTVSGCNTRVG
DLMGSGTISGPAKQAFGSLLELTWNGKEPVSLAGGGTRAFIEDGDELTLAGWCQGDGYRV
GFGTCVGEILPARGR*
>GCA_000452945.1_BPMSHR5858_genomic_03744_1 group_2488
MHLDTRLRLPDPDAFYEALIDMHRDLPDSESQLVNAKLVLLLANQIGDLDVLREAMALAR
SGAAVLEGAPGAPALQ*
>GCA_000452945.1_BPMSHR5858_genomic_03748_1 group_2489
MSSSESLEGGCACGAIRYRIVGTPNDAGFCHCRLCQRTTGAALIAWATVPIGAFEYVKGE
PRVFVSSAWGERRFCPQCGAQLEYRRSDAPSTVEVNYATLDDPSAIAPKAHTWYASRIPG
LEVAGGLPVRDDGDH*
>GCA_000452945.1_BPMSHR5858_genomic_03750_1 mmgC_2
MAAATFHWDDPLLLDQQLTDDERMVRDAAHAYAQGKLAPRVTEAFRHETTDAAIFREMGE
IGLLGPTIPEQYGGPGLDYVSYGLIAREVERVDSGYRSMMSVQSSLVMVPIFEFGSDAQK
EKYLPKLATGEWIGCFGLTEPNHGSDPGSMVTRARKVLGGYSLSGSKMWITNAPIADVFV
VWAKLDEDGRDEIRGFILEKGCKGLSAPAIHGKVGLRASITGEIVLDEAFVPEENILPHV
KGLRGPFTCLNSARYGIAWGALGAAESCWHIARQYVLDRKQFGRPLAANQLIQKKLADMQ
TEITLGLQGVLRLGRMKDEGTAAVEITSIMKRNSCGKALDIARLARDMLGGNGISDEFGV
ARHLVNLEVVNTYEGTHDIHALILGRAQTGIQAFF*
>GCA_000452945.1_BPMSHR5858_genomic_03757_1 group_19972
MMRTTRWRAHTLVEVMIAMALGLLILLAAMSLYRVQRAAYSAAVDAARLRDAAQAALALI
SQQIQMAGFVPLDAYDARAVPGLFGCAAGRPVGADGQTACDPLASRSDGLVVRYVGDGVS
TWPTASGQPTDCLGQGVGAADTQPLIVNRFYARVSASTGEPELYCEGSGRPGIAQPLVEG
VERLSLRYRLYGAARWADASALSVDDWANVAVVSVCVQVRGRRTGRPARYVDCEGRVASA
PDTRARLAWRRYVAVRNRAGT*
>GCA_000452945.1_BPMSHR5858_genomic_03758_1 group_2490
MTCCRSMRGSTLLEAAIACALLAVASLGVVAALLAAIRSERDTAACEPAVLVADSWAEMA
RAGGPSGVDWDASMSGLRHGKLVATNAGADISNVRVEWERSAGLAASPLRCDGDAAAARD
AAKSCVTLAYASTQRR*
>GCA_000452945.1_BPMSHR5858_genomic_03759_1 group_19975
MRAERDRWATAGFVLVEVMAALMLVALAAMLTAPTLAGARMRDRVDARARVFGALLAYAR
GEAVRLGARVVLCRSDATAKCIAAGRPCGGGAADWSCGWAVAAADGERGTRLLRRIAPDA
RVAVTGTTVDVVFTPPAGQVIGGFRSFEFAPADASGAWRGERWRRCLRIAAGGRVRFAEG
GCGAST*
>GCA_000452945.1_BPMSHR5858_genomic_03762_1 ydfG
MIVFVTGASAGFGAAIARAFVQGGHRVVAAARRKERLDALAAELGSALLPIELDVRDRAA
VEAVAAALPTEFAAIDVLVNNAGLALGVEPAHRASLDEWQTMIDTNCTGLVTVTRALLPG
MVERGRGHIFNLGSVAGSYPYPGGNVYGATKAFVRQFSLNLRADLIGTPLRVTDIEPGLC
GGTEFSNVRYRGDDEKAANVYANVQPLTAEDIADTIYWIATRPAHVNINTIEMMPVAQAP
AGLAVHRG*
>GCA_000452945.1_BPMSHR5858_genomic_03766_1 group_2491
MKPRQSRTAAYPPRPPRERGTGRAFLLAALMHVLLALFLYHGVHWQNSTPAGAEAELWTS
VPDTSTPQPAPTPPVKVAPPPPPVKNEEADIALQQKRREQQAAAAREAQLEEQRRQQQLK
AQQLAAQQAAQLAAQKAAEREKQKQAEKLKQQQLAEQQQRKLEQQKLEQQKLEQQKKQEQ
LAAQKKADAEKAEKAEKAAKAAAAAKANAAAKAKLDKERQARLAQLQGIAGGGSGGGEGL
AKSGTGTGSGGNAASPGYADKVRRRVKPNIVWAGERDSLVTVVAIRCTPSGDVLSTSIRR
SSGNSGWDQAVISAIQASVPLPPDTNGRTPSEITITFKAAE*
>GCA_000452945.1_BPMSHR5858_genomic_03774_1 group_3468
MSAPAYRGAIRLVLFDVDGVLTDGRLHVTGDGECMKSFHAKDGIAVALLHAHGIRSGILS
GRHSEALAWRARQLGFDVCVSGCGDKRAGYARIKSEQALADHAIAYVGDDVNDLPVIDAV
GVSYAPADAHVLVRRRVDYVVSRAGGEGAAREVAEHVLLRSGLSLDDAYRPLLDEWARHA
VVQ*
>GCA_000452945.1_BPMSHR5858_genomic_03776_1 group_2675
MTIDIAGGRHRDALPKRDAGAGGGARMKIVQVLESSATGTLSMVCLIANRLAREGHDVHV
VYSLRPDTPANFHALFDATVKLRHIQMKGVPLLPMLLELRRTLVGIGPDVVHLHSSFAGF
LGRIGTLFALREAAFFYSPHCISFMRRDISAFKRAAFVALERIACAKRCLYVACSDSERD
AVRSHLRQPVVVVENAVGAMPAPAAAESDSAHETVLRIVTVGGIRVQKNPRLFAEIARRL
CGSRVRFAWIGDGDAALKAELRDAGVEVAGWLGRGEVLARLRRTDVYLSTSSWEGMPVSV
IEAMLLGLPVVASACAGNVDVVRHLQTGAIFRGADDAVELLASLERDAALRARLAHAARQ
EARERFGEARFFRQLARVYASRPARVS*
>GCA_000452945.1_BPMSHR5858_genomic_03777_1 group_3469
MKVFILHPGKANYPEIAAYRDFLGPRGFDVHDGDASAYARFDGRHDCILWCIMGFYRALP
PARFVIHDYRSLSVGRLAALKDTVKRAWNAKPALRIFQNERMRDAMAFRDGIPSLLLPMG
VPDWIFGIANEPAGAGPAGRFCYIGEMSRERGFHKVLAAYRAAPRTARDSLVLVGQPEPE
IYEAFRHVEGLVFVGRVPQRDALKIVQRSDYAVCYFPYHRPHRFQTPTKLLEYAALGKKI
ICNDAPSNVDAINALGIRSHVTGAAIFDELLPLAQVRAQRNDPARLKRLEWTAVIARSGV
LAHIEAAASMRGVR*
>GCA_000452945.1_BPMSHR5858_genomic_03779_1 group_3470
MIVVVVVDSMERYYFAVRLVKAVRKEFDFLFAASEPLAHLMALAAGFRSVYLRRGAHAPA
MLDAAAGIRSDASIEVLNGQMTPERARADAQAVFAAMSGVFRRHLVSQCLMWNGQQLVCR
AVAHACATHGVPTKFVEISNLPDKLFVDRLGVNALSSISRNPAVIDGLPMPTEGEHRRWF
ARYEAYKARPLPQSRTSWARKAMSAANYALKLATQGVARKRLDAARATNGARAPAQAKVL
STKELSALRYVFLPLQVSGDTQIKLHSDVDNLKAIRLAFEHAANENADLIVKLHPAERDV
AVIDEVVRMQRVYHFDLVTSPTTDLIKHAHSVVTINSTVGLEALLYGKPVVSLGRCFYKE
FDRARLLKYIHAFLIDGIDYFGRADIAPRAARNVFSMKH*
>GCA_000452945.1_BPMSHR5858_genomic_03781_1 wzc_1
MNPNPTGTSTPADSEGDTDFIAVLDILIEGRWLIAAIALGVFIVGVAYAVFSKPVYQADI
LIQVEDSPDTSAAKSLLGDVSSLFDVKSSAAAETQILASRLVVSRAVDNLKLFIDAKPKR
FPVIGHWLARRSEGLSDPGLAGFGGYAWGQERIDVATFDVPRAMEGDTFELTMLDARRYR
LDGGDLERGAVGVVGRLERIAAKGGPIALRVDAFAAKPGATFVLVRHSRLRTIEALQDNL
DVQERVKQSDVVVASLRDTDPDLVSRALNEIGRQYIAQNIQRKSAEAAQSLEFLNGQLPT
LKQQLTDSEARLTKLRDEHGSVDLTEEAKLVLAQSADAKTRLLELQQKRQELLSRFMPKH
PSVVAIDQQIAALDGYRGTAEQQIKRLPALQQQFVRLMLDVKVNTDLYTALLNNMQQLQL
VRAGKVGNVRLVDTAAVPEVPVKPKKALVALASLLLGVLAGCGTAVGRSMLFHGISDPNE
IERRLGLSVYATVPRSDRQRALTERAKHKARALSLLCVAHPDEPAVESLRSLRTALQFAM
LDAKNNVVVIAGPAPGVGKSFVSANLAAVLTMAGKRVLLIDGDIRKGHLNDYLGLARGKG
FSELIAGSAQPDDVLHRDVIAGLDFISTGAMPKNPAELLLNARVSTLIDTFSQRYDAVVI
DSPPVLAVADTGILAATAGTAFLVTLAGSTKLGEIAESAKRLAQNGVRLSGVVFNGINPR
LGQYGYGSKYGGYRYVAYEYGAKRDA*
>GCA_000452945.1_BPMSHR5858_genomic_03782_1 ptp_1
MIESVLIVCEGNLCRSPMAAALFAAALPGRAVTSAGLNALVGLPAAPLAQDVMRERGVDL
SAHRAQQLGPAHCASADLILVMDRGQRRLVEERYPLARGKVFRIGEHENFDVHDPYRQPR
FVFERSARLLELGVNGWLSRLRLV*
>GCA_000452945.1_BPMSHR5858_genomic_03783_1 group_19898
MGSLGEAGGIAPPKRTCGARVRRLMKRVTLCAAFAALSACGVAPGMRMKQPANVPVSSAA
ADAPAEAGRKPRGEQLRVPITDIDLSLIRTLRDAQQAPRRAADLVSPASGYTIGRGDVLQ
ITVWDHPELAAALGTQQQTAARAADAPAGFVVDQDGTLQYPYVGRIAVVGLKPEQVQARL
ARQLAQTFRDPQVTVRIASFRAKQVYIEGEVHTPGSQALNDIPMTLYDAVSRAGGFSASA
DQRRVTLVRDGVERRIDLSGAAQGVNPSRIVLRDGDLLRIPPRDESGVFVMGEVNRPVTA
LPMRNGRLTLSEALSQAGSLNATTADAAQLYVIRGSLDAKPHVYRLDASSPVAMVLANQF
ELEPKDIVYVDGNGLVRFSRVLSLLLPAVNAGLTAAVVTK*
>GCA_000452945.1_BPMSHR5858_genomic_03788_1 group_2832
MRRVSVPTLLFAASACFFSLNAFSVRATFAAYALLALLALLRAPTLFARQQRLPAIVAYW
LLSVSSAFLVSAFGDFYANFFVKFVLIQTYVALAFWLFASGVLAMRSLERTCETLIYVHA
TFFIVQLGCYLAFGHFIDFDSYIRESDSEALYATKALSDSLISIRALGLYSEPSFYAMTV
VPAGAILLLAKRRMTAATIVAFATALLSFSIAAILICALLGGVHFFAGRTSIRIKLVIAA
VALAIAPAMYGVYDKRVNQSADYDALGSRTLVLRELRERDALADVFGSGLFWDERNNVGK
THLRGYQVRDSSFYVYLLFATGVAGATAFVGALFMLFRRRGRRSLLPYLLPLLLFKFHAL
YGMLWLTLLMFVVVAGHAERLPERREPPGTGTGRLSATGASGP*

>GCA_000452945.1_BPMSHR5858_genomic_03789_1 group_3473
MTTRWMPACLFCACVVASSAAQVPPATPPGSIIAHSAAATRVFLGSPSLAILPDGTYVAS
HDMFGPAALPDEVRVFASRDKGQTWEKLSVVTGQYWSSLFVVDGALYLMGTDRSMGTPVI
RRSTDGGATWTTPSDPLTGRFDVSGRYITAPVPVLIERDRVWRAFEVIDGGDLRSLVLSA
PVDGDLLDTRNWRATKPLPSDHAWLDGNFLTWEEGSVVATAEREPAVVLRVNTKHGPEKV
ALVHMLADGRSLSFDPAHDFIDFPGGGKKFTVRFDARSGRYWSLANAVPNADGGANLERV
RNTLALTSSPDLRHWSVERVLLQSADAQRHGFQYADWQFDGDDIVAVVRMAFDDGEGGAH
SQHDSNYVTFVRVRDFRHAAADKRDGAKPD*
>GCA_000452945.1_BPMSHR5858_genomic_03790_1 group_3474
MATAPADGERGATNDAAPRSSPWRFRAAAASDAHACAPLVFESGVREFGYFLGEPPARCI
AFLAFAFASKHGRFSWRRHRVALGGDGTVSAVLAAHDGRATTFDDLHVAWMLLRFFGVAR
TIRMLLRGLVLESELPAPKRSQTLIAHCATEARVRGCGAFSALFDDAMRAGLLGVEPGRE
VVLDVLASNERARALYERLGFAATPRARPRSARLPGELASVRMRFGGASR*
>GCA_000452945.1_BPMSHR5858_genomic_03801_1 rmd
MVGSHLADFLLENTDWEIYGLCRWRSPLDNVSHLLPRINEKNRIRLVYGDLRDYLSIHEA
VKQSTPDFVFHLAAQSYPKTSFDSPLDTLETNVQGTANVLEALRKNNIDAVTHVCASSEV
FGRVPREKLPIDEECTFHPASPYAISKVGTDLIGRYYAEAYNMTVMTTRMFTHTGPRRGD
VFAESTFAKQIAMIERGLIPPVVKTGNLDSLRTFADVRDAVRAYYMLVTINPIPGAYYNI
GGTYSCTVGQMLDTLISMSTSKDVIRVETDPERLRPIDADLQVPNTRKFEAVTGWKPEIS
FEKTMEDLLNYWRARISAGEKFLTR*
>GCA_000452945.1_BPMSHR5858_genomic_03802_1 group_3476
MKVFLVGSTGYIGKTLFDACSRRWRTLGTSTRDGADIVFSLARAEAFPYEQVSAGDVVAV
AAAISSPDACAKDYETAFQVNVTGTLTLIRGVVARGARVIFFSSDTVYGASEQLLSEEAE
LTPAGAYGAMKRRVEAELGENAAVKVIRLSYVFSLRDRFTQYLLGCAKEGKRADIFKPFS
RCVVYLSDVVEGVVSLIERWDAIDERVINFVGPELVAREDFVEKIRNLAAPELDYGFSEP
EGDFFVNRPRIINVSSARFEKLLGRRPRSLGEAINLELSN*
>GCA_000452945.1_BPMSHR5858_genomic_03808_1 kpsT
MIRLTRVKKTYQTKSGERTVLDGVDVDIASGEKIGILGGNGAGKSTLIRIISGAERPTEG
RVYRGMSVSWPLAFGGAFQTSLTGIDNLRFICRIYDVSVEDKLPFVREFTELGRYLNEPV
KTYSAGMRAKLAFAISMAVDFDCFLIDEVTAVGDERFRQKCHVELFEKRNSRAMIFVSHD
PGFIRANCQRATVLHGGKLYEFPEMDEAYHFYREMQRV*
>GCA_000452945.1_BPMSHR5858_genomic_03814_1 manC1
MNIHPVILCGGSGSRLWPMSRGGYPKQYLKLAGDRSLVQQTALRLRNIPDTTAPIVVTNN
EQRFLVAEQLRQVDITPSSIVLEPVGRNTAPAIAIAALLAMHDSPEALLLVLPSDHVILN
EPAFITAAREAAAIAADKHLVTFGITPTQAHTGFGYIRRGPLLADGGQAFKVDAFVEKPD
ASTAERFVRDSGYYWNSGMFMLKAATYMEELHRHAPEIARQAEFALASAKRDHDFLRLDA
DAFFACPNVSIDYAVMERTERAAVIATADLGWSDIGSWSALADIADTDAQGNALVGDVFT
DSVKNSYIRAEHRMIAALGLDDIVIVETADAVLVAHRDKAQDVRKIVEWLNASGRHESVT
HRRVVRPWGSYEGIDQGERFQVKRIVVNPGAQLSLQMHHHRAEHWIVVKGTALVTNDGKE
IILTENQSTYIPLGATHRLTNPGKIPLELIEVQSGSYLGEDDIVRFEDTYGRTST*
>GCA_000452945.1_BPMSHR5858_genomic_03815_1 egtC
MCRWLAYTGNPIPLETVLFRAKHSLIDQSLHSRMGATTTNGDGFGIGWYGQPDEVPFIYR
CVSPAWNDRNLREAARAIRSPLFVAHVRSATDTPVQETNCHPFRRGRWLFAHNGLIRRYH
TLRRDLMMRIDPGLFASIEGSTDSEVMFHLALTFGLEHDPLAALERMAGAIEDAGAHHRV
DAPLNMTVCATDGEQVVAVRYSSEHDSRSLFHSTSFRHLHELYPHDPRIRAIGDDAFLVL
SEPLVDLHDAWEKIPESTAIIARRGDIRQVPFQPRRST*
>GCA_000452945.1_BPMSHR5858_genomic_03817_1 dedA
LETLLHFVSLVVHIDAFLGDFIRQYGAWVYLVLFLIVFCETGLVIFPFLPGDSLLFIAGA
FAATGEMTLAGLLVLLLVAAVGGNTVNYLIGRAIGPKVFNTHIPGLERFLDRAALQKTHD
FYERHGGKTLVLARFIPVVRTFAPFVAGASAMSVARFQLFNVIGALIWVLLLVFLGYFFG
NIPFIRHYLNVIVLVGIGAAVIPVALGALWKLLRRKSDARKTQANR*
>GCA_000452945.1_BPMSHR5858_genomic_03819_1 miaA
MSERNAASARTVACLLGPTASGKTAAALALAARRPIEIVSVDSALVYRGMDIGTAKPTRD
ERAAVPHHLIDIVDPADAYSAAEFRADALRLVAQIAARGRTPLLAGGTMLYYRALTQGLN
DLPAADPDVRATLDADAARDGWPALHARLAGIDPATAARLAPNDSQRIQRALEVYLLTGQ
PMSALLAAPPRDDDAAAGLRFVPVALEPSERAVLHVRIAARFDAMLEAGFIDEVERLRRR
DDLHLGLPSMRCVGYRQAWEYLDGCTDYRTMRDKGIFATRQLCKRQLTWLRAMPERIVVD
CCAPDATMRAVDALERVLDGRAPA*
>GCA_000452945.1_BPMSHR5858_genomic_03825_1 folK
MTVAYLGLGANLGDARQALKDAVVCLAQQHAITVLGKSSLYRTAPVDASGGDYYNCVVKI
DTVLAARALLTLCQKIEHHFGRERPYRNAPRTLDIDILLYGELTIDEPDLVVPHPRLTER
AFALVPLVEIEPALAIPARGRADGFLADVAHQRIEKVQTCQCLAMRAAAEKNRCR*
>GCA_000452945.1_BPMSHR5858_genomic_03868_1 group_2499
MLTAPLAFAVLSWLAASRRRRRRAALVWALGFACAVGVVALTKFVYAGWGIGIAAWRFTG
VSGHTMLGAAVYPLVATICVRDAYPRRALAAGLAFALAIGMSRVLLGFHSWSEIVSGWLL
GAAVAASTARCLHGIGGARASQAQARGMHAARNVPMARRAHGGGAVGAHPHAAAGAAAVR
RAARAPRSTLMFVAAACAIAVSCYGRSAPVSAWISDAAPKVAEWGRVWFDDAR*
>GCA_000452945.1_BPMSHR5858_genomic_03869_1 group_2500
LKVAIVHDWLVVYGGAERVLAQMIDCFAQADIYSLVDFLDDRSCLRGRPVHTSFIQKLPF
ARSKYRSYLPLFPLAIEQFDLSGYDLILSSSYAVAKGVLNGPDQLHASYVHSPVRYAWDL
QHQYLNEAGLARGVKSALARTLLHYIRNWDARSANGVDLLAANSRFVARRIRKTYRRDAT
VIYPPVDVDHLALRDTKDDFYLTASRLVPYKRIDLIVEAFSHMPSRRLVVIGDGPEAAKI
RALAGPNVTLLGYQPFDVLHDHLQRAKAFVFAAEEDFGISPVEAQACGTPVIAYGKGGVC
ESVRAAGAAPTGLFYAKQTCDALIDAIDRFEAMPAGTFDPHACRANAERFSAARFRSTFS
RFVLEGYAALQAEMGETMQDAPLEPGGAPDGAPVERDAAAPHGACRNETLARI*
>GCA_000452945.1_BPMSHR5858_genomic_03870_1 group_19907
MDWTCCEFRHLSSNELYMILRTRNAVLVVEDAHTHLDIDGKDEFAIHVFATDKRGEQPAI
AAYARLLPGDDIDPETTIDKILTSAAHRDDRTIDALIEHVLAAAHARWPDAPVRVQAPAP
REGFYNRFGFRKVDGPYLEHGAPYVGMLRAASAPSKAVRNLLDLVGNATGANPGTSAGAV
PASAARKPRKANAESTTEGERYAFAGRLPADSGMNR*
>GCA_000452945.1_BPMSHR5858_genomic_03872_1 group_2502
MKVLVINDFARKGGAEEVYRTSVDVLRAQPGVEVATFDERAFALTQGGAARAWNAPAARA
LAAVLEREAPQRVLVHNYHNLLSPSVVPVIARYKRRTGCRAYLTCHDYHLVFYNPNLLTY
PRGRATPLPLDALARGARLFERSSPRGALHDAIKKLHWHAIDALVQPADAFDLLLCPSPY
MRDALARRGIARTALLHNPVSTALTPREPKCAERERLDLAFVGRIDPEKGLDEFLELARA
SRFERIASVTVYGDGGQRALLERKYAALIAAGQLRFAGRLDHARLFVALREHDALVLPSV
WAENAPLVIVEAAMIGLPVLVHDIGSLSSFGDEIGNKIRYAHNGASLAHALDALRAHLRE
PRRRYDWSLYTRRHYADRLAALLELSAREARELAPQCD*
>GCA_000452945.1_BPMSHR5858_genomic_03879_1 pknD_1
MSASVLAERARAGNRRAPRALAPPLPGDGDELQGEHRYRLGAVLGEGGAGRVHEAIRLDT
SARVAIKLLREDAPRGARERARLRARFRRETALCARLSHPHIVALLDSGETPDGLLFAVF
EHVAGRTLRALLAADGALAAETTGALMAQVLDGLAHAHANGVVHRDLKPQNVMVTTRDGE
PCAKILDFGIGALLPDAHAADELTLTATTEVLGSPQYCAPEQLRGEPPTAKSDFYAWGLI
VIECLTGHPVMQGASLADVLYQHLSPVDVALPPAIAAHPLGDVLRDALNKDPRQRAESAQ
ALANRFRAIHFPALVGGLRYGARHGRRALAEPGVAAREPGGTIALDAPVGRRQVTALCCR
VSVVSTLARPADAAADDEALDAFHAHWLTRCSDIAVRYGGHVAGALGDTLLFYFGYPEGI
DRPAQRACRAALEMTRHAGQPAGAPRSSDGDARACAPAWRIDIAGAIHVGTAFATARGAS
GGVLASAALGLQRIAQPGGILLSDEAARALERHVDAAPTALTFAAPGAAPQPVRELLGER
YERGPFESLELGDATPIVGRDREQAALARAWRDAVRAAAGGRRARRRATLVVGEPGIGKS
RLVHALRELVRAQRGACAACVCLPEQTNHALFPILRFVRAHWQLDVGDPHALDRMLEAFD
GDRASARATLAAWLGLPGGADTLRWSGARQQQALFDVLCQLLGSLGGGGPVLLIVDDVQW
CDSATDDFLDALAHHPACAAVCVVLTSRPERLERWRRGAERLMLRRLSAAATRNLIVSLM
PGAGTQRAALDFLVKRTGGVPLYVEETVRALAEGGVAPGAGARLPDLAQAGRCPLPGSLR
ETLELALERADDALDTVQLAATIGLEVDARLLAEASPHAGAELDERLRRLIEGRVLYAQH
RIGGATYVFRHALLREAAYESMPAATRREYHRRVAQALLACVAQGDPAARSASIADHFAR
AHAFAAAVPHGIEAARRALARALHDDAIRYAGAVRGWLLHCDYPGQHEDAVSVDLTLAHA
QMARDGWGDPRVREHTDRVLSRVGELSDAKAAAGALWTIAVYHHVAGDRDAVRRIGGQLS
ALARASARADLGVAADTLHGMALWIDGHYTLALDAFDAALAAYDPRRDGDHRHVFGLDTR
AWAQSARACVLWGIDDDVARTLALARDAVQLATCTDHLPTIGLTMMYLARMQQCVGDREG
ARATADAVLRLSRCHGLNAVERYAAVVRAWCDGDRDAARANVDALRASGCMLGLTYYASV
AADIDAERGDARAALATLDECLALAESMDERHYVAELLLKKARCVRDAHGRRAADEAQAL
CARAAAVARRSGMARIERKAQAELRELQRELRSTTGEER*
>GCA_000452945.1_BPMSHR5858_genomic_03893_1 group_3485
METRSIAYDCGGARLTGYFADDAPNTKKPAVLIAHEAFGLNEHIRARARRLAELGYAAFA
LDMYGAEGFPMPEAIRRHIELMSTPGLMHARASAALSVLMEQPGVDRERVAAIGFCQGGI
SALELARGGAPIRCAVGFHPGLMRPAGSQDGPIRAKVLMMIGARDPDVPAADQAAFAAEM
QEKQVDWQLHLFGGVGHAYTNPDADGWNKPGYGYSRAADQRAWTMMLALFDEVLGGAAAV
GKA*
>GCA_000452945.1_BPMSHR5858_genomic_03899_1 group_19988
MTFACIAAARRFDAAAHVPFAIAGRQVGWLRRGDVARLARWPDVFEFPAGRVELAARYDS
VDARSMALASPIGALAAEGAIVGWRDEIYAIRNRFDDPPLAYIERAASRFFGTMTYAVHL
NGIVEYAPSAPLAMWLGRRSATKATDPGMLDNVVAGGIGWGLGIEDTIAKECWEEAGIAE
ELAARAIPGRVAHVLRSLPEGTQAEQIFVYDLPLPRDFVPHNQDGEVAEHLLAGVDEIVA
WLRAGEATVDASLAMLDSLLRHRAIAPEDARGIDVLFAPPPLGLG*
>GCA_000452945.1_BPMSHR5858_genomic_03906_1 lptA
MNESFPCLFPSGARRAARAMVAAALVAAPLAGLAPAAHAEKADRDKPINVEADNLTYDDL
KQVTVATGNVVITKGTILIKGDRVEVRQDPQGYQYATATASGKKHASFRQKREGLDEYID
GDAERIDYDGKQDLTTLTTNATVRRLQGVSTVADTVHGSVITYDGQRDFYTARGGKDVAG
PGNPGGRVRAMLTPKNSGPAPLSGGPAQLAPSNDMQGAPNP*
>GCA_000452945.1_BPMSHR5858_genomic_03908_1 rpoN_1
MKASLQLRLSQHLALTPQLQQSIRLLQLSTLELQQEVATAIAQNPLLENDDDWIASPLRV
AADGSVIAQTPASSAPEPMAGAPNGTAATNERTERDETAGVDEYDSYSADGGDASQWNLD
DFGRSPAASDDDDLPPMQIHETSTSLRDHLMAQLRVTQASPRDRALVTFLIESLDDDGYL
SASLEEVLADLPAELEVDVDELNAALALVHSFDPAGVGARSASECLKLQLLRLDPSPTRA
LALDIVSQHLELLAARDFTRLRKQLKATDDALRDAHALIRSLEPFPGAAYGKAEADYVVP
DIIVKKAGQNWQAELNPEVVPKLRINNLYANILRNSRGDPSAGSLKQQLQEARWLIKNIQ
QRFETILRVAQAIVERQKNFFAHGEIAMRPLVLREIADTLGLHESTVSRVTTGKYMLTPF
GTLEFKYFFGSHVSTDTGGAASSTAIRALIKQLIGAEDPKSPLSDSRIAELLAEQGFVVA
RRTVAKYREALKIPAVNLRKSL*
>GCA_000452945.1_BPMSHR5858_genomic_03922_1 rplY
MKVVAFERQQQGTGASRRLRNAGKTTGIVYGGEAAPQMIELDHNALWHALKKEAFHSSIL
DLEVAGKSQRVLLRDVQYHPFRQLVLHVDFQRIDPKKKLHTKAPLHFLNAETSPAVKLSS
AVVSHVVTEIEIECLPADLPEFLEVDLSKIEAGQSLHAKDIALPNGVALTAHVDAENPVI
ASATIPAGAVSDEAAAGEGETPAA*
>GCA_000452945.1_BPMSHR5858_genomic_03958_1 speB
MNETLYGDGAIRRPSIYGSSVENTYAGVLSFMRRNYTRDLGGVDVAVCGVPLDLATTFRS
GARLGPAAVRAASVQLAELRPYPWGFDPFDDLAVIDYGDCWFDAHNPLSIKPAIVEHART
ILQSGAAMLTLGGDHYITYPLLIAHAQRHGKPLSLIHFDAHCDTWADDAPDSLNHGTMFY
KAVNEGLIDPKTSVQVGIRTWNDDYLGIHVLDAAWVHEHGPRATAERIASIVGGRPAYLT
FDIDCLDPAFAPGTGTPVAGGLSSAQALAIVRGLGGVNLIGADVVEVAPAYDQSEITAIA
AAHVACDLLCLWRQRKTGAR*
>GCA_000452945.1_BPMSHR5858_genomic_03960_1 groS2
MSLRPLHDRVIVKRLDQETKTASGIVIPDSAAEKPDQGEIVAVGPGRRDADGKRVEPDVK
VGERVLFGKYAGQSVKVDGNELLVLREEDIVAVVHS*
>GCA_000452945.1_BPMSHR5858_genomic_03961_1 groL_2
MAAKEIIFHDGARAKLVEGVNLLANAVKVTLGPKGRNVVLERSFGSPVVTKDGVSVAKEI
ELADKVQNIGAQLVKEVASKTSDAAGDGTTTATVLAQAIVREGQKYVAAGLNPLDLKRGI
DKAVAAAVDELKKISKPTTTSKEIAQVATISANGEESIGQRIAEAIDRVGKEGVITVEDG
KSLADELDVVEGLQFDRGYLSPYFINHPERQLAVLDEPFILLHDKKISNIRDLLPVLEQV
AKAGRPLLIVAEDVEGEALATLVVNNIRGILKTVAVKAPGFGDRRKALLEDIAILTGGQV
ITEETGLTLEKATLQELGRAKRIEVGKENTTLIDGAGDKPNIDARVKQIRAQIAEATSDY
DREKLQERVAKLAGGVAVIKVGGATEVEVKEKKDRVDDALHATRAAVEEGIVPGGGVALI
RVKQAIAALAGANADQKAGISIVLRALEEPLRQIVANAGEEASVVVATVAAGQGNYGYNA
ATGEYGDLVESGVLDPTKVTRTALQNAASIAGLLLTTDATVHEAPKDAPPAAPAGVPGAG
GPGFDF*
>GCA_000452945.1_BPMSHR5858_genomic_03964_1 bclA
MDNFCRTNLPAPTDLPEFVFELPGLQYPARINCAAALLDDAVTRRGWGERVAIRTESGAA
WSYRALFELSNRIANLLVRDGGLVPGNRVLLHGTNHPFLAAAWFAIVKAGGVVVTTMPLL
RAGELSKVIAQAQVTHALCEAAVSAELRAAMAAAPGVAFARYYETDDAAAFEPLLHACPR
TFEPVDTRADEPCIVAFTSGTTGRPKATVHFHRDVMAICHCFPQHVLKPNADDVFCGSPP
LAFTFGLGALLLFPLSVGASVVLLQRAKPQRLLAAIGAHRVSILFTAPAAYRAMLDELGA
HDIASLRKCVCAGEALPVPTRNAWLARTGIRIIDGIGATEMLHIFASADETQAKEGAIGK
AVPGYRLAILDERGERLPPYHVGRLAVQGPTGCRYLNDARQRDYVRHGWNLTGDAAYLDE
DGYLFYQSRADDLIISLGYTISPAEVEEALLSHADVLECGVVGAPDGRGGTLVCAHVVPR
PGVHGCDALTAALQQHVKARIAPYKYPRRIEYHAAGLPRNDSGKLQRFKLRQAAEEDVQA
A*
>GCA_000452945.1_BPMSHR5858_genomic_03966_1 pqsC
MSYNVRMASLACNLPSRLLRNDDAAFDEVEPIPSEWWQFWGIESRYVIDPHAGESELAMA
ERTARQALARAGVDAAELDLVLFNITSPFVTAADGARRFAPRLSRTLRERLGARRALDAD
VEMECASFVLQMQLAANLIKQGRVKRALVCSSERMSAVVDYASKSSTTFGDGAAAAVLVA
DAHDPHGADWLNAVYRSDATHYALATMQWRYPKPPADRDDAERQAERFGAYFTLKPEAQE
QIARFMPEAIPDIVERLLRKTGLRTDDIDAMVFHQPSEILVRAWAQRLGLAPDQYVIRLA
DCACLVSAAVPLALYESIRRGVVKPGSQVVIAGAGAGWGFGAQLWRWGETVVTDAGSPVD
ATEREHAA*
>GCA_000452945.1_BPMSHR5858_genomic_03972_1 group_2510
MITSTMRRDTASRLVKASPSAVYAAFVASEAVAQWLPPEGAMMEIQVFEPRVGGRFQMTL
IFASAPGKSTANTDVVVGRFVERVPQQRIVQAFEFDSPDPVFAGAMTMRWELEAAAGGTA
VTVVAENVPPGISQTDHETGMNSSLTQLAAYVESHD*
>GCA_000452945.1_BPMSHR5858_genomic_03975_1 ahpC_2
MPIINSPIKPFKATAYHNGDFVQVSDETLKGKWSVVVFYPADFTFVCPTELGDLAERYAE
FQKLGVEIYAVSTDTHFTHKAWHDTSDTIAKIKYPMIGDPTLAISRNFDVLIEEEGLALR
GTFVINPQGEIKLCEIHDNGIGRDAGELLRKVQAAQYVAAHPGEVCPAKWTPGADTLTPS
LDLIGKI*
>GCA_000452945.1_BPMSHR5858_genomic_03976_1 gbpA_2
MKSLFDAPSSRPRARAALTLGAAATLTASFAALLAPMNADAHGAVGFPIARQYQCRLEGG
YWDPPNGSAIPHDDCRAAYRAGNNSAYPFTQWNEVSANPVGQGNDLVQLKAAVPDGLLCA
GGDTSKAGLDKAPASVWRKTQLTPNNGHIELQWENTTAHNPARMRVFISKPSYDPSRPLR
WDDLQQIYDAPAPAPAPVPANGAGHLPGSIQSFYKLDVTLPAGRTGDAVLYSYWQRIDAG
NEGFFNCSDVTIAADERASGFPWVAARAFVEPGIAPRAGQQVRFRVMINDARGAEVVDVR
QPITPYNAERSVWAKQIADQVNGRYGNIAKIGVRSGNTIYFDATNLDANKVWLQPNYSSA
LSVVGAK*
>GCA_000452945.1_BPMSHR5858_genomic_03983_1 group_2200
MQTLFDTGSQQDAAEFGAALAAPAVTGRYDELRGGAAALSSASLAPAWRAFFTQLGSVGF
ADLDRRADALQRRMRENGLAYHPHERAAGGGAVRPWSLDLLPLIIAPDDWAAIERGVLQR
VRLSNAILADLYGEQTILRRGLLPPALVTGHPGYLRPMCGAQAPGGTWLHVAAFDLARGP
DGAWRLMAQHTQGPAGLGYLLENRLIVSRLFPRAFRGLHVQRLAASYRALLQSMQALSPA
RKNSRIVLLTPGPHAATYFEHAYLARYLGLTLVEGGDLTARDQRVYLKTLRGLEPVHGIL
RRVDDEWLDPLELRPDSLLGVPGLMQAVRAGNVLLANLPGSGFLESPGILGFLPRLAQAL
LDETLTLPAVHSWWCGEQAACDEALPLLARGIVKPTFPASVQAGGAFEPVIGARLSPRQL
ADWRARIAAQPAHYTIQADLPLSQAPTWAAGAGMGDGGARIVPRPLLLRVFALADGARSW
RVLPGGLARVGTRDELFNAPMPHGGSSVDTWVMTEGAVDPTTLLQTHLGPDDLQERSRAI
ASRAAENLFWLGRYTERATNLTRLARAALERLRGEDDVETPVHLHVLDALCRDNGLLPAD
APPAVDAPRAFQQALTRSLTPRADRSTGIASCLFGMRGAAAAIRERLSREQWRLIDEATQ
LFDEGGDDAEPEEQLGNDALQRLERLNLLLSAITGAQTDNMTRDDGWRLLSIGRQIDRLE
FLCGVLGHAFDGGAIHKQEGFELVLELFDSGITFRSRFQRCFDVAPLLSLVVLDTDNPRS
LAWVAQALRGRLSKVERSEGYALSELADGIPDIAGWPLHVLCETDGDGRHAALLARLQAC
GKAAWDVSNRIGERYFSHVRDAGRSLWG*
>GCA_000452945.1_BPMSHR5858_genomic_03998_1 group_3489
MNNDALPGHSPDLLDFDEDFIKIDAAICEYDSVGYAPQRKGESAFQWASIETGCLALLKK
AKDVRVGIWHLRACIARRGLSGLADGVRSLADLMSAPVEELHPRALPDESPGETLLIHLG
WLAGPQFLHQLGSSRFEDRDATLNDLIGGRAAAIVEDRDYRLRANTLVHDIQDSLSRIRE
SIAAAEQELNVSRALDLLSVAASRLTQAQAGGADGASVESDAPVDAPAGASAPGAQQPMA
GPGGVLRSRQEVGAALDRIVEYFRVHEPSHPAPIFLSRIQRMLGAGFEEVMAELYPEAAS
LVAQLSRPQSSK*
>GCA_000452945.1_BPMSHR5858_genomic_04002_1 group_19779
VSDLQLYNKLSKRIRRHSLQEVVADHLVDLMNHAIRGARMRIADDSPAAHSVLNFGCPPM
QMAGATKINPVHAAAHICEVIRRFEPRVDPASTVVKPRTESRKRLAQTIYFDVSMKARED
GAELRASLALDYLSGYFSLADDR*

>GCA_000452945.1_BPMSHR5858_genomic_04019_1 group_3490
MTRRAWAILMCAAAAAMPLRAAATGDADAVNDATNAALDLADATPVAAPETARPWKFSVQ
DAVRWSDGRGAASSWRNQLSLEFQFAHALTPTLGVNFAMRFDRFDPLSSGAASHRDVTLV
KEAYASWKPSALLALDAGRINERIGAAIGYNPTDFFKAGAVSLDVPPDPDSRHTNRLGSV
AVRAQRVWDSGSIALMLSPRLASRSVPGDPFATSDLQRTNGVDRWMLVGSQRIAPAIQPQ
WVLYGESGQSPQFGQNLSVLLGNAVVAYAEWTGGWRPSLIGSATGGGNRDRAFRTRSAVG
FTWTLPVDLSLTAELQGNSGGASASRWRALQASDAYAWGRAVRTSIAAQELPSRHGLFLM
AAWRNVFVRRVDLTGFAQLDLGAGQQYWLELRRRFDKFDLAVQFLRQTGPAWSRFGAMPE
ASSVQVLGIFYQ*
>GCA_000452945.1_BPMSHR5858_genomic_04022_1 yknY
MAIVRVENVSKTYLLDSVKVTGLAGVSVDIEANRFTVLSGPSGSGKTTLLNMIGCIDRPD
AGRVIVAGQDTAALCDDALSDFRAREVGHVFQSFNLLPVLSAYENVEYSLLMARTPARER
ARRVAYLLDAVGLAGKGAHRPSQLSGGQRQRVAIARALAAGPSLVLADEPTANLDSVTGR
AIIALMREMQRESGVSFIVSSHDPQVLEAADDVVQIRDGRIVERRRIAQEAQG*
>GCA_000452945.1_BPMSHR5858_genomic_04039_1 pleD
MNLLSSLSDLVIERVSFGIFVLDREMRVLMWNRFMQDHSGVAAADVIGRNLFDSFPELPR
AWLSRKIESVFQLGSFAFSSWEQRPYLFRFDHDRPITGGVDFMQQDCTFVPIARAREVEA
VCVTVSDVTHVSVMQREREEAVAKLQEYANRDGLTGIANRRFFEARLRDEFSRWQRYGGN
LSVLLFDLDHFKTINDRFGHVVGDNVLRVMAQRVADVVRSQDTFGRFGGEEFALLLPCTS
LDEAMLVAEKIRQTIGSTPIEAEGVRVPVTASVGGAAAHAGALTHETLVNEADAALYRAK
RQGRNCSVAFG*
>GCA_000452945.1_BPMSHR5858_genomic_04049_1 noc
MKPSQFAKGFQARPDITTSEKRTALDRLNAIDGLVKSEAAAPAAPPSKSSKKNLPVTIVE
TPPTADAADESSQYRSWRIDNRYQPGQVIELPLKAIKPSPFNPRHFYLKSSIAELAVNLA
KQGQQQAIHVIPDYDNPGSYFVSDGGRRVRALKEANKESVKAIVVDLPIGIQSYKLGYDL
NVQRDSQTVFDNAIVWKRFLEEKYFQSQKELAEHLGLDESTVAVALSIGKLPEAVMQEMV
ARPDRFGSNMAYQVGRYHAARGTDSTLRLINKILSDDLSTRQVADIVKGRATTQESPKPA
SRQRYAQRHEIKLNGVSVGDLKSYGEDRLELRLRGLTREKRDEILQQLERMLLSN*
>GCA_000452945.1_BPMSHR5858_genomic_04052_1 xerC_2
MSLPILTSPPSPPDIPSDLFDRDTSDWLNHPDTAFDAWLAAQQFRRSSADVYRAQWGAFL
GWLAKRQKNLATVDTETIALFVGELPIKKTQRVRYLRLIERVLDHVRKNEFGSTNPARFI
AQDGEATWRTARDNEPTSFLAPAERAALLAYLFSPLPAAGAALWKERRDRALIAAFLGGG
LKTGEARALTISCVKLGSTMLTIPAAQPEFTRETHLASFAIALFDAWLAERQRCGIPGEL
VFPASQSGRPMHKATMLRAVDAIVDAADIAQSRTARASPQTLRNTFAAELFENGVEPEKV
GQWLGFAQPISSNRLHRAWKNWRESLASALPNAESLTDEISTEPSQA*
>GCA_000452945.1_BPMSHR5858_genomic_04054_1 sutR_1
MASSSGARRTTAPASPATGATPPRVGEQIQRLRNERKLTLDDLSRAAGVSKSMLSEIERD
KANPTIAVAWRLTNALGISLDELFAQPKAPETIRVDGPHDIPTLAGHDAQYQLRVWGPIE
LAGKFEWYELTLPANGALVSNPHEPGTREHLTVLNGAVEIEAAGTARRLKAGDTARYAAD
GPHAIRNAGKSEAKALLIVIHR*
>GCA_000452945.1_BPMSHR5858_genomic_04055_1 kbl_2
MRDAYLAHLRGTLEQIRADGFYKTEREIASPQAADIRLAGGTHVLNFCANNYLGLADDSR
LIAAAQAALEQDGFGMASVRFICGTQTVHKQLEAALSAFLKTDDCILYSSCFDANGGLFE
TLLDENDAIISDELNHASIIDGIRLSKARRFRYKNNDLADLEAKLREADAAGARFKLIAT
DGVFSMDGIIANLKGVCDLADRYGALVMVDDSHAVGFIGEHGRGTPEYCGVEGRVDIVTG
TLGKALGGASGGYVAARREIVELLRQRSRPYLFSNTLTPSIAAASLKVLELLASDEGARL
RERVRANGAHFRDKMSAAGFALVPGEHPIIPVMLGDAQVASKMADALLAEGVYVIGFSFP
VVPRGRARIRTQMSAAHTREQIDRAVDAFVRVGKTLGVI*
>GCA_000452945.1_BPMSHR5858_genomic_04063_1 group_3495
MNSLTKRALGAQVRFVAMLAVLIFASAGSLRYWQGWIYWFVFSGATTWLARHFLKHDPAL
VERRMRVGVRAERELSQKIILGVVSVASAGLIVAMGAEWRVLRMPFDWRPVALGNALVIA
GFAICFAALRANRFASSIVEVERDQPVISSGPYRFVRHPMYSGAMVIFFGSPIAAQSTWA
WPFAAVLAVGVVARLVDEERYLSAHLDGYRAYCERVRWRLVPHVW*
>GCA_000452945.1_BPMSHR5858_genomic_04065_1 group_2514
MRPARAGRAFRFAVSRFAVSRFAMPIFVVLFALAALIWGAIHMFEAIAARFGDAVAIGAA
TVAAVAIAAAIARSVRRRRDIAPNAREDGWTHVLQRAWGDLRISATKGLLWLSQDGADGR
YTLTDLSGCRAETLDGRWYLVVAVRDDRRAEWKLPMDDKRDALRWARVLTLAKRQRL*
>GCA_000452945.1_BPMSHR5858_genomic_04083_1 hpcB_1
MGKLSLAAKITHVPSMYLSELPGRHHGCRAEAIRGHQAIGERCRALGVDTIVVADTHWLV
NAGYHVNCNGHFAGVYTSNELPHFIRDMRYEYPGNPALGHLIAASANERGIGTRAHEIDS
LELEYGTLVPMRYMNADQRFKVVSIAGWCMWHALDESRRFGEALRHAIDASDANVAFLAS
GSLSHRFNDNGSPEEAIHMISREFYRQVDLRVVELWKQGDFATFCKMLPEYNAHCHGEGG
MHDTAMLLGLLGWDRYDKPVEIVTDYFASSGTGQINAIFPLP*
>GCA_000452945.1_BPMSHR5858_genomic_04086_1 hpcE_1
MELSLVERGERVATVPIGTVYGALLNDRGALAALGDAVHAPPYHKPPRASVLYVKPANTH
AGDGAAVVVPAGVDALEIGATLGVVFARRTTRVPAEQAFDFVKGFTLASDVSIPHPDYYR
PAVRFKCRDGFCPLGPAIVARASIGDADALAVTVRIDGERRYEASTANLIRPVARLIADV
SAFMSFDAGDVLLVGVGPHAPRAAAGSTIEIDAPGIGTLRHTLVAEDAQ*
>GCA_000452945.1_BPMSHR5858_genomic_04092_1 sasA_3
MLRSLVRLYLVVVVCASAAILFINTSFTRFFYERISSAARETLSTYAFVLNDYLERHPGA
SRQPALRELAKHGNEGFGFMSIDDARKLLSAEQQRDLDRGEIAISYDGKDYFMPLVDGTV
LHARPDEPPSLDIQIYAYLTVALATLLAVALWIHYHWRDLRKLEAAARAFGDGRLSTRAE
LSRKSNIYELSQQFNDMAERIEASIQQQREMMHGISHELKTPLARLEFGLALLSSPDAPE
RMRERQAALRRDVRELDELVTELLTIGRLEQGERHLALMQVMVAELIDSVAASVAHEIAD
RRLTLDVSVRGAPACHVCDAKLVARALLNLIRNGMRYASTTISLRASLGEAGALVLSVDD
DGPGIAAADRDRVFEPFHRLDSSRDRQTGGFGLGLTIVRSVALAHGGDVRLDDTPLGGAR
FAITLPSLEAPGREQDARGPAA*
>GCA_000452945.1_BPMSHR5858_genomic_04106_1 ctaE_1
MRPGAPFDAAGDMSRIASRIAAASAAPARVPGEAELWIFILGDMLVFAVFFTILSAHHVL
HPALFDAGRRQLNLAFGLVNTLALLTSSAFVALGLGEAKAARLAAAKRRFVVAAALGALF
VAIKCAEYAQKIAAGANPLVNDFYMYYFVFTGVHLLHVAVGLIALALMIRRCVRPAAEPR
DVGFLEGAAVYWHMVDVLWVVLFLLIYLI*
>GCA_000452945.1_BPMSHR5858_genomic_04107_1 group_19725
VNTYRLERASAWSGLVFLLVFGLGWLVLARFFPPISPDAAAAGVAAIYRARRLSLMLASV
LMMSSIVFLMPVSALFVLLIQKIERRVGILTLMMGFTLTTNLVVTFYTALSFSLAAFRAQ
RSPELVQFANDLGFLQFMGGISMFVMAWFVSAYAILVESPRENPVLPRWVGYVSLWMGLL
YLPELLIYFFRTGPFAWDGIVGFWIPAALFVVYFLFSPVVLVPAVRRHFR*
>GCA_000452945.1_BPMSHR5858_genomic_04108_1 treA
MVTPRHRPLHVENPFYRRLLNAPAWVALVAAAGIGCTSATLARADSSPAHASTAAAVASA
ASAPGAASIPPPPSQLYGDLFVAVQTAQIFADQKTFVDSTPNADPATIVQLYQQQKGQPG
FSLKAFVAQYFTPPSDESVTPPPNQTLREHIDWLWPKLTRTTTTAPPYSSLIALPKPYVV
PGGRFREGYYWDTYFTMLGLQEAGREDLVDNMLDNFAYLIDTVGHVPNGNRSYYVSRSQP
PFFAYMVTLAAKAEGNRVYQKYLPALRKEYAYWMQGERTTPRGQATRNVVAMPDGSVLNR
YWDASDTPRDESYLEDVKTAQQASGRPAAEVWRDLRAAAESGWDFSSRWFGDNRTLATIR
TTAIVPVDLNSLMFNLETTIVKGCAVTRDFACVAEFAGRAGKRAVAINRYLWNRNGYYGD
YDWKLGKPRDNLSAAALYPLFAGVAWPERAKQTAKNVQKALLKPGGLATTTYDTAQQWDA
PNGWAPLHWIALVGLRHYGEKSLADDIGTRFLADVKGVYAAQGKLVEKYIVEGVGTGGGG
GGEYPLQDGFGWTNGVTLKLLDLYGG*
>GCA_000452945.1_BPMSHR5858_genomic_04119_1 group_2516
MAGKDIHVILDLSRCIEHGTQIPGPAVRGSVRPDTFMILSDHSIAFSNTHFTVPADNKPV
QEFMKYRANGDGKVEFQTMVLDPINFSVLRKNQYDCEVNKGVKFFW*
>GCA_000452945.1_BPMSHR5858_genomic_04128_1 group_4024
MTTAIAAAIDHDRLADFIERKWNDEILHALTDYIAVPAKSPAFDPDWAKHGYIERVIVDA
AQWAERQPVKGLRVEVVRLAGRTPVIFFETPATRAGSTDTILLYGHLDKQPEFDGWRADL
GPWTPKFEGGKLYGRGGADDGYAIYASLAALGALDAQGIGRPRCVGLIETCEESGSYDLL
PYVDALRARLGDVSLVVCLDSGAGNYDQLWLTTSLRGLVSGDLQVEVLEEGVHSGVYGGI
APSSFRVMRQLFERLEDAATGNLLPPSFHCEVPASRLRETQAAASILGDAVWKGLPWACG
QDGKPVLPTTADPREALLNSTWRPSLSVTGAQGLPALENAGNVLRPRTAFKLSLRLPPLV
DAAQAVQQLKALLELDPPYNAKVTFKPDAGAATGWNAPDVAPWLGAALDDASRRHFGANC
AYMGLGGTIPLMNVLQEGFPCAQFMVCGVLGPKSNAHGPNEFLHVPYAKKLTAAVADVIA
AAH*
>GCA_000452945.1_BPMSHR5858_genomic_04130_1 cdhR_1
MHPHLVVALAYDRLCTFEFGCAVELFALPRPELGVDWYRFAVCASEPGPVRAAGGIKVQA
PYRLALMDRADTIVIPGWRDPDETPPEPLLKKLRAAHRRGARLCSICSGVFVLAAAGVLD
GATVTTHWRYAERLRARYPALRVNPDALYVDAGRIVTSAGSAAGLDMLMHLVRRDHGSAI
ANRVAQRLVLPPHRDGGQAQFVPRPLPSVGGDRLAKLIDWMRAHAGQPHTLASLAARAAM
SPRTLQRQFRDATGLGPYEWLIRERVGLAKEMIERDPALPVARVAALAGFGSEESLRRHF
RRIAATSPAAYRRSFDAG*
>GCA_000452945.1_BPMSHR5858_genomic_04133_1 group_19870
MTTYRDTHTTSFRIGDVVTLKTGGPRMTVTYAGPVVFDTGEWVICQWFDENGEFRQEMFP
NETVVLEPRTISAGLARMRSLSVRGGMQA*
>GCA_000452945.1_BPMSHR5858_genomic_04134_1 murR_2
MPADFDELASLIRAQFSELSPQFQVGAAFLLDHPDEVAVSSMRKVAQRANVQPASLVRLA
QQFGFPGWNELRDLFVARVRTRPQPLTQRARSLVKPNAKASLAADLLAAQRHNLEASAAQ
NARALAEAAKLIRKAAHVHVAGFRSCYPVAFGFVYGYRLFRPTVSLLTGVAGSLDMELRT
IAKNSVTVVVSFAPYSAEATRVAQAARAQGGRIVAITDSAVSPIALHADAQLIFTHDSPS
FFPSLVAAHALVETLVAQLLALEGGEAIAALEQAEAELHAKGAYVV*
>GCA_000452945.1_BPMSHR5858_genomic_04136_1 rssA
MTGARGFARRLNASLRGRLLMWLLPAACVVGVLASTGTYWGALRELDDLLDDQMRSMSKQ
IQVDDAGKLSIEGRDRLAKRHPADYDSDDVLLQVWRGGKLEFTTDPATQLPPPAAAGFVT
LDAGGQRWHTYVRESAGTTIRVAQPRQARWEAIAGIAVHLLWPVLSLLPLLAIGLWLGIG
YGLRPLGAIAAGLKRRHANRLDPLDITTVPTEAQPLASEINDLLARLDRSFTLQRHFIAD
AAHELRTPIMGLSIQTQLLQRTASADERARILAQIQAGTTRLSHLAEQLLTLARLEPEAR
SAPFAELDLAALGRSVVAERARIAAAQHVDLGALGTAPVRVAGNADTLRVLLNNLVDNAI
RYAGAGARVDVDARIDAGGTPVLEVRDNGPGIPAADRERVWERFYRGGGAQLVSASGSGL
GLSIVKRIAEQHHADVSLEAGADGRGLTVTVRFGAARQA*
>GCA_000452945.1_BPMSHR5858_genomic_04137_1 group_3502
MTERSMQTPAAELVKRNTTRWPNESEAYRRARDALLVEEIELRRHVERVAALRRALPPGG
EVTGGYRFEGERGPCDFAQLFGDRQTLVVYSYMFGPQRERPCPMCTSLLGAWNGEARDLE
QRVALAVVARSPLERLVAFKRERGWRDLKLYCDLDGRYSRDYHAIGADGDEVPAINVFTR
RDGTIRHFWSGEMGGWSADPGQDPRGAPDPMPLWTILDMTPEGRGIDWYPSLDYPD*
>GCA_000452945.1_BPMSHR5858_genomic_04142_1 group_19818
MTNQKPDDARPHDDEPKLEQLEAAVDHLHDSIQSQSIAAGAAKGILFSLIETLGALVGDP
DLPEHARSGYEALRDKANELKANLENR*
>GCA_000452945.1_BPMSHR5858_genomic_04144_1 group_19710
MDIQKIEARLEFLREAERLKSVLRSAHTSTGRAESTAEHSWRLCLMAITLADELPGLDML
KVLKMCVIHDLGEALRGDVPAIRADAHPDKSAHERADLLTLTRMLDAPLRDEILSLWDEY
ERAASQEAQAVKALDKLETILQHAQGENPPDFDYAFNLTYGSRYTSAAPLFRAVREIVDA
QTRSRIDARGEPPPAR*
>GCA_000452945.1_BPMSHR5858_genomic_04147_1 bamA_1
MTVLASSARAGFRVPGFVAALLFVCAATGRPAPAAAADTATVASAATAAASAAGAPASTT
LPPAATPTRVASSAVAASSPASSLASSSASAATPANAAAAWSWTAARRADAPDSSSVSNA
VSNSGSNVAGTAAAHGPHAPHTAPDAGAGAPSAPTTPTAATPAAAGASASMAAAPTPAAT
DTLVCMPDGGGPHRPAIGLVLSGGGARGYAHLGVLKVLEANRIPVDCIAATSMGAVVGGL
YATGMTAQDMQRRLSQVNLADIAFDVTERSDLPQKKREDERLYIDSLTIGFDSKGFKAPV
GLVQGNRLQALLANWTAAVPTNQPFDRLPIPFRAIATDLQTGQKVVLDHGSLPLAIRASM
ALPGLFSPAEIDGRALVDGGLVGNLPVDAARAMGADVVIAVDIGSPLRPLDALASPADVM
QQMIGILIRQNVAEQRKQLTANDILLQPDLGKQTFTDFQNANQAIAAGEAAAVAALPRLA
RYALSPEQYEAYRAAHARPQQPIRITSIEIRTNGASVPKQIVRNALRVKPGDVYDPQAVS
ADLLSLTTSGNFENVTQQIINEGDEHRLVIDAQEKYWGPNFLLFGLGMSSSSTDEGGFRL
HLGYRRPWLTPSGLEFRADTTLGSDMQSVHVELRQPLSNKTGYYVAPYADYKRRFTNLYD
SDTDIKITQYRFQTSRVGLDLGLPLARLGDFRIGLAYTHLTASPTYNVPLNWFLPDDAPS
PGTLFPSAYGHQISARARLVIDQLDDPTFPRKGYFVEARVERSLSKSNDTFFDSDASFTD
VYGRLMVAQRFGRHSVNISVEAGKSFGGTNFGNPLGYTLGGFQHLSAYAADQLSGNALLY
GQVTYMNQLATFNASPIKALYVGASAEVGNVWTPDTRIGSGSLKQSYTFFTSLTTAFGPV
YFGVALAPGGRRNIYFQLGRTY*
>GCA_000452945.1_BPMSHR5858_genomic_04153_1 rna
MLRTFARAACALAVASACNNASAYDYLLLATSWEPGFCATHSATECGTLSGSYASTNLSL
HGLWPNNYDGNQPFYCGVPQNEIDLDNAHQWCSMDAYPVSSSTLAALSTYMPGVASCLDK
HEWYKHGTCAQAPSADAYWNNATGMVSRLSTTSFNTFLQSNAGNTVTRTQLLNAFEGAFG
SNTRGAVSLKCVKVNYVSYFTEAWISVKTDATSQFPSAASLVTDAGTQGTCPTSGIFIAK
*
>GCA_000452945.1_BPMSHR5858_genomic_04158_1 menB_2
MSTPMMHDAAHDTAARDVLMRVVNRVALITLNRPAALNALSHDMVRELAAWLERCRDDRS
IVAVVLRGAGDKGFCAGGDVRALYRMAKAPGRDWLPFFVDEYRLDYAIHTFPKPVVALMD
GITMGGGMGLAQGAALRVSTERSKIAMPETRIGFVPDVGATHFLARLPVELSLYVGLTGA
TLSGADALAAKLADLSVPSSWLDTFEARLERADWSGDVLRALRAVFAPPCNVVPHAALDS
QMPWIVRHFDKRSGVERIVATLRQDLTRDELSREQRQWLQTTLGALASHSPTMLEVTREA
LLRGRQMTLAESFRMELGIVARAIEEGDFREGVRAHLVDKDRKPGWAPATLVEVQRERIV
HFLASPWKREAHPLADLGTVAG*
>GCA_000452945.1_BPMSHR5858_genomic_04168_1 ephD_2
MQPLSDEAPLALFESVHTETAVAAGDVTLAAKTWGDASRPAVVLVHGYPDNSEVWRRVAP
LLAKSYYVIAYDVRGAGLSTKPARTADYRLERLVDDFAAVIDALAPNRAVHVVGHDWGSI
QGWEFVTEPRLAGRILSYTSCSGPNLDHVGYWLRQQLARPSPASIKRLAGQLVRSWYVYL
FHLPLIPELNWRLWLGRAWPALMRRLEHTDVGVRPTQTEDGVHGVRLYRANFLRRVLAPR
ERYAHAPVQVVVPLRDKFVSPALSADIARWVPTYYRREVAERHWLPMSEPARFAALAQEL
IEAVETGVQPPALAHARRRSGTGPFVGKRVVITGAGSGIGRCAAVEFAKQGASIVAVDID
EQAAERTALLVRLLGAQADVRRVDVGSADDMEALANWVGDELGGADVVVNNAGIGMAGGI
LDTSAAHWERILRVNLWGVIHGSRLFAKQMAARGAGGHIVNTASAAAFGPSRDLPAYATT
KAAVLMLSECMRAELADHGIGVTAVCPGFAETGIMASTQYAGAKSDQDEARLRKRATKLY
QMRGLKPETVAKAMVDGVLQNQPVVAIGAEAHAMRFVGRFAPWLGRLIARVSMASH*
>GCA_000452945.1_BPMSHR5858_genomic_04169_1 group_3504
MNAPENHYLIKARHVKFDFAKTPIQWIEGDPESTHIINTLNLLFPEGELWFCRVYNKALP
LITDAALRADADGFLRQEAVHSRSHGGVLKHYYERHGIDTKPFTQRLNWLFTKLLGEQPL
GLKIGHTRFWLRQQLAVIASLEHFFGYLGNWVLHARGLDDERTDPTIVDLLRWHGAEEVE
HRTVAFDIYRHLGGNYVERSIHMTFVILLLLYFIMSGAKYMYERDPGAGRYPGFVLSWWR
GSRRKHLPSFWKTIAAALRYYRPSYTPHAEGSTDVALAYLARSPAAQAASHGGNWVAQKP
AS*
>GCA_000452945.1_BPMSHR5858_genomic_04171_1 group_2818
MKTEDLIGILTDAGVDLAVGVPDSLLKSFCGRLNDPDCPLRHLVASSEGGAVGIAIGHHL
ATGGLAAVYMQNSGIGNAINPLVSLADRAVYGIPLVLIVGWRAEISASGAQVHDEPQHVT
QGRITLPLLDALSIRHLVLERAGGENDALAPSIAQLIAGARQTSQPVALVVRKDAFDDAS
ASRPGAAAPHAGRMTREQAIALIVEHADAGAAIVSTTGVASRELYELRDRLGHSHARDFL
TVGGMGHASQIAVGIALARPAQKVICIDGDGALLMHMGGLAYCAGAPNLTHVVINNGVHD
SVGGQPTLAAHLRLSHIAASCGYAFSRSVATPIELESALHHASRLDGSAFIEVTCRPGYR
SDLGRPRTSPAENKRHFMAFLSRNGATHERDDHAQESGIQDAIQCARH*
>GCA_000452945.1_BPMSHR5858_genomic_04175_1 group_19799
MTAYAIVLLIGVVAGVVSGVIGTGASIMLLPPLVFYFGAKQAVPIMAIAAVLGNVSRAYV
WRRDIDWTAFFAYSIAAAPAAALGANTLWALNPQWVDCALGVFFLSMIPYRYIARRSTFA
LTAWQLAAAGAVVGYLTGIVFSTGPLTLPIFSAYGLLKGGLLATEAAASLAVYASKLVAF
GQLGGLPLDVAIKGALVGLSLSAGISVGKAVTLRLSTNAFQRLLDLVMLSAGTTLLWGAI
R*
>GCA_000452945.1_BPMSHR5858_genomic_04177_1 group_2518
MLNWISRWALRHAPTPEKSAASMLVTARMELFTAEQRVIDAKLQADYWRTRVSFLEEVQK
QGIDPWVSARAQKADDAPAAAMRGAAGPRLAAST*
>GCA_000452945.1_BPMSHR5858_genomic_04181_1 group_19863
MQIQFTNDAPEYSGRELTLAFMAMVDGEPVQCHITAEALEDHFGAASPRFEDMVGAFDTH
RPRIEAAARRLLSETRAQCVVLRSGYVRFYEANGR*
>GCA_000452945.1_BPMSHR5858_genomic_04182_1 group_19952
MVRTELRVVLAAIATFVMLAGIAVAIHGSLFDQDAALRYGAAAIALGVTTCAIALNVWPK
DEKK*
>GCA_000452945.1_BPMSHR5858_genomic_04211_1 group_19964
MTTAGMQATELTVHRDTVRAEWVDYNGHLRDAFYLLIFSFATDALLDAIGLDDAARRARG
RSVYTLEAHVNYLREIKEGTRVRVDARVLAHDAKRLHLYLEMFAAGIDGAVSASEQMLLH
VDTGGPKAAPFDADVAERIANLHALQRAHPAPAYAGRVIGLPARR*
>GCA_000452945.1_BPMSHR5858_genomic_04212_1 lcdH_2
MAVITKIDTFAAIGAGVIGSGWVARALANGLDVLAWDPAEGAEMQLRANVENAWPALERA
GLAPGASPARLHFVPTIEACVADADFVQESAPEREALKLELHERISRAAKPDAIIASSTS

GLLPTDFYARAHRPERCIVGHPFNPVYLLPLVEVLGGERTTPDTVDAALGIYRALGMRPL
RVRKEVPGFIADRLLEALWREALHLVDEGVATTGEIDDAIRFGAGIRWSFMGTFLTYTLA
GGEAGMRHFMQQFGPALELPWTKLVAPKLTDALIDRVVEGTAEQQGPRSIKALERYRDEC
ITEVIAAIAAVKARHGMRYED*
>GCA_000452945.1_BPMSHR5858_genomic_04213_1 kce_1
MNQEVIVTCAVTGAGDTVGKHPAIPVTPKQIADAAIEAAKAGATVAHCHVRDPLTGRGSR
DPRLYREVVDRIRSADVDVIINLTAGMGGDLEIGAGENPMRFGKGTDLVGGLTRLVHVEE
LLPEICTLDCGTLNFGDGDYIYVSTPAQLRAGAKRIQALGVKPELEIFDTGHLWFAKQLL
KEGLLDDPPLFQLCLGIPWGAPADTGTMKAMVDNLPAGAHWAGFGIGRMQMPMLAQAMLL
GGHVRVGLEDNLWLDRGVHATNGSLVERARQIVERLGARVLTPAEGRRKLGLPPRGERPL
ERRAMTEYV*
>GCA_000452945.1_BPMSHR5858_genomic_04215_1 cdhR_2
MPEEFHFLLMPGFSALGFMSAVEPLRVANRFRPNLYHWRIVSCDGAPVVASNGISLNADA
AFAQVEHAHTIFVIAGFDPLARYTRALGDWLRREYRQGATAGGIDTGSFVLAEAGLFDAS
HALTLHWEALAAFQERYPHLKTTQELFEIDARRITCAGGTASIDMMLDLIGRRHGAELAA
TISEQFVLSRIRQRSDRQRLEIAARYGVHNRKLIQVIGTMEHHIESPLSSDALAREVGIT
RRQLERLFSTMLDDTPARFHLRLRLERARELLQQTDMTVAAVCAACGFESPSHFSRTYRA
QFGTSPRSDRHTSR*
>GCA_000452945.1_BPMSHR5858_genomic_04217_1 group_2519
MSKLQHLAAAVCGALCVSAAHAAPVWITLSEPALRELRALDPAVTSRYSAALATGDAKRT
ETIHVAQVDDSLLESLSQAIRRTRGHGPGFFVHATFDEARASLQPSAAKQAAAIDYPITY
SQQVRNWISQLQASNIVSTIVSLSGFTNRYYTTTHGVAASDWIAQQWKQLAGSRTDVTVE
QFTHAGWPQKSVILTIKGSDPAAGVVVIGGHLDSTVGRMSENTRAPGADDDASGIASLTE
ALRVLLANRYQPKRTLKFIGYAAEEAGLLGSQAIAKQFRAQNVNVVGAFQLDMTNYKGDP
KDIYLIGDYTNATQNTYLANLAKAYLPELAVGTSQCGYACSDHASWNAQGYPASFPFEAD
QNDNPYIHSAYDTLERSDSQGNHALKFSKLALAYAAELGGGLSASAKR*
>GCA_000452945.1_BPMSHR5858_genomic_04218_1 group_2520
MKKSNVAVMGALALAIGIAPGASRAQSSVTLYGILDAGITYVSNTGGSHVFKFDDGVSYG
NRIGFKGTEDLGGGLKAVFVLESGFRLGNGQLGFGGAQFGRQAYVGLQNNWGTLSFGNQL
DMTEEMVYLNNISAWASGYAIHQGDFDRFNGDRLPNSVKFLSNDFSGLKFGAMYSFGNVA
GDFHRNSAWSAGANYTHGDLSVGAAYTHLNNPSGIYAFDPYAMIGTRTFLGQPTVSVDPA
TGKVTDLFANTPMNVDSQGTFGVGASYTIDKLTLSGNFTYSTIKGFGNTSHMQVYEGGGL
YQFTPALSFVAGYQHTRFEGHHWNQGSAGVHYLLSKRTDVYLSGDYLRASNGVDAVIGYS
FTPSTTQTQADIRIGMRHSF*
>GCA_000452945.1_BPMSHR5858_genomic_04219_1 group_3511
MTPGVTHFQMEQAMKRYESIARRLARRAAAASPAFAALAWCAAAAAATTTAAAAEPAACR
DVRMAGPGWTDIEATNALAGVVLKALGYRQSVSNLSVPITYQGLKKGQLDVFLGNWMPAQ
APLVKPFVDARAIDVLHANLSHAKFTLAVPDYVAAAGVHSFADLAKYAQRFGAKIYGIEP
GAPANQNISRMLADKALGPANWQLVESSETGMLTQVERAVRERQWIVFLGWEPHLMNTKF
HLVYLSGGDAYFGPDYGGATVNTVARADFASQCANLARLFRQMTFTVDLENGMIAAMLQG
KRSAVDAAQHALRANPSLVEAWLDGVRTASGAPGLPAVRAALDAQ*
>GCA_000452945.1_BPMSHR5858_genomic_04225_1 group_3512
MMRDAANFEPADSRVTRPAFWQALPERWTSDVEETLVCCHVRQETHDVKSFFFRSPQGRA
FSFEPGQFVTLELDIDGETINRCYTISSSPARPHTISITVKRVPGGKVSNWLHDNLQPGA
PLRVLGPAGEFTCARHPARKYLFLSAGSGITPLMSMSRAHHDLAEDRDIVFVHSARTPDD
IIFARELDLIASTHANFRTAFVCERLGARTNWHGVTGFLSLPLLKLVAPDFMEREIFTCG
PAPYMKAVRELLAEAGFDRSRYHEESFSFETLAQGEAPADAPAGEDAAARGAQVRQYTIA
FAKSHREIACGADQHVLDAARRAGVRLPASCTQGMCGTCKVKLVSGQVDMKHNGGIRQRE
IDQGMVLLCCSKPLSDLVVDK*
>GCA_000452945.1_BPMSHR5858_genomic_04226_1 group_19891
MKVSADVRALVARRKAGYSLEAPFYLSDEIFALDMDAIFRRHWIQVGVEPDVPEPGDYVT
VQLGGDSILIVRDDDMQVRAFHNVCRHRGARLCNEEKGSVGNIVCPYHSWTYNLTGQLMF
AEHMGEKFDRCKHSLKPVHLENLAGLLFVCLADEPPVDFATMRAAMEPYLLPHDLPNTKI
AAQIDIVEKGNWKLTMENNRECYHCVANHPELTISLYEYGFGYQPSPANAEGMAAFERTC
VERAAQWEALNLPSVEVERLTDVTGFRTQRLPLDRSGESQTLDAKVASKKLLGEFRQADL
GGLSFWTQPNSWHHFMSDHIVTFSVIPLSAGETLVRTKWLVHRDAKEGIDYDVKNLTAVW
NATNDQDRALVEFSQRGAASSAYEPGPYSPYTEGLVEKFCEWYVGRLAAHIGA*
>GCA_000452945.1_BPMSHR5858_genomic_04227_1 group_19837
MNANRKLASIAVLVSVGRHPVSGVPRYSRNDAAALEIGRALAAEHRARLDVIHAGDARNP
ALADYLALGAASVEVLACADGDDAAAALAERVRGHDLVLTGTRAEGAYDTGMLPYRVAAA
LGWPFVGAAVDVSVDAARATLRQFLPKGVRRRVDAALPAVVAVHPLANAQPRYAYARLRA
GEVRVTRAATRAAADAGQWTTRAVERKPVRLAAADKRSGHARMLSATTTESRGGNVVIEG
SSVEKAQVILAYLREHQLIDY*
>GCA_000452945.1_BPMSHR5858_genomic_04230_1 stcD
MRYPHLFKPLQLNQLTLRNRIVSTAHAEVYAEPGGLPGERYIRYYEEKAKGGVGLAVCGG
SSPVSIDSPQGWWKSVNLSTDKIVDPLARLAEAMHRHGAKIMIQATHMGRRSAFHGEHWP
HLMSPSGVREPVHRGNAKIIEVEEIRRIIGDFAAAAKRVKDAGMDGIEISAAHQHLIDQF
WSPRTNVRTDEWGGSLENRLRFGVEVLTAVREAVGRDFCVGLRMCGDEFHEDGLDHEQLK
EIAQAMSETGLIDYLGVIGSGADTHNTLANCMPPMALPPEPFVHLAAGIKSVVRIPVMHA
QSIRDAGQAERLLANGMIDLVGMTRAQIADPHMVIKIRDGREDEIKQCVGANYCIDRQYN
GLDVLCVQNAATSREATMPHVIEKSRGPKRKVVVVGAGPAGLEAARVARSRGHDVVLFEK
NAEVGGQIMLAAKAPQREQMAGIVRWFDMETKRLGVDRRLGVEADEKAIMAEKPDIIVLA
TGGRSFTDQVSGWGVAEGHAVGSWDILSGRVEPGKNVLVYDGVSTHAGAGVADFIASRGS
NVEIVTPDVKVADDCGGTTFPIFYRRLYAQGVIHTPNYWLDRVYEEDGKKIAVLRNEYTE
EQEERAVDQVVIENGITPNDALYWKLKPESVNRGQIDVHKLFAAEPQPCLAETLGNGRFL
LFRVGDCISMHNIHGAIYDALRLVKDF*
>GCA_000452945.1_BPMSHR5858_genomic_04232_1 group_3514
MSTLHQDSIIIDGLNISKFERSVFEDMQKGGVTAANCTVSVWENFTKTVDNIALMKKQIR
ENGELLTLVRTTDDILRAKREGRTGVILGFQNAHAFEDNLGYVEAFADMGVRVVQLCYNT
QNLVGTGCYERDGGLSDFGREVITEMNRVGIMVDLSHVGGNTSSEAIAFSKKPVCYSHCL
PSGLKEHPRNKSDAQLKEIADAGGFVGVTMFAPFLKRGIDATIDDYIEAIGYVVNLIGED
AVGIGTDFTQGYSVDFFDWLTHDKGRYRRLTNFGKVVNPEGIRTIGEFPNLTAAMERAGW
KASRIRKIMGENWVRVFKEVWGA*
>GCA_000452945.1_BPMSHR5858_genomic_04236_1 rafR
VKVNLKALSEALGLSRTTVSRALNGYDDVSEATRERVMQAAREFGYVADPTARRLATGRA
DAVGIVYPFGAGDLGDPRFAEVVAGVTERLGESGLDFFIVSARPNAELDTYRRLVDGRLV
DGLIVARTLVDDPRLRFLQEREFPFVAYGRTASAKPYTWFDFDNEAGMHAAAQRLIAFGH
RRIALVCAPQTLNFAAQRRAGFERALREAGIEPDAALIVECAFTRDGGYGAAQTLLALER
APSAIVVDNNIAGAGVFRAIVERGRRFVRDVSLIVYDGVSPDVSFPHHASAVLQPTGHAS
GRAIAELMLGAIAHPGDAAHRLVSPVIEPGDTDGPFSG*
>GCA_000452945.1_BPMSHR5858_genomic_04275_1 htpX_1
MSAYTFTLLFAVAIVAMTGTKLWLASRQVRFVAAHRARVPAQFRETIPLAAHQRAADYTI
ERTRLTMFEIVVASAVLVGLTLLGGVGALDSLLAGWLGHGYGQQVALVAAVLVISSAADL
PFEYYRQFGIEERFGFNRMTKRLFFTDLAKNALLGAALGLPLLFVVLWLMNQAGALWWLW
TWIVWVGFQMLVLVLYPTFIAPIFNKFEPLSDDALRARIEGLMKRCGFAAKGLFVMDGSR
RSAHGNAYFTGFGASKRIVFFDTLLARLTGGEIEAVLAHELGHFKRRHVMKRMLWTFALS
LALLALLGWLAQRAWFYTGLGVMPSLSGSNAGIALVLFFLSMPVFLFFVTPLGSLSSRKH
EFEADAFAASQTDARDLVNALVKLYEDNASTLTPDPVYTAFYYSHPPASQRIDRLLQHS*
>GCA_000452945.1_BPMSHR5858_genomic_04276_1 rsgA
MKRAPTKQPAKPAARGGERAQGRVIAAHGRHYIVAPADGGPMLQCFPRGKKSEVAVGDRV
AYERTSADQGVIVEIGERRNLLYRSDQFKSKLFAANLDQLLIVLATEPYFSEDLLGRALI
AAEANELKPIVVLNKIDVEAALPVARERLAPYRALGYDVLELSVKGAPDDARTQLAPRLA
GHSTILLGQSGMGKSTLVNLLVPDAEAATREISAALNSGRHTTTFTRLYPLQDGGALIDS
PGFQEFGLYHLTEGRLERAFPEFRPLLAHCRFYNCHHLHEPGCAILEALADGRIAPTRHA
LYAQLVHEASQIVR*
>GCA_000452945.1_BPMSHR5858_genomic_04279_1 nudL
LIRRPIIDPEVLPIETTGADLPVVRPSSMTPAGLRERFAEPPEWTQEPAEAQLAHGVDPR
SAAVLVPLVVRERGLTVLLTQRADHLNDHAGQISFPGGRREPDDRDANATALREAREEIA
LAHERVELLGALPDYLTGTGFCVTPVVGLVHPPFTVQPDTLEVAEIFEVPLDFLMNPAHH
QIRVLRWEGGERRFFAMPYPRGPVGGQYFIWGATAGMLRNLYRFLSA*
>GCA_000452945.1_BPMSHR5858_genomic_04280_1 group_2523
MNAYTLQSIRYNDAPDALRRRFALLHQRVDPDDAFAPDGPPPVIHDPRLDAMSFYASTRS
NGEVVSYAAVVRKTIAHAGKRFSIGGLSWVATDPAHRRAGLGARTVSAATQWMAASGLDI
GVFTCDPPLARFYARAGDWPIAAHVTLIGSRAAGALRSDTLNKVVLMRLFTAKARAAEAE
LSRATIDLDLPPGQFL*
>GCA_000452945.1_BPMSHR5858_genomic_04288_1 rnpA
LRPWRRTAHFVIYGKPTGRDARLGLVIGKKYAARAVTRNLVKRLAREAFRTRRAEFAGWD
ILLRLHARFDKKAMPSAASAPLAALCAGEIRELLDRAAREVARRNGAKPASE*
>GCA_000452945.1_BPMSHR5858_genomic_04289_1 yidC
MDIKRTVLWVIFFMSAVMLFDNWQRSHGRPSMFFPNVTQTNTASNATNGNGASGASAAAA
ANALPAAATGAAPATTAPAAQAQLVRFSTDVYNGEIDTRGGTLAKLTLTKAGDGKQPDLS
VTLFDHTANHTYLARTGLLGGDFPNHNDVYAQVAGPTSLAADQNTLKLSFESPVKGGVKV
VKTYTFTRGSYVIGVDTKIENVGAAPVTPSVYMELVRDNSSVETPMFSHTFLGPAVYTDQ
KHFQKITFGDIDKNKADYVTSADNGWIAMVQHYFASAWIPQSGAKRDIYVEKIDPTLYRV
GVKQPVAAIAPGQSADVSARLFAGPEEERMLEGIAPGLELVKDYGWVTIIAKPLFWLLEK
IHGFVGNWGWAIVLLTLLIKAVFFPLSAASYKSMARMKEITPRMQALRERFKSDPQKMNA
ALMELYKTEKVNPFGGCLPVVIQIPVFISLYWVLLASVEMRGAPWVLWIHDLSQRDPYFI
LPVLMAVSMFVQTKLNPTPPDPVQAKMMMFMPIAFSVMFFFFPAGLVLYYVVNNVLSIAQ
QYYITRTLGGAAAKKKAS*
>GCA_000452945.1_BPMSHR5858_genomic_04296_1 group_2524
MQDLSTELAPELKPELKPELKLDDPFVDAVHTEFVQLLDAAARADDAHFLQAYDVWMDHC
RQHFEREERWMASTKFGPQYCHAADHNEVLKVAADVREKLARDAQFELGRRLLVEVAEWF
DQHVRTMDSMMVSHLKMLNFAMVDVPTEA*
>GCA_000452945.1_BPMSHR5858_genomic_04297_1 nsrR_2
MRLSEYTDYTLRVLMYCASNPDRLITIAELAERHDVSKNHLTKIVNSLARQGTLETTRGR
GGGVRLALPPERIGIGDVIRASETDFRLVECFDRNTNTCMLAAKCRLERLLDSALRAYFK
ELDGATLADLVGAPPKSGLSAAEKTHEFAIAMPPKSIARARS*
>GCA_000452945.1_BPMSHR5858_genomic_04301_1 sufS_1
MGDMLRSATMMASAFPALAQRVNGAPLAYLDNAATTHVPQPVLAAMRGFDERDRANIHRG
VHTLSQRATDAYERARDTLARFVGANGEHLLVFTSGATDALNLVANGLSVAGHTQAVLQE
GDEIIVSALEHHANLVPWQMAARRCGAKLRILHPDPQGSLHVRDLERLLGPRTRVFAVTA
CSNATGERPPYEALLAAARAGGALTVLDAAQAVGHDVPDLSALACDFAAFSGHKMYGPMG
TGALVGRRDALERLVPLRFGGDMVSWVGETDATFDALPARLEGGTPNVAGAVGIAAAADY
IDAIGRARIDAHVRALRDHAAAGLAALDGVTVLAPRTSSAIVSFVVDGVHPHDIGTLLDE
RGIAVRTGFHCAQPLLERLGCGPTTRASFALYNTHDEVERLVAGVAQALKVLR*
>GCA_000452945.1_BPMSHR5858_genomic_04303_1 sufC
MNRSEPLLQVFDLSVEVAKRRVLQSVSLAVPAGALVVLMGANGSGKSTLGMTLAGHPAYR
ATHGHVRFGGQDLLAMSVQERARAGLFLSFQAPPDIPGVKNNLFVRTALNAVREARGDTA
LDALDFLGDARAAAAQVGLSDAMLNRSMNEGFSGGERKRNELLQLALLRPRLAMLDEIDS
GMDVDGVRAAVALIGRLREQGTAFVIVSHYLQMIEALAPDTVLLLDRGRIAESGDLALAR
DIAAKGFARSDALAQA*
>GCA_000452945.1_BPMSHR5858_genomic_04305_1 group_3930
VKIPLATPQPSRQPTGLPVLRLGFRPFYLGGAYFGIVSIALWLASLRGHAVAGLSPAMSG
LAWHVHEMVFGFSAAIIVGFLLTAIRAWTSRETPHGAPLAALWLLWAAGRLLVWAGPEPL
AAVVDSAFLPIAAILLLRVLLAARNHRNVFLTVALFLFGALNALFHGWAAHGRLDLALQA
AYAAVGFVMLFVVVIAGRIVPTFTMNAIPGFTVKRWKWVETLAAPATVLALCADAARLPG
AIVAAVAFAAAALHATRIVGWRSWRVGARPILWILHVAYAWVPVGFAMLALAALDVAPHS
LAIHALTVGVIGGAIVAMITRTALGHTGRPLRAGPAEIACYWLLIAAALVRVFAPWIAPD
ATRVWIDVAGACWVAAFAVYALRYTGYLTAPRIDGKAG*
>GCA_000452945.1_BPMSHR5858_genomic_04307_1 hemN_2
MDMTTQSNDGADTTAPAPQHDVSAFADVQISEALIRRFDRQGPRYTSYPTADRFSDAFDE
RAYREHLSRRASAERNPPLSVYLHLPFCESLCYFCACNKIITQDHTRTSAYVDYLIREME
LVAPDLGRDRRTTQLHLGGGSPTFFAIDELARLMRALREHFDFAPHAELGVEIDPRTVNE
RTLQSLAALGFNRTSFGVQDFDPSVQEAVHRIQPLPMVERALEASRAAGFESVNIDLIYG
LPRQTPASFSRTLDEVIRLSPERIAVYNYAHLPSRFKAQRLIVEAQLPPAEDRLRIFIES
TRRLLDAGYVYIGLDHFAKPNDELGNALRERSLHRNFQGYTTQAECDLVGFGVSAIGKVG
ASYSQSTRSLKTYYRQLDAGRLPIERGFALTADDLLRREVIMTVMCSTPVDFAEIGHRHG
IDFARYFAPELAQLEPYRDAGLLTIDAQRIAVTPKGRMFVRAIGMVFDAYLGRSAAASYS
KLI*
>GCA_000452945.1_BPMSHR5858_genomic_04308_1 glxR_4
MKESKIPVQDFLARLPLFSVLSRGELDNLARGTMRVQVPRGQTVFSRGDPCGGFHMIVYG
QIKLSSFSPLGIEKIVRLLGPGDSFGEAVMFMDKPYAVTAKALSDTLLLHVTKAVVTEEL
DHNPAFARRMLASLSMRLHQLVVDVETYSLRSGTQRVISYLLEQTDAPVESGMQIRLETG
KKAIASRLNLTPEHFSRVLRDLCTRELVVVNGRDIFIPDVDRLRSEMQY*
>GCA_000452945.1_BPMSHR5858_genomic_04309_1 group_3520
MQRKLELDHSPALSVPLRFFVSAPLFALAAAGLLLWAGPDTLASRWSPAALALTHLCALG
MLATTMVGAMIQIMSVAAGIRIAAPVASATAIHACLTLGTLLLAAAFACAAPVLFAPAAL
LLGAAFAWFLLVCAIGFARERRRMPAGAAAVLRTVRLAWLALAATVALGASLGFALGGHL
PVFSVELVDLHAAWGLAGWVGLLTIGISYQLIPMFQATEPYPRAVARGLGPGVFAALVAA
SAVTFGPAGRAGFAADAIRLALYVLYAAFAAWTFHLLRTRKRPEADTTTLFWRTATASVA
GAGALWIANAGMGGRDIAVTTGVLLMMGGAWSAINGMLYKIIPFLLWYHAQKSSETAIPG
MPKVKEMLPERVARPQFFAHVAALLLLVAASVAPAPFARPAAVVLGVSAGWLALNMLGAW
RRYARVRRRVASVLHLRA*
>GCA_000452945.1_BPMSHR5858_genomic_04314_1 group_3521
MNHETKTLPIVYSCSGCSNVAQLANHVAVRLDRGGDAEMSCIAGVGGDVPSLLKIAHSGR
PILAIDGCPLVCAKKSLERHGIAPDAHLQLGEHGVRKRFHEDFAPGDASRILAIAKAEAA
RLAGSRPAPEPELAQAGEDAGAALERKR*
>GCA_000452945.1_BPMSHR5858_genomic_04316_1 group_19744
MTTTLQANPQAARAAIVLSDDERQPCEIWTRVMGYHRPVSSFNVGKQGEFHERKYFRERA
TDEAALRAAA*
>GCA_000452945.1_BPMSHR5858_genomic_04317_1 group_3522
LSETMHPCAAPNPASRDKPLIDVESSINEYLDRQDWRVNANANQGYSLGGLILNVSGKVI
ANYWLSHVYPSAIGEAHRNADLHIHDLDVLSGYCAGWSLRTLLNEGLNGVPGKVESGPPK
HMSSAVGQIVNFLGTLQNEWAGAQAFSSFDTYMAPFVRRDALTYAEVRQSVQELIYNLNV
PSRWGTQTPFTNLTFDWICPEDLREQVPVIAGEEMPFTYGDLQPEMDMINQAYIEVMQAG
DAAGRVFTFPIPTYNITADFDWHSPNAERLFEMTARYGLPYFQNFINSELKPNMIRSMCC
RLQLDLRELMKRGNGLFGSAEQTGSIGVVTVNCARLGYLHAGDERALFARLDTLLDYGKE
SLEIKREVIQHHMNNGLFPYTKRYLGTLRNHFSTLGVNGINEMIRNFTRDAHDLTTDWGH
AFALRLLDHVRARIVEYQEETGHMYNLEATPAEGTTYRFAKEDRRRYPDILQAGTPQMPY
YTNSSQLPVGFTDDPFEALERQDDLQRKYTGGTVLHLYMTEPLSSADACRTLVKRALTRF
SLPYLTVTPTFSICPTHGYLAGSHPFCPKCDEALLRRKLSQSQPMEA*
>GCA_000452945.1_BPMSHR5858_genomic_04322_1 alkB
MAIQQPVSADWTDTKRYWWLLGALTITLPLVAALLALSTGWHVFWWFGPLFAFGVIPVLD
LLIGVDSENPPESIVPRLESERYYRYVVSLATFVVYVAFAGAIWIVATQPLAWYDYLGFA
FSLGAATGISINTAHELGHKTSPLERWLAKITLAPVAYGHFFVEHNRGHHVRVATPDDPA
SARYGESFWAFLPRTVTGSIASAWRLEKARLARSGRSAWSWRNEVLQAWAMTVALWSAAI
AFGGAKVVPFLLIQAAYGASLLEVVNYVEHYGLGRRQLPTGRYERCTPQHSWNSNHVVTN
LFLYQLQRHADHHANPTRSYQALRHFDDSPQLPAGYAAMILLAYVPPLWFWVMNPRVVAH
YGADMAQSNIKPSIRQKVLAELAIANAP*
>GCA_000452945.1_BPMSHR5858_genomic_04324_1 cysC_4
MNMKGGFPPEAVAPVIWMTGLPGAGKTTTANALAERLLEDGAKAIVLDGDALRSGLCADL
GFSDADRMENIRRFAHVAKLFQREGYVVIVATISPLQAHRDLARSIVGEGFFETYVATPF
DVCRSRDPKGMYARAEQGRLMQFTGVSGVYEPPVAPDISIDTTEHSVAFSLAEVMAQLAR
VSALPAPLDAAASAPVAASAPAAAGG*
>GCA_000452945.1_BPMSHR5858_genomic_04325_1 anoR_4
MRAAMGNWAEDLLAGLDSARSEEEAFRSVETAAAALDFEYCAYGLRVPWPLSRPRIETRS
NFPEQWKRRYVEAGFLDVDPILAHGRRSQQPVVLAETLFASAHQMWVEAQSFGLRFGWAQ
SSFDAYGGMGMLALVRSREPVTAAELDAKEYRMRWLVRTAHAALGRMMLPKLMADPERGL
TEREVEVLKWAADGKTSGEISKILAISVDTVNFHVKNAILKLRTANKTAAVVRAAMLGLL
S*
>GCA_000452945.1_BPMSHR5858_genomic_04329_1 mngR
MSTPAYQEIKDFILARIHAGEWAEGDQVPSENELAREFKVARMTVNRALRELTAEQVLTR
IQGAGTFVARPKYESTLVAIRSISDEVAARGHRHHARVLDLVTMKADEALADEMQMPPHT
RLFRSVMLHYENDEPVQLEERWVNPAVAADYADQDFSAITPNQYLMRVAPLQRVEYRIEA
AMPDRAMREALEMADTEPCLVLHRRTWSQGVVASVVNLWHPGSRYRFTGHF*
>GCA_000452945.1_BPMSHR5858_genomic_04396_1 group_3530
MRQRTAKRLPPDADKLVGLSLALFASGSRVEDRFWEAKLDALLAKIVRNGNQTTLDAALD
HLQQNHPDAYGALADMAETHSESLSVEHDGKPYEALLVAIPVLAWTRYMIPSGALKNEIS

DVLRTHLQAHVLAQGTLVAMAPFLYSIDQLPRHHVETYRLAQQLAHAALGNHSVKLNYGD
LPETSPILADPRFLLAVVAAPAGAPLFRWQEEEHGSRIERGQCLEQWAAQGGVNLSAALP
GCEFECLLPDAYYSACRDADERIRPHTVRTAIRYLFDTIGAAPQELRAVLAGFGERRIDE
YRVAFTRRGSNDVIYGVVWPLYGRENGEASVDEATLEAEAPADGPLEEIASLLKEAGVTD
IRRHAGRFEPEYCDDCGVPLYADPLGEIVHAEMPEDATPAQPHFH*
>GCA_000452945.1_BPMSHR5858_genomic_04400_1 group_3531
MTLSITSNFDAGAIDVVSCERADAIRLRVRGDNRSEFAQWFYFRLTGARGERCVMTFENA
NDCAYPAGWRDYRAVASYDRVNWFRVPTSYDGQMLTIDHTPEFDSIHYAYFEPYSEERHS
EFLGAVQQMPQASVVELGRTVEGRPMSLVVLGTPDEAGAAKKKVWIIARQHPGESMAEWF
IEGLVKRLVGWGDWSGDPVARKLYDHATFYVVPNMNPDGSVHGNLRTNAAGANLNREWME
PDAERSPEVLVVRDAIHAIGCDLFFDIHGDEDLPYVFAAGSEMLPGFTEQQRVEQSAFID
SFKRASPDFQDEHGYPPGKYREDALKLASKYIGHRFGCLSLTLEMPFKDNANLPDEHVGW
NGARSASLGAAMLGAILEHVRAFA*
>GCA_000452945.1_BPMSHR5858_genomic_04407_1 group_2530
MATEHDPQSSQADYAAFEGTRAVGDAQRFDVDALAAWLAKHVGGFAGPLAVEQFKGGQSN
PTFKLVTPARSYVLRAKPAPAAKLLPSAHAIEREYRVMAALAGTGVPVAPMLARCDDESV
IGRAFYVMAFVDGRVLWDPSLPGMTPAERGRHYDEMNRVIAALHSIDPQAVGLADYGKPG
NYLARQIARWSKQYLASETEPIDAMRRLIDWLPQHLRPESGRDARVSIVHGDYRLDNLIF
DAHAPRVLAVLDWELSTLGDPLADFAYHCMAWHVAPERFRGIAGLDLRSLGIPDEAQYVA
RYCERTGLTMPENWNFYLAYNMFRIAAILQGIMKRVADGTASSAQALDAGRRARPMAELA
WRYAQHAR*
>GCA_000452945.1_BPMSHR5858_genomic_04408_1 bbsG_1
MHFDYSAKVEALRARLGAFFDEHIYPNERAFYEEIARNRRAGDAWRPVELIETLKAKARA
AGLWNLFLPDSARGAGLTNLEYAPLCEIMGRVPWAPEVFNCNAPDTGNMETLERYGTDAH
KAAWLEPLLDGVIRSAFLMTEPEVASSDATNIRTRIERDGEHYVINGHKWWSSGAGDPRC
KLYIVMGKTDPDAPRHAQQSMMLVPSDAHGVTVHRPLNVFGYDDAPHGHMEITLENVRVP
ASNLLLGEGRGFEIAQGRLGPGRIHHCMRLVGLAERALELMCRRASERIAFGKPVAAQTV
TQERIAEARCMIEQARLLTLKTAYMMDTVGNKGARGEIAMIKVVAPNMACQVIDWAIQAH
GGGGVSDDFPLAYAYASARTLRFADGPDEVHRNAIAKLELARHAPRSA*
>GCA_000452945.1_BPMSHR5858_genomic_04410_1 group_2531
MMNRSQQTFAGFAAACAFAAGSAAQAWAAPADAIVYPAGNGSDQADALQAALDALQTGQR
LVLAPGQYVVGRSLLVKNAQVVVSGYGATLVATNDADQTIEMRGRDSTLVGVTLVGTGAT
RLTTPESTKVDVTGAGVQVLDVEVRGGASAGIFVFGGTDVAVVGNTVRDTLADGIHTTYG
SRNVLVRDNTVQNTGDDMVAVVSYKGDGKLSSNVLIQNNTLLGNYWGRGATVVGGADVTI
TGNTVRNVQKAAGILVGQEDPANTYDARNVIVSNNTISDIEWPDPDNTRPPAYMAAIDVN
TWSGKATSVSVTDNRISRARYAGVRALGNVCQLRVSRNALASIDGTPIALQQAPSCAAGQ
IVCASNTLDGAALASPAGCSATDGLTITGANVARMPQVRAYLRQTAAPSAAKAGSAAASG
E*
>GCA_000452945.1_BPMSHR5858_genomic_04411_1 group_19827
MTARVHALPVVASADALHARVGAAPFASGWIDIDQTRVDRFADATGDHQWIHVDPERAAR
ESPFGGPIAHGFLTLSLIPALMTDALRFEQRMGVNYGLNRVRFLKPVPVGSRVRALFAVK
GTGPAKQGGVQATWAVSVEVELGGQPVPACAAEFITLHYF*
>GCA_000452945.1_BPMSHR5858_genomic_04413_1 acrC
MNFTFSEEQQQFAHALRRYLDERYGFDARRAIIHSDAGVCDEQWRAFAELGLTALPVPEA
HGGFGGGAVDMLVATQELGRALVVEPYWATAVGIEALRVAGSADGEDAALLARAAQGDAK
LAVAFHEPHARHDLFAIETLARPDGDGFALTGVKSVVRHGAQADVLIAPARLPNGAIGLF
AIARGAAGADIVDYRTIDGQRAATVRFSDTPARALVGGERDAAALERIADYGVVLLCGEA
IGALDALNHATLDYTKTREQFGVPIARFQALQHRMVDMLIHAEQARSLTYLAAVRYASDD
ANARRKAVSAAKARVGQAARFVGQQAVQLHGGMGVTDEVAAAHLFKRLSIIETTLGDVDH
HLARFASLPDFALLQDA*
>GCA_000452945.1_BPMSHR5858_genomic_04414_1 group_19833
MDLDYSPADDAFRAEIRAWLEANLPRALRDKVLNHKRLNRDDFAGWHRLLGTRGWSAPAW
PAEYGGPGWNATQRHIWEEECARIGAPPVLPFGVSMVAPVLMKYGSEAQKRRYLPRILDG
TDWWCQGYSEPGSGSDLASLRTRAERRGDHYLVNGQKTWTTLGQHADMMFCLVRTDPDAK
KQEGISFLLIDMKSPGITVRPIVTLDEDREVNEVFFEDVKVPVENLVGDENRGWTYAKYL
LGHERTGIARVGASKRELDFLKRIASRERKNGKPLIADPVFAAKIAALEIELMALEVTVL
RVVSSEASGRGPGPEASMLKIKGTEIQQALTELMVDAIGPLAAPFDAAFLEGERERSIAG
DDDAAPLAAYYFNYRKTSIYGGSNEIQKNIIAQMILGL*
>GCA_000452945.1_BPMSHR5858_genomic_04415_1 group_19836
MKVLDLQCPHGHRFEGWFASADEFEAQLSRKLVECPVCGATEVSRMPSAPRLNLSGATQA
QPADPRALQAQAMRALREVLAKTENVGERFAEEARRIHYNEAPARSIRGVTTPEDARALA
EEGIDVMPLPVPAALKEPLQ*
>GCA_000452945.1_BPMSHR5858_genomic_04416_1 act
MADLPNHDATLTETCIESESVYDGAFLKVKRDTVRLPDGKHATREYVTHPGAVMVIPLFD
DGRVLMESQYRYPIGKVMAEFPAGKLDPNEGALACAVRELREETGYTAREYVFLTRIHPI
ISYSTEFIDIYLARGLTAGERKLDDGEFLETFTATLPDLLEWVRTGRISDVKTIIGTMWL
EKAVSGAWPLGDVIRP*
>GCA_000452945.1_BPMSHR5858_genomic_04418_1 nuoN
MQNAPMNVLLPDALVMAAIIVAWLNDTFVGAAGRRLTYLIAVVASAAAGVWFALQALDPQ
QYYFFSRMVVVDSFASAMKAVVSIGFAVTLVYSRKYLEDRDLFRGDVFLLGMFSLLGQLV
MISGNHFLTLYLGLELMSLSLYAVIALRRDAAQSSEAAMKYYVLGALASGFVLYGISMLY
GATGSLELNEVLKAVASGRINDAVLLFGVIFIVAGVAFKLGAVPFHMWVPDVYQGAPTAM
TLFVGGGPKVAAFAWGLRFLVMGLLPLAVDWQEMLVILAALSLIVGNLTGIVQRNIKRML
AYSAISNMGFVLLGLLAGIVNGNPSAAASAYSSAMFYTIVYLVTTLGSFGVVMLLARRDF
EAETLDDFKGLNKRSPVFAFVMMVMMFSLAGIPPTVGFYAKLAVLEATVNAGLTWLAVLA
VVTSLFGAFYYLRIVKLMYFDAPQDTTPIAGDACKRAILVLNGLAVVVLGIVPGPLMTIC
LQAISHTLPL*
>GCA_000452945.1_BPMSHR5858_genomic_04419_1 nuoM
MHSFPILSTAIWLPIVFGLVVLAVGSDKHPGAARWIALIGSLAGLAVTIPLITGFDSSTA
ALQFVEQSTWIERFNISYHLGVDGISMWFVVLTALITVIVVIAGWEVITENVSQYLAAFL
ILSGIMIGVFSAADGLLFYVFFEATLIPMYIIIGVWGGPNRVYAAFKFFLYTLAGSLLML
VALIYLYTQTHSFDLATWQNAKIAMTPQVLLFIAFFLAFAVKVPMWPVHTWLPDAHVEAP
TGGSVVLAAIMLKLGAYGFLRFSLPITPDASHFFAPVVISLSLIAVIYIGLVAMVQADMK
KLVAYSSIAHMGFVTLGFFIFNQLGVEGAIVQMISHGFVSGAMFLCIGVLYDRLHSRQIA
DYGGVVNVMPKFAAFAMLFSMANCGLPGTSGFVGEFMVILAAVQFNFWIAFGAAFTLILG
AAYTLWMYKRVYFGAVANDHVAKLKDIGRREFTMLAVLAAFTLLMGLYPKPFTDVMHVSV
ENLLSHVAQSKLPLAQ*
>GCA_000452945.1_BPMSHR5858_genomic_04420_1 nuoL
MSTTLNENLLLAIPLAPLAGSLIAGLFGNAVGRKGAHRVTILGVALAFVLSALVFFQVLD
GASFNATVYEWMNVGSLKLEVGFLVDTLTAMMMVVVTFVSLMVHIYTIGYMSEEDGYQRF
FSYISLFTFSMLMLVMSNNFLQLFFGWEAVGLVSYLLIGFYFTRESAIYANMKAFLVNRV
GDFGFLLGIGLILAYAGSMNYGEVFAKRAELASLHFPGTQWGLLTVACICLFIGAMGKSA
QFPLHVWLPDSMEGPTPISALIHAATMVTAGIFMVTRMSPLFELSDAALSFITVIGAITA
LFMGFLGIVQNDIKRVVAYSTLSQLGYMTVALGVSAYPVAVFHLMTHAFFKALLFLGAGS
VIIGMHHDQDMRNMGGLRKYMPITWITSLVGSLALIGTPFFSGFYSKDSIIDAVKLSHLP
GSGFAYFAVVASVFVTALYSFRMYFLVFHGEERFRHPKTFGNNHGAESAAHHGHDDHGHA
HEPHETPWVVWVPLVLLAIPSVIIGAIAIGPMLYGDFFQHGVAFDKVIFIGQNHPALAEM
AEEFHGWTAMGLHSVSGLPVWLALAGVVVAWFLYLKRPDLPAAIRRAFGPIYTLLDNKYY
MDKINEVVFAKGSIAIGRGLWKEGDVVVIDGLVNGSARFIGWFAGVIRFLQSGYIYHYAF
AMIIGMLGLLTLFVTLGGK*
>GCA_000452945.1_BPMSHR5858_genomic_04434_1 tpiA
MNTTPGFGASTTRQTMSKQRIKRVIGNWKMHGRLSGNEALLDEVAQGAQAVPEHVGIGVC
VPFPYLAQARARLHGGRVAWGAQDVSAHEQGAFTGEVAAGMVAEFGAAYAIVGHSERRAY
HGESNEIVAAKAQRALAAGLTPVVCVGETLAEREAGATEQVVGVQLDAVLAVLSQDEAVR
IVVAYEPVWAIGTGRSATAEQAQQVHAFLRGRLAAKGAERVSLLYGGSVKADNAAELFGQ
PDIDGGLIGGASLKSGDFLAICRAAQ*
>GCA_000452945.1_BPMSHR5858_genomic_04435_1 ppsC
MKAIEITEFGAPDVLKLTERPQPEPKRGEVLIKVAASGVNRPDVFQRKGAYAPPPGASDL
PGLEVAGEIVGGDLADSAANPFGLKLGDRVCALLAGGGYAQYVAAPLAQCLPVPKGLTDV
DAASLPETFFTVWSNVFDRAQLGAGEGGAQETFLVQGGSSGIGVTAIQIAHALGFRVFAT
AGTDDKCRACEALGAERAINYKSEDFVEVVKSLTHDRGVDVILDMVAGAYVPRELAALAD
GGRLVVIALLGGAKAEINVAEILRRRLTITGSTLRPRPVEFKAAIAAQLKARVWPLIEDG
RVKPVIYRVLPAAQAADAHALMESSEHVGKIVLDWGANA*
>GCA_000452945.1_BPMSHR5858_genomic_04437_1 rpsO
MSVADIKKSEVVAQFARGANDTGSPEVQVALLTARITELTGHFKTHAKDHHSRRGLLRMV
SRRRKLLDYLKGKDADRYRALIEKLGLRK*
>GCA_000452945.1_BPMSHR5858_genomic_04438_1 group_19965
MSSYWMRFAAAASAALTLALPMPFAAAAATGEPIRIALVEGMSGPFANAGAAVERNLRFG
IERVNAQGGVRLRDGAHPLELVVLDSKGSVEEALVQLRAATDKGIGFAAQGNGSAVAAAL
VAALDKHNARDPEHRALFLNYSADDPALTGRDCSFWHFRFDAHAGMRMDALADVLARDRA
VKKVYLLNQDYSFGHDVSALARAALKARRPDIAIVGDEFHPIGRVKDFSPYAAKIRASGA
DAVVTGNWGNDLTLLVRATREQGLAAKFYTFYGNSLDAPAALGDAGVKRVLAVADWHPNA
GGARSDAFYRAFRARFPAAQDDYPVRRMSEMIEMLAAAMTRAGSADPVAVAKALEGMRYD
DGFHPAQMRAADHQLIQPLYVIEMERAGAPGVRFDNAGSGYGFATVLEAPPTQGAAPAEC
RMKRP*
>GCA_000452945.1_BPMSHR5858_genomic_04484_1 yhdE
MKLHIVAVGHKMPGWIASGFDEYAKRMPPELRIELREVKPELRSGSRTADSVMAAEQQRI
EAALPKNARVVALDERGRDWTTMQLAQALPAWQQDGRDVAFVIGGADGLAPALKSRAELL
LRVSSLTLPHGMVRVLLAEQLYRAWSITQNHPYHRA*
>GCA_000452945.1_BPMSHR5858_genomic_04486_1 rng
MNEEILINITPQETRVALVQQGAVQELHVERTLSRGRVGNVYLGKVVRVLPGMQSAFIDI
GLERAAFLHVADIWQPRAGDAPAAATHQPIEKIVFEGQTLMVQVIKDPIGTKGARLSTQI
SIAGRTLVYLPQEPHIGISQKIESEAEREAVRARLTAVIPSDEKGGYIVRTIAEDATSDE
LAADVAYLRKTWATIVAQGQRVPPTSLLYQDLDLAQRVLRDFANDDTTRIQVDSRETHQR
LVEFAAEFTPAVSPKLHHYTGERPLFDLYNIEAEIQRALSRRVDLKSGGYLMIDQTEAMT
TIDVNTGGYVGARNFDDTIFKTNLEAAHTIARQLRLRNLGGIIIIDFIDMENAEHRDAVL
AELKKALSRDRTRVTVNTFSQLGLVEMTRKRTRESLAHVLCEPCPTCQGKGQVKTSRTVC
YDILREILRESRQFNPREFRVIASQQVIDLFLDEESQHLAMLIDFIGKPVSLQVESNLSQ
EQYDIVLM*
>GCA_000452945.1_BPMSHR5858_genomic_04495_1 group_3542
MAHAPRLPYPKLASKPFNALLHLSETVWGGSLGKPLVDLVFLRVSQINGCAYCIDMHWRD
LVKLGVNERHLNAVAGWREAPFFDARERAALQWAESVARIPHTDPSDEAFAQLQAHFSDA
EIAELGFAIATINAWNLLNVSFRNPIPETV*
>GCA_000452945.1_BPMSHR5858_genomic_04496_1 group_358
MSGGFPRSAFRLPRPVAAAAPVVLATLALSGCALFRAPQASAPVVEAVAVPVEPASEPIA
APEPASAPEPAETAPPRKPHREAAPPRKPARVAPPAPAPAPAPAPLVTTRAIERSQVHAL
LDSEVRRGGKVIGRAVDMTADANGAPREMLVNLQGFMGVGDRKVSFPWKLFRFTPGGRHE
PVILDMPATAQLQPADRPKAVPLTGSTQAGAEPGQMRIIDADVERPNGAKVGRVVDVLIG
RDAQPQAVVLDVGGLVDPDRRTIAANWSALRFAPKDKSLRALLDLNDAQLKASPPYAGDK
PILAVSPAAGGAPAAAPATARAGAKR*
>GCA_000452945.1_BPMSHR5858_genomic_04498_1 group_2538
MRRASPAPCLNEMETIKAFNEARRQIQESVLELFKGLSLRERLLQGVLMAVQAASGACLA
YGIGRALHTEQAVWAAITAIAVTQHNYADTINLSRDQFIGAMVGGLIGFAGAATGAGHFV
AYAITIVTAIICCWCLNVGSAARLGAITATIVLLFPGEGPLWDIPLVRLGEVTLGTACAM
AVGWTMSRIERRWFAKR*
>GCA_000452945.1_BPMSHR5858_genomic_04499_1 group_2539
MKAHRTLRISMLAAGALVSSLAAAQPHGPGGPGDERHGPPPGKHVGRDHGKHGDDGARPG
EHRGDENAAWRRGERLPDEYRDRQYVIDDWRGYHLSPPPRGYHWVGIGGDYLLVRISTGV
ILQIGP*
>GCA_000452945.1_BPMSHR5858_genomic_04518_1 group_3546
MSQRILSPLVASCAAAALLSAFAAHAAPPIKGSVLGGNDGQLQYTVKVDSKQFGTMQETR
KVRSGETDDYNWKSVPPSGAVPMPDACPNADTLPRDANGAMVRQAQVRLAPAVDSKGVAN
VQLSFQASAPSGTKKVTVNGKALQCPNVVAVSQVKWLSIPTGGSKSITMRDGTKITVSIK
R*
>GCA_000452945.1_BPMSHR5858_genomic_04520_1 metC
MTASISKRALQTRVVQPDDRIPAGFESFVVPVARASTVVFPDLATMRNLDWRQDGQWRYG
LHATPTSLALAQRLAEIEGGAHALLQPSGLCAITNVYFGLVKEGDDVLIPDNAYGPNGDF
GKWLAKDFGITARFYDPMIGAGIADLIQPNTRIVWLEAPGSVTMEVPDVRAITAVARARG
IVTAIDNTYSAGLAFKPFEHGVDISVQALTKYQSGGSDVLMGATITADGELHAKLKLARM
RLGIGVSADDCSLVLRGLPSMQARFDAHSRSALSLARWLKTRAEIATVLHPQLPDCPGHD
TFMRDFTGAGGLFSVVFDERYDAAQVDRFVEALELFAIGWSWGGACSLAMPYDVQSMRTG
KWPHRGTLVRFYVGLEDEADLRADIERALTSALA*
>GCA_000452945.1_BPMSHR5858_genomic_04522_1 bktB
MQREVVVVSGVRTAIGDFGGSLKDFAPTELGARVVREVLSRAQVSGDEVGHVVFGNVVHT
EPKDMYLARVAAINGGVAQHAPALTVNRLCGSGLQAIVSAAQNVLLGDADIAVAGGAENM
SRAPYSVPAARFGQRMGDAKLVDMMIGALNDPFQSIHMGVTAENVAAKYGITRDAQDALA
LESHRRASHATKSGYFKDQILPIEIASRKGTVVFDADEHVRHDASLDDFTKLKPVFAKEN
GTVTAGNASGINDAAAAVVLMERGVADKRGAKPLARLVSYAHAGVDPAYMGIGPVPATKK
ALERAGITVADLDVIEANEAFAAQACAVSNALGLDPAKVNPNGSGISLGHPIGATGALIT
VKALYELQRIGGRYALVTMCIGGGQGIAAVFERL*
>GCA_000452945.1_BPMSHR5858_genomic_04523_1 kdgK_1
MPATFPDVLALGEAMVEFNQARPGEPNYLQGFGGDTSNFCIAAARQGARAGFVSAVGGDH
FGRLLLDLWRGERVDTSYVRVDDGAPTGVYFVSHGEHGHAFDYLRAGSAASRYAIRDLPL
DALAAARFVHLSGVSLAISTSACDAAFAAIAHARANGARVSFDTNLRLKLWPLPRARAVM
TEALRHTDVCLPSWDDVTALTGLDERDAIVDALLALGPRVVALKLGREGAYVATPGERRV
VPGFVVDALDATGAGDCFGGAFVARLAAGDDPFSAARYANAAAALSTQGFGAVAPIPSRD
AVERLLGA*
>GCA_000452945.1_BPMSHR5858_genomic_04524_1 rimO
MVSLGCPKALVDSEQIITQLRAEGYEISGTYDGADLVVVNTCGFIDEAVQESLDAIGEAL
AENGKVIVTGCLGAKKSASGSGLIAEVHPKVLAVTGPHAVGEVMQAVHSHLPKPHDPFVD
LVPAAGIKLTPRHYAYLKISEGCNHRCSFCIIPSMRGELVSRPVAEVMLEAENLFKSGVK
ELLVISQDTSAYGVDVKYRTGFWNGRPLKTRMTELVGALGELAAQYGAWVRLHYVYPYPH
VDEIIPMMAQGPLKGHVLPYLDVPFQHAHPEVLKRMKRPANAERVLERVQKWREICPDLT
IRSTFIAGFPGETDAQFETLLDFIREAELDRVGCFAYSPVEGASANALDGALPDDVREAR
RARFMEVAEEVSAARIARKIGKTLKVLIDEVNAEGGIGRTAADAPEIDGVVYVEPAAKAS
KRYKVGEFVSVKITGADGHDLWGEV*
>GCA_000452945.1_BPMSHR5858_genomic_04531_1 group_19821
MTRSSSLNARAGKSNRDDYSPSASPADASAGDSLDDDLTSDTLVAPARADTGADAPRTVD
VPLALAGERLDKALAQLFPEFSRSRLQQWIESDRVRVSGAPAKIRQPVPLGATIELLPEL
LPEQLAFTPEPVPLSIVYEDDTIVVVDKPAGLVVHPAAGNWSGTLLNGLLHRYADAAGLP
RAGIVHRLDKETSGLMVVARTLAAQTDLVRQLQARTVKRRYFALAWGRMPDEGTIDAPIG
RDPRERTRMAVVTGAAGKPARTHFRTVDTTLWARQPVSAIHCDLETGRTHQIRVHCAHVG
HPLLGDPVYGRARGKRSVAPLPGGFARQALHAWRLGLVHPASGRSMQWRAPLPADLAALV
DALGFGADDAEFGDDDAVYDDGYDEGMPYADDDED*
>GCA_000452945.1_BPMSHR5858_genomic_04532_1 bamD
MMHSTKRVAKPAAAWAALAAAAALVAGCHGLPQKSDETATWSNNKLYSEAQDALTGGDWG
KCAKYFEALQGRDPFGHFAQQAQINVAYCNWKDNETAAADQAVDRFIQLHPDHPDVAYAY
YLKGMIHFNDDLGLFGRFSGQDMSERDPQALRESYDAFKVVVDRYPKSKYAPDAAARMRY
IVNALASHEVHAADYYYRRGAYVAAINRAQLAIKEYKNAPAIEDALHIMMLSYAKLNQPQ
LADDTKRVLASTFPDSPYVTGHARPGSKKSWWQF*
>GCA_000452945.1_BPMSHR5858_genomic_04534_1 group_19854
MQTRSAETQQELYNRLVSLQELHQQLAREIEQKEGRSDIDDLTLHRLKKEKLLAKDKIIM
LQSQLEPDKRA*
>GCA_000452945.1_BPMSHR5858_genomic_04535_1 group_2542
MIRSFPTTLPTSRFSVVAFAVALAATLSCAPVAGVAQTPAPAATSPAAAADAGAFAAPAN
FDARQKVLNQRTAENDYRYAVAQHNCYSTFFVNHCLDKARETMRAERANISQEQLALDDE
QRAVRAQERDRQLALKEAERQAQAPQRAADEQANAAAYEGKQQENALKRAQRDAQAPQRA
ANQQAYDAKQAEFQRKLDAAHAQAAKDAQQRAENAARYEQKQRDAAQHKADVDERQRQAA
EKAKQKQQ*
>GCA_000452945.1_BPMSHR5858_genomic_04540_1 imuB
LLGIHLPRLPLDVCAPPSPDGGNGCAVLEQGVVLIADAPARANGVRAGMKRGGVLTLAPG
TRLVERDPAREADALRAVALALLRFSPCVALDDEATLVVDVGASLRLFGGLPLLCRQVRA
TLAALGYAARVAAAPTGRGAWLLARAPGKPGTRRRVVRAASLARALDALPCELLPDARPY
AGWFDGLGCRTLADLRRLPRAGLTRRCGPALVAALDRAYGDAVEPLAWMPVPPVFDARLE
LPERVEYAQAVLFVARRLVVQLCGWLAARQLSLSAMVFALEHERGRQAVPPTPLELAFAE
PARDETHFMRVLGERLARVALPAAVIAVRVTATRVESVAPPADDLFPEPGGTREARARLV
ELLSARLGADNVLRAAPVADYRPEAANRWLPLDAPTGKPVAPGVAPPRPAWLLAPPQPLA
MRGNRPFYRTPLRFVSSIERIEAGWFDGQLVERDYRIAQDDYGACYWIYQERGNRAAGQC

WFLHGLFG*
>GCA_000452945.1_BPMSHR5858_genomic_04542_1 ecfA2
MLSAQDLKLTFNPGTPIETRALRGLSLDIPDGQFVTVIGSNGAGKSTFLNSVSGDQPVDS
GRIAIDGVDVTRKPAWDRAPLVARVFQDPMAGTCEALTIEENMSLAMARGARRGFLPALN
RASRELFRDKLRLLNLGLENRLTDRIGLLSGGQRQAVSLLMASLRPSRILLLDEHTAALD
PKTAAFVLELTARIVDESKLTTMMVTHSMRQALDYGDRTVMLHQGQVVLDVAGDERRGLD
VPDLLQLFEKTRHEKLDDDALLLG*
>GCA_000452945.1_BPMSHR5858_genomic_04546_1 metQ_1
MKRRTLLKVVSAVAAGAAALSVSVGAQAQDKVIKVGTVAGPDAQVWQVVQKVAKEKQGLD
VKVIEFNDYVQPNAALDSGDLDANSFQHQPYLDSQVKQRGYKIVSAGLTYISPIGVYSKK
FKSLKELPAGAKVALPNDPSNENRALLLLQTQGVIKLKAGAGTGGSNATVLDVAENPKKL
KLTELDAAQLPRVLSDVDAAVINTNYALAANLQPTKDAIALESLTSPYANLIAVRAKDKD
QPWVKKLVKAYQSPEVKEFITKQFKGSMVASF*
>GCA_000452945.1_BPMSHR5858_genomic_04555_1 ctaE
MTGQNESPYYFVPHPSQHPISAAVGLLIMLGSFALWVNGEPWAPYTALVGLLWLLFVLYH
WFGDAIAESEGGMYGKRVDKSYRWSMSWFIFSEVMFFGAFFGALFYAREIAMHQLGSLDY
KLIWPDFSAVWPNEGPGALVGHFKTMGPWPIPTLNTALLLSSGATLTVSHHALREDHRRK
AIAWLAATLVLGVCFLFLQGFEYFHAYNELNLTLGSGVYGSTFFLLTGFHGFHVFLGGTM
LAVVLARMIRGHFKPDHHFAFEGAAWYWHFVDVVWLGLYVVVYWL*
>GCA_000452945.1_BPMSHR5858_genomic_04563_1 trmL
MFNVVLVEPEIPPNTGNVIRLCANTGARLHLIEPLGFPLDDAKMRRAGLDYHEYAQMRVH
RDWDAFVAAEAPDPARMFAFTTRGSGRFHDRAFEPGDWFVFGAETRGLAPALVDRFAPEQ
RVRLPMRPGNRSLNLSNTVAVVVFEAWRQAGFEGGA*
>GCA_000452945.1_BPMSHR5858_genomic_04566_1 secB
MSDVENQPFFNIQRIYLKDLSLEQPNSPAIFLEQEMPAVEVEVDVKAERLAENVYEIVVA
GTVTAKVREKVAFLVEAKQAGIFDIRNIPAEQIDPLCGIACPTILFPYLRSNIADSITRA
GFPPIHLAEINFQALYEQRLAEISQQQQQGGAPNGTTLN*
>GCA_000452945.1_BPMSHR5858_genomic_04567_1 grxC
VNKVVMYSTQVCPYCMQAERLLKLRGVEHIEKVLIDKEPERRAEMMERTGRRTVPQIYIG
DTHVGGYDDLSKLDREGGLKPLLEAA*
>GCA_000452945.1_BPMSHR5858_genomic_04579_1 ubiK
MKQPSDVFNDLQARIGDLLKNSPAKDVERNVKAMLTQGFSKLDLVTREEFDTQAQVLART
RARLEELEKRVAELEQKLADVQS*
>GCA_000452945.1_BPMSHR5858_genomic_04580_1 comM
MSLAVVRSRAPAAGRAPEVTVEVHLANGLPSFSIVGLPDLEVRESRERVRAALQNCGFEF
PVRRITVNLAPADLPKESGRFDLPIALGILAASGQLPAGALDGREFAGELSLTGALRPMR
GAFAMACGTARGQQADDDGSPAQAESPPARATVPNAAELYLPLPSAAEAALVPGVTVYGA
ADLPALCAHLADTPDGRLAPVAAPRLDALPAAATADLADVIGQAGAKRALEVAAAGGHHM
LMIGPPGAGKSMLAARLPALLPPLTDDEALTSAAILSSSRAGFSPAQWRRRPFRAPHHSS
SSAALVGGRNPPQPGEITLAHLGVLFLDELPEFDRHVLETLREPLEAGRITISRAALQAD
FPAACQLIAAMNPCPCGWRGDPGGRCRCSPDIAARYLRKLSGPLLDRIDIQIEIPALTPA
ELAERPAAAGESSETVAARVALARERQFARQRKTNHMLTGRETDTLCRPDSAGETLLRDA
GERFRWSARAYYRVLKVARTIADLAGDDAPSAAHIAEAIRYRRAFAPA*
>GCA_000452945.1_BPMSHR5858_genomic_04583_1 group_4109
MPPGVRARRAAVVLGALASLAACSPAYDWRTLHNDAGYTIDLPAKPTVEEHPVAIGGVPM
KMRMQAAHVGGAVFAVGTVMLPDERDATQHAVLDYLRTGLARNLRARPDERAVLVPLAAG
GQAAGVELRMSGAPAGAAGAPGADRQADRASGAHAPKTVIARLVARGRHVYEVIVIADAP
LASEPLEQFFGSLKLD*
>GCA_000452945.1_BPMSHR5858_genomic_04584_1 resA_1
MSPAPAPTSRRAGPLRYVAIAVAAVAIAVAGYFAFNGKSSAPEATFTLLSGQKLSTAADL
KGKVYLVNFWATSCATCMQEMPQMVDTYNRFKGRGLEFVAVAMNYDPPMYVANYAQTRQL
PFKVALDDGSVAKQFGNVQLTPTTFVIGKDGKILKRYVGAPQFAELDQLLQKALDTSA*
>GCA_000452945.1_BPMSHR5858_genomic_04585_1 dctD_1
MANRLQVIYIEDDELVRRASVQSLQLAGFDVVGFGSVEAAEKAIVGDATGVIVSDIRLPG
ASGLELLAQCRERTPDVPVVLVTGHGDISMAVQAMRDGAYDFIEKPFAAERLTETVRRAL
ERRALVLENHALRRELAGQGVVAPRIIGRSPAIEQVRRLIANVAPTDASVLINGDTGAGK
ELIARSLHELSPRRDKPFIAVNCGALPEPMFESEMFGYEPGAFTGAAKRRIGKLEYASGG
TLFLDEIESMPLALQVKLLRVLQDGVLERLGSNQPIRVNCRVVAAAKGDMSEHVAAGTFR
RDLLYRLNVVTIALPPLAERREDIVPLFEHFMLDAAVRYGRPAPLLTDRQRASLMQRDWP
GNVRELRNAADRFVLGVTEGIVGDAGPETDEHAEQSLKERVEQFERAVIAETLNRTGGAV
ATTADKLHVGKATLYEKMKRYGLSAKGETDR*
>GCA_000452945.1_BPMSHR5858_genomic_04586_1 dctB_1
MNAETAGRAAAPAPGGAGPDATRRAWPGPAAFSGGGYATIDDPYSPEAVTVKRRLLVVLA
LAAMLAAACALTWTLTWQSGIADLRDSAAARVDRTTNALKSTLDRYESLPYLLGGHPFVQ
DVLADPANRRQVNRANAYLEDMNRHAHATATYVITASGLCVAASNWRGPDSFTGIGYQFR
PYFIDAVKGGTGRFFGIGTISRDPGYYISQPVRRDGKIIGVAVVKLNLEWFQGADASEPL
VVADDHGVIFLSSVPQWKYHTLKPLPGPIATSIHETRQYAHYPVTPLPLRVERVLGPDAQ
IVRMGAARRAPRYLATRRLLGEPDWQLVTLAPIEPVDADARNATIVTAFGFVSLSLLAFY
WRTRRARVKEMLRSRVLLQSAYAELNRRVEERTADLSQANARLKKEVGERIRAEQELRAA
HDELVQASKLAALGQMAAGITHELNQPLAALRSFSDNTRVLLDRGEQAAARENLEAIAAL
TERMGKITNQLKLFVGRAKPRNERAHVARALRNVLALLKERMKDVELVITLRDETRDPAA
AARFDPSRDEPALIARCEDLRLEQVLINLLGNALDAVAAVDAPRIDVAIDATAATLAITV
RDNGPGIAPESLARLFEPFFTTKEMGRGLGLGLAISSSIARDAGGSLTARNAPAGGAEFV
LTLRRARTHHPDTSAAS*
>GCA_000452945.1_BPMSHR5858_genomic_04588_1 actP_1
MLRQSTARCARLMPAALFACAALAPAAARAMGAAAAPMPEKVVPNPVAIGMFFVFVFATL
ALTRWAARRTRSARDFYTAGGGITGLQNGLAIAGDYMSAASFLGLSGMVFMFGFDGLIYS
IGFLVGWPFVMFLIAEPLRNLGKFTFVDVVAYRFAQRPIRLLTSASALTIVVQMVGAGKL
IQLLFGLSYGVAELIVGVLMVVYVFFGGMTATTWVQVIKAVLLLAGATLLAALALGEFGF
SLDEMFRRAVAVHPGALGIMGPGKLIRDPANALSLGIALMFGTAGFPHILMRFFTVPNAK
EARKSVLYATGFIGYFYLLTFVIGFSAIVLLAQHPEFFRHDAAGAFNLTRDLVGGSNMVA
VKLAQAVGGNWFYGFIAAVTFATILAVVAGLTLAGATTISHDLYAQMWARGKPHERREMR
ISRAATLALSAVAIGLSILFEHVNVAFMVGLVAAVAASANFPVLATSIFWRGMTTRGAVA
GGGLGLASAVALTVLSKSVWVDVLHHAHAPVFLDNPALVSVPLAFAGIVIGSLTDRSERA
RRERDAFARQEFYAQTGALATRAATH*
>GCA_000452945.1_BPMSHR5858_genomic_04589_1 yjcH
METSVIETVVACRDYQQLVRARRRFSFTLTALMIATYYGFILLVALAPHTLAAPLYAGAT
ISVGVVAGVAIIIVAIGLTAGYVLRANRSFDRAVDALFNRS*
>GCA_000452945.1_BPMSHR5858_genomic_04591_1 Acd
MDFRHTEDRRMLADSLNRFIAEQYAFPVRDRIALSPAGYSDDMWRRFAELGAIGALFPQE
CGGFGGAGFDIALVFECLGRGLVVEPFVGALMAGRALAQAGGRAHLERLAALIEGRAIGA
FAHAEPDTHYEPHTVRTRAARDGEGWVLDGAKAVVDCGEHATFFVVSARIAGGDDDAAGL
SLFVVPRTAPGVTVRGYRKIDGGRAADMRLERAALADDACVGKPGEAADAIERAIGFGLL
ALAAEALGAMDVAKAHTLEYLRTRRQFGVPIGSFQALQHRMADLLLEIEQARSAVINAAA
QLDAPRVARERALSAAKYSVGRIGALVAEESIQLHGGIGMTWELPLAHYAKRLVMIDHQL
GDEDHHLARYAVLSRQ*
>GCA_000452945.1_BPMSHR5858_genomic_04595_1 smc_2
VRLSSIKLAGFKSFVDPTHFQVPGQLVGVVGPNGCGKSNIIDAVRWVLGESRASELRGES
MQDVIFNGSTTRKPGSRASVELIFDNSDGRAAGQWGQYGEIAVKRVLTRDGTSSYYINNL
PARRRDIQDIFLGTGLGPRAYAIIGQGMIARIIEAKPEELRVFLEEAAGVSKYKERRRET
ENRLHDTRENLTRVEDIVRELGANLEKLEAQAVVATKYKELVADGEEKQRLLWLLRKNEA
AAEQDRQRRAIGDAQIELDAQTAKLREVEAQLETLRVAHYSASDAMQGAQGALYEANAEV
SRLEAQIKFIVESRNRVQAQIAALVAQQEQWRAQADKAQGDLEAAEEARAVADEKAAIAE
DDAAAKHDALPALEARWRDAQTGLNDERGRIAQTEQALKLEAAHQRNADQQLQQLQQRHE
RLKVEAGGLDAPDEAQLEELRMQLAEHEAMLAEAQARLADAQEALPRLDAQRRAAHERVQ
AESAQIHQLEARLAALKQLQENVQTQGKIQPWLDKHELGALPRLWKKLHVEAGWETALEA
VLRERLAALEVSNLDWVKAFATDAPPAKLAFYAPPAAGEPLAAPGALRPLLPLVRIDDAG
LRAVLNDWLGTVFVADDLAQALAARMQLPQGGAFVVKAGHVVTRSGVQLYAADSEQAGML
ARAQEIENLTRQVRAQALLSDEAKAAAIRAEAAHTQASQALTEVRAQAERATQRVHALQM
DVLKLTQAHERYTQRSTQIREELEEIGAQIEEQRALRAESEANFERHDAELAELQARFED
NQLAFESLDETLTNARQEARERERAATDARFAARQSANRIDELKRSIQVAHEQAERVAAS
LEDARAELETINEQTAHTGLQDALEVRAAKEQALGAARAELDDLTAKLRAADEARLAAER
SLQPLRDRITELQLKEQAARMTGEQFAEQLATAEVDEAALKEKLMPDMKPSYLQGEVTRI
NNAINALGPVNMAALDELAAASERKVFLDAQSADLTNAIETLEDAIRKIDQETRALLQAT
FDEVNRHFSDLFPRLFGGGQAKLIMTGDEILDAGVQVMAQPPGKKNATIHLLSGGEKALT
ATALVFAMFQLNPAPFCLLDEVDAPLDDANTERFANLVRAMSDKTQFLFISHNKIAMEMA
QQLIGVTMQEQGVSRIVAVDMETAAGFAQN*
>GCA_000452945.1_BPMSHR5858_genomic_04596_1 group_2548
LQENERRSAGWPTLAILVGASVWGLIWYPLRLLAALGVTGTVASAATSIVACAFVLIVRR
KSIATVRRHWLLPALGVAAGVTNLGFVWGTIHGEVLRVMLLFYLTPAWTAICAHFILRER
LTAAGAALAALSLGGAVLMLWSPRLGVPLPSNPAEWAGLAAGMSFAMSNVLVVKASRELP
RMKPEMRTATLFAGAATMGVAAAFVEGLPPLPAGGELGRAALLVVGIGVTVAANNMLVQH
GLARVPANRASIIMLFEIVVTALTAWLFANEVPSAREWAGGLCIVVATLLSSRVHRARPA
EDEPREPRDGARAMV*
>GCA_000452945.1_BPMSHR5858_genomic_04599_1 yffB
MAGAKAAAGTVVYGIPNCDTVKKARTWLESHGVEFAFHDFKKAGVSAPLVKGWLADVPLD
ALVNRRGTTWRALTDEQKAAAESEAGAIALMIDKPSVIKRPVVVVDGRVKALGFSADEYA
GLFA*
>GCA_000452945.1_BPMSHR5858_genomic_04601_1 prmB
MTTSPTPFKTVRDLVRHAVSRFSQAKLAFGHGSDNAFDEAVYLVLHTLHLPLDTLEPFLD
ARLAPDEIDAVLAVIERRATERVPAAYLTREAWMHGHRFYVDERVIVPRSFIGELLDDGL
QPYVEDPELVGSVLELCTGSGCLAILAALAFPNASVDAVDLSADALAVAKINRDNYGLDE
RIALYHGDLYAPLPQFKWIDPAQRYDVIIANPPYVNAGSMAELPAEYRHEPEMALAGGAD
GMDIVRRIIGEARRWLQDDGVLVVEIGNERANVEAAFGGLELVWLPTSAGDGSVFLIHAS
ELPAVAG*
>GCA_000452945.1_BPMSHR5858_genomic_04606_1 radA
MAKQKTVFVCTECGGQTPKWQGQCPSCHAWNTLVESVESAPSAHRFQSLAKRAPVQRLAD
IEAADVPRFSTGIGEFDRVLGGGLVAGGVVLIGGDPGIGKSTLLLQSLAQIASERPALYI
SGEESGAQIALRAQRLALLEGGASAADLKLLAEIQLEKIQATIDAERPDVAVIDSIQTIY
SEALTSAPGSVAQVRECAAQLTRIAKQSGTAIIMVGHVTKEGNLAGPRVLEHIVDTVLYF
EGDTHSSFRLVRAFKNRFGAVNELGVFAMTERGLRGVANPSALFLSQHTEVVPGSCVLVT
QEGTRPLLVEVQALVDTANVPNPRRLAVGLEQNRLAMLLAVLHRHAGIACFDQDVFLNAV
GGVKITEPAADLAVLLAIHSSMRNKPLPKGLIVFGEVGLAGEIRPSPRGQERLREAAKLG
FTTALIPKANAPKQPIDGLRVHAVERIEQAIDQIRVLE*
>GCA_000452945.1_BPMSHR5858_genomic_04608_1 lplT_2
MAAQFFSSLADSALLIAAIALLKDLHAPNWMIPLLKLFFVLSYVVLAAFVGAFADSRPKG
HVMFITNSIKVVGCLIMLFGAHPLIAYGIVGFGAAAYSPAKYGILTELLPPERLVAANGW
IEGTTVGSIILGTVLGGALISPKIASHVIAHTPPSINTPAEAAMLVIMTIYVTAALFNLR
IPDTGARYPRQQHGPLKLVTDFADCFMALWRDKLGQISLAVTTLFWGAGATLQFIVLKWA
EVSLGMSLSEGAILQAVVALGVAAGAMIAASRIPLRKSLTVLPVGIVMGIAVMLMAFYTR
ELFPAHWALHLGRLHLPVYLLVAYVFLMFVGALSGFFVVPMNALLQHRGHVLLSAGHSIA
VQNFNENLSVLVMLCVYAALVWLDVPVGIVIVLFGTFVCLTMWLVMRLHVANQRRFDSVA
LIGESKH*
>GCA_000452945.1_BPMSHR5858_genomic_04609_1 thiD_2
MTRPIPNALTIAGSDSGGGAGIQADLKTFSALGAYGASVITALTAQNTRGVTGVHAPDAG
FVSAQLDAVFDDIRIDAVKIGMLANAAIVRAVAAALTRHAPRHVVLDTVMISKSAHALLA
PDAVDALRNELLPLADLVTPNLPEAAALLGVAAATTEDDMVRQGETLVARGAKAVLMKGG
HLGAERESPDWLIHAGGAQRFAGARLPVANTHGTGCTLSSAIAALLPQRATLAEAIADAK
IYLTGAIGASGQLDVGHGVGPVHHFYRWW*
>GCA_000452945.1_BPMSHR5858_genomic_04617_1 group_19733
MFKHILVPTDGSELSKKAIDGAIDLAQAVGARVTAYACLPQYPYSPFSEVVIEPPADFRE
RSEREARMHLDEVEAAARRAGVACESATSVHPSPYLGIIEAAERGGCDVIFMASHGRRGL
GSLLIGSETQRVLTHTKIPVIVYR*
>GCA_000452945.1_BPMSHR5858_genomic_04619_1 group_3557
MPGTPPAFPKAILFDAYGTLFDVHAVVAAAEQLFPGHGDALSQLWRRKQIEYSQLRTLAD
PAGGRYRPFWELTLDALRFAARTLGLALSAAAEKRLMDEYACLSTYPDTVPTLRALRARS
TPPKLAILSNGNPQMLDIAVKSAGMSGLFDRVLSVDAVRAYKPSPAAYALGTAAFDAAPA
DIAFVSSNGWDVAGAGWFGYRTFWLNRTGAPLEELGAAPAGTGAGMAELLAFLDAPRAAA
KAGGRARPAPRVSRLGAPPT*
>GCA_000452945.1_BPMSHR5858_genomic_04621_1 group_4073
MRYVFCYGTLRAGEINDIGRAAATHGIAAPRLVGAVAVAGRLYDFGNYPGMVAGGGRDLV
WGDVYAIDERLVPVLDEIEEVYPGVEGLFVRQRASVELGGRRYDCLYYPVGAHAIADKPR
IESGDWVQYRRARTA*
>GCA_000452945.1_BPMSHR5858_genomic_04623_1 nimR_2
MSPSVPLFAQPPAATIDVPSEFAPTRSHPMRVRARAIAAGMRVPPHAHAWAQLAYASRGV
LRLATAGSTWMVPPSRAIWVPPRIAHEVVIVEDAYLRTLYVDESAVPGGLDACRVVEVSG
LLRELIVALEARSLNRTRERLLAELVLDELTRAEPLPLAVPMPTEKRLRALCEAVLAHPA
QGESLEHWAAGVGASTRTIARLFKQELGVSFSQWRQQALLARAIPLLNQGRPLSHIANEL
GYQSQSAFSAMFRRAFGASPRAFIQRGDMHAASELASTDDGDAEPLL*
>GCA_000452945.1_BPMSHR5858_genomic_04701_1 norR_1
MDVAARHLIYYSHRPDADLQRCFDQHGWQVDVVDSPREMRRSAARDVIAGGLLDFSCGIG
AAELRELEASLKTPNVGWIATTRRGQMEDDAVRRLVRDYCFDYVTVPYECDRIVESVGHA
YGMVTLSEGLAPAAATVRNEGEMVGTCEAMLALFKMIRKVASTDAPVFISGESGTGKELT
AVAIHERSSRAGAPFVAINCGAIPPTLLQAELFGYERGAFTGANQRKIGRIEAANGGTLF
LDEIGDLPFESQASLLRFLQEHKVERVGGHQSIPVDVRIISATHVDMQIALRNGRFREDL
YHRLCVLKLEEPPLRERGKDIEILARHMLERFKGDAHRRLRGFTPDAIAALHNYAWPGNV
RELINRVRRAIVMSEGRMISAADLELSGYAEVAPMSLEEARESAERHAIEVALLRHRGRL
ADAARELGVSRVTLYRLLCAYGMRDDDGARAKRDEGLRRAC*
>GCA_000452945.1_BPMSHR5858_genomic_04706_1 group_3565
MPATPPAPLDRTETVFRFLAEPTSVNFGGKVHGGALMKWIDETAYACAAIWSSRYCVTVS
VGNIRFQRPILVGNLVELKARVVATGRTSMHIHVSVHAGDPKGGVLRQTTDCLVVFVAVD
ENGNPVPVPPFVPVNDEQKRLAQYAMDVRAALDKIVELKPEEVAKGEV*
>GCA_000452945.1_BPMSHR5858_genomic_04708_1 norM_2
MSPTGFTRAAAAPPPTLSRHAADTARLAAPLAIAQLSQMAMSVTDTVLLGSLGPDALAAG
GLGANLFFVVVTLLQGVLTSVSVSVAHARGAMAEDRVPHIYWTGFALSLLLAVPAFALLS
FAQPLLLAFGEPAALARNVGEYAAVLRFAAPGSLIGVGLMRSFLPAIGAAKRLLWVSLAG
VGVNAFLNYGLIHGAFGLPRLGFLGSATATTITIWLTAITLVALLHGRSTFRHFVAATRP
RLPLMGELFGIGWPVAITYGVESTLFLATGLTVGVLGESSLAAHQIALNVASVAFMVPLA
IGQAANVRVGYWAGAGAPVAARHAGFVALGLGVAFMSLSGLVLIVAPHAIVGLYLKLDDP
ANARTVVLATSLLGIAAVFQIVDGMQTVGSGCLRGLKDTRVPMLAATLGYWGIGFPTGYW
FAFHAGLGARGLWWGLAAGLASVAMLMTWRFHRKSAALGVRADARGQA*
>GCA_000452945.1_BPMSHR5858_genomic_04709_1 group_3567
MPNLDFTLTGEFVELHNLLKLTGLADSGGSAKMIVASGAVKVDGAVELRKTCKVRAGQVV
LLGDTRIAVHGAS*
>GCA_000452945.1_BPMSHR5858_genomic_04722_1 group_19887
VSAIAGNPIAIRAAAAADVPAMLALMRELAEFEQLTHLFVATEADLADALFGARPSAEAL
VAVNGGTLVGYALYFHNYSTFVGRRGLYLEDLYVQPSQRGTGLGTRMLRRLAAIAVERRC
GRFEWTVLDWNQPAIDFYRKMGADVMPDWRVVRVAGDALARLADGAA*
>GCA_000452945.1_BPMSHR5858_genomic_04724_1 moeA_2
MTTHTTPTPSATSGHTVALSVDEARALALRFAERVAQVERVALRDALGRVLAEDVASPFD
IPAYDNAAMDGYAFDGAALAAPAADGVLALAVAGRALAGHPFDGPRIAPHACVRIMTGAQ
MPAGCDTVIPQEKVDATGDVVRFPAAAVARGEHCRRAGEDLARGARALAAGRVVRAADLG
LLASLGIAEVAVRRRVRVAFFSTGDELQTPGEPPREGGLYDSNRATLAGMLARIGVETLD
LGIVRDDPAALEGALTTAAAQADAVITSGGVSVGDADFTRALLATLGDVTFASVAMRPGR
PLACGRIRASGREERPALFFGLPGNPVAVAATFIVIVRDALFAMMGAEAQPLPRYPAVCD
APINKRAGRTEYLRGRAARDASGVWRVAPAGSQSSASLKSLSDANCFIVLAHDAARVDAG
TTVDILPFDGAI*
>GCA_000452945.1_BPMSHR5858_genomic_04725_1 mobA
MPSHAAPACPITGLLLAGGRGTRMGGVDKGLQPLRGEPLALHVLRRLAPQVDALVISANR
HFDAYAALGAPFLAPIVADAHADFAGPLAGLAAGMRAARTPLVLCVPCDSPFLPDDLAAR
LAAALDAQHADIAFATTLDAHGGIAPQPVFALVRTALADDLAAYLAAGERKMRAWYARHK
SVEVPFGDERAFYNANSLRDLAGLERA*
>GCA_000452945.1_BPMSHR5858_genomic_04727_1 rne
MKRMLFNATQQEELRVAIVDGQKLIDIDIETAGREQRKGNIYKGVVTRIEPSLEACFVNY
GEDRHGFLPFKEVARQYFKEGVDMRSARIQDALREGQELIVQVEKEERGNKGAALTTFIS
LAGRYLVLMPNNPRGGGVSRRIEGDERQELRETMAQLQIPDGMSMIARTAGIGRSAEELQ
WDLNYLLQLWRAIEAASQSGHPGQPMLIYLESSLVIRAIRDYFQPDIGEILIDTTEIHDQ
ARAFMDIVMPDNVGKVKRYHDDVPLFSRFQIEHQIETAYSRTVPLPSGGAIVIDHTEALV

AIDVNSARATKGADIEETATRTNLEAADEVARQLRLRDLGGLIVIDFIDMESAKSQREVE
QRLKDALKHDRARVQMGKISRFGLMELSRQRLRPALSEGSHVTCPRCNGTGHIRDTESSA
LQVLRIIQEEAMKENTAAIHCQVPVEVTAFLLNEKRQEINKIESRFKVGVVLIPNKHLDT
PHYKLERLRHDDARLDDPRASWKMAEEAARELEAETGYSKRAEETKPRQEAAVKGITPER
PAPSPAPQRQSAPEAAAPVAAAPASGGGFMKWLKGLFGAQPAAVPAPAPAAQETAARPAR
ERAERGERGEKAERGERGGDRNRNRRGGAAQQAGGRDQAAAGGGRAQAPRAEREGKETRE
PREGREARGGRDNRESREARETRETREPREPREQREPREAREQREARERAETVDAAEAPR
GERRERGERRKPAPHVATLETVNRGESAHAETAEKPLYAPGAEAGVDADSNARDGEERRR
RRRGRRGGRREREDEVAGAVQAAEGSEGVTAEAEALEHAEHAQQRVAPVANQAPEHAAQA
GAGVAAAAAATVAAEAIAHVESGAKVEPQPAAEASAGDTEQAEAVPARSIEAPAQAASTG
QAEAVQAAPAAPAHVTPATADVAREPAAPAETASAETTPAEIPPAEAAPAEAPAAEASEV
AASAVASAPAAAAEPAPAAEPHRPAPVSAAPVSAAPAEADAAREPASVAEPHVPAAAAPP
PPASVATTPGASLDTALAAAGLVWVNTDTDKLRAAQAAAAQIVQPVRVPRERKPLPPVDA
TPMQQVETTHR*
>GCA_000452945.1_BPMSHR5858_genomic_04729_1 ppaX
MARQQFDLIVFDWDGTLMDSTAHIAQCIQAACRDLGVPAPSDEAARYVIGLGLRDALAVT
APSVDPADYPRLAERYRFHYLVKDQRTELFGGVREMLEELRDTGYLLAVATGKGRVGLNR
ALDQAKLTSLFDATRCADETFSKPHPAMLQELSRELGQDLARTVMIGDTTHDLQMAASAG
AAGVGVAYGAHSADALAALSPRFVAADVAALAGWLREHA*
>GCA_000452945.1_BPMSHR5858_genomic_04730_1 group_19903
MTGDGDARFVCAADALVDGGEGVRLDATLRGEPAVVFFVRYAGRAYGYLNRCAHVPMELD
WIEGQFFESSGLYLMCATHGAIYEPETGKCVGGPCRGARLRAVRVDERDTPDGRAVFWLP
DGDLRPAPSDH*
>GCA_000452945.1_BPMSHR5858_genomic_04732_1 rsmI_2
MTAGTLYLIPNTLGDGDAAMLAAVLPAAVQARAASLGYYIGENAKTTRAFLKKIGTERPI
QEIEIRELNVKTPASEIDRLLAPVLAGADAGLVSEAGCPAVADPGALLVRRAHERGVKVV
PLVGPSSILLALMASGLNGQSFAFHGYLPVDAAARAKRLRELEQQSRHANQTQIFIETPY
RNHAMLDALVATCAPSTLVCVAADLTLATETIASRPAAQWKKAPVPDLHKRPAIFLLLAS
*
>GCA_000452945.1_BPMSHR5858_genomic_04736_1 plsX
MTVKLTIDCMGGDHGPSVTVPAAVKFVRSHPDAHLMLVGIESAIRAQLKKCKALGEPALS
VVPATEVVAMDDPVEVALRKKKDSSMRVALNHVKEGAAQACISAGNTGALMAVSRYVLKT
LPGIERPAIAFALPNPTGYTMMLDLGANVDCEPQHLLQFAEMGHALVAALEGKERPTIGL
LNIGEEVIKGNETIKRAGELLRASTLNFRGNVEGNDIYKGTVDVIVCDGFVGNVALKTSE
GLAQMLADIIKEEFSRSLLSKLMAILALPVLLRFKKRVDHRQYNGAALLGLRSLVIKSHG
SADAYAFEWAIKRGYDAVKNGVLERLSRAMAENAAPLGESGRDANGAGQASPSAGQPAEP
SAALSSKT*
>GCA_000452945.1_BPMSHR5858_genomic_04738_1 fabD_2
MKFAFVFPGQGSQSVGMLNAFADVAVVRETLDEASDALGQDIGKLIADGPADELNLTTNT
QPVMLTAAYACYRAWQQAGGAQPSIVAGHSLGEYTALVAAGAIAFRDALPLVRFRAQAMQ
TAVPVGVGGMAAILGLDDDTVRAVCAEASATGVVEAVNFNAPAQVVIAGTKAGIEKACEI
AKEKGAKRALPLPVSAPFHSSLLKPASDKLREYLAGVDVKAPKISVVNNIDVAVVSDPAA
IKDALVRQAAGPVRWVECVQHIAREGVTHVIECGPGKVLASLTKRIDGNLVGASVFDPAS
LDEALKLAAA*
>GCA_000452945.1_BPMSHR5858_genomic_04741_1 fabF
VSRRRVVVTGLGLISPVGNNVADGWANLVAGKSGIVNITKFDATNFSTRFAGEVKGFNVE
EYIPAKEARHMDTFIHYGIAAGMQAMQDSGFQVTDENAERVGVVVGSGIGGLPMIEVTQT
ELLNRGPRRISPFFVPASIINMISGHLSIKFGIKGPNLAIVTACTTGLHCIGEAARLIEY
GDADVMIAGGAEATVSPLGIGGFAAARALSQRNDDPATASRPWDKDRDGFVLGEGAGVMV
LEEYEHAKARGAKIYAEVAGYGMSGDAYHMTAPLENGDGARRCMLAALRNAGVNADQVNY
LNAHGTSTPLGDLAETVAIKRAFGDHSKDIVVNSTKSMTGHLLGGAGGLESVFTVLAVHN
QVSPPTINIFNQDPECDLDYCANEAREMKIDVALKNSFGFGGTNGTLVFKRA*
>GCA_000452945.1_BPMSHR5858_genomic_04747_1 group_3574
VSAAVRFTLYGRGWCHLCDEMRDALAPVAAECSASVDYVDIDADPALVARYDEDVPVLLL
DGVEVCRHRFDEARVRGALARRGA*
>GCA_000452945.1_BPMSHR5858_genomic_04748_1 lepA
MDHIRNFSIIAHIDHGKSTLADRIIQLCGGLSDREMESQVLDSMDLERERGITIKAQTAA
LTYRARDGKVYNLNLIDTPGHVDFSYEVSRSLSACEGALLVVDASQGVEAQTVANCYTAI
ELGVEVVPVLNKIDLPAANPENAIAEIEDVIGIDAMDAVRCSAKTGLGVEDVLESLIAKV
PPPKGDPDAPLQALIIDSWFDNYVGVVMLVRIVNGTLRPKERIKLMATDAQYAVEHVGVF
TPKSRNLESLSAGQVGFIISGIKELTAAKVGDTVTHATKPAPEPLPGFKEVKPQVFAGLY
PVEANQYDALRESLEKLKLNDASLQYEPEVSQALGFGFRCGFLGLLHMEIVQERLEREFD
MDLITTAPTVVYEVVQSDGTTIMVENPAKMPEPARIAEIREPIVTVNLYMPQDYVGSVIT
LCEQKRGTQINMQYHGRQVQLTYEIPMAEIVLDFFDRLKSVSRGYASMDYEFKEYRTSDV
VKVDMLINGDKVDALSIIVHRSQSQYRGREVAAKMREIIPRQMYDVAIQAAIGAHIIARE
NIKALRKNVLAKCYGGDITRKKKLLEKQKEGKKRMKQVGSVEIPQEAFLAILRVEDK*
>GCA_000452945.1_BPMSHR5858_genomic_04751_1 era
MIAIVGRPNVGKSTLMNALVGQKISITSRKAQTTRHRITGIHTLDDAQYIFVDTPGFQTR
HSTALNRSLNRAVTSTLTSVDAILFVIEAGRFGPDDQKVLDLIPPGTPTLLVANKLDRVS
DKDTLYPFFQKMGGLREFAEIVPLSAKHPEDIQRLMNTIKPYLPEGDAIYGEDDLTDRSS
RFLAAEILREKVFRWTGDELPYTSTVVIDKFEEEGRLTRVFATILVERDSHKAMVIGKKG
AKLKQISTEARQDMEKLFDGPVYLETFVKVRSGWADNEAGLRAYGYE*
>GCA_000452945.1_BPMSHR5858_genomic_04754_1 acpS
MAIYGIGTDLAQVSRIAAVLERTGGRFAEKVLGPDELRVFHARRARSEARGIAFLATRFS
AKEAFSKAIGLGMHWPMTWRALQTLNRPSGEPYVVASGELAAWLDARGITARVTVSDERD
YAVTFVVAEAPDDVAAARSGAAS*
>GCA_000452945.1_BPMSHR5858_genomic_04756_1 atoC_2
MPHALIVEDDPNSLSGLTALLGADGFSVDTATSLAEARGALARSIPDVVLVDLNLPDGSG
FDLLQNLPQQQPNGSLPVIVLTGNATVESAIEGLRHGVWDYLLKPVNIPRLRSLLARIPR
PYELIEEVQALRATLRRLGRFGSLVGRSDAIQHVYDMIEHNARTETAVLLWGEPGTGKEA
AARTLHELSRRRKGPFIAFDCRAAVQSPRNIETNGRAIESMLLGRERGAYTGVERRDPGL
FEQASGGTLLLGEITALPLSLQEALLRALDSQNFMRVGGTTEIASDFRLIATTRRPAREA
VANGTLREDLWLRLDAASIGLPPLRERDDDVVVMAEAFVDDLNRDARTAGLNITDKRLAP
GFIRECLAYDWPGNVRELQERVRIAYDQSGDFIETLRAGDAYPQPGAALNGSSVQVRVGT
PLADVEDLLIRATLDAVGGTRHRAATLLGISPKTLYNKLQRMKIG*
>GCA_000452945.1_BPMSHR5858_genomic_04765_1 dmlR_17
MRDLNDLYYFVQVVDHGGFAPAGRALNLPKSKLSRRIARLEERIGMRLIQRSTRRFAVTD
IGQTYYAHCRAMLVEADAADEAVALVRTEPRGIVRMTCPVALLDALVGDMIAAFMAECPL
VEIHLEETNRRVDVVGEGIDIALRVRPPPLEDSDLALRVLAERGQCLVASPRLVERAGAA
PRVPADLAALPSIALGLPQDEHVWNLRGPDGAHAAIHHRPRLVTRGMLALRAAAVAGVGV
VQLPTMMVREQIARGELRQLLPGWAPRREIVHAVFASRRGLLPSVRALLDFLAQRFAALE
PD*
>GCA_000452945.1_BPMSHR5858_genomic_04768_1 group_3579
MNLLYWLEPWEPSPTVVIAVLVAGVLFARGARKAKVSASRRVAFWFGLVALYVALHTRLD
YFFEHEFFMHRAQHLVLHHLGPFFIALSYPGAAIRAGIPFAWRQRFVRPALAQPFVRATL
DVLFNPVVAVVLFVGLIYFWLLSPIHFVAMLDWRLYRVMNWSMVIDGLLFWWLVVDPRPA
PPARLAPGRRILMVVAAIPPQIVLGALIFFTPHELYPIYSICGRAFTWLSPLRDQQIGGL
LLWIPGSMMSVIGALIALRHWMRLSARGRLSDERAARAGLARRQAARASH*
>GCA_000452945.1_BPMSHR5858_genomic_04769_1 group_19726
MTLKPTLLAALAALSLSLNAHAAGAASISAQNAWVRWLPNKLPAAGYVTLVNASDKPIDL
VEVDSPEYGMTMLHQTVSNGSTQKMEMVDKLTIPARGKVDIAPGGYHFMLEEPKRAIKPG
DTVHLRMQFSDGEKLDAPFAVKSPAAAQ*
>GCA_000452945.1_BPMSHR5858_genomic_04773_1 btuD_4
LEALTPAQRNAHNAKLASYAERPLAFLFRYIRMHPVAHVVVLVSVLAAVGCALGSQYAIK
HLIDVLAAGRHHPGPLWGAFALLVGLIAADNLLWRVGGWVAAHAFVAVTGDLRRDLFQYL
SGHSPTYYAEKQPGTLASRITATSNAIYTSENTMAWNVLPPCIAVMGAILMIIVVNPLMA
LGLLSCSGVLAIVLFKLAKLGSARHHRFAAKAAAVDGELVDVIGNMGLVRAFGMTLREQK
RFSATVKAEMDARQQSLLYLEKLRLLHAVITAMLSAGLLGWALWLWDQGRATSGDIVLVS
SLGFTILHGTRDLAVALVDVTQHVARLAEAVKTLLEPHGMPDRSDAIALAPRGGRVDFER
VTFAYPHRRPILDHFDLHIEPGQRVGLIGKSGAGKSTVLALLQRFYDIQQGAIRIDGQDL
GTITQESLRQSIALVPQDISLFHRSIYENIAYGRPDATHEEVLAAAREARCTEFIEAMPE
GFETIVGDRGVKLSGGQRQRIAIARAILKNAPILLLDEATSALDSASEEAIQAALDRLMM
GRTVIAIAHRLSTLRNFDRIIVMSAGKVIDDGSPGVLRERPGLYRDLLAKQHGRHLAAPA
PAHDHSDDDDSSSDDERVA*
>GCA_000452945.1_BPMSHR5858_genomic_04774_1 bshA_2
MRIAQIAPLHEAVPPKLYGGTERVVSYLTEALVEMGHDVTLFASGDSQTSAKLEAVWPQA
LRLDPTIRDVMAPHMLLLEEVRRRADEFDVLHFHIDYYPFPLFSRQPVPHLTTLHGRLDL
PELQPIFNTFSDVPVVSISDNQRIPLSQANWLQTTYHGLPENLLKPIPDVKPSYLAFLGR
ISPEKRVDTAIRIAEQAGMPIKIAAKLDKADRAYYEEKIKPLFSLPHVEYIGEISEAEKA
DFLGNAHALLFPIDWPEPFGLVMIEAMACGTPVIAFKRGSVPEVIENGVSGFVVEDELSA
VAAVKRLNTLPRERVRAAFDARFTSKVMAQNYVKGYEELLRRKRRTVLREVNAG*
>GCA_000452945.1_BPMSHR5858_genomic_04777_1 hcnC_3
MHSKVVVVGGGVIGSATAYFLRTLDPGIDVTVIERDPTYARASSALSAASIRQQFSTPLS
IRMSLFGIEFLRSLGERLALDGERPSIDLHEGGYLFLATPAGVATLRENHALQTSLGAQI
RYLPRDALAATFPWLAVDDLAAGCLGERGEGWFDGYGLVQALRKKARALGAQYVSADVTG
ARLDGRRVTRLLTADGRAFDCDALVNAAGPWARTVAALVGVDLPVRARRRSIFNVTSPAV
LPRCPLVIDPSGVYFRPEGASFICGAAPPPARDFDDLPLDDVDHALFDELIWPTLAHRVP
QFEALRVVRSWSGYYEYNVLDQNAIIGPHPAVDNCIFANGFSGHGLQQGPATGRGVAELI
VHGRYTSLDLSPLGFARVLEGRPIVEKNVV*
>GCA_000452945.1_BPMSHR5858_genomic_04780_1 pdxY
MKNVLSIQSHVIYGHAGNSAAVFPMQRLGVNVWPLNTVQLSNHMQYGHWAGSAIDAAKME
QLVDGIAAIGALKRCDAVLSGFLGSPAQARAAVEIVRTVKATNPNAWYFCDPAMGQTGGI
RPEPGVEEFIVAELPELADGMAPNHSELQKLAGQRIETVAEAVAACRLIIRRGPQVILVK
HLHDRNSPADRFNMLVVTETEAWIGQRPLYAFPRHPVGVGDLTSAIFVARRLRGDSVRAA
FEHTLAAVHAVVKATYDARRYELELVAAQDEIARPSEWFGAWVTGAD*
>GCA_000452945.1_BPMSHR5858_genomic_04787_1 group_2568
MPRAKPRRFPMTMFRFRILPAAMLGAALALAGCDDRQSEQTVQRFKDFFNAIKPAPLFLK
GLTPGVTTEAEIRGQMGRPETERVYTDGSKRLEYPRGPMGNETYMVDIDASGRFVAATQV
LTAANFAKIRPGMTQDEVRRLLGKPTEVARYPLKPETVWSWRWLEDGVNQDAFFNVHFGP
DGLVYTTSRSDILKGR*
>GCA_000452945.1_BPMSHR5858_genomic_04788_1 hisQ_3
VFLYGFGPLIWAGTVQTIELSVLSLAAAVLLGLVGAVAKLSHNRVLRAIATGYTTLIRSV
PDLVLMLLLFYSIQIWLNQFTDLMSWDQIDIDPFVAGVLTLGFIYGAYFTETFRGAFLAV
PRGQLEAGSAYGMNGMRVFARIMFPQMMRFALPGIGNNWQVLVKATALVSIIGLADVVKA
AQDAGKSTFNMFFFILVAALIYLAITTASNLVLGQLEKRYSIGVRHAEL*
>GCA_000452945.1_BPMSHR5858_genomic_04800_1 group_3583
MATERFDWIDPYPARTLPRLRQVLDAPLTHGRWYALVDMGFSPTLREALAALNPDGAVIA
LYEGVYDGDGLLEISPCVTPLPDDAAHRAEVCAGLLRETDGRPMLSVLRAAHGLDALVAH
LRGQMEARAADDEAFLVRLADTRCVPTWAGVLTPAQRARFFAGIDAWWRFDRAASLVELD
VAPAGSETATGADDGEPYRLDGEQIAALRHASKIDTLLFHVRQRPESFGRLTATPSQAYA
CVSEAWANEDVPPSAGARQALDALEAAGWLVPAEVSA*
>GCA_000452945.1_BPMSHR5858_genomic_04801_1 group_2827
MSVITQFTSLFSAANCLYCLDGPGEIASLQIERWVGRETLSENFQWDVYALATDPGLDLD
AMLGQRVTIRTALANGTSAVRSGLVAQAECIGYDAGLARYCLQLVPWLAALAHGRDERTF
VRQGVADVLGAVFAPYGAIARWRFTADANQRIAELGARDYRVQYRTHTHFDFVRHVLAEA
GLGFCFVEEADAPAGHTMLIFDDSTQLPEDETSARMGGVPQRLSGTATEPDDVIVGIGQS
LSLTADRVTLISSDYRGNQSTSATASLGNPTGSRELYDDVGPEAFDSLREADDAARRHAD
AIVSAARFWTGYGTLRTARAGRALRIAGATWRMSRGGAPAPDAFVLTRVDQIGINNLPAT
VMERVERSLGPLPPANLDARVLAQAKAGGYANRFDAAPRDQAWRPTLEDGTGQRLNPVPT
ALGAQTAIVVGPQGETRPGSTGPVHTDAQGRLRLRYHWQADGDAGTYPTRAMQRLASEGH
GLQQTPRIGHEVLVQFVNGLVHRPIVLGGLFNGRGEGGEAPTPGGEAGQPLDESVYAQAS
DHAASAQGNLAGGHSPAWHGAGGGAKRHDHAGALSGFKSQEFDGQGYNQLVADDTDRMGR
MQMATTHAATQLNIGHLRHQADNYLGSFRGQGVELRSDAYGALRGARGVLISSYAPTGPS
QPAGDASALQSLLAQQATLAKLLDKSADTHKTLPFAAQRGAQRAGQSSMNGDAAPLDALG
KSFATTVGADGYAQASADASRRATGNALPHTGDAVLGVEAKGGQGVLAGQSLQWVAGETL
TIGSGNDTNLAVNRTLRVHSGQGIGWLAGANGANGAQGVGMSVIAGKDDLALQAQHDLMA
LRARDDLRVASIGADVEIAAKQTVRLAVSGGAHLTIEGGNVTFGCPGSLVVHAAQHTFVG
PDQLAPALPQFPEKVCVECLKHAMQSGSALAGKAI*
>GCA_000452945.1_BPMSHR5858_genomic_04925_1 group_2582
VGRAVSLKALAAASREDLAGILAGAPERAAAWVAAAAQNGIVEAQAVYGQYLLDGRGVAR
DAAAALEWFKHAARADHPMAMNMVGRCYEFGWGTAASAAVAVYWYREAARAGLDWGMYNY
ATMLALGNGVDEDRAAALAWFEKAAALGHAKSINLIGGFHEDGWVAAADRDAAFDCYRRA
AAAGDFRGQFNYARLLAERGRVAEALEWLARVPRTATPAFVAKMRAYLAASPIDAFRALA
ARQEKSAT*
>GCA_000452945.1_BPMSHR5858_genomic_04926_1 group_2583
MMLHIPGVLTKEQVAQCRDILDAADWTDGNATSGAQSALAKRNRQLPEGSPAARAAGDAI
QDALARNALFFSAALPLKVFPPLFNRYAGGDAFGTHVDNAIRLLRGTDFRVRSDLSATLF
LEEPEHYDGGELCVEDTYGVHRAKLPAGDMMLYPASSLHHVTPVTRGARVASFFWIQSMV
RDDADRTLLYQLDTQIQRLTAEKGGRDASVIALTGIYHNLLRRWADA*
>GCA_000452945.1_BPMSHR5858_genomic_04937_1 kdgR_2
MNARKPRPAGPADEPQDLDDADRYRAPALDKGLDILELLSERKDGLTRTEITKELGRNAS
EIYRMLERLVARQYVIRSPGGDRYSLSLKLYALAHRHPPMHRLIAEALPPMQRFADAAEQ
SCHLSVYDRGNLLVIAQVDGPGIWGMSVKLGSRVGLVDTASGHALLAFQSAEQRAHMLTE
HTKVKGEQPLAARNLDARVDAIRAAGHVRQDSRQLFGVTDITYPILGPAGHAIAVLTCPY
IRRIDAYVAPPIDEVVALLRATAAGLSMVGEAAV*
>GCA_000452945.1_BPMSHR5858_genomic_04944_1 group_3594
MEARLAGKTALVTAAAQGIGRASAERLAREGARVIATDIRIDALRDGPFDARVLDVRDGA
AIDALADAIGPVDVLFNCAGFVHAGSVLDATEDEWDFGFDLNVKSMYRTIRAFLPAMLAR
ERGSIINMASAASSVKGVPNRFVYGATKAAVIGLTKAVAADFVTHGIRCNAICPGTVESP
SLDARIAEQACARGEPIDAVRAAFVARQPMARIGRPQEIAALVAYLASDEASFTTGAIHL
IDGGWSN*
>GCA_000452945.1_BPMSHR5858_genomic_04955_1 phnY_2
MTTMPLRDHAAYRRETLRWCGQRAARARTLDVFDPYSGLRVGTVPLATVDDVRRAFDYAV
AYRPALSRYERSQILERAAARLTARAEEASTLISLESGLSKQDSRYEIGRVADVFKFASI
EALRDDAQSYSCDLTPHGKSRRVFSQRQPLDGVIVAITPFNHPMNQVAHKIAPAIATNNR
VIVKPSEKVPLSALYLADVLYEAGLPEPMLQVLTGDPREIADELLTHPSATLITFTGGVA
IGKSIAAKAGYRRIVLELGGNDPLIVLDDADLERAAALAAQGSYKNSGQRCTAVKRILVH
KRVAPRFTELLVEHTRAWTYGDPFDPANRMGTVIDEAAAALFEARVDEAVAQGARLLTGN
ARRGALYAPTVLDRVDASMTLVREETFGPVSPIIAFDTIDDAIRISNGTAFGLSSGVCTD
RADAIVRFVNELNVGTVNVWEVPGYRLELTPFGGIKDSGLGYKEGVQEAMKSFTNLKTFS
LPWA*
>GCA_000452945.1_BPMSHR5858_genomic_04957_1 phnV_3
MSADFPARAAPVPRRELRWTDAASRVASRALVALAAAACCWLFVLPVVVVALSSVAGHWS
GTILPAGYSLRWFERLGRPEFDALAASLEIGLGVAALGTALGVALALALEGRTRRGVGAL
VDALVMLPNGVPSVVLGLAVLIAYHAKPVDLSSSPAIVVFVQLALVLPFCYRCAAAALRP
ELAVLREAAASLGAPPAMVLRRVLLPQLVPAIRASLALGFALSLGELGATLTVYPPGFAT
VPIVVIGAVERGYYLPASALSLLLLGASLVALIAIAARAPRPAR*
>GCA_000452945.1_BPMSHR5858_genomic_04959_1 phnT
VDTASLTHPGAFDAARPHAAPRAGACGGVRIEHLSVRYGARTVLDDLSLEIGAGELLAVL
GKSGCGKTTLLRFIAGFVKADGLTGTLAVAGRDLTHAPPHKRNLGLLFQNYALFPHLTVF
ENVAFGLRARRMSSADIARRVADALKLVQLGDAGHHLPAQLSGGMQQRVALARALVIEPD
VLLLDEPLSALDANLRASVRSELKALHERLPNLTVVCVTHDRDDALVLADRALLMREGRV
AQLGTPQQLYDAPADGFVARYLGAANLLPPNVVFAFGDPRHDVRDKVACVRPERLAVRPL
GEGRLHGTIASVEWHGAALSLTVMLDAACDEPVLVTMQRGRGPAPERGARVSLDCEADDV
VLISP*
>GCA_000452945.1_BPMSHR5858_genomic_04962_1 gudP
MPQPIHTDAAPRAAVAAARAASPARPRSNARYRILALLAFGTMINYLDRTVLGVAAPGLT
KELGIGAAAMGVMFSAFSWTYVLAQVPGGLLLDRFGSKLTYYWSMTLWSLFTLLQGFVHG
VAPLLACRLGLGLSEAPCFPTNSRVVATWFPQKERAMATGTYTVGEYIGLAFLSPVLFAL
MGAFGWRSLFWVVGGVGIVFGFVWWKFYHEPHDHPRANAAELAYIEEGGGLVQATAGGGS
RAKFNWRVAGRLLKKRQLAGICLGQFAGNSTLVFFLTWFPTYLATERHMGWLKIGFFAVM
PFIAASVGVMFGGLFSDWLLRRGRSANVARKLPIIAGLLLASTIVLANYVRSNEAVIAIM
SVAFFAQGMAALGWTLVSDIAPEGLLGATGGIFNLAANLAGIVTPLVVGFIVAATGSFVG
ALAFIGVIALIGALSYIFVVGDIKRIEL*
>GCA_000452945.1_BPMSHR5858_genomic_04968_1 rhaS_6
MMTLLAVPPEKLSAASAASTAASTAAGPTPAAGNGACASPPSSGAALEGDDDGFAAMLAH
FAQLEPVFDALPDVAFFVKDVRARYALVNRTLAQRCGYKDKRDLYGKAADEVFPRRFGRS
YVEQDLTIIAGGRQLTDQLELHLYSGRQPGWCLTCKEPLRNAQGRIVGIAGLSRDLRAHE

SSHPAYSRIADVVRYIHAHYVQPLNLKHLAGMAGMSVAQLERYFHKVFHLTPRQMLLKAR
LDAATALLVTHDKVTDVAALCGYTDHSAFTRQFKATVGVTPTEYRLMLQEGRGA*
>GCA_000452945.1_BPMSHR5858_genomic_04972_1 group_3596
MKHIHIIDSHTGGEPTRVVVSGFPALGGGTMAERLAVLAREHDRYRAACILEPRGSDVLV
GALLCEPVSAGAAAGVIFFNNAGYLGMCGHGTIGLVRTLHHMGRIGPGVHRIETPVGDVE
ATLHDDLSVSVRNVLAYRHAKDVVVDVPGHGAVTGDVAWGGNWFFLVSDHGQRVAGENVA
ALAAYASAVRAALERAGVTGRDGAPIDHIELFADDPEYDSRSFVLCPGHAYDRSPCGTGT
SAKLACLAADGKLAAGVTWRQASVIGSVFSASYAAAEGGVVPTIRGSAHLSAEATLVIED
DDPFGWGIAS*
>GCA_000452945.1_BPMSHR5858_genomic_04974_1 yveA_2
MPAQGNAQRTAPLARPAPTDAGHGKFKKQLSLTDLTFIGLGAIFGSGWLFAASHVSMIAG
PAGILSWLLGGFAVLLLGIVYCELGAALPRAGGVVRYPVFSHGPLLGYLMGAITLIAFSS
LITIEVVAARQYAAAWFAGLTVEGSGNPTTLGWLVQAALLCFFFYLNYSSVKTFAKANNV
ISVFKFVVPLAVIVVLFAFFKPANLTLHGFAPFGMPGVEMAVSAGGIIFAYLGLTPIVSV
ASEVRDPQRTIPIALILSIVLSTLIYVLLQLAFLGGIPSAMLAGGWHDIGKAFSLPYRDI
ALALGVGWLAYMVVADAMISPSGCGNIYMNATPRVVYGWAKTGTFFKVFARVDEASGIPR
AGLWLTFALALFWTLPFPSWEALINTVSAALVLSYAVAPVSVAALRRTAPGLPRPFRASA
FAFTGPASFVIAALIVYWSGWRTVSWLLGLQIAMFAVYLACRRRVPTAHLSLAEQVRSCA
WLIAFYALMIVASYFGGFGGTGQLAHPYDTFAVAAVALVIYYWGAHTGVPSAKLQLDDDE
N*
>GCA_000452945.1_BPMSHR5858_genomic_04984_1 pdhR
MTFQPVESYTRVRLSDVVSDQIKSLISAGKLLPGQKLPTERELAAQLNVSRPSLREALVR
LEADGFIGSVGRGGFVVSDITAPVVSSPLADLLLQNPEASHDVLELRHGLETISTAYAAE
RATPGDLAKIKEAFDTLAGHSLKHEPGRLAEVDANFHLAIADATHNVALIHVMHGIHGLL
QESMHKSHRLVNHADAMERALLEQHTEIYEAIAAKDPERARAAAERHLRYVRALYEQAA*
>GCA_000452945.1_BPMSHR5858_genomic_05093_1 lip_1
MARTMRSRAVAAAVAFAVSAAPAAGIGTFLSLAGTQAAAAATSAVDNYAATQYPIILVHG
LTGTDKYANVLEYWYGIREDLQAHGAQVYVANLSGFQSDDGPNGRGEQLLAYVQQVLAAT
GASKVNLIGHSQGGLTSRYVAAVAPQLVASVTTIGTPHRGSEFADFVQTVLAYDPTGLSS
TVIAAFVNVFGILTSSTNNTNQDALAALKTLTTSQAATYNQNYPSAGLGAPGSCQTGAQT
ETVGGNTHLLYSWGGTAIQPTLSILGVTGATDTSTIPVVDPANVLDASTLAMLGSGTVMI
NRGSGANDGLVSRCSSLYGQVLGTSYKWNHLDEINQLLGVRGAYAEDPVAVIRTHANRLK
TQGV*
>GCA_000452945.1_BPMSHR5858_genomic_05094_1 lifO
MKGEGSVARRVALYCVAGAVAAGAVWYVVGAPAKRDAGASPASPELAASAPVAVAARADA
GSAASGLPASLAGSSAPRLPLDAHGHLAKARAVRDFFDYCLSAQSELSAQALDALVTREI
AAQLDGTLAQGEALDVWKRYRAYLDQLAQLPDGGAPGNKLDLAALQLALDQRASIASRAL
GDWAEPFFGAEQQRQRYDLARLRIVRDASLTDAQKAARLAALEQQLPLDQRAEQAREKQQ
NDAVARIAQLQKSGASPDEMRAQLAQSLGPEAAERVAKMQQANDAWRAKYDEYAAQRAQI
DAQYLSPQDRDTQIAQLRQRYFTQPGEALRAASLDRGGAATQTQ*
>GCA_000452945.1_BPMSHR5858_genomic_05095_1 gabR_3
VILSDWLAARLERHSAEPMYRQLLRLMQQAILSGELTPGTKLPSSRTLATDLAIARNTVL
HVYDQLTTEGYVLTTTGSGTYVADTRPDAAAMRAQAGAPAHPSDETRDAQQRAHGALSAR
GAQLIRHAGVSRRQWGAFMPGVPDVSEFPTRTWSRLQARLWKEANPELLTYAPGGGYRPL
RRALSDYLRVARSVKCSPDQIIITTGIHQSIDLSVRLLADVGDRAWVEEPCYWGVRSVLQ
AAGLALAPVPVDQEGLAPRAQDLKRPPRLVLVTPSHQYPLGMVMSLARRRMLLEYARQHQ
CWIIEDDYDSEFRYGSRPLASLQGLDDAGRVIYVGSLGKMLFPSLRLGYMVVPEHLVETF
RTGLSELYREGQLMQQAVLAEFIMDGYLTSHVRRMRALYGERRQLLIDAIHSRFGDALPV
MGDEAGLHLVIGLPNGCDDRAITQTAFDAGVIVRPLTTYYNHADTAREGLLLGYACVPNE
RIAPAFETLAQIIEAHLNRRAKQRAA*
>GCA_000452945.1_BPMSHR5858_genomic_05098_1 group_3610
LNTMTIATQSAPPTQSLRRELKAAEARRRTMALLLVAPLAIFLLLVFVVPIGALLTRAVQ
NPEIATALPRTVAALSNWDRKAAPADAAYAALASDLTQVADGEAMGALARRLNTEIPGYR
SLVAKTARAMPLTGDANAPLAPAQTRAKLIELDERWGDAAYWQAIAKNGGRYSPFYLLAA
LDHKQDGFGSIVPADPDQSIYLAVFGRTLLIGFAVTLFALALGYPLAYWISTLPERRANL
AMILVLIPFWTSVLVRVAAWIVLLQSEGLVNRALIDTGLISQPLALLFNRVGVYISMTHI
LLPFMILPLYSVMKSIPPSYQRAAVSLGSHPFAAFWRVYVPQTYPGVGAGALLVFILAIG
YYITPALLGGPNDQMVSYYVAYFTNVTINWGMACALGGLLLAATLVLYAIYGRFTRSQLS
LG*
>GCA_000452945.1_BPMSHR5858_genomic_05099_1 ydcV_2
MKLLAKPLFAPHMSGLERAWYFALRGLVVLTLLYLILPVLAIVPLSFSSSTFLVYPIPGF
SMRWYENLLASDEWRMAAKNSFIVAPAATLVATVLGTLAAIGLTKSDFRGKGLLMAVLLS
PMIVPVVVVGVGMYLFFAPLGLANTYTGLICAHAALGVPFVVTTVAATLQGFNQNLVRAS
LSLGANPVTTFFRVTLPVIAPGVMSGALFAFATSFDEVVVTLFLAGADQTTLPRQMFTGI
RENISPTIAALATILIVFSTCLLLALEWLRGRHAKRAVS*
>GCA_000452945.1_BPMSHR5858_genomic_05106_1 group_3613
MQSWGKTLARCAIAGLFTATAFAGTPPVGPASVPSTHAVAARIIDDLPARHAAGLDAATR
AQVERGRYVARLGDCVACHTGDKSKPLAGGLALRTPFGELHSTNITPDPQTGIGRYTFAQ
FDRAMRKGVAADGRRLYPAMPYPSYAKVAAEDMRALYAYLMKGVAPVRQANRALALRFPF
NQRWGLALWNWAFLDDTPFQPDATRSAEWNRGAYIVQGLGHCGACHTPRGLGFQEKAMSG
AGSAGRYFLAGETVEGWRALSLRALWTPQDTAEMLKTGRNRYGAVSGNMVDVVQHSTQYM
SDGDLLAVGAYLESLPAAGRDKPMQVPQPPAPAIARAGPPAADISSRVPADLYTSRGGLA
YLQFCNDCHRSDGAGVRDVFPPLAGNAALLSEDPSTLVHIMLTGWRSAQTASHARVLTMP
AFAQLSDREIAEILNFVRRSWGRADVRAIGAGTVGRMREQLRAGTGDGARFRTPRLADML
AESNANQLTLGARLNIETRQMLPKNVGNALNCASCHLNAGTVADGSPYVGVSAFFPGYAP
RAGRVITLAERINGCFLRSMNGKPLPPDSEQMQAMVAYFDWMRRETRPGDKVEGRGVGKV
SHRIVPAVDNGKRIYAAQCALCHGSNGEGVKDDAGRWVYPPLWGDESFNIGAGMARTYTA
AGFVQRNMPIAFHDRFPLGQGGLTDQEAVDVAEYFTHQPRPDFAGKVKDWPKDKKPDDAR
Y*
>GCA_000452945.1_BPMSHR5858_genomic_05110_1 citB
MAHNLHKTLKEFDSGSGKGKFYSLPKLGKELKTKIERLPVSIRIVLESVLRNYDGKKITE
EHIEQLANWKPNAKRVDEIPFVVSRVVLQDFTGVPLLADIAAMRGVAKRAGKNPKKIEPL
VPVDLVVDHSVQIDYFRQKDALDLNMKLEFQRNNERYQFMKWGMQAFDTFKVVPPGVGIV
HQVNLEYLARGVHKKKDDGDTVYYPDTLVGTDSHTTMINGIGVVGWGVGGIEAEAGMLGQ
PVYFLTPDVVGVELKGKLREGVTATDLVLTITEMLRKEKVVGKFVEFFGEGTKTLALPDR
ATIANMAPEYGATMGFFPVDEKTIDYFKGTGRTKAEIAAFENYFKAQELFGIPKAGEIDY
TKTLTLDLSTVAPSLAGPKRPQDRIEIGNVKSTFTDLFSKPVAENGFAKKADDLTAEYRT
SNGVAVKNGDVLIAAITSCTNTSNPSVLLAAGLLAKKAVEAGLTVAPHIKTSLAPGSRIV
TEYLTKTGLLPYLAKLGFEVAAYGCTTCIGNAGDLTPELNEAITKNDIVAAAVLSGNRNF
EARIHPNIRANFLASPPLVVAYAIAGNITKDLMTEPVGQGKGGRDVYLGDIWPSSDEVQA
LLKFALDPEKFEKNYSHLTKKGDLWSKIEGESGQVYDWPKSTYIAEPPFFGSDFSMEPAA
SIATVKGARALGIFGDSVTTDHISPAGSIKEDSPAGKWLKANGVQKADFNSYGSRRGNHD
VMMRGTFANVRIKNLMIPAKADGTRVEGGLTIHQPSGEQLSIYDAAMKYIDADTPTVVFA
GEEYGTGSSRDWAAKGTQLLGVKAVIARSFERIHRSNLVGMGVLPLQFKGSDSIQSLGIT
GDETYDIEGLGDDFKPQQDVTLVIHRRNGETKRVPVLLRIDTPIEVDYYKHGGILPFVLR
SLLAA*
>GCA_000452945.1_BPMSHR5858_genomic_05114_1 mdh_2
MAKPAKRVAVTGAAGQIAYSLLFRIANGDLLGKDQPVILQLLDLPQAQAAVKGVVMELDD
CAFPLLAGVVITDDPKVAFKDADVALLVGARPRSKGMERKDLLSANAEIFTVQGAALNEV
ASRDVKVLVVGNPANTNAYIAMKSAPDLPKKNFTAMLRLDHNRALSQLAAKSGKPVASIE
KLAVWGNHSPTMYPDFRFATAEGESLLKLINDDVWNRDTFIPTVGKRGAAIIEARGLSSA
ASAANAAIDHVRDWVLGTNGKWVTMGIPSDGSYGIPEDIIYGVPVICENGEYKRVEGLEI
DAFSREKMDGTLAELLEERDGVAHLLK*
>GCA_000452945.1_BPMSHR5858_genomic_05115_1 frlR
MTSNQASAPNPNGQAGSTQPGGEAAAAPSPTFSPLYRQIKALITQSLEAGEWKPGEIIPS
EVELAGRYKVSQGTVRKAIDELAADNLLVRRQGKGTFVATHSEERAQFRFLRLLADDGAE
HPHVSRLLECRRMRAPAEIARQLDLKPADPVVQVRRVLEFDGVDTVLDEIWLPGAMFRGL
TFERLSEYKGPLYAMFESEFGTRMIRASEKIRAIAAEPAVADVLRVPPGFPLLSVERVSY
TYGDRPVEVRRGWYVTTGYYYQNDLS*
>GCA_000452945.1_BPMSHR5858_genomic_05121_1 gltA
MTPSDVKATLSFSDNSPSVELPIYKGTMGPDVIDIRKLYGQTGKFTYDPGFMSTASCNSA
ITYIDGDKGELLYRGYPIDNLAQNADFLESCYLLLKGELPNAAQKKEFVATVTKHTMVHE
QMQFFFRGFRRDAHPMAILVAAVGALSAFYHDSLDINDPRHREVSAIRMIAKLPTLVAMA
YKYSIGQPFVYPRNDLSYSANFMRMMFANPCEEYQVNDVLVRALDRILILHADHEQNAST
STVRLAGSSGANPFACIAAGIACLWGPAHGGANEAALNMLEQIGTPENIPEFIKQVKDKN
SGVKLMGFGHRVYKNYDPRAKLMRETCYEVLNELGLHDDPLFKLAMQLEKIALEDEYFVS
RKLYPNVDFYSGIVQRALGIPTSMFTCIFAMARTVGWIAQWNEMIADPEQKIGRPRQLFI
GETPRTAKPIEQR*
>GCA_000452945.1_BPMSHR5858_genomic_05128_1 group_19971
MTRYEMKRQGGTQAEGPPVDFGRMHEEFRRAEDADLRALVDENLRRAEREWRRIQPAIDR
SAGDGDAALGAARPCACEAPGACGQPDLPDVPDPPIASDPADAPLAPSSGAHARAASCGH
ATKTLIPGYPVLDADRPSVGEFVAFVADMRNSSARFSEAAAFPHVRCGLQRIFYETSALI
PAIAHTAARAGGQATELLGDGLLILFHIERGKREQVIARACAAAARCIDSSIACVNDLLW
QRHRLPPLKIGIGLAYGPAIVKVVDANGRHPKVIGQCIWDATKLSGGENAIGLSRPLQDF
LRAKGFNERFEVIHA*
>GCA_000452945.1_BPMSHR5858_genomic_05131_1 group_2598
VSVNASVFRRIAAAFALAGCVLGLAGCNTVAGVGEDLNAAGRAIKRAAD*
>GCA_000452945.1_BPMSHR5858_genomic_05135_1 asd
MVGSVLMQRMQEEGDFDLIEPVFFSTSNAGGKAPSFAKNETTLKDATSIDDLKKCDAIIT
CQGGDYTSDVFPKLRAAGWNGYWIDAASSLRMKDDAVIILDPVNLNVIKDALVNGTKNFI
GGNCTVSLMLMALGGLFRENLVDWMTAMTYQAASGAGAQNMRELLAQMGALNGAVAAQLA
DPASAILDIDRRVLAAMNGDAMPTSQFGVPLAGSLIPWIDKDLGNGMSKEEWKGGAETNK
ILGKPAMGEPGSIPVDGLCVRIGAMRCHSQALTIKLKKDVPLDEINGMLASANDWVKVVP
NEREASMRDLSPAKVTGTLSVPVGRLRKLAMGGEYLSAFTVGDQLLWGAAEPLRRMLRIL
LDK*
>GCA_000452945.1_BPMSHR5858_genomic_05137_1 truA
MRIALGIQYDGAAFCGWQSQPHGKTVQDALERSLAEFAQTSLHTTVAGRTDTGVHGLGQV
VHFDTDLDRADFSWVRGTNAFLPPTVAVQWAKPMPDTFHARFAAFERTYYYALYVHPVRS
PMLAGRAGWVHTPLDVDAMREAAAHLVGEHDFSAFRSSECQAKSPVKHLYQIGIRPDGDF
IHFRFRANAFLHHMVRNLMGCLVAVGRGRYPSSWLAEVLESRDRDCAAPTFMPEGLYLAH
VGYPAEFAVPPAQLGSVPWSSVWADLDGRT*
>GCA_000452945.1_BPMSHR5858_genomic_05138_1 trpF
MMTDTPDATLPADAPPNDAPAAGARRDTGGDGSRDAPPPRTRIKLCGLSNAADIDWAAEL
GADAIGLVFYPKSPRAVTLEQAAALAARVPPFVSVVGLFVNPTEDEIARVANAVPLTLLQ
FHGDETPGQCASLAAAARLPWLRALRVGVATQPADLVESALHYSAARGLLFDTLVEHYGG
SGKVFDWSLIPAELARRAVLSGGLNAQNVGDAIRQVRPFAVDVSSGIEAPGAKGVKDRAR
MAAFVRAVREADAG*
>GCA_000452945.1_BPMSHR5858_genomic_05140_1 group_19729
MRDLIEEPAGGAASDAQATHAACALPAGIELHNRDFLHEAASLPDASIDLIVADPPYGLG
KDYGNDSDKRSGDEHLAWTREWLELAVPKLKSTGSMYVFCTWQYAPEIFSFLKTKLTMVN
EIIWDRRVPSMGGTTRRFTSVHDNIGFFAVSKGYYFDLDPVRIPYDADTKKARSRKLFEG
SKWLELGYNPKDVWSVSRLHRQHAERVDHPTQKPLEIIERMVLASCPPGGRVLDPFMGSG
TTAVACARQRRGFVGYEINESYCAIARERVASLAAVSAPADA*
>GCA_000452945.1_BPMSHR5858_genomic_05143_1 folC
MRRFDSKLDSMSTFPTLDAWLSHLESAHPVGIDMGLARIGQVKDALKLAFACPVITVGGT
NGKGSTCAFIETILVRAGYKVGCHTSPHLLRFNERARLDGRIVDDEELLPHFEAVEAART
SLPEAVSLTYFEFTTLAIMHLFASRGLDAVILEVGLGGRLDAVNVIDADCAIVTSIDVDH
IEYLGDTREKIAFEKAGIFRPGKPAICGDPAPPQTLVDHAGAIGADLWLVGRDFRFSTQP
GSERQQWTYAGRDKRYPALAYPALRGANQLLNASAALAALEALRERLPVSAQDIRLGLAN
VELPGRFQVLPGKPLVLLDVAHNPHAAAVLAQNLDSMGYYPYTHAVFGAMADKDLAGIVE
RLKGAIDHWHLTDLPLPRAAAADVLERVLRGAGVEHGAQHNITRHAGPADAFLDALKSAS
DNDRIVVFGSFYTVAGVMPVVDRRHDH*
>GCA_000452945.1_BPMSHR5858_genomic_05148_1 group_19896
MFQAGIGVNFEAALRDVSEYWSPRVVGRVNDQYVKVAKLKGEFTWHKHADEDEMFYVVYG
RLRIQFEGRDDVVLNPGDFCIVPKNTMHNPVAEAECGIALIETVTTRHTGDTPSPLAKSI
DAQLAG*
>GCA_000452945.1_BPMSHR5858_genomic_05150_1 gmd
MTQARKAIITGITGQDGAYLAKLLLDKGYEVVGTYRRTSSVNFWRIAELGVDKHPGLTLV
EHDLTDLGSSLRLVERTQPHELYNLAAQSFVGVSFDQPATTAEVTGLGALHLLEAIRVVN
PKIRYYQASTSEMFGKVQAIPQIESTPFYPRSPYGVAKLFAHWTTLNYRESYGIFGSSGI
LFNHESPLRGREFVTRKITDTVAKIKLGKASRLELGNLGAKRDWGFALEYVEGMWRMLQA
DEPDTYVLATNRTETVRDFVRMAFAAAGYQLEWSGKGEDERGIDTATGNVLVSVNPKFYR
PAEVDLLIGCADKARDKLGWVPKTTLEALCRMMVEADLTRNRHHETF*
>GCA_000452945.1_BPMSHR5858_genomic_05175_1 group_19966
MRHIKPQAALVATTNTQIGAQPMLGISVGIGFRLDQPSILVHEAAVWEALKAAAPSLPLY
EAALPKQRAEWLLAGHSVHAVGAGARARDVDWTAWVELDGVRKVVSCATSLGDEQAQSGY
ARIAVDHRHAAAGGARENPFGVASGTPPLQQLRTFGVGPAPLAAMGAINPDWPERAQWMP
TRPGTVDAMAQDGTHMGWPAEVDLRFFQQAAPDQWARGECWTPGARFELSGFGPRGEGFA
GELPRLAPVALVTRNGRPGIERLSFKQQTAWFLPDRGIGVLWWNGAVALDFLLDDSPTML
VTAFKDEAERIDVDALMKFADQRADLNCIDPLQQADHELMPAITRGWTWEMILDTEDHPR
FAPAPRGYEEVRARVEQNRRELVEARDASERLSAFEEANRNAKLPGAPRGGENWRTRLRQ
AKTPELANVTIRDADLSSLRFDGWKFDDVRFERCTLDRSEWTNCRLNQVHAVDCSFADVK
MSDGWWKGGKIQRCNLERSAWLNVEIERISLDECRLDDLKVAGGSWSMLSVQGRGGVRGD
VQDIQWNSVSWSEVSAPGWTWTRVRADDLAIVECAMAGLAVSQCTLAKPSILLTDLSASV
WQRSMLTFAVLSHGTSINGARLTDCVFKSSSLQELRADRVQVDHCSFMQLNAQHLHAQQS
HWSRTVLDGANVMHAQLTGTSFDRCSLKEAMFYGADMRQTRMRDCNLVRVRTSWIHPPEA
GAWRGNLSAGQLDVPRRV*
>GCA_000452945.1_BPMSHR5858_genomic_05177_1 group_19825
MSMREMAVAHDALAYEGDDDSASRRALTRIMKGGARGSDADAGAHALPADLSAREWVRES
ALPAAATAATLLARIEGERTAAGEWRVALRPDVVLRAKKAVSCVVAPRAGDLVQICREGE
RCWVLAVLERDAAGDETNDRTTDEVTLDFGDAHVALRARDVRVEARDRLSLEAAQLASRA
QVVTQAAAERQTHVSGTDATHAGSTLVHTERHLAMHAKSAVVTAASLLKIDAGQIHMG*
>GCA_000452945.1_BPMSHR5858_genomic_05180_1 group_19713
MSSLPVGPVAWSDGMLIETQHFQQLERHLAHQASLRLGQTSNHGWGFTLLDLDQDGLGLG
RLGLRHARGVFQDGTAFSLPSDDPLPPPLETELAQAGDIACLALQAARTGGPEMAFGDVE
LASRYRAVSTEVPDLAVGLDAPGTPRRLTIETGQLVTRLCWKSQLRSDEVALPIARVAGR
NASRTVSLDPRFIPPLLDTRAHLVLRSLIDELQSTLRVRLASTSAQRVLSTGGGVADLIE
LLLRQAIAEYRMRLANLDAFDPLPPAMLYHELVGLLGRLSVLPGVDEELADRELGYDHDD
LQTSFEPLAMMLRQALARVIETPVLPLRFEDRGDQVHICIVDKQWNLKKLIFAFSAAMPA
EKLRQLLPQQTKLGAVEQIQKLVDLQLPGARLNALPNPPRQIPYYAQSTYFEVESTDPFW
KQTLAGSAMALRIVGDFPDLRFEAWGLRDGKVA*
>GCA_000452945.1_BPMSHR5858_genomic_05181_1 group_3619
MSLLRNLSSTLRRVSMASNMLDRAAANPDLVTRVPTLSSLSSAAMTSSAVSTTGTTNTVA
SGGAAAGAPSFAPPAADAFPQADGAPRNPAVLQFPVPGGAPAPADARVAAPVVYSAQGEQ
AAIMKAGLQQASWNNPFVSHALPAVLQLQRHLAAGPLNQAAIRTQLGLEVRLYRERLAAS
GCEWEQIRDASYLLCTYLDETVNDAAREHAQVVYDGERSLLVEFHDDAWGGEDAFADLSR
WMKTEPPPIPLLSFYELILSLGWQGRYRVLDRGDVLLQDLRSQLHALIWHHVPPEPLGTE
LVAPAKRRRSWWTAGRAAAVALGVLVLAYGAISFWLDSQGRPIRNALAAWMPPTRTINIA
ETLPPPLPQILTEGWLTAYKHPQGWLLVFKSDGAFDVGKANVRADFMHNIERLGLAFAPW
PGDLEVIGHTDSRPIRTSEFPDNQALSEARARNVADELRKTALPGGARAPENAVQRNIEY
SGRGDAQPIDTAKTAAAYERNRRVDVLWKVIPDGAQQSGRSLNLQQPEKPGQVPMRPAMP
EGVEIAPDGQLPYATSTTMPATRPTTEGRQP*
>GCA_000452945.1_BPMSHR5858_genomic_05184_1 group_3620
MNKQIVAGIVVGLMSMSSHAQLGQLFQSVKEQVTQAATSQVNQGVRSATDEAVQATSTRT
RKAIDSVRSPSSSAAATSTSTSPSASAGTGDAALSEARK*
>GCA_000452945.1_BPMSHR5858_genomic_05207_1 clpX
MADKKGSNSEKLLYCSFCGKSQHEVKKLIAGPSVFICDECIDLCNEIIRDEAAAAGVEAS
LSKSDLPSPQEIRDILDQYVIGQERAKKILAVAVYNHYKRLKHLDKKDDVELSKSNILLI
GPTGSGKTLLAQTLARLLNVPFVIADATTLTEAGYVGEDVENIIQKLLQNCNYEVEKAQR
GIVYIDEIDKISRKSDNPSITRDVSGEGVQQALLKLVEGTMASVPPQGGRKHPNQDFIQV
DTTNILFICGGAFDGLEKVITDRTEKTGIGFGATVKSKQERDAGEVLREVEPEDLIKFGL
IPELIGRLPVVATLGKLDEAALMKILVEPKNALVKQYQKLFAMERVELEIRPDALQAVAR
KAIRRKTGARGLRSIIEQALLDVMYELPTLKGVSKVIIDDNVIEGDGKPLLIYEDTPKVA
GSN*
>GCA_000452945.1_BPMSHR5858_genomic_05208_1 lon_2
MSGTQLLPSERTALPLLPLRDVVVFPHMVIPLFVGRPKSIKALEVAMEGGKHIMLVAQKT
AAKDEPTEKDMYDVGCIANILQMLKLPDGTVKVLVEGLQRAQALSIEEQETQFSCEVMPL
EPDHADSAETEALRRAIVSQFDQYVKLNKKIPPEILTSLSGIDEAGRLADTIAAHLPLKL

DQKQHILEMFPVIERLEHLLAQLEAEIDILQVEKRIRGRVKRQMEKSQREYYLNEQVKAI
QKELGEGEEGADLEELEKRINAARMPKEAKKKADAELKKLKLMSPMSAEATVVRNYIDTL
IALPWRKKSKVNNDLSNAEKVLDEDHYGLEKVKERILEYLAVQQRVDKVKAPILCLVGPP
GVGKTSLGQSIARATNRKFVRMALGGVRDEAEIRGHRRTYIGSMPGKILQSLSKVGVRNP
LFLLDEVDKMGMDFRGDPSSALLEVLDPEQNHTFADHYVEVDFDLSDVMFVATSNSLNIP
PPLLDRMEVIRLSGYTEDEKVSIAQRYLLPKQKKNNGLKEGEIDVAEQAIRDIIRYYTRE
AGVRSLEREVSKICRKVVKMLLLKKASGAIKVDGENLDTFLGVRKYDFGLAAKENQVGQV
TGLAWTEVGGDLLTIEAAVMPGKGNVIRTGSLGDVMKESVEAARSVVRSRSRRLGIKDEA
FEKQDIHIHVPEGATPKDGPSAGIAMTTALVSVLTGIPVRADVAMTGEITLRGEVLPIGG
LKEKLLAAHRGGIKLVLIPEENVKDLADIPDNVKNAIEIVPVRWIDKVLELALERLPNAL
PEEEAKAAPVAEPAKDAGSTEVVKH*
>GCA_000452945.1_BPMSHR5858_genomic_05214_1 PSD2
MTNRTNATHATNATNATNAAREAAAERRRLGGWLPAEEAALARYRKTLAAHAREHADGVP
MTEAVRELAALAEGEPPLRMALTRAIDEARSRGYELGYETIGELMAVIDFVMTYAPPFSE
SGLVVCPLNALLDWPLCMPSGYAFFRDRAVNAQLKRVLNVWCGFLGGPHSRAHLNASEPD
GWFSAEAVKRIGMSQFHCDPQLPYWGFASWNDFFTRRFLPDARPVAAPDDPAVIVSACEA
APYHVAHDLKLLDTFWIKSQPYSLRDIFTAARGELARGFVGGSLYQAFLSAYDYHRWHAP
VAGRIAHAYLVDGTYYSSTEAEGADPAGLNDSQGYMTAVATRAVIVIDADAPEIGRVACV
FVGMGEVSSCMIDALPGQHVEKGEEIGYFQYGGSTYCMLFEPGVIGSFVAHPRYHEGAPA
VQVNAEVAAVAAK*
>GCA_000452945.1_BPMSHR5858_genomic_05215_1 sdhB_2
MTMADTRILDIYRYDPDDGRAPRMQRYELAPRAEDRMLLDVLGRVKAQDETLSYRRSCRE
GVCGSDAMNINGRNGLACVTSMQALPREIVLRPLPGLPVVRDLIVDMTSFFNQYHSVRPY
LVNEGVPPERERLQTPQEREQLDGLYECILCACCSSACPSYWWNPDKFVGPAGLLQAYRF
IVDSRDDASAARLDDLEDPYRLFRCRTIMNCVDVCPKGLNPARAIGQIRTMLARRAV*
>GCA_000452945.1_BPMSHR5858_genomic_05216_1 group_19873
MEPNARTMYEYGGYEIRVWASRNERGAWVSDVEIYRDGRRVLAAVPETVSPEWGTEEECV
RDAIGQGRDLIDRGDVARCRP*
>GCA_000452945.1_BPMSHR5858_genomic_05218_1 moxC_2
MTRKHIHFGVLIQGAGANMNAWKHPSVPPDASVNFDFYVDRARRAENAGIAFAFIADSAY
VTPKSAPHFLNRFEPISLLSALAVLTSKIGLVGTMSSSYSEPYNVARQFASLDLISGGRA
GWNVVTSSIEGTGKNYGRPHPDHAQRYAIAAEHLDVVQGLWDSWDDDALVRDRATGRFFD
PDKLHRLDHRGRFFSVEGPLNIRRSPQGQPVIFQAGSSDDGIDLAGRSADAVFSNGSTFD
EARVFYRRVKAAAAAAGRNPDHVKVFPGIGPIVGATQQEADDKYRQVRDLLSPREALAYL
SHFFQQHDFSVYPLDGPFPDIGTLGSDGFQSTTDNIKRLARERKLTLREVAYEVSTRRSN
IGTSEAFIGTPEAVASEMIRWVDEGAADGFMLGLPVTGFGLDDFVDHVLPVLSARGYFDP
VRRGATLRDHLGLPYKESRYARDAQVAAI*
>GCA_000452945.1_BPMSHR5858_genomic_05220_1 yhjX
MFAPETHRRDGSPRFFNPMMIAVVAGALVMSAAIGVRQTFGLFIGPFSFDHGLPVTTIAF
AIALHNLVWGAAQPFAGAAADRYGAGPLVAIGAVVFALGLAIAAVVPTGPMLVLGIGVLV
GIGISCTSFGVVLTAVGRGAPAEKRSMAMGIASAGGSLGQVALVPIAQWFTSHSGTMVSL
FVLAGCMIAIAPLGVLLDKNTRGSHVVAHETATISLKETLSYAVRHRGYCLLTLGFFTCG
FQLAFIGTHLPNYLLLCHMPAGLGATALALIGLFNMAGSWACGWLGGRYRQQHVLGWLYL
IRGAAIALFFLGPKSNASVVVFAAIMGLTWLGTVPLTSGLVAKVFGTRHLGTLFGVCFLS
HQVGSFLGSWLGGYVFDATGSYSLIWGATALAGLFAALLHFPINDAPAHGGAAVARA*
>GCA_000452945.1_BPMSHR5858_genomic_05221_1 rutF
MASPWTERATRAASDAHPAFEPPAPERARRDATRAEHRASTPNARASASNAAASPTPAPA
PDALAQHFKTAMRRLTSTVSIVATRDACARYGMAATAVSALSTEPPAILVCINRAATLHA
PLMRVRRFSLNLLHERQLELIAPFSGKLDHDARFAHGEWCDAHGVPMLAGAQATLWCAVD
GDFSYGSHTIVIGRVDSVSTAGPVAPLLWQDGAPCAARALAPCLHDAQ*
>GCA_000452945.1_BPMSHR5858_genomic_05226_1 bkdA2
MTMIQALRSAMDVMLERDGNVVVFGQDVGYFGGVFRCTEGLQNKYGKSRVFDAPINEGGI
VGAAVGMGAYGLRPVCEIQFADYFYPASDQIVSEAARLRYRSAAEFIAPLTIRMPCGGGI
YGGQTHSQSPEAMFTQVCGLRTVMPSNPYDAKGLLISAIENDDPVIFLEPKRLYNGPFDG
HHERPVTPWSKHPASLVPDGYYTVPLDSAAIVRAGGDVTVLTYGTTVHVSLAAAEETGID
AEVIDLRSLWPLDLDTIVESVRRTGRCVVVHEATRTCGFGAELIALVQEHCFHWLEAPVE
RVTGWDTPYPHAQEWAYFPGPNRVGDALRRAMEN*
>GCA_000452945.1_BPMSHR5858_genomic_05230_1 leuO
MIQNLRQLDLNLLLVFDALMQEQNLSRAAIRLHMSQPAVSNALTRLRQQLGEPLFTRTAR
GMTPTAQARTLYEPVRQALYLLQFGLGPQADFEPDTHHLFKLSMNDYGQTVLLPDLLAHI
KSRAPNVMLSVQSDDAGSIPAQLTTGTLDLAIDYLHFDNPELCYEPLHEEHLVVIGRAGH
PAFAGGLALRGYEESGHVSIQPRDRRGSPLEIVLGSARVRRVVHLQVPHYLTIPALVAKS
DLLGTIPRRLAERFADVYALQIAPLPIDIAPIQVSLIWHRQQDAQPGLRWLREQVVLTHR
AIIGCGEHAPA*
>GCA_000452945.1_BPMSHR5858_genomic_05232_1 choD
MKQQSYDYDYVVVGSGFGGSVSALRLSEKGYRVLVIEQGRRWTPENLPESTWNLSRWQWR
PALGLRGFFSMRFFRHVVVLHGNAVGGGSITYANTLLVPPDKVWREGTWAGLEDWERVMP
AHYATAKRMLGVVTNRRMDAADFRLKDMAKLIGVEKSFYPTEVGVFFGDDADAPGTRYAD
PYFGGAGPERTSCIGCGGCMVGCRHGAKNTLDRNYLYLAERLGAQVREQTKVVDVRPLDA
RADGAAGYAVEAVSLAAGARGAKSRLTCRGVVFAASSLGTQDLLMRLKEKGSLPRLSDAL
GKRVRTNAESLIGVRFPKSRVDLSKGVAIGSGIYIDEHTHIEATRYPSGSDTMGLLTTVL
TRGAPGGLRVLVWLGALAKLVLTRPLSAWRMIDPRGFARETMIFLCMQTLEGHLTMRLKR
RWFWPFSKQLATSGAKIPAYIPAANDFAQKAARALGGVPMTSLTEILLNVPMTAHCMGGA
AMARDARDGVCDGRSRVFGYRNMYVCDGSVLGANLGVNPSLTITALAEHAMSHVPAAREQ
RWDSTAETPVAA*
>GCA_000452945.1_BPMSHR5858_genomic_05241_1 IMPDH_2
VNAAEICTREVVACRRTDTVLDAAHLMRDRHVGDLIVVDDAGHAHAPVGMLTDRDIVLSL
IAKGVDPAALFVGEIMSAPAAVVHEHDSLWTIAQRMRLTGVRRMPVVNADGALVGMVSVD
DLLASVAALFDEIASISRRQIHFEEKARD*
>GCA_000452945.1_BPMSHR5858_genomic_05242_1 group_3627
MSDPELLTYQPIAVALPGPAEAGAPAVIDLPPPDLGAGLPLMSALSLRASTREFSSAALA
PTTLGELLWAADGVNRPTTGGRTAPSAHAFNEIDIYAALPDGVYRYDAALHRLLLKHAVD
ARNLTGYQDFVGAAPLDLIYVVRMSRLLSMPKPLRETFSAVAAGAIAQNVALYCASTGLG
CVVRGWINHRLLAEALSLNEDELPILAQTVGKPAAHP*
>GCA_000452945.1_BPMSHR5858_genomic_05261_1 rhsA
VNRRIVSSFLERCAAWLAAAKPHAVAVLHRLRHPTRRGVLLAVAALPALCVLYVLALIPF
TPSIGDIRKARVDAPAQILSADGKLLAEFKPSNREWVPLADISPKMVDALISTEDHRFYE
HHGLDWRRTAGAALRTFSGDRQGGSTITQQLARNLYPDEIGRAPTLTRKLKEAITALKIE
AVYSKAQILETYLNTVPFLYNAYGVEMAARTYFDKSADQLDALDAATLVGMLKGNSYYNP
VLNPERALARRNTVLAQMVKYGRLSPAAYASLQKKPLRIDFERQKEPPGPAPHFAQQLRK
WLIAWADRNDYNIYSDGLVVRTTIDSRLQTYATQALARQTNQLQGVANGMWNAGSGCAPG
NPLFRAFVRETPEFRAALDGGATGDAALKRLLADRGFARALCKAKADVQAGFLAIDPRNG
QIKAWVGSRDFTAEPFDHVQQARRQPGSTFKPFVYGAAFAAGATPDDTFVDQPVEIPLAG
GEIWRPDDDAPPTNKPMTLRDAIAYSRNRITAQLMMKVGPQKVARLARAMGVRDSALDAV
PSLALGTSPVTLKEMVSAYATIANVGEYVEPRMVTRIEGRNGEVLAEFASATPERALDAA
AARTLIDVMRGVVERGTGAAIRSRYGIRADVAGKTGTTQGDTDGWFILMQPELVAGAWVG
FDDGRVTLGGDWGQGARSALPIVGDFYQRAIRARLVDTRERFATEAPPSAFDTFRNKLGD
WYRYLFEKPEPPRKAAPPKAPRAPLEEVMPASEVEAASAAVARAASEAAAVASAASTASA
APPASGVPFAPGGASAPALPPLLPPMPQSAPLPPAAQPGSSLPNDNAPMSPTPTPDAPAA
GGSN*
>GCA_000452945.1_BPMSHR5858_genomic_05270_1 group_3628
MIDERWARAARELRCDAGFWSLRVVDERVDEHAVRNDVAQPLSSARDRGAMLVAWCGAGA
GYAATADLSPAGLQAALDVAAARAKASAALSLIDHRAVARPDASGGYVSPHADDALPGRA
EWLERLARECAAAAVDARIVERTAAVMLVQTDQLYMTSDGVRIEQRFRHVMPQLSVVAHA
RGDTQVRTLGNAGTLAQGGVDVLARYGFEGAGARVADEALQLLAAPNCPSGRRDLLLMPD
QMMLQIHESIGHPLELDRILGDERNFAGWSFVKPEMFGAYPYGSPLLNVTFDPTLPGEAA
SYAFDDDGGAARKQYLIRNGVLERPLGGALSQARAGLPGVANARASSWNRPPIDRMANLN
VEPGAHTFGELVAGIERGILMRTNTSWSIDDHRNKFQFGCEFGQLIENGELTQVVKRPNY
RGISASFWRSLRAVGDASTFGVYGTPYCGKGEPAQIVRVGHASPACVFADVDVFGGA*
>GCA_000452945.1_BPMSHR5858_genomic_05271_1 group_286
VAPVRRFGSAAASEIGSQNDPGVRSKAIDALTRAALAADMLDDPRTATSALDVSIQKQAL
NLLTNLQKKYKLSYLFITQDLAVMRAMAHRVRVMKEGRVVEAGDTPDVPDAPSHPCTRSL
LASSMPAARRSPRENEDD*
>GCA_000452945.1_BPMSHR5858_genomic_05275_1 cdhR_7
MSGVGPIVDALNLANEIDGRLLYRWQVCSWDGRAVPLAGGAQWHAHAAFNDAIVCDWLIV
VSERFQQFADYRLFLASLARVGQRTPLVTGIHHGVWWLAMAGQLSGYRVSVNWETYQQFA
EQFERSIVTQQIFEIDRDRATCAGGQASVDFMLAMIGRDQGPELAERIADALGTGPLRSG
EARQRIPFVTAPGERHPRLNDALLLMEANIEDPLTTDEIAGLVGVSRRQLERLFRQYLGA
MPSKYYLNLRLLKARTQLQRTSKSVVQVSLACGFSSAAHFSNAYRERFGVTPREDRRAWL
EKQHGGVEPPRAGAMVERPSGRE*
>GCA_000452945.1_BPMSHR5858_genomic_05281_1 astE
MTSSADSGRDAAWLDDFLALTLAGDAPPAEAGECAARAVRWRWLGDGLLRLEPADAAQRM
QSVLVSAGVHGDETAPIELLSTLVRDIARGALPLRCRLLVALGNPGAMRAGERYLDDDLN
RLFGGRHAQLAASREAPRAAQLEAAAALFFSTAGRARGARWHIDMHTAIRASVFEQFALL
PHTGEPPTRTMFEWLGEAQIAAVLLHTTKGSTFSHFTAQACGALACTLELGKVMPFGAND
LSRFAPADAAVRRLVSGRRDAPRGALPRAFTVVDQITKQSDALELFVANDVPNFTPFARG
TLLARDGDYRYAVRHEQERIVFPNPSVKPGLRAGLLVIETTRDTHAALA*
>GCA_000452945.1_BPMSHR5858_genomic_05295_1 group_3630
MSMIDIDPPSFQSPRPIAGDDGNRRTVLYGDYTVFSVFQPVFSVSHRRAIGYHASLRAHD
SQGNQVPSHEVFTQAARRGDLLELGRLAESLHLGNFHTFDSHDEWLFLSLHPAALMDTVY
GDALLANLKALGLPPQRVVLEVPEQAGGETPRFAAIVDGLRKAGFLIALGGFGAKHSNID
RVWHLHPDIVTLDRGILAQASEHSHLERVLPGLVSLLHESGQLVLMGGITTEREALIALE
CDVDFVQGQYFAGPSVEPVQPQAAAGVMDTLSAALRLRVAERTRAQQARLAPYVAALLQA
SALLREGRDLTEASKELLELPEAARCFLLDASGRQIGDNVLPRVHASQRAKRFSPLLHSE
GASWERRPYFIEAMRVPGRAHFTPPYLSINEAHLCVTASIATPAVQGMQVLCVDINWEAA
AHRD*
>GCA_000452945.1_BPMSHR5858_genomic_05297_1 group_2756
MATICVYGAGAVGCYLGGRLLAGGADVALIGRARIGDELRAHGLALSDYRGRDVRVPPSA
IAFSTADAALATAELVLVTVKSAATPAVGAALARIARPGAVVISFQNGVRNADVLRAALP
GATVLAGMVPFNVIARGEGRFHQGTAGALQAEAAPALRRFAAAFERAGLPLVPCEDMRAV
QWAKLLLNLNNAINALANRPLKEELAARSYRRCLALAQTEALRVLSAAQIRPARLTPLPA
AWVPRVLSVPDPLFARLGRKMLEIDPLARSSMSDDLAAKRATEIDWINGEIVRLAGQLGC
AAPVNARLCALVHEAERAAARPVWGGDALLAELREAARAG*
>GCA_000452945.1_BPMSHR5858_genomic_05308_1 tgt
MTTEGPSHDTHDTSAGIRPHGGLKFELLTTDGRARRGRVTLNHGVVETPIFMPVGTYGTV
KAVQPRELDEMRAQIILGNTFHLWLRPGLETIGAHGGLHRFMGWSKPILTDSGGFQVFSL
GDLRKITEEGVTFASPINGDRLFLSPEVSMQIQKTLNSDIVMQFDECTPYATNGVPTTHR
EAAESMRMSLRWAKRSLDEFERLGNPNALFGIVQGGMYEDLRDESLAGLSALGFHGLAIG
GLSVGEPKEDMMRVLEHVAPRLPADKPHYLMGVGTPEDLVAGVAAGVDMFDCVMPTRNAR
NGWLFTRFGDVKIRNATHKNSLKPLDETCGCYTCRNFSRGYLHHLHRVGEILGAQLNTIH
NLHYYLDLMREIREAIETHTFEAFQKRFAQDRARGVD*
>GCA_000452945.1_BPMSHR5858_genomic_05311_1 secF
MEFFRIRKDIPFMRHALVFNVVSLVTFLAAVFFLFHRGLHLSVEFTGGTVIEVQYQQAAQ
LEPVRATLGTLGYADAQVQNFGTSRNVLIRLPLKQGLTSAQQSDQVMAALKAQNADVALQ
RVEFVGPQVGRELATDGLLALACVVIGIVIYLSFRFEWKYAVAGIIANLHDVVIILGFFA
FFQWEFSLSVLAAVLAVLGYSVNESVVIFDRIRETFRRERKMTVQEVINHAITSTMSRTI
ITHTSTEMMVLSMFFFGGPTLHYFALALTVGIMFGIYSSVFVAGSLAMWLGIKREDLVKE
KKSAHDPNDPNAGAQV*
>GCA_000452945.1_BPMSHR5858_genomic_05331_1 gltA_1
MLSRRYLAMWCALALFAAAAVLAARHTISWLWIVPVAALVALGLYDLNQDRHAILRNYPL
WGHLRFLFEFIRPEIRQYFVEDDTDEKPFSRAQRSVVYQRAKNVADNRPYGTELDVKAVA
HEWISHSLAPTKLDGHDFRIRVGATRKQPYDISIFNISAMSFGALSANAIRALNLGAKQG
GFAHDTGEGSLSKYHREHGGDIIWEIGSGYFGCRRDDGTFDPEKFAKQAREPQVKMIEVK
LSQGAKPGHGGVLPAAKITPEIAETRGVPMGKDCVSPAAHSAFSTPRGLLEFVDRLRELS
GGKPTGFKLCVGHPWEFFGIAKAMLETGIVPDFIVVDGAEGGTGAAPLEFTDHVGVPLQE
GLLLVHNTLVGIGLREQVKLGASGKIITAFDIARTLAIGADWVNSARGFMFAVGCIQAQH
CHTDRCPTGVATQDPVRQRALVVPDKADRVYNFHRNTLHALQELVQAAGLSHPSELRAHH
IVQRVAPHEVRLMSQLLKYLEPGALLNGGHCGFSLYETWWPLARGDSFSPGEGAYARVGT
GAPARA*
>GCA_000452945.1_BPMSHR5858_genomic_05333_1 pepP
MNQPTEPALALDVYRQRRDRVLASLRAQGGGVAIVPTAPEVPRNRDSDYPYRHDSYFYYL
TGFAEPDALLVLDASAAGDAPRSILFCRAKNPEREIWEGFHFGPEAARDAFGFDAAFPYD
ALDAEMPRIVADAPALHYRFGVSAAFDARLNGWLDAVRARARAGVAAPGAAFDLGPLLDD
MRLVKDAHEQATMRRAADISALAHRRAMAACRPGIREYELEAELLYTFRRHGAQSPAYGS
IVATGANACVLHYPAGNAVVADGELVLIDAACELDGYASDITRTFPANGRFSGPQRALYD
IVLAAQEAAIAATRAGTQFDAPHDAAVRVLAQGMLDTGLVPKTRFASVDDVIAERAYTRF
YMHRTGHWLGMDVHDCGDYRERGAPRDDDGALPSRVLHPGMALTIEPGLYVRPGEDVPQA
FWNIGIRIEDDAFVTPTGCELITRGVPVAADEIEALMRDARPAPRPQP*
>GCA_000452945.1_BPMSHR5858_genomic_05338_1 ruvA
MRILGIDPGLRVTGFGVIDVSGHQLAYVASGVIKTPTADLPTRLGTIYDGVSTLIREHTP
DQAAIEKVFVNVNPQSTLLLGQARGAAICGLVSGGLPVAEYTALQLKQAVVGYGRATKEQ
MQEMVARLLSLSGRPGTDAADALGMAICHAHGGNTLNTLGGIAPALAKKGLRVRRGRLVG
*
>GCA_000452945.1_BPMSHR5858_genomic_05342_1 group_3635
MPRPLLHFVAAATFPFVAALAACGGDGTAPALAPTPAAVDAGFVSGHAPNPGPNAGDSPA
APAPTPGKWKNARAGDMLEVTLGELRPTQGALGYDQIYYKLGRYERQPDKKFDDFCADDG
LGGIAPDGYTALSALRDAASYACATSAAARDRSVLNPVVIGPNGDALYVTDGHHGLSTYY
ETPDGGAALRVHVVVKDNLADYSAAAFWQQMQNRGYTRLKDGDGRPIAPDRLPAGLGLKR
GMQDDRYRSLVYFTRDIGYAKPAQATDYLEFYWADWLRQQPQAFPLAGYDLTRAGATDPD
PATADTGYLNAVWNASARMVASTDPVIDGKTGADLGRADQVNGGKKYGKGEFDKLRQPIT
ADKPGKIAYALDYKSRRGLR*
>GCA_000452945.1_BPMSHR5858_genomic_05343_1 pspA
MPTTQILFIRHGETAWNRIKRIQGHIDIPLADTGLAQARQLAERLAREARGGARIDAVYT
SDLSRARQTAQPTADALGLPLVPRQALRERAYGVFQGHDSTEIEARFPDAFAQWQTRDPG
FEPEGGESHRAFYHRVLHEVERIVAAHPGGRIACVAHGGVLDCVYRFANDLPLDAPRKHA
LLNTSVNIVDYDDGRARVVRWADVDHLSGASDDDGYRKVL*
>GCA_000452945.1_BPMSHR5858_genomic_05345_1 tyrS
MSTDPTSKPAFPITDEVRHALAVTKRGVDELLIEEEFAQKLAKSAATDKPLRIKLGLDPT
APDIHLGHTVVLNKMRQLQDLGHTVIFLIGDFTSLIGDPSGRNATRPPLTREQIESNAKT
YFEQAALVLDREKTEIRYNSEWSMPLGADGMIKLASRYTVARMLEREDFTKRFQGGIPIS
IHEFLYPLMQGYDSVALNADLELGGTDQKFNLLVGRELQKQYGQEQQCILTMPLLEGLDG
VEKMSKSKGNYVGISEKPTDMFGKLMSISDVLMWRYFELLSFRSLDEIARFRGEAEGGRN
PRDFKVMLAQEIVARFHSQADAERALEDFNHRAKGGVPDDIPAVTLAGAPLAIGQLLKQA
GLVPSTSEALRNIEQGGVKIDGATVSDKALKVDAGEFVVQVGKRRFARVTLTA*
>GCA_000452945.1_BPMSHR5858_genomic_05356_1 yafJ
MCQLFGMNCAEPTDVTFSFTGFAARGGLTDHHADGWGIAFFEDKACRLFIDQQSSSTSPI
AEMVKRYPIKSKNTIAHIRKATQGHILLENCHPFMRELWGRHWIFAHNGDLQSYVPDLEG
GVYQPVGTTDSEKAFCKLMQGLREAFPSAQPPLPELFEQIGELTRDITRHGVFNFLMSNG
QALFAHCSTRLHYLVRRWPFSTAHLIDEDISIDFAKYTTPEDRVAVIATQPLTDDEVWTA
LEPGELVMFQCGDVAATMRIPVPEAVLEKLRNPALDASASAPRRSPALAASGAGDDLGDD
PVDF*
>GCA_000452945.1_BPMSHR5858_genomic_05357_1 group_3637
MLSPHEFATLMLVRHAPDQIDMDRAELDALLERQLVMLEHSAEGRRRAHLTAMGRSLLDA
FDPDDHPRRRPAKIHTCGESGR*
>GCA_000452945.1_BPMSHR5858_genomic_05363_1 glnQ_2
MISIKNVSKWYGQFQVLTDCTTEVKKGEVVVVCGPSGSGKSTLIKTVNGLEPFQQGEILV
NGQSVGDKKTNLSKLRSKVGMVFQHFELFPHLSITENLTLAQIKVLGRGKDEATEKGMKL
LDRVGLKAHAHKYPGQLSGGQQQRVAIARALSMDPIAMLFDEPTSALDPEMINEVLDVMV
ELAQEGMTMMVVTHEMGFAKKVAHRVIFMDKGLIVEDDRKEEFFANPKSDRAKDFLAKIL
H*
>GCA_000452945.1_BPMSHR5858_genomic_05370_1 purB
MSDTRPDTLFALTALSPLDGRYASKTEALRDWLSEAAFMRHRVTVEVHWLIALSRAGFAE
VPRFSEAAEQFLLQLVERFTAHDAARIKEIERVTNHDVKAVEYWLKESVKGQVELEQASE
FIHFACTSEDINNTSHGMMLAGARQHVILPALRTVHGRLVALAHAHAEQPMLSRTHGQPA
SPTTLGKEIANVAARLARAIERIEKVEILGKMNGAVGNFNAHLSAYPEFDWEAFSRDVIE
HRLKLTFNPYTIQIEPHDYMAELFDAVSRANTILLDLDRDVWGYISLGYFKQKTKAGEIG
SSTMPHKVNPIDFENSEGNLGLANATLRHLADKLPVSRWQRDLTDSTVLRNIGVAFGYSL
LAYDALIRGLDKLEVNPQRLNDDLDNCWEVLAEPVQTVMRRYGIENPYEQLKELTRGKGI

TRDALQQFIGTLAIAEDAKARLLAMTPASYIGKAVELAKRIA*
>GCA_000452945.1_BPMSHR5858_genomic_05372_1 gntT
MGAVQGSMLLIYTVIAIAVLILMIARYKVYPFLVLIIVSLGLGLAVGMPMDKIVKSFEGG
TGGTLGHIAIVVGLGTMLGKMMAESGGAERIATTLIDWFGEKNIHWAMMFVAIIVGLPVF
FEVGFVLLIPIAFNVAKRTGKSLLVVGLPMVAGLSVVHGLIPPHPAALLAVQQYGADIGK
TIAYGLIVGVPTAIVAGPLFALTISKFVKLPEHNPLASQFVETRAAGETRALPSFGITLF
TILLPVVLMLVGSWADLVFAPKSLPNSLLRFAGNSDVALLIAVLVSFFTFGKLQGFNREQ
IQKFCGDCLAPIAGITLIVGAGGGFGGILRDSGISQQIVATATHAHLSPLLLGWFVAALI
RLATGSATVAMTTACGIVAPIAAASGVTVRPELLVLATGSGSLIFSHVNDGGFWLIKEYF
GMTVGQTFKTWTLLETIISVLGLTFTLALSAVL*
>GCA_000452945.1_BPMSHR5858_genomic_05379_1 lptE
VIRRSFLMLVGSAVLLSACGFKLRGQQDYAFKRLYIAGAPPAAAARLTRLVEAGSDTKIV
KTPDEADVVLNLAESRGQGTLTLDRFGTVQEYQLNYSLNYTLFGKDGTLLIPPSLIALNR
AMTYSAKYTNAKAQESEILFADMQNDAIDQLMRRLAIVRTLHPEPGEGVPAVAPRAPLPP
PPL*
>GCA_000452945.1_BPMSHR5858_genomic_05385_1 fur
MTNPTDLKNIGLKATLPRLKILEIFQQSPVRHLTAEDVYRNLLHEELDIGLATVYRVLTQ
FEQAGLLSRSNFESGKAVFELNEGSHHDHLVCLDCGRVEEFFDAEIETRQQSIAKERGFK
LQEHSLAMYGTCTTENCPYRKH*
>GCA_000452945.1_BPMSHR5858_genomic_05391_1 cynT
MQDIIEGFLKFRRDAFAPRTALFKRLAASQQPRALFISCSDSRLVPELVTQREPGELFVI
RNAGNIVPSFGPELGGVSASVEYAVAVLEVADVVVCGHSDCGAMRAIASCACLDHLPAVR
SWLRYADAAKRVNEAREHAGEQARVDSMVRENVVAQLANLNTHPSVRLALEQGRLTLHGW
VYDIETGSIDALDGATGRFVALAEHPRVHATPPRVQRAA*
>GCA_000452945.1_BPMSHR5858_genomic_05393_1 sotB
MNSVTAPSPALTRGMTLLFACACGIVIGNLYYSQPLLAAIAASFGRQPAELGYLVTLTQL
GYAASLLVVVPLGDVVNRHTLIVRLLVANVVALVAVASSTRYGFFVAANMCVGFVTCSTQ
LLVPFAASLADDGARGRAIGTVMSGLLLGILLARVASGAIADWLGWRAVYAIAAAMVLLL
AGVLAAKLPKDRRHARVDYAALMQSLAALVRAQPLIALRSAYGALVFACFSLLWTGLTFL
LSQPPYGYTEGRIGLFGIVGAVGALAAASAGRLVDRGHGNAATGLFAAAVLASFAAIAAG
EQSLAALIVGILVLDIGVQGLHISNQSVIYALAGDARSRVTTIYLTSYFIGGALGSGAAG
AAFGIDGWRGVCIAGAILSGSLVALWGASQRVGARHAAR*
>GCA_000452945.1_BPMSHR5858_genomic_05395_1 tktA
MPPVPRFLDSFSGLDMTTSSPASTTLMANAIRALAMDAVQQANSGHPGMPMGMAEIGVAL
WSRHLKHNPTNPHWADRDRFVLSNGHGSMLLYSLLHLTGYDLPIEELKNFRQLHSKTPGH
PEYGITPGVETTTGPLGQGLANAVGMALGEALLAAEFNRDDAKIVDHHTYVFVGDGCLME
GISHEACSLAGTLKLNKLIAFYDDNGISIDGDVVNWFHDDTPKRFEAYGWNVIPNVNGHD
VDAVDAAIAKAKSADKPTLICCKTVIGKGAATKAGGHDVHGAALGAEEIAKTRDALGWKW
APFVIPQEVYAAWDAKEAGKRREGDWNAAFAQYRAKYPAEAAEFERRMAGKLPADWAEKA
AAIVAGANERGETVATRKASQQTIEGLAAVLPELLGGSADLTGSNLTNWKASKAVRANAH
GAGIQWGNHINYGVREFGMSAAINGLVLHGGYKPFGGTFLTFSDYSRNALRVAALMKVPS
IFVFTHDSIGLGEDGPTHQSVEHVASLRLIPNLDVWRPADTVETAVAWTHAVSHQHPSCL
VFSRQNLAFNARTDAQLANIEKGGYVLRDWDEEIVARKIILIATGSEVELAMKAVEPLAQ
QGIAARVVSMPSTDVFDRQDAEYRERVLPHGVRRVAIEAGVTDFWRKYVGLEGGVVGIDT
FGESAPAGVLFKHFGFTIEHVVETAKAVLA*
>GCA_000452945.1_BPMSHR5858_genomic_05409_1 araG
VAAALRFDNIGKVFPGVRALDGISFDVQAGQVHGLMGENGAGKSTLLKILGGEYQPDSGS
VLVDGRAMRFPSAAASIAAGVAVIHQELQYVPDLTVAENLLLGRLPSALGWVRKRDAQRF
VRERLAAMGVDLDAQAKLRRLSIAQRQMVEICKALLRNARVIALDEPTSSLSHRETEVLF
KLVDDLRRDGRALIYISHRMDEIYRLCDACTIFRDGRQVASHASLANVPRETLVRQMVGR
EISDIYHYAPRALGDVRLSARALEGDALRAGASFDVRAGEIVGFFGLVGAGRSELMRVIY
GAQRRTGGALTLDGEPLDIRSTRDAIRRGIVLCPEDRKEEGIVAHASVAENINISCRRHG
LRAGLFLDRKREAETADRFIKLLKIKTPNRRQKIRFLSGGNQQKAILARWLAEPDLKVVI
LDEPTRGIDVGAKHEIYGVIYELAKRGCAIVMVSSELPEVLGVSDRIVVMREGRIAGELA
RGEANEEAVLNLALPQGATAHAA*
>GCA_000452945.1_BPMSHR5858_genomic_05411_1 ucpA_3
VNRLAGKVAIVTGAGRGIGAAIARAFVREGAAVAIAELDAALAEESADAIARDTAGARVL
AVPTDVAQAESVAAALARTERAFGPLDVLVNNAGVNVFGDPLALTDEDWRRCFAIDLDGV
WNGCRAALPGMVERGRGSIVNIASTHAFKIIPGCFPYPVAKHGVLGLTRALGIEYAPRNV
RVNAIAPGYIETQLMHDWWSAQPDPQAARRETLALQPMKRIGRPDEVAMTAVFLASDEAP
FINASCITIDGGRSVLYHD*
>GCA_000452945.1_BPMSHR5858_genomic_05412_1 dgoA
MSMDAEALRFPAPYAPHRALMRAFGACPMIAILRGVTPADAAAHGAALYEAGFRIVEVPL
NSPDPFDSIAALRTALPADAIVGAGTVLRAGHVDEVVRAGGELIVMPHADGDVVRRAKTL
GIGCAPGVATPTEAFAALANGADVLKMFPAEQLGTPAVKAWRAVIDARVPLVPVGGITPE
NMGPYLAAGASGFGLGSALYAPGQTSAVTAVRAHAFINGWRIARHGAAR*
>GCA_000452945.1_BPMSHR5858_genomic_05414_1 kdgR_5
MNKTPMTLALDGATADRPDTPGMPDATLADSIGATSTPLDLAAQQAGTQTLLRGLAILEA
VAGGARDMRAIGAALGTTRSTTHRLVSSLVQARYLRQVQGGYLLGPKLIELGTIALEQMP
LTAVARPHLEALAQATLDTIHLGVRDGDDVLYIDKIPGTRGLEMRSRIGHRMPLASTGIG
KAMMLDLDPDTWRSLFDASKRALAGVNFKPDRRPDVGTFLQRMAHYAAGGYTFDLEENET
SIRCVAAPVRDASGAIVAALSVASTIPYMSLDRMNELVPLVQRGARAISADLGWSAPQAT
RRIKR*
>GCA_000452945.1_BPMSHR5858_genomic_05419_1 rnb
VNVFFEESGSFKAGSVLSRQGDAFQVELPGGRRAKVRAKDVLIEFEKPAAGELMQQADEA
AQQIDLDFLWECAPAEEFAYAALADEYFGASYGPVERAALVLRLHGAPVYFRRKGRGQYQ
RAPEEQLKMALAGLERKRQQALVQAGYEEELKAGRLPDAFAGKALGLLTKPDKNSIEYKA
LDAAALARGVSPARLMLDCGGIPSARALHEARFLAEYFPHGTGFPAVAVGKLPEDLPRAD
VNAFSIDDITTTEIDDAFSVEHLSDGRVRIGVHIAAPALGIVRGDAVDAIARARLSTVYM
PGDKITMLPDDVVDVFTLKEGDYRPALSLYIIVKRDTQEIVANETRAEYVYVKSNLRHNT
LDELVTEDALAAGTGDYPHKDDIAALWPLAQALFERRQVARAGYGLKREVQRNTDYNFYV
EGEHVSITPRRRGSPLDLIVSELAILANSTWGAFLHDHSVPGIYRTQRAFGMPSGPKRTR
MQTSAAPHEGLGVPQYAWSTSPLRRYVDLVNQWQLLACVQHGVTAKLAAPFKQKDADLYA
VVQGFDDTYAAYADHQRRMEYFWCLRWIKQEGRKQVSATVVKGELVRLDEVPLLLHVPAL
GVHARGTRVLLDVMSVDELTIEASVRLVSVLDAPMVSSGEPADEDEDADAADETLLDAAD
ESAQGEAEALAEAGGAASGNGESSANGEEQSK*
>GCA_000452945.1_BPMSHR5858_genomic_05420_1 group_19915
VILYLHGFRSSPQSFKARALAARLDELGRADEWRCPVLSVAPLAAIAAAEAEVAGVPAQR
VTVIGSSLGGYYATWLAEKHGWRAVLLNPATQPQRDLRRHLGEQPLWHGGGTIVVEPHHL
DELNALHVAAISRPERYYLFAATGDEVLDYREMLARYPGARTRVIEGSDHGISEFADYIE
DVLAFCDAG*
>GCA_000452945.1_BPMSHR5858_genomic_05425_1 accB_1
MDLRKLKTLIDLVSESGISELEVTEGEGKVRIVKNAPPVYVQPSAGFAPQVSAPAPVVTA
PSEAAGAAAAAAAAAAAAAAPQGHVVTSPMVGTFYRAPSPGADPFVQVGDTVKEGQTLCIIE
AMKLLNEIESDKAGVIKEILAENGQAVEYGQPLFVIG*
>GCA_000452945.1_BPMSHR5858_genomic_05427_1 prmA
MSYRELVAELPREHAEALSDALVELGALSVSVEDADADTPDEQPLFGEPGLVPERTAWQH
SRVIALVDATQDPAVLLAAAANEAGLAQPPRFELREVEEQDWVRLTQSQFEPIHIGEKIW
VVPSWHDAPQPDALVLELDPGLAFGTGSHPTTRLCMEWLEQTVQPGQTVLDYGCGSGILA
ILAKKCGAGRVTGIDIDPQAVEAARHNSERNRADVTYGLPDDCPDGEFDIVVANILSNPL
KLMASMLASKVKPGGRIALSGVLARQADEVASVYARYIDIAVWREHEGWVCLAGTRRESH
*
>GCA_000452945.1_BPMSHR5858_genomic_05441_1 group_2617
LESNFLSATVLLILITDPLGNIPLVIAALRDVPRERRVKVILREVAIAFVILLFFMVVGD
RFLRMMSLTDLSLRIGGGIVLFLIALRMIFPHPDGALGSDPRAGGEPFIVPLAIPALAGP
SALATVMLLTSQAPGKTLEWAGALTVTMLVCAVTLVLAERIQQWLGERTVIAFERLMGLV
LVAISVEMMMAGIRAFVHQL*
>GCA_000452945.1_BPMSHR5858_genomic_05445_1 proB
MRSIIADSKRLVVKVGSSLVTNDGRGLDHDAIGRWAAQIAALRGAGKEVVLVSSGAIAEG
MQRLGWSKRPREIDELQAAAAVGQMGLAQVYESRFAEHGIRTAQILLTHADLADRERYLN
ARSTLLTLLRLGVVPIINENDTVVTDEIKFGDNDTLGALVANLIEGDTLVILTDQPGLFT
ADPRKDPGATLVAEASAGAPELEAMAGGAGSSIGRGGMLTKILAAKRAAHSGANTVIASG
RERDVLVRLAAGEAIGTQLIARTARMAARKQWMADHLQVRGHVVIDAGAVDKLTAGGKSL
LPIGVVAVQGVFARGEVIACVDDTGREVARGITNYSSAETKLIQRKPSGEIETVLGYMLE
PELIHRDNLVLV*
>GCA_000452945.1_BPMSHR5858_genomic_05447_1 rpmA
MAHKKAGGSSRNGRDSESKRLGVKVYGGQAINAGGIIVRQRGTRMHAGENVGMGKDHTLF
ALVDGHVKFTTKGAAKKHTVVVVPAAA*
>GCA_000452945.1_BPMSHR5858_genomic_05457_1 mutT
MSGEVTNAAAPAATGRKVTEVAVGVLVRPDGRYLLAQRLIGKPYEGYWEFPGGKLEAGES
VEAALARELHEELGIAVTECHRWHMLEHDYPHAYVRLYFCKVTGWTGEPHSREGQAFVWQ
HLPVDVAPLLPAALPVLELLAREQGDAQQ*
>GCA_000452945.1_BPMSHR5858_genomic_05468_1 group_4306
LIGATLCASVTLMTAACGGSDGSGAPGASIFKSPHGSGSSSTTGTSGTTTSGTGGTSTGS
TQTTMALQMADFILAQQDSNGAIPDEAGSDTANTDSDMEYALIGLAAAYGATHDAKYLNG
LEKGIAWLAAREEMTDPAWRGSWRYVYSMTPPYDPIPTSPGAGIADVRGVDATSALFVYL
LYLDRQLTGSSALVTQYGANARAALDFVLAKNINPSGYSGSSWQLPVGSTTWQFWPYEYA
ADQGDVYLGMNGGALLFPDNPNYAAKAAFLRNNVPSQFYMTGAQRYSVGRDTGAPLDSDL
GIDTIFPQGYLPWVFGANSQSIGSIQWMINQTAADGSIRSPSTDPAYALSNVILLLGAPT
QGMQAPSTTLPWIVNNVLDPQTYGIHDYPGSPDQETNVSGFAVAALLGTKAFP*
>GCA_000452945.1_BPMSHR5858_genomic_05471_1 lpxK
MSARPGLLARAEARLTREWQRRGALAWALAPFACAFGAIAALRRAAYARGWKARVDCGVP
VVVVGNVTVGGTGKTPTVIALVDALRAAGFTPGVVSRGYGAKIAAPTAVTAASPPQQAGD
EPLLIARRTLAPVWVCPDRVAAVRALKAAHPEVDVVVSDDGLQHYRLARAVEIVVFDHRL
GGNGFLLPAGPLREPLSRRRDATLVNDPYSRALPPWPDTFALSLAPGDAWHLDRPSRRKP
LAQFAGERVLAAAGIGAPERFFATLRAAGVAPATRALPDHYAFATNPFVDDHFDAILITE
KDAVKLGTSWRDARIWVVPVEAALDPRLIALVVEKLRGRTSA*
>GCA_000452945.1_BPMSHR5858_genomic_05476_1 group_19800
MTRVLDYFSSLVAEDEGLPLTEAALSLAQDVYPDVDLQGVLAELDALAARLARRLPDGAG
ASDKLALLNDFFFRELGFACNHNDYYDPDNSYLNMVLKRRRGIPITLAVLYLELAEQVGV
PARGVSFPGHFLLRIALPDGDSILDPTTGRSLSETELVEMLEPYVAHAEGGTASVLRALL
QPATEREVVARMLNNLKAVYLQTERWQRLLGVQERLVILLPGQLDEVRDRGFAYARLDYL
RPALEDLETYLGERPDAADATIVEAQVSELRQRMQRDDDD*
>GCA_000452945.1_BPMSHR5858_genomic_05478_1 rpsT
MANSAQARKRARQAAKANSHNSALRSKFRTAIKAVRKAIDAGDQAKAAELFKAATKTIDT
IADKKIVHKNKAARHKSRLSAAVKGLQAQAAQ*
>GCA_000452945.1_BPMSHR5858_genomic_05479_1 group_2620
MAETLPTEYFIQGITKDGKKFRPSDWSERLAGVMSCYGPGASGPNARLQYSRYVRPTLLG
DLKCVIVDARLRDVEPMAFDFVMNFAKDNNLVVTEACELPDYGKTQE*
>GCA_000452945.1_BPMSHR5858_genomic_05480_1 argF
MTAKTIRHYLQFKDFSLEDYEYVLERTGILKRKFKNYETYHPLHDRTLAMIFEKSSTRTR
LSFEAGIFQLGGHAVFMSTRDTQLGRGEPVEDSAQVISRMVDIIMIRTFEQDIIQRFAEN
SRVPVINGLTNEYHPCQVLADIFTYYEHRGPIRGKTVAWVGDANNMLYTWIQAARILGFK
LRLSTPPGYALDTKLVDAESAPFYQVFDDPNEACKGADLVTTDVWTSMGFEAENEARKQA
FADWCVDEEMMSHAHPDALFMHCLPAHRGEEVTAGVIDGPQSVVWDEAENRLHVQKALME
FLLLGRLNH*
>GCA_000452945.1_BPMSHR5858_genomic_05481_1 murB
MPMSRPDSAVSLLPDYSLRAHNTFGFDARARVAARIGSPGQFASLARDPRVAGLDRLVLG
GGSNVVFTRDFDGLVLLDEIRGRALVREDDGAWYVEAGGGENWHAFVEWTLAEGMPGLEN
LALIPGTVGAAPIQNIGAYGLEMKEHFASLRAVDLATGELVEFDAARCAFGYRDSFFKRD
GRGRFAIVAVTFRLPKAWTPRIGYADVARELAARGIDARAARARDVFDAVVAIRRAKLPD
PLALGNAGSFFKNPVIDAQAFAALRAREPDIVSYPQPDGRVKLAAGWLIDRCGWKGRALG
AAAVHERQALVLVNLGGASGADVLALAHAIRRDVLGRFGVELEMEPVCL*
>GCA_000452945.1_BPMSHR5858_genomic_05483_1 plsY
MQILLATVAAYLIGSVSFAVVVSAAMGLADPRSYGSKNPGATNVLRSGNKKAAILTLVGD
AFKGWLAVWLVKRFGIGGEIGVALAAIAVFLGHLYPVFFRFQGGKGVATAAGVLLAVHPV
LGLATALTWLIVAFFFRYSSLAALVAAVFAPIFDVFLFGTHDNPVAWAVLAMSVLLIWRH
RSNISKLLAGEESRIGQKKKTGV*
>GCA_000452945.1_BPMSHR5858_genomic_05484_1 ybaK
MSKSKHVSETPATQFLRRHGVAFGEHVYEYVEHGGTSESARQLGVDEHAVVKTLVMEDEH
AKPLIILMHGDRTVSTKNLARQIGAKRVEPCKPEVANRHSGYLVGGTSPFGTKKAMPVYV
ESTILDLPSIYLNGGRRGYLVSLAPAVLATLLQARPVQCASVD*
>GCA_000452945.1_BPMSHR5858_genomic_05488_1 queG
MDRNPELAIADARPSQDGRAAPFRLDDAQLAELASRIKAWGRELGFGAIGISDTDLSEAE
AGLAAWLEAGCHGEMDYMAKHGMKRARPAELVAGTRRVISARLAYLPAGTLDGAPDAQGA
RRDWRAREAARIADPQAAVVSVYARGRDYHKVLRNRLQTLAERIEAEIGAFGHRVFTDSA
PVLEVELAQKAGVGWRGKHTLLLQRDAGSFFFLGEIYVDVPLPADAQTSPDAAPETPGAH
CGSCTRCLGACPTGAIVAPYHVDARRCISYLTIELHGSIPEPLRPLIGNRVYGCDDCQLV
CPWNKFAQAAPVADFDVRHGLDRASLVELFEWTAEQFDERMQGSAIRRIGYERWLRNLAV
GLGNALRAAPGGIGPDARAAIVAALRARLDDPCVSALVREHVEWALRAA*
>GCA_000452945.1_BPMSHR5858_genomic_05492_1 group_3941
MTDSIKAILKPHVRDIGNLTVRRTLPALAARTVGPFIFFDHMGPAAQPAGAGLDVRAHPH
IGLATVTYLFDGAIMHRDSLGSAQKIVPGDVNWMTAGRGIVHSERTPDEERARGQTVHGI
QTWVALPREHEAREPSFEHHAGATLPKLERDGVALTVIAGDAFGARSPVTTFSRTLYVAA
VFAAGGALALDADHDERAVYLVDGDLTIDGAPLEPAQMAVLAPGARVALASAGGARAMLL
GGDKLDGERFIEWNFVASSRDTIERAKRAWAAQQMGTVPGETEWIPFPEHRAH*
>GCA_000452945.1_BPMSHR5858_genomic_05496_1 group_3652
MFWEKKLAQWANEVREKADIPARLVLWNGQQLDFGTFAAPQVTLKVNSASALPLLLEPSL
DNLGEAYVKGKIDIEGKLADIINIGYSLARSTVTSASKLARVKRYFNHSKHSDRKAIQYH
YDVSNEFYQLWLDENMVYSCAYFENGDEDLGTAQIKKIDHILTKIGVQPGQRLLDIGCGW
GALVLRAAGKFGARCVGVTLSQNQFDLATERVKRAGLENQIEIRLQDYREIDGQFDRITS
VGMFEHVGRKNLPLYFSRIRELLADDGIAMNHGITSTDAESGETALGGGEFIDRYVFPDG
ELPHISLALEAAQRGGLEAVDVESLRRHYARTLDIWTENFEAKAEEARKLVDDEKFRIWR
VYLAGCAYAFEHDDVSIFQIVCRRAGQSAKTLPWSRRYMYEHALPR*
>GCA_000452945.1_BPMSHR5858_genomic_05499_1 C1-hpah
MKHLQASPPNFDTAAFRQALGQFATGVTVVTTRAPSGQLIGITASSFNSVSLDPPLVLWS
LAHKSASMPVFRTNSHYVVNVLSASQHDLCMRFATLKGNRFEGVSHAQGDSGMPVLDGAL
AWFECHNRSRYDEGDHVIFVGEVERCGVHPDAAALAPLVFQSGGFHGLTRL*
>GCA_000452945.1_BPMSHR5858_genomic_05501_1 kynB
MDTIWDISPPIAPATPVWPGDTPVGIERVWRIEAGSPVNVARVTLSPHTGAHADAPLHYD
ADGAPIGAVPLDAYLGRCRVIHCIGARSAVTPEHVRAALAGAPPRVLLRTYGQAPQHAWD
SAFCAVAPETIDLLAAHGVRLVGIDTPSLDPQESKTMDAHRRIRAHRMAILEGLVLDEIA
AGDYELIALPLKFATLDASPVRAVLRALPDAPR*
>GCA_000452945.1_BPMSHR5858_genomic_05503_1 kynA
MQPPGDDAAPRCPFAGAHAPDAPHVPEAAGDDAQAGWHRAQLDFSQSMSYGDYLSLDPIL
DAQHPRSPDHNEMLFIIQHQTSELWMKLALYELRAALASIRDDALPPAFKMLARVSRVLE
QLVQAWNVLATMTPSEYSAMRPYLGASSGFQSYQYRELEFILGNKNAQMLRPHAHRPAIH
AHLEASLQAPSLYDEVIRLLARRGFPIAPERLDADWTQPTRHDRTVEAAWLAVYREPNAH
WELYEMAEELVDLEDAFRQWRFRHVTTVERIIGFKQGTGGTSGAPYLRKMLDVVLFPELW
HVRTTL*
>GCA_000452945.1_BPMSHR5858_genomic_05508_1 gltC_4
MTFDLRQLRAFTTIAASGSLGRAADALHVTQPALSRILKRLEEQIGAPLFERHSKGVQLT
AIGEALLPHATLLQHEAEHAREEIDALRGLAKGTIKVGAVGSIASLVLPVALGRVLDRWP
NLRVVIVEGVWDRLVDALLTHEIDIALSTRVPDTDEVVAIAECRWDDVSHVVAAPDHPLR
AAARAPLTLADTRAARWAIPPRGTAPFEQMRAAFDAHGLAPPDIAVETRSVTALKSLVAH
AGFLSWMAEPMYRAERRARTIDTLAVEGVAATRTLTAFRRRHGILPGPAARLLEALALLT
REPF*
>GCA_000452945.1_BPMSHR5858_genomic_05518_1 sugB
MSDLTVSTPRGAFGAPRAALGALKRALPGLLAWLVALALFFPIFWMAITAFKTEQQAYSS
ALFFVPTLDSFREVFARSNYFAFAWNSVLISAGVTVICLLLAVPAAYAMAFFPGRRTQKV
LLWMLSTKMMPSVGVLVPIYLMWKNAGLLDTVSGLVIVYTLINLPIAVWMTFTYFNEIPK
DILEAGRIDGASTWQEIVYLLMPMALPGLASTALLLVILSWNEAFWSINLSSSNAAPLTV
FIASYSSPEGLFWAKLSAASLLAVAPILVVGWLSQKQLVRGLTFGAVK*
>GCA_000452945.1_BPMSHR5858_genomic_05524_1 zur
MASTHSLAARLSRADAFAAEHGLAWTPLRRQVYERVLAAGRPIGAYDLLTELEPQRGRVP
PTTVYRALEFLVEHGFIHRIESKNAFIACCEIGKPHEGQFLICEACGDTVEIPGGDLAKQ
LSSSPPAHGFEVHRQVVELTGLCGQCKTKHAHHG*
>GCA_000452945.1_BPMSHR5858_genomic_05526_1 mntB_1

MTHPMHALELDHVTLALGGRTILRDVSFAVESGEFVGVLGPNGAGKTTLMRAVLGLVPAH
SGAIRVAGEPVARGNPAIGYMPQIRSALAGRRVRGRDFVAMAADGHRWGLPHASAAVAAD
VDRVLDLVGASALAKRPLSELSGGERQRLLLAQCLLGTPTLLLLDEPLISLDPNHQRGVV
ELVRRVQRELGITVLFSAHELNPLLNALDRVLYLGNGVAALGTVDEVITKPVLSRLYGSP
IDVMRVNGRIFVMSGDVEVEKHDHGHEDEHDHGHEHGHAHSHGGGHSHDV*
>GCA_000452945.1_BPMSHR5858_genomic_05528_1 group_19842
METTTLECVSSHACHGGEQRFYRHESKTIGLPMKFSAYLPPQAAHGRVPALFYLAGLTCT
EETFAIKAGAQRFAAQHGIALVAPDTSPRGAGVPGEADAWDFGVGAGFYVDATEAPWSAH
YRMESYVSGELREIVAAELPIDAARLGIFGHSMGGHGALVLALRHPQLYRSVSAFAPIAA
PTRCPWGEKAFSGYLGADREAWKRHDASELVARADAKRFAEGILIDQGLADPFLPTQLHP
DVFEAACRAAGQPLTLRRHAGYDHGYYFISTFIADHLAHHARVLGR*
>GCA_000452945.1_BPMSHR5858_genomic_05532_1 uacT
MQSNTVHPCDEVLPSGKLLTLGLQHVLVMYAGAVAVPLIVGGALKLPKDQIAFLISADLF
ACGIATLIQTLGVWLFGIRLPVIMGCTFAAVGPMIAIGTNPGLGILDIFGSTIAAGAIGI
VLAPMIGKLLRFFPPVVVGTVISVIGLSLMEVGINWAAGGVGNPNYGDPVYLGLSFIVLA
LILAINKFGRGFVANISVLLGMIAGFAIAFAAGRVNTDGVAAAPWVGFVAPFHFGLPHFD
PLSIATMVIVMFVTFIESTGMFLAVGDMVERPVDQETLVRGLRVDGLGTLIGGIFNSFPH
TSFSQNVGLIGVTGVKSRFVCATGGMILVLLGLFPKMAQLVASVPPFVLGGAGIVMFGMV
AANGVKVLSKVDFVRNHHNLFIVAVSVGLGLVPVVSPNFFSKLPAAFAPILHSGILLASV
SAVVLNIVFNGMKGEKDARCDIRRAGRDFDGQPADLH*
>GCA_000452945.1_BPMSHR5858_genomic_05617_1 secY
LANSPSLAKPGRSTAKFGDLRRRAMFLLLALIVYRIGAHIPVPGIDPDQLAKLFQSQAGG
ILGMFNMFSGGALSRFTIFALGIMPYISASIIMQLMAIVSPQLEALKKEGQAGQRKITQY
TRYFTMVLATFQAFGIAAALENQPGLVTDPGAVFRLTTVVTLVTGTMFLMWLGEQITERG
LGNGISIIIFGGIAAGFPNAVGGLFELVRTGSMSIISAIIIVVLIAAVTYLVVFIERGQR
KILVNYAKRQVGNKIYGGQSSHLPLKLNMSGVIPPIFASSIILFPATILSWFSSGERKGW
MSDTLHNVAEALKPGQPVYVLLYTLAIVFFCFFYTALVFNSRETADNLKKSGAFVPGIRP
GDQTARYIDRILTRLTLAGAIYIVFVCLLPEFLVLRWNVPFYFGGTSLLIIVVVTMDFMA
QVQSYVMSQQYESLLKKANFKGGNIPMR*
>GCA_000452945.1_BPMSHR5858_genomic_05625_1 dsbD_2
MFNRIPRHAQSRFCFLIAVVAMLGVLFGTSLAARAADDFLDPAVAFKFSASEAPGQVDVH
FKIADGYYMYRERFAFAVKSGSATLGEPQLPAGHVKFDPTFQKNVETYRGDLTIHLPIKQ
ASGPFELAVTSQGCADEGICYPPAEHVARIEGAALGAAGTAPAAAGAGADTSAADGGSWY
ERVTSADYARSLLEGHGFLTIVALYFVAGMVLSLLPCSYPMIPILSAIIVGEGARATRAR
AFALSLTYVIGMALVYTALGVAAALVGQSLGAWLQNPWVLGAFALLLTVFALLLIGGVDI
TLPQRWQNGAAQTSGPRKGGRFAAVATMGALSALVVGACMTAPLFAVLAFIAHTGNALFG
GAALFSMGIGLGVPLLVIGIGAGTLLPRAGAWMDGVKVFFGVLLLAAALWIVWPVLNAAS
QLGLGALWLLIAAAALGLFTPHSGSSSVWRRLGRGLGAALAIWAATLLVGLAAGSTDPLR
PLAVLAARAAPSNGAAGAGAGAHEGPAFAPVRSIAELDEIVKTSTRPVMLDFYADWCVSC
KEMEHLTFTDARVGARLSQMHLVRADVTANSPDDQALLKRFGLFGPPGIIVFDANGQERG
RVVGYQSADRFLRSLDRMSLPAAWSAS*
>GCA_000452945.1_BPMSHR5858_genomic_05629_1 ccs1
MSVTTSGLQSRTGPSAVRRAVELLSSMRFAIALLVVLSIASIIGTVLTQDDPYPNYVNQF
GPFWADIFRSLSLYTVYSAWWFMLILIFLVVSISLCVIRNAPKMLADAKSWKDKVREGSL
RAFHHKAEFAVAGDRARATQTVAAFVAKAGYRHVVREAGGATLVAAKRGAMTKFGYIAAH
LAIVVICVGGLLDSNLPIKFQMWMFGKSPVNTSATISEIGAEHRLSASNPTFRGYAWVPE
GQYVSTAILNQPTGSLIQDLPFSIQLEKFIVDYYSTGMPKLFASDIVVVDRETGKRIPAR
VEVNKPFTYKGVSIYQSSFQDGGSLMEMTAYPMTGASAATFPFKGTIGNSVPLAAAGADG
ETVEFSDFRAINVENVTDANGKTDARGVAANRSIKELFDERLGSGAKTSKPVELHNIGPS
VQYKVRGKDGQAREFNNYMLPVDMGGERVFLAGVRASPNDPFRYLRIPADSRDSVGEWMR
LRAALEDPAVRIEAARRFARHSLGTGDAALRDRLEDSAQRVLTLFAGADGSIGRGADGNP
IGGFQSVATFIDRSVPKDEKEKAAALLLRMLEGSMWEVWQIARERAGEPPVQQAQDTVRF
VQNAINALSDSFLYGAPVYLQLDSFKQVQASVFQLTRAPGKNLVYLGSLLLVAGIFAMFY
VRERRLWFWLKDTGAGVSVLMAMSTARKTFDFEKEFVQTRDAAGAALGAPAFRAPDGAGT
KDSTR*
>GCA_000452945.1_BPMSHR5858_genomic_05631_1 msrP
MLIKKTLRAALAGDDIPRSEITPRAVFEHRRRILQATGAAAAGGLVGAHGLALAAYASPD
AAARKLAAPANPKFVVPEKVTSFKDITTYNNFYEFGTDKSDPARRAGTLRPHPWRVSVEG
EVRNPKVYDIDALLKLAPLEERVYRLRCVEGWSMVIPWIGFPLAELIKRVEPTANAKYVQ
FVTLADPSQMPGLSAPILDWPYSEGLRMDEAMNPLTLLTMGVYGQVLPNQNGAPVRVIVP
WKYGFKSAKSIVKIRFVDRQPPTSWNTYAPNEYGFYSNVNPNVDHPRWSQATERRIGEDG
FFTPKRKTLMFNGYGDWVASMYRGMDLKKYF*
>GCA_000452945.1_BPMSHR5858_genomic_05632_1 msrQ
MQSSTFAGSASPGAGARGRAPRRDAGRAGGAPGWLAPVKIAAFVAALYPLARIVFFGVTD
RLGANPIEFVTRSTGLWTLVFLCITLAVTPLRRLTGINALLRLRRMLGLYAFFYAVLHFT
TYIWFDKWFDVVEIAKDVVKRPFITVGFAAFVLLIPLAATSPRAIARRLGRHWQTLHKAI
YAIAAFGILHFWWMRAGKHDLALPKLYGAIVLVLLGWRVLVWARQRWGAAAAGGRAS*
>GCA_000452945.1_BPMSHR5858_genomic_05633_1 lysA_2
MTQAAFDYVDGTLHAERVSTIDLAERFGTPLFVYSRAALTAAYEAYAKACAGRRASIHVA
VKANSNLGVLNVFARLGAGFDIVSGGELARVLAAGGRARDVVFSGVGKSADEMRVALEAG
VKCFNVESIPELDRLNAVAASLGKRAPVSLRVNPDVDPKTHPYISTGLKANKFGIAFDDA
RATYRAAAALPNLEVVGIDCHIGSQITELSPYLDAIDKLLDLVERIEADGVQIHHVDVGG
GLGITYDDETPPDIGAYVRAVLARIDARGHGQREIWFEPGRSLTGNAGILLTRVEFLKQG
EEKNFAIVDAAMNDLARPAMYQAFHAIVPVAPRTDVAAAVYDIVGPVCESGDWLGRERSL
AIAPGDLLAIRSAGAYGFTMSSNYNTRPRAAEVIVDGANAHLVRPRETVESLFEHETILP
EGR*
>GCA_000452945.1_BPMSHR5858_genomic_05636_1 mrcA_2
MQSTNPTSPPPPPEPRRRPLWLKLVLGVAGVCVALAVSIGLVLGYALVVASPNMPSLDAL
TDYRPKVPLRIYTADHVLIGEFGEERRDIVRLQDVPDSLKKAVLAIEDARFYEHGGVDLT
GIIRAGIVALTNGHATQGASTITMQVARNFFLSSEKTYTRKIYEMLLAYRIERALTKDQI
LEVYMNQIYLGQRAYGFAAAAHAYFNKDLKDITLAEAAMLAGLPKAPSAYNPVANPKRAK
VRQQYILQRMLELNFITREQYDEALAEPLVVKSPSREYSVHAEYVAEMVRQMMYAQYKDE
AYTRGLNVVTTIDSADQTLAYRALRKGLMTYDRRHGYRGPEGFIELPANADDREQAIDDA
LVDHPDNGELIAAVVTEASARQIRVVFPNGSDATVSGDDLRFAASSLSSNAQPNKRVRPG
AIVRVFKNDDGSWAIGQLPQIEGAFISIVPQDGAIRSLIGGFDFNKNKFNHVTQAWRQPG
SSFKPFIYSASLDKGLGPATIINDGPLYFSAAETGGQPWEPKNYGGGFEGPMSMRVALQR
SRNLVSIRILNHIGPKYAQQYITRFGFDADRHPAYLPMALGAGLVTPLQMAGAFSVFANG
GYRINPYLIAEVTDPSGNVVARAQPLVAEQNAPRAIDPRNAYVMNSLLQSVAQRGTGARS
NVLKRTDLAGKTGTTNDSRDAWFGGYQHTLAAIAWIGYDSPRSLGDRETGGGLALPVWID
YMGPALKNVPEYKMPMPDGVTTLGGELYFDQFMPGRGFVATVGVNLAAAPPNSASDVPGH
VDEQEKQDIMNLFRGGN*
>GCA_000452945.1_BPMSHR5858_genomic_05644_1 ugpB
MKYKMLVRSLAFGGALWFGAQQAACAATEIQFWHAMEAALGERVNEIAAQFNASQSDYKI
VPVFKGTYDQALAAGIAAYRSGNAPAILQVYEVGTATMMQAKKAVLPVSDVFRQAGVPLD
EKAFVPTIASYYSDARTGRLVSMPFNSSTPVLYYNKDAFRKAGLDPNQPPKTWADVKADA
EKLKKAGYACGYTTGWQGWIQLENYSAWHGLPFATRNNGFDGADATLEFNKPQQIAHIQF
LQVMAKDGTFTYVGRKDEASAKFYSGDCAIMTTSSGALATIHKYAKFDFGTGMMPYDAGV
KGAPQNAIIGGASLWVLAGKDPATYKGVAKFLAYLSSPAVAAKWHEDTGYLPVTTAAYDL
AREQGFYAKHPGADTAIKQMMNKPPLPYTKGLRLGNMPQIRTIVDEELEQVWAQKKTPKA
ALDSAAARGDELLRRFEKSGG*
>GCA_000452945.1_BPMSHR5858_genomic_05645_1 ugpA
MTSHSRFGASVLPYLLVAPQLAITAIFFLWPAGVALWQSTQMQDAFGTSSEFVGFANFAH
LFADPLYVDSFRTTLVFSASVTVCGLVVSLLLAACADRVVRGARAYRTLLIWPYAVAPTI
AAVLWAFLFNPSIGVITYALARFGIVWNHALNGGQAMLLVVIASVWKQVSYNFLFFYAGL
QAIPRSLIEAAAIDGAGPVRRFLHIALPLLSPTSFFLLVVNLVYAFFDTFPVIDAATAGG
PGQSTKTLIYKIYAEGFQGLDIGSSGAQSVVLMVIVVALTVVQFRFVERRVQYA*
>GCA_000452945.1_BPMSHR5858_genomic_05653_1 alsT
MEGFVHALIDGINGILWNYVLIALLLGAGAWFTLRFRMIQLRALFLSMRLVGSKGEPGSI
SSFQAFATGLASRVGTGNIAGVAVAMTVGGPGAIFWMWMTALVGMSSAFVEATLAQIFKV
SHHDGTYRGGPAYYIQIGLRSRGFGVLFSLSLILAFGFVFNAVQANAIAEAFNTSFGVSR
AAVGLALVALTAPIIFGGIRRIAHVAQVIVPVMAIGYLALAVYAVATHVALVPDMIVLIV
KSAFGLEQAAGGLSGYAVSQAVSIGVKRGLFSNEAGMGSAPNAAATASTRHPVTQGLIQM
LGVFVDTIVICSATAFVILLSGQYEPGTSMAGAALTQRAISSHVGDWGGIYMAVAIFFLA
FSSVIGNYAYAEGNVGFVTSRRGALLVFRLAVLGMVMFGSVGQLPLVWAMADTSMGLMAL
INLVAILLLGKYALAAWRDYQRQRAAGVADPVFTRQTIPALAKVLPDDVWGEHGPLSQGD
KLAAGRASGADAVRAAGPVGSAR*
>GCA_000452945.1_BPMSHR5858_genomic_05657_1 group_19909
MDIQVNQQTFSLPDGATVADALAAYGARPPFAVAVNGAFVARTQHAARALAAGDKLDIVH
PVAGG*
>GCA_000452945.1_BPMSHR5858_genomic_05659_1 thiE_2
MSAALPDAFWPPADELTEAAERIRATLGAWPRPAVRTRICLAPPEQPRAADLWVAIAGDA
GAHAAQIARLNAAGAQAIVIDDASATLHTGAARHALASRAPLADDWIAALAAFLDCGFAA
SDALVLALAWRDGDEARGGDPWPVDPARFPRVLGLPAAPEPAFAPCPQRLGLYPVLPSAE
WVERVLDCGVRTVQLRVKDASPDTLRAEIERAVAAGRRHPDARVFINDHWRLALDAGAYG
VHLGQEDLETADLGAIARAGARLGLSSHGYYEMLVALQFKPSYLALGPVFATATKAVAAP
PQGLARLARYVRFAGPQAPLVAIGGIAPDTLGAVLAAGVGSAAVVSAITAAADYREAIVA
LQQNFGR*
>GCA_000452945.1_BPMSHR5858_genomic_05662_1 mlaD
MKKTALDFWVGLFVVLGFLALLFLALKVGNMSSLSFQPTYAVKMKFDNIGGLKPRAAVKS
AGVVVGRVKTIGFDTNTYQALVTIELDKQYPFPKDSSAKILTSGLLGEQYIGLDPGGDTE
MLKAGDTITMTQSAIVLENLIGQFLYSKAADAGGAKPGAAASGAPAPAPVAASAASAASG
AAAQ*
>GCA_000452945.1_BPMSHR5858_genomic_05665_1 group_19787
VSRFDSGATLTHASAKAALAQGLARIEAGATAVDCGALAQFDSSALAVLLAWQRAARARG
VTLDIENLPPKLASLAQAYGIDTLLSGRH*
>GCA_000452945.1_BPMSHR5858_genomic_05681_1 group_3665
MNHDPNCLFCKIAAGEIPSTKVHEDDEFVAFRDIRPAADTHVLVIPRKHLPTLSAVTADD
APLLGRMMVLVARLAERLGCAYTGGETGFRTVINTGPGGGQEVYHLHAHILAGPRPWRRM
G*
>GCA_000452945.1_BPMSHR5858_genomic_05683_1 tatB
MLDLGLSKMALIGVVALVVLGPERLPRVARTAGALFGRAQRYINDVKAEVSREIELDALR
TMKTDFEQAARNVENTIHDNLREHERDLNAAWNSAVSPGGSAAADAPDGPSAASGEPSWR
TIAAAPAKRRNWRVKKAVTPVWYKRATMRRTQVQSGAARVARHRPASLRRPARFL*
>GCA_000452945.1_BPMSHR5858_genomic_05688_1 petB
MATDNKEISTTGLLGWVDRRFPLTSTWKQHLSEYYAPKNFNVWYFFGSLALLVLVNQIVT
GIFLTMNYKPDSTLAFASVEYIMREVPWGWLIRYLHSTGASMFFVVVYLHMFRGLLYGSY
RKPRELVWIFGCAIFLCLMAEAFFGYLLPWGQMSFWGAQVIVNLFSAIPFVGPDLSLWIR
GDYVVSDVTLNRFFAFHVIAIPLVLLLLVVAHLIALHEVGSNNPDGIEIKEKKDEKGVPL
DGIPFHPYYSVHDFFGVCVFLMVFALIVFFMPEMGGYFLEANNFIPANPLQTPPEIAPVW
YFTAFYAMLRATTDPFKIVLMIVIAALGLLALVRARGKWKIALPVLAAALVVFMYFTESK
FWGVVVMGGAVVTLFFLPWLDRSPVKSIRYRPFFHKVFFGIFVVVFLTLAFLGTRPPSPA
ATLIAQVCALVYFAFFLGMPFWTPLGTFKQPPERVRFKSH*
>GCA_000452945.1_BPMSHR5858_genomic_05691_1 sspB
MQEISTKPYLLRALYEWCTDNGYTPHIAVRVDKSTRVPRQFVRDGEIVLNISFEATSQLQ
MGNEWIEFTARFSGKAHKLEVPVANVLAIYARENGQGMAFQVDVAAEMGEADGSDALAVA
EEGDDAGARHASADGAAEAAPEPGDAAEELPKSDGDGDGDGAKGSRPRLKIVK*
>GCA_000452945.1_BPMSHR5858_genomic_05737_1 pykA
MHRATKIVATIGPASSSPEVLSQMMHAGLDVVRLNFSHGSADDHRERAEMVREAARRVGR
EIAIMADLQGPKIRVGKFEAGKTTLTPGQPFILDATCELGNDERVGLDYKDLPRDLKPGD
VLLLNDGLIVLKVDRVLGDEIHTTVKVGGDLSNNKGINRQGGGLSAPALTAKDMEDIRTA
MSLGADLVAVSFPKNATDMEMARQLANIAGAPYGIKPKMIAKIERAEAIPALQEILDASD
GIMVARGDLAVEVGNAAVPALQKRMIRMARESNKLVITATQMMESMIHAPVPTRAEVSDV
ANAVLDGTDAVMLSAETATGKYPAVTIEAMAAVCVEAEKSEHVELDKDFLDRTFTRIDQS
IAMGALFTAYHLGAKAIVALTESGATALWMSRHYTHVPIFALTPRVGSERAMALFRNVTP
LHVDFNSDRDSALQQALELIVRRGYVAHGDMVVLTVGEPMGQAGGTNTLKIVRVGEHY*
>GCA_000452945.1_BPMSHR5858_genomic_05739_1 purC
MSTLYESTLRSLPLLGRGKVRDNYALGNDKLLIVTTDRLSAFDVIMGEPIPNKGRVLNQM
ANFWFDRLAHIVPNHLTGVAPETVVAADEVEQVKGRAVVVKRLEPILVEAVVRGYLAGSG
WKDYQATGKVCGVELPAGLSNAQKLPEPIFTPAAKAEMGHHDENISFEETERRIGTELAA
TIRDISIRLYKEAADYAATRGIIIADTKFEFGLDEHGELFLMDEALTADSSRFWPADEYR
VGTNPPSFDKQFVRDWLEAQNWNKAPPAPKLPDDVVAKTSAKYQEALERITGKTLD*
>GCA_000452945.1_BPMSHR5858_genomic_05740_1 purE
MSEVQTAHTHSAPLVGVLMGSSSDWDVMKHAVAILQEFDVPYEARVVSAHRMPDEMFDYA
QQARERGLRAIIAGAGGAAHLPGMLAAKTTVPVLGVPVASKYLKGVDSLHSIVQMPKGVP
VATFAIGEAGAANAALFAVSILSGTSADYANRLAAFRVRQNEAAHAMALPPL*
>GCA_000452945.1_BPMSHR5858_genomic_05748_1 htrA
MTTRILARGAVAVAVAAALSAGYVAGTRRAEPQIITPAVAALMPAEAAAKTGIPDFSGLV
ETYGPAVVNISAKHVVQRAAQRRAAPQLPIDPDDPFYQFFRHFYGQIPGMGGGRQPQPDD
QPSTSLGSGFIISADGYILTNAHVIDGANVVTVKLTDKREYKAKVVGADKQSDVAVLKID
ASGLPIVKIGDPAQSKVGQWVVAIGSPYGFDNTVTSGIISAKSRALPDENYTPFIQTDVP
VNPGNSGGPLFNLNGEVIGINSMIYSQTGGFQGLSFAIPINEAMKVKDELVKTGHVSRGR
LGVAVQGLNQTLASSFGLQKPDGALVSSVDPKGPAAKAGLQPGDVILAVDGVPVQDSSTL
PAQIAGMKPGTKADLQIWRDKSRKTVSVTLASLADDQAKAGADEPVEQGRLGVAVRPLSP
RERNGSSLTHGLVVQQSAGPAASAGIQPGDVILAVNGRPVTSAEQLRDAVKRAGNSLALL
IQRDDAQIFVPVDLG*
>GCA_000452945.1_BPMSHR5858_genomic_05749_1 group_3673
MRQQSVSQYLIVAALAFGLAGVARAQPGLPDVQHHGDVGFVSGGVGLDESTAFQRSESSW
PLALRFTGAGGEYLSDVHVSVLDAQGGEVLKTDARGPYMLVRLKPGRYTVRASYNGSEQT
HAVTVPAKGGTKAAFSWKAQ*
>GCA_000452945.1_BPMSHR5858_genomic_05750_1 group_3674
MSDAAGHRIEIEPNRHRVRVIHHGITYADSLAAYTLREPGAPEVQYLPRDDVNMSRLVPS
DRVALEHCAHKGPATYFHLHTEDGVIENAAWSYEEPSGIASAIRQYVAFDAACVDRIDVT
S*
>GCA_000452945.1_BPMSHR5858_genomic_05760_1 clpB_2
MRIDKLTTKFQEALSDAQSLAAGRDNQYIEPVHVLAALVAQQDGSARSLLSRAGVHIQAL
QGALNEAISRLPQVTGTGGDIQVGRELAGLLNQADKEAQKLNDTFVASEMFLLAVADDKG
EAGKLARQHGLTRRALESAIAAVRGGSQVHSQDAESQREALKKYTVDLTERARAGKLDPV
IGRDDEIRRSIQILQRRTKNNPVLIGEPGVGKTAIVEGLAQRIVNGEVPETLKNKRVLSL
DMAALLAGAKYRGEFEERLKAVLNDIAKDEGRTIVFIDEIHTMVGAGKAEGAMDAGNMLK
PALSRGELHCIGATTLDEYRKYIEKDAALERRFQKVLVDEPSVEATIAILRGLQEKYELH
HGVEITDPAIVAAAELSHRYITDRFLPDKAIDLIDEAASRIKMEIDSKPEEMDKLDRRLI
QLKIEREAVKKEPDEASQKRLALIEEEIERLGREYADLDEIWTAEKAAVQGSAQLKEEID
KVRADIARLQREGKLEKVAELQYGKLPQLEARLKQVTQAEESEQHNPTRPRLLRTQVGAE
EIAEVVSRATGIPVSRMMQGEREKLLHIEEKLHERVVGQNEAIDAVADAIRRSRAGLADP
NRPYGSFLFLGPTGVGKTELCKALAGFLFDSEEHLIRIDMSEFMEKHSVARLIGAPPGYV
GYEEGGYLTEAVRRKPYSVILLDEIEKAHPDVFNVLLQVLDDGRMTDGQGRTVDFKNTVI
VMTSNLGSQLIQSMSGSSQEEIKDAVWVEVKQHFRPEFLNRIDDVVVFHSLDRGNIQAIA
KIQLARLHDRLAKLDMALDVSEAALEQIGKVGYDPLFGARPLKRAIQQEIENPVAKLILA
GRFGPKDVIPVDMQDGRFVFERVVH*
>GCA_000452945.1_BPMSHR5858_genomic_05763_1 group_19812
MSNAPKQPSPRALTIGQVAALVGVSTHTLRYYEQAGLLRSISRTPSGHRLYAPADLDWLR
FVMRLKATGMPIAGMQAFAELRAQGDATFGERRALLAAHRDAVRAQIGALQTNLDSIVEK
IAYYERAERDAVRQARTTTQSQPDKDERWSNAFRKTATRAAGTS*
>GCA_000452945.1_BPMSHR5858_genomic_05764_1 group_2642
MEQRISEDRYTRGWNKLKEIDGKVGERVVAALAPIAPDFGRLLVEFGFGDIYSRPQLDLK
SREIATIASLATLGHAQPQLKVHIEAALNVGCTRDEIVEVFMQMALYAGFPAALNALFAA
HDVFGQRDAAHA*
>GCA_000452945.1_BPMSHR5858_genomic_05773_1 ruvB_2
MTYQVLARKWRPKDFASLVGQEHVVRALTHALDGGRLHHAYLFTGTRGVGKTTLSRIFAK
ALNCETGVTSQPCGVCRACREIDEGRFVDYVEMDAASNRGVDEMAALLERAVYAPVDARF
KVYMIDEVHMLTNHAFNAMLKTLEEPPPHVKFILATTDPQKIPVTVLSRCLQFNLKQMPA
GHIVSHLERILGEERIAFEPQALRLLARAAQGSMRDALSLTDQAIAYSANEVTETAVSGM
LGTLDQTYMVRLLDALAAADGPQILAVADEMALRSLSFSAALQDLASLLHRIAWAQFAPA
SVLDEWPEAADLRRFAELLSPEQVQLYYQIATVGRGELGLAPDEYAGFTMTLLRMLAFEP
ATTGGGAPAGGVPARVAGAAPKSGARAAAAVGASAVPAVTGATGATGAALAPKAAAARAA
TRAEAPPAAPARPATAERGDDASDGDAPVPAKANARASADSRCDERDAQPPADSGSASGA
SSSAPPDTAFESAPRAAAPSAATPAAVPDARAPAAASAEDAVEAAAPAAPESRPPTPAAG
APVSRATGAAAALDVLRNAGMRVSSGRGAGAARPAAGPAASPTKPAPRVQVSVPTPRARA

ATGDAPPNGAARAEARAESRGAPPPWEDIPPDDYVPLSMDDGFIPPDDGFVPVFDGGPDD
VRVAPKPAPAPPVDTRPLPEPIPLDAIGYDGEWPTLAASLPLKGVAYQLAFNSELTAADG
GTLKLAVPVPQYADAAQVAKLKAALADALGKPVDVAVEVGPARRTAAALAAAERAKRQRE
AEQEMSADPFVRQLLREFGASIVEGSIRPVGAEAGAADGASPTLH*
>GCA_000452945.1_BPMSHR5858_genomic_05775_1 recR
MSIKPPSALSELVEALRALPGVGPKSAQRIAYHLMQHDREGAERLGRSLLFATEHLRHCE
KCNTFTEAQICEVCSDPERDPALLCVVETPADQIMLEQTMTYRGLYFVLMGRLSPLDGIG
PKEIHFDRLVRRASDGIVKEVVLATNFTNEGEATAHYLGQTLKARGLAVTRLARGVPVGG
ELEYVDAGTIARAMLDRRTL*
>GCA_000452945.1_BPMSHR5858_genomic_05780_1 rpoS
MPKSKRHDPQAESEKISRAKQASVERTGASADEDEDAADNERDYESRDADPDESGEGRGD
AQPDLDDFRALLQAELTADTIQHYLNRISVKPLLTVEEEQRYSRLAKAGEFEARQVMIER
NLRLVVSIAKGYLNRGVPLLDLIEEGNLGLMHAIEKFDPTRGFRFSTYATWWIRQSIERA
IMNQARTVRLPVHVIRELNQVLRAKRHLEKNSMSTGEAAERREASIDDIAYLTGKTAEEV
TDILALNEHTASLDAPLDLDPASSLLDLLPDDQSQSPDAEVQHRELETLTRAWLSRLSDK
HRHVIERRFGLNHIEPATLEELADEMGLTRERVRQIQQEALVRLKRFFASNGVRKDAVL*
>GCA_000452945.1_BPMSHR5858_genomic_05783_1 rlmD
VSEAVPLSTRRASSAGDAPGRAPVLDIDSLDMEARGVGRVVDENGEPGKVIFVEGALPGE
RVTYSSFRRKPTYEQAQVVDILRPSVMRTQPKCAFFGTCGGCSMQHLDMRAQVAIKQRVL
EDNLWHLAKLRAEAMFAPIHGPSWGYRYRARLTVRHVAKKGGVLVGFHEKKSSYVADMTS
CEVLPPHVSAMLVPLRRLVEQLSIRDRMPQIELAVGARVTALVLRVLEPINAADEALLRE
FADTHRVQFWLQPKGPDTVAPFYPLDAQLDYTLPEFGIRMPFKPTDFTQVNHQINRVLVG
RALRLLAPERGDRVLDLFCGIGNFTLPLARLAREVVGIEGSDALTARALENAKENGVDGH
TSFACRNLFEVTADDLRALGAFDKFLVDPPREGALAVSKALAEIAQSGAGPLPARIVYVS
CNPSTLARDAGLLVHEAGYRLKGAGVVNMFPHTSHVESIALFERD*
>GCA_000452945.1_BPMSHR5858_genomic_05784_1 yccA
MNDYPYNFGRGGAIGTAETRNRVLRNTYWLLALSMVPTVLGAWVGVATGFSLFAATSPMM
SLLAFFAIAFGFMFAIERTKNSSAGVFVLLGFTFFMGLMLSRLLSFILGFSNGPSLIMLA
FGGTGVIFAAMATVATVSKRDFSGLGKWLFVGVIVLLLAMVANMFLQLPALMLTVSVLAI
VIFSAYMLFDVQRVVNGGETNYISATLAIYLDLYNVFTNLLALLGIFGGNRN*
>GCA_000452945.1_BPMSHR5858_genomic_05785_1 ndk
MALERTLSIIKPDAVAKNVIGQIYSRFENAGLKIVAARMAHLSRADAEKFYAVHAERPFF
KDLVDFMISGPVMIQVLEGEDAILKNRDLMGATDPKKAEKGTIRADFADSIDANAVHGSD
APETARAEVAFFFPEMNVYSR*
>GCA_000452945.1_BPMSHR5858_genomic_05797_1 group_3679
MDLAGSLLLALALMLIIEGAFPFVFPSAWRDTFRRIAERPPHHIRIGGFIVMALGLVLLL
LVT*
>GCA_000452945.1_BPMSHR5858_genomic_05799_1 purA
MSASAVNVTPGRNVVVVGTQWGDEGKGKIVDWLTDHAQGVVRFQGGHNAGHTLIIGGKKT
ILRLIPSGIMREGVACYIGNGVVLSPEALFKEIGELEEAGLSVRERLFISEATTLILPYH
IAIDQAREARRGAGKIGTTGRGIGPAYEDKVGRRALRVQDLFDARTFADRLRENLDFHNF
VLTQYLGGAAVDFQATLDTMLGYADRLKPMVTDVSRRLYEENHAGRNLLFEGAQGTLLDI
DHGTYPFVTSSNCVAGAAAAGAGVGPQKLDYILGITKAYCTRVGSGPFPSELYDADNPSR
QDQIGITLANVGKEFGSVTGRPRRTGWLDAAALRRSIQINGVSGLCMTKLDVLDGLDEVK
LCVGYKIDGEDVDLLPRGAAEVARCEPVYETFGGWKESTVGIDSWDALPANARAYLTRVQ
EVAGVPIDMVSTGPDRDETILLRHPFKV*
>GCA_000452945.1_BPMSHR5858_genomic_05806_1 group_3680
VTRVFTLDSLHGDDLKFHRLYGEEALGRMFDFRIEALADNHSLSLKELLGKPVTVRIRQQ
DESERHLNGIVARAALVGRRAERHYGYQLIVRPWLWLATRRSDCRIFQNKTVPEIVQDVL
GTYGFPIENHLTDTYAPRDYCVQYNETDAAFVSRLMEFEGIYYYFKHAAQTHTLMLCDAM
ASHVALPGYEHIPFIARDRTAIADEEHIDSWLPAQEVSIGKHETSDYDYTKPRADLSAQK
IDPRGHDHDGFASFEWPGGYRDDEPGAHYSRVRLEEQQAEHERALARTDVRGIAPGYLFT
LEHCPRADQNREYLIVRCQYRFQENAYATDSGNEAVVHESQTLVQPSSLPYRSPRATPRP
RTNGPQTATVVGPKGEEIWTDQYGRVKLQFRWDRYGQSDQNSSCWVRVSSPWAGGGFGGV
QIPRIGDEVVVDFLNGDPDQPIVTGRVYNGEKMPPWGLPGSATQSGLLSRSSPGGTAEHS
NAFRFEDKKGAEQLWMHAERNFDAETEQDHTLSVGHDHSHSVGNNEMMSVANDRQRSVGQ
NETVNIGKHRVAQIGGNETHGVAGNRTRQVGQNEAVTIGANREATIGGNHVETVAKDKTE
TIGQGKTLNVTQHYQTTSKSMKTAVAEHHAEEIGSRTSTIEKAHELNVGGSQSVNVGASH
TMSVKNNVHVGAGDQISLVCGDASITLKKDGTIAINGVTVESSASGSHSVRGKTVTSSAT
GEHTVEGTILKLNP*
>GCA_000452945.1_BPMSHR5858_genomic_05808_1 group_395
MTTNPKPEPAGVDAGSHANSNAGRRAGAAANVEANAEPDAMPNALRGAAADVAPSAAPSA
TPNATPSATSQAAAARRDAWWRRLRAAPHGYDLFHALRWLDALSPEHAPSGYASRPRDEP
VRFGQAPSLAFAAAMLADVRDDGPRPRVAIHGFGLFGPNGPLPSHLTEYAYERATQHDDP
TFAAFADLFHHRLILLFYRAWADAQPTVSLDRPARARFDRYVASLIGPCDALLADSPRGS
RPDSSFASSSDSLPDSPPDSSPDSRSDSRSDSPSDAHLDPHPDSPRGAGALAPHAKYHQA
GHLVRHTRNPEGLVQILRRYFGVRVRIVEHVPHWVMLDRAQRCAIRATRPTQPLGGAVLG
RAVRDAQSRFRIVLGPLTLDEYRQFLPGGVHARQLAQWVREYVGIEFDWDVQLELARGEV
PTLALGSRHGLGRTAWLGERLGPGPARDLVLGYDERMRASRARSGAADQAAAAGAFDRQP
A*
>GCA_000452945.1_BPMSHR5858_genomic_05814_1 group_3856
MQSSDPIESLLAGVAPDAPCGANLEYEQDFLRLQESATPRPEQQYGDTVIPAEAPDWSAV
ERLALELTARTKDLRVAAHLARSWTELRGMPGYADGLRLVAGMLDRWWDDVHPRLDADGD
HDPAPRANALAEIAGAHDCARAARRQALFDGGPSVRDAERVFDGRDGGEHGYPGGRERLI
ADLVRARDGGQPTLQAAFAALDALDAIRARVASALGGEWVPDASDFEKALRRIVRDGLPP
PAAAPAATNGAANVANAAHGGAANGVTNSAANGAASGGAPAFAANGRAWRDAELTSRDDV
QLGLEKMCRYFELHEPSHPAPILLRRAQRLLSLDFYEIIRDLAPESLPKLDLLSGQRNE*
>GCA_000452945.1_BPMSHR5858_genomic_05815_1 pknD_2
MTDPFSRDNEQETVTRGGGAEFAVRPLPLGHRLGELQLDEVLGVGGFGIVYRAFDRTLRR
AVAIKEYMPSMLATRGGDYTVSLRSVRFAQAFDAGRSAFLNEARLLAQFDHPGLVKVLHF
WESHGTAYMVMPFYEGRTLKQLLDGGAQIGETQLRHIVAALLGALDTLHRAQCFHRDIAL
DNILIRPDGNPILLDFGAARKRIGDLVDDSAMMIKPGYAPIEQYTDDPAFSQGPWTDLYA
LGAVMHAMVTGELPPAAVVRSIQDTYRPLAARELTAREPYSPAFLAAIDHALQLKIADRP
ESVAEFAAELGLREFDRPPYVAPMPAGAPQPREARDAGAAEDGARRAAASGSPEARRRGD
ADARPDGERARQPEQPPPARQSAAHRLGDSAQADGEGAGDESGAAAARWQGDDERADERP
DESTDEPAARAERAMAAQSGAQPAQTPPPPQASERGAAATRSDADSGAPSPRPWAADGRD
GEPSGPRVDHMQSQSQSQSQSPPSRSREHAGERDPLPARPGVADASRQAAPASRDGGAPP
PEASGPRVSDAPVSRDAAGADAEAAMSAAAAGAGAPARAGASNASASSGAPAVCAASTAS
TASAASRSTAPAGPPGNARARLRDRRTIAYAGGALAVLVALGIGVAHLVKPSGRTEATAA
NALAPLSPAAPPAPGGVSALAQRGEPPPALAPPPAPADTAVREPAPSVATTTAGASATPT
TSPGAAPPPSAQIANHDAGAASAHGAQASNLAPNGSLDATETVVANPPAGAEPPVSPSVA
QEAAAAAEQRPPDMQETIPVRLHVRPWGDVYVSGVKRGASPPLRTLSLAPGVYQIEIRNG
TLPPLRRTVKIDPGSKPVSIDYAFE*
>GCA_000452945.1_BPMSHR5858_genomic_05818_1 group_3681
MNTSSDSFSAGSGGFVPPNPGGAHPAAAPAAGAAAFQPRPGRWAASGTNPLVAAANPLLN
LVPQIRSTVHHPNPAWLREHLVVEIRQFEERAQQAGVASEAIIGARYCLCTALDEAAALT
PWGGSVWSSHSLLVSFHNETWGGEKFFHLLERLSQQPRQHLDLLELLYFCLALGFEGRYR
VLDNGRAQLDAVRRQLAQTIRSVRGEFDPALSPHWRDVVTRDVTRRFTVPLWVCVALALL
GGFGVFAGLRIALAGHSDRLFASIDALHVPKLQPAPPAPHPAPAPRVAKFLEPEIAAGLV
SVRDEADRSVIVLRGDGLFGSGSTSVIDRYMPVLTRVADALNQVQGNVRVSGYTDDTPVH
TARFASNWDLSRERAQAVRSLIAARLDRPERITAEGRGTLDPVAPNDSPANRARNRRVEI
TLMLAPGSDAARATKEAP*
>GCA_000452945.1_BPMSHR5858_genomic_05820_1 group_4063
MRPRFSVSVVLGCALLLAGCGATERSVAVPYSITLDVAPDVNPDLNRKPSPIVLKVFQLK
TASAFESADFFSLQDKPQSVLGADLLGVDRIILRPGDARTLHYRGNVDAGAIGIVAEYRV
LEKNRWRMTVPLPRAKQLNLYRFWQTSPGEMKLQVAVKNGGIGLGGDSRVRQ*
>GCA_000452945.1_BPMSHR5858_genomic_05838_1 gcvA_9
MNPSDLRDLRLPSLDGLLAFVAAARHGTFERAAEELCVTGSAVAKRVAALEQLLGVQLLV
RSGKALTLTAAGNEYLHQVHAPLHLLAAVPLHRRNAGKTERLRICAPPTFARQILVARLD
AFTLAHPGVELEVMLASPFPDAPPTGADLWIRGGGPIVSGATVLMVDALTPLAAPSLVER
FAPLRRPADLAALPLLCTPLEPWTPWFRVAGLDWPEPANGPKLLDLGLLLEAAVSGQGVA
LGSPALARPWLAAGALVPLFPRIVRPAAPYYLMPYAPRAAAGAFAAWLAEVCERAVRDGA
EFLSRHG*
>GCA_000452945.1_BPMSHR5858_genomic_05839_1 lldD
MSVPITCIEDLRVLAKKRIPRMFYDYVDTGSYTESTYRANEADFRKIRLRQRVGVDISNR
NLRTTMAGQDVAMPVALAPTGLVGMMRADGEILAARAARHFGVPFTLSTMSICSIEDIVA
HVGGPFWFQLYMMRDRAFIERLIERASAAGCPALVLTMDLQIAGQRHKDVKNGLSAPPRI
TLPNLLDMMRKPGWCLGMARTRRRHFGNIVGHVKGVTDMWSLDSWTREQFDPTIGWRDAE
WVRRRWNGKLIVKGVLDADDALRAADAGADAIVVSNHGGRQLDGAMSSIEALPAIVEAAG
KRVEVWLDGGVRTGQDVLKAVALGARGTMIGRAFLYGVAALGEQGARRALELIARELDTT
MALCGCTDIRSVNADVLARR*
>GCA_000452945.1_BPMSHR5858_genomic_05840_1 group_3934
MKLLRYGPVGQEKPGLLDADGRIRDLSSLIDDVADGALSDASLARLAAADPASLPLVAGE
PRIGACVGRIGKFVCIGLNYADHAAEAGLAVPAEPVVFGKWTSAVSGPHDGIEIPRGSVK
TDWEVELGVVIGRACKNVDEADALSHVAGYCVVNDVSEREWQIERGGQWDKGKGFDTFGP
IGPWLVTRDEIADPQALDLWLDVDGRRYQSGNTRTMVFTVARLIAYLSRCMSLQPGDVIS
TGTPPGVGMGVKPSPVYLKPGQTVRCGVAGLGEQQQRTRAAA*
>GCA_000452945.1_BPMSHR5858_genomic_05842_1 mdlY
MTSHRDTYVRSASVRAIADAVNGGRVRALEQAEAALERLALVEPLAAMTHVAPEIARRAA
ANVDAQIASGRRLPLAGVPFVIKDNLFTADMPTRAASPALLGFDATRNATAVQQLLDAGA
VPLGKANMHELAFGITSANGYFGAVRNPHDPARIAGGSSGGTACAVAAGIAFGLGTDTGG
SVRIPAAFCGVAGLRPTPRRYSTEGVLALSPTRDTVGPIAHGIDDLLLLDAVLTGDAGEP
APVALAGLRVGVPDAPYFDGLDPAVAELTERALAALTDAGARLIPLAAHALHEADEACGF
PIVGFEALAYWRRFSAERLDLPFAEFVARLGSDDVRHLFRMLIDAPPSSDAYAAAIDVLQ
STVRWYDQAFHRTGVDCIALPTVQCDAPLVTDSGHDLPSDAAYALFTRVIRQTGPATLGG
VPSVSIPAGLSKHTGMPVGLMLEGPAGNDRRLLAIAREVERLLARAP*
>GCA_000452945.1_BPMSHR5858_genomic_05851_1 group_2707
MLHDLVARFGPLIVFVNVLAAAIGLPVPAMPTLVLFGAMATLHPDALGAQLAPVLALAVL
AALIGDTVWYVAGRHFGGRALKTLCKLSLSRDSCVKKTERFFGRWGVRVLAVARFIPGLS
LISVPMAGALGTRYRVFVGYDGLGALLWAGCGVAIGFAFAKQIDWLFAGANQLGRAVLVV
IVALLAVYTAVRWMRRRALIRQLANARIDVDELERLLRAEPAPVVFDARSPEHRKLDPYA
IPGAQFADERDLRDIVAHYPATQKFVIYCSCPNEVSAALMAQRLKQAGFADALALRGGLD
AWRDTGRQLIELAPQPPSEAPVLAAAPKTA*
>GCA_000452945.1_BPMSHR5858_genomic_05852_1 oprD
MTQHRSMKRVARAVVLAWGLPMLAGVSIAGAARADDAAPAAPAQAPLAATLAQQATTPNA
LVNADATQAVLPQPPMSSQAKSKGFIADSHFDVLLRNYADVLDAKGGPHRHAWVQGVMAN
FESGYTGGLIGFGVDASIYAALKLDGGAGGGNMVHVAKGGGGSNQLAWAFPGIYDVKARI
SNTVIKYGLQAVDNPFMEQHDNRALPPTFLGATIVSNEFKNVMLEAGSFTKTNARGHTTL
TNLTTQYGGTRVDRFTYVGGTWDYSQTGSASFYAGQADDVWRQYYGSVKQSYGSPQTVKW
TGFGNFYSTHDTGDARQGKIDNNAYSLSLAAQHGPHELLLGYQQVLGDQFFDYLNETNGI
FLVDSMDVDYNAPHEKSLQLRYTFYGKEAGLPGFKAMVWGVTGWGADGSASAALDPTRSS
IYWKDGAPVQGRHHEFGFIPSYTIQSGKLKDTKITFIAMWHVGSAHYSDSTNQEYRLVVN
VPVKVF*
>GCA_000452945.1_BPMSHR5858_genomic_05853_1 btuD_16
MSAATDLISSTSSFVSASAGAPGGGETSGADAPGADARGGGASPHADALLSVRDLKVHFR
VPLGGYPWSPKGTLRAVDGVSFDVKRGETVGLVGESGCGKSTLARALIGLAPVTAGTVQW
RGVPIVSSARRDARRIRREMQMIFQDPLASLDPRMTIEQVVAEPLVTHQPGIGRDEVRRR
VLTMLERVGLSAHHLLRYPHEFSGGQCQRVGIARALIGEPHLVICDEPVSALDVSIQAQI
VNLLRDLQRELSLSYLFVAHDLAVVKAISQRVLVMYLGRVMEFGTRREVYGAPQHPYTRA
LLAAAPTPEPARERARRPLLLASEMPSPLDPPSGCAFRTRCPDAIDACAREVPQPEVRGV
TGARVACIRAGTGGAAR*
>GCA_000452945.1_BPMSHR5858_genomic_05860_1 group_19722
MKNKTTRLWRMLSDIRRAQRCTALRAAAAAADVDALLTQHDDELRDDDEATQDDASNETR
LRTDLRH*
>GCA_000452945.1_BPMSHR5858_genomic_05873_1 bpaC_2
MGQGSQATGDNSTATGQNAAASGESSTATGQGSQAAGSNSTANGAGAAATGHQSTALGAG
AQAAGSQGVASGWNANASGTQGVAIGGNASAQGNQSIALGANASATGDHSIALGAGSQAT
GSNSVALGQGSIADRDNSVSVGSDGGERNVTNVANGWHDTDAVNFRQLREVARYAYSGIA
AATALAMIPDVDAGKTFSIGVGTGGYLGYQAVAVGASARLGQNLKVRVGAGISAASTTWG
AGASYSW*
>GCA_000452945.1_BPMSHR5858_genomic_05878_1 htrE_2
MPVVAAEFDGTRTGEQSRPDAMGEAPSGAEPGTRGRQALLVASHASPSAPTTVSFNSRLL
MGGGIDLSRFERGNPVVAGIYPVDVTVNGERRGRMDVEFRDVRGGDSAAPCFTRAMLDRL
GVEGDIVAKRLDAARDATGAASGQTPTLSASACIGLRDALPDATYTLDSADLTLDLTIPQ
VDMRKTARGYVDPSRWDNGVNAGLLQYNLSGYASENKFFGNHASSLFLGLQAGANIGAWR
VRQRSNLMWANRSAGMSWRSLETYVQRDITALRSQITLGDSYTTGEIFESFGVRGMQLAS
DDRMLPVSLQSYAPTVRGVADTNARVAVRQRGNVIYEASVPPGPFEFDDLPPTGYGGDLD
VTITESDGRTKHFTVPFASVRQLLRPGMQRFNFTVGQYRDALSNGKPWVAQLTYQRGMTN
LLTGYAGLLSSTGYASGLIGVALNTPIGAFAFDVTSARTHLPGQGARNGFSSHLSYSKMV
PSTGTHFSMAAYRYSTANYYSLADAVTARYGYNAEERAWRNDYRARTRLQLNVNQRIGDR
SSAYVSSSLLSYWDGRGRDIQFQAGFSSVFKRVSYTVYAQRSRSSDDRTVTQVGVNLSIP
LGSGAYTKRNAFSSLTTSLSRASNGDSSIQANLSGSTAHVVPIDYGVNVSRSVTGDSNFA
SLGVYGTYRSPFGTYSGNASVDNRARQASFGANGAVVLHRGGVTLSPPLGSAAALVEAKG
ARGGKLINGQGATIDRFGYAVIPSLMPYRANTVAIDPSELPDDVELANTSEEVVPRNNSI
VFVKMETKRGRPVFAATETEDGKPLPMGSELFDVEGKSLGGVGQGGMAFLRGLQGAGNVV
AKWGAGASEQCTMPYLVPADQAQAKQSRAIVRIRLRCEPRLSADASQTPDGDGGTRND*
>GCA_000452945.1_BPMSHR5858_genomic_05879_1 sfaA
MAEHGMIEMNGLKFRRGAARRWPSIAAAAMFAASAACGAPPSPTVTIKFKGSYNPVTCTV
VEGNENKTVHLPTISTSSLANAGETAGATGFDISVLCDGYAGNVLAYFENGPTTNENGNL
DVEDPSDGQSASGVQVQLLNGDGSPIKVGDISTMKTVSVEASTPTPIPFFARYYATGKSG
AGKVRTYVTFVIQML*
>GCA_000452945.1_BPMSHR5858_genomic_05883_1 ompA_5
MAARESQARSADAQLATHSDFNALLSREFKPKTEQAREAVEHAVKTLAEQALANSVTLSD
DAYKSIEAIIGEIDRKLSEQINLILHHDDFQQLESAWRGLHHLVTNTETDEKLKIRFMDV
SKDDLRRTMKRYKGVAWDQSPFFKQIYEEEYGQLGGEPYGCLVADYYFDHTPPDVDLLSS
IGKVAAAAHAPFITGASPSVLQMDSWQELANPRDLTKIFTQNLEYAPWNSLRNSEDARYI
GLAMPRFLARLPYGIRTNPVDEFDFEESTDGSDHRKYVWANAAYAMAVNINRSFKHYGWC
TLIRGVESGGVVENLPCHTFPTDDGGIDMKCPTEIAISDRREAELAKNGFIPLIHRKNTD
YAAFIGAQSLQKPAEYYDPDATANANLSARLPYLFACSRFAHYLKCIVRDKIGSFKERED
MQQWLNEWIMNYVDADPANSSQETKARRPLAAAEVVVEDVEGNPGYYQAKFFLRPHFQLE
GLTVSLRLVAKLPSVKEAA*
>GCA_000452945.1_BPMSHR5858_genomic_05900_1 group_3687
MSKNGSPLKSDRSRHASAHHEAPDVPTPLVSLARYEGRDASPPRQHAIGAAGCVYFVQAN
RFEAWDARIGIPNDGIIFAAGPDGSLRMENHCARADCRVNGEAVAPGACRHLERGDVIAV
AAAELTIQDGASAREPSAPRQPPVDLIALGNALPRSEPAHFADLAELANLPAAAHPHGRA
SAGAAAPLVILDEEALSDAAPPAFDSTNHDPLAHLLTEYRRALIHQERSYAHVLKSIERP
QIAVPLPDDPFLDAPGRYRRGSLLGDLLGSRQNIDYVLDEMNPFRAEALFADEARHDVLL
LLAPRRRKKAHFPQTALLARHEHHLMSVDSHFPTLGAETDIQQEEPNDESI*
>GCA_000452945.1_BPMSHR5858_genomic_05901_1 clpV1_1
MTVSRQALFGKLGYSLFRSIESATAFCKLRGNPYVELVHWLHQMLQAADGDMQRIARHCD
IDLGELERDMVRALGELPAGASSISDFSHHIETAIERAWVLATLDFGDRRIRGAWLLAAL
LKTPELRRVLLSISSAFGRIRVDELDAMLPAWIDGSPEANETPYDNSDFGAAIPGEASSA
MQQKGGGSPLEQFCLDLTARARAGQIDPVVGRELEIRTMIDVLLRRRQNNPLLTGEAGVG
KTAVVEGLARAIAAGDVPPKLADARLLSLDVGALLAGASMKGEFEARLKGVLEAAAKATL
PVILFVDEIHTLIGAGGQAGTGDAANLLKPALARGTLRTIGATTWAEYKRHIEKDPALTR
RFQVLQVPEPDEAAAIDMVRGLTRTFSRHHGVVVLDEAIRAAVKLSHRYIPSRQLPDKAI
SLLDTACARVALSQHAPPRELQDVRRRLEAARVRLELLDGEQRIGLGDEAALAQSRAEIA
ALEVEEQSIDACWRQQIEAAQTLLSAREAAGDAAASAIDALRELEHALSNLQGDHAFVFP
EVNEAIVAEIMSDWTGIPVGRMVTDEVSAVRALPDTLAARVIGQRDALHQIGDRVQTARA
GLTDPKKPLGVFLLAGPSGVGKTETALALAEALYGGEQNLITINMSEYQEAHTVSGLKGA
PPGYVGYGEGGVLTEAVRRRPYSVVLLDEIEKAHPDVHEMFFQVFDKGYMEDGDGRYIDF
RNTTILLTSNAGSDLVSGLCMDEACAPDADGLCAALASELLKTFPAAFLGRVTLVPYRPL
AQETLSRIVRLHLDRVVTRMAENQAVALRYSDNVVDYVVNRCLVQETGARLLIGFIEQHV
LPLLAKLWLDSLASKHVLTRVDLDVADPAAAPTEAFALRAS*
>GCA_000452945.1_BPMSHR5858_genomic_05911_1 group_2026
MKIDLIVADDHPALIAGIEHALSKSSAISVAGTASNSTELVDLLARERCDILITDYAMPG
GSHGDGLGLLSFLRRHYPDLKIIVFTAIDNPALIASMARLGVRAVVNKMDHIDNLIASIQ
AVYAGATYIPAGVDAFVSSRPAGDEHTPEIERQLSAREL EIVRLYVSGLSVNEIAAYLHR

SKQAVSTQKMNAMRKLGIERDAGLFRYAYETGLMAAGVPSASPVSAQAVH*
>GCA_000452945.1_BPMSHR5858_genomic_05913_1 nicT_2
MTPPSSVAARPASDPLNDGYRKVTFRLIPLIFVCYLFNYLDRVNVGFAKLQMLSDLNMSE
TVYGLGAGIFFIGYVLAGVPSNLVLRRVGARAWIAIIMVVWGALSSALLFVKTPASFYTL
RFFTGIAEAGFFPGMVLYLTQWFSAEHRGRAIALFMSAIPISGVIGGPLSGWMLDHFSAG
QGGLAGWQWLFLLQGAPTIVLGVAVFALLEDRIDDARWLSADEKAALARAVATTAADAAI
ASAGRPNTLRDILLNGAVWKFGIVYFSIQMGVYAINFWLPSIIRSLGVGGSGAIGWLSAI
PYLFASVAMIAVGKSADARQERRWHLSVPMAVGVAGLLMAAQSGNAIVPAMIGLVLATAG
ALTALAMFWPLPTLVLTGSAAAGGVALINSLGQIAGFVSPYIVGWIKDATQRTDLALYIL
SSTMVVGIVLVMRTHARGAGR*
>GCA_000452945.1_BPMSHR5858_genomic_05915_1 ttuE
MRNESHTARERRRSTKIVATLGPSSSTETAIEALARAGADVFRLNFSHGTHADHALRHAA
VRAIEARIGHPIGVLLDLQGPKLRVGQFASGRAQIAKGRPFVFDRDPAPGDARRVSLPHP
EIFDAARPGHLLLVDDGKLRFRVDAVSSARIETTALLDGIVSDRKGVSVPDATLAIPALS
AKDRDDLEFGLSLGVDWVALSFVQTAQDVRDARALIGARAAIVAKIEKPQAVANIAEIVD
AADAVMVARGDLGVEMSLEDVPSVQKQIIRLARAAGKPVIVATQMLESMTLAPTPTRAEA
SDVAAAVYDGADAVMLSAESASGQYPVEAVDFMRKIISTTEADPIQPQLMKAIVTAHAPN
ATDAIGAAIEVVSDTLRLGVAVTYTASGATAIRLSRLRPSTAILSLTPRAEIARRLTLAW
GVRSRVAAEATGIENIVEIAIEAAHAEGFATDECPIVVAAGVPFGHPGSTNLMRIVWPTE
MTRQ*
>GCA_000452945.1_BPMSHR5858_genomic_05917_1 group_3691
MTASTLDLPRDCEHALRAASPPNIVDLLLRAARLHPHTGVRFIAAESEHKGAFVTYPELL
DEARRILGGLRARGYRSGMKVALLLEHASDFIPAFWACALGGFVPCPLVPIRNDPERWAK
HLAHVDTLLDHPLLVTTEALNNDLPGGASAVNLNALRASLPDASTHVAQPSDPAVFVLTS
GSTGNAKAVVLTHGNLLASMAGKNDRQQLAGADVTLNWISFDHVAALLEAHLLPLYVGAV
QLHVEAAAVLTDPLRFLRLVSRYRVTMTFSPNFLFGQLNAALEAMGDEALAAWRGAVDLS
SLRHVVSGGEAIVVATGQRFLDLLAPCGLARDALWPAFGMTETCAGSVYSREFPEGDAGR
EFASLGLPVAGLQMRIADDRNNVLPEGEAGEFQVRGPMIFQRYHNNAEATRAAFTSDGWF
RTGDLGRIERGRLWLVGRSKDSIIVNGVNYFSHELETTLEALDGVKPSFVAAFPTRGAGD
ESEQLVVTFTPSFPLDDEDALYRLVIAIRNSTILLWGFRPALILPLPEDEFPKTSLGKTQ
RAIMRKRLEAGSYDGYKARVADLANRQMGGYVAPDGQTEAAVAAIFARMFQVAPEAISAT
ASFFDLGGTSLDILKLKRHVEQRLGVIDLPIVTILQNPSVRALAARLAPGERVTAGEYDP
VVPLQLTGGKTPLFCVHPGVGEVLVFVNLAKYFVNERPFYALRARGFNEGETYFSSFDEM
VNTYVDAIRKRQPHGPYAVAGYSYGGAVAFEIAKVLEAQGERVDFVGSFNLPPHIKYRMD
ELDEVEGAVNLAFFLSLIDKQQSLTLPPQLRAAMPEQDPLAYLIDHAPPGRLVELDLDLA
KFRAWAGLAQSLLTLGRSYAPSGSVRAMSIFYAIPLRGTKDDWLNKELRRWDEFTRAPNR
YIDVAGEHYTLMGPAHVATFQAVLRAELDRALGGK*
>GCA_000452945.1_BPMSHR5858_genomic_05918_1 group_3692
MTVARNDGRGDARQRERAKRKVLITGATGQVARPVAEALAAQDEVWAIGRFGDAGVEHAL
RAKGVKTWRWDMERDTLDSLPDDFTHVLHAAVRRGEDGDFDRAIEINTVASGRLMTHCRH
AESFIYVSSGSLYARQALDHPYAETDPLNGVAHWLPAYPIVKISCEGVVRALSATLGLPT
VIARLNVAYGPYGHGGVPVLLYRQLLAGKPVPVPREGQNWASLIHTDDLMRQVPLLWEVA
SSPALVLNWGGDDAVGIRDCLQYVADLTGVDARFERSDVTRETYAFDNTKRRALIGDCTV
HWKDGVRRTIETHFPGAVKTQPAAQR*
>GCA_000452945.1_BPMSHR5858_genomic_05920_1 mmcR
MTEVDAGQTPHAAILQMVTGKWHAQALYVAAELGIADLLAQRERTADELAAETQTHPEAL
YRVLRALGSLGVFVEEDGRRFRNSPLGDTLRRDAPGSMRGFVRFAGMDAGWRAWGQLMYS
VRTGRSAFEHVVGGPGFAYIDSHPEIAAIVNDAMTSMSELEAPAVARAYDFSQARTIVDV
GGGHGFLLATLLRANPHANGVLFELPHACEGARQLFAKHGLTERVKVIAGDASHSIDAHG
DVFVMKHVICDWDDEQATRIMTNCAEAMRPGGKLLLVEAVLTPPGEPHFAKLHDLEMLIM
SSGGHGRTADGYGRLYAAAGLMMTAVRPTQGMNSVIEGVKP*
>GCA_000452945.1_BPMSHR5858_genomic_05922_1 group_1973
VDSSSNALYTRQIHAYVKALERGDVEAICVLFTPDAQIVSPFLGRMQPAPFFAKVAAASG
ESAIALTDVCVSASGARRAMGYLVYDWGLKDGSSVRFECVDVFEFDANGLIERMVIVYDT
HPIRDVVGDKYA*
>GCA_000452945.1_BPMSHR5858_genomic_05923_1 group_3694
MDGFLIQRPSRLQTLGAGTFALLTLGALVATAPHADLRLPPIEPFLPMCALTVFSTSLIA
AFLLGAQFIVMRRPILGTLAGAYAFAAITVALQLLAFPGVFAPTGLLGARHESAAWLWTF
WHGGFPLFVVLAMLTRYRLAKRLSNTRHVGRWLATLAGVPIAVAIALGLVALRADLPAPF
GPSLDNALLAADPVAIVIWTINLGAIAAALFVGRMRTMLDIWLVVVLLAGFTDTSLNLLS
TARFTAGWYVARLFSMIAPGILVCVLVWEVTVLYRRLFDAHASLMRTASHDALTMVYNRS
YFNEQFERAFHYARRHNTPLALVMVDVDHFKQYNDTFGHLQGDTCLAAVAAALSGVAGRA
NDFVARYGGEEFAVVLPGADERGAATIAERARQAVERLRIAADTAPGYVTVSAGYAASSG
NPFETPGALIAAADAALYRAKMLGRNVSAAPHHDMQDTAMRD*
>GCA_000452945.1_BPMSHR5858_genomic_05927_1 rclA
MTRRFDAIIIGTGQSGPPLAARLSAAGMKVAVVERGRFGGTCVNTGCIPTKTLIASAYAA
HLARRADEYGVTLGGPVAVDMKKVKARKDQVAARSNRGVEQWVRGLENGAVYHGHARFES
ARTVRVGEALLEAERIFIDVGGRALIPPIPGLDQVPYFTNSTMMDVDFLPEHLIVLGGSY
VGLEFGQMYRRFGARVTIVEKGPRLIAREDEDVSHAVREILEGEGIDVRLDADCLSVRRD
GGGIAVGLDCAAGAREVRGSHLLLAVGRVPNTDDLGLEHAGVETDARGYIRVDEQLRTNV
AGIWAMGDCNGRGAFTHTSYNDYEIVAANLLDGDPRKVSDRIPAYAMFIDPPLARVGLTQ
AEAARSGRRLLVGTRPMTRVGRAVEKGESLGFMKVIVDADSHAILGASILGVTGDEVVHS
LLDAMYAKAPYTTISRAMHIHPTVSELVPTLLQDLHPPA*
>GCA_000452945.1_BPMSHR5858_genomic_05930_1 rbsA_4
MQRSDPPRPLLEMRGISKTFPAVRALAGVSLTVHPGEVHSLMGENGAGKSTLMKILSGAY
QADPGGEILIDGRPISIDGPLAAREAGVAVIYQELCLAPNLSVAENIHVGRELRRGNGRR
GTIDRAAMARGCQDVLERLGADFGPNTLVGTLSIAEQQLVEIARAVHTRARILVMDEPTT
PLSSRETDNLFRLIRQLRAEGLAIIYISHRMAEIYELSDRVSVLRDGAYVGTLERDALSA
ERLVGMMVGRDISGFYKKAHAPYDPGNLLLSVRDIADGARVRGCSLDLHAGEVLGIAGLV
GAGRTELARLIFGAEPRVRGEVTLAGKAFAAHSPREAIDAGLVYLTEDRKRQGLFLDMSV
RENINISVCGRDARLGALDLARGAQRARDAIAALSIRVPHANVNVGALSGGNQQKVLLSR
LLETKPRVLILDEPTRGVDIGAKSEIYRIINELARAGVGVIVISSELPEIIGVADRVLVM
REGRIAGELGGRTDAPITQEAIIALATGSRTEASAAH*
>GCA_000452945.1_BPMSHR5858_genomic_05931_1 nagK
MRSPHLGQGSNSANVRRYNERLLLKTLRRAGSASKADLARLANLTGTAVGSIIASLAEAK
LIEFAGRRAEGQRGQPASLIRLDPRGAFGIGVRLDRMRIETALVNFAGDVIGRRSHDTLL
PQPAAVLEIVREDVAAMQQLLCAHERARLTGVGVAQPYNLGSWLREIDLPADAFGAWAEV
DFARELDRAIELPVFSENDGNAAAIAELFYGCGRQRDDFVYLFLGPAIGGGIAVDGDCLR
GVTGNAGDFAMMPVPPSRLPSVPKPSGAWDLLITRASLNGLARHLRHRGETLESRADLEA
CIARGGQAVDEWIDDCVDALAPALRAVLAVLESPVVVLDADTDAGLIDTLIARLHTALAA
LAPEARGTPVLVRGTFGPDAGAIGAATLPMFFNFSPRAGILRGAGVDSQEARYAAF*
>GCA_000452945.1_BPMSHR5858_genomic_05933_1 cga
MSARHFGFKQAVLVGSLAALAGAAALPANTSAAEAFGSPGAAPVRGPAAKSFLGTAVNGA
SRVYFTGYRGILSEVYYPVLDTPESVDLQFLVGDAGKTFVDEEKQQAYSAAQTDTRTMSW
QVTTGNSSHNWQIRKIVFADPNHNAIVQRVTFTALDGRKVGDFNLYMLSKPYLDNAGANN
TAQTVSGSGGTALVANHNSRYSALVASRPWKVVNGVGMTSNGFVGQSDGWTDLFGGTADK
TMDWTFSSATNGNVAQMGWLDLGDPSATSVSFDVVLGFGGTQDQALGDANTVLGSDLAGE
QQQYDTGWHNYAAGLSSQNGAADDGYYLAAMTLKTMQDKSNGAMIAGIGTPWGETQGDAN
AGGYHLVWPRDLFKFANALTTAGDTSSATSVVNYLFNTLQQTTDCGTAEYNAPGCAQGYS
RVGRFPQNAWVNGWPYWQGTQMDEQAMPIILAWRLGPSVFNPLWPKIKLTADYIVNTGPW
TYQERWEENAGYSPSTIAAEIAGLVTAADIATQNGDTASAARYLAAADYWQENVAAWTYT
SSGAFGNGSYYIRINPAGRAGTGTDRASFAPTAGPDTPQTLSVKNGGGSHDARRVVDGGF
FELVRMGVKRATDPTIANTISVYDSVLGQTLNVANAPALSPNAWFRYNFDGYGEHNDGGD
FDGTGAGRLWPIFTAERGMVEIARQGSGGAGSAYFSTLKQLTTPEGFVPEQVWSNSTTLP
DGWAVTTPAGYTPGQPTKSMAPLNWAMGEYISLLASIQAGRIVDVPSVVCARYNNCAAPP
QSGQVPVAVNVNASTQLGQQMYVTGNVAALGNWNTDLGIPVDPASYPVWRNTVNLPAAQA
IQYKYYRKNADGSVTWENRSGNRQLQTPASGTLALNDQVSW*
>GCA_000452945.1_BPMSHR5858_genomic_05935_1 dbpA
MTYPPTATPFSALALTPATLANLAQLGYTDMTPIQAASLPIALAGHDLIAQAKTGSGKTA
AFSLALLARLDARRFDVQALVLCPTRELADQVAQEIRRLARAEENVKVLTLCGGTPMRPQ
TQSLEHGAHIIVGTPGRIMDHLDRGHLTLDALNTLVLDEADRMLDMGFFDDIAKVARQCP
PTRQTLLFSATYPDGIAKLSQQFLRHPKRVKLEERHDDSKIRQRFYEVTEDERLHAVGLL
LNHYRPVSTIAFCNTKQQCRDLLDVLHAQGFHALALHGELEQRERDQVLIRFANRSCSVL
VATDVAARGLDIAQLEAVINVDVTPDPEVYVHRIGRTGRADQDGWALSLASMNEMGRVGG
IEQALAREVEWHPLAELKAGGDAPLLPPMETLQILGGRKDKIRPGDVLGALTGDAGFSGA
QIGKINVTEFSTYVAVERGIARDALRKLNAGKIKGKRVKVRLMDE*
>GCA_000452945.1_BPMSHR5858_genomic_05938_1 hipO_1
MHSAPSEHPVIPGIAAIAPELIDVRRRIHAHPELAFEETLTSDLVAGLLAGWGYDVHRGL
GKTGLVGVLREGQGARTLGLRADMDALPIAEATGLPYASRRANKMHACGHDGHTAMLLCA
ARHLAATRRFSGTLNLIFQPAEENFGGAKAMMDDGLFERFPCDAIFAIHNMPGRAAGDMA
FRTGAAMASSDRVTITLRGVGGHGAMPHFARDPMSAAGSIIVALQTIVAREIDAQQSAVI
TVGSVQAGETFNVIPETVVMKLSVRALNADVRALLAQRIEALAKGQAQSFGIEADVQYDY
GYPVLVNHDEPTAFAIDVARRMLGEARVEAQTAPLMGSEDFAYMLEARPGCYAFIGNGVG
SKGGCMVHNPGYDFNDDILAIGASYWVRLAEAWLAA*
>GCA_000452945.1_BPMSHR5858_genomic_05940_1 group_19953
MYNVTPRHAALWRSLLRDALHAFARAGGPADVTLLGEPSGEPDGELLPLWRRGDLLLSQT
CGYPYRMLGVGDAVHLIATPVFDAAGCDGPRYCSMLAVSAGAHARGATTLAACRGLRAAC
NGDDSHSGMNALRHAIAPHARGGRFFSAVTRTGSHLSALRALDAGAADLAAIDCVTLAYV
RDALPARLRRVRIIGATAAAPGLPFIASRALAGEHVAALRDALDAAVAIDAERARALRLR
GFARLAPDAYAEIERFEREAVALGYPALA*
>GCA_000452945.1_BPMSHR5858_genomic_05943_1 gltI_1
MMKPLRPLLSIAVGCALAAAAHADDAGPTLKKIRDTGVVTLGVRESSVPFSYYDQQQRTI
GYSQEIALKIVDEIKKTLNRPNLTVREIPITSQNRIPLVQNGTVDLECGSTTHTKERANQ
VSFSNSIFQYGMRLIVKKSSGVKDFPDLAGKTVATTAGTTEERLLRQWNAEKGMAMQIIS
AKDHADAFLNVKSGRAVAFFMDEPLLYGAKAKEANPGDYVITGNSPVSEAYGCMLRKDDP
GFKQLADRVIARMQRSGEAEALYVKWFNRPIPPKGVNLDYPLSADMKQLFANPNDKALD*
>GCA_000452945.1_BPMSHR5858_genomic_05944_1 dadA_2
MRICVLGAGVVGLTSAYCLAREGHDVTVLEARPDAVLDASFANGGQLSYSYVAPLADPAV
LGKLPAWLARRDSALRFVPRLDVDQWLWCAAFLAACRTARARRTAAEMLELGALSRIALH
ALVERETLDFDYARNGKLVVYRDRREFDRARRAMDHLIAAGARQQALDTAACVALEPALA
PASGLLAGGIHTPSEEVGDCRRFGIELARVLRERLRVAIHYETPVDTLRTEGATIVAART
PRGEIAADAFVLSAGNGSAPMLRRLGMRLPIYPLTGYSLTLPARHDATPRVSVTDLHRKI
VYAPLGARLRIAGMVEIAGLRSSGAARRLALLAQQAQEIFPHAGDYAHGETWCGHRPATP
DGKPLIGATPFRNLWLNTGHGALGFTLACGSARVLADLIADRPPALDTRAYALHR*
>GCA_000452945.1_BPMSHR5858_genomic_05948_1 group_19792
MVRKRIRAVFAAALVGLAAHAAHAQYTTDWIANTYGTIASHVGNNARSMWVSPEGVIYTA
SFWDENAGGVAIYQNGKTLGSIGTHAEFQGGAITGNATSIFAAMQYSTPQGSGTVGRYNR
ATLQRDLTIPVSVWNAVSRADVITGLATAGTLLYASDYFGNRVRVFTTEGVWQRDIGIAN
PGALALDDAGNLWVAQKNAAKIVEFGPTGALMNTIQMASASRPASLYFDASKRQLMIGDQ
GPDMNVKLYAIAGMPKQVGTFGVQGGYLDTTTGIRGQVGDRRFTRVVGIGKDAAGTLYVL
NNPWGGGWDLGRNGATDIHAYDALGNALWKLQALNFEAIAAPDPTTDGALFYSGMNVYSG
TAGGTFVANTVDPFTYPSDPRLDMNDYQRGQHFGQLVSVGGHKILVASGQNPGNFNFYHF
NAASGYIAIPDASLPGKGFNTSLQVTAGFSIDNKGDVWAGLNGTNAISHYPLAGIDASGK
PSWGAPTSIPTPASVQPTTRILYLSDSDTMILAQGIAGSWDWTAMNGRIEVYHGWSAGNV
TQPNPVIALTSPNPKSIASAGNYLFVGYVHTVPNIDVFDLNTGQLVATLTNSNTGMMDVG
NDVDSMYGLRAYLRSTGEYVITKDNYNGSSIVVYRWRP*
>GCA_000452945.1_BPMSHR5858_genomic_05954_1 qorA_2
MQAIEIRETGGPDVLEYVTRAAREPGRGELLVELAAAGVNFIDLYRREGRYPMPLPGTPG
EEGAGRVLAVGPDVTRFKPGDRVAWTTVMGSYATRVVVPEDKAIAVPDGIDLRTAAAALV
QGMTAHYLVNDSYRAREGDVVLVHAAAGGVGLLLTQWLKRRGVRVIGTVSTAGKAALARS
NGADDVVLHGAFDDLAAEVRRLTDGRGVHAVYDGIGAATFDASLASLRTRGTLVIFGGAS
GPVPPVDVMRLLWGGSLTLTRPYLEHFRASVDEFLWRAGAVFDGILDGSLKIRIGGAYPL
ADARRAHADLAARRTTGKLLLVP*
>GCA_000452945.1_BPMSHR5858_genomic_05965_1 group_2657
MIFFRIELIMRAFKLFRRGPPDIERAETAARRDFEPSCAGVDVEGNAGDETGACLAEAQP
PRAPQAGPQGATDGGDAVFGVIGAAFAADTPGERQGGTAAGASAVCGTVAAQDLLHTLYE
QYCRVLDDPRASLASDDAARLAIERTRDAGPQIDPQRSADGADSIDALLSRARVLDDAFG
TFAPGEAPDPADAEPVPEVLRLFAPAEYHAASARRPAGLPPALARREHQTLAIDSPLPAP
VATSEHGA*
>GCA_000452945.1_BPMSHR5858_genomic_05966_1 group_199
MTTPAFPAPPGARTSARPHDTPLDARLAAAQAAVRARPVDAASRWALFRLLCVTGQGERA
VGQLQAYARFAAQHAHIAHAYRDLIRAERFRARVMAGQARPGFVFDAPPWIDDLLEALRA
QAAGARDEADRARSRALDRAPFVAGRGPDAPFDWIADSDSRFGPVCEVVTAGHDRWLPFS
DLAGWRLARPGVLLALVWAPCVLTLADGSVAHGFMPARYPGSELADGGGAPGASAAAHEA
DAHESAAREADALRLGSRTAWRDVGRTGVFALGRKTWSTSAGDAGLFELHACEFGARARA
GAALGGAAREARGGEAADGCA*
>GCA_000452945.1_BPMSHR5858_genomic_05975_1 group_19893
MVMVNSSAGGMNAATSINLTPPGMPMPYPNNAPRANAVPNVSHIFVGGGPVHNLATVIPA
SNGDSGGSMGGVASQTVSGSSRNAQGASKTLVAGMPITRMTDPTQQNNNNTIGSGSSPSQ
TIVLNL*
>GCA_000452945.1_BPMSHR5858_genomic_05976_1 group_1975
MTKRIHAVLLAALLIAALAPPCQARSSTDWLANTFATDAMHVGNAARSMWVAPEGVIYTA
SMWDENEGGVALYQNGRSIGSIGAHNEFQGGAITGNAASIFAALQFNRNGGSGMVGRYDR
TTRRRDLLIPVSATTTERWADVVTGLATSGSTLYASDFPGNRVRIYTVAGVWRRDIDISE
PGALAADSAGNVWVAQKRAGAIAEFDSGGRLLNTIRMAAASRPSALYFDAATQRLMVGDE
GPDMNIKIYDVSRTPALVGTFGVRGGYLDSASGIKGQVGDKRFTRVAGLGKDAAGNLYVL
NNPWGGTWDLGRDGATDIHAYSPSGTLQWKLQSVNFEGIAAPDPGTDGAFFYSGTNVYAG
TAGGTFVANTVDPIDYPTDPRIDGRDPARGEHFGQLAAIGSNRILAAGSQNPDVFHFFHF
NAAHGYIAIPDGSIPGAAFGTTARIRAGFCLDSRGGVWAGLDKTNAIWHYPLTGFDANGK
PAWGPATVTPIPNSIKPLTRIVYLPESDTMILARGRPVGTDWTAIGTRIEVYHGWLAGNT
AAPNPVIGLSSANPKSITAAGNYLFVGYVHTQPNIDAFNLSTGALAATLINSSPEQIYVG
NDVDSMYGLRSYLRSNGEYVITKDNYNAAGIVVYRWTP*
>GCA_000452945.1_BPMSHR5858_genomic_05980_1 metZ_2
MTEPASADWRLETLAVHGGYRPDPTTRAAAVPIYQTVAFAFDDTQHGADLFDLKVQGNIY
SRIMNPTNDVLEQRVAALEGGIGALALASGQAAVTYSILTIAEAGDNIVSSSTLYGGTYN
LFAHTLPQYGISTRFADPREPAAFDALIDARTKAIFAESVGNPLGNITDIAALAEIAHRH
GLPLIVDNTVPTPYLLRPFEHGADIVVHSLTKYLGGHGTSLGGAIVDSGKFPWAKHAERF
RRLNEPDVSYHGVVYTDAFGAAAYIGRARVVPLRNTGAALSPFNAFQILQGIETLALRIE
RIGENALKIAQHLARHEKVEWVDYSGLPNHRDHALVERYLSGRAPGILTFGVKGGRAAGA
AFQDALKLFTRLVNIGDAKSLATHPASTTHRQLSPDELAKAGVKEETVRLSIGIEHIDDL
LADLDQALAKV*
>GCA_000452945.1_BPMSHR5858_genomic_05981_1 group_3699
MSVIGLDHYNLRAPRPLLDTLRDFYIDVVGLRLGARPPFRSHGYWLYAGAQAVLHLSQAG
PDETRRANVVNTFDHVAFSCDDLPGTLARLQRFGIRYSSADVPLTRQHQLFFDDPAGNGV
ELNFAVRDDG*
>GCA_000452945.1_BPMSHR5858_genomic_05985_1 pxpB_2
MTIRYTFGGDEFIFVEISESMSLDAFFKGTAITRELRRRNVPGVTEICPANASYQVRYDP
DVIEPDTLFALLKTIEAEVGEAALNIDTRIVEVPVLYNDPWKHETLMRFRERHQAPESTD
LEYAARVNGKRDIDEFIAAHAGSPWFVSMVGFVAGLPFMYQMVERERQLQVPKYLRPRTD
TPKLTVGHGGCFGCIYSVRGAGGYQMFGVTPAPIFDPAQRLDYLRDFMVFFRPGDIVKFK
PIDRAEYDAAVAAVEAGTFTLRVKPVRFALEAFLRDPHAYNRSLVEVLDAN*
>GCA_000452945.1_BPMSHR5858_genomic_05993_1 group_3700
VNAGFVQDRINQAVWRSRGALRGYGRASGWTDPGEAAAIAWIAERVRDEPILDVGVGGGR
TVPLLKTISADYTAVDYTPELVEICRRNHPGTRVHRMDARDMSAFADESFSLVVFSYNGI
DAVDHDGRMAILKEFARVLRPGGFTLFSTHNLAGPSYREGPSRLARLPRRSANPFAIALD
AARIAYSLPVGAFNYFRNSRFNRVFDGYAMRVCAAHKFGILIMYTDFATQQRQLADVGLK
LRAAFGGSSGEPVRRDQDLGGEAWLHFVASKPCG*
>GCA_000452945.1_BPMSHR5858_genomic_05996_1 tsr
MKLRRKIPLAFATALVLTSASAFYGIHALNRSLDTYGTTVRQNVANERMVSATLVAFKLQ
VQEWKDTLLRGKDPAKLDKYWRAFQQREQTVDALAAELKAKLPDGESRGLIEQFASAHAE
MGQGYRKGFEAFRASGFDPSAGDQAVAGVDRAPAVLLEKAARDIAADSARVSADAASDAA
HATAISIAATFALFALSLAGGVWFGGTVTRPLERALACVRRVATGDLSTPIDARGRDEIA
ELLAALKDMQASLSHVVRDVRHNADGVATASAQIASGNLDLSSRTEEQAASLEETAASMD
ELTSTVRRNAEHAQHACAVAAGASTKAARGGDVMRQVVDTMRGIADSSGKVAEIIAVIDG
IAFQTNILALNAAVEAARAGEQGRGFAVVAGEVRTLAQRSATAAREIKTLIEQSTERVGA
GSALVDDAGRIIGEIVDSVRQVTGIVSEIAAASNEQSVGIEQVNRAVAQMDNVTQQNAAL
VEEASAAAHALAEQAHALHGAVAVFSLHGERAGERGCAHAGQPAVEAAHDSPRTPLPVVA
PA*

>GCA_000452945.1_BPMSHR5858_genomic_06000_1 acnM
MNTANRKSLPGTSLDYFDARAAVDAIRPGAYDALPYTSRVLAENLVRRCDPAILADSLTQ
LVERKRERDFPWFPARVVCHDILGQTALVDLAGLRDAIAERGGDPAKVNPVVPVQLIVDH
SLAVECGGFDPDAFAKNRAIEDRRNEDRFHFIEWTKKAFENVDVIPPGNGIMHQINLEKM
SPVLQAQDGVAYPDTCVGTDSHTPHVDALGVIAIGVGGLEAENVMLGRASWMRLPDIVGV
ELTGKRQPGITATDVVLALTEFLRREKVVGAYLEFRGEGAKSLTLGDRATISNMAPEYGA
TAAMFFIDEQTIDYLRLTGRSDEQVKLVETYAKAAGLWADSLAHAQYERTLTFDLSSVVR
NMAGPSNPHKRLPTSDLAARGIAGKWEEKPGEMPDGAVIIAAITSCTNTSNPRNVIAAAL
LARNANARGLARKPWVKSSLAPGSKAVELYLKAANLLPELEKLGFGIVAFACTTCNGMSG
ALDPKIQREIVDRDLYATAVLSGNRNFDGRIHPYAKQAFLASPPLVVAYAIAGTIRFDIE
KDALGHDKDGKPVTLKDIWPTDEEIDAIVASSVKPEQFRTVYEPMFARAADAGERAAPLY
DWRPRSTYIRRPPYWEGALAGERTLKGMRALAVLGDNITTDHLSPSNAILPTSAAGEYLA
KMGLPEEDFNSYATHRGDHLTAQRATFANPTLVNEMAIVDGEVKKGSLARVEPDGKVMRM
WEAIETYMSRKQPLVVIAGADYGQGSSRDWAAKGVRLAGVEAIVAEGFERIHRTNLIGMG
VLPLEFKPGTNRATLAIDGTETFDVIGERTPRADLTLVIHRSSGERVAVPVTCRLDTAEE
VSIYDAGGVLQRFAQDFLESSRAA*
>GCA_000452945.1_BPMSHR5858_genomic_06009_1 hcaR_1
MDRLELRHVRAFLCVARHLHFARAADELDIAPPALTRLVQDAERLLGVRLFHRTRRSVAL
SAAGAAYLGEAQAALAHLARGRERAALAERGEIGRIELGYVSSAAYAGVLQRTVGAFRDA
HPRIRVDVREVPMSEVAARLDAGTLDAAYVRPPLPLPDGVSTHTVHRDVFVVAVHGQSPL
AARVSIRPAALAGERFVVPEQALGTLEAARRGRFEPIVDARPGALVAVLAHVSVSGGVAI
VPQALVGCVSLPGVAYRPIEGKPIASEVAVAYRRHEKAPAVRAFVRFVRQAAL*
>GCA_000452945.1_BPMSHR5858_genomic_06016_1 group_2659
MIAFLARRAARRTPTPTTASHSPISRSRGAGRHLFAALAVALIGLPHAHAEIRCGWLQNP
TPGNWWLDDRAGSWTLGTMGGPETEGMDVIPDMAGKQYVETNGSHGYACACLTVVTDKQE
RRIVKVLKAKQLPLSRCRRDKRLKEPT*
>GCA_000452945.1_BPMSHR5858_genomic_06017_1 kdpB_3
MTKLFAPAAPITTTLLVEGMHCGGCTSRVEQALAQVPGVTGAVADLAAGTATVAAASAID
TARLVAALDAAGYRATVATAPAATGNADARHGRARDEDDDAAAAPHTAVVTLTIGGMTCG
GCARRVEQALAAVRGVADAKVDLATTSAKASVARDVDSQTLVAAVERAGYRANVVRDARA
EAAPKPAACPFEDAARSAAPAAAFAVDESSAASPERVATQSFELDIAGMTCASCVGRVEK
ALAQVPGVARATVNLATEKAAVDADADAHVDTARLIDAVKRAGYRASPVSDPASALAPSP
EIAAARTAIELDIAGMTCASCVGRVEKALAQVPGVARATVNLATEKATVDADADAHVDTA
RLIDAVKRAGYRASPAIAACAPASRATATADAAATRPASPSADDRKLAEARRERALVIAS
AVLTTPLALPMFAAPFGVDAALPAWLQLALASIVQFGFGARFYRAAWHALKARAGNMDLL
VALGTSAAYGLSIWLMLRDPGHAVHLYFEASAVIVTLVRFGKWLEARAKRQTTDAIRALN
ALRPDRARIVEHGVERDVPLAQVRVGTVVRVLPGERVPVDGRIEAGVTHVDESLITGESL
PVPKGPGERVTAGSINGEGALTVATTAIGAETTLARIIRLVESAQAEKAPIQRLVDRVSA
VFVPAIVAIAFATFAGWLVAGAGVETAILNAVAVLVIACPCALGLATPAAIMAGTGVAAR
HGVLIKDAQALELAQRARIVAFDKTGTLTQGRPTVTAFDAIGIPRGDALALAAAVQRASA
HPLARAVVAAFDADADARRSSLAAAHADTPRAVAGRGVEARVDARLLALGSTRWRDELGI
AVPDGVARRAAALEAAGNTVSWLMRADAPREALALVAFGDTVKPNARRAIERLAARGIRS
ALVTGDNRGSATAVAASLGIDEVHAQVLPDDKARVVAQLKATAGDGAVAMVGDGINDAPA
LAAADVGIAMATGTDVAMHTAGITLMRGDPALVADAVDISRRTYRKIQQNLFWAFVYNLV
GIPLAALGWLNPMIAGAAMAFSSVSVVTNALLLRRWKGDAR*
>GCA_000452945.1_BPMSHR5858_genomic_00046_1 group_1740
MATHPTGTRDEWLAARLELLDAEKAHTRRGDALARQRAALPWVRVDKAYRFDTDEGGASL
ADLFRGRSQLLVYHFMFGPDYAAGCPSCSALADGFDGFVVHLAHHDVTLAAVSRAPLAKL
RAYRTRMGWRFAWASSLDGDFNYDFNVSFTDAQQRDGSVEYNYRRGGHAMDVTPAPAPVA
GFAADCGTDARTYARDRPGMSAFVLEDGAVYHTYSTYARGLDGLWGMYQWLDRAPKGRNE
TGVWWRRHDEYDGR*
>GCA_000452945.1_BPMSHR5858_genomic_00047_1 group_276
MDSLITAAARALAAGDPLGALNRVALRDDAPALALRGIAMAQLGEFVRAKALVRRAARAF
SPNEALARARCVVAEAEIALASRELGWPEKALDAARATLEAHGDSVNAAHARYLAIRRLL
LIGRVDEAARTLAALDAATHPPALRAAHELIVAGIALRRIETKPARAALARAERAARDAG
IPALAAEVERAFRVLDTPAARLVARGQTRALRLDEVEAWRASASLVVDACRRVVHDARAT
VPLARRPVLFALARALGEAWPGDAPREALVARAFRVKHADESHRARLRVEIGRLRALLGE
LADIRATQRGFVLIPRRAPDVAVLTRPHEDAHAAVLALLADGESWSSSALALALGASQRT
VQRSLDALAAAGNAQSFGCGRARRWTTPPVPGFATTLLLPAPLPGD*
>GCA_000452945.1_BPMSHR5858_genomic_00078_1 rluF
MRTKLTVKNPRPATPGRAPVRSGSLTARKVARPDPKAAGAKPAAAKPAAKSASAAKPAAP
RGAANAAPKRAPGPSHPAAASEGKRVAKPRVAHDAGRTGGERAPAKRATAPGAPGAPGAP
SAPRTRRTDAKPARRTDERPAGRAGNRPAGRDERAPRDSDARAFDAGTRGKDRAPREGAR
PGARGATGAKFGGAAHRSDDADRRTPRATRADSRARDAAPSSFAGKTTTAGKHAPQRADD
RYGAAGKRTSPRPERTERPARFGERPATRASASGERRPTARAATGSRLKLAQPIKRGGGE
LGESARGGEHGEHGERGKRIERGDETGLVRLSKRMSELGLCSRREADEWIEKGWVLVDGE
RIDTLGTKVRADQRIEIDSNARAAQAAQVTILLHKPVGYVSGQAEDGYAPAATLVTRENH
WSGDRSPLRFSPQHLRALAPAGRLDIDSTGLLVLTQNGRVAKQLIGEQSDIDKEYLVRVR
FGERTADIERHFPAESLAKLRHGLELDGVPLKPAMVSWQNGEQLRFVLREGKKRQIRRMC
ELVGLEVIGLKRVRMGRVMLGALPQGEWRYLGPDESF*
>GCA_000452945.1_BPMSHR5858_genomic_00121_1 group_1389
MSDARAVSIRGTAAVRDAAARRRTARGIVCAACAAAAVSAHAQTPSPLGEWQYSAGVPLD
KLFNPNPQTWQISVGAAATLQPRYDGSNQYRPMAGPTFDARYRDLWFVSTGEGIGVNVLR
GPNWRATLSAGYDLGRREADDRGHLTGIGNINPAAVIKLSADYVISHAFPLVLRADVRRS
VGGANGWVADLAAYMPLPGSSETFYWFAGPTVTFADSRYMNSWFGVNDAQAARSGHPRYA
PSAGVKSFGGGMTLVWFVTKHWFVTADGAIEQLVGSAARSPLTQRSTNAVVDVSINYQF*
>GCA_000452945.1_BPMSHR5858_genomic_00124_1 group_1746
MFFHNKRLQYTVRVAETNPGLANLLLEQFGGPQGELAAAMRYFTQAIAEEDAGRKDMLYD
IATEELSHLEIIGSIVAMLNRGAKGELAEAVDEQAQLYRKLNGGGNDSHVTQLLYGAGAP
LVNSGAVPWTAAYVDTIGEPTADLRSNIAAEARAKIIYERLINVTDDPGIRDALGFLMTR
EVSHQKSFEKALYSITPNFPPGKLPAVPEYADLYYAMSDGGGARRKSPWNDGHDLRLVEH
PPVPADGGDGKATVALDGDESTTLEAMRHRLKSDPSSDPTTGADLGSGGPDKPRDFPT*
>GCA_000452945.1_BPMSHR5858_genomic_00126_1 rpoN_2
MSATLALQMRQHLALTPRLQQSLRLLQLSSLEFQQELRQALDTNPFLEDVQSPDDDAAEA
APKPGETPAADANARADDGYAERDEGPFATDASPPAGQDVPLTAELSARGSSRRSDDASD
LEPGDWMTAEPTLHEHLHDALRLCQLTRRDRTLARMIIDALDDDGYLRQALPELAAAADP
LLHPAEQELLVALRLVQSLDQPGIGARTLSECLLLQLDAMPADTPGVECAKEIAAHHLER
LARRETAEMQRRIGCDTHTLRIACALVRRLDPRPGNQYGSTAGNYVVPDVIVRQVRNDWL
VTINPAVMPRARIHRRYAELFAQSSGSNQSPLGQQLQEARWLIRNAQKRFDTIRRVGECI
VERQRDFFRYGEIALKPLVLRDIADELGLHESTISRATGNKYMSTPHGTFEFKHFFPRKL
EAAGKGACSAAVARVLIRDMIAAEQAIDPLSDVALAQRLAGRGIVLARRTVTKYRQAMKI
PPAELRRRAPL*
>GCA_000452945.1_BPMSHR5858_genomic_00127_1 group_3057
MPAKSQAQQRAAGAALSAKRGRTKASELKRPSKSMYSSMSEKELEKLAGTKTKDKPKHKS
KS*
>GCA_000452945.1_BPMSHR5858_genomic_00128_1 group_2229
MLRYALIFFIIAIVAAVLGFGGIAAGAAEIAKILFYIFVVIFLVTLVLGVARR*
>GCA_000452945.1_BPMSHR5858_genomic_00129_1 katE
MTLPTDLPQRLVEIEHLLSDCFASPSTTVLHVDDSSGRLTIQVSWVAARAASSLLEARCA
LNLVFEPNVLRRYGRLDAAARARVRERLAALARQTVDAARPVEGGEADACNATLPVTDAL
LAAAPRRERLARTRRARAHSRCGRRARRPRHERDSSPMADSDRNAPHETPRAAQPDDARA
DRKSQDLEPYRSKPQGEALRTNQGVRIADDQNTLRAGARGPSLLEDFIMREKITHFDHER
IPERVVHARGSAAHGVFRVYESMAEYTKAAFLQDPAKETPVYVRFSTVQGPRGSADTVRD
VRGFAVKFYTDEGNYDLVGNNMPVFFIQDAIKFPDFVHAVKPEAPNEMPTGASAHDTFWD
FVSLVPESAHMVAWLMSDRAIPRSFRTMEGFGVHTFRFVNAQGVARFVKLHWRPVLGSYS
LLWDEAQKIAGHDPDFHRRDLWEAIERGDYPEYELGVQIIEARDQHAFGFDLLDPTKLIP
EETVPVKLIGKMTLNRNPDNFFAETEQVAFHPGHLVPGIDFTDDPLLQGRLFSYTDTQLS
RLGGPNFHELPINRPVCPFANLQRDAMHRQTIDVGQASYEPNSLNGGWPKETDPAGRDGG
FDSYPEAVEGAKVRVRSGSFADHFSQAALFYQSMSDVERRHIRDAYRFELGKVTRPEIRA
RVVDEILARFDAGLAAEVAEGLGLPPPPASAPAIVAQPLSPALSLLTRGKRSVRSRKVAL
VATPGADTRLIERVRRALTDARAVPVLVAPTLARIGELTPEATLAGMPSVMFDAVFVCGG
DGDGRDLVHSSDARHFLQEAFKHLKAIAAVGSGRQLLGAAHLPEQGDGVCVGHAADLDQV
VAKFFDALSEHRVWSREPLAQGVPA*
>GCA_000452945.1_BPMSHR5858_genomic_00132_1 lig
MAGKLSVYRSRRRFGETPEPEGARRSPRRTKGAGRAGDESKRRARGDAAASSRGERAARR
GRAQQPAPARARGRSALHYVIQEHHARRLHYDFRLELDGTLKSWAIPKGPSLDPSVKRLA
VHVEDHPLEYAAFEGEIPEGHYGAGSVIIWDRGAWTPAGGVAAARAGYRAGKLSFTLDGD
KLHGGWALVRSRGLRDSGGKHEQWLLIKERDGDARDEADYDVLRARPGSVLGGARSARAR
RGRAAEPTERDAAAPKRARASVPKGRAEAKTSGRAPRTVSNGTAGAGKGASKRASKRASK
REGTNTGNGARSGKGNGNGKGAGKGVATPRRKRAAAPQSSDVDSADTDSPAARGLALPAG
LAARFGLSGAKRAALPRTLRPQLATLVDAPPPGKDWLYEAKFDGYRVLIRIDRGAASRPI
AVYTREGLDWSAKFAAQLDALARLPVDRGWLDGEAVVLDRAGVPDFQALQNALGAGRSND
VTLFLFDVPFLNGFDLRGVPLERRRALLAALVAAHASDVLRYSESLAFEAADLLTGACDA
GLEGLIGKRRDGRYVEGRSRSWIKLKCRRRQEFVIGGYTEPAGRRSGFGALLLGVYARAP
GDEGDEGERRTRGAAETRDERRRASHASRTSRAPDAGAARTPQARPLALRYVGRVGTGFG
ERTLRALARTLREHETGRMPFADVPRERSGTPVHWVKPVLVAECEFAEWTGDGIVRQASF
VGLREDKPARTIVREQPQSMETEAMNEHADEHTDEHTDARAERRTGGRTRRKARDGGARD
APPLARHPKRGDAGSSARKGARDGEAGKRAAAGSSPSSSPSSSTSTSPSISASGRTRGGG
RSASRDRAGDADEGANEGANDHAPRERAGAPKVAGVRVSHPGRVIDPHTGTRKLDLVEYW
EWIAPWLLPHLKGRPVSLVRAPADIGGELFFQKHADKLEIPNVALHAGLDPGHEALITID
DVKGLVGAAQMGTIELHTWNAHVSNIEKPDRAVFDLDPDPSLDWRAVIEAAQLTRALLDE
LGLVSFCKTSGGKGLHVVVPLARHAGWDDVKAFAHAVARHMAATLPERFTATMGPRNRKG
KIFVDYLRNNRGASTIAAYSVRARPGLGVSVPIAWDELPDTTGAAQWTLANLHERLDALK
HDPWRAYAHVRQRVTAALRKRLSDG*
>GCA_000452945.1_BPMSHR5858_genomic_00150_1 group_972
MINILVASEDASRLAHLARLVGDAGRYRVTRTVGRAAQIVQRTDGLDAFDILMIDGAALD
TAELAAIEKLSRLHPGLTCLLVTTDASSQTLLDAMRAGVRDVLRWPLEPRALDDALKRAA
AQCAQRDTPDTRIVSFMSCKGGAGTSFIAGNIAYEIAEGSKRRTLLIDLNQQFADAAFLV
SDQTPPSSVAQLCGQLERLDGAFLDASVVRVTDTFHVLAGAGDPIKAADIREDALEWILG
VAAPRYDFVIFDLGVSLNAVSMVALDRSDHIEVVLQPSMPHVRAARRLQELLVSLGCPPE
RIQLVLNRQTRASERARAALEEVLSTRAAHVIADDPATVNEAVDQGVPLSRLSRNCGVAR
SLQAFARQLAEGEQRPRRDSARDAPLLARFFSRGAAPKLKSM*
>GCA_000452945.1_BPMSHR5858_genomic_00154_1 group_3061
MNLDYFTLTRRLVLAASLAASLGGCMSSTPVWDGRFGDSVRAVMQAQIIDPHAAEHARSA
PGVDGRAAAAALDNYDKSFQQTEPTPNAFVIGVGGGKTGGQ*
>GCA_000452945.1_BPMSHR5858_genomic_00156_1 group_2232
MKNSRAFTMLAIAILAGLAAVAFASRWLVQTSSSAVTPVAVAATDVSLGEPLGPNQLRIV
NWPTASVPPGAFTDLKPLEGRVVRTSLARGEPVLGTKLAPVGTKGGLSAVIAPGHRAITV
RVNDVVGVAGFALPGNYVDVIVNTQEQSKADPQQSISKIVLEKILVLAVAQQVSRDDTAP
KVVNAVTLEVSPEQAEKLDLARSVGTLSLVLRNQVDKDALSTGGATKNTLLGQPAAAPAP
IAAPAAQPRPVRTRVVAHARAAAGRDCVGVLSGVTGSVECF*
>GCA_000452945.1_BPMSHR5858_genomic_00157_1 group_841
VKANTGIYIVDDLGRQRIAPRDLLNRWLLPLDRNVDQLTFASGVRLEMPFDVWPVFSSNL
TPAQFSDDAFLRRLGSKLHVGPLPAGRYRDVYDANCAALALSSDASTFDYLLQHLHGPTR
MPFLACYPGDLLRLVAASVRYRGDAPTVTPQALHEAWRSFFGSAPDEPPFNPPDADRDW*
>GCA_000452945.1_BPMSHR5858_genomic_00174_1 fumA
MTVIKQEDLIQSIADSLQYISYYHPLDYIQALGRAYELEQSPAAKDAIAQILTNSRMCAE
GKRPICQDTGIVTIFVKVGMDVRWASENGAATMSVTDMINEGVRRGYTHPDNVLRASIVN
PPEGARKNTKDNTPAVIHYEIVPGDKVDVQVAAKGGGSENKSKFVMLNPSDSIVDWVLKT
VPTMGAGWCPPGMLGIGIGGTAEKAMLMAKESLMEPIDIQEIIARGPKDWVEELRVELHE
KVNALGIGAQGLGGLATVLDVKILAAPTHAASKPVAMIPNCAATRHAHFVLDGSGAAKLE
APSLDAWPKVQWEPNTETSKRVDLNTLTPEEVASWKPGQTLLLSGKMLTGRDAAHKRIAD
MLAKGEPLPVDFKNRVIYYVGPVDPVRDEVVGPAGPTTATRMDKFTETMLAQTGLISMVG
KAERGPVAIDAIKKHKAAYLMAVGGAAYLVSKAIRGAKVLAFEDLGMEAIYEFDVQDMPV
TVAVDSQGTSVHQTGPKEWQAKIGKIPVASA*
>GCA_000452945.1_BPMSHR5858_genomic_00195_1 uppP_2
VSLWFLVFLSVLQGVTELFPVSSLGHTLLVPALFGMHIDKHAPQLLPFLVALHLGTALAL
LWYFRERWIALIAGFFASLNGRKNDEGHLMWALIIGTIPTGLVGLLLEKRIERVFHDLRI
VAAALIINGVLLWLGDRIQRARAHRPPEKLTFKQAFFVGLAQVGALIPGFSRSGLTMIAG
NAAGLTADKAAEFSFLLGTPIIFAAGLLELPKLFHAPDQLADALLGGVLTAIAAYLSVRF
LMRYFEGRGRLASFGLYCVLAGLFCLGWFMFHAQPV*
>GCA_000452945.1_BPMSHR5858_genomic_00196_1 yeaD
MPSFQNQDILELIDGASRARIAPEAGGRLLSWSIGDTSIVFWPDAADWSNPAKIRGGNPL
LFPFLGRHRVEGRIGFWRDGEGAVRELPMHGFARDLPFRAQTDADGRGVTLTLDSSDRTR
AGYPFEFRFAARYRLVDGHTLDVTLSTTNLGDTPLPHYAGHHFYFALPHVKRASAVLELP
PTHRRRQLDDGSISAPEPGAPRYTLDDPAIVDRFHCLDGIPAEPVRVLMPGRRHAIEIDL
NRPGSAPWYAVTTWTEAPGSDFYCVEPWLGLPDAIHNGLGLQHVAPGATQTAALRIRVTP
LG*
>GCA_000452945.1_BPMSHR5858_genomic_00197_1 nlpD
MLLKTTRRLVVASLAALLAACGSAPVGPGFYRVERGDTLSKIARSNRQSVANIARWNQIT
NPDAIEVGQVLRVAPPPGTASTSGTTGASRGPSRQAPSSSATSPASSVKPASSISLVWPA
AGSVIRSFDGSKSKGIDIANTAGTPVIAAAAGTVVYAGNGLRGYGNLLIVKHDADFLTTY
AHNRALLVKEGQTVAQGQKIAEMGDTDNDRVALHFELRYGGRSIDPARYLPSR*
>GCA_000452945.1_BPMSHR5858_genomic_00198_1 group_19934
MRKPQMLRMIVSPTTDTTMDLLLIATGLSVIGIGALVAFARRVDPLSGRLHGRLRKR*
>GCA_000452945.1_BPMSHR5858_genomic_00199_1 acsA_2
MLPHAAHYAELVERFAWRVPAHYNIGVDACDKWADGSGRLALIHEHAHGVVARYTFDELR
SSSNRLANSFARAGVKRGDRIGILLAQGPETAIAHLAAYKLGAIAVPLFTLFGADALEFR
LGDSGAVALVTDRAGYEKIAPLHASLPSLATIYCIDGAPDLAEPGVLAFDAALAAESDTF
RPADTSADDPALIIYTSGTTGKPKGTLHAHRVLLGHLPGVEMSQNLFPARARLFWTPADW
AWIGGLLDVLLPSLHHGVPVLARRFEKFGGAAAFDLLARHGVTHAFLPPTALKLMRAAVA
RPRERYALALESVASGGESLGAELVSWGRDAFGVTINEFYGQTECNVVLSSCSALFEPRA
GTIGKAAPGHRVAIVDDAGNALPPGVTGNIGVRAPDPVMFIGYWRRPEATREKFAGDFLL
TGDLGIADADGFIRFVGRNDDVITSAGYRIGPGPIEDCLLEHPAVRMAAVVGVPDAVRTE
IVKAFVVLNAGYEGSAALARELQTHVKTRLAAHEYPRALAFVDSLPMTATGKIIRRALRD
I*
>GCA_000452945.1_BPMSHR5858_genomic_00207_1 group_224
MHAQCFFTLNGQRVSNLTCAGFGGVMAFSGNKQFVDKPDATAVANAGPIPKGRYYIVKRD
TGGRLGPIRDLALDLWSNSDRSTWFALYRTDAKIDDETYVNGVKRSAFRLHPNGRWGVSD
GCITVTTQVQFDQLRAYLMKQPTAKIPGTDIEYYGTVDVR*
>GCA_000452945.1_BPMSHR5858_genomic_00209_1 group_19678
MNWHCYVARELAHLEVAIGVLEKARKEFVHHTAVCDPAYWRLRLDTIRERLDEDPTLERQ
MSELLARIDRLEARNPRRESADFLA*
>GCA_000452945.1_BPMSHR5858_genomic_00211_1 group_19652
MPNCADIAAADHARRRALAMRRYRAGAALCAAVALVDAVWLHAGGYALQWSGVAVAARAI
AVCAAFSLVLRGVARVPRYASVAARFRYGEAARTFAWLALLAAFVAAAGILSYLCVTLKP
PLIDGALIRLDARLGFDWPAVYERIRAHPALQRALAFAYDSAQWQLIGIPVWLGLTRRDD
DLSEFIALFMLTSVMVLLISTPFPAASAFLHFRVADPIALAGVSDFVRLRDGTLRVIDPA
ALQGMVSMPSFHTVLGVLFAYALRRGRVVAPLALLFNGWMIASTPTQGGHYFVDVLAGLL
VACAAIAALRYAADARAGTGDTARSGRPRPSMAGPAADGEAALSARAARASRPSGDA*
>GCA_000452945.1_BPMSHR5858_genomic_00215_1 phoR
MNIIWARFLVSLVLLALVSVIVGAIAGPVPGLVFALAMLVVQGFVSTFHTQRLWRLLDAP
VYGEVPSAPGIWGEIYYRLHKLAKQWHAQVRQVEQQHSRFIQAIQASPNGVAMLDDRDQI
EWCNAIAEVHFGLDAKRDLRQHVTHLVRHPDFVRYLNGQHYDETLVMRGMGGNRQNVLAV
QVFPYGENRKLLLTQDITELERTDAMRRDFVANVSHELKTPLTVLSGFLETMRELPLSDD
ERARYLELMEQQASRMRHIVTDLLVLAKLEGESKPPADRAVDMRTVLDHLRDDAQTLSGG
HHEITFKVDETLAVTGAQTEIFSALGNLVTNAIRYTPEGGTIRVEWRRDGAQAVFSVADS
GLGIPAAELPRLTERFYRVDRSRSRDTGGTGLGLAIVKHVLQRHDAQLSIQSEEGRGSTF
TARFPAHRTISRRHAA*
>GCA_000452945.1_BPMSHR5858_genomic_00216_1 phoB_2
MPSNILVIEDEPAISELISVNLQHAGHCPIRAYNAEQAQSLISDVLPDLVLLDWMLPGKS
GIAFARDLRNNERTKHIPIIMLTARGDEQDKVLGLEIGADDYVTKPFSPKELMARIKAVL
RRRAPQLTEDVVSINGLRLDPATHRVAAQSDGSEIKLDLGPTEFRLLHFFMTHPERVHSR
TQLLDQVWGDHVFVEERTVDVHIKRLRAALKPAGCDAMIETVRGSGYRLAKHA*
>GCA_000452945.1_BPMSHR5858_genomic_00217_1 phoU
MSDKHLSSQFDADLNAVSSKVLEMGGLVEAQITEAMYALNEFDRAAAERVIEAEDRLNHM
EVEIDEECGNIIARRQPAARDLRLLMSISKTITNLERAGDEAEKIAKRVRRLVDEPAARG
VNIAEIKVSGDMAVTILRRALDAFARLDTVSAAQIVKDDKEIDQEFRAFVRKLVSYMQED

PRMISVGLDYLFIAKAIERIGDHAKNIAEFIIYIVKGTDVRHQPRDLLDREANS*
>GCA_000452945.1_BPMSHR5858_genomic_00221_1 pstS
MKLMQTAFAGLAGALFAVAAHADITGAGSTFAMPIYTKWAADYQQSGGAKVNYQGIGSSG
GLKQIVAKTVDFAGSDAPLKDDELAKEGLFQFPTVVGGVVPVINVPGVKAGELTLSGAVL
GDIYLGKIKKWNDPAIAALNPKVKLPDTDIAVVRRADGSGTSFIWTNYLSKVNDEWKSKV
GEGTTVNWPTGTGGKGNDGVAAFVQRLPGAIGYVEWAYAKKNNMVYTALKNSTGTVVEPK
TETFKAAAAGANWSKSFYQILTNQPGKEAWPVVGATFVLLHAKQDKPAQGAETLKFFDWA
FKNGAKAADDLDYISLPASVETEIRKQWKVKVTDAAGKAVAE*
>GCA_000452945.1_BPMSHR5858_genomic_00222_1 glmM
MGRRYFGTDGIRGKVGDAPITPDFVLRLGYAAGKVLASAPGRAASGARPTVLIGKDTRVS
GYMLEAALEAGFSAAGVDVMLAGPMPTPGVAYLTRALRLSAGVVISASHNPYHDNGIKFF
SADGNKLPDEIEAEIEAWLDKPLDCAASDGLGKARRLDDAAGRYIEFCKSTFPAAFDLRG
MKLVVDCAHGAAYQVAPHVFHELGADVIPIGVAPNGFNINDGVGATAPDALMRAVRANHA
DLGIALDGDADRLLVVDHTGRLYNGDELLYVLVKDRIATNGQVEGAVGTLMTNFAVEVAL
KEAGVQFVRAAVGDRYVLEQLRERGWQLGAEGSGHILSLDRHSTGDGIVSALLVLAALKR
SGKTLAQMLEGVTLFPQKLINVRMKPGADWKGSEAIRRAIDSAEQALSGSGRVLIRASGT
EPVLRVMVEARQATDANRHAEAIADAVKQATA*
>GCA_000452945.1_BPMSHR5858_genomic_00224_1 ftsH
LNNNMFSKAAVWLVIALVLFTVFKQFDKPRVQEGVSYSQFMDDAKNGKVKNVVVQGRNLT
VTPADGQKYQIVSPGDIWMVGDLMKYGVQVSGKADDEPNALVSALYYLGPTLLIIVFWFY
MMRQMQGGGKGGAFSFGKSRARLIDENNNAVNFSDVAGCDEAKEEVSELVDFLRDPQKFQ
KLGGRIPRGVLLVGPPGTGKTLLARAIAGEAKVPFFSISGSDFVEMFVGVGAARVRDMFE
QAKKHAPCIVFIDEIDAVGRHRGAGMGGGNDEREQTLNQMLVEMDGFEANSGVIVIAATN
RSDVLDKALLRPGRFDRQVYVGLPDIRGREQIMRVHLRKVPIANDVDAAVIARGTPGFSG
ADLANLVNEAALFAARRGKRIVEMQDFEDAKDKIFMGPERKSAVIREEAKRATAYHESGH
AVIAKLLPKADPVHKVTIIPRGRALGVTWQLPEHDNETYSKDYLLDRLAILFGGRVAEEL
FLNLVSTGASDDFNKATQTARAMVARFGMTDALGPMVYVDDENDGGPFGRGFTRTISEAT
QQKVDAEIRRVLDDQYGLARRLLEENRDKVEAMTAALMEWETIDADQINDIMEGRPPRSP
KSVPPAGDPSSGGSSGAEVKPGNATAPA*
>GCA_000452945.1_BPMSHR5858_genomic_00230_1 leuE_2
MFGHALGITDLWTYVFGVVFIILLPGPNSMYVLSLAAQRGVKAGYRAACGVFVGDTVLMV
LSAAGVASLLKANPLLFSVVKYGGAAYLLYIGVGMLRGAWRKLARPAADAGADVRRAVDG
ERPFRKALVVSLLNPKAILFFISFFIQFVDPAYAHPALSFVVLGAIAQLASFVYLSTLIF
TGARLADHFRRRRKLAAGASGGVGGLFVGFSVKLALATMS*
>GCA_000452945.1_BPMSHR5858_genomic_00231_1 carA_2
VLPSFSPALLALADGTVFRGYSIGAAGHTIGEVVFNTAITGYQEILTDPSYARQIVTLTY
PHIGNVGVNAEDVEATKVHAAGLIVRDLPALASNFRMARTLGDYLRDEGVVAIAGIDTRK
LTRILRDKGAQNGCILTGSDDETKALALARSFPGLAGMDLAKVVSTTKPFEWKQTEWRLG
SGYGMQETPRYRVVAFDFGVKYNILRMLAERGCHVSVLPAQSSAADALALNPDGIFLSNG
PGDPQPCDYAIAATREFIERGVPTFGICLGHQIMGLALGAKTLKMKTGHHGANHPVKDLA
NGRVVITSQNHGFAVDADSLPANARATHVSLFDGTLQGFELTDKPAFCFQGHPEASPGPH
DIGYLFDRFTALMDAAKARSA*
>GCA_000452945.1_BPMSHR5858_genomic_00233_1 group_660
MSTILAPHSGAAPTLPLIDSGHRRRGWRGWRGRLIAKAAMQILRPVEPCVQHGLPAPTTF
PPNGGARRFGLVHYGIMIPDLPAPHRFMATAIILGSAGMRVFDADFAAAPGDGPLQTATL
VHGTAAAVDRHFTRYSIPRDVESRADGSLLRLGDDLELSGRYPEFRLESRRDGFAADLTL
TATGDVTWFAKGLFYEHIALVTRYRGTIEHGGATTEVAGLCTYEYARGTSPYLLRNRFLP
QAAKLPWDVFSYQVIDLDADTQLLLAHVQVDQYPALTSAYLRVIGQRAHRLDADVHFRVT
STQARPAVGPDGHTMWLPETFAWRVRDRHGRTLFAIDATVDTPMLFGIGTGYVGGYRWEG
ERDGRPASGRGYIEYVDRRR*
>GCA_000452945.1_BPMSHR5858_genomic_00236_1 gloB_1
MNELEYVPVPAFDDNYIWLVSDGRQAVAVDPGEAAPVRRHLDARGWRLTAILLTHHHRDH
VGGVAELLAGAREGEPIAVFGPAHEAIEHITRRVAGGERVRIGAPALEFDVIDVPGHTRG
HIAYFQPAGPGNAAPHLFCGDTLFSCGCGRLFEGTPAQMLASLDALAALPGDTRVHCAHE
YTLSNIQFALACEPGNAALAAWRDEARARRARGEPTLPTTIAHECAVNPFLRADSASIRA
TLEARLHEAAPDRLAAFTLMREWKNRFR*
>GCA_000452945.1_BPMSHR5858_genomic_00240_1 group_19029
MLSLAKIAALFVLTAVAEIVGCYLPWLVLKAGKPAWLLAPAALSLALFAWLLTLHPAAAA
RTYAAYGGVYIAVALAWLRIVDGVPLSRWDVAGAALALAGMSVIALQPRG*
>GCA_000452945.1_BPMSHR5858_genomic_00241_1 proP_9
LTATPAPSSSSSAPTEGALPAAAHEITVVDQGLLKRAVGAMALGNAMEWFDFGVYSYIAV
TLGQVFFPSSSPSAQLLATFGTFAAAFLVRPLGGMVFGPLGDRIGRQRVLAMTMIMMAVG
TFAIGLIPSYDSIGLLAPVLLLVARLVQGFSTGGEYGGAATFIAEFSTDKRRGFMGSFLE
FGTLIGYVMGAGVVALLTASLSHDALLSWGWRVPFLIAGPLGLIGLYIRMRLEETPAFKR
QAEAREAQDKAVPKAHFRRQLARHWRALLLCVGLVLIFNVTDYMALSYLPSYLSSTLHFD
EAHGLVLILIVMVLMMPMTLATGRLSDAVGRKPVMLAGCVGLFALAIPALLLIRTGETAL
VFGGLLILGALLSCFTGVMPSALPALFPTEIRYGALAIGFNVSVSLFGGTTPLAAAWLVD
ATGNLMMPAYYLMGAAVIGAISVLALPESARQPLKGSPPAVASHREAHALAREIKRREAA
ERDDSGYPSAAALRA*
>GCA_000452945.1_BPMSHR5858_genomic_00249_1 madC
MTLDDVLASLFPSGHDVAADAAGLLAGHGMLAGRALDVIGVAHGAALGVDGALALAERVL
ATIRRGGETPILVLVDGDSQRMSKRDELLGLNECLAHLAKCLLHADALGHPTIGLLYGHA
AAGAFIATALATRTLVALPGAQPEVMDLQAISRVTKLPVDVLREKARSTAVFAPGPENFA
KTGAVDAVLDPARPLAEQLAAVLARPADRCDRRDRYGAERGGRPVAADVAQRVYELARAG
R*
>GCA_000452945.1_BPMSHR5858_genomic_00265_1 sir_2
MYRYTEFDKAFVRNRAAQFRDQLERWRQGALSEDAFRPLRLQNGWYVQRHAPMLRVAVPY
GELSSAQIRMLARIARDYDVPEPARYRAACDAQARLGTVRLPTRNAHFTTRTNVQFNWIP
LEKAADVMDLLATVDMHGIQTSGNCIRNISCDERAGVAPDEIADPRPFAEVMRQWTTLHP
EFAFLPRKFKIAITGAADDRAATDWHDVGLRLVRDAAGELGFRVSVGGGMGRTPMTATLV
RAFVPWRHVMNYIEAIVRVYNRYGRRDNRYKARIKILVKAEGQRYIDDVDEEFRQIVEHD
GGPHTIEQAEFDRVCASFVPPVPPPCAAGRHAKTDIDADMDAALARMRTLAGAQPAFARW
LERNVAPHRDPARRIVTLSFKRRHQAPGDASPEQLDALAHLAERFSAGEARVTHAQNVVL
PWVRAIDLHALWEAARDAGLASANVRLLTDMIACPGGDFCALANARSIPIADAIAERFDD
LDELHDIGDVDLHVSGCINSCGHHHSGHIGILGVDKDGDEWYQVTLGGSDGTAANGPAQP
GKVIGPSFSADEIVDVVDALVATYLRVRRTEAGRVERFIDAYRRVGPEPFKAAANAARHP
ETSVA*
>GCA_000452945.1_BPMSHR5858_genomic_00278_1 group_4416
MSYSLEGRILEVCDCKVLCPCWIGEDPDNGTCRATVAYHYDKGMIDDVDVAGLTVAFAAF
VPGNILQGNWRVIIFIDERATDAQFDALASVYRGERGGPLADFSKLFGDIVAIERAPIDF
DVRDGRGTLKVGNTTYAELEPYLGPTGEPTKLVESIFSTIPGSPAYVGKAGTFRMRERKL
GIELDLSGQNAIQGFFTFRS*
>GCA_000452945.1_BPMSHR5858_genomic_00279_1 dgcM
VSTVFTAHESPRVIGTIARLYTACLLIGFALVPAFLIAYVRFFIDPRLVFENHQFHELAI
AAATLEGLFVTYVTWRCYLASGEPLLRWLTLGFLGFALIYSLHGAFTGMAHQNIWLFLLY
GPASRLVMSILLLVGQLSYFRDADPPERRTHLSMWLPWIAGYVVVCFAVAAIAYSPIAGY
LWTRALMEGGAMIVSLVNVALLLVRRIRTPLMLVYGISVTSFALSSIAFILGKPWNHMWW
LAHAVFAGGFFMLSFGVVQALMTTRSFATVYSQQDLMARLAESMARTEGVLQELKRTNQK
LEHLATTDPLTGASNRRKFIEQMQSEIARAKRDGTAFSLLALDLDNFKTINDNYGHQVGD
VVLRNFVRQCLAAIRPYDHIARVGGEEFMVLLPRMLADTASGIAERVRTTIANASFGIDG
KHTPVTVSIGVSQYGRDGDTVDEILSAADKLLYRAKNEGRNRVVAS*
>GCA_000452945.1_BPMSHR5858_genomic_00303_1 group_1754
VTVRRSLLPRSPLAAVRRACAAMLIPVASLALSLGASAAHAQDTAICYNCPPEWADWAAQ
IAAIKQRTGIRVPFDNKNSGQAIAQLIAEQKSPVADVVYLGVSSAFQAKDKGVVAPYKPA
HWSDIPANLKDPQGYWFAIHSGTLGFFVNKDALDGKPVPRTWADLLKPEYKGMVGYLDPS
SAFVGYAGAVAVNQALGGSFDDFRPALDWFRKLKANQPIVPKQTAYARVLSGEIPILLDY
DFDAYRAKYKDHANVEFVIPKEGTISVPYVMSLVKGALHEANGRKVLDFVLSDEGQKLWA
NAYLRPVRAQALGADVAAQFLPASEYARAKSVDFGKLAAGQQAFGKQYLQAMQ*
>GCA_000452945.1_BPMSHR5858_genomic_00304_1 group_3082
MSDLTLPSRWRIALLAPALAVFVAFWLLPMVSLVRVSAGGGFVDAYLALLSNARYMKSLA
STVALSAAVTLATLALSTIAGLLLARRAFAGKRALIALLTFPLAFPGVVVGFMVIMLAGR
QGLIGMLSQKLAGDRWVFAYSVSGLFVGYLYFSIPRVIVTVIAAASKLDPSLEEAARSLG
ASRWHVLRDIVLPALAPGLVAAGAICFATAMGAFGTAFTLATDLDVLPMTIYTEFTLNAN
IATAAGLSIVLGIVTWAVLALARNLTGQATAASA*
>GCA_000452945.1_BPMSHR5858_genomic_00306_1 group_597
MTPPPKDAAALARPAAAAGPHAPPAVRRARRAPLDWLAAAQWAVTLALCAFLIVPVAMSV
LAGLTVNYFRGPSSGLTLRWLGEVWAQYHGSVFLSLEVALATLAVTLVAGVPAGYALARS
RSRVSRLIEEALVLPVALPGLASALALLSVYGGFAAFRTSLWFIVVGHVVFTLPFMVRAV
AAVAARADLRTLEEGAASLGASFATRFATIVLPNLRPGIVAGALAVLTLSIGEFNLTWML
HTPDTKTLPVGLADTYASLRLEIGSAYTILFLLMTLPLLVAMQRLGVEPPGARDRTRDRT
CKRR*
>GCA_000452945.1_BPMSHR5858_genomic_00311_1 group_975
MSSQPRHSAGTPGHYSRSLLLLLATIAGVSVANIYYNQPLLDAFRASFPGSASWIGVVPT
ATQLGYATGMLVLAPLGDRFDRRTLILLQIAGLSAALVVAAAAPTLGVLAAASLAIGILA
TIAQQAVPFAAEIAPPAARGQAVGTVMSGLLLGILLARTAAGFVAEYFGWRAVFAASVAA
LAALAAVIVARLPRSSPTSTLPYGKLLASMWQLVRELRGLREASMTGGAIFAAFSAFWPV
LTLLLAGAPFHLGPQAAGLFGIVGAAGALAAPYAGRFADKRGPRAIISLAIALIAASFAI
FALSGASLIGLVIGVIVLDVGVQAAQISNQSRIYALKPDARSRVNTVFMVCYFIGGAIGS
SAGVAAWRATGWLGMCAVGLLFSIVAAIVHFRGGAGAR*
>GCA_000452945.1_BPMSHR5858_genomic_00312_1 yedA
MTPFDRFLAFVARHPPRLPQSRSGRVALALVFIYFAWGSTYLALHVALESFPPLLLSGLR
NLLAGVGLFVFAMRRRPVRPTLVEIRNAALVGTMLVTMSSGLIALGMRTVSSGSAAVMVA
TVPLFATVIASVAGRRVTGGEWGAVALGMVGIVVLNSGGPSSPGSTLGSITVLAGALFWA
GGAHLAARLALPHDLFLSTALQIGLGGAASTCIAWVLGERIGHVAWVPGAAFVYLMLAGT
MAAYVAYGYLIRHTSPIIASSCMYVNPVVAVALGALLLGEPVTAATVIATIAILGSVGLS
FVFDPARRRAQ*
>GCA_000452945.1_BPMSHR5858_genomic_00313_1 yedY1
MSENDRKKRENDIARIDRASLLIDVRKSLALPSRRLFGKRMLTLGGLAMLTGCTLRDDAS
VDDFLEKVSRANDRVQAWLFGPTRLAPTYAEADITRPFPFNAFYGIDEVPRVDADGFRLA
VSGLVTGQRSWSLPALYALPHAEQITRHICVEGWSAIGRWGGTPFREFLRRVGADTSAKY
VGFKCADDYYESIDMATALHPQTLLAFEYDGKRLPPEYGFPMKLRMPTKLGYKNPKHIVE
IFVTNAYPGGYWVDQGYNWFGGS*
>GCA_000452945.1_BPMSHR5858_genomic_00323_1 group_3085
MPKTQGFSPESLISQFTRCDAGWCVRYHDVTLASAFQPVISITHKRVVGYEALARATDAN
GAPISPDTLFARAQARGETILLDRLTRCLHVANFAAQDTGACWLFLNVLPQMFDAGIAPG
AFIEALCAHFALPPTRVVLEVIEQPSRNEVALARTIDMIQHGDFLIAVDDFGTGFSNFDR
IWQIKPDIVKLDRSILERSLAASDAHRIIHHLVTMLHHAGTMVLAEGVENEDALQILMDA
DVDFVQGFCFGRPAPSLEHARRAAPARIEAAWRRFAERTKARCGDVVHPGFDTIERIMLT
GAADYAQTRNLRDAAQRLLTSPIVRRVFVAEANGEQVEPSVTIETPDAPSATARRLAPLL
PELHCNWSRRAYFQRAIAAPGRVALMGPHFSLTDGRDCYTAAVAIHLGTTLKVFCVDFDF
SSYGDGR*
>GCA_000452945.1_BPMSHR5858_genomic_00325_1 dadD
MSVVDERFAWTGGLADRLPEPADFGLALAEGFARRIGMLSRRVGASAAAARWAARAAFAA
SRATAAGHVCVALRALAQRYDEPYDDVRAALAASGVTAFDGIVRGGECPLVVDRDGRLYL
ARYFEYETRLANALVERSRCGGAPAGGADALSPDALGERLVRYFGPQKERGVDWQRVAAL
IALTGRVTIVSGGPGTGKTTTVVGVIACLLDAHPELRIALAAPTGKAAQRMQEALHARAG
SLPAELAARLPRTSCTLHRLLGGGPGGRFAHHRDNPLPYDLVVVDEASMIDVALAAHLLD
ALAPNARLVLLGDKDQLAAVEAGAVFAELSARPAFSPATCATIARALGVGEAEFVAALPD
GFVVQAVGAERVADARAAAADMDVAARGTAGAAAAAARATAAAAARMTTAAAAARGGAPE
SAAQRAPAPHAGRRGGGRASNRTVDDAQGSLFAFDDDAFGVGTSGGEPGAGAQGAGRRDG
GAPSLDGPHAGRRDGGASPGVAHSRRDAFDARARSGPADVSDLENAARAYAPAGAAGPAD
SPADASFVEPAAWIEADELAWLDNAVFSFSEGASANAEADGRRSAAATATDAGARTAPPV
AREATAPTPRETARSDGSAVAPLTDCVVWLERNYRFGLDSPIGRLSLAIRRGAVQDALDA
LSTQDDAAARFSDDGGATLSAATVEQLARGFAGYAAALRDALATPEPDALPLFDVLNRFR
VLCATRTGARGAEEVNARMAAEVRRAVRVPLALGAHWFAGRPVMVTRNDYALGLFNGDIG
IALPGARGVLRVWFRGADGRARAVSPAALPPHDTAFALTVHKSQGSEFDDAALILPASFN
RVLSRELVYTAITRARARVQVIGSRAVLALAIATRTARDSGLAARIADALRARQEGAR*
>GCA_000452945.1_BPMSHR5858_genomic_00328_1 yccS
MRYSVEIRKFFYSQYFFGGLRIALGVSLPAVLSLAVLHDRELGFTIATGALGACVVDMPG
PLKYKHNEMLACSVIGFFSALATGLATPNIFALWLTIVPLTFVLSLIVVYGNRWPQISFA
TLFMMVMTLEEKFTPLQALVNASWILAGGLWYTYWATIVSRWQARRIEQQALAESLFACA
SYLLARADFYDLDADLDECYRNLVARQITAVETQETARDIVLRNLPKLRRGKLDPGRTMM
FNLFINSVDLHEMFVGAHTDYPLVRSTFGGSDLMIFYRDLIRKAAADLEEIGFAVLENRL
PHSRISVKAELRAIEYEIELMRKKQFAATHPEAYAAVLATFRRIWSATRLIDKMRRNLSG
SADAQRTELKIDKALTRFLQKRRMSPLLIFSNLNMGSPSLRHALRVTIAVAVGFWLGRLL
PLTNAYWIVMTSIIILKPGYSLTKQRNAQRIVGTLIGCAASLALIYTVHEPHVLIAIMFG
SMVMSYSLLLFNYAASVVFTSSYVLLLFHLLAPGNMRIIGERAIDTVVGCMIAIAASRLF
PYWEYRAIGKLVADALATTRKYFEATWRAGRGKAAPAVPAADGAAAVPAVAAVVDAVAPG
LDGDYQYRLARKNVHIAFANLGQAFQRMMLEPKAHQKFVPELNDLLVQTHVLGAQITAAA
PLLRHACEQGNPPALERGMAAVREHLEQAEAGVPPPANQAETSKQLTRELDSMVVDAERS
DAGADLAHDLKVLAHQCKQMLASSLLIRKDASVIRLPE*
>GCA_000452945.1_BPMSHR5858_genomic_00329_1 proY_2
MKNLQRHLSARHIRFLALGSAIGTGLFYGSASAIRLAGPAVILAYIVGGAAVYMVMRALG
EMAVREPVPGSFGHYATENLGPFAGFVTGWTYTLEMVIVAIADITAFGIYMGFWFPHVPQ
WIWVLAIVAIICGLNLCHVKVFGELEFWLSIVKVGAIVAMIGGGAAILFTGMRLGSGGAP
SLANLWSHGGFLPNGVGGLVASLSVVIFAYGGIEVIGMSAGEAKEPERVIPRAINAVPMR
ILLFYVLTMIVLMSIFPWTGVGANGSPFVQIFSALGVRSAAEILNLVVISAAISAINSDI
FGAGRMMYAMALQGQAPRALAAMSARGVPWVTVLVMAGALLVGVVLNYLMPKDVFLMVAA
SATFATVWVWLMILLSQVAMRRRLGAAEVAALKFKVPFWPVGPALTIAFMAFVIAMLGYF
DDTRVALFVGAAWLAILALVFYTRIRPRLAAVAR*
>GCA_000452945.1_BPMSHR5858_genomic_00331_1 futA2
MFVKQARPLLRTLLRPLAFAFALAAVAPAAHAANEVNLYTTREPKLIQPLVDAFTKQSGI
AVNTVFVKDGLLERVKAEGARSPADVLMTVDIGNLLDLVDGGLTQPVRSASLDGAIPANL
RGAGGNWYALSLRDRVLYVDKNMQLDAITYESLADPKWKGKVCVRSGQHPYNTALVAAMI
AHDGEAAAEQWLRGVKANLARKATGGDRDVARDILGGICDIGLANAYYAGHMKHAQPGTD
ARKWGDAIKVVRPTFANAKSGGTHVNVSGAAVAKHAPNRDNAVKLLEYLASPPAQALYAQ
ANYEYPVRAGVALDPVIAGFGPLKVDPLPLVEIAKYRKRASQLVDKVGFDN*
>GCA_000452945.1_BPMSHR5858_genomic_00332_1 group_1138
LTTEADARVARRAHAPVWARAGGRARLSRIAAHGWLAGAAAIAAVVGTPLVALACAALGA
DLSHWRHLAEFVLPQAAANTLALLAGVGAIVTCIGTGAAWLVTAYDFPGRRTLAWALLLP
LAVPSYIVAFAYLDLLHPIGPVQSALRWALGFDSPREFRLPELRSLPGAIVVLGFVLYPY
VYLCTRAMFVTQSASLVEAARTLGAGRIETFVRVVLPLARPAIAVGASLALLETLNDVGA
SEFLGVQTLTVSVYTTWVTRSDLASAAQIALAMLAIVLGTLALERYGRRRERYAHGRRMR
AIAPRRLRGAAAAGAAVLGWLPVAVGFGAPAAYLVHETAQRVHLVGGVSAQLLASLGNTL
AVALTATVVTLGGGIVVAWATRRTRDSARFQPGRVCAPLALADRAIAAVLGRDALVLMGS
AAALVIAYAIRFLAIPAGGIEAGLARIPASLEQASRTLGESAGMTLRRVHLPLLRPALGA
SALLVFVDAMKELPATLLLRPLNFDTLATWLYAEAARGTYEEGAVAALAIVAAGLAPVIV
LARTQLKSGS*
>GCA_000452945.1_BPMSHR5858_genomic_00333_1 fbpC
MPAMLLELNAVTLAYDTPRGRRTIVDRLSLALERGAIGCLLGASGCGKTTVLRAIAGFEP
VRAGEIVLAGAALASRDADVPPEKRRVGMMFQDYALFPHLSAQQNVEFGLRRMPKGPRRA
RAAQMLELVGLADAANAYPHELSGGQQQRIALARALAPSPALLLLDEPFSNLDVDTRERL
AFEVRDILKAAGHTALFVTHNQAEAFALADRIGVIDDGRLAQWDTPYNLYHRPANAFVAD
FVRRDALADERRRALERGR*
>GCA_000452945.1_BPMSHR5858_genomic_00337_1 ywnA
MNTSSRFAFAFAVHVLALLSLQEGVPLSSEVIAGSVNTNPALIRRLLSMLAAAGLTASQLGA
GGGALLAREPSRITLLDVYRAVDDAQLFALHREAPNPACLVGRHIQGVLTDYIGDAQRAM
EASLATRTLADVTADVLRSEAAGQPPRGQQAGG*
>GCA_000452945.1_BPMSHR5858_genomic_00339_1 group_844
MPALPNPSRRHALKILAGAPMLPLSGVALPALLSGCGGDDDVAAAPPAAGFAAAAFASMA
APTLADPAAMATTTVGSTLSVSFTDGSTRNYKLAYRPFFVTGDRVPDGKGGTILAGGYYD
INGQPIIDRSVPGQERQFYSDCPDGSSLLTLKNANVAGVKGNTVFAVVQFEYTTRDLASA
SQYGRLPSPIAVLTLDQDPATGALKLVKYHNVDTSAAHGLWITCGASLSPWNTHLSSEEY
EPDATKAAGDAQFRAFSKNTFGSETAANPYHYGHLPEITVNPDGTGSVRKHYCLGRISHE
LVQVMPDQRTVLMGDDATNGGLFMFIADKAADLSAGTLYVAKWAQTSSAGAGAATLTWLN
LGHATSAEIEALANQLKAADIMDVATKDPGDASFTKINFSGTFNWIRIKPGMTKAAAFLE
THRYAAYAGGSMGFTKLEGTTVNAKDKVLYSAMSYIQKSMVRGNAAASPDIALDRAINAG
AVYALNLRGGQRDTSGAAIASEWVPVDMSAPAALVGEDLAAADALGNTAHPDKIANPDNL

KFSEKLRTLFIGEDSGMHVNNFLWAYNVDTQRLSRVLSCPAGAESTGLHAVDEINGWTYV
MSNFQHVGDWESPLHDKVKGVLDPLVRANYKDRFGASVGYLTAEPAAIQLKKA*
>GCA_000452945.1_BPMSHR5858_genomic_00371_1 farB_1
MAATAPASPSRSAEPAPLSGGTLALLTIGLALGTFMEVLDTSIANVAVPTISGSLGVATS
EGTWVISSYSVASAIAVPLTGWLARRVGEVRLFTLSVLAFTIASALCGLAENFETLIAFR
LLQGLVSGPMVPLSQTILMRSYPPARRGLALGLWAMTVIVAPIFGPLLGGWISDNYTWPW
IFYINLPIGVFSAACAFFLLRGRETKTTKQRIDAIGLALLVIGVSCLQMMLDLGKDRDWF
NSTFITSLALIAVVSLAFMLVWESTEKEPVVDLSLFKDRNFALGAMIISFGFMAFFGSVV
IFPLWLQTVMGYTAGLAGLATAPVGILALVLSPMIGRNMHRLDLRMVASFAFVVFAVVSI
WNSMFTLDVPFNHVILPRLVQGIGVACFFVPMTTITLSSIPDERLASASGLSNFLRTLSG
AIGTAVSSTFWENDAIYHHARLAESVNVYAQSTLDYQGALARLGVMGDVSTAQINQIVTQ
QGFMMATNDFFHISALAFVALAALVWVTKPKKGAGPAIGH*
>GCA_000452945.1_BPMSHR5858_genomic_00372_1 gltC_1
MELRALRYFVEVVRQQSFTVAAEQMHVTQPTISKMVKALEDEIGSPLLLRDGRQMVLTDA
GRIVYQRGQDVLAAQAQLQAELNDLGTLGRGELTIGIPPLGGSLFTPAIAAFKERYPRIE
LKLFEQGARMIEEALVAGELELGGVLEPVDPAVFDVLPMVRAPLWLVAPHASRWDGDATV
PLADLADEPFVFYAESLALHDAVLDACRKVGFTPQIVSRSGHWDFMAALVHAGVGIALLP
EPYCRRLDTARFTCRPIVEPEITWAVAIGWLKRGYLSHAARAWLDVARETGPVDPGDDLA
FGSLRAR*
>GCA_000452945.1_BPMSHR5858_genomic_00381_1 fliR
MFSVTYAQLNGWLTAFLWPFVRMLALVAIAPVTGHRSTPVRVKIGLAGFMALVVAPTLPP
MPPMPVATVFSAQGVWIIVNQFLIGAALGFTMQIVFAAIEAAGDIIGLSMGLGFATFFDP
HSSGATPVMGRFLNAVAILAFLAFDGHLQVFAALVDSFRLVPVSADLLRAAGWQTLVAFG
AAIFEMGLLLALPVVAALLIANLALGILNRAAPQIGIFQVGFPVTMLVGLLLVQLMAPNL
IPFVGRLFDTGVDFVGRVAAGMH*
>GCA_000452945.1_BPMSHR5858_genomic_00382_1 cueR
VGNLDIAEVAQRSGVPASALRYYEEKGLIASTGRRGLRRTFDARVLERLALIALGRAAGF
SLDEVASMFDAQGRPNIDRALLVAKADELDGAIRRLTSMRDGLRHAAACTAPSHMECPKF
RRILRAASDARRARKKAPPPGG*
>GCA_000452945.1_BPMSHR5858_genomic_00384_1 group_499
VRHAPIAASPRVPGERAIGWAAHYLIGVAFAALLLALRGVGWAGEPTLAPALAVGIGSVA
APLFVMQPAMGAGIAASRTPRPGAARFHSIVAHAVFGAGLYGAGCAANRLGVPALPGIG*
>GCA_000452945.1_BPMSHR5858_genomic_00385_1 group_4430
MRTTLRALSLAAVAAGLSFGFAAQPAFADDGGKITIMVGGITKLIYLPARLTQELGYFKA
EGLDVELQSQPAGVDAENELLAGAVQGVVGFYDHTIDLQSKGKDVKAIAVLGQVPGEVEM
VSTKAAGAIKSMADVKGKTLGVTGLGSSTSFLTQYLAQQHGIAANQYTMLPVGADASFIA
AVKQGRIDAGMTTEPTVSVLEKNGDAKVLVDLRTLDGTRAALGGTYPAASLYVQSAWADT
HKAQAAKLAHAFARTMQFIHTHSAEEIAAKMPADYQKDKALYVSALKASLPMYTPDGRMP
ADGPATVLKVLSAFNPSVKGKHIDLSKTYTNDFVSAK*
>GCA_000452945.1_BPMSHR5858_genomic_00392_1 group_6856
MQLHFESDLDYQLEAIEAVCDLFRGQEACRAEFSVTAQAARRRAGPQISLGMAESGLGVG
NRLTLDAHTFAENLARVQVRNGLPPSGAPSSNDFTVEMETGTGKTYVYLRTIFELHRRFD
FTKFVIVVPSVAIKEGVHKTLQITEQHFRRLYAGVPFDYFVYDSAKLGEVRSFASKSIVQ
IMIVTVAAINKKDVNTLYKDSEHTGGEKPIDLIRATHPIVIVDEPQSVDGGLEGRGKEAL
EAMRPLCTLRYSATHADKYQMLYRLDAIDAYERKLVKQIEIASATVEDAHNKPFVRVISI
GSRRGAIAARVELDVATAAGDVERQTVSVSDGDDLERVARRAVYANFSIGEIHAARGAEY
LVLRYPGGDAFLSVGDAYGDVDTHAIQREMIRRTIREHLDKELRLTPLGVKVLSLFFVDA
VDKYRKYDRHGQPFKGDYALLFEDEYRRAAKLPEYRALFDGVDAGLAAEAVHDGYFSIDK
KGGWTDTSDRSAGSRENAERAYGLIMKDKERLLSFDTPLKFIFSHSALKEGWDNPNVFQI
CTLRDIRSERERRQTIGRGLRLAVNQRGERVRGFDVNTLTVIAGESYEQFAENLQKEIEA
DTGIRFGIVETHQFAALPVPADDGSVQPLGVERSAALWMHLRDAGYLDARGRVQDTLRAA
LKLRALPLPDEFDALRALIVDMLRKLAGRLDVRNADERRHIALRRDAHGKAVYLGDAFRA
LWDRIRHRTTYRVNFDNARLIERCVAALKAAPAVTRARLQWRKADIAIDASGVEATETED
AGAIAIDEGEVDLPDLLTELQDRTQLTRRTIAKVLIESGRLDEFPLNPQRFIALVAAALE
RCKRDALVDGIEYRLLGEAQVHALSLFESEPLTGYLSSMRRDAVKSIHEDVPCDTPAERA
FVESLEQDDAVRLYAKLPGWFKIPTPLGSYSPDWAVLIAEGDGPRLYFVVESKSGVADGD
LRADERRRVQCGAAHFRALEAVADNPARYVRARRADDLPTAAADARDAA*
>GCA_000452945.1_BPMSHR5858_genomic_00395_1 livF_4
MQLEVKNLVVHRANKPVLHGVSLAVTPGRVTALVGANGAGKSTLVMSIAGALPATSGDVL
LGGAPLGALRPEAVRRLGVAVVPEGHHVLGDLTVRDNLRAAGAFLSARRLNGAIDRALAI
FPELEPKLDARANDLSGGQKQMVCVSQALIGEPHTLLIDELSLGLAPTVTKRLAQTVARI
ANDGVAVLLIEQFTTIALALATNAYVLERGRVAFAGSAQTLRERPEILHGSYLASKGSGA
NAA*
>GCA_000452945.1_BPMSHR5858_genomic_00402_1 moaF_1
MNDKPQDWKNYEDFAAGIDTNRLPATDALVGRALTFELPGGAFAANFVDGQTLSWRRGET
GGTDWYEAIEVAPDTFFVDVTFKSRPAEALTLVFNTATRRALGILSRIRSRDEAGAAPRV
AQDFLVGTIAGGQADAGVSGILGIAGAEPAETRDLIGTRTLNVYSPNHTYEHTYLSSARY
CWQCLVGEQRGHGDVDLATTYKFADDLYVFTFREFLIPVASVFVFNFAAGRSTGKFLGET
GDGAIANRPAGSFIRKLSQAVYPDDAQPI*
>GCA_000452945.1_BPMSHR5858_genomic_00405_1 group_4432
MKTQLKKGALDMCVLAMLARRDSYAYELVSTLAETMEISEGTIYPLMRRLQAEAWVTTYL
VESTSGPSRKYYSITDAGRASLRQMEDEWRSFVDEVNGVLALHGTRADGEERQ*
>GCA_000452945.1_BPMSHR5858_genomic_00408_1 group_4433
MTQPTARHLILDLLVAKDGEPLQVREAIIACRLFGLTENSVRVALARLAAESLVEGAERG
SYRLGPSAVELSEELTAWRTVESRLRRWSGAYLMVSCISLGRVDRTALARRERALEMLGF
REWDKSLYVRPDNIAGNVDSVRRRLTKLGVEPAAAVFVAMGFDAPQEARIRALWDGDALS
DTYRTLGAQLEDWLARAPELDLDVAAREAFLLGGKAIRQVVFDPLLPEPFVDVAARAAFI
ECARRYVGAGHRIWRALLDSEHPSAPGAARDAIVGHLH*
>GCA_000452945.1_BPMSHR5858_genomic_00417_1 group_4434
MSWSKAACALFALSFLGCAGASGTAGSIDAGTQVCPVRAQSPLQYVDVFDGAPSDLATLI
PDDAGERTGHWELGYVYDAGRYVTIRCKYADGRTKDVRLAERVKRCDYRIDGKKTLQVTC
AK*
>GCA_000452945.1_BPMSHR5858_genomic_00420_1 sigR_1
MPSADLDDPLRELLPRLRRFALWLTRDVHSADDLVQSTLERALSRWSTRRDASALRSWLF
TILYRQFVDATRRAKRYAWLLGRIKDDDEPQWPSAEDHALARSMLDAFGRLSTEQRSLLL
LVAVEGFSYQEVAELLDIPIGTVMSRLSRARQALRALGDGEISAPSLRLMK*
>GCA_000452945.1_BPMSHR5858_genomic_00421_1 srpA
MEHPSTPPGDGPSRGFWRWALIGGAVAAVAATFGYVGGWLAPHRLTPQRLVDALQRNGGL
HPGYRRNHAKGVCVTGYFEGNGAASAYSTVPFFAAVRTPVVGRFALPGGNPYAPDSSVPI
RSLALRLSAPDGEQWRTGMNSMPVFPVATPRAFYEQTVAGRPDPRTGKPDPAKLKAFFDA
HPETAAFRAWVKTAKPSASYVTETYYSLNAFYLVDAAGRRQAVRWRVAPERMADAGESAK
AGDPNVLREDLARRIGSGPQRWHLLLTLAEPGDPVDDATRAWPATRTTIDAGTLVLDRVE
AQDSGLCRDVNYDPTILPDGIRASGDPLLAARSAAYADSYLRRTSEEAGMPGAAHPLSEK
R*
>GCA_000452945.1_BPMSHR5858_genomic_00422_1 yceJ_1
MMKTTFSPLARLLHWVMAAMIVSMLFVGAGMVTTVSGRHAALVAMHKPLGVAVLVLACVR
VVVRLSSRPPALPADLPGWQKFAAYGSHLVLYALMIAMPLVGWAMLSAGGYPVTLGGGVR
LPSIVPADAVWFAWLRHAHRWLAYLFFATFLAHFAAALYHGMIRRDGVLRAMVGR*
>GCA_000452945.1_BPMSHR5858_genomic_00490_1 mdlC
MPPSIHRTSSINSTPAVTAAGARRASGPGGVLSADVARVASARRHTMPEIGARRGVDGDR
AAPAPRESFRQRLETVSSRAPTPPEADSRAANARTTGAADGAGATGAAAGPSHGATSLDA
ARAYLAELAGDRGGALTTLVAQLNGHDRRALDHLALTVHALHLSVDDASHAKTFAGIGDA
LASFLAAAARANAKPGSTPRDLCVEGNKGYRKLCECVGALLRSDGIANALFGRARERGDA
AADTLVSERLGGRMDGHVRTNGMVNRHGGDAQRRMADAAQRFALGARHATNLVDMVGTAL
ELLGRTDQLLNDVSLKRPAAEPGGPRAPGGGPAAPAGPASGAAQQPAAPVVVNNHNENNV
NVNLDGLERLVGDLGKMMGALLERMGAYDPSHTRPAAPDANGPAPVRTGGHTTVVKSPVS
VARTSNNGAPGSPAADGTRGDAPMPPTTPALPATPATPTTPTTPPAGGPSMDERTPATNT
LERGAGNGGASPTGAWRRGDDGGWRMARGNPVVRTTEGATRIDPFSRVGARAASGAGAAR
AWRDAAADATQRSLASGTSGGAADAIAPRAGAGAAAGGDAARVARRMPDVRAGGPEHDAP
GERAAGPNGSNGSAVGQPIAGHAPAGGPDAADFSDASGTGGTDARAPGMPPATGSWSRDP
ASGRWLRAGGGVPLRSTEGFGRIDPFAGVQRNASATGASAVSSPSFASASMSVGSTGALP
AGSSIASPDTSGIAPAVASGGALDVASNVASNVTSNVTSNVTSNVTSNVASGVTSGVTSG
VTSGVAPDGTAWTRPSALAPRNIDPRRMNRAQLERAGLYPLVDDALPPSSQSAPADAGDA
GDAGDAGDAGDAAARRASRPDTGAPQPGAAKTFPSVAPAHRIVTTEGAARVGWNDSIANG
TKR*
>GCA_000452945.1_BPMSHR5858_genomic_00574_1 eamB
MNYLTLSALPAGMLFALVTSITPGPNNTMLLASGVNFGFRRTLPHMFGISAGVAILMLSV
GFGLGEAFKRIPILYTLLETASVVYLLYLAWRIGTSGEVRAHNGKSRPMTFVEAIAFQWV
NPKAWMMVLTAATTIQLSADYGRNAVWMALVFVFVGLPCISVWAAFGQGLRGFLSNPRWL
RVFNLTMATLLVLSLYPMFAKVSGH*
>GCA_000452945.1_BPMSHR5858_genomic_00578_1 catD_2
MPFASVDGVRLHYRIDRAARADAPWLVFSNSLGADLSMWAPQIGPMTAHFNLLRYDTRGH
GHSDAPAGSYTIEQLAGDVIGLLDHVGVARAHFCGISMGGLTGAALAARHAKRVDRVVLS
NTSAKIGSPEIWAPRARKARAEGMAALADAVLPRWFTAAFFEREPRLIDVIRDTFNHTDK
DGYAANCDALNAADLRDEAKGIAVPTLVVTGAHDLSTPPEQGRALAASIAGAKHVEFDCA
HISNIECADGFNRTLIDFLTA*
>GCA_000452945.1_BPMSHR5858_genomic_00579_1 pcaB
MLEDSARLTALVCGSEPLNRIWSPHATLQRMLDVEAALARALAANGVIPANAVRPIEAAC
AASELDAQALMRDAALGGNLAIPLVRQLTARVKVRDAEAAKYVHWGATSQDIIDTAAVLQ
LRDTLDWLDPLLRDTCTALATLAAAHRTTPMIGRTWLQQALPITLGLKFAQWLDALLRHR
ERIVALRERALALQFGGAAGTLASLRDAAPAVARSLADDLRLALPAVPWHTQRDRIAETA
ACFGMLIGTLGKIARDISLSMQTEIGELAEPAAAGKGGSSTMPHKRNPVGCAAVLTAAVR
APGLVSTVLAGMVQEHERALGGWQAEWDALPELARLAGGALAQVAQIVEGLNVDVARLAD
NLDATRGLVLGEAVMLALGDRIGRLDAHHLVERASKQAVRSGRPLYDVLAADPDVAAHLT
PQALRQLLDPAHYVGEAHAYVDAVLALHASRH*
>GCA_000452945.1_BPMSHR5858_genomic_00581_1 pcaI
MVNKIFDSLQSAVADVHDGATIMIGGFGTAGMPAELIDALIAQGARDLTIVNNNAGNGET
GLAALLKARRVRKIICSFPRQADSQVFDALYRAGEIELELVPQGNLAERIRAAGAGIGGF
FSPTGYGTKLAEGKETRVIDGKHYVFETPIHADFALVKAYRGDRWGNLVYRKTARNFGPV
MAMAAKTSIVQVSQVVPLGALNPEHIVTPGIFVQRIVEVPQAAHAAELAAERAASAA*
>GCA_000452945.1_BPMSHR5858_genomic_00584_1 folE_3
MAKGMTAGKAAAPAAHQAARRTSNKASKTASSAATHEPAGAPANRPSRDEAEAAVRVLLR
WAGDDPQREGLVDTPARVVRAYEEFFAGYALEPRDILARTFSEVDGYDEMIVLKDIRFES
YCEHHMVPIIGRAHVAYLPNHRVVGISKLARLVDAFAKRLQIQEKMTVQIADTLFDVLQP
KGVGVILEAAHQCISTRGIHKPGVEMVTSRMLGTFRTDPATRREFLSIVANPSSVNLTNT
*
>GCA_000452945.1_BPMSHR5858_genomic_00586_1 group_4461
MSKSARNWGLEQIVAELRESREKLHRTRHPRGIRELPSRDAVCTIVSGLRASMFPTHYGA
PDLTDETVDYYVGHTLESTLRLLAEQIRRALRFLPEHAQTPLAVLEERAFEIAREFGTQL
PRIRALLVSDIQAAYTGDPAAQHITEILLCYPGVLAMMHHRLAHALYRLGVPLLARFINE
IAHSATGIDIHPGAQIGPSFFIDHGTGVVIGETAIIGERVRVYQAVTLGAKSFPADGDGM
LVKGNARHPIVEDDVVIYAGATILGRVTIGKGSVIGGNVWLTHSVPPGSSVAQGKIRESD
KDQAEKD*
>GCA_000452945.1_BPMSHR5858_genomic_00588_1 group_4462
MNPFERKTSAAGGRARPRSIELYNDATHDSTVDTDGKNREAARIASGGPISPDQMTTSNA
SLVNAMPETPDGFAGFDSRPGGNRPQFALRAGYAVVDKGFDAAALRDAQMADAFSPPYRP
PRRPGRIVELAPVAR*
>GCA_000452945.1_BPMSHR5858_genomic_00590_1 group_746
MQITGEMLIGAAAVRGSEGTMRAYAPAQGVELEPTFGAGGAADVDRACRLANAAFDPFRQ
APLETRARFLEAIAERIVGLGDPLIERAHAESALPVARLEGERARTVGQLRLFAAIVRDG
RWLSATLDSAQPERKPLPRADLRLQKIPVGPVAVFGASNFPLAFSVAGGDTASAFAAGCP
VVAKAHPAHLGTSELVGRAIRQAVADCGLHEGVFSLVVGAGNAIGEALVAHPAIRAVGFT
GSRAGGLALMGVAARRHEPIPVFAEMSSINPFFVLPGALRARGAQIAQGFVESLTLGVGQ
FCTNPGLVVALEGPDLKAFVDAAAQALSQKGAQTMLTSGIASSYESAVAARRAAAGVSEV
ARGVRSDARNAALPALFTTTHTQFVQNPQLEAEIFGPTSLVVACRDIDEMIALAEHVEGQ
LSATLHLEDDDVDLARKLLPTLERRAGRIVANGYPTGVEVAYAMVHGGPFPATSDPRSTS
VGALAIERFLRPVCYQDLPAALLPEALADANPLGLWRLRDGQLGKA*
>GCA_000452945.1_BPMSHR5858_genomic_00593_1 fnr
MLTPVARPAAAAAPVHASSTWAPRQAAHCSTCAMRHLCMPQGLAPEALARLESVICTARP
VRRGETLFREGDTFDNLYAVRSGSLKTIATRHDGREQVTGLHLAGEALGLDGICDDAHPR
TAVALEDSSVCVIPYSALKTLCSEAGTMQLRMHKLMSEQIVRETSQTMVLGSLNAEERVA
AFLLDVSSRYMKRGYSPNEFNLRMTREDIGSYLGMTLETVSRTLSKFHKRGLIEMQGRMV
RIVDFDGLHRL*
>GCA_000452945.1_BPMSHR5858_genomic_00596_1 group_847
MARRKLERWSALDLAEGLQLAHAVAALQTLGVVDAMAEPTTAQTLSAACDLDPELLRGVL
EYAASRTNLIRKTGDRFSASEHYTAQARFLLNLYAGAYADNASRLATLLRRPALAGTTVD
LVAHARAFEASEAGGGALAAIISQLHLNHVLDLGCGAGTLLHALAADDREFVGWGLDRNP
SMCKAARMRARQAGIAARVKVFQGDGRHPGASIPPRVLARVRNVVASQFVNEMFRGGTSR
AQAWLRRMRRLLPDRLLVIADYYGRLGHGFQTPHRETLLHDYAQLISGQGVPPPNADAWL
AMYRATGCRPLHVIEDRAATTRFVHLVAL*
>GCA_000452945.1_BPMSHR5858_genomic_00598_1 group_4465
MNARTIHALIAASTKDPQLIERWRRHPDLLRGAGIAPGDLDWRALEHFVGLSLKVRHNGL
RGDFPLSFRLLSVVGLEIEVFAAYASDCAESGHVFAATPQERARDLIAFLDRWLDPAQRN
HVLLWDILRHEYALACLRQAGDAPLADARPAARHRARATAGGIPTLNGSVVQHVMRCNPD
DLAAALHQKEPPLDAVPAVDTYCCYWRSAETGEIRLLRVDTLGYCAIEAIDGRRSAADIC
HRLLGNRRLLPQFLSSLDQLAGIGIVRFARPLGCAAA*
>GCA_000452945.1_BPMSHR5858_genomic_00599_1 group_4466
MKAFDRDAGAPRIGLAYGPGMAEFAARHAHLVDYIEVPFELLRFSPTIAELQQTIPFVLH
CASLSVAGFVPPDDSTVDAIERTAVRTGTPWIGEHLAYISADPIGEALGGTHEPTSLSYT
LCPQLSDETVRRVADNLAVLAPRFPVPLIVENSPQYFSIPGSTMGMTDFIRAIADRCDVG
LLLDLSHFLITAYNTGADVHREFARLPLERVVEVHLSGMSIQSGAAWDDHSLPASPILFE
LLERLLAVARPRALTFEYNWSPYFPLSVLTSHIERARALMGLA*
>GCA_000452945.1_BPMSHR5858_genomic_00601_1 group_4467
MNGLPLSLPRVDSTDSLFADPLAFLARARAQCGDVFVLREHGPIFSRASDCSGVIAVFGE
HRLRQVLTETDTFALPMSAAAKMALPKNLVNLNRGLHSMREPEHGRHKRLLTGTANRALF
DAHRSGMQTDLSRFCEMLNGDGRISVVSRMRELTAKLASRLFLGPQCEEDAELTFLLSAY
FTLRREASSPNAHDPLQYRDALIGAGRQLDRALRARIRRYRKAPAGECAGLLQRLATAGQ
PGSPALSEDEIVGHANVMFVSSTEPVAISLAWLLLVLSQLPDLRRALRAESAARASSPAS
PCGAPLLERVINETLRLLTPNALMVRATTQAVSLQGVALPARCEIVVCPFLVHREANAFA
RPHAFSPSRWETARPSPYEYFPFGAGGHFCAGRNLALSLIGEVLSTLLSRFDFVLDIEQF
IDWRIHIMLMPKGDPTLTAHPVDERRDTPSPKWRGPVAELFHFAPGLS*
>GCA_000452945.1_BPMSHR5858_genomic_00604_1 yqjH
MTQPTTERAVVRVRHTLKFRLLQVKRVVDVTPHLKRVTLAGDELDDFESASFDDHVKVFF
PPPGEDAPARPEFGPNGIVFPQDRPRPVARDFTPRRYDRAARELDLDFVLHHPGPASQWA
AQARVGQTLGIGGPRGSFVIPKDFDWHLLIGDDTALPAVARRLEELPAGTRVAAVLEVAD
PSAQIAFETDANLYVVWRHRGDAASGERGASRLVDALRELPLPAGEGYVWAAGEAAAMRG
VREHLCRERGVAKSRIRASAYWKHGAIAVHESLED*
>GCA_000452945.1_BPMSHR5858_genomic_00606_1 ybjG
MQNFNLAVFSALNAGSTPPLGAVRFGVFAADWLVYAVPALLLLTWVMGAHPTRRQAIEAG
VGACVALALAQVIGHFWYSPRPFVLGIGTQLIPHAPDGSFPSDHTTFVCSVAAGLLIARA
TRTAGAALAVIAAVIGWGRVYVGVHWPLDIVGGALVGTAGALAAHLYGRPVTALLDRVGE
SVHAVAFTRLHERPIRID*
>GCA_000452945.1_BPMSHR5858_genomic_00607_1 adc
MKPSQVRSKAFAMPLTSPAFPMGPYRFVNREFLIITYRTDMDRLREIVPEPLEVKEPLVH
YEFIRMPDSTGFGDYTESGQVIPVEYKGQPGGYTLAMYLNDHPPIAGGRELWGFPKKLAQ
PTLQTHIDTLLGTLDYGPVRVATGTMGYKHQELDLEEQAKRLAGANFLLKIIPHVDGSAR
VCELVRYYLQDIEMKGAWTGPASLQLAPHALAPVADLPVLEIVEARHLLADLTLGLGEVV
YDYLAQ*
>GCA_000452945.1_BPMSHR5858_genomic_00625_1 clsB_1
MSGETRHRLAQLRQMFLQERGSASRLAFTSGNAVRLCDGGAAFFPALIERIDAARERVAL
ETYIFCDDAVGRAVSDALVRAAARGVHVRVITDGIGTERLPLFGTWPDAGVEHRIYNPYL
FGRFGFSRTHRKLAVIDDACAFCGGINIVDDFEQNGERLPHPRWDFAVELAGPAVADVRA
AFEVQWHRIAAGHKRYAQYLPHDVDGPAFAARFRRWMRSHRWVKAGALRVVTEPSVAFVA
RDNIVNRRAIEKAYLAAIGRARQSILLANPYFMPGRKLRRALAGAARRGVDVRILLGRNE
FASLDMAVPFLYHALLRSGVRVAEYDKTMLHGKVAVIDDHWATVGSSNLDALSLMLNNEA
NVVLVRHREETAALRDAIAAAFADGREIDPALYAARPAAERFLNWFAYNAYRSAMKLLTV
GGYD*
>GCA_000452945.1_BPMSHR5858_genomic_00626_1 group_4473

MRKGEQTRAAILEAALDLASRDGLEGLTIGLLAERMQMSKSGVFAHFGSRDDLQVEVVRE
YHHRFENEVFFPSLREPRGLPRLRAMLARWIEKRIQEVTTGCIYISGAVEYDDRPDSAVR
EQLIASVKAWRAALLRAISQAKEEGHLRADTDPNLMLFELYSFTLGLHHDARFLHQPDAV
RLTWAALEKTIVSYQSESR*
>GCA_000452945.1_BPMSHR5858_genomic_00628_1 fadN
VSNFNIRKVAVLGAGVMGAQIAAHLVNARVPVLLFDLPAKEGPKNGIALKAIENLKKLSP
APFGVKDDAKYLEAANYEDDIAKLAECDLVIEAIAERMDWKHDLYKKVAPHIASHAIFAT
NTSGLSITTLSDGFADELKARFCGVHFFNPPRYMHLVELIPTAHTRPQILDQLETFLTSV
VGKGVVRAKDTPNFIANRVGIFSILAVITEAEKFGLRFDEVDDLTGSRLGRAKSATFRTA
DVVGLDTMAHVIKTMQDNLADDPFFPVYKTPAVLAKLVEQGALGQKTGGGFYKKEGKAIK
VLDAKTGAYVDSGAKADETVGRILKRPPAERLKLLRETQHPHAQFLWSIFRDVFHYIAFH
LESIADNARDVDLAIRWGFGWNEGPFEGWQAAGWKQVAEWVQEDIAAGRALANVPLPAWV
LDGVVAEKGGVHTAEGSWAPAAQRFAPRPDLAVYDKQVFRAPVFGEAGRDPKTFGKTLFE
TDSVRAWVDDRAGEDDVVIVSFKSKMNTIGPGVIDGLAQAIELAEKDYKAVVIWQPSSLK
LGNPGGPFSAGANLEEAMPAFMMGGAKGIEPFVKKFQQGMLRVKYANVPVVAAVSGLALG
GGCELMLHSAKRVVHVESYIGLVEVGVGLVPAGGGLKEAALRAADAATAAGMTSELLKFL
TKPFENAAMAKVSMSAHEARAMGYVKPSDPIVFNVFELLDVAKKEARGLAAAGYRAPLKA
KEIPVAGRSAIATIKAQLVNMRDGRFISDHDFLIASRIAEAVCGGDVEAGSTVDEEWLLA
LERRAFVELLGTQKTQERIMCMLQTGKPVRN*
>GCA_000452945.1_BPMSHR5858_genomic_00634_1 atm1_2
MRRYPASSEPAPAATGPRNDWQTIRSLLPYLSAYKWRVAFALACLIGAKVANLGVPVVMK
RIVDSLAAVQHLTALGRAEQSATVVLAGGVGLLVIAYALARLSTSLFTELREILFSKVTE
SAVRRLALQVFRHLHGLSLRFHLERQTGGMSRDIERGTRGIQQLISYSLYSILPTLVEVG
LVLGFFVVKYEAYYAYVTFAALITYIVFTVKVTNWRTHLRRTMNELDSRANSRAIDSLIN
YETVKYFGNEEWEAQRYDENLKRYRKAAIRSQNSLSFLNFGQQAIIGTGLVFILWRATQG
VLAGRLTLGDLVLINTFMLQLYIPLNFLGVVYRELKQSLTDMDRMFGLLSAAREVDDAPG
AGALKVSGARVRFERVDFSYEPSRQILHDVDFTIEAGSTTAVVGHSGSGKSTLSRLLFRF
YDLDRATGGAIMIDGQDIRDVKQETLRASIGIVPQDTVLFNDTIYYNIAYGRPSATRDEV
IAAARAAHIHAFIESLPKGYDTPVGERGLKLSGGEKQRVAIARTILKNPPILVFDEATSA
LDSRSERAIQHELDQIARHRTTLVIAHRLSTVVHADQIIVMDHGRIVERGTHAQLLRADG
LYAQMWALQQQRAAADGAAAEEV*
>GCA_000452945.1_BPMSHR5858_genomic_00638_1 surA_2
MKKTLRFAAAASGLVASLITVAPSASAQALRAQGASLADEVVAVVNNDVITGRELDQRVG
LIARRLQQQKAPVPPTDQLRAQVLNQMVLERIQVQRAKDDGIVVDNATVQATLGRLAQAN
GMQLDQYKARIEAQGVPWDLFVRDARTELMLSKLREKEVDSKITVSDAEVASYIASQRGP
NAGSQQDLRLEHIFVKAPANAPQADIDVAQKKAEGLLQQALASGANFERLAKNQSEADDA
KKGGDLGFKSPASLPSDVVDAVSKLRPGEVNPTLIRVPDGFEIVRLVERRASQNPAASPK
IVQTHVRHILLRVGEGKSESQARQQLIDIRRQIESGGDFEKFARTYSQDGSASQGGDLGW
ISPGETVPEFERAMNTLQDGQVSNPVRTEYGYHLIQVLGRRDAEGSVQQQMDIARQAIGQ
RKAEQAYSDWLRELRDSSYVQIKLPVAQ*
>GCA_000452945.1_BPMSHR5858_genomic_00642_1 gloA
MRLLHTMLRVGDLDRSIKFYTELLGMKLLRRQDYPDGKFTLAFVGYGDERDHTVIELTHN
WDTKSYELGTGFGHLALEVEDAYKACEQIKAQGGKVTREAGPMKHGTTVIAFVEDPDGYK
IEFIQRNAH*
>GCA_000452945.1_BPMSHR5858_genomic_00651_1 ybeY
MKSSRSSKPARAEAAQPASPRLSLFDAKGKVKTVEAQALRIDFADGRSLMFDLSGSSGDA
AVAIVAQHADPSLRATITLRPEHYDSVTVEVGADENPDFAHDETDGAGDESDDEDAQGRD
PELDLAVQYGDEIGDAQRKSLPKRKVIAEWLEPAIFSDAQFTVRFVGADEGRALNHSYRH
KDYATNVLTFAYGEEPDGVTVADLVLCCPVVEKEAREQGKTLVAHYAHLLVHGALHAQGY
DHERGEEDAAEMEALEIDILAKLGFPNPYR*
>GCA_000452945.1_BPMSHR5858_genomic_00654_1 dmlR_15
MDRLGDIRLFVEAAELGSLSAAGRKLNLTPAAASARLAKLEAQVSTRLFERSTRRLRLTD
EGRLYLNCCRQALQALDDADAMLQEGRNVVSGKVRLSSTADFGRHQLLDWLDEFNTRHPE
VTFAVTASDSSSNLWQDEIDLAIRFAAPPDGALIARRLAANRRVLCASPAFVDRYGMPKD
PLDLARFPCNVITVASGPLNVWRFTRGDDTQTYTVPLANAHETNDGGLTREWTIRGYGIA
LKSIWDIAADVRAGRLRVLLPDWRHHDAPLHAIYHSKRYMAPRVRVLLDFLVERFAREEK
ALEDLLNACR*
>GCA_000452945.1_BPMSHR5858_genomic_00655_1 ydhP_2
MPLPLLALAISAFAIGTTEFVIMGLLPDIARELAVTLPSAGLLVSGYALGVAAGAPLLAV
LTSRMPRKAALQLLMAIFIAGNVLCALAQSYGMLMVARVVTSFAHGSFFGIGAVVAASLV
SADKRASAIALMFTGLTLANVLGVPFGTFVGQTLGWRATFWIVAALGVASFAGVAALVPK
RRDAAGAALSRELRVLRQPQVWLALAMTVLGFGGVFVVFTYIAPILEDVTGFAPRAVSLV
LVLFGAGLTIGNTIGGRLADRALMPSLVAILVALIAVMAVFAKTSHMPAAAAVTVFVWGI
AAFATVPPLQSRVVEKAAHAPNLASTLNIGAFNVGNAGGAWLGGVALSHGVPLDALPWVA
AVVTLAALAVTALAARLDTRVSSGAATAAQ*
>GCA_000452945.1_BPMSHR5858_genomic_00656_1 group_4481
MHSPLALVRDVSSQDAAAAADADASHAPFDALERLVGVNLARLRAERQLSLDALARLSGV
SRAMLAQIESARSVPSIKVLCKVAAALKVSVAAFLRRHAVNGFEHLSAERAVRVVSSNGR
FSARALYPEGEPAVAEFHELRIAPLHTEHGAPRAPGTSVNLVVSEGTLEVSIHERRQLLA
TGDAIVFDADQPYTLRNPGDTEARAFRVTVNAEVPPRWHVPA*
>GCA_000452945.1_BPMSHR5858_genomic_00660_1 group_4482
MKTFLFVEARPIECAGIGYIRALGYAPVLFTSMQSRNKVLFDDLDLNLFDAVHHVDTLDA
RAMRERIDALRLPVGAVLGCYDDVMIPASELADALGLPHPSLAGLRRAYGKERVRDTLRG
RGYRQPAYQVLSAERAAPRPDIGFPCVVKPLRDAGAYGVSLCANEADYRAAVERFRSGDG
VSMLGSRHREFLAEAFVEGPFYGAELLHNGGRWHVLGINRIFVSPRDSLCMTGISHPSDL
PAADADAIADEIVEWVNLLGLAGGALNVEFILAETGPVLVEINLRIAGARAVRQIALTTG
IDMVEHLIDFVCGIDRPIAPREPARYGFVADAFVFAPAAGVVRGVAFAPNGAHYVASGFR
KLPLESARAAKNFGSVIGYVLAHGRTCDEAMRHARALADGVRVELG*
>GCA_000452945.1_BPMSHR5858_genomic_00662_1 group_4483
MTTSDRLQPSFVEVKNTSTLSGTIDLPASKSSSTRALLTAALTPGISTIRNVATGFNSNA
MKHNCERLGASFSSEGDATVVKGVDVMHVDREIVFDPGNSGVVLRLLMGVAGYLPDTRFV
TQYRYSLGVRSQAEMVAALRRLNVECEAVGPEARLPISMRSTRALGKHTEVSCKKSSQFL
SGLLYLGAIGERDLEIDVVDHITAPSMVHTTINNLAHAGVAVEYDAAFRRFFVPGRDRFK
PSEFTVGADPASTAAILALCGSLASDVTLNGFFEEELGSGAVIRYLTDTGTLIDELPGNR
IRIRGGASIRAQDFDGSLAPDAVPALAGRAAFAEGTSTFYNIEHIRYKESDRISDFRREL
DKLGVRSEEKLDQLIIHGNPRGYRGGAVVDGHYDHGLIMALTTIGLHCEHPVLIKEPHHV
GQTYPDYFADIGSIGANVDELIYPNVAAARA*
>GCA_000452945.1_BPMSHR5858_genomic_00675_1 adhB_3
MKRKSLFALSAVAIVAAAALVPVLWPGNDTLHGNAAVAATPADQAALIKKGEYLARVGDC
IACHTVRGGKPFAGGLPMATPFGTMYTPNITPDDQAGIGKWTSDDFYRAMHTGRSKDGSL
LYPGFPFASYTKVTRADSDAIYAYLRSVAPVSTPSRPHELRFPFNNRNLLIGWRTLFFKE
GEYKPDPTKSVEWNRGAYLVEGLGHCSMCHTSINMMGGPVSSAAFAGGLIPLQNWYAPSL
TNDKELGLGDWHVQELSDLLQAGVSHKGAVFGPMADVVHNSLQYMTDEDTRAMSTYLKSI
PQKAEAPKNMQYEPSKQFGTALLEQGKKIYADNCATCHGPQGEGKPTAYPPLAQNRSIMM
ESAVNPIRMVLNGGYPPSTFKNPRPYGMPPFAQSLSNQEVAAVVTYIRMSWGNNGSPVSP
QQVSDLRSAPLD*
>GCA_000452945.1_BPMSHR5858_genomic_00677_1 yebZ
MKLDSLWFGQAALAALGDVAFAVALGSAFIGAWLANDGARSVIAPSHPAWRQSLRSLAVA
AAVLVLADAGWLVYQAASMSGAGLRGAFGAMPTVLAQTHVGHAWAVACGGALLLLAVALA
RPSGPLAGALLALATLVVAAGKASLGHAADSGAFSAAVGVQTVHVAATAVWGGLVIAGGM
AVLPALGSSIARGAMIRIAQRLSGASIAALAFVIVTGALNTVRGTGGDFAALDGSTWGRV
LLLKLALVALALVLAALNRFSALPRLRRTASTEDAHTFRNVLHLEALAMIGVFIAAAVLS
STAPIAAG*
>GCA_000452945.1_BPMSHR5858_genomic_00679_1 group_3123
MLYTGSCHCGKVRFEVEGEIDGACACNCSMCARKGSLLWFVPRDALRLQTPDEDIATYLF
NKHVIRHRFCPSCGIHPFAEGTDPKGNAMAAVNLRCVDGVDLDALSVRHFDGRAL*
>GCA_000452945.1_BPMSHR5858_genomic_00681_1 group_1406
MSNKPAPDAPPAEQPCAVSRRGFLKFAGVSGLASAASGFAATRAAAAAPDGTPEQIHLTW
GDVDANEVVVSWASLAAATNPRVRFAGPNEAWRTVHGVQRTYTDGLNGEVVFTYHARLRG
LKPGAVYRYEVTADNDANAAQPFAARFETAPRGRAPFRWTSYGDLATPNTGWVLSSPGSR
FAVQAVERFQPLFHLLNGDLCYANLNPAHQPAVWRDFGNNNQTSAANRPWMPCPGNHEIE
FHNGAQGLDSYLARYTLPENGTRFAGRWYSFRYGAVLFVSLDADDVVYQDAAAFVAGPNP
LVPAASTGNEAIAPGTSLYVRGYSRGEQTRWLEQTLRRASRDRDIDWIVVQMHQDALSSS
KTGNGSDKGIREAWLPLFDRYGVDLVLCGHDHDYERSFPVRGCNHRAGVDAATGEVVDTL
QPRPVVTTDPADGRFDTSHGTIHMILGGGGTSAPLDVYGENPATGLPQARVFTKPNRPVP
GAAPNTFVRHGADAVEDAIWSARRDTGTGYGIAVFDYEPGEHGGRSTITVNYYHAPGADQ
HPTAEYELFETIVLSKPRRA*
>GCA_000452945.1_BPMSHR5858_genomic_00700_1 caaA
MMRSRTNGAIDTPRARRAQHRLLWACMLAGLVGAPALWMSQMLVSETLSSVACDPLGVPH
ALPQWPQFAGRLALFAALALLLGIACAALAWRGLRRSAAAAPRAARDADAAPPGPPAPAS
PRTTDRAPAHAPSRVPLDAANREPNAPGEPDELGELGELGEPDLPDEPDDAASKTRFIAV
CGLLAGVGFALGLIFTGLAVIFVEPCHPWR*
>GCA_000452945.1_BPMSHR5858_genomic_00713_1 group_3127
MRISTNWIKHLVLALLLGVTLSSYAAPFKFTVDGNIKRTNQPGRRAYVFSEAALMALPQH
SITTSTSWTPKATFTGPRLSDILKAVGATGTQIEFHCVDEYTFTIPTSDADKYGVILART
MNGKVLDNNNYGPLWIMYPRDQFPDELKTPISEAKFAWQIIGLTVK*
>GCA_000452945.1_BPMSHR5858_genomic_00718_1 group_747
MRGCRNALEAALSERGDAHDLVDAYEYAEKVIEDLADGIDYFQELVRHLMQAASAHTQWN
EFVDFLDRFQRDYSKQLTADNATLNRQAIRQKLEKLRAVPEAKFNRMDAQLRDIAHWMAK
EYTGSSLYDWLLGRIEDIVDAACESKQPGFLKAMETYLKRVNGLALQSMMLRSGQTRHAY
LAAVQKVAESPKLEQDRLLKLVGQFLAPAEVRVLDPASFKLKTATQRRKAVTVSIRPRPT
REERLAAAMARAEAEVFSIPNEDVAQGLRADLRLFQHPIRVSAMPTASARDVIRAMQAVE
AARSAIGGDLIVKKLPTRLENEFYGANDYEIDLKK*
>GCA_000452945.1_BPMSHR5858_genomic_00725_1 hmp_1
MTQMTAEQMARVKATAPVLAEHGATITKHFYQRMFGRHPELKNVFNQTHQKTGSQPETLA
KAVYAYAANIDNLGALGGAVSRIAHKHASLNIRPEHYPIVGENLLASIVEVLGDAVDADT
LEAWRIAYGQLAAILIGAEANLYENAAWSGFRPFKVAKKVRESDEITSFYLTPADGGAAP
GFEPGQYISVKRFVGDMGVDQPRQYSLSDAPHGKWLRISVKREAGHSEAVPAGKVSTLMH
DGVDVDSVVEVTAPMGDFTLNRHAATPVVLISGGVGITPMMSMASTLVAAGSEREVRFLH
ACRAANVHAFRDWLNDTTDAHPNVKRAVFYEVVGPNDRVGVDHDHEGRITPAALERHALV
PDADYYICGPIAFMKQQRDALVALGVAPERVQTEIFGSGALE*
>GCA_000452945.1_BPMSHR5858_genomic_00728_1 group_4489
MNAPAELSADLRAQRQREVVQALMAVLPTHCLLHRDEDTAAYECDALSAYRRLPLAVALP
ETESQVQRIVQICRRMNVPIVPRGAGTSLSGGALPIGHGVVLSLARFTRIVEVDSYARTA
TVQPGVRNLAISEAAAPYGLYYAPDPSSQIACTIGGNVSENSGGVHCLKYGLTCHNVLRV
RGVTIDGEIVEFGSLAPDVPGLDLLAVTIGSEGMFIVVTEVTVRLIPKPQCAQLVMASFD
DVVKGGEAVAAIIASGIIPAGLEMMDKPATQAVEAFTHAGYDLDAAAILLCESDGTQEEV
AEEIVRMTAVLREHGATRIQVSRSEAERLRFWSGRKNAFPAAGRISPDYYCMDGTVPRRA
IGPLLARIEQMETKYRLRCINVFHAGDGNMHPLILFNANDVDELHRAEAFGADILETCVE
FGGTVTGEHGVGVEKLNSMCVQFSREECDAFFAIKRAFDPPELLNPDKGIPTRARCAEYG
RMHVRGGLLPHPDLPRF*
>GCA_000452945.1_BPMSHR5858_genomic_00729_1 group_2281
MEEDDIVAGWAARIRDAAASGRALRIRGGGTKDWYGQALDGEILDTRAHHGIVSYDPAEL
VVTARAGTSLAELEATLAERGQMLPFEPPHFGRGATLGGAVAAGLAGPRRATTGAPRDFV
LGVAILNGRGDRLRFGGQVVKNVAGYDVSRLMAGSLGTLGLMLELSVKVLPVPAAELTLK
FDMSATDAVRKLNEWAGRPFPLSASAWRYGTLVLRLSGAEAAVKSAKTVLGGEAVDAVEA
ERFWEGVREQNDPFFSSLAPGHALWRLSLPSITEPMHLPGTQMMEWGGAQRWWITDADAQ
TVRMSAKQAGGHATLFRASESYDRSAGVFTPLPAPLMKIHRGLKTAFDPARIFNRGRLYP
DL*
>GCA_000452945.1_BPMSHR5858_genomic_00731_1 yggS
MSDLAARLASVHRRIDEAARAAGRAPGSVTLLAVSKTFPADAVRAAYAAGQRAFGENYVQ
ESIDKISTLADLRAELEWHFIGPLQSNKTRPVAERFDWVHTIDRLKIAQRLSEQRPAHLP
PLNVCVQVNISGEASKSGVAPADAAGLARAVAALPALRLRGLMAIPEPETDPEAKRAPHR
ALHALFEQLRAGGLALDTLSMGMSDDLEAAVAEGATIVRIGTAIFGARDYAH*
>GCA_000452945.1_BPMSHR5858_genomic_00734_1 ptxB
MKLIRSLVACVAALACAAAAHAQTLNFGIISTDSSAVLKRRWQPLIDDMNRKTGLDVKAF
FATDYAGIIEGMRFNKVQLGYFGNASALEAVERSNGEVFAKLRHANGETGYYSVLITNAN
SRFRTLDDVFRNPKDVTLGFGDPNSTSGTLVPAYYLFAPRGIAPRTAFRTVLPSSHEGNL
LAVVNDKIDIATNNTDVLGMLKKQHPDQYARVRVLWTSPQIASDPLVWRKDLPDATKQKL
RDFFYNYAKTDPHEQAVMAGIRNLVGFAPSSDAQLIPVRKIVLFQQKQKIETDMALSADA
RRAQLAALDARLAALDASGAADASSAQADKR*
>GCA_000452945.1_BPMSHR5858_genomic_00739_1 phnJ
MNAPDHCAAAAPDTHAAEGYNFAYLDEQTKRMIRRALLKAVAVPGYQVPFASREMPLPFG
WGTGGIQVTAALIGRDDTLKVIDQGSDETTNAVNIRRFFARTAGVATTTKTAQATIIQTR
HRVPETPLTERQILVYQVPMPEPLFRLEPRATECAKLHAHADYGLMRVKLYEDIVHHGSI
ATTYDYPVIVNGRYLSSPSPIPKFDNPKLHMNPALQLFGAGRERRIYAIAPYTPVASLDF
DDHPFEVQRWDAACALCGSRESFLDEMIVDDHGTRRFVCSDSDYCSDRRGGAQARADGGD
EGNGGGDGEMPAPLAAEQPLGETT*
>GCA_000452945.1_BPMSHR5858_genomic_00742_1 phnG
MSASAVSSLHAARRDWLAVLAKIPRDELERALDRVLDGAPAPAFDWLRAPEIGLAMVRGR
IGGTGDPFNLGEASVTRAALRLRGDRTNGTTGTTGADGMNGANGANRPHGATVGIAYVLG
RDKRRAELVALADALLQTPERHERLRAALIEPFRARLAQARAARSGEIATTRVEFFTMVR
GE*
>GCA_000452945.1_BPMSHR5858_genomic_00743_1 dasR
MRGISIETQRNDNAMTSVDHLLQDSPLERGSGVSVWRQIEQILAAEIAANGFGEDGRLPS
EGELAKRFDVNRHTVRRAMLGLAASGLVSVEQGRGTFVQSGAIDYTIGRRTRFTENLNRH
NHAAVGKMLSAARVKADPATAKALGLRAGAYVYRIESIHEADGVPLTYARSAYPAARFAE
LPAALERTERVTDALAEYGVADYLRKWSRIGSVLPEAAIARRLGINKQQPVLWVENVDVD
VDGTPIKYGTTYFAADRVQLLIENDNVPGHAR*
>GCA_000452945.1_BPMSHR5858_genomic_00744_1 group_515
MRAELAARAGAARFALYYAPPRASAWWRAGCAWLGRDPEDGATLAPPQPGALRRPVDALT
VAPRRYGWHATLVAPFALRPGIAPDELVAAAAAWARACAPLAIDVDVAALGAFVALRPAH
AHGDDALRALAARALRALAPLRRPPADAERAKRAAAPLSERQRALLAEWGYPYVLDEYRF
HMTLSDPLDDPAERAALIDWWKAAAAALGPLPVDHAAIFVEARPGTPFMLWKRLPFGQRA
AATDAAATTDAADAADAADAAESGEARR*
>GCA_000452945.1_BPMSHR5858_genomic_00746_1 ubiA
MLARFPLYLRLIRMDKPIGSLLLLWPTLNALWIASDGHPAPSLVVIFALGTLLMRSAGCA
INDYADRDFDRHVKRTAERPLTSGKIRAWEAIAIAVGLALVSFLLILPLNGLTKELSVVA
VFVAATYPFMKRFFAIPQAYLGIAFGFGIPMAFAAVQDTVPMIAWAMLAANVFWSVAYDT
AYAMVDRDDDLKIGMRTSAITFGRHDVLAIMLCYAAMLGIYVWLGAALHFGWPYWAGWAA
AAGCSIYHYTLIKDRERMACFAAFRHNNWLGGVLFAGIAAHYALAVR*
>GCA_000452945.1_BPMSHR5858_genomic_00752_1 oxyR_1
MTLTELKYIVAVARERHFGRAAEACFVSQPTLSVAIKKLEDELNVQIFERGASEVSVTPV
GDQIVTQAQRVLEQTFAIKEIAKQGKDPLIGPFRLGVIYTIGPYLLPTLVKQMIRRVPQM
PLMLQENYTLKLLELLKQGEIDAAIMALPFPETGLMVRALYDEPFVVALPAGHPWEKRRK
IDAADLKQETMLLLGNGHCFRDHVLGVCPELMHFSQTADGIQKTFEGSSLETIRHMVASG
VGITVLPRMSVAEVGPHASGPDADLLSYVPFEDPVPDRRVVLVWRKSFTRMPAIDAISEA
IAACGLPGVTKLDIPATVN*
>GCA_000452945.1_BPMSHR5858_genomic_00777_1 psdht
MSTLPNRPAHLTINGEPIPTLDEIAAQHFALTPWVMRTPVFERADLPSLDNTHVNFKFEL
LQASGSFKARGAFTNLLALGEDRKSVGVTCVSAGNHAAAVAYAAMRLGTSAKVVMFDTAS
PARVALCRHYRAEVVMASSPAEAFEIVRRIEAEEGRFFVHPFNGYHTILGTATLGYEWAT
QTPDLDAVIVPIGGGGLAAGTSTALRLANPRVHVYGVEPEGADAMSRSFAANHPVRMGAM
HTIADSLMAPHTEEYSYELCRRHVDRIVTVSDDALRAAMLYLFAHLKVAVEPACAAATAA
LLGPLRESLQGRRVGVLLSGTNIDSTALAAHLEHARLHRAAQP*
>GCA_000452945.1_BPMSHR5858_genomic_00778_1 group_6167
MRASHATRIGCSLALSVAVAALLPGCATSPQPASGDARCTQAGGTRELQADLLFGRDVAG
RGPITDAQRARFVADAVTPRFPDGLTTWDTRGQWRDRETGRIVREESFVIRIVAPDSPAT
LDSLAAIRRAYAERFHQQSVGLVLARVCASF*
>GCA_000452945.1_BPMSHR5858_genomic_00780_1 group_4495
MSLFDSISRTIKGLLNDAADSVQDPSRDARQIVRELDDSIGRAENSLIEIEAQVATQRSK
RDTADERVKKYEDGAKRALQSGDEALAREALAAQANAEAERDALARELDTLEPSVDKLKG
QIADMRARRNDLNARSNILQAKQEIAQAKDVAATALGGIGGKNLSEDFQKLEDKVALSNA
RSDARLNSADTNSGRALDDKLAALNKGPSVEDRLEALKKQLNTPAQ*
>GCA_000452945.1_BPMSHR5858_genomic_00782_1 mscL
MSIIKEFKEFAVKGNVMDLAIGVIIGGAFSKIVDSVVKDLIMPVIGVLTGGLDFSNKFVL
LGQIPASFKGNPESFKDLQAAGVATFGYGSFITVLINFIILAFIIFLMVKFINKLRKPEE
AAPAATPEDVLLLREIRDSLKQR*

>GCA_000452945.1_BPMSHR5858_genomic_00783_1 group_4497
MRMPHFLQLFLRRVFMSFPKKRRRAQPDGQESFLAVLRHAKPFAQLDTKIARARAQDEPV
LYAHVLPGLDVLLCSVRGAQPPYPAIAELRKRCIESIANALEQPLDGLENGGYWYEANGF
GFLVFASRARARILPEFGAGVKAGLRSGIGQPNAGDTTPRSL*
>GCA_000452945.1_BPMSHR5858_genomic_00784_1 glxR_2
MMGGNFNRYSNERKGEHVDVGFCGPGLMGAPMIRHLLTAGHRVHVWNRTRAKADALAGCG
AQVVDTPAELAARAQTVMLCVLDAQAVGDVVFGASGVLAGDAATRRVRRIVDHSSIAPAA
TRAYAARAAALGVSWIDAPVSGGVPGAEAGTLAVMAGGPQADVDAVRVLIGAYAARVTHL
GDAGAGQTAKLCNQAIVTATVTAIAEAVGLAQASGIDAARLAEALAGGWADSVLLQTFVP
RMTRAGHPPIGALKTFQKDVDAIADAARDAGAVMPVASTVQQVLRLGAAMGLAQADFAAF
IDIVRPQAKRGA*
>GCA_000452945.1_BPMSHR5858_genomic_00796_1 rbsA_5
MSGDMSGGACDLGRAGAAARRSPETAGAPVAPPGARGWTPARRVLLRGGPWLALALVFAG
FSVSSPLFVTFANLGNVLQQSAVTGLLAFGLTIVMIGGGADAIKGGLDLSIAANLGLCAA
VFAALTRDGHGDALTLAATCATGVAVGALNAWAVAWLRIPPLLATLATMNVCAGLELVLT
ENAPISSPSPLAAALVGTGPFGAPWLAYALVGVAAALAALAHRTRYGLALHALGAHRAAA
LAAGVDARRYVASSYVASGLSAALAALASCALLSGSAPGAGDYLLPVVAAVLLGVVFSRR
LVPTIPGTLVAVLFVGLLANGFQLNSVSSYWVSGVEGALILFVVAAVALLRRRRSQGAFD
A*
>GCA_000452945.1_BPMSHR5858_genomic_00802_1 ygcS
VSTLDLDAPISTPIRTASDVSRLINASGARANHARVIVLLALGGVFLDAYDLTTLSYGIE
DVTREFGLTPALSGLVSASIMIGTILGSLLGGWFTDKVGRYRVFMADMLCFVVAAIVAGL
APNVEVLIAARFVMGLGVGIDLPVAMAFLAEFSKFGGRGNKASRLAAWCPMWYAASSVCF
LIVFGLYFALPAEHARWLWRASLIFGAAPALAIIAVRGRYMNESPLWAANQGKLRDAARI
LRESYGIRAHAADDTPRAAPSQPPVSFRVLFRQPYLPRTLVASAMNLCIPFEYTAIAFFL
PTILTQFLGAGVFETIAATLALNVLFALTGGLLGMRLAYRLPSRRVAIAGFALQAAALVT
LALLGHPQTALGIGAAVLMLGLWLFAEGFGPGAQMMIYPALSYPASIRGTGLGFGRALTG
IGSAFALFVLPILNARLGAGMFWIVAIAAFVPIVFLAAIRFEPTARDVDVDIDAGSRAQG
DRARVRAAA*
>GCA_000452945.1_BPMSHR5858_genomic_00805_1 czcS
MIRRLLPRTLRARLTALIILSTAATLALSGVALYSALHNRLVGMSSYEMSATLAAMRTHL
ANVANVDDIPRKSDLWIDQLHGHQNLDLAIYDTDGRLRFATRGFVAPRPALGAPQTRVPA
SAAPAGATFSYLADDAPLRGGNPRTARIVVQYDGKNDHALLRAYAYTVVVIEVLAVVLTA
ALAYGIAMLGLSPLRRLVARAEQMSSSRLAQPLPELDTSGELKEMEHAFNAMLKRLDESF
VRLSQFSSNLAHDMRTPLTNLLAEAQVALSKPRTADEYRDVIESSIDEYQRLSRMIEDML
FLARSDNAQSHLAIRTLDAAAQAERVAGYYEPMAEDAEVSIVVRGKAEVRADALLYHRAL
SNLISNALNHAPRGSTITIECAQAADAATISVSDTGRGIEAPHRERIFERFYRVDPARHN
SASGTGLGLAIVRSIMENHGGTCGVDSEPHVRTTFWLKFPAHAA*
>GCA_000452945.1_BPMSHR5858_genomic_00812_1 group_1780
MLQLLSVAIDKPDTANFILGQTHFIKSVEDIHEALVGAVPGIRFGLAFCEASGKRLVRHS
GTGAALTELACRNALAIGAGHCFLVFLGDGFYPLNVLNAIKAVPEVCRIFCATANPTEVV
VAQSDQGRSILGVVDGFSPLGIETDEDVRWRKDLLRNIGYKA*
>GCA_000452945.1_BPMSHR5858_genomic_00817_1 proQ
MGFEQLASLRDQLAKAAAQKRAAKQPQKQPQKHSQKRDAKPAASPSSTDPVVLTIAKLQK
RFPQAFPRNPAPKVPLKVGILEDLVGQASALGLSEAELRDAMRTWCQGNRYWTCLVEGAV
RIDLAGGEAGRVSAADAKRARMLKGRRPAKPRAQPSRPEQAAQQQQTSACPPQGAQRTD*
>GCA_000452945.1_BPMSHR5858_genomic_00845_1 bepE
MNAGAPHDSGAARARFTDVFVRRPVLSLVFSLLILLVGLRALMSLPVRQYPALESATISV
ATDYPGASQELMQGFVTTPIAQSIATADGIEYITSTTTQGKSVVKARLRLNANADRAMTE
VMAKVQQVKYKLPADAYDSVITKLTDAPTAVMYLGFASDALSIAQITDYVVRVAQPLVTT
VPGVASAEILGGQNLAMRVWLDATRLAAHGLSAGDVAAAIRANNVQAAPGQVKGSLTIAD
ISANTDLTDVGAFNEMVVKSAPNGGGLVRLKDVATVEIGGENYNSSALMNGRRAVYIAVN
ATPAGNPLEIVRGVGAVLPAMERNKPASVQIANMFDGARFVNASIAEVRSTLTEAIAIVV
VVIFLFLGSFRAVIIPVLTIPLSLVGSAALMLAAGFSLNLLTLLAMVLAIGLVVDDAIVV
VENIHRHIEQGEPPVRAALIGAREIASPVLVMMATLISVYAPIGLMGGLTGSLFKEFAFT
LAGAVLVSGIVALTLSPMLGSLLLTSRMSEGRVAKAIERTLERVTHAYRRGLHASLAARP
AILLIGAGVLAGIVLLFSGVKRELAPQEDQGSIMVAVKAPQYANLDYMERYAPDIERVFR
TLPEADTSFILNSYGGSNLGFAGVNLVDWDKRTRSAAALQALIQARGDGIRGERVFAFQL
PALPASSGGLPIQMVLRTPSGFADLYAQMERIKAAAQKSGLFAVLDSDLTFDSRAVGVTI
DRNQANTLGVTMKDIADTLAVLVGENYVNRFNFKGRSYDVIPQVSRSERLSSEMLSRYYV
KTGSGQMIPLSTVIQVTNGSQANALSQFNQMNAATFSAVPAPGVTMGDAVAFLEAQKLPA
GFSIDWLGESRQYVQEGNRLAVTFGFALIVIFLVLAAQFESLRDPLVILVSVPLSICGAL
APLYLGFATLNIYTQIGLVTLIGLISKHGILMVSFANDLQRHEGLDRRAAIERAAAVRLR
PILMTTAAMVAGLVPLVFAAGAGAASRFAIGITVSTGMLVGTLFTLFVLPTVYTFVAKDH
GAAMRSERAQLLAATR*
>GCA_000452945.1_BPMSHR5858_genomic_00858_1 group_1127
MDNQRHKTLRSGMSARAARQLPGNRRLARLAGAAAAACAAGFGASTAHAQSSVVLYGLID
TSITYANNQRTHGAGSPGSPGWAVTSGALNASRWGLRGREDLGDGVSAIFALENGFSGAS
GALSQKGVDMFGRQAWIGLKSKEGGALTLGRQYDLILDFVTPLGASGPGWGGNLAVHPYD
NDDSNRNIRINNAVKYTSPTYRGWTLGAMYGFSNTAGPFGNNAAWSAGLSYANGPLKLGA
GYLRINRNPNAANANGALSTTDGSATITGGSQQIWAVAGRYAFGPHSIGAAWSHSATDRV
SGVLQGGSIAKLDGNSLVFDNFTLDGRYVVTPRLSLAAAYTYTMGRFDARSGETRPKWNH
MVAQADYAFSIRTDAYLEAVYQRVSGGNGIPAFNATIWTLTPSANGNQVVVALGLRHRF*
>GCA_000452945.1_BPMSHR5858_genomic_00860_1 mbtH_1
MDDELYFVVRNNEGQYSVWMDGRSLPAGWETVGEPATKQQCLQRIEQLWTDMVPASVREH
LNQHSGPGIDYAVR*
>GCA_000452945.1_BPMSHR5858_genomic_00861_1 group_1414
MPYDEILPIETLVRPKVSDFREHYLEKERPVKIARALDAWPAMQKWSLDYFENRFGDETI
GVESFQPDERGPGNNSPQGYVKHLRFQELKLKELIRILRTKPDHMYYMASHPFRKSFPNL
RADLAPHPYVQGHIEHIPGAHMDSYLWIGPAGTHTPIHTDPMPNFLTQIVGRKMVYLFPP
DQASKNLYIGQFERETFSPVDLEKPDLERYPNYRHCTPYRAIIEPGETLHIPRNWGHCVI
SMDISISISTFFITYPQLFRLVPEFFTQYVKRAFEGWRWKGMENERAGLNPPPR*
>GCA_000452945.1_BPMSHR5858_genomic_00862_1 mrsA
VSVGEAMDTEVRAAAQAVCLAFNENPEAVEPRVQAAIAAAAARINRYPFDAEPRVMRKLA
EHFSCPEDNLMLVRGIDECFDRISAEFSSMRFVTAWPGFDGYRARIAVSGLRHFEIGLTD
DLLLDPNDLAQVSRDDCVVLANPSNPTGQALSAGELEQLRQRAGKLLIDETYVDYSSFRA
RGLAYGENELVFRSFSKSYGLAGLRLGALFGPSELIAAMKRKQWFCNVGTLDLHALEAAL
DNDRAREAHIAKTLAQRRRVADALRGLGYRVASSEANFVLVENAAGERTLRFLRERGIQV
KDAGQFGLHHHIRISIGREEDNDRLLAALAEYSDHS*
>GCA_000452945.1_BPMSHR5858_genomic_00868_1 yecS_2
MPFSESVADAPSAVEAGYALSTRNVRIYLAYRVLTLAIIDRAIFVIFLMHKGFDAYQIGI
LQGVFFLANIVTEVPAGMFGDAFGRKRSVLLGLVAYCAYALGVIASDGFVPFVCLYALLG
VALALVYGSDTALLYDSLVVDGRVAHFNRVQLRANALGLISGAFAVLAGGLLQKVSWNAV
YAAYFAIYAAALGAWCLAIEPPEASAELSAGRKDVTKELFAFIRGHWRSVALPILGFTVF
AACTTPFFTFSQALFKENGFSVEHITWFFFAAQMLIGFVYLVMQRTMSFLGFYPVVLAST
LLTAIVLAAMFFNVAAVDFTGFFLVMIINPIVTVVANEYFNKRLPSRIRASFLSLIGLCM
SLTIAVMYFLYSYLAQYLAIYKVMASTSLIAACACAIFVVARCVDTKEQA*
>GCA_000452945.1_BPMSHR5858_genomic_00876_1 quiP_1
MASRTNRLPRWLKILPGVILLGALLVAAGAALFLRASLPRLDGDVRAPTLGGPMTIERDA
AGVPTVAARDRFDAAYGIGYLHAQDRFFQMDYLRRTGAGELAELLGPAALDFDREHRLFR
FRARAAAAFAQLPPDERRLLERYTQGVNDGLAALRARPFEYALLGEPPARWRPEDSLLVI
WAMYFQVQGTLASRDIARNWLTAHATQQQRAFLLPSSSGFDAPLDAPRIDEAPAPLPDAA
PDWFRAAGDGAAKRASLDFRSSVGSNNWAIAGSRSARGAAIVGDDMHLVLGLPNTWYRAA
FTYPGGAAPVRRAVGVTLAGLPAIVAGSNGHVAWGLTVGYADCLDLVPLERDGDDSRAFR
MSGARQIARRYVESIRVRGGASVSLTVLETTVGPVREIGGRPYAVHWVAQSPGAVNLGLA
RLADAVDVDGAMRVANTLGIPAENIVVGDRAGRIGWTIAGALPDRRAPRGGEGAAWRSLL
PPDAYPRVVDPSGGQLWTANSRQLAGGAYRLIGDGGTDLGARATQLRDGLTALGRTDEQA
AYRIALDDRALFIAQWRDRALRVLDDAALAGHPSRAEFRRLLEHGWTGRASVDSVGYTLA
RGFLYRLYDVTFDGLNARLKQVDAGADYELANLRWPAVVARLLDAQPPGWLPAGASSWRD
VQLIAIDRTIAALTADGAPLARASWGARNTLRIAHPFAGSLPLLGGWLTAPAAQMPGDSH
MPRVAAPDFGQSERMVVSPGHEEFGIFNMPGGQSGHPLSPFFLAGHDAWVRAEPTPFLPG
VARHTLRFAP*
>GCA_000452945.1_BPMSHR5858_genomic_00879_1 group_2286
MSDSVSIAKYMHRDCLHCPAPLVGTSFKPCHFADIVDNGLQDGFFEVHAENYMGKGGPPH
RALSALRETYPVSVHGVCLSIGGPSPLDAHHLARFRDVVARYEPVLVSEHLAWSSHGGAF
YNDLLPLPYTKATLDRVCAHVDQIQEAIARPLLLENPSTYVGFASSTMSETEFIAAVAGR
TGCGLLLDINNAFVSATNHGCSAADYLADFPLEQVGEIHLAGHSEQRDDEDEPLLIDSHD
RPVAGPVWALYASVIARTGPVPTLVEWDSQLPDWPVLRAQALEARRIMDAAVASPVEVAD
HAL*
>GCA_000452945.1_BPMSHR5858_genomic_00882_1 nrsf_1
LSPWRSNTPPCRSATLPVLRIFAMETRDLVTWLATDLTPVDPGAVLQRLYHALLVGFAGS
TALLVALYGIRSDMPQLILTTMFWVRFAFPLAVVFSATRLAERLGRPGARLCSVGLATAL
PVAAMLLATGGVLIATPPGYRLQLVLGTTWRTTVVDIVLLSLPPLAAALHAMKQLAPTRL
VLAGAGAGLLAGAQALVVYSLYCAEMSMPFWGVWYMLSLMLTMALGAMLAPIWLRW*
>GCA_000452945.1_BPMSHR5858_genomic_00883_1 nrsf_2
MTIAFGVRPDLSAVMRTPIFWAKLAFPICVAAGAIMATVRLARPGATVGAGRLVVGVGIA
AVWIAGLYAVATAAFGDRLAVVLGQTWRVCPFNIVLLSAPGFVAMFVAIRGLAPTRLRTA
GAAAGLAASAIATIAYCFHCPEMSPAFWGVWYLLGMALPAALGALVAPIWLRW*
>GCA_000452945.1_BPMSHR5858_genomic_00887_1 group_1416
MEPVSRNELTTRLADGLVTLLDVRPYDEFAQGHLPGALNIPLAALEARLSELPADTEIVA
YCRGPYRVLAFEAVTALRARGFKAARLEGGFPEWKAAGLVVETGAAD*
>GCA_000452945.1_BPMSHR5858_genomic_00915_1 group_818
LSTLIVLLPPREPAVPLQEWQWPELPFALVDKSGHTQRAGRAALALLPQAATTVLIVAAR
DLLMLEQALPPLKGPRLKQALPNIIEDQLIQDPQGCHIAVDPAALDGGRRVLAVIDRAWF
KFIVDAFTAAGHRHLRAVPVTRCLPPATRRDAAAAAETEAVADVALDRSAEHAAAADAPG
SGHAGATANAPAPAESIVAVALGLAATERAPSLAEEPAALLPQAPSAPRVELALARGALG
EGFAAPVSSAVATLEALANGTPLELYELGEPGAEPQLASVRPLDDKRLLPGAAIWPFDAL
VRRALDSRFDLCQFEFEFAPWRFDRATFMRLRLPLALAATTLAIAVIGANAHWWKLSRER
DALAAQITETLLSTFPKTTTVLDPAAQMTRQLDRLRIAAGELSPNDFLSLASGLARSLGP
LPPNGIASLDYHDRRLDVGFKPETKVDPDFAQRLARNGLTGEIDSSTGKWTIRSRS*
>GCA_000452945.1_BPMSHR5858_genomic_00921_1 group_4514
MNALSIRILSLALFAVLCATATYWVVTLSAHQAPLPAAAARTPVRTEDAAALFGGQLTRS
PVQDIHLFGILALQRGAAAIVGIVGIGGDAPHAVSLGSEIAQGAKLAEVRDRSIIVERNG
ARSEIFLPANTPSPAIYVR*
>GCA_000452945.1_BPMSHR5858_genomic_00923_1 hxcR
VTQALAQGAPDQPAAGAPSPIAARLLPYGFAKAGQILIAHQHADTLEVWISERTSSAALA
EVARNFGAISLQRMPADELAQAINHAYARQDGSAAQIVGEVEGEVDLSRLMQDIPEVEDL
LESEDDAPIIRMINALLTQAAREQASDIHIEPFENASVVRFRVDGTLRDVVRPKKALHGA
LISRIKIMAQLDIAEKRLPQDGRITLRVGGRPVDVRVSTLPTGHGERAVLRLLEKDAQRL
NLEALGMGRDTLVQFDKLISRPHGIVLVTGPTGSGKTTTLYASMSRLETATTNIMTVEDP
IEYDLSGIGQTQVNERIGMTFARALRSILRQDPDIIMIGEIRDLETAQIAVQASLTGHLV
LATLHTNDAASAVTRLTDMGVEPYLLASSLLGVLAQRLVRQLCPACKEERHEGGRAVWHP
VGCDKCGHSGYTGRRGVYELLVIDDSIRSLIHRNAADAEILATGRANGMRTLRDDAERWL
AAGATSLEEVLRVTGGA*
>GCA_000452945.1_BPMSHR5858_genomic_00929_1 group_4516
MKSAFSFLPNWPLAPDAIFWAGFALFAAGLCGELCYRAWRLPRITGYAVIGLVAGSFGFG
VIDASTDDTSRLLVNVALGLLLFELGSRLDLRWIRRNPWLIASSLAEATLTFVLVLAVLL
LLKVPGMIALVLAAIAISTSPAMVIQLKTELRAEGQVSQRLITLSALNSVYAVVLTKLVT
SWLHQEAYGNVFATILQPIYLLAGSFIVAYLFARACNYLFRHVAATMRDEHSFVALFGLV
VLAIAVAQVLKLSTMLTLLLAGIIVKNLEARPQLWPEHFGTAGWLLTVILFVLTLTSFEG
QDIAAGGLIAGALIATRFLAKLVGVLAFAKPSGLGVKQGIALGVSLVPMSALAYLLVDDT
YQLYPNFDPRLRAVVMCSIVVLQLIGPLVVYRSLSAVGERRDAS*
>GCA_000452945.1_BPMSHR5858_genomic_00933_1 namA_1
MSALFSPFTLRGVTLPNRIVISPMCQYSAENGEARAWHMIHLGHLALSGAGLLCIEATAV
EPDGRISAADLGLWSDATEAALEPVLAAIRKHSPIRVAMQLSHAGRKASSDVPWKGGQLI
AVDEGGWRPHAPSGVPHREHEPTPLALDAAGLERVKRAFVESAKRAARLGIDAIEVHAAH
GYLLHQFLSPLANHRDDAYGGSLENRMRFPLEVFDAVRAAFPDDRPVGVRVSATDWVSGG
WELDDTLAFAHELKRRDCDWIDVSSGGVSPLQKIPLSPGYQVPFAQAVKRAVGLPTIAVG
LISDPAHANRVIEAGDADFVAMARAMLYDPRWPWHAAAQLGATVSAPPQYWRSQPREYKA
LFGDASFGQR*
>GCA_000452945.1_BPMSHR5858_genomic_00934_1 group_4518
MSSRRIVVRRSGVHGKGVFAAVPIKAGERVVEYKGERISWKEALRRHPHDPDDPNHTFYF
ALEEGGVIDGKINGNSARWINHSCAPNCEAEEVGGRVYIHALRDIDEQEELFYDYGLVID
ARLTKKLKREYACHCGAATCRGTLLATSEEDEGKKKKAKKGGKDKKKKK*
>GCA_000452945.1_BPMSHR5858_genomic_00940_1 lrp_6
MLELDHFDLALLDVLQRFGRATHQQLGEEVPLSPSQIGRRLQRLEAAGVIDGYRVVLRPE
RLGLGVTAFTSLKLKHHGDSIIEQFQQQIDVLPEVLECHAVVGDADYLLRIVAPDLNALS
SFVMKKLMRVPGVDSVRSNIVLTTFKRNGPLPLGHLASGAPAA*
>GCA_000452945.1_BPMSHR5858_genomic_00941_1 group_4519
MARNIEIKARAREFEQLRERAAELATDAPLFYRQQDFFYDVPRGRLKLRRFEDGTPAELI
FYQRDDQDGPKASYYTRSPVTNADAMHALLATALTTRGIVTKERHVYLAGRTRIHLDRVD
GLGDFVELEVVLGPDDDEDGGHAEAHAVFAKLGVPSDDLVAVAYVDLLNAQAEPTA*
>GCA_000452945.1_BPMSHR5858_genomic_00942_1 yofA
MDLTLLRAFVAVAREGNLTRAAAQLHLTQPAVSLQIKNLQDLLGIALFVRTSRGLVLTRD
GQALLPHAERALDAAADVKRAAAALRHEVRGRLRIGTILDPEFLRLGGFLKRLVEAHPQI
ETALRHGMSGWVLDEVRARELDVGYYIGRPEEDDPRDAERFHTVTLTQFRYRVLAPAGWK
ERVQRARTWRELAALPWIWTPAASAHQRLLSRRFADAGAQPVKVAEVDQETSMLDLVKSG
VGLTLARDSIALREAHAHALTIVEHVAVPAELTFVTLADRRDEPAIAAALRLIDEQWAI*
>GCA_000452945.1_BPMSHR5858_genomic_00943_1 bauC_2
MTGSTHSNDSRVRALAHFIGGRALDGASDRYGDVFDPALGTVTARVPLASGAEVDAAVAA
AAAAFPAWSETSPLKRARVMFKFKELLDRHHDELAELITREHGKVFSDAKGEVMRGIEVV
EFACGIPNLLKTDFTDQIGGGIDNWNLRQPLGVVAGITPFNFPMMVPCWMFPVAIACGNT
FVLKPSERDPSASIRLAELLKEAGLPDGVFNVVHGDKTAVDALIAHPDVAALSFVGSTPI
AEYIHTQAARRGKRVQALGGAKNHLVVMPDANLDQAVDALVGAAYGSAGERCMAISVAVA
VGGVADALVERLAERAKALKIGNGMNADVEMGPLVTAAHRAKVSAYIDAGVAAGAKLVVD
GRRHVVAGGENGFFLGGTLFDDVTTDMSIYREEIFGPVLAVVRVPDFASAVELINAHEFA
NGVSCFTSDGGIARAFARKIQVGMVGINVPIPVPMAWHSFGGWKRSLFGDHHAYGEEGVR
FYTRYKSVMQRWPDSIAKGAEFTMPVAK*
>GCA_000452945.1_BPMSHR5858_genomic_00954_1 group_4522
MDREAKPVLAVQNLSVRYGKVEALHRAALTVGAGQIVTVIGPNGAGKSTLLNAVMGALPT
NGHAAGTVLYRGDDVDALPVEQRVALGMCLVPEKRELFGTMSVEDNLVLGAYRRKMAGDA
SFLDQLEPVFSLFPRLKERRKQAAGTLSGGERQMLAVGRALMGKPDLLMLDEPSLGLAPL
IVKEIFHIISALRGTGVAMLLIEQNARAALQISDYGYVLETGEIALEGPAGELAQNPQVI
ETYLGLARKAV*
>GCA_000452945.1_BPMSHR5858_genomic_00962_1 atpH_2
MQAFIANIQGLTAIGIGIIIGLGAIGACIGIALMGGKYIEACARQPELINPLQTKMFLLA
GLIDAAFLIGVGVAMLFAFANPLLSKLAG*
>GCA_000452945.1_BPMSHR5858_genomic_00966_1 atpG_1
MAGMKEIRGKIKSVQNTRKITKAMEMVAASKMRRAQERMRAARPYAEKVRAIAAHMSRAN
PEYRHPFMVANDGVKTAGMILVTTDKGLCGGLNTNVLRASLQKFKELEEQGQKVEATAIG
GKGLGFLNRFGAKVISQVVHLGDTPHLDKLIGAVKTQLDLYSEGKLSAVYLAYTRFVNTM
KQETVIEQLLPLSSEHFDTNDGTPATSWDYIYEPDAQAVVDELLVRYVEALVYQAVAENM
ASEQSARMVAMKAASDNAKTVISELQLSYNKSRQAAITKELSEIVGGAAAV*
>GCA_000452945.1_BPMSHR5858_genomic_00968_1 atpC_1
MATIKVDVVSAEEQIFSGQAKFVALPGEAGELGILPGHTPLITRIRPGAVRIESESGDEE
FVFVAGGILEVQPGAVTVLADTAIRGKDLDAAKAEEARKRAEETLQNAKSDIDLAKAQSE
LATAMAQLEAIQRLAKIRGKH*
>GCA_000452945.1_BPMSHR5858_genomic_00972_1 hemE
VAQTLLNDTFLRALLREPTDYTPIWLMRQAGRYLPEYNATRARAGSFLGLAKQPDYATEV
TLQPLERFPLDAAILFSDILTIPDAMGLGLDFAAGEGPKFAHPVRTEADVAKLAVPDIGA
TLGYVTDAVREIRRALTDGEGRQRVPLIGFSGSPWTLACYMVEGGGSDDFRTVKSMAYAR
PDLMHRILDINAQAVAAYLNAQIEAGAQAVMIFDTWGGALADGAYQRFSLDYVRRVLAQL
KREHDGARVPAIAFTKGGGLWLEELAATGVDAVGLDWTVNLGRARERVAGRVALQGNLDP
TILFAPPEAIRAEARAVLDSYGNHPGHVFNLGHGISQFTPPEHVAELVDEVHRHSRAIRS
GAGS*
>GCA_000452945.1_BPMSHR5858_genomic_00975_1 putA
MASTTLGVKVDDLLRARLKDAASRLERTPHWLIKQAIFAYLERIEHGQLPPELSGHTGVA
DLSDSHADGDDDGAPHPFLEFAQSVQPQSVLRAAITAAYRRPEPECVPFLIGEARLPASL

ASDVQAMAAGLVEALREKSSGGGVEGLIHEFSLSSQEGVALMCLAEALLRIPDRATRDAL
IRDKISKGDWRSHVGHAPSLFVNAATWGLMITGKLVTTNSEAGLSSALTRLIGKGGEPLI
RKGVDMAMRLMGEQFVTGETISEALANSRKYEARGFRYSYDMLGEAATTEEDALRYYASY
EQAIHAIGKAAGGRGIYEGPGISIKLSALHARYSRSQQERTMNELLPRVRSLALLARRYD
IGLNIDAEEADRLELSLDLLEALCFDPELAGWNGIGFVVQAYQKRCPFVIDYLVDLARRS
RHRLMIRLVKGAYWDSEIKRAQVDGLEGYPVYTRKIYTDVSYLACAKKLLAAPDAVYPQF
ATHNAYTLAAIYHLAGQNYYPGQYEFQCLHGMGEPLYEEVTGRDKLNRPCRVYAPVGTHE
TLLAYLVRRLLENGANTSFVNRIADKTVSVKELVADPVEDALKVVPLGAPHAKIALPREL
YGASRVNSMGLDLSNEHRLASLSSALLASAHHPWRAAPMLADDALAHAAPRDVRNPADLR
DVVGTVSEATPDEVSAALAHAVAAAPIWQATPVDARADCLARAADLLEAQMHTLMGLIVR
EAGKSLPNAIAEIREAVDFLRYYSAQIRDEFSNDTHRPLGPVVCISPWNFPLAIFMGQVA
AALAAGNTVLAKPAEQTPLIAAQAVRLLRDAGVPAGAVQLLPGNGETVGAALVADPRTRA
VMFTGSTEVARLINKTLSSRLDPDGRPIPLIAETGGQNAMIVDSSALAEQVVADVLQSSF
DSAGQRCSALRVLCLQDDVADRTLTMLKGAMRELALGNPDRLSVDVGPVIDADAKRTIDT
HIAAMKDKGHAVTQLEMPDACAQGTFVPPTIIEIGNVDELKREVFGPVLHVVRYRRSGLD
KLLEQIRATGYGLTLGIHTRIDETIAHVISRAHVGNIYVNRNVIGAVVGVQPFGGEGLSG
TGPKAGGALYLQRLLATRPAGLPRSLEAALVADAPQEGETRDNPSAALTALRDWLIEQRE
PALAARCDGYLSHVLAGATAVLSGPTGERNTYTLGARGTVLCIAATASGARAQFAAALAT
GNRALFAGAAGEALAAALPAQLKAHAGVRKQADAAFDAVLFEGDSDELLALVKEVAQRPG
PIVSVQGAAAGAFENGGDEDYALERLLTERSVSVNTAAAGGNANLMTIG*
>GCA_000452945.1_BPMSHR5858_genomic_00977_1 aidB_2
MNLIAALERAARATPDKPFLRDDGATLTYRQMQQRSRRAAEVLASLGVAPGERVAAMCLN
TPAFFDLLLGAWRIGAVVVPVNHKLQAPEVDHILRHSASRAVLFDAALAPVLANVEHDAL
RLATEGEAAGAAPFDRLVADAAGIAPATPDDAALAEILYTSGTTGRPKGCMHSHRTVALA
AATSALALSMTERERTLMAMPVWHASPLNNWFGGTLYVGGTVVLMREYQPLRFLQTVEAE
QVTLYFGAPVSYTLPLDTIADFASFDLTSVRAWLYGGGPIGATLARRLMRAYRSEAFYQV
YGMTETGPAGTVLHPFEQAMKPGSIGCQGTPGVDVRVVTAAGGLARPGEIGEIRLRADSM
MLGYLDDPGATRAAFDDDGWYRSGDVARVDDDGYLFVVDRIKDMIVTGGENVYSKEVEDV
LNAHPAVAEAAVVGRAHPEWGETVIAHVVLREAAGAARAGQAESAAGVDADALRAFCATR
LAAYKIPREYVFAERLPRTPTGKLQKYLLRARGGA*
>GCA_000452945.1_BPMSHR5858_genomic_00984_1 group_5664
MAQSASRYSPLALIVLAMLTEAPMHAYRIQQLVKLRGKDEVVNVKQRNSLYQTIERLQRD
ALIAVRETERDGAFPERTVYEVTDAGRDTARMWLREQLARPAREYPSFPAALSVLPLLSV
EDARRQFEARVAALEAELARLDETQNAALAMQIPRLFLLDGELMRVTLEAELDWVRSVIG
HLKVGALTWSEAWLREVAARFAQADSPDSADSD*
>GCA_000452945.1_BPMSHR5858_genomic_00986_1 kdpB_1
MTEATRAENRPRTPEADRGGAQSGDARTSARAAGCCSHHHPHGEVETQANLAVQSSNRDS
SDHAHAPHSHANGDHPHDHDHDHDHDHDHDHDRDRDRDHDGAACCAPAPVAFAPLPGARKAA
GGRVRSAFRIMQMDCPTEETLIRKKLGAMSEVAALEFNLMQRMLAVEHVPGAEAGIAAAI
RSLGMTPEQADAGASGRGALPAPADAPRPWWPLAVAGVAAAASEAATWLQLPVWLAAALA
LAAVATCGLGTYRKGWIALTNGNLNINALMSIAVTGAMAIGQWPEAAMVMVLFTVAELIE
ARSLDRARNAIQSLMRLAPDTVTLRQPDGTWQPVDAAQVALGAIVRVKPGERIGLDGEIV
AGRSTVNQAPITGESLPVEKTEGDAVYAGTINEAGSFEYRVTAVASNTTLARIIHAVEEA
QGAKAPTQRFVDSFARVYTPIVFAIALVVAIAPPLVLDGAWRDWIYRALVLLVIACPCAL
VISTPVTIVSGLAAAARRGILVKGGVYLEQGRRLAWLALDKTGTITRGKPVQTDFEMRAA
DVDAALVRGLAASLAARSDHPVSQAVAAASAAQAGAGGVPRAKPASFADVADFEAIPGRG
VRGKIDGVPYWLGNHRLVEELDCCTSALEARLDELERQGKTVVMLIDGARVLGLFAVADT
VKDTSRAAVAELHALGIKTAMLTGDNPHTAQAIAQQVGIDDARGNQLPQDKLAAVEALAA
GGRAVGMVGDGINDAPALARADIGFAMGAMGTDTAIETADVALMDDDLRKIPAFVRLSRA
THRVLVQNIAFALAVKAVFVGLTVAGMGTMWMAVFADAGASLIVVGNGLRLLRRGQ*
>GCA_000452945.1_BPMSHR5858_genomic_00990_1 yhdG_3
MQTESNGRPGAGGGAARPALQQTLGTWQLWGIAVGLVISGEYFGWSYGWASAGTLGFVVT
ALFVAAMYTTFIFSFTELTTSIPHAGGPFAYARRAFGPAGGYLAGVATLVEFVFAPPAIA
LAIGAYLHVQFPGLEPKHAAMGAYLVFMALNIVGVQIAATFELVVTLLAIFELLVFMGVV
SPGFAWSNFVKGGWAGADHFSAGAFHGMFAAIPFAIWFFLAIEGVAMAAEEAKHPKRSIP
IAYVAGILTLVALAIGVMVFAGGAGDWTKLANINDPLPQAMKYIVGANSGWMHMLVWLGL
FGLVASFHGIILGYSRQIFALAREGYLPEWLAKVHPRFKTPYRAILAGGVVGIAAIYSDE
LIQFGGQTLTANIVTMSVFGAIVMYIVSMAALFKLRRVQPRMERPFRAPLYPFFPAFALV
AALVCLGTMVYFNALVASIFVAFVALGYGYFLATRAQREAAPADALLEE*
>GCA_000452945.1_BPMSHR5858_genomic_01003_1 gcvT
MRRRFAELPFCFRARAQGPAALPGTPMTVLKTTPLHAAHRALNARMVDFGGWDMPVNYGS
QIEEHQAVRTDAGMFDVSHMCVVDFTGPRVRAFFEHAIANNVAKLQTPGKALYSCLLNPQ
GGVIDDLIVYYFTEEFFRVVVNAGTAEKDIAWFNQLNEQGGFGLTIAPRRDFAIVAAQGP
NARAKVWDTVPCARAATSELKPFNAAQVAGTPFGDLTVARTGYTGEDGFEIIVPATHVEA
LWNALAERGVRPCGLGARDTLRLEAGMNLYGQDMDESVSPLDAGLAWTVDLSAPRAFVGR
DALEAHGSRAAFVGLILQKENGRAGGVLRAHQKVATPHGEGEITSGTFSPSMQESIAFAR
VPKDVAIGDTVHVQIRDKQLPARVVKLPFVRNGKVLAA*
>GCA_000452945.1_BPMSHR5858_genomic_01006_1 group_983
MGYHFATAALGPLLLAQGRHVRRVTPRLAEADGPRGGADGAGPPLRLLVLGDSAAAGVGV
ASQSDALAGQLVRALAPFHRVEWRLLARTGATTQDLLDWLDAEPDARFDAVVTSLGVNDV
TDGVPPARWLDAQAALVERLTARFGAAHVVLTAVPPMHRFPALPQPLRWYLGLRAARLNA
ALSRWAAGRPACEFLRVDMPLTRDAMASDGFHPGTQACAAWAGQAAAAIRRRVAAATRLQ
AAVG*
>GCA_000452945.1_BPMSHR5858_genomic_01008_1 rep
MSAGLNSAQSEAVRYLDGPCLVLAGAGSGKTRVITQKIAHLIEAKGFEPRHIAAVTFTNK
AAAEMRERVGKLLEGKTLTTPGKEGRKVPVNQLTVCTFHSLGVQILRQEAEHVGLKPQFS
IMDSDDCFGMIQEQIGTTDKGLIRKIQTIISLWKNGLVAPEEAMTIAANEDEHQAALVYR
NYVATLHAYQAVDFDDLIRLPTELFARNEPVRDRWQNRLRYLLIDEYQDTNACQYELLKL
LAGPRAAFTAVGDDDQAIYGWRGATLENLAQLGKDFPKLHVIKLEQNYRSTVRILTAANN
VIANNPKLFEKKLWSEHGMGDSITVTPCNDEEHEAESVVFRLSAHKFERRTQFRDYAILY
RGNFQARVFEQVLRRERIPYVLSGGQSFFDKAEIKDLCAYLRLIANADDDPAFIRAVTTP
RRGIGNTTLEALGAFAGQAKVSLFEAVYMGGIEARLSARQVEPLRIFCDFIQRLTDRADK
EPATVVLDDMMEAIHYEAYLYDAYDERQAQTKWQNVLEFLEWLKRKGTKPEAEQAVDGEA
EGFHNADGLADTGKNLLGLIQTVALMSMLEGKEEDPDAVRLSTVHASKGLEYPHVFLVGV
EEGIMPHSGGSDDGPIDDERIEEERRLMYVAITRAQRSLHLNWCKKRKRARETIVCEQSR
FIAEMGLDEAPPPTPEEAPMTPKDRLASLKALLQK*
>GCA_000452945.1_BPMSHR5858_genomic_01020_1 group_985
MSPFMLSSIRSRILVACLAIVIGSLVINTALNYFVANRYNRESISQNLSAVLTGHEAGIA
DWVASKTQMIVSVEDAAISPDPIPALKQIAAAGGFTNVYVGYADKTAKFSDPTGIPPDYD
PTGRPWYKQAAQAGKPVVTPPYVDVGTGKLVVAFAAPIMRDGALKGVVSGDVAMDSVIAN
VKAIHPTPGSFGMLVDRSGHIVAHADSKLTLKPVTDLSDDLSLDALAAASADENAAPIDA
HVAGAAKLMRARAVPGTDWLTVVALDKSDATAGMHSLLLVSIGTLVALAAVAALIVGAIT
GVAFRGLARIRDAMESIGSGTGDLTQRLPDSGRDEVAQIARSFNAFVSKLQEVMRVIRDA
SESVRHAAGEIASGNHDLSRRTESAAASLQQTAASIEEITSTVTQSAGAARQANDIATNA
ASVASRGGTVVSDVVSTMHEIEGASGKIADIIGVIDGIAFQTNILALNAAVEAARAGEEG
RGFAVVAGEVRSLAQRSAQAAKEIKALIDSSVTSVSTGATLVQQAGQTMSDIVGTVSNVT
TIMREISNAADEQTRGIQEVNRAVAQLDEMVQQNAALVEQSAAAASALQTQAVELADAVG
QFKVA*
>GCA_000452945.1_BPMSHR5858_genomic_01027_1 group_4531
MSVETWLAFAGASIVLLMIPGPTILLVIGDSLANRHRSSWSTVAGVAAGDTTAMTISLAG
AGALLAMSAAAFTALKLVGGGYLVYLGIRSILHARRIGAMDVRIERRSAKRRFLSAWAVT
ALNPKSIVFFVAFVPQFISPHETFARQCAILLPTFVCLAAINAACYARIARFAAARLTGA
AAQRRFGYGGGAALIGAGALTLGMSRS*
>GCA_000452945.1_BPMSHR5858_genomic_01035_1 vioA
MSALPLRAVEPWIDERIFVTQPHLAPLAEFLPYLEAIWESKVLTNGGPFHQQLEKALCDY
LGVRHLALFTNGTLALVTALQALRITGEVITTPYSFVATAHSLLWNGIKPVFVDIDPSTL
NLDPAKIEAAITPQTTAIMPVHCYGRPCDVAAIQKIADNYNLKVIYDAAHAFGVKTPDGS
VLEHGDMSILSFHATKVFNTFEGGAIVCPDAKTKQHIDHLKNFGFVDEVTVVAAGINGKM
SEINAAFGLLQLKHIDDALARRAKIDAAYRDALARVPGIRCLPPVDAPVANHSYFPILVG
DDYPMSRDALYQRLRDHQIYARRYFYPLISDFPMYRGLPSAQRGNLPVAADAAARVLCLP
IYPALSDASLERIVSLMAGAHA*
>GCA_000452945.1_BPMSHR5858_genomic_01037_1 group_4535
MDQMSLIEHVLSITREIDHAVQMADWTEAARLTEERSPYLMSITAEQTPASLALVRQLQA
IDTALLANAKAAQAELQVEYRTAMDRLSQAGAYQAAAQL*
>GCA_000452945.1_BPMSHR5858_genomic_01038_1 fliD_2
MSTPVTSTTQQQTNSALQQAAQSIISGSTGNSSMDVNSLVTALVNAKTAGQSAALSTSIA
TDQTTLSALGTLKAALTALQAGIGSLSDGTLTQKFTATATGTGLTATTGAGAVAGSYSVA
VTQIATSQTLSSGAFNATQQLGTGTLTLSVGGKSTSISIDSTNNTLSGIAAAINSASNNP
GVTATIVTGTDGAHLVLRSASTGAANVINVGVSNLSGDNGLSSLAVTSTASTTGGQSTIR
SGGSVAWSQSTSAQDAEFTVGGIAASSASNAVSGAIAGVTLNLTQAAVGATQTLNVTTDT
TAQATAITNVVNLYNTVITTMSSLSSLSGAGTSSQSAGPLLGDSTLNMIRNSLARVVGAG
VTTGGSTTSLASIGIKFADGSSSSQTDGALTIDTAKLNAALQNSPSTVAALFNSTNGIGA
QLNTTIQNYVQTGGVFDTRSNALNQDLKSLAQQQTRLASYASQLTSQYNAQFTALNTLMA
QMNSNSNYLTQLFGGSNSSGAMANNK*
>GCA_000452945.1_BPMSHR5858_genomic_01040_1 rpsU_1
MTTILLKENEPFEVAIRRFRRAIEKNGLIAELRERQAYEKPTAVRKRKKAAAVKRLHKRL
RSQMLPKKLH*
>GCA_000452945.1_BPMSHR5858_genomic_01043_1 yigL
MYKAIATDLDGTLLNSDHQLDPYTIDTVRRLAGSGVPFVIATGRHYADVAGIRDVLGIRP
YLITSNGARVHGPDDTRIHAQDVPADAVRQLVRPELVGTHGRVIVNLFTNDGWLIDRDAP
QLLAFHQDSGFRYEIVDMLAHDGADIAKVLYIGEPEDLAVVSGNLARRFGDALYVTYSLP
DCLEVMTANVSKGRALRVVLERLGVDPAHCVAFGDNMNDIDLLETAGYPFMMNNANPDLA
LRLPKVPRIGNNFEAGVARHLRALFALEDSIAH*
>GCA_000452945.1_BPMSHR5858_genomic_01045_1 group_19856
MSQYAKIKAQIVELQAQAEEVRRQEIAAVVAELQGKIAEYGLTAQDLGFAERARRGRPPK
KGPLPAKYRDPKSGNTWSGRGKPPHWIVGKNRERFLIG*
>GCA_000452945.1_BPMSHR5858_genomic_01047_1 flhD_1
MSATSEMLAEINEVNLSYLLLAQRLLREDKAMGMFRMGISEELADVLVNLTLAQTVKLAA
SNQMLCRFRFDDHALLSSLADKGRSSAVSHAHSAILMAGQPVESLR*
>GCA_000452945.1_BPMSHR5858_genomic_01055_1 cheR
MMPARAASRLDAPEPQERSGDAARDFEFTSADFTKIRALIHRRAGISLSEHKRDMAYSRL
ARRLRARGLNRFCDYLELLEREDDPSEWEAFTNSLTTNLTAFFRESHHFPILAEFVKRRE
QPVSVWCSAASTGEEPYSIAMTLVEALGESAARSASVIATDLDTQVLAKADAGVYAFEQV
KHLSPERLKRFFLKGTGAQAGRVKVRPELRAMIRFAQLNLTDADYGLTKPFDAIFCRNVM
IYFDKPTQSQVLARFEPLMKPGGLLFAGHSENFTYVTQAFRLRGQTVYELTRDGQSAAPR
VRARIAAAAAEEELR*
>GCA_000452945.1_BPMSHR5858_genomic_01059_1 cheZ
VNEPIDMAVAGANAGERLADAADLASDRILARIGQLTRTLRDSMRELGLDKHVEKAAEVV
PDARDRLRYVVAMTEQAAERVLTAIEVAKPVQERLQQEAAALDARWAKWYETPIERAEVR
GLMDDTRAFLRALPDATGATNAQLLEIMLAQDFQDLTGQVIKKIMDMVYLIEQQLLNVLV
ENIAPERREQFAATAAALAAEQAGGAMTPESLLNGPQINPEGKTDVVQDQAQVDDLLASL
GF*
>GCA_000452945.1_BPMSHR5858_genomic_01060_1 group_4544
VRSSRFRRLAATAAGFMCAWLAAPAHAGLGGAPMTPPAADTAATVRSIQRSIRAAGGAST
AAVGYTVRETTLGSGTVVREYVSTAGTVFALSWQGPVAPNLSDLLGAYFPQYEAGVKASR
AAHGPRAPAAIDTGGLVIHTGGHMGSYSGQAWLPQALPAGFSTNDIQ*
>GCA_000452945.1_BPMSHR5858_genomic_01066_1 group_4545
LDKRITDQAEGLRRLLAGRASRVVAVTGGPSGVGCTSTVANLAAALTALGKDVLVVDERA
NVRSIAATLCGSWLRDGEPVRHELGFAVCEASRLARVGYSDAQLDALGDGAADIVLIDAQ
LDANGALSALARDAHDVLVVTRVSASAITEAYACMKRLHYAHARAQFRVLTNHVQSAADA
RAAYENLAGVASRYLRVSLSNAGCVAADSLIERARGLAHTVVDAFPSAAAARDYRQIAAD
LLYWPMRPGPGIGRVRTGSSAFERGAAHAA*
>GCA_000452945.1_BPMSHR5858_genomic_01067_1 fliA
MYNAQGKISQDEVLTQYAPLVRRLGLQLVAKMPASVDLDDLIQAGMIGLLDAASRYKEDQ
GAQFETYATQRIRGAMLDELRSNDWLPRSLRKTSREVEHAVHQVEQRLGRSANETEIAAH
LQMPLDEYQSMLQDLHGSQLVYYEDFDRSADDEPFLDRYRADHSDPLSALLDEHLRSALV
DAIERLPEREKLLMSLYYERGLNLREIGAVLEVSESRVCQLHSQAVARLRAKLREQAWVG
TES*
>GCA_000452945.1_BPMSHR5858_genomic_01069_1 group_4547
MTVILTWIINALALLIITYLVPSIHIKSFGTALIVAVVLGLINAVIRPVLILLTLPVTIV
TLGLFILVVNALCFWFASSLLKGFEVSGFWSAFFGSILYSIVSWLLSALIFGQRNIG*
>GCA_000452945.1_BPMSHR5858_genomic_01070_1 metF
MNPIELSFEFFPPKTAEGVDKLRATRAQLAPLKPKFVSVTFGAGGSTQQGTLDTVLDMQK
AGLEAAPHLSCIGSSKDSLRAILDQYRSHGIRHIVALRGDLPSGMGEVGELRYASELVSF
IRAEHGDWFRIEVAAYPEYHPQARSPKADLENFARKVKAGANAAITQYFYNADAYFRFVD
DAAKLGVDVPVVPGIMPITNFSQLMRFSEMCGAEVPRWVARRLESFGDDKDSIRAFGADV
VTGLCRRLIDAGVPGLHFYTLNGAAATKTVCERLGL*
>GCA_000452945.1_BPMSHR5858_genomic_01080_1 group_4549
MSAAPTMPSREIYVSTDVEADGPIPGPHSMLSFASAAFTEDKELIATFSANLELLPGAAA
HPVQAAWWKTQPEAWAACRRDLQAPEAALAAYVEWVEALPGKPVFVAMPAGFDFTFMFWY
MMRFTGRCPFSWSALDIKTLAFAITGLPYRKAIKPRFPKRWFDDYPHTHVALDDAIEQGA
LFCNMLADLRALQTAKASAPPVEEADESAQNAADGPRN*
>GCA_000452945.1_BPMSHR5858_genomic_01089_1 group_4550
MQKGHSETDLHRGLGQRQMRLIALGAAIGVGLFLGSANAIKMAGPGIILSYIIGGAVIFV
IMRALGEMAVHEPVAGSFSRYARNYLGPLAGYLTGWNYWFLWLVTCIAEITAVGIYMGIW
FPDVPRWIWALAALATMGSVNLLTVKAYGEFEFWFALIKVVTIVVMIAVGIAMIAFGIGN
HGVPMGISNLWAHGGFFPNGAQGVLMSMQMVMFAYLGVEMIGLTAGEAANPQKTIPGAIN
SVFWRILLFYGGALFVILSIYPWNEIGTQGSPFVMTFERLGIKTAAGIINFVVLTAALSS
CNGGIFSTGRMLYNLAQQGHAPRTFGETTASGVPRRAILVSMAALLVGVLLNYLVPEKVF
VWVTSISTFGAIWTWGVILVTQMKFRASLAPAEERALRFRMPFWPYASWVALAFLVLVIG
LMAYFPDTRIALYVGPAWLVLLVVLYFVLKLEPEGGKAAGLERRYGRG*
>GCA_000452945.1_BPMSHR5858_genomic_01091_1 group_4551
MKSTQILASCALVGAVAALAGCAGSPANGPQAESPSGRIIYVASPRSPHDVESCLTSRVS
QARVAQRNPTTVLIGPYSADSDWTVTLAPTASSGTTIGVYRPRSGDGDPEESELRFHIAR
CAV*
>GCA_000452945.1_BPMSHR5858_genomic_01093_1 group_4552
MVTVTFRFYEELNDFLARPLRRREFAHACMRGASVKHAIEALGVPHTEVELILVNGESTP
FSHVLEEGDRVAVYPSFEAIDIRPLLRVRAAPLRVTRFIADAHLGGLAQLLRLAGFDTLY
DNHYPDELIETIAAREARIVLTRDRELLKRRTITHGCYVRALKPQAQLQELFDRLDLAGS
ARPFRLCLSCNAPLRRIDPAEAAGRAPQGVLQRHTRFVTCDVCRRVFWEGSHWRRMRALI
EHVSQPKPPPG*
>GCA_000452945.1_BPMSHR5858_genomic_01103_1 group_67
MLELSAEQVAGLAQIDERGFVERVRQDLVKENPAFADDDSLSSRLWTAYRAARALGIERD
ENVVAFLRLEAYAPGFYEKPATKAWLTRPGRSADARFHDYLRVIKWRIEHPDGGLEHGGI
GISGNRSGGGGAWADLGARWRRLVGRRDGGGNGESVG*
>GCA_000452945.1_BPMSHR5858_genomic_01111_1 arcA
MSQAIPQVGVHSEVGKLRKVLVCSPGLAHQRLTPSNCDELLFDDVMWVNQAKRDHFDFVS
KMRERGVEVLEMHNLLTETVQNPAALKWILDRKITPDNVGIGLVDEVRAWLEGLEPRALA
EFLIGGVAASDIAGAERSKVLTLFRDYLGKSSFVLPPLPNMMFTRDTSCWIYGGVTLNPM
HWPARRQETLLVAAVYKFHPAFTDAKFDVWYGDPDRDHGMATLEGGDVMPIGRGVVLVGM
GERTSRQAVGQLAQALFAKGAAERVIVAGLPNSRASMHLDTVFSFCDRDLVTVFPEVVNR
IVPFTLRPGGDARYGIDIEREDKPFVDVVAQALGLKSLRVVETGGNDFAAEREQWDDGNN
MVCIEPGVVVGYDRNTYTNTLLRKAGVEVITIGSSELGRGRGGGHCMTCPVLRDPVDY*
>GCA_000452945.1_BPMSHR5858_genomic_01151_1 fpuB
MSARMRATRFSTFAPAAIAHYLEHVWLGAPDACDEPALAPDGAEPADLISAASLALRRDA
LVGAMVALYGGDARAHARALVSQWSKYYFNLAASAGFAAALLLGRPLDMAPQRMRVALRG
GMPVALLFEADALRPAQAEPASRYAALVDHLRATIDTLAALAKLSPRVLWANAGNLLDYL
FEQCAHAPRAGADAAWLFGPVDSRGEANPLRLPVRRVKPCSARLPDPFRARRVCCLRNEI
PGEDQLCGSCPLLLTMCDDALAKQDAAH*
>GCA_000452945.1_BPMSHR5858_genomic_01154_1 group_2759
MTQLSMPAAAACPTLGDLRVEPGLPTVVSPRDGADIALHEAAPLLREIADDVLERAGGVL
FTGFRVASIETFQRFAADFGDPLIGYEFASTPRSQVEGAVYTSTEYPPHRSIPLHNEQSY
TREWPLRIWFHCALAARTGGATPIADSRAVYRALDPALVARFAERELLYVRNFGQGLDLP
WQQAFGTGDPREVERICAARGIDCEWRDGDDGEPLLRTRERCQAVARHPRTGELVWFNQA
NLFHLSALDEDMQEALVDAVGIENVPRNVYYGDGAPLEPDALAEIRAVLDGQRIVFPWQT
GDVLMLDNMLSAHARDPFEGPRKVVVAMARSYREKRGA*
>GCA_000452945.1_BPMSHR5858_genomic_01155_1 mbtH_2
MTQAQQPASAPADADDVVYAVVVNDEEQYSIWPTFRAVPAGWREAGVRGPKAECLAHIEA

VWTDMRPASLRRHLDAARAS*
>GCA_000452945.1_BPMSHR5858_genomic_01156_1 group_2305
MAEVLARPAPAKPLPILASNGAFAPAAAPVLRAPSRPAARTPGAREHGALVDVLVANRPM
LVKLARGFVGCASRAEDVVHDVFVKLVDFPNQDAIRQPVAYVTRMVRNASIDACRRQTLE
NTYHADEDDGLDVPSPELSPEAALVVRDTLRHVYDALAQLPARSRAAFEMVRLREETLQS
AARALNVSQTLVHFMVRDAERHCVACVDASERGLACPAFCGARARTVKKCVRDSSIE*
>GCA_000452945.1_BPMSHR5858_genomic_01160_1 eltD_1
MTNRNDETQQMTAIVCHAPEDYRVERVAKPRANARELVIRIGACGICASDCKCHAGAKMF
WGGPSPWVKAPVIPGHEFFGYVEALGEGAAEHFGVALGDRVIAEQIVPCGTCRYCKSGQY
WMCEVHHIFGFQREVADGGMAEYMRIPSGAIVHPVPLGISLEDAAIIEPLACAIHTVNRG
DIQLDDVVVIAGAGPLGLMMTQVVKLKTPRRLVVVDPVEARRALARAYGADVTIDPARED
APAIVRALTGGYGCDVYIETTGVPAGVTQGMALIRKLGRFVEFSVFGKDTALDWSIIGDR
KELDVRGAHLGPYCYPVAIDLLARGLVTSKGIVTHGFSLDEWDEAIRVANSLDSIKVLMK
PRG*
>GCA_000452945.1_BPMSHR5858_genomic_01167_1 group_1097
VTLLEAGGRGRAYLREAPASVDRALAHARTLADARDAVTRAVWGAYVAVLDEAATGRRLF
LPDPLHSVRLYYRTDEDGRVDVDPQAANLLRGASIDWNLDYLIEFACTQFGPLGETPFAS
VRAVPPGCALVVDADGRCAIERAWLPRASAGDDVRASCAAALDEVYSRFARSHPNVCAAL
SGGVDSSAGAILLRKALGAQAPLAAVHLFSPSSPDCYERDLAARVADSIGARLICIDIDR
HLPFSERIVPTPPATLSQDMLFLGIDRAVSRAIGASSVLLEGQGGDLLFKAVPDANAVLD
ALRGKGWAFALRTAEKLAVLHNDSIPRVLLMAAKIALRRRLFGQDAAGTRQTMSKLFASH
APRAAASPTRMRRAGAPLDESVAMLDRFVSIMTPVTDAAYTNRLNPYLAQPVVEASFGLR
SYDSFDHRNDRIVLREIAAAHTPVDVLWRRTKGSFGIGFVKGIVSHYDAFRELIRDGVLM
RSGQLDEAELEHALKAVRVGQNAAAISLALVGCVEIFCASWQNFMTNRSAAIC*
>GCA_000452945.1_BPMSHR5858_genomic_01176_1 acrE
MKYEWARTRRLSAALAVAAFVAAGCGKHESEHDAAAPREASVVTVKKTSVPLSVELPGRL
DAYRQAEVRARVAGIVTARTYEEGQEVKRGAVLFRIDPAPFKAARDAAAGALEKARAAHL
AALDKRRRYDELVRDRAVSERDHTEALADERQAKAAVASARAELARAQLQLDYATVTAPI
DGRARRALVTEGALVGQDQATPLTTVEQLDPIYVNFSQPAADVESLRRAVKSGRAAGIAQ
QDVEVTLVRPDGSTYARKGKLLFADLAVDPSTDTVAMRALFPNPERELLPGAYVRIALDR
AVARDAILVPRDALLRTADSATVKVVGQNGKIRDVTVEAAQMKGRDWIVTRGLAGGERVV
VVDAAQFEAGTTVKALERGAAAQPASAPGRRST*
>GCA_000452945.1_BPMSHR5858_genomic_01181_1 mltF
MKFKPFATWLLGLSLVAAHAAAHADRLDDIRKAGVLRVATFDSNPPFGFVDPRTNRIVGL
DVDYAQALADRLGVKLKLQPTNPANRIPFLTSGKVDLVLANFTITDERAKQVDFSVPYFA
SGQQFLAKKGVLKSADQLNRLRIGADKGTTNEVTLREKYPQATIVAYDDTPFAFAALRAG
NVQAITQDGPKLIGLLANVPDKQNYEVPAFSISNDYMGVGAPKGEPRLVAFVNETLTDLE
ASGRAGRIYDAWFGPNTKTPLARIFKIGDKR*
>GCA_000452945.1_BPMSHR5858_genomic_01182_1 group_1420
MNERRAAFARRRERGAVVLWFVLFLPVLLLFGAFAIDLPRVAAARNELQNAADAAALAGA
ASLEAGAGAPAWAAAASAAAAALSLNASDGAALSSGDVQTGYWNVTGVPAGLEPTTLAPG
EYDVPAVQATVTRAPNQNGGPLSLLMGGLLGLVGTPAAATAVAVAGAPATVGAGGLFPMV
IDQCVLDQYWDARAGAPRVDPTTGAPYEFQVGNGRTYGGTCYAGQWTTFLVNANDVPTVR
GLMAHGNPTPLSIGDSIWIEPGVKTALYYDVPVGVTVVVPVATQISSKTYVPVVAFAAFY
VDASDGANLKAITGHFVGGYKIPASASGIGPAYGAYVAPRLAY*
>GCA_000452945.1_BPMSHR5858_genomic_01241_1 tolC
METEMFETENLGASSKFRHAGFHLTTKSFRLRANVAYVTVCAVLAAFGTGSESARAFCLD
EAYQHAISNDPKFLQARAEYDAARQKFPQARAQMLPQVSAQLEWGRYGTHANLFGIDVSG
NSTAAYGAAQLSQALFNMPYLYDMSRAKEFEESARQQLEVAKQELIMRVANACFDLLSAR
EKLQLADDEVGALTRLESDTRRMAQLGMKTIGDTAEIEARRSLAQSDEALARTDVEARRA
RYETLLGSAIDFTRWPRLAMHGTSPRIPTGDYQPQDNPSYQQAYRDLRVARLASKRINAE
HLPSVDLFATYSRGLNPNLRGLTDKNDFHQSAVGVQVTIPIFSGGSVHYRKIEADHVATQ
YQNRLREVEQQLSTDHRETLAALQSIGTRIRALQQSLQAARLAYDSSMKAHQVGYSTTYE
TLNLRTDISNIRQKLFESYLDALKLQLKLKGILGTLDEQSLVAVDSFLASNAAPADQKSE
*
>GCA_000452945.1_BPMSHR5858_genomic_01245_1 srmB
MHATPARPAAGALDAFHPVVAAWFSDTFAAPTGAQALAWPHIKAGRSTLVAAPTGSGKTL
TAFLSALDDLVRDALAHDGALPDETLVVYVSPLKALSNDIHVNLEAPLAGIAAEIARRGL
PVPQIRTAVRTGDTTQAERVAHRKHAPHILVTTPESLYVLLSSESGRRMLASTRTVIVDE
IHALANAKRGSHLALSLERLDALVGRKLPRIGLSATQKPVEAVARFLVGGRDDAPLDCAI
VDTGHTRTRDLALELPPVPLGPVMANDVWERLYDRIAELAAAHRTTLVFVNTRRMAERAA
RHLADRLGADAIAAHHGSLAKEHRFDAEQRLKHGKLKLLVATASLELGIDIGDVELVCQL
GSPRGIAPFLQRVGRSGHHIGGVPKGRLFPLSRDELVECAALLDCVRRGELDVLRIPRAP
LDVLAQQLVAEVACREWDEHALYRCVTQAAPYATLARETFDEVLKMLAEGFTSQRGARAA
YLHRDVVAHTVRGRRGARLAAVTSGGTIPDNADYAVLLEPQGVNIGTVNEDFAVESLAGD
VFQLGNQSYRIIRIETGRVRVEDAHGQAPNIPFWLGEAPARSDELSAGVARLRARLDALL
ARGDAAGAGAALPVNAASAPAPEREAAAAGQGARARAAAPHGTPALARRALPAALPAAPP
AVCARAAPSAAAPSTAIPSAAAPTVAAMPVPPEAELRAEPRRLAPALAWLTGKLDLDAEA
ARQIADYLARARAALGALPTQDTLVMERFFDESGGMQLVIHSPYGSRINRAWGLALRKRF
CRSFNFELQAAATDDALILSLSHAHSFALDEVWRYLRASSAEHVLIQALLDAPLFAARWR
WNATTSLALPRFAGGRKVAPQLQRMKSEDLLATVFPDQVACAENIVGEREVPKHPLVDQT
LDDCLHDAMDTEGWLALLRRIEAGDVALVARDLPAPSPLAAEILNAKPYAFLDDAPLEER
RTQAVLARRWSDPESADDLGALDAGAIEAVRDEAWPLVREPDEMHDALVGLACIADDEIA
SHDGWRACVAQLAAGGRAARLALAGGRGLWIAAERVACMRALYGADVALDPPLAPPPGFD
APWEPEAALVDVIRARLTGFGPLTAAALTAPLALPAASVAIALAALEREGYVMRGRFTPG
ARDDEWCERHLLARIHRYTVKRLRREIEPVELADFMRFLFHWQRVAPDTRGLGGDALAAV
VHQLEGFEAAASAWEDEILPARLADYVAGGLDEPCRAGKLTWTRLGAPAKAASAPVKTTP
VVLLPRRELAIWQALRDPAAQPALSARAERVRATLAAYGAMFFDELAAHARLLPVELERA
LGELVAAGLVNADSFAGLRALLRPAAQRGAQRPSHARRGGALIGGMDDAGRWALVQRLPL
APLDFGEGGSEDEAANDDDAPSGDGTPGAGPEAARRGGVREGGAEADARIARAMRATPPQ
AGRHAIAPAALEHVARTLLRRYGVAFWRLLACEAEWMPSWRELLPVYRRLEARGEIRGGR
FVAGLAGEQFALPDAIPILRELRRRPAEGLMLCVSGADPLNLTGTLLPGERVPALAGNRI
LFRDGVPIASLVGGQFAFARELDALAQDRARARLARHR*
>GCA_000452945.1_BPMSHR5858_genomic_01248_1 group_2317
MRVMVMVKATNESEAGKLPTREQFEAMGRFNEALVKAGVILAADGLHPSAKGKRVRFSGS
ARSVIDGPFAQTGELVAGFWLWKVASMDEAVEWVKRCPNPMESDSEIEIRPLYELEDFGD
AFTPELREREARLRDEIDEQGKKAP*
>GCA_000452945.1_BPMSHR5858_genomic_01360_1 acdA
MQFTEEHEAISRTVKRFIEQEVNPHVDEWEEAGIFPAHEVFGKLGALGMLGLSKPAAYGG
GGLDYSYELVMAEALGVCAAGGVPLAIGVQTNMATPALARFGSDALRDEFLAPAIAGRQV
SCIGVSEPGAGSDVASIRTSARRDGDDYVISGTKLWITNGTQADWMCCLANTSDGPPHRN
KSLIVVPLKSKGVHIEKKIRKIGMHSSDTAQIFFDDVRVPRRNLIGEEGQGFTYQMLQFQ
EERLYGAAAALVVLDRSIDETIDYTRARKIFGRPVLDHQVVHYRLAELKTEVEALRALTY
RATELYVQGGDVTTLASMAKLKAGRLAREVTDSCLQFWGGMGFAWESSISRTYRDTRLFS
IGGGADEVMLGIICKKLGTLPRE*
>GCA_000452945.1_BPMSHR5858_genomic_01361_1 paaF_3
MTGTTFSTPAPLHVAQADGVRFATLNRPARRNALSDDLVAALDAECERAAGDAAVRALVL
RGAGGMFCAGGDFGGFSAMMREPAPAGEPDPIAAANRRFGALLEKLAALPVPTLAVVEGA
AAGGGCGLAAACDRVLIASDAQLSLPETSLGLPPAQIAPFIVARAGAARGCWLMLTAGRL
SAADALDAGLADEIAGADGIDALLRRALARVLACEPAALRATKRIVADALRRERGAALDA
ASGEFAAALRSGAVAEGLAALSGKRAPRWASDAPLPPEAR*
>GCA_000452945.1_BPMSHR5858_genomic_01362_1 accA1_2
MSTRRFTRLLIANRGEIAVRIARTARRLGIATIAVHSDADARSPHVAACDDAVAIGGATP
AESYLSIDKLIDAALTSGAQAIHPGYGFLSEHAAFARRAAEAGLVFVGPRADAIDAMGDK
ARARRRMAAAGIAVVPGYDGDDQREARLLSEAERIGFPVMLKAAAGGGGRGMRRVDARDA
LPGALKRAVSEAQQAFGDGRMIVERAVIEPRHIEVQVFADAHDRIVHLGERDCSVQRRHQ
KIVEEAPSPAVDAALRERLGATAVAVAREIGYVGAGTVEFLLDREGRFYFMEMNTRLQVE
HPVTELITGQDLVEWQLRVAQGDALPLAQQDIRLDGHAIEVRLCAEDPADDFLPRTGSVL
TWRPGRHARCDHALADGLAISPFYDSMLGKLIAHGGSRAEAIDRLAHALDDTVLLGVPTN
RAFLARVLRHPVFADGRAVSTAFVAAHFPDNDSRRCAPTQAVWAIAAWLSTLAADRADAL
PGAWRGWRNGAPLPVPYRLSLANANAGADHPDARRGIVTAERQRAVVHAQGGSPIVLDAA
RPPCPGEPSTVSIDGRAHACCFAIAHGRLWLQVDGVDHAFAIHNREGRASAAAADSDGVL
RAPMNGRVIAVDIDEGATVSAGQTVMVLEAMKMEHAITAPFAGRIASLGARPGEQVSPGQ
VLARLEPQAG*
>GCA_000452945.1_BPMSHR5858_genomic_01363_1 kce_4
MNHPVVITCALNGIFTDPKQFNVPVTPEQMAREAKGAYDAGASCMHVHFRRQEPDKGHLP
SWDPRLAAEIAQAIREACPGVIFNQSTGIVGPNVDGPLACIRAIRPEIAACNAGSLNYLK
VKADGAWAWPPMLFDNPVEKVASFLTTMAETGAVPEFECFDTGIVRCIDMYARAGLFEGR
SNYNFVMGVESGMPAEPELLPILTRLLRADSTWQVTAIGRANIWALHRRTAELGGQLRTG
LEDTFYLPDGARATSNAQLVDAIAAIAREAGREIASPRDARRILGTRAQAAPHP*
>GCA_000452945.1_BPMSHR5858_genomic_01370_1 menE_3
MTTLRIFDARVHVLPDVVAKLVWKNYARDAWKIRYELLAEQSIAFLKAHGVARAAGICYA
GQPGIAPFLNDFVAQLASRHPAFLVPFGTVHPHDRDAARETARALDELGFAGLKIHCHLL
KIAPDDDALAPIFEALAERGKVLNLHSGAFAKNNAQMDEIRRFCNVERFRRAMRRTPELK
VVVPHIGYDEVQAYLDLLDEFPNLHFDTAMAFGGHRIATGEAVPDVRPLRETRYPRGAHP
RLPAPWKPALEQLVPQMRDRPDRFLYGSDFPLIPYDWDVEIEQLKRYLPADVLRKVLWDN
ARALFDGDAHRASDANDAGAHEDASAAAPVSAGQPATEPTR*
>GCA_000452945.1_BPMSHR5858_genomic_01372_1 menI
MQENLDADYFNRFGADYLPGLLGVVVKQVEKNHVLAELTIRQAVLAHGILHAGTIVSLAD
TACGYGCLASLPEGASSFSTLELKSNFMGTARDGVIVCDARAIHSGRTTQVWDATVRHRE
TDKLLATFRCTQMILWPTK*
>GCA_000452945.1_BPMSHR5858_genomic_01378_1 group_4586
MSSDILYTDSEISSSPSAFSTRFAYDSDWVAANNSINTSLIFKHNLKCIPYPLCLFFSPD
QEAVYPLIWNYYGPNAGNPVSIKLDQTKVTLSIGSGIPLHGYFDPQTGGWTYWRSGFFRV
TIPSQ*
>GCA_000452945.1_BPMSHR5858_genomic_01379_1 group_4587
MNSHIAAVDLKKAYRLINHGPTVLVSAHHDGVDNVMAAAWACALDFSPPKLTVVLDKAAK
TRELIEQSGRFVIQVPTAAQLQLTQEVGSHSLASRPDKLRRAGVELFSIDGHDAPFVAGC
SAWLACTLVREPHNQQAYDLFIGEVDAAWSDTRAFKNGRWQFEQADPAWRSLHHVAGGHF
YAIGEALSAREADRHDAEAAL*
>GCA_000452945.1_BPMSHR5858_genomic_01380_1 group_4588
MKLLLVGSTGLVGRHVLELALADPRVDGVTALARRALPAHPKLHALRVDFDRLPEDAPWW
RADAAICTLGTTMRAAGSRPAFRRVDHDYPLEVARIARRHGTPTYVLNSALGADPSSRFF
YNRVKGELERDLAGLGFASFTAVRPGLIGGQRDAFRAGERAAVLALTLFGPLLPRGWRLN
PAQRIAGALLDAALNPAPGLHVVASDQLV*
>GCA_000452945.1_BPMSHR5858_genomic_01391_1 group_1325
MSSPARHSPPQTITKDSLGCTSSGFLTDIERETALLRFYRKHTTGLRLEQVSTPASGRGV
LIGISLSGGHRRKILRGRRSVTHDFRADSVYVRDFSEDYRADMMSDFDFALVELSPSFID
GLAERGRPARIAGVAPTLAHDDRLLGELGRALALALQSGDAADAMLVDQLGIAIGTHAMH
AYGGLRADEPKQRRRLSAPLERRAKEMLAAGATSVDEIARACRVSRGYFINAFSATTGKT
PHQWLIEQRIEAAKHLLAHGDWTLARIAEHCGFSSQSHFTQSFAKAIGLPPGAWRRRARA
*
>GCA_000452945.1_BPMSHR5858_genomic_01392_1 rhaR_10
MGRPADTLASIPPSPTMNCTDLAPTSLREAFRGATIETIDARVEGYASVVLTRVRHAAHG
FGFVEFMPPADGYLVGMSLTPPAARAPRAAPDAAADADGDADAPHRAACVAVQILQDDEP
FRADLLQPFDMLFHALPRRTLAELAADLRMGGVRELASPAGGRDAVLQSLVGVLLAASAG
SSARSPLLAGHVARAMQIHIVQQYGVAAPSAPAKGGLAGWQLERAKAILTENIAGEVPIS
QVASACGLSRSYFIKAFRQTVGTTPHRWLLEHKIERVKRSLLSQSAPIADIAHQCGFSDQ
AHLTRVFTSMIGTPPAAWRRVNRP*
>GCA_000452945.1_BPMSHR5858_genomic_01414_1 group_2334
MYPWIWLWAPNFYFPLSGGVDQTIQPDLFDAINSRAGNGEIERKAFQVASYGRQLGLIIE
VLLDMAKKQEALSPEAKQSMERLEAIQGEIEQIKREAPTSIAADLAARLLQLKTQNPAEF
VRLRAQLLGIVKDGA*
>GCA_000452945.1_BPMSHR5858_genomic_01417_1 group_1708
MSFFTQFEGSRLYIGPVASGDAGWLAYLLPRSSPSTEEIDLDLALKQLPGSFLFSFTAPL
LSTTPLLDAFIGAVDQVIARSPADRGFLWLMDPTAPTISSVPLMGIAADGSYVSNALQAP
IFETLSLNVRAGVTLQLDGDTLRLGTANGIFFSGASAPGAPQASQIPAGTLPLSGAARGC
IRFPVFLQRQFLHDGWNWGFQFQIPHDDPVRSAIGEWLPLAFGHLPNPGDMLGFDASYDP
SDPWNAHAIAADRSVLVFTGQNQDGTATVLSSCYATTYGELLTLAPVAGPLDMQPDAARF
LFNAGAPDDKGNHDFQIAPEGDFVIHAHGPSDGGTYDLICGLQGTEFIAFQPFVDGGYAG
DRLRFTARSPAYAPRYPFTEVSPVGAPVDVNAPLLDATYTSSWGTVARAPGAPGAAAYVA
QPKGAALYGRDALIHSLGEKTLFGPMTPSVTLPSSAFPVVPYAGSSAGDGVTALTAEQMA
DFERQVVSPTRRRRIADGGAGVASSKAHALKLADMATATPYNTTTPSGLLVTVDAAGQWS
KILLGQNLSPTLRQMSFCNPGAELQQAFATSDLFLVAANAAHLGAFSADGSGACGPSATF
ANRINVGDWSFEAAVGQNNRYGDYADVMIVKGRRGPLWDPQNPAASLVASPDMWTQKDVF
AAPSDLVPNPAPPPDETVAAPNPDELTALSYWLQRYFQDALDQAEPEYFAKFNTIAKDPA
WTGILFLRMKISSIPDDLVGIVAGVQAPELFNAHHFGIEISQIANDPAAPDIALTTSSSM
FGLIYYVDPAYIAPPPGIDPRPVPPTAGTDYQFRVLTLKALFENTAVQRFQSYAQLTLNS
LFDMPVVSMGEGGNPYATLVLRGSYQDNGGRAVYSLGTTATSTFYFDNNLFTKIEVDSAQ
MSTRDSGANGQIVSWFALSGFLDFAVVQANDSSGAFDILSFGSHAGEDQLRRGLAYSNLG
VRMTFPVADPATIAFAFSASEIRFDMQTSTPRPDSLFVNFALGLQGLVQGSAQTSPEADG
FLNVVTDATLTGVNGDRWYGLAYQLNMGTPGNLAGTIGLTSGLLTAWAPVSADAGRYRAS
LGIQLPGTGGGANLISLQNVLRLSIGQMWLTYDRDKRSFLLMLTEIALKFMGLLKIPPSG
STLFYLFGNPQSEGKASGLGWYAMYNNTPASRSKSARLLGKARGWTE*
>GCA_000452945.1_BPMSHR5858_genomic_01419_1 group_19871
MPLARRDRALCAFVAAPLLWGLLAGVAAAQALAQPAASTVSVCAAVITGSSGTTVDVSYA
GLPGNQPKLYGDFVALWQSSVVPWSIAPAARLGIPTDAELGTVVMTGVSISATPYTVGYA
VGPEVDDVCASALLAADGSTGVVDTVTLQLASVGATSLTFRYHTLSGYLPATAGNWVGLW
RGRASPYNAAASVARVKVAADVTDDSVVMEGVTLSPGEIYTAIYFMGEPLTTAAVLLNFV
APAR*
>GCA_000452945.1_BPMSHR5858_genomic_01422_1 mauG
MLLPGAPPARVVDTIGNGTPQVSSKIDASAARFVPDPTLVALGRRIFFDTRLSEPRGMSC
AGCHDPGRAFAPTLSAAALAGPGVPEGSRPGRFSRRNAPSLLYVRYVPRRHFYQDDDAPA
PSPFGGLFSDGRADTLAEQIRGPLFDPNEMNNRSPAALLRKVDATELAPALAARFGDGVR
LDPAQLVRALGASVEAYLQSDEMAPFTSRFDAYLRQRTPLDAQQMRGLALFKNPDKGNCM
SCHTLSDTSSRPERSLFTDFGYDAIAVPRNRALPANRDPRHFDNGLCDTARRLRWPEPGQ
WCGYLRTPSLRNVALKQTFMHNGVFTSLRDAVAFYNTRSTDPRHWYHGAATFDDVPPAYR
GNINVNSTPMNRRPGTPPALTEAEIDDLVAFLGTLTDARYAAGAPPHLKIHDSQAFTIAP
*
>GCA_000452945.1_BPMSHR5858_genomic_01432_1 group_4601
MTSNLHDKPDSPCIGVCSTLFDEVCKGCGRTALEVANWVFMTDAEKQAVWERITREGTAM
RFQYDKL*
>GCA_000452945.1_BPMSHR5858_genomic_01438_1 group_4604
MLISLVLGSIVGAVLGLTGAGGGILAVPALVVGMGWPMQQATPVALVAVAGSAALGALEG
FRRGLVRYRAAFLMAAAGVPLTTLGVRVAHRLPQRVLLALFALVMLVVAVRLLRQALRHA
AGEPACSPLCVGRVNPDTGRLVWSWPAGIALASTGALTGLMTGLLGVGGGFVIVPMLRKF
TNVTMHGIVATSLMVIALVGTGGVIATIASGTPAPLDVTLWFTATTALGMAAGRWVSRYL
SARHVQLGFAAALVCVAAGLLAKAALGF*
>GCA_000452945.1_BPMSHR5858_genomic_01441_1 norR_2
MNQRDAIPIVPISPPDVGVLVSYLEQDPQPMIVLDPSYRILAANAAYQRQFGIAGEAHVG
RRCYQVSHHYDMPCDQAGEHCPMKQAGESRGLNRVLHIHHTPRGPEHVDVELRPIFDARG
AVIAYVERLTTVRSASAKPSGEGLVGGSDAFNAALSALQRVAPSMLPVLLLGESGTGKEL
FARALHEASARAMGPFVVVDCSGIAETLFESELFGYEKGAFTGASARKPGLVETAQGGTL
FLDEIGDVPLSMQVKLLRLIESGTFRRVGGVEALCADFRLVAATHKPLKAMIGDGRFRPD
LYYRISAYPISLPAVRERPGDMPLLVDSILRRIAALGPVAGQHFVVAPDALARLEAYAWP
GNIRELRNVLDRACLLTDDGVIRVEHLPDEVAGGARIEPGAPAKLSDDELARIARAFDGT
RRALAERVGMSERTLYRRLRALGIATREA*
>GCA_000452945.1_BPMSHR5858_genomic_01445_1 group_676
MRLKIGELAKRCGLTVRTLHHYDAIGLLKPSARADNGYRLYDRNDIARLHQIQALRRFGL
ALADVGAYLAQPGTPLASIVSRQIAMLDRQIEQAARLRERLVQLHRELADGAQPELADWL
KTLELMTMYDKYFSDEELARLPMYRNRQAPDPEWTALVDAVRALMDAGVAPEDERVRALS
VRWMTMLVRDTNGDPRIFAKLNLMHDREPSMQTHVGISPALRDYVLRAFSETKMRIYERY
LSPEETRFMRANYGKRAMQWPQLMADVRDAIEAGVTPSSPQGRALAQRWLELFRHYAGDD
PATQLKFREALANEPELMTGTWADDALLGFVREAMQHLAPARR*
>GCA_000452945.1_BPMSHR5858_genomic_01446_1 czcA_1
MRVSIPSLPSLAPRALAAAAAIAFAGALAGCALFAPSDAPPAMPSPAHYGATPLPERTAA
AQGLAQRFDVGAQPVPDWWKRYRSPALDALVDEGLRNSPTLAAAEKSLSAAREQLRAQIG
SSLLPTIDAGGQATRERALGMPAFGPQTVLYNVFAGQLQASYTLDLFGAARFANRALSKR

VDVNAYQLESARRALAANIVTAAVTSAALAEQVATTERLVALAREQASDAAARHRLGAAS
RTDMLNTRQSAAALEASLPALRQQAQSARHALAVLLGRTPDAAPANLALADLHLPQDVPV
VVPSTLLQARPDIRAAEAGLQAAAAEVGVATAQMFPQLSLSASMGHAGFKWPAMLSGAGA
IWSVGASLTQPLFHGGALLAQRRAARASYDAAVDQYKQTVLAAFQDVADKLAALENDASA
LDASNRSAVAARGAFDDTAARVRLGALPPSAARASEQQYRNARLDEIRATGTRLVDTARL
FQAMGTPIDGGPDARRVGGMSGSNDVSDVTGSSGVGGAHRVIDIHEAGAAQRADDAKRTG
DARMLRANGKTEAAQASVTASAAADAPAAPAPAAAASGANAIRIQARTPPTAGPSAGPSA
GAIDAASATGATSTARPANGRDTTVESNPTHSPS*
>GCA_000452945.1_BPMSHR5858_genomic_01449_1 group_19341
MLDFPAMKPIRLTREQSKDLTRERLLSAAHAIFTKKGYVAASVEDIASAAGYTRGAFYSN
FRSKAELLIELLKRDHEEAEADLQKIFESGGTREQMEAHALEYYSQFFRNNPAFLLWGEA
KLQATRDAKFRARFNEFVKEKRDRFTHYILTFAERVGTPLLLPADVLALGLMSLCDGVQS
YHAADPRHVTGDAAQQVLAGFFARVVLARAPD*
>GCA_000452945.1_BPMSHR5858_genomic_01452_1 cmpR_1
MTPDQLITFAAVAEHRNISRAALALHLSQPAVSGQLRQLQDEFGEPLYQRDGRGVRLTPV
GEALARHASQLRDTFAQARAYRDAVRGLERGTLRIGASTTPASYLLPYLLAAFQPLAPEV
AIQTMSGNTSDVVASLASLDIALIEGPPGDALPAGTTVHEWREDEIVAIVPASHPLAAPG
APGRARATLAALAAYPLVLREAGSGVRQLVERAFARAATPLAVAFEIAGVEAVKEAVRAG
MGVGFVSAMSLRHDDAALAMVSIAPEPLTRHFSILVPHGATPSRVATRFLELCIAQARAA
*
>GCA_000452945.1_BPMSHR5858_genomic_01453_1 group_1806
MLDRPETVSYLLASALFFALAVPLAAQRIAPNRFYGVRTRATLRNPALWYRRNRVFGVAL
MITSATFIVVAEYCRLRGVRLPDAALLAAFVAEIAVPIAVCFAAGRPPETPETPSRRAGD
GRRR*
>GCA_000452945.1_BPMSHR5858_genomic_01455_1 kdgK
MQVPLDVVTYGEAMAMFVAAEPGPLDSATHFTRRIAGADLNVAIGLARLGFRVGWMSRVG
ADSFGRYVLDTLAREQVDASCVTVDARYPTGFQLKSRATDGADPSVEYFRKGSAASRLSL
DDYVPDYVLGARHLHLTGVAPALSDSSRELAFHLARTTRAAGKTVSFDPNLRPTLWPSPE
AMARTLNALAEHADWVLPGLAEGRQLTGLDTPADIARFYLERGARGVIVKLGAAGAYFRV
ADGREGEIAAERVEHVVDTVGAGDGFAVGVVSALLEGRAIDEAVARGNRIGALAIQVIGD
SEGLPTRAALDRIENLSNARDRLEAPLAR*
>GCA_000452945.1_BPMSHR5858_genomic_01535_1 ompA_2
MNSTNVFGGIFVGMLSVALLGCASVEKPAAYGEPSPMFIFASNEGFVAPSKMAFPQGIPI
DPDALKKLQLGMSKDQVRDALGDRQYNGEDLPIWHRWDYLFQLRAPDGAVHQCQYQVRFL
GGLAYRGYWADSACRDIAARYADAAGLGVIAVAPCNPSDVAAPLPERVELPTDTLFAFDK
GGFEDISADGRRQLGDLVASIKAKIFSINHLIVTGYTDRLGSDEHNARLSSERARTVADY
MIAEGIPAAKITAVGRGAADPVVVCNNGEQPELIRCLQKNRRVEIRIKQK*
>GCA_000452945.1_BPMSHR5858_genomic_01536_1 group_2343
MAPRGREPIVPLPPAASGAPGQRAEHAAGLMTLEKMIALVGERDPRVLVSELSARAAHER
AAKAWAGHYPQLSVEYALGREGNWNAQGGPTSLSSVNTDNFHGGQVNLSWELYAWGATQK
RYEEAYALAKAEEDRYRRAWDLIVMEAISTYVNMYRLRKQLEITRQTLLGHSKLLKVVED
RVGAKLAPESRRSDALLRVQEIMADREDILSQLNEVDTTYRTLTGLEPPADLAEPQLAKE
YEAKLACLSFDDNLKLAYSIHPQIQANLRTIEAAAYAVEGMKAERLPKFNLYARYNNNRL
FSTQGSTLPFTGRGYEIGVNVVWTLFGSAYGNSIAERIFLKEKSEADNERAVREVERNIH
IARDALAAAKGRTAILDETLRLSDEVLNQRQHSFSVQAMSDEAVNGLSNAYYSRARAATA
LLNLHMREVVASYTLVESSGLMNEVFPAPLSVRGPIGPAPVRNFFPGRVAGMNPEDGMPL
ERNVMPDLLTRKVVLTDGMSTTAEPQSGTAPTPASSAGGVPASASDAATARSLEVPPVML
SPAREAPLDNGSPQPPAGDAQAPAAATGTTP*
>GCA_000452945.1_BPMSHR5858_genomic_01547_1 vgrG1_2
VSRVFTLDSPHGDGLKFHTLDGSDELGRLFEFRIEALIDSHSLSLKDLLGKPVTVRIEQQ
GRSTRYLNGIVARASLAGRRAERYYGYELLVRPWLWLATRRSDCRIFQNKTVPEIVQDVL
STYGFPIENHLAESYVPREYCVQYNGTDAAFVSRLMEFEGIYYWFRHAEDTHTLMLSDAM
SSHTALPGYETIPYIARDRTVIADVEHIDGWLPARQVSVGKHQTTDYDYTKPRADLSSQK
VDPRGHDHDSFASFEWPGGYRDDAPGAHYSRVRLEEQQAEHERASADTDVRGAAPGYLFT
LAHCPRADRNREYLIVRCQYRLQENAYASDQGTEAVVHQTMMLVRPSSLPYRSPRETPRP
RTNGPQTATVVGPPGEEIWTDQYGRVKLQFRWDRYGQSNQDSSCWVRVSSPWAGGGFGGV
QIPRVGDEVVVDFLNGDPDEPIVTGRVYNGERMPPWGLPGSATQSGLLSRSSPGGTTEHA
NAFRFEDKKGAEQLWMHAERNFDAETEADHSLSIEHDHTHTVGNDETMQVRNNRQRSVGQ
DETVNIGRHRVAQIKGNETQGGVGDRTRTVGKKETVTIVSDREATIGGNHMETVVKDKTE
SIGEGKTLNVVKHYQTNSKSMKTTVAEHHVEEIGERTSTIAHAHTLTVRGEHAVSVGANH
TTNVQHQVLVNAGDQRALVCGKSSVLMKSDGTILIQGVNVESSGSDSHSVRGKKVTTSAS
VEHTVEGTVLKLNP*
>GCA_000452945.1_BPMSHR5858_genomic_01552_1 shlB
MTTATRFAVLPITALITLAAQAQQTPATIDQAAAARANAEQNQQVQQRRDAQRRDATVQA
PGVRSDVPSPEAYPVLPAETPCFRIDRFALDVPDSLPAAAKVQGASALPMDRFAFARDWL
AHYAGQCVGKQGVDLIVKGLSQAILTRGYITTRVLVPGQDLSTGTLKLALIPGVIRHVRF
EDDKLRGTWKTAFPTRDGDVLNLRDIEQGLEQMKRVSSQDVSMRIAPGDRPGESDVVLDA
KRGKPWTVVASIDNSGTRATGKLQGNVSLGIDNPLGLNDIFNVGFNQDLEFGDKRFGSHG
WNAFYSIPWGYWTGTLSAYTSTYFQPLAAVNQTFVASGNMKTVDFRLNRVLARSRNDVFG
AQVRLTRRFGDSFIEGTTIPSSHQNATFLEFVLNDRHYFGSSQFDGSLAYRQGLGWLGST
DSMFAVEGGQTYRFKMVVLDANLSTPFVIGTQPFKYVTTFHGQYTGNTVSYLDSVTIGSR
YTVRGFDGERLLAASRGFYWRNELQAPIARTGLSAYAGLDYGRVWGPEPVALVGTQLAGA
VIGVKGSVSTRFGAYAYDLFAGTPVYKPSGFETARVTLGFQLTAQF*
>GCA_000452945.1_BPMSHR5858_genomic_01562_1 frc_1
MSGNVSKRLRASSGSEIGRAAAGASDAAGAAGTAGAFADAAHPLPPAGGLCMSSEGGVPP
GTAIADATATASDARTAFAANTKDAKDAKDAKDAKDAKDAKDAKDAKDAKDAKDTKDTKD
TKDTTSRPSLAAANHATGPVRPGALAGLKIIDLSRVLGGPYCTQALADHGARVIKIEPPA
GDETRGWGPPFLDDDAWYFTGVNRNKEGIVLDLSRDEGRAILWRLLEDADVLVENFKPGT
LARWGMDYERDLRPRFARLVHCAITGFGADGPLGGLPGYDAVIQAMAGLMSVNGDAGGDA
TRIGLPIVDMVTGLNALAGILLALAERERSGVGQSIDIALYDCGVSLLHPHLPNWFGSGR
TPARSGNAHPNIAPYDSYRTASAPIFLAIGNDRQFARLCAHLGVPALASDPRFIDNRSRC
AHRAELKASLETRLAGRDGETLARELMAEGVPCGPVRTIDALACDPHTLHRGLVVEMGRY
RGAASPIKLSRTPATYRTAPPTLGRDTRAVLDALGIDAATQARLADAGVLKEAKPMAT*
>GCA_000452945.1_BPMSHR5858_genomic_01563_1 abaF_4
MTFQHDERAERAPVGQPARAALAAFVGTTIEWYDFYLYATASALIFGKLFFPSGDAFFST
LASFGTFAVGFFARPFGGLVFGHLGDRVGRKKALVATLVIMAVGTVGIGLLPTYAQAGAV
APVALVALRIAQGIAIGGEWGGAVLMASEHASADRRTFVASFAQLGSPAGLILALLAFRL
VAGIEHDAFLAWGWRLPFLASAALLVVGLAIRAGVDESPEFARVKAQRRTAALPVADVVR
DAWRTLLLCLGANVIGVAGAWFVNTFMLNYTTQTLGLDRALILDCLFVVAFIQLGTQLAS
GWFAQRIGTGRFLTAAAALAIASPYPMFALVSTGRPVAIVVGIAIAVMCMSSSYAVMAGF
MSTAFDVRVRYSAISLSYQLCAALAGGLTPLAGTLLAHRFAGQWWPLAAFYSALAVVSLA
CIATLERGKRVGVGERRGALAKP*
>GCA_000452945.1_BPMSHR5858_genomic_01572_1 group_4617
MSKTVADVMQLAKDEDVKFVDFRFTDTRGKEQHVSVPVSAFGEDKFESGHAFDGSSIAGW
KGIEASDMLLMPDPNAAFIDPFYEESTLVLTCDVVEPADGKGYERDPRSLAKRAEAYLKS
TGIGDTAYFGPEPEFFIFDSVQWNTDMSGCFVKINSEEAPWSSAKEFEGGNTGHRPGVKG
GYFPVAPVDQFQDMRSEMCLLLEQIGIPVEVHHHEVAGQGQNEIGTKFSTLVERADWTQW
AKYIIHNVAHSYGKTATFMPKPVVGDNGSGMHVHQSIWKDGQNLFAGNGYAGLSETALFY
IGGIIKHARALNAITNPTTNSYKRLVPHFEAPVKLAYSARNRSASIRIPHVSNPKGRRIE
TRFPDPMANPYLCFSALMMAGLDGIQNKIHPGEAADKNLYDLPPEEDAKIPTVCAGLDQA
LEALDKDREFLTRGGVFTDAMIDAYLGLKEQELAKFRMTTHPIEFEMYYSL*
>GCA_000452945.1_BPMSHR5858_genomic_01573_1 glnL
MVLKNLIKARAGHPERLSDDDRLVRSGLLTGLEALPTVVLVLDRRTLRIAFANPSAEAML
DISRRQLSQMPWGEIFPNASELATTITAIGEERFHATHLDTVLDRPGREPLHVHAIVGFL
EGAPEFVLVELFENERQSRTDREERIHDLTAVNKQLIRNLAHEIKNPLGGIRGAAQLLEF
ELGERERDELREYTQVIIKESDRLQTLVDRLLEPHRHPHIVGDVNIHEVCERVRAVMLAE
FPRGLTIERDYDVSVPDLRGDKEQLIQALLNIVRNAAQALRERIAQGDARIELRTRVARK
VTIAKRLYRLALDLRVIDNGPGIPDDIRDRIFYPLVSGRDDGSGLGLTLAQTFIQQHDGM
IEVDSRPGRTEFQILLPLDH*
>GCA_000452945.1_BPMSHR5858_genomic_01577_1 narX
MAPALPDSPAPWRHRLSTRIVALSAIAWALVLAMVGGTLWLSWQLEGAGAAINDAGSLRM
RATRTYVELSGVGTEPRRQLAVELDAIDGTLARLRRGDPSRPLFLPNTTVIQHQLDIVTD
RWNAVLRPLAAGALTNAPRADAAAAYLVALPGFVDEADLLVREIESDNARKTTWLRLSQI
GLGLLACSGTVAVIYLLFLWIILPVMQLRDGLKRMAAREFAVRLPVQTRDEFGDLAHGFN
RMASELQEVYAGLEERVQQKTAQLAAQNRELSALYEITAFLNRPQAVDEMCAGFLSRAIA
QFDADAGSIRVTDPTGEKLHLVIAEGLSTELTELERCMPVDDCFCGTVTRADTAVLHDLR
GSAARAPTPCSREGFAAVAVFKVMTQDAVLGSFSLHYRDPSRLPDAERRVLETLGRHLGI
ALDHVRLSASARQLAVAEERNLVAQGLHDSIAQSLNFVNLQTQMLGDAIAHDNLADAREI
VPMLKHGVEQGYADVRELLLNFRTRLTQGELKAAIEETIARFEKQTRIACALNYRETGGA
PLPPDEQLQVLFILQEALSNVRKHAHARHVAVDVENGAAFRLSVADDGCGYDPAALAARA
DTHVGLTIMRERAARLRATLTLTGAPQRGARVELVLPADARQAA*
>GCA_000452945.1_BPMSHR5858_genomic_01588_1 tmoT
MNSPVTTSQETVFVVDDDEAVRDSLRWLLEANGYRVQCFSSAEQFLDAYQPAQQAGQIAC
LILDVRMSGMSGLELQERLIAENAALPIIFVTGHGDVPMAVSTMKKGAMDFIEKPFDEAE
LRKLVERMLEKARNESKSVQEQRAASERLSKLTAREQQVLERIIAGRLNKQIADDLGISI
KTVEAHRANIMEKLNVNTVADLLRLALSKKQA*
>GCA_000452945.1_BPMSHR5858_genomic_01591_1 aceF
MSQAIEVKVPDIGDYKDVPVIEVLVKPGDAVEPEQSLVTLESDKATMDVPSPSAGTVKEV
KVKVGDAVSQGSLIVLLDGAQAAAQPAQANGAATSAAQPAAAPAAAPAPAAAAGGGTVDV
KVPDIGDYKDVPVIEIAVKIGDTVEKEQSLVTLESDKATMDVPSPAAGVVKDIKVKVGDA
VSEGSLIVVLEASGGAAASAPQAAAPAPAPAAPAPAPAPQAAPAAAPAPAQAPAPAASGE
YRASHASPSVRKFARELGVDVSRVTGTGPKSRITKDDVTAFVKGVMTGQRAAPGAAAAPA
GGGELNLLPWPKVDFSKFGPFEAKPLSRIKKISGANLHRNWVMIPHVTNNDEADITELEA
LRVQLNKEHEKAGVKFTMLAFVIKAVVAALKKFPTFNASLDGDNLVFKQYYHIGFAADTP
NGLVVPVIRDADKKGLVDIAKEMAELSKAAREGKLKPDQMQGGCFSISSLGGIGGTHFTP
IINAPEVAILGLSRGQMKPVWDGKQFVPRLTLPLSLSYDHRVIDGAEAARFNAYLGALLA
DFRRIIL*
>GCA_000452945.1_BPMSHR5858_genomic_01594_1 pbpG_2
MKAQLFAPLRMMQSVALGTAVSVAVSLMVATAAVAPADAFAATAKTAQSAKGKKSAAKKS
LRAASSSAEPRAKGARKRVTYVSNGRHRSGVRRVAFQPRPPSVGQAFGLHETPDSLALRS
SVAYVVDQNTSEPLFDKNSHAVVPIASITKLMTAMVVLDSKSPLTDQIDVTDEDRDYEKG
TGSRLSVGSVLSREDMLHIALMASENRAAAALSRYYPGGRPAFIAAMNAKAKSLGMTDTH
FENSTGLSSQNVSSARDLVKMVNAAYQYPLIRKFSTDRSYEVNTGKRTLAYNSTNALVRN
PSWDIGLQKTGFINEAGECLVMQTTIHGRPIVMVLLDSFGKYSRFADASRVRNWLDNGGG
ERLTAANTSTGGT*
>GCA_000452945.1_BPMSHR5858_genomic_01596_1 lutA_4
MRVGLFVTCVVDLMRPEIGFSALKLLKQAGCDVVVPPAQTCCGQPAYNSGERAIARDLAE
KTLAEFESLDYVVVPSGSCGGMIRVHYGELFRDDPELMARYGRLRTKVYELTDFLVNVAR
VALEPGAFKGPVTYHDSCSGLRELGVKAQPRALLAQRGVAVAEMKDCEHCCGFGGTFAVK
YGDISTAIVDEKCANVRATGAGAVVLGDLGCMLNIEGRLRRTGDRDTRVLHVAQVLAGDV
*
>GCA_000452945.1_BPMSHR5858_genomic_01598_1 group_1813
MRKTISMYWPLAIVVPLAAAAYLYLSEAAPRRAPAVARDTSAELARAVSYGFVDDASQPL
PAASHVAPLGDPKAL*
>GCA_000452945.1_BPMSHR5858_genomic_01605_1 hscA_4
MAPAPHQRRLAVGIDLGTTNSLVAAVRNSIPEALPDDAGRVLLPSVVRYLDKGGRRIGHA
AKEEAAIDPRNTIVSVKRFMGRGKAEVEGAANAPYEFVDAPGMVQIRTVDGVKSPVEVSA
EILATLRQRAEDTLGDDLVGAVITVPAYFDDAQRQATKDAARLAGLNVLRLLNEPTAAAI
AYGLDNGAEGLYAVYDLGGGTFDLSILKLTKGVFEVLAAGGDSALGGDDFDHLLFEHVLA
QAGLEVAALAPEDVRLLLDRVRGAKEALSAAPQARVDVKLSTGEKLAQTITRDTFAALVE
PLVQRTLGPTRKALRDAQVSAADIKGVVLVGGATRMPVIRDAVAKYFGQPPLVNLDPDQV
VALGAAIQADLLAGNRSGGDDWLLLDVIPLSLGVETMGGLVEKIIPRNSTIPVARAQEFT
TFKDGQTAMAIHVVQGERELVSDCRSLARFELRGIPPMTAGAARIRVTYQVDADGLLSVF
AREQHSGVEASVVVKPSYGLGDDDIARMLEDSFKTAEVDMRARALREAQVEAQRLVEATE
AALVADGDLLDASERATVDALVASLRALAPGDDADAIDTATKALAEGTDEFAARRMDKSI
KRALAGRKLDEI*
>GCA_000452945.1_BPMSHR5858_genomic_01614_1 lolD_3
MNDRVFEQTMNQNHQDGGARECVLEARGVTKTFVQGGFNVQVLDNAQVSVRRGEKLAIVG
ASGSGKSTLLHVLGGLDEPSAGQVSLLGKPFTQLAERERNELRNRALGFVYQFHHLLPEF
TALDNVAMPLRIRRMSTEEARRHAREMLEQVGLGARAKHRPGELSGGERQRVAIARALVT
KPACVLADEPTGNLDGSTADHVFHLMLELSRTLDTSFVIVTHDPDLAARCDRILRLRDGV
LHEEPAVPV*
>GCA_000452945.1_BPMSHR5858_genomic_01615_1 yjjV
MWIDTHCHLDAPEFDADREVVAGAARAAGVSRIVIPSVGRTNFTTVRALAHATRGAVYAL
GIHPMYTPHARDEDLDRLRMEVEASLDDPRFVGIGEIGLDYFVPGLDERRQQFFYDAQLR
LAREFDLPVVCHVRKSQDRVLAGLRRAGVRRGIAHAFNGSFQQADAYLAQGLKLGFGGNV
TFERARQIRRLAGQLPLDALVVETDAPDIPPEWRDRQRNTPDQVPRIGAVIAELRGIGVD
ALSLSTSANALAALPRLALSSA*
>GCA_000452945.1_BPMSHR5858_genomic_01619_1 kdsA_2
MNLAGFEVGLDKPFFLIAGTCVVESEQMTIDTAGRLKEICATLGVPFIYKSSYDKANRSS
GKSFRGLGMDEGLRILAEVKRQLNVPVLTDVHEIDEIAPVAAVVDVLQTPAFLCRQTDFI
RACAQSGKPVNIKKGQFLAPHDMKNVIDKARDAARDAGLSEDRFMACERGVSFGYNNLVS
DMRSLAIMRETGAPVVFDATHSVQLPGGQGTSSGGQREFVPVLARAALATGVAGLFMETH
PNPAEAKSDGPNAVPLGRMAALLETLVTLDRAVKRVPFLENDFN*
>GCA_000452945.1_BPMSHR5858_genomic_01621_1 ftsB
MRLVTVVLIALLIVIQYPLWWGHGGWLRVHELRQQLNDQLQKNADAKLRNERIAGEVQDL
QGGTSAIEERARYEMGMVKDGEVFVQFVSPNSPAPAAQGNASSATSTRGDVSGAPMRVVP
NPVSRAKPERRHGGDKAHKPAHG*
>GCA_000452945.1_BPMSHR5858_genomic_01622_1 group_19947
MKPILVRAAPLAALIAAASLSGCVAPPSTTVLSRLPEQAAGTTAHTLTPDERKRYDEIDR
QVLREQNAAIAAEDAARAWSYYSPPPVTYYGGYYGGGWGRGWGSGFGYGYPGWWW*
>GCA_000452945.1_BPMSHR5858_genomic_01624_1 yrdA
VTIYKLGENAPSIHESVFVADSATIVGKVVLEENASVWFGATIRGDNEPITVGAGSNVQE
GAVLHTDPGCPLTIAPNVTVGHQAMLHGCTIGEGSLIGIQAVILNRAVIGRNCLVGAGAV
ITEGKAFPDNSLILGAPAKVVRTLSDEDIARMHMNTKSYAMRRAYFKEQLVRIG*
>GCA_000452945.1_BPMSHR5858_genomic_01627_1 group_4624
MPTMNADLHCHSNVSDGQLAPADVARRAHAGGVTLWALTDHDEVGGQRAAREAAEALGMR
YLHGVEISVTWASRTVHIVGLNIDPRNPALIDGLHRTRNGRAARAVAIGDALAALGIEAA
YEGALAYVSNPDLISRTHFARFLVDKGYAASMSDVFDRYLGDGKPGYVAHRWAKLSDAVA
WIRQAGGEAVVAHPGRYAYTSVEFDAFFGEFIELGGVAIEVVTGSHTPDQYREYADVARR
FGFEASRGSDFHAPGEGRTEFGSLPPLPSDLKPVWERWL*
>GCA_000452945.1_BPMSHR5858_genomic_01630_1 group_4625
MFPDRIFSGMRPTGSLHLGHYHGVLKNWVKLQSEYPCFFCVVDWHALTTHYETPEVIEKN
VWDVLIDWLASGIDPAQATLFIQSKVPEHAELALLLGMSTPLGWLERVPTYKEQIEKLKD
KDLSTYGFLGYPVLMAADILLYRGSLVPVGEDQVPHVEMTREIARRFNYLYGREPGFEEK
ALEAAKKLGGKRAKLYHELRNAYQQEGDDEALEQARAMLQESQSLSMSDRERLFGYLEGA
RKIILVEPQALLTEASRMPGLDGQKMSKSYGNTIGLREDAETITKKVRTMPTDPARVRRS
DPGDPDKCPVWQLHQVYTDETTHEWVQQGCRSAGIGCLDCKQPVVEGILREQQPMLERAQ
KYMDDPSLLRAIVADGCDKARKHATETMRDVREAMGLSYS*
>GCA_000452945.1_BPMSHR5858_genomic_01638_1 mutS
MATQIDASSEAAAATAAAQHTPMMQQYLRIKSEHPDTLVFYRMGDFYELFFEDAEKAARL
LDLTLTQRGASAGTPIKMAGVPHHAVEQYLAKLVKFGESAAICEQIGDPATSKGPVERKV
VRVVTPGTLTDAALLSDKSDVFLLALCVGHNKRGVASNIGLAWLNLASGALRLAELAPDQ
LGAALERIRPAEILAADGTIESVPAGMGAITRVPAWHFDIASGTQRLCDQLEVASLDGFG
AQALTSANGAAGALLIYAAATQGQQLRHVRSLKVENESEYIGLDPSTRRNLELTETLRGT
ESPTLYSLLDTCCTAMGSRLLRHWLHHPPRASVAAQARHQAIGALLDAPPNAGLDSLRSA
LRQIADVERITGRLALLSARPRDLSSLRDTFAALPALRERVAEIASNAAALGRLEAALEP
PPGCLDLLTRAIAAEPAAMVRDGGVIARGYDAELDELRDISENCGQFLIDLETRERARTG
ISNLRVEYNKVHGFYIEVTRGQTDKVPDDYRRRQTLKNAERYITPELKTFEDKALSAQER
ALARERALYDGVLQALLPHIEGCQRVASGLAELDLLAAFAERARTLDWVAPEFTDEIGIE
IDQGRHPVVEAQVEQFIANDCALNPERKLLLITGPNMGGKSTFMRQTALIALMAYVGSYV
PAKAARFGPIDRIFTRIGAADDLAGGRSTFMVEMTEAAAILNDATPHSLVLMDEIGRGTS
TFDGLALAWAIARHLLSHNRCYTLFATHYFELTQLPAEFPHAANVHLSAVEHGHGIVFLH
AVEEGPANQSYGLQVAQLAGVPAPVIRAARKHLAHLEQQSAAQATPQLDLFAAPPVVDEP
ECNEPPAATPHPALERLLELDPDDLKPRDALDLLYELHTLARSGPADAQR*
>GCA_000452945.1_BPMSHR5858_genomic_01639_1 group_1436
MKKTFAAIVSAGLLASSSAYADTLCTSGAITKIFIDNNGVMEVTVGTLAYLNGNKDVYSV
LTSAFVANKQVYIYAPNCQPGTSMGGFAVR*

>GCA_000452945.1_BPMSHR5858_genomic_01640_1 suhB
MHPMLNIAVKAARRAGQIINRASLDLDLIEIRKKQQNDFVTEVDKAAEDAIIETLKTAYP
DHAILAEESGESGDASEFQWIIDPLDGTTNFIHGFPYYCVSIALAHKGVVTQAVVYDPNH
NDLFTATRGRGAYLNDRRIRVGRRDRLADALIGTGFPFREKDGLDAYSRLFTEMTQACTG
LRRPGAAALDLANVAAGRLDGFFEQGINVWDVAAGSLLITEAGGLVGNYTGDADFLHRRE
IVAANPKVYAQMIPILSRYTHTRPASE*
>GCA_000452945.1_BPMSHR5858_genomic_01641_1 trmJ
VLVEPSHPGNVGAAARALKTMGFSRLVLVAPRVPRVQSDPEALAMASGADDVLASAHVVP
TLADALNGVQWSLALTARSREYGPPRLAPRAAAASALQQVRTGDIALVFGNERTGLSNEH
VERCSAIAHIPANPAYSSLNLAQAVQVLAYELRVALLEDDEAAVPPDAALGTLAQSDEIE
RMFVHLENALIALDFLDPRNPKKLMPRLRRLFARTGLEREEVNIVRGIAKHILLKAGGAN
ARAGASGEGNGNGNGNGNGESATGDADYSGRDVSRDD*
>GCA_000452945.1_BPMSHR5858_genomic_01645_1 ppiA
MKRLLLALGSAALLATAPAFAQSAATHPVVQLKTTQGDIRVELYPEKAPKTVANFLDYVK
AGQYNGTIFHRVIKGFMIQGGGYKTNFDEKPTRAPIPLESKNGLKNATGTIAMARTSDPN
SATAQFFINTVDNAGLDYPNPDGNGYAVFGKVVSGLDVVKKIEATPTTVRGPMRDVPEKP
IVIESATIVSK*
>GCA_000452945.1_BPMSHR5858_genomic_01647_1 cysS
MESLRIYNTLARDKQDFVPRQPGEVRMYVCGITVYDYCHIGHARMVVVFDIVQRWLRARG
YRVTYVRNITDIDDKIIRRAVENGETIQSLTRRFTDAMNADFDALGVERPDLEPRATEFI
PQMLGMIEKLEANGYAYQAKDGDVNYSVRKFANYGRLSGKSLEDLRAGERVAANDAKEDP
LDFVLWKRAKPQEPAGASWESKYGAGRPGWHIECSAMGCTLLGAHFDIHGGGQDLQFPHH
ENEIAQSEGATGQTFVNYWMHNGFVQVDSEKMSKSLGNFFTIREVLEKFDAEVVRFFIVR
THYRSPLNYSDVHLDDARASLTRLYTALKDATPDAQPLDWSEAHAQRFAAAMNDDFNTAV
AVAVLFELATEVNRTREPALARQLRLLAGLLGLLGREPREFLQHAAGAARTGALEPHEIE
ARIAARVAAKQAKNYAEADRIRAELLEAGIALEDKPGGSTEWRRV*
>GCA_000452945.1_BPMSHR5858_genomic_01656_1 mdtC_2
MSRNLFAVFIRYPVATCLMTAGILFAGVAAYFHLPVAPLPQVEFPTIQVSAVLPGADPVS
VASTLAQPLETQFSKIPYVTQMTSQSTLSSTSIVLQFSLERSIDAAANDVQSAIDAAAAQ
LPADLPSPPTFQKVNPADSPIMLLSAISSTLPLTTIDDYVETRLTKSLSQIDGVGSVSIG
GQQKPSIRIQLDPVKLASRGLSSEDVRRALSGLSGVNPKGVFNGTTRSYTIYTNGQLTEP
AQWNDAIVAYRDGTPVRIRDIGQAVLGPEDNTLAAWIDGRRAISVGIYKKPGANTVSTVD
KIRARLPELEASLPPSLKIAVLADRTQTIRASLLDIELTLLLNVVLVVVVIYAFLGSVRT
TIIPAVTVPVSLFGACALMWVCGYSLDNISLMAMTIAVGFVVDDAIVMVENIARHVEAGE
RPLQAALKGLSETSFTIASISLSLVAVLLPLLLMSGIIGRMFREFAVTLSMTIIVSAFVS
LTLTPMMASYLLRAHRHDAGRPPRPGLFERAFARTAAAYERALDVALRHRFVTLCAFFAS
VAASVFLYVGIPKGFFPQQDTGVITGISEAAQTISVEDMARHSMALAAIIRADPAVEHCQ
MAVGGSAYAGTTVNNGRWYITLKPRDQRDATADEVIRRLRPQFAKVPGVRMYLQAAQDVI
IGARLARTQYQLTLQSADVGALTTWAPRLLARLSGLPQLRDVASDQQVNGSALSVAIDRD
QAARYGLTPEAIDGTLYDAFGSRQVAQYFTQLSTYKVIMETLPSLQRDPGTLDRIYMKAP
SGALVPLSSVARWTTDTVQPLSVNHQSHFPSVTISFNLAPGVSLGEATAAIEAARASLRM
PPAVVGSFQGTAQAFQSTLATMPMLILSALIVAYLVLGALYGSFIHPWTILSTLPSAGVG
AIATLWLFKYDFNLIALIGVILLIGIVKKNGIMMVDFAIAATRERNMTSLDAIRSACLLR
LRPIMMTTMTALFGALPLMLTPGMGSELRQPLGYAMVGGLLVSQVLTLFTTPVIYLYLDT
LSNRLKARLRPLGADMPSNAHVEGATQD*
>GCA_000452945.1_BPMSHR5858_genomic_01662_1 lgrD_3
MNQSYDVVETALRLRAGLAEHLGAAGPDVDEPLVDHVGGALPPALLVDVVRACGGMHATR
VDVLGCATLRELAAWIERTGGAAREPRAAPDAAPNGARPEAPLAPEDADDAPASPAQEDL
WLAAQVQEEGAAYHLSIALRFDAPVDDASMRRALVEVIARHPSLRTTFRRDAQGLRQCVR
AAAHAEVDWSEAVLPAPDEAGERESGLRDALTRFAQPRFDLHAGPLLRARLHRGAPDCDV
VQIVVHHIVFDAWSRMVVSREIVDAYHALRAGRAIAWSAAAASPGAFARGQRDALTPQRV
ARLREYWRAQLAGVPPVLDLPTDFTRPAVRDGIGATALRMLPAALAERLAAVGARHGASL
FMTLMAAWQILLWRCSRQHEFAIGTPMAARDAGAFESTVGYLVNTVVLRASVRADEPFGG
LLARVAQTVLDAHEHKALPLREVIELAGADRNPSHTPLFQAMFEFHNERPGAEAAQSAPV
RAVPHDVGAAKYDLSLEIAYRHDGLACTIEYATALFARSSIDGLLAQYEALLEQIAAMPD
APVGSLSCLPDGERERVTRTWNATAFERTGSPLIHARFDARVRACPAAIALRTDAATMTF
AALGERVDALASRLLERTGGEPERIAICLERSFDMVSAILATLKAGCAYVPIDPQLPADR
VAFMLSDSAAALLLTIEPIRRERLASFDIDTLCLDAPAPPRAAPPRAAPAVDPHAAAYVL
YTSGSTGKPKGVAVTHANVTNLLDVMEASYPVGAHDRYLLKTNYAFDVSVPELFGWFVGD
GSLAILAPQAEGSPDLIVEALLRHGVTHLNFTPSLLRQFVTEAAADARFARGHRLRHVFV
VGEELTSALANDAWHALRPATIYNMYGPTETTVFATGYAHTAPIPNGRVPIGRALGNMRV
YVLDERMRPMPIGMPGDLYIAGDGVARGYLNRDELTAERFLPDPFTPGGRIYMTGDLARW
TRDGMLEFLGRTDQQIKIRGYRVELDEIASALNAHPLVGEAAVILKREPDGDARLVAYVV
PAEGAAAAPSRDERARLRDALVGALEQRLPDYMVPADYAFAHALPKGITGKLDRKALEAL
PVERPARDAALGAVAPRNDTQRALCAIWQAVLRVPALGVTDNFFAVGGDSILSIQAAARA
REQGIVFSARDIFRHQTVELLAAHVRWQQAPDEARRASAGDMPLLPIHHWFFEVDSTHVD
HYHQSRLLDVPAGVDAVFVRAWLAALVARHDALRLRLRDTPAGWRAHFAEAPHADASARL
MVCDVAWAGAPDAALDAFFADARRGVRLADGPLFVAALVAAGATGRGRLLLIGHHAVVDG
VSWRVIVADLRRTFAQWPALRPDAPGPASDAYQAWARALVAAADSPALHAERAHWLRVLR
APAPALRLDRAAGGDATRRATRIGAVSLNAADTRALLADTHRAYRTQTIELLLAGLLLAF
RRWQGHDALRIALEGHGREADALAHYGAERGLPDVGETVGWFTSYYPQWLTLAGAASAPA
PAADSVAAAIMAVKTQYRATPHHGIGYGILRYVAGDPELAAEAAAHAPEIVFNYLGQFDV
ANDEAAGIAVLDLSSRDDIAAARPREHALGIDGGVKDGCLAFEIDYSGAAFDDESIAAFG
AHFMHALREIAAHCKACAAWRPTPEDFPLAAVTQAELDVWHARHPSLETLYPATAIQRGM
VFHSLLPKQASAYTNQVHARVGGGTFDAARLRHAWQTVLERHAALRTAFVGFEREQPLQI
VLAHAACPWRDIDHRHLAPEARDAAFAALLAADKAAPFDFGRAPLMRFHLVRDDDAHYRF
IWTYHHAVLDGWSVQLVWSDLLRVYEALAAGRPAALPAAVQFDAVLAWRQRHASDADKRY
WREQIGARTQRTVLDIEQAGLAAASPAAPAVVERSLDEAASERLATAARACRVTLASVLQ
AAWALLLARHSGETAPVFGVTSSGRAIDVPGVETIVGPLIATVPARVDVDAALPLGDWLR
AVHERHVEREAHAQLELVDILRESGVRGGQPLFDSLLVVENYPLAEPSAARILGLRDYGY
AEDTHYGLTVSATPGPRLRVEINFDRSRYHIEDIEAMADNLKAVLTRVPDDLARPAGDVL
GGADAARAQAGAEGEPRLSRVSRQGELRLGRHLVPHLVEDHAAATPERRALVYNERAYSY
GELNRAANRIANRLMQAFPDLGTDALVGVRVSRSDRLVLTVLAIWKIGAAYIPIDPVLPG
QRMREMLELAGAKALVVDAAVAAAEPAVAGVPRIAFDDLVQDDPRLEDNPDVHLSGNDLS
YVLFTSGSTGKPKGAMIEHIGMLNNIANKALDLEMDEDSRVAQNASMSFDVSVWQMFIAL
TKGGTTFVYDERAVNDIAGLIRRMAADGVTILEVVPTYLIAVVEYLEEHPECVRPASLRF
LIVNGETVDATLIRRWFALFPATKLINAYGPTEASDDITHHIMSPGDEIVNPVPVGRALA
NFDLYIVDDELRPVPIGTRGEIVATGVGIGRGYIGMAGATAQAFVKSPFPDRYKGRLYRT
GDLGEMREDGVLMFHGRKDRQVKIRGMRIELDEVEASLRAIAAVRQAVVLAIRPENREAF
LCACVVPLDGAREEIVDALKAKLPPYMVPSVFRFERELPQLPSGKVDRNRLREQCLNETP
RASEHALAPRSPLERRLAEVFGEVLGHDAFGVLDDFFALGGDSFKAIRIAAKYGPPLEVT
DIYDHPTIEALAAHLERAPRAERSIVQMAGDPATAKAALVCIANAAGGPVNFVEMSRAMA
EQAGELAVFAVKLPRNAVDSDAAMLAEVTRLANAVCDDLLAASGLPIIVFAQCNGSALAI
AVARELTRRSADLRALCMGGALMRTASGKRDARTDDEILGFLGGIGSTLPGRPDERAFFL
HEFRYDSWLADVYYNHLVDEASRGALTPLDIPVWCLVGTDDPLVSQYETRHRDWLRIGRS
VKLAEFAGIGHYLLRDCPHALARTLGQAWEAVSRRSVEA*
>GCA_000452945.1_BPMSHR5858_genomic_01667_1 aptA
MSASAPASRSSERRPVRVLRVLARRHVDSTRMVRPRDFDTALVAQVPGMSDIGDGERYVP
LCVDWRDARLFLSRWDDDCAMTDVPFLYQRQRRTARQLLDVPFEQLEAPGRAARMTPIFI
FSVGRCGSTLLSRLLAAVGEQAVSEPDVLTSVAHFDDAAERAAALPARERIVQSCVAAFE
PACGPAPIIKLRARCNRAVDVFLNAMPHARYVFMCRNRDDWVRSSSRAFGDSGEALAELL
KASVEAFDRMHAARVDPLLVWYEDLLADPLAALRRILRARDDLDAHRAAVERALRADAQE
GSGLSRASLAARTGDAGALAAFDARWREIRPEALLREHGLARLR*
>GCA_000452945.1_BPMSHR5858_genomic_01670_1 oatA_4
MPVDLLVTGSGSLAEAVLFDLVAMPTLAAPMTIVVAARNAQRLAWLEQAAHARACALDTH
HRVACRAIDWEAPGDLARLLAEAAPKVVFHAASLQSMWEIARGEDAWAAEIRASGYGATL
PLQLALALRVARAIGERAPASRFVNACYPDASNEILARLGHDVTCGIGNVATLAATYAAR
LGVEAPRRLRMLAHHRQVAAAIAGQGRLHPPRAWIDGAAAIAPDALPDVRLPAGASLNLV
TGATGARLIAALLGRVERYAGHAPGPLGLCGGYPVTICGGRLELDLPPGLTLEEAVASNR
EWAREEGVSIDAHGNVRIASGRAALAADGGAGGEGSESSESSESSESVALPLSYRIDEIE
DVARHLMRVKADLQRRAA*
>GCA_000452945.1_BPMSHR5858_genomic_01682_1 group_760
LPRRLALANNEAHAQALGQLDREMARLNLDGSALADRVRAALTLTESGYPSLARLASKLF
LSERTLKRRLQAHGTPYRNMLEAARYRDACRLLTRTDRCVADVSTRLGYVNPSAFTRAFR
RWAGIAPSRYRDDR*
>GCA_000452945.1_BPMSHR5858_genomic_01685_1 group_4635
MPTLAKSLLLTLMACFAFSGEAGAAQKTFCYFHHNCPADGKLPKGKPPEQYTKRQCKNIG
GKSWGDGPTSCENVNK*
>GCA_000452945.1_BPMSHR5858_genomic_01689_1 group_4636
MSTYLELKAKADEILRQAEDARRSELETVLKEVRTRVAEYGLTPEQVFGARRGRRGQQVH
ASPTVPKYRDPKTGATWSGRGREPGWIKGKKRERFLIQQ*
>GCA_000452945.1_BPMSHR5858_genomic_01691_1 uvrA_2
LSSSNLIRIRGARQHNLKHIDLDLRTGEMTVVTGPSGSGKSSLVFDTLYAEGQRRYVETF
SAYARQFLDRMDRPQVERVDGVPPAIAIDQTNPVRSSRSTVGTMTELNDHLKLLYARAAE
LFDRRTARLVRHDTPETIYADLFARTQAHDPRIAITFAVELPENASDEEVEQWLSASGYT
RVQARRDVASPTGPRKVLDVVADRFRLHQIDKARAIEAIEASLKRGGGRVSIYVLAQAPE
GADGGAAQSQVWRYSTGLHDPDSDLRYADPQAALFSFNSAYGACETCRGFGRVIGVDLGL
VIPDARKTLRGGAVKPMQTPAWKECQDDLMRYAAKANIRRDTPWAELTPAEQDWVINGSP
DWNGKWQSQWYGVKRFFGYLESKAYKMHIRVLLSKYRSYTPCAVCGGARLKTEALLWRLG
TKANADAVLASADRFMPRGVDWTRAQLEALPGLTVHDLMLLPIERIRRFFDDISLPSALL
DDALKLLLAEVRTRLKYLCDVGLGYLTLDRQSRTLSGGEVQRINLTTALGTSLTKTLFVL
DEPSIGLHPRDLNRIVEAMRRLRDAGNTLVVVEHDPSVMLAADRLIDMGPGPGERGGTIV
YDGTPGAIGSARTLTGEYLGGRKHVAHASNWARRPVDARTPRIVLEGASEHNLRDVTVEI
PLQRLVCVTGVSGSGKSTLLQDVLYPAMARHFGKATESPGAYRRLAGAEQVGDVVFVDQS
PIGKTTRSNPASYVGAFDEIRKLFAKAPLALQRGYSAGTFSFNSGDGRCPSCGGSGFEHI
EMQFLSDVYLRCPDCDGSRYRAEILEVRIERGGRALSVADVLDLTVSEAAACFAADADVL
RVLQPIVDVGLEYVKLGQPVPTLSGGEAQRLKLAGFLAESAKARDGRRVVTEEARVARAR
LFMFDEPTTGLHFDDIAKLMQAFGKLLASGHSLIVIEHNLDVIRAADWLIDLGPEGGDGG
GLVLCAGTPDDVKACAGSHTGAALLQYERAMNVGADADASGDAGVPLQAALNAARARRAI
EGENVVRIVNAREHNLKALDVDIPHGKFNVVTGVSGSGKSTLAFDILFHEGQRRYLESLN
AYARSIVQPAGRPEVDAVYGIPPTVAIEQRLSRGGRKSTVATTSEVWHFLRLLYVKLGLQ
HCIHDGTPVTSQTVESIAAQLLRDHRGEHVGLLAPLVVNRKGVYTDLAKWAKARGNTHLR
VDGEFVTVDPWPKLDRFREHTIELPVADLVVSPDNEAELRQRLDETLELGKGVMHLLAPL
DGLRHAMDNRLCTAGVGAIKVLSVKRACPVCGTSYPELDPRMFSYNSKHGWCTTCVGTGL
ALTREQRAAYDDTVLAEDGRGREQTLPSDEQEPEGVGDEPCPDCGGTRLNPSARAVTFGR
HSIVDVAQWTVTDTRRWIEGLQLAGRDAQIARDIVGEIGSRLAFLEEVGLGYLSLDRAAP
SLSGGEAQRIRLAAQLGSNLQGVCYVLDEPTIGLHPRDNQILLSALRKLGDKGNTLVVVE
HDEDTIRRADHIIDIGPGAGRRGGTLVAQGGVADLSAQSASVTGRLLAQPMTHPLQPRRS
VNPPGKRNGHAVPQAWLTVHRARLHNLRDVTVGIPLARLVAVTGVSGSGKSTLARDVLMT
NLLDAVGRSVLSSPAARRARKAAQQDAPRAAKSRASVLARSAPRPVLNVTHAWQGCESIS
GWEHIDRVLEVDQTPIGKTPRSCPATYIGVWDAIRRLFADTLEARARGYTASRFSFNTGE
GRCPACEGQGVRTIGMSFLPDVKVPCDVCHGQRFNPETLAVTWRGRNIGDVLTMEIDEAV
EFFAPLSNIAHPLQLMKDVGLGYLTLGQPSPTLSGGEAQRIKLVTELSKVRDDVTRRGQK
APHTLYVLDEPTVGLHMADVAKLIRVLHRLVDAGHSVVVIEHDLDVIAEADWIVDLGPEG
GAGGGTIVAAAPPEGLVEVSASHTGRALGAVLARTGADRDEPLRSGTARAG*
>GCA_000452945.1_BPMSHR5858_genomic_01709_1 group_4639
MKDIDSPDDPPACVLIDRRIAELGDWRGDTLSRMRGLIRDADPDVVEEWKWRGTPVWSHD
GIICTGESYRSVVKLTFAKGASLDDPAGLFNSSLDGQVRRAIDIREGETLDADAFKALVR
AAIALNQSSGNARTQAKAKRAKPDAK*
>GCA_000452945.1_BPMSHR5858_genomic_01716_1 group_4640
MADPEFVSLVEMGLAMAMAAGRSIVPGASLTPRGRELVERIACAKAAAVQRI*
>GCA_000452945.1_BPMSHR5858_genomic_01729_1 codA
MKLINATLRKRSGLFSIVLDGGAIASVTPQPARVDAPHAPRADVIDADGKLVIPPLVEPH
IHLDAVLTAGEPEWNMSGTLFEGIERWAQRKATITHEDTKARAHAAIGMLRDHGIQHVRT
HVDVTDPSLAALKAMLEVKDEARGLIDLQIVAFPQEGIESFGGGRALMERAIEVGADVVG
GIPHFENTREQGVSSIRFLMDLAERSGCLVDVHCDETDDPNSRFLEVLAEEARVRGIGAR
VTASHTTAMGSYDNAYCSKLFRLLKRSQINFISCPTESIHLQGRFDTFPKRRGLTRVAEL
DRAGMNVCFGQDSIRDPWYPLGNGNILRALDAGLHICHMMGYQDLARSLDFVTEHSARAM
HLGERYGIEPGRPANLVVLDASDDYEALRRQAKALLSIRGGEVIMRRVPERIAYPAAR*
>GCA_000452945.1_BPMSHR5858_genomic_01738_1 group_588
MQRCLAHPARLLAATLLCVALAACGADDTNDSAAPPPARPNPTPDAPASTPAAAPTYYQT
KTPYRPQQDASRYEAPPAGYAPVYTEMLARHGSRGLSGFKYDGAIYDMLQKADAEGALTP
LGRQLKADTSAMMKANALLGYGVAGISTPGYGNLTQAGIREHQQLAARLLQRLPALFDTV
AGTTGGAARKIVVVHSGQDRAVDSSAYFSGALVAARPALAPAITSPSAIAGYPQGAPVAQ
AVGANRFLLYFHALKPSVDLVVSTDDPYYATYRDSLAYQAYGGSAAVAAKLDAIMRAPRT
SQVARTVLAALVTPGFAAKLGTAGYTFSNTGTYTFASADAKFANTLKGDGKTTIKSAADA
ANVLYNLLQVAPAMTAETGGVTMERYIAAPQAEYLAYLQDAEDFYEKGPGVAETNPITYR
MAQALVDDFFAEIDAIARGDLTNAAKLRFTHAEVVIPFASILKLKDVFAPVPQAQTYTYA
NNPWRGETVAPMAANLQWDVYRNGARLIVKMLYNERETDFQPACDGARIAPGSRFYDYAG
LRRCYGYS*
>GCA_000452945.1_BPMSHR5858_genomic_01742_1 group_4641
MRTADVACALAVGFASAAALADAPAMPSGGMLVAANGMTLYTFDKDAPNAGKSLCNGPCA
ANWPPYKASATDRPAGGYTIIKRDDGTLQWAYQGKPLYFFAKDAAKGDKKGDGFKDVWHA
VKE*
>GCA_000452945.1_BPMSHR5858_genomic_01743_1 group_77
MKRKSLAMAVAGIGIGAAVPPALAQSSVTLYGIVDSGITYTNNQKGHSAWKATSGNEQAA
RWGLVGREDLGGGTSAIFRLENGFDIENGTASQGGRLFGRRTYIGLANDRWGAFTMGRQY
NAVQETLAPLHIATVLGRYAHHPFDADDLDSTFRTNNSVQYQTPDWGGVRAIATYGFSNS
TSFARNRTWSVGASYARGPLQLAAAYVRVDYPAIDATGAIASDNYYGFVKGVARQQIWGA
GGAYAWGDATFGLLYTSALFSLQTGGAERFNNFEGSVRYRVTPAVQVALGETYTQVLASH
AAAHDAHYLQTSAGIQYFLSKRTDVYLNVSYQRASSNAVAWIKGLDGASSTRTQIVAVTG
IRHKF*
>GCA_000452945.1_BPMSHR5858_genomic_01744_1 group_4642
MSRTGFMPLSTLATMAVAAAISTPAAFAESAPVSGNGGIPSFYRWTDKVPQTRGQMLRTE
VLTPRQGLAAAGQNLRILYTSTDGIDGHTPIVVSGALFVPKGTPPQGGWPLMAWAHGTVG
SADICAPSYAGHSERDVRYLSHWLAQGYAIVATDYQGLGASGPHPYGLTRPLAYSVLDSI
RAVQNGGFELSPKRIVVFGQSQGGRAAFATAVYAKSYAPELDIGGVVATGTPYATIRHRA
DDAEHSKLHRSVIHNILRLNATPKLLDPSFATSDYLTERATPAFEFSQHACLHAIKQKIA
SDGLTFENSFKRSPQPVLDWIRREAAYPTLKSAIPIFIGTGGKDRNVPVSDQAALVKDAC
EAGDRIEWHVYPALDHSETVNGSLSDSTPFVARAFAGEHIDGNCASLPSP*
>GCA_000452945.1_BPMSHR5858_genomic_01751_1 group_4643
MNSPLPPAPAAFLPPISCLQDDDRPWLPMDASLPGLAIKYLHINVADDEMTVLLKMPVGL
ALPRHRHDGAVFVYTLQGEWRYREYDWIARPGSAVLEPAGSHHTPEALASETGAVVTFNV
MRGDLVLLDDAGRETARENCRVALLRERRFARGAPDAAAPFVTR*
>GCA_000452945.1_BPMSHR5858_genomic_01753_1 group_1126
MSDPSRENPPTDAHASSLVTHDEPVPRGRRAPRPLWTLVPRAAGYGVVFFVIWMVLTIMF
PNVFTRSSERAVVNNEVTLVTSPVEGVVTEQHVTAGKPFDANQPLATVQNPNVDRALLID
LTGKKLDNQQREDAARAELAGDESQLASTEHDLQRYQSVAQKEHAATIRALEARLAVARA
QVDQQEDIVNRNQAMQWAGAVSEAYTSASRYQLSILSNAKAAAAAELEHAVANGDASRSK
VYASATDGPAASLSQRGRLLGADIAQRKAEIAQFDAYGQSVDKLIAAEQQRLDRLSRIEI
RSGEPGVVEDVLAPPGTRVAAGATLIRASNCARSRVVAVFPRSLSDDLLPGTHLNVRMDG
VPAVLPASIAEVLPRASEGEQARYFVPFPPIEKNEIYVIAKLDEPLAPLSRRASARPDAR
CAMGRWARVSLDRGWLASNVSGLANVDPNWTAGARSALARGGQWLRDAGVQARRRLEDFA
TNAARRLGELAADGRRWLNERASAARRWLDDLKSAA*
>GCA_000452945.1_BPMSHR5858_genomic_01757_1 group_1104
MHVPFLRRASVGARLAALSSALFALLFAAFVWALTHAASGQIADQVHARIDEKDRSIAAM
IALFDEALTAEASRAMTLFASFLPAGYALDAARTIDVAGVATPALTAGGQTLNLDFSIPD
QFLQKSGAIATIFARRGDDFVRVTTSLKKQDGSRAIGTLLDRKGPAYAPLVAGRTYTGLA
TLFGKRYITQYKPIADASGAIVGALFVGIDVGAEMRLVENGIRQLKIGEHGYYFVLDASD
GPARGTLLVHPARAGQRADDAAAPYAQMLAAKEGQLSYTSTDAAAGDDGPRAKFVSFVTV
PQWQWLVGGIAIDDEVMAGMRATRNRFAAIGCAFVLAFAALFVAVVKRVVSRPLDAAAHA
SERFAAGDLSVRIGAHGKHDKHDKHGKHDARDGAARPPASGRSDEIGRLVRAVDGIGDGL
ARIVAQVRRGAADIAHGTVTIAAGSSDMAARIATQASSVEQTAASMEQITAAVQQNADHA
AQASALATGASSAATTGGAAVQRVVATMGDIQGVARRIAEITGVIEGIAFQTNILALNAA
VEAARAGEHGRGFAVVASEVRALAQRSAAAAKEIDALVGESATTAEHGFRIAEDARAAMQ
DIVARVDQVRAIIAEISAASREQSGGIEQVNLAVTQIGAATQQNATLIADAERAAAALRD
EAAQLAHAVSVFRLAADEGVLDAR*
>GCA_000452945.1_BPMSHR5858_genomic_01761_1 group_4644

MTSTAADSAIPPGAPSARTRVRRAAYRGHYDRPTLEAILDDAYVCHVAFADDSGVHCIPT
ACWREGDHLYIHGSNGSRMLKLAAAGAQVCVTVTHLDGLVLARSAFNHSMNYRSAVIYGE
FEVVPDARKAAVLDTFVERIAPGRSRDARPGDASELAATTVLRIALDEVATKVRTGGPKD
DEADLGLPVWAGVLPMREAHFAPLTDAAPAGVPDYVRRWQDAVPLRETQDEEARRAAAR*
>GCA_000452945.1_BPMSHR5858_genomic_01769_1 group_4645
MAALFEPFQLKDVTLRNRIAVPPMCQYVAEDGVPNEWHHVHLAGIARGGAGLVIAEATAV
SPEGRITPGCTGLWSDGQAAAFARSVAAIKAAGAVPGIQLAHAGRKASANRPWEGDDHIA
DGDPRGWRTIAPSAVPYGAHLPKVPREMTLDDIARVKADYVAAAKRALDAGFEWLELHFA
HGYLAQSFFSVHSNRRADAYGGSAENRGRFLVETLAAVREVWPERLPLTARFGVIEYDGR
DEQTLAESIELAKRFKLEGLDLLSVTIGFSTPAARIPWGPAFLAPVAERVRREAQLPVAS
AWGIDAPALAERVVSEGQLDLVMVGRAHLADPHWPYRAALELGVDRAPWTLPAPYAHWLE
RYRAA*
>GCA_000452945.1_BPMSHR5858_genomic_01781_1 group_608
VKRVGYGLLLLAFTSLTFAGERYVEIWNPPEARAGATGGSSAGSTAGTRRDPSARKPSAA
AGAKHKRVAPRVVEARMHHPPTAVKIAPKPRAIEDDAPRATPAPASPSFPGTPDIPRLFT
PDGNVLRVGTRGYHADVSR*
>GCA_000452945.1_BPMSHR5858_genomic_01783_1 ycbX
MPVISELFVYPIKSCAGIATVRAQLLVTGLEYDRNWMVTDPTGAMLTQRTHPRLALVRTA
IGERELVVTAAGMPELRTPLAASALAGAERLAATVWRDTVSALDTGTHAARWFSEFLGAP
ARLARFAPDARRVVGAKWTGPFTSYAQFADGFPLLVVGQSSLDDLNVRLRRKGASAVPMN
RFRPNVVLAGLDAYEEDYVDYLDVQTGGGGVRLSLVKLCTRCPVPTIDQRTGAPDPAWPN
EPTDTMSLYRGSKQFGGALTFGKNAIVLNGDGAFLEVGQSVDAEIAFGE*
>GCA_000452945.1_BPMSHR5858_genomic_01792_1 paaG
MSYQTIRIEIDQAAQIATITLDRPDKLNSFTRDMHRELQSALDDVQAANARALVLTGAGR
GFCAGQDLADLDFTPGAMTDLGEVIEAHFNPLVRRLQALPLPVIAAVNGTAAGAGANLAF
ACDLVIAAKSSSFIQSFVKIGLVPDSGGTWFLPQRVGFARALGLALTGDKLSAEQAERWG
LVWRVVDDAELAGAAAQLARELAQQPTRAIAAIKQAMRASLTHTLDQQLDLERDLQRELG
QSYDYAEGVRAFIEKRAPRFEGR*
>GCA_000452945.1_BPMSHR5858_genomic_01795_1 mltA
MGFSRRLAGWAAAVAAAALLAACVGSPVRQGARPAGAAIVPGQIAAARLTPVAWQQVPGW
QDDSLIGATIALRQNCARLARQANWQRACAAAMRLDDLDVGSARTFFETYFTPFQFANND
GTLDGLVTGYYEPLLHGSRVRRGPYQYALYRWPAGYRAGASMPARAQLMRSGALSGNELV
WVDDPIEAFFLQVQGSGRVVLDDGTVMRVGYGGTNNQPYRSIGKWLLDHGELGAGQATMQ
GIKAWARANPSRVDALLDTNPRFVFFREMPSQEDVPHGGADGPVGALGVPLTPERSIAVD
PSSIPLGTPVFLQTTRPMTNAPLNRLVFAQDVGTAIKGGVRADYFWGLGDDAGDQAGRMK
QNGRMWLLFPNS*
>GCA_000452945.1_BPMSHR5858_genomic_01804_1 ygiF
MAIEQEIKLALPAAQVDAAQRFLDARAGAGGREITLVNVYFDTPALALARAKSALRLRRA
PQGWLQTFKTVGTAAGGLHRRHEWEMPVAGEALEIDALCAACDVPSAVDALRAAAPQLIA
LFRTDFTRTLWHIEHAGATIEAALDRGEIVAQVSGDVRREPICEIELELVDGDAQALATL
ADQVAAALPGVTPDNLSKAQRGYRLRGGDLRTD*
>GCA_000452945.1_BPMSHR5858_genomic_01808_1 dsbC
MKKTIRIAALALAAATATLGCTAQADQSTDKLKAALQSRLGADAPIKSVTKSPIAGLYEV
NLGSQIVYSDASGDYVLLGELVNTKTHKNLTAERLAEINKIDFASLPLSNAIKVVKGNGA
RKIAVFSDPNCPYCKKLETTLQSVDNVTVYTFLYPVLSPDSTVKSKSIWCASDRVKSWQS
WMLEHRAPTSAANCDTTALDKNLALGRGMNVTGTPTVFLADGTRLPGAVSADELNQALAG
VK*
>GCA_000452945.1_BPMSHR5858_genomic_01814_1 nuoB_2
MANHPLTLEKDGFIVTTLDAAMAAAQKNSLWYMTFGLACCAVEMMHAAGARYDMDRFGMI
PRASPRQCDLMIVAGTLTNKMAPAMRRVYDQMAEPRYVVSMGSCANGGGYYHYGYSVVRG
CDRIVPVDVYVPGCPPTAEALVYGLMQLQRKVAERSTHSRPKLFARP*
>GCA_000452945.1_BPMSHR5858_genomic_01818_1 group_5332
MNQLIDTLGRRAVLIGVALGLAACAGGGPPGEAPATLPRIVAHRGGAADAPENTLDAIRA
AVANRADAIWLTVQLSRDGVPVLYRPADLSALTRSSGPVAGHTAAQLAQMNAGWQFRDAG
GRYPYRARPVGIPTLRDALRAIPPAMPIVLDMKAVPAAPQAKAVADVLTSEAAWPRVTIY
STDAAYQTAFASYPQARLFESRDATRGRLVDVLLGGACERAPEAPATAPIWTGFEMHRNM
TVSERFTLGEGVSPVKATLWTPATVACFRRRADVRILAIAVNDADDYRTAACLGLDAVLA
DSPREMAEIRSALRARPLRCETGAR*
>GCA_000452945.1_BPMSHR5858_genomic_01820_1 ettA
MAQYVFTMNRVGKIVPPKRQILKDISLSFFPGAKIGVLGLNGSGKSTLIKIMAGIDKDIE
GEATPMPNLNIGYLPQEPQLDPGKTVREAVEEGLGDLFQANKKLEEIYAAYAEPDADFDA
LAAEQAKYEAILASSDGGSPEQQLEVAADALRLPPWDAKIEHLSGGEKRRVALCKLLLEK
PDMLLLDEPTNHLDAESVDWLEQFLVRFPGTVVAVTHDRYFLDNAAEWILELDRGHGIPW
KGNYSSWLDQKEERLKQEEASESARQKAIKKELEWVRQNPKGRQAKSKARIARFEELNSQ
EYQKRNETQEIFIPVGDRLGNEVIEFKNVSKSFGDRLLIDNLNFKIPAGAIVGIIGPNGA
GKSTLFKMLTGKEQPDSGEVVLGPTVKLAYVDQSRDALDGSKTVFEEISGGADVLTVGRY
ETPSRAYIGRFNFKGGDQQKIVGNLSGGERGRLHLAKTLISGGNVLLLDEPSNDLDVETL
RALEDALLEFAGSVMVISHDRWFLDRIATHILAFEGDSQVTFFDGNYQEYEADKRARLGE
EAAKPKRLRYKPISR*
>GCA_000452945.1_BPMSHR5858_genomic_01828_1 prmC_1
MNTTKPSPATAAELLRASPLDALDARILLAHALGWSRTQLITRADEPLDAAARARYLALQ
ARRAAGEPIAQLTGAREFFGLEFDITPDVLIPRPETELLVETALDAIDGIASPCVLDLGT
GSGAIAVSIASERPDARVWALERSVAALDVARRNARKLLDPARAGGPLRFLESDWYAALD
PGLRFHVVVSNPPYIARHDPHLAEGDLRFEPRGALTDENDGLAAIRTIVAGAHAFVAPGG
ALWLEHGYDQAAAVRALLDAAGFADVESRADLASIERASGGRLPG*
>GCA_000452945.1_BPMSHR5858_genomic_01839_1 group_1621
MSTKTYEIRPNQSIELLKELHILTRDGKMNQDSRRKLKQVYHLFQFIEPLLADVEREKGG
VTLVDHGAGKSYLGFILYDLFFKARARADSHIYGIETREELVARSTELAARLGFGGMSFL
NLSVADSIASPTLPETVDVVTALHACDTATDDAIRFALAKRARHIVLVPCCQAEVAGVLR
RNKGKSLASALTEVWRHPLHTREFGSQITNVLRCLQLEAHGYQVSVTELVGWEHSMKNEL
IIAQYKNLPRRKPGERLDEILGMFGLAQLRERFFAPDTGTAGGAHAEPAAADAEPPRA*
>GCA_000452945.1_BPMSHR5858_genomic_01843_1 group_18276
MWINNRRLRYLSLLGALALAACGGDDGGTGSAVSLGAAHSPSNGANGANGTNGGAATPAA
VDKSTKPVEMPDTVVPVNYKLWFRPNADLNQFSGRADVEIKVLKPVNAIVVAGHRIQFTN
GKTTLQPGNVQLVATPQDKGDFYQLRPASGQIAPGNYSLHMEWQGIINFKSYDDPVNHTG
GSCGNDPYPGCSAAEGIFRVDLKSTDGTTSGAILTQGETNLSRQWFPGWDEPAFRPTYEV
TAEVPQAWRVVSNAAELPSVNVGGGYKLVSFEKTPPMPSYLLFFGGGLFDVLEDDFSSPL
PNSNGLHLRIFTPPGMREWARPAMQRTKQALDYYYRYTGIALPLKKFDTVAANDAFKDQK
GLNFGGMENWGSILEFADDILPEPGKPMSRYGNQVLTHEVAHQWFGDLVTTDWWDDVWLN
ESFARFFETKTTIQFFPDEFNWLDHIKSKYRVINKDISQDAFPIQPNFNGWASNDFVISA
SSFVYNKGGMVLKMLEGYLGEQTLRKGLQQYLNDYAFGNGTPKRLWDALSAASGQQVGPI
GDSFVRQTGVPLLALDTQCDLTKNQNVVTLTQSPFPNKNKYPGAQWTIPVTLAYGDGLVN
RKTLALKDTQTQIRLDGCSAVVANPTGFDYYVTNYSDAAWSALLTQINASTDPVLLLNLK
SEAALLVASNLAPPSRATSISSINSPAAMKLRQVPSILEMPKERPQLRYQGKFTPRQQRT
E*
>GCA_000452945.1_BPMSHR5858_genomic_01847_1 hlyB
MSLLDRLSFGIGSKLPMILQTEAAECGLACLAMIAGFHGHHVDLATMRTRFPVSLKGAGL
GRVIDIAQRLDLGTRALKLDLEQLGQLRVPCMLHWNFNHFVVLKEISGKHAIVHDPAQGV
RKLSLEEVSRSFTGVALELWPTGSFKPREASPAVKLRQLLGPVSGLSRSLGQILVLAVAL
EVFTLVSPFFLQWVIDEVIVSADRDLLTVLALGFGLLLLMQQATSAIRAWALMYLGTTLN
VQWRANVFTHLLNLPVQYFERRHLGDVVSRFGSIDTIQQTMTTSFLSAVIDGLMTIVTLA
MMFVYSRTLAFVALGTMALYALLRWLWYRPLRRATEDQIIHTAKQQSHFLETVRGVKTIK
LFNRQTERRSSWLTLLVEQINASLHVQKLQLLYQQLNGLLFGLEGLVIIWLGARLVMDGE
FTVGVLMAFNAYKGQFDSRVGSLIDKFFEVKMLQLQGERLSDIVFAKGEADAGVRHVPGE
TENLCASIEAENLVFRYAEGEPTVLDGVSLKIEPGESVALIGASGCGKTTLVNVLLGILE
PTGGKIRIGGVDVERLGLDRLRSLVGTVLQDDVLFAGSIADNISFFDPDADPKWVAECAQ
LAAVHADIVAMPMGYNTLVGDMGTVLSGGQKQRVLLARALYKRPKILVLDEATSHLDLQR
ERQVNAAVSALKMTRVIVAHRPETIASASRVIMLDAGKVALDKPTAALAAAPRPPAAPAA
APASAEAQAAPAAACAAPAAPPAQAATQPATGR*
>GCA_000452945.1_BPMSHR5858_genomic_01851_1 group_2363
MQRFLNVLTHPRTLTIVGFAALAAVLFIAADALQIGLAWAAVALGVALALWLATKLWRRW
RVRRANRQLGDMLEQQAETGKMSAAVAEPAKRAELDVLRTRLADTVKTIKTSKIGQVSGG
SALYELPWYIVIGNPAAGKSSAVINSGLQFPFADKNSAVIHGIGGTRNCDWFFTTEGILL
DTAGRYSVHEEDRSEWLGFLDLLKRYRPKAPINGIIVTASIAELTGNRPEFAINLAKNLR
QRVQELTEKLEVFAPVYVMFTKADLITGFTEFFSSNDRQEYDRVWGATLPYEPDEKRDVV
ALFDQRFEELYDGLKEISVAQLSVSRGNKLSPGQLSFPLEFSTIKPALRSFLATLFEDNP
FQYKPIFRGFYFTSALQEGETNSAAAQRIAHRFGLDSQSLPKPHSAFSKNGFFLRDLFSK
VIFADRQTVRQFASPTKTRMRYATFFGFVAALALALGGWTWSTIGNQQLVSNVQADLDNV
TRLQQNRNDLQSRLQAMDILEDRIEQLEQFRRDKPLSVSLGLYQGNRLEQHLLTEYYNGV
RQILLAPVSDNLASFLKDVNAHPDQLAPMTRAPESGATPAGPMPVSTHPAAPGAAPSPAV
AQGRQQGGLYNDASPTNVEDAYNALKTYLMLSDKRHVEQAHLTDQVARFWRGWLETNRGN
MPRDEMIKSAERMITFYLSRVSDNDWPMIEANLALVDQTRENLRRVVRGMPARQRVYEEI
KARASTRFAPMTIARIVGEGNTGLVAGSYAIPGTFTRDAWFQYVQPAIRDAATKELQAKD
WVLNTASADDLTLEGSPEQIQKTLVAMYKTEYAQHWQKFMQGIAVQGFGNFTQAVDAMNK
LGDPQDSPIRKVLETAYDQTSWDNPSLANATIKKAQTGVVNWVKQWFTRQPGGQIAANVD
INGNPADVPMGPIGQEFVGLGRIVATHDGTSMLKGYMDTLSKVRTRFNVIKNQGDPGPGA
RQLMQQTLDGNGSELADSLKFVDEQMLTGLTDSQRKSLRPLLVRPLMQAFAVVIQPASVE
VNKVWNAQVYQPFQNSLANKYPFAASAKVEAGAGEIAQVFGPDGSIAKFVGTTLGPLAVR
RGDTLSARTWGDMGLALTPDFTGGFARWVAPLAGGAATSAAASSEPQTVFQVLPQPSSGT
TEYTIAIDGQQLRYRNTPPQWTNFVWPNPQGSPGATLSATTFDGRTLQLVNEPGRYGLEK
LINSAQRKRRPDGTFDLTWTQGSVSVSVTMRIISTSQPTGGGDQPQQQSLRGLRLPSSVA
DASAGGAANATAQPGSGAAAAQVAVTAASASNAQGAQ*
>GCA_000452945.1_BPMSHR5858_genomic_01853_1 group_2201
MTKPRLPLLRARRFALALAALALGGAARADNVGPATVTPIDAGGVPVKSVPLAAPVTAAP
VAATAGAGATVSSPPPANATPGQVVAGGKVADEATKAAVLQRLRDTYGAANVVDQIEIGN
VATPPNWSANVQKLIGPQLKQISKGQLKIDGTQIDVKGEVHNEAQRQQLASDMANALNPT
YTIKNGLRVSASEQGLLDQTLANRTIEFETGSATLTPQGRAILDQMAGALAKMSNRTVEI
IGHTDNSGNRTSNIALSQARADAVKGYLVTKGIASQQLTTTGVGPDQPIAPNDSADGRAR
NRRIEFRAGQ*
>GCA_000452945.1_BPMSHR5858_genomic_01884_1 group_4656
MTPFRLFFVRIAATLALGLFSALHAHATCSVVSAAAASFGTVTSFAVARQPQSTSTTSSG
LSCSGALLGLFVIGDQINASITSANGGKLVGPTGDAVPYTVFADQNYSIKLDLGVTYNWA
SGQLLNLLGIFGGPAQTLPMYFRTVQGSNVAAGTYTDTLTIAWNWDYCSGIGVLGICLGR
DRGSGTAVVPVTITVTNDCMIAAPDVNFGAAPTVASFAPVTGSVSLTCTKGMVYTVGLSS
GANPHASGRRQMANGANRLQYDIFGPGAAAVWGQSANRAGSGAAADGRSAQQFPYTARLY
PDQPTPAVGTYTDSVIVDVRY*
>GCA_000452945.1_BPMSHR5858_genomic_01887_1 oprM_4
MRRALTRWRLSFALGALGALAALASACTVGPDFHPPRADVPAQWHDVTATSTSSTSSASS
AASPPAARSGAPNAGAPPRSLPTVEADPDPRWWRTFGDPVLDRLVERAAQDNLDVQAAVL
RIAEARAQVRAAAAQGLPDVRASASYQREQLGLKGFLEDGGVDAQLERLGAPGSPLERIA
PGAGAALRRGAEHTLDQLTSPVNLWQAGFDASWELDLFGRVRRSVEAAGAQTQAAIEGRN
DALLSLEAEVAQTYLQLRGAQALRTLAVALADEQRDVVALTRDQARKGLSSELDVQSADA
RLAQLRAQLPQFDQQIALLRNGLSYLLGDAPGALDDALREPRALPELPPTVPIGLPSTLA
RRRPDIRRAEAQLHAATAGVGVAVAQFYPDVSLTGQVGTRATHARELAHWSHLFYAFGPS
VSLPIFEGGALVANLRLSQAQQAQAAFGYRRAVLLALRDVDNALAAYRTDQTRRDALRDA
AAAERTTFELARDSYRHGLVSFIDVLDAERELSQARQQYVQGSLQLTTDLVSLYKALGGG
WQPVPAAQTPAPAALDAPTQDTPLGATAPARR*
>GCA_000452945.1_BPMSHR5858_genomic_01890_1 fabV
MIIKPRVRGFICVTTHPAGCAASVREQIAYVARRGPIERGPKKVLVIGASTGYGLAARIA
AAFGAGAATLGVFFERAPADAKPGTAGWYNSAAFHDEAAARGLQATSVNGDAFSDEIKHK
TIDAIRRDLGQVDLVVYSVAAPRRTHPKTGVTHQSTLKPIGHAVRLRGIDTDNEAIKETL
LQPATPDEIADTVAVMGGEDWRMWIDALDAAGVLADGAKTTAFTYLGEQVTHDIYWNGSI
GEAKKDLDRTVLALRGKLAARGGDARVSVLKAVVTQASSAIPMMPLYLSLLFKVMKARGT
HEGCIEQVDGLLRDSLYGAQPHVDAEGRLRADRLELDPAVQARVLELWDQVTDDNLYTLT
DFAGYKAEFLRLFGFGIDGVDYDAPVEPNVRIPNLIE*
>GCA_000452945.1_BPMSHR5858_genomic_01894_1 nagZ_2
MRKKSLVHLPVALAVAAALGACSGDDDATLESRADAIVERMTTRQKVGQKLMMAFRYWCP
DGQPACTAGMTEFPDAARDALRENGIGGVILFSNNLTGIEQTRRLIDGIRAAPAADSPLG
LMIGIDEEGGNVFRLPRVEATAFAGNMALGAAYEATRDDRLAYDMGRVLAAEIAAVGFNV
NFAPDVDVNSNPLNPVINVRAFGDDPATIGLLGRRMVQGMKSERVIGTFKHFPGHGDTDT
DSHYGLPVVIKSRADAYAIDLAPYRQAIEAGEAPDMIMTAHIQYPSLDDTRVATRTGEQM
IAPATMSRRIQHDILRGEFGYQGVTITDALDMKGIAGFFDEDDAVVKVFQADVDIALMPV
EFRTAADAGRLAALVDRVAAAVDSGRIDRAEFDRSVRRIVLTKLRHGIVASDRGRPIDEL
ASIGGPAHRAIERDIAQKSITVLRNENGALPLQAAGRRIFILTPWGEQAEAMRRRFVELG
HPLVTGAKLSAITWAEQQQAIDAADVVIVGTLSTGVTPVEHNGDPNARVSPPAPSAVRMR
QAAPANGEEEGSVIFDHVERADAAKDIGARPSVLAAIAAPSEAQQMRDAMDYAKARRKTV
IHVTMRAPYDVISYDDVADATLATYAYYGYEGGLRGPSLPAAVDAMLGVARPVGRLPVAI
HALNADGSTGPLRYARGFGLQY*
>GCA_000452945.1_BPMSHR5858_genomic_01900_1 alaS_1
MKAAEIREKFLKFFESKGHTIVRSSSLVPGNDPTLLFTNSGMVQFKDVFLGAETRPYSRA
TTAQRSVRAGGKHNDLENVGYTARHHTFFEMLGNFSFGDYFKRDAIHYAWELLTSVYKLP
ADKLWVTVYHDDDEAYDIWAKEVGVPAERIIRIGDNKGARYASDNFWQMGDTGPCGPCSE
IFYDHGPDVWGGPPGSPEEDGDRYIEIWNLVFMQFNRDAQGNMTRLPKPCVDTGMGLERI
AAVLQHVHSNYEIDLFQQLIKASARETGVADLANNSLKVIADHIRACSFLIVDGVIPGNE
GRGYVLRRIVRRAIRHGYKLGRKAPFFHKLVADLVAEMGAAYPELKEAEPRVTDVLRQEE
ERFFETIEHGMSILEAALAELDAAGGKTLDGELAFKLHDTYGFPLDLTADVCRERGVTVD
EPAFDDAMARQREQARAAGKFKATQGLEYTGAKTTFHGYEEIAFDDAKVVALYVEGASVG
EVKAGESAVVVLDHTPFYAESGGQVGDQGVLANAATRFAVGDTLKVQADVIGHHGELEQG
TLKVGDVVRAEIDAARRARTARNHSATHLMHKALRDVLGSHVQQKGSLVDADKTRFDFAH
NAPLTDDEIRRVEAIVNEQVLANAPGIVRVMPYDDAVKGGAMALFGEKYGDEVRVLDLGF
SRELCGGTHVHRTGDIGLFKIVAEGGVAAGIRRVEAITGDNAVRYVQALDARVNAAAAAL
KAQPSELLQRIGQVQDQVKSLEKELGALKSKLASSQGDELAQQAVEVGGVHVLAATLDGA
DAKTLRETVDKLKDKLKSAAIVLAAVDGGKVSLIAGVTADASKKVKAGELVNFVAQQVGG
KGGGRPDMAQAGGTEPAKLPAALAGVKGWVEARL*
>GCA_000452945.1_BPMSHR5858_genomic_01901_1 gltR
MDLAALAIFRAVVRENGVTRAAAKLNRVQSNVTTRIKQLEEQLGAPLFVRDGRRLVLTPA
GETLLPYAERLLALADEARHAVTENRPHGRLRLGAMESTAASRLPALLARYHQAWPDVTL
ELVTGTTGKLIGLVRDFELDAALVATPAGSEALGEPFETASAFTEELVLLTPRGHPRVRT
PRDVRLPTLVAFERGCAYRGCVERWYAAEGLKPERVLELGSYHAIVACVAAGAGVAVAPR
SVLAVQREPDSLGVHPIDGMPPIDTMLVWRQGHFSAALAALRDALRDAAAAANATDVAAH
EAAKVAPRRTRRGAARETLPA*
>GCA_000452945.1_BPMSHR5858_genomic_01903_1 degQ_2
VSRQTLGRALIHVAALGAVLAGFACLQPTPLAAGTLQSPAKAKRIAQIGAAGPVDFPTLV
ERYGPAVVSVSVPAQDPQMSASGLEALDPDDPFFAYFKSAATQPALSPENGPRAMAGAGS
GFIVGADGIILTTAYVVGQASEATVRLIDRREFKARVLAVDDSSDVAVLQIDATKLPTVR
LGDSSRVRTGEPVLTIGTPDGSANTVTTGIVSATARMLPDGGRFPFFQTDVTGNLDNSGG
PVFNRAGEVIGIDVQIYGSGERNPGVTFAIPIDMAMKVRAQVLQAQRQARQQAQPPMQQA
QQAQQAPPAAAQNALGVDAQDVGPGLAAAFGLPRPAGALVNAVEPGSPAAAVGLKPGDVI
VQIGDRPLGRSAELAGDLAALPPGASAPITLIRNRMPMTVMLGSGAAASAPTGATASPGN
AAAGRSETGGADRLGLTMHPLTDDERRSTGLPVGMVVDAVRGPAANAGIRPGDVVLELDD
TLIETPDMVPALEAKAGKVVAVLIQRGSERRFVSVKAR*
>GCA_000452945.1_BPMSHR5858_genomic_01913_1 iolE
MSAPDIRIGINPLSWMNDDLPSLGGETPLAVALTEGREIGYEGFELGNKFPREPQALKAL
LAQYDLSLVSGWYSGRLAERGVQEEIDAVGPHLELLAKNGATVMVYGEVAGTIQGSPAPL
YQRPRFVDDAQWDAYAERVDAFARYTRAQGVRLGYHHHMGAYVESPADVARLMASTSDAV
GLLFDAGHMTFGGGDPVAELAKHIGRVCHVHCKDVRPAVIRLARNRNWSFLDAVIAGAFT
VPGDGSIDFPTIVDMLKRHGYRGWLVVEAEQDPAVAPSYAYAQKGYRTLRALVDAPLPPV
GHKEAA*
>GCA_000452945.1_BPMSHR5858_genomic_01916_1 frcA
MSTPASHDTILALENVSKYFGKVIALSGVTLRVKRGEVHCLLGDNGAGKSTLIKTLAGVH
PPSDGQYLVDGKPVLFDSPKDALDLGIATVYQDLALVPLLSVARNFFMGREPQKKLFGLF
NVMDLDSSARIARAKLAEMGINVRDAHQPIGTMSGGERQCLAIARAIHFGARVLILDEPT
AALGVKQSLNVLKLIHAARAKGISVIFITHNVHHAYPIGDSFTLLNRGRSLGTFTKDTIS
KNEVLDMMAGGAEMQKMIAELDGATI*
>GCA_000452945.1_BPMSHR5858_genomic_01917_1 xylH
MGVVGKHMSAHAHEKPQPGAASDERLRGEAWFQRFLNRPEFAAISGAVLVFAVFAFGAGG
SGMFNLDGIMNWSQVAAYLGILAVGACLLMVAGEFDLSIGSMIGFAGMMVAIPTVYFHWP
IWLAVLFAFAGSMALGALNGYLVMRTKLPSFIVTLAFLFILRGLTLALSIMFADRTIVSG

LGDYAHADPVVHFLFQGVAFHELFRSLAHAGVGRLLDNGEPLVPGVPKVLLWWLAVTAVG
AFVLAKTRYGNWILAAGGDAHAAKNVGVPVRRVKISLFVLTAFCACLFAVLQVADIGSAA
ADRGLQKEFEAIIAAVIGGTLLTGGYGSVIGAAFGALIFGVVQIGITYTNISSDWFRVFL
GVMLLVAVLFNHYVRRRVSQS*
>GCA_000452945.1_BPMSHR5858_genomic_01918_1 group_4666
MRLCTGKAVLRACVAAIAVAAGVGGAAPAAQAAGARFALVSHAPDSDSWWNTIKNAIKQA
DEDFDVTTDYRNPPNGDIADMARLIEQSAARDYDGVITTIADYDVLKNSLRKVTAKKIPL
VTINSGTEEQSAQLGAIMHVGQPEYVAGHAAGEKAKAAGVKRFLCVNHIATNSVSFDRCR
GFADAIGADYKSSTIDSGQDPTEIQSKVSAYLRNHPNTQAILTLGPVPAAASLKAVQQMG
LANKLFFATFDFSDDIAKAIQSGAIKFAIDQQPYLQGYIPVAVLAIAKQNKTTDPAKIRQ
ILEANPKFQARLSTYGLQPSYGPKNIRSGPGFITKENLEKVIKYAGQYR*
>GCA_000452945.1_BPMSHR5858_genomic_01926_1 group_4668
MRFDRTQRRRLLGRRFLVEWIAIGCLGVAVILAGVAGRMTASVDRIIYDRLLTLRKLPIE
SNIVIVEIDNRSLDTLGRWPWQRSVHAEFLGALAKTHPAAVVYDVLFTEPSADDRALAKV
LDLVPTFLPVLLSPEEPDGTRTVNPPVPVLAQHVAGAGHINLEVDPDGIVRSVALFESDG
LHRWPQLMVPVFHALENGQFKLAQPAPIARRAHDIAQDDNGESRYFVPFSRASQNYPAIS
FVDVLAGRVNADELRGKIVIVGATASGLYDRFATPISGEFGPLPGVYIHASVLDTLMTGR
AISPVSGWLVFSASLLPLVVLLAGFLMLSPWRSLLLTLSLAALAVASSAALLYEARLWLS
PAPAIFGLIAAYPIWNWRRLEMTMSYLRYELQRLADEPHLLPEAPRERNEFGGDVLERQM
ALMAQAAQRVQDMKRFVWDSLDSMPEPVVVTDLAGTVLIANHAAKRYCARLDAPAPEGRP
LRVAFGELSFVKTVDGNAEQDVAIRTHWPAALDPTRDYENDVMERGIEVRDRSGLDHLLR
YAPCTNAQGNVTGWIAGLVDVTALHAAERQREEALHLLSHDMRSPQSSILALVEIERQRA
ESEHMRGLLARIERYAQRALMLADEFVQLARAESQTYQLEAVSFADLLIDASDEVWPQAQ
AKRIRIDTSFGDEPCWIGADRSLMTRALVNLLNNAVKYSPTDTVITCTLSVEPGAKRVHC
TIRDQGYGISLEDQRHLFERFKRFHANERPEVAGSGLGMAFVKTVVVRHGGCIGVDSTPG
IGTAITVSLPLLDEPAA*
>GCA_000452945.1_BPMSHR5858_genomic_01928_1 gtaB_2
MLKVTKAVFPVAGLGTRFLPATKASPKEMLPVVDKPLIQYAVEEAIAAGITEMIFVTGRS
KRAIEDHFDKSYEIEAELEARGKEKLLDLVRSIKPSHVDCFYVRQPEALGLGHAVLCAEK
LVADNPFAVILADDLLDGDPPVMKQMVDVFDHYHSSVIGVEEIPPEDTKSYGIVDGKEWE
DSIIKMSAIVEKPAPEVAPSNLGVVGRYVLKPRIFDHLRALKPGAGGELQLTDAIQALLA
DEQVLAYKYRGTRFDCGSKLGYLKATVEFALRHPEVKTEFETYLRSRSETQHG*
>GCA_000452945.1_BPMSHR5858_genomic_01930_1 uvrD
MPDLLANLNPEQLAAVTLPNEPALILAGAGSGKTRVLITRIAWLIRQGYAAPATVLAVTF
TNKAAREMMARLSAMMPIDTRGMWIGTFHGLCNRMLRAHYRDAGLPQTFQILDTADQLSA
IKRLMKAANVDDEKYPPKNVQYFINNAKEQGLRPEKVDATDSFNRKFVELYQAYEQQCQR
EGVVDFPELLLRCYELLAHNPPLRAHYQARFKHILVDEFQDTNKLQYAWLKLLAGGHNAI
FAVGDDDQSIYAFRGANVGNMRDFEHEFRVRNLIKLEQNYRSHGNILDAANHLIANNAHR
LGKNLRTDAGHGEPVRVYEAATDAQEAAWIVEEIRSLVNAGLARSEVAVLYRSNAQSRSI
EHTLMTAGIAYRVYGGLRFFERQEVKHALAYLRLIDNPNDDTAFARVVNFPTRGIGARSI
EQLADAARLYDCSMAAAIPYVAGKAGSSLASFANLIAKMRAETARMSLPETVEHVVRASG
LADFYQGEREGQDRLENLQELVNAATAFVSEEGYGLDTPARSIPLHARASAAPELAAGGA
DPADAVLAPANLADPAQNPDAMTPLAGFLSHASLEAGDNQAQAGQDAVQLMTVHAAKGLE
FTAVFITGLEEGLFPHENSALESDGLEEERRLMYVAITRAKERLYLSFAQSRMLHGQTRY
NIRSRFFDELPQHVLKWLTPKVEAGARWGGRSDNAGWGRDWFARPGGGARQGVVDAAVSA
PLPAFANQQREAQNGLRIGQQVFHTKFGEGTVTALEGSGTDAKAQVKFKRHGEKWLALAV
AKLQAVE*
>GCA_000452945.1_BPMSHR5858_genomic_01931_1 mtnN
MNQQTFFDRPLGILAALPEELGDLVAAMRDDGPVDTVTLGRRDYHVGVAHGVRCVVTLAR
VGKVAAAATASALIHRFDVRAIVFTGVAGGVASDVSVGDVVVADALLQHDLDASPLFPRF
EVPLLGLTRFAADAALAAQLKAACARFVDDEGETLNARFRLHGARVHAGLIVSGDRFVSS
EREVSALRDALPDALAVEMEGAALAQVCHEYGVPCAIVRTISDTADDHASTSFTTFLTDI
AGAYSSGILKRFLAAHARANAA*
>GCA_000452945.1_BPMSHR5858_genomic_01939_1 group_4671
MDTLQNMRVFVRVVDAGSFTAAAQQLNSTTAYASRAVSDLEAHLRTRLLNRTTRRIALTE
AGERYLQRCEQILAYVEQAEAEASDAHARPSGKLKVHCMTSLGQHYTVPAVSRYRQRYPE
VQVDLTLAQRLPDLLDEGFDVSLVVGRELPDSGLVSQRLGETFSVVCASRGYIEQHGAPQ
RPQDLAGHVCLGMVAPGLYWDEWKLVGPQGEQSVTLAPPPFRVNVAEALAAGIREGMGIG
VLPLYSAIAGLRNGDFVRVMPEYRSHVMNIYALYASRQYLDAKIRTWVDFLRDELPSILE
ADEAALERFTVQT*
>GCA_000452945.1_BPMSHR5858_genomic_01942_1 group_6048
MTSTSAFVDACRHAIGADHVLTDSHDTAPFLTDWRRRYQGAACAVLRPANTEEVAALVKL
ALEHRVALVPQGGNTGLAGGATPDAGGKQAVLSLARLNRVRALDPHNNTITVEAGVILAD
AQARARDADRLFALSLAAEGSCTIGGNLATNAGGTAVLRYGNARELCLGLEVVTAQGEVW
DGLRGLRKDNTGYDLRDLFIGAEGTLGIITAAVMKLHPLPAAKVTALAALESPHAALDFL
ALAQRAAGPLLTGFELMSDFCMKLVGKHYPQLRYPFDAAHPQTVLLELSDNESEAHARAL
FETMMEQAFEAGLVVDAVVAENLAQSRAFWDLREHIPLAQADEGLNIKHDIAVPISAIAH
FIDETDAAIQAAAPGARMVTFGHLGDGNLHYNVQMPEGGDPKAFLAEHQAPINRIVYDNV
HKHRGTISAEHGIGQLKIDDAQRYKAAVEIRLMRALKTALDPLNLMNPGKVLH*
>GCA_000452945.1_BPMSHR5858_genomic_01947_1 group_4672
MRVSDILKVKGNTLFTVTPDTPLRSAVDTMAEHDIGSLVVMEYGDLVGILTFREIILRLK
ANGGSIGEVLVRTVMDEPLTCTPETDIDEVRRMMLERHARYMPVLDKKVLMGVISFYDVA
KTVVEAQSFENRMLKAYIRDWPEPDSEERKPLAS*
>GCA_000452945.1_BPMSHR5858_genomic_01954_1 ompW_4
MRKALCTALGVAALTPLAAHAQSAGSNVVTLGWFHVMPQQSSTPLTTNVAPTPINTPLRL
PPSFTSAGTGLSTSGADTVGLTVSHFLTDHIAVTSVAGVPPVFKVSGHGTITPPGPAGAL
GTQNIGLAQVNPIVKSVRQWSPAVVFQYYFAQATAKFRPFVGIGVSYNWFSDLQLNTNFI
KQTQDNLGAILAAGAGKPGVTQVSAKASSSWQPVFNAGLSYNITDHWGLVASVTYIPLKT
TSTVTIKAADGTTLAESKSDLKADPIISYVAVSYKF*
>GCA_000452945.1_BPMSHR5858_genomic_01956_1 group_4674
MSDAFSKGVAMAIAVAPLFAACSGGGGGTPAPIAVPQCSGSSCGVQGPPSSTAANTSLCP
ADANIGSSTYLGGAGGGEIVSLNINATTMTYTLKWLESPVPLATGTVTPTRAGTTITGSV
AHPPAGTLPTAEQTRCAFVLLPGSGTAPATNSTYSTAADFNQANPPMILIGFGVAGGGIP
GATIQYSGLTIIPGVLQNIGQVPQRHFDFYPFLGFANTTTDLSKLPGTYNALVYHTVPSG
NYAAKAIASNETFDANGACTSTSASGCMTTGNPWTASGNGYFNSTQAPQILPQTQLPLIG
ATGKSAVAHMVLGQLNGATVPVVVRTGNVNLGTPPLHTDAQVDDESGIAVLGLAQAIASG
GIDGGYAGADSNFKYTATVIKGTTGTFVNPSTQQAETGFTLDYGQSTPGLLGVTTTDTSA
PGFVIASGGLYAALVQGTVNGGITQSSAIAGQTPSAPYFGVGAQVSK*
>GCA_000452945.1_BPMSHR5858_genomic_01958_1 fldP
MSNLVIVYHSGYGHTKKLAESVLAGAREAGANARLVAVGDLDDAGWAALDAADAIVFGAP
TYMGGPSADFKKFADATSKAWFGQKWKDKIAAGFTNSASMNGDKFSTIQYFVTLAMQHGM
VWVGTGLLPANTKAATRDDLNYAGGFTGLLAQSPSDASPDEAPAGGDLGTAKAFGARVAT
ATARWLAGNGR*
>GCA_000452945.1_BPMSHR5858_genomic_01962_1 group_2166
MNAPLDAGQRASLEAALQSVTLDDKYTLERGRAYMSGIQALVRLPMLQQERDRAAGLNTA
GFISGYRGSPLGGLDLSLWKAKRHLAAHRIVFQPGINEDLAATAVWGSQQVNLYPGAKYD
GVFSMWYGKGPGVDRTGDVFKHANSAGSAPHGGVLVLAGDDHAAKSSTLAHQSEHIFKAC
GLPVLFPSNVQEYLDFGLHGWAMSRYSGLWVALKCVTDVVESSASVDIDPHRTQIVLPTD
FVMPDGGLNIRWPDPPLVQEARLLDYKWYAALAYVRANKLDRIEIDSPHARFGIVTGGKA
YLDVRQALVDLGLDDDTCARIGIRLYKVGCVWPLEAQGAQAFARGLDEILVVEEKRQILE
YAIKEELYNWPDGQRPRVFGKFDEKDGAGGEWSVPMGNWLLPAHYELSPAIIAKAIATRL
EKFELPSDVRARIAARIAVIEAKETALAKPHVSAERKPWFCSGCPHNTSTTVPEGSRAIA
GIGCHYMTVWMDRNTSTFSQMGGEGVPWIGQAPFTDEKHVFANLGDGTYFHSGLLAIRAA
IASKANITYKILYNDAVAMTGGQPVDGVLTVPQITHQLAAEGAKRIVIVTDEPWKYDGHT
SQLAPGVTIHHRDQLDDVQRTLREIEGTTILIYDQTCATEKRRRRKRGAYPDPAKRVVIN
EAVCEGCGDCSVQSNCLSVEPRETEFGTKRQINQSTCNKDFSCVKGFCPSFVTVEGGRLR
KPKAVSADASALPPMPEPVLPAIDRAYGVLITGVGGTGVVTIGALLGMAAHLEGKGVTVL
DVTGLAQKGGAVMSHVQIAHAPADIHATRIAMGEADLVIGCDAIVTAGDECTSRMRHGTT
RVVVNSAQTPTAEFIKNPKWTFPGASAELDIRAAAGDAVDLVDANHFAVALLGDAIYTNP
FVLGYAWQRGWLPLAYASLVRAIELNGVQIEQNRAAFEWGRRAAFDPASVRAAAADDAHA
GATVIALHTKKAVDALIATRVEHLNAYQNAAYAARFSAVVSQARAAERALGADTAQELLT
EAVARNLCKLMAYKDEYEVARLHAAPAFVAKLAEQFEGDWKLSFHLAPPLFAKKDARGRP
VKQQYGPWLMPVFRVLAKLKFLRGTAFDVFGRTEERRIERALIGEYEALVQELTGSRLTR
DTLPLAVELASLPDGIRGYGHVKENNLRAVRAKWAQLLAQWRSPEGGQTRNVA*
>GCA_000452945.1_BPMSHR5858_genomic_01970_1 paaD
MPAFPASALGQPTAADAREGRRPVTGRAAAPGDDPLVERAWSALEAVPDPEIPVVSIREL
GILRDVRRASDGALEVVITPTYSGCPAMQQIAEDIGAALRQAGIAPHRIATVLAPAWTTD
WITADAREKLRAYGIAPPVGQCGEARGAHAAAPRVVRFMPKPPAAPPCPRCGSAHTERLA
QFASTACKALYRCLDCREPFDYFKPY*
>GCA_000452945.1_BPMSHR5858_genomic_01971_1 paaC
MTTTPQHLAYVLRLADNALILGQRNAEWCGHGPVLEEDIALTNMSLDLIGQARLLYTHAA
ELERRLTGAARTEDDYAYFRAEREFANYTFVELPHYGPLCGTAHAELDYAVTVVRNFLYS
TLMLHLWTALEASGDAQLAAIAAKSVKETRYHEHHAREWLIRFGDGTDESHRRAQAALDH
LMPYTREFFATDAIDDAAAAAGIGPAIASIEPAWRADIEAALAEATLALPTAGQYVPTGK
RGEHSEHMGYLLAELQSVARQHPGATW*
>GCA_000452945.1_BPMSHR5858_genomic_01983_1 catC
MLFHVEMTVNLPVDIDPVKAAALKAEEKEIAQRLQREGVWRHLWRVAGRYANVSVFDVES
PAHLHDVLSQLPLFPYMQIDVRALCRHPSSIHDDDR*
>GCA_000452945.1_BPMSHR5858_genomic_01995_1 group_2372
MLDDKERELRNSKRRALALLLAAAGVFAATLFAPRGFWSDGVKAVAEASMVGALADWFAV
VALFRRVPIPFVSRHTEIIPQNKDKIADNLAAFVHEKFLDPASIVALIKRHDPAARLAQW
LATPRNANVLGGYAARLVAFGLDMTDDARIQSFVKDAFHAVLERIDLSQSAGAILDTLTK
DGRHQALLDDGIAQIVGFLRDPDNRASIAAYIVDWLKYQFPKMEKLLPTNWLGEHGAELI
SNVVTRVLTQIAEDPEHRLRRSFDDAAARLVTRLKSDPAFIAKGEEIKRYLRDGDAFNRY
VKDMWDQLRAWLKADLARDDSVVHRRATALGGWLGERLAQSPELRDSMNEHVERAASEMA
PEFAEFLTRHISDTVKNWDAREMSRQIELNIGKDLQYIRINGTLVGGLIGLGLYAVSSAA
RWAGALPY*
>GCA_000452945.1_BPMSHR5858_genomic_02016_1 atm1_3
MVQKTTHPGGRAFTDVLAFTFRHWRAQPVRIAGIAALALATALGDVLTPLFAGRLVDALS
HGVAAADKAAAWSRALHAFGVLIGIGLACTLLRQAVFRNIIVLTLKMMSEIAANAFHRVQ
RFSTDWHANSFAGSTVRKITRGMWAIDLLNDTLLVALLPSIVMLTGATLLLGLHWPVMGA
VVGGGSVVFIAVTVAMSLRYVAPAARLGNLWDTRMGGALADAVSCNPVVKAFGAETREET
RLDGVVSKWRARTRRAWRRGTLNGGVQGAMLAAMQAAILGAALWLWAHDRASIGDITFAL
TTFLVLQGYLRDIGMHIRNLQRSVNDMEELVSLERQPLGIDDKPGAKPISIARGEIRFEH
VTFGYSGAHAKPLYEDFSIRIAPGERVGLVGHSGSGKTSFIKLIQRLYDVDGGRITIDGQ
DIAEVTQASLRSQIAIVQQEPLLFHRTLAENIAYARPGATRDEIERAARLASAHAFIDAL
PQGYDTLVGERGIKLSGGERQRVAIARAFLADAPVLILDEATSSLDSESEVLIQQAMERL
MVGRTTLVVAHRLSTVRALDRLIVLDKGRVIEEGSHDALIRLEGGVYRRLFERQALELAK
GLIEQTGGTVADWLRDEQDEHGGRDERNERGERDKPGALDGHDARRTHGVLDESRDGGQA
DNSAMAMEK*
>GCA_000452945.1_BPMSHR5858_genomic_02019_1 group_2373
MPLIHATAEMAPDAPNYVVELEAGKHRLQGDENEHEGGGNVGPAPYELVLSGLAACTVAT
LRMYMQRKAWPAARLLARAELHVDREGAQYVRRTIVVDGALDGAQRTRLAEIAEKTPVTL
FVKRGTRIETTLK*
>GCA_000452945.1_BPMSHR5858_genomic_02020_1 nodD2
MQPDHRLDLNLFRVLDAVYVQGGISAAARALHLTQPAVTHALNRLRAHFGDPLFVRQGNK
VVPTERTREVIADVQRHLKGLQDAARAQTRFSPGALDMSFVVGVRDVLESIALPQLVETL
AREAPGVRLVSRRVPVADIGRELGSGNLDLAIERRVPAGPRIAAEHVADDTLVVAMRRDH
ALAGAPLRRADYFAAGHVTVSPLGEPHTLDVLLGADGRFRRIQLVCQHYFAACQIVASGD
LLLTLPRSYALRMTRLLPLAAQPLPLRLKPYPILAYWHESRELDRAHQWFRQRVIEVVRA
ASPHPAGA*
>GCA_000452945.1_BPMSHR5858_genomic_02024_1 fabG_5
MSAPNLFDLSGKIAVVTGGSRGIGAEVAKLLGAFGAHVVVTSRRLDDCRAIADEIVAAGG
SAEAAACHIGELEQIDALYASLEGRHGRLDVLVNNAATNPYYGPIVDTDLGAFQKTVDVN
IRGYFFMSSRGAKLMGKNGGGSIVNVASVNGVIPGYWQGIYSITKAAVISMTKAFAIECA
ASGVRCNALLPGLTDTKFASALTQTPEVLKQVLAHVPMRRAAQPTEMAGAVLYLASAASS
YTTGTVLNVDGGYLSV*
>GCA_000452945.1_BPMSHR5858_genomic_02028_1 btuB_3
MDDHRRIAPPFARRLHPLSLLLAASLAHGETGAPPAERRSDAPPATALAPIFVTANPLGA
SALSSPTASLSGDALTLRRTDSLGDTLNGLPGVSTTTYGPLVGRPIIRGMDGDRIRLLQN
GVAAYDASSLSYDHAVPQDPLSVERIEIVRGPAALLYGGNAVGGVVNTIDNRIPREAITG
VSGALDASYGGANNARAGAALVEGGNGRFAFHLDAFGRETDALRIPGHAHSARQRALDGE
DASEPYGKLPNSDGRRYGGAAGGSYTWADGYVGASYSGYESNYGSVAETDARLQMRQERV
ALASEVRNLRGPFSQLKFDFGYTNYQHKEIEDGVTGTTFRNHGYEARVEARHRKLGPFEG
ALGVQVGQNTFSALGGEALAPTTRTTSVALFGLEQWQATDALKLSAGARIEHVRLDPSAN
GDDKFGFARSRDFNAGSVSAGALYQLAPAWSLAGNVSYTERAPTFYELYANGPHGATGQY
LIGRPDAQKEKAVSTDLALRYASGPNRGSIGVFYSRLRNYLAEYDTGRLVDDDGVPVAPG
ADDALREAVYRGVRAEFYGVELEGRWRAFERRGHRVDLELSADYTHARNADTGEPLPRIA
PLRATLAADYGYGPFGARAQLTHAWAQHRVPEHDLATDGYTSLGVVLTYKLRVGATNWLA
YLRGDNLTNQDIRYASSVVRNIAPQGGRSVSIGMRTTF*
>GCA_000452945.1_BPMSHR5858_genomic_02029_1 group_4680
MKRTIRKSWAALVPLLVAGSVCAAAPLCQTQKLGAHTSKMCVEQTPFKHDYYTLWVDDSP
IFMLPDDYVEKVALTHTVPEDGAIEFPLSKQGTPTVTISGGCAPVSETQGKGADAVNLET
GRVCSFNWGKEPVVKDLRFSFE*
>GCA_000452945.1_BPMSHR5858_genomic_02031_1 group_4681
MRKTLVMKVAIATVLGSLALAGCTTTPDKPANASSNASTREAIDARVNATLSRLYSTVPG
SRELVSKSRGVLVFPNVLQAGFIVGGQSGNGALRVGGSTLGYYNTSSLSVGLQAGAQSKA
LIFLFMSQDALDKFRNSDGWAAGADASVALVKVGANGAVDTSTVTAPVEVIVLTNAGLMG
DLSVNGTKVTRLKL*
>GCA_000452945.1_BPMSHR5858_genomic_02040_1 amiD_2
MAEEQQAKIINEGRRGFLARSSAWPVGMLMLPLAGCGDGDGVVPSAHAQDTPPPSGAGYT
IDRSIQSPNQDSRVRTLVLHYTAQTLADSIASLTSPQRQVSAHYLVPDAADDGRRFKVFE
LVPQARRAWHAGVSYWQGDRMLNAGSVGIEIVNLGFPPEDENLPLMNRRWYPYPDAQVAV
FGALAADVVAQHQVLPHKVVGHSDVAPGRKTDPGPLFPWKKLYDQYKIGAWPEAEAVDYY
RVNRPFAGDVASLQSKLLAYGYDTPQTGTLDAQTVNVVSAFQMHFRPSRYDGVPDVETVA
ILDALLEKYFNRGRAINKQTLPGATAPTGEKGSDAWPLAPAPSR*
>GCA_000452945.1_BPMSHR5858_genomic_02050_1 wzc_2
MVNTQAKHPYADLAVKTDEEDVVLGQMIQVILDDIWLLLGIALVVVALAGLYCYVAKPLY
SADAQVRVEASDNTSQALTQTQTGAMINSGPPTPPTDAEIEIIKSRGVVAPVVEQFKLNA
SVTPNTLPILGAIAARLATPGHPGKPWLGLSSYAWGGEEASIDSIDVTPVLEGKQLTLTA
GADGGYALADPDGAVLVRGKVGEREQGGGVTINVSKLVARPGTRFTVVRQNDLDAITAFQ
SAIQVAEQGKQTGVIQISLEGKDPEQTAQIANALAQSYLHQHVTSKQAEATKMLEFLKNE
EPRLKSDLERAEAELTQYQRTSGSINASDEAKVYLEGSVQYEQQVAAQRLQLAALAQRYT
DEHPLVVAAKQQLGQLEAERAKYDGKFRGLPATEVKAVALQRNAKVAEDIYVLLLNRVQE
LSVQKAGTGGNIRLVDAALRPGVPVKPKKVLILSAATLLGLILGTSVVFLRRNLFHGIED
PDRVERAFNLPLYGLVPMSAEQARFDAADKGNRVRPILACARPKDLSVESLRSLRTAMQF
ALMDAKNRVIVLTGPTPGIGKSFLAVNLAALVAHSGKRVLLIDADMRRGSLDRHFGTGGR
RGLSELLSDQVALEEAIRETSVPGLSFIPSGARPPNPSELLMSPRLSQYLDGLAKRYDMV
IVDSPPILAVTDATIFGELAGSTFLVLRSGMHTEGEIGDAIKRLRTAGVQLQGGIFNGVP
ARTRGYGRGYAAVHEYLSA*
>GCA_000452945.1_BPMSHR5858_genomic_02052_1 group_4683
MIATKVHVHLFYGADPRTYRQGDDFGCLYGYHHAQSEEFTLTYSRDAREGRLVSIGRRAL
KYALGFDLMHAWRNRAALLDSDVIWTHTEQEHLAAALVLKLSGARGAKPLLLAQSVWLLD
KWPGYGRVRRWAYRKLIGRADVLTTLARNNAELCRRYLRRDALQVYYGLNTQDFPLREPQ
AWSPRRPLRIAAIGNDRDRDWKTFVAAFGGDSRYDVRLATRRRIARALHAPNVEIAPASG
IVKQRELYDWADLIVVPLRPNHHASGITVMLEAAAVGKPMIVTDVGGLRDYFPHGTATYV
QPFDALALRRAADALAARPDQALARAQAAACRLRERDLTTQQFAAQHVRITRDLLTRRRA
HAAAEAALSFADSRSRSNG*
>GCA_000452945.1_BPMSHR5858_genomic_02053_1 group_4684
MSATTLPVPAAGGRARRRKLLSDPKHWAAQAGFWSVTALLIAAHQGTLLTYGFPCLAIAT
GLWLYFMSPARYVGFMWWMWFLSPEVRRLADWSKGAYTPTSLIQIAPLAVTMIAGLGLLR
HYRVLAERRGLPVLLILFGLMYAFLVGVVSSGPLAATYDLANWLYPVLIGFHIAVHARDY
PEYRKVILSTFVWGMLVMGVYGMVQFFVMPKWDALWMIGSQMNSQGDPVPYGVRVFSTMN
SSGPFAFAMMGALVFMIAATERVRWLAGAAGFISFCLCLVRSTWGGWVVALAIQLAKSSN
RVRVRIIGSAIVLAGLCVPLLSVGPVADRLGQRLQSITNLKDDRSYDDRNKFYATFAQTA
FTDVAGEGLGATGASTKLSSDSGSLGKYGSFDSGVMNVPFVLGWPGTLLYLLGLVWLLGR
ALRASLRLRRDKFVAACLSLSLSVFAMLVFTNSFIGTGGLLLFMSIFSILSAAHWQKLQR
ASVARFPGADQ*

>GCA_000452945.1_BPMSHR5858_genomic_02055_1 mshA_2
MKHTETSIKSLQIGMHWFPERAGGLDRMYYSLVGALPSAGVAVRGVVAGSERVAADTGGA
IRGFGPATSSLPRRMIAARHALRDVMRIERPDVVSSHFALYTFPGLDVTRGIPQVSHFQG
PWADESHVEGADSLGQKVKHRLEQAVYARSSRLIVLSHAFGQILTSRYNVDPARVRVVPG
CVDTAQFDLPMTPADARRKLQLPQDRPIVLAVRRLVRRMGLEDLIDAVKTVRRRHPDVLL
LIAGKGRLEGELRKRIDDAELGENVKLLGFVPDNHLAALYRAATLSVVPTVALEGFGLIT
VESLASGTPVLVTPVGGLPEAVAGLSEALVLPEVGAAAIADGLAAALSGSLVLPDADACR
RYARAHFDNTVIARRVAEVYEEAIRAAV*
>GCA_000452945.1_BPMSHR5858_genomic_02056_1 group_6726
MTRLAAPSLSLRRYGAVQASDVHDFHQIVLGVDGAMTMTVDGVDERIDRRCAWLIPAGAR
HDYAGLADNSQLVLDLPPGSLAVPERLFERARTMAIDPGLTTLVGELASRIARGPLADAA
SARRLHWQAAARLCGALVGELGGAPSQPAVGLDFARIDGWLRAHLAQPLRIADLAAHCGF
GMRRFHQLFCEAFGETPHRYLQRLRLDAAVMLLGDPKHSLTDIASEVGFADQSAFTHAFT
KRFGLAPGRWRSDRH*
>GCA_000452945.1_BPMSHR5858_genomic_02062_1 pphA_2
MDTAPLIRRHTANHAGRDFVVGDLHGCVDPLRALLHTVRFDPARDRLFSVGDLVDRGAQS
EAALALLERPWFHCVLGNHEDVLCSVADGKLPMSVWQRIGGEWAADLPAATLARHARRLR
ELPLVRVVGEGDARFNVLHAEFFGSDADLDRGDYAPDVITRLLWGRELAYGRTDPAARQA
GLSTTYCGHTPMHDVVRIGAQTFVDTGAFDPSGRLTLLDAKSGESWSMSVAAARHQGAAD
VPLP*
>GCA_000452945.1_BPMSHR5858_genomic_02066_1 lgrD_1
MARHIVQAIQSHAFATPYKCALSDSRGDLGYADLDSFSTRFAMRLQDLGCRPGDRVVMLA
SRRALLVAAIIGVFKAGCVHVPLDPRMPADRLRYILHDVAPTLVIADEDLIDAIEHALPA
AAPIVPVTELERLLDDADSPRLDALVQPLPLPPLDERAIAYCIYTSGSTGRPKGVLINHR
SIADFFEGTRAVYDVTAQSRCASFSPLNFDVYLMDMLFPLAQGASLYVHDDVNAPDLLFD
AIRVHDVTHFSAWGMMLGLIAQAEEFEAAPLPHLKTILTGTDVPDVKTVQRWLRKNAGVQ
VINAYGPTEATCAATAHVIREIEPERRTLYPIGKPLEHVRALLVDEGGNRITAPGVPGEL
MIGGTQVMQGYWNLPEETAARLVRLDGVPFYRTGDVCAYLADGSLYYMGRKDNEVKIGGY
RIHLSEIQRVINSVPHVYGSEVVLLESRYGETLLAAGVLLERGAPLDADCKADEIRQRLA
AELPAYMVPRHVKVLEQFPQLSSGKTDRKALLSILQQRINESNQEEVNP*
>GCA_000452945.1_BPMSHR5858_genomic_02081_1 mdlB_1
MANVGKAWNIDDLRKMARKRVPKYFFDYLDGGANSETTMRANENDFARWRLRQKVLTGAQ
SRVAGLGATYLGAEHRLPILLGPVGFAGMYWSRGEIAAGRAADDAGIGQCLSTFSICSLE
DVAAARSGPLYFQLYMFRDRDLTEDILARCRLANVDAVFLTVDTCYIPIRERDARNGFRA
DTRLSARGVWSMLKRPGWCVGALSHGVPRIGNVLRYPELGASLLEQSAAVGRMIETRLSW
ADVKWLRARWPGRIVIKGILDPDDARRAVDEGVDGILISNHGGRQLDPAPSVMDVLPEIA
EAVGKRTEILMDGGVRRGADVIKALALGASAVSIGRAYVYGLGAAGEKGVSRCLELLKGE
MLPALNMMGFESIAELRAAGKSALRMAVPVHGATAVVEGAEGGARRLTESVL*
>GCA_000452945.1_BPMSHR5858_genomic_02082_1 group_4688
VSVESVRQFLAEKAPDIDVIALSESSSTMTLSAAWDIKPAQIAKTLAMKVGDAHVLLVSC
GDSRLDNQKIKAALGGKAKMLSAEETVAVTGHPVGGVCPFGLPAPLPVYCDVMLKSYEFV
VPAAGSTHAALRIDPVRLAELVEAEWVDVCK*
>GCA_000452945.1_BPMSHR5858_genomic_02084_1 group_870
MSTPSAAAKFDPARAVEYAEQSRITLAGYDACHDLAACMLASAIPAEAHAAQVLVAGAGG
TAREIAALAGLAPGWRFTAVDPSRPMLDLASANVAAAGLDARVTTHLGYVDDLPADARFD
GATLIGVLHHVAGDGAKAALLRSIARRLKPGAPLVVAGNYRRYAEHPRLLSAWAQRWRMH
GATPEEVARKLATILRGADPPASEDAVHALLDAAGFREPVRFFASLFWGAWIAMRGA*
>GCA_000452945.1_BPMSHR5858_genomic_02119_1 rsmH
MGNEFQHRTVLLDEAVDALVTRPDGVYVDGTFGRGGHSRAVLARLGYAGRLIAFDKDPRA
IETAESIEDARFEIVHDSFAAMKGALDARGIGRVSGVLLDLGVSSPQVDDPARGFSFRAN
GPLDMRMDPTRGESAAEWLARASVQELTEVIRDYGEERFAFQIAKAIVARRAESDRLGPL
DSTGELAQIVGHVVKTREKGKDPATRTFQAIRIHVNQELADLQVVLEAALSLLEQGGRLV
VISFHSLEDRIVKRFLQAHASAPAVDRRLPIRAADLPSPPLKLLGRMFPNDAEVAANPRA
RSAVMRIAERVAP*
>GCA_000452945.1_BPMSHR5858_genomic_02125_1 murD
MFGDRQRPMVLVLGLGESGLAIARWCARHGCRLRVADTRETPPNLAALTAAGVDFEFVGG
AFSPALVDGGIELVALSPGLSPLAEDLAPLVAAARERGIPVWGELEFFAQALAALGANGY
APKVIAITGTNGKTTTTSLAGLLCERAGKKVAVAGNISPAMLDKLTEAIDAAALPDVWVL
ELSSFQLDTAHTFAPDAATILNITQDHLDWHGGFAAYAAAKGRVFGPRTVRVLNRDDAEV
MRFAPPAAAADAPRAVTFGLNEPAADGDYGLLRENGIAWLVEAIDRDGADAPAAPSRRRK
QEAANPPDIALKRLMPADALRIRGLHNAANALAAYALARAIGLPAAPLLHGLREYRGEPH
RVEVIATLDGVDYVDDSKGTNVGATVAALDGLAQRAVLIAGGDGKGQDFEPLAAPVARWC
RAVMLIGRDAPALREALADTGVPLADHATLEAAVRAASALAQPGDAVLLSPACASLDMFR
NYAHRADVFRSAVEDIALEKGTTL*
>GCA_000452945.1_BPMSHR5858_genomic_02127_1 murG
MTSTQRTLMVMAGGTGGHVFPGLAVAHRMQAQGWRVVWLGNPAGMEATLVPRHGIPMEYV
RFGGLRGKGLATKFALPFNLLRACAQSLRALRRVKPDVVLGMGGYITFPAGLVTVLTGRP
LVLHEQNSIAGLTNKVLAKLAKRVLVAFPGALPNAEWTGNPIRTELARTEPPQARYAARS
GKLRLLVVGGSLGAAALNEVVPRALALLAPDERPQVVHQAGAKHIDTLKENYEAAGLSCG
SDVALVPFIDDMASAYANADLVICRSGAMTVAEIAAVGVAALFVPFPHAVDDHQTTNAEF
LAEQGAAVLVQQRDLSAELLADWLRGQSRDSLAAMAERSRSLAKPDATDEVARVCAAVAG
ANLEGKQ*
>GCA_000452945.1_BPMSHR5858_genomic_02129_1 ddl
MSGIDPKRFGKVAVLLGGDSAEREVSLNSGRLVLQGLRDAGIDAHPFDPAQRPLAALKDE
GFVRAFNALHGGYGENGQIQGALDFYGIRYTGSGVLGSALGLDKFRTKLVWQQTGIPTPP
FETVMRGDDYAARAQDIVAKLGVPLFVKPASEGSSVAVEKVKSADALPAALEEAAKHDKI
VIVEKSIEGGGEYTACIAADLDLPLIRIVPAGEFYDYHAKYIANDTQYLIPCGLDAAKEA
EFKRIARRAFDVLGCTDWGRADFMLDAAGNPYFLEVNTAPGMTDHSLPPKAARAVGIGYS
ELVVKVLSLTLD*
>GCA_000452945.1_BPMSHR5858_genomic_02131_1 ftsA
MSKDYKDLLVALDIGTSKVVAVVAELKGEGHYEVIGLGQSESKGLKKGVVVNIEATVQSI
QRALEEAELMADCKITNVFTGIAGSHIRSFNSSGMVAIKEKEVTQTDVARVIETAKAINI
PTDQQVLHILTQEFIIDGQEDVREPIGMSGIRLEVKVHIVTGAVSAAQNIVKCVRRCGLE
VNDLILQPLASSLAVLTEDEKDLGVVLVDIGGGTTDIAIFAEGAIRHTAVIPIAGDQITS
DIAMALRTPTPDAEDIKVGYGIAKQALADPDEMVEVPGLGERGPRTLSRQALAAVIEPRV
EELFSLVQQVVRESGYEELLSSGVVLTGGAAMMPGMVELGEDIFLKPVRIGAPEYAGGLS
DVVRNPRYSTAMGLLVEGSAQRMRGRKVAVQSGSAGQIFSRMKEWFLSNF*
>GCA_000452945.1_BPMSHR5858_genomic_02144_1 group_180
MTKRTRRTHSAAFKAKVVLAAVKGERTLAELAHQFDVHPNQITEWKRQLQERAADVFGSP
GTASSELPVDLKSLHAKIGQLTLENDFLSGALDKAGLLSAKK*
>GCA_000452945.1_BPMSHR5858_genomic_02170_1 group_4699
MALTLINKKASPDGTPPARPAGWRDRAAAWARGPTLARDISIVLAFKLVLLIALKYAFFN
HPQAAHMSLPPAVVAAQLLSATPSAPTEGDRHDK*
>GCA_000452945.1_BPMSHR5858_genomic_02175_1 chb
MNRISHSLCAALLAAATLLPTASRAQLPARPTAGAAAPATAAPVRPASTPAELAARLANG
LAVRVAVDNNHAASAGVPCADLGADWASCATGRLILQNRGHSPLTDGGWKLYLHSIRRLL
RIDRPGFTLRHLTGDLYELTPQPGTVRLAQGERIELPFVAEYWLRRYSDVIPRPYVVVDG
AAPAVLRYDDTDDELRYVETLPADAQNNSPGNAPPAAAQPVANRALPSVKRQRALPGALD
LRGVELTLPELPSAQVAALRERAGTLGLDGARVPVWGVVAPRRLPADIAVPGGYRLAIGP
RGAFIEGADRAGLYYGVQTLFSLVPAGGATVPAMLIEDAPRFTHRGMHVDLARNFKPPAT
LRRLIDQMSAYKLNRLHLHLSDDEGWRIEIPGLPELTDVGARRCHDPSETRCLLPQLGSG
PDDRSGGGYLTRDDYVALLRYAAERFVEVIPEIDMPAHSRAAVVSMEARYRRLHAAGRER
EANAYRLLDAQDTSNLLTVQFYDRRSDLNPCMPGALNFASKVIREIASMHADAQAPLRIW
HFGGDEAKNILLGAGFQPLDGADPGKGRVDLAAQDKPWARSPACTALLRRGEIKSIDELP
TRFAKQVSAIVNANGIGTMAAWQDGIKHASGPREFSTRHVMVSLWDTIFWGASDSARDLS
AKGYRTVLALPDYLYFDFPYTRNPRERGYYWGSQATDEYKVFSLAPENLPQNAEVFGDRD
GNPFEVTSAGAAPSIEGIQGQAWGEVMRNGQLLEYMVYPRLLALAERAWHKADWELPYAA
GVRYKLGDTHHVDTAALERDWAGFATVLKQRELPKLERAGIGYRKPTFTLTGE*
>GCA_000452945.1_BPMSHR5858_genomic_02183_1 group_2687
MMPDTALPDTPDTPARTFPAAAHADAADLPTLAAQPVRERLAAFGAPPAALDALLRAGAT
RLLDSDMIIAQPRAHAAARAAERATTPLRYEQSAQTEAVLEQIAHEGLAPHFKPFMRGFS
HVMSALFLTPEQDAKVQAYRAAGHHLRFLMSDGGGPALAGWRSTVRVEHGRTRLAIDKVW
GIEAHRESIGLVAVRAPGAFMPAAYLVWPEQYRTLERAACGEPFLEGGLQLGNVRGEIET
TPEDRLRIGGPNVFNKYLTIVRPYFVRALMAHVGWLAREGRLALSPAHESVRRFVADAAR
AQSRSCVYGADVVQRVLALKFAANELLGDLVRGGAVRRFDDQRDLLALSKMEGSAYRCYY
EIRMGTKRA*
>GCA_000452945.1_BPMSHR5858_genomic_02187_1 group_4701
MQPYEDAARSQRWSMPVDALMMGDYEYRETALTNLYHKAKAAQWDVNRDIDWNVTLDPDN
PLGMPDGMLLVFGTDLWHKLGAAERAEMRHHWQGWLLSQILHGEQAALICASKLASAEQS
LPARLCAAAQIMDEARHVEAYAKLVNEKISVRYPMSRSLEGLLKDTVTSSALDITNLGMQ
VLVEGIALSTFQTVVAFSADPFVKDLFARIQRDEARHFAVGRLTLARVYSDMTSAEMKER
EEFVSEGASVLYEHLCADDIWEPLGYSKQACAQMVRESEVSRALRRSIFRRLVPTIREIG
LLTPRVRRTFEQLDVIDYEDMPLALDM*
>GCA_000452945.1_BPMSHR5858_genomic_02188_1 group_4702
MRPDQANHWIGIDAAAYELPDAAIEIGAWAARHAWPRAQVDAILESGCRFFHAAPQTSEV
ELASRAIGALVDSTGIAPSSIDLMVHVHTQTFSMPAPPRSLLYEVAHAHGIRPLWSGSIA
QLNCASIAAAIRMIDALFYAYPQAQSALVVSADRAYDEATRLRQFSGIQGDGAGCLLVTR
NSRRNRVGGVALHNHAAWYPGSDSANAVERAMISTEWLHTHRIVDAACALTGRPIAHYER
ILPINTDLRGWHALCRALRVPAERLFGANVARCGHVCCSDFAINLADDGLAALDAGRHVM
GVMQSNVGAFAAVALHPLEAASSSTPAEAYEH*
>GCA_000452945.1_BPMSHR5858_genomic_02193_1 group_1839
MGLATELLNRPATRYRGTDIDGAILEPEGLWVRGRNLNDLIGTVSFEDALWHLWFERMPT
PAEGAALRARLAQYGAHFAHGNPSTRAAAAVAQTGVDMVFAAATGLLRDIERPPFDAGEA
PGWGADFDTLLGCLAGAPYLMRAALGAPRVDDPGTSHAARVLRAAGASGGGSPHAERVID
ALLVAWHGGFGYVTPTVLVPRCAIGTGVSLSQAIAAGFMSSGPNHVGAARSAMQWLRALA
RRLDDEPDALDAVVRRAIGRVLDESGALLAGFGHPLFEADPRPPRIRALFAEWGFGGRHV
DMFDVACDEARKRKNLKPNIDFATGAALLDLGFTEPQWGIGVGLSARIAAMAAHAVERRR
RPAFGANSATARRMLAAVPVGWL*
>GCA_000452945.1_BPMSHR5858_genomic_02200_1 group_4704
MSSGIAPPHSVNYRKKYATEAHALSAIRPWRCAWVIARQWIGIAIAFALPIALVARLTGG
TSLAHAFAALGAPQRLGVAAALGAAYVYLACKQHALGIVMHDATHFRLFESRRVNELVGN
WLCAFPIGMVTSCYRRSHLPHHLFTNKPNDPYWARLVEDAHYAFPMSRAAFGRILLGDVF
GANLRAWWPTLRSWTGWSSVLDNREKLLTPSERRQFVAFWIGALALAAYFGVLSYFLLLW
VLPMFTLSLAFIRLRVIAEHDLEKAGHELERTRHVDGGWFERLAIAPLNINYHVAHHLFP
SVPLYNLPKMHALLMQEPAFREHAQLWRSYLGRKHGMVRSLLT*
>GCA_000452945.1_BPMSHR5858_genomic_02205_1 cheY_1
MALPVLVVDDSTLARKLLIKSLPSDWDVEITQASNGAEALLHYRAGKGAVIFLDLTMPVM
DGFQVLEHIREEGLDAFVIVVSADIQEGAVDRVKASGAIGFVAKPVSTERIVPILKGYGL
YE*
>GCA_000452945.1_BPMSHR5858_genomic_02209_1 group_4706
MTETIVVRMAHSDDYPNILALQKANQIGNLTEAERKNGFLSAEIPEERIGALASDTGIVV
AYDQAEFAGFFCISRLASWRGNAIIDALVASFEQSEEYASHRISETMCLFGPMCLNQVAR
GKGILEKMIGLAIESAKTQFENAISFIAVDNARSVNAVAKLGYRPVRRFTSGEREYHALI
AEFA*
>GCA_000452945.1_BPMSHR5858_genomic_02212_1 leuS_1
MSAPHADGARHAPPYRPQALEPRWQSAWRDARLFDADPQPGRPKWFIVELPPFATGQLHM
GHARNYVLADADARFRRMQGYNVLYTTGFDTFGLPTELAAREAGCAPERLARECSEAMAA
QFVRLGLGHDSRRITQYHVPAYYRWVQWVFLRLFEAGHCFRRDAPVAWCPQCEVTLAASL
VDDGRCWRCKRVVSTVVQPQWFVRESTFADEMLDGLDRLAGWPDEVKKIHRDWIGRREGL
RLRLPVRGRAHGLALHLDDAGWLPGVRFVAVGRRHPLAAAALRDGVQPGETILLADVAQA
AGAAGAVELPIIVEDAPGDAARAGRPDAVAVDRALAERHALAAPSPGAGGAPAVCDLDAM
LAAGQAERVVGYRLQDWNIARNRYWGPPVPIVHCAACGPVAVPEHALPVRLPEDVDLSRP
GNPLEHHARFRDVRCPRCGASARRDAQTLEAYSSPWWYHWMCKRVDVDDPFNRDDANAWL
PVDLMIGGADQVRTCFFHVRMIARALRGMGIADVDEPVTTLLALGMVKQDDRKMSKSAGN
AVSLASIIERYGADALRLAIIGAAAPENDVNWSDDLVRRQRAFVARLWAFVHRVAETEAG
IAAAHDAGDPPAAGRPDALQRRMQDWLATGGRRIAADFRRHDYHLALKNLAFLFERLQAF
DAAMRRRAPPRAEDVAILAGATRQLVLFLGPFAPHVAEELWQALGGAGLVAGAAWPGADV
VRSSAATAKREAFA*
>GCA_000452945.1_BPMSHR5858_genomic_02219_1 group_4707
MGMLDDVQLKAVHASLAPVVERLSEASATLASSAVPLLGVPPATAPPAAVPVAAAPPAGV
ELATQLSAHAAPVALAAPLVSLSIPALSQPVRT*
>GCA_000452945.1_BPMSHR5858_genomic_02224_1 group_4709
MDSDIMHIGGRIRRLRRELKKTLLEVATEANLSVGFLSQVERNLTGISISSLVNVAKALR
APLSALIDQPRQDQPDSHEGSRESYAIRPTQQRYERLSTTFAGSLLNAVKVSMMEGYSSE
WVAHAGDEFVFVLSGHVRYTVGKQDYPLGPGDSLHFDAHERHRVVNVGHGPVELIAVGTL
PLFGDGQPAFPSLPGERKPPRTRAARKRIEPPPTPSAQQDARHDEAAAPNAVTTHDGAPR
AAKKNTR*
>GCA_000452945.1_BPMSHR5858_genomic_02230_1 teaD
MYKKIFAALDGSRSASVALDEAIKIAETFDATVTAVCVVEHKPRLVDVDTAYVEPQDDDG
SATDIATRVLADAKARLAQHRVEGSVRAIDSYGEDIAEVLMRTAAECDADLIVMGTRGRH
GLRRLLLGSVAESLLRIADRPVLLRRHDEAAA*
>GCA_000452945.1_BPMSHR5858_genomic_02231_1 group_10070
MKFKTLLVHVDDSRQSGARTEFALDLARRWDAHVIGLYVVCQDLMRPLFRRDESLRLAAN
EAQHAARRDAAQARFTAAGERAGVNFEWRAPAGPPTDVATLHARHADLVVLGQPEPGDAA
AYIEPHFVDDVVMSGGVPAIVLPFTGAMRTFGENVLLAWDGSREAARAVADALPLLKRAR
FVTVESVLRRRATGAGLAPAGIDVAAYLGRHGIQASFSASAHVPGVDTGAALLNRAADLH
ADLLVMGAYGHTRARERVLGGVTRTMLESMTVPVLLSH*
>GCA_000452945.1_BPMSHR5858_genomic_02235_1 group_2775
MPIHAAHAPRRASRLSAAPMPPGSAFVAFEVRFVWHFPQAVPASLGPWAVFASVLVTQLG
MPVPAVPMLIVGGTMAAMGQASYASMLVAAVAATVLADSMWFFAGRARGRRLLNALVRFS
LSLDTTLRIARKVFEKHGAPLLVLAKFLPGLGLVSAPLLGTTAVAVWVFLFWDVAGASLW
ASVWLFGGAALHDEIVRLMQWVSASGGTLFDAFAAIFVTFLLYRWAMRMRFRRWLAKIRI
SPQQLDAMLKSAAPPIVFDARPRAVREKEAYRIAGAYPLDLDSPDPLHPDLMTRPIVVYC
VCPNEATAKRIVSQLQRKRIRHALALKGGLDAWEKHGYPVEPLPADFDAARYAAPPPVER
IAPVEPGGRDGGYPMRAGLTD*
>GCA_000452945.1_BPMSHR5858_genomic_02241_1 group_4712
MATYKELLAQLDDLKQQAKRARALELPDVLVGLRKKIVEYGLSQKDLFPPRLGRPKKADA
LPKPRYRDPDTGATWTGRGRAPAWIAGQDRERFRID*
>GCA_000452945.1_BPMSHR5858_genomic_02251_1 dinG_1
VSYVVAVRAMCEFTARAGDLDLRFTPAPSALEGIAGHATVVSRRGTGYEKEIALSGEHRG
LTVRGRADGYDPALNRLEEIKTHRGDLDRMPANQRALHWAQARVYGHLLCESRGLAELSV
ALVYFDVGTLRETVLTETHTAATLKAGFVEQCERFADWAAREDAHRAARDAALRALAFPH
ARFRSGQRELAVAAYRAARDGRVLLAQAPTGIGKTLGALFPLLKACGQAHLDRLFFLTAK
TPGRALALDAADTLRAAAGGVLPLRVLELVARDKACEHPDAACHGESCPLARGFYDRLAA
ARAAALEHGRLDRAAVRAAALAHDVCPYYLAQELARWCDVVVGDYNYYYDGSALLHALTL
QNQWRVGVLVDEAHNLLDRARGMYSAALDQRELAGVRKLAPAPLAKALERLNREWNALNR
EQAGAYVVHPDVPQRVLSAAQNLIGCFSEFAAQAPLALDDAVLRFCFDAIHFVALAEQFG
THSIFDATRVRAGAPRRGGHASILCVRNVIPADFLAPRHGAARATVMFSGTLSPFDFYRD
TLGLPGDTGTLDVEGPFRAEQLTVKVASHVSTRWRDRERSLEPIVELIAEQYAARPGNYL
GFLSSFEYLQQVVARLRERHPALPVWEQAPGMDEAARDAFLARFEPNGRGIGFAVLGGAF
SEGVDLAGDRLIGAFVATLGLPQVNGVNEEMRRAMDARFGSGYDYIYLFPGLRKVVQAAG
RVIRTEHDEGVLHLIDDRFHRGEVRRLLPRWWRLS*
>GCA_000452945.1_BPMSHR5858_genomic_02258_1 yajO
MALRSLGTSTIQVSPLVFGGNVFGWTADENTSFSLLDALADAGINFIDTADVYSAWVPGN
QGGESETIIGKWLKRSGKRDQVVIATKVGLLETRAGLSRENILKAADDSLRRLQTDYIDL
YFSHRDLADTAPLEETLGAYQTLIEQGKVRIIGASNYSGARLREAAELSRRTGLPAYQVI
QPEYNLYDREAYERDLEPAATELRLGVVTYYALASGFLSGKYRSEADLKKSARGRRVEQY
LNPRGLRILAALDAVAAKHGSTQTSVALAWQMARPSVTAPIASATSLEQLSALGAAIRLQ
LDAHDIRQLDDASAP*
>GCA_000452945.1_BPMSHR5858_genomic_02261_1 malT_1
MTYDPALSDADAAPSYARALQACVGSLQALARPTATVFYRIDASGEPVDFELFGMTAAMH
RAYVSRYRPLDPLHPSRCASQPGAVVTLASQLPDERRDTSSYWTGFLRRHGVVDVVEVLL
RDGGAVLAAFSLLRLAGDGRYSVPEIAALRAVQPVVEAALLPPLRAVRGIRRIACDVRLT
HREEQIARLVRDGRSNKAIARDLALGQPTVKTHLLRMFRKLGVSNRTELVGALFL*
>GCA_000452945.1_BPMSHR5858_genomic_02263_1 spuC_1
MSYNDSRFWHPMLHPNDMKRREPIRIVRGDGCHVYDERGRQLVDGVAGLWNVNVGHNRAE
VKEAIVRQLDELEYFQLFDGITHPRAEELSSKLIDMMEPEGMRRVLYSSGGSDSVETALK
IARQYWKVRGQADRTKFISLKQGYHGTHFGGASVNGNTVFRRNYEPNLPGCFHVETPWLY

RNPFTQDPEALGRICAELLEREILFQSPDTVAAFIAEPIQGAGGVIVPPANYWPLVREVC
DRYGVLLIADEVVTGFGRSGSLFGSRGWGVKPDIMCLAKGISSGYVPLGATAVNARIEDA
FAANADFGGAIMHGYTYSGHPVACAAALASLDIVLREDLPANAAKQGAHLIEALRPFVER
FDALGEVRGKGLMVALDLVADKATRAPIDPMSGYANAVAEVARENGVLVRPVGTKIILSP
PLVIQREQIDRIVAGLEAGFAATPFPGGR*
>GCA_000452945.1_BPMSHR5858_genomic_02264_1 steT
MADWSGYSNAAAGSAAGGAGGRPALAAGAVGFPTALASAVGLIMASPVILTATSGFGMGG
WAFAAAMLIAFVMMQAQATTFAEAAAMLPTAGSVYDYLSCGLGRFWAITGTISAYFLVHV
FAGTAETILSGIMALVNFESLNAAFERHDSAWLVGVGLVATFAFTNIIGIKVFSKLEIVL
TVGMWLSLTIFGVLGLVAAPAVQLDGWFGRSEIGASAPAVLSLVGMAMFMFVGCEFVTPL
APEMKTPGKTIPRAMALGLAGVAVCMFVYGAAIKRQVPNVAVTADGLTHLLDTPGAIPAF
ALQVLGRFGRAWFGIAFLCAGAATINTLMAGLPRILYGMAVDGALPKCFAYLHPRFRTPV
VGIVAAAVVPIAHAWIIHGDLGSIMHLVLAATCAWGTAYLLVTLSVVMLRIRRPDLPRPY
RSPWFPLPQIVSSVGIVLAIWYIAPPGTNSRDIYVPFGAMLGLTALYALFWTIVVQRKHP
FRPVPVEEVLRNECVR*
>GCA_000452945.1_BPMSHR5858_genomic_02267_1 group_1841
MHIRTWIAAGVAGAAVAPACAQSGVTLYGTVDTGVIYSTNQQVTHADGSTSGGHAWQMGG
GNLVPSRWGLQGTEALGGGLKAVFALEQQFLSASGQALQGGAAFSRQAWVGLRHDRYGTL
GVGRQYDSYTDTLGAYVSSNSWATPYGSHLGDVDNLNAAFNFNNAIRFTSADFHGLTFGG
TFGFGGQAGDFSARRGYALAVAYNRAPVSFGIGYLNLRQPLDAALGGANGYIGDFACGNA
GAMYCLLQDAVSMKAFGVGGSIAFGEATVALTYTHTRLGDSAYFATAARAQRASIAFDIA
ELNATYAFTPMLRGGIAYVFNRAKADSRGVTRFHQLNVGVNYSLSKRTALYAVAIGQIAS
GRGLGSDANGDPVNYAQIPVLANSNSNRQLAMMAGVKVDF*
>GCA_000452945.1_BPMSHR5858_genomic_02268_1 rhtB_1
MSLSSLALFAAALLVAAGSPGPSIAALVARVLTNGVRDVLPFLAAMWLGEVIWLTCAVAG
LTALAHTFAFGFLVLKFAGIAYLVFLAWKMWFAPADEPSGQLPRGQSPLRMFLTGIAVTL
GNPKIMIFYVALLPTMVDLSRAGAVAWLELTLTLLVVLITVDCAWAWLAARARKLLTSRR
AMKAANRASAAAMAGAAVAIAMR*
>GCA_000452945.1_BPMSHR5858_genomic_02269_1 group_2401
MTIGSSTIDYSDGCSHPIDHAPHSPVTARRSAAQPCPIGIRAARADEAPLMAAIEAAAAR
RFDDIGMPHIASSPPTDLADLRERIDDDRALVAFDAEGLRIVGFAIYRMLGASRLYLEEV
DVAPEQAGRRIGSALIEAVAARARAAGARQVVLSTFRHVPWNAPYYRRLGFVELDGNTLD
AALTAIRATHVAHGLDESQRVFMARMVHE*
>GCA_000452945.1_BPMSHR5858_genomic_02270_1 nemA
MPSLFDPLTIGDLTLANRIIMAPLTRARAGDTRTPNALMARYYAERASAGLIISEATSVT
PQGVGYASTPGIWSPEQVDGWRLVTDAVHAAGGRIFLQLWHVGRVSDPVFLDGALPVAPS
AIAPGGHVSLVRPQRPYVTPRALELDEIPGVVAAFRRGAENARAAGFDGVEVHGANGYLL
DQFLQDSANRRTDAYGGSIENRARLLLEVVDAAIDVWSAARVGVHLAPRGDAHTMGDSDP
AATFGHVARELGRRRIAFLFARESFGGDAIGQQLKAAFGGPFIVNENFTLDSAQAALDAG
QADAVAWGKLFIANPDLPRRFKLNAPLNEPNAATFYAQGEVGYTDYPALESAA*
>GCA_000452945.1_BPMSHR5858_genomic_02273_1 group_15303
MNKTLIVAAVAASFATVAHAQSSVTLYGVLDAGITYQSNVATPSGSGKSLWSVGAGVDQS
RFGLRGSEDLGGGLKAIFTLESGFNIGNGRFNNGGGMFNRQAFVGLSSNYGTVTLGRQYD
ATQDYLSPLSATGTWGGTYFAHPLNNDRLNTNGDVAVNNTVKFTSANYAGLQFGGTYSFS
NNSQFANNRAYSAGASYQFQGLKVGAAYSQANNAGANTTGATDPLTGFNIGGTNAASIQG
RSRVYGAGASYAYGPLQGGLLWTQSRLDNLANGAPTIRADNYEANVKYNLTPALGLGVAY
TYTNAKANGESTHWNQVGVQADYALSKRTDVYAQAVYQRSSKNANASIYNGDLSTPFSTS
INQTAATVGLRHRF*
>GCA_000452945.1_BPMSHR5858_genomic_02279_1 anoI_1
MRTFVHGDGRLPSDLAADLGLYRHRVFVEQLGWKLPSASEGFERDQYDRDDTVYVFARDD
DGEICGCARLLPTTRPYLLKELFPTLVAQDMPLPQSAAVWELSRFAANAEDPAGGGNPAW
AVRPMLAAVVECAARLGAKQLIGVTFLSMERLFRRIGVHAHRAGPAQQIDGRMVVACWID
LDAQTLAALDLDPLLCAPPAEAA*
>GCA_000452945.1_BPMSHR5858_genomic_02291_1 fabG3
MSAPVQAIALAGRRVLVTGGARGLGAAFVEALVAAGAKVAFGDVLDAEGRALAQRLRDAG
HDAHFIALDLAAPASVAAFVDEGARVLGGLDGLVNNAAITNSGGKRATELSIDTWDAVMN
VNARGVWLACNAALPHLARSGRGAIVNLASDTALWGAPKLLAYVASKGAVIAMTHALARE
FGEQGVTVNAVAPGLTEVEATAYVPAERHEYYLQGRALPRAQVPADVTGPVLFLLSDAAR
FVTGQLLPVNGGFVMH*
>GCA_000452945.1_BPMSHR5858_genomic_02292_1 group_4719
MADADIERKSWDQPAGESFAQWMDGRVARFETRRYDWDALKFQADFDPKYRRAQMRYVGT
GGTGVAKDANTVPAGGFTFSTMVIPAGNVGPSHIHIDVEEIFFVLRGRMKVICERDGETW
EAVLGERDLISVPPGVYRTEINVGEEDALMCVMLGSPKPITPTYPPDSPLAKIKR*
>GCA_000452945.1_BPMSHR5858_genomic_02304_1 group_4720
MLKLLVPVGQSQRSLQAVRHAAFLYHERCASEVVLVNVQPPLESTRLEAFHSLAKLRQIE
HDFSEGELAQAIGILKDADVKYSVAMKVGPIADAIAACATENGCDEIVVVAPRRDPVHAL
THCFHRSILDRLVRIASVPVTAVR*
>GCA_000452945.1_BPMSHR5858_genomic_02305_1 soxG
MPERLVPSPRSPSMPFDPIVDMPAHTLAEAIRRKDVSCVETMRAYLAHIERVNADVNAIV
ALREPDALLAEAAQKDAALARGEYAGWLHGMPQAPKDLAMTKGIVTTLGSPIFRTMTPSV
DAIVVERMRAAGAVFIGKTNTPEFGLGSHTFNEVYGVTRNPYDLSKSAGGSSGGAAAALA
ARMLPVADGSDFGGSLRNPAAFCNVYGFRPSQGRVPRWPSVDVFVQQLGTEGPMGRTVVD
VAQLLAIQAGYDRRDPLSLAEDPRRFTQSLDADMRGKRIAWVGDWNGYLAMEAGVLELCE
SSFDTLRAIGCDIDAALPAFPAERIWRSWLAHRHLLSGGNLLMHYREPSRRALLKPEAIY
EAQGLFALGAADVYEASAARSAWYQALIAFFERFDYIAAPSAQVFAFDVELRWPQAIAGR
AMDTYHRWMETVVPWTLAGCPVISVPVGFNAAGLPMGMQLIGRPRDDFAVLQLARAYEKE
RDWVSARPPALSGARND*
>GCA_000452945.1_BPMSHR5858_genomic_02306_1 group_4721
MKLDETDLKILKILQENGRLSNAELADRVALSAAPCWKRLRRLEADGFISGYQAILDRRA
LGYEIVAFVNISLDSHTEKTCRAFEAGVMAMPEVVACHNTTGQHDYLVQIVAKNFDTFSS
FVLNRLRSLPGVKELLSTISMRELKATSRLPAGELRITKQ*
>GCA_000452945.1_BPMSHR5858_genomic_02312_1 group_4723
MPRDNRLSRMLHVLIHMDRHVERATSEAIAKMLGTNPVVVRRMMAGLRDQGYVRSEKGHG
GGWELVCDLDAITLLDVYRAIGEPPLFSIGPHAEQSDCLVEKAVDARMTEALGEAQALLL
AHFGAITVGDLARDFDKRHARHKAGKRGNCAAA*
>GCA_000452945.1_BPMSHR5858_genomic_02316_1 proP_1
MTNKKPLSLTQIVLATSVGNALEWFDIAIYAFFAVHIAKNFFPTANETASMLLTFGSFGA
SYLVRPIGGMVLGAYADKRGRKAALMMSVGLMMIGTAIIAVIPPHASIGLLAPAGVFIAR
LIQGFSAGGEFGASTAMLIEHAPERRGLLASWQFTTQGLATLLASTFGFALAKLMPASEL
SAWGWRMPFFFGLLIGPAGLYLRRFLEDAADYTEAEHTATPVRDVLTRQKALLLTSIGAL
TVSTAVNYLLQYVPTFAIRELHLDASTGFAASIVAGLMLTLVTPFAGHLSDKIGRVKQMS
TAALLLFVTGYPAFAYVVSHVSVAALFGLVAWLALLKAVYFGALPALMSEIFPTSTRVTG
ISISYNIGVTVFGGFTPAIVIWLSSATGSKAAPSFYMMFTAVISLAALAAVSRGKEPLSS
AGTAA*
>GCA_000452945.1_BPMSHR5858_genomic_02317_1 ephD_1
MATVRDTLTVRSGDVRLAVYLSGPRNAPPVILVHGYPDSARVWERVREHLDKRFRVIAYD
VRGAGASDAPKRRADYKLSVLAQDLLAVADVTCGDRPFHLVGHDWGSIQCWEAATDPAVK
GRLASYTSISGPCLDHVFRAPLNLRQTLKSWYIAFFHLPLVPELVWRLGGAKLWPLWLRR
TERIVADADPDQLRNGLNGLKLYRANFIERARRPRERYAQAPVQIIVPRYDRYVTPALTR
GLDRWLGAHRRDEIAASHWTVIRDAALIAARIGRFAAAHEPKRHAIAAGGAKSGGQPRAK
QRTNQQASAGRRLKSVS*
>GCA_000452945.1_BPMSHR5858_genomic_02318_1 group_4726
MTTPHMMPVRRDIRFALPADRAHDWHVQGAPVTHFMNALSLLFPAGERFFMDSVRNYRDE
IHDPELKKQVAGFIGQEAMHTREHIEYNDLLQSAGLPAHKLDKRLWALLGWGKKVLPNSV
QLAITIALEHYTAILANQLLSGYTHRIDGSVEGYKQMWLWHAMEETEHKAVSYDVWNTVM
KPGVKTYLLRTGTMLATTLIFWAVVFDFHVRLMFAHRMRHGKLKGMWRLVKYLYSPKNGV
FPSIALEWLDYFRPGFHPWDHDNRVYLQGLDTLLANIDAANERYADQVAPRRVPLHSLPE
HQHA*
>GCA_000452945.1_BPMSHR5858_genomic_02321_1 group_4727
MLSHLPPRVRGLWLAAALAALLYGLSLGHAPYSGQPAAKAALGVLLLAAALRHPLARERA
WLGAALAASTLGDVLLALASWPPSFVAGLGAFLLAHLAYCALFAPWRVAPRGARAVALGA
VWLAAPALYAAFLPHLAALAAPVAVYVAVLGAMASFALCARTPGPQVAAGALVFVASDAL
IGIDRFLGAFSGIDYFIWFLYAIAQLTMAFGVLRRK*
>GCA_000452945.1_BPMSHR5858_genomic_02325_1 group_610
MPRVHRAFASLQTHYKKLDTAQKLYLNCALALLLYGVTYPLHAWWNFVGLALLSLAFWFA
GMVCDALTLYKRIYATHIGKGLLLLAFTGFTNFAIALANQRINAFAGVDPSKFPHTEIFV
AILSIPFIALLILMMLYTASVLFLPVYGVFFIAMDAKAKAFFSAGLMKQDDSEYPVITGI
FRFVSFFVFAGVVLFFAKRMLGDYDRFMTSTAERSVYQLEMYEKSPCDVPAGARVAFLDD
GNIVYALRVGAELKFFPQRCKRSGQ*
>GCA_000452945.1_BPMSHR5858_genomic_02326_1 bmr3_1
MSPDSSAPPAHADAGRPAASVRLTIAALLLVLLLSALDQTIVSTALPTIVGDLGGLEKLS
WVVTAYLLSSTVVLPLYGKLGDLYGRKIVLQAAIVLFVVGSALCGLARDMTQLIVLRALQ
GLGGGGLMVVTMAAIADLIPPDERGRYQGMFGGVYGIATVIGPLLGGFLVQHLSWRWIFY
INLPLGALAFAVIGAAFRPHTAYVRRRIDYAGAAFLATALTCVVLFTSQGGTILPWSSPQ
LWFTLALGLVATAGFVYEERLAAEPILPLELFRHRTFVLCCLIGFVVGLALFGSVTFLPL
YLQVVKGSTPSEAGMQVLPMMGGLLATSIGSGRLITTLGKYRLFPIAGTFVACAAMLLLA
TLSLDTPLRAMYLYMALLGGGLGMVMPVIVLAVQNTVEFRHLGVATSGATLFRSIGGSLG
VAAFGALFSNGLHARLAATLPPDTELPPSLGPSAVRQLPGAVRSAYLHAFASSLHTVYLC
AAAVIAIAFALAWLVEDAPLRERA*
>GCA_000452945.1_BPMSHR5858_genomic_02328_1 group_2405
MARIIPDDWKNLAATGAAERERETLATLERTLPDAYTVYHGVHWTRADHNFSVFGEAHFV
IVSPAGRVLVIEQKAGFLRETPKGLVKVYLQTERNVSIQLARTLENLHRRLTAALGAGAY
GVEELLYCPDYTIRNPAIAGVAPARIVDATRRHALAARIVDALPADEPPFPSAAKVHHFL
ADELSLTPDTSALVGQADTLVTRLSGGLAAWARRLEFEPFRLRVIATAGAGKTQLAVQAM
RGAISAGKRVLYVCFNRPLADHVARIAPQGVKIANYHQLCDWIARDGGHVPDFGARDAFA
QLEARFAQTPIAERWRFDTLIVDEGQDFQPSWAHALERLVAPGGAWWWLEDPLQNLYMRE
RVPLPGWVTLKETTNYRSPRDILDYVRDVVGSVEPLAAELVSGSPFGGSEISISAYDDAH
AQSPAQACIDATKRAITHALALGFRKQDIAVLSFRGREGSALATLDQLGPHRVKSFTGKY
DLFGNPEYREGDVLLDSLHRFKGQAAPCVILTEVDFDAFDERAARKLFVGATRATMKLIV
VASKRAAQRIAPSA*
>GCA_000452945.1_BPMSHR5858_genomic_02331_1 hcnC_1
MTTTYPLYKTLARADEPFPAQADVVIAGAGIMGCAAAYYLGLRGIKAAVFDKSRIAGQQS
TRAWGFVRQQGREAAEVPLMMAGMRLWEGLERELGADLEWRQGGCLYLADNDDDWASFRQ
WTDVARAHGLDTRTLARAEIDAHVSGLAAPALGGLYTASDGQAEPRRVAAAFAARAIEAG
ARFFEGCGVTAIDTSGGAVAGVVTERGVVKARRVICAAGATSFRLLDGVGIRLPQQAVRG
TCMRTNPLPAVSASTIWGHGLGIRQRANGAINLADDMQVDVDVTLGHLRGLSLFLPELWA
RREKFRFHLNGAAWRDLKVRAAGGAAVLEPRDPNPQPNPAHAPRALAKLKAVFPALRDAR
IVEAWAGLIDVLPDGIPVIDAPRAPDGLAIATGFCGHGFAMGPIVGSLLAEWVDTGATSL
DLSAFRAARFVDGTMRRPASML*
>GCA_000452945.1_BPMSHR5858_genomic_02332_1 kefC_1
LASPSEYRKSLRFRLRSARTPWKAPRPRTLFTRPAASPLRTLTLRLALVVGLCALAFLVL
YLDRDGLRDSTKSEPMSIADLVYFTMVTVATVGYGDIVPVTARARLIDAFFIVPIRIGIW
FIFLGTAYQFVIQRVIEEYRMKRLQKNLRDHVVICGYGLSGSIAVRELLESGVDPATLIV
IDSQEQALEAAAALGVTGLCGDPAHEDLLQQAQVRSAKAVIISVTDDPTAILLTLSVRSI
APDTKIVVRIQENLYQRQLRQAGADVIVSSTKIGALLLADAVESRYIVPFVNDMLSTRGR
ATLVERPAAAHEIGCMSNAVAGALVVGLDRGGRILSFYEDPPCRIEAGDTLVVIQSTRAM
ERAGIELAD*
>GCA_000452945.1_BPMSHR5858_genomic_02362_1 ycjP
MGRLIARGGTVAIGGAIALVAVFPLAWALLNSFKRLLDIVTPTPRFIFTPTLDNYRQVLS
SPEVLIGLGNSVAIVGAAVALGAALGVPAAYAIARYPVRGKRDIQFFLLSLRFLPPVAVA
IPLIAIWVDLGLYDTKLSMIVTYLLVTLSTITWLSIPVFRRLPREIEEAAALDGYGPYAV
FWRVALPVCASTLLGGIVFSFVLVWNELMIALALTSSRAATLPVVASAFTSLGQEVPWGV
INASTVLLALPPLVFVGMLSRLLNSMLKGR*
>GCA_000452945.1_BPMSHR5858_genomic_02363_1 group_675
LKALVLEKARELALRDIDLPLEVGPADVKIRVHTVGVCGSDVHYYMHGGIGPFRVDAPMV
LGHEASGTVVEVGRDVTYLRVGERVCMEPGVPRFDSKATLHGLYNLDPSVRFWATPPVHG
CLAPYVVHPAAFTYRLPPNVSFAEGAIVEPLAIGLQAAKKAAMKPGDIAVVVGAGTIGAM
TALAALAGGAARVILADVVKEKLALFAANRAVTTVDASTQSLADAVAHATDGWGADVVFE
ASGNASAYAGIVDLLCPNGCLVLVGMPLAPVPLDVVSLQAKEARIESVFRYANVFPRALA
LIASGAIDVKPFISRTFPFSDGLRAFEAAASGQPHDVKIQIEMD*
>GCA_000452945.1_BPMSHR5858_genomic_02367_1 dacA
MTTEPLSTTVARGSRRAACALLRTPSRLFAALFVSAACAAPNGARAAPAPAPAIQAASWL
VVDADSGKTLGAHNVDARREPASLTKLMTAYVAFDALERDALRWDDTVTVAADDIASVGR
DEARMYLTPGQRVRVRDLMRGLIVASANDAALVLAKRIGGSPAGFATMMNDTARRLGMRD
SHFVSPSGITTPDHYSTAHDLSILAQRLNRDFPAFYTFSSQRHFAYGAFAKTNKNRLLGA
DPSVDGMKTGHTNAAGWCMVVTAKRPVGGARARHRVIAVLLGEPTEKQRLADARALLDQG
FASLGADAPAANAVRRANARPGEHRAQARARML*
>GCA_000452945.1_BPMSHR5858_genomic_02370_1 napA_3
VTSASVKTVCPYCGVGCGMVLHVEDGEVVKVSGDAEHPTNLGRLCTKGSSAHVALRNAGR
LERAFVRDARELDPAPVPIADAIAETARRLRATLDAHGPDALAFYVSGQMSLEAQYLVNK
LAKGYVRTPHIESNSRLCMASAGSGYKLSLGADGPPGSYQDFERADLFFVIGSNMADCHP
ILFLRMMERVKAGAKLVVVDPRRTATAQKADLFLKIRPGTDLALLNGLLHLLHANGRTDA
DFIAAHTEGWDAMPGFLADYTPQRVAALTGLAEADIRLAAQWIGDAREWTSCWTMGLNQS
THGTWNTNAICNLHLATGKICRPGSGPFSLTGQPNAMGGREMGYMGPGLPGQRSALDGDD
RRFVEEIWGLPAGTIRPDAGGGTVDLFARMAAGDVKACWIICTNPVATVANRKNVIAGLQ
AADVVIVQDAFLDTETNRYADILLPGALWAEAEGVMINSERNLTLMRQALEPPGDAWPDW
RIVAAVARGMGYGDAFDYGSAAEVFDEIRRFANPNTGYDLRGASHAALRATPLQWPCPPE
DGRDRHPIRYLNDGASRPRVTLPDGRVPPLAFPTPSGKARFFARPHVAPAEMPTPEFPIV
LNTGRLQHQWHTMTKTGKVAMLNKLDARPFVEIHPDDAAALSIAANDAVEIRSARGRAVL
PAVVTDRVQPGACFAPMHWNDVYGDDLCVNAVTNDAVDPVSLQPELKFCAVALARVDAPD
SADAGAHAGAHAAPLHETGSREAQAQAAPRAHEAACADAAPEEHDMSDIDTFAALLGVPA
ATSAPQLTDAQRVYVAGFVSGLRSAEGRRAGGVPMLPASAPFDAKARLWLDGLLAGLFSR
APLGAAARVDARAAAAAAVPAPSGAADAVSAPSGVRIVRTRPKVVLLWASQTGNVESLTE
RYATQLMDSGFEIRVACMSDYPAASLAKAQYALLMTSTFGDGDAPDNGQAFWTALAAAGA
PRADGLRYAVLAFGDRNYDPFCGHGRRLDARLAELGATRLVERVDCDAEFQPAADAWLER
AIARIKEEDAALHAVPADGMIPSGAIPTKTRPAASRLVANLRLNEPGAAKDTRCVSLSTD
GAAIEYETGDALGVWPTNCPELVDELLALTRLKPDAPVTVAGMGELRLADALAKHLDITR
PHPDALAFVASRSRAGDALARLLGDDRKTDLKHWLWGQQLADVLHEFPLELSAAELVGML
KRLQPRLYSIASSPSAHRGEIHLTVSAVRYGNGRRQRKGVASTFLADRAADAAVPVFVQK
SAHFRPPASGDAPIVMVGPGTGIAPFRGFLHERRARGAKGRNWLFFGEQHADTDFYYRDE
LAQMRADGFLTRLDVAFSRDQTEKVYVQDRMLEQGAALWAWLEEGAHFYVCGDASRMAKD
VDAALKTIVARHGGMTDEQAADYVARLAKDKRYARDVY*
>GCA_000452945.1_BPMSHR5858_genomic_02373_1 nrtP
MSDKATRIELLDWRTVPMRAFHLTWIAFFVCFFAWFACAPLMPLIAREFRLSADQVANVN
IAAVAATIAVRLVVGPLCDRFGPRRVYAGLLACGAIPVFAVTLSHDYLTFLLCRLGIGAI
GASFVITQYHTSVMFAPNVVGTANAAAAGWGNAGAGAAQAAMPLVVAAALLLGLPDACAW
RAALAVPGAAMLVMAIVYWRGTQDCPQGDFVALRARGVTIDSGKKGGFASFAAACGNYRV
WMLFVTYGACFGVEVFMHNIAALYYVNHFALSLKDAGLAAGIFGLLALFARALGGWLSDR
VAARRGLDVRATLLFALILGEGLGLLWFAHAGSVPVALVAMVAFGLFTHMACGATYALVP
FIDRRALGGVAGIIGAGGNAGAVAAGFLMKGVGDIQQTLTLLGLFVTVSAFCALAVRFSA
EHKAREAALRERALAASGFTH*
>GCA_000452945.1_BPMSHR5858_genomic_02378_1 nrtA
MNTARIKAAAADMPERARLRIGFVALSDAAPLVAAQRLELGARYGLTLELCRQPSWASIR
DKLLSGELDAAHALYGLVYGVQLGIGGPRADLAVPMVLNRNGQAITFSNRLADAYRASGE
LKAALATLGRRPVFAQTFPTGTHAMWLYHWLASHGVDPLHDVRSVVIPPPEMVGALAAGE
LDGLCVGEPWNAVAEARGAGRTVAATSEVWPDHPEKALACRREFVALYPNAARLLARTLL
DACAWLDDPAHRMQAARWLAQPDTIGVPVEQIAPRLLGDYGAGPFAQPPAPIRFHAHGTA
NRPAASDGLWFLSQYRRWGMLSGDVDDAAIANGVAHTALYDEAVVLGGARRGE*
>GCA_000452945.1_BPMSHR5858_genomic_02381_1 group_2413
MQTRLDTRKVFTYLFAFFIAFLASCSDHDDISGGGEPPASALAFRLDRGGQINAFYRQDK
IAAHLLMRASTKPRLLVVFPAGNSGTGLWFGDTPQPVNWSQDTPIAPVTTQDAAGRPLYG
IGVDVSADVSNLSIRQALLSSVRFLRDYNGGATVPSDIVVQPSVSDNVATWQRNRVDGAP
GYALRVTVLDGGTVAADGTGRLMLQSPSGAPTLRLRVEAFSGETPLQPITRANLFTSAVN
PDPQSQDVFEFLSFSDKLLAGSWQYDTYFGRDTLISVRMLMPVLQPDVIEAGLGSVLDRL
SPDGKVAHEEGIGEFALIDNLKHGRPNDPTPTYDYKMIDGDYLLAPIMEDWLIEDPRGQA
RAAAYLAQKGDDGVTNGSRFVANLLRVARTAQPFAQQALAANLIHLRPGEIVGNWRDSTD
GIGGGVYPYDVNVALVPAALRATSNFLARGLLDPYLSADQRAVLASAGAAADVWERAAPP
YFQVAVTPNDARAKLAAYAPQAGVPAGTVPNTALQFDAIALDANGNPIPIMHSDGGFVLL

FGNPSPALLARIVTDVMRPFPVGLVTDVGMLIANPAYADPTLWPRFTNSAYHGTVIWSWQ
QAMWVAGLDRQLARTDLPGDVRTLLTQARQQIWQVIANAKDMRLTEMWSWSYANGQYRTE
AFGTRSTDVTEANAAQLWSTTYLAVRDPQLRAGKYWWEARAAGGDAPRRGSTAAAENAAN
GLNAVKGAAS*
>GCA_000452945.1_BPMSHR5858_genomic_02384_1 abaF_3
LTRGIPPSPPARQPRRAAGAAFVGTMIEWYDFYIYANAAALVFGELFFPSHDPFTSTMAS
FATFAVGFFARPLGGLFFGHLGDRIGRKKALMATLALMGAATVCVGLLPTYERVGLLAPV
LLVLLRIVQGIAVGGEWGGAVLMAGEHAPYGRRTFFASFAQLGSPAGLILSLVAFRAVAS
MEHGAFMSWGWRLPFLASIALLAVGVFVRLKVDESPEFAHEKAMSRTVARPLAEVLRTSR
AMLLFCLCANTIGIAGLYFTNTFMIAFTTQHVGVTKSLILDCLFIVAIIQFVTQPIAAWL
GGKLGDARFLKLAALFAMASPYPMFALVQTGRLAPMVIGIAIATVFLAGFYSVIAGFVSG
AFPTRVRYSAISISYQMCGAIAGGLTPLVGTWLAHRFAGQWWPLAVFYTCLAALSLGGVI
ALDARRHRRADVADAVLTR*
>GCA_000452945.1_BPMSHR5858_genomic_02393_1 group_1344
MTSRRLASRTAVAASLSALMLAACGGDDSANAPTAGGAAPLTPAVASPAGPTGSTPGATT
APAPSSTSAGQLSVDKMAFAQTHVVPSGGLSWTLPNASASLRPISRRDALVLVAIGQADA
VQPVLEAWKDGAKLGALALSPPSALPPTESGGRAYANDRWSAVVPAAWMVPGVSFSVSAS
NYTSSVAQAPVFGTDADVQLTILPFYLFGADDTNSPPLSTTQAPDAATQQEIFAKWPTAE
LKVRTHPAGRFSLATVVVGPRADRTGAAQPAYPVTALDQQKDGYGVMSAMLTLITNMRTA
NGDGPLNNQYYAPLIALNSNGQFANLGGGLGGVGSGAAVGDHRYTGIFIHEQGHAFGLNH
AGDEYAKGAYPYAGGSLSGSVWGYDPNHREFLDVLVPTTASSYAKCASSHQLDAQGRCYK
QDPMQGGAGDQSSGYKFATFSDYNTGRMQAWIASRVLADPASSTGYSKWDSAAQARAPYT
PTTDNNGLYGVNQNLPVQAGVPVHTIVVSFSKAGSAGASYIYPPFSYTGNLIATFDPTSA
ADRQAITVDKGTYPWYCKGTGCDYTLRVTYADGSRTYRVLQGGFRAWWTPTVYDANATNP
LSGSSFRVWAINVPGDKRIGKIELLDTPMVWNGMPANPTVLLSR*
>GCA_000452945.1_BPMSHR5858_genomic_02396_1 group_262
MSRFHVHLVVFFAGLAAVCWIGAGYAVSNPVALAVTLVIAAGYLAGALELRRYRQATSTL
AQAVAALSEPPAALGAWLERLHPSLRHAVRVRVEGERVALPGPALTPYLVGLLVLLGMLG
TLIGMVMTLRGTGAALESSTDLQAIRASLAAPVKGLGFAFGTSIAGVATSAMLGLLSALC
RRERLDAAQALDAKIATTLRVHSHAHQRDETFRLLQRQADLMPTLVERLQAMMHSLEQQS
AASAERQIAGQQAFLGKAEETYARLASSVGQSLTDSVAESARVAGSALQPVMETTMAGLA
RETAALHDALTQAVQRQLDGLSAGFETTAAHVADVWRHALADHQRSSDALAQRLHGSIDR
IVESFDRRSADLLDGVRARLDATASSVSDAWRGALAQQEQANEAHAERNRQALETAAATF
ERHSAALLRTIGESHSALQATLESRDEQRLATWTDSLGSIAAKLGTEWAQTSAQAANRQQ
TICDALAHTARDLSAQATAFEQHTAALLRTMSESHSALQATLESRDEQRLATWTDSLGSI
AAKLGTEWAQTSAQAANRQQAICDALAHTSRDLSAQATAFEQHTAALLRTMSESHSALQA
TLESRDEQRLATWTDSLGSIAAKLGTEWAQTSAQAANRQQTICDALAHTSRDLSAQATAF
EQHTAALLRAMSESHSALQATLESRDEQRLATWTDSLGSIAAKLGTEWEQTSAQAANRQQ
AIYDALAHTARDLSAHTQAHASATIAEISQLVQAASEAPRVAAEVVAELRQKLSDSMVRD
TAMLEERNRMLATLETLLDAVNHASSEQRAAVDALVATSSALLQRVGTQFTDEVGTQTDR
LGGVAAQITGSAVEIASLGDALGAAVQSFGESNDKLVAHLQRIEAALDKSLARSDEQLAY
YVAQAREVIDLSMMSQKQIIEELQRVGGERASAGAAAA*
>GCA_000452945.1_BPMSHR5858_genomic_02399_1 group_175
LRAEPALLDYFRETWSKLSADSQLRASLEQVPKNAGPLNSSSLVHRSLSLMRELSPEYLR
QFLSYVDALSWLQQMNGDEAATGDEAARGAVGGGGGASAKKNTRKRSR*
>GCA_000452945.1_BPMSHR5858_genomic_02403_1 dltA_5
MNHRLPHVERGDADEPSIETAAGEQPPLALPASVAQRRLWFVENGDVRASTYNVPAAFTL
TGPLDDAVLERALAFMQQRHPALRSRFRTRDGELRIELAPQPAPLARQDLGALDADVRAR
TAERLCANHANRRFDLERDAPIRCLLLRLGENEHVLAVNVHHIVFDDWSIRIFFRELGAV
YGALLAGATPDLPALDYAAAVAASVPAAARHAAARQYWARAMSGAPTLHKLPTDRPRPAE
PRMRGAVHKHVFARRHAEGIRALCRRAGVTPYMLGVAAFAALLHRYSGEDEIVIGSPFAN
RVTQAQQSLIGFFINLIPLRVRFDAGVNFLDLLAQVRETSFDAFEHAVLPFDQIVDAIRP
PRSSSHAPVFQIMFDYLKSGGMLELDGVGVTGSLVHTGTAKYDLTVSMEEGPDELAAIVE
YDTDLFDAGTIARLGGHFERLLENVLASPAAPIAEGSLLPADELRQVRRFTRPDEPYAHI
PFSPMPQRIREAARRAPHAVAIVHGDARMTYETLDRRSDALARALRARGVGRGSRVASLQ
SYSEKIVVAYLGILKAGAAYLPLDPADPRRLEKIEDAAPAMIVTARRDLEDVPQALRART
LTIDDPIECGKAPDAVNDAVNDVATDVVTDATARDAELDFATLAEADPAYVIYTSGSTGK
PKGVEVSHGSLNVSYHGWHRAYRFGKPGHPVTLQLAGMTFDLGIGDVSRTLACGGTLVMP
PRDGLLDAGRLHALMRAERVSFGDFPPVILRELIRHCNETGDRLDMLDTLVCGADVWFGH
ELHAARALCGPHARVLGSYGVTEAAIDSSYFDPDLHALAPDSVVPLGRPLPSCELLIVDP
LLQMTPIGVPGELLVAGPAVATRYLNNDALTAQKFLRGRVDEHGRVIAGDGQTRFYRTGD
ICRFLEDGTIDFLGRRDNQIKIRGFRVELGEVEGVLAAHPDVRQCAVVVRDEASGDPSLA
AFVVSDAPIAALRGYLRGRLPAYMLPAAIERLGDMPLTASGKIDRNRLKAWPLSAPDVPP
PDAATDVERRLLALWENLLSARVPSVHENFFQCGGHSLAAARLASSISQAFDISIGVSSV
FNHPSVAEQARLVEALASARAPRDVRQTAHADAAEPAGDDGLLSYSQQSLWLTAKRTPDD
FSYNIPVTWRLDGPLDAHALERAINDVVARHDALRTVFSSDVRTVVGPHRESSQEPTQRV
LDTLTIALRRVAVAPDDAASLPARLREAHSRAFDLNAGPLLRAVLFEIAPTHHVLDVTIH
HIVIDGPSFGLFWRDLQTAYRARVAGEAPGWQRPARRHADFVSRQRQALRGEAAARQLAY
WREQLRGLPAALPLPDAVLAAHAPGSARSLTFEMPDDVAAGLAALARRTNGSPFIVYLAL
FAAALRQQTGEADFAIGTPLSLRPHEGFADVLGFFANTMPLRMRLHGLDTFERVLRYVRE
QCLALYENGDVPFEYLVQALKPARAARRNPVFQTIFSCEFDDERLQLTGVDAHALALDAY
TAKLDLEMAINVSGGRVVCHLMSRPGSFDADALSSIRHHFLRTACSATRAHAEREAAEAR
ATHEVHEAHEAHEVHEVHEVHEAHEAHEVHEVHEAHEVHEVHEAHEMHDVHEARAQLADD
APGARRPSADDLFGLFARSAARHAQRVALDSPMLRASYAQLAERVSAAARALAAHGVRRG
DRVGIFVGHHPHNVTAMLAIARVGAAFVPMDPEHKPQWNRHIVDDAALTALVGGAWTADA
ARGFGLPVVDLDAPPPPASELADAPAAGGAHPDDCAYVIYTSGSTGRPKGVAVSHASVCH
NVRAMAEIMRIGPQSRMAQYVSPVFDVVLGEIFPALAAGAAIVFAERRRPLPGQALVDWL
DAQRVSHVWIVPSALAMVPEAALPALEVLIVAGEACPRELAQRWAAGRRLLNGYGPTEAA
IVVSLTDYHAQRERLILRPMGGARLHVLDEALRPAAAGAAGELFIGGACVAQGYLGQPAR
TAQAFVADPFDAEPGARMYRTGDVVRRLDDGAIQFIGRVDRQVKIRGFRIELDAVRAALM
EVPGVQAAEALAQPDASGQPLLVGYVVARRAKAELLDALRGKVPDAMVPSTLVFLDALPT
GSTGKTDLKALKALKTGDAARPAAAAADMPRAASQGRTLHRVREIWRTLLERDDIGDDEN
FFDAGGHSLRAVALHQRITEAFGDVIALTDLFEHPTIGALAAHLDAFAPRDGEAADDAAG
AAARAPADGVLDTDAIAVIGLAGRFPDAPDLDRFWERLLAGYEAGRTLSDAELDAHGVPA
ELYRNPHFVRRFKELEGKAEFDAGFFGYSPREAQVMDPQQRIFLELAWQALEQAGYGDRG
RVRSVGVFASAAFNYYLVQNVMPNAERLRLEPGQWLIGNDKDFIATRTAYKLNLLGPALS
VGTACSSSLMAVHLACASLRNGEAQMALAGAVALDPDQVGYLYAEGGIMSPDGRCRPFDA
AAAGTAGGSGGGVVLLKRLDAALRDGDTVYAVIKGSAANNDGADKVSYTAPSVAGQTAVI
RDALRAARVSADSIGYVEAHGTGTPLGDPIEVRALAQAFAEAAAPGALANGRCGIGSIKG
NIGHLDAAAGIAGFIKAVLALHREAIPPSINCETPNARIGFDKTPFSVVREARAWPRTAT
PRRAGVSSFGVGGTNVHVVLEEAPRVRAGESAEPSRWQLLPVSARSPSALREQWRQLRDA
LAHARPRVQDVAHTLQVGRTAFEHRGFAVVDAAADAPAQLDAAGSPPAFERRAAPPVVFM
FPGQGSQYPGMGAALYRSGGVFQAEVDRCAQLLRAHLDRDVRSLMFDAGASLLRETRYTQ
PALFAIEYALARQWLAWGVTPHAMIGHSVGELVAAAVGETLALPDALALVVARADAMQRQ
PPGAMLAVLADARELAALNGPGCEIAAINGPEQYVLAGDAARIAALEDACIAAGVACQRL
ATSHAFHSSAMDGAAREIDRASERIVRRAARIPLISNRSGRWLNEQDLRDAGYWGEHVRQ
PVQFHAGVRTLLDALDAPIFVEVGPGRALGNLIGGWAGLGPQRIVATLPHARERRDDMAA
ALRGVGTLWAQGVDVNWAGLHAPGAARRIALPTYPFERTRHWIERPAGARAAPAREADGV
PMRRGEDAADGSLTVSFALHERLWFLDEHRIFDGAPVLPGTACIELVRRAYSLVRPGAAM
TMRDVYFPTALILSTDESRNVRVVFRPPERVSDGAAAGGDLAFVLESNDANGPAGWTPHA
SGRIGDDPPACAAPASLAALAALDAPAALREQWGLTALDDVAALSAQAFADYGARWRGVD
ALWLGERAGLARLRLPAAGSGDLPDFALHPAMLDVATAFLPACLRPRDASVPFRYESIRM
HWPLRADCYSFAVETAPNVYDVTLFAWDEAARRADVLVAIGGFARREPAHRARDVAQWCR
TVSWRDAPAARALPPERWLVFGDEWFALAPAGSVLVREDDAFRAHGDNGYGVRPGEKADC
DRLIARLAEQGGVPAHVVYGWAQTDVDRAFAGLAALLQALGAHPADFRVSLVTKGARSAR
TLDACAAAAPAGLLKAVRWEYPRIVCRHIDIDDASDATIDALRAELSSEPATPPGAPPEL
PSSIALAGARREAPGFAALPDVARDDVLRDGGAYLITGGASGIGLELAAHIASRRRDVRL
ALLSRSPHDENAARFAALDEAAASVLRLTADVAHAAQLADALRTVRARFGRIDGVIHAAG
VEASGLLETGTPDAWRRVMAAKVHGARHLFDQLAGDPPDFIVLCSSLAAVVGGLGQADYA
AANGYMDALAQHWRQRGVAAIAIDWDTWSDTGMAFDHAARTRRSNDRPGALPGLANREGR
ALFDLALAHDASRIVISKRGFEQDRRDAPTRARRAAAPGDAQAALVALWQELLGVEQVGV
DDDFFDLGGHSLLATQLISRVRDQYARSPTLGEFLEEPTIARILRAIDHTGGDTAGDMSG
DTAGDAPDVDETLRYCVVPMVKAGSGAPFFCIPGMGGNITQLLPLANALGADRPVIGLQY
LGLDGKHAPHASVEAIAAHYVRCIRSVQPAGPYFLGGHSLGGKIAYEVARRLHAQGDAIG
LVAMFDSAAPPYSFVAHQDDFAIASMILGVFAYYAGKMEMMAGIDDARLRDAPRERLLAF
MGERLAQFGVIQSQSDTSAIRGLFNVYRAAADFSARYAPPHEHLPLPILLVKATEPMPDG
IKLPEIRETPAWGWENFTRLPVRTCEVAGNHYSCLMDGYVERIADALRDALASARQAIEA
*
>GCA_000452945.1_BPMSHR5858_genomic_02405_1 group_4738
MSQPVSEPTRVLDPASAHSASIELWLIERIACYLQRAPHEIRPDTPLDEIGLDSVYALTL
CGEIEEVLHLTVEPTIAWDYPTVAEIARELDARLAQRSAQGEAHAQ*
>GCA_000452945.1_BPMSHR5858_genomic_02435_1 hemS
MMNTAAPAASPARALAPDELRDAFLHLKETRKLRNRDVAQLLGVSEGEALAAFAGERVVR
LESSFVELFEEMPRLGGVMALTRNAAAVHEKDGAFEQMSHDGPVGLALGAIDLRIFYRNW
AAGFAVYEPTAHGVMKSLQFFDAQGDAVHKVYLRKHSDHAAFDAFVSRWRMPVQSPAFAV
EPAPPAHVERPDGEIDAAGLRAAWDAMTDTHQFHGVVRRHGVSRTQALRLAGASRAHRVA
TDAARRVLERAAQTRLPIMVFVGNRGMIQIHTGAVTNIRRMGTWINVLDEDFNLHLREDL
VASAWVVRKPTSDGAVTSVELFDAAGDNIAMLFGARKPGQPELAGWRELAGALPRLDTAD
AADAATVAHAADVPVATDAGAAR*
>GCA_000452945.1_BPMSHR5858_genomic_02442_1 cysW
VQRSFLERSLLRRFGAAPAGGLLALFFVLPLAALVSAAFAEGGRAFAAVLHDPLVGDAIG
RSLALAVGTGTLSTCVGVPLALSFAEQPPARRRWLLALLGVPLAFSGLVIAYGFILTFGR
AGFVTTLLAGLGADAAKVGGVIYTTFGLVVAYAYYLIPRVALMLFPAFANLERRPLEAAM
SLGAKPWRAWLDVAWRELWPSVMAAWCLVTAIALGTYGTALALAGTQINILPLLMYLKLS
DGQTDFSQAAVLSIVLTALCTCVLAMGEYVGRRHR*
>GCA_000452945.1_BPMSHR5858_genomic_02445_1 lsrR
MTRTAVHDAARRAAPLDHDMTNEELTQLAKCYYVDGLTQEELAQKFAISRPKVGRLLKRA
IEEGIVEIRVRHHPRAVQDLEQELVTRFGIQRAIISVDHKDQDSQRELLAGLVASYLDRV
LADGAIVAVGMGRNVSAVSRHAVSTTQRNCSFVSAIGGSYRGGETMNADHICRRLAARFG
GESETLYAPALVNDPQLFTALLENDVVRQSLDKARRASIALVGIGDILEDSNMVRMGWFT
PEEMAEAKRAGAVGDIMGYDFIDIHGRPATTMLHGRVIGLTLEDLKRIPNVIATASEPTK
ATGILGALRAGVINTLATTQSIAQTVLSLAQATETASAA*
>GCA_000452945.1_BPMSHR5858_genomic_02455_1 group_4742
MENAFNERGVMVTRNGLSAAGQIFALREIRGVEVRTVRKNKLVPCAISLTGAAAAIAGGA
LGSSALLVAGVMLVVVGYLAWTTQDVTHRLIVDMPDGKREAIMSVDREFVERVAHAVDAA
RAAGAST*
>GCA_000452945.1_BPMSHR5858_genomic_02456_1 group_4743
MVEDTVFSGLRTLLTHEHAFPVQSCRVSIEMQRPWGRPYRLVEWTMHLDAPARRRIVPAE
STEAEIAEAVASHVPGRLYEGGAALY*
>GCA_000452945.1_BPMSHR5858_genomic_02459_1 group_4744
MQTQTHPLISPAVGTARHITSFHYGPRGGKKVYIQASLHADELPGMLVATLLRRKLAALE
AAGRLRDEIVVVPVANPIGLAQHVFGDHLGRFELGSMQNFNRNFHDLAALVIPRIEGRLT
HDAAANLAAVRGAMREALAEQKPRTELESQRLALQRLSYDADIVLDLHCDCDAVMHIYTN
PDLWDDVEPLSRYLGAKASLLALNSVGNPFDEIHSFCWSELRGRFGERHPIPNGTISVTV
ELRSERDVSYELAEHDAQALVEYLTLRGAIDGTPAPQPPLEFAATPLAGTDPLVAPVSGV
IVFHTPVGVWIEAGQDVADIVDPLTDRVVTLKSSVSGVLYARQIARFATAGMEVARIAGA
TPIRTGSLLSA*
>GCA_000452945.1_BPMSHR5858_genomic_02471_1 group_4746
VRSSPRSSLARLARAAVFPVALALAASGMHAAFAQTTPLPPGAKQPSEFPRVKLRAGMYV
IDAAIAANDADREQGLMYRSRLAPNEGMLFVFNENAVHCFWMKNTLIPLSIAFIRADGTI
TDIDEMKAETTDNHCPRNNGVYALEMSSGWFAAKGVKPGMKIDGLPKPQ*
>GCA_000452945.1_BPMSHR5858_genomic_02474_1 mco
MLRRHFLSSALAAAAASLFARGAFAAGQAMDGMQGMEGMEGMDDMPDMKPAPAHARHGKP
APGPASALAAADALPAGAPLAALRVLANESREPGVFRATLVAQPVARALLPGAAPTTLWQ
FGADTQGPAVGPLIDVREGDAVEIRFVNRLPQPSTIHWHGLPVPPDQDGNPSDLVAPGAT
RVYRFTLPKGSAGTYWYHPHPHMATAEQVFRGLAGPIVVRAADDPLAGWPERHLFVSDLK
LARDGAIAPNDMMDWMNGRQGQFALVNGARRPRISLTGDERWRVWNGCSARYLRIAFDDG
RPFAHVGTDGGLFDAPREVASLLLAPGERAELVVRAGDRASHAVLTALEYDRGKMAMAMS
EAAHGSLPPDPALPLADVAFEPAAPRALPARLRAVPALGEPVARKEVVFGEQMDMAAMMR
AGAHGRPAGMRFMVNGATFEPHRATLTSRRGEIESWTIRNETDMDHPFHLHGTQFQVVER
EIDGETTPEPYRAWRDTVNVRKGERVRILTTQTERGERMFHCHILEHEDLGMMGTLKVV*
>GCA_000452945.1_BPMSHR5858_genomic_02480_1 group_2421
MNKLPAITLAFWVMKICATTLGETGGDLLSMTLDVGYAYSSLILFSFFIVTLAAQLAATR
YRPAIYWAVIVATSTAGTTMSDFMDRTLGLGYAAGSGILVAILLSIFAVWRLQGESLAVT
EIRTRKVEVLYWIAILFSNTLGTALGDFLADSSGLGFAGGAFLIGGLLAAIVLAAYFTNV
SKVFLFWAAFVLTRPFGATVGDLLTKPVAKGGLGFGTVGSSAILAGVLIALVIYAMFAEA
KRVKQARAPRMDWKQPALEE*
>GCA_000452945.1_BPMSHR5858_genomic_02486_1 purN_2
MKKLVILISGRGSNMEAIVRACAREGWPAEVAAVISNRLGAAGLEFAASHGIATAVVDHR
AFDGRDSFDAALAAEIDRFAPDLVVLAGFMRILTPAFVAKYEGRMLNIHPSLLPSFKGIH
THQQALDAGVALHGASVHFVIPELDSGAIVAQAAVPVVAGDDADALAARVLAAEHTLYPR
AVRWFVEGKLRLDAGRAIVAPDEARWLFADAIDTSTSEGV*
>GCA_000452945.1_BPMSHR5858_genomic_02487_1 rsmB_2
MKLHGFLIGQTEALLAEVLKFAGPADVTTSRFFRAHPKLGHAERGVIAEAVFAVLRRRME
YAHLAESGTGAPARRLALLGLMQTVGRSALKPFVSDAEWAWLDHVAKIDPASLPLRVRTN
LPDWIHQALAERLGADELAQFAAAVNYPAPLDLRANALKATRDQVIDSLRAAGIDAGATP
FAPFGVRVVGKPALTKLKLFEEGLIEVQDEGSQLLCSLVAPRRGEMIVDFCAGAGGKTLA
LGAMMRSTGRLYAFDVSEKRLAKLKPRLARSGLSNVNPVLIDSEHDAKIKRLAGKIDRVL
VDAPCSGLGTLRRNPDLKWRQSRETVDELAPKQASILASAARLVKKGGRLVYATCSVLEA
ENEAIVSAFLAAHPAFELVPASRVLAEQRIALDTGDYLSLWPHRHATDGFFAAVLERRAQ
*
>GCA_000452945.1_BPMSHR5858_genomic_02490_1 nadC
MQSAIKSVEYDRPLAAGAACGVGEAWAKVPDALAPDERDALKARIKALLVREKAVLVAHY
YVDADLQALADETGGCVADSLEMARFGRDHDAHTLVVAGVRFMGETAKILSPGKRVLMPD
LDATCSLDLGCPVDEFSQFCDAHPERTVVVYANTSAAVKARADWMVTSSIGLEIVADLHA
RGEKIIWAPDRHLGGYIQKKTGADMLMWQGSCLVHDEFKGIELDLLRHEYPDAKILVHPE
SPEGVVALADVVGSTTQLIDAAVKLDAQRFIVATDLGILHKMRLAAPGKTFIEAPTAGNS
ATCKSCAHCPWMAMNALSNLADVLERGHNEIFVEAAIAQRARMPIDRMLDFAARHKQRVQ
ASGDLQRDQALFANVGAA*
>GCA_000452945.1_BPMSHR5858_genomic_02494_1 rpmB
MARVCQVTGKAPMSGNNVSHANNKTKRRFLPNLQNRRFWVESENRWVRLRVSNAGLRLID
KNGIDSVLADLRARGEA*
>GCA_000452945.1_BPMSHR5858_genomic_02497_1 ispH_2
MSSTDTLSGQVAAADAEILLAQPRGFCAGVDRAIEIVERAIAMHGAPIYVRHEIVHNKYV
VEDLKKKGAIFVEELEEVPSGNTVIFSAHGVSKAVRDEAAVRGLRIYDATCPLVTKVHVE
VAKMRQEGVDIVMIGHKGHPEVEGTMGQVERGMHLVESVEDVRRLELPDPERVALVTQTT
LSVDDAAEIIGALKAKFPAIREPKKQDICYATQNRQDAVKFMAPQCDVVIVVGSPNSSNS
SRLREVAEKRGVAAYMVDAPEQIDPAWVAGKRRIGVTAGASAPEVLAQAVIARLRELGVT
NVRALEGIEENVSFPLPRGLNLSSAG*
>GCA_000452945.1_BPMSHR5858_genomic_02498_1 group_4749
MHLELVYAGPVDAAVALFAAAADGCDGGAAAQAAAAPAATPGDVIVAGAGRPAANFVAKP
ARIKRARIDVVAAVDIGTLARRTAVPRAASPSRRAPSTSDAVRAIADTTRRAIPSEAFDV
PQPMPASAREAGDGMRGGVGGAKAGAWTLYDFKEAKHTVLKS*
>GCA_000452945.1_BPMSHR5858_genomic_02503_1 group_4750
MSLDYGFVKAKIKAVAGLKAAARANETQYHVHLTLALPDGDWDVAINVGTNDADDLLKYK
LVYDFHHPVTQTLAAAAEGYTDLTGGDALPALDYVRSDVLNETGGWRMSDVMDGTEHPEP
IPSVLRLVDAAHQQNLDLYVFGRTYVEGNGIHDTHMNQGSSGPHFLHRPGVDTNDHNDVW
QDGALLVDLGGPQWAAYFAAFEQQAVPTDGLGNPLPGAGPIS*
>GCA_000452945.1_BPMSHR5858_genomic_02505_1 argD_2
MPLNDYPIDSLMYITNRPDIVFTHGKGSWLYDHTGKRYLDFIQGWAVNCLGHCNEGLVQA
LQAQAEKLLNPSPAFYNEPMAKLAGLLTQHSVFDKVFFANSGAEANEGAIKLARKWGRKF
RNGAYEIITFDHSFHGRTLATMSASGKAGWDTIYAPQVPGFPKAEINDINSVERLINDKT
VAVMLEPIQGEGGVIPATREFMQQLRALTKQHDLLLIVDEVQSGCGRAGTLFAYELAGVE
PDIMTLGKGIGGGVPLAALLSKADVAVFEAGDQGGTYNGNPLMSAAGHAVISQLVAPGFL
EGVRARGEYLKHKLLELSEERGFEGERGEGLLRALLLGKDIGPQIVEKARDMQPDGLLLN
AARPNLLRFMPALNVTNDEIDQMMAMLRSVLDSL*
>GCA_000452945.1_BPMSHR5858_genomic_02507_1 group_4753
VGTVQCAHGGYNAFLRINRYIMIATRILRRPRVLIVGCGDVGMRCAAQLRARRENLRILA

LTSRRSRCVELRAAGVVPVVGDLDARATLKRIARVAPVVLHLAPPQATGDVDRRTQALVA
ALASPRRPSRQLAPAYGRLRAWRTAARSARPPFQASAIVPDALPRPVVVYASTSGVYGDC
GGARVDETRAVRPANPRARRRVSAERQLRRATARGALSARIVRIPGIYAANRLPLARLEK
GTPALVEADDVYTNHIHADDLASILLRAAVRGKPARVVHASDDTELKMGDYFERVARAFG
LRSPPRIARAEAERQLEPMLLSFMRESRRLANARMKRELRIALRYPSVDDFLRTVSAPRP
LK*
>GCA_000452945.1_BPMSHR5858_genomic_02508_1 group_5385
MTIDNPIEISSVMLAFAALGAIIIGGLLTAMHLKRKYHPNLIGALIGALLCFLLIEALPA
IT*
>GCA_000452945.1_BPMSHR5858_genomic_02516_1 fucA
MVDTLKMGKAELVALAQRRLDNEIKDSGWSAREKLALTCRILFDDGHDSGLAGQITCRAD
APHTYYTQRLGLGFDEIAESNLLQVDQDLEVIGGQGIPNPANRFHTWIYRARPDVNCIIH
THPTHVAALSMLETPLVVSHMDTCPLHGDCAFLKDWPGVPVGNEEGEIISNALGDKRAIL
LSHHGQLVVGKTIEEACMLAILIERAARLQLLAMAAGAIAPIPPALAADAHDWISTPKRD
AVTFAYYARRALRRHPDCLD*
>GCA_000452945.1_BPMSHR5858_genomic_02517_1 group_4755
MTIRLKLLRKQKGWTLDVLAEATGLTKSYLSKVERGLSVPSIAVALKLSKALQVDVEQLF
SEGRDRELITVTRASERTSMGRASSERVRTYESIAAGVAPKKLLPFIVHPPRDYVSSAFR
EHEGEEMLFVHRGSVEIEFPNETIKLKTGDSVYFNALIPHRMRSVGATSAEVLVVVSNDE
GVQEDAQ*
>GCA_000452945.1_BPMSHR5858_genomic_02526_1 moaE
MATVRVQADDFDVSAEIAALRARNPKVGAIACFVGTVRDLNEGEQVDALELEHYPGMTEK
ALAAIAEDAIARWPGIDVAIVHRVGKLYPLDQIVFVATVSAHRGDAFQSCEFVMDYLKTQ
APFWKKEKTPAGERWVDARSADDAALARWGVESGNAREKRST*
>GCA_000452945.1_BPMSHR5858_genomic_02533_1 arnT_2
MNDTPTRLPLDRAAPVTLNRAALLLLVAALAVIWFAPLGLRHLIPSDEGRYAEMAREMFV
TGDWITPRYNGYKYFEKPPLQTWLNALTFAWFGIGEWQARLYTALASFAGVLFAGYAGAR
VFNPRAGFFAALVLASSPYWNLMGHFNALDMGLSLWMTVTLCALLLAQRPGIGKRAERGW
MWLCWASMALAVLSKGLIGLILPGAVLVLYTLASRDWALWKRLHLVSGLVIFFAIATPWF
VLVQSRNPEFFNFFFIVQQFRRYLTPEQNRPGAFYYFAPVLAVGFLPWLSVSVQSVRHAL
RIPRQPNGFAPMLMLATWSVFIFLFFSASHSKLISYVLPVAPALALMIGAYLPLVSREQF
RRHLFGYLVFLVAAAFGTLFLARLGDARTPNSQYLEFQKWVYVALAAAFALTLVAVWLNR
RGRAGVAAAAVAFGAGWLLLGTIGGTGHDAFGRYSSGALLAPAIREELAKLPPDTPFYSV
ETLDHTLPFYVGHTMIMVERKDELTFGISVEPNKWVPTVAEWEQRWRADRHALALMSPDR
YKQLSEAGLPMRVVARDNRRVVVEKPQS*
>GCA_000452945.1_BPMSHR5858_genomic_02541_1 group_1109
MPLPTPPSKLGSLLPPDEYKAKARPAKAAKQSASGGPAHAAAADYSPASVAEPMVVAANT
TTPLRAVAPAADASAGAPRARRAKQTAALLQPVPAAPAPVARSSKTDAAKPAAAAPAAPR
PAAKTRGRKDAEVEMQKLFVLDTNVLMHDPSSLFRFEEHDVYLPMMTLEELDNHKKGMSE
VARNARQVSRTLDSLVADAGPISAGIPLARLGSREALGRLYFQTTLTSIEPVEGLPQGKA
DNQILGVVRALQRERPDRQVVLVSKDINMRVKAHALGLPAEDYFNDQVLEDKDLLYSGVR
ELPQDFWTKHAKGMESWQDTKTGTTYYRVTGPLVASMLVNEFAYLEPQNGEPAFHAIVRE
LNGKTALLQTLRDYSHHKNNVWGITARNREQNFALNLLMNPEIDFVTLLGQAGTGKTLVA
LAAGLAQVLDDKRYNEIIVTRATVPVGEDIGFLPGTEEEKMQPWMGAFDDNLEVLQKTDD
AAGEWGRAATQELIRSRLKVKSMNFMRGRTFVDKYVIIDEAQNLTPKQMKTLVTRAGPGT
KIICLGNIAQIDTPYLTEGSSGLTYVVDRFKGWGHSGHVTLARGERSRLADYASDIL*
>GCA_000452945.1_BPMSHR5858_genomic_02550_1 sigR_2
MDCNETRALLGADVDHELSAADAWRIARHVGGCGACRLERERLVALRRAMRQAEYHRAPG
ALRARIAAGLPLAAAPFAQAPVQDLPASDVPVSDVPVSDVPVSDMPVSDVPVSDMPVSDM
PVSDVPVSDVPVSDMPVSDVPVSDAPEQDTPSQDMRAPNKPAVDRPPEAKPRFGADARGG
ARAGRWFARPGSRGPTLDRPGPGPRAAALPGLGWGVALTVALAAAAGFALDARRAATEHA
VDEIVASHVRAGLSSRDIDVISTDRHTVKPWFNGRLDFAPPVVDLSASGFALAGGRLDYV
GQRRVAVLVYRYRQHVIDVYVWPSGEGGARPYATVSQGYALDRWEAAGMTWWAVTDAEPS
ALAAFRTALDARVAAPRTE*
>GCA_000452945.1_BPMSHR5858_genomic_02553_1 cbaB
MRHFFFDPRALRGRARASAPSWPRRRWLFAASGAALAALAGVDASRVRAAAPRVIKVHAR
RFVFTPDRIALAPHESVVFELTAQDTVMGFSIPQYGVRADVPPGAVVRLAAQAGGPGTVQ
FLCDIFCGSGHETMNGVLVVG*
>GCA_000452945.1_BPMSHR5858_genomic_02561_1 psiF
MKIRSLMAALLVGGVLATPAFAANSQQDKMKACNAQAAGKTGDERKAFMKDCLAAKPAKK
MSQQEKMKACNTQAADKKGDERKAFMKDCLSAKPAA*
>GCA_000452945.1_BPMSHR5858_genomic_02563_1 group_19096
MHKRNLVLKAAAALVVGSLALAGCTTTPDKPANAATNASKRQAIDASVDATLSRLYSTVR
GSRELVAKSRGVLVFPDVIQAGLIIGGQTGNGALRVGGATVGYYNTSSLSVGLQAGAQSK
AIVFLFMTQDALDKFRNSDGWAAGADASVALVKMGANGAIDTTTATAPVEVIVLTNAGLM
GDVSISGTKVTKLKI*
>GCA_000452945.1_BPMSHR5858_genomic_02564_1 bshA_1
MRIAQIAPLTESVPPKLYGGTERVVSYITEALVDLGHDVTLFASGDSITRAKLDAVWPRA
LRLDASIRDRIAPHMLLMETVARRARDFDVLHFHMDYYSFSIFKRQDTPFVTTLHGRLDL
PEQQPVFDTFDTAPVISISNAQRHPMPQAKWLTTVYHGLPETLYTPQPVEQSYLAFLGRI
SPEKRVDTAIRIAQRCGMRIRIAAKIDAADEEYFEREIKPLLALPHVEYIGEIADHEKAA
FLSGAHALLFPIDWPEPFGLVMIEAMACGTPVIAFNRGAVPEVIDEGVSGFIVEDEIGAA
AAVNRLHMLSRERVRARFDERFTSRRMAQQYVDVYQSLIRAQKRSRFKVIDSAT*
>GCA_000452945.1_BPMSHR5858_genomic_02565_1 group_2424
MPPLTALAIAHYRSLRELIVPLATLNVITGPNGSGKSSLYRALRLLADTAQGRVIPSLAR
EGGLPSTLWAGPERFSRGMLAGDVPVTGTVRNAPVSLKLGFACEDFGYAIDLGLPPLNSA
TAFGLDPVIKRECIWSGPMPRPSTLLVDRRGAQLRTRDETGEWQTVPQPIASFDSMMTEF
SDPRGAPEMIAMRERIRSWRFYDHFRTDALAPARLAQVGTHTPVLSDDGGDLAAALQTIC
EIGDAAALDAAIGDAFPGARLEIDNPGKHGRFEVLMRQPGLLRPLRAAELSDGTLRYLLL
AAALLTPRPPALLVLNEPETSLHPDLLPALGRLIAQAAQRSQVIVVSHAARLVASLEREP
GSHSIVLEKELGATRIVDAHALEVPAWKWPAR*
>GCA_000452945.1_BPMSHR5858_genomic_02569_1 ycfH
MFVDSHCHINFEGLAERLPQVLENMREHGVTHALCVSVDLETLPAVLAIARDHDNVFASV
GVHPDHEDAREPTVAELVELAAHPKVVAIGETGLDYYRLEGRSIADMQWQRERFRTHIRA
AHATGKPLIVHTRASSEDTLRIMAEERASVPGGVMHCFTEPWPVAEAALAQNFHISLSGI
VTFKSAADVQEVARRVPLERLLIETDSPYLAPVPYRGKRNEPAYVSYVGRFIAAQREMTD
EALGAATSRNFFRLFKIARTGEASPVNQG*
>GCA_000452945.1_BPMSHR5858_genomic_02572_1 tmk
MARGKFITFEGIDGAGKTTHLQWFCDRLQERLGPTGRHVVVTREPGGTQLGETLREILLN
QPMDLETEALLMFAGRREHLALVIEPALARGDWVVSDRFTDATFAYQGGGRGLPRDKLEA
LERWVQGGFQPDLTVLFDVQPQVASARRGAARMPDKFESESDAFFARTRAEYLRRAHEAP
HRFAIVDSSESIPQIRKQLEGVLAAL*
>GCA_000452945.1_BPMSHR5858_genomic_02573_1 mltG
MSLLKKCAALAALAVVLLGVACAGGAYYWATRPLALAAPILDVTIKPRSSVRSVAQQLVH
GGVGVEPRLFVAMTRVLFLSSRLKSGNYEFKTGVSPYEVLQKVARGDVNEYVVTVTEGWT
FRRMRAELDANAALAHASAGMSDAALLRAIGAPAEVVARGTGEGLFFPDTYLFDKGTSDL
NVYRRAYRLMQARLADAWTARRPGLPFKTPYEALTVASLVEKETGHASDRAFVSGVFANR
LRAGMPLQTDPSVIYGMGDAYTGRLRKRDLQTDTPYNTYTRRGLPPTPIALPGEAALYAA
VNPAATSALYFVARGDGTSVFSDTLGDHNKAVDKYIRGQ*
>GCA_000452945.1_BPMSHR5858_genomic_02574_1 ygfZ
MSTPIASPAAQAIAPTAPPTLPRPSADDFAAVLERGAFAALEQFGIVDVTGADAATFLHG
QLTNDVEHLDAASARLAGYCSPKGRLLASFLAWRAGHDVRLLVSKDVQPAVQKRLSMFVL
RAKAKLADASGALVAIGFAGDVRAALSGIFDALPDGIHTKVDAPAGALMRLPDAAGRARY
LWIATRAEFDARLPALEAALPRVSAAVWDWLDVRAGEPRITQPAVEQFVPQMVNFDVIGG
VNFRKGCYPGQEVVARSQYRGTIKRRTALAHVAAGTDAAHAGVELYHSDDPGQPCGMIVN
AAAAPEGGVDALVEIKLAALGNGSVHLASASGPTLAFLPLPYALPVEA*
>GCA_000452945.1_BPMSHR5858_genomic_02577_1 nlhH
MPLNPKIAQVLDMIERAKRPDYHEQTPAQARAAYEKSAPILDVAPAPMFSVEDLRVPSRD
GGAFGARLYLPVEPSLAEPLPVLVYYHGGGFTVGSVNTHDALCRMFARDAQCAVLSVDYR
LAPEHKFPTAVDDAEDALVWLHAHASRFGIDSARLAVGGDSAGGTLATVCAVLARDRGIA
LALQLLIYPGTVGHQQTESHARLAKGYLLSADTIQWFFGHYVRDASDRDDWRFAPLDGTR
GVPSFERVAPAWIATAQYDPLSDEGEAYADKLRAAGNRVTLVAYAGMIHEFFKMGGYVPE
VRLAHADAVGALRAAFDGD*
>GCA_000452945.1_BPMSHR5858_genomic_02578_1 curA_1
MSQVNRRVLLVSRPEREARVENFELVETPLAPLAEGEVRVRNHFLSIDPYMRGRMNAGRS
YAEPQPLGEVMGGGTAGEVVESRNPAFAPGDRVIGAYGWQEYGTSAGKELRKVDTTRVPL
SAYLGAAGMPGVTAWYGLNRIIRPRAGETLVVSAASGAVGSVVGQLAKLAGCRAVGIAGG
ADKCRYVVDTLGFDACVDYKAGRLADDLAAAAPDGVDGCFENVGGAVLDATLARMNPFGR
IAMCGMIAAYDGAPAPLANPALILRERLLVQGFIVSEHFDVWPEALAQLASLVANRQLHY
RETIAQGLERAPDALLGLLKGRNFGKQLVALV*
>GCA_000452945.1_BPMSHR5858_genomic_02579_1 group_19053
MSDACATAGREPAIESPFVDALGVRLVSAKDGASEIVLPLAEQHMNTWNIAHGGVTMTLA
DVALAMAARSLTDDGVGVVTVEMKVNFMQPGRGELRAYGRVLHRSTTMAYCEGEVRDSAG
NFVAKALGTFKYMKRLAVGRDIARQRTRTDPGAHPGPSDA*
>GCA_000452945.1_BPMSHR5858_genomic_02604_1 pgdA
MTTNMSDAAGAPAASFAPERRAFLARAGGGLGLAAAGLALGAAAAPGRALAAGATATADT
GAASPAGGSPRRSPADEPETAHGAFWPNGARLVISISMQFEAGGQPPTGADGPFPPVVFP
PQVPVDLASATWFAYGYREGIPRMLDLWDRHGVKVTSHMIGEAVRRRPDLAREIVARGHE
AAGHGPRWSAQYALPRDEERRFLIAAREMVETATGARPVGYNCNWLRRGPNTLPLLQELG
YLYHIDDVSRDEPFIEQVNGQDFVVVPYTLRNNDILLIEGRNYSPGQFLEQIKLDFDQLY
DEAATRRRMMSISAHDRISGTPQMVRAWDAFLRYAQSHPGVAFMRKDDIARHALRSPLTL
REPETL*
>GCA_000452945.1_BPMSHR5858_genomic_02612_1 thrS
MVSIRLPDGSVRQYEHPVTVAEVAASIGPGLAKAALGGKLDGELVDTSALIDRDASLAIV
TDKDADGLDIIRHSTAHLLAYAVKELHPDAQVTIGPVIDNGFYYDFSYHRPFTPEDLEAI
EKRMQELAKRDEPVTRRVVSRDEAVSYFRSIGEKYKAEIIESIPASDEIKLYSHGSFTDL
CRGPHVPSTGKLKVFKLMKVAGAYWRGDSKNEQLQRIYGTAWTRKEDQDAYLHMLEEAEK
RDHRKLGKQLDLFHIQEEAPGMVFWHPKGWTLWQQVEQYMRRRLDAAGYLEIKTPMIMDR
SLWEASGHWQNYRENMFTTESEKRDYAIKPMNCPGHVQVFKHGLRSYRDLPLRYAEFGSC
HRNEASGALHGLMRVRGFVQDDAHIFCTEDQINSEAIAFNKLAMSVYEDFGFDRIDIKLS
LRPEQRMGSDETWDHAEEGLRNALKACGLEWEELPGEGAFYGPKIEYHIKDALGRSWQCG
TLQLDMMLPERLGAEYVAEDNSRRRPVMLHRAIVGSMERFLGILIEHHAGAMPVWLAPAH
AVVLNIAESQAEYARTVAQSLQKQGLRVSADLRNEKISYKIREHTLEKVPYLLVVGDKER
EAQTVAVRARGGVDLGVMPVEAFVERLREDIQAFK*
>GCA_000452945.1_BPMSHR5858_genomic_02613_1 infC
VRLVGVDGEPIGIVKLAEAFRKSEELDVDLVEIAPQAAPPVCRLMDYGKFKYQEAKKQHE
AKLKQKVIQVKEVKFRPGTDDGDYNVKLRNLVRFLEEGDKTKITLRFRGREMAHQEIGMR
MLERLRGDLDEVGQVEQMPKMEGRQMIMVLSPKKKK*
>GCA_000452945.1_BPMSHR5858_genomic_02615_1 rplT
MPRVKRGVTARARHKKIINLAKGYRGRRNNVYRIAKQAVMRAGQYAYRDRRNKKRVFRAL
WITRINAAVRQHDMTYSVFINGLKKASIELDRKVLADMAVFDKAAFAAIVKQVKAAVAA*
>GCA_000452945.1_BPMSHR5858_genomic_02621_1 oatA_1
MVSHLDILFFWGSSRIDTLQRAFFFWGGVDLFFCISGYVITGNLLRTLPSGRQGGFGAFA
IPFWIRRAWRILPSAWLWLAAPLILSVVANRTGYLGTPKGNIIDALSAVAQVANFHFWAC
YAFPATTCGIDQVYWSLSLEEQCYILLPVLLYFANRRTIAPILCVVVLIQVFLPRPILSL
LWFVRTDALALGALLALLQHRDWYAKVEPRPLGRGLISAIALAGMCIVLAALAGYLARIP
FHTGLIALMSCVMVWIASYGRGYTMRDGAVKRVLVYFGSRSYAIYLIHVPVYRLTREIWE
RIVITPSTIGGSFTLHFAFTAIPIVLVLAELNYRFVEAPLRVYGARLADRWAHARNVENR
KSIASVP*
>GCA_000452945.1_BPMSHR5858_genomic_02625_1 group_18989
MTDLRLTKRFAALLATGAFVAGCSSPSPTKIDYKSDSKSKEVSLAVPPNMLDETPDQRSL
PPQGGATSLSSLQQVQQAAPPTDSVAPAVPGMHIQRDGTESWLVIDDKAPAQIWPQVRRF
WQEQGFLLVVEQRDKGVMETDWNETHPQINDGLIRGVISKAMGNSYVTAERNKYRTRLDA
APNGGTYVFISQKGMREALTGANNDSSKWEPKPNDPGLETEYLKRLMAVIAQNDARVKAA
GVPITDDAAPSRAKASSNKSKEAAAAIAAKNVASASTSSTPSGASVPDVPPEVTLAEPYD
RSWLHVGLALDRANFTVDDRDRTKGLYYVRYVDPKDLSVEEQGFWSQLFHGKKEKKAKQY
LVNVKAVTEGETRIAVVDDKGNVDASAPARQIMSLLVNQLR*
>GCA_000452945.1_BPMSHR5858_genomic_02630_1 group_18935
MATYSNEAVLDALRRVQYRQVPWARRPGVFEYLRALGLMDTVRQKTVAPAPGFHAPVDIA
VLTDNGRVEFTRLERDEKQPDWETRRIEDYALSEASAVSILESRL*
>GCA_000452945.1_BPMSHR5858_genomic_02636_1 rimP
VQLTELIETTVTGLGYELVDLERTGRGMVCVYIDQPAGITIDDCEKVTRQLQHVLTVENI
DYERLEVSSPGLDRPLKKLADFTRFAGSEAVITLKKPLDGRKTYRGILHAPNGETIGLEF
ERKKGEAAMLDFTLADIDKARLIPHVDFRSRKQ*
>GCA_000452945.1_BPMSHR5858_genomic_02637_1 nusA
MSREVLMLVDALAREKNVDKDVVLGALEAALASASKKLFDEGAEIRVHIDRESGEHETFR
RWLVVPDEAGLQEPDREILLFEAREQKPDVEVGDYIEESVPSIEFGRIGAQAAKQVILQK
VRDAEREQILNDYLERGEKIMTGTVKRLDKGNFIVESGRVEALLRRDQLIPKENLRVGDR
VRAYIAKVDRTARGPQIELSRTAPEFLMKLFEMEVPEIEQGLLEIKAAARDPGVRAKIGV
VAYDKRIDPIGTCVGIRGSRVQAVRNELGGENIDIVLWSEDPAQFVIGALAPAAVQSIVV
DEEKHSMDVVVDENELAVAIGRSGQNVRLASELTGWQINIMTPDESAQKQNEERDALRGL
FMARLDVDEEVADILIDEGFTSLEEIAYVPLNEMLEIEAFDEDTVHELRNRSRDALLTMA
IANEEKVETAALDLKSLDGVTPELLAKLAEQGVQTRDDLAELAVDELVDMTGMEEEAAKA
LIMKAREHWFQ*
>GCA_000452945.1_BPMSHR5858_genomic_02638_1 infB
MASNNVAQFAAELKMPAGVLLEQLQAAGVQKASEDDALSETDKARLLDHLRKSHGATDGD
KRKITLTRRHTSEIKQADATGKARTIQVEVRKKRTFVKRDDVSETGADQAQAQTDEQAEA
ELKRREEEARREAELLEKQAQELRERQERLEREEAERRAREEAAEAERRRAEEEAAAKRA
AAAQAEAAQQAAAAREQAQRAQSEPAEQSAQDEARAAAERAAQREAAKKAEDAAREAADK
ARAEQEEIRKRREAAEAEARAIREMMNTPRRAQVKAVEPPKPAEPPAAKAAEAKGTLHKP
AKPAGEAAAARPAAKKPASGAPAPAAAPAGDRTKKPGTGKSGWQDDAAKRRGIKTRGDSS
GGVDRGWRGGPKGRGKHQDSASSFQAPTEPIVREVHVPETISVADLAHKMSIKASEVIKV
MMKMGQMVTINQVLDQETAMIVVEELGHRALAAKLDDPEALLVEGEIGSDAEQLPRPPVV
TVMGHVDHGKTSLLDYIRRAKVAAGEAGGITQHIGAYHVETPRGVVTFLDTPGHEAFTAM
RARGAKATDIVILVVAADDGVMPQTKEAISHAKAGGVPIVVAINKIDKPEANPDRVKQEL
VAEGVVPEEYGGDSPFVPVSAKTGAGIDDLLENVLLQAEVLELKAPVESPAKGIVIEAKL
DKGKGPVATVLVQSGTLSRGDVVLAGTAYGRVRAMLDENGKPTKEAGPSIPVEIQGLSEV
PGAGEEVIVLPDERKAREIALFRQGKFRDVKLAKQQAAKLESMLEQMGEGEVQNLPLIIK
ADVQGSQEALVQSLLKLSTDEVRVQIVHSAVGGISESDVNLATASKAVIIGFNTRADAQA
RKLAEANGIDIRYYNIIYDAVDEVKAAMSGMLAPEKREVVTGMVEVRQVFKVPKVGTVAG
CMVTDGVVKRSSSVRVLRNNVVIFTGELDSLKRFKDDVKEVKQGFECGMSLKNFNDIVEG
DQFEVFEVTEVARTL*
>GCA_000452945.1_BPMSHR5858_genomic_02640_1 truB
VQTMTTVSPRPRMARRALDGVLLLDKPVGLSSNDALMRAKRLYQAKKAGHTGTLDPLASG
LLPLCFGEATKFSQDLLEADKTYEATMRLGVRTTTGDAEGDVLDTRDVSCDEAAVRAALA
RFVGEIVQVPPMYSALKRDGKPLYEYARAGQTVEREGRTVTIRALALVSCALPDVTFRVT
CSKGTYVRTLAEDIGEALGCGAHLTMLRRTGVGPLTLEHAVTLDALDAATQDERDARLAP
VDALLSTFPCVKLDAALATRFLHGQRLKLSELAARPDAAEGGRVRVYDADDRLLGVARAS
EGVLAPERLVVTGA*
>GCA_000452945.1_BPMSHR5858_genomic_02641_1 group_19538
MAQPHAPLPPLAGGQLILGTIAVSLAVFMNVLDTSIANVAIPTISGDLGVSSDQGTWVIT
SFAVANAISVPLTGWLTDRIGQVRLFLASIILFVISSWMCGLAPTLPFLLASRVLQGAVA
GPMIPLSQALLLSSYPRAKAPMALALWSMTTLIAPVAGPILGGWISDNYSWPWIFYVNIP
VGIAAAAVTWMIYRSRESAVRRAPIDGVGLALLVIWVGSLQIMLDKGKDLDWFASTTIVV
LALTALIAFAFFVVWELTAEHPVVDLSLFRMRNFSGGTIALSVGYGLYFGNLVLLPLWLQ
TQIGYTATDAGLVMAPVGFFAILLSPLTGKFLSRTDPRYIATAAFLTFALCFWMRSRYTT
GVDEWSLMAPTFVQGIAMAGFFIPLVSITLSGLPGHRIPAASGLSNFVRIMCGGIGTSIF
QTAWDHRNNFHHAQLVEQANPYNPTFNQAVTQMGQLGFTRDQAHGLINNMATQQAAQLGV
NDLFYISAAIFVLLIALIWITKPERAGGGDSSAAASAAH*
>GCA_000452945.1_BPMSHR5858_genomic_02650_1 group_19192
MTPYRDITDEEWQRVAPLLPELRPRKELRGRPLADTRAVLNGVLWVIYSGANWSAMPRRF
PAYQTCHRRFKAWHESGALKRVLIELFGTQSDDLCELMAMRMRTHARSKAAQARSEAAST
TSSPVPRHDHTLEHVV*
>GCA_000452945.1_BPMSHR5858_genomic_02651_1 group_19847
MFQMTRSLRSITVAGALLVCAASAFAQADSPVGMWQTIDDHTGKPKALVQISADSNGELS
GKVVKGLGENDTPDRRCTACTDERKDQLIKGMTIIKAMKKDGEGWDGGNILDPENGKVYK
CKMKLEDGGQKLVVRGYIGVSLLGRSQTWMRSQ*
>GCA_000452945.1_BPMSHR5858_genomic_02661_1 group_19121

MAHDIQFADGATPFSQTQQFHDIKNAAHEHLLTRIEELGAEFGRWSRQAINQFVDLEIDS
FVRLRRIPLNENEVRAIAEALTKELAGFGPIEDLLADPHVEDILINGYSDVYVSKHGILT
KLPVRFTDNAHLLRIVRRILAPIGRRLDESNPMVDARLPDGGRVNVVIEPLSIDGPVVSI
RKFRKDPLKPADLLANGTFNDEIGALLEAAVVARCNILVSGGTSSGKTSLLNALAFHIPG
IERVVTIEDTAELSLNHPHVVRLESRPGGFDGTGVVSIRDLLRNTLRMRPDRIIVGEVRG
GEVLEMMQAMNTGHDGSMGTIHASSPRECLYRLEMLAGFAGFQGTESSLRRQIANAVDFI
VQIGRLSNGRRRILSVTEVTGLSDNIIATQELYRYEPRVNADGDEIDAWESLGIHPHSPK
LARFRQALSGGGFGGGEPYGRGGGFNV*
>GCA_000452945.1_BPMSHR5858_genomic_02665_1 group_92
MTTHDLTQTAAPRAGRGRALCAAALALACAAGAARAQSPQRDAPSASEIGHATQAWFALQ
ASNREAAPAQPMIGEAASLAYARYIESFRTKIPAFYGSAAGMSGMSGGGGGGGGAGYSLA
PASGSQ*
>GCA_000452945.1_BPMSHR5858_genomic_02671_1 group_6551
MSHTLPLHPSSSSRTRFARRRLATASAALSLSLSLMLGHGAAHAQVNAEASAAVESSAQE
AALPAELAKVAKLPVEQQARWLRTAARQGTLEKLDDATLTALFKSLDPQTVPDYVAAGPI
GHPSYEFTMLRQERISGKWSDTPDHMLVKVTRSPLRVYAKWLPDGAHSGQEVIYDSAKRA
DEMYGHLGGLLGKVPMWTAVDGTLARAQSNHQVRDLGTEFVANLYLTEAKKYREAGALKP
THVEAKTVKGVRVVALTYETPGGRPQFYAKKETLGLDLRQPYFRTVESYDNDGRVFEKIV
FEKITPKSLDETAFDPKNPDYKF*
>GCA_000452945.1_BPMSHR5858_genomic_02677_1 group_1015
MDLRSKHIDAPPQHAAHFAPLDEPPAPRARRTALDALAGLSIAGLLIPEAVAYAGLANLP
PQAGLIALVVGLVVYAIAGSSRFAIVSATSSSAVVLAATVMSEAGPGAAAQLMLAAALVA
TTGILFILAGAARLGGMSDFIARPVLRGFTFGLALTIVIKQLPKMLDVPIHHGDTLRVAL
DLLLGIAGCNVRSAALGATALAILFALGRRTRVPATLVVIVLGIAAGYWIDWHRYGIAVV
GTIDLQNLAFGMPVLGRSGWMQTAEFGFALMLILYAESYGSIRNFALKHGDTVSPNRDLV
ALGCANLVSGLLHGMPVGAGYSATSANEAAGAQTRMAGLFAAAVIALIAWLLLPQLARIP
EPVLAAIVIFAVSHSLHPEVFRPYWTWHRDRIVVIAALAAVIVLGVLHGLLAAIGVSLLL
TLRQLSEPNVSVLGRLRGSHDFVDVSMHEDAKPIPGVLIVRPEAQLFFANAERVLTMARR
LARDAQPPVHTVMLSLEESPDVDGTTIEALKTFGAECDARGWRLALVRLKPNVLRVLQRA
ADGGLRADALSELSVDESLQSLTAGELPRA*
>GCA_000452945.1_BPMSHR5858_genomic_02678_1 group_650
MASFKSSLRRGLRVSGCALWLALSKLDAAHAAAGPDASPDAWPSAALSDRAAVEVDASQL
LPTPQLVRWQVDLDRRGLRSTGSPAHERYIDVLRRRLARAGVQQLHTEATTLARWAVRHW
RLDVADGAPRERIDVAGYLPYSGDTGRDGIVAPIRYLAAGQTPDADVAGKIAVVEWPALP
LTGAFFRERALRVFDPDNAFAPSAPYVRTSFMLGTLTAMLDRLQAAGAAGVVMIADTSSA
EATRLYAPYDGRLRRVPGLFVDRATGAKLASLAERRATLRLSLDAGVERVHTRNLIGIIP
GMSDELTVVNSHTDGTNGIEDNGPNAIVAIAQYLSRLPRAALPRTVMILLSSGHFAGGVG
AEDFIARHARDGLVARIACVVTIEHLGAQEWLPSEQGALAPTGRTEPTALFMPAVPALVD
AADALVRRANAAPAFVMPPLNPNGDGSANDALWPGEGQYFWGRARVPTINLITGPTYLLN
YGVSTAEKIDYARLRREIAATTQMLLDLSRVPFDALRAVPPQTRAAAP*
>GCA_000452945.1_BPMSHR5858_genomic_02679_1 plcC
VANSLSRIEHIVHLMLENRSFDQMLGFLYTDDGNRSPTGQPFDGLTGNESNPDDLGRPVG
VYRIRATDPHPYLMPGADPGEGFQNTNYQLFCDDHPAPGAVPTNQGFVVNFKSAIATDQS
RHDKDALPGTTPEQIMGMYTPELLPVLSGLAKGYAVCDRWFASAPTMTMPNRAFALAATS
QGHLDDHVKIFTCPSIFGRLSDQGVDWAIFGYNRDPLTRHDFPDTQNADDSHFGHFRDFQ
ARAASGTLPAFTFLEPSWDASGNSQHPNYDVAAGEQLIHDVYYALRNGPGWNSTLFIVTY
DEHGGNYDHVAPPSGATPPGDGTVGEFGFDFTRFGVRVPAVLVSPLIAAGTVFRSAAGTI
DHTSVLKTIGERFGTAPLTARDRAAPSLGDALTLASPRAASDDPLAGVTVPVSRVSHPNA
AMPSKLDKLQAARIAALPLRNEKGYYEEADTPLASSAELSNFIRDRGAAWSQHRQRQQQR
RQQQQRQQPRSKPRR*
>GCA_000452945.1_BPMSHR5858_genomic_02685_1 amiD_1
MKFPLRHTLLASFSMLAAACTTPTLVNRGDYYADTSLPARSVDSRIRFLVMHYTEIGEAP
SLRVLTQENVSAHYVVPDAPRTERGAPVVYQLVPESERAWHAGVSAWQGATELNGVSIGI
ENVNRGPIDTPHGRMWAPYPPAQVDAIVRLAKDIVARYRIPPTRVVGHSDIAPQRKIDPG
PLFPWRALAKAGVGAWPDDETVAARLAGRPPKSLVDVRELQLKLARYGYDVATDGVLDDR
TRRVFSAFQMHFRPTDYAGNPDAETDAIAQALLDKYMPSQPAPERDAAALDAASGVAAQ*
>GCA_000452945.1_BPMSHR5858_genomic_02690_1 fliY_1
MKIASLKKLLVTGLIGASFAAATAHAADLLDEVKQRGTLRIGLEGTFPPFNSKNPQGELV
GFDVDIAKAIAAKLGVKPEFVTTEWSGIIAGLQAGKFDVIVNQVGITDPRKQTLDFSPAY
TFSSAQLIQRKDDARQFKSLEDLKGKKLGVALGTNYMDMAKSVPGIDVKTYPGAPEYLRD
LAAGRLDAALNDRLMLAYLTKNAQLPLRPGANVGSANPSGIPFKKGNPKFAKAIADAMTQ
LEADGTFTKISDKWFGIDVTKPVK*
>GCA_000452945.1_BPMSHR5858_genomic_02694_1 nudC_1
MKFCSVCGHEVIARIPPGDNRERFVCDQCGTIHYQNPRNVVGTIPVWGDQVLLCRRAIEP
RYGYWTLPAGFMEMGETTAEAAARETLEEAGARVEVQNLFTLLNVPHVHQVHLFYLARLV
DPSFEAGEESLEVRLFDEADIPWDEIAFPTVSQTLRFFFADREAGDYGVHTGDIFRSLRN
G*
>GCA_000452945.1_BPMSHR5858_genomic_02701_1 group_2706
MTTTHSRAHFAARRAAGRAARAVAPWLVPLVLVLAWEFAARAGALSSRVLPEPLAVVRAA
WALVESGDMWANVKVSTWRALVGFAIGGGVGLALGLATGLSKAAEVALDSTIQMIRNIPA
LAMIPLVILWFGIDEKAKLFLVALGVFFPVYINTYHGIRSVDAGLVEMARSYGVRGFALY
RDVILPGALPSILVGVRFALGLMWVMLIVAETISAQSGIGYMTMNAREFLQTDVVVVGIL
LYAVLGKLADVLAKWLERATLRWLPAHQPGAHA*
>GCA_000452945.1_BPMSHR5858_genomic_02719_1 rplK
MAKKIVGFIKLQIPAGKANPSPPVGPALGQRGLNIMEFCKAFNAQTQGMEPGLPVPVVIT
AYADKSFTFVMKTPPATVLIKKAAKVDKGSSKPHTDKVGKITRAQAEEIAKTKMPDLTAA
DLDAAVRTIAGSARSMGITVEGV*
>GCA_000452945.1_BPMSHR5858_genomic_02732_1 group_4772
MSGEKTPTRRVLAVLLVFRWLPVRWRTPRLCLWLLSHGPLPIDPCLPPLAWAQRCVQRGD
AVIRRRGRRATEPGDLQARSVYGSAVALGYYDLADVASPRTLQPVADSTWTREQLERLRQ
IGVGHGAALREYAGDYFYD*
>GCA_000452945.1_BPMSHR5858_genomic_02734_1 group_4773
LPTLDILTRQFPGHRASLKRTILATALACFNDHGLEATTIEMIRERCDTSVGNIYHHFGN
KDGLIAALFLCALEDQAQLLADYLARARTAREGVAALVHSYVDWVSAQPGFARFQFMART
AVASGPRAQELAERNRARNRRVLAWFAHAPQRDEMVQWPAELLPSLIVGQSENYCRAWLS
GRVKAPPVKYRDELATAAWRSVSK*
>GCA_000452945.1_BPMSHR5858_genomic_02742_1 tyrB_1
MFEHIDAYPGDPILTLNENFQQDPRTRKVNLSIGIYFDDDGRIPVMGAVREAEAIVQRES
GPKPYLPMTGLAAYRDAVQAVVFGEGSDARAAGRIATVQTLGGSGALKVGADFIKRYFPH
AQVWISDPSWENHRFIFERAGFTVNTYPYYDEATGGLKFDAMLAAIDALPHKSVVLLHAC
CHNPTGVDLDDAQWLKLIDVLQARELLPFVDMAYQGFGAGLAADAFAIRELARRGMPAFV
ANSFSKNFSLYGERVGALSVICEDAAAAARVLGQLAGAVRSNYSNPPTCGAKIVAQVLTT
PALRAQWEEELAAMCRRIAKMRSAIHDGLRAHVPAEALTRYVKQRGMFTYTGLTEAQVEC
LREQHGVYVLRSGRMCVAGLNEANVGIVADAIGAVIASGV*
>GCA_000452945.1_BPMSHR5858_genomic_02838_1 group_4779
MNKFGKHVVVAAALAALAAGAQAAGVVGNPKDGASKAAMCIGCHGIDGYRVAYPEVYRVP
LLGGQNQVYLENALKAYRKKDRHFPSMNAIAESLTDQDIADLASYYAAQKPDSKNNPYK*
>GCA_000452945.1_BPMSHR5858_genomic_02841_1 ydcO
MSSQPPASNLPRVAARANFFTDTSPSALVAGFVAMMTGYTSSLVLMFQAGRAAHLSDAQI
SSWIWALSIGMAVTTIGLSLRYRAPIVVAWSTPGAALLVASLPGVAYSDAIGAFVVCALL
LAAVGASGLFDTLMRRIPSGIAAALLAGILFEIGIEIFRAAQFQTALVLAMFFTYLIVKR
AAPRYAIVATLAAGVAAAGALGLLDFGRFHVALARPVFTAPSFSIPAIVSIGIPLFVVAM
ASQNVPGIAVLRADGYATPSSPLIATTGIASVVLAPFGSHGINLAAITAAICTGPEAHDD
RAKRYTAAVWCGTFYLVAGVFGATIAALFGALPKALVVSVAALALFGSIMSGLTNAIQDA
RQREAALVTFMVTASGLTLLSIGSAFWGLVAGVLTQAILNARRIA*
>GCA_000452945.1_BPMSHR5858_genomic_02843_1 group_4781
MQVQPYLFFGGRCDEALKFYGNALGARVNFLARYRDAPPNPERPTPPEMADKVMHANFQI
GDSVLMCSDGDCTSGRQQAHDGYSLSLDPKTVEDGKRMFDALLADGGAVTMPFEKTFWAL
GFGMLRDKFGVHWMINVEDPDMKK*
>GCA_000452945.1_BPMSHR5858_genomic_02846_1 speE_2
MTRLIKRASAEARAFRRAKPSPFNGSEKLKPPRAKRARAQFGDDFDLGGVAVLDEPRKPR
FAPVTFSEERGVRYLHFGTEWVQGAMRLSKPYHIELEYAQQMMAWLLFLETPKRVVQLGL
GTGALTKFAHRFLPRAHVEAVELNPAVIVAARSMFALPPDDARLSIHEADAWDFVNDPKN
RGAIGALQIDLYDATARGPVLDSVAFYRAVRACLADAGIATINLFGDHPSFVRNMKHLNA
AFDQRVIALPEVHDGNRIALAFSGPPLSISFAQLTERAKLIEAKLRLPARSWVKGLKASF
GERAGVFTI*
>GCA_000452945.1_BPMSHR5858_genomic_02849_1 htpG
MTQQTMSFQAEVKQLLHLMIHSLYSNKEIFLRELVSNASDAADKLRFEALENNALYESDP
NLRIRLSFDKAARTITIDDNGIGMSRDEAIANLGTIARSGTKEFFSKLSGDQQKDAALIG
QFGVGFYSGFIVADRITVETRRAGLPASEGVRWESAGEGDFQVDTIERAARGTTITLHLR
EGEDELLSSYRLKSIVQKYSDHVALPILMKKEEWDQEKGEMVEKDEDETINQASALWTRA
KSEVTDEQYKQFYQHVAHDHQDPLAWTHNRVEGRSEYTQLLFVPSHAPFDLWNRDYRGGL
KLYVKRVFIMDDAEQLLPQYLRFIKGVVDSSDLPLNVSREILQESRDVKAIREGVTKRAL
SMLEELANAEDDAGKEKYKTFWSAFGQVLKEGVGEDHANRERVAKLLRFASTHGDTDAQD
VALADYVARMKPEQTKLYYVTADTWQAAKNSPHLEVFRKKGVEVLLLTDRVDEWMLSFLH
EFDGKPLASVARGDLDLGALNDDEKKAQEETGEAMKPVVDKMKETLGEKVKDVRVTFRLT
DSPSCLVADDNDMSGYLQRMLKAAGQSAPSFQPILEINPEHPLVKALKADGADFGDWCHL
LFDQALLAEGGALEDPASFVKRTNALLLSRAA*
>GCA_000452945.1_BPMSHR5858_genomic_02854_1 group_4784
MANVYDKLKELGIELPVAGAPAAAYVMSAQSGNTVYLSGHIAKKDGKVWAGKLGLDVTTE
QGKLAARAVAIDLLATLHAHTGDLNKVKRIVKVMSLVNSTLEFTEQHIVTNGASELIAEV
FGDAGKHARSAFGVAQIPLGACVEIELIAELA*
>GCA_000452945.1_BPMSHR5858_genomic_02858_1 aphA_1
MLTYFHPDQSLHHPRTYFSRGRMRMPQEVPERAARLVAAAFAMGFPVREPDDFGIAPIAA
VHDTHYLRFLETVHREWKAMPEDWGDEAMSNIFVREPNALRGVLAQAARHLADGSCPVGE
HTWRAAYWSAQSALAAAAAVRDGAPAAYALCRPPGHHARVDAAGGFCYLNNAAIAAQALR
ARHARVAVLDTDMHHGQGIQEIFYARRDVLYVSIHGDPTNFYPAVAGFDDERGAGEGLGY
NVNLPMPHGSSEAAFFERVDDALRELRRFAPDALVLSLGFDVYRDDPQSQVAVTTDGFGR
LGHLIGALRLPTVIVQEGGYHIESLEANARSFFGGFGALRD*
>GCA_000452945.1_BPMSHR5858_genomic_02863_1 yahB
MRHSPEALLAFAEAANLGSFSAAARKLGKRQSTVSEAIANLEIDLGVTLFDRSTRQPTLT
DAARALLPEVQRVLEASEAIDQVAARLAGGEEARLTLVVSDTYQSSRYEETLAALDRRFP
TLEFECQIAEHDDVLDLIQQGRAQLGLMTARSTYPPDIGAATIAERSEIGLYVGPQHALA
AYGDAEVPLAALHDTRELRLNTYVAPRGERVETGLVAGRRCWLAPSYLQLLEMAVAGFGW
AELPRWMVEHFARERLVELNARGWPRRVPVDAVWSRGQPLGPAGSWLLETMLAA*
>GCA_000452945.1_BPMSHR5858_genomic_02867_1 uup
MSLYSITGAQLAFGHVALLDHADFSLEAGERVGLIGRNGAGKSSLLKIVADLAKPDDGLV
TRQQHLVTVYVPQEPEFAAGQTVFDAVASGLTDTRALLAEYDEIAHRLADTPEGAEHDAL
LARMNALQSSLDATDAWNWRTRVATTLAQIGLDGDTKIDALSGGMQKRAALARALVVQPD
VLLLDEPTNHLDFDGIRWLEELLVAQRAGLFFITHDRAFLDRVATRIVELDRGRLLSYPG
NFSAYQTRKAQQLEVERVENEKFDKLLAQEEVWIRKGVEARRTRSVGRVARLEQMRRERA
ERRNAQGNVRLDVAQGEKSGKIVAELTDVTKRYGERTVVERFSATVMRGDKIGFVGPNGA
GKTTLLKLILGELKPDAGTVRTGTNLQVAYFDQMRAQLDQEKSLADTISPGSEWVEIGGV
RKHVMSYLGDFLFAPERARSPVKSLSGGERNRLLLARLFARPANVLVLDEPTNDLDIPTL
ELLEELLTDYDGTVLLVSHDRAFLDNVVTSVIASEGNGLWREYVGGFTDWQIQRERAQRI
AHEEAAKRAAKEPASAKDDAPKGAAGRNAQRTVKLSFNEQRELDALPARIAALEAEQKAI
AAQLEDGSIFSKDPQEGTRLTERFAALDDELLAALERWEALEAKRKPS*
>GCA_000452945.1_BPMSHR5858_genomic_02870_1 group_1857
VAQVESPRQATAGSAEQAAGKLGGLLSLAFLLGLMTVMAAFGWIALREGTHRFLLPFVNG
NATRQIADAIASVRAHPSLEGIRQVSEEIWMMSLPTSVTRFSHSRLMEQGIYYTTMPRVN
QVLIAIHVLFSAFCVTFGSLQFWPSFRKRFMRAHRLIGAVYVATVPISTVSALAYLALTP
PHHLYAHLIGWIALWIFGVLTLIAIAMAVRALKARRIFEHQAWMALSFGCLLVAPLLRID
WVLLAPLFPHIDQETLNLVTMGVMLPQAQLITYALIVVNRQYARPMKQRTPAPLASRAGA
WFLRSQPGLLASTAVWGAVNVWAYGLGHGTAGLDAAARMLPADLLTREQEALHAYPGIAW
LMALSLTAAFPAAVLSLGARLRAASASVAARLDATAACLGLAAGAASVFLGWHIGIAPDN
HLFSGGTMYTVNGLVIAGFSLMLAATARRRQHAIAKESLVFLLCMLPFPALYFATLEAVG
RIRLPAAYLAAGQGFVIPVGFSSSLLFLAAFHVIFGQATREHN*
>GCA_000452945.1_BPMSHR5858_genomic_02894_1 group_4788
MEILPGRKRIGPATGGRRADQREVACAGWPLLRRNVELCRDPALP*
>GCA_000452945.1_BPMSHR5858_genomic_02946_1 group_18948
MLNELETLSQNIGRLIALNQRHQTERLALEEQVAQLRNEAQTLHAELEQLRDERNALAAE
RDTLSAKIDDAQVKLNAILEKLPRTKNVPDAENQLDLLAPQANDEGEGATGHGEHA*
>GCA_000452945.1_BPMSHR5858_genomic_02948_1 group_19892
MRYWLMKSEPDEASIDDLAAAPDQTLPWTGVRNYQARNFMRDTMQIGDGVLFYHSSCPEP
GIAGLARVSSTPYPDPTQFDSHSPYHDPKSTREAPRWVLVDVRFVRKSPLVPLAALREHE
ALANMRVLAKGNRLSITPVTPNEWRFITQRLMK*
>GCA_000452945.1_BPMSHR5858_genomic_02949_1 group_19530
MTKKSALAVAVTLAAALPIALALAPSAARAQSMGQMQPPAGVLSLSAQASTDVPQDVVDI
TLFYEQQAKDPGTLTAELNKRADTALAQARGVTGVTARTGEFSVSPSVDRDGKISAWRGR
TEVVLESHDFAAASKLAGQLSPMMQVGNVSFSLSPEAQRAAEQKLTSEAIKAFRARAEEA
TRAFGYSNYSIREVNVGSGRNVQPYPRMFAMAAPAMDSAKMSAPIAVEGGKTTVTVNVNG
SVQMK*
>GCA_000452945.1_BPMSHR5858_genomic_02954_1 group_614
VVEANASSIPVLTDVLVPGNPALARPPAAGPAQAGEPAAAAIRAGEGHDADRRDEATATE
PAAAVAAAAAAAAAAPAARDAALAPADEMRAPAPAEPSFALPAPDPLAVELPTAYPADAT
YADELPAAHPAPDLTPPAALPSSVPELAQAAPSFAADLEARQIAERLKSRMTRYLTGEGR
GLIEARCRDALHDHAGWLVGQIAREVALALETEVAGWVTEEVGAALARRNAPARSDTTQ*
>GCA_000452945.1_BPMSHR5858_genomic_02955_1 group_19301
MTRIDFHTNVGDSLAYACRLLRKAYQAGQPVVVCAEPARLRALDERLWTFSPLDFIPHCA
IDNAHAASTPIVLTADLERAPHHGILLNLGAQVPAQFARFERLLEVVGNAPDELAAGRDR
YRFYRDRGYALNNHKQGG*
>GCA_000452945.1_BPMSHR5858_genomic_02960_1 sumT_1
MGKVYLIGAGPGAADLITVRGARLLEQADVVLHDALVEPAMLDYAPRARRIAVGKRCGQR
SSAQHFINKQLVDAAREHAVVVRLKGGDPMLFGRADEEMRALEAAGIDYEVVPGITAALA
SAAALKRSLTLRGVARSVAFATKSRAPGSDEIREQVSADSLVYYMGRDNAPGIARQLIDA
GRAGATPVAIVEACSTPRERMLTLTLADMAAGAAQAWLDPAQPSLLMIGDAFAERAPRLA
DEPPGDAGGKRGMRDAA*
>GCA_000452945.1_BPMSHR5858_genomic_02963_1 group_19505
MTTNQRESTVSLRRACASLAALALLAACTSTPEAIQRKTGWMRSEVADSYIYAYPLVLMD
VAKEAATADGAPVNTLRHAQALPAPGAANPPLASVDMLDSTAWLDVADEPVIVALPDGRG
RYLDARALDMWTNVLWSTGPSANPRSGASRARTLAFVGAGWEGTLPEGVTRVDVPARNVW
LDVRIQTSGGRDLAAARKLQRAIRVEPLSVYTGDARGARADAGAASAEPASGAAASPVAQ
VAALDPPAFFSRVARALQDNPPPAADAHAQEILADIGVTAGAPVQWKGDKLIGAENGAAE
ARERLTALPPNALSANGWSWLGDSVGNYGQDYALRAYAASTQFGAGTRDDETVAVVKTDS
AGRPLNGAHRYVIRFAPNALPPARAFWSLAPYTPDGAVPELGRARRSIGERDRLRRNRDG
SLEIVVSAAPPGKGYASNWLPAPRADFELALRLYAPKRQAADGTWQPPAVVRK*
>GCA_000452945.1_BPMSHR5858_genomic_02966_1 group_19593
MKRNSKQFEIGAWLGACALAFAGAASAAAVQDRDHDSRPVDAKRVLLVSIDGLHEQDLAR
CIGANTCPNLALLAKSGVTYTNARTPGLSDSFPGLAALVTGGSPKSAGLFYDVSYDRTLY
APSDATCSGKQGWNVVFDETTGIDAMNGGALTHLDGGGAFNPQAIPHARVNGQCVSVYPH
DYVKTNTVFEVVKEHLRGSHTAWADKHAWGYDWVNGPSGKGVDDLARTEINSIDPATGTP
YTDIYTHTEKFDDYHVQAIVNQIDGKNSTGTAAAPVPTLFGTNFQTLSVAQKATVASGGG
YLDASFTPGPEVANAIAYVDGALGRIVAELRQRGLYDSTVVIVTAKHGQSPTDHTKLVKH
GDTLTALLEANGFVDPNGNFGQNNTASGNPNDGTGLVGTGFVQTDDVGLVWLRDPRQLSA
AVATLKANLGCNAPGICADGPQAYILYGPSVAERFGNPALGRTPDIVVQPNPGVIYTSSK
KKDEEHGGNAPDDSHLGLLVSYAGLRQGRTIDAPVLTTQVAPTILRSLGLEPRLLHAVAL
EGTRVLPGLGLER*
>GCA_000452945.1_BPMSHR5858_genomic_02968_1 group_19769
MAMKKTDLEKNKALKLTHAMKQTTSARFGKGAEDAAADRRERRRLEQAQGLVPFACKLNI
ELVARLNERAAAHPDGMTGLLTELLQRGLDETAK*
>GCA_000452945.1_BPMSHR5858_genomic_02974_1 cydB_2
MDVTVIWAAIIALGLFMYVVLDGFDLGIGIVFPFFPAEQERDLMMNTVAPVWDGNETWLV
LGGAGLFAVFPIVYSTVLSALYLPLVFMLVCLIFRGVSFEIRAKARRTKHLWDLAFIGGS
TGATFFQGVALGAFLQGIPVDGGRFAGDAFGWLTPFSLLTGLGLIATYALLGCCWLVAKT
EGDLQRRLHRVVWPLTVVLLGFIAVVSLWTPLQDTNIAQRWFATGLFWRLLPVPFLVAGC
AALMRRAVRERHDTTPFALALALVFLGYVGLLVSLWPYAIPQSVTLWEAAAPRSSQTFTL
IGAAVIIPVILAYTTMGYWVFRGKVRHEDQHYYHH*
>GCA_000452945.1_BPMSHR5858_genomic_02978_1 group_2744

MNAAPAPRYVELDFFRGLVLLVIVVDHIGGSMLSRVTLHTYALCDAAEVFVFLGGFATAI
AYNSLATRHTEAAARQRFIKRAFEIYRAFLLTAGLMLFITAALNAFAIDAPNMPTNDLDG
LMHAPLAALRDILLLRRQPYLASVLPMYAFFALLVPLALPVARGRWWWLLVAVSAAIWLA
ARRIAGYLPTVDGVPWDFNPFAWQFLFVLGIVARCQPIYPVLAKRPVGWFALAAAIAVVA
AGAYYRLRIEPFPTDPSIKQNLGALRLANFIAIAWLAAKLVHLGWMHRIARAMPWIGTIG
RQGLLCFVAGTGISLLVDSLLYAATDGYLDLRLGLAADAAAIGLLYVVAKLYAPLVARAS
DFVRQSPLLQPLRPFRLPLRLPLRRPKR*
>GCA_000452945.1_BPMSHR5858_genomic_02982_1 nlpD_2
MSKSEIDRSVAWLSGVAAAFVVAGCATTSPVTPTDTLAAAASSASAAQPAAAPPGDAAQH
PEHAAAPAPIATRYVVRRGDTLSAIAQANGCTVRELQAWNRMGRRTRIGIGQVLRIAPPG
AENAVASEAADGAGATRAAGNAAGAARAADPASGAGGAPVADAPQPASAPESAADRVADH
RVVQETKRHAQSIALAWPAKGAVVETFQPGRNRGIRIVGRAGDPVRAAASGRVMYAGTGL
NGYGTLILVQHNADFLTAYAHNRKVLVKTGDVVQQGEQIAEMGTGDSTRAGMLFEVRRDG
KPVNPMPYLASRQQG*
>GCA_000452945.1_BPMSHR5858_genomic_02989_1 dppB_1
MSLPAPPLETPRTLPARRPGVRWALRALRWTLTLAVTFAGLLALTFVIGRKVPIDPVLAV
LGDRASAAAYAAERIALGLDKPLATQFLIYARDVLHGNLGMSLLTSNPVLDDIRRVFPAT
LELATLATLIGVALGVPLGVAAAVKHNRPIDHVARFVGLIGNSVPVFWLGLMGLLLFYAR
LHWVAGPGRLDPVYDGMVDPRTGSLLLDAALAGEWDVLRNALSHIALPAAILGYYSVAYL
SRMTRSFMLDQLNQEYIVTARAKGLAERRVIWRHAFGNIAVPLLTVIALTYSNLLEGSVL
TEIVFAWPGIGSYLTGALLNADMNAVLGCTLVIGVMFITINLLTDALYRVFDPRAR*
>GCA_000452945.1_BPMSHR5858_genomic_03017_1 yphA_1
MKQTFLAPQKDLLLLLSRILLVILFVIFGWEKLLNFSGTAQFMAAEGTPLPSIAAVVAVA
MEFFVGIALLLGFCTRPLALLLGLYTLGTAFIGHHYWSMPAAEQMNMMIHFYKNIAIAGG
LLALCAAGPGRYSLDRG*
>GCA_000452945.1_BPMSHR5858_genomic_03020_1 hisQ_1
MLFQGFGPLLWAGTIETVKLAVLSLAASLVLGLAGAAAKLSSNRALASVGTFYTTLIRAV
PDLVLMLLLFYGIQILLNDVTDMLGAEQIDIDPFVAGIVTLGFIYGAYFTETFRGAFLAV
PRGQLEAGFAYGMGAWRVFARIMFAQMMRFALPGIGNNWQVLVKATALVSIIGLADVVKA
AQDAGKSTLNFFFFTLAAGAIYLAITTLSNLVLMYLEKRYSAGVRRVAL*
>GCA_000452945.1_BPMSHR5858_genomic_03051_1 clcB_3
MSIAAPHKRDFASNARLPRIALLALAIGVLGTFAAFALLSLIHLFTNLFFFQRFSFAEHS
PALNTLGPWAAAVPVAGGIVVGLIARFGSEKIRGHGIPEAIEAILFGKSRMSPKVAVLKP
LASGIVIGSGGPFGAEGPIIMTGGAIGSLIAQFVKVTAAERKTLLVAGATAGMTAVFGTP
VAAVLLAVELLLFEWRPRSFLPVALACAVAGFARAAFFGVGPLFPVETAAPTAAALGSCA
LAGLLSGALACGLSVSLYKIEDRFGKLPVHWMWWPAIGGLAVGIGGLIEPRALGVGYDVI
GDLLHGHIALQIALAILVVKAVIWVIALGSGTSGGVLAPLLMLGAGLGTLLGPVLPGGEP
ALWPLVCMAATLGATLGAPLTAIVFAFGLTHDANALLPLLTATLVAHGFATVAMKRSIMT
EKIARRGYHIYREYGVDPLERHYVDEVMTRSAVTIDADLSVEVVRARYFGATQAHRAYPV
VRDGVLIGMLDRATLDGSPAAHDGDGQADDARAAKMRAADVQMTDMRVADMRVADVLPRR
APLFSLADETCRLVATRLAVHQLERLPVVADPDTMRVVGIVSRSDLVKPALRHFDDEHKR
ERFRRAHPAAFVKRRFAPARKTG*
>GCA_000452945.1_BPMSHR5858_genomic_03057_1 group_2453
MIVDRLISEILFGFTGLMGIINPFGIAFLFLERTESLTEAQRAALARKVAINAFIVLLAA
FFVGTPILHFFGISLEALRIGGGLAVAVAGWQMLNEPDVPTGGDTAAQPVEPRNAMARAF
FPLTIPLTTGPGSIATAIALNANRTHKLSEFVLSSIVSVAVSALAALVIWQTYSRAAFLA
RYLGTEGTKVAMRVSAFLLLCVGVQIMLTGFSEFLRPIADQVK*
>GCA_000452945.1_BPMSHR5858_genomic_03107_1 efeU_1
MLSTAVIVFREVLEAALVVSIVLAATKGVPGRAWWVSAGLLGGVVGAAFIAAFADVISAW
ASGMGQEVFNAGVMFVATIMLAWHSIWMGKHGREMAQQLSQVGRAVAAGSRPLTGLAIVV
GVAVLREGSEAVLFLYGIAAGDPGQAPQMIAGGALGVLGGVGLGAGMYAGLLQIPLQRLF
SVTNALIVLLAAGMASQGTGFLVSAGWLPSWGDTVWDTSWLLKESSVVGKMLHTLVGYTA
RPAGIQIVAYVATLLVIVLLARRVARKQAIVARPTRAA*
>GCA_000452945.1_BPMSHR5858_genomic_03108_1 group_4818
MKTLQTISSLALALAFAVSAPHAFAADDVYSLTLKNHRFSPADLTIPAGKKVKVTIKNLD
ATPAEFESDDFKAEKVVPAGKQVDVYIGPLKAGTYEFHDEYHEAESKAKLTVK*
>GCA_000452945.1_BPMSHR5858_genomic_03109_1 group_19782
MAPAGGSAAGAMPADPGGASLGTCPSAPVESASAVLGAVMESGTVAYISPKIPISRALLD
GLRANGVPLENRVRFLGPCLGGQCAQWAGHRCGLIDAIVKEPAVLAPPEAGLPKCGIRST
CRWYAQHASAACMQCPVVIYEPHAG*
>GCA_000452945.1_BPMSHR5858_genomic_03112_1 group_4819
MAQLPEDKPVSRARRAIAAGLALLPVVAFARKTATKPVVQVWKTPTCGCCEDWLSHLKNN
GFDVVVHNVEDTSDVRQQAGMPERFASCHTGIVQGYALEGHVPAREIKRLLREKPNAVGL
AVPAMPIGAPGMDGPVYNGRHMPYDVLLIKRGGQSSVYQSYS*
>GCA_000452945.1_BPMSHR5858_genomic_03118_1 group_1869
MNSSLRHCPLMDCNALPRGLDGATHRQMLDQHALDDGEHTGAVPGA*
>GCA_000452945.1_BPMSHR5858_genomic_03121_1 glmS_3
MCGIVGAVAQRNIVPILIEGLRRLEYRGYDSCGVATVVDGEARRERSVSRVADLEAHVRT
AGLAGTTGIAHTRWATHGAPATCNAHPIFSRDHIALVHNGIIENHEALRTQLAGEHYEFD
GQTDTEVVAHLIHSKYRGDLLAAVRDATAQLRGAYAIAVFSKTEPNRLIGARVGSPLVVG
LKDGECFLASDALALAGITDQFIFLDEGDIVELTPRGARIVDRDGRPVVRAVQTVSAAQG
AAELGEYRHFMQKEIFEQPRAVAATIPDAGLFDPAVFGHDARRAFEQIDNVLILACGTSH
YAGLTARRWLETVARIPAQVEIASEYRYSDAVAMPNTLIVVISQSGETADTLAALKYAQA
LGHIDTLAICNVPTSAMMRQAGLRFLTRAGPEIGVASTKAFTTQLVALFVLAVTLGRMRG
YVDDAQLARCTQQLRRLPGALEDVLALEPRVKRWADEFACHENALFLGRGLHYPIALEGA
LKLKEISYIHAEAYPAGELKHGPLALVTNTMPVATIAPNDALLEKLKSNMQEVRARGGQL
YVFADADTRIDDGDGISVMRMPDYYGLLSPILHVVPLQLLAYHAACNRGTDIDKPRNLAK
SVTVE*
>GCA_000452945.1_BPMSHR5858_genomic_03125_1 group_477
MHAFRCRRGPPVPLAIVALACACVFGAPRARAAAAHADAPVAASIASFEPAGGPAAAADA
ADTAVMPAPPASFAPPAAPPVPRVATLASSASTVPTTPTTPTATTATTVPPAPIASAPAA
TPLSAPPLAPPAAAPLVIVPRRFDDTALVKVDPEGMPDGWPLAEVAPNQENQPALPEALV
AHVQAPAQRHTELSLAYSMAHLTNGYGDWYGVHLRGLYQAGGRTIIGELAQLHRFNESTQ
LGAITYIQDMGPDWFGGIGFSGTTSGTILPSARVDLSLNRKLLSNRSLVLSAGAGYAWNR
NGHRDQLYHLGAIWYAVPKWIFEAGWNYTVDSPGSVKAPAYYGAITYGEVGKSLIVLRGS
FGREAYQATGNNAQITDFRSHEVSLKWRYWVTRKWGTQVEFDYYHNPYYSRIGGEVGMFY
RW*
>GCA_000452945.1_BPMSHR5858_genomic_03128_1 sacA
MKKILLAFGTRPEAIKMAPLVRMLQTRADVDARVCVTAQHRQMLDQVLALFDIAPAYDLN
VMRQSQTLADVTTGILQTIGAVFDDFDPDIVLVHGDTTTTLAVSLAAFYRYLPIGHVEAG
LRSGDIWSPWPEELNRRVTDAVSSWHFAPTERAQHNLFSEGVPTQGVVLTGNTVIDALQD
VKRMLDADAPLAREIAAQFPFLGHDERVVLITGHRRESFGEPFAHFCDALRTLARRYPGV
RFVYPLHLNPNVQGPAHALLDGLPNVHLIAPQEYLSFVFLMSRAHFIITDSGGIQEEGPA
LGKPVLVTRDTTERPEAIQAGTARLVGTDTERIVGEASRLLDDDDAYDEMSRATNPYGDG
HASERIAHALLNMPHAADTARFPLAAIEAPCNALALGLRALRSA*
>GCA_000452945.1_BPMSHR5858_genomic_03138_1 ggt_1
MTDRIRPRPFRSRAGAIAAAFFAVAAAGCANAPGPGAQNAAAPASAASTPPAAPGAEAPA
APAELSSWVDKPGWTAQRDMIAAAHPLAAQAGQAMLKAGGTAVDAAIAAQMVLMLVEPQA
SGIGGGAFLLHANGKTIEAYDGRETAPAAATDRLFLDANGKPLPRAALGARTVGAPGALR
MLELAHRAHGKLPWRRLFQPAIRLAEQGFPLGPRLAAALANEPSLARDPAARAYFYDRNG
APKPAGTTLKNPQLAATLRLVAAHGANAFYTGAIARDIVATVRQAPNPGVLSMQDLAQYR
AKTREPLCADYRKWTVCGMPPPSAGGLVVAQILGLLEAQPDWRQIGAQKPVRNAVGVEPT
PFAAHLFSEAGRLAYADRAQYVADPDFVAPPGGSWKRLVERRYVAERARLIGDASAGEAS
AGSPSGLPPATAADRGAEPPSGSQITVVDRYGNAVSMASTLDERFGSRLMVRGFLLNSQL
LDFSPASAEHGKPVANRVQPGKRARSTLAPELVFEKGSSRLMMALGSAGGASTANDVAKT
LVGMIDWGMTMQQAIALPNFGSRNGPTELEQGRVSDALAGALKARGHDVRVVELGSSLQG
IQRINVEGQSVWFGGSDPRRDGIVAGD*
>GCA_000452945.1_BPMSHR5858_genomic_03146_1 dacC
MRLSSLGLTSLASSTAIAVAARNVALGIVLPAALVSTIAVAQAKPAAKAKHAAAPAAAAA
PTGAPATYVPGAVPPPGVNARSWVLVDASSNQVLASGNADERVEPASLTKLMTAYLVFEA
LDAKKITMEQIVTPSEAVRRVGRDESRMFIEANKPVSVHDLVYGMIIQSGNDAAIALAEL
VGGSEAQFVNMMNAEAQRLGMKNTHFADVNGMPDPNHYTTAGDLAKLSTHLIRDYPDYYN
IFSVKEFTYNNIKQPNRNRLLWLDPTVDGLKTGHTQAAGYCLIASAKRALPGAADATRRL
VTVMMGETKESDRVQDSMKMLNYGYTAFDSVRLYKASQPIDTPRVYKGKSNNVQVGVKKD
QFITVPRGLADKVKPEVALNAPLIAPLADGQVVGSVKLVADGKTVAEFPVVALQPVPEAG
LLGRIWDSILLMFSKKK*
>GCA_000452945.1_BPMSHR5858_genomic_03150_1 group_1200
MHAHFMSGSLSRRGRPALALAVALVMASASGCAGTPAALANIRYDLGPAQPAASSGTGPA
LKVLDVSAPDALNTDRFVYRLAYSDAQRIAAYRDSKWTAPPAQLLTQRLRGALSGRGAVL
AADDGVRAPVLKVELSEFEQVFDGRSESHAAVTARVTLTQEGKVLGQRTFVSRAPASTPD
AAGGAQALATASDALVSQLVAWLGVQAYAAVQ*
>GCA_000452945.1_BPMSHR5858_genomic_03159_1 group_4821
MPASFERKLITRDALAAMRASLPAPVVFTNGVFDILHRGHVSYLADAKALGACLIVGVNS
DASVRMLGKGDDRPINVQEDRMALLAALECVDWVVGFDEKTPVSLIEAVHPDILVKGGDY
DMDALPESALVRGWGGRALAIPFEHDRSTTALLKKVRAQSR*
>GCA_000452945.1_BPMSHR5858_genomic_03160_1 group_4822
MNTMLYPELYKSLEAVRWDMEKDIPWDKFDASLLTDEQAKTIKMNAITEWSALPATEMFL
RDNHHDSDFSAFMSVWFFEEQKHSLVLMEYLRRFKPELMPTEEELHAVRFEFDPAPPLET
LMLHFCGEIRLNHWYRRAAEWHTEPVIKHIYETISRDEARHGGAYLRYMKKALNNCGDVA
RAAFSKIGVLMASARRTEKPLHPTNLHVNQALFPRDTVQSRLPDPEWLERWLDEQIRFDG
EWEKKVVERILHNLSVLFERTFATAQELNRYRKEVTARLQSESGPAAAQPA*
>GCA_000452945.1_BPMSHR5858_genomic_03164_1 group_4823
MKLASLKDGTRDGQLIVVSRDLHTAAIADAIAPTLQRVLDDWAFYAPQLRDLYDALNHGR
ARNAFAFEPADCMAPLPRAFQWADGSAYVNHVELVRRARGAEMPPEFWTDPLMYQGGSDD
FLGPRDDIVCASEAWGIDFEAEVAVITADVPMGAAPDEALKAVRLVTLVNDVSLRNLIPA
ELAKGFGFFQSKPASAFAPVAVTPDELGEHWREGRLHRPMLVHWNGKKVGQPDAGVDMVF
HFGQLIAHAAKTRNVRAGSIVGSGTVSNKDAKRGYCCIAEKRCLETIEHGAPQTEFMRYG
DRVKIEMVDEAGKSIFGAIEQAVAPVDAAA*
>GCA_000452945.1_BPMSHR5858_genomic_03165_1 kdgR_3
MEGIYFTMVNPHGGPRREPPPPHSSRHAVNKSASPPSDAPDDDLADDHDTGEEKVRSGIQ
SIEVGFRLLDVLTGEPRAMMLRDLAQRAAMSPAKAHRYLVSFQRLGLVSQDPVSGRYELG
GFALQMGLARLARVDGVKLARIALTALRDQLDQTVGIAVWGNQGPTIVHWMESSHPAKAS
LKLGDVMPLLGSATGLVFAAYLPRGKTAAMIERELADTRRAAHYTGPRTRAEVDAVLADV
RAHRAARVEGMLLPTINAFGMPVFDAVGELALAIIALGHEGAFDIRWGGEVDTALRTCAQ
KLSYELGYSEGARDA*
>GCA_000452945.1_BPMSHR5858_genomic_03169_1 metH_2
MTDHTMRLAGLEPFNVTSGTLFINVGERTNVTGSKAFARMILNGQFDEALAVARQQVENG
AQVIDVNMDEAMLDSKAAMVRFLNLIASEPDIARVPIMIDSSKWDVIEAGLKCVQGKAIV
NSISLKEGKEAFVHHAKLIRRYGAAAVVMAFDEAGQADTLARKTEICKRSYDVLVNEVGF
APEDIVFDPNIFAVATGIDEHNNYAVDFIEATRWIKRNLPYAKVSGGVSNVSFSFRGNDP
VREAIHTVFLYHAIQAGMDMGIVNAGQLGVYAELDAELRERVEDVILNRRDDSTDRLLEI
ADKFKTGAAKKEENLEWRSQPVEKRLAHALVHGITNFIVEDTEEARAKIAAAGGRPINVI
EGPLMDGMNIVGDLFGQGKMFLPQVVKSARVMKQAVAHLIPFIEEEKRLLAQAGGDVRAK
GKIVIATVKGDVHDIGKNIVSVVLQCNNFEVVNMGVMVPCNEILAKAKVEGADIVGLSGL
ITPSLEEMAYVASEMQRDDYFRVKKIPLLIGGATTSRVHTAVKIAPHYEGPVVYVPDASR
SVSVASNLLSDEGATKYLDELKADYERIRAQHANKKALPLVTLAEARANKTKIDWARYTS
VKPKFIGRRMFKNYDLNELANYIDWGPFFQTWDLAGPYPAILNDEIVGESARRVFSDAKS
MLARLIQGRWLTANGVIALLPANTVGDDDIEIYTDDTRAEVLLTWHNLRQQSVRPVVDGV
MRPNRSLADFIAPKDSGVADYIGMFAVTAGIGVDAKEKQFLADHDDYSAIMLKALADRLA
EAFAEAMHARVRRELWGYAATEQLDNDALIAEQYAGIRPAPGYPACPDHLVKRAMFEALH
TDEIGMSVTDSLAMLPAASVSGFYLAHPDSTYFSVGKIGQDQLEDFARRTALSKADAERA
LAPLL*
>GCA_000452945.1_BPMSHR5858_genomic_03171_1 argS
MLPAQKHTLETLLENSVKQVVQASKGDADAAFVLPAIALERPKVAAHGDVACNVALQLAK
PLGANPRQLAEQIVAALTAQPEAAGLVDAAEIAGPGFINLRLTPASKQAVIGAVLAQGRA
FGASERDHGKRVLLEFVSANPTGPLHVGHGRQAALGDALANVLASQGYAVHREFYYNDAG
VQIGNLAISTQARARGLKPGDAGWPEAAYNGEYIADIARDYLNGETVAASDGEPVTGKRD
VEDLEAIRKFAVTYLRREQDMDLKAFGVKFDQYYLESSLYTEGRVEKTVDALIAAGMTYE
QEGALWLRTTDEGDDKDRVMRKTDGTYTYFVPDVAYHVTKWERGFTKVINIQGSDHHGTI
ARVRAGLQGLHIGIPKGYPDYVLHKMVTVMRDGQEVKISKRAGSYVTVRDLIEWSGGATP
GSEGSPELLDEATITRGRDAVRFFLISRKADTEFVFDIDLALKQNDENPVYYVQYAHARI
CSVINEWKSRYGATDALLPGADLSPLDSKQAMALMQKLAEYPDVLAHAAGELAPHAVAFY
LRELASEFHSFYNAERVLVDEQAPRTARVALLAATRQVLENGLAMLGVSAPSKM*
>GCA_000452945.1_BPMSHR5858_genomic_03176_1 group_4826
MLVVDKRFQRLPMSDAPMRLPCHCGTLRQATRAITSLYDAHLARHGIRITQFTILAALHG
GDALPTGELATRLLLDQTTLSRTLATLQSNQLVRAESGDDQRVRLWSLTPKGTALFKAAR
VDWEAAQNDVYRRIGKRNMQALDADIFALAKALSN*
>GCA_000452945.1_BPMSHR5858_genomic_03180_1 group_4827
MNALEHQLDYPFADALPEPGGFVEVAPGVRWLRMPLPFALDHINLWLLRDEIDGQAGWTI
VDCGIASEQIKAHWERIFDAHLDGLPVLRVIVTHCHPDHFGLANWLCDGGARARWNVRLW
MTLGEYLFGCVMAAGNGSNAGGPAAADHFARHGLADPAALDKLRNRRGYYADLVPAVPPR
YRRLREGDALAIGAREWRVVTGYGHSPEHCALHCAADGMLISGDMVLPRISTNVSVFDLE
PEANPLALYLESLGRYEAMAEGTLVLPSHGLPFRGVRTRIAQLRAHHDARLAEVRAACRE
RPMSAADIVPIMFKRGGLDIHQMTFALGEAIAHLNLLWLAGELARETGADGVLRFRHAG*
>GCA_000452945.1_BPMSHR5858_genomic_03183_1 thlA
MTAVDQDPIVIASAARTPIAGFQGEFASLAAPQLGAAAIAAALERAGLQPEQIDEAVMGC
VLPAGQGQAPARQAALGAKLPLSVGCTTINKMCGSGMRAAMFAHDMLVAGSVDVIVAGGM
ESMTNAPYLLPKARAGMRMGHGQVLDHMFLDGLEDAYDKGRLMGTFAEECAGEYAFSRDA
QDAFAIESLARAKRANEDGSFAWEIAPVTVAGKKGDAVIARDEQPFKANPEKIPTLKPAF
SKTGTVTAANSSSISDGAAALVMMRASTANRLGLAPLARVVGHSTFAQAPSKFTTAPVGA
IRRLFEKNGWRAAEVDLYEINEAFAVVTMAAMKEHGLPHEKVNVNGGACALGHPIGASGA
RILVTLIGALRARGAKRGVASLCIGGGEATAMGIELI*
>GCA_000452945.1_BPMSHR5858_genomic_03184_1 csgA
MKTVLIVGASRGIGREFVGQYLHDGWRVIATARDAAALAALDALGARALALDVAQPDDIA
AFDARLGGAALDAAVVVSGVYGPRTAGVEPIGVEDFDAVMHTNVLGPMLLMPILLPRVEA
SRGVLAVLSSKMGSIGEATGTTGWLYRASKAAVNDALRIASLQARHAACIALHPGWVRTD
MGGAQAALDPQASVAGMRRVIAQAAGDRERANGRFFQYDGVELGW*
>GCA_000452945.1_BPMSHR5858_genomic_03187_1 bioA
LSTHATDDWVARSLRAVWHPCTQMKHHERLPLIPVARGERAWLFDRAGNRYLDAISSWWV
NLFGHANPRINAALKQQLDTLEHAMLAGCTHESAIELAERLAARTQHTLGHAFFASDGAS
AVEIALKMSFHAWRNRGRGDKREFVCIANGYHGETIGALGVTDVALFKDAYDPLIRAAHV
VASPDARGARDGETAADVAARALGDVRRLFAERGEKIAALIVEPLVQCAAGFAMHDPSYV
RGLRALCDAHDVHLIADEIAVGCGRTGTFFACEQAHVWPDFLCLSKGISGGYLPLSLVLS
RDAIFDAFYDDDVTRGFLHSHSYTGNPLACRAAVATLELFDTDDVLAENARKSALMREAL
APLASHPHVRHLRERGTIFAFDVALEGDAARTFPRRFFEHALARETLLRPIGTTVYLMPP
YILDEGEIEWLASRTRATLDATLSEAHA*
>GCA_000452945.1_BPMSHR5858_genomic_03188_1 bioF
MNPLATLEQGLADIDAQGLRRCRRVADTACGAHMTVDGRAIIGFASNDYLGLAAHPRLVE
AFAEGARRYGSGSGGSHLLGGHSRAHATLEDELAAFSGGFSDAPRALYFSTGYMANLAAL
TALAGRGATIFSDALNHASLIDGARLSRANVQIYPHGDADALDARLRACDAPTKLIVSDT
VFSMDGDVAPLARLVALAETHGAWLVVDDAHGFGVLGPQGRGALAAHGLRSPNLVYVGTL
GKAAGVAGAFVVAHETVIEWLVQRARSYIFTTAAPPSVACAVSASLAVIASDEGDARRAH
LGALIKRTRAILRATHWQPVDSHTAVQPLVIGSNEATLAAMAALDAQGLWVPAIRPPTVP
AGTSRLRISLSAAHSFDDLARLEAALVTPIGAAA*
>GCA_000452945.1_BPMSHR5858_genomic_03189_1 bioD1
MSAPLSLFVTGTDTEIGKTFVSAALLHGFARAGLRAAAMKPVAAGAYERDGAWRNEDADQ
LDAAANVALPAAIRTPFLLKAPAAPHIVAAREGVALDIGTIVDAHRRACEMADVIVVEGV
GGFRVPLADTRDTADLAVALGLPVVLVVGVRLGCISHALLTAEAIAARGLPLAGWVANRI
DPAMPFADDNVDTLRAWLEREHRAPLLGALAHMSPPSPDAASHALDVNLLLNALRAAAPR
*
>GCA_000452945.1_BPMSHR5858_genomic_03190_1 bioB
MTEAQTACATTETPVAAPAAPRWRVADVIALYELPFNDLLFRAQQTHREHFDANAIQLST
LLSIKTGGCEEDCGYCSQSAHHDTGLKAEKLMEVDAVLAAARTAKENGATRFCMGAAWRN
PKDRHIEPIKEMIRGVKDMGLETCVTLGMLEEHQAKALAEAGLDYYNHNLDTSPEFYGQI
ISTRTYQDRLDTLERVRDAGINVCCGGIIGMGESRRERAGLIAQLANMNPYPESVPINNL
VAIEGTPLENAQALDPFEFVRTIAVARITMPKAMVRLSAGREQLDDAMQALCFLAGANSM
FYGDVLLTTGNPRAEADRKLLARLGMWASEASQLSA*
>GCA_000452945.1_BPMSHR5858_genomic_03192_1 group_442

MRTFAMKLDVVRRACAALGGAAVLAAAPALAAPPAPDGGRAPDLDAVLLHESMTVSADGV
TRTVTYRERMVRRDGHVWIERVPPSGAKRAHAGGAHAHEGLDAARTPPSRGGAHPAAAAA
DAHAGHRHFDFDSAARHVTNDGGRIGVEYVDAAQRTVVAVPPAEYETTGFDGSWDNAFYI
TPPSQLKRLAAQSKPGPAPGTRWYEQTVGAPRAQGTNRVLWSDALQAPLVVEYRSADGHA
SRKLTLTPAPRANALPWRQLQSYTRKAYVDYLD*
>GCA_000452945.1_BPMSHR5858_genomic_03196_1 group_4832
MTIAQQERPVAASPYRFEAEMTSGLERVAAHHHDLTLTSVFQPIFSLSHMRAVGYEALLR
AHDALDRPVSPLDVFGEAARHGEATQVDRLAQALHLENFARFGAEREWLFLNVHPGVLTD
PFQAAALFANLRRLDIAPRRIVLEVLEQRAEDIERLADAVREFREHGFLIALDDFGAGHS
NIERIWQLNPDIVKLDRIMLSHAQHRAGLTAILHGLVTLLHEAGKLVLIEGIETEHEAQI
ALACDADFVQGHFFGRPAPGLPDSAAASACIGELTERYRQQTEAAERRDAQRLAPYQRAF
ERAAERLAAGEPLDEVCWNFLALDHAARCFLLDERGRQAGRNVVLRADRASHETRFLPLA
DAQGANWLRRPYFRAAIAEPGRVQATRPYLSINEALPCVTLSIAVRVNDALCVLCGDIDW
IDADAAAGA*
>GCA_000452945.1_BPMSHR5858_genomic_03202_1 sbp_1
MKHNKGIGRRLAAGVAAALLATAAHADTALLNVSYDVTRELYKDINASFVAAYKQKTGET
VSVRQSHGASSAQALSVLQGLQADVVTMNQPNDIDLLAERGQLLPKNWRTRLPNGSSPYS
TTMVFLVRHGNPKQIKDWSDLAKPGVQVIIANPKTSGNGRYAYLAAWGYQKLKGATDQQA
LDFEKAIFRNVPVLDSGGRGATTTFTQRGIGDVLVTFENEVALIDSGAAGASFDAVYPSA
SILAEPPVAIVDKVVDKKGTRRAAQAYLDYLYSPAAQEIVAQHHLRPRDPNVLKQHASEF
KPLKTFTVEQIFGNWANAQKTHFADGGTFDKIIVDRK*
>GCA_000452945.1_BPMSHR5858_genomic_03209_1 group_2464
MNETSMMRRKALGALCSLALAGAPVGSRAATPFYSLPLSSKDWHKGFRLTDLHGHTKTPQ
DYRGNVLLLFFGFLSCPSICSTTMLELTQAKERVGAQKDKVKILFVTLDPNRDTSPTIAR
WLASFGDDNIGLRDSEAHVRKAATALNLKYERVEGDVPGAYTIDHGVQTYVFDPQGRLRL
IARAGIEPEYVAKDIVQLLSGR*
>GCA_000452945.1_BPMSHR5858_genomic_03212_1 group_4834
MNSLPTSTVISEASPARSEAWRGLRASLPVMLGFVPFALVLGAQAAQKGLGIVEVPLLTG
LNFGGGSEFTAIRLWTSPPHILLIVAMSFLVNCRHILVGAAFAPYLRHLSRRQTFPALFF
MCDESWAMALADARREGRGRVSLPYYLGVAAGLYLTWISCTALGAALGPTIGDVERYGFD
MAFTAVFLVLLRGMWKGARASRPWLVSLVVAAATYLLVPGAWYVAAGALAGLVAAVVCGE
PA*
>GCA_000452945.1_BPMSHR5858_genomic_03213_1 lrp_3
MKLDSIDRRILRALQRDGRQQNVELAKEVGLSPSPCLRRVRLLEEAGVIERYVAVLNPAK
VGRGLTVFTRVWLKEQDADTVAHFAEAVQHLPQVVECHLMAGDCDFLLRVVAADIDDYRQ
FQMQHLTRIRGVQSVKTEIPLQKVKLTSEVPV*
>GCA_000452945.1_BPMSHR5858_genomic_03214_1 group_773
MRQRLTAPAVVVVAALCVAPLVAACSDDASHANAGAGTSPAAQPGGAANAFNGGATTAGN
ASASAPLATPVIHYPPEDAADDASGAAAASAQSASHPG*
>GCA_000452945.1_BPMSHR5858_genomic_03216_1 group_1873
MQHTEQPRVGCGAAIVRDGRILLIKRKRAPEAGCWGLPGGKVDWLEPVERAVCREIEEEL
GIALERATLLCVVDHIDAANGEHWVAPVYLAHAFSGEPRVVEPDRHEALGWFALDDLPQP
LTHATRIALEQVTRAA*
>GCA_000452945.1_BPMSHR5858_genomic_03221_1 group_4836
MATPAPIDKLPDSARRVALLLRERGHAKAIVMLEETGKTSAEAAAGLGCSIAQIAKSILF
RRRADGAPVLVVASGANRVDERKVAAQVGEVGRADAKFVRDNTGYAIGGVCPIGHLVEPV
VLIDRDLLDLDSLWAAAGHPHAVFNLSPQELVSLTGAPVADVAQRGDA*
>GCA_000452945.1_BPMSHR5858_genomic_03249_1 group_1205
MKIVRLLLVLLWCATLPLTGLAASGLAGDCPMQQAMSMSEDGAMPSMQDCESMRSSVAGK
SGKAGKAANVFCKVMVQCQFGSLYHPMPAADVARPASPGRPLVFHYAKSLTVREPGGLWR
PPRTA*
>GCA_000452945.1_BPMSHR5858_genomic_03252_1 lemA
MRFHSSAPRAAATWIRLIRAALLAAFALTLSGCGYNAIQVQDEQVKASWSEVLNQYQRRA
DLVPNLVNTVKGYANQERDVLTRVTEARAQVGSIRATPELLSDPQAFAKFDAAQGQLTSS
LSRLLVVSENYPQLKSDANFRDLQAQLEGTENRITVARNRYIRSVQAYNTTIRSFPSNLT
AMAFGYKEKPNFSVENEAQIAKPPRVDFGTSSAPAPAPAAGASN*
>GCA_000452945.1_BPMSHR5858_genomic_03253_1 silP
MHDAHVHHDRSGDTNERRDPHRPHETKTPGETAVDPVCGMKVPIDTPRRYDYQGRTYRFC
SDKCATAFRASPQRFVGTPKQRAGASAAKPGTVYTCPMHPQIRANAPGACPICGMALEPL
TPDAAEDGNPELRSMTRRFELGLVLAVPLVLMTMGAMVLPFDIGAGIDALLARWPRPFGM
PASWSQWVQAALATPVVLWGGWPFLVRGWKSFVTRRLNMFSLIGLGVCVAYLFSVFALLF
PDTLPQAFRSGHEIPLYFEASAVIVTLVLLGQVLELRARSRTSSAIRDLLQLAPHTAVRV
NPDGSEQTVPLDAVVVGDTLRVKPGSKVPVDGIVVAGHSSVDESMITGEPVPAEKTEGSP
VTGATVNQTGTFVMRAQKIGTDTLLARIVQMVAEAGRSRAPIQKLADQVSGGFVLAVIVL
AALAFAAWAAFGPAPALANALVVAISVLIIACPCALGLATPVSIIVGVGRGAREGVLIKD
AEALELMEKVDTVVVDKTGTLTEGRPRVQTIVALGGQSERALLGDAASLEGASEHPLAQA
IVEHAARMGAERKPVASFDSVPGKGVKGTLDARTVALGNAHLMADLAIDCTAADADSNRL
REAGQTVMYVAIDGQLAGYIGVADPIKETTPDAVQLLKASGTRIVMLTGDNPVTANAVAR
TLSLDGVKAGVLPEDKYRHVQALQQPGHVVAMAGDGVNDAPALAQANVGIAMGTGTDVAM
NSARIVLVKGDLRGIARARALSVATMKNIRQNLFFAFVYNAVGIPVAAGVLYPWFGITLS
PIFASAAMAASSVSVIGNALRLRGAKT*
>GCA_000452945.1_BPMSHR5858_genomic_03255_1 group_4841
MNRAIRMRAAVAAAAFSASLPAGAAGPAIDPGRYLYVEGGSAHGVLTVKGNAFEIETIGG
NCHTCALSGTFDGRVGIARDGENVCRIAVSGGHGDLRLDTSGSDACRDFCGMRASFDGEY
RRPGAACTDRARDVRTERSHRQYAAHDYDAARTTLKALLAECGGFMGWIELDRAKSDLAL
TEYHRGDRAQCVAVLSDTIAYRAQQDHSDAFGLPPCDADNYKSTGDAILHNLALCQAPAK
R*
>GCA_000452945.1_BPMSHR5858_genomic_03260_1 mntH_1
MSTPTDISPPIALERDAVLDQAHVGDIKGAFGTIAHHDTAPRSGWWARVRTLLAILGPGL
IVMVGDNDAGAFGTYTQAGQNYGTSLLWTLLLLVPVLFVNQEMVLRLGAVTGVGHARLIF
ERFGKFWGAFSVIDLFILNALTIVTEFIGITFVLDFFGVSKVAGVCIAAALTMAAVSTGD
FRRFERFAVALCVLSLLLVPVLVSIHPPVHQMARDLFVPTWPAHAKRSDVMLLVIGIVGT
TVAPWQLFFQQSYVIDKRITPRFIKYEKADLWIGIVLVMIGAIAMISFSAALYAGRPEAG
GFTDAGGVIAGLAKYAGPTSAMLFAIALLDACIIGAAAVSLATAYAIGDVFRIRHSLHRG
VSDAKGFYLVYFGIVAAAAALVLMPGSPLGLLTEAVQTLAGVLLPSATVFLLLLCNDRAV
LGPWVNSARLNLFTGAVVWVLVMLSIILTASVMYPDMSGEAMLEVLAGGTLFAALGLAAT
FVLRRRAGGADRANVAHVDRAARDTWRMPPLDALPAPRVTLSTRIWMGVLRGYLVLAVGL
VIVKVVQMAFFR*
>GCA_000452945.1_BPMSHR5858_genomic_03271_1 group_4844
MSTLPACPQCAMENTYPDGEHYICADCGHEWPIAGDANAADDDAAAVVKDANGNVLADGD
AVVLIKDLKVKGSSITLKMGTKVKSIRLVGGDHEVDCKMDAGNFMLKACYLKKV*
>GCA_000452945.1_BPMSHR5858_genomic_03276_1 rutG
MADSYFPRWRAQLRDDAARVIAPDERLPWPQTFAMGVQHVVAMFGSTVLAPLLMGFDPNL
CILMSGIGTLLFFVLVGGRVPSYLGSSFAFIGLVIAVTGYAGGGANPNVPVALGGIVACG
VVYALIGLVVAAVGTRWIETLMPPVVTGAIVAVIGLNLAPIAVKGVSASTFDATLALVTV
LCVGGVAVFARGMLQRLLILVGVLIAYVIYAVLTNGMGLGKPIDFAIVAQAAWFGVPTLH
RPVFDPHAMLLIAPVAVILVAENLGHIKAVSAMTGRNLDRYVGRAFIGDGLATVVSASMG
GTGVTTYAENIGVMAVTRIYSTLVFVAAALIAIALGFSPKFGAVIQTIPGPVLGGVSIVV
FGLIAVTGARIWVVNRVDFSDNRNLIVAAVTLVLGAGDFSLKLGGFALGGIGTATFGAIL
LYALLRKEKEPGPMV*
>GCA_000452945.1_BPMSHR5858_genomic_03277_1 group_4846
MRISSTQLYSLNVDMMNDQQAQIAQLYQQVSSGISLTTPADNPLAAAQAVQLSATSATLA
QYTQNQTIVQTALQTEDTTLTSVNDVLNAAYQALMHAGDGGLSDSDRAALAAQIQGSRDH
LLTLANTADGAGNYLFAGFQPTTQPFSNKPGGGVTYAGDYGARAVQIADTRTVSQGDNGA
NVFMSVPFLGSLPVPAAGASNTGTGTIGAVSITNPSDPTNTHQFTITFGGTAAAPTYTVT
DNSVTPPTTTAAQAYSSGQGINLGGQTVAVSGKPAVGDTFTVTPAPQAGTDVFATLDTVI
AALKSPVGNSQTASTALTNTMATASTKLMNTMTNVLTVQASVGGRLQEVKAMQAVTTTNT
LQTTNSLSNLTDTNLPAAISQFLQLQNSLSAAQKAFVQMQNLSLFQYLNP*
>GCA_000452945.1_BPMSHR5858_genomic_03281_1 flgI
MLALAFAPAAARAERLKDLAQIQGVRDNPLIGYGLVVGLDGTGDQTMQTPFTTQTLANML
ANLGISINNGSANGGGSSAMTNMQLKNVAAVMVTATLPPFARPGEAIDVTVSSLGNAKSL
RGGTLLLTPLKGADGQVYALAQGNMAVGGAGASANGSRVQVNQLAAGRIAGGAIVERSVP
NAVAQMNGVLQLQLNDMDYGTAQRIVSAVNSSFGAGTATALDGRTIQLTAPADSAQQVAF
MARLQNLEVSPERAAAKVILNARTGSIVMNQMVTLQNCAVAHGNLSVVVNTQPVVSQPGP
FSNGQTVVAQQSQIQLKQDNGSLRMVTAGANLADVVKALNSLGATPADLMSILQAMKAAG
ALRADLEII*
>GCA_000452945.1_BPMSHR5858_genomic_03288_1 flgB
MLDKLDAEFAFGRQALDVRAYRQELLSSNIANADTPGYRARDVDFSSTLARALKQDGAAS
AGNAAQLPLAQPAGVASGMSMATTSGAHLAGNVKLIPTGGPSDDYGRAQYRIPQQPALDG
NTVDLDTERVQFADNAVHYQAGMTVMTQQIKTMIAAITSNSSS*
>GCA_000452945.1_BPMSHR5858_genomic_03289_1 flgA
MTTDAGRATRTRFAVALALAGWMGAALAQQAGDGGMIVIPGRGESAQTALANANAASARG
ANGAGAAGPGGADRTASPGGWHAPAAAQAAAAWSGRAGAASGAESAAESGDVRAATEPRT
GLIVIEPGPAESNGRIPAAAPSGWHPSAAMPRAESNGGANPVPPRTGSNRSAGSASASAG
WTAPPGAASQGNPAANVAPAPRVSAASRAPATTDMQRVAPVIAQDEGAAPAGRPANARAA
AQSWPARDPSVAAGGVIPVSLRAQPAPRTLPVRSAPIRAAATSAAGAQPAAASATAAGAV
PAAQQDGESIRRAALAFLQQQAAGLPGKTTVTVAPAFPRGLAACTTLEPFLPSGARLWGR
TTVGVRCAGERPWTIYLQAKLAVQATYYVAARQIAPGETLTAADLVARDGDLTLLPLAVI
TDAQQAVGATALMRVAAGLPLRRDLLKSAASVSIGQTVRVVASGQGFTISAEGSVLNNAA
PGQQVRVRMAAGQIVTAIVKDAATVEIPL*
>GCA_000452945.1_BPMSHR5858_genomic_03294_1 group_4848
MKLVIGDKNYSSWSMRPWVLMKHFGIPFDEVMIGLRLDDTAERIRAHSPSGKVPCLVDDE
GGSVWDSLAIVETLAERFPQHALWPRDPAARAHARSVSAEMHAGFPALRSEMPLNVRESH
PGRGATPAALADVARIDELWRTCVAASGGPFLFGAFSIADAMYAPVVLRFKTYAPPLSPE
ASAYAARVAELPAVREWIDGARGETRVIDIYGPSR*
>GCA_000452945.1_BPMSHR5858_genomic_03301_1 pxpA_1
MEIDLNADLGEGCGSDEALLDLVTSANIACGWHAGGAQAMRDCVRWAVEKGVSIGAHPSF
HDPENFGRKEMDLPASEIYAGVLYQLGALSAIAQAEGGRIAHVKPHGALYNQAAREPEIA
DAVVSAIHDFDPSLAVFGLAKSGFVDAARQAGLVAVEEVFADRGYRADGSLVPRSQPGAL
VDDENEMLARTLEMVRGQRVRAVTGEWVPLNAQTVCLHGDGPHALAFAKRIRDALEAAGI
DVHAPGALHAGERA*
>GCA_000452945.1_BPMSHR5858_genomic_03304_1 group_878
MPEAVAAREAARRSRNGVRRAATHAARAPRRWRARSLASAVLAGACATGGLAWPPAGALA
AGSVAAAPLPQAPIPAPGMSLPGFHAPPPSTSNGTVASGAVRTQPARMPFYVATKGKVTI
YVLGTLHVGDPADYPANQPFRRPILAALAASPTLALELSPDDLLESQDDVSKYGVCNYAC
LPRLLPPPLWQKLANRLRGNPAALAGIRNMRPWLASLVVETYDSLSAGLQTEYGTEAQLQ
NVFLRKKGGKVVGLETLAEQMRAFTGLTLAQQREMLAQDMVQTPAQNAADVRALHRLWRI
GDADAIAAWANAKTERLARARSIADSIDNKIVYERNRRFVARMTAIAAPNRPLFVAIGSL
HLGGPKGVLELLRQQGYRVDAG*
>GCA_000452945.1_BPMSHR5858_genomic_03309_1 dppA_2
MEHNRLLRALRATAIAGVAAASFGIAGSAFAQIPNKTLVYCSEGSPAGFDSAQFTTGVDF
TASTFPIYNRLVEFERGGTKVEPGLAEKWDISADGKVYTFHLRHGVKFHTTDFFKPTREF
NADDVAFTFERMLDPNQAFRKAYPVSFPYFTDMGLDKLIVKIEKLDPYTVRFTLKEPNAP
FIQNLAMEFASILSAEYADQLMKAGKAADINQKPIGTGPFIFRSYTKDATIRFDGNPDYW
KKGAVKISKLIFSITPDPGVRVQKIKRNECQVMSYPRPADIATLKADSNVDMPSLPGFNL
GYLAYNVQHKPVDKLEVRQALDMAINKKAILESVYQGAGQAASAPMPPTQWSYDKNLKAA
AYDPAKAKALLAKAGYPNGFPITLWAMPVQRPYNPNAKLMAEMIQADWAKIGVQAKIVTY
EWGEYIKRAHAGEHDTMLIGWNGDNGDPDNWLGTLLGCEAVKGNNFSEWCYKPFDELIQK
GRVTTSQDGRTKIYMQAQQIFAQQLPFSPIANSTVYQPVRKNIVDMRIEPLGYARFDGVS
VK*
>GCA_000452945.1_BPMSHR5858_genomic_03310_1 proP_3
MASIAASTKTAGQVLNRRAVVAAVIGNALEWYDFTVFSFMTVVIAQLFFPTSSEYSSLLL
TTATFGVAFFMRPIGGIVLGLYADRAGRKAALSLVILLMTFGIFLIAVAPPYAAIGIGGP
LLIVLGRLLQGFSAGGEFGSATALLIEAAPLSRRGYYGSWQMASQAAALLFGSLVGAAVT
RGLSTEALHAWGWRVPFLIGLVIGPVGFYVRRHLVDSEAFLQARERGEHVTLGEVFARHG
RAVVCGLGAVIALTVTVYVLISYLPTFAVKQLKLPYAQSFYAVIIGNLLLTALSPVAGAW
SDRIGRKGLSLWALGLTLVMIYPLFAWLDASPSIARLIAVQAMLSVTLAGYYGPFGAMIA
ELFPANVRSTGLSLAYNVAVMVFGGFGQFIVTWLIKTTGTPLAPTYYVMAGLALSLVALA
FVPTERDALD*
>GCA_000452945.1_BPMSHR5858_genomic_03313_1 lon_1
MTTAMVKQELAVASFSTVYDIERIETALNDLNEGASDALRATYEKMLKTGNLRFCVKPTR
MPAFDSLAQALPNFAEPLDDVRKQVALCLETDDRLELMPILLLGEPGIGKTHFAKALAKM
LGTAYHYVPMSSLTAGWVLSGASSQWKNAKPGKVFDALVNGCYSNPVIAIDEIDKAGSDA
QYDPLGALYALLEHDTASAFVDEFAEVPIDAGNVIWIATANDASAIPEPIMNRMNVYEIE
PPDASGARRIAQTIYGEIRNAHAWGQRFPAALGDDALDVLAAMPPRTMRRVLLHAFGAAR
LDGRDAVAPRDIRTDESAGRRRPIGF*
>GCA_000452945.1_BPMSHR5858_genomic_03315_1 group_4853
MHVPQTAPLAEPLPVADEITAVSWNLHKGRSPLGFTAWEAMRSWVASTHADVYFLQEAMA
RRMPRPVLAAGFGAPMAEPVDDIWHCQATEIARALDWQIALGPNVFKPSWRHGNAILSPH
PLDLGGRWDISAHRFERRGLLVARATLAGGAPVTLLCAHLALTRAARLRQMHWIAHWIER
NARTGPLVLAGDFNDWRNDSIPLFGEIGLSEVATLLGESGRTFPAFSPALALDKMFVRGM
TPLEWRAPGDETAWLSDHLPYVARLRLDPA*
>GCA_000452945.1_BPMSHR5858_genomic_03317_1 fpr_1
MSKFDTATVLSVHHWTDTLFSFTCTRDQALRFNNGEFTMVGLEVDGKPLTRAYSIVSPNY
EEHLEFFSIKVQNGPLTSRLQHLKVGDPVLIGKKPTGTLVADNLLPGKTLWLLSTGTGLA
PFMSIIRDPDIYERFEKVVLTHTCRLKGELAYMDYIKHDLPGHEYLGDVIREKLVYYPTV
TREEFENEGRITDLIASGKLFTDLDVPPFSPERDRVMLCGSTAMLKDTTELLKKAGLVEG
KNSAPGHYVIERAFVD*
>GCA_000452945.1_BPMSHR5858_genomic_03321_1 pucA
MESVDLEVLRTSARWLDEGRRALLVTVVRTWGSSPRPEGAMLAVRDDGLVVGSVSGGCIE
DDLIARVQASGVAHWTRPEAVKYGVTAEEAHRFGLPCGGTIELVLEPLTPASGIAALCDA
LEHGRLVTRTLALATGEATLASASATDGLHFDGARLVTVHGPRYRMLVIGAGQLSRYLCQ
IAAGLDYQVTVCDPREAYTDAWDVPGTRIVRTMPDDTVLDMRLDRRSAVIALTHDPKLDD
LALMEALKTPAFYVGALGSRRNSQARRERLREFDLSAAELARLRGPAGIYIGSRTPPEIA
VSILAEVTAAKNGVSLPTILQVEGAKAAREIEANSGVACGM*
>GCA_000452945.1_BPMSHR5858_genomic_03326_1 fliS
MFTPGHAGASAYARVGVETGVMGASPHRLIAMLYQGARQAIAHARLHLQQGNVAARGEAI
GKAIRIVESGLQQALNLEVGGDIASRLDSLYTYMCRRLLQANVDASEPMLIEVDGLLATL
EDAWTGIAPEVARMNAQAVTEQAR*
>GCA_000452945.1_BPMSHR5858_genomic_03333_1 group_445
MPPLPLIGMLLGAASTMIKSSAGAPGASSGDASSAAPFAQALKQSVQTQRQTGANADAPN
AGHAANAATADTAAASSPASGAGDRTTASAGDTKASGKPDDQDDDKHAAQPASADAAALA
AAASLQAQLQARPDAQAAQAAQAAAGATAAQAGAGDAAHPPATSAASAAAGDTDASAPAG
AQQPIGDALNAALSKLTGAGRTGTAAVQGAQPAGRNGDASAPLAANGAAFDKLLAGAKAP
TAQAAPTDASGANPATALANAAANAAPPDASGALAALQDAADSARATLAASSAPAALQQA
APAALAANASAAAASAAPSLAPPVGTPDWTDALSQKVVFLSNAHQQSAELTLNPPDLGPL
QVVLRVADNHAHALFVSQHAQVRDAVEAALPKLREAMEAGGLGLGSASVSDGGFASAQQQ
QTPQRQSSDGSATRRAFGASTADAALDELAAASSGGAARRAVGMVDTFA*
>GCA_000452945.1_BPMSHR5858_genomic_03334_1 alkJ_1
MQYDYIIVGGGSGGASLAGRLADACPDATIALIEAGGHTERNLLVNMPVGIAALVPFKLG
TNYGYETVPQPGLGGRRGYQPRGRGLGGSSAINAMIYTRGHPLDYDEWEQLGCTGWGWRD
VLPYFRRAEGNARGANEWHGADGPLTVSDLRFRNPFSERFIAAAHEAGYPLNDDFNGEHQ
EGVGFYQVTHRDGSRCSVARAYVYGRTRPNLHVIVDATVLRVVFDGKRATGVEFARAGRT
EQLAARAEVILSAGAFNTPQLLMCSGVGPAAQLRRHGVALVHDAPDVGENLIDHIDFIIN
KRVNSSELVGICMRGIAKMTPALFSYLSGRRGMMTSNVAEAGGFIKSEPGLDRPDLQLHF
CTALVDDHNRNMHWGFGYSLHVCALRPKSRGNVALASGDARVAPLIDPRFFSDERDLDLL
VTGAKAMRRILCAAPLASQGGRELYTDPGDTDAQLRAAIVAHADTIYHPVGTCRMGTDAR
AVVDPQLRVKGVDGLRVVDASVMPTLIGGNTNAPTVMIAERAADFIVAARNGQAAPTRER
VAATHGG*
>GCA_000452945.1_BPMSHR5858_genomic_03339_1 group_18951
MHVGSIVCTTHIAVPKGARGIVQRILGDMAMVTWYAGVPGESKELNTEPFFLEDLIDTGE
SVLPAGAALH*
>GCA_000452945.1_BPMSHR5858_genomic_03342_1 group_4856
MSTYFTIADFILLIPMALAGALFLGAVPCATEFRHNVLRVLGAMLGVAVAVLLVEGLPAL
I*
>GCA_000452945.1_BPMSHR5858_genomic_03343_1 gpr_1
MARWGTSGRKDPASFNVSTGIQMAYEAASERYADMQYRVSGKSGLKLPALSLGLWHNFGD
TTPISTQREILRTAFDLGITHFDLANNYGPPYGSAETNFGRLLREDFKPYRDELLISTKA

GWDMWPGPYGSGGGSRKYVLASLDQSLRRMGLDYVDIFYSHRFDAHTPLEETASALASAV
QQGKALYVGVSSYSAASTREIAKLLAEYKVPLLIHQPAYNMLNRWIERELLDALDETGSG
CIAFTPLAQGLLTSKYLNGVPADARINKPGGGSLKEAHLSAENLEHVRKLNEIAQRRGQS
LAQMALAWVLRDSRVTSALIGASRAEQVRENVAALAHLAFSDDEIAEIDRYATEGGINLW
EKPSTDQAI*
>GCA_000452945.1_BPMSHR5858_genomic_03357_1 regA
MSDKNFLVIDDNEVFAGTLARGLERRGYAVRQAHNKDEALKLAGAEKFEFITVDLHLGND
SGLSLIAPLCDLQPDARILVLTGYASIATAVQAVKDGADNYLAKPANVESILAALQTNAS
EVQAEEALENPVVLSVDRLEWEHIQRVLAENNNNISATARALNMHRRTLQRKLAKKPVRQ
*
>GCA_000452945.1_BPMSHR5858_genomic_03361_1 gph_1
LSARRQPRRPASPAASEQAPERPRRRRARPQAGGPVWLFDLDNTLHHASHAIFPAINRAM
TQYIIDTLKVERAHADHLRTHYTRRYGAALLGLARHHPIDPHDFLKVVHTFADLPSMVRA
ERGLARLVAALPGRKIVLTNAPETYARAVLRELKIDRLFERVIAIEQMRDRRAWRAKPDA
TMLRRAMRAAHARLPDAILVEDTRGHLKRYKRLGIRTVWITGHLPGHLPSYGRPHYVDRR
IGSLKSLRLGTRSGRRKCSRLTRMTRQ*
>GCA_000452945.1_BPMSHR5858_genomic_03365_1 ampG
MTKPPHDAPALNAHEDHPGWRAFLNTHMLICVFLGFTSGLPLFTLVYLVQAWLRSEGVNL
KEIGLFALIQFPYTWKFLWAPLMDRFIPRLPGWRPGRRRGWMFVTQLLVAGAIAALGFVS
PRESIWTVAALTTLVAFFGASADIVIDAYRRELLRDTEQGLGNAVHVNAYKVAALIPGSL
SLILSDHLPWDAVFALTAAFMLPGILMTLVVREPDVHGAPPKNLSEAVIMPFREFIGRDG
WRGALLVLGFIFLYKLGDTMATTLSTSFFLDLGFNKTQIGVIAKTTAFWASLAGGIIGGV
WLVKIGIARGLWIFGVLQIVSTLGFAWLAKAGPDPVLMAAIYGFETFSTGLTLAAFTAYI
ASTTDPRYTATQFALFTSLASVPRTLASASSGFIVAQIGWFDYFLVCAALGIPGMLLLFK
IAPWNGSMSGNEAARAG*
>GCA_000452945.1_BPMSHR5858_genomic_03367_1 rhaS_3
MSFEPLFADGGARLQRRILALFGQLAPNEGYTHAAIDGVRFIRSNRPVPRMPVLYDPSIV
VVCQGRKRGYVGERSFVYDAQQYLVLSVPLPFECETFATPDEPFLAISIRIDLAVIAELA
ILLDETHGASVSEPRGIYSTPLDAPLADAVLRLLEALASPADTRVLGPAIMREIGYRVLT
GAQGDAIRAALAQQHHFGRVARALRRIHADLSGELDVETLASEAGMSLAVFHAQFKSVTA
TSPMQYVKATRLHHARLMMVQDGLNAGAAAARVGYASASQFSREFKRLFGRSPSDEVRWV
QAGGRQPVALEAGE*
>GCA_000452945.1_BPMSHR5858_genomic_03372_1 gatA_1
MHAKSLTELRAALDAKECSAVELAQHYLKRIDAARDLNAFVHVDAELTLAQAKAADAALA
NGEAGPLAGLPIAHKDVFVTRGWRSTAGSKMLANYASPFDATVVARLSAAGMVTLGKTNM
DEFAMGSSNENSAFGPVKNPWDTSAVPGGSSGGSSAAVAARLAPAATGTDTGGSIRQPAS
FAGVTGIKPTYGRVSRYGMIAFASSLDQGGPMARSAADCALLLNAMAGFDERDSTSLERA
DEDYTRHLGKAWAAGGDAGKPLAGLRIGLPAEYFGAGLADDVRAAIDAALKTYEALGATL
VPVSLPKTELSIPVYYVIAPAEASSNLSRFDGVRYGHRAAEYRDLLDMYKKSRAEGFGPE
VKRRILVGTYVLSHGYYDAYYLQAQKIRRIIAQDFQEAFKSCDVIMGPASPTVAWDIGAK
GDDPVQMYLADIYTLSVSLAGLPGMSVPCGFGAGANAKRPVGLQIIGNYFDEARMLQVAD
AFQRATDWHVQEPAGV*
>GCA_000452945.1_BPMSHR5858_genomic_03374_1 mreB_1
MFGFLRSYFSNDLAIDLGTANTLIYMRGKGIVLDEPSVVSIRQEGGPNGKKTIQAVGKEA
KQMLGKVPGNIEAIRPMKDGVIADFTVTEQMIKQFIKTAHESRMFSPSPRIIICVPCGST
QVERRAIKEAAHGAGASQVYLIEEPMAAAIGAGLPVSEATGSMVVDIGGGTTEVGVISLG
GIVYKGSVRVGGDKFDEAIVNYIRRNYGMLIGEQTAEAIKKEIGSAFPGSEVKEMEVKGR
NLSEGIPRSFTISSNEILEALTDPLNQIVSSVKIALEQTPPELGADIAERGMMLTGGGAL
LRDLDRLLAEETGLPVLVAEDPLTCVVRGSGMALERMDKLGSIFSYE*
>GCA_000452945.1_BPMSHR5858_genomic_03375_1 group_774
MEYSPPPLFKQGPSALARLIFFVVLAIALLVSDARFNTLEIVRGVLGTVLYPLQRAALVP
RDLIVGAAELATSSTALRHENQQLRDRNLKLSLQANQAGQLAADNAHLRAVLELRQQIAA
QATPVEIQYDTGDPFSQKIIIGHGSQAGIQDGAPVVNEDGVIGQVTRVFPLQSEVTLVTD
RDLAIPVQVLRTGLRSVIYGTPKGDALDLRFVPTSADLLAGDELVTSGLDGVYPPGLPVA
KVVRVDKLADTAFAHVVCAPIAAVRGAREMLVLHYRNDIPPPPVTDDAASDAKAAKGKKA
TKGADKSAKAADKGADKDKGAKPAAAPPVPARSRPAGPAQPAAPLKPATAPSPGAPR*
>GCA_000452945.1_BPMSHR5858_genomic_03385_1 group_4863
MATILVVDDEMGIRELLSEILSDEGHVVDVAENAQAARDYRLRQAPDLVLLDIWMPDTDG
VTLLKEWASQGQLTMPVIMMSGHATIDTAVEATKIGALDFLEKPITLQKLLKSVEHGLAR
GAAPLPASAAAKPAAGAAVASAAALPTLGDDPAVALAGQTTAAIPFDIPLREARDAFERA
YFEYHLARENGSMTRVAEKTGLERTHLYRKLKQLGVELGKKPAEGAA*
>GCA_000452945.1_BPMSHR5858_genomic_03395_1 topB
MSKALIIAEKPSVANDIARALGGFTKHDEYYESDEYVLSSAVGHLLEIAAPEEYEVKRGK
WSFAHLPVIPPHFDLNPIAKSESRLKVLTKLIKRKDVDRLINACDAGREGELIFRLIAQH
AKAKQPVQRLWLQSMTPAAIRDGFAHLRTDMDMQPLADAARCRSEADWLVGINGTRAMTA
FNSKGGGFFLTTVGRVQTPTLSIVVEREEKIRRFVPRDYWEVRAEFVCAGGFYEGRWFDP
KFKKDEFDPEKRDSRLWSLPAAETIVAACRDHLGTVTEESKPSTQLSPLLYDLTSLQREA
NSRFGFSAKNTLGLAQALYEKHKVLTYPRTDARALPEDYLGTVKSTLEMLKESNNYLPHA
KQVLDKNWVKPNKRIFDNSKISDHFAIIPTLQAPKSLSEPEQKLYDLVVKRFLAVFFPAA
EFKVTTRITEVAGHHFKTEGKVLVEPGWLQVYGRDAEGADANLVPVQKGEKVKTDKIAAH
GLTTKPPARYSEATLLSAMEGAGKLVEDDELREAMAAKGLGTPATRAAIIEGLLGEKYLV
REGRELIPTAKAFQLMTLLRGLGVKELTAPELTGEWEYKLSQMERGNLQRDAFMQEIARM
TQTIVKRAKEYDSDTIPGDYATLETPCPNCGGQVKENYRRFACTKCEFSISKIPGSRQFE
IAEVEELLRKKEIGPLSGFRSKMGRPFSAILKLTFDDETKNYKLEFDFGQEQGGEEGEAP
DFSAQEPVGACPKCKGRVFEHGMSYVCEHAVANPKTCDFRSGKVILQQEITREQMAKLLE
NGRTDLLPNFKSSRTGRNFKAYLVKQPDGKIGFEFEKKEPKPAAAKKTAARSSAAADDAA
ADSGEKAEKKAAPARKTAARKTPARKTGS*
>GCA_000452945.1_BPMSHR5858_genomic_03399_1 group_4865
MDENTVRDLLDRLLAPWVRSLSLTPVSISEESVTLRLPFSGEFRHSGGVICGQVFMAAAD
TAMVVAISAALGEFRPMTTVALNTHFMRPVRKGDVLVTARVLRMGRNLVFGEIELFDEDG
KMAVHATSTYALVS*
>GCA_000452945.1_BPMSHR5858_genomic_03412_1 group_6434
MRRIDLLSALETFVCAVEEGSLSQAARRLAKTPSAVTKAIAGLEATLGVQLLERTTRRLA
LTESGRLYLATATDVLRRLADGARQLSERDDEPRGLLRVTAAHSFGHAILALLCAPFAQA
FPRVRIELSLSDHYVDIVGEGFDLALRMGNYDLPSQIVKPIGSNRGLLCASPGYLARRGR
PTAPADLAQHDCVIYRHPTLANTWTFERGGVRARLEPKGPLVTDDYGLALAAALDGAGVL
PCPQWSVVDALEAGRLVPLLVDWRFESASFGEDRICAAYPSSRRGSTKIRRLVERVQARL
ADSEARVARVLGDALCADLARR*
>GCA_000452945.1_BPMSHR5858_genomic_03413_1 group_502
MKIERYSKGLTVISLTAALWSLGMPVAPAAPASDARAQTPCGRGYGMGPGMMGGGYGRGM
MGGYGMGPGMMRGDGMGPGMMMGFGGWPSGLDLTSEQRAKINRIQDDTRKAHWALMGDMM
DQQAKLRDLYDAPKRDGAAIDETSRAIGSLRQKMIDASVDARKKMEAVLTSKQLDKLRAY
EKQADDMSW*
>GCA_000452945.1_BPMSHR5858_genomic_03417_1 alkA
LSAAVEPAAVRPDSVVLELPFKSPYDWPRVLRFFAGRAIPGVEAVGDGAYRRTVDHHGAI
GTLTVRKHPRKRCLVALVEGDAARHADAAFAARLSTMFDLQADPAAIGAHLARDAWFAPL
VGAAPGLRVPGAWSGFELIVRAIVGQQVSVKAATTIVGRLVERAGERLAPHAPGAIGWRF
PEPAALAACDVSRIGMPGKRAAALQGVARAVAAGDVPLEAYASDPAGVRAALLALPGIGP
WTVEYVAMRAWRDADAWPATDLVLMQAIVARDPALDRPASQRLRADAWRPWRAYAAMHLW
NEIADRAGSARGG*
>GCA_000452945.1_BPMSHR5858_genomic_03418_1 ada
MKQPPYPTDDARWAALVARDVDADGAFCYAVKTTGVFCRPSCASRLPRRENVSFFADTGA
ARAAGYRPCKRCRPEGLPRELDIVNRACAVLDAHRQDRFTLAQLSDAVHVSPFHLQRLFK
RVVGVSPRQYQAAQRGAALRDALQSGAAVTRAAVDAGFNSPSRLYESVPRELGMSPSAFR
RKGAGLKIDYATADTRLGVVLVAATSKGICKIAFGDAAAPLVDELRTGFANAQIAESPER
IAPFIAQIDAYLNGTRRRFELPLDLAATEFQQRVWDALRRIPYGETRSYTQIAEALGAPR
AVRAVASACASNPVALAIPCHRVVQKGGSLAGYRWGLPRKAALLDAEAQRAAGESVESAV
DHAA*
>GCA_000452945.1_BPMSHR5858_genomic_03423_1 speG
MQIQNEQHTLALRPLERQDLRFVHELNNDAKIMRYWFEEPYETFTELSQLYDQHVHDQRE
RRFVAFDSDGELVGLVELIELDYIHRRGEFQIIIAPNRQGRGFATRATRLAVEYAFKVLN
LRKLYLIVDKSNVAAIRVYEKCGFKHEAELIEEFFGNGQYHNALRMCIFQRDYFAAQPGA
ELHARE*
>GCA_000452945.1_BPMSHR5858_genomic_03441_1 group_4873
MLSALPPQPIPLCVTLLAVVAASIDLASRRIPNRLIALGLAGALVAQCALPGPHTGFAAW
LAGAATGAGLLLPFYLVRGMAAGDVKLMLAIGAWVGPTLALHIALATFVVGGAWALAVVA
GRGRFRKLLANLRHLFAGAALLRVASAARAAPEIDSVGMLPYGVAIAIGTLGVLFAAAA*
>GCA_000452945.1_BPMSHR5858_genomic_03445_1 group_4874
MSRSGGWLTSDRVLAYGGATLILSIALLGAWGWTTSGFTANTTVRPGIDFTVFWSGSHVM
LHGAPASVYDYPAFSRAEAAQLGAYVNRSFLPWVYPPTLLVLVTPLALLRYLPSLLLFCA
LGLLAYAKSVGALTGLRDRLAKPRLASFVILSFPGVLVAAVIGQNSLLTAACAAFATRLL
AKRPALAGLCISLLAVKPQMALLFPLLLVATRAWRACFAAAAGTLVFAAASVAMCGTASV
HAFLSGTTMLRELVLEQGTQYWLASPAPFSAMRLVGAPLRVAYAVQIGVGALAAAAALDV
WLRTRDTRLRAAALAVATLLATPYLWHYELAWLGIALFGVLACALDTGWLPGEQPVFVAG
WLLPFFEFFNRATHFPQIGPVVLLAVLLVVMRRARTMPGVAR*
>GCA_000452945.1_BPMSHR5858_genomic_03447_1 arnC_2
MRDQIDIPLISLVVPFYNEGDAVTRFFAEVMPLMEAIESIRFEIVCVNDGSRDDTLEQLV
AVGAREPRVRVIDLTRNFGKEAALTAGLDEANGDAVIPIDADLQDPPSLIPVMIDHWRDG
AEVVAAKRSNRACDTFAKRTAAALYYRVHNALSEVKLPVNVGDFRLMDRQVVNALRSLPE
RRRFMKGLFAWVGYRTVIVEYQREARCAGHSKFSGWKLWNFALEGITSFSTVPLRSWTYI
GLGIAALAFLYGGFIVARTLWLGNPVPGYASLISVMLFIGGIELVGIGVVGEYIGRIYYE
SKERPIYLVRRRYQARTKVSALPVGAAATRVAHGARAEFARRRAMPRARADSR*
>GCA_000452945.1_BPMSHR5858_genomic_03448_1 group_4876
MSPDAYLQMAATEATHWWFRARREILQALLAGLGLPAGARILEIGSGTGGNLGMLEAFGA
VSALEMDDTARAIANRRTGWRFDIRAGRCPDDVPFGDQRFDLICFFDCLEHIEDDAAALR
RISDYLAPGGAIVVTVPAYQWLWSGHDTFLHHYRRYDRAALERCARAAGCTIVRSSYFNT
LLFPLAVVARTIDRLLHRARSSGDALGPRAVNAALYRIFRTERGWLPKHSLPFGVSLFAV
LTKCSEPR*
>GCA_000452945.1_BPMSHR5858_genomic_03449_1 group_4877
MSAVADIGMRARIVRFGVSGAACTALHAAIAGVLMGAFAATAVQANAIAFVCATGASYLL
NTLWSFSAPLRWRNVARFVAVSVVGLMLTMAISHGVLALGLAAAWSIAAVVALVPPLTFA
MHRLWTYR*
>GCA_000452945.1_BPMSHR5858_genomic_03458_1 group_4878
MPTLEEKAAFAERLKFALQRSPEKISGATALALHFNLRHRGEHPISPQTAHKWLTGRTIP
TADKLRTLADWLRVDLHWLHYGPPPGAPGGKTPAPPRRDEKYAPTQETIELASKIEALSP
HHRYLMQELIEHFYGDPTKRR*
>GCA_000452945.1_BPMSHR5858_genomic_03463_1 group_691
MTLIVVFLLSCLSLVIWLVLLFGRGGFWRARAARRLPPDARGAAAAAGWPAVAAVVPARN
EADVIGEAVRSLVEQAYEGAFHLIVVDDHSTDGTAEAARAAAAAVGCADRLTVLAAQPLP
AGWSGKVWAQSQGIAAVRSLGLPADYLLLTDADIGHPPDAVAQLVTRAQAEQRDLVSLMV
RLRCDSFWEKALIPAFVFFFAKLYPFSWINDPRNRTAGAAGGCMLVRRDALEEAGGIESI
RGALIDDCSLAAQIKHRGAGRHPIRLDLADRSVSLRPYDSWRDIWNMIARTAFTQLRYSP
VLLLGTLVGMTILYLVPPVAALAYGARAWPAWLAWASMCTAYAPMLSYYRRSPWWAPALP
LVALFYVGATFASAVRYWRGKGGQWKARVQAPVRDR*
>GCA_000452945.1_BPMSHR5858_genomic_03465_1 auaH
MTDTQRDLVLVTGASGFVGSAVARAARRQGYRVRVLVRPTSPRTNVADLDAEIATGDMRD
EASMRAALRGVRHLLHVAADYRLWAPDPLEIERANLEGAVATMRAALAEGVERIVYTSSV
ATLKVTPSGASADESSPLAAGQAIGVYKRSKVLAERAVERMIAEDGLPAVIVNPSTPIGP
RDVKPTPTGRIIVEAALGKIPAFVDTGLNLVHVDDVAQGHLLALERGRVGERYILGGENL
PLQAMLADIAQSTGRKPPTIALPRWPLYPIALGAEAVAKLTKREPFVTVDGLRMSKNKMY
FTSAKAERELGYRARPYREGIRDALDWFRQAGYLR*
>GCA_000452945.1_BPMSHR5858_genomic_03484_1 potA_4
MKSTPSIDTAQLARPPVGTPAGTNEFVRIENVVKKFADSTAVDNVSLSIAKNELFALLGS
SGCGKSTLLRMLAGLETATSGRIFVDGEDLATLPPYRRPVNMMFQSYALFPHMSVESNVA
FGLKQEGTPKHEIRERVADALNLVQMSRYARRKPHQLSGGQQQRVALARSLVKRPKLLLL
DEPMSALDKKIRQKTQLELVNIIEKVDVTCVMVTHDQEEAMTMASRLAVMSEGRIVQIGT
PSQVYEFPNSRFSAEFIGSTNLFEGVVVEDEPDHIFVESEDLEARMYVSHGVTGPLGMPV
GISVRPERVRVSRERPSAPHNWARGVVTDVAYMGSYSLYHVRLPSGKTVVSNLSSSHLMQ
DGAPAYNDDVFVSWSPASGVVLTQ*
>GCA_000452945.1_BPMSHR5858_genomic_03485_1 potH_2
MRTSPSSPLAAAAPAARSAFSARAARVAARFTPSGRSVAIGVPFLWLALFFALPFVLVLK
ISFADQVMGIPPYTALMSVKDGAVRFALQLGHYAFLLQDSLYVATYVSSLKMAAVSTLLC
LLIGYPMAYYIARCAPATRNLLMMGVMLPFWTSFLIRVYAWIGILKDDGLLNHALIAIGL
IHSPLRLYHTDAGVYIGMVYSYLPFMVMPLYAHLVKMDLTLLEAAYDLGAKPWVAFTRIT
LPLSKNGVIAGSLLVFIPAVGEYVIPELLGGADTLMIGRVMWDEFFNNMDWPMASAVTVA
MVLLLLVPMAVFQYYQVKELEDAK*
>GCA_000452945.1_BPMSHR5858_genomic_03486_1 ydcV_3
MIKPNKPLCAGVLTLGFLFLYIPIVSLVVYSFNESKLVTVWSGFSLKWYAALWRDDELLT
SAWLSLKIGLLTAFASVFIGTWAGFVLARFGRFRGFTLYTGMINAPLVIPEVIQGISLLL
LFVALEQMAGWPKGRGMVTIWIGHVMLCVSYVAIIVQSRVKELNKALEEAALDLGATPLK
VFFVITLPLISQALLSGWLLSFTLSIDDLVLSAFLSGPGSTTLPLVVFSRVRLGLNPEMN
ALATLFITAVTIGVIAVNQMMLSRERRRERGMRQAFAPPAPCDSPESGPAGGAPRAARAA
VTRKPLGAA*
>GCA_000452945.1_BPMSHR5858_genomic_03491_1 group_1495
MNDGFLGMVRLAGVALQSVCFAIAVGVLLGDQWLARAASPWQAQVGRRLARTLRLAALGM
LLSSMLAFWVHCALMSETALWDAWPAVRSMLAGTGYGRAWLAGAALMLAVVGLSFARSGG
DARVPFALWLALAGVALARSNGGHPVDAGLFSAPVWIDWLHLLAISAWVGLVIVATYVVV
PRLADARGGERRNGAAFVQSLSDAATFALVVLLSTGAYNGWRGVGAPANLIGSAYGQVLL
LKLGLVAFAAALGGHNRFFEMPALLAAFKAPPDARAFVRSLRRFGVVLQVESVVLLGVIA
AAAVLASSPLPGTL*
>GCA_000452945.1_BPMSHR5858_genomic_03492_1 yobA_2
MKRIAHPKRAGLAALVAACAAIASAAAFAHGKLESAVPAPGGTVDTAPDALRLTFNESLE
SAFSTVKIADAGGAPVAAPKAKVDAANPRVLTLAVPKLAPGAYTVQWTVMTADAHKTKGS
YAFKVK*
>GCA_000452945.1_BPMSHR5858_genomic_03494_1 ftsP
MVSRRTFLGGSGAALLGAALVSKAGAASLPEAPTMAKAATQPPLAPPNGRPYTPVVTLNG
WTLPWRMKNGWKEFHLIAEPVVRELAPGMNAHLWGYNGQTPGPTIEAVEGEKVRVFVTNR
LPEHTTVHWHGMLLPCGMDGVGGLTQPHIPPGKTFVYEFQLEKAGTFMYHPHADEMVQMA
MGMMGTFIVHPKDRGAMPADRDFVFLMSAYDIDPGSFTPRVNEMTDFNMWTWNARVFPGI
DPLPVRAGDRVRIRFGNLTMTNHPIHLHGYAFEVAGTDGGWIAPAARWPEVTVDVAVGQM
RAIEFTANRPGDWAFHCHKSHHTMNAMGHQVPNLIGVPQQDLAKRINKLVPDYMAMGSTG
GAMGTMEMPLPENTLPMMTGTGPFGPLEMGGMFTVVKVRQGLGRNDYRDPGWFRHPKGTV
AYEYAGALPDD*
>GCA_000452945.1_BPMSHR5858_genomic_03500_1 vgrG1_1
VFGGYDTGQMSGQYTYTLELRSWLSFLAENRNSRIFQDLSVPQIVEQVFQGHQRNGYRFE
LEGTYEPREYCVQFQETDLNFVKRLLEDEGIYFWVEHEPDRHVVVISDTQRFEDLPLPND
TLEYLPDGEESRAIQGREGVQRLQRTRRIKSNNVALRDFDYHAPSKQLDSDAQVEQQSLG
GIPLEYYDYAAGYRDPEQGERLARLRLEAIQADAHALGGEANARALAVGRAFTLVGHPAL
SRNRRYYVTNSELTFIQDGPDSTSQGRNVAVKFRALVDDQPFRPLLTTPRPEVPGIQSAT
VVGPEMSEVHTDKLGRIRVHFHWDRYKTTEADASCWIRVSQAWAGKGWGVIAMPRVGQEV
LVTYVDGDLDRPLVTGIVYNGENPTPYDLPKDIRYTGLVSRSIKRAGGYQNASQITFDDQ
RGAERVMIHAERDMQQTVERNSSTSIAQDLNLSVKGTSTSVVGISISFTGISVSYTGLSV
SFTGVSASFTGVSTSFTGVSTSFTGVSTSFTGVDTGFKGVSTSFTGVDTSFKGVSTSFTG
VSTSLTGSSNSVTGVSNSMTGISSSWTDVSMSTTGQSQSITGVSLSYTGTSNSMTGTSTS
VTGTSTSITGTSMSNTGSSTSITGTSMSTTGSSTSVTGSSVSTTGSSVSTTGSSVSTTGS
SVSTTGFSFSYTGVSYSDTGIDLKKVGMQVKS*
>GCA_000452945.1_BPMSHR5858_genomic_03504_1 rimM
MAGHDSGNAKRGRSPSFGVFVRKPVERASAKGTSDGAVDSQAIRIDAAQSWPDDAVEVGA
VVDAYGLKGWVKLAAHAGAGRGGDALLKARDWWLQKGAERKFARVTQAKLHGDTVVAHPD
GSVDRDTALALRGARVFVRRGDFPALAADEFYWVDLIGLDVVNEAGVALGKIADMIDNGV
HSIMRVEYPATGKDGRPKTGERLIPFVGVYVKAVEQAAGRVVVDWEADY*
>GCA_000452945.1_BPMSHR5858_genomic_03508_1 group_4880
MSERYDVEYPVMPPAPRKGRGAVGNLQGRYETVEREAVDDGWTRDDDGEPRAPLRTQVFE
ERARTILTRNASPDIPFNVSLNPYRGCEHGCIYCFARPTHSYLGLSPGLDFESRIYAKVN
AAELLARELAKPRYVPEPIALGVNTDAYQPVERERRITRQVIQVMHDHGQPFAAITKSSL
IERDLDLLAPMAERRQVMAAVTITTLDPELARALEPRAATPSRRLRTIRALRDAGVPVGV
SIAPMIPFVTEPDLERVLEACADAGATHASYIVLRLPWEVAPLFTEWLAAHFPDRAERVM
ARVRDMRGGKDYDADFSRRMKGEGMWAELLKQRFRMAVKRCGLNERARGILDFSQFCAPR

RSKPPPPVRPRAAAQTGDQPQLSLF*
>GCA_000452945.1_BPMSHR5858_genomic_03517_1 group_4881
MDAAAPTSPRSADASRAAPQPTRVRTADGLELACYRWPSTAPSCAAPRATVALVHGLAEH
AGRYQAFAERLNAAGIEVVAIDLRGHGRSPGERAWAERFDRYLDDADALVASAARENTPL
FLMGHSMGGAIAALYAIERAAARHASLAGLILSSPALAPGRDVPQWMLAMSRFISRVWPR
FPALKIDAALLSRDPAVVAANRADPLVHHGSVPARTGAEILDAMRRIEAGRAALRIPVLV
YHGTADKLTEPDGSRDFGAHVGSPDRTLTLYEGNYHETMNDLERERVIGALIDWIAARVP
ARG*
>GCA_000452945.1_BPMSHR5858_genomic_03520_1 cysM
MAYKTIEDTIGNTPLVQLVRLPDDEIRARNNVILAKLEGNNPAGSVKDRPALSMIRKAEE
RGTIKPGDTLIEATSGNTGIALAMAAAIRGYKMVLIMPEDLSIERRQSMAAYGAEIVLTP
VKGGMEFARDLADQMQREGKGVILDQFANPDNPLAHYEGTGPEIWRETEGRVTHFVSAMG
TTGTIMGTSQYLKERNPTVEIVGAQPEEGSRIPGIRKWPEAYLPKIFDRSRIDRVENVSQ
AASEAMARKLAAVEGIFCGISSGGACEVALRVARQAENATIVFVVCDRGDRYLSTGVFPA
*
>GCA_000452945.1_BPMSHR5858_genomic_03530_1 aroA_1
VSGFYFDKLVIFGVGLIGGSLALALREGASGGRREVVGVGRSSASIGRALALRVIDRAAA
LDDDAQLRDALAGADLVLLAAPVAQTGPLLARIAPFLDASTIVTDAGSTKSDVVAAARAA
LGARIGQFVPGHPIAGRESSGVDAALADLYVGRNVVLCALPENAPDALARVEAMWRAARA
DVRAMSAEQHDRVFAAVSHLPHVLSFALVEQILGESDAELKFSYAAGGFRDFTRIAASSP
EMWRDVCLANRAALLDELDAYTAVLARLRAAIDAGDGAALEAVFARSRAARAAWRERGAK
PAPAVRSDTSGTGSHMEHLDLGPFSHAQGTVRLPGSKSISNRVLLLAALAEGETTVTNLL
DSDDTRVMLDALTKLGVKLSRDGGTCVVGGTRGAFTAKTADLFLGNAGTAVRPLTAALAV
NGGEYRIHGVPRMHERPIGDLVDGLRQIGARIDYEGNEGFPPLRIRPATISVDAPIHVRG
DVSSQFLTALLMTLPLVKAKDGASVVEIDGELISKPYIEITIKLMARFGVTVERDGWQRF
TVPAGVRYRSPGTIMVEGDASSASYFLAAGALGGGPLRVEGVGRASIQGDVGFANALMQM
GANVTMGDDWIEVRGIGHDHGRLAPIDMDFNLIPDAAMTIAVAALFADGPSTLRNIGSWR
VKETDRIAAMATELRKVGATVEEGADYLVVTPPAQLTPNASIDTYDDHRMAMCFSLVSLG
GVPVRINDPKCVGKTFPDYFDRFLALANA*
>GCA_000452945.1_BPMSHR5858_genomic_03531_1 pheA
MDDELNSRLKPLRERIDAIDTQLIALLNQRAAVALEVGEVKKHFNAPVFRPEREQQVIAR
LQDMSAGPLASEHISAIWREIMAASRDLEQTIHVAFLGPVGTYSEQAMFDYFGQSIEGLP
CPSIDEVFRSVEAGAATFGVVPVENSSEGAVSRTLDLLLHTQLLIGGELSLPIHHNLLTQ
TGKLDGVKRVCAHAQALAQCQQWLASNAPHLERQAVASNAEAARLAADDATVAAIAGDRA
ATHYGLQIAYALIQDDPHNRTRFAVIGQEPAGPSGHDQTSLIVSVKNEPGAVFKLLEPLA
RHGVSMTRFESRPARVGTWEYYFYIDIEGHRDDAAVQGALAELGRKAAFLKILGSYPRAR
*
>GCA_000452945.1_BPMSHR5858_genomic_03537_1 mupP
MSPSSPSFAAPLSDAPRLDACEAVLFDLDGTLADTAPDLAAAVNKMQRSRGIAQTPLDAL
RPLASAGARGLIGGAFGIAPADAEFDALRDEFLANYAADLCVHTTLFPGIGALLDDLDAR
GVRWGIVTNKAARFTDPLVALLGLAARAACVVSGDTASHPKPHPAPLLYAADRLSLAPER
IVYVGDDLRDIQAGSAAGMPTVAAAYGYCGDGAAPADWRAQHLVETTDDLQRLLRVLRYN
A*
>GCA_000452945.1_BPMSHR5858_genomic_03539_1 group_2474
MRIHELALFLRAWLRNPRQVGALAPSGPALAALMTAQIPGESARVIELGAGTGVFTRELL
SRGVAADRLVLVEADPAFAATLRRRFPALRVMNIDAAELGITHGLFGDARASTVVSGLPL
VAMPVEQAIAIVHGAFAMHLGPGGAFYQFTYLPRCPIPVRRLAAIGIRAERIGIAWANMP
PAAVYRLQRCADFSGSPSSLGRPWPSARPPSTAA*
>GCA_000452945.1_BPMSHR5858_genomic_03540_1 group_19751
MKLLTQLAVASAVAGALAAAATAAFAQNSPGMPGGILNDQFRLNEHPQMQFAASAPTKKY
QSGKKSDLRRKGDMGDPNGCNLKCPMD*
>GCA_000452945.1_BPMSHR5858_genomic_03542_1 nuoE
MSPHSVAPDALVRAHAQPGRSLVAILHAIQDDAGYVPPACVEPLAKALNLSRAEVHGVLT
YYHHFRTAPPAHVTIRLCRAEACRSMGGEALVAHAQARAGCRIDGGHGDRVALESVYCLG
LCAQSPSLTINDEPHAKMSPARFDALFDAAVRTKEPA*
>GCA_000452945.1_BPMSHR5858_genomic_03543_1 hndC
MSTRIHVPRDSAALAVGADALARAIEAQAAARGIAIELVRNGSRGLLWLEPLVEVSTPAG
RIGYANLTAADVPALFDAGWLDGAAHPACVGIVDEIPYLKRQQRLTFARIGITDPLSIDD
YVAHGGLEGLRNALALDGAAACETLVESGLRGRGGAAFPAGVKWRTVREAAAAQKYVVCN
ADEGDSGTFSDRLVMESDPYCLIEGMIIAGIATGATVGHIYVRSEYPHAIAALGEAIARA
RAAGWLGASVLGTAHAFELHVAKGAGAYVCGEETALLESLEGKRGVVRAKPPLPALSGLF
AKPTVINNVITLATVPIIFARGAAFYRDYGMGRSRGTLPFQLAGNVKRGGLVELAFGVTL
RELLFDFGGGTASGRPARAAQVGGPLGTYLPEGQWDIPLDYEAYAAVGAVVGHGGIVLHD
DTSNLAELAEYAMHFCALESCGKCTPCRIGSTRGVETIARIRAGDTSERQVRLLRDLCDT
MTAGSLCAMGGMTPYPVLSALDHFPEDFGLAAHASHAAPAA*
>GCA_000452945.1_BPMSHR5858_genomic_03544_1 napA_1
MTSSAFDSSRQGCGSGQCACRRAAQTRRADPFDDTDYGTPTRHADTDVTLDIDGRAVTVP
AGTSVMRAAIEAGINVPKLCATDSLEPFGSCRLCLVEIEGRRGYPASCTTPVEAGMKVRT
QSDRLQALRRNVMELYISDHPLDCLTCAANGDCELQDMAGAVGLREVRYGFDGKNHLSDA
KDESNPYFSYDPSKCIVCNRCVRACEETQGTFALTIASRGFESRVAASAGEAFMDSECVS
CGACVAACPTATLIEKSVARLGQPEHEIVTTCAYCGVGCALKAEMKGEQVARMTPHKNGL
ANEGHACVKGRFAWGYATHKDRITKPMIREKITDAWREVSWDEAIGYAAARFRRIQDEHG
RDSIGGITSSRCTNEETYLVQKLVRAAFGNNNVDTCARVCHSPTGYGLKTTLGESAGTQT
FASVGSADVIVVIGANPTDGHPVFGSRLKRRVREGAQLIVVDPRRIDLVDGPHVKAVHHL
QLRPGTNVALVNALAHVIVTEGLVDEAFVAERCEPHAFDVWRAFAARPENSPEATADITG
VPADAVRAAARLYATGGRAAIFYGLGVTEHAQGSTMVMGIANLAMATGNLGIEGAGVNPL
RGQNNVQGSCDMGSFPHELPGYRHIGDAAVRARFDEAWSTTLQPEPGLRIPNMFDAALDG
SFKGLYCQGEDIVQSDPNTQHVAAALASLDCLVVQDIFLNETAKYAHVFLPGATFLEKDG
TFTNAERRISRVRRAMRPLSGYADWEVTLMLSRALGYDMHYAHPSEIMDEIARLTPTFAG
VSYALLDELGSVQWPCNDAAPQGTPTMHVDQFVRGKGKFVITQYIASPEKVTPRYPLILT
TGRILSQYNVGAQTRRTENVRWHGEDRLEIHPHDANDRGIRTGDWVGVASRAGQTVLRAL
VTERMQPGVVYTTFHFPESGANVITTDSSDWATNCPEYKVTAVQVAPVAQPSEWQRAYTR
FRAEQLALLEQRTAERAPAIATGK*
>GCA_000452945.1_BPMSHR5858_genomic_03547_1 cpoB_1
MESAFDRAFAAHRAGRLDDAEHGYRAALAANPADADALHLFGVLRHQQGRHEEAADLVGR
AVGLRPNDAALQLNLGNALKALGRLDDAIERFRNALTLAPAFPLAHYNLGNAYAAQERHD
DAVDAFKRALALTPGDASIHNNLGNALNALGRHDDALEAFRRALELRPGHAGAHNNLGMA
LAALGDTDAAIAHFRAAIAAEPHFVAAHFNLGNALDAIGQHAQAQHAFEAALALQPRFAL
ALFGLANTLAARGRHRDALPHYERAVGLDPSFVLAWLNLGTAHHALGAHEMALRAFDQAL
RLDPSFTLAQMHRAVTLLTLRDFARGLPAYEARHALPGAAPLGPLPRWQGEPIANRTLLV
RAEQGFGDTLQFVRLVPLARARCARLILQVQPALLPLIAPMAARWRVSVVSADAARTPAA
DLVCPLLSLPFALGLEYDAIPSRTPYLDVPDAARRRFRGSLGGQAKRKFGIAWSGSAPVQ
DNRALPLDALAPLFALAGIDWIVLQPTLSDSEHAALDAHPDAARIHRLDGLTDFAATAAL
VDRLDGVVAIDTAVAHLAGALGKPLWLMLPVAADWRWSTGDDSAWYPRARLVRQSEPGRW
DDVVATVAGAIAHG*
>GCA_000452945.1_BPMSHR5858_genomic_03548_1 group_19150
MSTASPAIQQESKARTVFRVVSGNFLEMYDFMVYGYYASAIAKTYFPSGNAFASLMLSLS
VFGAGFLMRPVGAIVLGAYIDHHGRRKGLILTLALMALGTLTVATIPGYTTIGVLAPILV
LLGRLLQGFSAGVELGGVSVYLSEIATKGNKGFYTSWQSGSQQVAVVFAAFVGVLLNRAL
PVEQMTSWGWRIPFLIGCLIVPFLFLIRRSLKETDEFLAKRRRPSMGEIMKSMLENWGVV
LAGMGMVIMTTVSFYMITAYTPTFGKEVLHLSAIDALVVTVCVGLSNLVWLPLSGALSDR
IGRRPVLIAFTALTILTAYPAMQWLVGSPSFLRLLGVELWLSFLYGSYNGAMVVALTEVM
PADVRTAGFSLAYSLATTIGGFTPAISTLLIHETGNKAAPGLWLGLAAICGLIATLVLYR
SPEARNQYKTT*
>GCA_000452945.1_BPMSHR5858_genomic_03549_1 carA_1
MSENTSARADQSGSSSAATPPEHTGQAPSSSSVTDKPPAGATVHVGAFAAQIAAAQDIGD
PGSAPSGAIAEPGAPGIAAGGPGVEPAAVSAGPGAHAAAPSGATGAGASAAGAAAAGASA
KAGSPPPGFGAPPDFEASRPPPPGAATPAPPAYLKQSDTPWSVFGRIIAARARRLFDRAG
QRITQRTLRIGVSARIFHPEVGAPGLRGKTLQYLEESIAHWVMSRDVLVFMIPTVGHQGM
LHPSNIRLRDYAKHLDGLLLQGGADVSPQTYAASDARPEWPGDRVRDMYELELLHEFVES
GKPVLGVCRGCQLINVAFGGSLYQDIASDVPTAGAHVSEHYDQHRHSIRFPDGSTLANMF
PGRREAIVNSIHHQAIRDIGRDLNIEAVSAEDGIIEGIRYRRAPFVVGVQWHPEFHRAGG
PELLDCTPLLDTFLRAARETRL*
>GCA_000452945.1_BPMSHR5858_genomic_03551_1 dsbB
MNNLTLSLRRERRLLVLLALVCLALLAGALYLQYVKNEDPCPLCIIQRYFFVLIAVFAFI
GAGMASGAGVAVTEALIVLSAAAGVGTAARHLYVQLNPGFSCGFDALQPVVDSLPPARWL
PGVFKVAGLCETVYPPIFGILLPGWALIAFVLIAVPVAVSLLRHRGRLR*
>GCA_000452945.1_BPMSHR5858_genomic_03553_1 group_2475
MTPTFKDKIARAPKAELHIHIEGSLEPELIFELAQRNGVKLAYESIDALRAAYAFTDLQS
FLDIYYAGASVLLTEQDFYDMTAAYVERALADHVVHAEIFFDPQTHTERGVPIETVVAGI
DRALADAEKRGFSSKLILCFLRHLSEESALATFDAALPLFDRYARRLIGVGLDSSERGNP
PSKFARVFAKARERGLKLVAHAGEEGPPEYVTQALDVLKIDRVNHGVRSAEDAALVARLA
GEKIALTVCPLSNLKLCVFDDLTKHTLKTLLDRGVAVTINSDDPAYFGGYVNENYFATVD
ALQLGEADVYTVIRNGFEASFVSADERAALIAKLDAHWHAA*
>GCA_000452945.1_BPMSHR5858_genomic_03554_1 adeP_1
MDLYQGTAAPAKRSALERHFGIRESGSRLRTEIVAGVTTFLTAMYIIVVNPGILSQAGVP
FAAALTATVIVSFLGSCAMGLYARNPVLVAPGMGMNALFTFVMVHGGKMPWQTALGCVFW
AGVLFATLAAFNARKLVVDAIPANLRHAVSCGIGLFISLIGLVNAKFIVGDPVTIVHATS
LNPVIVTFLAGLAVTTVLVVRRVTGALMLGIVATTLLAIPIGRVWGDGSAYWPAAIATKT
LVNWNGLVAAPDFSALGQLDLLGSLKVAYWPFIFVMLFTAFFDALSTFMAISEAGNLRDR
DGNPRNIRQSMIVDAFSAVVSAPLGTSPANAYIESAAGISAGGRTGLVAVVAGLCFVPFL
FLSPLLSLVPAIATAPALVLVGVFMMESITKIEWHRFDEAIPAFLAMILIPLTYSITEGI
AYGFLAFVVLKLFTGRRGDVKPAMWVVAALSLLLVAQL*
>GCA_000452945.1_BPMSHR5858_genomic_03565_1 group_15429
MKRVAFALAALAVLANAAPGAAQEGRLPQPLRATDVASLYASGSGAPLVVEVWSLDCGYC
RENAAHLVEWQRRHPQVRVALVALDSLDEHGQELAGALAQMKLPAAVRQYANAEPMPERL
RAALDPRWRGELPRTLWIGEDGTRRAKSGLLAPAVLDAWLQRREP*
>GCA_000452945.1_BPMSHR5858_genomic_03580_1 gmk
MTDSNRDGAAAHSLHAGVYPGNLFMVVAPSGAGKSTLVNALLSKDPEIRLSISYTTRKPR
PGEQDGQHYHFTTVEDFRERHARHEFLESAEVHGNYYGTSRVWIEEQMKIGHDVLLEIDW
QGAQQVKKQFRNAVGIFILPPSLAALEERLKKRGQDEPNVITRRLLAAGSEIAHAAEAQY
VVINETFEHALAELECIVAATRLRFTSQYARHAELFVELGIHLPHAE*
>GCA_000452945.1_BPMSHR5858_genomic_03586_1 intA_2
MHTQQWTNTLATYAYPIIGALRPADIGTDLVRKVLEPIWLTKNETASRLRGRSFPRFFSG
TPESVVPLI*
>GCA_000452945.1_BPMSHR5858_genomic_03595_1 minE
MSILSFLLGEKKKSAAVAKERLQLIIAHERVGGRPPADYLPALQKELVAVISKYVKISND
DIRVSLERQDDLEVLEVKIEIPQA*
>GCA_000452945.1_BPMSHR5858_genomic_03596_1 minD
MAKIIVVTSGKGGVGKTTTSASFASGLALRGHKTAVIDFDVGLRNLDLIMGCERRVVYDL
VNVIQGEANLNQALIKDKKCENLYILPASQTRDKDALTREGVEKVINDLIAMDFEFIVCD
SPAGIESGALHAMYFADEALIVTNPEVSSVRDSDRILGILSSKTKRATEGKEPIKEHLLI
TRYNPKRVTEGEMLSLDDISEILRIKLIGVVPESEAVLHASNQGLPAVHLDGTDVAEAYK
DIVARFLGEDKPLRFTDYQKPGLLQRLFGSK*
>GCA_000452945.1_BPMSHR5858_genomic_03597_1 minC
MSLKKSPFFELRSGSVDTLLFIVKTADLDALRAELVKRFEATPEFFADDVVAIDVRRLAD
HERVPLDDIRGMLNDVRMRAIGVVAQPEQHAWAASAGLPLLEARDRRAPSSKAADEAPVQ
QAEPAAPAAGQAALFEQAGPTLADAGAPPESPAPAVAAQSATLVVDRPLRSGQQIYAKGD
LVVLGPVSYGAEVIAEGNIHIYAPLRGRALAGVHGNHDARIFCTCLEPELISIAGIYRTT
ENPLPADVLGKSVQIRLEQEKLMIEPLRLT*
>GCA_000452945.1_BPMSHR5858_genomic_03599_1 group_884
MAIGVKNSASNTVEVSINHWGTDGDTKPFKMAPGASDSWNRTDLRGFVMYVQLGGSATPY
YVLSTSNIVIYDDKVTDSGQTLLPANKRFG*
>GCA_000452945.1_BPMSHR5858_genomic_03605_1 ftsK
MAKAPYTAQAQALPHRMSRLLTEIRWILQVALLAFLLMALLSYSRRDPSWTHAAQVDHIA
NWAGRVGAWTADIMLLLFGLSAYWLIVLLGRRVAANYRRITRHDALPGEPEKPAGWLAEG
FAFVLVLLASDGIEALRMWSLKVQLPRAPGGVVGETVARGVAHALGFTGGTLALLIALAI
GLSLYFRFSWLSVAERVGDAIINAFTLAKLRREAERDRKLGEAAAVKREGKVEEERVRIE
EHEPVTIVPPVVTPAKSERAEKERQQPLFTDLPGDSTLPAIALLDPAPTSQETISADTLE
FTSRLIEKKLKDFGVEVSVVAAYPGPVVTRYEIEPATGVKGSQIVNLSKDLARSLSLVSI
RVVETIPGKNFMALELPNQRRQTVRLSEILGSEVYADAPSMLTLGLGKDIGGKPVCADLA
KMPHLLVAGTTGSGKSVGINAMILSLLYKASAEQVRLILIDPKMLEMSVYEGIPHLLCPV
VTDMRQAGHALNWTVAEMERRYKLMSKLGVRNLAGYNNKIEDAKKREEKIPNPFSLTPDD
PEPLGRLPNIVVVIDELADLMMVVGKKVEELIARIAQKARAAGIHLILATQRPSVDVITG
LIKANVPTRIAFQVSSKIDSRTILDQMGAESLLGQGDMLYLPPGSGLPVRVHGAFVSDDE
VHRVVEKLKEHGEPNYIEGLLEGGTIDGDEGSAAGTGEANGESDPLYDQAVEIVIKNRRA
SISLVQRHLRIGYNRAARLLEQMEQSGLVSAMSSSGNREILTPARDAE*
>GCA_000452945.1_BPMSHR5858_genomic_03608_1 group_4890
VHPRLTLAIAVLEAIATLAFAISGFVEARKNRLDSVGTFVVALATAFGGSTMRDVLLERR
PFYWVEHQAYVIAIFGLSLVAPLVLRMASRLSAERLLLAADAIGLGIFSISGTSIALDAQ
MPRFVAAMMGVTTGVFGGVIRDVLCNDIPLILRDSRPYATCAFAGCWVYLLLVWLQFDSV
YSVLAATAFILVARLVTYQLDIRLPH*
>GCA_000452945.1_BPMSHR5858_genomic_03609_1 msmX_1
MASLSIRDVYKTYPNGVPVLKGVDIDIEDGQFLILVGGSGCGKSTLLNMIAGLETVTKGE
IRIGDKVVNDLSPKDRDIAMVFQSYALYPSMTVRENISFGLNIRKVPKNEQKQIVDRVAA
MLQIEHLLERKPGQLSGGQRQRVAMGRALARDPALFLFDEPLSNLDAKLRIEMRAEIKLL
HQRLGTTIVYVTHDQIEAMTLGDRIAVMKDGVVQQFGAPQDIYDSPSNLFVAGFIGAPPM
NFINGKLVEQGSGVGIELDTGAMRGVLNLPFDAKRMNGHVGRDVILGLRPERITDARSAH
NGEGARLQPVDVTVDVTEPTGPDTHVFAQVNGKRIVSRVHPAANPQPQQKLSLLFDVSKA
VLFDPSTEARIA*
>GCA_000452945.1_BPMSHR5858_genomic_03611_1 melD_1
VAAPLSGNGTSAAAAHRTSPLSAFADRWIPKLVLAPSVAIALIFIYGFIAVTGYLSLTNS
RLLPNYDFDGFGRYSDLFGNDVWWTSAANLGWFGVPFIGICVVLGLFLAILLDQKIRNEG
ALRAVFLYPMALSFIVTGTAWQWILNPGLGLEKVMHDWGFTNFSFGWLDDPDKAIFCVVI
AAVWQSTGFVMALFLAGLRGVDGEIFKAAQVDGATLPTIYRKIVIPSMRPVFFSVLLILC
HITIKTFDLVVALTAGGPGTSSSLPAMFMYTYSFNRGQLGVGAASSVMMLATVVAVLVPL
MYLESRSTRNAA*
>GCA_000452945.1_BPMSHR5858_genomic_03612_1 group_4893
MKIRAIMGALGAAGLLFGAAAAQAAENVTVLHWWTSGGESKAVGVLKDDLQKQGYVWKDF
AVAGGAGAAAMTALKTKVISGDAPSAAQIKGPLIQEWADQGVLVNIDAAAGDWKQNLPPE
IDKIIKYKGNTVAAPFSVHRVNWLYINKAALDKIGAKPPATWPEFFQVADKLKAAGIQPV
AMGGQPWQDLTLWEDVVLSQGADFYRKALVELDQKTLTSDKMLEVFNTVRKIQGYFDSGR
NGRDWNLATAMVINGRAGMQFMGDWAKGEFEAAGKKPGKDYICAAVPGTANAYTFNVDSF
VFFQQKGQKAATPGQIALAKTIMTPAFQEQFSLLKGSIPVRLGVKMDKFDDCAKKSYADE
QTAIKSGGYVPSLAHGMAQGDATAGAISDVVTKFMNSQQDAKSALAALARAAKVK*
>GCA_000452945.1_BPMSHR5858_genomic_03616_1 fliY_2
MKAFKPMFTSALVAALLGVAAASAHAADLLDSVKQAGVLKIALEGTYPPFDYRNGDGQLE
GFDVDVAKEVAARLGVKPQFVTTEWSGILAGLQAGKFDVIVNQVGITPARRQALDFSQPY
VYSAAQLIQRADDKHDYGAPGALQGKRIGVTLGTNFADLAKTLQGVDVKTYPGAPEKLRD
VAAGRIDVSMDDRLMLPYLIKNAHMPLRPGATINGGETQMGIPFRKGNPKFAKALDDALA
SMRADGTLKKISVHWFGVDTSVPSAQ*
>GCA_000452945.1_BPMSHR5858_genomic_03619_1 dcyD_1
MNTTTLSTKLNAFARLNLIDAPTPLQYLPRLSAHVGRDIHVKRDDCTPLAMGGNKLRKLE
FLAADALGQNADVLVTAGAIQSNHVRQTAALAAQLGLGCVALLENPIAAARDDYLQSGNR
LLLDLFDVRAHVVDGLDDVDRQLEAAARRLRDEGRRPYVIPIGGSNPLGALGYVRAGLEL
AQQIRAAERDFAAVVLASGSAGTHAGLAFALAHALPGLPVIGVTVSRTDAQQRPKVRHLL
DGMSGLLDVALPAGARIDLWDDYFAPRYGEPNRAGIDALRLLARTEGLLLDPVYTGKAMA
GLIDGVARGRFDGNGPVLFVHTGGAPALFAYQDACRASR*
>GCA_000452945.1_BPMSHR5858_genomic_03622_1 group_4897
MQRDRHTRWLLTSGDIRKDAPFFGTLGVLIGVAQLVGFRCFHKADWGTKLLFEHIVFDTL
IVALIVVWLARIPTEWLRIERKREFPMLDKLIAHVARRAASFAVTAASVVAGVAAVAALS
GSPVHAVKFAFFCAYLLSIGEAVLNPLIAPGQSRLNGVAKALLIGMPLGFQLTG*
>GCA_000452945.1_BPMSHR5858_genomic_03623_1 group_4898
MTAFLLIWSPKKWPWPELPEMAARVKAGEAVADVWGCGFSRSILPGDRVFLHRVAQEPKG
VFGSGYVTRGPYEVPDPQTKRGYRLCIDFVYDWLVDGHSEVVIPRDALRVHPFSVQTWDA
QTSGTSIKPVAEGALEKRWAELTGKRKPPKRA*
>GCA_000452945.1_BPMSHR5858_genomic_03631_1 group_7037
MNAWLSWRPSERHAQMLRAVLRRGTRVSHHLFSVVGCCAVAVALALWLLPNVRGTLAAKV

MPFVSAAVQAGPARLLSGHPLPTFGPANADESESVNADAAPSTTAAPADAGAMLDAARNS
PSPVSLAKLIPTQRVAADARDDRVLASNREQALVATYLARRYRVAQEPVGQLVKAAFQTG
RDVGLDPLLILAVMAIESGFNPYAESGVGAQGLMQVMSKVHSDKFEYFGGTDAALQPVVN
IQVGALVLKDCIARGGSLAGGLRLYVGATTPDDGGYGAKVIGERDRLRDVARGRKVSIYA
SQQPAQGAVTVATVSTSSGATQKRVRTTLDGAHPLTIKAAAAPKAPQQDDVSADTAASSK
HETASPELGT*
>GCA_000452945.1_BPMSHR5858_genomic_03633_1 group_5695
MSFAPTSMLSDGFFLSLSLCLDIGLVNVAMLSLTLSHGFRPGFWLGVGSCVGDLVYAALA
LAGMAVLLRFEAVRWVVWLGGGAVLLFLAWKMACEALFPEPARNIDDGAPAALARASTSR
NFARGMLLAMSSPSAILWFAAVGGALIAKAGATTPASASVFLSGFFLGGLAWTLFLCTLA
SQGRKRAGAGLMRACHVVSALLFAYFAYSVIVGGYRDLILNAAA*
>GCA_000452945.1_BPMSHR5858_genomic_03655_1 group_4901
MTSTIHSLAVARGASPASSTASAASASPAAPRFALGLPPRPALLSRRAWLALVALCIAVG
VGVPLAALVAPEASAFHLSSYAMTLAGKLMCYAIAALALDLVWGYCGILSLGHGLFFALG
GYAIGMYLMRAIGRDGKYGSDLPDFMVFLDWHRLPWYWSGTEHLAYALALVVLVPAALAW
VFGFFTFRSRVKGVYLSIITQATTFAAMLLFYRNETGFGGNNGFTDFKRIAGFAITSPGT
RTVLLLTTFAALVASFVAARAIVTSKLGRVVTAIRDGETRVMFLGYSPLAYKLFVWVVSA
VLCAIAGALYVPQVGIINPSEMSPGNSIEMAIWVAVGGRGTLIGPIVGAFAVNGAKSVFT
AYFAEYWLFFLGLIFVLVPLVLPRGIMGLVETVMRKATSR*
>GCA_000452945.1_BPMSHR5858_genomic_03661_1 ureC
MTLRLSRRAYAEMYGPTTGDRIRLADTELLIEVERDHTLYGEEVKFGGGKVIRDGMGQSQ
LPAADVADTVITNAVILDHWGIVKADIAIKHGRIAAIGKAGNPDIQPGVTIAIGAATEII
AGEGLIVTAGGIDTHIHFISPQQIDEALASGVTTMIGGGTGPATGTNATTCTPGPWHMER
MLQAADGWPINLGFLGKGNASRPQPLVEQIEAGAIGLKLHEDWGTTPAAIDNCLTVADDT
DTQVAIHTDTLNEAGFVEATVAAFKGRTIHTYHTEGAGGGHAPDILKVCGEANVLPSSTN
PTRPYTINTLDEHLDMLMVCHHLDPSIAEDLAFAESRIRRETIAAEDILHDLGALSMLSS
DSQAMGRVGEVIIRTWQTAHKMKVQRGALTGDGARNDNFRAKRYVAKYTINPALTHGIAH
EVGSIEPGKWADLVLWEPAFFGVKPAMIVKGGMIAVAQMGDPNASIPTPQPVHYREMFAT
RGGALARTSLTFVSQLALDAGISARYGLAKRLVPVRGCRTVTKRDMIHNAWQPAIRVDPE
TYDVVADGALLTCEPAAVLPMAQRYFLF*
>GCA_000452945.1_BPMSHR5858_genomic_03664_1 ureG
MNAPRFAAPARRTKKLPPLRVGIGGPVGSGKTTLLEMLCKGMRERYDLVAITNDIYTKED
QRLLTIAGALPEARIMGVETGGCPHTAIREDASINLEAVERMLVRFPDADIVFIESGGDN
LAATFSPELSDLTIYVIDVAGGEKIPRKGGPGITKSDLLVINKTDLAPLVGANLEVMASD
TRKMRGERPYVMCNLKALDGVADVIAFIENKGLLTV*
>GCA_000452945.1_BPMSHR5858_genomic_03674_1 pglF
MTRAFCPKASWLSLGAFLFDLMAVVCAWLIAYVVRFNGAMPPEFLKGAFVALAWVVPLYG
LLFHVFGLYRGLWVFASLPDLLRISKAVVGGGLIVMIGAVMLQPVPIIPRSVLVLSPMLL
FLAMGGARALYRATKEFYLYGGLIGQGKPVLILGAGSAGASLARELSRSGEWRLAGLLDD
DPAKHGREVYGYKVLGSIGEVADWAEAAKAEYAIIAIPSASVEAQRRFATLCVRAGVRAM
VLPSLTALMPGQGILSQIRQIDLEDLLGREAVTIDTPHVEALLRGRVVMVTGAGGSIGSE
LCRQILKFQPAQLIAFDLSEYAMYRLTEELRERFPDLPVVPIIGDAKDSLLLDQVMSRYA
PHIVFHAAAYKHVPLMEELNAWQALRNNVLGTYRVARAAIRHDVRHFVLISTDKAVNPTN
VMGASKRLAEMACQALQQTSARTQFETVRFGNVLGSAGSVIPKFQQQIAKGGPVTVTHPE
ITRFFMTIPEASQLVLQASSMGQGGEIFILDMGEPVKIVDLARDLIRLYGFTEEQIRIEF
SGLRPGEKLYEELLADDETTTRTPHPKLRTAKAREVPDHLLDELLPWLMQHRVLSDDEVR
RDLRRWVPEYQPAVGPTLQSVPTGNGLVAGIGREAGR*
>GCA_000452945.1_BPMSHR5858_genomic_03686_1 rmlC
MAIQVTATALPEVKIIEPKVFGDARGFFFESFNAREFTELVEPGIEFVQDNHSRSARGVL
RGLHYQIQHAQGKLVRVVDGEVFDVAVDVRRSSPNFGKWVGVRLSAENKRQLWVPAGFAH
GFVVLSESADFLYKTTDYWYPEFERSILWSDKQIGIEWPIDREPLVAAKDAAGKAISEAE
VYA*
>GCA_000452945.1_BPMSHR5858_genomic_03691_1 pyrC'
MKIHIKGGTLIDPAAGTQRQADVFVAAGKVAAIGAAPADFNAAKTIDATGLIVAPGFVDL
SARLREPGYEHKATLESEMAAAVAGGVTSLVCPPDTDPVLDEPGLVEMLKFRARNRNQAH
VYPLGALTVGLKGQVITEMVELTEAGCIGFTQANVPVTDTQVLLRALQYASTYGYTVWLR
PLDAFLAKGGVAASGPVASRLGLSGVPVAAETIALHTLFELMRVTGARVHVARLSSAAGV
ALVRAAKAEGLPVTCDVGANHLHLIDVDIGYFDAQFRLDPPLRAERDREAIRAALADGTI
DAICSDHTPVDDDEKLLPFAEATPGATGLELLLSLTVKWAREAGVPLARALAAITSAPAD
VLKLPAGRIGEGAPADLCVFDPNAHWRVEPRALKSQGHNTPFLGYELPARVCATLVAGQV
AFERR*
>GCA_000452945.1_BPMSHR5858_genomic_03697_1 rubA
MEYKSWMCLICGWIYDEEAGLPEEGIAPGTRWEDVPINWTCPECGARKEDFEMVQI*
>GCA_000452945.1_BPMSHR5858_genomic_03699_1 groL_1
MAAKDVVFGDSARAKMVEGVNILANAVKVTLGPKGRNVVLERSFGGPTVTKDGVSVAKEI
ELKDKLQNMGAQMVKEVASKTSDNAGDGTTTATVLAQSIVREGMKYVASGMNPMDLKRGI
DKAVAAAVEELKKISKPCTTNKEIAQVGAISANSDSSIGDRIAEAMDKVGKEGVITVEDG
KSLADELDVVEGMQFDRGYLSPYFINNPDKQVAVLENPFVLLHDKKVSNIRDLLPVLEQV
AKAGRPLLIIAEDVEGEALATLVVNNIRGILKTVAVKAPGFGDRRKAMLEDIAILTGGQV
IAEETGLTLEKATLAELGQAKRIEVGKENTTIIDGAGEAVNIEARVKQIRTQIEEATSDY
DREKLQERVAKLAGGVAVIKVGAATEVEMKEKKARVEDALHATRAAVEEGIVPGGGVALI
RARTAIAGLTGVNADQNAGIKIVLRAMEEPLRQIVTNGGEEASVVVAAVAAGKGNYGYNA
ATGEYVDMVEAGVVDPTKVTRTALQNAASVAGLLLTTDAAVAELPKEDAPMPGGMPGGMG
GMGMDM*
>GCA_000452945.1_BPMSHR5858_genomic_03701_1 group_27
MKQNFKLTSIALSAAVAFGVAAATPAVAAGLDTLPVSASVTDLGGAGSGSAMLALAGGAQ
HGSIHASALGSADASISNVALDNLTGTLANAASTLQNAAANNPLSGVANNAVGTLTNLAN
NNPLPGALSNAVSTLQNAAANNPLSGVASNAVGTLTNLANNNPLPGALSNAVSTLQNAAT
NNPLSGVASNAVGTLTNLANNNPLPGALSNAVSTLQNAATNNPLSGVAGNAVGTLTNLAN
NNPLPGALSNAVSTLQNAAANNPLSGVASNAVGTLTNLANNNPLPGALSNAVSTLQNAAT
NNPLSGVASNAVGTLTNLANNNPLPVALSNAVSTLQNAAANNPLSGVASNAVGTLTNLAN
NNPLPGALSNAVSTLQNAAANNPLSGVASNAVGTLTNLANNNPLPGALSNVAGALAGAAS
NGSGALTGAAGNVTGALAGAGSTAAGAISGAMSNNPLPGVVNNVVGTLANAIGSNPITPI
TSVASGLANALSVANPAALTAAANTVAGTLARTANGTPVAGAIGGLVAALPVANPAGALT
SAANNAASTIATVAGTNPAAAIGGVAGALTGAAGTGVATASQLGSVGSALMGSGAASAGK
VLTSGSAAFGSAAASAGSLLTTGAAATSSVVNSLGSSVGAVVASLPNLSVSSSKSTAAAS
NPLAPVSSTVATLVGALPK*
>GCA_000452945.1_BPMSHR5858_genomic_03702_1 yybR
MRPKRFDSKSGCSVEVTLSVIGGLWKPVILFHLLKEKKRFMELTRLMPNTTQRMLTLQLR
ELEADGIVARHVYPQVPPKVEYELTPFGKTLAPVLISLREWGDAYRTHQLGAGSASSAEP
ARGAASCDAPARPRSRAAGAMKAA*
>GCA_000452945.1_BPMSHR5858_genomic_03713_1 group_4907
MTVNPTLAFIATCIGFGFIPGPALLQTVSLTLQHGRRAGVLSALGIHLGAFFQICAVAIG
AVVVLDTSPWLYHALRIAGGGYLIWLGIQRIRTREANDGGDGAPSTVPKNVISSSALIEA
SNPKSALFYLSFLLQFVDPSASLDVGWQLFLLGAGANLLFSLADLTCIALAHPLRKRAAP
GGTAMTIGRYLAGGLFIALGVAAIVER*
>GCA_000452945.1_BPMSHR5858_genomic_03729_1 pucD
MNQQAEPFLSVAADAERDAFRQVHVSRPHESAHLHVSGRATYIDDIPLVAGTLHAALGLS
AKPHARIASMRLDAVRAAAGVVAVLTAADIPGTNDCGPIVHDDPVLADGVVQYVGQPMFV
VVATSHDAARRAARLAQIEYEERPAILTAQAARAADSYVLPPMRLARGDAAARIAHAAHR
DAGELTLGGQEQFYLEGQIAYAVPKEDGAMHVYSSTQHPSEMQHLVAHALGLASHDVLVE
CRRMGGGFGGKESQSGLFACCAALAAWKLQRPVKLRPDRDDDMMITGKRHDFHYRYDVGY
DDAGVIEGVSVDMTSRCGFSADLSGPVMTRAVCHFDNAYWLPDVSIVGRCGKTNTQSNTA
FRGFGGPQGAFAIEYIVDSVARALGRDPLDVRRANLYGKTERNVTPYGQTIEDNVLPELI
AELEATSAYRARRAATRAFNAASPVLKKGIALTPVKFGIAFNVTHFNQAGALVHIYTDGS
ILVNHGGTEMGQGLNTKVAQVVAHELGVDFARIRVTATDTSKVANTSATAASTGSDLNGK
AAQDAARQLRERLAAFAAEHVGGGAARAADVRFAHDEVRVGEVAVPFEELVAHAYRARVQ
LWSDGFYATPKLHWDQAALRGRPFFYYAYGAAVSEVVVDTLTGEMRVLRADALHDVGASL
NPALDIGQVEGAFIQGMGWLTTEELWWNDGGRLMTHAPSTYKIPTVNDCPPDFRVRLFDN
RNAEDSIHRSKAVGEPPLLLPFSVFFAIRDAIAAVGDYAVNPPLAAPATGESILRAVRAV
RAARAGA*
>GCA_000452945.1_BPMSHR5858_genomic_03735_1 pucG
MSTTDHSPIPCPVVVPLDAILPEEPLLMMGAGPVPIPAAVAKANTIVINHLGATMAKIIE
QVKEMARYVFQTRTKWVLGVAGPGSAAMEMAVSNLAWAGTKVLSIKNGFFSARMAEMAMR
VGADVATIDVPDRSVASLDEIADAIARERPEIVTIVHGETSNTVWNRHLKEIAAIAKAAG
ALVVVDAVCTLSTMPLDMDAWGIDAVITGGQKGLSSIPGVSLIAFSDQAWERMKSRPEPN
THWCLDMALAENFWHNAGYHYTAPVSGVLALHEALRLVCAETLENRYARHLRCSLALQAG
ITGMGLELYAPEACRLNSVVGIAAPRGLTPGLICGHISKQYQVEISGSFGLPIVRIGQMG
EQCREHNLFRTLHAFGRTMVDLKVAVDLPAGVAALEQALSGRAA*
>GCA_000452945.1_BPMSHR5858_genomic_03736_1 group_5744
MNQIQTMRVFVCVAELQSFRRAARKLGVSNALVTRSIAMLETHLNTRLIHRTTRNLSLTE
AGIRYLDGCRALLEEFDQLEASVERAVHEPVGTLRVAVSGLLSPGRITPLVDSYRRQYPK
VRVQLTLTEGALDVIEAGYDAGLVTGGRLDGNPALIGHPLMPNTLALCAAPSYIERHGAP
HRPDDLVRHACIALPADQHASTWRFVDPDHFTHVVSLQPVYTVNSASMVRAATLAGTGIS
VLPESYIADALESGELVRILGDYRIDDPDTQLSIVYPNRQFLPARTRSFVEHALYHLGGQ
KTETNGHYFMREAGTAKRPDVVTGLQ*
>GCA_000452945.1_BPMSHR5858_genomic_03737_1 ldhA
VRVILFSSRQYDHDSFDAANQSFGYRLHFQPSHLDAETAILARGYEVVCPFVNDTLDTAV
LETLAAGGTRVIALRSAGFNHVDLAAAARLGLTVVRVPAYSPHAVAEHAVALILALNRRL
PRAVARTREGDFSLNGLLGFDLFGKTVGVIGTGLIGSVFARIMTGFGMRVLAHSLPPHDD
ALIALGVRYVPLDALLAEADIVSLHCPLMPATHHMIDADALARMKPGAMLINTSRGGLVD
TQALIDALKTGQLGHLGLDVYEEESGLFFEDHSDRPLQDDVLARLLTFPNVIVTSHQAFF
TREALAEIAHTTLSNIRAWQDGAPQNVVDAGRPG*
>GCA_000452945.1_BPMSHR5858_genomic_03738_1 bluB
MRFDEPAIAAVYRAIFERRDMRHFTSAPVDPAVLARLLRAAHHAPSVGFMQPWRFIRVTD
PALREQIHALVDAERRATADALGERHDEFMRLKVEGVRECGELLVVALADGRERHVFGRR
TLPEMDLASAACAIQNMWLAARAEGLGMGWVSLFDVDALRRLLAMPDGATPIAVLCLGHV
DAFYAKPMLEQERWATRMPLEACLYENGWDAPPTSARNT*
>GCA_000452945.1_BPMSHR5858_genomic_03740_1 ycaD
MKTILTRDFLALILSVAVVGLGTGATLPLTALALTEAGHGTRIVGILTAAQAGGGLAVVP
FVTAITKRLGARQVIVASVVVLAAATALMQFTSNLVVWGVLRVVCGAALMLLFTIGEAWV
NQLADDATRGRVVAIYATNFTLFQMAGPVLVSQIAGMTHVRFALSGALFLLALPSLASIR
KTPIADEPHHDAHDRWTRVMPKMPALVVGTAFFALFDTLALSLLPIFAMARGVASEAAVL
FAAILLFGDTAMQFPIGWLADKLGRERVHLGAGCVVLALLPLLPAVVTTPWLCWPLLFVL
GAAAGSVYTLSLVACGERFRGSALVTASSLVSASWSAASFGGPLVAGALMEQFGGDALIG
VLIVSAIAFVGAALWERRALPMQAARRGR*
>GCA_000452945.1_BPMSHR5858_genomic_03742_1 hmgA
MTLDFSKPGEAGYQSGFANEFATEALPGALPHARNSPQRAPYGLYAEQFSGTAFTAPRGH
NRRSWLYRIRPAAVHRPFELVSGERRIVAEFGDSDDVPPTPPNQLRWDPLPMPAQPTDFV
DGWVTMAGNGSAAAMSGCAIHLYAANRSMRERFFYSADGELLIVPQEGRLFIMTELGRLD
VEPFEIAVIPRGVRFAVALPDGRARGYVCENFGALLRLPDLGPIGSNGLANPRDFLTPHA
SYEDREGAFELVAKLNGRLWRADIDHSPFDVVAWHGNYAPYKYDLRHFNTIGSISYDHPD
PSIFLVLQSQSDTPGVDAIDFVIFPPRWLAAEDTFRPPWFHRNVASEFMGLVHGVYDAKA
EGFVPGGASLHNCMSGHGPDADTFEKASSIDTSKPNKVGDTMAFMFETRTLIRPTRFALD
TAQLQANYFECWQGLKKHFNPEQR*
>GCA_000452945.1_BPMSHR5858_genomic_03743_1 pcaK_1
MSAAANPARVLEIERVIDDTHRPAFHAMLLALCGLCLVIDGFDAQAMGYVAPSVIAEWGV
KKQALGPVFSASLFGMLLGALGLSVLADRIGRRPVLIGATLFFALAMLATPFATSIPILI
ALRFVTGLGLGCIMPNAMALVGECSPSAHRVKRMMIVSCGFTLGAALGGFVSAALIPAFG
WRAVFFVGGAVPLALAAAMAASLPESPQLLVLRGRHDAARAWLAKFAPRLAVPPDTRLVV
REAGPRGAPVAELFRSGRARVTLLLWAINFMNLIDLYFLSNWLPTVMRDAGYASGTAVIV
GTVLQTGGVIGTLSLGWFIERHGFARVLFACFACATIAIGLIGSVAHAFVWLLAAVFVGG
FCVVGGQPAVNALAGHYYPTSLRSTGIGWSLGVGRVGSVLGPLVGGQLIALGWSNDALFH
AAAVPVLCSAVFVIGLASVTRRRGTAAPNVA*
>GCA_000452945.1_BPMSHR5858_genomic_03747_1 group_4917
MVPPTIPELVARAGKLPFLCDHLALADGAQGAYARLRDLRLESVYEPIYDVSMPGAPQST
SLSDAIERYGDELGFQAVTHAGGAPFDPIGAVADDQELVALDRLARSLHAINFFGAQRHG
LLFLRVHERLLKSVKYDHGKHFSTVLKGFGLPPERIVIELPAAAVAHKTFLGYLTKSYQH
YGFKVADKLPDPGRILAVESDMARPDYIKMDAAIALREGMVKALVGYAQRVRIPLIFDHV
VDETQFELLRQYDVRLVQGPVFSQPAAV*
>GCA_000452945.1_BPMSHR5858_genomic_03761_1 glyA
MFDRAQSTIANVDPEIWQAIQQENVRQEEHIELIASENYTSPAVMAAQGSQLTNKYAEGY
PGKRYYGGCEYVDIVEQLAIDRVKALFGAEAANVQPNSGSQANQGVFFAMLKPGDTIMGM
SLAHGGHLTHGSPVNMSGKWFNVVSYGLNEAEDIDYEAAEQLAHEHKPKLIVAGASAFAL
KIDFERLAKIAKAVGAYLMVDMAHYAGLIAAGVYPNPVPHADFVTTTTHKSLRGPRGGVI
LMKAEYEKQINSAIFPGIQGGPLMHVIAAKAVAFKEALSPEFKEYQQKVVENARVLAQTL
VKRGLRIVSGRTESHVMLVDLRAKNITGKAAEAALGNAHITVNKNAIPNDPEKPFVTSGV
RLGSPAMTTRGFGPQEAELVGNLIADVLEHPEDAATIERVRAQVAELTKRFPVYR*
>GCA_000452945.1_BPMSHR5858_genomic_03771_1 flhD_2
MEQQKDVFEEIAAFNQRYLRLVRRWLCEDAERARTVLGISGELAIRLAAMTPTQLEQLAD
SGELVCRLRADAMPGRA*
>GCA_000452945.1_BPMSHR5858_genomic_03773_1 kdsD_2
MAHDGSQMNHHNYLDSARQVFDIESRALASLSARVGDSFGDAVDAILRSSGRVVVCGMGK
SGIIGRKIAATFASTGTPSFFMHPGEAYHGDLGMVTSADTFLAISYSGETDEVIKLIPFL
KSNRNYLVALTGNARSTLAQAAHSHLDAGVEQEACPLQLAPTSSTTAALAMGDALAVTLM
KARGFRPENFARFHPGGSLGRRLLSKVDDEMTVDGLPFVDERAPAIDVLQAMTRGRLGLA
IVRRETGFGIVTDGDVRRAIEAYGDTLFRRAASDLMSADPAMVPLGTRVEDALLMMEARR
INALLVFDGEDVVGVFKK*
>GCA_000452945.1_BPMSHR5858_genomic_03775_1 kdsA_1
MNVAISPGVTAGNSLPFVLFGGINVLESLDFTLDVCGEYVAVTRKLGIPFVFKASFDKAN
RSSIHSYRGVGLDEGLKIFAEVKARFGVPVITDVHEVEQAAPVAEIADVLQVPAFLARQT
DLVVAIAKAGKPVNVKKPQFMSPTQLKHVVSKCGEVGNDRVMLCERGSSFGYDNLVVDML
GFRQMAETTGGCPVIFDVTHSLQCRDPLGDASGGRRRQVLDLARAGIAVGIAGLFLEAHP
DPDRARCDGPSALPLHQLEGFLSQMKAIDDLVKRMPALEIR*
>GCA_000452945.1_BPMSHR5858_genomic_03780_1 group_695
MIRVRDSLISLCGVVAGQAALFGCISLIGRFQGPEALGHFNYLLALATFAGTLLACRYEL
ACVSDNPADSFNAFVNVMVLAGAAVSIGCVAIAVAGRTDLYIVAAYGFAAFAQLAAGGYL
NSLRRYGWIAASRIAVNGSFLVGVLAATFTPTFRARSGSIDAFRAYAAITAAVALLMLIA
VALHGRRRGYSFAVTRAFFVEHSRFAKYILPSTLCGSVLTYALAIVIPHWFDAQSAGYFA
AAYRLGFFPVSLIAQSLGGVFRRDAMAAMAGDDAHMLVPKVFGAYARALALLAAGYALGG
ALLFGPLVELFFGARWNGAIAFFLCMIPLFAFQMIYVPLSQIFLAARAQRIDFLFQLGCG
AVLAGALVTAKLAGLPVLASVRLFSFSGAVLMIAGIALTFSVMSGSVSRRRAGA*
>GCA_000452945.1_BPMSHR5858_genomic_03784_1 pglC
MPAIFPRVVDIAVIVFGAFLPILIEASGGPHAEIFDGTLVAFAAALSLSVFPACGIYEAS
RRRSPVHLISRTALAWLVVQGGTVVLLYVLHRAQILSSAWFAYWTVTTGIGLLIFRAVTL
AIFGLLARASRQVKAATLDQVGHVAQRMRTSGIAKRIVKRAFDVAAASCLIVVLSPALAV
IAFLVKRDGGPAVFGHVRIGRDGRPFKCLKFRSMVMNADAVLKALLERDPHARAEWEREF
KLKNDVRITPIGRFLRRSSLDELPQLMNVVRGEMSLVGPRPVVEAELARYGEDVRYYLAA
KPGMTGLWQVSGRNDTSYATRVSLDVSYVKEWSLRRDLVILLKTVNVVLRGSGAY*
>GCA_000452945.1_BPMSHR5858_genomic_03792_1 lpxC_2
MSVPTGWATSQGTLARALTIDGHGLHTGRRVGVRILPARPEDGVTGIAFRRVEQGRTLAT
LPVDPALRRAQPLCTMLRNADGVGVRTVEHLLASLLACEIDHAIVELDAEEVPILDGSAT
PWVDAIRTCGRVALDAPKRFIRVLRTVVVTEGEGEQRREMRIEPAPRYELSVRNDLRGFG
EMHWGGALTPAAFATEIAPSRSYGRVKWAVPAILAGYLRGVPILRGARPSCTASIVGKRV
LGGMRLPDEFVRHRVLDLVGDLALAGAPLLARVSAVRPSHEMNFRLVDALLAEPGAWQWA
DFSEA*
>GCA_000452945.1_BPMSHR5858_genomic_03796_1 group_4926
MERHGVNGRSLLPRDPAYYLDARRHIPPAVPGKPTGYNLYERACHDIRYRMANALYAHRF
PHYKSHRPRNGLQEYAGLAYRAVQQHVRDREAENVTRDLLERKRRYYLFPLQLNSDSQIV
DHSPFGGICDAIAIVLHSFAENAPDDSWLVIKNHPLDTGLIGYRQFATALATELGIEKRM
AFIDAGHLPTLLDQCRGVVVINSTVGLSAVHHRRPLVALGTAIYSMPGLTWQGSLADFWT
EAGSPDMNLYQAFLDYVMHHTQINGDFYTRTGIEMSVAGAVSRLEAVS*
>GCA_000452945.1_BPMSHR5858_genomic_03799_1 gmhA_2
MENRELTYITNSIAEAQRVMAAMLADERLLATVQKVADACIASIAQGGKVLLAGNGGSAA
DAQHIAGEFVSRFAFDRPGLPAVALTTDTSILTAIGNDYGYEKLFSRQVQALGNKGDVLI
GYSTSGKSPNILAAFREAKAKGMTCVGFTGNRGGEMRELCDLLLEVPSADTPKIQEGHLV
LGHIVCGLVEHSIFGKQ*
>GCA_000452945.1_BPMSHR5858_genomic_03803_1 group_5621

MAQRKKYSVYGSCQAPALAKMLNSCPTFARDWELVEMEPCFVASEEQIDRHLAETIPKLD
LFLYQPVSEGYRGEKYSSVFLRNSMPPGGSALSVQYMHWEGYHPTVNSPYGLPPHPEGYV
DALIAGAVVMDVDKETYLRHLEEIGASLRIDIDEIESWCVDELKSREVGENDGGKQIDIS
VTDFILANCRQKRLFYTMNHPTAALMREIAARCMLALGYTYSDISFDQNLDPLDVTKMSL
YPIYRDCFDFSELNRMNEYQVLYKKKAYEPYLLEQFEWFERSPKADVSAFFDRVAANRRW
VRTALRRAFES*
>GCA_000452945.1_BPMSHR5858_genomic_03810_1 group_1508
MLESVIAVSNPQAALARARERYSDYSPAIDTPTALGYVFNSWRFLLETGDGQVRLRIQSE
FEGTDLEHRQLVLRYLLTDARASSISVDELLPASSPYSVDAAELVDNPSSVVSKRATMES
FDAPAQESSWFWRVWTIGRLRLTPLFLVTVVLPTAIATLYYGLIASDVYISESQFILRTP
KRTSEPGLGALLQGVSLSKTSDDTSVVQDYIKSRDALVVLEQKMPLRDAFGGRYADIFSR
FPGVIGRDGFEYFYRYFKNHVAVDVGTATAVTTLRVQAYTSKDAYLINKYLLEMAEKRVN
ELNDRSRQDSLRYALAEVAGAEAKVKAASTALSKFRSKASLFDPDRQSNIQLELVAKLRG
ELIAKQAQLSQLRMLSPQNPQIPSLAASIGALEASIASEKGGVAGEKNSLSDRSVEYQRL
ALEQSFGEKLLASALTSLEQARADAERKQIYLERVAEPNEPDVAMEPKRVRNIFACFILG
LAAWGVLSMLVAGIREHQE*
>GCA_000452945.1_BPMSHR5858_genomic_03812_1 group_4930
MEFPRYYKKLASLGIKSVFLIHDLIPLTHAEYTRPGVEHTHRRRIHTALGYASGLIANSR
STLESLAAEATRAALPLPPCAIAHLASGVEPQPPRQRLLDAPYFVMLGTIEPRKNHWFIL
HVWRRLIEQLGNAAPKLVVIGRRGWECENVIDMLERCASLPGTVIEEANCSDERLHAWLQ
HARALLFPSFVEGYGMPLVEALGLGVPVLASDLDVFREVAADIPDYLDPLDGPAWAARIR
DYARDDSHERTAQLARIEHFREPTWVDHFERVDAFLDTLH*
>GCA_000452945.1_BPMSHR5858_genomic_03820_1 group_505
MTEGEHAIRQLLETWFVASRRGDLTTVLDLIADDAIFMVPGKPPFDKAAFAAASRDAHPA
RASDAAPRIDGRYRIDELRVLGDWAYLRNFIEIDVTPPAGEPVRRAAHTLTILRKSEGRR
RLVRDANLLASAA*
>GCA_000452945.1_BPMSHR5858_genomic_03821_1 purM
MNPPKSAPDAQGLSYRDAGVDIDAGDALVDKIKPFAKKTLRDGVLGGIGGFGALFEVPKK
YREPVLVSGTDGVGTKLKLAFHLNKHDTVGQDLVAMSVNDILVQGAEPLFFLDYFACGKL
DVETAATVVKGIATGCELAGCALIGGETAEMPGMYPDGEYDLAGFAVGAVEKSKIIDGST
IAEGDVVLGLASSGIHSNGFSLVRKIIERANPDLSADFHGRSLADALMAPTRIYVKPLLA
LMEKIAVKGMAHITGGGLVENIPRVLRDGLTAELDQHAWPLPPLFQWLRQHGGVADAEMH
RVFNCGIGMAVIVSAADADDALRQLADAGEQVWKIGTVRASREGEAQTVVV*
>GCA_000452945.1_BPMSHR5858_genomic_03824_1 pcnB
VIKKLIRKLLGQDARTAQDAPQDPAEPRSAAPAAPAGRAKPARGAAKKPRNGAPTIVHAD
VHGIDPALISKNAIRVTDTLQQAGFRAFIVGGAVRDLLLGIAPKDFDVATDATPTEVQRL
FRRARLIGRRFQIVHVQFGQELIEVSTFRALVDAPQEPPADAAPAKRLKRDELDRRTHAV
DASGRVLRDNVWGEQHEDAARRDFTINAMYYDPASQTVLDYHDGMADMRARLLRMIGDPA
TRYREDPVRMLRVVRFAAKLGFDIEEKTREPIKELADLINNVPAARLFDEMLKLLLSGHA
LACLKQLRKQGLHHGLLPLLDVVLEQPQGEKFVTLALNNTDARVRAGKPVSPGFLFATLL
WHDMRQRFEQYSADGEFPVPALHRAMDDVIDMQTEKLAIHKRYSADMREIWGLQLRLEKR
SGRSALRLLEHQRFRAGYDFLLLRCESGELDPAVGQWWTDFIDGDAATREALLTQGGSRD
KSPRKRRRRGGRSRRTADSAEGAEGAARGESDAND*
>GCA_000452945.1_BPMSHR5858_genomic_03838_1 glnE
MTDASDLLSLSYSHYLARAAAARPALAERIAAWAAAPVTRAALDARLDELLAQGGQPPSE
DALKKALRQLRGEAFGAVAERDLAGRADVAEVTGTMTDLAEAAIQRALALLAAELEAQYG
EPRGPSGERLALGVVGMGKLGGRELNVSSDIDLIFVYEDDGETAGGARGPISVHEFFTRL
GRRLIGVLSEATADGYVFRVDMRLRPNGDSGPLVCSLGMLEEYFYVQGREWERYAWIKGR
LVTERASAAARRLAQQLDAIVKPFVYRRYLDFGVIGAIRSLHEQIRQEARRRATMRPDKA
DDIKLGRGGIREIEFSAQVFQLIRGGQDAGFRVQPTLAVLRHASASGLITEEVRAGLTHA
YLFLRTLEHRLQYRNDAQTHAMPVDPAERAALAASLGFADYAALIDRLDQHRAFVEAQFD
QVFADKADGGARREDDQAAGCIWSGALADDGADEALVARLAELGFADPAAVLARLQAVWR
SSRYAGLPESSRVRFDRVAHRALEAAPGIDAAHRDETVVRCFDLLETVGRRGAYLALLTE
YPAALRRVLSVLGATRWGGGYLIRHPQLLDELLDDEAIDSPFDWPAFKDALRRRLAVADG
AEHQMDLLRHAHQAEVFRILLLDLAGRLSVEHVSDRLSELADAMLDVTIEVVWSQLAKRH
RDTPCFAAIAYGKLGGKELGYASDLDLIFLYDDPDERAADVYTTFARRLITWLTTATGAG
TLFDIDLRLRPNGEAGLLVTDLDAFRRYQLREGDAANTAWVWEHQALTRARYSAGDARIG
AAFEAIRVQVLTTPRDAAVLAREIVEMREKVLAGHPNTTERFDLKHDRGGMVDIEFAVQY
WVLLHAARHPEMIRNTGNIALLREVSRFGLMSEEEAETVGAAYRTYRKLQHRLRLDGMEK
ARVEPERVAAERQAVAALWARVFGA*
>GCA_000452945.1_BPMSHR5858_genomic_03844_1 queF
MNPEHSPLGKATVYANQYDASLLFPIPRAGAREQIGIGAPLPFFGTDIWNAYELSWLNAR
GKPQIAIATFYVPAESPNIVESKSFKLYLGSFAQTAFESADAVRDALKRDVSAACGASVT
VRLATPAEFRKLQMDELDGLSLDRLDLDAHVYETDPSFLTASHGEAPVEETLVTDLLKSN
CPVTGQPDWGSVQIHYVGAPIDHAGLLRYIISFRNHTGFHEQCVERIFVDILRACQPVKL
AVYARYTRRGGLDINPFRTNYNQPMPDNARTARQ*
>GCA_000452945.1_BPMSHR5858_genomic_03846_1 dan
MHSHPEVADTLIVGAQLYDGTGAPSVERDVAIRGDRIVAIGNLTNWLAEQVVEANGRALA
PGFVDVHTHDDTHVIDAPQMLPKISQGVTTVIVGNCGISASPVALMGDPPDPMNLLGARA
AFRYPTFAAYVDALGAAKPAVNVAALVGHTALRSNQMDRLDRAATGAEIAAMRAQLEEAL
AHGALGLSSGLAYGSAFAAPTEEVMALAEPLANAGAVYTTHMRTEFDAILEAMDEAYRVG
RHARVPVVISHLKCAGPSNWGRSAEVLASLERARRIQPVGCDCYPYSRSSSTLDIKQATG
DIDITITWSDPHPEMAGKLLRTIAAEWRVSEQDAARRLQPAGAVYHNMSEDDVRRILSHP
ATMVGSDGLPNDPLPHPRLWGAFPRVLGYYARDERLISLEDAVRKMTALSARRFGLARRG
EVRVGYHADLVLFDAARVRDAATFEQPQQPAHGIEAVWVNGALSYRDGAPTGVRAGRFVA
RGERAPATPGDAF*
>GCA_000452945.1_BPMSHR5858_genomic_03871_1 algA_2
MTDLATKPTPAASHGTLPRILPVILAGGSGTRLWPLSREHHPKQLIDLIADESPLSATAR
RLTGIANAELADTLLLVCGEQHRFTSAAQVSARGLDARILLEPAARNTAPALTLAALEAL
AHHGDPVLAAMPADHAVSDTRAFQETIALAARYAQDGAIVTLGVLPRRAETGYGYIKVGA
PLAGANGATGGYAIDSFVEKPHQELAQQYLQSGDYWWNSGIFVVRASTWLAAIRALQPDM
HAACEAAWRGGTSEGAFFKADAAAFAACPSDSIDYAVMERLTGARALGIPGVIVPLTAGW
SDVGSWDAIWELLPKDEAGNVARGEVLFEGTEDSFARSEGRLVACVGLKDVVVVETPDAV
LVANKHRVQDVKTVVARLKSGRHEQVRDHRKVQRPWGYYDSIDRGERFQVKRIVVDPGKQ
LSLQMHYHRAEHWIVVRGTAKVTRGDETFLLSENESTYIAVGAVHRLENPGRIPLEIIEV
QSGDYLGEDDIVRFDDRYGRASAQHAPDAAKPADAELAPAAGPARALGAQ*
>GCA_000452945.1_BPMSHR5858_genomic_03873_1 rfbX
MTTVRKNFLLLMTLQISTYVVPLVTLPLLTRVLGPEAYGQLSFVLAVITYFINFTNYSFD
LTATPRIALAHDDLAARSKIFWATFAAQAAITVAGFAILLALTFAIERFAIDRTLLVIGF
GMTVGAALTPGWYFQGIQKLRIYSMTLFACRALSVPAMYLCVRSPADLIDAVAINAAVPL
VCGVLVFGYLIVRREVVFVRIGAADVAASLKGGWQVFLASTSVAFYASTNTVLLGFVSGN
VAAGYFAAGDKLIRSAISLLQPLKAAAYPHVSYLMHHAREDAMSFLRKLLVVQGAMVLAI
SLAIFALAPLAVHILYGSTYEPTVGVLRCMALIPFMAGMTDLFGVQTMLPLGMKTAFSRV
LISSGVLNVALLPPLAKCFAEQGAAAAVLIAETAVAATLAYVLYRKRVSIFARPAPRPAS
RG*
>GCA_000452945.1_BPMSHR5858_genomic_03874_1 pdeB
MAFVPSGPSCDVNARCARFVRSAVAVALAVALLGAALLSVERFGAARSRASAAIPASIGA
RLSGCAFIAVRIGAALSRRVPSPAHAAAPGVARPDFDAIERVFGPIRMRERRAHGATCTA
DAPDASHAGWIEPARVCSAAAHAAPGRGAEGAPNTMPGPSEAGAAAVLQSQPYLDMIDST
FREPAAIVLALSLLGMIALALLPVFRIRRLALRLGEAQGALEMSEARARAALVAVGDGVI
FTGRAGRVECLNPAAERLIGMLADDCRGRPLVSTLRLSRTATSASSGAFAASAGLDAPDS
PGSAASPDAPLGDLEDGGSCDATLYRADGGAIAVRATASSIAPPPGHARRACGSGRVLVL
KSLATEHELVRRLAWQTTHDPLTGLASRAEFERHVGFALAADVREPVALLFVDLDRFRIV
NDTCGYAAGDAMLAALAARLVSCAASADVVARLGGDEFCVLLDARDEASAVSAAERLRAS
VDGFVFVWDGQPFSVTASVGVALLGGPGRAPRVEDAVRLAGIACDVAKARGRNRVQLADP
HDHELAHHISDVSWCARVRQALEYDDFRLYVQPIVDTATQGATGLPRARRGELLLRMGAL
GEREGVAPPGLFIRAAERYGLVTDIDRWVVRTVLDALARTRSRRFSEYAINLSGISIGDE
RFLDYVLEQFARTRVAPALICFEITETAAIANLAGALRFMHELKALGCRFALDDFGSGMA
SLSYLKQLPVEYLKIDGSFVTGIANDAASLDIVASINDIGHAMNCKTIAEYVDSAATLQK
LAALGVDYAQGYYIGRPVPWCEAACA*
>GCA_000452945.1_BPMSHR5858_genomic_03875_1 novR
VTNRTPNQPFARPARLAQAEWQARVQLAAAYRIFDHLGWSELIYNHISLRVPGEDTHFLI
NPFGLHYREVCASNLVKVDVDGHSVGHADWPFNPAGVTFHAAIHAALPDAHCVMHVHTTP
TMAVCATRRGLSFSNFYAAQLHGRVAYHDFEGITVRPDEGRRIVDSAAGRPVLLLRNHGP
VAIGRTLAHAFSLMWLLHRACEVQVATHALGEPLEIDAPILDACARDSLNLDPRHGAGQD
AFDALQRIVDRIDPGYRS*
>GCA_000452945.1_BPMSHR5858_genomic_03887_1 group_725
VAKTSGTADEILNLPSGGGSVSGDGGDFSVDLNTGTATLKFDLAVPAGPNGITPPHTLQY
SAGAGDGAFGIGWSLGLMTIRRRITPATGAAEPAPPGACSLVGVGELVDMGAPRFRPIVD
ATGLLIEFTGASWTATDKTDTQYTLGTSANARIGGGALPAAWLVDRCADSAGNAIAYTWL
DVGGARVPQAIAWGTYRLDFVYEARPDVLVDGSYGAPVTLDKRCARIELHATTEAPSLVR
SWTLLYDDDGGRGRSLLATIREQGHAADGSILAAPERTFAYSSPGAPALVPVTGWTTPLS
DPNTDLVDLNGDGLPDIVHLGHGMPTMHPNLGGGQFGAPRPLARTAAPLRLSAPSVAFAD
MSGNGSVDVLVLDAPFSGYYPLAAPGGSAPAGFGRPIVFERAPAVSPSDVRLRFVDLNGD
GITDILLDTGRGWLAYLRDGPASWSDAPRVLPPARTPPVSLADPHVHLADMTGDGLMDIV
RVTGGGLTYWPARADGGWDAAIAMTPSPAFGRDFDPHRLALYDVDGDGCADLVYVGLQSV
TLWRNVGATRLAAPVAIPGTPVALPGSYRIVDLLGSGTAGVHFQLPSIAGASRQSYLDLG
GGVKPYLLTDLAHGAAQSTHIGYRTSTEYARDDAHAGAPWRTYHPFPIQCVARTDQTDHG
TGATTSTRYAYHDARYDPATRTFLGFGRVDSEQLGDASCPTLRVETTFHLGLDPADPARP
LTGDEALKLGALRRKPLVTATYGLDGSPLEHRPYSITRHAYDALLVASTADNGKRIAVPY
CTTSTEERWERQSAAVSARTVDYLAITSEGDVTSQRTRAQRAGIAAPDQDVTTTTTLATG
GKNLRLPARVTQTAPGGDIVSESICFYDGDAFAGLPEGQATRGLVTRIEDRVFDDAFVAS
VWGDTPPDLTQYGYHRLPGEAGSWWKTRRAHARGANASGPTLATKGPLGALQTLQYDASG
QRVVKVIDALGNAVAATTDARVFQTASLTDANGHRTADAFDALGRVVATIGPLDTPALPT
IAFTYTVGAISTVTSATRGTHGGADAVPALTWIDGTGNVLGKGTPAAAPGEWTVTHAVRR
NARGLVAASFLPYAATGANWQPPPADTGATTSTYDALGRLVQLTRPDGLVVTSRREGDTL
ITSETWPGGAALDVERQVYDAAGQLVSVSRNAGDHWVEQRYAYVPSGKVRDVTLPGGAHV
AFTLDLLGRVFAQQSPDTGRTVFLLDASDNQRARTNAAGQIVRTEVDAMNRATNVYHDAE
PAPRVRYEYADASGAPPADGIVANRYTRLWRITDEIGTVDFEYDEAGRKTATTRTVAATG
QRFVTRRAYDALGRLARATLPASAPGGAARTIAYGYAADGRLVSASGVVKDAAYDRFGRL
TSIDYENGASTLIDYAANAGGIARVRVLDAARNVLRDTTLTRGGGLVQTLASAHAGDDSV
DFGYDPLRRLTSAHYRQGATAADAHDWTFDDAFNATAATDAGVLTYEPGTHRLASVGGAA
VAFDAAGRMTSGRFGAATFDAADHLSAVTLPDATHIAHTYDYQGRRVRSTKNGAQTYFSP
IEDIEFQGDTAIVWITFGKQRIAADIGGALTFVHPNALGVMDLITDSGGAYGTRVRQTPF
GYARAADGAAPAGGVAAVALALGAADATGLVCHGLRWYDPRVGQFISPDPVVTSVYTVGA
WNPYVYCLGNPILLADPNGCSFLSVLEIIGVAILAAACVVGAIFTGGATLVALGVLSANI
GGWLLAGVALGSLGGAIAGELAAQKAGGNLWAGAFLGAFLGGATSLIGGALGGAAAAGID
TLIGGTKTFLSFVAAGAIQGTLAGAGTGLAIGYAGGKGNAESMLIAMAKGAAWGAVLGTL
LGAGIGAIAGTGISGAAKPDNFLNIGAFGQKFADFTSTSTAINSADNAAGVTESLAQLSM
PNGFNAGNLLGLLPNLVTTNEQAAGWFSIPIGWLGPGVLNDAGFAGLVDTSMALDQAGFS
YAHQISLLLGAAPYFIDYAATMAQIVDANGVNNFEMAFNNAFGSGSPSNTG*
>GCA_000452945.1_BPMSHR5858_genomic_03892_1 yajR
MNPFFGAQREWQDVPVPARAYARLDNLHLPHADAPDRRAALRFDLIDADWLDDQAGADNP
LFDLGRPHRMRAAWKRWFAAARGGDADDPFRAAPLYVLEVSVPFDAREWRKRLGLTREAM
ANDAHGEILAAFRRRWYELTVLREADARAGAMLGLLHRGIADEGLRVCCFAALAPAAHAA
GLTALGTLAGSPALTSASLKPLVRDVRATYLGVYDVGQANANALLGGIRGRDLHGVVRHG
VFPALYFDLGMPLQSQRHTAPSNVRFCFTAEPPIVLSHWHDDHLGGVALPTAAGCPTSQQ
MDWIVPDYSLATRHKALVTAIWAAGGSVHVFSASPAGRRLQAATASGHLLTLTRGTARDP
NNGGIVMQVSSAQQQARRFADPPRSWLLPGDCDYKHFPSTFTPTNVIALVAPHHGADLGK
GSVPVPPAAGGYLRLVYSFGRDNKYQHPRQGFVDAHHAAGWTHPVVPATGAITCVSGDDV
RTTNVHSPWPAKPRGVLVGWSKAPHSIPACTLGGGATNCTLPLAKV*
>GCA_000452945.1_BPMSHR5858_genomic_03897_1 group_4942
MEKPSEPTREPQAREPHLRSVRLTSDFSLPKLSAIEIGSYLLALFGLWLVLHLKLLGGLL
AGLLVYQLVHMIAPAIERHMSSQRARWVAVVLLSTAIVGGLAGLTIAVIEHFEHTVPNAQ
NLLGQVMQIVDQARTRTPEWISNLLPVDAEQMRAKAAGLMNRHMDQLQQGGKSVARGFGH
VLFGMIIGAMIAIGIDRHKLRQPLSTALVARISRFSDAFRRIVFAQIKISALNAFFTAIY
LLVALPIFHQRLPLSKTLVLVTFIAGLLPVVGNLISNTLIVAVSLSVGMGTAIASLVFLV
VIHKLEYFLNAKIIGGQIESRAWELLLAMLVMEAAFGVPGVIAAPIFYAYVKRELIYLRL
I*
>GCA_000452945.1_BPMSHR5858_genomic_03901_1 apt
MMSTSDAPLDPVEFIHSRIRTVPDWPQPGVMFRDITPLLQSAKALRVLVDLFVERYVDAK
LDYIAGLDARGFIIAPIVAYELSVGFVPIRKVGKLPYATQRESYALEYGTATVEIHEDAC
KPGDRVVIVDDLIATGGTMMAGKNLLERLGAVVVEGAAIVDLPDLGGSALLREAGLPLYT
VTEFPGH*
>GCA_000452945.1_BPMSHR5858_genomic_03904_1 kdsC
MSAPPATAAERASRVKLMIFDVDGVLTDGGLHFTAAGDAMKSFNTLDGHGVKLLGEAGIA
TAIITGRRSEALAARAKEMRIAHLHEGVENKLAVFGELLRALGLEPGECGYMGDDWPDLA
VMLRCGFAAAPANAHPDVIARAHWVAEARGGHGAVREVCDAILRAQHRYDALLAAACGA*
>GCA_000452945.1_BPMSHR5858_genomic_03911_1 ptsN
MNRLAKILPLENVVIDLSVTSKKRVFEQAGLIFENQNGIARSTVTDNLFARERLGSTGLG
EGVAIPHGRIKGLKHPLAAFVRLAEPIPFEAPDGQPVSLLIFLLVPEQATQAHLEILSEI
AQLLSDRDTRERLHTEPDRDELHRLLTQWQP*
>GCA_000452945.1_BPMSHR5858_genomic_03912_1 hprK
MDTSSINAQSIFDDNAAMLKLSWLTGHEGWERGFSADTVANATSSADLVGHLNLIHPNRI
QVLGEAEIDYYQRQTDEDRSRHMAELIALEPPFLVVAGGAAAPPELVLRCTRSSTPLFTT
PMSAAAVIDSLRLYMSRILAPRATLHGVFLDILGMGVLLTGDSGLGKSELGLELISRGHG
LVADDAVDFVRLGPDFVEGRCPPLLQNLLEVRGLGLLDIKTIFGETAVRRKMKLKLIVQL
VRRPDGEFQRLPLESQTVDVLGLPISKVTIQVAAGRNLAVLVEAAVRNTILQLRGIDTLR
DFMDRQRLAMQDPDSQFPGKLV*
>GCA_000452945.1_BPMSHR5858_genomic_03918_1 lolB
MPRFAQPRARGVALVVAAAAAVLLGACATQPPRAPAAAPDARQTEQNRAYTGRFAVQYQD
PLGNPRNVYGNFDWQEQGTNVSLELRSPLGQTLAIVKSAPGGASLELPNRQPQFAPDVGE
LMQRALGFALPLDGLRYWLLPTPSPTTPATTVRDPQDGARIKEIRQDGWTIDYVAYADAP
AVGVKRINLARQTPPLDIKLVLDR*
>GCA_000452945.1_BPMSHR5858_genomic_03929_1 nagL
MKLYSYFRSSAAFRVRIALHLKKLPFDYVPVHLLRDGGQQLTDDYRALNPDALVPTLVDG
DAPLQQSLAIVEYLDEAYPAVPLLPQAPVDRAYVRAIALQVACEIHPLNNLRVLKYLKHT
LKVDDDAKDAWYRHWIEDGFKSLEARLSGDPRTGKLCFGDTPTLADLCIVPQVFNAQRFS
IGLERFPTIQRIHDHAMTLDAFKAAAPAAQPDAE*
>GCA_000452945.1_BPMSHR5858_genomic_03931_1 group_19987
MSATQPEWIAVCRLDDIPPLGARVLTREAGTPIALFRTASDAVFALLDRCPHKGGPLSQG
IVHGEQVTCPLHGFNVAFSSGCALPPDEGCTSVFSVKRDGDTVLLERTELATLGVDDAAQ
PAALASPGAAA*
>GCA_000452945.1_BPMSHR5858_genomic_03955_1 puuD_1
MMGAHPSHVVVEKYVTAVVDGADALAMLVPALGARQTPSDVLAAIDGLLLTGSYSNVEPH
RYGGEPSAPGTLHDPARDETALALARAAIDAGVPVLAICRGLQEMNVVFGGTLHQQVHAA
GGFDDHREDKRDTLDAQYAPAHEIRLAEGGLLRRLLGGAAHARVNSLHGQGIAQLGAGLA
IEALAPDGLIEAVRVPHARAFALGVQWHPEWRHANDALSTAIFRAFGDACRERMRIRMRH
DAGRLPTHAIGA*
>GCA_000452945.1_BPMSHR5858_genomic_03956_1 puuR_2
MSNEVATRLRHVRMKYGLSQRELAKRAGVTNGTISLIEQNRVSPSVGSLKKLLECIPMSL
AEFFTFEIDENRSVVSRRGEMPNLGNETIGFYLAGSGVKDRNMCIMREVYQPLADTGAEM
LTHAGHEGGVVVSGQVELTVDGTTWLLDPGDSYYFESRLPHRFRNPSAEHVCEVVSANSP
PTF*
>GCA_000452945.1_BPMSHR5858_genomic_03963_1 group_6436
MATIATEFHFAAQQFSIDVLLEKYAKGEERTVDDIYRRVARGVAQAEPPELRDEIEARFV
EHLQRGALGAGRIMSAAGSPIAATLINCFVQPVGDCIQGVDERGLPGIYVALLQAAETMR
RGGGVGYNFSAIRPRGAYVHSTGSSASGPCSYIDVFDASCRTVESAGARRGAQMAVLDCD
HPDLLEFIEAKHSKGRWNNFNISVAVTDAFMNAVEQDLPWQLVHRAEPSPALRAAGDLRR
RDDGMWVYAERRARAIWEKIMRSTYDVAEPGVVFISKMNDDNNLRAIESIRATNPCGEQP
LPAYGCCNLGPLNLTRFVIDPFAELTGGESAFDWRALAERTRTQVRFLDDVLDVTLWPLP
EQRHEAQQKRRIGIGFTGLGDTLVMLGLRYDSQQGRDFAARVGREMRDAAYRASVELARE
RGAFALFDANRYLEEGTFASRLPEDIREAIRRDGIRNSHLLSIAPTGTVSLAFADNASNG
IEPAFSWTYQRTKIMADGSRQTFAVEDYAYRLYRELGGDVGALPACFVSALQMSAHDHLA
MMAAVQPYIDTSISKTVNVPADYPFDDFESLYFDAWKSGLKGLATYRPNETLGAVLHTSE
PDDALEEADLDPLRIAIDHRPKGELPAVIEKVEYLTQLGKKSLYVAVSFMEVTGRIGGDE
VTIERPIEFFIPVGQRDESQQWITATMRSLSLAARGGFVARNLQDLRKVSWDRGQVRLGE
TQRLDGRRIPLWHDSEVAALSYAIQQILHRRGFLDAEGNQVPSRLLAKLPRGGADSHAVA

ALLAAEAGSAGGEGDEETAAARGAGAGELRTMFGRKCNSCGANAVIRKDGCDFCTACGEI
GACG*
>GCA_000452945.1_BPMSHR5858_genomic_03971_1 group_4957
MNVSQTCLLITALMPFVWTMCAKSSSRYDNHDPRGYLARLDGWRARAFAAHQNSWEAFAL
FTAALVVAWHNGANMQRVDQLAIVFVASRVLYGVLYLLNWATLRSLVWTVGLVCVVWLFF
AAP*
>GCA_000452945.1_BPMSHR5858_genomic_03977_1 group_4958
MHFPHAALNGATLADADAQTDAGTRDIDAARIVLTAIRRWLQPCADADDGARCRRQSWRD
VLAAAGLRADGLVHFDMLMRFLMCDTCHPLDTRCRCACGPMNDEARLLQALAHLQSTRSE
AALRALADWLPRCADSGVLKIARWFSIALLDAGLAIRPPAPPAAYMH*
>GCA_000452945.1_BPMSHR5858_genomic_03978_1 ydjA
MNATACAAPAGAAVADDPRGLLRGLLSRRSRWPLTEPAPGPRELDAIFDAALRAPDHGNL
RPWRFVVIRGAARHAFGDLLVELADARAPDAPPGSHAHRRAKALAAPLIVALGAALNRTS
KVPEVEQLLAVGAAAMNMLNAVHQLGYGGFWATGADSYEPAMRDALGFGADERLIGLLYI
GTPSAPASDARRPDRAAHVREWLGPAR*
>GCA_000452945.1_BPMSHR5858_genomic_03979_1 group_4959
MKPFDEMLQPGDTVRAPYERLKQWLDTQDPASLAQKAHDAEGVFRKTGITFAVYGDAEAA
ERLIPFDIVPRIISGSEWNRLSQGIEQRVMALNAFLDDIYHRQEIVRAGIVPKHLIAHND
AFLPEMIDFRPPGNVYTHIIGVDIVRTAENQFYVLEDNARTPSGVSYMLENRETMMQLFP
ELFQQVKVRPVETYPQLLRQSLAAVCPPGGNADNPTVAVLTPGIHNSAYYEHAFLADQMG
VHLVEGSDLQVIGDRVAMRTTEGFRPIDVLYRRLDDAFLDPLTFRPDSVLGVAGIMDVYR
AGNVTIANAPGTGIADDKAIYSYMPEIVEFYTGRRAMLENVPTWRCAEADSLEYVLAHLE
ELVVKEVHGSGGYGMLVGPAASKAERDAFAAKLRAKPSNYIAQPTLALSTTPILTERGLA
PRHVDLRPFVLVSDRIRITPGGLTRVALKEGSLVVNSSQGGGTKDTWVLAD*
>GCA_000452945.1_BPMSHR5858_genomic_03980_1 group_4960
MYRYIERAENTARIVDAGLRMALTRTSDAPAEWSSVLVSSGADDGYRQKYETYAADTVAD
YLLRDRDNPSSVLSCIECARSNARMVRTALTREAWESVNGAWLAIRRALAQPIRASALPA
ILDEIKRETALILGSFYSTMLRNEIFDFAQIGAFVERADNTARILDVKYHLLLPSVSHVG
TILDNYQWESILRCVAAHRSYRWVYDVQYKPMNIADYLILNGRMPRSLRYCYGRVVSSLE
HLAKDYGVTHGCHETAAKIKQSLEDNTVERVFKSGLHEFLTDFIAKNNSLGLEIAQAYNF
D*
>GCA_000452945.1_BPMSHR5858_genomic_03981_1 group_4961
MRLAIRHISRFRFDEHAPHVLQRLRLRPQSGPSQTVRAWQVTIDGVEPTLTYADGLGNRV
DLVRHERDKPEFAIVAAGVVETQDRAGILGHIDAYAPPWIHERTTDLTRAGEAIRALAEA
LPIEHKSLDALHWLMTEVHGRVAYEPRAQTAPRDAEAALRSGAGTSRDHAHVFVAAARAL
RIPARYVSGYLLTDGSMQRIAEALHQASGVAQQAMQVRRGAHDAPSAPAAPDDASRRTGV
AMQEAAVATQPQSGHAWAEAYVEGLGWVGFDPFMNRCPDERYVRIAAGLDYRDAMPVYGP
GAQPLGVEISVIQSPELV*
>GCA_000452945.1_BPMSHR5858_genomic_03991_1 group_4963
MIEAAQFVEAARARGFDWYAGVPCSYLTPFINYVLQDPALHYVSAANEGDAVALVAGATL
GGRRGIAMMQNSGLGNAVSPLTSLTWTFRLPQLLIVTWRGQPGVPDEPQHALMGPITPAM
LDTMEIPWETFPTQADAIGPALDRAIAHMDATGRPYALVMQKGSVAPHALEHASMPERRA
HAAARATPGAAAPSAWPTRHDALARIVARTPADSTVVLASTGFCGRELYALDDRPNQLYM
VGSMGCVTPLALGLALARPDLTVVALDGDGAALMRMGAFATLGAYGPPNLVHLLLDNGAH
ESTGGQATVSQYVSFAGIAAACGYATAIEGDDLGVLDAALDAARAGEQAVEHGGAHFARL
SIRTGVPDGLPRPTVTPVDVKTRLMRHIGAMPA*
>GCA_000452945.1_BPMSHR5858_genomic_03994_1 group_4964
MTRTGMILLSLGTAAFVALLAWQGLGAVAATLASAGWGLALVAAFHLVPLVIDATAIAVM
FRPGEPGSRLGEALRARWVGEAVNSLLPAGQIGGPVLMVRYLAQRGARLADAAAAITVST
TMQALAQMVFALVGIAAFSAYATHDAAAHLRTPALIATAVLGGCAALFYCAQRRGLFGRG
LRAASTLLGPRDWSSLATRADAIDHALDRLYRERAKVRATFVLSFVGWIVGTAEVWLALR
FLGHPVSWLDALLLESVGQAIRGAAFAIPGSLGAQEGGYLLLAPLVGLPPDAALALSLAK
RARELALGLPGLLYLHFSERNWQRRRAPQPIAD*
>GCA_000452945.1_BPMSHR5858_genomic_03996_1 group_6364
MDPRKSATHRIPSPPPRTWDARLARRLVTPLVGTPVTPNHLTTLRLLIGLAGAWCLTQPG
FGWINAGALLIVLSNFVDHTDGELARISGQSSKLGHFYDLASDAFVTVALFVSMGAGVVA
AGGQMAASPVLLGAVAGAAVALIFFLRMRIESFAGKAGTKQAFVGGFETEDVLYLLPVVT
LLDGVEPFLLAASIGAPLFAAWVVIDYWRIVRRGAGAGAPNSTEIQPSK*
>GCA_000452945.1_BPMSHR5858_genomic_04003_1 group_4965
MNFLDHYNDELRQLRDAGARFSKEHPQVASALGLHPDAVTDPFVERLLEGVAYLSARVQK
RLDRECAEFAQQALGRICPLYTASTPAISTFAFHPDLGSPDAFRGNTLPRGSLIAAHLPG
RKLPVMFSTAREVTLLPLRLASVECSRSITGLPSLLAQRLASSHAVLRFRFEIEGGASIA
ELAREGDGFKPLHLSVAGDLPRAYALHRALLADTTAWYALVSTNRGDEVLPLPMSGIRLS
GVDDEEALLPEAFGGLPGLRLLREYFAQPTRLLGVYVDALAAIAAKAPSARAFDLFFALR
DAPGDLVGDVDASQFRLFATPAINLYSKRFDPVPYDANQPEQWIPVDRMRPAAHHLWALT
EVFVCETNGRAHRARSVLETAGYEGHEGSIRYGMRREDALLVDGVRRDRFDPLASHDLIA
VSMVDDTLEPDDVATITGRALVADRDWRPSALLDAELQLLDPTAVKRIECLWPASAPRGK
PSIDACWEAVSHVGRNPLALRASQQQDVTARIVEQLLLAIDKDDALDRQRLESLRSVVLR
SRFVAAGRATPTALVRATQVEIDIAESLHADRGGWLFGRLLAQALAEAATLNDGIEIVVR
LDGEAASTHTNVATRDGRLA*
>GCA_000452945.1_BPMSHR5858_genomic_04013_1 mlaC_1
MKRHLFAFVAAAVVSVSAFAQTAPVEVVKNAVEGTVGAMKADPAARGGDMAKITQIVEAR
FLPATNFERTTRIAVGDAWKQASPQQQQELYKQFRILMTRTYAASLAQLGSQDAKFTFKA
AGAGGADALVRSTVTTPGDSQSVGYRLGKIGNDWKIYDIDMSGAWLIQVYQGQFKAQLAQ
GGIDGLIAFLTKHNARAN*
>GCA_000452945.1_BPMSHR5858_genomic_04014_1 group_4968
LHIPFERDGDLMDIGSYPHWLQDVVGTVRAARDRVRFHEVFSLMRDSRLAPRQLAAFFVN
GWPVVEQFPKYMSMNLLKANGTNSSGEEKARRYLIRNIRVELNHVEHWVNWAEASGVPRR
QLTDGDSPPAALALSHWCWKSSSADTLAASIAATNYAIEGVTGEWSADLCRSDVYEMGFP
EAVRGRAMRWLRLHSSYDDKHPWEALDIVATILGQSPSTEQVRDVAAGIERSFRYFEMSL
SCCLDA*
>GCA_000452945.1_BPMSHR5858_genomic_04015_1 group_4969
MREQALGMNDGGGQAAELRPALDVWPILLMVSVLVCVVIVSLAVVSFLRVYIGGESTWSK
SQKDAVIYLIRYAETGDERAYKLYEAAIDKPYRLSLARAALQRKPPDRASAREAIIASGI
DPTDATAAALLLPALSWVAKINYALQCWMQADVELAEFRSVASQLHALVSSGHRDDARVK
ALEYRAWEINERVSGLPDRFSKAFSDEFRSSVALLLVCYFVASMFLMYLALVWARKAALQ
RLSIQYALDRSEAFAEAALSSVAEAVITVGLAKNVDFINNAAEQLLGYPAAACVGAPVSS
VLMLLDKETGHAVDIVDEFWAREERTPIKREVHLLRRDHSKIVVQTTISEMGDRVHGHIG
YVLILRNMTREQQFLESLAWQATHDVLTGLVNRVEFERRLEIALADTASGKERADSMLLM
LDLDRFKEINDTCGHAAGDAMLREVTQRFQSCLDDDDVLARLGGDEFGVLLPHGAGSWPS
KNKAERLRRSLEDFVFLWEGERFTVSVSIGVLELCKAPRNLEMAVKLADIACYIAKERGR
NRIQLADPSDLQQARHVGDVQWSRRIKTALENDGFRLYVQPIVHTRPDSRTPERAEVLLR
MMDSAGRDISPAAFLPAAERYGLMGLIDRWVIRTVFRKLSALPAREYHEYNVNLSGASIS
DERFLEFVIAELLSSGLEPSVICFEITETIAVKNLELASRFMGELRTIGCRFALDDFGAG
MSSFRYLRSLPIDYLKIDGEFVSNMMSDRVSYGVVSAINEVAHSMRCSTVAEHVESEVEL
AMLRELGIDFCQGYFFAQPSPWREGGY*
>GCA_000452945.1_BPMSHR5858_genomic_04016_1 lip_2
MKSRFASAAAAVRGARKTLAKVWAAVLVAAMCAFAAPAAHADAATGALDDYAATRYPIIL
VHGLTGTDDYFGVVPYWYGMRPDLQQHGATVFVADLSGFQSDIGPNGRGEQLLAYVQQVL
AVTGAQKVNLIGHSQGGLTSRYVAAVAPELVASVTTIATPHRGSPFADFVLGALSLDPTG
LSTPIFGALLNVFGILTSHTHNTNQDAIAALNALSTPYATQFDAQFPSAGLGAPGACASG
APSETVGGNVHLLYSWSGSAYQPITLLGLTTGALDKSTIPLVDPANVLDPSTLVFLTAGN
IMALKQAGLNDGFTSTCSSLYGTVISTDYKWNHFDEINQLLGIRGAYAADPVAVLRTHAN
RLKLQGV*
>GCA_000452945.1_BPMSHR5858_genomic_04017_1 group_1078
MVEGLPRATPRAAGATLTFSVARPSAALMARLMRGEVHVWRARAEEAGSYLKRDCLSLSE
HERAARLRYGAHRELFVFARAMLRIVLGEYLSADPARLVFDVEPGGKPVLFGRDLEFSVS
HASGAVLLALARKRVGCDLESLGRKLDIDALAAASLGERERDVFARTPRRAQARLLLQCW
TRKEALLKAHGAGLRRDPRELEMPWPARRAARDECACGDVRWAVADIALGTGWRAAVAME
ERGADVHGFCLGW*
>GCA_000452945.1_BPMSHR5858_genomic_04018_1 group_4970
MSKRILQIATAILAAVPVTTGALGMMGIHDPLYASLGVALPADATLDGNLRFYAGVWFGL
GLGAFWTIPNIERNGVLFRALWTMIFVGGIGRLISLVSLGAPFAPFIGFTVLEIVGAPLF
VWWQSRVAATAG*
>GCA_000452945.1_BPMSHR5858_genomic_04021_1 group_5322
MHTFMLALRNLQRNRRRSITTLLAMVVGVCAVLLFGGFSKDITFGLQTDFVRRSGHLQIQ
RHGYYQYGSGNPVAYGIGGYARLIAQLRDDPVLAPMIAVITPTLQFGGIAGNFEAGVSRT
VLAQGAIADEQDRMQQWNDYGFPLTPKPYPLVGTAPDAAIVGNGVARVLHLCAPLRVPDC
GDGDVARQAAPQAASGASGADAPADVLALAAAEAHAQESGRGERAGAAHIEVLAANAYGA
PNVGGFTVAKAEQQGVKELDDVYLAMHLPRAQRLVYGGDAPRVTAIEIQLRHTAQLPAAR
ARLDRLFGGRFEGQPLDVLDFAALNPFYDQTNRMFSMIFGFVFVLIGAIVLFVVSNTMST
AILERTVEIGTLRSMGVRRGGIQALFVCEGALLGVVGASIGVLVALALAFVVNHSGLAWT
PPARIDSVALTVRVWGEWRLIALTFVGLAFVAGFSAWLPARHAARLSIVDALRYA*
>GCA_000452945.1_BPMSHR5858_genomic_04026_1 group_1598
MAQVAFLFPGQGAFYAGATRALSGAYPVVQRAMQAIDAVSVRRLRRPLLATVWDGHIGAE
DLLQHAPDLLQLAIYATSVSYFEALRARGVRPDVLVGHSFGEIAALTCAGAFSIETGAEI
VCDRIDALRAAAPADGRMAAVGASPDAVRAHLAACFRGAPDARAGRVRIAVENHVSQTVV
SGACEYVDEFRAYCARNNISAQVLNSPYAFHHEDLENARIEFGRRLKAYKNKALEYPVYS
PILNRYYTAADDLGACLARHLVLPVRFGDGVARLKAAGIGAFVECGALNALSRISVRALG
PGAVKTFPGASSAGGELAGLENVLAYFNGGSVMNDDIRASNLQLDFDEFWRNSGPGIVDK
IKGELKRFFDARGLQATPAPQIVADHGVAAGFAGGLTAGAADYAPKVAAGVSSAAAPVAA
PIVAAAPAAPLAVAPAAPVAQAAAVPAASARVPRDRLFVELVAIYAEAMEYPVEVFTETV
ELEAELGIDSVKQTEIIQRISARYGLPPLPANFRAGDFKAMGQIVDFVYEHQGEAALVTS
*
>GCA_000452945.1_BPMSHR5858_genomic_04027_1 group_1897
MLTSVLVRLVAWSVRRPIWVVVLSLVIAALSSVYVAHHFKINTDISKLVENDPKWAALGR
AIDDAFPQRNQTILAVVEAPAPEFAGAAADALAEGLRRETDAGRIGQVSEPAGGPLFEHD
GLLFLPEQDVATTTAQLASARPLINVLAKDPSIAGLATTLSTTLGVPLQSGQVKLSGMAK
LLSRSAATVDDVLAGKPAAFSWRALVDADAAREPARAFVTVQPVVNYGALKAGEQASRTI
RATAQALKLDERFGAAVRLTGEQPLADEEFASVQDGALVNGIATLAIVLVILWIALRSKR
MIASVFVTLFVGLVVTAALGLMMVGSLNMISVAFMVLFVGLGVDFAIQYGVKYREERHRD
PNLDHALVGAAHAMGMPLTLATAAVAASFFSFLPTAYRGVSELGLIAGVGMFVALFTTLT
LLPALLKLLAPPGERKPPGFPRLAPVDDYLDHHRKPILIGTLAVVIGALPLLAHLRFDFN
PLHLKDPRSESMATLLALKDSPEASVNDVSLLAPSLAAANAAAQRLGALPEVGRTTTLST
FIPDAQPQKLATIAAAARELLPALTQPAAAPVPDAQRVAALKRASNLLEYASEDYPGPGA
AAAKHLSESLAKLAAADAATRERAEHAFSVPLKIALNQLAMLLQPSEITRENLPPQIVRD
WIAPDGRALVQISPKVVKGADPGDDAMLRRFAKAVKAAEPGAIGGPISILHSADTIIRAF
LQAAALSVVSITVLLWITLRRFGDVLRTLVPLLVSGVVTLELCVLLGMPLNFANIIALPL
MLGVGVAFKVYFVMAWRAGQTGLLQSSLTHAVLFSAATTATAFGSLWLSHHPGTASMGRL
LALALSCTLIGAVVFQPVLMGKPRTKRVTNQSQGIDE*
>GCA_000452945.1_BPMSHR5858_genomic_04032_1 glnQ_3
MSMVEFRNVSKSFGHVPVLKDIDLRIDAGEVVVVVGPSGSGKSTMLRCINALEKITGGDL
LVDGQSVKGKAAVIHGIRLEAGMVFQQFNLFPQMTALENVMFGPIQVRGASRAQARDQAM
ALLAKVGLEARANHYPSELSGGQQQRVAIARALAIRPKLMLFDEPTSALDPELRHEVLKV
MRDLANEGMTMIVVTHEIGFAKQVGTRLLFMDQGGIAEDGDPKTLIDHPPTPRLKDFLKH
VS*
>GCA_000452945.1_BPMSHR5858_genomic_04033_1 glnP
VQFDWSAIWAALPDLMDGVRLTVFIAFFGLAGGFLVGMIAGMFRAYGPAVLNVLAQVYIE
LIRGTPIVVQVMFLYFALPLLAHVRIDGLSAAVVAITLNSGAYLAEVVRGALLSIPKGLT
EAGLAMGLSMPRVLVKIVGPLAFRRLIAPLGNQCIVSLKDTSLFIVIGVGELTRKGQEII
AGNFQAVEIWTAVAAIYLVLTGVMTLTLRFVEKRMRIL*
>GCA_000452945.1_BPMSHR5858_genomic_04034_1 glnH
MGRRSFIKTIVALALVGASAAGAHAKELVVGTDTSFMPFEFKQGDKYVGFDLDLWAEIAK
GAGWTYKIQPMDFAGLIPALQTQNIDVALSGMTIKEERRKAIDFSDPYYDSGLAAMVQAN
NTTIKSIDDLNGKVIAAKTGTATIDWIKAHLKPKEIRQFPNIDQAYLALEAGRVDAAMHD
TPNVLFFVNNEGKGRVKVAGAPVSGDKYGIGFPKGSPLVAKVNAELARMKADGRYAKIYK
KWFGSEPPKS*
>GCA_000452945.1_BPMSHR5858_genomic_04035_1 group_1516
MAGSGRPAAPVIAVAGMAFEARIARGAGVEAVYAARADRLERALAEAARAHGCAGIVSFG
TAGGLAPDLAPGALIVADSVDGPFGVLDTDAGWSARIVAALAGTPAASRARRGKVAAVGA
PVTGAHEKDALHRAKGALAVDMESHIAGAFAAARGVPFAVCRAIVDPAWRTLPKAATAGL
RDDGSTAILPILRELAKQPSQLGALLQVAGDARAARATLVQARQAFERAGAWRIG*
>GCA_000452945.1_BPMSHR5858_genomic_04037_1 hpnD
LAVSNPVVDDNETDAAAVTSGSSFYLAMRILPAEQRDAMFQVYAFCRAVDDIADEGGARA
ERAAQLDRWRADIDDCYAGKPRASLVPLAREIGKFGLHRDDFHAMIDGMAMDAAEDICAP
DEATLDLYCDRVASSAGRLSVRIFGMPDEPGRVLSHHLGRALQLTNILRDIDDDAAINRC
YLPRELLAREGIAIADPQTIARDPLLPRVCATLVERALHHFAQADAVMDASPRAQVRAPR
IMSGAYRLILEAAVARGFAPPRAPLRKPRARMLLLAARYALF*
>GCA_000452945.1_BPMSHR5858_genomic_04043_1 group_19453
MTYRIAVIVGSLRRESFNRALANAVASLSPADLSFEPVDIGSLPLYSQDYDADFPDVAKH
FKQTIEAAQGLLFVTPEYNRSMPGVLKNALDWGSRPWGSNSWSGKPAAVLGTSPGATGTA
LAQQHLRNVLAYLDVPTLGQPEMFIKHDASKIDAQGNIVNEDTRKFLQGFVDRYADWVHR
HAESA*
>GCA_000452945.1_BPMSHR5858_genomic_04044_1 group_4974
MQHARAAHDDPAYLAQLRHDLLRFARLQLRDADAAEDAVQEALAAAWAQAERFDGQSSHK
TWVFGILRNKLIDTIRARRRTINASALDAELDGEALLERELFADNGHWAPHAKPRPWPKP
ETILQQRQFWILFETCLDHLPEQIGRVFMMREFLDFAIDDICTELELKANHCSVLLYRAR
TRLRTCLTEKGLTTEDATGEM*
>GCA_000452945.1_BPMSHR5858_genomic_04045_1 group_19920
MLLGKCKDITRLLSDELDRPLSLDERLRVRVHLPACTGCRNYRMQIRMLREAARVAGGNN
GPRNE*
>GCA_000452945.1_BPMSHR5858_genomic_04048_1 soj_2
LAAEIIAVTQQKGGVGKSTIAMHLGAAFHEKGRRVLVIDADGQNTLVHWSSSSADSESGI
PFPVVNLAEAGSQIHREIKKFINDYDIIIVDCPPSITEKVSGVVLLAATIAVIPTSSSPA
DYWSSVGLVKLIQQAQVMNEDLRAVFLLNKTEEKRMLTRELKRALEELGFPLLKTQIPTR
EAYKQAMALGQTVLQMNDRGAKLAAAEIRACADEIVAMLP*
>GCA_000452945.1_BPMSHR5858_genomic_04050_1 group_4976
MATKRAKKTDVDVVSASSAELRKAVEAIAIQPKNGKITLLTRKLFNVLLAVAQQADDSGD
TYRALLSDIVANSAFDSNDTALVKEHLRRMVSVQVEWSTGTSSQKPGRKWGISTLIADAE
ILEDPATRRVWVEFSFAPKIKKKLLDPVQYARLSLQFQSQLRSSAGLALYEICVRYLTNP
SHLTMREPWEWWRPILSGTPDTEAGDEAKREYKYFKRDYLRPAIAEVNAVTNIFVELVEH
REGRRVAEIQFRVSERKQPMLALDEHPNVFDSTLVDRMVKIGIPLKEAQTLYADNEENRI
RAALQLTEQRVRSTTLPPVRSAPALFKDALKKGYAPPVDTVEAPPLGGSAKLAAAAANVG
QADDPKARLRSEYDAYRRKEAKQLYEEQGDAEREVARASFESDVLPALGSHLRDDWRRRG
LDSKLVETAFFDWLAQRTWGEPTDGDLLAFTLSQSRAA*
>GCA_000452945.1_BPMSHR5858_genomic_04053_1 group_1517
MTEQDLAALSFKAAAHDMETIVRHIAERYIRQGVPLTWRLLHAVEAEALADLGFASRHDP
MVRCLFDRPAELRFPETDDLVDFGQSNALPAVFSFAVAAYERAADARADTAAVPARSARA
SRAWGD*
>GCA_000452945.1_BPMSHR5858_genomic_04056_1 tdh_3
MKALAKLERGPGLTLTRVKKPEVGHNDVLIKIRRTAICGTDIHIWKWDDWAQKTIPVPMH
VGHEYVGEIVEMGQEVRGFSIGDRVSGEGHITCGFCRNCRAGRRHLCRNTVGVGVNREGA
FAEYLAIPAFNAFKIPPEISDDLAAIFDPFGNATHTALSFNLVGEDVLITGAGPIGVMAV
AIAKHVGARNVVITDINDYRLELARKMGATRAVNVSRESLRDVMADLHMTEGFDVGLEMS
GVPSAFTSLLESMNHGGKVALLGIPPAQTAIDWNQVIFKGLEIKGIYGREMFETWYKMVA
MLQSGLDLSPIITHRFAVDDYEKGFAAMLSGESGKVILDWADA*
>GCA_000452945.1_BPMSHR5858_genomic_04077_1 group_4979
MKKHRRLAGTTAISASLAACAGLASGHALAQSSVTLYGIMDAGIEYVNHAAPDGGGAFRM
KSGNKNTSRWGLRGVEDLGGGLKAVFRLESGIDLANGAFDDGPDSIFARRATVGLKGQWG
ELTLGRNFTPTYDYMLPFDPMGYAQNYSWATSSTATGGRKDGLFTRSSNAVRYDGAYGGL
RFGAMYGFGNVPGSMKTSSKYDFALGYESGPFAAVVTFDRQNGAADSVTPADPVNYVQGI
HAGVSYDFGRLKTMAGYRNYRRTYHTAAATQLSDMYWLGGSYDFTPAFSLTGALYHQNIK
GGTDADPTLVSLRAQYALSKRTVLYAAGGFAIAKHGQNISVSRDSVGYADTQLGVTVGMQ
QRF*
>GCA_000452945.1_BPMSHR5858_genomic_04081_1 hpcG
MLDSATIQLLARRLHVAERERRQIRQISLDYPEITIDDAYAIQRTWVDMKLAEGRVLKGH
KIGLTSKAMQSSSQIDEPDFGALLDDMFFEDGATIPTRRFIVPRVEVELAFVLGKPLKGP
DCTVFDVYDAVDYVVPALEIIDARSQSIDPDTKRPRKVFDTIADNAANAGVVLGGRPVRP

MDVDLRWVAAIMSRNGVVEETGVAAGVLNHPANGVAWLANRLARFDVSLDAGEIILGGSF
TRPCAAKSGDTFNVDYGSLGAIHCHFE*
>GCA_000452945.1_BPMSHR5858_genomic_04084_1 betB_1
MGIKHWIGGREVDSPETFTTFNPATGEPIAEVASGGAQEIDAAVRAAKDAFPKWAGTPAK
ERAKLMRRLGELIERNVPALADLETRDTGLPISQTRKQLIPRASENFHFFAEVCTRMNGR
SYPVDDQMLNYTLYQPVGVCALVSPWNVPFMTATWKTAPCLALGNTAVLKMSELSPLTAD
QLGRLALEAGIPAGVLNVVQGYGATAGDALVRHPDVRAVSFTGGTVTGKRIMERAGLKKY
SMELGGKSPVLIFDDADFDRALDASLFTIFSINGERCTAGSRIFVQRTIYDRFVAEFARR
ANNLIVGDPADESTQVGSMITRAHWEKVTGYVRLGVEEGARLVAGGPDKPANLPAHLANG
NFVRPSVFADVDNRMRIAQEEIFGPVACLIPFDGEEDGLRLANDTAYGLASYLWTRDVGR
AHRLARGIEAGMVFVNSQNVRDLRQPFGGVKESGTGREGGEYSFEVFAEIKNVCLSMGSH
HIPRWGV*
>GCA_000452945.1_BPMSHR5858_genomic_04093_1 dmoA_1
MTRQIRFNAFDMNCVGHQSPGLWAHPRDESWRYRELSYWTGLARLLERGKFDGLFIADVL
GIYDVYQGSGEAAIRQGAQVPVNDPILLVSAMAAATRHLGFGITCSLSYEHPYPFARRMS
TLDHLTGGRVGWNVVTSYLDSAARNLGLPRQANHDERYAIADEYLEVCYKLWEGSWEDGA
VVRDAARRVFAEPSKVHPIDHRGRYYDVPGIHLCEPSPQRTPVLYQAGASRRGKDFAAQH
AECIFVAAPSRTILKAFVADIRARVKRFGRDPRDVLIFNLHTVITGRTSADAHAKHADYR
RYASDEGALALMSGWTGIDLSKYALDEPLEHVESNAIQSAVEALSSADPTRKWTVREIAR
WGGIGGLGPVSVGDAREIADDLQSWVDETDVDGFNLAYAVTPETFVDVVELVVPELQRRG
AYQTDYAPGTLREKLHRAGPRLAAPHPAARYRVGGEPAAAGRLAAQQ*
>GCA_000452945.1_BPMSHR5858_genomic_04111_1 group_697
MTEQEAERIATHRHYKGGLYRVIGVARHSETEERLVVYEQLWPKERSLWVRPEAMFEETL
ADGTPRFRKLAD*
>GCA_000452945.1_BPMSHR5858_genomic_04114_1 atsK_1
MNDTVRRRRSEALSIRPLSGHIGAEVQGIQLGSQMAPNDIRFITQALLTHRVIFFRRQHH
LDDRAQELFAQAFGEIVKHPTMGGKTGSAILELHSHEGGRANSWHTDVTFGLRPPKLSVL
RALALPDAGGDTVWANTVAAYQHLPSSLQDLVDKLWAVHGNDFDYAASRVELLHDPVAKE
YRKKYAAQVIKTEHPVVQIHPETGEKSLLLGHYAQRFVQYDTHDSNRLYEILQAHITRLE
NTVRWHWAAGDVAIWDNRSTQHYAINDYGDATRVMRRVTVIGDIPVAVDGRKSVPHEA*
>GCA_000452945.1_BPMSHR5858_genomic_04120_1 group_1901
MDRRKFLSTSGMVIGAGAVQIVVPARTVYAATAASQLSNISSVAASSKLPNFANTKAILF
PGSDKTVEALLDNAKSKIKSGTFDGSLLATSIEKQNPITGLYVILETAAITVTESIPVVG
WLISGFLSIGLKYLNESLKEKTDYEKLIKSIVNKSIDENNYRLMKAAFEGVTQVYGNFSD
LAASLSTNKPKPDQKTQLISLFTNTISNCEQNAPTMLLKNLNDKAAYQGIPLFCSIANVE
LSAHADMLKNKVAFDLDDSFYKNNYAAMEKKAWRMHKKAREVVVELLDNVGDLSLDYPSV
WAKKNSIIRQIYLSGLGAFLENYRRRYEAKFLKKIKLRNSIEIYSNSYADQGVIEKTINA
DITRRSSLLSRLSGYSHIDRVQRIGVGYLSSSENSKGFFKEYESCTGRVNDRRDEKDIDS
KFPIDQFANSSRTGRELPVKIYSNAPKAGTYGLKIITNDGKNIDFGSMNGDLKKIYSFDV
PGYYISGLVVAESNAKRGSCAADSGLSIDFICSVHRHCSSIDYIAGFTDANIGIELDNSV
WSFENNSSSNVTGYADYLVGKNILSVPSGSVKFDLYNTTGKAISVTPVLMAAIPGRKDGE
SASLKVSIGNSVAQVKLSDKDSDKEDSYIQGIDGEKYLDAASGEKITVPVGRSSISIENV
DKKAGFKFASLLLIPG*
>GCA_000452945.1_BPMSHR5858_genomic_04143_1 group_534
MPRQADPASPRAPQAAAPASSSARCAAPGAAPPQSAQPPQSPQPAPSTPAFIAPEPDAQK
AVRGSQFVLKAGDAFVVSDALGDIAGRDDGLFVDDMRVLSQWRLTFGGRAPSLLSGATSA
DNASFTAHLTNRPLPPLGGRETPEGVIHIERMRVLADNVLHEALTLTNYGTCDAEVPLSV
SFGADFKDMFEVRGSRRERRGAVAPPCVEAGAVRLRYDGLDAVERSVRIGFEPKPDTLAV
DRADYTLTIAAQACVSLYLTVEARVGAAHAGGDAFARRPCVATGRGALRKALADVHRAMR
ERRRTIARVHTSNPLFNAWLDRSLADLGLLTTSLETGPYPYAGIPWFSTPFGRDAVITSL
QMLWLQPSLARGVLRFLAEHQARETSAFRDAEPGKIMHEFRKSEMAATGEVPFALYYGGV
DTTPLFVVLAGAYLEHTGDESLIDELWPALERAAQWVSTVCERNPRGLLDYQRTSARGLA
NQGWKDSQDSVFHADGRFPSGPIALVEVQAYACAAFDAMATFSRRRGHAADTVRYAQRAK
RLREQVEALFWMPEAGFYGIAVDGHGELCRVLASNAAHLLAFGLPEQSRGESVARVLGST
LFRTGWGVRTLAAGQPRFNPMAYHNGSIWPHDNALAARGLARYGDKRAVLDLLRALFEAA
VSFDMRLPELFCGFPRRRGEPPTAYPVACLPQAWAAGAPFMMLQACLGISVDAARGEVRV
ERPELPEGVDWLRVDDLRVGGDSVSLTFRRVEGQVVAAAEPGGARVVAVL*
>GCA_000452945.1_BPMSHR5858_genomic_04150_1 rstA_2
MSFRILLVEDDNRLSTLIAGYLRKHEYLVDTVLNGDDAVDAILTGRPDLVILDVNLPGKD
GFEICREAREHYDGVIIMVTARDEPFDELLGLEFGADDYVHKPVEPRILLARIKAQLRRA
PVRQADGALAQPESYTFGKFTIDRANRTVSLPDGTTPELTSAEFDLLWVLVCHAGEVVSR
DDLMLQLRGVEFDGLDRTIDGRISKLRRKLRDDASCPQRIKTIRSKGYQFSKNAWE*
>GCA_000452945.1_BPMSHR5858_genomic_04160_1 mmsB
MKIGFIGLGHMGAPMALNLLKAGHDVHAFDLSEDALRLVKDGGAQVAASPRDAAAGAAFV
VTMLPAAAHVRAVLTQENGVLAGLAAGAMVIDSSTIDPASAQAFGELVRGGGGAFVDAPV
SGGTGGAAAGTLTFMVGGDDADYARAKPVLAAMGKNIVHCGATGMGQVAKVCNNLVLGIT
MAGVSEAMALGASLGIDPKVLAGIINTSTGRCWSSDTYNPYPGVIETAPSSRGYTGGFGT
DLMLKDLGLANDAAKQARQPVYLGALAQQLYQTMSSRGDGKLDFSAVIRLYRPLGKDA*
>GCA_000452945.1_BPMSHR5858_genomic_04165_1 group_507
MPRSSPRPPPARGKLPARLARAAVLIGLLLACAAVAGAAAAARAPIVVIPVAGAIGPASA
DFVVRGLERAAREHAPLAIVQLDTPGGLDTSMRQIIKAIVASGVPVAAFVAPGGARAASA
GTYIVYASHVAAMAPGTNLGAASPVQLGIGGQAPAGGRREPAGAPGAAGAGATDTESIET
RKALHDAAAYIRGLAQLRGRNAEWAERAVREAVSLPANDALAQHVVDLLANDPADLAAKL
DGRTIVTATGAHRLDTAQAPLETLEPDWRSRLLATIADPSVALVLVTLGMYGLFFEFANP
GFVLPGVAGAISLLVGLFALQLLPISYAGLALVLLGIAFLIAETFLPTFGALGVGGIVAF
AIGALMLIDTDVPGYGVPWPVIASLAAAGAILVFGVSGFALRARRRPVVTGAEAIVGSIG
EVLDDGLVPDQPPQIGPSADPGAPLAPSAAGWARVHGERWRVASATPIAAGRRVRVTGRN
GLTLTVAPLYDNVVPPRGEPP*
>GCA_000452945.1_BPMSHR5858_genomic_04172_1 rdpA_1
VAYKTLSNALGTEVYGFDSPFPHDAEAVRALIEAWHAGGICRLRRQRLDMAEFVEFSRIF
GRPERALNQERKLTSREDLPELMIVSNILENGVSIGHLGSKEAYWHTDMCYTDVPPIASI
LYAIEVPAHGGNTEFMNMYRVHDALPASLRRQIAGLSIKHDRSYTAVGELRYGFDSVVDV
TTCPGSIHPIVRVHPVTQRPYLYLGRRLNAYVVGLPVGESEALLDELWRYTRLDGVTWTQ
RWEVGDIMIWDNRCTMHRRDAFDANARRLMWRTQIQADPARPL*
>GCA_000452945.1_BPMSHR5858_genomic_04173_1 phnY_1
LLNHRSAAFARVARALDARGKTSSGATLVVSHPYNRETVAELPLDSGDDARRKLQRAARF
RSALSRHERIAVFDKAIALLAAEKRDASILITLESGLCRKDTMYEVDRVINVLHAAIAEL
NRDDGQTFSCDNATSDERRKIFTVREPLRGVIVAITPFNHPMNQVAHKICPAIASNNRIV
LKPSEKTPLSALYLLDLFREAGLPEPMFDVVIGEPNALGAALVCDEHVELVAFTGSVAVG
KRIAQMAGYRRTVLELGGNDPLIVMEDADLERAAQLAAKGSYKNSGQRCTAVKRILVERT
VARPFTELLVEHSRRWQTGDPMDERVDIGTLIDDAAAIECARRVDEAREAGARVLLGHQR
DGAAYAPTVLERVGPALRLVQQETFGPVSPVITFCGLDEAVAIANSTRYGLSSGVCTNRL
DYITHLIAHLDVGTVNVWEVPGFRLESTPFGGVKDSGLGSKEGMQEAIKNFTNLKTYSLP
WDTLAHPHAA*
>GCA_000452945.1_BPMSHR5858_genomic_04176_1 group_4991
MDEASRLLFHPDPTRREGDDAIQLPRMTLHFALEPRAMMTWHARSHTEIRVLSARVWLTR
SFSVDDYWLKPGDVLHVPRGERVWLSADGPAAAEVSLTTAYVRRRSWIGGMVDGLLDVAA
ELTGPRPR*
>GCA_000452945.1_BPMSHR5858_genomic_04179_1 group_1521
MDQRIVPARGGPHAGGGHPAWMRVALAEQGVRRCGRGESNPRIVEYNGCTNLVGYDDKVS
WCSSFVNWCFSRVGIAATGSALARSWLEWGRALSEPAYGCVAVLMRDRPTSWKGHVGFYL
RHDDERVYLFGGNQRGAVREYAYARSRLLSYRWPDERGLS*
>GCA_000452945.1_BPMSHR5858_genomic_04187_1 group_1906
MAGGWPARAAVATGGAALGAAELAVTLARHYPADGLNRLYAAVLVALAAGVAVLLWALVA
RDGRQAAWRACAWLAVLTPCVWLCAGRC*
>GCA_000452945.1_BPMSHR5858_genomic_04188_1 fptA
MRYGVAGPERSRRGAAPVGRFGGRRWRSSLCLGVLIAAHPGAAALAGTAAKSDAQKNDMQ
RNENVREDTRRLAAGAPPAGGELKAISVSAPKDAADDPSVATVGKMPLALREIAQSVSVT
TRERIEQQNLFSLDDVMQQSAGVTVQPYVLLTTAYFVRGFKVDSFEFDGVPVVLGDMASA
PQDMSAYERAEILRGANGLLHGSGNPAATVNLVRKRPQHRFAASASASVGSWGRYRAQAD
IGGPLNPAGTVRGRLVAAYEDRGFFYDRAKQDTRSIYGIAEIDVARDTLVTVGAQYQSVA
SVPDMSGVPMARDGTSLGLPRSTFLDVGWGHFDWDTTRVFGSVERELGGGWKAKVSGEYQ
AVRSDLKYAGAYGAIDRATGAGGTLMGSAYQFSSYSRSVDANVQGPVRAFGLTHELLFGA
TYASSSSGQLTAPLLAGAGTPVNVYRWNPDGVPEPGVGPYRQDMQNDISQKGVYGLGRIK
LAQPVTLVLGGRLSWWNQDSLGAHYNAGRQFTPYGGLIWDVARDWSWYASYAEVFQPQTK
PTWDGGILTPVKGRTYETGIKGELAGGKLNVSLAAFRIDLDNNPRIDATRPCAGRSCYYV
NGGSVRSQGFEFEANGRITPWWSVWASYTFDTIRYAKDVANGGAFAAELTPRHLLRVWTN
YDLPWQERRWSVGGGVQVQSDFSAASSGVTMRQGGYALASVRLGYRYDRHWRAALNINNL
FDRTYYQSLSQPGWNNRYGEPRNVMLTVRGQF*
>GCA_000452945.1_BPMSHR5858_genomic_04190_1 irtA
MNAPLARMLRPFAAPLAAAVALQALAGVASLVPWLALGRIAEHWRAAADLDGAARAWLAA
AVAAGACWLGGQTLALHLTHRVDADLCDRLRARLADHLQRLPLNWFARTGSDRIARYVDQ
DVRALHQLVAHAPADITQLIVVPGAALACLLYLNPALLAFALVPLVLAAALFRWMRSARF
MPAFAERRAALEALMGDYAELARNPALVRQYPGAGIEAVALDSVAGFARAFSAWVGRIGG
LGACTQVLLGTPLLLAWVVFGAMGLVLARAPVGELCVFALLIWAVAAPVRALGHGADALR
DARAAAARLDALLALPGLPQPPDAGARPAPRDASIAIRGVCVEFDGAPVLHDIEATIEAG
TTTAIVGPSGAGKSTLLTLLARFTDPARGEVRLGGVDLRMLPPRVLRERIAVVFQHAAAL
DVSIAENIALYRPDASRDDIRAVARAACLADRIDALPGGYDCRYGRDVRFSGGELQRLAI
ARALLSHAPLLLLDEPASALDPQTGRALRNALHGGARTRVIVSHDLAAVRQADQILVMDR
GRIVERGTHRALLDARGLYARLWAERDRPAEEATR*
>GCA_000452945.1_BPMSHR5858_genomic_04208_1 group_4995
MHAWSAQHVPPQGGKVAVVTGANSGLGWQIAQTLAAKGAQVVMGCRDTAKGELAAHAIRT
RYPRARIEVEALDLADLASVCRFADAVADRHGRVDILCNNAGVMFLPLRHTRDGFEMQMG
TNHLGHFALTGLLLPALRASHRARVVTMSSGFNRLGKIRLDNMLAERGYNKYRAYCDSKL
ANLMFTLELQRRFDQACLPILSVAAHPGYAATHLQFAGPEMANSSLGTFAMRLSNRLVAQ
SADVGALPAIHAATAVDVDGGAYIGPAHLCETRGYPAEARIPRQARDVRMGKRLWEKSEQ
LTGVRYLDTPPPPGSRRRASRDDATFGAL*
>GCA_000452945.1_BPMSHR5858_genomic_04210_1 lip2
MLEPTIEAFVARVAAWYPDDAAARPIAEQRALYERFAAAWTPALPADVDTRDARLGAFGD
GREIRLRRYRSRCAAACGTVLFFHGGGFVVGSLDSHALMTARIAADTGLDVIAVDYRLAP
EHRAPAAFDDCLAVTLAALDGRLPFESIALPLRLAGDSAGGMLAASVATALRDGGRIGRD
GGGRAGGGVCAVALVYPMLGFEPQAPARDTHAHAPMLTLDDVHAYRRHYWGDAGAPPGAL
PLDARRFDGLPPVLAIGAEHDPLRDDARVYAERIRSAGGAARFWCGDGLVHGCWRALDTS
AQAARMHRAVCAFLLAPDG*
>GCA_000452945.1_BPMSHR5858_genomic_04222_1 purU_1
MSVPQRPHQFVLTLSCPSAAGQVAAVVGLLDRHRCYVDELTVFDDDLSARFFVRCVFHAT
DDADALRVDALRREFEPIAERFRMQWAIHDVAARPKVLIMVSKLEHCLADLLFRWKMGEL
KMDIVGIVSNHPDFAPLAAQHGLPFRHFPITADTKAQQEAQWLDVFETSGAELVILARYM
QVLSPEASARLANRAINIHHSFLPGFKGAKPYHQAHARGVKLIGATAHFVTDDLDEGPII
EQVVERVDHSYRPEQLLAVGRDVECITLARAVKAFIERRVFLNGDRTVVFQ*
>GCA_000452945.1_BPMSHR5858_genomic_04229_1 fadF
MNPAILITALLWLSVAGLAFAVAKRSAYWRLGRATTPGAFGLANLLAIPKRYFVDLHHVV
ARDPYIAKTHVATAGGALAALALVFVNYGLAIYSPWLDRLILLAALAMLVGVVFVWRRRH
AKAVPARLSRGPWNTLPWLLGAFALGLVLFMLVPAAAMSGAFALLCAALIGAGAFAMTVG
AAKGGPMKHAIAGLLHLAFHPRQERFAATRDAYTGDGGATPPTALKPSDVERDEYGVAKP
AEFRWNQLLSFDACVQCGKCEAACPAFASGQPLNPKKLIQDLVVGMAGGTDAAYAGSPTP
GIPVGAHGGDPQRPIVSSLIEADTLWSCTTCRACVQECPMLIEHVDAIVDMRRNQALVHG
AVPGKGAQVLANLRETGTMGGYDTAARYDWSVDLGAPVAQPGRPVDVLIVAGEGAFDMRY
QRTLRALVKVLNHAGVDYAVLGRAETDTGDVARRLGDEATFQQMARRLIDTLGALSFKQI
VTADPHVMHSLRNEYRALGGRYAVKHHTTFVAELAAAGKIKPQAADALREKRMTYHDPCY
LGRYNGETEAPRKLLKTIGIQIVEMERHGMRGRCCGGGGGAPLTDIPGRQRIPDIRIADA
RAVGADVVAVGCPNCTAMLEGVVGPRPDVLDVAELVAASLE*
>GCA_000452945.1_BPMSHR5858_genomic_04234_1 cdhR_3
MSPDRTASLSHFAFMPLPNFTMIAFTNAIEVLRMANYLSGQPLYRWSIISPQGGMVTASN
GLAVDTGPAECAGQPDIVFVCGGVDVQRATQPEHLAALRRFARAGVALGSLCTGTYALAK
AGLLAGYACAIHWENLSALKEEFPDTRFLKELFVIDRDRVTCTGGVAPLDMMLNLIASRI
GTARVTQIAEQFIVEHVRDTSAQQRMPLVARLGSANKSLFEVIALMENNIEEPLSREELA
RLANMSQRQLQRLFREHLGMTPTHYYLTLRLRRARELLLQTDMSIMHITMACGFQSACHF
SKSYRDAFGTAPTRERRKQVAPLAQPSMPGGAPAPSMMLHA*
>GCA_000452945.1_BPMSHR5858_genomic_04272_1 group_2701
MTRKTRIQPIEHPGEDADHGYDRPSKSQLKRDMHALQELGAALVELPKDAFKRMPMPEDL
ADAVREARRITDHEGKRRQLQYVGRVMRSLSADETAALRAALDAQRGVNKAETARLHWIE
RTREQLLANDDALTAFVREHPSADVQEGRTLIRNARKEAQQGKPPRYFRELFQWIKTASG
SASDAPDLSPDADDDDDEA*
>GCA_000452945.1_BPMSHR5858_genomic_04273_1 mog
MTTTKRNHPDELVIGLVSISDRASSGVYEDKGIPALAEWLAAALVSPWRAETRLVQDDAP
TISATLVELVDTLGCDLVLTTGGTGPARRDVTPEATLAVATKEMPGFGEQMRQISLNFVP
TAILSRQVAVIRETADHAALIVNLPGQPKSIKETLEGLRGADGKVEVPGIFAAVPYCIDL
IGGPYAQTNPAVCAAFRPKSAVRAPK*
>GCA_000452945.1_BPMSHR5858_genomic_04274_1 orn
MTDISAVAGQPALVRNELNLVWLDMEMTGLDPDTDRIIEIAVVVTNSTLDIAVEGPVLAI
HQSDETLAKMDDWNKNTHGRSGLIDRVRASSVTEADAAAQIAAFLAEHVPPGKSPMCGNS
ICQDRRFMARWMPELERFFHYRNLDVSTLKELCRRWQPAIYKGFQKRAMHTALADIHESI
DELKYYRERFLIPAAPAGETA*
>GCA_000452945.1_BPMSHR5858_genomic_04284_1 gyrB
MTEQHNSQPENSYGASSIQILEGLEAVRKRPGMYIGDTSDGTGLHHLVFEVLDNSIDEAL
AGYCNDIHVTIHADNSISVTDNGRGIPTDVKLNDKHEPKRSAAEIVMTELHAGGKFDQNS
YKVSGGLHGVGVSCVNALSSWLRLIVRRDGKKHFMEFHRGIAQNRVLEERDGEQVSPMQV
IGETENRGTEVHFMADPTIFGTVEYHYDILAKRIRELSFLNNGVRIRLTDLRSGKEDDFA
FAGGVKGFVEYINKTKSVLHPTIFHINGEKDSVGVEVAMQWNDSYNENVLCFTNNIPQRD
GGTHLTGLRAAMTRVINKYIADNEIAKKAKVETTGDDMREGLSCVLSVKVPEPKFSSQTK
DKLVSSEVRAPVEEVVAKALEEFLLETPNDAKIICGKIVEAARARDAARKAREMTRRKGV
LDGVGLPGKLADCQEKDPAKSEIYIVEGDSAGGSAKQGRDRKFQAILPLRGKVLNVEKAR
YDKLLSSEQIVTLVTALGCGIGKDDYNLDKLRYHRIIIMTDADVDGAHIRTLLLTFFYRQ
MPEMIEHGYVVYIAQPPLYKVKAGRDERYLKDDTELNAHMLRLALQGSELVPTENGTPISG
DALGELARSYLLARGVVERLSRLYDPAALEAVMDGVAIDLSSEASTEASAKALAAMLSDD
SKTEVRVVPAYDAVREQRSLRVERTHHGNVRVSVIDEEFQHTADYQQLVNTANTFKGLIG
AGAAIKRGERSSNVTDFKEAMKWLMADAERNLSKQRYKGLGEMNPEQLWETTMDPAVRRL
LRVQIEDAIAADGIFTTLMGDEVEPRRAFIENNALRAGNIDV*
>GCA_000452945.1_BPMSHR5858_genomic_04286_1 dnaA
MNDFWQHCSALLERELTPQQYVTWIKPLAPVAFDAAANTLSIAAPNRFKLDWVKSQFSGR
ISDLARDFWNAPIEVQFVLDPKAGQRSPAGATPLAPRAPLPSANPAPVAPGPASAPAVDA
HAPAPAGMNAATAAAVAAAQAAQAAQANAAALNADEAADLDLPSLTAHEAAAGRRTWRPG
AANANSEAADSMYERSKLNPVLTFDNFVTGKANQLARAAAIQVADNPGISYNPLFLYGGV
GLGKTHLIHAIGNQLLLDKPGARIRYIHAEQYVSDVVKAYQRKAFDDFKRYYHSLDLLLI
DDIQFFSGKSRTQEEFFYAFEALVANKAQVIITSDTYPKEISGIDDRLISRFDSGLTVAI
EPPELEMRVAILMRKAQSEGVSLSEDVAFFVAKHLRSNVRELEGALRKILAYSKFHGREI
TIELTKEALKDLLTVQNRQISVENIQKTVADFYNIKVADMYSKKRPANIARPRQIAMYLA
KELTQKSLPEIGELFGGRDHTTVLHAVRKIADERGKDAQLNHELHVLEQTLKG*
>GCA_000452945.1_BPMSHR5858_genomic_04287_1 rpmH
MKRTYQPSVTRRKRTHGFRVRMKTAGGRKVINARRAKGRKRLAI*
>GCA_000452945.1_BPMSHR5858_genomic_04298_1 group_784
MNEQYASIIDVRRIPHHKRHALIFGTFDALPGGEALQLINDHDPKPLYYQLEERAADSFE
WTYLEAGPSQWRVQIAKRGDALGHHDAGDSCCSGGGCGG*
>GCA_000452945.1_BPMSHR5858_genomic_04300_1 sufU
MNPIESPAVAADAQSTLYQELIVEHKRAPRRFGKLAEPTHEARGHNPQCGDDLKVQLRIE
GGRIGDIRFDGHGCAICIASASMMTEAVIGRDVDAARALQQRFRAVLTGQAEHDEAYLGK
LSSLVAVRRYPSRIKCALLGWHALAHALDTRVAAPGETEPHAS*
>GCA_000452945.1_BPMSHR5858_genomic_04306_1 tap_5
MSKHVFGPGMRLMARMKLPRKLVILAALFIVPLVAALYATLAVAWHAYATTQGERDGISI
LETTQDLLKAVQVRRGAGASVLAGNEAMRAKFDAASASAGRLAATLKQQVRGTDRFDIVT
AVDALASEYGKVAAGGLGTSAAALFGSHTDVIHAIFLFEKDLAVASRLGVDPDASNYHLI
DSVLFVLPGTAETVGVLRGKGAAALVGGALGEADKRELAVRAGSLAEQMSVIRHHLDRMK
DRLGAGVPDGVDLGAVAAFQRAAAALAGGDASIDAKAYFQRGTDAIDALYRVHGTLAQQL
RERLAARAHAEQVKVAAILALSVVLVACALYLFVAFAMSTHEDVAQLSDCMRAVGDGDLR
ARLAVRGGDEFAFIKRGFNALLDVWAQTLTETRRVADSVMVGSQQIAAGNLDLSGRTETQ
AASLEQTAASMEELTSTVRHNASNASHASTLSAEASERAADTGRIVTHAVSLMTRIHDSS
NRMAEIVTVIEGIAFQTNILALNAAVEAARAGEQGKGFAVVAGEVRALAHRSATAAKDVK

ELIHESIRHVTDGSSLFQRADDAIRSVNASIKSVDTVVSEIAKASEQQSDGIEQVNAAIT
QMDEVTQRNAALVEESAAASASLREQAEHMGKLVGRFVLA*
>GCA_000452945.1_BPMSHR5858_genomic_04310_1 group_5010
VSVSVVKTAFKNLVIKGRSLLPIVQGGMGVGVSAHRLAGTVASLGACGTISSVDLRRHHP
DLMARTGRSRDRALIDAANLEALDREIRAAKSLANGRGLVAVNVMRALSEYASYVRQSCE
SGAHAVVVGAGLPLDLPELTADFPDVALIPILSDARGIGLVLKKWMRKNRLPDAVVIENP
RYAAGHLGAPTTDSLNNPNFAFPTVLEGTFALFKELGIERERIPLIAAGGIHSHEQVRQL
FALGASAVQLGTPFAVTEEGDAHPNFKKVLVEAQPDDIVTFMSVAGLPARAVRTPWLTNY
LERERKLQRAAKPRKCLVGFDCLQQCGLRDGIEKHGQFCIDTRLAFALAGDIKRGLFFRG
SETLPFGHEIRCVRELIDYLLTGVKRAAAAAIAPATACAPMPALG*
>GCA_000452945.1_BPMSHR5858_genomic_04312_1 group_5011
MKIALGPVQYYWPRVATMQFYQAMSETPVDIVYLGETVCSRRHELRFADWIEIADMLAEA
GKEVVLSTQVLLESGRDLKTMREIAENGRYLVEANDMGAVRRSASRAFVAGPWLNVYNPP
TLAMLAGFGARRWVMPPEMSEAGLARMQAERPAHLETEVFAYGRIPLAFSARCFTARNRN
FPKDNCQYVCMAHPDGLPLHTREGEPFLVLNGISTQSARVYNLIGEIDALRALNVDVVRL
SPQSQHMIDVIDAYHGVLSGRTDAGDALAALHGRMPEASCNGYWHGKPGLEHVAR*
>GCA_000452945.1_BPMSHR5858_genomic_04319_1 group_699
MPAPKQPAASLARPLVRATAGVVIVALLACGIATGLARLGAPAPTHAVSRLAWHGVLMIP
VFFGAVIGVERAVALGARWAYLVPATAAAAGLSLLAGAPAMRAAALLTGAASIALAVSAT
VLRRQAALFLWVLAASVGCWWAGNAVWLLTGDPFAAVAWWLAFLILTIAAERLELTRVRP
LPRGASACFCGIVALLPAAAALASWQPLPALRLFGAGELLLAAWLVRYDVARRTVRQRGL
TRFIAVCLLSGYGWLAVSGALGLDGALAPGHPAHDAALHALTLGFVFSMILGHMPIIVPA
VARLRVAYRPFFYAPLALLHAAVALRVAGGLLDAFALTRAGGEIAALAIASFAATLAIGI
ALARRSASHRGAGKRAA*
>GCA_000452945.1_BPMSHR5858_genomic_04330_1 hutU_2
MNHPKHIDPRLDPTRVIRAPRGGEKTCKNWLAEAAYRMIQNNLDPEVAEHPHALVVYGGI
GRAARNWDCFDQILASLKDLNDDETLLVQSGKPVGVFRTHENAPRVLIANSNLVPHWATW
DHFNELDRKGLMMYGQMTAGSWIYIGSQGIVQGTYETFFAVANQHFNGDPSGRWILTGGL
GGMGGAQPLAATMAGFSMIAVECDESRIDFRLKTRYVDRKATTLDEALGMIEEAKRTGKP
VSVGLLGNAADVFTELVERGITPDCVTDQTSAHDPINGYLPQGWSVARWRDAQKVDPRSI
VQVAKQSMAVQVRAMLTLQARGAATLDYGNNIRQMALEMGVENAFDFPGFVPAYIRPLFC
EGKGPFRWVALSGDPEDIYKTDRKVKELIPDDAHLHNWLDMARERIAFQGLPARICWVGV
NDRYRLGQAFNEMVKTGELKAPIVIGRDHLDTGSVASPNRETEAMKDGSDAVSDWPLLNA
LLNTAGGASWVSLHHGGGVGMGFSQHAGVVIVADGTDAAHARLGRVLFNDPATGVMRHAD
AGYELAQRTANEAGLKLPMLGR*
>GCA_000452945.1_BPMSHR5858_genomic_04331_1 ves
MRAASSAMQASLIRADALVAAPWKNGGGVTREIAAHPLRERAGHGALDAFAWRVSVADVA
DAGPFSRFPGIDRTLVLLAGAGMTLVDAGGARHALRAPLDRVAFAGEAAIAAELHDGPTR
DFNLMTRRDAARGSVDVWRSGASHALRADTVLLFCASGAPVVDLGVDLGVGARVALGACD
TLRIDACAHGEFRCAIRGEGVLLAVGVTLLQDNAQGSIE*
>GCA_000452945.1_BPMSHR5858_genomic_04334_1 group_5013
MNTASQPPVFTLHRGTLPLLVSIPHAGTHLPDDIAATMTPVARHVDDCDWHLERLYDFAK
TLGASVLVPSHARYVVDLNRPPDDANLYPGQDTTGLVPVDTFDRAPLYAHGHEPTVTEIA
RRRERYWAPYHGALAGELQRLKDAHGRALLWEAHSIRSHVPRFFDGRLPDFNFGTSSGAS
AAPGLADKLAALVDRHGGYTAIANGRFKGGYITRHYGAPEQGVQAVQLELVQATYMDETR
PYSYDETRARRIAPLLEALVSAALEHH*
>GCA_000452945.1_BPMSHR5858_genomic_04403_1 ybdL_2
MMNAPHDMPTTPTLLSRLPEVGTTIFTVMSALAAEKGAVNLGQGFPDFDCDPRIVDAVAH
AMRDGRNQYPPMAGVAPLREAIADKIERLYGRRYDAATEITVTAGATQALLTAILCAVHP
GDEVIVIEPTYDSYLPSIELAGATPVFVTLNAPDYAIPFDKLAAAITPKTRAILINTPHN
PTGTVWREADMRALEDIVRGTGLLIVSDEVYEHMVYDGARHESVARYPELAARSIIVSSF
GKTFHVTGWKIGYVAAPAPLTAEFRKVHQFNVFTVNTPMQFGLADYLRDPEPYLTLPAFY
QQKRDFFRAGLANTRFALLPCAGTYFQCVDYSAISDLPEAEFAKWLTTEIGVAAIPVSAF
YHEPHESGVVRFCFAKREDTLATALERLARL*
>GCA_000452945.1_BPMSHR5858_genomic_04404_1 group_5027
MLRLCGFPVSNYYNKVKFVLLEHDIPFEASIVSVPIKDPAFLADTPLGKVPYLMTEHGSL
CESQAIIEYLAAIHPDKPIFPADPFAAAKAREITAVLELYLEWNVRELFPEAFFGGKVSD
GTKAHVEKRMPRALDGFKRLAKFSPYVLGDTFCIADIAAFIHLPVVALATKAIYGRDFVV
DAGIDWKAYVKRIEEARPAAKRVSDERKAFVAGLAKPG*
>GCA_000452945.1_BPMSHR5858_genomic_04406_1 gpmB
MSYQLPKRRRIYLMRHGDVTYFDASGRPFDQDAVPLNERGRMQAAAAGRVFAEQNVRFDR
VIASGLPRTIETAQRVLAETGQQLDIDIEPAWREIRGGFLADIPPAEQEAAFLRVLDGIV
AEHTRFLGGETIGELIDRVLPPLAALRADPSWDTVLLVLHGGVNCALLSHATVPGHRLFI
GTLSQSAGCINALDVGERDDDWVVRLVNYSPPEALHRDVRNTTMEVLYHQFLQSTRE*
>GCA_000452945.1_BPMSHR5858_genomic_04417_1 group_5029
MSASGWFIVLLALAGANLPFLNQRLFAVVPLKSAAAAKKSAWIRIGEMIVLYLIVGALGF
WLESRAGNRFEQGWQFYAITFSLFVVFAFPGFTFQYLVKRR*
>GCA_000452945.1_BPMSHR5858_genomic_04423_1 nuoI
MTAIQQFFKTFFLTELLKGLALTGRYTFKRKFTVQFPEEKTPISPRFRGLHALRRYENGE
ERCIACKLCEAVCPALAITIESETRADNTRRTTRYDIDLTKCIFCGFCEESCPVDSIVET
QILEYHGEKRGDLYFTKDMLLAVGDRYEKEIAAAKAADARYR*
>GCA_000452945.1_BPMSHR5858_genomic_04424_1 nuoH
MSLFDTINSGGAQLLGVAWPTVWALVRILVVAVVILLCVAYLILWERKLIGWMHVRLGPN
RVGPAGLLQPIADVLKLLLKEVIRPTAASRWLYLVAPVMTVVPAFAVWAVIPFQAGAVLA
NINAGLLYAMAISSIGVYAVILAGWASNSKYAFLGAMRAAAQMVSYEISMGFALVLVLMT
AGSLNLSEIVGSQQHGFFAGHGVNFLSWNWLPLLPVFVIYFISGIAETNRHPFDVVEGES
EIVAGHMIDYSGMAFALFFLAEYINMIVISALAATLFLGGWDAPFEFLSFIPGIFWLVLK
IFALLSVFIWARATFPRYRYDQIMRLGWKVFLPVCVFWVIVVGFWMMSPLNIWK*
>GCA_000452945.1_BPMSHR5858_genomic_04433_1 secG
MPYLKVLIIVVQLLSALGVIGLVLLQHGKGADMGAAFGSGASGSLFGATGSANFLSRTTA
VLATIFFVATLALTYLGSYKSAPSVGVLGAAPAPAASAPAASQTPAASAPAAVSAASAPG
QDVPK*
>GCA_000452945.1_BPMSHR5858_genomic_04449_1 kdpE_2
MSSAPATKEELDFALKVRRALDERAASLPAATADRLAAARRAALARKKPDAAIVLVPALA
GGAGTLELRPPGEPRSSLARRLARAWPLALLLAGLIGIAYWEDMQRTAELADIDAAMLSD
DLPLTAYLDHGFNAYLSHTH*
>GCA_000452945.1_BPMSHR5858_genomic_04451_1 group_1224
VIAPALPSEPPPPSVRRRLAALAYEGLLLFGVVFFAGLMFGVALQQRNGLDHRNLLAGWI
ALVVGVYFVWFWTHGGQTLPMKTWRLRVETARGAPLSAGRALARYALGWLWFLPPLALHP
LAGFPLPRTLAATAVWFALWALAARLHPSRQFPHDRLAGTRIVDAPRRG*
>GCA_000452945.1_BPMSHR5858_genomic_04452_1 group_2096
MGKKTSATSFFRDPAGLHAASAFLSGSAASDTLAPARPPAADTTRHDDNEPASHRYRTIW
LSDIHLGTSGCQAGYLLDFLKHNESEYLYLVGDIIDGWQLRKGWYWPQAHNDVVQKILRK
ARKGTQVVYIPGNHDEAARQFCDLAFGEIHVRGEAFHTTLAGKRLWIVHGDLFDGVIQHA
KWLAYLGDTLYTLILVLNRWFNRIRSRFGFQYWSLSQYLKHQVKNAVNFISSFEAVMTDE
ARRRGCDGVVCGHIHKAEIRDIDGVLYCNDGDWVESLSALVETMEGELKVVYWTVMRTPP
AALPRKTKAATA*
>GCA_000452945.1_BPMSHR5858_genomic_04482_1 nadD
MSRAATGRAAAPQGPTPNAEPAGLPALPRRIGILGGTFDPIHDGHLALARRFAHVLRLTE
LVLMPAGQPYQKQDVSAAEHRLAMTRAAAASLVLPGVAVSVATDEIEHAGPTYTVETLAR
WRERIGDRASLALLIGADQLVRLDTWRDWRRLFDFAHVCAATRPGFDFTAASPAVAAEIA
SRQASADVLQATPAGRLLIDTTLSLDVAATDIRAHLRACIARRAQVPDASAEHVSPAVWA
YILQHRLYHP*
>GCA_000452945.1_BPMSHR5858_genomic_04487_1 xanP
MQPAPLARPASHGADAAESAHDLVYGPDERPAPMIACVAALQHLLAILVPIVTPGLLICQ
ALGVSSRDTTLIVSMSLVISGIATFVQCRRFGPLGAGLLIVQGTSFNFVGPLIAGGSLMV
KQGTPVETVMAAIFGVVIAGSFVEMAVSRILPFVKRLITPLVTGVVVLLIGLTLIKVGLV
SMGGGYGAIAKGSFGSAQNLTLSGLVLGAIIVLNRVPIVWVRSTALVIALAIGYLAAAAM
GRLDFTGAREAALFQIPTPLHFGLGFSWALFVPMLIIYLVTSLEAIGDVTATSKISRQPV
EGPLWMRRIKGGVLVNGANSLLAGVFNTFPSSVFAQNNGVIQLTGIASRHVGIWIAGMLV
LLGLFPVVAGVLQAVPEPVLGGAAMVMFGAVAASGINILAGIRLDRRALLIIAVSLALGL
GVSQVPEILTNLPHALGNVLESGVATGGICALVMNWFLPEKQ*
>GCA_000452945.1_BPMSHR5858_genomic_04488_1 group_6139
MLVRECLECLRRLCVAGLLSAAAADGHAAVAAAQDPSIPYYSWYEVTLPESSGASCGNGT
PMRFYINRAQSDNLLYMMEPGGACWDHGTCTETATGAQAGLGAFNPDGIPHNYVNGTVQQ
SLKTSFLSPLMTRMDLAHILVGEPKVETQEWTQVFVPYCTGDIHMGSAVRSYASPSGDWR
IQHYSGLKNVRVIAQWLVDHGFGKPSRLLVYGTSAGGYGTLGNYATLRGTLQPQSHSSLL
DDAGTVFNTRFGADPAAYPSVGLYDKVRDEWGMTAPDGMITVNSRLTRYFDPGNMGSAYA
ALSATYPHDRFGYTTYRRDRIIAAYHYRPFVPAVIAAPDDATKDALSLAMFDRELSDLKQ
VLNPLPNFGYFIPWARDDFMGNHQVTAVSFTGSGIHENGIDADVGTFVADLLNQQDPADV
PVMKAYRTEQWSDFTFSTFLAWLDSIFNLTGEAGPISGHRS*
>GCA_000452945.1_BPMSHR5858_genomic_04489_1 group_1591
VKKSKALWRTAWLLAMIGVAVAGWRYVSPSQTAVAQSLKPGAASDERGGPPARARTDDAP
APAGPAERRLDLVALRKSLEGRDDAQAQVQRIVAFARFRDRIAAYGEGRNSMPAEERARL
AREILAELPEHVARNEIVPVQAEALSAALLADATSDSRTRDAAIQSMRQQWDAYAGRTVG
PSPASDPRYLAYARQSRDIVAQVQASVPDPDQQQAVIAQRLQALRVQLFDQASRSDTR*
>GCA_000452945.1_BPMSHR5858_genomic_04490_1 group_5044
MGSTLNPDDEPREPDERSPSSGDNRTLGPSDSSDSGSDVAGARRRPFDVDTELDNHALET
GDAELDSDTDRSGTGERASADGDSTFDDSADIEPDRTERVPRTKRGDD*
>GCA_000452945.1_BPMSHR5858_genomic_04492_1 group_5045
MFEPPLPSRQIDYEGFEIHVSPTPTHTDANRYAYAGYVCHPGADPRLPGHTVPFHADGEE
SFRSVDEALDEAVSIGRSIVDGTHPDLSVLSLVTHGY*
>GCA_000452945.1_BPMSHR5858_genomic_04493_1 group_889
MAVALFPNWLGQIQTRIVEFPDDTLHLGMPTPLQLNDALKHARLVWRRATPNG*
>GCA_000452945.1_BPMSHR5858_genomic_04502_1 mshA_6
MTPPVAPPAFTAQPAETAADVARPGVRRVRILFHINDFGRGGTETALLAWLNALDRSAFE
VGLSVTYPTADLTMWRAKAIPADVALQVLASERWMHALHQRERVRRLRGGEKLLHKASTH
ALIRPAVARRFLRLARGYDVVCDFDFSLRRIAGRGGMPWIGVSHYSFAARFGEKSAAYMA
RRVRQYERYTALAVLTPVMRREAESLFADTRVVVTALPNVIDPQALRARADEPAELPAGR
FIVSVARLDEGQKDHRTLLRAYAQVRASRADAPRLVLLGDGPDRGALEQLAAELGLRDAV
LFAGFCANPFPYVRAADMLILSSRYEGFGMVIGEAMALGTPVISADCPTGPRDLLDFGRA
GLLVPPGAADALADAIERMLADGALRASLVAHATRKIESFGPRAANARMQALAAQLLDRA
*
>GCA_000452945.1_BPMSHR5858_genomic_04503_1 group_13210
MTITTSRVERVLWIACPTLMFGVMFGHMNGVVNTTLALIGVGTLAGILSASRPPFRQWPL
VLPIVVWAAWSLASVGWSLYPRISLRAWFDEVLYPLVTFWGFWLFGSRVKRPEWPVLVVW
AACVLLALLSAFYWGHLQPPTPNTFPIRFYNRVGHTSTLVVFAMPLFATLLLRARWRAIG
AVGLVACGFVGLASLNRFFWPAAGATLVVAFYPLYRRRLGVSIVAIAILGATALGLLEYS
ARERDLSASTATASASSAARDITVAGQRVYVPGELNALGDTLSSDTRPKLWAFYTEQGKQ
HAWLGVGFGKPLPGHAYAADAPPLLLQIEPQALTHAHNLFLNTWLQTGYIGVALEALLLV
SLAAAFWRLRRDVPAVSAAGLALVVGMIAKNTVDDFMWQTTMLAFWSFAGFLLGCGERDA
CMARVSQNADAR*
>GCA_000452945.1_BPMSHR5858_genomic_04504_1 group_1526
MSRLHGRIHIFSRALPRLVLKPLRRKPAHLRRILIAHHLLLGDTLLLTPLVAKLREQHPD
AQIVLACPKAIAPLYAKRPFGVDALAFDPRDGVSVRRLLASGPYDLGIVAGDNRHSWLAL
GAGCRWIVAHAGDAPGWKNWPVDELVPYPEAPAAWADFAAALVDGPAPRLYRPSDWPAPQ
ASAPLPAALRERPYVVLHPGASTAVKRWPSERWRELASLIEAQGYQPVWSGGPSEVEFVA
QIGPDSAQPNLAGRLGLADLWHVLAGARAVVCPDTGIAHLARLVGVPTVALFGPGNAGIH
GAGRYWRDAPFIALTIADMPCRDQPSMFRRHVAWVRRCDRSAQTCVAWRGDHADCMGRLS
VDAVHRALQNVLALAPSSSSR*
>GCA_000452945.1_BPMSHR5858_genomic_04513_1 purT
MQIGQRLGTPLSPSATRVMLLGAGELGKEVIIALQRLGVEVIAVDRYPNAPGHQVAHRAH
VIDMTDPDALRALVDAERPHLVVPEIEAIATDALAAIEAAGVCEVIPTARATQLTMNREG
IRRLAAEELGLPTSPYAFAQSFDEFAAAVARIGFPCVVKPVMSSSGKGQSVVRSEADIEP
AWRYAMAGGRVNHGRVIVEGFIRFDYEITQLTVRAIDPASGQTRTSFCAPIGHLQVAGDY
VESWQPQPMSAKALERSRDIAHRVTSALGGRGIFGVELFVRGDDVWFSEVSPRPHDTGLV
TLASQRQSEFELHARAILGLPVEPALATPAASAVIYGGLDEAGIAFEGVRDALAVPGADL
RLFGKPESFAKRRMGVALATGANVDEARERAKRAAAAVRPVSAR*
>GCA_000452945.1_BPMSHR5858_genomic_04529_1 phaC_1
MQQLFESWLGAWRSFADPARAAAGDAPSPSPSPFAAFQPPQPFAFAMPAMPPMPPMPDWS
GAAASFAGLAPVASVPPARLQKLQADYSRDCLALIQQASAATPTVPELKDRRFSADAWKA
SPAHGFAAAWYLLNARYLQELADALETDPKTRERIRFTVQQWTAAASPSNFLALNPEAQK
NLVETQGESLRLGMMNLLADMQRGKISQTDESQFVVGKNLAVTPGAVVYENDLIQLIQYT
PTTATVFERPLLIVPPCINKFYILDLQPENSLVAHALSCGHQVFLVSWRNADASVAHKTW
DDYIDEGLLAAIDVVQQVSGREQINTLGFCVGGTMLATALAVLAARGEHPAASMTLLTSM
LDFSDTGILDVFVDEAHVQMREQTIGGKGGAPAGLMRGVEFANTFSFLRPNDLVWNYVVD
NYLKGRTPAPFDLLYWNGDSTSLPGPMYAWYLRNTYLENKLREPDALTVCGEPVDLSRID
VPTFIYGSREDHIVPWQTAYASTSLLTGPLKFVLGASGHIAGVINPPAKRKRSYWSYDAS
AKELPESANDWLDAAVEHPGSWWPVWIEWLDQYGGKKVKPRAHLGCARFPVIEPAPGRYV
LQRD*
>GCA_000452945.1_BPMSHR5858_genomic_04530_1 yfiH
MTMLRPLTLADCVQPDWQAPPRVRALVSTRNGGVSEPPYGAWGGDGGEVAGGMNLGRHTG
DDPAHVEANRARLLALAGQSRAAWLEQVHGTEVVRAEDVLAAGEGGAVRADASVAATPGA
VCVVMVADCLPVLLCDDAGRAVGAAHAGWRGLVAGIVEKTAERVAALAGGGADRLHAYLG
PAIGPGAFEVGADVRSAFLDAAPLADYAATERAFAARAPAPGKYLANLYALARLRLARVG
VTRVSGGAACTATERERFYSYRRDRVTGRMAALIWLAD*
>GCA_000452945.1_BPMSHR5858_genomic_04533_1 yoaA
LNSPLDAHPDARRDDPSAARSPTSGAATGAYTPSVKRIVELDAIFGEGGLFARALDNYRP
RASQIDMARAVAAAMEASGKLMPEPEIFETRKRPARRMQDGAAAGRDDACAAPPGDESAD
SGDNTLIVEAGTGTGKTYAYLVPAMLWGGKVIVSTGTKHLQDQLFQRDIPTVRDALAVPV
TVAMLKGRANYLCHYYLQRTADNGRLPSRQDTAYLQDILRFAKITRSGDKAELASVPETS
PVWSMVTSTRDNCLGQECPHYKDCFVMQARREAQQADIVVVNHHLFFADIMLRDTGMAEL
LPSANTIIFDEAHQLPETATLFFGETLSTTQLLELARDAVAEGLAHARDAVEWVKLGGAL
ERAARDVRLAFADDGIVRMSLAQLGDGHPLFGALDALDAALDALSTALAGQAERAESIAA
CQRRARELQDLLAGWVAPGAQDGAADAADAKPAAQTGEHGDPAEKVRWVEVFAHTVQLHE
TPLSVAPIFAKQRAGVPRAWIFTSATLSVRGDFTHYAAQMGLSSRRSMTLPSPFDYQAQG
LLYVPRNLPQPSSPAFTDAVFDAALPAIEASGGGVFVLCTTLRAVDRIAMKLRDTIDARG
WSTPLLVQGDASRTELLDRFRAYGNAILVGSQSFWEGVDVRGDALSLVVIDKLPFAPPDD
PVLAARLDALTKKGLSPFAVHQLPQAVITLKQGAGRLIRAETDRGVLMICDTRLVDKPYG
RRIWQSLPPFKRTREIDVVREFFAERREAEQA*
>GCA_000452945.1_BPMSHR5858_genomic_04536_1 yhgF
MTETVALKIVQRIADELSVQPRQVAAAVQLLDEGSTVPFIARYRKEVTGNLDDTQLRQLE
ERLLYLRELEERRATIIASIDEQGKLTDELRAAIDAADSKQTLEDLYLPYKPKRRTRAQI
AREAGLEPLAQALLANPLLDPQAEAAAYVNTDRGVADVKAALDGARDILSEQFGETAELL
GKLRDYLFERGVVSSAVVDGKQGEEGEKFRDYYDYSETIKTVPSHRALALFRGRNAGVLT
VKLGLGEELDAQVPHPGEAMIARHFGIANQNRPADKWLSDVCRWCWRVKVQPHIETELLT
QLRETAEHEAIRVFARNLKDLLLAAPAGPKAVIGLDPGLRTGVKVAVVDRTGKLLATDTI
YPHEPRRDWDGSLAKLARLAAQTQAELVSIGNGTASRETDKLASELIAKHPELKLQKIVV
SEAGASVYSASELAAKEFPELDVSLRGAVSIARRLQDPLAELVKIEPKAIGVGQYQHDVN
QRELARSLDAVVEDCVNAVGVDANTASAALLARVSGLNSTLARNIVDYRDANGPFPSREH
LRRVPRLGDKTFEQAAGFLRINGGENPLDRSSVHPEAYPVVERMLAKISKRIDDVLGNRD
ALAGLSPAEFVDERFGLPTVRDILSELEKPGRDPRPEFKTATFREGVEKVSDLAPGMVLE
GVVTNVAAFGAFVDIGVHQDGLVHVSAMSTKFIKDPHEIVKAGQVVKVKVLDVDVKRQRI
SLTMRLDDDAAPSAPGNRGGAERGAMRGGARAQRSREPEPAGAMAAAFAKLKQR*
>GCA_000452945.1_BPMSHR5858_genomic_04537_1 ynfM_2
MCGCAAFLGLYATQGILPQIAATFGTRIEQAALGITATTVAMAATAPFVGMLTRRLERRR
IITLAALLLAAPMLWTAHADSFAEFLAGRIATGVVIPVLFAVTVSYIGETWQDSTATEMT
SFFIAGTTLGGFGGRFIVNGVTAMSGWPRALDVLACAMLAAGVAIRLCLPRTRAAATAAA
PAAMSLRALGRNRPVLASYFIGACILFSQVTTFTYVGLYLAKPPYRFGTAAIGSIYAVFL
LAVVVTPMAARLSRRRPPADLLALAAGLGVAGALLTLHADVRVILLGLAIGSTGVFVGQA
AASAFVTRSTAEGRTFAVGLYLSFYYLGGSAGTVLPVPAWQRFGWAGCVALVVCAHLLAA
IAACTFWKPGGIDATRTRPSSITTSR*
>GCA_000452945.1_BPMSHR5858_genomic_04543_1 group_2543
MSLFSLLGALEIGLIFSLVALGVLISFRILNFPDLTVDGSFPLGGAVAATLIAAGHDPFT
STLAAIAAGACAGFVTGWLNVRLKIMDLLASILMMIALYSVNLRIMGRPNVPLITEPTLF
TVLQPDWMPDYVLRPALLAVVVVCAKLGLDWFFSSQLGLAMRATGANPRMARAQGIPTGR
ATLAGMALSNALVALAGALFAQTQGGSDISMGIGTIVIGLAAVIIGETLLPARRLFLTTL
AVVLGAILYRFFIALALNSEFVGLKAQDLNLVTALLVTVALVLPATRKKLFGRKTGGA*

>GCA_000452945.1_BPMSHR5858_genomic_04544_1 group_5053
MKRFKIVAAHSIAAGVAAFAMLGAGAAHAQTVKVLSIVDHPALDAIRDGVRAQLKAEGYG
DDKLKWEYQSAQGNTGTAAQIARKFIGDRPDVIVAIATPAAQAVVASTKTVPVVYSGVTD
PVAAQLVKGWGPTGTNVTGVSDQLPLDRQVALIKRVVPKAKTVGMVYNPGEANSVVVVKA
LKEILAKQGMTLKEAAAPRTVDIAPAAKSLIGKVDVIYTNTDNNVVSAYESLVKVANEAK
IPLVAGDTDSVKRGGIAALGINYGDLGRQTGKVVARILKGEKPGAIASETSSNLELFVNT
DAAAKQGVTLAPDLVKEAKTVIK*
>GCA_000452945.1_BPMSHR5858_genomic_04547_1 group_1843
MNRMSLNRKLWLALALVWVGLLGVGAWSAFETRTTMLAERKEGMANLVDSAAGVVKAYYA
LAQSGKMSDEDARRDALERLAAMRYGDSGYLFVMDSKPVVLMHPTLPKLVGTQVGDYLDP
DGKRLFVTILNAAKAGGHGFAEYRGRLPHSETAVPKISYVTRFAPWDWNISSGVFLKDID
TVYYETLVGHLAVVFVIGLVITAAMLVIIRNVRASLGGEPDEAASLAARIAAGDLTRPVA
VRAGDGTSMMAAMRDMQGRLRSTIGGIRRAAESIAAASHEIASGNHDLSQRTEQQAASLE
ETAASMEELTATVKQNAENARQASGLANNASEIALKGNDVVSRVIGTMGEINDSSRKIAD
IIGVIDGIAFQTNILALNAAVEAARAGEQGRGFAVVAGEVRSLAQRSATAAKEIKQLIDA
SVERVNNGSALVGQAGETMTEILQAVRRVTDIMGEIAAASEEQSSGISQVGRAVTQMDEM
TQQNAALVEEAAAAASSLQEQAARLRESVSAFQVGDGAAAGAGAAAGVVGGPNAAASSGR
VEPALA*
>GCA_000452945.1_BPMSHR5858_genomic_04548_1 group_5054
MYVIDIRYTAPLERIDDALERHRAYLQRHFDAGVFVACGPKVPRDGGVILAVRIERDRLD
AILETDPFVTEGLVTYTVTEFKTTRVAPGVNLPALPS*
>GCA_000452945.1_BPMSHR5858_genomic_04549_1 group_5055
MLKTLFARRRAQRGWLFACALAGAMLIAGCNRQPAFQNLDITGNSQFASDFALPDTAGKI
RTLGEFKGKAVVVFFGYTHCPDVCPTTMAELSQALQQLGPDGKRVQVLFVTVDPERDTAA
LLGQYVPAFNPAFIGLRPADEAQLKKVTKDFRVYYAKVPGKTPGSYTMDHTAASYVFDPN
GKLRLFVRDGQGPGPWVHDLKLLLD*
>GCA_000452945.1_BPMSHR5858_genomic_04551_1 ctaA
MYLLQLGLIGLCIALLPLSYVWVKADDDKFRKLVWITTFLTLDLVMFGGFTRLTDSGLGC
PDWPGCYGTSSPFIAHAAITAAHQAMPTGPVSMTKAWIEMIHRYFAMAIGVLIIAQTVIA
WAARLRRKPLHVSPWWPTSLLLLILVQGAFGAWTVTMKLQPVIVTIHLLLGLTLLGTLGW
LAARQTPLPAHEPGASRYRAAALAALVLLVVQIALGGWVSTNYAVLACTDFPTCNGAWIP
PMDFRNGFHLWRALGMTNDGDAITQDALVAIHWTHRTFAFVVVAYLAAFALKMRRFASLR
RPANGVLAVVVLQFVTGLTNIVLQWPLPVAVAHNGGAAILLLLVVMLNFRILSSRPGRVA
QPARDAAPA*
>GCA_000452945.1_BPMSHR5858_genomic_04558_1 ctaD_1
MSSIGHDVAAGHAHDDHAHETPHGWRRWLFATNHKDIGTLYLLFSFIMFLSGGVMALAIR
AELFEPGLQIMRPEFFNQLTTMHGLIMVFGAIMPAFVGFANWMIPLQIGASDMAFARMNN
FSFWLLPVAAVLLVGSFFSPGGATAAGWTLYAPLSTQMGPGMDFAIFAVHIMGASSIMGG
INIVVTILNMRAPGMTLMKMPMFAWTWLITAYLLIAVMPVLAGAITMVLFDRHFGTSFFN
AAGGGDPVMYQHIFWFFGHPEVYIMILPAFGIVSQVIPAFARKPLFGYSSMVYATASIAI
LSFMVWAHHMFVTGMPVTGQLFFMYATMLIAVPTGVKVFNWLATMWRGSLTFETPMLFAI
GFLFVFTMGGFTGLMLAMAPLDIQYHGTYFVVAHFHYVLVAGSLFALFAGWYYWAPKWTG
WMYNETRGKIHFWASMIFFNVTFFPMHFVGLAGMPRRYADYPAQFTDFNQLATIGAFGFG
LAQVYFLFAIVLPAYRGGGELERASDKPWDGATGLEWTVPSPAPFHTFEHPPTVE*
>GCA_000452945.1_BPMSHR5858_genomic_04561_1 tam_1
MARSERVYFGLFASVPDMSPASAKTSRPAYDPRRLRRIFDRRAAAFDAVSFLPREIAQRM
RERLDYIKVNPAGVLDAGCGTGDDLPLLRARFPQAPVFGVDASGGMLARAAARDTAETSW
RRFLPATLTKALGHRGPRVAQADFSALPFASGAFDLLWSNFALHWHARPDLVFPEWHRVL
RVDGLLMFSTLGPDTLRELRAACADAAAAAGEARAVARVIDFVDMHDLGDMLVESGFEIP
VMDQETLTVTYKSPDSLLADVRRLGAYPFERGAPGHASRRLRAALYDALEARRRDDGTIP
LTFEVIYGHAWKAAPRTTAEGFSIVRVQDIGKGRPKRS*
>GCA_000452945.1_BPMSHR5858_genomic_04562_1 group_1916
MANPAVLRCGIRAMAARAGIVLARFSAAALPNRCALCGNLSHRTICDCCDDAYWNEARLR
CPRCALPLPGARGAMRFHCGACAKAPPPFDATLALADYRAPLDSLALDLKFRAQLALGRE
FGERLARLATDALDGAPPLDVIAPVPLARRRLVERGYNQAWAIARPLARKLKVRADAALA
ARVADTAPQSRLAFDARRANVAAAFAVARPVAGLHVGVVDDVMTSGATLDALARTLKEAG
ARRVTNFVALRTAKD*
>GCA_000452945.1_BPMSHR5858_genomic_04570_1 group_5348
MRMKLKNIGLIAAGLATGVFATLQISASAQQAVTTAAAPLPLDQLRLFAEVFGQIKREYV
EPVDDKKLLTAAIKGMVSSLDPHSSYLDKTDYQELQEQTKGRFAGLGIEISQEDGLVKVI
SPIEDTPAFRAGIRPGDLITRINDRPVRGMTLDKAVKQMRGEPGTKVTLTIFRKSDDRTF
PVTVTRAVIRVQSVKMKLLDPGYAYIRITSFQERTTPDLAAKLQDIARQQPNLKGLILDL
RNNGGGLLQSAVGVAGAFLPPDSVVVSTNGQIPDSKQIYRDNYENYRLPSFDSDPLKNLP
AVFKTVPMIVLTNAYSASASEIVAGALQDSHRAVIMGKATFGKGSVQTVRPMTADSALRL
TTAYYYTPSGRSIQNKGILPDIPVDQYADGDPDDVLVTREVDYTNHLANTQDPNEKKELE
EREQRRMEQLRILEEQNDKKTPEQRQKDRERKPIEFGSADDFMMQQALNKLEGKPVEQSK
MIAADSTAKSAAAKAGAASAAKGASGAAAKPASAAKPASAAKPVSAPQPQ*
>GCA_000452945.1_BPMSHR5858_genomic_04574_1 group_5060
MAGILIIAHAPLATALRDCIAHIYGGLPARIGCIDVQAENDPAQVMAFAHAELARLKEEN
GALVLTDMYGATPANIAGQLAKVEGVRVLAGVNLPMLVRAVCYRTTPLDTLVDKALAGST
KGIHEIAAGTPPPRPSALGCGECAPIPPEPKAQTQPH*
>GCA_000452945.1_BPMSHR5858_genomic_04575_1 gshB
MDILFIADPLDRFKIYKDSTYAMMAEAARRGHAVYACEPNQLAWTGAGVEADVRRVTIAG
DTADLHRDRWYEAAARESRALPSFGAILMRKDPPFDMEYVTSTWLLELAERAGARVFNKP
QTIRDHSEKLAIGEFAQFVAPSLVTRDAARLRAFHAEHGDVILKPLDGMGGMGVFRVKAD
GMNLGSIIEMLSHDGARSVMVQKFIPEIKAGDKRILLIDGEPVPYSLARIPQGSEVRGNL
AAGGVGVAQPLTERDREIAAALGPVLAARGLLLVGLDVIGDWLTEVNVTSPTCFREIMEQ
TGFDVAGMFVDALERAVG*
>GCA_000452945.1_BPMSHR5858_genomic_04577_1 amtB
MRKILMSLLMAGSLLAGGIGAAMADDASSAPAAASAATASDTSAGAAASAPAASAAPAAP
AAPAASAPAAASAAAPASAAAAPAAPTAPFSVDSSKINSGDTAWMLTSTALVLFMTIPGL
ALFYGGMVRKKNVLATVMQSFAITALITVLWTVVGYSLAFTPGNGFIGGFSRVFLSGMNY
IHGDKATTLTVSHLASTIPESVYFVYQMTFAIITPALICGAFADRMKFSAMLVFMTLWSL
IVYVPIAHMVWEPTGWLSADGVLDFAGGTVVHINAGIAGLVSCLMLGKRVGYGREAMAPH
NLVLTLIGGSMLWVGWFGFNAGSAVAADGRAGFAMLTTQVATACAALGWMFAEWIAKGKP
SVLGIVSGAVAGLVAITPAAGFVGVAGALVIGIAAGVVCFWSATWLKHKLGYDDSLDAFG
VHGVGGILGALLTGVFAVKDIGGADGSLLLQAKGVAITLVYSGIVSFVLLKVIDIVIGLR
VTEEEEREGLDVILHGEHVE*
>GCA_000452945.1_BPMSHR5858_genomic_04587_1 dctA_1
VKKPFYKVLYVQVIFAIVVGVILGHYYPSLAVDMKPLGDGFIKLIKMVIGPIIFCTVVTG
IAGMQDMKKVGRVGGKALLYFEIVSTCALVLGLAATHILRPGVGFNIDPATLNGKEVASY
AAKAHGQSSVDFLMHIIPNTMIDAFAQGEILQILLIALLFGSVLAHLGERGRVVTDFIDG
ITRVLFGIVHIVTKLAPIGAFGAMAFTIGKYGVGSLVPLLKLIGTFYLTSVVFVLVVLGA
IARFTGFSIIRFVGYIKEELLIVLGTSSSEAALPQLMEKLEKAGCSRSVVGLVVPTGYSF
NLDGTNIYMTMAVLFIAQATNIELTWMQQLTLLAVAMLTSKGASGVTGAGFITLAATLAV
VPTIPLSGMVLILGIDRFMSECRALTNIVGNGVATVVVSAWEKELDRAKLRAALSGNGEA
AAGEAARV*
>GCA_000452945.1_BPMSHR5858_genomic_04590_1 uctC_1
MRNEPGARPLAGIKVLDFSRVLAGPWCAMVLADLGAEVIKIEHPRRGDDTRDWGLRVGDT
ETTYFNSVNRSKRSICVDLQTQAGRRIARELAAQADVVVHNFKAGGAEKLGLGYDALAAL
NPRLVHCAISGYDRSGPEAGRPGYDLVVQGEAGLMALNGEAGRPPLKFGVAVADLFTGMY
SAQAILAALYERHATGRGRRIEMALFDCGLMVTSYYGLEALLMGEDPPRYGNAHPSIVPY
GVFDAADGPIVITVGNNPQFARFCDVIGRPELAADARFATNIARSANRAALLPEILRELG
RRPRAALLAALAALADAGIPCGEVLGLREALTSERARSAGLVTRQPHPVAGEVDVLAPPY
RFDGERLPVRGAPPVLGADTDRVLGEWLGMSGREIAQLRSERVV*
>GCA_000452945.1_BPMSHR5858_genomic_04600_1 dapE_2
MSATLALTEQLIARASVTPDDQHCQQLMIERLAALGFECETIASHGVTNFWAVKRGTAGR
AGKLLAFAGHTDVVPTGPLEQWSSPPFVPTHRDGKLYGRGAADMKTSLAGFVVAAEEFVA
AHPQHRGSIGFLITSDEEGPATDGTVKVVEALAARGERLDYCIVGEPTSTATLGDVVKNG
RRGSMSGELVVKGVQGHIAYPHLAKNPIHLLAPALAELAAEQWDEGNEYFPPTTWQVSNL
RAGTGATNVIPGHADLLFNFRFSTASTVEGLQARVHAILDRHGLDYTLNWSVSGLPFLTP
RGELSNALDAAIRAETGVSPELSTTGGTSDGRFIARICPQVIEFGPPNASIHKIDEHIDV
RFVDPLKNVYRRVLEQLIA*
>GCA_000452945.1_BPMSHR5858_genomic_04603_1 gpx1
MSKLYSFNAQALSGGEVPLEQYRGKVLLIVNTASECGFTPQYAGLQQLYDRFRERGLVVL
GFPCNQFGKQEPGDASQIGAFCEKNYGVTFPMFGKIDVNGANAHPLYRYLTEEAPGILGL
KAIKWNFTKFLVNREGEIVKRYAPSTKPEDIAADVEKLL*
>GCA_000452945.1_BPMSHR5858_genomic_04634_1 yejE
LSRLAVSSRIDAARARVSPSPARRVWLRFKAQRLGYWSLVIFAIAFAASLAAPLWSNDKP
LVVRYDGQFYFPLFKSYPETTFGGDFPTPADYLDPYVRGKLDAPGNFVLYPPNRYYYDTL
NYFSKRPNPAPPSRENWLGTDAQGRDLLARLVYGFRVSVEFALILTFIGTLLGIAAGAVQ
GFFGGRIDIVGQRLIEIWSSMPELYLLIIFASVFEPSFVLLIVLLSLFGWIGLSDYVRAE
FLRNRTQDYVRAARAMGLTNWQIIWRHVLPNSLTPVITFLPFRMSGAILALTSLDFLGLG
VPPPTPSLGELLAQGKANLDAWWISLATFCVLVATLLLLTFMGDALRNALDTRIADTARA
AGRLQ*
>GCA_000452945.1_BPMSHR5858_genomic_04645_1 group_1919
MGIVKISDGMHESLRHASAALSRSINAQAEHWLRIGMLAELHPMLSYAEICRLLIETEIQ
GDEAAPSVDAARSA*
>GCA_000452945.1_BPMSHR5858_genomic_04648_1 cysN_1
MSIIENNEDLGVLRFITAGSVDDGKRALIGRLLYDGEAMEAEARRAPPRETRMRAGGAAP
DPASAPRTHVVPDLDAMAGGPSIAAGGHVGRDGGRTCAKPDRTRAASAHAPPRRLFSIGR
APATFGELVQGREPASGDDFLITLPITLSSTARFCRFRDSDRLYVFPASKKKSLKAAALF
LERFGILTGGVLQICSDVSEGKGLASSSSDIVATLRALAACFDIPLSPADMCAIIREIEP
TDGVMFDESVAFFHRRVELGKVMGRLPKICILAIDEGGTIDTVEYNCHRFEFSHEEADQY
AALLADVDAAISRSDVRQIGRAATLSAQMHQKRNPKRTLRQLEALMREVGAHGIVNCHSG
TFIGLCFDASGPDALDTIARAERTLRDALGQPISRFFTR*
>GCA_000452945.1_BPMSHR5858_genomic_04652_1 fdoH
MALQSLDIKRVSATTTPPPSAREPVTGSVAKLIDVSKCIGCKACQTACMEWNDLRDEVGT
NVGVYDNPADLTEHSWTVMRFAEYENPAGDLEWLIRKDGCMHCEDPGCLKACPSPGAIVQ
YNNGIVDFHEEHCIGCGYCVTGCPFNIPRISKQDNRAYKCTLCSDRVAVGQEPACVKTCP
TGAIVFGTKEDMKQHAAERVADLKGRGFENAGLYDPQGVGGTHVMYVLHHADKPSLYHGL
PDNPSISPMVKLWKGLAKPLAVAGIALSALVGFFHYTRVGPNDVTDDEEAAARDEARRIQ
EDAK*
>GCA_000452945.1_BPMSHR5858_genomic_04661_1 bglB
MHAKRLSIAVLSATLCALAHAAGNDAPSPDIASRDAYALRRAHALVRQMTLDEKLQLIHS
KYPMSDVPGGGAGFIQGIARLGIPDLNMVDSATGSGSTSQPSTTFPATIGLAASWDKRLS
YAFGAVIADQLRAQGFAMGLGGGTNLAREPRGGRLFEYLGEDPVLAGEMLAARTRGTQDR
KVIATIKHYVGNEQETNRMGGDDQIDERTLRELYLLPFEIAMKAARPGNVMCSYNRLNGD
YACENAHVLTDVLKNEWHFQGQVQSDWGAAHSTAKAINAGLDEEEDVGPTVFLTPALVKQ
ALANREIAPARLDDMVRRKLYAMIRTGVMDDPPRGGGTIDFAAANRFVQYAAEQSIVLLK
NQDRQLPLDAAGLKRIAVIGGHADAAVLAGGGSGNTRHPVTGAFPGCGGLTFPTTTGCNW
WPNPWLKLDVPIVQAIRDLAPGATVAFAGNSDRQSPFAAYTPQQIDAAADLARRSDVAIV
FVTQAAGEDFGELRSLALANPTNQDALVQAVAQANPRVIVVVESGNPVLMPWRDQVPAIV
QAWFPGEGGGNAIANVLFGKVNPSGKLPVTFPARDEDTPTWGADGTLAPNPVYSEKLKIG
YRWYDAHRIAPMFPFGHGLSYTHFSYSGLEVKQRPDAATTVSFALTNDGPVAGAEVPQVY
LGDLDDPQEPPKRLVGWDKVGLRAGETRRVRIVIPAEMRRVWDASRNGWALAKGGRIYVG
ASSRDIRLQQP*
>GCA_000452945.1_BPMSHR5858_genomic_04690_1 group_18945
MFAYIIEKLSNWFEAAERERREAYLASSSDIVQLEQRIRSLETNGYSL*
>GCA_000452945.1_BPMSHR5858_genomic_04692_1 rlmJ
MLSYRHGFHAGNHADVLKHTVVVQLLRYLNKKDKSYWYIDTHAGAGVYSLFDGYAAKTGE
FGTGIEKIWDATDLPEALSDYVDEVRALNRDGELRYYPGSPYLAWRLMREQDRMRLFELH
STEIDVLRHNFRDAGRRAMIYAGDGFEGIKALLPPPPRRALVLVDPSYEDKRDYAKTTAC
VEESLKRFATGCYAIWYPQVSRPESQKFPDHLKQLQPNNWLHLSLTVSRPPSDGFGLFGS
GMFILNPPYTLVDTMKEALPYLVEALGQDSGAKFAIEHRAN*
>GCA_000452945.1_BPMSHR5858_genomic_04696_1 dfrA
MTTLTLIVARARNGVIGRDNRLPWKLPEDLAFFKRTTMGAPIVMGRKTHESIGRPLPGRR
NIVVTRDAARRFDGCDTATSLDDALALAARDGAAEAFLIGGAQLYEEGLRHADKLIVTEI
DQDFEGDASFPAPDPAQWEAVSRDTHRAAQPNDFTYAFVVYRRRRAG*
>GCA_000452945.1_BPMSHR5858_genomic_04700_1 group_5072
MRIAACLARRRHSIAAALAASLAIACPYVTAAHENETYRIVDTRPEHVAARGRAERCARV
DDGAFARQRDHAPMPVPVATPRPLVDGTAVTLWDEIGPPAPMPVPTDARHTVRGDGANYT
RQ*
>GCA_000452945.1_BPMSHR5858_genomic_04704_1 group_2562
MPASDRHDHFPGLSHLGALLADPGRAAMLWALMDGSARPAGELTMIAGLSPSAASAHLAR
LTDGGLLALDVRGRHRYYRIATPDIAAAIEALANVAQAAAPQRPIPQPARTVPPDMRYAR
TCYDHMAGELAVQVYERLMSRGWLTASGGTLDATAAGAAQFAQWGIDIGGQRAKRRRFAC
TCPDWSERRPHLGGSLGAALLDSFCRRGWLEHAAKPRVLRVTPVGQREFDALLTGG*
>GCA_000452945.1_BPMSHR5858_genomic_04717_1 pncB2_2
MIITSLLDTDLYKFTMMQVVLHHFPAASVEYRFKCRTPGVDLVPYIDEIRAEVRSLCELR
FTDSELDYLRRLRFVKSDFVDFLALFHLNEKYISITPSQKGGGEIDIEIKGPWLHTILFE
IPVLAIVNEVYFRNTQRRPDYREGRGRLREKIKLLGAKPEFADCKIADYGTRRRFSKVWH
EEVLRTLQDGLGPQFAGTSNVYYAMTHEITPLGTMAHEYLQACQALGPRLRDSQTYGLEM
WAKEYRGDLGIALSDVYGMDAFLRDFDMYFCKLFDGARHDSGDPFDWGERLIKHYEENRC
DPRTKVLVFSDALDIPKVMQLYARFRGRCKLAFGVGTNLTNDLGYQPLQIVIKMVRCNGQ
PVAKLSDSPGKSMCDDKAYLAYLRQVFGITQPPDDDAGK*
>GCA_000452945.1_BPMSHR5858_genomic_04718_1 lutC_2
MDTSAARRNILARIRAAQGRPAELADVEREHAADYVERHPSGPRQPLGDDLVSRFVDEAL
RLATTVDTVASMRDAPAAAAAYLRRMGLGTQAIAWPTLDALDWAGAGLAVEFRKPVDGDR
IGITGCFCATAETGSLVLLSSPTTCASAALLPETHIALVSASRIVAGHEDAFALIRAERG
VLPRAVNFVSGPSRTGDIEQTIVLGAHGPSRVHAIVVRDA*
>GCA_000452945.1_BPMSHR5858_genomic_04721_1 group_5075
MTETLLLAAIVALAVALVVALVMLLRGGPRGGGLERLAEQIDDANDAHAHAAERLERELR
AEIVDGARHSRTELAGNFAQLQQTLAAQLTSVATVQNNQIDGFAQQLAKLVAGNTQQFDA
MRDTLQRQAQLAREEQGTALRHFGDTLNQQLTQLTQANDRRIGEVRATLETRLKEIEANN
AAKLEEMRRTVDEKLHATLEQRLGESFKLVSDRLEQVHRGLGEMQTLAAGVGDLKKVLTN
VKTRGTWGEVQLEALLEQMLTPDQYAKNVATVPKSGERVEFAIRLPGRHDEGAAGGAEPV
WLPIDAKFPREDYERLIDAQERADAAAVEEAARALEARIRAEARTIAEKYVAPPHTTDFA
LLFLPTEGLYAEILRRPGLTDLLQRDYRVTVAGPTTLTALLNSLQMGFRTLAIEKRSSEV
WQVLGAVKTEFGKFGDVLARTKSQLETVTRSIEAAEQRTRVMNRKLKQVEALPGEAANGL
LGVDGAEADES*
>GCA_000452945.1_BPMSHR5858_genomic_04731_1 group_5076
VLERVALAAIKEQRAARRWRIFFRFAFLAVLGVLAFAFLSVSGDGGKLASGRHTAVVTID
GEIAASTNANAEDINTALDSAFEDSGTVGVVLKINSPGGSPVQAGIVYDEIRRLRKKYPA
KPLYVVVSDMCASGGYYIAAAADKIYVDKASIVGSIGVLMDGFGFTGLMGKLGVERRLHT
SGENKGFFDPFSPETPKMDAHAQEMLDEIHAQFIKAVKDGRGARLHESPDIFSGLFWTGA
KSIELGLADDYGTTDTVARDVLKAPDLVDYTVKESLTDRVARRFGAALGKAALKAAVAGG
ELKLR*
>GCA_000452945.1_BPMSHR5858_genomic_04742_1 rpoE
LTEPFHAASAGTQHVVLRASACLRVACALFVPAVGAALYACAAPLVGGGGAAALAGLAAA
LVALRARAACVRHQPAALLIDAGAGVLAAFDRDGRRLARGQIAGCTQWSDLLVVLRVQGR
GRRVTPLVIPADAVDAAVFRALCVLGRRAARGALAAA*
>GCA_000452945.1_BPMSHR5858_genomic_04749_1 lepB
MNFALILFVLVVVTGIAWVLDKLVFLPRRKKAADTAVAEFDRQQERVGERFADENAPQTR
ARLRDEKLRQPWWLEYTASFFPVILVVFVVRSFVVEPFKIPSGSMVPTLLVGDFILVNKF
DYGLRLPITNQKITAGRPLERGDVVVFRYPKDESVDYIKRVIGLPGDTVAYQDKKLTING
QPVPEAPLPDYFDEERMNYAKQFEETLGTRKNAILNNPSVPPFVMGADDYPYRNNCQYDS
RGVVCKVPPGHYFMMGDNRDNSADSRYWGFVPDKNIVGRAFFIWMNFSDLKRIGSFN*
>GCA_000452945.1_BPMSHR5858_genomic_04752_1 recO
MGTNDGTSDVGMTDEADADPQPFAAPPATGAPAADKPARKPRRAAPRTSEFRIAEQPAFV
LHSYPYRETSLVIDVLTRDHGRIALVAKGAKRPHSALRGVLQTFQPLSLSWSGKSELRTL
TGAEWVGGMLPLAGDALLCGFYANELLVKFCAREDPHPPLFQHYLVTLTRLAHGEPPVQV
LRSFERVLLRETGYAMTLKRTVARRAVEPDKLYVFDPQRGVRDAGSDAPSHWPVIAGQTL
LDMEEDDYHRAQTVAQSKTLMRFLLNTYLGGTPLATRQILIDLQNL*
>GCA_000452945.1_BPMSHR5858_genomic_04755_1 nagZ
MKLPPGPVMLDVAGTTLTRDDARRLAHPHTGGVILFARHFESRAQLVALTEAIRAIRDGI
LIAVDHEGGRVQRFRTDGFTVLPAMRRLGELWGKDVLHATKAATALGYVLASELRACGID
MSFTPVLDLDYGRSKVIGDRAFHRDPRVVALLAKSVNHGLALAGMANCGKHFPGHGFAQA

DSHVALPTDDRPLDEILANDAAPYDWLGLSLSAVIPAHVIYTQVDSKPAGFSRVWLQDVL
RGRLRFAGAVFSDDLSMEAAREGGTLAQSAQAALEAGCDMVLVCNQPDAAERVLDELRTT
ASRESSRRIKQMRPRGKVLEWRKLMREPRYLNAQGLLRSTFA*
>GCA_000452945.1_BPMSHR5858_genomic_04757_1 efp
MKTAQELRVGNVVMIGNDAWVVSKTEYNKSGRNAAVVKMKLKNLLNGGGQESVYKADDKF
EVVVLDRKEVTYSYFADPMYVFMDADYNQYEVEAEMMGDALNYLEDGMACEVVFYNEKAI
SVELPTILVREITYTEPAVKGDTSSGKVLKNAKLATGFELQVPLFCNTGDKIEIDTRTNE
YRSRA*
>GCA_000452945.1_BPMSHR5858_genomic_04764_1 msrC
MFALSEALPSSKPALYETLAAQARALVETETDIVANAANFASLVYHSLDGLNWAGFYFFD
GRELVVGPFQGKPACVRIPLGKGVCGTAAQTRGTQVVHDVHAFAGHIACDSASQSEIVVP
LVARDGALIGVWDVDSPLVGRFDAEDATGMEALCRVFVEVAWENATRRGE*
>GCA_000452945.1_BPMSHR5858_genomic_04766_1 group_5083
MKQIVGVYSAPRSHWVGDGFPVRSLFSYDSHGAHLSPFLLLDYAGPARFEPTARRRGVGR
HPHRGFETVTIVYQGEVDHRDSTGAGGHIGPGDVQWMTAAGGILHEEFHSARFARDGGTL
EMVQLWVNLPAKDKMSEPGYQTLLSERIPSIELPDGAGRVRVIAGEFEGRRGPARTHTPL
DVWDVRVAQGGRVRLRADAGRTLALVVLRGTVLVNGEEVVREAQFVQFARDGSDVDIEAN
NDATLLWLSGEPIDEPIAGYGPFVMNSDEEIRQAIDDFNGGRFGTMSA*
>GCA_000452945.1_BPMSHR5858_genomic_04767_1 elaA
MSTFTTAASPIPAAAPSAPLEWRWKPFGALTPDEIYAILSARSDVFVVEQNCVYRDIDHA
DFDAWHLGAYDAQGRLACYLRVLLPDASEPDIRIGRVLTVRAFRAMGLGNAMLTVALERI
RAQWPDTPISLHAQAHLQKFYGAFGFAPSSDVHDEDGIAHVWMRVSA*
>GCA_000452945.1_BPMSHR5858_genomic_04771_1 otsB
MRTFARRARPPAAILFSESMQSIPLSLPLARTAFFFDFDGTLVDLAPTPDAIQVPPDVPV
LVDALRQLSHGAVAIVSGRGIDSIDAYLNLPGLPVAGLHGAERRDANGDTQRIGFDDPRL
LRIERELAALVDRHPGMLLEIKGAALALHFRNAPEREGVARAAAERLVADYADAYVLQPG
KMVFEIKPKGVDKGRAVAAFLNEPPFAGRMPVFAGDDLTDEQGFAVANANGGLSIKVGAG
DTTARARVDSVAALRAQLARWIAAGRAPA*
>GCA_000452945.1_BPMSHR5858_genomic_04782_1 group_2567
MIPFSVLDLAPIPAGADAAQALRHSVDLARHAERLGYRRYWLAEHHNMPGIASAATAVVI
GHVAGATRTIRVGSGGVMLPNHAPLVIAEQFGTLASLYPGRIDLGLGRAPGTDQTTARAL
RRDLIGSADSFPDDVVELQRYFAAPAAGQRVRAVPGAGLDVPIWLLGSSLFSAQLAAMLG
LPFAFASHFAPDYLMRALDVYRAQFRPSAALDKPYAMVGVNVFAADTDDDARRLFTSLQQ
QFLKLRRGTPGQLPPPVESLDALGATEQELANVAHALSFAAVGSRDTVHERLRRLIAQTG
ADELIVAAQIFDHGARVRSYEIAAQVRDALRDEAGV*
>GCA_000452945.1_BPMSHR5858_genomic_04783_1 group_1625
MTESLSIVGWVLLALVCASCGYAVLAACAPAPRVPRAAARDGFEPVSVLKPLCGSEPHLY
ENLATFCEQRHPRYQLLFGVASAADPAIAVVRRLQADYPDCDIELVIDARVYGSNLKVSN
LVNLAERARHGRIVIADSDIAVEPDYLTRVTAPLADPSVGVVTCLYHARSVGGFWTRIGA
QFVDAWFAPSVRITHLGGSSRFGFGATLALTRATLDAIGGFKALKDELADDYWLAELPRR
LGLRTVLSEVNVATDVAEPSFAPLWLRETRWLRTIRSLNPAGFAFLFITFTAPWLVIGAA
LAAWLGPASAAGATAAWAAAIGTLARLALHARGAAGWRAFWRDLPLVPVRDALLALEWLA
AAFGTQVVWRGARMTVVGGDARATVVEAGDGR*
>GCA_000452945.1_BPMSHR5858_genomic_04784_1 fom3
MKTLFLQAPSYDGFDGGAGSRYQAKREIRSFWYPTWLAQPAALVPGSRVVDAPADGLSVE
DTLKIAKDYDLVIIHTSTPSFPTDAMFAEDLKKMKPSMLVGMVGAKVAVDPHNSLTATQA
IDFVCREEFDYTCKDIAEGKPFAEILGMSYRAKDGSIEHNGPRPMIENMDELPFVAPVYK
RDLKIDNYFIGYLNYPYVSIYTGRGCRSKCTFCLWPQTVGGHRYRVRSVESVLAEVKWIR
DNMPEVKEIMFDDDTFTDFKPRVEEIARGLGKLGVTWSCNAKANVPYSTLKIMKENGLRL
LLVGYESGDDQILLNIKKGLRTDIARRFNEDCKKLGIKIHGTFILGLPGETKETIKKTIE
YAKEINPHTIQVSLAAPYPGTRLYNQAIENGWMEENKTINLVSKEGVQLAAIGYPHLPKE
EIYHQLEHFYREFYFRPSKIWEILREMLTSWDMMKRRLREGVEFFRFLRAHEA*
>GCA_000452945.1_BPMSHR5858_genomic_04793_1 group_194
MRSMVHLKRDECIAYRQSLDGADVLTPPKALDAGRVDYFAVVMPGCTEPEQVDGASFRVL
DRAAGRVRIALPSQDDGQCAWSR*
>GCA_000452945.1_BPMSHR5858_genomic_04798_1 group_195
MDARFSGVPGRVADAGQRVSGALHAAERRSALFAAWQENQIMNKCFVKLNGLIALFVFAI
LMCGGCSRTSKSSSDEAEMKHEVTSSSESEVMSLNLNALNYTEIPIGKFYVDGTWGGNVR
SRIGSGGLGILCCVALPKKWRPGLTVTVEWRDDEMYKNDPNSLTSRVVPVEKYEYFSDGF
LWVLFFPGGKIKAYASQWMPGFPGFPEGLQAPNVACPGHFTLLNSDPRCPAPDNRIKP*
>GCA_000452945.1_BPMSHR5858_genomic_04810_1 group_2569
MQLRTLLLALAIVAIAVATTYYIVTVDTDTPFATTEPAASAASAAGASEPTARLIAPGDG
APSTPIARDDAQSPLPDAWK*
>GCA_000452945.1_BPMSHR5858_genomic_04824_1 group_18983
MTRNARRTWQWISGLLAICAAALAALPARADVAAGPGAWSSQQTWGADTVNGGNLTGYFY
WPATQPTTPNGKRALVLVLHGCLQSASGDVIDNSRGAGFNWKSVADRYGAIILAPNATGN
VYGNHCWDYANTSPNRASGHVGVLLDLVSRFVGDAQYAIDPNQVYVAGLSSGGGMTMVLG
CIAPDIFAGIGINAGPPPGTTTLQIGYVPSGYTAQTAANQCKAWAGSKADQFSTQIAGAV
WGTSDYTVAQAYGPLDTAAFRLIYGGTFAQGSKVSIAGGGTNTPYSDSNGKVRTHEIVVS
GMAHAWPAGAGGDNANYVDATHVNYPAFVMDYWVKNNLRANGAPVQAGTPPTGLTVTNAT
QTSISLAWNPVANASSYNVYRNGNKVGSSTSTAYTDAGLIAGTAYSYTVTEIDPSLGESA
PSPAVSATTQPSFACSATTATNYAHVQAGRAHDSLGIAYATGSNQNMGLDNVFYTNTLAQ
TSAGYYIIGDCP*
>GCA_000452945.1_BPMSHR5858_genomic_04826_1 mak
MKPADTLEDVGRAQFVAHMARRPRRARRREAPGVADTPLWYKDAIIYQLHVKSFFDANND
GVGDFAGLIAKLDYIAELGVDTVWLLPFYPSPRRDDGYDIADYRGVHPDYGTLADVRRFI
REAHARGLRVITELVINHTSDQHPWFQRARRAKRGSVHRDYYVWSDTDLKYAGTRIIFLD
TETSNWTHDPVAGQYYWHRFYSHQPDLNFDNPAVVREVLQIMRFWLDLGIDGLRLDAVPY
LVEREGTSNENLPQTHAILKLIRATIDAEYPNRMLLAEANQWPEDVQEYFGDEDECHMAF
HFPLMPRIYMSIASEDRFPIIDIMRQTPALAPSNQWAVFLRNHDELTLEMVTDSERDLLW
QAYASERRARLNLGIRRRLAPLMERDRRRIELINSILLSMPGTPVIYYGDELGMGDNLHL
GDRDGVRTPMQWSSDRNGGFSRADPELLVLPPVMGTLYGYDAINVEAQTRDPHSLLNWTR
RILSTRRATRVFGRGAIRFLRPGNRKILAYLRELDGETPVLCVANLSRASQAVELDLSEF
AGCVPTEMTSDSPFPPIGQLPYLLTFPPYGFLWFALSEHGREPAWHQQYAEPLPEYLTLV
MRRGETQPGAALLDTLAQDALPSWLARRRWFASKERRVDSARFDALTPIPGEPFQYAEAR
VAVDGREERYVVPLASAWGSETPQPLFAQLALARVRRGHTVGLLTDAFALPSFAHGVLRQ
LRAGAAVPVAGGGRLEFRPEPDLARLDPGDAPGVRWFAAEQSNSSLVIGEAIVLKIVRKL
AAGVHPEAEMGRHLRRIGYRNVAPLVGEVVRIGADGAPHTVAILQEYVDNQGDAWTRSCD
FLKRAIEELALPAANDADAAAPSAEPEMIDGYATFAGIVGKRLGELHAALAQPSDDAAFA
PQRVSPAYVDGWIGDALGWFERAVGLLAERLDTLEGQTRAAAELLVAQRAAVAEALRALV
PRELDGCCIRIHGDFHLGQVLDVQGDALLIDFEGEPARPLGQRRAQSHPLRDVAGFLRSL
SYASAAAQFTIEKAPQQAAERKRALFERFGQAAADRFVAQYREALSAASREFVEPRYADR
LLALFLIEKASYELCYEAANRPDWLSVPANGLAALVARLIGGAAPQQEDAR*
>GCA_000452945.1_BPMSHR5858_genomic_04828_1 glgX
MPTPTPDRLEPGRSYPLGAHWDGLGVNFAVFSAHARRVQLCLFDPTGRREIARLDLPECT
DEIWHGYLPDAHPGTVYAFRADGAYEPQHGHRFNPHKLLLDPYARKLVGQFRWSDALFGY
RLHSSRADLSIDRRDSAPAMPKCVVVDEAVDWSTDRRPCVPWRDTVIYEAHVRGVSMRRA
ELRAPERGTFAALACPAFIDHLLALGVTSVELLPVHAFMHARALVNLGLCNYWGYDTAAF
FAPEPSYLATRRLDEMRIAIRQLHAAGIEVLLDVVYNHTCEGNQLGPTICWRGLDNASYY
RLAPDDPRRHIDETGCGNTLDLSHPRVLQMVMDSLRYWSNAFNVDGFRFDLCATLGREAA
GFDPGAGFFDALRQDPVLATRKLIAEPWDVGPGGYQLGRHPAGFAEWNDRFRDTVRRFWR
GDAGMRPELAARLAGSADLFNHHGRRTWSTVNFVAAHDGFTLADLVSYAAKHNDANGEDN
RDGRDDNCSANWGAEGPTEDRQVRAVRARVARSLLTTLFAALGTPMLLGGDEFGRTQQGN
NNAYCQDNELSWFDWSAAESADGRTMIHFVGRLAALRRAYPVLSEPRYPSGERDGAPGLH
EIDWFDERGDALTVPAWHDRERRALTMRRVGNGRAGRPEALLLMLNASSDAIVFHAPPPS
LVYRILLDSAAPDAAPRLMPADGVEIAAHAAVIAAASVPEAG*
>GCA_000452945.1_BPMSHR5858_genomic_04829_1 malQ
MNDSTATHAYALPFGATCVDARHTRFHLWAPACRSAAVERQDGETVAMTPDGGGGFDAIV
RCGPGTLYRYLLDGTLRVPDPASRFQPHGVHGPSEVVDPAAYRWRHDDWRGRPWRETVLY
EVHVGAFGGYARIARRLPALAALGITAIELMPVNAFPGARNWGYDGVLPFAPDASYGPPD
ALKSLVDAAHGLGLQVLLDVVYNHFGPDGNLLPRYAPAFFRRDRDTAWGPAIDFSCPQAG
AFFLENALYWLDEYRFDGLRIDAAHAIDDDAWLAGLARRVRAYAGDARHVHLVLENERNT
ASLLAGGRFDAQWNDDFHNSMHVLLTGEQAGYYRAYADAPIRHLARVLGEGFAYQGEPSP
LHGGAPRGEPSADLPPSAFVAFLQNHDQIGNRAFGERLRALVNDDALRAASALALLAPQI
PLLFMGEECGTTQPFQFFTDHRGALAAAVREGRRREFAAFPAFADPAHRDAIPDPNDPAT
FARSSLAAPGAEPPDANAWRRFYRGALAVRARFVTPWLDGARALGATVLARADGGHANAL
VARWRLGDGNTLAIALNLDARPAALAAPPDGKIVFETPPRARDALADARLAAHACIAWRS
GNVNGVARRGRAVDAKHMPAVKHANGVNGQDGAP*
>GCA_000452945.1_BPMSHR5858_genomic_04924_1 bfrD
MKSRSDELKLGKFTTLCSVLAASPAFAQDAAPPAASADHDKELAPIQIKGAAEHSYKADF
SSSVKFTAPLVDTPKSVTVIPQELIHNSGAATLTEALRTVPGITFGAGEGGNPLGDRPFI
RGYDTQGSMFVDGMRDTGATTREIFNTERIEITKGSDGAYGGRGGAGGSINLVTKAPHLG
TTAEASAGLGTDRYRRFTADGNWQFADHAAFRLNLMSHNNDVAGRDAVNNERWGVAPSIA
FGLGTPTRVSASYYHLSTDDLPDGGIPYFYTTANKPANVDTIYPANVDRHNFYGLVNRDF
RKTTSDIGTLRIEHDITPSLTVRNTTRYTESTQDYIWTQPDDSQGNVVNGRVWRRNNNRN
SAINSIANQTELFGEFRTGPFKHSFTTGIELSREWGKRDSYSVATGTGKICQQGIGAASG
YNCTSLGSPNPNDPWAGSITRNNDYAHARTVTKSLYGFDTVELSKHWQVNAGVRVDDYST
RFTDTRANGGKTYTRDDTLVNWQLGLVFKPARNGSVYASYATSSTPAGALLGEGSETQSL
TPGRGGVGANADQLAPEKNRSIELGTKWNVLDDKLSLTAALFQIDTTNARVTLPNNQYAM
VGNKRVQGLELGFAGQLTRAWQVFGGYTYMKSELRDNGRNTADNGHQFPNTPKHSFTLWT
NYDVTPKFTVGGGAFYMSKVFGDTANLRAVPSYWRFDAMAQYRINKKLDLQLNVNNLFNR
TYFDQAYPAHYASIAPGRSAFVTLNARY*
>GCA_000452945.1_BPMSHR5858_genomic_04938_1 rbsB_1
MTWINRGRTLAGCVALAGALAAAPAGADTGKVGLDLPLLTSPFWQSYNRYLLHYAKDMQI
DALAPVNSNGDPAQQITDMNTLLNLGAKGIVVGPLDSAAIGRALDAAAARNVPVVAVDVA
PTQGKVAMVVRADNRAYGEKACQYLGEHVRRGKVVQIMGDLASVNGRDRSEAFRACMKGY
PNLQVLEIPAAWKGDVAATALDSLLSANPDVKGIYLQAGGVYLSPTLQTLRRKQMLYPAG
DAKHVAIVSNDGIPQEYEAIRRGDIDATVSQPADLYARYGLFYIKAALAGRTFKPGPTDH
GSVIVQRAPGVLEDQLPAPLVTKANVDDKGLWGNTIK*
>GCA_000452945.1_BPMSHR5858_genomic_04939_1 rbsA_1
MNAIAAVTARACTDAGAPVVEARALTKRFGPHIALADVDLCVRAGESHALVGRNGAGKST
LVSILTGLRKPDAGVVRFAGEAAPPLADRDGWRARVACVYQHSTIIRELTVAENLFINRQ
PTKRGAIDWRALRRDARALLEHWRIDVSENARAGDLTVEARQLVEIARALSYGARFIVLD
EPTAQLDGDEIKRLFARIRELQRDGVTFLFISHHLQEVYDICQAVTVLRDARRIVSAPVA
ALPRDALIEAMTGERGALAVADAARRAPPPDDAPRALEVDGLAGDDYERVSFCVRRGEVV
GLAGATSSGRTSVAEAVAGLRAPRAGAVRVDGAALPPADVSAALARGVGCVPKDRHREGL
VLGESVAENASMTIAGALGRLGRFGLVPPARKHAFGARMIDALGIVAQGPRQPVSGLSGG
NQQKVVMARALANDPRVLVLIDPTAGVDVKSKEALLAVVERVRGGGAAVLVASGELDDLR
TCDRVLVMFRGRIVAEFQAGWQDNELIASIEGVDAHEV*
>GCA_000452945.1_BPMSHR5858_genomic_04954_1 phnZ
MALTLDDIRMLFDRHGDIAYSGEPVTQREHALQSAQLAEQAGANDALIAAALLHDLGHLL
NRRGETPTAHGIDDLHQYCVLPFLRPLFPDAVLEPIRLHVDAKRCLCAIDASYYARLSAD
SVRSLHLQGGVFSDAEAEAFLKRPYAPDAIRVRRWDDLAKMAGKPTPGLDHYMRAVARAA
IAK*
>GCA_000452945.1_BPMSHR5858_genomic_04956_1 phnA
MNRHIEVNGRVYRMPAEPTVAVCVDGCEYDYLEEAASAGVAPFLARMLREGSAFEADCVI
PSFTNPNNLSIVCGAPPSVHGICGNFFWDQNAQGGAGAAVMMNDPAYLRAGTLLAAAADA
GAKVAVVTAKDKLRRLLGWRMRGVCFSAEKANQANLGENGIADVLDFVGLPSPDVYSAAL
SEFVLAAGVRLAQSRCADLMYLSTTDYVQHKCAPGSEGANAFYAMMDRYLAALDALGWVI
GVTADHGMNAKHDPRTGEPNVVYLQDALDAWLGPRLARVILPITDPYVVHHGALGSFATI
YVEQGVDAGTVIGRLREFDGVELALDNATAAERFELPADRIGDVVVVGRRDVVLGTRARE
HDLSGLTVPLRSHGGLSEQRVPLLFNRRIEHVEPGRRLRNFDLFDLALNRTVQ*
>GCA_000452945.1_BPMSHR5858_genomic_04960_1 phnS
MTYTNFPRGGAWRRFALAAGAAALLQGAAAQAQAAAVVLYTADGLENLYRDVLPAFEKQE
GVKVNIVTAGSGEVVNRANVEKGSPKADVIVTLPPFIQQAGQFGLLQPYRSVNYKNVPAI
AKAEDGSWATFVNNYFSFAINPSVVKSQPKTFADLLHPDYSGKLAYSNPATAGDGMAVII
LTSALMGEDKAFDYLAKLERSVKFHTKGTGYLNVLLSRNEIAVANGDLQMDLDDAEHGGL
SIKPIFVAAKAGEPPTTFQLPYAIGLVKGGPNQDAGRKLIDYLMSADVQAKVPDMFGIPG
RTDVPLSGKNGEAVKRAIAGVKLIPVDWDAVMAKKPVWTERWKKEVIGDSGKQTEVVKPK
*
>GCA_000452945.1_BPMSHR5858_genomic_04964_1 aroQ_2
MNKPKVLVLNGPNLNLLGKREPRIYGSETLQDVERRCADAAHALGLALEFRQSNAEHQLI
DWLHDAREHVAGVVINPAAYTHTSVAIADALAALGKPIIEVHISNVHRREAFRHHSFVSA
LADGVICGCGTEGYVFALRRLATLIGGEGAR*
>GCA_000452945.1_BPMSHR5858_genomic_04965_1 hpd_1
MQRSIATVSLSGTLVEKLRAIRAAGFDGVEIFENDLLYFDGSPADVRAIAADLGLAIVLF
QPFRDFEGVPPERLARNLERAKRKFELMHALGANRMLVCSNMSPDAIGDDALLIDQLGAL
ARAAQAAGVVAAYEALAWGRNVKTYGHAWRLVDAVNHPSLGLALDSFHTLSLGDSPDGIA
RIPGERIAFVQIADAPKLAMDVLEWSRHYRSFPGQGDFDLAGFTARVIESGYAGPLSLEI
FNDGFRAAPTALTAADGYRSLLYLEETTRERLACDARRARRAGGAPGAGETRGERERHDA
HGERGEHRAHDKDDKPDDKRGGPDERESRAAHPRPAQPLFAPPPAPAHVGFQFIEFAVDA
AAAENVAGWLGKLRFRRAGRHRSKDVTLYQHGAASIVLNAERDSFADAFFQEHGLSLCAS
AFRVDDARLAFERAAGYGYAPFSGRVGPNERVLPSVRAPDGSLNYFVDEAPGAPTLYESD
FVLTDIDGPSEVGPLAGIDHVCLALPADALDTWVLFFKTAFGFEAERNWLVPDPYGLVRS
RAVRSPDGSVRIALNASVDRHTAVVRSLERYRGTGLNHVAFRADDIVAAIAEFAADGVPF
LRIPRNYYDDLATRYALPDETIDTLRRHHLLYDRDDAGGEFLHAYTELVDGRFSFEIVER
RGGYDGYGAANAAVRLAAQAQRRG*
>GCA_000452945.1_BPMSHR5858_genomic_04981_1 lutC_1
MSARDDILGRIRAALGSERARLAAQFPAPAQAKGAGASAAPNANAAFAPDASLSTASSAA
RGSAISLGDVDLVARFVAKAQQVSATCTHIATAAGAPAAVDAYLRDAGLADAPLVVAAAL
DALPWRAHRALPGVDLRRDGLVSVTPSFAAIAETGSVVCLSSSATPTSLNFVAATHVVLV
NRSAIVATMEDAWARVRATIATLPRAVNVITGPSRTADVEQTVQVGVHGPKRVLVLICDD
A*
>GCA_000452945.1_BPMSHR5858_genomic_04983_1 lutA_3
MDSSPQTAGAPGGTRETPRVGLFVTCLANLFRPSAALAAVQLLEAAGCAVDAPRAQTCCG
QPALNSGDFDDARGIARRTIEAFERYDYVVAPSGSCIRTLRHDYPDLLANDPAWRPRAER
LAARAHELTSFLVDVLGWAGPERAAVPECTLTYHDTCSGLRGLGIKAQPRALLARVEGVA
LTEMKDAEVCCGFGGTFCVKYPEISTKIVSDKTANVAASGATVLAGGDLGCLMNIGGRLK
REGSAVRVYHVAEVLAGMNVEPICGEGE*
>GCA_000452945.1_BPMSHR5858_genomic_04987_1 sfnC_1
MRPRDIAGRRGVDRQARHLGSIRMNDPRVLVEMTEQAGRPPRTLAELIAALRASAPERDR
AGGHAAREKRWIADAGLLTLAVPREFGGQEAGWPVIYHTIRALARVDSALAHLLGFQCLQ
IVSVDVWGSAAQRERYLRGTVEHDWWWGNAVNPLDTRLVARATGDGGYRLDGVKGFCSGT
RGSQRMTVSAHDSVTGKPVFAVVPTQREGIAVRDDWDPIGQRQTDSGSVAFDGVRVAPDE
VLHRSEAPPTPRATLRALVSQLVLTNLFVGIAEGALAEARDYVQRAGRPWLHSGVERAVD
DPYTLQRFGDMRVQTVSAEALADRAARALQGAWAKHEALTADARAEVALAISEAKIVAQR
AALDVSEALFDACGARATAAPLALDRFWRNARTHTLHDPLDYRLRDVGRYALTGALPDAS
LYT*
>GCA_000452945.1_BPMSHR5858_genomic_05041_1 group_1680
MARVLEIVLDFSLQDWQATRGARVRAARDLGAELARAWRICPPVKMRRGHERVTIEPCRF
VEAQPNDGSRWQTWIETTAQARRVLAARCRPFVPGVTVREHFDDYRGDVRVATPLADEQA
PIAAGSTPVPRDSAAAARDVLDGGDAPPSRGRASRAAAARTAARAAASTPVAPDADVSGA
ADASGSSRSSASGSASESAAESASDSVSAESTAPSAGAVPVGDAVAGTTSLESGSTARPA
PDFVVERRRGRWLDADGIEVELTLDDIVFAPAAASSDTIRAVASRVCELRLAVADPDDPD
DPDDPDARAAALRALFNAARELSGAWPASLSPISVLDRACAVDAPDANGAPAKAQSVDLS
NTRTQRAAFFALGCSVTGQWLGNEAGVRDTAEPEYVHQMRVALRRLRTLARLFPRYADAE
WKDAFSGDIRWLAGMLGAVRDWDVCVTSTLPALAAADGDEAAWAGTLDAARAQGDAARAE
LRQALGTARYTRLVFAWLEWLSLFPLGEDDPARGKAPSLKRHAAKRVSRLFGHLYGARRL
TALDAAARHRVRIDAKRLRYALEFFSSLASRRTREDTVRLLARVQNALGDANDATVALRC
LERLSAPPYQLGFARGYGAAAQRYAAEAAEQLLRGMRAPKIGGRKA*
>GCA_000452945.1_BPMSHR5858_genomic_05044_1 modC
MSLVVDIRKTLVTPERHFTLDVSFASNAQRIVLFGPSGAGKSLTLQAIAGLLAPDRGTIS
IGGDTLFDGAHGIDVPTQARRVAYLFQDYALFPHLNVRQNIAFGLKRGWRNPRAQELPDD
AAYWLRAFELEALAGHYPAQLSGGQKQRVALARALIAQPRILLLDEPFSALDVAMRQRMR
RELTDLQMRLDIPMVLITHDPDDVAAFGDQVVRLQEGRVLPDTPIAEWVTNMR*
>GCA_000452945.1_BPMSHR5858_genomic_05062_1 polA
MPEERNLEGKTLLLVDGSSYLYRAYHAMPDLRGPGGEPTGALYGIVNMLRRMRKDVSAEY
SACVFDAKGKTFRDDLYADYKAHRPPMPPDLALQIEPIHTAVRALGWPLLMIEGVEADDV

IGTLAKQAEQHGMNVIVSTGDKDLAQLVTDRVTLINTMTNEALDRDGVLAKFGVPPERIV
DYLSLIGDTVDNVPGVEKCGPKTAVKWLTQYGSLDGVVEHAGEIKGVVGDNLRRALDFLP
LARKLVTVETGCELAPHVESFDASLATDGEGRDALREIFAKYGFKTWLRELDSEPAAAAG
AAAAGAAPDPASGATAELPLATARNYATVQTWEQFDAWLATISAAELTAFDTETTSLDPM
RAQIVGLSFSVEPGHAAYVPVAHRGPDMPAQLPRDEVLAKLTPWLEDAGKKKLGQHLKYD
AQVLANYGIALNGIEHDTLLESYVLESHRTHDMDSLALRHLGVKTIKYEDVAGKGAQQIG
FDEVPLAQASEYAAEDADITLQLHHALYPQIAREPGLTRVYRDIELPVSLVLRKMERTGV
LIDGDRLSRQSGEIATRLVALEQEAYALAGGEFNLGSPKQIGQIFFERLQLPVVKKTPSG
APSTDEEVLQKLAEDYPLPKLLLEHRGLSKLKSTYTDKLPRMVNPDTGRVHTNYAQAVAV
TGRLASNDPNLQNIPVRTAEGRRIREAFVAPPGSKIVSADYSQIELRIMAHISEDESLLR
AFAHGEDIHRATAAEVFGVTPLEVTSDQRRIAKVINFGLIYGMSSFGLASNLGITRDAAK
LYIDRYFLRYPGVARYMEETRMRAKEKGYVETVFGRRLWLPEINGGNGPRRQAAERAAIN
APMQGTAADLIKLSMIAVDDWLERGGLRARMIMQVHDELVLEVPEDELSIVREKLPEMMC
GVAKLKVPLVAEVGVGENWEEAH*
>GCA_000452945.1_BPMSHR5858_genomic_05067_1 yeaW
MSNLSDALQLKSAHSQLPVTAYFDEALLAREIETLFKKGPRYVGHELMVPEAGDYFALPS
EDEGRVLVRNQASQIELLSNVCRHRQAIMLNGRGRTQNIVCPLHRWTYDLEGQLLGAPHF
PDKPCLNLHATPLQHWQGLLFEAEGRDVAHDLAQLGTKHHFDFSDYLFDHVEIHECNYNW
KTFIEVYLEDYHVVPFHPGLGSFVSCDDLKWEFGDWYSVQTVGVHNALAKPGSPTYQKWH
DQVLRYRNGVPPEFGAIWMVYYPGLMIEWYPHVLVVSWLIPRGPQKTTNIVEFYYPEEIA
LFEREFVEAERAAYMETAIEDDEIAWRMDAGRRALMERGESQVGPYQSPMEDGMQHFHEF
LRRQLGAI*
>GCA_000452945.1_BPMSHR5858_genomic_05076_1 dnaG
VIPHSFLQDLLNRVDIVDVVGRYVQLKKGGANFMGLCPFHNEKSPSFTVSPTKQFYHCFG
CGAHGTAIGFLMEHAGLTFPEAVQELAQSVGLTVPHEPSPMRGGGGSSDGHAPPAASKSV
TTALSDVMQTACDYYRKQLRGAPNAIQYLKRRGLTGEIALRFGLGYAPDGWQNLEAAFDD
YRHDSLVDAGLVIVSEKTDANGVARRYDRFRERIMFPIRNVKGQVIGFGGRVLDGGEPKY
LNSPETPLFNKGSELYGLFEARLAIREHKYALVVEGYMDVVALAQLGFPNAVATLGTACT
PIHVQKLLRQTDTVVFSFDGDAAGRRAARRALEACLPHAADNRTIRFLFLPPEHDPDSYV
REFGTDAFSEQVKRAMPLSQFLLNEATAGKELDQPEGRAKALFDAKPLLQALPANALRAQ
IMHMFADRLQIPFDEVAALAEVDARIAAPARQAPARNERRRVTESEKRALRNLVMHPRIA
AAIDDESVATLRALPRIGELFDEVIGHARALGDGAEFRLLSDVLRNSANAPTYDEIFREI
LVYDENVRDLMQQNPESDAAQEGLRERERIAGEELQAAVLKMRYDACCDRLNSLSKRSAP
SSEELAEIKDLHQNLADMKRRLGL*
>GCA_000452945.1_BPMSHR5858_genomic_05077_1 rpoD_2
MTKKLNEVSVDDDANQSDDQPTAASGKGEKAKARDRRAKEKALLKEAFATSTPGTAEELE
ERRTKLRALIKLGKERGFLTYAEINDHLPDNFTETEALEGIIGTFNDMGVAVYEQAPDAE
TLLLNDNAPAASSDDEVEEEAEVALSTVDSEFGRTTDPVRMYMREMGTVELLTREGEIEI
AKRIEDGLRHMVMAISACPTTIADILAMAERVANEESRVDELVDGLIDPNAADTDGFSEK
DAEEIENEDAEAEEDDEEEEEDDDGAAQATANAAQLEALKRASLEKFALISEWFDKMRRA
FEKEGYKSKSYLKAQETIQNELMSIRFTARTVERLCDTLRAQVDEVRQVERQILHIVVDK
CGMPRSEFIARFPGSETDLDWAEKVAAEGHSYGAILSRNIPAIREQQQRLLDLQARVVLP
LKDLKETNRQMAAGELKARQAKREMTEANLRLVISIAKKYTNRGLQFLDLIQEGNIGLMK
AVDKFEYRRGYKFSTYATWWIRQAITRSIADQARTIRIPVHMIETINKMNRISRQILQET
GLEPDPATLAEKMEMPEDKIRKIMKIAKEPISMETPIGDDDDSHLGDFIEDTNTVAPADA
ALHASMRDVVKDVLDSLTPREAKVLRMRFGIEMSTDHTLEEVGKQFDVTRERIRQIEAKA
LRKLRHPSRSDKLKSFLEGN*
>GCA_000452945.1_BPMSHR5858_genomic_05082_1 group_1934
MPNATDAQRRALRERRTIDYIPDHERHGKLSSQFTLWLGANLQVTAIVTGALAVVLGGDV
SWSLVGLALGQLLGAAVMALHGAQGPQLGLPQMISSRVQFGIYGAVIPLVLVCLMYVGFS
ASGTVLAGQALAQLFGVADPVGIFAFIAVVVVLAVFGYRTIHALGRVSSIVGVVAFVYMF
VRLLAGHDVGALLAIRHFSLSSFLLAISLSASWQIAYGPYVADYSRYLPRSTSARRTFWA
IGLGSVLGAQASMAFGVFAAALAGKQFAGHEVQFIVSLGASGATAALLYFAIAFGKLTIT
TLNAYGSVMSVATIVTGFSGRAQITQRARMAYVLAMVAAVAWLALVGRHAFLKDFSAFIL
FLLAFFTPWSAINLVDFYWVTRERYDVPALYDPDGRYGRWNVAGVSIYAIGVLVQMPFIA
TEFYTGPLVAKLGGTDISWIIGLVVPGVLYYVTARRQTAHVPERLILPDEPSLP*
>GCA_000452945.1_BPMSHR5858_genomic_05083_1 SIRT4
MTVSAVVSPQPQPPSSSPSPDAAPFADPRALDALHAFVERHPRLFVLTGAGISTDSGIPG
YRDRNGAWMRSQPIQYREFLDSDHARRRYWARSMLGWPVVGRAQPNASHHALARLGAAAR
IGRLVTQNVDGLHQRAGSVDVIELHGGIGGVTCLDCGAHHARAAIQHVLEAENPELLGAR
AQPAADGDAHLEWSALDHFRVPACPACGGLLKPAVVFFGENVPRERVAAAARALDDADAM
LVVGSSLMVYSGYRFCVWANDQRKPIVAINLGHTRADPLLTLKVEASCGQALAALIARLG
LAQRDAHGSPHV*
>GCA_000452945.1_BPMSHR5858_genomic_05085_1 gnd
MGKQAIGVVGLAVMGRNLALNIESRGYAVSVYNRSREKTDELIAEFPDRKLVPAHTLEAF
VASLETPRRILLMVKAGEATDATIAALKPLLDKGDVLIDGGNTHFTDTIRRNQELAQAGL
HFIGTGVSGGEEGALRGPSIMPGGQRDAYDLVEPILEQIAAKAPADGEPCVAYMGPDGAG
HYVKMVHNGIEYGDMQLIAESYAVLKQVAGLTNDELGAVYAEWNQGELDSYLIEITAKIF
GKKDDETGEHLVDVILDRAAQKGTGKWTSQNALDLGVPLPLITESVFARVLSSLKAQRVA
ASEVLSGPSPAPLEGDRAAFVESVRRALYLSKVISYAQGFAQLDTASKEYGWNLDLGTIA
KIFRAGCIIRARFLQKITDAYAKNAALANLLLDPYFQDIAANYQSALRDVVIAAVKAGVP
VPAFASAVAYFDSYRSARLPANLVQAQRDFFGAHTFERTDKPGSFHASWAE*
>GCA_000452945.1_BPMSHR5858_genomic_05088_1 group_2595
MAAVRDALTGRGRFIGLGAARGGRFATRGLALVPHRAAGDMTCLPLARLVPTGFGPDAVE
SVRRGPTLSGRSRTKVPAAVSMPPPRSELMEVRLRSILFVQRAGIGRTVRAALVAAGAAL
VGGCTVTPWTDSWQPTHVPSQPAPRASSGVPAGYYRVNSGDTLASIASAFGQRTLDIASW
NHMAPTDMVMPGQVLRVAPPPSTATFAPPPAAEPQPEAAASALAWPAHGTVTTPFGAGRN
HGIVITSTGGDRTVRAAAPGRVVYAGTGVAAYGPLVILKHENGLITAYGHNEKLLVNEGD
AVSAGQPVAEMATDASGRSTFEFEVRRNGKAVDPLGLLPRNGS*
>GCA_000452945.1_BPMSHR5858_genomic_05092_1 btuB_2
MTSTFLRHAPTARGDHARRRRRAITLTVPALAAGAFHLAPAVAQTSEAVHGHGTLGASGV
ASRADTDAASAKSDGAVREAASAAASRTTTGAAPDATTLPTIEIVAAPESTPLVVVTDPK
TPRQPLPASDGADYLKTIPGFASIRSGGTNGDPVLRGMFGSRLNILANGMPTLGACPGRM
DAPTSYIAPESYDKVTLVKGPQTVLYGPGASAGTVLFERVTPRFKTPGMRFDGSVVGGSF
GRNDQNVDVTAGTPDFYGRVSANHAHSQDYEDGNGRTVPSQWDKWNADAALGWTPDDNTR
LELTAGTGDGYARYAGRGMDGAHFRRETFGLKFDKKHIGDVLDRIEAQVFYNEADHVMDN
YTLRMPDPTSSMPMRMASEVRRRTLGARVAATLRLTDAFKLVTGVDAQSNRLDSRSAMGM
QNYGDKPWNPQANMWNAGAFGELTWYASDASRVIGGARIDYAAARDKRATTGGMKMSMRN
PTFDDLRSRVLPSGFVRYERDLASLPVTWYAGIGHAQRFPDYWELFSAKRGPNGSINAFS
AIKPEKTTQLDIGAQYKSDKLDAWVSAYAGYVQDFILFDYATGPMGQITQATNVNAQIMG
GEVGASWRPLAPWRFEGSLAYAWGRNVQSGAPLPQMPPLEARFGVEYTRGPWSAGGLWRV
VAPQHRYALNEGNVVGKDFGPSAGFGVLSLHAQYNVSKTVQISVGVDNVLDKAYAEHLNL
AGNAGFGYPANLPVTEPGRTAWVRLSTKL*
>GCA_000452945.1_BPMSHR5858_genomic_05096_1 group_5142
MKTDDVIVSFRGVRKTYDGETLVVKALDLDIRRGEFLTLLGPSGSGKTTCLMMLAGFEFP
TGGEIRLDGELLNNVPPHKRNIGMVFQNYALFPHLSVEQNVDYPLNVRRMPAAERAARVA
AALAMVQMERFAKRYPAQLSGGQQQRIALARALVFEPKLVLMDEPLGALDKQLREHMQYE
LKALHEKLGLTFVYVTHDQGEALTMSDRVAVFDKGIVQQLDTVDRLYESPCNEFVANFIG
DSNRLRGTVASVHGEFCEFRLEDGTRLVGLNAGNAQAGASGVACIRPERMSVAPGSGRAA
NGAANALAGEARSLIYFGDHVRMRCALPGQDECFVKVPLGTGALDAFAPGVPISLEFQPE
HLRVFA*
>GCA_000452945.1_BPMSHR5858_genomic_05097_1 group_5143
MHHTFNTPRRAVALLSLAAFCATAGAAELTVVNFGGANGDAQKAAFNQPFEKATGNKVTA
VEYNGEQAKVKAMVEAKHVNWDVVEVESGDLNRGCDEGLYEKLDWAKIAKKSDLIAESPQ
TCGVGFFVWSTALAYDADKLKRAPAGWADFWDVKKFPGKRGMRKGARYNLEFALMADGVA
PKDVYKVLATKAGQDRAFKKLDELKPNIQWWEAGAQPPQFLVAGDVVMSTAYNGRISAAQ
KEGKNLKVVWSGSIYDLDYWAIPKGTPNKALAEKYIAYTLSSKPQQDYAQHIAYGPANVA
AIKALDAKTLANLPNSPANGKNAVLQNITFWTDHGDELEQRFAAWASK*
>GCA_000452945.1_BPMSHR5858_genomic_05100_1 pcp_1
MNRTADFALSLSPLCKRLACVWPLALAAGIAHGATDWVDTHTKAFLNHAQIETLARGANA
ASLEVASGEATHVVVSLKLRNAEQLKAVARNVNDPHSAQYRQYITSAQFLANYAPTEAQV
KQVVAYLRKNGFVDIHVAPNRMLVSARGTAGTVKQAFNTSLVHFEYAGRAGFANASTAQV
PRALGDIVGSVLGLQNVARARPLTKIGAIAKPLALASGTATGHYPSEFPALYNATGVPTA
ANATVGIITIGGVSQALSDLQQFTSANSYPDVSTQTIQTNGSGGNYSDDQEGQGEWDLDS
QSIVGAAGGQLGQLIFYMADLDASGNTGLTQAFNQAVSDNAAKVINVSLGWCETDANADG
TLSAEEQIFTQAVAQGQTFAVSSGDEGVYECNNRGYPDGSNYTVSWPASSPHVLAIGGTT
LYTTSSGAFSNETVWNEGLDGNGKLWATGGGVSTILPNPSWQSGSHRKLPDISFDAAQST
GAYIYNYGQLQQIGGTSLSAPIFTGFWARLLSANGTGLGFPAARFYHSIPTHASLVRYDV
TSGNNGYSGYGYKASTGWDYPTGWGSINISNLNQLIQSGGFN*
>GCA_000452945.1_BPMSHR5858_genomic_05102_1 group_5360
MLPAFSMDVGRARSACAALHGAAGWLACLLPVAVAHAPAFWAWNALAAAACAAAGAGQAR
ASVHSRRSPLAGVAVLSGAQLADFASTYPDAVLALCSSASFVDDWVLHVRLFPFTAAAMV
GLSVAAARGERGRRTRWPKRIVDAAVALGAMMFAMNLAAMLLKTWAGLAGWPWGAHALVC
TMLVGMMLDHAAMRTMSDAIAAQRASHAIE*
>GCA_000452945.1_BPMSHR5858_genomic_05109_1 fhaB
MQRNEVVNRNVFRLVLNRVAGMPVPMPAAEVSRGRGKLGCGGVRAQRRGGAACAALLGVA
GPSLAFAAVVADPNGGAQRPGMATTANGTDLVNIVAPDATGLSHNKFNEFSPVGRGVVLN
NSVRPGESQIGGMAAQNPNLMQPATRALLEVTQQRSVLQGTLEAFGGKLDVLVANQHGVT
INGLTTLNVGRLGVTTGQVLPQAAGQLRLGVTQGDVLIDHGGIDTQGLDMFDVVSRSIAV
RGPIHDSSRAAGADVRLVAGATAYDPQTGHYEAIAADESKAPVQEGISGELLGAMHGRHI
VLVSTESGVGVRHDGPIKSANDIRVSANGEVTLGGPQRAAQEAVAGAQAVGGAGMQNVIA
GGTVSVCARGHVAIRGAVIAGQDVDLQGKSVKAGRMSAQRDALVTVADGVTLDGPVDAKR
HVWIGAHDDVVIREAAAGQNVVLLGRSVTAGRLDAQRDVLAAARDGVTIHEAAAAGQDVV
LQGSSARVGQMSAQRDVLVMAADGVTLDGPVSAQRAVWVETQGDVAGSEWIKAGRDVQIG
AAADLAGAVTAEEMQQLKAHGDAANRRRVKAGRNEPAGAAAERPAAAEQTVAVADAMREI
GVGGDRLSGLDAAPGTPFGAHPQAMFDDPAAQIARSARSTATAGGHAGSFMRVGDGHIAK
MTTSREAEIYENYRLALAGVIPDTVPPEEVDWRVGVTARQRQAMATFKGWAEMKGQRVVV
MQALGAKIAPEDKIELDVKIGASTVSRTELIGAGRTRWQALSKKVRLTAADLLRGSRSLV
GDDRGYTLAGRTSGGIALDARNSRNSVGRSSESLIREALDRSPDTRWRNAQHLLGQLQTI
REKMHALPLTFVASSVLIAIDKRKPENSVARLIDLAHPVQPFENEADYEKVNHRFEDGLD
KLIRLFQQVEK*
>GCA_000452945.1_BPMSHR5858_genomic_05112_1 group_5145
MKKLLIATVIGALSATMLAAAPSAFAQDSSATAKKATPKRPAPKRHLIPRSKKAQARAAA
KTDPVPEGAVKWACKDGLSYELAGDMKRDQIVTVHWANKNYKLPRQQTTTGADVFYDPAS
GMKLVVIPTKGMLFSDKDDSRLADECQTPEMAAGNGLAPTQSNELKPSN*
>GCA_000452945.1_BPMSHR5858_genomic_05118_1 sdhA
MAAIKTSLPRRKFDVVIVGAGGSGMRASLQLSRAGLSVCVLSKVFPTRSHTVAAQGGIGA
SLGNMSEDNWHYHFYDTIKGSDWLGDQDAIEFMCREAPNVVYELEHFGMPFDRNADGTIY
QRPFGGHTANYGEKPVQRACAAADRTGHALLHTLYQQNVAAKTQFFVEWMALDLIRDADG
DVLGVTALEMETGDVYILEGKTTLFATGGAGRIFAASTNAFINTGDGLGMAARSGIALQD
MEFWQFHPTGVAGAGVLITEGVRGEGGILRNANGERFMERYAPTLKDLAPRDFVSRSMDQ
EIKEGRGVGPNKDHVLLDLSHIGAETIMKRLPSIREIALKFANVDCIKEPIPVVPTIHYQ
MGGIPTNIHGQVVGTSRDHKEPINGFYAVGECSCVSVHGANRLGTNSLLDLVVFGRAAGN
HIVEHVKNQRDHKPLPADAADFSLSRLAKLEKSTSGEYTQDIANDIRATMQKHAGVFRTS
ALLKEGVEQMAGLKERAAAVHLKDKSKVFNTARVEALELANLIEVARATMVSAEARKESR
GAHAHSDYEHRDDENWLRHTLWYSEGDRLDYKPVQMKPLTVESVPPKARTF*
>GCA_000452945.1_BPMSHR5858_genomic_05125_1 aceE_1
MNDLSNGIRPVLATARRHDDTDPQETAEWLDALDGVVAHAGAERAQYLLAQLAEHAARRR
LAPPHAGATPYVNTIPVDEEPPYPGDLQLEERLAAVLRWNALAMVVRANRAYGELGGHIA
SYASAADLFEVGFNHFFRAAGDASGGDLVYFQPHSSPGVYARAFLEGFLSDAQLEHYRRE
IAGPGLCSYPHPWLMPDFWQFPTGSMGIGPINAIYQARFMRYLQNRGLLRTEGRKVWGFF
GDGEMDEPESIGALSLAAREGLDNLVFVINCNLQRLDGPVRGNGRIVDELESQFAGAGWN
VIKVLWGSDWDALFARDATGALARAFAQTVDGQFQTFSANDGAYNRARFFGQNDALAALV
AHLRDEDIDRLRRGGHDARKLYAAYDRAVRHEGRPTVILAKTMKGFGLGAIGQGRMTTHQ
QKKLDVEHLLAFRDRFRLPLTDEDVAQLRFYRPEKDSPEMRYLHARRAALGGCLPRRRAA
ATAALTVPSLPSWGQFALDGGREMSTTMAIVRMLGGLLKDAALGPRIVPIVADEARTFGM
ANLFRQVGIYSPLGQRYEPEDLGSMLYYREDTRGQILEEGISEAGAISSWIAAATSYSVH
DLPMLPFYIYYSMFGFQRIGDLIWAAADQRARGFLIGATSGKTTLGGEGLQHQDGTSHLA
ASTVPNCRAWDPAFAYEIAVIVDEGMREMVERQRDTFYYLTVTNENHAQPTLPADRLDAV
RGGILKGMYPLDAAALPAARVQLLGSGAILGEVQAAARLLRDDWRIDAAVWSVTSFTELQ
RDGIAAERAQRLFDADDANERSPTPHVTSALAATQGPVIAATDYARALPELIRAYVPRRY
VTLGTDGFGRSDTREALRAFFEVDRGSIVIAALRALADDGEVTRTVVRDAIARYGKRDAS
RTPPWER*
>GCA_000452945.1_BPMSHR5858_genomic_05130_1 leuC
MAQTLYDKLWNSHVVHTEEDGTALLYIDRQLLHEVTSPQAFEGLKLAQRPVWRISANLAV
SDHNVPTTDRSHGIADPVSKLQVDTLDANCDAYGITQFKMNDVRQGIVHIIGPEQGATLP
GMTIVCGDSHTSTHGAFGALAHGIGTSEVEHVLATQTLLQKKSKNMLVKVEGQLPRGCTA
KDIVLAIIGRIGTAGGTGYAIEFGGSTIRALTMEGRMTVCNMAIEAGARAGMVAVDDTTV
EYLKGRPFVPTGAEWDQAVEYWKTFRSDEGAQFDRVVELDAAQIVPQVTWGTSPEMVTSI
DGRVPDPEREKDPVKRDAMERALAYMALAPNTPIEAIKVDKIFIGSCTNARIEDIRAAAY
VVKKLNRRVAPNVRLAMVVPGSGLVKAQAEREGLDKVFTEAGFEWREPGCSMCLAMNADR
LEPGERCASTSNRNFEGRQGQGGRTHLVSPAMAAAAAIEGHFVDIRRLG*
>GCA_000452945.1_BPMSHR5858_genomic_05136_1 group_129
MPMTPRLLSSTTIFDGLPRARRAVALAVLALAGASAAYAAPDAASAASAEQAAPLTITVR
PGQSLNDIAVAVTQSHDPGVLARAGRALFDANPQAFMKRDPSRLKIGAQLMVPALDATGG
AAASGATAASAAASAASTSAVTGAAPHPSASASAPAAASRPAAASGASHAATASGAAAFA
ASGVAAGSASGANGASAPGVSAPGAASAHAPSAVAPAAPVASSGAGSAHVWSGSIQQAPA
GANGASAAGAEALLAPGAANRAPSGSGAQAAGASQARPSSLQQLLALKNRVLMELQKHGI
GKPAANVAPAPAAGSAQKPQASAAAGAAASATAAATTASETIASDAAAGSAASATPASAA
PAASVPAAAPRVSAPAHAAAPQAPFNTGAAVAIGAALAALIVGFALRRRKKGASADAIEP
GAFTPAPALDPDAAAAIADAPAGPNAAEDARHDEAEPDKAAAARRASSAAPLAEAKRPPA
QHGSEPPSADEDAAPAREPQAPAARPLTLKTPRPDEQGGAPQSAGDEPSGAKPAPYAWRQ
AFPSDAIAALDSLDMPLPPREAGAAADRPEQSQHAEPAGSRDAQSATNGLTAAPPPMGAE
NAREGQTHGEAGSATDFEGAGSFDNDARVAPQRPEGATPRPVTAPDAMGRDEAAARAAES
SPTESTRDAPRTPGAPAAESRLSTAGAGAAHSPMPDAAAAAEAAAPHAATPSDAAPHDDG
QTQAAPRPLDGGPQPPAVARHEAAPAHAEIPNAPLAPAAPAWAAPQPPGMSFAEPAQAEP
GDVPAAPDRAPAANPPPLGVAQFGALNLDFDLELPSGPSASLPAFTPEALAKIARNKLEL
ASEYVELGDLAGARTLLHEVIEADHADTRDEARVMLAKLADLS*
>GCA_000452945.1_BPMSHR5858_genomic_05139_1 trpB
MYNLPDDRGHFGPYGGVFVAETLIHALDELRAAYEKFKNDPEFVAEYNRELKHFVGRPSP
IYHAQRWSETLGGAQIYLKREDLNHTGAHKVNNVIGQALLAKRMGKRRVIAETGAGQHGV
ATATICARFGMECVVYMGAEDVRRQAANVYRMKLLGATVVPVESGSRTLKDALNEAMRDW
VTNIERTFYIIGTVAGPHPYPMMVRDFQRVIGDECKVQMPELAGRQPDAVIACVGGGSNA
MGIFYPYIDDAAVQLIGVEAAGDGLDTGHHAASLIAGSPGVLHGNRTYLLQDDDGQIIET
HSVSAGLDYPGVGPEHAWLKDSGRAQYVGITDEEALKAFHDCCRIEGIIPALESSHAIAY
GAKLAPTLPKDKILLVNLSGRGDKDMHTVAERSGITL*
>GCA_000452945.1_BPMSHR5858_genomic_05144_1 dedD
MGIFSFGKKEDDAPPRRGSRSGARAVRTERVERRPRRTERPDTDALLLDPTLPEKQRARR
RLVGAIALVVAAVIVLPMVLDSHPKPVTDDVQIDIPNRPSHQAAAANDEDASDVQAGVAH
DEPPASEPAAAPTPAVAAKAQAKPDAKEAKKPPTAAAKAPSAAQKPAAQPAQDTAAASVR
APAATATADGDASSPASPPGARFALQLGAFRDDATARNWATKLKAAGVPAYVEHRKQADG
STATLLRAGPFADRAAASAAIAKVREAGLTQ*
>GCA_000452945.1_BPMSHR5858_genomic_05149_1 tld
MRPSEPGARRALVTGIAGFTGRHLAARLEAHGYDVWGTVAPGTDVLGDPLLRRWRCVKAD
LLDVDSLHAAVADARPHAVVHLAARAHVAHGNPQDTYLVNVVGTRNLLASLAGLDARPHA
VLLASSANVYGNASTEVLDESAPPQPANDYAVSKLAMEYMAKLWLDRLPIVIARPFNYTG
VGQSEAYLLPKLVAHYARGEPRIALGNLDVSRDFSDVRDVVDAYARLIDAAPVGETFNVC
AERGYALKEVLAMLARIAGYVIDVSVDPRFVRASEVKKLVGSRRKLRAAIGDARRTPLDE
TLRWMYGDMRAALGAREDQPAS*
>GCA_000452945.1_BPMSHR5858_genomic_05152_1 group_3008
MAAGVAVCAATALAGAARAENVLPATTSWIGNTYGYGDGTWTQIDIRAIAVAPDGKVYTN
APWDESGAEASVYQDGKMLGFAGGTHGWGNAGGSAIAVNSRYAYVAIAVGNEKGRLVAPG
VWPDKGRQWYGISRREIGDVKQAAPFRGAKAGDARAKLAASFLMVNEVPTDTRAEVAGLA
ASDSVLYASDPLHDAVNVYDAQTMQKKGSWSAHEPGRIALAADGTLWLLTDTLNGPAKLA
HVRPDGRAIDDAPALPPDAQAVDVATDAKGRVLLADNGPRQQILIFSKSADGGYAMSGTL
GERGGIFSGVAGRPGPQRFNGLTGVGVDRAGNVYVATNGIGPRHGTIGAGLGSTLESYAP
DGRLRWQVQGLLFVDGAWIDPGRPNSVYTGNKRFELDLSKPAGKEWKYAGFLSNRFKYPD
DPVFNTDQWPGMPFARKVAGRTFLYLTDMHADHLKIYRFDPKRDGETAIPSGLLAGRARP

VDKVPNRPPGGDWIWRDANGNGRIDEGEFELNTTGKGMAGGWGWWIDTGGDIWRTSDTRG
IRRFRFGGLDKAGNPIYSYDKLTTYPSPAPFTELRRALYDASSDTLYVTGYTADAPRVGG
LSKEVGRVLVRFDKWSSGAPVMRYQVALPWRLDAKPIFTLIGLTQEGRYLFGVEPVGTVH
VYDKDNGRELGVIKPGPEVGRASGWVDVPFGISAYRRENGEYLVFVEEDARGKVMMYRWR
PQ*
>GCA_000452945.1_BPMSHR5858_genomic_05154_1 mshA_4
MNRDLAEHPLQRAATFQDVERAARDARSADAPPPAHRARDAARPLRVAIVHDWLVTYAGA
ERVLEQIVACFPDADLFGLVDFLDDRTFLRGKPVTTSFIQRLPYARTKYRSYLPLMPLAI
EQLDVSGYDLVISSSHAVAKGILTGPDQVHVSYVHSPIRYAWDLQHQYLEQSQLTRGVKS
ALARLILHYIRNWDVRTSNSVDRFVANSAFIARRIRKVYQRDAAVVFPPVDVDAFALSTR
KEDFYLTASRMVPYKKIDLIVDAFAQMPGRRLVVIGDGPDMRKIRAKAAPNVEIMGYQPF
AVLQDRMRRAKAFVFAAEEDFGISVVEAQACGTPVIAFGKGGALETVRDAASHERPTGVF
FDEQSVRAIVAAVDDFERAPARFKPEDCRANAERFSAAHFRRRFVAQIDALLPNAQARAK
LAGAMRPAGRASLAASGLRALVLDQSGVLGGAELSLLEIMKHLRESADVVLFDDGPFRAA
LDDAGVHVDVVGQRALAGVHKQGGVSPRAAGSLLALVREVARRARDADVIYANTQRAMVV
GALAGRLARKPVVWHLRDIVSDAHFGPKQRLAIKQCARLGVTRVIANSDASARAFLELTG
FERRAVQVVFNGISAEPFVALEPVRPAALRVRFGLPADAWIVGSFSRLAHWKGQHVLLEA
ARLYPDMHVALVGAPLFGEDEYAAELRGFVALHGLGERVHFLGFQRDVAACMKAVDVVAH
TSITPEPFGRVIVEGMLAKRPVVAARAGGVVEIVDDDVNGLLCEPGDAHALADALAALRT
DAVLCGRLVANGYDTAVNRFGTQIYVEQVERILVETARRR*
>GCA_000452945.1_BPMSHR5858_genomic_05156_1 group_2601
MDMHWIASGAVLLVMAVLSAYRDALKNEPINRSGLGRGGAPYSEEFE*
>GCA_000452945.1_BPMSHR5858_genomic_05179_1 group_5154
MGPFKSIAAALALGVLAGCSAFSSSKPEEPRQLHVTLVGGARLNVAPTGEPRPVQTCVYV
VTAADWLPTQGGDDSSCASRGQDSTVVADSRHVIAPNQLLQFSLNLPRSGELWLVADADY
ARRPANYGPLRVRVEGRGLIHMAVWLDRDGIYNALLPGPVPVGGAIAPAAVRPDEPPPER
KTKITYGTRRSRQ*
>GCA_000452945.1_BPMSHR5858_genomic_05198_1 group_5896
MTKTAFFTDERTFWHTGGAHALFFPVGGWVQPPSSAGYAESPDSKRRLLSLVHASGLAAK
LDMSSAPAATDDDLRRIHPAHYLDAFKRASDAGGGDLGELAPFGRGSYEIAALSAGLALA
AVDAVLAERTANAFSLSRPPGHHCLRDKPMGFCLLANIPIAIEAARAKHRVERVAVIDWD
VHHGNGTQSIYYDDPNTLTISLHQDRCFPPGYSGADERGAGAGAGSNVNVPLLAGAGDDA
YRYAFERIVLPALDAFRPELVIVASGLDANAVDPLARMQLHSDSYRYMTHALKQAAQRHC
GGRLVIVHEGGYSEAYVPFCGHAIVEALAGMRTDVADPMLELAIAQQPGERFNAFQRQLI
DEMATSFGY*
>GCA_000452945.1_BPMSHR5858_genomic_05199_1 group_903
MTIKTLRQEPALAGAASPADTPAFTSGTLAALVAFAAITPLLLLVAPAVAGQLGAQLGLS
ASRIGTYFFVELGAFSAATLPSYLWLGRIDARRIAWGATAVFCAGNLATAVWMPGFAPLL
ALRAATALGGGTLMVLCMTSAAASGNSDRVYGLWVVGQLIAGAAGLFLLPHLFDMVGLRA
LYAVLAALALCAAPLARRFPAVPRVRAQHAPRARAQAVRMAAALAIGGVLAFYVAIGGVW
TFASKAASAVGLDAQTSGNVLAIASLMGIAGAALASYLGGRAARRAMLLAGYGILAASLV
ALAAAPNANGYTLAIFGFKFAWTFVLPFMLASVAAVDATGRLIATLNLVIGSGLAAGPLA
AGLMLDGGGTLRALFSIAAAVSLVSLAAMLRVERDAR*
>GCA_000452945.1_BPMSHR5858_genomic_05201_1 group_5389
VAIRAGTRQDIDAQRDVPAAIVLDEAAWPPLAPRRRDFDGVTQTPGAASRELSLFLLDLY
AVASRASIGEFECRFFQLLSRYLPFDAAWTGVANHTPGGPVMHNSYLYRLPHAFFIDWKR
VRDCDPLAHRTLHGAYGRAAILSVVEPGLDGRFRDWCVKYGLAQLMCVCTLDRRFGLTTF
LSIYRHALNRPFTEEDARRYEDVIAHLAAALTINRAAQLTRLRSEAAASTVRAICDNFGV
LHHADAGFGAVLRSEWPDWAGSRLPPAFVEHLRRRASQPYSGDALCMRCMPVAGLFQLEV
RPRSLLDRLSPRELAAIRYYGEGRSHKEVAQQMEIAPATVRHYLRCAYRKLGMHDKSQIG
GVLGELDRAAEGAGAARDIDG*
>GCA_000452945.1_BPMSHR5858_genomic_05202_1 group_1243
MDRAPVRAAIVGIGAIGGLFAAGLARAGWDVSALARGATLDALRGRGLRIVGESGGETAV
ALRASDDAHALGEQDYVVVALKAHALPALASRIAPLVGPRTIIVAAMNGLPWWFFDGFGG
PLDGAALDAVDPGGVTAAALPPARAIGCVVHLSSATREPGVVVRGRGNRLIVGAPRAALR
EPGERVADALAAGGFDVERSADIRTDIWAKLWGNMNMNPLSALTGSAADRLLDDPHTNAL
ALRMMEEADAIGARLGLSAGMSGAERIAVTRQLGAFRTSMLQDLEAGRALEIGPILGVFP
ELGRRLGVPTPYCDAVLGLLRQRAFNAGL*
>GCA_000452945.1_BPMSHR5858_genomic_05204_1 glxK
MPQPSPIVVIAPDSFKGSLSAEQVANAIAAGVARARPDALVRRCPMADGGEGTLDALLAH
GGTRRTLRVPGASLAPRDAAVGLIDARTAVVETAEIVGITDPAGMSAPVAARSTRGLGSA
IRALLDDGVRTFYVALGGSSTNDAGAGLLAGLGMRAFDAQGREIEPTPERLAHVARVDFA
GLDARVAGASFVAMSDVDNPLTGAHGATAVFGPQKGVAPAQVAPLDAALDHFATLVEAAL
ALRPTAAAHARARDLPGAGAAGGLGFALHVLGARFEPGAEVVARQIGLDAALAGANWLLT
GEGRSDAQTLHGKAPFVACRHARAAGVPATLLSGAIDPAALPQLSEHFDGCFSPAPGPIT
LEAALRDAARLLENAAEQLARLRYGRP*
>GCA_000452945.1_BPMSHR5858_genomic_05217_1 glxR_1
MKLGFVGLGVMGQPMALNLARAGTELVVWNRTRERCEPLRAAGAQVADSAADVYRRARIV
ILMMATDAAIDAVLDRGKPAFASNVAQHTIVQMGTVSAEYSRGLEADIRAAGGRYVEAPV
SGSRQPAEAGRLVAMLAGEPAAVEEVRALLAPMCREIVATGQVPSGLLMKLAVNTFLIAM
VTGLAEAAHAARGFGLDMKQFQAVLDAGPMASSVSRVKIDKLVNEDFEVQASIVDVFKNS
RLATEAAHGAHLAAPLLEVCCELYRETEALGHGQADMAAVVHAIEARSVARYAGS*
>GCA_000452945.1_BPMSHR5858_genomic_05219_1 group_5157
MTNRMTNASTTLPTTNASPVPPLGERDAHVWYARTAACDTPALRERYRALLSAEERERLG
RFAFDHLKLEYLVTRALCRTVLSAYVDAVAPEQWRFRANAHGRPEIDAGDARPPLRFNLS
NARSIVACVITRTADAGIDVEERARSNDLDGIAASHFSASERAAFFALPPDARRTRFFEL
WTLKEAYIKARGVGLSIDLGEFSFALPAQPVRIAFDRHVDDDASHWQFALLDVGAEHQMA
LGIRDPHAAARPFDITMREIVPEPAASARCAAALD*
>GCA_000452945.1_BPMSHR5858_genomic_05224_1 lpdV
MSDTKTTTLLVIGGGPGGYVAAIRAGQLGVRTILVERDRLGGTCLNIGCIPSKALIHAAG
EFDKVRGFAGDSPLGIRTQAPAIDIARTVAWKDGIVKKLTGGVGALLKKNGVEVVHGDAR
VVDGKTVDVDTGGGARVRIQCEHLLLAAGSEPVELPAMPFGGNVISSTGALSPGRLPKRL
VVVGAGYIGLELGIAYRKLGVEVSVVEARERILPIYDAELTKPVAASLKRLGVRVLLGHK
VLGLNARGDAVCVQDDAHAQTELAADQVLVTVGRRPRTQGWGLETLQLDRAGAALKIDDM
CRTSMRNVWAIGDLTGEPMLAHRAMAQGEMVAEIVAGKKRHFMPAAIAAICFTDPEVVSV
GLAPDEAERTFGACVSASFPFAANGRALTLESADGFVRVVARRDDHLIVGWQAVGAGVSE
LAAAFSQSLEMGARLEDVGGTIHAHPTLGEAVMEAALRALGHALHI*
>GCA_000452945.1_BPMSHR5858_genomic_05225_1 bkdB
MGVHVIKMPDIGEGIAEVELGLWHVKVGDRVKEDQAIADVMTDKASVEIPSPVTGVVVAL
GGKEGDVLAVGSELVRLEVEGDGNHKAEPDGGARAAAAQPERVADTAHAHASAAAKSARG
EHGAGHGRDDARAASSGTSSGASHTQHAEREARGHRESSEYREDRSVSRGASQTDVERGP
ASPPPARRPGERPLASPAVRKRAWDLGVELRYVHGTGEAGRILHEDLDAYLQGRGAAAQR
ARGGQAAYVERHDEEAVPVIGLRRKIAQRMQDAKRRIPHFSYVEEIDVTELEALRAELNR
KYGDTRGRLTVLPLLARAMVIALREFPQINARYDDEAGVVTRHGAVHLGIATQSKAGLMV
PVVRHAEARDPWSIAAEVARLADAARAGRAERDELSGSTITITSLGALGGIASTPVINSP
EVGIVGVNRIVERPMFRGGAVVARKLMNLSSSFDHRVIDGMDAAEFIQAVRGLLEQPALL
FVE*
>GCA_000452945.1_BPMSHR5858_genomic_05227_1 bkdA1
MSQYGPLRLHVPEPTGRPGCKTDFSYLRLSPAGKVRKPPIDVAPADTSDLAYGLVRVLDE
HGRAVGPWAPDLDPDILRKGIRAMLKTRIFDARMQIAQRQKKISFYMQCLGEEAIAVAHT
LALERGDMCFPTYRQQGILMVREYPLVDMMCQLMSNERDPLKGRQLPVMYSTRDAGFFSI
SGNLATQFTQAVGWAMASAIKGDTRIASAWIGDGATAEADFHTALTFAHVYRAPVILNVV
NNQWAISTFQAIAGGEGATFAGRGVGCGIASLRVDGNDFLAVYAASRWAAERARRNLGPT
LIEWVTYRAGPHSTSDDPTKYRPGDDWTNFPLGDPLGRLKRHMIGIGVWSEQDHEDTKAA
LEAEVLAAQKEAERYGTLADEHVPNVASIFEDVYKEMPAHLRRQRQQLGV*
>GCA_000452945.1_BPMSHR5858_genomic_05228_1 group_5160
MKQQRIEIAEEFDAPVHQVFRFFSEHENLKSIFSPAKIRRISDGEPARNGVGSAREMRVP
GAPPFVETVTAYQENELIEYRITRGSPLRDHLGVMRFVPIDEHRTYFHYVITFEGKVPFV
APVVRHVLEGSIRRGLGKVATRRLWG*
>GCA_000452945.1_BPMSHR5858_genomic_05234_1 dksA_2
MAAFDEHQLQVLGALLRACRQALQADVRAGERQRADEPYANLAGPAPDEGDEANADLFVD
VDHALLGMKLAELRDVDAALQRIARRDYGCCADCGQPIAYERLLARPTALRCAPCQRAHE
RRFATQPRPSL*
>GCA_000452945.1_BPMSHR5858_genomic_05236_1 group_1665
MLYAWLEAQREFFRACQPWAGAAQRAGAGAGARPAGWPTWGPMARAADAVPAPPPFAIAS
IEIDGAAVPVDECVVDDTPFCALRKFSRRLRGDAAARPAVFLCTPLAGHHAVMLRETVET
ILATRDVYVTDWRNARDVPPEAGAFGLDDYVCTLERFVAGAAGDGLHMMAVCQATVPALA
AVALLAARDVHVTSVALLGGPVDTRSHPTLIDRFALEHDLDWFRRTAIDIVPPPYAGAGR
RVYPGFIQHAAIIVAHPQRRASLESRYWAAWMTGDFAAAARCLREMNEYGAVLDMTERYF
LDTIRVIFHERLLAQGRWSIGRRLVAPRRLTRTALCTIEGGRDDIAGAGQTHAAHALCDA
VPDAVRERITEPDCDHYDLFLGPRWRRTIHPALDAFWTRAEAARAAAGEPRPTAERAERR
IAH*
>GCA_000452945.1_BPMSHR5858_genomic_05237_1 nrdZ
MIGDMDDQPICDAVFADRYALPGEHSRAQACARVARALALAEPMATRSGAARRFYRNLLN
GAFGAGRIMARAGAAPDQTMASCFVHPIRAPAALTRFHPNLDEALDEARLALAMGADIGY
DFSAIPPAGARPDADHPDSPGVCAALDRFDRIGTQAGERDGRRRAQLAVLRCDHPDLLAF
AAAKHRHAGRHARWTTLELAVAATDAFMQAVEQDLPWTLRHTAPPRDAPGGALPGADGAW
TYASVPARHVWREIVSAARDGAGPGLVFVDAIDAADPLRGRERIDATSPCGAQPLPPYGS
AMLGAIDLSRFVRNPFGAGGEPRFDFAAFDTAVRIEVRLLDNALDVTRWPLAAHARESYQ
KRRIGVGVTGLADMLAMMRLRYDSPAAREMARYIASDLRHHAYAASAELAAERGAYPLCD
RHAHLDALRVGPPLPHAVCHAIERDGLRHSHLTSFAPVIGVSLAFGDHCSPGIEPARAWI
EHRPVRAGAAGAPGMRAENHAHRLFRSLRGERAALPDYFATDADVAPGERLAMRVALQPY
VDAGIDNTLTLANHYSLEEVNALLFAAWRAKLKSVAIRRADLARDAQPSDDDGV*
>GCA_000452945.1_BPMSHR5858_genomic_05238_1 group_1548
MRRSRPPKPAKPAAVRRRPHGAARAHAIRGNGPQRRAVRERAARRLSRDIDAALRRASTY
PHPAGPIVRIETHLSVVYLVGRFAYKRLKPFDFGFANFSELAARRRACEAELALNRPLAA
PIYLAAGPLVRRARGLRLFGAGTAVDHVVRMRRFDERMLFSRLLARGALDAADIDAAATR
LAAYHLHAPRDIPRRAYGSARELRRQLDDMLAPLERALGAALPASLRAWCVRRCDELAAH
LDARRADGYVRACHGDLHLNNVVKRGRDALMFDCIDFDDALRWIDVINDLSFLLMDLHAH
DRAALAHRLLNRWLDETGDFAGLAALPLYVAYRALVRALVATMRAGGDAAARAARIERAR
RYVDVAAHAARARRPCLLLCHGYSGSGKSVASRALADVSGAIRLSSDSERKRARPFAAVD
ARPLPASAYTPQQIDAQYERLRALARDVLRAGYTALVDATFLSHARRARFFALARELGVP
VYVLDFHASRACLERRVDARAAARDDHSDAGAAVLATQRASADPLDADERARTIGFDTDV
PLATLRSAGYWRPVLDALDAARVDAQATR*
>GCA_000452945.1_BPMSHR5858_genomic_05239_1 group_5162
METSPLKTTTPEAIPPFDPSASVEDKLRLAIHYAILAPSSHNTQPWRFMVGDGSVMLCAD
RLRALSVVDPYDRELLISCGAALLNLRVALSRFGFAYVIDMFPSPSDPDVIALVRLDPHG
YHDESLVPLFDAILERVTTRTPYANEAVPCDVQRALVDAGAAEGAEIACVDAPHALDEIA
ELIADADRLQFADPRFRRELANWVHPRRHDDGMPAFAAGVPALLDFATPVVASAIRTFDL
GGGLAAMHHKLVDGSPLIVGISTASDDRDAWVAAGQALERVLLVAAAAGLTASYLNQPIE
IDALRERLRPLLHVDAHPQLLLRIGRGPVVAHAPRRPLMDVVS*
>GCA_000452945.1_BPMSHR5858_genomic_05255_1 rsxB_2
MTQMALVIDLNVCVGCHACVTSCKEWNTSGDAGALSDLRPYDADPSGTFFNRVQTFEAGE
FPLADTIHFPKSCLHCEDPPCVPVCPTGASYKREEDGLVLVDYDKCIGCKYCTWACPYGA
RELDEVRKEMTKCTLCADRIHDETLPERDRKPACVLACPTSARLFGDIHDPNSTVSQAIR
ERGGYQLMPEWGTRPANHYLPRRTTTACGSGGGGGNDGSGGGGCGSGAACACRPADAKPA
PVSAGDALAHGQPNLAALAKRIRSE*
>GCA_000452945.1_BPMSHR5858_genomic_05259_1 icaB
LRHSGSRLTRFRLAAAACCLALASAARPAFAGANAPAAAEPPTDAQRPAILVYHRFSTSA
PPDSMTVRVSTFGAQLAFLRAHGYTFVPLRDVVAWAASPSAPLPDKAIAITVDDGHRSVY
ELLRPIVLRERLPVTLFIYPSAISNASYAMTWDELRALRATGRFDIESHTWWHPNFRTER
RRLAPDAFRRFAATQFAHSRARLEREIGGPVDLLAWPFGLYDDELTSLAAQAGYVAGFTL
DARKVRRGDAPLALPRLLIVDSCTPAVLARMLGERGDARADSHADLHVDSHAEQSR*
>GCA_000452945.1_BPMSHR5858_genomic_05260_1 group_211
MSTIDGRPRTRRARGVPPKRRHEDKCIPVVLAWTLGERVDVRPDSGIDLDIDLEIDSDID
SHAHTRGAIAMSAIDGRPRMPRAHRVPPKRRCEDKCMPAVLAWLPGRPARLRDRRCACNE
PYPSALGRSPRGGRAFDGCRRDARECVRRPAHGERSMTNPFAQWSRASACALAIVCFAPS
GAAFAAQATADVPTAGASIASAASTPAPGTPTSAAAAARSFVTSPAAIHGAVVDARTGKP
IPAAIVTIGGRPLRADDRGAFSADAAAADIAVRAPGYLATRARAEAGRPVTVALAPFRPK
AVYLSVFGITSKTLREAAVDLKGATAINALVVDMKGDRGITPYPSAARRASGAAARTPNA
PVVRDFAALVADLHRRGLYLIARIVVFKDDPLAAAHPEWTVRDADGNIWRDREKLRWIDP
SLRETWAHNLDVAEEAAKLGFDEIQFDYVRFPDARELRFSVPNTRANRTAAIAGFLRAAR
ERLAPYNVFVAADIFGYVCWNEDDTAIGQQIETLGGPLDYISPMLYPSGFTWGLPGCTQP
TADPGQIVRRSLVEARSRTGLPGVRFRPWLQAFRDYAFDRRDFAAAEIRAQVDAAEAADT
DGWMLWNARNRYDPRQLPK*
>GCA_000452945.1_BPMSHR5858_genomic_05266_1 group_19927
MKKILAVIAFLAVVGWLAATTTVLHAPSAQPCTDAWFDAIDKQFDITDNAGHGPDPGSGE
WLGVVERKAKLPESGQLTEQQRCEAIQRELSQRTYLVNRRLGLKLAL*
>GCA_000452945.1_BPMSHR5858_genomic_05268_1 aaaT
MERAATSPPPPDVAIRALRAADADALHALARQPSIDGRFASAPHASLDRRREWLEALIRE
QYALGAWSGETLVGWSELAPGRLRRSHTAAVAIAVHDAWHRRGVGGALMRETLRIADDWL
GLRRIEAQLYADDFVALAFHRRFGFEIEATQRGAALRGGVLVDGCLIARMKAPLPLAAAP
GGTPT*
>GCA_000452945.1_BPMSHR5858_genomic_05269_1 group_5556
MTDFAKPGRAAPIDWHAHFARLADEAEQLRRPGETVLLWFAGETSDFIRFNAGKVRQTGR
VVQGKLAVRLASGGRQASFAQTASGDWAADAPELAAALATLRESLRDAPDDPHLLFDTSS
WRQSSRRDGSLPDPDALAHVVADAARGLDFVGFYAGGSLARGFASSTGSRGWYEVENFDF
RWSLYDASGRAIKTVYAGESWSDAAFARKVDEAAARLPALGRAPKALAPGRYRAYLAPAA
VAEIVGLLGWDGFSARANASARSSLHRLHAGEAAFDPRVTIAEDFSLGVAPAFNADGYRR
DSVPLVAEGRSVGRLVSARTAREHGLSPNGAAADETPETLSIAGGALADADVLAALDTGL
YIGNLWYLNFSDKMNGRITGMTRFATFWVERGRIVAPVDAMRFDDSVYRLLGAELEQLGA
APELLLNDQSWSERPTGGAKLPGLLARSFELTL*
>GCA_000452945.1_BPMSHR5858_genomic_05284_1 desA
MNDSSHPIRVDEPMPHRKVIRSWITPFGERVIARSIVLLAVDYLLLFAAFAGALLAASGI
VKIVCGMAAGFITGRLFIIGHDACHQSLTPNHRLNRWLGRIAFLPSLTPYSLWEVGHNVV
HHGYTNLKGFDFVWAPLTPGEYAALSPARRLLDRIYRSGWAPGLYYLVEIWWLRMYFPAK
AYLGASRPIFRRDCLLVTGFAALWIGAVVWIAAATRQSMPLLLVTGVAVPFLFWCSMIGF
VVYVHHTDPRISWHANRAEWSRAAPFVSTTLHLTFPFGIGGLLHHIMEHTAHHVDMSIPL
YGLKDAQAKLEAMLPGRIVVQRFSWRWYFDTARRCKLYDTDRKRWTDYRGRATSDAHLRA
DAPMAEQRPIDAPHGPGGA*
>GCA_000452945.1_BPMSHR5858_genomic_05290_1 group_5170
MTSRLETVALPGGERVPKLGQGTWEMGERPAKRAAEIAALREGVDLGMTLIDTAEMYGDG
VTETLVGDALADVRERLFIVSKVLPHHASRGGVVAACEATLKRLRTDRVDLYLLHWRGSI
PLAETIAGFEALRDAGKIRYWGVSNFDVDDMEALVAEAGGAVCATDQILYNLARRGPEFD
LLPWLARREMPAMAYSPIDHMRLPKRTALDEIARERGVSPTRVALAWVLGQPNVLAIPKA
GSVEHVRDNRAALDLVLGEEELARLDAQFKSPRGKRPLEML*
>GCA_000452945.1_BPMSHR5858_genomic_05291_1 nagR_2
MKPSTEDRWSDLRPDPDNDTPLYLQLARKLGDAIHDNRWAAGEALPSERLLAESLGVSRI
TARKAIALLVEQGLIRRTQGAGNFIQPRYEDPLSRLSSFSEMLKRRGFTPSSTWLSRDIQ
PANRDEVIQLGLSPAASVTRLKRLRLADGIVMAVENSTFPATIVPDPQAIGDSLYSYLEQ
RGTPIVRALQHFRAVNATDEIARQMGIAPHDALLLITRIGFTHDQRAIELTDTYCRNDYY
DFVVELRK*
>GCA_000452945.1_BPMSHR5858_genomic_05296_1 tam_3
MDRTTLDAYDRHAADYAREWDEQPPPDDMYVLLERYFEPGPTADVGCGAGRDVAWLARRG
FDACGYDGSASLIAEARRSHPALRFEQALLPALAGAPSGAFRNVLCETVIMHLPASEVGA
AVARLVELLAPGGTLYLSWRVTAGDAQRDGRGRLYAAVDAERVHAALGDACDVLFEHEQT
SESSGKRVHRLVVRRRAAEAG*
>GCA_000452945.1_BPMSHR5858_genomic_05298_1 group_5172
MRTLVDQLAQYAAYHRDARNIATHLVGIPMIVVAVTVLLSRPTFGAWGAFALTPAALVAL
AAALFYLRLDLRFGVAMAALLALSLWAGRALAAQTTAFWLGTGLGLFFVGWVIQFVGHYF
EGRKPAFVDDLIGLIVGPLFVVAEVAFFAGLRGDVRDAVERRVGPVRVGARNARV*
>GCA_000452945.1_BPMSHR5858_genomic_05314_1 group_5173
MKVSFYRYMLAAFAAASVVVSGAALAQVDLAKSKVSAVSKQMNVPTEGVFKKFSAQIKFD
PAKAAQGSAQMTIDVASYDLGDQMYNDQVAGKDWFDAKAYPQASFVSTAIAPAGGNKYNV
SGKLTIKGKTVPVTVPVTVTQNGAAQVFDGVLPIKRSTFNVGTGEWKDTSVVADEVQIKF
HIVAAK*
>GCA_000452945.1_BPMSHR5858_genomic_05315_1 yceJ_2
MASFSSSASPAAAYTRTAIALHWLIALLIVCGFALGWVMTSIPGFTPAKLKYYSWHKWIG

VTVFAIAIVRVLWRATHVPPPMPAGMPAWQRAGAHAVHGLLYVLMLVIPVTGYLYSSAAN
IPVVYLGLVPLPRLIDPDPALKETLKTAHVLLNYGLLALVAAHVAAAVKHQWFDRDGVLS
RMLPFLK*
>GCA_000452945.1_BPMSHR5858_genomic_05329_1 nudJ
MKPEIWTPHVTVAAIVERDGRFLVIEEHTSSGLRVNQPAGHLEAGETLVEAVARETLEET
AHPFEPQALVGVYVAHYARPSSSGATYLRFAFCGRAGERIAHRALDDGIVRTLWLTADEL
RACEARHRSPAVMRCVDDYLAGRRVPLDFVHTHSVAPYRAAPDHQAVGK*
>GCA_000452945.1_BPMSHR5858_genomic_05332_1 yibF_1
MMKLIGSLSSPFVRKARIVLAEKKIDYKLELENVWAPDTNIHASNPIGKVPCLVMEDGAA
VFDSRVICEYVDTLSPVGKLIPPSGRERVEVRCWEALGDGVLDAAVAIRIEHTQREPQHR
SDAWIARQQRKVDDGLVAMSQGLAGKTWCVGNHYTLADIALGCTLGYLDFRMPEINWRER
HPNLDKHFAKLAQRQPFVDTVPQ*
>GCA_000452945.1_BPMSHR5858_genomic_05335_1 dusB
MPVLGSHVLRNNLFVAPMAGVTDRPFRQLCKRLGAGYAVSEMVASNAQLWKSEKTMRRAN
HAGEVEPIAVQIAGADPAMMAEAARHNVANGAQIIDINMGCPAKKVCNVAAGSALLQNEP
LVQRIVEAVVGAVGAGPDAVPVTLKIRTGWDRDHKNALRIATLAEAAGISMLTVHGRTRA
DLYRGDAEYDTIAAVKAAVRIPVVANGDITSPRKAKAVLDATRADALMIGRAAQGRPWLF
REIDHFLQSGELLPPPRIDEIRQVMNEHLEDHYAFYGEFTGVRTARKHIGWYTRGLSGAN
GFRHRMNTLETTREQLAAVNEFFDAQQALSDRLVYVDDEDGDRGEPDDSNQLAA*
>GCA_000452945.1_BPMSHR5858_genomic_05341_1 group_5177
MSAPLHDRPARSPLGHPFVERVRARVAGGVTHLTTGSGGPSLDYSSPPGDPGLFGPDAVC
WKVHADFTSMMTGGISALLLQALHPLALAGVWDHSTFRTDILGRLRRTATFIAGTTYGSR
ADALKLIERVKAIHAAVNGTGLDGRPYRASDPALLTWVHVAEVSSFLAAHLRYVNPLLPG
ADQDRYYAETARVAELLGATNVPKTRADVAAYFAAMRAELDAGERTHEVVRILMNAPVPK
PAMRPAAALMFNAGVDLLPAWAQQTLGLSSLAPVRRALVRPGVRIVAPVIRWALVNGVSK
RARRRVAAPPEPPDHGGT*
>GCA_000452945.1_BPMSHR5858_genomic_05347_1 yfcG_2
MIDLYYWTTPNGHKITMFLEEAGLPYRIVPVNIGRGEQFEPAFLRIAPNNRIPALVDHAP
ADGGAPLSIFESGAILLYLADKLGRFIPADLRGRNEALQWLFWQMGGLGPMAGQNHHFVQ
YAPEPLPYAIERYVKETSRLYGVLNKHLSDGRDYIAGEYSIADMASYPWIVPHERQRQRL
EDFPFLAAWFARVAERPGTVRAYARAKEINTAPTVDQQSRGVLFGQDASTVR*
>GCA_000452945.1_BPMSHR5858_genomic_05353_1 yhfA
MECKVSWMGQDGMAFAAQTGSGHLVTMDGAPEGGGNNLAPRPMEMVLVGTGGCTAYDVVL
ILKKSRQEVIGCSVTLQAERASEDPKVFTKVHFHFTVTGRNLNPATVERAINLSHDKYCS
ASIMIAKTAELSHSFEIVAA*
>GCA_000452945.1_BPMSHR5858_genomic_05355_1 pyrC
MNASVPASLTLARPDDWHLHVRDGAMLAAVLPHTARQFGRAIIMPNLKPPVTTTAQAQAY
RERILAAVPAGMTFEPLMTLYLTDNTPADEIRRARESGCVHGVKLYPAGATTNSDAGVTD
LLGKCAKTLEAMQEVGMPLLVHGEVTDPSIDLFDREKVFIDRVMEPLRRALPGLKVVFEH
ITTKDAADYVRDADAASGRIGATITAHHLLYNRNAMFFGGIRPHYYCLPVLKRETHRIAL
VEAATSGNPRFFLGTDSAPHAKGAKEAACGCAGCYTALHALELYAEAFDQAGALDKLEGF
ASFFGADFYGLPRSAETVTLRRETWELPREIDAGAGPVVPLRGGEAIGWRLV*
>GCA_000452945.1_BPMSHR5858_genomic_05359_1 group_5182
MSELFFDTDLLDEFDRLQKHMAHLFAGFPASLRATRVGAFPPVNIGSTDESVEIVAFAPG
LAPSEIDVSVDKGLLTIAGERKRVETDAPDDTRMYAHERFAGAFRRVIELPQDADPNNVS
ARYANGCLTIRVGKSEASKPRVISVQ*
>GCA_000452945.1_BPMSHR5858_genomic_05360_1 groS
MHVRPLYDRVIVKRIEQQRTTASGIVIPDSAAEKPEQGEIVALGNGRLLQDGTLRALQLE
VGDQVIFGEYAGQSIKVDGEDFLVMREEDVLGVLEAGERAARKAA*
>GCA_000452945.1_BPMSHR5858_genomic_05366_1 group_5184
MKFPKAMLMVAALSTFAGGAIAQETGTLKKIKDTGVIALGHRESSIPFSYYDQNQQVVGY
SRDFQMKVVDAVKKKLNLPNLQVKNIPVTSQNRIPLVQNGTVDIECGSTTNNLDRQKQAA
FSDTIFVIGTRLMTKKDSGIKDFADLKGKTVVTTAGTTSERLLREMNNKNQMGMSIISAK
DHGESFQTLETGRAVAFMMDDALLAGERAKAKQPGEWVIVGKPQSQEAYGCMMRKDDPAF
KKVVDDAIVQVEKSGEAAKIYSKWFENPIPPKGLNLNFPLSDEMKKLYANPNDKALD*
>GCA_000452945.1_BPMSHR5858_genomic_05367_1 gdhA
MSSQSQSPSVAQSIPSYLHADDLGPWGNYLQQVDRVAPYLGSLSRWIETLKRPKRILIVD
VPIELDNGTVAHFEGYRVQHNVSRGPGKGGVRYHQDVTLSEVMALSAWMSVKNAAVNVPY
GGAKGGIRVDPRKLSRGELERVTRRYTSEIGIIGPNTDIPAPDVNTNEQIMAWMMDTYS
MNQGQTATGVVTGKPISLGGSLGRKEATGRGVFVVGCEAAKKKGLEIEGARIAVQGFGNV
GGIAAKLFQEAGAKVIAVQDHTGTIHQPAGVDTVKLLEHVGRTGGVAGFEGAEPMPNDEF
WTVETDILIPAALENQITEKNASKIRTKIIVEGANGPTTTAADDILSANGVLVIPDVIAN
AGGVTVSYFEWVQDFSSFFWTEDEINHRLERVMREAFAGVWAVAEEHNVSVRTAAFIVAC
KRILMAREMRGLYP*
>GCA_000452945.1_BPMSHR5858_genomic_05371_1 gntK
MILIAMGVSGAGKSRIGEMLAQRLSCSYTDGDAFHSAANKEKMHHGIALTDEDRWPWLRS
IREAIEARQRAGETAVFTCSSLKRAYRDILRGNDRDVRFVYLKGSFDVLRERLKTRTGHF
FDPSLLQSQLDTLEEPGPDEAIEVSIELTPDEIVERVMRELDRERQQQQQQQ*
>GCA_000452945.1_BPMSHR5858_genomic_05374_1 edd
MATLHPTLASVTERVIARSRPTRQAYLARIDAAQGHFPARGALSCANLAHGFAGLEGSDK
FAIKAIREPNIGIVSAYNEMLSAHAPYKGFPDIIKAAARERGGVAQFAGGVPAMCDGVTQ
GNAGMELSLFSREVIAMSTAVALTHNMFDAALCLGICDKIVPGLLIGALQFGHLPTIFVP
AGPMTSGLSNDDKAKIRQQFATGQVGRDALLEAESAAYHGHGTCTFYGTANSNQMLMEVM
GLHLPSAAFVHPHTPLRDALTAEAARRVLELTAERGQYTPIGHVIDERAIVNGIVALLAT
GGSTNHTLHLVAIARAAGILIDWDDFDALSAAVPLLAKIYPNGKADVNHFHAAGGIAFLV
RNLLEGGLLHEDVTTVAGKGLAHYTKEPRLIDGKLTWVDGVGESADDKVLRPIAAPFQPD
GGLRLMQGRLGRGVIKISAVAPEHRKVTAPAIVFDSQEAVQAAFDAGELKRDFVAVVRFQ
GARANGMPELHRLTPLLGVLQDQGFRVALVTDGRMSGASGKVPAVIHVSPEALLSGPLGK
VCTGDTIVIDAQAGVLDVEIDEQAWAARAVAQPAHQAENEVGFGRELFGVFRAAAAPAEQ
GASVFGAWVGEAAHVTA*
>GCA_000452945.1_BPMSHR5858_genomic_05376_1 group_19482
MAMLWVKTFHIVLIASWFAGLFYLPRIYVNLAMETDPAARRRLLAMARKLLRFMTIIAVP
ALACGLWLWLVIGIGQGQGWVHAKLGVVLLLVVYHAYCGHLLKVFERGENRRSDKWYRVF
NELPVLGMLAAVALAVIKPF*
>GCA_000452945.1_BPMSHR5858_genomic_05377_1 proA
MDIDQYMTDVGRRARRASREIARASTAAKNAALEAVARAIGRDADALKAANARDVARAKD
KGHDAAFVDRLTLSDKALKTMIEGLRQVATLADPIGEMSNLKYRPSGIQVGQMRVPLGVI
GIIYESRPNVTIDAAALCLKSGNATILRGGSEALESNTALAKLIGEGLAAAGLPQDAVQV
VGTADRAAVGKLITMTEYVDVIVPRGGKSLIERLINEARVPMIKHLDGICHVYVDDRASV
AKALAVCDNAKTHRYGTCNTMETLLVARGIAPAVLTPLGRLYREKGVELRVDADARAVLE
AAGVGPLVDATDEDWRTEYLAPVLAIKIVDGIDAAIEHINEYGSHHTDAIVTEDHDRAMR
FLREVDSASVMVNASTRFADGFEFGLGAEIGISNDKLHARGPVGLEGLTSLKYVVLGHGE
GRE*
>GCA_000452945.1_BPMSHR5858_genomic_05383_1 dapB
MSSMKIAIAGASGRMGRMLIEAVLAAPDATLAGALDRTGSPQLGQDAGAFLGKQTGVALT
DDIERVCAEADYLIDFTRPEGTLAHLDAALRHDVKLVIGTTGFSEPQKAQLRAAGGKIAL
VFSANMSVGVNVTMKLLEFAAKQFAQGYDIEIIEAHHRHKVDAPSGTALMMGETIAAATG
RTLDDCAVYGRHGVTGERDPSTIGFSAIRGGDIVGDHTVLFAGIGERIEITHKSASRVSY
AQGALRAARFLAGHQAGFFDMQDVLGLR*
>GCA_000452945.1_BPMSHR5858_genomic_05390_1 cynR_1
MLLRHIRYFLAVAEQGNFTRAAESLHVSQPTLSQQIRQLEGVLGVQLFDRSARAVRLTEF
GTAYARHARAALQQLEAGRRAVHDVENLQSGSLRLAMTPTFSAYLIGWLIDGFNARYPNL
KLTIREMPQEQIEPLLADDELDVGIAFDATQSPEIETMPLWIETLALVVGRDHRHARRRK
PLSPDEAAGEPLILLSRAFATRGKVDQYFHRNGVTPRVAIEVNSISAMLEMVRGGRIATV
LPSAIAERRDDLCAIRLDPPLPQRTAALLMRKGGYRSAAMRAFVDHMLARRHEIGRREGA
RARDGVPSDGE*
>GCA_000452945.1_BPMSHR5858_genomic_05392_1 cynS
MTQSQHSQSAREALAERIVEAKTRKNLTFEQINEGTGLSVAFTTAALLGQHPLPADAARV
VAAKLDLDDDAQRLLQTIPVRGSIPGGVPTDPTIYRFYEIVQVYGSTLKALIHEQFGDGI
VSAINFKLDIKKVDDPEGGSRAVITLDGKYLPTKPF*
>GCA_000452945.1_BPMSHR5858_genomic_05396_1 speE_1
MPFVRALSESARARGFSALLPPSIIQTIGTARPAAPAALSTTLLFHPTPDVAYGFPNARR
LARIESEYQEIEVWETSLLGRLFTLDGRPMTSVGDEFVYHECMTHPAALAHPAPKKALVL
GGGDGGAARQLLRHPCIERIVVAELDAAVVDMARRFLGDVHQGALDDPRVELAIGDAARY
VDHACEHFDLAVFDLTPPDSPAAGLYTPAFYTRLKRILTPQGAVSLHLGSPLYHGARIAA
LLAGLRERFAVVAPLCAHVPLYGSPWLMAIASDTLDAASLFSHDIDARLAERHIEGLRFY
DAKLHSALFTLPHALRDTLGMHR*
>GCA_000452945.1_BPMSHR5858_genomic_05402_1 thiL
LIDRFFARRATGPHARAALGIGDDCALLAPEPGKLLAVSTDMLVEGRHFLADVDPRALGH
KTLAVNLSDLAAMGAAPRAFTLACALPRADADWLEAFSDGLFALAERHGCELIGGDTTSG
PLNLCVTVFGDVACGAALRRDAARDGDDVWVSGTLGDARAGLGVIRGEWRAGEREAAAFR
RALEWPQPRVALGVALAGIAHAALDVSDGLAGDLPHILERSNVRADVDVDAVPRSAALAT
LPADVQRRCMLEGGDDYELCFTAAPSARTAIDAAGARAGVAVTRIGTIRGLSAPTDARAV
TWRDASGAPLSLTLHGFDHFHAN*
>GCA_000452945.1_BPMSHR5858_genomic_05410_1 araF_1
MKRRTFITLAAAATVAAAGLPAQAAEPVKIGFLVKQPEEPWFQDEWKFAELAAKDKGFTL
VKIGAPSGEKVMSAIDNLAAQKAQGFIICTPDVKLGPGIVAKAKSHGLKMMTVDDRLVDG
AGKPIESVPHMGISAYDIGKQVGGGIAAEIKRRGWNMNEVGAIDITYEQLPTAHDRTTGA
TDALVAAGFPKANVIAAPQAKTDTENAFNAANIALTKNPKFKHWVAYGLNDEAVLGAVRA
AEGRGFKAADMIGIGIGGSDSALSEFKKPQPTGFFGTVIISPKRHGEETSELMYAWITQG
KAPPPLTLTTGMLATRENVAQVRETMGLAAK*
>GCA_000452945.1_BPMSHR5858_genomic_05422_1 group_5190
MSRKTLLDPRRVRDEIALAAARLIAEDGLDYAGAKRKAARQVLGDSRIAGEWLPDNDQIE
EELHEYLALFQSDTQPAELRRLRLVALAWMERLAAFNPYVAGAVLNGTANAHSDIHLQTF
CDNHKDVAIYLLNQNIQYDVSETRHFAGRGDIETLSFLWREARGAEPVGIHVALYTSDDL
RGAVKADARGRLSRADAQTLRALVDAAPSSPTESRTDE*
>GCA_000452945.1_BPMSHR5858_genomic_05423_1 resA_2
MNSKGIFAGIVVAAVAVAGGLAAGHWMRGNPASGPQAGTPAGAPAAGNPVDTLWAASYPD
VDGKPQRLAAFKGQKLVVNFWASWCGPCVEEMPELVKLSREYEKKGVRFVGIGVDSDQNV
KNFLKKVPVDYPIVVSGYAGADLARHFGNTAGALPFTVVIDETGKIRETKLGQIRPDELK
RTLDTL*
>GCA_000452945.1_BPMSHR5858_genomic_05430_1 adoK
LATLICGSIAYDNIMTFEGRFREHILPDQVHLINLSFLVPTMRREFGGCAGNIAYALNLL
GGDARMMGTLGAVDAQPYLDRMDALGLSREYVRVLPDTYSAQAMITTDLDNNQITAFHPG
AMMQSHVNHAGEAKDIKLAIVGPDGFQGMVQHTEELARAGVPFIFDPGQGLPLFDGATLR
RSIELATYIAVNDYEAKLVCDKTGWSEDEIASRVQALIITRGEHGATIRHRDGNEQIPAV
RAERVIDPTGCGDAFRGGLLYGIEHGFDWATTGRLASLMGSLKIAHQGPQTYAPTRAEID
ARFETAFGYSPK*
>GCA_000452945.1_BPMSHR5858_genomic_05431_1 pcp_2
MLTRKTLTLAAMLTATVTLAGCFTPPGSADVYSVGQAQREQTVRMGTVESVRAVRIQGDG
GGSALGTLGGGALGAVAGSAIGGGRGSVLTAIAGGIAGAVAGNAVGQGLSSANGVEITVR
LDNGDLRSITQAATPEVFRAGERVRLLSSGGVTRVTH*
>GCA_000452945.1_BPMSHR5858_genomic_05434_1 nrdB
MLNWDDEKTAVTPASGAQHNAMRDPAGMAVGLQAAPAAHQASSARDIFAGDFAVAPNASA
AVASEARVNVADKRIINGKTDVNQLVPFKYKWAWEKYLAGCANHWMPQEINMSRDIALWK
DPNGLTEDERRIVKRNLGFFVTADSLAANNIVLGTYRHITAPECRQFLLRQAFEEAIHTH
AYQYIVESLGLDEGEIFNAYHEVPSIRAKDEFLIPFIHTLTDPAFTTGTLEADQKLLKSL
IVFACIMEGLFFYVGFTQILALGRQNKMTGAAEQYQYILRDESMHCNFGIDLINQIKLEN
PHLWTAEFRAEIRELFKHAVDLEYRYAEDTMPRGVLGLNASMFKSYLRFISNRRCQQIGL
DPLFPNEENPFPWMSEMIDLKKERNFFETRVIEYQTGGALSWE*
>GCA_000452945.1_BPMSHR5858_genomic_05438_1 group_1052
MDIVLYALTVLLYGGLAVAGWRARRVGAARPLVASVPAVAPARESAPGGMGGLGRTILGA
ALLAHGVLLHMTIFPHDAMVFGFAFALSAMFWLGAGIYWIESFFFPLDGMRLLVLPLAGV
ASLLPLAFGGVRVLPYAAAPLFKVHFLIANIAYGLFAIAALHAVLMLMVERRLHALRHDG
LRDASGWVAGWLDTLPPLLTLEKLLFRLIGAGFVLLTLTLATGILFSEQIDARALKLDHK
TVFAILSWLMFGGLLVAHKASGWRGRGAARWVLASFVALLLAYVGSRFVLEVLLHRSVV*
>GCA_000452945.1_BPMSHR5858_genomic_05449_1 ispB
MIMSSTATPSLSAAHLLAPIASDMEQVNRVIRQSLASDVLLINQIAEYIIGAGGKRLRPA
LLLLVAGALGENTNQKHVLAAVVEFIHTATLLHDDVVDESELRRGRKTANALFGNAASVL
VGDYLYSRSFEMMVGVGKMRVMEILSEATTIISEGEVLQLLNMHDADVDEARYMQVIRYK
TAKLFEAAARLGAVLAGADAPTEAAAAEYGRRIGTAFQIMDDWLDYAGTAESMGKNAGDD
LREGKPTLPLIYLIERGTPEQSTLAREAIEHGGTDRFDTIFDAITRSGALDHTLECARQE
AQAAAAAISSFPSSNFKESLLELCSYSTSRQS*
>GCA_000452945.1_BPMSHR5858_genomic_05454_1 outO
MTPNPFLSGSPEHAAAAGPLAAFAALPTGMQLAFAIVLGLVVGSFLNVVVHRLPIMMKRA
WLAEIAEATGAPCADDSLPARYNLCVPRSACPHCGHALRAWENVPVLSYIALRGRCRHCR
TPIGARYPLIELASGALAAGALALFGPSGAALAAFGLCAALLAMSAIDMQTGFLPDSLTL
PLLWAGLCVNLWGTFASLRAAVIGAIAGYLFLWCILWLFKLLRGIEGIGYGDLKLLAALG
AWLGWEALPQVVLIAAVAGAAVGLVATWRGRMRFEEPLPFGPFLAAGGAATLFFGTPFYL
LLGG*
>GCA_000452945.1_BPMSHR5858_genomic_05458_1 group_5196
MDQLEQFLTRAEALLGRLEAILPPPPAAVDWAAAFAFRWRKRQGRGYLQPVAARSTISLD
DLRNIDRQKALIEQNTRQFVNGKPANNVLLTGARGTGKSSLIKACLNAYAADGLRLIEVD
KDDLHDLGDIVELISARPERFIVFCDDLSFEEGESGYKALKVALDGSVAAQSDNVLIYAT
SNRRHLLPEYMSDNETYKHLPDGEIHPGEVVEEKISLSERFGLWVSFYPFKQDDYLTIVG
HWLRHFGCDAAEVEAARGDALVWALERGSRSGRVAWQFARDRAGRKENV*
>GCA_000452945.1_BPMSHR5858_genomic_05469_1 group_5197
MAHTLPPLPYAEDALAPHISQETIQFHYGKHHQAYVTNLNNLIPGTEFENLPLEEIVKKS
SGGIFNNAAQIWNHTFFWNSLSPNGGGAPTGALGDAINAKWGSFDAFKEAFTKAAVGTFG
SGWAWLVKKADGSLDIVSTSNAATPLTTADKPLVTIDVWEHAYYIDYRNARPKFVEAFWN
IVNWDFAAKNFA*
>GCA_000452945.1_BPMSHR5858_genomic_05470_1 xseA
MHSESPFVGSGGAPAGDGVIPVSALNRAIGTMLERTFPLLWVAGEVSNFTRAASGHWYFS
IKDQQAQMRCVMFRGRAQHAEFTPREGDRIEVRALVTMYEPRGEVQLSVEAIRRTGQGRL
YEAFLRLKAQLEAEGLFAPERKRAPPTHPRAIGIVTSLQAAALRDVLTTLARRAPHVPVI
VYPAPVQGAGAAEKLAAMVATANARREVDVLIVCRGGGSIEDLWSFNDEALARAIAASDV
PVVCGVGHETDFTIADFAADVRAPTPTGAAELVSPQRALLLRELGERQGALARGLRRGLD
ARAQQLDWLARRLISPAERLQRQAMHVGQLAARLAAAGARPVRDARARFSLVQLRWQRAR
PDFAQARQVLAAFAQRLDATLQRRLERDTARVAACAARLEVLSPQRTLERGYAALIDAQT
GRAVRAPASLKPQRRLTVHLAEGSADVSLADVQPRLSDTF*
>GCA_000452945.1_BPMSHR5858_genomic_05482_1 group_5198
MPSFDVVSEANMIEVKNAVEQSNKEISTRFDFKGSDARVEQKERELTLYADDDFKLGQVK
DVLIGKMAKRNVDVRFLDYGKIEKIGGDKVKQVVTIKKGVSGDLAKKVVRIVKDSKIKVQ
ASIQGDAVRVSGAKRDDLQSTIALLRKEVTDTPLDFNNFRD*
>GCA_000452945.1_BPMSHR5858_genomic_05486_1 xerD_2
MARAVHACAADTAGGDAGRAPAQPADRPVDALAASPAWLDSRASIDLFCDAMWLEHGLSR
NTLDAYRRDLQLFAQWLAARHAASVDHASEPTLTEYIAARSDGKATSSNRRLSVFRRYYG
WAVREHRAAVDPTLRIASAKQAPRFPSTLSEAQVEALLAAPDVDTPLGLRDRTMLELMYA
SGLRVSELVTLKTVEVGLNEGVVRVTGKGSKERLVPFGEVAHGWIERYLRDARPALLGAR
AADALFVTARGDGMTRQQFWNIIKRHAQQADVRVHLSPHTLRHAFATHLLNHGADLRVVQ
LLLGHSDISTTQIYTHVARERLKTLHAAHHPRG*
>GCA_000452945.1_BPMSHR5858_genomic_05491_1 eamA_2
MAPKDLLLALVVILAWGVNFVVIKVGLHGVPPMLLGGLRFLLASVPAVFFVRRPRIPWRL
LVLYGSTILLGQFVFLFSAMYVGMPAGLASLVLQAQAFFTLFFARFALGERLRAQNLVGL
AIAAVGLVAIAVQGGRGMTLAGFALTIGAAALWALGNVVTKKVGKVDLVSLVVWASLVPP
VPFFALSYGFEGPQRIGAALTSLSGASIFAIVYLAFVATLLGYGLWSRLMSRYPAGQVAP
FSLLVPVVGLASSALLLDERLTHAQLAGAALVMAGLAVNVFGDRVVRRLFAAAS*
>GCA_000452945.1_BPMSHR5858_genomic_05497_1 group_536
MGDGSTRRKPAPPRQGDAQGDQFDLFGVEPLPLPERPSSTPSASPASSPSPATAADGKRQ
RAPRGRRRDASPDAAAAGDAPASDDIARTSAPAATDAPGTNDPAHGEDDEADARDSTRTK
PATAALPTPDSGMAGTVGDANDAGDRADPADPADPADPADPGVPDDPADPDDPPEPAATP
ALPGFDAPPAAAPKKRTRRRGVAPAAIADDVAAAARELPPHVRLGTSSWYFPGWKDIVYG
DDYAQSKLSREGLEAYAAHPLLKSVSLDRSFYAPLTVANYLRYAQQVPDDFRFVVKAPAL
VTDAVLRGARGEPTGPNPAFLNAQLAADEFVRPCLEGLGNKAGALVFQFPPLPDTLLADP
AALVDRLSTFLGALPALPAAPDGPRYAVEIRDASLLTPRFIRALAAAGVRYCVGLHAKMP
DPLRQAAALALLDGEPSGPLIVRWSLHGGFKYEQAKAKYEPFDQLVDEDPHTRTALAELA
ARYVLAGQPVLITVNNKAEGSAPLSCIALAKEIAAACARWRGEAA*
>GCA_000452945.1_BPMSHR5858_genomic_05502_1 kynU

MKTREEALALDRDDPLAPLREQFALPAGVIYLDGNSLGAQPRAAAARAQQVIGAEWGEGL
IRSWNTAGWFALPRRLGDRLAPLIGAADGEVAITDTISINLFKLLAAMLRHQARHAPKRR
VIVSERSNFPTDLYIAQGLIAQLDRDYELRLIDDPADLPDALDDETAVAMITHVNYRTGY
MHDMPSVTQTVRQAGALMLWDLAHSAGAVPVDLNGALADGAVGCTYKYLNGGPGSPAFVW
VPKRHQRAFEQPLSGWWGHRAPFAMQPAFEPDPGIARFLCGTQPIVSMSMVECGLDVFAQ
TDMHAIRRKSLALTDAFVALVESRCAGQPLKLVTPRAHHQRGSQASFEHPHGYEVMQALI
ARGVIGDYREPRILRFGFTPLYTRFVDVWDAVETLRDILDTEAWRAPEFATRAAVT*
>GCA_000452945.1_BPMSHR5858_genomic_05513_1 estB_2
MVLSSWSALGERVDAAIDAALAQRRLVGAVVLVARRGELAYRRAAGLADREAGVPMREDA
LFRFASVSKPIVSAAAMRAVAAGKLDLDASIARWLPAFTPALAGGRPARITARQLLSHTA
GLGYRFLETHAHGPYARAGVSDGMDCAGISLAENLRRIASVPLLYEPGTSWAYSLATDVL
GALIEAVCDQPLEDAVAEFVTTPLGMVDTRFYAHDAARLAAAYVDASDAAAPGGPRRMAA
LEIASPFPDTAGIRFEPARALDAHAFASGGAGMVGTASDVLALIEALRTGGDGWLPAARI
DEMARIQPGAEDLPTAPGYGFGLGFSVLRDPAAARSPESVGTWRWGGAYGHAWFVDRAAG
LSVVALTNTAYEGMSGRFVADLRDAVYGAGAAAQERAA*
>GCA_000452945.1_BPMSHR5858_genomic_05515_1 group_2624
MKTGRRHFVRSVASASAALAAAAWSPARAAIDAPTSPATALSLTPGRWSPNNVARLRAVL
AGHGASSPRYRPEHRPYAVFDWDNTSIMNDCEEALLMHQIDGLHYRLTPEQFSAILRQGV
PDGPFDAKLGYTSVDGKPVRMEDIAADVDADYRWLHANYRGLAGDKPLDEIHRSEQFRDF
RAKLYFMYDAICDTYPVEIGYKWIMYWYAGMTRDELQAMAFDSNVANLGDALRKVTYESS
RALPGKAGVIAATHFHGIRIHEEIRAVMDTLRSNGIDVYVSTASLDDVVRVFAGHPAFGY
GVPAENVIGMRLTMADGKYMNEYLPNWHFNYGPGKTVGIRRELESKKGYGPLLVFGDSDG
DAWMLRDFADTAVGVIVNRMKKGEIGIDSRKAAEQIGAKDARLVLQGRDENTGLMVADER
SIKYGKRDPKLLA*
>GCA_000452945.1_BPMSHR5858_genomic_05517_1 yieH
MTAHANDARVLICDCDGVLIDSETIAADVLVRELEARWPGVAARPIVLPLLGLRTERVLD
CASARAGRALADSDIEAIRRSVEAAAVQAPIVDGIDAALARIELTIACASNSRRHYVDAA
LRRTGLKRFFGERLFCADGVARPKPAPDVYLAAAQALGVAPSQCLVVEDSATGVTAASAA
GMTVLGFVGGGHASPRQVDALRAIGARRVFDDMRELPGLVSRWAETGVVEPH*
>GCA_000452945.1_BPMSHR5858_genomic_05519_1 ycjO
MRHLRLPLAHSASLVGAGPAADERARGKPARWLAAPSVAVLLLWMAIPLAMTIWFSLTRY
NLLNPDVKGFAGLENYRFLATDPSFLPAIWHTLALIGSVLAITVAGGVLMAVLFDRKFYG
QGIARLIAIAPFFVMPTVSALIWKNMILHPVYGLVARAMRALGMQPIDWFAQYPLTSIVI
IVAWQWLPFAFLILFTAIQSLDQEQKEAARIDGAGAFAMFFYITLPHLRRAIAVVVMMET
IFLLSIFAEIYTTTGGGPGDATTNLSYLIYALGLQQFDVGLASAGGILAVVLANVVSFFL
VRMLARNLKGEYEA*
>GCA_000452945.1_BPMSHR5858_genomic_05520_1 group_13488
MQRKTLTAAAARVAAFAALASSALAAQAATLTIATLNNPDMIELKKLSSAFEKANPDIRL
NWVILEENVLRQRATTDITTGSGQFDVMAIGTYEAPQWGKRGWLAPMSNLPADYDLNDVI
KTARDSLSYNGQLYALPFYVESSMTFYRKDLFAAKGLKMPEQPTYEQIAEFADKLTDRAN
GTYGICLRGKAGWGENMAYVSTVVNTFGGRWFDENWNAQLTSPEWKKAINFYVNLLKKNG
PPGASSNGFNENLTLTASGKCAMWIDATVAAGMLYNKQQSQVAEKIGFAAAPVAATPKGS
HWLWAWALAIPKTSKQQDAAKKFVTWATSKQYVEMVGKDEGWASVPPGTRQSTYQRAEYK
AAAPFSEFVLKAIQTADPTDPSLKKVPYTGVQYVGIPEFQSFGTVVGQAIAGAVAGQTSV
DQALAAGQAAAERAVRQAGYRK*
>GCA_000452945.1_BPMSHR5858_genomic_05606_1 rpsQ
MNDSVKTSLKRTLVGKVVSNKMDKTVTVLVEHRVKHPIYGKYVVRSKKYHAHDEANTYNE
GDLVEIQETRPVSKTKAWAVSRLVEAARVI*
>GCA_000452945.1_BPMSHR5858_genomic_05608_1 rplX
MNKIRKGDEVIVITGKDKGKRGVVLAVGAEHVTVEGINLVKKHVKPNPMKGTTGGVEAKT
MPLHISNVALVDANGKASRVGIKVEDGKKVRFLKTTGAVLSA*
>GCA_000452945.1_BPMSHR5858_genomic_05619_1 rpsM
MARIAGVNIPNHQHTEIGLTAIFGIGRTRARSICVASGVAFSKKVKDLTDADLEKLREEV
GKFVVEGDLRREVTMNIKRLMDLGCYRGVRHRKGLPLRGQRTRTNARTRKGPRRAAQALK
K*
>GCA_000452945.1_BPMSHR5858_genomic_05627_1 engB
MAFLLHQARFFTTVNHLRDLPPTVQPEVAFAGRSNAGKSTAINVLCNQKRLAFASKTPGR
TQHINYFSVGPAAEPVAHLVDLPGYGYAEVPGAAKAHWEQLLSSYLQTRPQLCGMILMMD
ARRPLTELDRRMIEWFAPTGKPIHSLLTKCDKLTRQESINALRATQKSLDAYRDAGYAGK
LTVQLFSALKRTGLDDAHALIESWVRPAAADEDRAAVAE*
>GCA_000452945.1_BPMSHR5858_genomic_05630_1 ccsA
MDLTQVSSSRAPRASGAAAPLLFDERPFLRRLSLVDWLFALALVAGAGFALARYHAHMDY
YDTAVLIGTVPALVALGWRWKPARLLMAAIAALSLVSLQLYHGDLARADSAFLLKYFLSS
QSAILWMSALFVLATLFYWIGMLSRSETGGAIGQRLTWIAVLMGFTGLMVRWYESYLIGA
DVGHIPVSNLYEVFVLFSLITALLYLYYEGHYGTRALGAFVLLVISAAVGFLMWYSIARD
AQQIQPLVPALQSWWMKIHVPANFIGYGSFALSAMVAVAYLSKERGVLADRLPALEVLDD
VMYKSIAVGFAFFTIATILGALWAAEAWGGYWSWDPKETWALIVWLNYAAWLHMRLMKGL
RGTVAAWWALTGLLVTTFAFLGVNMFLSGLHSYGKL*
>GCA_000452945.1_BPMSHR5858_genomic_05634_1 group_5219
MRIVLQMSAIVAALALAGCGQRGPLYMPVVPPLPPKPTEQTQPPPSDVTPDAETASARDD
GASGAADAPLTLSPDLSTQRTPKAAPASGASSAQ*
>GCA_000452945.1_BPMSHR5858_genomic_05642_1 aroB
MITVNVDLGERAYPIHIGADLIGRTELFAPHIAGASVTIVTNTTVEPLYGDTLRAALAPL
GKRVSTVVLPDGEAYKNWETLNLIFDGLLEQHADRKTTLIALGGGVIGDMTGFAAACYMR
GVPFIQVPTTLLSQVDSSVGGKTGINHPLGKNMIGAFYQPQAVIADIGALSTLPDRELAA
GVAEIVKTGAIADAAFFDWIEANVGALTRRDPDALAHAVKRSCEIKAGVVAADEREGGLR
AILNFGHTFGHAIEAGLGYGEWLHGEAVGCGMVMAADLSVRTGHLDEASRARLCRVVEAA
HLPTRAPDLGDARYVELMRVDKKAEAGAIKFILLKRFGETIITPAPDDAVLATLAATTR*
>GCA_000452945.1_BPMSHR5858_genomic_05643_1 group_5221
VSEIPGGGIPRETRDARVSNAGDGQAIPVAAPTTAALEAHLAPYAAHASRSRGRRHPEPP
PAARTEFQRDRDRIVHSTAFRRLEYKTQVFVNHEGDLFRTRLTHSLEVAQIARSVARNLR
LNEDLVEAISLAHDLGHTPFGHAGQDALNACMRDYGGFEHNLQSLAVVDELEEHYGAFNG
LNLCFETREGILKHCSRENARKLGELGERFLQSRQPSLEAQLANIADEIAYNNHDVDDGL
RSGLITIEQLAEVELWQCHYEAALAEYPHLEGRRLVHETVRRIINTLIVDLIDATTRNLA
RHGPTSLDDVRAAPPLVAHGEPIATQAAALKRFLYKNLYRHYRVMRMASKAQRVVTGLFN
AFTGDPRLLPPDYQAADAAHQPRLVAHYIAGMTDRFALKEYQRLFVMDEN*
>GCA_000452945.1_BPMSHR5858_genomic_05647_1 ugpC
MAALSLKGVRKSYGGAQYVLHGIDVDIADGEFVVLVGPSGCGKSTLLRMIAGLETVTEGE
IAIGGRVVNALEPKDRDIAMVFQNYALYPHMTVAQNMGYGLKIRGVERALIDARVQAAAQ
ILELGPLLARRPRELSGGQRQRVAMGRAIVREPSVFLFDEPLSNLDAKLRVQMRLEIQRL
HARLATTSVYVTHDQIEAMTLAQRVIVMNRGDAEQIGAPVDVYEKPATTFVASFIGSPAM
NLLHGRLSEDGAAFDVADGPRLPVAGAAGAGRGIAPGREWILGVRPEHMTPQPGEAFATL
AVDSCELLGADNLAHGRWGAHDVAVRLPHAMRPTRGETLPVALPARHLHFFDPATGKRAG
*
>GCA_000452945.1_BPMSHR5858_genomic_05652_1 gltD
MGKITGFLEYERRHEAYEAPLTRVKHYKEFVAALTDADAKIQGARCMDCGIPFCNNGCPV
NNIIPDFNDLVYRQEWRQAIDVLHSTNNFPEFTGRICPAPCEAACTLGIHDDPVGIKSIE
HAIIDKAWAQGWVAPQPAAHKTGKKVAVVGSGPAGLAAAQQLARAGHDVTVFEKSDRIGG
LLRYGIPDFKLEKWLIDRRMRQMEAEGVTFRTSVFVGRDPLPETIGNMAKETISPETLKD
EFDAVVIAGGSETPRDLPVPGRELAGIHFAMEFLPQQNRVNAGDKLADQLLAKGKHVVVI
GGGDTGSDCVGTSNRHGAKHVTQFELLPQPPEAENKPLVWPYWPIKLRTSSSHEEGCERD
WAVATKRFEGRNGKVEKLIAARVAWVDGKMQEVPDSEFEIKADLVLLAMGFTQPAAPVLE
AFGVAKDARGNARAGTEGDRAYYTSVDKVFAAGDMRRGQSLVVWAIREGRQCARSVDAYL
MGSSELPR*
>GCA_000452945.1_BPMSHR5858_genomic_05655_1 yddA
MTQSMQASAPAPDDPTQPQPSVSTWSLIKPYWVSEEWKVAWGLLVTIVAINLTLVWINVR
INSWSASFYNALQSKDVRDFPSLLITFAVLAFAFIILAVYGLYLRQMLGFRWRQWLTTRF
LNEWLGDRNFYRIERDRLADNPDQRIADDLQSLASTTLSLSLDLLSTVVTLISFATILWS
IAGAATISLGGHAITIPGYMVWVAMIYAIAGSYVMHRFGHPLVSINYQQQRVEADFRFSL
IRIRENAEQIAFYNGEPVETAHEQGLFQRIRENWWRVMRYTKRLTFVLNFYAQLASLFPI
AVAAPRYFAGAFTFGVLMQISRAFGTVSDSFSWFINSYGTLAEWRATVNRLREFKRVMGA
SHLKETMSPATEHGGINLHYVDARQITTSGLKLALPNGAALASIGDVAIEPGSRWLVRGP
SGSGKSTLMRALAGLWPFGDGAIDAPVDARMMFVPQQSYLPIGTLKAALAYPSATDTFSD
DACSEALRACGLADYVERLGETGHWTRILSPGEQQRLAGARVLLHKPDYLFLDEATSALD
ADNEARLYRLFDERLPKAAIVSIAHRESVASFHQQTLDVRRDEARVAA*
>GCA_000452945.1_BPMSHR5858_genomic_05661_1 mlaE
MISAIGRFVIGGLERTGYATRMFVRVVLEFFSLLRRPRLVTKQIHFLGNYSFVIIAVSGL
FVGFVLGLQGYYTLNRYGSEQALGLLVALSLVRELGPVVSALLFAGRAGTSLTAEIGLMK
AGEQLTALEMMAVDPLKNVIAPRMWAGVIAMPLLAAIFSAVGVLGGYVVGVLLIGVDPGA
FWSQMQGGVEVWADVGNGVIKSVVFGFAVTFIALFQGYEAKPTPEGVSHATTKTVVYASL
AVLGLDFLLTALMFS*
>GCA_000452945.1_BPMSHR5858_genomic_05671_1 hisD
MAIKIRKLDSAGEGFAAELRAVLAFEASEDDAIERAVAQILADVKARGDAAVLDYTNRFD
RLNAASVAALELPQSELEAALEGLEPKRRAALEAAAARVRGYHEKQKIECGSHSWQYTEA
DGTVLGQKVTPLDRVGLYVPGGKAAYPSSVLMNAIPARVAGVGEIVMVVPTPDGLKNDLV
LAAALLGGVDRVFTIGGAQAVAALAYGTQTVPAVDKICGPGNAYVASAKRRVFGTVGIDM
IAGPSEILVLCDGTTDPSWVAMDLFSQAEHDELAQSILLCPDEAFIERVEKAIGELLPTM
PRQDVIRASLEGRGALVKVRDMAEACRIANDIAPEHLEISALEPHQWGKQIRHAGAIFLG
RYTSESLGDYCAGPNHVLPTSRTARFSSPLGVYDFFKRSSLIEVSAEGAHTLGEIASELA
YGEGLQAHAKSAEYRMKGAGDRQKG*
>GCA_000452945.1_BPMSHR5858_genomic_05672_1 hisC2
MTTPEDIIRRDVLAMTGYPVPDATGFVKLDAMENPYSLPAPLAAELGERLAHVALNRYPA
PRPAALIDRLRAVTGVPAACDVLLGNGSDEIISMLAMACAKPGAKVLAPVPGFVMYELSA
KFAQLEFVGVPLRADLTLDIDAMLAALAEHRPALVYLAYPNNPTGTLYPDEDVERIIAAA
AASLVVIDEAYQPFAQRSWLPRAAQFDNVVVMRTMSKLGLAGIRLGYLVGLPAWLVQFDK
VRPPYNVNVLTQAAAEFLLERVDVLDAQAAQLRDARTALAHAIGALPGATVFPSAGNFLL
VRVPDAAAVFDVLLTERVLIKNVSKMHPLLANCVRVTVGSPEENARLLAALKLALP*
>GCA_000452945.1_BPMSHR5858_genomic_05677_1 hisF
MPRNIMALAKRIIPCLDVTAGRVVKGVNFVELRDAGDPVEIARRYDAQGADELTFLDITA
TSDGRDLILPIIEAVASQVFIPLTVGGGVRAVEDVRRLLNAGADKVSMNSSAVANPPLVR
DAADKYGSQCIVVAIDAKRVSADGEPPRWEVFTHGGRKGTGLDAVEWARKMAELGAGEIL
LTSMDRDGTKAGFDLALTRAVSDAVPVPVIASGGVGSLEHLAAGITEGHADAVLAASIFH
YGEHTVGEAKRFMAERGIAVRL*
>GCA_000452945.1_BPMSHR5858_genomic_05686_1 ybgI
MDRIELELYLNNTLEIGRFKDYSPNGLQVEGRRKIEKIATGVTASLAFLDRALEWGADAV
LVHHGYFWRHEAPQITGRKYQRLKRLLANDLNLFAFHLPLDAHPEFGNNAQLGERLGLIG
DARFGDQDLGWLATLPMPVSLEHFTAKVENTLGRAPLVLGDLDQQLRRVAWCTGGAQSFF
DAAIEAGADVFLTGEVSEHVTHAAAESGVAFVAAGHHATERFGIQALGQHLSERFDLEHI
FIDIHNPV*
>GCA_000452945.1_BPMSHR5858_genomic_05713_1 nanT
MATVGGQIAHAPMTGDEKRVIFASSLGTVFEWYDFYLAGSLAAYISKSFFSGVNPTAAFI
FTLLGFAAGFAVRPFGAIVFGRLGDLVGRKHTFLITIVIMGISTFVVGFLPGYASIGIAA
PVIFIAMRLLQGLALGGEYGGAATYVAEHAPSHRRGFYTSWIQTTATLGLFLSLLVILGV
RTAIGEEAFGSWGWRVPFVASILLLAVSVWIRLQLNESPVFLRIKAEGKTSKAPLTEAFG
QWKNLKIVILALIGLTAGQAVVWYTGQFYALFFLTQTLKVDGASANILIALALLIGTPFF
VFFGSLSDRIGRKPIILAGCLIAALTYFPLFKALTHFANPALEAATQKAPITVVANPDEC
SFQFNPVGTAKFTSSCDIAKSALAKSGLNYENVAAPAGTLAQIKIGDTAIATYDGKAADA
KDQAKAFDKTLASTLKAAGYPAKADPAQLNWPMTTVILAILVIYVTMVYGPIAAMLVEMF
PTRIRYTSMSLPYHIGNGWFGGFLPATAFAIVAAKGDIYSGLWYPIGVALVTFVIGLLFV
KETKDSNIYAQD*
>GCA_000452945.1_BPMSHR5858_genomic_05717_1 recX
MRKNREGADSQQAGHGARTASAGEPHAERETRATVSGEGRPSGRPATRASDDALVSFRRS
AACEAGEPFDPDESFDAHDAARPRASAGGGGAEGVRGVRDAGAGGGRAGDARAANRLRAP
AGLSSAAKREETAGDVYTRTSQAGPRSRRRASPFHSDSSPSASSETDGAARSSASQRPAR
TLKGRAVAYLSRREYSRAELARKLTPHADESDDVESLLDALERENWLSDSRFAESLVHRR
ASRVGSARIISELKRHAVGDALVESVGARLRESEFERAQSVWRKKFGAVPQTQAERAKQA
RFLAMRGFSSATIAKLLKGDADEFGGD*
>GCA_000452945.1_BPMSHR5858_genomic_05722_1 group_2638
MSDVLSVLPPLSLLALRRMGIARGLRRVWRVSRRRLRARGFTLIELMIVLAIVGVVAAYA
IPAYQDYLARSRVGEGLALAASARLAVAENAASGNGFSGGYVSPPATRNVDSIRVDDDSG
QIVVAFTTRVAAAGANTLVLVPSAPDQADTPTARVALSKGAVQAGAITWECFADGKASSS
LPAPGAGPLPTDAPTLAGKLAPPECRA*
>GCA_000452945.1_BPMSHR5858_genomic_05725_1 group_1958
MKTYSTYLALPLAASLLAGCAAFAPRDAAKLECTMPVAAYPENAKPLERRATVLVRAMIT
ASGNAENVTVTTSSRNAAADRAAVDTMSRIACSQTPACGGEPYPFTLTRPFVFEPRAKAP
Q*
>GCA_000452945.1_BPMSHR5858_genomic_05726_1 moaC
MSGSGFTHFDAAGHAHMVDVGDKQETRRIAVARGAIRMLPDTLALIRDGRAKKGDVLGVA
RIAAIQGAKRTADLIPLCHPLALTRVAVEFEFDDALPGVRCSAQVETLGRTGVEMEALTA
VQVGLLTVYDMCKAVDRGMTITDVSVREKRGGKSGDWKADEAGA*
>GCA_000452945.1_BPMSHR5858_genomic_05751_1 group_5232
MELNDALYIPLAPSVVWDALQDLALVRASLDHCESFSRLARGEYALALTVPLGPLRARYD
VRAHVVGERHDEPVHTRRTLNFRARADGIGALRGQIDVVLAPADDERGASRRAPTTRIEY
AVWATASGPLAELPGRQIQNALHELADDFFNEFCAVVQAKHGLAPNRARDGAPQRQHVFL
RPAAFTGIARRAHAQHLGGALTGRAASALHHRESNPVPLWAWAVMIFFVALLLYAARWLN
GG*
>GCA_000452945.1_BPMSHR5858_genomic_05752_1 ttgR_2
MARRTKEEALATRDRILDAAEHVFFEKGVSHTSLADIAQHAGVTRGAIYWHFASKSELFD
AMFDRVLLPIDELKAGTGEPHADPLGRIREILIWCLLGAARDPQLRRVFSILFMKCEYVA
DMGPLLQRNREGMRDALRNIEADLAQGVANGQLPADLDTWRATLMLHTLVSGFVRDMLML
PGEIDAERHAEKLVDGCFDMLRTSPAMRKDD*
>GCA_000452945.1_BPMSHR5858_genomic_05777_1 surE
MRILLSNDDGYLAPGLAALYEALRPLAEILVMAPEQNCSGASNSLTLSRPLSVSRSAATG
FYYVNGTPTDSVHVALTGMLDTKPDLVVSGINNGQNMGDDTLYSGTVAAATEGIMFGVPA
IAFSLVHKEWAHLGDAARVAAEIVRHYLDHPLPGQPLLNVNIPNLPYEELKGWRVTRLGK
RHPSQPVIRQTNPRGEPIYWIGAAGDALDASEGTDFHATASGYVSITPLQLDLTHTQMLA
ATRDWARAGSGAS*
>GCA_000452945.1_BPMSHR5858_genomic_05779_1 nlpD_1
MSMLRAMQNNRSKVPLSLAQRAICMAAFALLGACATRLDQAPVVDRSGSLGTTAATQPPA
PLPPAPPGYYRVKPGDTLYRIALENGQNYRDIAAWNNLTNPNQIEVDQLLRVSPTAGGAT
AAAPAAGAVVTPPLSSGPSNGAAAVPPIYGSGSTASAAPSAPAASASSEPSAPAANGNVS
FAWPVRGALLNTFDDSKNKGINIGGPAGEAVKAAADGRVVYAGNGLRGYGNLIIIKHDAT
YLTAYAHNRALMVKEGDAVTKGQKIAEMGNSDSDRVMLHFEVRRQGKPVDPLKYLPPQ*
>GCA_000452945.1_BPMSHR5858_genomic_05786_1 rlmN
MAGETIVNLLDLDAEGLVAYCGSLGEKAFRAKQLQRWIHQYNAADFDGMTDLAKSLREKL
KGRAVIGTPDILSDHVSADGTRKWLINVGNGNAVETVFIPEETRGTLCVSSQAGCAVNCR
FCSTGKQGFSRNLSTGEIVGQLRMAEFALRASLGRAPGPNGKAERVITNVVMMGMGEPLL
NYSAVVPAMRLMLDDNAYGLSRRRVTLSTSGVVPMMDRLGAELPVALAVSLHAPNDALRD
ELVPLNKKHPLRELMAACQRYLKVAPRDFITFEYCMLDGVNDTEAHARELLAVTRDVPCK
FNLIPFNPFPESGLVRSKTEQIKRFAQVLIDAGVVTTIRKTRGDDIDAACGQLAGAVKDR
TRLAERTGASKIIEVRAV*
>GCA_000452945.1_BPMSHR5858_genomic_05787_1 rodZ
MSEPQPSKSPETNAGEKPATAGGGMASVAAVGARLAELRRTKGWSVDDVSARLKVAAPKL
QALEAGDTSQLPGTTFALGLVRSYAKMLGVDPEPFAQALRREKGVPEVDLSMPASAGTDL
PRGRVSIPLGGASRGRPWLWAVAAVMVALVAAAMWRTGGDSSNWLARFKSSGAGEAASAA
HAQREGAASDLPASAAVSGTVMQAGALAASATQSVGGASAGDTAAASVPAAGAAPVVASA
AAALSASAPVAAAQPAQGAQQPAASAADGASAAVTAAPGQSVVALKVAQDCWFSVRDKNG
KELFSALVRAGETRQVAGDAPLKVTIGNKAGLESVAFDGKPVDPAKYSTARGNVARFALP
*
>GCA_000452945.1_BPMSHR5858_genomic_05794_1 hflX
LTSDNLINAALVGIDFGKTDFEASLEELSLLASSAGAHPAVTLTGRRSSPDAALFVGSGK
ADELRLACEANDVEIVIFNHALAPAQQRNLERVLNRRVIDRTSLILDIFAQRARSHEGKL
QVELAQLQYLATRLIRAWTHLERQKGGIGLRGPGETQLETDRRLIGERIKMLKSRLDKLR
RQHHTQRRQRARSGTLSVSLVGYTNAGKSTLFNALTKAQAYAADQLFATLDTTSRRVYLG
DEVGQIVVSDTVGFIRELPHQLVAAFRATLEETIHADLLLHVVDASSAVRLEQIEQVNDV
LREIGADSIRQILVFNKIDAVPELAARGGAVERDEYGNISRVFLSARTGQGLDSLRAAIA
EIATAEHLVGAAPLDGAPALPHEDHPVSEHGR*
>GCA_000452945.1_BPMSHR5858_genomic_05795_1 hflC

MRALLSINDPRWGRGGNGDKPRANESKRPNGRDDGPPDLDEMWRNFNRRLSGWFGGKGGG
NNGFRPDNGRAARVGVGIVAGVLIAIYLGSGIFIVQDGQTGVVLRLGQYKGSVGDGVHWR
LPYPFESHEIVDTAQVRSIEIGRNNVVRLANVKDASMLTRDADIVDVRFAVQYRVGSPTD
YLFHAVDPERSVSQAAQAAVREIVGARRADEVLAQDRDALRDALSKAIQRDLDRYRTGLV
VTGVTVQSVSPPEQVQAAVDDIAKARQDSEAAKNAAQAYASELLPRAQGDAAKMVDDAKS
YAERVVAQAEGDAERFKQVYAQYSKAPAVIRERMYLETMQEIYSNTTKVYVGNKAGNSVL
YLPLDKIVEAGRQRAAEAASAGVAPPASGAQAPAPAVSAPASAAAAASAPRAAAPAAASP
ASGSDSRSREALRSRSRTDDLQ*
>GCA_000452945.1_BPMSHR5858_genomic_05798_1 hisZ
MSTWLLPENIADVLPSEARKIEELRRRLLDRFRSYGYEMVMPPLLEYLESLLTSGGNELR
LRTFKLVDQVSGRTLGLRADMTPQVARIDAHLLNRQGVTRLCYAGPVLHTRPRGLHASRE
QLQIGAEIYGHAGLEADQEIQQLMLDALHLTGLKKIRLDLCHAGVLAALFARDAAAAERG
EALYEALAGKDVPRLNELTDDLGADTRAALRALPRLYGDASVLDDARRLLPALPEIARAL
DDLAHLAAQVKDAEVAIDLADLRGYAYHSGAMFAAYVDGVPNAVAHGGRYDHVGQAYGRA
RPATGFSLDLREIARISPVEARGAAILAPWKQDDALRAAVGALRDAGEVVIQALPGHDHV
LDEFACDRALVERDGAWVIEPR*
>GCA_000452945.1_BPMSHR5858_genomic_05801_1 kup
MTDTNHSSMRQHSLQSLAIAAIGVVFGDIGTSPLYSLKEAFSPAHGIPLTPSAILGVISL
LFWAIILVVGIKYVLFVMRADNNGEGGVLALMALSLRPLNPKSRITGLMMALGIFGACMF
YGDAVITPAISVMSAVEGLEVATPQLSHLVLPITIVILIALFWIQRHGTATVGKLFGPIM
VLWFVTIAALGIYHIARAPMIVSAINPYYAFSFMSEHVLLAYVVLGSVVLVLTGAEALYA
DMGHFGAKPIRLAAYVLVMPSLVLNYFGQGALLLLDPKAIENPFFLLAPQWAALPLVVLS
TVATVIASQAVISGAYSLTSQAIQLGYVPRMKILHTSELAIGQIYVPVVNWLLLFVILCI
VIGFKSSDNLAAAYGIAVTATMVITTILAAVVMVKVWNWNKLLVAMIIGVFLVIDLGFFG
ANLLKVEQGGWLPLGIGALLFFLLMTWYKGRHIVKERTAADGIPLAPFLQGLLAHPPHRV
SGTAIYLTGNDTLVPVSLLHNLKHNKVLHERTIFMTFVTRDIPYVKDHERVTVHDAGEGL
YIVKAEYGFNETPDVKAVLEEVARQRGMTFELMDTSFFLARETVVPTHLPGMSIWRERVF
AWMHQNAAKPTDFFAIPANRVVELGTKIEI*
>GCA_000452945.1_BPMSHR5858_genomic_05807_1 clpV1_2
MSDIGRVTLFGKLNTFLYETLEQATGFCRLRGNPYVELAHWLNQMLRRPDSDVHRILRRF
DIDAAAIDRGIVSALDRLPRGAGSVSDLSAHIDDAVERAWVYATLKYDAAQIRGAVLLLA
LVKTPQLRNVLYAIARDFERIVPDVLADELERIVAGSPEAPSPAARAAAGAIGDAGAAPS
AAREGSALARYAIDLTARARAGEIDPVVGRDGEIRQIVDILLRRRQNNPLLVGEAGVGKT
AVAEGFALRIVAGDVPPPLRDVELYLLDIGLLQAGASVKGEFESRLRGVIDEASSSERPV
ILFIDEVHTLVGAGGAAGTGDAANLLKPALARGLLRTIGATTWSEYKQYIEKDPALTRRF
QLVQVREPEEGAALAMLRGLAAKLEAHHRVLVLDDALQAAVTLSHRYIPARQLPDKAISL
LDTACARVAVSQHAVPAPIEDVRRRIDSLRVERELIARECALGAGDAQRLDAIDASIAGE
QTTLDALDARWQAERDALGKIVDWRASLLADDSSRVLDEAARADVQAKLSAALRALAELQ
GETPLVLPAVDTHAVAAVVSDWTGIPLGRMVRDEMQSVLKLAETLAERVVGQPHAVGLIA
ERIQTARARLDDPAKPHGVFLLCGPSGVGKTETALALAETLYGGEHNAITINMSEFQEAH
TVSTLKGAPPGYVGYGQGGVLTEAVRRRPYSVVLLDEIEKAHRDVHEIFFQVFDKGWMED
GEGRYIDFRNTVILLTSNVGSERVMQLCRDPQRLPDAQTLTDALRAPLREVFPAALLGRL
TVVPYYPLTDEMLARIVALQLARIERRIEAHHGIALRCADSATALIAERCRTIESGGRMV
DAILTHTVLPRISQEILRATIEGRALRAIDVSAEDGQFVYRFEEEGAT*
>GCA_000452945.1_BPMSHR5858_genomic_05812_1 group_5241
MNERAQTQADARAAAQPVVARDEFAALLQKEFKPKTAEARESVERAVRTLAQQALEHTVG
MTTDAYGSVKQIIAEIDRKLSEQINLILHHQEFQTLEGAWRGLHYLVTNTETDELLKIKA
LPASRNELARTLKRYKGVAWDQSPLFRKVYEEEYGQFGGEPFGCLVGDFHFNHSPPDVEM
LGELSKIAAAAHAPFIAGASPELMQMDSWQELANPRDLTKIFQNTEYAAWRSLRQSEDSR
YVGLAMPRFLARLPYGARTNPVDEFDFEEDTGAASHDRYTWANSAYAMAANINRSFKLYG
WCSSIRGVESGGAVQGLPCHTFPTDDGGVDQKCPTEIAISDRREAELAKNGFMPFVHRKN
SDFAAFIGAQSLYQPAEYHDPDATANARLSGRLPYLFACCRFAHYLKCIVRDKIGSFRER
DDMERWLNDWIMNYVDGDPANSSQETKARKPLAAAQVVVEEIDDNPGYYASKFFLRPHYQ
LEGLTVSLRLISKLPSAKAAGE*
>GCA_000452945.1_BPMSHR5858_genomic_05821_1 group_5242
MDFNSTYCFKLKQKVTGKPFPIIGTRIQRHGASPVHSTYNVNSRMKSRSSLYIFLGFFLA
GMGGCGTGATSPASTPTVAQATLDSARAAYDAGDYRRTIALLGGHAREIDGADVNTQVAA
HKLLAFSYCLTRRTTQCRAEFSRILDLNPRFDLSPAEKGHPIWGPAFEYARRKHALS*
>GCA_000452945.1_BPMSHR5858_genomic_05822_1 group_5243
MQLTVIEHAGEPVGTDGRNAVVFHAPGGTIGRDSDNHLVLRDDTRQISRLQALLQVADDA
CLLKNLSSVSTIEVNRVPIGYAQAQRLNMGDIIRIGPYLLRAEPDDATIERTVEAAATAA
AAAAPAASAAQAQAKGAGNKLWGLLHERFGLGKAQGAGEQSGARAAPARHHDSPASAAPR
DLNQLSTDPLDLFAQPHGDPDARAGAAREGEGRAPPTVTQPDHAPEWTQHVRVQPAQSAP
PAASRPGAPAARSGDIPAAGDASDMPSRVRASPAPAPATPETLLQAFFEGAGLDTAAEQH
HWSAEQLFVAGQLLALFANGTVELLSSRSILKREVKADMTMLLDRENNPLKLLPDGSAVL
RQMFGLPLPGFMTPQSAVSDAFQDLHAHQIGMVAGMRAALMDLLTRFSPQRLRERDAAPQ
WYEKRVPALYKARLWDRYATTHRDTLFAIEDDFASVFGKAFLSAYDAEVESYRGRCRR*
>GCA_000452945.1_BPMSHR5858_genomic_05837_1 abaF_1
MPTPTPGTPRPPRTPQPVRAALAAFIGTTVEFYDYYTYATAAALVLGDVFFPSDNPFVST
MASFATFFVGFVARPLSGAVFGHLGDRIGRKKMLVLTMFLMGAATTGIGLLPGYATIGVW
APVLLVLLRILQGIAVGGEWGGAVLMASEHASAGRRTLFASFPQMGSPAGLILSLLAFRY
VTSLDHARFVAWGWRLPFLASFVLLLIGVAVRLGVNESPEFERVRDSHAVARFPVAEVLR
TAWYPIVLAAAATTIGSAGFFFTNTFMISYVTTYLGMSKSFILDCLFAVTLIQLLSQPVS
ALLAQRFGETRFLVCAALLSMLTPYPMFVLVRTHDPVAIVAGIAFAVVALSAVYAVIAGF
MSAAFPTRVRYSGISIAYQLCALVAGGTTPLVGTMIAQRYRGEWLPLALFFSLLAFVSLV
GIVGLGRYRRNLHAAAPRAWAVNP*
>GCA_000452945.1_BPMSHR5858_genomic_05849_1 group_6091
LSFVIVLAALAFLMFAAYRGYSVILFAPVAALGAVLLTDPAAVAPVFSGIFMEKMVGFVK
LYFPVFLLGAVFGKVIELSGFSEAIVAAAIRYIGRSRANAVIVAVCALLTYGGVSLFVVV
FAVYPFAAELYRQSNIPKRLMPGAIALGAFSFTMDSLPGTPQIQNIIPTTFFKTTAWAAP
ALGTIGSVFIVVVGLSYLEWRRRSALARGEGYGTSLVNEPERVETKSLPHPILAVSPLVL
VGVANFALTKLIPVWYGGASYTVPPDVLPGVHTPITTPIKTVVAIWSVEAALLLGILLVV
ITAFKRVHERFATGTKAAVGGALLAAMNTASEYGFGGVIAALPGFIVVGDALKNIPNPLV
NAAVSVSSLAGITGSASGGMSIALAAMSDLFIKGAQAAQIPMDVLHRVVAMASGGMDTLP
HNGAVITLLAVTGLTHRESYRDIFAVTVIKTLAVFFVIAVYYLTGLV*
>GCA_000452945.1_BPMSHR5858_genomic_05855_1 oppC
MTPVTSSIELPTAADAPPRSRTPLGLAARRFVRNRAALASLAALAVIALMCFVGPLLLPN
DPAASDWSSISLAPTLANAHWFGTDELGRDLLVRTLIGGRVSLEVGLLGTFVSGLFGVAW
GAIAGFAGGRVDAAMMRVVDMMYAIPYLLIAILMMTLFGRSFILVVLTISAFSWLDMARV
VRGQTLSLRTREFVDAARAIGVTPAAIVRRHIVPNLLGVVVVYATVSVPAVVLTESVLSF
LGLGVQEPMTSWGVLIQDGAQKLEAMPWLLLAPAVMLCVTLYCVNFVGDGLRDALDPKDR
*
>GCA_000452945.1_BPMSHR5858_genomic_05857_1 oppA
MKHTHAFAAVLAALALTIAPSAPAVTVASNVTLADQQDLTRQVPAEVESLDPAHIESWTG
NTIGLDLFEGLARIDASGAVVPGVAQAWEHKAPDTWVFKLRRDAKWSNGQPVTAADFVYA
WQRLADPKTGSKYTILVEFVKNASAIIAGKQPPGDLGIRAIDPYTIEVKTEVPVSYFPEL
TAMAPLTPVNKDAVAKFGDAWTRPKNIVSNGPYTLVDWQPNNRIVMAKSDKYWNARNVVI
RKVTYLPIENDETALRMYQAGQIDYTYSIPAGGFGQISKQFGKELRPGLQLATYYYYLKN
SDPALKDKRVREALAMVLDREILTSKITQAGEVPMYGLMPKGVKGVQRPFMPDWASWPMA
RRVDYAKNLLKQAGHGDANPLTFTLTYNTNDLHKKVALFAASEWRTKLGVTAKLENVEFK
VLMKQRHDGKVQVARDGWFADYNDAMTFFDLIRCGSSQNTVGYCNPKVDSLVAEANQKLD
DGARAALLTQAHDLAMNDYPMVPLFQYSADRLVKSYVGGYTLTNYIDMRASQDMYLIKH*
>GCA_000452945.1_BPMSHR5858_genomic_05858_1 puuR_1
MVSSTENTRLANAGVALGSKIRALRQRLKRTLDETATAAGISKPFLSQVERGLASPSITS
LAGIAHALGVTVQYFVETPSEERSVCRGDQLRFFSFADSANLFARLTNVPEGRQLEAILV
RMPPGQKRSEVTTHAGEEFLYVIEGEVSLTLEGKSFVLHAGDSAHYQSTVPHSWVNTAKI
ESVVVWVGTPRLF*
>GCA_000452945.1_BPMSHR5858_genomic_05859_1 dtpA
MNTRVSQTRSFTTVFLIEMWERFGYYGMAALLVLFMVDKLGFTDGQANLTWGAFTALVYA
SPSIGGWIGDKVLGARRSMIIGAVVLCAGYLMLAIPNDQLAFMYTSLGVIVVGNGLFKSN
AANLVRRIYEGDNARIDSAFTIYYMAVNIGSTASMLATPWIKDHWGWHTAFAVCCAGMAL
GILNFVLMHRTLAHIGSQPDAQPVHWRRLAAVAAGGVALGLATMYVLGHKQLAVASVWVA
AFAILAIFAYMIAKSNRAERSGLIAALLLIAQVILFFIFYVQMQTSLTLFALRNVDPRFM
LFGTTLFTWSPAQFQALNPIWIMLLSPVLVAIYNGLGKRGRDLPVAGKYALGFGVVALGY
LAFSISGRFAVDGRVSSWFMVAGYGFYSLGELLVSGLGLAMIARYVPARMGGFMMGAYFV
ATGVSQYLGSVVANFAQMPSNDLPATASLPLYIELFTGLGWLAAAGMAIGLLMLPLMNKL
SREHHRCEAQRRETQALAAH*
>GCA_000452945.1_BPMSHR5858_genomic_05861_1 group_5250
LRTTKSPGISVIRQRLAVALPLALTLTLASCGGDDLTPAAQRWAMPGTELPLGPQGLAQS
VSTQTLAAGVAYYQIKRGAASAADFWTVNLGFYATQAAAQADAANLAAAGFATRVDASAG
TDLQGKVLGYWLSAGRYATQAEATAAAARIAQATQNRYKPGTRHTSLAGAPTTGPWIVNV
LAIDPSRAGAALSLALPGGDDLGAGGETVSAARARVNALAGVNGGFFTNINPFGAPLPPR
SPVGATVVDGRLVAAAIGRRPGLLLARDANGRQRATVVRNLATSITLTDAQGRAIAVQTL
NRPILGTVVNCGAQARTPTSEPAQDTVCTNYDDLVMYDSLYLRGGASNTLVDASYQGARY
ELVVDANGAVVAGHATLGAPPPPNGYVLQGLGASAAWLQAHATPGTRLAVSRRLSADGAD
LALASGMSLVEAGPTLSVPNLAQSAAQEGFAPTVGGVDAGEGAAANGNWYNGWYVARNGR
TAAGVAADGTILLVEIDGRQPALSVGTSIPETAAVMAWLGATSAVNLDGGGSSNMVVGGK
MVGHPSDAVGERGVGDTLMLLPG*
>GCA_000452945.1_BPMSHR5858_genomic_05862_1 pcpR_1
MNLRSIDLNLLVILDALLAERQVTRAGQRIGLTQPAVSNALGRLRYVFKDEILMRTPLGM
ELTPRAKALACPIRQILQQIEELFVPENQFDSFTSDRRFTLRMSDLTELLLLPPLLRNIR
GTAPHIRMNVMHLNADKTVEALDSGRLDMAVCAGLDHPDAISSQVLFQDRLVCVLSRRYP
DANRPLTLERFAALDFLNVSINPVDSSLIDTMLADMRFTRRIAFNVPHWLVVPSMLEALP
LAVIMSERHALSLGDSRLAIRELPLDLEPVTWTLYWHRRYDNSVAHEWLRNCITDVVDAI
QPRGVFEEASAVG*
>GCA_000452945.1_BPMSHR5858_genomic_05867_1 group_19411
MHSSQAYTADAPTRAEVDALAGATVIEFGANWCGICAGAQPAIRDAFSVHADVRHLKIED
GPGRPLGRSFGIKLWPTLVFLRDGTEVARVVRPTHAEQIETIGFAALV*
>GCA_000452945.1_BPMSHR5858_genomic_05871_1 group_531
MPIRQTHIWIFGFVIACVLAMSGPLTYTFENTFQQTVDIDRKKQGTVAEATAFDARSAEQ
PSAPDADRMHELPRIDAARSPTDHFDELPIHLAAFGGGEPSPSGPLPPGPLPDTSFAPFP
FQHGFDTAIHAGPPIPAPPESTLARTDAPPRDETPFEPGMPQPSETVDTQLSFYADMVAQ
AIERTAYDSIALAKLDSDTTAPPDMTPERHDALANMNAPMFYADMLTASIHVPPSNDRTL
SVPSLLTHQAIGIAWMELFARELPATATQAAPTRHAVSFAATLQAEPFDLAQWSEPDEAH
SDGPPLELDGQRTIAPPYPRAGTLPPGTRPTYTDSPPDTGIVQAAAMLATGFESAGKHAA
ALRGQPPGADECRMGARAIGLGRSGMERLAIEQLVSSARAAYDCARARIHGRATAFANAP
ASAQPSNHLDANGDPMRIAFLPRLASEPFQEAFSIPGDTAGIASAFAVADDQTGHRNRMA
AANSD*
>GCA_000452945.1_BPMSHR5858_genomic_05874_1 group_5252
MRNKVAGAIALSMLSLAYPRAEAAPIAFPCPIDARRDLTDTLACYRQTVENQPLNYTMTG
IVQLPGIEQRTYRLVSQSWSPDQLVRPDTWVHEVALYIPQDALPRRALVIANDGTRHPGE
GETPRTPNDFLPDTLADIARSTRTAVISVSDVPNQLLSYADNGKPKAEDDSVARSWTLFM
QSPHTRGAMPLHVPMAASIWRAMSLAERELTSLGIHRFVVSGISKRAWTTWLALIGDQRV
DAIAPFAIDLLSTRAALENMYRSYGGNWPLAFYPYYAEGIDRTLDSPAFGLLMRIEDPLS
YLGTRHGSRLAVPKYIVNASGDDFFVPDNSQRYFDKLPGAKALRMVPNSSHSDIRRATLD
SLVPFVKRIQHGKALPQVDAAPIETNGQTILRLRTSERPRQLLLWQATNPLARDFRYACG
IRYTSTPIAMNGARPQQLVLVPPTNGWRAYFVEATFDDGFVATSQTYILGDGYPSTPPPA
VGGSCRTLPGRDTHS*
>GCA_000452945.1_BPMSHR5858_genomic_05876_1 hifA
VKKKVVSFAVATLFGLVAAQSVYAQSIDQGQVKFTGELTANTCTIDAGSKDQIVPLPKVS
TSALAAPGDVAGSTAFEIKVSKCAADVKKVAAHFEMENMDPNTRTLKNEATGADAATNVT
IQLVNSDGTELPVGSTGSYFDVTGTDDARGATMIYGGQYYALDTTTAGKVESHTLFTLAY
E*
>GCA_000452945.1_BPMSHR5858_genomic_05877_1 yadV
VRLWKMCVAVAAACCLASTLSQAAIVITGTRVVYPESSREVNVRLNNVEQSPVLVQAWID
DGNAGASPDEIKVPFVLMPPVFRVEPKKGQTLRVMYTGDDLPKDRESVYWLNVLEIPPKP
TVEAEQNLLQLAFRTRIKLFFRPSALEDGNAAQARDQLKWKIVRNEKGVSVLRAENRSPY
YISISQATAKSGDKSVQFDPGMVPPFGSKEFVSKHDISTLSTHAEIAYKLLNDYGAEVAG
SATAE*
>GCA_000452945.1_BPMSHR5858_genomic_05880_1 group_910
MSIEMPAQPLRMLLQDTMLHVALLTQGARISRVHDWRARCIALVRQFEQALQDGGYRGGI
ANELGLAQCVFLDEITMRNLSTEQKEEWLRELLQFRFHSIRDGLSKVRARIDRLCRGHSA
NAALLEMYSLFHEFDLLGDKQGHALADARVSEARPTPGKKAVETAEVWKDGGAANEAAHG
RSWRIVSGLLAFAALMTMWLVIDTRLSHEVEQMEVTSGDEAAIHPASGD*
>GCA_000452945.1_BPMSHR5858_genomic_05884_1 hcp1_2
MSHDIFLKINGIDGEAEDATHKGEIEVLSWSWNVSQQSNMHLGSGGGAGKATIDDLQFEH
YIDRASPNLVQYCLLGKHIDEARLVVRKAGGSPLEYIKLTMSDVLVTQVSPAGVAQDESR
PRELVRLSFSRLKQEYVVQNPQGGSGGAITATFDIKKNAA*
>GCA_000452945.1_BPMSHR5858_genomic_05885_1 group_19974
MAFPFFFRREAARRTAVKRVVAAMVCASLAACASLPARKETADLIVKIKVSEGANPDEHQ
RPAPVMVRLYELKSAGAFENADFFTLQSDSRKVLGDDAIATDEFVMRPGDTRDIHRRADS
AGTSVGVLVGYRELGKSVWRAVHKLRPAPEDAWYRAFTPRTKIKLNVDVGQQTVTITELE
*
>GCA_000452945.1_BPMSHR5858_genomic_05892_1 rhsD
MSGISGLGGMLAGLETRHVTSLDVERRIDRYGNQPGNWAARHGVERALSGLLQRSVDAGS
GNKVCLAADEVDIKLPGRLPIMWARYHSSALKNAGLLGPGWRTAWEATLRREEDRLAYTD
DQGRVLSLPMPHRGSQVIVSSEQLHVAHLPDGRMVVADLTPHYSVFGEFDEAGVARLKYI
EDLHKQRIGCIWDDAGRLKRMRGTCGHELKMHYGSDGHRLTGIECVDGGPTGWLVHYGYD
DDGRLAEVRNRIGDVVRRFVYENGRIVREVGPLGQTTQYKWEVRRGAARVIERSTCRGAR
DRFNYDPDEGRCEVIDTFGHAAQWKFDRQGNISSHTDFAGSRYTFVYDSAGWPTQLNLPG
DRQVRITFDNLGRVTRETDPLGGERSTIYAFATREPVTVRESDGRLWVWRHDDRLRAIHH
QSPSHGSTQIAYSEHERGETHSYVTDRGTSITIERDRRGQVIGLGTAPDKATTYRRDAEG
RVVEIVDPVGAVTKIENDLLGRPLIVTLPGGRQERRVWNAAGQLVKITGPDGYSRLWHRD
SYGRVVQFVDEEGSVTAHEYDAHGRRIRTASGNGATQEFEWGPADRLVSIVDADRVTRTL
SYSMAGWLQQIATTAESMTRREIFVHDAVGRLIGRETEHAHYHYAYTQQGLLGTIDCSPT
EAGEAIGVAADTIRFEYNPFGLVTAEHGGAGVLHYEYDERGRTRAVTLPDGCVVKRERDE
DGRVALIGMTCREESRGIAAFRYDNLGRQVLRSQGELYLRSVYTSASELERCFAMSVVRT
PNNEMQAGEVRYWREFQYSIAGNITQVGDRFEGKTYCDYDRRGRLLRMLSDEVGIEYFTW
DAAGNLFDTAHVAASVVACPDHRMREYKGYQYEYDAWGNVICKRSAAAEQSFTWDAEGRL
MEVRSRQARVRFRYDALGRRTSKSIERHDADHARQANSSTEVVHFVWEGERLMQERSEKE
IRTFLYQPAADGFMSYAPLARVDQVRDADGNVQPMRLYHYHVDAAGTAVAMTDDAGALAW
AGRYSAWGRILPPTSLKVQIDQPLRFAGHYADDEVRLHLNGTRYYDPDTGRYLSPDRSAE
PGTSPYRYVSNPQTACNPTGRAVTLARPRAGKRLGSSVKRTFDLGEPAAGLVAKLDDVVD
GALIQLGQEV*
>GCA_000452945.1_BPMSHR5858_genomic_05894_1 group_2652
MKKKFSRFLAALVLLAATVDALAAGCNMLTAENIAWLTEQGKIARAHSSFPISFSSGMVD
VDPNLEIGGLIAEAKSIPSEELHFIWCSAPSGNVHFALNSSPLPSELGNSIYETGVPGVG
FRITQVRQSGSIGAIPRDTPWIEEKPGQDSSLNFGAGTVFRIELIKTSEALPSESTISLG
NLSRVYGDDNKTVVDFNAGSVKLRVLPICHVDQQEKNVDFGQFGPKDVSFDSGPTKDVKF
DVQCSGPTPPVSITATLAATPDSHDQSLIANAGDAMNLAIRLRDASTQQVLRPNDPSSEI
KVEPGGAMEHGFALEATVLRVGTAPPTAGTIDATSIITLTIL*
>GCA_000452945.1_BPMSHR5858_genomic_05899_1 group_5257
MTNRSESPQPSDPRDAMLRDALRAQSFEEIKARTAEAVRKRPSDTRERWLLFQLLCIDGA
WERALKQLQTWAGLEPQGEARAQLHRGLIRCEMFRADVFAGKRTPGFIDAQPAWVDTLLQ
ANAKLGAGDVAAADTLREAAFHCAPVTRGESAEMGSFAWLTDSDTRLGPIFEIAVAGGYR
WIPFEQLKSITFTPAGTLTDLVWRPVTALMLDATVLRGYAPTRYSGSENGATAIRLASET
TWRDIGTTNVVALGQRTWTTDQGDWGMLQIGGCRFTGESDDAAS*
>GCA_000452945.1_BPMSHR5858_genomic_05902_1 rcsC_4
MPGKLKELLIQPFKLNHRKALFDPRRMLAHQRLLTYGGAIGISAFILFCLLLMIRMEAMD
YVSNARSEFLIQQSKLTIKLQIRRTALMRAVMHAEGLWNRRIEPSTELLSQFDAHQGYAV
LHTPQGAPAALAVGIPAPGRSASAFAPYLALLETQLLMGDGQSPLPDDRVEGYSFSPDGT
YLSLLWSREDQAPEAFIESHSASGLIGDAMRGIRYLRDASTIVPLRADRRTVWLPPHPDP
LTGKPMIRIVQPAIDATGKLFAIYVRSLPLDMFYSNLNDAPAGSTLTIVDRKGNVILGSS
NQHGGDGGATASALGIRSLGDGFGGLVERYQDGMFVTSDQLPGTDWVLVHAYSWKTIAAA
LALRFSLVTGLVAGIIVMLWLFAIQFDRKGLIPNYRRAQRLLESEVLNRTILRTAPVGLG
LLSAQDGSVLLQNNAMLELSARTSDEPLPIRLWEAYRHAVGRSSDEDGALLQHDLAIERE
GANPVYLQANVVAARYRGANVLLCTLLDISLRKESELKLKEARIAADQANKAKSTFLATM
SHEIRTPLNAIIGNLELLRRSPLSEQQSERLHVVESCSNALLHIINDILDLSKVESGQMT

LENIPFDIHTLLQETVDAYAPLAESKGLALLSSIGEGVARHYTGDPTRIRQVLSNLIGNA
IKFTERGHISIDVHRAKAARDGSAIEIWVNDTGIGIRPDRIATLFDVYTQADASIHRRFG
GSGLGLPLCQRLVTIMGGSIEIDSLPDFGTTVTVALPLREARHATALSNGPAARRRNATA
LARPTLHVLVAEDHPANRALLRDQLEALDHRVDVVENGREALRAFGEHQYDIVLTDLGMP
ELDGYGFARFANKMRAGTPIVAMTAHATDEDYQRSKKVGIAEIVLKPLSLTTLEKVLQRH
AMRAPAEEIVTAAGPGPAHAEPRMRALPPAVAAALRTATLQSIDKIAQALAARNFETIRQ
ELHSIKGGFALTGNRLLIARCAATEELAEAEDLDTLRAAWRKLEASLLEALERL*
>GCA_000452945.1_BPMSHR5858_genomic_05912_1 group_5258
MNSYQEYKQHVKTLQATLDRERKAISAKVLQEIRECIAEFGFTPQEVFSSKKKREPKYYD
PVSGSTWTGLGREPEWIRGKDRKAFELSSHQRDDHCR*
>GCA_000452945.1_BPMSHR5858_genomic_05914_1 garK
MKIVIAPDSFKESLTAVETARQIEAGFREVFADADYRCLPIADGGEGTVDALVAAAGGER
RGASVADPLGRPTRAAFGILPDGTAVIEMAEASGLHLVAPDARDPRVTSSRGTGELIRAA
LDAGARRFIVGLGGSATNDGGAGMLQALGVHLSDADGRPLGPGGAALARLARIDAAGLDG
RLAECAFEVACDVDNPLVGPNGASAVFGPQKGATPEMVADLDRHLAHFAAIVARDVGPDV
AQLPGGGAAGGLGAAMSAFLGATLRPGVEIVTDALRLRDAIRGASLVVTGEGRIDGQTLR
GKAPIGVARIAAECGVPVVAIGGAVTHQAGALYAHGFDAVFASVGRACTIDEALRDAAAN
VRFAARNVAAAIRVGLKLGRHAPSRHLA*
>GCA_000452945.1_BPMSHR5858_genomic_05919_1 group_5259
MSNNMQTVRESYEAFHRRDLPGVLAALAPDVRWTHPDGMSPYGLGGTKHGHDEVIAFIRH
VPTHIAEMRLAPDEFIESGERIVVLGTRRVTAVNGRSATLKFVHVWRFENGRAVTFEDHF
DTAEMIRLITA*
>GCA_000452945.1_BPMSHR5858_genomic_05924_1 group_911
MKTNRLMAPLLVTCLMLATVAHAQTPPPLQDVNPNRHMNIAVAQASIRQAYDAISAAQAA
NHDRLGGHGDRAKRLLDEASVELKAAALTANGE*
>GCA_000452945.1_BPMSHR5858_genomic_05937_1 proP_7
METSALNFAGAPADARAAAKKRRLIAAAAVGNALEFYDFTVYSFFAILIGKLFFPVHSPF
GQLMLAVASFGVGFVTRPLGGLVIGVYADRAGRKRAMIATLLIMALGTATIAVAPTYAQI
GIAAPLLLVAARLLQGFASGGEVGASTTLLIEQAPQARRGFYASFQFSSQGLAALAGALT
GVLLTSTLSAAQLESWGWRVPFVIGTLFAPLGYWLRRTVDEAPAPASAPRGDAAPLPLAD
VLRHHGKAVFAGLGITIGGTSIHYIIVFYMAIYGVQVLHLPSWLSMSAGCIAGAILAVVT
PIGGHLSDRYGRNRIVWWTRVALMLAIYPAFVALNRWPGAASLLSIIAALASVHAINIGA
TGAMLGELFPRAVRATGGALVYSVGVAIFGGFAQFFVTWLIAATGNANAPAWYAIGCGAL
TLAALGSLDDKAGKPLD*
>GCA_000452945.1_BPMSHR5858_genomic_05939_1 tsaR_1
MKYHQLKAFVAVADEGSIRAAARRLNVSPAALTKAVKELELALGVSLVVRTARGVQLTRF
GQQLQVRASLIVAEMQRARDDIEQAQGATTGAVAAAITPAAAVTILPDAFRAFRRRFPLA
RVSIVEGFPGVALPKLHDGSLDFAVSVVVPELLAAEFEHAQLYASRHVIVARNAHPLAAA
TSLADLVDADWLMNPSPESSTQLMFNCFAEHGLPAPTRVVECPSYAIAHSLIQGSDLIAS
MPEQLLGVEWSRDRIAVLPIREALPAASVQVVTRRDSPLTPAAAMLVDCMRDAARRHGLR
*
>GCA_000452945.1_BPMSHR5858_genomic_05941_1 group_912
MAEYFDVDHAREIAALGERFAARTEWPTWLLIFVVYGGWLGVLLGVRAGQLPLAAATPLL
IALCAWHMSLQHELLHGHPTRSARVNMLLGYPPLAVWYPYTLYRDTHLEHHRDEDLTAPG
IDPESNYVPLAQWMRLPRWRRALWRARKCFVGRLVVGPPASVAAMGAATLRAWHGGDWRD
APMWAAHAVSVVALLAWLHGSIGVPWWYYLLAITWPAQSLAMIRSLYEHRAAEHPKARIA
INEAGLAMRMLYLNNNYHLVHHDLPTLPWYQLPRAYRMRRDAYARKCGGFVIRGGYWELL
KRHAWRVTDSPVHPFAGRGASDRASGGGATPGARVAVIEDCVQIGG*
>GCA_000452945.1_BPMSHR5858_genomic_05947_1 group_791
MEPITQAFAQQFSREWIDAWNAHDLDRVLSHYADAFEMSSPMIVQIAGEPSGRLCGKDQV
GAYWLKALRMIPALHFEWIATLAGIDSVAIHYRGANGRLALEVFHFGPGRRVVKAFAHYA
E*
>GCA_000452945.1_BPMSHR5858_genomic_05949_1 group_5262
MNRFVRLGAAMAVACALAACGGDDGGPAVAASSFAAAGANGATAGVADTAQADGAPRTGS
LTQYVDPLIGTLASNSPNPVPAGQAGSVVPAAGLPSGMIQWAPDTNTTPAPADSKEPGSP
AGYYYDLNAIQGFSVTHMSGAGCAGNNGEFPVMPTTDATKLAPTFSHANETAKPGYYSVL
LDSQIKVELTATLRTGFGRFAYPAGKPALLVIDATRTNTKTSTSGAITRVSANAISGSTV
GGGFCGNSVPVPVYFYATFDRPFAPASSISRGVAKLAFDSGATVRMKIGISYVSVDNAKA
NLDAENRTWDFDGVRALADAAWNDRLGAIRVSSSDADALKKFYTAFYHALWAPSVFSDVN
GQYIGFDKQVHTVAKGQAAQYSSFSGWDVYRSLIQLKAVLFPRETSDMIQSLVNDADQCG
AIPHWVNDNVEDGVMPGDAGSLMVSSAYAFGAREFDARGALAHMIRMANIPGTACAGVTT
NGGRASYLQTGYITSGEWGIASSTLEYTSSDFAISRFALALGDTATQKMLLGRSAYWQNL
LNASLNPPLIAARQSNGAWIAETPGSTDNYVEGNAEQYTWMVPYDPAGLFAQLGGNQAVV
PRLDKFFTVLNAGMSLPNFYMGNEPTFEVPWLYNWAGSPSGTQRVVRQIMQTAFSTKPDG
LPGNDDLGAVSGWYVWAALGLYPQVPGVAGFAIGSPQFDAIDVRLGSGRMLKIRAPGAPA
SGYVQSVAVNGRAQASPWIALDALDGGAVMHFKMGGAPSQWGAGEAPPSFGVPVARDVAD
SFNNRGISADGATNADGQGADFDGSLFSYSANALAQAGVRPGAPFTYGGASFVLGGASSL
DNAVAVGQTVMLPPGSAGTSVVVLGASNNGPSAGVARVSFADGTSAQVTLSFDDWTLNGG
STGATSAIAVTSAYRNAGNGQKDNVKTYIFAQKIPVPAGKVVTSVTLPRQVSAGKMHVFG
IGVAA*
>GCA_000452945.1_BPMSHR5858_genomic_05950_1 group_5263
MKCDRNDEIALAILGEILGIHRRYPEYTTDSDIRHEIEQIHAVQLPRIRAFVDAARPVEF
VLPAFPSKSPNPNKVLGRLPDMAEQLSLSFLNDLCERISGFYAPGAKLTICSDGRVFGDL
IRVDDRDITAYQHALGQLIAALRADRLSTYNLENFEAFAQRAANFDDMRRRLVDEFADPI
EAIRHRLMADEEGTLLYRAMTRFMFEDGFTPDYRGSKAALQKDSKTRALGVIQRSWAWGA
LLATRFPDAIRLSIHPQPAASLKIGVHMMPTRDNWLTPWHGVAVDLGGQFALMKRRDVEL
LGGRVVMRGGRPSHYAIERWRVAGDALAPLAGGGARPRADAGGALVISGEAV*
>GCA_000452945.1_BPMSHR5858_genomic_05953_1 asnO
MCGITGWIDFARDLRNEAPIVDTMTDTLARRGPDARGVWLDTHAALGHRRLSIIDPERGA
QPMLTPERGPRGLPRAVISYAGETYNFRELRAELGALGHRFDTHCDTEVVLRAYLEWGAA
FVDRLNGMYSIAIWDTARDELLLVRDRLGVKPLFYYPTADGVIFGSEPKAILAHPDVRAR
TSSEGLCDALLFLRTPGQVPFSGMREVKPGHVLRVRRGGLAEQRYWALEARPHTDDLPTT
IATIRALLDDIVSRQMISDVPLCALLSGGVDSSTIAALAQKRRRASGGGALSTFCVDFTG
HTRNFRADPIRPTADAPFALEVARHIGSDHHTIELDQAGLLDPQVREAVLRAWDLPFNFA
DLDVSLHRLFAEVRKHATVALSGEAADEIFGGYLWFSDPAARRAHTFPWLKLGAHRGLDP
RALFHRSFIDALRLGEYEAQLYRAALAEVPRLDGENAEDRRTRELHYLTLTRWLPVLLDK
KDRMGMASGLEGRVPFCDHRLVEYVFNIPWAMKTFSGQEKALLRAAAADLLPESVLNRKK
AAYPSIQAPGYDRGLIERLNRAIAAGRAPLEPFIDAGALRGLTEKTAAGSLSEFERILLE
SSSRLNDWLDLYRVELDGVPAG*
>GCA_000452945.1_BPMSHR5858_genomic_05955_1 hpcB_2
MGKIVGAGLISHAPVVMMPQAVRLRENGGRDFTLATGLARLRRDVFDAHDYDTVLVFDSH
WLTTTEAVVTAHARRAGRFTSDEMPNAIRQLPYDLAGDPALAHAIAEQARRRASWIDAVD
DPCLPLHYATLNPWTYLGRPDKRWLSVSVCQTATTEDFLRMGETVAQAIAQLERKVLLVA
SGGLSHAFWPLAELRGRMAGAASNIVTPAARAADERRIAWLEQGRHDRVIDAMPEFLRFA
PEANFGHYLMMAGALGARACAARARRFSEYENGIGTGHVHLWFSPADGGWARAETHNERR
TETDAASA*
>GCA_000452945.1_BPMSHR5858_genomic_05960_1 group_5618
MHPTTPVAAVALALACAACASGAAKQKEPLRLELAVRAAPTVNPDDRSRPAPIVVSVYEL
KTDGAFNAADFFTLQGPDKTVLADDLLARERFQLRPGERRVIRRDADPAAGALGVVAAYR
DLPNSVWRAAYPLPPARDAAWYRFSSPKLKLTIELDTHAIRITEPGK*
>GCA_000452945.1_BPMSHR5858_genomic_05968_1 group_6092
LLDYYNQELIYMRELGAEFARAHPKIARRLGMQAGEVADPYVERLIESFCFMAARMQIKL
DAEFPRFTGRLLEVLYPNYVAPTPSMAVARLYPSRTEGNLAQGFRVARGTAFAARVPASE
KTACRFRSGQDVTLYPLEIADARLTGIPPDIPALDRYVPAGAQVRGALRLRLRTTGNMRI
ADLRGLDRLPLYLAGDAQVASHLFELLHVAGVATLTAAPGEFAAPGRPPAAVTVDALAHE
GLGADEGLLPLTWAKFHGHNLLHEYFACPERFYFVALTGLEEGLRRIAGSEVEIVVLLEQ
SPERLADVVDASRFALFCTPVINLFPRHMDRIELSSGQTECHLVPARLAPLDYEVFSVEA
MYGQVAATSAELEFRPLYQTLNDDESNYGRYFSTRRERRLVSDFARRYGTRTPYVGTEMF
VSLVDQNEAPYSEDIRYLSVDALVTNRDLPSLVPRDGVRDLTAIESAPIDSVGLIRAPSA
PKAPYAEREIAWRLIRQLNFNYLPLDELDHRAGGQGLRGLLRLFLAGDEADSRRQVESLV
GVKTRPVTRKLPGAGPLVFGRGVECALSVDETGFSGVSPYLFGLVLEHYLARHVSINVFT
QTELNSIQRGRIARWPVRMGARGGA*
>GCA_000452945.1_BPMSHR5858_genomic_05973_1 group_5269
MSTRADRLRDAIRHGRAIRDTAIDAGDFDGHDWSGGVFERVRFIGVSMKRVRLDEAVFID
CLFRDVDMRQAGCARCTFDRCRLERVDLSASELRDCMMNGTHAAGVYFSGARASGLHCVK
SDLGDCGFDDARIESAVFSDTRLARAAFTRAAVRKAVFYRLDLTSAVFADAAFDDTVFAE
ANLAGQRLQGQRMHRCQFVGADLRHADFTGARLAGCNFQRAKLTGARLDGVDAPNTVFFE
ADAPDATCRDAALRGSIWVQADARRIDFTGSELDGAVFQRATCTGARFSRAKLEGADFSY
ADLTGAVFDEAGFARTAFHGATAPAIAWRDHPGAVACDAELSDAQAWSRQRDEQARREEC
*
>GCA_000452945.1_BPMSHR5858_genomic_05983_1 group_5270
MANMTQAAADGELDLSTLALPQAARMPAFSLTMAWWAVCSAVFYIVVGATLALDYGARNA
LIGMVLSVVSYGLVNAAISRYAIRTGLSVALFSRVLFGSAGAALATLIFFATAIYYAVFE
GSVIAVAAHGLMPALDYKWAALIVVCYSVPLVFGSVQHWLDKFNGVLLPFYLLGLLAAVA
LATHRYGYHAAWLDFGPKGGAPAHGWWHCFVYYMGVWVLMMFTFDYARFGREADADYHGR
WNFGMPFYLVTFVVNGAAGIYLVSTIPGLGALSEVSVVTALLKLMGVWGLLFVWVTQSRI
NTANYYLATVNMQAFFQKAAGLRAPKYVWALVVGAVVYALMMADIFSKLLQALAWQGVFI
VAWVGVALAHILSTRYEELLGSGIECRDTHVPAFNPGGLSAWFAGVFAGMMLTTATGFAQ
SLSAPVTFVVSWAVYRGMLASAKRTWFVRER*
>GCA_000452945.1_BPMSHR5858_genomic_05984_1 pxpC_2
MPTDTLNAIEVVKPGLATSVQDAGRHGYYHVGIPPSGALDQFSLRAANLLVGNDEDAAVL
ECTLLGPQLLFRRDALIAVTGAEMAPRIDGAAQPCNVALRIRAGGTLTFDYVKQGARAYL
AVAGGIDVPVVLGSRSTYTLGAIGGHEGRRLQKGDALPIGAAKRAAREGATLPDAMRTPL
AREAELRVLPGLYHHRLTDESARTFFEDTWVVAPEADRIGYRYRQGRALRFREREQPFGA
GADPSNIVDACYPIGSIQVPAGVEPIILHRDAVSGGGYATIGTVISADLDLIGQMQPNHQ
ARFVAVTMADALAARRDAQRRLARLRDALWPHGG*
>GCA_000452945.1_BPMSHR5858_genomic_05986_1 group_2027
MNDLDSTHASFYRPSRIGTVLVANRGEIAVRVIRAAHELGMRAVAAVSDADRDSLAARMA
DEAVHIGPSHAAKSYLDPAAMLAAARQCGADAIHPGYGFLSENADFAAQVEAAGLIFVGP
PARVIATMGDKARARETARRAGVPTVPGSEGVVASLDGAREVAARIGYPVMIKAAAGGGG
RGIRVAHDAAQLAAQLPLAQREAQAAFGDGGVYLERFIARARHIEVQILGDGENVIHLFE
RECSLQRRRQKILEEAPSPSLTPALRDALCASATRLARQVSYRGAGTLEYLFDDARGELY
FIEMNTRIQVEHPVTEAVTGVDLVREMLRIADGEPLRFAQGDIALRGAALECRINAEDPL
QDFRPNPGRIDALVWPAGPGVRIDSLLYPGYTVPPFYDSLLAKLIVHDESRPAALARAAR
ALRELRIDGVKTTAPLHRALLDDADVRAGRYHTNYLEAWMRDWLARLDARASAPRGLGEA
A*
>GCA_000452945.1_BPMSHR5858_genomic_05987_1 accB_2
MAQNEIVSPLPGTFYRRPSPDAAPFVDVGSTLAPGAVVGIVEVMKQFTEIETAAAGRVVE
ILVDDGEPVDAGQVLMRTEE*
>GCA_000452945.1_BPMSHR5858_genomic_05989_1 oxyR_2
MALTLRQLKYFVATAELGQISQAAIQLTISQSAVTSAIRELEDSLGTQLFVRTASGVALT
NTGRRFLNQAYTILSSVDEAMRIPNLESTLTGTLALAASYTVLGYFLPHHLMRLHTQYPR
LTIQLHELNRESIEEGLIAGRYDMAVLLTSNVSNPELVLEPVIHSARRLWVGAHHPLLRR
ESVTFADVAPEPFVMLTVDEAAYTALRYWNETPYRPNVILRTSSVEAVRSMVANGSGVAI
LSDMVYRPWSLEGRRIETVLLRDPVPPMSVGLAWRRNAEFSPAMHAVREYFRHTFMEPRM
GGGA*
>GCA_000452945.1_BPMSHR5858_genomic_05992_1 pip
LYPPIEPYAHGFLDTGDGHRVYWELCGNPNGKPAVFLHGGPGSGCSADHRRLFDPARYNV
LLFDQRGCGRSTPHASLENNTTWHLVDDIERLRAMIGVERWLVFGGSWGSALALAYAQTH
PARVAELVVRGIFTVRRSELLWYYQEGASWLFPDLWEDFIAPIPSAERADLIAAYRRRLT
GDDEAAKREAARAWSVWEGRTIALLPNAAHETYFGDAHFALAFARIENHYFVHQGFMEDG
QLLRDAHRLADIPGVIVQGRYDVATPARTAWELARAWPRASLEIVPDAGHAYDEPGILRA
LIAATDRFARER*
>GCA_000452945.1_BPMSHR5858_genomic_05994_1 group_5427
MNLHRSIVVPLVFSATFLAGCGSSVDGRFTSGMVAMGAVAGSTVAAYPIGADGTIGAQPL
AVTTSDASGAFSLPALDRWPALIRATHGAYVEEATGAAATLEGDGLEAVYASAPSTIVVS
PYSSAVVATARAAGGLTQANIAAAIATISAFVGDFDPQQTRPATMAAGAAAPAIEPGHQM
ALALGAESQSRTDAAADIATSIAAIVAQAAAGDTLDTCHAGAGDPRADGTLAAPVSSGCA
ITRGAARYAANARNTSGVTSLASLSAARTSRASPDQTAAATPDACADRAALLAQNLALFD
GRRDDVQANLVGGVTRDNWRTVTTRSTWGPTAAFYGTLATPAACADTDTFARELVIAAEN
YWIDQGINYCHHHIPGWTPPDDSAAAAPRYRNSSAGSTSGASSNGMTCTAQRSGTGAQIV
PGTAPSAGFSASEIQWNGVDCSDFTSWIYNFVGLTGVRLPTAIGSQACAANEGADALPTP
GVLLDIDSGNIDAMLPALKPGDLLYITQVDPLASGSDAGGYQLAHVVTWTGKRWSDLQAG
PDAARYALARLGQPGSRLGGDLAKYLPLADLATQNPWMIIDSHYAGPAYRPFVGWYRRSL
SNVRRIVGADAARRDPALAPYVIAPIASDARGDTLTLASPHANASAQQGWRMIYRQSGGT
PSCVRAGIAQ*
>GCA_000452945.1_BPMSHR5858_genomic_05997_1 norR_4
MSTLPETGVRPRVWAVGISRLRALFRDIADEYEERADLRIVARGYEEAITALATAGAERP
DAIVAAGSNGTYLKARVRVPVVLVNPTGFDVMHALARARRDAGRIALVSYGETPPEVRSF
AAAYGLDVVFASYQSAQEADTRVLEMRERGIETVVGPGLVTDLAANAGMGAVFVYSHASV
RAAVNTALEVAHATHAEALRRERLDNLLQHLRDGVVALDREGRIEAINERLALALGVSPS
AAVGRALDDLTPELRTLRAADGDTLATVRGVRYVVHRGPLASSGGAPGSVLTFQESRAVE
RLDRTLRSHQHAQQFTARYRLDDIVGTTPAIAYARDLARRYAKSDATVLILGESGTGKEM
IAQSLHALSARRKYPFVAVNCGAFPESLLESELFGYEEGAFTGARRGGKAGLFEAAHRGT
LFLDEIGEMPPSLQSRLLRVLQEREVVRVGSTEPVPVDIRVVAATHRPLFAAVEAGTFRA
DLYYRLNILNIGLPPLRERAADIPALAATLLAQAARREPRLAERLRDANDAARALDAANA
ALARYRWPGNVRELQNVIERIAVELADAGERGDAAVSADTLRAIAPEVFALDTLGGFGSF
GSAAGGGHGDASGGDIDADADARRPTLVQRRRRAEADEIRAALDACGGDRDRACARLGIS
KTTLWRKLAASEGKARGNKRGGGHGGDNDEGAGA*
>GCA_000452945.1_BPMSHR5858_genomic_06003_1 group_5274
MSQAQGHPVTPYGVAIHQAIADGDLAQMKSLRTQAQALLAQQGNLATALELLEVEIAKLE
RRK*
>GCA_000452945.1_BPMSHR5858_genomic_06005_1 group_5275
MATTGLFPVQLRVATPNLGAPVLWLNLLVNTVEKTASGFARITQTVYPPMHFRARVVGPF
HQMRIDPHAPQSVTLTLSGSPTGPVAPQVVILELNALLNEGWQSGTANYRYFYESRWHSI
EHAIVSKDNSRIPLDPPSEHVMPMYGVGLQEARASGDLSRMKALAQQAEQQLADHDVIAA
ELQKLEAEIARLEARR*
>GCA_000452945.1_BPMSHR5858_genomic_06006_1 pqqE
MSTTDARPARYLFDSDYQRFVPVHAVWEITLACDLKCLHCGSRAGHRRTNELSTAECLEV
IDALARLGTREVSLIGGEAYLRKDWTQLIRAIRSHGMYCAIQTGGRNLTPKRLAQAVDAG
LNGVGVSLDGLAPLHDKVRNVPGAFERALDTLRRARDAGIAVSVNTQIGAQTMEDLPALM
DTIIELGATHWQIQLTVAMGNAVDNDELLLQPYRLAELMPLLAKLYKDGVSRGLLMTVGN
NIGYYGPYEHLWRGFGDERVHWSGCAAGQNVIALEADGTVKGCPSLATVGFSGGNVRDMS
LEDIWRTSEGIHFGRLRSVDDLWGFCRTCYYADVCRGGCTWTSHSLLGKPGNNPYCHYRV
LELQKQGLRERIAKVQDAGPASFAVGRFDLVTERIADGEPVASVVRSGQVIELAWKNRGK
RSPEVGRVPPKLKMCRNCDGYVHAGEQTCPHCGGDIDAAARAHELDAQRRHALMNDLERL
LGLPASTFGGG*
>GCA_000452945.1_BPMSHR5858_genomic_06010_1 group_6016
MSAPVSVSDQAARLRRHFAQIVLPIWRGPGFNPALQLPFEAVAPDTHAPLPVTRYRAMAC
ARQLFIFSQAGDAQHAHALFAALCRHFRDPRHDGWFYSVDAQGAPLDRTKDLYTHAFVVF
ACAEYFAAFGNRDARELTQRTAALIVDRFAPRPGSALLDSARGEDFAAAAGGPLQNPLMH
LTEGWLAAGRAFGDTAFDDALLRTAQAVERTFVDPHTGCVAELPIGCADNRFEPGHQFEW
FYLVASAGARLAATGLPDALARAYAFAQRHGVDLDTGGVSAATDERGACVDGTQRIWAQT
EYLRALATHGGEPDALARQIARFAERFLHPRGWYECKTAQGEVSRADMPSTTPYHLATAY
ASLPAGT*
>GCA_000452945.1_BPMSHR5858_genomic_06012_1 group_9
MKRRTMRRACKRAVSAVSTGRGAMLVLVLMPVLVPVLVPVLVLMPMPVPMPVPMPMPMHA
ARARRRTRLSPRANRRIC*
>GCA_000452945.1_BPMSHR5858_genomic_06018_1 group_182
MSTNMKRLMTAALGAALAFGALSARAASFDCAHAANAAERAICGTPALGELDVRMAAYYE
MLQNARPADEGMAYREFRDALRDEQQRWRQRTRDACGARIDCLTNAYTARIAALRGVAAE
RLVLRMTGGSAASAGAADATYAIEGESITLANGESVRPAAPGSAMKRVTTLVARSAVATI
AGRPVEAVLLSDDPGGSGRFLYVATAQPGGGAPAVLLGDRVKPVSVSIERAATGGAVVVV
EYLDRPEGAPFAQAPTIKIVRRFALEQGRLVEQRG*
>GCA_000452945.1_BPMSHR5858_genomic_00064_1 fabZ
MSTEKINFDIHKILTLLPHRYPILLVDRVLELEPHKAIKALKNVTVNEPFFTGHFPKRPV
MPGVLIIEALAQAAALLTFAEAQPKDPENTLYYFVGIDNARFKRVVEPGDQLILNVTFER
YIRGIWKFKAVAEVDGKVAAEAELMCTVKTADAAP*

>GCA_000452945.1_BPMSHR5858_genomic_00076_1 map_1
MAITLKNEHDIAQMRIACRLASEVLDYITPFVVAGVTTGELDRLCHEYMTHVQGTVPAPL
NYQPPGYPPFPKAICTSVNDVICHGIPGEKTLKNGDALNIDITVIKNGYFGDTSRMFIIG
EGSILAKRLVQTTYECMWLGIDQVRPGAHLGDIGHAIQKHAEAQGYSVVREYCGHGIGTV
FHEDPQVVHYGRPGTGIELKPGMIFTIEPMINAGKRDIRTMPDQWTVKTRDRSLSAQWEH
TILVTDTGHEVLTVSAGTPARPTFAAPATAA*
>GCA_000452945.1_BPMSHR5858_genomic_00081_1 zipA
VLPAATTISAAPPAVVDRRIDCIVPIRLASPLAGDKILPAAQRLRRAGSKPVHIEGKPDG
GDAWELLQNGVRYEELRAAAQLANRSGPLNELEFSEFVTGVQQFADAIDGAPEFPDMMET
VSMARELDGFAAQCDAQLSINVMSDGAPWSANYVQAVASQDGLLLSRDGTRFVKLDAKQN
PVFMLQFGDTNFLRDDLTYKGGNLITLVLDVPVADEDILPFRLMCDYAKSLSERIGARVV
DDQRRPLPESTLLAIEQQLMKLYARLEEAGIPAGSPVTRRLFSQ*
>GCA_000452945.1_BPMSHR5858_genomic_00114_1 group_4395
MSRTKRMWREVEGAVEVTALAAIECALLVAILVALGFPHAGI*
>GCA_000452945.1_BPMSHR5858_genomic_00119_1 group_4397
MTIRIEPGDKREIAETLGRYFEARSIPFHIEEHPADVLHIGFHADGHPAAVTVTFDEDAF
AAFAQWDIEQKRLALRRLADEFAEMMAQRSPTAHAGDLHINRF*
>GCA_000452945.1_BPMSHR5858_genomic_00125_1 group_7231
MKPIRTVAALISAVALVAACGGDDTGTELGISNPQARFINAVPAGPSLDYYLNAQANATG
IAYKGVTRYRSVGSGSQTASYDVTGTSVTIASQTFNAANGHHYTTIAIPSTSTPISVIDD
PFAKGLLSDQARVRGFNASPNAQNVDIYVVPPGTNIAAQSPTLAGAAYQNAVPASGQDSI
YLSGGTYQVIVTLPGSKTPILTTPPVSIANNADWLLLTIPAGGVADVVPNDIHVLVAQGN
DADTSAQELGPQ*
>GCA_000452945.1_BPMSHR5858_genomic_00146_1 lrp_5
MSLDRTDMRILRELESDGRISNQDLATSVALSPSACLRRVKLMEEQGVIEGYRCVIAPKR
IGVEFEALVHVAMRPDIEDWHDSFVAAIQSWPEVVAAQVVTGESNYVLTVRARSLDHFSE
FVMNRLHRARGVMSTNSSIVLATLKRDGSVLDLASKPA*
>GCA_000452945.1_BPMSHR5858_genomic_00153_1 group_4403
MSRVTSSSGGARPCGRRRQRGVVSILVALMLAVLIGFVGLALDLGKLYVTRSELQNSADA
CALAAARDLTGAINLSVPEAAGITAGHLNYALFEQFPVQMQTNSNVTFSDSLSNPFQPKN
AIASPSSIKYVKCTTSRTGIVNWFIQTLNLVPGVTVANASVSATAVATVGAAQTTCAIPV
FICKAGTQTSPPVAGATYNIGDWLSAKTGSPPSFGAGNFGWSALDGSNSASSIANELTGN
YCALPATGSQVGTPGDKAATTNAYNTRFGIYANPYKNPSYGTPDFTGFAYDATTWPSQSN
AYSDFVSKRLTFASYQGDLITGINTGGSYNPSYYAAGADRRLALAPEVDCSVLLSGHSAP
VLSWDCVLMLDPMGSGGSATPVHLEYRGSSTASGSPCATQGTPGNGSSVGPQVPVLLQ*
>GCA_000452945.1_BPMSHR5858_genomic_00239_1 dnaQ
MRQIILDTETTGLNARAGDRIIEIGCVELLNRRLTGKNLHFYVNPERDSDPGALAVHGLT
TEFLSDKPKFAEIVDPLRDFVRDAELIIHNAPFDLGFLDAEFALLGLPSFTDHCAGVIDT
LVQAKQMFPGKRNSLDALCDRFGISNAHRTLHGALLDSELLAEVYLAMTRGQESLVIDML
DEAGDAHRNGDAPRMAFSGLDLPVLAATDAELAAHEAQLDALDKSAKGVCVWRKEAVGEA
*
>GCA_000452945.1_BPMSHR5858_genomic_00268_1 fabG_6
MADHSIKGKTVIIAGGAKNLGGLIARDLAAQGAQAVAIHYNSAASKGAAAETVAAIKAAG
ARAVALQADLTAAGAVEKLFVDTVAAIGRPDIAINTVGKVLKKPFVEITEAEYDEMAAVN
SKTAFFFLKEAGRHVNDNGKIVTLVTSLLGAFTPFYAAYAGMKAPVEHFTRAAAKEFGAR
GISVTAVGPGPMDTPFFYPAEGADAVAYHKTAAALSPFSKTGLTDIGDVVPFIRHLVSDG
WWITGQTILINGGYTTK*
>GCA_000452945.1_BPMSHR5858_genomic_00269_1 pgrR_3
MDRIDLFRVFARVVECANFTRAADTLGMPRSSVSAAIQELEGRVGARLLHRTTRKVTPTQ
DGAALYERCLRLIADVEETENLFRHSIVGPSGTLRVDMPGRIGRLIVAPALPEFLERYPQ
ISVELGVTDRAVNLIEERVDCVLRVGTLSDSGLVARAIGDLPLINVASPAYLARHGVPRT
PADLERHRPVNYASPSNGRVAPWEWVEGDEVKRVALRARVTVNSAEAYIACCLAGLGLIQ
IPAYDVNAHLRAGELVEVLPDHRAAPMPMTLLYPHRQHLSRRFQVFAEWLEALLRAKVL*
>GCA_000452945.1_BPMSHR5858_genomic_00273_1 decR_3
LAYSLDELDLRILAALQADASLSNLALAERALSSPPTCMRRVRKLTEAGIIERRIAVLDR
GRIGTSVTAIIEIGLDRQTAEDYDAFEAYACAQPEVTQCYRVSPGPDFVIVADLADMAAY
DEFARRLFTHASNVRNVRTFFSTHRAKFEANAPVRATPGKRAGG*
>GCA_000452945.1_BPMSHR5858_genomic_00281_1 group_663
MRAERANPRVLMSIRVGAGNLPHVRRLLHQVLGAALDIYTAVIDNQTGQASLQLELASTS
VSEAMTSIMRVLPEAEFGKIRLSPRAPSARAPRMPPRG*
>GCA_000452945.1_BPMSHR5858_genomic_00302_1 ccpA
MTPTIKDVAALAGFSIATVSRAINAPHTVHPATLEKIRAAIGALRFRPNPLGRQLRSDRT
QLIGVVLPTLANPVFAECLQGVDELATQAGFKLIVMSTEYDAARERHAIETLRAQRVEGL
MLTVADADAHPLLDELDRDGPLYVLMHNDTPHRPSVAVDNRRAAYDGVRMLIERGHRRVL
MLAGSLDASDRARLRVHGYAQALDERGLEPLPALELDFNAPALPHAMLAHLSARATRPTA
LFCSNDWLAMVVIRGLRDAHLAVPDDMSVLGFDGLAVGELLAPPLASVATPNREIGRAAW
RRLAERIAGKRHAQPALTLPHAVRDGATVAPPRDARDARAARIA*
>GCA_000452945.1_BPMSHR5858_genomic_00305_1 group_7239
MQTITMEQCKRIHGGDNLIDGTFNLLGGTFTINSSSLSTLLTNANTLVVNANNWLTNPKG
TLSSTKK*
>GCA_000452945.1_BPMSHR5858_genomic_00310_1 group_1394
MCRAPPLPASGSLFATAATRRVRLNQPARSPAVAGMLNKVPEVTLAFWIVKIMSTTVGET
GADYLAVHVGLGAAVTGALTACLLVGALALQWRARAYVPWIYWLNIVLVSVAGTQITDAL
TDGLGVSLHVSTPLFAAALAATFALWYRSERTLSIHTIVTPRREGFYWAAILLTFALGTA
AGDLATEALGLGFRLGIVAFGGAIALVAAAALGGMNRVLAFWLAYILTRPLGASLGDFLS
QAQAYGGLGLGTIRTSAVFLAVIVALVAGLSVAAARAGRLGARGR*
>GCA_000452945.1_BPMSHR5858_genomic_00316_1 group_1395
MKRNLIAFACVAGFALAAPFAHAQNDAMSKDGMGMSKDAMSKGAMSGDAMSKDAMSKDAM
SKDRMGKRHDHMAKGGMKAGPMSKDAMKEPMSSDKMSPMSN*
>GCA_000452945.1_BPMSHR5858_genomic_00330_1 group_7240
MNDPRSKKNPVFGVAILIVVVIALVIGTILYNAISQKHEYDEAHPHAASAASAASAQHG*
>GCA_000452945.1_BPMSHR5858_genomic_00334_1 decR_2
MDAIDRKLLELLQEDATLPIAELAQRVNLSQTPCWKRIQRLKETGAIRAQVALCDPRRLG
VGTTVFVAVRTNQHTEEWAARFTQVVRDMPEVVEVYRMSGETDYLLRVVVADIDDYDRIY
KKLIRSVPLFDVSSSFAMEQIKYSTALPVRDGGAG*
>GCA_000452945.1_BPMSHR5858_genomic_00335_1 group_4424
MNHAASSLPHAPDPHPKPRERAPSAPCRDRATPASTPIARFCTRLDAAFDSLRDETAPSR
HPAFARALRAALAEAAAAPGLLTPAQRESGAAGYRRHLLAADPAGRYAVVSLVWEPGQWS
PVHGHHAWCGYAVLEGKLTEIVYRWSDAACCAIDARRRARPMGAVSYVCAGRDAIHRLGH
AGAKTDAPAISLHVYGVAGEQVATHVNDVVAVDARAYA*
>GCA_000452945.1_BPMSHR5858_genomic_00336_1 group_4425
MTQGTLKIALFGATGMIGSRIAAEAARRGHQVTALSRNPAASGANVQAKAADLFDPASIA
AALAGQDVVASAYGPKQEEASKVVAVAKALVDGARKAGVKRVVVVGGAGTLEVAPGKQLV
DTEGFPDAYKAVALAHRDAYGYLSTVQDLDWTFFSPAALIAPGERTGRFRTGAGRLIVDE
QGNSKISAEDYAIAFVDEIEQGRFIRQAATAAY*
>GCA_000452945.1_BPMSHR5858_genomic_00404_1 group_4431
MSAANQNGSTEFDYIVIGGGSAGCVVTHRLVSAGHRVLLLEAGPPDNSFFVHTPATFVRV
IGTKRTWVYETEPQAHAAGRRMYVPQGRTLGGGSSVNAMVYIRGTPADYDGWRDAGCDGW
GWDDVLPFFRRAEHNHRLAGPLHGVDGPLHVSDSRFRHPLSHAFVQGAQEFGLPYNDDFN
GASQAGVGFYQTTTFEGRRGSTAATYLAAVKRDPLLTTETDAFVTRIVFENGAAVGVRYQ
ARDGEERIARARAEIVLCAGALASPKLLMLSGVGPAEQLLQHGIPVVHDSPEVGLNFQDH
LEVSLYGRAREPVSLAGQDRGLNALRHGIQYTLFHTGLLTSNVVESGGFVDTANGGRPDV
QFHVLPVLVGDVGREPLAGHGISINPCFLRPKSRGTVRLRSADPHAPILFDGNFLSHPDD
FAALMRGLSLAREIMRMPSMSKAIAGEMLPTDGGRVDLDAYVRSHAKTVYHPSGTCRMGG
DPDSVVDAQLRVRGVGGLRICDASVMPSLVSGNTNAPTIMIAERCAEFMLSPALAPTARA
GASTPQAAPQDASHAR*
>GCA_000452945.1_BPMSHR5858_genomic_00409_1 rhaR_6
MKKPLRGTGRPGVAILLPPRLLGGYVFLTQELLLLAGTMKARSLDVVNSRLFDIAILSHD
GRPVPTIGGLDVPATAALSDADAHEVVIVPAQFMPDAQIGALERVFIDWLERRYAAGALV
VGLNAAPLLAKAGLLDGRGATGLPSERTLFARHFPAVRYTPSKPLVVDGRLITVSGINPA
VDACAYVIDYCFGAGVSQRLLRVALTQSLPSYEHMAVWAAQYKRHGDAPVLAVQDSVERA
LAHPPTLAELAARAAMSERTLSRRFAAATGLTLRRYVAALRVELAAFLLRTSRMTLEHIA
DECGFASASALSHAFLAGQGCSPIQYRNRHRPDARREPAGGGGTAAGSGEGAAS*
>GCA_000452945.1_BPMSHR5858_genomic_00410_1 group_7246
MIPRTIFAEEHEQFRESVRRFIESEVMPHHERWEEQGYVDRDVWTKAAAAGYHCASMPEA
YGGVGADIRYSVVLFEEIARAGASGLGFGLHSEIVAPYILHYGSDALKARYLPKLASAEM
IGAIAMTEPGAGSDLQGVRTTAVRDGDHYVLNGSKIFITNGWHADVVIVVARTTPEGGSK
GTSLFVVDTGMAGFSKGKRLKKVGMKAQDTSELFFDGVRVPAGNLLGEENRGFVYLMQEL
PWERLQIAISAIASAEAALAWTLDYTRERRAFGRAVIDFQTSRHALAELKSEIQIGRVFV
DKCIELQLAGKLDAATASMAKYWTTDLQFKVIDRCVQLHGGYGYMWEYPIARAWADSRVQ
QIYGGTNEIMKELIARTL*
>GCA_000452945.1_BPMSHR5858_genomic_00426_1 hmp_2
MLRQLHSIPGLIAAALLIVVTLTGAALSIQPALERAAVAGPPRAPLDVATLAGRVSAQVP
GVEKLVRRPSGEIVAYYAGTDGPRASIVDPSTGAPLADYRPSPVVRWLTNLHRKFLLGDA
GRIATGVTAAAMLFIALSGLALLAHRMGGWRRLAGPIRGDGLQRLHNETARVALAGLVLS
ALTGLVLSLSTFGVIPERGAASALALDARPGIAAAAKAPIPIAQMAALRATDWSTLRQLS
FPAPDEPRGLIELKTTDGAGFVDPATGAALAYQPADGWQRLHALVKMLHTGEGLWWLGLV
LGASSLATPLLAVTGVMLWARRRRAQPKRAGDAPARDADTVLLVGSEGNSTWGFASALQA
ALAQAGRRVHVAPMNDGARGYPAARHVLVLSATYGDGDAPGNAQSFLPKLARLSAGQGAS
FAVLGFGDRQFPRFCGYARSVHDALVAKGLTPLLDLGTVDRQSEPEFRAWCARLGDALGL
ALDVRHAPTLPRTMPLTLVERDDYGGDPRAMTSVLRFAPAGRDGRTGRADRAGRDEHDER
DGRERRQEREERLEDAGRRAAWWTAFARRRLPSFETGDLLGVVPPGEASPRYYSLASASS
DGIVEICVRRHPHGVCSRYLTGLQPGDTIEAFVRPHARLRPHAGAAPVILIGAGTGIGPL
IGFIRHNAARRPMHLYFGARNANDGFPYRDELDGLVRDRRLRALTTAFSRAERGAYVQDR
LVADARNLRELVAHGAQIMVCGGRAMADGVARAWERILADSGSSVAQLKQQGRYVEDVY*
>GCA_000452945.1_BPMSHR5858_genomic_00427_1 apbE_2
MSKTSIEWSPGARLNRCRASGATMGTRYGAQFYAPPTADARAIAAALDAAVRAVDAQMSN
WKADSDLSRLNRATPGSWTPICANLAAVLVRAREIGRETDNAFNIGVGTLVDRWGFGPGA
AANRQADNEWAANRQAAGRHTVDRRTVARHMVDRHMVDRHTVDRHTADRQTADRQTKDGR
TPDRQPARPAGPANGLSGAIDARRRTSILRSPVPSPCRPIDELLEVDVARGRARRLADVA
FDLCGIAKGFGVDELARVLDRHGIGAWLVGIDGELRARGCKPDGSPWAIALEAPDYDRRG
AMGAIDLVDAAVATSGDYRHWADFGGERLSHTMDPRAGAPLRGDIASVTVVAPTCTDADA
YATALMVLGAQAGCAHAERHGLDALFVVRDGDALRTIGCGAFADAGPAG*
>GCA_000452945.1_BPMSHR5858_genomic_00428_1 gabD
MQFNDILAALDVDLSRWQGRALTARSPVDGATLATLAADAPADAERKIDAAHQAFLKWRT
VPAPVRGELVRVFGNVLREHKAALGRLVTLEAGKIASEGLGEVQEMIDICDFAVGLSRQL
YGLTIASERPGHRMMETWHPLGVCGVISAFNFPVAVWSWNAALAFVCGDPVVWKPSEKTP
LTAIACHTLFAKAVREFDKTHPGVVPEGLHQLVLGGRDVGEALCASPKVPLVSATGSVRM
GIEVAQVLSRRLARSILELGGNNGMIVAPSADLDLVVRAVTFAAVGTAGQRCTTLRRLIV
HRSVADQLLPRLEKAFASVKVGDPLEAGTLVGPLIDRASFDAMQKALGDAREQGGEVKGG
ERVELGRADAYYVRPALVRMPRQTAVVTRETFAPILYVLVYDDFDEALAIHNGVPQGLSS
AIFTNDMREAERFMSAAGSDCGIVNVNIGTSGAEIGGAFGGEKETGGGRESGSDAWKSYM
RRATNTINYSRELPLAQGVKFDV*
>GCA_000452945.1_BPMSHR5858_genomic_00430_1 group_7248
MNPNLAALLATVSDQPTIDELMRLMHVPTCFERCEPGVVTRAELAQALNMALFADLLERV
PTGRAYTRDVARDGGRVTFDHGALRTVRWPSCGALPPGEAAFTRILRPLGYRLNGTYALE
RLKMTGRSYAHLDAPDQIAQFFVSELHPERFSAAFQAAVTNVLGTSADPLTPQAVATLAE
LERDASLPLADACALLPVLVRCFGRQHATPALADYDTLLAESAEMAWIATEGNAFNHATD
RVPDVDALAAAQRALGRPIKPAVEVSRSGRVRQTAFRADPVARTFVDASGALVEREVPGS
FYEFITRDAVADAQTGRRALDLSFDAGNATGIFKMTAAA*
>GCA_000452945.1_BPMSHR5858_genomic_00431_1 gcvA_11
MRKGIPNLAALQVFEAAARHESFTRAADELALTQSAVSRQITSLEERLGVALFLRIKKRV
VLTPHGRHYAALVRRNLERIERDTLELMAQRGIGATLELAVVPTFASQWLIPRLPQFRAL
RPDITVNLSVRTGPFLFSDSTFDAAIYFGHALWPGTQGRLLFLEGDMLPVCSPALLGANG
RRITRDALLDLPLLHLSTRADAWRDWFRAQGLDNDLRAVRGPRYELFTMLTAAAHAGMGV
ALMPALLVADELASGRLCAPLDLRLPSDAGYYLVAPDAAADSEPFVALAEWLGSVSPARP
IS*
>GCA_000452945.1_BPMSHR5858_genomic_00433_1 puuB_1
MQNYYEATVTRTPYPALAERIDTDVCIVGGGLAGLSTALGLVERGVRDIALLDAQTVGFG
ASGRNGGFVFGGYSLDNADLLRTLGPESARALYRLTIDAVELIRARAARYGIDCELVDGG
VILANWFDDPARLCAPRKLMKEAFGVDWEPLDERALRTRLKTDRYRGGLFERNAFHFHPL
KYVLGVAAAAAQGGARIHEHSAARAIRRDGAGFVVATDGGAVRARRVVFAGGGYARGVYP
RIERAVLPIATYVIATEPLGARLADAIACEAAVYDTRFAFDYYRPLADTRILWGGRISVL
ERGPEAIARLLRRDLLRVYPQLRDVAVQFAWGGLMSYARHKMPQIGRDAHGVWHAVGFGG
HGMAPTTVAGEVLAAALAGERPVPDGFARFGLARTFGLAGLAAAQLTYSAYEARDALAAR
RCARAGQPFRQRL*
>GCA_000452945.1_BPMSHR5858_genomic_00434_1 nicS_2
MPPSDHAKMKQGSKAAAPDAEARRDEARRKYDPEQTKRNILDVATQEFSAMGLSGARVDA
IAERTNTTKRMLYYYFDSKEGLYEAVLEKVYGDIRTLERELNVGELEPREGLSRLVEFTF
DYHDRHRDFVRLVTIENIHGAKYIEQLKSFKNRNVSIIKTLEELLARGVESGVFRDDVDP
FDLHLLISSFCFHRVANRYTFGTAFGRDPSSPRLRARHRAMITDTVLRYVAR*
>GCA_000452945.1_BPMSHR5858_genomic_00436_1 group_4438
MSVSVRSYLSPTLVLLAFDWPDAQRRDDFLGFAIKRTPGFWSADGKTRAATSWLPNRLTF
DGPAAHTQGDAPTDQAPIQKFMWWDARIDPPDRGRSFRYDVYPVVGTPASLQVLEAEAGV
CAITLPAHVEDGVGTWFNRAVVSSQAFAKQVRALHLADGGAPSPDAALRLRTWLANDLQE
TFGLMLGADARRAVSAVYHLTDPLWAIPAFDAFGRAHGANALAIVYDAHETKRGSGEPPL
PSPNQPAVERLTGVATLAPRDKTRIMHDKFIVAGDETGPARVLTGSANFTAEGIAEQANV
LHTFESAALAELYNARARALASNPGIAETARLAPGWSEPVEIGSARVRVSFSPEPRGVRT
QIDTIVEAVRGAQHSVAFCLFMPTDAALRDACFAAGERGLMMFGLVNAISARGAQQADAD
RRDGKTLDAARLANLELYHRSRSNRDVIDAAYFSPATVPDGFETELRLFPGEPAPAYPPV
VVHHKFVVIDAEGANPIVYSGSANMSANSEHYNDENLLEIRDARIAGIYLAEFLRLYEHY
RARALAIASRRGDGGGSTAGNAGAHGGLALQPDSRWAKKYFVAGSPEEKARIALAAPVAA
R*
>GCA_000452945.1_BPMSHR5858_genomic_00439_1 exuT_2
MHQRTMGWVTVFLLFVVYGINYLDRVALSIVAPMVQRDLGIDAAQMGIVFSTFFVGYALF
NFIGGLASDRLGPKRVYVIAVGLWSIFCGMTAITVGFVSLLIVRLLFGMAEGPLCSAANK
MVNNWLPRESAATAMGLLSAGSPLGGALAGPIVGVLAAQLGWRPAFWIVCAIGLAWVLVW
IATTSDRPAPQASAMPAGGSGAAAAAARAAAAPRACASGGRTVDAAHASETADVPPLRDY
LKQPRILATGVAFFGYNYVLFFFLSWFPSYLVQAHHLNIREMSVATVVPWLVGTIGLACG
GVISDGIYKLTGNAMLSRRIVLVGCLLGAGVCVAIAGSVRSTQSAIALMSVSLFFLYATG
AIYWAIVQDIVAPGRVGAVSGCLHCMGSLSGVIGPAVTGFIVERSGSFVSAFVLAGAIAL
AGAVLSALFLRNRACDARVLRESPLL*
>GCA_000452945.1_BPMSHR5858_genomic_00442_1 group_4440
VKIERLNIIVTCPGRNFVTVKIVTDEGVHGLGDATLNGRELAVRAYLEDHVFPCLIGRDP
RDIEDIWQYLYRGAYWRRGPVTMTAIAAIDMALWDIKGKLAGMPVYQLLGGKSREGLMVY
GHANGRDHDEAIDAAHRHIAAGYRAIRVQSGVPGLAKVYGVGKTAGAYEPAEKGLPPEEP
WDTALYLRHTPELFRKVRDALGDAPHLLHDAHHRLTPIEAARLGRDLEPYRLFWLEDATP
AENQDAFRLIRSHTTTPLAVGEVFNSIWDCKDLIRDQLIDYIRATIVHAGGITHVRRIAD
YAAMYQVRTGFHGATDLSPVCMAAAVNFGLWAPNFGIQELMPHTALTDEVFPHNYRFDDG
YLVMDDVPGLGVDIDETLAANYPYERAYLPVARLRDGSMWNW*
>GCA_000452945.1_BPMSHR5858_genomic_00443_1 rspB
MLSVVVDQPHSMGVREMPTPEPAAGEVRVRVRYAGICGSDLHIFHGKNPFVSYPRVIGHE
FVGRIESVGAGVDASRIGEIVAVDPVISCGRCHACAIGRRNVCRSLTVLGVHRDGGFSEY
ACVPAANAHRIAPEIADTCAAIVEPFAVAANATARTGVLPSDVALIYGAGTVGLTLLQVL
KHVYGIRAFIADRLDERLALARKCGAAADEVIHAATETVPDALERRGVDGGPTLIFDAVC
HPSILEEAVRLAAPAARIGVLGFSSEPSSIVQAELTKKELTLCASRLNCAMFPQVIEWIA
DGRVHPEHIVTHTLDFRDVARAFELAERNPRESCKILLDFAAH*
>GCA_000452945.1_BPMSHR5858_genomic_00444_1 group_4441
MYRKGIVLEIQFPPQRLNDAAGDPYWIDLTLDEARRLHRQLSARLATEAGANQPLDTFSL
D*
>GCA_000452945.1_BPMSHR5858_genomic_00575_1 group_7279
MKPLLKAGDRIEGLHWVAEYRHRARLIRVLRERVEVGTYGAPSALFGDEEEMGASTPADH
RGREAALRAYLRHFAKVHDAEE*
>GCA_000452945.1_BPMSHR5858_genomic_00592_1 group_4463
MYQRILVAVDGSETARHAFDAALDIARTSGAELQPFYVVENAAIYYNVPGYDPSILRTQL
LQQGNELAKEFAEKMQAAGVKGALKIGEATSLADVPSLIVEGAKAFGADLLVLGTHGRRG
FKRLVLGSVAEQCVRHSALPVLLIPAAAHPEDRES*
>GCA_000452945.1_BPMSHR5858_genomic_00632_1 group_7281

MGLSDAPLLFNFEIESSEDLTYIPMIVRFNLDRFGLRISLEQWQMLPLEDRRLLARFPAD
EDVVIEPNFDHALFEMLRTHANVEPSWFQPDEHPAWRSVDAVPDALVAQSGLAGLPAPSA
SQWATLTPFQRYVLTKLSRKPKLNHDFVPAMREFGLAAH*
>GCA_000452945.1_BPMSHR5858_genomic_00648_1 lnt
MAEPISTRSQSSVSSTAAGRALPIWHYPAALLAGAANTLTFAPTPHGGWLQLVVFAWLFA
QLTRTTSWKHAAATGGAFGFGNFITGIWWLYISMHVYGEMAAPLAGGALVLFCLYLSIYP
AFSAGLWSLCAGHARNGKAVDARPFSPTWHGAFAFASAWAVGEWLRGTVLTGFPWLASGY
AQVDGPLAGYASIVGVYGIGWVLALVAALVVQAIVRAREGGGGHGRLRDNGGGNGRAAAG
AGAAPRASRVAAPALAALAALALGPLLALVPWTVPANAPLTVRLLQGNVKQDIKFEEAGI
KAAIEMYQKMITEKPADLVVTPETAIAVLIQELPEPFARAVRNFSDATDTAVLFGAVGGT
VTADGRIVDYTNSLYGVTPHSRDIYRYDKHHLVPFGEFIPWGFHWFVDLMKMPLGDFARG
APVQQPFIVRNQPVMADVCYEDIFGEEIAATIRDNPVSPGVLVNVTNLAWFGDTIALDQH
LQIARMRSLETGRPMLRATNTGMTAAIDAQGRVIGRLKPFTIGSLDVRIEGTAGRTPYVT
SGNATVLALSLLLLAFGFAFGPGLRRRG*
>GCA_000452945.1_BPMSHR5858_genomic_00650_1 chaC
VSRPAPARRAYPPSLGEARYLSDAELAASLGATLERWDRTSDLWLFGYGSLIWNPGMPAV
EALRAKVHGYHRGLYLWSRVNRGTPEQPGLVLALDRGGSCTGLAFRLAGRTAMPHLEALW
RREMAMGSYRPAWLPCALAGGERVNALAFVMRRDVPTYTGKLTDDVVKAVFGCASGRYGT
TLDYVSRTVEALRDSGMPDRALEALLARCR*
>GCA_000452945.1_BPMSHR5858_genomic_00665_1 glpF
MSPYIAEFIGTALLVLLGNGAVANVLLAKTKGKGADLIVIVMGWAMAVFVAVYVTASFSG
AHLNPIVTISLALAGKFAWAKVGGYVASQMFGGMAGAFLVWLAYRQHFAKEADPDLKLAV
FCTAPAIRSVTHNVLTEAICTFVLILGVLYLASPQVGLGALDALPVGLLVLGIGISLGGP
TGYAMSPARDLSPRIMHALLPIPGKRDSDWRYAWVPVLGPLVGGVLAANLYLYLHTTH*
>GCA_000452945.1_BPMSHR5858_genomic_00672_1 group_7283
MLADSNTTHEMPWRIEDIDLTRIDRQRAAANEDLLLLLCAASFIESGSDLYTSNLSQFFD
DDPEVSAWLNAEWEHEELQHGRALKAYIAYVWPEFDWDLAFANFFEEYSKTCSLEAFEKT
RALEMVARCVVETGTATLYRAINECSDEPVLKEITDNIRTDEVRHYKHFFRYFKKYNKVE
GNGRLAVLGALTRRVLEIKNEDSEIALRHVFAVRYPERAGDSAYCRERTARVNALVRRNL
SADMCVKMLLKPLELPAKIQPGLHYPLTKLTQHVFFR*
>GCA_000452945.1_BPMSHR5858_genomic_00673_1 group_7284
VTDPDRRTLERTTYDAWPRAARALFDGGDLAGKAGFTASLVVVDRGVDPPQPRTTLLSVG
ELYAPDARSLRFSLWPQSRAARALAAHPDATLAFVAGDAFHQARLRVDSSDSTEDGLARF
AATLVGGEAQRVGYARLTAGIAFELAEADAPAVLARWARQVAQLRR*
>GCA_000452945.1_BPMSHR5858_genomic_00682_1 sasA_12
MHGTFNLPLVALSLAIATLASYTALDLSALITLLDQPRMRRAWLAGGAAAMGTGIWAMHF
VGMLAFSLPIPLGYDFGHTFASLAIAVVVSYFALNAVTRAALTRERLAIGGVLMGLGIAG
MHYTGMSALRMQPAIAYDFTLFVASIAIAIGASTTALWIAHRLSNENEPRVLMKRIAAAG
VMGLAITGMHYTGMAAAHFGANAVCGAAGEISGAWLGATIALFTVTILSATLVVSRFDAR
TAFLRGMTDALEELVAKRTSELEGALRQYERTTHVLQRTRRKMEQEIDERKAAQARLEHE
KDEQRRLIRRLEETHVQLLQSEKLASIGQLAAGVAHEINNPIGFVNANLNTLRSWVQGLL
DVIAVQEALTGTLAADARAPLAAVARDIDLDYVRGDILALIDESIDGAMRVRRIVCDLRD
FSRPSGDAWAFADLHASLESTLNVVHNELKYKADIVREYGVLPLVECNAAQLSQVFMNLL
VNAAQSIGTHGTITIRTTHDGDTVSISIADTGAGIPEDAIGRIFDPFFTTKPVGQGTGLG
LSISHGIVEHHGGRIDVESHVGRGSTLTVTLPVRRKPEPAARTTAGDGANNVNDVHATNG
ASAARDAARAIERAAASAAPLTRGAAC*
>GCA_000452945.1_BPMSHR5858_genomic_00683_1 rssB_2
VNGTEESTTPIAPDAPQPHAADHARSAGSACAGADGGAPAAPVILIVDDEPSILSALKRL
LRTARYQVVTAESGAAALDVLAVGEVDLIISDMRMPCMTGAEFLARAQTLHPDTMRILLT
GYSEIDAVVSAINEGGVYRYLNKPWDDHDLLLTVRQALEQRRLRQETARLFALTQKQNEE
LVAFSTELEAQVHARTEEIRQTVMFLEDAQRDLKRNFMTMIQVGANMIELRCGVMGGESR
RIGELAKRLALALDMSELQAQDLYFAGLLHGIGKLSLPDELLRKSIDRMSAEESRLYHQH
PLRAQMVLTPVTQLNHVAHIIRHQYERFNGRGTPDGLAGDDIPLGSRILAIARDYEGLQR
GGVVNQRLQSEQAIALIKSQAGLRYDRQVVDRFVALVKDLATLGCNTPYARITGGQLLEG
MRLADDLRTSRGVLLMTKGSVVSAHQVALVRRYEAQESTPFAILIQTTTLAEAAASAGRE
C*
>GCA_000452945.1_BPMSHR5858_genomic_00687_1 group_10488
MKKILGDPASRRERRAFLGDVARLAGAGIIAGWTPIRPIAAHAQAAGAAPPNFPAGIALY
KQAFRNWSGEIAVADLWTAAPATPADVVAIVNWAADNGYRARPLGHMHNWSPLTVAANGA
HERTLLVDTTRHLCAVSVDPSTTPARVVAQAGVSLDTLLATLEQHGLGLTAAPAPGDITL
GGALAIGAHGTALPAANETRPPGHTYGSLSNAVLALTAVVYDTASSRYALRTFDRADPDI
GPFLAHVGRAFVVEATLQVGANQRLQCESFVDIPAAELFAAAGTRGRTVESFVQRSGRIE
AIWFPFTDYPWLKVWTVRPNRPSGARVVEEPYNYPFSDSISRELSDLVSRIVLNGEIQLA
PLFGKTQYTIAYLGLTNIFRPLTNLWGWSRSVLHYVRPTTLRVTANGYAVLTRRENVQRA
INEFVGAYRQRVAAYRAAGRYPMNGPIEIRVTGVDTPDDIGRGAVPPSLSAIRPRPDHPE
WNAAIWFDILTIPGTPDANRFYREIEQWMLSNYSGDYATVRPEWSKGWGYADTAAWSDDA
MLRTTIPDLFRQGLSSADDWDAALRTLERYDPRRVFSSPLLDRLMG*
>GCA_000452945.1_BPMSHR5858_genomic_00689_1 group_225
MVRMRGDGRAVDVSGFDVHPGARLHRVDDEANCERERRRVLEIDDGLHADAPGRVNLVHP
RDADDERREHGRREPHPDQADGAPSGVNGSPRPGATRPTSTPATSEGETYLGYATRIVAD
IRRMADAVRASRSVPRGLLRVNVMLGFGRTTITPLVSEFATPHPRVEVRLDVTDRPVDRV
DGGFDFAIRFGEPPDTRLNARRIMSNRRLLRASPVHLQRHGAPASLARCIVHRRNDDAHG
VWRFVRDGEAEGVKVRAAPSSNDGDIVPGWALDGHGILIRSEWDLAKYLESGRLRRVLPE
YALPSADLSACYPVRRNPSARARAPIDFLVARLGASAPAPVRTPEGGWRAGSRDVPPRVR
*
>GCA_000452945.1_BPMSHR5858_genomic_00690_1 selU
LNLQPASLAQIDDFDEIVDVRTPLEYAEDHIPGALNAPVLSNEERVIVGTMYKQVSPFEA
TRVGAAMVARNIAAHLDTTFADRPRNWRPLIYCWRGGKRSGSMTVWFNMIGWRARQLDGG
YKTYRRSIVDALGELPAGFRYVVLAGHTGSGKTRLLHALARAGAQTLDLEALAAHRGSLL
GALPSGEQPSQKAFDTALVSALRGFEHDRPVFIEAESRRIGAITLPLALLAGMHAADCVK
VETARDERVELLLQDYGHLFDQPAFFKIQLEKLVPLHGHARIRKWHAMLDEGRRGELFEE
LVDVHYDPAYTRSSGAHFARLPQAERFAFRPTARDVVEQARALLAHLDAAGARALDKRA*
>GCA_000452945.1_BPMSHR5858_genomic_00691_1 group_599
MVAGVVLWVALAVIFPTVVVPAAQHALERTIKQHDAGASAPGAPRGGTPGQSASPGGGEA
IEP*
>GCA_000452945.1_BPMSHR5858_genomic_00692_1 btuD_17
MPAPGERAPVIALRGVSMSFAKPSGEPLPVLADIDLAVREGEILGLLGRSGSGKSTLLRI
AAGLVKPTSGRVEYRGAPLDGPAAGIGVVFQTFALYPWLTVEENVELGLDAIRVPPDDAH
ARVTSAIELIGLDGFESAYPRELSGGMRQRVGFARALVGDPALLLMDEPFSALDVLTADT
LRTDFLDLWHARQLPIDAVLMVTHNIEEAVLMCDRILVLGAHPGRILAEVPIPLAYPRNR
LDPELQHIVDGIYATLTSRLGDALAAPNGKPGVPAQRLPNVSSHRLDSFVDAFVAHTRDG
CAELPRLASALAMPVGDLLPLAVALHVLEFAELRDGALRLTAAGRVYAQGDAGERKRLFR
EHLEHFVALAAHIRQVLDERAGHQAPRERFELELLDRLNPLDAANTLRVAIDWGRFAGLY
DYDDDARVFRRGEGGARG*
>GCA_000452945.1_BPMSHR5858_genomic_00693_1 cmpB
MMQHPDLLASGRRHLRTLGWRDAIAALLVLALIMLFGSGVRQMALPLDVARPEAISLSPL
ALPGYALRTVLRMLAALGASFVFTFAYATLAAKSRAAQAVLIPLLDVLQSVPILGYLSFT
VAFFLSIAPGTALGAELAAIFAIFTSQAWNMAFSFYQSLRTVPHDLDEASRSLRHTGWQR
FWRLEVPFAMPGLVWNAMLSMSGGWFFVVASEAISVGGLRIALPGIGAYVARAIAERDLA
AVGWAIAAMTIAIALYDQLLFRPMVAWAHKFRCDQTVSGPAPRSWLLDLLHRASLPQRLG
APLGAALHRAARARASLGVRLSTRGRAWAGRIARVAWLALCSAAAVYAGIALLRTTAPAL
SWSDLADVARNGALTLARVLVLIGVATVVWVPAGVLIGLRPRATALVQPVAQFLAAFPAN
LLFPLAVFAIVRFGLAPAIWLSPLMILGTQWYILFNVIAGASAFPDDLKEAAASLRVRTP
TWWREVMLPGILPYYVTGAITASGGAWNASIVSELVVWGHTTLDARGLGAYVARATASGS
YPRIALGVATMSLLVIAMNRLIWRPLYAFAERRSRLD*
>GCA_000452945.1_BPMSHR5858_genomic_00694_1 group_2829
MRTRATRCSRCAAIACAAAARDARSSGRTTRTSATRATERHLRGEWTLGQRVREALGERA
RSIGFTAYTGHVSAASHCDGDVERKWMRPALSAGYEHLFHATRLDRFFLPLREIAAPALH
DARLERAIGVIYPPETERDSHYFMSSITGQFDALFHLDETNPLEPLAPPGARQPRETPVS
AP*
>GCA_000452945.1_BPMSHR5858_genomic_00698_1 group_2278
MNRPLIRIALRPTGTLSFWKGFKWSLIVIVAIAVAIVARLVQIEIETSRLQARFLSELTR
DVGYSVDEGASNRIRFPDNGPYDLRLGYALLPSFQQRLLSRGFVVASQARVSDRMLSLAD
ERLFLPYGEKDQAGLSIVDSTGSPLFGVVYPHHAYVDFDTIPPLVVQSLLFIEDRYLLDP
SQPNRNPAIDWGRFSRALADQGLRFVNRHQATPGGSTLATQLEKFRHSPDGRTATPPEKL
RQIASASVRAYLNGPQTMAARHAIVVHYLNSVPLAARARVGEITGIGDGLAAWYGRDFDE
VNRLLAAPTTPENVAAQGVAFRQVLSLMIAQRAPSFFLNRGYPALQRLTDSYLRLLSTGG
VITPALRDAALAAHVERGAVPAAADTRSFVARKAVTSARAHLLGALGIDNVYQLDRLDLR
ATDTLNNGVQQAVAAGLARAATRDGAREAGLYGFEMLRSGDDPSKIQYSFTLYERRDGAN
LLRVQTDSVDQPFDINQGARLNLGSTAKLRTVVTYLQIVSELHARYANLSAAELAKVKPD
PTDALSRWALDYLAHTPERSLRAMLAAAVERKYSASPGETFYTGGGAQSFTNFDKSDNGK
ILTVHVAFQNSVNLVFVRLMRDIVHYEMIRSSGPSSSWLDDPAQRQRYLMRFVDSESRVY
VKRFYTRYAGKPDDDALAAMLGTVRKSPPRVATVLRSVAPEQPRAWFDAKMRAALHGTPA
AAKLSDDALDKLYTKYAIERFNLNDRGYIAGVHPLALWTLAYLRRHPAASLDDVQKASRD
ARIASYSWLFKTRYHATQDRRIKRMVELRAYDAIGESWRALGYPFDHLTPSYAAAIGASG
DQPAALAKLVGLIANHGEQVPNERISSLEFAHGTPYETRFVRAAAQPRPLLSPEIADQVH
MLLGDVVQKGTGRRLAQGLAFPDGQTLPVYGKTGTGDQRFNVYARGARLIESRKVNRSAT
FAFSLGDRFFGVLTAYAHEPYAARYDFTSAMAVQLLKSLAPALQPLMAAPAQPSANTGTA
PGADETNTDGASAQTAERG*
>GCA_000452945.1_BPMSHR5858_genomic_00702_1 ctaC
MNGANMSVKPAQRPERGFAKTRRRTAAYRPAATRAAGVARAAGAATPPLPAAPHAAAAHA
QAHAAQPAALPLAYVFDSAGPAPRPVLILGWALLALCTSVCLVIAVLPALALFLGRPASV
GLTERGGLGFVYVGSAISTALLLAALVYMLWVLAAVAKPPRPPAVTIAVTAYDWWWKADY
GGGPPDGFTTANELHVPVGEPVLIELRSADVIHAFWAPQLAGKTQAIPGQINRQWMQADR
PGVYRGQCTQFCGAQHAQMGFEIVAEPPDAYRRWYASQRRGAEAPRTADALRGQRIFADR
CAGCHAVRGTGAAGTQAPDLTHVGARRLLAAGALANTPDELRRWIADAQQVKPQSLMPSI
RLDPAQQRDLSAYLATLR*
>GCA_000452945.1_BPMSHR5858_genomic_00703_1 ydaP
MATVADFIVERLYDWGVRRLYGYPGDGINGMFGALNRADGKIEFVQVRHEEMAAFMACAH
AKFTGELGVCIATSGPGASHLVTGLYDARMDHMPVLAIAGQQARAALGGHYQQELDLSAL
FKDVAGAFVQTASVPAQARHLVDRAVRTALANRTVVALVLPNDLQELDYEPPKRRHGTVH
SGIGYTDPQIVPYEADLRRAADVLNAGSKVAMLVGAGALRATDEVIAVADRLGAGVAKAL
LGKAAVPDELPWVTGSIGLLGTEPSYRMMSECDTLLVVGSGFPYSEFLPKEGEARGVQID
LKADMLSLRYPMEVNLVGDSAQTLRRLLPLLNERADASWRAQIARWNAAWWNTLDARARV
NPGNGRGVNPQRAFTELSPRLPEYAIVTSDSGSCANWYARDLRMRRGMMCSLSGGLASMG
AAVPYAIAAKFAHPHRPVIALVGDGAMQMNNMAELITVAKYWKQWADPRWICMVLNNEDL
NQVTWEQRVMEGDPKFDASQQVPDVPYHRFAELIGLKGVYVDDTDQLGAAWEDVLAANRP
AVLEVKCDPEVPPLPPHVTLQQAKHFAQALMRGDPRERSVIVETAKQVLSGAFSGRADRT
RDEPDGDA*
>GCA_000452945.1_BPMSHR5858_genomic_00704_1 group_2279
MKPFSEPIIRCAKARAYRIPTDRPEADGTFAWTDTTIVVATLDAGDAYGIGYTYTDASAA
ALIERMLAPALDGRSAFDIGGALAALRRAVRNVGRGGVAATAISALDAALWDAKAKLLGL
PLAKLLGGVRDAVPVYGSGGFTSYDEHALRAQLEGWRAQGLHACKIKVGSAPRDDPARVR
IARDALGGGVALFVDANGAFAPRDALAFAQRIAECDVRWFEEPVSADDIDGLRFVRERLA
AGVDVAAGEYGYTLDDFRVLLASGALDVLQADATRCGGVSGFLEAATLADAFHRDLSAHC
APALHRHVCCAAPRVRHVEWFHDHARIERMLFDGAPTPRDGAIAPDMSRPGLGIELKAQD
AARFAV*
>GCA_000452945.1_BPMSHR5858_genomic_00706_1 mpdB
MKTRTNRPGGGGLARGRYAGYDVLAKRDTPSWNDATRRVLDARLAPREAPRYLDARQFAT
LDAVCARIVAQSEPRDARQADAEADAAAEAEVETATEAAAAAETAANEGASTRAPIPVAA
LIDERLLHDEGDGYRDARMPSYREAWRIGLAALDAHARAEHGRPFASLAGAERDAILRAA
QRGALDGPAWRGMSSALFFSKRVLSDVASAYYSHPLAWNEIGFGGPASPRGYVRMDFNRR
DPWEAPLAARDDGRADADADADADAASPRRLAQPGKRKGNRDAD*
>GCA_000452945.1_BPMSHR5858_genomic_00708_1 group_2280
MNDLLPHYEREIALLRRSMREFARRHPKIATRLGIADGQADDMHVERLIQSFALVGAQIG
AALDDEYPEFTEALVETVCPDYLRPFPACSIAQFDAPPLFGQPAEGVTIARGTTLESRIG
ANRFRTAYDVTLAPVSIVDARFRPASAAPSAARMPPQSTGIVSITFDALTAQPVLAALRA
RGLRVHLHGDAPLVAALADTLAMRAPAAFIELDGDGRWKPLSKVPLARVGFADAHALLDP
APGAAPFRLLMEYFAFPAKFDFVDIDVARIARAAGPCRRLSLHLPVVDVAMASQHARALD
TLGASNLRLFCTPIVNLFSQDAMPISLRDAEAVYPVTPQALKSNGIEVRSIDAVRIAREG
EPGASLDVTPYRSLLHGRRGGRDAPVYWVAQRERFAPGSPPAALRLVDADGANARLGADQ
LNVELTCTNGDYPQTLPIGEPDGDLLNEKDNLPGRIALLRRPTPARRFAREHGALWRMIA
AMTPHALLLQPSGLGALKALLVQHAARASSAAPQIDAIANLDHKVAVRWMAVKPMPTFVR
GIEIALTLNEAAFVTSGLKTFIDVMDGFFALHAPPNGFVQLVAYSRDTGRELHRCAPRGA
IAQLV*
>GCA_000452945.1_BPMSHR5858_genomic_00710_1 quiP_3
MTDTDERRMPPNRPAARVAGDLAGGRRAGATHGLAQTMASTRNRTRRHLKILSGLILCCA
LLALVLAAAYLRASLPQLDGELRAPVLGAPMTAERDALGVPTITARDRFDAAYGIGYLHA
QDRFFQMDMLRRSGAGELSELFGPIALGVDRDHRLYRFRALAAAAVAHLPPDDLRLLQRY
TQGVNDGLAALRARPFEYALLRMQPQRWRPEDSLLVIWAMYFDLQGRLASREFARHWLQA
HSTPEQLAFLLPPSSRFDAPLDVPGIAAAAAPVSATAPAWLRDARPATSVNGVSIDGQSA
VGSNNWALAGSRTASGAAIVGDDMHLSFGLPNIWYRAMFKLGGGAGLAARVVGVTLPGLP
AVVVGSNGRIAWGFTNGYADCLQLLPLERDARDPSRFRLGGAWQRARETVETIRVHGAAP
VELTVLDTTAGPVREIGGKAYAAHWIAQASGAANLAFAKMIEATGVEEAIGIAAGAGLPA
QNVVIGDSTGRIGWTVAGALPDRRAPGEAGARGDGAPSWRAALPPDAHPRIVDPAGGQLW
TANNRQLAGDAYRLIGDGGADLGARARQLRDGLTALERPDERAAYRLDLDDRALFMSLWR
DRALRVLDEAALAGHPARAEFRRLLASGWTGRASVDSVGYTLARGYLYRLYDVLFGGLDA
RLKQQQHDADYALANRRWPVVIARLLDEQPPGWLPADAASWRDVQLAAIDRTIDALSANG
APLADAAWGRRNTLRIAHPFARSVPLVGRWLTAPADPVPGDANMPRVAAPRFGQSERLVV
SPGREESGLFNMPGGQSGHPLSPFFLAGHEAWVKGEPLPLLPGPTEHTLRFVR*
>GCA_000452945.1_BPMSHR5858_genomic_00711_1 pgrR_8
MRADDFGAIDIGADVDTVADTDTDTDTDTDTDTDTDTDTDTDIDIDIDIDIEASVGGRPT
RVSPTIRRKRIVAKRLESRPSRALSCPVGDSKGDGMDRIQAMEVFTRVVDANSFTRAAET
LGMPRASVTTIIQNLEALLGVRLMHRTTRRLSLTPEGAAYYEHCIKIITEIAETDASFQV
GSRRPSGALRVHMPSSLGRRLVIPALPVFRQRYPDIVLDLGLSDRPVDPVEEGIDCMIRI
GPLEDSSMVARRIGMLKRVTCASPDYLKRFGEPAEIAELAAHHAVNFRASHGSRPVPWVF
VIDGKPVEVRLNGGVTVNDSDAYVTCGVEGFGMIQPTLFMVLPHLLDGTLQEVLPGFNPK
PKPISIVYPNSRHLSAKVRVFADWIAELFDSTPALEGGEDWRGSAAARTAPASRGQVAA*
>GCA_000452945.1_BPMSHR5858_genomic_00712_1 group_4486
MSRVEWRNEKIIVALGSLWILGFAAWAFLLYDLLGTSVKEGILEGPREGVFWTAAQYRNS
FSRFDRQLILYAAGENRDFDAVLLQLDSLEASFGFLERPSEVSAYWLSIPKARGDIAELS
RFMASLRRDVPALRARPDDTKRVLGELARQWPKVNALANYFRAIEMEQRDFTFHQLKEKR
RAIVMLGGVLGIILGALFLLLFYTVRTRGSLLEQQQAALDAERKASDRAFEMIAAKNAFL
GMVSHELRTPLQAICGSIEVLLARPQSEANMKTIKRLQNSAASLEAQVKDLTDYIKLRST
NRSVQSEIVEVAPLLADVLDPLRGRIRDKHLSTSLRVEPPALVVRSDRKLIQQIVSNLVE
NAIKYTNSGTIEISAALGGTSANRTMTITVRDTGVGIARHLLAKIFEPFFRVNDPGVRHV
DGIGMGLAVVQELVVALRGHVDVRSAVGEGSEFAVTLPVELPGRADALDDDAQPSPRAAH
RDLRALVVDDNENARETLGAMLATLGIRVDLRGTGKEGLRCFGECQHDIVVLDLELPDIS
GFEVAEQIRWATSSDAARKTTILGVSAYESALLKGDHAIFDAFIPKPIHLDTLGGIVSRL
RS*
>GCA_000452945.1_BPMSHR5858_genomic_00714_1 group_4487
MTQPDSANADLITATRVRDLLTRTGIAPRSHNTTIANILGLSFSVVTRKMKGQIPWNLSQ
LQDIASYFGVPPAILLDDKGVPPHPGVDMHDATLVIEARRFRCRAAIATKASSQADSDFV
ALQWQDSWVVYEKKHAPEGRTYAVDMIELRSSESSAYEARIAVVDDSPDVAETICEYFAE
KGVAAIAYYDSVSFRKALEVEDFDGYILDWLLGEETAAPLVRGIRASENADAPIFLLTGK
ISTGEASEDEIADIVSSFNARCEEKPVRLPILFAEVARALKIAYPPNAQAS*
>GCA_000452945.1_BPMSHR5858_genomic_00727_1 group_3129
VNHPAPPPRRPLPAAFVDALAARFGARMSVAQAVRAHHGRDESPFDPQLPDAVVFARSAD
DVRDAVLLCARYDVPLIPYGVGSSLEGHLLAVRGGVSLDLSEMNRVLSIDADDLTATVEP
GLTRRALNDALRDTGLFFPIDPGADASIGGMTATRASGTNAVRYGTMRENVLGLAAVLAD
GRVVKTGTRARKSSAGYDLTRLFVGSEGTLGVITEITLRLHPLPEAVSAAVCAFGSMSAA
ARTVIETIQIGVPIARVEFVDALAVRAINRNTQLGLAEAPTLFFEFHGTESGVREQAQQV
EALAEQNGGRGFEWATRPEDRTRLWSARHHAYFSMLQLKPGCRAVTTDVCVPISQLAACV
EETEADLNASSLPCPIVGHVGDGNFHVAILVDPTKPEELAEAEALNDRIVERALRMGGTC
TGEHGVGLHKMRFLAAEHGENAVDTMRAIKRALDPRNLMNPGKIFTIDA*
>GCA_000452945.1_BPMSHR5858_genomic_00745_1 phnN
MSAERLVYVMGPSGAGKDSLLAYARKHVREPRIAFAHRYITRKSDGHENHVELTRDEFAA
RAQLGFFALEWSSHGFRYGVGVEIDAWLAAGSVVVVSGSRAHLPAALERYPQMCVVHIDA

APHVLAERLATRGRETADEIRARLARSVRWAVPDGIALTAIDNSGTLDDAGRVLVALLEG
LARS*
>GCA_000452945.1_BPMSHR5858_genomic_00750_1 katG
MSNEAKCPFHQAAGNGTSNRDWWPNQLDLSILHRHSSLSDPMGKDFNYAQAFEKLDLAAV
KRDLHALMTTSQDWWPADFGHYGGLFIRMAWHSAGTYRTADGRGGAGEGQQRFAPLNSWP
DNANLDKARRLLWPIKQKYGRAISWADLLILTGNVALESMGFKTFGFAGGRADTWEPEDV
YWGSEKIWLELSGGPNSRYSGDRQLENPLAAVQMGLIYVNPEGPDGNPDPVAAARDIRDT
FARMAMNDEETVALIAGGHTFGKTHGAGPASNVGAEPEAAGIEAQGLGWKSAYRTGKGAD
AITSGLEVTWTTTPTQWSHNFFENLFGYEWELTKSPAGAHQWVAKGADAVIPDAFDPSKK
HRPTMLTTDLSLRFDPAYEKISRRFHENPEQFADAFARAWFKLTHRDMGPRARYLGPEVP
AEVLLWQDPIPAVDHPLIDAADAAELKAKVLASGLTVSQLVSTAWAAASTFRGSDKRGGA
NGARIRLAPQKDWEANQPEQLAAVLETLEAIRTAFNGAQRGGKQVSLADLIVLAGCAGVE
QAAKNAGHAVTVPFAPGRADASQEQTDVESMAVLEPVADGFRNYLKGKYRVPAEVLLVDK
AQLLTLSAPEMTVLLGGLRVLGANVGQSRHGVFTAREQALTNDFFVNLLDMGTEWKPTAA
DADVFEGRDRATGELKWTGTRVDLVFGSHSQLRALAEVYGSADAQEKFVRDFVAVWNKVM
NLDRFDLA*
>GCA_000452945.1_BPMSHR5858_genomic_00751_1 mfd_1
MMQPALSRARRNIAPRPAVIDRAKAAIRSLRPIAFAYLADRAHGG*
>GCA_000452945.1_BPMSHR5858_genomic_00762_1 group_980
MFRDTTLSSRDYEEMLARESRLVDLLSTKSPTMARANWRVTGDTLEEANSYPAFEADGAP
SRPALAVLTERGRGNKHGVSHAAIWNAATSAEEGASISCHVSDAKVLPDTLSVSLRVPGC
YATADDFADVIKAIVAAFHPAVIEASPDGYFEKQVFDDKPGVGWMLYLPKIITQQQVPEA
HALIPVPEAGKRQTGTIIVSVTDAPFSVDNPEHVAIANRIEIRLVDQDLLPAYVDI*
>GCA_000452945.1_BPMSHR5858_genomic_00768_1 ompR_2
MPLMETKNPSKILVVDDDPRLRDLLRRYLGEQGFNVYVAENATAMNKLWVRERFDLLVLD
LMLPGEDGLSICRRLRGSNDRTPIIMLTAKGEDVDRIVGLEMGADDYLPKPFNPRELVAR
IHAVLRRQAPAELPGAPSETTEVFEFGEFSLNLATRTLTKSGQEIPLTTGEFSVLKVFAR
HPRQPLSREKLMELARGREYEVFDRSLDVQISRLRKLIEPDPGSPRFIQTVWGLGYVFIP
DGAA*
>GCA_000452945.1_BPMSHR5858_genomic_00774_1 mfd_2
MKSKDGFYNTSFAAMRRAELVRHPAFANQPLVMSDNASPSQSPVALVKAGQRFAFDGAHG
SADALVIARYLAAHRAQVPLLAVICANAVDAQRLAQELAYFAPDARVRVLPDWETLPYDT
FSPHQDLVSERLATLHDLGEGRCDILLVPATTALYRMPPASFLAAYTFSFTQGERLDEAK
LKAQLTLAGYEHVSQVVRPGEYCVRGSLIDLFPMGSPLPYRIDLFDDQVDSIRAFDPDTQ
RSLYPVRDVRLLPGREFPFDEAARTAFRSRWRETFEGDPSRAPIYKDIGNGVPSAGIEYY
LPLFFDETATLFHYLPERAQLVFTGDLDAAIKRFTADTKQRHAFLSHDRERPILEPQRLF
LSDDDFYLLAKPFARIVLPAQPSGGWAAPLPNLALERHADAPLAAFAAYLETTKNRVLFT
VESAGRRETIAQLFAEHHLRPAGSDSFAAWLASDERFALGVAPLANGFAVPGEGYAIVTE
TELYGALGRRAGRRRQEQASNVDAMVRDLSELKVGDPVVHAQHGIGRYMGLVSMDLGEGD
TEFLHLEYAGDSKLYVPVAQLHVISRYSGADPDSAPLHALGSGQWERAKRRAAQQIRDTA
AELLNLYARRAAREGHAFGLDPRDYVKFADSFGFEETPDQAAAIAAVIGDMTSGKPMDRL
VCGDVGFGKTEVALRAAFIAVMGGKQVALLSPTTLLAEQHTQTFADRFADWPVRIVELSR
FKTAKEVNAAIAQINEGSVDIVIGTHKLLSSDVQFKRLGLVIIDEEHRFGVRQKEALKAL
RAEVDVLTLTATPIPRTLGMALEGLRDFSVIATAPQKRLAIKTFVRREEESVIREAMLRE
LKRGGQVYFLHNEVETIENRKAMLEELVPEARIVIAHGQMHERELERVMRDFVAQRANVL
LCTTIIETGIDVPSANTIIMHRADKFGLAQLHQLRGRVGRSHHQAYAYLLVHDPQALTKQ
AQRRLEAIQQMEELGSGFYLAMHDLEIRGTGEVLGDKQSGEIHEIGFQLYTEMLNDAVKA
LKNGKEPDLTAPLAATTEINLHAPAILPADYCGDVQERLSLYKRLANCEHGDAIDGIQEE
LIDRFGKLPPQAHALVETHRLRLAAKPLGIVKIDASEAAIGLQFVPNPPIDPMRIIDMVQ
KHRHIKLAGQDKLRIETRTPDLAVRVSTVKETLRALGQPQQPRAQAAAR*
>GCA_000452945.1_BPMSHR5858_genomic_00788_1 ecfG_1
VSLESDVVGWLPQLRRYARALTGDPAWADDLVQDTVERAFSRMRAFRPDSNLRAWLLTIL
RHLYIDQLRARREFAVDDDEGPWRTLEAPAGQIDALVLRDVQRMLYRLPLEQREVLLLVC
VEELSYQDASAVLGVPVGTVMSRLSRARERMRALLTDAPLRPPLSLTAARHHR*
>GCA_000452945.1_BPMSHR5858_genomic_00792_1 fecI
MPANTLRLQREIAALYTGHRGWLHGWLSKKLGCGERAADLVHDTFIRLLARDEPIDAAEP
RAFLTTIAQRVLYNHWRRERLERAYLDALAQWPEAVAPSPETRAVLFETLVEVDRMLDGL
PAIVRRAFLLAQLDGMTHAQIAAELRVSLATVKRYLVKAGAQCYFAIAA*
>GCA_000452945.1_BPMSHR5858_genomic_00794_1 btuB_4
MDSICNRRPRRAHAGVCADTANAPAGLPEAAQPAARLGAGRPRDAAAPRCARQATSHTSH
ASHTLHAPCMPQFAPRRVVARVAAAALFSACAAPGVSGAAEPAAATRGYDIPAGPLDAAL
TRFGREAGILLSFPGELTTGLRSPGLHGRADPAAALDRLLTGTGLVALRQPSGGYTLARL
PGPAAAGADAALAADTTLPTVAVRASGPHADSYRPPREAAGLRSDAPLAEVPQAVAIVAQ
QVLRDQRPRNLDDALANVSGITQGNTLGSTQDTVMKRGFGDNRDGSVMRNGMPIVQGRSL
NATTDSVEVLKGPASLLYGIMDPGGVINVVTKQPQLARRHAISALGSTYGGGRNGGELTF
DSTGAIGESRVAYRLIVDQTNEQYWRNYGEHRETLVAPSLAWYGRDTQVVLSYEYRRFLM
PFDRGTALDPRTNAPLAIPARRRLDEPFNDMRGESNLAQLAIDHQLAPDWKVHVGYSYNR
ETYDANQIRITAVDPLKGTLTRSNDATHGSRSTDSYGIAYVDGRVTLAGMRHDVQFGVDG
EYRQVYRADMLRQPIKTPFSYLNPTYGLVPPSTSVSASDSDQSDTLHTASLFFQDSIHLS
ERWMLVGGARWVRYSQLAGRGRPFQVNTNLSGTKWLPRAGIVYKWNDALSLYGSYTQSLK
PTSKIAPMAGGYVIDGSTAPEEGASWELGAKLDMPGGLAGTLAFFDIDKKHVLVSQYDDA
TNQTAWRTSGRARSRGIELDVSGRIGARWNVIASYAYIDAKTTEDPLYAGNRLWNVARHT
ASLAAVYDVGTVLGGDDLRIGAAGRYVGARPGDSANSFTLPAYATADAFATYDTRLGKQK
LQFQLNVKNLFNRTYYPSSVNRFFVSVGDARQVSLLTTLQF*
>GCA_000452945.1_BPMSHR5858_genomic_00795_1 rbsC_6
MRDATDRLSRFAALGALAAVLAFFAVAAPGFATLANVEHVLVNHFALLAIAALGMTLALS
IGAIDLSLGTAIDVASLAFVCALAHRAGCASAALAGLGAALALGAANALLVTRLGIAPFL
ATLGTLFIGQTIQQLATDGGQPIYLLSTAPPPAFDALAHGVWLGVAAPLWIVAALAGALH
VALERSTLGRQARALGAQPGVARHSGLRVGALAAGVFVASALVYGVAGLVLSSTVRSYAP
QAGNAYLLNAIGATFIGATLSEARRPNVAGTLLGVLLVSFVANGLLLIGWNFYWQQVASG
ALVFAVLALGSGVGRERRARDAGQSG*
>GCA_000452945.1_BPMSHR5858_genomic_00803_1 group_1778
MKAPIIAIAFGIALAACAPLRQDSRSGEAGPPIAAPNRGGGNTPPSGARRAANEVVDYAG
RPCVNQTLYVKTHPCIFPRLQR*
>GCA_000452945.1_BPMSHR5858_genomic_00806_1 czcR
MRILIVEDEPKTGMYLRKGLTEAGFIADWVEDGVTGLHQAETEEYDLIILDVMLPGHDGW
TVLERLRRAHSTPVLFLTARDDVGDRVKGLELGADDYVVKPFDFVELVARVRSILRRGQA
RESTVLRIADLELDLTRRKATRQGDVVLLTAKEFALLWLLMRREGEILPRATIASQVWDM
NFNSDTNVVDAAIRRLRSKIDDAYEPKLIHTVRGMGYVLEVRSASAPSR*
>GCA_000452945.1_BPMSHR5858_genomic_00809_1 czcC
MPKRIFAALVSAAIAASAVQAQAQPAPAPSAPDGPAAAPPDARAASVSVTPAVSPAPIST
STVSVAPPALSLDDALALAAANNPLLKSARSSADASVGALVQAGARPNPTVSFLQEGFGR
QERTTTAVVSQPIELGGKRRSRVDAASYGRESALASLDGRAAALRADVTGAFYGVLAAQR
QLQVADESAAIAARSADMADKRARAGKVSPVEATKARVAQTAAQIELSNAQTRVAAALAR
LANATGSADVRGRAASGDLDALPAAEPLPQLLGRLDDAPLARVAKADMLRTNAQIAIERA
KRVPDITVSAGVKHVVTGGAPDNQAVVGISIPLPLFDTNRGALLEAVHNAAKASADFDAE
RARLRLDLTQAYANYERATGEARRLKADVLPAARQSLDAMSRGYQLGKFGFLDVLDAQRT
LFDAQSRYVQALADAHQSYADIERVVGAPSRDASRPLAQLR*
>GCA_000452945.1_BPMSHR5858_genomic_00811_1 group_4500
MSLAAQDLIELLRLAQGGAGALTLHLLREIAGNLIAFGVADCRGGC*
>GCA_000452945.1_BPMSHR5858_genomic_00813_1 group_3139
MSMSVTLQECLRQKASRYEVIHHPYSHSNMEAAAAAHVPGDRLAKTVLLEDAQGYVAAVL
PTTHAVRLSELWAKTGRHLVLAKEVELRELFKDCDVGALPPVCMAYGMQTFLDDSLARQP
DVYFEAGDHEELVHMDRDEFLALMDKAERASFSHKIQGVVS*
>GCA_000452945.1_BPMSHR5858_genomic_00814_1 ald_2
MLIGVPKEIKVYEYRVGLTPAGARELALHGHQVLVQRGAGAAIGLQDDDYAAAGAMLADD
AADVYARADMIVKVKEPQPAECAMLRRGQILFAYLHLAPDPDQAAALVKSRAVCIAYETV
RAPGGGLPLLAPMSEVAGRMSIQVAAAHLESPGGGMGLLMAGVPGVPAAHVVVIGAGVVG
TCALQMAAGLGARVTVLDTNVDRLRELDLVFGNRIATLYSNTHAIDGAVRGADVVIGAVL
VPGASAPRLVTREMISKMRAGAVVVDVAIDQGGCIETSHPTTHAQPTYVVDGVVHYCVAN
MPGAVARTSTFALGNATLRHAIALANKGWKRALADDPYLRAGLNVCDGHVTHEAVARALG
LAYVHADEMLD*
>GCA_000452945.1_BPMSHR5858_genomic_00816_1 group_3140
VTSSKIQTLQKELGHLNLLLEEAKKKEKAEALAAIREQVKQFDITEVELLRAAGFVKDKP
KKLPAKYYDPSTGKSWTGRGACPKWLIGKNLDDYLIRPAPEPWWPGESA*
>GCA_000452945.1_BPMSHR5858_genomic_00818_1 mhpD
MTSLTERIDAAARHLVAARRSRVAGPLLPAACRPDDIEAALAIQRRVAERLGEPIGGWKC
ALPPPGRIVVAPILASSIRAAGGPFPVVADARAVRIEPEIAFVLDRDLPPRARPYDEGEV
RAAVREVRLVLEVLGCRYADPSRATVPELLADNQFNQGLCVGPIVSGGLRASLETIALEC
RGGVNRMLDGRHPDGHPLEPLRWLVNFLAARGESVRAGQIVTTGSYAGAIDAPLGQALTV
RFGELGGLAVEWVA*
>GCA_000452945.1_BPMSHR5858_genomic_00851_1 group_19960
MVQNRVDEIVQAIASETHTPAEAVSRLYEETLAEYSEGARVRDYLTVLVAKRVRETLRNR
TH*
>GCA_000452945.1_BPMSHR5858_genomic_00854_1 cheW_1
MNAILEAPHAARMSGQDAARDAQEFVTFRLGTEEYGIDILRVQEIRSYEEPTRIANAPAF
IKGVINLRGVIVPIIDLRLKFALDSAEYNTSTVVIVLNVAARTVGVVVDAVSDVLELAAA
DRRPAPEFGAAIDTGFITDLGSIAGENGNRMLILLDIERLIAAADVGLVG*
>GCA_000452945.1_BPMSHR5858_genomic_00859_1 group_930
MHVSTDTSAEAHSPAPNSLDMSDSDALRRIAEAVDGGAANIERIVPLARARERMPTRPRL
ERCGGGRVTAAHITLDSRARLDALLHALQRAIDQNADLRTCILGACLRRPMQVTLREVRL
RVHAATLDPDLDPAAQLAALSTGPGMRIDMQRPPWVLACIARIPGSGQWLLRLVAAPIAA
GFDALDALLRETVIHGDREPGPAPFHWTVETAVESCGGEPASLPTAGAVWPSNDVSRACD
PDAASCVEARIAAIASDLPGVVHGGPRDDLRALGRTPLQALRLARRIRDALGVTVPVESI
LASPTIVELAGYVEQLRSRDVRDGAAPVSIGEKPADADARAQAQADTDTAHTDCLIVIQA
GGAEQAPVFCIPGAGGSVASFVALASMLRADIPVYGLQPRGLDGLGPPDRSVEAAARRYA
RAILDAAPPGPPRIVGHSFGGWIALETARLLDGMGARCAPLVLLDSNPPPASQAWRAPSE
ADMLRTLVGLLEQAAGGAPSGIGDEEIARCAAAGEDARDALVHACMVRTALLPPRAPVEA
VRHLRRVFEAHSSTRYAPGGRYAGDATVIVANGDRDAGEMVPAFGWAALIERVEVAVTPG
NHMSMLAAPYVRHVALTMKTVWRMI*
>GCA_000452945.1_BPMSHR5858_genomic_00864_1 arnA_1
MNDMTREKVLLCIDNVAGLAVLKYLAGRPSVEVVGAIVHPQANATHLDEIVEVCARNGIA
PIDILEARARFDELIAPLAPDFIVSIYFDYILDDRFIELPAKDSINLHPGYLPYNKGFYY
YAWAVLDGTPAGVSIHRIVSAVDAGPIISQKRVLIDGTDTGDVIYDKHMDASVELFRTTW
PSIESGTYQLFRQRHKGTRKKITETNEAMEIDPNATYVARELIDLLRVFCFRGSGGCVMK
IDGKKYSIGIEFSELADNVNQIRGKRDNYLVG*
>GCA_000452945.1_BPMSHR5858_genomic_00865_1 argG_2
MKPKHILLAYSGGLDTSTALHFLKRHFDCRVTAYCANLGQKEDWERMKRRAAIAGADELV
IEDLRETFIGDFVFPALKANASYERDYLLGTPLARPAIVKGLIEYARKHDVDCLSHGCTQ
KGNDQVRFEMAAKILAPDLPTVAPWRIWSLQSREDLFAYCQQHGIPVESRPDNLLSHDEN
LVHITTEGDYLESVANAFDWRDANWITPPMQAPDAIETITLGFRRGVPVSVDGAALGPVE
LVERLNEVGARNGVGFQDIIENRINGLKVRGVFENPALTILHAAHRKLEKITLGRDVERL
RNLVSDDYGDIVYRGLWFSDERLCLQALIDESQKYVSGDVKVQLYKGSCTPCAVESEQSL
YSRELVTLHAGRAISGEDATGFLNTLGLRIGIEAARAGNTGAGA*
>GCA_000452945.1_BPMSHR5858_genomic_00866_1 argH_2
MSNPMANPWAGRFSQTMTDSLVAFNTSLPLETRLFEADIDGTAAHVEMLHATGLLETAEH
EALARALDEIRAAWRAGEIRLSPALEDIHMNLETLLVDKLGELGKKTHTARSRNDQQASA
QRLYFMRSTRELVDAIDALQRAILEHGERHDALVMPSYTHLQRAEFTYYAHWLATYVVML
ERDRSRFVDALARADQCPLGACASTGTSLPIDRRRSASRLGFREPTLHSIDSVSDRDYLV
EFCSHAANLMIHLSRLSEELISFTSQEFGFIALADGYCTGSSIMPQKKNPDVPELVRGKA
ASVIGNAMSLMALLKALPLGYNKDLQEDKTAWFAALDNCMSSLAILTELVRTMAPVPDRM
RQATLGGHIIATEYANYLVRKGMPFREAHRVVGELVKTADARGVDVSALPQAAFAEASPL
FGDDIGRVTVEDMVSRKNSYGSCGDQALGELLAQLRSMLDAHRPGR*
>GCA_000452945.1_BPMSHR5858_genomic_00867_1 pduX
MSEPMRSLIPAARTASGISYCSFGELLQGVLLEDDADFLVTLPIQKYSVSTFVPDPGSTE
IVTQPSGKTKAAALAKVILGRYGHNVGGTFYIDCDIPIGKGLSSSSADLLATARAIEVYM
GRELPLGELCRDMSGIEPTDGVMFAESVVYLQRKGVLWSRLGRLSGIQILSLDEGGTIDT
LEYHRRARAHRHNAEHRSEFNELLERIVAAFGARDLDEIGRVSTRSAYINQKINPKRHLA
SVHDVCQATRGLGLVTAHSGTCIGILYETGRAGHRENLARAADALSGYGNIKVYDTIQ*
>GCA_000452945.1_BPMSHR5858_genomic_00870_1 cysK2_1
MNDIMTQTFDRPAIPVYDSVSGSLDHPDLIRLAPGLVAAAFRLMKLVPAKYIIENAIASG
QLNPGMPVLETSSGTFAMGIGIVCAEKRIPFHIVSDAAIDERLQARLRQLGGRVQIVGAN
ATGSNVQVLRLEALQERLRENPGGFWPRQYDNPDNQHAYRAFAAQLIRTFGTNLTIVGTV
GSGASTCGTIRALREVDPSIPLVGVDTFGSVLFGLPVGPRALRGLGNSIYPNNLDHTCFD
QVHWVAPDAAFGSTRRLHRQHGLYCGPTSGAAFMVAEWLRAQRDDGRTIVFIAPDEGHRY
ADTVYDDAWLRGQGYAGADAAPAAAPVRAVSPNAASGPWAYVEWGRRTFEQASGRPRPAG
SALEQIRDVRPAPVA*
>GCA_000452945.1_BPMSHR5858_genomic_00871_1 ltaA
VKTFVFIESNTTGTGRLCLQKALLRGFDVLFVTSRPQLYPFLQEEMVVPLVADTADPQRI
ADALAPYAGIAGIFSTSEYYIETAATVATRLGLPAADPEAIRTCRDKGRLHRRLRDAGVG
VADTEIVSERTQLRDLAHGATYPRVLKPAFGSGSVGVRLVRTPAEMLAHGERMLDARGNE
RGIALARQVLVQSFVDGPEFSVEVVGLGAEHGHAVLGVTGKHLGPLPHFVEAGHDFPAPI
AAAQRDAIVAETLRALDAVGHRFGPAHVECRVSGGKVVVIEINPRLAGGMIPQAIEWATG
VDVLGAMIDLHAGTPPDLGPRRRGHAAIRFVLPARSGELRALSFEPDERFAGVRTRFMPL
KQLGQRIEPAGDFRDRLALVIASAADPDALAHALEDVDRCVTVAIGDAGAAGEGAGAGRL
RRTLHPEALAIVRKPAPRAERLAELDAFAAIDEAHLLMLVDAGICDRTRAATVLAELARQ
RDAKFAAIADAIAPRGTYALYEQLLIERVGIDAGGAVHTARSRNDINACVAKLRAREWFD
TCGGKLWRVRAAIVDKAQHTLDWPLPTYSQYQAAQPGSFGYYLWSVETALRRDQAALERL
DEELAVCPLGAGAGAGTDFPIRPGVSAALLGFARSFDSALDAVASRDLVLHFLAAIAIAS
TTLSRLAHDLQLWTMRETDFLALPDELSGGSSLMPQKKNPYLLEIVKGKLAHVAGALNAA
VFASQRTPFSNSVEIGTEMLAPCADAVQAFGESCDLLRLMVSGVTGDPAKMRAAAEAGLV
SATQVANALVRETDISFHAAHRQIGALITQALDAHEDPAAALDALVRQPGASIDEAAARL
AYGGGPGAAGAGLARSRALLRQSAERLWRRRAAWHAAHARRRGCVADLLAAAAA*
>GCA_000452945.1_BPMSHR5858_genomic_00873_1 group_7295
MMSAFSLTHAHLDFGHEGTRMTTFPISTARLVLRPFKRGDLAAFTAYRNHPDVARYQSWA
SYTDADAKAFFAEQQTLRFDTDGTWFQIAAEHQANGALVGDVAVHFFDEGRQAELGVTFD
VAAQRQGFALEAVSRVIELLFADLGKHRIVATVDALNVRAQRLLERQGFRREGHYRENIF
FKGAWGDEYGYALLNREWRALRQAAAGGTQ*
>GCA_000452945.1_BPMSHR5858_genomic_00878_1 group_4513
MSAKLSFHTALLVGAVASLVASGAHAAPLTKAEADAAVAAHKEKCFGVALKGQNDCAAGP
GTTCQGTSTVDFQGNSWKFVQGGTCTSIQVPGGGHGSLTPLKS*
>GCA_000452945.1_BPMSHR5858_genomic_00880_1 group_7296
MRSDAAASDYACAFAPGLTNPSIGAPDDAIAHGHKRVARRYDVYRNNVTVSLIEALAAIY
PAVQRIVGVEFFRAMARFHVRETLPVSPLLFEYGRDFPDFIAGYEYARTMPWLADTARIE
RAWLDAYHAADLSTLTADAFTAIPPDALAAVRLVPHPATRIVRSPYPAVSIFAMNRSDGP
VTSLHSAEAEDALVTRPAHDVIVSRLPLGGAVFLISLIDGASLGDAVAITLEDNPAFDLP
ASLRGMIEAGVFTAIHSGA*
>GCA_000452945.1_BPMSHR5858_genomic_00881_1 group_3146
MSDRHTANRAPLRALTSLVSRVRHGIQWLAHPTLAQFALRLALAVPFWKSGILKWNGFLR
LNDTAIDLFTSEFQLHLPGGPYPFPAPAVFAFLSGCGEVLFPVLLALGLGTRFAAAGLFV
MTCVVELTVPDGWPIHLTWAAMALGIAAWGPGRLSLDYLLGDRAPRS*
>GCA_000452945.1_BPMSHR5858_genomic_00885_1 sigF
LRALFVSGLAGHAPDYRSFLSELTRHLRGFLRKRIPQFDADIEDLVQEILLAVHNARHTY
RADEPLTAWVHAIARYKLMDFFRTRARREALHDPLDDHTDIFSEPDDDASDAHRDIGRLL
DHLPDKQRLSILHVKLQGLSVAEAAQLIGASESVVKVSIHRGLKALAARVRGVV*
>GCA_000452945.1_BPMSHR5858_genomic_00898_1 group_982
MKRYVLAVLAMVFLAGCGGGNGSDESGSAPANTGSAPASTIPGAGQYPPAASADPTYIGT
YAAVAGNNVVPVYFESGNSPTYNSNIPYIDVQICVPGTSQCVRVDHVQVDSGSTGLRIMK
SALNGLTLPLQANTSGAQISECMAYATSEVWGPVARSDVYVGGEFGGNQTLQVIDDSASA
SVPASCSSQGALTDTTSSFGGKGIIGINQLFSDGGSYYACTGTGCSQLDSSPVTVTNPVA
GFAQDNNGVSIQIQPISDLGATSASGQLIFGINTASNNRIAGAAILYTDKNGDLSVNTGT
RTMPGFIDSGTASYFFPDDGSIPICSDNRNWYCPPSPISVNVDVIAGDASVDRPSSYRLS
SYDSISADMEVAPTGESGSVFNDGASWYVLGLPFYINRRVYGSIQQSDSDPMYYAF*
>GCA_000452945.1_BPMSHR5858_genomic_00900_1 regX3
VVIYLIEDDEIQAQYYQSMLVEHGWQVKLLLDGERAFREIQRMPPDLIILDRRLPDLDGL
EVLMWVRKNYSNIPVLILTNAILESEVVAALEAGADDYVIKPPRKQEFVARVKALYRRAT
ETRTLSELIEIGPYRIQTSEKVVYFHHEAITLSPKEYEIIELLARNIGQIVSRDKMVYQV
WGRLPDEAAFRTLDTHIYRIRRKLELTRYNRLILRAVYMHGYRLEYMGINNMMAPAS*

>GCA_000452945.1_BPMSHR5858_genomic_00901_1 cysC_1
MDKLHAERNVEWQTTSVSRRDRESRQAHGALAVWLTGLPGAGKSTLAIEAERCLHQAGKR
TYVLDGDNLRHGLCRDLGFSADDRAENIRRAGEVARLFVDAGVIVLSAFISPFRFDRAKV
RALFDAGDFIEVYCDCDPRICEIRDVKGLYRRARAGLISEFTGISSPYEPPRHPDVYVNT
GKESLDACVDRIVRRVLHSLR*
>GCA_000452945.1_BPMSHR5858_genomic_00902_1 group_3148
MYGSGACPPRDLASAREWVARHPDDLAVLAACADVVLRAGQGGDIIDWLRALAVNSPNER
HRWRNNLAAVLFQQRQVSGALAELRGLTAECPEYVPGWLNYAAMLKAGAPDEYETALAQA
LRHHPRNAQLLAAFAPLRERAVAGAGRAMLLERWRSMATDGYRDEDSDAIAVALAHIDGR
AGQTDRRIAILLEAGARHTAQRRLAPLQESNRALADAIPGFYPTLRELRQPAHWVPWELA
LAYLGMQRWQEGFSCYESRLLWWGYSAGWTAAQKMRRWRGQDLVGATVLVTGEQGYGDQV
QFARFASRLKTLGAERVILGTNKPLQRLLRTLPGVDDVIALGDAIPGFDWHVPVCSLPHG
LGVTHTAELSAVPYLHVPQTLRRHWRARLDRALPTAGRAMRPRIGLCWAGKPTHVEDAAR
SIPIERLRSLIDRFATKVDWIRIQPGMSVQTPGMCCIDHAIGDFLELAALLDNLDLLITV
DSAPVHVAGALGVPAWLMNRHFGDWRWPRQETRSAWYGSVEVFHQATPDDWSGILDSIAE
RLAQGVVSKPDA*
>GCA_000452945.1_BPMSHR5858_genomic_00903_1 lapB_1
MIGMYEQSKAAKRRYRDGDFASRYFVGDGIDVGAGPDGLGRYRRQFAAMRSVREWDLADG
DAQLLAGVSDNTFDFLHSSHCLEHMRDAVEALHSWVRVVKPGGYLVICVPDEDMYEQGVY
PSRFNPDHKWTFTIQKPQGASWSPRSINLLEMLIPLSGVLEIERIQRVQEFFDDTLHGRD
QTLMPNVECAIELVLRKRGAAGAIMGDVRPAAAEGAAAMLPCARAGTGRFDAPAGREALM
LLDSGHRDAAERKALEALVREPADCTAMDVLERLYLDSARYAECADILEKLVAYAPNNAG
AWNNLGCCYGELRRFPQARAAFQRAVDANPEYVTAYGGLAAMHYALGDPASAEATYRAVL
ARSPEHVDSCVNLANCMLAQEKVEDARPYIERAGEKLLPDLNKPKDVALCAPREGLTVPY
RSRIENRPIFESMHARLGMSLLMLGRATEGLRLLEWRLGSAFVPGFDARRAQRMWRGEPL
RGKSLLLLWEQGYGDVLQGLRHVSRLAELAERIYLPADAAVRELFEASFSAWGQKVVVCD
WRDEPPAFDYYVSILSLHHRMRELGVACPEAVGPYLSCNPAEVERFRSYRLDENALAVGV
IYAGKSEMLLDAQRSLPESVVRDLTAPIAGVRFYSLQIGCRAADCAMLGKMNGVIDWSML
LSRFADTAAAMRALDLVVCVDTAAVHLAGALGVRAMLLNRFGSEYRWQLDRDDGVWYESV
SQVRQHAMNDWTNAVACVRSRLRQEALDLGRDAVVPGAQFSASEAGQPDVHRQLRAIEQA
LGNSQLDEARGLITGYLAQHADSPKAQCHAGVLELLSANPDAARAYLDRAIRLRPDYAVA
YSNRAATRKEGDAEREGDIIRSIACDPSFHAGWLNLARIRRPHDAALALQCARRAVQLAP
DDADAVLLRAELALDTGDFDEALSQFERLREQRPDAPESYANLIPALAALERRDDAIAAL
QRALELNSKHPGALNNGVQFYLRTQQYDKAMELAQRYVGAHGELASAHTMCGLVYHNLKA
YDRAEASLRRALEIEPHNAEALFALGTVLERVDRLAESEEVLRRALTIRRDYRVLVNLAV
TLNRQQRYEAARSLNDEALAIGGGKMKSELKPRMALMRKTFAGQPFRAELATVDMTFGPD
WNNALIALVQGDYVRGFAAYELRQEHQTVSGMTADEYRALLWRSEPLAGKRILLLPEQGY
GDVIQFLRYVPELKRKGATVLVGASAPLRRLLEVAPGVDEVVQLDDARRPAFDYVCPLMS
LAHRLDVSSAEDFLPSFPYLRVPEAASQTWRARFAADSRRKIGLVWAGRKIYPADRLRSV
PLPEMRDVLSDIPDVQWISLQVGDCASEAASWPAVLDVSAELTDFAETAALIESLDMVIT
VDTAVAHLAGALGKPVWMLNRISTDWRWGQEGQRCGWYPSMRIFRQQVHLSWAEPIGELA
DALEQWLDVERLNA*
>GCA_000452945.1_BPMSHR5858_genomic_00904_1 prsE
MSMKLIAAIFRPFLSLADAVLGYFERHAFEPASPEGRQSWRGSALGLEEASIAPARRAAA
LIPTVMLALLIVLVLWATFFKIDIIAAGQGKVIPSTTVQQLSTLEGGIVRELLVREGQIV
KKGQPLVRLDPVVAQGAVTEQAATREGLMASIARLQAEADGKATPLYPAGLKPEIVSEEE
HVRAQRAEALNSTIEVLQQQRAAKQAEAADYRGRIPQYVNNQHLLDDQIQRMLPLVGVGS
VAPNEITNLQRERGNLAAQIITTREGAAQASAQIAEASHKIEEKISTFRSEAREELARKQ
VQLQALEGTLSGKQDILDRTLIRSPVNGIVKTLYITTIGGVASPGKSVIDIVPTNDSLLI
EARIQPQDIAYIRVGDDAKVRITAFDSGALGSLDAKVELISPDSQADERSGSLYYKVQVR
THSSVVATQVGDLNILPGMVADVDVITGRRTIMSYILRPIVRGMSRAMSER*
>GCA_000452945.1_BPMSHR5858_genomic_00905_1 btuD_8
MSDTIPAIEHSAAESDDRTVERMGGPDAASELPLLARVHRMLAAAVTEAGRALGRGVSFD
ALLAGLTDGKDAMGQVDVIGLIEAAKAFGLVVESVDVPAASKLGTTSLIAIDTESAQDQV
LFFPSVEMPHERRAFRFNATGLCQYFEYRQDQVPTGVGVQWLRVDRVPYSATAGARDAGH
RPLPSKEHMAWLWDVLLAEKSAFRYALIASLGSTLIGLAMPFFSLVIFNTVAPNGAVATL
ITLMIGMLVAIAADFVLREVRAYIIDVTARRIDVVLSRRIYEHLLDIKLESWTGVAGTQA
DMLRGFATVQEVFTSGVLLLLIDLPFGLIALAMVFVVGGHLGWVPVFWVMVLLALLWRMQ
SSVHRSSQLSMQLGQERHGTLVETINALETVRAVGGERVLRRRWRDQTAMAAIASGTSRM
LTQFVAGVTTMSMQLESLMLLGIGAWMVGEKWLTLGGLIAANMLAGRALAPFLSIVSLGI
RLESARIAAGFISRFFQMPRTTRETQEFVARSDFTGVVTLNDVSFAYPAHKSGRATVANV
NLTVGAGERVAILGRMGSGKSTLLKLMAGVLSPMRGSVLYDGTDLAQLDLVTRRKVVGYL
QQNPVLVSGTLKENITLGCPDASDERVLRAAKIAGVDDFARTHPDGYSMRLGEHGRGLSE
GQKQVVGLAQLLLNDPYAILLDEPSSSLDLEAENRLVHALKTSLVDQALVVVTHRPAFLQ
LVSRIIVIDSGCVVLDGPRDDVLAKLSGRTQQTAGAKGMSSCP*
>GCA_000452945.1_BPMSHR5858_genomic_00906_1 group_7301
MNIHFISGLPRAGSTLLSVLLAQNPRIHATITSPLFSIATAMRQAMSTSSEHYQLMNERK
RLKLYAGLFDSYYEDAYKAGKTIVFDSNRSWLQLQALTSTVFPGARTIVCVRSPSWVIDS
LEQLFLANAAVIPQYFNDAFEARSLYSRTDALMRPDRMVGASWSNLREAFYGPQSEQLLI
VEYTALVADPEATLRTLYNFIGEPFFEHKYSELRIESQDDIDRFDYAMGVPGLHRVGGEV
RRRARATVLPPELYERLERMAFWREVNATNAKVLLGPPETPANRHATQTAE*
>GCA_000452945.1_BPMSHR5858_genomic_00907_1 group_7302
MGLLFEQSRLERVRTWAMANKQRFRPNGFGRQFALLQEFPDVPDAIWDIKRDVVAHFGLH
DLPQEPLYRDLCGVITEGGAVHPHRDSNQGMLVHTRFNVMVSRPDGGGVPMIDGMLVDVP
EGGIFRVDAGLLTHSCTPVIGARPRIILSFGFLSPLGRFSGLPFCISTP*
>GCA_000452945.1_BPMSHR5858_genomic_00908_1 group_59
VTVDTINPPPALTLSPLSDTFGSGTSGTNHDNITSATLPTFNGTAAAGSYVQLYDVTGGT
TVSMGSAVADSSGGWTTTLTSPLSGSASGVSHTLVAVGVDPAGNTSTVSGPDVVVIDNAA
AALPGPTMASASDTGASSSDGITSNTAPVFTGTGAEAGALVTIYANGTSVGHATADASGN
YTIQSNALGADGRYQITAQQVDIAGNTSPSSSVTAMTLDTSEPAPVNLHLVDDTFGQGTA
GTSSDNLTKDSRVTISGTASAGDVVTLMDGATSVGQVTADASGNWTIQTASLADGTHSLT
ASAVDLAGNTSPASSTLPVRVDTTTTLPSLTLSSSSDTFGAGTSGTNHDNITSATQPTIN
GTAEAGSYVQLYDVTGGTTVSVGEAVAGSNGTWTTQLVSPLSGSASGVSHTLVAVGVDPA
GNTSAVSGPDVLVIDTSTPSPSTPALTPADQFNGNPSTTLNARPTLTGTAEAGASVSLTD
SGVVVGVGVADSTGHWTIQTSALFAGGHTITATAVDIAGNSNVSGSSTIAVAANVPTPPA
PVLITPDDTSPIDSTNNDDLTRVTMPHFTGTTNQGYLVTLFVDGVSVGQGVAGGNGSWTI
QDGTSLADGAHQVTVRVSDPLTGATSAMGPGTTVTIDTTPPAAPVVQLASYSDSFGGGTT
GTNSDHLTNVTQPVLTGTADPNTVELLLEGTVTLGTATTNASGAWSFTIQNPLADGTHTF
AVTSYDQANNHSAATNLVVTVDTVAPAGTTPTLDPTSDSFSPINGIGNNHDNITNNLSPV
FTGVNQVAGGEVLLYDNGQLIGSATSLGAVGTTWTVSPGNLYPGTHFIQSRYIDPAGNTG
ALSSGVSVTIDNSEPAPTNLNLPASADSGQSNTDHITNVTMPPITGNANYQDVVVLTAVN
GTTTTTVGTATVGASGSWSIVPGSLADGTYTFSAVAVDVAGNTSNPGVPVQVVVDTHAAA
PSITLGTPYDTFGTGTSGTNSDELTRNTIPYMYGVAEPGARVTVVENGNTIGTVNADSST
GSYSIQIPPATVDGTYTFQAMQVDVAGNTSAYSAPNYVTIDTVAATPTLTALTPASDTFG
VGTAGNNHDNLTNASTIGIMGTAAEAGAALDLYQITVSGSITTSTSVAHTTAGAGGSYTF
ASFTAPGADGVYTYETREIDPAGNTSPMSPAFNVTIDRTVVAPVLTLQNDTQYSGSPNWS
TGTNSGGAGSANFGTTTDGLTKATQPIVSGTVENGSLVVVYRDGVSIGQATANSAGAWSF
TDPVSQADGTHAYAAYQVDRAGNTSGLGANLAVTVDTQIETARYLPAPTPAGASSNYYAQ
IGIPALANGLNAQEIYNVDAAGHTGVPVYFQIDPATAAKGKAGDIVVVDFTFTYANGTGT
ATVTTEASYTIQAQDLVVTAGNTPTLTTWVTPNTFFAGNNWLSSYTGVNQMTVQLQVRDQ
AGNVGFTNTPNVDTINGAMNASHVNIDLWTLATNTGTNTWLGNGVQVYRELGRPADQLGW
QVIVGDDFNNDGIHDLFMYAPTGGAANNNGRMYLVAGQADTVANGTGFVDLQNVGTTSDW
SENHGQGAATGGYVWTGVGNQSYTNWTSVADITGDGRGELAFDQTVGPFSQGNPEIVANI
LYSDAVGPGGFQQALTGLTRPEMTQFYVPVHGDSAYTSRGVSSNFAAEFGALATHYDFNG
DGRGDLLWADGGTGNMGNSTTVYGSVLVGFTPSSAVGLYDLRLPGSSMNQTASKGFDVYS
TDPLHPVGANSALTQAPNNVWAQPIGDLTGDGLNDIIVGQPNGTFNNLATNGRVAVIFGR
NDGQGVDINNLSAGQGFYITWNSAVNAGDVFGQQIKVMDLNSDGYDDVAISIRCSAAPNT
SSNDSFDFVIDGKWLANYVHTNLAAGGAVPTLSIDAIYNAMVNHTATPMGWIDGDPTNTN
FAGNTSYPLNVGTPVLKTDLNGDGYQDTVIRNGTQYYVLWGSPDYHAGTDTNYGTTFFNQ
PNHLSLLNNNVGGAPVQFADINGDGLWDMMISQPGNATDGTNGGLVEVVYGVNSQSGSAP
GSPLPFASSYTNSWTAGADTVTGSSGRDMLNGGAGNDTIVGNGGVDVLEGGGGNDMLVVN
GDNIAHFTTPGSYWDGGIMMGGEINTLQFDANNTANATLDLTQVSVGNNLSAHLHNIERF
DLNGNGNMLKFDLRSVLDTSAMGVFLDGGSSGETWSGSTWDPGYTGSHGNRTTPLGPNVP
MKQWVIDGNATNTLDFEPFTQMAQTGASTGLTGNWALVGHVTHNDVHGVSQNYDIYEFSG
QNAQIIVNHNIHLTHAGSGTTL*
>GCA_000452945.1_BPMSHR5858_genomic_00969_1 fadD3
MAADSGMGALIAPMHGLSYVRGPADVPLSDATIGQFLLDTVARFPDRAAVVFREQGVRRT
WREFADEVDALAAALIQLGIARGDRVGIWSPNRAEWLLTQFATARIGAVLVNVNPAYRLA
ELEYALNKVGCKLLIAAERFKTSAYAEMIAEIAPELATTRAGDVLCAARVPSLRTVVTMS
DAAHAGMLSFADVLARGRAALASARLDAIGATLDCRDPINIQFTSGTTGSPKGATLTHRN
VVNNARSIANVMRLTEADAMCIPVPLYHCFGMVLSVLACVSAGAKMVFPGAAFEPGATLA
AVSDERCTALQGVPTMFIAELDHPDFDRFDLSTLRTGIMAGSPCPIETMKRVVAKMHMSE
VTIAYGMTETSPVSFQSATTDSLEKRTTTVGRIQPHLEAKIVDATGAIVPVGETGELCTR
GYSVMLGYWDDEARTREAVVDGWMRTGDLATLDEEGFCNIVGRLKDMLIRGGENVYPREI
EEFLFRHPKIQSVQVFGVPDSKYGEEVCAWIVLRAGETMTDDELREFCSGQIAHYKVPRY
VRFVDELPMTVTGKVQKFVMRERMIDELGLSVQQTA*
>GCA_000452945.1_BPMSHR5858_genomic_00979_1 betI_1
MGVSFLSLFGSIRLAKMTSRHVQVSAPPDKSIAQPQPALRRRPRQRRAQASSAALQQAFV
RVLLERGYAKATIREIAAVAGVSVGTFYEYFGDKESLAALTIHHYVKTLATRMRAVVDAR
RGERGRAVAAALIDAQVDAIEQDPRVWAAMFAIERKVSPIDAYRRHYDDYVALWRDALAG
ASDAPADARLDVTARVAHALCYGWVSQSLLTQGAKVDVAQLRDELHRAAGAYLAAAMDGG
AGGGAAGG*
>GCA_000452945.1_BPMSHR5858_genomic_00982_1 group_671
MSERTTHDAARRTGASVGRGCGGGAPAGISPRMIRAHPFGYPKHLDRRRVAQAGCPKRRK
AVRRLHPPLAQPATALCRVPGYGALVQLWRRVRATLPGVVCFEAMRPAFYRCVAAHAPAR
RRRSTRTAASRR*
>GCA_000452945.1_BPMSHR5858_genomic_01021_1 rutD_2
MPFVTIDGQPLHYQIRGAGAPVLFGHSYLWDSSMWEPQLDALSKSYRVIAPDLWGHGRSG
PLPDGTRSLDDLARQMSELLDHLDIDTCSIVGLSVGGMWAVPLAHRAPQRIDRLVLMDTY
VGVEPDATRNQYFQMLEAIDAQGAIPAPLLDAIVPIFFRPGIDPASELPTGFRRALQAFT
TERLRDSVIPLGKITFGREDARAQLSALPADRTLVMCGANDVARPPEEADEIAALIGCEK
AFVPNAGHISNLENPAFVTQALSDWLGRGAARA*
>GCA_000452945.1_BPMSHR5858_genomic_01042_1 gspK
MGNDIFLIGIDGGGTGTRAVLADARGRELAQGSAGPSGLALGIEAAWRSIEAASAEACAR
AGVAFDWRHCVLGCGLAGVNHRAWLNAFRASAPLAALAIESDAYTTVVGAHGGAPGVIVA
LGTGSIAAALDASGACRIAGGYGFPSGDEASGAWLGLRALSYAQQALDGRAPLDALARAL
VAHTGAPDRDALIVWSCEANQTAYAQLAPIVLAHRDHPAAAHWIEQAGIEIGRMIDALDP
AAALPVALCGGLAQALASAVPEPHGARLVAPQADSAHGALRLAEREAARLGLTAPR*
>GCA_000452945.1_BPMSHR5858_genomic_01048_1 flhC
MASKSVVVEVKEITLAIELIELGARLQLLEAETNLSRDRLIKLYKELKGVSPPKGMLPFS
TDWFMTWQPNIHSSLFYNIYRFMREHGGCDSIQSIVKAYRLYLEHVGLSDDEAVLSLTRA
WTLVRFFDSGMLQMTPCTRCGGHFVAHAHDPHHGYVCGLCQPPSRAGKTRKAAAQGDQAA
IAA*
>GCA_000452945.1_BPMSHR5858_genomic_01052_1 cheA
MTLDITQFYQTFFDEADELLAQMEQLLLNLDVASPDPEDLAAIFRAAHSIKGGAATFGFT
ALTETTHILESLLDRARNHELTLTKEMVDAFLETKDVLSDQLADYRASAEPDAAAAAAIC
AKLERLKASASGAQAATAAAAAPAAPAAPAAPAAPVAAAEPVPAATAAPATSAMGLFGAP
EHVVEQAVEAAHPVAGAPEAAAGGDGPHLKITLTGVDDKDQELLTEELGNLGRIVGSTKT
GDALALWLETDVSSDDIVAVCCFVIDESQIAIGRGTAPGAGDAQAAPAARQPGEAAPEPA
AEQAAPEPAAAQPAAAPVADARPAAPAAAPSAPAPAAAPAAQAAASAAAQHQDDRKPRPA
AAASAEGSSIRVGVEKVDQLINLVGELVITQAMLAETTSAFDPALHDRLYNGMAQLERNA
RDLQEAVMSIRMMPMDYVFSRFPRLVRDLAGKLGKQVELVTFGQATELDKSLIERIIDPL
THLVRNSLDHGIETVEARRAAGKDAVGQLVLSAAHHGGNIVIEVSDDGAGLNRERILAKA
AKQGMQISENISDDEVWNLIFAPGFSTAEVVTDVSGRGVGMDVVKRNIQSMGGHVEISSQ
AGRGTTTRIVLPLTLAILDGMSVKVGNEIFILPLNFVMESLQPAADDIYTIGNGERVVRV
RGEYLPLVALHEVFSVDDARTDPTQGIVTIMQTEGRRFAMLIDELVGQQQVVVKNLETNY
RKVYGISAATILGDGSVALIVDVAALNRETRAAHGAHSSAALAAS*
>GCA_000452945.1_BPMSHR5858_genomic_01056_1 cheD
MSGLPIATNLYFDAHFHRHGVKLLPNEFYTTREDMVLVTVLGSCVAACLHDPIGRIGGMN
HFMLPDDGADPSAAASESMRYGAYAMEVLINELIKAGGRRERFEAKVFGGAAVLAGMTTI
NIGDRNADFVRRYLALERIRITAEDLQGVHPRKVAFMPHTGQAMVKKLRVQAPDVAAREA
ALAREAVDPHGERAPRVRPRVELFGTPAPKAQAKPRIELFGMRATQPATRKQEA*
>GCA_000452945.1_BPMSHR5858_genomic_01064_1 flhA_1
MNMPAGLLGKRAQLFAGTNLRALAGPILICMILGMMILPLPPLLLDLFFTFNIALSVMVL
LVSMYTMKPLDFAAFPSVLLFSTLLRLSLNVASTRVVLLEGHTGPDAAGQVIEAFGHFLV
GGNFAVGIVVFVILMIINFMVITKGAGRIAEVSARFTLDAMPGKQMAIDADLNAGLINEE
QAKKRRQSVAQEAEFYGSMDGASKFVRGDAIAGLIIMAINVIGGLIVGMAQHDMSFAAAG
KNYTLLTIGDGLVAQIPSLVISTAAGVIVSRVATDEDIGTQLTTQLFTNPRVLMITGSII
VLMGLIPNMPHFAFLLLGGGAIWLSRTMTKREAAKKAAGMVAEIAPPAALPADSHEATWE
DVTLIDPLGLEVGYRLIPLVDKNADGELLKRIKSIRKKFVQEIGFLPPVIHIRDNLELRP
NAYRIALKGVEVGTGEAYPGQWLAINPGQVTAALPGAPTQDPAFGLPAVWIDTALREQAQ
VYGYTVVDASTVVATHLNHLVVQHAAELLGRQEVQALIERTGKDAPSLVEDLVPKTISLT
TLQKVLQNLLDEGVPIRDMRTIIEAVSEQAGRITDPYDLTAAVRLSLGRAITQQWYPGAG
EMQVMGLDANLERVLSQALATGANPGLEPGLAHSLLIGTQDAMLRQQNMGLPPVLLVQHA
LRAMLARFLRRSLPQLKVLSYAEVPDTRTIKVVNVIGGTA*
>GCA_000452945.1_BPMSHR5858_genomic_01072_1 clcD
MASQWIDIPAGNDTFGGYLALPKRGKGPAIVIIQEIFGVNGHIRSVADQYAADGYVALAP
DVFWRTQPRIELGYEGADRDKGIELLQKTDVAQAVADIGAAAAALRARPEVAGKLAAIGY
CFGGRLAYLAAARQHVDAAVAYYGGGIQNHVDVAAQIAQPILFHYAGHDQSIPLDAVDKV
KAAFAGRANAEFHLYPDAQHGFNCSERASYDQRAAALAHGRTLTFLAERL*
>GCA_000452945.1_BPMSHR5858_genomic_01078_1 fieF
MSTFSNDDLASEKQVVARQSTFVSIVLNVILATLQVVVGVIAHSQALIADGVHSLSDLVS
DFVVLLANRHSGAKPDADHNYGHSRYETVASFFLGAILIAVGVGMLWRAGDRLVHLEDIP
PVHASALAVAVVVLLSKEGLFRYMLREAQRVRSAMLVANAWHARSDAASSLVVAIGIIGS
LAGVRLLDPIAAAIVGFMVARMGWTFGWDALQDLSDRALDESASAEIRALIVSTPGVRDV
HDLRTRKMGDSALVDAHILVDPMMSVSEGHYIAEAARSRALTDRRVLDALIHVDPEHDVA
RRTVLDLPPRDQLEGQIEATLEKAGLRASAINLHYLSTGLEIDITLAPDASRDRQGGHDP
AARLDIEALKRQFGARRLGMSRALASSDAAAVGRLS*
>GCA_000452945.1_BPMSHR5858_genomic_01105_1 yehZ
MKVPRFSRTLRAAAAALAVSLCAAAPGARAAGAVTVASKIDTEGNLLGNVISQVLKAHGI
AVVDKIGLGATPIVRRALTTGEIDIYPEYTGNAAFFFNKADDPAWKNAAQGYALARQLDY
AANRIVWLAPAPANNTWGVAVLNSIAQARHLKTFSDFGKWVAAGGKVKLAASAEFVNSAS
ALPSFEKAYGFKLKPEQMLVLSGGDTAATIKAAAQQTDGVNAAMVYGTDGGIAATGLAVL
DDDKHVQPVYAPTPIIREAVLNAHPQIADYLKPVFASLDLKTLQALNARIQINGEPAAGV
AASYLKAKGFVK*
>GCA_000452945.1_BPMSHR5858_genomic_01107_1 yehX
MIEIERVSRLFGDLAAVDDVSLAVARGTVTALVGASGSGKSTLLRMINRLIAPTSGTIRV
DGVDTASVPAETLRRGIGYVIQGHGLFPHWTVARNVATVPRLLGWRAARVDARVRELLEL
FELDYDTYARKLPHQLSGGQQQRVGVARALAAEPAILLMDEPFGALDPIIRGKAQDDLVA
LQRRLGITVVIVTHDIDEALRLGDQIAVMDAGRILQAGSPAEILGRPQPGVVERLVAGLD
RPLRLLALTRVDALAEPGAAPGEPISGALTLRDAISELLWRGARALPVSSADGAAARRIT
LDAIVAQAKRPA*
>GCA_000452945.1_BPMSHR5858_genomic_01109_1 group_7307
MRICIENIQPGVTREELQAFLFKYTGKTFTDIQFIGQSGPRPGALIDIDGANWSALAEIR
RRLHGMYWKRRRIGVWLFSFWDRSDAAEARRQREARTRQTRSTPMTTLRR*
>GCA_000452945.1_BPMSHR5858_genomic_01110_1 arcD_1
MAEVTQEQGVAAKSAANKLRLGALTALVVGSMIGGGIFSLPQNMAATAEAGAILIGWAIT
AIGMLTLAFVFQTLANRKPQLDGGVYAYAQAGFGDYMGFSSAWGYWISAWLGNVGYFVLL
FSTLGYFFPVFGEGNTPAAIAGASVLLWAVHFLVLRGIREAAFVNLVTTVAKIVPLVVFI
LIGFFAFKLDVFRTDIWGRANPSLGSVMNQVRNMMLVTVWVFIGIEGASIFSARAEKRSD
VGRATVIGFVGVLLALVLVNVLSLGIMTQPQLAKLQNPSMAAVLEHVVGHWGAVLISVGL
VVSLAGALLSWVLLCAEIMFSAAKDRTMPAFLRKENANHVPANALWLTNAMVQVFLLITL
FSQSTYLSLIYLATSMILIPYFWSAAYALLIAVRGETYEGDARARRKDLFVAALAVLYAV
WLLYAGGLKYLLLSALLYAPGALFFAKAKRELGKPVFTAAEKLIFAAAAAAALAAAYALH
SGLITL*
>GCA_000452945.1_BPMSHR5858_genomic_01112_1 arcB
MSFNLHNRNLLSLIHHNARELRYLLDLARDLKRAKYSGTEQPRLLRKNIALIFEKTSTRT
RCAFEVAAYDQGANVTYIDPASSQIGHKESMRDTARVLGRMYDAIEYRGFSQEIVEELAH

HAGVPVFNGLTDEYHPTQMLADVMTMREHSDKPLHDIRYAYLGDARNNMGNSLLLIGAKL
GMDVRIGAPKSLWPAADFIAQCEAFAAESGARLTLTEDPYEAVKGVDFIHTDVWVSMGEP
VEAWDERINALLPYQVNRKLIESTGNPRTRFMHCLPSFHNCETKVGKQIAERYPHLQDGI
EVTDEVFESPRCIAFEQAENRMHTIKAVLVSTLGGI*
>GCA_000452945.1_BPMSHR5858_genomic_01113_1 arcC1
MRIVIALGGNALLQRNQPMTEVQQRENVKIAVAQIAQIAPGNELVIAHGNGPQVGLLALQ
GAAYPAVAPYPLDVLGAQTEGMIGYLIEQEMGNLLPPDAPFATLLTQVEVDPADPAFEHP
TKPIGPVYSRDEAERLALEKGWHIAPDGDKFRRVVPSPRPRRIFEIRPVKWLLEKGTIVI
CAGGGGIPTRYDANGKLSGVEAVIDKDLCASLLARELSADLLVIATDVDGAYLDWGKPTQ
ALIEAAHPDELERLGFAAGSMGPKVQAAIEFARQTGHDAVIGSLADIVAIAEGRAGTRVS
VKVDGIRYRRPR*
>GCA_000452945.1_BPMSHR5858_genomic_01114_1 group_1794
MTGKRILVTGAGSGFGREVALRLAAKGHCVIAGVQIAPQITELSAEAARRGLALDAVKLD
VTCARERAQAARWDVDVLLNNAGAGEAGALVDLPVDIVRELFETNVFGPLELTQQVARGM
IARGRGRIVFVSSIAGLITGAYTGAYCASKHALEAIAEAMHLELAAHGVQIAVVNPGPYR
TGFNDRMMETPRRWLDPARDAATPETLTFPLEQFDPQEMIEKMVDVTEADGGLFRNLLPA
AAEQIVRDEQAHAWTRRQNERRGA*
>GCA_000452945.1_BPMSHR5858_genomic_01115_1 group_3175
MSPELTLERALRIVEAGVAHAARLSAAGAAVAIAVVDAGAHLAAFARMDDAFIGAIDLAT
RKAGTAARFRMPSASLGALSGAGGPVRSIEHSNGGLVTFGGGLPLVDAHGRCVGAVGVSG
GTVDDDAAIAQVCVDAFITLLNAAQGART*
>GCA_000452945.1_BPMSHR5858_genomic_01116_1 pgrR_1
MTMKRDPLGGLHVFMSVVTHGNFTRAAAALGLTPAAVSLAIGQLESELKVKLFSRSTRGV
SLTEAGQRYYDATAQPYRQVVDARDALNDARDEPEGLLRISALHLAKSVVTADVLAEFFR
RYPKVSVEIRYEDHLVDIVKERLDAGIRLADRLQPGMVGVKIAPALTCALVATPGYLAQH
GVPAKLADLARHMCIRFRFPKSGRLHKWPLRDGRRDVELDVAGRFVSTDTGAVIDAARAG
AGIAFVFMRERIEDDLRRGTLREVLPGACRTLPPMWLYYANRKHVPPKLRAFIAVLREHA
AADARR*
>GCA_000452945.1_BPMSHR5858_genomic_01117_1 yfcG_1
MSDLSEFPIMKKWPAQHPERIQLYSLPTPNGVKVSILLEETELPYEAHVVRFDTNDQMSP
EFLSLNPNNKIPAIIDPHGPGGKPLALFESGAILLYLAEKTGRFIPADPARRYETIQWLM
FQMGGIGPMFGQLGFFHKFAGREYEDKRPRERYVGEAKRLLGVLDGRLAHRRWIMDDAYT
IADIATFPWVRNLIGFYEARELVEFDRYPNVARALDAFVARPAVARGLNIPARP*
>GCA_000452945.1_BPMSHR5858_genomic_01119_1 group_382
MKLIGKQSGAALPLDETLPAPDAAPLRLDEWLPYQLFLPAQHVARVLAEFYGPRYGIGQP
AWRILATIVDRPGTNARQIGAALGMDAVSVSRGIAQLVGAGFARRDAARNDLRYACVMAT
PAGRAAFDDIAAVCVAVERRLLDVLTPEERATLAAALPKIGRESERIAIVGWQRLLDGA*
>GCA_000452945.1_BPMSHR5858_genomic_01120_1 braC_3
MNVRMGCLSLLAAVAFDAHAQSSVVRIGVAMPLTGPVAHLGKDVQNGAQLAVDELNRAPP
TIDGKPVKFALVVEDDQGDPRQAVQVAQRLVDARVAGVVGDLNSGPTIVAAKVYAAAGIA
QIAPAATHPAYTQLGYKTAFRLMATDNQQGASLAGLAAKLAKGRPIALIDDRGAYGQGLI
DQTEKTLRASGVTRIIRDYTTDTAVNFASILTRVKGAHAAVIVYGGADAQAGPMVRQMKA
LGIDAAFVGSDGVCTGQWTALSSGANEGQFCTQAGDPRARMAGYAAFERRFEARYGKVIV
FAPYGYDAVMLLADAMRRANSTEPAALLGALATTRYDGVIGRIRFSPQGDNLNGAVTVYR
VQRGALVPVSD*
>GCA_000452945.1_BPMSHR5858_genomic_01122_1 yycB
MNLLLSPAWRLAAVVLVGLNLRPALASIGPLLDTIQRASGLSDSAASLLTTLPILLMGAL
GLVAPWLARRVGVRAGVGLGVALIGLACAARVAAGAPAALLASALCAGLGIAAAQALLPA
FIKTQFAARAGSVMGFYSTSIMGGAVFASVATPFAAHAIGWTCALAGWALPAAAAALLWP
LATRATARPAHARPAARRAHGARAWLLAAFFGLGTGAYTLVLAWLPPYYVSLGWSPEAAG
GLLGGVTLAEVVAGLTVSALVERLPDRRPALFTALVSLFAGLIALIAAPLGLALPASLLL
GLGIGALFPLSLIVTLDHASSAAEAGALTGFVQGVGYLIAGLFPFVAGVIRQSVADLSPA
WGFMAAVCIAMAAMAARFAPPRPAIARSGARSGT*
>GCA_000452945.1_BPMSHR5858_genomic_01123_1 group_4106
MLSTRLRHLAAAALVLSASACASSPQAVPEVACRPDETMPKRQFRGTWIASVINLDWPSR
PSLPAAAQQAELSAWLDDAVRMNRNAVILQVRPTADAFWPSPFEPWSKYLTGAQGGDPGY
DPLAFAVAEAHRRNLELHAWFNPYRVAMDDRLDALVATHPARAHPDWVVRYGGKLYYNPG
VPAARAFVVDAIMDAAARYDIDAVHLDDYFYPYPVAGATFDDASAYAQYGTGFATLADWR
RDNVDRLVESLAQRIKAAKPWVKFGISPFAVWRNAATDPQGSRTSASVQTYDDLYADTRR
WVRERWIDYVVPQAYWARGFAPADYDEVVAWWANEVRGRDAHLYIGQAAYKVGTSNQSPG
WSDPHELSRHLAFNLTAPEVKGDVYFSAKDVRADRLGATTRLNRTWYSRPALVPTMPALG
GNAPPSAKALRAQRTPDGVRLQWQAGSAAAASYAVYRHALGRQDMCADSDARHLLATVRG
TQYVDVTARADRDYRYVVTALDRLWHESEPAYVAMPAAKPR*
>GCA_000452945.1_BPMSHR5858_genomic_01124_1 cobM
MTVYFIGAGPGDPELITVKGQRLVRRCPVILYAGSLVPAAVLDGHRAELVVNTAELDLDE
IVALLARAHGKGQDVARVHSGDPSLYGAIGEQIRRLAALGIPYEIVPGVTATAACAAALG
VELTLPGVAQTVILTRYAGKTTMPAGESLASLAAHRATLAIHLGVRHLARIVADVLPHYG
ADCPIAVIYRASWPDEARVTGTLADIVDKVSGRSIERTALILIGRVLSAEGFDESTLYAK
A*
>GCA_000452945.1_BPMSHR5858_genomic_01143_1 purN_1
MAKKKLVYIWSLRNAAADKAGQQVDYKGGTRYMKSVLESLVGALNDTELGDAYSLERVIY
DDDAGSPLDREKLAEYGFAYEPGKRWFYPPALRVQGRLVNDLLLAIPSEYRREPLASPAR
PAGKRAFEARLRETLDALGADLVVLDGLLVILDELVRPGAPYCRRIVNIHPGITRADSPY
ERRGAYATLDALFGARGQKVVDWKTGRTMPVEPLRMTGASFHYVDNGVDSGEVIHDVLNT
EIDPDDTILELRWNNFNRSLFPALHEGLAIMAGLDARVA*
>GCA_000452945.1_BPMSHR5858_genomic_01144_1 fhuA_1
MEWATSTRVRAIAAGVAFYAAAAGHAQAQAAQPGTDARQPGGEAKADTAAGGTLPAISVS
GAAERDASVGLVARRSMTGTKTDTPIIEIPQTINVVTAQQIEATGATDINQAFRYIPGFS
SYGSDNRSDWYAALRGFTPTVFVDGLQVPNTINLSSWRVDPYMIDSIAVLRGPTSVLYGQ
GDPGAIVDVQSKLANGERIRELGVQVGNYARKQLMFDIGDTIGKDGTLSYRIVGVGRDGN
AQTGPLADQRVSFAPSLKWQPNADTSLTLAATYLQDWGDTSSNFLPSRGTVLPNPNGTIS
DDLYTADANFDHYRKKQWSLGYQFEHKLNPVWTLRQNVRWMHLSLDDASVYGGGLDDADP
TMATMTRYAGLFQFNYSRFDVDNQAQAKFTTGPLSHTLLFGFDYNRQTTTDSEWLAKGPS
LNLYRPVYTPIPSDIFSGPNAYPRTDTKTTLNAFGLYVQDQIKWRRWVLTLGGRQDWTRT
SQDDIANAASFRQNDHAFSGRVGLTYLGDYGLAPYLSYSTSFNPQIGLKLAGGGLATPTK
GRQIEAGLRWQPPGKNLMLNAAVYQINQTNVAMSNPNDPTSSTFVQVGEVRSRGVELSAV
GNLSRELSVIAAYVYQDVKNVRANDNTLNKWPVDVPRPRQIASLWADWTWRTGPLTGFGV
GAGVRYMSAAAGAADNSLTVPSYTLFDAALHYELRNWRFALNATNLFNRRYVAGCQSDAV
CMYGNQRTVIATAKYNW*
>GCA_000452945.1_BPMSHR5858_genomic_01161_1 rbsC_2
MNMHDRTAAPSTSCAPAAASPARATGPARAAHALAALRRSTTFYPLVGLVAVCVAMVFAS
DSFLSAANLENVLRQVSINAIIGVGMTCVILTGGIDLSVGSVMALSGTLAAGLLVAGANG
AAALAAGIGVGVALGAANGLFVAFAGMPPIIVTLATMGIARGLALIYTGGYPIDGLPDWV
RFFGSGKVLGVQMPVLTMLAVYALAWLMLERMPFGRYVYAIGGNEHATRLSGVRVSRVKL
AVYTFAGLTSALAALVLTGRLMSGQPNAGGGFELDAIAAVVMGGTSIAGGRGSIVGTLVG
ALLLGVLNNGLNMIGVNPYVQNVIKGAIILLAIYIGRERRT*
>GCA_000452945.1_BPMSHR5858_genomic_01162_1 rbsB_2
MDEASGAARAPDEASEEAMDTILALTGITKRFPGVVALRGIDLRVARGEIHALLGENGAG
KSTLMKILCGIHPPDEGVIALDGEPRRFANHHDAIAAGVGIVFQEFSLIPELNAVDNLFL
GREWRGRLGLRERARMRRAAADIFARLDVAIDLSAPVRELSVAQQQFVEIGKALSLDARL
LILDEPTATLTPAEAAHLFGVMRELKRRGVAMIFISHHLDEIFEVCDRITVLRDGQYVGT
TEVARTDVGALVEMMVGRRIEQSFPPKPRLARDAAPVLEVDALQVRENGPVNRFALREGE
ILGFAGLVGSGRTSSALALIGAKPARVRRMRVRGRPVRVADPAAALAAGIGLLPESRKTQ
GLIPAFSIRHNIAINNLGKHRRLRWFVDAAAETRTTLELMQRLGVKAPTPHTRVDTLSGG
NQQKVVIARWLNHHTRILIFDEPTRGIDIGAKAEIYQLMRELSARGYSIVLISSELPEIV
GLCDRVAVFRQGRIEAMLEGEAIEPNTVMTYATSDVRGANHEHA*
>GCA_000452945.1_BPMSHR5858_genomic_01165_1 group_7315
MVRFLAKLLRSTIHGSHGVSLDAVSSTHGTPGFQTPDARVISRFGFN*
>GCA_000452945.1_BPMSHR5858_genomic_01168_1 atm1_1
MKLAALRSLADVIRLMLDKDTHARRPIVAGVLVEALTSAFMLLGPIALKLAVDEMARPRF
DAAKASVDIALFAVLWSASAISSIALLAYTGRIVHAVSNALLRRALHAQLPALASRASSD
SGYVQGLLERLPYSLQVIIEGVLWKIVPVAIQLAVALVLIALLVPLRYAAILFAVLAAYF
VFSHLSAEQYEKSAATTNEAAGALSAALGDVLANAPRVVYNGALAREIDYVGARAGARLD
VDWRRSWLLTRAAACQYGIIALGMAAMFVLCVRDIAAGRITLGDFMLLQTYVLQFALPLG
AYGFVLRQAGAALANVREALDIAPRPAPAAAIGAAPPPGRSGAPIAVRRLAFRRADRFAI
EPMSFDLPAGSYTAIVGHNGSGKSTLAKIVAGRLAPDEGVVAYGDVDLYRVASDARHRFA
LYVPQDVALLNRSLRENVRYYPSTLTDDDAARLLERLAFHKDGRPIDLDGEVGEGGARLS
GGQVQKVELVRLMGVDVPAIVLDETTSGLDPHSDALGIAMLRERLGQRTTLVLITHRIAN
VEAADQVLFLSGGRLVAAGPHRRLVDTCDEYRAFWRRQPEPADAKA*
>GCA_000452945.1_BPMSHR5858_genomic_01169_1 group_4559
MSALARLLPRRAPGWLAWACAALVVALAQPAWAVRCLTNSGQTSLTEPIGNVASYPTDAP
DGYVIWISAPRTTDGYCYKDLGGAGNLNIVDNIYFYANPDGKNPAAWGLEIGIRYRGVDH
FGAGSRPGTGVPTGFAVPPCSQTDFNAKRCPKIPVSITYQVVVRKKGAWVQPPGDVYTVF
QFDGEKGLNNTNPSFQYKLSGLQNLKPTPCMVDVTVTPEPGIVKFGQVQSSGNGFSPAVP
RKPFSLALTKKCSIAVRVDGYFETSQAVRDGLLVPQSDSNFGIGIEDRNGHAIAFNRQFV
LTQMPANVSYQNIVLDAVLKSFGAPKIGPFTGTATIRLFIY*
>GCA_000452945.1_BPMSHR5858_genomic_01170_1 yfcS
MTDMTHRFPCAARARRAAAALFIALAGAAHAAIVPDRTRVILNEGEQAAIVTLSNKSDAY
PYLVQSWLEDERGNRITTPLMVVPPLQRVEPKERNVLRIARMPGATLPADRESVFYLNIR
EVPPKTDTPNALQIALHTQMKLFYRPAGVQPARDEDPTLPMTLRVDAAARKLVFDNPTPY
HVTVVALAAGAHRAALPLDPVMVNPKSAASVPFSASAAGGLFVTHVDDYGGQVTVEYTCD
GLACRSVKR*
>GCA_000452945.1_BPMSHR5858_genomic_01171_1 papC
MLAAGSHARATEFNASFLSIDGRNDVDLSQFAQADYTLPGTYLLDVQVNDVFFGLQPIEF
VAHDDGQGARACVAPELVAQFGLKKSLVENLPRTMGGRCADLASLDGVTIRYQKGEGRLK
ITIAQAALEFADASYLPPERWSDGVDGAMLDYRVFANANHAFGRGAQQNNAVQAYGTIGA
NWGAWRFRGDYQAQTRAGGAVYAERAFRFNQLYAYRALPSIRSTLSFGEIYVDSDIFSTF
SMSGVAMKSDDRMLPPSMRGYAPLVTGVARTNAIVKVMQDSRVLYMTKVSPGAFALSNLN
TSVQGTLDVVVEEEDGTVQRFQVATASVPFLAREGQLRYKTAIGQPRTFGGAGITPWFGF
AEAAYGLPFDVTVYGGLIAASGYTSVAFGVGRDFGRFGALSADVTHARATLWWNGRTKRG
NSYRINYSKHVDALDADVRFFGYRFSERDYTNFQQFSGDPTASGLANGKQRYSAMLSKRF
GDTSTYFSYDQTTYWARPSDRRIGVTLTRAFSLGALKSVNLGFSAFRTQGAGGGGNQVSL
TATLPLGERQTLTSSVSAGEGGTSVNAGYLYDGANGRTYQLYGGTTDGRASANASLRQRT
PSYQLTAQASTVANAYASASLEVDGSFVATRYGVTAHANGNAGDTRLLVSTDGVPGVPLS
GSYARTNARGYAVIDGVSPYNVYDATVSVEKLGLDTDVTNPIQRTVLTDGAIGYIRFNAA
RGRNVFVTLTGDGGAPVPFGASVQDAATGKELGIVGEAGAAYLTQVQPRAKLVVRAGAKT
ICTPAALPDTLQLDGTPAPVACEAAAQHASR*
>GCA_000452945.1_BPMSHR5858_genomic_01172_1 fimA_1
MKKALLSAVACAALSTSAFAAGTGTLNFTGEIVAGACGIDAGSVDQTVRLGFVPANTFKA
AGDKSTPQNFDIKLVDCDTSVAKNAYFTFTGTSNATQPKLIATVGSATNVGIRLQSASGE
YLDNGAEQKGPVVLSNGTSVARFAAMYESTAASVTPGTADGVANFTVRYQ*
>GCA_000452945.1_BPMSHR5858_genomic_01174_1 acrB_1
VKAMMKPRARRRGARAARRPNGPRAWPLAVAAALVAGCTLAPRYERPAAPVPAHYWSTAT
GAPREAGPAAAGGPRAMPDARRGDAGRDARDARLDDWRAYFTDPALRALIDAALANNRDL
RIATLRIQEARGLYGVARADRLPSIDGSLGYERTRLYDPVLRESATSSLYRASVGVSAFE
IDLFGRVKSLSDAALAEYFATAEAQRAARISLIAEVASAYVTERALVDQLGLAERTLAAR
DAAYALTQRRYAAGTSTAIELRTAEMLVASARASKAALEREHTQAASALKLLAGDFMTAL
PADAPALDALAVARVSPGLSSDLLEQRPDIRQAEQRLVAANANIGAARAAFFPRIALTTD
VGSVSDAFSGLFSAGSSVWTFAPRLTLPIFAGGRNRANLDVADARKHIAVAEYEKTIQTA
FREVADALAARDQIDAQLAAQQAVYGADAERLRLAQRRYDSGVASYLELLDAQRSTFESG
QELIRLKQLRLTNAITLYRALGGGWSRAGCGGDGCA*
>GCA_000452945.1_BPMSHR5858_genomic_01177_1 ttgR_1
MARKTREESLNTKNRILDAAELVLLEKGVGQTAMADIAEAAGMSRGAVYGHFNGKIEVCV
AVCDRAFSRAVEGFDLSDERPALATLRLAASHYLHQCGEPGSMQRVLEILYMKCEQSEEN
APLMRRRALYELQTLRIAKALLRRAVAAGELDASLDVHLAGVYLLSLLEGIFGSMIWTTR
LRGDRWRDAEAMLDAGVDTLRASPALRVAEACAARGLPRANVH*
>GCA_000452945.1_BPMSHR5858_genomic_01180_1 gltJ_1
MDAWLAPKYLMWLWQGFLLTLGLSLAAAVTATLAGFALAIARHAKRASPRRAAAAYVVAF
RNTPLVVQLLFWYFGVAALLPEPAMAWLNAPHALDVGGLALRWPSFEFVAGWLGLSGYTS
AFVAEEFQAGLRGVPAGQHHAASALGLSPLQAFRFVVLPQAVRIALPPLFGQYMNLIKNS
SLTMAIGVAELSYASRQVETETFRTFAAFGVSTVLYVAAVALVEAAAQWTLRQRDRTFAG
R*
>GCA_000452945.1_BPMSHR5858_genomic_01199_1 group_2308
MTTRHLHNEARGYEPRRRSDEASGAAAHERDASPEAAAQHPEAPASGLKEARMNVADRKP
LPCPSDWTFELIEEYDTHISQVAEQYELDIYPIQLELISAEQMMDAYASVGMPVNYRHWS
FGKHFLSTEKSYRRGQMGLAYEIVINSNPCIAYLMEENTMTMQALVIAHAAYGHNSFFKG
NYLFRLWTDAHAIIDYLVYAKNYVAECEERYGLDRVEELLDSCHALMNYGVDRYKRPQKP
SLAKESELRREREAYLQSQVNELWRTLPAKKPEFADEEQEERYPPEPQENLLYFAEKNAP
FLEPWEREVIRIVRKIGQYFYPQRQTQVMNEGWATFWHYTLLNTMYNQGKLADGFMMEFL
HSHSNVIYQPPVTKPYYSGINPYALGFSMMSDIRRICEAPTEEDHKWFPELAGSPWLPAL
HYAMRNFKDESFIAQYLSPHLIREMRLFSVLDDDMRDALEVSAIHDDSGYQYVRQALSRQ
YDIHHREPNIQVWSVNTRGDRSLTLRHFMTDNRHLSNDSEEVLKHMARLWQFDVYLESVD
ETGTVRKRYECRYTAPEIRL*
>GCA_000452945.1_BPMSHR5858_genomic_01200_1 group_2309
VLHQIIDRRLAGKNKSIANRERFLRRVKNYIRRAVSDAVRDRSIKDIQSTQSITIPRKDI
AEPTFRHGPGGKRELVHPGNADYVRGDKIPRPPGGAGGGGSQASNEGEGQDDFVFELSRE
EFMQYFFDDLELPRLVKTHLLTVPSWKNVRAGWAAEGTPNNIDVVRSLRSALGRRIALGS
PLVNELRELEEKLAALKNEPGDHRVEIAQLEDAIHHLKGRIWRIPFIDPFDLRYVNRVKM
PQPSSQAVMFCLMDVSGSMDEQRKDLAKRFFILLYLFLKRNYERIEVVFIRHHTRAEEVD
EDTFFHSTESGGTVVSSALELMRKVMEERYSPTEWNIYGAQASDGDNWTDDSPKCRKILD
EDILTKVRYFAYIQVTPEEQNLWLEYAQLALSQPHLAMKKVESAADIYPVFRELFEKHVE
T*
>GCA_000452945.1_BPMSHR5858_genomic_01201_1 group_3179
MDIYSSFANRFEKTREEEFSLEEYLALCKNDPSAYAAAGERMLEAIGEPEQIDTRNDPRL
SRIFANKVIKVYPAFREFYGMEEVIEQVVAYFRHAAQGLEEKKQILYLLGPVGGGKSSIA
ERLKQLMERVPFYALKGSPVNESPLGLFDYDEDGPILEEQYGIPRRYLKSILSPWAVKRL
HEYNGDIRRFRVVRRYPSILRQIGIAKTEPGDENNQDISSLVGKVDIRKLEQYAQDDADA
YSYSGGLCLANQGLLEFVEMFKAPIKVLHPLLTATQEGNFKGTEGFGAIPFDGIILAHSN
ESEWKAFRNNRNNEALLDRIFVVKVPYCLRVSEEIKIYEKLIRNSSLAEAVCAPGTLKMM
AQLSVLTRLHEPENSSLFSKMQVYDGENLKDTDPKAKSYQEYRDYAGVDEGMTGVSTRFA
FKILSRVFNFDSTEVAANPVHLMYVLEQQIEREQFPPETEQKYLSFVKDLLASRYAEFIG
KEIQTAYLESYSEYGQNIFDRYVTYADFWIQDQEFRDHDTGESFDRAALNAELEKIEKPA
GISNPKDFRNEIVNFVLRARAANGGKNPAWVSYEKLRVVIEKKMFSNTEELLPVISFNAK
GSAEEQRKHEDFVNRMVAKGYTPKQVRLLCDWYLRVRKSS*
>GCA_000452945.1_BPMSHR5858_genomic_01233_1 group_754
MQELSPQEIAHVSGGHYDGGSLFDTLGNLIEIAGKGAETIGNTLTFGLLSPIGSAAYGVG
EVLTTEAVNLGNSLLNLFGIPAKHY*
>GCA_000452945.1_BPMSHR5858_genomic_01235_1 group_988
MGDKHRRRRARRDWRAPVSCWLGATLLACAWSAHAQDSGAARWRDGADGIGFFPGGDAPG
FDARAWGPVPGDARRAAASDARNGAAASAGSEATAAAPAADANAAPARKLTEERITLGER
VAPVADAARRVRADGDDGIGFADAPGGPPAGGATPSAACDDGACVPDGGDAGRAPRRPPA
GATPRFIAGVRYDRMPYELHPIDPERLPDLPEAQGPTLLEQLQGDDSNMIGVGWHYVLST
GRSTPVTTSTAALGIGSFANPGSAVSISNTNTPAFTFTHFFGEHVAAEIVAGIPPELTMR
GHGSIGLPFDKIFPGVQGRLPLVDLGNTQSNPLGTTRAWLGSAVFKYYLGKREDRLRPYV
GLGLSYTRFTNTNLNPVFAHKLASLGGLLSAGISLGDLQSLLTDSGALDRLLQAGANLIL
PNGVRATADVKSAWTPVFVVGANYQLTRQLSLSTALSYIPLKAAITVNINDTKGILASNT
TTLSANVLLCTMLLNFRF*
>GCA_000452945.1_BPMSHR5858_genomic_01236_1 cvaA_2
MPIRMNALREQKTPRKLTRGTTFGTVINGVVDVPISMRFFCYLSIMMFAMFIVALVRLTY
ANTESVMGMLTPRSGLIGVGAPPGWAVREVFVAKDQHVKAGQKLLSVTRDTSFVSQANNV
QGMRESIKRQRVEVGQQIDAAKLEYQSTIQQINQQIAAFDESRGLIDKQIQDQKRIVSEY
QERRDRVKQLLNEQVVTLEQYNQVNTQYLQASQAYQDLMLRRADLAKNAMKLRGDLETVQ
SKYDGSNAELKIKQEELNSKEYNIDESVNQVLYAPADGQIVRLDVVQGSVIDPPGTRVVE
ILPAKADGLIAELYIPSSKAGFVKPGQEVKLAYGSYPVEKFGTYRGKLLSVSPVAFTAKE
LNLPADNGAPQTYFKSWVELVDRKPAFEGKALSLKAGMTLRADIVLEKRTLLEWLFEPLY
RIRQRIFGTPA*
>GCA_000452945.1_BPMSHR5858_genomic_01237_1 group_4564
MTLMRLFLFNAFSYCFLLRRVVKRIALGRSPARGLGMMRWCAGVCLRLFPSIRGSIEMSV

RHMLSAHLNDDPVRIRGVADGIVRELFEAEVAGLMLRSHSIASMKTLIDRMECEGEGVLR
AALARGGPLMLAGLHFGNLMLYVSKLRLMIPDDRKMMIVMHKDAPGAFFDDAQRLVQEYG
AGKIELIDIEQRVHLRRLVAGLRKEAPVFLLFSDLHGRFGKTNACRLGGRWVRLAGGGVK
LAVEHDIPMLVSYATGVPFRDACKVHFVELDARPAPRMLGVADDASRVAAAHQRIADKLE
EALVKQPEQWHFWEHFTPYLMPSPLLEQNARAAAQTLRSA*
>GCA_000452945.1_BPMSHR5858_genomic_01238_1 apxIB
MWFNTNKVRPILQDEVTECGLACLAMIATWHGRDTTLRTLRNRYPVKIDSGLSFFDLMEI
ANDLGMRARGLQFEANEIDALKLPCILHWGMNHYVVLTKVCYGSVHIVDPALGETKLPLA
QALTQITGYALELNPDIGFKHVGTSDQIRLGDYLVGLTGIRKSILLGVAGGIAAQLLLLL
GPSYVQLTIDEALKKTDVDVLYLLTILFGIVFLFDTLYANLVGNIKSYLRNIVSQQLSAN
MVGHLARLPIGYYLFHNTGDSISRVSSVSEIGRFLVDGLIGGLLNVFTCVITLVLMLYYS
PVLAGISLAGMVLFALSRLAIQPQLQDAMSQILTRGAEADALLIENIRSAHSIKLLSSET
SRSSLWVNAFTQKLQAVRQQERLQLLFDAISKGIVHVEQLVIVTYGAWLVMHGQSTIGII
YAFIQYKNLFADKFVDSIQLYVRRKVLQVHMDRVADVLQTETEEPAPDAPVRSIEHFDGS
LSIRALTYTPKGGRRAILEDVNLDVPAGAKIAIVGRTGSGKTTLLNLICGLYPPAPGALF
VGGVDLSEVNLRLYRKHVAAVTQSDQLFRGTIRDNITNFAPAPRLGAMHEAARLACIERD
IDVLDNRYDTPLGDTQKFLSAGQMQRLLIARALYCEPRLLILDEFSSNLDQATTHEICRN
VLTLPCTIVLVTHDASILSMVDRIYEMRDGRLADITPARQPELEVQK*
>GCA_000452945.1_BPMSHR5858_genomic_01239_1 group_7328
MQAISEQQARAIRGGGSAYANSAAYTVSLANQSITPVVNGLLNEYQTQAVNQFKMALGSG
S*
>GCA_000452945.1_BPMSHR5858_genomic_01240_1 group_7329
MKVMTMEQAKVVNGGGPISSLGTLAAASGPLFTALGQFMPTLGGYLQNPKNFNSNGK*
>GCA_000452945.1_BPMSHR5858_genomic_01242_1 ggt_2
MNATNRFKSTVLAALVSSALAGFLPSAPAHAKGPQQRVVLDGSAVAVADKYSADAAEEIF
RQGGNAVDAAVAIAFTLAVTYPEAGNIGGGGFMTVYMGGKPYFLDYRERAPLAATKDMYL
DKDGNVVKGMSLYGDRAVGVPGTVAGMWEAQKRFGKLKWKQVIAPAIRYARDGFVVDEQL
AQRGVDASKEFGGKTNFDQYFSGLKAGATFKQPELAAVLARIANDGAKDFYDGKTADLIA
ASMKKGGGLITKQDLVQYKAVWRQPIQANWNGYRVITAPPPSSGGIGLVQLLTMKADRKA
DFDGVALNSPQYVHLIAEIEKRVFADRAQYLGDPDFYKVPVAQLTDAAYIAKRAAEVNPE
KPSDTKSVLPGLGTSMPEKAETTHFSVVDKWGNAVSNTYTINGYFGSGVVAEGTGIVLND
EMDDFSAKPGVANMFGVVGSDANAIEPKKRPLSSMSPTVLTKDGKVALVIGTPGGSRIFT
SIFQVITNLYDFKMPLKDAVGAMRFHHQLLPPNTIFWEPYHPIDGELAKGIEAKGYTLKG
QDFSGDIQAIRIDGKTPEAVSDPRGRGVSRVIR*
>GCA_000452945.1_BPMSHR5858_genomic_01247_1 group_718
MRRAPPIPILFAVACVAARAAGLPMPVLSRLPPGYEVMAARQGPDVDAGRISYLIVLHRP
ADSASEPSPRPLVIVEQQADGTFRLAARNDEVVLRANEGGQCDPFDPQDADENGLAVKGR
FFTVQNFVACGQHWSDYVTFRHDARTGRWLFANEIRTESFPLEGKPDRVRAIRADPRKPV
ALDAWRRGD*
>GCA_000452945.1_BPMSHR5858_genomic_01251_1 trmR
MNERQWSAVDDFFCSQLAPSDPVLDAALASSRAAGLPAINVTANQGKFLNLLATIRGARR
ILELGTLGGYSAIWLARALPPGGVLITLEANADHAALARENLGRAGLGGVASVVVGRAKD
SLERLIAERAEPFDLIFLDADKDSYPAYLPLMVALSRPGTVIVADNVVRQGRVADRRNQD
PDALGVREYFRLIAAHPRLTTTALQTVGSKGWDGFALTIVTA*
>GCA_000452945.1_BPMSHR5858_genomic_01252_1 group_3183
MPLGYMVIAGAVAVAIVMMSICAAILRDSRDDAFEHAQRAARNTLQMIERDVTRGFRLYD
RTLARIAAQFGDRRIAALPAGLRREILFDSAAAANEAGVMYVLDASGHVTLASTRGPLPA
ERFASYDFFAAQRDTPALGMYLSGPYRMSSRDGEPYVALSRRIERADGAFAGVVVLAVRI
AYFRQLLAAANVGPHGALVLMHTDGRILMRIPFDEKFVGLDLTGTSLYSRMQFGRSGEFF
DVEPIDHVKRFYLYRRLDDVPLVVQVAIADDDIYATWRLRAWRFGVLTGVFALIFVVMSA
YLAHSLWRKSTAEAALARLAGTDSLTGLYNRRALDETLEREWRRAVGGDRPLAILFVDID
HFKRYNDTYGHQVGDEVLATVSRCIAQQAARLGDVVARYGGEEFVVVLPDTPRGGALVVA
ERVRLGVRMLGIEHEGSEVGFVTVSVGVAVWHPDGASAPHIAAIVEAADQALYQAKADGR
NRVVDLGLG*
>GCA_000452945.1_BPMSHR5858_genomic_01253_1 group_1799
MAYAKSSSSVPTEFLYIGAAGEHYIMSECFRHNMEAFKLPIDKGFDLVVTRAYRHLSRLD
DVRKPVDPGLPDTPIYVQVKSRQATPVAPPRKKGERPVWEGHFPIKPADLDLICATPNSA
LACVLFIDTRGELMRSRTAYAWWMSSAYVNQLRDAGHFIAMPRKDALELWVRYVEPAPDS
GYKQNTYISLLKQCQIKGSEPGKKSTGFLLSEERFDFGKLGVE*
>GCA_000452945.1_BPMSHR5858_genomic_01256_1 pcaH
MDSSTLAPRDWPSHPAYIHPPYRSSVKRGPTQPLIPLKETLRNQHAPVYGADDLGALDHD
LTKNAAKNGEPLGERIVVTGRVLDEGGKPVRDTLVEVWQANAAGRYVHKVDQHDAPLDPN
FLGAGRCLTDANGRYRFLTIKPGAYPWGNHPNAWRPNHIHFSLFGDYFGSRLVTQMYFPG
DPLLALDPIFQGTPEDARERLISRFSIDTTEEGFALGYEFDIVLRGRDATPMER*
>GCA_000452945.1_BPMSHR5858_genomic_01260_1 gsiD
MNATVSSPPSAPASAGAPAAIRTPWSEFWRKFRKQRVALAAGAFVLLLVAVAIVGPHVVP
YDPENYFDYDALNAGPSAAHWFGVDSLGRDIFSRIVAGARISLAAGFFSVALGAAIGTFF
GLLAGYYEGWWDRATMRVADVLFAFPGILLAIGVVAILGNGMINVICAVAVFSVPAFARL
VRGNTLALKQMTYVEAARSIGASDRTIIMRHILPGTVSSVVVYFTMRIGTSIITAASLSF
LGLGAQPPTPEWGAMLNEARADMVNAPHVALFPSIAIFLTVLAFNLLGDGLRDALDPKLE
RR*
>GCA_000452945.1_BPMSHR5858_genomic_01261_1 gsiC
MLNFLVKRLFGLLPTLACVAVLVFLFVHLLPGDPARLAAGPEADDATVALVRADLGLDKP
LPAQFASFFARIAHGDFGVSTRSKRPVATEIGERFMPTLALTVVSMAWATLFGMAIGIAS
AVWRNRWPDRIGMTLAVSGISFPAFALGMLLMEVFSVKLGWLPVVPDGSWKSYVLPSVTL
GAAVAAVMARFTRASFVEVLNEDFVRTARAKGVREPMVVLKHCLRNAMIPVVTMMGLQFG
FLLGGSIVVEAVFNWPGLGRLLVDAVTMRDYPVIQAIVLLFSLEFILINLSVDVLYAVIN
PTIRFK*
>GCA_000452945.1_BPMSHR5858_genomic_01262_1 gsiB
MTQPHSFLPFSPRALLVACAGAIAMSSAPLALAERAAVMAVDSTFTTLDPYDSNDTLSQA
VSKSFYQGLFGFDKDMKLVNVLATGYEASPDAKVYTIRLRQGVKFQDGTDFNAAAVKANF
DRVTDPANKLKRYNLFRRIAKTEVVDPYTAKITLTEPFSAFINVLAHPSAVMISPAALKK
YGREIGLHPVGTGPFELVEWKQTDDLKVKKFAGYWKAGYPKIDAIDWKPVVDNNTRAALM
KTGEADFAFRIPFEQTAELKASPKVDVIERASIIQRYVSLNVMQKPFDNPKVRQALNYAV
NKEALAKVAFAGNATPATGIVPQGVDYAVKLGPWPYDPAKARALLKEAGYPSGFETTLWS
AYNNSTSQKAIQFVQQQLAQVGVKAQVQALEAGERVAKVESAQDPAKAPVRMYYIGWSAS
TGESDWAISPLLASSSIPPKLVNTAYYRNDAVDADLAKALETTDRAKKAELYADAQRRIW
ADAPWIALVQEKIVYARSKRLQGAYVMPDGSFNFDEIAIK*
>GCA_000452945.1_BPMSHR5858_genomic_01263_1 gsiA
MSASRAAPSLPDARVLAVDGLTVTFRREDAAFVAVRDLSFHVDRGETLAIVGESGSGKSV
TSLALMRLVEHGGGAIAGGAIALRRRGGAVLDLARATPSTLRTVRGADVAMIFQEPMTSL
NPVFTVGDQISEAIALHQHKSAGEARAETLRLLDLVRIPEARRVFARHPHQLSGGMRQRV
MIAMALSCRPALLIADEPTTALDVTIQAQILQLIRGLQDEMDMGVIFITHDMGVVAEVAD
RVLVMYRGEKVEEGACDAIFAAPSHPYTKALLAAVPRLGSMRGTDAPAKFPLLRFDPAAG
DALVVAGGDATAASGDAAREAVLFVDSDAAAASAASTVSTVSAVSADSAASAAPTACARP
AIDAGAPPLLRVRELVTRFPVKSGVFGRVSQYVHAVERVSFELRAGETLALVGESGCGKS
TTGRSLLRLVERVSGSIEFEGREIGALKGRELQALRRNIQFIFQDPFASLNPRLTVGFSI
MEPLLVHGVASGRQAQARVDWLLERVGLPADAARRYPHEFSGGQRQRIAIARALALNPKV
VVADESVSALDVSVQAQIVNLMLDLQRELGVAYLFISHDMAVVERISHRVAVMYFGQIVE
IGPRRAVFETPRHPYTKKLMSAVPIADPARRHAPRTLPADELPSPIRALGDEPEVAPLVA
VGPAHFVAEHRVGGAY*
>GCA_000452945.1_BPMSHR5858_genomic_01264_1 iaaA
MRRAAAPAAPFQENRPMTSPAIVAIHGGAGTILREAMTADAEAQYRAELAAILAAAQKVL
AEGGSALDAVSVAVRMLEDCPLFNAGRGAVYTADGTHELDAAIMDGATLAAGAICCATRV
RNPVLAARRVLEASEHVMFAGAGADAFAAAQGLELAPPGYFDTEPRRAQWLKARRAAGTM
LDHDAAAFAFGRGGGDGGGNDGGNGGGPGARAACAPEPLDPDRKHGTVGAVARDLHGHLA
AATSTGGITNKQPGRVGDTPIIGAGCYANDATCAVSATGTGEMFIRLATAYDVSAQIEYR
GASLAGAAYDVVMNKLPSIAGRGGIIAVDARGNLAMPFNTEGMYRGYARVGEAPVTAIYR
EDAA*
>GCA_000452945.1_BPMSHR5858_genomic_01266_1 betS
MPPSRSTPRTTLKPPVVVPSLAVIGALLAVCALRPNEAGALFGAAQQWIVERFDWFYVLA
ITTFLIFLVLIAASRFGNIKLGPDDAEPEFSFVSWTAMLFAAGMGIGLMYFGVGEPIQHY
LSPPTAAGGTPAAAREAMLMSFFHWGLHAWATYGVMGLVLAYFGFRYNLPLTLRSGLYPV
LREGVNGWIGHTVDAFALVGTVAGIAVTLGYGVLQLSAGLHTIAGWQTDTDTFRIGLVTV
VVALAGLSAASGLDKGVRRLSELNLMLAFALLAFVVVAGPTSFLLRAIGDNIGEYLSKLV
SLSFRTYAYEAPNDKGWFGGWTLLYWAWWVSWSPFVGMFIARISRGRTIRQFVIGVLLVP
TAFNLVWMTAFGNSAIWLDTHAAAGALAQTATNVDALLFRFFDYLPLTQLLSVAAIVLIA
VFFVTSADSGAFVIDAIATRGAPQSPVWQRLFWAAVLGVTASVLLVAGGLKALQALTLVA
ALPVALVMLALCYGLWRGLKADHAHYSQDMAPATSFWTGQHWRHRLSQILRHTSDADARQ
FIAQTVEPALRKVADELKAGGVDAHVARDDDDAVRLTVPAPAQRDFVYGARVSRKSAPAF
RIREAAEPEPQREHVCEVLTFFADGRLGYDIEYLRGDEIIADVLRQYERYVSLAADTRTH
LLNRAPEHAGPDGLSR*
>GCA_000452945.1_BPMSHR5858_genomic_01269_1 plcB
VAKILENVEHIVVVMFENRSFDTMLGWLYSPDAPPAHVLPPQSSPQFDGLRPEMSNPSKA
GKTIAVTREALNSVIPDPQETFVNVTSQILGPSDAGNPWPPMQGFVVNYETTQTTDPGHV
MQCHSVAQLPTLTALARAYAVSDAWFASVPSQTWPNRAFAHAGTSNGHVNNGAIPDPLEW
DVPTIFDVLSSIGASWAVYSDTTVTPSLTHTMFPSLWKDAYAERFRRFPDFVAACGNDAL
PQYAFIEPSFLFEPNDQHPPHDVRAGDKFLHDIWTAIRTSPGWPRTLLVVTYDEHGGCFD
HVLPPANATPPDNASNPGDEAFGFYRFGVRVPAVVVSPYIEAGTVFRSPTNVPYDHTSIL
ATLRDWLDIPPDRMLGSARIANAPTLAQLVTRDTPRTDLPTIAAPSTDFVHPPPDAPLND
VQKSLVTGTSRRDGHDPAAGVAPIATRQHALDFFKRRAAAKTA*
>GCA_000452945.1_BPMSHR5858_genomic_01270_1 group_7332
MKLSPGLVAELTVNLLLPWAAYRAALPYWGETGGLVASAVPPLVWSTVQLARFRRVDALS
ALVLFGIVLSIAAALAGGSARMLLMRESLASGAIGVAFLASLALRKPLVFHLARATLARE
AHDGAARLDALWAERPAFVASMRLMTLVWGCGLVAENLLRAWMVWNWPVERVLVVAPFVG
YGVYGALMMWTLWYRKVMRANALAGLPPGGALG*
>GCA_000452945.1_BPMSHR5858_genomic_01271_1 group_4568
MGFVEVERRRANVGKPNDIDGATLVSEVRSEPLVSAGGAPCDDAPLTLYGAYRQGFADYL
AGRPNPYRIGSRLAGIWHEGHAEAASNDALDFAVRVRFCWGARAAANPIAFFRRRR*
>GCA_000452945.1_BPMSHR5858_genomic_01273_1 group_6968
MTQRLLAAAWRRPVWLLWVPPAFWAGNVVLGRAFSTSFPPVSLAVGRWLVALVALAPFVW
RDVLRERRLLLRHWKLIWACGALGIAGYNALAYVALRTVPAASVAFLNSTLPLMVPLAAA
MLGVERITSRAIVGIVVSFGGVAWIVARGHWEDLVALRFDHGELIVILATANYAIYSVLL
RRKPAAIGPLAFLAATMAAGLAVLAPFWALELAGGARIPFDPKAIGAVLYIGVFASLIAF
ILWGHCVQVLGATVTGVSFHLVALFTAVLAALTLGEPVRSYHLVGIALILFGFFVATRAA
PGVRSAPNARSL*
>GCA_000452945.1_BPMSHR5858_genomic_01274_1 alaS_3
MNRLFCQDDPDTLLLSGTVVDTRPGAVRLSRHPFYPGGGGQLPDLGTVRWEGGVETVAGF
SEDGGGLWIEFANGVEIVGDVQSSVDAAFRRAMRELHTDLHVLNALVYRDFDGALVTGVQ
MNADRTARIDFDLPDADNDRLRALEAEMNDVVRRDLPVRTRYVPLADAQQASGLLRSRSV
APPPTADGLVRIVEIDGLDSQACGGTHLASTGGSSPVRILKIENKGRSNRRVRIGLEPR*
>GCA_000452945.1_BPMSHR5858_genomic_01278_1 mppJ
MQNVDRPTGSSLAGAAILLEIGAAYGIVDFIRHSNGFDVAALAERSGIGEPMIADYVDAM
CEAGLVTLAKAGERDIYRPADDFPDVVNDVGYLSWALLACAPLIVNAREFAADNALAQAR
HPRDGGLVARTSKWMGELSFYPQPERAILKLRPKKIVDLGSGSGGFLIRMLHQLDGARGV
GVDLSATATEQAKQAAEANGFAGRLEFLHRPIQSLVDDPAPLKGADVVHAGFVMHDLLPA
EERTLDALLETCREHASSGTVVIVDAVPLAPDRWERSFAAAFNHLHRNFMSRRLLSEDAW
REKFEQAGFGNVAVETLGHPGGRMILGSA*
>GCA_000452945.1_BPMSHR5858_genomic_01280_1 group_1801
MDISKIQHSAVALDASSDSSSGQQASGHTAARQTSSRGAGAALGGLKELQDLKRGDLIME
HGNMHAVMYAGGDKPLVHSTNGKFFGVLQQGPNYLTKKFGNEAPTKLPVYRMKNADSPIP
SEAARIGEQWAIRSGDTETTKTKTPFGENRLNDADEDKAVEWSPEATFRAARAALRSDEG
MPLSKQQGVSCSQFVTYAYQAAALKKGGDASGGAHEALDSVKQAGAHRQGAFKTAENKVQ
QQAGLVQGAHEQATQAMPHTLQVNAKHMTADVLAERLRERGGDFERVGYAVPYEKDGESG
VKLLPKDDPRATQKISNIE*
>GCA_000452945.1_BPMSHR5858_genomic_01281_1 acdS
MNLQKFSRYPLTFGPTPIQPLKRLSAHLGGKVELYAKRDDCNSGLAFGGNKTRKLEYLIP
DALAQGCDTLVSIGGIQSNQTRQVAAVAAHLGMKCVLVQENWVNYHDAVYDRVGNIQMSR
MMGADVRLVPDGFDIGFRKSWEDALADVRARGGKPYAIPAGCSDHPLGGLGFVGFAEEVR
AQEAELGFQFDYVVVCSVTGSTQAGMVVGFAADGRADRVIGVDASAKPAQTREQILRIAK
HTADRVELGRDITSADVVLDERFGGPEYGLPNEGTLEAIRLCAKLEGVLTDPVYEGKSMH
GMIEKVRLGEFPAGSKVLYAHLGGVPALNAYSFLFRDG*
>GCA_000452945.1_BPMSHR5858_genomic_01282_1 group_4572
MRATKKSSPKAAAPPAGVDIDRVDRAILRVLQRDASISNVALAAKVNLSPPACLRRVERL
KEMGLIRGIVALLEPKALNAGMLVMIGVVLDRSTPDSFGAFEQAAQKVSGCMECHVVTGE
FDYIMLIRTRDSDTFNRLHAEQLLYLPGVRQIRTFVVLKEILSTTSFPI*
>GCA_000452945.1_BPMSHR5858_genomic_01283_1 group_2164
MALPQDSVNMALFCDFENIALGVRDTKFEKFDIKPVLEKLLLKGSIVVKKAYCDWDRYKT
FKAAMHEASFELIEIPHVRQSGKNSADIRLVVDALDLCYTKAHVDTFVIISGDSDFSPLV
SKLRENAKRVIGVGVKNSTSDLLVANCDEFIFYDDLAREQQRALAKRDARKAEAGAKRTA
DDDKHRKHDGDARKAEAISLAVETFDALASERGESGKIWASVLKSAIKRRKPDFNESYYG
FRAFGNLLEEAQARGLLEIGRDDKSGTFVFRPIQTAAVEVMRPIEAAVDDASAGAKPSGK
KAHGKARSAKKRMPEQMPLIVETAAGAEGHAHADIEANAELAEASDSAPRGMRRERPADD
ARDTHPAHGAHEPHETRAAHEPARAGESAKVAEVAEAAEAQAADTAKPARSRARKTAVKK
AKGTQARAADASPVAAAEADAAGEPPHAAPHAFAPESATEAAPAKAPRKSAPRARRPRKT
AATSES*
>GCA_000452945.1_BPMSHR5858_genomic_01285_1 group_7334
MHATDDRLPRYQRLRDELAALIAARHWRLGEAIPTEQALAKRYDVAIGTVRKAVDLLVAD
GMLERFQGRGTFVRRASFDSSLFRFFRFQGRHGERRIPESRILRREVVEAPSAVAAALQI
PARASVIEMTRVRLIDGAPMLAEEIWLPYDRFAAFARLELASIGDLLYPVYEAHCNQIVA
SATETLTVEAIGPVHARLLRIEPGTPAVVIERLAYGYDRRPLEWRRSRGPASEFIYQAEI
R*
>GCA_000452945.1_BPMSHR5858_genomic_01287_1 group_1343
LSEQEWNAFAPAAAAPHAACGITAIDAHAHVFETGLPLAGRRRYAPDYDAPLDAYLAQLD
AHRVSHGVLVQPSFLGSDCRYLLRALARQPRRLRGVAVIDAGCAPAALDALDRAGVVGIR
LNLIGMPDPALDGPAWRATLERVAALRWHVELHAQAQRLERLIAPLLAHRVNIVVDHFGR
PEPARGIADAGFRDLLRAAATRRVWVKLSGAYRNWPFAPGEAHARARAAFDALAAEFGAE
RLVWGSDWPHTQFERTETFGRALAFARRWIPDDAMRRAIFVETPARLFRFGEQRGA*
>GCA_000452945.1_BPMSHR5858_genomic_01366_1 group_3197
MSSGQPPSVNEGASQPEGAIGSARPASLSSGSHGFARPPTILPRSAVRLDRRIAAVVTLL
PALGTLAAIALWIGGHGPGAVEWIVFALFYFATALGLEVGFHRHVTHKAFKAKPWVRTAL
IALGSMGAHGPVNWWASTHRRHHSTSDGDGDPHSPHLSGEGFGGRLRGLYHSHMGWLFVG
ESTRPAGWEKYVPDLYQDPLVFRQHMAYYRWVAIGLVLPPLVCGLASRSWTGVLLGFLWG
DMARIFAVSHCIWALNSLCHVIGRRDFHTTAHDRSRNSLLLAIPTFGQGWHNNHHAFPAS
AFTGLHWWQIDPGGLLVRVLERLHLVYDVHRPNAELIEKKRIAP*
>GCA_000452945.1_BPMSHR5858_genomic_01389_1 group_3201
MSSIAIQITHDFTCPWCARGEARERRGIARSGVRVPLVVTSRAFEVDPARPREGADRLAH
RMRRLGSRARAQRADAAARPIRRPARGGRANPMTFTASTR*
>GCA_000452945.1_BPMSHR5858_genomic_01390_1 group_4590
MNANLASSATACPVPQMARGLILLFAFSCGAIVANLYYAQPIIALIAPDLHMSGGMASLI
VSLTQIGYAAGLFFLVPLGDLVENKKLMVVTALTSIASLALAATVRAPGLFLAVSLIVGF
SSVAVQLLIPLAAHLAPSESRGRVVGTIMSGLLLGILLSRPVASFIADHFGWRAVFAFGA
ATMAVVTVLLMLTVPSRRPAHQATYMQLIRSLGRLVATQPALRERSLYQGLMFGSFSLFW
SAAPVELMHRHHLSQSAIALFSLVGAMGASSAPIAGRLADAGHTGRATVVALGLAAASFA
PALFVPGAGVAGLVATGILLDFAVQMNMVLGQREIYALDAHSRNRLNSIYMTSIFVGGAI
GSALASALYEHGGWTWIAIVGALFPLAALARFAFASRAAR*
>GCA_000452945.1_BPMSHR5858_genomic_01393_1 group_5279
MNHLRNLLGHLKHTDDFAAYRLPHDLIERACRYGSLVRVERGATLCASGARSDALYVVLD
GATRCLPHARLPLPCGIDHGPGTFTGELDLLNDVGCEVTVVASSACRLARIERATLRHLL
RREPALRDVLVRVFLLRRIGALASGAADTPLRERGAEPAFDVAIVGSGAAGMIAAAHAAS
LRLNTVVAEGRLPFGIDDEFASAAPPASPAPPARRQFGPFLTLRDHTVRLDCNAYPFRLV
THGGERLRAHAVVIASSLRPEYSPRRARVTLRANTEWLDGAVDVDDDGFVRTGRAASGTT
LAAAFESSQPGVFAIGAVRAGPRRSALGELTEGAVVVDAIRRFIRASAV*
>GCA_000452945.1_BPMSHR5858_genomic_01394_1 group_673
MKPNLPSLLTFNGGYVDTAGFLALQGLFTAHVTGNFVTLGATLVAGSTGAIAKLLALPVF
CGVVLAAGVARRLMLRAHAPALKILLGVQWLLLVAGAALAIRLGPFANGDAWPAIATGMA
LVMAMAIQNAVHRLHLPDALPTTLMTGTVTQLMLHLSERIAGDAAPVRGSPARLANMAWT
VVTFALGCAAAALAYLLGGAWAFALPPVLIAFALFLGDERKLEHAL*

>GCA_000452945.1_BPMSHR5858_genomic_01395_1 yhhW_2
MIEVRAANQRGRAEHGWLSSRHTFSFAEYYDPAQVGFSDLLVINDDRVAPGRGFGMHPHR
DMEILSYVLEGALEHRDTMGTGSVIVPGDVQLMSAGKGIAHSEFNHSADRPVHFLQIWIG
PSVRGVEPRYQQASVSDDDKRGRLRLIVSPQGQDGSLKIRQDARVYAGLFDGDASERFAL
PPQRFAYLHVARGSVTVNGVELHEGDGARIRDEQALQIANGKQAEVLLFDLRPIEVTAEW
A*
>GCA_000452945.1_BPMSHR5858_genomic_01396_1 dmlR_14
MQFDDLRIFVATVEAGSFTGAADKLLLSKQFVSRRVMALEASLGVRLLVRNTRKLAVTEL
GHELHARATRILADIAETEEAMSARRGDLRGTLKISAPMSFGIAHLSPLIAEFLVRHPAV
RLHVELSDRHVDLIGEGFDLALRIGALPDSTLIARRLGELRMVACCSPAYRRKHGAPATP
ADLHGRACLLYGQEGRTGWEFMIDGARRTFEVQGALLANNGDVVRDAAVAGLGIALLPHF
IVAHALDTGRLVAVLEPFAQAPLPINAVYPQHRQDMAACRTFLGFIEERLRERFAKPRR*
>GCA_000452945.1_BPMSHR5858_genomic_01397_1 group_4592
MFFCYVLFLIFYWSPAIVLQWGTYGMFDDAVPMWYKALRLIVPAYGLVVLARQPYYTPAH
LLYFVFGAWCVLVSGDLATSVNVLVFCATLVIMKGVRIDAAALRRVILAALPFVYGMGLL
EVFGVFVSAHTYGSELRVVSTFGGPNNAGIIFSSWALYFIIEFRNARRLRDLGHAAACVS
FVVLTGSLSAAVALAACAFVLNAYLIVPAVLVVVPTFLINDFFMLKLSYLMATLNGQGET
TGSFSDRVENVATIARVAGDNVVNLIVGSGLHSESDYLSVLGQFGVVGLILFICVLLTLP
RSLFMLLGLLQSLVTPFFFSFPSFPLIALLAAGEVAAEARKRAPRPAPG*
>GCA_000452945.1_BPMSHR5858_genomic_01398_1 group_4593
LPISITVGVPAYARPDELRELIESVLACDPPPDELLICEDASPQRDAIRQIVYAQRLALA
RSGCRLRYVENERNLGYDGNLRMLIAQASGDYLMLLGDDDQLYSNAIAETRRYVGAHPEV
RFVSRTYRRFSGHRDRGVLNETWLAKSDAVFDRGNSSPALIFRLCGFVGGLVVKTDWARE
LATSRYDGSLYYQYYLACHAYFGAGIGYIGKAIVAGRAQNPPLFGHAAAEAGVHVPGAYR
PAARARMWRGVLDISADVERALGVPLVGAVMREMAGRQSFHIFEMMPAQGRRATLELFVE
LRKLGLVRSPLPWLLASFVLTFGAAAASGFRMLRRLQFAVERGLRIKC*
>GCA_000452945.1_BPMSHR5858_genomic_01413_1 ilvB_2
MKASNAIARFLAAHDVRHCFELVGGMITHLLDSFAESGRFQIVSMHHEQAAAFAAEGVAR
RYMGRKVAVALGTSGPGATNLITGIGSCWLDSVPCVFITGQVNTHELKGERAIRQQGFQE
LDIVPMVRSITKHAVQVSDANELLPALHGALSAALSGRQGPVLIDIPNDVQRAEIADAAV
DRWLGEPLRIRDRCEPGDRDIDALRELCRSSSRPLVCFGGGARWAPSMPAWIAALEDARI
PYVSTLMGHERVPAGAHYFNMIGAYGNREANWAVQHCDLLIVIGARLDVRQTGADVGDFA
RHARVIQIDIDPAQLNNRIKVEQALCCDAESFFRRFAVRRDTFAGIRPDWADQLATLRTA
HARNEYPDWNISPGVVFARLNEAMAGKAVHYVCDVGNHQMWAAQGLRLGATQAVHYSGGM
GAMGFALPTAIGIACSGDGGGKTVVITGDGSLQVNVQELDTLSRLQLDVAIVVMNNYSLG
MVKNFQDMYFDGKNISTKQGYSCPSFVALARAYGIESMTVAAPAELQRAIDRVAGSRRPL
LVEIAMPEATECRPRLAFGRKLDDQYPKLD*
>GCA_000452945.1_BPMSHR5858_genomic_01415_1 group_674
MRSFRDTAGDSNAPALPSRPNLLLPLPSPAMKLNNPAGFFFTLFKKPTAANSPPDPASVM
AAPIPRENNIAIDDGRAVVRDSLSSAASVAATITGKGARCRERIRPEPTSANIRAAADFL
GAVRTAITNGTFDQPAERAAPGGLQAAREARRAPYGNCIGSRALRRVRRQDERRRREARR
DGIDRGGRVPQGRMRRLRSQEGRPRVDEARRAQCRQPPHADMPIGHGGRFAEYWRSPVCS
TALLTIAPVSMSTWST*
>GCA_000452945.1_BPMSHR5858_genomic_01416_1 group_4600
MAKELNAPSQQALVTSFTIKPEDVTGTQITYRYDTMPGNQPNAYGNTVFIWQTSQVFIPV
STQPAHSQIVQNNQPNGSSIFQGLSVSSESYLVAFAVGNSVKNIVAAVFVPSTDSHQPPY
DPVQAPSILVTNKGSTSVSFSYAMPAGMTPQADGDWVGLWQGQTEAVLYNMPPTWFVQLG
SNSNAGNFGINLSSGNIQRGVTYTLGYFKGGYAQSNPKQTTLAASTTFVG*
>GCA_000452945.1_BPMSHR5858_genomic_01418_1 group_487
VAGLNDVIERLSQGTQNVLTVHLGARITKVRDYLNALDEIRVFTANSQNFGHQSSSVNIL
RNLIRMGAPGPYTFALSASNSADYADLEEKIRLLIPQFRQVGVTFELGAGGRTADVTVVR
LDKALAPAQFAISGGFDDLENKTPPYHLLNVTNYVQLQPYAWNRGTNMVRIMPPGGTASE
YNLDELNPTTLLARRAFYLADPELTQSDREAIAQTPYANKARVIERLLERREAGEITLFP
VYGVTTKGSAYTSLYNAVTGALIAQNTHPAVKKTVMVQITTLTASEWEAFLFLMRDPAGQ
MVNKIRTTPDFRGWNEENKVKDRVQDLGSPNSVPTVEQLDEELSYLEDDQLLVVYIGKIP
APLFDALYASATLPPVLEGQNTAELMLNLGKPYFKITSNNSREADARFSYATLPLSSTGA
GTDATNSLDESFNGIYFTRPDNWYRDRPTYPPTQLPAMINAYVQPAGNARATYFAAQRTF
FHDELNDKLLRGLDLFVNLIGPAALEEHRLALAHAELDDANAPVHEAGAAARAPARIAHR
KPPTRVAGGDANGASELLEAFYEDLTSHTVDGVLDFLLAVTDGILNEFFRQVVIDVVFTI
TDTVTEINADKTEVTLTGKSKAFGAGNLTLAFSFTDNGGTIAGKMSGAFTDTVWAFPGAQ
WLSVANPSLALAIDSNAAVPVTGTVGATFTAGIAAKASLTLPSEPGRLLLQAEFLTPRPS
ITNIFQMLGGINIQALLPSQIQFFSDIEVQNLALRYSYANGVMEYIGVTLGTPENRSWQL
VPGVTVTGLSFSALTDYPGDLQRRSTRYVIGGRFDIAGGHAQLEARVPALRVTGGLIDGS
PPITLAAIVTEYLGADFAAAIPASVSSTAIEQLSFMVDQAQGAYSFSMDVSAQWPVPSAA
NALFTITGLNFAIDAVSRDINPPKADAGGNNGAGGTQTEIEGSFGGSLIVLPNSESPIGL
STTATYKTAAKAWTFDAQQTSGVVSLGALLVYYLGNTWQAPQGQEYAIDGLGLTITSSPT
DSTWAFTGKTADNWVVPFLDVSLAAKLRMGDAGAKAEVPGKFGRLDLEVIWQNIDLTVWF
DYNPKIKQYGITWGLLEGVVDGPDPTTQDWTATLGFKQNTTLGSMIETMVSWATGSKFGL
ESPWSFLNAIPLSNLALKYTFNQTTPSRNKVSFAVTIGPINLGFARIDSIDVGYQSTGED
RGVMVTLNGSFFWQSDPSTPLEWDASKPGTAPAPPGNGNKYLDLRLLAMGQHITLPCFAT
ADTVQKAIACMATLPDPKPGQIPAVRFDAQSAWLIGTDFGVLKIDSGQTGNNANALRVTN
DGSALAESSGYVLTLQAVFNDPHLYGLRIALDGAAAKVFKGLDFQIMYRQVSDTVGVYQA
EITLPDLMRHLTVGAYSLTLPVFGIAVYTNGDFQVDIGFPWNENFSRSFTIEAIIPPGIP
VLGSAGFYFGKLSSASTNRVPASSYGTFNPVLVFGFGMQVGFGKSIEYGILSAGFSVTVV
GILEGILAKWNPYQLTHSGREPSTQLQGDYYFWLRGTVGIVGRVYGSVDFAIVKANVDIT
VKLLLLQLTYESYVSITITVIASVDVSVSVKINLGLFKISISFSFSMRLKETFTIDNRGAA
PWLGDGRNVRGVLRLPVERRLSGFARAQARDSLLVSAPNWGNLRPGGVTDLSGYLVPGLT
AARDEWTPQGEPANQLSCWVALLLIESVPPAGQDAGASKLKAAGSAPDSSFEALAKMVLR
WAIAAVQGPMTPDEVDRCPVPATLLDWLADDVLVSTGDDPTPIPLDAVQAFLDTHFRFNL
RVPPTDQDASADTAYFPAPPQLRVMIPPYGNDYPGVQYTLGSYNALGENTLAELRAWFDQ
LAVQVEREQAANGAAARAFVEEAPLSMAGWMFSDYFLLLARQMVKAAQDALRDFKYALDA
NETPDDVVSWVNTTGQLNGLYTLNDVFGANALHALVAEKTLTIGVTSSISLAKTGQTFTS
LAKAFDDALPASAIASANAADAALLQPGATITYPGFDPYTSVAGDTLVSIAAHYQAKLND
LLADSDVLDAAGMLRIGASALMPYTAYTALATDTFASVAALPVYAGGFGAAALATANAGR
SVLLEGVKIEYPDKDAYTVQPRDTLGDVANAFGVTVSDLLATSAVLTQPGLLAPVASLTV
PAFRYTTQQGDDLAQVAARFGVTVSVLADQPANGTVAGLFDTGDTLDLPHLPQFPLAELL
AEAQRSGMLQHLSGIASSYTMHGLRFPTSGPTGTGGQWSIVPNEMGMWVHDVNGTLKLPP
QAGLYALTGQQFPLPALGADPFAATFDSVAGAGSSWLRFVDGNGGPTDRLTLSVTPGTPD
ATRIAQVTAAAKTRLVVPMDMLGAGKMYDTALATYPFTSALQWLSTNTVALPYGQPPAGV
QSLRVWQLPGALAALPDPATHAVNPRFALRVARYDDATGATETTGVDSYGWASTIGFTVR
RIPPVAGSPASVDTYEVVGASGAAIVVLEQLLSQVQADDSAYFGLSVGFAPDSATGGGEG
VQTGGAASVVFGIAQVNLSTETRPPAGAAFAALRETAGETPQLTLLNSPSEFVRLLWEAS
ITRSGGFFLYYYDRAAGGGLPDRIFNDRNEASLTLIVLYAKPAAVDDQDRVTNYMNAVVT
TDALDTGNAVLFAEAAPVPATVTSGAGETLASLAAQWYSDEADIAEANANVALRAGALVR
VSEGVYQAPPGGIALAQVASRFGTTVQALNDANPLWGGLPDPLPFPAAIRVPDLTLTAGT
SAHTASLADIAGWYGEPVDALASHNARVAQLFAAGVPLVIPGGPRVRSAAVQPGVQALAA
LRPAPPQVDGTSPDYGTELLLNNFSLLNQQVYGNVDFRPSDPPGLPAGPTTKAPEENGND
KVRTVVPADQVEAWNFSQALPYARFAKHVPQAPRAAVALPPASASPYFGVGGILQISFAW
QDYYGNVLSTPLSDPLAGDAAPYNDAPLLTGYTDPLVSLSQWPSIASNWQVLPGSGGANP
RLNIELSFDPSRYQGLLQASAATQTTITVVFTDALDAASVGELSRWQLVPGTVDSASLAA
DGKTVTLTVPALDDDLRYTVIAADIKAQASDMRYSGQASFDWPDNPVTRSSTVQQNASQD
LHVYTQLYYQLTDPAGVDLSAQSSLLADAHGAPGSVAYAPAAVDKLMDWLFGTAGAASSV
YAFVLDRSKFQSVAVPPAAGLPLDVDVPPQQVNTAQIFPLWTSFTMTRAHGPVLPGLETV
AGIRSASTRVAPLQDALGATGGTLGLVTFATGFEQALSTPGSVRLKVATGVDRTAPPATG
AASTVWAVRVGLAAGKAISYAIADAGNPAVFAPQPASNRLISRTQVPIYDYTTGKGISST
PSRTTDFTDVDLDTWCAQVFAAVDDVLTPQFTAPMQIVGELKSADYLQSILDGKKGLATV
AKLWMIPVFAGETSDPSAAREAFYQQLLVRLSAAYTTRAAVEFHANVTADVIEPAADQPP
RLFGPVTRNGPVFEAANVDGQALTTVFLLFSDPMDPVTAGNIENYALNSGTGVLTATVDR
GTVTLTLATDVQPGQTTVTVSNLKDATGRAVRPPLTRTVTTGSASLPASTLAFSSPKLTL
QAGDTRALTYLVNAPDSVRGAGGEIVSYVELDMTYQGSQIEHQIGALPGIEDYQASTWLS
FVVPDTDGPLAADLGNFAVPLVLRAFPASPAMTEQSGTPTHDLDTASLPLLKQWDYAFTY
SLPFHYPQDRIYGEVEFNLRTAPTLFASFPDAFAQLAEFITVFPKVNADLQTILAGIDAT
VDPMTDQQKIDDASIALQSFIQLVDELVDAAGGNTQGNGERRGGTGLTFQAPARLLTGDP
SLTFAFYEEEGSAEVGDTEGALVVTLVGAVPAGMGQPVVEIDPALYDAQPWQPPGDTQKA
GDVFHYVYKRKAGPGPEGSYLSAANGQNIPGRTVRLPALDILQRQDAWSTVWVERNRELV
PGKPSADAFVYTTPEVRFASPLYPTNDANAIIDVAAIPSGTPVKRSLQEHFDALFAYLLA
GDTLPQIVAQVEVTYGYALNAALDKIVLPVLMQAPLTVDVAGTGAGTIAKMTADWTAAIE
TWFSTYEPTGGGTLWMDLTLMSNLTGQPMPLLRMRRLMLSIAQVVPPLPCR*
>GCA_000452945.1_BPMSHR5858_genomic_01425_1 btuD_5
MTADVLIEARQLARRDAASGKTLLAPSDFALAAGERIALTGPSGSGKSVFLRALALLDPI
DGGELLWRGTRVPRPAIPRYRRSVAYVRQRPATAEGTVEDLLRYPYSLAAYRDARFDAAR
VARLAAQAGRGDDFLRKAASELSGGEAQIAALLRVLQLDPDVLLLDEPTSALDPDSTRAI
EALVAAWFDAAPAARAYLWISHDPAQAARVGRRRLTMRAGVLGDAATQDAR*
>GCA_000452945.1_BPMSHR5858_genomic_01435_1 potA_1
MNSQSGAAMAGAPSYRPQTGADGRAENFVQIIDVVKKFGETVAVRNVELSVRKGELFALL
GSSGCGKSTLLRMLAGLETITSGKILIDGEDLAQMPPYKRPVNMMFQSYALFPHMTVEAN
VAFGLKQEGVPKAELKERVRDALELVQMARYAGRKPHQLSGGQQQRVALARSLVKRPKLL
LLDEPMSALDKQIRQRTQIELVNILDKVGVTCIMVTHDQEEAMTMAGRLAVMSEGQIVQI
GTPHEVYEYPNCRFSAEFIGSTNLFEGATVEDEPDHVFIESPDLPCRLYVNHGITGPLGM
PVTVSVRPERIALTRKPADGAYNWGRGVVTNVAYMGGYSLYHVKLDSGKTVIANVSSLAI
ADLDSPGWGDEIYVRWSAAAGVVLTS*
>GCA_000452945.1_BPMSHR5858_genomic_01437_1 ydcV_1
MKPNRYLQVLMLAIGFGFLYIPIASLVVYSFNDSKLVTVWSGFSTRWYAALVEDDELIAA
AWLSLKIAVMTAFASVFIGTWAGFVLARMGRFRGFALYSGMINAPLVIPEVIQGISLLLL
FIELGKWFGWPAGRGMFTIWLGHVMLCISYVAIIVQSRVRELNPSLEEAALDLGATPFRV
FFSITLPLISQALVSGWLLSFTLSIDDLVLSAFLSGPGSTTLPLVVFSRVRLGLNPEMNA
LATLFIVAVTIGVVAANYVMLRQEKRRMAEAA*
>GCA_000452945.1_BPMSHR5858_genomic_01454_1 group_3209
MVDIVIVASAFGADAVRRAGHRAFVAAVADAGAAGFEVRRELFAADDDAVASALAALGRA
IAAAGLWSVYSTPEALYADDGDLNRDALAAALDQADALGARFVKFQLGGFAANPHAAELA
GATRGARARVLVENGQSRQGGALAQFTTLFAALRDARGPAPVGMTFDIGNWLWAGEAPLA
AARALAAHVEYVHCKAADGEGARRFAIAPPPGDRLCGDALALLPRTAPRGIEFPLDAARV
ADDAAARVAWLAAA*
>GCA_000452945.1_BPMSHR5858_genomic_01464_1 lrp_1
MSDSNTVLERVRAWCAATPRAVAVATADATMTYGELDRASDAVAAFLEAERIGAGSIVPI
EAMRTDDFVAGMLGIVKAGAAYCPIDHAYPEARKTHIVERTGSPLLLTAVSPRTPLACAR
APRTASIAALRRAGMPRSASPRTPRPNDAIYVIFTSGTTGVPKGVVVEHRSVDGLIAWHN
AQFGVDRTSRSTQIAALGFDAAHWEIWSPLCAGARLRFVDDDARRDANALVALLERERIT
HAFVPTVMARDVVAASEPGPSALRYLFTGGEKLNPVDTDRIRYRLIDYYGPTEATMWASF
HPVQSASLGLPPSIGTPVGGARIAIFDERLREAQSGAVGEIVISGPCLARGYLDDPRQTA
EKFLAHPSRPGERVYRTGDLGRRLPDGAIQFVGRLDDQVKIRGYLVEPGEVEIAIARQSG
VRRVAVVATSPADGAPRELVAFVVPADPAAPRRPLVGRLRAGVAASLPPFMVPGHFAIVD
ALPLSANGKTDKAALVAMHGRRAARADFAEVADAVERTVCESFADALGHADFGVDDSFFD
VGGHSLVAAAAVASLSARVGVALRLSDLYRRPSAAALAVDIRRRPSAGDPGALDLTPADV
LRRDAILPDDIAFDGAFDPQRLARPAHVLLTGATGFVGVHLLAQLLATTEAVIHCVVRAR
DAHDAERRVADKLRTYRLGVSERDRARIRCHAGDIAHDRLGMASADYDALSRCVDVVHHS
ASAVNFIKPYAAMKRDNVDGLVNVIRFAAAARVKALSLLSTISVYSWGHRITGKTVMRED
DDLDQNLDAVCADIGYVKSKWVMEKLADAARARGLPLITFRVGYATYHAQTGLSADYQWW
GRLVKTCIALRAVPELRELREGLSTVDYMTAAIAHIARNPAAPGKKFNLTHSGERNLSLE
DFFDRLERAFGFSFARVPFRDWFDRWKDDAATPLYPVLNLFRDPMHGGMCMVELYQHTYR
WEHANTSAFLAGSGVRPPEFDEPELRRYLVQSIGIAPACAAR*
>GCA_000452945.1_BPMSHR5858_genomic_01468_1 lysDH
MKIAIVGAGLIGHTIAHLLRETGDYEVVAFDRDADALAKLANEGIATQRVDSADAAAIRE
AVKGFDALVNALPYYLAVNVAAAAKAAGVHYFDLTEDVRATSAIRELAEGSNRAFMPQCG
LAPGFIGIAAHELVNGFTEVRDVKMRVGALPEYPTNALKYNLTWSVDGLINEYCQPCEAV
RDGRRQWVQPLEGLEHFSLDGIEYEAFNTSGGLGTLCETLEGKVETLDYKSVRYPGHREL
IQFLLEDLRLATDRDTLKSIMRRAVPSTKQDVVLVFVTVTGVKHGQLVQDVFTRKIFAKE
ICGMPMSAIQITTAGAMCAVLDLFREKKLPQSGFVRQEQVPLHAFLANRFGKLYEGGTLE
RMHALA*
>GCA_000452945.1_BPMSHR5858_genomic_01471_1 group_3211
MWYFSWILGIGVALGFGIINAMWLEANGNVPTGAPGDAPGAGGPATAPEGRRS*
>GCA_000452945.1_BPMSHR5858_genomic_01472_1 corA
MLINCAAYQDGRKLADIDIDDISDYVAKPECFVWVALKDPEPDEIERMGEEFGLHELALE
DARKGHQRPKIEEYGDALFAVLHTVEIDGQGELQVGELNVFVGPNYVLSIRSHTEQDFRA
VRARCEREPHLLKEGAAFVFYALMDQVVDRYFPTLEALGAELEALEDRIFEKTGTAAARA
LIEDLYRLKRRLVMLQQHTAPLVEPLAKLVGGRVPQVCAGMDHYFRDVYDHLLRIVRTIE
GRREMLVTAIQVNLGMISLAENEVTKRLGSFAALFAIPTMIAGIYGMNFNNIPELHWRFG
FYVCLGVMAAADLFLWRRFRKAGWL*
>GCA_000452945.1_BPMSHR5858_genomic_01474_1 group_7358
MKLRTWLLLGTSSAVLAACGWFHAGPGERDIDAAVRRALDAENRGALNSMLGQPLPVAAD
VASVKPLGDCTKLAERTYSCNVAIAWHDARSGEGDGALRTTLTFTEDNDGAWQTSHVDAA
LAAGVAKSLIDRISGAVAASGASQAQ*
>GCA_000452945.1_BPMSHR5858_genomic_01475_1 group_3212
VKLSILVAAACAAIALTSAAYAQTSQTYHFGEGQAGMSSGAQTAPARHAAKTPAHRHRKH
RAHRAVRSPRTYAHH*
>GCA_000452945.1_BPMSHR5858_genomic_01479_1 group_1430
MKLRNLSITAAAFALAFGLSACGQRDAGAPPAAAASASIAPAASVTLDASANASPSAASA
PAASNEAAPAATTAAAADTPNPGGEKVFKSVCFMCHQTGAAGAPIAGNKDDWAPRIAQGK
PTLYKHALEGFTGNKGTMPPRGGNPSLKDNEVEAAVDFMVAKAH*
>GCA_000452945.1_BPMSHR5858_genomic_01481_1 group_3213
MTHFFRNLPNEAARQIDALSRLLYDLREDRKRLLAAYGAADEAALFARIAAGEVDEHPAY
EHYLGAKTLADTRETIRGQLRALLLAQGA*
>GCA_000452945.1_BPMSHR5858_genomic_01482_1 nosL
MKRRFLSAAVRTSIAALAAAALVAACGHDAQTPPPAREITDATVSVLDGMSLKDYPGPKA
QIVYADGEPDFFCDTLGLFSVYLRPEHDRKVRALYVQDMGATDWQHPVGHWIDAKRAIYV
IGSKKPGAMGRTFASFAREADAARFAKAEGGKLYRFGEITPEMAATDGGVVKDQTM*
>GCA_000452945.1_BPMSHR5858_genomic_01483_1 nosY
MSTLPSRPLWAQIRVIAGKEFHDRIRNRWVLAISIVFAAFTLAISYFGAAAQGAVGFHGI
EPLIASLVSLAIYLLPLIALILGFDALVGERERGTLNLLLSYPLTRTELLVGKYVGLAAA
LTCATLAGFGAAGAVIAARMSAADWYQYAGFVLSAVLLGWVFLSLAVCASAFCASRTSAS
GVAIALWFCFVLVYDLLLLGALVLTGGAGLGPALPLLLLLNPADVFRVLNIFGPGDLRAM
FGAVSAFPAALASPLLLGPVMLAWIAAPLGLAAWRFNR*
>GCA_000452945.1_BPMSHR5858_genomic_01484_1 nosF
MIELTGVVKTFGPVRAVDGIDLAIAPGELFGLIGRNGAGKSTLFQLMLGLLAADAGNVRI
DGVSARDPRFRAVRRTLGYLPENVVLYDNLTGLETLRFFARLKGVDAAACAPLLDEVGLD
HAAARKVRGYSKGMKQRLGFAQALLGAPRLLFLDEPTNGLDPAGIRAFYRILHERRRAGT
TIVITSHILAEIQQRVDRLAIVRAGRLQAVGSVDALRGETNLPLTIELVVPDDDTREIEA
ALRRAPFGEPTMRDRRIAFACPRDAKMAALAALAPLGARIVDLRIHEPSLEDMFLGYAGC
EANA*
>GCA_000452945.1_BPMSHR5858_genomic_01485_1 nosZ
MPIPHALATLAAAALLMHAACAATLAVHPGERIGAALAAARPGDTVLVQHGRYEENLRID
KPLTLRGVGRPTIDGRLAGDVIRVAAPDVTISGFSIVDSGASLTAQNAGVYVAPGSDRTR
IERCELVYNLFGLWIERSADVRVTGNVIVGKRDLLSPRRGNGIQLYNTTGARIAGNTISY
TRDGIYVDVSHHARFEHNTIHDVRYGTHYMNSYYNVWDGNDVYHNRGGLAIMEARDQIVR
GNRVWGNTDHGIMLRTIQDSLIENNVVAGNQRGLFIYDAEYNTIRGNLVVDNKIGVHLWG
GSIHNDVTGNDFADNREQIRYVAASDVAWPGNYWSNYLGWDRRGRGIGDVPYKANDLVDR
LTWRVPSVKVLMNSPAVQALRVVARQFPLLAVPSVVDDAPRMRAAHAGWPQWVGKR*
>GCA_000452945.1_BPMSHR5858_genomic_01495_1 group_3215
MHLPNRTQVLVIGGGPAGATGAAFLAREGVEVTLVDKEVFPRYHIGESLLPSCLEILTLM
GARDTFDRHGFQRKPGAYFNWKGETWKLDFGELGGTYRYSYQVRREEFDHLLLQHARAVG
AQVHEGVSVREILFDDGRPCAALCVAQGAEEATTVEFDYLVDASGRNGLMSTRYLDNRKF
HEIFRNVAAWGYWEGLSWPDDCAPGSILVSSIPDGWWWAIPLADRPTSVGVVMHRDAFVA
ARRTGTLEQVYAQALALSPVMANLTEHARLVTPLKTEQDYSYTCDSFAGNGYFLSGDAAC
FLDPLLSTGVHLAMYSGMLAAASLASILRAEVTEREAAAYYRDSYRQAYLRFLVFVQTFY
EAHGKLGYYSKADELSHYMIEAGDIRRAFLNLVSGLEDIADAEQATSHLMGEMSRRIDQN
LALRKDKRALSSAIGSTQVEDNARFFDAIEGLPCLSANMALDGLYVSTRPRLGLQRVAAM
*
>GCA_000452945.1_BPMSHR5858_genomic_01548_1 group_759

MDAHADWMRFAKDPAARLMIWQTDETDAQLVQLYFQGQEETSCAVRTMRARFVNEARYAQ
ALTDELVAFYDSRREASYAQGLQADWQAPPRNDGASPVLHLLTVADSLMRHHPDIFPAMV
FVLEPAKVRDDAAWVQWLDGLLTTIAVAPQLGERVRFVVPRTDIAPFATLTQRHRAAVRV
VQGKYSMASVPRELLAESGERGPSGEFRRLFVMLTETIEGGSPARLEELRAAALKVAERE
QWFDQCVVVHLIAGAAYLKWRDRERAIEAYRSAADSGMRAVEAGHPAGHKLVANGLFGEA
SVHLTHKDFARSAYCYERAAASTSAAQDALMTVEAWRMSAVCWEKAGEREHALEAGFNAL
DAGLTIDESMRMNSNLRPVVEWMVSQVGAFDRRRDKLSEKVAALRGGS*
>GCA_000452945.1_BPMSHR5858_genomic_01549_1 group_437
VALPAVKHLDPVVGIDVHSVLVTPGTPPVFLPHPYVGFMLDRREYIDAALGVIGSIVFTF
TPLGKEAEAGLKWAKKQAKAELMSDPLIAEGVKLGKEAAPVAGDIAGAVGAGVGMGSMMG
RPIFVNGFLRATAGTHSFHVPGLHFPLGESFAPPPEDFEPSNDGESYMGSRTVLANNDPM
SFMALPAMSCWSVGLEPPPHNGAHTQRTALSMPSSVMLPIPVGRPVLVGGPPVMNMAALA
SSLFKAFRGSDWAKALADKLHLKPGFLRCNVLRAEPVDAITGEVVVQQHDFTVPGRLPLV
WNRYYTSHDTYVGAVGIGWQTPADIRLELTRHEGAVGVVAYFPDHATAFDAIPDAAGWPA
RVYDWQHGQALYRQDDRLVLRTRAGFEYEFALPAHWQHALERLAEDATLTLPVERMADLN
GNAWVFERGLDSSLTRLVEWKAEEATGRMIACAASASSRAGDHVSLLTALTLIDAGGRAH
PLVSYQHDRNVDLVAAIDAMGQPHRYAYTDGHRMISHTSARGISFYYSHRRHDDGVWRVD
HAWGDNGLFDYRFIYDPARHETRITDSLGHTTLLQSDERGMPVARIDPLGGVTSYRYDAQ
GRTNAETDPAGRTTAWEYDTFANLLMRTLPDGTALRTEYNTGHKPVCVAVPGGGQWRYTW
DERGNLLMQTTPSQASVRYAYDQYGQLIAHTGPRGTVTRFDYDRSGLLAVRTDALGHRTQ
YTHDALGNLVHVINALGQTSRYEYDNIGNLTRAIEPGGREVHCVYDADGNLVRYRDATGN
VTQLVYTALGHISKRQTPDGNVVEYRYDTEEQLVGVVNGRGEIYEIKRDELGRIVEETDY
WGQPRHYRYGAAGELLDITDPLGQTIEYRCDRLGRVVEKRMPDPLHPDGVRIDRFVYDQH
GDLVLAENPSSRVEFRYDADGRVIAERQGDDFTIASTYDASGNRTERKTRLVADSDVVEH
TVRYEYDALDAVIAIQIDDAAPIVLERDAVGQVCVEQLSPELRRELSYEAGGQLAKQTLL
GGTGLLFASEYAYDANDELVEKRDSRTGVERFKYDPVGRIVAHTDPTGRLRSYVYDPAGD
LLKTHIHERRTAGATEMTQTGTWVREGEFEGHYHAYDRAGNLIRRQDVGQDLTLRWDAAG
QLAEAVAVRPAIAGAGGGQVRIGTQYEYDAFRRRVGKLVLTRAAGSAELVLSRISRFFWD
GNTLVGQCTRGGGEGGGTAIPDAGDRMPAAQFIPVRDNRDDAPVSGFEHAYEWVYYPGMF
RPLAVVHCDLAATRASVPTTTNLLPLIGAVYFFQSDLNGAPVQMHAPGGRVVWEARYDPI
GRSEQLGLQLVEQPVRLQGQYFDAETGLNYNRHRYFDPNSGIFISQDPIRLSGGLNLYQY
APETNNWIDPLGCSGHRRRHEKMPPEGAPLTTGNLFRHAQDEGIPPIFSRKDDEFYHRLI
EIYAGTGVLNAYIRGIASHVEPKAGLILNEDGSGWKIGSLYINYPDGPCPDCRRLMPFIL
NDGSILYVTFPTLGLDGYSYGHFHGGVSGFFREGTPCNLRHPE*
>GCA_000452945.1_BPMSHR5858_genomic_01560_1 group_4616
MDLKQLRYFVAVAEELHFGRAAKRLFISQPALSFDIRKFEDELGVQLLARTSKSVALTNA
GEVLLGEARRLLLQAQEAQRLTIRSASGLAGRLKIGFVNSMLYRGLPSAVRRFEADYPNV
EVVLKEMNTHEQAQAILRAQIDVGYAHWGTFPPEIAVDTILSEPFLCCLPATHRLARRRR
VDLAALAHEPFILFPRDAAPHYHDLIIAQCVSAGFSPQIRHEARLWQTVLTMVEFEMGVA
LVPSVLREVRSERLRYLPLAGDALESRTLRLRRSNEREPVAECFGEYLDDEIRRLRGAAK
*
>GCA_000452945.1_BPMSHR5858_genomic_01561_1 caiA
MNMMKETVEANAPAAGAANIPDSRGINFFTSDPDCAALLRLHLGEARFRGFEPELRALGL
RASDELDRLASRADRNPPVLQPRTRRGEPLESIDKHPDYVELERVAYAQLGLAAMSHRDG
APPPLVKYALTFLFVQAEFGLCCPVSMTDSLTRTLRRFGSRELVERFLPRLASRDFDVLY
QGAMFMTEQAAGSDVGRIATRAVRGTSADGSATWRLFGDKWFCSNADADLAMVLARPEGA
PDGVKGLALFLLPKTLADGSRNRYRIVRLKDKLGSRSMASGEIALDGAEAYLIGEIGRGF
HQMADMINMSRLSNGVRAAGLMRRATTEALHVARHREAFGRKLIDMPLMQRQLVKMLVPT
EQARSLFMQIALLLAKADAGDAQAGGCVRILTPLVKFRACRDARRVAGDAMEVRGGTGYI
EEWSDARVLRDAHLGSIWEGTSNIVALDVTRAVKREGALAPLRAFLLGALDTARLPGASA
TRLRRALDRASDAIAHVARTGDDAFVRQAASALYHATTAILLASEGMRLAPDHRRLALAH
LVLRYKLSPVDPLATPRADADAEAEAEAGVLAALLHDRPVTLDDALRLLDDRSGDAA*
>GCA_000452945.1_BPMSHR5858_genomic_01564_1 group_7372
MARMIHCAKLGKEAEGLDFPPLPGELGKRLYESVSKQAWQDWLKQQTMLINENRLNMADP
RARQYLMKQTEKYFFGEGADQASGYVPPAQG*
>GCA_000452945.1_BPMSHR5858_genomic_01565_1 argA
MNSQTDLPAAQSGAAPQPSVDEAVHHAQFVDWMRSVAPYIHKFRNSTFVVGFGGEVVQHG
LLSALVSDIALLQAMGIQIVLVHGSRPQVEEQLHLHGVESAFSHGLRITDARALESAKEA
AGEVRLDIEAAISQGLPNTPMAHAHISVVSGNFVTARPVGILDGVDFAHTGVVRKIDAES
IRHSLASRKLVLLSPLGFSPTGEAFNLSMEDVASAAAIALRADKIIFLTESPGIVDEAGE
LVREMSLDAAAELLDSGDLQGDDGFFLKHAIRACRGGVARAHLIPQSLDGSVLLELFLHD
GVGTMISYENLESLREATPDDVGGILTLIEPLETDGTLVRRGRHQIERDIDHFSVIEHDG
VLFGCAALYPYPTEKIGEMACLTVAPEAQGSGDGERLLKRVEQRARARGLTRIFVLTTRT
EHWFLKRGFVKATVDDLPEDRRKLYNWQRKSLVLMKQL*
>GCA_000452945.1_BPMSHR5858_genomic_01568_1 group_6084
MLHVIAAALDGFVSDLQTWLYVDIVQPLLFKVGLMDYDEDTYDALYWVIVGALQVVAMYV
LLRPLEALRPVERWESRREARVDVIYTLIAKLGIFSLFFFFAMQPLFDSVQAWLRLHGIA
NVNLDYLWPGVTSRPIVAFAIYLIVLDFAGYWYHRWQHRIGIWWELHAVHHSQRQMSLWC
DDRNHLLDDVLQACFFAAIALVIGVSPSQFVVLTAVTNFMQSVQHANARLPFGRLAERLV
VSPTFHRRHHAVGYGHEGTKYGCNFGVLFPWWDMLFRTASWNRTPEPTGIREQYEGVSYG
DGFWAQHWLAFVRIARRLAPGKRRDVASA*
>GCA_000452945.1_BPMSHR5858_genomic_01575_1 xthA
MRIATWNVNSLNVRKQHVLDWLAQSDVDVLCLQELKIPDEKFPREALEAAGYRSWFAGQK
TYNGVAILARASLPFDETDIVRNIPGFEDAQQRLIAATIDGVRIVSAYFPNGQALDSDKF
VYKMQWLDALQAWLKDELQRYPKLALLGDYNIAPEDRDVHDPAKWEGQNLVSPQERAHFA
QLVALGFVDAFRRFEQPEKTFTWWDYRMLAFRRNAGLRIDHILLSPALAETCTSCTVDRV
PRTWEQPSDHTPVVAQLG*
>GCA_000452945.1_BPMSHR5858_genomic_01578_1 narV
MDYLNTFVYGIYPYIAAAIFLFGSLARFEREQYTWKSDSSQLLHRGALRTGNILFHVGVL
GLFFGHLVGLLTPVAVWDALGVTHQFKQLVAMSAGGVMGSLALAGLAILLHRRLTNARLA
AVTRRGDKVLLAWLLVTLVLGLSTIFESASHTDGHLMVQLMTWAQHIVTFRGDAASYVAD
APWLFKAHLFMGLSLFAIFPFTRLVHVWSGFASVTYVGRAWQLVRARR*
>GCA_000452945.1_BPMSHR5858_genomic_01581_1 narG_2
MSHFLDRLKFMSRVKSTFSDGHGAVVDEDRRWENGYRSRWQHDKIVRSTHGVNCTGSCSW
KVYVKNGLITWETQQTDYPRTRADLPNHEPRGCPRGASYSWYVYSAQRVKYPMIRGRLMQ
MWREARKTMDPIAAWESISQNPEKARRYKSVRGLGGFVRADWNTATEIIAAANAYTIKRY
GPDRVVGFSPIPAMSMVSYAAGARYLSLIGGACLSFYDWYCDLPPASPQVWGEQTDVPES
ADWYNSSYLLVWGSNVPQTRTPDAHFYTEVRYKGTKTVAISSDYGEMVKFGDIWLAPKQG
TDAALAMAMGHVVLKEFHASNQSAYFRDYVKQYTDMPMLVMLRERDGALVPDHFLRASHL
AASLSEANHPEWKTLAIDAATGDIVAPNGSIGFRWGEAAHNGGEKVGRWNLEMKDGGSGR
AIDPRLSLVDAHDEIVDVGFPYFGGEHEAVLARRVPAKRVALADGTSALVATVYDLQMAN
YGVDQGLGGPNVAASYDDDIPYTPAWQEKHTGVARHLVIQVAREFADNAGIRDNDWVEVF
NVNGTLTARAVVSQRVPAGMCLMYHAQEKIVNVPGAQTSGKRGGIHNSVTRTVTKPTHMI
GGYAQQAYGFNYYGTVGSNRDEYVIVRKMNRVDWLEEPLNEGAEQ*
>GCA_000452945.1_BPMSHR5858_genomic_01582_1 narK
MSTSLLVRWDPENPAFWQAKGRPVAWRNLAISIPALMLAFVVWSLWSVVVVNLDRAGFHF
SKNQLFWLTALPALSGATLRIFYSFLVPIFGGRRFTAISTATLLIPALGMGFALRDPGTG
YPTLLILALLCGFGGANFSSSMANISFFFPKAKKGLATGLNAGIGNLGVSVVQFVTPLVI
SAGLFGALAGDPQSVVAHGATTNLWLQNAGFVWVPFIVVATLAAWFGMNDIADAKASFAE
QAVIFRRLHNWLMCWLYVGTFGSFIGFSAGFALLTKALFPNVNPTAYAFIGPLAGALMRP
VGGWVSDRIGGARVTFWTFAAMIAAVGGVIATLPSGGAAGSFAGFLAMFIVLFALTGIGN
GSTFRMIPVIFLTERQRAAQGQDEAARKQAVLDAGKESAAVLGFSGAIGAYGGFFIPKSF
GTSLDMTGSAVPALVCFIAFYVSCVAITWWFYARRNASMPC*
>GCA_000452945.1_BPMSHR5858_genomic_01583_1 nasA_2
MAATSASPASLDKRAMPVLASSTIAFTLCFAVWMMFAILGIPLKKTLGLNDTEFGLIAAM
PVLTGSLIRVPLGIWTDRYGGRIVFFILMLVTVVPIWLISYATELWQFLVLGLFVGLAGG
SFSVGTPYVARWFPKARQGLAMGVFGAGNSGAAVNKFVAPALIAAAGTWTIVPRVYAVAM
LAMALLFWLFSATDPAHRSTNATSLRAQLRVLKNPRVWRYSQYYSVVFGGYVGLSLWLTQ
YYVGEYGFGIQSAAFLAACFSLPGGVLRAIGGWLSDRYGAYRTTWWVMWVCWVMFFLLSY
PPTDFTIRAAHGPLGFHLSLTPVAFTALLFAVGIAMAVGKASVFKFIADEFPNDIGAVSG
VVGLAGGLAGFALPILFGALVDLTGVRSTCFMLLYGAVSVSLVWMYFSFRAERAAHDQRV
LATRSA*
>GCA_000452945.1_BPMSHR5858_genomic_01587_1 folD
MTATLIDGNALSKTLRAQAAERAAALAARGHRPGLAVILVGDNPASEVYVRNKIKACEDN
GFFSLKDRYPATLSEPELLARIDELNRDPKIHGILVQLPLPAHIDSHKVIEAIAPEKDVD
GFHVANAGALLTGKPLFRPCTPYGVMKMFEAYKIPLQGANAVVIGRSNIVGKPMALLLLE
AGATVTICHSKTRELAAHTRAADIVVAAVGKRNVLTADMVKPGATVIDVGMNRNDEGKLC
GDVDFAGVSQVAGHITPVPGGVGPMTITMLLVNTIEAAERAAAAA*
>GCA_000452945.1_BPMSHR5858_genomic_01593_1 group_7376
MSLLTPEQIAAAQKANIESLFGLTTKAFEGVEKLIELNLQVVKSTLAESQENVQRALSVK
DAQELLALQASLTQPIAEKVLAYGRHVYEIASATQAEFAKVAEAQYEEQNRKVQALVDNV
AKNAPAGSETAVAALKSAINAANTTYETVQKAAKQAVEIAETNFNAAAAVATKAANTAAA
ASRRSGKTA*
>GCA_000452945.1_BPMSHR5858_genomic_01616_1 comEC
MRAKCGAFALGVVALQQQAALPGAAAWAGGALAFGLCVWLALVWRGDARARAAGFCACCC
AAALAGFGYAAARAQWRLADTLPAQWEGRDIVVTGAVRGLPSRDANGVRFLFDVDANDAR
IARFPATLSLGWYAFGRPGAQPPELVPGDRWRLRVRLKRPHGNANFGVRDAEAAWLARGI
RALGYVSAPRDARRLAGRASGVAAALDRLRARLRGRIAEALGDAAHRGIVVALAIGAQDD
IADDDRRILRDTGTSHLVAISGLHVGMVGGLCAWLAGGLWRRSGYVGRDWPLVVPAQKVA
ALGAIVGGAGYAALAGFNVPVQRAWWMLAAAGVAYLSARSLAPSSVLAAALGCVLLVDPW
AVTSAGFWLSFCAVAAILAVSSGWRAPRDLDETRGARHALGGVGHEGAPPSHRARWRAAC
GFAWGWAARAFGRLARRVRDAARAQFAVTIALAPLTVLWFAQIPLAGPLANAFAIPWVGS
LVTPIVLAGIVLPAPLDASAYALAHALVQALMRLLDATAGAGRTVWMLPAPDGFALAAAA
VGVAWALMPRGWPVRCAAPLAWLPLVAPAPLAPPDGAFRLTALDVGQGSAVLIETARHTL
LFDAGPGPEASNAGERIVVPFLRARGVRALDALVVSHADSDHAGGAPAVLGSIAVAQMTG
GLPPSNRLWRAARAAGVADALPCVAGQRWRWDGVEFDALWPAGGPRAGGATNAQSCVLRV
SAGGRAALLTGDVDARSERALVAGSRGALAAQVLVVPHHGSRTSSTEPFLDSVKPRIAIF
QVGYANRFHHPHPTVWARYAGRGIELPRTDRDGAVRVDVTSSGALAEPVRYRDAHRRYWM
GR*
>GCA_000452945.1_BPMSHR5858_genomic_01617_1 group_3234
LKDIIHFSHANGFPASTYRTIFAELADDYELRFIERIGHDRHFPVTRDWPHLVEQLLDDI
GRAYERPVWLVGHSLGGYLSLMAALKRPQWVRGVVMLDSPVIAGWRSSALRVSQWTGLDE
RLSPAAATRTRRVHWASRDDAWRHFRAKPAFARWDERMLSDYIDYGIPQTSAAGERTLAF
DRQIEYMIYRTLPHTLGPRLARGAPVPVGFLAGTRSREIRQIGLAATRRAVRGRIEWLEG
SHLFPMERPLETARALQCMINMLREAQGGAGARRG*
>GCA_000452945.1_BPMSHR5858_genomic_01635_1 roxA
MPKRPLPEPSGRVRERTRAPLSEAPTPLLGNLSPAQFMRRYWQKKPLLIRQAITGIAPPL
SRDALFELAADYDVESRLVTHFRNRWQLEHGPFEPEHLPSVKRREWTLLVQGLDLHDDRA
RALLERFRFVPDARLDDLMISYATDGGGVGPHFDSYDVFLLQVHGKRRWRIGAQQDLSLQ
EGLPLKILANFEPTDEWVLEPGDMLYLPPHIAHDGIALGECMTCSIGFRAPSAGELRAQF
LYHLAERGGLRTGARDNARYRDPAQPAVDSPAMLPAAMVKRVAATLAGIQWDEHDVGDFL
GCYLSEPKSNVVFEPPTPRLGEAAFVTQASRRGVRLDRKAALLYNARSYFINGDAHPLAT
AAKWLPELADTRRMEAKRFVTLSRDPAMTGLLHEWYCAGWIRVGDMG*
>GCA_000452945.1_BPMSHR5858_genomic_01637_1 group_4627
MRNADGTAPRAPLPSRIAASALACILAAALAGNARAAPASPRTGGDPYAFAVLSGVIASA
DDEPQARRLLDAIARDRSVSFIVYEGNLKGPKEACRDQLYEQRNVLFATSRVPLVLLPGQ
YDWAACGTRDAGAYDPVERLDFLRQNVFTEPASPGANPLPLTRESEVARFHPYRENVRWI
RDDTVFIGLNAPAPNNRYLTAGGRNGEFEDRAIANAFWIDHAAEYAKRRKARALVVFLEG
DPQFNRYERSERFGWLRFNRPRVRDGFREFKRALVKAAAVFRGSIVVVHPSGEPLADGFR
IDRPLRDDKGELVGNLTRVAIAPHARLTHWVRIGVDAAKQPMFNVSMQTVPKYLPQPPAL
PVVPHDDTPLPDMPEIPAQPVLPDSGASGAAPPGYGGWHGAHEGSFDHRAPSGATPGDAP
DIAVPGSMQGTR*
>GCA_000452945.1_BPMSHR5858_genomic_01642_1 dapH_2
MFTRLREDVATIRERDPAARSAWEVLTCYPGLHALVFHRLAHACWRSGWRWLGRFVSQLG
RFLTGIEIHPGATLGRRVFIDHGMGVVIGETAVIGDDCTIYQGVTLGGTSLARGAKRHPT
LERGVIVGAGAKVLGGFTVGAGAKIGSNAVVVKPVPAGGTAVGNPARVVVPAEAKRTPER
TAFCAYGITPNADDPMSLAIHGLIDHAAKEAQRVDEIVVALERLGAHLDALQGTDGARLD
LRRLSAAFEGRALDGGTAAER*
>GCA_000452945.1_BPMSHR5858_genomic_01659_1 tycA
MSVHDMTHPGELAPVRADRFGRRFATAAAAEIAAADSLTARFADVAARRGDALAVTCGDE
RWTYGALAARARRIAEAVRAAGEAGGEPVALLYPHGAPMIAAMFGVLGAGKFYVPLIADH
PLPHLQSIVRECGCRLVLAAPELAETAARLGVSARVIDDARLPPAHGPFDARGGDAVSYL
LFTSGTTGVPKGVMQCDRNVLHHAACYAASIGLDDDDRMTLLPYYGFDASVMDIFATLLT
GASLHLWDVRERGVDGIGEWLARERMTIWHSTPSVLRATFAAFARPAALRWVVLGGEAAT
GGDVALVARHGGPRCRLLNGLGPTECTTALQYVADPAADASVARLPVGRPVPGVEVELAD
ARGEACATEGELVIVSPFVALGYWGRAELSAERFRQTARPDGARRYRTGDLLRIDARGCY
EHLTRVDDQIKIRGLRVELGEIQATLAAHDDVLQAVVLPRLDEATQQQTIVAYVVPRAAS
ADVAALREYVASRLPAHMVPRAIVRVDAMPLLPNGKLNRRALPAPPRAEVAAGGRKAPRT
PLHRLLAACWADVLRRDAVGIDENFFELGGDSLLGAQLLSRVKRDLELDARLGDLFRHPT
VESLAEHLLACR*
>GCA_000452945.1_BPMSHR5858_genomic_01660_1 entS_3
MTDSMQDSLDEAARPARVSWLAALRGPFAYRTFAAIWVASLVGNIGGSIQTVAASWLMTS
MAPSPTMVSLVQTAFTLPIALFALLSGVAADAWDRRTVMLLSQALMFSVALCLVALAAAG
AMTPARLLVCMFVGGCAGAMFQPAWQSAVTEQVPARELSAAIALDSFSMNFARTAGPALG
GFIVASVSPNAAFVLSGLSYAGLIYALSRSIRGAAARPPVRERLATMLVQGVRYCGRARG
IRGTLIRSSLFGFLGSPVWALLPLFAKTQFGGEARTYGVLLASFGAGAASGALGGAAGRA
RLGREALVRLCTLTFAAGMLATAWSPCQAVAMLGLAVAGGSWVVVVSTYNLTIQTASPAW
VAGRSLSLFHSFIVGGLSIGSYLWGVAAQGSSINSAFAVSALMMAASACLAAWLPLPTHE
ALGERTHGEPRRT*
>GCA_000452945.1_BPMSHR5858_genomic_01661_1 group_3237
MSTTTVAIIGAGFCGATLATHLLRRPPVRPMRVLLINRSGAMARGVAYGTRALGHLLNVP
AGRMSAVAGDDDDFYRYASGRDPRVARGSFVPRRIYGDYLEARLTEAIEQAHAGIEFRSV
VGSAVRIAPVDGGARGAITMDGGAVIEADRVVLSSGNEMRRDPFIAESQRKFYDSHAYVR
DPWQPGALRGIAPDTPVLLVGSGLTMMDVVLDLRARGHAAPIHVVSRHGLMPLAHREMDA
PPSYDDRLAARMLARADVRHYVRAVRDAIRRGGDWRDVIGSLRAATPALWRQLPSDERRR
FLRHVRPYWDVHRHRCAPEPAARLQAEFERGGVAAVAGRVTGYSEHPNGVGVTVRRRGAA
VDERLEVGAVVNCTGPAPDFSARAGSLLGNLYADGLIVPDAIGMGFEIADDGAVLDRDGS
PSAWLRYVGPLLQARDWEATAVPELRQYVQRLADTLLAPRDERALT*
>GCA_000452945.1_BPMSHR5858_genomic_01663_1 group_7377
MTNHDHRIAFVHIPRTAGTTFGAYLDGLYAGRDVAKFYADDAHPVVNEKLDAFRRLAAGE
KASYALLRGHFVYGFDGGESGCRYVTLLREPVARLVSYYFYALKEPMNYLHTYLLQRRIG
LESFLMSDASIDLDNYQVRAVSGATFGSSRERVSGAHLALAKRNLVERFAAFGLTERFDA
SMRRFARQFGWAPAPRERRNEGAYGREVELGDACRAYVEEKNRFDIELYRFAQAHFESAA
ASGDDGRADRSADRPADHAVHTTRS*
>GCA_000452945.1_BPMSHR5858_genomic_01664_1 atsK_2
MQTLEIRPLSGTIGAQVRNRTLADVVESGRADEIRQALLRYKVLFFTNQPELSVETQIAF
GRLFGELETNFPSFTAKPDGQPEVTVFDGAVSTGRASIWHTDLSIAKTPSAMGILCVKET
PDSGGDTMWADLEAAYAALSPGMQAFLEGQRAVHDMMTPQYAQRPGAFQTRGRSDMDLSE
VFGAEHPVVRVHPETGRKCLFVNPFLTSHLVGFHSAESATILNYLYALMERPQYVVRWHW
SRGDVALWDNRCTMHTAVDDYGAGRRFARRVCVRGDVPYGVGERTPATAAVA*
>GCA_000452945.1_BPMSHR5858_genomic_01665_1 group_3238
MSNQLIQSGVDFSGSVFLASSNVRANAHRLNWRAKVLVSGYAADFKGKRVLDLASHDGRF
MHAALAHGATKVVGVEARDDHVDSARANLARGGHDRGRYDVICADLVEYLKSVDTGAFDT
ILCFGVLSHLIEHVDIVREVGRIAPGAFILDTWVARERWNLRERLRNRRVNAFVRRTQQG
GFGSVSGIARLRNWLNDVIPSEHSRTGNLVFLYEDAAAPGATVRQSGLMGWANRSLVEML
FDHYGLAHERVDWRAQGVADWSELEDYRLGRRESWVARARRA*
>GCA_000452945.1_BPMSHR5858_genomic_01666_1 group_7379
VKTFREKVRARWPHYDDRRGVFLPGATPDEMRRVACETQLDCKLTCRSHLHAYREYLPKL
YFEGNVADCDFDALPARYVIKPRCQTGVLMLMDDGVDLKTGRAAGHDEIRAFFARQWCYR
GLNNDVLVEEMIENRDGSPCVSQFKCLTFHGRVEYIHYVTVGYDLGCVAYDYDRDWNRVA
LYHSAQPVERGIARPDTFPMVIELAERLADYYRHWTGIEHVRVDLFDSDKGPVFGEYAGG
TNGGEGFTEEAQALLGALW*
>GCA_000452945.1_BPMSHR5858_genomic_01669_1 aptB
MRPAAHCVQALVERAAHDRRIWLLDGDLADSYGADRFAVAYPQRFLMAGIAEQSMVSTAA
GMAACGLRPWVFSFAAFLCYRAYDQIRVGLAQTGMPVTLVGSHSGGCGGPNGKSHLALND
IAVMASLPNVDVWAPACDRDTRFVVEAVMLTDRPAYVRYPREPWGELPGEPAECRWIGAR
RRIALVSCGLFSHVAHDAAEQLRALGVDAGALHLARLAPFPGEAFGTLVDGVDGIWVVDD
HTRFGGLADLLRANGYAVREPVFSWPANWSGDVGPTAELLRRYGLCAADVAARIAAELER

RGR*
>GCA_000452945.1_BPMSHR5858_genomic_01672_1 serC_2
MHRNQLNFSGGPGALPDTVLEQVRQAVVELPETGLSVLGMSHRSSWFSSLLAQAQADLRD
LLGIPDEYGVVFLQGGSSLQFSMIPMNFSRPGAAAPEYVTTGYWSRKAIGEASRVAAMRV
VWDGAASGYRTLPSLAALDWDARAPFRHYVSNETVEGLQFPDAADLPGSPLIADMSSDFM
SRPFDVRAYGMVYAHAQKNLGPAGVTVAIIRRALLERVPDTLPPMLDFRTHVEHRSNYNT
PPVFAIYVMALVLRWIRDEIGGVHAMRDINARKAAMLYATLDALNEVIDCHAHRAARSTM
NVAFRFRQPRLDTLFKEQSTEAGFCGLSGHRSIGGIRASLYNAVSEQAVSRLCTFLKDFA
IRHA*
>GCA_000452945.1_BPMSHR5858_genomic_01673_1 group_3240
MLSIIVTWRNRNELAGALPGLVETAGRLNGEVVIVNFGGDAASLARQLEGFEHAVQVAQW
REQRYFHKTKAQNLGAHAARHDMLFFCDCDIIVEPDAIVSLVRKLDAAPGAFATLKGVRE
TEQNARQAKNVVRFGYRLDVRIRNGRELTIVDNEEDAQDGTRQAPGLLFVRRRDFLAVNG
YNGRLHGWGWEDQDMISRLTLGAGLVRHGEAQRVPRPTNDQAWVANYQMKTRGESRDKMF
RQALAYYDADDFLGTYAQDVDLSAARAALACS*
>GCA_000452945.1_BPMSHR5858_genomic_01675_1 spuE_2
MKKVVGAMLLLAVGHAVSAELHIGSWPDYLPDTLIKKFQAETGIKTTLDTYASDAALTQK
LQSGGGGYDVVIAGDYYVPVLVKSGLLQKLDKNKLPNIANIKPEYRHPSFDPQRDYAMPY
TVVLTGFAYDSARVSGGKLDESWKSFFDPPAQLRGQIGDLDVEEELYMAASWYLGQDECT
ENPADAKRVLDVLQKQKPFVKTYSNDGTIDRLASKQIAVQHIWSGAAARAQDRLPSITFV
YPREGVRLFMDSLLIPAKAQNTASAYQFVNWMMRPENIAQVTNAVRYNNEIIGSERYIDA
ALLKNSAIKTPEQYKIRLRPYKICSPAAIQLRNKVWLKLKGNR*
>GCA_000452945.1_BPMSHR5858_genomic_01731_1 codB
MATRGEFSLSEVPAHERKGALSITMVLLSFTFFTGTMFAGGKIGVAFRVVDMLWVAVVGN
LLLAAYAAALAFVASRSGLNSVLMGRFCFGEVGSKLSDFLLGFAELGWYAWGTATVAIVL
VKLLGWPASVTTPLMVLFGFGFSITAIVGYRGMDALSRVSVPLMFALLVVSMWIATRDVG
GWPGLAKIAPTQPMSFAAAVTMVFGTFASGATQATNWTRLAKSGRAAVAASMIGFFVGNG
LMIVAGAYCAIVYQQSDIVEVMMLQGLSIAAVVMLCLNLWTIQGPTIYNVSAAACHLLRS
ERRRTLTLVGAAVGIVLAIGGMYEMLIPFLILLGSIIPPVGGVILADFWYRHRGRYPAIA
SARLPRFNIAGLAAYAIGAALAYASPWIAPLVGIAASSFCYIVFVQIAGRAVRAPSVQGE
*
>GCA_000452945.1_BPMSHR5858_genomic_01749_1 group_7384
MQARKPKSRALLGQLTDMDLRLLRVFRSVVQCGGMAAAELELNIGISTISRHVKDLETRL
GLVLCRRGRAGFTLTPEGQTVYEETLRLLASVEAFRSRVDGIHDRMGGELQIAVFDKTAT
NPHARLGDAIGRFHDEAPDVALNLHVASINEVERGIIDGSYQVGIIPAHRSSGSLVYADL
FDERMLLYCGASHALFNAPHTKLTWRQVRGYAFAGLGYHSPNMEVSHRAKLTRSATASDQ
ESIATLILSGRYLGFLPDHYAESFEAKGLMRPVAPQRFHYSCRFVSLLRRSPRPSRAALL
FQECLEIAHAGHRAKA*
>GCA_000452945.1_BPMSHR5858_genomic_01756_1 oprM_7
MHSPPFLSSPKRALPIALALGLAGCSMEPAYQRPDAPMPSAYPNGPAYATPGAPRANPGE
PAAAELGWRNFLADAQLQQLVALALANNRDLRVATLDIDEARALYRIQRAAQFPAIDASV
GLTSQRMSPALRAPGQSAAINSYDASVGLTHFEIDLFGRVRSLSHAAQEQYLATEEARRS
VHISLVAEVANTYLTLLADRALLALAQDTLRSQQDAADMIHRGKQAGAMAQLDEHRADTQ
VQTARVAAEQYTRQIAQDENALAVLIGGPLPAGVSRAAPLDDRALLAEFPAGLPSTLLER
RPDIMAAEHRLIAANANIGAARAAFFPRITLTGALGVASASLAGLFSGGVAWLFVPQLTL
PIFNAGSNQANLDLATVRRDINVAGYEHTIQDAFREVADNLAARATYEREVKAQEAMIRD
LAETKRLADMRFRNGVDDYFGVFDAQRQLFAAQQLLVTYKLAGLTSRVTLYKALGGGWVE
SAGAAAAQPRTGPAAPMARPAATPPQTTRPAPVARSQTVPPRTVQPPIAQPATAQPQTAQ
AAPVAQSQTIPLRTVQPPIAQSSTSQPQTAQAAPITQARTTPLRTAQPPAARQQAPQSAP
AAQPEAAPPQTAQPPIFQP*
>GCA_000452945.1_BPMSHR5858_genomic_01767_1 fabI_1
MRLQHKRGLIIGIANENSIAFGCARVMREQGAELALTYLNEKAEPYVRPLAQRLDSRLVV
PCDVREPGRLEDVFARIAQEWGQLDFVLHSIAYAPKEDLHRRVTDCSQAGFAMAMDVSCH
SFIRVARLAEPLMTNGGCLLTVTFYGAERAVEDYNLMGPVKAALEGSVRYLAAELGPRRI
RVHALSPGPLKTRAASGIDRFDALLERVRERTPGHRLVDIDDVGHVAAFLASDDAAALTG
NVEYIDGGYHVVG*
>GCA_000452945.1_BPMSHR5858_genomic_01779_1 dadX_1
MPRPIVARIHPDAVAHNLAIVRQKAPNARIWSVVKANAYGHGIEHVYPALAGADGIALLD
LDEAVRVRELGWTKPVLLLEGIFEAADVEIADRHRLTVAVHGDEQFKLLTAAKTTRPIDI
QLKMNSGMNRLGYRPHALRAAWERAAAAPSIGRIALMMHFANADEGEIDWQMSVFDAATQ
GLPGERTLSNSAAVLWHPRAHRDWVRPGTILYGASPTGASRHVADVPLRAAMTVTSRIIG
VQTLTPGETVGYGRCFAAEREMRIGVVACGYADGYPRHAPTGTPIAVDGVRTQVVGRVSM
DMLTVDLTPCPNAGVGSAVELWGEQVRVDDVAEAAGTIGYELMCALARRVPVTVDPSPAA
AALAQPVRTGSYGR*
>GCA_000452945.1_BPMSHR5858_genomic_01786_1 group_1442
MHHAPAGIAAVLFVAASTVCAQVPAGYPGNYQHVIDAAKQEGKLVVYSTTDTSLVRPLIK
DFESLYGVKVEYNDMNSTELYNRYVSESAAKSASADVLWNSAMDLQVKLVNDGLTAAYDS
PETPHMPQWAQYQKQAYGTTYEPLAIVYNKRLIAERDVPKTRADLIRLLQSQPERFKGKV
TTYDIEKSGVGFNYLTQDAHVNEKVTWELVRAIGATGPKLQSSTGAMMERISSGENLVGY
NIIGSYAYAKAKKDKSIGYVLPKDYTLVVSRLVAISKQAQHPNAAKLWVDYLLSRRGQTL
IANQANLYSIRADVSGETSMTSLAKELGDSLKPIQIGSGLLVYLDQSKRLAFLKQWQQAL
KR*
>GCA_000452945.1_BPMSHR5858_genomic_01793_1 paaI
MAARSPERSPTATAERSAEALARATAEAMYAADACSRSLGIEVLEVRPGYARARMPVRGD
FLNGHQICHGGLVFTLADSTFAFACNSYNINTVAAGCSIEFLRPVAGGDVLTAEATEQTL
NGRHGIYDIRVTNRAGETVAMFRGKSTQIKGTVIPVDR*
>GCA_000452945.1_BPMSHR5858_genomic_01815_1 group_4651
MPAYRRLHAPHPRLRDIVNCYFVELGRAGPMLYPATPSASITVFASGASVHTNGIAHAET
LMCRPQRSAVWATLLPGTTYVSVVFRPGQIRRVMPRGASAAYDDGSPLDAVFPRAAVHAL
VDGVRGAPTLGHAVAALEDWLMAQAHVAEEFDAGGILLPPACVDAPREALTAQFSRSPRQ
LERLFIDTFGASQREMRSLLRYARTLSRLIAADFALPALSDMAIDCGYYDHAQMTRAFRR
YAGMAPVQLAAAVRDARQASMTMYRYELADKRILLDA*
>GCA_000452945.1_BPMSHR5858_genomic_01816_1 bauD
MEGSQTGLGTGLKQRHVTMLSIAGVIGAGLFVGSGHAIAEAGPAALIAYAIAGLLVVLVM
RMLGEMAVAQPDSGSFSTYADRAIGHWAGFTIGWLYWWFWVLVIPIEATAAATILNAWFP
GVATWIFALGITSVLTVTNLFSVKNYGEFEFWFALIKVVAIVVFLAIGGAAIVGLLPGSN
VSGAARLVNAGGFMPNGVGAVLAAMLTTMFSFLGTEIVTIAAAESEHPERQIVRATNSVI
WRISLFYLGSIFVVAALVPWNDALLPAHGSYQRAMELIGVPHAKGIIDVIVLVSVASCLN
SALYTASRMIFSLSTRGDAPAFLRRTDASGTPRAAVLASTAFGFLTVIANYVMPEQVFGF
LLATSGAIALLVYLVIAISQLRMRSTLDATGERLPLRMWCFPWLTWAVIVFICGVLVVML
LREDHRMEVGATAVLALAVVVASRMNRRARGGDEPTPAASRVSANLR*
>GCA_000452945.1_BPMSHR5858_genomic_01819_1 group_1001
MPISLPRAARRIAVALAIAALPPAAAFAYSPTAPTAPTAHGEADLDRHDTYRNRDGDTVH
APAHSKSGRAPEGATARCRDGAYSFSRHRRGTCSGHGGVAAWL*
>GCA_000452945.1_BPMSHR5858_genomic_01822_1 rutB_2
MARAAVIVVDMQRGLLQRAKPAHRLDEVVAGINRLTAAARAAGAPVCFVQHDGGADDDVV
PGTPGWRLHAELTCADADWRIRKRASDAFHGTPLAAQLAAHGVDAVVICGYASEFCVDSA
ARRAALLGYRTTVVSDLHTTNDRAHLSAVQIVAHHNFIWSHCSFSGNGVAPRPLDEVIAA
EFA*
>GCA_000452945.1_BPMSHR5858_genomic_01835_1 rutE_2
MSVKPAPTSVSIHELIAGRWSPRAYSSEPVSAEHLHAVLEAARWAPSAYNAQPWRFIVFD
RSKDEVAFKRAFSTLVPFNQGWNAPAPVLIAVTAHTLTSKGEPSPTALYDAGAAAMSLVL
QAHALGLAAHQMSGFDVKAFRAAFAIPADVEPLAIISIGHYGDADKLDPVLRERERAVRT
RHPIGEVVYAHAWQKPFPSAD*
>GCA_000452945.1_BPMSHR5858_genomic_01836_1 group_4255
MNWAARRIGGRFHYGWLAAAVVFLILLAAAGTRATPSVLMVPLEHEFGWSRASISLAISV
NIALYGLTGPFAAAAMQRFGVRPTVLAALGVMGMGVALSSMMTATWQMVLVWGVMVGGAT
GVAALTLSATFVTRWFVARRGLVMGLLTASSATGQLVFLPMLAAIAQRHGWRPVVLTVAV
AAAIAVPLVALLLPERPSDVGLRPYGEPANAPRAEDGAKRNPIAVAFGMLATASKTRDFW
LLFFSFFICGASTNGYVGTHLIAMCSDYGMTQVQGASLLAAMGVFDLIGTTMSGWLSDRY
DSRVLLFWYYGLRGLSLIYLPHAFGIDFFGLPLFAVFYGLDWIATVPPTVRLATDVYGKD
AAPVVFGWIVAGHQLGAAFAALGAGMLRASLGTYTVASMISGGLCIAGALIVLRINRGTA
RVSASAA*
>GCA_000452945.1_BPMSHR5858_genomic_01838_1 group_7385
MTEVSPPDADILAATRHWLTRAVIGLNLCPFAKGVHVKRQIRYAISRARSLEAALTDLEN
ELRHLDAADPDEVDTTLVIFPNAFGNFLDYNDALWFADRLLRQLRLDGTLQIASFHPRYQ
FDGTEPDDIENYTNRAPYPILHLLRETSIERAVDAFPDAADIYERNQATMRRLGHAGWRD
WMAQRGDEEDSRENGNAGKPEGSSAAN*
>GCA_000452945.1_BPMSHR5858_genomic_01840_1 prmC_2
MRRRAFLPAGAKRGTVSSAARRAARRLNDSRFLVTAFPEFHWTDADGVEHVARWRSEAGS
PPPRRAVVADDRTTADVAYRLACEGTALVWQGDFQNARQLLQALARRIERKPKKARPAAT
PVDAFNLHRMAQAQRARTLGMLLIPLEADYSIALRRAPDLRAACEEAYGEGGEHSIASLR
ELLGIVGAHEWRKKGVPIAALGGERIHPHYGVFSPVRGEYVALVARAPLPSTSLAFEIGV
GTGVLAAVLASRGVGRIVATDQDKRALACAAENVARLGYARQIEIVEADLFPDGRAPLIV
CNPPWVPARPSSPLEYAVYDPDSRMLKGFLAGLAAHLAPGGEGWLILSDFAEHLGLRTRG
ELLGWIDAAGLVVVGRDDVKPAHPKASDPTDPLHRARAAEITSLWRLAARA*
>GCA_000452945.1_BPMSHR5858_genomic_01841_1 group_3261
MPLGASVCAEVCACRRFARSARRTIDPAMSDSTRPSRFAHVDAMRAVAVLFVMWTHYAEL
FDKLAGSQHGLDALQRSVNFGRIGVVIFFCISGMLIPSSLRGAPAEGTRRFVVRRFFRLY
PAFWLSLPLGYLVYWLLFGQRMDASGLLANVTMIPTAFGRDPVMGHYWTLETELYFYVLC
VLLFRIGALHRMRDLCAVCAGLCVLFVVTSALKIVPASALGQYKGMLYHLAIMFWGACFR
QAYEAPSKTFEWMRGQWPITYRAVTIALALLIVSISLLMAAANWRHGDFVHLSASFGYVL
GMAIFVVLATVLKIRLRLFAWLGEISYSIYLLHGIPLYLLLWACERYGITDLPLGFYMAL
PVLPAIALSWASHRLCEAPFVRFAHALTPKRRAAAAAAKV*
>GCA_000452945.1_BPMSHR5858_genomic_01842_1 group_7386
MHAWYRFRPFVGLWFAALAVLLALGGCAALTGRDPVRVSVVGIEPLVGQGLEMRFDVKLR
LQNPNDAPIDYDGVALDLELNGRPFASGVSDVRGTVPRFGEQVLSVPVTVSAFAAARQAF
GLADATVSGKLPYVLRGKLAGGMFGSVRFTDSGTLSLPASGGYGGGY*
>GCA_000452945.1_BPMSHR5858_genomic_01844_1 group_7387
MQELNQQEIDLVGGATVGGGLIGGLTGLIGAYGGALYDLGANLTGDLVGFATGVVQIGET
LVGGTLSNIGSIFTGTSH*
>GCA_000452945.1_BPMSHR5858_genomic_01845_1 group_7388
MQELNQQEIAMVGGASLLTNLVGAIGNEVLLLGQSAYNFGANLTDSAGAILTHTVDGVAS
ALTSTIGIIQGVLTGN*
>GCA_000452945.1_BPMSHR5858_genomic_01846_1 cvaA_1
MTPNTPLFRTAAQEAQRTQTLGEIVLIRPVSFAVLASAAASMALGVILLFTFGTYTRRTT
VDGVLTPDTGLVKVYAQQTGVVLKKNVVEGQHVTRGQVLYTVSTDLQSAAAGQTQAALIE
QAQQRKTSLQQELDKTRRLQQDERDTLQSKIASLRTELAGIDDQIAAQRTRASIAADAAS
RYAGLLAQDYISKDQAQQRQADLLDQRSKLNSLMRDRASTAQSLKEALNDLSGLSLKQQN
QLSQIDRSVIDVDRTLIESEAKREFVVTAPETGTATAVIAEPGQTADTSHPLASIVPTGA
HWQAYLFVPSAAVGFVHVGDRVLVRYQAYPYQKFGQYEASVVSIARTALSAAELATSGGP
AAQTASGTYYRITVALNSQNVMAYGRAQPLQAGMALQADVLQERRRLYEWVLEPLYSLTG
KL*
>GCA_000452945.1_BPMSHR5858_genomic_01849_1 group_609
MSAPDDSNSKAPPSLLSSDAGKGSGANHSRILANLEGRVSPQTENAPRSRKGPIALIALA
VVAIAGWGAWRLQQPSADSAEVAAAASAPAVAAAPASRAAPASAAQIAPASAPQPATIVN
DESAASAAKPAVAASVGDGSRLSRALANGASDVSTAGAAAGATAAAAAVAAAKPAKSAEK
RAAAKPDKQDAAHAKAEAAARKRHEKAQAADAAQAKKSRREAAAKKNEKDDPDADLLAAL
VARTKPADATKPADAAKPADAAKPSTSAAPKKAAAAAITSADTSLAARVKACSERGFFED
QLCRWRVCENHWGKDPACPTSATQNRQP*
>GCA_000452945.1_BPMSHR5858_genomic_01850_1 group_4655
LRVPDEKIAALLKGEQLVPPLPLPPEVFATREVEQVRPALKDASRDWNLLDTDFRTRLLL
VYKIMREQHGYEMALLEGYRSPERQNRLAQMGSNVTNAAAFQSYHQYGLAADNAFLRDGK
LVISEKDPWAMRGYQLYGQVAEEVGLTWGGRWKMMDLGHVEYHKPGFKLGRSSAR*
>GCA_000452945.1_BPMSHR5858_genomic_01852_1 group_7389
MTQTVQAQIAYFGKIPSRGDFVKSAHNPQLLQTLDHWIARAMELLADDPRWKIVYESAKP
MHFAFLGSKSRLAIAGHMIASHDASNRRFPFLAATALEVDRPLTFLARSPLAFARLWSRA
AAQMQSLLGPQEPPGALQALGDTQVPIDTGGPGSAHDGTFNDFVEHQTLFGLEQMLLASG
HPVKLRGAMLALGSLLRPVMQSGTSHIERGLTLPLPTDPFYRSLVAAFWLELVAPFVSRA
DFELAIFIGTIAERERLIIGFNGASAKTLHSVVDPQTYAEHNIDIDDPEWIDAHAQNDHG
ISKLVSYLDQPQLSLRVCIDTFREAFAGA*
>GCA_000452945.1_BPMSHR5858_genomic_01881_1 group_7396
MSTLQRVLFLLGVAAWPAAGRADTLLPRTQAFTVSAQIVAGCGVAGGGPASGLNFGTLDF
GAHPAVATGQVSAAVGGGALQIECSPGSTLKMTIDGGANASAGNTQRNLASGGARVAYRL
YSDPARTQAIAVGQAVSLPVSGTITLPIYGALTLPGGGAPAGTYTDTAQVTLSY*
>GCA_000452945.1_BPMSHR5858_genomic_01899_1 smtB_1
MGALSHIRVLDLSRVLAGPWCAQTLADLGADVIKVERPECGDDTRHWGPPYLKTPDGADT
REAAYYLAANRNKRSVTVDIATPEGQRIIRELAAQCDVVLENYKVGQLAKYGLDYASLAA
VKPGLVYCSVTGFGQTGPYAHRAGYDFIIQGMGGFMSITGERDGLPGGGPQKAGVAIADL
ATGLYSTIAILAALAHRDRTGEGQHIDMALLDVQVALLANMNTNFLASGKPPVRWGNAHP
NIVPYQTFETSDGWIIVAVGNDGQFRKFVEAGGRPELADDERFATNPARVRHRDTLVPIV
AAMTKTRTKREWLDALEALGVPCGPINDLAEVFDDEQVRARGMQVDLPHPSGASAKLVRN
PIRMSATPPDARSAPPLLGEHTDAVLRDMLGYGDAAIAALRDKRVV*
>GCA_000452945.1_BPMSHR5858_genomic_01902_1 group_68
MSSLPSFLAAASGPRGPSHDQRAACRAALAGAVALAVALGVGRFAFTPLLPLMLAGGELD
IRHGGWLASANYAGYFVGAMTCARIAVDPARMVRAGLAATVLLTFAMGLASPFWVWALVR
FVGGAVSAWTFVFASQWGLRRVVEHGAPAWGGVIYTGPGIGIVATGLIGFALAGRHAALG
WIGFAAASAVLTAFVWRAFGAAGAGTDGGQSRGGAKRAGDGVGVADAAGRAGERLAAGAT
GAAEGAEGAEAVADAVSAANVADTADTANSANSANSANSANSANSANSANSANSANSANS
ATATAERNSPVAGVAGQHGAMAAPAARAPHVESAAARRDAFWLVLLYGLPGFGYIITATF
LPVIARAALPAHSRWPDLFWPMFGAALIAGALLGARLPSGWDNRLLLGACCAVQALGVAL
GIVWPTAPGFALGSALVGLPFTAITLFAMREARRLRGERAAGLMGYATASYGVGQIAGPL
AAAPLAAHAGSFSPALWLAATMLAVGAAGFAAVALRAWRAKTRGGGVG*
>GCA_000452945.1_BPMSHR5858_genomic_01904_1 group_3267
MTDRRSHHDDDLLSLIDELEAESRAIDARMSHLDALVSWHDFLIFEAENLAKLQRSLADI
REQSGQIPHAPHDVRRFRLGLLWMGAVAAYEGAMHELFVNTLRVESLRAKALDRLRQLIR
DGASPHRDIKTADAQSVTRWFTLTTIADPLVAAKRFDTLYGIATITPDPAWCRRVLGIRN
AFAHRNGIGVDIREEALHELIDRLDDLASEFHSAFLHEAGKLTET*
>GCA_000452945.1_BPMSHR5858_genomic_01905_1 nudC_2
MTTADYRFCPRCARALAERADPVDEGGRLRRACPDATCGYVHWNNPVPVVAAIVEYEGRI
LLARNAAWPEGMFALITGFLEHGETPEAGIAREVREETSLEAESVTLVGVYEFIRKNELI
IAYHVRASGTIRLSPELLEYRLIEAPKLRPWRAGTGQAVADWMRARGLEFEFVDFPGAVA
NPG*
>GCA_000452945.1_BPMSHR5858_genomic_01906_1 group_4659
MADPQLITTAFDIPGYRIERSLGVARGIVVRSRSIVGTFGASIQTLFGGNISLYTSLCER
ARQDAYERMIDEARRMGGNAIVGMRYDATEIASGVTEVLCYGTAVQAVRAG*
>GCA_000452945.1_BPMSHR5858_genomic_01907_1 dhaT
VAYIYYLTHIHLGDDALAMLGAECARSGIARPLVVTDKGVAAAGLVDRALEALGLGVLPV
FDDTPSNPTEAAVLAAAQRYRDEGCDGLVAVGGGSAIDLAKGVAIAATHPAPLTRYATIE
GGSERITAAAAPLIAVPTTSGTGSEVARGAILILADGRKLGFHSWHLLPKAAICDPSLTL
GLPPGLTAATGMDAIAHCIETFLAPAFNPPADGIALDGLERAWAHIERATHDGGDRAARL
AMMSASMQGAMAFQKGLGCVHSLSHPLGGVKVDGKTSLHHGTLNAVVLPAVLRFNESAPS
VVAERRYARMRRVMNLPERADLSQALHDMTARLGLPTGLRQMGVDEQAFDHVIEGALADH
CHKTNPRIASADDYRRMLVESL*
>GCA_000452945.1_BPMSHR5858_genomic_01908_1 group_4661
MSSTKPVPLPRSAYRHFLPITTRWMDNDVYGHVNNVVYYSYFDTVVNEYLIRRGALDVER
GATIGLVVETQCNYFAPLVFPQRVEAGLRVARLGSSSVRYEIGLFAHGEASAAAQGHFVH
VYVDRATRRPVPLPEPLRAALAPLVAPAA*
>GCA_000452945.1_BPMSHR5858_genomic_01909_1 livH_5
VQSVVLDLLNGLSYGLLLFMLCAGLTLIFGMLGVLNFAHASFYMLGAYAGYAIAARTSFW
VALVAAPLAVGALGALCERTLLRRVRASGHLDELLLTFGLAYLIAEGVKLVWGLAPLAAP
VPAALDGAAFTVYGAVFSRYRAFMMLVSIAMLGALWAVLRASRTGLVVRAALTRRAAVEA
LGHDVPRVMTLVFAAGTALAALAGVIGAPLAVIEPAMAQTLGSVVFAVIVIGGLGSLAGA
LVASLVLGCAQTFAAASRASLSDAAQWAGIALPDAWRALTLAQFAPLIPYLLLIVVLACR
ARGLFGERADV*
>GCA_000452945.1_BPMSHR5858_genomic_01911_1 livF_6
MSALLELRGVRAGYGALPVLDGVDLRIAAGEAVALVGRNGSGRSTLAKAIMGMVRVEGTV
RIGGVDVAGAPTFAIARHRVGYVDEHRDVFALLSVEDNLRLGLRGASGAAARAALADARE

RYPMLAERWRTKAGVLSGGEQQMLALARALIGAPRVLIVDEPAEGLAPLAVEQVRACLAG
AKAQGVALLLIEQRMQLAPALTSRVAVMGRGRIVYDGAAKALDAGIVERWLSVG*
>GCA_000452945.1_BPMSHR5858_genomic_01915_1 iolC
MTSPTLSFAQNRPLDVICVGRVAVDLYAQQIGARLEDASSFAKYLGGSSGNIAFGCARLG
LKSAMLSRVGHDHMGRFVRETLAREGCDTSRLRTDTERLTALVLLGLKDRDTFPLIFHRE
NCADMALDEDDFDEAFIASSKALLITGTHFSTEQVMRASRRALDYARRNQVRTVLDIDYR
PVLWGLTGKADGETRFIANEGVTAHLQRILPLFDVVIGTEEEFRIAGGKDALLDALAMVR
AVTSAALVVKRGPLGCSIVDGDVPATLDALPVIRGVEVDVMNVLGAGDAFASGFLSQWLR
GRPFADAARAANACGALVVSRHGCSPAMPTPAELAYFLAEAKRDPARMRRPDLDATLARL
HRVTPARTPRDEVLGFAFDHRNQFFELARQTGASVARIDALKRLFVEAVAQTEAQLGLAG
RIGVLIDDRYGQDALNDATGRGWWVGRPVELPGSLPLEFDHGRSIGTTLASWPLEHVAKC
LVQYHPDEAAEPRIEQETQLRALYDAVQASGHELLLEVIPPRRADLPQAADTVYRALKRL
YNLGIHPEWWKLAPLDAAQWQAIDALIAERDPYCRGVVLLGLSAGVDELVGGFGAAAASA
TCRGFTVGRTIFHEPSRAWLAGEIDDAELIARVRRTFETLIGAWRAARTDARARGRHAGA
APRVAA*
>GCA_000452945.1_BPMSHR5858_genomic_01920_1 iolG
MTNAAPTIDVAVFGAGRIGRIHAGNLARRPGVRLKYVVDVNRDAAAALAAQYGAQAADVD
GALGDASIGASVICSSTDTHAELIVGAARAGKHVFCEKPVDLTLERARACEAAVARAGVV
CMIGFQRRFDPTFEALKRRIDAGEIGTPEMLVVTSRDPGAPPVDYIKHSGGIFKDMLIHD
FDIFRWILDDDADTLHATGSCLTDPAIAGAGDIDSAAVTIRTKRGRLCQINTSRRAAYGY
DQRFEVLGSAGMLQAGNVRPTEVTGYSARAVSTDLPEAFFLERYRAAYAREIAHFFAAVT
SGEPVRTTVADGVKALELAEAATLSWREARAVTLGERAA*
>GCA_000452945.1_BPMSHR5858_genomic_01922_1 tusA
MLIHKEVDARGLNCPLPILRAKKALADMESGQILKVLATDPGSQRDFAAFAKQTGNEIVE
SSTTQDKTFVFLMRRR*
>GCA_000452945.1_BPMSHR5858_genomic_01924_1 creB
MDFVLRGAMRIAVLDDDQAQTDFVSQTLTAAGHTCYAFKEGKALKKRLQRETFDLLVLDW
NVPDMSGEEVLKWVRANQVEHRLPIIFMTSRDDEAGITQILNAGADDYVVKPVSGPILRA
RIGSLLRRAYPVNAESSIREFDNYKFDANLKQAYVGDKPVSLTQKEFELALLLFQHLDRP
LSRAHILDLVWKQATDIPSRTMDTHISMLRTKLGLRPENGYRLAPIYGYGYRLERVMQGD
AE*
>GCA_000452945.1_BPMSHR5858_genomic_01927_1 group_7400
MKVLWTGAALAAALLSGCASVTTGVSASAAPGALAGARTYDFARTATQAGNADYRRVETL
VRGELAQRGFEEAAGTQARYRVTIAYATQPASVALTMPGCGGAQQPACVAVDGPAPFGLP
FAGPVYRHVLTLRFIERSSGGDAYRVSAAVQDRHPDALLAAPVLVRSALARVPFADGGGW
LVETKKRDADDAMPGVVSVKPARSN*
>GCA_000452945.1_BPMSHR5858_genomic_01943_1 group_3271
MSAPVAAKPLAAHAAALCLVALIALGVAWEAWLAPLRPGGSALILKAVPLALALPGVWRR
SVYTLQWASMLILIYLAEGIVRGMTDGGLSARLGWLEAALALGFFGAALAYVAPYKRAAK
QAAKQAARRAEQPEKHAPARARASDRTRP*
>GCA_000452945.1_BPMSHR5858_genomic_01949_1 aroC
MSGNTLGTLFTVTTFGESHGPAIGCVIDGCPPGMALTEADVQLELDRRKPGTSRHVTQRQ
EPDQVEILSGVFEGVTTGAPIALLIRNTDQRSKDYGNIAETFRPGHADYTYWQKYGVRDY
RGGGRSSARLTAPVVGAGAIAKKWLRERFGVEVRGYMSALGEIEIPFVDWSHVRENPFFA
PNADIVPQLEGYMDALRKDGDSIGARIDVVASGVPVGWGEPLFDRLDADIAHAMMGINAV
KGVEIGAGFASVAQRGSVHGDELTPDGFVGNHAGGVLGGISTGQDITVSIAIKPTSSIRT
PRRSITRAGEPAVVETFGRHDPCVGIRATPIAESMLALVLIDHALRHRAQCGDVSSATPR
IAARAPDAQ*
>GCA_000452945.1_BPMSHR5858_genomic_01972_1 paaB
MNNEWPIWEVFVRSKQGLDHKHCGSLHAADASMALRMARDVYTRRREGVSIWVVPSAAIT
ASDPNEKAEMFEPAGDKIYRHPTFYTLPDEVNHM*
>GCA_000452945.1_BPMSHR5858_genomic_01981_1 cyoC_1
MSYPSVSAAPRHAAHDHPPSHSVFGFWLYLMTDCVVFAALFAVFAVMAHQFAGGPTAVDL
FDIPGVALETALLLVSSITYGFAMIGAQRGRRGALFGWLAVTFVLGAAFLGLELREFSQM
IAEGAGPQRSAFLSSFFTLVGTHGLHVAIGLLWMAVLACQALRRPALTERDIRRLTCLSL
FWHFLDIVWICVFSFVYLASVI*
>GCA_000452945.1_BPMSHR5858_genomic_01982_1 cyoD_1
MADMHLSHDGEAHGTVGGYLAGFILSVLLTAASFGLVLSGALSSHASLIALAVLAFVQIV
VHLVYFLHMNGSSSQRWNVMAFSYTVLTAAILIVGTLWVMHNVGMNMMSR*
>GCA_000452945.1_BPMSHR5858_genomic_02000_1 group_172
MNAVRRFTALAGGMNPLHHDEALAETARFGGRIVSGTHYSARMMGMVATFAPFLGERRAA
LGLGLGLGLGLGLGLGLGLGLGLEFEFEFGFRKALAAGDTVTMTWRSLRSSRARS*
>GCA_000452945.1_BPMSHR5858_genomic_02001_1 sasA_13
MMNKTNRDNASDSLESGISVLLVDDQAFVGEVIRRSLVPESDIVLHTCTDAQRALAMAKE
VKPTVILQDLVMPDIDGLELVRAWRADAQTARVPIIVLSAKEEPAVKRAAFVAGANDYLV
KLPDTIELIARIRYHSNSYLMLRQRDEVLDFLSHDMRTPQTSILALLNVYRNEHGAMPPI
LERIETHARRALALADGFIHLTRAQSEKRPSETVSLNEVVLDAVDQMWEKAGSVGSRVRA
DVPPTECLCIGDRMMLTRAVANLVDNALKYGPGGSEVRCTLVEDGDGWLIGVEDEGAGIA
PDQRAAATESFVRLVPAGGAQRGGFGLGLAFVRATAAKHRGQILMRYTPRGFLAGLRIPK
DVSAA*
>GCA_000452945.1_BPMSHR5858_genomic_02008_1 hdl IVa
MSDGRGAFLSDVQERRFARGGSYGAVLRASPRRAPPAGARLAAAFGRARAFGVGPVRAAD
GGRCILKRHPTTQENIMQTLGVKALVFDVFGTVVDWRSGVIRDATPFLAKYGGAGADPAA
FADAWRAGYSPAMEEVRSGRRPFTRLDVLHRENLDALLPAFGIDRTSVADADLDALNLAW
HRLDPWPDSVAGLTRLKAHYIIAPLSNGNVILMIDMAKRAGLPWDAILGAEVAQAYKPTP
EAYLRTADILALRPDEVCLVAAHNGDLAAARRCGYRTAFVARAREHGPAQTTDLRAEQDW
DVVAADFIELAQRFGA*
>GCA_000452945.1_BPMSHR5858_genomic_02009_1 eriC
MKRLHAFEPVAALAHLCRWLALSAVVGALAGSASALFLWALDLATGTRIAHPWLLWLLPA
AGFATGWVYHRVGRSVERGNNLLIDEIHDPRAIVPTRMAPLVLVATVVTHLFGGSAGREG
TAVQMGGALADRVTQLVGLDRDARRIMLMGGIAAGFSSVFGTPLAGAVFGLEVLAIGRLR
YDALFACVAASIVADAVCRLWGVHHTVYVVPFVPAPSAAGVASAVAAGIAFGVVGRLFAA
ATHGLAALAKRRIAYPPLRPVAGGALVALAATLLHAPQYLGLGIPTIEAAFRAPLPVYDF
AGKFAFTVVTLASGFKGGEVTPLFYIGATLGNALSPLLALPAPVLAGLGFVAVFAGAANT
PIASTIMAIELFGAPIGVYAALACVVAYLFSGHAGIYRAQRIGHAKLPGPRGDVRVAKSA
AGQTTPR*
>GCA_000452945.1_BPMSHR5858_genomic_02010_1 group_8447
MGDVNRTGWPPRLPPRLKRIWYGAGGLRAGWAVLLYAAIVAAILLALGAAARIAHHPFRP
RGDLSTAGQMPFELALCAAALIATHVMSRLDRRSWLDYGLRARRAALHFGWGAFCSLAAV
SAIMGLLVATGGATIEYSGANRAAAFESAVTWACAFVLVALAEEIAFRGYAFFKLAQRTH
PVVAAALTSLAFGLSHVSNRNENIAGIVPVVIYGLVACLAIWRTGSLWWALGEHAMWDWS
ESFLFGAADSGLTAHDTLFRSHAIGPVWLSGGTVGPEASVLVFPALAALAYVAWRLPSRM
VSGPPLARGRRQAA*
>GCA_000452945.1_BPMSHR5858_genomic_02035_1 group_7405
MSYMLLIVEPRGQRSARTRAEGEALYERMRHFAGELQSRGVLIGAESLVSDDQSTRVQVR
NGEVRLVDGPYAEAKEMVGGFFLLDVGTQGEALAIAKDCPAAEWCSVEVREIGPCFR*
>GCA_000452945.1_BPMSHR5858_genomic_02051_1 group_3284
MKISVLVPTFRRPADLARCLIALQRQRVAPDEVIVVARPDDDVTHERLADPAVRGALALR
VVTVDVPGQVAALNRGLDAARGDVIAITDDDAAPRVDWIERISAIFAADPRVGAVGGRDW
VHEKGRLLDGEQPLVGRLTVSGKIVGNHHLGVGGAREVDTLKGANMSYRRTAIEGLRFDT
RLRGAGAQTHNDTSFSMCVKRAGWKLVYDPAVAVDHYPAERFDDDRRDAASMAALSNAAY
NLHLTLREHLSPVRREIAWWWWTFVGTRAYPGLVHVVLSAAAKNGASMRARWRAVRRGAR
DARRVSIAPHAGIAQ*
>GCA_000452945.1_BPMSHR5858_genomic_02085_1 glgB_3
MSECPHDPYAQHHSHCLPFGAQPCGAAGATTRTHFRVWAPAHAAVTLALDTAGGLHELPM
APAGDGWFETFADCGAHTRYRYRLDEALTIPDPASRSQPEGADGPSEVIDPRAFTWRHTF
WRGRPWEEIALYAIQPGAAGGYDGVRERLPQLAQLGVTALELLAAPQARDDSLPFAPIAA
YGGVDALKRLIDEAHGFGLAVLLDLDYARFGCGSDEMRRYAQPFFHTRDDPLQAPPLALD
HPQVCDFFCDNALYWLEEYRCDGLRIREADRIDSAWLREIADRVHAAMPTDRIVHLVLGS
ERHPSHLADTHFDAQWNGCGERALYRLTGRRDRAHDDGVSTHQSIHALARALTANGAVFQ
RAHPAGEGGMADVGLPLTSLVLSDGVVRDAREADLAALALSLLTPQIPLIFDEAAGDASR
RHFVQSALAVRAKLIAPRLIDVRPRRADMLRAADGVEADALVAAWRLGDGETLSIALNLS
PQALPFHGAPDGTIVFETPARARDRVDAGVLPPYALVAWLTGDVNQYALTHDVRRYADVA
PHSPGGA*
>GCA_000452945.1_BPMSHR5858_genomic_02091_1 group_7406
MKKVMLAAVLATAGVAAHAQSSVTLYGRLDAGIEYMNGLPSANGGSTSRWRAESGNWGTS
LWGLKGSEDIGGGNKIVFQLEGSFDTMNGNGPGAGSIWNRWATIGISNDTYGTLLLGREL
AIANGVWDFDPFGQSSWSTASLVRGRNWNKTSNNISYQSPKFYGFDVYGQYALSNATNWN
GNGSTSQGRSAGLQLTYTTSLFQIRGLYDEARDPANGKLDDVFNYSREYFAGVNVFLGQF
KLQAAYQASHADGGPVVNNGITTTQQVWGGVTWQATPAAALIAAVYHVNANHGGGNANIF
TVGGSYNLSKRTLLDVQVATVRNSKTANFGLNANPAGTDISTGNPKFGGSQTGVYAGIQH
LF*
>GCA_000452945.1_BPMSHR5858_genomic_02092_1 glnB_2
MKRITAIIKPFKLDEVREALAEVGLTGLTVTEVKGFGRQKGHTELYRGAEYVVDFLPKMK
IEVVVANNLVDQVIDAVIGAARTGKIGDGKIFVSDVERVIRIRTGEENEAAV*
>GCA_000452945.1_BPMSHR5858_genomic_02093_1 nadE_2
MKTRIALAQLNVTVGDFAGNVAKIVAAAQAAHDAGAHLLIAPELALSGYPPEDLLLRPAF
YAASDAALAALAAQLKPLAGLAVLVGHPLRAAGAGASSADGNANRPIERGAAPVDTYNAA
SLIVGGEVVGTYRKQDLPNTEVFDEKRYFATDAAPYVFELNGVRFGVVICEDVWHASAAQ
LAKAAGAQVLIVPNGSPYHMHKEAVRIDILRARIRETGLPMVYVNLVGGQDELVFDGGSF
VLDGAGELVAKMPQFEEGHAIVEFDGARALPAPIAPALPVEAQVYRALVLGVRDYIGKNG
FPGAIVGLSGGVDSALVLAVAVDALGAERVRAVMMPSRYTADISTTDAADMAKRVGVRYD
EIAIAPMFDAFRASLAAEFAGLAEDATEENIQARIRGTLLMALSNKFGSIVLTTGNKSEM
AVGYCTLYGDMAGGFAVIKDIAKTLVYRLCRYRNAAAEYGALDVIPERILTRAPSAELRE
NQTDQDSLPPYDVLDAIMRMYMEEDRPLAEIVAAGYAEADVKRVTRLIKINEYKRRQAPV
GIRVTHRAFGRDWRYPITSRFIESID*
>GCA_000452945.1_BPMSHR5858_genomic_02094_1 group_3291
MGILTSVLDASAAEWDALVARAPRPTPFLKHAFLSALHRAGCATDGTGWTPRFVTLADSA
TDALVAAAPVYAKRHSYGEYVFDWAWADAYQRNGLPYYPKLLCAVPFTPVQGSRLLAVDD
DARRRLAATLVALAEQSDVSSLHVLFPTGDEARALADMGMMLREGVQFHWLNDGYRDFDA
FLGTLEQKKRKNIRAERRKVADAGVTLRRVRGEDATDADWRFFARCYRQTYREHFSSPYL
NLEFFREIGATMPENLLLVIAERDGRPIASALAVYQRGEAGGGTLYGRYWGALEHVPCLH
FETAYYQLLEFCIEERLDTFEGGAQGEHKLARGFMPTVTHSAHWLAHPAFADAVGHFLAQ
ETNQIHAYVDELREHNPFRAGER*
>GCA_000452945.1_BPMSHR5858_genomic_02135_1 group_7409
MNRPPKQLFRPNDWSRPRAAADVLQRTDAFAALRAGVEQVAALKRDLVTLLPDYLANHVE
PGFIKDGTLTLFAAHNALAARLRQVEPRLLSELQARGWAVATLKVRVRPQPAPEPPRVKQ
ARMSTAGTDALRQLADTLEPSPLQTALARMAARHTKTPR*
>GCA_000452945.1_BPMSHR5858_genomic_02136_1 group_138
VTVAPIDPVTMQVDADFVRDAVDANTVMLVGSACNYPYGTIDPIGALSAIAVEKDVWLHV
DGCLGGWMLPWGEALGYPDIPAFDFRLPGVTSISADTHKFGYGPKGGSVLAWRDASFRRH
QYFLMTDWVGGVYGSPGLTGSRSGGLIAATWAALRGLGREGYLARAKAIFETAFDMQAAV
RAIPELRVLGKPTFCFAFTSDAFDIYHVNDFMRQRGWRLNGLRRPDALQMCVTGPQTQPG
VAEQFRCDLGAAADYARAHAQERPKTSGVYASDAAGVDLSDDARTRAFFTQVIDLFTDCP
L*
>GCA_000452945.1_BPMSHR5858_genomic_02165_1 group_4195
MQQIAVLLDGMIKKNAYLPRRIGGTMRVDEQPGYAIVDSGLPTDTFNLVIGKAHGQPDAA
AIARIAARFNDAGLPAAWWTCRELTDAAFARALGEAGFVADETSVGMLAELDALPPVAPP
PGFGVRQINAQHDVAHFGTLIGALFDPPDAFVDAFYARVAALDFDVAEPLKLFVGELDGR
PVSTTALYVDADTAHVFDVSTSAGQRKRGLASAIMHSALVHARERLGATRAALQASPDGL
NVYRRLGFRAVCEFQVYSNRAAVSPRGELGGAR*
>GCA_000452945.1_BPMSHR5858_genomic_02167_1 group_4698
MVAKGIDAIRALERDRFRAMVDGDGEALDALLSDKVCYVHTNGKRETKQQFIDAITAGRR
RYRQIEIQSQDVLPVGDATYVVAGRALIEMETNNGGLVFSIAYTAVQTHESGRWRLLAWQ
ATRCATDA*
>GCA_000452945.1_BPMSHR5858_genomic_02178_1 group_4700
LSNMLNEARESARVVAAQLADTRRVEALAQHLATHAPQVALTVARGSSDHAASYFASLTM
SRLGVPVASLPMSVATLQQAPLKVRGQLALAFSQSGKSPDLVNTMAALREAGALTVAAVN
VLPSPLAHACEHPLPLLAGPELSVAATKSYIAMLSIAAQLVAFWQRDAALASALRGLPDA
LEQAGRLDWSSAVDELRDVERMIVIGRGLGLAIAQEAALKLKETSGIQAEAFSSAEVRHG
PMELIERDYPLLVFAPPGPEQESLLQLARDMRARGARVLLAAPAGTPDATLPLARTAHAA
LDPIAAILTFYVMAAGLAPARGRDPDAPRHLHKITETH*
>GCA_000452945.1_BPMSHR5858_genomic_02185_1 group_3297
MIELAPIPEPAATALGAARLQLRALAYRRVAAAAGGVGELLEAAAQLGRLLGREALACAG
AAAGHAPIDDYVVCSTILSGLLMDDSPHHAAFVGAIASHGPRAADWLTHGYECASWGYVL
RETLGKAMHTGTRRLLLQIVDVDIHRFSYWLSHPQWGASGFGVCTLALDVAPGADWPLHL
GAAPPAHALAMLGRGFAAFGRERAGVPVAPPFFKEASRRALLKCLGDAPVHPDGHARFGH
AFGSDPWISLLLHRCAPGAPAAARAIVGSLALNGYFAIAEIGYDAAARFRLDDTA*
>GCA_000452945.1_BPMSHR5858_genomic_02186_1 group_1274
MTPFALRRYRLERAPRTPVLVRASSYGIPAATADDYARLAAYQPDSTAPVLLDHAQYVRF
MSELVDALWQALGAETRVFVRFRMPYNGALPANLLDPRVLIGSGLDTQMAFWSRVEAQPG
IGIVDRIRRDLDYPGDNTLAFACASERRLDALEAALLRDWPRYAIRLARAADAADAAGQP
ARPPARFA*
>GCA_000452945.1_BPMSHR5858_genomic_02194_1 COQ5_1
MLLKNLRPADDYDRFANEALAPWDALFIARIRELARALPPGTLADIGTATAVVPVRLAAD
PAMAQWRFVGVDLDPAMLDEGVPRIARLGLAGRVALKVGDALALPFDDASLSMVVSRATL
HHLPDKALSLAEMFRVLRPGGVGVVHDMRRDAPPALLERFTQMRAAANYPPTHLEEKVTL
DEARELVAAAGLAEHASVTSPPFGLGALGFEILLTKPGEPS*
>GCA_000452945.1_BPMSHR5858_genomic_02195_1 group_7410
MSEIRDSITFQNAAALKAQARLGPGVICGLRSERAPLPLVVSPHGDSALAADRDAALAWF
DARRAAFDALLLEHGGLLLRGFAVPDTHAFRALTDRYPPHAFGYIAGASPRKAIDGNVYE
STHLPAPYKLSLHQEKAYMSHYPRLIAFYCRQAAAVGGETPLSDMRAVTRRLPARTLERF
RGKGVMYRRNFSAKPMPAHFNQFYRRWQDAFMTDERAEVESLCRATQLEYEWLPDGSLTV
THVGPATVVHPRTGEEVWFNHASTQHINARVVHPTILRALQSFYKTRAALPYDIRYGDGT
PMPAEDLDPVYDAIDAEETAFRWREQDVLLLDNILVAHGRNPYSGQRDIQVAMMD*
>GCA_000452945.1_BPMSHR5858_genomic_02196_1 btuD_1
MSPVPFPAGEAGEALEGAQTAEAVEAAEAMEGGAAVARGATPGRAASALAARRIDVKAGD
FTFHATDIAFRTGALTAIVGPNGSGKSTLLEALFGFRRAQLEGATILGVPAARFMRDTRA
LRRFGAQLQRVEYAEHARVDEILAVHRALYRQQDAAVARALAIDELRAKPYSGLSKGQRQ
RLDLFIAFAHRPALVALDEPFTGLDRTMTRSVLGLLRGPLAGITIAMICHAGEELAIADD
VLWVRDGTLRYQGGKDALKRRLVGEFRALIHVDDDAQAERVRAVLARDAHVQRIVAPAPR
QIGAFGRAGLDAALRTLMEQAGIRHFEFAPTDEGDLLRACTEGATDA*
>GCA_000452945.1_BPMSHR5858_genomic_02199_1 dat_2
MQDVDPAYAFVRARESGARTYADTFDTVFAHGAGTVLTDTAGRRYLDFLSCAGTLATGHN
HPAVVERVRAFVDSGQAMQMLDMTTPVKHRFVERLFGILPPALARHARVQFCGPSGADAI
EAAIKLFKTATGRRSVIAFHGGYHGMTAGALALTGNLRAKHAVASLMPDVHVMPYPYAYR
CPFGLGAPHTAAASLHYIESMLSDPESGIAKPACVIVEAVQGEGGVIPAPPEWLAGLRAL
TARLDIALVIDEVQTGIGRTGAMFAFEHSGIRPDAVVLSKAIGGGFPLALVAYDERYDVW
QAGAHAGTFRGNQIAMAAGVACLDVIESEGLIAGAAAKEAHVRARLERLAARHPEIGDVR
GRGLMWGIELVDPDAAPDAAGARPAAPALARALKRYCFAHGLIVETGGRHGAVVRLLPPL
TVSAAELDLAFDTLDAGLAALGEHARAEFA*
>GCA_000452945.1_BPMSHR5858_genomic_02206_1 group_7411
MNDIALTEEHRDALQEISNIGMGRAGAALAKLLGAFVTLSVPDIKLVSARELLDELQRCE
RRADMPRPVRQSFQSDICGEALVLFGSDGRRELKELMGYDDATDDIEDEALSDIASLLVG
ACVHSVFEQLDRRLTFLRPTFVPPGSLASALGDERLGRWDVALLLEVHFTLEHGGFVAKL
VMLLPDAAIHKMKAALEQFLDAL*
>GCA_000452945.1_BPMSHR5858_genomic_02218_1 cefD
MTAPLKSLFLLDPSVTYLNHGAYGATPRPVFERHVQWQYELEREPVDFLSRRFAERMAQA
RAILAEYVDTERDNLVYVSNGTTGVNIVARSLPLGPGDELLTTDHEHGGIERLWRFTAQK
RGFEIVRHKVSLPVTTHARFVEDFWRDVTPRTRAILISQLTSPTALVFPVAAICARARAR
GILTIVDGSHVPGQLPLSLREMDPDFYVGILHKWVCAPKGSAFLYARPDVQPLVEPLVVS
WGWEPKNPGPSKFVEYHEWQGSRDISAFLSVPSAIAFQREHDWDGVRKRCIALASDAQRE
VAALTREPLYHPPGAHEWHGQMVCAQLPPQTDDIALLARLRNECGIDVSVDRFGGRPRIR
VSIQGYNGPDDVDRLLGSLKQLLRL*
>GCA_000452945.1_BPMSHR5858_genomic_02221_1 group_4708
MKTALTLKTAALALALAAPALSFAQPSSDTSITRAQVRDELIQLERAGYQPSKNRYPANL
QAAEARIAPMNGGADMSGYGGTTGGAMQSGQRVSPTSGGWDTPYRHR*
>GCA_000452945.1_BPMSHR5858_genomic_02232_1 group_4710

MYSIILVALDGSQTASHALDAALELAADAHARLVPVYVVDMPVFAFDTPGYDPSILVDAF
REEGRRVLDDAQARMTRRGVAGAPRLVEVEPPGEDVAERLERAAREIGASLIVMGTHGRR
GVRRLMLGSVAERLLRHARCPVLMIPARGAPAADANATHPTETA*
>GCA_000452945.1_BPMSHR5858_genomic_02233_1 group_3302
MSYKSILVHLDTSDRARARLETALTLARQFGAYLSAVFAVYTPEPTSFYVMAGTADYFAD
QQRRRDEKRAALERLFHAELKRADVEGQWIVADARANEAVPHYARYADLVIAGQTDPDDP
ETYVDDSFPDTLVLSAGRPVLLLPYAGMPSAIGTRVLVAWDGSREATRAVHDAAPFLALA
TKTTIVTVNGAAHEPPGARIPGADIALTLARHHANIDVRDLERARDASIGDVLLSHAYES
GTDLLVMGAYGHARWKELILGGVTRTIFASMTVPVLMSH*
>GCA_000452945.1_BPMSHR5858_genomic_02237_1 group_4711
MTIDTVSSRSIPLAWIAGVGASAGLGAALGRRFARGGYAVVLTGRTHERVAAVADEIRAQ
GGVAHVASGDIASEADVERLAIEVRTLGALRAAVFNAGNAVRAPTLELSVAQFEHAWRTN
VIGGFLFAKAALAQLLDADGGAPGDDTPRSLLFTGATASLRGRPPFAAFASAKAGLRSLA
QTLAREFGPRGVHVAHAVIDGGIDGERLRGAAPERVAQAGADGLLSPDAIADSYWQLHVQ
HRSAWTQELDLRPYNEAF*
>GCA_000452945.1_BPMSHR5858_genomic_02248_1 rbsK_2
METIAVIGSNMVDLVTYVTRMPADGETLEAPNFELGCGGKGANQAVAASKLGARVAMISK
LGDDLFAENTLRNFERFGVDTEHVRRVSGVSSGVAPIFVSPDSRNRILIVKGANRHLRPA
DIDAAAAKIEASRLVVLQLEIDIDTVYYAIDFAAARGIPVLLNPAPGVPDLDFARLAKLE
FLVPNETELALVSGMPTDTPDAVERAAGSLVERGVKHVIVTLGEKGSLLVSRAGAVRVPP
VAVDARDTTGAGDAYIGCFARHYVATADIPAAMRLASAYAAHSVTGLGTQKSYADAATFE
RFLQTIGFGA*
>GCA_000452945.1_BPMSHR5858_genomic_02254_1 group_420
VLTGSHTGTQPTGGRYDGICGVLGGLEVVRALNDAGVETARPIDVVVWPHEEGSRFAPPM
IASGVLAGVYSLDYGLSLAEAAGVTLGGALAQIGHAGDAPVGGAPLPAACDRAVAAAPHA
PRPMRITAPSGAIPARMRRRARHSPQLNRRLQQSSQTDSNNCDMLSYYE*
>GCA_000452945.1_BPMSHR5858_genomic_02265_1 group_388
MSGRASAIVRAAAIAGWRRRFAPAAAPGYRCGATLARVAADLGATLAAADGGDARRMAFD
DGVEWRAIERVDRQFLLHTVSVQLERTIAGPAAHGSASIAASGWVRRGPLAAAITRNADA
RFGALVPTLLAAPALNAPLAALDLTYCAIVACGSRWTLRIVPFGGSEVAGRMPPFRRYVR
LADTQRDALRAAFAGFEAALARMPHACRTRDAAGAPGGRAAAGACRRAASALSGGVHVVP
VAFARRRP*
>GCA_000452945.1_BPMSHR5858_genomic_02271_1 group_4715
MTFDADAIHKALASPVRRTILAWLKAPDKHFPASQAGPFPQGVCAGQIDALCGLSQSTVS
AHLATLERAGLVTSTRVGQWAFFKRNDAVIQAFLDSLQRGL*
>GCA_000452945.1_BPMSHR5858_genomic_02274_1 rnk
MRNRIYQLTELDVARLEKHAERNPRYQEMLDTLLERADIVEPNKIQANVVTMNSQIKLLD
ETAGQDRAWTIVYPDAANFEQGRLNVFSPVGMALLGSRCGERVKVTLPGGADATLKIVEI
VYQPEASGDYAH*
>GCA_000452945.1_BPMSHR5858_genomic_02275_1 group_4716
MKSFVYAAVAASILATPLASFAQADQQPPLTREQVRSELVQLEQNGYKPEAGETQYPANV
QAASQRMLPAQQTLTHADTSGYGVQPAGAAESGGRASRSPTSPAFGHSIYFGN*
>GCA_000452945.1_BPMSHR5858_genomic_02276_1 group_7415
MSLTVAQILRSKPDSGRTIHMVEKSDSVYNAIKLMAEKSIGALLVMDGANIAGIVTERDY
ARKVVLLDRSSKATRVEEIMTSKVRYVEPTQTSDECMALMTEHRMRHLPVLDDGKLVGLV
SIGDLVKSVIADQQFTIDQLEHYIHGIPAAAHPQ*
>GCA_000452945.1_BPMSHR5858_genomic_02277_1 ribBA_1
MKSSHPDSPQPDDGAAPECVTLVATATLPTRYGTFTSYAFRVAGGDAEHLALVMGDVAEQ
PSVLTRLHSECLTGDVFGSYRCDCGEQLDLSLRYIAAEGRGVLLYLRGHEGRGIGLSNKI
RAYALQEQGRDTVEANLDLGLPDDAREYDSAAAILRILKVTSVRLMSNNPKKFDTLARHG
IPVCERVALAVPVREENERYIRTKQLKFGHYYFDENE*
>GCA_000452945.1_BPMSHR5858_genomic_02280_1 group_3308
MNSPWLHRFRGSSTDGYVRLPMHAFAELRLVHVSSGIDSGLLVELRASDIDARIAGYTEW
ERPSSAGAAHLTVGWDWYIDGATGAFVIAWGDVRSNVMGVDGNGLDIGMDPTSAALSRRL
AQLNWPSAVTAAMLRRGDFAHAGPTLQ*
>GCA_000452945.1_BPMSHR5858_genomic_02281_1 anoR_1
MELRWQDAYLQFSAAENEQQLFQQIAAYTKRLGFEYCCYGIRVPLPISKPVVAIFDTYPN
GWMERYQEMNYLEVDPTVREGALSSNMIVWPEASASDATTLWSDARDHGLAVGVAQSSWA
SRGVFGLLTIARHTDRLTSAEINHLTLQANWLANMSHSLMSRFLVPKLAPESGVALTHRE
REVLCWTGEGKTACEIGQILSISERTVNFHVNNILDKLGATNKVQAVVKAIAMGLIDAP*
>GCA_000452945.1_BPMSHR5858_genomic_02282_1 mgtC
MTFEFALRLFTAFACGVAIGLERQIRQRTAGLRTITLVASGACLFVTLGVLTGNGIPGVT
QIAAYVVSGVGFLGGGVIMRDKGSIQGINTAATLWCSAAVGVLSGAGHYLPAFAGTGVVL
LTNTLLRGVSQAINATPVSNADLVREYQITVICLASDEVHIRTLLSNSMYAKPLSFQSLT
SEDVPREADAPERIKVTATLKLHPKDQQKLEQIASRMSMEKSISSVSWTAKEAEPLME*
>GCA_000452945.1_BPMSHR5858_genomic_02289_1 group_4718
MNETTVNFNPLLMPWRAPQPNNVAGKGQIEIPGRMPNLVWQTRKAEPTQYENDLGDALER
VFESGATELAEVVAGLNRIGFRAPDGAEWTAERFCAELAALAE*
>GCA_000452945.1_BPMSHR5858_genomic_02307_1 mdeA
MTYSSYHKKDIEGRPLHPETQMMSYGFDPFLSEGAVKPPVFLTSTFAFRSAEDGADFFDI
VSGRKPLPQGEAAGLVYSRFNHPNLEIVEDRLALLDGSEAAVVTSSGMSAISAIFLAFLR
PGDQLVQSVPLYGGTETLIAKYFREWGVGAHSIDNGLSPQAIGAALEAAAQQGPVRLCYV
ETPANPTNALIDLDGMRRELDAFEARHGYRPISVCDNTLLGPIFQKPSEHGVDMSVYSLT
KYVGGHSDLVGGGVTGRRDLVAKVRAVRSAFGSQLDPHSSWMLIRSMETVVLRMKQAART
ASAVAKWLATNPHQKVDVYHPELIVDDAYQAVYKRQCTGAGSTFAFVLNGGRAEAFRFIN
ALHLFKSAVSLGGTESLICHPASTTHSGVPEAARKAAGVSEGLIRVSIGLEHEEDLIADL
DQAFRRSGLSTGV*
>GCA_000452945.1_BPMSHR5858_genomic_02308_1 group_7417
MDTSTVDAVAIAAKLAKEISPHPPQWIIDLYAATLVQRHGFARTGDEIKQAQKAFFESRQ
AIVAALSTYRSPTD*
>GCA_000452945.1_BPMSHR5858_genomic_02327_1 trg
LRLFLVMNAFFSRFSIRTRIFSTLGLVAALLTITGVIGFFSMQNSNAALDDAYTRQLAAK
TALASASLNLAITRTTLDRVILHPEVPDAANTIAKAEQYLAASERGWRDYEALPRGDDEK
TLAAQANAARRALADEALKPMIDALKAGRHDEADRLMMTVAPPLSVAWTKATNALDDARA
AYGKAAYDDAEQMYGWLRLVLGGLIVFGLAACLGCAIGLHFAITQPLVRILGHLRRLSDG
DLTGELRWTSHDEMAELVSGLTTMQRSLSETVRKVTDGSESIATATRQIAAGNTDLSQRT
EEQAAALQQTAASMEQLTATVKQNADNAREAQNCADNATDIATKGATVVGEVVGTMAEID
QSSQKVADIIGTIEGIAFQTNILALNAAVEAARAGEQGRGFAVVAGEVRTLAQRSASAAK
EIKALIGESVERVATGSRLVGSAGDTMQEIQRAIARVTGIMTEISAASNEQRDGIEQVNR
AVSQMDQVSQQNAALVEQAAAAAASLEEQADGLRRTVGAFRVVA*
>GCA_000452945.1_BPMSHR5858_genomic_02330_1 group_7418
MANEAITTDSLAVAERVRELMTRHGIGKRQQTTELCRILDLSFSQGHRKLRGSSPWTLAQ
IKKVAEAYGEPAAQLFGAQALDPGMVGACARDAVLYAGIAEIACTAWIGGLLDAGARPEF
VAYEQQGRWRVLRHTGVLYQNAYDVHKIEIYPRRAESDRLLVAVIDADSVSADEVCRYLD
EQGFVTARFGSVDAFADALQGQAFDAVVTEWMFDGRNAADAIQAVRASDNPGAPIFVLTG
DLLTGRASEAEISEVIRVYDVVCYEKPARMAILSADLAKRLSRA*
>GCA_000452945.1_BPMSHR5858_genomic_02340_1 tap
MRSLSLNQKLTSMIVILWIGLLLIGATGAWQNRSSMIADRREQLAYLTAQAYNVSEHFYK
LSQQNAMPEADAKRAALEAIAAMRYGKDGYLSVNDSKPVVVMHPIKPELNGKDVSGFTDP
GGKHLFVEIVKAGNAEGGLGFVDYMWPKPGADKPIDKTSAVRHFAPWDWYLVTGMYMDDL
QAAVLASTGRWLAMTAVLGAAATILMVLVLRSVRSSLGGDLEAAVATAQRIAQGDLTAHV
DIRGGDRKSLLHALHTMQTALIDMVSRVRLGTENINVGASEIAAGNTDLSQRTEQQAAAL
VETASSMDEMTTNVKQNADSASQAAELADQAAQVATRGSSVVDDVVRTMDEITDASRKIG
DIIGVIDGIAFQTNILALNAAVEAARAGEQGRGFAVVAGEVRTLAQRSATAAKEIKSLIE
TSNTTVEHGATLVTRAGTTMTEIVQSVRRVNEILEEISHASREQSAGIDQVNRAVGEMDQ
VTQQNAALVEQAAAAAHSLKDQADALRSAVAQFALPG*
>GCA_000452945.1_BPMSHR5858_genomic_02341_1 penA_2
MRLNKKPKSSDPYVAVAKNPMLSSSLPMWRSKFVVILVFLAFGALIARAFWVQVANQDFY
VGQGQKRYQRTIELGAMRGSIVDRNGALLAVSLTTYEIWANPKQVDEAAHAPLAKLLDMP
LAEVRRRLTGDRSFVLLKRQVDPETAARIEKLDIDGVTQIADSKRFYPEGESAAHVVGFT
NVEDRGQEGVELAANGRLSGTAGQRQVIRDRLGRIVSDTGPLVPAQHGATIELTIDRRIQ
QLAYAQLKSTIIANNAQAGSVVVLDAQNGEILALANYPTFDPNDRARLTGQQLRNRAVID
TFEPGSTIKPVVVALSIDAGKVRPQTIIDTSPGAYKIGPNVIHDTSNHGAITVAQALQMS
SNIALAKLALNLPAETIWNKYQEYGIGRAPELTFPGVASGRLRPYKRWRPIEQATMAYGY
GLSTSLLQIAQVYTAYAGDGTIHPVSLLKASADPQSADAHRGHRVTTPATAAAIRSMLEM
AVGQGGTGRAARVDGYRVGGKTGTARKQVGASYVKGKYRALFAGMAPMSNPRLIIAVMID
EPAARSYYGGTVAGPVFSSVMSGSLQLLGVPPDAPIAADKPAGAGAKPQAKAKT*
>GCA_000452945.1_BPMSHR5858_genomic_02354_1 group_680
MTEVAPGALVTCGDWARAGSALVDAQRAKDDRPSALDGLSAGGMLTDHVAAVNEMVKGIV
GMTFPDQRMRQVRERDRPQPAWTETPR*
>GCA_000452945.1_BPMSHR5858_genomic_02355_1 group_7148
MAIQKITPFLWYSTQAEEAAAFYAGVFPDSRVARVTAVPGTGSTKLVEFVLFGQPFFAMS
HPRTESFNHAISFLVSCRDQAELDRYWSALLEGGGSPDDCGWLRDRFGVSWQIVPDDLIA
MMADPDPVKAARVAGAMMSMRKFDVAALRAAYAGAAG*
>GCA_000452945.1_BPMSHR5858_genomic_02356_1 zntB
MKPEMIIQTYGADTSGMVCGFRFAPDSAGEPVDAGVAAEWAAGRGARPDEFLWLHFNLAH
GASERWMRAHLDLPETFHEFLREGSHSTRIEQQDGVLRAVVNDVMFSLDLEASEIATMWA
YVDRRMMVTARAKPLRSIDTLRESVRQGERFRSPAELLIHLLRDQADLMMHIVRRASTDV
DRIEDRFLSERLSANRGQLSAMRRTLTRLQCMLAPEPGSIFRLLAKPPAWLLSADVQELR
ESTEEFSVVLADLAGLIERIKLLQEEIASRLDEQNNRTLFTLTLVTVIALPINIVAGFFG
MNVGGIPFSENKHGFWMMVLLVAGFTALAAWWTFWRRGDR*
>GCA_000452945.1_BPMSHR5858_genomic_02358_1 ybjJ_2
MSSDRPATRLATRLAFLVAGFGVACWAPLVPFAKARLGVDDAVLGALLLCVGLGSVIAML
ITGPLGTRYGSRPIIVAGGVALAAILPLLSVANTTPALGFALLAFGAALGSLDVAMNVHA
IEVEHAAARPLMSGFHALFSVGGIAGSSVMTFLLSMRVGAFASTLLCSAPMLAAILVARP
RLLRAARAAARPRFAMPRGVVPLLAMLAAITFLAEGALLDWSALLITDKGLVAAAQGGLG
YLLFSIAMTAGRLGGDAIAARVGDRSTMFWGGLLAVAGFVMLLAAPIAGVAMAGFLLIGL
GASNIVPVLFRRAGAQQAMPSALAVTLITITGYAGHLAGPAGMGFVARSVGLESAFWMLA
ALLGIVPLCARVVTANRT*
>GCA_000452945.1_BPMSHR5858_genomic_02359_1 group_4732
MKQAGIRIGTHAIALALGFALGIYALPILTAERGASETELRPVADRALYTGRFERSLTRG
DPLHWTDGKLSVTRAALAFEGRVAPGPDYKLYLVPGFVASKEAFLAAKPHARRVGELKTF
GDFVAPLAADVDIDAYTTVVIWCERFSQFIGAAQYR*
>GCA_000452945.1_BPMSHR5858_genomic_02360_1 group_4094
VNTIRLLGAAALVGACAPLAAAAATPVCKVPTLKVLAQKSLGLSVMEKSLPDYEKTSGTR
IEINYFGENDRRAKSRLDASTGAGSYQIYYVDEANVAEFASAGWIAPLLKYYPKEYDYDD
FLPGRRAVASYKGVAYFAPLIGGGDFLFYRRDLLDAAHLPVPKTLDELVAAVRKLNAPPK
LYGWVARGQRGSGMNVWRWAPFMLAQGGAWTDPHGQPAFNSPAAVQAAERYRDLFNYAPP
GAATYDWSNALEAFRSGKVAFMIESTPFADWMEDPSKSSVAGKVGYARPPAPLPSAAYGH
GLAISSVGAKDDCTRQAAGRFIAWATSKEQEQARLRNGVFSDYNRTSTIGSDYFRQHVKP
QILAGLNDTNPVTKATIWATPQWPDIGDNLGVALEEVFTGTQTDVRGALDDAAQYAKDAM
AHGARK*
>GCA_000452945.1_BPMSHR5858_genomic_02375_1 sumT_2
MNTMGKVTLLGAGPGDLDLLTMKAAKALAAADVLLLDDLVDPGIVALAPRARA*
>GCA_000452945.1_BPMSHR5858_genomic_02377_1 pdtaR
MGTTDQPSKLRVLLVTDTDKPIGELRDTLARLGYEMLDEVATPARLPAAVEEQRPDVVII
DTESPSRDTLEQLAVMNATAPRPVLMFSHDADQQLIRAAVGAGVSAYLIEGLSAERLAPI
LEVALARFSHDEQLRQRLAEVESELAERKLIDRAKRMLMDRRKLSEQDAYATLRRSAMDQ
GIRIVEAARRLLAAHSQS*
>GCA_000452945.1_BPMSHR5858_genomic_02379_1 folE_2
MHQTGNERACGAHDQPHAEIRRVRRFDPIAFEAAVRALLEASGIEIDSAHTGRTAQRVRE
LWQRRLLDGYEVDPAQALGAGFDDPREDMVIVRSIAVHGVCPHHLLPFRGVAHVAYLPGG
RLHGFGRIARMIDAISHRYTYQEWITHRIATALVTHGQARGAACLIEAEQLCLLMGENRR
GDERVITQCYAGEFERDVQARNEFLRGIHR*
>GCA_000452945.1_BPMSHR5858_genomic_02382_1 aphA_2
MLTYFHPDQLKHSPLTYLSRGKMRAPHEVPERAVRLLGAVTSLGFDVRAPEDFGAAPLAA
VHSEPYLRFLADAHREWRRIPEDWGPEVMSNIYVREPNPLRGVLAQAGYYLADGSCPIGE
HTYRAAYWSAQSALAGAAALGRGARDAYALCRPPGHHACRDAAGGFCYLNNAAIAAQALR
ARHARVAILDTDMHHGQGIQTLFYDRDDVLYVSIHGDPTNFYPAVTGYETERGAGRGEGF
NVNLPMPHGSPEAVFFAQLERARSRVERFAPDVLVFALGFDVYREDPQSKVAVTTEGFGR
LGALVGGLRVPTLIVQEGGYHLETLERNAAALFRSYEAAR*
>GCA_000452945.1_BPMSHR5858_genomic_02387_1 pgrR_7
MDRFQEMQVFVRIAERQSFSRASDDLRIPRATVTNLMKRMEARLGARLLERTTRTVRLTQ
DGEAYYRRCVRLIADLEEAEGAFRAAAPRGLLRVNLQGTLARYFVVPALPDFLARYPGIR
LHIGEDDRFVDLVREGVDCVLRSGNLQDSSMVGRRVAQLEQVTVASPGYLARHGEPAELA
ALAAHRAVDYVSSATGKPMPLEFTVDGRVTEVRLDAAISVAGVELYTGAAVAGLGIVQVP
RYRIADELADGRLRIVLGAYPPPPMPVSVLYPHSRQLSSRVRAFAQWLRERFDAAQAGAG
DGACGAPGSSGSAARAPSPARSEPAKALLSRRPRR*
>GCA_000452945.1_BPMSHR5858_genomic_02390_1 curA_2
MPQTKTTNRRIVLHSRPKGAPTPENFRTETAAVPVPAPGEVLLRTIWLSLDPYMRGRMSD
APSYAAPVELGGVMVGGTVSRVVSSSVPAFRDGDLVLSNAGWQDYAVSDGTDLMPLADVA
HPSYALGVLGMPGFTAYFGLLHIGEPKAGETVAVAAASGAVGAVVGQIAKLKGCRVVGIA
GGADKCRYVAGTLGFDACVDHHDPDFAKQLAAACPDGIDVYFENVGGAVFDAVLPLLNDH
ARVPVCGLIAHYNDSELPGGPNRLPLLTSSVLRKRIRMQGFIILDHYADVYPAFLKDMSE
WVAQGKVKPREDVVDGLDAAPRALIGLLGGKNFGKVVVRVGPDTL*
>GCA_000452945.1_BPMSHR5858_genomic_02391_1 yqhD
MLNFDFYNPTRIVFGEKTAARLNDLLPAAARVLVLYGGESARSNGTLDEVRAALGARDVR
EFGGIEPNPAYETLMRAVELARRERVDFLLAVGGGSVIDGTKFVAAAVPFEGDPWTILET
HGANVAAALPFGCVLTLPATGSEMNNGAVLTRRATRAKLAFRHPLVFPTFSILDPTKTYT
LPPRQVANGVVDAFTHIVEQYLTYPADGLAQDRFAEGLLQTLIEIGPKALAEPRDYATRA
NLMWVATLALNGLIGAGVPQDWATHMVGHELTARYDIDHARTLAVVLPSMLDARRDAKRA
KLLQYAARVWNIVDGPEDARIDAAIARTRAFFESLGVKTRLADYGVGADAIDGLIAQLEA
HGMTRLGERKDVTLDVSRRVLEASL*
>GCA_000452945.1_BPMSHR5858_genomic_02394_1 group_4736
VIQPTEVFKDNLAQLPAIDGIERIDLVDGGGAVVASIENKPGKQGSLAVYHYLREAFGTL
DAQAAAHGLAVFAEHTADARNRPGAHPNVDRLLAIAAGGEALRIDVLARG*
>GCA_000452945.1_BPMSHR5858_genomic_02395_1 group_766
MMQVPQRSALGGPTLIRLLARLAHADVPPSGQSLSDRLSQWLAWTDAIALSSALSMSPPA
AAAGARPAGDDPHGQGARVRASLAKAIARTGAFAAGRDGAARVAARGAPHDTPADFTLFR
HDYLSMQQAMEIDIGELRGRLRQTMAAQTPALARLAALDATMERALVARERSLFASVPKL
LGAYFERLREAEQQRLAEAEAKAHANAEADAHAEVARKTAAPAPHAWLDAFRQDMQSVLL
AELDIRFQPVDGLLAALRAS*
>GCA_000452945.1_BPMSHR5858_genomic_02397_1 pal_3
MSDEIDGGIEQTAPIWAAFGDLMSVLLGAFVLILVGVIGVQLELSSKLDAEVRQRQLESQ
RRKTLEQALAAPLAAGRVTLVNGRIGISGSVLFALNSDELQPAGRDLLKSLAGPLAAYLN
ANHEILMVSGFADDLQVRAGNRRFADNWELSAKRALTVTRAFIDEGVPASSVFAAAFGSE
QPVSPNSNDAGRAKNRRVEIAPVPRPTTGDGERRG*
>GCA_000452945.1_BPMSHR5858_genomic_02400_1 dhbF_2
MNMRDTSLRPLLPAQREIWLAEQLRPGTGVYNTAGYADIVGAVDHGALRDAFGHAMREAD
AVRATFSAHGDDAWQTPRDAPHADVPLVDVSGQADPAGVALAWMMRDMRTPPDFAAGPLA
RSALLRVGAARFFWYVCAHHIVADAYGTALVVQRTAQLYGARVRGAAPPPAWFGTLDALI
DEDRTYRESAAFADDRDYWRARLAAAPEPRSLSPVALPGQFAPAGDFHRQWGELDAATSD
GLRAMARAGAQGMPPLVAAMMAAYVHRFTHERDIVLGLPVMARLSRATRRTPGMVANVLP
LRFAFDRGTTFAALHAQCAAEMRASLRHQRYRGEAMLRDAQQRRPAERLHAQNVNVMAFE
RALAFGDCPARSHSLSNGPVDDFSITVYDDGARQPIRIAFDANAARYASGDLAAHRARFL
RLGVALANAAQRPIADAQWLTPAERRTLVGERGTPAPDAGEPFDTIAARFAARAAERPDA
IALVDRGRRITYGELNARANRLAHVLIEAGVGPEALVGLHMPRSAELVVGMLAILKAGGA
YVPLDPAYPASRIEFMVADARPMLSITTGEHAAQLPARTPTIVLDAADAQAALRRAPAHD
PVRPAPLDREHAAYVIYTSGSTGKPKGAVVSHRNVIRLLDGTRGWFDFGAAQTWTLFHSF
AFDFSVWECWGALLTGGRLVVVPYDVSRSPAEFLKLLVDERVTVLNQTPSAFRQLMQADE
AHADLSARLALRYVVFGGEALDARSLARWYERHADTAPRLVNMYGITETTVHVSYLALSR
AIAGMPANSLIGRPLPDLRVYVLDAALRPVPAGVPGEMYVAGAGLARGYLRRPSLTAQRF
IADPFGPPGTRMYRTGDVARWRADGGLDFIGRADEQVKVRGFRVELGEIAARLACDPSVA
QAQAVVRQDGPAHERLVAYVVPRAGATIDVCALRASLAAEMPEYMVPAAIVALDAMPLTP
NGKLDRAALPAPIVTGTSRRAPENRIEQQVCAMFAELLDAQTLGAEDNFFELGGDSLLAM
RAINKLRQTFDVELTIRDLFSAPTVAALSMRLDAQLAARRAHADGAELPAG*
>GCA_000452945.1_BPMSHR5858_genomic_02401_1 mbtH_3
MTHLLDDPDGSFQVLVNEEGQHSLWPAELPAPAGWRRTFGEAGRQACLDHIEAEWTDLRP

RSVAARCA*
>GCA_000452945.1_BPMSHR5858_genomic_02402_1 tetA
MDLHERTLDAPGAPAAAAHGAGSRLSREVWIILLTLAVDAVGLGVATPVLPDLLRAVGER
PAHVPVMLGSLITCFYFMQFVFGPIVGALSDAWGRRPILLIALGGGALSYAIGATARSYG
GLLAAHMLAGIAASSAAVATAYLADVTPPALRARRFALASSVLGLGLIAGPAFGGVLGAL
GPRVPFVAAGAIAALNFVSAACFLRESLPARRRVPVSWARANLFGSLRLVREDRVFRDLL
AAVCLGMLAYGIYLSCFVLANAARLGWGPRENGLALAGLGLGITLTQTLLLPRLVARLGE
HRTALVGFGAFVLAYGLYSRADSVALVAAALAVHALSLISDPSVRSLISVHAGPQRQGEY
QGALVCLTGLASSIAPLVGGNLFAFFTRAGASTYFPGAPFLAAAVLYALAMLAVWRAMAG
AARRRSASH*
>GCA_000452945.1_BPMSHR5858_genomic_02404_1 group_7426
MHRIEEEAFSAESRAEPDAGAALRALGWMMFGMEDLRLCADERRELAQLARYAQTLPIDG
FGGNGRHRCYAEAVLSPSARTLRWKPGIVGKDGTIEVEYEQETEFQPEYGGVRRRFARAS
DRVLQSSLIRKLIWFDFDLIGWAGASEPLQCGFHLVRLQALPGKPSRSTPDCLHRDGQPY
TAVHLVSRHDVTGGLNYIAPPRYSGRNVDELPADALTTFTLTEPLDSYIVDDAAVCHHVS
AVGCAPGAASGSRTVMLIDFSPIPLSS*
>GCA_000452945.1_BPMSHR5858_genomic_02407_1 group_7427
MSLNHAPFALASAFDRYLGDPSMPDNAFSYARCAQLDAHERFPDAVCAMLSQWGLHRYYV
PERFGGMLRDFEQPLQLIRMLARRDLTVAIGHGKTFLGAVCVWVGATHEQASRFARRIGD
GAIVSWGLTEREHGSDLLRSDTYAEADGDEYRIAGEKWLINNATRGDMISLLARTRREGG
PRGFDMLLVDKHALSQGRFRPLPKERTHGIRGADISGIAFDDARIPASARIGARGAGLET
VLKGLQLTRTLCASLSLGAGDHALRLALGFACERRLYGRTLIDLPNARRTLIDAYTDQLL
AEAAALVGARSIQALPDEMSVTSAVVKYLVPTRAEAMLDTLAELLGARAFLADTYAHGMF
QKVQRDNRIVSLFDGNTLVNLNAIVNAFQTLAKRYRGRDGADAGQALAAFDLGMPLPAFD
RGALSLVSRHGCRVVGALPALADGIAQRAACEPALSRLVAPVRRLAALTDDVHRRMLACP
PVRVHVPVEQFELAKRYALCHAAASALGLWWHTRHRVGEASAAPPWRDGLWLRAVLRRVR
VALGDAADFDDEAGDALWRALLRQHRDGLLFSLLPCELAEHRGEAA*
>GCA_000452945.1_BPMSHR5858_genomic_02408_1 group_4739
VPSNFAEHFENQVLKYGNRSQFVFLHGKPGDSAGEARLGYAQLDARARSIAAWLSACGAA
RRPVLLVYPEGLAFVAAFLGCLYAGAIAVPVPVPADARSVERTRRIVRDAGIALALTPTA
ADAHALANCLAAPGDAVSTVGALRCEATDALEALDASAWSRPALDASAIAFLQYTSGSTG
EPKGVVVSHRNLLHNEAQIQRVVASDADSVVTGWLPHFHDMGLIGMLLHPLYVGCDCVFM
APSTFLRRPVRWLEAATHYAATITVAPNFAYDLCARAVKREQLGGLDLSALKVALNGAEP
VRAKTIEAFSARFAPVGFDARAFSPCYGMAEVTLLASAGRPAAAPVLLDIDARALERHEA
SLAKPGAEALRVVGSGRAVDLDLRIVDPQTHAVAADGRIGEIWLRGDSVAQGYWRRPDAS
EAAFRARTAAGDGPYLRTGDLGFVRDGELFVTGRLKDLLIVNGRNLYPQDIEDFVRELHP
ALGAGAGAAFSIDADGERVVVVHEIKEARLDDGLSAAALSARIRTALARAFDIAAPSVVL
VPRGGVKRTTSGKVQRGLTRDAFLAARLRVLHEDIQVPVRSLRVAAGEGVAQDASRGAA*
>GCA_000452945.1_BPMSHR5858_genomic_02409_1 ecfG_2
MTGADIASTLPPLLPRLLAFARRLCGDTHDADDLVQRACLRALERPHQLQRDTSPLNWMM
AIVHSSWMSELRARHVRHRLQTDWNDDVIDALPGPTAADPQIHAMNAQIIRAVELLPETL
RAAMLLIAVEGLSYQEAAGVLDVPIGTVMSRLWRARQSIGAQFGARER*
>GCA_000452945.1_BPMSHR5858_genomic_02411_1 yigZ
MTYTLAATYTRELDIRKSRFIAYAIPVEDRDAAMAELQRLRAEHPGATHVCWALLAGGQS
GMSDDGEPSGTAGRPILEVLRHHDLDGVLGAVVRYFGGVKLGAGGLVRAYTDAIAAALID
AERVERIAYARLAIEIGYPDEARVRRWIEQEGHALVDSAYAMTVKLVIRLPAAAADAARD
MLFDLTQGRAGFPSVD*
>GCA_000452945.1_BPMSHR5858_genomic_02425_1 group_208
MSTKMDEDIKRWTAKRKSALVLDVIQGKTTVAEASRAYDLSPSEIENWVDDGKRGMENAL
RANPQDVKEQYERQEAYGEAMLE*
>GCA_000452945.1_BPMSHR5858_genomic_02431_1 group_2415
MHTLYLIAIVAEAMSGALMGMQRGMDRFGLALVGAVTALGGGTVRDVLLGRYPLTWVAHP
EYLLLTLAAATFASMTATHVARLKSLFTAVDALGLAAFSIIGCDVAAAVNGSPVVIVLAG
AITGVCGGMLRDLLCNEMPLVLRKELYASIALLTGGLYVGMKAIGAAQGFATVVALAVGF
ALRMLAVRRGWRLKAFQAADAG*
>GCA_000452945.1_BPMSHR5858_genomic_02433_1 hmuU_2
MSLAARRHSARPISAERGSTCASRALAALRRRRARWTLAALIAALAAASIAALCIGAYRL
SPAQLLGALTATDAQLAGDAALRQARAVLVDIRAPRVVLALFVGAGFGAAGAAMQALFRN
PLADPGLIGVSSGAALGASALIVLGPLFAARLALGLLPVAAFAGALAVAALVYRLAVARG
RLALPLLLLAGIAINALAGAGIGLLTYLADDAQLRSLTFWSLGSVGGAQWRTIAAVAPFA
LAGCVLLARERHALNALQLGEAEAMHLGVPVPRVKRRVLVGTAMCVGALVSCTGAIGFIG
LVAPHCVRLAAGADQRIVMPGAALFGALLAVAADLAARTLAAPAEIPLGILTALIGAPFF
LALLWKRRGALGG*
>GCA_000452945.1_BPMSHR5858_genomic_02437_1 group_7429
MNVQTLSGVLHAQELLFVSLIRVLPLETRQALADEFDRQIQLAETSRLEAPHDREAHDAF
LAHVRKLLIRLESMA*
>GCA_000452945.1_BPMSHR5858_genomic_02451_1 eltD_2
MPQTAEAPPQRAAERGETPALPATMRAVVCHGPQDYRLEQVPVPKPGPDEILTQVERVGI
CMGDIKTFRGAPSFWGDAVQPRYVKPPMIPGHEFVCRVVALGPGAERRGVKAGDRVISEQ
IVPCWSCRFCGHGQYWMCQKHDLYGFQNNVHGAMAEYMIFTKEAIVHRVPDSIPTDEAIL
IEPLSCSLHAADRANVGFDDVVVVAGAGTLGLGIIGAARLRHPKQLIVLDMKPERAALAR
RMGADDVWNPAEENVIEKIRAITGGYGCDIYIEATGHHRAVGQGLAMLRKLGRFVEFSVF
NDEASVDWSIIGDRKELDVLGSHLGPYMYPRAIEFIASRRIDVRGIVTHTFPLSRFADAF
AVMERGEQSLKVVLDPRG*
>GCA_000452945.1_BPMSHR5858_genomic_02488_1 group_1848
MMTNHLTRRLAAVAHDFGQPAMIWQAVAFFGSLALAWLLARHLRGRIDARRRAAGRTPGP
GALSLNRALFPLIGGVFMWLAETTFDELMPTSLLQLALVPLFGIGLIYILFFVARRVFAR
DGHTHAWLSIVEKLVSIVVWIAMALTVLGIQRDVLGWLDSVQFRVANARLTLLSIVSGAL
WVCVTLVVAMWLGSVLEERLMRARSLDANLRVVLARVGRALLIFAAVLIGLSLVGIDVTV
LGVFGGALGVGLGFGLQKIASNYVSGFIILLDRSLRLGDAINVGGLQGVVTQIRTRYTVV
RGLDGNETLIPNEKLITDVVQNQSSYLTRGYAKVAVQVAYTSDVERAMALLADAAKGVER
VLEDPAPTPYLVGFGADGIDLELGFWIADAAKGTAGVRSTVNRNIWRLFVAHGISIPFPQ
REVRVIGLPDGIAPAAGAAGPAAGRQPQDA*
>GCA_000452945.1_BPMSHR5858_genomic_02489_1 group_3336
LLNSLLDFLSNGLLRFSWWQIVLFTLAVTHVTIVGVTVYLHRCQAHRALDLHPIMSHFFR
AWLWMTTGMLTGQWAAIHRKHHAKCETEEDPHSPQTRGIWKVLLEGAELYRAEAKNEETM
RKYGHGTPNDWLERNVYSKYPILGVSLMMVIDVALFGLVGLTVWAVQMVWIPFWAAGVVN
GLGHFWGYRNFNAADASTNLFPWGIVIGGEELHNNHHTFATSAKLSNKWYEFDIGWMYIR
IMSAFGLAKVKKIAPTPRLAARKTVLDQETLQAVLSNRYEVMARYAKTLKRAYRQELAHL
KELGAREKYQLMRGARKWFHKDEAGLDEPQKRMLPEIFANSQKLHTFFQLRAELTAIWER
SNASREQLLTQLQDWCHRAEQSGIKALQEFATRLRRYA*
>GCA_000452945.1_BPMSHR5858_genomic_02492_1 nadB
MNFDVAIVGSGLAGLSVALNLAQTRRVALIAKRSMMEGASDYAQGGIAAVLDSADSVENH
VNDTLIAGGGLCDEAATRYIVEHGREAIEWLISQGVPFTKDDAAELGFHLTREGGHSHRR
IIHAADATGHAVLATLSERARAHPNITFFEDHHAIDLITSDRLGLPGLPGRHCVGLYALD
VQTGRTMTIEAPHTVLATGGAGKVYLYTTNPDTATGDGIAMAWRAGARVANMEFIQFHPT
CLFHPYAKSFLISEAVRGEGGLLKLPDGTRFMPAHDPRAELAPRDIVARAIDFEIKKRGI
DCVYLDISHQPEAFLREHFPTIYARCAEFGIDISKQPIPVVPAAHYTCGGVVTDLAGRTD
ITGLYAVGETSYTGLHGANRLASNSLLECLVIGRATAEAIEAAGYDSATHGPLPAWDESR
VADADEEVVVAHNWDELRRLMWNYVGIVRTDKRLERAQHRLKLLRDEIHEYYANFRVSRD
LLELRNLVDVASLIVDSARARHESRGLHYSRDWPHALPKALPTVLTPPRRASK*
>GCA_000452945.1_BPMSHR5858_genomic_02493_1 rpmG
MAKGARDKIKLESTAGTGHFYTTTKNKRNMPEKMEIMKFDPVARKHVAYKETKIK*
>GCA_000452945.1_BPMSHR5858_genomic_02504_1 ypeA
MSAGMSTAIVSDDIAIRPFARADTDAALAVWRDAFPSYSDASTPHRDPRRAIELKLATQP
ELFFVATAGGRVVGTVMAGYDGHRGWLYSLAVERGARRLGIGRALLAHAEAALAERGCPK
VNLQVLPGNDDACRFYEALGYHEEARISFGKRLATD*
>GCA_000452945.1_BPMSHR5858_genomic_02530_1 hom
MEPIKVGLLGFGTVGGGTFKVLRRNQEEIKRRAGRGIEIARVAVRNPAKALAALDGDANG
VSIGDDFNAVVDDPSIAIVAEMIGGTGLARELVLRAIANGKHVVTANKALLAVHGTEIFE
AARAKGVMVAFEAAVAGGIPIIKALREGLTANRIQYIAGIINGTTNYILSEMRERGLDFA
TALKAAQELGYAEADPTFDIEGVDAAHKATIMSAIAFGVPVQFDRAYVEGISRLAATDIK
YAEELGYRIKLLGITRRTERGIELRVHPTLIPAKRLLANVEGAMNAVVVHGDAVGTTLYY
GKGAGAEPTASAVVADLVDVTRLHTADPEHRVPHLAFQPDSLSNTPILPIEEVTSGYYLR
LRVADQTGVLADITRILAESGISIDALLQKESEQVDDANGETDIILITHETVEKNVNAAI
ARIESLATVVSKVTKLRMEALN*
>GCA_000452945.1_BPMSHR5858_genomic_02531_1 alaA
VKPIQKSNKLLNVCYDIRGPVLEHAKRLEDEGHRIIKLNIGNLAPFGFDAPDEIIQDMIR
NLPASSGYSDSKGVFAARKAIMHYTQQKGVHGVGLDDIYIGNGASELIVMATQALLNDGD
EVLLPAPDYPLWTAAVSLSGGTPVHYVCDESNRWMPDPDDIRRKITPNTKALVVINPNNP
TGALYSDALLVELIAIAREHGLIIFADEVYDKIVYDGKSHTALASLAEDVITVTFNSLSK
SYRSCGYRAGWMSVAGLTAENRRRAHDYLEGLGILSSMRLCANVPGQYAIQTALGGYQSI
NELIMPSGRLYKQRELAYDMLTSIPGVSCVKPEAALYMFPRLDPKLYPIENDQQFILELL
LRERVLLVQGTGFNWPTPDHFRVVFLPNVDDLTDSIERIARFLDGYRKRNSV*
>GCA_000452945.1_BPMSHR5858_genomic_02532_1 group_3340
MHQPGRGPRAAAPATAAARIAAGGPPRSGPDRRAARPLFRPFSTPRKTDLKLHQDPSNAL
NTVTGYGPDYVDINLQRHETSVIVLPGAPVVEWPVASFDALTPELFAMLLEPQPEVVVFG
SGARLRFPHPRLTAQLTAQRVGVETMDFQAACRTYNILMAEGRKVAAALLIER*
>GCA_000452945.1_BPMSHR5858_genomic_02534_1 arnF
MNPASLFCIVTGVMLNACAQLLLKAGVRAVGHFEFTRANIVPIGFKIATQLPIIGGLGCY
VLSVVVWIVGLSRVDVSVAYPMLSLGYVVNAFAAWYLFGEVLSVQRLVGIGIILIGVFVL
ARS*
>GCA_000452945.1_BPMSHR5858_genomic_02535_1 arnB_1
MSQTSVPFLPFVKPEIDEETIQGVADVLRSGWITTGPQCQRFEAALSEYCGGRPVRVFNS
GTATLEIGLRIAGVGPGDEVITTPASWVATSNVILETGATPVFVDIDPVTRNLDLGLVEK
AITPRTKAIIPVYLAGLPVDMDRLYAIARAHELRVIEDAAQALGSTWRGQRIGAIGDIVS
FSFHPNKNITSIEGGALVLNDEAEATLAQKYRLQGITRTGLDGMDCDVLGGKYNLTDVAA
RVGWGQLAHLERFNATRHALARRYFDAFANGPACALGVGLPVADFENTNWHMFQITLPLE
KLSISRGEFMEQLKQRGIGSGVHYPAIHLFTLYRARGFAQGMFPHAERFGASTVTLPLFT
QMSEAHVQRVVDAVNQICEQYGK*
>GCA_000452945.1_BPMSHR5858_genomic_02536_1 arnC
MTHPETRATHPEVSIVIPVYNEEAGLAALFARLYPALDALGTPYEVILVNDGSRDRSAAL
LADQFRVRPDTTRVVLLNGNYGQHMAILAGFEQSRGEIVITLDADLQNPPEEIGKLIAKM
REGYDYVGSIRLQRQDSLFRRKASAAMNRLRERITRIKMTDQGCMLRAYSRHIIDTINRC
GEVNTFIPALAYTFAQNPTEIEVAHEERFAGESKYSLYSLIRLNFDLVTGFSVVPLQWLS
FIGVILSLGSAALFVLLVVRRFIVGAEVQGVFTLFAITFFLLGVIIFALGLLGEYIGRIY
QQVRARPRYLIHTVLEARDGKPGVTLAAERREAAR*
>GCA_000452945.1_BPMSHR5858_genomic_02537_1 group_3341
MKPRAVVFAYHNVGVRCLQVLLARGVDVALVVTHEDSPTEHIWFGSVAAVAAEHGIAVIT
PADPAGADVRAALASAKPDFIFSFYYRHMLPVDLLALAARGAYNMHGSLLPKYRGRVPTN
WAVLNGETETGATLHEMAAKPDAGAIVGQTAVPILPDDTAAQVFDKVTVAAEQTLWRVLP
ALLAGEAPHLPNDLSQGSYYGGRKPEDGRIDWTQPAANVYNLIRAVAPPYPGAFTDLGGT
RFVIARARLAAPGTAAARAAVDLPPGLHVSDNALFGVCGDSRAVSILELRRPHDGGEAVV
SPAQFAQFIHSSRHS*
>GCA_000452945.1_BPMSHR5858_genomic_02538_1 arnA_2
MKAKKVLILGVNGFIGHHLSKRILETTDWEVFGMDMQTDRLGDLVKHERMHFFEGDITIN
KEWVEYHVKKCDVILPLVAIATPATYVKQPLRVFELDFEANLPIVRSAVKYGKHLVFPST
SEVYGMCADEQFDPDASALTYGPINKPRWIYACSKQLMDRVIWGYGMEGLNFTLFRPFNW
IGPGLDSIYTPKEGSSRVVTQFLGHIVRGENISLVDGGSQKRAFTYVDDGISALMKIIEN
SNGVATGKIYNIGNPNNNFSVRELANKMLELAAEFPEYADSAKRVKLVETTSGAYYGNGY
QDVQNRVPKIENTMQELGWAPQFTFDDALRQIFEAYRGHVADARALVEQHG*
>GCA_000452945.1_BPMSHR5858_genomic_02539_1 arnD
LARIVLKIDVDTLRGTREGVPNLARIFDRFAARATFLFSLGPDHTGWAMRRVLRPGFLKK
VSRTSVVEHYGVRQLLYGVLLPGPDIGRRARADMRAIHEAGFECGIHTWDHVYWQDNVRR
RPREWTVREMNKSRERFVEIFGAPPVTHGAAGWQMNDAAFEQIDAWGMRYASDGRGHSPY
LPVVAGKTLAHVQMPTTLPTLDEILGVDGIDTGNVAARVLEHTERNPHDQVFTLHAELEG
QKLAPVFEQLLAGWRAQGHTFATMGDYYATLDRAALPSYPVAWGEIPGRSGELIVQPD*
>GCA_000452945.1_BPMSHR5858_genomic_02540_1 bcp_2
VSVEVDRQVPDFTAPATGGEFSLSGIKGKKLVLYFYPKDNTPGCTTEGLQFRDLYPKFKK
AGAEIVGVSRDSLRSHENFKAKLELPFPLISDPDETLCALFGVMKLKKMYGKEVRGIERS
TFLIDGAGMLRHAWRGVKVPGHVDDVLSAVQAL*
>GCA_000452945.1_BPMSHR5858_genomic_02543_1 group_7442
MHSIQLALWVVAALVLVALVFDFMNGFHDAANSIATVVSTGVLKPQQAVVFAAAFNVIAY
FIFHLKVAQTVGKGTIDASIVDHYVVFGALFGAIGWNIITWYYGIPSSSSHALIGGLVGA
AVSKSGWGSLNVDGLMKTVAFIFISPLLGFILGSLFMLGVSWLYFRTAPSKVDRRFRRLQ
LLSASLYSLGHGGNDAQKTIGIIWMLLIASGYASAASDAPPAWVIGACYLSMGLGTLFGG
WRIVRTMGQKITKLKPVGGFCAETGGALTLFIASWMGIPVSTTHTITGAIVGVGATRKLS
AVRWGVAGNIVWAWVLTIPASALIAAAGWWIGHRVF*
>GCA_000452945.1_BPMSHR5858_genomic_02544_1 group_3342
MFGRFMPTEGKFFEIFNAHANYIVSGGRELELLIDNLADAEIHKQNVQSAEKAADKLTHE
AIDLLHKTFITPLDRDEIHKLITTMDDILDLMEDVATAVSLYDVRSVTSEASQLAHIVTQ
SAQHVQAAVALLSDMKQSGQILKACEEIDRWESEADRVLRAAMSKLFREEDDVKTLIKLK
AIYELLEQITDKCEDVANIIEGIVLENA*
>GCA_000452945.1_BPMSHR5858_genomic_02545_1 dnaB
MNAPKDPQIESLKVPPHSIEAEQSVLGGLLLDNGAWDRIADFLSQSDFYRYDHRIIFEHI
GRLIAATRPADVVTVYEALTTSGKAEDVGGLAYLNALAQNTPSAANIRRYAEIVRDRAVL
RRLVSVADEISADAFNPQGKEVRQLLDEAESKVFSIAEDGARGTQGFLEIGPLLTQVVER
IDTLYHTANPSDVTGTPTGFVDLDRMTSGMHGGELIIVAGRPSMGKTAFSMNIGEYVAVE
YGLPVAVFSMEMPGTQLVMRMLGSVGRLDQHRMRTGRLTDEDWPKLTHAVQKMSEAQLFI
DETGGLNPMELRSRARRLARQCGKLGLIIVDYLQLMTGSSQGENRATEISEISRSLKSLA
KELDVPVIALSQLNRGLEQRPNKRPVMSDLRESGAIEQDADVILFIYRDEVYNPDSPDKG
TAEIIIGKQRNGPIGPVRLTFLGQYTKFDNFAGAQNFYGE*
>GCA_000452945.1_BPMSHR5858_genomic_02546_1 rplI
MQIILLEKVANLGNLGDIVKVKDGYARNFLIPNRKARRATKDAIAEFEVRRAELEKVAAE
KLAAAQAVGEKLNGQTFEITQKSGVDGRLFGSVTNGDVAELLKKAGYEIEKAQVRMPEGP
LKMIGEHGVQVALHTDVVVDVTVNVIGDHA*
>GCA_000452945.1_BPMSHR5858_genomic_02547_1 group_7444
MARPTGKKFDKRRQQQNPLFKRKKFCRFTAAGVEQIDYKDTETLKDFIGENGKITPARLT
GTKAHYQRQLDTAIKRARFLALLPYTDQHKA*
>GCA_000452945.1_BPMSHR5858_genomic_02548_1 priB
VNRLQLTASVVEREPVRYTPAGVPIASATLQHRTEVVEAGIARQVELTIPAVAAGEASGR
LEACEMGVETLFTGFLAKKHRNARTLVFHITALQDIGKD*
>GCA_000452945.1_BPMSHR5858_genomic_02549_1 rpsF
MRHYEIVFIVHPDQSEQVPAMIERYKSTITSHGGQIHRVEDWGRRQLAYMIEKLAKAHYV
CMNIECDQTTLDELEHAFKFNDAVLRHLIVKMKKAETGPSPMMKEVQREEAKKAAAAQPT
EAQA*
>GCA_000452945.1_BPMSHR5858_genomic_02552_1 cpdA_2
MPNPIQPLKRRDFLRLAACGGGVAFASALPGWSFAANAGADFFFVQLSDAHWGFTGPAIN
PDARGTLPKAIEAVNALPVAPDFVMFTGDLTHTTDDPAERRARMRQFQSIVAQLRAKPLH
LMPGEHDASLDAGAAYREIFGDTHYAFDHKGVHFVVVDNVSDPAGRVGDAQIEWLARDLA
RQPKDARIVVFTHRPLFDLAPQWDWATRDGAKVVDVLMPYPNVTVFYGHIHQEHHAMTGH
IAHHAARSLMFPLPAPGSQDKRLPVPWDAAAPYRGLGWREVRVGDAARAPALTEMPVAAP
QPQPRA*
>GCA_000452945.1_BPMSHR5858_genomic_02554_1 dmlR_10
MDRYKQIETFVRVADAGSLAAAALEEGVSPVVLGRRIDALEKRLGVKLMYRSTRRLVVSE
DGAAFLERCRGLLTEWAQAENEVVAGRRAVGGHLIVSAPAAFGRKHVAPLAPAFVADKPD
LQMSFNLTDRVVDLVREGYDLSIRIGGAVDPNFVAVKLASNRRVVCGTPEYFRRHGRPRT
LDDLASHNCLAFNLQGGQNRGWYFRRNGKLVTMRVSGTLDCNDGELLHRWVSEGLGLGWR
STWEIAQQLARGELETVLDEYALPDYDILAVYPQQRYVPARVRYFIDYLRDVYARPGYWE
GAE*
>GCA_000452945.1_BPMSHR5858_genomic_02555_1 gcl
MARMRAVDAAVLVLEKEGVQTAFGVPGAAINPLYSALRRSGAIGHVLARHVEGASHMAEG
YTRAAPGNIGVCIGTSGPAGTDMITGLYSASADSIPILAITGQAPRARLHKEDFQAVDIE
SIAKPVTKWAVTVREPALVPRVFQQAFHLMRSGRPGPVLIDLPIDVQLAEIEFDIATYEP
LPVYKPRATRAQIEAALAMLNDAERPLIVSGGGVINAAAEELLVQFAETLGVPVIPTLMS
WGAIADDHPLMAGMVGLQTSHRYGNATLLASDFVLGIGNRWANRHTGSIEVYTKGRKFVH
VDIEPTQIGRVFGPDLGIVSDAKAALELFVEIAREWKAAGKLKNRGAWVADCQQRKRTLQ
RKTHFDDVPIKPQRVYEEMNKAFGRDTCYVSTIGLSQIAAAQFLHVFKARNWINCGQAGP

LGWTIPAALGVRAADPQRPIVALSGDYDFQFMIEELAAGAQFKLPYVHVVVNNAYLGLIR
QAQRAFDMDYCVQLAFDNVNAPEMNGYGVDHVAVAEGLGCKALRVFKPEEIAPALERARQ
MTAEFSVPVVVEVILERVTNISMGAEIDAINEFEDLADKAEHAPTAISLLD*
>GCA_000452945.1_BPMSHR5858_genomic_02556_1 hyi
MPKFAANLTMLFNEVPFLDRFAAAAEAGFSAVEFLFPYPYAKDELAERLDANRLRLVLHN
LPAGHWDQGERGIACLPDRIGEFQDGVGRAIDYARALKVPQLNCLVGIAPASVGRDTALA
TIVENLRFAAAELKKAGLRLLVEPCNAYDIPGFALNRSAEALDVIRAVGADNLFLQYDIY
HMQRMEGELAATIRRHIGAIAHVQIADNPGRHEPGTGEIHYAFLFDLLDELGYDGYVGCE
YKPRTTTAEGLGWLRRIARVAA*
>GCA_000452945.1_BPMSHR5858_genomic_02557_1 glxR_3
MATIGFIGLGIMGAHMARNLLKGGHRLVVNGAYPVPADLREQTQVVANSTAVAQAADIVI
AMVPDTPDVRNVLFAEDGVAKGLSAGKLVIDMSSISPLDTREFAKEINALGCDYLDAPVS
GGEVGARDATLTIMVGGPQRAFERAQPLFALMGRNVSLIGENGAGQTCKVANQIIVALNI
EAVAEALLFAARSGADPERVRRALMGGFASSRILELHGERMTKRAFEPGFRIDLHRKDLN
LALDGARRMNIALPHTASAQQLFSVCAANGGGAWDHSAIVRALEIMSNCEIGEPQSDKKV
A*
>GCA_000452945.1_BPMSHR5858_genomic_02559_1 kbp
MGLLSFIKEAGEKLLGKSDEQAATDPAAANQTAADAIKNYISAQGLDTSNLTVAFDGASR
TVTLTGSVADLDTKAKVKVAAGNVQGVAGVNDDDLQPDDPQVQYHDVVSGDNLWKIAEKY
YGDGSKNDAIFQANRPMLSSPDRIYPGQKLVIPPQS*
>GCA_000452945.1_BPMSHR5858_genomic_02566_1 group_3345
MLTIDDLWRIADAPLHSWLGTLVVSALVLLAVAGIHWIGARIVKPIARPYPFMSVIVSYI
DRPSFALLSLLALEFLWLQVDESLSHVGGLRTLAAVGTIAALTWLLMRLAAAVGDAIIRA
HPIDTADNLEARRIHTQTRVLARTVMVLIVIIGTGAALMTFPNVRQIGASLLASAGVAGL
VAGIAARPVLGNLIAGLQIALSQPIRLDDVVIIQGEWGRIEEISGTYVSVRLWDQRRLIV
PLQWFIENPFQNWTRSSAELIGTVFLYVDYRLPLEPLRAELARIVADAPQWDGRVQVLQV
TDATERSMQLRALVSARDSSLAWDLRCRVREGLIAFINAHYPHCLPRERTEWTTPDALAR
EPLRGERAPEPAASNAAFTAADPTTSEAGVPRAPH*
>GCA_000452945.1_BPMSHR5858_genomic_02567_1 group_4759
MLRAMFCAAAFAVPLTAAAFTAGDLDKLCAKTDVKSRASCAAYIEGAADGVYNTIDAIGG
TTGPRVGQYFCLPPDIKAQQMTDAVRKYIAENPKLADYNASTAVSLGLGKAFPCRSY*
>GCA_000452945.1_BPMSHR5858_genomic_02568_1 group_5586
MKSIKHLSKQRTMGARLRALALALAGGLGALAAAPAHAESLDGIVKAVKFDDIKDISKQL
KNGLDPNTIAPNGDPLLVIAAREKSDQVAAALAAAPNVELDKEDKAGENALMLAALNGDL
PLVKLLVDKGAEVSKKGWSPLHYAATNGNDEVVKYLVDKSAYIDAASPNGTTPLMMAARG
NHGSTVTLLLDQGADPALKNQLGITALDFAKHYNAPDATAILSKRTVRIGDAPAPNAQKS
AK*
>GCA_000452945.1_BPMSHR5858_genomic_02570_1 ywnH
MTQLAYRDATPADLPAIVAIYNSTVASRRVTADTEPVTVDSRRAWFDAHNPHARPLWVVE
EAGRVIAWLSFSDFYGRPAYGHTAEISIYLDEAARGSGLGSRLLEAALAKAPALGVHTAL
GFIFGHNEPSLRLFARYGFTAWGTLPRVAVLDGIERDLVIVGKRLADA*
>GCA_000452945.1_BPMSHR5858_genomic_02575_1 group_3347
MCLIVFDWQPDTTHGPVLTLAANRDEFFRRTSAPLAWWSDAPHVLAGRDLEGGGTWLGVA
RDGRFAALTNYRAPFDIRAGAPTRGKLVSGFLTGENVAPLDYLANVAEKAVFYNGFTLLA
GDVARGELAWYCNRPADAQPAPDAPVSVAPGVHGLSNARLDTPWPKLVGKRSALGALLTG
DAAAPLDALIEMMRDTREAADDALPHTGIPLERERALSAAFIETPEYGSRGTTVLRVARD
AEGRLRFDMKERCDDDGSHRIVRPGAFERAFTFDVDTGGHPRR*
>GCA_000452945.1_BPMSHR5858_genomic_02576_1 group_3348
MESIEIETGDWSRLGGDASRIRDAVFVREQRIPAELDLDDDDPHARHAVAYLVDGTAGAR
RAVATGRLLPTGAIGRVSVLADARGRGVGSRLLDALLAEARGRGDALVRLYAQQRAVAFY
LRIGFRIVGEPFVAAGVQHIEMAREP*
>GCA_000452945.1_BPMSHR5858_genomic_02580_1 group_4760
MIRHIVSWKLKESAEGATRAQNAQKLKEKLEACRGIVPGLLHLEVGLATPGLEATCDVVL
VSDFADKAALDAYQVHPVHEEVKKFVGAVRESRECIDYLVDDAR*
>GCA_000452945.1_BPMSHR5858_genomic_02581_1 group_3349
VFPKESTIRALIERWNRHYSTVLGIKSATERSERIAHDLYLVRNAGFGGVSPPPNLPGNL
VDKDDEIMACVEHYFLTRDWVANGKYPAWEARTLSGIYHLGKRIGVAPRHNKAKPVTPAS
PLQRALQLEGIKDGTIDRKLAGIQSPLVRKPPKY*
>GCA_000452945.1_BPMSHR5858_genomic_02605_1 group_3352
MPALFRRRPLAAIRAAAALSLAATLAPAHADTTGLIEPAHLSVDGSLPAAQRDAQILAAR
RYDTFWHNGDPALARAALADDFADRTPPPGRAPGLAGPLAASRTMRAAIPDLSCEIEQMI
VAGERVVVHLRFRGHFTGRFGPVQGKGQAIDFIATDIYRIERGRIAENWHIEDNLTLLRE
LGIAA*
>GCA_000452945.1_BPMSHR5858_genomic_02609_1 rutC_1
MAVTRHHVGPRLSEIAIHNGTVYLAGQIAEDTAQDIKGQTREVLAHVDRLLAEANSDKAH
LLSVQIFISDMANFPGMNEVWDAWVAQGATPPRATVEAKLANPACLVEVVVVAAQRG*
>GCA_000452945.1_BPMSHR5858_genomic_02620_1 group_19451
MWTPVQRVQRRPSRRAPGAWRAVHTAVSASRWFLRSCAGTAVTLAVLSAGCAESTPKSGN
GGAASTAIERTKADRLAREQQTARTVPTLASIKIDQPRPLTLRDSGQSGTLAFLREVDFR
IVGDLGFLVHELSATLVPTHPGAPTVFDDPTSFDIAVHRGTVTLDNTKLNALFGGYIFGY
RNAPLRNVRVSAGDGFLDIRGEMQRDGWVPFALKGKLEIRDGSTLVFHPTDVHVSGIEAG
PVMRAAKVQVSDLLKIDTPIVRLNGNDLVLNVDKLLPPPHLKINIVSLKLTSAGLDLVLD
DGTKAGFTMPENAPKHAMYLRGGDVKFMRTMPMNADIVIYPPRESSPDDAFVFDMYHYRD
QLVAGYFNFEPSGALSILMPSYRRLARPSAPSLGSAAARMNDSLIDAQQLSLGEARRQWE
AFAITPNGGAAQPNFRKVAAKHGGDVSMFGPRHISNGTTTIHLHNADFYIAGNIGFRVED
LVVQLVSKRAGEPVDLDDPNQYDIRILSGSVLAPWDAMSDLFNRHLLDYSPRSLNDLKLS
ADGGALRVRGGIKLWNQVPPGVWLPADMKGSLTLLDERHLAFTPTQVSVLGIPQAKLLRA
LGIELSSLAPLKRRGAELRGDSLVLDQYTVFPPPVLIGHMSQATVEPDGLRLTFRPAPNA
PVLRPPANLPGSYLWLEGGDTKMFNVLVLNMRILVEDTAKPRVRFDLYDYRDVVSRGSVR
MVHDGTLYVDVGKKDPLAR*
>GCA_000452945.1_BPMSHR5858_genomic_02624_1 group_3353
MNNKKSSPPTRRRPTRALVAAAAVTFAPVTAQSQLGSTVASPSARALVQASSAALPNGAV
TMRSYVDAGGTTIHEYATRTGLVFAYTWDGPTSPNLRKLLGERFPTFQSHAVTTDAHARA
SLHAGRVAQSDFVVEAGGRMRAYVGRAWLPGAVPPGVSIDDLR*
>GCA_000452945.1_BPMSHR5858_genomic_02634_1 scpB
MNTQEAKIVLETALICAQEPLKLGDLRKLFADGVSADTVRTLLEDLKQEWTGRGVELVAL
ASGWRFQSKPAMRTYLDRLHPEKPPKYSRAVLETLAIIAYRQPVTRGDIEEIRGVTVNTQ
VVKQLEDRGWIEVIGHRDVPGRPALYATTKQFLDDLGLKALDDLPALEEPAAHLEASLLA
QQAIDFPDDARGDGEAPAVAPETDAAAAQAGEVAEPASDSVVLRNEDTARADEPRQADAA
PTRADEPRQEGATDGLAARIGDPADNDAGRADVDVDAAPALGEADAPRGGDASGHAAREI
DSREMTEPADAGGEAAGTTASHTHAVAPYDHDAQSAADAVAAPHAGDVAADGERAESPSG
TNENANYVAQPDAEPADAAHAATREVAADPAAGRATKEDHGAIGGIPHDAEPVRAHAEEA
LDDTSDSLADAVRSASEAAPEHERQDDEDAPAKRRA*
>GCA_000452945.1_BPMSHR5858_genomic_02643_1 oprM_2
MKTFPLSACRTAVAVAVAALALAGCANYFGLKSDKAIAPAAQFESARSLPAQGGQWPSLD
WANQFGDPQLPKLIDEALEGNPTIAQAQARIAKASSYIESSRSTLLPKADAKYSWTRELY
SGNALFPPPYGGQWYSENNALASASWELDLWGKNRARLNAAVSQEKAAEADLQQARVTLA
TSVARTYNQLAQLYALRDIAGREISNRQTVGKITDGRVGAGLDTNVERQTARGNIATSQS
TLSDLDGQITNVRYQLAALLGKGPDRGLQIAAPVLSAGGAVALPENLPADLVSRRPDLVA
ARWQVEAAMHDVKEAKAEFFPDINLAAGFGFDAFGWGRFLNFASRQAQFGPAIHLPIFDG
GALRAQLKGRYADFDLSVANYNQTLINALNDVATQVASIRSVDTQMGDAQRALDASTRAY
ELAVIRYKAGLSPQLQVLNADSNRLAAEQTVTNLKMRRRDLQIGLVKALGGGFDAAGTRL
AAPAPASAPAAPAQHASN*
>GCA_000452945.1_BPMSHR5858_genomic_02654_1 group_312
MNINTISHGTLRTTLIAATVAAMLSLSACGGSGTISKGLDGSGSGSGGGNAISTTGGSGS
GGTSGAGGSGSGGSGSSGSTGGLSGGGGSTSGGGSTSGGGSTSGGTSTSSSINALGTIAG
NTGGIISGAGSTVSGLGTVVGSQTLPGVNPQTTQALGGVVQSLGGAVSALGSGVTSGIGQ
LGSSTNPIGTTVASTGGVVSQLGGAVTQTGNLVTSLGTGPLAPLAPLTSPLGGAVSTLGN
TITAGGTTLTTALSTGPVQQLTQTVSSAITPITSMVTGTTQTVGNVTGLGAPVNTLLGTI
GGGLNQAGALISKTGNNPVTTGLGQTVSATGNTITSVGGLLTGGSGATNPLAPITAAVGG
LTGTLSGVSGATSGTPLAPLTNVVSTVTGALSGATGGATSTSPLAPITGLVSTVTGALSG
ATGGTTGSTSPLAPVTGLVSTVTGALSGATGGTTSTSPLAPVTNLVSTATGALGGATAAP
ATSTTTSTTTSATTSATNPVASLTAPLTGTSSGTSGATNLLAPVTSLVGGLLGGIKK*
>GCA_000452945.1_BPMSHR5858_genomic_02663_1 group_371
MDPSRLGAIALVLGAIGVLMLAALAIMQAVLSRRTGRTLADALDQRAAALEAAAARVAAG
AAGAARAGMPEAAPDARRPRFAALLDRAGRFGMRLLDTRLGKQIVADEDRMLLEQCGYVD
AHTRGIFLSARIACAIALPAAVALVGGETVRTHLGAWVALSVIAGFMLPKTYVRRRAAAR
RQSVVDEMPLLVDMLRLLQGVGLSLDQSIQVVTNDFRGMLPVLSSELGIAQRQFVAGRTR
EQSLQRLATSFDNEDLRAIVRLLIQVDKHGGAVQEPLKQFGDRLREVRRAMLRERIGRLT
VKMTGVMILTLLPALFIVTAGPGMLAVTHALTAARR*
>GCA_000452945.1_BPMSHR5858_genomic_02693_1 aat
MVPWLGPDDPFPSVERALGAASGAPGLLAASADLLPSRLIDAYRRGIFPWYSDGQPVLWW
SPDPRMILVPAEFRISATFRKTLKRVLREPRWEIRVDCDFAGVMRACAQAPRRGQRGTWI
TAEIIDAYSSLHRAGDAHSIETWLDGRRVGGLYGVSFGRMFFGESMYAHASDASKIALAA
LVAHLREHRVEMIDCQQNTSHLASLGGREIARKTFVAHVRRAVAEPPIPWRFDKRVVAGL
LGQAASATAADAFDR*
>GCA_000452945.1_BPMSHR5858_genomic_02696_1 group_7452
MDIALDLQSIAVQEKTLVFPQFDAARAWALGSQLREFALARGHAVAIDVRTFGQPLFFAL
VDGATPDNVDWARRKGNVVAHFRRSSYAIGLRLQQAGATLADKHGLPAAEYASHGGAFPI
AVAGAGVIGSVTVSGLPQRGDHELVVEALCAQLGHAYATLALTGN*
>GCA_000452945.1_BPMSHR5858_genomic_02697_1 emrE
MQLPGYAWLAIAIVAEVVGTSALRAAEGFTRLWPTLVVALGYGTAFYCLSLTLKSMPVGI
VYAIWSGAGIVLITLVALVLYRQVPDWPAVVGLALIVAGVVVLNLFSKMQAH*
>GCA_000452945.1_BPMSHR5858_genomic_02698_1 menB_1
MNAFHSDSPSADVRAHVANRIGFLELNRPQALNALSTDMIRALHRALDAWRDDADVLAVV
IHSPHPRAFCAGGDVRFFHDAHRRGDQASIDTFFIEEYALNHAIFTYPKPYIALMHGVVM
GGGMGISQAARRTGGLRVVTDSTKMAMPETRIGLFPDVGMSWFLARTPGALGRYLAVTGA
TLDAASALYAGLADAYLPDAALPALLDALRTERLDDGAAVTARVAAEAARHAPTPAPDGS
ALACARASIDRHFAQPDVAAIAASLDAERDDAVRAAWADKARDAMRGGALSPLSMAVSLE
VVERARGATMADCLRRDLDLTRSTFARGDVIEGVRAVIVDKDHAPKWRYPAIADVTAQAV
AAMFESPWRAHEHPLRKLRD*
>GCA_000452945.1_BPMSHR5858_genomic_02699_1 group_4769
MAPRLHVALSMQKVILPFLSGFLAALFFREATLALLHTAGLIDATGFSTTPFAPLGIPEF
VANAIMSACWAILMAWLLRVSPERAAPWVPSLVFGGIVLTAARVFAIDPLRGIWPAGNML
PPLVVGFVANAVWGWGALVFMRAFMSDDGGSDDA*
>GCA_000452945.1_BPMSHR5858_genomic_02700_1 msuD
MNVFWFIPTHGDSRYLGTAEGARAADYDYFRQVAVAADTLGYDGVLLPTGRSCEDAWVVA
SSLIPATKRLKFLVAIRPGLSSPGLSARMASTFDRLSDGRLLINVVTGGDSAELEGDGLF
ADHDTRYAITDDFLHIWRGLLAESHENGGIDFDGEHLSAKGGKLLYPPVQRPHPPLWFGG
SSPAAHAIAADHIDTYLSWGEPPAAVEKKIADIRARAAARGREIKFGIRLHVIVRETQEE
AWRDADRLISRLDDDTIARAQQAFAKMDSEGQRRMAALHGGKRGSRQELEIYPNLWAGVG
LVRGGAGTALVGNPEQIAARMREYAALGIETFILSGYPHLEESYRFAELVFPLVKGGGNT
RRAGPLSGPFGEVVGNQYLPKASQS*
>GCA_000452945.1_BPMSHR5858_genomic_02714_1 rpoC
MKALLDLFKQVQQEEIFDAIKIGLASPDKIRSWSFGEVKKPETINYRTFKPERDGLFCAK
IFGPIKDYECLCGKYKRLKHRGVICEKCGVEVTLAKVRRERMGHIELASPVAHIWFLKSL
PSRLGMVLDMTLRDIERVLYFEAYVVIDPGMTPLKARQIMTEEDYYNKVEEYGDEFRAEM
GAEGVRELLRSINIDEQVETLRTELKNTGSEAKIKKYAKRLKVLEAFQRSGIKPDWMILE
VLPVLPPELRPLVPLDGGRFATSDLNDLYRRVINRNNRLKRLLELKAPEIIVRNEKRMLQ
EAVDSLLDNGRRGKAMTGANKRPLKSLADMIKGKGGRFRQNLLGKRVDYSGRSVIVVGPT
LKLHQCGLPKLMALELFKPFIFNKLEVMGVATTIKAAKKEVENQTPVVWDILEEVIREHP
VMLNRAPTLHRLGIQAFEPVLIEGKAIQLHPLVCAAFNADFDGDQMAVHVPLSLEAQMEA
RTLMLASNNVLFPANGDPSIVPSQDIVLGLYYATREAINGKGEGLSFTGVSEVIRAYENK
EVELASRVNVRITEMVRNEDTSEGAPQFVPKISLYATTVGRAILSEILPPGLPFSVLNKP
LKKKEISRLINTAFRKCGLRATVVFADQLMQSGFRLATRAGISICVDDMLVPTQKETIVG
DAAKKVKEYDRQYMSGLVTAQERYNNVVDIWSATSEAVGKAMMEQLSTEPVIDRDGNETR
QESFNSIYMMADSGARGSAVQIRQLAGMRGLMAKPDGSIIETPITANFREGLNVLQYFIS
THGARKGLADTALKTANSGYLTRRLVDVTQDLVVVEDDCGTSNGVAMKALVEGGEVVEAL
RDRILGRVAASDVVNPETQETLYEAGALLDETAVEDIERLGIDEVRVRTALTCETRYGLC
ASCYGRDLGRGSLVNVGEAVGVIAAQSIGEPGTQLTMRTFHIGGAASRAAVASSVEAKSN
GTVRFTASMRYVTNAKGEQIVISRSGEALITDDIGRERERHKIPYGATLLQLDGAAIKAG
TQLATWDPLTRPIITEYGGTVKFENVEEGVTVAKQIDDVTGLSTLVVIDVKRRGSQAAKS
VRPQVKLLDANGDEVKIPGTEHAVQIGFQVGALITVKDGQQVQVGEVLARIPTESQKTRD
ITGGLPRVAELFEARSPKDAGILAEVTGTVSFGKDTKGKQRLVITDLEGNQHEFLIAKEK
QVLVHDGQVVNKGEMIVDGPADPHDILRLQGIEALSRYIVDEVQDVYRLQGVKINDKHIE
VIVRQMLRRVQIVDNGDTRFIPGEQVERSDMLDENDRMIAEDKRPATYDNILLGITKASL
STDSFISAASFQETTRVLTEAAIMGKRDDLRGLKENVIVGRLIPAGTGLAFHKARKAKEQ
SDRERFDQIAAEEAFEFGTPSAPAEEPQHPAE*
>GCA_000452945.1_BPMSHR5858_genomic_02731_1 kdgR_1
MSYVVDAVDNALKLLNCVAEHPNLGVTRLASKLGINKSRTYRMLCTLELHHFVTRDARTS
TYVLGTQAFVIGIAAALQSALVRAAEHPMLALGQSVNETVMLRVREGFDTVCVARHATVR
DARMAGSVCNRRPLGVGAAGKVLLAFALDAVRDEYLARREKTAPGGSAVLADELDTIARR
GYAIGIGEAAASTVEIAVPVRDGRGATIASLSVAGPNLRMPHADTARYLEKLQACSRAIS
AEFEYM*
>GCA_000452945.1_BPMSHR5858_genomic_02747_1 group_3366
MKRSRSRFRASFAVAVALGVPSGFAHAQSLPGIPDPAAVGLTPGTGQNGAPTVSACFYNY
RTKVIAGFPTQIASGWIVTNQRNDPACIPPGGLPGPNWQEMTYFRDMPVGASMSACWSFS
WPSNWQPVGYTSDPSKCGYYPGGQIPGAQPNVVFLKRVQ*
>GCA_000452945.1_BPMSHR5858_genomic_02829_1 pgl_2
MTRRAHAPSPFAFPLRLSHWMKGFALALPLFGNPAFAQDASAGPAGGVYNLLVGTYTGSG
SDGIYVYRFDTDSGRVSPVSSAKAENPSYLVASRDGRHVYAVNELPGDAGPASVRGGVSA
FDFDAKTGALKFVNRVSAQGNDPCYLSLSPDGRYLVVANYSVASDPGGSFSVFPVEATGA
LGAAVLNVHHEGTGPVKGRQDGAHVHSTVFSPDGKYLFVQDLGADKLYSYRYTPDGSRGL
IGPTESRYTLAKAGSGPRHLVFGANGRFAYVTNELNASVDVYRYDDGRLVHVETVPMTAP
GFAGKVGGGALHLSPDGRFLYATNRGDANDIVIYAVNAADGKLTLVGRQSSLGKTPREFA
IDPSGKWLIVGNQDSDSVFVFRRDIASGRLEPNPARIRIDKPVDFKFVPVQ*
>GCA_000452945.1_BPMSHR5858_genomic_02830_1 group_7460
MPALIEDYALVGDGHTAALISRDGSVDWLCWPRFDSGACFAALVGTPEHGRWLVAPADDV
RVSATSRRYRGDTLILETDYECDEGAVTVIDFMPPGSGWSELVRIVVGRRGEMRMQMQLV
LRFDYGFSVPWVTRLPREAGVKAIAGPDTVVLRTPVPLSGENLHTVAEFTVKAEERVPFS
LSYAPSHLRLPPAREPLSMLARTENYWLEWSARCPLRGRYASAVRRSLITLKALAYEPTG
GIVAAPTTSLPEQLGGTRNWDYRYCWLRDATITLLALMRGGYYDEARAWRTWLGRVMAGS
PEQIQIMYGIAGERRLPEMELDWLPGYQDSRPVRIGNNAANQLQLDVFGEVMSALHIARV
GGLQADDTVWSVQTTLLAHLEKIWREPDEGIWETRGGRRHFTFSKVMAWVAFDRAIKSAE
MFRLSAPLERWRAIRDEIHADVCANGWNPRVRAFTQSYGSDALDASVLMLPLVGFLPPDD
PRIADTVEAIERGLMRDGLVLRYHTTEYDDGLPPGEGTFLACSFWLVDNLALLGRIDDAH
ALFRRLLSLTNDLGLLAEEYDPVTRRQVGNFPQAFSHVGLVHTAFNLMHHEEEMLRAAGQ
PAAADAVAGR*
>GCA_000452945.1_BPMSHR5858_genomic_02831_1 group_7461
MLYQFHEFQRAMLSPLTAWAQAASKSFANPSSPLSLVPGATRLAAGYELLYRLGKDYEKP
EFGIHQIVKDGHNIPIVEQTIIEKPFCRLLRFKRYADDSGAVGQLKDEPVVLVCAPLSGH
HATLLRDTVRTLLQDHKVYITDWIDARMVPAEVGSFHLDDYVGYIQKFIRHIGARNLHVI
SVCQPTVPVLAAISLMASRGEDTPLTMTMMGGPIDARKSPTSVNSLATNRSHSWFENNVI
HTVPANYPGEGRKVYPGFLQHTGFVAMNPERHAQSHWDFYQSLLRGDEEDAEAHRRFYDE
YNAVLDMAAEYYLDTIRIVFQEFRLAEGTWDIHGERVKPADIHSTALMTIEGELDDISGS
GQTHVAHELCTGIEQAHRRSLTAEKCGHYGIFSGRRWRTIIYPQLRDFIREHASQPKKPT
PQAGPTPNSPSTPSRSASGSETASAAPAKAAALKLSAKRTAAKTRAAKPPAAKRGAIKLA
APRTRKAA*
>GCA_000452945.1_BPMSHR5858_genomic_02832_1 nicS_1
MNQPKIKRDPEGTRRRILLAAAEEFATGGLFGARVDQIARRAETNERMLYYYFGSKEQLF
TAVLEYAFSALMEAERAIDLEGVAPVEAITRLAHFVWDYYRDHPDLLRLLNNENLHEARY
LQKSTRIREMISPIVKTLDGVLERGQKAGLFRTDIDSLRFYVTLSGLGYYMVSNRFTLAA
IFGRDFSAQHERAEMVKMNTELLLAFLLRR*
>GCA_000452945.1_BPMSHR5858_genomic_02839_1 group_3368
MRFEGSSHYVATDDLKLAVNAALTLERPLLIKGEPGTGKTMLAEEVAAALGMPLLQWHIK
STTKAQQGLYEYDAVSRLRDSQLGDERVKDIANYIVKGVLWQAFEAQRPSVLLIDEIDKA
DIEFPNDLLRELDRMEFHVYETRETVRARHRPLVIITSNNEKELPDAFLRRCFFHYIRFP
DPSTMQKIVEVHYPGIKQDLLRAALESFFELRGVSGLKKKPSTSELLDWLKLLLAEDIPP

DALRSKDQKQIVPPLAGALLKNEQDVSLFERLVYMNRHNR*
>GCA_000452945.1_BPMSHR5858_genomic_02861_1 cynR_5
MRAAAVRFLNDRLDWNLLRTFLVIMQERSVSRAAARLHVTQPAVSQALKRLEETLGHTLI
RRRGPHFEPTQAGAEVYRIASDIYGTISRLDTELDDAAADLTGTIRLLSVSRIESGVYDF
FLADFHRQYPRVDLQIEVMRSSDIISSLLQKTATAGLSLCRNPVDKLARASFLRQRYAIY
CGRHHRLFGQTQLTMNDLLAENFVSFTSDQIGDSLSPLTVFRDQKGFTGRIVAASPSLDE
VRRLIFAGYGIGCLPEHIVRDDLARQRLWRLPPDEGLIDVDIHLLWHRERKMNAAELAFL
EGFERAMQRYSFDERLAINP*
>GCA_000452945.1_BPMSHR5858_genomic_02862_1 group_5713
MHMQMTERVPAKTLTERFVHALTFEVTAIAICAPIVSWVLGLSLVHVGALTAAVSVIAMV
WNVTFNALFERIERRYRLTRTLAVRAAHAIAFELGLVAMALPLAAWWLEISLLEALLLDF
GILLFFLPYTFLFNLGYDRLRARWIAQRAMA*
>GCA_000452945.1_BPMSHR5858_genomic_02895_1 group_7466
MAGNLVIVCRDQDADAFDQLMQEYGSFQTRLSSTAWYLNMNIVPETLQEDILERVGKYTT
LYIFEATSVTYNTIDSNAAETLSTLFGE*
>GCA_000452945.1_BPMSHR5858_genomic_02898_1 group_3719
MYKKTSRLAIAAAVLALSFSAASRAAPPDMPPPGGLGDHHHMEGGPFMMMQRIHDKLNLS
AAQEQQWQAAVNTMKQNRDAMRKSHEALHQQFQAQQNQPILDLNALHAARQQAEQQNAQL
REQTASAWLAFYNGLNDQQKTTVSTALKQQFAKMDERREKMKERWEQRRAAKGASAPAPQ
Q*
>GCA_000452945.1_BPMSHR5858_genomic_02901_1 hisQ_2
MFLQGYGPLILSGTWQTVKLAVLSLALSFLLGLVGAAAKLSRNRVSNGIGTLYTTLIRGV
PDLVLMLLLFYSLQIWLNQLTDLMNWDQIDIDPFAAGVLVLGFIYGAYFTETFRGAFLSV
PRGQLEAGSAYGMTNWQVFARVMFPQMMRFALPGIGNNWQVLVKSTALVSIIGLADVVKA
SQDAGKGTLRFFFFTLLAGAIYLAITTISNFVLMWLEKRYSTGVRKADL*
>GCA_000452945.1_BPMSHR5858_genomic_02924_1 group_3381
MVKTFYITAAPVGAVPKFLDPLEPKFIPHALLELLPADAREATTQALEANGWEAVPAGGI
VREYGYDAPIDLTDYDGAQASASVQDALRNTGWTPCGTVWHRTQTSPSLAQPPLITRTTL
ERLSSVDLVRQIVLQLTTFGWTATEDGSLTWTHERIHSYLSPDFVERMRADKAAVLESLF
DNGWRVCGAGYWQPGKARSPYLPITADGIVDASREALREGAAVVHLHTRATDDQATLAIP
GLNTPIGIGSQRNHIVLDDYDRIVPTMLDLEPSAILNLSTSARGDRRASQSPLRRAHLKR
YGHAQLAPDVASFSPGPVVFQAGGGYDNPNAFLADQLAHFAEVGVRPEIEVFNHTIVENS
VTLYQSPLVKAGVPVLFMLVAAVDQYHRDPVSGDTSDDSLIDVPTRKAIAKLLQAGTDDA
HEKAVELAATQLRPTVDKLRDNFPSCKISLLLPGPFQALLVDVAIALDLDGIRVGLEDAL
NVFDARVPGGVRKACGTGDQVRWLRLELERRGIGIVDAEALRDELGMSRPDVALFRQAEA
ALAHYPADERLVSADTILDALRPIVDTYRKVEDRLATHLASAEALPADPAALAEHVLTAA
RSFGVTIRSFVEELDRYEDHEYLVARYIQVPQALNFARELLVPRGYSIDAYDRALEDYAR
PGKTVTREHASYSVRVDQFKPLPLRCLEYLVGIPCRYNGDYSNVVNLGLRQSPRYSATMA
LLYHALRELTLELRERSNASRKTCGPVWTVLETSANASEPPVRRDIAPDALTAAIDGVDW
VVLPSTPTTNYPLGLKLANGMAQLFHGFVAQIAADPTLRPSRQTHRDTPLRLLAITHSGR
RDDGETVIEASMLHNRFALNADPSGIYFSEESQLIYERLILPRLVDKPAKLAYNERQLVR
RDTAGFPLYQDGSRARRIKAEQIERLPFLKCFAHSSGIATAQQLDVQACRDGERLGLTAD
ELRAFFDRALLVSFGSAADIHLDWLGTSVVDVTAFNDVRSLAGTTSRHYLIQPGEHADVL
QHCLVHTQPADYRYDHATPVWQEGRQGKVVARLTGVFLLDDHARLDDGHSIRRYLAASPL
WLRQWIARFHDAPADAGAHAILRELQASMTDYRSSANQTTRRALA*
>GCA_000452945.1_BPMSHR5858_genomic_02935_1 cobP
VLVDCLTLWLTNLLCPADGPGVDDARCDAYVAELEAALRASRARVIVVSNEIGFGVVPLG
RQTRRFVDTLGRVNQRIAALATRVTVCVAGLPVTVKDEGEPRC*
>GCA_000452945.1_BPMSHR5858_genomic_02939_1 group_2445
MTDFESKLVLNIASALCSFVLIESKHYLIKWRSNGAQKRAQYSVWKPNIKPQKSNGHAWL
RSRWRKASRQSASEVRANQAKVFFWNSGQEPIVAADLSGKKPLTLQLAPAPITGVRIFES
VNAIGVKLASDHPFNRDKSRKIVTFDRIDPGHGFVVSLEFEAGHDPTPRMLGPILGASLD
YAGPIWAVDLACDGNVRRQGARARAARWISGIACAVGIGGQFVPVHGQAPLVTHSAPYWI
FFAVFAVSWTAFLMSSDICKQFSKRMPPSLHYWNEDNA*
>GCA_000452945.1_BPMSHR5858_genomic_02942_1 cobT
MTTSTFSLPSVDPLDDAALRATLARIIDHKTKPPGSLGRLEALALQLGMIQRTSAPSVAR
PAMIVFAGDHGIAAEGVSPYPQAVTAQMVANFVAGGAAINAFSRVAGLALEIVDAGVASP
LPASPQLVDAAIARGTRNFAQERAMSRDEALAALAAGAARVRHHAALGTNVIGFGEMGIA
NTSAAACMMSRLLDVPIDACVGRGTGLDDAGLAHKRAVLGRALERHAHAREPLDVLAAFG
GFEIAMIAGAFIEAAHARMAILVDGFIATSALLVADALAPVVREYCVFSHVSNEAGHRRM
LEHLGGEPLLALDLRLGEGTGAALALPLVRAAAAFVNEMASFESAGVANRDA*
>GCA_000452945.1_BPMSHR5858_genomic_02945_1 btuB_5
MRTPFVRAALAALSGLPCIALAQGDASASSASFASSVSYAPAAASPADADSALSTAPAAA
AASPASGAARGAEAVSADAASAVASGASSASPARAASAAQLAPVVVTARRGPQRLADAIP
QTTLFDAGDIADSGATDLPGLLALAPGAQIVRNGGPGASASLFLRGAQPTQSLVLIDGVR
VDSASLGQSQIAQLPLDQIERVEVVNGNVSSLYGSGAIGGVVQVFTKNGGNHPPRFDFSA
GYGSYHTQTQSAGASGRLDPDGRTTFSVSLARLKTDGFSALDPARKPQANPNANGYLNES
ASASLRHTFSDKWNAGVTYFQANGRNSYDDAYGSPADLNDLYSRVQQLSAFANGALADGW
TTRVSVSSGKDRNQSALNGAYTDHFNTDNRQYTWQNDFAIARGHKIQAGYERLDQSFESN
KYSAPERHVNSGWLGYTGRIGDSRFQANVRRDQYSDFGGANSYYLGYGFDVTEHWTVSAS
YSSAFRAPSFNDLYYPNSGNLSIRPERSHSVEAAVQYASDDVGVVRVTAFQTRYTNLIDY
RPGASGPYYVAQNVGRAKVQGVEGSWQGHVGGTDVRIAATLQNPVDETAGRDLNRRARRF
ASISANRSFGGWRVGGEWLASGARDDFGDRLGGYGIVNLSARYDITKSWYVSARIDNLFD
KDYELAYAYNTPRRGAYVTLGWQQR*
>GCA_000452945.1_BPMSHR5858_genomic_02975_1 cydA_2
METAFSALDLARVQFAFTVSFHIVFPALSIGLASFIAVLEYRWLKTGKAYYKSLCLFWSK
IFAVAFGMGVVSGVVMAYQFGTNWSGFSSFAGSVTGPLLMYEVMTAFFLEAGFLGIMLFG
WNRVSPRAHFGATLMVAIGTLISTFWILASNSWMQTPQGFEIVDGQVVPTDWFAIIFNPS
FPYRLAHMAIAAFVVAALVVAATAAWHLLKGRRDKAVKKMFSMSLWLLLVLAPLQAVIGD
QHGINTLRHQPAKLAAIEGLWETEPGDTALNLFGIPDMKAQTTRYAVSIPHLGSLILTHS
WDGEIRGLKSFPPQDRPNSTVVFWSFRIMVGLGLAMIGLAALAWLLRRRGRLYEAKWFHR
FALAMGPTGFVSLLAGWVTTEVGRQPWVVYGVMRTSHAVSPLSTQQVSVSLMTFVVVYFL
VFGTGVYYMLKLMRKGPALPGDTPDDAHDARPERTARRPLSAANQLIDAA*
>GCA_000452945.1_BPMSHR5858_genomic_02986_1 oppF
MSDAPMIDIRDVSIRFPTRGGHVDAVRGASLAVGAGEAFGLVGESGSGKSTLLRALTGLV
PLAGGTLSIDSRAVEGKLDRAFRRDVQMVFQDPYASLHPRFTVDETLREPLAIHRLPDAD
ARIARALAEVGLGPAFRFRYPHQLSGGQRQRVAIARALIVEPRVLLLDEPTSALDVSVQA
EILNLLKRLHRERELTMILVSHNLAVVGFLCSRVAVMRDGALVEQLGIEDVRAARVGSEY
TRTLLRATQGYRRAG*
>GCA_000452945.1_BPMSHR5858_genomic_02987_1 oppD_1
MPASSVSPNTPPLAEIEGLRIGFRTHDGALVEAVRDLSLTLAPGERLGIVGESGSGKSLT
GRALLGLLPAAAQWHARALRFEGRDLLALGARERRRLCGGAMGMILQDPKYSLNPVMTVA
TQMAEAFRLREPGLRGRALRERIVDALASVEIRDPARVADAYPHELSGGMGQRVMIAMMV
STGPRLLVADEPTSALDVAVSMQVLAVLDDMIARHRTGLLFISHDLPLVTSFCDRVAVMY
AGRVVETCAARDLVHAAHPYTRGLLAATPPLANPPDALPVLERDPAWLTQAAR*
>GCA_000452945.1_BPMSHR5858_genomic_02988_1 ddpC
MNASPDHSRTNPAATTLRAWLLSDAPASRRQATLGLAYRRWRRFAGNPLNLLGLAILAAL
VAIALVAPFVMPHDPLRQVLADRLLPPGSPSHWLGTDQLGRDILSRLIAGSRLTLGIAIL
VVAIVVPIGLAIGTTAGYCGGLVDSALMRITDVALAFPKIVLALAFAAALGPGVVNAVVA
ISITAWPAYARLARAETLRIANADFIHAARLQGASDLRIVLRYVMPLCLSSVIVRATLDM
AGIILTVAGLGFLGLGAQPPSPEWGFMVASGRNVLLDAWWVATLPGAAILVVSIAFNLLG
DGLRDVFDPRHGA*
>GCA_000452945.1_BPMSHR5858_genomic_02990_1 ddpA_1
MTLSMPKLVAALAAASAFAAVPAREAHAAPPKDLFVMATLLDEFISLDPGEIYELVPEEY
VANTYDRLVRVDLADPSKFNGDVAQSWSVSPDGLTFTFKLRAGLAFHSGNPLTADDVAWS
LQRAALLDKGPAAVLAGIGLTKANALANVKQLDALTVSVTTDRKYAPSFVLNVLGSWPAS
IVDKKLLLSHQQGDDFGNGWLKTHEAGSGAYQLVKWTAGDSIVLQRYDRYRLPLAMKRIV
LRHVPEAASQRLLLESGDVDAARDLSPDDLAAVANSGKAKVTPSPQATLLYLGLNTKNPT
LAKPEVQQALKWLVDYDGIQRHVVKTTYQVHQTFMPEGLLGALNANPYRLDIAKAKALLA
KAGVPNGFTVTMDVRNDYPYTEIAQAVQANFAQAGVKVQLIPGDNKQTLAKYRARQHDIY
IGEWSADYIDPHSNAQGFAWNPDNSDQSSYKMLAWRNAWNIPRLTAQTDAALAEPSAAKR
AQSYQAMQKEVLAKSPFVILFEKVAQVATRPGVSGLEVGPINDLVSYRNLKKQ*
>GCA_000452945.1_BPMSHR5858_genomic_02995_1 vgrG1
MKMSDIAGFLSLQNSRLLTIKTPLAGRSELVLSEFQCSEGLSVLFDMRLGLVSRDPTIEL
KQMIGQAVTISLQPSGAIVGGSPRHFHGYVTQFSHTGADGGLATYSATVQPWLWMLSRRV
DSRIFQDKSARDILDAVFSQYSALASYEFRAGRTLKPYSYCTQYRETDLNFVLRLMELEG
LFFYFEHTEDGHKLIIDDDSTRAKPVDGPPSLRYASGEILEDEAVVTQWAAQRQLMSGAV
SMKAYDYKVPAARRYVSGEADFNQGEVERYEIYDYVGLHGFDSTDRGEELARFRLESLAA
SGKTFSGTSTGRTLASGRYFELSAHYDHDNGPMQDRQFLLTNVRHHGVNNYQSSEGSGSY
HASFQCIRKKIPFRPPLAHARPVIPGPQTAIVVGPKGEQIHTDALGRVKIQFHWDRIGQR
NHGSSCWVRVSQPWAGGGFGSVQIPRIGDEVVVTFLDGNPDRPLIISSVYNAQNMPPWAL
PAGATQSGFLTRSHKGTAENANAIRFEDKLGEEEIWLHAERNQRIEVEHDESHTVGAKRT
KTVGADEIVTIGGAQTHTITGARTQTIGADHTQTIKGAHKQNVAGTHTQVIGGNASITTT
GPQPGAAGTVGDIEIQSSQGKIHLKAATEIIIEVGASVIHLKADGTIEISGPTHIGLNSK
S*
>GCA_000452945.1_BPMSHR5858_genomic_02996_1 group_4796
MRYSVNEGHLELPGQWLDRSVNALLPAMAEVTGSNLVLTRDELPYGVEFADYVDVQRAKY
RKELSGLQMQRDEPGVLDGRPCQFLAFTWNKDDLLIHQMAVIVLDAPLVLALTYTSPGRL
PDGVRDAIGAALASFRFHRGAQLPAS*
>GCA_000452945.1_BPMSHR5858_genomic_02998_1 group_3390
MSDFIKIAKVAAQIDWQWDGEWWVREASRLGFKHEKSSASLEVHADPDGNAWRARVKEGK
ILFIEITFEKFVDVESLSVSEYEDKVDEFFARFEAAVAELTPVLGKPVFSDGAAAKGFPD
DQEANWLALWHVKNARLMLEQKHESREFPFRLCFVIAPA*
>GCA_000452945.1_BPMSHR5858_genomic_02999_1 ssa1
MTRHKKRKTMKRSGAKLLAPVVVAAAAAVAARPGWAQAAPYPDPGRRGDPASWRTPEFTN
AWGLGAMHAEYAYAAGYTGANVAIGVLDSGYYAQHPELPDSRFVPVTAAGVSGVLNPNNN
NHGTLVSGVVGGARDGVGMHGVAPDATVYEGNTNAIDGFRFGVSDPKFPASDAKYFAEAY
DALAAKGVRIISNSWGSQPANENYSTLNKLTDAYKLHEAVRTATGRGTWLDAAAKVSRDG
VINNFSSGNTGYDNASLRGAYAYFHPELEGHWMTTTGYDQLSGQVYNQCGIAKWWCVMAP
TGVPSTSYSGGAAAPTGATYANFNGTSAAAPHASAALALIMERFPYMTSEQALSVLFTTA
QNMEPDPSRPDYTNNGLFSTVHPAKPGASGVPNAFGGWGLVDLRRAMNGPGQLLGTFDAA
LPAGTADVWSNDISDVALAARKREDDAEHRAWLDTLRTKGWEHGLPAGASDGDRIDYALG
VARETAYQAREYQGSLVKSGGGTLTLAGANTYRGPTTVDGGELRIDGSIAARAVVNPAGR
LTVNGRAADIAVNGGVATIAGTSANLSIDRQGRAAVTGTTADVRVASGFASLGGTSGNVA
VGALGVAAITGRTADVAVDGGRASLDGASGNVAVGNGGVVSGSGTVRTLTAAANGTVAPG
HSVGTLTVSGDVRFAPGSIYAVEVSPGGAGDRIVAGGRAQIDGGALALALENTPPPLTPE
QSRSVLGRRFEILNAAGGVAGRFDAPSGYLFVNPVLAYGPTTVSLTIDRNATPFASVART
ANERGVADALETADPGSAVYNSVLFAASAQAPQAALAQLTGEIYPAAYAALVNESRQVRE
TALERLWTARGAPGRAGAWARLLGAWGSARGGDVNGYTSSTGGFLAGADAALLDGVRAGG
FAGYSHTGVNLRNQPSSASFDSFHLGAYAGWQPGALGVRIGAAHAWHRGGVDRAVQYGAV
AENETTALHAETTQVFGEAGYRFALDGAATLEPFFGVAYVHLKNQGTTETGGAAALRVRQ
GNHDVTFSTLGVRGETRLGLTSRLQLTLQGSAGWQHALTDGQPSGTLAFATGSDTFTVSS
VPVAKDAAVLNVGAGLELGKNGWLRVGYSGSLASRQSEHAVQGSLHWKF*
>GCA_000452945.1_BPMSHR5858_genomic_03000_1 group_3391
MEAASRPGASSATRAWQDITIVDDLLPPDEHAQIHRFLAAGQWSHGWRSHNGAQTQPFWN
RHFAGATDPAQLDVGPGASDELARAAPLVHACWRRLSQTYLTQHALQSCYANGLPYGADG
ALHADSLAMGACTAVYYPHERWDPDWGGETVLFNKDRTDILCAVYPKPNRLLIFPGFVYH
VARGVSRACPTMRITLMFKTQWRP*
>GCA_000452945.1_BPMSHR5858_genomic_03003_1 group_3031
MLSSTTMSRSAGDFATAGDASSRTVRAARDALEGRRKGVAAALPFVGPAVIASIGYMDPG
NFATNIQAGAAYGYELLWVVLAANVIAMVFQAMSAKLGIVTGRNLAELCREHFPRPVVWG
MWATSEIAAMATDLAEFLGGALAFGLLLHLPLIAGMAATAVLTCAILTLEKRGFRPLEAA
IAALVGVIGASYLGELLIAPQDWHAAAYHLVVPQLRDGTALTLAVGIIGATIMPHTLYLH
SGLTQRRTTPRDARERRLLLRFSNREVAVALGVAGFVNIAMVMMASSAFHRAAPGMADIG
DAYHTLIPLLGPAAGALFLVALFASGVSSSVVGTLAGQVVMQGFLRRRVPIWVRRLVTIA
PAFAIVALGCDVTRAMVLSQVVLSLVLPLPMAALLLLSSRRALLGEHALRAPTLVAASAA
AAAIVALNVYLIWAAFN*
>GCA_000452945.1_BPMSHR5858_genomic_03004_1 yiaA
MNHHTNQLHPSTSFAFVAASWAALLAGVCAYLLGLWNAGMQLNEKGYYFTVLAYGLFAAV
SLQKSVRDRLEGVPVTGLYYGLAWASVILTIALLVIGLFNATLLLSEKGFYAMSFALALF
GSVAVQKNTRDLKAAGRGRAETEVAVDIAE*
>GCA_000452945.1_BPMSHR5858_genomic_03005_1 group_4797
MIVRPREHGFRMLFVWNGSVLKSILPQLALMSAVSIVALLTNGRILGEKVPLNPTPFTLA
GLALAIFAAFRNNASYDRYWEARKLWGGVLSAARALTSQALGYDALADGASFARATAGFV
YALKHQLRGTDPTHDLRRRLPADWLDPVLAAQFRPVAILHALRGRLAGRHRDGALTDAQL
WMLDAQLNELAAKLAGCERIASTPIPFPYHVLLHRTVYAYCVMLPFGLVDSIGIATPFVS
VFVSYTLIALDAIAGEIAEPFGDGPNHLALDALTRQIERSLFELAGLPLPDEMRAGPNHR
LS*
>GCA_000452945.1_BPMSHR5858_genomic_03006_1 ydeP
MKTKEPDARIEPYAHPAGGWGALKYVAINLFKEKVPGGNYRALLRQNQPDGFDCPGCAWP
DREHASTFEFCENGVKAVAAEATAKRVTPAFFAEHTVSALFDQSDYALEQHGRLTDPMVY
DAATDRYVPIAWNAAFELIANHLRALGEPNRAAFYTSGRASNEAAFLYQLLVRYYGTNNF
PDCSNMCHEATSRGLPATVGVGKGTVTLDDFEQADTLLIFGQNPATNHPRMMGELRACAK
RGATIVSINPLKERGLERFASPQHPAEMLTMSSTPIASTFVQPRVGGDLALIKGVAKRVL
ELDDAARERGGARVLDVDFIAAHTAGFDTFAADLRAQDWAALVGESGVPREQIDALARIY
VRGERVIATWGMGLTQHKHSVATVHMLSNLMLMRGNIGRPGAGLCPVRGHSNVQGNRTVG
IEEKPSDAFLERLGRVFDFAPPRSHGYDVVETIEAMLDGRIGVFIGLGGNFAMATPDTPR
TWQGLRRCGLTVHITTKLNRSHLVHGREALILPTLGRTEIDLQNGIAQGVSVEDSMCMVH
ASYGMNPPASPNLLSEVAIVARLGHALFGGDKIDWLGYMNDYAKIRDAIEASIEGFDDYN
ARIARPGGFHLRVASREREWLTPSGRANFIVHALPADTPIQRARARHGKRLMTLMTTRSH
DQYNTTIYALDDRYRGVFGERRVVFAHPDDLAMLGFEAGERVDLETVWDDGIERRVAGFL
LVAYDIPRGCLGAYYPETNPLVPLDSVGDVCNTPTSKSIPVLMHRSAGDAAHAA*
>GCA_000452945.1_BPMSHR5858_genomic_03007_1 hisC_1
VKRYEQLADDIEAMIRRGVYRPGERIPSVRQASVQHRLSVTTVVRAYLVLESRGAIESRP
QSGYFVRARQDAPALALHASAPVAVSSAVDVSRLVLSTLRSIARDDAVPLGSPYPDATPF
PAQRIARHAHAIARRRARWGVIDDLPPGNPELIRQIARRYLEGGVAVDPGEIVVTIGATE
AINLCLQAVAKPGDTIAVESPTFYAMLHAIERLGMRAIEVATHPGEGIDIDALARILATE
RIAACMVMPNFQNPLGFLMPDERKRALVELLARHDVPAIENDVYHELYYGAARPSTLKAF
DARGLVLHCASFSKSLSPAYRVGWAMPGRYRDQVEKLKFLNTLATPAIDQLAIAEYLRHD
GYDHHLRRIRKLYAQQASMMAALVRRFFPDGTRLSQPKGGYVLWIELPDGVDAMALYRLA
LAQKITLGPGHMFSTTLAYRHCIRLNYSYAWSPQIEAAVKTLGRLAAVCAKR*
>GCA_000452945.1_BPMSHR5858_genomic_03010_1 mgtB
MTQRTPSKHKQRGFLHYGARGDGDAGRPMRIEYASAQPLADTLAHLRTSECGLTTPDARQ
RLLGDGPNEIAHDKPPHWSRQLLAALNNPFVYVLLVLAAISLLTDVYFADPGDRDYVKIT
ILLSMVTISVLLRFVQEFRSLRAAEKLKAMVRTTATVQRRADDASAPIRHDVPMREVVVG
DIVHLSAGDMIPADVRLISSRDLFISQAVLTGEALPVEKYDTLGAVAQKSADPASRSSGA
AVRLERAEHGGSLLDLANVCFMGTNVVSGTATAVVVATGGDTYFGALAKNVVSHKRIETS
FDRGVSSVSWLLIRFMLVMVPVVFMINGLTKGDWLSALTFALAVAVGLTPEMLPMIVSAN
LARGAIAMARRKVVVKRLNSVQNFGAMDVLCTDKTGTLTQDKIILEHHLDVSGRKNEDVL
RLGWLNSFHQSGQKNLIDIAIVERANQLGEWIKPQGYRKIDELPFDFVRRRLSVVVEDPR
GTHLLVCKGALEEMLAVSTHVQDDEGVHPLDFVARKRLLAQANAYNEDGFRVLIVATRTI
ARGDEREQYRTADERDLVVRGFLTFLDPPKESAAPALAALRENGVAVKVLTGDNPTVTLN
VCRQVGLAPGKPLLGTEIEALHDTALEKAVEHTTVFAKLTPLQKARIVKALQANGHTVGF
LGDGINDAPALRDADVGISVDSGADIAKETADIILLEKSLMVLEEGVIKGRETFGNILKY
LNMTASSNFGNVFSVLVASAFLPWEPMLAVQMLVLNLIYDTSQMLLPWDKMDPEFLKKPR
KWEANNIGRFMLWVGPTSSVFDITTYVLMWTVFGAGALYHAQGGAAGQLVMNSGWFVESL
VSQTLVVHLLRTQKIPFLQSTASLPVLLSTTIAIAIGCWLPFSPFSEALGFIRLPGSYWL
WLAATMIGYIALAQIVKTLYVRRYKRWF*
>GCA_000452945.1_BPMSHR5858_genomic_03012_1 bprV
MNKKRSNPEQMRLGQVRAVAGILSMSVLVPLAGCGGGGDGGGSGTPSAAAQPTPAPAPAP
APAPAPSSGSSQSTNSSTSTAACPVTQAASTAAGETLVTRTVSHEAPVDHLIVKLQRTAA
ASASGARIMAAANDAARLDSVIQRVMSQWSAKSGAVRSYAQNIAPTNAVQVERTMSDGAA
LLALGQKMSADNAGALAQTFAADPDVAYAEPDRRVFARTVATDPDYAQQWNYFDPAAGIN
LPDAWNVTNGLPSVVTAVLDTGYRPHPDIIANLLPGYDFISDINTGNNGHGRGPDATDPG
DWVTQQELTDPSSPFYQCASAPSNSSWHGTQVAGIIGAAANNGIGIAGVSWYGKILPVRV
LGKCGGTTSDIADAMRWAAGIPVAGAPTNLTPAKVINLSLGGTGPCGDTFQQAINDVIAR
GTTVVVSAGNDGQATTLDRPANCKGVISVGATDSTGQRAWYSNFGSDITLSAPGSNILST
SNAGTTVPTTDAYGTHSGTSLAAPQVAGVASLMLAVNPNLTPAQIAQKLASTARPSPATA

SCLARAPGAGIVDAGTVVASATK*
>GCA_000452945.1_BPMSHR5858_genomic_03014_1 fdhS
MNNDTNPHARRSIDADEPMGGGLTRRQWLQRSLALTALGLAGSLTLRALADAPSAAPLDA
FMTLSEALTGKTGLNRAVGERLLRALQKGSFELGDRLPKLAGALAANALAPDQEALALRI
LEAWYLGIVENVVVTYEEALMFGVVSDTLVIRSYCPNKPGFWADKPIERQA*
>GCA_000452945.1_BPMSHR5858_genomic_03015_1 fdhL
MADTQQADVVVVGSGVAGAIVAHQLAMAGKSVILLEAGPRMPRWEIVERFRNQPDKMDFM
APYPSSPWAPHPEYGPPNDYLVLKGEHKFNSQYIRAVGGTTWHWAASAWRFIPNDFKMKT
VYGVGRDWPIQYDDIEHDYQRAEEELGVWGPGAEEDLLSPRKQPYPMPPLPLSYNERTIK
TALNNHDPKYHVVTEPVARNSRPYDGRPTCCGNNNCMPICPIGAMYNGIVHVEKAEQAGA
KLIENAVVHKLEVGPQKKIVAARYKDPKGVEHRVEGKYFVLAANGIETPKLMLMSTSHDF
PNGVGNSSDMVGRNLMDHPGTGVSFYASEKLWPGRGPQEMTSLIGFRDGAFRATEAAKKI
HLSNLSRVDQETQKIFKAGKLLKPAELDAQIRDRSARYVQFDCFHEILPQPENRIVPSRT
ATDAIGIPRPEITYAIDDYVKRGAAHTREVYATAAQVLGGTDVAFNDEFAPNNHITGSTI
MGADPRDSVVDKDCRTFDHPNLFVSSSATMPTVGTVNVTLTIAALALRISDQLKKEI*
>GCA_000452945.1_BPMSHR5858_genomic_03016_1 group_3395
MLKRTLSFMLAGCLALPGLSSAADAAASGAPASAASSVPAASAAPAAVRAADATLVERGR
YLAVAGDCMACHTAKGGKPFAGGLPMRAPLLGTIYTTNITPDKETGIGDWSFADFERAVR
HGVAKNGDNLYPAMPYVSYTKVTDDDVKALYAYFMHGVEPVRQPPRRNDIPWYLSMRWPL
KIWNLLFLKEGVYQPKPERGVEWNRGAYLVQGLAHCGTCHTPRAVTLQEKSLDETGGSFL
AGSVLSGWDGYNITSDPNAGIGGWSQPQLIQYLRTGSVPGLAQAAGPMAEAVEHSFSRMS
DADIGAIATYVRTVPAVADGAAKARSAWGKPAEDGIRLRGVALASTGIDPARLYLGNCAS
CHQMQGKGTPDGYYPQLFHNSTVGAPNPTNLVQVILNGVARKAGGEDVGMPAFRHELSDA
QIAALANYLTVQFGNPAAKVSEQDVAKLRAAQ*
>GCA_000452945.1_BPMSHR5858_genomic_03019_1 amj
MDTQLWIICGLTFVIHVIATLAYAVRIAGVRTRRIAVSFSLFGIIALVSRTANSFQGPFI
AKRVELDISRHLGEGLLADFRLFLLSATVASVVGALLIPTFQRYFSRAVHHFQANRSVSR
LLLRVFSRDGVGYIRGGARLPSPRNVTQLATGTGVSWQVIVLNVVAMAIWTVGVFASLYA
GYLKPELRVTCANLSSIINGFATVAMAVIIDPQVSVMTDDVIEGRLCENHFRRAITTLAG
ARVLGTLCAQAALVPAALVIVRVAELI*
>GCA_000452945.1_BPMSHR5858_genomic_03021_1 hisM_1
VIQIIGQYWQNYLFSDGYRLSGLAITLWLLVVSLGLGFGLAVPLAIARASRNRAVSGAVW
LYTYVFRGTPLYVQLLLCYTGIYSLSVVHDHAWLDSFFRSAMNCTLLAFTLNECAYATEI
FAGAIKSTAPGEIEAGLAYGMSRFKLYTRVILPSALRRALPSYSNEVILMLHATTLAFTA
TVPDILKVTRDVNSATYASFQAYGIAAALYAVVVFALIWLFRRMEARWLAYLRPQGH*
>GCA_000452945.1_BPMSHR5858_genomic_03022_1 hisP_1
MDKLTVDDLHKKFGDNEVLKGISLRAKAGDVISIIGSSGSGKSTFLRCINFLEQPCSGAI
TLGGETIRVTRDRRGAPRIGDPKQLQRMRVKLAMVFQHFNLWAHMTVLENVTEAPVSVLG
LSRAEADARAREYLRKVGLAPGVEAKYPSHLSGGQQQRVAIARALAMEPEVMLFDEPTSA
LDPELVGEVLKVMQALAEEGRTMIVVTHEMGFARNVSNHVIFLHQGRIEEEGHPQDVLVH
PKSERLSQFLSGRLK*
>GCA_000452945.1_BPMSHR5858_genomic_03023_1 murA_1
MSNLIVHGGAPLRGEITPSANKNAVLPILCATLLTDRPLRLVGVPDITDVRKILDIFRTL
GSDVSIDYASGVLDLHHRATAFDPAVHRLPEEMRSSIMLVPPLLARFGVARLENDVKGCT
LGVREIDPHVEVFERFGARIERTSDSLIVRADGPLTPNHHWLDYASVTTTENFVLCAASA
NGTSTLVNAASEPHVQEFCRFLAMLGVPIEGIGTSHLSVQGGRALAGGEYRFNEDFHEIA
TFLALGAITGGDIAVRNGSPEQFPLIDRTFAKFGVQVTHENGWSHALRDGPLKVKQPFTR
NILTKVEAAPWPYLPVDLLPIFIALGVQAQGSVMFWNKVYDGAMGWTGELSKFGAHVFLS
DPHRLITFGGLPLSPARVESPYIIRVAIALLMVAASIDGRSEILNAQPIRRAHPHFVENL
RSVGANVEWTSGE*
>GCA_000452945.1_BPMSHR5858_genomic_03024_1 group_7477
MQTDDTPIYEGGCTCGAVRYRMTSRPLIVHCCHCRWCQRETGTAFALNALIESDRLLLLR
GEVDIVDTPSNSGKGQKIARCPRCRIAVWSHYAGGGSAVSFVRVGTLDAPERLSPDIHIF
TSSKQPWVILPPAARAVPEYYSSDEIWSAESLARRTALRAKRDR*
>GCA_000452945.1_BPMSHR5858_genomic_03025_1 group_1862
MRKAFRHAVALSLLSWLVAPRVHAQYDPGATVGLGQGMIANNGVIGNLAIGSQVLERDAR
GDADRARAASPRTAPAALTYVGDAGVHRQVQADIVNFLAHGDASKRDEAKRVVEQGELLE
TFDRLMSRPAFDRRNLGDVFAAYLVMSWEVVNGADAHAQPAGVDAVRAKSSAALARDGRV
RAMSDADKQRLAETLAYLAMIAVTAQRDLRRSGNTAALEQLRAGVRLSTRKMIGVDVKGL
RLTENGLAR*
>GCA_000452945.1_BPMSHR5858_genomic_03026_1 group_7478
VKYLPLIALIVATSASAADTGVQNVGQSQKSAQDVSACIAKTWADKSQQQVVTQNVLANG
LAADVYVPGQQPPNGAAAVVRPAFSGNAKTWVGFRAGSSGAADGAAAGDINSCL*
>GCA_000452945.1_BPMSHR5858_genomic_03027_1 entS_1
MNRLPIAFRRLALSNLAAQSAEQIGLAVAPLVAVLSLGASASGTGLLQTAQTLPFLLLSI
PLGVWADRHSRKRIMAAAEGLRALALAATLALLAAHALTLPSLAMLGFVGACGTVAYSVA
APALIPALVGRAALARANGAIELARSAAYSAGPAIGAALVDRIGAQWAYGFAAALSAHAV
LLIVRLPDVRAPAAPRKRFVAELLDGARFVRRDRHLRPMIVTAVFFNVGFFTLQAVYVPY
AVRHLALGASQIGMTFAAYGIGMMGGAALASAIARRVRFGVVLTIGPVGGLVASLVMAVS
IFAPSFWLAASSFFLLGAGPILWSVASTTLRQAITPPAMLGRVCAINGTATYGARPLGAL
VGAVVAARFGIDACIWASVGGFVIQAAVIVRSAVSRLERIPDVSDAAPRYAPLPD*
>GCA_000452945.1_BPMSHR5858_genomic_03029_1 btsS
MRRVRGLAGLVLFNCIVGLSSWASRRDDAVLPYLVLANGIGLSALLLSLCADKLTRGKLA
VLPKVLIVAPVSVLAGFEIAASTIGHAPHLIGRASVREWLGYGSSFVVTGIACAFVSLFM
QAARMRASLETQRREAAEARQAETAARLALLQAQIEPHFLFNTLANVQSLIERDPARAST
MLDSLNRYLRASLGRTRKAASTLGEELELIEALLQIASIRLGERLRYAIDVPAPLRELAF
SPLLLQPLVENALLHGIEPSLDGGEIRVCGRRNGKLLELSVIDTGVGLGNGETRLHGGVG
LANVVARLTSLYGARGRVSVGATAGATRGVTATLQIPID*
>GCA_000452945.1_BPMSHR5858_genomic_03030_1 ypdB
MPTALIADDEPNLSAELAERLAELWSELEIVATPRNGVDTLAALNAARPDIAFLDIRMPG
IDSLKLASLVPDVHVVFVTAYDEYAIEAFERSAIDYLLKPVIDERLLRCIAKLQRGGFAP
PGAPPPAGARRPQEPTPIHWLTVGLKDTTRLVSVDDVLYFQALDKYTEVVTAGERHVIRT
PLKELLQRLDPDRFARVHRGVIVAYAAIDRIERDLLGRLRIHLRGHRDVLPVSRAYAGLF
KHM*
>GCA_000452945.1_BPMSHR5858_genomic_03031_1 katB
MISRHSMRWLSRALVAVSASGALAAHASTLTRDNGAPVGDNQNSQTAGANGPVLLQDGHL
IQKLQRFDRERIPERVVHARGTGAHGVFVATRDISDLTRAKVFEPGTQTPVFVRFSSVIH
GGTSPETLRDPRGFATKFYTAEGNWDLVGNNLPVFFIRDAMKFPDMVHSLKPAPDTNIQD
PDRFFDFFSHQPEATHMITRVYSDAGTPASYREMDGNSVHAYKFVNAHGGVTYVKFHWKS
LQGQKNLTAAQAEAIQGKDFNHMTRDLIAAIDAGRYPKWDLYVQTLKPDQLDQFAFDPLD
ATKVWPGVPEVKIGTMTLNRNPGNVFQETEQAAFAPSNLVPGIEPSEDRLLQGRLFAYAD
TQLHRVGVNGAQLPVNRPRAPVANYNRDGAMNGGAARGTVNYEPGAQAALAADPAFAASR
APLAGSTQQARIAKTRNFDQAGAFYRSLSASERANLVANLAGDLKQVRNDGVKYTMLSYF
QKADAEYGRKVTAALGADQGRVDALTAKLAD*
>GCA_000452945.1_BPMSHR5858_genomic_03033_1 group_7479
MFCDFYFPLIATLASALAPPGLRAKIREARVLSASADLARLTQDGCGDAERAYRIQLHAD
MQLAMALEFDAEESYRRSRKLMHAPKDEIRLLSFRNTGWQALFRRRLGTAMACFANVTSE
PAVDPCRRIEGLFGTMCVWFELGHLDEAGYLLDDIELAIERLLASHEDFALWRELAQTMR
ADLDLQRLMRHAPELSDHVYWQSARLSDPVRPPVERERLASRPIAEQDVRSPLLRARLDF
RTALRRLAGGQADALNDLMAHLDWADANGLTSYQNSARQEIALAGLIGNAPHVAEMVLAP
LANEIRVGHHHGHLEILYCLAKVNQAQGRTHHSQQLYSRYAMTAVHCARKGACDLARNSP
QGHAPHVADDISARLPAKYRRAYHFLLSNLERSDLAIGEIAEVIGVTVRSLQNTFKASLG
ATPSEIIRQERMKRIHRELQGEDARFGQRVLDSGNRWGVPNRSTLLNAYKRQFNEAPSQT
LHRKHPYKPGEDVASGVLASAAHPAAARQQHEASVSTAD*
>GCA_000452945.1_BPMSHR5858_genomic_03034_1 kshA
MNGTQRFPMDIPFGWFCIGYSDEVAVGDVKPLRYFGKDLVMFRGESGEIGVLDAYCPHLG
AHMGHGGTVEGDSLRCPFHHWAYRKDGFCTSIPYAKEMPLIAKREPIAVPYPVVERNGAI
WVWHHPQQIAPTYEVLEHEEFVAPGWSKPVRRFWSFASNPQEIAENGVDTAHFKCIHHMD
AMPEGEVTYEGHIRRSRTTGETSMIFPDGVTRTVETGTNVINSGAGQKISRFTGFVTISL
MEMATPVEADQVELRFCFAFPECPEGSPEWQGAMNTIDYFSGQLGIEGGDIPIWSHKIHR
EKPILCDGDGAIMRFRRYFEQFYAPRVVPVTASA*
>GCA_000452945.1_BPMSHR5858_genomic_03035_1 group_7481
MGDYLAQDARIERITVRDPEARTWFDRGLLWAYGFNFEAAVDCFQEAARIDPDCVLAYWG
IAYASGCNYNKQWKVFHPRMIARCMKLARDAIRRAQACRAGASALEMALVRAIECRFQAE
GAHDEALLRRWNDDYARAMRDVYLSCPDNLDVAALFADALINRTPWKLWDMSSGKTADGA
DTDEAIAVLERALAQVDANGLAPHPGLLHVYIHTIEMSPTPEKALRAADALRDLAPDVGH
LLHMASHIDILCGHYHDAIVANDRAIAANQRVLDRHPQWLEFRLYCVHDIHFKIYAAMML
GRFSAAWQGVLELEALITEALLRVEQPPMAYLLEGFLSVRVHVLIRFGKWREILEQPFPA
NAALYCNTTAMLHYARGIAYANLKLAGPAAQARRAFSIAQAALHEHRYVTNNTCADLLKI
AACVLDGEIAYHEDRFDEAFAHLRRAVELDDGLEYMEPWGWMMPTRHPLGALLLAQGHVE
EAERVYRADLGLDSTIYRSLQHPGNVWSLQGYVSCLRRLGKHETADALQPALDVARARAD
VEIGVSCFCAAPSKRGCCH*
>GCA_000452945.1_BPMSHR5858_genomic_03036_1 pksF
MRSSERDEILVTGVGVVSAIGQGRQPFIDALLDGRHAFGVMRRPGRQLPDDAGAGPAPAF
FGAEIDSIALPPMLDAHRRTASYSAQAAVACVREAWDDAALGDVDPLRIGLVVGGSNVQQ
REQALVHDTYRGRYAFLRPSYGLSFMDSDLCGLCTELFGIRGVAFTVGGASASGQLAVLQ
ACRLVESGSVDACIAVGALMDLSYWECQALRSLGAMGSDRFADEPDAACRPFDQDRDGFI
YGECSGAIVVERAARMRRAHVSPYAKWLGGCIRMDGTREPAPSLDGEMAVIEGALRDAGI
GADSIDYVNPHGSGSTLGDATELDALGRSGLAHARINTTKSLVGHGLSAAGIVELIATLL
QMRARRLHPSRNLDRPIDERFDWVRSTCGSDIRRALKLSIGFGGINTALCLERC*
>GCA_000452945.1_BPMSHR5858_genomic_03037_1 acpK
MDLQQVFDIIVRHTKEVIPGLDRHAFRTTDALKDLGANSLDRSEIVIMTLESLSLRIPLV
EFAGARNIGELAEVLYAKL*
>GCA_000452945.1_BPMSHR5858_genomic_03038_1 pksI
MSAHERVVEVREVADCVVQVVMQDRENKNTFSDALIRGLGEAFDAIRQNGRCKAAILTGY
GNYFASGGTQESLMYLSEGKGTFAEIAASASRNGANLYSLALNCPVPVIAAMQGHGIGGG
FVMGLFADFVVLGKECLYTTNFMKYGFTPGMGATLVLPHKLGLPLAQNMLMTARSFYGDE
LQKLGVPFPVVPRAEVAACALEIAKNLAEKPRLSLVTLKDHLVAELRSRLPAVIEQELAM
HEKTFGQPEVKERIRSLFGQ*
>GCA_000452945.1_BPMSHR5858_genomic_03039_1 pksH
MDMLSRPADYRTIRVRYDGDICFLQLHRPDAQNAINNRMIAECMDVLDRCEHAAKIVVLQ
GLPDVFCFGADFSDIAEKPDALIDSDAIYGLWHRLATGPYITIAHVQGKVNAGGIGFVAA
SDIAICDENVPFSLSELLFGLIPACVLPFLARRIGTPKAHYMTLMTQPVTAQQAFSWGLV
DAIGANTDTLVRKHLLRLRCLNKASVARYKSYAATLDDTLAAARPGAVRVSIEMFADPEN
RRKIARYVDTGKFPWEAD*
>GCA_000452945.1_BPMSHR5858_genomic_03040_1 pksG
MTAVGIEALNVYAGVASLDVSRLAEHRKLDMARFQNLLMREKSVALPYEDPITYGVNAAK
PIVDALTPDERDRIEMLITCTESAFDFGKSMSTYFHHHLGLKRNCRLFEVKNACYSGVAG
LQTAINFILAQVSPGAKALVIATDLSRFIVEEGGEALSADWSFAEPSSGAGAVAMLVSDT
PHVFRIDVGANGYYGYEVMDTCRPTTDSEAGNSDLSLLSYLDCCENAFLEYQKRVCDVDY
ASTFGFLAFHTPFGGMVKGAHRNLMRKTSRCSTQEIEQDFQRRAGPGLVYCQRLGNIMGA
TAMLSVASTIDNGEYRAPQRVGVFSYGSGCCSEFFSGIVDEEGQRRLRGMRIGEQLDRRY
ALSIDEYEHVLKESRVVRFGTRNAKIDDGFIPAARRAHGRETLFLSRINEYHREYEWIC*
>GCA_000452945.1_BPMSHR5858_genomic_03043_1 gerN_2
MNSIFVFLSQALLIVAVPPLLMNYLRVGAVVPLVVVQIVFGVVIGPSGLGRLSPETYTAL
FPPDSLTFLSHVSAIALFFFAIVTGLHLDATGLRGHGRKLGFISTASMLAPMSLGVLAGI
AISARYPQALDPHASVAEFVAGIGICCGVTALPVLAAILRETNLTSRRLGQFSLALAAVN
DGMLWILLAAFLSVTGANAANGGRLFITSALIVLYFVSILFVVRPLLLVLANRLRTSVDT
QILIACAVAVASAAATDRLGLHYLLGAFIAGAIIPGAWRDALLERMQPVTVNVLTPFFFI
STGLRVLIDVDAPGFLQITLIITLCTAIGKIAGTTLAARMTGESWGFCLGLGFLAQAKGL
MELVVATILLDSHVISRTVFSALILMALISTAFAMPLLRLKPVRGAALSA*
>GCA_000452945.1_BPMSHR5858_genomic_03050_1 stp_2
MTSEISGKRRWYALVVLCLGVLMIVLDTTIVNVALPSIAADLRFSENALVWVVNAYMLTF
GGCLLLGGRLGDLYGHRRLFLAGVTVFTAASLACGLAQTQLVLIAARAAQGIGGAVVSSV
ALSLIMTLFVEPGDRAKAMGVYGFVYAGGGSLGVLLGGLLTSALSWHWVFLVNLPLGAVV
YLLSSALLPKARAQATRAPLDIAGAVTVTAALMLAVYAVVNGNAAGWLSARTLSWLGGAA
ALFAAFVAIEARVAHPLMPLALCMKRNVAIANLIGILWAAAMFAWFFLAALYLQRVLGYG
PLQIGLAFLPANLIMGVFSLIVSALLVVRFGIRVPIAAGLLAAAAGLMLFARAPVGGDFV
ADVLPGMILLGTGAGIAFNPVLLAAMSGVDPADSGLASGIVNTSFMMGGALGLAVLASVA
TARTDALLAARTAVPAAMAGGYHAAFMLGAAFAIVAALIGMRLRAAPNPAAMQGLQGATQ
*
>GCA_000452945.1_BPMSHR5858_genomic_03052_1 group_4805
MTKPARELRKTDFEQLSEFRYQMRRFERFSERAAQSEGITPLQYLLLLHIKGYPERDWAT
VGELAERLQAQHHGVVALVTRCEALGLVRRRPSDADRRQVEVHLQAAGERVLARLAELHR
AELKSLRGAFQVPQIDY*
>GCA_000452945.1_BPMSHR5858_genomic_03053_1 IMPDH_1
MTYNFRLVARPIGAPFHLRYGSPTLSSSLAFAHRLARGLVRWLWRFVPIPVTLSRRERVR
SCAGALVGIGCVGVTMRALPGLPGDVPLLVAPMGASAVLLFAVPASPLAQPWSLIGGNLV
AATVGVACARWIADPVLAASVAIAGAIGAMFALRCVHPPSGAVALTAVVGGPAVHALGFR
FVAEPIALQSAALLGAALVYHALTGHRYPHGGAARPHADSNAHAAAAPLHARFVRADLEA
ALKNRSEWLDVAPEDLESLLRETELRAYARTFDELSCAEIMSRRPISIAPDTPLPAAMTL
LERHRIKALPVVDADARVVGIVTRADLSKAAPYATPGFLRNLSARLPRSLVGPAFVARAV
MSTRVHAVRTTTPIAELVPLFADHGHHHIPVVDADHQLAGIVTQADLIAGLYRQSQVRLA
A*
>GCA_000452945.1_BPMSHR5858_genomic_03054_1 dmlR_11
MDMLDNMRLFVRVVEAGSFTAVAKEIDATTAQVSRAVSNLEAHVQTRLLHRTTRHLGLTE
SGQRYFERAKSILAEIDYANAEARNALLRPSGKLRIHAMTGLGQSHVVSSIVRYQEDNPD
VSVELTLAQRMPNLVEEGYDVSIVSASHLPDSGYVAQTCGTSCSVLVASREYLARHGTPT
TPDDLAKHVCLRLDTPASPGGEWRLERDDGDETVYALPPAPFQANVPDALSVAVRAGRGI
GSIALYTALDDIREGRLVRVLPNYRLDTLSVYAVYATRRYLDAKIRTFLDHLRTTLAPAL
ENDLKELNRLNGGVGKKALREVA*
>GCA_000452945.1_BPMSHR5858_genomic_03055_1 chaA
MPISSSALPRWTLVAPLAAWLVLALSRVVPAEGFAIAFVAAALAGAVFSAVHHAEVVAHR
VGEPFGTLVLAIAVTVIEVALIVSVMLSAGPEKAGLARDTVFAAVMIVCNGIVGLCLLVG
GWRHGEQDFQGRGATKALAVLASLSVLSLVMPNYLSAAPGPRLSTSQLAFAGVSSLVLYG
VFVFVQTVRHRDYFLIDGSADESVHAVPPSGRTALTSIAFLFVSLVAVVLLAKLLSPAVE
RTVLRLGAPEAAVGIVIAALVLLPEGLAAVGAARANRLQTSMNLALGSALASIGLTIPTV
AAVFIWTGAPLTLGIGAMETVLLSLTLLVSTLTLSLGRTTVLHGVVHLSLFAAYLFLSIT
H*
>GCA_000452945.1_BPMSHR5858_genomic_03059_1 ycgM
MTDYVFAPPAVPSVEIAGCAARFAVRRVFCVGRNYADHAREMGADPNREPPFFFTKPADA
IVPASGTVPYPTLTSELHHEIELVIAIGRGGRAIAADAALDHVWGYGVGVDLTRRDLQAE
AKKAGRPWDWAKGFDASGPVTALHPAASIGHPQRGRIWLAVNGALRQQGDLADMIWPIPD
VIAYASRAVELRAGDLIFTGTPAGVAALQPGERVTGGVDGVASFEFVVGEKPAA*
>GCA_000452945.1_BPMSHR5858_genomic_03062_1 ackA
VDREILQNRPFDSLAIGDSASLVRTASRDDIDLFAAMSGDVNPAHLDAAFAAHDLFGHVV
VHGMWTGALVSAVLGTKLPGPGTIYLDQTLEFRHPVAPGDTITATVTVAEKRPEKRIVLL
DTRCTNQDGHVVLRGTATVIAPDTPITWERVPGPEVAVRRHDRYETFVRDAREQPPLRTA
IVHPCSPDALRAAIDARDANLIEPILIGPQAKIRAIADAAGVDLAGVPIEAVEHSHAAAA
RAVEMGGAGEVAALMKGSLHTDELLAAVIGPASRLRTERRISHVYAMDVPAYPKPLLITD
AAINIAPTLEQKRDICQNAVDLLHVLGVERPRVAVLAAVETVNPKMPSTLDAAALTVMAA
RGQIAGAIVDGPLAFDNAVSAVAAMSKGIASPVAGDADVLLVPDLEAGNMLAKQLMYFAG
ADAAGLVLGARLPIILTSRADSLRVRIASAALAKLVAERTRAGALAS*
>GCA_000452945.1_BPMSHR5858_genomic_03069_1 group_7487
MLYAILTPKAEAPLGYYDSSVTPTPEDMADFLAKTMGFDDRDEWIEAYGVEKLGYAPVH*
>GCA_000452945.1_BPMSHR5858_genomic_03070_1 deoC
MELSLNRNQLTEAALKALHLIDLTSLNDDDTEERIASLAASADTPVGTPAALCVYPRFVG
AAHAALARHGLVLPVATVTNFPHGAADPDAAARETADALARGADEIDVVFPYRALIDGDE
RVGRELVAQCRAAAGAQCLKVILETGELRDAAAIRRASEIAIEEGADFLKTSTGKVAVNA
TLDAAAVMLATIRVAGRMVGFKAAGGVRTAQDAAQYLSLAQRELSPGYLTSATFRFGASG
LLGNLLETLGHRAGATRGAY*
>GCA_000452945.1_BPMSHR5858_genomic_03071_1 psuT
VDILRSLCGIVVLLGVGYTLSVNRRAISARTVVAALATQLAIGALVLFVPVGRDALAGTA
HAVNSVLEMGQHGVAFLFGGLVGDKMFALFGDGGFAFALRVLPMIVFVTSLIAVLYYIGV
MKWLIRIVGTAMAKLLGVSRIEACSAVATIFLGQSEMPAFVKPFVRRMSGTEVFAVMSSG
MASVAGSVLAGYAGLGVKMEYLLAASFMAIPGGLLFGKMLCPTTEPSRVAVDSLEFDEKR
AANVIEAAASGAGVGMRIAVNVGTMLIAFIGLIALLNAMVELAVGWLGFAGVTLQSLLGA
LFSPLAWLIGVPWRDAPVAGSFIGQKLILNEFVAYGALSPYLKDAAQVVAAGLPVLAPKT

IAIVSFALCGFANFSSIAILTGGFTAVEPGMRSEVARYGLRALAAATLSNLMSATIAGLF
LSLS*
>GCA_000452945.1_BPMSHR5858_genomic_03073_1 yfcG_4
MSAHRPPAAPIRLHRSPLSGHCHRVELFLSLLGVPVEFVEVDMRGGAHRRPEFLALNPFG
QVPVIEDGECVLFDSNAILVYLAKRYGDPSWLPDDPQGAAAVQRWLSLAAGPIAYGAAAA
RLVTLFGAPLDHAGAIGIARRLFDVIEPELAGKRFATGEHPTIADVAAYAYIAHAPEGGV
SLDRYPNLRAWLARIEALPGFVPMPRSHAGPRAA*
>GCA_000452945.1_BPMSHR5858_genomic_03074_1 dmlR_8
MADIRDVNLNRLAIFVAVVEAGSLTAAAERLGLAKTMVSAHVQRLEAEIGASLVVRTTRR
LNVTEAGRAFYDACRDILRATETALAAVAGDAGPLRGTLRVSAPVDYGALVVAPALVALR
DAHPALEIELQCDDRVVDLVRDNLDVAIRLGRLADSNHRAAKLGGFERWLVASPAFVARH
GAPASPEALGALPYVMLSTVPRTTLELANAHGERASVRCRRAFASNTATACRAAVLAGGG
FGVATSFSSADDIAAGRLVRLLPGWSFPPGDIHAVFPSASHPSAKVRALIDMLKTRLAAP
C*
>GCA_000452945.1_BPMSHR5858_genomic_03075_1 group_4812
MSNSEMENSLELDKKGRPYGGIAPEARAAERRDALIRAGTRIFGTVGFRRATVRAICQEA
KLNDRYFYAAFDGTEALLRCTYEHHAEQLRASVKRAVDALGGSRDGLEARVDAGLAAFFA
FLRDACAARVLLLEVMGVSAETDATYQQNLREFGRMIISLAGGVSEDADTRADQRIIGIA
LVGALTNAGAAWLLTGYRDPEEKMVRNCRKVLLGTLQFLSGR*
>GCA_000452945.1_BPMSHR5858_genomic_03076_1 sadH_2
MRNFSGKVAAITGAGSGMGRSLAVELARRGCHLALGDVNERGLAQTAAICAAHGVKITSQ
RLDVASRDAVFDWARRTRDAHGKVNLVFNNAGVSLAAPAETARIEDFEWIMGINFWGVVH
GTQAFLPYLRESGDGHVINTSSLFGLVAMPTQSAYNATKFAVRGFTEALRMELELEGAPV
SVTCVHPGGVATNIATASRIDSSIAALTGQDIETHRRAANRLINVTTPEAAARQIIAGVE
RNARRVLVGADARRIDKLARLFGSAYQVFVLRFVRKSRERNLARQPAAARPTPSSSISKD
PA*
>GCA_000452945.1_BPMSHR5858_genomic_03077_1 ethA
MSSTTDWNAARALDERAPQAAAGGHARAKTDFDVLIVGAGLSGIGAAYYLRERCPDATYA
ILEGRETMGGTWDLFRYPGVRSDSDMFTLGFSFRPWHSDKAISDGQTILDYIRDTARAYG
IDKTIRFGHKVTAASWDSTSARWTVRVQRATGDARHDEAVYTCRFLYMCSGYYDYEEGHA
PSWPDMDAYQGRIVHPQHWPSDLDYKDKRVVVIGSGATAVTLVPSMAADARHVTMLQRSP
TYIVSLPARDAIANALRRWLPSGLAHRLVRLKNVLLTMYFYNLARRRPDATKKFIVRAAG
KQLGPQFDVNKHLTPRYKPWDQRVCLVPNGDLFKAIRAGRASIATDEIERFTPGGLRLKS
GQRLEADVIVTATGLKLKMLGGAAVTVDGKPVDLSQTVSYKGMMYSGVPNLASSFGYTNA
SWTLKAELIAQYVCRLLEHMRTHGYDVCVPRLAPGSMELEPVVDLTSGYIQRAAGVLPKQ
GTKKPWKSYQNYARDFATFRFGPLADGAMQFERRRPALAERDDTRQPAYEGR*
>GCA_000452945.1_BPMSHR5858_genomic_03078_1 rutD_1
MIAKLFHFLVYAFVALFAFTQYTAYRIRKMFPPEGKFVDVGADRLHYVEYGEGPPIVFVH
GLCGQLRNFAYLDLQRLAKSHRVILVDRPGSGRSTRGPRSSANVYAQARTIAMFIATLGL
DKPVVVGHSLGGAISLALALNHPQSVSRIALIAPLTHTETEPPGPFRGLALRSSLVRRFV
SLTLGLPVAMLQNRKAVELVFAPEAVPHDFGVKGGGLLGLQPHAFYSASSDLVAAPEDLP
DMESRYASLSVPVDVLYGRGDEILNWQRHGEALKKRLDAVNLTVVDGGHMLPVTQPAQTT
DWLLDVAGKPAAAEPEHTAESVES*
>GCA_000452945.1_BPMSHR5858_genomic_03079_1 cph2
MRKRTTSFATRFLQPWTQSLMSPTGVIVCGMLLLTLIWLFCARVLYQSHRDAYLHAVDNA
HNLVLLLERDIARNVELYDLSLQAVVDGVNDPRIMALDPAIRAKVLFDRAATGKYLGTIS
VMNEHGDIVLDSHFPHPPAVANFSYRDYFVYQRDHPKGGLYIGEPYAARLRHGAPTIALS
RRITRADGSFGGVVVGTLGIDYFRSLLDGLSVGPDGTAAVFETNGLMITRLPFDPKMVGR
SIADSALYAHVREHDAGVFTGVASIDGVRRLYVYKRLQGLPIVVNVSPAERHVFMQWRIR
AQRLGVLMLVFGIAIVGGTALLARELRWRRHAEMRLQRLARTDALTGLGNRRAFDENLRS
EWARALRTGRPLSLLFVDIDQFKDYNDHYGHQAGDDVLREVGGCLAVNVRRAADDVARYG
GEEFVITLPDTDAKSAAAIAEYIRRAVYDLDIEHVRSPYARVTVSIGLVTSHEHIAHSDA
TLVKMADAALYQAKSTGRNRVCGAQHA*
>GCA_000452945.1_BPMSHR5858_genomic_03080_1 group_7490
MRVGRPVPALPQPVCDYCGAKALLARFGDDAYPYREDHGELWICPPCDAWIGVFPRSRRH
VPLGRLANAELRHAKSELHAALEPLVAAKMRRDGCNAFEARAKGIRWLATQLGLDAASST
IHTLDLDACRNALRLVEQFMSRKSPPAHS*
>GCA_000452945.1_BPMSHR5858_genomic_03081_1 group_3397
MQRTQHVNRLFSKRFALSPLPLAIAVALSPLLAHAAPDWVPTRTDAFLIARAPAAAASQT
LAKTAPSYALNMIGTPELADTNVTPLELSQPLRVTIVLKSRNEAQLDSLVREVNQPGSAN
YRKYLTPEQFKARFAPTDAQVRAVVAHLKANGFGDITVSANNKLIFAQGNASNAEHGFHT
TLKRFSYRGKAVYANDSAALVPASLSPVVESVLGLQSAAVPHRLIHRGTPADARIQTDAI
TKNATASGTQTGHQPTDFAQIYNASGLPAATNTTVGIITWGDMTQTIADLNTFTRNAGLP
NVNTAVVAGSSGTLADDGDPGEWDLDSQTIIGTSGGVKQLIFYAAVNGDSNDSGLTNATL
TAAYNKAVTDNVAKVINVSLGEDEAAANSDGSLAANDAVFKQAVAQGQIFSVSSGDAGVY
QWSTSPYGAPGYVGTYSGGKVTTKINLAKYSVSSPASSPYVVAVGGTTLSTSGTTTWAGE
TVWNEGLAYADVSSSGQPLDNAVRLWATGGGVSGYEAAPSWQTAALGSSVTKRVVPDVAF
DAAQSTGAYLVINGQPNQLVGGTSLASPIFVGGWARVESANGNGLGLPTSAFYQGLPGNP
SLVHDVTSGNNGYNSYGYSAKSGWDDDTGFGSLDFAKVSASSVK*
>GCA_000452945.1_BPMSHR5858_genomic_03083_1 group_7491
MQSTLDTVANAQERPPRSATPFSIGLAGIAAGLLTLWASRGAAALTGTERSLLACAAIIA
TIGAYELFVARVHLRPSAGLANRALRPLSVARVSLRLAALASVYAGIALVYWLLPEYHGA
FYQPFWSLVGMLAPYVALAAPFYFGWMDRRMRDVDDAYLCWGRWLLRGQRPADWRPIREM
LAGWAVKAFFLPLMIVYLSTDADHIGASFASAASAPFSLATFRFMYDLSYTMDLMFGALG
YLCTFRLLDSHVRSTEPTTLGWLAALICYQPFWSLIGNNYIHYEGSMFWDNWLLSVPAIR
FAWGVVIVALLLCYALSTISFGLRFSNLTNRGIITSGPYRFTKHPAYLAKNLSYWMISVP
FVEPLGWRLAIMHSAALVAVNLLYWLRAKTEERHLMRDPDYRAYAEWIARHGMFARLGRM
LGRERRA*
>GCA_000452945.1_BPMSHR5858_genomic_03084_1 group_6863
MPARKTRRFDPQFDSPPDHYAPMPRLSKLHGLWAATFVIAVASDVLSGAIRYYTSLAGVA
ALGYLPKALMVACLGLAIVQRPKASHVLVAAYLAAQSCVSLANGVSLSAVGFWIWTIAPM
LFAIAMPPQALDALESPRMLGAFVALALLCIGGVLLNYFVKLPWVGGSVDIGGVSVQLAK
SSYVGTSSRLPGFGRSSATTGLMIGLLTTWIFPRLRSRLALAALLALAAAGIWATTNKTT
LVALALVVALHCLLRTPSLRTVCIWTSAVTIALPIAGWIVTLASTQDVGSSGSLSSMQDR
FINTWPLLIEGLLREHLIWFGIGPGGFGSAVGYYTADFGFNVGYADNMALYTVANFGLIG
AALIVAAFARLILALPANDRPVWLMLCFLLVSGVTTDICETIGCLLFLGLTLRSINLHAP
RRARVGSARIAFERLRYGAGVIAPNAR*
>GCA_000452945.1_BPMSHR5858_genomic_03086_1 etk
MDLKAQPIPAAPHAKRNDGLASLGWSVVRHRRLFATIVGVFFALGVLYALLATPQYRAEA
LLRIQTKPGASISALSDVSGTISNGPSANDESEVLTSRAIVGAAIEQIGANLEIRTLDHF
PLIGRLLASGHANDDKLASPLLGLSGFAWGGEKLKLGEFSLPDAALGEKFRLVAGDDGQW
SLYDKKDRLLARGALGKSVPFAVALDGERAQPGRIRVDTLRARSGIAFSIVKEPTQLVYD
DVMKKMKTVVSNRDSTLEEPSMMKLVYQADTPQRVQEMVNAIVRVYLDQDIKYRAEKAQR
NLDSLHARLPGLKKDLEKAEDALNRYRTTTGTIDVDQQGIALINRLNSLSEHQTVLQLAL
DNVKARFLPGSTTYQTVQTQLDQVKKEIQQTTTAANKLPTAQREFVRLSRQVSVATQLYT
SVLTNAQQLEIAVASTTPGVSVVDWANKPYKKAWPQRGIVVLGALLGGLFAGLAAAYLLA
RHRDELSGPQSIADVSDIPCVAVVAPSAESLALGDDRPGNARARVKPLAAQRPNDPGIEA
LRALRTSLRAALAHEGRGGGKVLVFAGPTARVGGSFVASNLAYLFADSNASVLFVDADMR
GGSHNPLGVGRNGGAIGLANVLEGGQPLEKAIVKLGKGKLSVMTPGTLTGSNPGELFERA
EFPQLLAALRTRYDFVIVDAPPILPYSDTLSIAAQDCDALLLVSRGRTTRASELETALQR
LDSVGAKIAGHVFNAYVAPPRPPRTSLRDDWREIVARWMKPRRPARPAQAKQTFRLAKGM
AAKASRN*
>GCA_000452945.1_BPMSHR5858_genomic_03087_1 group_4184
MKRLAAVAMSVTQSISIPIHYPAATAALLLLLLTGCGGGGDQSKVNAAASPANNLVVPAP
GTASPGTPAPAPGAPAPAETASVLPFFGVNGHYVDGGVYASVPLATQASHLAGLGMNVYR
QDVYIPDHVDTLASTVIPGLGSGITVLPMIQAHPWADPSLNGQPPTEASAYAYAYKLAAY
AAKKLAGIPMVEFGNEYDIDSHNAPIQGDGINVSDYDNSTFPIWRGALRGSLDGWRSVDT
NRTTKLIANATSGALHFGFLDGLMTGTQPDGTTGHPKITPDVIQWHWYSNGGDFENALGK
TGRYNVLARLKDRYNLPIVVTEIGVNTDNSDTQIAAYIAKTIPELVAAKAAYNVIGFNWY
ELYDDRSGAYGLLTNSAQEKPRYGLMRAAIAGAVPN*
>GCA_000452945.1_BPMSHR5858_genomic_03088_1 group_1334
MKGIRLVTLAATGLFVLMSSVAHAAGGGHGGGMGGMGGAGGNAGGMSAGHMSRAGMTNTN
GFDSGDRDKGIARASDRASTHANLADTQAGRTRSHATTHTTHQHHLGKHLAVGKAS*
>GCA_000452945.1_BPMSHR5858_genomic_03089_1 frmR
MSHTIREKQKLLNRVRRIKGQVEAIERALEEECGCGEVLQRITSCRGAMNGLLAVVLEDH
IRTHLVDADTGDNHEGSASEQLIDVVHSYFR*
>GCA_000452945.1_BPMSHR5858_genomic_03090_1 zitB
MSEFENVAFGAGHDHVFLGAAHGQNERRTWTVIALCTAMMVAEIVGGTLFGSLALVADGL
HMSTHAGAMLIAALAYTYARRHASDARFVFGTGKLGDLAGFTSAIVLAMIALLIAYEAIA
RFLSPVPIHFDEAIPIAVLGLLVNLASVWLLSGDHHGHGHHHGHAAHHAHGDEAQTISTG
AGVFAVSIFEDGVPPVFRISPAAKQACIAGARATITTIRADGSHQTFDMAERGDRLESID
AIPEPHAFTAIVRLHGSEHSLTFVEHEHAHHHGHGGSKDVTARDHNIRSAYIHVIADAAV
SVLAIVGLVLARAFGWMWMDPLAGIVGALVIANWSYGLMRDTSRILLDVSPDTRLADNVR
RSIESGGDRVTDLHVWRLGPGHMSAVVSVATGDPARDSRFYHQLLKRFTSLSHVTVEVHC
RA*
>GCA_000452945.1_BPMSHR5858_genomic_03091_1 group_7493
MTISPRDEQNRSVDLWFAYKVPKLTKDTDSDSATGYEYVYYDREIGSVQKSPNRMNNPKG
ALFYTLDSIFGDPGSTTGWILYNDEMPADADRSNNATLGHTKGVIAFDIASNSALWLLHS
WPKYTSPSAPGVPTPLYGQTFLCVSLDLETAGKIAAQMALHQQPQVYLPRTAGLDHASPL
YTLTQPLNASAPGDSDSLDFKTRGGAPFKVIAKNRKWGKDFWNDLVGPTLKADMDVETWI
RGKIPPILDSDGVHKTYDIKFIDLRKLGAPWAWPETQDHAKWGITTTDDWVCVGDINRMV
TQEKRGGGTIAFQDPQLWKALSQTDLIVPPPGKTEAQARAMIHRTHQPAD*
>GCA_000452945.1_BPMSHR5858_genomic_03092_1 group_4815
MNAHLPAGALVPLVTRHTDIAIAAPLRGTTTLPPVAWERIGQRAPVRIAPGARAPDDPLP
RADIVVITWTSAEWFALDHVFVDSAHTGDYNDYAWKQAWLPYTRGASPYAADAKSGALWG
LFQMVRIVDRSGRPWNVLLFKSNAHLAHSPWLDGLSAMLRCIVEDARPDRIYTIGTAGGA
RHDQRLGDTVLANAALLELQRPQNATSPEGGNMYRCPTWYPSTALVGEVESQLLFRMSEI
VTPQSLAALFDELKARHPDDPGLGELTLADLLNDAIRPECLRTPAIRPLKDAPLLTTDFY
YIAEGNDAHAYSCLEMDDAIIAQQANRLGVRFACVRNISDPIVRRRTDRGTPISEAVRAD
WSGLIYSTFGLQTSYNGALATWATIAGEGSAAYNPSREHPPADEADPLEVQLAFQVRSCG
TCSFFWPADPKKRTYGPYTAFDFDTTVPYPASANGRSGAVRWLSGRTRPPAFPNGEVIDG
CRKAPIMTIGINPNLTAFLPGQTGAAWCYPDFSSDGDTDAWAKYAWYYRYRTVYQEKLDL
DFVRRFMLPERRVIAARGGEVTGAARIDDNPAWSITVRYDGDAADTTIPIPGEPGDFPYV
LLFDTYRPHNRFAAGDVLAARVSVPEGIQVEVLQQPQSYYLQMVPVLERFERTLRDGGHP
GASLHVGEDVCQLDMVACASPHWKPGFLGGSDASVTAIVDNCVSRNAWAIKQMVQTRPAL
LYIVSESSWNMFHAALGAHVRRDPPLSSHPADKDYTLLKETTDPEHPAYVEFDVTIDGMR
YAHRTRLVITPHFSYNSFFLQQYRMSTQDWHAFGAAQPGCVAALTPQNGFTLVLPTQAYP
DDYVAIQLPADASAANAARAWLASQFPDAARTLGTYFVDAHASMASVLDELYANHTLTWH
DTDSGGYLSRNEGSCRFCVNRHWQFPNECRYDKTHEPPPPAGFLAKVARHLVATGKPAAE
NATTGAPL*
>GCA_000452945.1_BPMSHR5858_genomic_03093_1 group_7494
MSTTDVSQDMRVAPDSVRVWRGFRLPSLGLDQFLGKLGTVFVPATVKMQIDAGLRAYAPT
VPAGLPGKPATVPDETAILFWASPETYWNGFTRLAVRTYTLTHGGVYLTQDNQSRADFPV
RFDGALGVDQPVYLYDAPADWMHGAIRHLVAARPPAVDPASFRSNVAKALGDIQRSVPLA
GALACAGNDYVVYWELGPVAAGAAQAPSGVPRLQTALTDWSETFASAPTFLPIGLWDEWA
GMDVRSGSSFNMQFAREASR*
>GCA_000452945.1_BPMSHR5858_genomic_03094_1 group_4816
MNTPQESFPVAPNAVYVWRGFRSPTLDYEQFAQFLGSVFVPACVLLQPPVGLRAYLPTMM
PQAGKPAAVPDQTALMFWATPASHDRAMKALAVRIYQNLHGDVYDMTRSKLPEVPVALDP
HADALVAEQPYFLIDRAADWMLGSAKHFVGARRANLRAADFLAAVHVWATSFQIEPPNGV
DGALVCCGEDYVAAWIHGTYPQPRPCPALDQLAALTTPVLRMAPRPLALPAGLWDDWSGL
DLTRDTCINLQFPRPRTSRAAPREPEA*
>GCA_000452945.1_BPMSHR5858_genomic_03095_1 group_7495
MIPIEHFEVPHLALWKSCVAEVLAHALGVEHRPPAGIDTDHPLMRATDRYCRAMLENRPL
GAPAAGSDDEQAVQTYLSYLHHRRAHAHIAGDAQIERDIERQTQQYKFGNPLWQQMYIQY
FKYYWQYPYHKGGEPRYRSWQAADAGNGDLRYGVIEWKLSARARVAIVGDIGTGTDVAAA
VLVAALKFAPDAILHLGDVYFSGTRFETDHRLIGLVREVLRGNGRRVPFFTVPGNHEYFT
GAVSFLHALDSGELVDTPAQRQQASYFCLRTADDGWQFLGLDTGYHGHYLNVAAAAQQAT
LERLHIGRVETAGEGASPHWPTDRNPYFRHASLADLPVRDTTSPVDQVTVRTDEAVWHLD
KLASFPGRSILLSHHQLYSALDVCGIAQHRDASGTPDPADFNREWINTGLWRQFGPTFGD
RVAAWIWGHEHNLGIFADAYRPPDWPTEGDEAVRVFKTLPKGRCAGHSAIPVQAGEAPYA
QKYPVALKQADLQLDLTDGWYNRGFQILELAGAGKPARLSYFQVAGADPTPLPLFVEPIS
*
>GCA_000452945.1_BPMSHR5858_genomic_03096_1 group_2456
MSSLYSIRPDSGSEKVALERVWDGDSPLASGYTSFVALPRAGGPDLIAYNKATQKTDVYA
LGDAAPWISPVDSKIELAGGPWDSVASFVLGNVRYLLTYRADDGTFGFFALGADRSASPP
YKFALPRNTPTRGFTNVAPFTSLGQQYILGYDFDTGRVANFSVAVTPSSAGGAPPLLALN
VWYHEWAKGWTQFSFFQLGGANFFFKINTAKLNVNIDHIQDNPALGTVEVGSYLQDQLPD
ALKITNAACVPWAHGEPTLVTYIAASGATAVYRIHADCLGWTRIGAATTSAGASHVMPYR
VGDQSYALFYQG*
>GCA_000452945.1_BPMSHR5858_genomic_03097_1 group_2757
MNPNTVPPAARDVAWIKHALQTAIELEYSTLPLYLSAMFSLEVQNYTAYNAIRSIAMEEM
VHMAIAANLLAALGGSPQFKPIQIAYPTTGLPGGAEPDLHVGLARYSKPQLKNFLRIETP
GFLLRQLGRDEGYPTIAAFYESIRSAIARNANAVRAAVQAGGPANQVGDDIGFTTVTYKP
GVDPVESFDAGIGEILQQGEGSGRGDVFAGNPFEDEESHYARFAELYYGARYQAPSPPVD
FNPKNEPLFFGGPAIAAPVVVNTLAVPRDGYARILALDPERDTVESDLNAFDSGFSKILS
TLDAVWNGPSSTSWKTLGAAVHGMVDLRVLSCFNIMRHAVPAAAVSQLAELYPDEIGFLR
TYTDLSKPVFYGPRFVNVNGKQAS*
>GCA_000452945.1_BPMSHR5858_genomic_03098_1 mtnB
MSTASAAGIDPAEWKRRCDLAACYRLVAMHGWDDLIFTHISARVPGSDHHFLINPYGMMF
EEITASSLVKVDPAGNKVDDSPYPVNRAGFVIHSAVHAAREDVQCVLHTHTRAGVAVSAQ
RGGVRAVSQQSTFVLPSLAYHDYEGVALRDDEKPRLQADLGGATFLMLRNHGLLTVGPTI
ADAFLAMYLFETACQIQLAAQSGGELIEIPPEIVATSAEAAAVQTGGLGGAFVWPALLRK
LERRGGGYQC*
>GCA_000452945.1_BPMSHR5858_genomic_03099_1 group_3398
VRAHPGARILIVGQAPGARVHASGIPWDDASGKRLRAWLNVDDATFYDESLFAIVPMGFC
YPGRGASGDNPPRPECAPLWLDKLLAELPDIRLTLLIGQYAQRRFLGARRKSTLTETVRA
WPEYAPEFIPLPHPSPRNQGWLKQHPWFDADVLPALRDRVAELLPTPPGSRRTKR*
>GCA_000452945.1_BPMSHR5858_genomic_03100_1 pepX_2
MRFHRTEPRRAWIAVLAAAAMLAACGGDDGAGVPGASALAQVQQEGAAPSAAGAPQLAAR
FSPSGVPYASLSSGGRYRPVIQNGQVQPSLSGGTIAEEAWVETPVDSDGDGAKDRIHVRI
VRPSETASGARTPVIVLASPYYNGLADSPNHNVDVELDGTPHPAASASARIMAAAPQTRI
WQQLDAAAAGRSWIEGYFVPRGFTVVYADSLGTAGSDGCPTILTRDESIAMASVIRWLGR
GAAAKDANGKPLVATWSTGHVGMYGVSYDGTLPKMVASLRTRGLDAIVPVAGLTDMYGYY
RSGGLVRAPDGYQGEDVDVYIKALLTNPHPERCTHLIDEALQKEDRKTGDYSAFWAAREI
PSALAVAPALVAQGLADDNVRTDQSTSWYLAMRRQGVPTQLWLHRAHHTDPTRVPAMADA
WTGQVNRWFTRYLLGYDNGVERSPGSVIEQADGTLLKEASWPARGASSVTYFAGGDGAGT
GTLLTRPTGGPLAKFTDDARIMALALANANTGEHRSRFETAPVASATRLSGTATARVRLT
FSATANVTALLIDRAPDGSATIITRAWTDPRNRLSSWFSEPVLPGMPYDLRLAFMPRDYR
LEAGHRLGLVVLSSDNEATLRPTPGTELTLDPAGTSVTVPLLPA*
>GCA_000452945.1_BPMSHR5858_genomic_03126_1 group_686
MTTFNVMMPVAGVLALPFLPLLNELVRRSDVDALPIGEGPCVDYPRRAAQWLDALRASAR
GAPPAPDDAPPPWQALGLRVEHVDALRLARGDRRDGVLYADHTISLGAGAHAAYAFAEER
LEMRAGATLGGFAYAPNVDIDNTLVRGAVVGQAVRVHGCGGFANLHGTPIVFGRLAHAPG
RDAALTPRRAMSLSGRLAGVPHRVLGGRYLVARDIRLPSHTILHGTLIVDGRLTLGEGCV
LIGSVKAHSVVLECDALLNGAVFARDDVLLDEASCIDGVVSAGGLLRLTGARIGAADHPV
SACARDVSVVGHACVHGDLVAWRSGWYHAS*
>GCA_000452945.1_BPMSHR5858_genomic_03135_1 group_1870
MSATRSIRCLRGAVGMKLTGEPAARAHCHCTACRDFHGVSMLSATAWPAERAIVVEGDVA
SFAHPARQMSRTFCRACGETVFGTNRLGMRVVPNAIVARAAGGALPDGLRPTMHLFYRHR
IVDVRDALPKYLDGWDGPTDAA*
>GCA_000452945.1_BPMSHR5858_genomic_03156_1 birA
MNVATPPSMPATGGPLIDRARVDAHLAPAAREWSIEIVDTTGSTNADLGARLKALPRRRD
ALAAPIARVAYEQTAGRGRQGRPWFAKPGDALLCSVACVLPRPVGALAGLSLAVGAALAE
AFAARPAATGDAPRGDADGARRIALKWPNDLLVATERDGETAIVGKLAGVLIETVWNTID
ATAVVIGFGVNVRTADAAAAEVDALRARDATLAGGLPPVALAAVRAGATLTDTFAAALNA
LAVALPAFAADGLAPFAARWHALHAYAGREVVLLERGAELARGVAVGIDETGQLLLDTPA
GRQAIAAGDVSLRTPGAAR*

>GCA_000452945.1_BPMSHR5858_genomic_03157_1 coaX
MSGMCLLIDAGNSRIKWALADTARHFVTSGAFEHASDAPDWSTLPAPCGAWISNVAGDAA
AARIDALIEARWPALPRTVVRASAAQCGVTNGYAEPARLGSDRWAGLIGAHAAFADEHLL
IATFGTATTLEALRADGHFAGGLIAPGWALMMRSLGMHTAQLPTVSIDAATNLLDELAEN
DAHAPFAIDTPHALSAGCLQAQAGLIERAWRDLEKAWQAPVRLVLSGGAADAIVRALTVP
HTRHDTLVLTGLALIAHSA*
>GCA_000452945.1_BPMSHR5858_genomic_03170_1 group_7501
MITFKSKAAQDLDVLKDFAVYVLGLVGKQLGERGVITSDELDGAIAKLEDAVSQAKQARA
EHAGHFHEDEPDHAHHEVPPSLAQRVAPFLTMLREAKAQNADIHWGF*
>GCA_000452945.1_BPMSHR5858_genomic_03207_1 dapA_1
MQSRFEGIWLPIVTPFHARETGTVDHAALARLARHYSDEGIAGFVAGATTGEGALLNAGE
QEAIFATLRDAAPARPIVLGITASATHAAAERARALAALRPDGLLVTPPVYVRPTQAGVQ
RHIEAVVEAADLPVLVYNIPYRTGVNVELDTLKALSRDARVAGVKECGGTMERMLRLVHE
TPLRILSGDDNQNFAALCAGAHGAIASAAHVLPAWHVRVYALLREGRLDDARRIAVALQP
LVADLFAEPNPAPVKALLAAQGWCDNALRLPFLPVEEALAARLVAHWESLRQSEAAGA*
>GCA_000452945.1_BPMSHR5858_genomic_03208_1 group_2040
MKKREENSLSVSHSRRAFLVWVPASLLVTACGGDNGSSAPASIKKAVAALTGPGGATWTG
MGKGGILIQSPYSGAGNFELLAGDEYGMLHCLRRTNDNGAQWMVAVGDGMGSKNSVSGYC
HTYSTNYTNIEAAVVDDGAVKTYFRSNSTPWSAHKTIPIGDAQGFPAYLQSNRNGAQQLE
LVVGMRRGGMVHYWRDDKDGFTWKQSSIFGSGQVKGVSVMQGNYIGPDNNANLELVAWVD
DRLESWCCEGRAWRKICNIADGGVGGAPALLQSNYGTKGNFEVLVPLATGGIAHYWRNND
AEGIPWSFTGTIGTGAYVAAGLLQSQNGVKGDFEAIGLRASGKADVFRRGDTLVWENSAT
VSPFSEGAAADVGVSSHIVDVNVTGINSVLLKNGSVLMFGYYKGGSTNKTIPACIWNPVN
DQITAIPSFRNNFCAGQTAMPDGKVLIVGGHIGDTLKDVVVFDPDNHTATLVATMTKGRW
YPSTATLPDGQVFIISGTETAGWNTSVNDTWQTYANNALTAPEDVISPFSPYYPKSQTQI
DLYPFLFVLPDGKLLVHARNTTRFFDIGTRSWSATLYKTVSDNSRTYPFMGGTAVLPLRP
SENYRVKVVVAGGADKSAQIAIGDDSQYDNSVPGMTSCEMLDLGDAAPAWKAIAPLNEGR
VMCDLVTLPDGKLFIVGGNKTGKADYGRGPTYRPELYDPQTNTWTLLASTRIARGYHATA
LLLPDGRIAITGKDGDYQGSGLQYAETRVEIFSPPYLFKGPRPAIQSAPASINHGGSFTL
GLSSGTSPEDIGSIVIVACGSATHQINFSHRIVELVFAVSGGTLTVNAPPNANIAPPGYY
MMFVLSKLGVPSVSSIVHVAAGSAQAASATPEALTRQPRVATVSIEQAGDAPRCAPLTVA
EKQSINGMG*
>GCA_000452945.1_BPMSHR5858_genomic_03232_1 group_1486
MNTTRLTVKTRLRILVAIVVLVLAAAGGIGLAGVARVGDALTHVFEDRAKTLQRISSVDE
LVTQARYAVGDAVLDPSAAKTEAVAAAAARDVAAIDRLLADFRATPRAGAEREQAERLAA
DWATLRDKGIRPAVDLLKANNLSEAQWVVTQTLDPVAQRVKGEAAQLRQNQLEAAQRAYD
RARGVARLVQWAVAGCVLAGVAAVAILCAAMARALARQLGGEPDYAASVANEIASGNLAL
DLHAADAAAGSVLHAMHAMRARLAATIGTLQRAADAIAHATSDIAHGNHDLSRRTDEHAA
GLQQTASSMEQLAATVRTNADHARQASALALAASTQAEQGDAAARDAIARMEALSVRSQQ
IRSISSTIEGIAFQTNLLALNAAVEAARAGQAGRGFAVVAQEVRGLAHSSAQAAKEISAL
IADVTGEVELGADTVKAAGSTIVGMLASVRQVATLVDEISHASDEQRSGIDQVSRAVAQM
DEMTQHNALLVQKAAGAAGALADEAAALREAAAIFRVVA*
>GCA_000452945.1_BPMSHR5858_genomic_03233_1 degA
MGTTIRDVAQAANVSIGTVSRALKNQPGLSEATRARIVEIAHRMNYDPTQLRPRIKRLTF
LLHRQHNNFATTPFFSHVLHGVEDACRERGIVPSLLTTGPTDDVIRQMRPHAPDAIAVAG
FMEPETLEALAATGRPLVLIDIRAPGLRSINVDNAGGAALAMHHLFASGRRRVAFIGGSL
AHYSIAQRALGYRRAFFDAGMLFDPSLELSIDTSLGPDAGAANAMRQLLDAGGALPDAVF
AYNDAAALAAMRVCVERGVRVPEDIAFVGFDDIPAAAHAAPPLTTLAIDKEALGRQGVEL
LLADSPAQAELSLPVQLVARASSAGAARAPRTLTSVLGS*
>GCA_000452945.1_BPMSHR5858_genomic_03235_1 scrK
MSAGFPQFVSAGDILTDMVRTGGAHWTSRPGGAGWNVARAVARLGAPSACAGAVGRDCFS
DELWRESEAAGLDLRFLQRVERAPLLAIVHETQPPAYFFIGENSADLAFDPQRLPDGWRE
HVKWAHFGCISLVREPLASTLVALAAELREAGVKISFDPNYRNLMTDSYRPTLARMAELA
DLIKVSDEDLRHLFAGDDEASAIARLRALNPRAALLVTRGPDTATLYAGGETIDARPPRI
EVADTVGAGDATIGGLLFSLMTVPARGWRAHLSFALAAGAAACRHTGAHAPTLDEVVALV
AA*
>GCA_000452945.1_BPMSHR5858_genomic_03236_1 group_4838
MDGGNTYTKGLYTAKAHTRKTDNGRFQGYVILSRDEGGAPENTLYEVEATSANEAEAFDE
AKALAHRILGDLEL*
>GCA_000452945.1_BPMSHR5858_genomic_03237_1 group_7504
MLRWLLTTFIAVMVLTRAWPWLAKLGVGRLPGDVTLRLGSRVYPFPFMSTLVIMGIVSVI
ARLW*
>GCA_000452945.1_BPMSHR5858_genomic_03238_1 group_1307
MNPKAHEPASLPVPGPDALAQSDALAASLRAEIAAAGGWIPFSRYMERVLYAPGLGYYSG
GAQKFGRRGDDGSDFVTAPELSPLFAQTLARPVAQALAASGTRRVMEFGAGTGQLAAGLL
NALAALGVELDEYAIVDLSGELRARQRETLDEQASGAAARVRWLDALPERFEGVIVGNEV
LDAMPVQLVAKHAHGWCERGVSLGDAGAFAFADRPLARAEDAARLAALDADEGYVTETHD
AAAAFVGTVCAMLARGAALFIDYGFPRHEYYHRQRAQGTLMCHYRHRAHGDPFVYPGLQD
ITAHVEFSAVYEAGVGAGAELLGYTSQARFLLNAGITDVLAEIDPSDAQRFLPAANAVQK
LISEAEMGELFKVIAFSRGIDGTLDAFARGDRSHTL*
>GCA_000452945.1_BPMSHR5858_genomic_03241_1 ttcA
MNAPHTPHLNEAEAAAAVEANAAELGRRALTRREQKEAYENNKLFKRLVRQVGQAIGDYN
MIEHGDKVMVCLSGGKDSYALLDILLRLRERAPIDFDIVAVNLDQKQPGFPEHVLPEYLT
KIGVPFHIENQDTYSIVKRLVPEGKTTCSLCSRLRRGILYRVAGELGATKIALGHHRDDI
VQTLLLNMFYGGKLKGMPPKLQSDDGKNIVIRPLAYAKETDLEKYAELREFPIIPCNLCG
SQPNLKRAEMKALIRDWDKRFPGRVDNMFNALANVVPSHLMDARLFPFAGLRATGEADPN
GDIAFDEDPCGTDASAPGGAKSVSIVQFDDL*
>GCA_000452945.1_BPMSHR5858_genomic_03243_1 glmS_1
MCGIVGAVAQRNIVPVLIEGLRRLEYRGYDSCGVAVLDAGAPAPGTPKRARSVARVADLD
AQVRESHLEGETGIAHTRWATHGAPVTHNAHPIFSSDALALVHNGIIENFEPLREALRAK
GYAFVSQTDTEVIAHLIHSLYHGNLFAAVQQAVKQLHGAYAIAVTHKDEPHTVVGARQGS
PLVVGFGERENFLASDALALAGSTDRFTFLEEGDVCELKLEGVTVVDRDGARAARDVRVV
AAYGGAVELGPYRHFMQKEIFEQPRAIADTIAQTDAFDPPLFGERASSVFAGIDSLLILA
CGTSYYSGLTAKYWLESIAKIPTQVEIASEYRYRESVPNPRALVVVISQSGETADTLAAL
KHAQALGHAHTLAICNVATSAMVRQTELQFLTHAGTEIGVASTKAFTTQLVALFVLAATL
GKLRGHVDAAREADYLKQLRHLPAALNSVLALEPQIIAWSDEFARKENALFLGRGLHYPI
SLEGALKLKEISYIHAEAYPAGELKHGPLALVTEAMPVVTVAPNDTLLEKLKSNMQEVRA
RGGELYVFADADTHIVNDEGLHVIRMPEHYGPLSPILHVVPLQLLAYHTACARGTDVDKP
RNLAKSVTVE*
>GCA_000452945.1_BPMSHR5858_genomic_03244_1 group_4839
MQRLEHVLRRVKAGTGAPIDFSGTWKNELGSTMRIEQSGDSVSGTYESAVSENGGATSGQ
LSGYVDGDLIAFVVHWDQFQAITAWVGQGGPGASSDRINTLWQMTQQVEAGEEWASINAG
ADIFVKT*
>GCA_000452945.1_BPMSHR5858_genomic_03245_1 group_3410
MKQWIAALMTLAAAAAPAFAAEAAAGPNTPAPPRDARPPAGDALTEAVVRQVDLATGMIT
LKHGALDNVGMPPMTMTFKAKDIAMAKRVRAGDSVRVRVENVDGVMTVVKLEK*
>GCA_000452945.1_BPMSHR5858_genomic_03246_1 cusA
MIAHVIRWSIRNRLLVLLATALVAAWGVVSLNRTPLDALPDLSDTQVIVKASYPGKAPRV
VEDQVTYPLTTTLLGVPGAKTIRAYSSFGDAFVYVLFDDRTDQYWARSRVLEYLNQVQGR
LPQGASVALGPDATGVGWVYEYALVDRSGRRDLGELRALNDWFLKFELKAVPDVAEVASV
GGMVRQYQVVLDPDRLRAFGITQAAVADALGKANSESGGSVVEMAESEYMVRASGYLRSL
DDFRNVVLRTSESGTPVLLGDVARVQIGPEMRRGIAELNGEGEVAGGVIVMRSGKNALST
IEAVKAKLAELRRSLPAGVELVTTYDRSQLIGRAVDNLKDKLIEEFVVVGLVCALFLFHL
RSAFVAILSLPLGVLAAFIVMRHQGVNANLMSLGGIAIAIGAMIDAAVVMIENAHKHLES
HEHAHPGAPLSSAARWELIAASAAEVGPALFFSLLIVTLSFVPVFALEGQEGKLFAPLAF
TKTYTIAAAAGLSVTLVPVLMGYLIRGRIPREASNPLNRLLVRLYRPLLEATLARPWRAI
AIAAAALVLTAIPMSRLGGEFMPPLDEGDLLYMPTALPGISAQKAAELLQQTDRLIKTVP
EVATVFGKSGRADTATDPAPLEMFETTIRFRPRGEWRPGMTPGRLVDELDRVVKVPGLSN
VWVPPIRNRLDMLSTGIKTPVGVKIAGPELAQIDRIAAQVEAAVKRVPGVTSALAERLNG
GRYVDVDIDRRAAARYGLSVGDVQAVVASAIGGENVGEVIAGRERFPINIRYPREVRDSL
EKLRALPIVTERGAQILLRDVAAVTIADGPPMIRSENARLSGYVYVDIRGVDLKTAVGAM
QRAVAQQVALPPGYSIAWSGQFEYLERAAATLRTVIPVTLAVIFVLLFLTFDSAADALLL
MTTVPFALVGGLWFVWALGHAVSVATAVGFIALAGVAAEFGVVMLLYLKRAYERRIAAGE
PPNEATLADAIREGAVLRVRPKAMTVAVVLAGLVPIMIGHGSGSEVMQRIAAPMVGGMVT
APLLSMFVIPAAWLLLQRRRMRGAARAASQAPVAGAAGADFSSIHTGESR*
>GCA_000452945.1_BPMSHR5858_genomic_03247_1 group_9254
MNHIRLTRAALAVLAAVALSAAGYFAGARFGAHVDVAHRAHATNTTNAVDGADAKPGRKV
LYWRDPMVPNQHFDKPGPSPFMDMPLEPVYADDGGGAPGVRIDPGLQQNLGIRYAKVRRR
DVDEGFDAVGTTQFDESRTAVVQSRVTGYVDRLYASAPLQRIAKGAPIASLFVPDWLAPQ
EEYLALKRGAMDEALLAASRARMRALSIPDSVIANLDRTGRAQTHIVLASPETGVVSELN
VRDGAMVTPGQTLAKIAGLSTLWAVIDVPEALASGVRPGMRVDATFEGDPQRRVSGAIRE
ILPGVNATTRTLQARLELDNRALALTPGMLMRARVGASHAASRLVVPSDAVIATGKRSVV
IVRTADDRLRPADVTVGRDAGDETEVLAGLEEGETVVASGQFLIDSEASLKSVLPRLEAD
ASGALAARATRPPARSSPPAAQAGGAAPVYETTGKVEKITAADITFSHQPVPALGWGAMT
MSFDKPAPAAFANVKAGDTVRFAFEASGDGYRLTKVEPVGGAR*
>GCA_000452945.1_BPMSHR5858_genomic_03248_1 group_7506
MYSMLRAPHGRRARLHARTLVAALTFANAAAAFAQSAPLTLDAALQSAADRSAAMQAAQA
SVQASAELAVKAGQLPDPMLKAGIDNLPVDGPQRFTIGQESMTMRRIGIAQEWVSGDKRR
LRTALANEVVGRERAGYLAQLADVRRQTAIAWLDAAYAKKALALQRELADEMNRELAATQ
AAYRGAKATAADALQARALLAQARDRQANARQAFDTALIGLSRWVAAPALDVAGEPPAPV
SFVPSLPPDELRRVQPTLIAADEDVEVADADVALAASERRPNWTWEIAYQQRGAPYANMV
SIGVTIPLPLDRRDRQDRDVAQKRALATKARLMYEDARRQVDADIRTQSATLESGRARLA
ELGRALLPAADERVRLADAAYRAGAGSLADAFAARRARLDARLQVLDLERDVARAWAQLD
YQIVPSAMLAAAQ*
>GCA_000452945.1_BPMSHR5858_genomic_03250_1 group_7507
MDIARIARHLSMTRWRVAAAFPRRTLGKIERAIEASHEKHVGQLRFAVEGALHASMLLRG
ITARERAIDVFSELRVWDTEHNNGVLIYLLLADHDVEIVADRGIHARVDGDEWERVCRAM
EAQFRRGRFEAGSIHGIEAVTALLARHYPSRRAPGDAWPSTPVVL*
>GCA_000452945.1_BPMSHR5858_genomic_03254_1 group_7508
MKLEVKDMSCGGCANAITQAIQTADSSARVSVDVASKVVDVGSALGIERVVAIIEAAGFH
PAVLAA*
>GCA_000452945.1_BPMSHR5858_genomic_03256_1 group_7509
MFQMLPSMTFGRRLSVWWSCMWRQTLASAPVWILGVAIVGLSISRTHSAAGRPPSGGAAA
LAVATFFVCLVVCLPIAGYMVRGGFAAHALTAPERLAFRQALMVGLTTFGWAVLAALPIS
VATMPLRHAGYPLAGQAIGWVLNVAAGLYIVLPRQARRLRLLAGEAA*
>GCA_000452945.1_BPMSHR5858_genomic_03257_1 argG_1
MTTILENLPAGQKVGIAFSGGLDTSAALHWMRIKGAVPYAYTANLGQPDEDDYDAIPKRA
IQYGAEGARLIDCRAQLVAEGIAALQCGAFHISTAGVTYFNTTPIGRAVTGTMLVAAMKE
DGVNIWGDGSTYKGNDIERFYRYGLLVNPDLKIYKPWLDQQFIDELGGRAEMSEFMRQAG
FEYKMSAEKAYSTDSNLLGATHEAKDLESLESGIKIVNPIMGVAFWRDDVKIDKEEVTIR
FEEGRPVALNGVEYKDAVALLLEANRIGGRHGLGMSDQIENRIIEAKSRGIYEAPGLALL
YIAYERLVTGIHNEDTIEQYRENGRRLGRLLYQGRWFDPQAIMLRETAQRWVARAVTGEV
TVELRRGNDYSIIGTRSPNLTYQPERLSMEKVQSMFSPRDRIGQLTMRNLDITDTRDKLR
IYSQVGLLAAGESSALPKLKEDESGN*
>GCA_000452945.1_BPMSHR5858_genomic_03258_1 garB
MEFDYDLFVIGAGSGGVRLSRVAASYGARVGIAEEERIGGTCVLRGCIPKKLLVYASHYS
HDVEDAAGYGWTFDIGLFSWPTLIEAKDREIARLSGIYVDLLNKSGVEIHTGRATLVGAH
TVDVAGRRVTARHIAVATGSRPVLPPIPGIEHAITSREALELPELPQRIAIVGGGYIAVE
FAGIFNGLGVDVDLFYRGEQILRGFDDDVRHALHGEMTKQGVAIHTRATIEAIERGGDGG
LTLRLAEGAYGPYDAVLYATGRVANGDGLGLEAVGVARDANGAIEVDAYSATTVPSIHAI
GDVTARPQLTPVATRDGMLLAANLFGGKRIAADHRYVPSAVFSQPEIATVGLTEAQARAE
LGALDIYKTSFRALRHTLSGRDEKTFMKLVVARDSQRVVGAHMVGRDAAEIIQGIAIAIR
AGATKAQFDETVGIHPTAAEEFVTLRQKEPDDA*
>GCA_000452945.1_BPMSHR5858_genomic_03263_1 group_3411
MLFQFLAVRFVHGPLFARRRRAAAARAPSALAMRPTVRARLPFIPRPAGCAIVAATAGAV
ALGALLSASGQQVGLNQAIRVSWWFS*
>GCA_000452945.1_BPMSHR5858_genomic_03282_1 flgH
MKQVRFLPQPARRARAAAVRRPGAAALAAAALALAGCAQIPREPITQQPMSAMPPMPPAM
QAPGSIYNPGYAGRPLFEDQRPRNVGDILTIVIAENINATKSSGANTNRQGNTSFDVPTA
GFLGGLFNKANLSAQGANKFAATGGASAANTFNGTITVTVTNVLPNGNLVVSGEKQMLIN
QGNEFVRFSGIVNPNTISGQNSVYSTQVADARIEYSAKGYINEAETMGWLQRFFLNIAPW
*
>GCA_000452945.1_BPMSHR5858_genomic_03283_1 flgG
VNRSLYIAATGMNAQQAQMDVISNNLANVSTNGFKGSRAVFEDLLYQTVRQPGANSTQQT
ELPSGLQLGTGVQQVATERLYTQGSLQQTGNSKDVAINGQGFFQVLMPDGTNAYTRDGSF
QTNAQGQLVTSSGYQVIPAITIPQNANSLTIGSDGVVSVTQPGSNNAVQIGALQIATFIN
PAGLDAKGENLFAETTSSGAPNVSQPGLNGAGTLKQGYVESSNVNVVQELVNMIQTQRAY
EINSKAVTTSDQMLQTVTQMSR*
>GCA_000452945.1_BPMSHR5858_genomic_03295_1 group_5796
MQGQTIAMLGGTGFIGSRLVNVLVGAGAHVRIGARRREHARHLATLPVDIVELSAFDVRE
LARFVAGAQAAVNLVGVLRGGHGKPYGEGFERLHVALPAALAAACIEARVPRMLHVSALG
ADPHAPSMYLRSKGDGEIALRAQAAAGALDVTIFRPSIVFGPGDAFVNTFARLQRIFPVL
PLAMPDALMQPVHVGDVAQAIANACVRDATRGRTYELGGPRTYRLEEIVRYAGRLIGRPA
RIVRLPDALARLQARVFEMLPGEPLVSRDNLASLSVPSVMTGPIAPELGIAPASLEHIAP
TYVGDAALRSRFSDRRARR*
>GCA_000452945.1_BPMSHR5858_genomic_03296_1 slt
MSTSLFRVYRAVAAPLVAIALAAVAAACAAQTADEASSNDDRVFVQLREAARRNDPVRAA
QLAALIPNYPAPSYLEYFQIKPQLFDSAGHARLDAPDAPVLSFLSRYDGQAIADRMRNDY
LLVLGARHDWRAFDEQYKRFVLDDDTQVKCYALESRASRGENVADMARELLVEPKYYGDA
CVDLITALATNKQFSSDDVWAQVRLAYEQNYTTLGGKIADALGPRPVGFDQVTSAPPLFL
ARGVGSDATSRQLALVAITRMARNDPEAAAGQLASLASTLSAAEQAIGWGEIGYQATVKR
LPQAASWYRKSMDAPLSNPAYEWRVRAALLAGDWPMVRRSIEQMPERLRDDTAWIYWRGR
ALKESGDTLKANQEFERIAGQFNFYGQLAGEELGQKTTIPPRTKVTDAEIDAMSKVPGFA
LAQRFYALNLRLEGNREWNWPLRGMTDRQLLAAAEYGKRVDLLDRTVNTADRTTAEHDFS
LRYPSPYRDIVERYARTNGLDVEWAYGLIRQESRFITNARSTVGAGGLMQLMPATAQLVA
KKLGLGTVSRAQMHDIDTNVQLGTWYLSDIYQKFDDSAVLATAGYNAGPGRPAQWRQVLT
RPVEGAIFAETIPFNETREYVKNVLSNETYYAALFEKKPQSLKARLGFIAP*
>GCA_000452945.1_BPMSHR5858_genomic_03307_1 dppC
MSNMQNAMPSQAAPASGRVLALREFWANFSRNRGAVGAGIVVLALVAVALLAPVLAPHSP
IEQYRDYVKIPPAWLAGGNWQFVLGTDEAGRDILSRLMYGARMSFWIGFVSVVLALIPGV
VLGLVAAFFQKWADTPVMRVMDVLLALPSLLLAVAVVAIIGPGLTNTMFAIAIVALPAYV
RLTRASALGELQKEYVTASRVAGAGTPRLMFSQVLPNCTAPLIVQATLGFSSAILDAAAL
GFLGLGVQPPTAEWGAMLASARDYIDNAWWIVTMPGLSILISVLAINLLGDGLRDALDPK
LKRMA*
>GCA_000452945.1_BPMSHR5858_genomic_03311_1 group_7512
MKTSRRSFLITSVGAVSALALTREAFSADLPMLSETDPTAVALGYKADATKVDKAKFPKY
AAGEACAGCMLYQGKKGSASGPCGAFPGKQVAAKGWCNAFTKMA*
>GCA_000452945.1_BPMSHR5858_genomic_03322_1 group_7513
VNAPESIEKHSGVVVPYPGSPRKLDAFELAERRRRSRRATFIKWLRKVHGWVGLWGAALG
LLFGVTGVLLNHRAPPLKISSGEPQVSQLQLALPSPAPATPHAMAAWIRRELAFDGRLGR
VRKEPAQPVAWGGRRVEQPEHWQFGLFGPRSNVQVEYWKGNGYVSVKRTDNAFLTALNNL
HRGVGMNLFWVLLMDTIAGSMVLLSLTGVLLWTELNKRRTVGVVLVAGSVAAALAAGLT*
>GCA_000452945.1_BPMSHR5858_genomic_03345_1 ectD
MSRTSVLQSFIDNNDADHPMSSLHPELIHTQVQTLRERGFVVAPGLVAPERCAQLKTIAE
RQLREAAQPLEFEADLRYPGAPESRHAPGGHTVRRLLDAYARDAAFAERATAPEIGAWMR
AYFDETPVLSRAHHNCVMTKHPAYGSLTGWHRDVRYWSFERPDLVSVWLALGPETDDNGA
LWLVPGSHGAEFGPERFDEAKFFRGDVPANRRLIEQAVCPALAAGDVVFFHCNTLHSAGQ
NRSDQVKFSLVFTYHGDSNRPVPGSRSASKPEVRF*
>GCA_000452945.1_BPMSHR5858_genomic_03346_1 group_3421
MPFRLTSRLHRTPKSAPSAPHRIVRALRHHSARSSVLAERARQHQPRLDGLRAWLHAFLS
LMSAHAAARRTALRSLLQKPALRLRAPSLHRASVAPRRINRPRRLATSTGWFVFAAR*
>GCA_000452945.1_BPMSHR5858_genomic_03347_1 metK
VANDYLFTSESVSEGHPDKVADQISDAILDAILAQDKYSRVAAETLCNTGLVVLAGEITT
TANIDYIQIARDTIKRIGYDNTDYGIDYRGCAVLVAYDKQSPDIAQGVDRAHDNNLDQGA
GDQGLMFGYACDETPELMPLPIHLSHRLVERQANLRRDGRLPWLRPDAKSQVTVRYVDGK
PHSIDTVVLSTQHAPEIDLPALREAVIEEVIKPTLPADLIKGDIKFLVNPTGRFVIGGPQ
GDCGLTGRKIIVDTYGGAAPHGGGAFSGKDPSKVDRSAAYAGRYVAKNIVAAGLASRALI
QVSYAIGVAEPTSVMVNTFGTGRVSDETITKLVREHFDLRPKGIIQMLDLLRPIYEKTAA

YGHFGREEPEFSWEAADKALALAEAAGVEPAVQVA*
>GCA_000452945.1_BPMSHR5858_genomic_03348_1 lpxL
MLGRLGTRLAVAFLKFLAVLPYGLTARLGDGLGWLLYRIPSRRKRIVHINLQLCFPDWSD
ARREEIAGRHFRHAIRSYVERSVQWFGSAKKLEKLIQVDSAIDLTDPNLPPTLFLGLHFV
GIEAGSIWLNRSLQRRCGSLYQPFSNPTLEAIAKAARGRFDAEMVGRADSARVVLRWLRD
RKPVMLGADMDYGLRNSTFVPFFGVPACTLTAVGRLARTGHAQVVPFIGEVLPDYQGYRL
KVFKPWENYPTGDDDLDARRMNAFLEEQIPLMPEQYYWVHKRFKTRPPGEPGVY*
>GCA_000452945.1_BPMSHR5858_genomic_03350_1 group_4104
MNDREVADYLLANPEFFAQHAELLATIKLANPHGKAAVSLQERQMEMLREKNKHLERRLA
ELLRYGHENDGLAAKFNRWTARVIAERDPYALPRAIAGGLADVFDVPQTALRLWDVAETY
AQADFARNVGEEVRLFTNSLATPYCGANTGFEAAQWLTPVAAPPAADDGEAPAASGAAES
VALIALRASAEGEGAGAFGLLVLGSPDPRRFHDGMGTDFLTQIGTLASAALTRLLPH*
>GCA_000452945.1_BPMSHR5858_genomic_03354_1 dksA_1
MTKKLLTEAEILKMSDKDYMNEDQLAFFKNRLEQLQAEILRNAGQTTENLRETVIVPDPA
DRATIEEEHALELRTRDRERKLLKKVQQSLARIESGDYGWCEETGEPIGIPRLLARPTAT
LSLEAQERRELRQKLFGD*
>GCA_000452945.1_BPMSHR5858_genomic_03359_1 group_1877
MTEPAATPGLISKRCPRCGSAFDCGARTRPFTCWCAAMPAMPAGAQPATGARCLCPACLA
DEIARRVAASTAGEGPARPERED*
>GCA_000452945.1_BPMSHR5858_genomic_03368_1 yahK
MSTTYAYAATSATTPLAPFEFARRALRDLDVQIEILYCGVCHSDLHQARNEWKNTTYPVV
PGHEIVGRVTAVGAQVTRFEAGELVGVGCLVDSCRTCPSCAQGLEQYCENGFVGTYNGRD
RATGDITYGGYSTQIVVDEAFVLRVPDTLDPAGAAPLLCAGITTYSPLRQWGAGPGKKVG
IVGLGGLGHMGVKLARAMGAHVVLFTTSPSKIEDGKRLGAHEVVISKDDAQMNAHLNSFD
FILNTVAAQHDLNPFIGLLKLDGTMTLVGAPEHDHPSPRVFNLIYKRRRLAGSLIGGIAE
TQEMLDFCAQHGITSDIEVIPMSQINAAYERMLTSDVKYRFVIDIGSIRPQRAA*
>GCA_000452945.1_BPMSHR5858_genomic_03380_1 garL
MSTLTHSLKARLRDGDEPLFGLWLTLASEAATEALAHAGFDWLCIDMEHAPNDSRDVAAQ
LRALAAAHLPSEPVVRVPAREPWFVKRALDAGARTLMFPGVETADEAAHAVRLTRFQAPD
APDGLRGVAGIVRAAAYGMRRDYVQTANAQIATIVQIESARGVDEAERIAATPGVDCVFV
GPADLSASLGHLGDTKHPDVAAALEHVLAAGRRAGVPVGIFAADTAGARQSLEAGFRVVA
LSADVVWLLRATRQALQEVRG*
>GCA_000452945.1_BPMSHR5858_genomic_03382_1 queE_1
MTYTVKEIFYTLQGEGANAGRPAVFCRFAGCNLWTGREADRDGAVCRFCDTDFVGTDGEN
GGKFKDAASLAARIASLWPAGEAYRFVVCTGGEPMLQLDQPLVDALHAAGFTIAIETNGS
LPVLESIDWICVSPKADAPLVVTKGNELKVVVPQDNQRLADYAKLDFEHFLVQPMDGPSR
ELNTRLAIDWCKRHPRWRLSMQTHKYLNIP*
>GCA_000452945.1_BPMSHR5858_genomic_03387_1 group_7518
VTIKRFFPLRLAAVLWVALTLCLAAVPAARADTIAVQRASLQSDGSGWSLDARFDFELNS
SLEDAVNKGVPLYFTTDFELSRARWYWFDEQPVSVSQTIRLSFQPLTREYRVSTGGLQLG
FPSLKEALAVVKHVTSWHVIDKGQVRPGETYTASVRMQLDTALMPKPFQVDAVNNRDWNL
VSDWKRFIFTVAERAK*
>GCA_000452945.1_BPMSHR5858_genomic_03389_1 htpX_2
MFNWVKTAMLMAAITALFIVIGGMIGGSRGMTIALLIALGMNFFSYWFSDKMVLRMYNAQ
EVDEATAPQFYRMVRELATRANLPMPRVYLIDENQPNAFATGRNPEHAAVAATTGILRVL
SEREMRGVMAHELAHVKHRDILISTISATMAGAISALANFAMFFGGRDENGRPANPIAGI
AVALLAPIAGALIQMAISRAREFEADRGGAQISGDPQALASALDKIHRYASGIPFQTAEE
HPATAQMMIMNPLSGGGLQNLFSTHPATEERIARLMDMARTGRFD*
>GCA_000452945.1_BPMSHR5858_genomic_03398_1 group_4864
MFDQVVFAGGGNRCWWQAGFWDVVRPALDLRPRVIAGISAGAATACMLYTRDSRWVMRYY
EEALRHNRRNVHWGNLLRAEPVFPHYRIYRQALLDIYGEPFAKLAGAPDIRIGVSHVPRW
LGARSAVAAGLIAYNIEKYVRKTLHPTLGRSLGFRPEFVRAQDCARVDELADLVLQSSCT
PPFTPVLRRDGRPVLDGGMVDNVPVGALDASPGAVLVMVTRRYPRPQTFVVDGGGQRRLY
VQPSSKVPISSWDYTSPHQMQHAYDLGRRDGERFVGAMPQCLDEDGAVR*
>GCA_000452945.1_BPMSHR5858_genomic_03414_1 group_3429
MEEAYARRALLLHLGSVLQILSRLEGAPDNQSIEALLAANGMLADVPLVEYVVQDMTVRE
FSARALRAFCLWPQLLLEDPLDCAALALPVRKHLFAGNDAGWKAYAATLREVAPGFDTAR
SYAAEPHEAEFDLADERDAETGDGPVERDDEADDDGRLRERAAASVESATPAGAPVRRES
YE*
>GCA_000452945.1_BPMSHR5858_genomic_03422_1 group_7520
MLKRTGLFLALTGGIVAFSVAQANGDASLKPQQEIQLTKNAWGCLSKDNLDSVLNHERDG
KAQAKQQYFDDYRCLSVPEGQRFRVVSVDKGDVQFVSAENSDQQGLWTDARFIKQ*
>GCA_000452945.1_BPMSHR5858_genomic_03424_1 group_4871
MEAIVIEQVILGVFLVLPLLIVAVLYSDELWQEHRLQHPRDEHTPHIDWRHPWRILRRGH
*
>GCA_000452945.1_BPMSHR5858_genomic_03427_1 group_1029
MPAPPPHQYAMEQDGTYGYEPALSEDDVRSGKATKPLVMMRYVGRRDGTYVLLLIDPDNE
NYATRVTCQAPCNFAKLQTMSGTMVLKTETIRVAPNSLIGAMLDDALSGQLRPYGQTVTM
PRPTAVPSADIPAMASAQAAVQSTARPADQSAPQDSSPQSDLQRTSFDCSKVGSIPEYLI
CHDPELAASDRELANIYQQAKDAVPDKAAFAARTRRQWNFRQRNCRDKPCLVSWYAYQKE
TLTKIAQTGDVNAQ*
>GCA_000452945.1_BPMSHR5858_genomic_03460_1 group_7523
MTSSDIATITGWHAHVYFDASSRDAAWAFRETIERRFGASVQLGRFHERLVGPHPTWSYQ
IAFEPTQFADVVPWLVLHHGALDVFLHPNTGDELRDHRDAAAWIGRSYTLNLDALTP*
>GCA_000452945.1_BPMSHR5858_genomic_03468_1 group_1210
MSVDAMASPALAGRSIEVDFFRGVVLLTIVVDHIGASVLSRVTLHAFALCDAAEVFVFLG
GFATASAFLSVSARHGAGAARRRFMRRAAQIYRAFLATSTLMLVVSAVLDHYGIDAPNMA
LDDISVLLASPLTVLVELLTFKRQPYLASVLPMYVLFALATPAIVPLARAKPWWLVFGSV
LMWGCAPWLAAQLLDTESFRWSFNPFAWQLMFVFGVLMRCWPLHRDAATRAGGTAITAVA
LAVVLACAYYKLGSGLPLPEGEFKRHLAWPRVMNFAAFAWLMAELVRYGWIARLARAARP
VVAVGQRGMLCFVAGAAISLTLDSVLHGMHGGARMMQLGAGLAADACALALMLSIARSDL
LLQRLRRASAAAAAT*
>GCA_000452945.1_BPMSHR5858_genomic_03483_1 group_3435
MSVSHLRHAVARAALVALAGASALALPAAHAAGAELNVYNWSDYIAKDTIPNFEKQSGIH
VKYDNYDSDDTLQAKLLAGSSGYDIVVPTSNYMAKQIQAGVYQKLDKSKLPNLSNLDPTL
MKMIADADPGNQYGVPWAFGTDGLGYNVQAVKKALGENAPVDSWALVLDPANVSKLKGCG
VSFLDQAVDVFAATLQYMGRNPNSTNPADYQAAFEVLKKVRPYITQFNSSGYINDLANND
VCVALGWSGDVGIAHRRSAEAKRPYDIKFSNPKEGGLLWFDVMVIPKDAPHAEAALKWIN
YIEDPKVNAAITNEVFYPSANRAARQFVVPAVAQDPTVYPPEDVLKKMTLMRPMPADILR
LENRLWAQLKTGH*
>GCA_000452945.1_BPMSHR5858_genomic_03488_1 group_7524
MAKRMPPDFHGPPMTDTFARRADALARRSYYEATVRRPADDPALAEALDVDVCVIGAGFA
GLSTALDCRARGLSVAVLDARRPGWGASGRNGGQLIAGFAKDGEIERQLGVEGARAAWAL
SLDAIALVNERIARHGIDCDLARGYLSVATHPRRVPELRQWMEGATARWGHPSLEWLDAR
AVRARIASPRYVAGVHDPLSGHLHPLKYCLGLADAARREGARLFSHTPALEVVRGARPIV
RTPAGEVRCRHVVACCNAGPGGVLPAATAARIAPVASYIVATEPLGADRANALIPRREAA
CDNNFFLDYFRLSADHRMLFGGRASSAGASPDALVDAMRKRMTGAFPQLADTRIEFAWGG
FVDVTRNRAPDFGAIDPNYFYLQGFSGHGVALAGIAGRIVAQAIAGDSRAFDLFARLRHR
RFPGGPAWRQPALALGMLYHRVRELLR*
>GCA_000452945.1_BPMSHR5858_genomic_03489_1 puuB_2
MLKFINQPHAASYYAATVNDTTRHPPLEGTIDADVCVIGAGLTGISAALNLAERGHSVAV
LEASKVGWAASGRNGGQLIGGFACGIDAFEPYLNADEIQLVWDMGLETLAIAKERIAKHA
IDCAFVPGYLSAANKPRDVDALRRSRDEAARRFGYTRLRYVERDALAQYVQSSRYLGGLF
DPDSGHLHPLNYTLGLARAAVASGARIHEDSAVTRIASEAGGHVVSTARGAVRARFVVLA
CNAFLGALAPALSRKIMPVGTYVIATEPLGEARARALMPAQAAICDSRFALDYFRPTPDA
RLLWGGKVSYSTLEPRKLADAMRRDMLKTFPQLADVTIEHAWGGFVDITMNRAPHFGRLT
PTVYFAQGFSGHGVNTTGLAGKLIAEAIDGQAARFDVFGKIRHRDFPGGAALRMPALVLA
MAWYRLRDLL*
>GCA_000452945.1_BPMSHR5858_genomic_03490_1 group_491
MKLFFDMKIGTRLIVLILAALAALCAIAGFGIYESRRVYTAASYSTVNTVPSFVVLDDAQ
RAFDSMLLLVNQQVFSTTADQAKALEPRIAQARREVDAQFAKYETLLSNDKDKALLAADR
ARVSQLDAVRENVLALSRDGRKQEAGELMGTRMTELAQQTNAALAAHRAFNVDLGQAGSN
EAKDIIDRAVTLEASAAAVVLVLVLWLGVLISRSITKPMGNAVKFARTVADGDLTTHIGA
ASKDETGQLLKALGDMNDSLKRVVERVRMGSDAVATASGQIAAGNLDLSSRTEEQAASLQ
ETAASMEELTSTVRQNAENAQQASGLASNASDVALRGSSVVDRVVDTMNDISERSSKIAE
IIGIIEGIAFQTNILALNAAVEAARAGEQGRGFAVVAGEVRGLAQRSSSAAKEIKELISA
SVQKIRDGSALAGEAGRTMTEVTQAVARVTDIMGEIAAASAEQSRGIEQVNLAITQMDEV
TQQNAALVEEAAAASHSLEEQGHELKQAVAFFRFDGSAAGGTADTAAPRHRAAPPPHAAA
TAPAAHATPAPHAPHASHASHTIRTAHATPAPRAFDAVKPAASAAPAAAAVSAPALSGAR
ATAADANADWETF*
>GCA_000452945.1_BPMSHR5858_genomic_03493_1 group_563
MKNVLATIAIGCALAVSNAAHAAGEMSEMSEMSGMSGMDMQGSAPQAGAAHVGMSHGEVK
KVDTAAGKLTIKHGPLENLGMGAMTMVFKVKDPAMLSQVKAGDTIDFVADEVDGALTVVK
LQKP*
>GCA_000452945.1_BPMSHR5858_genomic_03497_1 polX
MLDSAPMSYWRKLFVIVLLVLSLPVQTLAAVSMNCDPAPADDGGAPAHAWLQASAENGHG
MHDAAAADDGHRHDGRGGAHHDHAHACSTCLSCCVGAGLPAASVAAASFDAARTRLFLPR
DTRAVSFLTGGIERPPRTSLV*
>GCA_000452945.1_BPMSHR5858_genomic_03503_1 trmD
MDEATQSAIQFDVVTLFPEMFRALTDWGITSRAVKQGRFGLRTWNPRDFTTDNYRTVDDR
PYGGGPGMVMLAKPLEAAIGAAKAAQAAQGVATSRVVMMSPQGAPLTHERVARMAAEPGV
VLLCGRYEAIDQRLIDRCVDEELSLGDFVLSGGELPAMALMDAVVRLLPGVLNDAQSAVQ
DSFADGLLDCPHYTRPEEYEGVRVPDVLLGGHHAEIERWRRQEALRNTIAKRPDLIARAR
REKLLSRADEAWLASLAKEAKQAS*
>GCA_000452945.1_BPMSHR5858_genomic_03521_1 group_3437
MLKKLLATAALCAAAAHGWAAVDVNAANDDALRGIRGIGPAKAKAIVDERSAHGPFKDAA
DLARRVKGMGGKTVERLQAEGLAIGAARAPAAAGAPQKAAAAGKK*
>GCA_000452945.1_BPMSHR5858_genomic_03522_1 hldD
MTLIVTGAAGFIGANIVKALNERGETRIIAVDNLTRADKFKNLVDCEIDDYLDKTEFVER
FARGDFGKVRAVFHEGACSDTMETDGRYMMDNNFRYSRAVLDACLAQGTQFLYASSAAIY
GGSSRFVEAREFEAPLNVYGYSKFLFDQVIRRVMPSAKSQIAGFRYFNVYGPRESHKGRM
ASVAFHNFNQFRAEGKVKLFGEYNGYGPGEQTRDFVSVEDVAKVNLHFFDHPQKSGIFNL
GTGRAQPFNDIATTVVNTLRALEGQPALTLAEQVEQGLVEYVPFPDALRGKYQCFTQADQ
TKLRAAGYDAPFLTVQEGVDRYVRWLFGQL*
>GCA_000452945.1_BPMSHR5858_genomic_03523_1 rfaE
MSSPSLPSADKRASLAATARAVSREQIAKSRVLVVGDVMLDRYWFGNVNRISPEAPVPVV
HVQRQEERLGGAANVARNAVTLGGQAGLLCVVGRDEPGERIVELLGESGVTPYLERDPSL
PTTIKLRVLAQQQQLLRVDFEASPTHEVLLAGLARFDALLPSYDVVLMSDYAKGGLTHVT
TMIAKARAAGKPVLVDPKGADWARYRGASLITPNRAELREVVGRWTSEDDLRARVTKLRE
ALAFDALLLTRSEEGMTLFSHEGELHVSALAREVYDVSGAGDTVIATVATMLGAGLPLVE
AVVLANRAAGIVVGKLGTATVEYDELFH*
>GCA_000452945.1_BPMSHR5858_genomic_03524_1 rkpK_2
MKITIIGTGYVGLVTGACLAEIGHDVFCLDVDPRKIDILNNGGMPIHEPGLQEIIARTRA
AGRIAFSTDIEASVAHGEIQFIAVGTPPDEDGSADLQYVLEAARNIGRHMTGYKVIVDKS
TVPVGTAQRVRAVIDEALAARGLAGSAEHRFSVVSNPEFLKEGAAVDDFMRPDRIIIGVD
DDAAGAIAREKMKKLYAPFNRNHERTIYMDVRSAEFSKYAANAMLATRISFMNEMSNLAD
RVGADIEAVRRGIGSDPRIGYHFLYAGVGYGGSCFPKDVQALIRTASENGQPLRILEAVE
DVNHAQKNVLLDKIEKCYGADLAGRTFAVWGLAFKPNTDDMREAPSRRLIASLLARGATV
RAYDPVALDEARRVFALDLHDGADALARLAFVDSADDALAGADALVIVTEWKEFKSPDFA
HLKSVLKAPVIFDGRNLYEPDAMAELGIDYHAIGRPYVEPFSSERG*
>GCA_000452945.1_BPMSHR5858_genomic_03525_1 lapB_2
MDLDFWWLLAIPVAFALGWMASRYDLNKLLSESANLPRSYFRGLNFLLNEQPDKAIDAFI
EVAKLDPETVELHFALGNLFRRRGETDRAIRVHQNLLSRNDLPVSERDHALFELGQDFLK
AGLLDRAEETFHMLAEGDYALDAQRALLTIYEIEKDWTKSIDTAARIEAMGAPRLTTEIS
QFHCELAQDALQRKSAELAAEHLRDALAVNPQNARAAILSGDTAAAAGDHRAAIEHWRRI
ETQNPAYLPLVADKLMKSYAALDRAAEGAELLTGYAQRYPSNDLLDVVYQYVAQLRGNDV
AHALARSQMEKAPNLSGMLHLLDAQIAAADEPRRSELEMMRALVKQRTKNLPRYTCQNCG
FRARLFYWQCPGCSGWETYAPRRVEPAVA*
>GCA_000452945.1_BPMSHR5858_genomic_03526_1 lapA
MKFIVWLIRVLVFVLLLVLALANTQSATLNFLAGYNWQAPLILIGLAFFGVGLIAGLLSA
LPAIFRMRMENGRLKRDLRAVRENPAVVDQPPMPPVI*
>GCA_000452945.1_BPMSHR5858_genomic_03527_1 ihfB
MTKSELVAQLASRFPQLVLKDADFAVKTMLDAMSDALSKGHRIEIRGFGSFGLNRRPARV
GRNPKSGEKVQVPEKHVPHFKPGKELRERVDGRAGEPLKNDEPEDAQ*
>GCA_000452945.1_BPMSHR5858_genomic_03528_1 rpsA
MSDLQTSNPNTESFAALFEESLTRQDMRAGEVISAEVVRVDHNFVVVNAGLKSEAYIPIE
EFLNDQGEVEVQAGDFVSVAIDALENGYGDTILSRDKAKRLASWLSLEKALDNNELVTGT
ITGKVKGGMTVMVNGIRAFLPGSLVDTRPVKDTTPYEGKTLEFRVIKLDRKRNNVVLSRR
AVIEATQGEERAKLLETLKEGAIVNGVVKNITDYGAFVDLGGIDGLLHITDIAWRRVRHP
SEVLSVGQEVTAKILKFDQEKNRVSLGIKQLGDDPWEGISRRYPSGTRLFGKVTNITDYG
AFVEVESGIEGLVHVSEMDWTNKNVAPSKVVQLGDEVEVMVLEIDEDRRRISLGMKQCKP
NPWDDFSRNFKKGDKITGAIKSITDFGVFIGLPGGIDGLVHLSDLSWSEAGEEAVRKYKK
GDEVEAIVLGIDVEKERISLGIKQLEGDPFSNYVAMNDKGSIVDGVVKSVDAKGAVITLT
PDVEGYLRASEISQDRVEDARNVLKEGEKVNAMVINIDRKSRGINLSIKAKDSAEQQEAM
RGLQADSSAAATGTTNLGALLKAKLDGQNQ*
>GCA_000452945.1_BPMSHR5858_genomic_03529_1 cmk
MKSTRPFHPTPVITIDGPTASGKGTVAALVAAHLGFHLLDSGALYRLAALASIRYQVEPD
DADALASLVDGLHITFREGCAQLDGVDVSDEIRAEAVGNRASAIAVHASVRAALVARQRA
FRKTPGLVADGRDMGTVIFPDAVLKVFLTASVEARAARRHKQLMQKGFSANMDNLLQDLR
ERDARDSNRAAAPLKPAADAKPLDTSALTIEQSVEQVLAWYRELGQPA*
>GCA_000452945.1_BPMSHR5858_genomic_03532_1 serC_1
MAVCATAQAAISFSRAAFGRFEAAGSLPRFFTMRVFNFSAGPAAMPEEVLRQAAAEMLDW
NGSGMSVMEMSHRGKEFMSIHEAALADLRELLGVPANYRILFLQGGGIAENAIVPMNLLG
SRTTADFVITGSWSQKSFNEAKKYCVPHLAATGKTDAGFTRVPAFSEWRMSDDPAYVHLC
TNETIDGVETFDIPDLGNVPLVADVSSHILSRPIEIEKYGAIFGGAQKNIGMAGVTLVIV
REDLLDRALSICPSAFEWKTVALNNSLYNTPPTYAIYIAGLVFKWLKAQGGLAAIEARNI
EKSKLLYDAIDASGFYLNKVEKPARSRMNVPFFLADESRNEDFLAGAKERGLLQLKGHKS
VGGMRASIYNAVPLAGVQALVEYMKDFERRCA*
>GCA_000452945.1_BPMSHR5858_genomic_03533_1 group_4885
MMQKRFKQLVLLAAMVPAFAMAQSLSNQTSAPAAAAPIDADKKAAIKDLLDAIDAPKLVS
AIANSAEMQSKQLVPAILSDALSENKTLNDKQKQAAVPTLQKNAVPKLVDGAGKVFGTQQ
FTNDAMQAQYDAYAKYYSTSEIKDLTTFYKSPTGRKFIQVQDQVGRDVVNGLMQKYMPQA
IKATRDQADKEVAAVKPGK*
>GCA_000452945.1_BPMSHR5858_genomic_03534_1 gyrA
MDQFAKETLPTSLEEEMRKSYLDYAMSVIVGRALPDVRDGLKPVHRRVLFAMHELNNDWN
RPYKKSARIVGDVIGKYHPHGDTAVYDTIVRMAQDFSLRYMLVDGQGNFGSVDGDNAAAM
RYTEIRMAKIGHELLADIDKETVDFGPNYDGSEMQPLILPSRIPNLLINGSSGIAVGMAT
NIPPHNLNEVVDACLYLLNNPQASVDELIEIIPAPDFPTAGIIYGVAGVRDGYRTGRGRV
VMRAATHFEEIDRGQRMAIIVDELPYQVNKRSLLERIAELVNEKKLEGISDIRDESDKSG
MRVVIELKRGEVPEVVLNNLYKATQLQDTFGMNMVALVDNQPKLLNLREILQCFLSHRRE
VLTRRRTVYELRKARERGHVLEGLAVALANIDEFIAIIKAAPTPPIAKQELMAKSWDSSLV
RDMLTRAETENAAAGGRDAYRPEGLSPAFGMQGDGLYRLSDTQAQEILQMRLQRLTGLEQ
DKIIGEYRDVMAQIADLLDILARPERITTLIGDELTQVKAEFGDARRSKIELNATELNTE
DLITPQDMVVTMSHAGYVKSQPLSEYRAQKRGGRGKQATQMKEDDWIETLFIANTHDYIL
CFSNRGRVYWVKVYEVPQGSRNSRGRPIVNMFPLQDGEKINVVLPVKEFSADKYVFMTTA
LGTVKKTPLEAFSRPMKKGIIAVGLDDGDYLIGAAITDGAHDVMLFSDAGKAVRFDENDV
RPMGREARGVRGMQLDDAQQVIAMLVAGSEEQSVLTATENGYGKRTPITEYTRHGRGTKG
MIAIQTSERNGRVVAATLVDPEDQIMLITTAGVLIRTRVSEIREMGRATQGVTLISLDEG
TKLSGLQQIAEADADADDDAVEGGDA*
>GCA_000452945.1_BPMSHR5858_genomic_03535_1 group_6066
MNKLSKLAFIAATAVMAASASAQSVPASRQAVNDNWVNGTGEWVWMNGTNELCWRDAFWT
PATANAKCDGALVAQAPAPAPVAPVAPAITSQKITYQADTLFDFDKAVLKPAGKQKLDEL
AAKIQGMNVEVVVATGYTDRIGSDKYNDRLSLRRAQAVKSYLVSKGVPANKVYTEGKGKR
NPVTGNTCKQKNRKQLIACLAPDRRVEVEVVGTQEVQKTTVPAQ*
>GCA_000452945.1_BPMSHR5858_genomic_03536_1 ubiG
MRPASPAAAFLATSPARLYTRGLAGAPPPSFPHPLADMTNADPHELQKFSDLAHKWWDPN
AEFKPLHDLNPVRLSWIDAHAHLPGKRVVDIGCGGGILSESMASLGAQVKGIDLATEALG
VADLHSLESGVSVDYEAIAAEALAAREPGAYDVVTCMEMLEHVPSPANIVAACATLVKPG

GWVFFSTLNRNLKSYLLAVIGAEYIAQMLPKGTHDYARFIRPSELARFVREAGLQMVEIK
GIAYHPLAKRFALSNDTDVNYLVACRRGA*
>GCA_000452945.1_BPMSHR5858_genomic_03550_1 cnbH
VPIRFVNPIEPDTVMTTFAPFPPLARLAADLAAGVTTSRALVEAALERIADPAGQGALAF
TEVDAPRARAAADAHDRLRAAGTVLSPLAGIPVSVKDLFDVAGQVTRAGSRVLADAPPAA
DDAVAVARLKRAGAVLVGRTNMSEFAFSGLGLNPHYGTPRSPYRRDVPGDERIAGGSSSG
AAASVADGMAAVALATDTGGSIRIPAALCGLTGFKPTASRIPKQGGVPLSTTLDSFGPIG
VSVACCALVDRILAGLEPHVCAARPLEGVRLGVLTNYVTDGMDAHVATSLDAALKHLEAA
GAIVSDARFAPLDTLPGINRIGFASIEAHAWHRPLVARHRDAYDPRVLARILKGEPASAA
DYLDLVAARAALIDEAARTVWQRFDALVAPTVPIAPPRIAELERDDDAFARTNALVLRNP
SAFNFLDACALSLPCHPRDTAPVGLMLAAAPHHDDALLAIGQAVEAVLATIR*
>GCA_000452945.1_BPMSHR5858_genomic_03552_1 group_3439
MQTWLTDLQHLLARGDAAVLVTVARVDGSAPRDAGTKMLVTRDAARHTIGGGHLEWKAID
IARQLLKEGAHTPHARRLERLALGPSLGQCCGGAVVLAFERLDIAELGWVTSLAKRVAAG
APTVRSVSFGPDADTVMLSEPEPGATRPDCLLWDMGGAPLLTETIAPHAFPVVLFGAGHV
GAALVRVLGTLPCHVRWVDERDAQFPPPEALAGIRNLSLDANDAPDEAIDEAGPGAYFVV
MTHNHARDLELAHRILARGDYAYFGMIGSRTKRMQFEHRLAARGIDPLQIARMQCPIGVP
GIVDKAPEAIAIAVAAQILQTVDARRANGRPAASPGASAC*
>GCA_000452945.1_BPMSHR5858_genomic_03556_1 mneA
VTEAFLISTGAVALAEIGDKTQLLSLVLAARYRKPLPIIAGVLAATLINHGFAGALGEWL
GVYLTPAVMRWALALSFIGMGLWILVPDKLDADEANANRSRLGVFGATFVAFFLAEMGDK
TQIATVALAARFQDYVGVVAGTTLGMMLANVPAILLGDRFAHRLPTKLVHGIAAVLFIVL
GALAFLHGGA*
>GCA_000452945.1_BPMSHR5858_genomic_03557_1 group_3441
MKRIPFIRGAALVAGALIVLLSGCTSYVTTQVTAFSNWSGNDATRTYAFARTAEQQNSLE
QSTYEQIVGNELSTYSFRRVPTKDARYLVRLAYGIKSDWVSVPQPVYYDPWFGPGPYWRG
GPWGPFGPWGPFPAGYVNQSYQIFQRSLEIRIAERATGTEVYNVAARNAGEGSSLLQAMP
YLARSALADFPLGNGTVRTVRLPVDKNGAPAPMPANERAVPEVPAMPASGAAVAPAR*
>GCA_000452945.1_BPMSHR5858_genomic_03558_1 pepN
MSDIAAPNAEIRRSDYTPPAFLIDTVSLEFDLEPARTIVTNTMRVRRNPDAAPAPHFELM
GEALVLIGARVDGKPHDAVRVHEHGLSVENVPDAFELTIENACAPESNTTLSGLYVSSGN
FFTQCEAEGFRRITYFVDRPDVMASYTVTLRADQAAYPVLLSNGNLVDAGDLPNGRHFAK
WEDPFKKPSYLFALVAGKLVKLEETIKSASGKDKLLQVWVEPQDLGKTRHAMDSLIHSIR
WDERRFGLELDLDRFMIVAVGDFNMGAMENKGLNIFNTKYVLANPETATDVDFANVESVV
GHEYFHNWTGNRVTCRDWFQLSLKEGLTVFRDQEFSADMSAGAEDDAAARAVKRIEDVRV
LRQLQFAEDAGPMAHPVRPERYVEINNFYTMTVYEKGAEVVRMYQTLFGRDGFRKGMDLY
FRRHDGQAVTCDDFRHAMADANGRDLALFERWYSQAGTPRVTVRTAYDAAAKRYAVTLRQ
GYGDAAPAARDTQKGPLLIPFAIGLIGADGRDLPLRLEGEAAASGTTRVLELTEAEATFT
FVDIDAAPLPSLLRNFSAPVIVEYDYRDDELAFLLAHDSDPFNRWEAGQRLATRALLTLA
SRAAAQQPLTLDDAFAAAFKRVLTDDTLSPAFRELALTLPSEAYLADQMTQADPAAVHRA
RQFVRRQLATALRAEWLSVYERHQTPGAYAPTPGDAGRRALKNLALAYLAELDEPADAIR
LATAQYDAANNMTDRACALVALLSAAAASADAARAADRALDDFYRRFENEALVIDKWFSM
QATRRGTPEHPTLDIVRKLLAHPAFNLKNPNRARSLIFGFCSANPAQFHAADGSGYAFWA
DQVLALDALNPQVAARLARALELWRRFTPSLREKMRDALERVAANAQSRDVREIVEKALA
*
>GCA_000452945.1_BPMSHR5858_genomic_03559_1 metE
MTTAHILGFPRIGAQRELKFALERYWRDGASADAERALVDTGRALRAEHWRIERDAGLDC
VTVGDFAWYDHVLTTLAHVGGLPRRFGFDARALTLADYFAAARGNAAQPAMEMTKWFDTN
YHYLVPEYSPATTFGPGVEWLFDEVREARALGYRAKAALVGPLTLLWLGKARDGLVERLA
LLPRLVPAYRALLARLREAGVDWVQIDEPIFSLDLPDAWRDAARPTYEALAPGAPKLLVA
TYFDDASEHAALLKALPVAGLHVDLVRADAQLDAFVADYPADKVLSCGIVDGRNVWRNDL
DRSLARLAPVRDALGERLWVATSCSLLHVPVDLAHEPRLDEELKTWLAFAAQKTREVAAL
RDALVKGRAAVAAEFDDAAVAAAARRTSARIHNPLVKRRVAALTDADARRASAYSVRAAA
QRARFGLPLLPTTTIGSFPQTPEIRRARAAFKQGVLDHLGYLEAMREQVRIAIDKQLAYG
LDVLVHGEAERNDMVEYFGELLWGFAITSNGWVQSYGSRCVKPPLVYGDVYLPEPMTVGW
ASYAQSLSAKPVKGMLTGPVTMLQWSFVRDDQPRATTALQIALALRQETLDLEKAGIGMI
QIDEPALREGLPLKARERAAYLDWAVRAFGIAASGVADDTQIHTHMCYSEFGDILPSIAA
LDADVISIETTRSNMELLDAFETFDYPNEIGPGVYDIHSPRVPDADEIERLILLALERIP
AQRLWVNPDCGLKTREWRQVDAALAAMVDAAKRVRQKVEEAAPA*
>GCA_000452945.1_BPMSHR5858_genomic_03560_1 metR
MIRSPLEFRHLQTLVALRDTGNLSRAAQVLCLTQSALSHQLKALETHFGLPLFVRKSAPL
TFTAAGKRLLALAEQVVPAIEEAERDIARLALGTGGALRIAVECHTCFDWLMPAMDAFRQ
RWPEVELDIVSGFHADPIGLLHQDRADLAIVAEADADEAVDYHPLFRFQIVGLVGNDHAL
AHQPSLTAADFADETLITYPVPDEMLDIVRQVLKPAGIEPKRRTSELTIAILQLVASRRG
VAALPLWAVATYLDKRYVSARPVLRADGSTLSGELYAATLPAFSTRAYAADFVATMRETS
AASLPEIELL*
>GCA_000452945.1_BPMSHR5858_genomic_03561_1 group_7535
MSITRRTTLSKYLIEQQRETHNLPADLRLLIEVVARACKAISYNVSKGALGDALGTAGSE
NVQGEVQKKLDILSNEILLDANEWGGNLAAMASEEMETFFPIPANYPRGEYLLVFDPLDG
SSNIDVNVSIGTIFSVLRCPDGQQATEQSFLQPGTEQVAAGYAVYGPQTVFVLTTGNGVN
CFTLDREVGSWVLTQSNLRIPEDTREYAINASNARHWYEPVKRYIDELNAGAEGPRGENF
NMRWIASMVADVHRILNRGGIFMYPADKRTPDRPGKLRLMYEANPMSFIVEQAGGAATTG
LKRILDVQPTGLHQRVPVILGSKNEVERVARYHEQAQS*
>GCA_000452945.1_BPMSHR5858_genomic_03564_1 group_4886
MAHGLIMWLIIGAIAGWLAGLLVKGGGFGLLVDIIVGIVGAVIGGWLAGILGIHIGGGFI
SSVIVAVIGAVILLFVIRLFKRA*
>GCA_000452945.1_BPMSHR5858_genomic_03566_1 group_5743
MVAVRHRLAGVALAAFACGALAHDAQPAAGASSAHAAHMAMDAATAGAAQKTPLATGAAF
DARHRLWVAWVEGQHVVVAHSDDAGRTLAPPVTVNALPEPIYTSAENRPKLAASPDGRAI
YVTWSMPLDAPYTGMVRFSRSLDGGATWSVPVTVHRDRQAITHRFDMLTIDPQGRIFVAW
IDKRDLVAAQRAGRSYDGAAIYYAVSTDGGATFEPERKVSDHTCECCRIAMTVDSDGRVQ
AAWRNVFPGQIRDHAVAVLHASADRAVEPVRATFSNWHVEACPDHGPALAITADGVRHLA
WFGVVDGKADVFYSRLDARGAPLGAPFAFGDDPAAPDEQASHAALVARRDTLWLAWKAFD
GDTMRIRLRRSDDGGAHWSAPRTLAWTSGASDNPQMLADRGRVYLSWRTRNDGYRLIDVE
GQR*
>GCA_000452945.1_BPMSHR5858_genomic_03567_1 btuB_1
MFHQLAAPGARRRLAAACAAALAWPAAHAASTAAAVPADSTPAAAAEMTASGKTLDTVKV
TAQRAAFAPDTPGVVEALTREQIDSHVNVTTEDALKYAPNLMVRRRYIGDRNSVFAGRDF
NELQSARGLVYADGILLSNLLGSSYSYPPRWSLIQPDDIARVDVLYGPFSALYPGNAIGS
TVQITTHKPQRLEASVSTQFFTQRYRDGYGFADSFGGNHQSARIADRVGRFWYALSLDRL
ENDSQPMQYASPNGAFDPRLGAPVPVTGAVSDIGPNGRPRTIVGAQTIERTEQLNETLRF
GYAFTDHVDATVTLGHWENHYRQHGDTFLRDAAGNPVYGGNVSFGGRNYTVSPGAFAPQT
GDQENWLYGLGLDARLASGWKLSATASAYEVSRDVLRSASGAPPGAWDGGPGTVSHGDGT
GWRTVDLRAESPDVRGHRFSFGYHFDTYFLRNATYNTADWQNAVPTTLVNRYRGNTRTQA
LYAQDAWRVAPGWLATLGLRYERWDAYGGELGGATATLGYAERGATAFSPKLSLEWQPAS
AWRLRLSFATGTRFPTVAELFQGTISNNAIVNNNPNLQPEKAIDWDFTAERDVGFGVVRA
SVFQSDLRNSIYSQTTLAGASTYTNVSNVDRVRVRGVELAFSGQDVALKGLDVDANVSAT
NAQTLADAANPNYVGARWPRIPRMRANLLASYRFDEHWMTSVGVRYSGRQYNALDNSDVN
PGVYGGTSSFMVVDLKARYRFDRHWLASFGIDNVTDRRYYVFHPYPGRTFYGELKWSL*
>GCA_000452945.1_BPMSHR5858_genomic_03568_1 group_7536
MSLVWLALVLNVLSPVVGYARAAANAADSPFALELCSAAGAQRVVVDSGGEPRRGDASLA
HVAHCVYCPGFAANLALGSSTPALPGFVRAFVYASRVERPAVFFRRGVRIAQPRAPPETV
PV*
>GCA_000452945.1_BPMSHR5858_genomic_03569_1 group_7537
MDERPTRIPPPEKVMSPDPEPVAVEFLAELPEHVRAFFDEQHRLCTPK*
>GCA_000452945.1_BPMSHR5858_genomic_03570_1 group_3442
MKQSLRASLAAAVLGCIVLSAPRIAAADGDDTALTNLVALASQRLALAEPVAHWKWLNGK
PISDPPREAALLADVEQRATANGVDPAYARAFFDDQIAASKQVQNALFATWRATHGPEGP
APDLATSTRPQLDRLTQSLIAALARVAPLRDAPDCPSRLARSVSNWKTLTRYDSGREDAL
GTALSHVCTAGSTSAVG*
>GCA_000452945.1_BPMSHR5858_genomic_03572_1 cspG_1
MDTGIVKWFNDAKGFGFITSDNGGEDLFAHFSEIRMEGFKTLKENQRVSFEVKTGPKGKQ
AANIQAL*
>GCA_000452945.1_BPMSHR5858_genomic_03573_1 group_3445
MLNGIKRFAHRLGIARRSNSGRAPAASTTFATEFDATWHGDHWQNLLASPLDARHYVMED
WAQPVMPEPDEQAHASPPGRRKRWYEQ*
>GCA_000452945.1_BPMSHR5858_genomic_03574_1 group_7538
MKKTLIVAALSGVFATAAHAQSSVTLYGLIDAGITYTNNQGGHSAWSQSTGSVNGSRWGL
RGAEDLGGGLKAIFVLENGFGINNGTLKQNGREFGRQAFVGLSHEQYGALTLGRQYDSVV
DYLGPLSLTGTQFGGTQFAHPFDNDNLNNSFRINNAVKYTSVNWAGLKFGALYGFSNNNQ
FANNRAYSAGVSYSYAGFNIGAGYLQLNNNFGPTVSNASGAVALDNTFVGKRQRVFGGGL
NYTFGPATAGFVFTQSRVNRATAIGAGASGVSSGIALDGTFMRFNNYEVNARYAITPAWT
VAGSYTYTAGFIENHHPGWNQFNLQTAYALSKRTDVYLQGVYQKVNNDGTGLGAYINGIG
GMSSTEKQIAVTAGLRHRF*
>GCA_000452945.1_BPMSHR5858_genomic_03575_1 greB
MNKAFVKESDGDDDDLDFAQAAIPPGAKNYITPAGHKRLRDELLHLIDEARPDVVKLVSW
AASNGDRSENGDYIYGKRRLREIDRRIRFLTKRLDLAEVVDASRQENADQVFFGATVDYA
TQDGNEHTVTIVGVDEVDLDRGHVSWISPVARALLKARVGDTVTLLTPVGPEPIDVLDVR
YPPPGAA*
>GCA_000452945.1_BPMSHR5858_genomic_03593_1 group_7539
MKRTLFALALVALTLAGCVVVPARPVYYRPAPVVIY*
>GCA_000452945.1_BPMSHR5858_genomic_03606_1 trxB
MSTPKHAKVLILGSGPAGYTAAVYAARANLSPLLITGIAQGGQLMTTTDVENWPADADGV
QGPELMQRFLAHAQRFNTEIVFDHIHTAKLHEKPIRLIGDSGEYTCDSLIIATGASAQYL
GLQSEEAFMGRGVSACATCDGFFYRGQNVAVVGGGNTAVEEALYLTGIAKKVTVIHRRDK
FRAEPILVDRLLEKEKEGAVEIKWDHVLDEVTGDDSGVSGVRIKHVTTGATEDVAVQGLF
IAIGHKPNTDIFKGQLEMKDGYIITNSGLSGNATGTSVPGVFAAGDVQDHIYRQAITSAG
TGCMAALDAQRYLESLHDHK*
>GCA_000452945.1_BPMSHR5858_genomic_03613_1 zwf
MHTDSSFTFVLFGGTGDLSMRKILPALFEAHRANMLSEAGRIVAVARHAADREGYLQWVE
EHVKPHAAKAAGGAFDEAVWRSFLERIVYVKLDLGRAEDYALLRDTVGGLSGIRVFYLAT
GPSLFVPICKALAAVGLNEGARIVLEKPLGYDLRSSNAINDAVGEIFAEDQIYRIDHYLG
KEPVQNLLALRFGNALFEPLWRREWVESIQITIAEELGVEARGDFYDNTGALRDMVQNHL
LQLLSIVAMEPPHSMDSDSVRDEKLRVLRALKPVDPRDIGKVAVRGQYHAGVIKGAQVPA
YATEPGVKADSQTETFVALKVEIENWRWAGVPFFLRTGKRLADRVAEIVVNFRPVPHSAL
GPTALRAGANRLVIRLQPNESIRLYCLAKQPGEGMNLASVHLDLAFDQFFKEGQMEAYQR
LLLDVINGRLALFVRRDEQEAAWRWVEPILNEWARTTKPPKPYAAGTWGPAAASAMLAQH
GTCWLEEEN*
>GCA_000452945.1_BPMSHR5858_genomic_03620_1 group_1061
MPPVTQTDPPGETTVKDKRFILRPGFYDGAQGPFYRGDSVAVEGLLSCFAPLRDAVDVEY
IDAPRPRHALVALIGEAHQSAPVIVLGAGRTPKAPSIAVREHDGTRFIDSPADIRRYLSS
QYGVAHAS*
>GCA_000452945.1_BPMSHR5858_genomic_03621_1 bcr_1
MPHVARRRPDGRLILLLGALAACGPIATDMYLPSLPAIAGGFGVTAAAAQRTLTSFMAGF
SIGMLLYGPLSDTYGRRPVLLGGIALFTLASVGCFVAGSIDMLIAVRFLQAFGAGAASVL
ARAIARDAHEPADAAKVLSMVAIVTAIGPLLAPLIGGQVLRFSGWRGVFVVLALFGAVCA
TAAYLRVPETWPREKRASSAVLNSFAAYGRILVDPVAWGHMLCGGMAFASMFAYITATPF
VYIDYFHVSPQHYGLLFGLNVIGIMLGNFLNARLVGRVGSLKIIAGASLLSGAASFAVAF
FALTGLGGLWSIVASLFFVVGVVGILSANCTTDLMLRYPHNAGAAAAVFGAMQLALGALA
SAAIGALANGAPFAMGVTIGATGLLCLGGRMLVLRWHGRPVKGALGRVDAA*
>GCA_000452945.1_BPMSHR5858_genomic_03630_1 group_7541
MNTAADSLLRLASRVDAIEPFYVMEIVKEAAVLERAGRDIIHMSIGEPDFTAPEPVVDAA
AAALRRGVTQYTSALGIAPLREAIAAHYARAHGLSIAPERIVVTAGASAALLLACLALVG
RDDEVLMPDPSYPCNRHFVATAEGRAVLVPSGPETRFQLTADDVKTRWGERTRGVLLASP
SNPTGTSLEPAELGRIIGAVRARGGFSIVDEIYQGLSYDGAPVSALSFGDDVVTVNSFSK
YFNMTGWRLGWLVVPPALVGTFEKLAQNLFICPSALAQHAALACFEPDTLAIYEARRAEF
RRRRDFIVPAIESLGFKVPVMPDGAFYVYAQCGGVAHPAAGDSAALVNAMLHDAGVVLVP
GMDFGVHAPRDYIRLSYATAYSRLEEAVERLGKLFGR*
>GCA_000452945.1_BPMSHR5858_genomic_03634_1 thadh
MSVQSASDLILPTFDDVTDAAARLAGVAHRTPVLTSSTADARAGATIFFKCENFQRMGAF
KFRGAYNAISHFDAEQRRAGVLTYSSGNHAQAIALAARLAGIRATIVMPHDAPAAKVAAT
KGYGGEVITYDRYTESREEIGARLAQERGMTLVPPYDHPHVIAGQGTAAKELIDEVGELD
MLVAPLGGGGLIAGTALSAAALSAGCAVIGVEPEAGNDAQQSLERGEVVRIPVPRTIADG
AASTHVGAYNFPVIQRLVKRIVTASDAQLVDTMRFFAERMKLVVEPTGCLGAAAVLDGVL
PVAGRRIGVILSGGNVDLARYGEFLRG*
>GCA_000452945.1_BPMSHR5858_genomic_03635_1 group_4899
VKKLIPLIAVAALGLGAASAARAHVSIGVGIGVPIAPAYPVYAAPPPVYYAPPPPPVVYA
PAPVYYGPPAVVVGGGYYGRPYWRHYHGYYGRPYWRY*
>GCA_000452945.1_BPMSHR5858_genomic_03637_1 group_7543
MFGEIARFLLNTVFTLFGAALLLRVWLQAVRVPPYNPVTQAVLQATNWLVLPLRHIVPGV
RGIDWASVVAAVVTSLVYVALMVTMAGVDALSIIPTILVVALLTVVKWALNLVLWLTILM
ALLSWLNPRSPAMAILYQLTAPLLNPLRRLIPNLGGIDLSPILLFVIVQVLIMIVTRAAV
SLTLFGI*
>GCA_000452945.1_BPMSHR5858_genomic_03638_1 group_7544
MDGIFLRFYVHENHRLHWKPLWEWLLEEANRMGIAGGSAFRAMAGFGQHRVLHEDRFFEL
QGSLAIEVEFVVTEDEAKRLIERVSREKVRVCYATIPARLGVIDNLGDGAPGAAPPAA*
>GCA_000452945.1_BPMSHR5858_genomic_03639_1 crcB
LFYSIVAIFVGAGLGALLRWFLSIGLNALLPEVPLGTLASNLIGGYLIGIAVVAFATRAG
LPPEWRLFVITGFMGGLTTFSTYSVEVMTHAVQGEFGWALAVAALHLIGSFTLTGLGMWT
ARAWLAPA*
>GCA_000452945.1_BPMSHR5858_genomic_03640_1 group_4900
MKNNSSLSILIAAACIQAFAATASLAQGPAHPPSYVEGTRVPKGFARPPFHTNPARFSAT
TVSGLAPATVRHAYGFDSIANQGDGMVVAIVDAYDDPKIESDLGVFSKNFSLPPCTTSNG
CFKKLYASGSKPSPNAGWALEMSLDVEWVHAIAPKAKIVLVEAASNSFNDLMTAVDVAVG
AGASVVSMSFGGSEFSSETSFDSHFGAPSNVTFVASSGDSGNGTEYPAASPYVVAVGGTT
LSADASGNYVGETAWSGSGGGVSAYELEPVGQTLWPIPYAGQRGVPDVAYDANPNSGFAV
YDSVTYQGQSGWFVVGGTSAGAPQWAALFAIANSMRTAAGKAKLAGAYNQLYTVGKTAYG
SDYHDVTSGTNGSCGMICTASGGYDYVTGLGSPQALNLVQALVAQP*
>GCA_000452945.1_BPMSHR5858_genomic_03641_1 tam_2
MTQNIYDDPDFFEGYGQLPRSRDGLDGAPEWPSLRAMLPNVAGRRVVDLGCGYGWFCRWA
AEQGAARVLGIDVSARMLDRAREMTTSPAVRYERADLERLSLPSASFELAYSSLAFHYVE
HLPALFGAIGRALVPDGRLVFSIEHPIYMASRRPEWRIDENGHRFWPVDGYRIEGRRTTD
WFAAGVVKYHRTLGTLLNQLIAAGFRIDHLDEWGPSDEQVAAQPALAQERERPMFALVGA
RR*
>GCA_000452945.1_BPMSHR5858_genomic_03642_1 SIRT5
MSDSRRPPSNPSPAMRDGAPPVDAERIDAAVRALSRADALLVTAGAGIGIDSGLPDFRGA
EGLWRAYPGLGHVGYAFHEIASPRAFRERPRLAWGFYGHRLAMYRATVPHEGFGILRRWI
GAMRHGGFVFTSNVDGQFQKAGFDPERIVEVHGSIHAMQCMSSCSAYTWDAESFAPVVDE
PACSLVGALPRCPRCGGLARPNILMFDDADWLGARYDAQHEALRAWLARAGRVAVVEVGA
GTAVPTVRAFSERTGDDVIRINVREAHARRDDVIGLPGGALETLRALDAAWRRD*
>GCA_000452945.1_BPMSHR5858_genomic_03644_1 trmB
MIHDDDPNAPGAPHDATAAPASATRAAPAAGDDDANPLHLRRIRSFVTRAGRVSTGQRRA
IDELGPRFVIPYGSAQPDWDAIFGRRAPRVLEIGFGMGASTAEIAALRPGDDFIGVEVHE
PGVGALLKLIGEQQLSNIRIIQHDAVEVLAQMIAPDSLDGVHIFFPDPWHKARHHKRRLI
QPPFVAQLAAHLKPGAYLHCATDWQNYAEQMLEVLSADPSLENTAQDYAPRPGYRPVTKF
ERRGLRLGHGVWDLVFRKKHAG*
>GCA_000452945.1_BPMSHR5858_genomic_03645_1 uppP_1
MDWILICKALALGIVEGLTEFLPVSSTGHLIVAGSFLRFHPEQAKTFDVVIQFGAILAVC
WEYRRRIIDVVTGLPAQREARRFTMNVVIATVPAVALALLFEKTIKSVLFAPVPVAVALV
VGGAAILWVEGRQRERSEPARVQSIDALTPFDALKVGLAQCCALIPGMSRSGSTIIGGML
FGLERRVATEFSFFLAIPVIFGATLYETAKDWRAFNVDSVGLFAIGLVAAFVSAFACVRW
LLRYVASHDFTAFAWYRIAFGLFVLLVGYSGWIEWT*
>GCA_000452945.1_BPMSHR5858_genomic_03646_1 group_7547
MTDTSAPDRRRFLIAACAAAGMTVSLAACASTFPFIPDHYTFSQGDVQRAVARKFPYQKT
VAQVIDVALANPVVGLLPDQNRVAVRLDARFESPFLRAPVNGKFTVSGQLAYDAPSRSVV
LKAPAVDSVNVDGDAQLYTQQISAAAGLLATQLLTNYPIYTFKPEQLQFAGVNYEPGTIT
ILTNGIRVAIVEK*
>GCA_000452945.1_BPMSHR5858_genomic_03648_1 hemN_1

MTQPIPSEALFRPDLLAKYTANGPRYTSYPTAPQFRDNFDPADYIRAAADPGASASDLSL
YFHIPFCATACFYCGCNKIATRNRKRARPYLDQLKHEIALHAALFDPERTVTQLHWGGGT
PTFLSNAETTELMAATREYFTLASDRTGEFSIEIDPRTASPATLVHLRTIGFNRVSLGVQ
DFDPDVQRAINRVQPLKMTSDLVRAARATGYHSIGIDLIYGLPHQTVSSFARTLDTIVEL
EPDRLSVFGYAHLPHLFKMQRQIDAAALPPPATRIALLGLAIERLTRAGYVYIGMDHFAR
PGDELVRAQRNGTLQRNFQGYSTRADADLIGIGASSIGKVGDVYAQNAKDLPRYGAALAD
QRLPIARGVKLSADDRLRRDVITHLMCNLVLPFSHVEAAYGIRFANKFARELDTLRGFER
DGLLSIGADRLTIHPAGRLYVRNIAMVFDAYLSGASEHRYSKTV*
>GCA_000452945.1_BPMSHR5858_genomic_03665_1 waaA
MLRAIYRGLWWLVAPLAVLRLVWRSRKERGYREHIGERFGFGPGRALARDVDEATPIVWV
HAVSVGETRAAQPLIDALLRARPDAHVLLTHMTPSGRATGEQVFGERVSRCYLPYDLPRA
VRRFLRAWRPSLGLVMETEVWPTLIDECRRADVPLVLTNARMSARSFGRAAKFGAAARDV
FGGFSRVLAQSPADAQRLTSLGARNVAVLGNLKFDMTTPPELAARGRAWRAAIGERPVWV
AASTRDGEEALVLDAFVSLKTPGALLILVPRHPQRFAEVAALVERRGLRHVRRTEWAHDA
AAAAAGRPAASALPADVSVLLGDSMGELGAYYAAADVAFIGGSLLPLGGQNLIEACAVGV
PVLIGPHVFNFTQATADAVTAGACAQVRDPADLACTLDALFADHARRTAMGAAGAAFAAR
HRGATARTVDVLNALLPAPRVAPAANDEGDAGGLDAGAALD*
>GCA_000452945.1_BPMSHR5858_genomic_03690_1 group_4905
MRALRKARLVLHLLRGMATVALRFERATPERRRDLTRRWTLKMLQLCGMRLVVHNDAARL
DARALVVGNHVSWLDIYVINAWRPTPFVSKAEVRAWPVVGWLAERLGTVFIQREKRSEAK
RIMHELSERLGNGELMCVFPEGTTSDGLGLLPFHSNLFQAVVSADCSVQPICLMYEDARG
RQSVAPAYVGDLPLGKSLDMVLANGPLTAHLYVCEPIAATGDRREMAALARASVADALAR
MQAGVERPLDADLARLAAAAYPTPAEPAGEPVAVAEPEVGADAISRREG*
>GCA_000452945.1_BPMSHR5858_genomic_03700_1 groS3
MNLRPLHDRVIVKRLDQETKTASGIVIPDAAAEKPDQGEVLAIGPGKRDDKGAPIALDVK
VGDRVLFGKYAGQTVKVDGQELLVMREEDIMAVVNAK*
>GCA_000452945.1_BPMSHR5858_genomic_03705_1 group_4906
MRNRKFVPFVVALAFAAAAATPALAVTVVRVDGQPMNPNGEPFSATSAPLETTLSKGSIS
ANCVATFNGTITPAGIVNITSTTFTGTNSLCGLIKGSASGTNPWTGQADSATQLTINNAQ
VNVTLLGQCGPSKVVTSWTDANSSLTFSNAALAPDCKVTGTVVTSPKFHVQ*
>GCA_000452945.1_BPMSHR5858_genomic_03712_1 ttr
MSHAIGVRRIGPDEAAACVDALSDVLIDCVEGGASVNFMAPLARDKARAFWREAAEGVAR
GERTLFVAEDADGRIVGTVQMITRQPENQPHRADVAKMLVHRDARRRGVAQRLLAALDDA
ARAAGKTVLVLDTVTGGDAERLYARAGWQRVGVVPDYALMPDGAPCATTFFYKQI*
>GCA_000452945.1_BPMSHR5858_genomic_03716_1 group_4908
MSLPHIDPLYSLSGLFVGFLVGLTGVGGGSLMTPILVLLFNVHPATAVGTDLLYAALTKA
TGTFVHGLKGTVEWRITGRLAAGSVPAAAITLWFLHAHGMHSQETSRMIQFVLGAALLLT
SLSLLLRPQLAAFAAKRTRALPPNPTRTLAATVLTGAVLGVLVSLTSVGAGAIGVTVLLL
LYPALSTTRIVGSDIAHAVPLTLVAGVGHWMLGSVDWAMLVSLLIGSVPGIVVGSHLSAR
APEGLLRRVLAATLVAVGAKLVLS*
>GCA_000452945.1_BPMSHR5858_genomic_03719_1 group_4909
MKSIVSKAVAGMIVAALACGNAFAQASGAAAEASAPAAAQNASKAAAKTARKSNRKLGYA
VRKAISKVNGVNVSNITVRTKGGVVTLEGSVPDASQIDKAVEAAKGVAGVVSVNNKLSVQ
PQ*
>GCA_000452945.1_BPMSHR5858_genomic_03739_1 ydhP_1
MPLPLFALAVAAFGIGTTEFVIMGLLPNVARDLGVSIPAAGMLVSGYALGVTIGAPILAV
VTAKMPRKAALLALIGVFIVGNLFCAIAPGYATLMVARVVTAFCHGAFFGIGSVVASNLV
APNKRAQAIALMFTGLTLANVLGVPLGTALGQAFGWRATFWAVTGIGALAAAALAFCVPK
RLEMPAAGIAREFGVLRNPQVLMVLGISVLASASLFTVFTYIAPILEDVTGFTPHDVTLV
LLLFGLGLTVGGTVGGKLADWRRIPSLVATLASIGVVLAAFAGTMRTPLPALVTIFVWGV
LAFAIVPPLQILIVDRASHAPNLASTLNQGAFNLGNALGAWLGGTAIHAGVPLAKLPWAG
AALAMAALALTLWSASLERRAAPGPAGCA*
>GCA_000452945.1_BPMSHR5858_genomic_03756_1 group_7550
MMRVMVGVDVERLPMFRARPTPCDAPRRREPRMRGSRGVALPGVLAVTASLIVMSFAWFE
IAKTEVRRMTSVASRSIAFRAADAALEACVIALESGAAPHPSAGDGTTSTREPGGWREPG
AFDGIGAFRPYAGWPGAAQAPSCLIEAWRLPARPDVRAFLVTARGVGASNDTAEWLQLQV
ALDGGRVERRWRRVVGRPA*
>GCA_000452945.1_BPMSHR5858_genomic_03778_1 cysE
MRTTLKRTLDAIAEDRRRNAFFKTRCVVVFFRIANYLACRSRLTLALGAPVIALYIVVCD
WIMGIEIPVKTKIGKGLTLYHGTGLVINGYCEIGERCVIRHGVTIGNTLRPDGTYSGVPS
VGDDVEFGAHSIALGEIRIGHRARVGAGAVLLRDVPDGGVAVGVPARVLENGKEQSAA*
>GCA_000452945.1_BPMSHR5858_genomic_03787_1 group_3472
MWSAISDLGDAALTLPLSAACCAWLLRASPERRYAVSWLALLAAGMLVVGLTKILYAGCG
VQIRAIGFRVVSGHTMLASAVWPMALLLGLQWLRSNAALAAGLALAALIGTARVFDEAHT
VSEVVAGWALGTLVTVSFVRWQRAPAMPARLWPYASASLLAVMAIAYGLHAPIQAAIERY
SPFLCRSFPW*
>GCA_000452945.1_BPMSHR5858_genomic_03791_1 kbl_1
MALGEHLRQQLAAKALKRQLERATDAAAAPGVPGTPAAQTASARSRFESMPQYQQVRIMR
EMGEKLRVDSPFFRVHDGVAGATTQIGGREYLNYANYNYLGLAGDPAVSARAKDAIDRYG
TSASASRMVAGERPVQRELERGLASFYETDDCVAFVSGHATNVTVIGSLFGPGDLIVHDA
LAHNSIVQGAQLSGAKRLGFAHNDWRALDALLARVRREYRHVLIAIEGLYSMDGDLPDLA
RFVEIKQRHGAFLMVDEAHSLGVLGRHGKGIREHCGVPSADVDLWMGTLSKVLAGCGGFI
AGCQPLIDILRHLAPGFLYSVGLSPVLAAASLAALERLQAEPQRIATLRERGLQFLTQAR
EAGLNTGTSEGYAVVPVITGSSLKAAQWANALFDEGINVQPIFYPAVEEKAARLRFFICS
THEPEQIERTIAALVRLAR*
>GCA_000452945.1_BPMSHR5858_genomic_03797_1 gmhB
VKNRALFLDRDGVINRDDGYVFEIEKFVFLDGIFELAGAAKALGYLSIVVTNQAGIGRGY
YSEDDFFRLSDWMKGVFATEGAPIDGVYFCPTHPEHGIGRYKVESRFRKPNPGMILAAQH
DFDLDLGASLLVGDKESDIQAGSTAGVGTTLLICDRDASRVATTASAVVRNPRDVIPFLT
GPGPDAGSF*
>GCA_000452945.1_BPMSHR5858_genomic_03798_1 hddC
MREAIILAGGFGTRLRTVVSDVPKPMAPIAGRPFLEILLTRLSEKKFSRVVLSVGFMAEK
IMSHFGDRFAGIDLAYSVESDPLGTGGALKATLPYCEGDHAFVFNGDTYLDLEVDELDDG
WQTGGFPTIVARQVPDTGRYGRLVVDGGRVTGFAEKGVSGPGLINAGCYVLPKDILAGET
AEAFSFETDFMSSAVQSRRFDVFVTRGQFIDIGVPEDFYRAQDELSGICK*
>GCA_000452945.1_BPMSHR5858_genomic_03806_1 group_778
MRERTVDYSFCGMVRSRERLARQIVFHHIPKTAGSSFNQILRTLYRDDEVCNAALDDELD
EVMADETRRYELFVGHFSFDALHRHFGGATRLTFLRDPVQRCISQYHNWHDASRYSDAWI
GRSDTNPDVIKALKMTSEMSLGEFVSSDNLVISDSAQNMMTRYLAPSVEWKKERGYYDAE
LVEKAKRNLVEYFHFFGLTEQFDRSLVLLAHTLGIRPWERSDALLTNRNPKKASFDSVYN
TTPEEGGVLRDYNLMDIELYEFAVKEFNRRFDAGYQKLVECAFEYLADKDTRDMGNAGDF
YTFDMTNAVGARGLHFLESTRLPCGANVLGRWTGLEPRAVWEIPLRAGRDSHVVIEVDYI
DSVSQEALAPEHFTLNGMPARQHAFSAEGSIQRLRLVFSAGAALAGRMLHTLKLTTPLVR
AEDGTRDVGVLLLRLQSYSV*
>GCA_000452945.1_BPMSHR5858_genomic_03818_1 mutL
MSEFTDSAAGRSSTPPADAPSPASRPLRAIQPLPDQLISQIAAGEVVERPASVVKELGEN
ALDAGAGTLRILLDEGGVKRISITDDGCGIPADELPLALMRHATSKIRSLAELEAVATLG
FRGEALASIASVAEMFITSRTEDAAHATRIDAQTGVVGPAAGTRGTTIEVRELYFSTPAR
RKFLKSEQTEFGHCLEMIRRAALARPDVAISVLHNGRAVEHWNASEPAARVAKILGDGFA
TAHLPLDERAGPLAVYGCAGLPTASRGRADQQYFFVNGRFVRDKLLTHAVRAAYEDVLHG
DRYPSYVLFLDLPPEAVDVNVHPSKIEVRFRDSRSIHQFVFHAVQRALARHAGASPETTA
GGHAAHLAPIVPASADSAAAPGASFVRSGPTAGAGVGQPASGNTWLRQSRMTQGTLPVAQ
PLALYDALFGRKDTGAGTPHGATLALEAHDAHDAPDGANAPGAPLYATMPGDDATPAFSP
AGAAGLPMHDEQPLGFAVGQIHGIYVLAQNARGLVIVDMHAAHERILYEQFKRALADRTI
AVQTLLIPVSMTATPVEVGTAEEERETLDALGFDLAVLSPTTLAIRAVPALLKDADLQAL
ARAVLADLHAFGGSRVLTERQHELLGTLACHHAVRANRRLTLDEMNALLRQMEATERADQ
CNHGRPTWYQLTLGDLDKLFMRGQ*
>GCA_000452945.1_BPMSHR5858_genomic_03828_1 menF
MTDEAGAVFALFDDCDSTAAARSSRLYSGFSHERACADPAGLDAVCAAVADDARRGLHAV
VLGDYEFGRDLQLGKRGGGALRFLLFAECRMLSREEADAWLASRDGGSAEPSAAGVAHVT
KSVARDAFDAAIAAVHDALRAGDSYQINYTYRLHFDAFGAPLALYRRLRARQPVRYGALV
ALPGGAWIVSCSPELFVEKTGALLRARPMKGTAPRSDDPREDAAAARFLANDAKNRAENV
MIVDLLRNDLARIARTGSVTVPALFSVEPYASVWQMTSTVEAGIVEGATFADVLRALFPC
GSITGAPKHRTMQLIDALETTPRGLYTGAIGWLDAEARADGAEPAAGAFAPERAASSSPG
TASDGACAGETASKQASKRAGATSPAGACGDFCLSVAIRTLTLDAPSAGGERRGTMGVGA
GIVLDSVAADEYAECELKARFLTDAEPGFQLFETMLATRDAGVRHLDRHVARLRASADAL
GFAFDEAAARQRIGARCVQLGDGEHRLRVALAKDGVLDIVAAPLAPLAGAAVAVLLAPEH
GFAPMRSGDFLLAHKTTRRADYDRAWKAAQACGAFDMLFFNERGELTEGGRTSVFVKLAG
RWFTPPLSSGVLPGVMRGALLDDPAWQASERVMTLDDVLRADALMLTNALRGAMPAQLVR
AAGAAAPR*
>GCA_000452945.1_BPMSHR5858_genomic_03829_1 dnaJ
MAKRDYYEVLGVAKNASDDEIKKAYRKLAMKYHPDRNPDSKDAEEHFKEAKEAYEMLSDG
QKRAAYDQYGHAGVDPNVGAAGAQGFGGFADAFGDIFGDIFGQAAGGGRARGGPQVYRGA
DLRYSMEITLEQAAHGYDTQIRVPSWAACGVCHGSGAKPGTKPETCPTCHGQGTVRMSQG
FFSIQQTCPKCHGTGTYIPEPCVHCHGSGKVKETKTLEVKIPAGIDDGMRIRSAGNGEPG
INGGPSGDLYVEIHIKPHAVFERDGDDLHCQMPIPFTTAALGGEIEVPTLAGRASFTVPE
GTQSGKTFRLRGKGIKGLRSSIAGDLYVHVQVETPVKLTDQQRDLLKQFEKSLAEGGPRH
SPQSKSWFDRVKSFFE*
>GCA_000452945.1_BPMSHR5858_genomic_03830_1 dnaK
MGKIIGIDLGTTNSCVAIMEGNQVKVIENSEGARTTPSIIAYMDDNEVLVGAPAKRQSVT
NPKNTLFAVKRLIGRRFEEKEVQKDIGLMPYAIIKADNGDAWVEAHGEKLAPPQVSAEVL
RKMKKTAEDYLGEPVTEAVITVPAYFNDSQRQATKDAGRIAGLEVKRIINEPTAAALAFG
LDKAEKGDRKIAVYDLGGGTFDVSIIEIADVDGEMQFEVLSTNGDTFLGGEDFDQRIIDY
IIGEFKKEQGVDLSKDVLALQRLKEAAEKAKIELSSSQQTEINLPYITADASGPKHLNLK
VTRAKLEALVEDLVERTIEPCRTAIKDAGVKVSDIDDVILVGGQTRMPKVQEKVKEFFGK
EPRRDVNPDEAVAVGAAIQGQVLSGDRKDVLLLDVTPLSLGIETLGGVMTKMINKNTTIP
TKHAQVYSTADDNQGAVTIKVFQGEREMAAGNKLLGEFNLEGIPPAPRGVPQIEVTFDID
ANGILHVGAKDKATGKENKITIKANSGLSEAEIEKMVKDAEANAAEDHKLRELAESRNQG
DALVHSTKKALTEYGDKLEAGEKEKIEAALKELEDVLKNASSDKAAIDAKVEAVATASQK
LGEKMYADMQAQQAGAAGAAGAAAEGASAQGGAQPADDVVDADFKEVKKD*
>GCA_000452945.1_BPMSHR5858_genomic_03832_1 grpE
MENTQENPTDQTTEETGREAQAAEPAAQAAENAAPAAEAALAEAQAKIAELQESFLRAKA
ETENVRRRAQDDVAKAHKFAIEGFAENLLPVLDSLEAAVGDTSGDLAKVREGVELTLRQL
TSALEKGRVAALNPVGEKFDPHLHQAISMVPADQEPNTVVAVLQKGYTIADRVLRPALVT
VAQPK*
>GCA_000452945.1_BPMSHR5858_genomic_03833_1 hslR
MNFKISTEPGARLRIDKWLWAARFFKTRSLAADAVEKGRVRIGGAAVKPAKEVRVGDRVD
VTIERVVWEIDVLGVCDVRGPASVAQTLYAETEAGRAKRLAELERRRHFREPSAELHGRP
TKRDRRIIDRFSNGS*
>GCA_000452945.1_BPMSHR5858_genomic_03834_1 hemH
MRFDLEPPSHASAAHRVAVLLVNLGTPDAPTPRAVRRYLAQFLSDPRVVEIPQLVWQVIL
RTLILPLRGRASAKKYAAVWLPEGSPLRVYTERQVESVKPLFAANGYRVIVDYAMRYGTP
SIADVLAQLKRAGAERVLLLPMYPQYSSSTTATAFDAAFAALGRMRNQPEVRTVRHYADH
PAYIHALAEQVRQYWAAHGRPAFDAGDKLVLSFHGVPKRTLDLGDPYHDQCQQTAALLMS
ALGLTPFECRVTFQSRFGKAEWLQPYTAPTLKELGAAGVRRADVFCPGFTADCLETIEEI
GIEVRDEFVHGGGKEFHRIPCLNASPAWIAALGEIAAENLQGWPVRVAMAPEAVS*
>GCA_000452945.1_BPMSHR5858_genomic_03835_1 hrcA
MLDPRARTLLKTLIERYIADGQPVGSRTLSRYSGLELSPATIRNVMSDLEELGLVSSPHT
SAGRVPTPRGYRLFVDTMLTVESPIDSDAVTRLVQTTLQAGEPQQRVVAAAASVLSNLSQ
FAGVVLTPRRSHVFKQIEFLRLSDKRILLIIVTPEGDVQNRMIATQRDYAPAQLTEASNY
INAHFAGLSFDEVRRRLREEIDQLRGDMTALMHAAVTASTEEPDDEETVLISGERNLLEV
ADLSSDMARLRKLFDVFDQKTSLLQLLDVSSHAQGVQIFIGGESTLVPIDEMSVVTAPYE
VNGKIVGTLGVIGPTRMAYNRVIPIVDITARLLSLTLSQQ*
>GCA_000452945.1_BPMSHR5858_genomic_03836_1 nadK
MKIGHQFHTVALVGRSNTPGIAEPLASLAACIAKRGFEVVFEADTAQAIGSAGYPALTPA
EIGARADVAVVLGGDGTMLGMGRQLAPYKTPLIGINHGRLGFITDIPASDMREVVPMMLA
GSYEREERTLLEARIVRNGEPIYHALAFNDVVVNRSGFSGMAELRVSVDGRFMYNQRSDG
LIVATPTGSTAYALSSQGPILHPQLQGIVLVPIAPHALSNRPIVLPDDSKIAIQIIGGRD
VNVNFDMQSFTALELNDTIEVRRSKHTVPFLHPVGYSYYATLRKKLHWNEHPSSEEDDDA
*
>GCA_000452945.1_BPMSHR5858_genomic_03837_1 recN
MLRHLSIRDFVIVAALDLEFDSGFTVFSGETGAGKSILIDALALALGERADASVVRTGSG
RADISAEFTPHDRVARWLDEHAFDADDTVMLRRVVDANGRSRAFINGTSATLAQLREVGE
MLVDIHGQHAHQLLMRADAQRELFDTHAGLAADAAAVARGYRAWRDATHAIDAAQAHERE
RQLEREKLAWQLAELDKLAPQPGEWDEITVEHKRLTHSANLIDGVQGALGAISESDDAML
TQLGAIVSRLRSLAEYDPALNDALASLEPAEIQLQEASYSLSHYAQRLDLDPDRLAQVET
RLDALHSTARKFRLPPETLHDEHEARRAQLAELDAAADLSALQAVADKAKQAYLADAQKL
SKARAQAAKALGVAVTTGMQELSMAGGSFEVALVPLAEGGAHGLEQVEFRVAGHAGVPLR
PLAKVASGGELARISLALAVIASAASPTPTLIFDEVDTGIGGGVAEVVGRLLHQLGQMRQ
VLCVTHLPQVAARGDHHFQVAKGEDGEGGTVSTVVPLDRASRIEEVARMLGGLEITATTR
KHAKEMLTA*
>GCA_000452945.1_BPMSHR5858_genomic_03839_1 group_3479
MSDRQDSAATTEAGPPQHHHPVLRRVFKGVLAIAIAAYFIAAAAFLGLRYLLLPRIDEYR
PRIEAFVSQKLHAELRIGRLAPHWSGMQPGVDISRLTIRGRDGRVALSVPHATAALSWRS
LARFAPTLSSLVVDDPDLLTERRADGSLFVAGVAVPTTKTGADDTFSAWLLKQEAIVLRG
GTLRWRDAQHDAPELALTGIRIAVLNDGIVHRIALQAPANGTLFHGPLDFRARFVHQALA
PIGKPSNWTGDAYLSTGPVDLPTLARYANIRVTAYAGRIDNAIWAHFGDGHLYRAGGELR
GYDVALRVRPTQPRLDIPVARFGWDVAIDPKRDYTLHLSRLHAELGQPPLADGTPLSRAL
ALHTLTARYRVPNVDEGQLLSVSGDRVDLGILAEFIRGLPLPARLRNELVRFDPRGLVAN
YAIEVERAKPASPEFVDEERRSGTAPIIRYRFQGDLQGISFEAQEPPPGLSPHGHPRAGI
PGVENLWGHVDADEKGGAARFDTVDAAVTVPGVFDEPRLAFDKLRGRASWTITPAPGERH
ARVDVAVPEFRVENADAAIAVAGSYANPGHGRGSLDLKADFERAAVARITRYLPTSLSDH
LRLYLGHALQAGQVTKGATIVAKGPLETFPYEHDPKAGVFHIVAPFAGGRFEPTPQPPRT
LANGTPNVWPALDGIDGVFELEQNRLRFDIDRARYKGVALAKVTGRIDDLGNPTHSPLVI
EGRAHGPLADLLDYANHSAIAGMTGHIGNLVRAQGPATLALKLTIPQHVAHPHVGVDGAL
GFAGNALEADGVPPVTRLRGNVRFTQYTASVDRLTARFLGGDVRARGALAENGRYAFDID
GRLSLDAARGLNLRGAAAAALERVVGDAPYRLAVRGAKGGLPDITANSDLTGVALEFPAP
FAKPAGTPMPFSFVLAPEPQASGKPLERADLALGPLAATYLLDVERGRPVRAVRGALGMN
RMPDLPQEGVSAAIDVHELDADAWQAFAQGFGKSPAAREQPAASPVDLASFAPKRFALHF
GTLKLLKRNWENVIVGASHIDDIWQANIASNQVSGYLSWAPGGGPTGAGVLSARLAKLVI
PESAEHDLVGRAIDLPTPTHHAMPAIDLVVDQVVARGHDIGRLEVDARNVDENGIPVWQL
DKLELANPAAKLTATANWRTSRRALAHGADEEDAPRRTVFDFTLAIDDAGALLERVGLPR
TVENGRGTLSGKVAWRGGPTAIDYPTLNGRLSLDLAHGQILKVDPGAAKLLGVLSLQGLA
RFLTLDFRDVIGKGLPFETITGTGRIDNGIARTQDFEMKTSPAKVTVTGSVDLAHETQDL
NAHVAPKVSASAAAVGAAIINPFLGLGVLAANLALSQTLAHAFAMDYTITGSWAHPHIER
VSGDRGKMGFAPAAVEH*
>GCA_000452945.1_BPMSHR5858_genomic_03840_1 nit1
MTDRHASASPFPVAALQMVSTPDRERNLAEAGRLIADAAASGARLVLLPEYFCFMGHQDT
DKLALAEAYCDGPIQRFLAERAKAHGIWVIGGTLPLKAPEPSRVLNTTLVFDPQGREAAR
YDKIHLFNFEKGDESFDEARTIRPGDAVRTFDAPFGRVGLSVCYDLRFPELYRRMGDCAM
IVVPSAFTYTTGRAHWETLLRARAVENQCYVLAAAQGGKHENGRRTWGRSMLVDPWGEIV
AVRDEGAGVVAGEIDPARIADVRQSLPAWRHRVLA*
>GCA_000452945.1_BPMSHR5858_genomic_03841_1 tldD
MNIIEPGIPQGDFLQGARNLSLAKDILLTPYGLDEALLTRTLADIFTHRVDYADLYFQAT
RSEAWSLEEGIVKSGSFSIDQGVGVRAVAGERTAFAYSDDLSPEAIRQAAVATKAIAAAG
GGKQKIRAATALKGVSGRDLYLPADPLASLDATAKVKLLERVEQMARGRDPRIKQVMAGL
AGEYDVVLVARSDGALAADIRPLVRVSVTVIAEHNGRREIGTGGGGGRFDYGYFTDDVLA
RYVDDAVHAALVNLDARPAPAGAMTVVLGPGWPGVLLHEAIGHGLEGDFNRKGSSAFAGR
IGEQVAAKGVTVVDDGTLPNRRGSLNIDDEGNPTQCTTLIEDGILKGYIQDTLNARLMKM
PVTGNARRESYAALPMPRMTNTYMLNGDKDPREIIESVKHGLYAVNFGGGQVDITNGKFV
FSASEAYMIENGKVTYPVKGATLIGSGPESLKYVSMIGNDMRLDTGVGVCGKEGQSVPVG
VGQPTLRLDNMTVGGTAS*
>GCA_000452945.1_BPMSHR5858_genomic_03842_1 aroG_1
MPPHNTDDVRIRELKELTPPAHLIREFALGEAVSELIYNARQAMHRILHGMDDRLIVIIG
PCSIHDTKAALEYAGRLVQERERFASELEIVMRVYFEKPRTTVGWKGLINDPHLDNSFKI
NDGLRTARELLLQINEMGLPAGTEYLDMISPQYIADLISWGAIGARTTESQVHRELASGL
SCPVGFKNGTDGNVKIAVDAIKAASQPHHFLSVTKGGHSAIVSTAGNEDCHVILRGGKAP
NYDADSVNAACADIGKAGLAARLMIDASHANSSKKHENQIPVCADIGRQIAAGDERIVGV

MVESHLVEGRQDLKEGCPLTYGQSITDACINWDDSVKVLEGLAEAVKARRVARGSGN*
>GCA_000452945.1_BPMSHR5858_genomic_03845_1 yabJ
MKRYGVGEAKGTGGQVMPFARAVEADGWLYVSGQTPMVNGEVVEGGIVTQSKQAIENVIA
ILKEAGYGLEHVVRCGVWLDDARDFASFNKVFISYFGEHPPARACVQSSMVIDCKVEVDC
IAYKAPAK*
>GCA_000452945.1_BPMSHR5858_genomic_03847_1 hexR_2
MTPFAEQPAFDIVARIAECAPELRDAERKVAALILGDLAQAAHASIGALAARADVSVATV
TRFAKAVGCRDVRELKLRLAQAAAVGQRFLAPAAADAQRDETPVSRVYEDVRIALEHNHR
LLRQTSFDAAAAALAAANMIYVYGQGGGSTALADELRFRLVRFGRPVASYQDALLQRMVA
ATAAPGCVVVALSVSGRVPEQLESCALAKRYGATLVALTAPASPLAQLADHLVPVVAFET
DFIFKPSTSRYAMLMALDVLVTEVALRLGDDCRELLRRMKHALDMHRGGGERQPLGD*
>GCA_000452945.1_BPMSHR5858_genomic_03848_1 group_4934
MKVTNYQEATIDPFGKGLGNVPSASVPLSDAGRLEWNLLAEDVSLPAAVLYADRIEHNLK
WMQAFVAEYGVKLAPHGKTTMAPQLFRRQLDTGAWGITLATAHQTRAAYHGGVRRVLLAN
QLVGRHNMAMIAELLTDPDFEFFCLVDSVDGVEQLARFFGAAGKPLNVLLELGVPGGRTG
VRDDAQRDAVLAAIARHAGVLELAGIELYEGVLKEEGEIRAFLRRAVKLVHELASAGRFA
RAPALLSGAGSAWYDVVADEFAQAAAAGVIDVVLRPGCYLTHDVGIYRKAQTDIFARNPI
ARKMGEGLLPALQLWAYVQSIPEPERAIVALGKRDAAFDAGLPEPARHFRPGAGGLPREI
APGEGWEVTGMMDQHAYLKIPAGADLKVGDMVSFDISHPCLTFDKWRQLLVVDPQYRVTE
VIETYF*
>GCA_000452945.1_BPMSHR5858_genomic_03849_1 yhbO
MAAKKILFLTGDFAEDYETMVPFQALLAIGHHVDAVCPGKRAGEKVKTAIHDFEGDQTYT
EKPGHQFTLNATFDEADATGYDALAIAGGRAPEYLRLDPKVIALVRAFAEAKKPIAAICH
AAQLLAAADVIRGKRISAYPACAPEVRLAGGEFADIPVDAAVTDAPFVTAPAWPAHPAWL
SQFLALLGTRVEL*
>GCA_000452945.1_BPMSHR5858_genomic_03850_1 group_7559
MEEQDRIAILQQFGRTYRAFMSAFEAHVGQPLPRWRIMVALEQHGGASSQKRLVELLRVD
PGALTRQLKTLETLGWIEREADARDNRVTNVKLTQAGRAAFEACLPRRNAFLHDTMASLP
DDVLNALSGALSMLEARVAEAGAK*
>GCA_000452945.1_BPMSHR5858_genomic_03851_1 bmr3_2
MAVHTAAHHSSGQVLPFRESLLAMLGISFVTMLVALDQTVVGTALPTIVAELRGFDLYAW
VATSYLLASVITVPIFGRLGDYYGRKPFVIASLVVFTGASVLCAMAQDMLSLVLARGLQG
IGGGMLVGTAFACIPDLFPDSVVRLRWQVLMSSAFGIANAVGPSLGGVLTQNYGWRAVFY
VNLPVGLLSIFFVWRYLPHMRHVEHTRKMRLDWPGALLIAVTLGAMQLFVEWLPKYGIGG
WPTLLLVAALAGGVALWRWEKRCAQPILPFDMFSNRALSTLFILAILAGFAMFSLLFYAP
LLFQGGFGMSPQQAGVVITPLVVFITIGSIMNGRVITRIRRPNAMLYVGFLLFALACAGV
VASTHATPGWALMALMVAGGVGLGFVLPNLTVFAQQTAGREHLGIATALLQSLRMVGGML
GTALTGTLVNQLYANGVRDALAADGALRWQARLADPQVLIDRAAQSRVVDELVRAGHHGA
HLLEAARESLVGAIHIGIGVGAFVALAAVWQSRCVPPVELRRKLEPAVLAE*
>GCA_000452945.1_BPMSHR5858_genomic_03854_1 group_3483
MLTLRKIGVGMLVCALATGIAHAETVKLVANLQPSSEVPPTTSRGAGALDATYDTATRTL
RWHATYRDLTGPATAAHFHGPAPVGQNAGVQVPIPKDALASPIVGEKALTDEQVGDLMAG
KWYFNVHTKAHPAGEIRGQVLPAN*
>GCA_000452945.1_BPMSHR5858_genomic_03855_1 group_3484
LFGGRLAGGGGRSYAWAFDYQEGLGRYAAAGFAWYNEGHIPNHHRDGQAVQLWGRLPLRE
NRFVISAGVGPYRYFDTVAATEGRGYSNSHGWGMLMSVRAAYYTSHRWLAQLQVNRTQVF
SGPNTTSLMFGIGYQLDAPDTPGPRDRAAGRAGRVTANEVTAMLGETILNSRRSPTALGG
SLEYRRGIAKYVDWTATYLYEGAKQSVRRNGIASQVWLTRAFLDDKIALSAGAGAYLTLN
ERDIRGRPGEGDGRVSGIVSISASYRFDDRWLARVTWNRVVTRYDRDTDVIQAGLGYRF*
>GCA_000452945.1_BPMSHR5858_genomic_03856_1 mshA_1
MNLSSPLSNPSPRVRRDRQDFDVTTPSVSQAAGVLRDAPPRAAFAGDHAGVAALPRPVAI
NGKFTSQRLTGVQRVAHEFTSALARLLPGERNPTLVVPRDHASDTLPPTVARRVVPRLRG
ALWEQLALPFATRGQTLVSLCNIGPLFKRNQVVMIHDVAVLDFPQGYSLKFRLWYRFAFW
MLKRRARHILTVSRFSKERIVARLGVAPTDVSTIVSGVDHFGRIEGDPSVLDRLGLAYDG
YVLIVGSLAPGKNLARTLEAIARLERMRPELRFVIAGGSNVRIFGASALGERASAGNVTW
AGYVSDGELKALYENAGCFVFPSLYEGFGLPPLEAMYCGCPVIVSREASLPEACGDAALY
CDAHDAIDIAATIAQLMGDAELRRELREKGRARASRYRWDAAAKQLIGVLRALD*
>GCA_000452945.1_BPMSHR5858_genomic_03857_1 group_4936
MFKPLAWASAAALALSGCALAPGPALDSSRMNDNLSAPTDSTVYDVKLITPQLVYTLKQA
DEADTRAKEAGLAQSLPAASADYRVGPDDVLGIVVWDHPELTRGGGNGTGADTSPLADIG
TLQASGGLGGVLPQQVSAFGTNGQGEVDSPGQRVAANGTIFFPTLGRVRVEGMSPVQIAA
LLARRLDKQIRNPQIDVRVMQYRSQRVQVTGDVKNPGQLSLTGSHLRVVDAINRAGGGNP
DADLQRVLLTRGDQVMTIDVNRILNRGDLRQNIVLQANDIVHVPDRTQNRVFVMGEVPKP
QTVYMNQGQLSLADALSAAGSIDPMGANPRQVIVIRHPNPPLAQAPGVQSGLQEGLKKVS
YAPERNKPEVFRLDMTQVDALMLATEFDMKPLDVVYVGTAPAARFNRMLSQILPTAESFY
LIWSVSRGR*
>GCA_000452945.1_BPMSHR5858_genomic_03858_1 wcaJ_1
MVGLLNKLLDVALVAVGALLAQMLYDGGLFDVSDAQRTAIALLCALTLLVFAAIGVYDGA
REQMSYRTLSRVLLGWLGVAAAAAAFTLLLQRDADVSLAWLGRTMLMSGAVLLLGRASIH
GVLNGLRRTGAKPRAVAIVGSEVYGRAVIEQLRASSLHGYEAVCVFDDSARAADAELRVG
GVPVVSDLREFKRRVRAGAIKEIWLALPLSHERRIQRIVREFRHDFVNLRFLPDVRGITF
FNRSVAQVAGMPAINLATSPLSIPQLWPKFIFDRLFALAVLIPLSPILAALALAVKLSSP
GPVLFRQKRKGVDGREFEILKFRTMRVHAEETGVVRQASRNDSRITKVGAFLRRTSLDEL
PQFFNVLFGQMSVVGPRPHAIEHDDLYKELVDGYMYRYRVRPGITGWAQVNGYRGETRKV
EKMAARVKFDLFYMQNWTFWFDIKIILITLVKGFVGRNAF*
>GCA_000452945.1_BPMSHR5858_genomic_03859_1 group_7562
MLHQNNGFHANALLGSLADDSLRALAPHLELVKIKSAQLLCETDEPMRHLYFPTTAMMSV
LYLMEDGAMVEVAAVGSEGVVGVSTLADYACGGASGRIEVRSGGYAYRVPTQVFRREFDR
SIDTFQLMLRYWQAAMTQIARGALCNRHHSVSEQLSRWLLLAHDRVEGDELAVTQQTIAN
MLGVRREGITEAAGKLQEAGLIRQRRGHITVLDREGLEARACECYGVIRGEFNRLILEAR
RDAHAPQPIPAEGRPLRVAGYHGAVRSAA*
>GCA_000452945.1_BPMSHR5858_genomic_03860_1 group_7563
VTYREYKEPATRAGGIIELPHRTDANRFLASLGVAERATLASHLQLVHVKSGQVLCEPGM
ALEHVYLPVTTVISIQYASSDGMTLEIAEIGNEGIVSPQLVGTDGASPCRVITCRDGFSY
RLGARVLSNLLEESAQTRQAIFRCTHLLMAQAKQISFCSRHHVLKNQLCRWFLLAYDRSR
SVEIQVTHSMLAQMLGVRRETVTDAAGDIQKMGLIRQYRSSIILVDLPGLDAQSCGCHAI
IREEMKKIQAAPTSASGTLTELRS*
>GCA_000452945.1_BPMSHR5858_genomic_03861_1 group_7564
MKNELRTIIKDVAHLEASIDHIADGDDLYEAGLSSLNTIQLMLAIEKHFNIEIPDEMLNR
QLFQSIDSLAGAVTQLQRAEQSA*
>GCA_000452945.1_BPMSHR5858_genomic_03862_1 group_7565
MKRFDAETGNARVAAVESLLADDHTLNEAARRVAQVAARFADAVDRDARFPTEAVDAMRE
ERLLGALVPIELGGRGASLAAVAAACRIIGQACSSAAMIYAMHQTQVASIVDHALSSDWH
RRFAEQLAEREWLLASATSEEEVGGNLRNSRCAIEADGDMFTLAKLAPTISYGAQADAIL
VTARRDRDAPAAEQVLVTLLKENATLTRRSGWDTFGMRGTCSEGFALDARGYCEQIFPVP
FAQIAERTMVPTSHILWAAVWTGIANDAFLRAHQFFRAQMQKQGGALPPSGRRIADALAL
LQAMQARIDGALRLHEHSASRSWSATMAQAAEINTLKTYVSTTALEVVKHATMICGMASY
KNGTPYTLGRHIRDLHSAPLMISNDRIEANTANLLLALRPATLEREI*
>GCA_000452945.1_BPMSHR5858_genomic_03863_1 group_7566
MTQPSVRAAETANAPIDRSGVNKLRDELVEAGLLISTGIQGLFGRSERFERVVDALDAYI
TRLGADQHAEVLRFPPAMSRPEFERSEYLKSFPQLAGTVHSFCGNEHDHQRVLQCLDRGD
DWTESQKPTYVVMTPAACYPVYPVLAQRGALPEDGRIVDAFSYCFRHEPSLDPTRMQLFR
MREYIRVGTPEQILAFRQDWIERGTRMIDALQLPNSIDLANDPFFGRGGKIVASSQRDQN
LKFELLIPIEYDARLTACLSFNYHMDHFGLLWGIKTADAAVAHTGCVGFGLERLTLGLFR
HHGLDPHAWPQAVRDVLWGHR*
>GCA_000452945.1_BPMSHR5858_genomic_03864_1 group_4937
MSSILLDAPGNGASLEDIFTDIRHRVRHYRPHALHGPQMVWKQTNCYVDLWIEVLGWWGL
NPYAALPFTITLDFEGDQFSFFKFPFEDLETLYGIVVQEHAIFEPVDVHVEHQVARGNLM
LVEVDAWYLPDTRGNSYQSEHTKTTIGIDAIDRERRRVGYFHNSGYYLAEGVDYDGLFGK
HGPAHLVPYVERAKRQFKPLEERELCDASLALLARHLKRRPASNPIAAFREQLARDAKTI
ASQPISYFHAYSFNTLRQLGANFELFGRYLRWLEASGCGGPIDTLVAACDTIASEAMVLE
FRLARASARGKEERGESSLDILERAHASLIDGVARIAPPGRVEPNPPFGTLYVPPRPVTA
VR*
>GCA_000452945.1_BPMSHR5858_genomic_03865_1 group_4938
MKSAPDRVARGAAQWTLIATPAGAIARPSELGEAGWCAASVPGTVAQALAAARRFDPAHP
YPLGDSDYWYRTTLHGAGPRIVRLNGLATIAEVWLDDTLLLCSDNMYVAHDLPVTLGGAH
RLALCFRSLDRHLAEHPPRGRARWRTRLVDTPALRGVRATFLGRMPGWFPAIEPVGPWRP
IDIVNPAGAPTIVHDTLRATLDGRDGVLDATLEFAAPLPRTARAQLVCGEHAAPLEATGP
RTARATLRIANVTPWWPHTHGEPALYDVGVAIGGATIALAKTGFRTLAVERGADGRGFAL
SVNGTPLFARGACWTSADPVGLHADAPAYRRALALARDAGCNMIRVGGTMIYEADAFYAL
CDELGLLVWQDFMLANFDYPSNDPRFAESLKREAEQFLGRHMARPSIAVLCGGSEIAQQA
AMVGLAPDERRVPATEQWLAELCAAHRPDAAYVSDSPHGGVLPFAPREGVTHYYGVGAYL
RPPEDARRAGVRFASECLAFANVPCDATLASIGSPAAHEPAWKRAVPRDPGAPWDFDDVR
DHYLRALYGVEPARLRSIDPARYLTLSRAVVADLVGETLAEWRRVGSSCAGALVWQFQDV
MPGAGWGLVDAHGRPKSAWHALRRVSQPRQILLTDEGLNGLDVHVLNDAPAPLEARIELV
ALRDGKTPIARAARTVHVAAHAGQCVNSADLLGRFFDFTYAYRFGPREHDVVIASLYASD
GALLSQAFHFPERTAPTVFERGDIGLEASAAYRDGRWCVQVQTRTFARYVHVCAPGLLPD
IDWFHLAPGAAARIEFAADPHSPAPDHRPPEADAAHCAPPAIEVRALNSNKTIRPRIEN*
>GCA_000452945.1_BPMSHR5858_genomic_03867_1 zraR_1
MTTSAIIGGPSRGSIRQRPLIYWTHQPSTLLRKELARRDWKISVVTSANQVRTTAGEVTG
GILDLSAVSIDEMNAVAAVCASLRNVAWVALVEAQQANVPNVRTLLRDYCFDYITLPASH
QRIADAVGHAYGMECLFAPSTAPIASADQGMIGTCDAMLQLFDTVRRIARTDAPVLVSGE
TGTGKELTAAAIHQHSARRAGPFVAVNCGAIPPHLLQSELFGYERGAFTGASTRKIGHVE
SANGGTLLLDEIGDLPLESQASLLRFLQERTIHRLGGSEPVPVDVRIVSATHVDLRGAMN
EGRFRADLYHRLCVLRVDQPPLRARGKDIELLARHMLERFRADARHRVRGFSTDAISALY
KHDWPGNVRELINRVRRALVMAEGRLITAHDLELDHCTEVAPLSVAAIRKSIEREVIELA
LLRNRGRLAGTARELGISRATLYRWMEAYGIERPRSAASPAN*
>GCA_000452945.1_BPMSHR5858_genomic_03883_1 group_4941
MSSSDLIQPRQVLVSIASATATYTALSPHVPGFLHVVAQGEAPLHVGNIHLAHDPGFVGG
VLLEVVGIAGRPSPTVKQYRTPAYSFESCYHREIFIRGANGIIPVKVGLFRREAAQAAEA
ALDAD*
>GCA_000452945.1_BPMSHR5858_genomic_03890_1 group_7567
MIAPDPESTFRGLPLTPEQNAEVMHYIKVRTQRGLPWDTPELEAMIRDMMLPPEDEAADS
DFLSDEARAVAERAMSSVEDEMDPIEAREEWRAAMEAENMKGLRR*
>GCA_000452945.1_BPMSHR5858_genomic_03909_1 group_4945
MTTFRIEAACLPIFVADARGEQVSVGGFAVDDAAAGKRPPAGDGLQRLAQIHRRDLEGFA
IFGNRAARDHEPLFR*
>GCA_000452945.1_BPMSHR5858_genomic_03917_1 group_4948
MTLPLKLSQKRLAAARGPRAVPVRRAIGVALVAAWALAALPAHAQDDAGDDAPQAAFASA
LPEEQKDLPNVALTSQIVYQVLAAEVALQRSLPAPAYQTYLALARDTRDPRMAQRATEIA
LAAQSPADALTAANLWREYSPGSQRAAQVDAALLVLGGKPAEAQPMLSQELARATGENRG
QAIIALQALLARGPNRVGGLTVLQDLLKNDMGRPEARLAIARQQLATDDKDGATQSLKEA
LRIKPDYLPAALMLSQMGPGERAAGIASFEKFVQQNPKSRDGRLALAQLYLADDRLDDAQ
KQFDAMRRNDSSDPTPLMAIALIKIQQKHLDDATTYLKQYVKVAQKKPGADVGQAYVYLA
QIALDQNNEALAAQWLDKVDEASQQYVPAQVTRAQLLQKQGKADEARKLLANLQASDPRD
AAVIARTDASILFTSKRYKEAADRLAQAVEDFPDDPDLRYDYAMASEKIGQYTTMEQQLR
LLMRAQPDNPQAYNALGYSLADRNLRLQEASKLIEKANSLAPNDAFIMDSLGWVKYRLGD
TAGATAILKRAYDLQPNAEIGAHLGEVLWRSGSRDEARAAWRAAQKLEPDNDTLVQTLKR
LQVNGL*
>GCA_000452945.1_BPMSHR5858_genomic_03950_1 catD_1
MTVIVVANPKGGVGKSTLSTNLAGYFAAQGAWVALADLDRQQSAHAWLDLRPAGLPAIET
WALDPDSPSKPPRGLEYAIVDTPAGLHGNRMNVALEFADKVIVPLQPSMFDILATQQFLE
RLASEKAVKKGAIKVGIVGMRVDARTRSADQLHRFVEGLDLPVLGYLRDTQNYVQLAAHG
LTLWDVAKSRVDKDLEQWGPIVEWAQQ*
>GCA_000452945.1_BPMSHR5858_genomic_03952_1 cobQ
MIANANANANANANANANANANAARPRGTLMIQGTTSDAGKSTLVAGLCRLARRTGARVA
PFKPQNMALNSAVTANGGEIGRAQALQALAAGIEAHTDLNPVLLKPTGDRGAQVIIHGTA
RANLDARAYHDYKPTAMRAVLESYGRLRGAYDAVIVEGAGSPAEINLREGDIANMGFAEA
VDCPVVLVADIDRGGVFAHLVGTLACLSDSERARVRGFVINRFRGDPALLEPGLRWLEAR
TGKPVLGVLPYLHGLTLDAEDMLPASARTSAARRDAGVLRIVVPALPRISNHTDFDALRA
HPRVDFTYWKRGPVPDADLLILPGSKNVLADLAWLRDAGWDALIKRHLRYGGKVIGICGG
MQMLGRTLADPHGVEGAAGATSAGLGLLDYATTLTPEKTLVNAAGRLAFGGDARVAGYEI
HMGRTEGPALASPALMLAGRGGERPDGALSADGQILATYLHGLFDTPHACAALLEWAGLD
GAEALDYPALREASLERLADTFAEHLDLDRVFAAFA*
>GCA_000452945.1_BPMSHR5858_genomic_03962_1 group_4956
MQGFKRKIVYVACFELIAIAITTTGFALLTGQAPAHASIAAVASSAIAVLWNLVYNTAFE
WWEARQSRRGRGLLRRIAHAAGFEAGLVVMLVPLFAWWLDVSLWQAFVLDLGLIAFFLAY
TFAFNLAFDRVFGLPASAQPAPSEASPG*
>GCA_000452945.1_BPMSHR5858_genomic_03968_1 pqsE
MQHGTPGRIDSRLAIVGTADVPLYVIVNDEAATLVEGGLSGMTELVWRQLHELLSDYGGI
RHLRYWLITHSHYDHCSLLGTLAPRMPWLHIVGSAATADALQSPSACRTIRALDAQACAA
WEPVAAAAELSDIPLYPLNPGRSLDIGEGMRMRALALPGHSACLLGYHCPQLDLLFVSDA
LGEYHAPTQWLPLVFQDLAAYRLSLDAIERQHASRIALGHHGIVGGGIARHAVRHARDGL
AARDDEARAARGDANATRALARQWTERYAARSANVVPRALHLKSMERMIDLFQRAA*
>GCA_000452945.1_BPMSHR5858_genomic_03988_1 cdhR_6
MRLIVYILVFPDLNLLDLSGPLQVLSSANELSREAGADAPYDIRVVARGGRSVRTSTGIS
LSTQPLAPIDAPADTVIVAGGAGVDAAARCADTLAWLRAHARQARRVVSICNGAFLLAAC
GLLDGRRAVTHWQCCDALARRHPRVRVEHAPIFVQDGPIWTSAGVTAGIDLCLRLVSNDC
GHTLALALARYLVVFLVRPGSQAQFSESIEMQSASGQFADLHAWIRRHLRADLSVPTLAA
RVNMSERSFVRHYRNAFGTTPAKAVERIRVETARNLLGETALPVKQIAQRCGFGSVATLR
RSFARAFDTSLHEYRERFRQA*
>GCA_000452945.1_BPMSHR5858_genomic_03989_1 inhA_2
MSATARRPLKTGFLLAPGVAVMDLFGAHAVFGFAPRSELHLLWKTREPISALPPFPIAAT
TTFADCPDELDAFVVGAVPADVIADDEVIAFVRRQASRARYVIGICGGVLLLGAAGLLNG
RRATTNFHVLDALAELGARPVGGGEVVIDGNLYTAGPATGGFEAALLVLAELRGAEAAKH
VELTIEYHPRAPFGVGTPALAGPALTNEVLAAHAWFFDPCKDAARAAYGRGR*
>GCA_000452945.1_BPMSHR5858_genomic_03992_1 prpB_2
MNAREPNFTESRSARLRRMLTSPSLEFLMEAHNGLSARIVREAGFKGIWASGLAISAQFG
VRDNNEASWTQVVDVLEFMADASDLPILLDGDTGYGNFNNVRRLVRKLEQRGIAGVCIED
KQFPKTNSFIDGERQPLAEIDEFCGKIKAGKDSQSDPDFSIVARVEALIAGWGMDEALRR
ANAYAQAGADAILIHSKLSRPDEILQFAREWSGRAPLVIVPTKYYSTPTDVFRQAGISTV
IWANHLIRASASAMQAVAREIQDSETLVNVEERVASVNEIFRLQDADEYSAAERIYLSSS
ARASSAALVLAASRGNGLEAVTEDKPKVMLPVAGKPLLRWLVDGFKKQGVNDITVVGGYR
ADAIDTSGVKLVVNERHAQTGELASLACAAERLTGDTIISYGDLLFRSYILRDLAESEAQ
FSVVVDSSQTQPSNQSVRDFAFCSAADDRGLFGQKAYLRRVSSDASEAAPHGRWIGLLNV
RGAGVARLKAMLGTLQARDDFDALDLPALLNALVDAGEQIEVQYVHGHWRGVNDLDDFRR
AGDFAHGQTPYAEQNAGSGSAR*
>GCA_000452945.1_BPMSHR5858_genomic_04020_1 group_7574
VRRLFRQWWAGLSFCAAALAHAQTAPDPQVLLAQSDAIRTPAKSFVLTATLTEYRAGKQV
DGNTLSIYSKPDAPGGAFRTLVRFVAPARDTGKLMLKNGNDLWFYDPANQASIRISPDQR
LLGQAANGDVVTVNLAHDYRAALKGAEDVMDGDRQTRHCYKLSLTGNAPGLTYHRIEMWL
DASNSRPVKAKFYSESDRLLKSAFYRRYTAELGVERPTETVIIDGLDPNWVTVMRFSGYA
WRDIPDAWLQRDYLPRFQAQ*
>GCA_000452945.1_BPMSHR5858_genomic_04029_1 group_3999
MRLPEFNRDTATDEQKAVLDEILSEPRGNLNGPFLGWIESPALAQHAQRLGAFCRYRTGL
PPRLSELAILVTAARWRSQAEWHIHHPIALEAGVPAAAAEAIRTGAAPSFDDADDALIFR
FATELYETKRVSDATYDAAVARFGHRVVVNLVALLGYYALVAMTLNVFDMRADGQTALPF
AE*
>GCA_000452945.1_BPMSHR5858_genomic_04030_1 group_4971
MTTVFHRAPRASLPIAVAGDGIEIVDSTGTRYIDACGGAAVSCLGHSNQRVIDAIKRQAQ
QLPYAHTSFFTTQPAEALADRLVAAAPAGLEHVYFVSGGSEAIEAALKLARQYFVEKGEP
ARRHFIARRQSYHGNTLGALAIGGNAWRREPFVPLLIEAHHVSPCYAYREQRADETEEAF
AQRLADELESRILALGPDTVAAFVAETVVGATAGAVPPVREYFRKIRAVCDKYGVLLILD
EIMSGMGRTGYLFACEEDGVAPDLLTIAKGLGAGYQPIGATLVSDAIYRTIVDGSGFFQH
GHTYVGHASACAAALEVQRVIDEERLLENVKARGEQLRASLAARSAEQPHIGDVRGRGLF
VGVELVRDRDTKAPFDPRLKLNALVKREAMQRGLMVYPMGGTVDGHLGDHVLLAPPFICT
APQIDTIVERLGDAIDAALASIGAAALR*
>GCA_000452945.1_BPMSHR5858_genomic_04031_1 group_4972

MKFLTPVVVTGSPHDIGRQLGELARPAMDAYIAQSGAWRAVARWRGHPFVAELRRAAEAA
FPALVAELDGIAAGLGWHAQDVFVWNCRGELIHNAPDGCTTLAACTPRERMIAHNEDGDP
HLRGKCMLVDVRPDGKPGFVGFYYPGSLPGHTFAASRAGLVQTINNVRIRRPAAGVPRMI
AARAVLDAASLDDALRVLRDTPHASGFHHMLGCAGDARIVSVEASVARTSVLAVEGVYGH
ANHLIHPGGDAEAQIVTASSADRQARLAQLLPPLRGAADASALWCVLADRAGGGLPIYRD
DPDDPDDENTLATALFSIGDDGVAFEVRERGRVRFQQFVPRDAQRPPVHRHERGHA*
>GCA_000452945.1_BPMSHR5858_genomic_04036_1 shc
MNDMTEMHTLDATAAPAGLDAAVARATDALLAAQQADGHWVYELEADSTIPAEYVLLVHY
LGEAPNVELEQKIARYLRRIQQPDGGWPLFTDGAPNISASVKAYFALKVIGDDENAEHMQ
RARRAIHAMGGAEMSNVFTRIQLALYGVVPWYAVPMMPVEIMLLPQWFPFHLSKVSYWAR
TVIVPLLVLNAKRPVAKNPRGVRIDELFKGAPVSTGLLPKQPHQSAGWFAFFRAVDGVLR
LVDGLFPRYTRERAIRQAVAFVDERLNGEDGLGAIYPAMANAVMMYAALGYPEDHPNRAI
ARRSIEKLLVVGEQEAYCQPCLSPVWDTSLAAHALLETGDARAREAAVRGLDWLVPRQIL
DVRGDWISRRPHVRPGGWAFQYANAHYPDVDDTAVVAMAMDRVAKLDRTDAYRESIARAR
EWVVGMQSSDGGWGAFEPENTQYYLNNIPFSDHGALLDPPTADVSGRCLSMLAQFGETSA
SSEPARRALDYMLKEQEPDGSWYGRWGMNYIYGTWTALCSLNAAGLGHDDPRVKRAAQWL
LSIQNADGGWGEDGDSYKLDYRGYERAPSTSSQTAWALLGLMAAGAVDNPAVARGVDYLL
GTQREHGLWDETRFTATGFPRVFYLRYHGYRKFFPLWALARYRNLKRANATRVTVGM*
>GCA_000452945.1_BPMSHR5858_genomic_04042_1 group_1898
MTPERPDASRSTSPDNDDWEDDASFAPGAPEHDFAVRRVTLIVMLVAAIVLPCVYVAVMA
YNDMRTREATARDVTMRTVRVAEEHALKVFDLSETLDARIVDLVQDLDDAAVRNAEADIH
DALNTIGGGYPQVASVSIFGASGMLLANSLYYPAPYASIAERDDFVGIRDGKVIEHISRV
MPGPLKAANPGPVFNTGVARRHSDGTFAGMVSIALKSSYFNAFYRDLLGGADAPMSMALT
RTDGAVLASYPPASRGAPDDAASQADSANPFGGAQGDQHAGIVRVKRDGDSEIVAYRQVG
SYPVFVSCAYRTSAIRHAWYEHLSVLFISMFAPSAALWGVIWLSLKRLRAEEQAWERWQA
EASMRRSIESAYRQSRKMETLGNLVGSVAHDFNNLLMIISSNVQIARRRGVQHLDTELAA
IERALKNGQSLTRQLLGVARKQPLRSETIDIAHWLASCRELLKTSLGSKSTLAVDTQLDL
WPVRVDVAELELAMINLAVNARDAMPQGGRFKISAHNITFRREDGFPLTGDFVQISVEDT
GLGMSPEVLARAFEPLFTTKAKGMGTGLGLPQVFAFCERSGGLATIDSEVGAGTSVRLYL
PRTTDAVPPVDAPNAVDNLAASTPAGLRILLVEDNGEVAAGTEALLELLGHQVTHAPNAD
DALRMLNAAGAEPGAPGTFDLVISDIHMPGTINGIDLAEAIERGDMKLPVILVTGYAEEF
DRTRRVHARLLSKPFDIALLDDLLQKIQHERDARLAGHED*
>GCA_000452945.1_BPMSHR5858_genomic_04057_1 yceI_4
MKPVRWAGRFACAIALAGVAASCTPLRVVTHSVSTAEAAVPAGRYTLDPHHWSIVFDVDH
FKYSRFTMRFDRANAQLDWRPGGGLADSGVSASIDAASVDTRVPLLDKLVAGADALDAAR
NPQIRFDSTRFARTSATQGTLTGNLTIRGATHPVTLAVTFNGYGRNPLTKQDTLGFSASG
TFSRAQFGVTSWYPAVGDDVRVRIEAEFVKEGEAPAQ*
>GCA_000452945.1_BPMSHR5858_genomic_04064_1 gstB
MLKLLGKAPSINVRKVLWTCAELKLAYEREDWGAGFRSTQTPEFLALNPNGLVPVLVDGD
FVLWESNAIIRYLANRYGGAHLYPVEPHARARVDQWLDWQTTALNRAWSYAFLALVRDAP
DHGDTARIRASCESWARHMAILDAQLGTTGAFVTGADYTLADVALGLSINRWLRTPFDKP
DFPAVSAYFDRLAARPGFAEHVDNGLP*
>GCA_000452945.1_BPMSHR5858_genomic_04082_1 hpcD
MPHIIVEYTANIRDDARIPVLLKSINETLIAQGGVFPTGGIRSRAIELQDYCVADGTAAD
AFVHVTLKIGAGRDDATKKAACDALFDAIKAHFEALYARRYLALSMELAEFSETGSYKHN
NIHARYKRGA*
>GCA_000452945.1_BPMSHR5858_genomic_04091_1 rstA_1
MNEVPLKYRVLLIEDDDRLAQLIREYLDSYEFSVTVVRRGDLAVGAVREHQPALVILDLM
LPHLDGMEVCRRIRAFSNVPVLILTARSDVYDQVAGLETGADDYVTKPIEPRVLVARARA
LLRRVQPGASVEAPASETLVFGELAISPPNRTVTWRGEPVDLKTAEFNLLLILARAAGTV
LSRDDILKQLRGIEFDGIDRSVDSGISKLRRRFEDTSSEPHKIKTIWGRGYLFSPSAWDD
*
>GCA_000452945.1_BPMSHR5858_genomic_04095_1 soxC
MSAIYETAVQPARGATARDAHLIASDAEAIDAARALAARLAEGAAERDRERRLPYEEIDW
FSQSGLWAITVPKAYGGAGVSHVTLTEVVKIVAAADPSLGQLPQNHFGLVDVIALTGTDE
QKRLFFGEILKGKRFGNGFSEKGTKNVLDLKTKVIRDGDGYRVDGTKFYSTGALFAHYVP
VLGIDDARRGWLAYVPKGTPGLAVIDDWSGFGQRTTASGTVRLDNVRVPASHVFAAHRVS
DQPTLNGPLSQILQAAIDAGIAHAALDDTLRFVRERSRPWIDSGVERAADDPLTIRETGR
LVIQLHAADALLERAARTLDALAAQSELSEDDVARASVAVGEAKVLTTEIALLASEKLFE
LAGTQATLAEHGLDRHWRNARTHTLHDPVRWKYHLVGNYYLNGVAPARHAWN*
>GCA_000452945.1_BPMSHR5858_genomic_04096_1 group_7577
MRLTIRINGSDAPAQQFAVLWLDTDEQLWSREAHQGIDLPAWGKVKDVEGAVALCSADSG
EALCRLQGLSFSNVRRLSEDEEHGNAVLGGKNPQGAWRLQAVDSASIQPEHREFTIVR*
>GCA_000452945.1_BPMSHR5858_genomic_04097_1 group_4983
MNMNADGETAVGVVREFWRLMGANDFHAVAAVLAEDFVLDWPQSNERIRGARNFAVLNSE
YPAHGPWTFAIGRLIGGEGQAVSDVSVTDGVQQARAISFFTVVHRKITRIVEFWPEPYAA
PYDRSHFVERID*
>GCA_000452945.1_BPMSHR5858_genomic_04098_1 rhaS_1
MDQRHTRLESAAHAPPRPDAQTLAPREAARRELAALIERFAPADGAHPSAIPALSFFRCS
SPVDLGCSVTRAAFVFAAQGAKRVTVAGQAYEYDHQQCLVTSVDLPMLSQVTRASAGAPY
LCVKIALDVQRIAELSAEMRMPPPEAVPTGEGIVVGALSEPLFDAALRLVRLLDTPADIP
ILAPLIEKELLYRLLTSGLGARLRHIAVAGSQTYRIARAIEWLRHHYTEPLRVETLAQQV
NMSVSSLHHHFKHVTTLSPLQYQKQLRLHEARRLLLGQHGDVGSVALRVGYDSPSQFSRE
YSRLFGAPPLRDVVQRRRNGTGVQE*
>GCA_000452945.1_BPMSHR5858_genomic_04100_1 iorB
MTTDLDLPDAVRASRRTFLKAAGAAAAVGLTIGFEWLDASRPALGATAPAAPAGDAAGAF
APNAFLRVAPDNSVTVIAKHVEMGQGAYTGIATIVAEELDANWRDVRVESAPADASRYAN
LQFGKLQGTGGSSAMANSWMQLREAGAKARAMLVSAAAAQWNVPAAELTTGDGVVRHAAS
RREATYGSLATAAAALPVPEQVALKSPSDFRLIGKPVPRVDARAKSDGTAQFTIDVSMPG
MLVALLQRPPVFGGKVKSFDASAAKAVPGVVSVVQVERGVAVVAKGFWAAKQGRDALKVE
WDESGAEKRSSDAIMREYRQLAERPGASARRDGDAEHALAGAAKRLRASYEFPYLAHAPM
EPLDAVVKLSADRCEIWAGDQFQTIDQGNAARVAGLDPSQVQIHTLYAGGSFGRRANAWS
DYVVEAVSIAKALGANGTPVKLQWTREDDIHGGLYRPMYFHRLDAALSKDGKLVGWRHRI
VGQSILAGTPFEAVMVKNGIDQTSVEGAANLPYAIPNVSVELTTTRSAVPVLWWRVVGSS
HTAYAVEAFIDEAAHAAGKDPYLFRRDLLAHAPRLKGVLELAAEKAGWDPGKPLPKGRGR
GIAVAEAFKSYVAQVAEVSVDEHGAVKVERVVCALDCGTAINPDVIAAQMEGGIGFGLGA
VLHGAITLKDGKVEQNNFDGYRVLRIAEMPKVEVYIVPSGEAPTGVGEPGVAPIGPAVAN
AIFAATGKRHYALPFPAAGDATA*
>GCA_000452945.1_BPMSHR5858_genomic_04101_1 group_3498
MDCKYLYIDNIKINFVRRIIEIDGDAISLTPREFDVVEFLLDNMNKVVSRQAIQEAVWGR
ELAVSSRTLDTHVSRIRSKLGLDFDKNMRIIPIYSIGYRLVLFGAATTRVEPHDEQHATH
AEQPRAGNAHAPAEPLAPIAPIARPAVAPHLERVAEYH*
>GCA_000452945.1_BPMSHR5858_genomic_04105_1 group_4984
MGAQTRMTIDSARARALTVKIMRVWAALVLATVGSWTESLRAASPAIVAAVVGLAALKVA
LIVFCYMEVGRAPRWLKLVCGAWIGIVFGIVAACHLLPRDVLRLIG*
>GCA_000452945.1_BPMSHR5858_genomic_04110_1 group_5321
MIVRRPRGNASRRAARHADSRYSGTSVSCVHSPPLVSAPLSIASARALHLAAQGLLTPPR
RKAVKADVLAAVRRMGQLQIDTIHVVARSPYLVLFSRLGAYAPQWLDEHLADAKLFEYWS
HEACFLPIEDFGLMRHKMLNPIGMGWKYAAEWHAQHRDAIDALLAHVRARGPVRSADFAR
GAGKGNGWWDWKPEKRHLEVLFSTGQLMVAERRNFQRVYDVAERVLPDWDDARDLPPRAA
VVPRLVGNTCRALGIVRADWIADYYRLPKRSYRDELHALADAGELLPVAVDGWHADAFVH
RELAPLVDAARDAALRPTVTTLLSPFDPVVWDRRRASALFDFDYTIECYTPAHKRRYGYF
CLPILHRGRLVGRVDAKAHRTQRVFELKAVHIEPGVRLGAGLAADVGRAIRKLADWHETP
VVEAGHAPKEIARAIGADRVAKPR*
>GCA_000452945.1_BPMSHR5858_genomic_04127_1 group_4985
MNWKAEERRADELAEQLDALGPIGVARFFSGASLRLDGVLFGFVHAGSLFLRVDDDTRAA
FERAGMRPFSYSGRTRTVVVGGYYETPADVLEDVGMLRDWCRDAHRAALRAGAGARRAKR
KA*
>GCA_000452945.1_BPMSHR5858_genomic_04141_1 cyc1_2
MHHERISLRAAMREMRALASAQVERFPAGQAEPLRFVAASGAPPEIAAAARHCAANMGNW
ISSVYWFHHESDRYGTARDF*
>GCA_000452945.1_BPMSHR5858_genomic_04145_1 cyc1_1
MTYMTYMTYMTYVTYVTYAACAACASCHGQNGQGAGTFPRLAGQHADYLRRQIDVFRNGT
RANAPVMSAVAHTLDGDPAKAVAAWLQSR*
>GCA_000452945.1_BPMSHR5858_genomic_04146_1 mreB_2
MAAPFLGRFFAHNVAVDPGTASTLIYVRNRGVVLNQPSVVCFRKRGGPAERARVEAVGEQ
AKALLGRSPEHLEAVRPLRHGVVANYHAAEQMMRQFVGMSHARSLFGRRVEFTICVPSNA
TAVEQRAIREAALAAGASRVSLISEPLAAALGAGLPVSEAVGSMVVDIGGGTTEVAVIAL
GGIVYREAIRVGGDQFDAAIVNHVRNLYGVLLGEHTAEHVKKTIGTASCRVPRESIHAIG
RSVADGLPRTIELSNHDIADALAAPLNQVVSAVKSALENAPPELITDIADRGLVLTGGGA
LLANLGKRLHDETGLAARVADDPLTCAVRGAGEAMGRGGFDAGDEDAGDFGMNR*
>GCA_000452945.1_BPMSHR5858_genomic_04157_1 tdeA
MTTRPIHRLARRAAALAGTLALLSGCTSASRNAPPPAPELPAQWAGPVARQASATPDGWW
RRFGDPMLDRLIDDALRTNNDLAAAAIRVYRAQLQAGLADTNLTPNVTLGATGNASRTLD
THQTRRTSGVTATLGYELDLWGKLAAQRDAARWELEATQADRDAARLALIGTTAALYWQL
AYLNQQIAFGDANIAYAERTLKLVQSRHAAGATSGLDLAQAERNLAALRADQTQLIQQRT
ENRNALAILFGRPPQRQAAERDALPDTPLPEVAAGVPAELLGRRPDLRAAEFRLRESLAN
VDVTRTSFYPTFTLTGSVGTASTSLERVLQNPVATLGLGLALPFIQWNTMQLQIKVSQSQ
YEEAVVNFRQSLYRALGEVENALAARVQLDAEADERGRALALARRAETLAKARFAAGATG
VQPWLDEQQRLRDAQSALARNQLGRLNNRMNLYQVLGGAG*
>GCA_000452945.1_BPMSHR5858_genomic_04174_1 group_3505
MSALPPLAAIRTFLVACRAGSFTAAADELCVTHSAVSRQIQTLESWLGTSLFEKDGQRMV
PSVHARAFAQELGGALDALTDVVQRYGKGSARQALRVSVPTTFGMRWLIPRLAAFRDVHP
DATIQVLTVTTQQQPSGGNCDVAIRRDDQVFNPESAIRFLSDHHTVVASPSLLSRTPLER
PEDLVRHTILETETRPRHWDDWFAEAKLSASRFTRRQRFDHFHVTFQGIVDDLGVGIGPV
VTLSRDLANGKIVAPFSDIRIAPHNYYAVTPIGIQKTALHREFEDWLVATAAAAETRAA*
>GCA_000452945.1_BPMSHR5858_genomic_04180_1 adeP_2
MESIKRYFGFAQAGTDLRTELLAGVTTFLTMAYIIFVNPAILGDAGMPKESVFVATCLVA
ALASLIMGLYANYPIACAPGMGLNAYFAYTVVKGMGFTWQAALGAVFISGCLFLLVTLFR
VREVIVNGIPKSLRIAITAGIGLFLGIISLKTAGVIVGSPATLVTLGDLHKPTTVLAIVG
FFAIVTLDYLRVRGAILIGIIGVTILSFFFGGNQFHGVFSAPPSIDATLFKLDIGAALSS
GILNVILVFFLVELFDATGTLMGVANRAGLLVEGKMHRLNKALLADSSAIVAGSLLGTSS
TTAYIESASGVQAGGRTGVTAITVAVLFIACLFIAPLAGVVPGYATAPALLYVSCLMLRD
MVDVPWDDATEAVPAALTALLMPFTYSIANGVAFGFIAYGGLKLLTGHAKQVKPVVWIIA
AIFLFRYFYLGAE*
>GCA_000452945.1_BPMSHR5858_genomic_04183_1 acoR
MPQSSVLPAAAGRVDLLAQAHARSLSVGLRASELPDFSPLSRAALRELIDGSQSLYTHAR
PVMDTLHTQIADTQSLVLLTDRDGVILHSIGDADFVEKANRVALCPGVSWAEDARGTNAI
GTALIAGQALAVHGAEHFLRANHVLTCSCAPIVDPFGRLLGALDVSGDPRGFSPHTLALV
KMSAQLIENHLFANQCADALRVRFHAHDEFVDSLFAGLVAFSPSGALIAANRSAQFQLGA
NLDALQQRGCDELFGLSFAKLAQHAAREPLACFTLTLGTGVRVRARCEFAHAQRTAVAVS
GRAAPPRATDAARDEEPDAITFATLDTGDARMTAVLQRVGKVRGRDLPVLVLGETGTGKE
WLARALHHASPRRDGPFVAVNCAALPDSLIEAELFGYEDGAFTGARKRGSPGKIAQADGG
TLFLDEIGDMPLAQQVRLMRVLQERAVMPLGGARAVPVDVRIVCATHRDLRAMIAAGTFR
EDLFYRINGLAVTLPPLAERSDLETLVARVLARLARSEPMPATVSAAVLAAFARCRWPGN
LRQMTNVLRTAGMLAEGEARIDVEHLPDDFWLDCDAHARDAAPPRTCADAAAGAAAAGAS
TLQDHQVALIDGALDRHRGNVSAAARELGLARNTVYRHLRRRRSRE*
>GCA_000452945.1_BPMSHR5858_genomic_04184_1 petF
MTEERSHWVRVEPLGAGFDAPESLSLLEAAGFAGVSLPRSCRNGTCRTCLCRLREGSVAY
RIEWPGVSAEEKAEGYILPCVAIAQSDLVIEVPHAE*
>GCA_000452945.1_BPMSHR5858_genomic_04185_1 group_508
MRDLLEHRRLVITITLLYLSQGIPIGIAMDAMPTLLRHDGAPLHALAFLPLVGLPWVVKF
LWAPVVDNRWSARIGRRRSWMLPMQCIVVLCLASLAQIGMTVATAGWAVGLLAVASLASA
TQDIANDGMAAEHFRGGALAKINAIQIAGVMIGFFGGGAGALTLIGLFGQRAAFLALAAV
PLASLACVLWLMPREAGRPGAPCAHDASLARFAKRPLSLSLLALALLSAMTAVSGFGLSK
LFLNDAGWTLEAIGRLGMAGGIVTVVLGCGGGAWLVRRLGLWRAFSAGVSCAGAGALLWF
AQANGWLASAPHWVWLCTVLGSLSTGITSVSIMTAGMRFAGDGDQAGTDVTAVQSTRDLG
ELIASSMMLTLTSKVGYSGGFLAGAVLAALAALIAARLRRREAGRPARLHVRAVSDAPMT
GPD*
>GCA_000452945.1_BPMSHR5858_genomic_04189_1 irtB
MTAWLKRWRCLPADTRRSLVRGVGWAVAAALLDGLCGVLLVPLIRAWFAGGALLHWAAAL
IALTLGHALVLYAAQLGGYRAGGALALGLVDRLVRHLPRIAAPQTVRDSDPEGLLRGPVM
QAMGIPAHLLGPLIGAVVTPLAVVAGLAAIDWRIALCLAAAAALLFALLNRSGARTLALE
EARAAGERDLAEQLQAFAAHQGLLRFAGRDGSGRAALQRALDERHRRARALMRRSLPLEL
GFAGAVQAVFVAMLVGGAAAVAAGRLDPATVVAVLVLLVRFIEPLAQLTQLDQALRGGWR
ALDAVLTVLHAPRFESPERGLRPQDASVAAVRVGYRSETGATLLDDVDLHCPAGGFVAIV
GPTGAGKSTLLGLLARLDDPSSGRVLFGNVDLRRLSETTLAGARTVVFQHSGLFRGSLGW
NLRMGRPDASGAALDAVLDALGLSRDVERLPHGLDTDVGPGGQLLSGGQRQRACIARGLL
AKAPLLLLDEPTASLDELSARRVRDCLVAMRGQRTRIVVTHHPALARAADEIVVLEAGRV
RARGTHAQLVAADPWYAEFARADSECEREPEVETPATATAGISAPARGADAGHR*
>GCA_000452945.1_BPMSHR5858_genomic_04191_1 dltA_2
MSTHPLPVVVAGSRFGQFYAAGLAASGAYRIAGILGQGSARTAALARRVGAPVFTDPGAL
PDDVRIACVAVGGAARGAQGAALARGLLERGIDVLIEHPLLPAEWDAVLRTAVRHGRRCL
LNSFYPHLPAVARFVDVAQRLSRIGRPLHVDLACSVQASYAALDVLASALRGVGPWSIEP
ARQASSPMRECTMVLAQTPVALRVHNEMAAAADGRMHLLFRIALMTAHGTWTLASPHGPL
AWEPALREPPADDDGLFPIFGAAAPPHGLPSMQLYEAEPPAWDAIHARHWPRAAVRAVDR
LASGDGLAASNQRSVEVTRVWQHLTAQLGFPEPPPGDASADTLEPLLERAA*
>GCA_000452945.1_BPMSHR5858_genomic_04195_1 dhbE
LPAAARNAASSDAPDWPDDFARRYRAAGYWRDETFYGALADRARRSPAATAVVDGERRVS
YRALLERIRRLAGGLRRLGLARGDTAVVHLPNGVRFIEACFALFQLGVRPVLALPAHRQH
EIGAFCRFTNARAYLGAARLGEFDCRPLAHALRASCPALAHIVVAGDDHAFLHFDSLYDA
PPVADCAARADDIACFQLSGGTTGTPKLIPRRHREYLYNVRACSDASGFGADTVYLAALP
MAHNFTLCCPGVIGALLAGGRVVATEHPEPECGFALIARERVTHTALVPPLALLWLDAQR
ERRADLSSLRVLQVGGARLMDHAAERVTPVLGCRLQQVFGMAEGLICCTRLDDPPARIAR
TQGRPVSPADEVRIVDEAGRAVAPGEIGELQVRGPYTIRGYYRLAEHHAAAFTADGFYRS
GDRVRRTEEGDLVVEGRDKDQINRGGEKVSAEEVENLLLAHSQIRDAALVAMSDPLLGER
TCAFVVARPPAPTSLALKRHLRDHGLAAFKIPDRIEFVPSFPETGIGKTSKKSLRDLLRR
RLEAARA*
>GCA_000452945.1_BPMSHR5858_genomic_04196_1 pikAV
MKPPPPHAGWVRELRLSPCPRAQLVCMPHAGGAASFFRGWAGALPWDLDLLALQYPGRED
RFGEPCARSIDALAGPAAEALVEYARQPLVLFGHSLGAALAYEVALRLEQRGAPPLYVAV
SSHPPPHRQRASSLHLQSDAALLNDVARLSGEHAALLDDPALRAIYLPMIRDDYRAIETY
RRERPPMLDAPLGVMLPLADPELDRDEAQAWQDVASRPIRVTTFDGDHFYLRHAYPALIA
HIAGQIDRSLRPLKEHS*
>GCA_000452945.1_BPMSHR5858_genomic_04197_1 pchB
MKTPDECTGLADIREAIDRLDADIIAALGARMQYVKAASRFKPTEASIAAPERVAAMLPD
RRRWAEQAGLDGAFVEALYAQVIAWYIEQQTRHWRRQRGLA*
>GCA_000452945.1_BPMSHR5858_genomic_04198_1 pchA
MSARLSALLDTFARAARRAADTGEPVLAAASFPLRPLNLFDLIASWDDGATPWCFVEAGD
EPGAMRAPGKPPALFGWDCALDLNEAGDARFARLDARWRALVRAAVVTGAQPPRIVGGFR
FDPLGPRYAHWHAFADASMTLAKLLVVREGEAYWAVCQRVVAAHDDPAALARECDARIGM
LASLAPAADDEAPRLLRVQALQAAAWQHEVRRAVDAIRGEAFSKVVLARDVLQQYARPVA
IGPLLRRLRRRDPRAHLFAIRRRDGCFVGATPERLARVAHGRAHTHALAGTIARGNDPAH
DRALGAQLMASAKERLEHALVVDAIRDALAPLSRAIDVPAQPSLLRLPRLQHLSTPIAAA
LNERATLLQVVAALHPTPAVAGHPRAAALEHIRAHEGFDRGWYAGPIGWIDAHGNGDFVV
ALRSALISAGACRLFAGCGIVAASEPAREYRETSLKLSGMQAAIEARDAAAAARTVAPIP
SAAH*
>GCA_000452945.1_BPMSHR5858_genomic_04199_1 group_4993
MGPMVWRGASDSITRWLAAPLIAAGLTGAASAAPVDMPDLQHAVRQAFSARLGKQAPAAP
AVRDDLIESVRSDPEAGWVLGTVTQVVPNDTPAYPVTKLFIARRASEGWAVGIEGTDAFY
TLAAASPAKLLAGDERAHLNAGRAPTAPPRKAVPAQTGLALPWQQGTAWYWTGGAHGWSG
DSRPFNSLDFSGGNGQVLAARDGTLYKSCERNGSAIVKVVHDNGYTSTYYHMVQLTQAGS
GTRVRQGQYLGRVGNGLPCGGQTTGPHVHFALSQGGSDVPVNGKTIGGWQFFEGSNAYSG
YAVRNQRRVSVQASLTNYGADDSGGPTEPSPPVKATVQSPGPVNLRSAPSLSASIVGTVA
NGAAVQLACYAYGDTVQGNWGATRLWYRLDSNRWVSDGFVYTGSNDPVVSACAN*
>GCA_000452945.1_BPMSHR5858_genomic_04201_1 livF_1
MLKLEKIHTHYGAIEALSGVSIEVKKGEIVTLIGSNGAGKTTLMMTVCGTPRASSGRVLF
EGEDITAMPTHEIMRRGLAISPEGRRVFPSLTVLENLKMGGFFANRHEIDEGVEHVYRLF

PRLKERSAQRAGTMSGGEQQMLAIGRALMSRPRLLLLDEPTLGLAPLVIAQIFDIIRTIR
AEGVTVFLVEQNAHKALQVADRGYVLETGRVVLADTGANLLENDRIRAAYLGG*
>GCA_000452945.1_BPMSHR5858_genomic_04202_1 lptB_1
MSATTEMLKVSALQMRFGGLLAVDGVDFDVRRDEVFAIIGPNGAGKTTVFNCIGGFYRPT
GGRVVLDGHSIGGLTSHLVARKGLVRTFQNIRLFQSLTVVENLLVAQHRQVKSGLLHGLF
ATPAYRRAEREALERAAMWLERMNLTAVANRPAGTLSYGHQRRVEIARCMITNPRLLMLD
EPAAGLNPQEKVELQHLIDRLRREFGISVLLIEHDMSLVMGVSDRILVMEHGRPITIGRP
DEVRNDPRVIKAYLGEE*
>GCA_000452945.1_BPMSHR5858_genomic_04203_1 group_3509
MNQAMTSVHTAAGVPAVQSLKHAVTAAVLTAILTVPILGLQLKLDGYQVVLEAHWRPVWI
AVAAVFLFQLFKPALMRAKAAVRLPAAPPLGVREQRAIVWALVALGFVWPFFGSRGAVDV
ATLALIYVILGLGLNIVVGFAGLLDLGYVGFYAVGGYTYALLNQHFGLSFWECLPLSALA
SATFGFVLGFPVLRLRGDYLAIVTLGFGEIIRLLANNLTSITGGPDGISGIPKPTVFGFE
MARSASVEGAKTFHELVGLEYSGEHVVIFLYLIALVLVCFTLFVTSRLIRMPMGRAWEAL
RDDEIACRSLGLNPTRIKLSAFTLGASFAGIGGAFFAARQGLVNPESFTFIESALILAIV
VLGGMGSQLGVILAAILLTLLPEVARGFAEYRMLIFGLVMVLMMMWRPQGLLPASRPHVE
LPK*
>GCA_000452945.1_BPMSHR5858_genomic_04204_1 livH_1
MNDFFPQFAQQLVNGLTLGAIYALIAIGYSMVYGIIGMINFAHGEIYMLGAYVGLVTLTA
IGTSAGYPLPVVLGAALVVSVVVTGLYGFAVERVAYRPLRGGPRLVPLISAIGMSIFLQN
YVQIGQGARDMSVPVLISGAFDIHLGGDFDVTVPYARLLIVCVTIVLMIALTLFIAHSRM
GRACRACAEDMRMANLLGIDTNRVISFTFVLGAMLAAVGGVLIGLTIGKLNPYIGFVAGI
KAFTAAVLGGIGSIPGAMLGGVLLGLAETFASGYMPAEYKDVVAFGLLVLILLFRPTGLL
GKSDIEKV*
>GCA_000452945.1_BPMSHR5858_genomic_04220_1 betC
MPDTAEPTDIQPNILVLMADQLTPFALRAYGHRATRTPTIDRLAAEGVVFDAAYCASPLC
APSRFALMAGKLPSALGAYDNAAELPAQTLTFAHYLRAAGYRTMLSGKMHFCGPDQLHGF
EERLTTDIYPADFGWVPDWTRPAERPSWYHNMSSVLDAGPCVRTNQLDFDDDATFAARQK
IFDVARERAAGRDARPFCMVVSLTHPHDPYAITREYWDLYRDEDIDLPAVRMDFDASDPH
SRRLRAVCEVDRTPPEDLQIRRARRAYYGATSYVDAQFGALLATLEQCGLADDTIVIVTA
DHGDMLGERGLWYKMTFFEGACRVPLIVHAPRRFPAARVPAAVSHVDLLPTLVELATGER
RADWPDAVDGRSLVPHLRGEGGHDEAFGEYLAEGAIAPIVMMRRGSHKYIHSPADPDQLF
DLRNDPRELDNLANTPAAAKHVAAFRMERVARWDLDALHQQVLASQRRRRFHFEATTQGR
IRSWDWQPFQDASQRYMRNHLELDALEAAARFPRPHA*
>GCA_000452945.1_BPMSHR5858_genomic_04221_1 gcvA_3
MTRPDRLPPMQTLSAFEAAARLASFTAAARELGSTQPAVSQRVVQLEEDLGTPLFERGRR
GVTLTEDGARLFAAVRQSLDALRAATADIRSRRANGTFTLVTDFGFATYWLMPRLDDLKR
AMPGVDVRVVTSQDIDPQREHADVAILFGAGDWPGCTSTRLFQEHVTPVCSPAFRTAHAD
IARPADLLRAPLLHVQPTRPERWLAWRDWFDAHGLAAPPEPHGLTFNSYSLVIQAALMNQ
GVALGWAPLVDTPIAAGQLVRLVDAPVVTPRGYYLVLPPARPEARAVPLFRRWLLGACA*
>GCA_000452945.1_BPMSHR5858_genomic_04224_1 group_7583
MKFLTKLIGCGVLAMMLAAVAPAQADTKPTLKIGYVEGWDDSVATSNVAARIIETKLGYP
VQLVPVAAGVMWQGVARGDLDATLSAWLPVTQGAYWEQFKTKVVNLGTNFPDAKIGLIVP
AYVKAKTIADLNAQKNDFAGRIVGIDAGAGVMRKTDDAIKAYGLNYSLMPSSGSAMTAEL
ARSIHSNKAVVVTGWAPHWMFAKWKLRFLDDPKKVYGEAEHVDNVVNPALETKAQPVVAF
LKKFQWKPGEIDGVMLAIQNGSKPAAAADAWIAAHGDRVGEWLGAER*
>GCA_000452945.1_BPMSHR5858_genomic_04250_1 group_6691
VIDSIRSSSTGPRDAPGKRLAVGISLFARDGQAIWENGIHQNIAFLAMMLKQSDRIGPVY
FLNGGDAKALPAGLDLGGLDVPLVQPRDVSHALDVVIEMGAQLPVDWLKHMKALGKKLVV
CFVGHTYSGLTETPMFNRPSGHIFNGGPFDEVWILPKSEKIDAPLLRTLTRAPVHVVPHI
WSPYFIERRARELEAAGLRFGYAPGRAKWRLSILEPNISVVKTCHYPMLAADEFHRAQPD
AVEHLFVVNSLHMKEHPTFVHFANSLDLVREHKATFEPRIDLPSFLAQHGDAVISHQWEN
PQNYLYYDVLYGGYPLVHNSPMLGDAGYYYPEFDSAAGGRALQHAWRHHDAHLDDYRRHA
DALLQGVSIDNRRNLDAYVARLFA*
>GCA_000452945.1_BPMSHR5858_genomic_04251_1 dpgD_2
MSETTARDVPNGGKRLVVGVSLFVRGAGQSLWENGIFQNCLLLVLLLRQSPLVAEAVMVN
GGTSDADPQLMLGEWNVPLVSMDEAMTRCDVLIEMSAQFNAGWLAEYQRRGGKVVTMRVG
NDYVIDIERAIFDKPSGFLFSGARYDGVWTIPEFENSCGHYLESGLRSPVTIVPHIWHPL
LFDKARATLGPGLEFGYKPGRPRWRVTMFEPNICMVKTSIVPMLVAEEAYRAQPDFVEFV
RVCNTLHMKEHTTFVHFARSLDIVDHGLTTFEGRFAVYEFMAAYGDAVVSHTWENAQNYL
YYELLYGDYPLVHNSPYLGRAGYFYPDFDCQEGGRQLLKAFREHDANLDAYREQSRHVLD
SVSIYNAQNVAAYTEAIAALYR*
>GCA_000452945.1_BPMSHR5858_genomic_04253_1 accA1_1
MFDKILIANRGEIACRVAATCRRLGIASVAVYSDADAHAKHVAACDEAVHIGGAAAADSY
LRIERIIDAARATGAQAIHPGYGFLSENEDFARACEDAGIAFVGPPVEAIAAMGSKAAAK
ALMHAAAVPLVPGYHGDDQSPELLQREADRIGYPVLLKASAGGGGKGMRVVERAADFAAA
LASCQREAAASFGNDRVLIEKYLQRPRHVEVQVFGDTYGNAVYLFDRDCSVQRRHQKVLE
EAPAPGLADALRQAMGEAAVAAARAVGYVGAGTVEFIMTGDAFYFMEMNTRLQVEHPVTE
MVTGLDLVEWQLRVAAGEPLPLRQPELHVRGHAIEARLYAEHPARGFLPSTGRLKHLRFP
AGAEFAAGAAVRIDSGVREGDAITPFYDPMIAKLIVHADTRAQALATLGRALRECEVVGL
HTNAEFLQRIVASRPFADADLDTGLIERHRDALFAPRPAPAAALGLACAALFAREQAGAG
GVASPWAALPNWRLNAGYRRTIEWRALDDDTALEVRYLEHGGAQSLEIGGVAQPFAWSRG
AGPLDFDVTLGGVRASGRVHVDGDTFHVFCQGVARAFEWRNLLAHAADSTEGEGRLTAPM
PGKVIAVLVEPGRKVEQGDPLIVMEAMKMEHTIGAPAAGVVAEVLYAVGDQVADGAQLLV
MGPA*
>GCA_000452945.1_BPMSHR5858_genomic_04255_1 pccB
MPIIESKLNPRSDEFRANAAALEALVADLREKIDRLALGGGQAARDKHLSRGKLLPRDRI
AQLLDPGTPFLELSQLAAYGMYNDDAPGAGLITGIGRIAGRECVIVCNDATVKGGTYYPV
TVKKHVRAQEIAAENRLPCVYLVDSGGANLPNQDDVFPDRDHFGRIFFNQANLSAAGIAQ
IAVVMGSCTAGGAYVPAMSDESIIVRNQGTIFLGGPPLVKAATGEEVSAEDLGGGDVHTR
LSGVADHLAQNDAHALAIARAIVGNLGARGGAAPAMREPLPPRYDARSVYGVIPVDTRKP
FDVREVIARIVDDSAFDEFKARYGTTLVTGFAHIWGQPVGIVANNGILFSESATKGAHFI
ELCCQRKIALVFLQNITGFMVGRKYENEGIARHGAKMVTAVSTAKVPKFTVIIGGSFGAG
NYGMCGRAFGPRFLWMWPNARISVMGGEQAASVLATVRREGIEAKGGEWSAQDEEAFKQP
IRDQYERQGHPYYASARLWDDGVIDPAQTRDVLGLALAAARNAPIEDTRFGVFRM*
>GCA_000452945.1_BPMSHR5858_genomic_04257_1 group_5001
MTATQKADRADAPRAPAGRKSQQRVQDILRAGREVFAEKGYEHATAAEIAQRVGVSEATV
FSYFRGKRELCARVIADWYDEIVDAFETGMPRDGSVRQQFAFVVRTHLRLMLVNGTSLCA
LVLSEGRAKQHALADELTALQRRYTAPLMDVLARGQATGQVRADMPLSLLRSMVFGPIEH
VLWDAILGHRKLDTETTAEQLIDLLWTGLQPPAPENAALVRFRSDVADAMRRLDQACGRA
PRKAAAGRDAQKPA*
>GCA_000452945.1_BPMSHR5858_genomic_04258_1 norB
MDTSETVLRHRQDDRVSDVLKWTLLVVAVLTFALLAWATVVTYRSAPPRPARLVAPDGSV
VMTARDITDGKASFQRADLMDYGSIYGMGSYFGEDYTAEYLLRLGQLTNDNIALSRFGQP
FAALDQDSRNSVQAAMRQQLKGIDLSRTDALVPAEVARAIRSLRVEISRALTHHDFIKGW
TQAYSLDERHAAKTADFLLYAALTTVARRPGSDVSWTQNWPYEPLVGNAPTTNTFGWTWA
SYCFTFLCFGIVLFVYHRYIRVDDLGTMDPTLVRFRALTRSQRKVGKYFLVVAAVLLQIA
AGAILGHYYTDRTSFYGIHFDDVLPFNFWRSIHIQAPIVWIGLSWIGAGLFLAPAISGRE
ARGQGPLVDGLFWATVVIVAGALIGDYLGIMGVIENGWFWFGNQGLSYLELGRFWQVGFF
IGLVFWSALILRGLWPGLASIKQAARQFWTGRIRLEHLIWASTTNIALLYAFGMIPLFGV
NPSFTITDFWRWWVVHLWVEQSFEFFAAAISAYLLMGLGLVSRQLAERSVYLELILIFLG
GVLGTGHHIYWCRRPRHVGSRRQHVLVHRGAAARAAHPRRDGAPQPDPETAAVPLRSRVH
VCAGLRVLELRRSGRVRRRHAQRAARQLLRARHLPHAEPRAHVAVRRVRPARDRIDLFLP
AACRRRAIEIQRETRPPGVLAV*
>GCA_000452945.1_BPMSHR5858_genomic_04259_1 group_698
MNPAVDPSPLSRAAQALALASIGGALALGACTSRPSGERYAPSRMDFNPGPAIVTQTRHT
GPFASGSALSFDAALQPLDAARDQAIRLDTTHTVIRIAPGIAFAAWTFGNQVPGPTVHVK
VGDRVRLSMTNRSDEPAPGGLQLTAPMMHSMDFHAAMVSPTDKYRSIAPGQTMHFEFTPN
YPGVFMYRCGTPMVLEHIASGMYGVVVVAPRDGYPTRADREYVIVQSEFYTKPDPQHRSV
GTDALHVLDGERLRRKAPTYTVFNGRYNGMVTQPLIAKPGERVRLYVLNAGPSDTSSFHV
VGAIFDRVWLDGNPDNQLRGMQTVLLGSSGSAIAAFVVPEAGAYVMVDHQFANASQGAVG
VIDAGAHEESTIEHHNIPASATPTDAEAIQGKLDFESKCLACHTLGHGAKLGPALLGVTQ
RRSDAWLRRWLASPEAMVASDADARALRAHYPITMPDQNLSDSEIRRYVRYFHWADEASK
QRDHAMP*
>GCA_000452945.1_BPMSHR5858_genomic_04260_1 lysA_1
MDHPDADQEHERQLIACGREQYEWVVRAREAAASSLADWDARAEALLDGVRGMPAHFDHA
AARYAARTGEPLTLDHARAFCAATMLRAVEHPAGTDDWHLLCRLDEIPDIGDYVTGYAFQ
ESMTAVRASEHEVRVFLNVCPHRQTQLFQAAGSFDPHRHVRCPYHGWTFDVDGRCVNAPG
ARRCEFGADFREQDFSLSRVPARIEDGGILVRKAVSRVGRRMAALYERVYFAFPDVKRPF
VPDRFRVLLRIGFLAADIRHARDSSILSALRAELRQHLLTYPDLPVRYASLAKRHGEPTV
DAVDAVDAVDAVDLGPRLAGGPLRETVATWVYGDAELHALERDTLVLPAWHFVGHVAELR
GGERTLTLDVSGERAYVCRARDGTIHAGRLVDIDRRTDDPARLRADGALAPIDVDVWQGL
IFVRLAAGRGGVADIWEHARMLEPYRIGQMAPLAGKGWYDFAVEADAKILWENFLEMYHF
PAAHQAMIRLFELSSRSDLLTLREPASTRLSPIEARYRDALKRGATHGADEEARQRAMAD
GDAALARPLQLTVFGSIPETAQTATMLGITVFPDHIQAMSFIPVGPRECRVKIRSYGHAA
EGALDIARQCNVEQLQIELVEEDIQLNHRSQIAASTRSYERRGVLHDMEVSVADFQQLLR
RVLPVTRCVRRPPVGALARANQALRLRDRDERGAPRRACADAARRARAPAAASAFAAPPP
VSRQRNAETLEALDTPAFVYDERTLRDALSSMHTLLGDSGCRLLYPLKPFSHAGALECIG
PHVDGFAASSPFEAAFSRMLLGRAGSVHLATPALRDRDLGTIVDNCDYVALNSLSQWRRL
HDALAGRLSCGLRINPQYSVVADARYDPCRPHSKLGVPLDAIERVVASDPRALHGLQGIH
FHTNCDSTDFSRLAETIEHLEKRLARTLDGLAWINLGGGYIADGARYVDRFRACAKRLRE
HFGVALFIEPGAAIVRAAGTIIASVVDLFDSGGMPIAVLDTTVNHYPEVFEYQFQPDVDG
HHPAHAHAYLLAGASCLAGDVFGTYRFDAPLRVGSRVAFVNAGAYTLVKAHRFNGINLPS
VYRRGEDGALAAVQRFSYEDFLRQNGGLDHAYP*
>GCA_000452945.1_BPMSHR5858_genomic_04261_1 group_1220
MHIHEKAFSVSGPLASASLDHLRELTRQQLGPDETPVRFVVTRSDEHGFDCEIGVLSDGE
MPESMRADGLFRCERRTRGGGGEFNAVLIVPTGVGAEIGGHDGDAGPVAMLLSSICDRLI
THPNVVNAADINELPANGLYVEGSVICRLLLGQIGLQPVRANRVLAVVGSNECPTFVNAA
INAVNAARAAYGLDCPRVLHLDPGIRLTSTYASSGRAAGTVENLDALVALLAEHRDTFDA
VAIASHIHVPDACRTGYYAGTLGLVNPWGGVEAMLTHTLSTLFSVPSAHSPMYEDPRFAV
RDYGIVDPRIAPEVVSLTFLQCILKGLRRAPRIVELARAGNAAGTLSVSDVSCIVIPQGC
VGLPVLAALAQRIPVIAVRENRNVMRNDLRSLPWAPGQLRIVDNYWEAAGVLASMRAGLN
PDSVRRPLASVDVVRASAAHAHGASSIATAANIA*
>GCA_000452945.1_BPMSHR5858_genomic_04265_1 ydjE
MASTQSAPRATVPATTIDAGVISARLDRLPPTRSVWKLVALLSLGFFFELYDLLYSGYVA
PGLVKGGILSATTRGLFGTTGVASFIAALFAGLFIGTIACGFLADRFGRRAVFTWSLLWY
TAANVVMAFQDTAGGLNFWRFVVGLGLGVEMVTIGTYISELVPKQIRGRAFACEQAVGFV
AVPVVALLAYLLVPHAPFGLDGWRWVVLIGAHGAIFVWWIRRQLPESPRWLAQQGRLDEA
ERVLAALEAKVEAEYGRPLPPPAPAEPIAPRGRFADMWVPPYRRRTVMMTIFNVFQTVGF
YGFANWVPTLLIKQGITVTTSLMYSSVIALAAPIGPLIGLAIADRFERKTVIVAMAGAAM
IAGLLFSHASAAWLLVALGVCLTLANNIMSYSFHAYQAELFPTAIRARAVGFVYSWSRFS
AIFTSFAIAAVLKGFGTPGVFVFIAGAMAIVMASIGLMGPRTKGVALEAISR*
>GCA_000452945.1_BPMSHR5858_genomic_04277_1 group_5004
MRLKAPDLATAQHWANVLGVAGVGCELHNCYAPGALGGLPIDACQPEIWLDDERDEALAR
RLIEAASAGPAAGAAPWRCARCGETLDAQFTACWHCGASRDPRDG*
>GCA_000452945.1_BPMSHR5858_genomic_04318_1 group_1724
MPLVRSTNHSPLAPPDGRHAALWSTGFRSFYLGGAGFGAIAMLAWLGALAGHDSAGRAAP
ASGLFWHAHEMVFGFGGAIVSGFLLTSLRTWTSTNPAQGPSLAALWLLWLAGRVLMWTGP
GWAAAAVDIALLPAIAYALVRALIGAKNRHGIFLPVALGMLAALNALFHLWIRQGHADWA
LRCVDLAAGLLVLFIVIIGGRIIPSITATARPELPVRQWRLVESSVMSVTLAALLADATG
APAPAIAALAAAAAALHAARLVGWRFWAVGRRPMLSILHVAYASIAAGFALTALSAYGFV
AHSLALHTFTTGAIGCAIVGMITRTALEHTGREPVAKLAERICYGLLIVATLARVAGPAL
APAATAQWLAVAGVGWTAALLVYVVRYAGLLMSPAPSAPGPDARSGG*
>GCA_000452945.1_BPMSHR5858_genomic_04321_1 group_3523
MRSTRRLWTWLGLVCLASFAVLLWLGRDIYLAAPPIPNTVVSENGRVIFTGEQIKRGQQV
WLASGGQQLGTVWGHGSYVAPDWSADWLHREALELRDAWARQQFGAPYEKLPVDQQGMLD
ARLKHEMRANRYDAANDRLTLSDTRAAAALRVASHYEALFGADPSLDTLRDQYAMTAGAV
PGAADRTALASFFFWSAWSATTDRPGETGTTYTNNWPHEPLVGNTPTAANGMWSIASVII
LLGAIAAMVWYHTAHGEHDAPLAVPTNDPLFNVKPTPSMRATKKYFYVVIGLLLTQVFMG
AITAHYAVEGHSFFGIPLADVMPWVVSRTIHTQFGVLWIATAWLATGLYISPLLSGREPK
LQKLGVDVLFWALIFIVVGSTVTGWLGSLKHLGTNFSFWIGNQGLAYTSMGRVWQILLFV
GLLFWLFLMGRALMPALKNKATEGRGLIGMVFLSAACIGLFYASSLAWGQNTHYSMIEYW
RWWLVHLWVEGFFEVFATAVIALIFARLGLIRFESANRAIVMETIVFLFGGILGTLHHLY
FTGATTSIIAVGAVFSACEVVPLALVGFEGWQTYRKSSAAPWIQSYKWAILCFVAVGLWN
TVGAGLLGFSINPPISLYYVQGLNMTPAHGHAALFGVYGMLGIGLMLFCLRGMSERNAWR
DTLLKPAFWLLNVGLFMMVFLSILPSGIYEAYASVTQGLWYARSPEIVHAPIMNALVWLR
VPGDVVFAFGVLYLGWFALRLLRRPAPSQPDERAVGQAAQRT*
>GCA_000452945.1_BPMSHR5858_genomic_04335_1 puuD_2
MNDPLGNGPRPVVGICADRKTVGLHVAHVAGEKYVNAVVDGAHALAIVLPALGERQPADE
LLALVDGLLLTGSYSNVEPARYGGPTSAPGTLHDAARDATALPLVRAAIDAGVPVLAICR
GMQELNVAYGGTLHQQVHALSGHADHREDLSAPVDVQYGPAHPVRLVPGGLLRRLAGAET
VEVNSLHAQGIERLGDGLTVEAHAPDGLVEAIGVRGARAFALGVQWHPEWRFDGNALSRQ
ILAAFGAACRARRRATAGRAPRA*
>GCA_000452945.1_BPMSHR5858_genomic_04384_1 scoB
MAWNRDQMAARAAKELQDGFYVNLGIGLPTLVANHVPAGVEVWLQSENGLLGIGPSPTED
EVDADLINAGKQTVTTLPGSSIFSSADSFAMIRGGHINLAILGAMQVSQRGDLANWMIPG
KMIKGMGGAMDLVAGVKRVVVLMEHVAKGDQHKILEQCTLPLTGVGVVDLIITDLGVIEV
TPNGLKVLELADGVTVDEIQTKTGAPLDVSALA*
>GCA_000452945.1_BPMSHR5858_genomic_04386_1 group_7594
MPAIDYKTAFHLAPIGLVLSRDRVIEDCNDELAAIFRCARADLIGRSFEVLYPSSDEFER
IGERISPVMIAHGSYADDRIMKRAGGELFWCHVTGRALDRTAPLAAGVWTFEDLSATRRV
AVELTPREREIAAQLVTGKTSKQIGRVLDISSRTVDIYRARLMRKYGTGNTAELLQRLLG
Q*
>GCA_000452945.1_BPMSHR5858_genomic_04387_1 group_3528
MINAKTGFLFFIFVMSLIGCRPAFAQTNTPVTGIDLGGEIIEIIKQNPQDPTQAINNYMS
GVAATNVRWVRIHLDWSAIQPASTWIFTSDPNPGRSSGLTWGATDIVVRAARNYGIRILG
VITSTPRWATHAQCPSYYRQSTDDAYKLCAPNLTSYTNFARAAAKHYGNDSYGGKIDAWE
IWNEPNCGPNFIPHDPVLYTRLVKSAYPAIKQANPSASVYAGGSSGCLTSPNNGTGALPS
SGVAGLVNSTDARYPAPTQWEPRDWLAVMYANGARGYFDGLAHHPHCHSDDWQLPGDQCP
STSVNSVYPDYSNPFNIMWHTFTSPSYGWPAPTGKTFASYTGTSLRELMNANGDGGKPIV
ITEFGVATTASDGATDFTGQLGGSAQQYANADFQTIAPAYLTQANQEREYKALIAWIANQ
PYGQYGPVYAYCYSDMALSSPYSANIYEPYFGLVTITASTATSGTTGAPKIAGAYQAASG
QYYPAWPNFGAAATAAANHTPGYAPVGPGW*
>GCA_000452945.1_BPMSHR5858_genomic_04389_1 group_3529
MSDYTPVFEMSMPIRWGDMDAFGHVNNTVYFRYMEEARVTWFGKLGIGGGNGEGQGPVVV
NASMEFLRQLHYPGDVIARLSAAKPGRSSFDTAFELTRSDDPERVYARGAARCVWVDYAL
GKSVPLPDLLRNTIESALQAKVD*
>GCA_000452945.1_BPMSHR5858_genomic_04390_1 gno
MGRSINLEGKVALITGASSGLGQRFAQVLSQAGAKVVLASRRVERLKELRAEIEAAGGAA
HVVSLDVTDYQSIRAAVAHAETEAGTIDILVNNSGVSTMQKLVDVSPADFEYVFDTNTRG
AFFVAQEVAKRMMMRAGSGNAKPACRIINIASVAGLRPFSQIGLYAMSKAAVVHMTRAMA
LEWGRHGINVNAICPGYIDTEINHYLWETEQGQKLQSMLPRRRVGKPQDLDGLLLLLAAD
ESQFINGSIVSADDGLGLA*
>GCA_000452945.1_BPMSHR5858_genomic_04391_1 group_7595
MTPASLIEQYGPRESMEYDVVIVGGGPAGLSAAIRLKQLAAEKGAEIGVCVLEKGSEIGA
HILSGAVMDPRAITELFPDWKEKGAPLNVDVTEDRFLFLTEKSSFQVPNWALPDNFKNHG
NYVISLGNVTRWLGQQAEALGVEIFPGFPAAEILYNDDGSVKGVATGNMGVGKDGEPTEN
FQLGMELHAKYTLFCEGARGHLGRQLMDKFKLNANSDPQAYGIGIKELWEIDPVKHKPGL
VIHTAGWPLSSDTYGGSFLYHMDNNQVVVGFVVGLAYSNPYLSPFEEFQRYKTHPAIRPF
LEGGKRVSYGARAITAGGLLSLPKTVFPGGALVGDDAGFLNASRIKGSHAAIKTGMLAAE
AAFDAVQAGRQADELTAYPDAFKTSWLHTELYRARNFKQWMGKGLYLGTLMVGVEQKLMN
GNVPWTLHHQHGDHEMLKPASQCQPIDYPKPDGKLTFDRLSSVFISNTNHEENQPAHLTL
KDASVPVNVNLRTYAGPEARFCPAAVYEFVKNDDGSDRLVINAQNCVHCKTCDIKDPTQN
IVWVTPEGGGGPNYPNM*
>GCA_000452945.1_BPMSHR5858_genomic_04405_1 paaF_2
MSAELLASRPPESESTLVLTLSNPGARNALHPDMYAAGLEALNTAERDPSIRAVVLTGAD
RFFCAGGNLNRLLDNRAKAPSVQAASIDLLGEWITAINTVTKPVIAAVEGAAAGAGFSLA
LACDLLVAADDAKFVMSYARVALTPDGGGSWFLARALPRALAAEILFEGKPAAATRLHQL

GVVNRLTKPGTTRDEAIAWADQLGTMSPNALARIKSLIADAQTQPLAAHLAAERDHFVAS
LHHADALEGITAFLDKRAPIYKR*
>GCA_000452945.1_BPMSHR5858_genomic_04440_1 can_2
MNRPKSMLVANIAWARETSERSPDFFDALSRGQNPRVLWIGCADSRVPAEAITQSAPGEL
FVHRNIANIFQPDDDNCASVLEYAVKVLKVDHVIVCGHYGCGGVRASLLPPSPELPHVNR
RIAPLCALAGRHRAELDGVPPDAAADRLAELNVLEQVRLLRSSPIIRDAAPAPLVHGWIF
SLADGRLKELASGYPAAQRPPQAERAQAAAA*
>GCA_000452945.1_BPMSHR5858_genomic_04441_1 bicA
MKNLRASFSTFPRDFVAGTVVFLVALPLCLGIANASGVEPFAGLVSGIVGGLVVAVLSGS
PLSVSGPAAGLVVIVVDGIAQLGSFQAFLLAVLLSGVIQFGFGLLKAGRFAAYVPSPVIK
GMLAAIGILLIVKQVPFALGLSGDGDAPGHAVFASSAIALASLALLAVWETRAMRRFAFV
RLVPAPLAVVILGIGATVLLGFVSPNFAPPAEHRVALPELASFAALGEALKTVDLGPNFA
YLLSPDVWRIAITIAVVASLETLLSLEAVEQIDPKRRPSQPNRELKAQGVGNLVAGAIGG
LPITSVIVRSSVNVNAGAQSRMSAIVHGVMLVVSVFALTWLLNLIPLASLAAILIHTGFK
LAKPALFTSVAKQGPGAFLPFAVTIAGVLAIDLLAGIALGLACSVFAVACANLRSPATLA
QHDDHYLLSFRKDVSFLGKVQIKQYLAQIPDRAVVIIDATRADYIDHDVREMIDAFVAEA
PRREITVDYRRQVQHVRGAGLRWFFRNPAAQQAPQ*
>GCA_000452945.1_BPMSHR5858_genomic_04442_1 leuA_2
MTDKLIIFDTTLRDGEQSPGASMTKEEKIRIAKQLERMKVDVIEAGFAASSNGDFDAIQT
IASQVKDSTICSLARANDKDIQRAADALKPANSFRIHTFIATSPLHMEKKLRMTPDQVFE
QARLAVRFARKFTDNIEFSPEDGSRSDMDFLCRVLEAVIAEGATTINIADTVGYGVPELY
GNLVKTLRERIPNSDKAIFSVHCHNDLGMAVANSLAGVKIGGARQVECTINGLGERAGNT
SLEEIVMAVKTRKDYFGLDLGIDTTQIVPASKLVSQITGFVVQPNKAVVGANAFAHASGI
HQDGVLKARDTYEIMRAEDVGWTANKIVLGKLSGRNAFKQRLQELGVSLDSEAELNAAFA
RFKDLADRKAEIFDEDIIAIVTEEESALAQEHEHYKFVSLAQRSETGERPQAKVVFAVDG
DEVAGEASGNGPVDATFNAIETEVGSGAELLLYSVNAITTGTQAQGEVTVRLSKSGRIVN
GVGTDPDIVAASAKAYIAALNKLYSNADKLNPQRA*
>GCA_000452945.1_BPMSHR5858_genomic_04443_1 group_7597
MAAFKPRRPRNGNAPWPRSFRHKKSVAQDVAPVESRRAARQQFLRTRGIYLLPNAFTTAA
LFCGFFAVVQAMNVRFETAAIAIFAAMVLDGMDGRVARMTHTQSAFGEQFDSLSDMVSFG
VAPALVMYEWVLKDLGRWGWLAAFVYCSGAALRLARFNTNIGVVDKRFFQGLPSPAAAAL
IAGFVWLATDNRVPMKLGWLPWVAFVLTIYAGVTMVSNAPFYSGKALDVRHRVPFAAILL
VVVAFVLVSSDPPLMLFGLFVLYGLSGYVFWAYMAVRGRANPARSSQREH*
>GCA_000452945.1_BPMSHR5858_genomic_04444_1 psd
MNYPHPIIAREGWPFIAIAAVVALLIHAVGGFGLAWPFWLLLVFVVQFFRDPPRAIPTQA
NAVLCPADGRIVAVETAHDPYADREALKISVFMNVFNVHSQRSPVDGAVQKVEYFPGAFL
NAALDKASAENERNAVVIQTGAGHTVTAVQIAGLVARRILCYVRAGEPLSRGQRYGFIRF
GSRVDVYLPKGSRARVSIGEKVSASSTILAELPEQP*
>GCA_000452945.1_BPMSHR5858_genomic_04445_1 ilvC_1
MKVFYDKDADLSLIKGKQVTIIGYGSQGHAHALNLKDSGVNVTVGLRRGGASWSKAENAG
LAVKEVAEAVKGADVVMMLLPDEQIAAVYAQEVHANIKEGAALAFAHGFNVHYGQVIPRA
DLDVIMVAPKAPGHTVRGTYAQGGGVPHLIAVAQDKSGAARDIALSYAAANGGGRAGIIE
TNFREETETDLFGEQAVLCGGTVELIKAGFETLVEAGYAPEMAYFECLHELKLIVDLIYE
GGIANMNYSISNNAEYGEYVTGPRVVTEETKKAMKQCLTDIQTGEYAKSFILENKAGAPT
LQSRRRLTAEHQIEQVGSKLRAMMPWIAKNKLVDQSKN*
>GCA_000452945.1_BPMSHR5858_genomic_04446_1 ilvH
MRHIISVLLENEPGALSRVVGLFSARGYNIETLTVAPTEDQSLSRLTIVSIGSDDVIEQI
TKHLNRLIEVVKVVDLTDGAHIERELMLIKVRAVGKEREEMKRMADIFRGRIIDVTEKTY
TIELTGASDKLDAFIQALDAGSILETVRTGSSGIGRGERILKV*
>GCA_000452945.1_BPMSHR5858_genomic_04447_1 ilvI
MNMPSAEFSTSESLSPQKTGSIGATVLMKALADENVEFIWGYPGGSVLYIYDELYKQDKI
QHVLVRHEQAAVHAADAYSRSTGKVGVCLVTSGPGVTNAVTGIATAYMDSIPLVVISGQV
PTAAIGQDAFQECDTVGITRPCVKHNFLVKDVRELAQTVKKAFYIARTGRPGPVLIDIPK
DISKTPCEYEPVKNVSLRSYNPVTKGHSGQIRKAVSLLLSAKRPYIYTGGGIILADASRE
LNQFADLLGYPVTNTLMGLGGYRSSDRKFLGMLGMHGTYEANMAMQHCDVLIAIGARFDD
RVIGDPAHFASRPRKIIHIDIDPSSISKRVKVDIPIVGDVKEVLKELIEQLQTAEHGPDA
DALAQWWKEIEGWRGTDCLKYDRASEIIKPQYVVEKLWELTDGNAFVCSDVGQHQMWAAQ
FYRFNKPRRWINSGGLGTMGFGLPAAMGVKMAHPDDDVVCITGEGSIQMCIQELSTCKQY
DTPVKIISLNNRYLGMVRQWQQIEYSKRYSHSYMDALPDFVKLAQAYGHIGMRIEKSADV
EPALKEALSLKDRTVFLDFQTDPTENVWPMVQAGKGITEMLLGSEDL*
>GCA_000452945.1_BPMSHR5858_genomic_04448_1 group_5036
MASDKELADFLAGVERRAFKQAAYAVRDDDASLDIVQDAMIRLAEKYGDRPAAELPLLFQ
RILQNAIHDYFRRQKVRNTWVSLFSSLNNTDDDEFDPLETLESADGDGVESGETRLEREQ
VLALIDDEIQKLPARQREAFLMRYWEDMDVAETAAAMGCSEGSVKTHCSRATHALALALK
AKGITL*
>GCA_000452945.1_BPMSHR5858_genomic_04453_1 mgtA
MKIMIVTDAWEPQVNGVVRTLKSTARELTALGHRVELVTPLEFRTVPCPTYPEIRLSILP
YRRLRERLDAFEPHALHIATEGPLGLAARRYARARKLPFTTAYHTRFPEYVQARFGVPLA
ATYRFLRWFHGASLAVMAPTPVVKDDLERFGFDNVVLWTRGVDLDIFRPIESKVLNTARP
IFLYVGRVAIEKNVEAFLKLDLPGSKWVAGEGPALAELKSRYPEANYLGVLTQAELAKVY
AAADVFVFPSRTDTFGLVLLEALACGTPVAAYPVTGPVDVLGDGGAGAMNDDLREACLEA
LKIDRRHARAWAERFSWRAASEQFASHLKPLQKSACPHTEGAAV*
>GCA_000452945.1_BPMSHR5858_genomic_04454_1 group_3536
MRAKAPPASPSRKQAAHAGAARPAHVRLAPSETHEPAEPHEPLGPDDPLALPHNPYKGNR
GLMRAWYALKNSYNGFRVAIREESAFRQELTLAAIMLPIAAFVPVEPASRALLIGTVLLV
LIVELLNSSVETAIDRISLERHELSKRAKDLGSAAVTVALCTCLSTWGLILGPLVWRRLA
G*
>GCA_000452945.1_BPMSHR5858_genomic_04456_1 group_5038
MNSVCVYCGSANGAKPVYADAARAFGRALANAGLTLVYGGGRVGLMGVIADEVIASGGRA
IGIITELLYDKEVGHTGLTELHVVPDMHHRKKMMAELADAFVAMPGGAGTLEEFFEVYTW
AQLGYHRKPVALYNVDAFYQPLIALLEHTVDEGFMQRTYFDALCIDAAPDALIDQLVRYR
PPVRDKWTFVSTQEA*
>GCA_000452945.1_BPMSHR5858_genomic_04457_1 ygfF
MTNTQRKVVLITGASRGIGRASALLAAARGWSVGINYARDAAAAEATADAVRAAGAQACV
VRGDVAHEADVIHMFDAVQSAFGRLDALVNNAGIVAPSLPLADMDIARLKRVFDTNVLGA
YLCAREAARRLSTDRGGGGGAIVNVSSIAARLGSPNEYVDYAGSKGAVDTLTLGLAKELG
PHGVRVNAVRPGLIATEIHASGGQPGRAERLGAQTPLGRAGDADEVAETIVWLLSDAASY
VTGALLDVGGGR*
>GCA_000452945.1_BPMSHR5858_genomic_04458_1 arnT_1
MQGNASPGGRRTTASAQRAGSSDNHSRSPSVALAAGAPGTVGAAGERAIGSRGAAATRSA
LTGWRAWFVAAAVLCAYLLPGILGHDPWKQDETYTFGIIQHMLESGDFVVPTNAGQPFLE
KPPLYDWVATGFAWLFSRFLPLHDAARLASALFAALAFGFTARAARVAMGAARWLELPVI
GTVALCAGSLVVIKHSHDLMTDVALMAGAAMGFCGLLELVIRHAGDAFGARAERPPANRF
AAPLFGLGVGIALMSKGLFVPLVFGATLAATLVLYPACRSRAFFRSLAIAALVCAPFALI
WPTALFLRSESLFLVWFWENNVGRFFGFSVPTLGAENDKPLFIWRALLTVGFPVAPLALV
ALARGLWRDWRAPRVALPLTFAGIGMVVLHISATSRQLYILPFIAPLALVAAQAIPRLPQ
RLHAAWDYASRLLFGAAAALVWIVWSLMSDHGGPRAGLQWLGRWLPLDWTMPIEPALVLS
ALAITIGWVGLLPSLRLAGKWRGALSWAMGALVAWGLVYTLLLPWLDVAKSYRSVFEDLN
RRLALEWNDGDCMASVDLGESEAPMLYYFSGVLHQPVAQPNASACTWLIVQGTRANPPVL
DAEWKPFWAGARPGDEQEMLRVYVRTPAAARPAHP*
>GCA_000452945.1_BPMSHR5858_genomic_04459_1 proP_6
MKDPLESACPPAPGYGAGADGAACAAAMAGMAGVDGVDGVDGVDGADAARADARRDVAAV
ALGNAIEFFDFGAYATFAVMIGRTFFPTQRPFVSLLLSVSVFGLGFVVRPLGALVIGAYA
DRAGRKPAMTLTLAMMAVGTGAIAVLPGYETIGVAAPILLVVTRLIQGLAWGGEAGPATT
YILEAAPPERRAAYACWQVATQGFAAVAAGLAGYALTLALPEAELDAWGWRVPFALGLLV
LPVGVYIRRRLSDTIDPRHAYCTTREILRELNARHRRPIAIGLMILLGSTITQYFLNYMT
TFALTELHLPGGVAMLATFVTGAALAVGALAGGSLSDRVGRRAILIWPRVLLLLLLFPAL
QLIVSRPTPAVFLATLTVSLGLHGMSGAALIVLIAESFPQRVRSTGFSIVYAVAVSLFGG
TAQSIVTWLIGTTGNPMAPSGYLLVANAICIAAGWLAAETRPGGRGTR*
>GCA_000452945.1_BPMSHR5858_genomic_04460_1 aam_2
MSELLDLSACSLAERFKNRSVTPVDYAHALLDHIARWEPHLNALCRFDPARVRDEALAST
RRWAAGAPLSEIDGVPVTIKELIATQGDRVAQGSAAAADAPPATVDAPAAARLREAGAVV
IGKTTVPDYAMLSSGLSSLYGITRNPWRLDLNPGGSSSGAAAAAAAGYGPLHVGTDIGGS
IRLPAGWCGIVGFKPSNGRIPIEPYYTGRCAGPMTRTIDDAALLMRFLSRPDWRDATSLP
PETIDWSIAPAHVKGMRIGLMLDAGCGLAPDPQIVAAVEAAAERFAAHGARIVAAPPVLS
RPMLEGLDRFWRARLWADLERLPDAARARILPYIAEWAGQGASVSGVEAVRGFGATFEMR
AAAARLFQSVDAVLSPTNVVTGFPAEWAGPTDDPARPFEHIAFTVPWNMGEQPALSINCG
FARDGMPIGLQIVGPRFADRLVLQLGRAYEAWRGEMPRWPAPPR*
>GCA_000452945.1_BPMSHR5858_genomic_04479_1 yebC
MAGHSKWANIKHKKAAADAKRGKIWTRLIKEIQVAARLGGGDVNSNPRLRLAVDKAADAN
MPKDNVKRAIDRGVGGADGANYEEIRYEGYGIGGAAIIVDTLTDNRTRTVAEVRHAFSKF
GGNMGTDGSVAFMFDHVGQFLFAPGTSEDALMEAALEAGANDVNTNDDGSIEVLCDWQEF
SKVKDALEAAGFKAELAEVTMKPQNEVDFTGEDAVKMQKLLDALENLDDVQEVYTNAVVV
EE*
>GCA_000452945.1_BPMSHR5858_genomic_04480_1 purD
MKLLVVGSGGREHALAWKLAQSPRVQLVYVAPGNGGTAQDERLKNVDLSSLDDLADFAES
EGVAFTLVGPEAPLAAGIVNHFRARGLKIFGPTKEAAQLESSKDFAKAFMKRHGIPTADY
ETFTDAAAAHAYLDSKGAPIVVKADGLAAGKGVVVAMTLEEAHAAVDMMLSGNKLGDAGA
RVVIEAFLDGEEASFIVMVDGKHALALASSQDHKRLLDEDRGPNTGGMGAYSPAPIVTPQ
MHARVMREIIMPTVRGMEKDGIRFTGFLYAGLMIDRDGNPRTLEFNCRMGDPETQPIMAR
LKSDFSKVVEQAIAGTLDTVELDWDRRTALGVVLAAHGYPDAPRKGDRINGIPAETAHAV
TFHAGTTFDGDKLVTSGGRVLCVVGLADSVRGAQQAAYETINQINFEGMQYRRDIGYRAL
NRKTV*
>GCA_000452945.1_BPMSHR5858_genomic_04481_1 hemF
MTDSTYDVNRVRAYLQGLQMRIADALGAFDGTPLAADTWRRGPGERLRGGGCTRILEAGG
FFERAGIGFSDVAGDALPPSASASRPQLAGRGFEALGVSLVLHPRNPYCPTVHMNVRMLI
ATKPGEAPVFWFGGGMDLTPIYGFEEDARHFHRTCRAALEPFGAELYPRFKKWCDDYFFL
KHRNEARGIGGIFFDDFSELGFERSFEMLQSVGDAFLPSYLPIVERRRDTPYGERERAFQ
AYRRGRYVEFNLVFDRGTLFGLQSGGRTESILLSMPPTAGWRYDWHPDPGTPEARLQSEF
LVPRDWA*
>GCA_000452945.1_BPMSHR5858_genomic_04494_1 sigJ
MTETSRPAPSDEPFAALRPRLFSIAYRMLGTRADAEDVLQDAWLRWHQAERDTLDSPEAW
LVTVVTRLSIDRLRAAKAQRDAYVGWWLPEPLVDIDERTPEAAAEFADDLSVALLWVLER
LSPEERAAFLLRQAFERDYAEIAQLLDKSEAACRQLVHRASTRVQQEHRRFDVSRERHRQ
LVERFARAAASGERNAIRALLADDVELVGDGGGKVPSFFKVLRGALRIANLYWAVGRRHA
GQIEYRLAQINGEPGLLRFIDGRIESAQAFATDGARIVAIYAVRNPDKLAHIPNRPA*
>GCA_000452945.1_BPMSHR5858_genomic_04500_1 group_5047
MAEPTLGVALIAYNAAARLAECLSALSFADDVIVVDGGSTDATVDIAKAHGARVIVAADW
PGFGPQKNRAVSALDTDWILSIDTDEIVTPELAASIRAAIRAPRAPVYSLDRLSSFCGTW
VRHSGWYPDWLPRLFKRGAARFSDDLVHERLVFDGPSAKLDGKLLHYSYEDFETVLRKLD
AYSSAGAEQRRAAGKRGGLAKAFGRGAWAFVRTYVLRCGFLDGRAGFMIAVFNAETVYYR
FLKLGLARQRKARD*
>GCA_000452945.1_BPMSHR5858_genomic_04501_1 rfaQ
VLLTTPLVRSLKARWPDAQIDMIVFRGTEGVLEHNPDIRRVITVAPRARPKERIADALRI
WRKYDLACAAISSDRARFYAFFAGRRRIGLVDPGRLTRLTRFILNGIVLDEHRDVHTVTS
NLSLADALDVPRCADVVAPGIGDDPAARARFDAKLYGTPALSRGEPYIVLHPYPMFRYKQ
WHEDGWVDLVRWARSSGFAVALSGGPAQAERDYAARIARAAGEPVLNMTGELSFGESAEM
FRHARLFIGPDTGATHVAAACGVPTIALFGPSNPTRWGPWPAHWPAGDEPWPLRGSGRRG
NVYLLQGEGDCVPCKLEGCDRHLDSWSRCLSELPSARVIGVAAAMLGFDAANAPADVATP
VDVSRLAARK*
>GCA_000452945.1_BPMSHR5858_genomic_04505_1 group_3543
MSGARPAERAPTEREDRRPMKIGLSCNALKYGGGLERYAIDLARGLADAGIRPSVFARSF
DSSVPEYRRVESHRINVSFLPGKCRDAWFSWRLRAARRASPVDVLIGCNRVDSSEIAICG
GTHLGFLDAIGRVPTFADRRQIALERRQYAHARFVIAHSMLMRDELLHFYGLGDDKVRVL
FPPVDAARFTPVHAVRRAELRTRFGFADDEIVLLFPSSSHERKGLPLIEAVLRDAGPGIV
VAVAGRPPERTSERLRYIGYVKDIEDGYRAADFTILASKYEPFGLVGVESVMCGTPVILP
SNIGCCDAIAPSAKLVFAPGDTAGLRAALDEAVRRTKAGAARVGSGAAAHAAIRYDPSIA
RHVAQLLDLAVEAAGDGPR*
>GCA_000452945.1_BPMSHR5858_genomic_04506_1 msbA
LSVKPTLSKPIGGQDASSPAVVMRRLWPYVKPLVWVLVAGVLAMAAVAATEAGIPALLKP
LLDHGFGSKGDMTTKLYVPAAVVGLALARAIAQYASGYLLQYVSNRILLDLRIQMFERMI
HTGVSFFQRETASTVINAVVFEVNQVLSVLMGVTITLVRDSLTVVFLLGYLFYLNWRLTL
IVAILLPCIGWLVGKINRRLRRLNREHQTLTNQLAYIVEETVGGYKVVKVHNGEPYEIGR
FNELSRKLRGYSMRMTVSGGLAQPLTQFLASIALAVVLTIAVVQSANDQTTVGGFVAFVT
AMLLIISPLKHLMDVNQPLQRGMTAAELIFGLIDEPREPEGGGKPLARASGAIEFSHVSF
SYGMSRDGRQTLDDVSFTVAPGEMVALAGPSGSGKTTLVNLLPRFFDPSSGSVRVDGVAL
PEYSLRDLRNQIAMVSQDVVLFNDTIAANVAYGQAPERDRVEAALRAANLWETVTAMPDG
IDTLVGDNGMRLSGGQRQRLAIARAIYKDAPILILDEATSALDSESERHVQAALETLMKG
RTTLVIAHRLSTIERADRILVLEGGKIVESGSHRELLEQGGLYAHLHRIQFQQDAG*
>GCA_000452945.1_BPMSHR5858_genomic_04507_1 dnaE
MSDPRFVHLRVHSEFSIADGIVRLDDIVKSAAEDGQGALALTDLGNAFGLVRFYKEARDA
GIKPIAGCDVWITNHDDRDKPSRLLLLVKDKRGYLNLCELLSKAWLTNQYRGRAELDASW
LEGELAEGLLALSGAQQGDIGLALAAGNEAAARRHAQRWARVFPGGFYIELQRYGQPGGE
AYIQQAVTIAAELKLPVVATHPLQYMTADDFTAHEARVCISEGDILANPRRQKRFTTEQF
FRTQGDMAALFADLPSALANTVEIAKRCNLTLELGKPKLPRFPTPDGMSLDDYLVQLSQE
GLDKRLVQLYPDEAEREAQRGKYNQRLDFECGTIKKMGFPGYFLIVADFINWAKNNGVPV
GPGRGSGAGSLVAYSLGITDLDPLRYNLLFERFLNPERVSMPDFDIDFCQHGRDRVIQYV
KEKYGADAVSQIATFGTMAAKAAVRDIGRVLDLGYMFTDGVAKLIPFKPGKHVTIADAMK
EEPLLQERYDNEDEVHQLLDLAQRVEGLTRNVGMHAGGVLIAPGKLTDFCPLYTQGDEGG
VVSQYDKDDVEAVGLVKFDFLGLTTLTILDWAERYIRRLDPSKENWSLAQVPLDDPTSFQ
ILKKANTVAVFQLESRGMQGMLKDAQPDRFEDIIALVSLYRPGPMDLIPSFCARKHGREK
VDYPDPRVEPVLKETYGIMVYQEQVMQMAQIIGGYSLGGADLLRRAMGKKKPEEMAKHRE
IFAEGAAKNGLTREKSDEIFDLMEKFAGYGFNKSHAAAYALLAYYTAWLKAHHPAEFMAA
NMTLAMDDTDKVKILFDDCLVNGLAVLPPDINRSNYRFEPVAEADGKRSRTIRYGLGAIK
GSGQNAIEEILRAREEKPFADLFDFCERIDRRVVNRRTIEAMIRAGAFDSLHENRAQLLA
SVPLAMEAAEQAAANALQAGLFDIGGVPAHQHALVDEPAWDDKRRLQEEKGALGFYLSGH
LFDAYRDEVRRFVRQKLGELKEGRDKVVAGVIASLRTQMTQRGKMVIALLDDGTGQCEVT
VFNEQFDANRALFKEDELLIVQGQARNDAFTGGIRFTAESVMDLERARSRYAQAVRMTMN
GNADAAALRRVLEAHVAKPDETPPAAAPAPRGGRDGGRRAQAAIPNGLAVRIAYSNARAQ
GEMRLGDAWRVKPSDALLADLRAAFGGSVVEIVY*
>GCA_000452945.1_BPMSHR5858_genomic_04508_1 group_5048
MTTVNLAAYRFVSLDSIEQWRPLVAARCNTLGLRGTILLAPEGINLFIAGPREATDAFVD
YIRHDPLFEGKFADLPFKESLSDSQPFRRMLVRLKREIITMKKPAIKPELGRAPSVDART
LKAWLDQGHDDAGRPVVMLDTRNAFEVDVGTFDRALDYRIDKFSEFPAVIEANRADLEGK
TIVSFCTGGIRCEKAAIHMKDVGIENVYQLEGGILKYFEEVGGAHYHGDCFVFDYRTALN
PQLAPTADVTCFACRAVVPADAQQSPLYVPGKCCPACHPGDSGTPGRRAEPGAEPARAV*
>GCA_000452945.1_BPMSHR5858_genomic_04510_1 rhlE_1
MSDSVAKPVDATFDQFGLAAEILKAIAEQGYTTPTPIQAKAIPVVLSGRDVMGAAQTGTG
KTASFSLPIIQRLLPQANTSASPARHPVRALILTPTRELADQVAANVHAYAKHTPLRSAV
VFGGVDMNPQMAELRRGVEILIATPGRLLDHVQQKTANLGQVQILVLDEADRMLDMGFLP
DLQRILNLLPKERQTLLFSATFSPEIKKLASTYLRNPQTIEVARSNAAASTVTQIVYDVA
EGDKQAAVVKLIRDRSLKQVIVFCNSKIGASRLARQIERDGIIAAAIHGDRSQSERMQAL
DAFKRGEIEALVATDVAARGLDIAELPAVINFDLPFNAEDYVHRIGRTGRAGASGDALSL
CSPNERKQLADIEKLIKRTLSLETLALDLPRHRHDDRGGRRERDRDERRGAPAGRRSAGG
ERTHHPRREAPIDDFFLKPYVPSPSANQPDEAKPVQPEKKAPKQPLAALLGGFGMPRKTS
SS*
>GCA_000452945.1_BPMSHR5858_genomic_04511_1 group_3545
MNRKTRAAIGVAALCAAAGAAHAARLTVEEIDADARETVAYRCANAPKPVRVSYWRAGNG
QSFALVPVNGVRLLFVDTVSASGARYQAGRYVWWTKGRDANLYDEIAGERAPPVLADCSE
VRKPRGKG*
>GCA_000452945.1_BPMSHR5858_genomic_04512_1 group_5050
MKWMLIVALCASTAGCGLAAAPCRVASAGLKIVPLVGHVAAAPTDACAGVIDPD*
>GCA_000452945.1_BPMSHR5858_genomic_04514_1 group_1039
MFNSSAAARMRLRIARSLRAPLGALTVAALLAACTMPTHPDSGAPAPDPFNPATIQLIDD
TSWELVGWRNADGSARDIPHGDNGEPIKLALSTQTGVRRAAGFSGCNRYMGTYDVKNGVL
AFGPLAGTRMACVSPARAALERDYLAALGHISKAGVQMRAPQQLVIITDADATLTFARRD
GK*
>GCA_000452945.1_BPMSHR5858_genomic_04527_1 phaB_1

MSQRIAYVTGGMGGIGTSICQRLHKDGFRVVAGCGPNSPRRVKWLEDQKALGFDFYASEG
NVGDWDSTKQAFDKVKAEVGEIDVLVNNAGITRDVVFRKMTREDWQAVIDTNLTSLFNVT
KQVIDGMVERGWGRIINISSVNGQKGQFGQTNYSTAKAGIHGFTMSLAQEVATKGVTVNT
VSPGYIGTDMVKAIRPDVLEKIVATIPVRRLGSPDEIGSIVAWLASEESGFSTGADFSLN
GGLHMG*
>GCA_000452945.1_BPMSHR5858_genomic_04538_1 group_3549
MNSITTTPNHLLDIVYPVLEGMAPGALQRALADAFDAESASLAQIEHAFNTLTSGKPPQE
ALRMFFASWSKTNNSAASVSGLANRVTLLARSDPGSPASARLYDVCASLQRITDEDLGAL
GGALHADLFYNMATTICGDDAWLLKSNCVASAQAFKDWTDRQRLRNRDLMQGLLTTLVHE
VYTHGEVEFIHPLYKAWVEHRMNVPAERVRFVVAWVTVHTGGTESAHFAHAVRAIDTFCD
AMDVRIDPASSTALFREYLRRKADVMRTCAAAL*
>GCA_000452945.1_BPMSHR5858_genomic_04539_1 dnaE2
MDAASGILPDYAELFCRSNFSFLHGASSAEELVERAAKQGYRGIAITDECSLAGAPRMHV
AAKAVGLPLVVGAYFGVTPDDAAPGHDPGPGAFGLVLLAQNREGYGNLSELISWRRMNAP
KGTYRLTPRMLAAPPRALAHLRGVPDCFAILVPTYPARADVLDAQLAWFDALFGERARLG
LVQLQRALDGAHREQVRAAGERRGMHIVALGDVTMHIRSCKPLQDTMTAIRLGMPIAECG
HALAPNGEQHLRTRQRIAQLFPADALAQTCRMLDACHFSLDDLRYEYPHEIVPAGHTPTS
YLAQETWAGARRRYPDGVPDTVRQRIEFELALIADLKYEPYFLTVYDIVKYARNKDILCQ
GRGSAANSVVCYCLGVTEVNPQQSTLLFERFLSRERGEPPDIDVDFEHQRREEVIQYLYE
KYGHDRAALAAAVSTYRPRGALRETGKALGVDPMLVERVAKEHRWFDGSRDLLARFASVG
LDPEVPLIRTWAEIAARLLNFPRHLSQHSGGFVVSRGKLTRLVPVENAAMEGRRVIQWDK
DDLEALGLMKVDVLALGMLSALHRAFDMITAWRGPPLPDGRPFRLEHIPQDDEATYDMIC
RADTVGVFQIESRAQMSMLPRLRPRGYYDLVVQVSIVRPGPIQGGAVHPYLERRRIAAGE
AHGEITYPSEALERVLERTLGIPIFQEQVMQIAIVAAGFTPGEADALRRAMAAWKRKGDL
GKYHERIVAGMLERGYSREFAEQIFEQIKGFGEYGFPESHAASFAKLAYASSWLKRHEPA
IFLAALLNSQPMGFYPPAQLVQDAKRHGVTVLPIDATKSGWEASLEAQPGAAPPDGRPAV
RLGLSLVRGLGEEAARRIGAARAAGPFASVDELARRACLERRDLEALAAANAFATLAGNR
RDALWQAVAAAPERGLLAAAPIDEAVRPALGAPTEADDVFADYRTIGLTLNRHPVALLRP
ALDARRLSSAAALRDRRNGRLARACGLVTARQMPGTAKGVLFVTLEDETGCVNVIVRPEL
LERQRRETLDSQLLAVSGVWQCESDVRHLVAQYLEDLTPLIAGLRTESREFH*
>GCA_000452945.1_BPMSHR5858_genomic_04554_1 group_5057
MHILVPIAFVLIIASMVSALYFMMHDRGHTKRMVWSLAMRVGLSISLFLFILFANWMGWI
HSTGIPIGR*
>GCA_000452945.1_BPMSHR5858_genomic_04556_1 group_7609
VVSGTGNKSTFVQTIKAVMWSFFGVRKRRDLEADATQLNPLHVLIAALIGAALFVGVLVL
IVHAVVG*
>GCA_000452945.1_BPMSHR5858_genomic_04559_1 pal_1
MEILGKEAMKTIKRALTGVLACSALLLSGAALAVGDSPGGPRVNEINLQPPVTKIAEELY
DLHTMMLILCTVIFVGVFGVMFYSIFAHRKSKGHKAANFHESTTVEIIWTIVPFIIVVLM
ALPATKAVVAMKDTTNADLTVKVTGYQWKWGYDYVKGPGEGISFLSTLSTPRTEVNGRQP
ISDTYLQEVDHPLVVPVNKKIRVITTANDVVHSWYVPAFGVKQDAIPGFVRDTWFKAEKV
GTYRGFCTELCGKEHAYMPVVVEVLSDDDYAKWVGTQKAKLAAGAVDPNKVYTRAELMAH
GEEVYKANCAACHQPNGKGVGAFPALDGGKIVNGPIAGHLEQVLKGKGAMPSWASLSDLD
IASVITYERNSWGNHKGDSLQPKQVADARNGKLPEDAQQADGGAAANAASGAAAQTQAQA
PALPAAIYFETGKSELPADAKDAIAAAAEYVKAHPDAKLALSGFTDKTGSADANAELAKR
RAQVVRDALKTAGVAEDRIILKKPETITGGADAKEARRVEIGPAA*
>GCA_000452945.1_BPMSHR5858_genomic_04564_1 ymdB
MPNIGPTSVEACVVDITTLTLDAIVNAANASLLGGGGVDGAIHRAAGPELVKECATLGGC
ATGDAKLTRGYRLPAKYVIHTVGPVWRGGGHGEAELLASCYRRSLEVAAGAGCASIAFPA
ISCGVYRFPPADATAIAVRTVAGALAGELADARFERVVFACFSSDMLDFYRTALVRF*
>GCA_000452945.1_BPMSHR5858_genomic_04581_1 group_5062
MTIVVRAARRTDWPALRALYLHARRDTFAWLPAERFAAGDFDEHTRGEALLVAQARDEGI
VGFVSVWEPERFVHHLYVAGTRLREGIGAALLRALPGWPAARYRLKCLVRNERALAFYRA
HGFVEIDSGVSADGEYRLLGTREAATAARPGKRD*
>GCA_000452945.1_BPMSHR5858_genomic_04605_1 group_5065
VSEPIGAPWGGGCRIVEWIGGDGRIARRVAAVNVTEAEVYAMIRRPVEGRRYLMSDDEQM
PRDTLPRR*
>GCA_000452945.1_BPMSHR5858_genomic_04607_1 dadX_2
MPRPISATIHTAALANNLSVVRRHAAQSKVWAIVKANAYGHGLARVFPGLRGTDGFGLLD
LDEAVKLRELGWAGPILLLEGFFRSTDIDVIDRYSLTTAVHNDEQMRMLETARLSKPVNV
QLKMNSGMNRLGYTPEKYRAAWERARACPGIGQITLMTHFSDADGERGVAEQMATFERGA
QGIAGARSFANSAAVLWHPSAHFDWVRPGIMLYGASPSGRAADIADRGLKPTMTLASELI
AVQTLAKGQAVGYGSMFVAEDTMRIGVVACGYADGYPRIAPEGTPVVVDGVRTRIVGRVS
MDMLTVDLTPVPQAGVGARVELWGETLPIDDVAARCMTVGYELMCAVAPRVPVRAE*
>GCA_000452945.1_BPMSHR5858_genomic_04610_1 group_1530
MSAAASPVPPVALRDAAVRDLAWLLSSASLLAPSAGAPLAQPWASVADAARTAAWLAALD
AQPAPLHDALAHARPVRLGRYAECLLGFFVGHAPSLRLVAANLPLRSNGRTLGECDFLIE
TARGERLHWELAVKCYLCVATAGTASLADFVGPNLADRFDLKRAKLVDHQLRLTARDEFA
SLGHAGPWRAQMFVKGWLFYRADDGANAAGAVADPPEIGADHPRGFWITREAWPKFALAA
ARRDVAWIVLPRLAWLAPRAFDDADVACGRCVPLASDAVIDEVARQSGPLLVAALARDAS
GGWRETARGFIAPDDWPRRASAFAAR*
>GCA_000452945.1_BPMSHR5858_genomic_04614_1 group_7610
MSDITAQFEQAQIDVKQLTERPGNLTLLRLYALFKQATDGDVHGDKPGFTDIVGKYKYDA
WDALKGTSQDDAKKQYVELVESLKNGTAS*
>GCA_000452945.1_BPMSHR5858_genomic_04620_1 aceB
MTTTLKLPQGMAITGEIKPGYEAILTPEALELVAKLHRQFELRRRELLAARVERTQRLDA
GERPDFLAETKSIREGDWKIAPLPADLQCRRVEITGPVERKMIINALNSGADSYMTDFED
SNAPSWTNQIDGQINLKDAVRRTISLEQNGKSYKLNDKIATLIVRPRGWHLDEKHVTVDG
QRVSGGVFDFALFLFHNAQELIARGSGPYFYLPKMESHLEARLWNDIFVAAQEALGVPRG
TIRATVLIETILAAFEMDEILYELREHSSGLNAGRWDYIFSAIKKFKNDRDFCLADRAKI
TMTVPFMRAYALLLLKTCHKRNAPAIGGMSALIPIKNDPEANDRAMAGVRADKARDAGDG
YDGGWVAHPGLVSLAMEEFVKVLGDKPNQIGKQRDDVQIEGKNLLDFQPEAPITEAGLRN
NINVGIHYLGSWLAGNGCVPIHNLMEDAATAEISRSQVWQWIRSPKGTLDDGRKVTAELV
REYAKVELANVKQVVGGATAPYDRAAAIFEQMSTSENFTEFLTLPLYEEI*
>GCA_000452945.1_BPMSHR5858_genomic_04622_1 group_7611
MMFATTDLCDAHEDRLAAGTLRVLEPVFRPFGGVRRFAGPAATLKLFEDNSLVRTALEQD
GAGRVLVVDGGGSLRCALVGGNLGKLAEKNGWAGIVVNGCVRDSDELAECRVGVLALAAH
PRKSDKRGAGVSDAPVDVRGTRIVPGDWIYADADGVLVSDDALLE*
>GCA_000452945.1_BPMSHR5858_genomic_04624_1 group_7612
MSDSAAAPLVYLLRPAASGDFEFAEALTHGNMNAYYQRHGLRWRADLFFASWRDSENFIL
ELDSVSIGLLRITEEGDSLHIRDVQIAPGHRQRGAGTYLLETSHRFAKARGLRETQLRVF
VDNPAARLYLRMGYRLAGPRLAQFGSIRHMARPVS*
>GCA_000452945.1_BPMSHR5858_genomic_04630_1 gltX
MTRPVRTRFAPSPTGFIHLGNIRSALYPWAFARKMKGTFVLRIEDTDLERSTEASVDAIL
EGMAWLGLDYDEGPYYQMQRMDRYREVLAQMLEKDLVYPCYMSTEELDALRERQRAAGEK
PRYDGTWRPEPGKVLPEPPAGVTPVLRFRNPLTGSVVWDDAVKGRVEISNEELDDLVIAR
PDGTPTYNFCVVVDDLDMGITHVIRGDDHVNNTPRQINILRALGGEVPVYAHLPTVLNEQ
GEKMSKRHGAMSVMGYRDAGYLPEAVLNYLARLGWSHGDAEIFSREQFVEWFDLEHLGKS
PAQYDHNKLNWLNNHYIKEADDARLAELAKPFFAALGIDADTIARGPDLVGVMGLMKDRA
STVKEIAENSTMFYRAPAPDAQALAQHVTDAVRPALAEFAAALKTAEWTKEAIAAALKAV
LGAHKLKMPQLAMPVRLLVAGTTHTPSIDAVLLLFGRDVVVSRLAAALA*
>GCA_000452945.1_BPMSHR5858_genomic_04631_1 group_3558
MPPEIESMSSPRLSRRAFTLACASAALAACTSTGTKPTTGAATPTNPTQPREKPIRVGVA
LGGGAARGFAHVGVLKALEARGIPVELVAGTSAGSVVGALYASGMNALQINKLALTMDEA
SISDWATPFRSRGLLQGIALQNFINKTLNNRPIEKMAKPLGIVATDLKSGQPILFQRGNT
GLAVRASCSVPSVFEPVRIGDREYVDGGLVSPVPASFARKMGADFVIAVDISSRPDGAAT
GNPIEMLLQTFTIMGQTIKTYELDKYADVVIRPNLAAMGGSDFNQRNAAILAGEEAVARI
MPELQRKLAAARSATAAA*
>GCA_000452945.1_BPMSHR5858_genomic_04632_1 group_7613
MQHRYLTQACTRVVAGMFVGILIAAAPGAFADEVSSSSQNAINSAQSEPASSSSNLQQVN
VPAPAKSQGGAKAFLSGMAGKAGDVVVGALNMIGVRYRWGGNTPDSGLDCSGFVRYVFQD
TLGMSLPRRAEEMSRVGEKVRMSELKPGDLVFFNTMRRTFSHVGIYIGDNKFVHSPSTGS
TIRVDDLDSGYWEKRFTGARRIETQFSTKPDDLRQRVKATIGDGGVNASASGSN*
>GCA_000452945.1_BPMSHR5858_genomic_04633_1 yejF
MSAAQIARGAGAPPSPSTPPHEPLLSIERLRVRFGGTLAVDDVSLAIGRGERVALVGESG
SGKSVTALAILRLLRDADVSGAIRFAGETLADKSEREMRALRGADIAMIFQEPMTALNPL
FTIGEQIGETIELHDGVSKRDARARAIALLARTGIDRPDERVDSYPHQLSGGQRQRAMIA
MALACRPRLLLADEPTTALDVTIRAQIVELLLELQRDAAAERGMAVLLITHDLNLVRRFA
ERVAVMEKGRLVECGPVERIFDAPEHPYTQRLLHSRPQRTVSPVLPIAPVVLDARGVTVQ
FRQKLPGFAGWLRSGRFTAVSDASVSVRQGETLGIVGESGSGKSTLAMALLGLQRVAHGD
IEFEGRSLASFRGRERVALRANMQVVFQDPFSSLSPRQTIGRIVSEGLELHRPRLTADAR
RDKVIAVLREVGLDRTVLHRYPHEFSGGQRQRIAIARALVLEPRVLVLDEPTSALDVSIQ
QQVLKLLSNLQRKYNLGYVFISHDLEVIGAMAHRVAVMQGGAIVETGDVERIFTEPVHPY
TQKLLKAVR*
>GCA_000452945.1_BPMSHR5858_genomic_04636_1 group_2554
MTIGSPRARRPRSPQRAAPSEQAARAAAPRRAARARAALARFARRAAAGVALAFVAAPAA
HAVYAIAQYGEPKYPAGFAHFDYVNPDAPKGGTLVLANPNRLTTFDKFNPFTMRGNPAPG
IDLLFESLTTGSADEPASAYGLLADDIAVAPDGLSVTFHLNPRARFSNGEPVTAADVKYS
FDTLKSPKAAPQYPAYYADIARAVIVDAATVRFEFRRKNRELPLIAGGIPVFSRKWGVRA
DGSRIAFDQIAFEQPIGSGPYLIERYDSGRTITYRRNPAYWGAALPVRIGTNNFERIVYK
LYGDGVARLEAFKAGEYDVLVEYIARNWARRDVGKRFDSGELVKREFRQHNGAGMQGFFM
NLRRPLFQDVRVRHALDLAFDFEWLNRQLFYGAYTRLNSYFADTDLQATGTPSAGELALL
APLRAQLDPAVFGPMTVQPSTDLPASLRANLLKARALLAEAGWTYRDGALRNAKGEPFVF
EILDDSGSAFEPVVAAYIRNLAKLGIVAKYRTADFALLQKRLDAFDYDMTTVRYPGVQVP
GAEQVARFASRYADEPGSDNLTGLKSPAVDAILKALTQAETRDELLDATHALDRVLMHGY
YAVPQWYSAVHRIAFKRTLAYPSVLPLYYSAEGWVASTWWARPEHGASAR*
>GCA_000452945.1_BPMSHR5858_genomic_04637_1 fabI_2
MGFLDGKRILLTGLLSNRSIAYGIAKACKREGAELAFTYVGDRFKDRITEFAAEFGSELV
FPCDVADDAQIDALFASLKTHWDSLDGLVHSIGFAPREAIAGDFLDGLTRENFRIAHDIS
AYSFPALAKAALPMLSDDASLLTLSYLGAERAIPNYNTMGLAKAALEASVRYLAVSLGAK
GVRVNAISAGPIKTLAASGIKSFGKILDFVESNSPLKRNVTIEQVGNAGAFLLSDLASGV
TAEVMHVDSGFNAVVGGMAGLEE*
>GCA_000452945.1_BPMSHR5858_genomic_04638_1 group_7615
MKSLRAASLAAGFAALLASGAALAGVSVGINVGVPAPVYVAPAPVYAPPPPPVVYQPAPV
YVPAPVYGPAPSIVIGWHGDRYWDGRRYWGRDEWYRHRGRDWDRGHWDNGRHNGWRH*
>GCA_000452945.1_BPMSHR5858_genomic_04639_1 napA_2
MAVEKTACILCSRNCGLVVDVTDGKFAKIRGDDDHPVSKGYLCQKAARLTYYQHHDDRLT
HPLRREPDGSFVRVTWDEALDDIAQRLVALRERHGGDAFAFVGGGGQGNHVGGAYSRQLL
AAMRSRYAYNALGQEKTGDFWLNGRLFGRQDCHTTEDIEHADYVLVIGANPYQAHGIPNA
RDTLRDLKKDPARTLVVVDPRRSETAKFADLHLQVRPGADAYLMSAMLAIMLRDGLFDRA
FIAQRCTGFEFVERVLRDVPIAEYARRADVPLEDIERVAHGFATARAACLRIDLGIQHTL
HTTLNGYLEKLLFLLTGHFGKRGGNNLHSFLLPVIGHTDERAIRHGKPLKRTAHHRMFPI
AGIYPPNILPDEIERAGEKRVRAAFVDSANPVVTYADTNAYERAFGKLELLVVIDVAMTE
TARLAHYVLPAASQFEKWEATGFNLEFPANAFHLRHPLLPPLGESLPEPEIYTRLLEKMG
ELPRRFPLLERIARFEPRASKHLAYLGALGIVLAANKKWQRHAASIVYRTLGRALPGDAA
ASAPLLPLAMLYAQRHYAAVRRAGFRGNRATLGTALFRAILERRSGTLISVHEFDEMWRF
IRHPDARVHLHIPEMIDELRALATETPPGADYPFVLMAGERRAYNANQIYRDPTWRKVDP
HGSLRIHPDDAAALGVADGGKLVCASATGSIEVVAELDDSVRRGMVTLPHGYGMRYKGGP
PVGPELNRLTSGAHCDPLSRTPYHKYVPVHLRVVDTPVAAPATLEAESA*
>GCA_000452945.1_BPMSHR5858_genomic_04640_1 fpr_2
MSANAHETVLSVHHWNDTLFSFKTTRAPGLRFKTGQFVMIGLEIDGRPLMRAYSVVSAHY
DDHLEFYSIKVPDGPLTSRLQHLRAGDKLLVAGKPTGSLIIDNLRPGKHLYLLSTGTGLA
PFISVIRDPDYYGAFDKIVLMHGVRWKSELGYFDHITTELPENAYFGDLVRDKLIYYPSV
TRETFERQGRLTELIESGKLFDDVGLPPLDPAVDRAMVCGGPSMLADLVEMLERRGFVEG
TSHAPGDYVIERAFVEK*
>GCA_000452945.1_BPMSHR5858_genomic_04643_1 group_5068
MGVIPAGPNYATYGRTFRNTRGIRIQYRRLPMPLPCESTRIDTDRPPEATALPASRTGRL
GRRTASGGPKPASGARSITGTPIIGEGPLNNDRYSNGRYSILPSTSIEQSFETETNMRYS
PKHKEASRARLVEAGAALAKRNGFANTGMDALTAAAGVTTGAFYSQFRSKPEFLYAIVEH
EMSKVLATVENRSKDDLVATLRSYLSTAHADHPELGCPVPTLGAEIARADVATRRMFEDL
IKRFQASLATALHDDEAAWTLGCAAIGAVLVARAMASTERRSEVLRSVLAYTMRSFDADA
RKKK*
>GCA_000452945.1_BPMSHR5858_genomic_04644_1 map_2
MARSRQVPIRSKAEIAMARRAGTMAAQVLSMIAPHVKAGVTTNALDRLCREYIVDVLRAR
PANIGYHGYPKTICASVNHVVCHGIPSDHALQDGDIVNIDVALDHDGWYGDTSRMYCVGE
PAAPARRLVDATYEAMMAGIAAVRPGATLGDVGHAIQSVAHREGFSIVREYCGHGIGRIY
HDEPQVLHYGRPGAGLRLAPGMIFTIEPMLNAGRAETRQLADGWTVVTQDRSWSAQWEHM
VVVTEDGFDILTPWPDGEPARSPAATGAAGAA*
>GCA_000452945.1_BPMSHR5858_genomic_04646_1 cysK2
MTGIFVFIESNTTGTGELLVRKALQRGLTPYFLTANRGKYPFLDAIRVVTISLDTSDADR
IHRFVSSLDGVAGVMSSSEYFIEVASEVARRLGLPTANTEATRVCRDKKHLARTLAEHGI
DVPRTHALALDADADADADADADADADAVALSALDGLAYPVVVKPRMGSGSVGVRLCASVDE
VAEHCAALRRAGTRAALVQAYVEGDEYSVETLTVARSTQIVGIVRKRLGREPHFVEIGHD
YPAPLSSPQRERIERTVLRALEALGYAFGPAHTELRVRGDTVTIIEINPRLAGGLIPVLL
GEVFDVDLLDHVLDMWLGVAAFADLTAKRYGAIRFALPAREGVLRGPLALPADIAARPEL
RHFHPIAQPGDALRLEGSFRDRIAAVVCAGDHRESVEALAERAVAGLSIDIGDDARAAAL
NESNESNGANAATPGLPPRLQAIVYGDGASEAPLAELDHLFDLNEAHLVMLGATRIVAPE
RVRPLLDAHRRLRRAGYAPLLARPRPRGLYMLVEAYLIETLGEDVGGVLQTGRSRNDINA
ATTKLHLRDATSRAFDALWRLRRSLVFKASANVDCAFPIYSQYQPALPGTLAHQLLAFDG
ALAHETHALFALFQHIDVCPLGAGAGGGTTLPIDPEFVCRLLGFEQPAPNSLDAVANRSG
VVHFLSAMNAIGLVLSRLAQDLQIWTTAEFALVSLPAALTGGSSMLPQKKNPFLVEFVKS
RAGVPFGALASCSAALGKTPYTNSFEAGSPMNGLIAQACAAIEDAAAVAVLLIDGLEAAQ
ARIDAHLRDTGVVAMAVAESLVVRRSIDFRSAHTRVAQAVRDSAAQGRSSHDALAALDPD
FVSRAPLEWARSHRFGGGPGAADLNHGVARACRALADDEAVFRRKQDVWREAEQMRRLAA
QQLAGD*
>GCA_000452945.1_BPMSHR5858_genomic_04650_1 fdoG_1
MLQLSRRQFLKLSATTLAGSSLALMGFSPAEALAEVRQYKLARTVETRNTCPYCSVGCGI
LMYGLGDGAKNATSSIVHIEGDPDHPVNRGTLCPKGASLIDFIHSPSRLTQPEYRAAGSD
KWQPISWSDALDRIAKLMKADRDANFVETTDDGMKVNRWLTTGMLAASAGSNEVGYLTHK
TVRSMGMLAFDNQARV*
>GCA_000452945.1_BPMSHR5858_genomic_04651_1 fdoG_2
MTNHWVDIKNADVILVMGGNAAEAHPCGFKWVTEAKAHRNARLVVVDPRFTRTASVADYY
APIRTGTDIAFLGGVIHYLLTNDKIQHEYVKHYTDFSFIVREDFAFDDGIYSGYDADKHA
YPDKSTWDYERGDDGFVKVDETLAHPRCVYNLLKQHYARYTPEMVEKICGTPKDKFLKVC
EMLATTAVPGRAGTVLYALGWTHHSVGAQMIRTGAMVQLLLGNIGIAGGGMNALRGHSNI
QGLTDLGLMSNLLPGYMTLPMQAEQDFDAYIQKRAQQPLRPNQLSYWKNYRAFHVSFMKA
WWGDAASAENHWGYDYLPKLDKQYDLLQTIELMHAGKMNGYICQGFNPLAAAPSKRKTSE
ALAKLKWLVIMDPLATETSEFWKNHGEFNDVDSSKIQTEVFRLPTSCFAEERGSLVNSGR
VLQWHWQGAEPPGQAKSDLEIMSGIFLRMRDMYRKDGGKYPDPIVNLSWPYANPESPTPE
ELAMEFNGRALADLPDPKDPTKTLVKKGEQLAGFAQLKDDGTTASGCWIFCGAWTQAGNQ
MARRDNADPTGIGQTLNWAWAWPANRRILYNRASCDVNGKPFDPSRKLIGWNGKTWTGAD
VPDYKLDEPPETGMGPFIMNPEGVARFFARAGMNEGPFPEHYEPFETPLAANPLHPGNPR
ALNNPAARVFPDDRASFGKVDQFPHVATTYRLTEHFHYWTKHARLNAIVQPQQFVEIGED
LAKEIGVAHGEQVKVSSNRGHIVAVALVTKRIKPLMVDGRKVQTVGVPLHWGFKGLTKPG
YLANTLTPSVGDGNSQTPEFKSFLVKVEKA*
>GCA_000452945.1_BPMSHR5858_genomic_04653_1 fdoI
MKHDDPNLIVRYTANERSNHWITAISFVLLALSGLALFHPSMFWMTALFGGGQWTRILHP
FVGLVMFASFAVMVVRYWRHNLLDAGDRQWLRQMDDVLANREDKLPEVGRYNAGQKLLFF
VMVACLLLLLVSGIVIWRRYFSLYFPIGVIRAAAVVHAAAAFALIVGIVVHVYAALWVKG
SIGAMVRGTVTVGWARKHHPKWFRESVK*
>GCA_000452945.1_BPMSHR5858_genomic_04654_1 fdhE
VTDPTQRILDPSEIASLDHSAIPRVRMPERGAVFAARAARLRKLADLNPIAGYLRLMAAV
ADAQHELLQTFVPNAPEKAAAERAQRHSMPLMPALGGERDPRWRDLLQSLLDRVERAGLV
NPPLAKLIDGMRLREAAALDAQADALVAQRFAEIEPASAPFLMAALQVVWTALASRIAPA
DVPYLEQPGLCPVCGAQPVASIVRVGGQYQGYRFLQCGLCSTEWHMVRTKCSHCDSTKGI
AYHGIEGGSEAIKAESCDACHTYRKIGYQEKDYDVEPLADDLASLTLDLLMGEAGYRRAS
PNPLLWPDLPAEAGAAAGDERT*
>GCA_000452945.1_BPMSHR5858_genomic_04656_1 selB

MIVGTAGHIDHGKTTLVRALTGVDTDRLKEEKARGISIELGYAYTPLPNGDVLGFIDVPG
HEKLVHTMAAGACGIDFALVVIAADDGVMPQTREHLSILQLLGVAHGALALTKCDRVDAA
RVARVRDEIRAWLAASPLADAPVFETRASEPDDAGVAALNAHLRDTALAWRARRDDGLFR
LAVDRVFTLAGQGTVVTGTAVAGRVRTGDSLAVARTGETVRVRSIHAQNRATDVGHAGER
CALNLAGIDKAALARGDAIVDARLATLSPRIDVELTLTADADLTITHWTPLHVHLGTLHR
VAHVALLEGETLGPGRRARAQLNFTEPVFAAPGDRFIVRDAQATRTVGGGRVLDPFGPAR
KRRTRARRAWLDALAAWLDEGRLDALLDEAPLGIARATLMHLTGLPAQAWALPADAVSVA
APGKHADEARVLARGHWDALRTRVLDALAAFHQRSPDEQGPDVARLRRIAAPLADDALWR
ALTDALIAEGAIVRSGPWLHLPSHAVSFDAAEEALAGRLLPLVAAGRYDPPWVRGHAAAT
LMAEDAVRALLRKLARRGDVHQVVRDLFYHRDVIAELARLIARLAGEHGGGLDAATFRDA
TGLGRKRAIQILEFFDRVGYTRFHRDLHWLRADSRLLAGSR*
>GCA_000452945.1_BPMSHR5858_genomic_04660_1 group_7617
MATTAEEDVLLHILHNVYSVTGGEVTPSSVSVPTVELKLQFDDNTHRISGFGRIAWGKNK
PPKVTSEFVDLLIHGDFGYFPPARKQRDANFPVKTLQINLVGYPNLHWNPRWGIGPQLPI
IFHQNAVIPVGSQTGYAAYSYGVPSDPESEISVTDATIRQLLL*
>GCA_000452945.1_BPMSHR5858_genomic_04662_1 entS_2
LTSVAPPSATSEAVSHRNTAVLVVFTALTNLADGVTKVALPLMATALTHSPARISGVSLT
LTLPWLLVALHVGVLVDRFDRRTLLWLANAARMAAMALLIALLPSGRVTLPVLYASGLTL
GLAEVVALTSAAALIPDAVAPSSRERANAWIAGAETVCNEFCGPLTGGMLVAAGTAIALG
AVAVGYFGGGVALFFLIGRFRVAHAPHGRPPPVRLQIAEGLGCLWHQPLLRLMAVALTVL
CMCWGAWLALMPLFATTVLGLDSRGYGVTVSALGVGGFVGALSVTLLNRRFGRRTVMLTD
LLGTFAMMAVPVLSTNLWAVAASAFAGGLGGTLWTVNARTISQHLVPGPLLGRYNAAARL
FSWGAMPIGAGLAGAIAELLGMRAAFAALAVAALLLIVPFLRVASAQALRIGPERRH*
>GCA_000452945.1_BPMSHR5858_genomic_04689_1 hmrR
VNIGEAAKVSGVSAKMIRYYESVGLLNPAKRTDSGYRVYHDDEIHLLRFIRQARRLGFLV
DDIRKLLALWQDRSRASAEVKAIALEHVAELDGRIAELTDMRNTLVNLAAHCSGDDRPDC
PILERLADPCCDPK*
>GCA_000452945.1_BPMSHR5858_genomic_04691_1 group_1535
LCVIATTAVAGSDSARAPLILDTQSGIHDGKSGIVLQNAPLSRAPIVAPAQPAPMTELPS
GSTMGSTAPMVVIPYIEVPGSAVPLKAPPQRQ*
>GCA_000452945.1_BPMSHR5858_genomic_04693_1 gpr_2
MNYRRLGRSGLQVSELSIGSWVTYGNQVDRRAARESLAAARDAGVNFFDNAEVYAGGKSE
EIMGHALKSLGWPRISYVVSTKFFWGLAEAPNQYHTLNRKYLLNAIDASLQRLQLDYVDL
VFCHRPDPNTPIEETVWAMSDIIARGKALYWGTSEWSADEIRAAYEIADRHHLRKPTMEQ
PQYNLFHRKRVEQEYRRLYEDFGMGLTTWSPLASGLLTGKYRHGVPAGSRAEIHGYDWLR
AQLTDPAKNRVVEALGEIANDLGCTVGQLALAWVLKNPRVSTVITGASRVEQIIENMRSL
DVTTQITPDMKQQIEQIVGDAYQ*
>GCA_000452945.1_BPMSHR5858_genomic_04697_1 zraR_2
VPDFRPPSDCAPPGHAAQRSASPADAAARQLVYLSRTPDTTLVAHLRARRWNVHVARSAN
EAARRVKPNQPQAGIADLDGFAPRELPTLEAVLRQQQVGWIALAGDTRINDPDVRRLIRQ
YCFDYMQGLPPHETIDYLVGHAYGMVALCDLDVTAGAAATGDEMVGACDAMQQLFRTIRK
VAATDATVFISGESGTGKELTALAIHERSERRKAPFVAINCGAIPNHLLQSELFGYERGA
FTGASQRKVGRVEAADGGTLFLDEIGDMPLESQASMLRFLQEGKIERLGGHESIPVDVRI
ISATHVDLDAAMREGRFRDDLYHRLCVLKLDEPPLRARGKDIEILAHHILHQFRSDGARR
IHGFTSCAIEAMYNYHWPGNVRELINRIRRAIVMSDSRQLSAADLDLAPFAARQATTLAE
ARERAERRTIEASLLRHRNRLTEAAAELGVSRATLYRLMVSHGLRELSWGAQRAGASDVD
DEAGPA*
>GCA_000452945.1_BPMSHR5858_genomic_04702_1 thyA_2
MKQYLDLVRTILDTGTWQSNRTGIRTIGIPGAMLRFDLQQGFPAVTTKKLAFKSAIGELV
GFLRATRSAADFRALGCKVWDANANENAQWLANPYRRGVDDLGDVYGVQWRRWPGYKVLD
AHADAQIADATSRGFRAVARFDEDGAQKVLLHKAIDQLRDCLDTIVRDPSSRRILFHGWN
PAVLDEIALPACHLLYQFLPNVERREISLCLYIRSNDVGLGTPFNLAEGAALLTLVGRLT
GYTPRWFTYFIGDAHIYENQLGMLKQQLEREPFDSPRLEIAERVPEYAKTGQYEPQWLER
IEPSDFTLVGYRHHEPLSAPMAV*
>GCA_000452945.1_BPMSHR5858_genomic_04710_1 group_7620
MSKIRAMLIGAAVAAMATSAFAQTGGVATQGSAGGSADVLGQKAGASAQLNAGGDVSAAS
PEAAENAVKHSAKKHVKHAKAQAKAKADEAAELGADAKTKASGAIEGAAGTAGNAVGGVT
NAAGNVVGGATNAVGGVAGAAGGLAKGVAGGANGSVSVNGGMKAGAGE*
>GCA_000452945.1_BPMSHR5858_genomic_04711_1 group_5073
MGQITLTLKDDTVVSLRKDYDAFVRVSLKLDPQFVTPSFEDFLRAKLLDSMVPLTEHAVQ
RLLQGGQYAWAKRTLDKEFPDVVSILIRQAGDFGFGFASRSEWTPDELAKACRDWAAAVV
SGAQGDAALVDPLAAQIKSAVTDIQTLEERMQTPAWRLAESLRQRVYEAKLACEMSVGSV
AREKLGELRGLLRLGIAHGSFQKQEAQQIMEYLRLLKPEIFIEEPYDIFTRIAAWLRSMF
MDATPARGAQQNASQPPGQRRS*
>GCA_000452945.1_BPMSHR5858_genomic_04713_1 group_19445
MKNRYFAALATCAALLPIASTARGEEVASVNTNFRLTGSDRVVVEAYDDPLVSGVTCYVS
RARTGGIKGTLGIAEDPTEASIACRQVAPIRFTEPLRQQTDVFSERLSFIFKTLHVVRVV
DKKRNTLVYLTYSDRIATGSPKNSVTAVPVPAGTPIPLR*
>GCA_000452945.1_BPMSHR5858_genomic_04726_1 group_3569
MSRRIIPVTPVSAAPDLSGAPLTPTGTLRDALARPLRDLRISVTDRCNFRCVYCMPRTVF
GKDYPFLPHSALLTLEEIERLARLFVAHGVEKIRITGGEPLLRKNLEFLIERLANMTTVD
GRPLDITLTTNGSLLMRKAKSLRDAGLSRVTVSLDALDDTLFKRMNDADFASADVLDGIF
AAHAAGLAPVKVNMVVKRGTNDAEIVPMARRFKGTGVVLRFIEYMDVGMSNGWNMAQVLP
SADVIARIAEHFPLVPLERHTPSETAQRWGYADGSGEIGVISSVTRAFCGDCTRARLSTE
GRLYLCLFASSGHDLRSLVRGGATDEQIAALIGHIWQARTDRYSQLRGSAQATDEAAGEA
KRVEMSYIGG*
>GCA_000452945.1_BPMSHR5858_genomic_04750_1 rnc
MPLSQLESRLRYEFRNAELLRQALTHRSHSATHNERLEFLGDSVLNCAVAALLFQRFGKL
DEGDLSRVRANLVKQQSLYEIAQALNISDGLRLGEGELRSGGFRRPSILADAFEAIIGAV
FLDGGFEAAQGVIKRLYVPILDHIDPRTLGKDAKTLLQEYLQGHKIALPTYTVVATHGAA
HNQQFEVECTVPKLDIKVSGSGASRRAAEQAAAKKALDEVTAVAPMLAAKPKRSKSARGA
KHVEPEIVPGVKGVQEALDLRSPERKERGANRVAGGEAKGVESKAVDAKHGESRAAQLRA
DDAKAGETKAGEARASADAKAGAHTHAAAMPAVPEQADDAARSPATTPVAVIRAAHVEHG
LDKGEPRASKPAEKTAAATDKPPEKASDKPSPEKTSEKTPDKSHEKQLDKSSEPVAEKAL
DKTADKPDAAARLPAETADRPPRARDASSSAEPDLAAAPVRVADAGH*
>GCA_000452945.1_BPMSHR5858_genomic_04772_1 otsA_2
MSRLIIVSNRVAPISEGEPAAGGLAIGVYDALKETGGMWFGWSGEVIASGVPQIRIEEHG
PVTFATIGLSRRDYDQYYRGFSNATLWPAFHYRADLIQFDRHEFEGYSRVNVWLAQQLVP
LLRDDDVIWVHDYHLIPFARALRTAGVKNRIGFFLHIPFPAAQVLVNVPPHRELVEALCS
FDLLGFQTDTDLRAFFEYVEYEAGGTVDRAAHPARVDAFGRTLRAAAYPIGVYPDEIAAL
AKAGENGRAVKSLAASLRGRQLVMSVDRLDYSKGLVERFRAFEKLLEHETSFRNHVSFLQ
IAPSTRADLRAYQDIRRQLEGESGRINGRFAELDWAPILYIHRQYERPVLAALYRLARVG
YVTPLRDGMNLVAKEYVSAQDPDDPGVLVLSRFAGAACELTGALIVNPIDIGGMADALSR
ALSMPLAERRARHADMLARLRENNVSVWRDHFLRDLGR*
>GCA_000452945.1_BPMSHR5858_genomic_04778_1 braC_6
MRLRSSLPSLALAAAFACGATGAARAADETPVRIGFAAPLTGVNAGYGKDLQNGVQLALD
DARAQKITIAGKPARLELVVEDDQADPRIGVQAAQSLVDKKVSVVVGHFNSGTTIPASVV
YDKAGIPVIDPAATNPVITSRGLANVFMVIATDGQNAGNAGRYAVDVTKAKRIAIVDDRT
AFGQGEADEFDKAVKAAGAAIVAREYTSNQAVDFRAQITSLKSKNVDLVFFGGLDSLAAN
FIKQMKQLGLNAQFVGGGGVKDAEFIKIAGPAAEGAMAWEYGRPLDQLPQGKDFESRFKK
RFGVDVLSYAQFGYDATWAAIKAMQAAGTTDPSGYRPALKKIDFEGITGRISFGGDGSLK
SGMSTLYQVKSGAWQTIVTKGG*
>GCA_000452945.1_BPMSHR5858_genomic_04792_1 group_265
MSIPTDVLQALFGGWSQHDRFLWITTPLGANALVAESLHGWEALDDGGFRLQLTALAEHP
SLPLAQLIGAPILIEWQAQEGRDARRPFHGHVIAAELVGYNGGLARVRLVVEPWLTLLRQ
RVDSYNYLDASVVEISEQVFRRYARGVIAPAWRWALADAAKYPKRSLTAQAGESDFDFLE
RLWAEEGIFYWFEHEGDARASSLGKHTLVLADSNQRFAPDEPEIVGFHQTSDDDPQGCIQ
HFMHARRWRIGSVARASWDHRSLSTRPTGARANGAVAPGEDRDVAGPYAYQTGAIGDRRA
QQQLDAQRVAALRSEGRSTRRDLRPGLRFAIAHHPTLGVSDAFICLRVEHSARANVDATV
RSAIEQRLGAIASIAGAAPVPHGPASALNAALGADTHHGGSLMRDDAVYRNRFVALPAEQ
TYRPLAASGHGARMHPVAVMPGAQTAIVVGAGDPVHTDRDHRIRIQHHAQRGENAASRDD
HPHAANAPANRGAGTWTRMMTPVGGDNWGGVSVPRVGQEVWTEWLEGQPDRPVAVAALYN
GRGNAHAQHNAQAGGPSGSTGNAAAWFAGNAHAAVLTGFKTQDMSMSQQGTGGYRQFMLD
DTAGQSSARLYTTDRNSGLTLGHVKQTQDNRRQADRGYGAELATDAAGALRGGAGLLIST
APGVSQMDASAPSQVLAQHRQTLQSLAALAHKQGAEPGGAVPEAAGGAGAASASTGAAAD
KPLPAVDGIEQSREAIGATREGSGGDTAGGGGGGAIAWSKPHLVAHGEAGLAAMSAKSHV
WVSGTETVLSAGQDVQLTAKGKTSVVANHGISLYTQGAAGDGRPVAGRGIALHAASGAVS
VQAQNAGKLSASAQKAVTLASAQGSASVQAQQRVLLSAAKAYLKMEGNDIVVGAPGRADF
KAAAHQLTGPKSAGASAQMPKSEPKLCEYKTRAADVAHEGTMKSAA*
>GCA_000452945.1_BPMSHR5858_genomic_04795_1 group_3582
MHRIFRNSVIGVLWCGCVAQAGAMSRGPVPLDVKRIDGKPAACIPTSDEGQAEMRLGFIG
VSRATGPVSPDVIYWQVEVSDNVPPVYLKRGDCIVYGQSIPGATVLTPPKRFDVGKWYNF
AVMPGGKAEGPIYRGTFCISGKQGDSGRVVLRSENKDACSSGR*
>GCA_000452945.1_BPMSHR5858_genomic_04807_1 group_3778
MRTIRMASIDARLFIFGLLLFLSGLAHADTPSAAQPSPYGIVSVDIDGTRLCMEVGGAAV
APLAGVDVGECRQSADQQWSLVPNGQNFQVVAKHSLQCLTIANAATSPGAPLVQYPCNGG
SNQQWSVSPSRSGYKLVSAHDALCASAAGASPGSRMVQQRCDEHASQTLYVSAPPELVGA
LVGMNGGLCIDGGAAGAQAVQRTCDDAPGQQWRIRPNGSAYKIHLASTNLCLGTRDASRG
TGAAIESQRCANVASQLWTIRAASEVDGRNGYANGYWQFVSANSGQCIVVQNASTADGAN
LIQYPCGQGNAGSNTMWRVNRTSRSSWTGAIALPLVPSAAAHLPDGKILMWSADSTISFG
GGGVDGNTYTTVFDPIAQKATDAVLTSLGHDMFCPGLNLLADGKIFVNGGVSSRKTSVYD
PATHAWSPSNLMNIARGYQSSVTLSDGAVFTLGGSWNVATPDFRLGDKYGEVWRPDSGWS
ILSNVPDIIGPDPAGAYRGDNHMWLVAARDRWVFYAGPDATLRWIDTTGNGRIVEAGKRG
DDAYSISGNAVMYDVGKVLTVGGAPAYDNGVASASAYVIDISAGPQVPPVVRKVQPLAYS
RGFVNSVVLPNGQVVAIGGQAVTIPFSDDQSVLVPELWDPSTEAFTRLAPMTVPRNYHSE
ALLLPDGRVMASGGGLCGSGCNTNHPNVQILTPPYLLNADGTAASRPVIAAAPEQAANGS
TIAVSTDAPIRSFALVRMSSSTHSVNTDQRRIPLTFRQSSGGDGGYAYTVAIPADAGVAI
PGQYMLFALNAGGVPSVAKTIRIGA*
>GCA_000452945.1_BPMSHR5858_genomic_04812_1 group_5089
MTAELDARVRALTLRAQQLFDDGRPEQGALLAAQALALAPDDALALKLVGVAECMRGDHA
AGLVYLERACLSAPGDANLHYNVAVAHECTGSHERAALSYRHCLRLQPDHADALWNYGEY
LRLNGHFEAAARCFEALQAQACRYPSMHHRMAVVYTHLHRFDDARRHFALAMDENIDTRV
TRWERAHLRLGTRDFARGWPDYDTRFDIGYLINVHCHPFPIRLWQGEPLAGKTLLVHGEQ
GLGDEIMFASIVPDIVRQAARVVLACAPSLVSLFQRAFPSAIVRAHRAGVAPARVDDLGA
IDYQSPIGSLPRWLRASEASFGTGAPYLAADPARVAWFGARLRALAPRADRALKVGLTWG
SNPAAAVPSAARRATRKSMPLRLLAPLARVPDVQYVSVQNAELGEQAATVPELDLIDFSS
ALRDFADTAALVANLDVVVSVDTSVAHLAGALGKTTYTLLMRNCDWRYGFEGERCVWYES
MTLLRQTTQDDWLPVVDRVIDALARYRKQ*
>GCA_000452945.1_BPMSHR5858_genomic_04814_1 ehaG
MNKTYRVSWSASRGAWMVAPETARRKGKGHSLTIVCAIASGLLLAAPAWADTVSPSGTDN
VYGVDATDPGVSTNQGNTAYGAQAGAKVTGSYNTAIGYQAGQNVNAIDTVSIGKQATASA
NDAIAIGTNTKASGPADIYMGLNAGAGAGSTTSPDGTVTLGIRNMGLGESAGSYVTGQNN
TGIGYQSGMNVTGDQNVGLGQQAGQFVTGTGNSAMGHLAGSTVSGSYNAAFGEYAGTNTS
GGANAAFGFYAGRYINGTNNTALGAYDLPVVNGTWYGSYVTGSNNLGAGHNSGAYVSGAS
NVGLGDGAGTFVTGSNNVAIGTAAGSGAYTSGPSGATLNAALVASNTVSIGTRATASQSD
AIAIGKGATASGAQSISIGTGNVVSGKGSGAIGDPSTVSGAGSYSIGNNNTVANSNTFVL
GNGVTTTQDNSVVLGNQSTDRAAVAVSSETINGTTYNYAGVASPANGVVSIGGVGTERQL
INVAAGQVSATSTDAINGSQLYATNQAVIAEDAKVNSLGGGVASALGGNAAYNATTGAIT
APSYAVYGTTQNSVGGAIDALQALAPLQYTSGPGVTTPNAPGSAPTNTVTLVGAGGPGAN
TTPVTLTNVAPGKLSATSTDAVNGSQLYATNQQVANLVSSVNNGGVGPVQYSDPSAPTTP
NGGKPSQDLTLVGAASGPVALHNVAPGTASTDAVNVGQLGAVTTGLGGGAAIDPKTGAVT
APSYTVYNADGTTSNVGNVGAAIDAINSTGIKYFHANSTKPDSQALGADSVAIGPNAVAN
NAGDVALGSGAVTSQAGGTLSETINGVTYSFAGTTPIGTVSVGAPGVERTITNVAAGRIG
QSSTDAINGSQLYGTNQSIEALTDKMNSLGNTVANTLGSGASYNPQTGAVNGPANSGGVV
TPTVIQEAANKWVSANPSTYVAPVATGTNGMAVGSGAVSTGQNSVALGTNASDGGRSNVV
SVGAPGAERQVTNVAAGTQATDAVNLGQMNGALAQQTDSFNQRLGAVQQDVDNVARAAYG
GIAAATALTMIPEVDKDKTIAVGIGGGTYRGYQAVALGATARITENIKVRAGVGMSSGGT
TAGIGASMQW*
>GCA_000452945.1_BPMSHR5858_genomic_04815_1 phoP
VVLIYLIEDDEVQARCYAAILQHAGYSVRVLPDGERALREIQRAAPDLIVLDRRLPDIDG
LEIIAWVRERCAPLPILVLTNAVLETDLVEALEAGADDYLIKPPREREFVARVNALRRRA
SISKQFEGTIEIGGYRIATVERAVYLAGERIPLSPKEYEIVELLARHVGQVVPRKTIISR
VWGRAAEEEVSRSLDTHIYRIRRKMNLSRRNGAMLRVIYTHGFRLDDFSHV*
>GCA_000452945.1_BPMSHR5858_genomic_04817_1 group_3586
MKLSVLFMAALPLIAIAASAHAQPRHPAVYSPAAGVLCDRYVCADDQGISRALTERYLGK
RVAAKAFSQGDFDPTEFTFANGVFCDVKERLCRDDRYYGADGKRSGAVSRRYTELLFGRR
SGG*
>GCA_000452945.1_BPMSHR5858_genomic_04818_1 group_5090
MKLVFPACLAFVAGIVCFARFAASGSHVDAFGRLHEPFGLLPLGWALTLLGGTGLLIGAA
QAVVRRSARRR*
>GCA_000452945.1_BPMSHR5858_genomic_04820_1 gltC_2
MAAEWMGCDRFRANSMKVSLRHLRAFVAVAREGSFTAAAGALCLTQSTLTKTIREFEQAL
GVHVFERNTRRIALTPYGAAFLVDAVRILNDLDLALDGVNQQLLGNSGTVRVASGLAFAS
TVMPCVIRALHAQEPGIHVDLLDDTSGGVIRRIESGEVDVGIGSYVGMAGNALSARRLLS
ARLGVLFPVGYPKIPSRPTLASLNEIPILRDSDESSIAAAFRQQAPDLWERMRRKVSVTN
LDLQLALVREGIGACIVSALAASHPSARGMPYRLLVEPEMTRDVYVFTRRDVSLLPAAAR
FVDILMNVLPSIDFAEGVELACDGA*
>GCA_000452945.1_BPMSHR5858_genomic_04823_1 group_5091
MDRIFMTRTEALEFLLKAHQTAVDKVGHPFRKPTPADHAAVEALDRLLLDVRARRIDTFQ
INASSAQVIVTD*
>GCA_000452945.1_BPMSHR5858_genomic_04825_1 glgE
METRPSFAPNLYFCDARLVGPLHAWPPLFERVAAWGFDHVLIGGLWAASRRGYPRHVADP
DRPAESFATSLDATSALARLSDSAREHGLRVAVEVVVDRVAREHPLHDAHRDWYVVDERD
DALIDPRTAALAHDVAHANVGSAAALDALADWWRARLGALADAGAAAFLVDAPQRMPAHW
WAALLRALRQARADLPVIAGVPGREREALAQLACAGFDAAFSSLRWWDLRAPWFVEEHRL
LRRVGAPIAFPDAFDGPRLAHDWRQAAPETIERAHRRALWTAAALGTGWLVPMGFERGVA
VELMAREPRADAYRAALDSAPFDLSGAIAEANALRRATPALRGNGEIAQLTGADAPATVL
LRGARTALEYDDEAALIAVNPDLAHPAAIAPCAALAGVPGGFTRFAPFADGRRPRMGALE
PFALAAGACTLLRAQRARPVTTAPAEDRRGNRPGTRASVTAALAGERIAIERIEPVVDDG
RFAVKRVIGERLAVRAAIFADGHARLAAAVQWRAADENGWHEARCAAEGNDAWRADIPLE
RLGRHLFRVIAWRDDWATLVDEIGKKHAAGQAVALELEEARRLAADVLTRAPEANPAALA
VLREFAAALDAAPPDQRLALIGAPHVADAFAALRERAFATRDAPVFPVDVERRAARFASW
YEMFPRSASDDVRRHGTFDDVVAHLPRIRDMGFDVLYFPPIHPIGTTARKGRNNSLQAAP
DDVGSPYAIGSPAGGHTAVHPQLGSLDAFRRLVAAARAHDLEIALDFAVQCSPDHPWLTE
HPGWFAWRPDGSLRYAENPPKRYQDIVNPDFYARDAMPALWIALRDVVLFWIDAGVRIFR
VDNPHTKPLPFWAWMIADVRARHPDTVFLSEAFTRPSMMYRLAKLGFSQSYTYFTWRESK
REFIDYLTELADGPAREYFRPNFFVNTPDINPRHLQQAPRTQFVIRAALAATLSGLWGMY
SGFELCESDALPDSEEYRDAEKYELRARDWRRPGHIGDEIARLNRARRDNPALQTHLGIR
FAHAPNDAVLVFSKATPAHDNVVVVAISLDPWHPQATDFTLDAALYRGWGIADGERLVAV
DQTADHVETWHGRRHYVALDPHVRPFAIWRVAPAAGVARGARDDARDVPAQEVHER*
>GCA_000452945.1_BPMSHR5858_genomic_04827_1 glgB_2
MTDAPFDRADIDALLDARHPDPFACLGPHRVGDATVVRTLLPGALRVRAIAAGGGVLGEL
RQVDPAGCFAGALPDGRERGERPRYRLSIDWPDARQDVEDAYAFGTLLDEDALARFAAGD
PRAALACLGARALDMDGVPGVRFAVWAPGASRVSVVGDFNGWDARRHPMRLRRPWGVWEL
FVPRIGAGERYKFALRARDGAALPLKADPCACRTEAPPRTASIVADLDALERFGWHDDAW
LRARASLDLAHAPVSIYEVHPESWLRVAAEGNRSATWDELAQRLIPYAAGMGFSHVELTP
IAEYPFGGSWGYQSLSPFAPSARFGPPEGFARFVEHAHAAGLGVIVDWVPAHFPDDPHGL
GKFDGTALFEHADPREGWHPDWHTHVFNVGRREVGAFLIASALAWAHRYHVDGIRVDAVA
SMLYRDYSRAAGEWVPNVYGGRENLESIAFLKHFNDTLHGPAAPPGVATFAEESTAWPGV
TAPTAEHGLGFDFKWNMGWMHDTLAYLREDPIHRRHHHDRLTFGLVYAFSERFVLPLSHD
EVVHGKGSLAAKMPGDAWQRLANLRAYFGFMWAHPGKKLLFMGGEFAQWGEFAHDATPQW
DLLDAPAHRGVQRLVRDLNRLHAAEPALHALDDRPAGFAWLVGDDRNNSVFAFVRRDDAG
RMLVAVCNFTPVPRTDYRLGLPAPGRWAEVLNTDGAAYGGTDAGNGGALQADEIPAHGER
WSAALRLPPLATLWLRPA*
>GCA_000452945.1_BPMSHR5858_genomic_04882_1 anr_1
MLQAQTQNAYTPAQQRVTVPLRAAKRDDAAKRTAARCSSCAMRSVCMPVDLNTHEFAKLD
ALICSTRQVKRGDTLFRTNDGFQSIYAVRTGSFKTVVMHRDGQEHVTGFQIVGETLGLDG
VCAGHHNSDAVALEDSTVCIIPFAQLEAVCREVKPMQHHVHRLMSGEIVRESSQMMLLGT

MTAEQRVAAFLLNISERFQKRGYSAAEFNLRMTREEIGCYLGMKLETVSRMLSKFQRDKL
IAPRGKQIRIIDPVGLARV*
>GCA_000452945.1_BPMSHR5858_genomic_04927_1 group_763
MPVTNAIESLHMQLRKIIKARGHFPSDEAALKLIWLALRNVVAKWTGSRHDWKSAMMQFA
LLYPERFNMGV*
>GCA_000452945.1_BPMSHR5858_genomic_04945_1 group_5109
MKLLRYGAKSREKPGLLDAQGRIRDLSGVIDDVAGDALGPDALARLRAIDPASLPLVDGA
PRLGACVGRVGKFVCIGLNYSDHAAESGMDVPSEPVVFGKWTSAICGPDDDVELPPGSTK
TDWEVELGVVIGTGGRDIDEARALAHVAGYCIVNDVSERAYQLERGGTWDKGKGCDTFGP
LGPWLVTADEVPDPHRLKLWLDVDGRRYQHGSTATMIFRVPFLISYLSRFMSLQPGDVIS
TGTPPGVGLGQKPPVYLRAGQVMTVGIEGLGEQRQRVVQR*
>GCA_000452945.1_BPMSHR5858_genomic_04946_1 tsaC1_1
VNLNLNDKVVIVTGGGSGIGGAITLALAEEGAIPVVFGRSPLADAFAERLRALQSRARFV
RVDLVDGARCREAVDETVAAFGRIDGLVNNAGINDGVGLDAGRGAFVASLESNLIHYYVI
AHYCLPHLKASRGAIVNVSSKTAITGQGGTSGYCASKGAQLSLTREWAASLADDGVRVNA
VIPAEVMTPLYANWLARFDDPDAKLQSITRKIPLGKRMTSAEEIAWTTVFLLSERASHTT
GQWLFVDGGYTHLDRALT*
>GCA_000452945.1_BPMSHR5858_genomic_04969_1 hcnB
MTNEILTVDVAIVGAGPAGLAAARAAARSGCTVAIVDDNPRAGGQVWRHGPHAAPGALAR
ASFGVLRQPNVRHLAATRVVAAPRPGALLVEDEQRALVLGYRKLILCCGARELLLPFAGW
TLPGVTGAGGLQALVKGGAPVRGERVVIAGSGPLLIASLATARAAGARVVALVEQAPPGA
LARFALSLAATPSKLAQAARLTRGFVGTRYLTGAVVREAHGDARVRAVTIEQGGARETLD
CERLACGFGLVPNLTLALALGCAVRERAIAVDDAQRTSVEHVYAAGECTGVGGVELARIE
GELAGLAAGGAHAAPHGRARMAALLRRRARWRRFAARVARTFALRDAARALPPDDTLLCR
CEDVTIGAVRAHASARDAKLLTRCAMGACQGRVCGAAAHAYFGWDDAPPRPPFSPARIDT
LLAASDDALL*
>GCA_000452945.1_BPMSHR5858_genomic_04973_1 yjhH
VSRHAIQWSGVFPAVSTQFKPDFSLDVDATHRVVKNLVNDGVSGLVVCGTVGENTSLGTA
EKLAVIEAARDAAGGKVPVIAGVAEFTTEFARNTVREAQRVGVDGVMVMPALVYSAKPHE
TAAHFRAVASSIDLPVMVYNNPPIYRNDVTPDVLIALQDCENIVCFKDSSGDTRRFVDLR
NAVGDRFVLFAGLDDVVVESIAVGAQGWVSGMSNAFPKEGETLFRLAKQKRFDEALALYR
WFMPLLHLDARPDLVQCIKLCEELVGRGSALTRPPRLALQGDTLAHVKAIVAKALETRPT
LPDVGL*
>GCA_000452945.1_BPMSHR5858_genomic_04977_1 proP_10
MPASELSAVSSAPPRALTGRDYKTLGLAALGGALEFYDFIIFVFFAPAIGQLFFPHDIPD
WLRQLQTFGIFAAGYLARPLGGIVMAHFGDLVGRKRMFTLSVLLMSVPTLLMGLLPTYDS
VGILAPVVLLLLRVLQGAAVGGEVPGAWVFVSEHVPPQRIGYACGTLTAGLTIGILLGSL
VATAVNSRFSTAEVAAFAWRIPFLLGGVFGLFSVYLRRWLHETPVFAEMKARKTLAAEIP
LKAVIRDHGRAVIVSMLITWMLSAAIVVVILMTPTLLQKQFHIAPATALFANSIATLCLT
AGCITAGSLADRFGAKAVLSIGGVALAACYYAMYTQIAVDAARLVPLYGLAGFTVGTIGA
VPFVLVKSFPAVVRFSGISFSYNVAYAVFGGLTPVIVSLLMKSHSLAPAHYVAAICVLGA
VAMLFSKDAE*
>GCA_000452945.1_BPMSHR5858_genomic_04978_1 mdtD_2
MDTHLPHPALSRAALVRIVSTVSAGFVITQLDVTIVNVALARIGIDLHTSVAGLQWIVDA
YTLALAGLMLSAGALGDRFGARRLFAAGLALFAVASFVCGIAANAATLIAARALQGLAAA
AMLPNSLALLNHACAHDPRLRARAVGWWTASGAISIAAGPVIGGVLIAQFGWRSIFFVNL
PLCAAGLAATRLWIDSDDTRAAAASSASSASSASSAADGAHHSHPAADRRASGPNACAPA
PKRAAGAGTRGLDLPGQCLAVVALTLFTGAVIDWHPALVAIALAAGAAFVFVESRSAHPM
MPLALFRRRTFSVAVLFGVCMNLSYYGIIFVLSLYLQRVRHNTPLEAGLAFLPLTGGFLL
SNVASGWATAHYGPRRPMIAGALIGATGFALLSAVRADTPIAMLVVPFLLIPGGMGLAVP
AMTTTVLASVERARAATASAVLNTARQAGGAIGVAAFGALASGARPPDIVSGLHASAYVS
AALFMFASAMAAFVHGAPQPAVGKPANANAR*
>GCA_000452945.1_BPMSHR5858_genomic_04979_1 yphA_2
MTRSVDSGVIFFARLLLAVLFLWGGAMKLMGYGEFVGYLKGLGVPFTQVAAPVIVALEAI
GGVLLIVGYQVKPLALVLALYTLAAALIGHNFWDATSSALQRDMVVHFWKNVAIAGGFLL
LYVTGAGGASIDGARRPSSSYGSLR*
>GCA_000452945.1_BPMSHR5858_genomic_04980_1 glcA
VQVWNQVYLPLGGVGWSALAAGAPIILFFVSLAVLRLKGHVAGALTLALALVIAIAVYGM
PAERAFASAAYGFAYGLWPIAWIIVTAVFLYKIVVKSGQFDVIRSSIVALTDDQRLQMLL
IGFSFGAFLEGAAGFGAPVAITAALLVGLGFNPLYAAGLCLIADTAPVAFGALGIPVIVA
GQVSGLDPMAVGAMAGRQLPFLSFFLPFWLVFVMDGVKGVRETWPAALVAGGSFALVQFF
TSNYIGPELPDVTSALASLVALASFLKVWRPLRAREAARAQRLVSAGGGALALGGMPGGL
GGARATGGREPSPYSFAQIARAWSPFVVLTVMVTIWSMKPFKALFAKGGALAFTTLQFHV
AHLDKLTQKMAPVVAHPTPVPAVFNWDLLAATGTAILLSAIVSMAILNVPARTGVRTFFE
VLKDLKRPVLSIGLVLAFAFVENYSGMSTTLALLLAGTGAAFPFFSPLLGWIGVFLTGSD
TSSNALFCSLQATTAQQIGVPSTLLVAANTTGGVAGKMISPQSIAVACAAVGLVGHEADL
FRFTLRHSLFFALVVGVMTCVQAYWLTGMIVH*
>GCA_000452945.1_BPMSHR5858_genomic_04982_1 lutB_1
MSATPLNAPATFKARASAALHDAQLQEALGMLGEGWIEKRRRVVSLLPEFEMLRERAKAV
KDHVLANLDTYLERYEAAVTRAGGVVHWANDAADAREIVLRICRDANAKTITRGKSMVGE
EIGLPAALEGAGMEVVETDLGEYIVQLAHEGPSHIVFPALHKTIGQVTELFAGNHSSAQY
DGPRETVADVVGEARQVLREKFFVADVGITGANYLVAETGSNVICTNEGNGDLTSTLARV
HIVTAGIERVVPTLDDVALFVRLLGRSATGQEITSYTTFSTGPRRGDDLDGPQEYHVVLV
DNGRTRMLAGEFRDMLRCIRCGACMNHCPVYGAVGGHAYGWVYPGPMGAVLTPLMQGIEH
DTDLPNACTLNGRCAQVCPVKIPLPTLLKKLRGMQFDARLTAPGTRFLLRAFGAIARRPA
LYHWLTRVGARALRALGGKRGSIARVPFASGWTDSRELPKPPPRTFFETWDATRTNQPEV
RR*
>GCA_000452945.1_BPMSHR5858_genomic_04985_1 group_3597
MSPDHPWNSLEIVKLAAEVLTPVSVAAFGWLVSHRLKQLELVQWTNQKLIEKRLALYDTI
APLLNRLLCFYTWVGPWKDISPDDVIRAKRELDQTIHIYRHLFDDDVYRAYQAYLDALFD
THSGAGRDARIKSTIASPDGDRIGNGTYRWNPAWAERFTDANVASRDEVRARYQRIMDGL
RISLGVAR*
>GCA_000452945.1_BPMSHR5858_genomic_04986_1 group_7639
MKARPMLEYALHPLEDRLVHVDELCRADPVPRGWARCPLCLEALYVVQLRDRSHARRFAH
LAGEFARCPLVNDALPNPLAVHVGPPLDEHGRQARATFFRHWQRHLHTIRQTASAFGIAR
FMRAIELADVLKLWAWPTLAQRDIPYILLVLTEFIAAPRGERKQAAWSRFWFDASVQRVD
DLRKPRGVLPRLFRLRYRLPRMSKFPNARHLIDCQPVLMNDIGPSDAALGTPRADIAMFE
AFAARFARRPAE*
>GCA_000452945.1_BPMSHR5858_genomic_05028_1 group_163
MKARSPAVLAAPAHCADTPRTPDMPDMPDMPDTRGTPAHDASPRPEHKPHGKMLHPVVMM
IWVLAAAVALTWLVDSGRYARNGKLVVPGTYQVVPKATSLATLVAPAVSKSTPQLAMPAS
LVSAFVAVPGGLVKNAPLIVMVMFVGGMFGVMRRTGVVDAGIDRLLQLTGNDVYLLTPLL
MILIGLGSTLLGFISEYLVIIPMVMVIAKRLGLSNLFAVALVALAAKIGYIASVTNPLAL
AVAQPLVGVPLFSGVALRAAVFAVYLALGILYLLHHVKRSGYRRERAKALAHAHGVARLS
VRHQATLALLAAAVAMLVFGTRELKWGNVELAAFYAFVAIAAAAIGGLDSRSAADAFVDG
MKSMMLAALLMGLAASVELLLQNSLVLDTLIHLFTRLANGQSPVWVANGLMAVQMVLDVF
IPSVSGKAAVSMPIIGPIAQLSGVSGQTSVLAFVLGGGLTNLVTPTSGMLLAYLATARVD
FGAWLRFVLPLFAVLLALSCGVLAFAVQIGY*
>GCA_000452945.1_BPMSHR5858_genomic_05040_1 group_5128
MPFTRIALREGKSAEYRWALSEGVHCALQHAFAVPVDDIFMTVTEHSADNFFYGRDYLGI
ARSDDLVMIQITANNTRTLEQKRELYRLIADQLAQRPGVRREDVFVSLVEVLKEDWSFGN
GIAQYVS*
>GCA_000452945.1_BPMSHR5858_genomic_05042_1 mopA
MTADSLSSRSDARAARESLALSGELWLHAGGQTLGGAARIALLAAIGETGSITRAAKAVG
LSYKGAWDAIDTMNNLAGEPLVLRATGGKGGGGTTLTPRATALIAAFRAIEREHRRFIDA
ASAAVEGFEVNWKLIGRIGMKTSARNQLFGKVLAVKHGAVNDEVVLALPGEHTITAVVTH
ESVQELGLAPGVDACALVKASWVVLAVEDGSPLRLSARNQLQGVVETVTRGAVNSEVLLA
LDGGMTLAAIVTNDSVDALGLAQGVSAVAAFKASSVILAVNG*
>GCA_000452945.1_BPMSHR5858_genomic_05046_1 modA
MSAPVRSARRLFLRLAMLSALVAGFAAPAAHAADELVVSAAASLTNAFKAVGDAYEKQHP
GTKILFNFGASDVLMQQIAKGAPADVFASADQKAMDRAAAERVIADGTRRDFAANSLVLI
VPADSRATFGSLRDLTAPAVKRVAFGDPASVPVGRYTEGALKAAGVWNDVGAKGVLAANV
RQSLDYVARGEVDAGFVFGTDAAVMPERVKVALTVPTQTPITYPIAVVKDSRHAAAAQGF
VAFVLSPAGQAVLAKYGFKPAAGGAN*
>GCA_000452945.1_BPMSHR5858_genomic_05047_1 group_4201
MFDLPPRLWISITALGGAGLTLPLAVTIAVWLALGYSLRRAAAWLAVLGTAIGAVALTKI
AFLGWGIGVRAWDFTGFSGHAMLSTSVYPVAMFLALARARPAVRVAGVVVGLAAGVAVGL
SRVALDAHSPSEAVTGCVIGALAALSFIAGSWRAMPHRWSAPAVATSLVAVTIALHGITV
PSHRWVTAVALQLSGHERPYVRARWKANPNYRPASRPTQQQTLGAPAAVERA*
>GCA_000452945.1_BPMSHR5858_genomic_05048_1 group_5131
MTKSLTKFWLGGVKRMLHMPNARTSASALKAAQKMTADWPFAETMAAAAAASVASVAASA
DTVPPAARESRVRPRAAAWAGGEWIRADHPLPPAFGRFVKHLEYGLYVPSGAKTLGLPLV
VMLHGCKQDMNQFSQGTRMNLLADRYGFAVLYPEQSLSAHAHGCWHWYEDTVHGGRGEAQ
AVVALVDALVAERGFDASRVYAAGMSAGAGLVSLLALHFPRRFAAVALHSGPAFGDAHSG
ITAMDVMRRGLYRDPAAVVDALVEPGAHPGMPAFIVHGDDDRVVVPKNADELAVQFLRLN
GLADADGEPAGVERFETRAADVRTLDYRRDGESVVRLCRVHGLAHAWAGGDDSVPFHSAA
GPDASELIWSFFASRERAAVAS*
>GCA_000452945.1_BPMSHR5858_genomic_05049_1 group_3604
MYLLSRLFLFLTKSPEQRAKERADAYLAGASDLYDLEFRMRKLDREAALDRA*
>GCA_000452945.1_BPMSHR5858_genomic_05050_1 ohrB
MNILYKTAATSTGGRDGRATSHDQKLDVKLSAPRELGGAGAEGTNPEQLFAAGYSACFLS
AMKFVAGQNKQTLPADTTVTAEVGIGPNEEGGFALDVELRVALPGLDAAAAKTLVDRAHH
VCPYSNATRNNVAVRLVVA*
>GCA_000452945.1_BPMSHR5858_genomic_05051_1 ohrR
MNDSPTFPLTLDDQLCFALYSTSLAMTKAYKPLLDKLALTYPQYLAMLVLWETDDMAVKD
IAARLSLDPATVTPLLKRLEALGYIERVRDTNDERIVHARLTADGAALKHKARAVPAELF
CAMRQTPDFLIRLREDLTRLRQALTDADGA*
>GCA_000452945.1_BPMSHR5858_genomic_05053_1 group_7645
MQLIEVSAKTGYVWFRQGIWLFRRNPLAFVTLFFTYLLAMMLVSLVPVIGAALPLLLIPG
IAVGFMAACRDTIAGKQVLPTILIDGFRSYGPTVTQRLLALGGLYIVSMAAVFACSALGD
GGTLLKIMFGLGAENLGPEALDSPGVRIAVLIAAALYAPVAMMFWFAPVLTAWHDVPPVK
ALFFSVVSCWRNKGAFTVYGLLWFALALGVSFGLAALMQALGASAYALTVMMPASIVITA
MLYCSFYATYRGCFGVQEPGAQNPPNASGR*
>GCA_000452945.1_BPMSHR5858_genomic_05054_1 thrB
MAVFTAVSDADLALWMRHYDLGDVVAFRGIPSGIENSNFFLTTTRGEYVLTIFENLTAGQ
LPFYVDLMSHLAKHGVPVPAPVARDDGTLFGELHGKPAAIVTKLEGAAQLAPGVEHCVEV
GQMLARMHLAGRDYPRHQPNLRSLPWWRDTVPAIAPFVTGEQRALLEGELAHQAAFFASD
DYAALPEGPCHCDLFRDNALFAHAEPDTGHSVRLGGFFDFYFAGCDKWLFDVAVTVNDWC
VDLPTGALDAARADALLRAYQTVRPFTAGERRHWGDMLRAGAYRFWVSRLYDFHLPRAAQ
MLKPHDPGHFERILRERIAHAGALPETHACN*
>GCA_000452945.1_BPMSHR5858_genomic_05056_1 amn
MKNDPNRRSRTRTPGSPCVEAFDDPIAAVARLSEIYETNTAFLRDAFARYRGNEAFDEHV
RACYPFVRIRTDVNTHIDSRRSYGFVAGPGVFETTVTRPDLFANYYREQLRLLAKNHHVR
IEVGVSAQPIPVHFAFSEGIHLEGDLDRDRLVAMRDVFDTPDLAYLDDRIVNGTYEPAPG
EPHPLALFTAARVDFSLHRLRHYTATLPTHFQNYVLYTNYQFYIDEFVKLGRTMMSASDD
PEVRAYRSEYTAFVEPGDVITYNANLGEDASEGTPPPRLPQMPAYHLKRADGSGITMVNI
GVGPSNAKTITDHIAVLRPHAWVMLGHCAGLRNTQRLGDYVLAHGYVREDHVLDADLPLW
VPIPALAEVQVALERAVAQVTQLEGAELKRVMRTGTVASVDNRNWELRDHREPVQRLSQS
RAIALDMESATIAANGFRFRVPYGTLLCVSDKPLHGELKLPGMADQFYRGQVDQHLQIGV
KAMEILRTNGLDRLHSRKLRSFAEVAFQ*
>GCA_000452945.1_BPMSHR5858_genomic_05058_1 group_3605
MKNFVAKIYIVALGSLIVTSSHSQQIRTHGAVTQITLPEHGSQASSIDFKNAIPIPLPKP
IMAPPTMLDSLMYNAKPSLGVPGYSPGGAGDGKQRPMRLVAPMDLNQIGIEPQEFGSSNQ
PYTTSQVNAYGDNTQYYYPFRPAGKLFFNIGSASYLCSASLIKRGVVVTAAHCVANFGQR
QFYSNWVFVPDYSNGSAPYGVWSAASATVMTSYYDGTDSCAVSGIVCQDDVAVILLAPQG
GSYAGNSAGWYGYGWNGYGYNPSGLALIDQLGYPVALDSGVFMERNESQGYVSSGYSNNT
IIGSLMTGGSSGGPWLVNLGMPPTLSGTTFGNSGAHNTVVGVTSWGFTDLAVKQQGASPF
TSSNIVTLVNTVCSGTPAAC*
>GCA_000452945.1_BPMSHR5858_genomic_05059_1 group_5135
MNKILVSACLAGLPVRYDGSSKTVEDALLHTWRDEGRLVVVCPELAAGFGTPRRPAEIRL
HRGGGDVLAGAASIWDDTGADVTARFIDGAHHALRQAIAHGCRYALLADGSPSCGSGFIY
DGTFSGNRHDGSGVAAALLEQHGIRVFAPSKLKALASLIERDG*
>GCA_000452945.1_BPMSHR5858_genomic_05060_1 group_1241
MKRSLIAFALTGAGIALSGAAHAVNVDVNIGTPAPVVVAPAPAPVIVAPAPAIVVGWHGD
RYWDGHRYWERREWEEHHRHRGFCPPGHAKKGEC*
>GCA_000452945.1_BPMSHR5858_genomic_05061_1 group_7648
MNKRKVIPSLRSLADQERATAKKARASWQMFTIMAEFIEATEYLSEIRPAVSIYGSARLK
PDSPHYKLTVQIARKLSDAGFAVISGGGPGIMEAANKGAHAGKAPSVGLNIELPHEQAGN
HFQDISLRFRHFFTRKVTFVKNSDAVIVMPGGFGTLDELSEVLTLIQTKKSRLVPIVLVG
SAFWSGLLQWFRDQMIPMGLINPDDMNLMKVIDDPDQVLEAVLAFYEDRGEETEESHPPR
PEEDRMFYL*
>GCA_000452945.1_BPMSHR5858_genomic_05063_1 ndh
MHRFIIVGGGAGGLELATRLGDRYGARGDRPARALVTLVDRYPTHIWKPLLHEVAAGSMD
PFTQELEYAAQARWHGFEFHQGELIALDRASRRVTLAPVADNDGAELLPLRELEYDTLVI
AIGSTTHYFGVQGAQQHSIALDTVAEAERFRKRLIAACMRAEHQPAEPAVRTAEGGPEQA
VEPTDAAQVSASPRIQVAIVGGGATGVELSAELRNTAQVLSAYGLHKLDPRHDVGIVLIE
SGPRILPALQERVSTATAELLEKLGVRLLLGERVTEVAPGVVRTASGKSVRADLTVWAAG
IKAPSVLAKLDGLEVNKLGQLAVRRTLQTETDPDVFAFGDCAGCEWPGHERNVPPRAQAA
HQQASFMLRALANRLEGRPLPEFTYRDFGSLVSLGHFSAVGNLMGGLIGGNMLIEGLFAR
FMYMSLYRLHIAALHGYPRMVLDTVAHWLRRSTLPRVKLH*
>GCA_000452945.1_BPMSHR5858_genomic_05064_1 group_7649
MNEPTRQTTGDLSERRGPNRRRRAPRSPARQRGSLAIIAAIAIGVVIAALGAVDLGNLFY
QRRALQSIADLAALAAAQTMDDGCAKPAATAQSAALGNGFDSTASGQSMTVVCGRWDVKD
NVGPSFFAGSASGAAAGSDAQLNAVQVTVTRAVPYYFLGAQRTIAATSTAEATNVGAYSI
GTTLAQLQGGVANALLNGLLGANLNLSVLSYQGLANARIRIKDLMAAANVGTVSALLSTQ
TTVPQLANWMLSALSQTSVANADLQTSIGALQTIVSANIPGGRTFTIGNTANSAGIFSIG
LSNPQAALDATFSPFDALLVAAEIATGQTAFSLANGLNIGGLNANLQVQIIQPPVLGIGE
AGIDPVTKTWRTIARTAQVRLYLNIGLGTANLPLGLLGALLPVQVNLPLSLQIAPGQAWL
QSASCTASPSTCASAIGVQTGLTNLCIGDTPANMSASLPFTCSTPATLVNVANLVTIKSL
VSFPADVPASQTPTLTFYGTTGGYQSTNSNGVGSVLGNALSGLGASLQQTQISLFGISLP
LGPIQTALNSFLGGVLPPLLSGLDAAIVPLLQLLGVQVGESTIHDMSLTCGVSQLVY*
>GCA_000452945.1_BPMSHR5858_genomic_05065_1 group_3607
MLKPEVDSLVPHVPFSRRKFVQAALGGTFAAAVLPVSAQTITTDAAGLDVDTVQIRSGDA
SVPAYRAQPDGKSNLPVIVVIHEVFGVHAHIADICRRFAKLGYLAIAPDLYARQGDPSKH
ASIQELIDQVVSKVPDRQVIEDLDATVRWAGKNGGDLSRLGVTGFCWGGRQTWLFAEHNP
HVRAAVAWYGKVAGETNEMTPFNPDDHAAQLKAPTLGLYGGKDDSISQRSLARMRERLAA
AGTQAARESEIVVYPDAGHAFFADYRPSYVKADADDGWKRAIAWFRHHGVM*
>GCA_000452945.1_BPMSHR5858_genomic_05066_1 sseB
MPHTHYTTLISADNLAERLAAAPGSVLVFDCRFDLADTDLGEQAYAAGHIPGAHYLHLDR
DLSGAKTGTNGRHPLPARDALVDTLKAHGLRQNQQIVAYDAQGGMYAARLWWLLRWLGHD
SVAVLDGGLQAWQSEGHAVSQDTPPKSMGDFRAGAPLATVVDAQTVLANLATKDRLVIDA
RAADRYRGENETIDRVGGHIPGALNRFFKDNLTADGRFKSGHELREAFSPLLGATEPKKV
ILQCGSGVTACHNALAMEIAGLHDPALYAGSWSEWSADPSRPVATGANP*
>GCA_000452945.1_BPMSHR5858_genomic_05068_1 xseB
MAKTATPGACASDPGSGPLPENYEMALAELEALVARMEGGTLSLEDSLAAYRRGAALVAF
CQQQLEKAEQQVRVLDGASLKPLSAGTAAADGEDDDL*
>GCA_000452945.1_BPMSHR5858_genomic_05069_1 ispA
MTFDQWMRSVLERVETALGHYLPAGTVAPAQLHEAMRYAVLGGGKRVRPLLCHAAGELTG
ASEAARDAASAALEMIHVYSLVHDDMPCMDDDALRRGKPTVHVKYDEPTALLVGDALQSQ
AFVALTDAAALPADRQAALVRELALASGSIGMAGGQAIDLASVGVALTRDALEKMHRMKT
GALLRASVRMGALAGETPSGDAMRALDAYAAAVGLAFQVVDDILDVTTDSATLGKTAGKD
AANDKPTYVSIIGLDASRALAAQLRADAHAALEPFGARAQRLAELADLVVNRVS*
>GCA_000452945.1_BPMSHR5858_genomic_05070_1 dxs
MYDLLKTIDDPADLRRLDRRQLQPLADELRAFVLDSVSKTGGHLSSNLGTVELTIALHYV
FNTPDDRIVWDVGHQTYPHKILTGRRDGMKTLRQFDGISGFPRRSESEYDTFGTAHSSTS
ISAALGMAIGSKLNGDDRFSIAVIGDGAMTAGMAFEAMNNAGVSEDAKLLVILNDNDMSI
SPPVGALNRHLARLMSGRFYAAARAGVERVLSVAPPVLELARKLEEHAKGMVVPATLFEE
FGFNYIGPIDGHDLDSLIPTLQNIKELRGPQFLHVVTKKGQGYKLAEADPVLYHGPGKFN

PAEGIKPSTTPAKKTYTQVFGEWLCDAAELDARVVGITPAMREGSGMVEFEKRFPERYYD
VGIAEQHAVTFAGGLATEGLKPVVAIYSTFLQRAYDQLIHDVALQNLPVVFAIDRAGLVG
ADGATHAGAYDLAFLRCIPNMTVMAASDENECRQMLHTALQQPNPTAVRYPRGAGTGVAT
VKAFTEIPLGKGEVRRRTSQPDGKRIAILAFGTMVAPSLAAADALDATVANMRFVKPIDA
ELVQALARTHDYLVTVEEGCVMGGAGSACVEAMMESGAVRPVLQLGLPDRFVDHGDPAKL
LSLCGLDGDGIAKSIRERFLSHAADVASPAKRVA*
>GCA_000452945.1_BPMSHR5858_genomic_05071_1 folE2
MNLMNPEFAMPDVQSTVDTRQMPIQRVGVRAVRHPLTVRTAEGETQATVGTWNLDVHLPA
DQKGTHMSRFVALLEERGGPLTADAFRTMLATMLEKLEARAGRIEVSFPYFVNKTAPVSG
VRSLLDYEVTLTGDVRDGLTRVFAKVLVPVTSLCPCSKKISQYGAHNQRSHVTIDAELAA
DVPVEDLIRIAEEEASCELWGLLKRPDEKFVTERAYENPKFVEDLVRDVARRLDADERIV
AYVLEAENFESIHNHSAYALIERDKRRGA*
>GCA_000452945.1_BPMSHR5858_genomic_05072_1 tsaD
MRPPHTIMLVLGIESSCDETGLALYDTERGLLAHALHSQIAMHREYGGVVPELASRDHIR
RALPLLEEVLAASGARRDDIDAIAFTQGPGLAGALLVGASIANALAFAWDKPTIGIHHLE
GHLLSPLLVAEPPPFPFVALLVSGGHTQLMRVSDVGVYETLGETLDDAAGEAFDKTAKLL
GLGYPGGPEVSRLAEAGTPGAVVLPRPMLHSGDLDFSFSGLKTAVLTQMKKLEAAHAGGA
VLERAKADLARGFVDAAVDVLVAKSLAALKATRLKRLVVAGGVGANRQLRAALSAAAQKR
GFDVHYPDLALCTDNGAMIALAGALRLARWPSQASRDYAFTVKPRWDLASLAR*
>GCA_000452945.1_BPMSHR5858_genomic_05073_1 group_7653
MEKFDIAVIGAGAAGMMTAAVAGQLGRRVVLLDHSPRLAEKIRISGGGRCNFTNLYAGPD
NYLSANPHFCRSALARYTPRDFLGLLKNHRVTWHEKHKGQLFCDHSSDAIIDVLKSECDA
GGVAWRRPVTVDAVRHAQADGFALDTSAGAIGAQALVVATGGLSIPKIGATDFAYRIAKQ
FGHKLVDTRPALVPLTFSADAWAPFAALSGVSLEARVSTGAKRGGGEFVEDILLTHRGLS
GPGILQISSYWRSGEPIHVDLAPQIDAANELIRAKAGSRRQIGTLLAEWVPARLAHAWLE
AERVHADARLADLPDKTLRRIGETLSRWTLTPSGTEGYRKAEVTKGGVDTRELSSATMMS
ERVPGLFFVGEALDVTGWLGGYNFQWAWASGVAAGRAAADGTRRLTG*
>GCA_000452945.1_BPMSHR5858_genomic_05074_1 rpsU_2
MTIIRVKENEPFEVAMRRFKRTIEKNGLLTELRAREFYEKPTAERKRKKAAAVKRHYKRI
RSQMLPKKLY*
>GCA_000452945.1_BPMSHR5858_genomic_05075_1 yqeY
MSLRDQISEDMKAAMRAKESERLATIRLLLAAIKQREVDERVTLDDAGVTAVVDKMIKQR
KDSISQFEAAGRTDLVEKEQAEVAVLSAYLPAQLSDAEVAAEVQAAVAQTGAAGPQDMGK
VMGVLKGKLAGRADMTAVSALVKAALSK*
>GCA_000452945.1_BPMSHR5858_genomic_05084_1 group_5138
MPIDSASRQWAPAAERNREPILAVLKRVLPARGAVLEIASGTGQHAVHFAAALPDLVWQP
TDVDAAARESIAAWAADAALPNLRAPLALDVCVEPWPLAAADAIVCVNMIHIAPWAAVCA
LFVGAARALPDGGVLYLYGPYRRGGAHTTESNAQFDAQLRRRNPAWGVRDLEAVVELGGA
SGLALDEVVEMPANNLSVVFRKRA*
>GCA_000452945.1_BPMSHR5858_genomic_05086_1 group_3609
MKSLIATVAVAAALAVPAVSFAQQSNGPVTRAQVRAELIQLEQAGYNPASDRVNYPEEIQ
AAEAKIHGQQNVAAQADTSGYGAQPAAAAESGSPNKVKRIADGAPVYKGR*
>GCA_000452945.1_BPMSHR5858_genomic_05089_1 group_5140
MQKAAFVAAYRLVGCGLILSATCHSIARRWNAPTFRLDNFLSYFTQLSSLYTAIVLLAGL
WLATKPPSRRYESARGAVVLYMAITGIVYELLLAHLDAVHHATPYYTNWILHRVIPIAVF
LDWLYVAPRVRIDWSQLARWLAFPVAYLGYTLVRGALIDWYPYPFVDPRAHGYLMVAAYS
GAIAAGSIGFAALIVLLGNRTGAPAPQAEHA*
>GCA_000452945.1_BPMSHR5858_genomic_05090_1 yodB
MNRILAGQERYNRPAVFFHWAIFLLVALAYFAIEVRGPKGTDSRVFWSNVHYLAGSLVLA
LAVLRIVWRFAGGAPRELPQPAWLALFAKLAHLALYAFIVAQPLLGIMTLNFGGKPVTLA
GIDWSFTLFAPNAALRSTVKEAHELIGNVFYYVIGLHALAALWHHFVKRDATLRRMTF*
>GCA_000452945.1_BPMSHR5858_genomic_05091_1 group_1936
MPNRFRKLTAWLGMLAIALAIVAPLVSQASARSAAPSSGAVICGDEHRAQPAEAGIAHDG
PVHDGVAHHALHLDACGYCAFFAHSPAIGAAAPSLFAIHVAAVAPPIPSGAAAPSLERHT
HAYPRAPPTDA*
>GCA_000452945.1_BPMSHR5858_genomic_05105_1 group_1545
MTWLADIVLVVHALLVLFIVGGLAAIWTGAALGWRWVRARTFRLAHLLAIGIVAALAALG
IDCPLTVLEDRLRTGAPDAQGFVQRWVGRLLYHDLPAWAFTAAYVLFALLVWLTWRRIPA
RDK*
>GCA_000452945.1_BPMSHR5858_genomic_05147_1 metZ
MDDSLNFDTLAVRSGTLRSEFNEHSEALFLTSSFCYASAAEAAERFKHSEDYYTYSRFTN
PTVTMFQDRLAALEGGEACIATASGMAAITSVVMAALQAGDHLVSSRSLFGSTLGLFGQI
FAKFGIETTFVDPADLDAWRAAVRPETKMFFLETPSNPMTELADIEAVGRIAKAAHALFV
VDNCFCSPVLQQPLKLGADVVMHSATKFLDGQGRVLGGALVGSKAFIMEKVFPFVRTAGP
TLSAFNAWVLLKGMETLSLRVNRQSENALEIARWLEAHPAVKRVFYPGLESHPQHALAKR
QQKSGGSVVSFELNGDTPEQQRANAWRVIDGTKIVSITANLGDTRTTITHPATTTHSRIA
PEAREAAGITEGLIRLAVGLEDPADIRDDLARGLAG*
>GCA_000452945.1_BPMSHR5858_genomic_05151_1 group_1940
MTRDTFVAPPAHTGRIVQLDGLRAFAVLAVFFQHALKAPLWIGVDLFFVLSGLLITGILL
ERKARGESYFGYFYARRVRRIVPPYLLLLVVSSILFGVEWARHWPWYAFFSTNIGLSLGG
IGHESLNVLWSLAVEEQFYIFWPFVVLLVPERALARVAAALIVAAPLLRAIATPWFDSFW
PIYYLTPFRMDLLAAGALLAVALRRDRRALEPYYGAAIAAACAALAVLAWLHLSFPRFRA
ANTPLSNATLYSVSLVLCTSIVVIALRGRGLAQRVLTHPALVYVGTVSYTIYLIHLSVLY
ALRPLHPNRYLTAALAFGVTLAYASASWYGFERRLTRGAPRQALAGAARASA*
>GCA_000452945.1_BPMSHR5858_genomic_05153_1 group_6967
MLKNVAINFFGLVLPTFVSLVTVPAYIKALGVERYGVVSLVWTLIGYFGILDLGMSMAAQ
NHISKALASGDAKESARVFWSAFWLNLATGIAGGLLIYFGAFMYTAYFTKVSAELQHEVY
LALPWLALAIPLANVSWVFAGAINGAERFGVFNTNQTIGTFLFQLMPLFAAWLVAPTLQV
VLAAAVGARFIAAVMLGAASMKVLGIRRIAPPQLGTAKGLFSFGGWMLIASTTGMIADTL
DRVMLGAGLGAKYVTYYTVPQNLVTRLNMLPNALVRTLFPRLSALGRDHADTLVKQSLEF
LNGVFTPIAIAAIFALGPFLALWVGRDLAELSAPVGRVLVISVWLVGQASVTRILIQSQV
NPARAAAAGLVQMPFFVGALWFGIHHFGLIGAAVVVAARALVDYGVLLYLSAIRMRAIVL
DMVAHLAFLLASLYVAHALPGLAAAIAACAAALALNVGWSITMTPGMRALARSALMRLNP
RKSV*
>GCA_000452945.1_BPMSHR5858_genomic_05155_1 group_5512
MNASSMASGARPSATDHKEHLIDALRGFAALVVAYFHCRQVVWVGMQHFHRTYGHALEPG
AIVGYLTFPFAWGSAGVPIFFVISGYCIHRNAALKLAADPSYRLNAPNFWTRRFARIYPV
LLAALVATLVFDSISLQIEPVSHKIRDIGLASFVVNLLSLQGVAGYTYGSNGALWTLSLE
VQFYAIYPLLFAARRRFGMVPVVVAIALVNVASAWLLERHDLQFFTSYWLSWTVGAWIAD
ARAQRGACAAPAPSRRWYAAAALGIALGCVAFHFGQYGAFQLWAAGFACYLHAALARPAP
AFAPMRALSWFGDFSYSLYLIHLPFFVCLASVLYRSEPQLSIWPSFAFIAAVIPVAYLFY
RMFELPAMRWSTSLKPKRAVAASARQQMPV*
>GCA_000452945.1_BPMSHR5858_genomic_05157_1 group_5150
MLKVTKAVFPVAGLGTRFLPATKASPKEMLPVVDKPLIQYAVEEAINAGITEMIFVTGRS
KRAIEDHFDKSFEIESELEARGKEKLLELVRGIKPSHVDCFYVRQPAALGLGHAVLCAEK
LVHGEPFAVILADDLLHGDQPVLKQLVDVFNHYHSSVIGVETIEREDSRSYGIVEGREWE
EDIIKLSGIIEKPAPEHAPSNLGVVGRYVLMPSIFEHLRKIKPGAGGELQLTDAVQSLLT
EEQVLAYRYYGTRFDCGSKLGYLKATVELALRHPEVGREFEAYLRNCLPALAAVA*
>GCA_000452945.1_BPMSHR5858_genomic_05158_1 oatA_2
MRAVKHVRGLDGLRAVAVTLVFLSHRAHFGAVDVGQIGVWTFFLISGFLIVGELHRSRLA
IERGASSRRYAFWLFAGKRALRILPVYYLLLAALAIAHRSVYQRGVDLGLRWHAVFLSDY
WIGVVKEGWPGSTSHFWSLAVEQQFYLIAPIVLLAARASRHLTICAAAVALAAAAHLLLY
RLHVSPVAVYAFSPWNFALIALGGIGGILRSGGIAAPEGVAGALAPWIGTAGAAFCATLA
LWAAPLPATAGGLADIGLGLSLGILLFWIVSHPEHPAVSLLEWAPLAYLGTISYGFYLFH
NLIPERIGVVPGLYERLRVPSAWQELLPEILQFALAALLAHLSWQYLEKRVLDYKKPLDA
ALRRRFALSERPAAR*
>GCA_000452945.1_BPMSHR5858_genomic_05159_1 group_7658
MATSAKKVAKKAAAPAKKVAAKKAAPAKKVVAKKAVAKAPAAPTPLKDKFTKASLATHIA
ERAAVEVKAVKAVLAALENVVLGSIHKKGAGEFTLPGLLKITAQAVPAKKKRFGKDPFTG
AERWFPAKPASVRVKARALKKLKDAAA*
>GCA_000452945.1_BPMSHR5858_genomic_05161_1 group_1242
MKKLALSTLSLALLGAAGAAQAQSSVTLYGVIDTSITYVHGNDGKANNAWLMGSGNLQGS
RWGLKGTEDLGAGLKAIFQLENGFNSNDGTLGQGKRMFGRQAFVGLQHDQYGTLTLGRQY
DPLVDLVQPVTADNYFGSLFATPGDVDNNDNSLRVNNTVKYTSPVFAGFQFEALYGFSGI
AGSPGQGQTWSAAAAYNNGPIGVAAGYFYTSNPSPTAGSRSTWSGSSDAIFDGAINSGYT
SAKSIGIAQVAGQYVFGPVTVGLGYSNAQYKPDGFSGFSSTEKYNTGRGFVTYQATPALL
LGLGYAYTKASGDTDAKYHQVSIGADYALSKRTDVYLAGAYQHASGTQRVDATTTQTAQA
SIGSYGVNGTKSQEMVALGLRHKF*
>GCA_000452945.1_BPMSHR5858_genomic_05164_1 doeB
MTDTPRFRNPIRSEVDLDANGKHAGYLRVPHSVHRSAYGWIPVPIVSIRNGSGPVVLVMA
GNHGDEYEGQIVVSRIVREIDAAHVRGQLIFLPMANFPAADAGTRVSPIDGGNLNRSFPG
DPAGAPTAVIADYIEHTLLPRAQYLIDLHSGGSSLLYQGGNLLALEPRDAAERARVRALL
AAFGLPRAFLHAPNPVHASSAARRQGALSILTELGGGGMSDPALIRDARRGLLHFLGFVG
TLRGPLVPDSPPATTRFMRVSGAHHYVYAYDKGLYEPLVELGARVRAGQPAALIHFPDTP
LREPVTQCFASDGEVVCKRVPAQVQRGDCLFQLAEDDAPA*
>GCA_000452945.1_BPMSHR5858_genomic_05197_1 group_5155
MQLPKRFAAACLAAAFLCGAPPAVSKTTSESLLERLVALIADALPSSAKQAPAGQVVEAA
FSPDGGAEALVLKVIGTSRASLRLAGYSFTSPKVVRALLDAHRRGVDVAIVVDNDGNRSK
ASKQALNLLVNAKVPTRTIDRYAIHHDKYIVVDGRHVETGSFNYSAAAAARNSENVLVVW
NNPALAAQYLKHWQNRFDQGTAYRSAY*
>GCA_000452945.1_BPMSHR5858_genomic_05203_1 marR
MARSKVGLDQFMTYRMHVLNKLSDRGIGELYRMKLGISLPEARIIASVGAFGPFSIMELA
RRANLDKSQASRAAEALIRRGLVKRGPSDDDGRVVLIALTPAGAALNRKIMPIARKWNET
LLDGLSESEKAAFSRALDKIIAGARERED*
>GCA_000452945.1_BPMSHR5858_genomic_05222_1 group_3624
METGIGLARRLSRVARGAARRALSILTAGAGRAGYAVDARVASAGPLYIGAGWFLIGAAA
ARPGAAEFALFDDRFATTARHRKEET*
>GCA_000452945.1_BPMSHR5858_genomic_05223_1 group_7665
MARQSTSVPFELSSAELARTRVGIVDGKRISLGVQGDALRGFVLERRCKSPGEPVSTQRV
GLRDPAAVAAFVEHDPYVVQLGIDYRALLDVHRAADDAGSHGAFAVHDARYARPASEAGG
AFRPAEHAGAAPAASGVPTASAASVAQPEFAVECEHDGALLALMRRICASCGATQCFYHW
FVVDEDTGEFTAHDLLIGGAPAWAQRYVHQHWYLNDPAVAHARDNTQPLRGSALAELRSD
HWLNHYAQTQGLGSNVFFPAHRRDVSTFGLLHVAAPLPAPHGEDALWRNRRVLRGLANEM
LEWRVVRRRRELAQELSLAAQDVLALRLVARGGGARHVAEELRLDERAVYQLFTAINRKM
DSKHIKSSATKAKRLGLLAEGYISK*
>GCA_000452945.1_BPMSHR5858_genomic_05229_1 group_7666
MTQDVTVFRARYRASVHPRYNAWLHGGFVLAYGCAAIAFFLKDVARVSAVQWASVPAAFL
LFNWLEFTVHRHVGHVKRRFGAMFYRRHTGDHHSFFANEQIAYREARDWRVILFPAWLIV
VFSLGLFALHAVLGRFDANVAGLFASTMLGGYLLYECVHACEHLPDAHPLAKWPWIRQMR
VLHRVHHRDALMRSCNFDVIWPLMDWLHGTLRWSEAPGADMSTRMRHEIDIACAGADVLA
YASDASRWPEWHPSSLRVDGPAGSLAAGSRFDEDVRAAGRVDRLHWDVVRNEPGVMWQAR
AANTAGSLRILLTYECSGGPRGARFVRTLHYHSPNPLLRLANALVMRGRVERESAYSLTL
LKRRLEAARDA*
>GCA_000452945.1_BPMSHR5858_genomic_05243_1 hspA
MSNVTRYDPFSLEPVSDLFQGLFRPLRGIADADDEKLASMKIDVTENDQAYAVKAELPGV
DKNDINVQIDGNTVSISAKVERNKELKEGERVIRRERYSGEFARTFSLANELDRDAASAQ
YQDGVLSLTLPKKASSEKKKLTIG*
>GCA_000452945.1_BPMSHR5858_genomic_05244_1 group_1549
MLPGHFPDMGDMMGDMMNAKSIAAAALAVAQAGLAHAQSSVTLYGALDAGVLWQNTSAAS
FSPTAPNTGAVVRFKDGGVYSSLWGIKGTEDLGGGYRVNFRLQGAFDGGTGKFGLSDAPG
AVAAFNQIATLGVSGPFGTFNAGRQIVPMIYAMADTDVRAARFFGSVLTAWLGLNTAAGW
PGNSTNAPIGALYDSDALVYQSPAFGGASLALEYAPGGVPGGVQRGARASIVLKYASNAL
NLAAVYYNGHDTNPAPGASATGLDNNRFVYVGAQYKLKDVAVSASYGNGRNPARGDRVNL
DMFAFGLGYRVSPFFRVSSGVYYLKDRNRSANRSTSIVLGADYELSKHTTLYAQGGRVDN
RGTMAQMLAYGQPVAPGLATTAAMLGMRHSF*
>GCA_000452945.1_BPMSHR5858_genomic_05245_1 menE_2
VHATDQHSALNAAPSAYVYPLLVGQLLTHARGADAGRQIVYRGDVRHSYAQFRERIGRLA
GALAALGVGADATVAVMDWDSHRYLESYFAIPMMGATLFTVNVRLSPRQIAHTLRDARAS
VVVAHADFLPLLEQVRDALGDSTCVIVACDGGAMPATSLPLAGEYERLVAAADPDYPFAD
FDENARAVLFYTTGTTGDPKGVCYSHRQIVLHALATAAALGAARDGQRLHRDDVYMPITP
MFHVMAWGMPYIAVMLGLKIVLPGRYRAHALLDLRQAERVTFSHCVPAVLQMLLDEARLR
ACELSGWKMIVGGSALPASLCRAARARRIDVFAGYGMSETGPVVALAQLRGTEGGVDVHA
DEDAGIDADVGAGIDVDTDVDADINANAGANASDSSVTTAAAHADDETAKRCAAGWPPPL
VELRVVDAAMRDVPRDGRTPGEIVLRGPALALGYRGNPQASAALWAGGYLHTQDVAVMHG
DGAIRIVDRIKDVIKTGGEWVSSLELEGLLNDIAGIRESAVVGVPDARWGERPAAVVVCE
RDAPLAAQDVRARLLEHVAANRISRYAVPECERIVFVDALPKTSVGKIDKKSLRRLVERI
D*
>GCA_000452945.1_BPMSHR5858_genomic_05246_1 malT_2
MPPTAPGAERAATEIVLKTVAPREPPHLLARPRLGAGDAALAGCRAVLVQAPAGFGKTSL
LAQWRREHLSRGTAVAWLSADERDDPRRFLLALVQAVRTACARPTFARTLTDGPATPARE
LEGVTGWLAEIAQFSLDVLLIVDEAERLPDAGRRALTYVLHNAPQNLRIVVAARHGFDAH
AGDLIAYGQCRLIGADALRFRLDETIALVGERIGARVDADTCARLFELCDGWPLGLQLAL
AAMERAPDLRLSVEPAAAVPGALRDRLVDALIAKLAPDDERFLTGISIADMLHPALCRAL
TGDAHAPERLARLLRETPMFTASDDGPWCRLHTLVRDVLRMRLAQWRDDERNRLHDRASA
WLAAHGMIEEAARHAHAAGRHQAAFELAERCLHDAVKQGHLQAMLDWLELLPEAELERRP
RLRLAAAWALALGERHPEAARQVEQVLANPGASVELRYECALILSAAAYYADDPDRFVAL
FEPWARAAPVGTPWLAQVHANRLGARAILLGEPAQARRFQSAAPRADCRGEHGYVVRWGE
YMTGMSYLCEGQLHLAEEVLFPALASAEIDLGRRHPLTCMLAALCATLAYEADRIEQAAA
LVANRLDVLERGGTPETVALAYRTAARVAAARGAEHRALDLLESLHALGVARRLPRLCVA
SLAEQARMHAVYYRAQTCEALVQRIDDIMRREAPRRGPLWRRSAALAQASAHAAASVAAR
RWPQALDALAQAGGLACEMKLGRERIESMALRAFALEQTGADGRALLAEAMNLARTFHLT
RTLADAHPAVADWARRALDQASVAGGGARAAPRVVRPAQRPKAGSPRAVPSVVLTPKERE
ILELLGRNLSNKEIALAMTVGEETVKWHLKNLFGKLDASSRKHAVRRALVLGLLESAE*
>GCA_000452945.1_BPMSHR5858_genomic_05247_1 group_479
MYRHFLVPVDDTDAGIDTVAYAVEFARSIGARITFVQTRIEFEAADAARRIGETERKEQT
QRTDDAGKSGECGRSDESVRPAPTPDAAARPAAEATPAARAPELPIAKAEAAARAQGVPC
DSVRAAGATTADALAGAMLAHDCDLLCVGPALGDAAAAPPHACVADRLAARGIAVLTCAF
RRTPAAARAIAALYAAHREAAGALGAWLAQLRAAIAAGRALDADAAHAIANGLSHLRDGR
QPKAARRLYAALRGATGALDAELGELERQRLRNARMLSGLLEAIHAGIAREAPPVRLEHA
LSAYAQCVCEHAGRGEGVIVPAAQRYLADDDWRAIDASLALIASGPAAATRGA*
>GCA_000452945.1_BPMSHR5858_genomic_05248_1 group_5164
MAHAIRFHETGGPEVLRREETQVGAPGPGEVRVRHAAVGLNFADIYFRTGLYPAKLPSGI
GVEAAGIVTAVGGNVAGLAVGERVTYTGSPLGAYSTERVMPADSLIRLPDAIGFDTAAAM
TMRGLTSAYLMRRIHAWQAGDTFLLHAAAGGVGLIVCQWAKLLGLNVIGTVSSDEKADIA
RAHGCDHVIVHRREDVAKRVRELTNGVGVAVVIDSVGKSTFAASLDSLKRRGLLVGVGTA
SGPFPPIDAMQLAIKGSLFFTRPALADYIADPAEKAALAGELFDHVAAGRIRIEINQRYE
LNDAVQAHRDLEAGRTTGSSIFVVDGD*
>GCA_000452945.1_BPMSHR5858_genomic_05250_1 kce_3
MQFFDDSLHPENMDKVVITVAPYGPEWMPEDFPEDIPVTMAEQVQKAVDCYNAGATVLHL
HVRELDGRGSKRLSKFNELIAGVRAAVPDMIIQVGGSISFAPEDDGQAAKWLSDDTRHML
AELDPKPDQVTVAINTTQMNIMELLYPEYLAGTSLANSALHAAYSEMTVPAGPAWVEEHL
RRLQAAGIQPHFQLTGIHALETLERLVRKGTYTGPLNLTWIGIGGGFDGPNPFNFFNFIQ
RVPDGCTLTAESLLKNVLPFNTMALAMGLHPRCGIEDTIIDQHGERMTSVRQIEQTVRVA
RELGREIASGKQARDIYRIGVQYRDADETLAANGMAPNRQPGSRRERRFAA*
>GCA_000452945.1_BPMSHR5858_genomic_05251_1 exuT_1
LHIHPADVDDGALAAPIPRPYAWTVFALAFALLLSDYMSRQVLNAVFPLLKAAWGLSDTR
LGSLSGIVALMVGTLTFPLSVLADRWGRVRSITLMAALWSVATLGCAAATSYGELLVARA
FVGVGEAAYGSVGVALILGLFPPHLRSTLTGAFMAGGAFGSVLGMALGGAVATQLGWRAS
FGAMACLGFALVVCFRLVVTERRIAGSGARAARSGRERTPALAHASLRTVLAELFSTVSV
VCAYVGSGLQLFIMGAVIAWMPSFLNRYYALPPDKAAAAAAGFVLLGGSGMIACGIVTDR
LSRACPERKWAMAIVYCAISFALLAIGFRLSPGAPQLGLLALGMFVVAGTSGPAGAMVAN
LTPPSIHSPAFATLTLVNNLLGLAPGPLVTGFIADRIGLVGALQIAPLVALASAFAFAVG
RFRYGDGLRRLNPLSPSAAAE*
>GCA_000452945.1_BPMSHR5858_genomic_05252_1 group_5167
MNDLSSTVLIVPGLRGHVPDHWQTLLEHRLPNARSVPPLERDGLSCAARMEALDAALRRI
AGPVILVAHSAGVMTTVHWARLHRREIKGALLAAPADLETPLPDGYPQPDALRRHGWTPI
PRKRLPFPSIVAASTNDPLARIERAAALADAWGSRLVDLGAAGHLNPASGYGEWPGALAL
IAELEDARPRAAQCGEAR*

>GCA_000452945.1_BPMSHR5858_genomic_05253_1 group_1084
MSAGACHRSRDRGNRGDQLGDARPCKLTVECWFDFVCPWCWIGKRHLHAAVDALRALRPD
VRAEIVWHPYRLLPDVPRGGLPYDAFYRARLGGGDAVDARRAQVRQAGREAHLEFAFERI
RTMPNTHAAHAFVDAAADAAHAARVDVDAMIERIFAAFFAHGRDIGDVATLECIALESGL
PQHTLPSRLGRLPPRSPRPLGVAHRDIAGVPAFRFGGRETLSGARSPAVLLHAMLRSLCA
*
>GCA_000452945.1_BPMSHR5858_genomic_05265_1 gatA_2
MKRRQFLHTLGALGPTHALLASAPASAGQTALDERAGASAPASRAVRESLERIARIDRDG
PRLNAIIELNPDAEAIAHALDAERAAGVARGPLHGVTVALKDNIATGDRMSTTAGSLALD
GVRATRDAHLVARLRRAGAVIVAKANLSEWANFRSTRSTSGWSARGGLSRNPYALDRTTS
GSSSGSAAAVAAGLVAVSVGTETDGSIVSPAAINGCVGLKPTLGRVSRDGIVPVSHTQDT
AGPIARTVRDAARLLGALAGGDARDSATASAPAPADYVAALDANALRGARLGIARAYFTG
HDEVDVQIERAIAEMKRLGAVVIDPVDLPKADYEEDEKVVLLHEFKHGLPSWLRTFAPHA
RVRTLADVIAFNARQHAREMPYFGQELLLHAQEAGGLDAAAYRDALARCGRRARDEGLAR
VLREHRLDALVAPTEGTAWLIDLINGDSGGDGFSTPAAVAGFPHLTVPAGLVRGLPVGVS
FVGAPWSEARLLALGYAFEQATQWRREPRFVERSNVPAAR*
>GCA_000452945.1_BPMSHR5858_genomic_05287_1 cyaA_2
VPLQIGARDVAFPFLNSLSFWMTAVAALLMNVSLVVGEFAQTGWLAYPPLSELQFSPGVG
VDYYLWALQISGVGTLITAINFFVTILRMRAPGMTLMKMPVFTWTALCSNVLIMATFPIL
TVALALLALDRYLGMHFFTNDAGGNAMVYLNLIWAWGHPEVYILVLPAFGIYSKVIATFS
KKPLFGYKTMVYASCVIMVLAFLVWLHHFFTMGSGANVNAFFGIMTMVIAIPTGVKIFNW
LFTMYRGRIEFTAPVLWTIGFMVTFTLGGMTGVMMAIPGADFVLHNSLFLIAHFHNAIIG
GVVFGYLAGFHYWFPKAFGFKLDEKLGKRAFWCWFIGFYVSFVPLYVLGFMGMTRRLNHY
DNPAWHPWLIVAACGVALIALGVLHQVAQVWVAVRNRNAPGYRDTTGDPWDGRTLEWATA
SPPPVYNFAVIPIVHALDELAYRKEHGIGVGKNAVYQDIHMPSNTSAGLFVGMFSLVLGF
ALVWHIWWLAIASFVGIVAMVVLYSAGENDGYYIAADTVREIEERRAGARAPARPAEVEL
EAN*
>GCA_000452945.1_BPMSHR5858_genomic_05294_1 hsrA_4
MPPAPPAAAPSEKSLTVMLWLVATGFFMQTLDATIVNTALPSMAASLGESPLRMQSVVIA
YSLTMAVMIPVSGWLADTLGTRRVFFSAILIFTLGSLLCANAHTLPLLVAFRVIQGVGGA
MLLPVGRLAVLRTFPAERYLPALSFVAIPGLIGPLIGPTLGGWLVKIASWHWIFLINVPV
GIAGCIATFYSMPDSRNPAAGRFDLKGYLLLTIGMIAISLSLDGLADLGMQHAMVLVLLI
LSLACFVAYGLYAVRAPQPIFSLELFGIHTFSVGLLGNLFARIGSGAMPYLIPLLLQVSL
GYGAFEAGLMMLPVAAAGMFSKRIITVLITRHGYRKVLLANTIMVGLMMASFALVSDAMP
TWLKIAQLALFGGFNSMQFTAMNTLTLKDLGTGGASSGNSLFSLVQMLSMSLGVTVAGAL
LATFTGMMRSVTPTHTLPAFHATFVCVGLITAASAWIFAQLSPDVRGAVRRKTDPSERA*
>GCA_000452945.1_BPMSHR5858_genomic_05299_1 group_4111
MHPLFEPYRPQLDAHPWFGALPAELRDALTAAALPRRLVTGQVLFRRGDAPCGLYAVLSG
ALSVGAVDASGKEALLTVAEPTTWFGEIALFDELPRTHDAIALEDALLMHVPQAALQQLL
DETPRYWRYFALLMAQKLRMSFMNVEALTLMPAAQRLAARLLMIAEGYGGIGTRHRRIRL
SQERLGAMLSLTRQTANQLLKDLAARGVVRLHVGEIEVVDLDALRAASGAANRARSDGAG
GAGGAGGTNEA*
>GCA_000452945.1_BPMSHR5858_genomic_05321_1 group_3632
MFESLTTLIKKWRASRSASHQLDALLATADANAPYAERAEWLIELAHWLHRDGSMTAAAP
AQPESGERTFAAHARLRYVFHVLDRNPAWKTNVARLLRGVLRESDGISLLCDAGMPVHSG
FFGALVERIDSSLIPPAPNRRELAALFTLMFPAPADAQWIDALPDDLLARLADLFAFDVT
DKERHQPGSFSRDLLAALHNLTCQISSTGLSQTVRSRLSDEDARTPLEAQPFYRLTRAML
AVETAQAALDDGGDASKLLHEVNYLRVLLDECRVSIDDVFSHLYRNGVSVDIVFQVERMR
MRILRAETLLNAWMARDDLRGLARLTAELVDANQNSQSVSHLVRSNFALFARKLVETNAD
TGEHYISRGRAEYLKMLRMAAGGGLVTVATVCIKFGITGAHLQSMFEGLLAGINYAASFM
LMHFLHFTLATKQPAMTAPTIARELDGTGHDEGVKQFVSSVIALIRTQAAAICGNVLVVL
PVCFAVQMFASGVLHANLISPDKAHATLKSFSLFGPTPIYAALTGVLLWASSLLAGWADN
WFVLHRVGDALAYNRRLRITLGVAGAAKLAHFCRSNVAGVVGNVGLGLMLGLVPAIVSAF
AFSFEVRHVTLSAGSIGVAIGVLGKETLRSAELWWAVAGVFSMAILNVAVSFALAFTMAV
RSRSLRPTKVRALVAAIGRAVLARPASLVWPPRGAGGAAGESRAH*
>GCA_000452945.1_BPMSHR5858_genomic_05324_1 group_7669
MTTANNESVIAWFDIPSVDFDRAIRFYETVLQTALQREVIGGVPTAVFVHPEAATGGAVV
YDPQRLKPGADGTVVYLNANGPVTAALERAKRAGGIVQGAVIELPNNYGYIGYLIDTEGN
RVGLHSLKCS*
>GCA_000452945.1_BPMSHR5858_genomic_05325_1 group_267
VLGIEFAVGVGVGVGVGVGFGFGFGFGFGFGFGFGFGFGFGFGFGFGFGFGFGNIETRAISR
RNDRAHRFAYENGACPWSRAALSSRHSNACPRSRSPAMTRRADRLFQIAELLRGRRLTTA
QQLADWLSVSPRTVYRDVRDLQLSGVPIEGEAGIGYRLNRAASLPPLTFTAEELAALAAG
ARMLETWGGARFAGGARSALAKIASAMPADKRATLDRLAVFAPSFHVDRAFSEKVDALHE
AVDGRRIVSFAYCDRLGAQTTRHVWPLGLVYWGGRWTIGAWCELRGDFRTFDIARMSEVA
VHGPFPDQEGKRLADYLRKANAPVR*
>GCA_000452945.1_BPMSHR5858_genomic_05327_1 group_7670
MEIINHTVINVIIFVLAVYVGYHVVWNVTPALHTPLMAVTNAISAIVIVGAMLASALTLG
GVGKAFGTIAVALAAVNVFGGFLVTRRMLEMFKKKEPRQKADHAKEGAR*
>GCA_000452945.1_BPMSHR5858_genomic_05330_1 mnmA
MTKRRVVVGMSGGVDSSVTAWLLKEQGYDVVGLFMKNWEDDDDGEYCSTRQDWIDVVSVA
DLIGIDVEAVNFAAEYKDRVFAEFLREYSAGRTPNPDVLCNAEIKFKAFLDHAMSLGAQT
IATGHYARVRERDGRFELLKAFDHTKDQSYFLHRLNQAQLSKTMFPLGEIPKTRVREIAA
QIGLPNAKKKDSTGICFIGERPFRDFLNRYLPTKPGPMKTPDGKTVGKHIGLAFYTFGQR
KGIGLGGSKDGSGEPWFVAAKDIASNTLYVVQGHDHPWLRSRELVAGNVSWVAGEPPADG
ARCGAKTRYRQADAPCAFGRAAQAGDERFSLVFDEPQWAVTPGQSAVLYDGDVCLGGGII
ESAATGRAGTAPAGRAPALVEAR*
>GCA_000452945.1_BPMSHR5858_genomic_05340_1 ruvB
MIETDKLAAERIIAATPASSHEEAFERALRPRQLDEYVGQEKVRDQLEIFIEAAKRRSEA
LDHVLLFGPPGLGKTTLAHIIAREMGVNLRQTSGPVLERAGDLAALLTNLEANDVLFIDE
IHRLSPVVEEILYPALEDYQIDIMIGEGPAARSVKLDLQPFTLVGATTRAGMLTNPLRDR
FGIVARLEFYDAEQLSRIVRRSAALLNAQIDPAGALEIAKRSRGTPRIANRLLRRVRDYA
EVKADGNITAAVADAALAMLDVDPVGFDLMDRKLLEAILHKFDGGPVGVDNLAAAIGEER
DTIEDVLEPYLIQQGFLQRTPRGRVATLLTYRHFGLSAPDAANPVRNLWDTPDAEC*
>GCA_000452945.1_BPMSHR5858_genomic_05358_1 group_5181
MSDRTELATRYEPAGTAVTTPTDTEAAAAQARTAPAVDVFENRHGITLYADLPGVPREKL
DVRVQDGNLTIEAEAVVPTPAGLRLRHAEIRHPRYTRSFALSPDFDVSRIDAQLRDGVLK
LTIPRRDEAKPRRIEVSAG*
>GCA_000452945.1_BPMSHR5858_genomic_05365_1 gltJ_2
MSYHWNWGIFLSPVSTGEPTTYLGWLMSGFWVTVKVSLVAWVIALVVGSLFGVLRTVPNK
WLSAIGTVYVSIFRNIPLIVQFFVWYLVVPELLPVSIGTWIKQLPPGTQFFTASIVCLGL
FTGARVCEQVRSGINALPKGQRAAGLAMGFTQWQTYRYVLLPVAYRIIVPPLTSEFLNIF
KNSAVASTIGLLDLSAQARQLVDYTAQTYESFIAVTLAYVVINLIVMTFMRWVESKSRLP
GYIGGK*
>GCA_000452945.1_BPMSHR5858_genomic_05369_1 nasA_1
MDTLRTSLRSGNWRSLLACFLYFDTGFTVWVLYGPLAPFISKDIAMTAAQQGFLVAVPVL
AAAILRVTLGNLYQSADGRRIALMGVLLSALPAAVLPFVPGTPSYTLLLVLGVFLGIGGA
SFAVALPMAGSSYPPKVQGLVLGLAAAGNIGAVLDGFMFPHIAAALGWKFSTAAALPLLA
IAGFALFAWADDRAEKSGSAARAFGAFVITLGGLVALVLAVHAGLFGAGKAGVLLLPVMG
ALLAIAVLPGRYRAVLAERDTWVIMLIYSITFGGFVGMSSYVTLLLTTLYQMPKIEAGLF
MSLLAFLGAIVRPFGGHLADRITGVRALMAILAAIAAADFAFAAVMPPLAGGIALLVCLY
VAFGLGNGSTFQLVPHRWQGKTGLLSGIVGAAGGIGGFYLPVVMGIAKESTGSYQLGFAT
FGALATCALVALAMLRAQWLRWAAPELAAAEPSAGAGMPIGGAARVGD*
>GCA_000452945.1_BPMSHR5858_genomic_05375_1 hexR_1
MLPRIETIRAQLRPSERKLADFVLAAPREVLDLSMTELAARAGASQPTIARFCQALGCSG
FREFKIRLAQSVAPGVASVYRDVEPDERAPGIIGKVFDRTIGALIEVRNSLSATSVAEAI
ELLTRASRIEFYGAGGSGIAAQDIQHKFFRLGVPSVAYSDPHTFSMSAALLGANDVVVAI
SNTGRTRDIVDAARAALACDAKVISITHSHSPLAQLSTVSLASNVAEETDVFSPMTSRMS
HLAIGDILAVGVALRRGPELVDRLGRAKEAITRRRIGADAKAQKETGKDATKVAASGVAG
GAAESAQQAARAQAPQVGEKTRIARDIEDVKD*
>GCA_000452945.1_BPMSHR5858_genomic_05398_1 rsmE
MSGAGTTAAVPRFFVDAALRADATLALPADVARHAHVLRLQPGDALALFDGTGGQYRARL
VEIDKRGALARIDAFEPAEAEPPYRVTLAQGIAGGDKMDWVIEKAVELGVAAVAPLSTAR
GVVRLSGERADKRVSHWRGVVRASCEQCGRNRLPEIAPVRGFDAWLDALPATPADGELRL
LLSPRASVPFASLPAAPPAGDVTLLIGPEGGLAPDEEQAARARGFAALSLGPRVLRTETA
GAAVLAALAARWGGW*
>GCA_000452945.1_BPMSHR5858_genomic_05403_1 group_8687
MPTDPTPRPADSADSASQPGATPAPASSPAPRRDSPQDPQRIARRRATVRFMFSHPVHIV
SLGFGSGLAPFMPGTFGTLFGWLTFVALNRYLTVPEWWALIVAGFVAGIWMTGFTAKKMG
IADPGPAVWDEIVAIWLVMLLVTPATFVEQLWAFVVFRFFDMVKPPPIRYFDRNLKGGFG
IMFDDLIAALMTLFVIALWRSFGVQ*
>GCA_000452945.1_BPMSHR5858_genomic_05404_1 pncC
MPTDSVVHQLAIRVGNKLRDEHLTLVTAESCTGGMIATAITDISGSSGWFERGFVTYSNA
AKIEMIGVPADLIDKHGAVSEPVARAMVEGALRNSRAQVALAVTGIAGPGGGSEHKPVGT
VSFGWSNRLHTSVETLVFKGDREQIRVQAAAHALRGLVKLIDEQER*
>GCA_000452945.1_BPMSHR5858_genomic_05405_1 pyrF
MSSTPTFIESLRTAWQRTNSLLCVGLDPEPTKFPAHLENQPDAIFDFCREIVDATAPFAS
AFKPQIAYFAAHRAEDQLERLIAHIHLQHPGLPVILDAKRGDIGSTAEQYAREAFERYRA
DAVTVNPYMGFDSIEPYLEYEDKGVVVLCRTSNPGGSDLQFLETGGRPLYQVVADLAASQ
WNAKTGQLALVVGATFPREIEIVRGIVGDMPLLIPGIGAQGGDVAATVAAGRTADGAGMM
INSSRAILYASRGDDFADAAARTAERTRDAINAHR*
>GCA_000452945.1_BPMSHR5858_genomic_05436_1 ampD
MSEAGRFAVGADGWVDGARREPSPNFEARPDGAAISLVVIHNISLPPGEFGGDAIVALFQ
NRLDCDAHPYYAAHLRGVRVSAHFLVRRDGALIQFVSCDARAWHAGASSFFGRERCNDFS
IGIELEGADDVAFDARQYATLAALARALAARYPIDAFAGHEHVAPGRKTDPGPHFDWQRF
ADDGGFPPRYFPYRKH*
>GCA_000452945.1_BPMSHR5858_genomic_05439_1 ffh
MLDNLTQRMARVVKTLRGEARLTEANTQEMLREVRLALLEADVALPVVRDFIAKVKEKAL
GEDVIGSLSPGQALVGVVQKELTAVIGGDYEGKAAELNLAVAPPAIILMAGLQGAGKTTT
VGKLAKLLREKYKKKVLTVSCDVYRPAAIAQLKTVSEQVGADFFPSTPDQKPVDIANAAV
DWAKRHYHDVLLVDTAGRLGIDEAMMQEIAALHTALKPVETLFVVDAMLGQDAVNTAKAF
NDALPLTGVVLTKLDGDSRGGAALSVRHVTGKPIKFVGVAEKLDGLEIFHPDRMANRILG
MGDILALVEEAQRGVDIQAAEKLANKVKKGGDFDLNDFRAQISQMKNMGGLSSLMDKLPA
QFQQAAAGADMSQAEKQIRRMEGIINSMTPAERAKPEIIKATRKRRIAAGAGVPVQEVNR
MLNQYDQMRTMMKKLKGGNLQKMMRGIKGMMPGMR*
>GCA_000452945.1_BPMSHR5858_genomic_05440_1 hpt
MNREEALHIFQHSEEIVSAGEVNASIGRMAGEIRGQIGEEFPLVLSVMGGAAVFTGMLLP
HLDFPLEFDYIHLTRYRNTTKGGPEMHWRVAPRESVKDRIVLVLDDILDEGETMAAIRDR
ILEMGAKRFMSAVLCEKTLAKAKPLHPDFCGFAVPDRYVFGCGMDAKGYWRNLPTIRALT
ADV*
>GCA_000452945.1_BPMSHR5858_genomic_05460_1 secA
MTTGFLQKIFGSRNQRLVKQYQKTVAAINALETQIETLTDDQLRGKTGEFRQRIAVGESL
DKLLPEAFAVCREASRRVLKMRHFDVQMIGGMVLHYGKIAEMRTGEGKTLVATLAAYLNA
LAGRGVHVVTVNDYLAQRDAEWMGRLYNFLGLSVGINLSGMEHDQKQAAYAADITYGTNN
EFGFDYLRDNMVYETDSRVQRPLNFAVVDEVDSILIDEARTPLIISGQAEDHTELYVRMN
ALPPLLERQIGEEKADGTGVEKPGDYTLDEKGRQVFLTESGHEKAERMLAEWGLIGDGES
LYAPQNITLMHHVYAALRAHTLFHRDQHYVVQNDEVIIVDEFTGRLMPGRRWSDGLHQAV
EAKEHVKIQSENQTLASITFQNYFRMYAKLSGMTGTADTEAYEFNEIYGLETVVIPTNRP
PKRIDKQDQIYKTAKERYDAVIRDIRECHERGQPVLVGTTSIENSELLSHLLKQAGLPHE
VLNAKQHAREAAIVAEAGRPKRITIATNMAGRGTDIVLGGNVEKQAAFIEADESIPADEK
ARRIQQLHDEWETLHEQVKTAGGLHIIGTERHESRRIDNQLRGRAGRQGDPGSSRFYLSL
EDPLLRIFAGDRVRAIMDRLKMPEGEAIEAGIVTRSIESAQRKVEARNFDIRKQLLEYDD
VSNDQRKVIYQQRNELLEAHDIAETIGAMRHGVISEVVRQFVPAGSIEEQWDLPELEETL
RNDWQLDLAIQEMVNESSSINADEILDAVTTAADEHYEAKVALVGRESFSAFERSIMLQT
LDRLWREHLAALDHLRQGIHLRGYAQKNPKQEYKREAFELFAAMLDAVKQEVTRIVMNVQ
IQSPEQLEEAAEQIEEQGGQLGNVEFQHADFAAAAAAATAGGAVVADATAEMVSHAMSHS
GPAGEVPRVGRNDPCPCGSGKKYKHCHGKLN*
>GCA_000452945.1_BPMSHR5858_genomic_05461_1 group_3649
MTLALLTVVALVAALLRFTKRRRASRALFVATVAAFFAIGCGLVPAWLLRALQAPYAARP
AIEWGERNAIVMLGLGTEKIAATGAVEPGTFSYSRVVEAASLYRDCRRARADADCKILVS
GGDARRNGVPEASVYRDALVGVGIDAADVLSEPRSMNTWQNAQFTRVALDGYRADRVLLV
TSGIHLRRSMLYFAHFGVAAIAVRAEYLQAVTFPLPLAYNFAIADLALHEYLGIARYRLY
NALGWNPARTRPGDA*
>GCA_000452945.1_BPMSHR5858_genomic_05466_1 group_2619
MKLPYTKSKSFAASLARAVVAFALAGIGAAAFAATFDLSPEQPGRVRGARDDAVIRALPA
SFKFVEPDTLTVGIAPSQPPISAYATDARTVVGFDADLAQLLADSLGRKLKIVALAWPDW
PLALESGRVDAVLSNVTVTEERKAKFDFSTYRKDQVGFYVKSDSKIASIRQPRDVAGLRV
ITDAGTNQEKILLEWDRENVAHGLKPVTVQYYDDHAVKSVALQAGRADAIFSVNSVLAYQ
AAQQHRTKLVGAVSGGWPRTADIAVTTRRGSGLAAPFTLAINGFIRNGTYRRVLERWSLG
SEAIDASRTNPPGLPKT*
>GCA_000452945.1_BPMSHR5858_genomic_05467_1 group_7678
MTDSIIDTAPHDARAKPLIDALIDEYATRYHAYRPDSRASAQAELARYPAELFAPPEGAF
ILLIRGGETIGGGAFKRYDAQTAELKRIWTRADLRRQGLARRVVRELELRAARQGYRRVY
LTTGFRQPEAWALYLDTGYAALFDRDVDPQIYVHLPFGKDLIEPGRIGALDDLRAPVPAA
PVE*
>GCA_000452945.1_BPMSHR5858_genomic_05489_1 tsaE
MPEQPSHAHAAPPLPAPLAERTLALADEAATIALGERLAHALDAMRGARAAAHAFDGLQI
QLYGDLGAGKTTLVRAMLRGLGHAGRVKSPTYTLVEPYALARSDGELEVYHFDLYRFSDP
AEWADAGFREYFNSGAICIVEWPQRAGALLGVPDLVFSLDVAGEGRLLTARAYSASGKAC
LERC*
>GCA_000452945.1_BPMSHR5858_genomic_05493_1 cnoX
MDTTLATFERDVIEASLAAPVLVDFWAPWCGPCKTLGPLLEKLEQEYEGRWKLVKVNVDE
NQELAAHFQTRSIPHVIAFAGGQPVDQFVGVLPEGQLRAFLDRLVPAPDEAERYAAQTAL
AEERIDDALSHLENALALNPGYDDARLDLVELLLALNRIDDARAEAARLSPQTTGGADAR
YQAIKTRFDALDATADLPPTDALEARIASHPADLDARFDLAQSLIARRAYEGALEQLLEI
VLRDRTFGDDVGRKTMISVFELAADQPELVSAWRRKLSAALN*
>GCA_000452945.1_BPMSHR5858_genomic_05498_1 msrA2
MSEATSEVATLGGGCFWCQEAVFLDVDGVTAVESGYAGGHTRNPGYRDVCDGDTGHAEVV
SVSFDPARIGYREILEIFFATHDPTQLNRQGNDVGTQYRSVIFTHSDAQRDAALELIRTL
ERDKLYDDPIVTQVEPLNGNYWPAEAYHQNYYARNPGQGYCAVVIGPKLAKFRQKFAHRL
KSRGA*
>GCA_000452945.1_BPMSHR5858_genomic_05516_1 malK_2
MASVLLRNIGKRYDDNEVLRNVNLDIADGEFVVFVGPSGCGKSTLMRMIAGLEEISGGEL
LIDGAKVNDVPSAKRGIAMVFQSYALYPHMTLYDNMAFGLKLAGAKKQEIDDAVKQAARI
LHIDHLLDRKPRQLSGGQRQRVAIGRAITRKPKVFLFDEPLSNLDAALRVKMRLEFARLH
DELKTTMIYVTHDQVEAMTLADKIVVLSAGSVQQVGAPNALYHAPANQFVAGFIGSPKMN
FLAGVVESASADGALVRFESGETQRAAVDASTLRAGERVTVGIRPEHLHVGVTGGDGIVA
RTMAVESLGDAAYLYAESAVAPDGLIARIPPLDTYRAGEMLRVGAQPEHCHLFDADGRAF
KRKPRHALAA*
>GCA_000452945.1_BPMSHR5858_genomic_05522_1 group_11379
MAAIVAAGELLAEFVAAKRGQGFDAPGEFAGPFPSGAPAIFADQAARMGASVAYAGCVGR
DAFGDAIVARLASHGVDVAHIRRAARPTGIACVAYRADGGRQFVFSIEGSASALLEPADV
VPSMFAGCRYFHVMGSSLTSENAIAAVKRGVIEAARVGAKISFDPNVRAEMLAFAPMRAA
LREMLDACHLFLPSEADLPFFCGPQPAERAIAGLLATHPLLERVVLKRGAAGSAAFDRTG
SVAAPAYETAEVDPTGAGDCFGGTFVACAIAGLPVASALARANAAGALAVSRIGPMEGNS
MPADIDRFLAERGVRCAA*
>GCA_000452945.1_BPMSHR5858_genomic_05525_1 psaA
MSLAAAPRMFASCSSRSPSAPLALRASPLLRAARALAAGVAALALASAAFAQNATLKIVA
AENFYGDVAKQIGGARVAVSSVLSNPDQDPHLFEASPKVARELQHADLVIYNGADYDPWM
AKLLAASKNAKRATVVVAELVGKKAGDNPHLWYDPATMPAAARALAAALGSADPAHKAEY
DANLAKFVASMKPIDAKVAGLRARYKGVPVTATEPVFGYMADAIGLDMRNPRFQLATMNN
TEASPADIAAFENDLKRRQVRVLIYNSQAVEPMTKRMLKLAQDARVPTVSVTETQPAGKT
FQQWMLAQLDALGAALGKRP*
>GCA_000452945.1_BPMSHR5858_genomic_05537_1 group_5206
MTRRSTKDNLGLRRPTIPVAYLRHLLTVFEERGIDARDARAGTGLRDDVLGEPDARIAPS
QWGRLVLNAIRLSGDEGIGFELGLRLRMSVHGFLGYAALTAADLRASLLVNARYFRMRNR
QYRLHYAERDDGATLSLHGVQASPVLQHHAMFEIVLTGIAQNVGTVAGRADPPIELQFAW
PEPDYYARYRDRLPPVRFGCAANAVWVSRDALDWPLPLADDVAHRQALAQVEREYADVRH

EEGDLVARVRAELTLTAHGYPGPDALARRLLMSTRTLRRRLLDAGCGYRALVDEARLRDA
RRLLRASDLDLKTIAARLQFNDPANFTRAFRKWTGLTPSAYREAGGGDER*
>GCA_000452945.1_BPMSHR5858_genomic_05540_1 benM_4
MELRHLRYFVAVAEEQNFTRAAERLHIAQPPLSRQIQQLEEALGVMLFERHARPLKMTDA
GRFFYSHAVQLLAQTAELEAMTKRVGKIERALSVGFVGSTLYGMLPRIIRRFRAEHGEVE
LSLHEMSTMDQIQALKDGRIDVGFGRIRHEDARVRRVVLREERMIVALPDGHPLEAAKPV
LSLHDLVNETMIIFPKSPRPSYADQVLAAFHDRALKPRRIYETRELQIALGLVAAGEGVS
VVPHSVYGLKRDDVGYKELDDPNLVSPIIMSTRMLDESEDIRAMLELIYRLYDEFKMTYL
PPHG*
>GCA_000452945.1_BPMSHR5858_genomic_05543_1 cbdB
MKTIDLADIQAFLYRESRLLDDKAWDAWLDCYRADAVFWMPSWDDADALVTDPQREISLI
YYPNRQGLEDRVFRIKTERSSATVPDTRTSHNIANVERESADGDVHTVRFNWHTLSYRYK
TVSSYFGMSRYAIDFSGDAPKIVSKYVVLKNDYINQLIDIYHI*
>GCA_000452945.1_BPMSHR5858_genomic_05545_1 group_3658
MNTHHTPGRFSGKVTIVTGAAQGIGRGVARRAAREGATAVLVDRADLVHEAAADIVAAGG
RALAVVADLETYAGAEAMARAAVDAFGSIDVLVNNVGGTIWTRPFDRYEAAQIEAEIRRS
LFPTLWCCRAVLPGMIARGGGAIVNVSSIATRSVHRVPYAAAKGGVNALTASLAFEHAAH
GVRVNAVAAGGTDAPPRKVPRNTEPQGDDDARWYRAIVDQTLSSSLMHRYGTIDEQVAAI
LFLASDEASYITGSVLPVGGGDQG*
>GCA_000452945.1_BPMSHR5858_genomic_05546_1 aes_3
MQGRAANRHAAFESFECSVHAPAQAARGARAVIRRAIEEQAMLAGMMLIGSAVTFGVWAG
RNPLVVLNAAARRPTFVGRFGLTYGAGPRRALDVYLPAAREPRPAGGGAPLVVFFYGGSW
QRGRRGDYRFVGEALASRGCVVAIPDYRLYPDAVFPGFVEDAAAAVRWARDHAAALGADP
RRIHVAGHSAGAQIATLLATDSRFLRAHGLDKRDLAGVVGLAGPYDFLPLEDATLKRIFP
EPVRDASQPIRFVDGREPPMLLASGLRDATVKPGNTVRFASRVAAAGGAVQVRLYPGIGH
ALLVGALGLPMRRFLPVLDDVAAFVRAAPRAPA*
>GCA_000452945.1_BPMSHR5858_genomic_05549_1 group_2625
MIRAARSALAARAGRAFDLPTLGRTLAVCAPTIALYGATGDARWLLASIATVSIAIAVER
VGIAPLGALAQSAAIVAGFLSLSCALSAWPAFVAGCATLAAAAVALSRCGARLRSLGNFV
FIPSLYLTCEAAAAHHSATRLVPYLGAAMLPAIALSGVQAWRNAPAAHRLHALARWRGAR
ELGRPAASSDAAQAIVAVALAVAAAALLVEWRHIDNGQWAIWSAASVVTGDAATARVKLR
DRGLGALAGVPLGLAAGHALPHGALVYTLATLASVLTLVAFRHYATGFGARCACIACASW
VAQQSTAIAAERVVNVLAGGLIGIACVLATHWFATRPLTRRHNARANRRA*
>GCA_000452945.1_BPMSHR5858_genomic_05551_1 lysP
MIAIGGSIGTGLFVASGASISQAGPGGAMLAYLLIGLMVYCLMMSLGEMAAFMPVSGSFA
TYGAKYVEEGFGFALGWNYWYNWAVTIAVELVAAQLVMHYWFPDVPGVWWSAAFLGVMFL
LNVLTVRGFGEAEYWFALIKVVTVIAFIGVGLLMIFGILKGGPSNGWSNFTIGDAPFAGG
LPAMMGVAMIAGFSFQGTELIGVAAGESENPRTTIPRAVRQVFWRILLFYVLAIFVIGVL
IPYTDPNLLKSDVTDVGVSPFTLVFRHAGLAFAAGVMNAVILTAVLSAGNSGMYASTRML
HNLATEGRAPKLFAKLSAGGVPRNALYATTAVGALCFLSSLYGDKTVYLWLLNTSGMTGF
IAWLGIALSHYRFRKGYVRQGFSVDQLPYQSSLYPYGPIFAFALCIVIALGQDYQAFLAN
RIDWIGVLATYVGIPLFLVVWLGYRLVHKTRVVRYEDMDIASWVAEHERAAAHEAREAAA
VRVPSSPEAAS*
>GCA_000452945.1_BPMSHR5858_genomic_05552_1 group_1954
VNAIVRRDIRFALPPDRICDWHTEGVGVTHLFNALSLLFPAGERFFMDSVRNYRDRIEEP
NLKRDIAGFIGQEAMHTREHVEFNDLLQAAGLPAHKLDKRLWTVLGWFKKVLPHSMQLAI
TMALEHYTAMFTGLVLANRITTSEVDGYRQMWMWHSMEETEHKAVAYDVWRTVMKPSLAS
YLLRTGTMLFVTLVFWATLFDFNTRLLISHRRRIGGRLGIGKLVKYLWGPRNGIFAQIAR
EWIDYFRPGFHPWDHDNSVHLERLDDLIAEIDASNARQAAKAAPRRVPLHPVPQAV*
>GCA_000452945.1_BPMSHR5858_genomic_05553_1 group_2626
MKLTFLGATETVTGSKYLLEDAGTRVLIDCGLFQGTKKLRLRNWSPLPVPADSIDAVVLT
HAHLDHSGYLPVLVREGFRGPIYCTHGTAALCEIMLIDSARLLEEEADFANRHGYSKHHP
ALPLYTTDDAKRTLERLAPRDFDAPSALAGDLHFRLLPAGHILGAASVVVCWRHTLLAFS
GDLGRAHDPIMRAPVPPVHADYLVVESTYGDRLHDASDPQTELADVFAKTFARGGVVVMP
CFTVGRAQEILYYIAQLKQTGRMPHVPVFLDSPMATSVTEIYRRRLSDHRLTMSDADAIG
HAAVMVRTVEQSKAIADHHGPMVIIAGSGMATGGRVLHHLARYAPDSRNTVLLVGYQAAG
TRGAALAAHEPTLKIHGEYVRVRAQVQMISSLSAHADYGEILGWLGAQSSAPARTFVTHG
EPAAADALRRRIEETLHWRCDVPSYLESVELEDAPTRAA*
>GCA_000452945.1_BPMSHR5858_genomic_05556_1 adhA
MRAMVFDGGAPRLREAELPDPVPAAGQLLIDVHACGVCRTDLHVVDGDLRDPKRPVIPGH
EIVGTVAALGALAKGFTIGERVGVPWLGRTCGHCAYCASGRENLCDAPGFTGYTIDGGYA
QRTVADARFCVHVPSRYDDIHAAPLLCAGLIGYRTLKMAGSARRIGLYGFGAAAHLVAQV
ARFQGRTVYAFTRPGDAAAQRLARRLGAAWAGGSDEAAPEPLDAALIFASVGALVPAALA
AVVKGGAVVCGGIHMSDIPSFSYDLLWGERRIVSVANLTRDDAAEFMRIANDVPLDVEAT
PYPLAQANQALDDLRAGRLAGAAVLTMR*
>GCA_000452945.1_BPMSHR5858_genomic_05559_1 group_13555
MGIGMQIAYFGFAGSAQLEAEASIQLIRLERFSARLAGCHLAIEALGAGAASPTYDVRLD
LITRGYELKPIPHMTGTDPLDAIRRAFDAAERELAAALA*
>GCA_000452945.1_BPMSHR5858_genomic_05560_1 group_2737
MNPPPPPRAKRRRTRAASASAPAPGMRPTPRRTRTAPKRRAPLSSTVERLVSSPTYRQAD
EDLAFLQRPEMCGVRLQLDYWKTEETLQRFGICDTVVVYGSTRIASPAVARARLADAQHR
LAERPNDPERRHAVSVAVRLLERSHYYGVARDLGRLVGETGRSPHPRRLTIITGGGPGIM
EAANRGAHERGAPSIGLNITLPREQFPNPYVTPELCFRFHYFAIRKLHLLERAKAAVFFP
GGYGTCDELFEVLTLLQTRKIAPLPVVLVSRAFWRSAVDFGFLVDEGMIDPCDAALFRFC
ETADEIWAAIGGPHGPA*
>GCA_000452945.1_BPMSHR5858_genomic_05564_1 cysC_2
MSASSPSRARPGARPHDGAPAPVRDVRSAPRRDASAPENLFWQAGRVAADARSTQFGRPP
LTLWLTGLSGAGKSTLAGELERLLLRDRWPCAVLDGDNLRHRLNRDLGFGAEDRHENIRR
TAEVARLMNGVGLVVIASLISPFRDDRAMAREIVGSDWFSEVYVSTPLDECERRDPKGLY
RKARGGLLPNFTGVSAPYEPPVSADLAIDTTRLSVDAAAAMLRAHLIARCAGRHVD*
>GCA_000452945.1_BPMSHR5858_genomic_05570_1 group_2627
VRQRIVFVAAIVGFLASLAAAWVYSIQEPPQPPVFKPASNPYPRGVYATGIVESDQSSGA
NVNLYPEVTGTVTRIFVREGDAVKAGDALLAIDDAVQRATAAQLAAQADAAQALLDEIRA
QPRRETLDVAAAQVEAAAASLKTAQDQYDKQQRAFAIDPRAVSRDALDNALNGVKLARAN
LGVLTRQYRLTRAGAWVYDVRNQEKQAVALRKAADAAAALLARYTLRAPADGVVLAMNAA
VGSYLSPQGMYDTYTQGATPALVLGSPANHLQVRCYVDEILISRLPAGKMPSAHMSVRGT
AMEADLAFVRVQPYVTPKIQLSDQRAERVDVRVLPVIFRVVPRKDLRLFPGQIVDVYIAS
G*
>GCA_000452945.1_BPMSHR5858_genomic_05573_1 group_1250
MTHPDRKAIASTVAVFASQIGLIAAVRGLFVADGSAPYFVLFAIVAGVLGFVDLYYRPIF
APAFARQRVAFVHRRRGAARDVS*
>GCA_000452945.1_BPMSHR5858_genomic_05576_1 oprM_5
MRNDARHDERHQRRRGIARRAARHAAALALCAAVSGCLSLAPDYARKPPPVPAAYPAYPG
EPGAGSRTLEQLGWRAYFVDPRLQALIETALANNRDLRIAAQRVEQARAAYGIRRADQYP
TIAANAAYLRFREPGGLLLPSPIIGELYSASLSEVQWEIDFWGRVRSLKAAALDTFLASD
ASRRAVTVSLIGQVADAYLALCAYDERLALTNETIASRRDSLRIFRRRYEEGAISKLDLT
QSEILLQQAQTLGSQLQQARDVQQHALDLLVGTNAAARTPASLDDASIAPDLAPGLPSEL
LANRPDVVAAEYQLRAAHANIGAARAAFFPRIALTSSIGTGSTELHNLFASGTAAWNFIP
NLSVPIFDAGRNIGNLKLANARRDEALAQYEKTIQTAFRDVADALAARHWLADQVRIAQE
TLATQAERARLAKLRYDSGAARFLEVLDAQRDLLNAQQQLVQTRRALLASRVALYAALGG
RLDEPARDGAAPRPSPEPSAKQGPLS*
>GCA_000452945.1_BPMSHR5858_genomic_05577_1 group_5211
MIQVTRKQIVAAAAVVVVALGAYYGWTLLRHQGPGDGFASGNGRIEATEIDVATKLPGRI
DAILVDEGDFVKAGQPLANMQIQVLRAQYDEATAQRQRALNTAAGVQSQVAQRKSDKAAA
QAMVVLRESELDAAERRLARSQTLSREGASSLQELDDDRARARSAQAAVSAAAAQVAAAQ
AAIEATEAQLVAAHSAVTAADATVARVQADIDDSQLASPRDGRVQYRVAQPGEVLPAGGK
VLNLVDLSDVYMTFFLPETVVGRVALGADARIILDAAPNYVIPATVSFVSSTAQFTPKTV
ETANERQKLMFRVKARIDRELLQKHLKLVKTGLPGVAWVRVDPSKPWPAQLMVKVPQ*
>GCA_000452945.1_BPMSHR5858_genomic_05578_1 btuD_15
VSAPSVVRFSDVSLRYGKTVALERITLEIPAGLTIGLIGPDGVGKSSLLALAAGARALQT
GAVDALGGDMRSRRHRERVCRRIAYMPQGLGKNLYPTLSVEENLQFFARLFGHDADERRR
RIDALTRSTGLFAFLSRPAGKLSGGMKQKLGLCCALIHDPDLLILDEPTTGVDPLARAQF
WDLIARIRSERPAMSVIVATAYMDEAQRFDWLIAMDAGRVLATGAPAELLARTGCDSLEA
AFIALLPEDERRGHKPVTLEPLRADAQTGTAIEARGLTMRFGDFTAVDHVSFRIRRGEIF
GFLGSNGCGKSTTMKMLTGLLPATEGTAQLFGKTVDPKDINTRRRVGYMSQAFSLYSELT
VRQNLVLHARLFGVPAAEIDARVDEMARRFGLADIYDMLPDSLPLGMRQRLSLAVAMVHK
PELLILDEPTSGVDPVARDSFWQLMIDLARRDRVTIFISTHFMNEAQRCDRISLMHAGRV
LASDAPAALVRARGAATLEEAFIGYLVDASAQEAQDAQEAQGGAGERAAGESLAGRGAAT
DGDADGAHARPAAYANGAAQHRAPDADAAGTDVAAETGARDDARTAHAAGHSAGNDVRAT
VGSEVGSEVGSDVGSEVGSEVASEAGLAASAESGSERGVAHGAERAREPAADAGAAIDVE
PGASAESATAAVAPARHAESAATSPRAGNAPAGTPFAAAPAEPPHRAFSAQRALSYMWRE
MLELRRDPVRATLALIGSLVLMCVIGIGISLDVEDLTYAVLDRDQTELSHDYALNLSGSR
YFVERPPIADYAALDRRMRDGELSLAIEIPPNFARDVERGAPAQIGMWIDGAMPQRAETI
RGYAIGMHTMWLADKARHRLGVTLAPRAEVVTRYRYNPDVKSLPAMIPAVMPLLLLMLPA
MLTALAVVRERELGSILNLYVTPVTRTEFLIGKQVPYVVLAMLNFLLMTMLARIAFDVPV
KGSFMTLLLAVLIFNVVATGIGLLASTFTRSQVAAIVMTIIGTMIPTVQFAGLLTPLSSL
EGTGRLIGLVYPATYMLSISRGVFNKALSLHDLYSQFWPLAACVPVILGATILLLKKQER
*
>GCA_000452945.1_BPMSHR5858_genomic_05579_1 yhhJ
MRRLAIVYRLGVKELWSLVRDPILLVLIAYTFTASIYSAATARPDTLHMAPIAIIDEDAS
PLSARIASAFFPPQFALPRMVTLAEADRGMDAGDFTFVLDIPPNFQRDVLAGRPAQIQLN
VDATRMSQAFTGNGYVQQIVSGEIDDFARRYRANTAPPVDIAMHMRFNPNLDQTWFGSLM
EVINNVTLLSMVLTGAALIREREHGTIEHLLVMPVTPGEIMLAKVWSMGLVVACAAAFSL
AFVVRGALHVPIEGSVALFVIGMAIHLFATTSMGIFLATLVRSMPQFGMLLVLVLLPLQL
LSGSLTPRESMPFAVQNIMLAAPTTHFVEIGQAILYRGAGIETVWRQFLILAVIGTALFL
LSLKRFRKTIGQMA*
>GCA_000452945.1_BPMSHR5858_genomic_05580_1 rcsC_9
MHVEVFAQHPSCPGWHGEMARRIRAFDWSATELGPLERWPASLVAAVRTVLASPVPLVML
WGASGYMLYNDAYAAFAGGRHPYLLGQPVELGWPEVADFNRNVMRTCLAGGALSYRDKAL
VLLRNGRPEEVWMDLYYSPLAGDDGAPAGVLAVVVETTERVLAARHRDRAKAALHETNAE
LRRLTETLEQRVADAITERASIEEQLRQAQKMEAIGSLTGGIAHDFNNVLQVVSSNLQML
SAELGEDARAQRRIASAANAVARGAQLASHLLAFARRQPLAPAVLEPRGLIDGMREMLHR
ALGESVHVDADLRDDVWNVLADRHQFENALLNLAINARDAMRGEGTLKITACNIVLEAGR
HARRNGLPPGEYVAFDIADNGVGMTPDILERVFEPFFTTKPDGQGTGLGLSMVFGFVKQS
GGHTSIESAPGRGTTVRLLLPRCREAPAGEPAAQRRLPAEGGRETILVVEDDADVRLAVV
DMLAQLGYKVITASDGEAGLLALESGAPIDLLFTDVIMPGRIKGGELARRAAQRVPPVPV
LFTSGYTRDEIFHAGRLDPGVMLLGKPYRRDELAHKVRSVLDAHASARRAHGARHAQADP
RKKTAAARVQR*
>GCA_000452945.1_BPMSHR5858_genomic_05581_1 tar_3
MPTLSFRQKLWLPLVISLIALLLVSISSAWLSYQTRMEERRNDLTNVAHVGLSIVREYAA
LAQRGVLTDAQARKEALERLRSVRYGSDGYFIVIDSSPRMIMHPIKPDTVGKDLRHVADA
DGRHHYLTFASVAQSPQGGFVDYVFAHPNAHPAKAVDKLGYVIRYAPWDWIISTGAYIDD
IQAAFAKSLCLSAGVFALLAALLALSVVYTNRGIERMIGGDPRIAAHMAGVIASGDLATP
FATRREDRSSLMFAISQMRDALANAVAQIRASAGYVATTAGEIADSNMDLSSHTECQATA
LQQAASSMQRLTERVLDTADRARAASELAGSAARITDRGGDMVVRVVAAMNDIRAESRKM
VDIIGVIEGIAFQTNILALNAAVEAARAGNEGRGFAVVASEVRVLAQRSASAAKDIRALI
GRAAERVGNGAELVEATGATIGEAQHAICRVTGIVQDIAAAATDQSQGLEQINAAVSQMD
SVTRRNATLVEHAAIAAQSLNEQSRCLQDAVAAFRTEALGC*
>GCA_000452945.1_BPMSHR5858_genomic_05584_1 group_2630
MAQYMKAAVVHAFGEPLRIEEVPVPTPGAGQILVNVKASGVCHTDLHAADGDWPVKPTLP
FIPGHEGVGFVAAVGEGVRHVKEGDRVGVPWLYTACGHCEYCQTGWETLCHEQQNTGYSV
NGSYAEYVLADPNYVGHLPSNVAFDEIAPILCAGVTVYKGIRVTDTRPGQWIAISGIGGL
GHVAVQYAKAMGLHVVAVDVAPQKLELARKLGAAFVVDASKDDPAAVIQKEIGGVHGVLV
TAVSRGAFAQALGMVRRGGTVSLNGLPPGDFPLPIFSTVLNGITVRGSIVGTRRDLQESL
DFAAEGLVRAHIHRDKLEHINGVFSALREGKVDGRIVLTGQ*
>GCA_000452945.1_BPMSHR5858_genomic_05585_1 atpG_2
MSDKLAAIEARTDTARQLQTVIGAMQGVAAARAHEAQQRLPGIRASAATVGAAIGDALSA
GARSGDTAAGARAAPRARLVVVLCSEQGFVGAYNAQLIEHATRPEPSVSREYMMVGSRGA
MLADAGGVPLVWRTPMAARADDVVHLANRITDALYAHLAKRGAQPVSIVHAMPGGAARLE
VTERRLLPFDYARFDTAPRAQPPLVHLPPATLLAELAQAYVFVELCEAAMLAFAAENEAR
TRAMIAARESVERALGDLLQAYRIARQDEITADIVELAASGL*
>GCA_000452945.1_BPMSHR5858_genomic_05597_1 rplC
MSLGLVGRKVGMTRIFTAEGDSIPVTVLDVSDNRVTQIKTVETDGYTAVQVAFGSRRASR
VTKPLAGHLAKAGVEAGEILKEFRIEADKAAELSNGAVIGPDLFEVGQKVDVQGVSIGKG
YAGTIKRYNFGSGRASHGNSRSHNVPGSIGMAQDPGRVFPGKRMTGHMGDETVTVQNLEI
ARIDADRKLLLVKGAVPGAKGGKVFVTPAVKTRAVKGAK*
>GCA_000452945.1_BPMSHR5858_genomic_05654_1 thpR
MTMAGDRLRCFVALTLDRASRDALAALPVAAGARRTPRDQLHVTIAFLGAIERAKSELLG
ARIPALAAAGAVPPVDVERVVCWPSTAHARLVVAELAPQPSLLALGERVGGALRALGLPP
DSRAFKPHVTIARFPRDAHRVTIDGACGAERRSPFALRFETLTLYESVLTRMGAEHRVLA
AAALPA*
>GCA_000452945.1_BPMSHR5858_genomic_05714_1 qseC_3
MATPVRSAHASRRASAAAPDTDAARDARYENPFAPPDEADAAEGTRPRSLFGEILDWMLA
PLLLLWPMSIAVTYLVAKTIANGPFDRALETDAYVLARQITPINGVAELRLPQATLDFLR
ADNVDSLYFQVLGTRGELVAGEADLPLPRDDDRPAPGVVVFRDDLLRGNDVRVAYTSVAL
QGATGSQPVLVQVGETLDKRNALANDIIKGVILPQFVILPLAILLVWFGLSRGLAPLTAL
QAHIRGRRPDDLSPVEARRAPPEIEPLVTSFNDLLARLEQNITLQKRFIADAAHQMKTPL
AGLRTQAEFALRHEVNADVARSLEQIATSSEQAARLVTQLLALARAENRATGLTFEPVEI
ASLARQAVRDWVQAALAKQMDLGYEGPDTDAPLRIDGQPVMLREMLGNLIDNAIRYTPAG
GRITVRVHAERAAGAVHLEVEDTGPGIPPNERERVVERFYRILGREGDGSGLGLAIVREI
VAQHGGTLTIDDNVYQTSPRLAGTLVRVSIGLRQEAPE*
>GCA_000452945.1_BPMSHR5858_genomic_05716_1 recA
MEESKKGSGLTAEKSKALAAALAQIEKQFGKGSIMRLGDGEAVEDIQVVSTGSLGLDIAL
GVGGLPRGRVVEIYGPESSGKTTLTLQVIAEMQKLGGTAAFIDAEHALDVQYASKLGVNV
PELLISQPDTGEQALEIVDALVRSGSIDMIVIDSVAALVPKAEIEGEMGDALPGLQARLM
SQALRKLTGTIKRTNCLVIFINQIRMKIGVMFGNPETTTGGNALKFYSSVRLDIRRIGSI
KKNDEVIGNETRVKVVKNKVSPPFREAIFDILYGEGISRQGEIIDLGVQAKIVDKAGAWY
SYSGEKIGQGKDNAREFLRENPEIAREIENRIRESLGVAAMPQGAGSEAEIMDEEE*
>GCA_000452945.1_BPMSHR5858_genomic_05718_1 group_7705
MPLSEPVPRQLRHRRAIRAEAYERADGLWDIEACLTDHKPRDVSLASGVRPNGLPIHELW
LRVTIDRELNIVDVDASSDWVPYPGQCEAAKPAYRVLIGLNLFRHFRRDVSRLLSGVAGC
SHLTELCAVLPTAAIQAFAGDVWNVQESLGDGANEGDRPAGQPPFQLGRCHALRFDGEAV
RQFYPRWYGASRPAREGADSAKE*
>GCA_000452945.1_BPMSHR5858_genomic_05719_1 sucC
MKIHEYQGKEILRKFGVAVPRGKPAFSVDEAVKVAQELGGPVWVVKAQIHAGGRGKGGGV
KVAKSLEQVREYSNQILGMQLKTHQTGPEGQKVNRLLIEEGADIKKELYVGIVIDRVSQK
VVVMASSEGGMDIEEVAEKTPEAIHKVAVEPSVGLQDAEADDLAKKIGVPDASIPQAREI
LKGLYKSFWETDASLAEINPLVLTGDGKVIALDAKFNFDSNALFRHPEIVAYRDLDEEDP
AEIEASKFDLAYISLDGNIGCLVNGAGLAMATMDTIKLFGGEPANFLDVGGGATTEKVTE
AFKLMLKNPGLKAILVNIFGGIMRCDVIAEGVIAGSKAVNLNVPLVVRMKGTNEDLGKKM
LAESGLPIISADSMEEAAQKVVAAAAGK*
>GCA_000452945.1_BPMSHR5858_genomic_05720_1 sucD
MSILINKDTKVITQGITGKTGQFHTRACREYANGREAFVAGVNPKKAGDNFEGIPIYASV
KEAKAETGATVSVIYVPPAGAAAAIWEAVEADLDLAICITEGIPVRDMIEVKDRMRREGR
KTLLLGPNCPGTITPDELKIGIMPGHIHRKGRIGVVSRSGTLTYEAVAQLTALGLGQSSA
VGIGGDPINGLKHIDVMKMFNDDPDTDAVVMIGEIGGPDEANAAQWIKDNMKKPVVGFIA
GVTAPPGKRMGHAGALISGGADTAEAKLEIMEACGITVTRNPSEMGRLLKAAL*
>GCA_000452945.1_BPMSHR5858_genomic_05721_1 group_7708
MFEFFASLHWGAVAQIVVIDILLGGDNAVVIALACRNLPAQQRVRGVLWGTAGAIVLRVV
LIAFAVLLLDVPLLKFAGGVLLLWIGVRLMAPAEDAHENVKPADKLWEAVKTIVIADAVM
SLDNVIAIAGAAEQADPGHRIALVIFGLLVSIPIIVWGSTLVLKLLDRFPIFITFGAALL
GWIAGGLMVNDPAGDRWPLLDTPAAIYGASVAGALFVVIAGYVLKKRRTASSATHSR*
>GCA_000452945.1_BPMSHR5858_genomic_05723_1 group_3669
MPSSFLRSLSLIALAVALILPYAITNHTYPIPTFYSEFAAFALYWGLGASVVLLVKAERA
EQPFAAPMALVAPLGFGAVLLAQIALLPLRQPSMNWLAMGYLLGALVAMQAGYALARVNM
VDMVARMIAGATIVGGVVAVACQFVQLFHLETTFSPFVVSYGVTVERRPYGNMAQANHLA
TYIAFALAGALYLVQTRRMPALAWAALSALLSVGLALTVSRGPWLQVGVMVVAGFWMAFA
QARRDPAASRARAWAIPVVLGVLFVAVNVAVRWANVHYHLGLAESAADRMRDAGQIAPRL

ALWKYGLTMFREHPLLGVGWGEFPIHQFELARRLGGVEIANNSHDIFIDLLAKSGLLGLG
VLFVALVAWFVRALRVPHTESRVFGFALVGIVLMHALVEYPQQYTFFLLPVMFVIGLLET
KPLRALPGRAAFVLFAALSVAGLASLYPILRDYQRAEVLYYGTNPAEQYRAHQSFLFGAW
GDYGAATLLAISRENLPAKLAAHESAIALLPGETVLRRYAVLQALDGRETDALDTIERLR
VFAEELHDWPVQLAALYKLLDDQPSLKSFKAALVAKYGTPAANLSADDEEDDGDD*
>GCA_000452945.1_BPMSHR5858_genomic_05724_1 group_4090
MKKRIHRIAAAGILVAGVLSLSAAHAQLGDFLKQGADAGNGGAGGIAGALGNLGGGGGAA
SAGSLLTPGSTGNVAGLLQFCVKNDYLGDGGASSIKDALMSKLGAGVTSDSTYASGASGI
LDAGNGRTLDLSGGQSFKQQLTKQVCDKVLSQAKALL*
>GCA_000452945.1_BPMSHR5858_genomic_05727_1 group_1563
MRVKSSFAVLLCAALALPPGGHAQSRGDAPPLESARAAGAEDAAARARDALSTVPSGIAP
GVFGMYGGAQSRLADPASGTPSLRAPLRSLQLPDLGDGSGGSLTPQAERRLGERVMREVR
RDPDYLDDWLVRDYLNSVAAKLSAAAAAQFIGGYMPDFELFAMRDPQINAFSLPGGFIGI
NSGLVAATQTESELASVIGHEMGHVLQRHIARMIGASEKSGYAALATMLFGVLAGILARS
GDLGSAIAMGGQAFAVDSQLRFSRSAEREADRVGFQLLAGAGYDPYGMPGFFERLERASV
GDAGVPAYARTHPLTGERIADMDDRARRAPYRQPRQSAEYGFVRARLRMLQNRAPTDYAN
EARRMRAELDDRVAPNVAANWYGIALGEMLGGRYDDADRALAAARDAFARTAAREGEAAR
TSPSLDVLAAEIARRAGRGDDAVRLAAAAQARWPGSHAAIAAHLQALLAARRYGQAQALA
QAEANAAPRQPDWWNYLAQASLGRGDALTQRRALAEKFALEGAWPSAIRQLREARDLKSA
GFYEQSIISARLHEFEARYKEEREEDKDDRRG*
>GCA_000452945.1_BPMSHR5858_genomic_05728_1 group_5762
MDDIVKQALAKWPNVPHCTGWLLLDRRGGWRMRDDAAQAAGALGSPVRHPALVDFIARNY
ERDEHGQWFFQNGPQRVYVELAYTPWIVRLAAAPAGGGCEPARVALTDHTGRAFEPARAW
LDDAGGVLFADTSTPPRVAALHDHDLALFADRADLDDDGAHGVLHLGGGVDLPLEPIRRD
DVARRFGFVASPAARAGERAAGGKPTD*
>GCA_000452945.1_BPMSHR5858_genomic_05729_1 group_3670
MSTALPPLPAADETARPAASSFYRAPLWLPTSHAQTIVPALFARRPDVAYRRERWDTPDG
DFIDVDWLAPPAGAAPGPLAPLAVLFHGLEGSSDSHYARVLMAAARARGWHGVVPHFRSC
GGEMNRMPRFYHLADSAEVDWILRRLAKSHRGPLVVVGVSLGGNVLLRWLGERRGDTSIV
AAAAAVSTPIDVHAGGRALSQGFSMVYTRSFLKTLKWKALVKLEQYPGLFDKQALLAART
MHDFDDIVTAPLHGFADANDYWTQATTRPLLAAIDVPTLILNARNDPFLPASALPGPREV
SRAVVLDQPEHGGHAGFMTGPFPGRIDWLPARVFEFCSRFTDHG*
>GCA_000452945.1_BPMSHR5858_genomic_05730_1 group_1959
MPRFARLFEAAADTLNAYYQAVADANLDALMVLWIDEDFASCIWSDGTHLHGLEQIRSGF
GARLSTQPVTIEPLDIRVYDSLGTVVYTVAEAHQQADLTAEPQMVFTTYVMIHERGEWRI
AHIHASPIPEQTATQFAAKIRHAQGPLH*
>GCA_000452945.1_BPMSHR5858_genomic_05731_1 rfaF_2
MRRALVIAPNWIGDALMAQPLFALLKKLHPRIVIDAVAPAWVAPVLERMPEIHDVHATEL
AHGKLQMLHRWQLASDLRELGYDAAYVLPNSLKSALIPWLAGIPLRIGYTGEHRYALLNV
RHANPGKARETRAPMAQHYAALAYAPGAKLPESFQTLPPPRLDADLNETARVSARFNLDT
RKPLIVFCPGAEYGPAKRWPPEHFAALAQSVSQSFPYTQIVALGSPKDAAAAQAIAERAP
NVRSLCGQTSLTEACALIARANAVVTNDSGLMHVAAALRRPLVALYGSTDPRHTPPLSEL
AKVQWLHLECSPCFERECPLGHLKCLRELSPEQVFGDLRGMLVGQR*
>GCA_000452945.1_BPMSHR5858_genomic_05738_1 cbbA
MPLVSMRQLLDHAAENGYGLPAFNVNNLEQVQAIMAAADEVGAPVIMQASAGARKYAGEA
FLRHLIEAAVESYPHIPVVMHQDHGQSPAVCMAAIRSGFTSVMMDGSLEADGKTVASYEY
NVDVSRKVVEMAHSIGVTVEAELGVLGSLETMKGDKEDGHGAEGTMTREQLLTDPEQAAD
FVKLTQCDALAIAIGTSHGAYKFTKKPTGDILSIQRIKEIHARIPNTHLVMHGSSSVPQD
LLAEIREFGGDMKETYGVPVEEIQEGIKHGVRKVNIDTDLRLAITGAIRRYMFENPGKFD
PRDYLKPAREAAKKICVARYLAFGCEGQAAKIKPVPLEKIAEKYKAGDLAQVVR*
>GCA_000452945.1_BPMSHR5858_genomic_05743_1 group_7709
MNPLQRQQQPAGAARRRFWRGAQVALASAAFALLAACGGGDDNGSSQPSAGVNMQVVSFG
DSLSDVGTYSPQILIGFGGGRFTTNPGQVWTQDVAAYYGGTLTPAFEGGFGVPLQAAGGL
GYAQGGSRVTQQPGIGHADASVANADYAQATTVPVATQVQQYLQQHGSFNANQIVLVNGG
ANDIFYQVQVAQAQGNTPAAQLAAAQQIGLAAQQLAGVVQQIVAAGATHVFVSNVPDIGG
TPLAASTGQQAALTQLSTIFNSTLVAALKALNVDPAKAVLIDAFTWQDGIAANYQGNGFS
VANTGTACNLQSMIAAATKAGVANPTAFGSSLFCSPQMYTVANADQTYMFADTVHPTTRL
HALFAQYVEQQIAKTGVGK*
>GCA_000452945.1_BPMSHR5858_genomic_05747_1 sasA_10
VRSIRHQLLIWLLAIVVAGVGLAGWMIYRQALAAANELFDYQLQQIAAALPSEPFSQVLG
SQTNGDEGIVIQIWNRNGVLMYFSHPRAPIAPRAELGFSTERTERGAWRVYGAIVGDNVV
QLAQPLSVRNRLAASVALRTLWPLIVLLPFLGAAVWMIVGRGLAPLQRVTRAVEARRPEA
LDPLPDSPLPLEVRPLVRALNGLLARLSAALDTQKAFVADAAHELRTPLAAVQIQAQLVA
RAQDDVSRREAIVDLQSGVTRATRLAEQLLALARAEPGDATVRERVDLRAIVEECVAAHA
PLAQRRDIDLGFERVEDASVDADPAALRVMFNNLLDNAVKYTPAGGRIDVSLTRGEGARA
CVQIGDSGPGIPAAERERVFDRFYRDTSARARTDVAGSGLGLAIVKRVAAQQRASVTLGE
AAAGGLLVSVSLPCAGPA*
>GCA_000452945.1_BPMSHR5858_genomic_05754_1 acrB_2
MAKFFIDRPIFAWVIAIILMLAGVAAIFTLPIAQYPTIAPPSIQITANYPGASAKTVEDT
VTQVIEQQMSGLDNFLYMSSTSDDSGNATITITFAPGTNPDIAQVQVQNKLSLATPILPQ
VVQQLGLSVTKSSSSFLLVLAFNSEDGSMNKYDLANYVASHVKDPISRINGVGTVTLFGS
QYAMRIWLDPTKLTNYGLTPVDVTSAISAQNVQIAGGQLGGTPAVPGTVLQATITEATLL
QTPEQFGNILLKVNQDGSQVRLKDVAQIGLGGETYNFDTKYNGQPTAALGIQLATNANAL
ATAKAVRAKIDEMSAYFPHGLVVKYPYDTTPFVRLSIEEVVKTLLEGIVLVFLVMYLFLQ
NLRATIIPTIAVPVVLLGTFAIMSMVGFSINVLSMFGLVLAIGLLVDDAIVVVENVERVM
AEEGLPPKEATRKAMGQITGALVGVALVLSAVFVPVAFSGGSVGAIYRQFSLTIVSAMVL
SVLVALILTPALCATILKPIPQGHHEEKKGFFGWFNRTFNSSRDKYHVGVHHVIKRSGRW
LIIYLAVIVAVGLLFVRLPKSFLPDEDQGLMFVIVQTPSGSTQETTARTLANISDYLLTQ
EKDIVESAFTVNGFSFAGRGQNSGLVFVKLKDYSQRQSSDQKVQALIGRMFGRYAGYKDA
LVIPFNPPSIPELGTAAGFDFELTDNAGLGHDALMAARNQLLGMAAKDPTLRGVRPNGLN
DTPQYKVDIDREKANALGVTADAIDQTFSIAWASKYVNNFLDTDGRIKKVYVQSDAPFRM
TPEDMNIWYVRNGSGGMVPFSAFATGHWTYGSPKLERYNGISAMEIQGQAAPGKSTGQAM
TAMETLAKKLPTGIGYSWTGLSFQEIQSGSQAPILYAISILVVFLCLAALYESWSIPFSV
IMVVPLGVIGALLAATLRGLENDVFFQVGLLTTVGLSAKNAILIVEFARELQQTEKMGPI
EAALEAARLRLRPILMTSLAFILGVMPLAISNGAGSASQHAIGTGVIGGMITATFLAIFM
IPMFFVKVRAVFSGEKEDADEALRLAHEHMHRDDKPEHGDDAGKKD*
>GCA_000452945.1_BPMSHR5858_genomic_05762_1 group_7710
MLKKLLMLVVALSLSLTAALAAAVEVNTADQAALESVKGLGPVKSKAIIDERAKNGPFKD
ADDLANRVKGLGAKSVAHLEAAGLTIGGSTTPPTGAKAKTGTAKPAATATPAPAPAPAPT
TSSATPAPSAATPAPATATPSTAASAPAAKKSRSSKKQDKAAAASAAAQASAPAAASTTK
AKGSKKSKKNKAATPASAASGA*
>GCA_000452945.1_BPMSHR5858_genomic_05765_1 norM_1
MFGDIRRIAGLAWPVLVGQLAIIAFGVIDTAMVGRFSALDLAALGLGMSIYVSIYIGLTG
ILSALQPIAGQLYGARRYAEIGEEVRQAMWLAVLLAAPGFLLLHFPHPLLRLAHAPAELT
DRTVAYLRILSFGLPASLLFRIYNALTNAAGKPRLAMILQVGALAVKVPLNLWFIFGGLG
VPALGGPGCGLASTLINWALAIVGFTLLARLDVFEPLAIFARFTRPVWHRQKALLVLGVP
MGLSYLIEVTSYTFMALFIAQFGTTTLAGHQIAGNIGAVLYMTPLSIGVASSTLVARALG
AGRFDEARALGRHGVALACGLACAYGIVVFALRPLIVSGYTPNPAVAAAAMPLVAIVTVY
HFFDALQITTAFVLRAHKVAVVPTAIYAIALWGVGLGGGYLLGFDVGGAVPAQFTGARGF
WIASAASLMLAGGGLALYLRHIGKRETGGAR*
>GCA_000452945.1_BPMSHR5858_genomic_05766_1 group_7711
MKPVVRLTASATRALPRWLLLTLCIVYAAFGLFGRDPWKNEDAAGFGVMWTMAQGGLHDW
LLPNLVGKFVTSDGPLGYWLGGLAIRALPWVDASNASRVYTGVLFCVACAFVWYAAYLLG
RRAEIQPFKYAFGGEPEPRDYGRTLADGALLVLLACFGLAERGHETTPQLAQFACIATFV
YGLVRAIDKPTQGALWWGLALGLVALSGNPVLVAALALGTLALYLVTPEIRCVQLPAIGL
PLAVAVFAIWPLAAYIAFPDDANWFFNQWLHGSLMRFSGPPTTVLAYAAKNLPLFTWPAW
PLAIWAWVSWAGLRRRPHIAIPLSVAAPLLALVILQSQQTNRMYMLLLPALAVIATFALP
TLKRGAINAIDWFAVLSFTILGTFVWLVWLASLTGFPHPLARNLGRLVPGYEPHFKVLSF
VCAVAATACWLMLVRWRISRQPKVLWRSVVLSSAGTTLMWVLLMTLWLPIVNYSRTYRDV
AQQIAAHLPSDYECISPVRLGDAQIATFAYFGDMHFSFTDDCDVILRQDRADFGEPSSIS
QYVWRLVWEGRRVADRDERFRLYERIERPKTPVKRRPPRKAR*
>GCA_000452945.1_BPMSHR5858_genomic_05767_1 rpmE2
MKQGIHPDYREVVFQDMSNGFKFITRSTIQTRETIEFEGKTYPLAKIEVSSESHSFYTGQ
QKIMDTAGRVEKFKNKFGARASGKAAK*
>GCA_000452945.1_BPMSHR5858_genomic_05768_1 mtfA
MLSKLNQWLDARRRDRALRSHPIPDVLWRETLERLPFLSYLSDEDRARLRETTSLFLAQK
SFSTAHGLELTDAIVVGIAAQACVPVLNLGLGLYRGWVGIVVYPGEFVIRKTVEDEDGVV
HEVEHDASGEAWEGGPVILSWEDVQMADGSDAYNVVIHEFAHKIDMMSGAADGHPPLFRR
WHAPALDAERWADVFDHAYDQFCDRVDAVPERAWARFERDSLIDPYAADHPSEFFAVCSE
ALFVKPRPFEAEFPELYRLLARYYRQDPARTGCLAGSAAHV*
>GCA_000452945.1_BPMSHR5858_genomic_05769_1 farB_2
MSAAPGRPPLWSAANLRGDFFPWVLAIVTGLDYFDNAAFSFFASYIAGGINASPDELVWA
SSAYAVTAVLGILQQQWWVDRLGHRRYVAGCMLMFSLGAMAAAAADTSLQLAFARGFQGY
FIGPMMGACRILIQVSFAPKDRPPATRAFLIMLLLGSALAPIAGGLLVAHFTWRALFACT
VPAGILFAALAFVALPDSGHTPPDERGGAHFWPYVIFALAQGALQIVMQQVRFQLFAGSP
LLVLLAAGGLAALAWFGHHQWHHPAPLVRLHAFRERTFRVGLLLYLFYYYETTGYSYLIS
RFLETGLGYPVENAGRLVGTMSLISASALFVYLRYAKLLTHKKWIIVPGFALAAFAALWM
TRMSPEVGEAALVAPLLMRGLLLLFIVLPVANLTFRVFAIDEYSHGYRLKNIVRQLTISF
ATASVIIVEQHRLAVHQTRLVEQANVYNPLFRQTLDTLTRGFAAAGHAFSDAHALALVTV
SRTIAQQASFLASLDGFYFLAGVAICGGLFAAWQKEID*
>GCA_000452945.1_BPMSHR5858_genomic_05770_1 tipA
MSDAPPTLLLTVRDAAERLGVTPRTLKYYEERGLVTPSRSEGRYRLYDAADLERFERILR
LRALGFSLHGITEMLKRPLEPAGDGRRRYSDASLRDIHADLSQQIATLDARIAAAQRELK
EVRALKQELQHDIDYVERRLAGENPDELIAQRRAAARARKTGGTSA*
>GCA_000452945.1_BPMSHR5858_genomic_05771_1 rho
MHLSELKSLHVSELIEMANGLEIENANRLRKQELMFAILKKRAKTGETIFGDGTLEVLPD
GFGFLRSPEMSYLASTDDIYISPSQIRRFNLHTGDTIEGEVRTPKDGERYFALVKVDKVN
GQPPEASKHKIMFENLTPLHPNKPLSLEREMRGEENVTGRIIDMIAPIGKGQRGLLVASP
KSGKTVMLQHIAHAIKQNHPDVILFVLLIDERPEEVTEMQRSVAGEVIASTFDEPATRHV
QVAEMVIEKAKRLVEMKHDVVILLDSITRLARAYNTVIPASGKVLTGGVDANALQRPKRF
FGAARNIEEGGSLTIIGTALIETGSRMDDVIYEEFKGTGNMEVHLERRLAEKRVYPSINL
NKSGTRREEMLIKPEILQKIWVLRKFIHDMDEVEAMEFLLDKIRQTKNNAEFFDLMRRGG
*
>GCA_000452945.1_BPMSHR5858_genomic_05772_1 trxA
MSEQIKHISDASFEQDVVKSDKPVLLDFWAEWCGPCKMIAPILDEVAKDYGDKLQIAKIN
VDENQATPAKFGVRGIPTLILFKNGAVAAQKVGALSKSQLTAFLDSHL*
>GCA_000452945.1_BPMSHR5858_genomic_05782_1 group_5237
MRLISWNVQWGRDANGTVDLARTVDAARGLGDFDVLCAQELTRGFHALPGGPSGDQYAEL
AALLPGYAVFDAIGVDLPPLEPGAPRRQFGNALATRLPVERALRHALPWPASAAAPSMPR
CALEVTLRAPFGPLRVVVTHLEYYSAQQRLAQVDALRRLHREAAAHARASAPPETPASPF
APSARAADAIVCGDFNSAYRSDAYQRFLAPFPDAPRFVDAWLARHPGKTPPMTAGVYDTA
QWSEGPMTCDFAFVTDTLVRRLSRCEIDGTVRASDHQPIVLELSDA*
>GCA_000452945.1_BPMSHR5858_genomic_05816_1 group_123
MSSEAMSTMQTTPACDGEPPAWYGKIPGAGDFVNHRLSHELAGWWERWLQQGMAAMRQRG
DDELARYYTVAPVWNFLIPAGAGAQCVQPGCLAPSCDRVGRYYPVIATLPMRTADYWSAL
PDAADAFYWQVGSALLDAIRHARAPVQLEQALAKVRLVPGAAAGSAWFGGAGDGPGERDW
RGGARDAARESDAGWAIGGAGDIDGAGVRGAGDEGDGGDRGEVGGMSGVGAADGGDARNG
ALAQPMPSPPATPATPAWPGLSQYFDPHGATSFWWTNRADGSPLRTHAHTGAPDSRLFLR
LFGGVQHGV*
>GCA_000452945.1_BPMSHR5858_genomic_05827_1 fabG_3
MAVEKVALITAAGKGMGAAIARELASAGYRVALMSPSGSAVALAGELDGLGLRGSVTEEA
DLDRLVRETLARYGRIDAVVNNTGHPPKGELLSIADASWHDALDLILLNVARLMRRVTPV
FQAQGGGAAVNISSFAAVAPEQPMPVSSVLRAALGAWTRLYAERYAAENIRMNAVLPGFV
DSWPETPGIVARIPAGRFGRTREIAKTVAFLLSDGAGYVTGQSIRVDGGIVRAM*
>GCA_000452945.1_BPMSHR5858_genomic_05830_1 dsdA
MPVGRSLSLDPNLLAQLQSHSPTLWLNPHQGMPLPDFAPTAADLADADARLRRCAGLLAE
LFAELRPSGGLITSPLQPAEPLKRAARAGHAQAGAWYVKRDDALPVAGSIKARGGFHEVL
ALAESIAERHGLAGADTDRRALASGAARARFARHTVMVGSTGNLGLSIGMLASALGFRTV
VHMSADAKAWKKARLRTRGVEVVEHAGDYAKAVDAGRRQAAGMPCCHFVDDEGSRMLFLG
YATAAAELAAQLAQAGRPVDARHPLFVHLPCGVGGAPGGIVYGLKALYGEHVHAFVAEPT
ASPCVLVQLAGDAAHPRSVYDIGLDNRTEADGLAVAQASPLAAALLRAQAAGAFTVDDRQ
LFAHLLDARERLGIDLEPSAAAAFGGPAWIAGSDAGRAYLRGRGIDPDAATHVIWATGGS
LVPAQEHRRFQAHARAQRQVGGAGA*
>GCA_000452945.1_BPMSHR5858_genomic_05831_1 gcvA_10
MDIKLDTPLLGALRCFEAAGRLGSFTKAADALHLTQSAISQQIRQLEARLGYLLFVRHAR
GLALTPKGGALHEATVAAFASLGRTLRQLSAPDAPLQVSCLPSFALQWLMPRLTEFHRLH
PQTPVRLRAEFHAIDRETMSADDIDIAIRWDPFSYGTLHVEPLFDESLVAAAAPAYLDRH
PPRATAKWVSSVVPLHDASAWPEAPRYVEWRTWLDAVVPGYAGAPRGPEFNLASLAIAAA
VAGQGVVVCRQALVDDEIARGRLVNVFARRVPAPAQYKAICARPEDARVKAFVGWLRAES
DSYAARRQTQARP*
>GCA_000452945.1_BPMSHR5858_genomic_05834_1 gloB_2
VILRQFLHCDPVGISYLLGCGGKSTGVVVDAVAEPGTYLQAAADAGMKIDYVIDTHVHAD
HLSTGPALADAAGAAYVLSAKADVVLPFKGVEDGDEIRVGNVVIKVLETPGHTPEHISLL
VTDRTRAEEPWLALTGHTLMIGDLGRTELAESAEAGARNLFRSVRTLKSLPDYLPLLPGA
FAGSACGRSLSATPLSTIGFEKRHNAAFGIDDEAAFVRFMLSDMPPAPPEAARLRAANSG
RNPDEI*
>GCA_000452945.1_BPMSHR5858_genomic_05835_1 yfcJ
MSKSTMPAGGVAIRLGLKENWRQFALLVLINAFVGGMVGIERTVVPLIGSETFHIQSTTL
ITSFIVSFGLVKAVANLISGQLADTWGRKRVLVAGWLLGLPVPFMIIAAPNWEWVIAANV
LLGLSQGFAWSMTVIMKVDLVGPKARGLAVGLNEFAGYFAVGLTAFLTGYLASRHGLRPA
PIYLGVAYAIAGLTLSILVVRDTRDHVCLEAGKPKEATSLSFHDVFLLASLKDRNLFAAS
QAGLINNLNDGMSWGIFPLFFTGLGLGVERIGILKAAYPIVWGVFQVVTGPLSDRWGRKG
LIVAGMWVQAAGLVLTASMGEFRWWLVASVLLGLGTAMVYPSLIAAVSDASDPRWRARAL
SVYRFWRDLGYAIGALSAGLIADRFGFADAILSIAAVTFLSGAVVAIVMHARH*
>GCA_000452945.1_BPMSHR5858_genomic_05836_1 iscS_1
MTHIYLDYNASTPIDPAVADAMRPLLPEAFGNPSSGHWASVPAKVALDRARAQVATLIGA
APDEIVFTSGGSEANNLALKGTVLASSNAGGHVITQSTEHPAILKPLQFLERLGTKVTCV
PVDGNGRVDPDAIRRAITPRTTLITVMHANNETGTIQPIEDIGTIAREHGIRFHTDAAQS
IGKIPADVDALGVDLLSIAGHKLYAPKGIGALYVRKGVGLEPLIHGAGHENGRRAGTESA
LLAAGFGEACALARDLAAMDGIRALRDRLWHALQDRFDDRVRLNGHLEYRLPNTVNVSFV
GRVGADVLAALDGVAASTGSACHAGRMTLSPVLAAMGIAEQVGMGAIRFSLGRHTTADEI
DTVVAKLSAIARG*
>GCA_000452945.1_BPMSHR5858_genomic_05854_1 oppD_2
MALLEVKDLGVRFTRREGEPVDAVSGVSFSLEAGRTLGIVGESGSGKSQTVMALLGLLAG
NGSTSGEALFDGENLLRMNTRQLNAVRGDRIGMIFQDPMTSLNPFLTIERQMTESLQLHR
KMSRKHALKRAIDALEAVRIPDAARRIRMYPHEFSGGMRQRVMIAMALLLEPEILIADEP
TTALDVTVQAQIIELLRELNRERGTSIVLITHDMGVVAGLADDVMVMYAGRTVEYASADA
IFSAPSHPYTIGLLNALPRLDAPDDAPLVAIPGNPPLPGQGLLGCAFAKRCAHASERCGV
ERPDLIAYSDAGAVRACHRPIAEITGGLR*
>GCA_000452945.1_BPMSHR5858_genomic_05866_1 group_19493
MQQSIQHITVEARGRGLVEFTPQVRAFVEVQSVSTGLLTVFCRHTSASLLIQENADPSVQ
RDIERYFAALAPEDDARYEHDTEGADDMPAHLRTALTQVQLSIPVEHGRMVLGTWQGIYL
FEHRRAPHRRDVVLHLIGE*
>GCA_000452945.1_BPMSHR5858_genomic_05869_1 group_410
MTARPRTRRHARAHADAGANANADADADADADADANADANADANADADADADADADANVN
MKADARCDTRKHAQRHCTHAFVAHDPRLMTRDVAENRRVWLLKRNCALSPRQSLISMGLL
AALTLAIAMPFAIAGAWAVFACAIGEIAAMCGCFLRYARHAVDYDCVALTEQRLEVIQCD
GAQLRRYDWNPLWVAVDLDAADARDPTIRIRHGSETVLVGRHVTLARRRHVAEELNAALA
ASAGDFGDTDFAPSVRQRQPEPLAAHR*
>GCA_000452945.1_BPMSHR5858_genomic_05886_1 group_5255
MSWHNKVVWNEGLFLLPQLFQQQERYFEYFAHKRAAVLSPFFWGFSRYEIDQESLSFGKL
VFKSGAGIFPDGTPFDVPGHTPPPPPLTIAAEHQDQVIYLAVPLRLPNTEETAFDEQAGS
LARYSAFEIELRDSNAIGQGPKPVQLANMRLRLLPEKELTQSWIGIALTRVKTLHADGSV
ALYDGDHIPPVSQYGANPLLREWATQLHGLAKLRADALATRLSGSDGRAGAAAEVADYLL
LQVLNRYEPLLEHICRIREMPPVTLYRELSMLAGELSTFVRPQTRRPRPTPGYDHAQLYA
SIRPLVDEVHYLLNQVLIRGAQPIPLTEQPHGIRVATMLPSELAGYSSLVLAVGAQMSPD
VLQQQFASQTKISHPQRLPELIRSHLPGMTMIPLPVPPRQIPFNSSYIYYELSRTGPFWE
QIAQQGGLAMHIAGHFPELKLELWGVRHK*

>GCA_000452945.1_BPMSHR5858_genomic_05889_1 group_3686
MNIRNADTYTFDKLPSRHESSTQALERAIASNCTTLRTRIREYREIVAFRRQPHSKKLAR
ALWTAAWRLPRVDEDWVAALSSRGNLATIAGVLGEWLGTHAMPVGRVAAIDPPGGGDEIP
EPRAAYCMRCVVEFGQKVVDARAPIDLDLAASHLVDAALSIGANLLIDVLLRRARVRIRH
PSSAGGDGA*
>GCA_000452945.1_BPMSHR5858_genomic_05895_1 group_7717
MNATPPAAALRYADLLEPVSPDAPCGPDLEYDPAFVMLQSAIAPRKDAQYGEFVEAPQPA
NWAEAERDCLALLLRTKDIRLVVILMRCRIRQSGAEGLRDGLALLNELLARYGEALHPVP
FFEGERDPVVYANAIATLADPDATLADIREIPLPKASGLQLQLRDIEKALAVTRVKDALA
PESASRLLKEWWNRRDKTIAALAQAQRIVADLIASTRESLGDDAPDLSGIAKLLHPFAQA
QLESPYSANAAQPQGDAKPATGDAAHARAADTSTQAGDTDTQAPAAMPIAPAQPPMDRWG
ALAAIQATRLWFEQNEPSSPVIVLLRQSERMVGKRFSEIANAIPAELLAQWDAIDV*
>GCA_000452945.1_BPMSHR5858_genomic_05896_1 group_7718
MTAHPPAARPIDAHSPLHPDALRAWFDPQAPWRAGFLSLLRAIAARDTRMPLPGTACLPK
EEAFRIGQRPSMAFAPREIASLDVQRGRLDIQLFGLGLWGPQGPLPLHMTELAYNRAESY
QDHAIAHFSNLFHHRALALFYRAWASSQATVSLDRAEHETFSFYIGSLMGTDPEEAARTH
PPTHARYAACAHLVREARNPDGVAATLSHYFGVPIAVDEYVFHWIRIAPPERCLLGARAA
STVMGEGALLGDMVADCQHKFRLVIGPLDLDQYLRLTPHGNDLPTLVDWVRAFVGHEYDW
EIKLLVKPRAAPPARADTTHRLGYSTWLGESRDDRPVVGMVFEPEKYCS*
>GCA_000452945.1_BPMSHR5858_genomic_05897_1 group_7719
MDPQFLDHYNRELTYMRELSAEFAAQHPKIARRLGMQGIEVADPYVERLIEAFCFMSART
QLKLEAEFPRFTQRLLEVAYPNYVAPTPSMAVARLRPSLREGDFSKGFKVPRHSMLRSSI
PPGEQTACEFRTGQDITLWPIEIAGATLTAVPPDLPDLQRSLLPHTKLRGALRLRVRTVG
EIRFSQITGLDRLSLYIGGDERIASHLFELIHASSVASVVRAPGAARGEGAVVAKNAVDF
EGLSPDQSLLPLVWNTFHGHNLLHEYFTCRQRFYFFALTQLNAGLSRIDGKEAEIVLLLD
RLPDELVTHVEAARFLLFCAPIVNLFPKRTDRVEINRAQTAFHLIPDRTRPLDYEVFSVS
RVFGQKAETSTEVTFNPLYQTLHSDIGNYGRYFSILREPRTTSTNARKYGTRTPYVGTEV
YVSLVDQAEAPYADDIRYLSVDAWVTNRDLPRLIPRNGVNDLTMQDSVPIEGVSLVHPPS
APREPYATGETAWRLIRQLSFNYMPLAELDHRDGGQALRNMLRLFVGTSEREQATQIDSL
VGARTEPVVRRLPGHGLLVYGRGVRCELTVDESGFSGLSPYLFGLVLEQYLTRHVSINVF
TETELRSMQRGLVTRWKPRMGGRGAV*
>GCA_000452945.1_BPMSHR5858_genomic_05898_1 group_5256
MRHPEGRHRAAYLPSLLDRLQDDAPHALSESPDAYAPSADEMRRIVQRDLSLLLNTSNLD
DEVDARRYPLVSASVVNYGVPPLAGSYLHDRNRETIERLVRTAIAHFEPRLIADSLAIRP
LAAQPGASYNKLTFEISALVQWSPYPLELRIQSTFDLELNRVTLEKTTLNGN*
>GCA_000452945.1_BPMSHR5858_genomic_05916_1 cdaR
MMIDARLATEIVVRTVEVMPFDVNVMDAHGVILASSDAARVGEIHAGAQLALARARAVEI
DAAMAANLSGARPGINLPLSVRGKVCGVIGVTGEPDAVRGFGELLRVTAELILEREQLTV
ELRRNARHREEFVLQLLRRTAAAAELEAWADRLGIDMRLAHAAIVCRLSGVDTEPEAGVG
QIEAMQSQLAQEWPQLLSAKTSYRELVMFDAVDARAGRDGVLATHAGKRLAALEGTLRKV
ATQPFSLALGIALDGIDGFERSYQCALATLRVGAARTPQRAAYSYYDFVLPVLLSALSQG
WQARQLRLPLTRLLARERRSGALLDTLKAWYAHDGHPSATAQALGIHRNTLDYRLQQIRD
ATGLDLGAIDDRLWLYIALQTMAGSDDGDDDGLDGDAGLGDERPAS*
>GCA_000452945.1_BPMSHR5858_genomic_05926_1 group_7722
VKPRSTNMSIRLARLIALSMLATLVAGVSGCVVEPVQPAAHWHPAHYDVYGYWHPGHWGP
*
>GCA_000452945.1_BPMSHR5858_genomic_05946_1 yueD
MTTSPSPLPAVRAIVTGHTRGLGEALAEQLLRRDIAVLGLSRGRHPSLASHAPERLVEAE
LDLSEPARVQAWLGGDTLGAFVAGASRVLLFNNAGTVEPIGPLDTQDTAAIARAVGLNVA
TPLMLASALAKLAPDALERRIAHISSGAARNAYAGWSVYCATKAALDHHARAVALDANRA
LRICSVAPGVVDTGMQATIRATSDERLPSRERFAQLKSSGALSTPDDAARQLIDYVLSDD
FGSTPTADIRHLG*
>GCA_000452945.1_BPMSHR5858_genomic_05956_1 group_7723
MSEVGIRNLNHLRVFMAIVEKGSFTAAAECLSMSKSLVSEYLSRLEAEIDTQLVMRSTRR
IAPTDAGNKLYCASQAFVSGLYDVIGSIRCLRHESTGLLRVAAPSGFSTTHLSSIAATFI
HQHPQIELEIVCNDDEIDLVGERVDLAFETGWPKKKGFRMKMLGAFDQVLVASPEYSRRH
AVPRHPDDLPGSHWIGHGGLANLSYSVFGNEERSVRIQTPGRLKVKSVLLAHQMALAGAG
ISAFPDYLVAEDLREGRLHRLLPTWTMPKGGIYAFRTAPRQASVRERLFLAAVQAYLAGL
CGERARAGAVPT*
>GCA_000452945.1_BPMSHR5858_genomic_05958_1 group_3697
VTSETVHHPDIPAPAPTFDSVAATFARREPAPAPAGEPPAARLAAIRLARNPLLEAARVL
LRALADMPERLDRDDIPQLRLLLEQEVRLFQRLCEQANIRRDHMLGARYCLCTALDEAAM
QTSWAQSASGNLGTWISEGLATSFHEDRQGGDKVYLLIGRLMNSPHEHIDLLEVIYRILS
LGFEGRYRYEADGQRKHETVRQRLYNEIASQRGPVSVALSPHWQPGPRSRSAPFRDFPAW
VTAAVLSLIALGLFGCFKYALSTRSADVQQRIAAIARMAPPAAPAELRLATLLAGEIAAG
TLSVEENARRSSVTFRGDAMFAPGAAGVNPAMGPLIRKIAAEIAKVPGKVTVRGYTDNQP
IKSRQFASNEALSEERATQVMQMLQSAGVPASRLEALGKGGAEPIGDNRTPQGRALNRRV
EITVAR*
>GCA_000452945.1_BPMSHR5858_genomic_05961_1 group_5266
MAQDIFLKIDGINGESLDDSHKDEIEVLNWNWEIQQESTMHTGSGGGAGKASVKDLTFEH
AIDRASPNLMKYALTGKHVDQAVLVMRKAGGNPLEYLKLTMSDVIITRVRPSGSRDDTER
SRETVSLSFAKVKQEYVVQNAQGGSGGAVTTSFDIKGNKEA*
>GCA_000452945.1_BPMSHR5858_genomic_05962_1 group_5267
MKKQQAQTAAAGVQPQADSDFAQLLAQEFKPKTEQAREAVEYAVRTLAEQALAQSATISD
DAYKSIAAIIAQIDHKLSEQINLILHHADFQKLESAWRGLHHLVSNTETDERLKIRFMDI
SKEELRRTMRRYKGQSWDQSPLFKQIYEEEYGQLGGEPYGCLVADYYFDHTPPDVDLLGS
ISKVAASAHTPFLSGASPSVLQMESWQELANPRDLTKIFTQNLEYASWNALRNMDDARYI
GLAMPRFLSRLPYGVLTNPVDEFDFEEDTNGANHRRYAWTNAAYAMGVNINRSFRLYGWC
SLIRGVESGGTVENLPCHTFPTDDGGIDIKCPTEIAISDRREAELSKNGFIPLVHRKNTD
HATFIGAQSLHKPAEYDDSDATANANLSARLPYLFACSRFAHYLKCIVRDKVGAFKERED
MQRWLNEWIMNYVDADPANSSQDTKARRPLAAAEVVVEQAQGNPGYYQAKFFLRPHFQLE
GLTVSLRLVAKLPSIKEAA*
>GCA_000452945.1_BPMSHR5858_genomic_05963_1 group_5268
MSASSSSQKFIARNRAPRVQIEYDVEVYGSEKKVELPFVMGVLADLSGKPVAPLPAVADR
RFLDIDIDNFDERMKAIKPRVAFAVDNTLSGDGQLMVDMTFESIEDFSPAAIARMVGPLR
QLLEARTQLANLQTYMDGKSGAETLVNQLLQDPALLRSLAAAPKPQLAGGPDAGNARGAK
DAPDTANHDSDAA*
>GCA_000452945.1_BPMSHR5858_genomic_05978_1 kefF
MNVLLVYAHPEPKSLNGSLKNFALRHLEEAGHAVQVSDLYAMQWKAALDADDNTARRPDT
RFDPSLDSKHAFENGTQAADIAREQDKLRWADALVLQFPLWWFSMPAILKGWVERVYAFG
FAYGVGEHSDTHWGDRYGEGSMKGKRAMLIVTTGGWASHYGPRGINGPIDDLLYPIHHGI
LYYPGFDVLPPFLVHRTNRIDETRFATIRDALGQRLDTLWTTAPIAFRAQNAGDYEIPQL
TLRADIAPGRSGFAAHIA*
>GCA_000452945.1_BPMSHR5858_genomic_00082_1 group_2226
MQRRYQGQPVGRGGAAGWQERNERGLRGGERGMQYDDEALLRPDLDVEGDYRMLAEDIGP
EDWGSDWADWDDRNPYDERRRGYEGGRREREGGRGPQRMRGVGAQGNVGEARGGYAGPRG
YGDDGWYQGGGRDDDGDLRKFRGQSRLVAGSRSPYRGGEREERAGPEPWRAGGERSAYRD
ELWSPYRTREYDRSRDYRDLLHGGGGQREMLHRQGPKGYTRSDERIREDICERLAYAFDL
DVSDVSVEVNGGRVELQGTVPERWMKHEIEDIADNCMCVRDVDNRVRVLPRERGETTGTA
GMAAPPEGQPMRPEHRTSTMAGGAGGAGGQSPGGSVAGGQPSGALSGTGGHDPRH*
>GCA_000452945.1_BPMSHR5858_genomic_00084_1 group_4388
MNTLNTLQPSAKVPTNERGGSAFADRQAAVAAVELLLPAFSAALQSDFVGDSGCLHIVVM
DPALGPSLASFDDAILYEFSLPDPQYWDADYRAYARAKARLSWETGRDGHLLQANEPHRL
RAGDTTLWGSVALDGIVVGVSGAQPWFDEAFAGCIAHCLRAIAKHRAHATPDTLAL*
>GCA_000452945.1_BPMSHR5858_genomic_00085_1 group_7225
VSQLSSDPISGASLAGRTAFVTGGGRGLGAAICGELARAGAHVVVADLDEARARGVAERL
ARDGARAVPLGVDVGDETSVRHAFDAARESLGDIDVIVNNAAIDVTAAIDDVSVDAWRQV
LTTNLFGPYLLCHAAVPMMKARGRGHIVNIASTAAKRAWPNASAYHATKWGLLGLSHALH
AELRPSGVHVSAIVAGGMRTPFLLDRFPDIDADTLQPPEHVAAAVRFVLTQPPGTVIPEV
MVLPVKETSWP*
>GCA_000452945.1_BPMSHR5858_genomic_00086_1 hisB_1
MAVTDARARGAVLLDKDGTLLENVPYNVDPARMRLARGAARALATFGELGMPIAVVSNQP
GVALGRFPETALGAARHRLAELFAQQGAMLAGFFYCPHHPAGTVVRYACDCACRKPQPGL
LFRAAAQLGVSLDASWMIGDILDDVEAGRAAGCRTILVDCGNETEWRYGGLRTPDFIASR
LDDAAWIVARECAATAAPGEAPQEGRR*
>GCA_000452945.1_BPMSHR5858_genomic_00088_1 novN_2
MAKSRPPLYTLGINAAFHDSAACLVRDGVVVAAAEEERFTRIKHGKRPVPFSTWELPYHA
IDYCLAEAGITLAEVAHVAYSYDPWQQIERDGREPTFTLPLSPSAHARANGHASPWEPLF
LSYIVNAPRQLADGAPHHLRRRFRGVTPDGPFRWHFVEHHLSHEASAFLAAPFSHCAVLT
MDGRGEKATTSYGVFDGHVYRRVGQVDLPHSLGLLYERVTRHLGFLHSSDEYKVMALASF
GKPAFAHALRRLVRYVGDGRYETPALDLRELLGPPRERGARVEPRHCDIAHSLQAVLEEV
VLEIVDWLAGATGEKRLAMAGGVALNCVMNAKIRDRGRFEDVWVQPAAGDAGTALGAALW
TDFSERGERGGWRMDHAFLGPAYDERAIEACLIDAKLPYRRLANVAAETAALLAANRIVG
WFQGRMEFGPRALGARSILASPIDPGMQQRLNTIKDREDFRPVAPVVIEEKAYRWFTGGK
HERLRAPFMLFVYDVAPGAQARIPAVRHVDGTARVQTVNRAQHPLYYDLLNEFDALTGVP
VLVNTSFNTRGEPIVRTPRDALECFWTSPLDALVIGPFLLEKPL*
>GCA_000452945.1_BPMSHR5858_genomic_00089_1 rfaF_1
MNAPHWRPPPVVSIVVPTYRRPELLERCLGALASQVFDPGTYEIVVVDDDAAGSARPVVD
ALTVRMGGLPAIRYVSAPRTQGPAGARNAGWREAAGPVIAFTDDDTIADPLWLRNGCSAL
LAQPNASAAAGRIEVPLAPCPTDYERDAGGLAHAEFATANCFVRRAALERVGGFDERFTR
AWREDADLMFALRERAGPIVDARTATIVHPVRPARWGVSIAQQSKVFFDALLYKKHRDVY
RRHIRSVPPWHYYAAVLALLGACVALALGLHAAAAACAAAWAGITAAFCWRRLRGTAHTP
SHVAEMIVTSIAIPPVSLYWRLRGALHFRVLFL*
>GCA_000452945.1_BPMSHR5858_genomic_00092_1 luxO
MSQHSSDGRRLRGNSCLIQALLHKIEKVAATRASVLIVGESGVGKDIVARLLHDMSPRRR
GPFVPVNCGAIPADIAESQLFGHEKGSFTGAAAQHIGFFEAARGGTIFLDEVAEMSRDLQ
VKLLRALESNTITRVGGSEPIALDVRVVAATHHNPAEAVQEGRFREDLFYRLAVFALRVP
ALRHREDDIEGIAQELVDTLNSRHSTRKRLSGQALKTLRTYSWPGNVRELRNAIERAYIL
SDEQIEQLPVRRLAPRDDVRRNAMTLPLGMTLAHSQQRFIAASLKYFDGDKPRTAKALGI
SLKTLYNRLALMREHDGETAQH*
>GCA_000452945.1_BPMSHR5858_genomic_00093_1 atoC_1
MPYILIVEDDADTRDMLCALAHTQQLACDAAATLEEARALIATHTPDLVLCDLVLPDGNG
MDLFDELPKGAHVEMVLTTGHASLETAIDALRRGATDYLVKPLNMQRLNSIFARVPRTGV
LHEEIAALRSELQRLGRFGRMLGRSPVMQTMYDAIGRVAGTEASVLLMGESGTGKELAAQ
TIHDLSLRRKGPFLAVNCGAIAANLVESEMFGHDRGSFTGADRQHKGFFERADGGTLFLD
EITEMPVESQVKLLRVLETGRLTRLGSARELDVDVRIVAATNLDPEAAMADGKLRPDLYH
RINVFPITLPPLRARGDDIPMLAEAFLRQLGEDSGRKMRFSASALAALAEYDWPGNVREL
RNFVQRACIFNDGEVIDTLPPPIMDEVASEPGDDGDAVTVPFGTSLEEVDRRIILGTLAQ
CGGVKTHAAEVLDISLKTIYNRLAQLEAEGDDKTKR*
>GCA_000452945.1_BPMSHR5858_genomic_00095_1 group_4390
MTSTHRWLVISPHLDDGVFGCGQLLAQSPGSVVVTVFAGVPPPRTPAPPWDRRAGFATGA
DAMRARRAEDARALRVLDATPVWLDFLDDQYGAPAAPDDIAKQLAATLDAHPGFDVAAPA
GLFHRDHLATNRAALAVLARGPRARRWLFYEDALYRRVDDLMAKRLADWRRQGWIARVAA
PLAHARAGRAAKAEAVRAYASQVALFDAHALVDLHVSETYWRLQCETSSA*
>GCA_000452945.1_BPMSHR5858_genomic_00096_1 rcsC_5
MANFVRGHAKNYAAVTGFIAIGTLLQAALVRFGGADMPLAPYFPFIAAAAWATSLAGGIA
ATAASAALAWLLFLNDPAAYSAPLAGRLMQLATFAVVSCAICAIVVALKGSRRANDALRR
REAGARHYYETLLQSLTQGVVVSDRHGMLTYMNPSAAALIGCDATAVRGLPFRDVYRAYD
GRGARVRASALEHVLRSGRSASVEGHWLAPHDGQRVPIAEAASPLVDSAGDMTGAVLIVR
DISIERERSAASQMLRRLVEASPDAIVGVGSNRRITSWNPAARRMFGYDEHAARGCDVAM
LIAPRWLRNHPLPESVPDMREPIENVDVLCVRQDGGRMRATMSASPVFDERDACIALSLT
LRDTGEQRRRERRVHRSLRGARDARRQADTSNRLKDELLATVSHELRTPLNVIYGWIEVL
RNPVDGALQQQAIDAIDRSARSLSRMVGDILDASSLATGKLRLDAMPVDLVRIVNDVTGA
LGAAAQADGIALSVECALATCIVSGDGERLRQMLSNLLSNAFKFTPRGGRVTVSLERHDA
QAKLTVADTGQGISPEFLPHVFEAFRRAEGAPASPRRGLGLGLSIVRHIAELHGGKVEAA
SAGKNRGTTFAITLPAGWQPMGALAWAAQAEGGARMTLDAQRILLVDDDATSRASLAAAL
KTLGAEVVVASSGREAIGKVDAMRPTVVLSDLAMPDGDGYWLLDTLRHVDADADADTRTR
AANDAPVLAVTAHAGRDSERRVLAAGFDGYLCKPVDLQTLARAILRATGSG*
>GCA_000452945.1_BPMSHR5858_genomic_00097_1 galE1
MKDPERKRILVTGGAGFLGSHLCERLVAEGHDVLCVDNFYTGAKDNIAHLLDAPNFELMR
HDVTFPLYVEVDEIYNLACPASPVHYQRDPVQTTKTSVHGAINMLGLAKRLKARILQAST
SEVYGDPASHPQRESYWGHVNPVGIRACYDEGKRCAETLFVDYHRQYGVDIRIARIFNTY
GPRMHPADGRVVSNFIMQALADAPLTVYGDGRQTRAFCYVDDLIDALIRLMAAPGPMPEP
MNLGNAEEVSMLQIAREVVRVTGASVAIEFRPLPADDPRQRCPDLGFARERLGWRATTTL
ADGLAATVRYFIQRQAAHHAPGNLVHSTRVASHMLARVNAQGRSRVSA*
>GCA_000452945.1_BPMSHR5858_genomic_00099_1 group_839
MKEMQSRVPRYTQPHSPHHAAKRFGRLATAVLIALAAACLALPQAAAAAGDGTGGAGAAT
GGANGSANGSANGPNDMTGANTANRATDMNGASSSHSTSAGTKLRDTAITTKVKAALLAT
NDLSSGDIHVKTRRGAVQLAGTVPDERQRTLAVDVTRQVDGVKTVRDKLTVQPK*
>GCA_000452945.1_BPMSHR5858_genomic_00100_1 hcaB
MKCTLKPIGEQTIVITGATSGIGLVTARKAAKKGAKLVLVARNDTALEALCEEIRQHGGL
AVAVAADVSRYEDVQRAAAKAVETFGGFDTWINNAGVTIFGSALSVPLDDQRRLFDTNYW
GVVHGSLVASEHFRRKSDFRGGAIINMGSEASDAPVPLQSAYAASKHAIKGFTDSLRIEL
EADNVPVSVTLVKPAAVDTMFVMHAKNYMNVEAKLPPPIYDPDIVADAILFAAEHARRTL
FVGGAAKLASSGAYHAPRLFDRLATSLFSRGQRTMRPARPRDDNALYEPTRPLHEREGMD
GVVLRSCAYNTVMQRPRVAGALALTAAALVVAALARARRDVAS*
>GCA_000452945.1_BPMSHR5858_genomic_00102_1 group_1387
MAMIVAGRFTTFEQAEAGARHLYARDFGPDDVSVFFLNPSGQHARFPIGGDVYADSAARP
GGSGAAYGAALGVIVGLVVGFVVYMFGWRYWVVWPAAALVGGYVGAFGGALRRMQGRQIE
GLGTLRQSDTGVMLAAHVTGETAGAVEAVLRASGAANVERVDGEWVDGEWADFDPARPPR
TRPDVSSR*
>GCA_000452945.1_BPMSHR5858_genomic_00103_1 ydaD
MSTAEASSPQGRTPQPGREREMAAKPRDEAAGYVGSGRLDGKVALVTGGDSGIGRAVAVG
FAKEGADVAIVYLNESDDAAHTKHLIEQTGRRCETIALDIGERANAHVALRRAVERFGRL
DVLVNNAGEQHVQTDIGQISEAQLTHTFRTNIFAMFFCTQAALTHMKAGARIVNTASVTA
YHGNPVLIDYSATKGAIVSFTRSLALNLASRGIHVNAVAPGPIWTPLIQSTFDDEQRARF
GANVPLKRPGQPDELIGCYVLLASDGADYMTGQTLHPNGGTIVNG*
>GCA_000452945.1_BPMSHR5858_genomic_00105_1 group_4393
MAKKQLGEGVRELLYQALETELGGAKVYEAALSVVTDEALREEWQKYRDETLHHQEVLRS
VFDALGLDPDEKTRGRAAVAGLGKALVEVIAYARGGGNPLVAQIVAAECVVLAETKDHLN
WELIGLAAGELSGDAARVLKDAHDEVEPDEDHHLYHTRGWTRELWMDSMDFKAALPPPEE
VRKVDTAIGAARAEHARGRMAR*
>GCA_000452945.1_BPMSHR5858_genomic_00106_1 group_4394
MKALCWHGKHTVRYETVADPAIEDPRDAIVEVRACSIGGADLHLYDAVIPGLKDGDVLGR
ECVGEVVEIGAGVSRRRVGERVVVPFAIACGECDQCRRGNWSVCETTNPHRALADKVFGH
ATGGLLGCGHLAGGYPGGQAQYVRVPHADVAPATIPDGVDDERALFVGDSLATGWQAAAQ
CEIEPGDVVAVWGAGAVGLFAAMSARLLGAAEVIAIDRVPERLALAQKLGATPLDFERLG
VADTLAERTRGKGPDKCIDAVGLEAQAGEALDAVTDRFRPTVAAGADQPHVLREMIYACR
PGGVLSVPGVYGGLVDKLPMGALMHKGLTLRAGQTHVCRWTAGLLERIADGRLDPSCVIT
HRATLEEGPAMYETFGARRDGCIRPVLRPRH*
>GCA_000452945.1_BPMSHR5858_genomic_00107_1 mntH_4
MDELRSQRRAERAARNEHAGGAPPGAAARRAGNACDKPDAHGAHDAHDAYGSCDEPGADG
RRRARAAPWYRRLGPGLLTGASDDDPSGIATYAQAGSQFGFAMLWLTVVTLPLMVAVQLA
SAHVGRGDREGLIAAIRRYYGAPLAKTLVVLVALVNVANVGADLSAMGDATALLVGGKSY
WFALAYGAGMFALQVRLPYPRYARYLKWLSIALFAYVIEVALVDVDWAAVAHALIAPRIA
FSHEYATTAVAVLGTTISPYLFVWQAALEVEEIAQERRAGTSAGASRKSSTSRGQGRRIL
IDTWVGMLASNVVAFCIMLTASAVFFAHGVHDIATTRQAAKTLEPIAGRAAFLLFACGIV
ASGLLAIPAFASSAAYGFAEAWRFRASMNAPLARAPLFYAVIGACLAVGVALCFSPINPI
KALYWSAVLNGVAAVPMLAMVMRLASRRDVVGRLTNGIVLNTLGWSATALMAVAVGVMVF
DLVS*
>GCA_000452945.1_BPMSHR5858_genomic_00108_1 tmung
MARDARAAHSPRRRRAADDDDAPQTLDACRRCGLWEHATQAVPGAGPPRAAIMLVGEQPG
NQEDREGAPFVGPAGRVLERALAAAGLDRASVYVTNAVKHFKWAPRGKRRMHKTPAQQEI
DACRYWLERELDALGPRVVVALGAIALKAVLDDRHARLQSAMGVVQSAGGRYVIATYHPS
FVLRVPDPDARKRAYAAIVEACRIAHELAQRTDGAPRP*
>GCA_000452945.1_BPMSHR5858_genomic_00109_1 group_1745
MAFKTYAGGAGALALLAALAGAHPRTGDAAHALAALMGAAAPAPRADFDTDALTGLGIAS
LSVLADVAEHEARRTLGVDLAQTLDAAWAALRGYAGSDWHTRGGDERARIARHRRASA*

>GCA_000452945.1_BPMSHR5858_genomic_00110_1 group_7228
MTYERTESGEDVVLWRSLDGERAGHRRVLVVDDYRDAAEALRVLLDARGFECRVVDDPFE
VCDTARDWQPFAVVLDIAMPGLDGLEIARRLRRERATAHMLLVACSAFSARHDLERAHDA
GFDAHCAKPLTPHRLLRFLEAASAGTHDD*
>GCA_000452945.1_BPMSHR5858_genomic_00111_1 group_7229
MNTPQQGTTRIIGKGRASAAGPGPDVMAASTLDGDKVYTTDGDDVGKIKEIMLDVRTGRV
AYAVLSSGGLLGIGDKLLAIPWSALTLDTDRKCFLVAVSSERIKNAPGFDKSHWPTMADP
NWAASLHQYYGSAPYWSDIEEELGIDAPPHEASPDRPERGGRH*
>GCA_000452945.1_BPMSHR5858_genomic_00112_1 hrp1
MQRINEIMSRDVVHVAPSDSIRHAAELMARFDIGALPVCQNSRLIGMITDRDLAVRAVSA
GKAPDTKVHEIASGPIEWCFDDDQVDNVQKYMADAQVRRMPVVDHDKRLVGMLSIGDIAT
RTDGASRDEVANTLEGVSQPRRS*
>GCA_000452945.1_BPMSHR5858_genomic_00116_1 group_3056
METPKSTVHYESNVCGGSFESVLDRFGNWKREPLVYRPERRMFEGKDSVRRLGDEAFDSP
DKARRALIRSCAPRDRFALAAPICDDDHQMWLVMAAFEA*
>GCA_000452945.1_BPMSHR5858_genomic_00120_1 group_1388
MTIRRISTKLAGVAAMALTAACSQNGPYGGSPRAGSTGGTITDGVNQPPAAVTPSDGARY
GSPGYHGAQPMAPGGDVGDGNGAGTGSMPAPGAPGSSD*
>GCA_000452945.1_BPMSHR5858_genomic_00134_1 group_3059
MATRKTSPRKQAPRKTARRPAASGRKRARAAQEKSGRYWSRDVTEHSDALDLEHDVFKSD
NPEAIAQSLKRSALDSTRRKANPFQSAMSMLNFYVNRAGRNLPKTRRTTLERAKKKLREA
FGRKP*
>GCA_000452945.1_BPMSHR5858_genomic_00137_1 group_739
MKKTDTREKNAAAKPDSPSNIGECIRLPCSGQFYVLERALSKPRVSGAFDGIEAALDAQT
AKFAERNAAAARAFAQLRVRCVREQRAALQIDGVTVGATLSSLTRSFLRPPLLAPEADVA
EARFAHVLLVELTLHPVGKDEVDAYLYVYRELADDPLDHGLREHCTEIETPAFVEPFVQP
DATRIERMSMRMMAASRGEVRRKTIDAYDVGATTSSLGLHRTIAGTMTLAVAHGGGTRRF
DVSPHRQRVRAGTSRLPLDKVIHWASERAVGFSRSSTATATSSAFLSQFARPIARLLDKT
PSSVLIERRAFAEAIDEYARLTGLAWVAGRGAPQAWKTVDDVLDALGDTFVVDGPALDGS
GAPLDRSRPFAEVCYRSRAPSIPGLKSTDAVRLKVGSRTCKIVLPSAVGHMGGAKDAQRR
GLGEILNRSRAFRAVFDGGRVLFCSEGAYGSDDLALATTQLASIFRSVAALGDVHSEKGK
THEHATSFQAQSCFHVIETERALAHPDSALICDDSTVEWADYIELDSAAPRIRWMHAKVQ
KVASEAAKRARQDKTTLPPHVSPTVAVRHSPSLSASDLEEVVGQAIKNLARLRLRTSDPE
FSGRHNTWMSETCALPSKSRIARLRRLGGLRRKADIEARFDAAAIDPHAVYEVAIVVPNY
SKTQVESELAKIGAGDAQPSVLQMFWLLSGFMHACLEVGAKPLVFMHA*
>GCA_000452945.1_BPMSHR5858_genomic_00138_1 catM
MDQSFQVNLNRLRYFHAILEHGSIRAAAERLNTAPSVVSRQLQLLESEIGAPLFERRQHG
MIATEAATLLVEYTHACRIGLEQFHESLHGELRGRVEIALTVGYVEDLMDAVITGFLHRH
PHVAVELNVRTADGVVDDVLRKRAHIGLTYNPSLPPEIRECLSVAHPTFLAVAPDHPLAK
RGEPLELRDAVRYPLALMPAPYYGVGQIVHTVAHAENIPLAPRLVTNSVPAMRRFVRSGN
GVAFLTRLLVEDDLERGEIVALRMRNPLFNSPCGKVLVHNERALPRAANEILARIVRDLP
SFKS*
>GCA_000452945.1_BPMSHR5858_genomic_00139_1 ttuB_3
MREQNASGNTRSRIYRKVDIRILGFLAVCYITAYIDRINISFAKLAMQTDLGWSEAVYGL
GAGIFFLGYMAFEVPSNLWLTRIGARLTLSRIMVLWGLTSAAMMFATDVTTFYVLRFLLG
VFEAGFSPGMLFCLTRWYPQARMARAMAMLLCAGPVGAVIGGPVSTLIMTQLAGAHGLAG
WQWLFVLEGLPAVLLGIVLFFTLSERPADARWLSENEKAILLSELAAAPRAASRHTAFRG
VLKDPRIYAMAFSYFAIICGLYAVSFWLPTLLRGAGVKNLLHVGALSALPYLAAIVAMQV
IGRRSDRRGERRVHSALALLLSAAALTGTVLAGDRFVVAMVALVAATALMYSSYAVFWAI
PADYLKGDAAAGGIALINSIGLFGGFVSPTIIGWIKTATGSLQAGLMAMVALFVAGAIVL
LANRVDANKWSAVAS*
>GCA_000452945.1_BPMSHR5858_genomic_00177_1 group_3065
MIVCVCKSVSDRKIRASLAEGVNTFEELQFELGVATCCGKCEETVREIMAEQGVCASRCG
VERPAAVPVAVTFYERKAA*
>GCA_000452945.1_BPMSHR5858_genomic_00191_1 tyrB_3
MSLFSAVELAPRDPILGLNEAFNADTRPTKVNLGVGVYTNEDGKIPLLRAVRDAEKARVE
AGLPRGYLPIDGIAAYDASVQKLLLGDDSPLIAAGRVVTAQALGGTGALKIGADFLRTLN
PKAKVAISDPSWENHRALFDMAGFEVVAYPYYDAKTNGVNFDGMLAALNGYEPGTIVVLH
ACCHNPTGVDLNDAQWAQVVEVVKARRLVPFLDIAYQGFGESIEADAAAVRLFAAANLNV
FVSSSFSKSFSLYGERVGALSIITDSKDEAARVLSQLKRVIRTNYSNPPTHGGAIVAAVL
ASPELRASWVQELGEMRDRIRAMRNGLVERLKAAGIERDFSFINAQRGMFSYSGLTSAQV
DRLREEFGIYAVSTGRICVAALNTRNLDVVANAIAAVLK*
>GCA_000452945.1_BPMSHR5858_genomic_00243_1 group_1393
MHERPPITAETTWFAARDGRLLTGTFYAPGAPSASNGRSVLVSGALGVPRGFYERYARFL
ASDGFTVLTYDYRGIGDSRKPPAEADGASLRDWGEADLPGALDHLAARAPSHRLLAVAHS
VGGQLFGLAHNNGRAAALLSICCPSGYWRLWPWPHRLWLAPLWFCVVPALGGVLSHLPVR
RAGLGFAADLPAGVMREWARWCRHADYLVDDEGHPLREHFRGYRGRILAYTVSDDWMAPP
AAVQALHRFYSTAQVEHREIAPDRRRHAIGHFGYFRDDNRALWPLGAAWLREQ*
>GCA_000452945.1_BPMSHR5858_genomic_00245_1 pld
MTCRVRFASGLSFFCALLCAVSAFECARAAQLPSRDAALGAYFDADAGSTDIEVDLIHRA
RTRVLVAGGASVPAAVAAALCAARGRGVDVRVVLDRTGRATRYSGAAFLANAGIDVVLSD
RVPNAHGPFVVVDDAVVLGRASFAQTRDARAAGNFDVFYRAPQLAQSYATAFWRLYGLAR
R*
>GCA_000452945.1_BPMSHR5858_genomic_00248_1 mdcG
MSSRAPAADAPLVRHTLVRVRAQAWPALLAAHGGIGTLPFVRDWAARGWPLIVRRRSPCD
AGVPLGLPLPPSAGKRRVALAAAREQIDSACALPTLADVGRQAPPAWRPTLARLRALADA
HRVDCRVFGSLAWQTLTGLRYLSDESDLDVLLMPCGASSSASLFRFSSSFASVSVSPSSP
SVSIPQSSRAWPVPALAGASSSAAVTGAEASAPMTAQMEASASRLSRRSGSSSMPSRRDA
IAPFLAALAEIDADAPMRIDGELLCADGAGVNWRELHAGGRAVAMKTATGVELVAPRTFL
EAWR*
>GCA_000452945.1_BPMSHR5858_genomic_00271_1 group_1752
MTTIDLNAGALAPVADEIDAVDLRVTGALARELNGVLVRNGPNPLRGRFDGGDVLSWWPQ
DAMLHAISFDDGRATRYRNRWARTRRWARVHDPAREPSLVDTNPNVNVLAHAGEILALAE
GGAPLAITAGLDSIGAARRHPGLAHGMAAHPKVDPHTGELIAFRADWNRPWLRYLVADAA
GAQTVDTEIALPAPSMMHDIAITATHSIVFDLNVAYDFSMLSRGHRMPLRWHDARGARIG
VVPRRGGDARWFDIAPCFIQHVVNAYDLDRPAIVLDVVRYPWFLRVARDGRGFDDNPPGV
LWRYVIDLVTGTVAEQPLDDAGIELPRIDARRTGRRHRFLYAAEQPTPVELRGIVRYDLD
GGSTQRYRVPPGDQNSEPVFVPRPGAAGEDDGWLLVCVYRHATDTSDVVILDGRSIGDGP
IATVHLPRRVPAGFHGAWLPAGA*
>GCA_000452945.1_BPMSHR5858_genomic_00275_1 gabR_4
MARRARIIEIPSLGMLERTAGNLSRQLAQALRDAVRRGEVMPGDALPSTRLLAASLRIAR
GTVIDAYEQLIAEGFLESRGGVGTRVAPALAEPRGARRRAAAAARSALPPPAAEYARVAR
EFAPLPPAPFAISVPAGATAPDDVWRRLGNRLRARGPAAPAGYSDPLGVRALREAIAGYV
RKSRSVHCAPDQIIVTSGAQQGLYLACQVLLGAHDRAWVENPAYRGLTAILECTGRRDAM
VRVPVDAEGIDVDAGVRLARDARAAFVTPSHQYPLGMPMSMARRAALLAWARASGAWVVE
DDYDSELRYEGYPFPSLQGLDPARVVYLGTFSKILFPSLRLGYLIVPDELVDALRGARVL
MDRHAPTADQHVLAAFIAGGHFDRHIRRVRGVYAEQRAQLIDTVGRLLSGDLAWLQPGDQ
GMHAVLWLAAGVDDLRVAAMAAQAGVAVRPVSPMFAPGTARPGLVLGFGGFGREQMDAAA
RRLAEVIAAASGSAVPRAGAGRRRRDGAGGTDA*
>GCA_000452945.1_BPMSHR5858_genomic_00277_1 group_4415
VLNATSPRRVFVPLAIASIVAAWGALWTWDSSPYARFLHPVDWSATSLAGLCRLVPAGGA
IVPAALHAAAWLLMIVAMMLPTVLPLLRTFERLTAARPERNRLMLLVVLGYLAAWSAFGL
LAQAGDAGLHALARRAPWLAWHGWLVGAAVLALAGAYQFSPLKYRCVDKCRSPLMFVSEQ
WRGGHPRRDSFALGLRHGLFCVGCCWMLMCVMFALGAGSVGWMLAIGAAMAAEKNLPHGR
RLSAPLGVALLGWAASIAVAGMGAA*
>GCA_000452945.1_BPMSHR5858_genomic_00309_1 group_1755
MIKRKLIAFAAAPALALAAFGIGTVSPAGVAPSMAEAATVSKLGDLSPFRAIAVDVAQRV
DKGDLAGAKARIKDLETSWDNAEPSLKPRAAADWHAVDHAIDRALTALRADRPQAGACRQ
ALADLLAVIDRPGGAR*
>GCA_000452945.1_BPMSHR5858_genomic_00315_1 group_2243
MSISFSVSLRAVAAALSLAAAGAAQAAADGACMARSPDGRQALVELYTSEGCSSCPPADD
WLARLAARRAALRVVPLALHVDYWDGLGWADRFAQHRFTERQHALAARGGGRFVYTPEVA
VDGRELRDWRDADAFGRRVVATAAEPARVGIALAAKRRAGALDVALSVTPRVGAPRALDA
YLALYENGVESQVRAGENRGATLRHERVVRQWIGPLAATADAHASLDVRRALPLPANLRA
ADAARYGVAAFVEDPATGDVLQALDLPLCG*
>GCA_000452945.1_BPMSHR5858_genomic_00327_1 recC
MLHLFYSNRHETLADALLDDLAFASDGAGPWAAQQVIVPSAALRRRLELDLAKRHGVCAN
VRFSYLAQWLWAQIGEVLTVPAHSPFAPDRLVWRCYRLLSDHGAAPWRASPRLAAYLAAA
DAPMRHELAHRIATVFDHYLTYRPEWLAQWQDGESIFALEGAPRHGGEAARDDEAWQAAL
WRALLAELSDSGMPPAHRFLREARAFDPDVIARARAKWPDAVSVFALPTMPPLHIALLRE
LSRWIDVRIYALNPCREFWFDIVTAARAESLDAADRLDYQEVGHPLLAEWGRQTQAQLHM
LHELTESAAASDASRYADNPAPTWLARVQNAILDLQPEAALGEPPAERGVEVHVCHSLAR
QLEVLHDRLLAWFDADPSLAPSDVLVAMPDLATAGPLIDAVFGTPAGADAARIPYRITGL
PPSQANPVARVLLDWLALPERDVGAPELVEWLRADAVAARYGIDATALETVQAWLAASGA
RRGLAPDAAEPPNPAVPAPRHTFADALTRLFLGYAMPDGAAPVSTWLPIAGAEGGEAELL
GRLARFTDDLDGFARRIARDLTPQGWGDVFADALARFFDSGPAFADALSGVRDALDAMLA
AMAEGAAGAALPAAVVRAGLAAALDDPARGGVPWGGVTFSSLTSLRGLPYRIVCLIGMDD
GVPPSLARADEFDLMAVFAKLGDRQRRDDERNLFLDLLLAARERLLIAYTGRSIRDNAPL
PPAALVDELLDHLAAASAGPDAPPDAVDAARRAFVIEHPLQPFSAEYFAPGGALFTYDAS
RAMLARLLAEGARAQARPFFAAPLPAEPEQPVAFADFERFWRHPARSLLRARLGIVLLDA
QAELVDTEPFELDYAGSDALAERVLPLLIETADAAAFEHALRVADASPEVPGGATGAVRR
DHALDALAQLAGNVRRALADGATRVPFAFDVAPAWPAAGDARWFGEHDAALAAAATREPL
RLHGALARLTAAGQVIYRYARPGARDYLAAWLAHLVYCAIDPGGPRRTLWFGSGDTFEFA
PVAAPLEQLAPLAALFRAGRRMPLAFFPRSAWALVSESESKAAGVWINDRVAGEADDASL
ALAWRGANPSLDEPFGALAHLVFDPLVEHLRRAS*
>GCA_000452945.1_BPMSHR5858_genomic_00338_1 group_1756
MTTEIAHATAHLRRAGRRPLRAVALAAIAAAALAGCKKPEEASAPPDNAASAAAHAARQT
TQKLDQVASFVNQQIEAAKQGVASAASAVPPVTASSVSAAAQGAASAVLGQAATAAGTSL
EAAGRKLQQWSQTAALGTSGAAAASGAAASQ*
>GCA_000452945.1_BPMSHR5858_genomic_00342_1 mdtB
MNPSRVFILRPVGTALLMAAIMLAGLVALRFLPLAALPEVDYPTIQVQTFYPGASPEVMT
SSVTAPLERQFGQMPSLNQMSSQSSAGASVITLQFSLDLPLDIAEQEVQAAINAAGNLLP
SDLPAPPIYAKVNPADAPVITLAVTSKTLPLTQVQDLADTRLAMKISQVSGVGLVSLSGG
NRPAVRIQANPLALASYGLNLDDLRTTISNLNVNTPKGNFDGPTRAYTINANDQLTSADQ
YNDAVVAYKNGRPVMLTDVAKIVAGSENTKLGAWVDAEPAIILNVQRQPGANVIQTVDNV
KAILPKLQESLPAALDVQIVTDRTTMIRAAVRDVQFELGLAVALVVLVMYLFLANVYATI
IPSLSVPLSLIGTLAVMYLSGFSLNNLSLMALTIATGFVVDDAIVMIENIARYVEEGDSA
LEAALKGSKQIGFTIISLTVSLIAVLIPLLFMGDVVGRLFHEFAITLAVTIVISAVVSLT
LVPMMCAKLLRHTPPPESHRFEAKVHGLIERVIERYGVALQWVLDRQRATLVVAVLTLAL
TALLYAVIPKGFFPTQDTGVIQAITQAPQSVSYGAMAERQQALAAEILKHPDVVSLTSFI
GVDGANITLNSGRMLINLKPRDERSESASDVIRSLQRQVANVTGISLYMQPVQDLTIDST
VSPTQYQFMLTSPNPDEFATWVPKLVDRLKKEPSLADVATDLQNSGKSVYIEIDRTSAAR
FGITPATVDNALYDAYGQRIVSTIFTQSNQYRVILESEPQMQHYTDSLNGIYLPSAGGGQ
VPLSAIATFRERPAPLLVSHLSQFPATTISFNLAPGASLGEAVKAIDAAERELGLPASFQ
TRFQGAALAFQASLSNQLFLILAAIVTMYIVLGVLYESYIHPITILSTLPSAGVGALLAL
MITGHDLDIIGIIGIVLLIGIVKKNAIMMIDFALEAERVEGKPPREAIYQACLLRFRPIL
MTTLAALLGAVPLIVGSGAGSELRQPLGIAIAGGLIVSQVLTLFTTPVIYLGFDALARRL
RGFFERSGGPAAHPDA*
>GCA_000452945.1_BPMSHR5858_genomic_00343_1 mdtC_1
MNLARPFITRPVATTLLALGIALAGLFAFVKLPVSPLPQVDFPTILVQASLPGASPETVA
TSVTSPLERHLGSIADVAEMTSMSSVGNARIVLQFNLNRDIDGAARDVQAAINAARADLP
ASLKSNPTYRKVNPADSPIMVVSLTSKTASPAKLYDAASTVLQQSLSQIDGIGQVSLSGS
ANPAVRVELEPQALFHYGIGLEDVRAALASANANSPKGAIEAGPHRYQLYTNDQATKAAQ
YKDLVIAYRNHAAVSLSDVSSVVDSVEDLRNLGLMNGERAVLVILYRSPGANIIDTIERV
KAALPQLTAALPADIQVTPVLDRSRTIRASLADTEHTLIIAVSLVVMVVFLFLRNWRATL
IPSVAVPISIVGTFGAMYLLGFSLNNLSLMALIVATGFVVDDAIVVLENIARHIENGTPR
LQAAFDGAREVGFTVLSISLSLVAVFLPILLMGGIVGRLFREFALTLSLAIAVSLVVSLT
LTPMMCARLLPEAHAPRDEGRVARWLERGFEWMQRGYERTLSWALRHPFTILMTLVATIA
LNIALYIVVPKGFFPQQDTGLMIGGIQADQTTSFQAMKLRFTEMMRIIRANPNVANVAGF
TGGAQTNSGFMFVALKDKPQRKLSADQVIQQLRPQLAEVAGARTFLQAAQDIRAGGRQSN
AQYQFTLLGDSTAELYKWGPILTEALQKRPELADVNSDQQQGGLEAMVTIDRATAARLGI
KPAQIDNTLYDAFGQRQVSTIYNPLNQYHVVMEVAPQYWQSPEMLKQIYISTSGGSASGV
QTTNAAAGTYVATTARASTAGAAAQSAAAIAADSARNQALNSIASSGKSSASSGAAVSTS
KSTMVPLSAIASFGPSTTPLAVNHQGLFVATTISFNLPPGVSLSKATQVIYQTMAEVGVP
PTIQGSFQGTAQAFQESLKDQPILILAALAAVYIVLGILYESYIHPVTILSTLPSAGVGA
LLGLLLFKTEFSIIALIGVILLIGIVKKNAIMMVDFAIDASRQGKSSFDAIHEACLLRFR
PIMMTTMAALLGALPLAFGRGDGAEMRAPLGIAIAGGLIVSQMLTLYTTPVVYLYMDRLR
VWAEKRRDRRSSGGAAIAGE*
>GCA_000452945.1_BPMSHR5858_genomic_00344_1 yhfK
MSMKVLLIGASGRTGRALADLLLKQQDFEVTALVRRPDFALPGAKVVVADLTGDFSSAFN
GITHAIYAAGSAESKGATEEEQIDRDAVARAADYAKARNVQKLVVISSLSAYWPQSSPEF
LRHYSQMKREGDERVIASGIDYVILRPGPLTDDPGVGKIALTEERLDDAPPVARQDVAWA
AIEAIKLGISKKTIGFVGGSVPIEQALRA*
>GCA_000452945.1_BPMSHR5858_genomic_00345_1 bcp_1
MNRKLASGAAAALFALYAVAAHAVLKVGDAAPDFSAQASLGGKTYTYSLADALKKGPVVL
YFYPAAFTTGCTIEAHAFAEAIDAYKRYGATVIGVSADDIGTLTKFSVSECRSKFPVAAD
PDAKIIREYDAKLPAISKANRVSYVISPDGKVLYEYTSLSPDKHVENTLNAVKAWAASHR
QP*
>GCA_000452945.1_BPMSHR5858_genomic_00346_1 group_1758
MEAIRNKTPRKGSWRAALHQLRRFAWSGGALMPARAGTPRADDTVRSWLEQTVGAVDFLA
HVDRDLRFLYVSDASLRFIGYHRDYLHTLTLRELIPEQDTATLDGLLARASQTGDVEKAT
LCLVKSLTYPLDVEVRAVRSRHHGIDGFAIAAFDVSSWRAIEARLTYELHHDPMTGLDNL
SALLPALMRAQQIADEGGGCTALLLLDLDDYQRINRALGYDAGDALLRDTAQRLQRLATR
GERLARVASDKFAVMLSAPTRVAALDAAEALARQLQTAVQRPYAYDGQAVHLSASVGIAI
YPDARATSKHAQHHSPLLRRADRALAQAKAAGGNALAFHQPTDDPADAERLKLEADLYNG
VRNGEFSLHFQPITKSLSGAVVGVEALIRWHHPVHGLVAPATFIPLAESIGLINYLGNWV
LKAACMQLVSWDRQGIALQYVAVNVSPQQFRDPRFTQSVRDAIKLTGIDPRRIVLEITES
LLMHDPNHAKTLLEEVTELGIRFAVDDFGTGYSSLAYLQSFPLAKLKIDRSFVENLLTSR
NDRAIVSAVVGLAQTLELELVAEGVETEAQRTLLTEMGCNHIQGWLVCQALPSDELAQRF
EAQELHLHATA*
>GCA_000452945.1_BPMSHR5858_genomic_00347_1 group_2246
MPITAQLPRTALLDKLWARMNERGDFPLLSDALRATMAAMKNDELDFTALVRVVLSDFAL
TQKVLRLANSAMYMAFGGNITTVTRALMVLGMDAVGHLVVGLKLVDHFHHSAPRRIDAKL
ELNRALLSGCVARKLTEHVDLRAGEQAVVCTLMRQVGKLLVVCYLDAEWEEIRRRAAEHD
DDEAACRAVLGVGYDEIGLEAASRWRLPELIRAGMTRFDANDGAPHEVQWLRAVSHCSTD
VATALTSTHAQQRDARIAALAHRFHGVLGTDADELADIAASLSREETSDTVMREIVELRA
NADKIARGASTPQACLEAGLADLRALPSEHVLGPVLALASESVLAGLAFTRTVMFVRHDD
GVFAARLGFGPNVDAALERLRFHETFEPDVFHLAITNSVGIFIEHARDPKMTKRLPAWYL
DAFDDARAFVLLPVIAERNAVALLYGDWALGQPARKITPQEMGVLNELAAELGRFFCGAQ
GDDR*
>GCA_000452945.1_BPMSHR5858_genomic_00348_1 cbaA
MSTERLFSIRNRWLTVSVGLTLFAVVASVLIGFVWLPSVQKDAQFQGIWNAICSAAGVPR
AWLPSQQEPVPSAVRTSTVVVTPQMMAGADSLSIGRGATLAMRCTMCHGARGMSDANSPN
LAGQYAAVIYKQLKDFQTGARTNAVMAPMVVNLSDQDMRDLAAFYSYLPRTATALAAAPP
RIVASGAPMRNIAPCASCHGGMDNKVGGPRLEGEPEIYINAQLQAFASGARHNDISEQMR
NVARQMTPAEIAAAARYYAGRP*
>GCA_000452945.1_BPMSHR5858_genomic_00350_1 group_3087
MSKQPMPDRLDGAQLAARIERRWAILAVGLMIVLVATMVYTGLHWAMMPPSRVETIDPTT
LHVSGEFVESNLGTAREPDGSVTVRVIGQQYSFTPQCILVPANTPVTFRATSADVVHGFL
ITDTNINSMLEPGYVATFRTTFDKPRDHLMPCHEYCGTGHEGMWAHVKVIDRAEFDKLAS
SARRLSCVKE*
>GCA_000452945.1_BPMSHR5858_genomic_00351_1 group_3088
MSDTSQRHGGDPEPHDDAVDEIVRSGPRGALALAGTATAIVIALWFAFYFLVFLPRGVIQ
*
>GCA_000452945.1_BPMSHR5858_genomic_00352_1 adhB_1
MEASVHKHLIVKFRSACANAALGLLAATLAASAVHAQPAADAATLKRGEYLARAGDCVAC
HTVPGEKLFAGGRAMPTPFGTLYSPNITPDNDAGIGKWTADEFYTMMHTGRSRDGSLLYP
AMPFAAYTKVTRADSDAIFAYLRSVPPVKLANRPHDMRFPYNNRQLLIGWRTLYFKEGEY

QPDHTKSAEWNRGAYLVQGLGHCSMCHTAINALGGSSEANAFEGGLIPMQAWYAPSLTSN
KEAGLGDWSIADITGLLQAGASHRGAVYGPMAEVVYDSLQYLSDDDVRAMSVYLKSLPQR
GSDAESAPKTTMAASEQARLSKLGAKIYDAQCASCHGKTGRGKTPAFPPLAGNQSIQMTS
AVNPIRMVLNGGYAPGTAKNPEPYGMPPFAQSLSDDEVAAVTTFIRTAWGNRGAPVSAKE
ANALRSAPLY*
>GCA_000452945.1_BPMSHR5858_genomic_00353_1 group_1759
VTSRPTAVLAVLCGLLSQTALSQEPQTERAPDTMEARVLACAACHGRQGEGTSNDYFPRL
SGKPAGYLYNQLVAFRDGRRKYPPMNYLLAYLPDAYLRKIADHFASRRPPFPTIAAPAAA
PAVLERGRLLVKSGDPSHKIPACASCHGASLTGMEPAVPGLLGLHAEYLSAQLGAWRYGT
RTSIAPDCMQQVASRLTDTDVTAISAWLASQPAPANPSPAPAGSLDMPLACGSEPR*
>GCA_000452945.1_BPMSHR5858_genomic_00354_1 group_1141
MVRHAFASTPAPSRAASVSILHPRALASIALASAVALTAAGCGERAEPRRDGATPVLSAS
FRPKSGFVVPVSIAGKTYHFLVDTGASHTAIDNRLAQSITRPSTDEQIPIAYRTMLEKGL
TTADGVLPRERVRLWQPLPIALGSYQVPSFYPWLGLDLSLLSQVLGTQIDGIVGIEIFRQ
LSWVADNRSGTLTVWRHPPAAQRFAHCVPYQDSFGQSPAVSVDFRDRWTMFRFDTGARYS
IASAPTLAYLASHKAAMPLGGTVPSMSANGVGESRDHFVSGLSFDGRPVGRLRVAEGGGD
MLGMNFLARLDRYMFVPSTMEFCYDAGRFTQDDPQPLRTIAIRFVDGRVELFHNRPEDLR
RYGLENGDVLVEIDGKRVEAPAIDDVRDRLSTAPAGSLDIVVERGGSRRAVRI*
>GCA_000452945.1_BPMSHR5858_genomic_00355_1 group_3090
MGELADESRIFLSTLQPERRERGLAMTAVALSVVLFATLAPFVRIQMPQVWAFIPIYQSA
IVICDLITAGLLLGQFGILRTPSLVVLAGGYLFTACMASVHMLTFPGLFLPSGLLDAGPQ
TTAWLYMFWHSGFPLCVIFYTLASRRERAEPLYAGRLALPVIVCIGTALVATGAFAALAT
SGRELLPPIMANNHYTPLMRGVVTTVWLMTLAALILLTSRRPHSVLDLWLIVVLCAWLLD
IALSAVLNHGRFDLGFYAGRVYGLVASSFVLLALLLENGKLYARTVHALQGERIEHRRVQ
EKTVQLNEANELLEQRVAERTRELQAANDELRREIVERERAEEALARSREELREIAAISS
TAREQERGRIARELHDELAQTLAMLRLDLERVSTTGNELAGSFMEMRGLLDGAVAATRRI
ASDLRPLMLDDLGLVPAIQWLVQAFQQRHGIDCTLIVDPPELDLVEPQASTTFRILQESL
TNVGRHARASHVDIRLIADDREVMLSVRDNGVGFDTENPRKPNSFGLVGLRERAYLVQGT
LSVQTAPGQGTAIEVHIPLVRAREVTVLQSGELRE*
>GCA_000452945.1_BPMSHR5858_genomic_00356_1 ssuA_2
MTADDTPRTALSPSAHDDTRRSLLKAAGAVALAAPLPGGLAALSASGAAHAQRSAKTLRI
GYQKYGNFVVLKARGSLEKRLADRHVSVQWIEFPGGPQLLEGLNAGAIDVGTVGETPPVF
ALAAGVDFVYVGSEPPAPLGEAIVVPHDSPIRAVADLRGRKIALNKGSNVHYLLVNALRR
AKIDYRDITPVYLAPADARAAFAQRGVDAWVIWDPYLAAIERQTNARAIANGEGLVNNIQ
YYLASRTFAEAEPQLVRALLDEVGAVDRWARANVATVAAQLSPLVGLDTATLERALKRAS
YGVQPIDDAALAYQQRIADTFTALKLIPRRIDVAAARWQAA*
>GCA_000452945.1_BPMSHR5858_genomic_00357_1 group_7242
MQFEELSDDEWARIEGLLAAQPVRNTRCGRPRAQARAVANAVLWVLTTGECWAALPRRYP
SSPTCRRRFDEWSADGTLAALIERLRGAGRQVELRERIGAIDAKPHARARRRFQHNAFWS
SPESWCAPAVAVVGGVIEAGRDDCVPEHGEGNAN*
>GCA_000452945.1_BPMSHR5858_genomic_00358_1 serA
MAAYAYRAGISGLRYDSTTTLGNMTSVLLLENIHDSASRHFADANVDIERRKGALAGAEL
RRALQQHQLIGIRSATHLLRDEIDAARHLLAIGCFCIGTSQVDLPAAAHHGIPVFNAPFS
NTRSVAELVIAEAILLLRRVPEKSVLAHAGEWAKGAGGSFEARGKTIAIVGYGNIGAQVG
VLAEALGMRVVYYDVQAKLSLGSAQPARSLGEAIALADVVTLHVPAHASTHNMIDASVLA
QFKSNAILINASRGTVVDIDALRDALVQKRLSGAAIDVFPREPKTNADRFESVLQGLPNV
ILTPHIGGSTQEAQENIGEEVSSKLVAFLTRGDTAGAVNFPQVSPGECTSAARLLNVHGN
TPGVLAVLNTLLAQDGANIVAQHLQTRGDIGYVVTDLDRVPSDAFLDTLSGNRAFTRSRL
LRSA*
>GCA_000452945.1_BPMSHR5858_genomic_00359_1 group_7244
MATSVPDPTRQLNKATVSSAYALFMLMLAEERGIADADLLAHTGVTRTKLEEPNARITPL
QQAAIVFNLLGMTNDPSIAIEIGLRSSLTKSGLIGFGLMSCATLGEAIQLGIRYLPTRVP
FFSVRFTEFEHTVQIDILEAFPLGRLRQFAVENFMVETAILFNSLLTPSHDKTMKANAEL
CFEWPEPPYFARYRDRLPRCHFGAPANQIRCEAALLDEPIKTANAHTAQMIVQQCEAELA
RLGYAESIVERVRNLLIRGSHGYPSLDALARELHLSERTLKRKLSDYGTTYSALLDEIRL
RDALRLLEGTTLTVEEIAARVGYTDRANFSRAFRRWTGTSPSDRRRT*
>GCA_000452945.1_BPMSHR5858_genomic_00360_1 group_7245
MNKTLRFKNDAEKVAHVRNEVNAASDALRAKYPLLDNQNLIGAAVMAICVATLIGSAYLY
AIGAIAWYVALPIATLATSLIHELEHDLIHLMYFKKTPWAYHAMMTLCWLTRPGTINPWT
RRRMHLHHHKVSGGESDLEEYGITNGERWGLKRLLMLADGMLAVALRPLGMRRKVLQYVA
AQPAQARGDRVRLRIEQLMSYMPIGHVYYVLWHAFIAYHAGLFALHALGYHPAVPALVQQ
TMHVVDFLAVTWLGPNFVRSFCINFISSNMHYYGDIDSRNVVQQTQVLNPWWLIPVQLFC
FNFGSTHAIHHFVVRDPFYIRQLTAKRAHAAMRAVGVRFNDIGTFRRANRWNETRAA*
>GCA_000452945.1_BPMSHR5858_genomic_00362_1 ypdF
MIDRLKYSFTDLARYDASVVDTHAGLSRLIDALQLDALVITSQDEYISEWLPRCNNPRYA
LSGFDGSVGSGVFLSAAAARRIGVPQFVLFVDGRYHLQAEKQCDPALVQVEKLGLNVAMW
PAIGDWLAAHAADIKRAGYDAPRLSVAQRARLLGQTERAGVQWTSLAGREIDGAISLPGW
RVERPVFELPRSATGVSIAANVATLNSRISEHLGDPRAKAAFITCAADDLSYLLNSRGYH
LPYASSHVGFLFVVGEAVALFLPEGFDRCPVRIDSYPALHVIRNDAAALQRFLAQFDVEY
VCYGFEAVNCALPDTVRSVWPDARHVDHNPVEAMRASKTPEVLDQFRDAFARSSEAIAEA
MRWAKAGEPGKRHSEHDLARVISEAYAARAAVALSFTTIAANGANSASAHYTAVSPDVEL
TEGELVLLDSGAYFEGGFATDCTRVVLRRTHADTRPQPWQREIYTVALKACIKGLVTRFP
QAATGADVDAAVRGVCRAHGYDYNHGTGHGIGIHVHEGGVRFNVGSQYGLVPNAVISVEP
GIYLPGKGGVRIENIVVIHPSEREPGMMEFENLVTVGYDWDLIDVDLLTDDERAYLRDYE
RLCVERGTQVTDCPLL*
>GCA_000452945.1_BPMSHR5858_genomic_00425_1 group_4436
MKHALPIAALAGALAAPSLAHARQVTFETQLADYGGDGAYVVLYVTDRAGRYQGTLWMAG
RQTKYYRHLRDWYRATGGSARLDGITGASVGAGRTLRATLDVADALLDAGYEIHIDTAVE
DMNENASEVVVPLNAAGAGKAVKGRGYVKSFRYSL*
>GCA_000452945.1_BPMSHR5858_genomic_00429_1 lysN_2
MTLAAAVDYLFAAPFADPQGSPIRELFKHVGKPGMISFAGGYPSAELFDREGIAAAFADA
ARDDPVACLQYGDTAGQPGLRAALADWMTRARGVACDASQVLVTTGSQQGFDLLVRALIE
PGDVALVEAPAYPAALQALRLAGAKLVPVRTDADGVDPDALAAQLAAWPAAERRPKLLYT
VPTFANPTGATLPPARRARLAAVAADARLVLVEDDPYADLRFAGEPVKPVLAHDGASDWT
VYLGSLSKIVAPGLRIGWAVAPAAVLRRMTVAKQTSDLCTAPIAQEAIRRYLASGRLAAH
LRTIAQTYGSRCRALVSALAQFVPDDVEWREPSGGMFVWARLGGGRDASEVLKRALEHNV
MYVPGAAFYARDADAGSLRLSFAAPGEAEIFEGVRRLRAALAACRQRT*
>GCA_000452945.1_BPMSHR5858_genomic_00432_1 group_714
MTIRIDDIVPLSIARARFSELADDARDGGEKIITKNGAAYVALIDARRLDHYHRLERERA
ECACAALLHGGGDDGRDSDDRGNGGDAAGDAARSGGRDSPDGRGAAIAAGRETGAANATD
ATGAPGTMGAMAAPCDSDGADALAPDALAPDALAPDAPSAERRAGPD*
>GCA_000452945.1_BPMSHR5858_genomic_00437_1 glpR_1
MLAEQRHRYILSELGRAGALSVAQLVNALNVSRETIRRDLNALAARGLVITTHGGALAAD
RHEPSVSERDASHADAKRVIGRRAAEFVPDGASVLLDSGSTVHAVALALADRHRLSVYTN
DWRIAGVLGRRNGNRVTLLGGELSDDEDATFGLDTIHQLAQYHVDFAFVGAGGITPDGTL
TDYSRVVAEVRGRMLAAAAAPIVVADHSKFGRVTPVRINGSDAARFLVTERAPDKAMRRA
LTARGIELVVCG*
>GCA_000452945.1_BPMSHR5858_genomic_00440_1 por
MVQTGRFLVMPLLSSDHCRALPPEVSRPRYDRRALRTGIVHLGLGAFHRAHQACYTETLV
ERGDLRWGIAGVELRRRHTVERLAAQDHLYSVTERAGDAARTRVVGAVHRTLFAPQALAT
LLGLIADPSVSIVSLTVTEKGYYRRPGGGGLDLDDPAIRRDLAQPHAPSTTLGVLAAGIR
LRAAHAPLSVLSCDNMPSNGDTLRALLAQYAEQTDGALARRIRCDVAFPNTMVDRIVPAA
TPESLDWVQSRIGVRDEAAIVCEPFAQWVFEDRFAGARPRWEDAGALVAADVRPYEKMKL
RLLNGSHSAIAYAGQLRGRRTVSDAMADPLIDALARGVMTRELLATLDVPAGYDVRAYCA
SLIERFRNPALAHRTAQIATDGTQKVPLRWLPALAESAAAGVERPFLERALAMWLHYVEV
ARDESGRPLVLEDPGAQALAARLHGAPGATDAVRAALGLIASRDAARWPEALTARVGAHL
ETVRTRGTDALLRPLLDA*
>GCA_000452945.1_BPMSHR5858_genomic_00441_1 rspR_1
MSDKEPAEDVVVAALNGFVLDAERSYTQQVHALLRNAIVRGALPPRTALSEAVISATIDV
SRTPVREALAQLADERLVQIYRKVGTLVAPISVSMLEEGRFARGTLECANHVDLAQKITP
AQLAELARLVDAQRDAVAAHDVDTFFTLDETMHGRMFEFAGRPHVWAMLQPMKRQFDRVR
WLLLDRVEDHARRALHEHEQILAHIAARNMALLGASVAVHVDRVNAHLPEVRERAPHYFV
D*
>GCA_000452945.1_BPMSHR5858_genomic_00445_1 group_1367
MPASSGPLRMLRSASAAAAAAFVALWSTGFIVARAIKPYADPNLYLLARFCGTALLYAGI
ALAARAPWPAPREWGRHLLAGALLQGFYLSASYWAVAHGLSAGVMALLGALQPLATAALA
APLFGERLAARGWFGMALGIAGVALVLAPKLGDAPAAAGDAPAWAVVAIAIAAVGSITAG
SLYQKTSLAKADLRIAVALQNLGAALVAGAAAVALGETRWHGAPALWFSLAWGIGMLSGV
AVMLLMAMLRRGEAARATSLLFLAPPLAALQGYALFGETLVPLQLAGFALALAGVTLARR
N*
>GCA_000452945.1_BPMSHR5858_genomic_00446_1 nadE_1
MSRPDQAARRRAIAAELHVSPTFDARDEAERRIGFVADYLRTAGLRACVLGISGGIDSST
AGRLAQLAVERLRASGYDARFVAMRLPYGAQHDEADARRALAFVRADETLTVDVKPAADA
MLAALAAGGLAYLDHAQQDFVLGNIKARERMIAQYAVAGARNGVVIGTDHAAESVMGFFT
KFGDGGADVLPLAGLTKRRVRALARMLGADEPLVLKTPTADLETLRPQRPDEHAYGITYE
QIDDFLEGKPMDDAVAETVLRFYDATRHKRALPYTMFDWPGHPA*
>GCA_000452945.1_BPMSHR5858_genomic_00448_1 group_7251
MVPHRTRIYVAATVDTKGAEAHFVKDRIADAGLAAVVVDLSTRVPGLAADIGADAVAAHH
PDGAAAVFCGDRGRAIAAMAVAFEHYIKSRDDVAALIGIGGSGGTALVTPAMQALPIGMP
KLMISTMASGDVSAYIGSSDIAMLYSVADIAGLNRISRQVLANGAHMIAGAVRDMQPPHA
DLKPALGLTMFGVTTPCIQAVTSRLDARFDCIVFHATGKGGPAMEKLADSGLLDGVLDLT
TTEVCDLLMGGVLACGADRFDLIARSRVPYVGSCGALDMVNFGHIDTVPPRYAQRLLYKH
NPQVTLMRTTPDENRRIGEWIGAKLNACDGPVRFLIPEGGVSALDAPGQAFWNPQADEAL
FDALEATVVQTERRRLVRVPAHINDARFADAAVEHFLSLHAAHRN*
>GCA_000452945.1_BPMSHR5858_genomic_00449_1 group_4442
MKRLSRVEILEKLRATIARGEPIVGGGAGTGLSAKCEEAGGIDLIVIYNSGRYRMAGRGS
LAGLLAYGNANEIVVDMAREVLPVVKSTPVLAGVNGTDPFCNFDAFLDELTRLGFSGVQN
FPTVGLIDGNFRANLEETGMGYALEVDMIRLAHRKGLLTTPYVFSEADAIAMTEAGADIV
VAHLGLTTGGTIGANTARTLADCVPLVRSWADAAKSVRDDVIVLCHGGPIASPEDAAYIL
RACPQCHGFYGASSMERLPAETALTDMTRRFKEIDRHAAS*
>GCA_000452945.1_BPMSHR5858_genomic_00450_1 group_7252
MNHTSFASSVIDAPIERVWAFFRDFDGLPAFHPAIVESRLEPGPDAYTVGAVRYLTLADG
YVREKLLKLDEPNHALEYAIVESTMPVRDYVAGVQLVPVTDSGKTFAQWWANFTTQGVEL
QPVAESISEHVFAAGFRALADKLRAR*
>GCA_000452945.1_BPMSHR5858_genomic_00451_1 group_1609
MKDLQSIRGQLRATFLSVLLGAVIAASAQAGAATGKVPGDFGPPQGEPLHATLVSLPNVP
PPITRRYPAKVIVDLEVVEKVLPIADGVNYTFWTFGGAVPGNFIRVRQGDTVEFHLKNRP
DSKMPHNIDLHAVTGPGGGATSSFTAPGHESRFTFKALNEGLFVYHCATAPVGMHVANGM
YGLILVEPPEGLPKVDREYYVMQGDFYTNGKYREKGLQSFDMDKAIDERPTYVVFNGAEG
ALTGERAMRARTDETVRLFVGNGGPNLVSSFHVIGAVFDKVRADGSNVTQNDVQTTLIPA
GGAATIEFHTRVPGNYTFVDHSIFRAFNKGALAILKVDGPENKAIYSGKELDAPYSGDAV
TAAAGSGPAGAPGAAAGVKGGVKGGASPVRTSAAPLSLPAQVKAGGTVFASTCAVCHQSA
GTGLPGVFPPLAGSDFLAANPKRAIGILLHGLNGKVTVNGHDYDSAMPPMAQLTDDEVAN
VLTYVLNSWNNPGGRISADEVSKARAKAPAVAEAAH*
>GCA_000452945.1_BPMSHR5858_genomic_00452_1 group_3099
MHAPSSRHACRAASAVRLARVAIVALTLAFAAAGAHAARFVRLPGGDFESALPQDVPGRS
TPVHIDAFELQDTPVTVRAFAGFLRAHPEWRRDRVAHVFAGPAYLADRADPLHPASPTPP
DAPVTGVSWFAARAYCASEGARLPTWLEWEYAAAADATRPDARNDPLWRRQILSWYEQPA
ARVLPSVGGAPNVYGLRNLHGLIWEWVDDFNALFISGDSRTQGDPDQQRFCGAGAISIVR
RDSYAVLMRVALLSSLTGADSTGSLGFRCARSISGDRP*
>GCA_000452945.1_BPMSHR5858_genomic_00618_1 ubiJ
MTLAAKPFAAAVNHLLARESWARDRLIPYAGKTARLEISPVTLVLLVQPDGYLAAVEAHD
ARRFDVSIALSSGADGGAFDAAAAFLQGGQAAVMKHVKIDGDAEFATQIAKLAEHLRWEP
EEDLSSVVGDAAAHRIATVVRSAGARARRTGRNVLDSIAEYWLDENPQVVRKSALADFDA
GLARARDTLARVEKRIERLEQKIDVRAGGSSRRAQ*
>GCA_000452945.1_BPMSHR5858_genomic_00621_1 group_848
MPIYAYRCEACGYAKDVLQKMSDAPLSQCPECGQDAFRKQVTAAGFQLKGSGWYVTDFRG
GSSGSSAPAAGGEKAGGEKAGGETAAAPAAADSTASAPAASTPAPAAAPASST*
>GCA_000452945.1_BPMSHR5858_genomic_00684_1 group_3124
MTRRSTFSLAGACAAALFAFATAASAQNAPAPTQPRPEAAADTRLHEADQAFVADATKAV
STQRDAARIADSRSSDRAVKAFAARVAADNEKLSQALRAASPRGVDVPHNDPDPAVLDGI
KNLRGDAFDKAYIEQVALDGNRKALSVFQAEIASGRNEQLKNAARKGLPVIQAHYQAAQE
LASRKHLTSDASK*
>GCA_000452945.1_BPMSHR5858_genomic_00695_1 group_3125
MQIHEVRIPIGKVELNGLLAAPEQASGIVVFAHGSGSSRLSPRNQEVAAVLQRAGLATLL
FDLLTLEEQRRDAVTAEYRFAISFLARRLVSALDWLRERPDVGALPVGLFGASTGAAAAL
IAANARGRVVRAVVSRGGRPDLAGDALPRVRVPTLLVVGERDDEVLRLNRVAAGWLIGES
KLVVVPGATHLFEEPGTLDEVARVAADWFVAHLGEGRPSPEGARR*
>GCA_000452945.1_BPMSHR5858_genomic_00696_1 pncB2_1
MRAKPACTRALLADLYEFTMLQSYFDCGMNDTATFELFVRRLPAHRNFMMAAGLEQALDY
LANLAFDADEIEALAGMGLFGDGFLASLASLRFTGTVRAMREGTVFFADEPILQVTAPLR
EAQLIESRVMNLLHYETIVASKAARAVLAAPGRALVDFGLRRAHGAEAALLSARASYVAG
FAGTATLLAGLRYGIPTYGTMAHSYVQAHDDEALAFEHFARSFPRNAMLLVDTYDTEAAA
GKVIEVARRLARDGVEVKGVRLDSGDLAQHAMRVRAQLDRAGLGRVTLFASGNLDEYRLA
ELIGQGAPIDGFGVGTRMNTSADAPYADCAYKLTEYAGVPRRKRSEGKATWPGRKQVFRR
YRDDGLLDGDCLALHDDAQAGVALLEPVMIGGERVAPSPSLADVRAHARAQLDTLPAALR
DLARAAEYPVAVAGTLAALAQRLDARAASGA*
>GCA_000452945.1_BPMSHR5858_genomic_00697_1 group_3126
MDDYFRDRVDAGRQLAAHLADYANRDDVVVLALPRGGVPVAFEVAKALRVTLDVLIVRKL
GAPGNPELAMGAIATGGVVLLEQSVLRAMRVTERELADTIERERDELARREAAYRGDRAP
PALDGRTVIVVDDGIATGSTMLAALEAVRAHRPARLVAAVPVASPSSVRKVEAAADATVC
VMRPDGLMGIGQFYRNFEQTSDDEVRALLARAREWARPNAPDAPDAPDAPPAAAGGDD*
>GCA_000452945.1_BPMSHR5858_genomic_00699_1 adhB_2
MALIAARGPRLRFSAAAATAATASALVMAALLGGCERDAGDAGGAGGARAAAAPPPASAA
SAAQAAGARAAARIASDASRADPAAIARGRYLARAGDCAACHDAADHTPYAGGQPVNSPF
GPIYASNITPDPDAGIGRYSLRQFADALRLGKAADGRRLYPAMPYPSFAKLDDSDVAALY
AYFMHGVQPSDKRAPATRLPFPFNQRWGLAIWSALFGNRERFVPNPQRPAAWNRGAYLVQ
GLGHCGACHTPRGPAYDERGYDERSAAYLTGGVNDHWFAPNLTGAPLDGLGRWSERDIAA
FLRTGHARHGAAFASMAPVVEASTALLSDDDLHAIAVYLKSLPAQRTPAIALVPGRARQP
PARAADDGTQQRPGAGVYFAFCARCHGADGAGTPDRGPALAGNSLVLAPDPTSAIRIVVE
GSAEPRVAGRDARRMPGMRGRLTSTEIAQVVSFVRGAWGNQAAPVSDREVESLRSAVHR*
>GCA_000452945.1_BPMSHR5858_genomic_00705_1 group_849
MTRHSRHAAQREAQRMMKASRELNRSAILLALSVLADSGVEHYRGSFRNRAMLAPLAASA
LSLAAAAHGHADGAPRRHPARDAVHLGAAAAAVAGVGFHVYNVLKRPGHLSWHNLFYGAP
LGAPVALLLSGLLGAAGERLRACPEQAPRLCGLPAGRALAALVAAGLAGTVGEVALLHFR
GAFHHRAMVAPLVVPPVAALLVAHAALAPARPNRWFSRAWLKATAALGIAGVGFHAYGVA
RQMGGWRNWAQNLLAGPPLPAPPGFSALALAGLSAVSLAEREAGA*
>GCA_000452945.1_BPMSHR5858_genomic_00754_1 queA
MLTLSDFDFDLPPELIAQTALPERSASRLLEVDNTNPSAPPRLVDRRFAELPACVAPGDL
LVFNDTKVLKARFFGRKASGGKIEVLIERVTGERTALAQIRASKSPPPGTTLTLADAFDV
TVGERVEPFFTLHFPDDCLVLIERHGRLPLPPYIEHAPDAADETRYQTVFAANPGAVAAP
TAGLHFDDAVLAALEARGVERATLTLHVGAGTFQPVRVENLAEHRMHSESYELTDALVEK
IAATRARGGRVIAVGTTSMRALEAAARDAQAAGRPLAATRAETDIFITPGYRFRVVDRLV
TNFHLPKSTLLMLVSAFAGIETIRAAYRHAIDARYRFFSYGDAMLLTRRDDAAEATHGGA
*
>GCA_000452945.1_BPMSHR5858_genomic_00790_1 group_278
MDAAGSFASFAATLACALSAWLAPSAGIAQTAWPSDAPRVAGASGPPRTSGLARTPAAVT
DAPGAATAGALAASPPAATRYRLDPRHSGVTFRVDNFWHAHLTMRFARMRAELSDVGDDR
LASRVDVTVDAASLGANVPFVAALVKGSAMLDVARYPEIRFVGTRFERTGATTGVLTGEL
TIRAMTRPITLTVRFDAGRQLEPDTGARAEREGDESDESGESNPAERRAGQGRRAAELRD
GRPRDAGPGVADSRDAMRRETDSGSAHRGAEPPERIARAAPDAARPLAFVADGHFSRTAF
GLSRWLPAVGDDVRLRIRAEFVRARADP*
>GCA_000452945.1_BPMSHR5858_genomic_00791_1 group_7287
MIRNVLIALVCAAPVAALAEPPKAAGGMIVDDDGMTLYTFDRDTMPGKSACTGGCTANWP
AALADAYDKPGGDLGFIAAAGGKHQWTYKGRPLYRFSGDAKPGQHTGDGFGGMWHVARP*
>GCA_000452945.1_BPMSHR5858_genomic_00793_1 fecR
MSGAKPAGGPPAVPERVARRAVQWWVELGAAPASDATRARLARWRAEHPAHDAAWRHIEA
VNGRLGHVASALSARAARAALLPPRSRGRRAALKALAVLLFAGGGAWTGQAYLPWRAWGA

DVRTALGERRTITLADGTTVVLNTDTAIDIRFDDATRGVRLLRGEIMVTSGRDAARAVPR
ALVIATAHGDVRPVGTRFSVRERGDATRVDVFDGAVRIAPRRAVGRAPLITAGHGADFTR
DALAGPPRVLGASAGAWVDGILVASNLRLADLLAEIDRYRPGRVRCDAAVANLRVSGTYP
LDDTDRILDTLRETLPVDVEYLSRYWVTVVAARD*
>GCA_000452945.1_BPMSHR5858_genomic_00849_1 hsrA_2
MRAGDFRVTAAVVASALFMQNLDSTVVATALPSMARELGVNVVFLSSAITSYLVALTVFI
PVSGWIAEHFGAKRVFIAAIAIFTAASVMCAAANGLATLVAARILQGAGGALMVPVGRLI
LYRGVSRHEMLAATTWLTMPALVGPLLGPPLGGFLTDALSWRAVFWINVPVGVAGAALAA
RLVPASAGERRAPADARGMLLVGAALAALMLGVETAGRDVLPAGAPALCLGAGVALGGLA
IRHCRRVAHPVVDLSLLGIPTFHAATIAGSLFRAGAGALPFLVPLTLQVGFGASASRSGA
ITLASALGSLVMRPMTHAALHRAPMRTVLIAGSVSFAAVLAACATLSPAWPDAAVFALLL
VGGLSRSLSFASLGALVFSDVPSERLSAATSFQGTAQQLMRAVGVAVAAGALHLAMLIGG
RGQASRTDFACAFAAIALVVLASVPMFAALPADAGEGLAGPARRPRP*
>GCA_000452945.1_BPMSHR5858_genomic_00899_1 group_853
MSTKVLTIIVGTLAMVAAVYPDLAAAALGGEPVNYEAGSQVPPHIYDVRPESASQQPYTV
QRDLIQNRTEVHQYIDGNGKVFAVAWSGSAKPNLQTLLGGKLFSTMIRSQSASHHGSRHS
LNQWNDDLVVQSQSHGHLFRGRAWIPSMLPQGVSPEVIQ*
>GCA_000452945.1_BPMSHR5858_genomic_00911_1 grsB
MVVGLLAILKAGGAYVPLDPSHPPERLRRMLDDTNPVAVLVDDIGADALASFESHVAARS
PRVHLSRDIAQWRACSPANPSTPRERAARRLAYVIYTSGSSGEPKGVMNEHRGVVNRLWW
MQQTYALDERDAVLQKTPFSFDVSVWEFFWPLMSGARLVIAKPEGHKDPAYLSELIDRER
VTTLHFVPSMLQAFLEDEGAARGCGSVKRVMCSGEALPPSLVKRFYRCLPDARLHNLYGP
TEAAVDVTAWACDAEEGGASVPIGRPIANTRIYILDGYGQPVPRGVAGELYIGGVQVARG
YLNRPELTRERFVDDPFVAGGRLYKTGDLARWRTDGSLEYLGRNDFQVKIRGFRIELGEI
EAQLAKVAGVREAVVLAREAAPDTKRHAASNENSASNSGEKRLVAYYTGDADVVALRAQA
AQHLPSYMVPSAYVRLDAWPLTPNGKLDRHALPAPADDAYARAEYEAPQGAKEEALAAIW
KDLLPVERISRHDNFFELGGHSLLAITLIEHMRRANLHADVTTLFTAPTLADLAVRVGAS
RGDTYVPPNLIPGDAANGTPESAEVEELRL*
>GCA_000452945.1_BPMSHR5858_genomic_00983_1 group_1154
VSVRRHAGDGDVHVGVSLADLPAYGADSVERVVYGIVREMGEPIAAEHGIGALKRPFPGY
ARSTAEIAVMRAMKAAFDPLGMLNPGKAL*
>GCA_000452945.1_BPMSHR5858_genomic_00987_1 group_1155
MDWFKWLAGGRHRGYGGRRGHHGGEAGGGHGHGGGHGERGDHGSHGYGGHGWGRQGVQRG
GGWNDARDGAPLRGEAGAPAQHAACAKCGAINGAEAKFCAQCGASQTPPVCGACRAPLAA
SARFCPQCGTAVSVQRG*
>GCA_000452945.1_BPMSHR5858_genomic_01007_1 group_816
MLALKLTLVPLFLLLVSIAGRRWGPSIAGWLAGLPVVAGPILFLVAVERGPAFGAHAALL
SLSAIAASEAFSFAYAWTCRRHRWPLALAAGLAAWAAAASALARLPATPPAATAVAFAAT
CFGQSCLPRGATLAPRAPLSHADLAGRLAAGAALALAVTSLAGALGPAWSGLLAVFPLLG
SVLAVSSHRAHGPDFVVPLLRGMVFGRFSFAAFCLCVALTLPRQPALQAFAEAAALSVAV
QGATKRLAERGPRPTALAQTATPD*
>GCA_000452945.1_BPMSHR5858_genomic_01022_1 group_586
MPKQSAAIVVIAAFSCGAMESSRYIISDVSRNQSSLPSRHVPDSTRTAGESGRSARAPDA
PIQSTSRNVRNRNASMRRATRALLRSLDIRHFFVVGRARLPLSLRTTTIIACCESLVATH
KKSSRLASFRHRGAADAIGPAACAVANALSCRDTCLRSHAQAAAVPANRAARDGLSSRQS
VAAAPNPRIVPAGAALSGRMRRRIACMREHAGAMPRPPAAAHASVPFVPSARFARFAPFA
PSALPIRCFT*
>GCA_000452945.1_BPMSHR5858_genomic_01106_1 yehY
MTPRTAGGAAAPAPAPRPRAMPAWAARVDKVGVLIAALVAYAAFVLPFVTLRANRIAAGA
ELAPAAVFPALHAYALDALWAAGALFALVHSRAAWRAAVGVGLVFALGVAIGAAPAHLVT
PDTPLARVSPAAGAWLLLFAFAVLIADALARIALAPAMRLVALAAASAALAAFIHGGFWD
GLSVMQEYAVRADTFRNEAIRHLALVAGSVAAAVALGVPLGIGCTRSAALRGALLPLLNV
VQTIPSIALYGLLMAPLAILAARVPLAARLGVSGIGVAPALIALFLYALLPIVSSVVVGF
AQVPAAVVEAALAMGMTGRERLVAIELPLALPVVLSGVRIVLVQNIGLTAVAALIGGGGF
GTFIFQGIGQSATDLVLLGALPTIALALVTAVLFEAATDLAKGARR*
>GCA_000452945.1_BPMSHR5858_genomic_01127_1 cobL
MTAWLTVVGIGDDGYAGLGKPARRALADAAVVMGGERHLAMLPARHRAQRAPWPKPFDLA
PLFARRPARVCVLASGDPMLFGVGALLARALAPNEWRVLPAPSSLSLAAARTGWPLQDVD
AVSLVGRPLASLHRHLYPGRRLFVLSADGATPAAIAGALVARGFGPSAMSVFEHLGGPLE
RRLDARADAWRDARAAALNVVAIECRACADAPRLALTPGLPDDAYRHDGQITKRDLRALA
LARLAPAPGELLWDVGAGCGSIGIEWMRAHPSCRTIAIEAHPDRQRFIEHNRDALGVPAL
ELVAGRAPDALAGLRAPDAVFVGGGVSREGVLDACWAGLKPGGRFVAHAVTLQGEAALVA
WRERHGGTLTRVSVAHAQPLGGFDAWRQALPVTLYDARKPRGAQPEHA*
>GCA_000452945.1_BPMSHR5858_genomic_01129_1 cobH
MLDYLRDGRAIYRESFATIRAQADLTGVPPDLEKLAVRVIHACGMVDIVRDLRFSAGAGE
AGRAALAAGAPILCDARMVAEGITRARLPASNPVICTLGDPEVPHLARELGNTRSAAALE
LWRAHLAGSVVAIGNAPTALFHLLDMIDAGAPRPALILGFPVGFVGAAESKALLAADSRG
VPFVAIAGRRGGSAMAAAAVNALASEAE*
>GCA_000452945.1_BPMSHR5858_genomic_01130_1 cobI
MTRKTGRLFGLGVGPGDPELLTLKALRMLQAAPVVAYFVAKGKKGNAFGIVEAHLSDAQT
HLPLVYPVTTETLPPPLCYETVIADFYDTAAALVAAHLDAGRDVAVICEGDPFFYGSYMY
LHDRLAPRYDAEVVPGVCSMLGGAAVLGVPLVYRNQSLAVLSGVLPEDELRQRLASADAA
VVMKLGRNFDKVRRVLDALGLARRALYVERATMAAQRIVPLADVDPMASPYFSLLVVPGE
KWQG*
>GCA_000452945.1_BPMSHR5858_genomic_01132_1 chiA
MNFSMLSRIVPRALAAGCLFAAAGASQAAGVYAPYVDVTLYPTPLVDQIGVQQGIQQFML
AFVVSGGNQCTPSWGGVQPIGNGATGDLLDKIATSVTAYRAKGGDVAVSFGGAAGQPLMQ
ACSSVAALKGAYQTVIDTYSLTHVDFDIEGASQQDSAAVARNFQAVAQLQADYAAKGKPL
HVTLTLPAMPTGLVQDGLNVLNAALANNVTLDAVNIMTMDYGPSGIDMGAAAISAAQGLY
SQLDTAYKSAGKPQTDAQLKQLVGVTPMIGVNDVAGEIFTLANAQSVQTTAANNNYGFVG
IWSITRDKACDGSSQYASPICSGVAQQPYAFSSVFKQLGGHWGAGVTQDPNYGGGSDGGG
KPQPGAPWSATQVYTAGATVTYQGTTYQAQWWTQGDIPGQASVWKPVGGNVPAWSSTTAY
PGGACVTYQGAKYCAKWWTQGDVPSAGGPWTRA*
>GCA_000452945.1_BPMSHR5858_genomic_01133_1 group_3177
MKRFLSCVALVCFALPAAHAQTQTYHFEEGGTTPTGVTSQPASAAAQARTQRAKPHRHPV
RRNKHHRTHRRARSAQNDRFYSQP*
>GCA_000452945.1_BPMSHR5858_genomic_01134_1 estB_1
MLDETPTIEIETGPNPAFAVILMHGLGADANDFVPLVPELRIANGPAVRFVFPNAPEIAV
TANNGYVMRAWYDILSFEGVNRQVDEAGIDASCASVRGLIAEQNRRGIATSRIFVAGFSQ
GGAMAYSAGLTHPDALAGLIVLSGYVPSPGFIDARLADANRTTPIFAAHGTDDDILPIRL
GEAARDFARDKGASVDWHAYPMPHSVCIEEIDALRRWLHARIAALQAA*
>GCA_000452945.1_BPMSHR5858_genomic_01137_1 cobN
MHLLRTTPGGFVDDTAGVVRIDQRPADIVILSSADTTLSLLASAVPTLGEGFPSVRLANV
TFLRQPASVDFYVDDVLRHARVVVIDHLGGEAYWPYGIEQAVALAARAGQKLAMFSGDLQ
EDPNLIAKSTVAPELCRQWWRYLREGGPANAQALLRSIAHHALGVGDEPEPPRPLPAAAL
YHPAQASPSLEHWRARWREHAPVVAILFYKAHWQAANTAVFDALAAALEREGLNPLPIAV
TSLKDAMSRAVVDALCADANVSLVLNTTAFAAGALDAAEPEVLAGDAPVLQVILSGGNRD
AWLADPHGLNARDIAMHVALPEVDGRIVTRAVSFKGLAYRCPHTEVDVVRYQPDDERIAF
VAELSRRWCRLRTLENAHKRVALVLANYPASEGRIGNGVGLDTPASALAVLAMLRDEGYR
VGELPDDGDALLARITAGVTNDPATRALRPAFQSYPLDDYLRRFAQLPQAARDALNARWG
PPEADPALRQRRFPIAGWRAGHVFVGVQPSRSRGDDDYANYHDADLVPPHAYLAFYFWLR
DAFRIDALVHVGKHGNLEWLPGKSVALSDACWPDLILGPMPHLYPFIVNDPGEGSQAKRR
AQAVIVDHLMPPLTRAENYGPLQDLERQVDEYYDALMVDARRAKVLRETILMTIVEHKLH
EELSIAPPAGRDAEDALLTRVDAWLCELKEAQIRDGLHTFGSSPRGRQRRDTLAALARFP
SGDGRGERAGLIGALARDLALGEAFDPLASDWAAPWTGPRPAALGAMSDEPWRHAGDTRE
RLEALASRLIERRCGDGSPAGEGGGAPSGASRFAAPEYAAAHWPHAHAVLERIVRDVLPC
LDACGDEELRQLRRGLEGRFVPPGPSGSPSRGRPDVLPTGRNFYSVDTRAVPTQAAWTIG
LKSAQQLVERHLQEHGDYPRAVGLSVWGTATMRTGGDDVAQALALLGVRPKWAHGSHRVT
DFEILPIEIFDRPRIDVTLRVSGFFRDAFANVMHLFDAAVQAVAELDEPEHLNPVRARVR
READALVARGIPADEARRRAGWRVFGARPGGYGAGLQALIDGRRWQTDADLAHAYRNWGG
YAYAQNSAGEAAHDAFGARLATIDAVVQNQDNREHDLLDSNDYYQFQGGMAAAVRHASGR
QPGLYHGDHSNPAAPRIGTLREEIARVIRSRVVNPKWIDGVKRHGYKGAAEIAATVDYLY
GYDATARVVSDHQYALVADAYLHDADTRAFLERHNPHALHGICERLVEAMQRGLWQAPGA
HRDAIEGYLLASEQRLEGGRR*
>GCA_000452945.1_BPMSHR5858_genomic_01138_1 yjiA_1
MQMRKIPVTIVTGFLGSGKTTLLRHILQHAGGRRIVVIVNEFGELGIDGEILKGCGVGCD
EAGREVDGRLYELANGCLCCTVQEEFYPVMEQLVERRAHIDHVLIETSGLALPKPLVQAF
NWPSIKNGFTVDAVLTVVDAPATARGQFAEHPAAVDAQRRADPNLDHESPLHELFADQLS
SADLVIVNKTDLLDDAALAAVEATIRDEVPPQVKLVRARRGELDLATLLGLNAASEETIH
LRHDHHGSADDADHHHDEFDSVVVEARVSSREAALGALGALVEAHAIYRVKGFAALPGVP
MRLVVQGVGRRFDSYFDRRWRDGEAPASRFVLIGEDLDAAALQRAFDAALAAHGQAA*
>GCA_000452945.1_BPMSHR5858_genomic_01139_1 hoxN
MLESFLRLFNDSPAELRSKIVGIYAMLIAFNVGAWAWAFAAFHGQPVLLGTALLAYTFGL
RHAVDADHIAAIDNVTRKLMHEKKNPLGAGLFFSLGHSSVVILMTVAVALTAATLAERFE
GMKAWGGAIGTSVSAFFLLVLAFANLLILISVYRTFRAVRRGEPLVEQDLDILLNQRGFF
ARIFRPLFAIVSRSWHLYPIGFLFGLGFDTATEIALFGISATQAHGGLSFWSVMALPVLF
TAGMTLVDTTDGIMMMGAYRWAFVRPIRKIYYNMTITFVSVLVAIVIGGIEALALIGGKL
ALKGGVWDFAAMAAEHFGVLGYFVIGLFAASWIVSALIYRIRRYDDIDVTLSA*
>GCA_000452945.1_BPMSHR5858_genomic_01141_1 cobO_1
MKTDSESHQRMAERRRAGHEKKQAAATVEKGLLIVHTGNGKGKSTAAFGMAVRMLGHGMR
TGVVQFIKGALHTSERDFLGANAQCDFVTMGDGYTWNTQNRDADIATARRGWDEARRMIE
SGDYRMVILDELNTVLKYEYLPLDEVLATLAARDPALHVVVTGRHAPDALVDAADLVTEM
RLVKHPYKEQGVKAQRGVEF*
>GCA_000452945.1_BPMSHR5858_genomic_01142_1 cobB
MPACPALFVAATASGQGKTSVTAALARHHRRLGRRVRVFKTGPDFLDPMLLARASGARVY
SLDLGMVGERGCRALLARAAAEADVILIEGAMGLFDGTPSSADLAQAFGVPVAAVIAAQS
MAQTFGALAFGLAHFRPGLPFHGVLANRVGSERHAQLLRGALPGSIRWLGHLPRDARIAL
PERHLGLHQPDDVADLDARLDRAADAIAHTALAALPPVVEFACGDAANAEADAGDGGATI
PARDASRTAHRTATAARVAAGRGALAGATMRAAHGERAARSADAAAHETVDDALLDAAKR
AGTGDGALSPSPDAGGAIGAIGAIRSTETTEAVPSLVGCRIAIARDAAFSFIYPANLDQL
EALGAELAFFSPLADEPVPDGSHALYLPGGYPELHAARLAANATAARSIAAHVAAGRRVV
AECGGMLYLSDALTDADGVTTPMLGLVPGHATMQRRFAALGMQTLATRFGAMTGHTFHYS
RLATPLAPVAAATRPDDGAAGEPVYRHGPIVATYLHAYWPSNPRAAAALFAGDAP*
>GCA_000452945.1_BPMSHR5858_genomic_01145_1 pvdA
MQRDTVFDLIGVGFGPSNLALAVRLAEAGEPAAFAHCYVERQPAFGWHRGMLLDDCRMQI
SFLKDLVTLRDPKSRFTFINYLYEHGRLSEFVNLKNFYPTRVEFHDYLSWVANAFDERVH
YGETVTAIEPVAAPGAPGEIDALRVFSKGADGRERQRVARALSVGVGGAPSVPDAFAALG
PECVVHSSSYLTSIERLVGAPGAHARKRIAVIGAGQSAAEVFVDLARRFPHVEASLVIRS
GALKPADDSPFVNEIFNPAFTDVVYAQPASGRRSLIERFRDTNYAVVDRPLIEQIYEMLY
LQRVAGEAPGAPRHRLLANTAIESAARTAHGQVELALRDRLTGDARTERFDGLVLATGYR
RDTHLPLLEGLAPHLGDALAAGDVARDYRLATPAHFKPRIYLQGCCEDSHGLSDTLLSVL
ARRADEIAASLADAADTTEYEEADPAAGRARPRHENNGVSDGRLAIAL*
>GCA_000452945.1_BPMSHR5858_genomic_01147_1 lgrB_1
MTSFPTALHHRIHALARLDPDAPALASFAPDTVRLTRGELDERAARLAAQLRAAGVGAEV
PVGVCVARSCDLFVALLAVMKAGGAFVALDPRHPAARLDWVARDAGLAHGIVDASADAAM
RARFARCFDVGGVAAADPAAPREHGGDVHPRAAAYMIYTSGSTGTPKAVVVEHGPLAAHG
DALAESLPIGPDDRVLHFASVNFDVAIEAWLVPLAVGGSVVISDPPPFTPDAAHALISRE
RVTNTTLPPAYLREFAAVCAREGVPPSLRVLLFGGEAMSQDAFDEIRRVFPAIRLVNGYG
PTETVISPMLWPVAPGTTPALDAGNGYASLPIGWPIGRRVARVERADGTVARGEAGELLL
GGACLARGYHGRAALTAERFLPDPAGEPGARIYRTGDLARERADGAFDYLGRIDDQVQVR
GVRVEPAEIAACLLTHPGVRDAGVLAETAGGRTQLIACVALAAPPRETPSGGDARGDAPP
DDDALRAHVAAHLPAAWLPHRFVRFDKLPYTLNGKLDRVALRDAVAARPAEAAADVDAPR
TDTERRVAALWQRLLNDPAPIGRVDRFFARGGDSLAAMQLQTAIRLDWRVNLRLDTLFDD
APLAELAARIDAAEREARQPAAIGTTARRAIAADAIDARATGQMGRTGAAGEATEAAGAL
ERPASFAQQRFWVLARTQDAGAAYHVSFHWDIDGALDLGTLQRALDMLIARHEAWRTTLV
ENDDGVVLQRIHAALPVRIAAVDLRGEAGASRAARLAELTERHAGAPFDLSDGPLVRALL
VTLADGAQRFLLTTHHAVSDGWSSRCAFAELTAAYAALAEGRAPELPALPIQYADYAQWQ
RDALDAHETARELAYWRGALDGAPAPLALPLDRPRAAERDYRGGRLARRLSPAASDAVRA
AARRLHASPFTVLLAAFDAWLFRLTGERDLVVAAPIAQRARPETAPLVGLFLNTLALRAR
VSPAQSFESLVASVRDAAFGAFAHQDVPFDKVIDAVKPPVRRGDEWLRVKFAQQFDLECR
ASLPGAAVRMAPGLDTAARFDYALDFTDDPRGIEFVAAYARDGIDEATAHAWLDSFAALV
EDAVREPRRPIAALAVAQAGAPCALIAGRPLATAPDVLALFAREAAEHPHRVALADADTR
LTFAELDDASNRVALALRRDAARDEAPGAETPVAICIERSARFVVAFLGVLKSGAYAVPL
DPASPRERIAAALAACGARRMLAAGALDALGEFDGVAVQDIDACAHDASLANAAAPRVPP
QPEQAAYLIFTSGSTGAPKGVVVPHRALADYVAGMLDELAFAPHASMAMVSTVAADLGHT
ALFGALCSGRTLHLLPAQAAFDPDRFAHEMATREVGVLKIVPSHLHALLDAQRAADVLPA
HALVTGGEALPWALVERIAALKPDCRVINHYGPTEASVGALVCDTSAPAQADLRAAAASS
PGEAARGVPLGRPLPNAHACVLDAYGSSVPVGAIGELYLGGPGLARGYLGRAAASAERFV
PHPHVAGARVYRTGDRVRLRADGRLDFLGRLDDQVKIRGYRVEPGEVSAALRALPGVAQA
ETLALEHEGRLRLAAFATPEAGARIAADALRDALAARLPDYMVPAALVVLDALPVTANGK
IDRAALRARAAAPAPATAGDEDAPQGPIEATLAEVWRDVLKAARVGRHDNFFELGGDSIL
VLQVIARARKRGVKFTPKQLFDGPTLAELARVAVAIEAAAPASGAAHGVAIGANAAAARR
DEAVLTPAQLRFFALDIPRRGHWNQSIALDVAGAFDFDAFARAFDALLTHHPVFRERFAP
TGDGGGWQRSAAPRAFDTLPLAAAAARDEADALAQFDALQATLDLTHGPLACAFAAVLPS
GATKLYLAIHHAIVDGVSWRVLLDDLDAAYRAACERRAVRLGPTGASASEWAARLARAAR
DPAGPFAGELPYWAALAAPHDDLRPDRPDAAATNAHADVVIQTLDAALTREVLTDANAAY
RTQAVELLIAALVAALGQHTGAAACRLELEGHGREALFDELDASRTLGWLTSHYPVAFAV
EATPAATLASVKDALRAVPNKGLGFGVLRHYGDDATRAALARVARPRVTFNYLGQFDAPR
DAALVPRFGGAGRERDPAGPLGNALAIHAYVDANGERALKVHWVYGATQFDRATIDALAA
RFDAALRALAAACRARVAERGAGATPGDYPLARAGGLTQAALDRLPFDARAIDDIYPLSP
MQQGILFHSLFAPERATYVNQLVATLVDPDVERLRAAFDAAVPRHDILRTGFAAHEAAPM
QIVHRHACMPVEIVDWRGAHASPAALDTALDAWLAADRARGFDLAAPPLMRVTLIRTDDA
DWRLVWTRHHLLLDGWSTARLFADVLRDYIEPPRANPFAAPARTRYRDFIAWLARRDAQA
DRAFWLGRLARLDEPTHVAERARGASAAAHEAAGRANWRATLPAADTARIGEAARRMKVT
VNTIVQGAWALALQRITHRRAVAFGATVAGRPHALPDVDTVLGLFINTLPVITAPLPQLA
ARDWLASLQRDNAAALEHAHTPLYEIQQWAGLGGALFDTLVVFENYPVDEAWQGRDARAL
QKRDLRNIEATDFAVTLVIEAGDTLAIDYGYDPARIGPARVEALHRAFAACIAGLVDHPD
APLGTISCASADDLALIARANATELDWPAAQRAPLFAQFEAAARARPDAIALECFASSDG
GDGARAQMRYGELDAKADRVAAALAASGVRPDSVVALCVERSFDMVVALVGTMKARAAYL
PVDPDYPAERIAYLLGDAKPPVVITQAHLRARVDAALAGADAAVVTVDELLARAAGAEPE
AERVAAAADVAPGQLAYLIYTSGSTGQPKGAGNTHGALANRIAWMQRAYRLAPDDVVLHK
TPFGFDVSVWEFVWPLAVGAKLAIAAPGDHRDPARLVAAIDAHRVTTLHFVPSMLAAFVA
YLDDFGAAARCASVRTIVASGEALAPELVARVAALLPHAQLHNLYGPTEAAIDVSHWRCT
ADDAAADAVPIGHPIANLRLHVLDAALHPAPVGATGELYLGGAGLARGYLGRAALTAERF
VPDPFVPGARLYRTGDLARRRADGALDYLGRLDTQVKLRGQRIELGEIEALLRATDGVRD
AVVIVRDERLVGYVACATPAGFDAAAQIERLRARLPAYMVPAQLVALDALPVTPNGKCDR
RALPAPVFDARVVDAPRTATERALAAIWQRVLTLPELGRDDDFFALGGHSLLAAQANAQA
NLQWSLTLPLRTIFDERTLARCAAAIDRARDAGRERDAAGAIDALLGELEAQ*
>GCA_000452945.1_BPMSHR5858_genomic_01148_1 yojI
MTDATGRLDAPPSTDSRPAVKHARRLIVALLKRSRGAFAIALAACVLNGASSVLLVATLS
RALSEPQAADASLAWRFGLCAVIALVTRIVTGVLFARLSQDTMAQLREHVARRVAAAELQ
DVERIGAAPVQSVLTDDATNVSMLFFALPNLVMHGSIVFGCLGYLAWLSWPVCLLAVAAI
LVGSLGYHAGDKRAIAWLEDAGRSQDKLFGYLGALFSGAKELKLHQARSRQFVDGQLGAA
IAEVRDHRRRAFSAYAIGVGWIVFLFYVFLGVAAFWPALGVRGDGPAASGYVVVFLFMLL
PLDGLLNNVPTLNAARVSLERIEKLMAQFGALRTAPPPDENASHEPFDTLALRGVTHSYF
HERDERMFRVGPIDLTLKPGELVFIVGGNGSGKTTLAKVLTGLYEPEGGVIELDGKPIGL
AERAAYRERFSAVFNDFHLFDALLGIVDPNDAARAQADARANALVAKLSLDHKVQVVDGA
FSTRALSTGQRKRLALVVAYLEDRPLYLFDEWAADQDPAFKAVFYEQLLPELRARGKTVI
VISHDDRYFPIADRVLKLDNGIIVSETFGARHAALANGTEG*
>GCA_000452945.1_BPMSHR5858_genomic_01149_1 group_4240
MSPTALLVPTFTHMLQSLSAWLDKAAAHQRALGAEPDDVLTKRLAADMFVLAAQIRFTAF
QAQEPVYRLRGDALPESLLEVRRAGWQADAAPGSLRDAQACLADARALLAGLAPHALDEG
AERPIALELPNGAVFDMTGEQYLRDWALPQFYFHAIAAYAILRHHGVALGKPDYVSHMFA
YLRPGTMPQDR*
>GCA_000452945.1_BPMSHR5858_genomic_01158_1 golD_1
MSVAFDFHGRSFLVTGASSGIGRAAAVALRGCGARVVAAARNARELERLAHETGCEPLEL
DVGCDASVRAALSGERMRDAFDGLINCAGVTSLAAAIDTTADEFDRVMAVNARGAMLVAR
HVARAMIRAGRGGSIVNVSSQAALVALPSHLAYCASKAALDAMTRVLCVELGPHGIRVNS

VNPTVTLTPMAERAWSDPHASGPMLAAIPLGRFARVADVVAPILFLSSDAAAMVSGVALP
VDGGYTAR*
>GCA_000452945.1_BPMSHR5858_genomic_01166_1 group_4558
MVPARHCHVAVFDQAIVALDIRRSRYFLYDAACARDFADHYLDFKPIDAPRALKPLIADR
ILVAESPAAARSRIADYRGWSFDAFDSGIWGSRVLGKRDAARFEWLPFWRIVRSAASLKM
RGFHALAALDRLDARGAPACGDDASRTVERLLRASIWSPFRITCVQMSLALATHLRRQNV
PAQLVIGVRPMPFVAHAWVEVDGRVCGDEPDLKKSYGEIYRTPRCDASAAPFGLAA*
>GCA_000452945.1_BPMSHR5858_genomic_01183_1 group_7319
MKRLARLVHDERGVVSLEFVLAFPFMMLVLFGIVDTSLLLCDKAVITNASREAARAGVVV
RVPQLAATDITNIALSYAQGSLVSGGTVGAPVVNVDQSAGTSPGSPLKVTVSYTYQGLVL
GSALSALTGPITLTAVTVMNYE*
>GCA_000452945.1_BPMSHR5858_genomic_01184_1 bepA_1
MPVALLLAALLAGCASHGVFQPRPVLSQRGTVGDGDLSVAESALAAGDAELAATLFERAL
KADPRSLPAQVGLGDAMYQTGELARAGVLYAQAAAAAPDDPRAQLGLARVALRERHLDDA
LARYAKLNAARPDNIAAAAGLGTVLDLLGRHDDAQRVYRAALARHPDAHALKTDLGLSLV
LANRAREGANVLLDIAGLPNAPAQAREDLALAYGMLGNDDAAKRILVHDLPAASAEDNLR
FYRNVRERRAAGAGGGGGAPGPAGAAGATNAANAANAAAGPTMTTAANAPRAAHAAAAPQ
APYAGRAIADAGLAPVRVEVVK*
>GCA_000452945.1_BPMSHR5858_genomic_01185_1 group_1161
MTPAALSITVLLALVAVAALAWLLTRDVAVRRRVAQRARASAARAAAVYAPVDDGARSVP
RRVAQRVASLGERVPVVDATQRMKLAARLASAGFREHRAVSVMAGLKLVVGACAGLAALA
LGAHVPRVGDYVVLRVLMMAGAFVIGMMLPEHALGFVLARRQKTIAAYLPDALDLLVICT
NAGNSLAVAIRRVASELKTICPPLADEFSVCADELQIGGDAAFALNAMAARIGLPSMRAL
VTTLVQSQQYGTPITQSLRMLSRTERAMQMIALEEKAAKLAPKMTLPMMLFVMPTVALIA
AGPAVIRLIAVFHK*
>GCA_000452945.1_BPMSHR5858_genomic_01187_1 group_190
MFGRRPAASFAPGDAGGAHEFAPDAAPAGAAPAGGHASAPDASGAARAREPAGASGASAP
GGVPQSGVAPSGHRPASRDDHIDRDDGHDRAPAQDHHEALIRSETFKTIRAVVFSSMNMS
AALMMSRAEVREGIEQAAADVIARERLTVTLAEQTLIVDEILNDMFGVGPIEPLLADERV
TDILVNGPDQVYVERAGKLELTPLKFRDNAHVINVAQRIAAAVGRRVDESSPMVDARLAD
GSRVNVVLPPIAIRGASISIRKFAKRNITLARMAQQGNLSQAMVEVLKIASACRLNIVIS
GGTGSGKTTLLNALSHFIDSHERIVTIEDAAELQLQQPHVVSLETRPENSEGLGGVSQRD
LVRNALRMRPDRIILGETRGPEAFDVLQAMNTGHDGSMTTIHANTPRDAITRLESMVMMA
NGNLPLVSIRRQIASAVHMILQIERMRDGVRRVTRVTEIAGMEGDVVITQDLFAFRYDAS
AFQEQVHGMFESSSLRPAFAQRAAYYGLEGALLGALQP*
>GCA_000452945.1_BPMSHR5858_genomic_01189_1 group_2307
MAIRPEHTTRRRASRAVAACAALAALAGCMSGHPPYGMPDASTIGYDARTGLARAPDCAA
LEQRSQMIDAGRARPGVSFGCATYGNLAAMLARPADLVAPLPYAGADAALGASAVRRYDE
GRATPLNPTSTTTSVTH*
>GCA_000452945.1_BPMSHR5858_genomic_01190_1 sctC_2
MWSVIDAIGKRRQAASASPRARAFAAARRAAAAALWWLALALGMAVAPRFARAADLGPVL
AVPAGGGEMVKLPAPAVAVFVADPDVADVHVPTPQAVFVLGKKAGTTTLFALGANNRTIL
RETVVVDVDTPSLQRILDARFPQLRLTLAGAPGSLMVSGRVPSAADADAVVQTLKPYLRQ
QEALVNRLTLARPIQVHLRVRITEVDRNITQQLGINWSALGASGNFVGGLFNGRTLFDTA
SKAFDLSPSGAFSVVGGFHTSRYSIDGVLDALDQEGLITMLAEPNLTAISGQTASFLAGG
EFPIPVAQDTTGAITIQFKPYGVSLDFTPTVLADNRISLKVRPEVSEIDPTNSVTTGSIK
VPALTVRRVDTTVELSSGQSFAIGGLLQSKSSDVLAELPGLARLPVLGKLFSSRNYLNDK
TEVVVIVTPYIVQPANPGELRDALDDVTRPSSDIEFVLQRSLGIDPLGGDAPRLAGPAGF
VY*
>GCA_000452945.1_BPMSHR5858_genomic_01192_1 group_1421
MILPLKLVASWTLASLALADLRTRRLATFAVALVGALYAALALVGAPGDGGFASHAALGA
AAFALGAAMFRAGWIAGGDVKLAAVVFLWAGPAHAWPVAFAIGVGGLAVGAVCIAAGRAP
RVLAWFAPARGVPYGVALAAGGLLAVWAPAACRLPGCLG*
>GCA_000452945.1_BPMSHR5858_genomic_01193_1 group_7320
MLRFIQSLWRDERGVSALEYSVLAGIVVVAVAAAGAIFGGGSGLPELFQNMITKVTSVQT
NGH*
>GCA_000452945.1_BPMSHR5858_genomic_01194_1 ssuC_2
MSAAPRIVPPVRDEYELALAPAASIDAIAPAAGIAAARAGDPAAGAGERGRAAPLARLYA
RLGVRRLRDAQWLRKTALALALIAAWEIVARVVDDDLLLPSFGATALAFAHGVANGELQA
KIAVSLSMLLRGYALGVALAFALTALAATTRLGRDLLTMLTAMFNPLPSIALLPLALLWF
GLGTGSLLFVLVHAVLWPLALSTFAGFQAVPATLRMVGRNYGLTGVRHVVAILVPAALPS
ILAGLKVAWAFAWRTLIAAELVFGASAGGGGLGWYIFQNRNELYTDRVFAGLAAVIAIGL
VVEHGVFDALERVSVKRWGAHTL*
>GCA_000452945.1_BPMSHR5858_genomic_01195_1 nrtD_1
MAAEPSTLFSNPADAHACATSGKLVRVDRVTLEYRHRERVVRATQQVSFDVYGADRFVLL
GPSGCGKSTLLKAIAGFVAPVEGAITLAGEPVRGPGADRIVVFQEFDQLAPWKTVRENVA
FALRAARRLSRRDAAERSRAALDKVGLAAFADAYPHTLSGGMKQRVAIARALAMEPRVLL
MDEPFAALDALTRRRMQHELLRLWDDARFTLLFVTHSIEEALVIGNRVLLLSPHPGRVRA
ELNSHHFDERSVGRADFQRTAERIQRLLFDESGARA*
>GCA_000452945.1_BPMSHR5858_genomic_01196_1 group_60
MCWMHGIHGIHGIHGIHGIHGIHGIHGIHGIHGIHGIHGIHGMRWTRGRRGSSARHD
MATRAAAPAVDAATRRDDDVAPMRRQCGAHDIDIENASLCRAATHLNLDP*
>GCA_000452945.1_BPMSHR5858_genomic_01198_1 kgtP_1
MNDQTAVAAAARQDVRRRVLAIVGASSGNLVEWFDFYIYSFCALYFAPAFFPSGNTTTQL
LNTAGVFAAGFLMRPIGGWLFGRIADKHGRRAAMMISVLMMCGGSLVIAVLPTYAQIGAL
APLLLLVARLFQGLSVGGEYGTSATYMSEVALQGRRGFFASFQYVTLIGGQLCALLVLVI
LQQTLSSDALKAWGWRIPFVVGAAAALISLYLRKSLDETSTSESRKAKDAGTIRGVWQHK
GAFLTVVGFTAGGSLIFYTFTTYMQKYLVNTAGMHAKTASNVMTAALFVYMLMQPVFGAL
SDKIGRRMSMILFGTGAVIGTVPLMHALGGVTSPLAAFGLIVVALAIVSFYTSISGLIKA
EMFPPEVRAMGVGLSYAVANAIFGGSAEYVALWFKSVGSESSFYWYVTVLCAISLLVSWR
MRDPSREGYLRNEP*
>GCA_000452945.1_BPMSHR5858_genomic_01203_1 rbsK_1
MTAAVERAAGGASPARAGRVMVVGSLNMDLVVRSARLPRPGETLAGRSFAQAAGGKGGNQ
AVAAARLGAQVAMLGRVGADAHGAALRAGLVAEGIDCVALSVSATATTGVALIVVDDASQ
NAIVIVAGSNGEVTPESIVGHEAAIAAADVLICQLETPAAAVRAALAAGRRLGKAVVLNP
APAAGPLPADWLPLVDYLIPNEIEAAALTGLPVRDPASAETAARALAAAGARNVIVTLGG
QGVLALTADGDARHYPAPRVAPVDTTAAGDTFIGGFAARLAARDAPHDAIRFAQRAAALS
VTRAGAQPSIPTLAELDAFAPGPA*
>GCA_000452945.1_BPMSHR5858_genomic_01204_1 rbsR
MATIKDVAAIAGVSFTTVSHVVNNSRPVSADVRAKVERAIRELNYVPSAVARSLKARSTA
TIGLVVPNSTNPYFAELARGIEDQCALSGYCVFLCNSDDDPEKQRNYLRVLQEKRIDGLI
IASAGDDAVLAQTLADTREPLVVVDRNIEGLPADLVQIDHERGAYLATRHLLELGHAKIG
CITGAVSTAVSAMRVHGFIRAMAERGIDIVPGAIAESDFSCMGGYHAAVRLFDSVRPSAI
FAGNDLMGLGALRAAAERGIRVPADCSIIGFDDIELSRYTYPALSTVGQSVRALGEMAAQ
TLIERIGGGASGAPQRRRVVSPRLVLRESTTVYVEPADAGYRG*
>GCA_000452945.1_BPMSHR5858_genomic_01205_1 group_2310
MNHRSIDRNADAPSGPAGTVAAKPAGVRLGLSNYVGLALALLALIALFSVLSSHFLTYDT
FSTIANQIPDLVVLSVGMTFVLIIAGIDLSVGSVLALAASVVSVAALRGQWPPLPAALAG
VAAAAATGTLTGAVTVGWRIPSFIVSLGVLEAARGVAYQLTNSRTAYIGDAFDFLSNPIA
LGISPAFLIAVAVMIVAQFVLARTVFGRYLVGIGTNEEAVRLAGVNPRPYKIIVFALMGA
LAGLAALFQISRLEAADPNAGAGLELQVIAAVVIGGTSLMGGRGSVVSTFFGVLIISVLA
AGLAQIGANEPTKRIITGAVIVVAVVLDTYRSRRARG*
>GCA_000452945.1_BPMSHR5858_genomic_01207_1 purR
MRSLCGQPYAGSFFRRIPFAAQMFETRCRFGLRKRLLVAKATLDPFGTRNHCQESRMDVS
LRRRVLAVAAVCVAVAAAGPFSVARAETAHKPKVALVMKSLANEFFLTMENGAKEYQKHN
ASQFDLVTNGIKDETDTASQIRIVEQMIVSKVDAIVLAPADSKALVPVVKKAVDAGIIVV
NIDNRLDPDVLKSKGLNVPFVGPDNRKGARMVGDYLAKRLKAGDAVGIVEGVPTTTNAQQ
RTAGFQDAMKAAGAKVVSVQSGEWEIDKGNAVAAAMLNEYPNLKALLCGNDNMAIGAVSA
VRAAGRQGKVYVVGYDNINAIKPMLKDGRVLATADQYAAKQAVFGIDTALKALAEHRKQA
DMTGVVATPVDLVTKP*
>GCA_000452945.1_BPMSHR5858_genomic_01208_1 ssuR
MNFQQLRFVREAVRQNMNLTEVANVLYTSQSGVSKQIKDLEDELGVDIFIRRGKRLTGLT
EPGKAVHQLIERMLLDAENLRRVARQFADQDSGHLVVATTHTQARYALPKVIRQFTEVYP
KVHLALRQGSPQQIAQMILNGEADLGISTEALDRFPDIVTFPCYSWHHVVVVPKGHPLVG
RENVTLDDIAEYPIITYDQDFTGRSHIDQAFANAGAVPDVVLTAIDADVIKTYVELGMGI
GVVAAMAYDAQRDTGLVALDTQHLFEASTTRVGLRRGAFLRAYAYRLIEMFAPHLSEGEI
AGQLREAA*
>GCA_000452945.1_BPMSHR5858_genomic_01209_1 cysA
MGITVRNLHKRFGEFAALDDVSLDFPAGELVALLGPSGCGKTTLLRVIAGLEHADSGQVV
LQGLDVASVGARERQVGFVFQHYALFRHMTVFENVAFGLRVKPRRERPSEAAIRAKVHEL
LSLVQLDWLAQRYPSELSGGQRQRIALARALAVEPKVLLLDEPFGALDAKVRKELRGWLR
RLHDDLHISTIFVTHDQEEALEVADRIVVLNHGRVEQVGSPQAVYDHPRSAFVYEFLGAA
NRLDGTVSGNGFVAHGAAQAIAVDADFAGPARAYVRPHDLELAAPHARAQGIAADVRRVV
PLGGSVRVELAARSGEVLEAELDRNAWRALALDVGDALTAVPRAVRVFPAR*
>GCA_000452945.1_BPMSHR5858_genomic_01210_1 cysW_1
MSRDTARSRAGAARADAGPEFGARFGLESGLESGDDARPRARAPARARRLDPVSEPRAVR
WLLTGAALAFLALFLVVPLAAVFFEALRKGVDFYLESLADPDAWSAIKLTLVVAAIAVPL
NLVFGVCASWAIAKFEFKGKAVLTTLIDLPFSVSPVISGLVYVLLFGAQGWLGPWLQAHD
VQIIFAVPGIVLATIFVTFPFVARELIPLMQAQGADEEEAARVLGASGWQIFRRVTLPNV
KWGLLYGVILCNARAMGEFGAVSVVSGHIRGQTDTMPLHVEILYNEYNFAAAFAVASVLA
LLALVTLALKLFAERRLSAELAHGRDDASAPAAHPGAAVTSSIS*
>GCA_000452945.1_BPMSHR5858_genomic_01211_1 cysT
MTTFTFRKPSALPGFGVTLGITLAYLSLVVLIPLAATFLKTATLTWDQFVAATTSPRVLA
SYRLTFSAALGGALVNAVFGFLVAWVLVRYTFPLKRLVDAIVDLPFALPTSVAGISLAAV
YAQNGWIGRYLAPLGVKVAFTPFGVLVALTFIGLPFVVRTVQPVLEDFEREQEEAAACLG
ASRWLTFRRVVLPAVTPALLTGFALAFARALGEYGSVIFIAGNVPMKSEITSLLIITKLE
QYDYAGATALAVVMLVVSFVMLLLINTLQWYLQRRTSRGASGPTPAAHAATALGGAQ*
>GCA_000452945.1_BPMSHR5858_genomic_01212_1 sbp_2
MVKRNTGLAGGARRLIASLALGAAAALGALTPALADTTFLNVSYDPTRELYQDVNQAFGK
EWKARTGETVNFKQSHGGSGAQARSVLDGLQADVVTLALAYDIDALANKGLVSKDWQKRL
PDNASPYTSTIVFLVRKGNPKGIKDWDDLVKPGVSIVTPNPKTSGGARWNYLAAWAYAQH
QPGGTAQTAKDFVTKLYRNAGVLDSGARGATTSFVQRGIGDVLIAWENEAFLSIKEFGAD
KFEIVVPSASILAEPPVAVVDKVVDKKGTRKLADAYLNFLYSRQGQEIAARNYYRPRSRD
VPAALTKQFPKLKLYTVDDTFGGWTQAQKTHFADGGVFDSIYKPQ*
>GCA_000452945.1_BPMSHR5858_genomic_01213_1 lexA
MIKLTARQQQVFDLIRRAIERSGFPPTRAEIAAELGFSSPNAAEEHLRALARKGVIELAA
GASRGIRLLGIDDAPHQLTLPHAALMQLSLPLVGRVAAGSPILAQEHISQHYACDPALFS
SKPDYLLKVRGLSMRDAGILDGDLLAVQKRTEAKDGQIIVARLGDDVTVKRLKRRPGGVE
LIAENPDYENIFVKAGSAEFALEGIAVGLIRPGEF*
>GCA_000452945.1_BPMSHR5858_genomic_01214_1 group_7324
MERLARLLPFRQFGRLRHLRGRTPCAPLVAAHSRDHADTQPSLLPSALPAFARVSLNGSL
HHAEPARRAPVRVYHGPSRFIMVGTVDAICRMIDRCIAEERSLTQQGL*
>GCA_000452945.1_BPMSHR5858_genomic_01215_1 group_1275
LNSIARPRGRFLKPVAIGIALIVVAAVALYAYASPYLALRQMKQAIDARDAQAISVHVDF
PALRISLKQQLTDELMRRIDAQKRDNPLAILGAMIGSALVGPLVDAYATPDGVAALMSGL
PPHARPGQRPPELNPPSDPASAAAAGTGAPPGNAGVPGSASGAAEQTASAAAASTSGASG
TSGTPPGAPSATPGPSPQTSAGYRGFDEFVVTYRRDDGSARYAAVFRRSGIFGWKLSAID
LHDD*
>GCA_000452945.1_BPMSHR5858_genomic_01216_1 group_4560
MASYKKILLCYDGTLEGRKALRCGADLALELKAETHLLSVVDMRSSIAQSAGLLTDVACS
RFEETAREILQEGVDWLTERGVHAQGHFAFGYPIDEIANLATKLNVDLVVVGHRCRSGLS
RWWMGAGNTQLLDRVSCSILVACASPAEQKAESAKEGAATIGH*
>GCA_000452945.1_BPMSHR5858_genomic_01217_1 lnrL
MSVAPIDFQQVEKRYDDKLVVDGLSFHVQPGECFGLLGPNGAGKTTTLKMLLGITHPDAG
SISLCGEPVPSRARHARQRVGVVPQFDNLDPDFTVRENLLVFARYFGLTAHAARALVPPL
LEFAKLESKADAKVGELSGGMKRRLTLARALVNDPDVLVLDEPTTGLDPQARHLMWERLR
SLLARGKTILLTTHFMEEAERLCHRLCVIEEGRKIAEGAPRMLIEAEIGCDVIEIYGPDP
VQLRDELAPFAERTEISGETLFCYVDNPEPIHARLKGRTGLRYLHRPANLEDVFLRLTGR
EMLD*
>GCA_000452945.1_BPMSHR5858_genomic_01218_1 yadH_1
MDTRHYTPSPTLAARRRESAFALALPANATNWIAVWRRNYLVWQKLAIASMFGNLADPMI
YLFGLGLGLGLMVGHVEGVSYIAFLAAGTVGSSVMISASFESMYSGFSRMHVQRTWEAIM
HTPLSLGDIVLGEIVWAASKAMLSGAAIMLVAGVLGYAKFPSMLVALPVIALAGIAFASL
AMIVTALAPSYDFFMFYQTLALTPMLLLSGVFFPLAQLPEFAQHIAHALPLSNAVDLIRP
AMLDRPATDVARHVAILAAYALGGFFVCARLFRRRMMR*
>GCA_000452945.1_BPMSHR5858_genomic_01219_1 group_7327
MTQHKVHPRLEQALTRGDLAIRQANSTRATAVLNALGKMIVEASATIGVEASIDIAQGER
VYDPVNGVWPQKMLVSFDGPVEDADPEELRTVYLLADDPGTMFRVEWHRADGKLGRQEGG
PLATVAFLTDVEMPWSDDDE*
>GCA_000452945.1_BPMSHR5858_genomic_01220_1 group_3181
MSSSRSYTPHPALGLATFVVLAVAGLFYVKWFPYYHKAFAAAEHHSIGQSILMGAAAHAP
QPSLQAALDYAWAYGKAIWQAMVLGLLLGSAVQALLPAHWVARALGGTGFGSVAAGGLLA
LPGMMCTCCAAPVVAGLRERDASPGGALAFWLGNTVLNPAALVFMGFVLGWHWSALRLVL
GVAMVFGVGYLVNRLAGAQPRVVDDALRAKLVAEQAAVGNAFVRWMKIFARMTVRLVPEY
LVLVLLLGAARAWLFPHIGPDIGNGVGWIVAFAIAGMLFVIPTAGEVPIIQAMLSLGMGV
GPAGALLMTLPPVSVPSLAMLARSFKPATLALVAALVVAFGVVGGLAAVALGF*
>GCA_000452945.1_BPMSHR5858_genomic_01221_1 tas
MEYRTLGDSGVKVSLIGLGTMTWGEQNTESDAHAQLDYAIDQGVTLIDTAEMYPVPPKAQ
TQGRTETYLGTWLAKHPALREKLTIATKIAGPARQPHNPTHIRGAGNQFDRKNLTEALDG
SLARLRTDYVDLYQLHWPDRSTTTFGRPAYPWIDDPYTVPIEETLAVLGDFVKAGKVRHV
GVSNETPWGVAQFLRAAEKLGLPRIVSIQNPYSLLNRTFENGLSEFSHRERIGLLAYSPL
AFGWLSGKYEGGARPEGARITRFERFQRYSKPHAVQATSRYVALARRHGLSPAQLALAFV
NSRPFVTSNLIGATSLAQLRENVASVDVKLAPEILAEIDALHEQQPNPAP*
>GCA_000452945.1_BPMSHR5858_genomic_01222_1 group_4562
MNPLSMSGLELLRAAAAGDLPAASIAETIPMRPESVELGYVKMIARADGRHLNPLGGVHG
GFAATVLDSVTGCAVHTMLDPGVGYGTVDLHVKMLRPVPRDVDLVAEGRVIHLSRNLGVA
EGTLKTPDDKVVAHASATCFIQRPA*
>GCA_000452945.1_BPMSHR5858_genomic_01223_1 group_4563
MKWDDVGSMPCSVARTLAVLGDRWTMLILRNAFLGHRRFEAFQTQLGLTRHVLAERLARL
VDEGIFTKRAYQDRPPRFEYRLSQKGLDLYPVLLALAAWGDRWKDDGNGPPVVLRHRRCA
HVMHPVMVCSECGEPIDPREVEPMPGPGWTKQQADKVD*
>GCA_000452945.1_BPMSHR5858_genomic_01225_1 gltT
MKKRRNITSYIVIAMILGIAVGYGCHSAFPDPAIAKEIAGYVSLLSDVFLRLIKMIIAPL
VFATLTVGIAHMGDTGAVGRVGVKALGWFFIASFTSLLLGLLAATLLQPGSHLSLPLPAT
DAAVNLKTSAFTLKDFVIHLVPKSIAEAMANNEILQIVVFSIFFGTALSALGDAGKRLTG
VIEDLAQVMLKVTGAVMWFAPVAVFAALASTITTEGLGILLTFAKFMGSFYIALALLWGV
LTLAGLVFLGKRTFTLIRLIREPFLLSFATASSEAAYPKLLDALDRFGVNRKISSFVLPI
GYSFNLDGSMMYCTFAVLFIAQVYGVHLPLGTQITMLLLLMLTSKGMAGVPRASLVVIAA
TLNQFHLPEAGLLLIMGVDMFLDMGRSATNAVGNSIAAAVVAKWEGQLDAPRDDADSDSD
SGDASRGSPRITQRV*
>GCA_000452945.1_BPMSHR5858_genomic_01226_1 dusA
MMDWTDRHCRSFHRTISRHAWLYTEMVTTGALIHGDVARHLAFTPDEAPVALQLGGSEPA
DLAHSAKLGERWGYDEINLNCGCPSERVQRGAFGACLMNEPQLVADCVKAMRDAVSIPVT
VKHRIGVDAVEDYAFVRDFVGTVAAAGCDVFVVHARNAILKGLSPKENREIPPLKYDYAY
RLKRDFPQIEIVINGGVTTLDEVEQHLRHVDGVMLGREAYHNPYVLAGIDARFYGASAPA
PTREEVEAKLVEYCAAELARGTYLGAIVRHALGLYRGVAGARGWRRVLSDNKRLGRGDLS
IFDEARTHLTHPEEFFEKKALQS*
>GCA_000452945.1_BPMSHR5858_genomic_01243_1 group_1798
MASHAAHHASGWAAGLVAAALVAQAGAAGPWHVYSLAAFAAGVAGGTAPDWLEIAWWRRT
RRLWVTHRTITHWGVGWAALLAFAYRSLGHRHPWAPPLFGFACGGLVHLIADWPNPLGVP
WLWKRHSLNWWNSGHCDLIVVAAAWAGALWLAQAAWGHAAPLAHWLGWLRTA*
>GCA_000452945.1_BPMSHR5858_genomic_01244_1 group_989
MTDDITTPSPAPRPLPGFGPLRWIVRHWRGGYSLAVSFWLNQAPIYALRTICSQVLMTRV
FELLDPGQGPGDNDALANALAAVPPVVLADAALSLLNVWWIVGVWRAASRHIAATGRRFW
ARAAQVMVALAAARIVYGIAVIPHSIESLQLLVIHPGSLLTDPVGTLRNWLF*
>GCA_000452945.1_BPMSHR5858_genomic_01246_1 yjeH
MNAPAEPGATTVPHRAGSLTILQGAALYIGAVLGTGVIALPALAAEVAGPASLLAWAALV
MLSIPLAATFAALGARYPDAGGVSTYVRKAFGPKAAAVVGWCFYFAVPAGAPAAAMFGGA

YVAAVTGGAHATMIASAAVLIALVAAANAFGVTVSGRMQLVLSALLVALLVAAVLASAPH
ARAANLHPFAPHGWLAVGEAAALLVWSFAGWEAITHLAAEFRRPARDLPRSTGIAVAVVG
LLYLSVAAASVLVLGPAAGESGAPLAALIARGIGGNAQLIATAAALLLTLGTMNAYFAGA
AKLGAALGRDGALPRWFAQGSQTGGVPRRSLAVIAALAGCALVAAVATGMGPKPLVLATS
GNFVMVYVLGAAAATRLLPRGGLAHRASIVALLSALLLLATTGWYLLWPLLLTASAIGYL
FITGRRAAAKR*
>GCA_000452945.1_BPMSHR5858_genomic_01249_1 aroG_2
MQNIDNPQRDRETGFADATQDTTRIDDVRIGAVRPLISPALLQDELPVPSAVQALVEASR
DAIGDVLHGRDDRLLAIVGPCSIHDHDQALDYARRLKSAADALRDDLLIVMRVYFEKPRT
TVGWKGYINDPRLDGSFRINEGLRAARRLLIDINALGLPAGTEFLDLLSPQYIADLIAWG
AIGARTTESQSHRQLASGLSCPIGFKNGTDGGVQVAADAIVAARASHAFMGMTKMGMAAI
FETRGNDAAHVILRGGKRGPNYDRASVDEACAVLRAAGQREQVMIDCSHANSNKSHLRQV
DVAEDLARQLSDGERRITGVMVESNLEAGRQDLKPGVPLQYGVSITDACLSWAQTEPVLH
TLAQAVRRRRAA*
>GCA_000452945.1_BPMSHR5858_genomic_01250_1 group_858
MITFGEPLIALFDGSAVRNIQVGAPCYPTSRRYAAIDHGGAPAGA*
>GCA_000452945.1_BPMSHR5858_genomic_01255_1 gbpR_2
MHNRIADGRVKFRHLQCFLAVAQFGSVQKAAHSLSITQPAVSKTIAELESILGVRLFERG
RQGARPTREGQLFMPHASACVLALRQGVGLLAREGGGAAATLEIGMLPTVAASLAPAVLK
TLAAEWPRAVVRITTVANAELLERLKAGAIECAIGRLSEPERMVGLSFEHLYNEPLVAVV
RAGHPLAASASPAAQLARYPVVLPPYGTMIRQAAEQLLSACGAPPLESFVEVLSVSVARA
LALENDAVWFVPRYAAEFDLAAGTLTRLALPVDGADEPVGLILRTDAQPSPVARALIDAV
RAIARRRLAGAGTTSGRARGARGATGKGRGRRDA*
>GCA_000452945.1_BPMSHR5858_genomic_01257_1 pcaG
MTTLKQTPSQTVGPYFAYGLCPQQYNYDLKSLFTAEIATPDAAGEPILLVGRVLDGDGHP
VGDALLEFTHVDGDGRYPASRDEAAQRGFSGFARVGTGTHAQNRFVVRTVKPARTSDTDA
PHIDVTVMMRGMLAHAFTRIYFDDEAAANAADAALASVPAVRRATLIATRDAQANGVTVY
RFDIHMQGERETVFFDV*
>GCA_000452945.1_BPMSHR5858_genomic_01258_1 bapA
MMKVLISTDIEGVAGVVSVEQTRAGNPEYERARRWMTAEANAAVEGAFAGGATRVWVNDS
HGGFRNLLPDGIDARARVVLGKPRPLGMMAGLEAGPDLVFMIGYHAKAQTRGLLAHTINS
SAFARVVLDGRDVGEAGLYGALAHEYGAHVALLSGDDVFVDETRPLFPAAQFVTVKEAAG
FASGASMTPAAACGAIGVAARQAVEQAREGARAARPHAPRAARCELQAQSVAAADLFCQW
PSLERVDALTLRFFAPSIEHVVRTLNCLSAMSFMLR*
>GCA_000452945.1_BPMSHR5858_genomic_01272_1 ileS_2
MSEYKETLNLLQTPFPMKGDLPRREPALVERWAAQRVYARMRAAAAGRPPFVLHDGPPYA
NGDIHIGHAVNKVLKDIVLKSRFAAGYDAQWIPGWDCHGMPIEHRIEQLHGRGLPPAAVQ
RLCREYAFEQTERQRRDFLRLGLLGDWPHAFRTMDFRTEANELRFLERIRARGLLYRGQK
PVNWCVDCQSALAEAELEYARKTSVAIHAGLRVRDPVDFASRFRRRPVLDKPAMLVVWTT
TPWTIPGNAAAGVRADAPYGLYDTPGALIVVAQPLADALFASLGVDHALCALARGRELVG
LALGQPFFAGRDVPVVEAEFVTLDAGTGIVHLAPAHGAEDAELCRRLGIAGENVVDGAGR
FAADLPEIGGLPLADGIARIVAKLRADGTLVREAAFEHAYPHCWRHKTPILFRSTPQWFI
GMDIECEQGEADPGRAAVTEEAGATGEARKVGKAEEAEEAGPVKTLRASARDAIADVPFY
PPSARQRMEAMIDGRPDWCVSRQRTWGVPLPYFVRRDDRSLHPRSARLVEAVAARVERGG
IAAWSRLRPAELGVDENAYEKLSDTLDVWFDSGSIHATVYRDAARAGTGGYPADLYLEGA
DQHRGWFGASLMTGCAADGRAPFRAILTHGFVVDGAGRKMSKSLGNTVSPQRIADTRGAD
ILRLWIASTDYAAEMSISDEILERVVETYRRMRNTLRFLLQNVADFDPRDDAVPAGQLLD
VDRYALARCREFVDACRSAYARYDFLAVTRLAHGYCAEELGGFYLDALKDRLYASVADGV
ERRAAQTALHSVLANLLISIAPILSFTAEEAWTVFAGDERDSVFLHTWDEHAPPPDDAAL
ARWAHVRALRPHVTKALEEARGAALIGRSSEAELVVRAPRDVLDALAPLHGELAAVFIVA
GVTLETADQIAVAVARTPLARCERCWRHEPSVAAHASGDALCARCRHALSRRARAERSEP
RAAVGSR*
>GCA_000452945.1_BPMSHR5858_genomic_01275_1 group_604
MREVTDDGQQWDGRAAAFAGVPPGGGAPAAFATPQTIGRCVEALRAIVGERLAVGRAVRE
QHGKDISFHPGAPPDAVAFVRTTAEVAEITRACHRLRVPLIPFGTGTSCEGHIAALRGGV
CIDLSGMNRILRVSAEDLDCTVEAGVTRKQLNAHLRDTGLFFPIDPGADASIGGMCSTRA
SGTNAVRYGTMRESVLALEVVLPNGDVVSVGSRARKSAAGYDLARLFVGAEGTLGTITGV
TLRLHPRPDRLSAAVCSFPDLKAAVDSVIGVIRSGVPIARVELLDALQIVACNRYAKLSL
DERPTLFFEFAGFGAAVDEQIDWVGEICAGYGGGEFRWASRPEERSALWAARHDAWWAAL
ALRPGCEGMPTDSCVPVSRLAEAVLAARADVDALGLVATICGHVGDGNFHVCIAIDPADA
DELARATELNRRLALRAIALGGTCTGEHGVGYGKIGYLEREHPTALQLMAAVKRALDPRG
IMNPGKVLHHDRYPPFD*
>GCA_000452945.1_BPMSHR5858_genomic_01276_1 group_3186
MAAYLLGRMAISDPSKYARYKALTPDIIARYGGRFLTRGGEKLTLEGPADERRIVLVEFA
DADAAARFYHSADYQHAKSLREGAADMQLMIVGGFERK*
>GCA_000452945.1_BPMSHR5858_genomic_01277_1 group_7333
MPITVDELVQKVKSHRCYSHPVFLNWAKVDPEPKVVGALFHQIQHFCASTRPGWNFPRAL
AEHGLQKQSELMTEIVDSESGHGPELATMAGYIVNRAAGGAVIPDLYDQAAVEGALKRYS
DELLGSLPGYDDETGLTTQVRRAINVFERRKLVDVESTYRNLGTALALEMISNRQLIPGE
KHCLVDSGLYGASLEVPEMHYLLEHWGEVGAEQQHEENARAAVKPALESEHAALVIEGAQ
EFLDALASVWDLLDASLLESGYVKKAA*
>GCA_000452945.1_BPMSHR5858_genomic_01279_1 rhaS_9
VAKLDHRNQSRYWHSPGISGVDLLLADFTTHDYAPHVHDSLVVAVTEVGGSVFKSRGQTR
LAEPNAVLVFNPCEPHSGRMGGSSRWRYRSFYLAEAGLARVLTLLGMAQPRFFTSNVLDD
PQLVEQFLTLHRAMDEQDDLLRQQELLVSSFGTLFSRHGLQAGLGAGPGFGTKAGLPALK
PALDLMNDCFDHALTLEQIAGAAGLTSFQLITAFNRVIGLTPHAYLNQLRLRAALRELQA
GHSLADAALTSGFYDQSALCNHFKRTFGMTPMQYTRALAPGKRALAPIGI*
>GCA_000452945.1_BPMSHR5858_genomic_01284_1 abaQ
VDETSIRERAVMAKIARRLMPLLVIMFLIAFIDRQNVGFAKLQMVHALGMTETSYGLGAS
LFFIGYLLFEVPSTLALHRFGARLWLARIMMTWGVITVLLGFTHSTGVFYVFRFLLGVAE
AGFYPGVIYYLTLWFPQSHRTRVLGLFTLGSALANMLGSLAGGLLLSLDGRLGLAGWQWV
FVATGLPAIAVALVAWRYLPESVERATFLSDDEKRVALAALKREASPEAVSESPWRALVD
PRVLMFALAYMLMSTSLYGVTYWLPTLLKSNGVPSSLNGLLNMIPWAIAALLLLWLPARL
KRERIVLKAMAAVAGVGVAGFALSLALPGLPLRFAALVLGGACIPLLYPCFWSLPPRFFS
GARAAASIAAINSIGNLGGFFAQNLMPYVGKAAGSAGAPMLVPVVCLAALGVGMLAAASM
AGRRGPAGMAA*
>GCA_000452945.1_BPMSHR5858_genomic_01286_1 naiP_4
VFTWFAAVSPRERRTFWACFGGWALDALDMQMFSLVIPAIVAEWSLSRTQAGFVGGATLV
ASALGGWVAGMLSDRFGRVRTLQWTIACFSVGTFACAFATSYPQFVTLKTLQGIGFGGEW
AAGAVLMAETIRAEHRGKAMGAVQSAWALGWGSAVLLYALAFSLLPAELAWRAMFAIGLL
PALLIVYVRRGIAEPSARALNPSGPCDPPEPPNSPDGAPPSGASPAVRASAPLVGIFSRG
TRRMTLIGALLGVGAHGGYYALMTWLPTYLKTERHLSVLGTGGYLAVIIVAFFCGCLASA
QLLDRIGRRGTILAFAACCVGTVILYLFAPLGNAAMLALGFALGFFAAGIPASMGALFNE
MYPRGTRGTGVGFCYNFGRVASAGFPVLVGHMSAGMPLGTAIGIGAAVAYSIVVVAVLML
PETRGRTLDDCASADADAARAGAGRIGAPAAPGARHR*
>GCA_000452945.1_BPMSHR5858_genomic_01290_1 acr3
MNPSNVAPPENAAAQPAINFFERYLTVWVALCIVAGIALGQALPDLFRQIGRMEYAQVNL
PVGLLIWVMIIPMLVKVDFGALHEVRRHVKGIGVTLVVNWLVKPFSMAFLGWLFIKQFFA
PMLPAAQLDSYLAGLILLAAAPCTAMVFVWSRLTGGDPLFTLSQVALNDSIMVIAFAPLV
GLLLGMSAITVPWATLLTSVVLYIVIPVILAQLWRKRLLANGQAALDAAMAKIGPWSIAA
LLATLVLLFAFQGEAILAQPLVIALLAVPILIQVFFNSALAYWLNRAVGEKHDIACPSAL
IGASNFFELAVAAAISLFGFHSGAALATVVGVLIEVPVMLLVVRIVNRTKGWYERT*
>GCA_000452945.1_BPMSHR5858_genomic_01291_1 arsC
MTTNILILCTHNSARSVLAEGMLDHWAAKLGKDVRAHSAGSAPSGRLNPFALEALRGAGV
DVGGYRSKSWDEFVGDGAPVMRAVITVCDAAASETCPYWPGSPVKVHWGYADPSNAPGGD
AGKRLAFELTRQAIGYRMLQLLALPIDRMSDAELQTALTDISRN*
>GCA_000452945.1_BPMSHR5858_genomic_01293_1 group_4573
MEIDQTVAALAALAHTSRLAVFRTLVQAGPQGLPAGRIAALLDVPPSSLSFHLKELAHAQ
LVTSRQEGRFVFYCANFATMNGLLAYLTENCCGGNPCSPAAACSTARGHRS*
>GCA_000452945.1_BPMSHR5858_genomic_01294_1 group_3188
MRHWVHQSRGPVPIGSDAHKQMFCNMLLDTHNPYKPAVIDWPALRPDALRRLTSLPIWDI
AVQTEGRASVRVATFASTVRDPLLRAALELDAAEEARHKVVLSKLVQAYGIRLAPEPAYP
APKDAEWAWMVTGFSECIDSFFAFGLFRSAQRSGYFPPELVETFEPVIQEEGRHILFFAN
WFGWYWRNLSWWRRPWLFARVAAVWAYLIWERIGIARGIDADGVARDANFPANTAATVGN
ALNPRELIELCLVENARRMAGYDTRLLRPTIVPRLARFALRFIRK*
>GCA_000452945.1_BPMSHR5858_genomic_01295_1 group_7335
MNRSIRYKGYEVAPAAARLPNGLFAANLTIEKAGPAPARAYSFDAIDFFFDEEHALAYAF
RWGRLWVDSHC*
>GCA_000452945.1_BPMSHR5858_genomic_01296_1 group_4574
MERTLTAFDPHRRPFIVGIGGTTRAASSTERALGFALRGAQAAGARTRLFDGPFLHTLPH
YAPERRALTDAQRELIDAVRAADALVIATPGYHGGVSGLVKNALDTLEELRADERPYLDA
RAVGCIVTAYGWQAAGTVLTSLRAIVHALRGWPTPFGATVNTLETRFETAGSCSDSKVAA
QLETVGAQAAEFALAFASHRAATRVAGGETLAPVLEAAAG*
>GCA_000452945.1_BPMSHR5858_genomic_01297_1 ousV
MDAPKVVVEGLCKVFGSNPRQALDMLAAGATKDEVFARTGQVVGVHNVSFDVREGEIFVL
MGLSGSGKSTLIRLVNRLVEPSAGKVMIDGRDVAAVRRAELTALRRTDMSMVFQSFALMP
QRTVLSNAAFGLEVAGMGRKDRERRAMDVLEQVGLAQFAHKLPAELSGGMQQRVGLARAL
AVNPSLMIMDEAFSALDPLKRKEMQNVLLQLQKEQRRTIMFVSHDLEEALRIGSRIAIME
GGRLVQVGTPQEIIANPADDYVRAFFEGIDTSRYLTAGDLMLTGAVPTLSKLDAKHVAAS
LNGSAEYAFVLDEARKIRGFVTRDALNGATPNVRQVESIPRDASLDHVVERCVAHPHALP
VVDDDGCYCGSVDRAVLLKAITRSRGSHV*
>GCA_000452945.1_BPMSHR5858_genomic_01298_1 gbuB
MSELIPLGSWVDQSVHYLLDHDAKTFDAIGRAIEGLAALVEHGLQAVPMWLMMACFIGVG
LWRVGWRFAFFTAASLFLIFATGFWDQTVVTLGLTLSSTIVSLVLGIPLGIWAAKSKWVA
AIVRPILDLMQTMPAFVYLIPAAMLFGLGRVPGILSTVIFAMPPAVRLTSLGIRHVNREI
VEAGQAFGCTPWQLLYKVQFPNALPSIMQGVNQTIMMALSMVIIASMVGAGGLGNDVLAS
IQRLDIGLGFESGLSVVLLAIILDRITESFGRAPGTARAPLFSGLRQLFGPRSRSPVVQA
*
>GCA_000452945.1_BPMSHR5858_genomic_01299_1 group_7338
VTSAAAAAPAACAEPVPSVAHFGFLTLPNFSMIAFTSAVEVLRMANYVARADHYRWSIFS
LDGAPVRASNGIAVRPTQPLDVDDPPDVVIVCGGIRIREAVDERVRDALGALAARDVPLG
GICTGAYALMACGLLDGYRCAVHWENLSALHAEFPRVRFADELFAVDRDRLTCTGGTAPL
DLMLNLVGARLGQPLAAQVSEQFILERIRGATDPQPIPVDARVGFSRAELIEVVRLMEAN
IEEPLSLEELARLVRLSQRHLQRMFKIYLNVSPTHYYLTLRLKRARDLLRTTDASIARVT
AVCGFHSPCHFSKAYRAQFGHAPSHERRVSAR*
>GCA_000452945.1_BPMSHR5858_genomic_01300_1 group_3189
MKRNLIAAACGLAIAAAPFASARAGDAPTCKAVRFADVGWTDIAATTGLASTMLAGLGYA
PTKTIASVPITFAGIKSKQIDVFLGYWSPTMDPMIAPFTKAGTIKVLAAPNLTGAKYTLA
VPDYVYQGGLKSFADIQKYADKLNGRIYGIEPGNDGNALIKKMIDGNQFGLGKFKLVESS
EAGMLVEVNRAIRDKQWIVFLGWEPHPMNVQMKIDYLSGGDDVFGPNYGEAKVLTATPPD
YAARCPNVAKFVSNLQFTTSIENHVMLPIMNKEDPNKAAAEWLKANPQSLDKWLAGVTTF
DGKPGLPAVKHYLGIQ*
>GCA_000452945.1_BPMSHR5858_genomic_01301_1 group_2321
MRWAILFIFVACAAFTHRRGKVRHGFFRQLSDHSTFTAPLNCFAYAFSNVPNTPYIDDAH
FPELAILKREWRTFRDEALALRDASRIKASGEYNDIGFNSFFRKGWKRFYLKWYDRPHPS
ALAHCPRSLAILSKLPSVKAAMFAQLPPGGKLGLHRDPYAGSLRYHLGLSTPNDDECAIV
VDGDPYAWRDGEAVMFDETYLHWAENRTGHDRIILFCDIERPMKYRWAQAVNGAIGGVLM
RAAASPNETGDRTGGLNRLFRYLYAVRRVGKRLKAWNRTVYYALKWALFGGILIAILWR*
>GCA_000452945.1_BPMSHR5858_genomic_01302_1 group_7339
MAAFYDFGARKLRDLYDRRLHCGAVLDTATLFPDAARFTHAWRAIRDEALAVSRDLNRIP
RFHEIMREQAAISANDARDWRMFIMQAYGVRFAQNLGRCPTLASIVAASPDVLSASFSFL
APGKHIPPHRGPFRGILRGYLVLSMPMRANGTPAAVLKIDGRDHRLRDGQFLLWDDTFEH
EVWNESDAVRIVLLLDIRRRDLPPALALLSGALIRLVRVAIRLRGGAIPV*
>GCA_000452945.1_BPMSHR5858_genomic_01303_1 denD
MKVLITGGAGFLGQRLAKQLLARGELTGPNGAPRRIDELVLLDVVQAHDFDDARVTARVG
DIADRAVLEAAIDARTHAVFHLAAIVSGQAEADFDLGMRINLDASRLLLDVCRARGHRPR
VVFTSSVAVYGGALPELVRDDTALEPQSSYGAQKAIAELLLSDYTRRGFVDGRVLRLPTI
SVRPGRPNAAASSFASGIVREPLNGEQAVCPVPGGTRLWLLSPRRAIDALIAGCELDGAA
LGNRRAINLPGLSVTVDDMIDALREVAGIEAVKLIRRAEDERVVKIVGSWPGRWDTSRAE
ALGLAGDASFVDVIRGYLEDERR*
>GCA_000452945.1_BPMSHR5858_genomic_01304_1 otnI
MPRFAANLTMMYNEHAFLERFAAAARDGFKAVEYLFPYEFPAAELKARLDAHGLVQALFN
APPGDWAAGERGIASLPGREDEFRRAIDTALDYARVIGNDTLHVMAGLIAPTQDRARHRD
VYLRNLAHAAEAARAQNVTIVIEPINPRDMPGFFLNRQDDAQAICREVGAPNLLVQFDCY
HCQIVEGDLAMKLKRDIAGIGHIQIAGVPQRHEPDVGEIHYPYLFELIDVLGYDGWIGCE
YRPKAGTSEGLGWLAPYL*
>GCA_000452945.1_BPMSHR5858_genomic_01305_1 nicT_3
MIPSQAAAARPLEAAGQAELDLTYRKVFWRIVPFLMLCYVVAYLDRVNVGFAKLQMSQDL
AFSETVFGLGAGIFFLGYFLFELPSNMLMHRIGARIWIARIMITWGLLSALFAFVKTPTQ
FYVLRFLLGLAEAGFYPGVILYLTYWFPTHRRAKIIAVFMSAIPVSGIFGNPLSGWIMER
FHGGSGFHGWQWMFVIEAVPAIAIGIATILYLDNNIRGAKWLTEREKKLLADEIEAQPHE
RDKHAHSLSTVLRDPRTWWMSLIYFTFVTGQYGLTFWMPTLVKSTGITDTLRIGLLSAIP
FAFAIVAMNLFGRSADKRRERRWHLIVPALMGTVGFSVAAAYSHNTTVSILFLSIAAGGV
LTCAPLFWSLPTAFLAGTGAAAGIAIINSIGNLAGFASPYLIGYLKDATSSTSAGMYVLA
AMLVIGALAVWLTPAKLVNR*
>GCA_000452945.1_BPMSHR5858_genomic_01306_1 otnC
MSPIETKLREEICEIGASLYARGHAVGSAGNISAKLDDGWLITPTDACLGRLDPAGIAKV
DPAGNAVSGSKPSKTLALHRGIYARNAEARGIVHTHSTHLVALTLAGVWSEADVLPPITP
YYVMKVGHIPLIRYRRPGDPAVAAEVAALADRVRGVLLERLGPVVWGPSVAHASYALEEL
EETARLWLITHPKPEPLGEPALAELRDAFGARW*
>GCA_000452945.1_BPMSHR5858_genomic_01307_1 otnK
MSTDQALRPLLGCIADDFTGATDLANMLVKSGMRTVQTIGVPAAGAAVPADAIVVALKSR
TIPAADAVAQSLAALDWLRAQGCRQFFFKYCSTFDSTDAGNIGPVADALLDALGGERAFT
IACPAFPENGRTVYRGHLFVGDALLGESGMENHPLTPMKDANLVRVLQRQTPSKVGLIRH
DAIALGTCAVRETIDTLRREGVRIAIADALTDRDLYVLGEACADLPLITGGSGVALGLPS
NFRLGGLLPERGDAAALPAIGGASAVLAGSASKATHAQVAAWRAERPALRIDPFAAARGE
PVVDQALAFARAHLPQPVLIYASAAPDEVKQVQQALGIEAAGHLVEATLAAIARGLREMG
VRKFVVAGGETSGAVVQALGVKALRIGAQIDPGVPATATTEGSPCGTTEGSPCGATEGSP
RGTPDAQPLGLALKSGNFGSVDFFEKALRALEGAA*
>GCA_000452945.1_BPMSHR5858_genomic_01308_1 ltnD
MSRNVGVIGLGAMGLGVARSLLRAGFRVHACDVREDVLAAFAAEGGVRCATPAELGALCG
VVVTLVVNAQQTDAVLFGEQGAAAAMPRGGVVISSATVAPDFAAQLGARLAAAGLLMLDA
PVSGGAARAASGEMTMMTSGPAAAYEACGDVLAAIAGKVYRLGDAHGAGSKVKIINQLLA
GVHIAAAAEAMALGLREGVDPDALYDVITHSAGNSWMFENRVPHILNGDYTPLSAVDIFV
KDLGLVLDTARRSQFPLPLSATAHQMFMSASSAGHGGEDDSAVIKTFPGIALPARR*
>GCA_000452945.1_BPMSHR5858_genomic_01309_1 lldR_2
MFEKIPTRAMSDTVAQQLLKQIEIGGFAGTGKLPTEAVLAQEFGVSRTVIREAISRLKNE
GVVEPRQGSGVYIARHAAIRPLRIDYAEAVEASSLPHLLAVRRAIEAEVAAEAALCRTDE
DMEDIDAALAKIDDAVAEGRDGVAEDVAFHRTIASVTGNPYFLKTLSFLNQYLEAGVKVT
RGNEATREDFSRQVREEHAAIADAIRARDPSAAGNAARAHMYNAARRLEQAGIC*
>GCA_000452945.1_BPMSHR5858_genomic_01311_1 group_3191
MSNTPPELLKAADIEAMEAVRSVHVLNANAVRLKKPLSEKTGITQFGFLIVTLMPGHESS
ECHRHLYEEECVYILSGYGEATLGERAYPVGPGDFLGFARRGPAHVLKNTGDVPLVFIAA
GQRLEHDVCEYPRVGKRLFVAGEMEGYVDWEEGPREMLGD*
>GCA_000452945.1_BPMSHR5858_genomic_01312_1 group_2322
MRANRNNGLTAGALPPNSAAPDDAAKAAPPVAQSDRGAGPLAGLTALKADRAQCMSARPS
IRSTTKAPAVMSHARADTSRELTPVEARIAQTFVRPEHLRAIAAAARAGSFAVSFRASGQ
LSLKRLANGAPAKGHDILEKTIKPGSVQEVYGADSERRLAQIERADIDGYVGHWGRDKLL
GLYMGPAANERDTPMLFAALRSTPSGHRYYPVDHDDLEGSLALLKRTQGPSGTAGSFWRS
LPYTGDYDTHDMINMAGHRAPVPSGSRDEHRIMRSLNVAIEAIDPIRKASSGSHRLIQHG
PQHNYVAHMRTNEQGAVIDEAVARASFPLAMCDRGTWSVIETPEALAAFYRQHGVVLKEK
WREKRANTFTA*
>GCA_000452945.1_BPMSHR5858_genomic_01313_1 group_4577
MSSERYIQLIQDLCATVGLANVDNVLEARTIEVEGFDVRLDCFDSDAEAMYANFHFGTVT
AGRTLTVFRLMLEANLLVYAQDQAQLGIDTNTGGVILIIRLPFSLGIDGQGLADLLAHYA
EHGRYWRQSIIESSDEMFEGIASGEYFWLRA*
>GCA_000452945.1_BPMSHR5858_genomic_01315_1 group_4578
MYEPLEPFAQDFNEVRTLLDDPAGGDRLRELCAAIEAVAERLGAAPAATELDRSNLAKLY

RGFLATSRVLAGLQETARAPAS*
>GCA_000452945.1_BPMSHR5858_genomic_01316_1 group_6821
MKLLRILTGVHAGAQLQLAPGTHRIGSDDGADIRLTDWHAADLLLHVKDDVVRAQFVETD
AERAPSASDSSETVLFVDLAPMQFDATVLCVGPDSSAWPSDFDLLSTLILRRAPQPLWRH
RYARTATACIALGSVMLAVAAISTSQTSRAAPAPNVGYRAQRIVDALAAAHIDGLQARPI
GDAVVVTGMVAHAADAATVRTLLASIASSGIVSRYDIAQQVSHNIEDSLGVPGAQVEYAG
KGRFVVKGPPGRHEALEAAVARVRSDLDPNVTGVTVAAPESAAETAAENSASYAEMVAAD
GVQYAQTSDGVKHIYVADTPAQAGDNVPMRGNARSATRDVPPLPPNMPD*
>GCA_000452945.1_BPMSHR5858_genomic_01317_1 group_2104
VNRSFRSSNHLSSPAQLQQSADAQQRAADRQAARNSALTAALYRGTTNFTYSKDPAPRGR
LPSVSKTNALARKRKITSRLRRNAAGNGDGDDVDGALVPHFEAEPGGDRDRGGRGGREQP
RDDPMDIETCRRLPRMQAAESRAAAAPRGRFDAVADLHAAPGQEGERADAIARAWESEMF
GLRVFSPNAPRTAEILELSLDFLLIQRRIGPIPVSTLARLRESPAPTPVACGRDRLPGET
APDDRRRFNLLFPLLWLQAGMPRTAIRLDCAIAKLLAIRNALPDNPLATDDAQAEPTPAP
PASSSE*
>GCA_000452945.1_BPMSHR5858_genomic_01318_1 spaQ_3
MEIDSLIRFCTQAMLLCLSVSLPPVIVAALVGLGVSFVQAITSLQDQTLPQVLKLIAVTI
TIMVAAPTGCAAILHFANQMMQLAVPQ*
>GCA_000452945.1_BPMSHR5858_genomic_01319_1 spaP_3
MVQFSDITGLLLVVIAISLLPFIAMVVTSYAKIVVVLGLLRNALGVQQVPPNMVLNGIAI
LVSLYVMAPIGMQAAKALDEQQLASQSSQAIIQALGSAREPFRSFLEKHTPEREKRFFIR
SASVIWPKEEASLLNERDLIVLAPAFALSELTDAFKIGFLLYIVFIIVDLVIANVLLALG
LNQITPTNVAIPFKLLLFVAMDGWSTLIHGLVMTYK*
>GCA_000452945.1_BPMSHR5858_genomic_01320_1 group_2323
MASPFSVDPNADPPCGVATRQPGNRSNAFDGPANRSAPPQSTARQNAATGIAPLSPVRVE
PAVARASRTVFDARLEALAASVSGLASWHVSVVDNAHVFIDRLVDAAVIELSIGPSGGMG
GFPLFIAIDLDAHPDLGIAAYPERDAVCAHREASESVALRQAVSGILLEPLVKCLDRFGL
CSPSVVSITRPRPVDRAGDWRDQPAVELTLALDERRVACAISLPMRGYDIVDALLRTQPT
PRKPAMSIPGALIIGARPLAVDTLNVLEAGDVLLRGLFPHFNATLLSTGTRATESLRAVA
AWGARGHVRLHAAVQVDVRSFVISKELSMSEEVDQASAGLGVVTTDEPTRIGELELPVQF
EIDTVSLPIDQLSALEPGYVIELPVAVTDARLRLVVHGQTVGYGELVAVGEHLGVRIIRM
AHRHGTVQ*
>GCA_000452945.1_BPMSHR5858_genomic_01321_1 group_7343
MLAREIGAFCSDGAIVCAGNWEAQLTLDPELLPRTTLHVALSRFIIRIRFDVPDGETRDL
LLTHRSLLERELDEVMRCWGEARDIQLSVW*
>GCA_000452945.1_BPMSHR5858_genomic_01322_1 ssaV_2
MFKSLKLPAGGEIGIVALVIAIISLMILPLPPVAIDLLLGINITISVTLLMVTMYVPDIA
ALSAFPSLLLFTTLFRLSLNIASTKSILLHAEAGNIIESFGQLVVGGNLVVGLVVFAIIT
TVQFIVIAKGSERVAEVGARFTLDALPGKQMSIDADLRSGLLSTEEARKKRATLAVESQL
HGGMDGAMKFVKGDAIAGLIITMINIVAGIAVGVAYHGMSAGDAANRFSVLSVGDAMVSQ
IPSLLLSVAAGVMITRVTDERAPKRRSLGDEISHQLGSSARALYFAAFLLLGFAAVPGFP
AALFVLLAAALAFAGYRLSRKGSPSSGAARGQEQEALRAMQRSGAKADVPPILPRAPQFA
CAVGVRIAPDLASGLALPRLDEALDLERARLQDELGLPFPGVTMWIHPTLSAATFEVLIH
DVPHLSVTLPARKAMLPQQRHLSPAHAALTDEQKARHRDLIERSEAGPPIDASGQATHWI
DEPAVTAKDSVWRAEQIIAYASVAAIRTHAPLFVGIQEVQWILDQLGADSPGLVAEVLKI
LPSTRIADVLRRLLEEQISIRNVRTIMESLVAWGAKEKDMLMLTEYVRGDLSRFLAHRAA
QGERVLSAVLFDLQAEQHIRQAIKQTPTGNFLALPPDDASRLVDRIQSLVGANARTDIAL
VTSMDIRRYVRRMIERRIDWLAVYSYQELGEHVELRPVGRVSLQG*
>GCA_000452945.1_BPMSHR5858_genomic_01323_1 yscU_2
MSDEKTEEPTDKKLRDARRDGEVSRSTDLSDAVSMSAAILLLVAAADHFGDAMRALVNGA
LAFVSADHSLVEMTARLYQFGGIALSAVMPLLFVAALAGIGGSVLQVGLQISLKPVMPNL
GALNPAEGLKKLFSPRSAIESIKMIVKAVIVFCVAWKTIVWLFPLIAGALYQSPPELSRI
FREILAKWLMVVAGLCLLMGAADVKLQRFMFMQKMKMTKDEVKRESKNDEGDPLLKGERK
RLARELAAAPPQHQVAHANFVVVNPTHYAVAVRYAPDEHPLPRVIAKGLDEAAIALRRAA
QDANIPIIGNPPVARALFRIGVEEPVPEELFEIVAAILRWIDAIGPRRNERA*
>GCA_000452945.1_BPMSHR5858_genomic_01324_1 group_7344
MKADTPDYMQCRPEIVGGLIETVSTALLRTFPKVTADPYDIELILDALRVMRPNVVEIQM
LDGVLYMACGRWDDAIRILSEVCATAPHFGYGKALLAFCLSTKGDDAWKQHAAEALESSP
NDDTLVLVKALQAREDMVKAVQANRNGGPFVPPQSVCELNAELERSAPQAEQPIAERVEV
AQYGATLRA*
>GCA_000452945.1_BPMSHR5858_genomic_01325_1 group_7345
MNMDVSRTALQLSLEQLTNGTAGNTPAHLAEKFSALMRKAPMEPVQAPRSAGLETVSRLA
AAQDMELQRSVSDVADVERRAPMLSLSEVNVETIRLTYEIASTQLDMEAKMSVVNASKTA
IETLMKNQ*
>GCA_000452945.1_BPMSHR5858_genomic_01326_1 group_7346
MIFGRVPLHRLVVRLLGVLALSVLLAGCKKELYTGLSEQDVNEMMVALLENGVDASKDSA
DGGKSWKLNVDGDQLVHAMEVLRTRGLPRSKFDDLGNLFKKDGLVSTPTEERIRFIYGMS
QELSSTLSKIDGVLVARVQIVLPNNDPLAQTAKPSSAAVFIKYRPSADITALIPQIKTLV
MHSVEGLTYDQVSVTAVAADAVDLARSRPAAVIMPPWLVGLLAFGAMSIAAAGLFVVSRR
PWTRAGAHEAGSAAPWRKRVAELAAHLRRRQRAN*
>GCA_000452945.1_BPMSHR5858_genomic_01327_1 group_7347
MTAASSPSPHQRAAALLERYRRNFANAVCWVHPSWTCALLGIDDSRCAVWRAALERAGEA
TRIPCSQAIACAAGVARPSLDMLLEPALRRPARGSTVAARIPNLALLDVLPHELGLAVLR
MRALCLRSAEVRVVIDKRTRSRILAWTGIHPDHFMQDPRSIGGADAAWLKAAVGMPPLAS
HDALTLSIEGWMLLQCDTGAPPSSDTITLMRLALPRSLATPHWLAELAPRLDAFGGEHLF
ARLPRLLPEYAWLFG*
>GCA_000452945.1_BPMSHR5858_genomic_01329_1 yscN_2
MSADEPLRTAEFSHLADAIEREILAVSGVVRTGRVLEVIGTLIKVSGLDVALGELCELRT
REGRLLQRAEVVGFTRDVALLSPFSQLEQISRTTQVIGLGRPLSIPVGDALLGRVIDGLG
EPLDGGPPLASETLQPLIAAPPEPMSRRMIDAAMPTGVRVVDAMMALGEGQRMGIFAPAG
VGKSTLLGMFARGAACDVNVIALIGERGREVREFIELILGQAGMARSVVVCATSDRSSME
RAKAAYAATAIAEYFRDRGRRVLLMMDSLTRFARAGREIGLAAGEPPARRGFPPSVFAEL
PRLLERAGMGRTGSITALYTVLAEDESGSDPVAEEVRGILDGHMILSREVAAKNQYPAID
VLGSLSRVMPLVVDDGHMRAAARVRELLAKHREVEVLLQIGEYKAGANPLADEAIAKADA
IRSFFSQRTDDYAAAEDTIARLYALSGAN*
>GCA_000452945.1_BPMSHR5858_genomic_01331_1 group_3194
LTPALFALQGWAGALLDDVTLLAICSVRLYIVMSVFPPTADGLSQGVVRNALVILFGSYV
AYGQPAGFVQTLHGTPLIVTGLREAVIGVVLGIAASTVFWAIEGAGCYIDDLTGYNNVQM
TNPARGEQSTPTATLLTQVATVAFWTFGGMTFLLGTLYESYRWWPITLTFPRMPALIEAF
AVSQTDSLMQMVTKLAAPMMLVLLLIDFAFGFAAKSASKLDLMGLSQPVKGAVTVLMLAL
FVGTFVDQAREQVTLSTLSKQLREWSHAMSRG*
>GCA_000452945.1_BPMSHR5858_genomic_01332_1 group_7348
MFSMLYFPMVSALSLLGADAPTHLHSHLKLILGGEFNAALERSSEWAETTVASERTSWDL
QLHADLQLVLGFEVEAEENYRRAQRKIRGSNSKIRIATCRNAAWQALFRYRVTTALACFS
RICDEPGIEAGGLVEARFGIACALYEMGRIDDAFDAIDSMEKIAEQQSDEMRAHWKDLIA
VLRFDLVVQSELRRAAAFVDHVYWQSAQSMSRVDRAHGVSEAAVSVETPLLRGRVAYLLQ
LRCAAAGNRDAVAELARCLDAAGEQGFVDFRYTLRLEIALALLAGDAPNLAQFVLEPISD
TLHGAESSRRYREYFYCAAKVHLAQDHTQESLALYRRYALIAMRCLREDALIGRQFLVGQ
ELKQLPQSDDVTVRLPLKYRRAYHYILQNLNRSDLSVREIAAEIGVTERALQNAFKIYLG
LSPRELIRSRRMERIRTELVDFTLTGERNVKEAARKWGVQNGSTLVIAYRKEYDETPSET
LAR*
>GCA_000452945.1_BPMSHR5858_genomic_01333_1 sctC_8
MKAKALLCAAWCVAIFGTIGIGAANAMPVRWRSTVVHVVLEGKDLKDVLRDFAASQGVAT
SIAGNVQGTVTGRFDLSPQRFLDTLAATFGFVWFYDGNILSITNANDMTRQVLPLDHASI
GELRSALRQIGMDDKRFPIFYDEVSGTVLVSGPAQYVQIVTDIAQRLDTLSGRRNGSTIR
IFKLKHAWAADRNVQIDGNTVTMPGIATVLANLYRVRGRAGTGQSSAAPGVQRVQPMGDV
SGSAYGGNRLMPPLPPSMMGGGRGDAFAKGASIDGESGGAASAAPGGAPRTEEASAERDE
LPIIQPDPMTNSVLIRDLPDRLAQYAPLIQQLDIRPRLIEIQAHIMEIDNDLLREIGVDW
RAHNSHGDIQTGTGNTAQNDYSGGQINPFFGTTTLAGNAVVNAAPAGVSVTAVVGDAARY
LMARVSALQSTSKVKIDASPLVATLDNSEAVMDNTTRFFVKVSGYASAELYSVSTGVSLR
VLPMIVQDGSETRIKMNVHVTDGQLTGDQVDNLPVITSSEINTQAFVGQGQSLLIAGYST
DKRANGVAGVPWLSKIPLLGALFRYHSDSQNHMERVFLLSPRIIDPGT*
>GCA_000452945.1_BPMSHR5858_genomic_01335_1 group_2324
MAISPTIATNRQVVGSVGSENVPGPAAEALQSVESQLHQLMARQLQGGVAMAQRQGSDIR
AAQYAPLDDYTRMYDQADRDRWADLAALGQGVSLERPLAAMQILSGERQPSAAQTAAALQ
FVNDNPSLKTAMQNTGGLKPDGSVDPGKVADLLKEVEANLAKADEDVKAYLNDHPDADRD
ALDTVRAAALLEAYEPIVGQSAGYQSSGKNHSYSTGGKNNGGGLTTKQQVADLQNNGGFS
AALKSAAKAWSTSGAFDNLDRAGDDKATEKVDQIFSANNLHHFITKDAPTSDAAEHAFLE
KASLENITAGTDISKLNQDIFANPQDYSPQQKAAVMVKLMETLVDVQAGGADKLRDVTAT
VSALKQDIMMLANDPETNAYLHQMVPPEMRDLNATFEQAGGLEGDTGAAGDMSVADNANR
VSARNIVDQTKDALDEAEKVAKRLGKLGNEGAGEATDATQSIADGAETAADGAVQGGVEE
VGGAATSAVDGAEGAITGVADGVEAADGAVTGVAEGVEGALAGAEGAVVGAEAGAGVAEG
VMGAVAGGMAAAAPVLAVGAAIAGIAGIVLAIVEAVKKAQNRKAFAENVNPTLNQFGIPL
PT*
>GCA_000452945.1_BPMSHR5858_genomic_01336_1 group_7350
MRRVLLDRSDEVMRFVADRTGESRYDDYATIGLERDGRIVAGVVYQGHGGPNVLMHFALD
GSRHLATPAFVCAVFTYPFKVLRCNRVTGLVRTDNHEAQRLDEHLGFVREGVMRAGASDG
TDFILYGMLKAECRFLSGKYVEALRVELAGSIDGVLASVHS*
>GCA_000452945.1_BPMSHR5858_genomic_01337_1 gbpA_1
MPVIIRSSGASIHQSDLSSRRSEADASDPAAAGPAREPDRVAGAESRSFVQPAPLNKTAD
KVKGGTPAIPLSLQVLGLYQRDKVCAQPVRPDHASLPESTRAEGLPRLDGKLPKRIETAP
FARALVQGNAGSGHNSRDVFLNVTGGVSGKAAPSNADHLTDVLLSDCRPGESLAGTPDDN
AFMIGGHSEIIDVATRPDGLSRADKARQMAIVMFGVTDPANDQIAFAERKLTALENAETR
HLAMSLFTGEEGREALSGWIADFDPGGAARLQRMHQLMYAAAVVLKQNGRAAARLLRNYC
GMSTERPDLQLKRQSYSMQRVLVSARGTSLDILGIPPSITGREHHAEQKSVYRNALAIAQ
YLHEKLDAAYKPDSLEDMLAAAQIVSAEALGGSGDRTEREYHFSRVVDNLGSIYQEERLS
IDALACAANLISIDRDNGAKALGAVAMLSSSEQADVMDALEQAGELVRLLPLGIRRDIAT
VADLDKVLNANAEGPGARASVSSVDPMGQASAQKAIGKLRAALAVAPDIGNIPIGAAARA
AIPMQQKIAYFKWRNGLADERKARLVFERLYKLNKYADRAIPRGKNLTGRLRRIGDDIAG
AFGHRKNPVGMLEEGLLLGGFLREDVFNEKIAPAFHSLIEKYRESGAGAGAGAGAGVAKA
RSEVVSRVFEKLKIDGFSSKIEIKSDDLKKWVALAVAGGRNDAVKRSLEFESIRRDLNNG
FSVDVLERWAAEAGLDDGNFRRQLNDLRSVSNNGYGEIPELGDRTSVFSAFSEIVARYRA
GKSLKFSSGGVGGLALGDSQSIGPLVSVSPTLGAGGGRVATVEIGGASGRGGLIAVTRAN
SQKLKVGASVFAGPQFIHVVRAGGSANLELVSAEQTREKGAIVRFVPRAGSLGGGEAWRG
WAQDCLSIIGNSADSADLLERLAEEYVNAGEIGIGMTEKEENVISSSAAVGAGVKASSSG
GDFRGGAGVSASIGVKREWRRHKAVRELDNKMGNHKFDQSSSTSVTWNAGLNASESYQLN
PTPDFSTSASASASAQAFSVGGTLFSAKEGASVAIRLRDGMVDSELAILERTRNRGRRAVEL
IARDYPLWVKAMGKTDAEGEAALNRFFASELGRGVNRPLRPEVPETYVASYRMTKEAASA
LNHYRAQIDLEQMLGTSKNRVSELRRRYAEVIENSASWRPSSIQHYVETSSAKSVGLHNV
FGVEAKKSSSTSTVQVLAHQAASEYVDPVALARHPVSVGSVQAGGDAPSDVGGRSTHGSA
VKTGE*
>GCA_000452945.1_BPMSHR5858_genomic_01341_1 group_3195
VTSGALRKLDQRFAPSASWADAGDVLARADALSVARRDDGRVECTARFAGVRAVGCADGP
DLRVALARAFVRARFGDAVDTPPEKPHQVRGGKIEPYDAGEPIPQHGDASATGNAGDIRS
VPRM*
>GCA_000452945.1_BPMSHR5858_genomic_01342_1 clsB_2
MMPPDTPTPAGAPPNAPTAGDALRDVAPQTGARALGAAAARAQPRTRELRIATYNIHGGL
GAWTASAAERIAVVLDELRADVIALQEVPLGGVRGADVLAHLRACTGMHAAEGPTIDTPA
RRYGNAVLSRFPIRAARMLDLSFGNREPRGALDADIECGFGVLRVVATHLGLSATERSAQ
VARLLAAFDTRALPVILLGDINEWFVRGRALRALVTHFRRAPAPRTFPTLCPLFSLDRIW
VHPGEWLVDVAAHRSARARRASDHYPLVARIRAAPETDADIAQAPASASRTPPERPCEPT
DSTEPASTTGAPARSPATTPRCASSPSRRASSSP*
>GCA_000452945.1_BPMSHR5858_genomic_01345_1 group_1424
MSDCSFHSRRPRRARRAAGVCAALAVLAATLWAAPACAQPEPPPASGAVTETAPDSPTRA
FERADRKMMDAMRAAPYTGDVDRDFVAHMAPHHQGAIDMAQVELRSGKDPALKQLARRII
AAQRDEIAFMERWQKQHGAGR*
>GCA_000452945.1_BPMSHR5858_genomic_01346_1 appB
MMEIDALLLSRLQFAWVIALHILLPALTVGLACYVATLEALAWATGRDVYRRLSAFWLRI
FAVSFGMGVVSGIVMPFQFGTNWSRFVAATSNVIGGFMAYEVLTAFFLESAFLGVLLFGR
RRVPQWAHALAAALVALGTLLSSFWILAVNSWMQTPSGYRFVDGRFYPESIVAVLLSPSF
PYRLAHTVAAC*
>GCA_000452945.1_BPMSHR5858_genomic_01350_1 group_4582
MRETNDKQQRGNVTQRHDTPAAPGGNRVTQAFERFAARVTQWAGSPVAFGVAVAITIGWL
ACGPLVHFSDAWQLVINTGTTIITFLMVFLLQRNQNRDSVALHLKLDELVAATKSASNRL
IGIEDASEDELKRLAHAYLELALRARTLRETSGEPARRDADDDSDGDGDDAASEAQERPT
PAD*
>GCA_000452945.1_BPMSHR5858_genomic_01388_1 group_4589
MDKLLALQTFAVVADTGGFSKAARQLGTATSSVTRLMDALEASLGVTLLTRTTRHVTLTD
AGATYLEQVGALLSQLVEADESVADAGGEPVGTLRVSVPVTYARIVLGPHLSAFLDAYPR
VSLDLAVSDAYLDLATDRIDVAVRIGEPSRDPGLVVRGLAPNRRIVVASRDYLDRMGTPA
APGDLAAHRCLRFPYRQGRQSWTFSRDGHSEKIDVGGRLTANSLDILREGVLSGQGVALV
PMWLVHDDIVAGRLLPLFESWCVTPPGGDAQVYAAYLPNRRHSRKVQTFIAFLAQRMPGA
PGR*
>GCA_000452945.1_BPMSHR5858_genomic_01421_1 group_7354
MGQELKTRNIFWRIVLLGIVSLSMVTMTSNVCVAAPGFPAIVAHRGGTGDAPENTVYAIS
KALQNSADAVWITLQLSSDGVPVLYRPTDLKVLTSGSGPISALSAAQLARLDAAYYYNPK
DGYPLRGKGYGIPSLEEVLKTFKDTFFYLDIKSPDADPNRMATALSQVLERTGALSRVRI
YSTEAKYTAAVQAMPHFETRDETRTALANVTMNHTCQLSPKFGSWYGYELRRDVTLIEET
TLGVSPPSPSRLVWNLEAKTCFIDTGKGSILLIGVNSADDYATAASLGAAAVLVDSPAQA
RHWPTQVSMP*
>GCA_000452945.1_BPMSHR5858_genomic_01424_1 fetB
MNTPLQDLSLVDVGIAAALVAINGAISVALSLGLGRKLAWAAARTVVQLLAIGYVLGWVF
GHPRWYVVLPLVALMTLIAGFAGAARGKRTYAGQRADSVLSIWASSWFVAAIGLFVVIRI
HPWYAPQYAIPILGMILGNTLTGVSLGVERMMEELTARRDRVETALALGATRWEAAQDAA
RQAVRAGMLPTLNQMAVVGVVSLPGMMTGQVLAGQSPLQAVRYQIVIMFLIAAASALGTV
GAVLSTYRRLFSAEHRFLASRLVEREKTANGG*
>GCA_000452945.1_BPMSHR5858_genomic_01456_1 group_7355
MSATLAIRRWWTIMPIVFITYSLAYLDRANYGFAAAAGINQDLGISKGLSSLIGALFFLG
YFFFQIPGAIYAERRSVKTLVFWSLVLWGACASLTGVVSNIPSLMAIRFLLGVVEAAVMP
AMLIFISNWFTKRERSRANTFLILGNPVTVLWMSVVSGYLVHEFGWRHMFVAEGLPAIVW
ALCWWCLVQDKPAQAKWLTGDERRALDAALAAEQAALKPVRNYRDAFRSPAVVKLCAQYF
CWSIGVYGFVLWLPSIVKNGSALGMVATGWLSALPYLAATIAMLAASWASDRLGSRKGFV
WPFLLIGAAAFAASYTLGSTHFWLSYALLVVAGAAMYAPYGPFFAIVPELLPKNVAGGAM
ALINSMGALGSFVGSYAVGYLNGATGSPAASYAFMSAALVAAVVLTLSVKQPGATPPLAH
PLQGK*
>GCA_000452945.1_BPMSHR5858_genomic_01457_1 group_1808
MKHRIVVYKPLPDDVLAALRARADVVLAEGADALARALPDADGALGASLRITPELLDRAP
RLRAWSTISVGFDNFDVADLTRRGIVLAHTPDVLTEATADTVFALILASARRVVELAEYV
KAGQWRQSIGEALYGTDVNGKTLGIVGLGRIGTALARRAALGFRMPVLYTSRSAHPQAEA
QFGARRVELDELLATADFVCLQVPLSPQTRHLIGARELAKMKRDAILVNASRGPVVDEAA
LIDALRAGAIRAAGLDVFEHEPLAADSPLLSMRNVVALPHIGSATRETRHAMARCAAENV
IAALDGTLARNIVNRDVLQRTPSTP*
>GCA_000452945.1_BPMSHR5858_genomic_01458_1 group_4606
VSDPRTQTSRPATISDVAREAGTGKTSVSRYLNGETNVLSADLRQRIETAIERLNYRPNQ
MARGLKRGRNRLLGLLAADLTNPYTVEVLRGVEAACHALGYMPLICHAANELEMERRYLQ
LLTTYRVEGMIVNALGADENALKSLRGGGIPTVLVDRSVEGFAADMIGLDNAAAIETGLR
HLFARGFQSLHFVVQPFEHVSSRRLREAAFRTQVAASGRAVHPTIVLEPKEPESTQRALA
ELDARIDAAAAESVRPAIFAANAPVALAIARHLHARFGDAWQKRAALIAIDDPDWAELIG
VTTIRQPTYDIGYRAVEFVHERIDGGTGDFHESLLPGELHVRASTAS*
>GCA_000452945.1_BPMSHR5858_genomic_01459_1 group_4607
MTVAPVTLAALVIATLVVALLPIGLYRFLRQPRALDRRDALIGVVVYTVFATLLERGLYA
LLLSQPSAVTWLSKPVTFAVFGTAVAALFEEVGRYLGLRWLAKRYGPSDGDARGLANGIG
HGGAQAWFVGVVVLAQWVYLAWLANRGELNAQLQTMPADLALRIQVMLATMSPVSVAVSA
LERIAAFVVQLALSVLMWRGIRANRRWILPAMIVLHPVIDLPSLVYNGGGVPFEWMVALY
LLLTAGFAGALMKLCRPARAAR*
>GCA_000452945.1_BPMSHR5858_genomic_01460_1 group_7356
MDDYQYDCASAEFGELARVICDLFPEQTRFAERADEAGRLLSVHWLAMRFGATPRKTTLD
VRFTSAALRRYLALKPMQRARSHAVLRAYVEATLGSLEERHAAGETVPRETTLELGDEFA

*
>GCA_000452945.1_BPMSHR5858_genomic_01461_1 group_2338
MIELKNVDLSNDPDALRVVKDETVQVEFAAQTGELTSLEGPNRYAAGDALVTGSTGDRWV
VSRERFDAKYAPVAAGGAHGEPGAYRNRPAVVLAKRMDAPFSIARSAGGDTLRGDAGDWV
MQYAPGDYGVVQAKRFAQVYRAAD*
>GCA_000452945.1_BPMSHR5858_genomic_01462_1 group_756
MPMRHAAARAVLAAAFTITLAAAPAGARAAPPLPALRADANRVSVSGLSSGAYMALQYQV
AYSASVVGVGVIAGGPYYCAAGSVANTDLCRGLVPNMVPDSGRLVAAAQGFAASGQIDPL
ANLQRARIYLFSGTKDTLVRQSAVDATWSFFWLVGVPVTNIVYVADVPAGHAFVTPSAGN
ACDANAAPFISHCTVGQSGYDQAGALLDAIYGPLAPPAATPTGRAIAFDQREFAPASSGL
AAQGYAYVPRTCDANAGCKVHVVFHGCLQSAAVVRDMTTYDNWADANGIVVLYPQVAKTS
TPNDPQGCWDWFAYTGQNYAWKSGAQMRAVRAMIERVTSAP*
>GCA_000452945.1_BPMSHR5858_genomic_01463_1 group_992
MKKISLTFAAALALAATLAHAQTGASAPAAAASAPAAAARHEARVEERITYLHNQLKITP
EQESQWKTFADTMRDNGATMARLYRERMENKTASALDDMKQYAELTQANADGAKKLADAF
APLYASFPAEQKALADSTFRKWLHPEPGKARARKHPGKADGQPASSAGGAQ*
>GCA_000452945.1_BPMSHR5858_genomic_01466_1 group_862
MTTSPTYSRPERLLQIASWGIAVAFALFLDMLGSLVIRDMAFAPRGGPPTLARYADTQAD
AALRATQRELQREQGALTDKAGAFSIARNRAQQEYDQAKESLRNWIATRSATGDASRNPD
VLARTRRLDELQQAVAGWQRQQDQIADQLDALRKRQDDVGARLAQSRAQAEWRYEQASRR
YELVVFAWRLAFALPILVFAVWLFVRYRKVRYWPFVYGFGMFALTAFFVELVPYLPSFGG
YVRVIVGIVLTVFAGVYMLRAFQRYVERKREEMRRSRHERAQAIGYEKAIAAYQKKLCPS
CDKPWNLGGDGATFCIHCGLKLFDQCGGCGTRNFAFFPFCSGCGMPVSRAEGERRRAGGA
*
>GCA_000452945.1_BPMSHR5858_genomic_01470_1 group_13213
VTYLVFFCGHAGTGKTTLAKRLIRPLMKTTGEAFCLLDKDTLYGRYSSAAMGALTHDPND
RDSPLYLKHLRDPEYQGLLDTARENLELGISAIVVGPLSREVRDRRLFDRAWLGIGPDVE
LRIVWVHTSEETAHARIVARGNPNDAYKLAHWDEYRQRRFIPTGEQCEGLLMFDNTAPSD
ADIDALRDHIAPPPARGVSTVPPLPA*
>GCA_000452945.1_BPMSHR5858_genomic_01473_1 group_993
MYYLLTYELVDDYLDRRGAYRADHLALAKAATARGELLLAGALADPADEAVLVFDAPSPA
VAEAFAQADPYVANGLVKRWRVRPWTVVIGKLAAPA*
>GCA_000452945.1_BPMSHR5858_genomic_01477_1 mscK
MIRLATFARRVALIGLLHLLCAALPAAAADLASGASVPPAPAISRDEALAELKRVQAALD
RIKQQASAATTYKQLDALDESTQALSADVDKLTAALVPTRAQLQAQLDVLGPPPAPGAAP
ETPAVARQRADLNARKTQLDAALKQAADEKESLANLTQQYSKLRRSLLRDQLVLRSSSIL
GPRFWAPLFRPSDDDRQELGAFSAQIGQVLAAAWEPGRRLGTALLLAAALAVWTLGRRLA
ERALAWFSLTKLPETRVRRSALATSIALSSVVTTGIAVQFVYLALTRDYALSSALQDFAD
ELAKLALTCALIAGLGRALLCTHHPSWRLPALADEVALAMRPFPAILAALLLVAGALEEL
NRTADTSLQVTLFGRGIVSLVVVVTIGASLLRANRARTALAAAGEPPEARSTLAGLIHAG
VTLTVVAALAALLIGYISVARFLTYELVWFEIVIGSVYLLTRLTRDAYASAFAANLSSGR
LIKHLFGLNDSHLDQARTVLTGISTSVLLLVAVIALLTGGFGTTPNDLLVSVISMFGGDK
LQRLNIVPARILNAALALAIGIYLLRSVRRWLDNELLPAIGLDQGMRASLVTLFSNLGYV
ALVLLALSLLGVKWDNLAWIVSALSVGIGFGLQEIVKNFVSGLILLTERPVKVGDMISIS
GVEGDIRRINVRATEIQLSDRSTVIVPNSQLISQNLRNVTMGNSTQGVATLMLTFPLNTD
PEQVRDLLLDAYREHASVLEKPAPSVTFSQLTPDGITLSVTGYVGSPRIAAATKSDLLFE
ILKRLRAAGIALSNPQTLMVQNLPPPSSE*
>GCA_000452945.1_BPMSHR5858_genomic_01478_1 group_2340
MKALFRILLIAFFVVAARPGAASIYEQELPPELFSQPDLCKWTDCASVLPGANAFSARKG
SPPYVEAYADDGHARALKGYVFLSTDIADIQGYSGKPIVTLIGMDTKGTITGVRVLKHSE
PILLVGIPESKLLAFLRQYVGKFAGARFEIGNGDAGAASLDAISGATVTVIAENQLISRC
AVAIASQVGIVKETVLPQAKLLPSDARETWQALVERGAIAHLAIRPDDVGAPDTGAPYLD
LYYGYLNAPAIGKSILGEHEYAQLMSRLKPGEHALFVVANGTESFKGSGFVRGGIYDRIQ
VRQGIHAFTFKDSDYLNLYALRAAGAPSFNETGIFIVRSTRFSAAYPWKFVFLGHRSDQE
TGAKTFSAFASPYWLPGEFLAGGRPRVDEDAPVWRKAWAGKRLEIGLFVGWIAAVMLFFA
TRERWVRRAKRADKRWVSWPKTASWLIAIGFAGWHELAQPSITQVLTLVHALASGWNWGL
FLSDPFIFVFWIAIAVTVLVWGRGLFCGWLCPFGSLSELLYKIARATGLKSLQRKLPPHW
HNRLRKLKYAVFAVLLVVSFFSMPWAEKLAEIEPFKTTFLVGVLNRAWPFVLFWTAVVGA
SIFVERPFCKYLCPLGASLALPGRLRVIKLKRKPECTSCHACAVGCGSQAIDPAGRIDPM
ECLLCLDCMVMYYDAHSCPPLSKERKRREKAGLPLTPVGKDGRYIPIERV*
>GCA_000452945.1_BPMSHR5858_genomic_01480_1 group_1431
MLHIELKERIETEFETARVESVTLRRDALDVRLANGVELTLRIANAREYAMDWRWGEAAM
RIDTAPRHAPPEAAHPNHLHAPDGRVLADPVTEFGGEPWRNVGALIERLAAAPLLGHEPP
SAHGLPSDARAIAS*
>GCA_000452945.1_BPMSHR5858_genomic_01487_1 apbE
LPGLKKLVGSSLALAVVVTLMAWLALRSPQVYVQGTYVFGTRVQLALYGVPLDRAQQATN
AVFAHFQAMQRGLHAWQPSEITRLNRSIAAGEPFRASPATAEILRAARRLSLDTQGLFEP
GIGRLIRLWGFQSDQFRVAPPSPDAVRRELARGARIADLAISPDGVVTSANRAVAIDLGG
FAKGWALDDAAAILRRQGIANALIDVGGNLLALGSKGGAAWRVGVQDPRKPGTLATLELR
DGEAIGTSGDYERFFQAEGVRYCHLIDPRSGFPAVQSEAVTVLVAPGPHAGALSDGASKP
PFIAGRAAMPLARRLGVQAVLIVDAQGRVWATDAMAARARFADPALRAARLD*
>GCA_000452945.1_BPMSHR5858_genomic_01488_1 group_864
MTNATHEPREPHELHEPRESHTPCIDLQDVSLQRGGRAILQRVRVTIGYGELVAMVGLNG
AGKTSLLSLVATLAAPSGGRIFVDGFDALAQPARVRERIGVVFQESALEPRLTARENLWF
IARCQGLTGRAAHTRVDALLEALGLRAHAATLVRRLSGGERRRLELARALVAQPRVLLLD
EPTLGLDVAARDAFWAEIKRLVADGRTVLCSTHHTDEACDADRVVVLDRGELIALGSWRA
MRAPVPGVIRLRVPDIDAARRWLAEHGYPFEVDAQGIAVAGADPQAVLPTLLRQMPCRVL
GADIAAPGLKDVVGHWIAARRGGDAAPRFAGRPA*
>GCA_000452945.1_BPMSHR5858_genomic_01489_1 trg_2
MSAVAAIVWFDLKRFWASPLRILFGLTQPLLYLFVLGAALRSGTYAEIGGYQAYIFPGVV
GLSLMFTAISAAVGIVHDRQTGLLNALLVSPVRRVDIALGKIGAGALLAWLQALLLLPFS
PAIGIGLTAPRLALLVAAMAFAALAFSALGLALALPFRSVIVFPVVSNTLLLPMFFLSGG
LYPLDLAPDWIRAAAAFDPAAYGVDLMRGVLSAQYAIAPSRSIAALAACLVVAGTLVVRG
VGRQHA*
>GCA_000452945.1_BPMSHR5858_genomic_01492_1 group_2063
MMATVDEDDIGTASGRDEGDWVPNRFCLRNAWFPLAHTFEIGERASRWQIYSQPCYLWRA
RGRIHASRRHPDLPAAPAMPAAPDSPFEPPERYPVVERFGYAWIWYGDPERASDALVPDV
PFLPREGGLPERMQGNIRLDCCTPLLVENLLDLTHADYLHANLLGDEQSEEDRVDVRFTS
ETVTMIRQCTNKSIAPIMRWFGGVRAKYQDVHVVIHVHVRSSVAVAYGRYMPGIDLPIFH
PCVPESRDRCRLSFALNMTRTPWLLRALMPLTPYIVLPQDNRMIGPQSTRYRDAGERRDL
YSRFDRAGLRYRLLLQQLARRQRDGDFSYAPDALPGQDARGILGMPD*
>GCA_000452945.1_BPMSHR5858_genomic_01493_1 group_1165
MDTIVPFLSDSTARWLPLALSEQVSGKAPLAVMCMGQAIVLFRDAAGAVRAMEDRCAHRR
APLSLGRVTPEGRLQCGYHGWTYDGASGTCVAIPNLSPHERVPAHYAARAYETLERDGYV
WARAARGVDTREPAAAAARATRCAQRFAGCATVSLAYDDYVAALADGPHLLMRIAGLRIT
DYVIADPEPRGGRLAMERGVTWAARRHDHRFGAHYPWTLRLTPPQDGAPASVEIDTRDGA
PMLWASVAIAPAARGATNLLWRGGVAADTGGAGPALFRLWARLRRSPFAMLEHVDGHALA
KLDASCSRAWPRAQHARVIPVESDRLPCEGSR*
>GCA_000452945.1_BPMSHR5858_genomic_01494_1 gerN_1
MADWILQLCVVVVVAFTLGALAQRLGQGRVVGELLAGLLLGPSAFGALTVAGHGALIGPQ
TAALLSRLGELGLVFLMFQTGMHITWTTAHGRSSGLVATVVAGFGMALPFATGCAIAVAA
SGSFQPQASHLAYVLFCGIALSVSALPVMMRIIADARIAGRACATLSILAATLTDSLGWL
MLAAVGTLATSGLSPASTVHAMGVLVAFVVVSISLVRPIVLGVLDRTRRAGSGSAMLICA
LCYVLLSSWAAARLGFHGAFGALVAAANLAGRHDLRQLWDARFAGLADLVLIPLFFVSMG
FQASFSALDTGAAWGWLAAFLVGGVATKFAGCYAGARLCGIDRAEATLVGVLMNCRGTVE
LMVLAIGLQLRIIPASLYTVLLIATLTMTWLSAMSTHRLAKRRPLRSPPAGASDAPAGFR
ATQAQAAGGDARLPGLPEASRPSR*
>GCA_000452945.1_BPMSHR5858_genomic_01497_1 mdh_1
MSNLSSAERTDSFFIPCVTLIGPGCARETGARAKSLGAKKALIVTDAGLHKMGVSEIVAG
HIREAGLQAAIFPGAQPNPTDVNVHDGVELYRREGCDFIVSLGGGSSHDCAKGIGPVTAG
GGHIRDYEGIDKSTVPMTPLISINTTAGTAAEMTRFCIITNSSNHVKMAIVDWRCTPLIA
IDDPSLMVAMPPALTAATGMDALTHAVEAYVFTAATPITDACAEKAIALIGEWLPKAVAN
GESMQARAAMCYAQYLAGMAFNNASLGYVHAMAHQLGGFYNLPHGSATRSCCRTCASST*
>GCA_000452945.1_BPMSHR5858_genomic_01499_1 group_2703
MDSAAEYGKVMQAIAANESAAVTYVQDSRLSATEVQTTHPIWSGVQNGDRLGVSWSGGDF
QHQRIFVYEGNGNAGKWKKAEAADIPLKQGGAPSILNQTPLMPLMPQQQVRETSASAKSV
ADVVSVSLYRRVTGNIDIYTVKVGSRIQGIYGVGMTAPGWIQADGVGTDIVVFQVRASGT
PGLFYSGVASATVIRE*
>GCA_000452945.1_BPMSHR5858_genomic_01500_1 group_4608
MRRKKSSIWLPYFDLLSLTLRARPRQKVAKPKPSVTRPTSAAKTRSTKRTRAAHAGRTSA
DNARGTWLRSFHSARPAAGRLINHLAYALYLPAVPAAAASMPAVVMLHGCKQTAESFATG
TRICDLAQRAGFAVLLPEQAKTSHSHRCWNWHGDSSQSEAPAVASLVDAIVRQYGFDRER
IYLAGLSAGAGLAAGLAMRYPELFAAVGLHSGPVFGAPSSTLAAMSLMRGGSREDPLRVI
ENCVDVSDHPGMPALIVHGEHDTVVAKQNAMQLGLEFARINRLIDGQGTLRVGEQHVYSR
KGVDYTDYLKSGRLVVRVCIVHGLRHAWSGGDPREAFHSATGPDATAMFWHFFRPRRRKR
AQ*
>GCA_000452945.1_BPMSHR5858_genomic_01501_1 group_7362
MSLSTDVAFFRLLADSYHRLVGKSLVPEGMSAAEGAAWLYEAAPFGMLAHDASADPVFVY
GNRRAQAIFEYDWDELTTLPSRLSAEPVERRERRSFLDRVARDGFVSDYRGVRVTKSGKR
FWIEHATVWQLTDSAGHYRGQAAMIPEVRPIGESA*
>GCA_000452945.1_BPMSHR5858_genomic_01502_1 fimA_2
MKAKGMQYGLTAVLLSMLSGGAAYSQTATTGTINFTGSITDVPCEIDTAATSSNVTMAKV
FANDFSGVGSTTGTTAFKIVLKNCGASTSGATVRFMGTTDSANPAALQTTAGGAGGVALQ
LVDDTGTPISIGSSSKAYTIAEGDNTFNFAARYIATSATVTGGAANATAVFALTYK*
>GCA_000452945.1_BPMSHR5858_genomic_01506_1 bpaC_1
VLVAAIFPAATFVSRTGVVTPAPPRVIADFDGSSYDTTLSTEVCRPARFALTVLRLVDSD
VTLSAVPSSDVDNDAMPLSAVASPVDALVDSDARSPTAPSSDVDSDATPLAAVESPAAVL
VDSDANWPTLTASVALTPAATPVSVRGPWPPAKSTVRPGELAPTLIAPVEASCVTTPTPS
DSMLVESEAMLVAVDVESEVMPVAVDVDNEVNWDMLTASVGLTPAARPLSNTPPVAFDIV
NTLPFMPPVRK*
>GCA_000452945.1_BPMSHR5858_genomic_01510_1 group_3216
MPIVFVAFCVLLISLWYIDILTGWLIAAFALLIEAMLIGIGFFVFVMVRISHRRRREREA
KTALCRANQPDDADVPFPVGFDGVKTVHCMFGSGQVGGFRVSCYNTNCPLMRGATETERL
PGQAERA*
>GCA_000452945.1_BPMSHR5858_genomic_01511_1 group_4610
MSNVALHTRKVVVADDHPIVLRAVTDYVNSLPGFHVVASVSSGDALLSAMREQEVNLVVT
DFTMHQANDDKDGLRLISHLMRAYERTPIIVFTMLTNSGVISQLCRMGVAGLVGKEEEIA
ELGRVCVSVARGVSQSLSPGMAHRLAAVGSIRPGEAAFNALTPKELEVVRLFTGGMSLTD
IARTLNRSLGTVSTQKRSAMRKLHVDTNVDLINCAREQGLL*
>GCA_000452945.1_BPMSHR5858_genomic_01512_1 rcsC_1
MQGLLQELDGSPLRKFYSLESNLKRERRVFTIVIVLLVCAALSIAAMTVTGLFQTAFRQE
EQSARIHEKEVVDVFLQRRMMLTTASLVLQLRMNGAPSALNVPAPNACTPMAHNVRDDAI
LRESCDYTVQLLANSGQTPSVEMVTADGSVGYGYLFPTGDLSALRSSTPSELVSAVLERY
GKRGLDPLEAARKKRILWFAVGRGGRGEELHMIGASVVFKDERLYALVLTSVDLHSLVSP
IERAGRVQQPVVVDSEGVPLVNADDAETVRKVDGRLAGQQDGLYHWIPGFGWALRRPAPF
SGFGHMTYLLPLDLQLRSMRYELSLVGGATLVLIVLLFVAFRYWNYRFLTRIYEEASRAL
ESEMLNHLLVHATPVGLCIVRRATLEIVVANPIARTMLGLRLSDRHLPQELLSAFESSLA
EQDTQSDDARIFQFPFTLSRAGHAAVHIEITYAPAMLNAREVFFCAITDMTAHHQAEILL
REAKLTSDAAAKAKVAFFASMSHEIRTPLASLVGNIELIARGPLAPEQQARVKAMETSAR
GLMQIVNDVLDFSKIDVGELSLMEEWSNIAELLDRLALSHAPLATQQGLKFYMVFDRSLP
ARLYFDPIRVSQIVNNLLSNALKFTPSGKIVLRAGWRAGALEISVTDSGIGIPDDLKHRL
FLPFTQGDSNRLRQARGTGLGLSICARLCELMKGRIDLESTVGVGTRIAVTLPLGVSEAD
SSDAYWTLPYRRVAVLGRAQENLEWLANLFDPGVTAVTAFSRPAEPIDAHAHDFLMVTDE
FSPAEVLPWWRRPDSIVWVGQAGPLVPRRRDDGGVEISMYSLAGLKSATHMLAAGRTALA
EAGHEPPGAEAGMTVLIAEDNLLNRSLLLDQLTTLGVRVIEAKNGEEALALLLKEPVDVV
MTDIDMPMMDGFQLLAEMRRLGMTMPVYAVSASARPEDVAEGRARGFTDYLAKPVSLERL
ETVVRACCSAPAGARADEDAQDELPGLPDVPPAYASAFVAQAGSEIAEFDAILRERALPK
LRRWLHGVSGGIAVLGPSALHEQCQELRAYARESGEWNREIELQALAIRNALERMVAALT
SA*
>GCA_000452945.1_BPMSHR5858_genomic_01513_1 dosP
MEAGITRRRGAFFAREHEHGLSLDEADLERAFEAGEFVMDYQPIVSIRSGAVTGAEATIR
WDHPEWGALPQRVVHAAADRLGAASRIAGHAFGEACRQLARWKQQGAGVSALSMRLSGAQ
LGAESVFERLALSIEAFDISSSQFTLEVPEMSAPEESLSLLDRLKRLRQKGYGIVLGDFG
AHHTAMSTLMVLPVTGVKFGESFTERLPGSPTAEAILSSVSRLAHDLGFTLTVSGVENGR
QFELLRRFRDIELQGAFLFEPMRAEVWHERVNPRALSQLNLREPPF*
>GCA_000452945.1_BPMSHR5858_genomic_01515_1 group_7364
MIAYQRFKELKIELEARLADERRAAKEAVLTEIRACVDEFGFSLDDVFPPDGGHGRRKVR
AKYYDPASGATWSGVGREPVWIRGKDREQFELRCSGGVQESS*
>GCA_000452945.1_BPMSHR5858_genomic_01516_1 group_1700
MATELDPPDSFDPFDSIDSYDLFDSLDERQTNFTLSCAMDWGQNIKSPTTDILGKLIIGY
LGKSTKTKNNRFSIVWGPAVYVDPLSGYAANVTAIFKNTSNDKDYRLAFSGTNPNSQFAD
IVIEDNDVKNLTPFSKYVSDCPPDALIANGTAKALDITLQTLPSNSSASLTDFLKNLPKG
STLTVTGHSLGGVLASTFALYFSQSSGYPFKNVFCHSFAAATAGNVAFVAYSDGSLTGKM
KRLWHRYDVIPSSWNLSTMNRIKYIYEPNLFPTEDFIRKIRKIVYYFSNKHIYYAQPGSL
DHSQQFELEGAYHSGFKVFTDEAGWQHTFGYLAPLSLTANDLNEGEPWQPSLGTLA*
>GCA_000452945.1_BPMSHR5858_genomic_01551_1 cdiA2
MVVQENGRASHGSGSRDASAGVVPAWLSLSPFALRHVALAVLVAAGVVPIWVNAQVVAGG
AHAPSVIQTQNGLQQVNINRPGSSGVSMNTYNQFDVPKPGIILNNSPINVQTQLGGIIGG
NPNFQAGDAARLIVNQVNSNNPSFIRGKVEIGGAAAQLVIANQAGLVVDGGGFLNTSRAT
LTTGNPNFGPDGSLTGFNVNQGLISVVGAGLDTANVDQVDLLARAVQINAKAYAKTLNVV
AGSNQVDYNTLNATPIAANGPAPTIAIDVSQLGGMYANRVFLASSENGVGVANAGDIAAQ
AGDLTLQANGRLVLSGHTNAAGNMSLSASGGIQNSGVTYGKQSVTITTGADLTNSGALTA
QQNLTANVGSLNSTGTLGAGINVDSTVGTSGDLNVTSSGQLTATGTNSAAGNATFTSSGV
NLSNSATAANGNLALSATAGDVNLAGSTVSAKGAVNAQASGTVVNDRGNLSSGAGMTLGG
GSLSNQGGRANSQGPLSVQMAGTVSNQNGMLSSQSTADVRGSAIQNNAGLIQSAGKQTIA
GASIDNSAGRLISLNADGLSVTATGALTNAAGANVSGDPGGVIGGKGDVTVQGNTVTNSG
SMSADATLHVIGQSVDNGNGALHAGQTTTVDAGNHLSNAGGRVEGQSAVLNGATLDNSQG
TVNAATVSLNGTTLLNHGGTVTQTGTGPMTVAITDTLDNSNNGLIQTRSTDLSLTSTTLI
NDSPVDLSERSEWGQHGLGVEGWRAADSRSRYSVGKFLYGTGSERFVDGFARRAVQCNPV
E*
>GCA_000452945.1_BPMSHR5858_genomic_01576_1 narL
MTIRVLLIDDHTLFRSGVKLLLQRQPDFEVVDEASDGVEGVKRAKQHQPDVILLDLNMPG
LSGLETLQLLVQDVPSAHVVILTVSEDAEELTTALRDGACGFLVKNIDAETLVAGIRKAA
AGEPVIAESMTAKWIASLRAPVPAARPAPAAALAPAAHASVDPDQLTPREHDIVRALARG
ASNKEIAREFDVAESTVKIHVQNVLKKLNLSSRVQIAVYAVERGLTAPHPAEA*
>GCA_000452945.1_BPMSHR5858_genomic_01597_1 lutB_2
MQVQSMHFKARAGEKLADARLQENLRRLSTKFVSGRAEAVTQIDFVATRAQLKARRDRAL
ERLDVWLEIFEREAARRGTTVLYAQTTQDAAKLVADIARRHDVKKVIKTKSMVTEEANLN
AVLARLGVQSIETDLGEYILQINGNEPPSHIIAPVLHRDKEEIADLFAKTHGRARLTDIS
EMTREAREVLRPHFLSADMGVTGGNFLIAETGSVALVTNEGNEGMCTVLPRVHVAVTGIE
KVLPTLEDLATAMRLLPRSATGQKTSNYFSLLTGVRGDGDEDGPEHAYVVLVDGGRSGLI
GGEFQEMLRCIRCGACMNHCPVYQKVGGQAYGWVYPGPMGSVLTPSYVGLANALDLPQAA
TLCGECDSVCPVGIPISGLLRKLREKQVERRLRPWRERAALAAWGFFARRPALYALATKL
AVRVLERAGGSAGGVAGMLGRMPFAGGWARGWTATRDIPAPAGRTFRELYAASRTHLDPS
STR*
>GCA_000452945.1_BPMSHR5858_genomic_01658_1 cysC_3
LSIAAVNSLRRTIAPSHRVLDACGSGAACLLASRYGAKQVVALAAAEATLVGARAGANGP
GERISFVDAEQAAALVARQGRFDVVLGLAGGGRDGLDPHYGARLDAFARRFGTANVAVVP
NGVRYDAQLVEWREAERLEADVAARQRALEARYDLTLGPVLDQVAAGAAPRGRVRADALR
ALCARLPFLSCRPGDGGDAGTSGTSGTAAPEHVELTAAHGGRADGVLWTQELIHDGIVIE
RIQGCSWLDERVDLEAGDAIAVPVADMARAAPLDGGEAPPQLAPAPRASGAPAPLVFWLT
GISGAGKTTIATRFKQRADADAWPTVVLDGDTLRGGLNADLGFCDADRAENVRRIAEVAA
LMADTGLVVVVSCISPRRSFRETAREIVGAERFVEVFVDTPPAVAQARDPKGLYRRARAG
LIASFTGIDSDYEPPPDPQLRIDTTTHDVEDATQMLRRYYVDARLSGARAGTAECAA*
>GCA_000452945.1_BPMSHR5858_genomic_01684_1 group_4634
MLLLRHAILTALAATACASAHAQTADDAPVAAPACAIAIVTGVGGAATNLREYLAARERD
RYRYLADHPLDCRVSEDGRASGCTGLAYLRRERVSVYDDGDDTVQNVVARVDLDHGTYPA
IIAVQKRDVRCEQ*

>GCA_000452945.1_BPMSHR5858_genomic_01730_1 group_7381
MSLTISEILQLPGLEELQLRAGERSVQRPVRWYYVAENEGIADWVMGGELVFVTGINHPR
DEANLLQLIREGAKSRIAGMVILTGEAFIRRIPDSVVALAEQLEIVLIEQPYLLKMVIVT
QLIGTALARHENTLRSQRDIVNQLLTGDYPSIDIAAHRARNLQLALDRPRRVVALRLAGV
PALFEGRDPAAAEALLQDARQTVQRGLDDWLRDEEGALPVVEQGELFVLLLPCDDPRFRK
QKLALGALRDALNRQTGPLALFVGISSTVGAARHYCRGLAEARQALGVAEGMRAGQGLCD
YSELGVLKLLAAIPDPTLIDGFVKETLGNLLDSNRKHPTMLIETLEALLQENGNAIKAAE
QLSIHRNTLNHRLRRIETQSGQSLADPYFRLNASVALLAWRMSDTQRQEF*
>GCA_000452945.1_BPMSHR5858_genomic_01745_1 group_3022
MMKFLTREIVVLAALDWALLAGCAPVQAPPSSSSAPPASSCCGPITASGQTLRHTLDASD
VEHLWLPRQHVDWSTGKPDPTAEGFKSHCSAFAAAMGARLDVYMLRPPEHSQILLANAQA
AWLASDSGRAAGWRELHEAYEAQAAANRGELVVAAFQSADPKMPGHMAIIRPSLKSNVQL
ADEGPEIIQAGAVNRLDWNVRDGFARHPGAWPNGIKYFAHVVPAK*
>GCA_000452945.1_BPMSHR5858_genomic_01854_1 group_1310
MPTNLTELLTPISEASPCGDDLLFSAEFDAIQHARKFDDPSLDQGEWITDIKEADWGFVV
EQASALLRERTKDLRLAVWLTEALAIEDGIGGLTQGYTLLTSLCRQFWDHVHPLPDGDDA
EYRLGNVAWLAGRTVELLRAVPLTDGAANAFSELDWDVAQHVAQAIRRDPEQASEIARGK
PSVEQIEASKRMTPVAFYARLLGELKAFQAALDALEQELDGRAGDAAPSFRQARDAYETV
YRLAERFAKEQGYSADAQPQAVAPPAPSERAEPSFKTPLHSEEPVPRHAHAPSAPAPIVI
AGIQNRAQAVEQLRAVAKYFRGTEPHSPVAYLADKAAEWADMPLHQWLASVVKDDGSLAH
IRELLGLKPDDNA*
>GCA_000452945.1_BPMSHR5858_genomic_01855_1 group_2364
MSTPLKTLIAKLNPVCRKATERAASHCFARGHYEVDLEHLFLALLDESTGDVPLVLRASG
VDPHALRADLERELERLKTGNTRTPVFSVHLSELFEQAWLIASLDSQIGRIRSGHLLLAL
LTGPDLAQFAQRMSSQFARVRVDDLKHKFDEIAAGSSEAEPRHADADVAVPDGAAASGDA
PRGPSKTPALDTYTTNLTQRAREGKIDPVIGRDAEIRQAIDILMRRRQNNPIMTGEAGVG
KTAVVEGLALRIAADDVPPPLRGVALHVLDMGLLQAGASVKGEFENRLKSVIDEVKKSAH
PIILFIDEAHTIIGAGGQAGQNDAANLLKPALARGELRTIAATTWSEYKKYFEKDAALAR
RFQVVKIEEPSEPLAAAMLRGMAALMERHFNVRVLDDAITEAVRLSHRYISGRQLPDKAI
SVLDTACAKVALAHSSTPAAIDDAKKRIERIDAEIAALEREAASGAAHDARLAELREARD
ADLKALAEDAARYEEERALVTEIGALRAELDAARESSADGKPVDVDATRAKLAERVDALR
ARQGNQPMVPLQVDGHVVAEIVASWTGIPLGRMVKDEIETVLNLRDLLGARVIGQDHALG
AIAQRVRTATANLEDPNKPRGVFMFVGPSGVGKTETALALADVLYGGERKLITINMSEYQ
EAHSVSGLKGSPPGYVGYGEGGVLTEAVRRNPYSVVLLDEVEKAHPDVLEMFFQVFDKGA
MDDAEGREIDFRNTLIILTSNVGSSAVMQACLNKAPQELPDAETLAETLRPQLYKTFKPA
FLGRMKVIPYYPISDDVLAEIIELKLERIRRRIEANHKAAFEWDESLVDAVLARCTEVDS
GARNVDHILNGTLLPEIAEHVLSRIADGEAIVRIAARAAETGEFEYTVE*
>GCA_000452945.1_BPMSHR5858_genomic_01856_1 group_7390
MQTAQRRIDPGVVERLLDEPHRFEFFQAVRLLETWFAKHDAPRHAASRPGEIVARRIGFR
NTLSLGFPPSEIERAQSYDAAGAALKDDVRRNAALEAGALSRVELTPAFFGLLGGQGALP
LHYTEQIIAREQLKRDHAAHAFFDVFSNRATALFYSAWKKYRLPLHYELDRDERYLPLLL
ALAGVSNDDARSTVQQGRGALLDEAIAGYALAARHRPVSAAYLQRTLSEYFRVPIRVEQF
VGKWYDVPPDQLSVLGDVNVTLGATALVGERVWQRDMRARLVVGPLTKRDYEAFLPGEER
AIALERMLTLLAGVTLEYEVSLVLRRAEVGASVLGAGARLGWDAFLCTREAGRDRADARY
ELHVIH*
>GCA_000452945.1_BPMSHR5858_genomic_01857_1 group_7391
MEELLPYYERELSFLRRYSLEFANRYPKIAARLAQTGEQCEDPHVERMIESFALLGARIN
KKLDDEYPEFTEALVEVLYPHYLRPFPSCSIAQFGAPHAFAELTEPRIVERGTELKSRAI
RGVQCRFRTAYDVTLAPIRLADARYGSVAVAPAAARLPGNATGVISIAFESTSPQLDLAA
LKLGKLRAHLHGEQSFVAALADCLFAHALACYVEPDRGGRWIALGKTPFEHAGFDETDAL
IDYPARSHPAYRLLTEYFAFPDKFNFVDFDLGALARRAGPCRQLTLHIALRDVRGDSHVA
RLLDLLSASHFRLFCTPVVNLFRQHGEPIRVDHRAVSYPVIGDARRAFAYEVYSIDSVHL
VRQRADRETLVEFRPFYSLHHGEAAQAGHYWFARRNDDVARKSPGYETEISVVDVDFEPG
APQTDTLSLDLTCTNRDLPAALATGLDGGDLFIDGGEQPGPIALLRRPTPSARFERGHAA
HWRLVSHLALNHVSLAEEGLAALKEALVLYDLRRSAVSTRHIDGIVGIEQKSAVQWLPGK
PFATFVRGVEIRLAIDDEHFVGSSLATFVRMIDAFFGLYVHLNSFVQLIVVSRRTGEEIM
RCKPRSGESILA*
>GCA_000452945.1_BPMSHR5858_genomic_01858_1 group_7392
MKRFEPSFLDKLFDDEPHVPAPAAMRQLSLEELKNTVARDVEAILNTRIALTEEDLAELP
ECQRSVLTYGLNDFAGLSLASHYDRTFICKSIQQAIARHEPRLQQVQVAFELNEQSTNAL
YFAIQALLVVHPAEEPVTFDAMLQPSTLQYSVTRSRAAKV*
>GCA_000452945.1_BPMSHR5858_genomic_01859_1 hcp1_5
MLHMHLKFGSPAIKGESADKDHEGWIELKSWDHSIVQPRSATASTAGGHTATRCEHGDMV
FTKEIDSSSPLLYQHASGGTTFDEVTIDFLRADGEGQRVKYLEIKLKYVIISSIAPSVHT
EGLPVETFSLKYAAVQWKQTQQKIGGNQGGNTQGAWSLTKNDKTYAV*
>GCA_000452945.1_BPMSHR5858_genomic_01860_1 group_12194
MNQQTAAAQSTGVQAGTESSLLDEIVEKSKVAKSESEHARAKDLIGELVSQVLDGTVVVS
DNLSATIDARVAELDRLISSQLSAVMHAAEFQRLESTWRGLDYLVKESNTGSTIKIKALH
APKRDLVRDFKNATEFDQSALFKKVYEEEFGTFGGSPFGVLVGDYEISRQPEDQYFIEQM
SHVAAAAHAPFIASAAPELLGLESFADLGKPRDLGKVFDTVEYAKWKSFRDSEDSRYVGL
TLPRFLGRLPFNPKDGAIAESFNFVEDVDGTDHDKYLWCNASWAFAARLTAAFDDFGWCA
AIRGVEGGGLVEDLPTHTFKTDDGEIALKCPTEIAITDRREKELSDLGFIPLVHCKNSDY
AAFFAAQSVQKPKKYSTDSANANAVLSAQLQYIFSVSRIAHYLKAMMRDKIGSFASAQNV
ETFLNRWISQYVLLDDDATQEQKAQFPLREASVQVSEIPGKPGAYRSVAFLRPHFQLDEL
SISLRLVADLPKPANS*
>GCA_000452945.1_BPMSHR5858_genomic_01861_1 group_7394
MAKKESIQKRLQKVRPPRVQLTYEVERGDAIEIKELPFVVGVVGDLAGQSEIEQPKLRDR
KFVNIDRDNFDDVMKAIEPRATFQVENRLSEAGGKFAVDLKFRSINDFNPDEVVGQVEPL
RRLLEARSKLADLRNKLAGNDKLEDLLSEVLSNTQQLEALKKSADGGDKQGE*
>GCA_000452945.1_BPMSHR5858_genomic_01862_1 group_11195
MKDRLFAKLSGVVLACGIIAGCASQPTPPTTEAFNKSLADADAVAKTGDQERAIGLYQQL
AKSDPTREEPWSRIAQIQFQQGHYGQAIVAAQEALQRDKTDRQAKSVLAVAGLRIATESL
GELRQDSSLAGDAKSDAQALAKQLRDTLGEAALFPPEQQATKPVVKKRRIVRRAKPVHEA
PRAAESETAAAPATPPAAPAQPAATPAPAPAKAAGGDPFSALR*
>GCA_000452945.1_BPMSHR5858_genomic_01863_1 group_7395
MNPHVTRYALPLAACILLGGCAAAVPLLGSAGSAALQMVGVGKPDVPDSQKPPRNLGLTL
YAAQNLNAANDNRPLALVVRLYALKDPTSFQQAPFDSFTDPTKEKTALGADLLSVREITL
IPGQHYNATEKVSREAQAFGVVALFRDPALQRWKLTFDPAKSEKSGIIIGLHNCAMTVTG
GTVIPSQQGMPTQRLDLLSSVNCG*
>GCA_000452945.1_BPMSHR5858_genomic_01864_1 group_3889
MSYSAKVLWGEGLFLRPQHFQRQDAYHEARLFESIQAIQPYNWGVRSVRFDRDALGSNVL
RASELSLVFPDGALYSAPQADELPPPIALDTLPEGINEFTFYLALHPLRETGSNYADDRD
AGFVSRFVSEQASVADHFTDAAEADITFLKTNVKLIAHSEPRDQLLSIPLVRVRRTATSG
FEIDDGFVPPCLAIEASPILHQRLRQLIDALQAKVNALYGFHREPTKNIIEFRSGDIASF
WLLHTASAAFAALAHLHQHSALHPERLFQELLRLAGQLMTFSKGYVLADLPAYRHDDPGP
GFARLDSILRELLETVISTRYFAITLEEVRPSFHVGRLDSGKIDDKTAFYLAVSADMPSV
ELVDAVPARFKVGAPDDVDKLVLSAMPGVRLAYTPQVPPAIPVRPGACYFALDARSPLYE
RMLQAQSAMIYAPSGINDLKFELIAVTS*
>GCA_000452945.1_BPMSHR5858_genomic_01865_1 group_2365
MSYAPSLFGGNTPPPAPQTASSTDAGFQARSLVDLLYDGFFMLFLLKNGREPNDASEFGT
RIQEFLSEFERGAKKLNIAADDVYAAKFAFCAAIDESVLSSPFKIRADWERRPLQLVLFG
EQLAGEKFYQYLEECRAQGAARLQSLEVFHMCLLLGFQGKYLLEGPEKLAYLTARLGDEI
ANMKGKRAPFAPHWPLPDQISHRLKREVPLWAIGAVFALVGLLAFVGLNTYLKDSTVRAL
APYSQVIKVGPESANLTISLP*
>GCA_000452945.1_BPMSHR5858_genomic_01910_1 btuD_13
MSSARAVAVRSRLPEALAWLAFVALLVVPALAWPRGWMLGYLAQTAAMIVLALSYNLLLG
TTGLLSFGHAAFAGLGAFVAAHGFNRYGLPLPLVPLAGGAAGAAFGALFGFVATRRAGTA
FAMITLGIGELVAAAAWSVPEWFGGAGGVSIDRAAGPALAGWEFGKPMHAYALIAAWCVV
AAVAMRVLVRTRLARLANAVRDNPARVAALGTDPCRVRFAMAIAAAFFAGVAGALTLIDV
EVATAESVGMLRSGAVLFATVIGGTTAFFGPVVGAVLLTFFSVFVASVSRAWLLYLGLLF
VAVVLAAPGGASGRWRAWRDARRHGGARFDARRALPGAGAALAGGVAIVFAAELAYALQF
AQNDARRAVIGPWRFDAGAPLGWGIAALAAALGMGLAHAARRWGRGAPRTNGPAFADAID
TCGAACGSAELKARDAGDVDGSTNAASAANETNAVSIASTTSTTSMTSATSATSATSATS
ATSATSATSARHAARVSDNAPSPVFEVAAEPVGGGVHAASAPPALRMADVSKSFGATPVL
VGIDLTIRPGERHALIGPNGAGKSTLFNLIAGALAPTRGRIALNGIELGRRRPHAVARLG
LARSFQQTSVFARMSVYDNLRCAALHAPRARRRWRRGPAAAAAIDRAADAALASIGLAAH
RDALAGSLSYAQQRALDVGLALASGASVFLFDEPTAGMSREQARRTIELIRRATAGKTVL
MIEHDMDAVFGFADRVSVLVRGELVATGSPAAIRANAAVRAAYLGEAFER*
>GCA_000452945.1_BPMSHR5858_genomic_01912_1 iolB
MSLLVKGAGDGMSIARVTPESAHWKHVGFAAYRLKAGEAIALAEAAREMCIVVLTGTVDI
EAGDGTCWTALGSRDSVFDGVAPYALYLPPKLSVIVRASRDAELGVASAPAAGRYPARLI
KPSQMRRSVRGSGANTRYVCDILPQTEPAEALLVVEVRTPSGHSSSYPPHKHDTDNVPVE
SSLEETYYHRVHPPQGFVFQRVYTDSRDLDESMAAGDHDVVLVPRGYHPVVVPYGYESYY
LNVMAGPTRVWHFKNDPAHEWMLNGGR*
>GCA_000452945.1_BPMSHR5858_genomic_02007_1 cheB_1
MNIGIVNDLPLAVEALRRTIALRPEHRVLWVATDGAQALDFCVAQPPDLVLMDLVMPRID
GVSATRSIMERSPCAILIVTANVGANASYVYEAMGAGALDAVDTPTLEQGGSADPSQPLL
AKIDQIGRLLATRMPLAAPAAAAPAPQGALPPLVAIGASAGGPTALTALLRRLPDDFPAA
LVIVQHVDQAFAIGMAQWLDGYSPLPVRIARQGSVPQAGEVLLAATNDHLHLTSRGVLAY
TRRPEETPYRPSVDVFFHSVVDHWKGEAIGVLLTGMGRDGALGLKAMRTKGHYTIAQDEA
TSAVYGMPKAAAAIGAASAVLPLERIADQLISLVQRNRQRWR*
>GCA_000452945.1_BPMSHR5858_genomic_02072_1 lgrE_2
MSAREAAAESPWLIVHEARNARVRLFCFHYAGATASIFRTWPGGLPDWIEVVAVQLPGRE
YRLGEPLIEHAEPIVEALADIVPPLLDLPYAFFGHSMGALIAFDLAHLLRARGFAQPLLF
VASGRSAPRFRWRDAGIQTLPDEAFIAAVRDYNGTPEALIADPALRELWLPRLRADLTIS
ATYRYVGPPPLDCPMLVLHGSNDGLVSDAGLSGWLAETIGAVRYVGFSGGHFFMHGEEAG
VLAEVRRELERVLARAAQDEPNGMREPAARMSHLTEESGT*
>GCA_000452945.1_BPMSHR5858_genomic_02204_1 zraS
VCWTRFDVLARRAAPAASRSIPPEPNRAMSESTVTQMSAAIVEQLNCGVFSVDHDMRILA
WNGFMQYHTGRSRDDAIGQDLFALFPELPARWLRKKLESVFVLGVAMYTSWEHRPYLFRF
DHSRPITGTIDAMRQNCSFVPLDDGRGGVAAVGVTIVDATDICIAHEVLGNREKRLTDAL
AELTARHAELSDLNRQLAHAHQQLLQSEKLAAIGQLAAGIAHEINNPVGFVLSNVNTLGG
YLGALIAHAYAVERLVAERQPQLAPALAELARGADLDYLCEDAPALVDESKEGLARVRKI
VVDLRDFSRVDSVHQWEWVDIHHCIESTLNIVRNEVKYAADLVREYASLPMVRCIPSQVN
QVVLNLVVNAAQSYSSMRADGGHGDVPRGTITIRTGVDAGDCPRIWFEVIDAGCGIAPEN
LKRIFDPFFTTKPVGKGTGLGLSVAYGIVSAHGGEIAVSSAFGAGTTFRVTLPVERTGEP
QPEAAAVSHSTPVASGIAS*
>GCA_000452945.1_BPMSHR5858_genomic_02207_1 mtgA_1
MNGLASTLARRAAQAAVPRWVWPFARPAGWSLGSAFGSAFGSAFGSAFGSAFGSAFGSAF
AAFGLALAPLLAAPPVRPRARRNPRPCGSPRRAALPRARTTASRTLAVAMLAAPLAAHAL
PAYDDVRRDWRSTDWLLLARDGTPLQRTRVELTERRGDWIALADVSPAFREAIVVSEDKR
FYAHSGVDWRGIAGAAWGNLWNERTRGASTVTMQLAGLLSDTPRRSGQRSLPQKAAQAMN
ALRLERGWRKDQILEAYLNLVPFRGETVGLDAMSHALFGKAPSGLDVRESAVAAALVRAP
NASAAKVAERACRILRDLRAAQPCESLDGYVRFVTSAPPGAARDGREALAPHFARRVAAE
VKPGAGARVRSTLDAPLQRFARDTLTRALAELNAPAHRRHVQDGAVVVIDNATGEIRAWV
GSSGALSAARDVDAVLAPRQAGSTLKPFLYAQALDERRLTAASLLDDAPIDLAAGGGLYI
PQNYDKHFKGWVSVRTALGGSLNVPAVRTLVLVTPHRFARTLTALGLPLAEEGDYYGFSL
ALGSADVTLLSLANAYRALANGGVARATLDVPGGSGSRNGNGNGNDAVTSAPFAPAVSKS
KVGASAVFPRAVSTKAEPAKPVPTAALSPTSASTSTAGKPAAHAAAGTRVFSEAAAYIVT
DILADNNARVRTFGFDSPLATRFFSAVKTGTSKDMRDNWALGYTSRYTVGVWVGNADGSP
MWDVSGVTGAAPVWAAIVGYLHRDLASRAPRPPAGVEQRRVAFERDVEPARPEWFIAGTA
LDTVRLAAPATAAAGGADRPRAPLSIGAPTDGTIFAIDPDIPPKHQRIWFERAAGRAARF
SWRLDGKVIGRGERLAWMPWPGSHRLELVDARGNVADAVGFEVRGAFAKTAAPPKP*
>GCA_000452945.1_BPMSHR5858_genomic_02208_1 group_338
MKHDDKHNKPGHPNPSILRLLARVGAPAALGAAAALSLHADAARTVNVSPQGTVAEVRQA
VVKFDEAMVAFGSASAPDPARLACADPAAARGHGRWLDEKTWAYDFENDLPPGVRCTVAL
NDTLRSAAGHAVTGPRRFAFQTGGPFPVAVRPGAREIEERQVFVVKLNGPADERSALASI
WCEAAGIGNRIPVAPADAPTRAALLDHFHWKKDAARVLTLACAQALPAGAKMQLVYGRGV
ASPSGIANDTERRYDFTVRAPFAASFSCERENAKAPCTPLRPLTLSFNAPVARRAAGEIR
LRGPHGAVEPFFKPDDRAEEVTSVQFAAPLPAQAALTIELPPALRDVTGRTLSNADLFPL
ATRTAPMPPLAKFSSGTFGIVERFAEPDSPALVPVTLRNVEADLRIAGLNAGGAQFSNLK
VENDSEIRRWMQLVERFDGRAMSVESIDKLRPGLLARGQHPVYVPLAAGERAPKPQHRQI
DIRSLSLLAGEPGVRTLTLPKADPKALRPFEIVGVPIDKPGFHVLELASPALGRSLLAKP
AKMYVRTAVLVTNLGVHLKLGRENSVVWVTTLDKGKPVPNAQVRVSDCNGDEIAAGRTNA
QGLVTIDAPLEPKRACDSSNGDGDYFVSARVDDPKTGPDMAFVRSSWNRGIESWRFNVPT
DMSDTPTVRAHTVFDRTLVRAGETVSMKHFVREETLRGLAFPPRYPSRATIRHLGSGQTY
RVPLAWAADHTADTRFALPAAAKLGEYSVELEDGPEDAPSASYYGGSFRVEAFRLPVFKG
SIGVRDAKASPLVGAKDAPLAVQIDYVSGGGASNLPVQVSALVKRAEPPFAERYPDFGFE
PYRPQTQDATADDEDTQDGENASRDTDPDATKLIADKIALTLDRTGSGALTLKGLPAVDA
PKRVALEATFADPNGEVQTIRGDAMLWPAAVVAGIQAGHWVSVGQRVPVQALVVDLQGRP
RASAAVEIKGVARVTTSSRKRMVGGFYAYDNQSDTRELGVLCSGKTDAQGRLACDATLSQ
AGNVQLIAVAKDGDGRASNASTSVWVTREDDLWFGGENTDRIDVIPEKASYEPGDTARFQ
VRMPFRHATALVAVERGGVMQTRVVELNGKNPTVDLKVGDTWGPNVYVSVLALRGRLRDV
PWYSFFTWGWKAPLEWARAFWREGRRYEAPSALVDLSKPAFRYGLGEIKVGTGAHRLGVA
VTTDAARYPVRGTAHARVKVTLPDGKPAPAGTQIALAAVDEALLELMPNRSWDLLDAMLQ
RRAYGVETATAQMEIVGRRHFGRKAVPAGGGGGMAPTRELFDTLLLWNPRVTLDANGSAS
VDVPLNDALTRFRIVAIAATGAERFGTGSATIRSTQDLQLISGLPPLVREGDAFRAQVTV
RNTTERKMDVVVTPRVPGIDAAPRKISLAPDSAQEIAWDVTVPETALDAAGALNWRIEAA
EQGGKRAADALALAQKVVPALPVTVQQATLAQVDGTLSVPVAPPAGAMPDARGAPRGGIA
VSLQSTLADGLPGVRRWFERYPYRCLEQQASRAIGLRDAAQWQALTARMPVYLDRDGLAS
YFPPSSDDAHSGSPPLSAYLLVLADEASRADARFALPEDVRTQLEAGLARFVEGRIERDT
WAPRQDRDLRKLAAIEALSRYGAAQGRMLGSIEIAPNQWPTSAVLDYHAILTRVKDIARR
DEKRAQAEQILRARLAYQGTQLVFSTARGDDLWWLMTSNETNAARLALAFAGEAGWKDEM
PRVAAGLLALQKNGAWQTTTANALGLLALERFSRTYERAPVAGATKIALGDDTRSIAWSQ
PAGAGGATVATGATGAAATAGAALASGASASAAAKPAATQSRTPPPSSGTPPPSAATRAA
AAHSVTLPWPRGARTPGTLSIVHEGSGRPWATIESLAAVPVRAPFAAGYRIAKTVTPVSP
AVSGALTRGDVLRVRLDIDAQSDMTWVVVNDPIPAGATILGSGLGRDSEAATQGEKSPDG
AWPAFVERDFDGYRAYYDYLPKGKLTVEYTVRVNNVGTFGLPPTRVEALYAPSVYGLWPN
PPMTVKPAVASKP*
>GCA_000452945.1_BPMSHR5858_genomic_02260_1 budC_2
MNGVDSRRAAPPRGGLAADFRGRTALVTGAAQGIGAAIADAFAAAGASVAIADVQGDAAA
AFAARLAARGAADGQTVRAYRVDAARRDELFGLVAQAEADSGRLDIVVHNAAYFPLTPFG
AIAPDVLERTLAVNLSALFWLTQAALPAFERVGRGRVLATSSVTGPRVAYPGLAHYAASK
AGVNGFIRAAALELARRNVTVNGVEPGMIRTPAAGNLGDAAHGERIARGVPLGRLGEPED
IAAAMLFLASDAAGYITGQTIVVDGGATLPETAAALG*
>GCA_000452945.1_BPMSHR5858_genomic_02324_1 bar
MKATDAPTSSAIVRDATEHDLEAIQAIYAHHVLTGVASFEETPPSVDDLRARREAVRRHG
LPYLVAELDGVVAGYAYATPYRPRSAYRYAIEDSIYVNEACRGRGIGRVLLAALIARCEA
GPWRQMIAVIADGGRGGSTSLHRALGFEPIGTLRGVGFKHGRWIDTALMQRVLGDGAQTP
PSNADASH*
>GCA_000452945.1_BPMSHR5858_genomic_02334_1 group_2408
MSEADREQGKARRTQVMGDAFVERAMSDLDGFSRPLQDWLNEHAWGSTWRRGGIDLKTRS
LCTCAMLAALGRGHELKGHIRGALNNGASLIEIREVLLHSALYAGAPAAVEAFRHAREVI
DALALDVPDDGA*
>GCA_000452945.1_BPMSHR5858_genomic_02357_1 glpR_2
MNDAIPLARRDAIAGRLALGQPVQAAALAAEFHVSEDAIRRDLRALAAEGRCRRVYGGAL
PVTPACAPMAARIDAARERKAALARTAASLIERGELLFLDSGSTALALVEYLPEDAELTI
ATNSIDIAAAVLRRADLSLIMIGGAVDQAVGGCVDASAVQSVARMNIDRGFLGACALSPQ
RGLAAFGLADATFKRAVVAASARCVVLATTDKLAARAPHRVAALDEIDCIVVEHDLPRED
RAALSSAGASILAANPPAQP*
>GCA_000452945.1_BPMSHR5858_genomic_02366_1 xylB_5
VTYLGIDLGTSEVKAILTDADSAPLAAGGAPLAVERPHPHWSEQSPQAWWHATLDAIAAV
RAMHPRGFAALRGIGLSGQMHGATLVDRAGQVLRPAILWNDTRAAAECVELEALVPESRA
ITGNMAMPGFTAPKLLWLAKYEPAVFRAAHKVLLPKDYVAWRLSGEFVSDMSDASGTLWL
DVGRRDWSERMLAATELSREQMPRLVEGSAAAAQLRDALRREWGVAGPVTIAGGAGDNAA
SAIGMGVANAGSGFLSLGTSGVLFAGNDRFAPNPGDAVHAFCHCLPERWHQMSVILSAAA
SLDWFAKTHDVAVDTLPALAERADASRAPLFLPYLSGERTPHNDARARGVFFGLASEHRA
GELAYAVMEGVAFAMADGYAALRRAGTTLDDASFIGGGSKSAFWAQLCANATGLAMHRHA

SGAVGAALGAARLARLAATRDALDDVCVAPAIAETYRPDPAATGLLGERLERYRRLYRAL
KAEFAG*
>GCA_000452945.1_BPMSHR5858_genomic_02383_1 hyuC_2
VWEIDGRRLWNSLTELARIGGTARGGVRRLALTEADRRGRERFAQWCRDAGMAVRVDAIG
NLFARRAGADDTQPAVLVGSHLDTQPEGGRFDGAYGVLAGLELVRTLNDRGIVTGKPIEI
VSWTNEEGARFTPAMLGSAVFTGALPLDAALAARDADGIALAEALEACGYRDALRSAWMR
GAHGARGLEPALAPLHERVDAYFEAHIEQGPVLERHGATIGVVTGGQAIRWLDATVTGAA
AHAGTTPMPYRKDALFASAELALAIEALVARHAPDALATIGQSAIDNASRNTIAERVTFS
IDLRHPDDARLDAIERELRRVCAQIAGRRGVAVDISRHWSSPATPFDPACIALVEAAARG
GGYAHERIVSGAGHDAVHLARCVPSAMVFIPCVGGLSHNAAERALPEHVEAGANVLLGAV
LARAGVRERAARSARSPRPAPAMQAAHSGTDAGTFAQPH*
>GCA_000452945.1_BPMSHR5858_genomic_02385_1 argP
MSNIRSHVSRVLMDRLDWNLLRTYLTIMQERSISRAAARLHLTQPAVSQALKRLEDSLGR
TLIQRRGAQFQPTRAGEEVYRIATDIYGHMSRLDNELDDRGGELIGSVRLLCVSRVESPV
YDEFLAEFRRTYPRVDLHVEVMRSADILSSLLQKTATCGLALCRTPVDKIELRSFLRQRY
AMFCGRHHRLFGRQALKIEDLLAENFVSFTSDQIGDALAPLTVFRDQRGFTGRIVATSPS
LDEIRRLVFAGYGIGCLPEHIVRDDLARQRLWRLPPEEGLLDVEIFLMWNRERRMNAAEN
VFIDAFQRYVQRYPMAERLGMLH*
>GCA_000452945.1_BPMSHR5858_genomic_02388_1 fabG_7
MNATPLTRVALVTGASRGIGAAVARRLARDGFSVAVNYASSGAEADALVAGLRAGGAAAL
AVRADIGRADDVRRMFDAVERQLGRVDVLVNNAGILKTGPLADATDDAFDRIFDINVRGT
FNTLREAAKRLADGGRIINFSSTTLALKLPGYGLYNATKGAVEALSHVFAKELRGRRVSV
NVVAPGPVATSLFLEGKTDEQIQGYAKMPPLERLGEPEDIAGVVSFLAGPDGGWVNAQVL
RANGGLA*
>GCA_000452945.1_BPMSHR5858_genomic_02389_1 frc_2
MFDLAASSHTEPSGHARAAAYLRDIWQAANGDPEWLRTLAFGGAGVLPSAFPVTDFASAA
IGAASVALAEFVHRATGRLPGVRVDRRYASIWFGTSLRPRGWNMPPLWDAIAGDYRTADG
WIRLHTNAPHHRAAALAVLGAPPERQAVARAVGQWRGDALESAVVEHGGCAAVMRTEDEW
ARHPQGRAVRAEPLMIHDDVSIDGPRPAWPMSAERPLRGVRVLDLTRILAGPVATRFLAG
FGAQVLRIDPIGWDEPGTVPEVVLGKRCARANLACADGRALLRRLIGEADVMVHGYRPGA
LDRLGFDADERRRINPGLVDVSLDAYGWSGPWRGRRGFDSLVQTSAGIAAAGMREAGADR
PVPLPVQALDHGTGYLLAAAAIRGLVRRLATGAGTSTRASLARTAVLLTSGGLQEAGRQP
IAPETAGDLDAWIEDTSWGPAQRVRAPVTIECAPVQWPHPARALGTSPAEWA*
>GCA_000452945.1_BPMSHR5858_genomic_02412_1 group_398
MPMSARTNAAFTALLAAALFGATTPLAKTLLGSLAPFMVAGLFYLGSGVGLGAFMLMRRL
ARGAGAGASPAGHARLPLAELPWLAGAVAAGGAAGPALLMLGLATPPAATSALLLNLEGV
FTALIAWAVFRENVDAQIFAGMAAIVAGGVLLSWHPGAAGVPLGALLVAAACACWAIDNN
LTRKVSTHDAAAIACVKGLVAGTVNLGIALALGARLPAAADSAAAMLTGFAGYGVSLVLF
VVALRNLGTARTGAYFSVAPLFGVGLSLALWPEWPPLSFWAAAALMALGIWLHLRERHEH
PHTHEALEHSHRHRHDTHHQHAHDFDWDGTEPHTHAHRHTPITHTHAHFPDIHHRHSH*
>GCA_000452945.1_BPMSHR5858_genomic_02432_1 hmuV
MLNADHLHVARDGRAILNDLSIRIAPGCVTALLGRNGAGKSTLLGVLAGDLPAGGLARGA
TVRGGVALNGEPLHAIDAPRLARLRAVLPQASRPAFAFSAREIVLLGRYPHARRAGALTH
ADGEIASQALALAGATALDARDVTTLSGGELARVQFARVLAQLWPPPGAAQPPRYLLLDE
PTAALDLAHQHQLLDTVRRLSRDWNLGVLTIVHDPNLAARHADRIAMLADGAIVAQGAPA
DVLRPEPIARCYGFRVRLVDAGDGVAPVIVPA*
>GCA_000452945.1_BPMSHR5858_genomic_02529_1 thrC
MNYISTRGAGIGERHTFSDILLGGLAKDGGLYLPSEYPRVSADELARWRALPYADLAFEI
LAKFCDDIPADDLRAITRRTYTADVYRHTRHGGNAADITPLTTLGAENGAPVSLLELSNG
PTLAFKDMAMQLLGNLFEYTLAAHGETLNILGATSGDTGSAAEYAMRGKRGVRVFMLSPH
RKMSAFQTAQMYSLQDPNIFNLAVKGVFDDCQDIVKAVSNDHAFKAQQKIGTVNSINWAR
VVAQVVYYFKGYFAATRTNDERVSFTVPSGNFGNVCAGHIARMMGLPIEKLVVATNENDV
LDEFFRTGAYRVRSAEHTYHTSSPSMDISKASNFERFVFDLLGRDPARVVQLFRDVEQKG
GFDLAASGDFARVAEFGFVSGRSTHADRLATIRDVFERYRTTIDTHTADGLKVAREHLQP
GVPMIVLETAQPIKFGETIREALGQEPSRPAAFDGLEALPQRFEVVDASAQQVKDFIAAH
TGA*
>GCA_000452945.1_BPMSHR5858_genomic_02560_1 ansA
MNAPTSSHSPHVSNPAPRALPRIAVLATGGTIAGAAPDAAQTAGYQAGALGVDRLLAAVP
ALAQIATIEAEQVASIDSKDLSPALWTTLAERIDALAANDAIDGIVITHGTDTLEETAYL
LHLTVKTAKPVVMTAAMRPATALSSDGPLNLLNAVAVAGSAAARGQGVLVAFNNRIHSAR
DVVKTSTYAVDAFHSPELGALGWVQDGRIEFARRATRPHTLDSRFAIGAAWPQVEVVASY
AGASRVIVDACVAAGARGLVVAGTGNGSIHAALQAALADAAAKGVAIVRASRVGSGHVMR
NGAANDDALGFVSAGSLNPYKARVLLMLALAAGVADVRELQQVFDTY*
>GCA_000452945.1_BPMSHR5858_genomic_02562_1 group_1013
MAWRQHRWFRRWLIVIVFWAVPVAIVAVREIREEMAYNYADLKLALTTWQLTDAQRAAGA
AKVCRGEPDEARAAGCPADVLAANARRQQEARDEFAVRKRTLASYLWHAFVGYWIVPAAF
LFACGVVIGLVRRALRRPPIKPPAKPPLEPPLEPPVSHH*
>GCA_000452945.1_BPMSHR5858_genomic_02571_1 holB
MIYPWQTDDWNRLQALRAHWPHALLLYGQAGIGKLAFARHLAKGFLCETPRENGEPCGAC
AACTWFEQGNHPDYRIVVPEALAAELPGAADEPKAADADEGGKKARTPSKEIKIEQVRAL
VDFCGIASHRGGARVVVLYPAEALNVAAANALLKTLEEPPSGVVFLLASARVDRLLPTIV
SRCRQWPMAVPAPDAAAAWLAAQQVADANALLAEAGGAPLAALALANDENRPLRDWTLAQ
LAAGAACDPFACGETLQKLPVPVVLGWLQRWLYDLLAERAAGAPRYFPAQRAALARCAAA
ADANAFARFLKTVTRQRAVENHPLSARLVFEELFLGYRELFASP*
>GCA_000452945.1_BPMSHR5858_genomic_02596_1 lolD_2
VCKRVADTAGELTILAGIDLTVRAGESVAIVGASGSGKSTLLGLLAGLDSATEGTVRLFG
QDLGRLDEDARAALRNGAVGFVFQSFQLMPHLTALENVMLPLELQGGVGAREAAQRARAL
LEEVGLGARTGHYPKLLSGGEQQRVALARAFVTRPALLFADEPTGSLDAATGEAVIDLMF
ELNRAHGATLVLVTHDTELARRCGATIVLEAGRLRGADRARQEAG*
>GCA_000452945.1_BPMSHR5858_genomic_02597_1 tesA
MKRTVHWKAGATLAVVATFAGPASAATNAAPAPNADGGKPVIVVLGDSLSAEYGLPRDTG
WVALMRARLAAERIDYSVANASISGDTTSGGRARLPAVLARLKPAVVVVELGANDALRGV
PLAATEGNLREIVAQARQARAQVLLVGMYVPSNYGPDYTQKFHGVYTGLSKEMRVPLVPF
LLAGIENKPDMFQADQIHPTQAAQRQLLDNVWPALKPLLRTSSR*
>GCA_000452945.1_BPMSHR5858_genomic_02598_1 ppiD
MLDFFRNHQRLMMFFLLLIVLPGLGIVGIQGFRGFFDESANVAAVNGHKITRAEFDGTLR
QQVDQARQVLGAQFDAKAFDTPERRQQLLDGLIQQRALADETQRLHLSASDGAVRQTLLS
DPVIASLKKADGSFDAERYTQMLAMQGMTPDQYQERVRYNLALQQIPASIVSSAFTPKSV
ARRLTELAEQQREVQPMVLKSADYAAKVQPTDAQISAYYDAHKQAFATRETATIQYLVYS
QAAAAAAAQPSDVDIKKYYDDNIAHYRTDAQVRVSHIFIAAAKDASAAEKAAAKAKAEQL
LAEVKAHPDRFAQIAEKNSQDAPSAAKGGDLGFITRGSTAGGAAFDDAAFALKKDEISGV
VQGDFGFHILKATDVKPAVVKRLAEVKDSIATDLRQQFAAKAFADNAEGFTSTVYEKAKS
LQPAADKYKLTIQTATVSPQPNPALPPDSPLNNAKFLAAVFAADSVKNGNNTQAIDIGNN
TLIAARVTNHQPSTVPALDAIKDQVRAKVVADEAARLAQQAGEAKLAELRKSKSTAGFAA
PDKISRTQAHGLPPAAVSAIYKVDPKTLPAYVGVDLGNDGYAIFRVNSVVAAAPADDQRL
AAAQQQLAQVYAQSETQAYLASLRARSKVKLYGSTAGGAQDSSN*
>GCA_000452945.1_BPMSHR5858_genomic_02635_1 group_1851
LTDIHDIDSSESAHAVATARADDAPEQSAADAGGEDRPRRGLRRGPRSLIARRRAAAKSK
HSDAPESADAAPAADAGAGADVAKAPARAPRGKDAAAKPPRKTAGKREGAARQGAQPKRG
AQQAAAAVAPSAEAGQDDVFAYVISPAFDADNNAPGGGVRAPMLRRGRQTQPKRVLSPDD
DAPKLHKVLAEAGMGSRREMEELIIAGRVSVNGEPAHIGQRIMPTDQVRINGKPVKRKLP
SKPPRVLLYHKPTGEIVSHADPEGRPSVFDRLPPMKTAKWLAVGRLDFNTEGLLMLTTSG
DLANRFMHPRYSVEREYAVRVVGELSEASRQRLLHGVELDDGPANFLRIRDGGGEGTNHW
YHVALAEGRNREVRRMFEAVGLMVSRLIRTRHGPIPLPRGLKRGRWEELDEAQVRRLMST
VGLKAPTEDKGGKRGGPAERRQPDPMQTSMGFINREPVLTTHGQLDQPRRGRRGPAGGGF
GAGLGGGYAGLPGYGGASRQGGRDVDGNRASYGGAGANKRGAGKGGRNPNGNRAEGGARG
GPRTPQQRNRSRSR*
>GCA_000452945.1_BPMSHR5858_genomic_02662_1 group_590
MSSAALWALALALLCVAGAFALWRRGEANRERAHAARYIDSRLEPGARASAQPKMPAAAE
PKRAAPMPAAGAAGGARAEKPAEGLARWRERAADAWLNVSNRAGVSEIRAPLAALAATTA
VATLWAGLRGGLLAACAALVAGATLAVFWLVSRMQKRRLRIVRQLPSFLDGIVRLVTLGN
SVPAAFQATLQTTEAPLRGCLDHVSRMLRSGVEIDRAMVSIAALYRIKEFELVGSVLRLS
VKYGGRADVMLDRMAVFMRDLEQAERELVAMSAETRLSAWVLGALPVGIGSFVIATNPKY
FSAMWLDPTGRQLVYLAFILQIAGGYWLYRLARLR*
>GCA_000452945.1_BPMSHR5858_genomic_02702_1 mopII
MTGTTLAATYGPISGADLEAELAQPRIADGDAQDAAVYERDGGAHAPPDGDRADVRRAAG
AGDASVRLTRVSKRYGERAVLADVDLSIGRGSFVSIVGRSGCGKSTLLRLVAELETPSAG
TLVKRGDGGGALDTRIMYQEARLLPWKTVLQNVMLGLGRRAKDDARAVLDEVGLLARAND
WPAQLSGGQRQRVALARALVHRPQLLLLDEPLGALDALTRIEMHALIERLWREHRFTALL
VTHDVQEAVALADRVLLIEAGRIAFDQRVPLDRPRARASAAFAALEDRVLQRVLTGSDAA
PAAPNAAGPEGASRGRAAPASGLRWAV*
>GCA_000452945.1_BPMSHR5858_genomic_02704_1 group_7453
MFPTLSPEAIEALKWIDQFGSGRPLPVGFRLALEELLNDGFVYQSGPDRVDITDDGKAYL
SEAYD*
>GCA_000452945.1_BPMSHR5858_genomic_02705_1 group_7454
MEPKGRDIGDFGEPYKGYDIEVKTEQVWAGEKAHYRVLQGGAVMIDWRLVHIDGLLLSER
RVIEQTLAVARSAVDRELLGAGVV*
>GCA_000452945.1_BPMSHR5858_genomic_02729_1 group_8
LSTAVNGGGTKYFHANSTQPDSQALGADSVAVGPAAIAAGASGIAIGNAANAAANGAVAI
GQAAVAKGGLAVSIGVSNTASGDGAVAIGDPNVATGTGAVALGADNSANGQGAVALGNAN
IATGTGSLAFGNTSTAAAAGAVALGAGAIANNANDVALGSASVTAAANPVASALIAGQAY
SLAGGAPASVVSVGAPGAERQITNVAAGRISATSTDAVNGSQMNAMTQALESLSTSTASA
LSTAQSGLGSLSTGLSSTQSSVSSLSTGLSTTSGNVASLSSGLGTMQSGIASLSTGLSTT
NSSLASLSTAVSGGGVRTSSLGDTSAGNGANASGGNGTAVGGAASASGTDATALGQASNA
SGNHSTALGQASSASGSGSTAVGQGAGAPGDGASAFGQGALASGTDSTALGAHSTAAAPN
SAAIGANSVASAPNSVSFGSRGHERRLTNVAPGIDGTDAANMNQLWGVQSSVDQAARRAY
SGVAAATALTMIPEVDPGKTIAVGIGAGSYQGYSASAIGVSVRFSDNLKAKLGVGISAQG
STYGAGVSYQW*
>GCA_000452945.1_BPMSHR5858_genomic_02749_1 mauG_2
VTGGKRSGAARRAGAAFAARGPALAGAPGARNARRIVSTLAIAAVTRAAGGALAACASAI
AFASGAAAPGALDAVRAAHPASSLSPARTPGAGGAAHVKPVQEALRAKAASPSPSLLPSP
PSPTTSLLPGAPPQRVVATIGRGTPQVASKVDPTAAAFHPDPALAALGKRVFFDPALSEP
RGTSCASCHDPGRAFAPTLARAALAGPRVPQGSRPGHFSRRNAPSLLYVRYVPRRHFYQD
DDALAPAPFGGLFSDGRADTLAEQLRGPLFDPDEMNNASPAALTRKIGGTALGAALAERF
GPSVRRDPERMVRALGEAMQAYLQSDEMAPFSSRYDAYVMQRAPLTPQERRGLALFRNPD
KGNCMSCHTLSDTASRPERSLFTDFGYDAIAVPRNRALRANRDPRHFDNGLCDTAAKLRW
PEPAQWCGYLRTPGLRNVAIKESFMHNGVFDTLRDAVAFYNTRSTDPKRWYHGRDTFDDV
PAAYRGNINVNSTPMNRRAGTPPAMTDADVDDIVAFLRTLTDARYVGLMPAAPDGKAARP
*
>GCA_000452945.1_BPMSHR5858_genomic_02750_1 group_7455
MSNQSIGAPDSTAVRVALWRALHARIDAPPHVLDDEIGLALAAPDDDWRSRPDMDPQASR
GYRASIVGRARFVEDLVDEQADRGVAQYVVLGAGLDTFAQRRAKLASHLRVFEIDQPGTQ
AWKRQRLIALGYGVPQWLRLVPVDFETSGAWRAQLSAAGFDDSRPAVVSSTGVSMYLTRE
AIAGTLSQIATLAPGSTLAMTFLLPLELIDDPAERAQHAAVYERARAAGTPFVSFFSPSD
MLALAREAGFREARHVSTDDLVRRYFESRPDGLRPASGEAFLVAST*
>GCA_000452945.1_BPMSHR5858_genomic_02751_1 group_2436
MEHLVRIVNDTDRQILAWLRSQVGDERVERAAQHMGRVRKPYLSAVCRYLGVWPPISLRY
PPRHGAVDHAVGDRYLTLIRQHLAAHTAGR*
>GCA_000452945.1_BPMSHR5858_genomic_02752_1 group_4775
MAKKFPLHPKHPERNCWGCDRYCSVDALSCGNGSDRTQHPIELLGDDWYEWEQENEAAQQ
PIQSVPAAAR*
>GCA_000452945.1_BPMSHR5858_genomic_02753_1 mshD
MREVEARLRASGTQAVFGALTDDGGLIGIAGVRREALAQLAHKAVLWGVFVAPAHRGNGL
ARRLIDAALTHASRDWGVEQVNLCVNTENEPAKALYASLGFETFGVERRAMRVGDRFYDE
QHMVRVRR*
>GCA_000452945.1_BPMSHR5858_genomic_02754_1 group_7456
MSLLFAFLSGLLFGAGLLLSGMANPAKVQGFLDLAGNWDPSLAFVMAGAIAVGSIAFAFA
KRRERSLLGLPMQIPPRGAVTPRLVAGSAAFGVGWGLVGFCPGPAVVSLGLGGAKAFGFV
AAMLIGMAAFECGERFRAKR*
>GCA_000452945.1_BPMSHR5858_genomic_02755_1 group_1471
VAVDLVHFTPGLSFAGGLLIGAAAALLVFFDGRIAGISGIVGGLLAGGRSDAGWRAAFLV
GLIAAAVAASALDWFSMPAVDAGWARVIAGGVLVGIGTRYAGGCTSGHGVCGISRGALRS
IVATAVFMAAGIATVFVLRHVVGG*
>GCA_000452945.1_BPMSHR5858_genomic_02756_1 bigR
MSAACSPIDLSGMRASADAACALLKVLSNPDRLLMMCELAQCGELCVSDIEARLDIHQPT
LSQQLGVLREHALVDTRRDGKNIYYALTSEPAVAIMAVLYEQFCATHQKGKRRGR*
>GCA_000452945.1_BPMSHR5858_genomic_02758_1 yceJ_3
MTPSTRTPDRYDLLSRILHWSVAALVLAQFALGWTMPDVHGTAPPVGLVAWHVALGTALI
VVIAARLLWSIVRRSPPAHQQGALLSFAASATHKLLYVALIAVPLLGWLNANGRTWQLKL
AGAIPLPTIAAPHSLGAAIGEWHSLSATLFIVLIGMHALAALVHRFVMKDGAFERMI*
>GCA_000452945.1_BPMSHR5858_genomic_02759_1 sufS_2
MSAAPVALPDAAPPAGLPDPATLARLASEFLAALPGQPAAPNAGAGSGAVGGVPSALPAA
APMLASVSNPAPPGSPLAGPGGTGTGVPGIGVPPGKVPGANLVPAPTHVLSLGNRTPALV
GHAAAQNGWPDSAVAIAPALEPRAGGVALGVPPVPEPDAVRRAGDASAAAAPSPWSYYFV
EPASDDWWRDAARTPIDVPRDGVASPRAFGLPDENAWRDLLSIGWPAADRHRASRYFVDD
AQPTNAHAPGAGAHPPFDIAAIRRDFPILAERVNGKPLVWFDNAATTHKPQAVIDRLAHF
YAHENSNIHRAAHALAARATDAYEHARATVQRFIGAASPDEIVFVRGATEAINLIAKTWG
VGNVGEGDEIVVSHLEHHANIVPWQQLAASVGAALRVIPVDDAGQVLLGEYRKLLNDRTK
IVSVTQVSNALGTVVPVKEIVELAHRAGAKVLVDGAQSISHMRVDVQALDADFFVFSGHK
IYGPTGIGVVYGKRALLDGMPPWQGGGNMIADVTFERTVFQPPPNRFEAGTGNIADAVGL
GAALDYVARIGIERIARYEHDLLAYAAGVLAPVPGVRLIGTARDKASVLSFVLKGYETEE
VGRALNAAGIAVRSGHHCAQPILRRFGLEATVRASLAFYNTRDEVDAMVDVVRELAARRI
*
>GCA_000452945.1_BPMSHR5858_genomic_02760_1 mmpI
MSTTVSAQTALGDNAARQLANATKTVPQLCTITPRWLTHLLQWVPVEAGIYRLNQVKNPE
AVRAACTAREDEGALPRTFVPYEEEPREYFLSAVSTVLDVHTRISDLYSSPHDQIKEQLR
LTIETIKELQESQLINHPDYGLLANVAEAQRVFPLTGAPTPDDLDELLTKVWKEPAFFLT
HPLAIAAFGRECTRRGVPPPTVSLFGSQFLTWRGIPLIPSDKVPVADGKTKILLLRVGDK
RQGVVGLYQPGIAGEQGPGLSVRFMGINNQAIASYLISLYCSLAVHSPDALAVLDDVEIG
KYHDYPDTYR*
>GCA_000452945.1_BPMSHR5858_genomic_02761_1 sutR_2
MTALVQRFGAAIRQLREARAWSQEQLAEHAGLNRSYVGEIERGTAIASIVTVDKLARAFG
VPIARLLSPAGDAGPPPHGFGAPPASSAPLSSTR*
>GCA_000452945.1_BPMSHR5858_genomic_02762_1 dapH_3
VAVFDIDDIVQSLQTVRQRWREVQRRSLEPGGRELPAREALAGIVETFKGVLFPMRLGPP
DLRQESENFYVSHALDDALHALLAQARLELRYKGRHDADAPAEAAIDAKADAAVRAFAAR
LPDIRALLDSDVLAAFHGDPAAGSVDEVLLCYPGVLAMIHHRLAHALYRLELPLLARIVA
EHAHAQTGIDIHPGAQIGGGFFIDHGTGVVIGETAVIGERVRVYQAVTLGAKRFPRDASG
HLEKGLARHPIVEDDVVVYAGATILGRVTIGKGAVIGGNVWITQDVPPGSHVTQAVTRSD
PARPADAAASSPRPAGAHDATLSAAQALR*
>GCA_000452945.1_BPMSHR5858_genomic_02763_1 glpE_2
MTIELEARATHAVLDAAREAALAAGLPYAGGVSPRDAWALVAAGHARLVDVRTAEERTFV
GHVPDSLHVPWATGTSLTRNPRFVRELEAKTGKDAVVLLLCRSGNRSALAAQAAAKAGFT
QVFNVLEGFEGDLDDAGHRGTTNGWRLHGLPWRQS*
>GCA_000452945.1_BPMSHR5858_genomic_02764_1 group_2032
MPHASWPEQADPHQFLEELDSAASVGWVDAQNARTHDAPWLDEAHYRALVERFTRALLPR
ERPVIPQRWQDWAYDVWQDEQHPKGLWRRTRWTSWRSGHADWQTLIDLDALGEAQGVQWV
FDDQLILEPDGDRALIVLSDGGADAVVVREFDIAQCRFVDDGFSIEAAGKHSVEWIDRDT
IYVGWDDGGATVTRSGYPREVRRWTRGTPLSSAPVVFRGARGDISVDAQYDPLDRHHAIE
QAINFYDANTYRLAEDGAWARYDVPPHVEVGYWSGWLLLQPRLDWTCGGARYAGGSLLAI
REDAFVAGERAFAALFEPNERTSACGWTHTRRYVLVSWLDDVLTRTMLWLPERQDDGAWR
WHARPFPARGLAQVDVSPVEPTFDDEVYVSVDDYLKPPEYSLANLASDDLSAWTLLDRWP
TQFDASELTVRREHARSRDGTLVPYTLVGPRDVLDNAARAPRPCLLNGYGGFAIALTPDY
DPLLGIGWLEKGGIAVFAHIRGGGEFGTQWHESARQTQRQRSFDDFIAVAEKLVADGVTS
AAQLGIRGGSNGGLLVAACMIQRPDLFGAVVSDVPLLDMQRYALLHAGASWLDEFGDPDD
PAHASALAAYSPYHRVARDIAYPPALFTTSTSDDRVHPAHARKMVARMQAQGHRNVWLIE
KTDGGHGSADAIDTAEHEAIGYVFLWTHLSRGAHDARE*
>GCA_000452945.1_BPMSHR5858_genomic_02828_1 ftsK_1

VQRQALASESGEAAQGVAGAAAAGDSRETTQVSPAGARPDKAAPSAAGANPIAPLPGASA
ITAHEDAPTSAAPDAATPVIAAMDSAMPNAVAPASAIASNAGMSPASASAAAPRMASAPA
SAAVPNTHPPLPRAAAAVPGVASIGVAAPGVIVTNAATALPAAPGRIASPAGASAVAPGA
MTPNAASTDVAPAAAPASDVSPNVVPAPAVETNASVPPASASPAARHVNAPMVASTGAAA
PAPSIPSSLPPSTVTSNAERRAATTAPTAAPAGLAPNPVAASSFVAPTTSAAPGQFAPAA
TAPADNAPAAAEAPPGRVPNPPAGAGFVTPTSPTPGPLPPAAETPAATATPTAPPPGLAP
NPPAGAGFAATPEAVAHPFGNPSAPAPGAIPESPATAPSVALTANGTEAPGAPQAFAPSP
VPAMPAAPAAADPASAAPAAEPVRPSRPPAPNAFEFHAPAASNVELPTLDLLEPASDTIE
AISDEHLAQTGQIIEQRLQEFKVPVTVVGASAGPVITRFEIEPALGVRGSQIVGLMKDLS
RGLGLTSIRVVETIPGKTCMGLELPNAKRQMIRLSEILASRQYQHSASQLTIAMGKDITG
NPVVTDLAKAPHMLVAGTTGSGKSVAINAMILSLLYKATPEDVRLIMIDPKMLELSVYEG
IPHLLAPVVTDMKLAANALNWCVGEMEKRYRLMSALGVRNLASFNQKIRDAAAKEKKLGN
PFSLTPEDPEPLSTLPLIVVVIDELADLMMVAGKKIEELIARLAQKARAAGIHLILATQR
PSVDVITGLIKANIPTRVAFQVSSKIDSRTILDQMGAESLLGQGDMLFLPPGTGYPQRVH
GAFVADEEVHRIVEYLKQFGEPQYEEGILDGPSAEGGTQDLFGEAPDAEADPLYDEAVAF
VVRTRRASISSVQRQLRIGYNRAARLVEQMEAAGLVSPMGINGSREVLAPPLPE*
>GCA_000452945.1_BPMSHR5858_genomic_02904_1 group_4789
MRYELYYWPDIQGRGEFVRLALEAAEAEYVDIARESSRKGMGVPAMTRLLESDAMACLPF
APPFLKAGEQIVGQTANILLFLGARHGLAPRDDAGRLWAHQLQLTVADFVVEIHDTHHPI
AGGLYYEDQKAEAAMRAEDFIEHRLPKFLGYFSRVLAHNPHKHGYMVGAHLSYVDLSMFQ
LIEGLRYAFPKAFARVAKQHAALIELGDRVAQHPPVARYLASARRIPFNEQGIFRHYAEL
DR*
>GCA_000452945.1_BPMSHR5858_genomic_02905_1 group_2443
MNSTPDTFQRPALRELTPLEARVLGVLIEKQHTVPDTYPLSLNALTAGCNQKTARSPVMN
VSEAEVLTAIDGLKRLSLASEGSSSRVPRFEHNMNRVLGIPSQAAALLTMLLLRGPQTAA
ELRLNSARLHGFADISSVEAFLDELAARTPPLVVKLPRAPGARESRWMHLMCGDVAPDEA
PTYGAHEDAVPPSEFEALKAEQKALTAELAQLRAFVEYMANELGIDVGKLTRGA*
>GCA_000452945.1_BPMSHR5858_genomic_02908_1 sdaB
MNVSVFDLFKIGIGPSSSHTVGPMIAACRFASHVEDANLLGFVRRVRAELYGSLGATGKG
HGTDKAVLLGLEGHLPDSVDPDLIEPRLKAIRAEKSLALLGKRGIRFDEKDDIGFYRKLM
SGTSVVHPNGMRFQAFDEHGQLLVEKEYYSVGGGFVVNRDGDRVNGVRAAADVPYPFRTG
DDLMRACRASGLSIAELTFRNECALRPADEVRAGLVAIWRTMAACVERGCKADGELPGPM
RVKRRAAELYTRLRDRSEESLRDPLSMLDWVNLYAMAVNEENAAGGRVVTAPTNGAAGVI
PAVLHYYVKFVPGSNEQGIVDFLLTAAAIGIIYKETASISGAEVGCQGEVGVACSMAAAA
LAAVMGGTPGQVENAAEIGMEHNLGMTCDPVGGLVQIPCIERNAMGAIKALNASRLALKG
DGQHYVSLDSVIKTMRETGADMKTKYKETSRGGLAVNVIEC*
>GCA_000452945.1_BPMSHR5858_genomic_02909_1 soxB
MSRYSIFSLFRNGLSYHENWEKQWKSPEPKREYDVVIVGGGGHGLATAYYLAKEHGVTNV
AILEKGWIGGGNTARNTTIVRSNYLWDESAGLYEKAMKLWEGLAQDLNYNVMFSQRGVMN
LAHTLQDVRDTERRVNANRLNGVDAEFLTPAQIKEIEPTINLNSRYPVLGASIQRRGGVA
RHDAVAWGFARGADRAGVDIIQNCQVTGIRREGGAVVGVDTVKGFIKAKKVAVVAAGNTT
TLADMAGVRLPLESHPLQALVSEPIKPVVNTVVMSNAVHAYISQSDKGDLVIGAGIDQYT
GFGQRGSFQIIEGTLEAIVEMFPVFSRVRMNRQWGGIVDVSPDACPIISKTDVKGLYFNC
GWGTGGFKATPGSGWVFAHTIARDEPHALNAPFALDRFYTGHLIDEHGAAAVAH*
>GCA_000452945.1_BPMSHR5858_genomic_02910_1 soxD
MLLIECPWCGPRAESEFSCGGEADIARPLDTDKLSDREWGDYVFMRKNPRGKHREQWLHT
QGCRRWFVAQRDTVTYEFQGYETFGTPAAHAAKDTSK*
>GCA_000452945.1_BPMSHR5858_genomic_02913_1 group_4790
MKPVEEPFAALDPARSRGRGWSVFVDALKVPARIGIHAHEHAAPQPVVIDARLAYRREPS
EASGDGWIDYDAYCARIASFLARKPHTRLLETLALEIAVLSFDEWPALDALTLALHKPKI
RPGTKRVGVELDWTRGDYRAWRAASEPNGASSG*
>GCA_000452945.1_BPMSHR5858_genomic_02914_1 cspG_2
MDTGTVKWFNETKGFGFITPDSGGDDLFAHFSEIRSEGYKTLAENQKVSFETKRGPKGLQ
AANIKPL*
>GCA_000452945.1_BPMSHR5858_genomic_02915_1 group_7470
MTSMTYRGYAVQSFAHRLPDGYFSANLQLENARLGDDDGRYSFYALDYFHSEDEAVCYSA
CWARNWIDTRG*
>GCA_000452945.1_BPMSHR5858_genomic_02917_1 group_4792
VFASFEPTHTGFVAEIDGCRCSIEGAPSPIAERIDWRWTIAQPTPENPDGSDPYQYEVLA
TGETVTPLQAEQQIVAWLEAHPPEDA*
>GCA_000452945.1_BPMSHR5858_genomic_02918_1 ghrB_1
MKPRLLVLIPLRDHAHAHIADVFDILYALTPDQRRRAIAERGAAVQAVLTNGTTGLFAEE
IACLPTLEFVAALGAGYENIALDDARSRGITVVNGAGTNDDCVADHAFALLLAAVRAVPK
LDAACRAGVWRDALPMRPNMSGKKLGIVGLGNIGQKIARRAAGFDLEIGYHNRRPREGAA
YRYFAGLDALAHWADFLVVATPGGAATHHLIDARILDALGERGFLVNVSRGSVVDTAALA
DALREGRVAGAGLDVYEGEPQPPRALAALDSVVLTPHVGGWSPEALERSVRQFLENAARH
FAGRPVLTPI*
>GCA_000452945.1_BPMSHR5858_genomic_02920_1 sasA_14
MASGIGRFGKRASRLIDFGLWAAAALLLLTAGVGALGTARLSDNEADIHRASEVISGLER
LLALARDMETSQRGYILTGKTDYLQPYKAALPEIRKQFDALTTLVQDDAVQQQRLATLRM
LLEKKQHELAATIELRKTEGFDAAQAIVSSDAGKIFMDRARVMVNEMEATVQQQLRTLRD
RARATRDIAIGVGLASGLLTLAVCVSFGYVMRRLLLVEAAAAHDLFEQRELLHVTLASIG
DGVIATDVGGRITFFNRVAEEMTALCADAVVGQPIERVLMFRHATTQRVIDNPARVAVER
RRTASLPAHTRFVGRSGGELPVDGNAAPTFDANGKLVGAVLIVRDVSERERAEERFRLAV
EAAPTAMIMVGRDGRIVLVNSQAEKLFGYARGDMIGHGIDMLVPERFRAGHWAHRDAFFG
NPDARPMGEGRDLFGLRRDGSEFPVEIGLNPITTSEGAFVLCSIADITARKRAELELRRR
SEELARSNQDLEQFAYVASHDLQEPLRAVAGPLQLLQRRYQGQLDARADEFIGHAVDGAT
RMQSLIDDLLSYSRVGRLEDARRPVDCEHVLDEALRNLSAAIRESGAQVTHDPLPTVHGI
EAQLVLLFQNLIGNAIKFRGKDRVARIHLEAHTRDDAWLLRVRDNGIGIDPQYFERIFLI
FQRLHTRREYPGTGLGLALCKRIVEHHGGRIFVESEPGQGTTFSFTLSRPGADETRWPRD
PRE*
>GCA_000452945.1_BPMSHR5858_genomic_02921_1 rcp1
MIADERGNFVEILLVEDSPTDVMMTKEALEYYKVLNPLNVVEDGVAAMAYLRREGPYAGA
RRPGLIILDLNLPRKSGREVLQELKLDPELSTIPVVILTTSKSEEDVARSYGLHANCYIT
KPVDFAKFTDVVHSINEFWLSVVTLPPAKS*
>GCA_000452945.1_BPMSHR5858_genomic_02922_1 rssB_3
MSSPSKNVEILLVEDSVTDALLIGEALIDVTEFAPHVVNAELLSAALECTKRAHFDVVLL
DLGLPDAHGLDTFRQFHRQVPDVPVLVLTGLDDLSVGLQAIQEGAQDYLLKKRVETPLLS
RAIRYAMERHRVAAALAASEERFQLAVSGASAGLWDWNPKTGAMYLSPHFKKIMGYEDHE
LPDEITGHRESIHPDDLKRVLDALKAHLEHRDTYDVEYRVRTRSGDYRWIQSRGQALWNS
AGEPYRMVGWIMDVTDRKRDEDALRVSREELRRLSAHIQSIREEEKTRIARELHDDLGQQ
LTALKMAVTLVDNELKDAGWASRTSRLPNVYALIGQLLDTVRRIAADLRPVMLDDLGLMA
AIEWLTHEFVERYRIPVTSRIEADDIAFTRGSATEVFRIVQEALTNVARHSEATEVTLDI
IRDEPNCIVRIADNGCGTARDTRPNRHSLGLLGMRERAARLGGEIRIDTAPGRGFALTAT
LPLAVIEAADSA*
>GCA_000452945.1_BPMSHR5858_genomic_02923_1 naiP_3
MPAAPAVPSRAARASAAARLERLPFSGYHKLIFVIIAIAFFFDSVDLGTMTFVLGSIKRE
FGLSTATAGLVASASFFGMVLGAAVAGLLADRFGRRPVFQWSMVLWGAASYLCSTAQSVD
ALIVYRVLLGIGMGMEFPVAQTLLSEFVPAAKRGRLIALMDGFWPLGFITAGIVSYFVLP
LFGWRTVFALLALPAVFVLVVRRLVPESPRWLEHRGRLAEAGRVLAHVEAKVMKSARVAS
LPAPSRLAESACAQGRGAFGEIWSAAYRRRTTMVWLLWFFALLGFYGLTSWLGALLQQAG
FAVTQSVFYTVLISLGGIPGFLCAAWLVERWGRKPTCIASLVGGGAMAYVYGQSALFGGS
VTLLVGTGLAMQFFLFGMWAVLYTYTPELYGTGARATGSGFASAIGRVGSLIGPYAVGIV
LPVFGQGGVFTLGALSFVVAALAVGVLGIETQGLALEALASTHAADDDARDPAGAAAN*
>GCA_000452945.1_BPMSHR5858_genomic_02927_1 betB_2
MSVYGLQRLYIAGAHADATSGKTFDTFDPATGELLARVQQASADDVDRAVASAREGQREW
AAMTAMQRSRILRRAVELLRERNDALAELEMRDTGKPIAETRAVDIVTGADVIEYYAGLA
TAIEGLQVPLRPESFVYTRREPLGVCAGIGAWNYPIQIACWKSAPALAAGNAMIFKPSEV
TPLSALKLAEIYTEAGVPAGVFNVVQGDGSVGALLSAHPGIAKVSFTGGVETGKKVMSLA
GASSLKEVTMELGGKSPLIVFDDADLDRAADIAVTANFFSAGQVCTNGTRVFVQQAVKDA
FVERVLARVARIRVGKPSDSDTNFGPLASAAQLDKVLGYIDSGKAEGAKLLAGGARLVND
HFASGQYVAPTVFGDCRDDMRIVREEIFGPVMSILSFETEDEAIARANATDYGLAAGVVT
ENLSRAHRAIHRLEAGICWINTWGESPAEMPVGGYKQSGVGRENGITTLEHYTRIKSVQV
ELGRYQPVF*
>GCA_000452945.1_BPMSHR5858_genomic_02928_1 betI_2
MPKLGMREIRRAQLIDATLRSIDEAGLPGTTLASVAQRANISTGIVSHYFGDKDGLLEAT
MRHVLRDLWAATTRRRAAASDAPRARLRAVVAANFDDTQISAPVMKTWLAFWSQSMHEPT
LRRLQRVNTRRLHSNLCAEFAKTLPRARAREAASGLAALIDGLWLRGALAGEPLDTKAAL
KLANDYIDQLLAPRV*
>GCA_000452945.1_BPMSHR5858_genomic_02941_1 cobS
MNPLAELRYFVVALGYFTRVPVPRAISGQAVDLAHAARYFPLVGVCVGALAAAVHLAASR
VWPPGVAALASIAATLVATGALHEDGLADSCDAFGGGYTREDVLRIMRDSRIGTFGAAAL
VVALALKWQALAAMPPARAAWTMIAAHAASRTLAASLLLTLDYARAEGKAQPVAHRMRAG
AFAFAAALGLPWLFWPDWRAGALALAAMLVLRAGIARYLDARIGGYTGDCLGFAQQVFEL
SIYLAMLGWMSS*
>GCA_000452945.1_BPMSHR5858_genomic_02994_1 group_1365
MIIGLSDNGILLYSNTEYAKTMSKSPAFWEPYCHRSEAEFHLSNFAARVLGRHAGVVFFD
GIDAARLKQAIPDAHGDTLDLVAAVVRPPLMPIEPAETLDALTLEQLSREWGRRIAASAP
SARPPPPAQGVAHARRFIDAPKTAFDAAAVGDTFGVLAGAVSIVAIVGGGIALAPVLGAA
AGAASLLLLAEDSSMLRYELKGDEVRKKKLETSWHCKLVETVGPLLVLPELAVGGTRALA
SLPTAARQAGEAAEETAQAAQRLATQRHAFDAFKRASLDNPDRAARRTGMHAMRTETSGL
AAGVRRARQKMLAARRELLLLRTIEAPAYLASVYGAGVYGMDPPEGGSRSTDWLRHRMDA
QSSGDPDHPAHLLIPRHPAVDGSNGAPAMQLHVAVGHRPEAAT*
>GCA_000452945.1_BPMSHR5858_genomic_02997_1 group_771
MGEVQSSAPSDSQQKLAALAQRGSNADAVQTVSNMGLAINAAQVTAAGTSAWAAGTFQCF
AGRVIAPLGGAMLGGALAEALGADRPVTWVLDKMGLPAVAKPGKAPARVGHKIVHENAFI
GALTGLLAGIAVGVAIAAAAAAIVATGGAAAVAIAAAGPFVVGFVSGAVGGFVGAAVAKG
IGHTGSVTGAIAHGSPNVSFEGAPVARVTDPVTCSKDPGMPPPQIAQGSLTVSVNGLPLA
RIGHKITCSAVIQEGCTTISADETTGTFGKIDANVSLLEQLVLTATDVIMMRSATKEGGL
LDGVLRELLGEPIDMATGDYADYRTDFTWPHVLPLTLSRAYAGRQPVEGLLGDRWISNWS
QRLRYRRPADGPATVTFFDADGQQLVYPVPHEPFNAINFWAPHYALHGSRARAVVFDERS
QQSLIFEPAHAEDDVARLTRIEDRNGNTIDFEYNALGRLCTVRHSGGMTLWVTCDSRGLL
QSVSEQPGGEGELVRYRFDGKRLTDVHSRFQGEFHFGYTDEGWLNHWRDSGATQVALRYD
ERGRVIATRTNTALYDDRFEYDDEARLTTYIDALGHRHQRWFDAQNRLIRSRDPLGRVMC
ASYDENGWLASRTDPLGRVSTYRHDCRGRPLQVTDAFGRVSRYGWNGAGQLVEQQDHNGK
VQWHYSAEGNLVALRARSGETRFRYDARGLLVGRTDPDGAVHAWRYDGAGRPERWTDPLG
RHTYLEHDRYGRLIARIDAAGHRTTYGYERGPSNPRELLASITYPDGAIARFQYDSEGLL
TEAVNPLGQRTRYAWGAFDLLASVTDPGQATTRYHRDGAARLIGVTNAAGQHWKFERDPA
GQLIAQTDWSGRRTRYVLNPMGQVTEKHLPDGVAIRYEYDSQDRLVALTGPRRRHVFAWT
ASGLLTRAEVWTRNDEAGEWRRDDRLDLEYDDACRLVEESQHGRAVGYEYDPMGRTRSMA
TPSGRTLWQYDAAGQLGSMESNGHRFHFDYDGLGLETLRRYTPTQAHVARHPQWVEPYSE
GYAQQQDHDGRWRLTKQACATWAELRERGAARTRHYEWDAAGRCVGMHEARRGLPIAQDR
WRYDARGQVVDAHYERTETRSGRERYEYDALGNVSTRQIDAGEAMTHVYHGDQLVSAGPS
RYEYDARGRMIARTEGRDGFRPRTWRYQWDDFDQLEQVLTPEGERWRYRYDAFGRRISKA
CLSTPKAGRPARIDYLWCGSRLIEAWRAYGERDGTQYEIQRWHYRPGTHSVLAQERLKYD
DKPDLQNSEWFALACDPNGAPHTLYSSDGRIMWSARRELWGRVADDPDRDTVRHAVREQL
RTSLLTGDAFDPPDCELRFPGQWADEESGLHYNFHRYYDPATGQYLSPDPLGLAGGLRTH
AYVHDPLQWVDPLGLQGYQGSGKPEFIGNRKLPQNDLPWIKYQKRVTGRPYEETWRVGDH
NVNLDGKRAGYTVEAKWTGKNSAAWESSPYNPEHEFYNESKILDQAGRLLEFNDASGGSG
VRYAVSNAEAQQHFTQLFEQHFPTEMQDGTLSVWHVPGNGM*
>GCA_000452945.1_BPMSHR5858_genomic_03002_1 oxdD
MTNLSRRKMLAGTAGALAAAGIAVSAKAASFGNPDSPAEGAVNARNPQSLTDPGPQNPAL
MKEFPSFQSPPATDINGMPIFWASFNNAHKRIQNGGWAREVTQEDFAISETISGVNMRLA
RGGIREMHWHQQAEWAFMLDGRCRITVLDEEGRPSVQDVKTGDLWYFPPGLPHSLQGLGV
DGAEFLLAFDNGRASEFNTLLVTDWIAHTPPDVLALNFGVPADAFRRIPLDNLWIFQGDD
PGPLAAAQRASASSRGAPKHPFIFSMGDMKPNVKTRGGEVRIVDSTNFAVSKTIAAALVT
VKPGGMRELHWHPNADEWQYYIQGDARMTVFDTGPKAQTADFRAGDVGYVKKSLGHYVQN
TGTTDLVFLEIFKADRYAEVSLSDWLAHTPPQLVEAHLHIAPDVIARFPRNRPDVVPA*
>GCA_000452945.1_BPMSHR5858_genomic_03009_1 group_3393
MKRMLPLAAASALFAPAGAHADHPFTSDDTNTQGDGNWQYELNVERTSKQPDIGRQQLWN
TTLTRGFGERVDLYVQAPYTHVQTRSDEDGSGLGDLEIGAKWRVLERGPLSIALKPRFTM
PTSDDARGLGNGRASAGAMLLAQYDVARLQVLVNAGLMYQPNRQGNLASIWQACGAIVYR
ATDKLRLGVDIGVSRNPERGAGAHPAYVIAGAIYTPRDWLDVDLGYRRGLNDQIYDHALM
AGLTVRW*
>GCA_000452945.1_BPMSHR5858_genomic_03011_1 group_7476
MEADDSSRLPLAGVAPGGACAASAAPASDSDSPLIQPNARDAMRRETRVRR*
>GCA_000452945.1_BPMSHR5858_genomic_03028_1 group_1478
MFPTLHLPTYVYVLFCAFVYIGVKRCFARQVTPLLPLLSPIGFAMLGATSLHARFPRAGV
GADAAALASLAAGAALGWLHASRWRLRFGVEHSRLAVRLPGDASLLVTLLLTFAAETYMH
YAVAAAKPWAATHAFALLPFAAWGLLVGMPLGRAINVVARCVRYTNGPRDKDGAYTLDSN
*
>GCA_000452945.1_BPMSHR5858_genomic_03032_1 group_2452
LRRAWVGNRVVFFILMQASDPSSGYLSDVTFPDRFHRELSPTWLNYASVLGGARPKELGR
PFRYLDLGCGFAHSTVINAAAFPHAEFHACDFNPAHIEAAARRASRLGIGNVAFHEASFD
ALLDRDLPPFDFIVMHGIYSWVDAGMRRVIRQLLSRRLADAGLVYLSYNCQPGWAAEAPL
RKLMLELAQAADGGIEARTGSAIAGMRKLGTPSLRYFRDNPAAAEALAALANDPLDYLAH
EFLNGTWKIHYSVDVVDEMAEAGLAYAGSATLADNHPMLLIDRQAADAIAALPNARLRHL
AEDFAVNRRFRRDVFVRGARASTAPAEALRHLDEIAIGCTTEIDRIDTRVTIPRGAISFQ
PDFIADLRALLRHGAMRIGEIVARLGAARRNSREIRQNLLFLVASGTLTPFAQPGGPTDT
GARRAASPAAAAALAGSVDDAAPAFVPSELLGNGLAVSPDEAAQALRWIAGEAMPRPERL
ARVGVLRGA*
>GCA_000452945.1_BPMSHR5858_genomic_03044_1 tnaA_1
MNGLKERVGSGYVAIPILVALRGARAPGAAGDRDAAWTERIAAAARLDADGAAALREWLN
DAAPAAGADDARLACGDAIPTDILEPYRFEPDARAFFDDAFRECLNRWSRRLLEQDGATW
LSDVVLVPFLVAAARARESAPSCEARSIRALADAFGADLRRLLDASRVLDDGDIGDIGAM
RASLECVTSFRDGLLHADRRLSNGAPAACRPQVCAAAYEDLEAAVLRRFDASANARAGRR
PRYLAFVGGEDRDLPERLVERIVRQGEPRNGGGTGVEPPRIVIPRADAAPADDSAQPDPA
AAFGVDAIDDVLPVCVLSGVDDAAAIRRLRALLVRGAHAAVVLACHGLPAASAALPDAWA
AARPARMLRLLGFGAARDAQAFLVDMAADGLFARTPPIGYPRASRARRATLGEFEARDYR
VRPAAPHDLPALQALELACWPAALRMPEATLAARVGRHAAGQFVLELDAREAAAGPRRLA
GVIYSQRIASVRALDGVDADTVDRLHEDGGPVIQLLAVNVDPACQSRRLGDQLLEFMLQR
CAALADVESLVAVTLCRDFHKHASMPIDDYLRLRNAFGFLADPILRFHELHGARIERPMP
GYRPRDARNAGFGVLVSYDLARRARNEAGAPAASDAPATPDGAPARADGGHAGVAASPRP
ERDAATAADPDALDAFIEAEIRRIVGGGAELAYARDLPLMQLGLDSVGLLELAEALALRC
GVALPATFLFQHNTPARIVAYFDASRHAPPRAGCDTGCETAGAAAAAARPSSRGCPATEP
GAAREPAQAAPFAPDGIAIVGIACRLPGGLDTPEAFWDALKAGACVVGELPGDRWTWPAD
IDPGARHRGIDRGGFLDDIRSFDAGLFRLSPKEVATMDPQQRILLELAWEAIERAGHCAD
AVAGSRTGVYVGASGSDYRLLLERAGTGVDAHVATGASMAVIANRISYTYDLRGPSIQVD
TACSSSLVALHQAVQALRAGECDQALVGGVNVICHPGNTIAYYKAGMLSPQGRCKTLDDA
ADGYVRSEGAVMLMLRRLEQAVADGDPIHAVIRGSACNHGGLTGGLTVPHPDRQADLLRA
AWAAARVSADDIGYLEMHGTGTRLGDPIEVRGLADAFGARDDAAARGTCGIGSVKSNLGH
LEAAAGLAGVLKTVLALKHREVPATLHFSRLNAQISLARTPFAVVDTHRAWPARGGARRL
AGVSSFGSGGANAHVVLEEYPSEAPPRAAAGDALFVLSAHSREQLAEYARRVLAYCERRL
QSGDDAPAAAAVAHALQRRQAMAWRLAFVAASLEEAVRRLRAFAAGAAQPGTFVGGGAPK
TSVADFVNQNPDVQQVVSAWLRERQLAKLARYWADGVRIANWSALYDARPACVPLPCYPF
ARERHWIAARPAEASEATAAAAPPAAPDDAYRAAPRLEAESRDAAGASFRVRLDGDASFL
IDHRLRGRKILPGVVHFELAHAAWAALARADAPAIEFRDLAWTHPVDVATPERVLGVRLR
RVAARPGAHAYEAYSPPDAAGGGERVHARGTVLDVAGPPEPALDLDALRARFDGEPGAHE
RDAHDCHRAFERMGFGYGPAHRGLRGLRWRGGARGATEVLAHIVLPDCIADARERHRLPP
GLLDAAVQAAMAAAIGRDALGASPACVPFSLDRLVCAGSCPARAWVWARRRDGARAALAP
VDLDVCDDRGRVWASFRALTFRPLREAAGRDVAAPRLFRPRWAARPLSGERASAGAADVA
HWLVLCGFDEGAALRRDLATLRARLPDTSIVAIDSDAATLESRFADSAGQLFELFRELAL
SGATPRAAVQVLVPADGPERAFAALSALVRGARLEHPSWSVQLLSLERATQAADAAARAL
ENRGDDADFVRYRRARRESLVFEALPAGRDEPPRPWKEAGVYLITGGAGGLGLAFAKDIA
AHVRRATIVLAGRAAAPDAALSASLGAIARPAGVDIRYRSVDVGDAPAVAALVGDLLREH
GRLSGVIHAAGVTRDALLVRKPRAEFDAVLRPKVAGAAALYAATQDIDLDFLVSFSSIVG
VTGNLGQTDYGAANAFLDALAGLRAQMGGERRARVLSIAWPLWRDGGMGREPEVAAHFER
AFALAPMDTAAGIDAFHRALASDAAYVVVANGDADWTPERAIARALSARARPGAPEPAER

AHGPAREAAAAACEAVAVSASASSLSPSPSPSSASASVSAECAPADDGAARRHAVVAYLT
RQIGAVLGHAPDSLDIDAPFTSYGMDSILALDTTRAIETDLGSLSKTLFFEHENVRQLSD
YLLDEHADRLAACAWFAQAGEPARAAGPALAAGGEPTRETAAPSAAADTTADTTADTTVG
IAIETSTETPTAATTQAGAAPEGRYRRVAKAALPADGQLAAAVAAIGGAAATKGVALFEI
WPELFVDSAGHGYCHLLVDGGVLFAAQHAGDARHRAALFAALLAYCDRHGYAFGYLDLSE
GRKPDLEAQCGLLAAPVGVVQIVEAIASFSLAGGRMRRLRYMVERFRKAGACRVVEYRAP
DPDVAREIRRVICAWSDAKKVVNNVDIVLGEMASGSLHERYRVFLTYLDDVLQNVIMIAR
DGDGYLMDQEYYVADMPLGGTEYAVTEILAALAAEGRERFSLGLTWGLFDTGEGSSDPAA
DAFLASTQTQLRRIFERGAANRQYKSKYGTRDHAVYLYRRPGKPEPAIVGCLSQFYRKGL
THHEVRRLAGLADAPVPVAAPATVPQVTPAPAPVATPTADERAYDVTRIDAATIRVDLVS
DSWAHVDYPFMRARAATLDRHAPPARGGDPGRAVAELLGFAQCLLTTSGRAAEHLFFRAR
RSARTRVPQNLLFESTLHNLVKSGFEPVELPDARALDPDSRDLFRGGVDLAALDRELQAH
ADATAMVMLELCNNASGGYPVALAQIRAIAAACRRHGVPLVMDVTRIVKNAELIRRGEAG
YAQRGLWEIVREIADHADAVVGSLCKDFGLGAGGLLAARDARVVANAAGIARLEGGLPGP
AELRRIAAAFDDRAYLEREIGRQLDFARDLHIELERCAVPVVQPGAGHCVLVRVDQLAPP
GGSAPSRGAYLRLLAERYGVRGGLHLVGNLRDSHLNACVRLALPLGFDDPRGPGALAAAL
AAARDGRDHALDDLMRAPRARAAHGGRCADGIAIIGLSGRYPDAPTLDAFWRNLVSGRRS
ISEIPAERWDWRDHYERDPDTAVAHGKSYGKWGGFLDGFSAFDPLFFQIAPREAEFIDPQ
ERLFLEACWHALEDAGCPPSALTRAQRAKAGVFGGMTKQGFNLYGAGGAQPYQSTSLAAL
VNRVSHCFDFNGPAVAFDSHCASALVAIHEACQYLRREPEGIAIAGAVNLNLHPSNYQQL
SKMQVLASGAESASFASGGLGYVPGEGVGAVVLKDYRRALEDGDPIYGVIRGSAVNQNGR
MNRFGMPSQKQQEAVVRAALAQAGVDPRSITYVEASAHGSAVGDAIEMAALTRVFGARER
ADGRYRIGSVKPNIGHGEAVSGMSQLTKVLLSLRHGQLPPTLVCGAPNPDIDFDALPFEL
NTSLTDWARARVDSERVPRRAGITSTGASGLNAHLVLEEHAAPAVPAQAGPGEADARAHV
FVLSARDRARLDDYARDWIAFLNDDPQRDLAAIAYTLQVGREPMACRLAVVAADCRDLAG
KLARWREAAHADCDDVFHGEARAAAGKPHREAARDAREPRDVARAWVGGAVVDWAARHAG
ARPARVAGLPGYPFERRSYWPGAAAALAAAPATARATAASDASEAARALEAREAHEATRA
TRMTRAARESEARQAPEAPSMSEATEATEVAEVAEVAEVAEARDARPGTVADDAAARLEA
AFLPRFIELVADVFRLPAGELDADRPLDEYGINSFLIKVLNVRFADIVGRVSSTLPFEYR
TAGEMARHFLTAHRDACAAWVAFDGAASPGAADASSAPPVPAASAPAASAASAASATPAT
QATQATQATQATGPTGEPSRASAGVPSGASSSIKRPGATWDEPIAIVGVSGRYPQARDLD
AFWDNLMRGRDSITEIPPERWPLDGFYDEDRERAIGASRSYAKWGGFIDGFAEFDPQFFN
LSPREASNMDPQERIFLQACWEALEDAAYTRARIAREHGGRLGVFAGITRAEFCLYGAGN
LKQGKAPFTSFCSLVNRVSYFLDANGPSIPIDTMCSSSLVAVHEACDKLRLGECEVALAG
GVNLSLHPYMYVSLSAQRMLSSDGRCKSFGLGGNGYVPGEGVGVIVLKPLSRALADGDRI
HATIRATSINHGGKTNGYTVPNPIAQQNVIRSALDRAGVHARAVSYVEAHGTGTELGDPI
EIAGLSGAFRRDTSDRGFCAIGSVKSNIGHLEAASGLAGLTKVLLQMKHGLLVPSLHASE
LNPNIDFPASPFVVNRETRAWERPVIDGREHPRIAGVSSFGAGGTNAHVILEEPPRQASP
ARAPTPAGAPALIVLSAKKPEQLRRYASELLARLRDADYRARVDADGLRSLAYTLQVGRE
AMDERLAVIADSVQALEGKLRQFVDGKTDIQDLHVSRVGRSAHHVI*
>GCA_000452945.1_BPMSHR5858_genomic_03061_1 phaC_2
MDTRHAPESGAPDAPLPAHPPASYAPESPYRIFDLAKEASVAKLTSGLSPASLQLALADW
LIHLAAAPGKRAELATLALRHAALLGQYLLEAATGRTPAAPAQPSPGDRRFRAGAWQLEP
YRFWHQSFLLAEQWWRAATRDVPGVSPHHEDVVAFSARQMLDTFAPANYVATNPEIAQRT
ALTGGANLAQGVWNYLDDVRRLITKQPPAGAEQFELGRNLATTPGRVVFRNHLIELLQYS
PTTPDVYAQPVLIVPAWIMKYYILDLSAHNSLIRYLVGEGHTVFCISWRNVDASDRDLSL
DDYRKLGVMDALDTIGAIVPGEKIHATGYCLGGTLLSIAAAAMANTGDDRLASITLLAAQ
TDFAEPGELQLFIDDSEIHFLESMMWERGYLGAHQMAGSFQLLMSNDLIWSRVIHDYLLG
ERTPMIDLMAWNADSTRMPYRMHSEYLRHLFLDNDLATNRYVIDGQTVSVHNIRAPFFVV
GTEHDHIAPWRSVYKIHYLSGSDVTFVLTAGGHNAGIVSEPGHAKRHYRMKMTAAAAPSI
SPDEWLAGATDFEGSWWPAWHAWLARHSSPQRVAPPPLGKPGARTLGDAPGTYVFQK*
>GCA_000452945.1_BPMSHR5858_genomic_03064_1 pfkB
MPEIVTLTPNPAIDVATCVERVTDTRKLRCGPARRDPGGGGINVARVLTRLGADCSAVYL
AGGGTGLALRGLLADEGVRAHGIDIAGETRENFSVLETSTGREFRFVLPGPALAAHEWPR
CVEALGRLADASRYLVMSGSLPPGMPDDCYARLARRASARGVRTVVDTSGPALAAALDAG
VYLVKPSLGELRALTGLPLEDDGARLAAARAIVAGGRVQIVALTLGDAGALVVSRDDAVR
LPGVKVAVRSAIGAGDSFVAGLVAALNRGANVADAARHALAAASASLLSTGTALGTKEDI
ARIYRELPGTIDA*
>GCA_000452945.1_BPMSHR5858_genomic_03065_1 group_1863
MKSDAALKQDVEQELYWDPSLDARQIDVSVRDRIVTLRGFVPSWAQKYAAQKAAQRVAGA
RALVLELKVLPSEAPRDDEVLAAAIVSALAWQEGLAGQEIHVDVDEGCVTLTGEVAFGYQ
RQAAETVVSRMRGVIGVVNRIDVRSRDAGATVRERVADALARRAQREAAHIEVDERDGIV
TLTGTVDSLIERRAACGAAWSVKGVHEVIDHLSVA*
>GCA_000452945.1_BPMSHR5858_genomic_03067_1 cdd
MTHHALIEAAKAAREKAYAPYSNFKVGAALVTNDGKVFHGCNVENASYGLCNCAERTALF
SALAAGYRPGEFAAIAVVGETHGPIAPCGACRQVMIELGKPTLEVVLTNMQGDVRVTSAG
DLLPDAFYLA*
>GCA_000452945.1_BPMSHR5858_genomic_03068_1 deoA
MTFLPQEFIRKVRDRAPLDTADVARFVQGVTAGDVTEGQIAAFAMAVYFNELPLSARIAL
TLAQRDSGDVLDWRGARLNGPVVDKHSTGGVGDLTSLVIGPMVAACGGYVPMISGRGLGH
TGGTLDKLEAIPGYDVAPSVDMLRRVVRDTGLAIVGQTAQLAPADKRIYAVRDVTATVES
ISLITASILSKKLAAGVGALAMDVKVGSGAFMPSAEQSAELARSIVDVGNGAGMRTAATL
TDMNQALAPCAGNAIEVRCAIDFLTGAARPARLEAVSFALAAQMLTMGGLAADAHDARRR
LRAVLESGAAAERFARMVAALGGPADLVERPERHLPRAAAAAPVAAARAGWIERIDARAL
GLAVVGLGGGRAKIGDTLDYSVGLSALAELGERVEAGQPLATVHARDADSAAQATDAVRR
AYRIGAEPPAQTRVVHAVIE*
>GCA_000452945.1_BPMSHR5858_genomic_03082_1 group_237
MQKLFNKSAIAIGISTLLVLSGCGSVDGPSLSASGVKPATVGNTGTSGTSGTSGTSGTSG
TSGTSGTSGTSGTSGTSGTSGTSGTSGTSGTSGTSGTSGTSGTSGTSGTSGTSGTSGTSG
TSGTSGTSGTSGTSGTSGTSGTSGTSGTSGTSGTSGTSGTSGTSGTSGTSGTSGTSGTSG
TSGTSGTSGTSGTSGTSGTSGTASSPLGNILAQTGNLITATGTVVSATGNQISGTSVPGV
NGATTTNLGNAVSSLGNGVQTLGNGTAAGLGTLGTSTNPLGPTLTSTSGVVSNVGNAVSS
LGGVVTSIGTGPLAPLAPVTSPLGGAVGTLGSTVTQVGSGLNNVLTSAPVQQLETGVSSI
INPISNVVSGTTQTIGTATGLGAPVNNLLTTVGNGLNAAGAKVSGSTNNQVVQAVGGVVS
QLGNTVTSVGGLLTGGTTNPLAPITGALSGGATNPLAPITGIAGSLSGTLGANAGLNAGA
TQPGTTTNPLAPVTGLLGGLSGSAGGNAAAGLTAVLAPVTNVVGSLTAGVNGTVAAPSPA
PTTGSTTTAPATGVVGSLTGGTTASGTSTTNLLAPVTNLIGGLLGGVSGK*
>GCA_000452945.1_BPMSHR5858_genomic_03085_1 group_7492
MKRSTPNPRRRLLVALAGTMLTVPGLPRAATRIVLDAARTNADATRGYIDLPGFAASRKN
TGIAVHAIDDTSLLDAVKDAGFSFVRADLFWEAVDTPHGWDFSRYDTLVAELSERGLGAL
FILGGIHRVHSPNRPPTTPAQLAAFRRYVFNAAQRYKGKPVRFEVWNEQDHKTYWHGEPS
PVAYRNVLKLAVQAVKAADPDAIVATGGVQQVNRTFIRAVGDIASPAQPAPDAVSVHPYR
QTEPETVFADYAALRRDLSSYRRPPQIWATEWSYPSYGYNYVADIGDGHSSVARARQANY
AVRLLLANWISQIGLTSYYDIRDDGRDRTNMEHNFGLLDADDAPLPAYIAVRRLFAFTAD
AVRARYFLDPDARYVVLKLSAARATKYALWCYGDGNAISVDLSRLPAGATVTNMYGAALD
AGNRRRLDVPEARGPVFITIEA*
>GCA_000452945.1_BPMSHR5858_genomic_03102_1 copR
MKVLIVEDEPKTGMYLKRGLTEAGFVCDWTQDGISGLHQLQTENYDLAILDVMLPGCDGW
TVVRELRRTHRTPVLFLTARDHVDDRVKGLELGGDDYLAKPFDFAELLARVRSLLRRGQA
TDAPVLRVANLELDLIQRKARRQGNTILLTAKEFLLLWLLMRRQGEILPRSLIASQIWDI
NFDSDSNVVDAAIKRVRAKVDQDYEPKLIHTVRGMGYVLEQRGEPCESCA*
>GCA_000452945.1_BPMSHR5858_genomic_03103_1 rcsC_8
MMRRLAPRTMRARLAIKFALSTSVLLAISGLLLYETLTRTFAYSSERAMETTLAGVVAHI
GEARSIAELREKGTKFYYSFHSDEKVDFAIYDAQGTAYMQSGWYVPYGGVQQAKPGLDPA
IVDDRRNAKRYLVADARLTGNPGLPLRVAVQYDFRREQSLLRTCAISIVAIILVGTAVAA
TVAYWIAAYALMPLRRFAAQADQISSSRLAQPLPETEMAGELRELAVSFNRMLVRLNASF
TRLKEYSSDLAHDIRTPLTNLLAEAQVALSQRRSVDEYRQVIESSVEEYQRLARMVDDML
FLARADGHQNKLDIEPIDAYAMADRIVRYYETVAEDRGIGIAVEGRASFSGDSILVQRAL
SNLLSNAIRHAPDHSTVVVRCEQSEQDVVIAVTDTGRGIEAVHLPRIFDRFYRVDPSRYD
SASGTGLGLAIVRSIMTEHDGDCWVESMPNVFTTFFLRFPREHAAGCGDCPPAGHDSRWH
LEQT*
>GCA_000452945.1_BPMSHR5858_genomic_03104_1 group_3399
MKATFLTVALAAAFVSPAAFADAPAGKTRAQVYQELVEAKANGLDYVTDASYPEISPLYA
PRFANKAKPQQPAHAVADTRSDAKAGSSGNLSEAAQDEHCVGPRTFCNPYAGS*
>GCA_000452945.1_BPMSHR5858_genomic_03105_1 bla_2
MKFRHALSSAFVLLGCIAASAHAKTICTAIADAGTGKLLLQDGDCGRRASPASTFKIAIS
LMGYDAGFLRNEHDPVLPYRDSYIAWGGEAWKQPTDPTRWLKYSVVWYSQQVAHHLGAQR
FAQYAKAFGYGNADVSGDPGQNNGLDRAWIGSSLQISPLEQLEFLGKMLDRKLPVSPTAV
DMTERIVESTTLADGTVVHGKTGVSYPLLADGTRDWARGSGWFVGWIVRGKQTLVFARLT
QDERKQPVSAGIRTREAFLRDLPRLLAAR*
>GCA_000452945.1_BPMSHR5858_genomic_03113_1 group_772
VCERARRPCHEGEQFTEAQIVYALKQAEPGTTVAKVCRKLRISDATFYSRRTKYGGLSPS
ELHWVKPLEDENAKLERLVADLSLDKAMLRNVPSKSSETFSAP*
>GCA_000452945.1_BPMSHR5858_genomic_03149_1 group_1110
MTPAARRQRRHSTLARQAFAVYAALVVYASLYPFTGWRSLGIGPFDFLLAPLPRYLTAFD
VVSNVLGYLPFGALAVLALYPLRGVPAALAATLLGALLSGAMEALQTYLPTRVSSNLDLA
ANALGALVGAAAAAPAATALIERGVVRRVRFAWFEREASTPLFLSALWAGAILFPSPFLF
GIGDWPSELWERADVTMRGALLAWAPDAWNVPAWPERLDGLLSDSAWETLLAALGLFAAL
AVVSLAMRERAPRVRLVVGFAACALALKAAATFMQSYTGLVLDWATPGALRGIAAGLVAA
LVALRLPAAWRAALAAAALAAALVLVNVLPVNPFFDFALSGWQQGRYVHFNSIARWLAWI
WPYAALVWLAGRAERAWLARRGAGASRRASGKRRRSL*
>GCA_000452945.1_BPMSHR5858_genomic_03155_1 esiB
MMNAIKKLLPTHAAPPPDGARRPPPRRRLLHARLSPVGMLAVAAALAAGVAVTERIERPA
SGAPAVSRAQLDEWRAMVEQASEPNALGRLRALAQRGSADAQAALGLALAGSRDLALRDE
GWRWLETAAHATPPENAPTSAGERDARLALGKAWLLGAGGAARDYPRALAMLRPLAAAGD
PNAAYYVGLVYRSGYGTPADATEAARWFELAARHDIPAAQFMLANAYRDGSGVRRDEARA
LALYRQAADHELPEAVQALAIAYRNGELGLPRDADAFHSQWIEAAHALKHPALAP*
>GCA_000452945.1_BPMSHR5858_genomic_03191_1 cutC
MPASSILLEVIATTIGDARAAARAGADRIELVTAISEGGLTPSVGVIEAVVAAVPIPVNV
IVRPHSRSFSYDASELAAIARDVRAAVAAGANGIVFGMLDAHGDIDLDALRRIADAADGR
DLTFHRAFDVARDLNAAFDTLLRVPSVTSVLTSGGHPSVLDARDVVARMVRRAAGAACTV
LAGAGLTVDALGDFVRATGVRAVHFGSGVRERGEVLAPIDERLVAKARAALDAATRG*
>GCA_000452945.1_BPMSHR5858_genomic_03305_1 ddpF
MNAVHETRESAARNDVVLVADQLAKHYAVRRGMFGHGTVKALNGVSFSLARGKTLAVVGE
SGCGKSTLARQLTMIEPPSAGHLSIDGRDVAGADRETVAALRRRVQMVFQNPFASLNPRK
TVEQTLAEPLAINTRLTAAERAQRIAQIMRTVGLRPEHAKRYPHMFSGGQRQRVAIARAM
ILDPQIVVADEPVSALDVSIQAQILNLFMDLQEQFKTSYVFISHNLSVVEHIADDVMVMY
FGSVAELGDKKTIYARPRHPYTRALMSATPAIFEADRKIQIKLQGEMPSPLNPPSGCAFH
QRCPYAVERCRVEEPKLRDVDGRLVACHRAEEVGEANA*
>GCA_000452945.1_BPMSHR5858_genomic_03324_1 group_185
MTGIDTAAAAILASRIDSLLDAIQPAAGGAAAPQVGTSGAPPASAAPRPAASGPPPASAQ
TALSEVGLTLDAISRYGGAATPAVVGAAPLLPAAPTAVAAQAASLDPLLATPDARAGNPA
AATAPGAQAAQAAPQAALPASPAAALSLALVQAVAQSGLFYESHLAQWLAGQRTTAELAR
EPQARLASGAEPDAQAGAQPGQMGDALDDALADLLAVRLPLPQGGRDAPARGAARPPGQP
AGGADEGAAASGRAPTNAAAPARSVDAYAALADADGKPGAPSAQTALARAGMPPTGADTP
ASIAASLHAATLPIVRQQLDLLATDQFRWIGEAWPGARLDWTIEPDEQHGRERPAPDADF
PDARGWRTRLTLALPSLGTIDAELVLNGEQLAARLRASEAGAARLAPHGEALRARLQALG
LQVSGLSIRAIDGVPDSFDAAAARAAASAYARAAAGGEAVAKDGFGGAAGATGGAGGAGV
EHRASSSSPVETDDDWEFAQ*
>GCA_000452945.1_BPMSHR5858_genomic_03328_1 fliF
MDSQANSLINPDARSSLAGASPQAAAAAGALPGAAAGGADFGLGGFAERIPGLSRMKTNP
RLPFLIGAALAIAAIVALVLWSRAPDYRVLYSNLSDRDGGAIIAALQQANVPYKFADAGG
AILVPANQVHETRLKLAAMGLPKGGSVGFELMDNQKFGISQFAEQVNYQRALEGELQRTV
ESINAVRAARVHLAIPKPSVFVRDREAPSASVLVDLYPGRVLDEGQVLAVTRMVSSSVPD
MPAKNVTIVDQDGNLLTQTASATGLDASQLKYVQQIERNTQKRIDAILAPIFGAGNARSQ
VSADVDFSKIEQTSESYGPNGTPQQSAIRSQQTSSSTELAQSGASGVPGALSNTPPQPAS
APIVASNGQPAGPAATPVSDRKDSTTNYELDKTVRHVEQSMGTIKRLSVAVVVNYQPSTD
AKGRVTMQPLAADKLAQVQQLVKDAMGYDEKRGDSVNVVNSAFSAAADPFANLPWWRQPD
MIELGKDIAKWLGVAAAAAALYFMFVRPALRRAFPPPAEPAAAAVPALDGPDDMLALDGL
PSPDKKQLAEEDEEHPALLAFENERNRYERNLDYARTIARQDPKIVATVVKNWVSDER*
>GCA_000452945.1_BPMSHR5858_genomic_03337_1 group_324
MLGRRARRRSTARAAGRRATGGQPAGGADEGAAASGRASMLHVEGVVYRSVCATPARPVE
LTLAYRQDNDNPLFGAPKDMPRKSLTASR*
>GCA_000452945.1_BPMSHR5858_genomic_03379_1 group_3024
MSGKRRGAPLSTRTPRVLRGAVIGSLAAAALGAAAGGALAQAPGAPSRPDPQRETEAAVA
DYNAGNLRAALVQFHDAAARGNRLAAFDYAMMLINGEGVTANVPEGLRWLRRAADAGMTQ
AQYVYGRMLDDGEFVARDPAAAHDWFLKAAQQGHVQAELALANQFLDGRGTPRDNRQAFA
WYKRAADAGEPVSQYVTASFYERGGDGVARNLDIARAYYAAAAAQGDDAAALKYRELTAA
QKAGPGGASAPKATGPSY*
>GCA_000452945.1_BPMSHR5858_genomic_03386_1 group_3425
MRVRRATSGKSLLIRVIVSTVAITALLLLVLLAAASANTEFFDRYYSWLYSANIIVALVF
MLVVLGLVGMIVARLRKGKFGTRLLAKLAVFFTLVGVVPGGIIYIVSYQFVSRSIESWFD
VNVETALTAGLNLGRGMLDTSLADLQTKARLMSEQLSSADANTNGTTLTLLRLRDQFGVQ
DATIVEPGRGTGASPDVHIIAQASGNFAALIPDDMPTPLMLSQAREHGVFAAIEGEVDGD
PRARGSKGALRLRVVQRIPDSSTSLLQPAERFLQLTQPVSPTLARNADAVQRAYREYQEK
SLGRSGLRKMYIGTLTLALFLATFIAMMIALALGNQLARPLFLLAQGTKEVAEGDYTPKR
EIKTRDELGFLTQSFNAMTRQLSEARLAVEKNRVALEHSKTYLESILANLTAGVFVLDRQ
FRLTTANRGAERIFRQPFNSLIGTTLDQIGVVAEFGAMVRKAFADREAAGRGGGEDRGHW
QQQFAVEVPGETEPLTLLVRGTRLVSTAAAPGQPDDPQTSGYVVVFDDISDVISAQRSVA
WGEVARRLAHEIKNPLTPIQLSAERLQMKLSDKLSQPDADVLKRGATTIVNQVAAMKRMV
DDFRDYARTPPAVLVNLQLNDLVSEVLTLYGVGEGKSTIVVEQTPLPVIRGDATQLRQVI
HNLLQNAQDSVADVEHPRVLLETKTVEYGDPDANGKTRVAVRLTVSDNGPGFPARILTRA
FEPYVTTKAKGTGLGLATVKKIVDEHGARIDLRNRMHGDVVEGAQVSILFLQLADDTAGA
GGGTNGGTAPAKTRATVQTKAA*
>GCA_000452945.1_BPMSHR5858_genomic_03471_1 mcl1_2
MKTSNEPVWRSLLYVPAHVPRFVAKAAGCDADALILDLEDSVPPEWKAPARDGLADTVAG
LAAVGHEVLVRANGPLEPLVADLRAALAAGVHGIVLPKVRGASHIEAIDELLDALEAEHG
IAAGRTRIVAIIETPGAFQAMERIARASPRIAAMMLGGGDFALHCGAEARADVLRVPKQL
LVIAARAARVQPLGLVGGLDEIRDLDTFERLARESADLGYAGATCIHPAQVEALNRAFRP
SDAAVRAAREIVAAYDDARANGRGALRVGDRMVDAPGVAHARNVLARADAIARRPRRCAA
A*
>GCA_000452945.1_BPMSHR5858_genomic_03545_1 group_616
MESRHLIDMANQIGAFFASMPDHDEAVAGIADHIRRFWEPRMRRALLAALDDPQDEAARD
VAPIVREAIDTHRASLEPAPVAPHAA*
>GCA_000452945.1_BPMSHR5858_genomic_03546_1 oxyL
MAVVPLSTPPVLIVGAGPTGLAAALCLARARVPVRIVDKAARPARYSRAIGIQARTLELL
EQQRVVDRFVELGHRARLAILYSAGQRIAELDFDPLQTRYPYLLFLDQTVTERLLTEHLA
TFDVHVERGVTLTHCAQPEGALDVRLRHGDGRDERLQPSYVVAADGARSTVRHLLDVDFV
GHAFEQTFMLADLELGADWPDDEIHLFTTGEGLAGLFPMGGGRYRLVADRPPRNGALDDE
REPSLELCREVVRARVTPDLKVGDLAWSSYFHLHSRMVARLRVGRVFFAGDAAHVHSPAG
AQGMNTGIQEAFNLGWKLARVLAGNAPERLLDTYHAERHPIERDVLRQTSFATQVVEADR
GPLKLLRDHIVPILASFGPLRDAARRTVSELAIQYRKSPLTLERVLDGGPRAGERAPDVL
VHVLDGPLGKAPGVARLFDLHDPAHFTLLVLEPREAIDDTLPSDPGEDGRALAGALERIM
PGAVRCWCATDAEGEGAPLLADAYGRARPVFYLLRPDGYVAARGRPATDANALLRHCETW
FSGMRLET*
>GCA_000452945.1_BPMSHR5858_genomic_03555_1 guaD
MTQTAFRARLLRFDGDPAQSDDALAYDEDGLLIVENGRVVAAGAHAALAARLAPGATLVE
MRDKLIAPGFIDTHVHYPQTEMIASPAPGLLPWLDRYTFPTERRFADPAHARDVAEFFLD
TLLACGTTTALVYCTVHKQSADALFGASEARGLRMIAGKVLMDRHCPEFLRDTAQSGYDD
SAELIARWHGHGRQSYALTPRFAPTSTHAQIEACGALARLHPDVFIQSHVAENLDELRWA
AELFPERRSYLDVYDHYGLLRRRAVYGHCIHLDDDDRRRFAETGAIAAHCPTSNLFLGSG
LFDFERANARHMAVTLATDVGGGTSFSMLQTMNEAHKIARMTGHHLSATRMFWLATAGAA
HALDLADTIGTLAPHAEADFVVLDPAATPLLARRTARAESLEELLFALALLGDDRAVYRT
YAAGRCVHRRDIADAG*
>GCA_000452945.1_BPMSHR5858_genomic_03563_1 group_1211
MGVGYAVAEMIENGGVIIGGSAGIANDLAEGRLAPGFDDRDRIASPYEHRLVEHRLVWAR
PASHAPEIRPSIDSNLALGEQAPELER*
>GCA_000452945.1_BPMSHR5858_genomic_03571_1 dinG_2

MSASPPPHPAIDTPLAFVDLETTGGSAAEHRITEIGVVVVNANGVSTWTTLVDPQQPIPP
FIQQLTGITDAMVRGAPTFADIAGALFERLDGKLFVAHNASFDRGFLRAEFERAGIAFNP
DVLCTVRLSRALFPRESRHGLDALIERHALAPSARHRALADADLIWQFWQKLHAVIPAEQ
LSEQIVRTTRRFRLAGALTEAHLESAPAGCGVYALFGDGDAPLYVGRSVRVRQRLRALLT
GERRSSKETRLAQLVRRVEWRETGGELGALLAEADWIASLAPSFNRRSDRGATGDAHWPF
GGPVAFEERGESRVFHVIDQWRYVGAASSIERAATLAADARAAGEGAGSAAPAVRRILQT
HLARGLQLIPIPLAGAAPAAA*
>GCA_000452945.1_BPMSHR5858_genomic_03592_1 clcA
MQRPSLFPTLTRRTQRIWRQYGVFWLGAVVVGLTAVMYARLIDWGYEAFRSLGQGRPWLP
FLLTPAIAALSVWLTRRFFRGAEGSGIPQVIATLHAQPSAFGSRLLTLRILFGKVLISLL
GIAGGFTIGREGPTVQIGASLMFNLRRFYPRSNALIERQLVLAGAAAGLSAAFNTPLAGV
VFAIEELSRSFSARASGVLITAIIIAGVIALGLNGNYTYFGTIEIGQHFPKMLAAAVLLT
ALVTGVAGGLFGWLLLNTGRWLPARLLALYRERPIVFAAACGLAIAAVGVASGGTTFGSG
YAEARGLLDGREQLSVLYPFLKMISMVASYLPGIPGGIFAPSLSIGAGFGNLLHFVFSGM
HLPMLIALAMVGYLAAVTQSPITSFVIVMEMINGHALVISLMATALIASRVSRLFVPPLY
ETLAERYLKPPQPKQAATAPAHTGIAAVKMPEDAESAPADPLDTLRDEAANEANTNERDA
GDAGGHTGAGAAAHADDAPCGAADAARAGAPPDASPASAPGAAGASNTRANDPTPPSGDD
ENREPPQRPAQ*
>GCA_000452945.1_BPMSHR5858_genomic_03636_1 group_2996
MSLSATPRIGFIGAGRLARCVARRFARAGYDVVAVASRSSASAAALAAQIDAERAARQAG
SNPAPHSALNSVPNITPDVAANGAPNASSKTASKVPSSAASQAASSVAPASAPDAGRSRA
RCAALDSPQAVVDAADLIFVTTPDDALGRIAAELRFAPARAGEQALVHCSGASGVDLLDP
ARAQGAATGGFHPLYLFGGDDADLARIDGCSVTIEADGALKDALMALAAALGCHPLSIPA
GGRMLYHAAANYAASFALCNLAECVELWRSLGFAEDDALRALLPMLAGTIETARDKGLAN
ALAGPVSRGDVGIVERQLALLESRGGDHAAFYAFHTRRAVALARRRASPPPSLDALERAL
DASLARSLDLARPACDEP*
>GCA_000452945.1_BPMSHR5858_genomic_03698_1 thiD_1
MSSNTPPIVLTFGLSDPTGGSGLQADLMTLASMGCHGVSVLTGYTVRDSAACDEVTGLDP
DTVAAQARMLLEDMPVAAFKIGAATRAEVVSAIAEVVADYDGVPLVLAPDFTLDDEHVLA
ADDLRESIADLLAPQTTLLVADHATLIALAQPDGDAEAPNLDAALSHLLGQGCEYILATE
TGSHRLVNTLYGEEGQIRQDLWERTPHRLMGITDTLGAAIAALLANGQEPPEAVREAQEY
LYQAARDAFRPGMGAYLPDRFFWARSNEDESPAPAGIAKRTN*
>GCA_000452945.1_BPMSHR5858_genomic_03731_1 yhhS
MSADSADSVPSPRSAFATTLQIVSVVSFTFICYLTIGLPLAVLPGFVHDELGFSAIVAGA
AISVQYFATLASRPLAGRCADTLGPKRTVLRGLAACGASGALLLSAFAFARWPAASIVLL
VASRLVLGIGESLVGTGAILWGIGRVGTAHNARVISWNGIATYGALAIGAPVGVAISHAL
IPAVLGMLVIALAALGYYLARLITPVPLVHGERMSYASVLTRVLPHGLGLALGSAGFGSI
ATFITLYYAARHWPNAALSLTVFGTLFIGARLLFANTIKTHGGFRVAIVSFAFECAGLLM
LWLAPVPHVALVGAALTGFGFALIFPALGVEAVALVPPASRGAALSAYSVFLDLSLGITG
PLAGYVAGAFGYPQVFLCAAVAAAAGVALSTVLYQRQARLSGSGAAA*
>GCA_000452945.1_BPMSHR5858_genomic_03763_1 ybgC
MIEPSRQPDAAPDFAWPVRVYYEDTDAGGVVYYANYLKFFERARTEWLRACGIDQSRIAD
ETGAIFIVRHAALDYRAPARLDDALQIVCRIARIGRASVEFEQQAWRGNTLLVAADIRIG
CVDRSAFKPTAIPPPVFAALERGPAASDRASRVNGACVSAV*
>GCA_000452945.1_BPMSHR5858_genomic_03852_1 group_7560
MEPLTLVAGPPLSAEDLDTLRRAKLALESPSLTIKLTSVVGAPVEKMIGKLPGFATDKIN
DATQLALRKCLHIALRTLGKSGGVASAGETPGKPSNLLHKLAVATTGAAGGAFGLFALPV
ELPVTTTLMFRSICDIARSEGEDLSSAETQLQCLAVLGMGGGFGPLPGAGKDDGQADGDA
GGERDKKEEDADIGYFVLRGALAQAVSKASSEIASKGFATHGSAALLRLVQSIASRFSVQ
VTEQIAAKSIPAIGAVLGATVNTLFIDHFQQAAHGHFAVRRLERKYGQPAVKAAYQAIDA
SAVR*
>GCA_000452945.1_BPMSHR5858_genomic_03888_1 group_207
MALPQSYRELTGEVATQMRALRKTQPDLMSAFAALAAAGTKEGALGKKTRELVALGIAIA
SRCDDCIGFHVQTLVKLGTTREEFEDLLATAVYMGGGPSMMYAAHALRAVEEFGGE*
>GCA_000452945.1_BPMSHR5858_genomic_03954_1 puuA_2
MQDIDEFLKKHRITEIEAIIPDMAGIARGKIIPRSKFETGESMRLPQAVMIQTVTGDYPE
DGSLTGVTDPDMVCLPDASTIRLIPWAIDPTAQVIHDCVHFDGTPVEISPRRVLRRVLEL
YKAKGWKPVIAPELEFYLVDMNKDPDLPLAPPIGRTGRPETGRQSYSIEAVNEFDPLFED
IYEYCEVQDLEVDTLIHEVGAAQMEINFLHGDPLNLADRVFLFKRTVREAALRHHMYATF
MSKPMENEPGSAMHVHQSIVDEETGRNLFTGDDGGPTDMFYAYIAGLQKYTPALMPIFAP
YINSYRRLSRFMAAPINVQWGYDNRTVGFRVPHSSPVSRRVENRIPGVDCNPYLAIAGTL
AAGYLGMTRRLEPTEPLASDGYSLPYQLPRNLEEGLTLMSACEPLAEILGEKFVKAYLAL
KETEYEAFFRVISSWERKHLLLHV*
>GCA_000452945.1_BPMSHR5858_genomic_03965_1 group_2508
MKILGAGIAVPGGAAHALEFANDYALKAARQALAKARCLPSELDLIVSLSVSPNRMADAA
PIAGPRLAHPVQRDLRASNAAVFDLLDADWTLALDLVQSHCRQLGYRRALVVKAEALADV
EGAQAGGYADGAGAIVLAPGRGDRYHASYADLDAPSLATLAGVGARHAHETGAVARFEGG
FDPAAGSFRRVPRNAGAAMREVVGDVRSKVDAPISELFRESWLRGWLEDEAGFAGEPRVD
VIDGDAGVPAAFQLPAWLAARAGGGAAAGVRGDAQVVAALTLDAFKPRIACIAMEV*
>GCA_000452945.1_BPMSHR5858_genomic_04194_1 pchR
MTSISLASSRRKPAVLMPGNHPGVVHVDAGMKLLSGTLCSESRDWYEEPLERGLKLVLVQ
SGQLRCRVPGQPEQCIKGPGLCLIANDGEFTTQQIYSWDAPLRYTIVQLGIDALDRGLSL
LPEKLLAPTGGDPRIVSCPASKALQALAAQIATCPFEGAVRDYYLTGKAFELTALSAQLV
STQRAPTSIDVRITSSDIERVYAASGILTRDLQAPPTLDALCARVGMNPRKLTAGFRKVF
GASVYAYLQEYRLRTAHEMLCAEDANVSTVAYRVGYSPAHFSIAFRKRYGISPSDIRSAP
NAHGDADPACAAI*
>GCA_000452945.1_BPMSHR5858_genomic_04200_1 group_1522
MRARQPNASIDAGKRDIGRARVEVDKEAAVKNEYADYMKRNEFRDM*
>GCA_000452945.1_BPMSHR5858_genomic_04248_1 group_617
MQLKQGDLIAAADAIDAWRAAEPASADALACRAHWLRLLGRFDEAAAALEPALAATPPCA
AAWAERARLDRLAGQAERAHAAFDAAHRADPAATAWLAEWIELLHPLHRPALALPVAQAL
CEHAPDSAQSWFLLGLTHHYAGDYAAAAAAYRRADALDPAYPMLRNNLAALRYQTGMTAE
ALALAEAAIRAEPDNQMAWCNCSNAWLALREPARALIAGERACALGPNYAIAQLARANAL
KELQRWPDALAAAAHAHRSAPDDPVMQWSLAMLQLLHGDYANGWANHEARWNGSRELGDR
PRPSPQQQWRGEPLAGKTLMLWGEQGFGDALQFARFAPIIAEQATRAGAQVVFACFAGLE
PLFARSFAGAPMRIVRHDAPQLPAFDHHLPVGSAPLLLGVRPDTIPAAGGYLRADPARAA
QWAARRPADGRLRVGLVWSGSRTHQRNPLRAIDPAACAHAWRDLTGVALHSLQIDGAADV
ATMRAAGLDVIDHTAELPSFDDTAAYLSSLDLVVTVCTSVAHLAGALGRPTRLLLDVNPH
WVWMIDREDSPWYGSLRLYRQPRYRDWTTVLDRVRDELAALAAARA*
>GCA_000452945.1_BPMSHR5858_genomic_04252_1 group_7585
MTPPVDLNGLRIGITIGLREPNESLWINGIKQNALFLAKLLMKSSRGYRVTIVNTTDVPI
TDALPWDRRIFDTRSFADVKDSLDVLIELGGQIDGEQTAYLKARGVKLVSYCCGFEYIHA
TQSILFGRRMWDTLFINRGYDEIWAIPQIAPSSLPFLQSLRRSPGRVVPFVWDPMFLGER
TRLLPERGEYRPSGKRAKRLTVMEPNHDVVKFCVYPMLIIDEAYRREPDAIAFAHVTNAE
RLAHDSPEFVMLMNYLEIVRASKASFVGRFDTPTFLAEHTDVVVSHQWENPLNYFYFDVC
WQGYPLVHNAHLVPDIGYYYPDNDVQTGAEVLMRVLREHDDDWEGYAARQRSLLRRYTSK
NPTLVAEYDTLLANLVNDRR*
>GCA_000452945.1_BPMSHR5858_genomic_04256_1 mmgC_4
MSNLPGVNFMLGEDIEMLRDSIAHFAAKEIAPRAAEIDRTDQFPMDLWRKFGELGVLGMT
VAEEYGGANLGYTAHMVAMEEISRASASVGLSYGAHSNLCVNQIHRNGTDAQKRRYLPKL
ISGEHIGALAMSEPNAGSDVVGMKLRAEPRGARYVLNGTKMWITNGPDCDTLVVYAKTEP
EAGARGITAFIVEKGMKGFSVAQKLDKLGMRGSHTGELVFEDVEVPEENILGELNGGVKV
LMSGLDYERAVLAGGPTGIMAACLDAVVPYVHDRKQFGQPIGEFQLIQGKIADMYTTFQA
CRAYLYAVGRHLDAGGSAHPRQVRKDCAGVILYTAEKATWMAGEAIQILGGNGYINEYPV
GRLWRDAKLYEIGAGTSEIRRMLIGRELFAETA*
>GCA_000452945.1_BPMSHR5858_genomic_04264_1 group_1909
MTARQLTPALASRFANLALSHLTREYPNKLTHSLAGPQDVLGPRALHPIFYGSYDWHSCV
HGYWLVTRLLDRFPDLPEGPRIVATVDAHFTAGNVAGEAAYLNLPHNRGFERPYGWGWLL
ALSAQLASMKSDAGRRWSATLAPLTDLFVERFAEFLPKATYPLRVGTHFNTAFALSLALD
FARQTGHAALEALIVDTARRWHLRDANCQAWEPSGDEFLSPALMEAELMRRVLPAAEFLA
WFDAFLPSLAAREPATLFTPATVTDRTDGKIAHLDGLNLSRAWCQRALARALPDGDPRRA
ALADAADAHLASALEHVAGDYMGEHWLASFALLALEA*
>GCA_000452945.1_BPMSHR5858_genomic_04266_1 group_7587
VSDNAYTNAFGQPIGAPVPGWQPRPAPPRTPAQGRFCRIEPVDPGRHADDLHAAYATAAD
GRDWTYMSAGPFADAESYRAYLRKAAETDDPLHHAIVDLATGKAVGTFALMRADRANGAI
EVGFVAYSPLLQKTCAGTEALFLLMRRVFDELGYRRFEWKCDSLNAPSRAAALRYGFTYE
GLFRQAVVYKGRSRDTAWYSIVDGEWPALRSAFEQWLAAGNFDAHGRQRQSLSTLIAAAR
RR*
>GCA_000452945.1_BPMSHR5858_genomic_04268_1 qseB_1
MRVLLVEDDPWLGEAVRAQIEKDGHPTDCVASLAHARRHVAVAPYDLILLDLLLPDGRGL
DFLRELRAGGSTVPVIILTALDQVANRIAGLNAGADDYLIKPFDLSELSARVSAVARRYS
GNPNPLVKLGELAVDIAARSVTRAGARVDLTASEWALLETLIQHPGMILSKPQLEERLYS
FSDEVGSNTIEVHVSRLRKKLGAELIETVRGLGYRLRRD*
>GCA_000452945.1_BPMSHR5858_genomic_04269_1 group_1910
MKTSRLIKAVALAALASLVLGGALGIRAAMADDGDDCRAPLADWKPRDAVRALAQQKGWR
VDKLKADDGCYEIKGHDADGKRFKAKLDPVTLDVVRMKREGERKRDHDDDDDHGRAPDAR
APAGGPPAGAPPGGVLKPGSKPDVQIR*
>GCA_000452945.1_BPMSHR5858_genomic_04291_1 mnmE
MLATDSDPIVAIATASGRGGIGVVRLSLGRAGEAAARALSDALCGARLMPRHASYVPFLD
GAGEPLDRGIALYFPAPHSYTGEHVIELQGHGGPIVLQLLLQRCLDAGRAHGLRLAEPGE
FTRRAFLNDKLDLAQAEAVADLIEASTEAAARSAGRSLDGAFSRDIHALVDDVIALRMLV
EATLDFPEEEIDFLEAADARGKLAHIRERLAHVLGDARQGALLREGLSVVLAGQPNVGKS
SLLNALAGAELAIVTPIAGTTRDKVAQTIQIEGIPLHIIDTAGLRETEDEVEKIGIARTW
GEIERADVVLHLLDARSGLGPGDEAIAARFPDGVPVVRVLNKTDLTGAPASVTRTGGGAA
RADVCEVRLSAKRGDGIDLLRGELLRIAGWQAGAESVYLARERHLIALRAAQAHLARAAE
HAEQNAQALDLFAEELRLAQERLNSITGEFTSDDLLGVIFSRFCIGK*
>GCA_000452945.1_BPMSHR5858_genomic_04326_1 group_3989
MQDPQPDALPPEPFVAPAADGFPVRGFAWRHRAPAAGRPVTVINCATSVRCRYYFRFAAY
LFSQGSDVLVYDYRGIGESRPASLAGFHATWLDWGRLDCDAVLQYAARTCAGQPVDVVAH
SVGGVVLGLAASNPIVRRALTVGAQYAYWRDYAGSHRLRMLAKWHVAMPMLARVFGYVPA
KRLGWMEDTPRGVALSWSRSRPRFEDAYVRAPIRETPDARRDLVERFTRLTAPMLAIGLS
DDEFGSVEAVERLLGYYTRSAVTHLRIAPEQIGVASIGHFAFFHSRFEQTLWPIALGWLK
TGALAPETPGRVHRQPCAPEPARAARDEASGRTAAR*
>GCA_000452945.1_BPMSHR5858_genomic_04385_1 scoA
MNKVYPSAAAALEGVVHDGQTFAVGGFGLCGIPEALIAALRDSAVKGITCISNNAGVDGF
GLGLLLETRQIKKMISSYVGENKEFERQYLSGELELEFTPQGTLAEKLRAGGAGIPAFFT
QTGYGTVIAEGKETREFDGKHYVLEPSLTADVALVKAWKADKSGNLVYRRTARNFNPMCA
MAGRITIAEVEEIVENGELDPDAIHTPGIFVQRLVLNATPEKRIEQRVVRAKGD*
>GCA_000452945.1_BPMSHR5858_genomic_04450_1 group_5037
VSQKRGLAVFFGCVIALVVAYAATYPRFHPQPAAVAAASAPVIASAAIALSSGEFPPLPP
SSPLSWGRLTPQQHVALAPFASQWDSFSDERKRKWLKIAARFPRMSPEAQKRLQERMTEW
VRMTPEQRRVARENYLVSKDLSAQAREKAWKAYQQLSPEQKEKLAAAERRRRPTVVSAPP
TGKTDRDINRLVNAHDRHPASAPTASIAPSAASAPAAAPVPASPATTGAVTPPTAPTPVS
PSEAPSLFNGS*
>GCA_000452945.1_BPMSHR5858_genomic_04455_1 group_7600
MEAKPPRRTRERILELSLKLFNEIGEPNVTTTTIAEEMEISPGNLYYHFRNKDDIINSIF
AQFEQQIERRLRFPEDHRPTIDETWSYLQYMADFMWTYRFLYRDLNDLLARNRTLETHFK
QIISHKVRFAREMCELLAADGELVATPAEIEVIATNMAVIATYWLSYQYVMHPRKYNDQD
AIREELHQVSMHVISVIAPYLRGRSRQIFDDLVSGKLPKREFHDYLPPRDGSPRKDTKQ*
>GCA_000452945.1_BPMSHR5858_genomic_04461_1 group_3537
MTAPHASDGTREAAARERAALDSRAREVLDFWFGAPDDPAFGTARPMWFGGAPALDAQLR
ERFGPLADAAKAGELDAWAGTPLGALALIVVLDQFPRNIHRGTPLAFAADRAALAHAKAL
VASGGDRALPTGHHRAFAYLPFEHDESPDSQREAVRLCAQIKDEAGCAGYHDYALRHAVV
IERFGRFPHRNAILGRASTDAEAAFLKEPGSSF*
>GCA_000452945.1_BPMSHR5858_genomic_04462_1 alaS_2
MATQALFREDAYLTRCDALVTAVDEAGIHLDRTVFYPLGGGQAGDSGTLTLAGGAALAVA
DTRKAKFDGATPDDAVHVLAPGQAALAASVKPGDALVAEIDWAPRHRRMRLHTASHLLCA
VLPYPVDGCSVTADYVRIDFATVEPIDREHVQARLAELVGGAHPVTTEWITDDEMVARPE
LVRTMSVKPPMGLGRVRLLRIEGVDLQPCGGTHVRNTAEIGALRVAKLEKKSARTRRLVL
ELA*
>GCA_000452945.1_BPMSHR5858_genomic_04464_1 group_1317
LTAAGKARPAGRRFGLVDLLRQAGRMTLRDWRAGELTLVALALVLAVAALTSVGFLADRL
RQGFERDARRMLGADFVVRADHPVDPAFAHEARTLGLATATTAVFPSMVGAAKPADSADS
AGSAKAGTPAPSAAADGAGSSPPVRLAAVKAVSAGYPLRGAVEIAPEANAPGRAAGSIPA
SGTVWVDPSLLDALHLKVGDALRVGGRTFTIGAAITRELDRGFSFVNFSPRVMLRADELE
STGLTGYGSRVTYRLLVAGDDAAVERFARYAHARVDGGKLRGVALESLQEGQPQVRQTLD
RARHFLTLVALLTALLAAVAIAMAAQRYMRRHLDGCAAMRCLGASRRTLSALFAIEFALL
GLAGGAAGAALGYVGHLVLLRALGSLIDVVLPPPSAWPAAIGVGTGLVLLLGFALPPLAP
LTRVPPVRVLRREWGDAGRVAWVGYAMGIALFAALLIAAAGNLTLGAIVAGGFAGSLVLF
ALIARLALVALARLVRDGRVAAGLGWRHALASLERRGAASALQITALALGLMCLLLIAIT
RNDLVAGWRQSTPPDAPNQFLIDIQPDQRDDVVRYLAANGIADPALSPMVRGRLVAVNGK
PVNPDDYKRDDARRLVDREFNLSYTTRLPEDNRLVAGDWFGASGVPQISIEAGLAKTLGV
KTGDTLRFDVTGLTVDAPVTSVRKLDWGTFRVNFFVLMPPAALRDFPATYITSFHLPAAR
ERVLDGLIARYPNLTAIDVGPILAQLERVLLQVIGAVQFLFGFTLAAGVLVLYTALAGSR
DERVREAALMRALGASRKQVVAVQRAEFVVVGVLAGAMAAAGALVVGAVLASRVFDFQLA
PDPWLVPAGIAAGVACAGAAGWLGLRGVLRRPALQSLRDA*
>GCA_000452945.1_BPMSHR5858_genomic_04465_1 group_7603
MVEAISLGAGLAWASGLRLYLTVLIAGVFARAGFIHLPDTLAVLTSPWVIGSAAVLAVAE
FLADKVPAFDSLWDAVHTFIRIPAGAVLAAASLGHADPALLVVAGLAGGTLAGSAHIAKA
GTRALINLSPEPISNWIASTTEDGLTVGGLVLAFFVPLLFLALMIGFLAFSAWALPRLWR
GVSGGFRGMANHMVSRLNSIGSKRD*
>GCA_000452945.1_BPMSHR5858_genomic_04466_1 gdx
MPWLWLLVAGLLEVAWAAGLKSSEGFTRLGPSVFTIVTALGSFGLLAVAMRELPLGTAYA
VWTGIGAVGAFIFGIVMLGEAVTLARIASASLIVAGLVGLKLSSAG*
>GCA_000452945.1_BPMSHR5858_genomic_04470_1 kdpC
MKSLFRPLIVVFVVLVAVTGLAYPAVMTVFGQAVFPAQANGSLIEKGGRVVGSALIGQQF
DAPQYFWGRLSATSPMPYNAAGSGGSNLGPLNPALKDQVKSRLDALKAAGTDLSQPVPVD
LVTASASGLDPEISPAAADYQVARVARARKMADADVRRLVADHTSGRQFGVLGEPRVNVL
KLNLALDAAQAAH*
>GCA_000452945.1_BPMSHR5858_genomic_04471_1 kdpB_2
MTQHSATRSMFDPALVRPAIVDSFKKLTPRTQLRNPVMFCVYVGSILATILWIAALTGQA
EAPAGFILAVALWLWFTVLFANFAEAIAEGRSKAQAASLRSAKKDVMAKKLNEPHPKSPV
RIMTASDLRRGDVVLVEAGDTIPADGEVIEGVASVDESAITGESAPVIRESGGDFSSVTG
GTRVLSDWIVVKVTANPGEAFLDRMIAMVEGAKRKKTPNEIALTILLVALTIVMLLATAT
LLPFSMFSVDAMHSGYVVTITALVALLVCLIPTTIGGLLSAIGVAGMSRMMQANVIATSG
RAVEAAGDVDVLLLDKTGTITLGNRQASAFLPAPEVTEATLADAAQLSSLADETPEGRSI
VVLAKERFNIRQRDMGALHAVFLSFSAHTRMSGVDLSPEREIRKGAADAVKKYVESHGGR
FPAEVTNAVTDVARRGSTPLVVAEKRGALARVLGVIELKDIVKGGIKERFAELRKMGIKT
VMVTGDNRLTAAAIAAEAGVDDFLAEATPETKLSTIREHQAAGRLVAMTGDGTNDAPALA
QADVAVAMNTGTQAAKEAGNMVDLDSNPTKLIEIVEIGKQMLMTRGSLTTFSIANDIAKY
FAIIPAAFATTYPQLNALNVMHLATPASAIMSAVIFNALIIVLLIPLALKGVKYRPLGAA
TLLRRNLLVYGLGGILVPFVGIKLIDVVLAALGWV*
>GCA_000452945.1_BPMSHR5858_genomic_04472_1 kdpA
MNANNLLQAAIFIVVLIAAAVPVARYLTRVMDGSSAVVRVFGPLERALYRLAGIDPRTEM
SWKQYALATVAFNALGVLFLYALLRVQGWLPGNPQGFGPMTVDGALNTAVSFVTNTNWQD
YTPEQTVSYLAQMLGLTVQNFLSAATGIVVVLALIRGFARHTAQTIGNFWVDVTRVTLYV
LVPMAAIIAALLMSQGVIQNTKAYQDVPTLQTTSYAAPRLDAQGNPVKDAKGNPVTVQTS
VKAQTLAMGPVASQEAIKMLGTNGGGFFNGNSSHPYENPTPFSNFLEIFAILIIPAALCL
VFGNTIGDRRQGVAVLAAMTVALAAAIGIETSAEQGGTPVLASLNVDQAASPLQAGGNME
GKETRFGIAQTGLFVVATTAASCGAVDAMHDSLTPVGGLVPMLLMQLGEVIFGGVGSGLY
GMLVFALLAVFVAGLMIGRTPEYVGKKIESYEMKMVSIVVLLTPLLVLVGTSIAVLADAG
RAGIANPGPHGFSEILYAFSSAANNNGSAFGGLSVNTPFYNWMTAIAMWFGRFGTIVPVL
AIAGSLAAKKRIAATSGTLPTHGPLFVVLLLGTVLLVGALTYMPALALGPGVEHLMLFVG
AH*
>GCA_000452945.1_BPMSHR5858_genomic_04473_1 group_5040
MSWMLWLAGASTFLLFAYLVYALLRAEDIE*
>GCA_000452945.1_BPMSHR5858_genomic_04476_1 mgsA
MSTPRIALIAHDAKKDDIVALAGAYRATLAQCRLVATGTTGGRIAQAHGLDVERKLSGPL

GGDLQIGAELADGRVDIVIFLRDPMTAQPHDPDITALVRACDVHDVPVATNVATARVLLD
DLARRLTANA*
>GCA_000452945.1_BPMSHR5858_genomic_04477_1 fabG_4
MDLGIAGRTALVCAASKGLGRGCAQALAAEGARLVIVARTRETLEAAAADIRAATGADVT
AVACDITTPEGRAAALAACPQPDILVTNAGGPPPGDFRDFTHDDWIRALEANMLTPIELI
RASVDGMIARRFGRVVNITSSAVKAPIDVLALSNGARSGLTGFVAGLARKVAEHNVTINN
LLPGLFDTDRIATTLAAAAQAQGATVDELRARRTKEIPAKRLGTPDEFGAACAFLCSVHA
GYITGQNWLLDGGAYPGTF*
>GCA_000452945.1_BPMSHR5858_genomic_04483_1 rsfS
MDIRKLQRVIVDALEDVKAQDIKVFNTSHLTELFDRVIVASGTSNRQTKALASNVRDKVK
EAGGDIVSSEGEDTGEWVLVDCGDAVVHILQPALRQYYNLEEIWGDKPVRMKLGGPAAHA
RGARAADGADDDESDEHEAPAARPARKATAARRR*
>GCA_000452945.1_BPMSHR5858_genomic_04491_1 group_7607
MRPEFSRSIWRAVAVGVTTAGLAMAAASCSKQSDSNSGSSSTGGATGGVQPAPMAPSGSS
GTAPGGASSGG*
>GCA_000452945.1_BPMSHR5858_genomic_04509_1 gluQ
MTRYRGRFAPSPTGPLHFGSLVGALASWLDARAWGGAWLVRIEDIDGPRTVPGAAEDMLA
TLRGFGFIADEPPVWQSARVAHYEAALARLTAAGLVYPCGCSRKEIADSLRAAHERHTTL
AYPGTCRTGLHGKPARAWRLRVPDGAAAVVAFDDRWQRAQTQNLATEVGDFVLKRADGQW
AYQLAVVVDDGDANITHVVRGADLLDSTARQIYLQRCLGLPTPRYLHVPVVLDANGEKLS
KQTGAAALDPAAPLPALAAAARHLGLALDGAACASLDAFQAAAIAAWDARFGPNARG*
>GCA_000452945.1_BPMSHR5858_genomic_04521_1 group_2541
LKLITAIRGAALAAALATAAGLAFAQSSSARAAAAPGAADAAPAAAAAQADAPPPAVTLK
PNPAYARLPRYEGTVGGRPVVVRLGPKTDEPGVHGECQYLDTGAVILLAGDRDGDTLEIE
ESDDGTNITGVWIGRFDPSGGLSADRMNADQSDPLPVALHPAGAQSASAAVGAPQAGARG
QSVGGVSNLRTGE*
>GCA_000452945.1_BPMSHR5858_genomic_04571_1 moeB
MNDEQLLRYSRHILVDEIGIEAQQRFLDAHAIVVGAGGLGSPAAMYLAAAGVGTITLVDA
DTVDLTNLQRQILHATDSVGRKKVESGRDALARLNPDVKVNAVAERVDDAWLNAHVPHAS
VVLDCTDNFATRHAINRACVAHRVPLVSGAALRFDGQISTFDFREPGSPCYACVFPEDQP
FEEVACSTMGVFAPTVGIIGAMQAAEALRVIGGIGATLVGRLMMLDSLRMEWNTMKIARQ
PDCPVCGAGGAH*
>GCA_000452945.1_BPMSHR5858_genomic_04572_1 ptsI
VSFTLHGIPVSRGIAIGRAYLIAPAALDVAHYLIEAERIEAEIERFRTALGAVRRELDVL
RADLTDDTPTEVAAFIDVHAMILGDAMLVQETIDLIRTRRYNVEWALTEQLDVLAGHFDD
IEDEYLRERKADIEQVVERVLKALAGAPSAAQALDRAAGNGRDEMIVVAHDIAPADMMQF
KTQSFQAFVTDLGGRTSHTAIVARSLGIPAAVGVQHASALIRQDDLIIVDGDQGIVIVDP
APIVLEEYSYRQSEKALEQRKLQRLKFSPAQTLCGTKIDLLANIELPDDAKAAVDAGAVG
VGLFRTEFLFMSKVRMPEEEEQFAAYKRAVELMHGMPVTIRTIDVGADKPLDVHDEGYET
APNPALGLRAIRWSLSEPQMFLTQLRAILRASAFGQVKILVPMLAHAQEIDQTLDLINEA
KRQLDAAGLAYDPNVRVGAMIEIPAAAIALPLFLKRVDFLSIGTNDLIQYTLAIDRADNA
VAHLYDPLHPAVLHLIAFTLREAKRAGVPVSVCGEMAGDPALTRLLLGMGLTEFSMHPSQ
LLVVKQEILRAHLKALEKPTADVLASFEPEEVQAALARLASAEPRADVAA*
>GCA_000452945.1_BPMSHR5858_genomic_04611_1 udg
MAWTEAALEELGLAPAWVRRDAARAADVNEAAAHAAGEAAVAAVARGAQPASPDAPREPA
RRIARDSGRDGGGDGGQDGGQDGASGDAGSASPNDAAHAMARVAGAEASAADAVADEHAA
LARQAGDGRGQTRQAAAESGARAAADAPAPAAAPASRTRDATIARGASPDEPGVAGVVGV
VGVADEPSVAGGARRARTSGGGADVPARALDWTDAAEATRPAAAPAPAALTGGDAGAAAS
DEDMSWFDLEPVHEPVLPDVAARPAATTPSVAELGWDELRARVADCERCRLCEKRTNTVF
GVGDEHADWMLVGEAPGENEDKQGEPFVGQAGKLLDNMLRALALKRGENVYIANVIKCRP
PGNRNPEPDEVARCEPYLQRQVALVKPKLIVALGRFAAQTLLKTDGSIASMRGRVHQYEG
VPVIVTYHPAYLLRSLQDKAKAWSDLCLANDTYRSAAPAADPP*
>GCA_000452945.1_BPMSHR5858_genomic_04612_1 rimI
MSGVLMTDRYLAPMTDADLDEVVEIERAAYEFPWSRGNFEDSLRNGYFGVCMRHVTGVLV
GYCVLMPVVDEMHLLNLCVAPVAQRSGVGLALLREAVRIARAERLDGVLLEVRPSNPRAI
RLYERFGFVSVGRRRNYYPAKHRSREDAIVMRLALTKEGGAHGMD*
>GCA_000452945.1_BPMSHR5858_genomic_04615_1 rhlE_2
MTSSHSLSPLNAIADEALGLPAAAAQPAASDACDAASDEPTFASLGLSPEIVSALQAAGY
VKPTPVQQRAIPAGIAGRDLLVSSPTGSGKTAAFMLPAIERFAQLQKAQAQQPRAPREGQ
PGERRGRRPQPVARPGLLVLTPTRELAMQVTTAASTYGKHLRRLRTVSILGGVAYGQQLM
LLAKNPEILVATPGRLLDHLERGRIDLSELKMLVLDEADRMLDMGFIDDIDTIVAATPAT
RQTMLFSATLDGKIGSLTNRLLKDPERIEITQKIESRSNIAQTVHYVDDRDHKDRLLDHL
LRDVALDQAIIFTATKIDADQLAGRLADAGFQSAALHGDLPQGARNRTIRALRERRVRVL
VATDVAARGIDIPGITHVFNYDLPKFAEDYVHRIGRTGRAGRSGTAVSLVHHAEQGALKR
IERFVRAPLPVNVVEGFEPRKAPPPRNDRGNGRGRPGGNGNGGRRFGGKPGGGHGGGRSG
GWSGKPGGGNREAYGARRGDGQRNGPRRGNSAA*
>GCA_000452945.1_BPMSHR5858_genomic_04642_1 group_2555
MTFVVMEGCIRCKHTDCVAVCPVDCFHEGPNFLVIDPDECIDCALCEPECPIDAIRAAAE
LPDDQRHFVALNAELARHPNWPRIIGKKPALPDYAAWADVQGKLAQLDRNGA*
>GCA_000452945.1_BPMSHR5858_genomic_04663_1 thcB
MTTHSASTPPALPAAQCPFRPVSDDADAHPAGCPASAGAAAFDPFDDAYQQDPPEYVRWA
REREPVFYSPKLGYWVVARYDDIKTIFRDNRTFSPSIALEKITPTGPDANAALASYGYAM
GRTLVNEDEPAHMARRRLLIEPFTPDALRAHEPMVRRLAREYVDRFVDDGRADLVDQMLW
EVPLTVALHFLGVPEADMDTLRAYSIAHTVNTWGRPRPDEQVAVAHAVGQFWQYAGKVLE
QMREDPSGPGWMQFGIRMQREHPDIVTDSYLHSMMMAGIVAAHETTANATANALKLLLEH
PRAWRELCDDPALIPNAVDECLRHSGSIVAWRRIATRDATVGGVAIPAGAKLLIVTSSAN
RDERRFADPDLFDIHRDNPADHLTFGYGSHQCMGKHLARMEMQIFIDELSRRLPHMRLAE
QRFTYVPNTSFRGPEHLWVEWAPAENPQRRGHGARRCALAVRIDESNARRAGRALRVDSV
MPVAERVLRVRLVAPDGAPLPRWSAGAHIDVECGDTGLSRQYSLCGDPAERGVYEIAVLR
EPASRGGSAWLHERLRPGMRVKIRGPRNHFRLDGHAARAILIAGGIGITPIGAIAREARA
RGIDYALHYCGCSRRTMAFVDELAALHGERLRLHVSDEGTRHDFSALDVAAGTQVYACGP
ARMLDALAACAAHWPAHALRVEHFAAAAPRVDPRTDQPFEVELKDSGLTLTVPANRTLLA
TLRAANVDVQSDCEEGLCGSCEVRVLAGEVDHRDSVLSADERALNTRMMACCSRARHGRL
VLEL*
>GCA_000452945.1_BPMSHR5858_genomic_04664_1 pcaK_4
MTARVDVRARLDDSRFDAYHWRIVAPCAAYRVMDGFDVQATGGVAPAVIREWGIAKEALS
PAFRAGWFGMLIGSLVFSALAAVFAAIFMAAFPIVGGQPALNALAAVYYPAPLRSTGIGW
RSGVGWIGSVVGRRAAAC*
>GCA_000452945.1_BPMSHR5858_genomic_04666_1 group_3561
MAFKILSCDGGGIRGLITALLIQDLDRRSGILAKADGFAGTSTGGLIALALARGVSISEV
VDVYRNRGSEIFQESGAWLEQRATIATDASFAAFAGPGLFACQYVNTGLKRIAQELLRGG
DLTELHRLTVINSSRLWDPALRSWSACTFSNASGNAYRHVALVDAALATSAAPSYFPPYE
VPSLGYFADGGLFANNPSMTAVSEALASRLGGELGNLRVLSFGTGISQAGIRPSAIGDPL
RWGVTTWFWPFESHDVPASALLGLTLDTTAMLATQQATQLLGGGYRRANFVLGETVGLDD
WRKASLLESETAAYMKTPAWEGVCQWVDANWR*
>GCA_000452945.1_BPMSHR5858_genomic_04667_1 group_3562
MSADEIRTRRRKLLAAALGAALAPALPALAATQRAQPLAGQSGYHGGLITPPVPVPDMPL
RTADGRATRLRALLAGRVTALQLFYTGCSSTCPIQGAVFHRVQTLLGPHPKPDIQLLSLS
ISPLEDTPQRMHAWLARFGARPGWIAAAPELKDVDALQAFFGGGRTGLDNHGTQVQMIDR
RGALVWRTYELPSPETIAALLGHA*
>GCA_000452945.1_BPMSHR5858_genomic_04668_1 group_2849
MPGEPNARRASLTGLALKLAFGFGFAFALAWPGHAARADDSAARGRALFRGDAPLQGRLP
MHPGDLPAPLVRCANCHAAGAGAAVPNSIAPRLTRAWLGDLQSRRGGPPSRYDRNAFCAL
LRTGLDPTYVLINVEMPRYRIGERDCTALWRFLNEAPHERR*
>GCA_000452945.1_BPMSHR5858_genomic_04669_1 group_2557
MHFFRFAWRHGRYWLCLLIASVLLPQGAAAAVTASYTDIVGADSGLCVSTAGNSSASGAG
VVQTSCAGLSNTTWSFVPVGNRYHIVLQGSGLCLNVPGGSLNSGTQLIQYACQGNGQTND
QWTVVAVGSSYRIVSASSGMCVNVSGASHASGAALIQYPCQGAGALNDQFNLYLPVVAAT
NVTAANSNLCVSVNGGSTAAGASIVQGTCSNQGATGWSLLPAGSGYHVVSQGTGQCLNVY
GGYTTSGAPIIQYPCQGDAQTNDQWTLVPVGSKYRLISVSSGMCLNVSGGSLSPGAPLIQ
YPCQGANALNDQFSLSLPQTFPVTLPSAWSPVIPLPVNPIGIANAPNGKLVMWSADQQLS
FQNDVGSKATQTQTAVFDPATNTATQYLETSAGSDMFCTGTAMLPDGRLLVNGGDSSPKT
TLYDWTTNTWSAAATMNIARGYQGDTLLSNGSVLTLGGSWSGGQGGKTAEVWTNGGAWTL
LPGVPETNIVGPDPQGIYRGDNHLWLFAQGNGTVFHAGPSSQMNWISTAGGGSIQSAGMR
GVDPFSINGTASLYDVGKILKAGGAKSYQQNGSVTTYASNSVYQIDITRGPNQPASVQRL
NGMTYQRAFANSVILPNGSIVMIGGQSVPMPFTDTTAIMVPEIWDPATQRFNLLKPMQTP
RTYHSTAILMADGRVFAGGGGQCGTGCAMNHLNAEILTPPYLLNADGTPAPRPVITNAPA
TAKLGATIAVSTQGPVASFVLMRLSSVTHTTNNDQRRIPLAIASSGGTSYQLAIPADPGV
VLPGYYMLFALNAQGVPSVSASIRIS*
>GCA_000452945.1_BPMSHR5858_genomic_04670_1 rssB_1
MSKRAKASSPSISIHSNGSAGEDDARPAHAAGDAPADAPVVLLVDDEPSVLSALKRVFRP
KHYRTLCADSGEAALDLLAAGEVDLIVSDMRMPRMSGAELLTRVGERQPDTMRILLTGYS
EIDSAVRAINEAGIYRYLTKPWDDHDLLLTVEQALEQQRLRRETARLDALTRKQNDALRA
FNAGLEAQVQARVEEIRQTMLFLEDAQRDLTHNFAAMAQVCANMIELRCGAPGGESQRVG
EAARDLALALGMSGWQALELFFAGLLHGIGKLSLPDELLHRPLDRLTPDESRLYYQHPLR
AQMVLTPVQQLGRVAHVIHHQYERFDGRGTPDGLAANAIPLGSRILAVARDFDALCSGSM
FTRPYTVQQAMAVLVSQSGLRYDARVVDRFVALSKEAAAVDRRASVSRINASQLAEGMRL
ADDLRNGRGILLMTKGSVVSSHQAAQVRRFEASETAPFVIFVDAAGGEPPRGMLD*
>GCA_000452945.1_BPMSHR5858_genomic_04671_1 group_3563
VTMLSSTIPAVHSTCAHARDAVREVHAALANCDAELVLFFCSSRFDLDALADEMRERFRG
TRVIGCTTAGEIGPAGYRNDSLVAVALPRALFTVETALLEDLQTFTIASGHACALDALHD
LERRAPRASGANSFALLLIDGLSVREEPVTRTLQGALGDIALVGGSAADDLRFERTAIFY
DGRFRDDCAALIVASTALPFRTFKTQHFRCGTERLVVTQADAERRTVSEINGLPAAEEYA
RLIGARVEDLSPGHFAAAPVVVLIDGTDYVRSIQKLNPDGSLTFYCAIEEGLVLRVARAL
DLVDNLQATFGDLRDSFGEPQLVLAWDCILRHLEMMQRGTRDTAAEVLKANHAVGFSTYG
EQYGGVHVNQTLTGIVFSRAPEPDRG*
>GCA_000452945.1_BPMSHR5858_genomic_04672_1 sasA_5
VAESSLVDDASADIPSLKKEIVRLNKIVRSLMDRAERSTIVCGSDFSLFQMAVTLEDQVR
HRTRELEAALHENQKIMHALQRTQALMAQEIEERKRTQAELETEREAQRHLIEQLAQAHG
QLLQSEKLASIGQLAAGVAHEINNPIGFVDSNLRTLKTWVRQLLDVMAIEDALIADCGDA
ALARLRAAHAEVDLDYLRGDIGTLIDESIEGASRVRRIVQDLRDFSRAGSEEWNFADVHE
GLEATLNVLRNELKYKAEVVKDYGELPAVECMPSQLNQVVMNLLMNAAQAIVEHGTITIR
TRREGDGVTIAIEDTGVGIPADRLAKIFDPFYTTKPVGKGTGLGLSVSYGIVEKHGGRIT
VDSEPGNGSRFTIWLPIVRQRSLQDVAAG*
>GCA_000452945.1_BPMSHR5858_genomic_04673_1 group_1533
MNEPNPYFQEVVDAHVDIERWLSGRAEHGLLPALLERFSPHFSMIAARGAALDRAGLDTL
FRQGHGQRSGLRIEIDELQQIGAWRDGAVIGYRETQIDGEGRSNTRRSTVVFERDAASRI
VWRHLHETLLGA*
>GCA_000452945.1_BPMSHR5858_genomic_04674_1 hsrA_3
MSARSRIAAVYLLGFSIDLANMFAINAAYPALQRALHASVAQLAWVGNVYMLGLTVVIPL
GAWLAHRFGERRLLTASLLVFALGSAGAASAPSIGALLASRLVQGLGGGLLIPVGQAMTY
RAYPREARARLTSIVMMVALLVPAASPALGGVIVDHASWRVVFWSMLALAASTCTLALAW
LPADVARDAPRTLDAASLGLSAAALVALLLGLAFASRRGAGAPAIAMLALACTAGVAYVR
HARRSSAPLLDLALLSQPLLRVGVVIYLCVPGIFTGVNLIASLFLQNALGLGATQVGALM
VPWAAASFAAIATTRRAFPAVGARPLLACGVVVDCIGIALLATPLAAHEAGRVLAYVAMG
FGASLCTSTSQTAAFLDVPADRMGEASALWNINRQLSFCLGVAVLGGMLNFLLAAGGGNA
PLAYRRCFAFGALLTLLPLPLVARLGSRRAAVSPAVSVDAQSNRT*
>GCA_000452945.1_BPMSHR5858_genomic_04675_1 dmlR_12
MVSLDRFDVFRAVVEAGSFTGAANVLNQARAAVSFHVKQLEAELGVTLLTRTTRRVELTE
AGERFYERCLRVLAEAEEAIDDARAEHGGMQGSLRVTSTVEYALRIVAPALSEFMSAHPA
LRVRLETHTSQADLVRDRFDVAIRLGRLEQFRDLPYRGVCIDTYDVRPVASPALLAAHGV
ARIDSPEALARLPQLGHSRLSQVASWLLHDRDGREHVYRPAAKPHLIADNASVLRAFAQQ
GGGVALLPEWLVRDDLASGTLIDALPGYRFAQQGVYALYPSARHVPQKIRAWIDFIKRYV
RRRAEG*
>GCA_000452945.1_BPMSHR5858_genomic_04676_1 group_891
MATPDVGFLARPELNALRDVDEPIVFAQAGLSGLSLFEEASYRGVRAAYRALA*
>GCA_000452945.1_BPMSHR5858_genomic_04677_1 cynR_3
MRFDITDLRLFVSVIEAGSITHGASRVNLALGSASGRIKQMEETAGVALLVRERLGVKPT
EAGHTLLRHARATLASMQRLTDDLGEFANGLRGTVRLLTNTNALTEFLPGPIARFLAQHR
NVSVELEEMLSHEIVAALVEGVADIGIVAAREDIGGLQRFPFATDRLCAVVPASQPEFAN
ARDVSFASLLDYDFVGYAAGAAIQTYLEAHARELHRRLRPRIHLRSFDAICRLVENGVGV
SVVPESAARRCRRTMSIKTLALRENWALREMRVCVADVDALPMYARQLLGHLVESGRAFA
RRR*
>GCA_000452945.1_BPMSHR5858_genomic_04678_1 mcl2
MSISIPHDAAAPAACDRLHSALRSWLFTPATRADRFSNASECGADVLIVDLEDSVAPADK
AYARDHARAWLSSGQAHARPASAVRINAPDTVAGHADLGMLIDAPRAPDFIVVPKVETQQ
RIAHLDALLRDAGRLPRYGFVAMIESAAGADAADAIVRACGSIGALMFGAADYASDVAAQ
PDALALQIARVRLAAACARGRIAAIDAPCFALRDAAALDAELAFAVRNGLRGKAAIHPAH
VAPINAAFAPSPQRIAWARRVIGAASQGAAVVDGRMVDEAVAREARGVLAAAR*
>GCA_000452945.1_BPMSHR5858_genomic_04679_1 mch
MSTTISAYRQIGDKRFREISGLYFEDFEPGNVYEHRPGRTITDVDNIWMTLLTMNTQQVH
FDAEYAAKTEWKKLLVDSTFTLALLTGMSVRTVSAKVVANLGWDKVRASAPVFAGDTLYA
ESTVLSRRESKSRPTQGIVTVETKGVNQHGVIVMSFERTMLIYKRGHSPEAGANY*
>GCA_000452945.1_BPMSHR5858_genomic_04680_1 group_3564
MRNDRDAREIYAGKLTSPQRAVSQVRSGSAIALGMAMSQPPALLAALAERAQAGDVDALR
VYYFHAEAFLRETLLRYSLMGRILPHCMFMGPPERELVKLGEADGGRKVVFFVPNSFSQS
PRLFAEHVDVDTMLVMVSPMDRNGYFTFGTNNDYTSSVARAARRLLVEVNPNMPRVFGNS
LLHVSEVDAIVESDQPLPELIPKPGGELDRVIGEQIAELVTEGACLQMGIGALPNAVCGA
LVGHRHLGIHTELLTPGLVELIRRGVVDNSRKTLDRGKSVFTFAMGDRAMYDFIHDNPAL
ESHPVDYVNDPCNIARNDNMISVNSTIEMDLTGACNSEHVNGHQYSASGGQLDFVRGAYA
SRGGKSILAFHSTAAGATVSKIVPQLTGPVTTPRTDTHLVVTEYGIANLKGLSSTERARA
LIRLAHPDFRDALSVHARRLHLI*
>GCA_000452945.1_BPMSHR5858_genomic_04681_1 poxB
MASTIADYLARTLAAAGVTKIWGVTGDSLNGLSDSLRRDGGIRWMHTRHEEVAAFAAGAE
AAQTGRLAVCAGSCGPGNLHLINGLFDCHRNHVPVLAIAAQIPSTEIGLGYFQETHPQEL
FRECSHYVELVTNASQFPRVLERAMRAAIGEHGVAVIVLPGDVALSAAPARTPSWFDFAS
PVTTPADADLERLAALLDGSDAVTLLCGSGCAGAHDEVVALADALAAPVVHALRGKQHIE
YDNPNSVGMTGLIGFSSGYHAMQSCDTLLMLGTDFPYRNFYPTRARIAQIDRRASALGRR
APLDVGLVGDVRATLSALLPRIRRKTRRGFVERALAHYAAARKGLDELARPSPAGRPIHP
QYLTRVVDELAAADAIFAADVGTPTLWAARYLTMNGRRRLHGSFNHGSMANALPQALGAQ
AAHPGRQVVSLSGDGGLSMLLGDLLSARQLDLPIKVIVYNNSSLGFVAMEMKAGGYLDTG
TDLHETDFAAIAKGAGLYSLRVTESEALPDALGKLLAHPGPALLDVVTDKHELAMPPKIE
WAQAKGFSLYMLRAVLSGRGDEVLELAQTNLR*
>GCA_000452945.1_BPMSHR5858_genomic_04682_1 argD_1
MHRFETIRQLESSARTYASSFEAVFESGAGALLRDQSGREIIDCLSCAGALPLGHNHPEV
NDAIRRFFQSGHVQQALDLSTPAKFEFVKTLFETLPDDWARRAKIQFCSPSGSDAIEAAM
KLTRFATGRQTIVAFSGAYHGMTRGALAAMGNLGVKSGLGLGAGDVHFAPYPYRFRCPFG
TDGTETDALSIQYLRNLLADPESGVSKPAAVVVEVVQGEGGCIPVSDAWLRALRQLTRAH
EIALIIDEVQTGFGRTGSMFAFERAGIRPDVLVLSKAIGGGFPLAVVVYDEVLDVWPRGK
HAGTFRGNQIAMVAGKATIEILRRDRLDAHAAQMGALLVDGLREIARAHPEFGDVRGRGL
MVGVEIVDPSSDGAPQDGELARAIKLEAFRNGLLVETGGRHGAVLRLLPPLIVNRADISA
ILERLDASIAQAKRGRGPRGGANHRSDERDGEAARPVAQGRSALPQAEPVKREAERRRGL
LKPAFLLLWLGETALDFGSALMSFALAAWIFQKTGSAERFSFAVLSAAIPALLLTPVAGA
LADRFDRRWVIAGCDVAMAIMIGALAWLLFRGALAPGHLYFFNATSAAIGCIRMPAYRAA
VTAIVPRERLTQASGFTGTSQALLRIAAPLIAGYVLADYGLKGIVGLDMLMIAAGSAAIF
AALLRAAHAIRGVVGAVDASLIEGVSASFAAAVRYFKSVPMFQGLAAYNMLQESLLLLSS
VMLTPLVLSTHSSSTLGLILTYGALGGLAGSLLLLVLRIEARLMSLALFADAVLAMLIAL
VGFSRSPAAWCAFVFLAYFASSASSACTSALWARKTPREMRGSIFALNTSMNLAAMSIVV
LVGGVLCQRLFEPALADGGAWAPTVGAWLGTGKGRGIGLLLVLCGAAGCVASLLALTCTR
LRHLDALVPDQTHDDARDLLRNGVAVT*
>GCA_000452945.1_BPMSHR5858_genomic_04699_1 group_1040
MTALPDASHPSSRVAWPAVEPGFAAASVRAMPIDAAARGIARLCRAVARAVRQCVARIGR
ARDDCEDCEDSDAHDDRHDRGSLDARSEPAPMPRMTTRAATRASASRPTESKRSRLAIRR
DRRRASQTAASIDRIASANPSGSSPAPRRAGNTDLQRRSVTILALRAEAGAASDSELSEA
APKHGSLGQPAGGHDMRRALRHGDALAHLIGIVQVVQIAGAVRATANVLVLRPPAGTLFE
SGIHACPGALQRPLAEAVKLAPAICSTFAAGRRRVAGGATHARWRTAAPRVPVRADRADD
ATAGVDARRMPATPGRAYCPSGRCGSRRAARASAIAKASASASGRRGSGDARDVGGPAAS
RIAANLLHARPASAGRVTAGAIGIIETKAAGNAGSRQDNHSPSPR*

>GCA_000452945.1_BPMSHR5858_genomic_04806_1 aqdA1
MTMERLYRGMDRATLDRAYDNTRAVADFPAVLADFRRRSAQLYARVGGRREVRYGERPRE
RFDWLSCGRAGAPRFVFIHGGYWQNCAKEDFAFVADGPLAAGFDVVLAEYTLAPEASMTQ
IVGEIARLLDRLSADRDGLGGAGRPLCLSGHSAGGHLSAVHRGHACVTSTLAISPLVDLE
PISLSWLNDKLRLSEREIADYSPLFHVGKGAPTIVAVGAHELPELVRHADDYAHACAAAG
EPVRRVHVPGRTHFSVLEELAQPDGALTRLLGAFVGGA*
>GCA_000452945.1_BPMSHR5858_genomic_04811_1 group_7621
MDHDPQLTERWLEIHAEIEALRRKAKAAAIMQIELLMTQYGISPGELKGVDGAPAHRKPK
ARYWNPETGQTWSGRGRMPKWLVGRELASHLIDGESALKRMKLLNRDAQDADEG*
>GCA_000452945.1_BPMSHR5858_genomic_04813_1 group_7622
MFKTIVLAACAAAVLGACTSATGPRFNAYVLSDAAGAKTYKVECHGLFEGPGVCRDKARE
ICADKDVHVLTGVGPLGHDDDVRTLAFQCGEPAQPVAQQPQPAPAAAPAAEPIRLNADAM
FAFDRADAASMTEQGRQQLSQLAQRLTDRHAQTVSIVGYTDRLGSDAYNRQLSQARAKTV
GDYLIAAGVPADSVHAEGRGASDPLVQCDQRERAALIACLAPNRRVEVVAAGLDGA*
>GCA_000452945.1_BPMSHR5858_genomic_04819_1 kdpE_1
MSAARKVLLVEDDEAQANWAKLVLTRGRFDVTHCQTGGQAIRAMTKEVPDAVVLDMRLPD
VHGLEVLVWIRRNFFDVPVIVLSNAMQEMQIVEAFSAGADDYVLKPAREAEFLARIAAMM
RRKGHAVSGTIEIGGIAIDLGWKAVRIDGEPVRMAPIEYEILEMMARHLGSVVQREVLVN
RIWGRSIDEAVSRSLDTHVYRIRHKLKVHGVSNVSLRSVYNLGYRLEWCRGRALAGAAPG
RDDALRLAEGGGEAMLPSLL*
>GCA_000452945.1_BPMSHR5858_genomic_04821_1 group_3587
MPHESPPSSKSRRLVLKLGGGTVLGTLLAACGGGGGADAAAAAPQQPPPAGPSSSANVAM
TLPADAPRIARDFAGLSIEKAALSYPLLSGENGNMVGLFNRLGAGVLRIGGNSSDASGWQ
RTGPDETSGVITPAAVDRLASFVQACRWRVIYGLNFVGNDPATIADEAAYAAQALGVQLA
GFEIGNEPDLYAQHGLAPNANTYPGFVSRWTTFANAIRAAVPDAVFTGPATAWNYQRYTV
PFASDAAGLVSLLTQHHYRNPDSATIEAMLSPDPSLAPMLQALQGAASARGIGFRLAETN
SYWGGGKPGVSDAHASALWVINFLFAVAQGGASGVNLHTGGGASYSAIKTNKTAGTVAAI
GPEYYGIYLFNQAAGGRLMQTRVDSAGTTLFAHAVAADGGGVRLILVNTDANSGYDVAVD
CSSVPNARAGIVTTLGGPSLGSLTGTQIDGATFALDGSGAPQGGRPVACVNGVLGVHVAS
ASALLVDFA*
>GCA_000452945.1_BPMSHR5858_genomic_04832_1 fldP_2
MAGVSAEFKAFEEATSGAVMTKGFLWRSKIAAGFTNSGAHAGDKLSMLMQLALFAARYGM
HWVNLGLPPANDSMAGSPAELNRLGFGLGAGAQSNTDQGPDAAPPEQSEQSEQSEQSE*
>GCA_000452945.1_BPMSHR5858_genomic_04833_1 group_7623
MEIDAISRALQIYFDVMYECDLEKFDLVFHPTSSLFTMKDGELSVRPFARYRSEIAARTP
PKSVSQPRMDAILQIAVLSPEIAFAQVRVRLFEKVFIDNLNLLKFDGRWMIVAKIFHHAR
TIAAA*
>GCA_000452945.1_BPMSHR5858_genomic_04834_1 gabR_1
MVETLAGLFDFEMDPQSGVALTRQLRDQLRRAIAAGSLASGRRLPSSRALALHLRVSRNT
VCAAIEQLAMEGYLDVSRGRRPVVAPMPSAGLVAIGSDRPVSDGRSGMSQWAQRIDRSDW
PFISEGRPTPFAPCVADARAFPHAIWGRCLRTATRYAVARTSTACNRPGLRRALLHHLVE
YRGVNADAHQVFLMPTAQAAIALVARVLLDPGDLAWVESPGYGGARAAFEAAGATVQGIA
LDQSGMAFETSTDTPRLIFVTPAHQHPTGLLMPPARRQALLRFAARVGARIIEDDYDSEF
HYEGRPVAALQGADNADSVFYVGTFSKSLHADIRVGYVVVPGHFVDVFAKAQRHTGQIVG
ATLQDALAEFIDDGHYAAHIRKTTRLYHARRDYLCDALKAVGSELTVSPPDGGMQVVARL
GPLRDDREICRRLAEAGVTARPLSPHYCAQTGAQGLFLGFAAWNECEIDAGVRILARVIR
EPEPSK*
>GCA_000452945.1_BPMSHR5858_genomic_04835_1 nolA
MRLKIGELAKKVGLSVRALHHYDAIGLLSPSQRTDGGARLYGRDDLIRLHRIEALKRFGY
SLPAIQASLDGPPAGAPLQILRRQIAALDAQAARAQRLSRHLQHLVAMVAAGGETAAIDW
LNVLELMNMYQKHLDDNELDTLLASGPDTVAPMDPSWVGLVDEVRDAMRRALPADGDAAQ
ALAWRWSRLMNRMTRNDPALAGKLMGIQLGEPRAQHIVGITPAMLTWIGEACAHARCTLF
AKYLNPAQIAEVRRRQLADTHMHAWLALVAELRAHMEAGVDAGAAPVQAIVARWQQLFRD
SFCGEDEGLEAKVRDALMREPDLQLGGGFDEALLVYLNKAHIAGRDIASGDDGPKPSALM
VAKQRAAHQLLDRPLVLDDPIALSILGAAEEQALRADLDRFRYPASLGMRSSVVVRSRLA
DDMRAEAIGRGVRQYVVLGAGLDTSAYRHPEAPGRLFEVDLPATQRWKQARLREAGIALP
RSLRFVPVDFEHVSLADGLARAGFDPGAPALFSWLGVTMYLDEAAVVETLRFIAGCAKGS
AVLFEYVTPLSGLPPMMRIAMEQLTAQLAARGEPWKCFFEPAALAEMLIGLGFGSIGAWS
PGELNRRYLADRADGLHIGATPARLILATV*
>GCA_000452945.1_BPMSHR5858_genomic_04963_1 aroE_2
MSGRSFLIGLIGAGIGGSLSPAMHEAEGCGQGLNYVYRRIDLDALGLTADALPELLAAAE
RMGFDGLNITHPCKQRVIEHLDELSADAAALGAVNTVCFAHGRRIGHNTDWSGFAKAFAR
GLPGASLERVVQLGAGGAGAAVAHAALTMGAAQLALFDVDPVRARSLADQLQARFPHARL
SAGGTLADALANATGLIHATPTGMLGHPGLPLPAHLLRSGLWVADIVYFPLETELIRAAR
ALGCRTLPGGGMAVYQAVDAFELFTGRAPDAERMFEHFRALVAR*
>GCA_000452945.1_BPMSHR5858_genomic_04970_1 hcnA
MIIYLDRRALEVADGTTVAAALALAGDDTARTSCTGAARAPFCGMGVCHECRVTIDGRRR
LACQTLCREGMQVERTR*
>GCA_000452945.1_BPMSHR5858_genomic_04971_1 hcnC_2
VTDSPTDVIVIGAGIVGAACAHEFALRGLRVVVVDDGGGGATAAGMGHLVAMDDNAAELA
LTHYSIGLWRALRDAMPEGCAYRNCGTLWLAADAHEMDLARAKQAALGARGVAGELIGRA
ALAALEPMLRTDLGGALKIPGDGILYAPAAAHWLLHRLPGVALRRARAVALDGARVALEG
GDVLRANAVVVANGVAARALVPELPLRPKKGHLLITDRYPARVSHQLVELGYAASAHASD
GTSVAFNVQPRPTGQLLIGSSRQFDTEDPRVEAPVLARMLRRALGYLPALAGLNGIRAWT
GFRAASPDGLPLLGEHPSRRGVWLALGHEGLGVTTAPGSARVLAALMFGERAAIDAEPYR
PARFLSTSKSAAAGELR*
>GCA_000452945.1_BPMSHR5858_genomic_04975_1 LhpG
MRLTGELLIGQSATFGQHGAFRAIEAATGAPLEPAFGGASLQDLDAACALADDAFDAYRE
TSLDARAAFLDAIGRHIVALGDALIERCAAESGLPRARIEGERARTVGQLALFAAVVRDG
GFIDARIDPARAQRKPLPRVDLRLRNIAIGPVAVFGASNFPLAFSVAGGDTASALAAGCP
VIVKAHGAHPGTSELVGRAIQAAVRECGLPAGVFSLLFDASREIGQALVADPRIKAVGFT
GSRRGGVALMRIAAARPEPVPVYAEMSSINPVVLMPGALAARHDAIAQQFVASLTLGAGQ
FCTNPGLILAIDGPALRAFEKAAAAAIDAAAAQTMLTPPIHASYREGVVRLASHRAVERL
GEGPAGGPFAARAALFATTADALLSHAELRDEVFGPASLVVRCADADALHRVLKSLEGQL
TIAAHLGERDAPLFAALRPTLERRAGRILVNGFGTGVEVGHAMVHGGPFPATSDPRTTSV
GARAIERFLRPVSYQDVPDALLPDALRDANPLRVPQRIDGALAPREARDA*
>GCA_000452945.1_BPMSHR5858_genomic_04976_1 ybiC
MHDRPDLARPDVVLSLDDVRALALRVLAHHGLSDAHARAIANVITQGQRDECHSHGVYRL
LVCARSLRLGKVDPQAVPTLRRLSPSIVAVDAHRGFSLLAFETGLPVLVEMARRHGIAAM
AINRCYHFSALWPEVEALAAHGLAGVAMNPSHGWVAPEGGTRPVFGTNPLAFAWPRPGGT
PFVFDFATSAIARGDLELHAKAGKPIPPHWAIDANGAPTTDPQAALQGAMRTFGGHKGSA
LAAMIELLAGALIGDMTSAESLAFDEGAGATPCHGELVIALDPKTFLGDALDAGLARGER
MFEAIVAQGARLPSQRRFDARARSAAHGVRIPARLHDEIVALLD*
>GCA_000452945.1_BPMSHR5858_genomic_05026_1 group_311
MLTFNRHTVALDERFACFVAQPLDAALPIVVDSPHSGIAYPPDFGAVAPRHAILTSWDAF
VDELWAGAPERGCALVGARFPRAYIDPNRAVTDIDAALLAEPWPEPLAPQSYTRRGMGLI
RRYALPGVPLYDRKLSLDEVRHRIDTYYLPYRRALADAAEPLYAKHGALWHVDCHSMKSR
GNAMNLDAGAARPDVVVSDRLGTSADPAFTRWTADWFAHAGYRVRINAPYQGGDLLNALA
APARRRHSIQIEFNRALYMNETAFDKHAGFAALKRTVDAYLDALAAHVRAQLPPPRGEGV
SR*
>GCA_000452945.1_BPMSHR5858_genomic_05055_1 group_7646
MKPLLFAAALTALALVGPARAADAAQAAADAKAQAAANEAAGLPDLTKINRPGAEVTSKV
DFNDVRRTPSFHEKSKNGTEVTEFRDRGKPVEINVKSNFGTRYQMSASPDTSPRPHDAGV
PVTRLPSVNLHY*
>GCA_000452945.1_BPMSHR5858_genomic_05081_1 group_1544
MLEQTAEHWGQSYFSETFNRREWQAHPLSIQRQYELQGNLMREEWLFSRYLNRKRVQRAA
SLGAGRAETEIALLELGAVEHFDLFDVSSVGIEYAKSIAEEKGFGHKVTCHVGPIGCAEL
NENTYDLITFVASLHHMEPLAETLERANRALTQHGIIWCANEYIGPDRFDYPVAHAAIAK
SFFQQIPPGLRNHWHRELQFPTPQEVAEVDPTEAPCSSKIEPTMRDMFPALEIMPLYGAF
AFMVFWGLNHDALYETPEGTELTRFILGMDKALTDAGILPTYFAHIIARKNNAPRQQIHR
PGRHSSALLYRYAQRAASILRHLKIR*
>GCA_000452945.1_BPMSHR5858_genomic_05107_1 group_567
MARPLNSALAKAFSRLRYPFDVHLGYARGCAAYAPSLRHSDEMTAERDASADLSTAHRRP
ANRMPALARAFDRRERPVGKSGRTGRTGAC*
>GCA_000452945.1_BPMSHR5858_genomic_05126_1 lrp_10
MSAPSRRLDRIDIGILNQLQQNARITNAELARAVNLSPTPCFNRVRALEKLGLIKQQVTL
LDPQPLGLRINVFIQVSLEKQAEDALRRFEEAIDKRPEVMECYLMSGDADYLLRVVVPSM
KSLERFILDQLTTIPGVSNIRSSFALKQVRYRTALPLPADGLTLADADDAPQEWV*
>GCA_000452945.1_BPMSHR5858_genomic_05163_1 group_7659
MTTHRFDVFGRQILVERTGGRWVPFYPGADGKRRPAHFVIPDFLEADELGQYLGDLFHES
ATPDNGDVKRLDLPEPD*
>GCA_000452945.1_BPMSHR5858_genomic_05194_1 group_7661
MGKANTIELTNNTSFTLVLHTIYANTGNWSGDYPPAYLRPNDTLIFTSTLDGKGDLNGSA
RFDILDTAVKRCPDATYVQLNWDNPVGADNGGSSSVVGATAQFFNVSGPTMSGDYGHPTF
SWSVNGA*
>GCA_000452945.1_BPMSHR5858_genomic_05210_1 group_7663
MKLPLEISIQGMPRSAALEEAIARHAAKLERYCAEIIRCRVSVILDDKHKHQGKPFDVHV
DVTIPGREIVSNRESDEDVYVALRDAFGNAERMLDDAASKRRDQTRRSA*
>GCA_000452945.1_BPMSHR5858_genomic_05211_1 group_7664
MSHRTRPFKAGSHEAVYPGLAQDPYAMRRKLPEPTVCPGCGAVFSAGRWQWIARPDGARE
RQCGACRRSAEKMPAGYLHIDGPFAAEHWTELLQLLRSREEQARTAHPMQRIMSIDTDGG
ATVITTTDVHLARNLGSALKSAYRGSLDLKYSPDAQLVRAHWRR*
>GCA_000452945.1_BPMSHR5858_genomic_05249_1 tauD_2
MRIEPLTCAIGAELVGVCLADAAHDDGLFAEIRAALLAHRVLFLRDQDITRAEHVAFARR
FGELEDHPVAGSDPDHPGLVRIYKTPEQPNEHYENAWHTDATWRQAPPLGCVLRCVACPE
VGGDTMWANMVLAYENLPEHVKAQIADLRARHSIEASFGAAMPIDGRLALKAQFPDAEHP
VVRTHPETGEKVLFVNAFTTHFTNYHTPARVRVGQDANPGAALLLGYLLSQAYIPEYQVR
WRWRKNSVAIWDNRSTQHYAVMDYLPCDRRMERTAIVGDVPY*
>GCA_000452945.1_BPMSHR5858_genomic_05256_1 soeC
MKPAFSVIFLTTLSGASQGLMIALFGVELAARLGWIASPPQAFFVASAALAVLLGSLGLF
ASFFHLGRPERAWRAVAMWRTSWLARECIALPCFLACTTAYGGAHWLALPHTLALGLLAA
VASGALFVCTGMIYICLRFLQEWATPLTMTNFVLLGCASGGTLATAGAAWFAPPLVPALA
ACACALTLAGGAARVASLARNARLRPRSTVQSATGIRAQQVVQRSRGFTAGAFNLREFFH
GKTPQTLRRVKWTFLICAFAVPFALLAIGGATAPTVMSFALACGAFAIQYAGLVAERWFF
FAEARHPQNVYYQSAA*
>GCA_000452945.1_BPMSHR5858_genomic_05285_1 cyoD_2
MAHSHPSHGDEAHGTVGGYIAGFVLSVLLTAASFGLVMGGVLPPREALVSLAVLAAVQIV
VHLVYFLHMNTSSSQRWNVMAFSYTVLTALILIFGTVWVMHNVGMNMMSR*
>GCA_000452945.1_BPMSHR5858_genomic_05300_1 srpC_2
MHASRFVIGRAWPVFVAFLRLGVTSFGGPIAHLGYFREEFVVRRRWLSERSYADLVGLCQ
FLPGPASSQVGIAIGLSRAGYAGALAAWAGFTLPSAIAMIAFAYGVAGAAGPALAPGALH
GLKIVSAAVVAQAVWGMARNLCPDARRVTIMTAATGVALVVPSTWTQLAVIAGAGCIGLL
HVDHARDGAHEPLPISLGRRAGACCLALFVALLVGLPLAARASGSHALGIADAFYRAGAL
VFGGGHVVLPLLQASVVAPGWVGGQTFLAGYGAAQAVPGPLFTFAAFLGASMKDAPSGWL
GGALALAAIFAPSFLLVAGAIPFWERWRTNARMQAALAGVNAAVVGLLLAALYQPVWTGA
ILAPRDFVAALVAFVALTFWRVPPWGVVIVAGVAGWVVFSV*
>GCA_000452945.1_BPMSHR5858_genomic_05346_1 anmK
VTSNRMQTGHPADGVYFGLMSGTSMDGVDGIAVRFEAGKPPAVLSEAFVGFADTLRDALF
ALQQPGDDEIEREALAANALAARYAVCCHELLRTAGLSPDDVRALGVHGQTVRHRPERGY
TRQLNNAALLAELTRIDVIADFRSRDVAAGGQGAPLVPAFHATVFGSPDETRVVCNLGGI
SNITILPAGGGPQGEGHARNDTVRGHDCGPANALIDAWVERHLKQPFDDGGRFAARGKVD
ETLLAALLDEPYFRQNAPKSTGRDLFNADWLDAKLAGFQHLAPENVQATLTALTAATVAD
EIARHAGDCRAVYVCGGGARNPVLLDALATALAARGLDAAVATTAALGVPPQQVESLAFA
WLAYRFNARAPGNVSTVTGAAGERVLGALYPR*
>GCA_000452945.1_BPMSHR5858_genomic_05433_1 group_5194
MRVSSNYTLRQNPLPADVYQDVRCEAGYSSAYADDACLR*
>GCA_000452945.1_BPMSHR5858_genomic_05435_1 group_1553
MQTTDNATSQFESAASRPFAGGAQGAPALASQTTYADYKVIRRNGSVVSFEPSKIAIAVT
KAFLAVNGGQGAASARVRELVEQLTQTVVRALVRSRPNGGTFHIEDIQDQVELALMRGGE
HNVARAYVLYREKRHLERVHAGEAEAAEASPAGGINVTDNGVTRALDLNALRALIVSACE
GLGDAVSPEPIVAETVKNLYDGVPMTQVYDSSILAARTMIEKDPAYSQVTARILLHTIRR
EILGEEVTQAEMTTRYAEYFPQFLKRGVEAELLDDKLLQFDLKRLGDALDASRDLQFGYL
GLQTLYDRYFLHVDGTRIEMPQAFFMRVAMGLALNEIDREARAIEFYNVLSSFDFMSSTP
TLFNSGTRRSQLSSCYLTTVADDLDGIYEALKDNALLSKFAGGLGNDWTRVRALGSHIKG
TNGKSQGVVPFLKVVNDTAVAVNQGGKRKGAVCAYLESWHLDIEEFLELRKNTGDDRRRT
HDMNTANWIPDLFMKRVMEGADWTLFSPSTCPDLHDKFGAEFEAAYVGYEEKVARGEIKL
FKKIPAQQLWRKMLGMLFETGHPWITFKDPCNVRSPQQHVGVVHSSNLCTEITLNTSDTE
IAVCNLGSVNLVAHLAKQADGSYALDEEKLKRTVSVAMRMLDNVIDINYYAVAKARNSNL
KHRPVGLGIMGFQDCLHLLRTPYASDEAVEFADRSMEAVCYYAYWASTELAEERGRYSSY
RGSLWDRGILPQDTLKLLTEARGGYVEVDTSATLDWPALRARIDSFGMRNSNCVAIAPTA
TISNIIGVSACIEPTFQNLYVKSNLSGEFTVVNEYLVRDLKERGLWDEVMVADLKYFDGM
LSRIDRIPADLRAIYATAFEVDPKWLVEAASRRQKWIDQAQSLNIYMAGASGKKLDEVYK
LAWLRGLKTTYYLRTMAATHVEKSTVAHGALNAVPTSGGTSGGAAGGAQGAAGGFGAAGG
DAPSGAMNAAPVEADGPVCTMRPGDPGFEECEACQ*
>GCA_000452945.1_BPMSHR5858_genomic_05462_1 group_1246
MTDRRTFVGGAAAFAAAWAAAPFGARAGIAVDLDPRQAGRVRAERDPAVVDAARRYRWVD
DRAFTVAIAPHAPPVATFATDARTVVGADPDYAQLVADALGRRLALVPVAWADWPLGLSA
GKYDAVISNVGVTEQRKRKFDFTTYRLGLHGFYVRAASPIARIAAPRDVAGLRIITASGT
SQERILLEWDRRNVAQRLKPVDVLYFDDDAASRVALLSGRADAELNPNATLAYQAARDGR
IRCVGNVNAGWPLKADVAIATRKGSGLADALTLATNGLIRNGKYRQALARWGLLSEALDR
SETNPAGLPSF*
>GCA_000452945.1_BPMSHR5858_genomic_05463_1 btuD_14
MSDITHAVGGAPPTSGLPAARADAPRFRIVPARHRARTAGTVLAAVLIALVLNSVLGNPQ
WGWPVFAEWFLSEPVLSGLARTLLLTALGALFGFALGVPLALARVSRSPLLAACAWAFIW
LFRSIPLIVLLLILNNLGYLYEHVRLGVPFTGITFAEVATTDLISPFCAAVLGLTLNHAA
FAAEGIRGGILSVDQGQLEAAAALGLPRARQATRIVLPQAMRAFLPVAFNDLIALAKGSS
MVYVLAMPELFYTVQVIYRRNLEVIPLLMVATVWYLIILTVLSIVQAQVERRYARGAVRE
PERSPAAALVGRLAGALPRRRVGAAGRSDAPGDARPASKAAGAPSAADAPGAAAPGASAS
FAASRAPGRARSPGAARTPASGRRGGEIAVQRVSKRFGAHKVLDDVSFVAPAGSVTVILG
QSGSGKSTLLRTINHLERVDGGFIDIDGELIGYRREGDTLYELKEKDVLRRRADVGMVFQ
NFNLFAHLTVLENIVEAPVASGRATRAEAERDARELLARVGLAAKAHAYPRQLSGGQQQR
VAIARALALKPKVLLFDEPTSALDPELVNEVLDVIKELARSGTTLVIVTHEIGFAREVAD
TVLFMEHGRIVEAGPPAQVLGAPAHPRTRAFLSKVL*
>GCA_000452945.1_BPMSHR5858_genomic_05464_1 moxC_1
MTQSTIPFGIMLQGAGSHMNAWRHPSNPPDASVNLDFIVRIARQAEANGIAFAFVADGLY
INEKSIPHFLNRFEPISLLSALATATSRIGLAGTVSTSYSEPFTVARQFASLDLLSGGRA
GWNVVTTPLEGTAKNYGTRHPEHDLRYEIADEYLSVAQGLWDSWDDDAFVRDRASGRFFD
RDKLHALDHAGRFFEVAGPLNIQRSPQGQPVIFQAGSSDAGISLAGKYADAVFTHAPSLD
ETRAFARRVKDSAIAHGRRADDVKIFPGIGPIVGRTAAEAEEKYRAIRELLTIDEALAYL
GRYFDHHDFTQYPLDAPFPELGDVGRNSFRSTTDRIKAEARAKGLTLRETALAVATPRPQ
FIGAAHDVADALIRWIDAGAGDGFILGFAVQAQGLDDFVELVIPALEARGRYRRTLVGST
LRDHLGLPRKESRRAAAAPA*
>GCA_000452945.1_BPMSHR5858_genomic_05465_1 group_905
MSYSLSLLDKSPIAEGATAADALRFTVALAQRAEALGYRRFWIAEHHGAPGLASSAPEIV
VSHVLAHTSSIRVGSGGVMLQHYSPFKVAETFKLLASLAPGRVDLGVGKAPGGLPLTTRA
LQSLHDKPRRPAFAEQLAELDAFLNWGVAEDHPLAGALALPVPPEAPERVLLGGSPESAA
LAARLGWQFCYAGHFNGDEANVERSLDAYRSATGRAPLLALYAVAAPTKGDAERLIGALR
IFKLKLASGQSVNLGSAQAAAEFARQSGATDYRIDELRPHVIAGAPDDVHRELDALSRRY
GLAEFVVDSPVTHYPARLASVELLAPQRQPAGT*
>GCA_000452945.1_BPMSHR5858_genomic_05509_1 mtlK
MSSPSTAAPAILHIGVGSFHRAHQAWYLHRVNEASPAAERWSLVVGDIRDDLRASREALA
AQHGVYTLETVTPQGERAYETIRSIARVLPWSMDLAALVDAGAAPACRIVSFTVTEGGYY
LDEHDRLDVANPDLAADLQGARLTIYGALAALLAERAARGAGPLTLQSCDNLRNNGARFR
AGMRAFLARRGLVELLAWFDANVACPSSMVDRITPRPTPDVRERVRAATGFDDACPVMGE
AFIQWVIEDRFAAGRPAWEKAGAELVDDVHPYEEAKIRILNATHSCIAWAGTLAGHSYIH
EGTRDAHIRRFAHAYVTDDVIPCLTPSPLDLARYRDVVLERFGNPHIRDTNQRVAADGFS
KIPGFIAPTLAESIARGVEPVSTAVLPALFLRFLQRWAQGALPYAYQDGVMDARVARAIA
GAADPVAALAGERPLWGALAGMPALERALRAGAARVDAWLASR*

>GCA_000452945.1_BPMSHR5858_genomic_05512_1 nepI
MSDCTTRPAGFARPSREAARLPLAGLLALATAGFITIVTEALPAGLLPLMGRDLRVSDAL
VGQLVTVYAAGSIVAAMPLVAATRGMRRRPLLLAALAGFVVANTATAASPYYAPVLVARC
VAGVSAGLLWALLAGYASRMVDARQRGRAIAIAMLGAPVAMSVGIPLGTALGAALGWRAT
FAGVTALTLALIAWVRASLPDAPGRPSGERLPVARVLRMPGVLPVLAVMFAYVLAHNILY
TYIAPFLASAGMGARIDATLFAFGAASFAGIGLTGVWIGNGLRRLALASIALFALASVLL
GVASGSPAVVYASVAVWGLTFGGAATVFQTASANAAGEAADVAQSMIVTVWNLAIAAGGV
AGGVLLERFGAGAMPWALVALLVPAWLGAWRARRHGFPAARAP*
>GCA_000452945.1_BPMSHR5858_genomic_05539_1 group_3657
MANLDMALVVGAAGGIGGEMVRALRGAGWRVRALARSAPPAGDGAADIDWRLGDASNPHD
VAEAARGCAAIVHAVNPPGYRRWSEWVLPMLENTIAAAKRERATIVLPGTVYNYGPDAGG
LLREDSPQHPLTRKGAIRVEMERRLRDATEAGARVLIVRAGDFFGPGARNNWFSQGLVKP
GRPVRTVRAPGRAGVGHQWAYLPDVAQAMVELLARRDALDAFSCFHLAGHWDADGTRMSA
AIRAAAARRSGVEPRVAAFPWWLVTLASPFVETLREMREMRYLWREAVRLDNARLVAALG
REPHTPLEAAVETTLDALGCLNGRASRAAPSVR*
>GCA_000452945.1_BPMSHR5858_genomic_05542_1 cbdA
MIPIDPDRRATPRPIDDFLVEDKARGDYRLHRSAFTDEMLFELEMKHIFEGNWIYLAHES
QLPNANDYYTTTIGRQPIVIARNRHGELNAFVNACTHRGAMLCRHKRGNRASYTCPFHGW
TFSNGGKLLKVKDPEGAGYPDCFNRDGSHDLKKVARFENYRGFLFGSLNPEVEPLAAHLG
DAARIIDMIVDQSADGLEVLRGSSTYTYEGNWKLTAENGADGYHVSAVHWNYAATVNHRK
TDAQHEDTIRAMDAGNWGRQGGGFYAFDHGHMLLWTRWANPEDRPNFDRRDEFAARCGGD
VADWMIRNSRNLCLYPNVYLMDQFGSQIRVLRPLAVDRTEITIYCIAPKGEAPDARARRI
RQYEDFFNASGMATPDDLEEFRACQQGYAGRAVEWNDMCRGASHWIEGPDEAARRIGIRP
LMSGVKTEDEGLYTVQHRYWIATMKQALAAERSGA*
>GCA_000452945.1_BPMSHR5858_genomic_05544_1 benC
MEHRIALQFEDGVTRFIACRDGETLADAAYRQQINLPLDCRDGACGTCRCRCESGDYALP
ASSYIEDALTANEAAQRYVLACQTRPRSDCVIDVPASSAACKTGAARHEGTLASVERLSA
STIHFSIDVDEPAKLAFLAGQYVNVEIPGLGATRSYSFSSRPGDARVSFLVRNVPGGRMS
RYLTDEAAPGQRIAFSGPHGSFYLRDAARPALFLAGGTGIAPFLSMLDVCASRDGAPPVR
LVYGVTRDADLVALERLGDVERRLAGFAYRTCVADEASAHPRKGYVSAHVEPEWLNGGDV
DIYLCGPAPMVDAVQAWLRERGVTPANLYFEKFSSSNAP*
>GCA_000452945.1_BPMSHR5858_genomic_05547_1 group_7682
MITSHHLQQASRLRGQVTALNRRLRREAQADPVQFSQLVVLGAIDRLGGDVTPSALASAE
RMRSSNLAALLRELEGGGLLQRHADPADGRKSRVSLTASGKRMLYGNRAKREEWLVRAMH
ACLTAGERELLAAAGPLLERLAQFDEAAGAASPAPSPRSSR*
>GCA_000452945.1_BPMSHR5858_genomic_05548_1 rutB_1
MNTALIGLDYIVDIMHPTGKIARSAAHAAQRDVVGRFNRALAAAKQKDWLRIGVKVGFEP
GYADLPAHSPMFGRAQAFGALDLSGAGTAFHPDLDAAAFQLVVVKPRVSAFYATRLEAAL
RARRIERVIVAGVSTAWAVQAAARDAHDRDYEVLVLEDACAAATEDEHQRSIDTLRGIAR
IVTLDDLAAL*
>GCA_000452945.1_BPMSHR5858_genomic_05554_1 phaB_2
MQAKRVAFVTGGMGGLGAAISRRLHDAGMAVAVSHSERNDHVSTWLMHERDAGRDFKAYA
VDVADFESCERCAEKVLADFGKVDVLINNAGITRDATFMKMTKGDWDAVMRTDLDAMFNV
TKQFIAGMVERRFGRIVNIGSVNGSRGAFGQANYASAKAGIHGFTKTLALETAKRGITVN
TVSPGYLATAMVEAVPQDVLEAKILPQIPVGRLGQPDEVAALIAFLCSDDAGFVTGADLA
INGGMHMS*
>GCA_000452945.1_BPMSHR5858_genomic_05557_1 group_7685
MHQDTLFDSLLAAARRRSITEGEVMHMLDDEIARLADGARIHDYLRVIAIRRVRERIVSH
ARAADEAHARRPGAR*
>GCA_000452945.1_BPMSHR5858_genomic_05558_1 group_7686
VRITVHLDTFASTDPAAYAILWIDTTERRWSREGHAGVELPAWGNVVCRGGTTRVTGADD
PHSLCVLEGLDLGAKQGPFEGETGAAHWYPHAHRAPVVGAWHVQCIDETVAPAEHELFTG
REAS*
>GCA_000452945.1_BPMSHR5858_genomic_05561_1 group_7687
MSYHDVYTESGYLAYGRAEAAPDTEADRPARPRPAAARADRTDALLALAREAGMLVTLDG
QIGRERYQSVAGSVASLMRFVTALEQSLLDGDQ*
>GCA_000452945.1_BPMSHR5858_genomic_05562_1 group_5210
MKTDRQIKQEVEDELTGDPMIDVAHFDVDVAGHIVTLTGHPSSYAEKLAAEKAANRVAGV
RAVVVDVQVRLPSDDVRTDEAIADAVRSTLHWTVGLHDAAVHVQVEAGWVTLSGQVDWPY
QSHVAARAISQMRGVTGVTNQIEVLGDISADEIAGGIRRAMQRHAEREANHIDVTVKDGT
VTLAGKVGSYAERAVARGAAWSARGVRAVVDDLVVE*
>GCA_000452945.1_BPMSHR5858_genomic_05563_1 group_906
MTHAATGAAIGTPIGAVRPRPVLMIPLRRCGSHALRLRLNLNSQFYSPYPLHIVDFMPLL
PLYGDLADDRAYFRLVADIVGLQAASMVKWPGVAFDPVEIFDALRHAPRSAHRIVWELLL
RAGEREGARVVMDKSLDSVHYADELMTLYPDMLFLNVVRDPRAQVASMNRAIIHDFDTLL
NAQAWVAAHRAADVLIARHPQRVLTIRYEDFLSDQAHTLQRVCAFFGIDFLPRMLDIANS
PEARHISRMSELWASNCFAPIAANADKFKQQLSTAEIATIETLAHEYMQRYGYQQMTDAT
AMPDAFAAAAARRRSDARRRHAWRELEQSNFRDFVLRRHRADYLETVRARLQRHASAQAD
SRADLRADSPAGAPGRRDTLTAAFDVTD*
>GCA_000452945.1_BPMSHR5858_genomic_05568_1 icaB_1
MSFRSRFTWRTAVFGPGFRPVSRIVATTASSGAPAVAAPPTDVAVLVYHRFSNVCGADPM
TVGVATFEAQLAHLRRLGYRFVPLRDVIGWLRGEPVALPSKAIALTIDEGHASIFDWART
VALRERVPITLFVYPSAIGEAPGALTWHQLRVLHKTGWFDVQSHAWWHPDLNAAHRPPSG
TFREATRAQFAQARARIAREIGNQVDLLAWPFGAFDGELGAAAREAGYVAGFTLEPSKIR
RDTPLLTLPRFLMVEECTPAVLRRLLSKSDAAREEAARR*
>GCA_000452945.1_BPMSHR5858_genomic_05569_1 arpC
MCTSHPASRPRGGRPRLAAIALGCAALVAGCAVGPDFIAPARPAVDAYVPDGAARTRFAA
SDATQTLAPDAPFPDAWWRLFGSREIDAAVDDAFAGSPTLARARATLARSEHALITGAGV
FYPQADAQAGASRQRYVPLRTGANLPASLFNLFTLSTTVSYALDLWGGERRHVEGLAAQA
DAQRHALAAARLTLAANVVNTMIARAAYADEIALTRELIALTDEQIRLTRAQVSAGTSAY
TAVLALQAARASLDASLPALEQKTGEADDLLATLAGRYPAEHVPPQVSLAAIALPARLPR
TVPSELVRRRPDILQAEAVLHAASANIGVASAALFPSVTLSASGGFDATRASALFGPAGR
AWSVGASITAPLFHGGALWYQRKAAVDAFDESNAAYRQTVLSAFAQVADTLRALDHDATA
LDAQARAMSAARDARRLATVQYEAGTAGYLQVLLADQQYRQARLAWVQASAQRLQDTVAL
YAALGGGWGGARQP*
>GCA_000452945.1_BPMSHR5858_genomic_05571_1 lolD_1
MAGLAIDARGLTKWFGEGEARTRALADVSVQARFGEMLLIVGPSGSGKTTLLSVMSGILR
PDAGSVIVDGVDLWAQDNDAIAEFRLNRIGFVFQDYHLFARLTTAENVAIPLILKRRDWN
RALADAHEYLDVVGLRNRASLPPVKLSGGEQQRVAIARAIVSQPDILILDEPTASLDGDT
GRTIIDFVKHKVLNDKRCIVIVTHDSRIFDYADRILRMEDGKLTAIERGGGL*
>GCA_000452945.1_BPMSHR5858_genomic_05572_1 group_7690
VNGILRLAFKLLVNDSAKFTALLVGITFAVFLMVEMTSLFAGILDKSSSTVINVGAKMWV
MDPGVQTIASSIGMPAYVLDAVRSVDGVKYAVPIYSGGALVKLADGTYQAVTVIGLDDAS
LLGRPTMKAGRIEDIYAENGFIAIDDAEFPKLKNPALGTTFELNDHRGVIVGIAKVASSG
LFGTPTLYTTYARATRYIPSTRYTTSYILVEPKSQRDIARIKREVAALGYRAYTKEEFMR
RISDFYKYETGVGTNILLMTAISFIVGLSISGQTFYAFIIENLEKFGALKAIGAKNRELV
AMILFQATFTALTGYGLGVGLCTALISVARLRLPSYAALITYGNLALAFGMVVVIAGLSS
YLGVRRVLRIEPFDIFRG*
>GCA_000452945.1_BPMSHR5858_genomic_05575_1 group_1072
MRNRNGDEQAATARHEAPARPPRCRLPADIRINQILDAALAEFSASGFAGARIDDIAARA
GMSKGGVYTHFGSKDEIFEALLARTLMPPLPAASSAVRAARRARRAGAGPAARTSGARRA
AQASTNGAGDADERPGPGEPREPGAIREPGAIREPGAAGSNAAAEFPDAPPPVPDSLAAI
VDLLTGDLYTWATSEPVIAMLRLLIAESRRAPRAVERWRKHVEQAFATAVGRLVRQGVDA
GELRDGIAARAPLLLVAPLTQACLRRALRAAPPSRREVAASRRDYARFIEALLT*
>GCA_000452945.1_BPMSHR5858_genomic_05582_1 group_7691
MRRIYFLLPDADTARAIVDELLLKRIEWRHIHVVANDRVPLENLPEASIAQSSDLLPALA
RGTAAGGVTGALAGLAAIAFPPAGLTIAGGAVVAFAIAGAGFGAWVGTMIGVSVPNTRLK
HFEDAVDNGELLMMIDVHRDRVEEIEALVKQHHPNAHLEGTDPTIPAFP*
>GCA_000452945.1_BPMSHR5858_genomic_05583_1 group_306
MRGAGRASPSRARRYENVSKEPNTHREAGAPVHRYRLQVRNMNGECRTYELRLSSLDDGY
TLEIDEWLNFPDMHLGDVTPVHVSRPVQFAVPAHEFYRRRRAFRTELIESVAAALRAGEV
VRMRDSPDEYPGGHAGNPWVNWIRADLRANPGGRPPPVGDDMQDLIH*
>GCA_000452945.1_BPMSHR5858_genomic_05588_1 atpE
MNNLIEVVSIAAAALAVSFGAIGPALAEGRAVGAAMDAIARQPDASGTVSRTLFVGLAMI
ETMAIYCLVVALLLLFANPFVK*
>GCA_000452945.1_BPMSHR5858_genomic_05589_1 atpB_2
MNASPLSTVPLFSVGPVGISAPVLATWAIMAVLVALALVARRRLSVDAPSRLQSVLELFV
ATVDAQIADTMQTPPARYRALIGTIFVLVFCANGASLVPGVEPPTAHLETDAALALVVLA
ATLYYGVRARGVRGYLATFAEPTWVMIPLNVVEQLTRTFSLIVRLFGNVMSGVFVIGIVL
SLAGLFVPIPLMALDLLTGTVQAYIFMVLSMVFIGAAVSGDETSAPRRGQENT*
>GCA_000452945.1_BPMSHR5858_genomic_05590_1 group_5214
MIEMWETARASFDGIAAAIGLALGFVAGACHFVSLGWNSRLFVAGRAGVALALQLARVAL
AVAVLVVLARAGFGMLVAGSAGFFAARAIAVRRAAALTGARR*
>GCA_000452945.1_BPMSHR5858_genomic_05637_1 group_907
MQRRSWWSVGRRFAAGIDVNRRTVRIAVLSRRARGGAVRLEALDAEPLPAGAFDEPADAD
WVAVAAAMAAALGRVPPMRPAGWVRRTMAGVMALPAGAFSSATLAWPRADAAAAVEPAVR
AAAERLTGLARDALAVDWCVNDAGAASELVFATAAHVHLDVRIEAAASAGVELVAIDGEP
PAALRAIRHVAAREVDSGEPYLGVWIGESGVHGWLVSADAVAAQIRYPAPEHADLADALR
ELARAGAPRCAFVAGDPEWLGGARISLADVGDLLGCIVLPFECAACGPLGAVDDGLAHSP
GFAVAFGLALRGVFE*
>GCA_000452945.1_BPMSHR5858_genomic_05663_1 group_5223
MQTIRISPAVLAITAAAALSGCATVQTPTKGDPFEGFNRTMYTFNDKVDQYALKPVARGY
QWAVPQPMRDSVTNFFSNIGDVYIAANNLVQLKIADGVGDIMRVVINTVFGVGGLFDVAT
LAKLPKHANDFGVTLGHYGVPSGPYLVLPLLGPSTVRDTAGLAVDYAGNPLTYVRPDGVS
WGLFGLNLVNTRANLLGAGDVLEAAAIDKYSFVRNAYLQRRQALIGGATGAQSNLPDYGS
EAPPPNYEMPEEGAAAPASGAASAPAAASAPTAASGAAPGAAEPASAAAPNPSSATNVPA
QQVVPPSPGGIRFPSIRLH*
>GCA_000452945.1_BPMSHR5858_genomic_05715_1 tctD_4
MRILIAEDDSILADGLTRSLRQSGYAVDHVRNGVEADTALSMQTFDLLILDLGLPRMSGL
EVLRRLRARNSNLPVLILTAADSVDERVKGLDLGADDYMAKPFALNELEARVRALTRRGA
GGGPTVVRHGSLSFDQVGRIAYANERVLELSARELGLLEVLLQRIGRLVSKEQLVDHLCE
WGEEVSNNAIEVYVHRLRKKIEPSGVRITTVRGLGYCLEKAAPAGASDAPNAAASPAAEA
VAGQPVR*
>GCA_000452945.1_BPMSHR5858_genomic_05736_1 pgk
MSQVKRLTDLIAEGKLSGKRVFIRADLNVPQDDHGNITEDTRVRASVPAIQAALDAGAAV
MVTSHLGRPTEGEFKPEDSLAPVAKRLAELLGRDVPLVANWVENGVEVAPGQVVLLENCR
VNKGEKKNSDELAQKMAKLCDVYVNDAFGTAHRAEATTHGIAKYAPVACAGPLLAAELDA
LGKALGNPKRPLVAIVAGSKVSTKLTILKSLAEKVDQLIVGGGIANTFMLAAGLAIGKSL
AEADLVGEAKAIIDEARKRGASVPIPTDVVVAKEFSPTAAATVKKVADIEADDMILDIGP
DTAKALAGQLEKAGTIVWNGPVGVFEFDQFGNGTKTLADAIASSAAFSIAGGGDTLAAIA
KYGIHDKVSYISTGGGAFLEFLEGKKLPAVEVLETRA*
>GCA_000452945.1_BPMSHR5858_genomic_05825_1 ubiC_2
VRDRAQHRVGARALETQWKPLTGYGSLPLGHILYDDPAIVRAPFECALLMPDDPLDAISR
RYARAAAPPRARRSAFVRNGATLVVSECFLPQFWSGFAQARLAGA*
>GCA_000452945.1_BPMSHR5858_genomic_05826_1 group_2647
LKLVEPGKPDVSYGLHKLKGSQASVGGKGGAMPFGEPRAARERVDALERWIGNGAPDN*
>GCA_000452945.1_BPMSHR5858_genomic_05828_1 gcvA_8
MPARKPGRLPPLNALRAFEVSARHLNFRAAADEIGVTQGAVAQQVRHLEDALGVALFERL
PRGLALTADGLAYFADVRRALNLIGDATDRLAKRRANVTISTTPSFASRWLIPRLASFGD
EHPGVDLRVIADSQFATFQGDGVDLAIRYGKPPFGKGLVTRLLFPVDIYAVCSPALLAPR
RARPRARSRRARAAARRA*
>GCA_000452945.1_BPMSHR5858_genomic_05832_1 group_3682
VITVITSFQLPAPLAREEARSLFLRAAPKHQGVVGLCANTMSCRKAEGRVYLWNARAQAE
TMYTDAWKASAYETFGAFPSITYFESPAVVDNATNAIFSDE*
>GCA_000452945.1_BPMSHR5858_genomic_05845_1 group_1566
MNGWSEQAGGPHAASPVLRPAACRARLDGVDTGARCDSGLPSRLARQRVSPRRSAAACTR
EHGAPGSTATSASGRNPRVEVPSRGSPRPAARHSKPDSSVLKRPSRNIGPAGSGIENGGP
ANSMHCPIAGNRRAGQKIAVQSSERRGPSLTRFEFNAQCHCAISVKYENNGDARHECRAN
LRSRPRTEAPLAIPLPPPLRQASKLREPAWQAVEERLFGENSGISFMPAPVRRDSIAARM
RKLSRKCRLHGFAVHVRWLFPPPGWPIRTDKWRYHALSARRRFSGIAVEHALAARIECET
ARRTRRGDTRHAPPKVSVFGSQGEAIRDWVSSFGEIARLPRQNRKRRSEWMRRRETFGSI
RQVRIDGRIARHPPERIGFIAESTHNPLRREIAGPP*
>GCA_000452945.1_BPMSHR5858_genomic_05848_1 group_512
VDKKEKVETNASISGGWVRLPADYGDVLRRAAESLFKTFEHSSVGTLIVDKDARVVWINQ
RYAARFGFADPQQAIGRDCEAVIPHSLMREVVATGRPILLDIMETGREPLVVTRLPLTDD
AGETVGAIGFALFDELKTLTPLFSRYMQVQQELIATQRSLAQARRAKYTFASFVGTSAVS
LETKRQGRRAAQVDSPVLLLGETGTGKELLAHAIHAASARALKPLVTVNVAAIPDALLET
EFFGAAPGAYTGADRKGRVGKFELADGGTLFLDEIGDMPVPLQGKLLRVLQDKEFEPVGS
NRIVRANVRIIAATSAELPALVAEGRFRADLYYRLNVLTIHAPPLRERASDIEALVYTML
EELAAQHGLAEHCELTDDALRLLCAYPWPGNVRELRNTLERALMLSDRALIDARALAPFI
GPARGAGGGVGAGGGAGAGGVGPAAVAIAAQTAMADTRAAASSYADAFAAWERQFLIDAL
AASNGKVTEAAARIGIGRATFYKKLATLGIDT*
>GCA_000452945.1_BPMSHR5858_genomic_05872_1 group_1567
VGVSRRQMSAPERSAGHKALTSRDDLKRCASMKSTRRNMLGRVTRATRRDSHASIAVPVF
RSVGRAIGAPVRPAILESHRSLRLRAKNGRSSDDSIQPRVQAGTSIKLHLGKDLFAIESQ
SQRAGLSTPHQKPTGFTKPLSIAQKNPRNAASRILDSLMRRAETPADMRHRRAGGKQALQ
LVACLPSEKAAGIPCRDESVQPCRQAVNKRVEVCGARRQGCIHHAGFMQAIRVSSLNAGR
RSYLKSRTAFRRQWIKSVSRPPA*
>GCA_000452945.1_BPMSHR5858_genomic_05875_1 group_789
MPNARASDYSIAIGNGYQSVSRTPTGTGFGFLLDATRSGEDAYSIGPGLSFAVRIGAVSL
TAVGKALHVRSADGRSGLTTPIGVIAGIPAASRLSLVARAFYTPRAVFNAALGAYTQATA
GVRWTYKPINLEAGWRYEALKGRNGTRDTRLTNGPYLAAGFSF*
>GCA_000452945.1_BPMSHR5858_genomic_05942_1 group_790
MSKKLIQIMAVAALSAAASLPAFAGDMNNALGGALGGVAGAAVGGAVGGSTGAVIGGAVG
GGAGGAVTSNRRERTGAIIGGALGGGAGTAAGNAMGGRGGGLIGAAVGGGAGAALGGNIS
RSNSYNDDYDRGYRRGKHHGKHHHRH*
>GCA_000452945.1_BPMSHR5858_genomic_05964_1 clpV1_3
MAISRQALFGKLGATLFRAIESATVFCKLRGNPYVELVHWLQQLLQQSDSDLHRIVRHAG
IERDALDRDIARALAALPAGAGSISDFSHHVETAIERAWVLATLRFGDRRIRGAWLVAAL
VDTPELRRVLLSISPAFARIPHDDALDDVLPAWTAGSPEAADAPYDHADSAPASPGEPSG
ATRAAPNGSPLERYCTDLTARARDGDIDPVIGRELEIRTMTDVLLRRRQNNPLLTGEAGV
GKTAVVEGLALAIANGDVPPKLADVRLMSVDVGALLAGAGMKGEFESRLKGVLEAAAKSV
APVILFVDEIHTLIGAGGQAGTGDAANLLKPALARGTIRTIGATTWAEYKRHIEKDPALT
RRFQVLQVPEPEEPAAVHMVRGVARAFARHHRVTVRDEAIRAAVALSHRYIPSRHLPDKA
ISLLDTACARVALSQHAAPGELQHVRQRLLAARAERDLLEQEARIGLDAGQSLAAVRERI
EALAAEEAAVDARWKAQADAARALLAAREAALAECHRESCAETRAGSLSESRTESRTESR
ARSHIDSSACAHSDVPAEMHAGSRTATRPDTHAEAHAAPASPPPAADTPHAGAAPGLREL
ERALAAAQGDAPLVFPEVDEAIVAQIVADWTGIPVGRMMTDEVAAVRALPATLEARVIGQ
PDALRQIGERVQTARAGLADPKKPLGVFLLAGPSGVGKTETALALAEALYGGEQSLITIN
MSEYQEAHTVSGLKGAPPGYVGYGEGGVLTEAVRRRPYSVVLLDEIEKAHRDVHELFFQV
FDKGYMEDGDGRYIDFRNTTILLTSNVGAELSASLCADASLAPDAAALRDALMPELLKVF
PAAFLGRVSVVPYRPLEARALARIVRLHLDRVVARMAERHRIALAYDDAVVDYVVGRCLV
QETGARLLIGFIEQHVLPRLSALWLDAFPSKAALARIDIGVADAAAPAARALVFRPGQAS
RAGPPNAPLTAVQAG*
>GCA_000452945.1_BPMSHR5858_genomic_05979_1 rhaR_7
MEWLHGHLREPSTRDAIARQSAMSVRTLARRFQEWTGTLPLQWLRPARVRRAQPLPQTTA
LSVERIATEAGFGLASAFRERFVRIVGTTPKRYRQAFRLQAG*

### S3 *B. pseudomallei* proteins from the core genome with a signal peptide (removed before counting cysteines) and even number of cysteines (263 proteins, fasta format).

>ABA47916.1 putative membrane protein [Burkholderia pseudomallei 1710b]
MTSSRRSCSARTGRRGFTRSSCVTRDAAAAASRAVSIRISRGIHMKRHAAWAGMALGGALGAAPALASAA
TLDGAALSAFWAVPFAGILLSIAVFPLIAPVFWHHHFGKIAAAWAVAFLVPFAATFGFGTAFGTLMHALF
EEYIPFIVLLSALYTVAGGICVRGNLHGTPRLNTGILALGTVLASVMGTTGAAMLLIRPLLRANDNRKHV
VHVVVFFIFLVANAGGSLSPLGDPPLFLGFLNGVDFFWTTIHLALPMLFICAILLTLFFVLDTYFYRKGG
EEGKPFLDPTPDSHGMSIEGKVNFVLLGAVIALVLMSGLWKPGIAFDLFGTHVALQNAVRDVALVAVALV
SLAVTPRSAREGNAFNWAPIEEVAKLFAGIFVTIAPVIVILRAGGDGVFAPIVHLVTGADGKPVDAMYFW
ATGVLSSFLDNAPTYLVFFNLAGGDAQTLMTTGATTLAAISAGAVFMGANSYIGNAPNFMVKAIAESRGV
RMPSFFAYLGWALAILVPVFLLTTWVFFSAQAAQ

>ABA48964.1 branched-chain amino acid ABC transporter, periplasmic amino acid-binding protein [Burkholderia pseudomallei 1710b]
MATAAARRAAGRVFCNRGAGGIGRKIAVEAAEGRVRCARARGAQQQEERAPRRFHVERETGAAACRFDRC
AMSPCPLDRVSRSVWRHTMKMNRWIEVAFAAGCLCTAGLASAQVKIGVTVSATGAAASLGIPEKNTVALL
PKEIGGKRVEYIVLDDASDTSRAVQNTRKLIDEDHVDAIVGSSITPNALAMIDVVAQGKTPAISLAASAH
IIAPMDAKRAWVFKTPQNDRLMADALAGYMAKHGVKTVGFIGFADAYGDGWYGVFSAAAAANGLRIVANE
RYNRTDASVTGQVLKTLGARPDAVLIAGAGTPAALPAKTLKARGYTGKVYQTHGVANNDFLRVCGKDCDG
ELLPAGPVLVADQLPDSNPAKQPALAYKAAYEKAYGAAAVSTFGGHVWDAGLMLQRAIPDALKKAQPGTP
AFREALRGALENVKDLPVSHGVINTTPADHNGFDTRARVIVQIVGDKWKLQAD

>ABA50111.1 ABC transporter, periplasmic substrate-binding protein [Burkholderia pseudomallei 1710b]
MACRFESGPRHQARPSARTPLRRAPAIRIRAASRRPRFRRAPPARLPSVREAALSRDSTARSFDAIHPPR
SRPPRLLSASCPPRVAERRPAAALQPRPLAFAAVISSSSTPPIIPSTKTFTRLRFPLHDAHHQRRRRARR
LFDRHRVARDQRAAHRPSGDARKDPRGDRRAALSPESARPAIAQRPHAIDRRRAADARESRVRRMPAGRR
RTRDAGRLQADRDVDRIRCGARAPCDRDTARAARGRADAHRRRRRRAPAARRARPRRPALRADAQRHAAS
PVGGGRQSPRRVRRRADADRARPSARADARGLARRIRSRAAARARLCAGTRRARARTAARARARLQCTRA
AARDARASERARDAPHRALLQQRLARDGRDSRAARRAPRGARRHVGARLRRPRGRRAARAAARERRDAES
RDRPRRVAAPCRAHRRQAPCATRADAAARGARRRDRRAAARCARCARGAHRLNRAFAFRRFPFASPSRRP
AVTVRRSLLPRSPLAAVRRACAAMLIPVASLALSLGASAAHAQDTAICYNCPPEWADWAAQIAAIKQRTG
IRVPFDNKNSGQAIAQLIAEQKSPVADVVYLGVSSAFQAKDKGVVAPYKPAHWSDIPANLKDPQGYWFAI
HSGTLGFFVNKDALDGKPVPRTWADLLKPEYKGMVGYLDPSSAFVGYAGAVAVNQALGGSFDDFRPALDW
FRKLKANQPIVPKQTAYARVLSGEIPILLDYDFDAYRAKYKDHANVEFVIPKEGTISVPYVMSLVKGAPH
EANGRKVLDFVLSDEGQKLWANAYLRPVRAQALGADVAAQFLPASEYARAKSVDFGKLAAGQQAFGKQYL
QAMQ

>ABA50268.1 family S13 unassigned peptidase [Burkholderia pseudomallei 1710b]
MANRPHALRFLHDHVAEAADVADQVRAELLAQRVDIDLDGVAADLLAPAVHLLLELALRQHRARTREQRF
EQRELARRQRDGRPVERHLMRRRIERQRAVPDHRFAAPRLAADQRAHPRGELVEIERLDEIVVGARVEAL
HAIGDRIARGEDQHRQREPARAQRAQHVDAVALGQPEIEQQQIVRLAAERGERRVAVLHPVDRAAVGLQR
LAHAFGDHPIVFNQQYPHPEKRPSHAFEMADIVAPPRAGTDLRAISALLARRFLYNVRLAPRARPLLTSP
SVLPHSSMNHHACRIAASPSPPPPAAVRARRTLARGAMLVAACAALAFAAPADARRKPKPPRYPAAVSAA
RNVLPASVLVALQRARVPASSLSVVVERIGDRTPAVAWNASRPMQPASTMKLVTTYAGLSLLGADYRWRT
SAYADGDVDENGTLHGTLYIKGTGDPKLVPEELIDLVNKIRRAGIVNVDGALVLDKTFFAPQTRDLPPLD
DDASAPYNVGPDPLLYAFKALSFTVTPTDSGAIAVDVVPALANLNVDNRLVEGQGSCDAARPTLVTDANG
ELTASFAGDYPPSCGPTTTNLAVLNHSTFFARGFLALWRQTGGSFSGTIAEGKVPGRARPVASHHGPVLS
SVVHDINKFSNNVMARNLFLTIGAVEHRPPATPAQSADTIRAFLAHSGLPTDGLALENGSGLSRDERVSA
LGLADMLQAANASPVAQAFVDSLPIAGVDGTMRNRLTNAPVGGNAHIKTGTLRDVRAIAGYVASADGSSY
VVVSFINDDRASAARAAHDALLEWVYEGPR

>ABA50277.1 aminopeptidase P [Burkholderia pseudomallei 1710b]
MSPIITGRCAFVDNRPSGLWMVCNNRVGRAGHATRTKQRRLPRGFPFRFAPCDRGARRRAARIGHANRAC
KAALGSSTRESGTRINAGMKRAVRLHISPRTAPSRIGRSLPPAVRAAACGCHREHAFARGPRARMQPRSR
IAARAASHCRTAPRPHERRAIRRARPARRRYNRVMNQPTEPALALDVYRQRRDRVLASLRAQGGGVAIVP
TAPEVPRNRDSDYPYRHDSYFYYLTGFAEPDALLVLDASAAGDAPRSILFCRAKNPEREIWEGFHFGPEA
ARDAFGFDAAFPYDALDAEMPRIVADAPALHYRFGVSAAFDARLNGWLDAVRARARAGVAAPGAAFDLGP
LLDDMRLVKDAHEQATMRRAADISALAHRRAMAACRPGIREYELEAELLYTFRRHGAQSPAYGSIVATGA
NACVLHYPAGNAVVADGELVLIDAACELDGYASDITRTFPANGRFSGPQRALYDIVLAAQEAAIAATRAG
TQFDAPHDAAVRVLAQGMLDTGLVPKTRFASVDDVIAERAYTRFYMHRTGHWLGMDVHDCGDYRERGAPR
DDDGALPSRVLHPGMALTIEPGLYVRPGEDVPQAFWNIGIRIEDDAFVTPTGCELITRGVPVAADEIEAL
MRDARPAPRPQP

>ABA50993.1 dipeptide ABC transporter, periplasmic didpeptide-binding protein [Burkholderia pseudomallei 1710b]
MVFSPACSSYESSSNIRLRRPRPALNRRPGGNMEHNRLLRALRATAIAGVAAASFGIAGSAFAQIPNKTL
VYCSEGSPAGFDSAQFTTGVDFTASTFPIYNRLVEFERGGTKVEPGLAEKWDISADGKVYTFHLRHGVKF
HTTDFFKPTREFNADDVAFTFERMLDPNQAFRKAYPVSFPYFTDMGLDKLIVKIEKLDPYTVRFTLKEPN
APFIQNLAMEFASILSAEYADQLMKAGKAADINQKPIGTGPFIFRSYTKDATIRFDGNPDYWKKGAVKIS
KLIFSITPDPGVRVQKIKRNECQVMSYPRPADIATLKADSNVDMPSLPGFNLGYLAYNVQHKPVDKLEVR
QALDMAINKKAILESVYQGAGQAASAPMPPTQWSYDKNLKAAAYDPAKAKALLAKAGYPNGFPITLWAMP
VQRPYNPNAKLMAEMIQADWAKIGVQAKIVTYEWGEYIKRAHAGEHDTMLIGWNGDNGDPDNWLGTLLGC
EAVKGNNFSEWCYKPFDELIQKGRVTTSQDGRTKIYMQAQQIFAQQLPFSPIANSTVYQPVRKNIVDMRI
EPLGYARFDGVSVK

>ABA51121.1 Periplasmic binding protein family [Burkholderia pseudomallei 1710b]
MRATSRTARTRRRRRTAARNGCSARRARSSSSRRCASSPAAHPDARRRTHAHVRDPRHRTPELVARRRGR
VAARAVAHGARGRARGPHRPERQRQDERAALRVPLRAAACRHGDARRAGRVACAAALERAAHRGAAAGGA
GRFRADGRGTRAHGPHAAQAPVRRGHRGRCARRRSRAARRRSVGAARAGLRVAVGRREAARAARARARPA
AGAAAARRTDQSSRRAPSARTACARARAARDDARDDPRSEPRRRVLRPAARARGRAHRRERRACRRADAR
AHRRRVRRRRDRRCAPGVGAAARHADLSGGCRTMLKRTAAERGAPALRAAFTLPFAAAAMAAAAIVATVA
TAAEANAYPVTVRSCGRDVTFERAPARAVSNDVNLTEMMIALGLQTRMAGYTGIAGWKTGNARLRAALAG
VPELATHYPSLEALVDARADFYFAGWNYGMHPGGPVTPASLERFGIRTYELTESCSLVMRRPPASLDDVY
RDLANLGRIFGVDARAAQVAGAMRARIDAVRRTLASAAGGGTPPRVFVYDSGTDKPMTAGALAMPTALIA
AAGGRNVMDDVPRGWTQVGWESVVARDPQAIVIVDYSAVTAEQKKQFLLSQPALAQVEAIRARRFIVIPY
DAATPGVENAAAVETIARGLHPRAFAKAAR

>ABA51731.1 Substrate binding domain of ABC-type glycine betaine transport system protein [Burkholderia pseudomallei 1710b]
MGLAAVPGCEPALHAQSPRTRRARGSRAFSSSARMTPGVTHFQMEQAMKRYESIARRLARRAAAASPAFA
ALAWCAAAAAATTTAAAAEPAACRDVRMAGPGWTDIEATNALAGVVLKALGYRQSVSNLSVPITYQGLKK
GQLDVFLGNWMPAQAPLVKPFVDARAIDVLHANLSHAKFTLAVPDYVAAAGVHSFADLAKYAQRFGAKIY
GIEPGAPANQNISRMLADKALGPANWQLVESSETGMLTQVERAVRERQWIVFLGWEPHLMNTKFHLVYLS
GGDAYFGPDYGGATVNTVARADFASQCANLARLFRQMTFTVDLENGMIAAMLQGKRSAVDAAQHALRANP
SLVEAWLDGVRTASGAPGLPAVRAALDAQ

>ABA51907.1 trehalase [Burkholderia pseudomallei 1710b]
MLARATTCCRHALPFARRVRSGRGARCRGGVVGAARVCARRARAARVPDASIRARCAAPISGEPLARTMT
AVFHRRGSALRAGVHAMPKKRRAARSAGLRGIGTKRPRRRPTRPAAGHRSRSAVPARRAGAAAPASPGAA
CLVRAGACATRRMPRPTVICAKCGKNRLPERRFSCESAAAAYDRQTRRHASGPRDQETVLPFISKTGGGD
MVTPRHRPLHVENPFYRRLLNAPAWVALVAAAGIGCTSATLARADSSPAHASTAAAVASAASAPGAASIP
PPPSQLYGDLFVAVQTAQIFADQKTFVDSTPNADPATIVQLYQQQKGQPGFSLKAFVAQYFTPPSDESVT
PPPNQTLREHIDWLWPKLTRTTTTAPPYSSLIALPKPYVVPGGRFREGYYWDTYFTMLGLQEAGREDLVD
NMLDNFAYLIDTVGHVPNGNRSYYVSRSQPPFFAYMVTLAAKAEGNRVYQKYLPALRKEYAYWMQGERTT
PRGQATRNVVAMPDGSVLNRYWDASDTPRDESYLEDVKTAQQASGRPAAEVWRDLRAAAESGWDFSSRWF
GDNRTLATIRTTAIVPVDLNSLMFNLETTIVKGCAVTRDFACVAEFAGRAGKRAVAINRYLWNRNGYYGD
YDWKLGKPRDNLSAAALYPLFAGVAWPERAKQTAKNVQKALLKPGGLATTTYDTAQQWDAPNGWAPLHWI
ALVGLRHYGEKSLADDIGTRFLADVKGVYAAQGKLVEKYIVEGVGTGGGGGGEYPLQDGFGWTNGVTLKL
LDLYGG

>ABA52048.1 spermidine/putrescine ABC transporter, periplasmic spermidine/putrescine-binding protein [Burkholderia pseudomallei 1710b]
MLRQGAARHRRARCVRARRADLARVPARASAGLRMNAKPRVARRASYNHHASSSVTRGNIMHHTFNTPRR
AVALLSLAAFCATAGAAELTVVNFGGANGDAQKAAFNQPFEKATGNKVTAVEYNGEQAKVKAMVEAKHVN
WDVVEVESGDLNRGCDEGLYEKLDWAKIAKKSDLIAESPQTCGVGFFVWSTALAYDADKLKRAPAGWADF
WDVKKFPGKRGMRKGARYNLEFALMADGVAPKDVYKVLATKAGQDRAFKKLDELKPNIQWWEAGAQPPQF
LVAGDVVMSTAYNGRISAAQKEGKNLKVVWSGSIYDLDYWAIPKGTPNKALAEKYIAYTLSSKPQQDYAQ
HIAYGPANVAAIKALDAKTLANLPNSPANGKNAVLQNITFWTDHGDELEQRFAAWASK

>ABA52198.1 beta-glucosidase [Burkholderia pseudomallei 1710b]
MRRPANVRYQTARAAQPTATSRISSASASASASASAATATATATATAAAAAAIPRCRPASILPLDSRSIR
SPKPDPATARRARARPAGGRSVAYARPNPRQSKTSSIDHPIVQKFTCSQSTYALSLNLKFCYIPTQPPLT
PDSSPAARGRSVTHDRVSSVTRLTAPVAARSRGVSRRAERQNQANIHGDKRMHAKRLSIAVLSATLCALA
HAAGNDAPSPDIASRDAYALRRAHALVRQMTLDEKLQLIHSKYPMSDVPGGGAGFIQGIARLGIPDLNMV
DSATGSGSTSQPSTTFPATIGLAASWDKRLSYAFGAVIADQLRAQGFAMGLGGGTNLAREPRGGRLFEYL
GEDPVLAGEMLAARTRGTQDRKVIATIKHYVGNEQETNRMGGDDQIDERTLRELYLLPFEIAMKAARPGN
VMCSYNRLNGDYACENAHVLTDVLKNEWHFQGQVQSDWGAAHSTAKAINAGLDEEEDVGPTVFLTPALVK
QALANREIAPARLDDMVRRKLYAMIRTGVMDDPPRGGGTIDFAAANRFVQYAAEQSIVLLKNQDRQLPLD
AAGLKRIAVIGGHADAAVLAGGGSGNTRHPVTGAFPGCGGLTFPTTTGCNWWPNPWLKLDVPIVQAIRDL
APGATVAFAGNSDRQSPFAAYTPQQIDAAADLARRSDVAIVFVTQAAGEDFGELRSLALANPTNQDALVQ
AVAQANPRVIVVVESGNPVLMPWRDQVPAIVQAWFPGEGGGNAIANVLFGKVNPSGKLPVTFPARDEDTP
TWGADGTLAPNPVYSEKLKIGYRWYDAHRIAPMFPFGHGLSYTHFSYSGLEVKQRPDAATTVSFALTNDG
PVAGAEVPQVYLGDLDDPQEPPKRLVGWDKVGLRAGETRRVRIVIPAEMRRVWDASRNGWALAKGGRIYV
GASSRDIRLQQP

>ABA53699.1 putative inner membrane glycosyltransferase [Burkholderia pseudomallei 1710b]
MAANTGNVRIHAHAFAAHASRCRVRDGFSRVDSPMNQNAAHEPAAVARLRRARLVARPFPSDGSPFALNQ
RRSVPTRYDVSTPKERPTGRNRKPAARRAFGRTARQAARRDGRYARHAAPVSFCKSVRGERSRRRCGHGD
TSMKSGRAMSLCVQCVLLLIVFACSLSARGTALGPASASLATAPFARTAQARAFAQPSLPHAEAVAFAFA
QAEPFDAERPAAAGRPAAADAPPAPGKGIVISLCAAYLWVFALLTLVYASRHYVFGLDRLFKPQRAPYRA
ITHADWPEITVFVAAHNEEAVVADCLTALLATTYPRERLTIVPVNDRSTDNTRALIDEVQARAPELIKPF
HRESGKPGKAAALKDALREIRGDIMVVFDADYLPRPGLLKELVAPFFDPEVGAVMGRVVPQNADRNLLAR
LLDLERAGGYQVNQQARNNLGLVPQYGGTVGGVRKSALDAVGGWRDDTLAEDTDMTYRLLLSNWRTVYLN
HAECYEEVPERWPVRARQLTRWAKGHNQTLFRYLIPLLRSPVTPRRCRLDGALLLGVFVMPALLALAWGI
ALALYLTNGIDSLVLGLLVSVFALFAFSTFGNFGVFFEIVVAARLDGRATRLRLVPVNVVGFCVTIAAVV
AALWGLALDALLRRELRWDKTERFRRPLNSGR

>ABN85841.1 conserved hypothetical protein [Burkholderia pseudomallei 668]
MCAFVAAPLLWGLLAGVAAAQALAQPAASTVSVCAAVITGSSGTTVDVSYAGLPGNQPKLYGDFVALWQS
SVVPWSIAPAARLGIPTDAELGTVVMTGVSISATPYTVGYAVGPEVDDVCASALLAADGSTGVVDTVTLQ
LASVGATSLTFRYHTLSGYLPATAGNWVGLWRGRASPYNAAASVARVKVAADVTDDSVVMEGVTLSPGEI
YTAIYFMGEPLTTAAVLLNFVAPAR

>ABN86005.1 glycine betaine/L-proline ABC transporter, periplasmic glycine betaine/L-proline-binding protein [Burkholderia pseudomallei 668]
MLTTSLLMRRSTMKRNLIAAACGLAIAAAPFASARAGDAPTCKAVRFADVGWTDIAATTGLASTMLAGLG
YAPTKTIASVPITFAGIKSKQIDVFLGYWSPTMDPMIAPFTKAGTIKVLAAPNLTGAKYTLAVPDYVYQG
GLKSFADIQKYADKLNGRIYGIEPGNDGNALIKKMIDGNQFGLGKFKLVESSEAGMLVEVNRAIRDKQWI
VFLGWEPHPMNVQMKIDYLSGGDDVFGPNYGEAKVLTATPPDYAARCPNVAKFVSNLQFTTSIENHVMLP
IMNKEDPNKAAAEWLKANPQSLDKWLAGVTTFDGKPGLPAVKHYLGIQ

>ABN91854.1 type I pilus protein [Burkholderia pseudomallei 1106a]
MLFLLGVAAWPAAGRADTLLPRTQAFTVSAQIVAGCGVAGGGPASGLNFGTLDFGAHPAVATGQVSAAVG
GGALQIECSPGSTLKMTIDGGANASAGNTQRNLASGGARVAYRLYSDPARTQAIAVGQAVSLPVSGTITL
PIYGALTLPGGGAPAGTYTDTAQVTLSY

>ABN92885.1 ABC transporter, quaternary amine uptake transporter (QAT) family, periplasmic substrate-binding protein [Burkholderia pseudomallei 1106a]
MQAARLSRNRLRNPSNREATMKSTTTFVFGAALAAACALPSQSFAQDSAACRNVRFADIGWTDITSTTAL
ASLLFDGLGYKPTTTIASVPISFAGLKNRQLDVSLGYWWPVQQHQLQPFLDSKSISVVEPPNLSGAKATL
AVPSYVYQAGLKSFDDIAKHRAELDGKIYGIEPGSSANATIQKMIDTNQYGLGGFKLVESSEAGMLVTVE
RAIRDKKWVVFLGWEPHPMNIQIGMNYLSGGDAAFGPNYGEARVYTLTSPDYMARCPNAGKLVGNLRFTT
QMENQLMQAVMNKVKPAEAAKAYIRKNPQVLDAWLAGVKTYDGKDGLAAVKAYLGL

>ABN94432.1 putative peptidoglycan-binding LysM/M23B peptidase [Burkholderia pseudomallei 1106a]
MTGVVSSMSKSEIDRSVAWLSGVAAAFVVAGCATTSPVTPTDTLAAAASSASAAQPAAAPPGDAAQHPEH
AAAPAPIATRYVVRRGDTLSAIAQANGCTVRELQAWNRMGRRTRIGIGQVLRIAPPGAENAVASEAADGA
GATRAAGNAAGAARAADPASGAGGAPVADAPQPASAPESAADRVADHRVVQETKRHAQSIALAWPAKGAV
VETFQPGRNRGIRIVGRAGDPVRAAASGRVMYAGTGLNGYGTLILVQHNADFLTAYAHNRKVLVKTGDVV
QQGEQIAEMGTGDSTRAGMLFEVRRDGKPVNPMPYLASRQQG

>ABN94719.1 putrescine ABC transporter, periplasmic putrescine-binding protein [Burkholderia pseudomallei 1106a]
MRRRSNWVFPNGIANRAGGARAPRAHLHHQGNNMSVSHLRHAVARAALVALAGASALALPAAHAAGAELN
VYNWSDYIAKDTIPNFEKQSGIHVKYDNYDSDDTLQAKLLAGSSGYDIVVPTSNYMAKQIQAGVYQKLDK
SKLPNLSNLDPTLMKMIADADPGNQYGVPWAFGTDGLGYNVQAVKKALGENAPVDSWALVLDPANVSKLK
GCGVSFLDQAVDVFAATLQYMGRNPNSTNPADYQAAFEVLKKVRPYITQFNSSGYINDLANNDVCVALGW
SGDVGIAHRRSAEAKRPYDIKFSNPKEGGLLWFDVMVIPKDAPHAEAALKWINYIEDPKVNAAITNEVFY
PSANRAARQFVVPAVAQDPTVYPPEDVLKKMTLMRPMPADILRLENRLWAQLKTGH

>EBA48499.1 putative amino acid ABC transporter, periplasmic amino acid-binding protein [Burkholderia pseudomallei 305]
MSAPVRAHTPTGKGNSMKRRSLLKFGSMAGVMALAGQSPIARAADSGKGPIKVGILHSLSGTMAISETSL
KDTALMTIADINKNGGVLGRPLQPVVVDPASNWPLFAEKARQLLTQEKVACVFGCWTSVSRKSVLPVFEE
LNGLLYYPVQYEGEEMSRNVFYTGAAPNQQAIPAVEYMMSAEGGGAKRFFLLGTDYVYPRTTNKILRAFL
KSKGVKDSDIQEVYTPFGHSDYQTIVANIKTFAQGGKTTVISTINGDSNVPFYKELGNQGLKATDVPVVA
FSVGEEELRGIDTKPLVGHLAAWNYFMSVKGPANAKFKEQFAAWVKSQNLPGGAKRVTNDPMEATFVGIH
MWKQAVEKAKSTDVDRVRTAMIGQSVAAPSGFTLTMDGNHHLHKPVMIGEIRGDGQFNVVWKTKTAIRAQ
PWSPFIAGNQGKPDVVGSIPEFLRRRRAALA

>EBA51467.1 putrescine ABC transporter, periplasmic putrescine-binding protein [Burkholderia pseudomallei 305]
MHGRAAGAAFGRAGARPPTRNDERAPPEFRVGDPFGPRGLFSGWTRADNPIKLRPPAASRRPAPCNRKVR
MCSVCATTAGNFCGGDGLVVAFGVFKGRRETTRRSGEADGDRIGSIRENGMKAKVAGRLTALALCAGATV
AAAKDTQLNVYNWSDYIAKDTIPNFEKQTGVKVRYDNYDSDDTLQAKLLTGNSGYDIVVPTSNYAGKQIQ
AGIFTPLDKSKLPNLKYLDAQLMALVAGADPGNKYVVPWAYGTTGLGYNVDKAQKVLGKVPLDNWDILFK
PENLSKLKTCGVSVLDAPDQMFAATLHYIGKDPMSTNPADYQAAMQVLKKIRPYITQFNSSGYINDMVGG

DICFAFGWSGDVVIAKHRALEAKKPYKLEYYVPKGGAPVWFDVMAIPKDAKNKDAALQWINYIEDPKVHA
SITNAVYYPSANAQARKYVRPDVANDPAVYPPPDVVKTLFLLKPLPPEIQRLQTRLWTELKSGR

>EDO83218.1 NAD-dependent glycerol-3-phosphate dehydrogenase family protein [Burkholderia pseudomallei 406e]
MLSMKVAVLGAGAWGTALAAHLAVRHDTLLWARDAALVAELAVRRENARYLGGVALPPGLRYEADLATAL
SHAQADDALCVIAAPVAGLRALCRAMRDARRVPAHFVWVCKGFEADTRRLPHQMVAEELPDHASYGVLSG
PSFAREVAQGLPVALTVASASAACRERTLAAFHHGAMRIYTGDDVVGVEVGGAVKNVLAIATGIADGLGL
GLNARAALVTRGLAEMSRLGVALGGRAETFTGLTGLGDLILTATGDLSRNRSVGLQLAAGRSLDDILAAL
GHVAEGVRCARAVLSIARERGVDMPITEAVCAVLFDGVAPRDAVSGLLRRDAKAE

>EDO83840.1 chitin-binding protein [Burkholderia pseudomallei 406e]
MTESLKESANASHFDIQNNRLISPHRGTRTVSFLAWTHLEERYMKSLFDAPSSRPRARAALTLGAAATLT
ASFAALLAPMNADAHGAVGFPIARQYQCRLEGGYWDPPNGSAIPHDDCRAAYRAGNNSAYPFTQWNEVSA
NPVGQGNDLVQLKAAVPDGLLCAGGDTSKAGLDKAPASVWRKTQLTPNNGHIELQWENTTAHNPARMRVF
ISKPSYDPSRPLRWDDLQQIYDAPAPAPAPVPANGAGHLPGSIQSFYKLDVTLPAGRTGDAVLYSYWQRI
DAGNEGFFNCSDVTIAADERASGFPWVAARAFVEPGIAPRAGQQVRFRVMINDARGAEVVDVRQPITPYN
AERSVWAKQIADQVNGRYGNIAKIGVRSGNTIYFDATNLDANKVWLQPNYSSALSVVGAK

>EDO85817.1 putative lipoprotein [Burkholderia pseudomallei 406e]
MRMPKQSAAIVVIAAFSCGAMESSRYIISDVSRNQSSLPSRHVPDSTRTAGESGRSARAPDAPIQSTSRN
VRNRNASMRRATRALLRSLDIRHFFVVGRARLPLSLRTTTIIACCESLVATHKKSSRLASFRHRGAADAI
GPAACAVANALSCRDTCLRSHAQAAAVPANRAARDGLSSRQSVAAAPNPRIVPAGAALSGRMRRRIACMR
EHAGAMPRPPAAAHASVPFVPSARFARFAPFAPSALPIRCFT

>EDO89205.1 class D beta-lactamase [Burkholderia pseudomallei Pasteur 52237]
MPAAGAGRTRADRLLADPFCFSGLARGAATIVLVARVLVRKLSVCHNASPITRARAMPFSDRPAFQAHAF
DPDRANAVCRAVAFVGRKRSRAASWLDDPSRFAMFKSQQFIAAIWLSPGCCIVFAARVAQFAAAAETGRM
GQQDGLDERLLGVCGVRAVEEDRRRDSGEQNRIRRANEWRPAIESSLRSFGPNDEPFPTLCGTTMKFRHA
LSSAFVLLGCIAASAHAKTICTAIADAGTGKLLLQDGDCGRRASPASTFKIAISLMGYDAGFLRNEHDPV
LPYRDSYIAWGGEAWKQPTDPTRWLKYSVVWYSQQVAHHLGAQRFAQYAKAFGYGNADVSGDPGQNNGLD
RAWIGSSLQISPLEQLEFLGKMLDRKLPVSPTAVDMTERIVESTTLADGTVVHGKTGVSYPLLADGTRDW
ARGSGWFVGWIVRGKQTLVFARLTQDERKQPVSAGIRTREAFLRDLPRLLAAR

>EDO95399.1 putative ABC transport system, exported substrate-binding protein [Burkholderia pseudomallei Pasteur 52237]
MGGISSNHGSFGIARYRRRCAACVRRVYERLRHQLFLGWCRHDHLAYVHLQRIEDRNYAKGKCYLDSRHW
TFVDSTTDGLGSRQEGCRTDCHAGEIRAVSREHSNDRNRMKKVVGAMLLLAAGHAVSAELHIGSWPDYLP
DTLIKKFQAETGVKTTLDTYASDAALTQKLQSGGGGYDVVIAGDYYVPVLVKSGLLQKLDKNKLPNIANI
KPEYRHPSFDPQRDYAMPYTVVLTGFAYDSARVSGGKLDESWKSFFDPPAQLRGQIGDLDVEEELYMAAS
WYLGQDECTENPADAKRVLDVLQKQKPFVKTYSNDGTIDRLASKQIAVQHIWSGAAARAQDRLPSITFVY
PREGVRLFMDSLLIPAKAQNTASAYQFVNWMMRPENIAQVTNAVRYNNEIIGSERYIDAALLKNSAIKTP
EQYKIRLRPYKICSPAAIQLRNKVWLKLKGNR

>EDS82086.1 putative phosphatidylethanolamine-binding protein [Burkholderia pseudomallei S13]
MGRRAVPRGSSRRLLSLALVCACALAPVGGSRALAEDAFMLTSDDLHPGGRVGAPHVFNQGDCKGGNRSP
QLAWRNPPPGTKGFAITMFDPDAPGRGWWHWAVVGIPASVSSVPADASASGFLRKLGAAEARNDFGTDGY
GGPCPPPGKPHRYVITVHALKATDLRVATGRPAQMFDHEIGTESIGSAQLVVTYGR

>EDU07436.1 conserved hypothetical protein [Burkholderia pseudomallei 1655]
MPQREAMSALARLLPRRAPGWLAWACAALVVALAQPAWAVRCLTNSGQTSLTEPIGNVASYPTDAPDGYV
IWISAPRTTDGYCYKDLGGAGNLNIVDNIYFYANPDGKNPAAWGLEIGIRYRGVDHFGAGSRPGTGVPTG
FAVPPCSQTDFNAKRCPKIPVSITYQVVVRKKGAWVQPPGDVYTVFQFDGEKGLNNTNPSFQYKLSGLQN
LKPTPCMVDVTVTPEPGIVKFGQVQSSGNGFSPAVPRKPFSLALTKKCSIAVRVDGYFETSQAVRDGLLV
PQSDSNFGIGIEDRNGHAIAFNRQFVLTQMPANVSYQNIVLDAVLKSFGAPKIGPFTGTATIRLFIY

>EEC34719.1 carboxymethylenebutenolidase [Burkholderia pseudomallei 576]
MGIVCGFRLGGAGGGARRDCIAPAAVIRIHVGDACAARARQREERSASMLKPEVDSLVPHVPFSRRKFVQ
AALGGTFAAAVLPVSAQTITTDAAGLDVDTVQIRSGDASVPAYRAQPDGKSNLPVIVVIHEVFGVHAHIA
DICRRFAKLGYLAIAPDLYARQGDPSKHASIQELIDQVVSKVPDRQVIEDLDATVRWAGKNGGDLSRLGV
TGFCWGGRQTWLFAEHNPHVRAAVAWYGKVAGETNEMTPFNPDDHAAQLKAPTLGLYGGKDDSISQRSLA
RMRERLAAAGTQAARESEIVVYPDAGHAFFADYRPSYVKADADDGWKRAIAWFRHHGVM

>EEH25189.1 bacterial extracellular solute-binding protein, family 3 [Burkholderia pseudomallei Pakistan 9]
MHCIVEPARLRLAAAIRTAAAASIGRRPPRRRAHADRATTPIARCTARFVTRPPFRQETTMMKPLRPLLS
IAVGCALAAAAHADDAGPTLKKIRDTGVVTLGVRESSVPFSYYDQQQRTIGYSQEIALKIVDEIKKTLNR
PNLTVREIPITSQNRIPLVQNGTVDLECGSTTHTKERANQVSFSNSIFQYGMRLIVKKSSGVKDFPDLAG
KTVATTAGTTEERLLRQWNAEKGMAMQIISAKDHADAFLNVKSGRAVAFFMDEPLLYGAKAKEANPGDYV
ITGNSPVSEAYGCMLRKDDPGFKQLADRVIARMQRSGEAEALYVKWFNRPIPPKGVNLDYPLSADMKQLF
ANPNDKALD

>EEH25224.1 putative glucan 1,4-alpha-glucosidase [Burkholderia pseudomallei Pakistan 9]
MLFCYKNHNRRIHNGDEKMSARHFGFRQAVLVGSLAALAGAAALPANTSAAEAFGSPGAAPVRGPAAKSF
LGTAVNGASRVYFTGYRGILSEVYYPVLDTPESVDLQFLVGDAGKTFVDEEKQQAYSAAQTDTRTMSWQV
TTGNSSHNWQIRKIVFADPNHNAIVQRVTFTALDGRKVGDFNLYMLSKPYLDNAGANNTAQTVSGSGGTA
LVANHNSRYSALVASRPWKVVNGVGMTSNGFVGQSDGWTDLFGGTADKTMDWTFSSATNGNVAQMGWLDL
GDPSATSVSFDVVLGFGGTQDQALGDANTVLGSDLAGEQQQYDTGWHNYAAGLSSQNGAADDGYYLAAMT
LKTMQDKSNGAMIAGIGTPWGETQGDANAGGYHLVWPRDLFKFANALTTAGDTSSATSVVNYLFNTLQQT
TDCGTAEYNAPGCAQGYSRVGRFPQNAWVNGWPYWQGTQMDEQAMPIILAWRLGPSVFNPLWPKIKLTAD
YIVNTGPWTYQERWEENAGYSPSTIAAEIAGLVTAADIATQNGDTASAARYLAAADYWQENVAAWTYTSS
GAFGNGSYYIRINPAGRAGTGTDRASFAPTAGPDTPQTLSVKNGGGSHDARRVVDGGFFELVRMGVKRAT
DPTIANTISVYDSVLGQTLNVANAPALSPNAWFRYNFDGYGEHNDGGDFDGTGAGRLWPIFTAERGMVEI
ARQGSGGAGSAYFSTLKQLTTPEGFVPEQVWSNSTTLPDGWAVTTPAGYTPGQPTKSMAPLNWAMGEYIS
LLASIQAGRIVDVPSVVCARYNNCAAPPQSGQVPVAVNVNASTQLGQQMYVTGNVAALGNWNTDLGIPVD
PASYPVWRNTVNLPAAQAIQYKYYRKNADGSVTWENRSGNRQLQTPASGTLALNDQVSW

>EEH28759.1 putative secretory lipase [Burkholderia pseudomallei Pakistan 9]
MKNPIRNTDVQPQHRRARRTLRVLRRAAHRLAMWDDTVEIASASAFRDRSQSARLPRRTLEEASLMSRTG
FMPLSTLATMAVAAAISTPAAFAESAPVSGNGGIPSFYRWTDKVPQTRGQMLRTEVLTPRQGLAAAGQNL
RILYTSTDGIDGHTPIVVSGALFVPKGTPPQGGWPLMAWAHGTVGSADICAPSYAGHSERDVRYLSHWLA
QGYAIVATDYQGLGASGPHPYGLTRPLAYSVLDSIRAVQNGGFELSPKRIVVFGQSQGGRAAFATAVYAK
SYAPELDIGGVVATGTPYSTIRHRADDAEHSKLHRSVIHNILRLNATPKLLDPSFATSDYLTERATPAFE
FSQHACLHAIKQKIASDGLTFENSFKRSPQPVLDWIRREAAYPTLKSAIPIFIGTGGKDRNVPVSDQAAL
VKDACEAGDRIEWHVYPALDHSETVNGSLSDSTPFVARAFAGEHIDGNCASLPSP

>ACQ98979.1 serine protease, subtilase family [Burkholderia pseudomallei MSHR346]
MTSRKWARSRASQAKHAIYAATFFAAAALSAHAAAAWVDTQTGAYPALAQQALAASQASAAATAAGKAID
TAPGEPVRVVVSLNLNDEARLDRFLRDLHTPGSAAYGRHLTPAEFAAQYAPTPQQVALVEAHLRRAGFRD
IEVAPNRLLISATGTAAAVKTAFNTRLKRFTLEGRRVYANQDAAQVPAELGRIVGAVLGLDNATLARTYN
RQAAVTGAVGGAKASLAARASDASAAASGAPVLTGHDPLEFSRIYRAGSTPTASQTTVGVIMAGDAAPVL
RDLDTFAAKAGLARVAATVTRTGPPGSDYSDNSGLSEWDMDSQAIVGAAGGAVKGLVLYAAPSMLLSDIT
SAYNRAVVDNVAKVINVSLGVCEADARASGTQAADDRIFKSAVAQGQTFVVAAGDAGAYECSVSRVSGGQ
GVPARSNYSVSEPATSPYVVAVGGTTLSTDRTTLAYAGEVAWNEGLQPIGVYDAYGSYDGTRRLWATGGG
YSRSEAAPAWQRSVLGASAKARALPDVAFDADGRSGAHVYVNGRTEQWGGTSLAAPIFTGIWARVQSDNG
NRLGFPLASLYRYAPANGAFAHDVKSGNNGSGGYGYKAGAGWDPVTGFGSLDIANFAAFVKQTADFAR

>EES27008.1 bacterial extracellular solute-binding protein [Burkholderia pseudomallei 1106b]
MERSPMQRASGRPFSSASSKERSMKRIAWLAAVLASLACAAARAADGNVLNIYNWAEYFAPDTIAGFEKE
TGIKVRLDVYDSNEALQTKLTTGNSGYDLVFPSNDFLARQIQAGLYRKLDKSRLPNLTNLDPAIVARAAE
VDPGNQYSVPYMQGTFGLGLNVAKVKQALGGPLPANTLELIFNPAYAAKLERCGIAFNDAGSEVFPLALR
YIGRDPNTTDPRDYEAALDMMKKIRPTIRQFIATPVMNDLATGDVCVVTGYSGAVLVAARRAAEAKNGQQ
IVYSLPSAGAPFWFDSMAIPKGAAHADHALRFIDYILRPDVVAKISNKVMYPNPNRVATPLVDRRLTANP
AIYPDAATMRTLWVKRPMPPQAIRMQTRYWTRFKTGY

>AFI68205.1 Gfo/Idh/MocA family oxidoreductase [Burkholderia pseudomallei 1026b]
MRRPSRSAPFRKQTGSFMSSTLKIGLMGFGFAGATFHAPVITHSGRTRLAAIATGQPERATAAYPEAAVV
PDLDALAARDDVECVVIATPNDTHVDLAKRALEAGKHVVVDKPVALSAADALVLARLAAARGRVFAPFHN
RRWDGDFLTVRQLVESGELGRIVCFESHFDRFRPHVRTRWREEAARGGGLLFDLGPHLIDQALALFGPPE
TVSATVKTRRDGGDAPDFVHLQLGYPDKDVVLHASALAAIEPARFTVLGTQGGYQKHGLDTQEDQLKAGL
TPDDVEFGGGNPPGVLRGLDGELEVERPVPTLDGQYAEFYRALAASIHDGAPFPVAPQDAVDVMTIIETA
MRSEREGRRLPFVREAV

>AFR15131.1 hypothetical protein BPC006_I1246 [Burkholderia pseudomallei BPC006]
MRAARRHCLTRKKNMNRKTRAAIGVAALCAAAGAAHAARLTVEEIDADARETVAYRCANAPKPVRVSYWR
AGNGQSFALVPVNGVRLLFVDTVSASGARYQAGRYVWWTKGRDANLYDEIAGERAPPVLADCSEVRKPRG
KG

>AFR15554.1 type I pilus protein [Burkholderia pseudomallei BPC006]
MSACAGVQAQTSPLTGTVNSQLVLTTGCAVDTGGGSVNSANFGTLDFGTQPSGFTGRLTSAAKGGGSTST
QVTCSPDVTSIQVTIDGGQNASKGATVGTGTRALANGASFVPYEVYADAGHSQQYVSGTAQSVAVPTPGA
AFELPLYGVVNKTNASALAAGTYTDVLNVTLGW

>AFR16106.1 hypothetical protein BPC006_I2236 [Burkholderia pseudomallei BPC006]
MRHASITRRFAQIVAIKGEVRMSQRILSPLVASCAAAALLSAFAAHAAPPIKGSVLGGNDGQLQYTVKVD
SKQFGTMQETRKIRSGETDDYNWKSVPPSGAVPMPDACPNADTLPRDANGAMVRQAQVRLAPAVDSKGVA
NVQLSFQASAPSGTKKVTVNGKALQCPNVVAVSQVKWLSIPTGGSKSITMRDGTKITVSIKR

>AFR16694.1 glycosyl transferase, group 2 family protein [Burkholderia pseudomallei BPC006]
MRVVESASKSERSWDRVRNRPRRSIASPSRLPPDRREARGMTESLSIVGWVLLALVCASCGYAVLAACAP
APRVPRAAARDGFEPVSVLKPLCGSEPHLYENLATFCEQRHPRYQLLFGVASAADPAIAVVRRLQADYPD
CDIELVIDARVYGSNLKVSNLVNLAERARHGRIVIADSDIAVEPDYLTRVTAPLADPSVGVVTCLYHARS
VGGFWTRIGAQFVDAWFAPSVRITHLGGSSRFGFGATLALTRATLDAIGGFKALKDELADDYWLAELPRR
LGLRTVLSEVNVATDVAEPSFAPLWLRETRWLRTIRSLNPAGFAFLFITFTAPWLVIGAALAAWLGPASA
AGATAAWAAAIGTLARLALHARGAAGWRAFWRDLPLVPVRDALLALEWLAAAFGTQVVWRGARMTVVGGD
ARATVVEAGDGR

>AFR17326.1 glutamate/aspartate ABC transporter, periplasmic glutamate/aspartate-binding protein [Burkholderia pseudomallei BPC006]
MEAASRRTRGRHRNGVARVRIRAPNAGLSAHVEMRMPPGKSVVNNHYFHKNYFAKLARVFAKEMTTMKFP
KAMLMVAALSTFAGGAIAQETGTLKKIKDTGVIALGHRESSIPFSYYDQNQQVVGYSRDFQMKVVDAVKK
KLNLPNLQVKNIPVTSQNRIPLVQNGTVDIECGSTTNNLDRQKQAAFSDTIFVIGTRLMTKKDSGIKDFA
DLKGKTVVTTAGTTSERLLREMNNKNQMGMSIISAKDHGESFQTLETGRAVAFMMDDALLAGERAKAKQP
GEWVIVGKPQSQEAYGCMMRKDDPAFKKVVDDAIVQVEKSGEAAKIYSKWFENPIPPKGLNLNFPLSDEM
KKLYANPNDKALD

>AFR18870.1 collagenase [Burkholderia pseudomallei BPC006]
MTIHDHQSNSRQQRANMKNSHNVVNRFIVAASIIIGVVLYSSAWANPQPMHTKQARMPRIPQNLPLSPDQ
AKYDLPLSKYDRATLMEPLRRKQSAKPDRRTRPGADCRDMSIMTQYHGTALADYIANLPDYECHYGLFSI
DRAMAAQIFNSENVWAVASRLTQEINRYDATNITLVNLLIYLRAAYFQYDAAQLADPIPGLVVWLRPYIL
QSLSGDALYLENSRAPSTANELMILITNMKDEAYYLPTLKDRIAFYTASATNPQAAAPLLQRSAAGGFTG
LLTVFFYAHQRSGAQPMLDSDATLPETLNRFVTANRAYLSNTSAAYQLADAARETYRFLRYPSQKPRVKK
MIQDMLASTTMTGPDNDLWLAAAEAADYGDPGNCADYGTCDYQKRLIEAVLTHRYSCNANVRILAQDMTV
PQFQSACQSVAQEEDYFHRMMKTGHVPVANDHNDTIEIVVFGDYDNYRKYASVIYGISTDNGGMYVEGDP
SAPGNQARFIAHEASWLRPEFKVWNLEHEFTHYLDGRYDMAGDFAASTAKPTVWWIEGLAEYISRKNDDQ
ESIDAVRTNAYRLSDVLQTTYSSGDYVTRAYRWGYMATRFMFERHRADVDAIVSRFRVGDYDGYADYVAY
MGNRYDSEFVDWARGATTTGEPPLPPTKAGH

>AFR20596.1 serine metalloprotease [Burkholderia pseudomallei BPC006]
MSFLAGSAAKQAATAQHRRLIPFNFVWGSMSMSILIRTASFKATVLCAALAGLVSAAQAETAAAPQVPGP
ADAVNQLIVKLRAVKTPPGATAAKAERADVQAVIDRVLAARNARAAGRAFGAAAASAPGNPDDPAAGIRI
KRDMSGGATVLSLQRHVSLAQAEALARDFAADGAIEYAEPDARMHPFVVPNDTRYSEQWGYFNPTAGANL
PKAWDRTTGSARVVVAVIDTGYRPHADLAANLLPGYDFISDIPSANDGNGRDSDASDPGDWVSAQEDGDP
SGPFYGCGASDSSWHGTHVAGTIGAVTNNGVGVAGISWVGKVLPVRVLGKCGGMLSDIADGMRWAAGLPV
PGAPSNPNPAKVLNLSLGGYGRTCSSTYQNAINEITSRGANVVVAAGNNGGSVSTTQPANCRGVIAVGAI
DSRGVRASFSNTGAAVKISAPGVGILSTLNAGKTSPGADSYASYSGTSMATPHVAGTVALMLAVNSTLSP
SQILQRLQSSARPFSSGSSCSTSTCGAGLLDAGNAVDAAAQ

>WP_004185598.1 MULTISPECIES: 16S rRNA pseudouridine(516) synthase [Burkholderia]
MDLESILFTQGFGSRRQCRALVEAGHVAVGGATCADAHASFDTTQLVFEVDGVAWPYRARAYVALNKPAG
YECSREPQHHSSVFALLPPQFAERGVQCVGRLDQDTTGLLLLSDDGQFVHAYTSPKRKVPKTYVATVRHP
LDDAQLDALRAGVLLHGEPKPVAALAAVARGERLLEMTIAEGKYHQVKRMVAAASNRVEALHRSRIGGFA
LPEDLAEGAWRWLDERDLAALRDTAETLSG

>WP_004185616.1 MULTISPECIES: right-handed parallel beta-helix repeat-containing protein [pseudomallei group]
MMNRSQQTFAGFAAACAFAAGSAAQAWAAPADAIVYPAGNGSDQADALQAALDALQTGQRLVLAPGQYVV
GRSLLVKNAQVVVSGYGATLVATNDADQTIEMRGRDSTLVGVTLVGTGATRLTTPESTKVDVTGAGVQVL
DVEVRGGASAGIFVFGGTDVAVVGNTVRDTLADGIHTTYGSRNVLVRDNTVQNTGDDMVAVVSYKGDGKL
SSNVLIQNNTLLGNYWGRGATVVGGADVTITGNTVRNVQKAAGILVGQEDPANTYDARNVIVSNNTISDI
EWPDPDNTRPPAYMAAIDVNTWSGKATSVSVTDNRISRARYAGVRALGNVCQLRVSRNALASIDGTPIAL
QQAPSCAAGQIVCASNTLDGAALASPAGCSATDGLTITGANVARMPQVRAYLRQTAAPSAAKAGSAAASG
E

>WP_004188402.1 MULTISPECIES: succinate dehydrogenase flavoprotein subunit [Burkholderia]
MAAIKTSLPRRKFDVVIVGAGGSGMRASLQLSRAGLSVCVLSKVFPTRSHTVAAQGGIGASLGNMSEDNW
HYHFYDTIKGSDWLGDQDAIEFMCREAPNVVYELEHFGMPFDRNADGTIYQRPFGGHTANYGEKPVQRAC
AAADRTGHALLHTLYQQNVAAKTQFFVEWMALDLIRDADGDVLGVTALEMETGDVYILEGKTTLFATGGA
GRIFAASTNAFINTGDGLGMAARSGIALQDMEFWQFHPTGVAGAGVLITEGVRGEGGILRNANGERFMER
YAPTLKDLAPRDFVSRSMDQEIKEGRGVGPNKDHVLLDLSHIGAETIMKRLPSIREIALKFANVDCIKEP
IPVVPTIHYQMGGIPTNIHGQVVGTSRDHKEPINGFYAVGECSCVSVHGANRLGTNSLLDLVVFGRAAGN
HIVEHVKNQRDHKPLPADAADFSLSRLAKLEKSTSGEYTQDIANDIRATMQKHAGVFRTSALLKEGVEQM
AGLKERAAAVHLKDKSKVFNTARVEALELANLIEVARATMVSAEARKESRGAHAHSDYEHRDDENWLRHT
LWYSEGDRLDYKPVQMKPLTVESVPPKARTF

>WP_004188537.1 MULTISPECIES: efflux transporter outer membrane subunit [Burkholderia]
MTMRNDARHDERHQRRRGIARRAARHAAALALCAAVSGCLSLAPDYARKPPPVPAAYPAYPGEPGAGSRT
LEQLGWRAYFVDPRLQALIETALANNRDLRIAAQRVEQARAAYGIRRADQYPTIAANAAYLRFREPGGLL
LPSPIIGELYSASLSEVQWEIDFWGRVRSLKAAALDTFLASDASRRAVTVSLIGQVADAYLALCAYDERL
ALTNETIASRRDSLRIFRRRYEEGAISKLDLTQSEILLQQAQTLGSQLQQARDVQQHALDLLVGTNAAAR
TPASLDDASIAPDLAPGLPSELLANRPDVVAAEYQLRAAHANIGAARAAFFPRIALTSSIGTGSTELHNL
FASGTAAWNFIPNLSVPIFDAGRNIGNLKLANARRDEALAQYEKTIQTAFRDVADALAARHWLADQVRIA
QETLATQAERARLAKLRYDSGAARFLEVLDAQRDLLNAQQQLVQTRRALLASRVALYAALGGRLDEPARD
GAAPRPSPEPSAKQGPLS

>WP_004189053.1 MULTISPECIES: DUF2501 domain-containing protein [Burkholderia]
MKKRIHRIAAAGILVAGVLSLSAAHAQLGDFLKQGADAGNGGAGGIAGALGNLGGGGGAASAGSLLTPGS
TGNVAGLLQFCVKNDYLGDGGASSIKDALMSKLGAGVTSDSTYASGASGILDAGNGRTLDLSGGQSFKQQ
LTKQVCDKVLSQAKALL

>WP_004189371.1 MULTISPECIES: hypothetical protein [Burkholderia]
MKLLTQLAVASAVAGALAAAATAAFAQNSPGMPGGILNDQFRLNEHPQMQFAASAPTKKYQSGKKSDLRR
KGDMGDPNGCNLKCPMD

>WP_004189720.1 MULTISPECIES: hypothetical protein [pseudomallei group]
MNRAIRMRAAVAAAAFSASLSAGAAGPAIDPGRYLYVEGGSAHGVLTVKGNAFEIETIGGNCHTCALSGT
FDGRVGIARDGENVCRIAVSGGHGDLRLDTSGSDACRDFCGMRASFDGEYRRPGAACTDRARDVRTERSH
RQYAAHDYDAARTTLKALLAECGGFMGWIELDRAKSDLALTEYHRGDRAQCVAVLSDTIAYRAQQDHSDA

FGLPPCDADNYKSTGDAILHNLALCQAPAKR

>WP_004189892.1 MULTISPECIES: OmpA family protein [Burkholderia]
MNKLSKLAFIAATAVMAASASAQSVPASRQAVNDNWVNGTGEWVWMNGTNELCWRDAFWTPATANAKCDG
ALVAQAPAPAPVAPVAPAITSQKITYQADTLFDFDKAVLKPAGKQKLDELAAKIQGMNVEVVVATGYTDR
IGSDKYNDRLSLRRAQAVKSYLVSKGVPANKVYTEGKGKRNPVTGNTCKQKNRKQLIACLAPDRRVEVEV
VGTQEVQKTTVPAQ

>WP_004189909.1 MULTISPECIES: 3-hydroxyacyl-CoA dehydrogenase [pseudomallei group]
MEIRDNVFLITGGASGLGAGTARLLTEAGGKVVLADLNQQAGEALARELGGVFVKCDVAREEDAQAAVAA
AAKLGTLRGLVNCAGIAPAAKTVGKDGPHPLELFAKTITVNLIGTFNMIRVAAAAMAANEPAQTGERGVI
VSTASVAAFDGQIGQAAYAASKAGVAGMTLPIARDLSRNAIRVMTIAPGIFETPMLLGMPQDVQDALGAM
VPFPPRLGKPAEYAMLVKQIFENPMLNGEVIRLDGAIRMQPK

>WP_004190112.1 MULTISPECIES: thiol:disulfide interchange protein DsbA/DsbL [Burkholderia]
MKKLLSSLFLSLSLVAGFAQASPSAPVAGKDFEVMKSPQPVSAPAGKVEVIEFFWYGCPHCYEFEPTIEA
WVKKQGDKIAFKRVPVAFRDDFVPHSKLFYALAALGVSEKVTPAVFNAIHKEKNYLLTPQAQADFLATQG
VDKKKFLDAYNSFSVQGQVKQSAELLKNYNIDGVPTIVVQGKYKTGPAYTNSLEGTAQVLDFLVKQVQDK
KL

>WP_004192320.1 MULTISPECIES: EamA family transporter [Burkholderia]
MNPVSLFCIVTGVMLNACAQLLLKAGVRAVGHFEFTRANIVPIGFKIATQLPIIGGLGCYVLSVVVWIVG
LSRVDVSVAYPMLSLGYVVNAFAAWYLFGEVLSVQRLVGIGIILIGVFVLARS

>WP_004192750.1 MULTISPECIES: phosphate starvation-inducible protein PsiF [Burkholderia]
MKIRSLMAALLVGGVLATPAFAANSQQDKMKACNAQAAGKTGDERKAFMKDCLAAKPAKKMSQQEKMKAC
NTQAADKKGDERKAFMKDCLSAKPAA

>WP_004193385.1 MULTISPECIES: fimbrial protein [Burkholderia]
MKKALLSAVACAALSTSAFAAGTGTLNFTGEIVAGACGIDAGSVDQTVRLGFVPANTFKAAGDKSTPQNF
DIKLVDCDTSVAKNAYFTFTGTSNATQPKLIATVGSATNVGIRLQSASGEYLDNGAEQKGPVVLSNGTSV
ARFAAMYESTAASVTPGTADGVANFTVRYQ

>WP_004193508.1 MULTISPECIES: glycine zipper 2TM domain-containing protein [Burkholderia]
MDNQNTTTQQRQRLHPLVATAAGAVIVASLAATAAITGVFPKASSTNAQNAQTQAALIASQPAVDTAAAA
SAALAAQAAQQAAQQPQQTVAQAAPKPTSHQTTRHHHKTSAAPQPPQYAQQPYPQQQQSTYCASCGTVAS
IVPVRTAGTSSGLGAIGGAAAGGLVGNQFGRGNGRTAMTIIGALGGGLAGNAVEKQVRSETDYQVQVQMQ
DGSTRTFTYHNPPPFGQGQRVRVENGTLVGA

>WP_004193933.1 MULTISPECIES: PHB depolymerase family esterase [Burkholderia]
MTRNARRTWQWISGLLAICAAALAALPARADVAAGPGAWSSQQTWGADTVNGGNLTGYFYWPATQPTTPN
GKRALVLVLHGCLQSASGDVIDNSRGAGFNWKSVADRYGAIILAPNATGNVYGNHCWDYANTSPNRASGH
VGVLLDLVSRFVGDAQYAIDPNQVYVAGLSSGGGMTMVLGCIAPDIFAGIGINAGPPPGTTTLQIGYVPS
GYTAQTAANQCKAWAGSKADQFSTQIAGAVWGTSDYTVAQAYGPLDTAAFRLIYGGTFAQGSKVSIAGGG
TNTPYSDSNGKVRTHEIVVSGMAHAWPAGAGGDNANYVDATHVNYPAFVMDYWVKNNLRANGAPVQAGTP
PTGLTVTNATQTSISLAWNPVANASSYNVYRNGNKVGSSTSTAYTDAGLIAGTAYSYTVTEIDPSLGESA
PSPAVSATTQPSFACSATTATNYAHVQAGRAHDSLGIAYATGSNQNMGLDNVFYTNTLAQTSAGYYIIGD
CP

>WP_004194152.1 MULTISPECIES: ribonuclease [Burkholderia]
MARKWLRNGALASVFAIVAMGNIGATPGSLVSAAYAREAATVDGAAAQVDTIPASRLPREAATTLSLIAA
GGPYPYGKDGAVFGNYERILPKMRRGYYHEYTVPTPRARNRGARRIVCGGPLRRVDNCYYTGDHYNSFKR
IVD

>WP_004194433.1 MULTISPECIES: polyisoprenoid-binding protein [pseudomallei group]
MNKQLMIAAGALAAALSFSAHAAPATYQFDPSHTYPSFEADHFGGLSVWRGKFDRSSGTVTLDRAAKTGT
VDVTTKVASIATGSQKLDEHLQTADFFDSAKYPDATYKGTIKFEGDRPAEVVGNLTLHGVTKPVTLKIDS
FKCMPHPMLKKEVCGVDAVGEFNRDDFGLDYGKQYGFKMKTKLLITAEAVKQ

>WP_004194773.1 MULTISPECIES: c-type cytochrome [Burkholderia]
MKKPQTALKTAAALALAAGFAIGTAHAANVAKGKELVESHNCAACHGAKLDNPINAEYPRLAGQHADYLV
WAMRQYQMGLTNPLLGRNNAIMQAQVQSLSIADMKDIAAYLESLQGSLVFKK

>WP_004196853.1 MULTISPECIES: BON domain-containing protein [Burkholderia]
MSNSRVRQTLVRTTLVIGLSAGLAATLQGCVLAVAGAAAGGGALVATDRRTLGAQTEDREIQVKALTQMN
NGLPDGSHVNVTVFNRRVLLTGEVPSDAAKQRAEEIVRAINNVNAIVNELAVGPASSLSDRANDSYLEGR
VKTAMIAEKGISANNYKVVCERGNVYLMGLVTTDEGSRGADVASRVPGVEKVVKVFQYIKPQDAQALTAA
TPASGASAAAAPATAPEGATVGAVPDSSVTATPLSAPAPVSNSSNVHPGNPKAGTP

>WP_004197153.1 MULTISPECIES: sensor histidine kinase [Burkholderia]
MSHSLRGRLLWWLLLPLAVFVAIAGAMSYDTARKTADLVQDGALVASARVIAEDVDWEGGALVANVPPAA
LELFASPAQDHVYYKVRTGGGRLLAGNPDLDGPAAPAASGAQPVLFDTALGGLAIRAVAYTRELYNAGNT
ETVTVVVGKTQTSRQMMIAAIWHPQLWRLALMLALAMALVYLGLTFELRPLMKLKEDVADRGPMELEPIR
TERLHFELRPIVDAINQCIAQLNLHAATQRRFIADAAHQLRTPIAVIDTQIQCARQRENGDAALAALLAS
MQRSSRRMADVTDKLLLLAHAEAASPARLAARVDIAAVVSGVLEEAIVLAERRRIDLGAELDDDLQVAGS
ESLLSALLMNLVDNAVRYAHEGGRVTVSARRDGDAVVLEVVDDGPGIPAEARPHVFKRFYRVARDEEGTG
LGLAIVEEIAQSHGGAVSLATGPGNRGVRMTVRLPAYRN

>WP_004197194.1 MULTISPECIES: iron-sulfur protein [Burkholderia]
MKTSRRSFLITSVGAVSALALTREAFSADLPMLSETDPTAVALGYKADATKVDKAKFPKYAAGEACAGCM
LYQGKKGSASGPCGAFPGKQVAAKGWCNAFTKMA

>WP_004198637.1 MULTISPECIES: flagellar biosynthesis protein FlhG [Burkholderia]
MDKRITDQAEGLRRLLAGRASRVVAVTGGPSGVGCTSTVANLAAALTALGKDVLVVDERANVRSIAATLC
GSWLRDGEPVRHELGFAVCEASRLARVGYSDAQLDALGDGAADIVLIDAQLDANGALSALARDAHDVLVV
TRVSASAITEAYACMKRLHYAHARAQFRVLTNHVQSAADARAAYENLAGVASRYLRVSLSNAGCVAADSL
IERARGLAHTVVDAFPSAAAARDYRQIAADLLYWPMRPGPGIGRVRTGSSAFERGAAHAA

>WP_004204600.1 MULTISPECIES: GNAT family N-acetyltransferase [pseudomallei group]
MSVAPAARRPGVGAALLDAALAAAARTAGQVALAAVDGRGAARAFCERHGFVAYGVEPRALDAANGCADA
LPMVKFPPLAPPHA

>WP_004521493.1 MULTISPECIES: DUF2282 domain-containing protein [Burkholderia]
MSAKLSFHTALLVGAVASLVASGAHAAPLTKAEADAAVAAHKEKCFGVALKGQNDCAAGPGTTCQGTSTV
DFQGNSWKFVQGGTCTSIQVPGGGHGSLTPLKS

>WP_004521540.1 MULTISPECIES: heme-binding protein [Burkholderia]
MSPELTLERALRIVEAGVAHAARLSAAGAAVAIAVVDAGAHLAAFARMDDAFIGAIDLATRKAGTAARFR
MPSASLGALSGAGGPVRSIEHSNGGLVTFGGGLPLVDAHGRCVGAVGVSGGTVDDDAAIAQVCVDAFITL
LNAAQGART

>WP_004522915.1 ammonium transporter [Burkholderia pseudomallei]
MRKILMSLLMAGSLLAGGIGAAMADDASSAPAAASAATASDTSAGAAASAPAASAAPAAPAAPAASAPAA
ASAAAPASAAAAPAAPTAPFSVDSSKINSGDTAWMLTSTALVLFMTIPGLALFYGGMVRKKNVLATVMQS
FAITALITVLWTVVGYSLAFTPGNGFIGGFSRVFLSGMNYIHGDKATTLTVSHLASTIPESVYFVYQMTF
AIITPALICGAFADRMKFSAMLVFMTLWSLIVYVPIAHMVWEPTGWLSADGVLDFAGGTVVHINAGIAGL
VSCLMLGKRVGYGREAMAPHNLVLTLIGGSMLWVGWFGFNAGSAVAADGRAGFAMLTTQVATACAALGWM
FAEWIAKGKPSVLGIVSGAVAGLVAITPAAGFVGVAGALVIGIAAGVVCFWSATWLKHKLGYDDSLDAFG
VHGVGGILGALLTGVFAVKDIGGADGSLLLQAKGVAITLVYSGIVSFVLLKVIDIVIGLRVTEEEEREGL
DVILHGEHVE

>WP_004523300.1 NAD(P)-dependent oxidoreductase [Burkholderia pseudomallei]
MEAIMKLGFVGLGVMGQPMALNLARAGTELVVWNRTRERCEPLRAAGAQVADSAADVYRRARIVILMMAT
DAAIDAVLDRGKPAFASNVAQHTIVQMGTVSAEYSRGLEADIRAAGGRYVEAPVSGSRQPAEAGRLVAML
AGEPAAVEEVRALLAPMCREIVATGQVPSGLLMKLAVNTFLIAMVTGLAEAAHAARGFGLDMKQFQAVLD
AGPMASSVSRVKIDKLVNEDFEVQASIVDVFKNSRLATEAAHGAHLAAPLLEVCCELYRETEALGHGQAD
MAAVVHAIEARSVARYAGS

>WP_004523794.1 DUF3455 domain-containing protein [Burkholderia pseudomallei]
MTAALLTSATMAAHAAPPPPPELVPRDAARDRAFAAAGIQVYSCEYDGNHRLAWTFQHPEATLYDAAGVA
VIKHGAGPAWEAQDGSRIVGEKLADAPSPNADSIAQLLLSTRASASGSLASVRYVQRLDTKGGTAPAAAC
TAEHQTGGSPYYARYVFWK

>WP_004523965.1 MULTISPECIES: hypothetical protein [Burkholderia]
MKTFSHALKRVVALVIAMPAAAGWASGQHRHGAADADELPRAFAAKEMGRSAAGARSYGSPVTSVFEEAK
GALPAFDAKRSGAAGAAGFAHELGAAERRQRLAEEDRAVQRLRDSRHDGRGRWPGGAMPPRVPGRPDIAI
EKPVPGVTCASGSDNSIGCGTGF

>WP_004524043.1 MULTISPECIES: hypothetical protein [Burkholderia]
MQRPSLIASSLLTASMFALSFAAHAQQTGIVRFTGMIVEPPCSFSVAGGAGTHAQLQPACPRPAKGSVTF
VDPQSGAALRTVTFTELSRAIALPAGQTGDARPIVAVVSYL

>WP_004524330.1 MULTISPECIES: helix-turn-helix transcriptional regulator [Burkholderia]
MFSMLYFPMVSALSLLGADAPTHLHSHLKLILGGEFNAALERSSEWAETTVASERTSWDLQLHADLQLVL
GFEVEAEENYRRAQRKIRGSNSKIRIATCRNAAWQALFRYRVTTALACFSRICDEPGIEAGGLVEARFGI
ACALYEMGRIDDAFDAIDSMEKIAEQQSDEMRAHWKDLIAVLRFDLVVQSELRRAAAFVDHVYWQSAQSM
SRVDRAHGVSEAAVSVETPLLRGRVAYLLQLRCAAAGNRDAVAELARCLDAAGEQGFVDFRYTLRLEIAL
ALLAGDAPNLAQFVLEPISDTLHGAESSRRYREYFYCAAKVHLAQDHTQESLALYRRYALIAMRCLREDA
LIGRQFLVGQELKQLPQSDDVTVRLPLKYRRAYHYILQNLNRSDLSVREIAAEIGVTERALQNAFKIYLG
LSPRELIRSRRMERIRTELVDFTLTGERNVKEAARKWGVQNGSTLVIAYRKEYDETPSETLAR

>WP_004524503.1 transporter substrate-binding domain-containing protein [Burkholderia pseudomallei]
MKMNQTLLALACCAGLAHAQTAPQPGAPASRLDDVLARGTLRVCTTGDYKPYTYRREDGAFEGIDIDMAA
SLAKSLGVKTAFVKTTWPTLTDDFVAKCDIAVGGISTTLERQKRVFFTQPYVSDGKTPIVRCADAERYQT
IAQIDQPQTRVIVNPGGTNERFAKQHLTRAKLTVFPDNVTIFKQILAGNADVMVTDASETLLQQKLNPGL
CSVHPDKPFQFGEKAYMVPRGDVVFQQYVDQWLHLARETGELRAISDKWLK

>WP_004525656.1 SGNH/GDSL hydrolase family protein [Burkholderia pseudomallei]
MTSRRWFSALSCCLALAASQPGSAAQADAPARWVASWATALQPIPDLAAPPPLYRAPDVAGRTVRQIVYP
TLAGKAVRIRVSNAYGKTPLAIGEMNIGRSAGGAAVAAGSSTAVTFGGRRETEVPPGQERDSDPVAYDVR
AGEPYALSLYLGSRQTMTVWHRVSNQVNYVSAPGNHTGDASPDAFRTRFTQSAWIAELAVEVQQPGAAAI
AAVGDSITDGLRSSLNRNRRWPDALAARLERAGAGDIGVANLGISGNRLLSDSRCYGIALERRFERDVLT
RAGVKVAVLLIGINDINFAAMPARSGLDCDAPHTRVDAQALIAGYRRVIAAAHARGVAVFGATLTPASLP
PAREAIRREVNEWIRTSGAFDGVVDFDAALRDPAKPSTLLRRYNSGDDIHPSDAGYAAMAEAVPLERLAA
AAGRR

>WP_004525898.1 flagellar basal body P-ring formation protein FlgA [Burkholderia pseudomallei]
MTTDAGRATRTRFAVALALAGWMGAALAQQAGDGGMIVIPGRGESAQTALANANAASARGANGAGAAGPG
GADRTASPGGWHAPAAAQAAAAWSGRAGAASGAESAAESGDVRAATEPRTGLIVIEPGPAESNGRIPAAA
PSGWHPSAAMPRAESNGGANPVPPRTGSNRSAGSASASAGWTAPPGAASQGNPAANVAPAPRVSAASRAP
ATTDMQRVAPVIAQDEGAAPAGRPANARAAAQSWPARDPSVAAGGVIPVSLRAQPAPRTLPVRSAPIRAA
ATSAAGAQPAAASATAAGAVPAAQQDGESIRRAALAFLQQQAAGLPGKTTVTVAPAFPRGLAACTTLEPF
LPSGARLWGRTTVGVRCAGERPWTIYLQAKLAVQATYYVAARQIAPGETLTAADLVARDGDLTLLPLAVI
TDAQQAVGATALMRVAAGLPLRRDLLKSAASVSIGQTVRVVASGQGFTISAEGSVLNNAAPGQQVRVRMA
AGQIVTAIVKDAATVEIPL

>WP_004526045.1 family 20 glycosylhydrolase [Burkholderia pseudomallei]
MNRISHSLCAALLAAATLLPTASRAQLPARPTAGAAAPATAAPVRPASTPAELAARLANGLAVRVAVDNN
HAASAGVPCADLGADWASCATGRLILQNRGHSPLTDGGWKLYLHSIRRLLRIDRPGFTLRHLTGDLYELT
PQPGTVRLAQGERIELPFVAEYWLRRYSDVIPRPYVVVDGAAPAVLRYDDTDDELRYVETLPADAQNNSP
GNAPPAAAQPVANRALPSVKRQRALPGALDLRGVELTLPELPSAQVAALRERAGTLGLDGARVPVWGVVA
PRRLPADIAVPGGYRLAIGPRGAFIEGADRAGLYYGVQTLFSLVPAGGATVPAMLIEDAPRFTHRGMHVD
LARNFKPPATLRRLIDQMSAYKLNRLHLHLSDDEGWRIEIPGLPELTDVGARRCHDPSETRCLLPQLGSG
PDDRSGGGYLTRDDYVALLRYAAERFVEVIPEIDMPAHSRAAVVSMEARYRRLHAAGREREANAYRLLDA
QDTSNLLTVQFYDRRSDLNPCMPGALNFASKVIREIASMHADAQAPLRIWHFGGDEAKNILLGAGFQPLD
GADPGKGRVDLAAQDKPWARSPACTALLRRGEIKSIDELPTRFAKQVSAIVNANGIGTMAAWQDGIKHAS
GPREFSTRHVMVSLWDTIFWGASDSARDLSAKGYRTVLALPDYLYFDFPYTRNPRERGYYWGSQATDEYK
VFSLAPENLPQNAEVFGDRDGNPFEVTSAGAAPSIEGIQGQAWGEVMRNGQLLEYMVYPRLLALAERAWH
KADWELPYAAGVRYKLGDTHHVDTAALERDWAGFATVLKQRELPKLERAGIGYRKPTFTLTGE

>WP_004526282.1 DUF192 domain-containing protein [Burkholderia pseudomallei]
MRSSPRSSLARLARAAVFPVALALAASGMHAAFAQTTPLPPGAKQPSEFPRVKLRAGMYVIDAAIAANDA
DREQGLMYRSRLAPNEGMLFVFNENAVHCFWMKNTLIPLSIAFIRADGTITDIDEMKAETTDNHCPRNNG
VYALEMSSGWFAAKGVKPGMKIDGLPKPQ

>WP_004526472.1 hypothetical protein [Burkholderia pseudomallei]
MDSFPASLRLLGALGKLTACAAGLSLALAAPAFADLLDQRAELVNKYVNEMHADPLVADCAAHGNFIAST
SSAFDHVEFAPNAFDSGSASITPWNDSFDEGKQRVKVDNIVTVEGLGVRASGGDPEPLKFRCGYVGTQML
AFSWNDPVPPAKPRVERATSSKKKLRAHSGKGKVKAKATGKSGKRAVSKKSSTTKKAAKKRSTRKS

>WP_004526830.1 cellulase family glycosylhydrolase [Burkholderia pseudomallei]
MINAKTGFLFFIFVMSLIGCRPAFAQTNTPVTGIDLGGEIIEIIKQNPQDPTQAINNYMSGVAATNVRWV
RIHLDWSAIQPASTWIFTSDPNPGRSSGLTWGATDIVVRAARNYGIRILGVITSTPRWATHAQCPSYYRQ
STDDAYKLCAPNLTSYTNFARAAAKHYGNDSYGGKIDAWEIWNEPNCGPNFIPHDPVLYTRLVKSAYPAI
KQANPSASVYAGGSSGCLTSPNNGTGALPSSGVAGLVNSTDARYPAPTQWEPRDWLAVMYANGARGYFDG
LAHHPHCHSDDWQLPGDQCPSTSVNSVYPDYSNPFNIMWHTFTSPSYGWPAPTGKTFASYTGTSLRELMN
ANGDGGKPIVITEFGVATTASDGATDFTGQLGGSAQQYANADFQTIAPAYLTQANQEREYKALIAWIANQ
PYGQYGPVYAYCYSDMALSSPYSANIYEPYFGLVTITASTATSGTTGAPKIAGAYQAASGQYYPAWPNFG
AAATAAANHTPGYAPVGPGW

>WP_004526889.1 dicarboxylate/amino acid:cation symporter [Burkholderia pseudomallei]
MKKRRNITSYIVIAMILGIAVGYGCHSAFPDPAIAKEIAGYVSLLSDVFLRLIKMIIAPLVFATLTVGIA
HMGDTGAVGRVGVKALGWFFIASFTSLLLGLLAATLLQPGSHLSLPLPATDAAVNLKTSAFTLKDFVIHL
VPKSIAEAMANNEILQIVVFSIFFGTALSALGDAGKRLTGVIEDLAQVMLKVTGAVMWFAPVAVFAALAS
TITTEGLGILLTFAKFMGSFYIALALLWGVLTLAGLVFLGKRTFTLIRLIREPFLLSFATASSEAAYPKL
LDALDRFGVNRKISSFVLPIGYSFNLDGSMMYCTFAVLFIAQVYGVHLPLGTQITMLLLLMLTSKGMAGV
PRASLVVIAATLNQFHLPEAGLLLIMGVDMFLDMGRSATNAVGNSIAAAVVAKWEGQLDAPRDDADSDSD
SGDASRGSPRITQRV

>WP_004527347.1 MULTISPECIES: hypothetical protein [Burkholderia]
MKKTFAAIVSAGLLASSSAYADTLCTSGAITKIFIDNNGVMEVTVGTLAYLNGNKDVYSVLTSAFVANKQ
VYIYAPNCQPGTSMGGFAVR

>WP_004527508.1 MULTISPECIES: UDP-glucose/GDP-mannose dehydrogenase family protein [Burkholderia]
MKITIIGTGYVGLVTGACLAEIGHDVFCLDVDPRKIDILNNGGMPIHEPGLQEIIARTRAAGRIAFSTDI
EASVAHGEIQFIAVGTPPDEDGSADLQYVLEAARNIGRHMTGYKVIVDKSTVPVGTAQRVRAVIDEALAA
RGLAGSAEHRFSVVSNPEFLKEGAAVDDFMRPDRIIIGVDDDAAGAIAREKMKKLYAPFNRNHERTIYMD
VRSAEFSKYAANAMLATRISFMNEMSNLADRVGADIEAVRRGIGSDPRIGYHFLYAGVGYGGSCFPKDVQ
ALIRTASENGQPLRILEAVEDVNHAQKNVLLDKIEKCYGADLAGRTFAVWGLAFKPNTDDMREAPSRRLI
ASLLARGATVRAYDPVALDEARRVFALDLHDGADALARLAFVDSADDALAGADALVIVTEWKEFKSPDFA
HLKSVLKAPVIFDGRNLYEPDAMAELGIDYHAIGRPYVEPFSSERG

>WP_004527659.1 hypothetical protein [Burkholderia pseudomallei]
MRRASVPTLLFAASACFFSLNAFSVRATFAAYALLALLALLRAPTLFARQQRLPAIVAYWLLSVSSAFLV
SAFGDFYANFFVKFVLIQTYVALAFWLFASGVLAMRSLERTCETLIYVHATFFIVQLGCYLAFGHFIDFD
SYIRESDSEALYATKALSDSLISIRALGLYSEPSFYAMTVVPAGAILLLAKRRMTAATIVAFATALLSFS

IAAILICALLGGVHFFAGRTSIRIKLVIAAVALAIAPAMYGVYDKRVNQSADYDALGSRTLVLRELRERD
ALADVFGSGLFWDERNNVGKTHLRGYQVRDSSFYVYLLFATGVAGATAFVGALFMLFRRRGRRSLLPYLL
PLLLFKFHALYGMLWLTLLMFVVVAGHAERLPERREPPGTGTGRLSATGASGP

>WP_004527768.1 MULTISPECIES: TlpA family protein disulfide reductase [Burkholderia]
MNSKGIFAGIVVAAVAVAGGLAAGHWMRGNPASGPQAGTPAGAPAAGNPVDTLWAASYPDVDGKPQRLAA
FKGQKLVVNFWASWCGPCVEEMPELVKLSREYEKKGVRFVGIGVDSDQNVKNFLKKVPVDYPIVVSGYAG
ADLARHFGNTAGALPFTVVIDETGKIRETKLGQIRPDELKRTLDTL

>WP_004528277.1 lipoprotein [Burkholderia pseudomallei]
MIRNVLIALVCAAPVAALAEPPKAAGGMIVDDDGMTLYTFDRDTMPGKSACTGGCTANWPAALADAYDKP
GGDLGFIAAAGGKHQWTYKGRPLYRFSGDAKPGQHTGDGFGGMWHVARP

>WP_004528537.1 D-alanyl-D-alanine carboxypeptidase [Burkholderia pseudomallei]
MTTEPLSTTVARGSRRAACALLRTPSRLFASLFVSAACAAPNGARAAPAPAPAIQAASWLVVDADSGKTL
GAHNVDARREPASLTKLMTAYVAFDALERDALRWDDTVTVAADDIASVGRDEARMYLTPGQRVRVRDLMR
GLIVASANDAALVLAKRIGGSPAGFATMMNDTARRLGMRDSHFVSPSGITTPDHYSTAHDLSILAQRLNR
DFPAFYTFSSQRHFAYGAFAKTNKNRLLGADPSVDGMKTGHTNAAGWCMVVTAKRPVGGARARHRVIAVL
LGEPTEKQRLADARALLDQGFASLGADAPAANAVRRANARPGEHRAQARARML

>WP_004529035.1 MULTISPECIES: trypsin-like serine protease [Burkholderia]
MISCKLKALESSKIRISEKSSRTYQGIQMKNFVAKIYIVALGSLIVTSSHSQQIRTHGAVTQITLPEHGS
QASSIDFKNAIPIPLPKPIMAPPTMLDSLMYNAKPSLGVPGYSPGGAGDGKQRPMRLVAPMDLNQIGIEP
QEFGSSNQPYTTSQVNAYGDNTQYYYPFRPAGKLFFNIGSASYLCSASLIKRGVVVTAAHCVANFGQRQF
YSNWVFVPDYSNGSAPYGVWSAASATVMTSYYDGTDSCAVSGIVCQDDVAVILLAPQGGSYAGNSAGWYG
YGWNGYGYNPSGLALIDQLGYPVALDSGVFMERNESQGYVSSGYSNNTIIGSLMTGGSSGGPWLVNLGMP
PTLSGTTFGNSGAHNTVVGVTSWGFTDLAVKQQGASPFTSSNIVTLVNTVCSGTPAAC

>WP_004529236.1 MULTISPECIES: DUF411 domain-containing protein [Burkholderia]
MAQLPEDKPVSRARRAIAAGLALLPVVAFARKTATKPVVQVWKTPTCGCCEDWLSHLKNNGFDVVVHNVE
DTSDVRQQAGMPERFASCHTGIVQGYALEGHVPAREIKRLLREKPNAVGLAVPAMPIGAPGMDGPVYNGR
HMPYDVLLIKRGGQSSVYQSYS

>WP_004529364.1 peptide ABC transporter substrate-binding protein [Burkholderia pseudomallei]
MRRALPFRYHYQSHTMKHTHAFAAVLAALALTIAPSAPAVTVASNVTLADQQDLTRQVPAEVESLDPAHI
ESWTGNTIGLDLFEGLARIDASGAVVPGVAQAWEHKAPDTWVFKLRRDAKWSNGQPVTAADFVYAWQRLA
DPKTGSKYTILVEFVKNASAIIAGKQPPGDLGIRAIDPYTIEVKTEVPVSYFPELTAMAPLTPVNKDAVA
KFGDAWTRPKNIVSNGPYTLVDWQPNNRIVMAKSDKYWNARNVVIRKVTYLPIENDETALRMYQAGQIDY
TYSIPAGGFGQISKQFGKELRPGLQLATYYYYLKNSDPALKDKRVREALAMVLDREILTSKITQAGEVPM
YGLMPKGVKGVQRPFMPDWASWPMARRVDYAKNLLKQAGHGDANPLTFTLTYNTNDLHKKVALFAASEWR
TKLGVTAKLENVEFKVLMKQRHDGKVQVARDGWFADYNDAMTFFDLIRCGSSQNTVGYCNPKVDSLVAEA
NQKLDDGARAALLTQAHDLAMNDYPMVPLFQYSADRLVKSYVGGYTLTNYIDMRASQDMYLIKH

>WP_004529401.1 MULTISPECIES: class I SAM-dependent methyltransferase [Burkholderia]
MSPDAYLQMAATEATHWWFRARREILQALLAGLGLPAGARILEIGSGTGGNLGMLEAFGAVSALEMDDTA
RAIANRRTGWRFDIRAGRCPDDVPFGDQRFDLICFFDCLEHIEDDAAALRRISDYLAPGGAIVVTVPAYQ
WLWSGHDTFLHHYRRYDRAALERCARAAGCTIVRSSYFNTLLFPLAVVARTIDRLLHRARSSGDALGPRA
VNAALYRIFRTERGWLPKHSLPFGVSLFAVLTKCSEPR

>WP_004529879.1 oxidase [Burkholderia pseudomallei]
MIIYLDRRALEVADGTTVAAALALAGDDTARTSCTGAARAPFCGMGVCHECRVTIDGRRRLACQTLCREG
MQVERTR

>WP_004529973.1 copper oxidase [Burkholderia pseudomallei]
MKMVSRRTFLGGSGAALLGAALVSKAGAASLPEAPTMAKAATQPPLAPPNGRPYTPVVTLNGWTLPWRMK
NGWKEFHLIAEPVVRELAPGMNAHLWGYNGQTPGPTIEAVEGEKVRVFVTNRLPEHTTVHWHGMLLPCGM
DGVGGLTQPHIPPGKTFVYEFQLEKAGTFMYHPHADEMVQMAMGMMGTFIVHPKDRGAMPADRDFVFLMS
AYDIDPGSFTPRVNEMTDFNMWTWNARVFPGIDPLPVRAGDRVRIRFGNLTMTNHPIHLHGYAFEVAGTD
GGWIAPAARWPEVTVDVAVGQMRAIEFTANRPGDWAFHCHKSHHTMNAMGHQVPNLIGVPQQDLAKRINK
LVPDYMAMGSTGGAMGTMEMPLPENTLPMMTGTGPFGPLEMGGMFTVVKVRQGLGRNDYRDPGWFRHPKG
TVAYEYAGALPDD

>WP_004530215.1 DUF4148 domain-containing protein [Burkholderia pseudomallei]
MKATFLTVALAAAFVSPAAFADAPAGKTRAQVYQELVEAKANGLDYVTDASYPEISPLYAPRFANKAKPQ
QPAHAVADTRSDAKAGSSGNLSEAAQDEHCVGPRTFCNPYAGS

>WP_004530759.1 alginate export family protein [Burkholderia pseudomallei]
MIARRAVALGGWALMSVLAAARAHAADAMDAPAAAAAAAAPGSRFARVSATSESAPSSAFEFASPAASAS
TSASALAVVPMSALTLASAPAPAPAPSAAPSPSPTSSPVQPGVRPGASCAKTRPAILFNRWQEDWSALAD
PCVPRRPLDALKYVPLFGRVDSYLSLGAGLRERLELNDAPLFGLGRARGDTYVLQRVQMHADLRIAGHVQ
AFVQLEDARPFGKDNVGPVDRNRVDLRQAFVTYVDAIGSGAFKARVGRQEMAFDLQRFVSVRDGPNVRQA
FDGIWADWEQGPWRLIGYATQPVQYRDDGAFDDVSNRNLTFSGVRIERQRVGPGDLSAYYSRYNRTQAQF
PDGAGGEHRDVFDVRYAGKRRNVDWDIEGMYQTGRVGAQRIEAWAVGSLAGYTFAGVGWMPRIGLQVDAA
SGDRRPRDGRIETFNPLFPNGYYFALAGYTGYTNLIHVKPSLTLKPSSALTLLAAVGLQWRATTADAVYA
QGATPVPGTAGRGGNWTGFYTQLRADWAVTANLAAALEVVHFQIGDALRAAGGRNADYVGAELKFGW

>WP_004531414.1 2-dehydropantoate 2-reductase [Burkholderia pseudomallei]
MDRAPVRAAIVGIGAIGGLFAAGLARAGWDVSALARGATLDALRGRGLRIVGESGGETAVALRASDDAHA
LGEQDYVVVALKAHALPALASRIAPLVGPRTIIVAAMNGLPWWFFDGFGGPLDGAALDAVDPGGVTAAAL
PPARAIGCVVHLSSATREPGVVVRGRGNRLIVGAPRAALREPGERVADALAAGGFDVERSADIRTDIWAK
LWGNMNMNPLSALTGSAADRLLDDPHTNALALRMMEEADAIGARLGLSAGMSGAERIAVTRQLGAFRTSM
LQDLEAGRALEIGPILGVFPELGRRLGVPTPYCDAVLGLLRQRAFNAGL

>WP_004531547.1 MULTISPECIES: ABC transporter substrate-binding protein [Burkholderia]
MLEIKHKLVQTAAACALAFAACAAHAADKPLKSIGITVGSLGNPYFVTVVKGAQAQAKEINPSAKVTAVS
ADYDLNKQFTQIDNFISAHVDMILLNAADPKAIEPAVRKAQAAGIAVIAVDVAASGADATVQTNNVKAGE
LACDYLAKKLNGKGNVIIENGPQVSAVVDRVNGCKAVLAKNPGLRLLSSDQDGKGSREGGMNAMQGYLTR
FPKVDGLFAINDPQAIGSDLAAKQLRRTGIVITSVDGAPDIETALTSDTQVQASSSQDPFAMAKKAVSVG
YGIMNGHKPANAMILLEPTLITRENVKRYKGWSAH

>WP_004532251.1 MULTISPECIES: urea ABC transporter permease subunit UrtB [Burkholderia]
MPLSRAARALAALAACAAFSFAAPRAALAVTAADVAALAGDDFDAKRAAIDRLAAGHDAAAAALLNALAN
GDALATDAGRILIQHGDAARDALTNAPAQAGDAQPVMLNNLLRTKIANALSGLDLASPDIDTRRRAIDAL
LKRPDAALKPMIDAARAKETDPVLKRRLDALWAIAALRDADPAKRLEAVRLVAARSDLDMIEQLRPLVAK
RPDGGDAEPDARVREAAQQGLGALYAIQRRGEIAGTLFAGLSLGSVLLLAALGLAITYGLIGVINMAHGE
FLMIGAYATYVVQTLVQRYLPGAFDWYPLAAIPVSFAAAAALGIVLERTVLRHLYGRPLETLLATFGVSL
ILIQATRMIFGAQNVQVVNPSWMSGGVTVMQNLILPYNRLAILAFALVVVGIAWAVLTKTRLGLFVRAVT
QNRRMAACVGVKTARVDSYAFAFGAGIAGLGGCALSQIGNVGPDLGQSYIVDSFMAVVLGGVGQIAGTVL
GGFGLGLVSKAIEPFWGAVLAKIAVLVMIVLFIQKRPQGMFALKGRSAEA

>WP_004532434.1 MULTISPECIES: class I SAM-dependent methyltransferase [Burkholderia]
MARRKLERWSALDLAEGLQLAHAVAALQTLGVVDAMAEPTTAQTLSAACDLDPELLRGVLEYAASRTNLI
RKTGDRFSASEHYTAQARFLLNLYAGAYADNASRLATLLRRPALAGTTVDLVAHARAFEASEAGGGALAA
IISQLHLNHVLDLGCGAGTLLHALAADDREFVGWGLDRNPSMCKAARMRARQAGIAARVKVFQGDGRHPG
ASIPPRVLARVRNVVASQFVNEMFRGGTSRAQAWLRRMRRLLPDRLLVIADYYGRLGHGFQTPHRETLLH
DYAQLISGQGVPPPNADAWLAMYRATGCRPLHVIEDRAATTRFVHLVAL

>WP_004533337.1 F0F1 ATP synthase subunit gamma [Burkholderia pseudomallei]
MSDKLAAIEARTDTARQLQTVIGAMQGVAAARAHEAQQRLPGIRASAATVGAAIGDALSAGARSGDTAAG
ARAAPRARLVVVLCSEQGFVGAYNAQLIEHATRPEPSVSREYMMVGSRGAMLADAGGVPLVWRTPMAARA
DDVVHLANRITDALYAHLAKRGAQPVSIVHAMPGGAARLEVTERRLLPFDYARFDTAPRAQPPLVHLPPA
TLLAELAQAYVFVELCEAAMLAFAAENEARTRAMIAARESVERALGDLLQAYRIARQDEITADIVELAAS
GL

>WP_004533800.1 D-threitol dehydrogenase [Burkholderia pseudomallei]
MTNLWDRAFDLTGRVALVTGGAAGIGHACARLLAQRGASVALVDRHPETAGIAATLEGGAARHSGMSLDL
RDCSAAQAGVALAASRFGGVDMLVNSAGVALLDKALDVGEAAWDATMAINVKASFFVAQAAARQMIAGAR
GGRIVNLASQASVVGLERHAAYCASKAAIVGMTKALALEWAPHGITVNAVSPTIVETALGKQAWAGEVGE
RAKREIPAGRFAQPDEIAALVLYLLSDAAAMMTGENVVIDGGYTVR

>WP_004533895.1 D-amino acid dehydrogenase [Burkholderia pseudomallei]
MRVVILGSGVVGVASAYYLARAGHEVTVIDREAGPALDTSFANAGQISPGYAAPWAAPGVPLKAVKWMFE
KHAPLAIRLDGTRFQLQWMWQMLRNCTTERYALNKGRMVRLAEYSRDCLQALRAETAIQYEGRTGGTLQV
FRTQQQLDGAAKDIAVLREANVPFELLSSDELKKAEPALAAVSHKLTGGLRLPGDETGDCQLFTTRLAAL
AEQLGVKFRFNTRIDALAVAGGKIAGVQCGGEMVRADAYVVALGSYSTNLVASLVKIPVYPLKGYSITAP
IVDAAKAPVSTVLDETYKIAITRFDDRIRVGGMAEIVGFDKRLRDARRGTLEMCVNDLFPGGGDTEKATF
WTGLRPMTPDGTPIVGRTPVPNLFLNTGHGTLGWTMSCGSGQLLADLMSGKKPVIRADDLSVHRYLSETD
GEHRPAYA

>WP_004534049.1 Bcr/CflA family multidrug efflux MFS transporter [Burkholderia pseudomallei]
MPHVARRRPDGRLILLLGALAACGPIATDMYLPSLPAIAGGFGVTAAAAQRTLTSFMAGFSIGMLLYGPL
SDTYGRRPVLLGGIALFTLASVGCFVAGSIDMLIAVRFLQAFGAGAASVLARAIARDAHEPADAAKVLSM
VAIVTAIGPLLAPLIGGQVLRFSGWRGVFVVLALFGAVCATAAYLRVPETWPREKRASSAVLNSFAAYGR
ILVDPVAWGHMLCGGMAFASMFAYITATPFVYIDYFHVSPQHYGLLFGLNVIGIMLGNFLNARLVGRVGS
LKIIAGASLLSGAASFAVAFFALTGLGGLWSIVASLFFVVGVVGILSANCTTDLMLRYPHNAGAAAAVFG
AMQLALGALASAAIGALANGAPFAMGVTIGATGLLCLGGRMLVLRWHGRPVKGALGRVDAA

>WP_004535146.1 TonB-dependent siderophore receptor [Burkholderia pseudomallei]
MLIAAHPGAAALAGTAAKSDAQKNDMQRNENVREDTRRLAAGAPPAGGELKAISVSAPKDAADDPSVATV
GKMPLALREIAQSVSVTTRERIEQQNLFSLDDVMQQSAGVTVQPYVLLTTAYFVRGFKVDSFEFDGVPVV
LGDMASAPQDMSAYERAEILRGANGLLHGSGNPAATVNLVRKRPQHRFAASASASVGSWGRYRAQADIGG
PLNPAGTVRGRLVAAYEDRGFFYDRAKQDTRSIYGIAEIDVARDTLVTVGAQYQSVASVPDMSGVPMARD
GTSLGLPRSTFLDVGWGHFDWDTTRVFGSVERELGGGWKAKVSGEYQAVRSDLKYAGAYGAIDRATGAGG
TLMGSAYQFSSYSRSVDANVQGPVRAFGLTHELLFGATYASSSSGQLTAPLLAGAGTPVNVYRWNPDGVP
EPGVGPYRQDMQNDISQKGVYGLGRIKLAQPVTLVLGGRLSWWNQDSLGAHYNAGRQFTPYGGLIWDVAR
DWSWYASYAEVFQPQTKPTWDGGILTPVKGRTYETGIKGELAGGKLNVSLAAFRIDLDNNPRIDATRPCA
GRSCYYVNGGSVRSQGFEFEANGRITPWWSVWASYTFDTIRYAKDVANGGAFAAELTPRHLLRVWTNYDL
PWQERRWSVGGGVQVQSDFSAASSGVTMRQGGYALASVRLGYRYDRHWRAALNINNLFDRTYYQSLSQPG
WNNRYGEPRNVMLTVRGQF

>WP_004535373.1 sn-glycerol-3-phosphate ABC transporter substrate-binding protein UgpB [Burkholderia pseudomallei]
MKYKMLVRSLAFGGALWFGAQQAACAATEIQFWHAMEAALGERVNEIAAQFNASQSDYKIVPVFKGTYDQ
ALAAGIAAYRSGNAPAILQVYEVGTATMMQAKKAVLPVSDVFRQAGVPLDEKAFVPTIASYYSDARTGRL
VSMPFNSSTPVLYYNKDAFRKAGLDPNQPPKTWADVKADAEKLKKAGYACGYTTGWQGWIQLENYSAWHG
LPFATRNNGFDGADATLEFNKPQQIAHIQFLQVMAKDGTFTYVGRKDEASAKFYSGDCAIMTTSSGALAT
IHKYAKFDFGTGMMPYDAGVKGAPQNAIIGGASLWVLAGKDPATYKGVAKFLAYLSSPAVAAKWHEDTGY
LPVTTAAYDLAREQGFYAKHPGADTAIKQMMNKPPLPYTKGLRLGNMPQIRTIVDEELEQVWAQKKTPKA
ALDSAAARGDELLRRFEKSGG

>WP_004535631.1 ABC transporter substrate-binding protein [Burkholderia pseudomallei]
MRFTHIAAAALVAAASVAAAKPLTVCTESSPDGFDVVQYNSLVTTNASADVVFNTLVSYDEATKKVVPAL
ADKWDVSADGLAYTFHLRPNVAFQTTDSFKPTRALDADDVVFTFGRMLDDANPWHKVAGASGFPHAQSMG
LAKLVKAVTKIDESTVKFELNEPNATFVPILTMGFASIYSAEYADQLLKAGKQADLNAKPVGTGPFVLKS
YTKDAVIRYEANPTYWGPKPKVERLIYAITPDPSVRAQKVKAGECQIALSPKPQDVAAAKADRALRVVET
PAFMTAFVALNTQKKPLDNDKVREALNLAFDRATYLKVVFDNTATPATNPYPPNTWSYAKSIAPYPHDPA
KAKQLLAQAGFPNGFSTTIWVRPTGSVLNPNPKAGAELLQADLAKIGVKADVRVIEWGELIKQAKLGQHD
LLFMGFAGDNGDPDNILTPQFSCNSVKSGLNFARYCDPKLDKLIADAKETADQAKRAKLYEAAQKIIHDE
ALWIPLGYPTAAAITRPNVAGYRVSPFGRQHFDAVSVQ

>WP_004535805.1 c-type cytochrome [Burkholderia pseudomallei]
MTIAKRVKRTMSAAAAAMAVVSCAMAATPAAHADAGDGLKVARSNACMGCHAVDRKLVGPSFQQIAERYK
NDKQAEPKLAKKVKDGGSGVWGAIPMPAHPRMSDADVRSVVQWVLAGAPSK

>WP_004536717.1 cytochrome c [Burkholderia pseudomallei]
MLKRTLSFMLAGCLALPGLSSAADAAASGAPASAASSVPAASAAPAAVRAADATLVERGRYLAVAGDCMA
CHTAKGGKPFAGGLPMRAPLLGTIYTTNITPDKETGIGDWSFADFERAVRHGVAKNGDNLYPAMPYVSYT
KVTDDDVKALYAYFMHGVEPVRQPPRRNDIPWYLSMRWPLKIWNLLFLKEGVYQPKPERGVEWNRGAYLV
QGLAHCGTCHTPRAVTLQEKSLDETGGSFLAGSVLSGWDGYNITSDPNAGIGGWSQPQLIQYLRTGSVPG
LAQAAGPMAEAVEHSFSRMSDADIGAIATYVRTVPAVADGAAKARSAWGKPAEDGIRLRGVALASTGIDP
ARLYLGNCASCHQMQGKGTPDGYYPQLFHNSTVGAPNPTNLVQVILNGVARKAGGEDVGMPAFRHELSDA
QIAALANYLTVQFGNPAAKVSEQDVAKLRAAQ

>WP_004537213.1 D-amino acid dehydrogenase [Burkholderia pseudomallei]
MRICVLGAGVVGLTSAYCLAREGHDVTVLEARPDAVLDASFANGGQLSYSYVAPLADPAVLGKLPAWLAR
RDSALRFVPRLDVDQWLWCAAFLAACRTARARRTAAEMLELGALSRIALHALVERETLDFDYARNGKLVV
YRDRREFDRARRAMDHLIAAGARQQALDTAACVALEPALAPASGLLAGGIHTPSEEVGDCRRFGIELARV
LRERLRVAIHYETPVDTLRTEGATIVAARTPRGEIAADAFVLSAGNGSAPMLRRLGMRLPIYPLTGYSLT
LPARHDATPRVSVTDLHRKIVYAPLGARLRIAGMVEIAGLRSSGAARRLALLAQQAQEIFPHAGDYAHGE
TWCGHRPATPDGKPLIGATPFRNLWLNTGHGALGFTLACGSARVLADLIADRPPALDTRAYALHR

>WP_004538457.1 MULTISPECIES: cytochrome c [Burkholderia]
MEASVHKHLIVKFRSACANAALGLLAATLAASAAHAQPAADAATLKRGEYLARAGDCVACHTVPGEKLFA
GGRAMPTPFGTLYSPNITPDNDAGIGKWTADEFYTMMHTGRSRDGSLLYPAMPFAAYTKVTRADSDAIFA
YLRSVPPVKLANRPHDMRFPYNNRQLLIGWRTLYFKEGEYQPDHTKSAEWNRGAYLVQGLGHCSMCHTAI
NALGGSSEANAFEGGLIPMQAWYAPSLTSNKEAGLGDWSIADITGLLQAGASHRGAVYGPMAEVVYDSLQ
YLSDDDVRAMSVYLKSLPQRGSDAESAPKTTMAASEQARLSKLGAKIYDAQCASCHGKTGRGKTPAFPPL
AGNQSIQMTSAVNPIRMVLNGGYAPGTAKNPEPYGMPPFAQSLSDDEVAAVTTFIRTAWGNRGAPVSAKE
ANALRSAPLY

>WP_004538458.1 MULTISPECIES: c-type cytochrome [Burkholderia]
MTVTSRPTAVLAVLCGLLSQTALSQEPQTERAPDTMEARVLACAACHGRQGEGTSNDYFPRLSGKPAGYL
YNQLVAFRDGRRKYPPMNYLLAYLPDAYLRKIADHFASRRPPFPTIAAPAAAPAVLERGRLLVKSGDPSH
KIPACASCHGASLTGMEPAVPGLLGLHAEYLSAQLGAWRYGTRTSIAPDCMQQVASRLTDTDVTAISAWL
ASQPAPANPSPAPAGSLDMPLACGSEPR

>WP_004538839.1 hypothetical protein [Burkholderia pseudomallei]
MNLHRSIVVPLVFSATFLAGCGSSVDGRFTSGMVAMGAVAGSTVAAYPIGADGTIGAQPLAVTTSDASGA
FSLPALDRWPALIRATHGAYVEEATGAAATLEGDGLEAVYASAPSTIVVSPYSSAVVATARAAGGLTQAN
IAAAIATISAFVGDFDPQQTRPATMAAGAAAPAIEPGHQMALALGAESQSRTDAAADIATSIAAIVAQAA
AGDTLDTCHAGAGDPRADGTLAAPVSSGCAITRGAARYAANARNTSGVTSLASLSAARTSRASPDQTAAA
TPDACADRAALLAQNLALFDGRRDDVQANLVGGVTRDNWRTVTTRSTWGPTAAFYGTLATPAACADTDTF
ARELVIAAENYWIDQGINYCHHHIPGWTPPDDSAAAAPRYRNSSAGSTSGASSNGMTCTAQRSGTGAQIV
PGTAPSAGFSASEIQWNGVDCSDFTSWIYNFVGLTGVRLPTAIGSQACAANEGADALPTPGVLLDIDSGN
IDAMLPALKPGDLLYITQVDPLASGSDAGGYQLAHVVTWTGKRWSDLQAGPDAARYALARLGQPGSRLGG
DLAKYLPLADLATQNPWMIIDSHYAGPAYRPFVGWYRRSLSNVRRIVGADAARRDPALAPYVIAPIASDA
RGDTLTLASPHANASAQQGWRMIYRQSGGTPSCVRAGIAQ

>WP_004540441.1 MULTISPECIES: molybdopterin-dependent oxidoreductase [pseudomallei group]
MLQLSRRQFLKLSATTLAGSSLALMGFSPAEALAEVRQYKLARTVETRNTCPYCSVGCGILMYGLGDGAK
NATSSIVHIEGDPDHPVNRGTLCPKGASLIDFIHSPSRLTQPEYRAAGSDKWQPISWSDALDRIAKLMKA
DRDANFVETTDDGMKVNRWLTTGMLAASAGSNEVGYLTHKTVRSMGMLAFDNQARV

>WP_004545891.1 branched-chain amino acid ABC transporter substrate-binding protein [Burkholderia pseudomallei]
MMLSRLTSMSLAAMLVAAGAAAHAETVKIAIAGPMSGSVAQYGDMVKAGALTAVEQVNAAGGAGGNKLEV
VMMDDACEPKQAVAVANKIVSQKIKFVIGHVCSGSTIPASDIYENEGVVMITPSATAPQLTEGKKRKFIF
RTIGRDDQQGPAAAQYIINKVKPKKVAVLHDKQSYGQGIASAVKKELDAAKVPVVIFEGINAGDTDYSAI

VTKLKSQGIDFVYFGGYHPEMGLLMRQAREQGVKAVFMGPEGVGNKDVTAIAGPASEGMLVTLPADFTID
PANAGVMKAFAQKKRDPNGTFQMPAYSGVKIIADSIAGAKSTDPAKVAAYMHQHTFDTPIGKVAYDAQGD
LKSFKFVVYKWHKDATKTAAN

>WP_004546221.1 multicopper oxidase family protein [Burkholderia pseudomallei]
MLRRHFLSSALAAAAASLFARGAFAAGQAMDGMQGMEGMEGMDDMPDMKPAPAHARHGKPAPGPASALAA
ADALPAGAPLAALRVLANESREPGVFRATLVAQPVARALLPGAAPTTLWQFGADTQGPAVGPLIDVREGD
AVEIRFVNRLPQPSTIHWHGLPVPPDQDGNPSDLVAPGATRVYRFTLPKGSAGTYWYHPHPHMATAEQVF
RGLAGPIVVRAADDPLAGWPERHLFVSDLKLARDGAIAPNDMMDWMNGRQGQFALVNGARRPRISLTGDE
RWRVWNGCSARYLRIAFDDGRPFAHVGTDGGLFDAPREVASLLLAPGERAELVVRAGDRASHAVLTALEY
DRGKMAMAMSEAAHGSLPPDPALPLADVAFEPAAPRALPARLRAVPALGEPVARKEVVFGEQMDMAAMMR
AGAHGRPAGMRFMVNGATFEPHRATLTSRRGEIESWTIRNETDMDHPFHLHGTQFQVVEREIDGETTPEP
YRAWRDTVNVRKGERVRILTTQTERGERMFHCHILEHEDLGMMGTLKVV

>WP_004546662.1 hypothetical protein [Burkholderia pseudomallei]
MKSLQIAVVALSLSAVMATAHAQPAAADAGRQPANRTEAVQAAGRVPAAHQDRQDPNACVGPVSFCNIYF
GS

>WP_004546872.1 DUF3472 domain-containing protein [Burkholderia pseudomallei]
MRRSELKFNCIAATILAAVAADATAAGACLNGSTIASTTRAPLVARQGSVFSSTLYDPAITSNNRTHNPV
MLTVKVTNNGRPVAGCDVAWQPRGAGGASGWLFPASASTDANGIASAWWVAGSGAAQTAVASIRRFDGTT
QGVAIGGSAQPHATRANSIHLNYEPASDWTAFRVDVTPEALAPTTYWEAIGWPGAYTGIQSIDGKQDGLV
LFSVWDVNGKSPQIIAKGPGVDCTQFGGEGTGYKCAKRHAPVAGRTYRFMASIAPVAGQNQTDYSVWFTD
TSTNARELIATLRYQKAVQSANYANSFVEDWATQGASCLGATQRAGQYGNVWALDRASAQWRTVKRASTS
AVYTPDHNEVCSNYQFSVVNGRFRMSTGGHAVGQPLNLPNGPKSFPLTLP

>WP_004547886.1 MULTISPECIES: DUF1853 family protein [Burkholderia]
MSAAASPVPPVALRDAAVRDLAWLLSSASLLAPSAGAPLAQPWASVADAARTAAWLAALDAQPAPLHDAL
AHARPVRLGRYAECLLGFFVGHAPSLRLVAANLPLRSNGRTLGECDFLIETARGERLHWELAVKCYLCVA
TAGTASLADFVGPNLADRFDLKRAKLVDHQLRLTARDEFASLGHAGPWRAQMFVKGWLFYRADDGANAAG
AVADPPEIGADHPRGFWITREAWPKFALAAARRDVAWIVLPRLAWLAPRAFDDADVACGRCVPLASDAVI
DEVARQSGPLLVAALARDASGGWRETARGFIAPDDWPRRASAFAAR

>WP_004548048.1 TonB family protein [Burkholderia pseudomallei]
MKTYSTYLALPLAASLLAGCAAFAPRDAAKLECTMPVAAYPENAKPLERRATVLVRAMITASGNAENVTV
TTSSRNAAADRAAVDTMSRIACSQTPACGGEPYPFTLTRPFVFEPRAKAPQ

>WP_004548157.1 M48 family metallopeptidase [Burkholderia pseudomallei]
MQAKLRVARAALTFVGGVALGVAGPVLGFELCMAEPMLVADNGAPPVPSAPAVATPAAPAAAPAAGNPQP
YALGDQQVRYGNAIVFRSLIPSPLLEQLTDNEYRQTVQDAAQRKRLLPPNNARVKRLRTIVMRLAPYAVK
WSERVKGWNWEIEVLRSRSIRAFCLPGGKVLVDSGLLERLRLTDDELGVLFAHEIAHALREHARSSLGEQ
QAASLGTGATPLPPLFGLSEPLPAPLGVVERFASVRYDPTDETEADVIGGDIAARAGFDPRAAITLWDKL
AVATRADKDNGFIHAHPYDTRRRNDLRKRLADLMPLYRKALVKNADARANAAGAGAAAGAKQRGAATANR

>WP_004548306.1 sulfatase-like hydrolase/transferase [Burkholderia pseudomallei]
MSTIKRIAAAAFAAAALSGGFGHAAHAAGQAKNVIFFLGDGMGPATVTASRLYKVGEAGQLTMEKLPRTA
RIKTFSNDAQTTDSAPSMAAYMTGVKMNNEVLSMSPDTRAIAPGSDANGNKTVNRCGVGNGTPAATLLEL
AKARGKAVGAITTTELTHATPAATYSHICHRDAQYDIAAQAVPGGAGYNAALGDGVDVLMGGGRNHWTPY
DPLANRRGRADGRNLLAELQAKGYAVVATKDQLAQAGAGKLIGLFSTTSHLEYELDRVAGKGEGATQPSL
AEMTAKAIDVLRKNPNGYFLMVEGGRIDHALHGTNAKRALEDTVAFDEAIRTALAKVDLSDTLIVVTADH
DHTMTINGYSKRGNPVLDISRNYRDGQPNKDADGNPYTTLVFGNGANRPNARVPVDSSTATNDAYLQEVG
VRMGSAGSETHGGGDVMLFADGAGAKAFKGTLDNTKVFGLVKAAFGF

>WP_004550304.1 MULTISPECIES: hypothetical protein [pseudomallei group]
MRAMFCAAAFAVPLTAAAFTAGDLDKLCAKTDVKSRASCAAYIEGAADGVYNTIDAIGGTTGPRVGQYFC
LPPDIKAQQMTDAVRKYIAENPKLADYNASTAVSLGLGKAFPCRSY

>WP_004550328.1 SCO family protein [Burkholderia pseudomallei]
MNETSMMRRKALGALCSLALAGAPVGSRAATPFYSLPLSSKDWHKGFRLTDLHGHTKTPQDYRGNVLLLF
FGFLSCPSICSTTMLELTQAKERVGAQKDKVKILFVTLDPNRDTSPTIARWLASFGDDNIGLRDSEAHVR
KAATALNLKYERVEGDVPGAYTIDHGVQTYVFDPQGRLRLIARAGIEPEYVAKDIVQLLSGR

>WP_004553586.1 S8 family serine peptidase [Burkholderia pseudomallei]
MNRTADFALSLSPLCKRLACVWPLALAAGIAHGATDWVDTHTKAFLNHAQIETLARGANAASLEVASGEA
THVVVSLKLRNAEQLKAVARNVNDPHSAQYRQYITSAQFLANYAPTEAQVKQVVAYLRKNGFVDIHVAPN
RMLVSARGTAGTVKQAFNTSLVHFEYAGRAGFANASTAQVPRALGDIVGSVLGLQNVARARPLTKIGAIA
KPLALASGTATGHYPSEFPALYNATGVPTAANATVGIITIGGVSQALSDLQQFTSANSYPDVSTQTIQTN
GSGGNYSDDQEGQGEWDLDSQSIVGAAGGQLGQLIFYMADLDASGNTGLTQAFNQAVSDNAAKVINVSLG
WCETDANADGTLSAEEQIFTQAVAQGQTFAVSSGDEGVYECNNRGYPDGSNYTVSWPASSPHVLAIGGTT
LYTTSSGAFSNETVWNEGLDGNGKLWATGGGVSTILPNPSWQSGSHRKLPDISFDAAQSTGAYIYNYGQL
QQIGGTSLSAPIFTGFWARLLSANGTGLGFPAARFYHSIPTHASLVRYDVTSGNNGYSGYGYKASTGWDY
PTGWGSINISNLNQLIQSGGFN

>WP_004553625.1 polysaccharide deacetylase family protein [Burkholderia pseudomallei]
MSFRSRFTWRTAVFGPGFRPVSRIVATTASSGAPAVAAPPTDVAVLVYHRFSNVCGADPMTVGVATFEAQ
LAHLRRLGYRFVPLRDVIGWLRGEPVALPSKAIALTIDEGHASIFDWARTVALRERVPITLFVYPSAIGE
APGALTWHQLRVLHKTGWFDVQSHAWWHPDLNAAHRPPSGTFREATRAQFAQARARIAREIGNQVDLLAW
PFGAFDGELGAAAREAGYVAGFTLEPSKIRRDTPLLTLPRFLMVEECTPAVLRRLLSKSDAAREEAARR

>WP_004554341.1 MULTISPECIES: nitrous oxide reductase accessory protein NosL [Burkholderia]
MKRRFLSAAVRTSIAALASAALVAACGHDAQTPPPAREITDATVSVLDGMSLKDYPGPKAQIVYADGEPD
FFCDTLGLFSVYLRPEHDRKVRALYVQDMGATDWQHPVGHWIDAKRAIYVIGSKKPGAMGRTFASFAREA
DAARFAKAEGGKLYRFGEITPEMAATDGGVVKDQTM

>WP_004554469.1 SCO family protein [Burkholderia pseudomallei]
MLAGAALVSLPACAAESREIAHRPWGAIEPAEAAPDIEVSLADGTRGRLVDLLRGKTSAIQFVFTGCSAT
CSLQGAIFQSLQAQLARHPIGGAQLLSISIDPANDTAVAMTAWLRRFGAQPGWQAAIPASGDLARLARLY
RDERNPADSHIDQAFIASRSARFVWKTDHLPTPESVHDALRYYASH

>WP_009918239.1 hypothetical protein [Burkholderia pseudomallei]
MRAATSNPIASNIARRTANHGYFAALYSCAMKIVRLLLVLLWCATLPLTGLAASGLAGDCPMQQAMSMSE
DGAMPSMQDCESMRSSVAGKSGKAGKAANVFCKVMVQCQFGSLYHPMPAADVARPASPGRPLVFHYAKSL
TVREPGGLWRPPRTA

>WP_009950543.1 ShlB/FhaC/HecB family hemolysin secretion/activation protein [Burkholderia pseudomallei]
MTTATRFAVLPITALITLAAQAQQTPATIDQAAAARANAEQNQQVQQRRDAQRRDATVQAPGVRSDVPSP
EAYPVLPAETPCFRIDRFALDVPDSLPAAAKVQGASALPMDRFAFARDWLAHYAGQCVGKQGVDLIVKGL
SQAILTRGYITTRVLVPGQDLSTGTLKLALIPGVIRHVRFEDDKLRGTWKTAFPTRDGDVLNLRDIEQGL
EQMKRVSSQDVSMRIAPGDRPGESDVVLDAKRGKPWTVVASIDNSGTRATGKLQGNVSLGIDNPLGLNDI
FNVGFNQDLEFGDKRFGSHGWNAFYSIPWGYWTGTLSAYTSTYFQPLAAVNQTFVASGNMKTVDFRLNRV
LARSRNDVFGAQVRLTRRFGDSFIEGTTIPSSHQNATFLEFVLNDRHYFGSSQFDGSLAYRQGLGWLGST
DSMFAVEGGQTYRFKMVVLDANLSTPFVIGTQPFKYVTTFHGQYTGNTVSYLDSVTIGSRYTVRGFDGER
LLAASRGFYWRNELQAPIARTGLSAYAGLDYGRVWGPEPVALVGTQLAGAVIGVKGSVSTRFGAYAYDLF
AGTPVYKPSGFETARVTLGFQLTAQF

>WP_009952932.1 SDR family oxidoreductase, partial [Burkholderia pseudomallei]
AEATADAVRAAGAQACVVRGDVAHEADVIHMFDAVQSAFGRLDALVNNAGIVAPSLPLADMDIARLKRVF
DTNVLGAYLCAREAARRLSTDRGGGGGAIVNVSSIAARLGSPNEYVDYAGSKGAVDTLTLGLAKELGPHG
VRVNAVRPGLIATEIHASGGQPGRAERLGAQTPLGRAGDADEVAETIVWLLSDAASYVTGALLDVGGGR

>WP_009956690.1 glycerophosphoryl diester phosphodiesterase [Burkholderia pseudomallei]
MGQELKTRNILWRIVLLGIVSLSMVTMTSNVCVAAPGFPAIVAHRGGTGDAPENTVYAISKALQNSADAV
WITLQLSSDGVPVLYRPTDLKVLTSGSGPISALSAAQLARLDAAYYYNPKDGYPLRGKGYGIPSLEEVLK
TFKDTFFYLDIKSPDADPNRMATALSQVLERTGALSRVRIYSTEAKYTAAVQAMPHFETRDETRTALANV
TMNHTCQLSPKFGSWYGYELRRDVTLIEETTLGVSPPSPSRLVWNLEAKTCFIDTGKGSILLIGVNSADD
YATAASLGAAAVLVDSPAQARHWPTQVSMP

>WP_009966818.1 hypothetical protein [Burkholderia pseudomallei]
MKNRYLAALATCAALLPIASTARGEEVASVNTNFRLTGSDRVVVEAYDDPLVSGVTCYVSRARTGGIKGT
LGIAEDPTEASIACRQVAPIRFTEPLRQQTDVFSERLSFIFKTLHVVRVVDKKRNTLVYLTYSDRIATGS
PKNSVTAVPVPAGTPIPLR

>WP_009966896.1 carbohydrate ABC transporter substrate-binding protein [Burkholderia pseudomallei]
MKIRAMMGALGAAGLLFGAAAAQAAENVTVLHWWTSGGESKAVGVLKDDLQKQGYVWKDFAVAGGAGAAA
MTALKTKVISGDAPSAAQIKGPLIQEWADQGVLVNIDAAAGDWKQNLPPEIDKIIKYKGNTVAAPFSVHR
VNWLYINKAALDKIGAKPPATWPEFFQVADKLKAAGIQPVAMGGQPWQDLTLWEDVVLSQGADFYRKALV
ELDQKTLTSDKMLEVFNTVRKIQGYFDSGRNGRDWNLATAMVINGRAGMQFMGDWAKGEFEAAGKKPGKD
YICAAVPGTANAYTFNVDSFVFFQQKGQKAATPGQIALAKTIMTPAFQEQFSLLKGSIPVRLGVKMDKFD
DCAKKSYADEQTAIKSGGYVPSLAHGMAQGDATAGAISDVVTKFMNSQQDAKSALAALARAAKVK

>WP_009972253.1 MFS transporter [Burkholderia pseudomallei]
MHPLRSILPSALFTAVGLLATDLYLPAVPSLPQQLGGSIESAQATLAAFSAALAVSQLVWGAAADRFGHR
RTLAFAVLLQLVAGAACALAPSMGALIGARLAQGFGVGAAMVIVPALVRQSFGDGGAVRALAWLGIVESA
VPGLAPLVGAALLVVADWRTSFWIIVALSAIAAPLVFRVIPTARAMRACAPANVGAHAGGYRRLLRSPVY
LGYALGHALCFAALLAFVASAPQVVEIWLGAGPSTFSLMQACGVAAFMLSAARSGKWSDALGLDRIIALG
ALLQFAASAAFLLLAYADWRSTPLVVASWMLFCGSLGLRGPASMARALAAEPAVAGRAAGLLMFFGLGGA
ALATQAVAPFLRLGLAPVAWMCAGFTLASGAVVLWGIAIRGRHRAAATEIA

>WP_009981055.1 ABC transporter substrate-binding protein [Burkholderia pseudomallei]
MKLKTLAHACLAVAAAWSVGAVQAADSVKIGFITDMSGLYADIDGQGGLEAIKMAVADFGGKVNGKPIEV
VYADHQNKADIAASKAREWMDRGGLDLLVGGTNSATALSMNQVAAEKKKVYINIGAGADTLTNEQCTPYT
VHYAYDTMALAKGTGSAVVKQGGKTWFFLTADYAFGKALEKNTADVVKANGGKVLGEVRHPLSASDFSSF
LLQAQSSKAQILGLANAGGDTVNAIKAAKEFGITKTMKLAALLMFINDVHALGLETTQGLVLTDSWYWNR
DQASRQWAQRYFAKMKKMPSSLQAADYSSVTTYLKAVQAAGSTDSDKVMAQLKKMKIDDFYAKGSIRTDG
SMIHDMYLMEVKKPSESKEPWDYYKVVATIPGEQAFTTKQETRCALWK

>WP_009981467.1 esterase family protein [Burkholderia pseudomallei]
MRASNHYAVMRRFVSPLVSPLAALLAALLAAALTAAAHAAPASSIVTRTFRSPALHRDWSYTVYLPAGYN
PEGARYPVLYLLHGNAGNANDWITQGRLQLTADALIERRDIAPVVIVMPQGGTDWYVDRKEKMQSAFLDD
LIPDVEAHYAVSNQRAGRAIGGVSMGGYGALRFAFLEPERFCGAMLLSPAIYANEPPASSAARYVGVFGD
RQFDPKVWHELNYPALWRGYFAQPLRLRMFIAAGDDDLSIQAESSALYTSLRRAQNPAALRIVDGAHTWD
VWRRLIGPALKYTLECVK

>WP_009981622.1 hypothetical protein [Burkholderia pseudomallei]
MIHIRLKRASFLLLLGAQAVSLGAAAGTLSGTVSSAGTPLAGAMVTVFDAAQARRDTVYTDHNGRYRITV
DFAGELRVRARTPYFKDAAQDLALAPDASKTLDFSLARQTVADELSASLPASAHLATLPWSSQDSRTAFI
SQCNYCHQVGNALTRTPRDEAAWGATVRRMEGYAALLTDRQARDITHTLYQGMNAHAVAAVEKYLYDDRL
APAKIREWAAGDGLTFIHDADVGFDDHLYGADEGHDKIWELDRKTGRLTEWKEPDVDLPVGGIFSGVQLP
IGVFSGKHGPHSLAQAPDGRFWITNALSSTLASFDPATKRFKLYELGHSHLYPHTLRIDRAGIVWFTIVA
SNEVARFDPKTGRFTIIHLPDGGIWRAMSHYLFPLVVKMAAWFPGQNLHLALTHHKWAFQGRDAFPFPYG
IDVNPVDGSIWYAKLYANKIGRIDPKTLAVTEFDTPLGGPRRLRFDPQGNLWIPAFDDGGLMRFDTRTHR
FETFKLPLLAHNEYEVPYALNVQPKTGDIWITSNMSDRIFRFVPSTQTFITYPLPTRVTWLRDMTFTQDG
AVCSSSSNLPAYGIEGGRASFICLYPDGERGGTAANAAGNANAAANLARVRAAAARRTPERTAPARAALA
RAGSNRDGAIQTAAIQADGTRTDRSAH

>WP_011203825.1 MULTISPECIES: chorismate mutase [pseudomallei group]
MKQSLRASLAAAVLGCIVLSAPRIAAADGDDTALTNLVALASQRLALAEPVAHWKWLNGKPISDPPREAA
LLADVEQRATANGVDPAYARAFFDDQIAASKQVQNALFATWRATHGPEGPAPDLATSTRPQLDRLTQSLI
AALARVAPLRDAPDCPSRLARSVSNWKTLTRYDSGREDALGTALSHVCTAGSTSAVG

>WP_011204325.1 MULTISPECIES: tRNA (adenosine(37)-N6)-threonylcarbamoyltransferase complex transferase subunit TsaD [pseudomallei group]
MLVLGIESSCDETGLALYDTERGLLAHALHSQIAMHREYGGVVPELASRDHIRRALPLLEEVLAASGARR
DDIDAIAFTQGPGLAGALLVGASIANALAFAWDKPTIGIHHLEGHLLSPLLVAEPPPFPFVALLVSGGHT
QLMRVSDVGVYETLGETLDDAAGEAFDKTAKLLGLGYPGGPEVSRLAEAGTPGAVVLPRPMLHSGDLDFS
FSGLKTAVLTQMKKLEAAHAGGAVLERAKADLARGFVDAAVDVLVAKSLAALKATRLKRLVVAGGVGANR
QLRAALSAAAQKRGFDVHYPDLALCTDNGAMIALAGALRLARWPSQASRDYAFTVKPRWDLASLAR

>WP_011204795.1 MULTISPECIES: prolyl oligopeptidase family serine peptidase [pseudomallei group]
MMTGWIAACVLSAAQADPLPSDAPSAPAAAPYAPRVSHAPRAAGALANAERFDYGDSGLPPVAANLNETI
IRIPVDAAGAITLEATVYKPDGPGPFPLVVFNHGKNPGDLRAQPRSRPLSFAREFVRRGYAVVAPNREGF
AGSGGTYIQEGCDVERNGVAQARDVAATIGYMSKLSYVDARHVVVAGTSHGGLVSLAYGTEAARGVRGII
NFSGGLRQDLCEGWQKNLVDAFDTYGSRTHVPSLWLYGENDSVWSPALVAQLRDAYMSHGASTLFVDFGR
YKDDAHRIIVDRDGVPIWWPPVASFLAQLSLPTSVRYAVANPHEPKASGYAAIESVDAVPFIDDAGRAAY
RRFLAQHPSRAFAVSSEGAWSWAEGGDDPMALALEGCRKQGAGACQLYAVDERVVWRDAGTQTADESTSA
AHALASR

>WP_011205039.1 OmpA family protein [Burkholderia pseudomallei]
MEKTGRGGGMNSTNVFGGIFVGMLSVALLGCASVEKPAAYGEPSPMFIFASNEGFVAPSKMAFPQGIPID
PDALKKLQLGMSKDQVRDALGDRQYNGEDLPIWHRWDYLFQLRAPDGAVHQCQYQVRFLGGLAYRGYWAD
SACRDIAARYADAAGLGVIAVAPCNPSDVAAPLPERVELPTDTLFAFDKGGFEDISADGRRQLGDLVASI
KAKIFSINHLIVTGYTDRLGSDEHNARLSSERARTVADYMIAEGIPAAKITAVGRGAADPVVVCNNGEQP
ELIRCLQKNRRVEIRIKQK

>WP_011205373.1 ABC transporter permease subunit [Burkholderia pseudomallei]
MASKERDVQRSFLERSLLRRFGAAPAGGLLALFFVLPLAALVSAAFAEGGRAFAAVLHDPLVGDAIGRSL
ALAVGTGTLSTCVGVPLALSFAEQPPARRRWLLALLGVPLAFSGLVIAYGFILTFGRAGFVTTLLAGLGA
DAAKVGGVIYTTFGLVVAYAYYLIPRVALMLFPAFANLERRPLEAAMSLGAKPWRAWLDVAWRELWPSVM
AAWCLVTAIALGTYGTALALAGTQINILPLLMYLKLSDGQTDFSQAAVLSIVLTALCTCVLAMGEYVGRR
HR

>WP_011852052.1 lipoprotein [Burkholderia pseudomallei]
MKSNSSLSILIAAACIQAFAATASLAQGPAHPPSYVEGTRVPKGFARPPFHTNPARFSATTVSGLAPATV
RHAYGFDSIANQGDGMVVAIVDAYDDPKIESDLGVFSKNFSLPPCTTSNGCFKKLYASGSKPSPNAGWAL
EMSLDVEWVHAIAPKAKIVLVEAASNSFNDLMTAVDVAVGAGASVVSMSFGGSEFSSETSFDSHFGAPSN
VTFVASSGDSGNGTEYPAASPYVVAVGGTTLSADASGNYVGETAWSGSGGGVSAYELEPVGQTLWPIPYA
GQRGVPDVAYDANPNSGFAVYDSVTYQGQSGWFVVGGTSAGAPQWAALFAIANSMRTAAGKAKLAGAYNQ
LYTVGKTAYGSDYHDVTSGTNGSCGMICTASGGYDYVTGLGSPQALNLVQALVAQP

>WP_017881844.1 MULTISPECIES: sugar ABC transporter substrate-binding protein [Burkholderia]
MQRKTLTAAAARVAALAALASSALAAQAATLTIATLNNPDMIELKKLSSAFEKANPDIRLNWVILEENVL
RQRATTDITTGSGQFDVMAIGTYEAPQWGKRGWLAPMSNLPADYDLNDVIKTARDSLSYNGQLYALPFYV
ESSMTFYRKDLFAAKGLKMPEQPTYEQIAEFADKLTDRANGTYGICLRGKAGWGENMAYVSTVVNTFGGR
WFDENWNAQLTSPEWKKAINFYVNLLKKNGPPGASSNGFNENLTLTASGKCAMWIDATVAAGMLYNKQQS
QVAEKIGFAAAPVAATPKGSHWLWAWALAIPKTSKQQDAAKKFVTWATSKQYVEMVGKDEGWASVPPGTR
QSTYQRAEYKAAAPFSEFVLKAIQTADPTDPSLKKVPYTGVQYVGIPEFQSFGTVVGQAIAGAVAGQTSV
DQALAAGQAAAERAVRQAGYRK

>EQA86889.1 DeoR faimly transcriptional regulator [Burkholderia pseudomallei MSHR338]
MQAAALAAEFHVSEDAIRRDLRALAAEGRCRRVYGGALPVTPACAPMAARIDAARERKAALARTAASLIE
RGELLFLDSGSTALALVEYLPEDAELTIATNSIDIAAAVLRRADLSLIMIGGAVDQAVGGCVDASAVQSV
ARMNIDRGFLGACALSPQRGLAAFGLADATFKRAVVAASARCVVLATTDKLAARAPHRVAALDEIDCIVV
EHDLPREDRAALSSAGASILAANPPAQP

>EQA87624.1 ABC transporter substrate-binding protein [Burkholderia pseudomallei MSHR338]
MGAATGAGAARADASQTLRFGLEAQYPPFESKAANGALQGFDIDVGNAVCNAAKLTCKWVETSFDGLIPA

LQGRKFDAINSAMNATDQRRQAIDFTTVIYRVPTQLIARADSGLEPTPASLKGKRVGVLQGSIQEAYANA
HWAGAGVQVVAYQDQNQAYADLTAGRLDGTLVLAPAGQRGFLSRPEAKGFAFVGPPVRDDKILGSGIAFG
LRKGDDALKARLNAAIDKLKADGAVKALGRKYFGDIDISAK

>WP_023360415.1 tetraacyldisaccharide 4'-kinase [Burkholderia pseudomallei]
MSARPGLLARAEARLTREWQRRGALAWALAPFACAFGAIASLRRAAYARGWKARVDCGVPVVVVGNVTVG
GTGKTPTVIALVDALRAAGFTPGVVSRGYGAKIAAPTAVTAASPPQQAGDEPLLIARRTLAPVWVCPDRV
AAVRALKAAHPEVDVVVSDDGLQHYRLARAVEIVVFDHRLGGNGFLLPAGPLREPLSRRRDATLVNDPYS
RALPPWPDTFALSLAPGDAWHLDRPSRRKPLAQFAGERVLAAAGIGAPERFFATLRAAGVAPATRALPDH
YAFATNPFVDDHFDAILITEKDAVKLGTSWRDARIWVVPVEAALDPRLIALVVEKLRGRTSA

>AHE31311.1 methyltransferase domain protein [Burkholderia pseudomallei NCTC 13178]
MQASDPSSGYLSDVTFPDRFHRELSPTWLNYASVLGGARPKELGRPFRYLDLGCGFAHSTVINAAAFPHA
EFHACDFNPAHIEAAARRASRLGIGNVAFHEASFDALLDRDLPPFDFIVMHGIYSWVDAGMRRVIRQLLS
RRLADAGLVYLSYNCQPGWAAEAPLRKLMLELAQAADGGIEARTGSAIAGMRKLGTPSLRYFRDNPAAAE
ALAALANDPLDYLAHEFLNGTWKIHYSVDVVDEMAEAGLAYAGSATLADNHPMLLIDRQAADAIAALPNA
RLRHLAEDFAVNRRFRRDVFVRGARASTAPAEALRHLDEIAIGCTTEIDRIDTRVTIPRGAISFQPDFIA
DLRALLRHGAMRIGEIVARLGAARRNSREIRQNLLFLVASGTLTPFAQPGGPTDTGARRAASPAAAAALA
GSVDDAAPAFVPSELLGNGLAVSPDEAAQALRWIAGEAMPRPERLARVGVLRGA

>AHK68005.1 bacterial extracellular solute-binding family protein [Burkholderia pseudomallei MSHR520]
MNTIRLLGAAALVGACAPLAAAAATPVCKVPTLKVLAQKSLGLSVMEKSLPDYEKTSGTRIEINYFGEND
RRAKSRLDASTGAGSYQIYYVDEANVAEFASAGWIAPLLKYYPKEYDYDDFLPGRRAVASYKGVAYFAPL
IGGGDFLFYRRDLLDAAHLPVPKTLDELVAAVRKLNAPPKLYGWVARGQRGSGMNVWRWAPFMLAQGGAW
TDPHGQPAFNSPAAVQAAERYRDLFKYAPPGAATYDWSNALEAFRSGKVAFMIESTPFADWMEDPSKSSV
AGKVGYARPPAPLPSAAYGHGLAISSVGAKDDCTRQAAGRFIAWATSKEQEQARLRNGVFSDYNRTSTIG
SDYFRQHVKPQILAGLNDTNPVTKATIWATPQWPDIGDNLGVALEEVFTGTQTDVRGALDDAAQYAKDAM
AHGARRAQVTRVTQVARGTRRRDRQMKARRRDAAPRPRRPQPPRPRRCLAARASASQETPMFNHGKTSLP
WLFLGPPLALMAVLGLVPTIAAINLALKNRVLRYADSDYVGLRNFVRLASDRRFLNAIEVSALWESVTVA
GAVAVGIVLAVFLFERVHGRWRHAAALILIMPVLLPRVSAAFIWKFMYAPLTGILGWLLGALGMTNVAWL
ADPRLALAAVALVDVWQWGLFFCVVVLKLLETLPPEPLEAARLDYASTWQVYAYIALPMLKAPIMSLVFI
KMVESLRSFDLIYVMTKGGPGIATETLDMYAYAQGIGLSGKVSYASSMAVLMMVATTLVFTFIWKRVDKW
DD

>WP_024428428.1 type II and III secretion system protein family protein [Burkholderia pseudomallei]
MDTKPQLSRPTPARSRAVRGAALAAACCAGLFTVFAAAAQDAVENAAIAKGTAAAPMALGRAPMQMTISM
APAPAAGAAAAAAAVPLRGPSCVGEMRESTSVSVPLGKSLLVPLDEPVRNRTIGNPAVAQATMVSPRTLY
LVGMSVGTTNMIVQGKSGACRIIDVAVGADAGGLQASLRQLMPRERDIRVSTAADTIVLAGNVSSAQAAQ
QAVAIAKAYSNRTNAGAAGSGGGARADVLNMLTVTSPQQVMLEVKVAEVSKTLINQMGSAVNIQGGFGSW
TGALVTSLLTGVGSGIAASKSNKLPLNGSIDAQNTDQPVKILAEPRLVTISGQEATFLAGGKIFIPVPQS
SGIGTNTITLQEEEFGVGLKFTPTVLSNGRISLKVAPEVSELSPTGVTLSAANVGNVSILPLITTRRAST
TVQMGDGETFAIGGLIKDNAQGTLKALPGIGELPVLGALFRSTSFQQDRTELVFIITPHLVQPLETADVP
LPTDSFTKPNEADVYATGNMEGRGGLRQHGAAPAAAPANAAPGAAAPTGAAPAAAVPAPAAPAPAAPAAP
AAAAAPAKDAAASPAPAKQAPIAAQAPAPQPAKPAASAPALANARPPATVQAPPPALPSAAALAQRQGAH
AAAPDTAHAGPAESAAATASPAPAAALAAPAAAGATAKRQGAQPPQQLANAGAIAPAADR

>WP_024428446.1 AAA family ATPase [Burkholderia pseudomallei]
MSVVDERFAWTGGLADRLPEPADFGLALAEGFARRIGMLSRRVGASAAAARWAARAAFAASRATAAGHVC
VALRALAQRYDEPYDDVRAALAASGVTAFDGIVRGGECPLVVDRDGRLYLARYFEYETRLANALVERSRC
GGAPAGGADALSPDALGERLVRYFGPQKERGVDWQRVAALIALTGRVTIVSGGPGTGKTTTVVGVIACLL
DAHPELRIALAAPTGKAAQRMQEALHARAGSLPAELAARLPRTSCTLHRLLGGGPGGRFAHHRDNPLPYD
LVVVDEASMIDVALAAHLLDALAPNARLVLLGDKDQLAAVEAGAVFAELSARPAFSPATCATIARALGVG
EAEFVAALPDGFVVQAVGAERVADARAAAADMDVAARGTAGAAAAAARATAAAAARMTTAAAAARGGAPE
SAAQRAPAPHAGRRGGGRASNRTVDDAQGSLFAFDDDAFGVGTSGGEPGAGAQGAGRRDGGAPSLDGPHA
GRRDGGASPGVAHSRRDAFDARARSGPADVSDLENAARAYAPAGAAGPADSPADASFVEPAAWIEADELA
WLDNAVFSFSEGASANAEADGRRSAAATATDAGARTAPPVAREATAPTPRETARSDGSAVAPLTDCVVWL
ERNYRFGLDSPIGRLSLAIRRGAVQDALDALSTQDDAAARFSDDGGATLSAATVEQLARGFAGYAAALRD
ALATPEPDALPLFDVLNRFRVLCATRTGARGAEEVNARMAAEVRRAVRVPLALGAHWFAGRPVMVTRNDY
ALGLFNGDIGIALPGARGVLRVWFRGADGRARAVSPAALPPHDTAFALTVHKSQGSEFDDAALILPASFN
RVLSRELVYTAITRARARVQVIGSRAVLALAIATRTARDSGLAARIADALRARQEGAR

>WP_024428496.1 hypothetical protein [Burkholderia pseudomallei]
MRTMIAAGALALLAGCAGGPAGFGSWGGPSFAQLQRGCGDVRDYGDDARSVYSAVFDAWVAKCHGKLTDA
RFCAFENELAQRHAALGTNADAAARGRWVSYLNDARARALSWRAAVDPSLRGG

>WP_024428498.1 FAD-binding protein [Burkholderia pseudomallei]
MKKILGDPASRRERRAFLGDVARLAGAGIIAGWTPIRPIAAHAQAAGAAPPNFPAGIALYKQAFRNWSGE
IAVADLWTAAPATPADVVAIVNWAADNGYRARPLGHMHNWSPLTVAANGAHERTLLVDTTRHLCAVSVDP
STTPARVVAQAGVSLDTLLATLEQHGLGLTAAPAPGDITLGGALAIGAHGTALPAANETRPPGHTYGSLS
NAVLALTAVVYDTASSRYALRTFDRADPDIGPFLAHVGRAFVVEATLQVGANQRLQCESFVDIPAAELFA
AAGTRGRTVESFVQRSGRIEAIWFPFTDYPWLKVWTVRPNRPSGARVVEEPYNYPFSDSISRELSDLVSR
IVLNGEIQLAPLFGKTQYTIAYLGLTNIFRPLTNLWGWSRSVLHYVRPTTLRVTANGYAVLTRRENVQRA
INEFVGAYRQRVAAYRAAGRYPMNGPIEIRVTGVDTPDDIGRGAVPPSLSAIRPRPDHPEWNAAIWFDIL
TIPGTPDANRFYREIEQWMLSNYSGDYATVRPEWSKGWGYADTAAWSDDAMLRTTIPDLFRQGLSSADDW
DAALRTLERYDPRRVFSSPLLDRLMG

>WP_024428546.1 alginate lyase family protein [Burkholderia pseudomallei]
MVRQTFPGRAQALRQRLSALAPALVAAAALAAAGPARAAMNFCAAPALQSSEATHAEPGVQALIKSVDAH
LNDEPKALPRVHTEGTLPHEGIYDQSAEALNDMELIRNAALAWRVTNQSRYLALVDRFLSTWVNTYRPSF
NPIDETRFESLILAYDMTASALPVKTRNAAAAFIAALGNGYVQQIDAQKRPLKGTWRNNWQSHRIKLIAL
AAFTLGDRRMMNAAQRLFVEHLADNIEPDGTTYDFLERDALHYAVYDLQPLATAALAARRFNRNWLRERA
PNGATLAAALDWLAPYARGEKTHEEFVHSPVPFDAKRREAGLPGYSGMWEPKNATELFHLAARLDGRYAG
IAQQLSPMPPAWLAACLPLPAR

>WP_024428578.1 metal-dependent hydrolase [Burkholderia pseudomallei]
MASHAAHHASGWAAGLVAAALVAQAGAAGPWHVYSLAAFAAGVAGGTAPDWLEIAWWRRTRRLWVTHRTI
THWGVGWAALLAFAYRSLGHRHPWAPPLFGFACGGLVHLIADWPNPLGVPWLWKRHSLNWWNSGHCDLIV
VAAAWAGALWLAQAAWGHAAPLAHWLGWLRTA

>WP_024428614.1 nitrous oxide reductase family maturation protein NosD [Burkholderia pseudomallei]
MPIPHALATLAAAALLMHAACAATLAVHPGERIGAALAAARPGDTVLVQHGRYEENLRIDKPLTLRGVGR
PTIDGRLAGDVIRVAAPDVTISGFSIVDSGASLTAQNAGVYVAPGSDRTRIERCELVYNLFGLWIERSAD
VRVTGNVIVGKRDLLSPRRGNGIQLYNTTGARIAGNTISYTRDGIYVDVSHHARFEHNTIHDVRYGTHYM
NSYYNVWDGNDVYHNRGGLAIMEARDQIVRGNRVWGNTDHGIMLRTIQDSLIENNVVAGNQRGLFIYDAE
YNTIRGNLVVDNKIGVHLWGGSIHNDVTGNDFADNREQIRYVAASDVAWPGNYWSNYLGWDRRGRGIGDV
PYKANDLVDRLTWRVPSVKVLMNSPAVQALRVVARQFPLLAVPSVVDDAPRMRAAHAGWPQWVGKR

>WP_024428630.1 autotransporter domain-containing protein [Burkholderia pseudomallei]
MTIGSKTKLALGIALAMPAAAWAYTFYALGDSLTDNGRVVRLTGILPNATSTIFRGGRSSNGPVWAEYLP
GLIGARFAPDDDYAINGALSGHGGYLNIIPTRPTWRTLPGFVDQVEQLIAAHPRLRRDDLVGVWIGTNDQ
DLTKASLNGIEPFLGVPRPANIAEMSAYTLTNLNAQLQRLIGAGARQFVILNLNDAGGTRPGYIDYNGKL
PEDLARFSRQGVNVHLFDVSALLNQMRRNPSAYGLNDTPDVQCRYVPSCSGGSIELQNTYLTADGTHMMT
SVHEYIARYLANQLNAPAAISTAPGLGLDVARAGALSALNAADGGPLGPARVAWSDRLSLFADVGYTRNF
HGAAGGMNAFDSDVEMFSIGADYKLSEASRAGALLSSGNANGSLAGGQGRIGLHAYRLGVYHAFERASLF
VRAYAGAGWSRYRLDRAAVLPGAVRASTSGFDFGALVKAGYLFALGGVRLGPVADVGYTQLVARGYTEDG
DPILAQNVGVQRLKGVSAGAGVRFAAPLAAIGLRAGELSAEAQLRHDAFGDRTLVTAQRYAPGLPIGTAV
DGASATYERLSVALTVNPAKRWRGKLVVQTDLGTAQRRSYTLLALLGGTF

>WP_024428767.1 ABC transporter substrate-binding protein [Burkholderia pseudomallei]
MKKLALCAALALAAGGAFAKEWKTVRIGVDASYPPFESTAPSGEIVGFDVDLAKEVCKRINAKCTWTPQD
LDGIIPALKAKKFDVIVSSLTVTDKRREQIDFSDKLYDAPARMIAKAGSPLAPSVESLKGKRVGVEQGST
QETFAKAHWEPKGVTIVPYQNQDQVYADLGSGRLDATLQDELQADYGFLRTPRGKGFAWAGTAVKDPRTL
GDGTAMGLRKEDADLKDAINRALASMHKDGTYDRLSRKYFPYSVYSAK

>WP_024428782.1 autotransporter domain-containing protein [Burkholderia pseudomallei]
MLMTRHKKRKTMKRSGAKLLAPVVVAAAAAVAARPGWAQAAPYPDPGRRGDPASWRTPEFTNAWGLGAMH
AEYAYAAGYTGANVAIGVLDSGYYAQHPELPDSRFVPVTAAGVSGVLNPNNNNHGTLVSGVVGGARDGVG
MHGVAPDATVYEGNTNAIDGFRFGVSDPKFPASDAKYFAEAYDALAAKGVRIISNSWGSQPANENYSTLN
KLTDAYKLHEAVRTATGRGTWLDAAAKVSRDGVINNFSSGNTGYDNASLRGAYAYFHPELEGHWMTTTGY
DQLSGQVYNQCGIAKWWCVMAPTGVPSTSYSGGAAAPTGATYANFNGTSAAAPHASAALALIMERFPYMT
SEQALSVLFTTAQNMEPDPSRPDYTNNGLFSTVHPAKPGASGVPNAFGGWGLVDLRRAMNGPGQLLGTFD
AALPAGTADVWSNDISDVALAARKREDDAEHRAWLDTLRTKGWEHGLPAGASDGDRIDYALGVARETAYQ
AREYQGSLVKSGGGTLTLAGANTYRGPTTVDGGELRIDGSIAARAVVNPAGRLTVNGRAADIAVNGGVAT
IAGTSANLSIDRQGRAAVTGTTADVRVASGFASLGGTSGNVAVGALGVAAITGRTADVAVDGGRASLDGA
SGNVAVGNGGVVSGSGTVRTLTAAANGTVAPGHSVGTLTVSGDVRFAPGSIYAVEVSPGGAGDRIVAGGR
AQIDGGALALALENTPPPLTPEQSRSVLGRRFEILNAAGGVAGRFDAPSGYLFVNPVLAYGPTTVSLTID
RNATPFASVARTANERGVADALETADPGSAVYNSVLFAASAQAPQAALAQLTGEIYPAAYAALVNESRQV
RETALERLWTARGAPGRAGAWARLLGAWGSARGGDVNGYTSSTGGFLAGADAALLDGVRAGGFAGYSHTG
VNLRNQPSSASFDSFHLGAYAGWQPGALGVRIGAAHAWHRGGVDRAVQYGAVAENETTALHAETTQVFGE
AGYRFALDGAATLEPFFGVAYVHLKNQGTTETGGAAALRVRQGNHDVTFSTLGVRGETRLGLTSRLQLTL
QGSAGWQHALTDGQPSGTLAFATGSDTFTVSSVPVAKDAAVLNVGAGLELGKNGWLRVGYSGSLASRQSE
HAVQGSLHWKF

>WP_024428812.1 AzlC family ABC transporter permease [Burkholderia pseudomallei]
MNSLPTSTVISEASPARSEAWRGLRASLPVMLGFVPFALVLGAQAAQKGLGIVEVPLLTGLNFGGGSEFT
AIRLWTSPPHILLIVAMSFLVNCRHILVGAAFAPYLRHLSRRQTFPALFFMCDESWAMALADARREGRGR
VSLPYYLGVAAGLYLTWISCTALGAALGPTIGDVERYGFDMAFTAVFLVLLRGMWKGARASRPWLVSLVV
AAATYLLVPGAWYVAAGALAGLVAAVVCGEPA

>WP_024428855.1 MFS transporter [Burkholderia pseudomallei]
MKTILTRDFLALILSVAVVGLGTGATLPLTALALTEAGHGTRIVGILTAAQAGGGLAVVPFVTAITKRLG
ARQVIVASVVVLAAATALMQFTSNLVVWGVLRVVCGAALMLLFTIGEAWVNQLADDATRGRVVAIYATNF
TLFQMAGPVLVSQIAGMTHVRFALSGALFLLALPSLASIRKTPIADEPHHDAHDRWTRVMPKMPALVVGT
AFFALFDTLALSLLPIFAMARGVASEAAVLFAAILLFGDTAMQFPIGWLADKLGRERVHLGAGCVVLALL
PLLPAVVTTPWLCWPLLFVLGAAAGSVYTLSLVACGERFRGSALVTASSLVSASWSAASFGGPLVAGALM
EQFGGDALIGVLIVSAIAFVGAALWERRALPMQAARRGR

>WP_024428898.1 NAD-dependent epimerase/dehydratase family protein [Burkholderia pseudomallei]
MLIWKTALSLTWKAGLVSPASAPAAAAHVDRVVLTGATGFIGGAVLVSLVNAGLLDRVVCIVRACDRAHA
LARLRAAALRSGLAPYWAERLSEANVIAGELDGALADADAAHIALASHVIHCAGVASLADARIVNETNVG
ATLRFARRFAGSRRLQRFVHVGAAFACGLRARGTIREDDTPARGREIDFAPYTRGKRDAEAQLRALGLPL
VVVRPSCVVGHTLLGTQPSASTFWMFRIVHAARRFTARPMARIDVIAVDDCARALMLLALKPSLAHDTYH
VSAGDEAPTVTQIVRAMDEAVGLDDEPRYALCSPAEFPSIARDVLGRRDAPRERVIRRALQSYAAFAELD
HVFDNARVRREIDFEPLPFVDYVNECMRTSRGIDVLAQMPRTAAR

>WP_024428916.1 Gfo/Idh/MocA family oxidoreductase [Burkholderia pseudomallei]
MSTHPLPVVVAGSRFGQFYAAGLAASGAYRIAGILGQGSARTAALARRVGAPVFTDPGALPDDVRIACVA
VGGAARGAQGAALARGLLERGIDVLIEHPLLPAEWDAVLRTAVRHGRRCLLNSFYPHLPAVARFVDVAQR
LSRIGRPLHVDLACSVQASYAALDVLASALRGVGPWSIEPARQASSPMRECTMVLAQTPVALRVHNEMAA
AADGRMHLLFRIALMTAHGTWTLASPHGPLAWEPALREPPADDDGLFPIFGAAAPPHGLPSMQLYEAEPP
AWDAIHARHWPRAAVRAVDRLASGDGLAASNQRSVEVTRVWQHLTAQLGFPEPPPGDASADTLEPLLERA
A

>WP_024428940.1 branched-chain amino acid ABC transporter substrate-binding protein [Burkholderia pseudomallei]
MSSYWMRFAAAASAALTLALPMPFAAAAATGEPIRIALVEGMSGPFANAGAAVERNLRFGIERVNAQGGV
RLRDGAHPLELVVLDSKGSVEEALVQLRAATDKGIGFAAQGNGSAVAAALVAALDKHNARDPEHRALFLN
YSADDPALTGRDCSFWHFRFDAHAGMRMDALADVLARDRAVKKVYLLNQDYSFGHDVSALARAALKARRP
DIAIVGDEFHPIGRVKDFSPYAAKIRASGADAVVTGNWGNDLTLLVRATREQGLAAKFYTFYGNSLDAPA
ALGDAGVKRVLAVADWHPNAGGARSDAFYRAFRARFPAAQDDYPVRRMSEMIEMLAAAMTRAGSADPVAV
AKALEGMRYDDGFHPAQMRAADHQLIQPLYVIEMERAGAPGVRFDNAGSGYGFATVLEAPPTQGAAPAEC
RMKRP

>WP_024429081.1 protein-disulfide reductase DsbD [Burkholderia pseudomallei]
MFNRIPRHAQSRFCFLIAVVAMLGVLFGTSLAARAADDFLDPAVAFKFSASEAPGQVDVHFKIADGYYMY
RERFAFAVKSGSATLGEPQLPAGHVKFDPTFQKNVETYRGDLTIHLPIKQASGPFELAVTSQGCADEGIC
YPPAEHVARIEGAALGAAGTAPAAAGAGADTSAADGGSWYERVTSADYARSLLEGHGFLTIVALYFVAGM
VLSLLPCSYPMIPILSAIIVGEGARATRARAFALSLTYVIGMALVYTALGVAAALVGQSLGAWLQNPWVL
GAFALLLTVFALLLIGGVDITLPQRWQNGAAQTSGPRKGGRFAAVATMGALSALVVGACMTAPLFAVLAF
IAHTGNALFGGAALFSMGIGLGVPLLVIGIGAGTLLPRAGAWMDGVKVFFGVLLLAAALWIVWPVLNAAS
QLGLGALWLLIAAAALGLFTPHSGSSSVWRRLGRGLGAALAIWAATLLVGLAAGSTDPLRPLAVLAARAA
PSNGAAGAGAGAGAHEGPAFAPVRSIAELDEIVKTSTRPVMLDFYADWCVSCKEMEHLTFTDARVGARLSQM
HLVRADVTANSPDDQALLKRFGLFGPPGIIVFDANGQERGRVVGYQSADRFLRSLDRMSLPAAWSAS

>WP_024429096.1 MBL fold metallo-hydrolase [Burkholderia pseudomallei]
MILRQFLHCDPVGISYLLGCGGKSTGVVVDAVAEPGTYLQAAADAGMKIDYVIDTHVHADHLSTGPALAD
AAGAAYVLSAKADVVLPFKGVEDGDEIRVGNVVIKVLETPGHTPEHISLLVTDRTRAEEPWLALTGHTLM
IGDLGRTELAESAEAGARNLFRSVRTLKSLPDYLPLLPGAFAGSACGRSLSATPLSTIGFEKRHNAAFGI
DDEAAFVRFMLSDMPPAPPEAARLRAANSGRNPDEI

>WP_024430699.1 membrane protein [Burkholderia pseudomallei]
MNVSQTCLLITALMPFLWTMCAKSSSRYDNHDPRGYLARLDGWRARAFAAHQNSWEAFALFTAALVVAWH
NGANMQRVDQLAIVFVASRVLYGVLYLLNWATLRSLVWTVGLVCVVWLFFAAP

>WP_028359473.1 Fe(3+) ABC transporter substrate-binding protein [Burkholderia pseudomallei]
MFVKQARPLLRTLLRTLAFAFALAAVAPAAHAANEVNLYTTREPKLIQPLVDAFTKQSGIAVNTVFVKDG
LLERVKAEGARSPADVLMTVDIGNLLDLVDGGLTQPVRSASLDGAIPANLRGAGGNWYALSLRDRVLYVD
KNMQLDAITYESLADPKWKGKVCVRSGQHPYNTALVAAMIAHDGEAAAEQWLRGVKANLARKATGGDRDV
ARDILGGICDIGLANAYYAGHMKHAQPGTDARKWGDAIKVVRPTFANAKSGGTHVNVSGAAVAKHAPNRD
NAVKLLEYLASPPAQALYAQANYEYPVRAGVALDPVIAGFGPLKVDPLPLVEIAKYRKRASQLVDKVGFD
N

>WP_029671417.1 multicopper oxidase domain-containing protein [Burkholderia pseudomallei]
MNPAVDPSPLSRAAQALALASIGGALALGACTSRPSGERYAPSRMDFNPGPAIVTQTRHTGPFASGSALS
FDAALQPLDAARDQAIRLDTTHTVIRIAPGIAFAAWTFGNQVPGPTVHVKVGDRVRLSMTNRSDEPAPGG
LQLTAPMMHSMDFHAAMVSPTDKYRSIAPGQTMHFEFTPNYPGVFMYRCGTPMVLEHIASGMYGVVVVAP
RDGYPTRADREYVIVQSEFYTKPDPQHRSVGTDALHVLDGERLRRKAPTYTVFNGRYNGMVTQPLIAKPG
ERVRLYVLNAGPSDTSSFHVVGAIFDRVWLDGNPDNQLRGMQTVLLGSSGSAIAAFVVPEAGAYVMVDHQ
FANASQGAVGVIDAGAHEESTIEHHNIPASATPTDAEAIQGKLDFESKCLACHTLGHGAKLGPALLGVTQ
RRSDAWLRRWLASPEAMVASDADARALRAHYPITMPDQNLSDSEIRRYVRYFHWADEASKQRDHAMP

>KGC51336.1 L-xylulose reductase [Burkholderia pseudomallei]
MTGASSGIGRAAAVALRGCGARVVAAARNARELERLAHETGCEPLELDVGCDASVRAALSGERMRDAFDG
LINCAGVTSLAAAIDTTADEFDRVMAVNARGAMLVARHVARAMIRAGRGGSIVNVSSQAALVALPSHLAY
CASKAALDAMTRVLCVELGPHGIRVNSVNPTVTLTPMAERAWSDPHASGPMLAAIPLGRFARVADVVAPI
LFLSSDAAAMVSGVALPVDGGYTAR

>KGC89431.1 spore Coat Protein U domain protein [Burkholderia pseudomallei]
MRIAATLALGLFSALHAHATCSVVSAAAASFGTVTSFAVARQPQSTSTTSSGLSCSGALLGLFVIGDQIN
ASITSANGGKLVGPTGDAVPYTVFADQNYSIKLDLGVTYNWASGQLLNLLGIFGGPAQTLPMYFRTVQGS
NVAAGTYTDTLTIAWNWDYCSGIGVLGICLGRDRGSGTAVVPVTITVTNDCMIAAPDVNFGAAPTVASFA
PVTGSVSLTCTKGMVYTVGLSSGANPHASGRRQMANGANRLQYDIFGPGAAAVWGQSANRAGSGAAADGR
SAQQFPYTARLYPDQPTPAVGTYTDSVIVDVRY

>KGD30356.1 hypothetical protein DO70_5810 [Burkholderia pseudomallei]
MIAFLARRAARRTPTPTTASHSPTSRSRGAGRHLFAALAVALIGLPHAHAEIRCGWLQNPTPGNWWLDDR
AGSWTLGTMGGPETEGMDVIPDMAGKQYVETNGSHGYACACLTVVTDKQERRIVKVLKAKQLPLSRCRRD

KRLKEPT

>KGD44734.1 hypothetical protein DP43_550 [Burkholderia pseudomallei]
MMKFLTREIVVLAALGWALLAGCAPVQAPPSSSSAPPASSCCGPITASGQTLRHTLDASDVEHLWLPRQH
VDWSTGKPDPTAEGFKSHCSAFAAAMGARLDVYMLRPPEHSQILLANAQAAWLASDSGRAAGWRELHEAY
EAQAAANRGELVVAAFQSADPKMPGHMAIIRPSLKSNVQLADEGPEIIQAGAVNRLDWNVRDGFARHPGA
WPNGIKYFAHVVPAK

>KGU91512.1 bacterial regulatory helix-turn-helix, lysR family protein [Burkholderia pseudomallei MSHR4032]
MAAIRTFLVACRAGSFTAAADELCVTHSAVSRQIQTLESWLGTSLFEKDGQRMVPSVHARAFAQELGGAL
DALTDVVQRYGKGSARQALRVSVPTTFGMRWLIPRLAAFRDVHPDATIQVLTVTTQQQPSGGNCDVAIRR
DDQVFNPESAIRFLSDHHTVVASPSLLSRTPLERPEDLVRHTILETETRPRHWDDWFAEAKLSASRFTRR
QRFDHFHVTFQGIVDDLGVGIGPVVTLSRDLANGKIVAPFSDIRIAPHNYYAVTPIGIQKTALHREFEDW
LVATAAAAETRAA

>KGV04506.1 beta-lactamase Toho-1 [Burkholderia pseudomallei MSHR4377]
MRRSLLVAAISTPLIGACAPLRGQAKNVAAAERQLRELESTFDGRLGFVALDTATGARIAHRGDERFPFC
STSKMMLCAAVLARSAGEPALLQRRIAYAKGDLIRYSPITEQHVGAGMSVAELCAATLQYSDNTAANLLI
ALLGGPQTVTAYARSIGDATFRLDRREPELNTALPGDERDTTTPAAMAASVHRLLVGDALGAAQRAQLNA
WMLGNKTGDARIRAGVPADWRVADKTGTGDYGTANDIGVAYPPNRAPIVFIVYTTMRNPNAQARDDVIAS
ATRIAARAFA

>KGV09504.1 lysine-arginine-ornithine-binding periplasmic family protein [Burkholderia pseudomallei MSHR4503]
MVTATALAAGSAFAADLKEIRFGVEASYAPFEYKTPDGKLAGFDIDIGNAVCAKLKVKCVWVENAFDGLI
PALQARKFDAINSDMTITEQRRKAIDFTDPIYTIPNQLIAKKGSGLLPTTASLKGKRIGVLQGTIQEAYA
KKRWAPAGVEVVPYQTQDLAYEDLKSGRLDATFQDSEAGAKGFLSKPQGAGFAFAGDHVSDAEILGTGVG
FGMRKNDAQLKSAVNQALKELKADGTIDGLAKKYFSVPVTLK

>KGV62168.1 traB family protein [Burkholderia pseudomallei ABCPW 91]
MPEAVAAREAARRSRNGVRRAATHAARAPRRWRARSLASAVLAGACATGGLAWPPASALAAGSVAAAPLP
QAPIPAPGMSLPGFHAPPPSTSNGTVASGAVRTQPARMPFYVATKGKVTIYVLGTLHVGDPADYPANQPF
RRPILAALAASPTLALELSPDDLLESQDDVSKYGVCNYACLPRLLPPPLWQKLANRLRGNPAALAGIRNM
RPWLASLVVETYDSLSAGLQTEYGTEAQLQNVFLRKKGGKVVGLETLAEQMRAFTGLTLAQQREMLAQDM
VQTPAQNAADVRALHRLWRIGDADAIAAWANAKTERLARARSIADSIDNKIVYERNRRFVARMTAIAAPN
RPLFVAIGSLHLGGPKGVLELLRQQGYRVDAG

>KGX12970.1 hypothetical protein X984_4408 [Burkholderia pseudomallei]
MLLLRHSILTALTATACASAHAQTADDAPVAAPACAIAIVTGVGGAATNLREYLAARERDRYRYLADHPL
DCRVSEDGRASGCTGLAYLRRERVSVYDDGDDTVQNVVARVDLDHGTYPAIIAVQKRDVRCEQ

>KGX72041.1 hypothetical protein Y026_6279 [Burkholderia pseudomallei TSV28]
MNHKKGDVARFENMQKGINMKRSRSRFRASFTVAVALGVPSGFAHAQSLPGIPDPAAVGLTPGTGQNGAP
TVSACFYNYRTKVIAGFPTQIASGWIVTNQRNDPACIPPGGLPGPNWQEMTYFRDMPVGASMSACWSFSW
PSNWQPVGYTSDPSKCGYYPGGQIPGAQPNVVFLKRVQ

>WP_038707916.1 Tat pathway signal protein [Burkholderia pseudomallei]
MDRRKFLSTSGMVIGAGAVQIVVPARTVYAATAASQLSNISSVAASSKLPNFANTKAILFPGSDKTVEAL
LDNAKSKIKSGTFDGSLLATSIEKQNPITGLYVILETAAITVTESIPVVGWLISGFLSIGLKYLNESLKE
KTDYEKLIKSIVNKSIDENNYRLMKAAFEGVTQVYGNFSDLAASLSTNKPKPDQKTQLISLFTNTISNCE
QNAPTMLLKNLNDKAAYQGIPLFCSIANVELSAHADMLKNKVAFDLDDSFYKNNYAAMEKKAWRMHKKAR
EVVVELLDNVGDLSLDYPSVWAKKNSIIRQIYLSGLGAFLENYRRRYEAKFLKKIKLRNSIEIYSNSYAD
QGVIEKTINADITRRSSLLSRLSGYSHIDRVQRIGVGYLSSSENSKGFFKEYESCTGRVNDRRDEKDIDS
KFPIDQFANSSRTGRELPVKIYSNAPKAGTYGLKIITNDGKNIDFGSMNGDLKKIYSFDVPGYYISGLVV
AESNAKRGSCAADSGLSIDFICSVHRHCSSIDYIAGFTDANIGIELDNSVWSFENNSSSNVTGYADYLVG
KNILSVPSGSVKFDLYNTTGKAISVTPVLMAAIPGRKDGESASLKVSIGNSVAQVKLSDKDSDKEDSYIQ
GIDGEKYLDAASGEKITVPVGRSSISIENVDKKAGFKFASLLLIPG

>WP_038708181.1 collagenase [Burkholderia pseudomallei]
MTEVFRKTRRWSAVAALSAFVGLAGAASANTQPMQPTQQKQARMPRLPQNLPVSPEQAEYNLPLSEQDRA
ALTRPSPLKQPAKRGKRSAPGADCRDMSVMTQYRGAALADYIANLPDYECHYGLFSVDKTLAAQIFSAEN
VHAVASRFVQDIYRYDASNLILVNLLIYLRSAYYQYDVSGIANPIPNLAVWLRPYIKQSLEGAALYRENA
RAPSTANELMKLITNMKDEAFYLPTLKARIAFYTASATNPQAAAPLLQPSAAGGFTGLLTVFFYAHQRSG
AQPMLDSDATLPETLNRFVTANRASLSNTSAAYQLADAARETFRFLRYPAQKPRVKKMIQDMLASTSMTG
ADSDLWLAAAEAVDYGDPGNCADYGTCDYKKRLTDAVLTHRYACNAGVRILAQDMTLPQLQSVCTSVAQQ
DDYFHRMMKTGRKPVAGDRNDTIELVIFDDYANYRKYASVIYGISTDNGGMYLEGDPSAPGNQARFIAHE
ASWLRPEFKVVWNLEHEFTHYLDGRYDMAGDFAASTAKPTVWWIEGLAEYLSRKNDNQESIDAARTGAYRF
SDVLGTLYSSSDYVARAYRWGYMATRFMFERHRADVDTIVSRFRVGDYDGYANYVAYIGNRYDGEFVDWA
RAATTAGEPPLPTKR

>WP_038714058.1 spore coat protein U domain-containing protein [Burkholderia pseudomallei]
MHKREVQVVSGAAFMLFFAAVSAGQLTGTMQVNLQVSRGCEVAGVAASGDLGRLDFGAQGPLWSDYLTAD
GRATSSGAVRVVCSPDVNGFLVSIDGGRNGDQSTRYLVKRGANGRVAGRIPYNVYRDAARSVPYVPLMPQ
SFFVDGGRDDVTLPVYGVVNGMTRAVPSGTYEDLLGITLDW

>WP_038724299.1 sugar ABC transporter substrate-binding protein [Burkholderia pseudomallei]
MRLCTGKAVLRACVAAIAVAAGVGSAAPAAQAAGARFALVSHAPDSDSWWNTIKNAIKQADEDFDVTTDY
RNPPNGDIADMARLIEQSAARDYDGVITTIADYDVLKNSLRKVTAKKIPLVTINSGTEEQSAQLGAIMHV
GQPEYVAGHAAGEKAKAAGVKRFLCVNHIATNSVSFDRCRGFADAIGADYKSSTIDSGQDPTEIQSKVSA
YLRNHPNTQAILTLGPVPAAASLKAVQQMGLANKLFFATFDFSDDIAKAIQSGAIKFAIDQQPYLQGYIP
VAVLAIAKQNKTTDPAKIRQILEANPKFQARLSTYGLQPSYGPKNIRSGPGFITKENLEKVIKYAGQYR

>WP_038726191.1 hypothetical protein [Burkholderia pseudomallei]
MKRTIRKSRAALVPLLVAGSVCAAAPLCQTQKLGAHTSKMCVEQTPFKHDYYTLWVDDSPIFMLPDDYVE
KVALTHTVPEDGAIEFPLSKQGTPTVTISGGCAPVSETQGKGADAVNLETGRVCSFNWGKEPVVKDLRFS
FE

>WP_038727163.1 hypothetical protein [Burkholderia pseudomallei]
MKSSVLFMAALPLVAIAASAHAQPRHPAVYSPAAGVLCDRYVCADDQGISRALTERYLGKRVAAKAFSQG
DFDPTEFTFANGVFCDVKERLCRDDRYYGADGKRSGAVSRRYTELLFGRRSGG

>WP_038729084.1 hypothetical protein [Burkholderia pseudomallei]
MRNRKFVSFVVALAFAAAAATPALAVTVVRVDGQPMNPNGEPFSATSAPLETTLSKGSISANCVATFNGT
ITPAGIVNITSTTFTGTNSLCGLIKGSASGTNPWTGQADSATQLTINNAQVNVTLLGQCGPSKVVTSWTD
ANSSLTFSNAALAPDCKVTGTVVTSPKFHVQ

>WP_038730428.1 peptidase [Burkholderia pseudomallei]
MQTSRKALPLALGLAIGLGAALPAWADSKAPSPQEESRRASLTRGVVAPAEQAGKTGQFRPGAVAVTLAS
PAFHAKKADAAAMAREYVTARAAQLGLDKAALANLVVASERADAAFTVVRFQQRAAGLPVYDSDIAVTVA
PDGRVLYVASKAVSGVAAVSSKTQAVDEQQALDRARAYLGVGGFVNVQSQLVAFVDGAGTHTAWKVSGRP
QDSLHGDWELIIDAGSGEVLRAQDKASYATDGSGLVFRPDPLSPTKSSYGSPGFKDNNDADSPQLSAARV
RVTLKDLTQTSGGYKLSGPYASCIDFDAPLDKACPVQASTTFDFTRSNLYFEAVNAYYHIDTFLRYVNLT
LGVKALPYQYAGGVQYDPHGQSGDDNSSYSPSSGRLSFGQGGVDDAEDADVVIHELGHGIHDWITNGGLS
QVEGLSEGTGDYLAAAYSRDFNQWSPSDAQYHWVFNWDGHNEFWAGRVTNYNVGRTYAQIRNAAIHTAGQ
YWASCNMVARDAIGGAAMDKAFLKGLSMTNGSTNQKAAAQAVLTAAAALGYSSAQLNAIGDAYNKSCTYG
VTVPQKL

>WP_038730764.1 cytochrome c4 [Burkholderia pseudomallei]
MNRLSKSLVVLEFAVAGLAGFMAEARAADAAKPDLDRGKAIATQVCAACHGVDGNSATGSFPKLAGQHPD
YLVKQLHDFKTQPGAKGPVRVNSVMVGFASALNDQDARNVAAYYGSQSAKPGAARNAATVPVGQKIYRGG
IAEKGVPACASCHGPTGQGIPVQYPRLSGQWADYTVAQLTAFQQGTRSNDAMHQIALRLTDSEVKAVADY
IAGLH

>WP_038740236.1 hypothetical protein [Burkholderia pseudomallei]
MISFKRKMFASPASRPARAGKFVLEACFAGALAGAAHAQGAPQPVVDWEIQVVRDGQTIDAFEQKTTVGQ
ARSDTHSLPIAPAAGCAATSAAASAADAPGSGLSRTITVAPLYVEDGAVALAIDAQETLADEGATPPAGA
PCTFALPRRIVASHPGLNARAAQWTDWTLAERNPQLVYRVRARVVED

>WP_038741088.1 hypothetical protein [Burkholderia pseudomallei]
MTTRARVSILPRMLVGAACAAAALGAHAQNNLNFLNDTPISYFSKADTASLAKAVQKVRDEGKDGETVDW
VNDGRGTKLAAKLTPSTTEQEGRTCREIKTEIEAKGQSMTLRPLYCKTAAGKWQLQKR

>WP_038741497.1 triacylglycerol lipase [Burkholderia pseudomallei]
MARTMRSRAVAAAVAFAMSAAPAAGIGTFLSLAGTQSAAAATSAVDNYAATQYPIILVHGLTGTDKYANV
LEYWYGIREDLQAHGAQVYVANLSGFQSDDGPNGRGEQLLAYVQQVLAATGASKVNLIGHSQGGLTSRYV
AAVAPQLVASVTTIGTPHRGSEFADFVQTVLAYDPTGLSSTVIAAFVNVFGILTSSTNNTNQDALAALKT
LTTSQAATYNQNYPSAGLGAPGSCQTGAQTETVGGNTHLLYSWGGTAIQPTLSILGVTGATDTSTIPVVD
PANVLDASTLAMLGSGTVMINRGSGANDGLVSRCSSLYGQVLGTSYKWNHLDEINQLLGVRGAYAEDPVA
VIRTHANRLKTQGV

>WP_038749854.1 TonB-dependent siderophore receptor [Burkholderia pseudomallei]
MKSRSDELKLGKFTTLCSVLAASPAFAQDAAPPAASADHDKELAPIQVKGAAEHSYKADFSSSVKFTAPL
VDTPKSVTVIPQELIHNSGAATLTEALRTVPGITFGAGEGGNPLGDRPFIRGYDTQGSMFVDGMRDTGAT
TREIFNTERIEITKGSDGAYGGRGGAGGSINLVTKAPHLGTTAEASAGLGTDRYRRFTADGNWQFADHAA
FRLNLMSHNNDVAGRDAVNNERWGVAPSIAFGLGTPTRVSASYYHLSTDDLPDGGIPYFYTTANKPANVD
TIYPANVDRHNFYGLVNRDFRKTTSDIGTLRIEHDITPSLTVRNTTRYTESTQDYIWTQPDDSQGNVVNG
RVWRRNNNRNSAINSIANQTELFGEFRTGPFKHSFTTGIELSREWGKRDSYSVATGTGKICQQGIGAASG
YNCTSLGSPNPNDPWAGSITRNNDYAHARTVTKSLYGFDTVELSKHWQVNAGVRVDDYSTRFTDTRANGG
KTYTRDDTLVNWQLGLVFKPARNGSVYASYATSSTPAGALLGEGSETQSLTPGRGGVGANADQLAPEKNR
SIELGTKWNVLDDKLSLTAALFQIDTTNARVTLPNNQYAMVGNKRVQGLELGFAGQLTRAWQVFGGYTYM
KSELRDNGRNTADNGHQFPNTPKHSFTLWTNYDVTPKFTVGGGAFYMSKVFGDTANLRAVPSYWRFDAMA
QYRINKKLDLQLNVNNLFNRTYFDQAYPAHYASIAPGRSAFVTLNARY

>WP_038750526.1 SCO family protein [Burkholderia pseudomallei]
MSADEIRTRRRKLLAAALSAALAPALPALAATQRAQPLAGQSGYHGGLITPPVPVPDMPLRTADGRATRL
RALLAGRVTALQLFYTGCSSTCPIQGAVFHRVQTLLGPHPKPDIQLLSLSISPLEDTPQRMHAWLARFGA
RPGWIAAAPELKDVDALQAFFGGGRTGLDNHGTQVQMIDRRGALVWRTYELPSPETIAALLGHA

>WP_038760383.1 fimbrial protein [Burkholderia pseudomallei]
MKKKFSRLLAALVLLAATVDALAAGCNMLTAENIAWLTEQGKIARAHSSFPISFSSGMVDVDPNLEIGGL
IAEAKSIPSEELHFIWCSAPSGNVHFALNSSPLPSELGNSIYETGVPGVGFRITQVRQSGSIGAIPRDTP
WIEEKPGQDSSLNFGAGTVFRIELIKTSEALPSESTISLGNLSRVYGDDNKTVVDFNAGSVKLRVLPICH
VDQQEKNVDFGQFGPKDVSFDSGPTKDVKFDVQCSGPTPPVSITATLAATPDSHDQSLIANAGDAMNLAI
RLRDASTQQVLRPNDPSSEIKVEPGGAMEHGFALEATVLRVGTAPPTAGTIDATSIITLTIL

>WP_038761248.1 membrane protein [Burkholderia pseudomallei]
MKRSLIAFALTGAGIALSSTAHAVNVDVNIGTPAPVVVAPAPAPVIVAPAPAIVVGWHGDRYWDGHRYWE
RREWEEHHRHRGFCPPGHAKKGEC

>WP_038762696.1 hypothetical protein [Burkholderia pseudomallei]
MTTIKAGDASGAGCAAAARARMTTRIMRRALAVLGGAAFAGWFGAAFAASSGTTVPPLDTTPWLASVQAP
KTPFFTRASDAQGVSFHFANSGHDVERGREVVVSETADGDMWQGGMVHATYIDAAGVETSGWLVRSHLRR
VKKPVPPPPTWDGRWRSSAGARRLIVHGERISYSFAGGGGTGGAQPRVEMLLRMRPVSDDEAVLSRMQAP
DGGMCDLAVRRLGDYLIVSARDCFIPGVSPEGILRKQR

>WP_038765499.1 type 1 fimbrial protein [Burkholderia pseudomallei]
MKKKVVSFAVATLFGLVAAQSAFAQSIDQGQVKFTGELTANTCTIDAGSKDQIVPLPKVSTSALAAPGDV
AGSTAFEIKVSKCAADVKKVAAHFEMENMDPNTRTLKNEATGADAATNVTIQLVNSDGTELPVGSTGSYF
DVTGTDDARGATMIYGGQYYALDTTTAGKVESHTLFTLAYE

>WP_038774138.1 MmgE/PrpD family protein [Burkholderia pseudomallei]
MNRRHFLAAAVAAGLPLAATLAPRGMRAQPNATQAAAPTLARQLAEYAAGLRYEDLDSATIDIVKSHLID
ALGCALAALDEPPVRIARDAARSAGGGGPSTIIGTAERTSPDLATFATGTALRYFDFNDAYAGREIGHPS
DNIAACVAVAEAQHASGRELILSIALAYEIACRLMDAAAISPRGWDHTCYSLPAAALAAGKLMRMPVEAL
TQAVNLSLNSHLALNQTRVQQLSNWKALADADAARNAVFSTQLARAGLTGPSPIFEGEAGFFRQVSGPFE
LETSRFGGRGEPFRIARCFVKYYPAQGFTQTAIPAALDVASQAGDLSRIRRIDVHTTRVGYVTAGSEPEK
WKPSTRETADHSLPYVVARAMLDGDIRTTSFSDAALRDPVLHALIAKIRVEEDPALTAGYPARAANRVTA
HCRDGAVYAKQVDDLPGSPTRPMRREDFEAKFVKNGGARLSEQRMRAALDRLWRLDELQDVAALPPLFVA
G

>WP_038775093.1 triacylglycerol lipase [Burkholderia pseudomallei]
MKSRFASAAAAVRGARKTLAKVWAAVLVAAMCTFAAPAAHADAATGALDDYAATRYPIILVHGLTGTDDY
FGVVPYWYGMRPDLQQHGATVFVADLSGFQSDIGPNGRGEQLLAYVQQVLAVTGAQKVNLIGHSQGGLTS
RYVAAVAPELVASVTTIATPHRGSPFADFVLGALSLDPTGLSTPIFGALLNVFGILTSHTHNTNQDAIAA
LNALSTPYATQFDAQFPSAGLGAPGACASGAPSETVGGNVHLLYSWSGSAYQPITLLGLTTGALDKSTIP
LVDPANVLDPSTLVFLTAGNIMALKQAGLNDGFTSTCSSLYGTVISTDYKWNHFDEINQLLGIRGAYAAD
PVAVLRTHANRLKLQGV

>WP_038778478.1 membrane protein [Burkholderia pseudomallei]
MKKYALALWLMLVSSMSAMAGAIIPSGATLVAGQYAISNNGNFAFGVDAASGKLYFHYRFPTQAWGASAM
SQMYGTQPEGALYVGKGDRLVMQTDGRLVFYSGNDVVWSNWYYGPAPIPGSYAVVEDFGVVAIYPPQGGY
PTFTWPFASVTPNSGALATVVGYPFGVNFPVLNGRQWLQIFNNVTYYLWNKSFAADSVAMQTDGNLVVYN
KGQPVWSTGTSGNPGAYMLVTPVGFVLGKQYGGILANVPYPVSVDQSSRASGPNDPKPTPAAPPPPSLNL
PINASWKCYYVKDWLACYAPG

>WP_038781359.1 ExeM/NucH family extracellular endonuclease [Burkholderia pseudomallei]
MRTPTPLLAALSLLSTVAAAPAFAAAASPVSTNCGGSATPIAEIQGAGAPSPLAGQNVSIEAVVTADFGG
ADGFGGFFVQQADPQRRNQPGVSEGLFVYSPKARARAGELVHVTGKVEEKYGQTQLTLSGAVAVCASGQS
VTPATLALPVDSPNAFAAYEGMLVRLPQTLTVTEVYELGRYGSVLLSNGRLRTPTSVVPPAQAKAVAEAN
ARNRLILDDGSNKQNPATVPYPAPALSAANTLRAGYTAADVEGVLELRYGAWRLQPVPGARTPTFGASAN
PRTAAPARDPRANLRVASFNVLNYFNGDGAGGGFDDPSNRGAKNYDEFVRQDAKIVSALKALDADVIGLM
EIENDGYGPLSAVRQLAAKLGENWRVVDPGSARLGGDAIAVALIYDSRKVKPIGNAATLAIDDKNRQPLA
QTFRPLGGSRAVTVAVNHLKSKNCPDATGDDLDQGDGQGCWNATRSRAAAKIADWLARNPTGAPSEGVLL
IGDLNSYTYEDPVRTLESRGYVNLVSSKIGSGAYSYVYNGEAGYLDHALATNALAWRVKAVHDWHINADE
PIALQYTLAYKSAEQQRTYYAPDAYRSSDHDPVLIDIALADEYAAAASRRGAEMAAAHSRRPWH

>WP_038783934.1 hypothetical protein [Burkholderia pseudomallei]
MGDRGLRGGRRRTIAHDTAAGVALPSLAALPPGVLPADLAAASSGALLSVGGGPSQQIDDHAFTSRNYTA
LVCSSGLRQEFTTPHCSR

>WP_038790363.1 sugar ABC transporter substrate-binding protein [Burkholderia pseudomallei]
MTWINRGRALAGCVALAAALAAAPAGADTGKVGLDLPLLTSPFWQSYNRYLLHYAKDMQIDALAPVNSNG
DPAQQITDMNTLLNLGAKGIVVGPLDSAAIGRALDAAAARNVPVVAVDVAPTQGKVAMVVRADNRAYGEK
ACQYLGEHVRRGKVVQIMGDLASVNGRDRSEAFRACMKGYPNLQVLEIPAAWKGDVAATALDSLLSANPD
VKGIYLQAGGVYLSPTLQTLRRKQMLYPAGDAKHVAIVSNDGIPQEYEAIRRGDIDATVSQPADLYARYG
LFYIKAALAGRTFKPGPTDHGSVIVQRAPGVLEDQLPAPLVTKANVDDKGLWGNTIK

>WP_038797268.1 PepSY domain-containing protein [Burkholderia pseudomallei]
MKTSRLIKAVALASLVLGGALGIRAAMADDGDDCRAPLADWKPRDAVRALAQQKGWRVDKLKADDGCYEI
KGHDADGKRFKAKLDPVTLDVVRMKREGERKRDHDDDDDHGRAPDARAPAGGPPAGAPPGGVLKPGSKPD
VQIR

>WP_041189005.1 c-type cytochrome [Burkholderia pseudomallei]
MESRVSSRRLFRPLLAVVLMSAAGLLSTAHAQTKPTEPAAAKAPLKAPDTMAERVRGCTACHGTHGQGTD
NDYFPRLAGKPAEYLYNQLVNFRDGRRKYPPMNYLLTYLSDDYLREIAQHFSDERPPYPAPTKPTVPAAV
VERGKQLALHGDPARKLPACVACHGNGLTGMQPAIPGLVGLHSDYLSAQIGAWRSGTRHAKAPDCMHDVA
SKLSDEDVTAVTAWLAAQPAPANPVPAPARSMKTPLACGSEPQ

>WP_041190807.1 TlpA family protein disulfide reductase [Burkholderia pseudomallei]
MKRVAFALAAFAVLANAAPGAAQEGRLPQPLRATDVASLYASGSGAPLVVEVWSLDCGYCRENAAHLVEW
QRRHPQVRVALVALDSLDEHGQELAGALAQMKLPAAVRQYANAEPMPERLRAALDPRWRGELPRTLWIGE
DGTRRAKSGLLAPAVLDAWLQRREP

>WP_043294325.1 hypothetical protein [Burkholderia pseudomallei]
MLKRTALFLALAGSIVAFSVAQANGDASLKPQQEIQLTKNAWGCLSKDNLDSVLNHERDGKAQAKQQYFD
DYRCLSVPEGQRFRVVSVDKGDVQFVSAENSDQQGLWTDARFIKQ

>WP_043299325.1 hypothetical protein [Burkholderia pseudomallei]
MKYLPLIALIAATSASAADTGVQNVGQSQKSAQDVSACIAKTWADKSQQQVVTQNVLANGLAADVYVPGQ
QPPNGAAAVVRPAFSGNAKTWVGFRAGSSGAADGAAAGDINSCL

>WP_043304483.1 hypothetical protein [Burkholderia pseudomallei]
MTMHGEPNARRASLMGLALKLAFGFAFALAWPGHAARADDSAARGRALFRGDAPLQGRLPMHPGDLPAPL
VRCANCHAAGAGAAVPNSIAPRLTRAWLGDLQSRRGGPPSRYDRNAFCALLRTGLDPTYVLINVEMPRYR
IGERDCTALWRFLNEAPHERR

>KIX45952.1 hypothetical protein SZ28_00940 [Burkholderia pseudomallei]
MRRALPIPILFAVACVAARAAGLPMPVLSRLPPGYEVMAARQGPDVDAGRISYLIVLHRPADSASEPSPR
PLVIVEQQADGTFRLAARNDEVVLRANEGGQCDPFDPQDADENGLAVKGRFFTVQNFVACGQHWSDYVTF
RHDARTGRWLFANEIRTESFPLEGKPDRVRAIRADPRKPVALDAWRRGD

>WP_045588731.1 hypothetical protein [Burkholderia pseudomallei]
MSGITGSIPVAQAGAMSRGPVPLDVKRIDGKPAACIPTSDEGQAEMRLGFIGVSRATGPVSPDVIYWQVE
VSDNVPPVYLKRGDCIVYGQSIPGATVLTPPKRFDIGKWYNFAVMPGGKAEGPIYRGTFCISGKQGDSGR
VVLRSENKDACSSGR

>WP_045597613.1 amidase [Burkholderia pseudomallei]
MKRRQFLHTLGALGPTHALFASAPASAGQTALDERAGASAPASRAVRESLERIARIDRDGPRLNAIIELN
PDAEAIAHALDAERAAGVARGPLHGVTVALKDNIATGDRMSTTAGSLALDGVRATRDAHLVARLRRAGAV
IVAKANLSEWANFRSTRSTSGWSARGGLSRNPYALDRTTSGSSSGSAAAVAAGLVAVSVGTETDGSIVSP
AAINGCVGLKPTLGRVSRDGIVPVSHTQDTAGPIARTVRDAARLLGALAGGDARDSATASAPAPADYVAA
LDANALRGARLGIARAYFTGHDEVDVQIERAIAEMKRLGAVVIDPVDLPKADYEEDEKVVLLHEFKHGLP
SWLRTFAPHARVRTLADVIAFNARQHAREMPYFGQELLLHAQEAGGLDAAAYRDALARCGRRARDEGLAR
VLREHRLDALVAPTEGTAWLIDLINGDSGGDGFSTPAAVAGFPHLTVPAGLVRGLPVGVSFVGAPWSEAR
LLALGYAFEQATQWRREPRFVERSNVPAAR

>WP_045889382.1 hypothetical protein [Burkholderia pseudomallei]
MRANRNNGLTAGALPPNSAAPDDAAKAAPPVAQSDRGAGPLAGLTALKADRAQCMSARPSIRSTTKAPAV
MSHARADTSRELTPVEARIAQTFVRPEHLRAIAAAARAGSFAVSFRASGQLSLKRLANGAPAKGHDILEK
TIKPGSVQEVYGADSERRLAQIERADIDGYVGHWGRDKLLGLYMGPAANERDTPMLFAALRSTPSGHRYY
PVDHDDLEGSLALLKRTQGPSGTAGSFWRSLPYTGDYDTHDMINMAGHRAPVPSGSRDEHRIMRSLNVAI
EAIDPIRKASSGSHRLIQHGPQHNYVAHMRTNEQGAVIDEAVARASFPLAMCDRGTWSVIETPEALAAFY
RQHGVVLKEKWREKRANTFTA

>CFK48476.1 oxidoreductase [Burkholderia pseudomallei]
MSERTTHDAARRTGASVGRGCGGGAPAGISPRMIRAHPFGYPKHLDRRRVAQAGCPKRRKAVRRLHPPLA
QPATALCRVPGYGALVQLWRRVRATLPGVVCFEAMRPAFYRCVAAHAPARRRRSTRTAASRR

>CFL10512.1 Pectinacetylesterase [Burkholderia pseudomallei]
MLVRECLECLRRLCVAGLLSAAAADGHAAVAAAXXXXXXXACAPRTTDPIEMEESMPVRECLECLRRLCV
AGLLSAAAADGHAAVAAAQDPSIPYYSWYEVTLPESSGASCGNGTPMRFYINRAQSDNLLYMMEPGGACW
DHGTCTETATGAQAGLGAFNPDGIPHNYVNGTVQQSLKTSFLSPLMTRMDLAHILVGEPKVETQEWTQVF
VPYCTGDIHMGSAVRSYASPSGDWRIQHYSGLKNVRVIAQWLVDHGFGKPSRLLVYGTSAGGYGTLGNYA
TLRGTLQPQSHSSLLDDAGTVFNTRFGADPAAYPSVGLYDKVRDEWGMTAPDGMITVNSRLTRYFDPGNM
GSAYAALSATYPHDRFGYTTYRRDRIIAAYHYRPFVPAVIAAPDDATKDALSLAMFDRELSDLKQVLNPL
PNFGYFIPWARDDFMGNHQVTAVSFTGSGIHENGIDADVGTFVADLLNQQDPADVPVMKAYRTEQWSDFT
FSTFLAWLDSIFNLTGEAGPISGHRS

>CFK95952.1 extracellular ligand-binding receptor [Burkholderia pseudomallei]
MNIKMQKLLPISAAAMLLAAAATNAAADQVVKIGHVAPLTGGIAHLGKDNENGARLAVXXXXXXNRLLPI
SAAAMLLAAAATNAAADQVVKIGHVAPLTGGIAHLGKDNENGARLAVEEINAKGLTIGGQKVTLQLDAQD
DAADPRTATQVAQKLVDDKVVAVVGHLNSGTSIPASKIYSDAGIVQISPSATNPAYTQQGFKTTYRVVAT
DAQQGPALANYARSKGVKSVAVVDDSTAYGQGLANEFEKKAKALGLKVVSHDATNDKAVDFRAILTKIKG
ENPDAVMYGGMDATGGPFAKQAKQLGLRAKIFAGDGVCTEKLADLAGDATDNVVCSEAGASLEKMPGGAA
FKAKYEKRFGQPIQIYAPFTYDAVYIIVDAMKRANSTDPAKIVAAMPKTNYTGVIGTTTFDSKGDLQHGV
ISLYDYKGGKKTFLDEVKM

>WP_050040327.1 LysR family transcriptional regulator [Burkholderia pseudomallei]
MDRFKQIETFAAVVAKGSLSAAAHAEGVAPAIIGRRLDALEERLGVKLLVRTTRKLTLTFEGSAFLEDCQ
RIINDMQNAEASVSAGGVKASGHLRVSAPAGFGRRHVAPLVPPFTGAHPDVSVTLDLSDRMVDLVNEGFD
CAVRLGELPDSSLVSLKLGENRRVCVASPAYLARAGMPGTLDDLARHNCLALAANANQQRGWSFVDGGKV
VSIRVSGTMECSDGAVLHEWCLEGHGLAWRSWWEVGADIAAGRLVSVLDAFAAPPVGIHAVFPQRRHLPL
RVRLFLDFLKHTYERPGYWG

>WP_053292838.1 M20/M25/M40 family metallo-hydrolase [Burkholderia pseudomallei]
MSKLQHLAAAVCGALCVSAARAAPVWITLSEPALRELRALDPAVTSRYSAALATGDAKRTETIHVAQVDD
SLLESLSQAIRRTRGHGPGFFVHATFDEARASLQPSAAKQAAAIDYPITYSQQVRNWISQLQASNIVSTI
VSLSGFTNRYYTTTHGVAASDWIAQQWKQLAGSRTDVTVEQFTHAGWPQKSVILTIKGSDPAAGVVVIGG
HLDSTVGRMSENTRAPGADDDASGIASLTEALRVLLANRYQPKRTLKFIGYAAEEAGLLGSQAIAKQFRA
QNVNVVGAFQLDMTNYKGDPKDIYLIGDYTNATQNTYLANLAKAYLPELAVGTSQCGYACSDHASWNAQG
YPASFPFEADQNDNPYIHSAYDTLERSDSQGNHALKFSKLALAYAAELGGGLSASAKR

>WP_053293009.1 glycoside hydrolase family 2 protein [Burkholderia pseudomallei]
MKSAPDRVARGAAQWTLIATPAGAIARPSELGEAGWCAASVPGTVAQALAAARRFDPAHPYPLGDSDYWY
RTTLHGAGPRIVRLNGLATIAEVWLDDTLLLCSDNMYVAHDLPVTLGGAHRLALCFRSLDRHLAEHPPRG
RARWRTRLVDTPALRGVRATFLGRMPGWFPAIEPVGPWRPIDIVNPAGAPTIVHDTLRATLDGRDGVLDA
TLEFAAPLPRTARAQLVCGEHAAPLEATGPRTARATLRIANVTPWWPHTHGEPALYDVGVAIGGATIALA
KTGFRTLAVERGADGRGFALSVNGTPLFARGACWTSADPVGLHADAPAYRRALALARDAGCNMIRVGGTM
IYEADAFYALCDELGLLVWQDFMLANFDYPSNDPRFAESLKREAEQFLGRHMARPSIAVLCGGSEIAQQA
AMVGLAPDERRVPATEQWLAELCAAHRPDAAYVSDSPHGGVLPFAPREGVTHYYGVGAYLRPPEDARRAG
VRFASECLAFANVPCDATLASIGSPAAHEPAWKRAVPRDPGAPWDFDDVRDHYLRTLYGVEPARLRSIDP
ARYLTLSRAVVADLVGETLAEWRRVGSSCAGALVWQFQDVMPGAGWGLVDAHGRPKSAWHALRRVSQPRQ
ILLTDEGLNGLDVHVLNDAPAPLEARIELVALRDGKTPIARAARTVHVAAHAGQCVNSADLLGRFFDFTY
AYRFGPREHDVVIASLYASDGALLSQAFHFPERTAPTVFERGDIGLEASAAYRDGRWCVQVQTRTFARYV
HVCAPGLLPDIDWFHLAPGAAARIEFAADPHSPAPDHRPPEADAAHCAPPAIEVRALNSNKTIRPRIEN

>WP_057050932.1 CoA transferase [Burkholderia pseudomallei]
MSGNVSKRLRASSGSEIRRAAAGASEAAGAAGTAGAFADAAHPLPPAGGLCMSSEGGVPPGTAIADATAT
ASDARTAFAANTKDAKDAKDAKDAKDAKDAKDAKDAKDAKDAKDTKDTKDTKDTTSRPSLAAANHATGPV
RPGALAGLKIIDLSRVLGGPYCTQALADHGARVIKIEPPAGDETRGWGPPFLDDDAWYFTGVNRNKEGIV
LDLSRDEGRAILWRLLEDADVLVENFKPGTLARWGMDYERDLRPRFARLVHCAITGFGADGPLGGLPGYD
AVIQAMAGLMSVNGDAGGDATRIGLPIVDMVTGLNALAGILLALAERERSGVGQSIDIALYDCGVSLLHP
HLPNWFGSGRTPARSGNAHPNIAPYDSYRTASAPIFLAIGNDRQFARLCAHLGVPALASDPRFIDNRSRC
AHRAELKASLETRLAGRDGETLARELMAEGVPCGPVRTIDALACDPHTLHRGLVVEMGRYRGAASPIKLS
RTPATYRTAPPTLGRDTRAVLDALGIDAATQARLADAGVLKEAKPMAT

>WP_060547731.1 hypothetical protein [Burkholderia pseudomallei]
MRTANVACALAVGFASAAALADAPAMPSGGMLVAANGMTLYTFDKDAPNAGKSLCNGPCAANWPPYKASA
TDRPAGGYTIIKRDDGTLQWAYQGKPLYFFAKDAAKGDKKGDGFKDVWHAVKE

>WP_063597677.1 type 1 fimbrial protein [Burkholderia pseudomallei]
MKAKGMQYDLTAVLLSMLSGGAAYSQTATTGTINFTGSITDVPCEIDTAATSSNVTMAKVFANDFSGVGS
TTGTTAFKIVLKNCGASTSGATVRFMGTTDSANPAALQTTAGGAGGVALQLVDDTGTPISIGSSSKAYTI
AEGDNTFNFAARYIATSATVTGGAANATAVFALTYK

>WP_065793661.1 sulfatase-like hydrolase/transferase [Burkholderia pseudomallei]
MKRNSKQFEIGAWLGACALAFASAASAAAVQDRDHDSRPVDAKRVLLVSIDGLHEQDLARCIGANTCPNL
ALLAKSGVTYTNARTPGLSDSFPGLAALVTGGSPKSAGLFYDVSYDRTLYAPSDATCSGKQGWNVVFDET
TGIDAMNGGALTHLDGGGAFNPQAIPHARVNGQCVSVYPHDYVKTNTVFEVVKEHLRGSHTAWADKHAWG
YDWVNGPSGKGVDDLARTEINSIDPATGTPYTDIYTHTEKFDDYHVQAIVNQIDGKNSTGTAAAPVPTLF
GTNFQTLSVAQKATVASGGGYLDASFTPGPEVANAIAYVDGALGRIVAELRQRGLYDSTVVIVTAKHGQS
PTDHTKLVKHGDTLTALLEANGFVDPNGNFGQNNTASGNPNDGTGLVGTGFVQTDDVGLVWLRDPRQLSA
AVATLKANLGCNAPGICADGPQAYILYGPSVAERFGNPALGRTPDIVVQPNPGVIYTSSKKKDEEHGGNA
PDDSHLGLLVSYAGLRQGRTIDAPVLTTQVAPTILRSLGLEPRLLHAVALEGTRVLPGLGLER

>OMR39540.1 sorbosone dehydrogenase [Burkholderia pseudomallei]
MRSLVALFHHVSSGPPALPRAASAACLVLATIVTVATAPAASAALPIDELRVPPGFRVQVLADDVPTARE
MAWSPRGILYVGSMNGRVHALVVRDGHVREHHVIASGLEMPVGVAYRSGALFVSAVSRILRLDRIDERLA
APPKPVVVTNALPTDRHHGWKFIAFGPDGKLYVPTGAPCNICVADRDRYAMIGRMNADGSGYEVYARGVR
NTVGFAWHPATRELWFTDNGRDLMGDDRPDDKLNRAPRAGLDFGYPFCHGGDVLDPQFGRGHTCSSYAPP
VLKLGAHVAALGMRFYTGGMFPPEYRDNIFIAEHGSWNRSRKVGYRVVRVIASPDGRAAREETFVHGWLR
PDESVWGRPADVLPLPDGSLLVSDDYAGAIYRITYDATH

>OMW33686.1 hypothetical protein AQ807_07070, partial [Burkholderia pseudomallei]
NSNSNSNSNSNSNSNSNSNSNSNSNSNSNGNGNSSRHRYHPRLRRTPYERSSNYPLEVPQGGSMSTNMKRLM
IAALGAALAFGALSARAASFDCAHAANAAERAICGTPALGELDVRMAAYYEMLQNARPADEGMAYREFRD
ALRDEQQRWRQRTRDACGARIDCLTNAYTARIAALRGVAAERLVLRMTGGSAASAGAADATYAIEGESIT
LANGESVRPAAPGSAMKRVTTLVARSAVATIAGRPVEAVLLSDDPGGSGRFLYVATAQPGGGAPAVLLGD
RVKPVSVSIERAATGGAVVVVEYLDRPEGAPFAQAPTIKIVRRFALEQGRLVEQRG

>ONC96520.1 UDP-N-acetylmuramate:L-alanyl-gamma-D-glutamyl-meso-diaminopimelate ligase [Burkholderia pseudomallei]
MRGGASAALSMFSGNRRDVRPAADFNYAHAARSSMHIHILGICGTFMGGLAVLARAAGHTVTGCDAGVYP
PMSTQLEAQGIQLIEGYGAEQIDLKPDLFVIGNVVSRGNPLMEAILDRGLPYVSGPQWLGEHVLAGKWVL
AVAGTHGKTTTSSMLAWVLEDAGLNPGFLIGGVPLNFGVSARLTDSSFFVIEADEYDTAFFDKRSKFVHY
RPRTAILNNLEFDHADIFPDLAAIETQFHHLVRTVPGVGRLVTNGRDDALERVLSRGCWSDVERFGVDGG
WQALPVENGVPVDGRFAVYWRSERVGAVDWQVQGEHNRMNALAAIAAARHVGVPPAQAAAALAAFRNVKR
RMEVRGSVDGVTVYDDFAHHPTAIETTIAGLRTRIGRENTRILAVLEPRSNTMKLGVMKAQLPASLADAD
LVFGYGAPSGRDALGWSLPDALAPLGGKARAFDDLHALVKAVTASARPGDHVLVMSNGGFGGVHQKLLDA
LSARGDAAPARSGA

>WP_076802983.1 MULTISPECIES: chitinase [pseudomallei group]
MERTMNFSMLSRIVPRALAAGCLFAAAGASQAAGVYAPYVDVTLYPTPLVDQIGVQQGIQQFMLAFVVSG
GNQCTPSWGGVQPIGNGATGDLLDKIATSVTAYRAKGGDVAVSFGGAAGQPLMQACSSVAALKGAYQTVI
DTYSLTHVDFDIEGASQQDSAAVARNFQAVAQLQADYAAKGKPLHVTLTLPAMPTGLVQDGLNVLNAALA
NNVTLDAVNIMTMDYGPSGIDMGAAAISAAQGLYSQLDTAYKSAGKPQTDAQLKQLVGVTPMIGVNDVAG
EIFTLANAQSVQTTAANNNYGFVGIWSITRDKACDGSSQYASPICSGVAQQPYAFSSVFKQLGGHWGAGV
TQDPNYGGGSDGGGKPQPGAPWSATQVYTAGATVTYQGTTYQAQWWTQGDIPGQASVWKPVGGNVPAWSS
TTAYPGGACVTYQGAKYCAKWWTQGDVPSAGGPWTRA

>WP_076819717.1 NAD(P)H-dependent oxidoreductase, partial [Burkholderia pseudomallei]
STERALGFALRGAQAAGARTRLFDGPFLHTLPHYAPERRALTDAQRELIDAVRAADALVIATPGYHGGVS
GLVKNALDTLEELRADERPYLDARAVGCIVTAYGWQAAGTVLTSLRAIVHALRGWPTPFGATVNTLETRF
ETAGSCSDSKVAAQLETVGAQAAEFALAFASHRAATRVAGGETLAPVLEAAAG

>WP_076848120.1 cytochrome P450 [Burkholderia pseudomallei]
MSKHRQAPGPRGGLVMGNLAAYKRNPITMLLRLHQQYGDVARNRLGPFVTHALAHPDHIQYVLQENHRNY
VRGRFYDNFKMFFGDGLLTTDGEFWRRHRRVVQPLFHKKQVDAHTAAVGDAALALAHRWSALPPGKALDV
VEEMMHLSLRMLGLMVFNTDVSSHAEAVGPAVRFGIEAMMPQGNLNDFIPRWAPTRFNRRIAHARRAIDT
IVAKIIADHREARCEPSDVISLLLNARDPDTGAPMTQQEVHDEVMTVFLAGHETTGAGLAWALYALAQHP
AVLRQLRDELDARLGGRAPTVQDFEQLPYLSQVVDEVLRVYPPIWGFTRDLVEDDEIGGYRIPARSSVFM
SPYVTHRHPAFWRNPDAFDPENFASDAPARHRFVYFPFGGGMRKCIGFQTALLQMRVLVAVVAQHFDLNA
LPGHPIELGATISLRPVHGIRLIVKPRERQQSHLARVREHESARALRPLGDTATTAGAACPMTAAREPGA
SAAAAVAPAALAVPAASAASAASAARPAPLAQPAAPAIVDSQPQTRAQVYEAVKQMTALGRQLFFDPSLS
GSGKLACASCHSPQHAFGPPNALPAQFGGDDLRQQGFRAVPTLKYLQKVPAFSEHYHESDDEGDESVDAG
PTGGLTWDGRVDSGAEQARAPLTSPFEMNGTPEKVARAVRAAPYAPAFRAAFGARVLDDDRATFEAVLQA
LGTFEQAPDVFYPYTSKYDAYLAGRARLTRAELHGLQVFNDEKKGNCASCHVSRRGLDGSPPQFSDFGLI
ALGVPRNRALAVNRNPNFYDLGACGPERRDLKGRDEFCGLFRTPTLRNVALKKTFFHNGVYHSLDDVLRF
YAERDTHPEKFYPVKRGVVQKFDDLPKRYWKNLNDEPPFERKRGDPPAMTDAEIRDVIAFLGTLTDGYDP
RAKPAGGAR

>WP_076852667.1 depolymerase [Burkholderia pseudomallei]
MRHAAARAVLAAAFTSTLAAAPAGARAAPPLPALRADANRVSVSGLSSGAYMALQYQVAYSASVVGVGVI
AGGPYYCAAGSVANTDLCRGLVPNMVPDSGRLVAAAQGFAASGQIDPLANLQRARIYLFSGTKDTLVRQS
AVDATWSFFWLVGVPVTNIVYVADVPAGHAFVTPSAGNACDANAAPFISHCTVGQSGYDQAGALLDAIYG
PLAPPAATPTGRAIAFDQREFAPASSGLAAQGYAYVPRTCDANAGCKVHVVFHGCLQSAAVVRDMTTYDN
WADANGIVVLYPQVAKTSTPNDPQGCWDWFAYTGQNYAWKSGAQMRAVRAMIERVTSAP

>WP_076887541.1 peptidoglycan DD-metalloendopeptidase family protein [Burkholderia pseudomallei]
MVWRGASDSITRWLAAPLIAAGLTGAASAAPVDMPDLQHAVRQAFSARLGKQAPAAPAVRDDLIESVRSD
PEAGWVLGTVTQVVPNDTPAYPVTKLFIARRASEGWAVGIEGTDAFYTLAAASPAKLLAGDERAHLNAGR
APTAPPRKAVPAQTGLALPWQQGTAWYWTGGAHGWSGDSRPFNSLDFSGGNGQVLAARDGTLYKSCERNG
SAIVKVVHDNGYTSTYYHMVQLTQAGSGTRVRQGQYLGRVGNGLPCGGQTTGPHVHFALSQGGSDVPVNG
KTIGGWQFFEGSNAYSGYAVRNQRRVSVQASLTNYGADDSGGPTEPSPPVKATVQSPGPVNLRSAPSLSA
SIVGTVANGAAVQLACYAYGDTVQGNWGATRLWYRLDSNRWVSDGFVYTGSNDPVVSACAN

>WP_076889967.1 hypothetical protein [Burkholderia pseudomallei]
MRNADGTAPRAPLPSRIAASVLACILAAALAGNARAAPASPRTGGDPYAFAVLSGVIASADDEPQARRLL
DAIARDRSVSFIVYEGNLKGPKEACRDQLYEQRNVLFATSRVPLVLLPGQYDWAACGTRDAGAYDPVERL
DFLRQNVFTEPASPGANPLPLTRESEVARFHPYRENVRWIRDDTVFIGLNAPAPNNRYLTAGGRNGEFED
RAIANAFWIDHAAEYAKRRKARALVVFLEGDPQFNRYERSERFGWLRFNRPRVRDGFREFKRALVKAAAV
FRGSIVVVHPSGEPLADGFRIDRPLRDDKGELVGNLTRVAIAPHARLTHWVRIGVDAAKQPMFNVSMQTV
PKYLPQPPALPVVPHDDTPLPDMPEIPAQPVLPDSGASGAAPPGYGGWHGAHEGSFDHRAPSGATPGDAP
DIAVPGSMQGTR

>WP_076902853.1 hypothetical protein [Burkholderia pseudomallei]
MKKLLIATVIGALSATMLASAPSAFAQDSSATAKKATPKRPAPKRHLIPRSKKAQARAAAKTDPVPEGAV
KWACKDGLSYELAGDMKRDQIVTVHWANKNYKLPRQQTTTGADVFYDPASGMKLVVIPTKGMLFSDKDDS
RLADECQTPEMAAGNGLAPTQSNELKPSN

>WP_076903047.1 cytochrome c1 [Burkholderia pseudomallei]
MKKLLSTLAMFIVSACLLTSASVRAEEGEFPLDRAPDNTENLVSLQHGAQLFVNYCLNCHSANLMRYNRL
TDLGISQKEIEKNLLFATDKVGNTMTVAMRPEDAKTWLGVAPPDLSVEARARSRDWLYTYLRTFYRDDTR
PTGWNNAVFPNVGMPHVLWQLQGERIAKFEEKTDEETGEKTHKLVGFQQVTPGTLAPADYDAAVGDLVAY
LNWMSEPAQQTRKRLGVWVLLFLGVLTFLAWRLNAAYWKDIK

>WP_080248664.1 c-type cytochrome [Burkholderia pseudomallei]
MSVKPAQRPERGFAKTRRRTAAYRPAATRAAGVARAAGAATPPLPAAPHAAAAHAQAHAAQPAALPLAYV
FDSAGPAPRPVLILGWALLALCTSVCLVIAVLPALALFLGRPASVGLTERGGLGFVYVGSAISTALLLAA
LVYMLWVLAAVAKPPRPPAVTIAVTAYDWWWKADYGGGPPDGFTTANELHVPVGEPVLIELRSADVIHAF
WAPQLAGKTQAIPGQINRQWMQADRPGVYRGQCTQFCGAQHAQMGFEIVAEPPDAYRRWYASQRRGAEAP
RTADALRGQRIFADRCAGCHAVRGTGAAGTQAPDLTHVGARRLLAAGALANTPDELRRWIADAQQVKPQS
LMPSIRLDPAQQRDLSAYLATLR

>WP_080248669.1 TonB-dependent siderophore receptor [Burkholderia pseudomallei]

MEQRKMEWATSTRVRAIAAGVAFYAAAAGHAQAQAAQPGTDARQPGGEAKADTAAGGTLPAISVSGAAER
DASVGLVARRSMTGTKTDTPIIEIPQTINVVTAQQIEATGATDINQAFRYIPGFSSYGSDNRSDWYAALR
GFTPTVFVDGLQVPNTINLSSWRVDPYMIDSIAVLRGPTSVLYGQGDPGAIVDVQSKLANGERIRELGVQ
VGNYARKQLMFDIGDTIGKDGTLSYRIVGVGRDGNAQTGPLADQRVSFAPSLKWQPNADTSLTLAATYLQ
DWGDTSSNFLPSRGTVLPNPNGTISDDLYTADANFDHYRKKQWSLGYQFEHKLNPVWTLRQNVRWMHLSL
DDASVYGGGLDDADPTMATMTRYAGLFQFNYSRFDVDNQAQAKFTTGPLSHTLLFGFDYNRQTTTDSEWL
AKGPSLNLYRPVYTPIPSDIFSGPNAYPRTDTKTTLNAFGLYVQDQIKWRRWVLTLGGRQDWTRTSQDDI
ANAASFRQNDHAFSGRVGLTYLGDYGLAPYLSYSTSFNPQIGLKLAGGGLATPTKGRQIEAGLRWQPPGK
NLMLNAAVYQINQTNVAMSNPNDPTSSTFVQVGEVRSRGVELSAVGNLSRELSVIAAYVYQDVKNVRAND
NTLNKWPVDVPRPRQIASLWADWTWRTGPLTGFGVGAGVRYMSAAAGAADNSLTVPSYTLFDAALHYELR
NWRFALNATNLFNRRYVAGCQSDAVCMYGNQRTVIATAKYNW

>WP_080248725.1 patatin-like phospholipase domain-containing protein [Burkholderia pseudomallei]
MTVLASSARAGFRVPGFVAALLFVCAATGRPAPAAAADTATVASAATAAASAAGAPASTTLPPAATPTRV
ASSAVAASSPASSLASSSASAATPANAAAAWSWTAARRADAPDSSSVSNAVSNSGSNVAGTAAAHGPHAP
HTAPDAGAGAPSAPTTPTAATPAAAGASASMAAAPTPAATDTLVCMPDGGGPHRPAIGLVLSGGGARGYA
HLGVLKVLEANRIPVDCIAATSMGAVVGGLYATGMTAQDMQRRLSQVNLADIAFDVTERSDLPQKKREDE
RLYIDSLTIGFDSKGFKAPVGLVQGNRLQALLANWTAAVPTNQPFDRLPIPFRAIATDLQTGQKVVLDHG
SLPLAIRASMALPGLFSPAEIDGRALVDGGLVGNLPVDAARAMGADVVIAVDIGSPLRPLDALASPADVM
QQMIGILIRQNVAEQRKQLTANDILLQPDLGKQTFTDFQNANQAIAAGEAAAVAALPRLARYALSPEQYE
AYRAAHARPQQPIRITSIEIRTNGASVPKQIVRNALRVKPGDVYDPQAVSADLLSLTTSGNFENVTQQII
NEGDEHRLVIDAQEKYWGPNFLLFGLGMSSSSTDEGGFRLHLGYRRPWLTPSGLEFRADTTLGSDMQSVH
VELRQPLSNKTGYYVAPYADYKRRFTNLYDSDTDIKITQYRFQTSRVGLDLGLPLARLGDFRIGLAYTHL
TASPTYNVPLNWFLPDDAPSPGTLFPSAYGHQISARARLVIDQLDDPTFPRKGYFVEARVERSLSKSNDT
FFDSDASFTDVYGRLMVAQRFGRHSVNISVEAGKSFGGTNFGNPLGYTLGGFQHLSAYAADQLSGNALLY
GQVTYMNQLATFNASPIKALYVGASAEVGNVWTPDTRIGSGSLKQSYTFFTSLTTAFGPVYFGVALAPGG
RRNIYFQLGRTY

>WP_080248756.1 PhoPQ-regulated protein [Burkholderia pseudomallei]
MRNKVAGAIALSMLSLAYPRAEAAPIAFPCPIDARRDLTDTLACYRQTVENQPLNYTMTGIVQLPGIEQR
TYRLVSQSWSPDQLVRPDTWVHEVALYIPQDALPRRALVIANDGTRHPGEGETPRTPNDFLPDTLADIAR
STRTAVISVSDVPNQLLSYADNGKPKAEDDSVARSWTLFMQSPHTRGAMPLHVPMAASIWRAMSLAEREL
TSLGIHRFVVSGISKRAWTTWLALIGDQRVDAIAPFAIDLLSTRAALENMYRSYGGNWPLAFYPYYAEGI
DRTLDSPAFGLLMRIEDPLSYLGTRHGSRLAVPKYIVNASGDDFFVPDNSQRYFDKLPGAKALRMVPNSS
HSDIRRATLDSLVPFVKRIQHGKALPQVDAAPIETNGQTILRLRTSERPRQLLLWQATNPLARDFRYACG
IRYTSTPIAMNGARPQQLVLVPPTNGWRAYFVEATFDDGFVATSQTYILGDGYPSTPPPAVGGSCRTLPG
RDTHS

>WP_080300428.1 glycoside hydrolase 68 family protein [Burkholderia pseudomallei]
MTYSKQAVSRRHKRLALSAAALTAAACMSAHAQSDGAGPAPTPHTQQAYDPESHFTMRWTRADMRQLVKQ
SHTAGADKNSLPPALTMPDIAQNFPLVDSNVWVWDTWPLADMRANQLSYKGWEVIFSLSADPHAGYTFDD
RHVHARIGFFYRRAGIPASQRPANGGWTWGGHLFPDGASAKVFGTAPMTNNAEWSGSARLTHGENVSLYY
TALSFNRSAPGGADITPPIAIITRADGHIHADDKHVWFSGFDDHKALLQPEGKMYQTGQQNTYYSFRDPF
VFTDPAHPGNTYMVFEGNTGGPRGARTCTEADLGYAPNDPYREDLNAVMNLGAVYQKANVGLAIATNPQL
TEWKFLPPLLSANCVDDQTERPQIYLKDGKYYLFTISHRTTMAAGIDGPDGVYGFVGNGIRSDFLPLNGG
SGLVLGNPTDFSAPAGAPYAQDPNQNPRAFQSYSHYVMPGGRVESFIDAIGARRGGTLAPTVKIDIHGDS
TTVDRAYGAGGLGGYGDIPANLPAVGAGHHD

>WP_080310460.1 DUF3761 domain-containing protein [Burkholderia pseudomallei]
MHSLPLGRRYATEFLQRFAAVKMPSGRRARRQRRTMRETMPISLPRAARRIAVALAIAALPPAAAFAYSP
TAPTAPTAHGEADLDRHDTYRNRDGDTVHAPAHSKSGRAPEGATARCRDGAYSFSRHRRGTCSGHGGVAA
WL

>ARK97515.1 galactose oxidase [Burkholderia pseudomallei]
MTEWIRVRSPAFKRKTQFKRFQWEKARDRSPFRYMEINMKKREEKSLSVSHSRRAFLVWVPASLLVTACG
GDNGSSAPASIKKAVAALTGPGGATWTGMGKGGILIQSPYSGAGNFELLAGDEYGMLHCLRRTNDNGAQW
MVAVGDGMGSKNSVSGYCHTYSTNYTNIEAAVVDDGAVKTYFRSNSTPWSAHKTIPIGDAQGFPAYLQSN
RNGAQQLELVVGMRRGGMVHYWRDDKDGFTWKQSSIFGSGQVKGVSVMQGNYIGPDNNANLELVAWVDDR
LESWCCEGRAWRKICNIADGGVGGAPALLQSNYGTKGNFEVLVPLATGGIAHYWRNNDAEGIPWSFTGTI
GTGAYVAAGLLQSQNGVKGDFEAIGLRASGKADVFRRGDTLVWENSATVSPFSEGAAADVGVSSHIVDVN
VTGINSVLLKNGSVLMFGYYKGGSTNKTIPACIWNPVNDQITAIPSFRNNFCAGQTAMPDGKVLIVGGHI
GDTLKDVVVFDPDNHTATLVATMTKGRWYPSTATLPDGQVFIISGTETAGWNTSVNDTWQTYANNALTAP
EDVISPFSPYYPKSQTQIDLYPFLFVLPDGKLLVHARNTTRFFDIGTRSWSATLYKTVSDNSRTYPFMGG
TAVLPLRPSENYRVKVVVAGGADKSAQIAIGDDSQYDNSVPGMTSCEMLDLGDAAPAWKAIAPLNEGRVM
CDLVTLPDGKLFIVGGNKTGKADYGRGPTYRPELYDPQTNTWTLLASTRIARGYHATALLLPDGRIAITG
KDGDYQGSGLQYAETRVEIFSPPYLFKGPRPAIQSAPASINHGGSFTLGLSSGTSPEDIGSIVIVACGSA
THQINFSHRIVELVFAVSGGTLTVNAPPNANIAPPGYYMMFVLSKLGVPSVSSIVHVAAGSAQAASATPE
ALTRQPRVATVSIEQAGDAPRCAPLTVAEKQSINGMG

>ARL94721.1 DNA protecting protein DprA [Burkholderia pseudomallei]
MIGQSPWARACGIRSISDDENFITALRSPQAALFEEFIMSPRALTRAELSGWLRLAGASGVPAAACAALL
GAFASLEALFAASHAELAALVGDAAAQSVLAPPAADFEQRVDAALAWLDEPGNALVTRHDPAYPGPLAEL
YDPPPLLYIKGRVALLHARAVAIVGSRGATPQGLADATRFARELSDAGLAIVSGLARGIDGAAHRGGLDG
ASGTVAVIGTGADLVYPACHHALAHEIAERGALVSEWPLGTPARSAHFPQRNRLIAALSGGVLIVEAALR
SGSLITARLANEMGRDVFAIPGSIHAPLSQGCHALIRDGAKLVETAADVLDEFGLDPARPVNSKRGAPAA
ASADADLDDDTRRLLDAIGYGPVPLELLAQRTSLPSGTLHRLLLQLELAGRVAALPGGRYTRIDGAQAEP
AQARGTSAVLHSGA

>WP_085538790.1 shikimate dehydrogenase [Burkholderia pseudomallei]
MSERSFLIGLIGAGIGGSLSPAMHEAEGCGQGLNYVYRRIDLDALGLTADALPELLAAAERMGFDGLNIT
HPCKQRVIEHLDELSADAAALGAVNTVCFAHGRRIGHNTDWSGFAKAFARGLPGASLERVVQLGAGGAGA
AVAHAALTMGAAQLALFDVDPVRARSLADQLQARFPHARLSAGGTLADALANATGLIHATPTGMLGHPGL
PLPAHLLRSGLWVADIVYFPLETELIRAARALGCRTLPGGGMAVYQAVDAFELFTGRAPDAERMFEHFRA
LVAR

>WP_085546935.1 Flp pilus assembly protein CpaB [Burkholderia pseudomallei]
MKNNRAFTMLAIAILAGLAAVAFASRWLVQTSSSAVTPVAVAATDVSLGEPLGPNQLRIVNWPTASVPPG
AFTDLKPLEGRVVRTSLARGEPVLGTKLAPVGTKGGLSAVIAPGHRAITVRVNDVVGVAGFALPGNYVDV
IVNTQEQSKADPQQSISKIVLEKILVLAVAQQVSRDDTAPKVVNAVTLEVSPEQAEKLDLARSVGTLSLV
LRNQIDKDALSTGGATKNTLLGQPAAAPAPIAAPAAQPRPVRTRVVAHARAAAGRDCVGVLSGVTGSVEC
F

>VBI96880.1 adenylylsulfate kinase [Burkholderia pseudomallei]
MTSMSLLAAVYSGRREVSIDGCHEVQNIAVAAALDGGQAAPSAHGAVQFTDRKLMDKLHAERNVEWQTTS
VSRRDRESRQAHGALAVWLTGLPGAGKSTLAIEAERCLHQAGKRTYVLDGDNLRHGLCRDLGFSADDRAE
NIRRAGEVARLFVDAGVIVLSAFISPFRFDRAKVRALFDAGDFIEVYCDCDPRICEIRDVKGLYRRARAG
LISEFTGISSPYEPPRHPDVYVNTGKESLDACVDRIVRRVLHSLR

>VCQ54129.1 phosphoserine phosphatase [Burkholderia pseudomallei]
MESLSGFVVFVQVAETRSFVAAARALGVSASAVGKRIARLEARLNVRLFHRSTRSIALTAEGARFLERCR
RVLAEIEEAEQELSRSAHAPSGRLRVSLPALSAPVLPVLADFMAAYPDIQLDLDFTDRLVDVIDEGFDAV
VRGGEPRDSRLSARRLGTFAQVVVGSPDYFARRGTPRTPADLAHHTCLLYRFPTTGKLERWPLRPAPGDA
DVEPPQSMICNNVETRVCFAIRGRGLACVPDFAVRDELASARLRTALDAYIERPQTFHVLWPSGRHASPK
LRAFVDFIVARSTWPPASRGRGAPMRTGAPPGMRPRRGAPRAATAHRTSSHLIASPRDTPRRCPCRRGPA
APPAARESVVKSAFESVRRGKTARAARVGRISTSGCPFSTGAPFYGQRCWAAPASAGGSAGGTRPARPRR
ANPPRRNSMKTGRRHFVRSVASASAALAAAAWSPARAAIDAPASPATALSLTPGRWSPNNVARLRAVLAG
HGASSPRYRPEHRPYAVFDWDNTSIMNDCEEALLMHQIDGLHYRLTPEQFSAILRQGVPDGPFDAKLGYT
SVDGKPVRMEDIAADVDADYRWLHANYRGLAGDKPLDEIHRSEQFRDFRAKLYFMYDAICDTYPVEIGYK
WIMYWYAGMTRDELQAMAFDSNVANLGDALRKVTYESSRALPGKAGVIAATHFHGIRIHEEIRAVMDTLR
SNGIDVYVSTASLDDVVRVFAGHPAFGYGVPAENVIGMRLTMADGKYMNEYLPNWHFNYGPGKTVGIRRE
LEAKKGYGPLLVFGDSDGDAWMLRDFADTAVGVIVNRMKKGEIGIDSRKAAEQIGAKDARLVLQGRDENT
GLMVADERSIKYGKRDPKLLA

>VBC72697.1 Uncharacterised protein [Burkholderia pseudomallei]
MKHDAAHHDERAISMNHSAITWAAVLALCAGFAVTNAHARDSAECRRLSDATGGADDNFRPPLTATVTGA
GRAYFHSAPASACIARRVFVVPGDTVTVYKPYRHWYQVTYVNGRTGEDVEGWIEAGRLRLGGHLGGGQ

>VCG67913.1 3-demethylubiquinone-9 3-methyltransferase [Burkholderia pseudomallei]
MRPASPAAAFLATSPACLYTRGLAGAPPPSFPHPLADMTNADPHELQKFSDLAHKWWDPNAEFKPLHDLN
PVRLSWIDAHAHLPGKRVVDIGCGGGILSESMASLGAQVKGIDLATEALGVADLHSLESGVSVDYEAIAA
EALAAREPGAYDVVTCMEMLEHVPSPANIVAACATLVKPGGWVFFSTLNRNLKSYLLAVIGAEYIAQMLP
KGTHDYARFIRPSELARFVREAGLQMVEIKGIAYHPLAKRFALSNDTDVNYLVACRRGA

>WP_122634654.1 mechanosensitive ion channel family protein [Burkholderia pseudomallei]
MIRLATFARRVALIGLLHLLCAALPAAAADLASGASVPPAPAISRDEALAELKRVQAALDRIKQQASAAT
TYKQLDALDESTQALSADVDKLTAALVPTRAQLQAQLDVLGPPPAPGAAPETPAVARQRADLNARKTQLD
AALKQAADEKESLANLTQQYSKLRRSLLRDQLVLRSSSILGPRFWAPLFRPSDDDRQELGAFSAQIGQVL
AAAWEPGRRLGTALLLAAALAVWTLGRRLAERALAWFSLTKLPETRVRRSALATSIALSSVVTTGIAVQF
VYLALTRDYALSSALQDFADELAKLALTCALIAGLGRALLCTHHPSWRLPALADEVALAMRPFPAILAAL
LLVAGALEELNRTADTSLQVTLFGRGIVSLVVVVTIGASLLRANRARTALAAAGEPPEARSTLAGLIHAG
VTLTVVAALAALLIGYISVARFLTYELVWFEIVIGSVYLLTRLTRDVYASAFAANLSSGRLIKHLFGLND
SHLDQARTVLTGISTSVLLLVAVIALLTGGFGTTPNDLLVSVISMFGGDKLQRLNIVPARILNAALALAI
GIYLLRSVRRWLDNELLPAIGLDQGMRASLVTLFSNLGYVALVLLALSLLGVKWDNLAWIVSALSVGIGF
GLQEIVKNFVSGLILLTERPVKVGDMISISGVEGDIRRINVRATEIQLSDRSTVIVPNSQLISQNLRNVT
MGNSTQGVATLMLTFPLNTDPEQVRDLLLDAYREHASVLEKPAPSVTFSQLTPDGITLSVTGYVGSPRIA
AATKSDLLFEILKRLRAAGIALSNPQTLMVQNLPPPSSE

>WP_122651768.1 metallophosphoesterase family protein [Burkholderia pseudomallei]
MSNKPAPDAPPAEQPCAVSRRGFLKFVGVSGLASAARGLAATRAAAAAPDGTPEQIHLTWGDVDANEVVV
SWASLAAATNPRVRFAGPNEAWRTVHGVQRTYTDGLNGEVVFTYHARLRGLKPGAVYRYEVTADNDANAA
QPFAARFETAPRGRAPFRWTSYGDLATPNTGWVLSSPQSRFAVQAVERFQPLFHLLNGDLCYANLNPAHQ
PAVWRDFGNNNQTSAANRPWMPCPGNHEIEFHNGAQGLDSYLARYTLPENGTRFAGRWYSFRVGAVLFVS
LDADDVVYQDAAAFVAGPNPLVPAASTGNEAIAPGTSLYVRGYSRGEQTRWLEQTLRRASRDRDIDWIVV
QMHQDALSSSKTGNGSDKGIREAWLPLFDRYGVDLVLCGHDHDYERSFPVRGCNHRAGVDAATGEVVDTL
QPRPVVTTDPADGRFDTSHGTIHMILGGGGTSAPLDVYGENPATGLPQARVFTKPNRPVPGAAPNTFVRH
GADAVEDAIWSARRDTGTGYGIAVFDYEPGEHGGRSTITVNYYHAPGADQHPTAEYELFETIVLSKPRRA

>WP_122821554.1 hypothetical protein [Burkholderia pseudomallei]
MPTLAKSLLLTLMACFALSGEAGAAQKTFCYFHHNCPADGKLPKGKPPEQYTKRQCKNIGGKSWGDGPTS
CENVNK

>WP_122826861.1 lytic transglycosylase domain-containing protein [Burkholderia pseudomallei]
MEKRFVIVAMFAAGAWSASAPARADCYDEAAKYQKVNPLVLRAIAWQESHNRPEALNKNANGSTDYGLMQ
INSIHLPTLSRYGITKDTLMEPCKSVYIAAWHLRRKMDKYGNTWQAIGAYHSETPSLRDKYARQIADILA
RWKLLPASAASATQDTQSAQR

>WP_122827599.1 cytochrome-c oxidase [Burkholderia pseudomallei]
MRRLRRRWLFAASGAALAALAGVDASRVRAAAPRVIKVHARRFVFTPDRIALAPHESVVFELTAQDTVMG
FSIPQYGVRADVPPGAVVRLAAQAGGPGTVQFLCDIFCGSGHETMNGVLVVG

>WP_122830792.1 peroxiredoxin [Burkholderia pseudomallei]
MNRKLASGAAAALFALHAVAAHAVLKVGDAAPDFSAQASLGGKTYTYSLADALKKGPVVLYFYPAAFTTG
CTIEAHAFAEAIDAYKRYGATVIGVSADDIGTLTKFSVSECRSKFPVAADPDAKIIREYDAKLPAISKAN
RVSYVISPDGKVLYEYTSLSPDKHVENTLNAVKAWAASHRQP

>WP_122831074.1 NAD(P)/FAD-dependent oxidoreductase [Burkholderia pseudomallei]
MEKFDIAVIGAGAAGMMTAAAAGQLGRRVVLLDHSPRLAEKIRISGGGRCNFTNLYAGPDNYLSANPHFC
RSALARYTPRDFLGLLKNHRVTWHEKHKGQLFCDHSSDAIIDVLKSECDAGGVAWRRPVTVDAVRHAQAD
GFALDTSAGAIGAQALVVATGGLSIPKIGATDFAYRIAKQFGHKLVDTRPALVPLTFSADAWAPFAALSG
VSLEARVSTGAKRGGGEFVEDILLTHRGLSGPGILQISSYWRSGEPIHVDLAPQIDAANELIRAKAGSRR
QIGTLLAEWVPARLAHAWLEAERVHADARLADLPDKTLRRIGETLSRWTLTPSGTEGYRKAEVTKGGVDT
RELSSATMMSERVPGLFFVGEALDVTGWLGGYNFQWAWASGVAAGRAAADGTRRLTG

>WP_139900217.1 c-type cytochrome [Burkholderia pseudomallei]
MKRKSLFALSAVAIVAASALVPVLWPGNDTLHGNAAVAATPADQAALIKKGEYLARVGDCIACHTVRGGK
PFAGGLPMATPFGTMYTPNITPDDQAGIGKWTSDDFYRAMHTGRSKDGSLLYPGFPFASYTKVTRADSDA
IYAYLRSVAPVSTPSRPHELRFPFNNRNLLIGWRTLFFKEGEYKPDPTKSVEWNRGAYLVEGLGHCSMCH
TSINMMGGPVSSAAFAGGLIPLQNWYAPSLTNDKELGLGDWHVQELSDLLQAGVSHKGAVFGPMADVVHN
SLQYMTDEDTRAMSTYLKSIPQKAEAPKNMQYEPSKQFGTALLEQGKKIYADNCATCHGPQGEGKPTAYP
PLAQNRSIMMESAVNPIRMVLNGGYPPSTFKNPRPYGMPPFAQSLSNQEVAAVVTYIRMSWGNNGSPVSP
QQVSDLRSAPLD

>WP_139910463.1 hypothetical protein [Burkholderia pseudomallei]
MNNLIEVVSIAAAAPAASADSQALDAALDCSSTGHAFVAPLVASGAVRSQPMHVEANSMNAFRTNRSLSA
YGFSVYVVLGYQANDPLFAHGDGEPIGDWAYGVVVRGSKQAVEAKVRAAGSQAVVKDAFPFLTAIVCTSP

>WP_139910601.1 phospholipase D family protein [Burkholderia pseudomallei]
MSKPPRRALVLCAALALSGCATHPPATTLERTVSHALPPDASTPLADALAVQARAHPGESGFVVLPRGDE
ALQMRIAVARAATKTLDIQYYIAAEDTTGKLLLGAALYAADRGVRVRMLVDALNFKDIDKLMAALDAHAN
LEVRVFNPFGAPRLGMFARTANVFTRIDNFTRRMHNKAMISDNQIAIVGGRNLGDEYFNASPTLQFRDLD
VLAAGPVTRAVSASFDAYWSSALTYPLPALNRRRYDAKDLDAARDALRAHWRANATPYNAKPLNATPLAA
QIARNELGLVWASAEFTADSPEKIAAPDDSYKSPPMQRLFALTRDAQREFLVLSPYFVPHDAGVNALGRL
TARGVRVAILTNSLAATDAIAVQAGYAPYRVPMLERGVELYEYKPDPGRSRIGMLGSRSRASLHAKAYVI
DRKILVIGSMNLDPRSAHLNTELALVIHSPRLANEIANLFDEVTKPTISYRVTLAPDTPGAAQTTGAGAP
AWPLVWTEIADGQVRTYSVDPNAGFYRNLLTGLCLLLPIDDQL

>WP_151269450.1 molecular chaperone [Burkholderia pseudomallei]
MPVRARARRAAAALFIALAGAAHAAIVPDRTRVILNEGEQAAIVTLSNKSDAYPYLVQSWLEDERGNRIT
TPLMVVPPLQRVEPKERNVLRIARMPGATLPADRESVFYLNIREVPPKTDTPNALQIALHTQMKLFYRPA
GVQPARDEDPTLPMTLRVDAAARKLVFDNPTPYHVTVVALAAGAHRAALPLDPVMVNPKSAASVPFSASA
AGGLFVTHVDDYGGQVTVEYTCDGLACRSVKR

>WP_151271354.1 DUF1223 domain-containing protein [Burkholderia pseudomallei]
MSTDMSISFSVSLRAVAAAFSLAAAGAAQAAADGACMARSPDGRQALVELYTSEGCSSCPPADDWLARLA
ARRAALRVVPLALHVDYWDGLGWADRFAQHRFTERQHALAARGGGRFVYTPEVAVDGRELRDWRDADAFG
RRVVATAAEPARVGIALAAKRRAGALDVALSVTPRVGAPRALDAYLALYENGVESQVRAGENRGATLRHE
RVVRQWIGPLAATADAHASLDVRRALPLPANLRAADAARYGVAAFVEDPATGDVLQALDLPLCG

>WP_151277731.1 cytochrome c [Burkholderia pseudomallei]
MNSAKHVVVAAALAALAAGAQAAGVVGNPKDGASKAAMCIGCHGIDGYRVAYPEVYRVPLLGGQNQVYLE
NALKAYRKKDRHFPSMNAIAESLTDQDIADLASYYAAQKPDSKNNPYK

>WP_155027764.1 ABC transporter substrate-binding protein [Burkholderia pseudomallei]
MKPLSLNMKRALASACIGAAGFLAQQAALAQSCGLANGKPATGAPIPIGAVVGKTGPDDFSSSARAAAAY
FKCVNANGGINGRPVQYLVEDDQWNPETASQVASKLVRDRKVLALAGNASFVECGANAKFYEQENVIAIA
GVGVPRECYFARNYAPLNMGPRLSMTEAALYAKQQYKATRMVCIAPNIPSLGAWSCEGPALWGKRNGVSV
DTIVMDPDSADPTSVVLQAASKNPQAILLGLPKGLMVPILSAAEQQNLGRRIHFVSAASGYDLGVPKAIG
PYWKGNFDVNLEFQPLDAQTPDNRNWLAVMDKYGDRKDPRDTFSQAGYLAARLVTDTLLKLPANQLDRAH
VTAALREVKDFRSDILCGPFYVGAGERHNANNAGRMAQSTGTGWKTVSTCQAVDDPQLADIRAAEKKMH

>WP_161788821.1 ABC transporter substrate-binding protein [Burkholderia pseudomallei]
MSIHAESDWASEPIDSFEAARQSKIVTFSRARVDMNVRMRCLSLLAAVAFDAHAQSSVVRIGVAMPLTGP
VAHLGKDVQNGAQLAVDELNRAPPTIDGKPVKFALVVEDDQGDPRQAVQVAQRLVDARVAGVVGDLNSGP
TIVAAKVYAAAGIAQIAPAATHPAYTQLGYKTAFRLMATDNQQGASLAGLAAKLAKGRPIALIDDRGAYG
QGLIDQTEKTLRASGVTRIIRDYTTDTAVNFASILTRVKGAHAAVIVYGGADAQAGPMVRQMKALGIDAA
FVGSDGVCTGQWTALSSGANEGQFCTQAGDPRARMAGYAAFERRFEARYGKVIVFAPYGYDAVMLLADAM
RRANSTEPAALLGALATTRYDGVIGRIRFSPQGDNLNGAVTVYRVQRGALVPVSD

```
>WP_162492307.1 hypothetical protein [Burkholderia pseudomallei]
MSSRVVACRRVSSRVVACRRVSSRVVACRRVSSRVVACRRVSSRSSRSSRSSRSSRVVAGRRGFAFGARA
ARRRRTRSSGAPTVEAIVFQKDSMKKILAVIAFLAVVGWLAATTTVLHAPSAQPCTDAWFDAIDKQFDIT
DNAGHGPDPGSGEWLGVVERKAKLPESGQLTEQQRCEAIQRELSQRTYLVNRRLGLKLAL
```

S4: gene ontology analysis (see excel file)

S5: virulence prediction of BpsDsbA substrates (see excel file)

S6: DsbA substrates epitope predictions (see excel file)
